# RLP/K enrichment sequencing; a novel method to identify receptor-like protein (*RLP*) and receptor-like kinase (*RLK*) genes

**DOI:** 10.1101/2020.02.26.966085

**Authors:** Xiao Lin, Miles Armstrong, Katie Baker, Doret Wouters, Richard G.F. Visser, Pieter J. Wolters, Ingo Hein, Vivianne G.A.A. Vleeshouwers

**Affiliations:** Plant Breeding, Wageningen University and Research, Droevendaalsesteeg 1, 6708 PB, Wageningen, The Netherlands; Cell and Molecular Sciences, The James Hutton Institute, DD2 5DA, Dundee, UK; School of Life Sciences, Division of Plant Sciences, University of Dundee at the James Hutton Institute, Dundee DD2 5DA, UK; The Sainsbury Laboratory, University of East Anglia, Norwich Research Park, NR4 7UH, Norwich UK

**Keywords:** genotyping by sequencing (GBS), pattern recognition receptor (PRR), potato, *Phytophthora infestans*, RLP/K enrichment sequencing (RLP/KSeq), RenSeq, receptor-like protein (RLP), receptor-like kinase (RLK), INF1, SCR74

## Abstract

- The identification of immune receptors in crop plants is time-consuming but important for disease control. Previously, resistance gene enrichment sequencing (RenSeq) was developed to accelerate mapping of nucleotide-binding domain and leucine-rich repeat containing (*NLR*) genes. However, resistances mediated by pattern recognition receptors (PRRs) remain less utilised.
- Here, our pipeline shows accelerated mapping of *PRRs.* Effectoromics leads to precise identification of plants with target PRRs, and subsequent RLP/K enrichment sequencing (RLP/KSeq) leads to detection of informative SNPs that are linked to the trait.
- Using *Phytophthora infestans* as a model, we identified *Solanum microdontum* plants that recognize the apoplastic effectors INF1 or SCR74. RLP/KSeq in a segregating *Solanum* population confirmed the localization of the INF1 receptor on chromosome 12, and lead to the rapid mapping of the response to SCR74 to chromosome 9. By using markers obtained from RLP/KSeq in conjunction with additional markers, we fine-mapped the SCR74 receptor to a 43-kbp *G-LecRK* locus.
- Our findings show that RLP/KSeq enables rapid mapping of *PRRs* and is especially beneficial for crop plants with large and complex genomes. This work will enable the elucidation and characterisation of the non-NLR plant immune receptors and ultimately facilitate informed resistance breeding.

## Introduction

To protect themselves against pathogens, plants have evolved two layers of defence (Jones & Dangl, 2006). The first layer of defence is formed by extracellular receptors on the plant cell surface that are often referred to as pattern recognition receptors (PRRs). These surface receptors typically represent receptor-like proteins (RLPs) and receptor-like kinases (RLKs), which can recognize apoplastic effectors, microbe-associated molecular patterns (MAMPs) from plant pathogens and danger-associated plant breakdown products (DAMPs). The second layer of defence is mounted upon recognition of cytoplasmic effectors by internal immune receptors that typically encode for resistance (*R*) genes of the nucleotide-binding domain and leucine-rich repeat (NLR) class. Stacking and pyramiding *R* genes and *PRRs* is believed to contribute to more durable plant disease resistance (Dangl *et al.*, 2013).

Potato is an important food crop. However, the global yield of potato is threatened by potato late blight, which is caused by the oomycete pathogen *Phytophthora infestans* that led to the great Irish famine in the mid-1840s (Haverkort *et al.*, 2008). Traditionally, breeding for late blight resistance in potato has relied on introducing *R* genes from wild *Solanum* species into potato cultivars (Vleeshouwers *et al.*, 2011a; Jo *et al.*, 2014). However, these NLRs are often quickly defeated by fast-evolving *P. infestans* isolates in the field (Wastie, 1991; Fry, 2008). Another, currently largely unexploited layer of immunity occurs at the surface of plant cells. This apoplastic immunity is believed to generally provide a broader spectrum of resistance and is based on RLP/RLK-mediated recognition of MAMPs or apoplastic effectors. Some MAMPs, like Nep1-like proteins (NLP), are conserved among different pathogen kingdoms (Gijzen & Nürnberger, 2006; Oome et al., 2014). Other examples of well characterized MAMPs are flagellin and elicitins, from bacteria and oomycetes, respectively (Felix *et al.*, 1999; Derevnina *et al.*, 2016). INF1 is a well-studied elicitin from *Phytophthora infestans* that triggers defence responses upon recognition by ELR, an RLP from *Solanum microdontum* residing on chromosome 12 (Du *et al.*, 2015). Other types of apoplastic effectors are extremely diverse and include small cysteine-rich proteins such as SCR74 from *P. infestans* (Liu *et al.*, 2005). Cloning and characterising plant surface immune receptors, including the receptor of SCR74, will further our understanding of plant immunity and help to engineer crops with more durable disease resistance.

Recent advances in sequencing technologies have facilitated whole genome sequencing and enabled genotyping by sequencing (GBS). This development has led to the emergence of several novel approaches for map-based cloning, such as genomic re-sequencing (Zou *et al.*, 2012; Zhu *et al.*, 2017), bulked segregant RNA-seq (Ramirez-Gonzalez *et al.*, 2015), Indel-seq (Singh *et al.*, 2017), and QTL-seq (Takagi *et al.*, 2013). In addition, when targeting certain types of gene families (e.g. *NLRs*), target enrichment sequencing significantly reduces the complexity of the genome prior to sequencing (Hodges *et al.*, 2007; Jupe *et al.*, 2013). *R* gene enrichment sequencing (RenSeq) aided the re-annotation and mapping of *NLR* genes in potato. All *NLR* genes from the potato reference genome DM1-3, v4.03 (doubled monoploid *S. tuberosum* group phureja clone) were predicted and an RNA bait library was generated to represent these NLRs (Jupe *et al.*, 2013). This work led to the accelerated genetic mapping of late blight *R* genes *Rpi-ber2*, *Rpi-rzc1, Rpi-ver1* from *S. berthaultii, S. ruiz-ceballosii* and *S. verrucosum*, respectively (Jupe *et al.*, 2013; Chen *et al.*, 2018). When combined with single-molecule real-time (SMRT) PacBio sequencing of larger DNA fragments, RenSeq generates a true sequence representation of full-length *NLR* genes which enabled the rapid cloning of *Rpi-amr3* from *S. americanum* (Witek *et al.*, 2016). RenSeq has also been successfully applied to other crops and has led to the cloning of two stem rust resistance genes *Sr22* and *Sr45* from hexaploid bread wheat (Steuernagel *et al.*, 2016). Furthermore, used as a diagnostic tool and referred to as dRenSeq, the methodology enables the identification of known functional NLRs in potatoes (Van Weymers *et al.*, 2016; Jiang *et al.*, 2018; Armstrong *et al.*, 2019). These successful advances in enrichment sequencing indicate that, with adaption and optimization, the sequence capture technology can be applied to other types of immune receptors, such as RLPs and RLKs. Consistent with other genome reduction technologies such as RenSeq, GenSeq and PenSeq (Jupe *et al.*, 2013, Strachan *et al.*, 2019, Thielliez *et al.*, 2019), we refer to this adaptation as RLP/KSeq.

In this study, we established a pipeline to accelerate the identification of surface receptors that perceive apoplastic effectors, by using the potato – *P. infestans* pathosystem as a proof of concept. We developed a pipeline (Fig. 1) that consists of two steps: 1) effectoromics, i.e. screening wild *Solanum* species to identify plants that recognize the apoplastic effectors INF1 and SCR74 and 2) RLP/KSeq to accelerate the genetic mapping of the underlying immune receptors through bulked segregant analysis (BSA). Ultimately, we fine-mapped the SCR74 receptor to a 43-kbp *G-LecRK* locus.

**Fig. 1.**
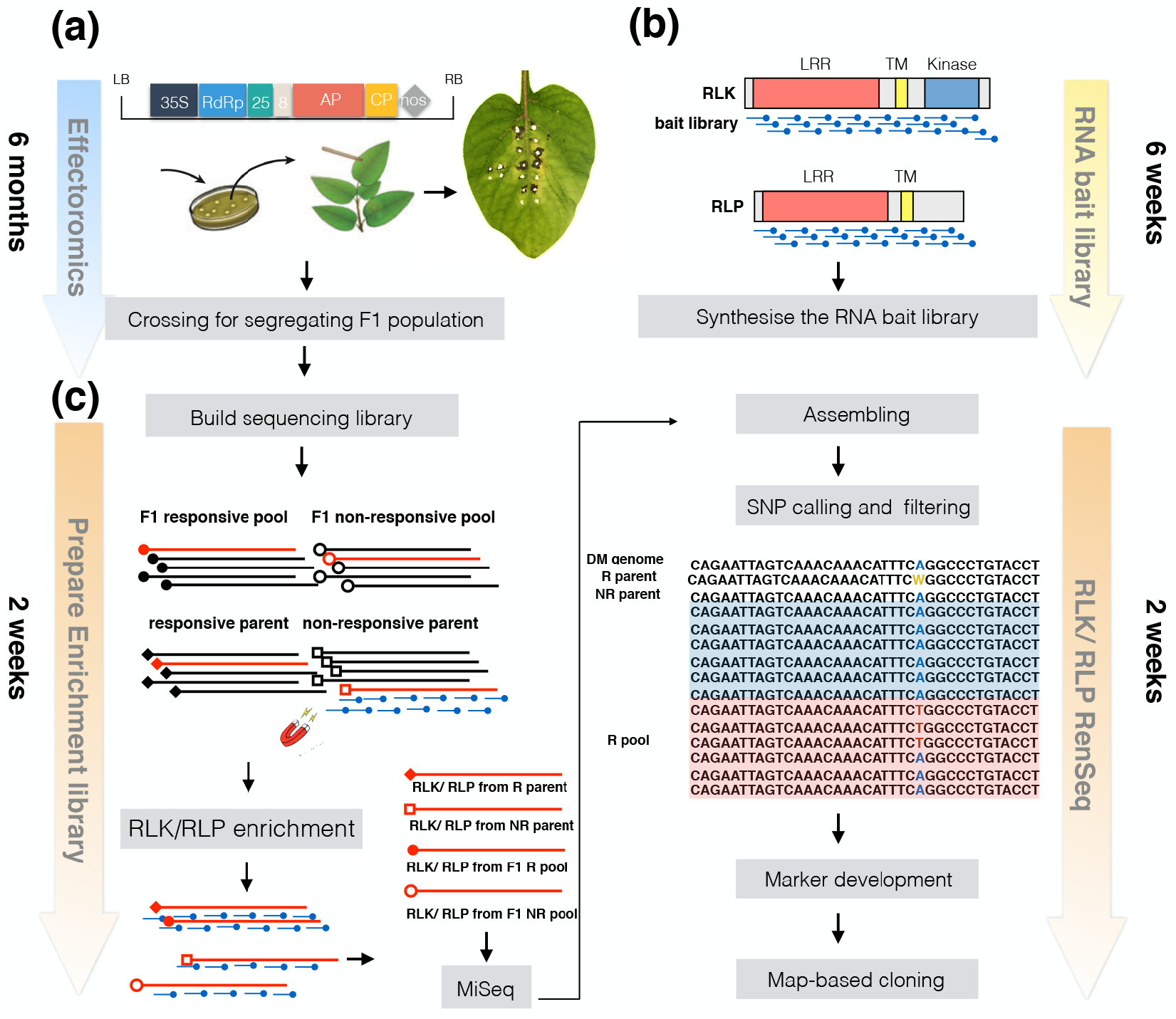
Overview of the effectoromics and RLP/KSeq pipeline for the fast identification and mapping of surface immune receptors. (a) Predicted *P. infestans* apoplastic effectors are cloned into the binary potato virus X (PVX) vector pGR106 and transformed into *Agrobacterium tumefaciens* for functional screening by PVX agroinfection. Agroinfected leaves are scored at 10-14 dpi for occurrence of cell death phenotypes. Responsive and non-responsive genotypes are crossed to create segregating F1 populations. (b) Prediction of the *RLP* and *RLK* genes from the reference genome enables the design and synthesis of the *RLP/RLK* bait library for bespoke target enrichment sequencing in the selected plant species. (c) An F1 population is screened for segregation of the recognition phenotype and pooled, based on their response pattern. Responsive and non-responsive pools as well as the respective parents are subjected to enrichment sequencing. RLP/KSeq derived reads are mapped to the reference genome, and SNPs linked to the recognition phenotype identified. Candidate markers are tested on the segregating population by SNP genotyping technologies such as LightScanner.

## Materials and methods

### Plant material

*Solanum* genotypes used in this study are listed in Fig. 2 and Table S1 (Vleeshouwers *et al.*, 2011b). These plants were maintained *in vitro* on MS20 medium at 25°C, as described by Du *et al.*, (2014). Top shoots of plants were cut and clonally propagated *in vitro* 2 weeks before transfer to soil in a climate-controlled greenhouse compartment at 22°C/18°C day/night regime under long day conditions. The F1 population 7026 was generated by crossing *Solanum microdontum* subsp. *gigantophyllum* (GIG362-6) with *Solanum microdontum* (MCD360-1). The plants were grown in a crossing greenhouse until flowering. Flowers from GIG362-6 were emasculated before they were fully opened and pollinated with pollen that was collected from MCD360-1. After 4-5 weeks, the ripe berries were removed from the plants. The seeds were collected and cleaned by water and dried on filter paper. Seeds were sown on MS20 medium or were soaked on filter paper after 3-4 months dormancy. Gibberellic acid (GA3) was used for breaking dormancy if necessary.

**Fig. 2.**
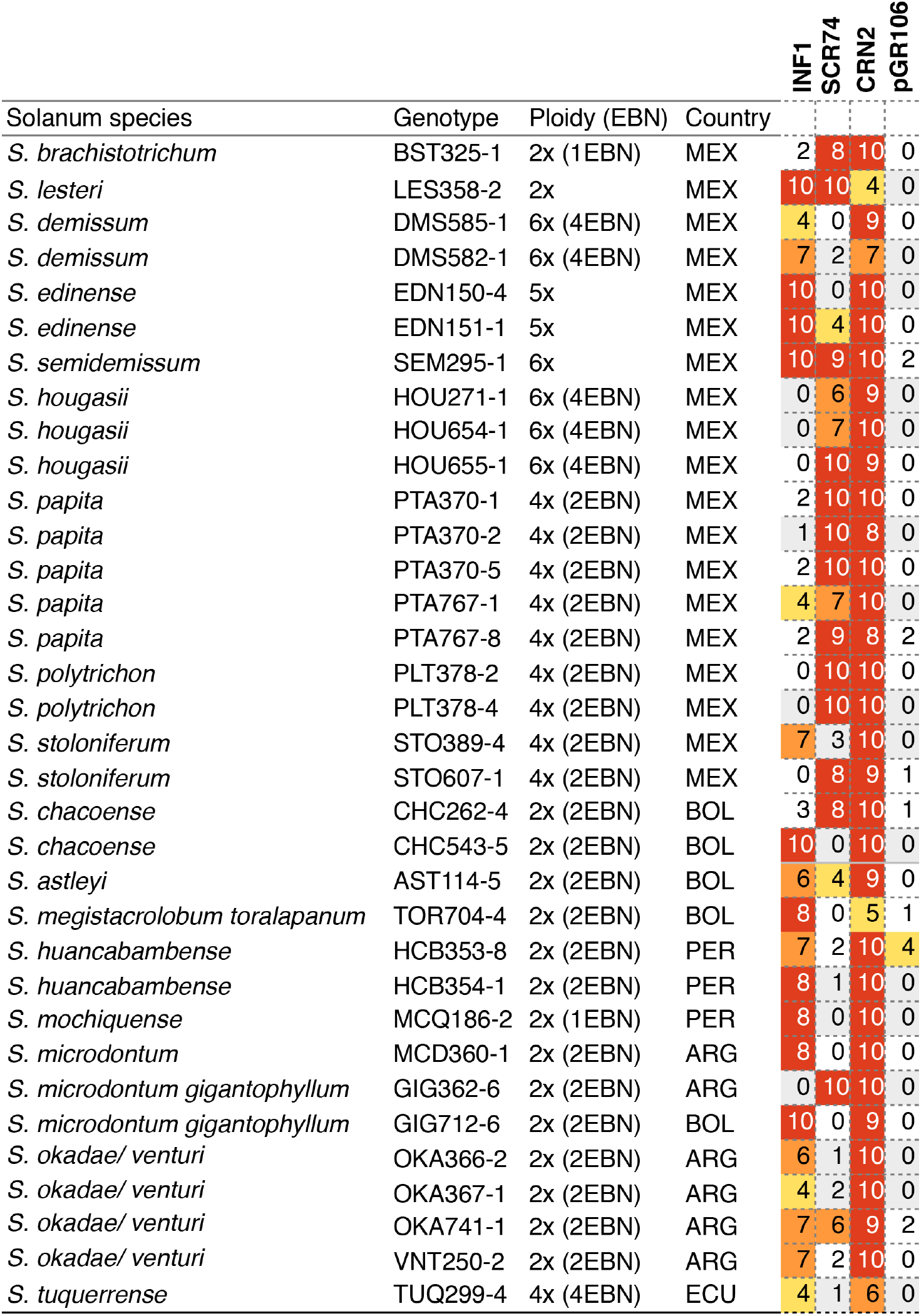
*Solanum* species show specific response to INF1 and SCR74 after PVX agro-infection. *Solanum* genotypes that response to either pGR106-INF1, pGR106-SCR74 or both are indicated. The empty vector pGR106 and the vector containing the CRN2 cell death-inducer, pGR106-CRN2, are included as negative and positive controls, respectively. The responses are scored from 0-10 and presented as a heat map ranging from no response (0-2, blank), weak response (3-4 yellow), medium response (5-6 orange), and strong response (7-10, red). Experiments were independently repeated at least 3 times. The ploidy level and endosperm balance number (EBN) are shown. The countries of origin are indicated by abbreviation: MEX=Mexico, BOL=Bolivia, PER=Peru, ARG=Argentina, and ECU=Ecuador.

### Cloning of effectors for PVX agroinfection

*Inf1* (XM_002900382.1) and *Scr74-B3b* (AY723717.1) were cloned into the potato virus X (PVX) vector pGR106, and electro-transformed into *A. tumefaciens* strain GV3101. Recombinant *A. tumefaciens* strain GV3101 carrying the effector constructs were grown for 2 days in LB medium at 28°C with kanamycin (50 μg/ml) for selection. From the suspension, 1 ml of *Agrobacterium* culture was plated out onto LBA plates supplemented with kanamycin (50 μg/ml) and incubated at 28°C for 2 more days. The *Agrobacterium* culture was collected from the Petri dishes with a plate spreader and used to inoculate 3-4 week-old plants through toothpick inoculation (Takken *et al.*, 2000; Du *et al.*, 2014). Per leaf, two spots were inoculated for each construct and three leaves were used per plant. In total, three replicated plants were used for each genotype. Cell death responses were scored two weeks post infection on a range from 0 (no response) to 10 (strong response).

### Design of customized RLP/ RLK enrichment library

In total, 444 *RLP* genes and 533 *RLK* genes were predicted from the DM genome by HMM, BLASTp, and InterPro, from both PGSC and iTAG annotations. Sequences of predicted RLK and RLP genes from DM are summarised in Note S1. The respective homologs of the 444 and 533 *RLK* genes from the *Solanum chacoense* (M6) genome (Leisner *et al.*, 2018) are in Note S2. Further, 18 known *RLP/RLK* genes from other *Solanaceae* species were included (Note S3). All *RLP* and *RLK* genes were included in the enrichment bait-library design and represented by 120bp fragments allowing for two times coverage (60bp overlap). Duplicated oligonucleotides were removed. Unique RNA oligonucleotides were synthesized to generate a customized MYbaits enrichment library (Arbor Biosciences Inc., MI, USA) comprising 57,020 probes (Note S4).

### Preparation of sequencing libraries and target capture

Genomic DNA was isolated from GIG362-6, MCD360-1 and the F1 progenies using the DNeasy Plant Mini Kit (Qiagen, Valencia, CA). Equal amounts of DNA were pooled from 30 responsive as well as 30 non-responsive progenies for the INF1 recognition phenotype and 29 responsive and 30 non-responsive individuals for the SCR74 response phenotype, respectively. DNA concentrations were measured using a Qubit fluorometer (Thermofisher, Dubuque, IA, USA). The NEBNext Ultra™ II FS DNA Library Prep Kit (New England Biolabs) was used for fragmentation/adaptor ligation and indexing of samples. The Bioanalyzer with a high sensitivity DNA chip was used for detecting the size of DNA after fragmentation. DNA from parents and pools was enriched for RLPs and RLKs with the customized MYbaits custom kit detailed above (Note S4) (Arbor Biosciences Inc., MI, USA), following a hybridization period of 24 hours. Post enrichment PCR was performed, and products were quantified by Qubit. Paired-end sequencing was performed on a single Illumina MiSeq lane for six individually indexed samples including the INF1 and SCR74 responsive and non-responsive bulks as well as the parents of the 7026 population, GIG362-6 and MCD360-1.

All RLP/RLK enriched Illumina MiSeq raw reads were deposited at NCBI Sequence Read Archive (SRA) under accession PRJNA396439.

### Read mapping and SNP calling

Paired-end Illumina MiSeq reads were quality and adapter trimmed with fastp (https://doi.org/10.1093/bioinformatics/bty560) to a minimum base quality of 20. The trimmed reads were then mapped to the DM (v4.03) or Solyntus (v1.0) reference genomes (https://www.plantbreeding.wur.nl/Solyntus/) using Bowtie2 (v2.2.1) (Langmead & Salzberg, 2012) in very-sensitive end-to-end mode. Discordant and mixed mappings were disabled and the maximum insert was set to 1000 bp. Two score-min parameters were used in different mapping runs: “L,−0.3,−0.3” and “L,−0.18,−0.18”, approximately equal to 5% and 3% mismatch rates respectively or “L,−0.54,−0.54 for the Solyntus reference (9% mismatch). The BAM files for the bulks were sorted, merged and indexed using SAMtools (v0.1.18; (Li *et al.*, 2009)), as were the BAM files for the parents. Pileup files were generated for the bulk and parents using SAMtools mpileup with default settings and piped into VarScan mpileup2snp (v2.3.7; (Koboldt *et al.*, 2012)) with --strand-filter 0 and --output-vcf 1 for variant calling.

### Diagnostic RLP/KSeq

A dRenSeq-type analysis was conducted to ascertain the presence and sequence integrity of the known functional target gene *ELR*. The mapping condition for the diagnostic analysis of the RLP/KSeq-derived reads was as described previously (Armstrong *et al.*, 2019), and adapted for RLP/KSeq. For this, the *ELR* sequence was used as reference (GenBank: MK388409.1).

### Read coverage and on target estimation

The percentage of reads on target was calculated as the proportion of reads mapping to a targeted *RLP*/*RLK* region in the DM reference (Note S1). The mean read coverage to *RLP*/*RLK* genes was calculated from the previously generated BAM files using BEDTools coverage (Table 1).

**Table 1.** RLP/KSeq read statistics: Enriched reads are mapped to the DM genome v4.03 at 3%, 5%, mismatch rates and the number of reads that map to target genes are specified. Avr. RD: Average read depth.

|  | PE reads | Total reads | Missr | Reads mapped | % | On target | % | Ave. RD |
| --- | --- | --- | --- | --- | --- | --- | --- | --- |
| MCD360-1 | 2284289 | 4568578 | 3 | 1757896 | 38.47797 | 870460 | 49.51715 | 54.26 |
|  |  |  | 5 | 2541388 | 55.62755 | 1189339 | 46.7988 | 72.88 |
| GIG362-6 | 2589095 | 5178190 | 3 | 2024698 | 39.1005 | 1020145 | 50.38505 | 64.03 |
|  |  |  | 5 | 2957160 | 57.10799 | 1397279 | 47.25071 | 86.25 |
| INF1 Responsiv | 2251593 | 4503186 | 3 | 1669192 | 37.06691 | 902139 | 54.04645 | 55.3 |
|  |  |  | 5 | 2450878 | 54.42542 | 1210312 | 49.38279 | 74.39 |
| INF1 Non-respr | 2345320 | 4690640 | 3 | 1741082 | 37.11822 | 952555 | 54.71052 | 59.12 |
|  |  |  | 5 | 2564836 | 54.67987 | 1282185 | 49.99092 | 79.74 |
| SCR74 Respon: | 2598380 | 5196760 | 3 | 1970580 | 37.9194 | 1057408 | 53.65973 | 65.75 |
|  |  |  | 5 | 2905738 | 55.91442 | 1421961 | 48.93631 | 88.58 |
| SCR74 Non-res | 2502842 | 5005684 | 3 | 1818864 | 36.33597 | 969584 | 53.30712 | 59.66 |
|  |  |  | 5 | 2691420 | 53.76728 | 1311170 | 48.71666 | 80.93 |

### SNP filtering

SNPs were filtered using custom Java code (Note S5) to retain informative SNPs present in both bulks and parents. SNPs were filtered based on the expected allele ratio for responsive/non-responsive bulks/plants (responsive: Rr; non-responsive: rr). To be retained, each SNP had a minimum read depth of 50 and alternate allele ratios reflecting the expected genotype: 0-10% or 90-100% alternate allele for non-responsive and 40-60% alternate allele for responsive bulks/plants. BEDTools intersect (v2.20.1; (Quinlan & Hall, 2010)) was used to extract SNPs present in both bulks and parents (informative SNPs) and to relate the informative SNP locations to annotated *RLP/RLK* genes. The number of parental, bulk and informative SNPs and variant genes were plotted in 1 Mb bins over each chromosome and visualized using R.

### High resolution melt (HRM) marker development and SSR markers

The BAM and VCF files for the filtered informative SNPs were visualized using Geneious R10 (Kearse *et al.*, 2012) (http://www.geneious.com). Primers were designed in Geneious R10 for the PCR products to contain the informative SNP(s) and a size between 80-150bp. Primers flanking the informative SNPs were manually selected on the conserved sequences of both parents, R and NR bulks. The HRM markers were tested on the parents and the F1 progenies with the following protocol for a 10 μL reaction mixture: (1 μL template (20 ng gDNA), 1 μL dNTP (5mM), 0.25 μl forward primer and 0.25 μl reverse primer (10 Mm), 1 μL LCGreen® Plus+ (BioFire), 2 μL 5x Phire Buffer, 0.06 μL Phire taq, 4.44 μL MQ water). Black 96-well microtiter PCR plates with white wells were used and 20 μl mineral oil was added to prevent evaporation. The protocol for PCR cycling is: 95°C for 3 min, (95°C for 10 s, 60°C for 15 s, 72°C for 30 s) with 40 cycles, then 72°C for 2 min followed by 94°C for 40 s. The LightScanner® System (Biofire) was used for measuring and analysing the melting curve. The primers used in this study are listed in Table S2. A further 78 SSR markers described in Milbourne *et al.*, (1998) were used in this study (Table S3).

### BAC library

A BAC library of plant GIG362-6 was generated by Bio S&T (Canada). A BAC clone that spans the mapping interval was isolated using molecular markers (Table S2) and subsequently sequenced using PacBio sequencing (INRA-CNRGV).

## Results

### A wide range of wild *Solanum* species respond to apoplastic effectors of *P. infestans*

To explore the recognition spectra of apoplastic effectors from *Phytophthora infestans*, transient effectoromics screens with INF1 elicitin and SCR74 were performed on a wide range of wild *Solanum* genotypes (Fig. 1a). In total, 100 *Solanum* genotypes were screened for responses to INF1 and SCR74 by PVX agro-infection. An empty vector and the general cell death inducing crinkling and necrosis-inducing protein (CRN2) were included as negative and positive controls, respectively. An overview of all tested plants, including responsive as well as non-responsive plants, is presented in Table S1. A set of 34 *Solanum* genotypes showed specific cell death responses to INF1, and/or SCR74 two weeks after agro-infection (Fig. 2). These responsive plants belong to 17 different wild *Solanum* species, vary in ploidy levels as well as endosperm balance numbers (EBN), and originate from different geographic origins (Fig. 2). In most cases, the specific effector responses were clear and highly reproducible (i.e. clear cell death phenotypes scores >7). In some cases, we observed more variability (cell death phenotypes scores ranging from 4 to 6), but these variations were also observed for the positive control CRN2 in some genotypes, which suggest that these plants were less amenable to the PVX-based transient expression system. As expected, response to INF1 elicitin was confirmed in MCD360-1 (Fig. 2), which is the source of the elicitin receptor ELR (Du *et al.*, 2015). In addition, other *Solanum* genotypes were also found to respond to INF1 (Fig. 2 and Table S1). Similarly, SCR74 was recognized in various plants including GIG362-6 (Fig. 2; Table S1). In conclusion, responses to INF1 and SCR74 are widely found in wild *Solanum* species, which suggests that surface receptors that recognize these effectors are present in these plants.

### Response to INF1 and SCR74 segregates independently in *Solanum microdontum*

To genetically map the gene encoding the immune receptor that recognizes SCR74, and to confirm the location of the INF1 receptor (ELR), a mapping population was developed (Fig. 1a). We crossed MCD360-1 with GIG362-6 and generated the F1 population 7026 (Fig. 3). From this population, 100 progenies were tested for responses to INF1 and SCR74 by PVX agroinfection. The population segregated for clear responses to INF1, with 53 responsive versus 41 non-responsive offspring clones, which is close to a 1:1 segregation (χ2=1.532, P=0.216). Reproducible segregation for responses to SCR74 was also observed at a near 1:1 ratio (χ2=0.36 P= 0.549), as 47 responsive versus 53 non-responsive offspring genotypes were identified. Importantly, the responses to SCR74 were independent of the responses to INF1. Both segregation ratios are consistent with two different dominant loci that mediate the responses to INF1 and SCR74, respectively.

**Fig. 3.**
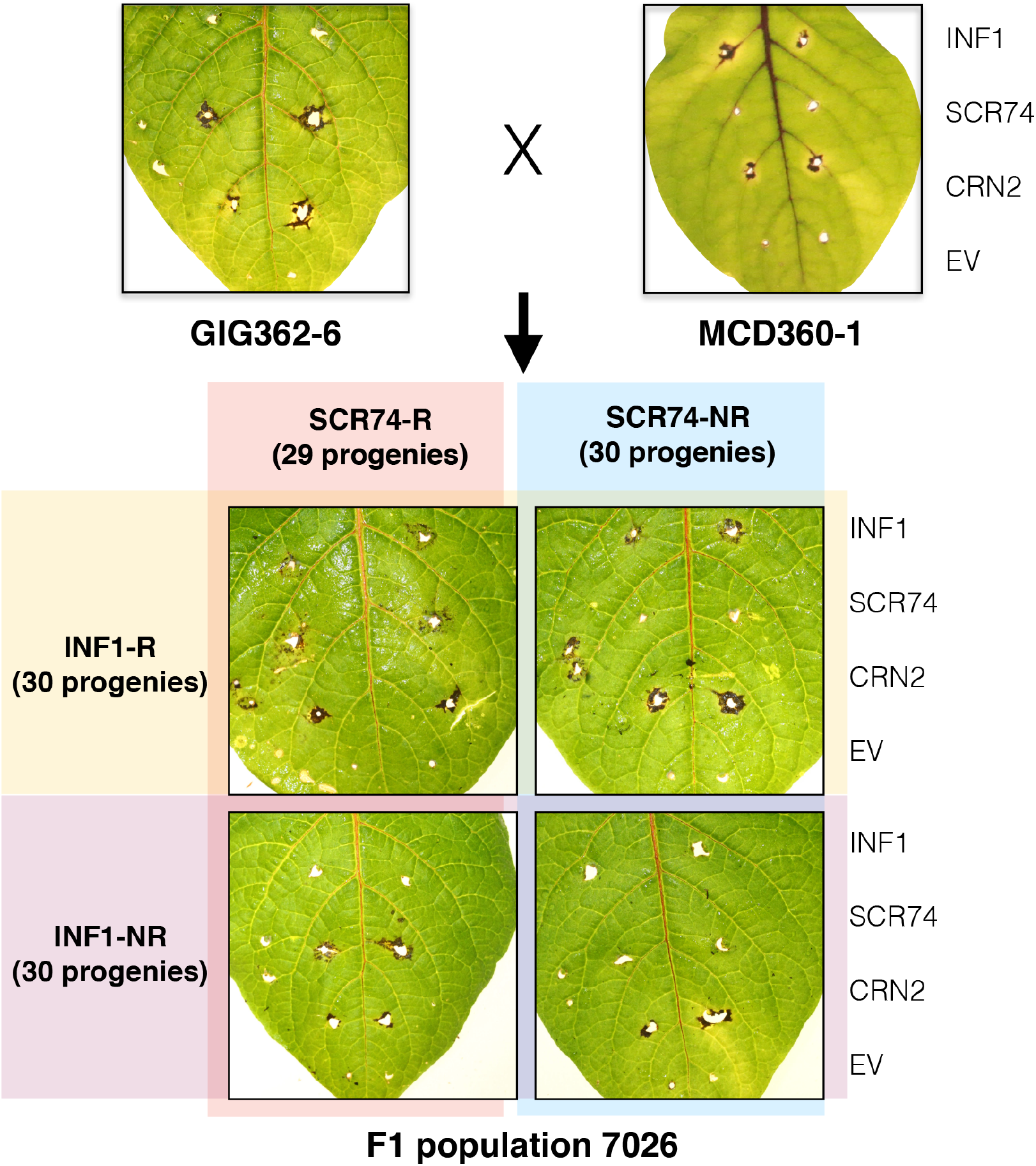
Independent segregation of responses to SCR74 and INF1 in F1 population 7026 of *S. microdontum*. *S. microdontum* ssp *gigantophyllum* GIG362-6 (SCR74 responsive) was crossed with MCD360-1 (INF1 responsive) and progeny plants were assessed for phenotypic respsponses to INF1 and SCR74 through agroinfected with pGR106-INF1 and pGR106-SCR74. The empty vector pGR106 and the vector containing the CRN2 cell death-inducer, pGR106-CRN2, were included as negative and positive controls, respectively. For INF1, 30 responsive (INF1-R) and 30 INF1 non-responsive (INF1-NR) progeny plants were identified. Similarly for SCR74, 29 and 30 SCR74 responsive (SCR74-R) and SCR74 non-responsive (SCR74-NR) progeny plants were selected for the RLP/KSeq, respectively. Representative images are shown at 14 days post infection (dpi).

### Designing the RLP/RLK bait library for target enrichment sequencing

For mapping the gene that confers recognition of SCR74, we developed an RLP/KSeq approach, based on adapting previously described RenSeq targeted enrichment technology to non-NLR genes (Jupe *et al.*, 2013) (Fig. 1b). Since the INF1 receptor ELR was originally cloned from MCD360-1 (Du *et al.*, 2015), we used this genotype and the segregating progeny as a positive control throughout this study.

To design a comprehensive bait library for *Solanum RLPs* and *RLKs*, we combined 301 *LRR-RLK* and 404 *LRR-RLP* genes previously predicted in potato (Andolfo *et al.*, 2013) with *de novo* identified genes. A combination of BLASTp, MEME and Pfam searches was utilized to predicted 533 *RLK* genes and 444 *RLP* genes from the potato reference genome DM1-3, v4.03 (Note S1), including 70 *RLK* with WAX or WAX-EGF domain, 38 *RLK* with malectin domain, 11 *RLK* with antifungi domain, 6 *RLK* with ANK repeat, 11 *LysM RLKs*, 24 *L-LecRK*, 103 *G-LecRK*, 1 *C-LecRK* and 22 other *RLK*s with TM. Additionally, 18 known *Solanaceae RLP/RLK* genes from were included (Note S3) alongside the *RLP/RLK* homologs from *Solanum chacoense* (M6) (Leisner *et al.*, 2018) Note S2.

A customized target enrichment RNA bait library with 2x coverage comprising 57,020 120-mer biotinylated RNA oligo probes was synthesized (MYbaits custom kit, Arbor Biosciences Inc., MI, USA) (Note S4). The long RNA baits can tolerate mismatches like SNPs and indels (Clark *et al.*, 2011) and were used for the mapping of the INF1 and SCR74 receptors (Fig. 1a and 1b).

### Bulked segregant analysis (BSA) and RLP/K enrichment

To map the genes that mediate response to INF1 and SCR74 using RLP/KSeq, we used a BSA approach. Normally, for mapping one gene, it would require two pools, i.e. responsive and non-responsive, plus the two parents (Fig. 1c). In this case, since we multiplex for two target genes, we composed four bulked pools. These comprised response to INF1 (INF-R: 30 plants), no response to INF1 (INF1-NR: 30 plants), response to SCR74 (SCR74-R: 29 plants), no response to SCR74 (SCR74-NR: 30 plants), progeny individuals, respectively (Fig. 3). DNA was isolated from each clone and then pooled prior to indexing. DNA from the parents GIG362-6 and MCD360-1 were individually indexed and included in the enrichment.

### Mapping reads to the reference genome and SNP calling

The *RLP/RLK* enriched DNA libraries from the bulks and parents were sequenced with Illumina 2×250 bp chemistry on a MiSeq platform (Fig. 1c). The number of raw reads that passed quality control ranged from 4503186 to 5196760 in different samples/pools (Table 1). High-quality paired-end reads were mapped to the potato reference genome (DM v4.03) using Bowtie2. To compensate for differences between the potato reference DM and *S. microdontum*, two mismatch rates of 3% and 5% were used for the read mapping (Table 1). The mapping rates ranged from 36% to 57%, with reads on target accounting for 46% to 55%, depending on the mismatch rate (Table 1). The resulting coverage of known *RLP/RLK* genes was calculated and ranged from 54 x to approximately 89 x. To enable the identification of informative SNPs whilst ensuring sufficient accuracy, a 5% mismatch rate was used for further analysis. SNPs were called by SAMtools and VarScan from different samples, and the output SNPs were filtered by a custom java script (Note S5; Chen *et al.*, 2018).

### Diagnostic analysis of RLP/KSeq-derived reads confirms presence and sequence integrity of INF1 receptor ELR

To validate our targeted enrichment sequencing approach and to confirm that RLP/KSeq specifically yields sequence representation of expected RLPs/RLKs, we used diagnostic mapping of enriched samples to previously characterised, functional gene sequences as a control. In line with dRenSeq (Van Weymers *et al.*, 2016 and Armstrong *et al.*, 2019), we refer to this approach as dRLP/KSeq. As a proof of concept, we assessed the sequence representation of the known INF1 receptor ELR that was expected to be present in the INF1 responsive parent of the population 7026, MCD360-1, as well as in the INF1 responsive bulk but expected to be absent in the non-responsive parent, GIG362-6, and the non-responsive bulk.

In line with this expectation, dRLP/KSeq revealed continuous coverage of ELR in the progenitor parent of the INF1 receptor and the responsive bulk. Indeed, a very similar nucleotide representation profile was observed for both samples and only the very 5’ and 3’ region of ELR are not resolved due to a lack of flanking sequences in the reference that prevented the mapping of RLP/KSeq-derived reads that extend from the gene into the 5’ and 3’ UTR, respectively (Fig. 4a). In contrast, functional ELR sequence representation in the non-responsive parent and bulk was very limited and discontinuous which is in accordance with the absence of the function receptor sequence in these samples. The partial coverage observed hints at the present of non-functional ELR homologs.

**Fig. 4.**
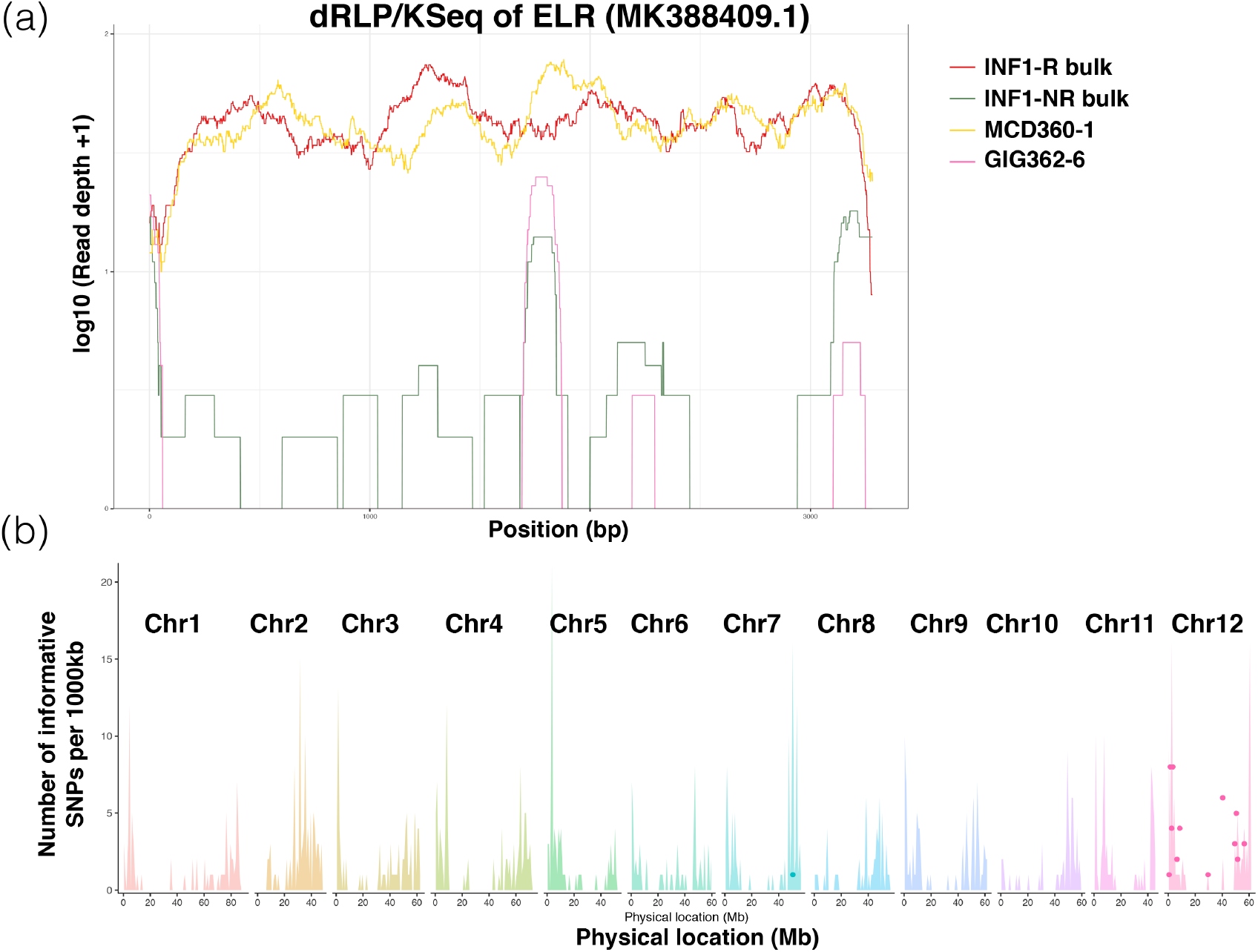
*ELR* was recovered and independently mapped to chromosome 12 by RLP/KSeq. (a) Diagnostic RLP/KSeq for ELR. The *x-axis* depicts the coding DNA sequence (CDS) of ELR from start to stop and the *y-axis* indicates the read coverage of functional ELR with RLP/Kseq derived reads mapped to the reference under highly stringent conditions on log scale. The yellow and red horizontal lines indicate full length ELR sequence from MCD360-1 and INF1 responsive bulk without any sequence polymorphisms. The green and pink lines show a low and discontinuous read-coverage from GIG362-6 and INF1 non-responsive bulk. (b) Mapping of ELR: The *X-axis* represent the physical positions of the 12 individual potato chromosomes and the *Y-axis* the number of RLP/RLKs or SNPs per 1Mb bin. The background color spikes represent the number and position of annotated RLP/RLKs and the colored dots depict the position of significant and linked SNPs in a 1MB bin. The peak in chromosome 12 indicates various SNPs that are linked with ELR, which confers response to INF1.

### *De novo* mapping of the INF1 response using unrelated potato reference genomes coincides with the physical position of the ELR receptor and identifies linked SNPs in closely related homologs

Following the successful dRLP/KSeq analysis, we assessed the suitability of using RLP/KSeq-derived reads for the mapping of receptors using ELR as an example. SNPs from the population parents alongside INF1 non-responsive and responsive bulks were called and filtered for the expected ratios of heterozygosity as described by Chen et al., 2018. In short, for a single dominant gene segregating in a diploid population, the allele frequencies were set at 0-10% or 90-100% for the INF1 non-responsive bulk and parent as well as 40-60% for INF1 responsive bulk and responsive parent. The SNPs from the bulks were independently validate through comparison to parental SNPs, and only the accordant SNPs at the correct ratio were maintained as informative SNPs (Table S4). Allowing for a 5% mismatch rate for the positioning of RLP/KSeq enriched reads and determined SNPs from *S. microdontum* against the *S. phureja* reference genome (DM), 99 SNPs passed the filter criteria in the bulks and 4,323 SNPs in the parents. Among those, 48 SNPs were shared in both bulks and parents (Table S4). The number of informative genic SNPs per 1 Mb interval was placed on the 12 chromosomes of potato. With the exception of one significant SNP on chromosome 7, the remaining 47 SNPs were positioned on Chr12. The SNPs were found to localize in two major locations on chromosome 12, one near the bottom and one at the top of chromosome 12 where ELR resides (Fig. 4b). The majority of SNPs were localized in two *RLP/RLK* loci that correspond to 19 polymorphic genes. Intriguingly, the sequence that is most similar to *ELR* in the DM reference genome has not been placed on any linkage group and is currently found in the unassembled chromosome 00. This highlights some remaining ambiguity within the DM reference genome which does not contain functional ELR.

Thus, we also mapped the reads to the recently released but largely unannotated potato genome Solyntus (v1.0) (Materials and Methods). Essentially, we observed a somewhat similar distribution of informative SNPs as seen in DM (Table S5). However, in Solyntus, the most similar sequence to *ELR* is 92.8% identical and spans the physical position between base pairs 2,491,817 and 2,495,104 on chromosome 12. Allowing for a 9% mismatch rate, we observed an informative SNP at position 2,493,665 in this gene (Table S5). In summary, despite the absence of a true *ELR* gene in DM and Solyntus, RLP/KSeq led to the correct mapping of the ELR locus on chromosome 12 and identified an informative SNP within the closest homologue of ELR in Solyntus.

### RLP/KSeq accelerates mapping of the SCR74 response gene on Chromosome 9

To map the single dominant gene that confers the response to SCR74, the same SNP filtering approach was performed as shown for ELR above. The SNPs that meet 0-5% or 95-100% allele frequency in the SCR74 non-responsive bulk and parent as well as 45-55% allele frequency in the SCR74 responsive bulk and parent were identified and then independently corroborated between bulks and the parental material. This resulted in the identification of 61 informative SNPs of which 60 could be placed on chromosome 9. The SNPs correspond to 16 polymorphic genes (Fig. 5a. Table S4).

**Fig. 5.**
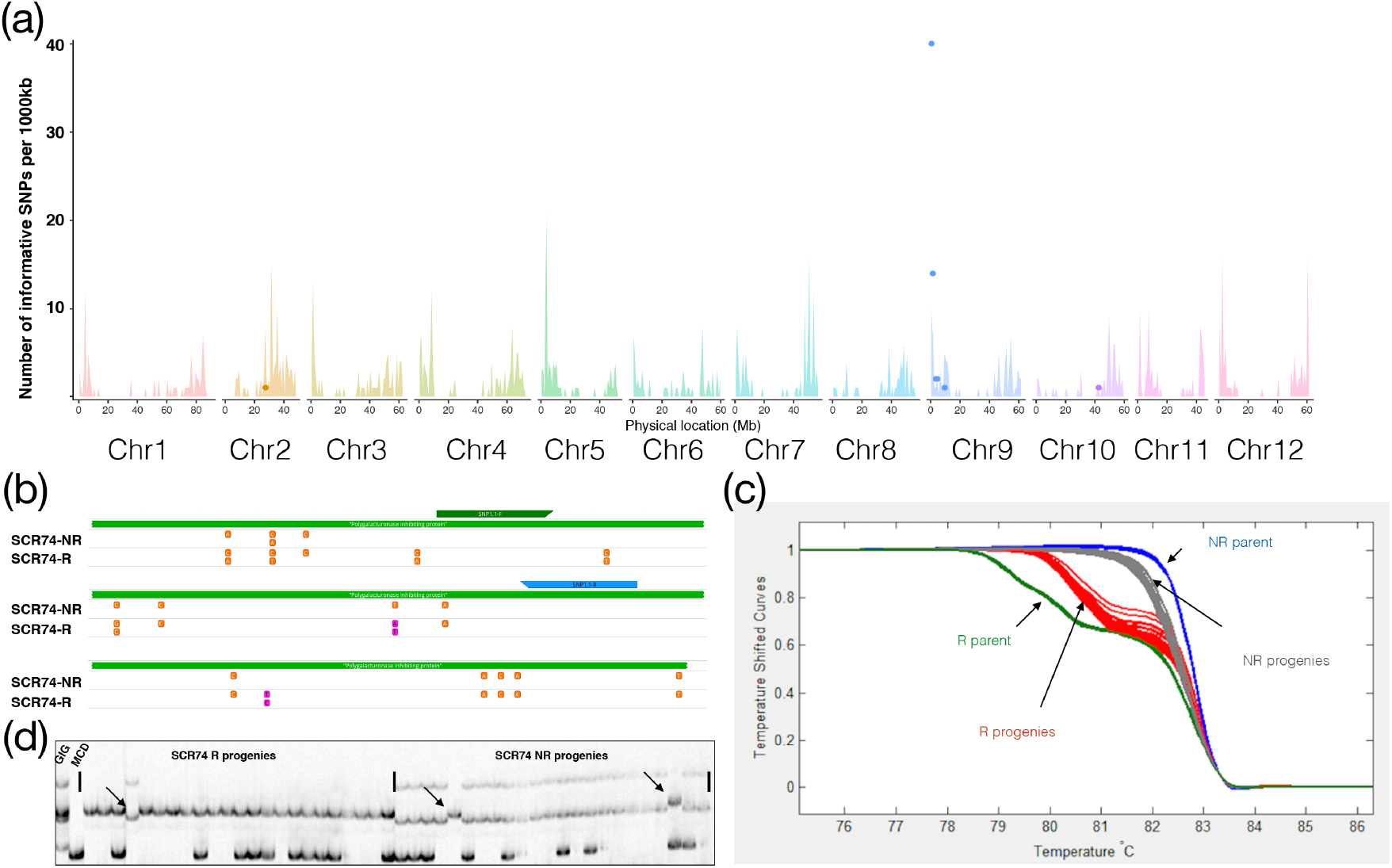
The gene conferring response to SCR74 was mapped to chromosome 9 by RLP/KSeq. (a) Mapping of SCR74: The *X-axis* represent the physical positions of the 12 individual potato chromosomes and the *Y-axis* the number of RLP/RLKs or SNPs per 1Mb bin. The background color spikes represent the number and position of annotated RLP/RLKs and the colored dots depict the position of significant and linked SNPs to SCR74 in a 1MB bin. The significant accumulation of SNPs on the top of chromosome 9 indicate the map position of SCR74 receptor. (b) Within the identified region on chromosome 9, a polygalacturonase inhibiting protein (PGIP, PGSC0003DMG400006492, green bar) contains 15 SNPs (orange). Two of these SNPs (pink) show a near 1:1 frequency in the SCR74-B3b responsive pool and, together with marker RLP/KSeq-snp1.1 (green and blue arrow), flank the interval. (c) Melting curves of the high-resolution melting (HRM) marker RLP/KSeq-snp1.1 tested on the mapping parents and progenies. (d) SSR marker STM1051 on chromosome 9 is linked with the SCR74 response. The mapping parents GIG362-6 and MCD360-1, as well as the responsive progenies and non-responsive progenies were tested with STM1051 and 3 recombination events (arrow) were found. This figure is reproduced from (Domazakis *et al.*, 2017).

One of the identified SNPs, RLP/KSeq-snp1.1 (A->T), corresponds to PGSC0003DMG400008492, a polygalacturonase inhibiting protein (PGIP) gene that resides at position 0.16 Mb on Chromosome 9 (Fig. 5b). This SNP displayed a 59% frequency in the responsive bulk and 0% or 100% frequency in the non-responsive bulk and was used to independently corroborate the mapping position of the receptor on potato linkage group 9. We converted the SNP to high resolution melting (HRM) marker, RLP/KSeq-snp1.1, and tested it on the mapping parents and the 56 progeny genotypes of the F1 population, our result indicates that the SCR74 receptor is linked to this marker which resides on chromosome 9 (Fig. 5c).

To further confirm our RLP/KSeq methods, 78 SSR markers dispersed over all 12 potato chromosomes were tested on 56 F1 progeny of population 7026 (Table S3). SSR marker STM1051 was found to be linked to the SCR74 responsive phenotype, and 3 recombination events were detected (Fig. 5d). This marker resides in position 6.15Mb of DM chromosome 9 and independently confirms the RLP/KSeq mapping analysis for the SCR74 response (Fig. 5c). Consequently, the SCR74 response gene was mapped to a 10.7cM region on potato chromosome 9 between RLP/KSeq-snp1.1 and STM1051, which spans a 5.99Mb physical distance based on the DM genome.

### Fine-mapping of candidate SCR74 response gene to a 43kbp *G-LecRK* locus

To fine-map the SCR74 response gene, we first genotyped 1500 progenies of population 7026 with flanking markers RLP/KSeq-snp1.1 and STM1051. As a result, 120 recombinants were identified. To further narrow down the mapping interval, we genotyped 500 additional progenies and developed more SNP markers for genes predicted to reside in this interval and for which RLP/KSeq had identified SNPs (Figure 6a). The latter included two *L-LecRK* genes, PGSC0003DMG400008822 and PGSC0003DMG400008897, and a *G-LecRK* gene, PGSC0003DMG400024259 (Table S4). By testing those RLP/KSeq markers and other SNP markers developed within this interval (Table S2), we located the candidate gene between marker S111 and S105 (Figure 6b). In DM, the mapping interval contains eight genes, including three receptor-like kinases with a G-type lectin domain (G-LecRK) genes, a putative reticulate-related 1 like gene, a serine/threonine-protein kinase ATG1c-like (autophagy-related protein) gene, a prenylated rab acceptor family gene and an uracil phosphoribosyltransferase encoding gene. Remarkably, of the markers, S55, was derived from the RLP/KSeq analysis and locates within a *G-LecRK* gene (PGSC0003DMG400024259). This marker displays perfect linkage and co-segregates with the SCR74 response (0 recombinants out of 2000 F1 progeny, Figure 6b). To obtain the physical representation of GIG362-6, a bacterial artificial chromosome (BAC) library was generated. A BAC clone that covers the mapping interval was isolated (Figure 6c), unlike in DM, only 2 *G-LecRK* genes are located in this region. The physical distance between the two flanking markers in the GIG362-6 responsiveness haplotype is 43-kbp.

**Fig. 6.**
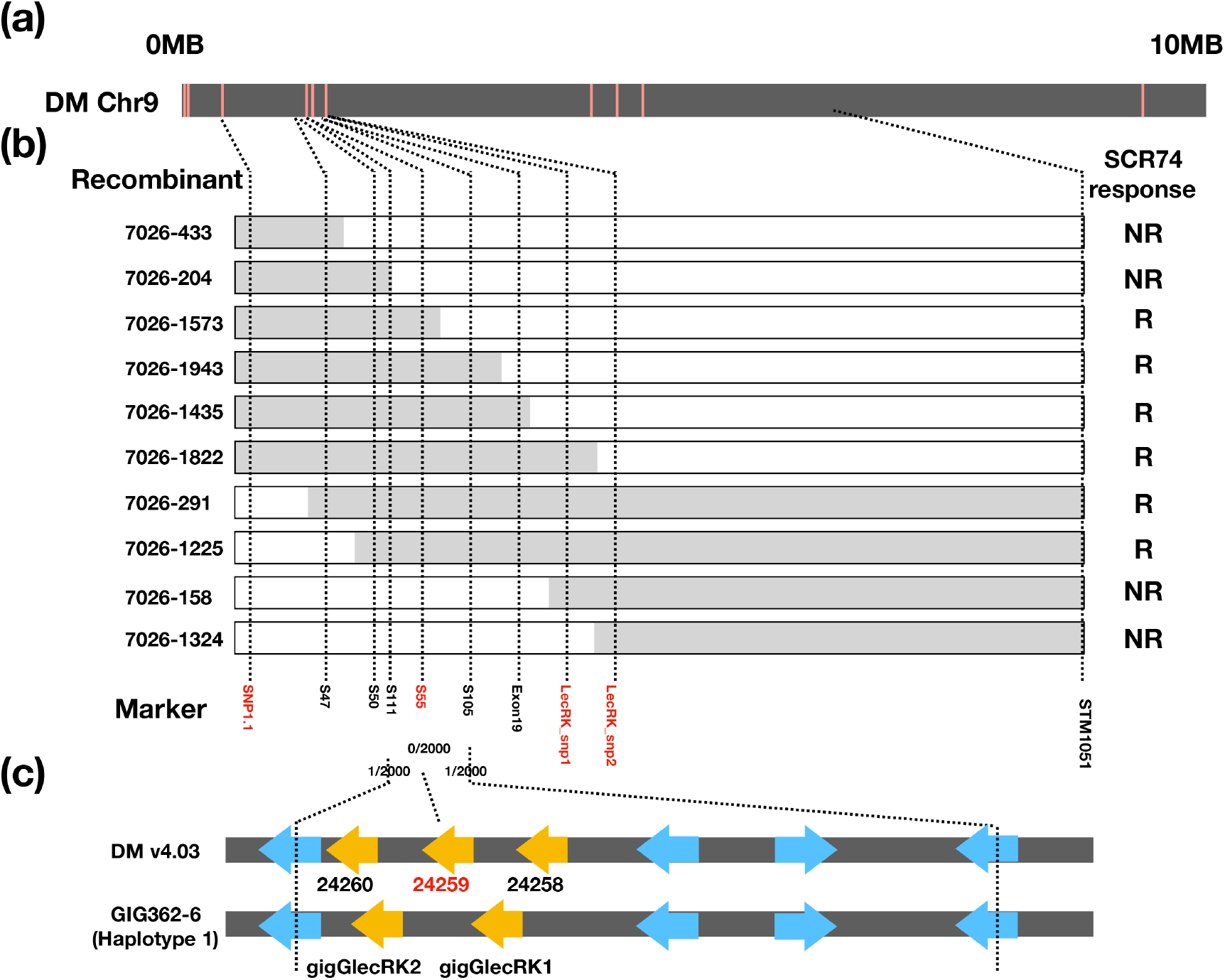
Fine-mapping the SCR74 response gene to a *G-LecRK* locus. (a) Graphical representation of the top 10MB of chromosome 9 of DM (black bar) and the location of informative SNPs (red lines). (b) Overview of recombinant screening, showing representative recombinants with their genotyping and phenotyping results abbreviated as (R) for response and (NR) for non-response to SCR74. The position of the responsive haplotype of GIG362-6 (grey bar), the non-responsive haplotype of MCD360-1 (white bar) and the markers (dotted lines) are indicated. RLP/KSeq-derived markers are marked in red. (c) The *G-LecRK* locus of DM and GIG362-6 based on the sequencing of a BAC clone that spans the SCR74-responsive interval on chromosome 9. The location of *G-LecRK* (yellow arrow) and other predicted genes (blue arrows) are marked.

## Discussion

In this paper, we present a workflow that combines RLP/KSeq with effectoromics of apoplastic effectors, to rapidly map plant surface immune receptors (Fig. 1). We used potato and *P. infestans* as a model system. We screened wild potato species that mount specific cell death response to the apoplastic effectors INF1 and/or SCR74 of *P. infestans*. *S. microdontum* MCD360-1, which responds to INF1, and GIG362-6, which responds to SCR74, were crossed in order to generate a population that segregates for both responses independently. In parallel, we designed bait libraries based on predicted *RLP* and *RLK* genes from the potato genome. We subjected pools of genomic DNA derived from responding versus non-responding genotypes to a BSA RLP/K enrichment sequencing approach, using a bespoke bait library to enrich for genomic DNA representing our target genes. This approach quickly led to the identification of SNPs that are linked to the phenotype and could be used as molecular markers to genetically map the genes encoding the putative RLP/RLK genes. Here, we have shown that RLP/KSeq can successfully identify informative SNPs in the ELR receptor that underpins INF1 responses, obtain full-length sequence representation of ELR in responsive parent and bulks through dRLP/KSeq analysis, and fine-mapped a novel receptor for SCR74 response to a 43kbp interval containing two *G-LecRK* genes.

With continuous advances of sequencing technology, genotyping by sequencing has already been applied to clone plant genes in multiple crops (Huang *et al.*, 2009; Austin *et al.*, 2011; Mascher *et al.*, 2014; Pandey *et al.*, 2017). However, when the genome size is large, or when certain types of genes are expected, targeted enrichment sequencing becomes a preferential option, as it can dramatically reduce the genome complexity (Hodges *et al.*, 2007). RenSeq and its descendants like dRenSeq, MutRenSeq, SMRT RenSeq and AgRenSeq have been demonstrated to be powerful tools to clone plant disease resistance genes (Steuernagel *et al.*, 2016; Van Weymers *et al.*, 2016; Witek *et al.*, 2016; Arora *et al.*, 2019; Armstrong *et al.*, 2019). However, they all target *NLR* genes. RLP/KSeq can complement the RenSeq toolbox by targeting additional types of plant immune receptors, including RLPs/RLKs that also function as effective immune receptors (Boutrot & Zipfel, 2017).

Effectoromics has proven to be a medium to high-throughput approach to identify plants carrying *R* genes as well as surface immune receptors (Vleeshouwers *et al.*, 2011a; Du *et al.*, 2015; Domazakis *et al.*, 2017). The specificity and robustness of effector responses enables the identification of multiple receptors in a single segregating population (Fig. 3). Another advantage of combining the enrichment sequencing with effectoromics is that targeted libraries can be used for PRR or NLR, depending on the matching effector response. Effectoromics was pioneered for the potato – late blight pathosystem and has been successfully applied in various other *Solanaceae*, such as *N. benthamiana*, tomato and pepper (Takken *et al.*, 2000; Oh *et al.*, 2009; Lee *et al.*, 2014). Beyond *Solanaceae*, the approach has been used in other plants such as sunflower (Gascuel *et al.*, 2016), as well as in various plant pathogens such as fungi, nematodes and insects (Catanzariti *et al.*, 2006; Sacco *et al.*, 2009; Hogenhout & Bos, 2011). This demonstrates the wide application of the effectoromics strategy for pathogens with well-characterized genomes. To summarize, our newly developed pipeline enables the rapid identification of plants carrying novel immune receptors and genetically map the genes responsible for the phenotype. This strategy is complementing the current RenSeq toolbox and will help to understand the first layer of the plant immune system, to ultimately achieve more durable disease resistance in plants.

## Acknowledgements

This work was supported by a NWO-VIDI grant 12378 (X.L. D.W., V.G.A.A.V.), Short Term Scientific Mission (STSM) of COST Action SUSTAIN FA1208 (X.L.), China Scholarship Council (CSC) (X.L), the Rural & Environment Science & Analytical Services Division of the Scottish Government (I.H.) and the Biotechnology and Biological Sciences Research Council (BBSRC) through awards BB/L008025/1 (I.H.), BB/K018299/1 (I.H.) and BB/S015663/1 (I.H.). We thank Dr. Hendrik Rietman for performing effectoromics screens, Isolde Bertram-Pereira for culturing *Solanum* plants, Henk Smid and Harm Wiegersma for help in the greenhouse. We thank Dr. Helene Berges and Caroline Callot from the French Plant Genomic Resource Center (INRA-CNRGV) for their help to sequence the BAC clones. We thank Prof. Dr. Evert Jacobsen for reviewing the manuscript.

## Author Contribution

L, V.G.A.A.V and I.H. planned and designed the research, X.L., K.B., M.A. and D.W. performed the experiment. X.L., M.A., V.G.A.A.V, I.H. R.G.F.V. and P.J.W. wrote the manuscript.

## Supplementary Information

**Note S1.** Fasta file of the 977 RLP RLK genes from DM genome used for generating the RLP/KSeq enrichment library

**Note S2.** Fasta file of the 977 RLP RLK genes from M6 genome used for generating the RLP/KSeq enrichment library

**Note S3.** 18 additional known RLP and RLK genes from Solanaceae species

**Note S4.** The 2x bait library used in this study

**Note S5.** Java script for calling the informative SNPs

---

**Table S1.** All tested Solanum genotypes for INF1 and SCR74 response by PVX agro-infection

**Table S2.** Primers used in this study

**Table S3.** SSR markers used in this study

**Table S4.** Summary of the SNP calling outputs under 5% mismatch criterial

**Table S5.** Using Solyntus genome for the mapping and SNP calling

