## Supplementary material for "RLP/K enrichment sequencing; a novel method to identify receptor-like protein (*RLP*) and receptor-like kinase (*RLK*) genes": Note S1

>PGSC0003DMG400009417 C-LecRK

AATTAACTACTCTGTCTGAACAACACTTTCTTCTTCAAAGAAATTCAACTAACCATTGGAGACTCCAAATGTGAACTCTTGACCTTTCTCTCTCTAAAAAAAACCCTATCCACCCACCCCATATCCACAATGCCCAACCCCACACTCAAAATCATTCTAGGTTTTGCTATACAGTGAGAACAAGTGATGATACTGATGGCTGTTAAACCTCTGTTCCCTTTGTTTCTACTTACTTGTTTTCTGCTTCACCTCTGTGCTTCCACTTCTGTAAGTAAAGAAAAGATTATAGTACCTTTTATTTAGAGAAAAGGGTGTCTGAATATGTTATCAGAATTCATGTTAGTGTGTGAATTTGGAGAAAGTTTTGATTTTTATTTTTTTGAATTTTTTTCAGGTCCCTAATAAGACACTTGATTCACAAATAGTTTCATCTTCTGGGAATCAAAGTCATGATGATGGTAATGTTTTGGAATTAATTAGATTATTGTTCTGTCTATTTCCTTGGAACTGATAGATTGGCTTGTGTTGATTAGATTAGGGCTCTTTTTAGAATCAGTAATGGAGTTGCATCTATAATCATCTTTTCAACATAATGTTCAACTACTATATGTTACTTTGTGTATAATATGTTGGGATTTCTTACTGGTGCTTCTTATGTAAATTGTAATGTGTGCATATCTATGTGTCATTTGCTAATTTAAACTAGCATAGTGAACTTTTGAGTTTGGAGAAAATAAAGGCTGGCGAATCCAATCTTGTTTATCAAGTTGCTATAGGTTTAAGGTCTACAACTTGGGGTATTTTGTGCCAAACTATTATTCATAAATCAAGATTTTGAGAACTACGTCACTTCCCAGCATCATTAAGCTCAGGCCATAAAAGACTGTCTCGCAACCAGTTATCCCTCGAAAAAATAAAAATAGAAGATGAACTTGCAACTATTTATTAATCAGCTTCCTACTTCTAAAGAAAATTAATTGACCAGCCTCTCGATTCCGGTTTACAAAGTTATGGATTCCCATTCCTCCAAGACAGTTGTTCTTGGAATATTCTTAGAACAATATTCTATTTGACTGATCGATTATCCTTCTAAGAAATCCTGGTGGTCATAGTCACCTGATTGAATTTAATTTATTCGGTCGACAAAGGAGCAGAAGTAGATTCTGCTCATTTCATGCATAAAGGCATGCTTAGGTTGCTCTTAAAGTGAAACTTTGGTTCTATTTGAGGGAGATAATATATATGCTTTTATGGTGGATTAATGATGGCCTTGATGAATAATGTTGTCCAGTTCCTTTCCTATTGAGGAACTCCTCCCTTGGAGTTTCTTTTGTTTTGATGATATGCATTTCACCATTTTTTTTCCTAAAAAAACTGCACTAGATGTCGGGAGGACTCATGAGGTGATGCATAGAGCTCACCTGGTAGTTGTCTAAGAGGAGGAGGTGGCCAGATGAAGCATGTGACACTTACAATTTCACCAGTTAGAGGTTTTGCTATTACATTAGGCAATGGAATGACGAGAATAACAGCGTGTTTTATGTTGATCAGCATGTAGAGCAGCAGTAATTGCGCTAGTATTCTAGCTTGATAGTACTACATCTTTCATTTGAGATAGCACATCATTTTCTTTTGAAATGGGCATAGACTGAAAGCAATAGGGAAAATAATGATGATCCAACTTTATCCTACTTTTGCCCCCTGCTATTCTTGAATATTGATGGTTTTTACAGGATATTTGCAGTCATGTGGAATGTGATTCCTACATACTCTATATGCCTCTTTAGTAAGACCTGTGCTTTCAAGTCTCAAGTTATCCCTGCAACTTATGGTGTTAGACATACTGGTACACGTGTAGAAATGTCATGCCAAGAACCATCTAGCTGTTCATTTAAGTTTCCACAATTACGAAGAACCTTTCTTTTCCATTCGCTCTGTAGGAAGAAGAACAAAGTTCTAGTTTTTATTGAACTCGCATGGGTTTTAAAAAAGACGTATATTTGTTGGTACATCAGTGAATAAGGCTGTGGATCATCTCAGAGAAAACTGACTAATTGAGTGAACACAATTAGGAGTCTTGGATGGAACAATTATGGAGAATGTTCATCTGTCTAAAGGGGATAAGATGGGAGAAACCAGGGAGCATTAAAGGAGTCCTGAGCAGTTGGAATAGAGATGGCAATGCTTCTGACAAGGAGAAGAGATGGAAACTTGTTCCAGCATGTATTTGGTGGACAATCTGGAAAGAGAGAAACAACCGGAGTTTTGAAAATTGGCAGAATAGCTTACAGAAGATCAAGATGGACTGCTTAGCTCTTTTCTATTTTTGGTGTAAACAAGATATATTAACTAGGACAGAAGACATCTTTGATGTGCTAGATTGTTTATAGGCAGCAGTTTAGACAATGACTATTTTATTCTGTAAGTTGTAACACTTTTGAAGGGGAACTACACTTTTGTTGTAGTATACCTGTGGATATATAGACTAAAGTTACCAATTTTTTTTTAAAAAAACAATTAGGAGTCCAAGCTGAACGAAAAGAGAAGAAAACACTGGAGCTTAATTTTTTTTTTAAAGATTCAGTGTTCTCTTCTCTTTTCGTTCAGCTTGGACAAGCTTAGTTTAATGCAAGTTTGGCTTTGTTTGTCAGAACTTGGTACTTTCATTTATCCATCATACTTGAATCTAGCACACCTTCCCTCTCAACAAAGGAAAAGTGAAAACTGAGAACACACTGTCGATGTCAAACCTCCACTAATCTGAGTGATATCACACAATCCATAACACGGAGGCCAAGAAACGACTACTCCTATCAGGTTCATAAGACACAGGGGTAGAGGGCAATGACCTGGTGTCAACATCGTTCTTGGGTCTTCTTATTTCGTAAGCTTAGTTTGAGATGGCAGTGGAGGCCAATTGTGGTGAACCGTGAATCTTTCAATCTTCAAATCTTGGAGACCTTATCTGAAATGAATTCTAATTTTGAAGCTGATCCCTCAATTGTAACCTTTGTAAAAAAAAAAAGTAAATAAATCCATTAATGAAGATAAAGATACTTTTAGGGTGAAAAGTCATCTTCAGTTGTGGTATTAGAAATAACTAGTTCAGAATACCCGAGCTTTATTGTCCATTATGAGGTATTTATTAGATATAAACAAGTTCATGATACCTGAGTTTTACCTACCTAGATATAAACAACTATTCCTACCTAGCACTTTCCAGTCCCCGCAATCTTACTAGTATTATTTACAACTGAAGTCGATGATCAAACTTAAGGAAACAACAGAAAGAAAAAACATTAAAACTACAAATATAGAAAGATGGTGAAAAGAATCAACAAATGTGTACAAGAGTTGGAGATCATGTGGCTAAATGGATGGGGTTAGTGCAGGGATGGCAAGTTTGAGCCACTTTTATAACATTAGGAACACATATAAACTGATAGAATAACATTAGGGGTATAGATGAACTGATAGTATAATGGAGGGTAAAATTAAACCTTTTCAATAGTACAGGAAGTGAGGGACAAATATGACCCTTTTCCTAATTTCTTTTCTTCTTCTTAGATAAATGAAGAAGTTAGATCAAGTATTATTAAAACACTAGTGCAACTGAAACTTATTTGGACCTCTCAACGCGTGGTTTTCTTAGTGTATGTTTTTTATTTTTAACTGGTCAAGTTATTATGTCTCATCTCTTTGGCACTGTTGTTTATTGTTTCGCAAGCAGGAAGATGTGATCCTGGTTGGTTTAGCAGCCCAAATGGGAAAAAGTGTTTCAAATATAATGCGACCTCTTTGCAATGGAATGAATCAGAGACTTACTGCAATGGTTTTGGAGGAAACCTGGCTGCCATAGCATCTTCAGAAGAACTGAACTTTGTCCAAAAGCTCTGTGGCAATGATAGTAACGGATGCTGGGTTGGCGGAAGGAGCATCAATACCACAGCTGGGTTGGGTTGGAAATGGTCTGATAATGCTTCTAGCTGGAATGAGAGTATTGCTCCTAAGACAACTTTGGACTCAAGTTGCAAAAATTTCTCATGCTATGGCTTTAAGTCGGTGGATTTGTGTACATCGATAACCAATGGCGCTATTATAGCTGAGAGATGCAATGTAACTCATGCTTCTATATGCACACTTGATGCTGGTAAGTGATTGCAGACCAATTTATTTCTAATTGTTTCTTTCACCCAGTGTAATTTTGCAAAGTAGGAGCAAAACCCTTTCCCTGATATAGTGTGGTTATACTATGTGATTAAAAGCTTCATAGTCCCCTTTTGTTTTTTCCATGAGAAGTGCATTTTAGCTAACCATTATACAGGCCTACATTGTTATCTGGATAAATCTATTACATTCCTTAATTGGTTTTTAAATAATCTTCTAAATTTTACATTAGGGAGCAAGTGTCGCCATATGCGTTGCCACAGGGAGTATCTCATTATTCTTGCAGTTGTGAGTGGGTTGATCCTTTGCACTACTTTGGCTATTGTGATATGGCTTCTTGTATTTCGGCGAAGTAAAAGACGCAAAAGATCATCGCGTGCATCATTAGCATTAGTACCCCCGTCTTGGAAAATTTTCAATCGTGATGAAGTAAAGTCCATGACAAAGAATTTCAGTGAAGCAAATCGTCTCCTTGGGGACGCCAAAACAGGTGGTACCTATAGTGGGCTTTTACCTGATGGTTCAAGGGTGGCTGTGAAAAGATTGAAGAGGTGCGGTATTCAGAGGAAGAAAGAGTTCTACTCTGAGATTGGAAGGGTTGCAAGACTTCATCATTCAAATCTGGTATCTATCAAAGGATGTTGCTATCATCATGGTGACCGCTATATTCTGTACGAGTTTATTATCAATGGACCCCTGGATAGGTGGCTACACCATATTCCTAGAGGAGGTAGGAGTTTGGACTGGAGCATGAGAATGAAGGTCGCCACCACCCTTGCTCAAGGGATTGCGTAAGTAGTGTTCCAACATTATCTTCATGTGCATTGTAGATTAGGAATAAATATTTTAACTTATCCTTGAGGTGGATTGATTTTAGTCAGAAATGTGAGTTTATTCTTTAAATTTTATCATACTTCCCTTCTTTGTCATGAGTATGAACTCATATATACATGTTTGGGTTGATGTCCAATGAAGATGTTATCGTGGAAGTCAAACAAGGGTGAAAATAGATGATAAGAATGCATTATCTATATTGTTAAGAAAAAAAGGTGTGTTATCTGTCTTGCAATATTCCTTTTCGTATTTTCTTTTTTCTTCCCTAGATTTTTCATTCAGAAGAAAGTCAAATACTGCACATCTTGACCAAGATACTTCTTGTTAACATATAATGCAATTAATAAATGAAAACCTTTTTAGTTTACAACATTCGTTCCAGTACTAAGGAGAATTCACTTCCTAGGGATCTACCTATCTGCTTTTCCCACTCCAGCTAGATCTTACCATTGCCAAACGTGTCTTTGCAGCACTGGCCTTCTGGCTTCTCGTGTTTCCTTTCAGGTTTGGGGTGGATCTTTCCTTTCTCAATTATCACATACTCGGTTAGGTCGCCCGTGTGTGTGTCTTTCAATCTAAGTGTTTGAGCAGGAACTCTCTTCATGAATCTCACTAAAATATTGTAGGAAAGCTATGATTGATTTTCTATTCGGGGATTCATTATTAACTATTATGACTCTATTTATTGGTGACAGAATTGTTGGACTTTTATCTAGGGAATATTTCCTTGACATGGAAATTATGGCAAATAAATTTTTCAGGGGTTTGTGGGACTACTCACCACCTAACCCCCAACAACAAATATACATGGAAGAAATATTGAAGTGTACAAAAAGAAGCCAATATATTATTCCAATTATTCACGATGATAGAGTGAAATAATTAGCATAAGATAGACCAAACCTTAGGCTTATCTACATCACCAGATTTAATATATGCAATGTATTATGTATGGTCCCTGGTAGTTTAACTTTTCATGGACTTCTTTGTTAAGAGGTATTATTTTCCATAATTTATTCAGTGTAAGTTGAAAAATATAGGCCACTTGAGTAAGCTGGGTTCATTTCATAATGTACATACTTTTAAAGATTTCAATAGTTTGCAAGTAGGTCTTTGTAATTTTTATATGCTTCAAGGAATTCAGACAGAAGTGGGTGAGAAAAGATGGAATACTTAAGGAAATGGATACCAGCGACATCTTAGTAAGAAAATAAAAACTGCATATTACTGTCCTGTTGTCTTCATGAGTTTTTGTTCTGAGTGTCTTGCCTTGTTCTAGGGCGGATGTCTATGGAAAGTGAAGTCATAGAAAATAGGATGTCTATGGAGGTCACTGTTATAAGCGAAGCAACCTAGATGCCTAATATATCAATGGATAGATACTTTAAGACCAGTCATGCCTATGTCATATTTGTTAAAAAAGGGAAAATAAACAGAGACAATTCAGGGTCTATTACTAGTTTATTACCTTTGTAATTTATGGAATTGAAGGATCTAGCAAGTATGTTAATCAACTGCATAAGGTGAAAATGACTGGTCTGGTTGTATTTTTAACCATGAAAGGTGGTGATAGTTGTTAAAACTCTGTTTTGTGTCACTGTGTCAGCTAGTGCACAAGAGGATGTGCGAGTGGCCAAAGCATGACATTGTTTGTTTGCTATCTCTATGATGGCAGTTTCTTTGCTTATATTGTTAGTGCTTACTTTCATCAAATGTGACAGGTTTCTTCATGACAAAATGAAGCCACAAGTTGTGCATAGGGATATCCGTGCAAGCAATGTTCTTCTTGACGAAGACTTTGGCGCCCATCTAATGGGAGTTGGTCTGTCTAAGTTTGTACCATGGGAAATTATGCATGAGAGGAGAGTCATGGCAGGTGGTACTCATGGGTATCTTGCTCCGGAGTTTGTTTATAGAAATGAGCTTACTACAAAGAGTGACGTCTACAGCTTTGGAGTCCTCTTGCTTGAAATAATTAGTGGACGAAGACCAGCACAGGCAGTTGATTCTGTTGGATGGCAAAGTATATTTGAATGGGCCACGCCTCTGGTGCAGGCTCATCGATATGTTGAACTCTTGGATCCTCTCATATCATCTTCTTCCTCTTCTCAGATTCCTGAAGTTGGAGTCATCCAAAAAGTTGTTGATCTTGTTTATTCTTGTACACAACATGTACCATCAATGCGTCCGAGGATGTCACATGTCGTCCATCAGCTACAACAATTAGCTCAGCCCCCTATTGTAAAGTAAGTGGATAAGAATTATAGTTCAGCATTTCTTTTGACTGAAACCAGCCTGCAATCTTTTATCCTAGTTGCAAGGGCTAATTGTATTAGGCATTATTTTTGAACTTGACATGCAAGGCCTTGAGGCTTGAACCAGTCCATGATGGTTGAGATGTAATACAACAACTAATCTGTATCACCAAAGCCAGTATGCTTTCATCTGCTGCTGCTTTGCATTGTTTACTTGTAGCACCTGTATTTCTAAAATTTGTTTCTGTTTCCTGTGTTAAAATATCACTATGCATAACAACCAATTCAATAAATGTCCATGAAATCTTG

>PGSC0003DMG400000179 G-LecRK

CAAATATTCTCTGAGCTTTTCAATGTTTCCTTAAAAACCAAAATCTTTTTTTCTTTAGCTAAGCAAAATTCATCACCATGATTCCACAGAAAGCAATTTTCATTTTACTTTTCATCACAATTTCTATGGCATTTGAGGTCAGTTCAGAAACAGCAAAATCAAGAATGCAAAATGTAAACATGACCCATGTAGCTGAACTCACTAAAGAGTCTTTTTCAGCTGAAAGAAAGTTGGGATCTTTAGAAGGGAGCATAAAGTTTGAATCTTTGAACAAATCTAGAGTTTTACTAAGTGGGAACTTCACTGTTTTGAGTGAAAACAAGACATTTGAAGTGGGGTTTTTCAAGACTAATGATGAGAGTAAATGGTATTTAGGGATTTGGTTTGCTTCAATTCCTACCCCTACTTATGTTTGGGTTGCTAATCGTGAAAAACCCATCAAGAATCCATTTTTAGCTACTTTGGAGATTAGTGAGGATGGGAGATTAGTTTTGAAGGAAGATTCAAGAACCATTGTATGGGAAACAACCAATTTGGAGAAAGCTAGTGATGTGAAGCTTTTGGACCAAGGGAATTTGGTTCTTGTTTCTAATGAAGGTGTTTTGGTATGGGAAAGTTTTGATTTTCCTACAGATACATGGCTTCCTGGGATGAACTTGACTGCAACAAAATGGCTGACATCTTGGAGAAGTACTAATGATCCATCACCTGGAAGGTACTCTTTAAGGCTCCAACCACATAGTTATGGTGAGATAGTTCTTTTTTATAATGGTACTTATCCATATTGGTCCACTGGGAAATGGAGTGAAAATGCATTTGTTGATGTACCAGAAATGACTGTTCCTTACATTTACAAGTTCAATTTTGAATCACCTTTTACCCCTATGGCATCATTTGGATACTCAGCGGTGACATTAGAGAATGGAATGCCTCCTCCTTTAACTAGATTCATGGTTGATTTCACTGGACAGGTTAAACAGTTCACTTGGATCCAACAGGCGCAAAGTTGGAACATGTTTTGGTCACAGCCAGAGAATATGTGTAGTACTTACGGTCTTTGTGGGAATTTGGGGTTCTGTAATAGTAAGTCATTGAATCCATGTAAGTGTTTACCAGGGTTCACCCCTTTGGATGGTGATTCATGGGATGCAGGGGATTTTTCGGGTGGCTGTCACCTCGAGAGTAATGAAGTTTGTGGTGAGAATGATAGGTTTGAAGAGGTTGGAATGGTTAAATATGATGGAGCAAGGGTAGTTTCAGTTTCTGGAACTAGAAATGAATGTGAGAGAGAGTGTTTAGGTAATTGTTCTTGTATTGGTTTGTATCACAATGAGAAGACTAATTTGTGCAAGAGTTTATATGGGTCATTGCTCAATCTTAGGAATCTTACTTCCGATGGCACTGTTGAGGATAAGCTATATGTTAGAGTTCAAGGAGGAGGGAATACTCGAAAGAATCAGATTCAAGTCAGATTACTTGTGATCGAAATGATCTGTGGATTTGTGGTAATTCTGTTAGTTGGAATTGGGACTTTCTTGGTAGTTAGGAGGAGAAGAGTAAGGAGGAAGAAAAAAGATGAGGAAGATGTATTTCCTATAATGAATTTGAAGGTATTTTCATACAAAGAGCTTAATGCTGCAACAAAGGGATTCTCGGAGAAGCTTGGTCATGGTGGTTTTGGAACTGTGTTCCTAGGGGAGCTATCGGATTCTTCACTCGTAGCTGTGAAGAGACTTGAAAGACCTGGTGGTGGTGAAAAGGAATTTCGAGCTGAGGTATGCACAATAGGGAACATTCAGCATGTTAATTTAGTGAGATTGAGAGGATTTTGCACTGAAAATTCTCATAGGCTTCTGGTTTATGAGTACATGTCAAAGGGTCCTCTTAGCGCATATTTAAGGCGAGATGGTCAGAATCTAAGTTGGGATGTTAGATTCCGAATTGCTGTAGGGACAGCAAGAGGTATTGCTTATTTACATGAAGAGTGTCGCAATTGTATAATACATTGCGACATTAAACCAGAGAATATTCTGCTACATGAAGATTTTTCAGCAAAGGTTTCTGATTTTGGACTTGCAAAGCTCCTTGGTAGGGACTTTAGTAGAGTGTTGGCAACCATGAGGGGTACCTGGGGCTATGTTGCACCCGAGTGGATCTCTGGATTAGCGATCACAACGAAAGCTGATGTATATAGCTATGGCATGACATTATTGGAACTGATAGGTGGTCGACGCAATGTAGAGTCACCACCTTCAGCTAAAGGCGAAGAGGGTGGAACAGAGGAGAAGTGGTTTTTCCCACCGTGGGCAGCACGTCAAATAGTCGAAGGAAACATTGCTGCAGTTATGGATGAAAGACTACATGGAACGTACAACCTCACGGAGGCAGAACGTGTGGGATTGATAGCTATTTGGTGCATTCAGGACGATGAATCAACGAGGCCTTCCATGGGAATGGTGGTGAAAATGTTGGAAGGAGTAGTCGAGGTGACAACGCCTCAACCTCCGAAGCTGCTCCAAGCATTAGTATCTGGTGAATCATTCCATGGGGTAGGAATTGATTCAAGTAACGGGACATCACGAGACGGAGGTTTCAGCTCGGGTTATAATCCTCAACGTTCAAGATATTCAAAAGACTCTCAAGCTTCAAGTTAGGTCTGCTTCTCAACTAGTATGCAAGTTGATTTTTCCCAGCTTCACTGAATGTAGAGTTCTTTTTTATTGTCTTCAAAATCCCACTGAGAATTAGTAGCAAAATGCTCATCATGGTATTTTGTAAAATGTATCATGAATATAGAAAATCTTGTTTACTTTGCTTAAAAATTGTCACTTTTTTTTTTTTTGTAAATTCCAAAGAACAAAAGCAAAACAAAACATACATTTCACTTTTGAAGGAATGTATGTTTGGATCAGAGGGTCGGGGCAGAGCTAGTAAATTACACTATTGATATATGGTTAAAATATTTTTTACGTATATATAGTAGACATTGAACCCTTTAACTTCTTCGTGTGTTTACTTCTTCATATTTTGAACTCTTTAATGAAAATCCTGACTTGCATAGGGTTATAGGAGAGGGCTTCCTAACCTTAGAATGAGACGGTAGTTGAAAATATCAAAAGTTATAATAACGGAATTATTTAATGGATATATTTTTATTGGTAGATATTTTGTATAGTGAACTTAAAATGAGATATAAGATATAGGTGTATAATATAAAGCATTAAAACATGTGTTTGGATTATACCAACCAAACATTATGCAAAATTTTATACTGAGATTATTTTTGTTACTATGGTAAACCAAATGACTATAATAAGAATATACCCTCTAACCTAATTTAATGGGAGAGTTAATTTTATTTTTTATAAATGAGTGTAACGTTTGTACCTGCCCAATTATAACCACACTTTACTTTCCTAAGTTTAATTTACATAACTGTCTTGACTACAACAATAACAACTAACCACTTGTCAATAGTTTCCTCTTTTGTAGCTAAATAATATAACAGAAATTGAAACAAGAAAGACCATATACTCACACAAACAAGCAAACAGAAAAAGGAGAAATGTTTTCCTAACTTGTCAACTTTAGTAACAAAATTTTCCCAAAATTAATTTTTTCAAAAGAGTGTAAAGATTGAGGGGGTTAAAGAAAATGGTCAAAGGAGAGCAGAGGTAGACAACTCTGCACTCTCACCAAAGAAGTTTGTTACCTTGGAAATGGAGACACCTGGCAGTTCAGGTGGGGAGGTGCCGATATATGGGGGCTCCTCGTCTATGGAGCCGACTGAGTTACACTCGGCTCCATCTACCATGCTGGCAGATTATCCGCCAGTACAAGGCTTTCAAGATTTGAAGAACTTGGTTAGCATGGAATCTGGGAAGTTATGGACTATTGCAGGTCCCATAGCCTTTAATATTCTGTGTAATTATGGGATTAATTCTTTCACAAGTATATTTGTTGGACATATTGGTGATGTAGAACTCTCTGCCGTTGCCATATCTTTATCGGTGATAGCAAATTTCTCATTCGGATTCTTGGTTAGGTTCTTTTTTTGTTCTTACTAATTCTCAAAGTTCTGTTCTTCTTTATATTATGTTATATGGGCATTGAGTATTCATATAGCCGACCCCAACTTGCTTGAAATAGCGGGATAGTTGTTGTTATTGTTGTGGTTGTTTGTAAAATGTAATACAACGAAGATATGTTGGATTTTGTCACTCTTTGATGAGACTTATGACTATAGATCACAAATAGTTTATTATGAAACAGAACTAGGATTTTGATTGTCAAATGATATAAAGTCTCAAAAAGGTTAGAATAAGCATTCTTTTAGGTTATTGGTTATGTTAGTTTTCTAATTCTTCATCTTTTTTTTGATGGATTATCAGTTAGGCATGGGTAGTGCACTTGAGACACTTTGTGGACAAGCATTTGGTGCAGGCCAAATAGAATTGTTAGGAGTTTACTTGCAGAGGTCATGGATAATACTTGTTGCTTCTTGCTTCTGCATAATGCCACTCTACATCTTTTCCACGCCGATACTGAAGTTATTAGGCCAAAGAGATGACATAGCAGAGTTAGCTGGAAAATTTTCGATACAAATCATACCTCAAATGTTCTCCCTTGCTATCAACTTCCCGACCCAAAAGTTCTTGCAAGCACAAAGCAATGTCGCGATCCTGGCATGGGTTGGGTTCATGGCTTTAGCCATGCATATCGGGGTGCTCTTTCTTTTTATCAAAGTTTTCCGATGGGGCGTTACTGGTGCTGCTGCAGCCTATGATGTTTCTGCTTGGGCTATTGCTCTGGCTCAAGTGGTTTATATTGTTGGCTGGTGCAAAGATAGTTGGAAGGGGTTGTCTTGGTTGGCCTTAAAGGAACTTTGGCCCTTCGTAAAGCTTTCTGTTGCATCAGCAGTGATGATTTGCTTGGAGATCTGGTATTTTATGACTATAATTGTTCTAACTGGTCATCTTGAGGACCCCGTCATCGCTGTTGGATCTCTTTCTATCTGGTACAATCTTGCAAATATGCACTTAAAATTTATAATGTGTTCATAATTGTTCTGTTTTTCAAGCTTAGATTCTTTCTTCTTCATTGTTTCAGTATGAACCTTAATGGATGGGAAGGCATGTTGTTCATTGGAATTAATGCAGCAATAAGGTATACTACTACTCTACTAGTTATAGTAGGACATTGTCAGTGCACAAAAGTTAAACTCTAACGAATAGTGTAACTGCTAACGGTTATTTTCATTTGTTGATATTTCAGTGTTCGCGTTTCCAACGAGCTTGGATCAGGCCATCCTAGGGCTGCAAAGTATTCCGTCTTCGTCACAGTAGCTGAATCTCTCATGATTGGGATATTCTGCATGGTACTAATCATATTAACCAAGGACCATTTTGCATTGCTTTTCACAAGCAGTGAGAAAATGCAAAAAGCTGTTTCTAAGTTGGCATATCTTCTTGGTGTTACAATGCTGCTTAATAGCGTTCAGCCAGTTATATCAGGTACCAAAATATGTGTGTTTCTTACTCAAACAATTCTCCACACACACATCAGTTGATGTGCCAATCAAATAGTAACTATTTGGTACAAATCTATGTTGCTTGGACTCTTCAAAAAATGTCAATGGGTGTGTGTCGGATTCTCGATTAGTAGAGTATTGTTGAAGAACCCGCACAATCAAAAGTGAAGAGTCCGCGCAACTTTAGGTACAAACTCAAAGAATAAATTAGAATGTTTCTAATATTGAAGGTGTTGCTGTTGGAGGAGGATGGCAAGCACTAGTGGCATACATTAATTTGGCTTGCTACTATGTCATTGGCCTCCCACTTGGGTTCCTCTTGGGCTATAAAACAAGTTTGGGAGTTCAGGTAACAACATAAATCTTCTTGTATTAATTTAGCTCCGGTTTGACCATAGATTAGTAGTTTAAAAACAAAGGTATACTAAATTGTTCTTTTACTTGATCTCAACTAGACATCTTTGCCCTCCAATTCTGAGTCTGCACAAGTAGATACGTAAACTTATATAAAGTTGAACAAGTAGACACATGTGTCCTATATGACATAATGAAAATCAGTTTGGGTGTGTCTCACGTGAATTGACACGTAGGACACATTTATCTACTTGTTCAACTTTATACAAGTTTAAGTGTCTACTTGTGCATACCGAAAGTTGTAGGGCATATATGTCAACTGAAGTCAAGTTAAAGAACGTTTATGTATAATTTTGAACATTGTTTGTTCATGAAATTTGGTCAGTTTTCTAAAAATATTCAATTGAGCTAGTTTTTGGATGAAATTTCACGCCTACTCACAAAACAAGTAAAATGCATATCCAAACATAACTTCATATTCCAAAAACTATTGTTCAACACAACTTCAAAAACTCACTTTTTCAAGTTTCAACTAAATCTATGGTCAAAACTCAAACTCAAACGTTAGCTTAGTTTTAAGTTTAAATTTTAGCTGGAAACTTTGATATAACGATAAAGTTATTTCGAATAAACTATTGATATTTGTATCCGGATAAATTGTCTACATTACACTAATTGTCTCTTCATTTTTTTTTTCTCTTTTCTGTAGGGAATATGGATGGGCATGATTTTTGGAACTTTTTTGCAAACAATAATTCTTTGTGTTATCGTATATAAAACCAATTGGAACGAAGAGG

>PGSC0003DMG400000915_ G-LecRK

ATGGAGTTCATCTTATTCTTTCTCATTTTATCTTTGTCCAGTATTCCTGTCTTTGGTTTTTTATCAAAAGAATTTGTCTATCCCAACTTTACTGCTTCCAATTTCCAATTTGTCGATAGCACCGGAAGTTTCTTGGTTTCTCGCAATGGAACATTCAAAGCTTCTATCTTCAATCCCGGTCCTGAACAAGTTAATTTCTATTTGTGTGTTATCCATGCTGAATCCAATACCATTATCTGGTCTGCAAATGGTGATGCACCTATTTCAACTTCAGGAATTATGAGGTTGACCAAGAAGGGTATTGGTATTACTGAAAAAGATGGCAGTTTTAAGTGGTCAACTCCACCATTAAAGTTAGCAGTATACGCGATGCAGCTAACTGAGACTGGAAATCTCCTTCTCCTTGATCAGTTTAATGGGACACTTTGGCAAAGCTTTGATCATCCGACAGATACAATTGTAATTGGACAGAAGTTGCGTGTTGGCATGATGCTTTCAAGTGCAATAGCAGGCGATGACTTGTCAAAGGGTCATTACAGGCTTTCTTTGTCTGCTTCCGATGCTATGTTACAGTGGCAAGGTCTGACATACTGGAAATTGTCCATGGAAACCAAAGCTTATACTAATTCAAATTATGCAGTGGAATACATGTCTGTCAATCAAACAGGTCTTTATCTTTTTGGTCAAGGTGGTTCTGTAGTTGTGATAATGGTGAACTTGTTACAATCCACTTTTAGAATTGCCAAATTGGATGGTTCAGGTCAGTTGATCGTTATTAGTTTTGTTGGCTCTGACATCAAACGAGATTTTGCTGGGCCAGTGGATGGGTGTAGAGTACCTTATGTTTGCGGAGGGCTTGGTGTATGCACCTCTGATGTATTATCAGATAATCCTAAATGTTCCTGTCCTGCTAACTTCAATCTTGGATCACATGATTCAAGCAGCTGTGTGCCTAGTGATAGTTCCTATTCTTTGCCAGTTTCTTGTAATTCCACCAACCACAGTAATCTGTCGAATTCTTCCTCCATGTCGTACATAGGACTTGGCTTTGGTATGGACTACTTCACTACTGATTTTACCAAGCCCTTCAGGGATGGGGTAAATTTGTCCCTTTGTCAAAATTTCTGCTCAGTAGACTGTTCATGCTTGGGGATATTTTATGCAAACTCGTCTGGTTCTTGTTACAAACTTGAAGATGAAATAGGGTCAATTATGTTAAGGACTAGTAGAAATGATGATCTGTTGGGATTTGTTAAGACCCTGGTTGGAGTTTCAACAACCTTAGGAAACAATGATAATTTTGATCAGGAAGCTGCAGACTTTCCTTTAGTTGCAACAGTGATCTTACCTTTCACAGGAATCTTCCTACTCATGGCGCTGGGTTTCATTATGTGGAGAAGATCTAGACCCCAAAGAATGGGAAAGATAGAGTCGAAAAAAAGCCAACCCAGTTCTCCTTCTTCAGAGGATCTGGATGCCTTCTCCATCCCAGGATTACCTGTACGGTTTGAGTACAAACAGCTTAAGGCTGCAACTGATAATTTCAAGACTCAGATCGGCACAGGAGGCTTTGGTGCTGTATATAAGGGTGTGCTCCCTGACAACACTCTGGTTGCAGTAAAGAAGATAATAAATTTAGGCATTCAAGGGCAGAGAGACTTTTGTACTGAGATTGCAATCATTGGCAATATTCACCACATAAACTTGGTCAAGTTGAAAGGTTTCTGTGCTCAAGAGAGACAACGCCTATTGGTTTATGAATATATGAACCGTGGATCGCTTGATCGCACTCTTTTTGGTCATGTACCTGTATTGGAATGGCAAGAACGGGTTGAAATAGCTATTGGTTCTGCCCGCGGACTTGCATACTTGCACAGTGGCTGTGAACAAAAGATTGTTCACTGTGATGTGAAGCCAGAGAACATTCTCTTGCACGATAATTTTCAGGTGAAAATATCTGATTTTGGCCTCTCCAAGCTGCTGAATCGTGAAGAATCCAGTCTATTCACAACAATGAGGGGGACTCGTGGCTACCTTGCTCCTGAATGGCTCACAAGTTCTGCAATCTCAGACAAAACGGATGTGTATAGCTTTGGAATGGTACTCCTTGAAATTGTAAGTGGAAGAAAAAACTGCTCTTTAAGAACACAAAGTCACAGCCTTGATGATAGTGCTACTGGAGATCACTCATCATCTTCATCTGCACAGGGACTTGTTTATTTCCCTTTATTTGCATTAGAGATGCATGAGCAAGGAAGGTACCTTGAACTTGCTGACCCAAAACTTGAAGGGCGGGTCAGTGGTGGAGATATTGAGAAGTTTGTGCGGATAGCATTGTGCTGTGTCCATGAGGAGCCATGTCTCAGGCCAACTATGGTTAGTGTTGTTAGCATGTTAGAGGGAGAGATACCACCGACTGCGCCAAGAATGGAATCTCTAAACTTTTTACGCTTCTATGGGCGACGGTTTTCAGAGGCATCAAACATGGAAGAAACTGGAGGCCAGATAGATGCCATGTTATATCCGCAGGCAAATACTTCTCATACAACCTCTCGGAGCGTCTCTAATGCTTACTTCTCTTATATTTCGTCCCAACAGATCTCTGGTCCAAGATAA

>PGSC0003DMG400001059 G-LecRK

AAAAATTTCATACTAGAACAATAGTTCTTTAATGGCATGCCTAAAAATCCTACTATTTTTCCTCTTCATGAACATTTCTTTTGCCCAAAAAAACACAATTTTTCAAGGCCAACACCTTAGAGTTGGAGAAAAATTGGAATCAGCAAACAAGGAATTCAGGCTTGAGTTTTTTAGCCTTGATGCCAATAAAACACAATATATTGGCATATTCTACAATTTACCCTCTAACATGACCTTATTCCCCGGTGATGATCGACCCGTTTGGGTTGCAAATCGCGATATTCCGATACAATATGCATCAGGTAACACTCTCACATTGGATGATGATGGAAAGTTGAAAATTTTTTATGGAGGGGACAGTTTTGTTATGTTAAGTTCTTATAATGTAACAACAAGAAATGCAAGTGCTACACTATTTGACAATGGCAATTTTGTGTTGGTGGAATTGAATACTAATGGTTCAGAGAATAAAACATTGTGGCAAAGTTTTGATTATCCTACTGATACACTATTACCTGGGATGAAACTTGGGAGAAATGTTAAAACAGGACAGATATGGTCACTGACTTCTTGGATAAGCAAAGATGTACCTGCCTCGGGGTCATTTACGATTGGTATAGACACGAATGACATTGACCAATTGATAATATGGTGGATGGGAAGTGTGTTTTGGACGAGTGGACGCTGGGGAAATGAGACGTTTGGCAATGTATCGCGAGTATCACACTATGAGTATGTTAACTTGAGTTTTGTTTCGACAGAGGATGAGAAATATGTGACTTATTCGGTCAGTGAGACTCGAACTTTGTCTAGATATGTAATGGATACATTTGGTTTTATTAAGGAAAGAGGAGCAGCAGGACCTTTTGGGGTGTGTTTTTACAAGCCTAGTGCTGGTTGTGTGACACAAGAATCGATTGGGTGCTCGATCAGAAACGATTCATGGTTCAAGAGGAGGCCAAATAGTGTCTCTGGTAATAGATTTAGGTTTGCAAACAACAACATGAGTTTATTTGATTGCAAGAAACAGTGTGAGAAAAACTGTTCTTGTGCTGCTTTTGCCTCTATCACAGCTACTGGTACTGGTTGTGATATTTGGAGTAACGTATCGATTTTGAATTCACTTGAATCAGATGTTTTCACTCTTGATGATGAAAGAGGTATATTCATATTTACCTCATTGTCTTTTTTATGTTGTTGTCGTGTGACCAGGAGGTCGCGGGCTCCAGCCTCTTGCAGATATGCAGGGTAAGGCTGCGTATAATAGACTCTTGTAGTCCAAGGAAGTGCATTGTTTCTATGTCTAGCAAGTAAAGTTAGTAAAGCGGACTCTAACGATGAATTGTGCAATGTGTATTCATAGATTTGGTGCCAATAAGAAATCGGAACAGCACAGATGTACCAAGCACAGTCTCACTTCCTCCAGATTTAGCACCAGAACCATCTCCTAATTCAACGTCAGGATCGTCTAAAGGAACTTGTATGTATTGTGTCTAAATCTTGCTCTTCAACTATTGTATATGATTGAAGGTGAAAACCTTTTTTTGATTGTTTCCTCCATATTATCCATTTAAGAAAGAACTATACATTTTAGGCACGTATTCTTTTGTAGTATCATCATTACACAATCTAGTCCTTGTTTCGAGTATATCCAGAGAAAGGTCATTTAGATCATCTGTGTTGATGTTGTTACTGCTCCTTTCCTGAATGTTCTGCACTGTTTTCCTTTTTACTTGTCATGTTCTCTTCACTGTCGTTTTTCCATTTCCATAACTACTTTTATTTGTTGCTCTTGAGCCGAGGGTCCTTTAGAAACAGTCTCTCTACCTCCATGTGGTAGGGGTAAGGTCTGCGCACACTCTACCCTCTCCAGACTCCACTTGTGGAATTTCACTCGTTATGTTGTTTTGTTGTTGATGCAGCAACAAAATGGTGGATATGGCTTATAGCTATAATCGGTTTGACACTATTCGTAGGACTTAGCTCCCTATGCTACTTTCTACACGGAAAAGGTGAGCACTCGAAGAATTTAGCTAGAAGCTTGCACCAGTAAAACTCAACTATGTTGCTCGGACTCTGCTAAAATGCTGCCACAGTACCCCTGTCGGATCCTCAAAAAAAAAAACTCACTACTTTTGAAGGATCTGACACACACCAAACAGCATTTTTTAAAAGTCCGAGCAATATAGAAATGCTCTATACTATGGGTTCTGATCCCACAACTTCCTCTTTCACTATCAGGGAAAGCAAAAGCAACAGCTTTGCTATTGCTTAATCAAACAGCAAATATATCTAAAAAGAGAAAAATAGACAAGAAAATGAGCCATGAAGTGCAGTTATACAGCCTCGAGAGCCTAGCAATCGCCACAGACAATTTTTCACCAGGAAATAAGCTTGGGGAAGGTGGATTCGGACCAGTTTACAAGGTAAACATGAAACCTTTTATATGAATATGATTCACGTAAGCATAAATACTTAAACGAGTCATTATTTTTCAGGGAGAAATGGCTGATGGGCAAGAAGTAGCTATAAAAAGACTTTCGACTAGTTCAGGACAAGGCCTATTGGAGTTCAAGAATGAAATTCTATTGATTGCAAAACTTCAACATACTAACCTTGTTAGGCTTCTAGGATACTGTACTCTACGCGAAGAACGGATTATAGTTTATGAATACATGCACAACAAAAGCCTAGACTTCTTCCTCTTTGGTAAATATCATGTTTTTCATCTTCAACAATTAGAACAACTTTGTCAAGAAATTTCATTTTTTACAATGATATGAACAGATAGTAACAAAAAGGAGCTACTAAATTGGGACACACGATTCAGAATCATCGAAGGGATTGCTCAGGGTATACTGTATCTGCATAAATATTCAAGGTTGAAAGTGATTCATAGAGATTTAAAAGCAAGCAACATCTTACTTGACGCGGAAATGAATCCAAAGATATCAGACTTTGGCATGGCGAGGATATTTGGAACACAAGAATCTGAAGCAAACACTAAAAGAATTGTTGGAACACAGTAAGTTAATAAGCAATGTTTTCATATACAAGTTTTTATACAATATTACTTGAAGTTCTGAATACATTTTTCGGATACTACTTCAGTGGCTATATGTCTCCAGAGTATGCCTTGAGAGGCATTGTTTCAACGAAAACGGATGTATTCAGCTTTGGAGTTTTACTGTTAGAAATTGTTAGTGGCAAGAAAAACAACAGCTGCTATGATTCCGAACATCCACTAAACCTCATAGGACTGGTAAGTTTCTAGACTATATTATGAACATATCTCGCGAAATGGAGAAACATATAACTATTAGTACTATGTTTTTAGGCATGGGAATTGTGGAGAGATGAGAGAGCTTTGGAGCTAATAGATGCAACACTAATTGAATCATGCTCGCGCGATGAAGTAATGAGATGCATTCATGTAGGACTTCTTTGTGTGCAAGATTATGCAAAAGATAGGCCTTCAATGTCAAATGTTGTTTCAATGCTTATGAATGATACGAAACAACCACCACCACCACCAGAACGACCAGGATTTTTCATCGAGAGAGGGGATCAACGTGCAGAGATCTCCGAAGAAGTAGTCAGATACTCGATAAACGGGCTATCAATATCAGAGTTGAGAGCAAGATAAGATACAAGATGGTCAATTGGGAAGTAACTTATTTAACATTCTTTTGATGAAGTGTAAAAAGTCACCAACTTCAAGTACATACTTATAGGACTAATTTCGAGTCCTCTTTCTTTGATACTATTACATTATGATGCATTTTGTGAGTAGAATATTTGAGACGGATCCAGAATTTAAGTCAAAGATGGTAAACGAGATAAGGCAAGGCGAAGTTATGGCAATGGAGACTTCACCATTACCTTCCCCCTCCCGCCTGATAGCAACGTCACCAGTTACATTTAAAATGTTAGAGAATACACTTGACACTCATCGATTATATAAGTCGAAAGAGAATATCAGAGGAGTTTCTGAACTATCAATATTCGTATATTAGAAAAAAGAGGTAACTGAATGTACATATATTTTCTCATTCTGAGCTTTACATGTTCCACAAATGGGCTGTGAATTAGTTTACACTCAATTAACTACAAAAATAAAGAACAATTTTGGTTTTCACCTAAACTCACTACAATTCTGATGATTTAGCTTCATCAATATTGTTATCTTTATCGCTAACAAAAACTAAACGATCACCATCTCTGATCAACTCTATTTCGTCGATTTCAGCTCCATCTTTGCTTACAACTTTGGCCTGAGAGATACCATATCTACTAGAACCAATTTGAAGTAGCTCTTTAAAACTCTGTGGGAGTAAAATGAGTTTTCCCCCGTTGTCTCCTTTCTCAGGGCAACACACTGTCACTCGGGGAGCATAAGTCTTGGTTGTCATGACACTTACATTACTGTCGTTTGCAGATAAAAGCACGTCGTGCTCATTGGTCTGCTTTGCTGACATTATCCCGAATAATGAGTTGTGGAAGTTATTAGACCTACGTCTAGGACGTGAACGCCCTAATGATCCGTCTAGTCCTAGAAATGAGACTCCATGGGATGCAGGGGTGATTGTTGGTTCACTTTTGAACCTCCCAAGAAATCTAACCCCTCGTTCCTCGGGGATAGGATCAACGGATCGAGTTCTCATCATTACCCTGGATTCAAAGAGTTCTTTGATGTCTTCATGTCCTTGTTGCTCAGCAAGGTCCCGGGGGGTCCAATTATGTTCGTCTGGTTGATCAACGTTAGCCCCTCGATCCAAGAGATACTTTACTATTTCTATGTTACCTTCACAAACAGCAACATGGAGAGCTGATGAACCATTGATCTTGGGAAGCGTGACATCCCCGCCATAACGGACAATCTCCTTGAGCAAACTCAAGTTGTTTTGTTCAACAGCAACGCAGGCGAAATGTCCCACATCACCAGCTGATAGTTTAGCACCGTTGTTAACGAGTAACTGGATAACTGACTCGTGCTTCCCCATGATAGCCTCCCACAATGGCACATTGCCTTCTGAATCTGTGTATACACAAGAAACAAAATCAAATTCTTATATCTGGCTGTGTCGGTGGACATGATAGGAAAAGGAGACAACTGATAGAGTGTGAAACTCGCAAAAAAAACGATAGTTGAGGTGTGTTTTTGACTATAAACTCTCTAATCTTTGGACAAAAACAAACTTTTGGCAACAACAGAATTTGAAAAAAGAAATGGAGCATACCTCTACTATTGGGATCAGCCCCGAAATCGAGCAGAAGAACCACACAGTTCTCAATGCCTTTAGACGCAGCGATA

>PGSC0003DMG400002017 G-LecRK

ATGAATGCGATTTGCGTAGCAACAGATACTATATCAATGAACCAACAAGTTGTAGATGGTGAAACATTAGTTTCAGCAGGAGGTATCTTTGAGCTTGGGTTCTTCAGTCCTAATGGATCCCAAGGGCGATATGTCGGGATATGGTACAAGGGATTACCAGTTGAAAAAGTTGCATGGGTTGCAAACCGCGAAAGAGCATTAATTGGATCATATGGAGTTCTTATGATTAATGCAAGTGATGGAAATTTAGTCCTTGTGGATGATAAAAATATGTCTGTCTGGTCTACAGAAATTAATCTGGTATCGAACAATAACACAATTGCTGTACTGACTGATCAGGGGAATCTTGTTCTCCGCGATAAGAACGAGTTGAATGAATCTGTACTGTTATGGGAGAGCTTCAATCATCCATCCGATACTCTCTTGCCTCGTATGAAGATTGGGCTGAATACTAGAACAGGGGAGCAAATAGTCGTCACGTCGTGGAAAGATGAGAAAAATCCATCACTGGGAAGTTTTAGCACTGCTGTGGTGGATCCTCATGAACTTTTACAAGTACTTATTTTGAATGGCTCATCGAAGCATTTCAGATCGGGGCAGTGGAACCGGAGAAGTTTCATAGGAGTGCCTAGCATGACTAAAGACTTCCATAATGGCTTTCAACTACTTATGGATAATAATGGGGAGGAACTTTATTTTACTTATGCTATACTCAATTACTCTGCTCCAACATTGATATACATTGATTCCTACGGAGCGATAATTCAAATGGATTGGGATGCAGAAGAGAATAAATGGTTAAAGGTGCAGGTAGTACCTAATGGTCAATGTGACTTGTATAACACATGTGGTCCTTTTGGAAGTTGCAAGGATAATGATTCACCAATATGCAAATGTTTGACGGGCTATGAACCAAAGTCTATCGGTGAATGGAACGAGGGGAATTGAACTAGTGGTTGTGTCCGGAGAGCACAATTCCAGTGTGAGGAGACGATCACTAGTACGAATTCTGATGATGAGAGAAAGAGAGGGGAAAAGAATTCTGATCACTTGTTTGGCATATGCTTATGTGGATGGAATAGGCTGCATGATTTGGGGAAAAGAACTTATTGACCTCCAAGAGTTTGCCCGTGGTGGAGAGCAACTATACATTCGCCTCGCACATTCCCAACTTGGTAGATACAGATACACTAATTCTATGTTGCTCGGACACTCCAAAAATGTTGTTGCACCCGTATAGGATCCTCCAAAAAACACACTATTATTGGAGGATCCTACATGCACACAACAATACTTTTGAAGAGACTGAGCAACATAGCACCAATTTTCTCTATAAACATCCCATTTTATTACTACTCGAACGAAACTTCACTTCATTTTGGATTTGTTTTAGTAGTTTGCCTATGTTATTTGTCTCTAATCATTTGTTGGATTCACATTCTCCTTTTATGACTGTAGTGTTCGGGCCAGCTTTAGTGCGCCTGAACTATTCTCAGAGACAGAGCTAGGATTTTGAGTTTATGGATTTTGGATTCTAGAAAAAACAAATTACTGGGTCTTGAATAAATTATTTATACTTATTAAATGGATTCTTAAACACAAAAAATATATGGTCTAGCCCAAAGCTACTGGATTCTACAGACTCTGTAGACAGGCTTGTAGCTCCGCCTTTTCCATCGAGTACTTGCTACCTCTCACCAATAGGGATACTGAGTAATTCTGCTCACTAAGGCATAAGTAGATGTGAAAAAACACCTAGTATGGTTTTGCTTCCATCGGAATCTGAACTTGAGACCTCATGTTTTAGTTTTTTTCTTATCGAGTTGCCTTTTGTCATGCAGATAAGAGGAGGATGCTAACGGGAGCCCATATTGCAATTATTGTCTCAGCGACATTCGGCCTGATTCTTATTGTTTGCACTATATATGTATTGAAGATCAGGAAAAAAGGTAAAAAAAAAAAAAAAACAGTCAATTTTTACAGTTGTTAGAGGCACAACTTTTCGCTATCTTTTGGAAAGACTTTAATTTTTACTAGATTTTATCAGGCATTGGGAGTTTAGTCCCTTTTCGCATAGCTTCAAAACACTCTTCAGCAACAAATTTGCTTCATAACAGTTTTACTCAAGGTAGAAGTACAGAATTACCAGTTTTCAGCTTCAAGTGCATAGAAGCTGCTACTAAGAACTTTTGCTATCGAAATAAACTTGGAGAGGGGGGATTTGGACCTGTATACAAGGTATATATATATATATCACTCGTGTGTGCTTTAATCATCGAAAAAGTAGAGCAAATTTCACATGGTTTATACTCATTTGCTTATTAGTTGAAGTCATGTGTGATTCATATATAGGGGGTATTAGCAAATGGACAAGAGGTTGCTATTAAACGACTTTCGAGGAGCTCAGGACAAGGCATTGAAGAGTTCAAGAATGAGGTTACACTGATTTCTGGACTTCAACACAGAAACTTAGTCAAACTGATGGGATTTTGCGTTCACGGGGAGGAAAAGTTGCTGATCTATGAATACATGTCCAACAGAAGCTTAGACGTTTTCATATTCGGTGAGCTAATTAACAAGAGTACTTAAACAGTACTTTTCTTGATGAATTATGATTAAATTACTCTTTGTGCTTGGCAGGAGGATCAAGATCTAATGATAAGGCGGCATTGAGTTGGAATCAACGATATAGTATCGTTGAAGGGATAGCTAGAGGTCTTCTTTATCTGCATCGTGATTCACGACTAAAAATCATTCACAGAGATCTTAAGACAAGCAATATTCTGTTAGACGAAGATATGAATCCCAAAATCTCAGATTTTGGAATGGCTAGGATTTTCGGAGGGAATCAAACTATAGCAAATACCAACAGGATTGCCGGTACCTAGTAAGTATTAATTGTTCATATATAGCAGTGAAATTCATCATAACTAAATTGTATATTTGTGTATCAATGCAGTGGTTACATGTCCCCTGAATATGCAATACAAGGATTTTTTTCCGAGAAGTCGGATGTCTTCAGTTTTGGTGTCATATTGTTAGAGATTATCAGCAGTAAAAGAAACAACAGCTATTGTCACTCTGAAGACTCTCTCAACCTCCCGGACCAGGTGAAAAAAGAGAAAAACTGAAAGAATTTCTCATTTTCTCATATACAGTCAAACCTCTTTATAATAACGTAATTTCTTGTCTAGCTGCTATAGCAAAAATGTAGTATAAAGAACATATAAATATAACATAATATGGAAGTTCGAGAAAAACTAGGCCGAGGATGATTGTTATCGAGAGGTCTGAATGGACTTGAATATTCTTCTATCCTATACTACTACTATTTTCACATGACAAAACTAATATCAATGGATTTTATAGGCATGGCAACTGTGGAATGAAGGCAAGATACTCGAGTTTCCGGATCCAAAGGTGTTTGGTTCACTTGAAACACCAACACAAGTTGTTAGATGCATCATTGTGGGATTGCTTTGTGTGCAAGAACATGCAAGTTACAGGCCAAATACAGCTACAATTGTTTCCATGCTCTGCAGTGACACACTTCTTCCTCCGCCTAAGCGGCCGGCATATAGCAGCAGAGGCTCTGATGATCCAAGCATTTCACCTTGCTCCAGAAATTCCATTTCTATTTCAAATGTTGAAGGACGTTAA

>PGSC0003DMG400003313 G-LecRK

TTCAAAATTACATATTTCAAAACTTCTGTATGAATTAGGATTTCATACTATAACATGAATATAAAATTTTCAATTCTTGTTTTACCTTTACTTGTTGTTTCATCTTATTGTTCAACAATCTTTAGTTTATCATCTGAAGGTTCACTTTCTGCTCCACAAGATTTAATTTCGTCTCGAAATGGAAAATTCACTGCTGGTTTCTACTCTGTTGGTGATAATGCATATATCTTTGCTATATGGTTTACTAAGTCATTGGCTGATGGAACCAACACAATTGTTTGGATGGCTAATCGAAATCAACCGATAAACGGACAAAAATCACATCTGTCGCTGTTTAAATCAGGCAACCTTGTACTGATTGACGCGAATCAGATCAATGTTTGGGAATCTGGTACACAATCATCAACTTCCTCTGTTGAGTTAAGATTGTTGGATAATGGAAATCTTGTTCTTGTAACTTATGAAGGTCAAAGATTATGGCAAAGCTTTGATTCACCAACTGATACACTTCTTCCTGAACAACCACTTACGAAAACCTCAAAGCTTGTTTCGCGTAGAAGCTTGACTAATTTCTCCTCTGGATTCTATCAGTTACATTTTAATGAAGATAATGTCCTGCATCTTGTCTTTGATGGAATAGAGATGACAAGTGTGTTTTGGCCAAGCCCGTGGTTAATCGTTTGGGATGCAGGTTAGTAGAGACAGATTCAGAATTTAAATTTGATTCGTTAAATCCTTAGTTTGTTGTCACTCAACTCACATATATTTTTGCATGAAAAATACTAGTTTTAGTTGAACTCATTATATGTTTCATTTGAGTGTATTGGTTAGTAAAGATGGATTCAGGATTTAAATTTAGTTGGTTCAATCTTTAGTACAACTCATTTTTTTTCGGTTTAACTGAACTCACTGCTGATCATGTCTGCAGGTCGATCCACTTACAATGACAGCAAAACAGCAGTTCTTGATCGTTTTGGGAATTTTACATCGAGTGATCGATTTGGATTCCAATCTGCTGATTATGGTGTGGAATTGCAAAGAAGATTCAGCCTTGACGTAGATGGTAACCTTCGATTGTACAGCCTAGATAAGCTGAGCAACATTTGGAAAGTTTCTTGGCAGTTATTTCCAGCAGCCTGCAGGATTCATGGAGTTTGTGGATTGAACAGTTTGTGTTCCTATGATCCTGTCTCTGGCAGAAAATGTTCTTGTGTACCAGGATACAGGATGAAAAATCCTAAAGATTGGTCCTATGGTTGTGAACCAGAATTCAGTATCTCTTGCAACGATACAAGCTCGTTGGACTTCATTCCACTTCGCCATGTTGAGTTTTACGGGTATGATATTGCATATTTTCGCAATAAAACTTTGCAAGAATGTAAAGATTTGTGCTTGAAACACTGTGATTGTAAAGGTTTTGAATACAAGTATGTTGAGGGTAATGGCACCTATGGTTGTTACCCGAAAACACTTCTGTTCAATGGCTATGTTCAAGCTAGTTGGCCTGATTTTGTTTACCTTAAAGTGCCTAAAGCACAACGTGCCAGGCAGGAGCAGTATGAAGGAAACCTTCAATGTGACGATAAAAAGGTGATGTTGGATAGAGCTTATAGAAGGAAAGAACATGGTGGATGGATGAAGCCATTCATTTGGTTAGTTGTTGTAGCTGGAGTCTTTGAGATTCTCTGTGTTCTCACTTACTTTATTAAAACGAGAAAAAGCTCGAATGAAACCACACAAGGCTACCTTCATCTTTCGACAAGATTCAAGAAATTCACCTATGCTGAGCTTAAAAAGGCCACTTCCAATTTCAGTGAAGAGATAGGTCGAGGAGGCGGTGGTATTGTGTACAAAGGGAAGTTGTCTGATGACAGAGTTGCAGCTATAAAGTCTCTCTATGGAGCCAGCTATCAAGGAGAAGCAGAATTTTTAGCTGAAGTGAGCACTATAGGCAACGTTAATCACATGAATTTGATCGAAATTTGGGGATATTGTGCTGAGGGAAAGCACAGGCTTTTAGTTTATGAGTACATGGAGTGTGGTTCTTTGTCAGATAATTTGAATGCCAACAATCTTGATTGGGAAAAAAGGTTTGAAATCGCGTTGGGAACGGCTAAAGGACTTGCATACTTGCATGAAGAATGCTTAGAATGGGTTTTGCATTGTGATGTAAAGCCTCAGAACATACTTTTAGATTCCAATTACAAGCCAAAAGTTGCTGACTTTGGGCTTTCAAAGATATTTAACAGAGGTGGCTTGGATCATTCAAACTTTTCCACGATAAGGGGGACGAGAGGATACATGGCTCCAGAATGGGTATTCAAGATGCCTATTACATCCAAAGTCGATGTCTATAGTTATGGCATTGTTTTATTAGAGATGATCACAGGTAAGAGTCCAGAAGTTTGTGCTGATGGTGACTCGGGGGATGATGATTCTATGGGACTAGGATTGTTGGTCACTTGGATTAGAGAAAAAATGCGCGAAAACAGTGAAAGAAAATCATGGATTCAACAAATTGTGGATCCTGCATTGAATGGTAAATTCGATTTGGAAAAAATGGAGATCTTATTAGAGTTAGCTTTGCAATGTTCGGAAGAAGATAGAGATGCAAGACCTACCATGTGTGAGGTGGTGGATAAAATGCTTCATCCAGAAAATTTCGAGTTAAAATTAGACATGTTAATGTAGTTTCTCAAGATGTAAAATCTTGCCAAAGCTGATATAGTACAAGCTTATATGTAATGTCTCTTGAGTAGCTAGTACTTTTGAATTCCAAAATGTAAATCCTTATAGAATGACATAGTTTCTTAACAATATTGTGCATTATAGCTACTGTTATGGTCTTGTGCATCAGTGCTAGATGGAAAGTGTGCAAGAGTACAT

>PGSC0003DMG400003982 G-LecRK

GAAAAGATAAAATCTATGGCTGCTTTGCTATGGTTATTAATCTTATCTGCATTTCATGCAGTGACACTAGCACAACAAAGACAGTTTAATATAACTCTTGGCTCTTCTCTAACACCTACTGCCAATTCATCATGGTTTTCACCGTCCAGGCGCTTTGCCTTTGGGTTTTATGAACAAACCAATGGCTATGCTGTTGGAATCTTAATTGTCGGTATGCCTAAGAAGACAGCAGTGTGGACAGCAAATAGGAACAGTCCTGTTGTTCCGAGCAACGCTGTCTTGCTCTTAACTAATGATGGAAGGCTCATTGTACAAGTAGGAGGACAAGAAATCACTGTCGTAAATCTCTCTGGTCAAGTCATTGCTTCAGCTTCCATGCTGGATACTGGCAACTTTGTGCTCTATAATTCTGATCACAGTATCATATGGCAAAGTTTTGACAATCCAACAAACACCCTTTTACCAGGTCAGCATATATCAGCTGGACAAGAGCTGTTCTCGAGTGCTTCAGTAGCAGATGATTCATACGGTATTTTCCGTCTCAAAATGCAAGATGATGGGAACCTCGTTCAGTACCCTGTCAATACACCAGATACTGCCCCATATGCTTACTACTCAACACGCACTTCTAGAGTAGGGAATACTGTTACACTGAATCTTGATGATGACGGCCTCCTCTACTTGCACAATTCCACTAAGAGGCTAAGAAATCTAACCAGAGGAGGCTACCCTAGAGAAAGGACCATCATCTATATGATGAAAATTGATGCTGATGGTATTTTTCGAGTCTATTCACATTCTCTGAACCAACAGAACAGCTCCGTGATATGGTCATCAACAAATGATGGCTGTAATCCAATAGGCCTTTGTGGGCTTAATGGGTTTTGCACCAATATAGATGATCAAGCTAAATGTTTGTGTCTTCCGGGATTTGATTTTGTAATGCCAGGAAATTGGAGCGCAGGCTGTGAAAGGAATTTCACCGCAGAAACCTGTCGGTTAAAGGAGAGCACTTCCAAATACTATGCTATGAGAACAGTTGAAAACACAGGATGGGAAGATAGTACTTATGTTGTTTTGGCCGGAACGACCAAGGAAGATTGAGACCAAGCCTGTCTACAGGATTGTAACTGTGAGGCTGCTCTGTTCAAAGACAGAGAGTGTAGAAAGCAAAGGCTGCCTCTGAGGTATGGAAGGAGAGAACTGGATAATTCTAATCTAGTATTGGTCAAGGTGGGCACTAATTTCATACCCAACAAAGGGGTGCCTAATCAGACGATTGAAGAAACAAAAGGCAAAAAGCTAAGGATAGACATCCTAATTGCCAGCATTACATTGGCTGTCTTTGCTTTGTTGGTGTTAGGCATCTCTGGATTTCTCATTCACAGAATTCATGTCTGGACTTACAGAAAAATCCAGGATAGTAGAAGTGTTCAGTTATGTGAGGATGTTGCTCCACGAGCATTTTCCTATGCAGAACTAGAGCAGGCAACCAGTGGCTTCAAAGAAGCGTTAGGGAGGGGGGGATTTGGAACAGTATTTAAAGGGATATTGGCAGAAGGTCAGAAAGTGATAGCTGTTAAGAGACTAGACAAAGAATTGGTAGAAGGGGAAACAGAGTTCCAAACTGAAATAAAAATCATTGGAAGGACACATCACCGAAATCTAGTCCGTCTCCTTGGTTATTGCCTGGAGGGATCAAGAAGGCTTCTGGTGTATGAGTACATGACCAATGGGTCACTTGCGGACATACTTTTTACACCAGAAAAACAGCCCATGTGGGAGGAAAGGTGTGGCTTTGCTAGAGATATTGCAAGGGGCCTCCTATATTTGCATGACGAGTGTGACACACAAATCATCCATTGTGATATAAAGCCTCAGAACATACTCATGGATGATCAGTACTGTGCAAAAATATCTGACTTTGGAATGGCCAAGCTCTTAAAGAAAGACCAAAACAAGAACTTACACAGGGATAAGAGGGACCAGAGGCTATGTTGCACCAGAATGGCACCGAAATCTGCCAGTCACTGTGAAAGCAGACGTTTACAGTTTTGGTGTTGTCCTATTGGACTTGATTTGCAGGCGGAAGTTTGTTGATATGAGCCTTGATGAAAATGAGGCAATTCTTGAATTCTGGGTTTACAATTGCTTTGATGCAGGAGAGCTAGATAAACTAGTCAGGGACGAGGAGGTTGATAGAAGACAATTCGAGAGAATGGTGAAATAAGCATTGGTGCATTCAAGAGGAACCATCACTTCGCCCTTCAATGAAGAAAGTTCTACTGATGCTGGAAGGAACAGTAGAAATTCCAGTGCCTCCTAGTCCTTCTTCTTTTCTAAGTGCCATATAGTTTTAAACTTCTTCACTTTTTGTTTCAAGTATCCCAACTTTCCCATAGAGAGCAGAGTACTTTATTATAATTGTATCAATTTTCTACATCATTCTCCACCTCTTCTAGCCATCAGTTATCTTCTGCTGTGGTGTATCTCAAGGCTTATGTTTCCATTTTGAGATTTGTCCATATCAACACTTCTCTTCAAATTTACTTGTAGTATTTGAATTGTAAATACTAATTT

>PGSC0003DMG400003999 G-LecRK

ATGGCTGCTTTGCTATGGTTATTAATATTATCTGCATTTCATGCAGTAGCACTAGCACAACAAAGGCAGTTCAATATAACTCTTGGCTCTTCTCTAACACCTACTACCAATTCATCATGGTTTTCGCCGTCCAGGCGCTTTGCCTTTGGGTTTTATGAACAAAACAATGGCTATGCTGTTGGAATCGTAATTGTCGGTATGCCTAAGAAGACAGCAGTGTGGACAGCAAATAGGAACAGTCCTGTTGTTCCGAGCAACGCTGTCTTGCTCTTAACTAGTGATGGAAGGCTCATTGTACAAGTAGGAGGCCAAGAAATCACTGTCGTAAATCTCTCTGGTCAAGTCATTGCTTCAGCTTCCATGCTGGATACTGGCAACTTTGTGCTCTATAATTCTGATCACAGTATAATATGGCAAAGTTTTGACAATCCAACAAACACACTTTTACCTGGTCAGCATATATCAGCTGGACAAGAGCTGTTCTCGAGTGCCTCGGAAGCAGATGATTCTTTCGGGATTTTCCGTCTCAAAATGCAAGATGATGGGTACCTTGTTCAGTATCCTGTCAATACACCAGATACTGCCCCATATGCTTACTACGCATCCTACATCCGTGGAGTAAGGAATAATTTTTCACTGAATCTTGATGATGACGGCCTCCTCTACTTGCTCAATTCCAACAATAGTCTAAAAAATCTAACCAGAGGAGGCTACCGTAGAGAAAGGACCATCATCTATATGATGAAAATTGATGCTGATGGTATTTTTCGAGTTTATTCACATTCTCTGAACCAACAGAACAGCTCCGTGATATGGCCATCAACAGATGATAGGTGTGATCCAAAAGGCCTTTGTGGGCTCAATGGTTTTTGCACCAATATAGATGTTCAAGCTAAATGTATGTGTCTTCCGGGATTTGACTTTGTAATGCCAGGAAATTGGAGCGCAGGCTGTGAAAGGAATTTCCCCACAGAAACCTGTCAGTTAAAGGAGAACACTTCCAAATACTATGACATGAGAACAGTTGAAAACACAGTATGGGAAGATAGTACTTATGTTGTTTTGGCCGGAACGACCAAGGAAGATTGTGAACAAGCCTGTCTACAGGATTGTAACTGTGAGGCTGCTCTGTTCAAAGACAGAGAATGTAGAAAGCAAAAGCTGCCTCTGAGGTATGGAAGGAGAGACTTGGGTAATTCTAATCTAGTATTGGTCAAGGTGGGCACTAGTTTCATATCCAAAGAAGGGGTGCCTAATCAGACGATTGAAGAAACAAAAGGCAAAAAGCTAAGGATAGACATCCTAATTGCCGCCATTACATTGGCTGTTTTTGCTTTGTTGGTGTTAGGCATCTCCGGATTCCTCATTCACAGAAATCATGTCTGGACTTACAGAAAAATCCAGGAGAGTAGAAGTGTTCAGTTATGTGAGGATGTTGCTCCACGAGCATTTTCCTATGCAGAACTAGAGCAGGCAACCAGTGGTTTCAAAGAAGCGTTAGGGAGGGGAGCATTTGGAACAGTATTTGCAGGAATTTTGGCAGAAGACCAGAAAGTGATAGCTGTTAAGAGACTAGACAAAGAATTAGTAGAAGGGGAAACAGAGTTCCAAACTGAAATAAAAATCATTGGAAAGACACATCACCGAAATCTAGTCCGCCTCCTTGGTTATTGCCTGGACGGATCAAGAAGGCTTTTGGTGTATGAGTACATGACCAATGGGTCACTTGCGGACGCACTTTTTACACCAGAAAAACAGCCAACGTGGGAGGAAAGGTGTGGCTTCGCTCGAGATATTGCAAGGGGCCTCCTATATTTGCATGACGAGTGTGAAACACAAATCATCCATTGTGATATTAAGCCTCAGAACATACTCATGAATGATCAGTTCTGTGCAAAAATATGTGACTTTGGAATGGCCAAGCTCTTAAAGAAAGACCAAACAAGAACCTACACAGGGATAAGAGGGACCAGAGGCTATGTTGCACCAGAATGGCACCGAAATCTGCCAGTGACAGTGAAAGCAGACGTTTACAGTTTTGGTGTCGTCCTATTGGAATTGATCTGTAGGCGGAAGTGTGTAGATTGGAGCCTTGATGAAAATGAGGCAATTCTTGAATACTGGGTCTACAATTGCTTTGATGCAGGAGAGCTAGATAAACTAGTAGGGGACGAGGAGGTTGATAGAAGACAATTCGAGAGAATGGTGAAAATAAGCATCTGGTGCATTCAAGAGGAACCATCACTTCGCCCTTCAATGAAGAAAGTTCTACTGATGCTGGAAGGGACAGTAGAAATTCCAGTGCCTCCTAGTCCTACTTCTTTTTTAAGTGCCATATAA

>PGSC0003DMG400004000 G-LecRK

ATGCCTAACAAGACAGCAGTGTGGACAGCAAACAGGAACAGTCCTGTTGTCCCGAGCAACGCTGTCTTGCTCTTAAATAATGATGGAAGGCTCATTGTGCAAGTAGGAAGCCAGGAAAACTCTGTCATCAATCCCTCTCGAGCCATTGCTTCAGCTTCCATGCTGGACACGGGAAACTTTGTGCTCTATGATTCTGATCGCAATGTCATTTGGCAAAGTTTTGACAATCCAACAAACACACTTTTACCTGGTCAGCATATATCAGCTGGACAAGAGCTGTTCTCGAGTGCCTCGGAAGCAGATGATTCTTTCGGGATTTTCCGTCTCAAAATGCAAGATGATGGGTACCTTGTTCAGTATCCTGTCAATACACCATATACTGCCCCATATGCTTACTACGAATCCCACATCAGTGGAGTAGGGAATAATGTTTCACTGAATCTTGATGATGACGGCCTCCTCTACTTGCTCAATTCCAACAATAGTCTAAAAAATCTAACCAGAGGAGGCTACCGTAGAGAAAGGACCATCATCTATATGATGAAAATTGATGCTGATGGTATTTTTCGAGTTTATTCACATTCTCTGAACCAACAGAACAGCTCACCGATTTGGTCATCAACAGATGATAGGTGTGATCCAAAAGGCCTTTGTGGGCTCAATGGTTTTTGCACCAATATAGATGCTCAAGCTAAATGTATGTGTCTTCCGGGATTTGACTTTGTAATGCCAGGAAATTGGAGCGCAGGCTGTGAAAGGAATTTCCCCACAGAAACCTGTCAGTTAAAGGAGAACACTTCCAAATACTATGACATGAGAACAGTTGAAAACACAGCATGGGAAGATAGTTCTTATGTTATTTTTGCCGTAACGACCAAGGGAGATTGTGAACAAGCCTGTCTACAGGATTGTAACTGTGAGGCTGCTCTGTTCAAAGACAGAGAATGTAGAAAGCAAANNNNNNNNNNNNNNNNNNNNNNNNNNNNNNNNNNNNNNNNNNNNNNNNNNNNNNNNNNNNNNNNNNNNNNNNNNNNNNNNNNNNNNNNNNNNNNNNNNNNNNNNNNNNNNNNNNNNNNNNNNNNNNNNNNNNNNNNNNNNNNNNNNNNNNNNNNNNNNNNNNNNNNNNNNNNNNNNNNNNNNNNNNNNNNNNNNNNNNNNNNNNNNNNNNNNNNNNNNNNNNNNNNNNNNNNNNNNNNNNNNNNNNNNNNNNNNNNNNNNNNNNNNNNNNNNNNNNNNNNNNNNNNNNNNNNNNNNNNNNNNNNNNNNNNNNNNNNNNNNNNNNNNNNNNNNNNNNNNNNNNNNNNNNNNNNNNNNNNNNNNNNNNNNNNNNNNNNNNNNNNNNNNNNNNNNNNNNNNNNNNNNNNNNNNNNNNNNNNNNNNNNNNNNNNNNNNNNNNNNNNNNNNNNNNNNNNNNNNNNNNNNNNNNNNNNNNNNNNNNNNNNNNNNNNNNNNNNNNNNNNNNNNNNNNNNNNNNNNNNNNNNNNNNNNNNNNNNNNNNNNNNNNNNNNNNNNNNNNNNNNNNNNNNNNNNNNNNNNNNNNNNNNNNNNNNNNNNNNNNNNNNNNNNNNNNNNNNNNNNNNNNNNNNNNNNNNNNNNNNNNNNNNNNNNNNNNNNNNNNNNNNNNNNNNNNNNNNNNNNNNNNNNNNNNNNNNNNNNNNNNNNNNNNNNNNNNNNNNNNNNNNNNNNNNNNNNNNNNNNNNNNNNNNNNNNNNNNNNNNNNNNNNNNNNNNNNNNNNNNNNNNNNNNNNNNNNNNNNNNNNNNNNNNNNNNNNNNNNNNNNNNNNNNNNNNNNNNNNNNNNNNNNNNNNNNNNNNNNNNNNNNNNNNNNNNNNNNNNNNNNNNNNNNNNNNNNNNNNNNNNNNNNNNNNNNNNNNNNNNNNNNNNNNNNNNNNNNNNNNNNNNNNNNNNNNNNNNNNNNNNNNNNNNNNNNNNNNNNNNNNNNNNNNNNNNNNNNNNNNNNNNNNNNNNNNNNNNNNNNNNNNNNNNNNNNNNNNNNNNNNNNNNNNNNNCAGAAAAATCCAGGAGAGTAGAAGTGTTCAGTTATGTGAGGATGTTGCTCCACGAGCATTTTCCTATGCAGAACTAGAGCAGGCAACCAGTGGCTTCAAAGAAGCGTTAGGGAGGGGAGCATTTGGAACAGTATTTAAAGGGATTTTGGCAGAAGGCCAGAAAGTGATAGCTGTTAAGAGACTAGACAAAGAATTGGTAGAAGGGGAAACAGAGTTCCAAACTGAAATAAAAATCATTGGAAAGACACATCACCGAAATCTAGTCCGCCTCCTTGGTTATTGCCTGGACGGATCAAGAAGGCTTCTGGTGTATGAGTACATGACCAATGGGTCACTCGCGGACGTACTTTTCACACCAGAAAAACAGCCAACGTGGGAGGAAAGGTGTGGCTTTGCTCGAGATATTGCAAGGGGCCTCCTATATTTGCATGACGAGTGTGAAAACACAAATCATTCATTGTGATATTAAGCCTCAGAACATACTCATGAATGATCAGTTCTGTAGCAAAAATATCTGACTTTGGAATGGCCAAGCTCTTAAAGAAAGACCAAACAAGAACCTACACAGGGATAAGAGGGACCAGAGGCTATGTTGCACCAGAATGGCACCGAAATCTTCCAGTCTCTGTGAAAGCAGACGTTTACAGTTTTGGTGTCGTCCTATTGGAATTGATTTGTAGGCGGAAGTGTGTTGATATGAACCTCGATGAAAATGAGGCAATTCTTGAATACTGGGTTTACAATTGCTTTGATGCAGGAGAGCTAGATAAACTAGTCAGGGACGAGGAGGTTGATAGAAGACAATTCGAGAGAATGGTGAAAATAAGCATCTGGTGCATTCAAGAGGAACCATCACTTCGCCCTTCAATGAAGAAAGTTCTACTGATGATGGAAGGAACAGTAGAAATTCCAGTGCCTCCTAGTCCTTCTTCTTTTCTAAGTGAGATATAG

>PGSC0003DMG400004928 G-LecRK

GGATCATTCACTATGGAAATTGGAATGGTATCAACCAGCAGAAAGCAAATGAAAAATTTTCTGCTTATGTTGCACATATTGTTATGTGGTTTTGTATGCTCAATTTCTAGTGCAAAAGATACCATCACTACAACTGACTTTCTAATAGAGGGTAAGACCATTGTTTCATGTGATGAAAGCTTTGAATTGGGATTTTTTTCCCCAGGTAGTTTCGCGAATAATTGGTATGTTGGAATATGGTACAAGCACGACAACGTGCCTGATAAGTCTGTTGTATGGGTTGCCAATAGAGAAACTCCACTCAACAATACATCAGGTGTTATGCTGAAAATCGTTGATCCAGGACGACTTGCTCTTGTTACTTCTGCTAATACGAGCATATGGTCTACTAATTCGTTGAGACCTTTGGCTGTGAAGAATCCAATTGCGCAGTTGTTGAATTCAGGGAATCTTATCATAAGAGATGCAAATGATACTAAGCCTGAGAACTTCCTGTGGCAGAGTTTTGATTATCCTACTAATACGCTCTTACCAGGCATGAAACTTGGTAAGAACTTTGTGACTGGACAGGAATTTTACCTGTCCTCATGGAAGAATGAATATGATCCGGCTCCAGGGAAATATACGTATCATTGTGATCCTACTGGATATCCTCAGAATGTCATGAGGAAAGGTAAAGTTAAGGTGTTCAGTGGCGGACCATGGAATGGACTCCGCTGGAGTGGTGTACCAGGATTGACAAAAAATACTATTTATACATTTAAGCTGGATTTTGATGAAAAGAAAGCAATTTACAGTTACGCGCTCGTGGATAAATCTGTTTTATCGAAATTAACCTTGAACAGTCATGGTATGCTTAAGCGTTGGACGTGGGATGAGAAGATAAAGGAATGGCATGTTTACCTGGCTTCACCAGCAGATGCCTGTGAAAATTATGGAGCATGTGGTGCCTATGGTAGTTGCAACATCATATTTTCTCCTGTTTGTCGTTGCTTAGACAAGTTTGTGCCTAAAGATCAGAAAAATTGGTCAGTGACAAAGTGGTCCGGTGGATGTGTTAGAAGGACACCCCTTAATTGCCAGAATGGAGATGGATTTCTCAAGTACTCTGATATAAAGCTACCTGATACACAGTATTCACGGTTTGATGCGAGAATGACACTCCGAGAATGCGAGAAAGTTTGCCTAAAGAATTGTTCTTGTATGGCTTACTCAAATCTTGATATACGCAATGGTGGGAGTGGCTGTTTGCTTTGGTATGGCGACTTAATTGACATTCGAGAATTACCTGGAGGACAGGACATTTACATTAGGATAGCTAACTCAGAGTTAGGTAAAATTTATGCAACAAATTCATTACTATTTGATAAATATGAAGCATTAGCTGAATGAAGTAAACTTCACAATTATTAGCTTTAGTTGTGTTTTGCTTATCTATTTGTTAAAAAGTTCACATAGCATATATTTAGTTTATTTTACCCTCCATTTTTTTTAGGCTCGAAGAAGAAAATGAAAGCACTTGTGCTGAGTTTATCACTGTTGAGCGGAGTGTCTGTCATTGGCCTGATCATATGGCTGTATATTTTGATAAAGACGAGGAAGAAGAAGAGGAAGATGATTCTGAAAGATGATCTAGAACTTCCACTGTTTTCCCTATCAACAGTAACTAAAGCGACCTCTAATTTCTCAGACAAGAACATGCTCGGAGAAGGTGGATTTGGATCTGTTTACAAGGTAGCATAACTTGTAGGTTTATTATATGTACTATAATTGACAGTAAACAGATAAGTAACACTTTGAATTATCACTGTCAAGACATCCTTAGCTTCAATATGAGTCAGCTGTATGTTATGATATTGTAGGGAATCCTGGAGGGAGGACAAGAAGTAGCTATGAAGAGGCTCTCAAAATCTTCCTCGCAAGGAGTCAACGAGTTCAAGAATGAAGTTAATTGTATTGCCAAACTTCAGCATCGAAACCTTGTGAAGCTTATAGGTTGCTGCGTTGAAGGAGAAGAGAAAATTCTGGTCTACGAATACATGCCCAATAGAAGTCTTGATTTCATCATATTTGGTATGCTTACTCTTGCTTAACACATAAAACCTCAAGTACTGCTCATAGTATAGCATTGAGCTTGCAGCTTTCACCATGATTTTGGAAAAGGTTGTTAATTTCAAATGGAAGTAGATTCTGTTTAACATACTAAAAGCAGTTAGAGCATCAGTTACTATACTAATTGTTGGAAGATCTGTTAACAGATGAAGACAAGAGCTCGATACTCGACTGGCCTAAGCGCTTCAACATCATCAATGGAATTGCTAGAGGACTTTTGTATCTCCATCAAGATTCTCGGCTAAGAATTATCCATAGAGACCTCAAAGCTAGTAACATTTTGCTAGACGCTGACATGAACCCCAAGATCTCGGATTTTGGCATAGCTAGAAGTGTGATAGGGAATGAGACGGGTGCTAATACACATCATGTGGTTGGGACACAGTAAGTGATTTCACACTACGACCTTTTAGCATTTATCTCATGTACTACTCATATAGTATCATCTGTGTAACACTATTTTCTGTTTGATATGTGATGCAGTGGCTACATGTCCCCAGAGTATCTAATACATGGTGTTTTCTCGATAAAATCAGATGTATTTAGCTTTGGCGTGTTGGTTCTAGAGATAATCAGTGGTCGAAGAAACAGAGGATTTTTCCATGAAAGTCACAGTATCAACCTTCTCGGACATGTAAGTTTTGTATGTTAAAAAGTACTTACATATTCAGATTTCAGACCACTCGTAAGGTAATTACAAGTGATAGTGTAACTAAACTGATAGTGTAAAAAGTCTTACTAAGTGTCGGTGTATATAACTTAAATCTATTAAACACGCTTCACTTGCTTGTACTGATCTGTTATTAGCAGGTTTGGAAACTTTATAAAGAAGGTAGACCATTGGAGCTGATTGATGTGCATATTACAGATTCATGTTATTTTACTGAATTACTACGGATAATTCATGTCGCTCTATTATGTGTACAACATAGTCCAGAAGACAGGCCAGACATGTCTGAAGTGGTTGTGATGTTGGCTAATGATGCCATTTTGCCTGAAGCTAAGGAACCTGGTTTTTACACAGAAAGCAAATTCCCTGATTCTGAATATTCTACAAGCATGTATTCGAAAAACGAAATCACTATTACACAGTTAGATCCCCGGTAAATTAAATTTCATATTCTTGTAGATCTTTAAATGTCTAGAAGCTTGAATAACAGAGGAGAACATTTTATGAGTTGTATCATATTAGGAAAGAATTTACCTAACTCTTGAATTAAAAAAAGACAGCCTGGAGCACTAAGCTCTCGACATTCATGGATTCAGGTCAGAATTGGACCACACGGGTCTATTGTACGACAGTTTTACAGGCACTTGAGTATTAGTTACAAATTGCCATTGTTGCGTATGTTACATGGTTTTGTTATACTTACAGATACGTATCCACTCTGGGTGTGTGTAAGAATCTAATATGAATACTACAGTTTTTTTTGAAATTTTAATCTACTCTCCA

>PGSC0003DMG400004930 G-LecRK

ATTAAAAGGCTTAGGCATCAAACACATAGATGGAATCTAAATACTTCATACTTTTCTTACTATCTTTCTACTCTTTTATACCGAAGATTAGTTCTACTACGACAAATGTAATCAATACTAATCAGTTCATAACAGATGGTGACACTATTGTTTCATCTGGTGGAACCTTTGAGCTGGGATTTTTCAGCCCCAATGGTTCCACAACGCGATATATTGGGATATGGTACAAGCAAATTCTTCCTTATGTCCAAACAATTGTATGGGTTGCAAACAGAGAGAAACCACTCACCAATACATCTTCAGTCGTTTTGAAGGTCAACAAGCCGGGAATACTTGCTCTACTTAATGACAAGAATGAAACTATATGGTCCACGAACACCTCACGATCAGTCCAAAATCCAGTTGCAGTGCTGTTGGATTCTGGTAATCTTGTCCTAAAAGATGCAAATGATGACAATCCAGAAAATTTCCTATGGCAGAGTTTCAATTTTCCAACTGATACTCACTTGCCTGATATGAAACTCGGAAAGAATTTTAAGACTGGTATTGAAGTTTACCTTTTAGCATGGAAGAACGATAATGATCCAACTCCAGGGGAATACAGTCTCCACATTGATCCTACTGGATATCCACAAGGCCTCATTAGACATGGGGCAAGGGTATCAGCTCGAGCAGGACCATGGAATGGTTTACGGTGGAGTGGAGCACCTGCACCACTACAAACACAAACCAGCATATATACTTTTCAATTTGTTTTCAATGAAGAGGAGGTTTACTATAGTTTTTCTCTCATTAACAACTCACTGTTAACAAGGTTAGTCCTGACTAACAATGGTTATATACAACGTTTGACGTGGGTGGATAGGACAAAGAGTTGGCATCTTTACCTCAATATTCCTCTGGATACTTGTGATACATATAGCTTATGTGGTGCATATGGGAGCTGTGTCATAGATAGTTCTCCTGTTTGTGGATGTTTGGACAAGTTTGAACCTAAATATCCACAAAATTGGCAAACGGGAGACTGGTCACAAGGGTGTGTTAGGAAGACACCTATTGATTGCAACAAGGAACATGGATTTCTAAAATATTCTGGAATCAAGTTGCCAGAAACTAACAACTCTCAGTACAATAAGACCATGACACTCGAAGGATGTAGGCAAGTATGCTCAACGAACTGTTCTTGCACAGCTTATTCGAGCCTAGATATAAGCAATGGAGATAAAGGTTGCTTGTTTTGGTCTGGGGAGTTGATTGATATCAGAGAGCTCTCTGGAAGAGGACAAGACATTTACATACGAATGGATTCTTCAGACATAGGTAAGAACTTATGTTAACGAAATTTCTAATGATGTGTTTTTGAACTTTTAAATTTTTTAACATATGTAGACCATATTATATATGCAAAAGTAGTGGCCTATATCAATTCGAAGAAGCTAATTTAGAAAATGGTTGTAGTATCTCAGGCAAGTTCAAACAGAAAGAAGACAGGGACAGTACTCGCGGTGAGTTTCTCGTTGTTGGTGGCAGTGATTCTGCTAGGCCTGATTCTGTTCATGTACATACGTAAAAAGAAGAAACTAAAACTCAAAGAAGATTTTGAGCTGCCACAGTTTCAATTGTCGTTAATAACAAGAGCCACAAATAACTTTTCGGTCAACAACCAGATTGGAGAAGGTGGATATGGACCTGTTTATAAGGTATAAACTCAATTTTTTTTTGGAATAGTGACCAGGATAGAGATGACTTTCATTCTGCAATACTCGCATCTTTGTAAAATATATTTCCTTCTGATATACTTTTTCCATTTCAAGGGAGTACTGGAAGAGGGACAAGAAATTGCTGTGAAGAGACTGTCAAGGACTTCCATGCAAGGAATTGATGAATTCAAGAATGAAGTTACCTATATTGCCAAGCTTCAGCATAGAAATCTTGTGAGACTTCTGGGTTGCTGCATTCAAGGGGAAGAAAAAATGTTGATCTATGAATACATGCCTAATAAAAGCCTGGACTCATACATATTTGGTTAGGATTTCTAACCTACTATAAAAGCAAAAATGAGTAATGAACCATGTTTTCAAATTAAAGCTGCTAAAAAAAAAGAAGTTTTTTATTTTTTTTTGTAGATCAAACAAAGAAAAAGTTACTTGATTGGTCAAGGCGTTTCGACATCATTAATGGAATTGCTCGTGGCTTATTGTATCTCCATCAAGATTCTAGACTACGTATTATCCATAGAGACCTTAAAGCAAGCAATGTTTTGTTAGATACAGAAATGAATCCAAAGATATCAGACTTTGGGATGGCTCGAAGTGTTGCAGGAAATGATATGGGAGCAAAAACATGCCATGTAGTTGGGACACAGTAAGAATCGATCATACAGAAGTACTCTCATATGAAATCCCCCTTCTTAAATTTCTCATGGTGTAGTTCTTTTTTTGGAACAGCGGTTACATGTCCCCTGAATATGCAGTAGATGGAATCTTCTCAGTAAAATCAGATGTGTTTAGCTTTGGCGTATTGGTGTTAGAGATTGTGAGTTGCAAGAAAAATAGAGGATTTGTCCATGAAGATCACAACCTTAACCTTCTCGGTCACGTAAGCACTCTATACATTCTTGTCTATATTACATATAGAATTTGTCATGATCTAAATTACTCAAACAGGCATGGAAGCTTTACAAGGAAGATAGGTCCTTGGAACTAATTGATGAGAAGCTAGCTGAATCTTGTCATATTTCTCAAGTTTTGAGGTCAATTCAAGTGGGACTATTATGTGTGCAACAATGTCCTGAAGATAGACCAAACATGTCTTCTGTGGTTCAAATGCTAGGTAATGAGAGTTTACTGGCAAAAGCTAAAGAACCAGGATTTTTCATGGAAAGAACGCTACATGATTCAGATAATTCAGGAATGCAGACAGGAAGTTCGAAAAATGAAATAACAATGACAATGTTAGATCCTCGGTAGAGTACTGTGAATGAGGCAACTACTGTTGATAAGCATGAGTAACAAATTATTTTAGAGGGCGTGATAACGACATTGAAGCGCCTGAGAAGCGCTCAAGCATCACCTCAGGGATTAAGCGTGCCTTTGAGTATTTAGAAAAATGGTTCATAATTGCCATGCTATTATAAGTATCTATTCAACTTGAATTTTCATTTGTATCTTATCTTAGAAAGAAATGTAAATTCTAGTAAGCAAAAATGTCCAAATTATATAGCCTATGATAATACATAAAATCGAATCAAACCGACAGATGCACGAGCTCTCCAATATCTGCATGGAAGAGTCCAATATGTCGTTGAAATTAAAGGTCACAAACTTTGGATTTCCATATTTGAGTAAAGAGAAGAAGATAGGGAAAAAGGCCATAAGCCCACATTGACCGTGCACGGCCTTTCATATCCATCTCCGCCCATAATAAGGGAAAGTAAACTAATCGGGTATGAATAGATAAGAAAGTGAGATTTGAATCCAACCCGGCC

>PGSC0003DMG400005044 G-LecRK

ATGTCACCAGAGTATGCAATGGGAGGACTATTTTCTGAGAAATCTGATGTCTACAGCTTTGGCGTATTGCTACTAGAGATTGTCAGTGGAAAGAGAAACAGCGGATTTTATGACCACGAAAGGCATCTCAACCTTCTCAGTTATGTAAGTGCAAAATCAGCAGGATCTCTTGCCACAAGTTATTCACACAAACTTAGAGTATTCATTTCAAAATGATTTCAGGCGTGGAAGCTACGGACTGAAAGCAACGGATTAGACTTAATGGATAAATCAGTTTTGGACTCAGATTCCTCTGCAACAGTGCTGAGATGCTTACATATTGGCCTGCTTTGTGTCCAGGATCATGCTGCTGATAGGCCATCAATGCCGTCTATCGTTCTTATGCTAAGTAGTGAGATGGATCTTCCTCAACCAAAGCAACCTACATTTATATTTCAAAGATGGTTGAATTCTAATACCCAATCGCAAATCAGCAAAACTCAGTCTGTTAATGATATTACTGTCTCTGTTGCAGAAGGACGATAA

>PGSC0003DMG400005689 G-LecRK

AAAAAAACCTGGTCAACAAAAGAAATCAAAGCCAACAGCATGCTTGTCTCTGTTTTGGTCATTCTATTTTGACTGTAAACATTCATGTGTGGATTAGAGAGCTTCTCTCATATGGCTACCTACAGAATTCTGATTGCTTTCATCCTCGTGTTATCCTGTCACTTTTTGCGTGCAGATTTCAACTCAATATTGAAGGGAGAAAGTTTAGCTGATTACCAACAGCTGGTCTCAGCAAATGGCTTCTTCAAAATGCAATTTTTCAGTCCAGGCAAGTCAAGGAACCGCTACTTGGGTATATTTTCCACTCGGCCTAGTCTTGATTTGTACTCTGATGTTAATGGAGGAGGAGATGAAAAGGCTTTGTGGATAGCTAACAGAGACGATCCGATAACCGATACATCTGGAAGTCTTATGATAGCTCCTGATGGCAGATTAATAATTTCTCATAAAGAAGGTAATGTTACTTTGTTCAGTGCTACACCAACAACAGCCACTAACTTAACTGCTATACTACTCGATAATGGGAATTTTGTGCTGAGGGAACTGAATACCAATAGCTCTGTAAATCGAACGCTATGGCAAAGCTTTGATTATCCAACTGATACACTACTGCCTGGAATGAAGCTTGGGATTAACTTGCGAACGGGGCATAAATGGTCATTGACTTCATGGGTAAATGATCAAGCACCTGCTTCAGGGTCTTTTACCTTTGGTCTTGACCCTAATGGTACAAATCAGCTGATGATCTTGTGAATGGGCAAGGTCTATTGGAAGAGTGGCCCGTGGCCTACTGGCCATCTCGCTCTTAAATATGTCTCAAATGAGGATGAGAAATACTTTATGTATACTGTTGAATTTGAATATCAAAGATACTCTGTGAGTCCATTTGGAATAATTCAGGATACTTTTAAAGGAAATGCAGTGTTTGGTAATTGTAGTAATGAGACTCCCCATGCTGGCTGCGTAAAGCAGGTATTGCCCCAGTGCAGGGCAGCTAAAAAGTACTGGTTTGAGCTTAGACAAGTTTACATGTTTGGAGACAGCGTTAAATTTGATGAGAACTATACCTTGAGCCTCTCAGATTGCAAGGCCAAGTGTGTGAATGATTGCTCGTGTGTTGCCTATGCCTCAGTCGATAGTGACACCGGAATTGGCTGTCAAATTTGGGGGAATGACACGAGTTTTGTGACAGCACAAAACAGTCTAGCCAGAGATGTTTTCTTTCTTACAAGAAAAGGTAATTCTTGAGTACAAAGTTATTCAATAGAGTGAAAGATCAAAAGACATGTTAATAATGTTGCATCTCATATTATGATTACTTTATCCAGCTCAAGATAAATTGTTTTGTGAGGGCATCACACTAATGTCACCAATCTGCATTACTCTCATTTGCATGTCATTCCAAAAATTACTAGTACTAGAACAAGAAATTTTCTCCGATCGCAATGCTGCAGGATTTGAGTTAATAAGAGTGTCATCTTCATTTCAAAAATACACTCGAGCAAATGAACAACCATACCTTACAGAATCTTGTTTTGGAGGCTATTAAATTATTATTTTTTGGTATTTCTAGAGTACCGACAATTATGATAGCATCATAGTGTATTCCCAAGGTAAATATGTGGAATTTTGGTCATAAACACTGACTTACAATTTTGACAGATAGGAAGCGGAAATGGTGGATATGGTTGACAATAGCAGTGTCCCTAATGGTATTTGCGTTTATTTGCTCCTTATTCTGTCTGATACGAAGAAAACTCAGAGCAAGAGGTGAGCAAAACATCAGCTCTAGCATTAGTCTAAGGTGACACTTATTTGTCTTTGAAATCTTCAATTTTGCATTTGTCACCTAGGCAAAGTCAGACAAATGGAGAAAATTTTATACGAGCTTGAAGATCCTAAGACCATTTCTGGCCAGTACAATACTAACAAGACTGCTAGACTAAGGAAGAAGTTCAGGCATGATAGACACATATTCAGCCTAGAGACTATGAATATGGCCACAAACAATTTCTCGTCTTCTAATAAGTTGGGACAGGGCGGTTATGGACCTGTTTATAAGGTAAAAGTTTAGAAGCTTTTTGTCTTAAGCTGATTTTTAGATTTGCAAGTAGCAAAAATTTCAGGGGATGTTGCTCGACGGACAAGAGATAGCCATCAAGAGACTTTCAAGAAGTTCAGGGCAAGGATTGGTTGAGTTTCAAAACGAAATTGTGCTGATTGCTAAACTCCAGCACACTAATCTTGTTAGGCTTTTAGGATGTTGCATTGAAGGGGAAGAGAAGATATTAGTTTATGAATACATGATGAACAAAAGCTTAGACTTCTTCCTATTTGGTATGGATTTCCAATTTCATGCCTAAATTGCTTAATTAATCATATTTGATAACTAATACCCCTTCCCATTTTGATTGAATAGATCCTAGCAGAAAAGATTCACTAAAATGGAATACGCGTTTGAACATTATTGAAGGAGTTGCTCAGGGACTACTTTATCTCCATAAGTATTCGAGGTTGAGGGTGATTCATAGAGATCTAAAAGCCAGCAATGTCTTGCTTGATGACAATATGAACCCCAAAATCTCAGATTTTGGTTTGGCCAGGATATTTGGAATGCAAGAATTTGAAGCAAACACGGAGCGAATTGTTGGTACCTAGTAAGTCCTTAATAGATATAATTATCTAACAATGGTGATTCACTGTGTCCTTTTTATTATGTGCTAAGTTGCTACCTTTTCAGTGGTTATATGTCTCCAGAATATGCCATGAATGGCATTGTGTCCATGAAGACAGATGTATTCAGTTTTGGAGTTCTAGTACTCGAGATTGAGTGGCAAAAGAAACAACAGCTGCTATCATTTAGAACGCCCACTAAACCTCATCGGATATGTAAGTTCCTCTCGTAAAATATCAATATAGCATGATCAAGAGGCGGATGGAGTGTTTATTCAATGAGTTCAACTAAACCCAATACTTTAGAAACGAAACATAAATATACAACATCAACATACCCATTGGAATCCCACAAGCCACAAGTTGGGTCTGGGAGGGCGGTGTGCACACAGACTTACCTACCTAGTAAAGGCTAGGCTGTTTTCAATAGACCCTCGACTCAAGACACGAAACATTGATATACATGTGAAAATCACTTGAATTATAAAATTCTGAAGTATAATGAGTCTAGTTCTAAAAACTTTAAAGTTCAACCTGTCAACTTTACATTGTTTTGGATCCACCCTGGCATGATCACTTAGTTGAACAAGGAAATTTCAAACTTGATATGCTCAATTTGTCTCAGGCATGGGAATTGTGGAAAGCAGGCAGCGTCCTTGAAGAACTAACAGATCCTGTATTGACAAATGAATCAACACCAACGAATGAAGTGATGAGATGCATTCATGTAGGATTACTCTGTGTTCAAGTCAATCCTATGGACAGGCCATCCATGTCAAATGTGGTTCTGATGCTCACAAATGATAGCCTACAACTATCCGTCCCCAAGCAACCAGCATTTTTCATTGAGACAGCTATGACTGAGACTGAAACTCGCGAAGAAGTTGGACATTGTTCCACAAATGGGCTATCAATTTCAGATATGGTAGCGAGATGACTTTCATCATTGCACCCCTCAAAATTGTAAAAAATGGTAATATGCAATCTCCCTTGTGATACAAAATAAAAACCACTGCTTGTCTCCTTCATCCAGATTTTACTTTTTCTAGCGTCGATATTAAGTATGAGTTGTTGACAAAATAAAGTGAGTAAGCTTGTTAGAGCCTTTCAAGATTATACTGTGTCTAAACAATTTTTATACATTTGTTTTGAATGCAAATGAGGATTCAGAGTATCAATTGTAACTGAAATCAGAACTCATAAATGTAGTAATTGTTACTAGTTTTGCAAAGATGATTAGTATCTGTACATTAGAAAATCAAATGGATGTTTCCCACGTTCCTTTGTTGCTTG

>PGSC0003DMG400007073 G-LecRK

CCAATCTCAAGAAGCAAATGTATCATCTAACAAACAAGCTAAAACAACAAACTAAACATGTCTTCTTCCTGGCCTACTACACTTCTTGCCTCTCTCTTTCTTTTCTCCCAAATATTTTCTTGCATTGCTCAAGTTCCAGCAGAAAACACCTTTAAATTCGTCAATGATGGCGAATTAGGAGATTATGTTGTAGAATACAGAGCAAATTATCGCGTTATTAGTATTTTCAATAACCCATTTCAGCTCTGTTTCTACAACACAACTCCTAATGCATGGACATTAGCATTGCGAATGGGCACTGTCCGTTCTGAATCACTCATGCGTTGGGTGTGGGAAGCAAATAGGGGAAACCCCGTTAAAGAAAATGCGACATTCACGTTTGGAACTAACGGAAATCTCGTATTGGCGGATGCTGACGGTCGAATCGCTTGGCAGACCAACACGGCCAATAAGGGGGTCACCGGGTTCAAATTGTTACCTAACGGTAACATGGTGCTACATGACTCCAAGGGTAAATTCGTCTGGCAGAGTTTCAACTATCCCACTGACACATTACTAGTGGGCCAAACTCTCCGATTGTCCGGCCCGAATAAGCTTGTGAGCCGGGCCTCGGTGAAAAAGAATGCAAACGGGCCTTATAGCTTGGTGGTGCAGCCGAGATTGTTCGCTGTCTACAAGGGAAACAAGGTAGACGTGGAATTAGCTTGGTTCGATTATGGGAATAACACGTTGGAATCGATAAAATTGAATAGTGGGAATCAAAGATTGAAATTGGATTATAGATTTGCAAAATCAACACAGAGAAGTTCACATGTGATGGCATTTACTAAATACAACACAACATTGACATATCTTAGACTTGAAATTGATGGAAATCTAAAGGCCTACACTTTTGTTAGAGATGAAGAAGCAGATGAATTCCGTTGGAAAGTAACTTACAGTAGGATTTGAAACGTCTAAATCTGTATACACATTATCCTTTACGAGCAAATTTCACGGGTTATATTATTATTGTTGTTGTTATATTGTTGTAAACGTAGCTCTATAAGATAATGATTAGTGTGTCCAGTACAATAAGTGGGTCCACAGCTAGTAGTCCAATAAATGAGTACTTCCTATTTAATGTTAATTATATTTTTCTTTTTCCCTTTTTGAAAAAATATTATGTTTTGGTTGAATTAAGA

>PGSC0003DMG400007455 G-LecRK

ATTTTTTAGAGGAATGAGGTGGAAAAAGGAAAAAAAAGAAATTGGGTCCTTTGTGATTTCTTGCATTGGCAGTAACCAACGTCCAAAGGCCAAATCTCATCGATTGTGACATTGTAACAAAATATACATAATTCATCAATCTTCATAAACCTATCATCATAATCATAATCAATACAACATTATTGTAAACAAAACAAATATTCCTTCTTGTATATACACACAATATTAAAAAATTATAAAGTTTTTTACATTCTTGATTGTGTACGTGGGAAATTGGGCAGCAAATTTGCAATTTTTTATCCTGTATCTAATGCTTTGTGGATGTTCTCACAATTTGATTCACCTTTTTGGAGTGGGGTTTCCTTAGAAAGGGTAAATTCTACATAATCTTGAATTTCTGTATCTTTTTCTTGATTTGGGTATCTGGATATTCGAATTATTCTGTTTGGTGTTGTGGGTGTTTCTTGAAACTGCATTTGATGATCTGGGGTACTTTTATTTTGGTCATAGTTGCTGGTGATATCATTATTTGGCTGCTTTGTTTAGAAAGATTGAATCTTTTTTACTCTGATAAGGTTCTGCAGTGTGTGGAGGGATTGTTCTTATGAATTATAGCTAAAAAGGAGATATTGTTCTGCTTTAGACCTGTCACTAAATTCTGTGTCCTGTGGCAACATTTTGATTTCGCGTAAAATCTGATATGTTTAAGGTTGAGGTTGAGGTTCTTGTTGTGTTGAAAATCCTAGCTGTTTAACTTGGGAGTGGTGTGTGTGTATACTATGAGTGAATTGGTTGGTGAGTTTGGATAAATTTTGGAATAAGAGGAAGTAATTTACTGCATATTAGGGATCTTAGTTTTATGTCAGCAAGTCTGCATTGTATAAGAAATGTTGTAAATGTAGCTAGCTTTGGTGGTGGTGAGATGTTAAAACTTTAATCATCCACCTCCAAGTTATAGAATTTGTAGTTCCATTATCTGTGACTTCTCTCCTGGCTCGTTACTCTCTTTCGACAAATGTAAATATAGTCTGTCCTAGCCTCACTACCAATCTTATAGACTTATAATGGATTCTTCACCATCTCATGAAGAATCTGCTCCTCCAACTGATGTGGCACCTCCAACGTTTAGTCCTCCGCCTACACCTTCTAGTCCATCTGATAGCTCACCACCACCTCCATCTTCTAGTCCACCTCCTAGTTCACCACCACCTGCATCTTCTAGTCCACCTCCTAGTTCACCTCCACGTGAATCAACTAGTCCACCACCTAGATCACCACCACCTCAATCTTCTAGTCCACCACCAGCCTCACCTCCACCTGAACCAACTCCTCCTTCTCCTGTCAGTTCACCTCCACAAACTTCATCACCTCCCCCTGCATTGTCACCTCCGCCACACGTACAGAATCCAGAACCTCCTCCAGCACCACAATCACATGGAAATTCATCCCCACCACAATCTCCTGAACCTAAAAAAGGTTCATCATCATCTTCACCACCTTCACCACCTTCATCCAACTCACCTGGGGATGGAGGTGGAGGTGGAAATTCACCACCATCCCCAACATCTAAGTCATCAGATAATTCGCCTCCTTCACCAGCCTCAGGCCCTCCTAAAGATTCGCCACCTTCACCAGCTGCATTATCTCCAAGTAACTCACCATCTTCTGATTCACCCTCAAACTCATCAACACCAACATCTGTAATACAATGGTCACCACCACCTCCATCCTCTAATGGTGCACAATCTTCTCTTAATCCTCCTTCATCACAAGTTCCTAGTAACCATAGTTCCGGAAACCGTTCAACAGGAAGTCCAAAGCCTGCTGGTGGCAACAGCAATGGTACTGCAGAGACAGTTTCCATTGGCGTTGTAATTGTTCTTTTTTTGGTTGGTATCATCGGAGCAGTTGCATGGTGCATACGGAAGCGAAAGAAGAAGAATTCTGTCCATTGTAGTGGTTATGTCAAGCCAATCTCTGTGTGCTCCTCTCCAAAATCAGGTAAATTTTCAGAAACTATAAAATCATTCCTTTTAGTTTGTTATTCAGTGCTCAAGATGCTTGCAAGAAAAATCAATGTAATATTTGTAGTTGAGACCTTTCAAGTGAAATCTGAGAAGATGTGAAAGTTGTGTACAGAGTCTGCTTTATTGAAGGTCAATGAATCAACACCTGAGAGAAATGGAACGAGCTCTAATTTCCTCAATTCTCCAGCAGACCCTGGCGGATTCAACAATGCAAAGACATGGTTTACTTATCAAGAACTTGTTGAGGCCACTAATGATTTTTCAGCGAATAATGAATTAGGGAAGGGTGGATTTGGTTCTGTATACAAAGGCTATCTAGCAGATGGAAGATATGTCGCGGTAAAGCAGCTAAATATTGGTGGGAGCCAGGGCGAACGGGAATTTAGAGCTGAAGTTGAAATCATTAGCCGGATACACCATCGTCATTTGGTTTCACTTGTAGGTTATTGCATTTCAGAGAACAAAAGGCTCCTCGTTTATGACTATGTCTCCAATAATACCCTTTACTTCCATCTTCATGGTAAGCATAACATTTGTAGATTTAATTATTTCATTTGTTATGTTTGGTCACTAAGTGAGCTGTTGATACTTGCAAGTTACAGCACAAGGGAGGCCTGTCATGAATTGGCCAACACGTGTTAAAATTGCTGTTGGCGCAGCTCGTGGAATAGCTTATCTGCATGAAGATTGTAATACTGTTCTCTTCCTCAATTCTTATGAGTTATCCATGTTTTATTCACATGATATCTTTCATTGGTTGAACAAACGTCCCCCTTATAGTTAAAAGAGAATGTACAGTTAAAAACTTTACCACAACTTGGAATTAGTTTTAATGATCCCTAAAGCTGTGGCGGGATCATGGATGGCAATCTAAGCTCCACACCTTCTTCCTACTCTTCCTAAAAGCGTACATTAACGAAAGGAAAATAAAAAAGAGAGAGTAAACCAAACATGTAGTAATAATTTTCAGCTCATAGAAGTAGAAGACCTTCCAGTTGGCATTTCCTTTTTGTAATGTAATGATATTACTTATGCAGTTTAGAATTGAGTTCTAAGGTCCTTTTTCATGCCATAAATTACCTCGTTATATAACTTTTTGGCATGAGTGTAAATTATCATATCTGCATTAAGATTGTAATTAGTGTTCTGTTCCTTAATCCTTTTATGTTATCCATGTTTATTTTTATTTCAGTTCGTTTGTATGGCTGAAAAAAATGTCTCCTCTTTCCTGCCCCAAAAGTTAGAGAGAAAGCAGGAGGTCAAAAATGTTACCATATGACGAATCGGATTGTGATATGAGATTGTGTGTTCCTATCATATCACTTACATTCTTTTTATTGATCCCTACTCCACTAGCTGAACAGCTGTGGTAAATTAAATGATGCAATCTGAGCTACTTCCAGAACCAACTTTGTCCTATTATCCCTACAAGAGGACTTGAAGTAAAAAGAGAAAATGGAACGTATAAATAGTTTAAGCTCGATGGAAGTAGAAAGATCTTAGTATTGGATTTCCCTTTGGTAATGTTATAATATAACTTCCGCAATTTAGAATTGTGTCCTCTGTTTTTATGCATCTCGCAAATTATCTCGATTATATGACCTTTTGGCAGTATAAATAATGAATTCTAATCACTTCTCTAATTAGTAACTTAAAAGATGCTTTTGTAGCTGAATGTACCAAATGAGGCTCAAAATCAAACTAAGACTGAACAAAGCATGATTGGTTCCATTGAGAGAACAGAAAAAATTGTATGACTCAGTTCTTTATACATTCTTGTATTGCACCATGTAGTTACATTCCCGAGAAAGAAATATTTTAATCCATGAGCAGGCATAGTCGCCTTGAACATGAGTATATCTTCCACACTGAAGTGAGCGCATCAGGCGCTGCATTTGTAGTGCATTATATGCTTTTCGATTGCCAATATTCACTCTACGACAAGCATTTGAATAAATCTTCACTGATCGAGCCAGGTTGAAAAGTGAGAGAAACATGTTCAACTGCTCTTGTGAAGCATATAGTGAAAAAGCAAGTGACAAACTGGTTTCTCCATATAACTAAGCTGAATGTGTTAATCATCACTGACTTATTATGTTATACTGTTGAATAAGTAATGAAAAGTTAAGAAAAGTAGACCAGGCGACGCGATTGTGATTTTCTATGTTCATTTCCTTCATTTTTGTTACTTGAGCTGCCTTTAATTATATAGTTAAAGTATGATACTCCTTTTTTATTTGGTTCTTAATTAATTCTTTTGGTGTGGATGACAGGTTATCCTCGTATTATTCACAGGGACATTAAGTCATCAAACATTCTTTTAGATGATAACTTTGAGGCCCATGTATGTCTAAGTCTTCGAACTCCTATATCCACATTCTTTGTCTCGCGAGTTGTCTTTTACAATTAATTTACTTTCTACAGGTTGCTGATTTTGGACTAGCTAAATTAGCTCAGGATGCAGAAAGTCATATCACAACACGTGTCGTGGGAACATTTGGGTATGTTTCAGTGTCAACTTTTATATTAAGTGGACAGCATAAGATAAATATTTTATTCCATCTTTTTCAATTTCACATATTTATTCTAATTCCTTAGATCTGACTCAGATACATGGCTCCTGAATATGCATCAACTGGCAAGCTGACTGAAAAATCTGATGTATATTCCTTTGGCGTGGTACTTCTGGAGCTAATTACTGGAAGGAAGTCAGTGGATACATCTCAGCCTTCAGGGCAAGAGAATCTAGTTGAGTGGGTATGTTGTAGCCAATCTATTATTTGTCTTTCTCGAAATCCATATTTTCATCAGGTCTCCTTTATTACCTCTTTGTTAGAAATTTTGACATTAATATCCAACCTAAGGCAATGAAAGTGCATATCAGCATGCAGAGAGAACTTAATTTTTAGTCCTTTTAGCTAGGATAACCTACAAAATTTCATTTGAGTCAGGATTTGGTTGTTCCTCCGTCTTTTCCCATACTGCACAAGAACTTGGTAGCTGATATGTGAAGAAACCGCCATTTTGCTACTGTACAGTAGTAGGCCTCTAGTTGGTCTGCTAGTGAAGTTCCTCTTCTTTATTTTTCCCTTTTTTACCATGAAATGATGCCTTATCCTGGGAAGAATGTGGTTGCAATGCTGTAAGCCTGAAACATTGTGTCTGCATTTTTTTTTAAATTTTACAACAATCAAACAGCTTGATCAAGTTATCCGCTTTAATTTCATTGATCCTTTAATGTTAGAAAACTGAAAAACCATTAACAGTAGCAGCCGAAAACTTGTATCTTTTATCATGCTGAATGTGTGAATTCCATTTACTATAGTTATTTTATTTTCGAACTCTTTCGTTTTCTTGTCGTCAGTGCCTAGTAGCAGTAGCCAGTTTACAAAGCTTTTAATGCCTTGTATTCTGGTACTAGTTTCCACTACTAAATTTGAACCAGGCTTAAAAATAACTACTCCATTATATTGTACTGTAACTTGATTTTTTTGTTGTCCAGGCACGACCTTTGCTTAGTCGAGCACTTCAAAAGGAGGAATTTGATCTGTTGGCAGATCCGTGCCTTGAAAAAAACTACGTTGGCATGGAGATGTTCCGAATGATTGAGGCTGCTGCATCTTGCGTGCGTCACTCATCTGCAAAGAGACCAGCAATGGGACAGGTAACTCTTGTTCTGTTATATTGAAAACCTATGCAAAAGACACCTTCAAGACTCTTCTTGTGAAGGGCATTTACAACATTCTGAATTCTGATCATGGTTATGCTTCGTAGATCATGAGAGCTTTCGATGGTATGGCTATACAAGATCTGTCAAACGGGATGAAAGTTGGTGAAAGTGCAATACATAGCGCTGCACTACAATCTGCAGAAATAAGTTGGTTTAGGAAAATGGCATTCGGTAAGCAAGATTTTAGTTCGGATTTTTTCAGCGAAAGTAATCAGAATAATAGAGAGTCAAATGAACATCCTTAGAGGATTTTATAGAACAAAAGGGCTTATTGAAGCTATGTATGCATCTGTTGATACATCAAAGGTACCTTATTAGCTTAGAGATGCCTTGGTTTAGCTTTCATATGTATAAGTTCCCTTTTGGCTTTGAGTAATTTTC

>PGSC0003DMG400008493 G-LecRK

TAAAAAGGTGTATATAGCTAGCGTCCGGCATGCTTTTATTTCAACTCTAATGGCTTTTGATTTTCAATCACAACAAGAAATCATGTACGTATAGATTATTATCTAAATTTTCATGTTTTTCCCATTTGGCTAAGTAGATTTTATAGTAGTTGATTTTTTTTTCCTATTTCCTTTTCATTAATTTGCAGAAATATTAGGTGCTACAAAGTTAAAATTTTGAATATTTGAGCAATGGGTTGGAATTGCTTTGGAGTTTTGCAATCATACAAAAACAATAGAGAGACTCAAATCCAAGGTAAGTACACTCATAACTCTTATTTCTGTTTTCTCAATTACATGCTTAAAACAAATCGTTCTTATAAAATTTAGAACCCAGAAAATTTAAAGATTTGACGCTTCTGCTCTACTTTTTGTAGTCCATCCAAATTTGAGAAGTACTTCACATAATTTTCATTTTCTTTTTGTGTTTTTCATTGTGTAGAGTTCACGACCAACAATGTGAGGGTTTTTTCCTATAACTCATTGAGATCAGCAACAGGTCACTTCCATCCATCTAACAAAATAGGAGGTGGTGGTTTTGGAGTTGTGTATAAGGTAATCATCTCGTTTATGGTGTCATGCCTTTTAATGGTCCTTGTGGATGCTAGTTATTAAGTAGTGAGTTTTAACTTATATACGTTCAAAGTTTAAAAGAATCTTTTCACAGTCAAGTCACTTAAAAGATAAATATAATAATTTATCCATTCGAAATGGGATTAGCTAGTAAGTAGATGCATTTAAGGGTGTAGCCTAATGATTAATAAAGTGGGATGAGACCAATGAGATTTTTAGGTTCAAATCTCAACGGAGGCCCAAAAAGGGTGTAGCCTAGATAGACACCTTCCAATGTAACTAAGCTTTGGTGGGTAGAGTACCGATGGAATAGTCGAGTTACATGTAATCTCATACGAACACCACCATCATAAAAAAAGAAAGTAAAATAAGATAGATTACCTGCTACATTAATAATATATATATGTTAAATCTTTTATGTATTTCCACTCTCTGTACTTGTGCAACTTTGTTCTTTAAATGTAGGGAGTTTTGAGAGATGGAACTTGTGTTGCCATTAAATGTCTCTCTGCTGAATCAAAACAAGGGACTAAAGAATTCTTAACAGAGATTAATATGATTTCAAATACCCAACATCCAAATCTTGTTCAGCTGATTGGTTGTTGTGTTGAGAGTGGCAATAGAATGTTGATATATGAATACTTGAAGAATAACAGCCTTGCCAGTGCTTTACTAGGTAATGTTCTCACAGATCAGTTTTAAAAAAAAACTTATTTTGAACCAAATGAACATGACATAGTTGAAGTTAGGAATGTTAAATGAGCGGATTGGACTGAATCTGGACGAGTTAAAATTGTTATTTCAGGTTCTAATGGTAAACGCGTTGCCCTGAATTGGCCTCAGAGAGTTGCTATATGTTTAGGTACAGCATCTGGTTTAGCATTTCTTCATGAAGAAGCCTCACCACCGATTGTTCATCGAGATATTAAGGCTAGTAATATCCTTATAGATGAAAACCTTCATCCTAAAATTGGAGATTTTGGTCTGGCTAAGCTTTTTCCTGAAAATGTCACTCATCTAAGTACGCGAGTAGCAGGAACAATGTGAGTAGTTACATTGAATTTCAGTTTCATAACATTCGATATTCCTCTCCTGTCTCGTCTCAGTATGGTATTCTGATGTGCAGAGGTTATCTTGCTCCGGAGTATGCCTTGTTTGGACAACTCACAAAAAAGGCTGATGTTTATAGTTTTGGAGTTCTTGTACTTGAAATCATAAGTGGAAGAAGCAGTAGTAAATCAGCATTTGGAGTGGAGCTCTTGGTTCTGGTGGAATGGGTAGGATTATACTTTCAGCTTTATGATCTTCAAGATTCATCAGTAATCAGCATTTTGAAAGTTTACGCGTTGCAGGTATGGAAGCTAAGAGAAGAAGGACAGCTTCTGGAAATCATTGATCCAGAGCTAACAGAGTATCCAGAAACTGAATTACTGCGGTTCATCAAAGTTGCTCTCTTTTGCACACAAGCAGCACCTACCCAAAGACCTAATATGAAACAAGTGATTGAGATGTTGTCTAAGGAAGTCAACCTCAACGAGAAGTCGTTAACTGAACCAGGAGTTTATAGACCACATAGCTCGAAACGATCAAGTTATGGTAGTCTGCAGAAATCATCCTCCGCGGGGAAGAATGGGGTGCAATCTGTAAATCCTTTTGTAACATCAACAAAGTTTGAGAGCTTTCAGAGTGTCACCCAAATGCTTCCTAGATAATGTTAACTGGTGTTCTTATTCTTCATACTTGTATAATGTCGTTATTCTAACGTGGGCTATCGAGGAGTAAATGTATTGTCTCTCATTAGACCTAGCTTTTTTCATACATTTTCTCATGTAATGTAAAGTTGGCAGTAGCTGATAAGGGAGAAAGAGAGACAATCACACAGAATGAGCTACTATGTAACTCCAAATTTCAGATATGAAGATGAAGTTTACAGCAGAGACTATGAACACAGTAAAGGATAGAAACTAGGAGAAACTTTTCTCTATGCACATATTAGCTCCTAAAACTAGAAATGAAGGTAAAAACTTCCTGATAGGAGTTATTCTCAAACTCATAAACATATAAAGGACGACTGCTTTACAAGATCAACACAGTAGCATTTTCATTGTGCAGCAGAGAATAAACCTTCAAGGTGCAGCAGTAGTAGCTTGAGGCTTTGCAGAGGAAGCAACAGAAGTTTCTATTCCTGTCGTTGCATCTTTTCCTGGTTCCGAAGATGGTTTTGCAGAAGGCTTCATATGGTCAGGGGGGTTGTGTTTGAGTTTCACAAGTATTTGACGCATTCTAGAGAGTTCAGTTGGCCTGCCAGGTTTGGGTCCTTGGATGACATTGATGAATGGCTGCTTCTCCTTATCCTTCTTTCCTTTCTTTTCTATGGTCGGGACCTCCTTTGGGATAAGCTCTCCAATTGTCCTCCAATAAGCATTATTAGCTTCAGTATGGAACTTCTCTTGGTTGGCCAAGAATAACTGGAAAAAAGAAAAAGATGACACATGAACTGCAACACATTGATTGACTAAGGAGGAACTTGCCGTACTGTGGAACTATGTTTAGAGATTTGACAAACCATAAAACCAAAGACATAGTTCTGTGCAGATTAGCTAATAAAACTCACCAAAGCACAAATACTCAGCATAGATTGTACTAAAGGTTATCAGCACATTTTGATCTCTTGTGGAAACCTTCCACTTTCAATTTTTTTAATTTACACCTCTTACTTAGTGTTAACTCTAAATCATATGAAATGTAAACAGATAAATGTAGAAACTGGTTTTCTCTGTGTGCCTGCCTTGGTTGAGTCGAAAGTACCAATACAATGACTTTTAAAAAACAAAAGTAACTTAAAAAAGAGAGTAAAAAGTACCATGATTAAAAATTGGATAGTACTTTTACCAAGTTAACTAGGTTAATCTCCCTCTCTCAACACTCATATCCTATACCGAAGATTTGAAAGCAACGATTAGGAAGAGAAGGTAACTAGATGCACGACACATCAGACCAGTGATAATTTTCCTTCCAGCTAAAGCAACTGTTTGGAAAACAAACACCCAGCACAAGCAAATTTATGCCAAACTAGTTAACAAACCATGACTTTAAGCAAAAATGCCTCGGAAAGATTTCTAATTTATATAGCTATAGGTCTCACCTTCTCCTTCTCCCGGTTAGCAGCTATGTTGTTCTCACATCTAAGTTTCCATTTCCTATAGTACTCTGCTTTGTATTCCTCTGCTTCTTTAATTATTTCCCTAACAATTTCTTTCTCCCTTTTCTCCTTCTCCTCCAGCCGTATGGCGTTCAAT

>PGSC0003DMG400008508 G-LecRK

TTTAAATTTATTTTTAGCAATCACTCATTCTACCAGCAAATTGTGCAGCCATAGAAATGAAGAATAGCAGTTCAATTCTGATACTTGTGCATTTTTTATGCTTTAGCCTCATCACTAATCTTTCCCTAGGAGGGGACACCATTTCTGTGAATGAATCTGTTTCTTCAGGCCAAACAATTATCTCTTCAGGTGGGAACTTTGAGCTTGGTTTCTTCAGACCAGGCGACTCTCGCTCCTACTACATAGGCATATGGTACAAGAAACTATACCCTCAAGCAGTAGTATGGGTAGCAAATAGGGATAAACCACTTGATAGTGCTGATGCTAACTTGATAATTAGTCAAGGTAACCTGGTGCTTCTCGATAGACTTCAAAACTCAATTTGGTCTGCACTTACTGAGAACATCAATCCAAATATTTCAGTTGCTGCAGTTCTTCGTGATGATGGGAATTTTATTTTGAGTGATGTGTCAAAGGCATCAATGCCCTTACTACTTTGGCAAAGTTTTGATCACCCAACACACACTTTTTTGCCTGGTGCTAAGATTGGATATGATAAACGGACACAAAGAAAACAGGTTTTGGTTTCATGGAAGAATTCGAGTGATCCGGCACCAGGAATGTATTCTCTAGAAATGGACCCAAAAAATGCTCAATATGTTATAAAATGGAATAGGACTACAGAGTATTGGGCAAGTGGTTCATGGGATGGCCAAAGATTCGGCTTAGTGCCTGAGATGAGCTTAAACTACATATATAATTACAGCTATATCGACAATGAGAATGAGTCATATTTTACATACTCCCTTTACAATAGTACAATCACATCAAGATTGATAATGGATGTCTCGGGACAAATTAAGCAACTGTCATGGTTGGATGGTAGCATTGATTGGAATTTATTCTGGACTCAACCAAGAGAATCTTGTCAGGTTTATGCTATATGTGGGGCATTTGGAGTTTGTGATGAAGCTAATGCAACCTGCAATTGTTTGAGTGGTTTCAAGCAAAGATCAGATGCAGAATGGAATTCAAATGATTATTCCAGTGGCTGTGTAACAGACGAAAAGGTGCAGTGTGATGCAATTACTGAAGACAAAGATAGTTTATGGATAACTTCAATTGTGAGAGTACCTGCTTCTCATAATACTAATATCACAGTTGGGACAGCCTCACAGTGTAGATCTGCATGTTTCAACGATTGCTCTTGCACTGCTTATACTTATGATGGTTCCGGTACCTGTTCCATCTGGACAGGGGATCTGTTCAATCTGGAACAACTCAGCACAACTGAGTCAAAAAGGACAATATTTGTCAAACGCGGCTCACCTGAAGGTTAGAATAGCATATTAACATAGCTTGATTCTTTATTTCATATAAGTACTATGTTATGGATAGAATTCCTTATTAACCATTAGCACTTAACAACAATGATTCTCCTTTCAGCTCAAACTAAAGCTAAGAAGTCGATGAAGCTAAAAGCAATCCTTTCGTCAATATCAGTTTTGATGTTTCTACTCATAGGCAGCATTAGCTACATTTACTATAAAAGAAGAATTGCAAAAAGAGCAGGTAGTTGCTGAATGAATCTTACTCTGAAAGATATTTTCTCTTATAGCATTTTGAACTTACTTTCAAAGGTCATTCTCAATCAGATAGAAGTAAAGGTATTCAAGGGGCACATAAATCTCACTGGCACAAAGCTGAAGGAGAAGCAAAAGTATTAATGAATGAAAACAGTGATGAAGCGATTGATGTTCCATATTTTCACTTGGAGACTATTTTGGAAGCTACAGACAACTTCTCAAATGCGAATAAGCTCGGACAAGGAGGATTCGGTCCTGTTTACAAGGTTCAAACTTCATTCCTACATCTATATGGACTGGCATGGCCATATTGACCTATAGTAATGAGATGAATTTTTTTTGCAGGGTATCTTTCCGGGCGAAAAAGAAATTGCAGTAAAAACGTTATCCAGTCAATCAGGACAAGGCATAGATGAATTTAAGAATGAAGTGACTCTAATTGCAAAGCTTCAACATCGAAATCTGGTGAGGCTATTGGGTTATTGCATCAATGCAACGGAACAAATATTACTTTATGAATATATGCCGAATAAAAGTTTAGATACATTCATATTTGGTGAGTTAGCCTTGTTAAGTTCTTTATTCTCAAAGTCATACATTTCTTATAATGATCTAATTTTATTGACGTACCAAACTTTGTCATGGTAACGATGCAAGATGGAACACTTTGTCAATTATTGGATTGGAAGAAACGTTATGACATCATACTGGGCATTGCACGGGGTCTATCTTATCTTCACCATGATTCACGACTAAGGATCATTCATAGAGATTTAAAAACCAGTAACATTTTATTAGATGAGGAGATGAACCCGAAAATTTCAGATTTTGGCTTGGCAAGGATTGTTGAAGGAAAAGTAACAGAAGCAAATACAAAGAAAGTAGTGGGGACCTAGTAAGTATTATATCTCTTCCTGTCATTAATTTCTTACAAAAGGAGCTTTGGATAAGATCTTATACTCAAATGTTTTGTTCACCTGATGAAATCTAGCGGCTATATGTCTCCTGAATATGCACTAGATGGATTGTTTTCCATCAAATCAGACGTATTCAGTTTTGGAGTAGTCGTACTTGAGATAATCAGTGGAAGAAGGAATACAGGATTTTATCAGTCAGAAGAAGCCTTAAACTTGTTGGGCTATGTAAGTGAAGAAACTTCCCATGACTTAATTATTATATCTTCTTCTTAGCTTTGTTTCTCTGATAGAAAAATTATCACAGGCATGGAAATTGTGGACAGAAAAAACTGAGATACAATTAATAGAGAAGTCACTACTTGAATCATGCAACAAAAGTGAAGCATTGAAGTGCATAAATATTGCACTCCTGTGTGTACAAGAAGATCCAAATCATCGTCCTAATATGTCAGATGTCATTTTAATGCTGGGAGGGGAAGGTACTAACCTTCCAACACCTAATAGACCAGCTTTTGTAATAAGGACACATGCTTCTAGTACATCTTCTTCTTCCTCCGATAAGAAATACATCGTATCCAACAATCAGGTGACGATTACAGTTGAAGAAGGACGTTAGCAGGAATAGTGAAGGCATTTAATTCGTATGGAAAGGGCTAGGGGTGGCACTAGCTAGGACTTAAAGATGAGCCATATTATACTTTTTTATCCTTGTTAACCTCTACTTTGGTCCTAAAAGAGTCCAGGGATAATTGTGTGTCGGTACTTGGCCAAATGTCGTCTTCTCTATAATAGTTAGGACAAAAGGTGGAAATGTTGGGAAAACTTTCAAGTGTTTGTACTCAAAACATATACAATTTTGCTTTAGAATGTAGTAGGTCTTAGCAATCAAGTACAAGAGTAATTTGATTATTCCTGATGCCTTCAAATGTACAAGCAATGTAATTTTTCTTTCTTATTCAATATCTTATTGGC

>PGSC0003DMG400008523 G-LecRK

CACACCCAAAAAATCAACTTGTTTGACTACTTTCATGGACTCTGAAGAGAAAAGTATACCTCCGTCAACACTTATTTTCTCTTATTGTTTCGTTCATCATCTTCTTTACCATGGAAATGAAGAACAATCATTCGTTTCTAGTACTTTTTTTGTGCTTTAGTCTCGGTACCAATCTTTCTATCGGAGAAGACACCATTTCTATGAATGAATCTGTTTCTTCAGGGCAAACAATTATCTCTTCAAGTGGGACCTTTGAGCTTGGTTTCTTCACACCAGGTAACTCTTTTAAATATTATCTAGGCATATGGTACAAGAATATTTCATCGCAAACTGTAGTTTGGGTAGCAAATAGGGAGACACCAGTTAGTGATGCTGCTCACTTGACAATTATTCAAGGCAACCTAGTGCTTCTCGATAAATTTCAAAGCTTAGTTTGGTCAACAAATATTTCAAGGAGTGTACCTCCAAAAAATTTAGTCATAGCAGTTCTTTGTGACGATGGAAATTTGATTTTGAGTGATTTGTCATCAAACTCATCAATACCGTTACTACTTTGGCAAAGTTTTGACTACCCAACACATGCATTTTTACCTGGTGGTAAGGTTGGATATGACAAACGCACACAAAGAAAACAGGTTCTGATTTCATGGAAGGATTTGAACGATCCGTCACCTGGATTGTTTTCCATGGAATTAGACCCACGGCGTGCCCAACTTGTTATAAAATGGAATAGGACAACACAGTATTGGGCAAGTAGTTCATGGAATGGCCGGACGTTCAGCTCAGTACCTGGGATGAGGTTAAACTACATATTCAATTACAGCTATATCGACAATGAGAATGAGTCATATTTTACATATTCCCTTTATAATAGTGAAATCCCATCAAAATTTATCATGGATGTCTCAGGACAAATTAAGCAACTCTTATGGTCGACTAGTTTGGATGATTGGTATCCATTCTGGGCTCAACCAACGGAACAATGTGATGTTTACGCCAATTGTGGGGCATTTGGAGTTTGCAACAATGTTAATTCATCCTGCAATTGCTTGAGCGGTTTCAAAGCAAGATCTGATGCAGAATGGAATTCAAATAATTATTCCAGTGGCTGTGTAAGAGTAAGAGACCAAGAGGTGCAGTGTAACGGAATTACTGAAGACAAGGATAGCTTTTGGATGAATTCAATAGTGTCATTACCTGCTTCTCAAGATACTAATATCACAGTTGCGGAAGCCTCACAGTGTAGATCTGCATGTTTCAACAATTGCTCTTGCACTGCTTATACCTATGATGTTTCTGGTGCATGTTCATTTTGGACCGGGGATCTGTTCAATCTACAACAACTCAGCAAAACTGAAACAGAAAAAACTATCTTTGTCAAAAGTGGCTCACCTGAAGGTTAGAATAACATAAAATAACTTAGCTTAATAAGTATCGAGGTCCACTTTTGCATTATGTTTTTCGAGAAAGATGTTAAACATATCTGACTTGTTTTTGGCTCTCTACTTTCAGATCAAACTAAGGCTAAGAAATCGATGAAGCTGGTAGTTCTACTTTCATCCACAACAGCTTTTATCGTTCTATTCATAGGAAGCTTTAGTTACATTTACTATAGAAGAAGAAGAAGAAGAATGACAAAAGGAACAGGTATTTACTGAATGAATCTCACTCTGGAAAATATTTCGTTCGTACCTTTTTTTAACTTAATTTGCTTTCTCAATCTTATTGAGGTAATGAGGTCATTCTGTTTTTCTTATTCCTTTTCCCTAAATGATACTCAAGGGACACAAATATCTCACTTGGACAAGATTGGAGGAGGAGAAATCATTGATGTTCCATACTTTTGTTTGGAGACCATTTTGGTAGCTACAGATAACTTCTCGAATGCAAATAAGCTCGGACAAGGAGGATTTGGTCCTGTTTACAAGGTTAAAATATAATTATTGCACCTCTGTGTACTGGTATGGCTTAATTGACCAATAGTAATGAGATGAACAACTTTTAACAGGGTATCTTTCCAGGCGGAAAAGAAATTGCAGTGAAAAGGTTATCTAGTCATTCAGGACAAGGCATAGATGAATTTAAGAATGAAGTGACTTTAATTGCAAAACTTCAACACAGAAATCTGGTCAGGCTTTTGGGCTATTGCATCAACGGAAAGGAACAAATTTTACTGTATGAATATATGCCAAATAAAAGTTTAGATACATTCATATTTGGTAAGTTTGCCAAGGTATGTTCTTTATTCTCAAGCACCTATACTATTGACAAGATTTAATCTTTGTCATGATATTTATGCAAGATGGAGAACTTTGCAAATTATTGGATTGGAAGAAGCGCTATGACATCATATTGGGTATTGGACGAGGTCTTGCTTATCTCCACCATGATTCACGACTAAGGATCATTCATAGAGATTTAAAAACCAGCAACATTTTACTAGATGAGGATATGAATCCCAAGATTGCAGACTTTGGCTTGGCGAGGATTGTTGAAGGAAGAAGAACAGAAGCAAAGACAGAGAAAATAGTTGGGACCTAGTAAGTATAACATCCTTCCTATTGATCATTTTTCCCTTTTGCTTTTGTAAAATACATCTAACACGTTTTCGACATGGTGTAATCAAGTGGCTATATGTCTCCTGAATATGCATCAGATGGTCTGTTTTCCCTCAAGTCAGATGTATTCAGTTTCGGAGTAGTCATACTTGAGATGATCAGTGGACGAAAGAACACAGGATTTTACCAATCAGAAGAAGCATTAAACTTGTTAGGTTATGTAAGTGCAGAAACTCAGACCTACATCATTTTTGTCCTTTTTGCAAGTTTTGATATCTAACTGGAAGTTATCACAGGCATGGAGAATGTGGATAGAAGAAAGGGCAATGCAATTAACAGAGAAGTCATTACTAGAATCATGCAATAGAAGTGAAGTGATGAAGTGTATCAATGTTGCACTCTTGTGCGTGCAAGAAGATTCAAATGATCGTCCTAAAATGTCAGATGTCATTGTAATGTTGGTAGGAGAAGGTACGAGTCTTCAGAGACCTAATAGACCAGCTTTTGTAATAAGAACACACGCGTCTAGCAGACCATCTTCTCCCTCAAAAGTCTCTAACAATCAAGTGACAATTACAGAAGAAGGAGGACGGTAGTAATGGTGTGAGATCATCTTAATTTTCATTTATATATGATACAATTGTAATGTTTTCCTGCAGACTTCTAAAGCACGAGCAGAAAAATAATCTATATTTTGTTAGTATCTTGATTGTAATTTGTAGCCATGACATTGTTATCCTTACACATATATTACATGCACTTTTC

>PGSC0003DMG400008524 G-LecRK

GTGAAGTTTCTCCACTTGCCATAATCTTGTCTCCAACCCCATTAAGAGAAAAAAAAAATTAAACAACAGAGCAAGAAAAGTTGGAGCTCAAGAAGGTGAATGATACTCAAAGAAGTGAAATAAAGGAAGAAAAATTGTTTGGATTTTCTAAAATCCAATTTAATTTAAGGTAAATTAGTTTTAGGTACTTTTAAACTTTATTTTGGAGTAAATATAATGTGTATGTTATTCATTTGTTTTTTGTGGCTTTTATGGTTCTTGAGTGATTGATTTTTGGATTTTCATCTTGGTTGTTGAATGTTATTTGCTTTTTGTGTGTGTGTGTGTGTGGATTTCATGGTTCTTGAGTTTAATCTGATTCAATAATGAATTCTTGAAAGTTAGTATAGTTTTTCATATTGAATTTACATGTGGAAACTACTTTTGATGTTGACTTTTATGTTATAAGCATTCTAGGTGGAAAAGTTATTTAAAAATTGTTATATATAACTTAAATCCTTTTGGTTTTTGTCATTTCAGGCATTTATGTTTGTTATTGGGTATATGTGTTATCAATTACTACAATTACCTAAAAAGAAAAGTTGTGTCTTACTACTCTCAACATCATCTCGTTGTCTAGCAAGATATGTCTAGTTCTCCCTCTACTTACCTATCCATTAGGAAAGTGGAACTTTCTTGACCTTCGAAAAGGTATAAAATGGTTTTATAAATATGGTATGGTATATGGACTAATTTTTTAGGTGCTACTTGCTTGCTTCAAGCACAATACTTTGATTTCTTTTTAAAGTTTTAGACAGAGGTGGAGCCAAGATTTAAAATTTATGAGTTTTAGATTCTTTTTATGTTACTGGGTTCTAAATTGATAAAATGTATAAATTCAAAAGATTTTAAAGACAAATACAAGATTTAAACTAAAACTACTGGATTCAGACTGAACCTGTAGCTGATATTCTAGCTCCGCCCCTAGACTTTTTTATTAGCATCCTCAAATACTAAAGTGGTCTTATTGCCCCTTCTTTTGTCACACTTTTTAGCTAGTTTTCTCAAAATGCATGAAAAGGAAGAGATGCTCTATTTCATCTTTTAAGTTCAATTCTTGAGGCTATATCTTGAATTCATTTTATTTTTTTTAAATAAATAAAATTGAGGGCCTAAGAAAGAGATTACAATGTGGGGAATCGAATACTCACACAGGCAGACAATCAACTGAGTTACGAGATTCTCCTTTATCTTGATTTCACATGATTTGCTATATTTTGTTGTGATTCAAGTTGTCTATCTATTTTAGCTTTAGGTTATCAGTTCTGTTCCTATTTCAGGCTAATTGCAAATGATTTGAGGCTTTGAAGGTTTACGAGGATGTTGGACTTAAACAAAATAGAACCTGAAGAAGGAAAGCTAGACTTGGATCTTCGTCGCGTTAACTATGATCTTGATGACAAATCTCATCTTAGATATGGTGATTGGGTCACTTTCAAGAGCCATTCATCTAAAAAAGTTAGAATCTCTCGACCTGAATCTCCGAGAGGTGAAAATTCAGATAGACAACCTCAGGATGCAGATTATCCCTCACTTAGCTATGAACTCGAAAGTGAGATTCTATCCAGGTTTCCTAGGTGGGAGTATTGGAAACTTAGCCTCGTAAACAAGAGATGTTCAATGCTTCTAAAGAGTGGTGAGATATTTGAGATTCGTAAAGAGAATGGATTTAAAGAACCTTCGGTATATATGTTAGCAAGTGGGGAGACTAATTGGTGGACGTTTGATAGAGAGTTTAATTCCCGGAGGAAAATCCCTGATTTACCATCAGATGTGTGCTTCACATTTGGCGATAAGGAATCGCTATGTGCAGGCACACATTTACTTGTTTCGGGTAGGGAAATCGATGGTCTTGTTATATGGAGGTTCGAATTAGCAACGAATTGTTGGTACAAAGGTCCTTCTATGGTTAATCCAAGATGTTTATTTGCATCAGCTACTTGTGGCACCTCTGCTTTTGTTGCTGGTGGTGTTGGTATTATGGCGAACAGTGAGGTCTATGACACAGCTGAGAAATACAACCCCGATAGCAGATCGTGGGACCCACTACCGAGGATGAAGAGGAAGAGGAAGCTTTGTTCAGGATGTTACATGGACAAGAAATTCTACGTGATCGGGGGGAGGAATGAGAACGGAGAGCTTACCTGTGGAGAATTCTTCGATGAGGGTAAGAACAAGTGGGAGCTGATTCCGGACATGTTGAAGGATGATCCAGTCCTAACATGCCACTCACCACCCCTCATTGCTGTCGTGAACAATGAGCTGTACTCACTCGAGGCATCTTCAAACCAGCTGAAGCTTTACCTGAAAAAAACCAACACTTGGAAAATGTTAGGACAAGTGCCAGTTAGAGCTGATTCAAACAGAGGATGGGGCATTGCTTTCAAGTCATTGGGAAATGAACTTCTTGTAATCGGAGCTGCTTCATCGTCAGCCTCTTATTCGGGTAATTCCATGGCTATATATACTTGCTGTCCAGATCCTGATGCAATTGAGTTGCAGTGGCAACCTCTTGACAGTGGTCGAAATCGACTTAGTAGCTTTATCTTAAACTGTTCTGTTATGGTAGCTTGAATAATGTCATTTATCTAGCAAACAACTTGTGTATTATGATCAAAATATATAAGTGGCATATGGCCACTGGTTTATGATATTTCTTGCATATATTCACTAATGTTTGCTCTGCTATGTTTGATGTAATTTGCATTTATTGTTGCATTATTATGATTTATATTGAGGTGTTGTTCTAA

>PGSC0003DMG400010240 G-LecRK

AAAACAATGTCTTCTTTGCTTGTAATAACACTTTGTTCCTTATTCATAGTAACATTAGCCAATGTTCCTCCTTCCAAAACCTTCAAGTACATAAATGAAGGCTCATTTGGCGAATACTCCGTTGAATATTTTGCAGATTATCGTGTGTTGGACATTGTCAACTTCCCTTTCACCCTTTGTTTCTTTAATTCTACTCCAAACACCTTCATACTTGGGCTACGAATGGGTCATCGTAGATCTGAGTCCATTATGCGATGGGTCTGGGAGGCTAATCGTGGTAAGCCTGTCCGCGAAAACGCCACTCTAACGTTTGGTACTAATGGGAATTTGGTACTAACTGATGTCGATGGCACGGTAGCTTGGCATACCAATACTGCCAACAAAGGTGTAACAGGGCTAAAACTTTTGCCTAATGGCAACTTGGTTTTACATGACAAAAATGATAGATTTGTTTGGCAAAGTTTTGATCACCCAACGGATACACTTTTGGTGGGCCAAAGTTTAAGGGCTAAAGGCCCATCTAAGCTTGTTAGTAGAAAGTCCAGTAAAGAACCTGTTGATGGGCCTTATAGTTTTGTCCTAGAAGGGAGACATTGGGCCATGTATTATATAAGTGTAAATTCACAAAAGCCTTTACCATATTTCAAATCAGATGATTTTGGTAATGGGAAAGGCTCTTTAGCCAATTTTGATTTCTATTGTGAGCCAGAATATGGTGAAAATTATGCTTTTGAGCTAGGATTTAGGATTAACATGACCAATTCTCCTAGCTCTGGTACTTCTATATTTACTAGGCCCAAATACAATAGTACATACACTATGTTTAGAGTTGATATGGATGGTAATTTAAGAATGTACACTTATGAACCAAATGTTGATTGGGGAGCATGGGAAGTCACATATCGACTTCTCGATAGGGACAACGATGGAAAAAGTGAATGCAAGCTACCGAGGAGGTGTGGATCATTAGGCGTTTGCGAGGATAGCCAGTGTGTGGCGTGTCCGAGCCCTCGAGGCTTGATTGGGTGGAGTAAGGGATGTGCACCACCAAAGTTGCCAGTATGTAGGGGAGGATCAAATGTGGACTATTATAAGGTGGAAGGAGTTGAACATTTTACTAGTGATTATAATGAAGGAGTTGGGCCTATGAAAATAAGTGAATGCAAAAATAGGTGCAACAAAGATTGCAAATGTGTGGGCATTTTCTACAAACAAGAGTCATCAAAGTGCTTGGTTGTACCTGAGCTAGGGGCTTTGGTTAAAGTGGAAAATTCCTCCCATGTTGGCTACATCAAGATCTCAAAGTAGATGAAGTTATTACTGATTTATATTTATATCAAGCAAGCATGTTATATTTTTTAAATTTCAGTAATAATAATGTAACTTTTGTCAAAATATAATAAATAATAAATGCAC

>PGSC0003DMG400011251 G-LecRK

ATGTTAATTTCTCCAAAAAGGAAATTTAATTTGGGATTTTTTAGTCCAAATGTTTCTAATAATCAAAGGTTTGTTGGTATATGGTATGTTGATGGTCCAAAAAATTCATTTGTTTGGGTTGCAAACAGAGATAAGCCAATTTTTGATAAAAATGGTGTTTTTACTATTGAGAAAAATGGGAATTTAGTGGTGAAAAATGGTCGTGGCGATTTAATGTGGACGAGTAATGTTGCTGCTATGAATAATAATTGTACAGCACATCTTTCTGATTATGGAAATCTTATGATGTTTAATGGCAATAACAGGGAGTTGTGGCAGAGTTTTCACCATCCTACAGATACATTTTTACCAGAAATGAGATTTTATTTGGATGAAGTTTTGAGGTCATGGACTAGTGAAAGTGATCCTTCACCTGGACGGTACGTAATCTTGTATCGTGTGATCATCTTATCTGAAATCTTTTATTTACTTAATTTTGTTTTGTACATGCCCCCTCACGTGTGGATGAATTTTTTATATAATGTGTGGTGGTGAGACTCTCTGATAGCATGTTGCGTTTGTGACATCTCATTTAAAAGCTTAATTGAGAATGTGCTTTTTTATTTTACGTAATTTTGTTTTGAACCCCTCACGTGTTTGACTCGTGGAAAGTTGAAAGACAGAAGGGTTGTTGGTCAGTCACGGACTATTTGATTTCATTATGACGATTCAATGTTAATGACCTCATTCAATATTTTTTTTTTTTGATCCTTCACTATCGTGGGCTGGATGGATAGCTAGTAGTTGTGAGTAACAGTCCCCTTTTTATAGATTAAAGGTGATGATTTTTTGATATTGGGTGGCAGTGAGACTCAAAACCTAAGAACTCAGCTTGCCCTGATATGGTAATGCATTTGTGCCATCTCATCCAAAAGATTAAGTTGAAAATATGTTTTTCTTCACTTAATTTTATTTTTAGCATGCCCCCTCACATGTTTCTTGAATCTTTGATATTGGGTGGTAGTGAGACTTGAAACCTAAGAACTCTACTTGCACTGATATGGTAATGCATTTGTGACATCTTATCGAAAAGATTAAGTTGAAAACATGTGTTTCATAACTTAGTTTTGTTTTTTAACACGACCTCTCATGTGCTTCTTGAATCCTTGATATTGGATGGCAGTGAGATTCGAACCTAAGAACTATGTCTGTTCGGATACAATGTTGCATTTGTGACATCCCAAAGTCAAAGAGCATATTTTTATTTACTTAATTACGTTTCCAACAGGTACTCATTAGGAGTAAATACTCGCGGATCACCACAAATTGTGATCTTTGATGGAGAGAAGAGACGTTGGCGGAGTGGATATTGGGATGGACGCATATTCACAGGCGTTACAGATATGAAGCCCGAGTATCTACATGGTTTCAAGCTTTACAATGAAGGAGATAAGCTATACTTCACTTACACTGTATCAGATCCTTCTGATTTGGTGAGGTTCCATATTAGTCCAACCGGATACGAGCTGGAACAAAGGTGGGATAAAGATAACGAAAACTGGAGTATAATACAGTCTCATCCATCAGGTGATTGTGACTTGTATAACTTATGTGGGAATTTCGCGAAATGTGATATCACATATTTGAAAAAATGCATTTGTTTAGTTGGATTTGTACCGAAAGATTTGGGGCAGTGGAATGCGAGGAACTGGTCAGAAGGGTGTGTTAGGAGGAAAGAAGTGGAATGCAGAGGAAATAACAGTGTCTTGAAGAGTGGTAGTGGAAAGAAGGACGGATTCTTTGAGATTGAGAAAATCAAGTTGCCGGATTTTGCTGATACTGCATATTTACAAAATATCGATGAGTGTCGAAGCATGTGCCTTGAAAATTGTTCATGTACTGCTTATGCTTTTGTTAGCGGAATCAATTGCATGATGTGGAGTGGAGATTTAGTCGATATGCAGCAGTTTCAGGAAGGCGGAAACACTCTCTATGTTCGCCTTGCTGATTCTGAATTTGGTAAAAATGTGTACTATTTTTTCTGTTGTTATTCATATAATCATTTTAGATGTTCTTATTATAAACTTTCTTGAAACAGCTGGAAGCAACAAGACAGTCAAAATTGTGGTAATATCCGTTATGGTAGCCGGGGCTTTTCTTGTATGTATGGTAGTTTTTCTACTATGCAAGTACAAAGCCAAAACGCGAGGTAAGATTGCAGAAAAGCTGGTGTTTTACGACAGTCATTTATTGTTATGCAATATACCTATATCTTCTTCAATGTCCCATTTTCAGTGTCCAATAAGATCAACCAAATGGAGACAAGCGTTCCAACTAGGAGCGGAGAGTTTTCTATGAACACATCAGGAGCAGGAGACTTGAGTATTGAAGGGCATCAAGGAAGTGGTTCAGAACTTATATTCTTCAGTTTCAGCGGTGTAGCAGCAGCTACCGATAACTTTTCTAACGAAAACAAGCTGGGACAAGGTGGATTTGGCCCTGTCTACAAGGTTGGTTTCCATTCAATTGACAAGCGTGTCTTACTTTTCTAGAGAAGTATGACATTATTGTGTTAAACTAACGTCGATGTTTGTATATTTAGGGAAAACTACTATGTGGCGTAGAAATTGCAGTGAAGAGACTTTCAAGAAAGTCAGGACAAGGTGTGGAGGAATTCAAGAATGAGATCAAGCTAATCGCCAAATTACAACATAGAAATCTTGTTAGACTTATGGGATGCTGCATTGAAGGAGAAGAAAAGATGCTTTTTTACGAGTACATGGCCAACAGAAGTTTAGATTCATTTCTATTTGGTTAGTACTCTAGATATGCTACAAATTATTTCATCGATAGAGAGATCGATCACTTACAATTATTTTGGTTGATAAATTTTTGACAGATCCTGTTAAGCAAGCTCAATTAGACTGGAGGAAACGCTTCAACATTATTGAAGGGATTGCACGAGGGCTATTGTATCTCCATAGAGATTCGCGACTCAGAATAATCCATAGGGATTTAAAGGCTAGTAACATTTTGCTAGATGAAGAGATGAACCCGAAAATTTCAGACTTTGGTATGGCTAGGATATTTGGTGGAAATGAAAATGAAGCAAACACAAATAGAGTAGTTGGGACATAGTGAGTATCATTTGATTTTTATACCTTTAAGAGTGAGTACCTTTAACTTGCTCAAAGAACTAACTCCATCAACGCGTTTCTTTTTTTGTTTTTACTTAAATCAGCGGATATATGGCTCCTGAATATGCAATGGAAGGTTTGTTCTCCGGAAAATCTGATGTTTATAGCTTTGGTGTACTATTACTGGAGATCATCTGTGGACGGAGGAACACAAGCTATCGTTCAAATGAACACTCTGGCATTATTGGTTATGTATGTCCAATAACTACTTCTCGATATCTGTTGTGAAGTTAAATCTTTTAACAACAATTCTACGTTTTTTCTTTTTAGGCATGGCAGAAGTGGGATGAAGGTACACCAATGGACTTAGTAGACCGTTCTATTTGGGACGAGTGTCAATATGATGAAGCATTGAGATGTATACAGTTAGCACTAATTTGTGTACAAGACATGGCAGTTCACAGACCGAGCATATCATCCATCGTGTTAATGTTGGAGACGGACAATATACGATTGCCCTTGCCTAGGCAACCTACATATACCTCAATGAGAAAACATGAAGATGCAGAAACATGGAATGAGAAGCAGGATTTTTCTGCAAACAATGTCACAATTAGTGTTATAGTTGGTAGATGA

>PGSC0003DMG400013383 G-LecRK

AGACTCCATAGCTGTAAAGCTTGCAAACGATGTCGCTCCATCCTTCTTCTTCATCCTCACTTCTCTTCTTCCTTCTCTTCTTCTTCTTGAACCCCTTTTTCTCTTCTTCACAAAAAATCCTCTCTTCATTCAATTCAAAATCCCCTCCTTGGAATCCAACCCAAAACCAAATCCTACTTTCACCCAACTCCACTTTCGCTGCTGGGTTTCTTCAATCTTCCCAAAATTCCTTCAATTTCTCCATTTGGTACTACAAAATCCCTGTCAAAACCATTGTTTGGTCAGCTAACCCAAATTCCCCACTTAATTCCTCAGCTACCCTTTTCATTTCTTCCTCCGGTGAACTCAAGCTAACCCCATCTACATCTTCCTCAGCTCCAAATTTATGGCCTTCAATTATTCGAAACACCTCCTCTGTTCTTTTCCTACAAGAAGATGGAAGCTTGGTTTATGGAAACTGGAACAGTTTTTTGAACCCAACTGATACTTATCTTCCGACCCAGAACATCACCGGAACAAATTTAACTTCCGCTAATGGAAAATTCCAGTTTGATGGATCGAATTCGAATACCCTTTTCTTCAATGGAAACGATAGTTATTTCACTTTTTCCCAAAATGCCCTTCAGAGGTTGGAGGAAACAGGGGAAGTTACTCAAGTTAATGGAAAATTTGTTTCGTCGGATTTCGGGGAGAAAGGGAAACTGAGAAGGATGAAATTGGATGAAGATGGGAATATGAGAATCTATAGTTTTGATCTTAGTTCAAGTTTAGCTAAAAATTGGACAATTGTTTGGCAAGCAGTTAATCAATTATGTACAATACATGGTACTTGTGGTACAAACTCTATATGCTTGTATGATACTTCTACAACTCAAACTTCTTGTGTTTGTCCACCTGGTTTTAGGAAAGACACTAGTAAATCTTGTGTGAGGAAAATTCCATTGATGACGAAGGATTCGAAGGCTAGTAAGTATTTACCTCTTGATTTTGTTAGTTTTACTGGTGTGGGTAACCAAACAGACCTTAAAGCTTTGAGCTTTTCGAGTTGTGAAAAAAATTGTTCAGATAAGAATGATTGTTTAGGTTTTTTGTTTAAGTATGATGGGACTGGATATTGTGTCCTTGTTTTGGAGAAATTGTTGTATGGATATTGGTCACCAGGTACTGAGTTTGTTATGTATTTGAGAGTTGATAGTAGAGAGAACGATATATCGAATTTCATAGGCATGACTAGTTTGATGGAGACTTCATGTCCAGTGAGAATTAGCCTTCCATTTCCACCAGAAGAGTCTAAGACAACCACTAGAAATATTGTGATCATTTCGACGATTTTTGCAGCTGAGTTGATTAGTGGAGTGTTCTTCTTTTGGGCATTTCTCAAGAAGTATATTAAGTATAGGGATATGGCTAGGACTTTTGGTCTTGAAGTCATGCCTGCGATTGGACCAAAAAGGTTTAGCTTTTCTGAGATCAAGAATGCAACTAATGATTTCACTGATAAAATTGGGAGGGGTGGATTTGGAGATGTTTACAAAGGGAAGTTGAGTGATGGACGTGTTGTTGCTGTGAAGTGCTTGAAGAATGTCAAAGGTGGTGATGCTGAGTTCTGGGCTGAGGTCACGATCATAGCACGTATGCACCATTTGAATTTGGTGAGGTTGTGGGGGTTTTGTGCTGAGAAGGGTAGAAGGATACTTGTGTATGAGTATGTACCTAATGGTTCATTGGGTGAATTCCTCTTCCAAAAGTCTCTGATCCAGTCACCAGATGGACAGAAACCTATACTCGATTGGAATATAAGGTACAGGATTGCGCTTGGTGTGGCACGAGCAATTGCCTATCTACACGAAGAGTGTTTGGAATGGGTATTGCATTGTGATATCAAGCCTGAAAATATTCTCTTAGGCGATGATTTCTGCCCAAAGGTGTCTGATTTTGGGCTAGCTAAGCTAAAGAAGAAAGAGGAGATGATGACCATGTCAAGATTTCGAGGCACGCCAGGATATGTAGCACCAGAATGGACCAAGGCTGATCCAATTACTCCAAAAGCCGATGTGTACAGCTTCGGATTGGTGCTGTTGGAGATAGTTTCAGGTACAAGGAACTTTGAGCATCATAACTCAAAGGTGGAAAGTGATCAATGGTTTTTCCCTGCTTGGGCATTTGATAAGGTGTTTAAGGATATGAATGTTGATGATATTTTGGACCCTCAAATCAAACAATCGTATGATTCAAGGGCACACTTTGACTTAGTGAACAGGATGGTGAAGACAGCAATGTGGTGCATCCAAGATCGGCCAGATGCACGACCATCAATGGGGAAAGTGGCTAAGATGTTGGAAGGGACGGTGGAGATCATCGAGCCAAAGAAGCCTACCATTTTCTTCTTATCAGAGGAATAATATATGGCTCAATCCTCTTCATAATTTCCTCAAAATGCTGCCTCAACAAGTTGCAGTAAGCATGTATTCCATGTTGTTTTGTATTGTAATAAAAGAAAGTAGATAGGAATTTGGATCATGTTGTGTTTTGTCTGTTTTCTTTTTCTTTTCTTTTTTCCAGTTTTGTAATCTCTCTTGTGTTGTAGTTGTAAAGTACATTTTGATCAATGAAGAAGCAAATGGTTGTTACATAGAAG

>PGSC0003DMG400013882 G-LecRK

TTCATTTTGGTCCAGAGTTTAAAGTTTGAGGAACCTAAACAGAGAAGATTACCTTGTTAGAAAGCATCAGTTATGTGTCAGAAGAAAGAGTTGTTGATAGTAATTGTTTCATTTCTTGTCCCATTTTGTAGTTCCATAGATACCATTTCCTTTAACCAATCTCTCAAAGATGGTGATTTGTTGATCTCTTCAGGGAAATCTTTTGCTTTGGGTTTCTTTGGGAATTCCCCTGATAAACGTTACGTCGGAATTTGGTACAACAATGTTCCTGAACTAACTGTTGTTTGGGTTGCCAACAGAGACAATCCTGTCAATGGCACATCTGGGATTCTCACCATCGATTCAACTGGAAATCTTGTTGTACATCTCGATGCGGAAACAAAAACTGTAGCCTGGAGAACAAATGTTTCTTCAGCTACAAAACGAGCAGACTCGTATACTGCTAAGCTGGAGGATTCAGGGAATTTTATATTGTTTCAAGACTCCAAAATGGATGTTACTGAGTGGCAGAGCTTTGATTATCCTACCAATACTTTGCTTCCTTCAATGAAATATGGAATAGACAAGAGGACGGGTTTGAACCGGTTTCTAACATCCTGGAAATCGTTGAACGATCCAGGCACTGGAGAGTATCGTTACACAATGGAGCTCAATGGGACACCTCAAGTTTTCTTGTACAAGAACTCTAGCAGGATTTGGCGGACAGGATCCTGGACGGGTCATGGTTGGAGTGGTGTACCAGAAATGAGCCCACGATTCATCTTCAGCCTTAGTTATGTGGACAATGATACCGAGGTTTCCTTGACATATGGCATACATGACAGTTCAATAATCTCAAGAATGGTCCTGAATGAATCCGGGATCGTGAACCGGTTAACCTGGCAAGAGAGTGAGCAAAAGTGGGTGCAATTTTGGTCTGCCCCTAAAGACCCTTGTGACAATTATGAACATTGTGGAGCATTTAGCAATTGCAATCTGTTCAATTTGGCTGAATTTGAATGCAGTTGTCTTCCAGGGTACGAGCCAAAGCTAAGCCGACAATGGTTCTTGAGAGATGGCTCTCATGGATGTTTGAGGAAGAAAAATGAAGAGGTGTGCAACAATGGTGAGGGATTTGTCACATTAAGTCATGTAAAGGTCCCCGACACAGGTGCAACTAGGATGAACAAGAGCATGGGATTGAAGGAATGTGAAGAGTTGTGTCTGAAGAATTGCTCCTGCACAGCCTATGCAAGTGCTAATATTAGTGCTGGAGGAAGTGGATGCATCACTTGGTATGGTGAGTTGATAGACATAAAGCAATTTACAGATGGGGCCCAAGATTTATATATCAGAGTCTCTGCATCTGATTTAGGTAAGGAGTTAACTTAATATATTATGTGAGGATTTAGGTAATTCAATATTATGTGGAGAAGAATGTGCAAGTTGATCATTTTTGGTTAGTAGTAGTACACAGTATCTTACCCTGGTGTTTACTGGAAGTTCTCATTTTTTGTTTGTTCAGCGCAATTCTCGAAGAATTCAAGAGGCCATAATAGGAAAAGAATGATAGGGATTCTAGCAGGGTCTGCTGCAGCAATAATTCTTGCACTCTCTTTGGCATGTTGTTTGGTGATCAATATAAGGAGAAATGGTAAGGATATTATTCTACATCATACTTTAGTCGAATATTTTCAGTATCTGTACAGCAGCTATAATGGTGAAGCAACTATAGCAACCTAGACCTTCTACTACCTTTCTATATCGTGCAGCAAAAACAATAACATTACTGCAAATTTCATTTGATCCTAATCGGTGTTTGCTGATGTCACAGATAATGAGAGAAGTGAAAGTATAGCATCCTATGACGGCATGGATGAAAGTAAACATACAGAAATCTCTATCTTTGATTTAACCACAATAACTAATGCCACTGATAAGTTCTCTGATGCTAACAAACTTGGTGAAGGAGGATTTGGTAGCGTATACAAGGTAACAACTCCCAAACATGATTGAATTCCTAACAATCAACATTATCTTTATAGGAGGCTGCAAATAGATTTAATAATACTTGTTAAACCAAGATTTTCTCCCTAAAAATAAGAAGAAGAAGCTATGTTTACCAAATAATAATCTGGAATCAATATCAGGGTCACCTGACAGATGGACAAGTAATAGCTGTCAAAAGACTTTCAGTAACTTCAGGACAAGGGACCGAGGAGTTCAAGAATGAAGTCACACTAATTGCAAGACTTCAGCACAGAAATTTGGTGAGGCTTTTAGGATGTTGCGTTCAACGAGGAGAGAAGATGTTGGTTTATGAATATATGCCAAACAAATCCTTGGACAGTTTTATTTTTGGTAAGGAACATCTCCAGAACCGGAATTTCTTCCATTGACCTCCAGTCTCTACATGCTTTTACATGTTTACTGATGATTAATGATTACTGCTTTAACGTGTGTGTGTGTGTCCCTAAAAGATTTCTTATGCACAAATTTTTAAACTGTGATCCAGATAAAACAAAAGGGTCTTTGTTGGATTGGGGAAAACGGTTTGAAATCATACATGGGATTGCACGAGGAATGTTATACCTTCACCAAGACTCTAGACTAAGAATTATTCATAGGGATCTAAAAGCAAGCAATGTTTTACTCGATGCCTCCATGCAACCAAAAATATCAGACTTTGGAATGGCCAGAATATTTGGAGGTGACCAAATGGAAGCAAACACAAATCGAGTTGTTGGAACATAGTAAGTATGTCAAGTTGTGTATAAACATAAATGAGTTATGTTAGATATGTGCATGTGATCTTTAAAATGATAAATCTTGACATTGCTTATCTATTGCAGTGGTTATATGTCACCTGAATATGCAATGGTAGGGCACTTTTCAGCAAAGTCTGATGTTTTCAGTTTCGGGGTTTTGTGTTTGGAAATTATCACTGGCAGAAAGAACAACAGTCATAAGGATCAGGAGAAATCTCGACATTTAGTTGGATATGTAAGTACAAACATATATATGCTGTTAAAATCGAGAAATTGCATTTATTATCTTACAGTTGCCTTAGACATAAGCTGTGTCATGGATGGTAGTACTGCAAATAATAATTTGGAAACAATTCCCTTTCAGGTTGTTAAAAGTGCTGAATAGATCGAGATTTTGTATTTCAGAATTTCAATTTTTGGTGTTATAGTCGCTTCTCTTATAGAGAAGAAATTGTCTGAAATTTTGATTGTGATGCATTGTGGTGTTTCTATTTCAGGTTTGGGATCTTTGGAAGGATGAAAAGGCTTTAGATGTTGTTGATCCATTGTTAGGTGGCTCATATGAAGCTTGTGAAGTTTTGAGATGCATCCACATTGGCCTTTTGTGTGTACAACCGTTTACAGATGACAGACCGACTATGTCAGAAGTGGTCTTCATGCTGTGCAATGAAACAAATCTTCCTTTCCCTAAACAACCTAGCCTTATATTCAGATCAGAGAATCATAGTTCAGTTACACACTCTTCATCAGCAAGCATAGCAACATCATCTGTCAACGATATGTCTATTTCAACAGTTCATGGTCGCTAGAGCTTATGCTAACTATTCAAGAACTATTTTGAAAGAAATACAGAATTCCTTTTTCTGTACTTTGCTTTCATAGTATTAGAGAGATTTATTGGTGGGTTATACACAACATTTAACATGCTAGTCCATTCTTGATTCAAACTAGTGGGGTTGGGCTTTAATAGCACTGGAACCAAAACTGACTTCTGATTCCCCATGGCCCATGTTACAGACTTAAAGGTAGGTGGGTGGCAAAGGATTGGTAGGAAAGAAGCATAAACACAGAGGGTTTCTATTTCATCTGTGAATTAAATGGTGCTCTTTGTCAGCTGTTTTTGATAATTGCCCCACTTCATGTTTTCTTTAACAGTCTTCTCTGTGTTCATCAATGCTCTAAAAGCAACAAAGGCTAATCCTGCTAGCCACACACAACTTTTCAATTTTTTTATTGATTTGCAGGATAGTAATTTGGTTGTGGAGTTTTGTCAATAAATGAGATATTTAACTATTATTAGTTGGTTATGTTTTTCCTCAAAAAGTTATGGATAAGAAAATACAAACATGTAAGTTATCTTATGTAAGATTTTAACTTTTGTTTCATAATTAGATCATGAGATAGACATGTATCAAATACAACAATAATATTATGATCATGAGTAAGAGCACTGTTGTCTCAATCTTTCTGTGATGAGTATCTGTAAAAGTAATATCAACTAGTCTGTTGAAAATCTTTTAGCATTAAATGCATTGAATCATTGAAGACTTATATCATACTTTGGGAAGTAATGAGCACTAGATACATTTGATACTCAGATAACAGAAATACATATACACGTCAGACAACTAACGAACATAAGGTGCGGTAATTGTCACCTCGTTGATCGAAGATGATGCTGTTTCTGTAGTAATCAAGTCTGAGTTACGGGTAATGAAGGGCTTCTTCGGGGAGGGAACCGTTGCTTCAATACTTAGCATGGACAGAATTGTTGACATGGTTGGTCTGTCTGCTGCGTTGAGTTGCAAACACAATAGTCCTACTAGAATGCAGCGTAATGCTTCCTCACTGGGACAAGAATCAACTATTGATGGATCTACCAGCTCAAAGGCCTTACCATCATTCCACAACTCCCAGGCCTGAAGCAGCAGTTCA

>PGSC0003DMG400013885 G-LecRK

GGGATGTGTACCAAAATACCATCTTCATTCCTTCCCTCTTCCTATCAAACAAGAGAACAAGTACACCAACAAATGTCTCAAACACCAACACTGCAGCAAAACAAAATACTCTTTTCTTAAATTCCATAATGAGAAACACCAAAGATTACTACTTTCACCAATTCTTGTTCTTGATTTTGATTTCCCAATTTCTTCATCCAATTCTTGCTATACCCACTGATACAATCACCCCAACACAGCCTTTAACCAAAGATCAAACCTTAGTATCTTCCGATCAACACTTTGAATTAGGCTTTTTCTCTCCCGGCGGTGCAAATTCAGACAAATGGTATATCGGTATTTGGTACAAAGAGATTCAAGACAGGACTATTGTCTGGGTTGCAAACAGAGCTAAACCCCTTTCAGCTTCATCAACATCAGTCCTAAAAATCACAGAAATTGGTACCCTTCTTCTCGTCGATGGCCAAACTGGAAATCCCGTCTGGTCTTCCGATCAAACTCCGGCGACCAACGTAGTAGCCCAACTCCTTGACTCCGGGAACTTTGTCATTCGACCGGAAAACGATGACAGAGAACAGAGCTACCTGTGGCAGAGCTTTGATTATCCAACAAACACTTTATTACCTGGGATGAAACTTGGGTGGGATTCAAAATCCGGGATGAATAGAAACATAACATCATGGAAATCAGCAATCGATCCAGCTCCCGGAGATTACACCTTCAAGATCAACACCAGTGGGTTCCCGGAAGTTTACTTAACAAATAAACAACAAATTATCTACAGGAGCGGCGCTTGGAATGGAATTAGATTCAGTGGGGTACCGGAAATGAAGGCAAGTGAGATTATCTCATTCGAATTTCAGTTCAAATCAGATGAAATTACTTACATGTTCAAATTACACAACAAAACTTTATACTCAAGATTGTCTGTGAGTCATACTGGTTTTCTTGAAAGATTCACTTGGATCCCAACAAGCAACATATGGAATCGATTTTGGTATGCTCCAAAAGATCAATGTGACGGATACACAGAGTGTGGGGTATCTGGAATTTGTGACACAAATATTTCACCAATCTGCAAATGTATGGTGGGATTCAAGCCTAAAAATCAAGTGGCCTGGGAATTGCGAGATGGGTCTAATGGTTGTGTTCGTTTTCATAAATTGGATTGCAAAACTGATAAATTTAACATATTGAGGAACATGAAGTTGCCAGATACGACGAATTCGTTTGTGGATACAAATATGAATTTGGATGAATGTGAAGCTATGTGTATGAAGAATTGTTCTTGCACGGCCTATACAAATTCGAATATTAGCGGAAGTGGGTCGGGTTGTGTGATTTGGAGCACCGAACTTGTTGACATGCGGCAGTATGCGGCGGCGGAAGGTGGCCAAGTTCTCTACGTTAGGGTGGCTACTTCTGATGCAGGTATCATCCTATTTTTCTACTTCTTTTTTCGCTTAGTACAATGATAACGAATTAACGATTAATAATCGATAAAAAATAGTTGTATTTAAATCGTATTAAAAAGAATCTGTACAATTAATTTTAAGTAATTTGGGGTTGGAATTTGATTGTTAGATAATCCACTTCTGTGTTAGGATCAAATTTTGGGGATGATTTGAGTCGACTCGGCTGCACGAGTGGAGTTTGTTGGAATTTTTTATTTACTTGTTAGGTTGGATGATATTTCAACACAGAGAAGTAGAGTAAAATAGAAAGGGTGATGAAATTTTTGGAAAATATTTTTGTGGAAGTTTGGAGAGTATTTCCATGATTGGAAATATGCAATTTAATACTTTTTATCTCAATTTATGTGCTACACTTTTTAGTTTGTCTACAAAGGAATGATGCATTTTTTTTGGTAAGTAAACATTTTATTTGTCAAGTAAACGTTACAAAACAACATTTTTATATTTAGTACTAACAATTTATTATCTTTAATTAGATGATTTATAGTCACACAAACATCTATGACTTGTTTAGACCACACATTTCAAAAATCACTCTTTCTTAAATTTTATATCAAATCAAATTACGTCACATAAATTGAGACAAATAAAGTAGTATAATGTTTTGCACATTTTTGTCGGAGAATTCAAGTTTGATGCAAAGTTAATTTATTTAATTCATTCGGCGAACTTGCTGTTCTTATGTTCTCTGAATTATTCCACTATATAAACTGTATATATTGTTTTGTGTTTGAATTTATTTCCATAAATTCAAAAGTCGAGGACAAGTTGAAAAATCTGTATGTCTCAAAATTGTTACTCATTTTTGCATGTCATCCGATCCAAATATCCTTGAATTATATTATTCTTTTGTCATTTTAATAAGAAGCATATTACACATTATAATAATTTTAATTTACTCTTTTCTGATTGAGTTCCAAAGTTGGCAAACTTAGCCCTTGATCAAAGATAAACCATGTGCATGCAGTTAGTTTTGCCTCTAATAGATGAAATTATAAAAATATATTTTACTTTTTTGAATTCTTCTTTTCTATTTTATTTTTTTGGGTATTGCTATTTTTTTAAAATTCCCCCCCCAAACGTCATGCAGAAATTAAATTAGAGTATTTATTTTGAAAATAAATACTCTATTCCCTTTAAACAACACGTATAGTTGTCTTTTAATATCCTTGGTCCCCTTTTTAAATTCCCTTTCACGTTACCTACTTTCGATCTAATATTTTATATAATCAACTAGATTTCAAAATAAAAATAAAAATTCATAATTGCTAGTTGAAAAAACAAGAAGTTAGTATAATAATTATTGAGTGCATTTAATCCCGAAAAGGACTTATAAACAACCTACTTTTCAAAGTTATTCTGCTAATGAGTTTGCCAATATATTCTTTGTGGTATGCACGTTTTCAATATAATTTTTCTTTTTCTATGCTTTGTGTATTTTACTTTGAAAAATCGAAAAGTATAAAAAAACCTAACCATATCTACCAATCCCCATCAACAATTTGCCTTCAATAGGGACTTAATTTTATTTATTTACTATATTTGGACTTTCTTTTTCTCATTTGATTTGATAGGGATGATTAGTATTTGGAAAATATGTTACACAGTCAAGTATCTCAAGGCTCTAAAAGTAATCAGCCACCTAATTCCACCTTCTTTCCAATCACCTATTCATAAATTTGAACAAACATTATAGATTTATTGTCACAATTCAGCTCGTAATTAACGTTCAGATTGAAATATTAATAATAAAAGGTAACAAAGTAATCAAAATGTGCATTTGCTTTTTCTTTTGTACTTTCCTCTTTAAATCTGCATGCTTAAGAGGAAACTTTTAATTATGGTGGGAAATAAAGTTTCCAAATAAAGCTTATTTAGCTCTAAATTATTTTCAATTAATTGGTCCCACAATAAATGGGTCCCACAAAGAGATACTCCTTTGCAAATCAAAACGGTGACGAAGCAATTTCCTGGTCGGTGTAATGGGCCCTTACATGGCCGTCGATCTTGACTTTTATTGGGCCATCCCGGCCCAGTTGTTTTCCCTTTTTTCTCCTTTTTAGAGAAAAAAAGCATATGTTTCCTAATGTAATACAAGAAAATGATTGGACACCAATTCCTTTGACACGTTTGGGAAAAGGTCATATACACGTTTGAAAAGTAAGGTTTGGTGGGGTAAATTAATTCAAATATTACAAAGGGATTAAGAAGTATATATTAAGCTAACTATTTTTTTTTACCAAGAATATGTCAAGGATGATATTAATAGTTTTTTTTGGACACAACTTTGTGAAGTAGGTGAATTAGATCATGTTTAGTGACACCATTGGACTGTAGTCTTTTCAATTCAAACTTTTCTATTGTCACCGAAAATGATTTTGAACAACACTATAGGTTTTCATACATAAATAGACATTCAAATTTGGCCTAAGTAAACACCTTAACTTTAGTATACATACCTAGACACCTCAAATTATTTCCTCCGTGTCAATTGAACTCTCCAACTTACAAAATGATAATCTAGACACCTTCAAAATTAATGTGTCAAGTCAGCATGGGTGTCCATGAGACATAGTGGGACGAGTTGGAGGCGTCTAGATATGCTCTTTTGAAGTTGGAGTGCTTCCTTTGTCAGCTGAGGTCAAGTTTGAGTGTTTCTTTATGTATTATGCTAGTAAAAAATGATGATCTTCTTTTTGCAGCACTAAACTCTTGAATAAAATTAAGCATGATTAATTAACTTGAAAAACTTTTAATTTCAGTGCAAATTGGAGGTGTGGGATCTGGAAATAGCTCTGGCAAGACAAAAATAGTTGCAATAGCTGCTGGGGTTACAGTTGGTATAGCTCTTGTGCTATTTGGACTAACCCTTTGCTTCTTATCAAAGAGAAGAAAACATCAAGGTTCAATCCGAACTAAATCCGTAAACAGAGGTATTATATGAATCTATCATGATAAATAAGATTGTCTAATATAAAATGCTTCATTTTCTAATGAATGTTTCTTATGATCCTAAACCAGGCACTAGCGAACGAAGTCAAGAACTTCTGATGAATGCAACTATTATTCCAAGTAAAAGAGAATTTTCTGGTGAAACTTCCACGGAAGAGTTTGAATTGCCATTATTTGATTTCAGCACCCTAGCTACGGCTACAGAAAATTTTTCTGATGCGACTAAGTTGGGCCAAGGTGGATTTGGTTGTGTTTACAAGGTAAAATCTCAGTTATATGTAAACCATAGTGGTATAATAAAGAAGAAAAAATGATCGACGAACTCAATAATATAATCTTTTGATCAGGCAATGCTGGTTGGTCAAGAAGTAGCAGTCAAAAGGCTCTCGAAGAATTCCGGACAAGGAGTAGAGGAATTTAAAAATGAGCTAAGATTAATTGCCAGGCTTCAACACAGAAATCTAGTCCGGCTTCTTGGCTGTTGTGTTGAAATGGAAGAGAAGATGCTGATCTATGAATACCTGGAGAATAAAAGTTTAGATTCAATTTTATTCAGTAAGTCTTTTTCGTTGTATCGTCAAGAGAGTAGTGTTTCACATCATTTACTTACATGCTTATATAATTTGTGAGTCACATTTGAATATGCTCCTCAGATAAACAGAAAAGCTCGCTGCTCGATTGGCAAAAGCGGTTCAATATTATTTGTGGCATTGCTCGGGGGCTTCTATACCTTCACCAAGATTCAAGATTTAGGATTATCCACAGAGACCTTAAAGCAAGCAATATTCTGCTTGATAAGGATTTGACCCCCAAAATATCAGATTTTGGCATGGCAAGAATATTTGGCGGAGATGAGACTGAAGGAAATACAAAAAGAGTAGTTGGAACCTAGTAAGCACTCAACATAATTTCTTCACATAATTCATAGAACCTTTAAAGCTGTTACTAAATCTGTTTTTTTGTTTTGATGTAGTGGTTACATGTCTCCGGAATATGCAATGGATGGCCTCTTCTCTGTTAAGTCCGATGTTTTCAGCTTTGGGGTTTTGGTGTTAGAAATAGTAACCGGGAAGAAGAACAGGGGATTCTATTATCAAAACAACCAACTCAACCTTCTTGGACATGTAAGTCCACAGATCATATGTTAGCTTCATTATTGGTAATACATTCATTATTTGATTCATTTCTGATAAACAAAGTAATCGATTATGCAGGCGTGGAGACAATGGAAAGAAGGGAGCGGCTCAGAGCTACTAGATCCATCTTTTGGAGAATCATTTTCTCCGTCTGAAGTAATGAGATGCATACAGGTTGGACTGTTGTGTGTCCAGGAACAAGCGGAAGACAGACCAAACATGGCCACTGTGGTTTTGATGTTAGGCAGTGAAAACGCGTCACTACCTCAGCCTAAAAACCCAGGATTTTGCCTTGGAAGAAGACCCGTTGATTCTGATTCATATTCGACTAATTATGAAGAAACTTGCACTGTCAATCAAGTTACAGTTACCATGATAGATCCCCGGTAGTTGATGGTGACGCCCTCAGAAAGTGTATATTACTACAAGAGGTTCACAATTTTCTCTTGTATACTTTCTCAGTTTTTGCTTTTTGGGTAAGTTTTGTGTAAGATAAAGAGTATAATTAGGTCTTTTTTTCTCATTACATACAATTGTGGGTTCAAAAATCTCTTTTTTCATCATCAATGTAATAGTCCAATCATTCCTTTTCCATATCCTTAATTGAAATCCATTGTAAATTACTGTGC

>PGSC0003DMG400013886 G-LecRK

GCCTCATCAGAACCTTCTTCTTTCTTTTTGTTTTCTATACCCTTTACCATTCTCCTTTGTAAAATTCCCAATTTTCTCCATTCTTGAATTGTGGCATCATCATTTCCATTTCCAAATTCTTGAATGGCCACTGGCTTTAGATTTCCCCCCTTTTGGTTCTTATTTCTTTTCATTCTTTCTTGTTTCTCCTCATTCTGTTTAGGTGGTGACAGAGTTACGCGAGGAGAGATTTATAAAGATGGTGATAACATCACTTCTAAAGGAGGGGATTTTGTGTTGGGGTTCTTTAGTCCTGCAAATACCAGTAAAAGGTATCTTGGTATATGGTATGTTGACGTACCAGTAAAGACATATATTTGGGTTGCCAACAGGAACAATCCTGTTCATGATAAAAATGGTACCTTTTCAATTAATGAAAATGGGAATTTGGTGGTTAAAGATGGACATGGGGATTTACTATGGTCGTCTAATGTTTCTGTTAAAACCACAAATTCTACGGCTTGTCTGAGAGACGAGGGCAATCTTGTCATCCTTAATAATGACAGGAATGCTACGAGGTTAAATTCTGAATTGTGGGAGAGCTTCTCGGATCCTACAGATACATTTTTGCCTGGAATGGAAGTTCTCATAGAAAGACAAGGCCAAGAGCAAAAAGTTTTCCGTTCTTGGACAAATGAGAGTGACCCTTCACCTGGGAGGTACTCAATGGGAGTTGATCCTCGTGGAACACCTCAGATTGTTATTTGGGATGGACCGAATAGACGATGGAGGAGTGGACATTTTGATGGAGCGGAATTCATTGGTGTTCCAGATGTGATTAGAACTACTTTTTTCTCTGGCTTCAGAATTCAAAACGAAGGAGACAATAAATTGTTACTCACTTACAGTGCCTCAAATACATCTTCTTTTGTGAGGTTTCAGATTACTGTGACTGGAAATGAGCTGCAACAGAGGTGGAACGAAGACCAGGGTGAATGGAACACGTTGCAATCTAGGCCTGTAGGTGGTTGTGACTTGTATAACTTCTGTGGGAATTTTGCAGAATGTGACAAAGATGTATGCCAATGTTTAAAAGGTTTTGTACCACGTGTTCAGGAGCAGTGGCATGCTGGGAACCGAACCGAAGGATGTGTTAGGAAGACAGAATTGGAATGCAGGAGAAATAGCAGTGTTTCAAGGAATGATAGTTCAAAAGATGATGGATTCTCGACTATTCGGAGAGTTAAATTACCTGACCATGCTAATGTTTCAGAAATAACCATAGACGAGTGCAAAATCAGGTGCCTGAATGATTGTTCCTGCAATGCTTATGCTTATGTCAGAGGAATCAATTGTATGATGTGGCGTAATGATTTAGTTGATATAGAGCATTTTCAGGAAGGTGGGAACACTCTTTATGTTCGTCTTCATCCTTCAGATATCGGTAAGCACAATACTTTCTGCTGAATTCATAAATCTCTTACCAGTTTTCTTCTCATTCGTGCAATTATACAAATCTAAAACCAAAGTAGTTTGTCTTTTAGTACAAAGATCAATTTCTTGTTTGTCTTTACTCTTTATTTGTAGCAAGCTCAAACAACAAGCAATATATCTTTTTCTTCCTTGTGTTTGAAGAATGATTAGCCTTTCTAATTTCTTGATTTCTCCTTTGAATATCTTTTGAAACAGGTAAAAAAAAGAAGACGATCATAATTGTTGTAATATCAATTCTAGCAGCTCTGGCTTTGGTTGTTATGGTGGCCATTTGGCTAGTATGCAAGTACAGGGCCAGAAAACGAGGTAAATTTTGATGCAGTAACATTTCTGACTTGTTAGTTGCATTTATTATATGTTCCATATTGTACTTATACCTTTGACTTGCAATTTTCAGAATCCAAGAGGACCAGTGAAATTCCCAAAAACCATTTAGTTAGGAGTGGAGAGTTCTCCACGGAATATTCTGGACCAGGCGACATCAGTGCTGAAGGGCATCAAGGAAATGGTTCAGAATTAGCATTTTTCAGCTTCAGCATGGTGGCCACAGCTACTGACGACTTCTCTCTTGCAAACAAGCTTGGGCAAGGGGGATTTGGCCCCGTTTATAAGGTAAATTTTCAATCTTGATAGATTATTCTGTTCCTTTAGGAAGTATATGGCCATGTTTTGGACTAAAGTTGATGTTTTTCAAATCTTTAGGGAAAGCTACCTTGTGGTCAAGAAGTTGCTGTAAAAAGGCTTTCACAAAAGTCTGGACAAGGCGATGAGGAGTTCAAGAATGAGATCACTCTAATAGCAAAATTGCAACACAGAAATCTTGTTAGACTTTTGGGTTGCTGTGTTGAAGGAGAAGAAAAAATGCTGATTTACGAGTACATGCCCAACAAAAGCTTAGATACATTTCTATTTGGTAAGTTCAGGAAATTCTATCAAAATAATTTCTGCTGCTCCCACAGATGTACTATAGCAACACATCTCATTGAGATAGACACTAACAACTGTATTGGCTGTTAAACTTTGACAGATACTGTTAGGAAATCCCAGTTAGACTGGAGAAAACGCTTCAACATTATCGAAGGAATTGCCCGAGGATTACTGTATCTCCATAGAGATTCAAGGCTTAGAATAATCCATAGGGATCTAAAGGCTAGTAACATTCTGTTGGATGAAGAGATGACCCCAAAAATTTCAGACTTTGGCATGGCCAGAATATTTGGAGGAAACCAAAATGAAGCAAATACAAATAGGGTCGTAGGAACATAGTAAGTATCATTTGATTTTAAATCTTTTAAGAGACTAATTTCAAAATTTCTTGTTCCTGGAGCTAAATCCATTTCATTTTGTACCTTGTTGCTAAACAACAGTGGATATATGGCTCCTGAGTATGCTATGGAGGGCCTGTTCTCTGGCAAGTCTGATGTCTACAGCTTCGGCATAATATTGCTTGAAATTATATGTGGACGTAGAAATACAAGCTTTCGGACAGATGAGCATTCAGGCATCATTGGATATGTAAGTCTGACTAATTTCTCCACCTATTTCAATCATTTTAGTCAAGAACAAGCTGTCTAAATAATGGGGCTAATTTCTTGTGTGATGAGCTATGACGGAGTGGTCATCTTTCTCATATTTTTCTGGTGGAAAATATTTCAGGCATGGGAGAAGTGGGAAGAAGGTAGACCAATGGACCTGGTGGATCGTTCAATTTGGGATGGTTGTCAACACAATGAAGCATTGAGATGTATACACTTGGCATTGCTTTGTGTTCAAGATTTGGCGGCTCACAGACCAAACATGTCTTCTGTGGTTTTGATGCTGGAAACAGACAATGTAAGGTTGCCCTTGCCTAGGCAACCTACCTATACTTCAATGAGAAGATCTGTGGATGAAGATATATGGCATGGGAATCAGGATGTAACATCTTCCAACAATGTCACAGTTAGTGTACTAATAGGTAGATGATTGGTTAATGCTTGTTCCTTCTAACATACCTGCTTCATTTAAGAGAATGGGATCAAGTAGTGTAACTATGCCAAGTTATACTTGGAACTATGTTCATAATTTTTTGTTAGCTAGGTACACTTTAGCACATCATGTGTATTGTAATTAGAAGGTGGTTCAGCTCTCAAAGCTGAAAGAGAGATATGTCACAACCTTGTTCAATTAGTATGGCTGAAGTTTTTTTCTTTACTGTTTGTAGATGGAGATGCTCCACCAATCGAGCAAACCCCACTTGTCATATCTAGTTTTTTTTTTTTTATGACAAAGGAAACCCGCAGCCGCTACCCTTTGGGTGCGCACAGGTAAAACCCCTGCTCCTATGCAATAGCTCGCAAACCACATAAATCAAGCATTTCAATCGGTCCGTATATATCACTGTATAATCAAGGAACAGAATTACATGCTTGTTATCTCCTATCACTTCAAGAAACCATTTTGCAGTTCTTCTAAAGCTTGAGAACTAGTATTTTGTCCATTGAAGCACAAGCAATAGTTATTACCAGGTTCTGCATAACTTGAACTATCTGTAGGGAGATCAGCTTGTGTTCCCTTTGAGACAGAAGGGGAAATGGAATTATTTACTATGTAATTAATCAGTGTCAATTTAGTGGTGGTTGACTTGTTGCATGTAACTCAAATATGGAAG

>PGSC0003DMG400013887 G-LecRK

AAAAATTTCACAAGTTACACGCTTTATATTTTGGTTAAGCACGCACGCCACGTAGTTAATGTGGTGCGTAAAAGTCACACCTTTGGAAAGGTTTGGTATGTAAAGAGATTCCCATGGCTGGGGAGTGTGTAGAAGTGATTTGGCTAAGAGTTCACAGATTAACTAGGTATTAGGCCTATTTTCTACAGTAAACTGTTTTTCCCGTAAACATTTTGCAGAGATGGTTGGTGTGTCATTGAGTCAATACCCGTAGGAAAAGGACAATCCATTAAATTCACATCAGAAATTGTCAACATGATACAAAATCCTGCCCTTATATTCTCAGAGCTAAACATATCAGCCCAAGTAATTTTTCAGTATTGCAGGTATCAAATGCACAAGAAGTACTCAGAAACATAGTAAGCTGTAGACGTAGAAGCAAAATGGCCACTTAACAAATCATTTGAGACATGGGTCCAGCAATTAATTGACACCCAATTGAAAAGTAAACGGATGAGAACATAAAAAAGATAAGTATCCAAGAACCTTATTTGACTAAAGGAAAAGAAAGGTCATCCTTGAGGTTCTCAGCTCTTACACTTCAAATGATAAGACTGTACCTTTTGGAAGTAAAGAAGTTAGAATGAGCATATCTATGCTTTTTCATCTTTTACTCTATTGTTTTTATGTAGTTTTTTCCAGTGCCAATACTTCAGACACCATTAGCAGTAGTGAGCCTGTGAGAGACTCAGAAACTGTATTTTCTAGAGGCAAAAGATTTAAACTGGGTTTTTTTAGCCCCAGGAATTCTGCAAAACGTTATGTAGGGATTATGTTTAACCTACCATCACCAACACCAACTGTAGTATGGGTAGCTAACAGAGATGAGCCTTTACACGATTCTAGTGGAATACTCACAATATCAGAAGATAGCAATCTTGTCATGTTGAATGGACAGAAGGAGATAATTTGGTCATCCAGTATTTCAAACTCTATGAAGAATTCCACAGCACAACTCTTGGACACTGGAAACTTAGTCTTGAAAGACAGCTCAAATGGGAGAGTTCTATGGGAAAGCTTTCAATATCCTACGGATTCTCTCTTACAGCTCATGAAAATGGGCACTGATAAGCGTACTAACACAACAACTCTCCTGAACTCATGGGGGAGTCCTGATGATCCATCCGTTGGGAGCTTTTCAGCTGGAATTCAACATCGATACATTCCTCAGTTTCTTATTTGGAAAAACAGTTCTCCTTACTGGCGGGGCGGTCCATGGAATAAACAGATATACATAGGATTACCAGAAATGAATTCTTTCTATCTCTATGGTATTGACCTAGTAATTGATAATGCTGGCACAGCATACCAAACTTATTCAGACCCAAATCAGTCTAGGATACTCTATTATTCCTTGAATTCAACAGGGTCTTACCAGGAGAAGGTTTGGGATCAAAGTAAGAAGGATTGGGTGGTAACATGGGCAAATCCCCGAACAGAGTGTGATTTTTATGCCAAGTGTGGGACATATGGAAGCTGTAATCCAATGAATTCTCCAATATGCAGTTGTATACAAGGTTTCAAACCAAAAAATGATGGAGAATGGAAGAAGGGAGATTGGTCTGGTGGATGCATCAGAAGTACTGCACTAGACTGTGAAAGGAACAAAACTGATATTGAGAAGGACAAGCAGGACGGGTTTCTGAAGCTGCAGACAATGGGAGTGCCAGATTTTGTAATTTCGGTACCTTCTGCAAAAAAAGACTGTGAGAGTGACTGTTTGAGTAACTGTTCCTGCACTGCATATTCATACTACAGAGGCATAGGTTGTATGCATTGGAATAGAAGCTTAGTTGATATTCAAGAATACTCCAGGGATGGGGCGGCTAGTTTGTTCATTCGCCTTGCCTACTCTGAACTTGGTAACATACTTCTTTACAACCTTGAGCTTTCAATTGTTTAGCTTGTTTTTTTTTTTTTGGAACTACAAGGAGTTGAAATTTAAAGTCTTCTTGTCTTTGAAGAATATAACCAAGACTCCCAATAAAATCTATAAAACCATCGAATTAGGCTCAAATTCTCAATTAATGTATTGCTTAGCTCTTTTCTATTTTTGGTGTAAACAAAGATATATTAACTAGGACAGAAGACATCTTTAATGTGCTAGATTGTTTATAGGCAGCAGTTTAGATTAAGACTGTTTTATTCTGTAAGTTGTACTCCCTCTGTCCCAATTTATGTGACTTACTTTCCTTTTTGGTCAGTCCCAAAAAGAATGACACATTTCTATATTAAGTAATAATTTAACTATAAAATGTTTATTTTACCCTTAATGAAATGATTTACAGCCACACAAATTTCTATCATTCATTTTGGACCACAAGTTTTAAAAGTCTTCCTTTCTTTCTTAAACTTCGTGCCGAGTCAAAGTACCTCACATAAATTGGGACGGAGGGAGTAATACTTTTGAAGGGGAACTGCACATTTTTTGTAGTGCACCTGTGTATATATAGACTAAAGTTACCAGTTTCAAAAAAAATAAAAATCAATTAGTTCTATCACTTCCATGGAAAGAAAGATATGCTCAAGAACAGTCTATGGCTCATACATGTTTGCAGCTGCAAATGACAAGAAAGACATTCCTGTAGCAGCTATTGCAACCACAGTCTCAATAGGCTCAATAACAGTTATCTTGTGTGGATATCTTTTCTGGAAATTGTTGGCTAAGCACAGAGGTAATTAATCAAATAAGATTTTTGAGAAGACAAACTAGAACTTCTCTGTCTGCAAATGGTTTGATTTCTGAATGTCCACTCTGAACCAGAAAGGAAGAGGAAAAATGAAGCACTCTTAAGAGAAGCATCTCCAAAATTTTACCAAGAAGGCATGAGTAAAGATGACATCAATCAAGTCAAAATTGAAGACATCACCTTGTACAGATTTGACATGTTAGCGAATGCCACTGACAGTTTTCACCCAGCTAGCAAGCTGGGGCAAGGAGGCTTTGGTCCAGTCTACAAAGTAATGTTTCTTGATGCATCTCTGCAGTGTTCTGACAAATGAACACAATATAAAATAAATTAGATCAATTATCTGCAGGGACAATTGCCAGATGGACAAGAAATTGCAGTAAAAAGGCTTTCACAGTCTTCTGGTCAGGGGCTACAGGAGTTTATGAATGAGGTCGTTGTGATTTCTAGACTACAACATCGTAATCTTGTTAGACTCCTTGGCTGTTGCACAGAGAGAGGGGAAAAGATGCTGATTTATGATTTCATGCCAAATAGAAGCTTAGATGCATATCTTTTTGGTCAGTGCAGAATTTTCTTTCATTGGTTTTGCCTGTCCTTCAACTTAAGTACTTCCAGAATCAAAAGAAAGCTGTATTTCTTTAAGGAAACAAAATCCATCATACATTTCTCGGTGATACCTTTTCAGGTTCACACCAGGAAAAGTTCCTTGATTGGAGTAAACGAGCCATCATTATTGAAGGAACTGGTCGAGGCCTCCTTTACCTTCACAGAGACTCAAGACTACGAATTATTCATAGAGATCTGAAGGCAAGCAACATCCTGTTGGATGAATACCTAAACCCAAAAATTTCGGATTTTGGCATGGCGAGGATTTTTGGAGGCAATCAAGATCAGGCCAGCACTATAAGAGTTGTTGGCACATAGTATGTTTGCACTGCTTCCTCCATTTTTTCATGGATGAACTGCCATCATTAGTCATTGAGTTTCTGTGTAATATCTTATGCAGTGGTTACATGGCCCCTGAATATGCAATGTACGGAAGATTCTCAGAAAAATCAGATGTCTACAGCTTCGGTGTGTTGATCTTGGAAATTGTCAGTGGAAGGAAGAACTCTAGTTTTTATGATGATGAAGATGAACTGACTCTACTTGCATATGTAAGTTCATTCTTGCAACTATATATAAAGCTTTAATGTTCTGCTAGCTCATCAAACCAAAAAAATAGAGGAGAAAAATTATGAAACTTCTGAAAATAACATTTTTGCTGTTTGACAATGTCTTTGTTCATTAGGCATGGAAGTTGTGGAATGAAAACAATATCATAAAATTGATAGACCCCAAAATATTTGATTCAAGCTTTGAAAAACAGATGGTGAGATGTGGACATATTGGATTATTATGTGTTCAAGAATATGCAGAAGATAGGCCAAATGTCTCCACAGTTCTGTCAATGCTCACTAGTGATATTGCTGAACTACCTACTCCTAAACAACCTGCATTTACCGGAGGACATGCTTCACCACAGCAAGTATCTTCTAAAAGCCAAGGGTCCGTGAATGCTGATACCATTACTATATTGGAACCACGATAGGATACAGAGTTTTCAACAGGCAGGCAAAGATCATTACTTTCCTTAAAATGGTAACAGTAGCATAGTATTTCTTGAATGTGCATAGGGTAAAACACGCTCCTTTGCAATAACTTGCAAAACACACAAAAGAGGTAAAAGTGCACTAGCTATGTGATGAATTAATTACAGAGTGAGCATGGAGAACATGAAACAGTTGATCAATTTTTAACTTTGCATTATGAAAATCCTTAGAAAGAAAGTAGACAAAAGGAAAAAAAGAATCGCAAATCCCCAATTAACAAGTTCAAATAATGGCAATCAAAGACTTGAAAAATCATACCTACTGTACTGATACACGAAACGGCGTCGGCGAAAAATAGCCACTCGTTTTACCACTGTTTTTCTTAGAAAATTACAGAGAAGTGAATTGGTGAATTAATCAATGTTGAGGAAAAGTTAACTCTGCACTATTTTAGGTCATTCACACAAGTTATTTGGGTGTTGGTTAAAGACTATAATTGTTTCATTTTTGTTTTAATTTTGCATAAACTAATGTGAAACTTACCTAAATCTCACTAGTTTAGGAGTTAATTACTTATATATATTTTAGTTTGTAATATTACGTGTGTTACCCCCAATTTTGGTGCACTAAGAAACATCCAGATACATGTATTTCTGAATACATGGGCCAAAACTAGGTGTAAGTTGTTCTAGATACATTGTATCCAAGTAGATTCACATGTATTTCGTATACATAAAAAAATTGGTTCGCCTCCTTCGCCTCTCTCCCATATCGCTCACCACTCTCCAATGTATTAGGGATATATAATAAATCTCGCTCACCTCTCTCCCTATTTTAGTCTATCCAGAA

>PGSC0003DMG400015476 G-LecRK

TCCAAGTATTGTTTTGGGGGGTTGGGGCTGGGGAAAACTGCTAGACTAAGATGAAAATTTTACAATGTTAACATTTCTCTCTCTGTTCCATTTGACAATCATTATCCATTCTTGTTCTTCATTGAGGTAAAAAACTTTTATTTATTTATTTATTTATTCCCATATTATATAATGCAAGTAACATTGTTCATGGAAATATCGTAGGGACATTGAAGAATCGTTGAGAAGAGCAGTTAACGGGAGATAGATAGATATTGAGGGAATTTTTATCATTTGTTAAAAATATGGATTCATCTCCTGGAACTGCTTCCTCCAATGATACTGATTCTAAAGAAACCCCTTCTTCAAATGGAACTCCGCCTTCATCTTCAGATTCAAAAAATGATTCTCCTTCGTCTTCGTCTTCGTCTTCGTCTTCATCTCCACCACCAAAAAATGAATCTCCACCTTCATCACCACCACCACAATCTCCTCCTCCATCTCCTAATGCATCACCACCTCCAAAAAATGATTCTCCTCCGCCTTCTTCACCACCACCATCCTCTTCTGAATCTAATTCATCACCATCAAAAGATAATAATAATAATAATCAATCCCCAAATCCTTCTTCTAATACACCTTCACAATCACCACCACCATCATCCACGTATAATACTCCATCAACCCCAAATAGTCCTAAAAATCCTCCATCATTTCTATTATTTTCTCCACCACCGCCTGCTTCTTCACGTGAATCGCCTGCACCTCCTCATGCACATTTGTCTCCTCCTCCACGCTCAGGATCTAATCAACCATCACCTGCAAACGATGATTCCTCAAGTACTGTTGGTATAGTAGCTGGAATTGCTGTCGGAGGCATCGTCATAATTGCATTAATCATTCTATGTGTATGGTGTAGTAAAAGGAAGAAGAAACAGAGATACTATATGGCTCCTGGACCTAAAGGTATGTTACTAATTACTAATAAAGTGTTGTTTAAAGCATAGTATTACTTTTTAATTTTCGATTAAATATGTTTTTGATTAGGTGGTGATCCATACTACAATAACAACAATCAATGGAATAGACGTCAGTCGATGGATCATGTAATGAAGGTACCACAAAGTGAAATGGGGACTCAGAGTGGTTGGGCTGGAGCTACTCCACATCAGCAGGGAGCTAGTTCAAGCGGCGAATTTGGCTCAGGTTATTCAGGCCAAGCAGTGCCATTGCCTTCTCAATCACCTAACATGGGACTTGGTGGTTTCTCCCAGAGCCAATTCAGTTATGAGGAGCTAGCTAAGGCCACAGATGGATTTTCTCAGGCCAATCTGTTAGGACAAGGTGGTTTCGGATACGTGCATAAAGGTGTTTTGAATGATGGAAGAGTGGTTGCCATCAAGAGTTTGAAAGCAGGAAGTGGACAAGGAGAACGAGAATTTCAAGCTGAGGTTGAGATCATTAGCAGAGTTCATCATCGTCATCTTGTGTCCCTCGTTGGATATTGCATTGCTAATGGACAACGCATGTTGGTCTATGAATATGTTGACAACAAAACCTTGGAGTTTCACCTTCATGGTACATTCTTTCCTATTCAATACTCACTACTTCTTCTACGATCATGTTAATTTTATTGTGAAATAAGAGAATCCTTTTCCTAGTTGATTCCCAAATTTCGATTCAATATATTTCTTAATTCTAGTTTTGATAAGGGTGAAATCAAATTCATTATTGCAGGAAAAGGTCAACCTGTCATGGATTGGGAGACCAGACTTAAAATTGCTTTGGGATCAGCTAAAGGACTTGCTTATCTCCATGAAGATTGTATGAACATTCTAATCTACACTTTCACCTCTCTCTGTCTCTCTTTATCTCTCTCTCTCATTTCTTGATATATACATGCAGGCCAATATAGAATTATCCACCGCGACATCAAGGCTGCAAATATTCTACTAGACCTCAATTATGAAGCCTTGGTAATTAAATGATCAGCCACATTTGATATGTCAATGAAGATATGAAGAGCAAGTTTTATGTTAAAACCTCGCGTTTTTGTTTAAGAAAGGTTTATTAACCATCACAATGCCTTTTAGATTTAAAGGAAAATTTTGGTTTATTAGGTGGCAGATTTTGGATTGGCTAAGCTTACTTCGGATAACAACACTCATGTATCAACTCGTGTTATGGGAACATTCGGGTGAGTCAAGCATAGCTACTTAGTCGTATAAAACTTGTATTCAACAATCAGAACCAATTATGTATTTTAACATACTTCAGATGTCTCTAAGCTAGAAAGTTACTTGCAGGTACTTAGCTCCTGAATATGCATCAAGTGGCAAACTAACAGAGAAGTCAGATGTGTTTTCATATGGAGTAATGCTGTTGGAGCTCATAACCGCAAAAAAACCTGTTGATCCTAGCAATATGATGGAGGACAGTTTAGTAGATTGGGTTTGTTCTAGATCCTCAGTTCACATTTGATTGCGCCAAGATGTGCTTAGACTTTTAACTTTTGTTTGAGGTATTGATCTAATATATGAATGTATACAGGCTAGGCCCCTTCTTACAAGAGCACTGGAAGAGGAAAAATATGATGGATTGGTAGACCCACGTCTTGAAGGGAACTTTGATACAGATGAGCTACATCGCATGATAGGTTGTGCTGCTTCTAGCATTCGCCACTCTGCTAAAAGACGTCCCAAAATGAGTCAGGTATGATTACCCTCTCTAGAAGATAACAAATACTGCATATTTACATTATGATTTAACATGTTGTAGCATATTTTGGTGGTTTGCTTATATGGACTCGAGAAATTCATATCTTTACATGATATCAGAGTAAGGTTCATCTCCGTTTGGACAAAATGACTTGTTTCACTTATATAAATATTATGTATTATGAACAGATTGTACGTGCATTGGAAGGCAATTCATCGTTGGAGGACTTGAACGATAATCCAAAACCAAGCAAGGTTCCTAGCTTTGCCGCGTCAAATGGGGCACCCACTCAGAGCTATGATACTGGTATGTATAATGCTGACATGATGAAGTTCAGGAAAATGATTATGCCAACCCAAGAATTCAGCAGCAGTCCAAGTGAGTAATGAATCAACCCTTCTTCCTCCTCCTCAAGCAACGATTCAACACGCGGGCAAGAGTAAAAAAAGAATGTACCTAAAGCTTGAGATATTTTTTCAAAACATATTCTTGGCTGGAACTTATAACAGAGATTTGGGATTACAAAAATTAATGTTTATGATGAACATTCACTACTCATATCTTGATTTGAATAACAAAGATTATAGGATCAGAATACAATTATACTTTTTTTGTTTCAATATGTACATACAATTTATTCCTACAAAAATGTTGAAGATCTTTATGTTTAAC

>PGSC0003DMG400017034 G-LecRK

GCTAACCACAAAACCACTACACTTCAATGTAATCCTTCTGTCTCTTAATTAATAGTACCGTTTATGTACTATTTGATGAATGTAAAAACATTTAAAAAGACTCACTTGATAGGTAAGTCACTATTTTAAGGAGTGGGGGGGACACTACTAATGTTGTAACATTATAATCATGCCAAACTTTTTATTTTCTTTCATCTTGTCTATACATTCCTAACAACAACAAAGTCCTGGAGCAAAAGCCTATTGATTCACCAGTTTGCTAACCTACTTTGTCGGCTATGTGCACTCTGTAACCAGAATAGAGCTCTGGACTTAAAACCACCCACTTTTTGAAGTTTGGAAGGCGGCGCCTAAACTAGTGAGTAACATTCCACACATGAAAATTTATGAGCTTTTTTATCTCATTTGCTGGTTGGCTCATGGATACTGCACCTCTGATATTCACATTGGTTATCAGGTAAGTCTTGCTATACCAACATCATATTGCAAAGGATTCATAGGCAGGGCTTTTCTAATGGAAACTGATCACAACCAAATTGGACCAAATTTTAGAGGTGGAATCAGTGTTGAAGCCAATAATGATAAATACACATGTTCACTTGATGTTTTTCTGGGAGATGTCAAGGTTTGGAGTTCTGGTCATTTGTCACAATTCTACACAATAGACAAATGTACACTTGAGCTAACTCAGTATGGAGACTTATTGTTGAAAGGGGGTCTAGATGACAGAATTGGATGGAAAGCTGGAACTTCAGGACAAGGTGTCAAGGTAATTAATTGTCAATCTTTAGATATAAGTTGTATATGCAAACAAAAGGGAAATAACATTTCTTAGTATATGTTTTTTGAGGTAAAACTATAAAAAGAACAAATTATTATGTGCAACAATGCTAATATATGACTTCTTCTGCAGAGACTGAACTTACTTGGGACAGGTAATTTGGTGTTAGTTGATGCTATGAACTTGATTAAATGGCAAAGTTTCAATTTTCCAACTAATATTATGCTTTGGGGTCAGAGACTAAGCTCAAGAACTCGGTTAACGTCTTTCCCTAGCAACTCTAGTTTGTCATATTCTTTTGAGATTCAGTATGACAAGATTGCATTGTACTTATACTCTGGGAAATCGAAGTACTCGTATTGGGAATATACACCTTTGGAATTAGATGGCCTGAATATCACATATGTTGAGTTAACTTCTAACGCGCTGGAGATATTCAACAACGAAAATCGTAGAATTGGAAGGATAACATCAGACAAGCCAGAGCCCTTGAGATTTTTAGCATTGGGAAATAGCACAGGGAATTTGGGATTCTACTATTACTCATCTGACAAAGGAAAGTTTGAAGCTTCATATCAAGCACTAAACACTAGCTGTGATCTTCCGTTGGCATGTAAACCTTACGGTATATGTACATTTTCAGAGAAGTGTTCTTGCATAAGACTAATAAAAAGAGGAGATGGATTGCTTTCTGATTGCACTGAAAATGTAAGAATTTGTGGAAGAAATGATTCCCAAATGCTAGAATTACAAGGCGTTACGAGTGTATTAAGAAGCAATCCTTATAAGATCAATGTGACCAAAGAAATATGTGCAAATTTGTGCTTGGATAACTGTACATGTGTTGCAGCATTACATTTTTCTTATGAGAATGATCCCGCGAAACCTGGAGAATGCTATGTGTATGGACTAGCTAGAGGAGTTAAACAAATCGAAAGAGATGGAAGATTGAGTTATATGGTGAAGGTACCAAAGGGAACTGATCAAGGTCATGATAAGCACTCTCATTGGAAGAAATGGATTCCAATTGTTGTAGGAGTGGTTGATGGAGTTGTATTATTACTTGTTTTAGGAGGAGTTGGATACTATGTAATCCGAAAGAGAAGAAAGTCGTGCGTAGATACTGGTCACAGCAATTGATATTGTATAGATAAAATTTTGATTTCAAGATAACATTTTCTACATCGAGATAGCAATTAACCGATTCTCATGAAAAATCAAGGATGAAGCTCGAGTGACAGATACTAAACTTGTAGCCAAAAACTTAAATTTTTGAACTTGGCTTGGCTTGATGAATTGCCTGAGATTGAGTAGAATTCGAAGAAATAGAAGTGTCAAAGGTATGCAAGCAATCTTTTCAGGGTTAAAATGTTGTTACTTCACAAGAGATTTTTCGATTCCTAAACTGACATGTTGAATTTGTTGATGTCTAAGGTTAATTAGAGAGAAACACTCTACGATATTCTATACTTCAAAAATTGACTTACAAAATGCGCGGCCAAATGTCC

>PGSC0003DMG400017106 G-LecRK

CGAAGTATTAAAAATCACAACAAACACATTATACTTACATCCCTATATAATCTCCTTTAACAATGGCTCCTAATTATGTGATGATAGTTGCTCTGATAACATTTATTTGCTTTTCACTATCTTTTGGTGATGATGCTATGTTACCAACGTCACACAACAATCTCATAGACAAACCTAAGTACTGCAATAAAAGTTGTGGAAATGTCAACATCCCTTTCCCATTTGGTATGACCCCAGATTGCGCTCTACGCAATCAGTTCCTTGTCACTTGTAATTACTCCTTTCATCCTCCCCAGCCTTTTTTAGGAAACAGCAGTATTGAAATCACAAGCATATCTCTCTCTGGGCAACTTAAGGTCTTACATTTCATATCCAGAGATTGTTATCGTGATGGGATAAGCACGAATAGAACCAAGCCATATATAGTATTACCATCGTTATTAACTGTGGACAACACTGCGAATAGCTTCATCGCCGTTGGCTGTGACACGTATGCAATTGTCCAGGGTTATTATACCAAGTTTGGAACTGAAATCGAGTATCTCCGCTACATAACAGGATGCATGTCCATGTGCTATAGTATTGATGAAGCTGATGATGATACATGTTCTGGGGTCGGTTGTTGCAAGACATCCATCCCTAAGGGAACATCGAATGTCAATATAACACTAAGCAGCTATAATAATTATAATGATACTAATATGACAAATGACCCTAGCTGCAGCTATGCTTTTGTGGTCGATCAAGCTAATTTTAACTTTTCTAAGAGTTACCTCAGCAGTTTACAGAACACGAAGAAGCTTCCTCTTGTGCTAGATTGGGCAGTTGGTGAAGACAAATGTGAAATAGCTAAGAGAAACTCGACTGCATATGCATGTAAGAGTAACAACAGCTATTGCCATGATGATTCAAATGGTTATCGATGTTCCTGCATGCAAGGATATGATGGGAATCCATACCTGAGTTGTCAAGGTATGTATATGTTTGAGAGTTTCCTAATATGATTTTAATTATATCAAATCTATCATTTTGGAAATGAATGTTTACGTTATCATACGTTTGTACTCTACGAGTCTGCAGATATCGACGAATGTATAGATCGGAAACCAGATTATAGATGTGCCAAGGATGCCATTTGCCAGAATACAGAAGGGAGTTATACATGCATATGCCCTCCAGATTTCAGTGGTAATGGTACAGTATGCAATCGTGACAATCCTCAGAGAATAACTCAAAATATTTATTTTGTTATAGGTACCTCTCTTAACTTCTTTCAAATCGATAACAATTTTATTTTCACGAACTATATACATCGACCTAAAATGTCTAATAAAGTTTAAATATTTTTATACAGGTGCTACAGCAGGGGTTGCTTTTGCGATTATTATTGTTTTTGGCTGGTCATACACTGCATTCCAAAGGAGAAAGATGTCAAAGATGAAGAAGAAGTTTTTCCAGGAAAATGGTGGATTAGTTCTACAACAGCGACTTACAAGGGAAGAGGGATCAAGTCATAATAATACAATAAAGATTTTCACAGCGGAAGAGCTAGAAAAGGCAACTAATGGCTTTGACAAAGATAGAGTTGTTGGTCACGGAGGTTTTGGAACTGTGTACAAAGGATATCTGAAAGATAATTGCATCGTTGCCATAAAAAATCTAAGGTAATTGATCGCAATCAGATCGAACAATTCATAAATGAAGTCCTTGTCCTTTCCCAAATCAACCATAGAAATGTGGTTAAGCTCTTAGGTTGTTGCCTCGAGACGGAAGTTCCACTATTGGTGTATGAATTTATCAGCAATGGAACACTTTCTGAACACTTACATGATAAATTTAAGGCGTCAACTCTTTCTTTGGATATTCGTCTGAAAGTTGCTGCAGAAACTGCTGGAGTACTTTCCTACTTGCACTCTGCAGCTTACCCTCCTATCATCCATAGAGATATAAAATCAGTCAACATTCTTTTAGACAAAAACTACACAGCCAAAGTGTCTGATTTTGGAGCATCCAGGTTGGTTCCTGCTGATCAAACAGAGTTATCTACATTGGTTCAAGGAACTCTAGGATATCTAGATCCTGAATACTTGCAAACAAATCAACTTAATGAAAAAAGTGATGTATACAGTTTCGGAGTAGTGCTTATAGAGCTCCTAACCGGTAGAAAGGCTTTATGTTTTGAAAGACCAGAAGAAGAGAGATGTCTAGCTCAATATTTCATATCTTCAGTGGAAAAGGGTCATTTGTTCGATATTCTAGACAGCAACATAGTCTGTGATGAGGTAAATGCAGGGCAATTGAAAAATGTTGCTGTGCTTGCCCAAAGGTGCTTAATTCTTAAAGGGGATGATCGACCTACTATGAAGGAAGTTGCAGCAGAATTGGAAGCTGGATCAAAATTGAAGCACTCGTGGGCTCAAACTGATCAGCAGTCCGAGGAGATTAAGTCCTTACTGCCTGGATTTGGATATGAATATTGTGAACACAGTATCCATATTGATAGTGTAACGAGCCATGTAACTTTACCATTTCCTGGAGGAAGGTGATCAATCATGCATGTTAATTTTGCTCAGAAGCACGAGCTCAAAACGGCCACGGCTTCAACATGAATTTCTTTTTTCTCTTGGGCATGTTAGAAATATGAACTAAGTTTGATAAGTTGCATCTCTTTTATTTGTCCAGACATAATAAATTTGAGTAACAAATTAATGTCAAGTGTAGTTATTTATTATGTATCCGAGTAAAAATCAACACATTTTGAGAGAATCTATGTATTATTTTATGC

>PGSC0003DMG400017277 G-LecRK

ATGATTGCAGAGCAGAGAGTGTATCTTCCTATCCTTCTCTTCATTTGTCTCCACTCCTTTTCATCCCGGTTTGCTTTCTCGCAACTTTTCTGGAACTATAATGTTAGTGTTTTGAATTCTACTACTGGACTTTCCAATTCTTGGATCAATTTGCCATATGTTAGAGTTTACAATACCTCCGAGCCTTCTGAAATGACGCCTATTATTCGGAGCAGAAAGGATAGCCCTCAGTTTCTCTTTGCCTTCTATTGCCCCCTTCTAACTAACGGTACCACAGAATGCCTTCTTGGTATCCTTTTGTATCACAACAGATTCAACGGGGTTTCAAGTGTTCCCCAGTTAGTTTGGTCTGCTAACAGGAACCATCCAGTGAAAGTCAATGCAACGTTGCAACTAGGCCAAAATGGCAACTTGGTCTTGACAGACTCTGATGGCACTCTTGTTTGGTCCACTAATACTACTGGAAAATCTGTTTCTGGCTTAAACATGACAGAAATGGGAAATCTTGTGCTCTTTGATAAGAGAAAACAAGCAATTTGGCAGTCTTTTGATCATCCAACAGATTGTTTGCTTCCGGGGCAGAAGTTGGTTTCTGGTAGGAAGCTGATAGCAAGCATTTCAGGATCCAATCAGTCCTTTTCTTTTACTGTTCTTAATGGACGCCTGGTGTCTTCCATAGATACCAACCCACCTCAGTATTATATCGCTTCAAGTGGGGATTATAGTCCTCTTTACGATTTTGACGGTCGAACCTTTACTGCTCTACAAGACCCTGATACATCCCAATTCATAAAGCTTGGGCCTGATGGACATTTAAGGTTATACCAGTCGGATGCATTTGATTGGAAACAGTTAGATGAGGTCATGAATTCATATCTGGGGAACTGTGGGTACCCAATGGCGTGTGGAAGATATAGCATTTGTACAAATGATGGGCAATGTAATTGTCCGGTAGAAGGAAACTTCTTCAGGCCAATAAATAGGAACCCGGATCTCGGGTGTTCACAGCTAACATCTATTTCTTGCAACTCTTCACATTATCATAGTCTTATAGAGCTCAGCGACACCACATATTTTGCATTTGAGATCAACTTTTATGCAAGTTCAAACATGTGGTTTGAGGGAACAAAGATGGAAAATTGCAAAGCGGCCTGTTTGAGTAACTGTTCCTGCAAAGCTGCTGTTTGGTCTAAAACTCTAAGGAAAAACTGCTTATTACTGAATGAAGTATTATCTCTTAAGGACAACTGGTCTGGTAGTGACAAGACAAAAGTTTTTCTCAAGGTGCAGAATTTCGCAAAGTCTCAGTATCAGCCGCCAATCGTTTCTCAAAGAAAGCAATCAAGACCTCTGAAAGTGATTGTAGCATCTGCTCTTGCTGCTTTAGTGGGAATAATTTTAAGTATTAGTGCATGGTTTGTTCTTTTCAAAAAGAGGACACTGTCTGTCAAGGCTGGGGATTTTCTGGATTTAGCACCAATCTTACCGGGAATCCTAACTCGATTCTCTTACAATGAGTTGAAAATAATTACACAAGATTTCAGCAGAAAGCTCGGGGAAGGAGGATTTGGCTCAGTATATGAAGGAACACTGAGTAATGGCAACAAAATAGCTGTGAAGCGTCTGAATGGTTTAGGTCAAGTAAAGGATTCATTCTTAACAGAGGTAAAGATAGTTGGTAGCATTCACCATGTCAATTTGGTAAAACTCATTGGCTTTTGTGCTGAAAAAGATCACTGGGTTCTGATCTATGAGTACATGGTAAATGGATCATTGGATAGGTGGATTTCTCATGAAAAGCAAGAAAATGGGCTTACATGGCTTACGAGGCAAAGGATAATATCAGATATTGCAAAAGGATTAGCGTATCTTCATGATGAATGCAATCATAAGATAATTCATTTGGACATCAAACCACACAACATCCTCTTAGATCAAAATTTCAATGCTAAGATATCTGATTTTGGGTTGTCGAAACTAATTGAGAAAGACAAAAGCAAAGTTGTTACTAGAATGAGAGGAACACCAGGGTATTTAGCTCCTGAATGGTTGAGGTCTGTAATAACCGAGAAAGTAGATGTATATGCCTTTGGAATTGTGCTCTTGGAAGTTCTTTGTGGGCGAAAGAATTTGGATTGGTCGCAGGCTGACAAAGATAATGTCCATTTGCTTAGTGTCTTTAAGAGAAAAGCAGAACAAGATCAGCTCATTGATATGGTTGACAAAAACAACGAGGATATGCAGATCCACAGGGAAGCTGTGATGAAAATGATGAGCCTCGCAGCATGGTGTGTGCAAGGTGATTTCAACAAGAGGCCTTCCATGTCATTGGTGGTTAAGGTACTGGAAGGTTTGGTGTCTGTTGAAACCAACTTGGATTTCAATTTCACAAACCTAACTGAGGCTGGGGCAGGCAACCAACAGATGGAAGTCCCTATCAGTTCAATACTGCCTTCAGTTTTATCTGGACCAAGGTAAACAATATGCTTGAGAATTATGACAATATTTCCAATTCTAGTGTAACTTCATAAATGTAATATTCAGTAACTTTGCATTAGTT

>PGSC0003DMG400017278 G-LecRK

AATTTATTCAAACTAAATAACTTATTTTGCTAAAAGTTACTGCAGTAACTAAAAGCAAACTTCAACAAAGCATTTCAGCAGACAATTCGAGTCAAGGTTTATTTTCTCTTACTGTTCTCAATGGAAGCTGGGCCACATACATAGATTCTGATCCACCTCAGTATTACTACACTTCAGAAAATCCCAATAGTCCTTATTGGAGTTTTGATGGTCATACCCTTACTGCTTTACAATATCCTCCTACATCGATAGCTCAATACATACAGCTTGGGCATGATGGACATTTACGCGTATACCAATTGGACAACCTTGAACAGAATTTGGTTTCTGGTCAGAAGCTGACAGCAAGCGTTTCAGCAGACAATTCGAGTCAAGGTTTATTTTCTCTTACTGTTCTCAATGGAAGCTGGGCCACATACATAGATTCTGATCCACCTCAGTATTACTACACTTCAGAAAATCCCAATAGTCCTTATTGGAGTTTTGATGGTCATACCCTTACTGCTTTACAATATCCTCCTACATCGATAGCTCAATACATACAGCTTGGGCATGATGGACATTTACGCGTATACCAATTGGACAACCTTGAAAAAAATATGCTCGGCCTTAATTGGAAAGAGGTAGGTGACATTTTGACACTCTCTGACTGTAGGTACCCAATGGTGTGTGGGAGATACAGCATTTGTGTAGATGGAGGACAATGTACTTGTCCAGTTGAAGTTAACTTCTTCAAGCCATTTTCTGAGAGGATGCCTAATCTCGGATGCACTGAGCTGACGCCCATTTCTTGTGACTCTTTGCAGTATCATAGTCTTGTAGAGCTAAAGAATACTGACTATTTTTCATTTCGGTTCAATGATGAACTAAACTCGAGCGTATTTTGGCTTAAAGGGAAAAAGATTGAAGATTGCCAAAGGGCATGTCTGAGTAACTGTTCTTGCAAAGCTGCTGTTTATGATCCGTATCTGAATGTGAGTCTACTACGAGGGAACTGTTTGTTACTGAATGAAGTCTTCTCTCTCATAGACAGTGCAGTAGGAAGCGGCACGACAGTTTTTCTTAAGGTGCAGAATTCCTCAAATGCACCAGTCATTTCCCCTAGACAGAAATTGCCTTTCAAAGTTATCATAGGATGTACTTTGATTGGGATAATTTTGATTCTTTCAGTTTGTTCTGCTCTTTCCAAAAAGAGATCGGAGAAACGCAAAGCTGGTGATTTTCTGGATCTAGAACCAATCTTACCGGGAATCCTAATTCGATTCTCTTACAATGAGCTGAAAATAATTACACAAGATTTCAGCATAAAGCTCGGGAAAGGAGGATTTGGCTCTGTATATGAAGGAACACTGAGAAATGGCACCAAAATAGCTGTGAAGCATCTGGATGGAGTAGGTCAAGTAAAGGAATCATTCTTAACAGAAGTAAAGACAGTCGGTGGCATTCACCATATCAATCTGGTAAAACTCCTTGGATATTGTGCCGAAAAAAGCCACAGGCTTCTAATCTATGAGTACATGGTGAATGGATCGTTGGATAGGTGGATCACGCAAGAAAATGGGCTTACATGGAACACAAGGCAGAGGATAATATCAGATATTGCGAAAGGATTAGCTTATCTACATGAGGATTGCAGCCATAAGATAATTCATTTGGACATTAAACCACAAAACATCCTTATAGATCAAAATTTCAATGCTAAGATATCTGATTTTGGTCTGTCGAAGCTAATTGAGAAAGACAAAAGCAAAGTTGTGACTAGAATGAGAGGAACGCCGGGGTATATAGCCCCTGAATGGTTGAGCTCAATAATCACTGAAAAAGTTGATGTGTATGCTTTTGGAATTGTGCTCTTGGAAACTCTCTGTGGACGAAAGAATTTGGACTGGTCCCAAACTGATGAAGAAGATGTCCATTTGCTAACTGTTTTTACGAGAAAAGCAGAACAAGAGCAGCTCACAGATATGGTTGACAGAAACGAGGATATGCAGCTCCACAGAGAAGCAGTGACAGAAATGATGAGCCTTGCTGCGTGGTGTCTACAGGGCGATTTCTCCAAGAGGCCTTCCATGTCATTGGTGGTTAAGGCACTCGAAGGTTTGGTGACTGTTGAAACCAACTTAGATTATAATTTCAGAAATGTACCTAAGGTTGGGGCAGGCAACCAACCGAGGGAAGCCGCTATCAGCTCAAAATTGCCATCAGTTTTATCGGGACCGAGGTAAACAGTATGTCTAATTCTAGCACAAATTCTTACGGTGTATTAGATTACAATTCATGAATGTAATATTCAGTAACTTTGCATTAGATTTTTTGAGTTTCATGTAATTATCATGTAGAAGCTCCTCAATTCATATGTGACAGGAGATATAATATTGCATTAGCTTTTCCTCATTTTTTGTAAGGGAAGTATCAAGACTCGAGAACAGCATGAATCGGATATCGAGGGCCAGATGCTCTGCC

>PGSC0003DMG400017282 G-LecRK

GAAAAAACTACTTGAATAAAAGGATGAATTTTAAGATTGATACTGATCAATCAAACAGAGCCTTCTTTCTGAAAAAAATAGTACAATGGTGCTTAATTATAGTGCCATTCTTTTCATGGCCAGTCTCCTCTTCATTTCTTCACAGCATCTAATTAGTTGTCAACCTCCTGAAACATTTATCACCAATTCCTCCATTTCCTGGATCAATAGCCGCGCCCGCTCCTATGTAGTGAATTCCACAGATGGAGCCACACTGACACCCATCCTATCGAGAGAAAGGAACGAGACTCTGTTTGTCTGTGGTTTCTTATGTGATTACAATGGTACTGCTTGCGTTTTTGGCGTCCTTTTATCCCCAAAGATCAGTAACAACAATTCTTATCTATATTACGCGAGATTAGTTTGGTCAGCAAATCGGAATAATCCAGTTAGAGCTAATGCAACCTTGCAACTTAGGCAAGATGGAGGTTTATTCTTGATGGACTCAGATGGCACTTTGATTTGGAGTACAACCACAAGCACAAGAGGAAAGGTTGTTACGGGATTCAAATTGACAGTACTGGGAAATATTGTTCTCTTTGACAAAAGTAACGACATCATTTGGCAGTCTTTTGATCATCCTACAGACACTTTGGTTCCTGGACAAATGACAAATGCTGGACAAATGCTGAGATCCAGTGTATCAGCAACAAATTGGAGTGAAGGTTTGTTCACATTTGAAGTTCAACTATTTGGCTTTTTAGCATACATTAAGTCAAATCCACGTCAACTCTATGTCTCAAGACTTGGTGGGGATAATGGTTCCCCGTTTCCTGGACTAATCTTTGATTCGAAAATAATTATTTTCAACGGGACATTAATCTTAAATTTTGCATTTTCAAATACTAGATTCATGAGGTTTGGGAATGATGGGCATTTAAGGGTGTATCACTGGAGAGGATCTTGGTTAGAGGAGGCTGATCTATTGACAGACTACATTGGTGAGTGTGGGTATCCAACCGTGTGTGGGAATTACTCTGTTTGTGCAAATGGGCAGTGCACTTGTCCTCAAACCGCGGTTGATCAGACTAACTTTCTCCAGCAAATAAATTACAGGCAGCCAAACCAGGGGTGTGTCCTTATAACACCTATCTCTTGTGAACATTCCCAATATCATATTTTGATGGAGCTTAAGTACATAGCTTATATTCCCCTCTATTTGCAATATGGTACCAACAAAACAGATTTGGAGAATTGCAAAACGTCTTGCTTAAGCAATTGTTCTTGCAAAGCAGCTCAATTTCAATTTAGTGGTCCTAATAACACAATGGGTCACTGCGCATTGTTGAATGATGTGTTTTCGCTCACATACGATTATGGAGCACCCAATAAGACCACTCTTTTCATCAAGGTGCAGAGCTCTTTTAACTTACAAGCCTCTCCATCACTTGTCTCTTCAGAGAAGAAATCAAAGCGTGTCTCAGCAATTGTAGGATCCACCATCGGAGCTTCATTTGGTTTGCTTCTCATGGTTTTAACTTGCTTTGCCTATATTTTCAGAAGGAGAAAAGGGCTCGAGGAAGATGAGGAGGAGTTCCTTGATCAAATACCAGGAATGCCTACTAGATTTTCCTACGAGGAACTAACTGTGATGACAGAAAATTTCAGTGAGAAACTTGGGGAAGGGGGATTTGGTTCTGTATTTGAAGGAACAATGAGTGATGGCACCAAAATAGCGGTGAAGCGTCTGCAAGGTTTTGATAATGTGAAGAAATCATTCTTAGCTGAGGTAGCAACAATCGGTAGCATTCAGCATGTCAATTTGGTAAAACTTGTTGGATTTTGTGCTGAGAAATCACACAGACTTCTGGTTTATGAATATATGGCCAATGGCTCTTTAGATAGGTGGATTTTTCACGGAAAGCAGGAACGATCTCTCACATGGGATATCAGGAAAAAGATCATATCAGACGTCGCCAAGGGCCTAGCTTACCTACATGAAGACTGTAACAACAAAATTATTCATTTGGATATTAAACCTCAGAACATTCTTCTTGATCATAATCTCAATGCAAAAGTATCAGATTTTGGGTTATCAAAGCTTGTGGGGAAAGATGAAAGTAAGATAGTTACAACAATGAGAGGAACACCAGGCTATTTAGCACCTGAATGGTTACATGAGGTCATTACTGAGAAAGTGGATGTTTATAGCTTTGGGGTTGTGATTTTGGAAATCATCTGCGGCCGGAAGAATTTGGATCGTCATCAAGATGAGGATGATATGCATTTGCTTAGTTTGTTCATGAGAAAAGCAGAGGAGAGACAACTCTTGGAAATGGTGGACAAGAACAGTGAAGACATGCAGCTACATAGAAAAGAGGCTGTGGAAATGATGAAGATTGCAGCCTGGTGTTTACAAAGTGATTATACTAAGAGGCCTTCCATGTCGTTGGTAGTTAAGGTGTTTCAGGGTTTAGTGGCTGCTGAAACCGACTTGCATTACAGCTTCACATTTCCAACAATGACGAGAAGAGTTGCCGGAACTAATCAAGAAAGGGAATCTGTGGTTGGTATCAGTTTGCCACTTCCATCACAGTTGTCAGGACCTAGGTAAATTCTTAGTAGTATTTATCATCAAGACTGAATAGCTAATACGAAAAGCCATTTTGTGTTTTTCATAATTCCTTATTATATCAATACTATTTCAGAAACATAATTTAGAAAAGTAGAACACACTGTCTACTAGAAAAATGTTAGTTTTCGTGCAGTCGAATTAACTTATTGAGTATTGATAAATGGATTGCCTCCTCCGTATTGATGAATTAAATTTGATTGCGC

>PGSC0003DMG400017284 G-LecRK

AAAAAAGGCACAACAGAAAGAACTCATATATTTTAGAAAAATATGTTTGTAGAAAAGAGACTGTATTTTCCTATCCTTCTCTTCATTTACCTTCTCTTTTGTTCGTCTCGATTTACTATATCACAATCTTTACAGGACAACAAGATCAGTCTTTCAAATTCTACAGCAGGGCTTGCCGGGCCTCTGTTGTTCGTCTCGATTTGCTATATCGCAACCTTTACAGGACTACAACATCACTCTTTCGAATTCTACTGCAGGGCTTGCCGGGCTTTCCTCTTTCTGGATCAACAGGCCCTCTCTTCTTGTTAATTCCACCGAGGACTTCTTTGGTTTCGTGACACCCATACTTCAGCGAAAAAATGCTGGCTCACGATTCCTCTGTGGCTTCTACTGCAATTACAATGCCACAGAATGCCTTCTTGGTATCCTTTTGTACCACAACGAGAACAATATGATAGATAAACCCCAGTTAGTTTGGTCTGCTAACAGGAACCATCCAGTGAAATTCAATGCAACCTTGCAACTAGGCCATGATGGCAACTTGGTCTTGACAGACTCCGATGGCACTCTTGTTTGGTCCACTGATACAATTGGGAAATCTGTTTCTGGCTTAAACTTGACAGAAATGGGAAATCTTGTTCTCTTTGATAAGAGAAAGCGCACAATTTGGCAGTCTTTTGATCATCCTACAGACTCTTTGCTTCCAGGGCAGAATTTGGTTTCCGGGCAGAAGCTCATAGCAAGCGTTTCAGCAACCAATCGGAGTCAAGGTTTGTTTTCTCTTACTGTTCTCAATGGAAGCTGGGCTGCTTACACAGATACTGTTCCGCCTCAATACTACTACATTTCATCCTATGCTGATAGTTCTTATTACAGTTTTGATGGTCAAACCTTTACTGCTTCACACAATCCTACTAACTCGACAGCTCAATTCATGAAGATTGGGCCTGATGGACATTTAAAGGTATACCAATGGTCTGTAGTTGATTGGAAAGAAGTATCTGACATTTTGTTGCCAAATGTAGAAAACTGTGGGTACCCAATGGTATGTGGAAGTTATAGCATTTGTACAAATAACGGGCAATGTACTTGTCCGCAACAGGAAAACTTCTTCAGACCATTTTCTGAGAGGAAACCAGATCTTGGATGTTCACAGCTGACTTCCATTTACTGCAACTCTTTGCAGTATCATAGTTTCATAGAGCTCAAGAATACTACATATTTTTCATTTAACATTGATCAGGAACTAAATTCGAGCAAATTGTGGCTTGGGCAGAAAAAGTTGGAAGATTGCAAAAGGGCCTGTCTGAGTAACTGTTCTTGCAAAGCTGCTGTTTTTGAATATAATTGGAATGGGGCTCGAAGAGGGAACTGTTTGTTACTGAATGAAGTTTTCTCTCTCACAGACAGCGAAGAAGGAAGAGGCAAGACAGTATTTATTAAGGTGCAAAATACCTCAGAGGCGCAGACTCGGTCTCTAATCATTCCTGGAGGAAAGAAATCAAGACCTTTCAAAGTGATAATAGGATCTACTCTTGCAGCTTTCTTCGGGATAATTTTAAGCATAGCTACTTGCTTTGTTATTTTCAGAAAGAGGGCACATGAGTCCAGAAAGGCTGCGGATATTTTGGATTTAGCACCAATCTTACCGGGAATTCTAACTCGATTCTCTTACAATGAACTGAAAATAACTACAGATGATTTCAGTAGAAAGCTCGGGGAAGGAGGATTTGGCTGTGTTTATGAAGGAACACTGAGAAATGGAACTAAAATAGCTGTGAAGAATCTGGATGGTGTAGGTCAAGTAAAGGAATCATTCTTAACAGAAGTAAAGGCAGTCGGTGGCATTCACCATATCAATCTTGTAAAACTCATTGGATTTTGTGCCGAAAAAAGCCACAGGCTTCTAATCTATGAGTACATGGTGAATGGATCGCTGGATAGGTGGATTACGCATGAAAATCAAGAAAATGGGCTTACATGGAACACAAGGCAGAGGATAATATCAGACATCGCCAAAGGGTTAGCATATCTACATGAGGATTGCAGCCATAAGATAATTCATTTGGACATCAAACCACAAAACATCCTTCTAGATCAATATCTCAATGCTAAGATATCAGATTTTGGGTTGTCGAAGCTAATTGAGAAAGACAAAAGCAAAGTTGTGACTAGAATGAGAGGAACACGGGGTTATTTAGCTCCTGAATGGTTGAGCTCGGTAATCACTGAGAAAGTTGATGTATATGCTTTTGGAATTGTCCTCTTGGAAATTCTCTGTGGGCGAAAAAATTTGGATTGGTCCCAAGCTGATGAAGAAGATGTACATTTGCTAAGTGTCTTTAGGCGAAAAGCGGAACAAGAGAAGCTCATCGATATGGTTGACAAAAATGAGGATATGCAGCTCCACAGGGAAGCAGTGACAGAAATGATGAGCCTTGCTGCATGGTGTCTACAGGGCGATTTTTCCAAGAGGCCTTCCATGTCATTGGTGGTTAAGGCATTGGAAGGTTTGGTGACTGTTGAAACCAACTTGAATTATGATTTCACACATGTACCTGAGGTTGGGGCAGGCAACCAAGAGAGGGAAGTCATTATCAGTTCAAAATTTCCTTCAATTTTATCGGGACCAAGGTAAACAATATGCTTAAGATTTTGTCTTTGAAGGAACCAACATATGGTTGCGTAATGTCAAATATCGCAACATTTTCAATTCTAGCATAACTTTTTTACCGTGTATGAGATTATAATTCATGAAATGTAATGTTCAGTAACTTCTATATTAAATTTTTGAGCTTCATGTTACTATCATGTAGAGGCTTCTCAATCCATGTGCGAAAATAAGTATAATATTGTACGTGGTGGTGCAGCTACTGAATTCAAACCAGATGAACTGTTCTTGGATCTTAA

>PGSC0003DMG400017286 G-LecRK

AATCAGGCAGATGTTATCTTCCCATTGGTTAGTTGAGAAATGTGATTTGGAAAACATTTCCTTCTGACTTGACGCGTAAATGATTCGCAAAATTGGGTATCACTGAACTCCACACATCTCAAAAGTATTCTTAGCCACAACATTATGAACTACTGGAAGATGCTAACCTACATGTAAAATACAGTTTGGATATTCTGTATGCTCATTTCTTTCTTCAACTATACACCACAAGATTCCATCTATTTTATTGGTTTTGATTAGGAAACAATGTTTGCAGAACAGAGACTGTGTTTTCCTATCCTTCTCTTCATTTATCTCCTCTTATGTTCATCTCAATTTGTTATATCACAATCTTCTCGGGAATTCAGCATCAGTCCTTCGAATTCTACTGCAGGGCTTGCCGGGCTTGCCGGGCTTGCCGGGCTTTCCTCTTTATGGATCAACAGGCCGTCTCTTAGGGTTAATTCCACGGCTGACGGCTTCTTTACTTTGACAACACCCATACTTCAGCGGGGAAATGCTGGCCCACGATTCCTCTGTGGCTTCTACTGCATTTACAATGCTACAGAATGCCTTCTTGGTATCCTTTTGTACCACAACAAATATGACGAGCCGAATGGTGTGATAGATAAACCCCAGTTAATTTGGTCTGCTAACAGGAACCATCCAGTGAAAGTCAATGCAACCTTGGAACTAGGACAAGATGGCAACTTGGTCTTGACAGACTCTGATGGCACTCTTGTTTGGTCCACTCATACAACTGGCAAATTTGTTTCAGGCGTAAACTTAACAGATATGGGAAATCTTGTGCTCTTTGATAAGAGAAAGCGCGCAATTTGGCAGTCTTTTGATCATCCTACAGACTCTTTGCTTCCAGGGCAGAATTTGGTTTCTGGGCAGAAGCTCATAGCAAGCGTTTCAGCAACCAATCGGAGTCAAGGTTTGCTTGCTCTTACTGTTCTCAATGGAAGCTGGGCCGCTTACACGGATACTGATCCGCCTCAATATTACTACATCTCTTACTATCTTGAGAGCCCTTATTACAGTTTTGATGGTCGGACCTTTAATGCTTTACAATCTCTTGATAAATCAGCAGCTCAATTCATGAAGATAGGGCCTGATGGACATTTAAAGGTATACCGATGGTTTATAGCTGATTGGAACGAAGTATCTGACATTTTCTCGCAATATGTAGAAAACTGTGGGTACCCAATGGTATGTGGAAGTTACAGCATTTGTACAAATAACGGGCAATGTACTTGTCCGCTACAGGAGAACTTCTTCAGGTCATTTTCTGAGAGGAAACCAGATCTTGGATGTTCACAGCTAACTTCCATTTACTGCAACTCTTCGCAGTATCATAGTTTCGTAGAGCTCAAGAATACTACATATTTTTCATTTAACATTGATCAGGAACTAAATTCGAGCAGATTATGGCTTGGGCAGGCAAAGTTGGAAGATTGCAAAAGGGCCTGCCTGAGTAACTGTTCTTGCAAAGCTGCTGTTTTTGAATATGGTTGGAATGGGACTCACAGAGGGAACTGTTTGTTACTGAATGAAGTTTTCTCTCTCAAAGACAGCGAAGAAGGAAGAGGCAATACAGTATTTCTTAAGGTGCAGAATTCCTCAAAGCCGCAGACTCAGTCTCTAATCATTCCTGGAGGAAAGAAATCAAGACCTTCCAAAGTGATAATAGGATCTACTCTTGCAGCTTTCTTTGGGATAATTTTAAGCATAGCTACTTGCTTTGTTATTTTCAGAAACAGGACCCGTGAGTCCAGAAAGGCTGCGGATATTTTGGATCTAGCACCAATCTTACCGGGAATCCTAACTCGATTCTCTTACAATGAGCTGAAAATAATTACAGATGATTTCAGTATAAAGCTCGGGGAAGGAGGATTTGGCTGTGTTTATGAAGGAACACTGAGAAATGGCACCAAAATAGCTGTGAAGAATCTGGATGGTGTAGGTCAAGTACAGGAATCATTCTTAACAGAAGTAAAGGCAGTCGGTGGCATTCACCATATCAATCTGGTAAAACTCATTGGATTTTGTGCCGAAAAAAGCCACAGGCTTCTAATCTATGAGTACATGGTGAATGGATCGCTGGATAGGTGGATTACGCATGAAAATCAAGAAAATGGGCTTACATGGAACACAAGGCAGAGGATAATATCAGATATCGCCAAAGCGTTAGCATATCTACATGAGGATTGTAGCCATAAGATAATTCATTTGGACATCAAACCACAAAACATCCTTCTAGATCAATATCTCAATGCTAAGATATCAGATTTTGGGTTGTCGAAGCTAATTGAGAAAGACAAAAGCAAAGTTGTGACTAGAATGAGAGGAACACGGGGTTATTTAGCTCCTGAATGGTTGAGCTCGGTAATCACTGAGAAAGTTGATGTTTATGCTTTTGGAATTGTCCTTCTGGAAATTCTCTGTGGGCGAAAGAATTTGGATTGGTCTCAACCTGATGAAGAAGATGTCCATTTGCTGAGTGTATTTAGGAGAAAAGCGGAACAAGAGCAGCTCATTGATATGGTTGACAAAAATGAGGATATGCAGCTCCACAGGGAAGCAGTGACAGAAATGATGAGCCTCGCTGCATGGTGTCTACAGGGCGATTTTTCCAAGAGGCCTTCCATGTCATTGGTGGTTAAGGCGTTGGAAGGTTTGGTGACTGTTGAAACCAACTTGAATTATGATTTCACACATGTACCTGAGGTTGGAGCAGGCAACCAAGAGAGGGAAGTCATTAGCAGTTCAAAATTTCCTTCAATCTTATCGGGACCAAGGTAAACAATATGCTTAAGATTTTGTCATTGAAGGAACCAACATATGGTTGCGTAATATCAAATATTGCAACATTTCCGATTCTAGCATAACTTTTTATAATTCATGAAATGTAATGTTCAGTAACTTCTATATTGAATTTTTGAGCTTCATGTTACTATCATGTAGAGGCTTCTCAATTCATGTGCGAAA

>PGSC0003DMG400017315 G-LecRK

CAACAATGTTTGCAGAAAAGAGACTGTATTTTCCTATCCTTCTCTTCATTTACCTTCTCTTTTGTTCGTCTCGATTTACTATATCACAATCTTTACAGGACAACAAGATCAGTCTTTCAAATTCTACAGCAGGGCTTGCCGGGCTTTCCTCTTTCTGGATAAACAGGCCATCTCTTATTGTTAATTCCACCACCGATGGCTTCTTTGGTTCAACAACACCCATACTTCAGCGGGGAAATGCTGGCCCACGATTCATCTGTGGATTCTACTGCAGGTACAATACCACAGAATGCCTTCTTGGTATCCTTTTGTACCACAACAAATACAACGAGGAGAACGGTATGATAGATAAACCCCAGTTAGTTTGGTCTGCTAACAGGAACCATCCAGTGAAAGTCAATGCAACCTTGCAACTAGGCCATGATGGCAACTTGGTCTTGACGGACTCTGATGGCACTCTTGTTTGGTCCACTAATACTACTGGAAAATCTGTTTCTGGCTTAAACTTAACAGAAATGGGAAATCTTGTGCTCTTTGATAAGAGAAAGCGCGCAATTTGGCAGTCTTTTGATCATCCTACGGATTCTTTGCTTCCAGGGCAGAATTTGGTTTCCGGACGGAAGCTCATAGCAAGCGTTTCAGCAACCAATCATAGTCAAGGTTTGCTTGCTCTTAGTGTTCTCAATGGAAGCTGGGCTGCTTACACGGATACTGTTCCGCCTCAATACTACTACACTTCATCCTATGCTGATAGTTCTTATTACAGTTTTGATGGCCAAAGCTTTACTGCTTTACACTATCCTACTAAATCAAAAGCTCAATTCATGAAGATTGGGCCTGATGGACATTTAAAGGTATACAAATGGTCTGTAGTTGATTGGAACGAAGTATCTGACATTTTGTCGCCATATGTGGATAACTGTGGGTACCCAATGGTATGTGGAAGTTATAGCATTTGTACAAATAACGGGCAATGTACTTGTCCGCCACACGAGAACTTCTTCAGACCATTTTCTGAGAGGAAACCAGATCTTGGATGTTCACAGCTGACTTCCATTTACTGCAACTCCTCGCATTATCATAGTTTTGTAGAGCTCAAGAATACTACATATTTTGTATTTCAGCTCAATCTTGAACTAAGTTCGAAGATATTTTGGCTTGAGGGGAAAAAGTTGGAAGATTGCAAAAGGGCTTGTCTGAGAAACTGTTCTTGCAAAGCTGCTGTTTTTGTATATGATTGGGAAGGGGCTCAAAGAGGGAACTGTTTGTTACTGAATGAAGTTTTCTCTCTCACAGACAGCGAAGAAGGAAGAGGCGAGACAGTATTTCTTAAGGTGCAGAATACCTCAAGGGCGCAGACTCAGTCTCTAATTATTCCTGGAGGAAAGAAATCAAGACCTTTGAAAGTGATAATAGGATCTACTCTTGCAGCTTTCTTTGGGATAATTTTAAGCATAGCTACTTGCTTTGTTATTTTCAGAAAAAGGACACATGAGTCCAGAAAGGCTGCGGATATTTTGGATCTAGCACCAATCTTACCGGGAATCCTAACTCGATTCTCTTACAATGAGCTGAAAATAATTACAGATGATTTCAGTAGAAAGCTCGGGGAAGGAGGATTTGGCTGTGTTTATGAAGGAAGACTGAGAAATGGCACTAAAGTAGCTGTGAAGCATCTGGATGGTGTAGGTCAAGTTAAGGAATCATTCTTAACAGAAGTAAAGGCAGTCGGTGGCATTCACCATATCAATCTGGTAAAACTCATTGGATTTTGTGCCGAAAAAAGCCACAGACTTCTAATCTACGAGTTCATGGTGAATGGATCGCTGGATAGGTGGATTACGCATGAAAATCAAGAAAATGGGCTTACATGGAACACAAGGCAGAGGATAATATCAGATATCGCCAAAGGGTTAGCGTATCTACATGAGGATTGTAGCCATAAGATAATTCATTTGGACATCAAACCGCAAAACATCCTTCTAGATCAGTATTTCAATGCTAAGATATCAGATTTTGGGTTGTCGAAGCTAATTGAGAAAGACAAAAGCAAAGTTGTGACAAGAATGAGAGGAACACGGGGTTATTTAGCCCCTGAATGGTTAAGCTCAGTAATTACTGAGAAAGTTGATGTGTATGCTTTTGGAATTGTGCTCTTGGAAATTCTCTGTGGGCGAAAGAATTTGGATTGGTCCCAAGCTGATGAAGAAGATGTCCATTTGCTAAGTGTCTTTAGGCGAAAAGCGGAACAAGAGCAGCTCATAGATATGGTTGACAAAAACAATGAAGATATGCAGCTCCACAGGAAAGCAGTGACAGAAATGATGAGCCTTGCTGCATGGTGTCTACAGGGCGATTTTTCCAAGAGGCCTTCCATGTCATTGGTGGTTAAGGCATTGGAAGGTTTGGTGACTGTTGAAACAAACTTGAATTATGATTTCACACATGTACCAGAGGTTGGGGCAGGCAACCAACAGAGGGAAGTCATTATCAGTTCAAAATTTCCTTTAATTTTATCGGGACCAAGGTAAACAATATGCTTAAGATTTTGGATTTGAAAGAACCAAATATGGTTGCTTAATGTCAAATATTGCAACATTTTCAATTCTAGCATAACTTTTTTATGGTGTATGAGATTA

>PGSC0003DMG400017318 G-LecRK

ATAAAATTATGACTTATATTTTCTTTCCTCGTGAAAATTTCTTCAGGCCATCTTCTGAGAGGAAACCAGATCTTGGATGTTCACAGCTGACTTCCATTTACTGCCCCTCTTTGCAGTACCATAGTTTCATAGAGCTCAAGGATACCGCATATTTTTCATTTCAGTATAATCATGAACCAAGTTCGATCATATTTTGGCTTGAGGAGAAAAAGTTGGAAGATTGCAAAAGGGCCTGTCTGAGAAACTGTTCATGCAAAGCTGCTGTTTTTGAATATAATTTGTATGGGAATCGAAGAGGGAGCTGTTTGTTACTGAATGAAGTCTTCTCTCTCATAGACAACAACGGACAAGGAAGAGGCAAGACAGTATTTCTTACGGTGCAGAATCAGTCTTCAATCATTCCTGGAAGAAAGAAATCAAGACCTTTCCAAGTGATTATAGGATCTACTCTTGCAGCTTTCTTCGGGATAATTTTAAGCATAGCTACTTGCTTTGTTATCTTCAAAAAGAGGATACATGAGTCCAGAAAGGCTGCGGGTTTTCTTGATCTGGAACCAATCTTACCAGGAATTCTAACTCGATTCTCTTACAATGAGCTGAAAGTAATTACAAAAGATTTCAGTAGTAAGCTCGGGGAAGGAGGATTTGGATCTGTTTATGAAGGAACACTGAGAAATGGCACCAAAATAGCTGTGAAGCATCTGGATGGTCTAGGTCAAGTAAAGGAATCATTCTTAACAGAAGTTAAGGCAGTCGGTGGCATTCACCATATCAATCTGGTAAAACTCATTGGATTTTGTGCAGAAAAAAGCCACAGGTTTCTTATCTATGAGTACATGGTAAATGGATCATTGGATAGGTGGATTACGCATGAGAATCAAGAGAATGGGCTTACATGGAACACGAGGCAGAGGATAATATCAGATATCGCCAAAGGCTTAGCGTATCTACATGAGGATTGCAGCCATAAGATAATTCCTTTGGACATCAAACCACAAAACATCCCTCTAGATCAATATCTCAATGCTAAGATATCAGATTTTGGGTTGTCGAAGCTAATTGAGAAAGACAAAAGCAAAGTTGTGACTAGAATGAGAGGAACACGGGGTTATTTAGCCCCTGAATGGTTGAGCTCAGTAATCATTGAGAAAGTTGATGTTTATGCTTTTGGAATTGTCCTCTTGGAAATTCTCTGTGTGCGAAAGAATTTGGATTGGACCCAAGCTGATGAAGAAGATGTCCATTTGCTAAGTGTATTTAGGAGAAAAGCTGAACAAGAGCAGCTCATGGATATGGTTGACAAAAACAACGAAGATATGCAGCTCCACAGGGAAGCAGTGACAGAAATGATGAGCCTAGCTGCATGGTGTCTACAGGGTGATTTTTCCAAGAGGCCTTCCATGTCATTGGTGGTTAAGTCATTGGAAGGTTTGGTGACTGTTGAAACCAACTTGGATTATGATTTCACACACGTACCTGAGGTCTGGGCAGGAAACCAACAGAGGGAAGCCATTATCAGTTCAAAAATTCCTTCAATTTTATCGGGACGAAGGTAAACAATATGCTTAAGGTTTTGTCTTTTTAGGAACCAATTTGTGGTTGCATAATGTCAAATATTGCAACATTTCCAATTCTAGCATAACTTAAGGTTTATGAGATTATAATTCATGAACTGTAATGTTCAGTAACTTCTATATTAAATTTTTGAGCTTCA

>PGSC0003DMG400017327 G-LecRK

AGAGACGCGTTCCCAAAAGAAAGGCATTATTAGAACACAGGAGGAGTTGCTAGCAAAGCAGATTTCAAACATTTTCTTTTGTTAGCTTTAGTAAGAAATGATTGCAGAGCAGAGATTGTATTTTCATCATGTCCTGGTCATCATTTATTTTCAGTTTTTTTCATTCAGGTTTGCTTCATCCCAGATTTCATGGTACCTAAATGGCAGTTATTTGAATTCTACTGCTGGACTTTCCACTTCATGGACCAATAGGCCATTTTGGTTTGATGCACAATCCACATATGGATTATCTCTCTTGACACCTATCCTTCTGAGTGGAATAAATGGAAATAAATACCACTTCTGTGGCTTCTACTGCAATAGCCAAAGTGTTGAATGCCTTCTCGGGATCTTCTTTGCATTCCCGGACAGCTCCTCTACTGACAACAAATTCATGGATCCCCAGTTGGTTTGGTCTGCTAATCGAGACCGTCCAGTGAATGCCAATGCAACCTTGCAACTAGGAAAAGATGGCAACTTGGTCTTGGAAGACTCTGATGGCACTTTTGTTTGGTCCACTAATACTACTGGAAAATCTGTTTCTGGCTTAAACATGACAGAAACGGGGAATCTCGTGCTCTTTGACAAAGCCAATCGCACAGTTTGGCAGTCGTTCGATCATCCTACAGATTCTTTGCTCCCGGGGCAGAGCTTGGTTTCTGGGAGGAAGCTCATAGCAAGCGTCTCAGCAACCAATTGGAGTCAAAGTTTGCTTTCTTTAACTATTATCAATGGAAGCTGGGCCACTTACATCGAATCTGATCCACCTCAGTTTTACTATGCTTCAAGTAATCCGGATACTAGTACTTATTTCAGTTTTGATGGTCAAACACTTACTGTTCAGCAAGGCCCTCCTAAATCACCAGCTCAATTCATGAAGCTTGGGCCTGATGGACATTTATGGGTATACCAATGGGACGAGAATGCATTAAATTGGAAAGTCATATCTGATGTTTTGTTTAATGTACATGCAGAGAACTGTGGGTACCCAACGGTGTGTGGAAGATATGGCATTTGTACAAGTAATAGGAAATGTAGATGTCCGCCAGAGCGAAATTTCTTCAGGCCATTAGATGAGAGGAATATAGATGATGGATGTTTACAGCTGACTTCCATTTACTGCAACTCTTCGCAGTATCATAGTTATGTGGAGCTCAAGAATACCACATATTTTGCATTTGAGTTCAGTGATGACCGAATTTCGGGTATATTACCGTTTGAGGGGAAAAAGTTGGAAGATTGCAAAGGGGCTTGTCTGAGAAACTGTTCCTGCAAAGCTGTTGTCTTCAGATATGATTCGGATGGGGCTACAACTGGGAGCTGTTTATTACTGAATGAGGTCTTTTCTCTCGTAGACACCGACGACGGAATGGACAAGAGAGTATTTCTTAAGGTGCAGAATTCCTCAAACACTCAGAATCAGTCTAAAACCATTTCTGGAGGAAAAAAATCAAGACCTCACATAGTGATAATAGGATCTACTCTTGCAGCTTTCTTTGGGATAATGTTAAGTATCACAGCTTTCTTTGTTATCTTTAAAAAGAGGACAAAAGATTCCAGAAAGGATGGGGGTTTTCTGGATCTCGAACCAATCTTTCCGGGAATGCTAACTCGGTTCTCTTACAATGAGCTAAGAATAATAACTGAAGATTTCAGCAGAAAGCTCGGGGAAGGAGGATTTGGCTCAGTATATGAAGGAACACTGAGTAATGGCACCAAAATAGCTGTAAAGCGTCTGGATGGTCTAGGTCATGTGATGGAATCATTCTTAACAGAAGTAAAGATAGTCGGTGGCATTCATCACATTAATCTGGTAAAACTCATTGGATTTTGTGCTGAAAAGACCCAAAGACTTCTGATCTATGAACACATGGTAAATGGATCACTAGATAGGTGGATTTATCATAAAAATGAAGAAAATGGGCTTAGATGGCATACAAGGCAAAGGATTATAACAGATATTGCCAAAGGATTAGCTTATCTTCATGACGAATGCAGCCAGAAGATAATTCATTTGGACATCAAACCACAAAATATCCTTCTAGATAAAAATTTCAATGCTAAGATATCTGATTTTGGGCTATCGAAACTGATCGATAAAGACGAAAGCAAAGTTGTAACTAGGATGAGAGGAACACCAGGGTATTTGGCTCCTGAATGGTTGAGGTCAGTTATCACTGAGAAAGTAGACGTGTATGCCTTTGGAATTGTGCTCTTGGAGCTGTTGTGTGGTCGAAAGAATTTGGATCGGTCCCAAGCTGATGAGGATGTCCATTTGCTAAGTGTTTTCGAAAGAAAATCGGAACAACAGCAGCTTATGGATATTGTTGACAAAAACAATGAGGATATGCAGATCCACAAAGAAGCAGTGACAGAAATGATGAGCATTGCTGCGTGGTGTCTACAGGGAGATTTCACCAAGAGGCCTTCCATGTCTTTGGTGGTCAAGGCATTGGAAGGTTTGGTGTCTGTTGAAACAAACTTGGATTATGATTTCACAAGCCTACCCGAGGTTGAGGATGACAACCAACAGAGGGAAGACACTATCAGTTTAATATTGCCTTCAATTTTATCTGGTCCAAGGTGAACAACTAGCCTAAGAGGTTGTAAGTTTTTTTTTCCTTTCCCAAGTCTATGTCTTTCTTTTGTGAAAATGTATTTCCATGCTTCATCACCAGATCCATCTTTCTCTTGTACATAAATTTGGCTAACAGGTAAACTATGTCATAATGTAAAAGGTAAACTTAATTAGCTTTGATAGATAACCAATTAACGTCTCGTGGCAGAGATGGATAGCATTTTCTTTCTAATTTGCATCTTTGTGGTCAATACATCTTCATATGCAGCACATTTTATATTTCTAAGAGTAAGGGGTCTACTTATTCTGTGTTATGTACAGAGAAAACTGCAG

>PGSC0003DMG400017330 G-LecRK

AAACAACCCAACATCTTAAATTTAAGTTCAATCATTTTTCATCTCAATGATGAACTAAAGGCCCTTTGCCTTCTTTGGAAAACCAAAAAATTGGTCTTTATGGCATTAAGCCAGTTTTTACCTTTCTTCTTCTTCATTCTTTCTTGTTCTTTTCATTTGATAACTAGCCAACCTTATGATTATCCATCTGCAAATCTTTCTACAACTTGGATCAACAGTGTTTCTGCAGACCATTCAGTGAATTTCACTGATGGCTCAACAATAAGAGCCATACTACTCAGAGGAACTTTTGGTCCAAAATATGCTTGTGGTTTCTACTGTAATGGAAATTGTGATACTTACCTCTTTGCCATATTCATTGTGCAAACCAACAGTGCTTCCCGAATTACTCTTCCCGCGATTGGGTTCCCACAAGTTGTTTGGTCTGCTAACAGGAATAAACCTGTTAGGATCAATTCTACTTTGGAACTTACAGCACAAGGAGATTTGGTGCTTAGAGATGCTGATGGTACTTTAGCTTGGTCAACTAACACTACTGGGAAGTCTATTGCTGGCTTAAACTTGACCGATGAGGGGAATCTTGTTCTGTTTGATTCGAAGAATGCAACTGTTTGGCAATCGTTTCATCACCCAACTGATGCTTTGGTTCCAGGGCAGAGGTTAGTACCAGGGATGAAGCTAACAGCAAGTGTTTCAACAACAAATTGGACTGAGAGAGGTTTGTTTTCTCTCTCTTTTACCGATGATGGTTTTGTTGCTTTGGTAGAGTCAAATCCTCCCCAAACATACTTTGACGTGACTATTGGAAGATTGAATGCAAGTAGAGGCTCAAATTATGTTGTGTATTTGAATGGTAGTCTAGCTTTACTCTCAAACTCTAGTAATTCCAGCAATTCACAGACACTGGTTTCATTTAGTCTAGCGACCTCAGCTCAATATATGAGACTGGAGTCTGATGGACACTTGAAAGTGTATGAGTGGCAAAGTGGGTGGACTGAGATGGATGATCTCCTCACAGGTTTTAACGGCGAGTGTTATTACCCCATGGCCTGTGGAAGATATGGCATTTGCTCAAGGGGGCAGTGCAGTTGTCCCAAATCGAGCTCTAATTCAACGACCTATTTCAGACAGATAGATGATAGGCAGGGTAATCTGGGTTGCTCTGAGGTCACAAGGCTGACATGTAATGCTTCAAGTAACCACAGATTTTTGGATCTTCAAGATGTGGACTATTTCACGTTTACGACAGATATCAGAAACACTAGTATGAATACCTGTAAAGATGCATGTTTGAGGAACTGTTCCTGTAAAGCTGCTTTGTTTCGCAGTGGTTTAAACTCATCCACGGGAGACTGCTACCTACCATCCGAGATCTTTTCACTAGCGAATAATGAGAAGGACAAGACAAGGTATGATTCCCATGCATTTATAAAAGTACAGGTTGAACCTGAAATAGCTGCTGCTAAAGAGAAAAGGCGTGTTAATGGTGCTATACTGGGCTCTGTTATAGGAGTTTCCATCTTAGGGATCATAGTTGGAATTGCAGTTTTCATATTCTGGAAGAAGAAGAGAAAGGCCAATGAAGATGAGGAGGATTATCTAGATCACGTACCAGGAATGCCGACTAGATTTTCATACGATGATCTAAAGGCTGCAACGGAGAACTTCACCAAGAAGCTTGGTCAAGGAGGATTTGGGTCAGTTTTTGAAGGGTGCTTAGCAGATGGCACAAAGATTGCAGTGAAATGTCTTGATGGAATAGGACAAGTCAAAGCGTCATTCTTAGCTGAGGTTGAAACTATTGGCAGCATACATCATGTAAACTTGGTGCAATTGATTGGCTTCTGCGCTGAGAAATCTCATAGGCTTTTAGTATACGAGTGCATGAGCAACGGATCACTAGAAAAATGGATCTACCATGGGAAACAGGAGCAAACTCTAGATTGGAACTGCAGGAGGAAGATTATTCAAGACATAGCCAAAGGATTAGCCTACCTTCACGAAGAATGCAGGCAAAAGATTCTGCATTTGGATATTAAGCCCCCAAACATACTATTGGATGAGAAGTACAACGCTAAACTTGCTGACTTTGGGCTTTCAAAGCTAATTGATCGGAATCAGAGCCAAGTTATGACTCAGATGAGAGGCACTCCTGGTTATTTGGCTCCTGAATGGCTTAGTGGAGTCATAACAGAGAAGGTAGATGTCTACAGCTTTGGCATTGTGCTCTTGGAAATTTTGAGTGGAAGAAGACATTTTGAAGCATCGGAGTCTGAAGACCAACAGGTAATGGTGAACTTATTCAGGAGAAAGGCAGAGGAAGGGAAGTTGGTGGATCTTATTGATAAGCACAGTGAAGATATGCAGTTTTACAAAGAAGAAGTAGTAAAGGCGATGCAGATTGCTGCCTGGTGTTTACAAAGCGATTTCACGAAGAGGCCATCAATGTCGATGGTGGTCAAGGCCATGGAGGGTGTTATAGATGTTGACAAGGATCTAGACTACAGCTTTAGACCACAAACTGTTTCCGCAATACCATATATCAGTTTTGCAGATTCAACTCCTTTATTGCCTTCGATCCTATCAGGTCCAAGGTGAGGAGGAAGTTCCTTGGAATTTTTGTTAATTTCTTTTTTTTATGCTATTGCAGCTTGCTCGCTGCTGTGTAAATTACTTCTCCAATGTTCACAAAGTTATGATTGTTTTCTTCAATTTATT

>PGSC0003DMG400017331 G-LecRK

TTTTAACTCATCTTCCTCTAGTGTTCTTAATGGCTTTATGCAAGATTATTCCTTTTGCCTTCATCCTTCTTTGTTGTTCTTGTTATCTGATATCGGGCCAACGTTTCGATTACCCCACTGCAAATCTTTCTACAACTTGGATCAACAGTGTTTCTGCACCCCATTCAGTGGATTTCACTGATGGCTCAAGAGTAAGGGCTATACTACTCAGAGGAACTTTTGGTCCAAAATATGCTTGTGGTTTCTACTGTAATGGCAATTGTGATACTTACCTCTTTGCCATTTTCATTGTGCAAACTAATAGCGCGTCCCAAATTACTTCTCCCGCGATTGGATTCCCACAAGTTGTTTGGTCTGCTAACCGGAACAATCCGGTTAAGATCAATTCAACTTTGCAGCTTACAGCACAAGGAGACTTGGTGCTCAGAAATGCTGATGGTACTTTGGCTTGGTCAACTAACACTGCTGGCAAATCTGTTGCTGGTTTAAGCTTGACCGATGAAGGGAATCTTGTTCTGTTTGATTTGAAGAATGCAACTGTTTGGCAATCGTTTGATCACCCGACAGATGCTTTAGTTCCAGGGCAGAAGTTAGTATCCGGGATGAAACTAACAGCAAGTGTTTCAACAACAAACTGGACCAAAGGAGGTTTGTTTTCCCTCTCTGCTATCGATAATGGTTTGGTTGCTTTCATAGAGTCAAATCCTCCCCAAACATACTTTGACGCGACTATTGGTGGATTGGATCCTAGTGGAGGATCCAATTATGTTAAGTATTTGGATGGAAGCTTAACTTTCTTCACAAATTCGAGTGGTACAACCGAGTTGGTTCTGGTTTCTATTACTCCAGCATCCTCTGCTCAGTATATGAAACTCGAGTCCAACGGACACTTGAAAGTGTACGAGTGGAAAAATCGATGGGGAGAGGTGGATGATCTCTTGACAGGCTTTCGTGGTGTGTGCAATTACCCCACAGTCTGTGGAAGATATGGCATTTGCACAATGGGACAATGTAGTTGTCCCATAAGCTCTAATTCAACTACCTATTTCAGACCGATAAACGTTAGGCTGCCTAATCTTGGTTGCTCTGAGGCCAAAAAGCTAACTTGTAACAACTTAAAGAAACACAGACTTTTGGAAGTTGAAGATGTGGACTATTTCGCATTTACTGCAGATATTAGTAACACAGATGTGAATACTTGTAAACGCGCATGTTTGAATAAATGTTCCTGTAAAGCTGCATTCTTTCGTAGTGGTTTAAATTCTTCCAGGGCCATAGGGGAATGCTACTTACCAACTGAGATCTTTTCACTAATGAATAATGAGAAGGACAAGACAAGGTATGACTCTGTCGCGTTTATAAAAGTACAGGTTTAAGCTAAACCAGCTGCTGCAATAGGGAAAAAATCGTGTTTTATGTACAAAAATGAATCTATGGGAGGACATGGATGGAAACTTAGCAAATTATGCACGGGATCTTGAATAAGTTCATGCTAGACATGAACTATATGAAGGTATTATTACATAAATAAATGGTTGGATAAGCCAAATTAGTGTTAAGAGTTCTCTGTATTCCTTCATTTCTCTTCTTCTCTGTATTACACTTTCATTTCTGTTTGTTATCTGTAATTTCAATTCAATAGACAATAATAAAGACA

>PGSC0003DMG400017946 G-LecRK

AGCTATGCAAAATTTCCACGCGACATATCAACATAGTAGCAAATAGATTATGAAGCTTCATATTCTGAACATCTCCATGAATCCATCTGCAAGTGAAATCAAATTCCATCTGCTATCTCATTTTAATGGAGGCCATAAACATACTTTACTTCTTGTCATCCATATGGATTCTGTCTCGTGTGAGTGCTCAACATACCATACCTGCAAATCAACCTCTAACAGATGGAAGCACCATTATTTCATCAGATGGCAAGTTTAAGTTGGAATTTTTTTCCCCTGGGACATCCGGGAAACGCTACTTAGGAATATTGTTTAATAAAGTTACTGTACAGACAGTGGTATGGGTTGCTAATAGAGACAATCCACTCAATGACACATCCAGTGCGCTAAATCTTACCAGACAAGGAATTCTTACTCTTCTAAATGGTTCTGGCAGAATCATTTGGTCCTCCAATTCCACTAGACATGGGAAAAATCCAGTAGCACAACTCCTTGATTCAGGCAATCTTGTTGTTCGAGATGCAGCCGAGGATAAGCCTCAGAACTACCTGTGGCAAAGTTTTGACAATCCAGGTGACCCAGCTTTACCTGGAATGAAGGTTGGGATTAATCTGAAAACTGGTTTTCATCGTTCCCTTTCGTCATGGAAGAGCACAAATGATCCTTCCAGGGGTCAGTTTACCTGGACATTCGATACTGGTGGATTCGTACAAACATTCATTATGAATGGTTCCATTGAACTCTACAGAGCTAGACAATGGAATGGTCGAGTCTTTCCTAATGCACCATCTAGTGACACTTCGTGGAATGGTTATAACTATACATACTTATCTGATCCTAATGAAATATCATTCATGTATGAACTTACAGACAGCTCTATCATTGCAAGGGTAGTGATGCAACTAAATGGATTTCTACAGCTTTCAGAATGGAATAATCAGACCCAAAATTGGGACGATGTTGTTAGTGCACCAGCTGATAACTGTGACATTTATGGCCAGTGTCATTCATATGGTTTGTGCAACAGCGGCAATTCCCCAATCTGCAGATGTTTGGATAAATTTGAACCCAAAGATCCAACAGAATGGGCAAGGGGAAATTGGTCAGGTGGCTGTGTTAGAAAGACAGCAATATTGAATTGCCAAAAGGAAGTTAAATTCTTCAAGTATCAAGGCATCAAGTTGCCAGACACTCGTTTTTCCTGGTACAGTGGAGGAGTGACTCTTAATGCATGCGAGGAATTGTGCTTGAGAAACTGCTCGTGTATGGCTTACGCAAATCCAGACATAACAGGGACAAATGAAGACTGCTTACTTTGGTTTGATGAGCTGATGGATATCAGAGAGTTTGGTGCTACTGGGCAAGATATCTATGTAAAGTTGGACTCTTCTGAGTTAGGTATAAATCTGTTTCTTTACTTTTTCATGTAAGGCTGTTAAGTATGCTACATTAGATTTTGTGAGCAAGTTGAGGTTTCTCTACATTATTTGAAAAACAAAACAATATCTTTTTTCCTTCTTATGAATTTATTTCTGTCATCAGGTAACTCCAGTACAGGGAAAGTAAAGAAACTGAGAATCAGCTTGCCACTGGCAGCATTAGGTCTGTTCCTAGTACTATGCTTAATCATGTACATAAGGCAAAAGAAGAAGAATCAGCAATATTTTAGAAAAGGTAAACCTCAGAGGTTTTGCTGCTTTTGAATTTGTGGTGCGTTTTTTCTGGAGTTACATTTGGTAAATATCATAATTTTTCTCTTTACGTCCAACTAAGTCCTTATTTTAACAGGAAGAAGACCAAGAATCTCTGAAATGTTGTGCACTAAAGAAAGGAAAGATAAAGAGCTAGATTTACCGTTGTTTGATTTTGAAACTATATCTCATACTACCAATAACTTTTCATTGAACAACAAGCTTGGAGAGGGAGGTTTTGGACCTGTTTACAAGGTAACGATTTTAACTTCAGCTCTTTCTGTTTCATCCAAAAGAACTGCCTAAAATGGAATTTCCGAACCTTTAAAAGGAAACACATGTTTCAGTAAGAAAATTGATATGTCATTCTCCTATTTCATCTAAACAGCAATGTCAAAATCTCGTTTATTGTTTGTCAGTAAAATATTCTGGATAAGCTGATTCAACTAATTTCCTGGATATCTAGGGTATGCTGGAAGATGGGCAAGAAATTGCAGTAAAGAGACTTTCAAGATACTCAGCACAGGGAACTGATGAGTTCAAGAATGAGGTTATCTTCATTGCTAAACTCCAGCATCGGAATCTTGTGAAGCTTCTTGGTTGCTGTATCCAAGCAGAGAAAAGATGTTGGTTTATGAGTACATGCCAAATAATAGCTTGGATTGGTTCCTTTTTGGTTGGTTCTCACTTCTCTCAACTCTTAATACATTTGGTTATCTGTAGTAATAGTTCAATCAGCTGCAGATGGATTTGGCCATCATCAGATGGAGTTGGTTATCACTTTTCTGAACTCTTGATTTTTGCAGATACAGATAGGAGGTCACTGCTTGATTGGCCTAAGTGCTTCCATATTATTAATGGAATTGCTCGCGGACTTCTCTATCTTCATCAAGACTCAAGATTGCGGATAATCCACAGAGATCTTAAACCTAGCAACGTTTTGCTAGACATTGACATGAACCCGAAGATATCTGACTTTGGCATGGCAAGAAGTTTTGGAGGAAATGAGACTGCAGCCATGACAAAAACAGTGGTTGGGACATAGTAAGAATCTTTAAACCGATTAAGTATTTGCAGTCTACTAAAAAGCTACTTACTTTGTTTATTTGACATTATCCTACAGTGGGTACATGTCCCCGGAGTATGCAGAAAAAGGAGAATTTTCAGTGAAATCAGATGTATATAGCTTTGGAGTTCTAGTACTGGAGATTTTAAGCAGGAAGAGAAATAGAGGGTTCTTGCATCCAGACCATGACCATAATCTTCTAGGACATGTAGGTGATCTTCTTTTCTATCTTATATGTTAAGTAAAACTAATAGCATCTAACTGCATGATAGTAAATGTTCAGGTGTGGATCCTTTTCAAAGAAGGCAGGGTTATGGAAGTAATTGATACACAACTAAGGCAGTCATGCAATCAGTCTGAAGTTCAAAGATCAGTTCATGTAGGTCTATTATGTGTGCAGCAGTGTCCAGAAGATAGGCCGAATATGGCATCAGTTGTGCTCATGTTGGGTAGTGATGTCGCACTACCCTTGCCTAAAGAGCCAGGCTTTTTCAATGGCAGGAACCGATTCACTGAAGCTGACAGTTCATCTAGCAAGCATGGCGAAACTTCAGTGAATGAATTAAGCATCACACAGTTGGATGCCAGATAGGAGATAAAATTTCAGGTCATGGATGTTTTCTGTATGCTATATTAACTTAATTGTTGCAGTCATATAGAACAAGAACTACTGGTTGTTCAACTAAATAGTTTTACTCAAGAATCTGTGGCATGGTATTAAGCATGACAACTTGACACTCAGTATGCTGTATATAAAGATAGGAGCAGCAGAGTTGTGAGAAGAGCTAAAAGCATTCTCTACATACAAAAGCACCTAGTACATTTGCTTATGAAAGAAGAACAACTTATTCTGTAAAAGACTTTCAGCCTCCCTTGGGGAAGTAAGAACAATTTTTCCTTTGTTGGCTTGTATGAGTCCACGACTAGTCAGGAGACCAGGTCCCTTGACTGTAGTAATAACAGGAAGTAATGTAAATGCTCAAAGTAAAGAACATAGATGTTTACCTGGAAACTTCCTTGCTCAAGAGAGGAAAAGCTACGACCTGTCACACGAGATTTCTCAAAACTCCAATAACTTCAAGTAACTTGTGAGTACTGACTCTAGTAATCACAAGGATTAAGATATTTATCCTACAAACCTTTAAACACCTCTATTACAAAACACAAAGCAATAACCCTATTGTCCACTTTATCAACTACCTCTAAGTAGAATAGACTTCAACAAACACCACTCAGCTAACTCAAGCCAAAAACTTAACTAGCACTAATATGGAATGGCATGATCTATCTTCCCGTATAACATAGGAATAGATTCTTAATAAAAGACCATAAGACACTAGACTCAAATAACAACTTCAGACTCTGCAAGACTAAATTGGGTCCCTCATTACGTGTCAGAGTAGTTCTTTCTTTAAAACGGGTTTTGCTTTTGAATAATCCTAATTGCAAAAGCTAGGTTGTTTCTTGTCAACAAACAACTTAGCAGCATGTCATTGGTCGAGGACTGTTGTCCTTTCTGTGCCTCTACTGCAGTGTCCACCAACCACTGCAGTTTCTACAGCTTTAGAACTGCACTTCATTGACTTGTCCTTTACGACAAAAACATAGAGGACTAGGTCCCTTTACTAGATCCTATTTCTTTATAGCTTGTTAAATCATCAAAATCCCAAAATACAACATTTCCCTCCTTTTTGATGATGATAAATTATTGCCCATGTCTACATATCTTTTGCTTGCACATTCCCCCTATCAACATGCGTCCCTTGTCGGAAAGACCTGATCCCTTGATCTGAACATGCTTACATGTCATCAGTGTATCTTCCTCCTTTTGACATTATAGAAAAGTAAGAACAGTAAATCAAGCAAAATCACATGCATTCTGGGTAAATTCATGGCCACTAGACAATACTAATATGAACAAACAGGCAGAAAGAAAGGAACGAGAAACAAACATTCTTGGTTTTCCAACTCCGCCTTCTCACTTTTCCATTGGTCAAACCAAAATTGAGTAGCAAACTTTGGTTGATAAGCGGCCAACTCCGCCTTCTCCACCGTAGAAAAAATCTATAATTCCAACTATCTTATACACCTTATCAATGAACTCTTAAGGATCCTCATCAACCTTGGAAACATGGAACTTTGAAGGATTCAACCTAGTTAAGTCCCTTATCCTTGTCACTGCCGTACCCTCATTTGGATTCATGGGAGCAACAACTTCTCCAATTTATCGGGTGGTCATAACTTGAGTCAACACTTGAAAAACGGCTCAAAATTCAGCATGAGTCACTTGCTTGGCCAAAGGATCAACCAGAACTTGAGGAACCTCGGGCTCCATATTTTTACATGTATTCCTTTTCACAAAAACTCTTCGAGGGGCCATATCCTGAAAAGCACCGAAC

>PGSC0003DMG400017949 G-LecRK

TATATATCAGCTATTTCATCAATAATCCCAAAGGAGAATCGTGCAATTTCCTTTAGTAAAACAAGGATGATGGAAAGCAGACATGTTCATTTGTTATTATTATCTTACATGTTTTGTGTTTTGATTCTTACTAGAGCTGCAGATAATATACGTAGAGATGAACCTTTAAGAGATGGAGAAACGATTGTTTCGTCAGGTGGAAATTTTGAGTTGGGATTTTTCTCCCCAACTCCAACATCGAGTAAACGATACCTGGGAATATGGTTCAACAAAGTATCTGTACAGACAGTGGTATGGGTTGCTAATAGAGACACTCCACTGAATGATAAAAATGGTATGCTCAATTTTACAACACAAGGAAATCTTACTCTTGTAAATGGTTCTGGCAGTGTTGTTTGGTCTTCCAATACCACAAGAGGAGGCGTGCAAAATCCAATAGCACAGCTTCTAGATTCAGGCAATCTTGTTGTTAGAGAAGATGCAAGTGAGAATTCGAATTATGTATGGCAGAGTTTTGACTATCCTGGTGACACAGCTTTGCCTGGAGTTAAGATTGGGATCGATCTTAAGACTGGTTTTCGTCGTTTCCTCTGGTCATGGAAGAGCAGAAATGACCCTTCTAATGGTCAATATAGTTTCATATTTGATCCTAATGGATTCCCACAGCCATTTCTCATGAACGGTTCCATCCAACGCTTCAGGGGCGGAGCATGGAATGGTCAAAGCTTTGCTAATTCACCATCTCTACTACCAAGTCCTTCATATAAATACATATTTGCATCCGATCCAGACCAAGTATACTTTACTTATGAGCTCATTGACACTTCTGCTATTGCAAGGGTAGTGATGCAACTACATGGGTTTCTAGAGCTTTCGACATGGAATAATCTAACTCAGAATTGGGATAACTTTGGTAGCGCACCAGCAGACAACTGTGACATTTATTCCCAATGTAACGCATATGGTTTGTGCAACAGTGGCAATTCTCCAATCTGCAGTTGTTTGGATAAATTTGAACCCAAAGATCCAACAGAATGGGTAAGGGGAAATTGGTCAGGTGGCTGTGTTAGAAAGACACCAGTATTGAATTGCCAAAAGGAAGTTAAATTCTTAAAGTATTCAGGGATCAAGTTGCCAGACACTCGATTTTCCTGGTACGGTCGAGGAGTAACTCTTAATTCATGTGAGGAATTGTGCTTGAGAAACTGTTCATGTGTGGCCTATGCAAAACTAGACACAACAGGGACAAATGAAGACTGCTTGCTTTGGTTTGATGAGCTGATGGACATCAGAGAGTTTGGTTCAAGTGGGCAAGATATCTATATAAAGTTGGACTCCTCTGAGACAGAGATAAGTACAGGTTGGTATGCTTCTTTACATTAGTCTTGAGTTTTCTAGACCAAATTGAACTGTCTATTAAAAATTTGAAAAGACTAAGATGAATTTATATGAATCATCAGGAAACTCCAGTAAAGAGATATTAAAGAAATTGAGAATCAGCTTGCCATTGGCAGCATTAAGTCTTCTCTCAGTGCTATGCTTGATCTTGTACATTAGGCATACGAAGAACAAGAAGAAGAAAGAGGAGGAGAATCAGAATCAACAACGTTTTAGTGAAGGTAAACCGGATGTTTTGCTACTTTGAAATTTGAAATAAGTGGCTTGAATTGATGGTAAATCCTAGAAATTCCTCTTAACATCCAATTAAGTCCTTATTTTGACAGGAAGCTCTGAAATGTTGTACATTAATAAGAGCAAAAATGATGATCTAGATTTACCATTGTTTGATTTTGCCACTATCTTGGAGGCTACCAATAACCTTTCTTTGAGCAACAAGCTTGGAGAGGGTGGTTTTGGACCTGTTTACAAGGTAACAATGTTAACTACAGCTCTTTCTTTTTCTCTAAACAAACTGCTTATCTTCAATATATTATATATGCTGGAACATTGACAAACTTTCGGAACCTTTAAAAGGAAACAAGTTCCCGATTAGAAAATATGTCTCCGATTCTCCAGAAATGTCATTAAAAGTTAATTCCACTTCAAACAAGCTATCTAAAAAAGTTTACTCAGAATTTTTCAATCGAAAAACTAGAGAAGCTAATTGCCTGAGGATATGTAGGGTGCGCTGAAAGATGGACAAGAAATAGCAGTAAAGAGACTTTCAAGATACTCAGCACAGGGAACGGATGAGTTCAAGAATGAGGTTATCTTCATTGCTAAACTCCAACATCGGAATCTTGTGAAGCTTCTTGGTTGTTGTATACAAGCAGAAGAAAAAATGTTGGTTTATGAGTACATGCCTAACAATAGCTTGGATTGGTTCCTTTTTGGTTGGTTCTCACTTCTCTCAACTCTTAATACATTTGGTTATCTGTAGTAATAGTTCAATCAGCTGCAGATGGATTTGGTTATCACTTCTCTGAACTTTTGATTTTTGCACATAAAGATAGGAGGTCACTGCTTGATTGGCCCAAGTGCTTCCATATTATAAATGGAATTGCTCGAGGACTTCTCTATCTTCATCAAGACTCAAGATTGCGAATAATCCACAGAGATCTTAAACCTAGCAATGTTTTGCTAGACATTGATATGAACCCGAAGATATCTGACTTTGGCATGGCAAGAAGTTTCGGAGGAAATGAGACTGGAGCCATGACAACAAGAGTGGTTGGAACATAGTAAAGATCTTTAAACCAAAGAGCTACTTACACTCTACTAAAAAGCTAACTTTCTTTATTTGTGATTTAAAAAAAAGAAGAGATACTTACTTTCTTTGTTTGACGTTATCCTACAGCGGCTACATGTCTCCGGAGTATGCAGAAGAAGGGAAATTCTCAGTAAAATCAGATGTTTTTAGCTTTGGAGTTGTAATTCTGGAGATTTTAAGCGGGACGAGAAATAGAGGGTTCATTCATCCAGACCATCACCATAATCTTCTAGGACATGTAGGAACTCATCTTATACTTACAGCGTAAAAATCATGTTAGTTAAAATTATAGCATCTAACTGCATGATGTGAAATGTATGTTCAGGTGTGGATCCACTTCAAAGAAGGCAGGGTTATGGAAGTAATAAATACACATCTAAAGGAGTTATGCAACAACCTATGTGAAGTGCAAAGATCAGTCCATGTAGGTCTATTGTGTGTGCAGCAGTGTCCAGAAGATAGGCCTAGCATGTCATCAGTTGTGCTGATGTTGTCTAGCGATGTTCCATTGCCCTCACCTAAAGAGCCTGGTTTTTTTACTAGCAGAAGCCGATTCGGTGAAGTTAATAGTTCATCTAGCAAGCTTGGCGAACATTCTGGAAATCAATTAAGCATCACACTATTGGATGCTAGATAGGATATATATATATATATTTTCATTGTTTGCTTGTAGAGGCTAGTGTCACGGACTTAGAGTCTCACTAATGTTCCGTGCGGTACTCGACAACTTACACAGGAATCTAATTACAGTGGCGGGACATGACACTAGGCTTTATTTACTTTATTCTTTACTTCACAGAGCTCAACACAGCTTCCAGTTGTACAACCAAACAACTTAAAGTGAATTTGTCAAATGTACAAGTGCTTTCTCAGTCTAATAATTCTTCATAATCTATATTGTCCTCACTCCTCAGATTGTGAAATGAGTTGAGACAATTTGTCATAATTGTACAAAGGGCATTGGACAGCTCTTTTTCAAAACAAGTACAGTTGATAGTATTTTAATTAACAACTTGACGGTGATCAACAGGTTGGTCAATC

>PGSC0003DMG400018452 G-LecRK

CTTCAACCATGTTAACAGTTAGAACTTGTCCTTCTTTAATTTCTCCCCCATTGATATCATGGCTCTCTATTTCTTACTTGTGTTGTTTATCTTTGGATCCGCTAGAGCTCAATCAAATCCACCTAGTACAATAAGATTAGGATCTTCTCTTCATCCCACTGGACAATCTGTAGCATGGTACTCAGCTTCAGGCCATTTCGCCTTCGGCTTTTATCCACAAGGAAGTGGTTACAAGGTGGGAATTTGGTTAGTGGGAGGCTCCAATAACACTATTGTGTGGACCGTGTCTCGTGATGATCCAGAAATATCTGCTGATTCTACCTTAGCTTTCATTGATGGAATTGTAGTTCTGAAAACAAATAACAGTGAGGATAAGACTGTTGCTAGTTCGTCAGAATCTGCTTCTTATGCCAACTTGCTTGATAACGGTAACTTTGTTCTTTACAATCAAGATCATGAAACTATCTATGAGAGCTTCAGTTTTCCTACTGATACAATTTTAGGTGGCCAGAAATTAACAAGTGGGGGTAAATTGGTTTCTAGTTTCTCCTCCACAAATCACTCCTCAGGGAGATTTCGCCTCTGGATGCAGGATGATGGGAACCTTGTTGCATACCCCAGAAACTTGTTTCGGCCAGTAGATGCCTATTGGTCTTCACAAATGTACTGTGATGGTTGCAGAAATCTCCTGCTTCTTAACAGCACTGGCATCTTATTTCTGATCAATGACACTGATTCTTCAGTACTTCGACAATTATACACTCCACTTGTACAGAATAACAGAGCTATTTATCGGGCCACTCTAGATTATGATGGGAATTTTCGACTCTATTCTCACGAATTCAACTCCAATGGGAATTCTAAAATGGTAATGGAGTGGGAGGCAATTGATGATATTTGTCTTGTGAAGGGTTTCTGTGGCCTTAATGGCTATTGTGTTGGAAGTAATCACAATGCATCATCTTGTATGTGTCTACCAGGGTCATACTTGAGAGAGAATGATCCAAGCTTTGGTGATTGTCAAAGGAACTTCACCAGAGGAAAGTGCATTAATGGGAAAGAAGATGTAAGTGTCTACAAGATTACTACCACAACCAATTTGACCTGGGAGGATCCTCCTTACTTTGTTACATCACTTCTAGGAAAAGAAGATTGTGGTAAATCTTGTTTGGAGGACTGTGATTGTGATGCTGCTCTTTTCGATGAAACTGGCCAGTGTATGAAACACAAGCTTCCTCTGAGGTATGTTAAAGGTGCTCCTGAAGGATCTCGCACTGCTTACTTCAAAGTCAGCAATCTAGCAGTGCAAAATCCAACAACGGATTCAGTGAAGCCACCATGGGTGGTGATCTTGGTTTTATCGATAAGTTTCGTTATGTATTCAGGTACTGCTCTTGCATTTTCAGGCTTTTATGTCTTCAAATTCCGGATAATAAAGTACAGGAAGCTACTTCAGACAGGAACCACAGGCTTAACAAAAGATTTCATACTCAAAACTTGCTCTTACAAGGAGCTAAAACAGGCAACGGATGGTTTCAAAGAGGAATTAGGAAAGGGTGCTTTTGGAGCAGTTTATAAAGGGAGCTTTGACAAAGGTAAAAACCTTGTGGCAGTGAAGAGGCTAGAGAAGGTAGTGGAAGAAGGTGAAAGGGAGTTCAGAGCAGAAATGAAGGTCATTGGTAGAACTCGACACAGAAACTTGATTCGTTTGCTAGGATATTGTGCTGAAGGATCAAAAAGAGCTCTTGTTTATGAATATATGAGCAATGGTTGCCTAGCAAATCTGTTGTTCCGAGGAGCTACTCGGCCGGATTGGAATCTAAGAGTAAATATTGCACTAGATGTGGCCAGGGGAATCCTCTACCTCCATGAAGAATGTGAAGCTCCAATTATCCACTGTGATATAAAGCCACAAAACATCTTACTCGACGAGTTGTTGACAGCTAAAATCTCAGATTTTGGTCTGGCCAAGCTGCTAATGCCTGATCAAACAAGAACATTCACTGGTGTAAGAGGGACTAGAGGGTATTTAGCACCAGAGTGGCAAACAAATGCCCCCATCTCTGTCAAGGTTGATGTCTTCAGCTATGGTATTGTATTGCTAGAAATCATTTGCTGCAGAAGAAACATTGAAGTCTGTGTGACTAAAATAGAAGAAATCCAGCTTTCTATATGGGCATATAGTTGCCTTGTTGAAGGTGAAGTAGACAAGCTTGTTGGAAGTGAAGAAGTAGACAAGAAAACCTTGGAAGGAATGATCTCTGTGGCGATTTGGTGCATTCAGGACGAACCAGCTCTTCGTCCTTCCATGAAAAAAGTATTACACATGTTGCAGGGAATTATTGATATTCCAATTCCTCCATGTCCAAGTCCCAGTAGTTAATTTCAGTAGTTGTAAGTGGTTTGGTTTATTATATTGTTGTTTTCTTCAATAACATCTTTTAATGGAAATTTTAGTGAGGATATATAGTTAGTTCAACATACATGAGTTCAGATTATGTTTTATTTTAATGAGATCATTTGCAGGCCCTATGGGTGTAATCTTGAAAGATATTACTCAAAAATGAGAATGTTTTGAAGTGTCAGAGTATCAAAATACTCAACTATTACACTAGGAGATTATCAAAATTATGCTTTCCATCATATGTTGGGAGTTGCGAAAATTTAGACATATATACAAGTATATGTGGGAATATTTAAGGTGTTGAGAGCCACTAGATTTGGAGCTTCTAATTAAGGTGGATAACACTTCAATATAGTAACATGATTGCTGATATTATGCAGGGAAAGGGTATTCCTTGATAGTATTGTGGTTAAAGGCTTAAAGCAAGGAAGACTCTTTATCAAACCAATGGTATTATATATTGCTGTAGAAAAGGAAATTAAATAGTGGATGCAATGGCTAACTGGAGCTTAAAGGGGCATACATCTATCAGTTCAATGATCTACTACAAGAAGCTAAAGCTGCATACAACAAATTGAACAAAATTCAGGTGCTTAGTCAAAGGTGCAGAGTATGCAACAAGATTCTTGTAAGAACTTCATGCATCAAGACATGATAAACCACTTCATATTTTGTATGTTCTTATAAAATCAGATAGAAAAAGTAGTCGAAAGATGATTTTATTAATTCAGTAGTCA

>PGSC0003DMG400018457 G-LecRK

GGAATACTACTGCCATTTAAAATTGGAAATGTCCAAGTTAGTATACAAAAGAATACTTAGCAAAGAGTATATCTATATTCTCATCCTTGTGTCATGGCTAACTTAACATTTACTTTCATGTTGTTGATGCTTCCATTTGTGTTTGCAGCTATTCAAGTTGGAGCTCAGCAACAGCCCTCCAACATAACCTTAGGTTCTTTTCTTTATCCGAACCGAAATCCAAGTTATTGGCCATCTTCTTCAGGCCTTTTCTCATTTGGTTTCTACCAAAAAGGGGATGGATTTCAAGTTGGCATATGGCTACCTGATAAAACTGTTGTTTGGACTGCAAATAGAGATGATCCTCCACTTCATTTGGATGCATATATCGAGTTCACAAAGGGAGGCAAACTTATACTATGGACAAAACAGAACAAAAAAATCATTGCAGAATCTCCACAGCCAGTAACTTCAGCATCCATGCTTGATTCTGGTAACTTTGTTATCTATAACAAAACAATTGTTGTTTGGGAAAGTTTTGATTATCCAACAGACACCATCCTGGCCGGTCAGCGTCTAACATTGGGCCAGAGCCTTGTCTCTAGTGTTACTGAAGTTGATCATTCAAGTGGAAGATTTTATCTTAGTATGCAGGCTGATGGGAACCTTGTTGCCTACCCAACCAATTATCTTAATGGACCTGAGGCTTCCTATTGGGGTTTTATGCTACAAAGCAGAGGTGAATTTGGTGAGGTTAGCTCAGTTTTCATAGGTCTTACTCCAACAGGCCAACTCTTTCGGAATGCTTCTAATGGTTTTCGTTTCCAAGTCAATACCATAGGGAATACCTCTAATCAATCATCAACCAACAAAGCAGCAGCAATTTATCGCGCAACTCTTGATCCTGATGGGATTTTTAGGCTTTATACACATAACTTTGACAACTCAAGCTTGGAAATCAATTGGTCAGCGCTGGAAAACCAATGCAAAGCCAGGGGATTCTGTGGTTTAAACAGTTATTGTACTGCAAAAAATGGCAGCAGCATAGGCGATTGTTCTTGTTTTCCTGGATTTTCATATGTCAACCGTGCAATGAAATTCCAAGGCTGCTATCGGGGATTCGTTTATGAAGAGTTTTGTGGAAGCAGCAGAGACTCAACATTGGCCTACAATGTCACGGCGATTCAACACTTGAAAGTTGGAGGATATCCATTTTCTCAGGTTTCAATGGTTGAGCAGGACTGTAGAAATTCTTGCTTGGATGATTGTACTTGTTGGGCTGCTCTATACATGAATGGAGTTTGCAGCAAATTTAAGCTTCCATTAATTGATTCGACAATTGAACAAAATGATGAGTCCATCAAGAGCTTCGTCAAGCAGAGCTATAACATCTCCCAAATGGCAGGTCACTCTACTCCCTATCCTGGAAAAACAGGGAATAGAAATGAAAGTCGGAAAGAGATAATATTGATTCTATCCCTGGTTCTTGGTTCTGTAGCATTTCTGTTCACTGTTTTTGCAATTTTCAGTTATATTTTCTACAGAAGCAGAGCGGACGAATACCAAAAGCTATTGGAAAATCCATATTTGGGGCCAAATGAGGAGTTCACTCTTCGATCATTTTCTTACAGTGAGCTTGAGAAAGCCACTGAACATTTCAAGGAAGAGATAAGACATGGTTCATTCGGGAACGTCTATAAAGGAATTCTATCAGAAGGTAACAGAACAGTTTCTGTAAAACGACTGGAGAAAATCGGGGATGAAGGAGAAAGGGAATTTAAAGCAGAAATGACTGCGATTGGGCAAGCTCGTCACAAAAACTTGGTTCATTTGCTTGGTTTTTGCTTGGAGGGATCAGAAAAAATTCTTGTGTATGAGTACATGAGCAACGGGTCACTCGCAGATATCATATTCAATTCAGAAACAAGACCTTCTTGGGAGCAAAGGATGAGACTTGCACTGGACATATCAAGGGGAATACTTTATCTGCATGAAAAGTGTGAGACTTGCATAATTCACTGCAACATAAAGCCACACAATATCCTAGTAGATGATTCGTGGACGGCTAAAATCTCTGAATTTGGGCTTGCAAAGTTTCTGGTGCCTGATCAAGTAGGAAACCAACTTCAGCTTAAAGGAACAAGAGGATATTCAGCACCAGAGTTGCAGAATAGTGCATTGATCTCCGATAAGGTAGATGTTTATAGCTACGGAGTAATGCTCTTGGAGATCATATGCTGTAGAAGTAATATGGATGTCAACGTCTCAACCGAGGATGAAATCTTTCTTCCTAATTGGGTACACAAATGCTTTGTCGAGAATGATATCAGAAAGCTAGTGGGAGATAAGGAAGTAGATCATGCGAAGTCCTTGGAGAGGATGGTGAAGGTTGGATTGTTATGCATTCACGATAATCCTGATCTGCGCCCTTCTATGAGGAATGTAGTTCTCATGTTAGAAGGTACTATGGATATACCATTTTTGGCATGGTATTAAGTCTTTTACACTTCGTTTGGCTGGTTGCTACATCTCATTCAAAATGTATCATATTATATTGTATTGTGTTGTGTACTTGCATTGTATTGTACTGTTTTCTTTTTATGAATACAAC

>PGSC0003DMG400019062 G-LecRK

ATGGTTCTGTGTGGTATTATTTATATAATTTATCTGAAAAGATCGGCAAGAAGTAAAGGTAACAATGTAGAAATTTCATCTGCTTTTTCTCAACTACTCCAACTTTTGGATATCGTAATCACTTCAATTGTTTTGATTATAGAAGCCAGAGATATTGTTTTGGGAATGCCTATGGAGTACTTGCCTCATAGAGATAGTTTTGATGAAGATTTAATCACTGAAGATGATAAGAAACGGATTGATGTCCCGTTTTTCAGCTTGAATAGCATACTAGTTGCTACTGAAAACTTCTCAAATGCAAGTAAGCTCGGGAGAGGTGGATTTGGACCTGTTTATAAGGTTTTGCAACTTTATTTTTCTTTCCTTCTTCTGTTCTAAGTTGTTTTACCATTCAGATTTTTTTCTTTCGTTTTCATTATATAACAGTCCGGAGATATATTTATGCAGGGTAAGTTTCTCGGAGGTACAGATATGGCAGTTAAAAGGTTGTCAAGTAATTCTGGACAAGGTGTTGAAGAATTTAAAACTGAAGTAATGTTGATTGCCAAACTGCAGCACAGAAATTTAGTTAGGCTTCTAGGCTATTGTGTTGAGAGAAATGAAAAGATTTTGTTATATGAATATATGCCCAACAAAAGCTTGGATACGTTTCTATTCGGTTAGTATACTTCCCTCCCTACCATGAAACTCGTTGTAGTAATAGTTTTATGATTGATTCTCTCTGAAGTTATCTTACAAACCTTGAAATTAGATCATGCATTCTGTCGATTACTGGACTGGAGGATTCGTTTTGAAATTATATTGGGTATTGCTCGAGGGTTACTTTATCTTCACCAAGACTCAAGGCTAAGGATCATTCATAGAGATCTAAAGACGAGCAACATATTGTTAGATGAAGAAATGAATGCCAAAATATCAGATTTTGGCTTGGCAAGGATTGTTGAAGGCAAAAATACAGAAGCCAATACTAACAAAGTCGTTGGAACATAGTAAGTCTCTTAGTTCTGCTTCACCTTTGATTTGGTGGGAAGAAGATAGATATATACACACCTAATGTATTGGTTTACACAAACTAGATGCTAAAATAACCTATTCTGGAACACTAAAAACAGTGGCTATATGTCTCCGGAATATGCATTGGAGGGACTCTTTTCAATCAAGTCTGATGTATTTGCCTTCGGAGTAGTCCTACTCGAGATCGTAAGTGGGAGGAGAAACATGGAATTTTTCGAAGATGTAAACCTCATTGGTCATGTAAGTACCAAAATTTCCCTCATTAATATACATCTCATTCTACTTAAAGCTTTCATTGACTCAAAATTAGCATTTCACAGGTATGGAGACTGTGGATGAAAGACAGTGCATTAGACATGATGGATCAAACCATAGTTGAGTCGTGCGATGAAAGTGAAGTGCTAAAGTGCGTTAATGTGGCACTCCTATGTGTTCAAGAAGATCCAGCTGAGCGTCCTGTAATGTCAAATGTTGTTTTTATGCTTGGAGGTGAAAGCATAACACTACCAAGACCAAATCAACCAGCTTTTATAGCACGAAGAAATAGTGCTGGTGCAAGTACATCTTCTTCGTCTTCTAAGCTATACAACACTTCCAACAATGGGTTGACAATAACTCAACGAGTAGGCCGATGA

>PGSC0003DMG400019237 G-LecRK

GCCAAGTGTAATTCAAGAAGTGGACAACTTTGTCAAAATTCAAAATTGTCCAAAATCTTGCAAGCCCCACTTTGTCTTCCAAGTAATTCAGTCACAAAAATGAAGTCTTGGTCATATTGACATGTTATCACATCACCAACACCCTTTTAATATTTTTAAAATTTTTTTTGCCTTATTTTGTGTAACCAAGAAATGAAAAAGGGCTAGTAGCTATGGAGAAATTTCTACCCTTTTTGCTTCTTTGCCTTTTAATTTTTCTTCGAAAAGTTAATGGACAAGGAGATATGATAACTACCACTAAGTTTATCAAAGATGGTGAAACTATTGTTTCTTCTGATGGAACATTTGAGTTGGGGTTTTTCAGTCCTGGAAATACTTCTACAAATCGTTATGTTGGAATTTGGTACAAGAAAATTTCTGTTATTACTCCTGTTTGGGTTGCTAATAGGTTGGTTCCACTTACTAATAAATCCGGGGTTTTGAAAGTTATTCAATCGGGCGGTGTTGCTCTTCAAAATGTTACAAATTCAACAATTTGGTCGACAAATTCATCGAGATTCGTGCAAAACCCTGTTGCCCAGTTGCTGGACACGGGGAATTTTGTGTTGAGGGATGCTAATGATCCGAACCCAGAGAATTTCCTTTGGCAGAGTTTTGATTACCCTACTGATACGCTGATTGCGAATATGAAACTTGGAAGAGATTTGATTACTGGGTTTGAAAGGTATCTTAGTTCATGGAAAAGCAGTGATGATCCTGCTCCAGGGGATTATACTTATCATTGTGATCCAACGGGTTATCCACAGGACGTTATGCGAAAAGGGGCTGATGTGATTTATAGAGCAGGGCCATGGAATGGTTTAAGGTGGAGTGGTGCACCTAATATGGTGAACAATTCAATCACTTCGTTTGGTTTGGTAATGAACAATAAGGAAATTTATTATAAATATGAACTCGTGAACAAATCTGTAGTTTCAGCTCTAGTAGTAAAGCCTAATGGGAACACTATGCGTTTGATTTGGATCGCGAAGACTCAGGGTTGGGTTAACTACCATTCTGCTGATGCAGATGACTGTGACACTTACAAATTATGTGGTGCATATGGGACTTGTAACATTTTGAGTGATCCTGTGTGTCATTGTTTAGATAAATTCGAGCCAAAGCATCAGGATGATTGGAATAGGGCAGATTGGACTAGTGGTTGTGTCCGGAAAACCCCATTGAATTGTACTGGAGATGGATTTATTATGTATTCTGGTGTTAAACTGCCTGATACACGAACTTCTTGGTTCAATGAGACCATGAGTTTAGATGAATGCAGGGCGGTTTGCTTGAGAAATTGTTCGTGTATGGGTTATACTAATCTGGATATACGCAATGGAGGAAGCGGGTGCTTGATATGGATTGAGGAGCTGATTGACATCAGACAGCTATCTCAATCAGGACAGGATATCTACATTAGGATGTCTGCTTCAGAAATAGGTAATATGCAATTTACTGAGGTTTTCTTTATTACACAATGCTTGTTATTAAGAATGATGTATTGGATAGTTCAATTCTGTAAGGCTGATTAACGAACGAAGAGTAGTGCATTTACAGGGAAATTAGGTGTAAACATCCTCAAGCAATGGATCCTTTTGTTGAGAATAAGACTTATATGACCATCTTGTTCTACAATAAAATAATTTGCACAGGATTTAAAAAGTTTATCGGAAATTCTAATTCAAACAACTACCTGAAGAAACTTTTGGATGGAAAATGTTGCAGGTTCTGCTGGATCAAGTTCAAAGGGAGATAAATCTGTTATACTTGCAGTAGCTTTGCCACTATTGTTTGCGTTGATTCTTCTAGGTTTAGGTGTTGGCCTGATTCTTTATAAACGGAGAAGGAGAGAGGATCCAGTGGTTATGACAAGGGGTAAGTGAACATAACAAGTTTTTATCTGTTGAAAAGCCGGAAGTAATTAATGGTTATTAGTTCTGTATTCCATTAGTATCTCTTTTCAAGCTCATCGTTTGAACAAAGTCTGCTTGGTTGTTCAAATTGCAGGGAGATTTAGTGGTCACAACAACAAGAACGATAACACTAATCAAAGTCACCATGAAGATTTTGAGCTACCACTTCTTGATTTGCTTACATTAATCAATGCTACTGACAACTTTTCAATTGCGAACAAGATTGGAGAGGGTGGCTTTGGACTAGTTTACAAGGTAGAAACATTTGATCCTTGAGAGCACATTGAAACCTTAAATCGATTTAACTTTTTGGTTGAAATGAAAGTAAGTTCTACTTTTACAGGGTGTGTTAGAAGGGGGGCAAGAAGTAGCAGTGAAGCGGCTATCAGAAACATCAAAGCAAGGAATTCACGAGTTCAAGAATGAAGTTAACTGCATTGCTAAACTTCAGCATCGAAATCTTGTAAAGCTTCTGGGATGCTGCGTTCAAGGAGAAGAGAAGATGTTGGTCTACGAATACTTGCAAAACAAGAGCCTTGATATATATATATTTGGTTAGTCTTTCACATCCAGTGTACATCCATTTCTTTCAAGTGAACTTCTGATGTGATTTCACTGAGCTTAGTTTCATGTTTAGGAGAAATTATAATAAGTTGAAAACATTTGGCAGATGAAGAAAGGAGCGCATTACTTGACTGGCCAAAGCGCTTCAACATTATCAATGGAATTGCTCGTGGACTGATGTACCTCCATCAAGATTCAAGACTTAGAATCATCCACAGAGACTTGAAAGCGAGCAATGTCTTACTTGATACTGAAATGAACCCGAAGATATCAGATTTTGGGATGGCTAGAAGTTTTGGAGGAGATGAGACTGGAGCCAATACACGCCGTGTTGTTGGAACATAGTAAGATTTAATCTTTAATTTTACATGTAGTTGTTACATTCACATGTGCATTTCCTGCATTATATTGTTACTTGTCCGCGTAGCTTGGGATTTTCAAAAGCAAAAAAGAAGATATTTAACTTGCCTTGTTCTACCCCGCTCTGTCCTGCATAAGAGCTCCTCTACCCTGCCCTATATAAGATTAAACCCACCCTATCCCATTGCCATCTCTAATTATATAGTCCTTTGTTATCGTTGTTGCAGTGGTTACATGTCCCCGGAATATGCAGTCGATGGGATATTCTCAGTGAAATCAGATGTCTTTAGCTTTGGAGTATTAGTACTAGAGATTGTGAGCGGAAAGAAGAACAGACGATTCGTTCATCCAGACCACCACCTCAACCTTCTTGGACATGTAAATACTATGTCCCAACTCCTATGTGTAAAAGGACTTTACTCAAAGTATCTGTTAATGAGTTCAAGAGTAGTATATAAACTATTGACATTGTCAATGTATATAACTTAAATCGTTTGTTAACGGATGAGTCATTATATGCAGACATATATGCTACATAAAGAAGGAAGATCATTGGAGCTAGTTGATCCAAAACTGGTGGACTCATGTAATATATCAGAAGTACTAAGATCAATCCATGTAGGTTTATTATGTGTTCAACAAAATCCAGAGGACAGGCCAAGTATGTCAACTGTGATTATGATGCTGAGCAATGAAGGCATTTTGCCATCAGCTAAACATCCTGGCTTCTTTACTGAAAGGAAAATTGGTGATGTTGATCAATTTTCTTGGAGTACACAAACACCTAGTTCTATAAATGAAGTTACCATCACATTGTTGGATGCTCGATAGGAAGAATGTCGAGTGATTTCTGAGATGATCACTCAACTAATAGTTACTATATCAGAAAGTCACTTTCTTTGTTAAAAGATCATTGGAATGTGATTTTCTGAGATAATTACTACTACTTAAGTAACCATCTGAGAACTCAACCAGAATTGTATTAAAATCTTCTCCTCTTTTCTATTTTATTTATTTAGTTAGAAATTTTTGTATTTCAACAACATACACAGTGTGATCCCACTTGTATTCTTTTGTAGTAACATATTTGTCTACATTGTACGTTAG

>PGSC0003DMG400019238 G-LecRK

ATGAATGCAATACAAAGGAATGATTTTTACTTGTTATTGTTATTGTACTTGTATTCGATCAACCATACTTTTGGTGCTACAGATAGTACTATAACAGCTACTCAGTTTCTGAAAGATGGCGATAAAATTGCTTCACATAGTGGAAGCTTTGAAATGGGGTTCTTCAGTCCAGGTGATTCAGAGAATCGCTATGTGGGTATATGGTACAAGAACATATCTGTTAGGACTGTGGTGTGGGTAGCTAACAGAGAAGCTCCTTTGACCGGTGAATCAGGTATCTTGAAAGTCATTGAACAAGGAATTCTTGTCCTTCTCAATGGTACTAACAATGTTGTGTGGTCAACTAATACCTCTAGATCTGTGCAGAATCCTGTTGCACAGTTGCTGGATTCGGGGAATCTTGTTTTAAAACAAGCTGATGATGATAACTCGGGAAATTTCCTCTGGCAGAGTTTTGATCACCCAAGTGATACACTATTACCAAGCATGAAGCTCGGTTGGAACTTTGTAACTGGCCGAGAGGTGTACTTGTCATCATGGAAGAACGAGGAGGATCCAGCTCCCGGTGATTATACATATCATTGTGATCCTTCTGGATATCCACAGAACATCCTGAAGAAGGGTTCAGATGTTGTGTATCGGTCTGGACCGTGGAATGGTCTTCATTTTAGTGGTGCATTAAGCTCAAGAGACAGTCCTTTGTATACATTTGGGATATTCTCAAGTAAGACGGAGGTGTATTTTGGATTTTATCTCACATCTTCGGTTATCACAAGGTTGACACTAAGTCAGAATGGTGCATTACAGCGTTGGACTTGGGGTGACCGTGCACAGGATTGGATCCCTTACCTCTCAATACCCACAGACAACTGCGATGTTTACAAACTGTGTGGCGCATATGGTAGCTGCAACAGTCAAAATTCTCCTGTATGTGGATGCTTAGACAAATTTGTGCCAAAACATAATGAAGATTGGCAAAAGGCAGACTGGTCAACTGGCTGTGTTAGGAGGACTGAACTAAACTGCCTCCAGGGAGATGTATTTTTGAAGTATTCGCATATTAAATTGCCAGACACACGGAATTCCTGGTCCAATGTGACTATGACACTAGAAGAATGCAAGACTATATGCTCTAGAAATTGCTCATGTATGGCTTACTCAAATATTGACATACGCAATGGAGGAAGTGGATGCTTATTGTGGTTCAAGGATCTGCTTGACATTCGACAGCTATCTAAAGAGGGACAAGATATCTATATAAGAATAGCTGCCTCTGAATTAGGTATGTTTGTTACTGAAACAGTTAACAATGATTTGTCATTTGTAACTTGAAAATAAAAACCTAAATTTATGGTTGCTGATACTTGGGGTTAAGCAGAGGTAAACTTAAATTTCAACTGTGCTTATGCTTATAAGTATCTAGTACTAGAACGAAATTTCAAAGTCTTGCGTGCTGTATATATATATACTAGTTTCTGAGCACATGCTTCGCACGTGTGTTTCATGTGAATTTTTATAGATACGTTAGAATGACTAAAATTTCATGAAAATTTATATATGTAAATTGTAATGAACAATAAAATGCAACAATTACAAAATAAGTATAAAGACTTACGACAATAAAACATTCATAGAAAACTAAATTAAGAAGTCTAATAGAGAAAAAAAAAAAAAGACAAAAAGAAACAGATTTAAGAAAAAATACAAGGTAGACATTATTCTTATAAGACTGATACATATATAGATGGAAAAATAAGTACAAAATTAGTACATAACTCTCCGAGTTCTTATCGATTCCTTCCATATCCTATCAAAATTCAAGAGTCTTCTTCATCTACTTTAACTATAGAATAAAAATAATAATAGCCTAATAATCTTGGTAGGGATTTCATGAGATAAAGAAGTTGAATTTATATAGATAGAATAGAAGGAATATCAAAAGATATCTTAAAGTGCTAGTTTAAATATTTAGAGGTTATGAATATGAAAAGATGAGAGTTATGAATATTAAGAGTATAAAAGTTGAATAAGTAATTAGAAAATAATAGTAAATAAATGAAGAATATGAAAAAGAAAATTAAAGTTAGCAAACCATGTAGAAAGAACAACTAAAAGTTCTTATCATGTAATTAAGCATCAATGAATTTAAATTTGTGAAGCTAGAGTCATTCTTTAATAATAATCAAGAGGACATTATCAATGTGTGCGTCTCCATTTTGCAGATTTGCACCCTTCAAATTATTAAATTTCGTTAGTTATCATAAAGGATAATACTTTATCCACATAAATTATATTAAAGGAAAAAGTTATGAGAAGAGGAAAAGTATAGCAATTTAGGTATAGACTTTTTTTAATTTTTTCTTTTATATATATAGATTATAGATGATAAATATTAATAAATTAGATATTTAATTAGGAATTAATCTTAGGAAGTCTAAAAGTCATTAATATTTTAATTAATTTTTTTTTAAAAAAAATTATAATTATTATATACATATAGATTATAGATGATAAATATTAATAAATTAGATATTTAATTAGGAATTAATCTTAGGAAGTCTAAAAGTCATTTAACATTTTAATTAAACTTATACAATCAAATATTTAAATATTTTATAACAATTTAATTAATAGAAGGGGTAAAACCGTAATTCAACTTTCAACTTTTTTTGTTCATGCTTATAGTAATATATGATATATATATATATATATATGTATAGCTAGGAACTAAAACTTTAGTAGAATAAAAGGCTACTACTTAGATTGAGGTGTTGAATTTTAAAGCTAAATCTATAAGCATAATGAGTTATTGCATCTGTCAGCTAAAAATCTAAGACGAAATAAAAGAGGAATTTAGTTGGAAAAACTCATTGAGAACTAAAAATATGATTTGCAGAATACCTCTACTAAATGTCAACAAATTAAAAATGACGTACCGTACTTCAAAGTCCAGCAGAATGTTGTAACTCAATATCAGTTATCAACTCCCCCAAGTGATCAACAGGTAGAGACAGTCAAATTTATTGATTACTTTGTTGGAATGGACCTTGCAGATAGTCTGGAGAAATCAGATGGTGAAAAAGGAAGAGCACTATGCTGGATTCTTCCATTATCCGTTGGTTTAGTTCTAGTAATCCTTAGCATATTGCTGATATGCCACAGAAGAAGGAAGAAGGCTTTAAAGCTTACAACGAAAGGTAAGCTGGATAGGTCACTTTTTTGCCTGATCGTTGAGCAAAATAGAGGAAGTGAAATTCAATTAGAAGTGAAGACGGTCTTTGCTGCTTTAACACAATTCAATCAATTTCCTTTTAGCTGCAGAAAAATCGAGATACAGCGGAAGTATCAAGCAGGATTACAACAGTGGTAGTTGCACTGAAGAATTTGAGATACCACTGTTTGACTTATCTACCATAACTAAGGCTACCGATAACTTTTCAGTAAACAGAAAGATTGGAGAGGGGGGCTTTGGTCCTGTTTATAAGGTAAAAACACTTGTTTCATGGGCAATTTTGTACAAGAAAAATACCAGCTTATCACACAAAGTCTAGATGACTGACAGAAGCTTCCTGGTCCTCCAGAGCAAAGCCTTCATCTTTAGTAGGCAAAATGTAATGAATTAAGTATATGCTGGAGCGTTTTTTGACCTAAACTATACTTAAACTGATTCTCATAATTTCTCCAGTCCTTTTTCGTGGGAAGCAGCTTACTTTGTACCTTATACTTTCAATTTAGATATTATTTATTGTTTTTGAGAAAAAAGATATAAAATATATCGGAAACAATCTCTCTACCTTCAAAAGGTAGTGGTAAGGTTGTGTACACACCAGTTACCACCCTCCCCAGACCCCACTTGCGGGATTACACCAAGTATATTGTTGTTGTACGTTATGTAAAATATTTAACTTGCAGATTTTGATGTTTAGGGAATACTTGAAAAAGGACGAGAAATAGCTGTCAAGCGGCTGTCTAAAACATCCACGCAAGGAGAAGATGAGTTCAAGAACGAAGTGGTATATATTGCCAAACTTCAGCATAGAAATCTTGTGAAGATTCTTGGATGTTGCATTGAAGGAGAAGAAAAAATGCTGATATATGAATACTTGTCAAACGGAAGTCTTGATTCATTTATATTTGGTAACAAATTCCCATACCAATATTGTGTATTCTGAATTATCTTCTCCCAGTTTCGAATCTCCCATCAAATAAGAAAAAAAAAAACATGTAAGGAGTGAACTAAAACCTCAGCTTTATTAGAGCCTAATAATAATGTACACAGTAGCCCACATTGATCTTTACGTTCAATACGTGAATGGACGGATTTAGTTTGATGTTTTAGAACTACTTTCTATCTATAGCTTAGTGAACCAACAAAATTTGGCAGGTGGCAGACAAAGCAGGATATTAGACTGGTCTGAGCGATTTCACATCATCAATGGGATTGCTCGGGGACTTGTGTATCTGCATCAAGATTCTCAACTGCGGATAATTCATAGAGACCTGAAAGCTAATAACATCTTGCTAGACAATGATATGAATGCAAAGATATCAGACTTTGGTATAGCAAGATGCCATGAAGAGGATGGTAATGAAGCCATGACAAACCGAGTTATTGGAACATAGTAAGAATCTATTACTGGGTAGTAGGAAATGCTTGAGATAGTTATGTTTTCCAGCATTAGTAACTATATTTTGTGTACATTTTCTGCAGCGGATACCTCTCACCAGAATATGCGCTATATGGGCAATACTCTGTAAAATCAGATGTGTATAGCTTTGGTATCTTAGTACTGGAAATTGTGAGTGGAAAAAGCAACAGACGATTCGCTCCTTCAAACCTTAACCGTAGTCTAATCGGGCATGTAAGATTTTTATGTATTTTTCTTATTTATACATTGGTGTTAGCAACTAACAAGTATCTTGCAAGGCTAGTGATGCTTATAAGCTTGTAAAATCTACAGGCATGGGAACTCAACAAGGAAGGTAGGTCAATAGAACTACTTGATGAACGCGTAGGTGATTCTTGTTCTACACCTCAAGAAGTGGTACGATCAATTGGTGTTGGTCTATTATGTGTCCAAGAACGTCCAGATGATCGTCCAAGTATGTCTTCCGTAGTTCTAATGTTAAACAATGAAGGTACGCTGCCTCAGGCTAAACTACTAGCTTTTTACATGGAAGGAGATGCATCAGATACTGAATTACTTTCTACCCAATCACATATTACAACGGATGACACCCCGTTAGAAATTAGATGA

>PGSC0003DMG400019241 G-LecRK

TTGATTTTTCACATTTTCCCAAATATTCCAAAAGGACTAAACAATTTCCATGAAACGCTTTAGCTTTTTCTATTTCTGTTCCTCTAATATAGTGTCATTTCTATTCATATTTTCCACTGCTTTAGACACTATATCGAACGATAGCCCCATCAAAGATGGACATACCATAGTTTCAGCTGGTGGAAACTTTGAGCTTGGATTTTTCAGCCCTGGAAATTCCAAGAATCGGTATATCGGTATATGGTACAACAATTTACCTAAAGGCAGAGAAGTAGTGTGGGTTGCCAATAGAGTGAACCCACTGAATGAAACTTCTGGCATATTAACAGTCAGTAGCAAAGGGATTGTACTACTCAATGGAAACCAAGATGTGATTTGGTCATCAAATTCATCGAAAAGTTTGATAAAACCAGTGGCTCAACTTTTGGACGCTGGAAATCTTGTCTTAAAAGATGACAGTTTGGTAAATCAAAAAGACTATGCTTGGCAGAGTTTTGATTATCCAGACAGTACTTTGTTACCTGGCATGAAATTAGGCCTTAACTTGGTCACGGGCAAGTATTGGACTATGTCGTCGTGGAAGAGCAGTGATGATCCTTCTCCAGGAGAATATTTGGATCGCCTCGACACCAGTGGTTATCCTCAATTCTTTGTGTGGGAAGGTCCTGCAATAAAGTTTAGCTCAGGAATATGGAATGGACATTTATTTGTTGGAGGGCCAAATTTGAAACCAAATCCTTATTATACATTTGAGTTTGTGAACAATGACAAGGAGATCTACTATAAATATGAGCTTATCAATACTTCTATTCCTACCAGGTTGGTCTTGAATCCAGCAGGCCTGCTTCAACGACTCTTATGGATCGAACGGAACCAGAACTGGTTCCTTTACTCCACTGGACAGATGGATAATTGTGACCGTTATGCCTTGTGTGGTCAATTTGCGCAGTGCAACATCAATGACTCACCTCCATGTGATTGTTTGAGAGGATTTCAGCCTAAAAATCAACAAGGATGGGATGCAGCAGATTGGTCTAGTGGGTGTGTACGAAGAACTCCGTTGACATGTGGGACAAGTGACAGGTTTCTGAAGTATTCTAGTGTGAAATTGCCAGATACACGCCATTCTTGGTTTGACAAAAGCATTGGCCTCGAGGAATGCCAAAGGCTGTGTCTGAAAAACTGTTCGTGTACAGCATATTCCAATCTTGATGTTAAAAATGGCGGGAGTGGATGCTTACTTTGGTTTAATGAACTCGTTGATATCAGAGAATATGCTGAACTTGATCAAGATCTCTATGTAAGGATGGCTGCATCAGAATTAGGCAAGTCTCCTTATGCTAAGATGAATCATACCACAGGGAATCTTAGTAACTCAATTGATTGATTGACTATTTGAACTTCCACCTTGTTAACGAGGGTTGGGTTCCCAACCTCGTAATCCCCTTTCCCATTTCCCATTTAGCCTTCACCTACCCCCAACCCCATTTCAAAAATAAGAATCATACCGGAAGGTTTTCACAAGTAATAATGAATTTTCACAGGTTCAGGCTATATGGGAAACACAAGAACATCAGTCATTGCAATTATATTAACAGTTTCAGCAATCATCCTTGTTGGTTTTCTATTTTGGTTTGCTATGCAGAGAAAGAAAGGGGAAAGAGGTTAGAAAATTTGTCCGTCGCGTTGAAAGTATAGTTATTTCATTCACCTCAATCTCTTTACAAAATAAAGTTCAATTTTACAGGAGTAGGAGAAGGAGAAGGAAAGGAAGATATGGAATCACCTTTGTTTGATGTGATGACTGTTTCTGCTGCAACTAATAATTTCTCATCTGCTAATATCATTGGAGAGGGTGGTTTTGGATCTGTTTATCGGGTAACTAGAATTATATTCTCTGTGTATCTTAGTCAAATATATGAATGATTCTCTTGTGACTTAGACTTCGCGTGGATTATGTGACAGGGAAAACTATCAACAGGACCAGAAATAGCAGTAAAGAAGCTCTCTAAACATTCCGGACAAGGTTTTGAAGAGTTGAAGAATGAAGTTGTTCTGATTTCCAAACTTCAGCATAGAAATCTAGTTAGGCTTCTAGGCTGTTGTCTTGAAGGGGAAGAAAGGATGCTAATTTATGAATATATGCCAAACAATAGCTTGGACTTCTTCATCTTTGGTTCGATTCTCTTGACGCATTTCATCAAAAGATTCATCTTATCATATCAGCATTGTTCTGCTAATGCAAAATGAACTGATGTTTCTGCAGATGAATGCAGGAAGAAACAACTTCCCTGGGAAAATCGTTTTCGTATTGCAATGGGAATATCAAGGGGCATTCTTTATCTTCACCAGGACTCCAGATTAAGGATCATACACAGAGATCTAAAGACCAGCAACATTTTACTGGACAGTGAATTGAATCCCAAAATTTCTGACTTTGGACTGGCTAGGATTATTGGTGGTGACCAAAACGAAGCAAGAACAAAGCGAGTAATAGGGACATAGTAAGTTCAATTAGATAACTCATTGATGCTTTCATCTTTTCCTCTGTTAGAGTTCCACATTGGTTGGGGAATTGACTGATTTATATGGACTTGGGCAATTCTCCCTCATAAACTAGATTTGAGTTGAGTTAGACTCAAGTGTCATATCCTTCCATGGTATCAGAGCCAGGTAAATCCCGATTGGGGCTCCTGGTCCATGTTGGTGGTTTGGTTATATGAAATTGGACAATCTTTCCCCTTATGAGCTAGCTTTTGGGGTTGAGTTAGGCATGTGTCAAATCTTTACAACTTCCATAAAATCATTTAACCCATATTAAACATTATATATGTCTTTGTGCAGTGGCTATATGTCTCCAGAATACGCAGTTGATGGGAAATTTTCAGTGAAATCAGATGTCTTCAGTCTTGGTGTTCTTCTACTAGAAATAGTCAGCGGAAGAAAGAACAGAACATTTCATCATCCAGATCACCATCATAGTCTTATAGGACATGTAAGCAAGTTACAAATCCAGACATTTGCTTTTGTCATTTCATACTACATTTGTATTTCCTACATGTCATTGAAAACTATACTTGATTTTGCTATAGGCTTGGTTATTATGGAACGAAGGAAAGGCTTTGGAACTAATCGATGATTGTTTAAAAGAATCGTTTGTGGAATCCCAAGTACTAAGATGTGTTCATGTGGCACTGTTATGTGTTCAACGACTAACAGATGAACGACCAACAATGTCATCAGTGGTTTTCATGTTGAGCCATGAAGAAGTGGCATTACCTCAACCAAAGGAGCCTGGTTTCTTCATAGAGAGAAGTATAGCTGAAACAGATCATTCAAATGAGAAAAGATGCATCAGTGACAATGTTTTGACATTAACAATTCTTCAACCAAGATAGCAATGGCGGAACCAGAAATTCAAGAACCTTCTATTGTATAAAGGTGATTCAACAACTAATGTATATACAATACAGAAAATCTGTTCATCCCTATATGTACAGTAAAATATTTTGACGAAGGGGCTTCAGTTGAAACCCCTCTAGACCATGCAGCTCCGCCCCTGATTGTTATTGGACGAGCATTTACTAACATGCTTACATAGAACAATTGCGTATATCTTTCTGATATTTGATTAAGACTCACGAGCCATCGGGACCATTGTGTCAAGTTGTGCAAATTGTCAGAACATGATGGAAATGGACATGAAAGTTGTTACACAGTATAATGACTTATAGTAGATCTGAATGGCTAAATGATAGCCACTGACTTTGTTGAGCAATTATTTTTCCTCATGACTACAAATCAATATCATGTCCTGGGGGAGTGTTTCTTTGCAATCACATTCATCATATCTTTTTCCACATCTCTATCACCAAACTCGAGTTTTGTGCAACTTTTTCCTGGTTTTTCTACCCAGTGTTCAGTACCAGCTTTGGGGGCTTTTTTTGAAAAATGCTAAACAGGGTAGAGCAAGGTGGGTCGATATCTTAGCAGGTCAAGGCTTGCCTCACCTTAACCCGCCCTATTGACATCCCTACCTACTTCAAGGTAAGCCAGCTTCAACAAGACGCCCATAAATCCCAGAATGCCTACCTTCCTAGTTCCTGGAACTATGTAAATCTTTGCATTTGATTCTCATTTGAACAGCATACAGATTTATATTCAGAGGTAGTTACAATAAGTAGAACAAGGAAATGTTCTCGAAAACACTCACAAAATAAATAATTCAAAAAAATGAAGTCATAGCCAACAGAAAAAAAGGCCGTTCAATCAAGCCAAAAAACAAATCCCTTGATGGTTTATACTCCTAAGGCATAAAGCAGTGCAATGGAAGCTCCTACAGTGATTGTCCAGAGATATATGGTTTGAGACAAAGAATTAGCACCACTGCATTGGTCATTAGAACCACTGAAGCATCCAGGTCTACTCACCTTAGGTTCTACTCCCCCTGTGGAAAAGCTACTCTTTGGTTTGCCATTGCTGAAGAGTACACACCAGAAATAGGGGCCACCACCACCGGAGCCACTCACAGCAGCACCAACCTCAGTGTGATTCTTGCTATAGAGAATATCCAAACTCTTGGTATTTCTTATAAGAATATCATTGAATGCTTCAGAAGGTTCGGCATACTTGGATTGACATGCGAGAAATCTTCCAGTTATTTGAGCAAGCGATGACGCCTGCACACCACAGTTAGGGGCAAATTCTTGGGCGAATTCAGATTCAGCAGGTTTCTTTCCATCTGGCCCTCCAACTACTTTACAATCACCTCCATATGCTTTTATATATTGCAGAGCCAAGCATGCCAAGCCAGGGTTACTGTATAAGGAGGATAATTTATTCGCAGTTCTATTACTATTAAGGGCAGATACCAGCTCATCAGCAGGATTATCAGTTACCTTGGCTGCAGTTTATCAAGCCAATATATGTGTTAGAGGTTGATCATAAAGGCAATAATAGTGAATCAAGATCGAGAAGCTATCTGCTTGAAGGAATACACATTGGGAATGACAAGCGTAAATTACATCTAAACCCAAGAGGCTTGAGCACATGTCAGTATGCTTTCGGTAATTGGAAGTACACGCAATTATCTAAGACTGACTAAAGGAAGAAGAGCCAGAGAAGGAGAGTAAACATCATTTCGGCTGTGACTATAATTCTAGCACACTCATGGAAAGATTATCAACAACCACCCTATTATCAAGAGATCTCTAACAAGAAAGATACAAAATTAGGGCTCCAACATAGTATTTGAAGGCATAAACTAAAGCATCTTGAGCACTGTCCTTCCATTAGTGGGCTAGAGTCCCACATTGGTTGGGAAAAGACTGGTGGCTTGTTTATATTAACTTAGACAACCCTCCCCTAAAAGCTAACTTTTGAGGTTGAGTTAGGTCCAGATATCATATCTTTACCTGGGCGGCAAAGGGTGTTGGAATGGGTTAGAGTCCTCATTGGTCTCCTTATATGGACTTAGACAACCCTTTCCTCATGAGATGCCTTTTGTGGCAAGGGGTGTTGGAATGGGTTAGAGTCCACATTGGTCTCCTTAGATGGACTTAGGCAATCCTCCCCTCATGAGATGGCTTTTGTGGTTGAATTAGGTCAGGTGTCATATCTTTACGGGTATCTTCTCATGTTACCACAAATTACAAAGACCGAAAAACCAATTTGAGCAGCTTCCCCTAACTATTCACACTATTCACAAAATCAGAATTTGCAGATTGACTAAATAGTTTCTAGTCAGTGCTATAAATGCAACATGCCAACAAAATCAGAATTTGCAGATTAACTAAAGAGTTTCTTGTCAATGCCATAAATGCAG

>PGSC0003DMG400019275 G-LecRK

CTTTTTGATGTAAAAATCATGAAGGCATTACTTAGGGAAAGTTTCTACTTATTCTTGTTCCTGTGTTTGTATTCAATCCATCAAAGCTTTGGTGCTACAGATACAATAACTACAACTCATTTTCTAAAAGATGGTGATGATGATATTACTTCACCTGGTGGAATATTTGAAATGGGGTTCTTCAGTCCAGATAATTCGGAAAATCGCTATGTCGGTATGTGGTACAAGAACGTATCTGATAGGACTGTGGTGTGGGTTGCCAATAGAGAAGCTCCTTTGAGGAGTGGATCAGGTATCTTGAAAATAATTAAGTCTCGACTTGTCCTTGTTGATGGAACTAACAATGTTGTGTGGTCAACTAATACCTCTAGATTGGTTCAGAATCCTGTCGCGCAGTTGCTAGATTCTGGAAATCTTGTTGTTAAAGAGGCTGGTGATGATAGCTCGGGGAATTTGATATGGCAGAGTTTTGATCACCCAACGGATACGCTATTAGCAGGCATGAAGCTTGGGAGGAATTTTGTAACCGGAAGAGAGGTGTACTTGTCATCATGGAAGAATGAGGAGGATCCAGCTCCCGGTGATTATACATTCCATTGTGATCCTACTGGTTACCCACAGACTATCGTGAAGAAGGGCTCAGATGTTGTGTATAGCTCTGGACCGTGGAATGGTCGATATTTCAGTGGAACACAAAACTCGAGACTAGGTACTTTCTATACATATGGAGTATATTCAAGTAAGACAGAGGTGTATTTTTTGTATAAGCTCACATCTTCAGTTCTTGTGAGGTTGACAGTAAGTCATAATGGAATTCTACAGATTTGGACTTTGGATGACAGAGAACAGGGTTGGGTGCCTTTGATCTTAGTACCAGCAGATAAATGTGAGGTTTATAATTCATGTGGCGCATATGGTAGCTGCAACAGTCAAGATTCTCCAGTATGTAGATGCTTAGATAAATTTGTGCCCAACAATAGTGATGCTTGGAAGAGGGAAGATTGGTCAGGTGGCTGTGTTCGGAAAACTAAACTAAATTGCCTTCATAGAGATGTATTTTTGAAGTATTCGCATATTAAATTGCCAGACACACGGAATTCCTGGTCCAATGTGACTATGACACTAGAAGAATGCAAGGATATCTGCTCAAAAAATTGCTCTTGTATGGCTTACTCAAATCCTGACATACGAGGAAGTGGATGCTTATTGTGGTTCGGGGATCTGCTTGACATTCGGAAGGGAACCAATGGACTGCAAGAGATCTACATAAGAATGGCTTCTTCTGAATCAGGTAGGCTTGTTGCTACTTAAAGATTTTTGATAGATTACCTAAAGATAGTAAAGGAATTTGGGGTTGCAAGCACTTTGAATAAAGCAGACTGTATTTTTGTGCCTATGCTTGTAAACATCTAGTAGTGTCAACTGCCAGCTCCTACCATATATAACCAGGAACTAACAAATTATAGTAGAACGGAAAGCGACTCCTTTAGAGTGAAGTTGGTCTGAACTAATAAGCTGAATCTACAAGCATATGACTGGTTAGACCCTGTTCAATTTTTTAGAACTAATTGGTAAGCACTTAGTTTTCTATAGAACAAGCACTTGATATGCAGTACTCCAAAGGCTTGCAGAACCCTTAATAACTCAGAATAAGAAATCATCTCTCCCATCCGCTTGCAGAACCTTTAATAACTCAGAATAAGAAATCATCTCTCGAATGGATTGGAGAAAGATTGAAATTCATTATTTGTCTCAGTTTAAATGGTTACTTGCAGATGATCAGGAGGAATCGGATGGAAAGAAAGGAAGAGTACTTTTCTGGATTCTGCCATTATCAGTTGGTCTGATTCTGGTATTCCTTAGTCTGCTAATCTACCATAGAAGAAGGAAGAAGGCTTTAGAGCTAAAAAACAAAGGTAAGCTGGATCCCCTTTTATTTTTCTTGACGTCTAAGCAAAATAGAGGAATAGACATTTAAATATTAGTGAAAAGAACTTCATAGTTGTCTGATATTCACAACTTAAACAACATTCACTTCATTTTCTATTTCTTAGAAGCAAAATTACATCATTTGGTTTTGGCAGGAAGGACAGGATGTGGGGGCAATTGCGCTGAAGAATTTGAAATACCACTGTTTGACTTATCTACCATAGCAAACGCTACCAATAACTTTTCAATTGACAGACAGATTGGAGAGGGAGGCTATGGACCTGTTTACAAGGTATACACACACTTAAACCCTTGTCCTTCAGAGTAAAGCTTAATCTTCAGACACAATTTAGGAGTTGTATGATCAAAGGTGTAATGACAAAAACCTTTGCTTTGTGAAACTAAAAGGAGACAGATACAATAGCTAAGGGAAGAAAGGTGGGGATATGGGGGATGTCCTTCTCAGAATGTGTCCTAGAACCACTATCTTAGCTTTTCCAATAAGTGTTGTCGACAATTAGCTCAAGTTAGTCAGTCTAGAGTCTTTAGAGGGGATATCCTTCTGGTGCGTGAACTTGTACATTTCTAAACATTCTCCGGATGTATTTTTATGAGTTAAATTATCTTTCTCTTCTGCATCTCAAGCCTTTATAGTGAGCCGACATTTTGATCAACTTAGATATTTAAAATATTAACATGCAGATATTGTGATGTTTAGGGCATACTTGAAGGAGGGGAAGTAGCTGTCAAGCGGTTATCTAGAACTTCCACACAAGGAGAAAATGAGTTCAAAAATGAAGTTGTATATATTGCCAAACTTCAGCACAGAAATCTGGTGAAGATTCTTGGCTGCTGCATTGAAGGGGAAGAAAAAATGTTGATCTATGAGTATTTGCCGAATGGGAGTCTTGATTCATTTATATTTGGTACAACTTCTCAATCCAATATTAAGAATTTGAATAATAATTGTACTCTCGTATTATTTTTCCATCGACAAAGAACTGATTTTGCTTTAGGCTTTAATAGTACCTTCCACTATACCTACTTAGAATGTTAATAACCCAACATTATTTGGCAGATGACGCACAAAGCAAGGTATTAGACTGGCCTAAGCGATTTCACATTATCAATGGTATAGCTCGGGGACTTATGTATCTGCATCAAGATTCTCAATTGAGAATAATCCACAGAGACCTGAAAGCTAACAATATTTTGTTAGACAAAGACATGAATCCAAAGATATCAGATTTCGGGATAGCAAAAATATGTGAAGAGAATGACATTGGAGCCAAGACAAACCGAGTTGTTGGAACATAGTAAGAACCTTCAACTGGGTAGTTAGGAATTTGAACCTAAGACGAGTTTTGAGATATTAAGCATGTTTTCCAGCACTAGTAATTCTACATTTTGTTTACATTTTTATGCAGTGGGTACCTCTCACCGGAATATGCATTGCACGGAAGATACTCAGTAAAATCAGATGTATTTAGCTTTGGTATCTTAATATTGGAAATTGTGAGTGGGAAAAGCAACAGAAGATTCTCTCATCCAGATCACAACCTTAATCTACTTGGACATGTAAGATTTGATTAGTTTTATTTTACTTATCCTTTGGTGTTATGTTATCTCCTAGTAACAGAGATCTGCAAATCTAGTCAAATGCTAATAAATCTTGTAATCTACAGGCATGGAAACTCTATAAAGAAGGTAGGTCAACAGAACTACTTGATGAATACCTAGGTGATTCTTGTTCAACAACCGAAGTGGAACGATCAATCTGTGTTGGTCTATTATGTGTCCAACAAAGTCCAGAAGACCGTCCAAATATGTCTTCTGCAGTTATGATGTTAAGCAATGAAGGTGAATTGCCACAGGCTAAACGGCCAGGATTTTACATAGAAAGAGATGCACCAGATGGTGAATTATATGCACAGAATACAGTGAGTGAGACCCCCATCACAATATTAGCTGCCCGATAAGAAAATATTTGTCCACTCGCTTCCACAACAAGCTTGCAGTTTTAATAACCTTTTCTTTAATTCTTTGTAGAAATGTGGTGAAAATTTTGTGTATGGCTTTGCCTTATTGGTGTATTAACTATGTCAAACTGACTCAAACATATGTAAAGTGCTGATTCGCTGTTGCTTCTCAATACAAGGAAGAGGTTAGCCAACAAATCAAGTTCAAGAACATTAGCCTCAAACCCAAACACAAGCTCCATAGAAAAAGATGGTGTTACATCAGTCCATAAAACTGTTCTTAACCTGGCGATATGATCTACGGATTTTAATCTATCGCAAAACTGTTTTAGTTTTAGTACTTTACTAATGGATTTTGTCACAACTTTCAATAAATCAAATTCAGGTTTTAAACTTTGATGAGACATTTCCTCAATCAAACTAGTATCATTTTAGTCATTACTTGTACTTAATTTCCTTTCTACTGTCATTACTATAATAACAACAACATGATCTGGAAAGGGTAGAATGTATGCATACCTTACCCTTACCTCGAGATAAAGAGGTTGCTTCTGATAGACCCTTGTCTCAAGGATAAACAAAATAACGACACTGAAGGAGCCATGGACTGTCATAGTATAATCTATAGCAAATCACAATTACTCCATGTCCAATCATTTAGAAGGGGAACGAGGAAGCAATAAGTTTCGGTACTTAACCCCTGTTCTCCAATGTGCACAGATTATATATAACGTTAATTGGAACAATAGCATTTCGTAATTGATGAAAGAAAAATATTCGTACCACGAGTTAGCCGCCGGTAGCGTTGGCCGATTCGGAGCACTGAATTTTGATTGGATTCGCCTGATTACCGAGCACTGAAAAAGGAAATGAAGGTACAACTTCAATTATACAACTTTACACCAATTTATGAATCAAAGATGAACTTCCTCATAGATACAGATGAATGCATATCGGCAGTAGAATTAGATGAAAATGAGAGAGGATTTGACAAAGCATATCATCCTAAAATACACTATAATTATACAATCTTAACACAACACATGAATAATTAAATGATAGAAAAATTAATAATTTCAATTTTTTGTGATATTCTCAAGCAAAGGGATCACAGTGCACATTTCAAGTAATTAAAGTTTAAATATGCGTAGTTTAAATATCACAAACTTGCTTGGACATAAAATTATTACGATTTCTAACAAGATATTAAAATATTATCTAATTTTTTCATGCTCAAAAGTCAAGAACTATTGACCTCCTATAATATTTCTTAATTCCACATCAGCCGCCGTTTAGTCTTAGTTATCCAGTCAAATCTATGAAGACCCCATTATATTCGAACTGCAGCAACATTTTCCTCATGTTTGATACTGCTCAGAACATATATGCCTTTTACTAAATTATTCGTAAAATATACTATTCAAAAGTCCACCAAAAGCAGCCATTAACTTTTTTAAAAGCAGCAGGACTTGAAGCAGCCTCAACTCTCATTAAAAAGCTAAGACTTTTCATAGTCATAGACCTTTCACTCCTTTCAAATAATACTCTTTCATATGTTAGAAAGCTCAAGTGATGGCTTTTTTAACACTAGTTTGTTCAGTTTCTTCTTGTAAGATTCACCGTCTGCAAACAAGTCAAAGTGTCATCACTATGCTACATTTGACATTGAAAAAAATTGGATCTCAAAGCATGAAGGCATTACTAAGGGAAAGTTTTTACTTATCTTTGCTCCTGTGTTTGTGTTTCATCCACCGAAGCTTTGGCGCTACAGATACAATAACTACAACTCATTTTCTGAAAGATGGTGATGATAATGTTGCTTCAACTGGTGGAATATTCGAAATGGGGTTCTTCAGTCCAGGTAATTCGAAAAATCGCTATGTGGGTATGTGGTATAAGAACATATCTGTTAGGACAGTGGTGTGGGTTGCCAACAGAGAAGCTCCTTTGACTGGTGGATCAGGTATCTTGAAAATAATCAAGCCAGGAATTCTTGTCCTTGTCAATGGTACTAACCATGTTGTGTGGTCAACTAATACCACTAGATCTGTGCAGAATCCTGTAGCGCAGTTGCTAGATTCTGGAAATCTTATTGTCAAAGAAGCTGGTGATGATAGCCCGGGGAATTTGATATGGCAGAGCTTTGATCACCCAACGGATACGCTATTAGCAGGCATGAAGCTTGGTAGGAACTTTGTAACTGGCAGAGAGGTGTACTTGTCATCATGGAAGAATGAGGAGGATCCAGCTCCTGGTGATTCTACATATCACTGCGATCCTTCTGGTTACCCGCAGAATATCCTGAAGAAGGGGTCGGATGTAGTATACCGCTCTGGACCATGGAATGGTCGCTCTTTCAGTGGGAACCGAAACTCAAGAGAAGGTCCTTCCTATACATTTGGAGTATATTCAAGTAAGACAGAGGTGTATTTTGGATATAACCTCACAACTTCAATTATCGTGAGGTTGACATTAAGTCACAATGGAGTCCTACAGGTTTGGACTTGGGGTGATGGAGACCAAGGTTGGATCCCTTTCCTCTTAATACCCGCTGATAATTGTGATATGTATAAGTTGTGTGGCGCATATGGCAGCTGCAACAGTCAAGATTCTCCAGAATGTGGATGCTTAGACAAATTTGTGCCCAACAATAGTGATGCTTGGAAGAGGAAAGACTGGTCAGGTGGCTGTGTTCGGAGAACTGAACTAAATTGCCTCCAAGAAGATGTATTTTTGAAGTATTCGCATATCAAATTGCCAGACACACGGAATTCCTGGTCCAATGTGACTATGACACTAGAAGAGTGCAGGAATACCTGCTCTAAAAATTGTTCTTGTATGGCTTACTCCAATTCTGACATACTCAATGAAGGAAGTGGATGCTTACTGTGGTTCAAGGATCTGCTTGACATTCGGCAGGGACCCAACGGAGGGCAAGATATCTACATTAGAATGGCTTCTTCTGAATCAGGTTGGCTTGTTGCAACTTAATATTTTTTGATAGATTGCCTGAGGATAGTAAAGGAGTTTAAGGTTGCAGGCACTTTAATAAAGCAGACTGTAGTTTAGTGCCTATGCTTACTATGATACTATCTACTGCCAATTCCTACCATATTTAGCCCGTAACTACCAAATAATAGCAGTACAGAAAGCTACTATTTTTTAGTGAAGTTGGGATGAATTTATAAGCCAAATCTGCAAGCATGATGACTATAGAACAAGCATTTGATATGACTTGCATTACACAAGGAATTGAAGAACCTTGACAACTCAACCTCAACAAACTAAAAAGGATGTGCTGTACTCCAAAGGCTTGCAGAGCCTTTTATAACTCAAAATAAAAATCATCTCCAATGGATAGGATAATTGCAATTTATTATTTATGTTTGTTGAATTGATACTTGCAGATAGTCTGGAGCAATCAGATGGAAAGAAAAGGAAAGTACTTTTCTGGATTCTACCATTATCAGTTTGTTTGATTCTGGTATTCCTTACTCTGCTGGTCTACCATAGAAGAAGGAAGAAGGCTTTAGAGCTCAAAAACAAAGGTAAGCTGAATACCCCTTAATTTTTTCTTGACATCTAAAGTCTAAGCAAAATAGAGGAAAAGACATTTAAATTATACTGAAAAGAACTTTCACTTCATTCGCTTTCTCTTATAGGCAAAATTATGTCATTTGGTTTTGGCAGGAAGGTCAGGATGCAGTGGAAATTACAAGATGAATTACAACCGTGGCAATTGCACTGAAGAATTTGAAATACCACTGTTCGACTTATCTACCATAGCAAAGGCTACCAATAACTTTTCAATTGACAGACAGATTGGAGAGGGGGGCTTTGGACCTGTTTACAAGGTATAAACACTTAAGACCTGGGTCTTCAGAGTAAAGCTTCATCTTTAAATACAATATAGGAGTTGTATGATCAAAGGTATAAGCCTTTGCTTTGTGAAACCAAAAGGAGACCTATACAATAGCTAAGGGAAGAAAGGTTGGGATATGGGGGATGTCCTTCTCAGAATGTATCCAAGAACCACTAGTCTAGCTTTTCAATAAGTGTTGTCCCAAGTTAGTCAATATAGAGCCTTTAGCATTTTCAAGAAAGTGCTGCAAGAGTAACCCATCACCCTTTGGGTGGAACTGCAGATTAAAGTGTTGAGGATATAGTTTGTTTAAGAAGAGAGTCATTTGTTCTTTGTAGTCATTTATTTGTTCATCTTTCCATGGTGACAAGTGGCAACTTGTTCCCGCATCAAAAGGGCGAATATTTCTTATAGACTTTAGTTTTAATTTTAAAGAAGATTATCATAGGGCTAAACTGAAAATATAATGAACTTATAATAGAGATTTGGCACATTTGACCTAAATTTGTTAACGGAGATTTTGATCAACTTAGGTATTGAAAATATTAACATGCAGAGATTGTGATGTTTAGGGCATACTTGAAGGACAGGAAATAGCCGTCAAGCGGTTGTCTAAAACTTCTACACAAGGAGAAAAAGAGTTCAAAAATGAAGTTTTATATATTGCTAAACTTCAGCAAAGGAATCTGGTGAAGATTCTTGGCTGTTGCATTGAAGGGGAAGAAAAAATGTTGATTTATGAGTACTTGCCAAATGGGAGTCTTGATTCATTTATATTTGGTACAACTTCTCAATCCAATATTATGCATTTGAATAATAGTTGCATTTCAGTGACTTGATGCCTATAGGCATACACTGATTTTCTTCCAACTTCAAAATCCCCATAAAGCAGGAAAAAAAATGTAAGGAGAGAACTAAGACCCCAGTTGTTTAAGAGCCTAATAATTACATACTTGGTAGTTCAAACTTCTCGTATTATGTTTCCATCAGCAAGGAACTGATTTTGTTTTTGGTTTTAATACTACATTCCACTAAACCTATTTAGTGTGTTAATGACCCAACATTATTTGGCAGATGACATACAAAGCAAGGTATTAGACTGGCCTAAACGATTTCACATTATCAATGGCATAGCTCGGGGACTTATGTATCTGCATCAAGATTCTCAATTGAGGATAATCCACAGAGATCTGAAAGCTAACAATATCTTATTAGACAAGGACATGAATCCAAAGATATCAGACTTTGGCATTGCAAAAATATGTGATGGGAATGACATTGGGGACAAGACAAACCAAGTTGTTGGAACACAGTAAGAACCTTATACTTGATAGCTAGAAATTTGAACCTAAGAAGAGTTTTGAGATATTAGGTACGTTTTCCAGCATTAGTAATTCTATTTTTTGTTTACATTTCTATGCAGTGGGTACCTCTCACCGGAATATGCATTGCACGGAAGATACTCAGTAAAATCAGATGTATTTAGCTTTGGTATCTTAATATTGGAAATTGTGAGTGGGAAAAGCAACAGAAGATTCTCTCATCCAGACCACAACCTTAATCTACTTGGACTTGTAAGATTTGATGAGTTTCATTTTACTTATCCTTTGGTCTTATGTTATCTTCTAGTTACAGATATCTGCAAATCTAGTCAGCATTAATAAATTGTGTAATCTACAGGCATGGAAACTCTATAAAGAAGGCAGGTCAACAGAACTACTTGATGAATACCTAGGGAATTCTTGTTCAACACCCGAAGTGGAACGATCAATCTGTGTTGGTCTATTATGTGTCCAACAATGTCCAGAAGATCGTCCAAGTATGTCTTTTGTGGTTATGATGTTAAACAATAAAGGTGTATTGCCACAGGCTAAACGGCCAGGTTTTTACATTGAAACAGATGCACCGGATGGTGAACTCTCGTCTAGCCATACTGAGACCCCCATCACAATATTGGTAGCCCGATAAGAAAATATTGTTTCACTTGCTTTCAACAAGCTTGCAGTTGTAAAAAATCTTTTCTTTTATTCTTTGTAGAAATGTTGTAAAAAAATATTTGGTATATCAGCTATTTCGAAC

>PGSC0003DMG400019276 G-LecRK

AAAAAATGAAGTCTTGGTAATCTCTGAGACAATCTTACTGAAAAGAATACCACAAGAGATGGAATATGTTAAATTCTTTTTAACAATTTTGAAGAAGAATGCAATGCAACAAAAGGTAAATATAACTATCATGGATATAACAATATTTCAATTTATTCTACTTCTCTTGTATTTTGTGTTTCTTCAAAATGTTAGTGGTCAAGGAGATACCATAACTACAACTCAATTCATTAAACATGGTGATACTATTGTTTCGTCTCATGGAACGTTCGAGTTAGGATTCTTCAGTCCTGGAGATACTTCTACAAATCACTATGTTGGAATATGGTACAAGAAAACCTCTGCTACTACTCCTGTTTGGGTTGCCAATAGACTAGCTCCTGTTGCTAATAAGTATGGGGTGTTGAAAGTTATTCAGCCAGGTCTGATTGTACTTCTGAATGATACTAATGCAACTGTTTGGTCCACAAACTCATCGAAATCTGTACAGAATCCTGTTGCTCAGTTGTTGGACACAGGGAATTTTGTGGTGAGGGATGCTAATGATCCGAATCCAGAGAATTTCCTCTGGCAAAGTTTTGATTATCCTAGTGATACGTTGTTGGAAGGTATGAAACTTGGGACGGACTTGGTGACTGGATTAGAAAGGTATCTTACTTCTTGGAGAAGTAGTGATGATCCTGCTCCGGGGGATTATACTTATCATTGTGATCCTACGGGTTATCCACAAAACCTTATGCGAAAAGGGGCTAATGTGACGTTTAGAGCAGGGCCGTGGAATGGTATCAGGTGGAGTGGTGCACCAAATCTGGTAAACAACTCTATCATTTCTTTTGGCATGGTTATGAATAGTAGGGAAATTTACTACAAGTATGAGATGGTGAACAAATCCGTTATTTCAACATTTGTTTTAGAGCCTTATGGGAAAGCTATGCGTATCATTTGGATTGGAAAGGTTCAGGGATGGGTAAATTACCACTCTGCAGTTGTGGATGACTGTGACACCTACAAATTATGTGGTGCATATGGGACTTGCAACATTTTGAGTGATCCTTTTTGTCAATGTTTAGATAATTCGAGCCAAAACATCCTGATAATTGGGAAAGGTCTGACTGGTCTAGTGGTTGTGTCCGGAAGATCCCACTGAATTGCACAGGAGATGGATTTATTAAGTATTCTGGTGTTAAACTGCCTGATACGCGAAATTCTTGGTTCAATGAGACAATGACACTAGATGAATGCAGGGTAGTTTGCTTGAGAAATTGTTCGTGTACGGGTTATACTAATCTGGATATACGCAAGGAGGAAGGGGGTGCTTGCTATGGATTTGGGAGCTGGTTGACATCAGACAGCTACCTGCATCAGGACAGGATATCTACATTAGGATGTCTGCTTCAGACATAGGTAATTAATAATTAATATGCGATTTCATAAATCATAACCATGTCAATTGTACTTCAATGCCAGAAATTCAACTCATATGAGGGACAAAGGTAATCCAAGTCCACATTATGGGAATAAGACTTCAAGGACCGTCCTGTTTTCCTCGACATAAATACAACAAATATAAAAAGAGAAAAAACTTCTAGCATTACTAGAAACAACTATTGATAATATTTCGATGGGAAATGTTGCAGGTTCTGCTGGATCAAAAGGAAAGATAGCTTTCATACTTGCAATAGCTTTGCCACTATTGGTTGCATTGATTCTGCTTGGTTTAGGTGTAGGCCTGATTCTTTGTAAACGAAAAAGGAGAAAGGATCCGGTGGTTACAACAACAGGTAAGTGGAAGTACCAATTCTGTCTATCGTAAAACCAGAAGCAACTAGTGGTCATTAGTTCTGTATACCACTAGTATTTCATTTCAAGCTCCTTATTTGAGCAAGGTCTACTTGTATGTTCAAATTGCAGGGAGATTAGGTGGTCACAGCAACAAGAATGATAACAGTAATCAAATTCACGAGGAAAATTTTGAGCTTCCACTTTTTGACTTATTAACGTTAACCAATGCTACCAACAACTTTTCATTTGCAAACAAGATTGGAGAGGGTGGCTTTGGACAAGTTTACAAGGTACAAACATTCCTCAGACCCCAAGTAATGATACTCGAGACTTAATTTCTTTGCTTTTCTTGAAAGGAATATTAACTCTTGGAGTCAGTACATACAGCAGTGTTTGCTAGTATGATGAGCGGAATGAGTGTTAAGTTGAGAGCCTTGTTTCAAGACCGATAATGTTTTATAAAATAAAGGTTTCATATTCAAAGAAAAGGAACATCAGCTGGAAAAGATTACCAACCATGAGATAAGCTGTGGAATGAAATGTGCAAGTAGATGCTGCCTGAACAAGAACTTGATTGCACTAGTAAGGAAAAAAATGACGATTCACTCCATCAAATTTCTAACGGGACTGCTACTTTTACAGGGTGTATTAGAAGGTGGACAAGAAGTAGCAGTGAAGCGGCTTTCAGAAACATCAGAGCAAGGACTTCACGAGTTCAAGAATGAAGTTAAGTGCATCGCAAAACTTCAGCATCGGAATCTTGTAAAGCTTCTAGGATGCTGCATTCAAGGAGAAGAGAAGATGCTGGTGTATGAGTACTTGCCAAACAAAAGCCTGGAATTATATATATTTGGTTAGTCTTTCACATCTACTGAACGGCCATTTCTTTCAATGCAATCAATTGTGATTTCACGAAGGTTGGTTTCATTGTTATAGAAACCTTTAGTAATAAGTTGAAAACATTTGACAGATGAAGAAAGGAGTGCATTACTTGATTGGCCTAAGCGCTTCAATATTATCAATGGAATTGCTCGAGGACTGATGTATCTTCATCAAGATTCTAGGCTCAGAATCATCCACAGAGACTTGAAAGCTAGCAATGTCTTACTTGATATTGAAATGAATCCAAAAATATCCGATTTTGGGATGGCTAGAAGTTTCAGAGGAGATGAGACAGGAGACAATACAAGCCGTGTTGTTGGAACATAGTAAGATTTAATATTTGATTATATATGTAGTCATGAACACATGGTGCATTATATAATCCTGTAGTTTCTTTATTGCAGTGGTTACATGTCCCCGGAATATGCAGTCGAGGGGATATTCTCAGTGAAATCAGATGTCTTTAGCTTTGGGGTATTAGTACTAGAGATTGTGAGTGGAAAGAAGAACCGACGATTTGTTCATCCAGACCACCACTTCAACCTTCTTGGACATGTAAGTACAGTAACTCCTAACGGTAAAAAGACTCAGAAAAAGTGTTTGTTAATGCATTTAAGTTTTATGTGCCGACAGTAAATTACATGTAAATCTCTATTATAAGAGCTTGTTAACTAACATACTATCACAAGTTAAACTACAATGCTAGAGTATAAAGTCTTTGTACTGTTAGTGTGTATAAGTTAAATCCTTTGTTGATGGAGGGATCGTTTTATGCAGGCATGGATGCTTCATAATGAAGAACGGACGTTGGAACTAGTAGATCCATATCTCGTAAATTCATACTATATATCAGAAGTACTAAGATCAGTCCATGTGGGATTGTTATGTGTTCAACAGAATCCAGAGGACAGGCCAAGCATGTCAACTGTGATTATGATGCTGAGCAATGAAGGCATTTTACCACTGCCTAAACATCCCGGTTTCTTCACTGAAAGGAAAGTTGAAGGTCATGATCAATTATCTTGGAGTACACAAACACCAAGTTCTATAAACGAAGTTACCATCACGCTGTTGAATGCTCGATAGAAGCATATCATGGCCATAGTCAGTGGCAAAGCCAGGATTTTCACTAAGGGTTTCAAAATATGAAAGAAGTAAACACACAAAGTCAAGACGATTCAACAACTATTATATATACATAAACAATTAATTGTAGACCTTTATATATATAGTGTAATTTTTTGGCGAAAGGGTTCGGATGAACCCCTTGCATCAACTTGGCTCCATCCTTGTCGATTGTGAGTGTCATTCATTTCTCAGAATTGTATTAAAATCCTCCCTTTTTTTCTTTAAGAAATTAGAAAGCAAAATCTGCAGGAATTGTATTGAAATATGTCTAAATTTTAAGGCTTTGAATTCAATTTCAG

>PGSC0003DMG400019278 G-LecRK

ATGAAAGGGAAACATTTTTTATCTTCTTGCTCCATTTTTCTTTCCATCTTGCTAATTTCCACAGCATTAGACACAATCACAACAGAAAAACCAATTAGAGATGGTGACACAATTGTTTCAGCTGGAGGGGTTTACGAGCTTGGATTTTTCAGCCCTGGAAATTCGAAGAATCGCTACGTTGGCATATGGTTCAAGAAGATAGCAACTAGAACTGTTGTCTGGGTTGCCAACAGAAACATTCCACTGAATGACAATTCAGGAGTGTTATTACTCAATCCCAATGGAATTCTTGTACTTGTTGACAATTCCAATGCCTCAATTTGGTCATCAAACTCATCAAGATTGTTAATGAATCCAAAAGCACGGCTCCTGGATTCCGGTAACCTTGTTGTCGGTGATGGAAATAATAGTGAACGAGAAGTTAATTTCGTGTGGCAGAGTTTTGATTATCCAGGAGATACTTTATTACCTGGGATGAAGCTTGGACGTAATCTGGTCACGGGCATGGATTGGTACATAGAGTCATGGAAGAGCACTGATGATCCTGCTGCGGGTGAATATATAGAACGTCTTGATTCTCATGGATACCCTCAATTTTTCGTGTGGAAAAATTCGTCTATAGTATATAGCACAGGGCCATGGAATGGTATCGCATTTAGTAGTAGTCCTAAAAATCAACCAGCTATATATTATGCTTTCGAGTTTGTTATTAAACAGAAGGAGATTTACTTTAAATACGAGCTTAATGAGTCCCTGCCCACCAGGGTGGTGATCAATCAGGCTGGAATGGTAGAACACCTAACATGGATCGAGCGTAATCACAGATGGATAGTCTACGTATCAGCACAATCTGATAATTGTGATCGTTTTGCTTTATGTGGTCCTTATGCAAGTTGCAACATCAATAACTCTCCTCCATGTGACTGCTTGCGAGGTTTCGAGCCTAGGTATTGTTAAAAATATGCCTTCACACGACTCCTCTGTCTGTTCCAATACCATGTTAATTCATTAATTATGTCTTCACAGGTATCCTGAACAATGGTATGCAGTGGACTGGTCTAATGGTTGTATAAGGAAAACTTCTTTGTCTTGTAACCAAGATGGTTTTCTTAAATTTAGGAATATCAAGATGCCGGATTCTAGACACTCCTGGTATAATGTAAGCATTAGTCTTGAAGAATGCAAGAAAATGTGCTTGGCTGATTGCAATTGTACAGCCTACTCAAATCTTGATATAAGAAATGGCGGAAGTGGATGCTTACTATGGTTCGGTGAACTCATTGATATTAGAGAGTACAACAAAAATGAGCAATACCTGTTTGTGAGAGTTGCTGCTTCAGAATTAGGTAATGACCCTACTCTATACCATTTTTGTGTTCTTTTGCACAATATAAGCATGACACCTTAGAGTTTAAGGAACTTTACTATCTGATGTCACTGATAGTAACATATGATAGTCGCGGAAGCAATTCATATGTCAATTGAAGACTGCTTATGGACTTTTTATTAAACAGATTCAGTCAGGAATCGGAGGGGAAAGAGGTCAGCCCTGATTGCGGTCATTTCAGCACTAGTAGCAACTTCTATCCTCATCTTTCTAGCTTGGTTTACCTTCCAAAGAAAGAACAAAAAAACAGATAAACATACTGGAGGTATTAGAAAATTGTTAACTGTGAATGCTTATTTACCTCCATCTAGCTAACATTGTTCGACTTTATCAAATGATTGAGGAAAAGTTCAATTATATCATGTCTGTGATTGGAAAATATATGTTGACAGGTTCAGAAGTTGGAAACAAGGACCTCGAGTTGCCATTGTTTGATTTAGTTACTGTTACTACTGCCACTGAAAATTTCTCTTCTGCGAATGTGATTGGTGAGGGTGGCTTTGGACAGGTTTACAAGGTAAATCAACTTTGATCTTCAACTAAAAGCTACAAGCATATTTGACCCTAGAGAGAAAATCGCAGGGTATTCTACCAGATGGACTAGAGATAGCAGTAAAGAAGCTATCAACGTATTCTGGACAAGGCGTTCAAGAGTTAAAAAATGAAATTGCTCTCATTTCCAAGCTGCAACATAGGAACCTTGTCAAGCTTTTGGGTTGCTGCCTTGAAGGAGAAGAAAAGATGCTAATCTATGAGTTTATGCCCAACTCTAGCTTGGACTGTTTCATTTTTGGTCTGCTTTTCGTGATTCATTAACTAAATATTACTACTTGATAAGTCCTCAATATCTCTGTTATCACTCAAGAATTGGTTTTACATTGTGCAGATCCAAGCAGAAAAGCTTCACTTGCATGGAAGAATCGCTTTGAAATTTCTGTGGGAATATCTCGAGGTCTTCTTTACCTTCACCAGGACTCAAGATTTAGAATTATTCATAGAGACCTCAAGACCAGCAACATTTTACTAGATGGAGACATGAATGCCAAAATTGCTGATTTTGGCCTTGCCAAAATTTTTGGCGGAGAACAAGTGGAAGGAAATACTAAGAGAGTGATAGGGACATAGTAAGTTCTACAAATCTTTATGATAGACATTCAATCAGTACTTGAACTTAAATTTGATGAGTTGATGCAGTGGATATATGTCACCTGAGTATGCTGTTGATGGGAAATATTCAGTAAAATCCGATGTATTTAGTATCGGCGTTATCATTCTAGAGCTAGTCAGTGGCAGAAGGAACAGGAAATTTCATCATTTGGAACATTATCACAATCTTTTGGGACATGTAAGCACATACAACACTAAACTTCGTTCCCTTAACTCACTTTTTCTATAGTGAATGCATTTGTATAAATGTGTTCTGTTTGTAATTTAGGCATGGTTACTTTGGACTGAAGACAAAGCGTTGGAACTGATGGACGAATGTTTGAAAGAATCATTTTCAGAATCTCAAGTGTTGAGATGCATCCAGGTTGGCTTGTTGTGCGCCCAAAAACACCCTGAGCATAGGCCTACAATGGCATCAGTAGTTTTCTGGTTGGGAAATGAAGGCCTGGTTCTTCCTCAACCAAAGCAGCCTGGATTTTTTATAGAAAGGAATTCAATGGATTCAACAGAATCAGCTCAATTTATCAGTTAGCTCGTTGTAATCGGATAAGGGCATTGTGAATTCAAGTTCAATCAACACATATAATATCTAATTAATGAGAAGAAAATCTGAAAGTGTGTACACACATTTCAGCACACGCTTTGTAGCACAATTAATCCGGATTCACAGGGGAAAGTTCGACTTTTTGGGAGAAACTAAAGTCTCCTCATCAAAAGTGACTCCATTCAAGAAGTTGAAGATTCTTCTTGTTTTTTCTCTCAGTTGAATATTAGATGGTGACTTTACAACTTTTTAAAATAAATGCATTAATATTAATAGAAAATAAAAAAGACATCAGTTCCTTTTTTCTTAGTTCTGCCATTACTAGTCCAAGATGTCACATCGTCATTAACACTAAATCATTTTAGGAAGACTTTAGGCAATTTACTCCAATACATTTCATGTGTTTACACCAAGAAAATAGATGCATGCCATATGTTTGTATGAATAGTTCTATAGCGTGCAATTGCAAACACCAGGTTAATGGTGGAAATGGAAAGTCAAAGGCATACAACTTTTGACAAGTTCAAACTTTGACATCTAATTCTTGGCAGCAGTTGTGATTGGCCTTTCATTCAAAAAAGTTCTTTTTTTCATTTTATAGTTGTGATTGGCCTTTCATTCAAAAAAGTTCTTTTTTTCATTTTATTTATATAAAACATTTTTGGACATGGTTGTTAATTCTAAACTGAGAAAAAAAAAGTTATCAGTATCTTTTCTGTGTCAAGGCTTTTTTGAACTTGTCAATCAAAAACATGAGTCAGAGTCTATCAAAAGGGCTACTGTTTTTATTCCTTTGCTCACATTTTTTCTTCATCTTACTAATTTCCGCGGCATTAGACACGATCACAACAGATGAATCCATTAGAGATGGTGACACAATTGTTTCAGCTGGTGGGATTTATGAGCTTGGATTTTTCAGTCCTGGAAATCCGAAGAATCGTTATGTTGGCATATGGTACAAGAAGATATCACCTAGAACTGTTGTCTGGGTTGCAAACAGAGATATTCCACTTAATGACACATCAGGAGTGTTAACACTCAAACCCAATGGGATTCTTGTACTTGTTGATAATTCCAATACCTCAATTTGGTCATCAAACTCATCAAGATCATTAAAGGATCCAAAAGCACGGATTCTGGATTCCGGGAACCTTGTTGTCAATGATGGAAATGAAAGAGACCTGAAAATTAATTTCGCGTGGCAGAGTTTTGATTATCCAGGAAATACTTTTATACCTGGCATGAAACTTGGACGTAATTTGGTCACGGGCATGGATTGGTACATGTCATCATGGAAGAGCATTGATGACCCTTCTCCTGGTGAATATATAAACCGTCTTGAATCTCATGGATACCCGCAATTGTTTGTGTGGAAAAATTCAACTATAGTGTCTAGCTCAGGGATATGGAAAGGTAATGCATTTACTGTTAGTGCAAATAGTAAACCAAATACACATTACACTTCCGAGTTTATTATTAATCAGCAGCAAATTTACTACCAATTCAAGCTTAAGAACGAGTCACTGCCCAGCAGGATGGTGCTTAACCCGGAAGGGCTGATAGAACACATAACATGGATTGAGAGCAGTCAAAGCTGGTTTCTGTACTCAACAGTACAGATTGATAGCTGTGGTCGATTTGCTTTATGCGGTCCTTATGCAAGTTGCAGCATCAATAACTCCCCTCCATGTGACTGCTTGCAAGGTTTCAATCCTAGGTATTGTTAAAATGAGTGATCATCAAACCTTAAACTATTAGAGATAACACACTTGATTACTTAGTTATGATTTTACACTCCCCTTCATGTGTGTGCTTGATTCTATTTCTTGGGCCAAGCACATAGATATTATATTGATTACTTAATTATGACTTCACAGGGTTCCTCAACAGTCTGCAGCAGATTGGGCTACTGGTTGTGTTAGGAGAACTTCTTTGGATTGTAAACAAGACGGTTTTCTTAAATTTACGGGCATCAAGATGCCTGATTCTAGAAACTCCTGGTTTAATGAGAGCATTAACCTTGAAGAATGTGAGAAATTATGCTTAGCTGATTGCAACTGTACAGCCTACTCAAATCTTGATGTTAGAAATGGCGGAAGTGGATGCTTACTATGGTTCGGAGATCTCATTGATATTCGAGAGTTGAGCCAAAATGAACAAAATCTGTTTGTGAGAGTTGCTGCTTCAGAAATAGGTAATGACCCTACTCTGCACCATTTTCATTTTGTTGTTCAATATAAATACACAGATCTTTGAAACTGACGAATCCCTGGTCAACAAATTACTGATAACCAAGTTTGCAGGCTTCATAACAAAATAAGACGTATATGTAAATTGAAGACTGCTTATGAACTAATTATTGAACATATTCAGACAGGAAGCGGAGGAGAAAGATGTCAGTCCTGATTGGTGTCATTTCAGCAGTGGTAGCAACATTTATCCTCAGCTTTTTAGCTTGGTTTTCCTTCCAAAGAAAGAAAAGAAGAATAGGTAAATATAGTTGAGTCACTCTTGTCTACCAACTTTATAAGAAAATTGTCAAGTCTAAATGCTGTGCTGGCTAGGTGAATTTTCTATCTAGCTGACATTGTTTGCCTTTACCAAATTCTCATTGATATCATGTTTGTGGTGGGCGAATGTATATTGACAGGTCCAGCTGTTGAAAATGAGGACATGGAGCTTCCATTGTTTGATTTAGTTACTGTTACTAGTGCCACTGGGAACTTTTCTGCTAAGAATGTGATCGGGAAGGGTGGATTTGGACCGGTTTACAAGGTAAATCAACTTTGATCCTCAACGTCTAACTTGAAATGGCATCTTTGACCCTTGACATAAATTTCAGGGTATCCTACCAAATGGACAAGAGATAGCAGTAAAGAGGCTATCAAAGCATTCTGGACAAGGCTTTCAAGAGTTAAAAAATGAATTCGTTCTCATTTCCAAGCTGCAACACAGGAACCTTGTTAAGCTTTTGGGTTGCTGCCTTGAAAGAGAAGAAAGGATGCTAATCTATGAGTTTATGCCCAATGCTAGCTTGGACTATTTCATTTTTGGTCTGCTTTTCATGATCCATTAATAAATAGTACTTGATATAAGTCCTCAATATCTCTGCATCACTCAATGATTAGTTCTACATTGTGCAGATCCAAGCAGAAAAACTTCACTTTCATGGAAGAACCGCTTTGAAATTGCTATAGGAATATCTCGAGGTCTTCTTTACCTTCACCAGGACTCAAGATTAAGAATTATTCACAGAGATCTCAAGACCAGCAACATTCTATTAGATACTGACATGAATGCCAAAATTTCTGACTTCGGCCTTGCCAAAATATTTGGTGGAGATCAAGTGGAAGGAGAAACTAAGAGTATAGTAGGGACATTGTAAGTTTCTACACATCTTTAAATCGTATAATCTTTATTATATTAGATCAGTACATGAGCATAAACTTATTGAATTGATGTAGTGGATATATGTCCCCGGAGTATATTGTTGATGGGAAGTATTCAGTAAAATCAGATGTATACAGCATCGGCGTAATCATTCTTGAAATAGTTAGTGGCAGAAGGAACAGGAAATTTCATCATTTGGAACATCTTCACAATCTTTTGGGACATGTAAGCATATACATTACACAACTTCATTCCCTTAATTCGCTCTTAGCTTTCTATTGCGAATACATTTGTATAAATGTGTTCTGTTTGTAATATAGGCATGGTTACTTTGGACTGAAGGCAACGCGTTGGAACTGATGGATGAATGTTTGAAAGAATCATTTTCAGAATCTCAAGTGTTGAGATGCATCCAGGTCGGTTTGTTATGCGTCCAAAAACTCCCAGAGGATAGACCTATAATGGCATCAGTGGTTTCTTGGTTGGGAAATGAAGGTCTGGTTCTTCCTCAACCAAAGCATCCAGGTTTTTTCACAGAGAGGAATCCAATGGAATCTACTGATGAAGAATGTCTAAGTAACAACGCGACGTTAACAGTTCTTGAACCAAGATAGACAAATAATCACTATTGGATGTTTTGAAATGTATTTACTTTGTCTTATTTGTCAAACAGTACTAGTGTATATATGATAGAATATACTAAAAGAGATAGTTTAATGCCAGTGTTCATATCAAGGCAGGGTAAATAATTGATATACACTTCTTGTCTTTGATAATTGAGCAATCAATGGACTAATTATTGGCAAAATTTGTAAGCACAAGTAGTTAGAGGTATCTATATCATGCATTACTTTTTTTGGGGACAATGAATATATTCATCAAACATAGTTGCTATACATGATACATAAGCCCTATTCTATATGAAGTATCTTTTTTTGTTTCCCACTCGGTGTCTGGATCCCACATTGGAGCTCTGGCTAAATTTGGATTGCGTATAGCAGGA

>PGSC0003DMG400021804 G-LecRK

TGAATGTCTGCGACTTCCCTGTTGATGACCCATTTTGGCCGTCTCCATTACCAAACTACTAGTCTTTGATTGTGTTGCAAAGAGCTCCAAAAATGGAAGAAATTCGCTAATAAATAAGCTGCATTTTATTTTTCCAACCATGAAGCTGCTACAAACAACTATTCATTTCTTGTTCGTTTTCTACTTATTTTCCATTTTCCAAATTTCATTAGGTGATACCATAACAACAACTCAATTTCTCAAAGATGGTGAACCCAGTATCACTTCATCCGGTGGAAGTTTTCAGATGGGGTTTTTCAGTCCAGGTAATTCCAAAAATCGTTATATAGGTATATGGTATAGCAACATATCTGTAACAACCGTTGTTTGGGTTGCCAATAGAGAAGCTCCTCTAGGAACTAATTCAGGTACCCTAAAAGTTATCAAACCAGGACTACTTGTTATAGTGAATGATAGTAACCACATCATATGGTCTACCAATACCTCGAGATCTGTGCAAAACCCTGTTGCAAAGTTGCTTGATTCTGGAAATCTTGTTGTTATTGATGCTGGTCATGGTGTTGGTGATGATATTAAAATTGGGGATTTTCTCTGGCAGAGTTTTGATTACCCAACTGATACCCTTTTGCCCGGGATGAAAATTGGGTGGAATTTTGTAACTGGGAAGGAACTGTATTTGTCGTCGTGGAAGAACGAGGAGGATCCAGCTCCGGGTGATTATACGTACCATTGTGATCCTTCTGGTTACTTGCAGAACATTTTGAAGAAAGGTTCAAAGGAGGTATACAGATCTGGGCCATGGAATGGCTTAAGGTTTAGCGGAGCAACAAGTTCAAGACAAAGTCCATTTTATACATTTGGTGTGATTTCAACTAAAACTGAGGTGTATTTTAGTTATCACCTTCTAGCTTCGGTTATTACAAGGTTTTGTTTGAATCCAAATGGTGCGTTACAACGATGGACTTGGGGCGATAGGAATAAAGGTTGGGCACTTTACCTTTCATTACCAACAGATAATTGTGATACTTATAAGTTGTGTGGAGGATATGGTAGTTGTAGTAGTTTGAATTCTCCGGTATGTGGATGTTTAGAAAAGTTCGAGCCTAAACATGTTGAAGATTGGGGGAAGGCAGATTGGTCAAGTGGTTGTGTTAGGAGGATTGAACTGAATTGCATCAAGGGAGATGGATTTTTGAAGTATTCGAAACTTAAATTACCAGACACGCGGAATTCATGGTTTAATGTGACTATGAACCTTGAAGAATGCCGGAAAGTTTGCTTGAGAAATTGCTCTTGTATGGCATACTCAAATCTTGACATACGCAATGGAGGAACCGGTTGCTTACTGTGGTCTGAAGATCTGCTTGATATTCGACAGCTAGCCAACGAAGGACAAGATATCTACATTAGAATGGCTGCCTCTGAATTAGGTAAGTTTATTGCTGAAAATATAAAATAAATAGTTTTGAGATTGATTTCGTAAAGACGGTAAATTTATGAGTTGCAGGAGCCTGGGATTAAAAAAAAAAGTGAATTTTCTTGTAGACACTGCCAATGACTACTATGAAATCCATAAATGAAAGTAGTTAATGGTAGGAGTACCGTTATCCTTTTTGTAGTGATGAATCTTTTATTTGTTCCTAAGAAAAGGAATTTGTCCATTTTTGGGGGTCTCAAAGGTGCGTGGAAACGCATATGAACTCTTGAATGGGAAAAGGATAAAACTCCACTTCCTTCTGAATGGGTAGTCAATCCTATGTGTCGATAGGAGCAGTTGACAATTGAATCCGATTTTGACCATTATTTTCACATCAAACATGTTGTACTATAACGACAAAGTAAAACTCACAGATTTAACAGAACAAAATACTATGAAATTAGTACAGCTACGTCTCTAGATTAAGAAAAAAAATGAGTTAGAGTTGAGCTATGCTGAACTGGTGACCACTGACCATGATGTCTAGACAAATAAAAGAGCTTTAATGTAGTCATGTCTAATAAGCTGACGGAGAAAGTCGAACTGATTACCTATATTTGGGTTACATACTTGCAGCTAGTCAGGTGAAATCAAATGGACACAAACGAAAATTACTCTCCTGGATTATCCCGTTATCGGCTGGAGTGATTTTGGTAATCCTAAGCCTAGTGATCTGGATTAGAAGAAGGAAGATAGCTTCAGAGAAAAAGAAAGGTAACCTCGACAAGTCATTATGTTCGGAATATAATTAGTAGATGAATTCAAATTCAAAGGAAAATCTTTCTTGTCTAGTATTTGCCAAACTGAATTTGGTATATCGGTCTTTTGACAGGATGCTGGGGGAACAATGGCAATTACAAGATGGATTACCTCAACGGAAACCTCAGTGAAGAATATGAGTTACCACTGTTCGACTTATCTACAATAGCTAAATCTACGAACAACTTTTCAGGGATGAGCAAAATTGGAGAAGGTGGATATGGACCTGTTTACAAGGTAATCACTTATAGCATTTGCACCTGTTTACATGAAATAAGCAATTCACAGCCCACATAATTGAGCATAGAGAGATTCGGGAAGACTGACAGTAAGGAATGATTTATCGACTTTATAGTTTTGGAAGAGTTTTATAAACATTAGCTAAACTTGATTGAATTTATAATAGTTGCTCAATTCTTCATAATTTGAGCTTCTACCTTTTGGTGGAAGTATAATTTGATTTTCATGTTCAGTGTTGAAAAAACTTAACATGTAGATATATGATGTTTAGGGCGTGCTTGAACACGGACAGGAAATAGCTGTGAAGCGGTTGTCTAGGACTTCCACACAAGGACAAGAGGAGTTCATGAATGAAGTGATGTATATTGTGAAACTTCAGCATAGAAATCTGGTGAAGATTCTCGGATGTTGCATCGAAGGAGAAGAAAAAATGTTGATCTATGAGTACATGCCAAATGGAAGTCTTGATTCATTTATATTTGGTAAGGATTTCTCAAAAAAAAAGAAAAAAATTGTGCTTAATAGTTACATACATCTCAGTTGATAGATGCCCTGTAGGAGTACCCAAATAATCCTCTTAGCCCAAAGTTATGTTTTATTGTTACTTTTCCGATTTATTTCGATGTTTAAATCCCACTCTCAGTTTTATCTGATTTGTGTGAATTAATGTTCCAACTATGAAACTTTTAACAGATGACACACGAAGCACGGTATTAGACTGGTCTAAGCGCTTTCACATAATCAATGGGATAGCTCGGGGACTTATGTATCTGCATCAAGATTCTCAGTTGAGGATCATTCACAGGGACCTGAAAGCTAACAACATTTTACTAGACAATGACATGAATCCAAAGATATCAGACTTCGGTATAGCAAGAAGTTGTGAAGACGACAAGTTTGGAGCCAAGACACACCGAGTAGTTGGAACATAGTAAGAATCTCAAGATGATATTTTACTGAATCATTCAAAACTGGAATAGCAGAAGAGCCTTGACATTATCATAGTATGTTAAAATCAAAAAATAAAATCAAAAAATTATCATAGTATGTTTACCCAGTATTAGTAACGCTAATTTCTGTTTACGTTTTATGCAGTGGCTACCTCTCACCGGAATATGCTGTACATGGTGTCTACTCGGTGAAGTCAGATGTATTTAGCTTTGGTGTCTTAGTTCTTGAAATTGTGAGTGGGAAAGGGAACAGAAAATTCTCTCATCCAGACCATAACCTTAACCTCCTCGGACATGTAAGGGTTGTGTTTTTCATTGTCCATCAGTAAATTATCTCGTAAACCTGGTCATGCTAACACGATAGTAAAATCTACAGGCATGGACACTCTATAAAGAAGGTAGATCAATGGAATTACTCGGGGATTTCCCCATTGATGTTCATTCTACACCTGAAGTTATACGATCAATCCATGTTGGTCTATTGTGTGTTCATCATCGTCCAGAAGATCGTCCAAGTATGTCTTCAGTGGTTATGATGTTAAACAATGAAGGTGTACTGCCACCTGCTAAACAGCCAGCTTTCTTCGTAGAAACAAATACACCAGATAATGAATTCTCTTCTAGTCAATATGCACATAGTACAGTGAATCAGATCACCATCACAACATTAGAAGCCAGATAGTAAATTCTTCATTGCAAATAGCAATCTATTATACCCTTCATTCTTTTCTTCTTTGTAACAACGTAGCAATCTTATCGTAATAGTGTACCCATCATATCAAAATCTTGGATATGCCTCTGCACGCGATCCTTCTCTGCCAGCCATAAACAGCTAATCATAGCAATGCAGGGGGTTAAATAGGGAAAAGATGTAAAAATCACAAAACAACTTTCAGGATCGGTATATATTCTCTCTCTTGTATGTGTTATTTTGCTTTTGTTCATATATAGTTTAGTATTGATTTATATGTATGTGTTTATCTATTAATTAAGAAGATGGAAGAATGTGTAGATCCAACACTAAAAGGATCATTTTGTAAGATATATAGACATGTTATTCAAAG

>PGSC0003DMG400021993 G-LecRK

AAAAGTATGAAAATTTTATGATGATGAATTGATGGAGGACAAAAAAGTTTATTCCTTAGCCCTTTTACTTGTGATTTTATTGATCACACTCATCGTTATCGCACGTATAACGTTGAAATATTCGGAGACATTTTTCCTCATTTTCGGAGCTGATGTTGCGTTGATAGTTGGTGTTTTTGCTGGTGTTTTGATCCGAAGGAAGTTCAACAACAGGAGGTTATTGCTAGAAAATCAGTTAGACTCAGATGGACGCGAGCTGAGGATCGAGTATAGTTTCCTAAGGAAAGTAGCTGGTGTTCCAACAAGATTTAAGCTGAAAGAGCTCGAAGAAGCAACCGATAACTTTGGATCGTTGGTAGGACGTGGATCATCAGCTTGTGTTTTCAAAGGTGTGCTTAGTGATGGTACATCAGTAGCTGTTAAAAGGATAGATGGAGAGGAGCGTGGCGATAAGGAATTCAAATCAGAAGTTGCAGCTATTGCTAGTGTTCAACATGTTAATCTTGTACGTTTACTTGGATATTGTAGTGTTCCTCCTTCAGGACCTCGATTTTTGGTTTATGAGTATATTGTTAATGGATCACTTGATAATTGGATTTTTCGAAAAAGGGGAATTCGAGGGTGTTTGTCGTTGGATTTGAGGTGTAGAGTTGGTCTAGATGTGGCTAAGGCACTTTCTTATTTGCATCATGATTGTAGATCGTGTATTTTACATCTCGATGTCAAGCCTGAGAATATACTTTTAGACGAGAATCATCGTGCTATTCTAGCTGATTTCGGGTTGTCTAAGTTGATGGGGAAAGATGAGAGTAGAGTTGTTACAACGATTAGAGGTACTCGGGGGTATTTAGCCCCCGAGTGGCTCCTGGAGAACGGGATTTCTGAGAAAAGTGATGTTTATAGCTACGGAATGGTGCTATTGGAGATGATTGGAGGGAGGAGAAATATTACTTTAGCTGAAAATGGGAAGAGTACCAATTCAAAGAGTAAGTTCAGCTATTTTCCTAAGATTGTGAGTGAGAAATTGGAGCAGGGGAAGATCATGGAAGTCCTGGACGAGAGGCTCGTCCATGAGGCAGCTACAGGAGGTGTAGCTGTAGAAATGCAAGTGAAAAGATTGGCGTGTGTGGCGTTGTCTTGCATCCAAGAACGGCCTAGTCTTAGGCCAACAATGGCACGAGTCGTTGAAATGTTGGAAGGTCGTGGGCCAGTAGAAGCGCCTAGACAAACGACAATGTTGATCCTTGATCTATTAGACGAGGGTGTGGATCATCATCCTGGGATACCTAGGGTTGCTGCAGCCATGGCAAGATCTAGTGATACTAATTCGCTTCGCTCTTATACAATGTCTATTCTCTCAGGGAGGTAGTACCTACATAATGCATAAGGGATTTCGTCTGCTTTCTGTCAAATTTTTCAATATCAATAGGTTGATTGTTTCTCGAGTGTTGTATCCTGAATCCGAAAGAGTGTACTTGAATTCTCCCCCTAACAAAAAAAAGTGTATGAATTTGCTTTTGTTTACGCCAAAATATA

>PGSC0003DMG400022391 G-LecRK

AAAGATAACATTCAAAGCATAACGAAAAGAGAAAGAAGTGTGAGCAAATAGTATTGCAGGCTGCAGCTATAGGTATTCTCCATTTTAGTTTATACAGAACTAATTCACATGCAAATTGGCAGAAGAAAAACTTCATAAGCAATGAAATTAAGCACAAGAGAGATACTTCTGTTTCTTTCTATATCAACTGCTTCAGGTATCATTAGCACCAATAAATTCCTGCGAGATTCAGAAACTTTAGTCTCCAATGACAAAAGATTCATATTCGGGTTCTTTAGTCCTGAAAATTCGACGAATCGTTATGTTGGTGTTATGTTTAATGTCCAACCACCAACTGTTGTATGGGTTGCCAACAGAGAGAAACCTTTACAGGATTCTAGTGGAAGAGTGACAATATCTGATGATGGAAATCTTGTAATCTTGAATTCACAGAATAGGAGTATATGGTCATCAAATATTTCACCAGCTGTGAGAAATTCCACAGCGCAGATCTTGGATACTGGAAACTTAGTTTTGAACGATAGCTTCAATGGGAGGGTTCTCTGGGAAAGTTTTCGGGATCCTTCAGATTGCTTCTTGCAGACCATGAAAATTGGCGTTGATGTAAGTACTAACACGACAAATCTGCTGAAATCATGGATAAGTCCTTCAGATCCATCTGTTGGGAGTTTCTCAGTTGGTATTCAACCTGAAACAGTTCCCCAGATTTCCATATGGAAGAATGGGAAACCTCATTGGCGTAGTGGTCCATGGAATAAACAGGTTTTCATTGGGGTACCAGACATGACTTCATTCTATCTCAATGGATTTGATCTAGTTAATGACAACAAGGGCACCGTGTACCTTACCTATTTATATGCAAATCAGATTGAGCTGACGTTTTTCACCTTGAACTCAACAGGGTTTTTGCAGCAGAAATATATGGATCCTAGTAAGAATGATTGGGAAGTAACATGGGAATTTCCTGCAACTGAGTGTGATTTTTATGGAAAATGTGGACCTTTTGGAAGCTGTGATCCTACAAGCTCACCAATCTGTTCTTGTTTAGAGGGATTTAGGCCGACAAGTGAAGAGGAATGGGGAAAAGGAAACTGGACTCGTGGATGCAACAGAAAGTCCATGTTAGAGAGCGAAAGAAACAGTTCTAACCTTGAGCAAGGGAAGCAAGATTGGTTTCTGAAGCTGCAGTCAATGAAAGTGCCGGATTCTGCTATTTGGGTACCTTTTGCAGATGAAGATTGTGTTAACGGTTGCTTGAGGAATACTTCATGCATAGCTTATTCATACTACACAGGCATAGGATGCATGCATTGGGAAGGAAGCTTACTTGATGTTCAGAAATTCTCCATCGGTGGGGCAGATTTATTCCTCCGCCTTTCATACTCTGAGCGTGGTAAGACCTGCATTTCAGAACTATAATATAAGCTATAAAACTCTGCTTGCCAATGACATATGTATAATGGACACTTATTTTGTAATGCAGATCAAAAGAGAGAATATAAAGTAGTCATTGCCATCATAGTCCCAGTAGGCTCAATAATTCTTGCCATTTTCGGATACATTTCCTGCAAATATGTAGCTAAGCGCAGAGGTAACAAGCTTATCTGAAACATTAGAAAATCCATAGTAGCTTTAATCATACGAATCACTTTATCTCTAATGTCCTCTTGAATCAGGATGGAAGAGAATGAGTAAGATCTTATCAAGTGAATCATCGCCAAACTATTACAAGGAAGACAAGATTACAGAGGACATCAATCAAGCTAAATTGGAAGAACTGCTTGTATACAACTTTGATATCTTAGCAAACGCAACTGAGAATTTTCATCTGTCCAGCAAGCTTGGACAGGGAGGTTTTGGTCCAGTTTACAAAGTAAGGATTTGTTTGATTGAGTGTCTTACAATTGTTTTTAAGAAAACATAACCAAGTATATGATATAATTGCAGGGGAAATTGCCAGATGGACAAGAGATTGCTGTGAAAAGGCTTTCACAGTCTTCTGGTCAGGGGCTGCAGGAGTTCATGAATGAGGTTGTGGTGATTTCAAAACTTCAACATCGTAATCTTGTTAGACTTTTTGGGTGCTGCATAGAAAGAGGGGAAAAGATTCTGGTTTATGAATACATGCCAAAAAGAAGCTTGGATGCCTATCTCTTTGGTTCGTTATGATTAGTTTTCTTTTATTTTCCATAGTCTAACTAAAACAAACTTTTCTTTAATTTAAACTTGTCAAATTCGCGATGTATTTTCAGGGTCACAACAACAAGAGGAAGAGTTCCTGGATTGGAGTAAACGTGTGATCATTATTGAGGGAATTGGTCGAGGCCTTCTTTACCTTCACAGGGATTCAAGACTAAGGATTATTCATAGGGATTTAAAGGCCAGCAACATTTTGTTGGATGAATACCTGAACCCCAAAATTTCAGATTTTGGGATGGCAAGGATTTTTGCAGGCAACCAAGATCAGGCCAACACAAGCAGAGTAGTTGGAACCTAGTAAGTTCTTGTATCTTTCCCTTTTATCTTTTCATGAATGAAAAGAACACAAAAAGTAAACATTCAAAATTATAACATGATGCAGTGGTTACATGGCACCTGAATATGCAATGGAAGGAAGATTCTCAGAAAAATCAGATGTTTATAGCTTTGGAGTATTGTTATTGGAAATTATAAGTGGAAGGAGGAACACTAGCTTTCACCAAGATGATGGTGCATTAAGCCTGCTAGCATTGGTAAGTTCTTATGAAAATTAAGCTCCAATTCATATGAAATGTCAACTTTTCATAGTTCAAGTTTCCAATGTTTTGTCAACGTGCTTGCTCTTCAGGCGTGGAAATGTTGGATTGGAAACAAGATTGTGGAATTGGTTGATCCCAAGATAACTGACCTGCACCTTGAAAAAGAAATGGTGAGATGTGTACAAGTTGGATTATTATGTGTACAAGAATATGCAGAAGACAGACCAAATGTCTCCACAATTTTGTCTATGCTCACCAAGATGTTTATAGCTTTGGAGTATTGTTATTGGAAATTATAAGTGGAAGGAGGAACACTAGCTTTCACCAAGATGATGGTGCATTAAGCCTGCTAGCATTGGTAAGTTCTTATGAAAATTAAGCTCCAATTCATATGAAATGTCAACTTTTCATAGTTCAAGTTTCCAATGTTTTGTCAACGTGCTTGCTCTTCAGGCGTGGAAATGTTGGATTGGAAACAAGATTGTGGAATTGGTTGATCCCAAGATAACTGACCTGCACCTTGAAAAAGAAATGGTGAGATGTGTACAAGTTGGATTATTATGTGTACAAGAATATGCAGAAGACAGACCAAATGTCTCCACAGTTTTGTCTATGCTCACCAGGGAAATTGATGATTTACCAAGTCCTAAACAACCTGCATTTACAACAAGACCGAGCTTTTCCAAAAAATGCACTTCTAAATCTCAAGGCTCCATTAACAATGTTACCGTTACTATTATGGAAGGACGATAAGGCATATACGATTTTTATTTAGGCATGATAGTATTCTTTTCTCTGTAATTATTACTGTTTCTGGTTCTTATTATAATGGTTCCACTCAATATTACCTCAGTACATTAAGAATCCAAGAGAAAAGTGTTAACCCTGCAAGTCACTTATGCTTGATAACAGAACCCATATCCAAGCAAGAAGAAGCAAAGCTAGACTTCAAACTCAGAGTTGTCAACAAGTAGCCTAAACAGGTGTTTGCAGAAATTTAGAGAATCCTATCCACGGATTGAGATCATTTAGAGTTAGCTTTGTATTACAGCTGACTCTGAAGATGAACAACTTTTCTAGACAAAGGCATTCTTACAGCGTGGTTGGAGGGTCACCAAATCAACAGGGAAGCCTGTAGGAAAGGGGAACATGCATCCAATTCACATCAGAAATTGCTGAAATGATGTGAAATCTTGCACCACTTTTGTTCTTGGAACTCAAGTCATGTCAGCCCAAGAATATTTGCAGTATCACAGGTAACAAAGGACAAGAAACCTTCTGAGATAGCATGTAAATTTTTCAAAATTCAGACAAAGATTACTCCTTTCCTTAGAATTGTTAATAAACCTATTATTCTGCTGATGCCTTTACTAAAT

>PGSC0003DMG400022426 G-LecRK

GAAATATCGTAACTAATAGAGTAGCATTTACATATAATCTTAGTCCAGGTAGGGGTGAATAGGATGGAATGTCACTTATGGAACTTGTTTTTTTGAAAGTCATTTTCCTGATTTTCCGAGAAACTTGTTATCCTAGTCAATTGAAAACAAAAGAGCTGGAATCATATTGAATAATTGGTCCCAGTTTGAGGAAAATCCTTACGGTAGTCTATCTAAGCACCACAGATTATTCTGAAGACTTGTAAATTCAAATCTGAATACCAATATTCCATAGCCATTGTTTTTATGAGCTAGCAACATGAAGTCTACTAGTGTCTTCTTGTTTGTATTCTTGCTGTATTTGAGTCACTTCAAATGTGATTCTAGGGACATCATAACTTCAAGGAACATATTAAAAAACAAAAGGAAAACTCTTGTGTCAGCTGGACAGATATTCGAGCTGGGACTCTTCAACACACGATCTAAAGACGGAAAACTTAGAAATTATGTTGGCATATGGTACCGGGTGAGTCCCAAAACTATTGTGTGGGTTGCCAATAGGGACAGACCAATTCCCGTATCAGATGAAATTTTGGCTATTGGCATTGATGTGGATGGTAATCTCAAAGTGTTGGACTCAAAGAAGAACTCTTACTTTTCAACAGGACTCGTTAGTGCATCTTCTTCTAAGCGTACAGCAAATCTCTATGACTCAGGAAATTTGGTTTTAATTGATGAATTGTCAGGCAAGAGATTGTGGCAAAGTTTTGACAACCCCACTGACACCTTCCTTCCTGGTATGAACATGGACAGTGCATTGAAGTTGACTGCTTGGAGGGTTACCCAAGAGGATCCCAGCACTGGAGATTACACGTTTCGGCAAAATACAGCAACCAAATTTGAGTACAACATACTATCCCAAAAAAGGGGCCTTCACTGGAAAGCCTCTGTTTCTGCTAAGCCCTTTAGTTCCAGGGAACTGCCCTCTCTTGTGGGATCAATGTTAACCAATTTCAGTTTCAATACAGAACAACAATATACCCCGAAATTGAACAAGTCACCGACTGATGATGTTTCATCTGTGCCAGACTTTAATTACACAAGGCTACTACTGAATTCCTCAGGGGAAATACAATTATATGGCTGGAGTAACGAGAGTACAGGGTGGTTACAAATATGGTCAGAACCACAAGATCCATGCGATGTGAATGAAAGTTGTGGGAAATTTGGAATTTGCAACAGTGGGTTTATGTCTCGGTGCAAGTGTTTGCCAGGATTTGACCCTGATTCTCCAGATGAATGGACGGTTGGGACTTGTACTGATGGTTGTTCAAGAAAGTCAGTTCGTGGTTGCAACAAGAAGTCAGAATTTGACACATTCTTAAACTTTTCAACTATCAATGGGAAACTTTGTTTGCATACCTAACAAGGTGTGGATCACCACCGACAAATCCTTTTAGTTGTATCCTCCATTTTATTCACCTACGCAAGTTTCAATTTCGTTTGGTTCAGAGCATGTAGGACAAAGAGGATGGAGTGGAGAGAAGGCAACATCCTTGAATCTCAAAGTCCTGATTATTTCTGTCAGTGTAGCTGCTGCTGGAACTATAACCATAACCATTTGCTATGTAATCTACCATAGAAAAAAGGTGGCAAGAAGGAAAGGTAACAAATAGCACAAGCTTTTATATAAATGATATATGTATTGTGAATTTGGTGCTAATTTTGAACTCACTTTATGTAATGAGAAGCCAAGAAAATAAGTTTGGGGAATGGGATAGAGTACTTGTCCGAGTGTGGAGGTTCATCTAAATATTTGGTAACCGAAGATGATAAGAAACGGATCGATATCCCATTTTTCAACTTGGAAAGCATACTAGTCGCTACGGACAACTTCTCAGATGCAAACAGGCTTGGTCAAGGTGGATTTGGCCCTGTTTACAAGGTGGTGGTGATTCTCACATGTACCATTTTTAAAATCATTTTTCTAACTACAGCTAATATTGTACGTACAGGGTAAGTTCCCTCAAGGACAAGAAATGGCAATTAAGAGGTTGTCAAGTCATTCTAGTCAAGGTGCAGAAGAATTTAAAAACGAAGTAATGTTAATTGCCAAACTCCAACACAGAAATCTAGTCAGACTTTTGGGCTACTGCATCGAGGAGAACGAAAAGATCTTATTATATGAATATATGCCCAACAAAAGCTTAGACACCCATATATTTAGTGAGTCTACTTACTTTTCTTCTCAGAACTCCATTATAGAATCTCTCCCGTTGGTTCAGTGAACCTCTAAATTTCATTATACATACAGATAGATACAAGGGTCAATATTTATTTAACTAAACCATTGTTCAACTAAAACTTTGCTTTGTTGGGGTCATTATTGTTACATTGGAAAACAGAAAACAAGGGAATTTCATTTTGGTGTGGAAAGATCTTTGTATTAGCAAAAGCTTAGGAATTAGAATGCTTAATTTCTGTAATTGGTGAAAAGTAAGATCTAACTGTGATTTAGTTGTTCTTTGAGTTCACTCTACACAAATACCTCGGGTACTACTAGAAGGGGATTAAGTTTCTTAAAGACACAATGTAAATTCAACTTTGACTTGATGGACTTGTTCCGGATAGTTCTCTTGAGACATTTATCAAACATATTGGCTGTGTTTGGTATGAAGGAAAATGTTTTCCTAGAAAAACAAATAGATTTTTTATTTATTTTCTCATGTTTGGTTGGTGAGGAGAAAAGATTTTTAGTGTCTGGTTTTTGAATCAAAAATATTTTTGAGAAATATCTTTTATTACTAGAGTAGAAAATAATTTTTGAAATTGAAAATATTTTTTAAAAANNNNNNNNNNNNNNNNNNNNNNNNNNNNNNNNNNNNNNNNNNNNNNNNNNNNNNNNNNNNNNNNNNNNNNNNNNNNNNNNNNNNNNNNNNNNNNNNNNNNNNNNNNNNNNNNNNNNNNNNNNNNNNNNNNNNNNNNNNNNNNNNNNNNNNNNNNNNNNNNNNNNNNNNNNNNNNNNNNNNNNNNNNNNNNNNNNNNNNNNNNNNNNNNNNNNNNNNNNNNNNNNNNNNNNNNNNNNNNNNNNNNNNNNNNNNNNNNNNNNNNNNNNNNNNNNNNNNNNNNNNNNNNNNNNNNNNNNNNNNNNNNNNNNNNNNNNNNNNNNNNNNNNNNNNNNNNNNNNNNNNNNNNNNNNNNNNNNNNNNNNNNNNNNNNNNNNNNNNNNNNNNNNNNNNNNNNNNNNNNNNNNNNNNNNNNNNNNNNNNNNNNNNNNNNNNNNNNNNNNNNNNNNNNNNNNNNNNNNNNNNNNNNNNNNNNNNNNNNNNNNNNNNNNNNNNNNNNNNNNNNNNNNNNNNNNNNNNNNNNNNNNNNNNNNNNNNNNNNNNNNNNNNNNNNNNNNNNNNNNNNNNNNNNNNNNNNNNNNNNNNNNNNNNNNNNNNNNNNNNNNNNNNNNNNNNNNNNNNNNNNNNNNNNNNNNNNNNNNNNNNNNNNNNNNNNNNNNNNNNNNNNNNNNNNNNNNNNNNNNNNNNNNNNNNNNNNNNNNNNNNNNNNNNNNNNNNNNNNNNNNNNNNNNNNNNNNNNNNNNNNNNNNNNNNNNNNNNNNNNNNNNNNNNNNNNNNNNNNNNNNNNNNNNNNNNNNNNNNNNNNNNNNNNNNNNNNNNNNNNNNNNNNNNNNNNNNNNNNNNNNNNNNNNNNNNNNNNNNNNNNNNNNNNNNNNNNNNNNNNNNNNNNNNNNNNNNNNNNNNNNNNNNNNNNNNNNNNNNNNNNNNNNNNNNNNNNNNNNNNNNNNNNNNNNNNNNNNNNNNNNNNNNNNNNNNNNNNNNNNNNNNNNNNNNNNNNNNNNNNNNNNNNNNNNNNNNNNNNNNNNNNNNNNNNNNNNNNNNNNNNNNNNNNNNNNNNNNNNNNNNNNNNNNNNNNNNNNNNNNNNNNNNNNNNNNNNNNNNNNNNNNNNNNNNNNNNNNNNNNNNNNNNNNNNNNNNNNNNNNNNNNNNNNNNNNNNNNNNNNNNNNNNNNNNNNNNNNNNNNNNNNNNNNNNNNNNNNNNNNNNNNNNNNNNNNNNNNNNNNNNNNNNNNNNNNNNNNNNNNNNNNNNNNNNNNNNNNNNNNNNNNNNNNNNNNNNNNNNNNNNNNNNNNNNNNNNNNNNNNNNNNNNNNNNNNNNNNNNNNNNNNNNNNNNNNNNNNNNNNNNNNNNNNNNNNNNNNNNNNNNNNNNNNNNNNNNNNNNNNNNNNNNNNNNNNNNNNNNNNNNNNNNNNNNNNNNNNNNNNNNNNNNNNNNNNNNNNNNNNNNNNNNNNNNNNNNNNNNNNNNNNNNNNNNNNNNNNNNNNNNNNNNNNNNNNNNNNNNNNNNNNNNNNNNNNNNNNNNNNNNNNNNNNNNNNNNNNNNNNNNNNNNNNNNNNNNNNNNNNNNNNNNNNNNNNNNNNNNNNNNNNAGGGGGGATGGGTGGGTCGGGGGAGAGTTTTGGAAATTGTTTTCCATAAATTTGAAGGAAAGTCATTTTCCGAAGTGTTGATTTGAAAAGCATTAAATCCAACCAAACATGAGAAAATTGGAAACACACCCATTGTAAGTAAACATATTTGATTTCTAAAAAACTGCTATATATGTAATGTCCCTTCAATTGTTCTTTTGATCATCAATTGGTTTCGTTTTTGCTAATGTTGTGATTTTCTACTTGTTCTGCAAGATCATAGTGTACGCCCATTGTTGAATTGGAATATCCGCTTTGAAATCATATTGGGCATTGCTCGTGGACTTCTTTATCTACACCACGACTCCAGATTACGTATCATTCATAGAGATCTAAAAACGAGCAATATTCTATTAGACGAAGAGATGAATGCCAAAATATCAGATTTCGGCTTGGCCAGGAATGTTGAAGGCAAAAGTATTGATGCTAAGACGCAAAAAGTTGCCGGAACTTAGTAAGTATATATATATATATATGCTTTCAGTTTAGTTTTCTGCATAATATAATCGCCAAAAAACTTATGAAATTACAAGTATATCGGAAATTGTGCAATCACCTGTTTATAGAGAGCAGCCTTGTTTCATGAACTAGCCCCTCTCAAATCCCCACTTGATCAGGAATTCTTCATTGACAGTGTCACCGGAGCTAGATGGATCTTAATTACCGGCGTGATATTATTGACTTTTTAGTCAAAATCATTCTTAAAATATAGAGAAAATACATTAAAAAAAACCCTGAATTTGTCCGCTAAATTTACTTTAGTACTTTTAGCTTTATAAATAATCATTTACTTCTAAATCTCATTTCAAGTGAATTAAAATCCACCCAAAAAGCTAACGTCGCAAATAAAAAAAAAGAAAAGGCTAGTGAAAACGTAAAAAATCAAAATAAATACTCTCTCTGTCCCTAATTACTTGTCCACTTTTCCTTTTATAGTTATTTTTAATTATTTGTCCATTTTGACAAATCAAGAAATGACAATTTTTTTTACCTATTATACCCTCAATTAATTACTTTGAAAAAAGTAGAACTTTTTGAAAATCTTAAATTTTAATTCATCTACTTCATAATTAATAAGGGTAACATGATAAACTTACTATGTCAATAATTATTTTCTTTAATAGGTGTGTCAATTCAAAAGTGGACAAGTAATTAGGGACAGAGGAAGTAAACAAATAGGCCTTTTTTTTCTTTCCAACCAATAAGGTTTCATCTTCCTCACACCCCACCACCCCACCACCTTTCTCCACCATAACCCCACCCTTACGCCTCGAACCCCCCACCCGCACCTCTCTTTCCCCTATTTCGAACCCTGCCGCCACCCACACTTGCCTCACCCCTTTCTTCTTCTTCTTCGAGGTCCCGTCCCTGCCACGTTCTTTCTTCTTTTTCAAAACACTCCACACATTCCTTCATTTCAAATTATTCTTGCTTATTCTTAGAAATGATGAATAGGAAAAAAAATGAAAAAACCAACAACTTATTATTGCTTCAAGATCTTACTTTTGGAAGGGTTGTTGATTGTGATGAACTTGAAGTTTTTTTAATAAAAATAATATAATTTCTTCATCTTAAATTATTTTTTTTAATTTGTACTCCTATTATTTATTTATTTTGATAATGTGCTTTAAAAATTGAATGTTTTCTAGTCTTGGATCCATAAAGTTTGATTTTATTTTTATAATCTTAGATTAATTCTCAATTTTTTTTCTATTTTTTACCTTTCTAAAATTGTATAAATAAAAAGAAACAAAAAAGAATTAAAAAAACTCAAAATTTAAATAAATTTCAAGGTGGCGCTGATGTGACAATGATGTGGCATTGATTTGGCAATGAAATGGAGCATGTGTAATACACTTGCCATTGTGAGACTGATATTGTAATATTAGATTTGAAAAAAAATCACTTCAAAAATGTTAAGGGGTATTTAAACGTCCATGTAGTTTAAGTGCTAAAGTAAGTTGTGCTGCCAAGTTTAAGGGGTTTTTTTTATGTGTTTTCTCTAAAATATACCCTAAACTTAAACTATATCCTTAAAATATTATTCTTATCAAGAAAACATTTTTCAAGTATTTAAAAGTGAATAATTTTCATCCTTCTACTTGAATTTATGAGAAATCTAACAAATCCCACAAAAATCAAGTTATTCCTTCATAATTATTATTCTTAGATACGCCAAAAATACTATCGTGAACTTTTATGGTAATTAAGTTTAGCATTATGCAAAATCTTTAAATTTGACAGTTTTTTTCATTTATTTTTCATGTAATTTCATATTTAAAATATTATTAGTTGGTATAAGTTTCTTCTCAATTGAATATACTAGAAGTGACGTTTGTCGGAAACTTCCCCTTCTAATAATGTAGAATGAAACTCAAATGCGAGACATGATCAATATTTTGATTACTTCAATATCAGATTGACCCAAATATTTATTGATCTAAATACAATTTGACTTTGAAAAAGGTAAAGGTAATTAAGCTTCAGCTTATTTCTGTTATTAAATGAGAAAAAGAGCTGATCATGTATCCTCACACCTTCCAAACATTTTTTGTTTCAGTAAGCAATGGTGTCTAAGATTCTTAATATTTATTTTCGAAACTTCTTATAGCAGGTAATGAAATTATGCAAGAAAACTTTTGTTGATAGTCACATGAATTGCTCATGTTTTGTAGGATCCCTTAGTTTTTTGACTATATCTAACATAAGGATGAAAATTGTTAACTTTCAAATATTTAGAGGATGTTTTTTTGCCAAAAAAGATATTTTAAGGATGTAGTTTAAGTTTGGGGTATAGCATAAGGATGTTTTTGACCAAAAACTTTGATATTGTTTCCCGGTAGTATTTTTCCCCAATTGAGTTGTGCTATCATGGGCCGAGCCTCTTTTACCTGAAAATTATGCGGTGAATAGAGGTTAAATGTTAATTATTATGAATATATATTTAATTTTGTATATTATTCACCTAACTAAAAAGAAAATTTGCAAATCCTCGCCACAGGCGTTTACTGATCATGTACTCATTCATCTACTCAAGGCCGTTTTTATCTCATATACATGGTGGTATACGTTATCTTGACTCATACATGATGGGATACACTATCTAAGATAATTAATCCTGGGATAACTTGTTCGTTTAATTGGATCAATTCTTGTCTTGTGATATACAGCGGCTATCTTGCACCAGAATATGCATTGGAAGGACTTTTTTCAATCAAGTCTGATGTTTTTGCTTTTGGAGTGGTTGTACTGGAAATCATTAGTGGAAACAAAAACTTGGATGTTCTTGAAGACACAAACCTCTTGGGTCATGTAAGTACCTCTCTATTTAGAACATGTAATCTTAAGACATAAACGAAATGTGAAGATTGACATATAAAGTAGGCCTTCTGTCCCAATTTATATGAATATTGGGCATCGTTGATAACTCGAAATTAAATATAATTACAGGCTTGGAAACTATGGATGGAAAACAAGGCATTGGACATAATGGACCAAAGTCTAGTGGATACATTCAATGAAATTGAAGTAGTAAAGTGCATAAACATAGCACTTTTATGTGTGCAAGAAGATCCAGGTGATAGGCCAATTATGTCAAATGTGATAATTATGCTTGGAAGTGATAGCATGCCTCTTCCAAGACCCAATCAACCAGCTTATGTCACAAGGAAAAATTACACAATAGGAGCAAGCACATCATCCTCCTCTTCTTATAATTATAATAATTACTATAATTATGATTCATTCACTGGCTCCAAAAATGAGTTGACCATCACTGATATGCAAGGCCGATAAATATTATATAGTAGTAGTAAGTTTTGCACGATATGGACATGTTAAATACATATTCTAAGGTTAGTGTGAATGACAATATTTATGTAGTAGGCGGATCTACGTAGAGGAGAGGGAATGCAAGGGCACCCGATCGAACACTCGAGAAGAACTTCATGTATAAATATCTCGAGTATATATGTGTATGTACTCCATTAAAATCTATGTGTGTTAGCATATATATATTGAATGTAAATATAGTAATTTGTATATTATGATAAATATAGGGCTAATAGGTTAGGATTAGCAGTACTATTCTCTTGTATGGAGCACACAATTGAAGCAGACATGTTGATGCTCAGCTTTGTGCTTATAGTAATTCTTGGATAATACTAATATTTGTACTGGTAATAAAATCCTTGTTAATCCTCTAAATGTTGCATTAGTTTTC

>PGSC0003DMG400022787 G-LecRK

CACACTTCGACTAATTCATAGAATACGTATCACCTCTCACCAGCAACAAAATCTTGATTTGTGGAAAAAAAAAAAACCAAAGTGATGATCACTAGTCTACAATATTTGCTTGTTCTTCATTTTTCTTACTTAGGACTTTGCTTATCAATATGTAACATTAATAGCACTCAATATCTAAAAGATCCAGCAACTTTATGGTGTCCTGATAACATGTTCAAGTTGGGATTTTTCAGTCCTGGAAATACTACGAATCGTTACGTTGCTATTTGGTACAACTTCTCAGAAACAACTGTTGTTTGGGTAGCAAATAGAGATAAAGTACCATTGAAAGATTCATTTGGTGTTTTCAAGATTTCTGAGGATGGAAATCTTGTTGTGATGAATGGAATGAAAGATGTTATTTGGACGTCGAACATTTCATCTACTTCACACGTTGTTACGAGTTCTGTTGCTCAGTTACGAGGAGGGAACCTTGTTTTACTTGACAACTCGAAAAATAGGAACATAATATGGCAAAGTTACCAACATCCTAGTGATACATTTTTGCCTAAAATGAAAATACCTATTAACTCTGTTACAACAGGGGAGAAAACAGGGCTAAGATCTTGGAAAAGTCCTTTGGATCCAAGTTTTGGTGATTTTTCGACTAGCATTACCGCGATTCTTCCTCAGCTTTTGATTTGGAAAGGGAAGAATCTATACTGGCGAAGTGGTCAATGGAATGGTCAGGTTTTTATTGGAATAAAGGGTATGCATAGAGTGCACACTGGTGGATACAATGTTGTTGATGATGAAGAGGGCACTGTCTATCTTACGGGTTTTGTTGAACGAAAAAAACCTACAAGGTTTGTCTTGGACTCGAGTGGGAACTTGGTACAATCATACTGGGATGGGATTGAGAGGATTTGGAAGTTTCCGTGGTCAGCTATCGAAAATGATTGTGATGTTTATGGAACTTGTGGTCCATTTGGGAGCTGCAATTCATTGTATTCGCCAATTTGTTCGTGTTTAAGAGGGTTCGAGCCAAAGAATATCGATGAATGGGAAAAGGAGAATTGGACTAGTGGTTGTGTGAGGAGGAAAAGTTTGCAATGTGATCAAGAAAACAACAAAACAAGTACAAGAGATGGATTTTTGAAGATGGGATACATGAAAGTGCCTGATTTCGCTGAATGGTTACCTCCTAAGCTTGAAGATGAATGCAGAAGCCAATGCTTGAGTAATTGTTCTTGCTTAGCTTATGCTTTCGACACAGGAATAGGCTACATGTCGTGGAGCGGATACTTGATTGACATTCAACAGTTTCAAGCCTCTGGCACGACACTTCACATCCGTGTGGCACATTCTGAACTCGGTATTGATTCATTGTTTCTTGATTTGCCACTGTTCCATTAATGTTTCCAGTAGTCTTACATGATTTGGTTTCGTCTGCAGATCATCACGGAGAGATTAGACTAATCGCAACTCTAGTGATTGTATCGTCTTTTGTTGTTTGCGTTGGTGTGTATCTTATATGGTTTTGGATGTCAAGGCAAAGAGGTAAAATATGGTTTCACTTTAAAAGGAACAATCATTTAGATGGAAAGAAGATCTGTTTGGGAGAAATTATTAGTAATGTTGGTTTGGATGAGTTGCCAATTTTCAAGTTTGAAGTGCTTGCAATGTCAACAAATCAGTTTCATGACAACAATAAGCTTGGACAAGGTGGTTTTGGTCCGGTATACAAGGTAAACCTGTTTTTTTCTTCGCGTGAACCAATATGTTCCAACTCCGGTTCTTGTAAATGCAACTGTCACGTAGTATCGTTTACCTGTTTATTGTTCGTAGGGAGTATTAGCAGACGGGGTGGAAATAGCTGTCAAGAGGCTTTCGACTGTCTCAGGACAAGGGATGGAAGAGTTCATGAATGAAGTGTTGTTGATATCTAGAGTCCAACATAGAAACCTTGTTAGATTATTGGGATGTTGCATAGAAAAAGAAGAGAAGATGTTGATCTATGAGTATTTGCCAAATAAAAGCTTGGATGTTTTTCTCTTTGGTTAGTCCTAAAAGAAACTCTTACTCTGCGATATATAGTCAGACCTTTTTATAACAGTCATTCTCTATAATAACATTCCACTATAACAACCAAGTTTCCTTTGGAACCGATTTTAATGTTATGTTTTTTATAATAGCATTCACTATAGCAAGCCAAAAAATCATCCAGACAAACGACATTGTTATAGAGAGGTCTGACTGTATGTTCATTATAGCTTTCCTCTTTTATCTCATGCAAAAGCTTAAGTTGTTATAGAAAGAAATTGTCTTTCGATGAATGTTTAGATCAAACGCATAAAGGGATGTTGGATTTGAGCAAACGATTGCACATTATCGAAGGGATAGGGCGAGGACTACTATATCTACATAGAGATTCAAGAATAAAGATAATCCATAGAGATCTAAAGCCAAGTAATATCTTGTTAGATAATGATTTCAATCCAAAGATTTCAGACTTTGGCATGGCTAGGATTTTTAAATGCAATCAAGATCAAGCAGAGACAAGGAAAGTTGCAGGAACATAGTAAGTATATATATAGCCATTCAATATGCTAAGTTTTCTTCTTGTCTAAAAGTTCCCACATTAAGCTAATTCGTTGATTTATCGTTGAATTTGCAATGCCTTAGTGGCTATATGGCTCCAGAGTACGCGATCAAAGGGATGTTTTCTGAGAAATCGGATGTTTTTAGCTTTGGAGTAGTGTTGTTGGAAATAATGAGTGGGCAAAAGGTTGCAAGTTTTTGGAATGAGGAGATCTCATTAAGCCTTCTTGGATATGTAAGTTCCTTTAAAACTCTCTTGTACTTAATTGTGAGAAGCAAAAATATATTGATAAAGAAGTTCATTAGAATCTAACTTTAGAACTTGATATTGAGTGTTATAGCCTTGTGAAATAAGGACATTTTGATAGATCGAATTGAATGTTATTAACGCTCAAAACTTTAGTACAACCTTTTCTGATCCCATTGATGGGAAGACCCATATTATACCAACATCTCTTTAGGTTCTTCGATCATCTAGAGGTGTATATTTCCATTCTGCTTGGATCCAGTATCATTAAGCTTCAAATCCTGAGTGCGTCTCATTGTTAATTATTTAATAAATCTAAATACCACTAAAACAATTTATTTATTTTTCAGGCATGGGAACTATGAAAAGAGAATGATTTATCAAATTTTATTGATCAAGTGATATGGGATCCCAACTTTGAAATTGTGTTGACAAAATGCATACAAATAGGTTTGTTGTGTGTTCAAGAATTTGCTAGAGATAGGCCAACAGTTTCAAATGTTCTTTCCATGTTTTCAAGTGAAGTTATAAGTCTTCCAACACCATTGAAGCCTGCTTTTGCTAATTTTTATGATGATAATGGAGTTGGTTCTGTAAATTATGTCACTTTGACAAACCTTGATGCTAGATAATTAATATGAAATATTGTGAAT

>PGSC0003DMG400024258 G-LecRK

ATGGTGTCATCGACTCAATTCAGGAGGTTTACATATACTGAACTTAAGAAAGCAACAAAAAACTTCAAGGTTGAGCTAGGAAGAGGAGGTTCAGGAGTTGTTTATAAAGGAGTTTTAGCAGACGGAAGAGCGGTAGCAGTCAAGAAACTTTCAAATGAGTTTCAAGAAGAATTTTGGGCAGAGATGACTACCATAGGAAGAATCAACCATATGCATTTGGTGAGGATGTGGGGATTTTGTTCTGAAGGTAGACGTCAGCTTTTGGTCTATGAATACGTCGAGAACTCATCACTAGACAGGCATCTTTCCAGGGCTGATATTCTTGAATGGAAACAAAGGTTTGCAGTGGCACTAGGGACAGCCAAAGGTCTGGCATATCTTCATCATGAATGTCTTGAATGGGTGATTCATTGTGATGTTAAGCCTGAAAATATACTTCTCGATAGTGAATTCGAGCCCAAGATTGCAGATTTTGGACTTGCAAAGCTGTCCCAAAGAGGCGGCCCTGGTTCAGATTTCTCCAAGATTCGCGGTACTAAAGGCTATATGGCTCCTGAATGGGCTTTGAACCAACCTATAACAGCAAAAGTTGATGTTTATGCCTTTGGGATTGTAATACTAGAAATGATCAAAGGAAGTAGGCTGTCAAGTTGGGTGGTGGATGATAGTGAATGTGATCATGAACAAGATTCACAGCTCGGAAAATTTGTCAGGATGGTGAAGAGAAAGATTCAAAGTGGAGAAGATTCTTGGGTGGAGAAACTTGTCGATCCACGACTAGAAGGCAAATTTGGCAAGAACCAGGCAGTGACACTGATTGAAATAGGCCTTTCTTGTGTAGAGCAAGACAGAAACAAAAGGCCTACAATGGCCTCAGTAGTCCAAACATTGCTGGATTGTGAAGATGAAACAACACAAATAACATAG

>PGSC0003DMG400024259 G-LecRK

ATGAAAAGTTTGAAACTTTACTATCTTTTTCCATTTTTAGTTTTTTCATTTCTTGTTTCTTCTTTCCCTTTGATAACATCATCAAGAATTTTGCTTAAAGGTGATTCTTTTTCAGTTAAAGATGATTCACATTTCATTACTTCTCCACAAAAAACATTCACTTGTGGTTTTTATCCTATAGAAAGTACTAGTAATGCTTATTATTTTGGAATTTGGTATACAAATTCAAGAGATAAAACTGTTGTTTGGAATGCTAAACAAGATAGGCCTGTTAATCTCAAAGGGTCAAAGTTGACATTAAGAAAGAATGGAGCTTTAGTTTTAACAGATGTTGATGATACAATTGTTTGGCAAACAAACACAACATCATTTGATGTTGAAAAGGCTGAGGTTCTTGAAACAGGAAACTTAGTATTGAAGAATCCTAAAGGTGATGTTTTGTGGCAGAGTTTTGATTTGCCTACTGATACTTTATTGCCTGATCAAAGATTTACCAAGAATCATAGATTAGTTCCTCCTTTGAGGAAAGGGAGTTTATCACCTGGATACTTTAGTTTGTACTTTGATAATGATAATGTATTGAGAATGATTTATAATGGTCCTTTAGTTTCGAGTATATACTGGCCTAATCGCGAAAGGAGTGCGCCCCAGAATGGTAGAACTAGTCAGGTGGATACTGTAGTTGCACATTTTGATGGAATGGGTAGATTTTTCTCAAGTGATATGTTGCAATTCAATGTTTCTGATATGGGGCTTGGGATATTGAGAAGATTGACTATAGATACTGATGGTAATTTGAGGATGTATAGTTTGGATAACTTGACAGGGTTGTGGAAGATGTCGTGGCAAGCTATTGCACAACCTTGTGCTGTGCATGGAATGTGTGGAAGATTTTCGATATGTACTTATGTATCGAAGCCTAAATGTACATGTCCTCCTGGATATGAGATTGTTAATGGAAGTGATTGGAGTAGAGGTTGTAGGCCAATGTTTAGATCAAGTTTGGAGTCTAAACAACATGTAAAGTTTGTGGAGATTCCTAATGCTGATTATTGGGGTTTTGATCTCAATGCCACTACTCCTTTGTCATTTGAATCTTGTAGGGCATTGTGCTTGGACGATCCACGTTGTCGTGCATTTGTTTATAGGTTAACTGGTGTGGGATCATGCTATACTAAAGGCATCCTTTTTAATGGATACAGGTCTCCTAGTTTCCCTGGAAGTGTATTCTTGAAACTCCCTATGAATCTAAGTGCATCAGAATCAGGTCATTTGATGCTTGAAGGTACTGATGCAAAATGTGGATTGAGTCCTGAAAACTCTCTTTTTGGTTCTCTTTCTATGTATGTATTGGATTTTAAGAAGGTGAAGTGGATTTATCTTTACGTGTTTTGTTTCACCCTTGGTGGAATAGAGATTCTGTTCTTCGTGTTAGGCTGGTGGGCGTTGTTTAGTAAACATGGGATTCCTGCTTCCATTGAGTATGGATATAAAATGGTGTCATCGACTCAATTCAGGAGGTTTACGTATACTGAACTTAAGAAAGCAACAAAAAACTTCAAGGTTGAGCTAGGAAGAGGAGGTTCAGGAGCTGTTTATAAAGGAGTTTTAGCAGACGGAAGAGCTGTAGCAGTCAAGAAACTTGCAAATGAGTTACAAGAAGAATTTTGGGCAGAGATGACTACCATAGGAAGAATCAACCATATGCATTTGGTGAGGATGTGGGGATTTTGTTCTGAAGGTAGACGTCAGCTTTTGGTCTATGAATACGTCGAGAACTCATCACTAGACAGGCATCTTTCCAGGGCTGATATTCTTGAATGGAAACAAAGGTTTGCAGTGGCACTAGGGACAGCCAAAGGTCTGGCATATCTTCATCATGAATGTCTTGAATGGGTGATTCATTGTGATGTTAAGCCTGAAAATATACTTCTCGATAGTGAATTCGAGCCCAAGATTGCAGATTTTGGACTTGCAAAGCTGTCCCAAAGAGGCGGCCCTGGTTCAGATTTCTCCAAGATTCGCGGTACTAAAGGCTACATGGCTCCTGAATGGGCTTTGAACCAACCTATAACAGCAAAAGTTGATGTTTATGCCTTTGGGATTGTAATACTAGAAATGATCAAAGGAAGTAGGCTGTCAAGTTGGGTGGTGGATGATAGTGAATGTGATCATGAACAAGATTCACAGCTCGGAAAATTTGTCAGGATGGTGAAGAGAAAGATTCAAAGTGGAGAAGATTCTTGGGTGGAGAAACTTGTCGATCAACGACTAGAAGGCAAATTTAGCAAGAACCAGGCAGTGACACTGATTGAAATAGGCATTTCTTGTGTTGAGCAGGACAGAAACAAAAGGCCTACAATGGCCTCAGTAGTCCAAACATTGCTGGATTGTGAAGATGAAACAACACAAATAACATAG

>PGSC0003DMG400024260 G-LecRK

ATGAAAAGTTTGAAACTTTACTATCTTTTTCCCTTTTTAGTTTTCTCATTTCTTGTTTTGTCTTTCCCTTTGATAACATCATCAAGAATTTTGCTTAAAGGTGATTCTTTTTCAGTTAAAGATGATTCACATTTCATTACTTCTCCACAAAAAACATTCACTTGTGGTTTTTATCCTATAGAAAGTACTAGTAATGCTTATTATTTTGGAATTTGGTATACAAATTCAAGAGATAAAACTGTTGTTTGGAATGCTAAACAAGATAGGCCTGTTAATCTCAAAGGGTCAAAGTTGACATTAAGAAAGAATGGAGCTTTAGTTTTAACAGATGTTGATGATACAATTGTTTGGCAAACAAACACAACATCATTTGATGTTGAAAAGGCTGAGGTTCTTGAAACAGGAAACTTAGTATTGAAGAATCCTAAAGGTGATGTTTTGTGGCAGAGTTTTGATTTGCCTACTGATACTTTGTTGCCTTATCAAAGATTTACCAAGAATCATAGATTAGTTCCTCCTTTGAGGAAAGGGAGTTTATCACCTGGATACTTTAGTTTGTACTTTGATAGTGATAATGTATTGAGAATGATTTATGATGGTCCTTTAGTTTCGAGTATATACTGGCCTAGTCGCGATAGGGATGTGTATGGGAATGGTAGAACTAGTCAGAATAGTAGTAGATTTGCTTATTTTGATGGAATGGGTAGATTTTTCTCAAGTGATATGTTGCAATTCAATGTTTCTGATATGGGGTTTGGGATATTGAGAAGATTGACTATAGATACTGATGGGAATTTGAGGATGTATAGTTTGGATAACTCGACAGGGTTATGGAAGATGTCGTGGCAAGCTATTGCACAACCTTGTATTGTGCATGGAATATGTGGAAGATTTTCGATATGTACTTATGTATCGAAGCCTAAATGTACATGTCCTCCTGGATATGAGATAGTTGATGGAAGTGATTGGAGTAGAGGTTGTAGGCCAATGTTTAGATCAAGAAGTTTGGAGTCTAGACATGTCAAGTTTGTGGAAGTTCCTAATGTTGATTATTGGGGTTTTGATCTCAATGCCACTACTCCTTTGTCATTTGAATCTTGTAGGGAATTGTGCTTGGGTGATCCACGTTGTCGTGCATTTGTCTATAGGCGTACTGGGGAGGGATCATGCTATACTAAAGGCATCCTTTTCAATGGATACAGGTCTCCTGGTTTCCCTGGAAGTGTGTTTTTGAAACTCCCTATGAATCTAAGTGCATCAGAATCAGGTCATTTGATGCTTGAAGGTACTGATGTAAAATGTGGATTGAGTCCCCAAAACGTTCTTTTTGGTTCTCCTTCTATGTATGTATTCGATTTTAAGAAGGTGAAGTGGATTTATCTTTACCTGTTTTGTTCCACCCTTGGTGGAATAGAGATTCTGTTCTTCGTGTTAGGCTGGTGGGCGTTGTTTAGTAAACATGGGATTCCTGCTTCCATTGAGGATGGATATAAAATGGTGTCATCGACTCAATTCAGGAGGTTTACGTATACTGAACTTAAGAAAGCAACGAAAAACTTCAAGGTTGAGCTAGGAAGAGGAGGTTCAGGAGCTGTTTATAAAGGAGTTTTAGCAGACGGAAGAGCGGTAGCAGTCAAGAAACTTTCAAATGAGTTTCAAGAAGAATTTTGGGCAGAGATGACTACCATAGGAAGAATCAACCATATGCATTTGGTGAGGATGTGGGGATTTTGTTCTGAAGGTAGACGTCAGCTTTTGGTCTATGAATATGTCGAGAACTCATCACTTGACAGGCATCTTTCCAGGGCTGATATTCTTGGATGGAAACAAAGGTTTGCAGTGGCACTAGGGACAGCCAAAGGTCTGGCATATCTTCATCATGAATGTCTTGAATGGGTGATTCATTGTGATGTTAAGCCTGAAAATATACTTCTCGATAGTGAATTCGAGCCCAAGATTGCAGATTTTGGACTTGCAAAGCTCTCCCAAAGAGGTGGCCCTGGTTCAGATTTCACAAAGATTCGCGGTACTAAAGGCTACATGGCTCCTGAATGGGCTTTGAACCAACCTATAACAGCAAAAGTTGATGTTTATGCCGTTGGGATTGTAATACTAGAAATGATCAAAGGAAGTAGGCTGTCAAGTTGGGTGGTGGATGATAGTGAATGTGATGACGAACAAGATTCACAGCTCGGAAAATTTGTCAGGATGGTGAAGAGGAAGATCCAAAGTGGAGAAGATTCTTGGGTGGAGAAACTTGTCGATCCACGACTAGAAGGTAAATTTAAGCAAGAACCAGGCAGTGACACTGATTGA

>PGSC0003DMG400024426 G-LecRK

CATTGTTTGGGAGAACGGCAGGGTTTAAACGACGTGTTTTGACATAAAGTGTGATCATCAAGACTTTATAAAAATGTCTTCATCATCTAAGGATGATCAATCATCCAATACACCATCATCGGATTCTTCATCATCTTCTTCGAATTCTCCACCGTCTGGCTCATCCTCAAATTCTTCGCCATCCCAAGACAATTCATCTTCATCCAATGATTCAAAATCCCAATCTCCTCCTCAAAGTAATTCATCATCTTCTCCGCCATCAGGGGATAATAATAACTCGTCTCCTTCCGGTGGTAACAACAACAATAATGACAATAACAACGACAACAAGGACAACAACAACAATAATGGTAACAACAACAACAACAATGATGGTAACAACAACAACAATGGTAACAATGATAACGACAACAATAACAATAAGAATGACAATAACAACAATAATGGTGGGAACAATAACAACAACAACAACAATGGTGGTAACAATAACAATAACAACAACAACAATGGTGGTAACAACAATAATGGTAACAAATCTTCACCACCAGGAAAATCGTCAAATTCAAATGGAGGTTCATCACCTCCTCCACCCCCACCACCACCACCTCCTCCACCACCTCCTCCTTCTCCTTTTTCTCCCAACTCCGGGTTTCTTTCTCCTCCAAAGCATAAGCCACCACCAAGCAAGTCCTTAGATAATAATAATAACAACAAGCCTAATGATAACGGGACAGCCATCATTGTAGGAGTTGCTGCTGCAGGAGGTCTATTACTTCTTGTCATGTTAGTTTTCCTTATATCATGTTGTAAACGCAAAAGGAGAAGGCCACGCGACCAAATGGGCTACTACAGGGACAATTCTCATGGAGGCAATAGTATGTTCTTTCTTCTCTTAACTTATACTGTATGTTTTATGTTGAGCATACTGTTTTCAAATAGAATTAGCTTCAAATGCAGTAGCTATGAAGAGAACTGGTAAATGTGCAGAGTACTGAATATGTTTAGGTTTAAATAGTGTTAGTATTGAGAATGTATATATGGTGCAGGTACTGACTACTATAACAGTGGACCATATGGGAATTGGCACAACAACAACAATATTCAATCAACGGATCATATGGTGAAAATGCCTAATCCTCAAGCACCGAGTACTAACGTAAGTTCAGAACTGAGTTGGCCAATAGCACCTCCACCACCACCACCAATGATGAGTAGCAGTGAATTAAGCTCAGCTGCTTTCTCTGGTCCGCAACAACCTCCATTGCCACCCCCACACCCATCAATGGCTCTTGGTTTTAACCAAAGTAGTTTTACATACGATGACTTATCTACCGCAACTGGAGGATTTGTTAAGTCTAATCTATTGGGACAAGGTGGTTTTGGCTTTGTACATAAAGGGGTCTTGCCTAATGGTAAGGATATAGCTGTCAAAAGTCTAAAAGCTAATAGTGGACAAGGTGAACGAGAATTTCAAGCTGAAGTTGAAATTATTAGTCGTGTACATCATCGACATCTTGTCTCTTTAGTCGGTTATTGCATTGCTGGATCTCAAAGGTTGTTGGTATATGAGTTTGTTCCTAATGGCACTCTTGAGTATCACCTTCATGGTATTAACTTAATTTCCTTTTTTTTATATCTAAATACTCATTCTACATATGTCGTATTTAACTCATCGATCCCTTATTTAATTAGGAACTGGTCGCCCTGTTATGGACTTCCCTACTAGGCTTAAAATTGCATTAGGATCTGCCAAAGGCTTCGCGTACCTTCATGAAGATTGTAAGTTCATATATACTATTCTCATTTAAGCACTTGTCTATATATTCTCATAAAAAATTAAACTTCTATTTCATATATAAATTACAGGCCACCCTCGTATTATTCACAGAGACATTAAAGCTGCTAATATTTTACTGGACCATAACTGTGAAGCTAAGGTATTTGGTAAAAATAGTTGAAACATTGATTCCTCATTCAGATTTCTAACAAATTTTATTATTACTCCCTTTGTAAGGTTGCTGATTTTGGATTAGCTAAACTTTCCAACGACACTAATACACATGTTTCTACGCGTATCATGGGCACTTTTGGGTGAGTTTAGTTTGATAATTCTTCAAATATCCAATGGATTTGCATTAATCGAAAGAGTATATTTATTTGTGTAGGTACTTAGCACCAGAATACGCTTCAAGTGGCAAGCTAACTGAAAAGTCTGACGTCTATTCTTATGGTGTTATGCTGTTGGAACTTATAACTGGGCGTCGTCCTATAGACATCAGCAGTGATGATGATACTTTAGTGGAATGGGTATACCATGATGATTACTTTACATCTTCAAAATTAATTCAGTAGGACATCGATCAGATTATATGTATATATATGTGCAGGCTAGGCCAATTCTGATTCGTGCAACAGAGGGTGGAAACTACGATGAATTAATAGATCCACGTTTGGAAGGAAATTTTGATGCCCAAGAGATGCTATGTATGGTAGCTTGTGCTGCTGCATCCATCAGACATTCAGCAAGAAGACGTCCCAAAATGAGTCAGGTTAATTTCTTCTTAATATTCGTCAAAAGCTTTTCCTAATTCACAATTTTAAAAAGTATAGTGGTCCAGCGATTGACATATATATATGTTGTAGATTGTACGTGCTTTAGAAGGTGATGTGTCTCTGGATGATTTAAATGAGGGAATGAAGAAGTCCCCGAGTACTGCAATGTTTGGTTCTAGTGAAAGTTCTGAATACGATGGAGGTTCATATGACCTTAAGAAATTCAAGAAATCTGGAATGTCGAGTCAAGAATTCACAAGCAGTGAACATGGTACCGGTGAATTTGTCCATTCCGGTGGTGAATCTCAAGAACATGGGCAGAAAAGGCACACCCCTTAAGGAAAATGGATCATACCTTCAACTTCTGATGGATCTATAAGTAATAATTACACACATTTGTTGGGCTTACAAGGATGCTTTAGGCATTAAAAACTAGTTTTATGGGGGAAATATAATTATGTATTTAAACAACAATTGTGAAGCTTTATTAGAGATTTGAATGCGGGTTGTAAAGAGTATCACATCCTCCTAACCATTCTGTGGTTGGCTTTGGTTGTCAGTTGAATCATTTAGTTTTGTTCGACTTTGGAAGTTGGGATATATACTAATTCTGAATTGTTTTCAATATATTTCGTCAATGTATCTGCTAATTACTAATCTTTTCTTTGTTTA

>PGSC0003DMG400024998 G-LecRK

GAAGAATTAGCATCAAAAATTTGAATCCATGAAATTTCTAGCTTTTTTATTCATTTACTTCAATTTTCTTTCAATTGTACTAGCATTAGACACCATAACTATTGATCAACCTATTAAAGATGGTGACACAATTGTTTCATCAGGTGGTGTTTATGAACTTGGATTTTTCAGGCCTGGAAATTCGACGAATCGATATGTTGGTATATGGTACAAGAAAATATCAACTGGAACTGTTGTTTGGGTTGCTAATAGAAATAATCCGTTGAGTGATAGTTCAGGTGTGTTGATGATCAATCCTGATGGAATCTTAGTACTTGTTAATAGTACTAATGTGACGATTTGGTCAACGAATTCATCAACAATCTTGAAGAATCCAATAGCAAGATTGTTAGATTCAGGTAATACTTGTTATGTTTCGTTTCTGGAGAGTGAAACTACATGAACTTTGACCAACATTTAACATGTATTTTTTACCATACTAGAAGAATTGCAAGTACTTTTCGTATACTTTTTAAAAATCTGTAAATTTTAATTTTGAAATATTGAGTTAATGTAATCTAATCTGATTTCGTAATCTTTTCATCAAGGAAGGAAATGAAAATAGAATTGAGATTTTCACCTGGGAGAGTTTTGATTATCCAGGGGATACTCTGTTGCCTGGAATGAAGCTGTGTCCAGAAATACTTGTCATGTTTAGTTTCTCGAGAGTTAAACTGAATGAACTTTGATCAACATTTTAACATGTTTATTTTTAACCATACTGATATGAGAAGAACTGCAATTTGTAGTACTTTTCGTATAATTTTTACTACTGTTGTTTCTTTTACTTTGGTTATCCTTTATATTTGTACTGTAACTACTTTTCTTTCTTCAATATTTTCATCTTGAAGTTCAATTATTGCTTTTCTTTCAAAGACGAGGGTTTATCAGAAACAAACTCTGTGTATGTTGTTGTCGTAGTAGTAGTACTTTTAGTATAGTTTTTTAACTATCTTAATTTTAATTTGAAAATACTGAGTTAATCTAATGTAATCTGATATCAGGTAATCTTGTCATCAGGGAAGAAAATGAAAACAGGCCTGAGTTTTTCACCTGGCAGAGTTTCGATTATCCAGGGGATACTCTGTTGCCTGGAATGAAGCTCGGGCGGAATTTGGTAACAGGCCTGGATTGGTACTCGTCGTCGTGGAAGAGTCCTGATGATCCTGCTATAGGTGAATTTGTTGAAAGGATGGATGTTCAGGGGTATCCACAGTTATTTGTATGGAAAGGTTCATCAATTACATTTAGTTCAGGTCCCTGGAATGGTTTAGCATTTAGTAGCAGTCCTAGTTTGCAACCGAATACGTATTTTACGTTCGGTTTTGTACTGAATCAGGAGGAAGTTTATTACAAATATGATCTTAAGAATGGATCACTGCTCACCAGGGTGGTACTAACCCCTGGAGGATTGATAAATCACTATACCTGGATCGATCGTACGCAGAGTTGGTTCCTTTATTTGACAGCACAGTTTGATAACTGTGATCGTTTCGCGTTGTGTGGTCCTTATGCAAGATGTGTCATTAACAACTCGCCCCCGTGTGACTGTCTGAGGGGTTTCGAGCCAAAGTATCCACAGGAATGGGATGCAGCTGACTGGTCTAGTGGATGTGTTAGGAGGACTCCGTTGGCGTGTCAGCAAGATGGATTTAGGAAGTTTACGGGGATTAAAGTGCCTGATACGCAGAAATCATGGTTTAACGAGAGCATTGGACTTGAGGAATGCGGAAAGTTGTGCTTGGCTGATTGTAACTGTACAGCCTACTCAAATATGGATGTTCGAGATGGTGGAAGTGGATGCTTACTCTGGTTCGGAGACCTCATCGATATCAGAGAACTTAGTCCAAATCAGCAGGACCTGTTTGTGAGAGTGGCTGCTTCAGAAGTAGGTAATAACTACTTACTGTTTTTCGTTTCTCTAACTTGATCTTGTTGAACGTGGCGAAAAACATGTATGTAAAATTTCAGCAAGTATGTATCTAGATAGTCAAAAGTCTCGATAGTTAGCTCGTGGAGTCTTCAGTCAATAGAAAAAACATCTATGAATGTGTGTTGGTGCATTGTTCTCTCTCACTCAAAGACGGAACGTTGCCTGTTTCTTATGGCCTACAATTTTTGTTGGGAACTCATGAATGTGAATCATATTGTTATGACAATGTTGGAATTTTCTGATTGATTATGTACGAACTTATCGACAGACAAGAAACGAAAGAAAAAGAAGTCAAGGCTGATTGCAATCGTATCAGCAGTAGCCGCGACTTGTATCCTCAGCCTTCTAGCTTGGTGTGCATTATTCCACAGAAGAAAAAAGAGTAAAGGTAAACAACTATTGTTCATCGTTATCAAAAGATCGTGTTGCTAACGTTCTATTTGTATCAGGCTAATGTATACATTTATACCAATTGATTGCTCATTTTACAGGTAGGGAAGTTGGAGCAGACGATATGGAGTTACCATTGTTCGATTTAGTAACTGTTGCTAACGCCACTAAGAACTTCTCTTCTGCTAATATTATTGGAGAGGGTGGCTTTGGACCTGTTTATAAGGTACAATAACTATCCCATCGTCGAATTTTAGCTAAAAAAAGGCATCTTTAACACTTTGTTTTGAAAAAAAATTCAGGGGAAACTACGAAATGGACCTGAGATAGCGGTGAAGAGGCTTTCGGAATACTCAGGACAAGGTCTTCAAGAGTTGAAAAATGAACTGATACTGATATCCAAACTACAACACAGGAATCTTGTCAAGCTTTTAGGTTGTTGCCTTGAAGGAGTAGAGAGGATGCTGATATATGAGTATATGCCAAACAATAGCTTGGACTATTTCATTTTTGGTCCGGTTCTCTCGACTAGTCACCTTTATACTATGTTGCTCGTATCCTCCAAAAAATGATGTTGCACCAGTGTCGGATCCTCAAAAGATGCACTACTTCTGGAGGATCCAACATGGGCGCGGAGGCATTTTTGGAGGACTTGAGCAACCTACTAATAAATGAATGGATGAGGATTCACTGACATTTTACATATTGCAGATCCAAGCAGAAAAGAATCACTTTCTTGGAGAAATCGCTATGAAATCGCGATGGGAATATCTCGAGGCCTTCTCTATCTTCATCAGGACTCAAGATTAAGAATTATACACAGGGACCTGAAGGCCAGCAACATTTTGTTGGATACTGACTTGAATCCAAGGATTTCTGACTTTGGTCTAGCTAAGATATTTGGTGCAGACCAAATGGAAGGAAAAACAAAGCGAGTAATCGGAACATAGTAAGTCTTATAAACAACCTCTATGATAGATGTTTTTGTTTGTTACAATCAACAAGTTGATCAACTTAAATTTGATGCATTGTGTAGCGGATACATGTCTCCGGAGTATGCTGTCGATGGGAAATATTCAGTGAAATCTGATGTCTTCAGCCTTGGCGTGCTTTTACTTGAAATTGTCAGTGGCAGAAAGAACAGGAAGTTTCATCATTTGAGTCATCATCATAATCTTTTGGGACATGTAAGTACATATAGCACTAAATTTTTACCCTTTAGCCTCTTTTTTTACTAAACACTCGACCCCGTCTGAGCTTACTCACATCACCAGTCCCAAGCTCCCATAAAGGAAGAGGGTTGCAGCCAGTATAAAACTAGTCAGTTCATGTCGAATCCTCACAATACAGTCGGAACTGAGGCATTGTAGTAGTTGTTATAGCCTGTTTTCCATTAAAAAAACTTGACCTGCCTGAGTTTACTCATATTACCGGTCCCAAACCCCCATAAAGGAAGAGGATCGCAGTAGGCTAGAAGCCATTATAAAACTAGTCAATTCATGACGAATCCTCATAATACAGTTGGGGCCGAGGTGTTGTAGTAGTAGTTGTTATAGCCTTACATTTCCATTCCCAAGCCCCTACAGAAGTAAGAGGGTCACAATAGGCTAGCAGCCAAGCATAAAACTAGTCAATTCATGACGAATCCTCATAATGCAGACGGGACTGAGGCGTTGTACTAGTCCACATAGCCTTTCTTTTACTAATGCAAGTGTATAATTCATGATGACTGAGGCATTGTAGTAGTCACCGTTGTAGCTTCTTTTTACTAAATGCATTTGTATAACTGTGTTATCATAAAAAATATTTCAGGCATGGTTACTATTGAATGAAGGCAAAGCATTGGAACTAATGGATGAATGTTTAAAAGACTCATATGTTGAATCACAAGTGTTGAGATGCATTCAAGTAAGTTTATTGTGTGTACAAAAACTCCCTGAAGACAGGCCTACAATGGCTTCAGTAGTTTTCTGGTTAAGCAATGATGGTGTGGAATTGCCTCAACCAAAGCAGCCTGGTTTCTTTATAGAGAGGGATTCAACTAATCAATCAAATGAATCAACTGATGAAAGATGTGTTACTGACAATGAACTTTCACTAACAATTCTTGAGGCAAGATAGACAAAATATGAACTTGCAGTCGTGTGACCAGGAGGTCAAGGGTTCAAGTCATGACAATAGCCTCTTGCGG

>PGSC0003DMG400025558 G-LecRK

AAAAAAAACTCAAATCTATGGCTGTTTTGCTATGGTTATTAATCTTATCTGCATTTCATGCAGTAACACTAGCACAACAAAGGCAGTTTAATATAACTCTTGGCTCTTCTCTAACACCTACTACCAATTCATCATGGTTTTCACCATCCAGGCGCTTTGCCTTTGGGTTTTATGAACAAACCAATGGCTATGCTGTTGGAATCTACATCGTCGGTATGCCTAACAAGACAGCAGTGTGGACAGCAAATAGGAACAGTCCTGTTGTCCCGAGTAACACTGTCTTGCTCTTAACTAATGATGGAAGGCTCATTGTACAAGTAGGAAGCCAGGAAAACTCTGTCATCAATTCCTCTCGAGCCATTGCTTCAGCTTCCCTGATGGACACGGGAAACTTTGTGCTCTATGATTCTGATCGCAATGTCATTTGGCAAAGTTTTGACAATCCAACAAACACACTTTTACCTGGTCAGCATATATCAGCTGGACAAGAGCTGTTCTCGAGTGCCTCGGAAGCAGATGACTCATTAGGGATTTTCCGTCTTTCCGTCTCATAATGCAAGTTGATGGGTACCTTGTTCAGTATCCTGTCAATACACCAGATACTGCCCCATATGCTTACTACGCATCCTACATCCGTGGAGTAAGGAATAATTTTTCACTGAATCTTGATGATGACGGCCTCCTCTACTTGCTCAATTCCAACAATAGTCTAAAAAATCTAACCAGAGGAGGCTGCCGTAGAGAAAGGACCATCATCTATATGATGAAAATTGATGCTGATGGTATTTTTCGAGTTTATTCACATTCTCTGAACCAACAAAACAGCTCACCGATTTGGTCATCAACAGATGATAGGTGTGATCCAAAAGGCCTTTGTGGGCTTAATGGATTTTGCACCAATATAGATGCTCAAGCTAAATGTATGTGTCTTCCTGGATTTAATTTTGTAATGCCAGGAAATTGGAGCGCTGGCTGTGAAAGGAATTTCACCACAGAAACCTGTCAGTTAAAGGAGAACACTTCCAAATACTATGACATGAGAACAGTTGAAAACACAGCATGGGAAGATAGTTCTTATGTTATTTTTGCCGTAACGACCAAGGGAGATTGTGAACAAGCCTGTCTACAGGATTGTAACTGTGAGGCTGCTTCGTTCAAAGACAAAGTGTGTAGAAAGCAAAGGCTGCCTCTGAGGTATGGAAGAAAAGACCTGGGTAATTCTAATCTTGCATTGGTCAAGGTGGGCATTAATGTGTTAGACAACAAAGAGTTGCCTGATCAGAAGATTGAAGAAACAAAAGGCAAAAAGCTAAGGATAGACATCCTAATTGCCAGCATTACATTGACTGTTTTTGCTTTGTTGGTTGATGTTATTTTAGTTTTGATGATTTGACAAACAAAGGGATCCTATAATGGTACCTGATACCTTGTGACAATTTGTTGAAAATGCCAGCTGGAAAAAGTATAGAAAGTTGGTGAGGTTGGTGCACGGCTTCCAAGCATGCCAGAATCCTCAGCCAATAGCATGGCTTCCTTGTGTGTGTCTATTAGATAAAAGGCTTTAATCAATAGACACATTTTTAGTTTTTGCATTCCTGAAATTCTCAACCACACAAGCAAAAATCACTCCTATCAAGAAGTATTATCCTCAAGAGCAACAGGGACCTGATAGAGCTGCTTGCTGCTTAGTTGTCTTGAGATTATATCTTCCGTGCTTATATTGTAAATCTATTCCTAATCTATAAAGGAATTAGATCGTTGCTTTGTTCATCTAAAAGTCTAAGCCAGTTAAGTGTAACTGGCTTAGTGGGCAGTGTTGTATCTACTTAGTCAAATTCTAAGTCATCTAGAGTTAGGTGGTTTAGTGGGCAGTGTTGTATCTGCTTAGTTGAAGTCTAAGTTATCTAGAGTTAGGTAGCTTAGTGGGTAGTGTTGTATCTGCTTAGTTGTTTTCTTTGTAGTAGAGTTACTACATAGAGGGGGAAGGGATTTAGAGGTAAATTCCTAGGTTGCAACTGCCTGTAATCTAAACTGTTGCTCCGTTTGAGTGAAATGGAGTTGGATTGAAAGCCTGTGAGGACAGGTCGTGGTTTTCACCGCCCTTGAACAAGGGGATTTCCACGTAAACTACTTGTGTTGTTTTCTTTATTGTTTACGCCTCATTTATTGTTCATACTGAGTCAAGGGACCTGGTCCGTTGACTGATAGTGGACGCATACATCTTATCATTGGTGTTAGGCATCTCTGGATTTCTCATTCACAGAATTCATGTCTGGAGTCTGGACTTACAGAAAAATCCAGGAGAGTAGAAGTGTTCAGTTGAGCGAGGATGTTGCTCCACGAGCATTTTCCTATGCAGAACTAGAGCAGGCAACCAGTGGTTTCAAGGCGTTAGGGAGGGGGGCATTTGGAACAGTATTTAAAGGGATTTTGGCAGAAGGCCAGAAAGTGATAGCTGTAAAGAGACTAGACAAAGAATTGGTAGAAGGGGAAACAGAGTTCCAAACTGAAATAAAAATCATTGGAAGGAAACATCACCGAAATCTAGTCCGCCTCATTGGTTATTGCCTGGACGGATCAAGAAGGCTTCTGGTGTATGAGTATATGACAAATGGGTCACTTGCGGGCATACTTTTTACACCAGAAAAACAGCCCATGTGGGAGGAAAGGTGTGGCTTTGCTCGAGATATTGCAAGGGGCCTCCTATATTTGCATGACGAGTGTGAAACACAAATCATCCATTGTGATATTAAGCCTCAGAACATACTCATGGATGATCAGTTCTGTGCAAAAATATCTGACTTTGGAATGGCCAAGCTCTTAAAGAAAGACCAAACAAGAACCTACACAGGGATAAGAGGGACCAGAGGCTATATTTTGTGCTAAGTGACTAGCAACCGTCGTTGTAGTATTTGAATTGTAAATACTAATTTAGGAGAGTTACAATTTTCTCCATAGTGTAAGCGAATCTGCAACAACCACAAC

>PGSC0003DMG400026154 G-LecRK

GTCTCACTGAAATGAAATAGAGGGCAAGAGCTACCTATTTGAATTTATGTTCTTCATTCAAATCATCAACAATTCCATAAAACAATGCAATTAAGTAATTTTCTGTTGTTGTTGTTGTCTGCAGCATTGATGATTTTACATCGATGTGATGCAAGGAGGAGCACCATTGATGGAAGCAGTAAATTAGTTGACAATGGCGAAACCCTCGTATCTGCTGGAGAAAATTTCGAAATGGGATTCTTTTCTGATGATGCCGGGTTAAACAAATATGTAGGCATATGGTATTACAAATTAAGTCCTAGAACTGTTGTATGGGTTGCTAATTGGAATAATTCGATTCAGGGTAAAAGAATAATGAATGAGGATAATTCGGTTGTCGTTGAAGATGGTAATCTCAAAGTTATAAGTAATGGGTATACTTATTTCTCCACACAATTAGGTAGTGGTTCTAACAGGAAAGTAGAGTTGTTGGATACAGGGAATTTAGTTTTAGTTGATGAAGGAGCTGAAATGTGGCAGAGTTTTCGCAATCCAACCGATACATTCCTACCAGGTATGAAAATGGATAGCAGTTTGAATTTGACTGATTCAAAATATGAGAACTACATATTCCGATTAGATCAAGCAAGTGACAAGGAATATGTTATAGTACTTCCAAAGCAGGGCAAAATTCTCTGGAAAGGGTCTGCGGAATCAGGAAAGTTAAGTTTCAGTGAGATGCCTGGTTATGTTGCCTACTTGTTATCCAACTCCACTAATAATTCATTGGAATCGATAGGAACATTTAACAAGTATAGGCTCTTAATGAACTCCTCGGGGGAAATACAATTCTACGGTTGGGATAAGGAGATCAGTGGATGGTCTTTGAGGTGGTCAGCGCCAAATGATAAGTGTGATTGGTATAAGTATTGTGGGAAATTTAGCATTTGTAACAGCAAACGTGAGCCAGTATGCAAATGTTTGCCCGGGTATAAGCTCAATCCTCCAGATAATTCAAAAGCAGGAGAATTTTCAGGTGGTTGTTCAAGTATGTCAGTTAGCTCTTGCAATGAAGACAATGTAGAAGTGCTCGACACGTTCTTGGATTTGAGGTCGATGAAATTCAAGAGTCCAGACAGGATATTCAGTAATAATAGTACCCGTGAGGACTGCAGGAGAATTTGTCTAGGCAACTGTAAGTGCCAGGCCTATACTTACCATGATTCTGTATGCCGGATTTGGGTCACTAGTCTCATGTATCTCCAGGAGAACTATGCTGGTGGCTTCAACATCTCTGTTCGTGTCAGCATTTCCGATATTGGTAATTTCTTTTTCGATTTAACTTACGTACACAGACAGTTTAAGCTAATGTGTGACCACAAATAGTACAGTAATTTGATTTTTGTGCATTGCAGAAGCAACAAGGAGAAATTGCAAGCCGTGTGGCATAAACCTAATACCCTATCCATTAAGCAGTGAACCAAATTGTGGTGACCCTTTGTATTATAGTTTCTCATGTGATGATTTAGTTGGTCAGGTAAGCTTCCTCACATCGAATGGCAAGTATACTGTCGTTAACTTTGATAAAGATAACAAAACATTTGTTATAGAAGCAGCTCATAAAGAATCTGTTGGTACTTGTGATGACAAAGGACCGGTGACAGGAATTTCTTGGTTCAACCAGTCATCACCTTTTAAGGTGATAAACTGGTGTTACAATCCCACAGAGAATCTTAATTCAGGACCTTTTAGTCGAGGGAAAGATTTAATATTGATTAGTTGGAAACCACCATTGGAGCCATTTTGCAAGACTTCAGAAGATTGCAATGATTGGCCAAATTCTAGTTGCAATATGACAAAACAAGGAGAAAGAAGATGTATATGCCAAACTGATTACAAATGGAATGGCTTGATTTTAAATTGTTCCTCGAGTTTAGGTATATGGATCAATGTTACTTACTACAAATGATTTTCTTGCCAATAGTTCAATGATACCTTTTCCTATGTTCTCCATTTGACCTTTTATATGACCAACACAAATTTCATTTCTTAAATTTGCTTCAGAACTTGGAACACAAGGATCATTCATTGCGAAGCTGGCATCCTCACGGAATCAGAGAACCCTTGTTATTTCCATCTCTGTAGTTCTTGGAGTTATAACTTTGTGCAGCATTAGTTATATAATTTATCAGAACACTAGAGTGACAAGAAGTAGA

>PGSC0003DMG400026155 G-LecRK

ATCTGATTGAACGAAAAAACCAAGTCTTACCATGACTATCAACATGATCTTAATTGATGCTGTGCATCTTCTGTTATCAAATTTGGTGCAACCATGCAACTGCAACATTATTCGATTCGTCTACTTTTTGTGATCCTAACATGTTTATCTATCAATATTCATCAGTCTCTTGGAGTTGAAATGATCTCAGTAAATCACTCTGTTTCTGGAAACCAAAGTATAGTCTCTTCGGGCGAAAACTTTGAATTAGGATTCTTTTTTCCTGGTAGTTCGAAAAATTACTATCTTGGTATATGGTATAAGAATGTAATGCCACAAACAATAGTATGGGTAGCAAACAGGGAGAAGCCACTTTCTGCTACTGATATGAATTCTGTTGAGTTGAAAGTTTTGGATGGTAATTTAGTGCTTATTACTGAATCCAAAAACTTGTTTTGGTCCACAAATATCAGCAACACAATTAGTTCAAATACACTAATGGCAATTCTTTCTGATGATGGTAATTTGATTTTGAGTGATGGATCAAATTCAACTACACCACTTTGGCAAAGCTTCGATAACCCGACTAATACATGGTTGCCGAATGCTAAGGTGAAGTATGACAAGCGTACAAATACAACAAAGATTCTTACTTCTTGGAAGAACTCAGAGGATCCATCTCCGGGGATTTTTTCCGTGGAAATGGATCAGAGTAATAAGCAGTTCTTGATAAAGTGGAACAGGACTGAGATGTACTCAGCTACTGGATCTTGGAATGGGCGTATTTTCAACATGATGCCTGAGATGAGTTTGAATTCTGATAGATACAGTTTTTCATATGTTGATAATGAAAATGAGAGCTATTTTACCTATTCCCTCCGTAACTCTTCCAAAATCAGATTAACACTAGATGTCTCTGGGCAAATCAGGCATCTGATTTGGGCCGAAAATTTAAAGGAATGGCAAATTTTTACATCTCAACCAAGACAACCCTGTGAGGTGTATGCTAGTTGTGGTGCCTTTAGCATTTGCAATAAGGAGTCCGCGACCTTCTGTAATTGCTTGACTGGTTTCACACCTAGATCTGATACAGAATGGGACTTGAATGATCATTCTGGTGGCTGTGTTAGGAAGGAAAGCTTGCAGTGTGGTGATGGTAAAATGAAGGGCGGTTTTTCGGAGAATCCCAAAGTGACATTGCCCGGTTATTCTCTAACTGTGCCGGCTGCTAGTTCTGAGGAATGCCAGTCGACTTGCTTGAGTAACTGTTCTTCATGTAATGCATATGCTTATGATAATAATGTGTGTTCAATTTGGAATGAAGTTGTGAATCTGAAGCAACTCTCACCAGGAGATGGTAGTGGAAGTGTGATTTATACCAGACTTGCTGTCTCTGATGAAGCACGCGGTGAAGGTAAGTTTAAAGTTAGGAGTTGTATTTGATAGTGTTTGCCATTTCTATGTATGCAATGCTCATCTCCAAATAGCTACCGATAGAGCTTTCTATGGATGTCTGTTTTTTCTTGTCTGATGTTTCAGTTCTTTACCTCATGTCTTTCAACAGTACGAGTCTATATATTCTCTAAATCTTAGTGCTCATAAGAATTAACTAGGCAATTATAAGTTTACGCAAACACCTTCTATGTTTTAAACAAACATCAGATCGCCACTGACCAAATTTAAGTTTACGCAACATCCTCTACGTTATAAACACACATAAGATAGTCACTGACCAAATGTATGGTTTTCATGATTTGATTAGTTACAACTAGCCAAAATGAGTTGAAGACCTTTCGCCAGTAAACATTACCAGTTCGCTGAGTTCCCACATCAGCTCTTTAAGGATGGGATGGTCTTTTCATATGGTTTTGGACAGTTCTTACCTCATAAACTAGCTTTTATAGTTGAGTTAGCCTCAAGGTCCATTTCTTATCAAGTTATTAGAATCAGACCCATCCCAAAATAGTCTATTTCTTGTCACAATTTCCTTATGTTTCTTCTTTCTGTCATCAGATGCAAATTCAGGGAAATTGTCCCTGAAGGTTAAAGTCATCATATCTGGCGTGGTAGCCGTTGCAGTTCTACTTCTATGTAGCTTTAGCTACATTTATCATAGGAGAATAATGTCAAAGAAAACAGGTACTTGTGTTGCTATTACAATCTCTGGAGTAGTTGTTTTACCACTTCCAGAAAAATTTCTCCTTCCATTTTTTTTTTCCGAATCAGATCAAGAGAGATATCTCAAAAGGCTGCTCTTTTTTTTATTTTTTTATTTTTTATCTTAGCAAGCCAACAAAGCACAGGAGGCAATCCAGTTCCACATTGGTTAAACAGAGAAAGAGAAGCACAAAACCTGATTAACGAAAATGATAAGCAAAGCATTGCCGTTCCATTCTTTAGTTTGGAAAACATACTAGCAGCCACAGATCATTTCTCAGATGTAAATAAACTTGGACAAGGAGGTTTTGGTCCTGTTTATAAGGTTAGCATTAATAATTCATTTTAACATGAGTTCAACTTTATGTAGTCATTTCATAGCATCCTAGTTTACACATTGAAAATCTTTCAGGGTATATTTTCAGATGGACAAGAAATTGCTGTGAAAAGGCTATCAACCCAATCTAGACAGGGCATTGAAGAATTCAGGAACGAAGTCATACTGATCTCGAAACTTCAACATAGAAATCTAGTCAGACTTTTAGGCTACTGTATCACAGGGTATGAACAGATTCTACTTTACGAGTATATGTCAAACAAAAGTCTAGACACCTTCATATTCGGTTAGTATATATATCTGAACTTATCTTATTTCTTAATCTCTCTATTTACTTGATCAATATTGTTAAATTTTCTCGTTGTTTACCAAAATAAGATCCAACACTTAGCAAGTCATTGAAATGGAGAAAGCGATTTGAGATCATATTAGGAATTTCTCGAGGGCTTCTCTATCTTCATGAAGACTCAAGGCTCAGAATCATTCATAGGGATCTGAAAACAAGTAACATTCTTTTGGATCAACAAATGAACCCAAAAATATCAGACTTTGGCTTGGCAAGGATAGTTGAAGAACAAACAACAGAAGCAAGTACCAAAAAAGTGGTTGGAACTTAGTAAGTAATACAAACATGACAATGTTATGAACTTTGCTTAATTCCATCTCACATCTGACTCATGATAATTCCATCATTAATTTAGCGGATATATGTCTCCTGAATATGCATTGGAAGGAGTATTCTCAATTAAGTCAGACGTATTCAGCTTAGGAGTAGTTATACTGGAGATAGTAACCGGAAGGAGAAATACAGGATTTTATCAATCTAAAGAAGCATCTAACCTTTTGGTCCATGTGAGTGCCTAGAGTACATGTTAATTTTTTCCAATATGTATGTTGGACTTCATATTCATTATTCTAAATCATGATATCACAGGCATGGAATTACTGGAAAGAAAAAAGAGCACTACATTTGCTGGATCATTCATTGCTCGAATCATGCAACCCGAAAGAAGCAATGACGTGCATAAACGTTGGGCTCTTATGTGTGCAAGAAGATCCAGGTGATCGTCCTTCCATGTCAAATGTTGTAATGATGCTTCGTAGTGATAGTACGTCTCTTCCAAAGCCTAATCAACCTGCTTTTGTGTCAAGGAGAAATGTGTCTAATTCAACTTCTGTTTCATCTGGTACACCACCTAGCTTCACCAAAATTGAGTTGACAATCACAGTTGAAGAAGGCCGATGATGTGATGAATTCATTAGTTTTTCAAATGAATTTAAACGCGAAGTTTATTTATTTTTTCACCTTAGATTAAAAAACAAAATTACAGGAATACATCATCTTTTTCTTTTTCTCAATTTTTTTGTTGTTTAGGCAGTAGAAGACTTAGGACCAGTACTGCTGATATTTATAGTGAAAACTGAAAGGTATATATGCTTAATATTGTGTACAACATAGAATATATTGTGTTGATAAAAAAAAGTTCATTTTCTGTTTCATTTTGGTCTCTTCTACCTTCTTAACTTGTCTTAAAAGGCCATTCAGTAAGAATGGAAGAACAAAAAAG

>PGSC0003DMG400026156 G-LecRK

TTTAAACAGGAGCAAATTACAAGGAGGTGTACTAGGAAAAATAATGGAGTTGGATTGTGTTGTAGCTATTTTAATTGTATTTTGTTGTTCTTTCGTTACGCTTGTGGCGAGTGAAGAAAGTATCAAGATGGGCAAATCAATCACTGGTAATCAGAAAATGATTTCAGGTGGTGGAAGTTTTGCCTTAGGCTTCTTCACTCCGCGTAATTCTACATTTACCTATCTTGGCATATGGTACAACACGATCCCCGAACAAACAGTCATTTGGATTGCTAATAGAGAATCTCGAATTCCTCAGAATTCTACTGTTGTTTTCACTATTGGTGATGATGGGAATTTAGTGATTTTTGACGTAAAAGATCAACTTATTTGGTCATCAAATGTTTCATCCATAACTTCAAATTCTACTGTAGGAGCACTACTAGATAATGGAAATCTAGTGCTCATACATGGTGAGTCTGATATATTGTGGCAGAGCTTTGATCATCCTACCGATACGTTTATGCCTGATATGAAACTTGGTTATAATAGAAAAACAGGACAACGGACTCTGATCAATTCGTGGACAAGTAATGAAGATCCTCGGCCTGGAAATTTCTCGTTTGGGATAAATCCTGAAGGGCGGACGCGCTTTTATATATTGAAGCAGAACAGTATTTATTTTAGGTTTGATGATAGTACTGAATACAGTGGAGGGAATATCGGAGGAATTGCTTGGTATATTTCTGTTGTTTCGGGAAATGATGGAGTTTTTGTTACGTATGGTTATACCAAAGGGTTGGTAACATTAAGGATTGTTTTGCATCCTAATGGTTACATTCAGTTGCTGGTATGGCGTCAAAATGCTAATAAGTGGATAGTTAAACTTCAGGCACCAGAATCTCTTTGTGAACGTTACGCGGAGTGTGGTCCATTTGGCAGCTGTGATGTTGGATCAAGTGGATTATGCAGATGCTTAACTGGATTTGAACCAAGATTTTCGACAGATTGGGAGAACGGGAAATGGAATGATGGTTGTGTTAGAAAAGTAGCATTGAGTTGTGATGGCGGAGACATATTCTTGAAACATGAAGAGGGTATGAAGTTTCCGGATCATTCTATCTCCTTAGGACATATGAGCATTAAGGACTGTGAAACGCGGTGCATTAGGAACTGTTCTTGTTCAGCTTATGCTTATTCGAATAGCACTTGTTTAATTTGGTTTGGAGATTTACTGGACCTTTCCCACAACTTTTCTGGTGGCAGAGTTCTCAATGTTCGAGTTCGTGGCTCCGAGCCAGGTATTTTCACTTCTATCCTTACTAATTCTACTAACAAACGATTCTCTACACGTCTTTGGTCAAGAAAACTAAATTTATTTGCTTTTTGTCTTCTGGATGCAGTTACTCATGGTGGATCTAGGAATTTGGCTCATAGACACAAAGTTTTCGTTGCGAGTATTGTTTCTGCAATATCTATCATCTTGGTTCTTATAAGTATTTTTTTCTTTATCTTCGAAAGAAAGCACTTGAGAAGACAAGGTATGTTTAGTGTACGTCTTAATTGTAAATACAAATACTGAATTCGAATTTCGTATGGGTTACACTGATTTGCTTTACTGATTGCTGTTTGAAATTAGGCTGGAAGAATGGAACTAAAGCTTTCATTAGGTCCGTGTCTGGTTTATATTCGGTTGAAAAAAGCGATATGAAGTTGCTGCAGTACAACCTTCAGAAAATAAGAGAGGCTACTAACAACTTTCATGGAGAACACAAACTTGGAGAAGGTGGATTTGGTCCTGTGTTTAAGGTGCTAAAACTAAATTCATTGTTGATATTTTACTGCTGCTACTTTGAGAGAGACGTCAACGTCTCATTTTCGTTTCATGACTATGGTGGATGTGGATTTACATATTCTGTTTCAGGGATTTTTAACTGAATTCGGAGACGTAGCCATCAAAAGGCTAAGCAAGAGGTCATCACAAGGACTGGAGGAGTTTATGAATGAACTGAAGCTAATCGCGAAACTGCAGCATAAGAATCTTGTCAGCCTCTTAGGATGCTGTGTTGAAGGGGAGGAGAAAATACTAATCTATGAGTATATGTCCAATTGCAGCTTGGACAAATTTCTGTTTGGTTCGTCTTCTACTCCGTTTTAAGACTTCACATAACTTAAATTTTTACTCCTCTCAATTCCCTTTGGTTAATGAGATGTTCAAATGTCTTTTTAGACCCTTCGTTGAAGGTTACACTCGATTGGGTTACACGTTTGGGGATAATAGAAGGAATTGCTCAAGGAATGCTTTACCTGCATAAGTACTCGAGATTAAAAGTCATCCACAGGGACTTAAAAGCAAGCAACATTCTTCTAGATCAAGAGATGATCCCAAAAATATCAGACTTTGGAATGGCAAGGATTTTCGGGACTGATCAAACACAGGCCAATACAAATCGAGTCGTTGGTACATAGTAAGTAATAACCCCTTATGATTTGTGGTTTCCAAGCTTTTATGTCATTTGTGCATATCTAATGGATACATATTTATACAAATATGGCATAGTGGCTATATGTCTCCAGAATACGTAGTATATGGCCAATTTTCAGAGAAATCAGATGTTTTCAGCTTCGGAGTATTGCTGTTAGAAATCCTGAGTGGTGAACGTAACTCAGACTTTTTAATGACTGAAATTTCCGCGTCCCTCTTAGGATGGGTAAGAACACTCAAGACTTTAATTAATTCTGCTTTCTTTCACTTTGTTTTCATGTTTTTTATTTATGGTGCAACTCATTTGAACTTTTTTAAGGCATGGAATAAGTGGAAAGAAGGAAAAATCTTGGAGTTGATTGATCCGTCCATAAGAGAAACTTGCGACAATAACAAAGCAACAAGATGTATTATAGTTGCCCTTCTTTGTGTTCAAGAAATCCCTGTTGATCGACCAACAATGTCTGATGTATCTTTCATGTTAAGCAACGAAACAACTCCTATCCCTGAACCAAAAGAACCTGCTTTTCGTAGCAGCTGGCAATCTCAGCAGTTAAGTGATGTATCCATAAATGAGATGACGATTACCTTGCCAGCACCGCGATGAAATGATTTAAGTTCTCCTACTTCTTGTGGTAAATTTTTCTTGTTAGCATAGAGCTGTCATAATTTTTTTTTGTAGAACTTGCATAAGTGTAGCTCTGGAAGAAATATAGACTAAATTTCTCTGTTTAATTGATGTTTTCATGTTTTGTTTTTTTTCTGTTCCTACAGTGTATCAAGTTTGTGAAAATGTTGTTTTGAACATTCACTTGGCTT

>PGSC0003DMG400026183 G-LecRK

AAAACAATGGAGTTGGATAGTTTCTTACCTCTTTTACTGTTAATTTTCTCATGTTCTTGCTTACCCGTTCTTCCTTGCCAAAATTGTGTCAAAGTTGGCAAATCAATCACTGGGAATCAGAAATTGGTTTCAGCTGATAGAAGTTTTGCCTTAGGATTTTTCACTCCATCAAATTCAACGTACACCTATCTTGGCTTATGGTACAACATAATCCCTCAACAAACAGTCGTTTGGGTTGCCAACAGAGAATCTCCAATCCCACAGAATGCTTCACCAGTTTTCACTATCGGCGATGATGGAAATTTAGTGATTTTCGATGAAAAAGGCAAACTCATCTGGTCATCAAATGTCTCATCAACAGCACCGGAATCATTATTAACTTCAAATTCTACTATCGGAATATTGTTAGATAACGGGAACCTTGTTCTTCGACACTACGGTGAGTCCACTTATTTGTGGCAGAGCTTCCAGTATCCTACAGATACGCTATTACGCCATATGAGATTTGGTTTCAATAAAATTAGTGGCTTACGAAATGTGATCACTTCATGGACAAGTAATGACGATGCTCGCCCTGGAAATTTCTCTTTTGGAATAGACCCTACTGAAGGGGTTATGCGTTTTGTCATTTGGAACCGAAACGGTATTTATTATAGATTTGACGATAGTTGCAGAGCGTATATACTAGCAACAGTGTGGTGTTTCACTGTTGTTACTGAAAATGATAATGTATATCTTAATTTAGCTTACTACAGAATCTTTCTGAGATTAAGAATCGTTCTGAATCCGAGCGGGCACCTTCAGGTGATGGTGTGGAATCGGAATGGAAGTAACAGATGGAATGTTGAATTTGAAGCACCACAATCTAAGTGTGACCTTTACGCACAGTGTGGTCCGTTTGGTATCTGTGGGATCAGATCAAGTGGATTATGCAGTTGTTTAGACGGATTTGAACCTAAATTTTCGACAGATTGGGCAAATGAGAAATGGATTGGAGGTTGTGTGAGAAAAATAGCGTTGGGGTGTGATGGTGGAGATGGATTTTTTAAGCACGAGAATATGAAATTGCCAGATCATGCTGTCTCTTTAGGAAATATGAGCACCGAAGAGTGTGAAACTCAGTGCATTAAGAACTGCTCCTGTTCAGCTTATGCTTGTACGAAAAACGCCTGTTTTATATGGCATGGAGATTTACTGGATCTTGGGCATGACACTACTAACGGCAGAGCTCTCTATGTTCGAGTTCACGTGTTAGAGCTAGGTATGTTTACTTGTATCATTGCTGGTTCTGTTAACAACAAGTGAAAGACTAGTATAATTTTCGTGTTGTGGATGCAGTTACTGATAATCTATCTGGGAATTCAGCTCGAAGATACAAAATTCTCACTGTGAAAATTGTTTCTGCAATTTTTGCAATCTTGCTTCTTGTTAGTACCTCTGCTTTTATCTTCAAAAGAAAGCGCTTGAAAAGACGAGGTGAATCTAGCATTTCTAGACTGTAATGATTTCAGATTTCATATTGTTTACAGTGCTTTGCCTTACTGATTGTTGTTTGAAACTAGATAAGATGGACGGAGATTCTGCTTTAGTTAGCTCCATGCCTAGTGGATCTTTAGTTGGGAAAGATGATATGAAGTTGGTACAGTACAGCCTGCAGAATATAAGAGACGCAACAAACAACTTCCATGAAAGTAACAAACTTGGAGAAGGTGGTTTTGGCCCCGTGTTTAAGGTGCAAAATTCAATATTTTGATGTTCGTATTTAATTTCATGAATATAGAGAATTGTGGCTTTAAACATTCTGTTTCAGGGATCTTTGGCTGAATTTGGAGACGTAGCCATCAAAAGGTTAAGCAGGAGGTCATCACAAGGACTGGAGGAGTTCATGAATGAGTTGAAGCTAATAGCGAATTTGCAGCACAAAAATCTAGTCAGCCTCTTAGGATGTTGCATTGAAGGGGAGGAGAAGATATTGATCTATGAGTATATGCCCAATTGCAGCCTGGACAAATTTATTTCTGGTTATACTCATTAGTATTATTTTCACGAGAATCATAACTTTGTTTTCCTCATTTCTGTTGGCTAATTGGATATTCATTTTTTTCTTTTAGACCCTTCGTTGAAGGTTACACTCGATTGGGATACACGTTTTGGGATAATAGAAGGAATTGCTCAAGGAATACTTTACCTGCATAAGTACTCGAGATTAAAAGTCATCCACAGGGACTTGAAGGTAAGCAACATTCTGCTAGATCAAGAGATGATCCCAAAAATATCAGATTTTGGAATGGCAAGGATTTTCGGGACTGATCAAACACAGGCCAGTACAAAACGAGTGGTTGGTACATAGTAAGTATAGTAGCCGCTTCTGTAACCAGTTTTTATGTTACTACCCCTTGTGTATAGCTACTTAAGACTGATTTACGCCAATATGGCATAGTGGCTATATGTCTCCAGAATACGTAGTATATGGCCAATTCTCGGAGAAATCAGATGTATTCAGCTTTGGAGTATTGCTGTTGGAAATCTTGACTGGTGAACGAAACTCAGACTTCTTCATGACTGAAGTTTGTGTGTCTCTATTAGGATGGGTAAGTACATTCGCGCATTTGAAAACTAAGTCCAAGTTTGTGTCAATTTCTTTTAACATTTCACATCATTTGGCATGTTAGGCATGGAAAATGTGGAAAGAAGGAAGAATGTTGGAGCTAATAGATCCATCAATAAGGGCGACCTGCGACAGTAACAAAACAACACGGTCTGTTCTGGTTGCTCTTCTTTGCGTTCAAGAAATTCCTACTGATCGACCAACAATGTCTGATATAGTTGTCATGTTAAGCAACGAAACAACTCCCATCCCTGAGCCAAAAGAACCGGCTTTTCGCAGCAGCTGGCGACATCAAAAGTTAAACGATGTTTCCATAAATGAGTTGACATTTACTTTGCCAGTACCTAGATGAGAGGATTTAGCTTTCCATATTTCTTATGGAATTAATCTTATTTTGAAAAGAACCATCTCAAAAATGTTTAGCACTGTAAATAAAAAATCAAAAACCAATTAAATTACTCCCTCTGTCCCAGTTTATATC

>PGSC0003DMG400026184 G-LecRK

ATGGCGAAGAAAACAGGTTTGTGTTGCTATTACAGCCGTTGCAATTCTACTTCTTTTTTTTGCTCTATGAATCAGATAAAGAGAGATATCTCAAAAGGCTGCTCTTGTTTTCTGATCTTAGCAAGCCAACAAAGCACAGGAGGCAATCCAGTGCCACATTGGTTCAACAGAGAAAAAGACGCACAAAACCTGATTAACGAAAATGATAAGCAAAACATCGCCGTTCCATTCTTTTGTTTGGAAAACATATTAGCAGCCACAGATGATTTCTCAGATGTAAATAAGCTTGGACAAGGAGGTTTTGGTCCTGTTTATAAGGTTAGCACTATTTTTTCACTTCAATAGTTCCGCTTTATGTAGTCATTTCATAGCATCCTAGTTTACACATTGAAAATCTTTCAGGGTATATTTTCAGATGGACAAGAAATTGCTGTGAAAAGGCTGTCAACCCAATCTAGACAGGGCATTGACGAATTCAGGAACGAAGTCATACTGATCTCGAAACTTCAACATAGAAATCTAGTCAGACTTTTAGGCTACTGTATCACAGGGTATGAACAGATTCTACTTTACAAGTATATGTCAAACAAAAGTCTAGACTCCTTCATATTCGGTTAGTATATATATCTGAACATATCTTATTTCCTAATCTCTCTATTTACTTGATCAATGTTGTTAAATTTTCTCATTGTTTACCAAAACAAGATCCAACACTTAGCAAGTCATTGAAATGGAAAAAGCGATTTGAGATCATATTAGGAATTTCTCGAGGGCTTCTCTATCTTCATGAAGACTCAAGGCTCAGAATCATTCATAGAGATCTGAAAACAAGTAACATTCTTTTGGATCAACAAATGAACCCAAAAATATCAGACTTTGGCTTGGCAAGGATAGTTGAAGAGCAAACGACAGAAACTAGTAATTAA

>PGSC0003DMG400026185 G-LecRK

ATGATTTCAGTAAATCACTCTGTTTCTGGAAATCAAAGTATCGTCTCTTCAGGCGAAAGCTTTGAATTAGGATTCTTTAAACCTGGTAGTTCGAAAAATTACTATCTTGGTATATGGTATAAGAATGTAATTACACAAACAATAGTATGGGTAGCAAACAGGGAGAAGCCACTTTCTGCTACTGATATGAATTCTGTTGAGTTGAAAGTTTTGGATGGTAATTTAGTGCTTATTACTGAATCCAAAAACTTGTTTTGGTCAACAAATATCAGCAACACAATTAGTTCAAATACACTAGTGGCAATTCTTTCTGATGATGGTAATTTGATGTTGAGTGATGGATCAAATTCAACTACACCACTTTGGCAAAGCTTTGATAACCCGACTGATACATGGTTACCGAATGCTAAGGTGAAGTATGACAAGCGTACAAATACAACGAAGATACTTACTTCTTGGAAGAACTCAGAGGATCCTTCTCCGGGGATTTTTTCCGTGGAGATGGATCAGAGTAATAAGCAGTTCTTGATAAAGTGGAACAGGACTGAGCTGTACTCAGCTTCTGGATCTTGGAATGGGCGTATTCTCAGCTTGATGCCTGAGATGAGTTTGAATTCTGATAGATACAGCTTTACATATGTTGATAATGAAAATGAGAGCTATTTTATGTATTCCCTTCGTAACTCTTCCAAAATTAGATTAACACTAGATGTCTCTGGGCAAATCAGGCATCTGATTTGGGCCGAAAATTTAATGGAATGGCAAATTTTTACATCTCAACCAAGACAACCGTGCGAGGTGTATGCTAGTTGTGGTGCCTTTAGCATTTGCAATAAGGAGTCCACGACCTTCTGTAATTGCTTGACTGGTTTCACACCTAAATCCAATACAGAATGGGACTTGAATGATCATTCCGGTGGCTGTGTAAGGAAGGAAAGCTTGCAGTGTGGTGATGGTAAAAAGGGCAGGTTTTTGATGAATCCCAACGTGACATTGCCCGAATATTCCATATCTGTTCCGGCTGCTAGTGCCGAGGAATGCCATTCTACTTGCTTGAGTAACTGTTCCTCATGTAATGCATATGCTTATGATAATAATGTGTGTTCAATTTGGAATGAAGTTGTGAATCTGAAGCAACTCTCACCAGGAGATGGTAGTGGAAGTGTGATTTATACCAGACTTGCTGTCTCTGACGAAGCATACGGTGAAGGTAAGTTTAAAGTTAGGAGTTGTACTTGATAGTGTTTGTCATTTCTATGTATGCAATGCTCATCTCCAAATAGTTACTGACAGCTTTCTATGGATGTTTGTTTTTTCTTGTCTAATGTTTCTTCAGTTCTTTACCTCATGGACTTCAACATTACGAGTTCTCTAAATCTTAGTGTTCACAAGAATTAAATAGGCAATTATAACTTCACATAAACAGACCTTACGTTTTAAACAAACATCAGATAGTCACTGACTAAATATAAGTTTACGCAACACCCTCTATGTTATAAACACACATAAGATAGTCACTGACCGAATGCATGGTTTTCATGTTTTCATTAGTTACAACTAGCCAATATGAGTTGAAGACCTTTAGCCAGTAAACTATTACCAGTTTGTTGAGTTCCCACATCGGCTCTTTAAGGATGGGATGGTCTCTTCACATGGTTTCGGACAGTCCTTGACTCATAAACTAGATTTTATAGTCGAGTTAGCCTCAAGGGTCCATGTCTTATCACATTATTAGAATCAGACCCATCCCGATTCTTTGTTTACCCGATATTCTGCCCCATAATATTGTTCACGCTCCAGTTGGCAGGCCTAGGCATGAGCGATGTTGAGTTCCCACATAGTCTCTTAAGGGATGAGATGTTTTTTTTATATGGTCTCGGACAATCATCATATCATGAAGCTAGCTTTTAGGGACAAAAGTTTATTTCTTGTCACAGTTTTCTTATGCTTCTTCTTTTCTGTCATCAGATGTAAAATCAGGGAAATTTTCCCTGAAGCTTAAAGTTATCATATCTGGCGTGGTAGCCGTTGCAATTCTACTTCTATGTAGCTTTAGCTACATTTATTGTAGTAGAATGATGGCGAAGAAAACAGGTTTGTGTTGCTATTACAGCCGTTGCAATTCTACTTCTTTTTTTTGCTCTATGAATCAGATAAAGAGAGATATCTCAAAAGGCTGCTCTTGTTTTCTGATCTTAGCAAGCCAACAAAGCACAGGAGGCAATCCCGTGCCACATTGGTTCAACAGAGAAAAAGAAGCACAAAACCTGATGAACGAAAATGATAAGCAAAACTTCGCCGTTCCATTCTTTTGTTTCGAAAACATATTAGCAGCCACAGATGATTTCTCAGATGTAAATAAGCTTGGGCATGGAGGTTTTGGTACTGTTTATAAGGTTAGCACTATTTATTCACTTCAATAGTTCCGCTTTATGTAGTCATTTCATAGCATCCTAGTTTACACATCGAAAATCTTTTAGGGTATATTTTCAGATGGACAGAAATTGCTGTGAAAAGGACTAATCAACCCAATCTAGACAGGGCATTGAAGAATTCAGGAACGAAGTCATACTGATCTCAAAACTTACTAACATAGAAATACTAGTCAGACTTTTAGGCTACTGTATCACACGGGTATGAACAGATTCTAACTTTACGAGTATATGTCAAACAAAAGTACTAGACACCTTCATATTCGGTTAGTATATATATCTGAATTTCTTAATCACTCTATTTACTTGATCAATGTTGCTAAATTTTCTCCATGGTCACCAAAACAAGATCCAACACTTAGCAAGTCATTGAAATGGAAAAAGCGATTAGAGATCATATTAGGAATTTCTCGAGGGCTTCTCTATCTTCATGAAGACTCAAGGCTCAGAATCATTCATAGAGATCTGAAAACAAGTAACATTCTTTTGGATCAAGAAATGAACCCAAAAATCTCAGACTTTGGCTTGGCAAGGATAGTTGAAGAACAAACAACAGAAGCAAATACCAATAAAGTGGTTGGAACTTAGTAAGTAATACAAACATGTTAATGTTATGAACTTTGCTTAATTGCATCGCACATCTGACTCGTGATTATTCCATCATTATTTGGTTGATTTAGCGGATATATGTCTCCTGAATATGCATTGGAAGGAGTATTCTCGATTAAGTCAGACGTATTCAGCTTAGGAGTAGTTATACTGGAGATAGTAACTGGAAGGAGAAACACAGGATTTTATCAATCTAAAGAAGCATCTAATCTTTTGGTCCATGTGAGTGCCTAGAGTACATGTTAATTTTTTCCAATATATATGTACGTTGAACTTCATATTCATTATTCTAAATCATGATATCACAGGCATGGAATTACTGGAAAGAAAAAAGAGCACTACATTTGCTGGATCATTCATTGCTCGAATCATGCAACCCGAAAGAAGCAATGACGTGCATAAACGTTGGGCTCTTATGTGTGCAAGAAGATCCAGGTGATCGTCCTACAATGTCAAATGTTGTGATGATGCTTCATAGTGAAAGTACGTCTCTTCCAAAGCCTAATCAACCGGCTTTTGTGTCAAGGAGAAATGTTTCTAATTCATCTTCAGTTTCATCTGGTAAACCACCTAGCTTCACCAAAATTGAGTTGACAATCACAGTGGAAGAAGGCCGATAATGTGATGAATTCATTAGTTTTTACGCCAATTTTCTCCGCATGAAATTATTTTTTCACCTTAAGAGAAGATAAAATACATCAAATCAACTTGATAGCAAAACTTTTTATCAGTTTAACTACATTGGTG

>PGSC0003DMG400026810 G-LecRK

CAAATTGGAAGAGACATTTTTCAAAGTGTCTGAAACCTTTTTCAGACCACTCCCTTTTACTCCCAGAAAAAAATTGCTTTGTTGATTTTCCGTCAGTCCCTTCTCTTGCTTTCCAGGGAAAGATAAAGAGTTGAAAATCTACAATGTGAGGCACTTAAATTTTGCCTCTTTACTTTACTGGTGAGTCATATCTCTTGTTAATTTAAGCTCACCACTCAAAACTTCCCCAAGATACTAACTTTCTGTTTCACTCTAGCTTTTTCACTGAGATATGTTGATTTTAAAGCTTTTTCACTTGTGGGGTTCTCATTTTCTTCAAATCTTGAAATACCCATTTAAGGATTTCAACTTTTGGAGTTATGTTAGACCGAGTACTTTTGCTCTACATTATCCTTAAAATTGGGACGTAATTGTAAAGATCCATTCTCTCTTGCATGTGCACTTAGATTTGTTCTCTTATCTTTCCTTAAAAATATTTTTTTCTCTTACCTAAATTGCTCTTTTTTTGCTTAGAAAAATTGAGATATATATGTGTGTATGTGGACTCATGGTTTCTTTGCAACGCGTATTTTGTCAAAGGCATGTTTGTTTCCAAGTCTATGCGTCTATATTGACTTTTAAATAGTTAATTTTTGGGGTATTAAACTACTCCCTTATTAGGTTTTCAACTTTGGATGCTTGCAAAACATTTTTTAGTGTCACTTTAAAGTATAGAAAAGTGGTTGAATATGCATAAGGACCTAAATAGGGTCCCATTTGTAGTTTTGAACCAGGATTTCAAACAAAAGCTGTTTTCTTAAGCCATTTTAAATTAATTTTCTTTGTGTACTGTTTATTTTATGGTCCATCTGTAGGTCATATTTTTTTTAGTTCAATCATAAGGTGTGGATCTTTTGTTTTCTCACATGGCAAGTTTCTTATAGGGAAGCATTTGCTGTTTCAAGAAAGTGTCTGCAGGCAATATTCTTCCAATTGATGTGGTTTCATTCTCAAAAAAAATATTCTTCCAATTGATGTGGTTTGGATGTCGTTTCTGAGGTTCAATGCTTCCCTTCTCTTTCAATCATTTTCCCACCTTTTGTTTGATCTTGGTGTAACCTTGAATCAATTGGTAGTTTGAGAGTTTTACCTTATAGTTGATAGTGATATTTCAATTCCTTTACAAAAGAACAGTTTCTTGTTCATTGTAAATCAGTGGAATGACTGAACATATTTTTTGTTGATATGCAGAAGCTGGAACCAGCTTTCTGTCTTTTCATCTGCTTCCGCAAGTAGTGACAGATATGGATAGAGGAGACAGAATAATTGCCTGTTATTCTAATTTGCTGCTTTCTGTCTCTATCGGGTTCTTGCTTTTTTCAGTTGTCAGTTCACAAATTCCTTTGGGTTCCAAACTAACAGTAGAAGAGAACAATTATTGGTTCTCATCCAATAAGGACTATGCAATTGGATTCTTGAACTTCTCCGACCAGTATAGCTTTGGTGTCCGTTTCAATGCAATTTATATACCTAGCAGTGAGCAAGCAGCCATTTGGACTGCAGGGTGGAATGTTAAAGTTAGTAGCAAAGCATATTTTGAGCTTTCCCTAACTGGGGAAATGATATTGTTTGATCCAGCAAAGGGAAAAATTGTTTGGCAAAGCAAAACTGGCAACACATCTGTTGAATCAGCTGTTCTTCTTGATGATGGCAACTTTGTCCTGTTAAATAGGAATAAGAGTGCTGTTTGGCAGAGTTTTGAAAGTCCATCTGACACAATGCTGTCTGGCCAGAGTTTATCTGTTGGCCAGTCTCTTCGAGCTTCTAGCAGGGGTTCTTTAACAAGTTATTACAGTCTTCATATGAATGTTTCTGGTGAGATGCAGCTTAGGTGGGAGACTAGTATCATTTACTGGACTGTTGGGGGTCCTAAATCTGCTGTACGAGCCATTCTTGGCTCTGACGGAATCCTCCAACTACTTGATCAGCAATCGCAAGCTGTTTGGTCAGTCTATGGTGAGGACCACAATGATTCTGATGTGAAGTTTCGGTTCCTCAGACTAGATTCTGATGGTAATCTTCGGATATACTCATGGGAAAATAATGCAACATCATGGAGAACAGTTTGGCAAGCTATAATTAATCAGTGTGATGTCTTCGCAACCTGTGCTACCAATGGTATTTGCACCTTGAATGCATCTGACTCTTATGTTTGTTGGTGTCCATTTAGATCTACAAGAGACTCAAACTCTGAATGTCTAATTCCATACAAACCAAGTTGTGAATCCGGTTCATCCATGATCCTTCATGAGCATATGTATTTGTACGGAATATACCCACCAAACAACACAGTTGTGCAAACCAATTTGCAGCAGTGTAGAACTCTGTGTCAGAAAGATCCATCCTGCCATGCTGCATCCTTTATCAACAATGGCACTCCACAATGCCACATGATGAACTCCCGGTATGTTGGTGGTCAATCAGATCCTTCTTTGGGTTCCGTTACTTTTGTCAAAACTTGTTCAGACCCAATTGCTGTTTTACCTCCGCCTGTACCTGCTCCGAGAAAGGTATCACAGAAAATTTGTGTTGCTTGTCTGTTAGAAGTTGCTGCTGCGTCAATCGTTGTGTTTGTCATGCTTCAATTCAGCATTGGAGTTTACCTTTTTAGGAGAAGGAAACATATGATGCAGAAATCTGCTTTACCCCATATAGTGCCTAATGCTACTGGCTGCATCGTGTTTTCTTACTCTGAAATCAAAGACCTGACTGATAATTTTAAGCAACAAATTGGGCAGAATGTGTTCAAAGGTCTACTTCCAGATAATAGGCTTGTTGCAGTTAAAGATCTCAATGCCTCTATTGATGAAAGGAGGTTCCGGGCTGCAGTTCTAAAAATAGGAAGCATATACCACAAAAATCTTTTGAGGCTGGATGGTTATTGCTGTGAGTCCGGCCGTAGGTTATTGGTTTATGAGCTTGCCAAATATGGTTCTGTTGACAAATGTTTAGAAGAACCTAGAATGTGCAAGAGATTGACGTGGAGAAAAAGAATGGATATATGTTTGTCAGTGGCAAGGGCCATATCTTACTTGCATACAGGATGTAGGGAGTTTATTAGTCATGGAAATCTTAAATGTGAAAATGTTGTCTTAGATGATGAATTAGAGGCCAAAGTTAGTGAATTTGGACTGAGGACTATTCAAGCTGAGGCTTCCAGTAGCGGGGGGTTAGCAGAGACAGATGTAAGAGATTTTGGCAAAGTGATGGTGGTATTAATCACTGGATGCCAAAATGCGGATGAAGCTTGTTATTGGTCCTATGAAAAATGGGTGAAAGCTGAAAAGGAAATGGCCATGGATCAGCGAATTGTGGTTGGCACCGATTCAGAAGAGTTAGAACGAGCATTAAGAATTGCATTTTGGTGCTTACAAGGTGATGAACGTATGAGGCCATCAATGGGAGAGGTAATCAAGGTACTGGAGGGAACTCTGACTGTCGATCCTCCACCACCTCCTTTTGCTCATAATCAACAGTGGCCACCTGATGATGAATCACCATCTGAATCATATTCAGCACTGTAGAAGTTCTAATTCGTATATTTCATTCCCATTAAGTCATCTGTTTCTGTACAGAACTTACCAGGCACCTTCGACTTGGCATGATGTAGAATGCCTAGTTTTTCCAGTATGGCATGCTTAATCTGAAATTCAAGACGAGCATCCCACTCTGCCTCTTCTGAAACTGTAACCTTGGTATGTAACCCTCCAAGTACAGTTTTGTGTTAGAAAATCCTGATTCAGAAATTCAGGTTTCTTTTTTAAGTTTTTTATTCTGTTCATGTTAATTATCACTGCTTATCACATGTGTTTACATTTGATATTTTCTTGGAGGTGACTAAACTTCCTTAACTCTGCCCAAAATGTCTCCTTGGCTTGTGCTATCAGTTGGCACTGGCTTTTTTACTAGGCAGTGGTCCAAGAGATCGTGGATGAGGTTTGAACCTGGACATCTCTATTCTTAGGATTACTCTGACCATTAAGTCGATCCTAACTTAAAATGGAAATGGAAGTTTGTGTCTCTTTTTCTTTACAGAGGTGGTTTCCATGCTATTTGCACCTCGACCATTTCACCAGGTACTTGGTATCTCCACCAACCTAGGTATCAGGTAACTCTATCTCCAAAGCTTAGACAAATGAAAAGAAATCACCTAGCATTTTGAATTTCGTTGATAGGTCACACTTGTGTTTTATATATGTTCTGTTTTGATTTTGGCTTGCTGCACCTTTGTTGCCTGGTAAGTAGGGTCATTTTCGGTGAACATCCATGATCAGTATTTCAGCAATGAACACAATTCATTATTTTTATTAGTGAAAGAGTTG

>PGSC0003DMG400027233 G-LecRK

CACAACCAAAAAGTTATGTTATGACTTTTGTGAGTTTGTCTTAAATCTAAATTTACAAACTCGAAAAAGTTATTGTAAGTAGCAGAGTGGCCTTGCTAGTACAATCCACTTATATGCTAGGTAGCAGAGTAGATTGCTACCGCGATCAACTGAAATTCAGCTCTCTATTGCTCCAAAGAAAACATTTTGACATGTGTAGTCTTGCTGTCAGGATCCACATAGAGATATCTTGGTTATCGTGACCAATTGAGATTCAAACACTTTGTTGCTCCAGTGAACAAAAAAACATTTTCATAGAGTAATCTTGCTGTCAGAATCTACTGAGATATCTTGGTTATCGTGACCAATTGATATTCAACACTTTTTTGCTCCTGTTCCATTTCATATAGTAAGTCTTGCTACCAAGATCAGCTGAGATTTAGCTTTTTGTTGCTCCGATTCATTTTCTTGTTTCTATTAACAAAAAAATATTTTGATAGAGTAATCTTGTTGTCAGGATCCACTGAGATATCTTGGTTATTGTGACCAATTGATATTCAACACTTTGTTGTTGCTCCAATTCCATTTCATATAGTAAGCCTTGCTACCAAGATCAATTGAGATTCAGCTTTTTGTTGCTCCGATTCGTTTTCTGTTCTCCTTTTTCAAGTAAGTAAAGTAGGTTGACTCGCTACTTTACCTTCTGTGTATAATGAGACGGAACAGAGTAAAAAATGCAACTTATAAGTATGGTATGCAAAAAGATTCTTGATTTCAAGAACTGATTTTTTTTTTCAAGTTTTCATTGTTGTTAAGAGCATGGGATCATCTTTTTTGTTATTCATTGTTACCCTTTTGTTATCTTGTTTTCTTGTACACTCTGGACCTCTTTCTCTTCAACCATTAACTCCAAATTTCACTGCTTCAAATTTCAAGTTCATTGATACTTCTGGATCATTCTTGTCATCACCAAATGGAACTTTTAAAGCAGCAATAACAAACACAAAATCTCAAGAAAGATCCTATTACTTTGTCATTGTTCACTCGGAATCACATGTAGTTGTTTGGTCAGCAAATCGCGATATGCCTGTATCTGATTCAGGTGAGTTGCACCTTTCTGTCGATGGCCTTGCTCTTTTTGATGATTCAGGTCACACCGTTTGGTCTGCGAAGCGTAGTAGTACTAGTAGTTCTGTTACTTCAATGCAGCTTCTTGAATCAGGAAACTTGGTGTTGGTTGATGCTTTTAATAATACTGTGTGGGAAAGTTTTGATTCCCCGACAGATACAATCGTTGTAGGCCAACGATTGCCTGTTGGGAAATCGTTGGTTAGTTCAGTGAACGAAGATGAAATTGCTAAAGGCGATTATAAACTTGTGGTTGTAGAGAATGATGCAATGTTACAATGGAATGGAATGACTTATTGGAAGTTAAGTATGGAACCTAAGGCTTTTACTGATGCATATACTCTGGTGGAGTATATGATGATCAGTAGCAACGGGTTGTTCCTTGTAGGGGCGAATGGGACGGAGAGAGTGATTCAGGTTATTTTGGATGAGGTGAAAGATCCTGACTTTAGGATTGCGAAATTGGAAGAGAATGGACATTTCGGTGTCAAACGATTCAGTAACGGAAATTGGATGTCTGAATTTGATAGTCCTATTGATAGTTGTCGTGTTGCTTTCACTTGCAAGAAGCTTGGAGTTTGTGATGAGGGGAGCTGTTCTTGCCCACCGGGATTTCGTGTTAGTTCTGAGGTGAATGGTAGTTGTGCTCCAGTTGATAGAAATCTGGTAATGCCAGTTTCTTGTAATGCATCTTTGAATATGAATGTAACTGAATTGGGAAATCGTGTTTCGTACTTGAGGCTAGAGAATGGTATGGACTATTTTGCTAATGATTTTATCGAACCTGTGAAGCGTGGTGTTAATGTATCTGCTTGTCAAGATCTTTGTTCTAAGAATTGTTCTTGCTTGAGTGTTTTTCATGATCAGTCGTCGGGGTCATGTTATATGATTGAAAATTTTCTTGGTTCGATTTTGAGGGGCTCGGATTCTGGTAACGGAAGAGGTAGATTGGGATACGTTAAGGTCATTTCTGAACCGTCATCATTTGATCCAAATGACAACTCAAGTGACAAGAGGTCAAGATTACCCGTTGTTGCTTTGGTTTTACTTCCTTCTTCTGGGCTTTTCCTGATCATTGTGATGATGGCTGGGATTATGTGGTTGATGAGAAGAAAAAGATTGATGCAGATTTCAGGGAAAGAGTTTGGACGGACGGATTCTTCTTTGTTTGCTGAGCTTGATAACATTTCCATACTGGGGTTGCCTGTGAAATTTGATCATGAGGAGATTCGGGTAGCTACGGAGTGTTTTAGAAATCAAATTGGTACTGGTGGATTTGGAACAGTATATAAAGGTACACTTTCTGATGGAGCTGTTGTTGCTGTCAAGAAGATGAATGCTCTCGGGGCTCACGGAAATAGGGAGTTTTGCACTGAGATCGCGATTATTGGGAGAGTTCACCATGTTAATTTGGTCAGTTTGAAAGGATTTTGTGCACATAGGGGAGAAAGGTTTCTGGTGTATGAATATATGAATCGAGGCTCATTGGATCGAACCCTTTTTGGACTTGGACCTGCCTTAGATTGGCATACCAGATATGAAATCGCGCTTGGAACAGCACGTGGGCTTGCTTACCTGCATGGTGGTTGTGAGCAGAAGATCATACATTGTGATGTGAAGCCAGAGAACATACTCCTCCATGACAACCTGCAAGTAAAAATATCAGATTTCGGGCTTTCCAAGCTTCTTAATTCTGAGCAGTCTAGTTGGTTTACCACGATGAGAGGAACTCGAGGCTACTTAGCTCCTGAATGGCTGACTAGCTCAGCCATAACGGAGAAATCAGATGTCTACAGCTACGGAATGGTGCTACTAGAAATAGTACGAGGGAAGAAGAACTCCTCGTTCCAACCACCAAATGACACTACTTCTCAAAGCGAAAGTAGTGAAAGGAACAGGCTTTCTCCATCGTCTCTTGCCTCAGCAAATCAGCCAATCTACTTCCCATTATTCGCGTTGGAAATGCATGAACAGAAAAAGTACCTGGAGCTAGTGGATCCTCGTGTTCTGGGGAGTGTCAAAAGTGAGGAAGTCGAGAAGCTAGTACGTGTGGCTTTGTGTTGTTTGCATGAAGAGCCAACTCTGAGACCAACAATGGCCAATGTTGTTGGCATGTTAGAAGGTGTGTTGCCTTTGGCTACGCCACAGGTTCAATCACTCAATTTTCTACGATTTTATGGTCGTAGATTCACTGAAGCATCAACTATCGATGGGGATCAAGAGGTTAATGTGTTTGAGTTACATCAACAAAGCAGGAACCGTAGCAGTACTACTAGTAGCTCTTACAATTCCTTCTCTTATATGTCTTCACAGCAAGTCTCAGGTCCAAGATAGCACTAGTAAATCAAACGTTGCTCAAACACTACAAAAATGTTGTTACATTCGTGTCTGGTCCTCCAAAAGTGCATAGCTTTGGGAAGATCCAACCTACTTTTGACATTATTTTTGGAGAGTCTAAACAACACGGAGTAAATCTGCTTTTTTAGTTCAATATGTTATGCTACTGTTTGAAAACTTTAGTTGTAATATGATGTTTGTACCTCTTCTGTGAGGCAGTAGAAAGAGTATAAATATAAGTTGTAATTTATTCAATTTC

>PGSC0003DMG400028461 G-LecRK

AAATACAACATCAGCTTCAAATCATCTTTACCCTATCCAGACAAGTAGTTAAGCAGATGGGAATAGATACATTTCGATGATCGTCAAGTCATTGATATAAATTAGAGATTTCACAGATGAACTATTCGTATGTATAAGAATTCTTTTCCCTTGAAATATGTTTTTTGGTCAATGAACATTTTAGTTTTTTTGTCTAAACAGAATCATGGCTTTACCCTCTGTCAAACATTTGTTGAAGCTTTTGATCATTGTGCTCCTTGAGAACCTTACTTTGGCTCAATTTCCTGATTATCACACAGTGAAAATACCCACAACGTGGATAAACAATAATGTATCGTCTATCAATGTTACAAAGCCCTTGACATGGCCTTTTTACTCATTCAATCAATCATCTGGACCTTTCATGCGAATTCTGATGCTCAAAGAGATAGGAAAGGACATGGGATATGCTTGTGGTTTTGCTTCAAATGGAGAGGATGATTCCTTTGTGTTTTCCATTGGAATTATACGTCTTCGTCAGGACTTTCTAGGAGCTCAGCGCTGGACTGACCTTGAGCTGGTCTGGTTTGCAAACAGAAACCATCCAATTCATGAAAATGGAACGCTGCAATTGTTGCAAGATGGAGATTTAGTGTTAAAAGATGTGGATGGAACTCTTGTATGGTCTACAGGCACCGCGAATAAGTTTGTTTCGGGCATTAAGATGATGGATACTGGGAACCTTGTGCTTCATGACATGTATAATCAGACTGTCTGGCAGTCTTTTGATCATCCAACAGATGCATTGCTTCCAGGCCAGAAATTAAGGTCTGGTCAAAGACTTGCGGCTAGAGTTTCTTCGTCCAATTGGAGTGAAGGTAACTACTATATCTCTGTAACCAATCAAGGCTTGTTTGCATTTTACATATCAGACAAGCCGCAGATGTATTTTAAGTTTTTGGTCAGTGGAGAAAGGGATAGCTTTGATGAAAGTTATGTCAAAGCAGTTAATGGTACTCTTGCCTTGTACATATCTTCAACCGAGCCGAATGAGCCAAATGCTGTCTTCTCAAGACCTTCAAGGATGAAATATTTGAGATATGATTATGATGGACATCTTCGAGCCTACACCGAGGGTAGTGATCAGACAAACGATCTTTTGGCAGATTTCATTGGGCTTTGCGATTACCCCACAGCTTGTGGCAGTTTGGAGCTTTGCTCAAATGGAATTTGTTCTTGTCCAGGTCCATTAATAAAAAGTAATGATCGACAAAACAACGGATGTGTGGAAGTTAGTCCGGTGGAATGTGATGGACAACGCTCCCACAGAATGGACCGTGCTTCGGATGTCTATTACTTTAACTATGTCGACACTGATGCAGCAGCTCTGAGAGGAACAGATGAGGAAAGTTGCCAAGAGTTGTGCTTGAAGAACTGTTCTTGTAAAGTAGTTCTATTCCGGTATTTCACTAACTTTTCAAGTGGTGACTGCTATTTACCCTCTCCTGTTCTGTCACTTATAATTGATGGAAAGGGAAGAAGTGTCTATGAATCATCTGCATTCATCAAATTACCAAATGATGCAGAAAAAGGTAAGTCTACAGCTACAAGAAGAACTGGCATTATAGCTGGATTAACTGGTGGAACAGTTCTTCTGATAGCATTAACTGTTGGCATCTTGATTGCTTTCAACAGAAAACACACATTACAAGAAAATAATGATGATTACTCAGGGGAGATATCTGGTCTTGTAAGATTTTCTTATGAACAGCTGAAGATGGCAACTTGGAATTTCCAGAAAAAGCTTGGTCAGGGAGGTTTTGGCTCAGTTTATGAAGGAGTTATTCGAGATGGTCAGAAAGTAGCAGTGAAGGTTCTTGATGGTTTTGGCCAAGGAAAGAGGGAATTCTTGGCAGAGGTCCAGACCATAGGAAGCATCCATCATGTTAATCTTGTTAGACTAATTGGAGTTTGTGCTGAGAAAGAGCACACAATATTAGTTTATGATTTCGTGAGTAATGGATCATTGGATAAATGGATTTTTGGCACATCCTCTACTCAATTTACCCTTGATTGGAAAATAAGAAGAAAAATCATCCATGATATAGCAAAAGGATTGGCTTATCTTCATGAAGAATGTATGCAAAGAATAGTCCACTTAGATGTTAAGCCGCAGAACATTCTGCTGGACGAGAATCTTTGTGCAAAAGTATCTGATTTTGGCTTGGCCAAGTTGGTGGATAAAGATCAGAGCCACATAGTGACCAGAATTAGGGGAACCCCTGGATACTTGGCTCCTGAATGGTGTAGTGCATTCATTACAGAAAAAGCAGATGTCTACAGCTTTGGAATTGTTGCAATGGAGATCCTCTGTGGACGCAAGAACGTGGACTATTCACACTCACTGGAACATCCACATTTGCTTAGTTTATTTATGGAGAAAGCCGAAAACAATCAGCTTATCGATATGATGGGAAACTACAGCGATGATCCACAATGCAATACATCAGAAGTCATCCGTATGATGAAGCTTGCTGTCTGGTGCTTACAGAATGATTTTACTCTGAGGCCTTCCATGTCAATGGTGGTTAAAGTAATCGAGGGGACAATGGACATTGAATCTCAGTTAGACTATAGTATTCCAAATTCACAAACAATTGCAGCAATCAAAAGAGTGGCAGACACCACGACTAACTTGTTTCCTTCAATTCTTTCAGGACCTAGGTAAGCATTTCTAAGCATGATGAAGAGTCAATAGAGAAAAGAAAATCCCCCCTCCTTTATCTTATTTCTTGTCTGATCCCATGATCATCATTGTTGTATAGCTATTTATTGGTTTCATTTGTGTTGTTTTTCTGTGACTCAGTAAGATACTTCAATTAATTAAGAACCATGGCGTGATTTCCA

>PGSC0003DMG400028617 G-LecRK

GTCACAGTAAAAGTTCTACAAACAATATCATCGAAACTTGGGAACAGATACTGGAATGTCAACCGAGCCTCTTGTACGCAGGAGAGGGATTTTAACATGACGGATCCTAATTATGATGAAGTTTTCAGCAATGTAAAATGTAATTGTTCATTCAATAATGGCTCAGTTTGCCATGTTGTAACAATGTAAGCTCTGCTTTGATTTTTAAAACTTCACATTTTACTCTTAATCAGAAAGTTTTACAGTTAAACAGATGCCATCTTCTGCATCATGTAATGAGAATTTCTTACACTTTTTTTGTTGATACAGCCAATTGAAGCGTCTTAATATGACAGGAATTCTGCCTCCAGAATTTGCAAATCTTACTCATCTGCAAGAATTGTGAGTATCAGTTATAGATTTTTCTCCCCTTTCCGATTACAGAAACTTTTATTCAATTGTTTTTATAAAATCGTAACAGTAGTAGCAAGTTAACTATTTGTTGCTTTTATACCCTTCGAGGTGGCCTAGTGGTCTGGGCTTGGGAATTCCATGTCGGAGGTCTCAAGTTCAAAACCCCTTGTCAAACGTAAACAAGGGGTTTGCCTTCTGGGTCGAGCTCGTCACACCAGGCTTGCCTAGTGCGGGTTACCTCTCCTGTGTGCATAAGAGTGGGGGTTTTACCTGTGGCTTAATCAATTTTTGCATTTCCAGAGATCTGACTCGTAATTACTTAAGTGGATCGATTCCCTCAAGCTATGGCCAGCTTCGCGTCACTAATTTGTAAGTTCTTTTCTGTTTAATCTTTTCAATGAAGAATTTTTGACATTTTGTTCAAACTCAGGTCACTCTTGGGTAACCGTATCAGTGGGCCGATTCCGAAGGAGTTAGGTGACATTTTCACACTGGAAGAACTGTAAGTTGAAGATATTGCATACTTCCTTTAATTTTCTACATATGTCACATTCATGAATTCCATATTTTCAGAGTTTTGGAAAATAATCTACTTGAAGGACCTCTTCCTCCGAACCTTGGTAGCTTGAACCGCCTCAGAAGAATGTAAGACATAGATTCAACGTCTAATATCTACTTATTGTTACATCTATTCCAAATTATTTGGCACTTTTCAATTCCCAGTTGTCAAACTAAATTGCATTTCTTTCACCGTTTTTATTTGGAAAGAACTGTCAAGTTCATGAGGTTAATGAAGTATATCCTACCCATGATAGAACGATTTTACTCACTAATGTATTGGTACCTAAAATAGTGGTGTCAACGAACCACTCAACAAGAGCAAAATACGCTCAAATTTATTTGTGCAAAAGAAGAAGAAGTTCAGAAAATTCGTAAGGAATAATCTGAGGAAATAGATTTATTTATAGCCATAGAAGTACTGTTTCTGACAGTTTGCAACCTTCCAGAATACCTTAAGAGAAGTACTATTTCGGAAAGTTTCCTTAAGATTGTTGTTTATTTGTAATAATAGGATACTTAGAACAATACTTTGCAGAACTTCGTAAGAAACATCACTTCATATTCCCTTCATTTCTCATTTATTATTATTATTATTATACCCAACACTTTTAATATGAAGAAAGAGTTAATCGAGTCTAATTTAGCTCCAAAAGTTGGTCAAATTGTATCAAACATAAGCAAAAAGTGTCGATCAAATTGGGATGAACAGTGTGATTTTTCAGTTATCATGGTTAATTGATGGATCATTTTGCAGGTTGCTTTCTGCCAATAATCTCAATGGAACTATACCTGAGAACTTCAGCAACCTTAAGAACATGAGAGATTTGTAAGCAAGAAAATGTTATAAGAACATTTGACTTTTGTGCTTCAATCTCTTAACTCTTTCGTCTGTTTGCAGTAGGATCGATGGGACTAGCATATCTGGAACCATACCAGATTTTATTGGCAATTGGACCGAAATGGATAGATTGTAAGTATGCATCTTGTAGTTGTAAATGTTGATGGAATATTCTTCTCCCGGAAGGATTCTAAACTTTAAAAAAAGCAAAAAATTGGTATCATACATTGTATTTAATAGCAAATATCATGAAACTTTTTGCAGAGACATTCAAGGAACGTCAATGGAGGGACCTATTCCTCCTACAATATCCCAGTTGAAAAACATGACACAGTTGTAAGTACACACGTCTCTCGTACTACCTGCTATAGCGGAAGTTCTCATGTGCATTGTTGTTTTCAGGACAATATCTGATTTGAGAGGAAAACAAATGCAATTCCCAAATCTTCAAGGCTTGAGGAATATGACAAGGCTGTGAGTAGTTGGAAATTTAGCTATTACATCTCTTCATTTTAATTTCGTATTCTCATGGTTCTTTAGTTTGACTCAGGACTTTGAGAAATTGCTCAATTTTTGGTCCGATTCCGAGATATGTAGGTGACATGCCTTTGAAACTTTTGTAAGTTTGTCATCTCCTCTTTCGATTTTTTTTTTTTGTCTATAATACAAACCTGGAATTACCCTATATTGTATAAAAGGAAAATCTTCTACATACCAATGCATTCGTCGTATTTGCATGATGGAGCAAAGATTTTCACTAATTGTTTGGATACCACAGTGGTTTGCTTTGCTGTCTAGTTTGATGCTAAACCTGCCTGCTTCTGATAACAGTTATTTAATTACTTGCAGGGACCTGAGTAACAATATGTTGAATGATACAATCCCAGATACATTTGAGCAGTTGAATTTTGATAACATGTAGGTACCTTGCAAGTGAAACTTTTACAAGTTGAAGCATAAATCTTGTACATGTTTGTTTTATTTCTTTTCTCTCGAAGTTTTCATTCGGTTATAACAAGCAAAAGTTTTATCTGCAATGTAGGTTGCTTGGTAACAACACATTAAGCGGAGCAATTCCCAGCTGGATGTTTAGTAAAAGAGAGAACATGTTGGTATTTGCTCAATTTATTCCGTTTTCCTCATCTTAGTAGTAAACTTTGGTTATTATTATAGTTAGCCACACTTCAAAGTTAACAGTTTTTTTCCCATAGCTTTTTGTTTACTGCAAGGACACATTTACAAGAATTTGATGGCGTCGACAACTACAATTTTTTGCAGTGATTTATCATACAACAATTTCACCCATACAACTACTCCAGGATGCCAGTCCTCTACCATGTGAGCATTCATTTATTATTGCTTTTTTTATTAGTGAGAATTTCGGTCCTTTAACTACATAACTTACTAACATTCTTTCTTTAATAGAAACTTAGCCGCCAGCTATTCAGATACAATGAACACGTTCAAAAACATGTAAGTCCTCCTAAATGTTGTCTTATGCATCAGGTGCTAGTTTGTTTGATGCCTTCTATCTATTGTCGGCTTTATCTATTCGAAATTGTAGGGATGCCTGGTGTTCCATGAAACCCCTCATTTGTCCCACAGAAGCGAAATGTAAGAGTTACATCCTATTAACCTTATTTTTTTTATACCTGAAAATTAATAAATGAAGTAGCAGTTTGTCAAAGGACCTCAGGGGTTATGTCTGGATCGTTTACCACTGAATCGACTCTTGGAGTGGCAGACTGTAGAGTTTGAAGTACAAAAGCATAGCCTTGGAGTTGAGAGATAATTTCACAACCCTTCCTCATGTTATGGTCTTTATGTCAATTGCAGATACTTCGCTATTCATAAACTGTGGAGGAGGTAGAACTACATTTGAGGGAAATGATTATGAGGAGGACAAGATCAGTTCGGGTCCATCATATTTTTCTTCGTCCTCCGATAAATGGGCATTCAGCACTTCAGGAGTGTACGTGGGTCGAGTAAATGCTAGCTATATTGCAACAAATACATTTTCACTAGATGTGTCCGGTCCTGACTTTTACAACACAGCACGCCTTGCCCCAAATTCTCTCAAATATTATGGGCTTTGTCTACGAGGAGGTAGCTACAGAGTGCGCCTTCACTTTGCTGAAATAATGTTTTCTAATGACTCAACATATAGCAGCCTTGGAAGGCGGATATTTGATGTAGCAATACAAGTAAGTTTGAGGCCTAACTTCTTTTGTAAATAATAAATATGTAATCTAAAATGTTGATTTATTCAGTTCAGATATGATTTGGTCACTTGATGTAGATATACTGAGTTCTGATATGATTGGCAACTTTCTGATTTAATATTGAATGTTGGAGATAATTTTGATTTCTTACACAGGGGAGAGTAGTTTTGAAGGATTTCAACATCATGGAAAAAGCTAATGGTGTTGGAAAAGGTCATACCGAGGACTATGATAATATTACTGTTAGTAGCACTCTGGAGATTCACTTGTATTGGACCGGGAAGGGGACTAACGCCATTCCAGAAAGAGGTGTATATGGACCTCTGATTTCAGCTATCACAGTGACACCAAGTCCGTTTATCTCTCTCTGCCTATAGACTTTACACTTTTCTTCCCTCCCACAATCTTATCCCATTTTCTTCTTGCAGACTTTAACACAGGCAACGGGTTACCTGTCGGAGCAGTTATTGGCATTGTACTTGCTTCAATTGTGGTGGTTCTGATAGTGTTATTTGCTCTGTGGAAGAAGGGCATTTTTGGAGGGAAAAATAATGAAGAAGAATTAGGTAAACATACCTCAGAATTAATTAATCACTCTCTATATTTGATATGGTGATACTTTTGTTGTGATAATCAGAATTATTTACGAGATTTCTAGCTGCGACATACAAATAATTATCTGCCTGTAATTGACCAGCATTGAGATGATTGTTCTGTTGGTGGATCGACATCATGACATATCATAGATAGTTAGCACACAACTAAATGCTCATGACTGCTTAAACTATATATCTTCGTCATACAGATGTTTTAGAAGATCTATATATACTGACGTTTGCAATATTCTTCTCGTTTACAGAACTCAGAGCCCTTGATTTACAAACAGGTCATTTTAGACTTAGACAAATTAAAGCTGCTACAAATAACTTTGATCCTGCCAACAAAATTGGTGAAGGAGGGTTTGGACCAGTTTACAAGGTATCAAATCTCTATCTCTCTCTACCCATACTGATTTTGCGCAAGGATAGCTTGTGGTTCTGTGGTGATATTTCCTACTGTGCGTAACGTTTATAGGAAAATAGCTGATATAATCCATATCCTACGTTCTTCCTTCCGTGTGTGATATGATCGTTGTGAATTAATCTTGCTATCTGTAGGGTGTACTCTCGGATGGTGCAGTGATAGCTGTTAAGCAGCTGTCTTCCAAGTCGAAGCAAGGAAATCGTGAATTTGTCAATGAGATAGGCATGATTTCAGCTCTACAACACCCAAACCTTGTCAAGCTTTACGGTTGTTGTATAGAAGGGAACCAGTTATTAGTGATATACGAATACATGGAAAATAACTGTCTTGCTCGAGCTCTCTTTGGTAATGCTTTGCAATAACATTTTGAAAATTATTAATCATGTAATGATAATTTTATGCTTATTAGTCCTCGTATTCTCCTTTCTTAGGCCGTGACGACCAGAGGTTGAACCTAGACTGGGCAACCAGAAAAAGGATATGCTCAGGAATAGCAAAAGGGTTAGCTTACCTTCATGAAGAATCAAGGTTGAAAATCGTACACCGAGATATCAAGTGTACCAATGTGCTTCTTGATAAGGATCTTAACGCTAAGATTTCCGATTTTGGATTAGCTAAATTAGATGAAGAAGAGAACACTCATATTAGCACCCGAATTGCTGGAACAGTGTGAGCCTTCTTATCATAACTATCTCTGTACTATCATTTGAATTTGTTCGTCATGGTCAGTTTGAAGTATCTTACATTTCTTTTTCCGTTTTTGAAGAGGTTATATGGCTCCCGAGTATGCAACGAGAGGCTACTTGACAGATAAAGCTGATGTTTATAGCTTCGGAGTCGTTGCATTGGAGATTGTCAGTGGGAAAAGCAACACAAACTACAGGCCAAAAGAGGAATTTGTTTACCTTCTTGATTGGGTAAACTATCTTTCTACTAATATCTGCAACATTATATCTTTAGTTTAAGACCACAAGATGCAAATGTCTTTGTTACTTTCTTAAACTCTGTACGGAGGTATCTTTGCATATTTAGTTTTTCTTTTCTCTAGCTAATTTGATGTTTGCTATGTATGTGAATGTTTGTTTTAGGCTTACGTCCTACAAGAACAGGGAAATCTGTTAGAACTAGTGGATTCACATCTCGGTTCCAATTACTCCAAAAAAGAGGCAATGCAGATGATAAATATTTCTCTTTTGTGCACAAATCTCTCTCCTACTCTCCGACCATCCATGTCTTCTGTGGTGAGTATGCTCGAAGGGAAACTCCCCGTGCAAGCACCAATAATCAAGCTGACTACATCAACCAACGAAATGAGATTTAAATCGTTTGAAAAAGTTTCACATGACAGCCAAACAACATACTCTCAGGAGTCTCAAGGACAAGCACCTTGGAATGATTCGTCAGGGTCAGGGTCTCTGCCTGCTAAAGATGAAAAGTTTAGTTCCTCCAGTAGACTTCTTCCAGATCTTTATGATATAAACATAGATTGATTTTGATTTTGATTTTCACACAGGATCAAATATTTATTTTCTTGTGTATACTACTTTGTAA

>PGSC0003DMG400029520 G-LecRK

CCCCACCCCACCCCCTCCTTTGAATTTTCTGAAAATTCCACATAAACACGGTTCCTTAATGTAACAAAGAACCTTAAAATTTGTAAGGACAAGTTATAATTTTCTATTGTTTTATACTTTGTACAAAGTCCTCTTGAGTTTTGTAATGGACTAATTATTTTTGTGTACCCGCTATGACCAAACCCAGAAAAATTCTTGAGAACTTTTTGAAACCCTTTTTCTCAAGTTCCAACAAAGGTAAGCAAAAATTCTTGTGAGGGTACCCTTAATCTTGATTTTTTTTTTTGTTGTTCTAATGGTGATTTTGGGTTTTTTTCTTGTTGTGAATTGACTAATCAAAAAAAGGGAAGAATGACGACTTAGAGAAAATTGCAGCACAAGAACAGAAGCAATTCTCTTTTGGAGTTCTTGTTTCTGCTACCAAAGATTTTCACCCAGATAACAAGCTTGGTGAAGGTGGATTTGGGCCTGTATTCAAGGTAAAAAATAAAAACAAACTGAAAATCAACAAAATTGAGTCTTTCTTGTGATTTGGGTGTTAACAAAAACTCAAGCTTTCTTGAAAAGATTGAGTCGTTTTGTGATTTGGGTGTTAACTGAAACTCAAACTTTCTTGAAAAAATTGAGTCTTTTTGTGATTCTTGAAGAAAATGAATCTTTTTGTGATTTGGGTATGAACAGAAACTCAAGCTTTCTTGCAAAAATTGAGTCTTTTTGTGATTTGGGTATGAACTGAAACTAAGGCTTTCTTGAAAAAATTGAGTCTTTTTTTGATTTAGGTATTGAACAGAAACTCAAGATTTCTTGAAAAATTGTGTCTTCGTTCTTTATAGGGGAAATTGAGTGATGGGAGGCAAATAGCTGTGAAGAAGCTTTCACATAGTTCAAGGCAAGGGATGAAGGAATTCAAGAATGAAGCTAAGTTGTTGGCTCGTGTACAACACAGAAATGTGGTGAATTTGTTAGGGTACTGTATACATGGGGTAGAGAAGTTGCTTGTCTATGAGTATGTTGCTAATGAGAGCCTTGACAAAATCCTCTTTAGTAAGTTTTATTACTACTAGTTTTTATCTTCTTAGCTCTACTAGATTTTGTATGTGCTTTTAAAATGAAATGAAATGACATTCATTGAGAGGCGTAGCCAGGATTTTTAGTTTATGTGTTCTAGGTCATAATTGTGTTTGTTTACTGAGTTCTGGATAGATTAGTCGTACTTATTAGATGAATTTTTTAATGCAATAACAAGGTTTGAGCCAAAGTTACCAGGATCTGCAGAACTAGTAGACACCTGTGGCTCCGCCCTTGTCTTCACTATCCAAAGATTAAAACAAATTGAACTGCAGATGGATGCTCTTGTCAGTCTTTTTAGGAGTGAGATACTGGAAATAGCCTTGGGATAAGCATTTTCATTTTGCTATTGGGTCCAGAAATTTGTTCTTTCAGTTACGTTGTCAGTAATCTAGTTCAATGATCTCTGTGTCAGTCCGTGAAACTACTAGAGACCGTCATATTGTTCTGTATCCGTGTGTTTTACTTGCTTAACAAACTCTAAGCTCAAATGTTCCTTGAACTCCAGTGATGGACCATAATGTTAAAATTAAATTTTCTCTTGACATCCAGCAGCTAACTTGTGGCTGTTCTTTCTGTCGTCTTTCTTCTTTTTTTGATAACTTTGGTGTTTGGGCCAGCATGTGTGCACCTCGACTATTTCATGGAATACTTTTATCCTCCTATCAATACATGTATCATGTAAATCTGTTCACCAAGGGTCAGACAGATGAAAAGAAACTACATAGCTTGTTTTTTTGCCTTTGCTAGGATTCGTACCCCACCCATTGTGTTCATTCCAGCTTCATCAACCTTAGGCGGCCGCAGTATGTTATTGAAACTTCTTCTGTCGGAAACAGCCTCCCATTCCCCACTAAGGTACATCATACACTCCCTAGACACCACGCGTGGGATTACACTGGGTATGTTGTAGTAGTATTTGTAGATATTGAAACTCCTTGTACTTCTCAAGAATTCTGTAACCCTGAAACATCATTCGACGACATTTGAGGACCTTAACTTCCTCGTGTAAAACTTATCAGTAGAAGCAATTTGTGATTCCAAGCCCGAATATAATATAATTCTTCAAGCAAGAAATTTTTCTCATAGTGGTCTTAGTAAAACCAGACTCTTTACTTTATGAATCTTTTTAACCTTTCCGAGTTTTGAGCAAGAAGAAAGATGTTTAGTAGTTTTAGAAGCTAGTACTGACTCTTATATTCTTGTATTAGAATCTGCTAACAGAGATGCACTCAGCTGGAAGCAACGGCATGACATAATTGTTGGCATTGCTAGAGGGTTGCTATATCTTCATGAAGGTTCACATACTTGCATTATCCACCGTGACATAAAAGCTAGCAACATCCTACTTGATGACAAATGGGTGCCCAAGATTGCTGATTTTGGAATGGCTAGACTCTACCCTGAAGATAAAACACATATTAATACCCGTGTAGCTGGTACCAAGTATGATTCTATCGAACCTTATAAACTGTGTTTAGTAATGACTATGTAGAAATTTCATTGAATAATATATTGCACAAGTACACTCACTTGATATTGCTAACAACTCTCTTTGATAATTTAACTTGGAAGCCAGTGGCTATATGGCGCCAGAGTACGTTATGTATGGACATCTATCTGTTAAGGCTGATGTATTCAGTTTCGGAGTTGTAGTTCTGGAGCTCATAAGTGGCCAGAAGAATTCAACCTTTAACAGAGATCCTGATTCTCGAAGCCTTCTTGAATGGGTAAGAACTTGATTTCTCTGCGCTTTTCAGTCTTGATATTCCTTGAAATTTCTGAATTACAAAAATAAGCACCGGTCAATGCCTTCGTTCCTGTTGAGTATAATAGGAGAGTCCTCGGAATCAATTCGAGCTTCCTATAATCATGTAACAGAAAAAACAATATAGCCATATGGTAGAATTTTTTAACTACTTGAAAAGTAAGATCGAACTACTTTTGTCTATGTTATAGTGCAAAACATAATCAAATCGGTAGGTGGAGGGAGAAGGACTTTACATTTAAGGTATAGGGATACTCTTGATGGTGTGAGGCCTTTTGGAGAAAACTTTTGTGGCTGTGACCCTAAGTGGATAGTAGCACACCATGTAAAGGTATTTTCTAGGCCGACTTAGCCTATAAGCAGATAGTAGAGCCAATTATTTGACGGGTTGAGTATGGAGATGAAGGCATTGAGTGAGTTTGCTGCTAGTCGTGGTAAATGCACAAGAAAAACACTATCAGTCTTGGTGATTGACCACACGGTACTAGGTCCGGGGAGTGTGTCGTTCTCGGGTGAATCGTGATTCGAGAGAATATTTGTGGAGATGCGAACAAAGTCTCATATTGGTAGTCGAAGTAGAAAAATAGCTATATATAAGGTGTTGGGATGGTCTTACGGTGTGTGGGGTAAAAACGTGCATGCTTGGCCCGAAACAACAGTATTAAACCATGTTTAAAGTGTCTTTGAGCGGGGTTAGCCTAACTTTGTACCAGATGGTGATCATGATTAAATGTTGTTATGGTTGTAGGCGTACAGGCTTTACAAGAAGGGTCGGAGCTGGGAGATTATGGACCCTGCACTGGCACAATCGGCTATCCCCGATGAGATAGCAATGTACATACAAATTGGCTTATTATGCATTCAAGCGGATCCACCACTACGTCCAACCATGCAACAGGTCGTGGTGATGCTATCAAAGAATCCTGGATCTCTTGATAGAGAACCAACTAGACCTGGATATCCCGGTTCAAGAATTAGATCTCACAGACCAGGTGGTGGTACGTCTCATACAGCTGGAACCTCGGGTGCATCCAATCCTTCTACATTCAGCTCCTCGTCTATTAGCAACACTGTTAGTGCAACTGCAAGTTCATCCACACCTACAAGACGGAGATCGGACCCTCATGGAAAACGTCCTGTGAGAGATTAGGCGCGTTGTTGTTGGGAGATTTTTTCTTATTCTTTTTGTGTATACATTCAATTTTTGTAGCATGTAAATTGGTTACAAATAGAGATCAATGAGTATGATGAATCTTAGTGACAGTTAACATGTAAAAGAAGTGTATGCAGTTACTCTCAGGATATAAAACTACGTGGCACAATATATAAATGTGTCCTTTAATTTGGCCTCGTTTTATAATTATACCCTCTAACTTTAGGTGTGCACAAGTAAACAACTTAACTTTTATAAAATTGAAGAAGTAGACATGAGTTTTACGTGGCATGATATACATTGGACACCATGTAGGATGAATATTGTCACGTCAGACATGTGTGACTATTTGTTTAATTTTATGCAAATTTAAGTGTTTACTTGTGCACACACAAAGTTGGAGAACATAAATGTAAGTTGAGGTCAAATTAGAAGGCATATTTATA

>PGSC0003DMG400030209 G-LecRK

TTTTGATATTTTCTTTCAATTCAAATTAATAAACCAAAACGCCCAATTCTTGGCCAAAAATGAAAATTCAACTCAGAAGGAGGCCAAAGTAGAAAAGTAAAGTCAAAGTCGACACTTCTTCTTCCCACCTTCCCCTGTTTTCTCCACCTCCACAACAATGGCACACAACTCATAAATCCACACCAAATACCTCATTCTTCTTCACCCATTTCCTCCAAAAATGAGACTACTATTCATACTTTTCACCATTCTTATCATCACCATAGAAGCTGCAGATATTCTTCCTAATTCTACCCTTTCAGCTTCCAACCCAAACTCCAAATGGTCATCTCCAAACAACACCTTCTCTCTTTCCTTCCTACAGCTCGACCCCACAAACCAGAGTTCCTATTTTGCTGCTATTTCTTACAACAATATACCCATATGGAAAGCCGGCGCCGATACCGTTAACGGTGGAGCAGTTGATATCTCTGGTGAACTACGGTTCCTCTCTAACGGTGACCTTCAGCTCATAAGTGGGTCTTCTGGTTCTGTTGTTTGGAGTTCAGGAACTGTAAATCGTGGTGTTTCTTCAGCTTCTCTTGATGATAATGGAAATTTTCGTCTCAAAAACGGTACGGTATCGAACATTTGGTCTACTTTTGATAACCCAACTGATACTATTGTTCCTGGACAGAATTTCACTAATAATCATGTTTTGAGATCTGGGTTGTATTCTTTTAGCCTTATGAATAATGGGAACTTGTCACTTTTGTGGAATGGTTCTATAGTGTATTATAATTCTGGGTTGAATTCTTCTGTAAATGTTAACTTGTCTTCGCCTAGTTTGGGAATGCAGCCTATTGGGATTCTTTCACTTTCTGATCCTAGTCTTTTGAACCCTGTGAATGTTGTTTATAGTAGTGATTATGCTGATGAAGGGAACATTTTAAGGTTTTTCAAGCTGGATAATGATGGAAATCTTAGGATTTATAGTTCTACACAGGGTAGTGGGACTCAAAATGTGAGATGGGCTGCTTTAACTGATCAGTGTCAGGTTTTTGGTTATTGTGGGAATTTCGGGGTTTGTAGTTATAATGAAACTGATCCAACTTGTGGATGTCCTTCGCAGAATTTTGAGCTTACTGATCCGAATGATAGTCGAAAAGGGTGTAGGAGGATAGTTGAACTTAGTAATTGTCCTTCCAATGCAACTATGTTGCAATTGGATAATGCTAAGTTCTTGACTTATCCTCCGGAATTATCTGAGCAGATTTTTTCTGCTGGTATTTCAGCATGTAGGTTTAACTGTCTTGTTAATGGTGCTTGTGTTGCTTCGACTTCTTTAGCAGATGGCACGGGGATGTGTTATATTAAACAGCCAAATTTTGTTAGTGGCTATCAGGCGCCGACCCTTCCTAGCACTTCATTTCTTAAGATATGTGGGCCTGCAATGCCTAATCCTACTGCAGTTTCAGACACTGTTCAGGAGAAGAATGGAGGGAGGGTTCCTGGCTGGGTTGTCGCAGTTGTGGTTGTGGCTAGTGTTTTGGGTTTGATCCTGTTGGAGGGTGGTTTGTGGTGTTGGTGCTTTAGGAACAGTTCTAAATTTGCATCACTGTCATCTCAATATGCACTTCTTGAATATGCCTCTGGTGCCCCTGTGCAGTTCACATTCAATCAGCTTCAGCGGGCAACCAAGGGGTTTAAGGAGAAGCTTGGTGCAGGAGGATTTGGGGCTGTTTATCGGGCAGTTCTGGCTAATAGGGCTGTTGCTGCAGTAAAGCAGCTGGAAGGAATTGAGCAAGGAGAGAAGCAGTTCAGAATGGAGGTTGCTACTATTAGTAGCACACACCATTTGAATTTGGTGAGATTAATTGGGTTTTGTTCTGAAGGGCGCCACAGGCTACTGGTCTATGAGTTCATGAAAAATGGTTCTCTTGATAAGTTCCTCTTTTCAGAAGATCATTCATCGGGAAGGCTTTTGAACTGGGAGCAACGGTATAATATTGCTTTGGGGACGGCAAGAGGTATCACATACCTTCACGAGGAATGCCGAGATTGTATAGTTCATTGTGACATAAAGCCTGAGAACATACTATTGGATGAGAACTACGTTGCCAAAGTATCAGATTTTGGCCTCGCCAAGCTCATTAACCCGAAGGACCACAGACATAGAACCTTGACAAGTCATGTGAGGGGTACTAGAGGATACTTGGCTCCCGAGTGGCTGGCCAATCTTCCTATAACTTCCAAATCTGATGTGTACAGTTATGGTATGGTATTACTGGAAATAGTGAGTGGTAAAAGGAACTTCGAAGTCTCTGAGGAGACTAACCAGAAGAAGTGCTCTTTGTGGGCATATGAAGAATTTGAAAGGGGTAATATGGAGGCCATTATGGACAAAAAGCTTTCCAATCAGGAGATGGATATGGAACAAGTTATGCGAGCAATTCAGGTGAGCTTCTGGTGCATTCAAGAGCAACCATCTCAAAGGCCAACGATGGGCAAGGTGGTTCAGATGTTAGAAGGCGTTTTTGAGATTGATCGGCCACCAGCACCTAAGGCTACTGAAGGGTCCTTTGCTGGAACCAGCTTAAACACCAGTAGCACAAGCGGTCTCTCCACCTTTGCAGCCTCGGCCCCAGCTCCCTCTTCATCATCATCATTCCAAACGGCAGGATTCCAGTCTTCAGCTTCTGCAAAGAATGTCGATAGGCAATCATCATCACTTCTACACTCCGAAATTAAGTGAGTCTAAACCATACACTTTAGATTCCCTACAAGAACTGATGGGAGGTTTAGTATCTCAAAGATCATTGTTTGACTTGTGTCTTACAATTCATATTGTATAATAGATAGGGTTTGCATATATATTGATATGTTTCATATTAGAAAATACCACTCACTTATTTTCCATATCCTATTATTCACCAGAATTCGACGACTCTGTCCTGTCCTGTTTGTGGTGCTTGTCCCATTTCTTTTGATAATGAAGCTAGGGGTGTGCATCAGTCTGGTTCGATTTTATGTATTATTAGTTCGATTTATCGATTTTCGGGTTGGGGATGATTTGGTCAGGGGGTTGATGTGAACTAGAAACATGGCTATAAAATAAT

>PGSC0003DMG400030225 G-LecRK

ATGTCTAGTAAAGCTGGTCATAATGCACCTGTATTTGAGAAGTACCAGCAGAAAGCAAAGAACTTTATGTGTTCATGTCTTGCAAAGGGTGACAGAAATACCCAGAAAACTCCTGGAGGTCTCATCTATAGGCAAAGATGGAACAATATGTATTTTGTTACTAGTGCTGCATTCCTTGGCACTACTTATTCTGACTATTTGGCCTCTGCTGGTAAATACCTCAAGTGTTCTTCTGGTTCTGTTTCTCCGAATGAGCTTCTCTCATTCTCTAAGTCTCATGTAGAAATTTTAACTTTGATAGAGTGCTACAAGCCTGTATTATGACGATTCATCATAAATTAAAGCATGGAACTAGCAATGTGTTCTTGGTGAACTAATCATCATGTTTGATGTTGTTCTAGGTGGACTACATTCTTGGTGACAACCCGAGAGCAACAAGTTACATGGTGGGATATGGAAACAATAACCCAAGACAGGTACACCACAGAGCCTCCTCCATTGTCTCGTTTAAGGTTAATCCTAGTTTTGTTATCTTTCGGGCAGGCTATCCTACCTGTTATAGTAGAAAGGCGAGTGATCCTAATCTCCTTACTGGTGCTCTTATTGGTGGACCTGATGCCTATGACAACTTTACTGATCAAAGAGATAACTATGAGCAAACTGAGCCTGCCACTTACAACAATGCTCCGTTGATCGGTGTGTTGACAAGACTTCATGCTGGTCACAACGGTTACAATCATCTCCTCCCAGGTACATTTTAGTGAATTAACATCTTTATATGAGTATTTTGAGGATTCATCATAAAATGACTAGTTTTAGCTAGCCATCCACCTACTACAACTTGCTTGGGATTGAGACGTAGTAGTTGTTGTAGTACTAAATTATTCGTATAACTCATTTAGTCTCAAGTTCTAACCTGTAATGGAGAAAATCCACTCAATGACACATCCAGTGCGCTAAATTTTACCAGACAAGGAATTCTTACTCTTCTAAATGGTTTTGGCAGAGTAATTTGGTCCTCCGATTCCACTAGACATGGGCAAAATCAAGTAGCACAACTCCTTGATTCAGGCAATCTTGTTGTTCGAGATGCAGCCAAGGATAAGCCTCAGAACTACCTGTGGCAAAGTTTTGACAATCCAGGTGACACAACTTTACCTGGAATGAAGGTTGGGATTAATCTGAAAACTAGTTTTCATCGTTCCCTTTCATCATGGAAGAGCACAAATGATCCTTCCATAGGTGAGTTTACCTGGACATTCGATACTGGTGGATTCCTGCAAACATTCATTATGAATGGTTTCATTGAACTCTACAGAGTTAGACCATGGAATGTCCGAGTCTTTCCTAATGCACCATCTCATGACATATCTTGGAATGGTTATAACTATACATACTTATCTAATCCTAATGAAATATTATTAATGTATGAACTGACATGTAGCTCTATCATTGCAAGGGTAGTGATGCAACTAAATAGATAA

>PGSC0003DMG400030553 G-LecRK

AAAAAATGTCTCCTCTCTTTGTTTTTCTCTTTGTACTCTCTTTTTTCTCTCTAAAAAATTTAGCAAAAGCTAACAATTTCACAAGCCAAAATAGTTCAATTATTGGTAACACAATTGGTGGAAACAAATGGATTTCAATTGCCTCAACAATAGTTTCTTCAAGTGGTAATTTTGAAATGGGATTCTTCAAACCAGGTAACAATTCCAATTATTACATAGGTATATGGTATAAAAAAATAAGTAAACAAACTATAATTTGGGTTGCAAATAGAGAAACACCAATTTCCATATATGAAATGGATTTTGCACAATTCAAGATTGATAATGGTAATTTAGTACTCATAAATGGTACTAAACACATAATTTGGTCCACAAATATCAATTTTACAACAAATTTCAACAATTCACAAGTTGTAGCAACACTATTTGATGATGGCAATCTAGTATTAAGTAATCAGGATATTACCAATTCGACGAATTATCTATGGACAAGTTTTGATCATCCCTCTCATACGTTCATGCCTGGTTCAAAATTCGGTTACAACAAACATACAAGATTTAAACAGGTACTTACTTCATGGAAAAACGTGAACGATCCATCACCAGGACCCTTTACACATGAAGTTTACATGGAAAATAAGTATATTGGACAAGGTGTTAATATGTGGAACCATAGTGTTGTTTATTGGAATAGTGGACCATGGACAGGTAACAATTTTACAGGGGTACCATATCAACCAAATCCTATGTTTAATTATACATATGTTAATAATGATGATGAGGTTTATTATATGTATAATTTCTTTAACCCTTCATTAATATCAAATTTTATTATGGATGTTAATGGACAAACAAAACAAGTATTATGGATGGATAGTACTAATGATTGGAATGTGTTTTACACTGATCCAAAACAAGTTTGTGATGTTTATTCTTATTGTGGAGGTTTTGGTATTTGTAATGAGGTAAACTCTACTTCAACATGTGATTGTTTGAATGGTTTTAAGCCTAAGTTTGAGAAAGATTGGAAATTGAGTAGTTTTTCAAGTGGATGTATGAGGAAAACAAGTTTGAATTGTGGTGATTTTGGTGAAAAGGATAGGTTTTGGATGTATAAGAATATGAGATTACCTACAAATAATGAGAGTTTGGGAGTTGGAAATGAAGTTGAGTGTGAAAGTGGTTGTTTGGAGGATTGTGATTGTGTTGGTTATGCTTATGGTAGTGGAAATATTGGGTGTTTGGTATGGAAAAGGGAGATGTTGAATTTGCAACAACTTTCTCAAGATAATGTTAATGGAAGCACAATTTATGTTAGGCTTGCAGCTTCTGAGTTTTCAAGTAACCAAGGTAATGACTTGCAATTTCTCCTATTTTTTTGAATTTTAGATGCATGATGTACTTTATTTTGTTTTTTTGGTTCTCGTTAGGTGTCTGATATCTGTTTTGGGATCTGACTAAGATGTATTCGCACCACTTTTAGGGATGGGGTGGGGGAAGGGGAAGCTTTTCCTACCGAAAACGACTTCAATCTCAAAACTTGAACTAGAGACTAGAGACATCTAGTTGAGGATAGATGAATACTTAACAGTTACCACTCCACCACAACCTTTAGTAATACATATCTACAGTTACCATTCCACCACAACCTTTATACATATATATATACTTCAAATTGTTACCTTTGTAGGTAGAGTGGATTTACCTAGTTTATAAAGTTTCTTTTGAGTCCAAATTTGAATTTCATTAATTACTTCTCCTCTTTTTTTTCTCTATTATCAAGAAAATGAAAACCAAAAACACATATTTTCTTTGTTTAACGATCCAGTATCTGAAATTTACTGGCTTGATTAATTTAGATTTACATCGGGGTAAGCCTAATTAAGGTGTTAAATCTCTTTCTATTAATTCATTATTGAATTCGAGACGTCTGGTTAAAAATGGAGGCCATCCCCTCACAATTAGTGACACATTAAAACATAACATTTTTTTTCTTTCTCTCTTTTGTTATTTGTCTATTTATGAACTTGTCATTAATCTTTTCATTTTACTTGGTAATAATGTGTTCTTTGCTCTCTTCATTAACAGATCAAAAGCAAACTTCAACAAAGCTGAAAATTGCAATTCCTATTGGTGTAATTGCTGCACTTCTTATTCTATCATGCTTTACCTACATATACTATAGAAAAAGAAGAAACTCAAAAGTCAAAGGTAACAACAAATGTCTCTTTTTTTTAATGACATTTTTCTCCTTTTTATTAATAAAAATATACAAAATTTAGTATTATCAAGACTCTAAACTCATACATTTTTTTAATTTTTAAATATAGAGAGTACTACAAAATTTCACAGACAGAATACAAAAGGAGAAGGACTAGAATTGATAGATATTCAAGATGATGATATTGAGGTTCCATTCTTTAGTTTTGAAAGTATATTAGTAGCAACAGATGACTTCTCTGAACAAAATAAGCTTGGTCAAGGGGGATTTGGTCCTGTTTACAAGGTATTTATGCAAATATCATTTTGTAATTTATTTTTATATGTGGCGCACAGTTTGAAATTCGATTTGTATAGGGATAGAGGTACAAACAGGTTTGACAGAACCCAATAATTTTGGTTCAGACCTTATATTTTTGTTTAAAAGATCGATGAATTAGTATAAATATTTTATCCAGATCCCAATAAGCTCTCTTTTCAAAAATCTGAAATCTATACACTCAAAATTCTAGTTTGCGGCATGTACTACTTTTCTTATTAGTTGTCATGTCTTTTTTACTGCCACATTTCCCTTTTTCAAACTACTTTGAAATGCTTTTCTTGAGTCTGAGCGTCTATCAGAAACAATTTCTCTATTTCCATGAGGTAGGAGTAAGGTCTATGTACCATCTACCCTCTCTATACTCCACTTATGAAATTACACTACTGAATATGTTAGTAGTATTATATTGTTTTTGCTGTGGTGGCCGAACCTGTGACGTGCACCTAACCAACATAATCACTTTTTAAATAAATATATTGTATCACAGTATTTATTTTTAAAAATAAATAATTTTTTTGTAGGGAATATTTTCTGGAGGAAGAGAAATTGCATTAAAAAGGTTATCAAGTCATTCTGGACAAGGCATAAATGAATTCAAGAATGAAGTAATGTTGATTGCAAGGCTTCAACATAGAAATCTTGTTAGACTTTTGGGTTATTGCATTCAATCAAGTGAAAAGATTTTACTCTATGAGTATATGGCAAACAAAAGTTTAGACACCTTCATATTTGGTTAGTATTTTTATTTTTTAACTAATATCTTTATACTGTCGATGTATATAAGTTAAATTTATTTGATTGTTTTTCTTGCATAAGATCGAAAACGAAGCGTATTATTGGATTGGAGAAAAAGATTTGAGATTATAGAGGGAATTGCTAAAGGACTTCTTTATCTACATCATGATTCAAGATTAAGGATCATTCATAGAGATCTCAAAACAAGTAACATATTATTAGATGAAGAATTGAATCCAAAAATATCAGATTTTGGATTAGCAAGAGTTGTTGAAGGGAAAATTACACAAGCAAATACAAATAAAGTGGTTGGAACCTAGTAAGTACTCATAACTCCCTATTGTGATGTGACCCTTCTCCAGAATCTGCTAATGTAGGATGTTTTGTGCATCCGAGTTCAGACTGTCCTTTTTAAAATAATTTGCCAATTGTGTTGATTTTCTTGTTCATTAATATGTTTTTGTTGATACTATTAATGCAGTGGTTATATGGCTCCGGAGTATGCAATAGACGGATTATTCTCAATTAAATCAGATGTATTTAGTTTTGGAATAGTTATACTCGAGATCATCAGTGGAAGAAGAAATACTGGATTTTTTAATCAAGAAGAAGCGTCAAATCTCTTAGGCCTTGTGAGTGAATAATTACTTAAATTCTTGCTTTTTTTTTGCAACACTGTCACGGAGTTTCAAACTTCATAGGTTAAACTCCAGTACATTGCCATGGAATTTCACCTATGAAATTTGAATCGATCTCCATGACATTTATTTTTCAATTACATGACTGAAAAGTGACTAGTCCGCGCTATTTATTACAAAACTTTATGTTAACATAAATATGTGAACTATAATTTAGGCATGGAGATTGTGGACAGAAGATAAGTCAATGGACTTAGTGGATCAATCACTACATAAATCATGCAACAAAGAGGAAGCAATTAAATTGATAAATATTGGGCTTTTATGTGTACAAGAAGATCCAAAAGATAGGCCCAATACATCAAATATAATAATGATGCTTGGTAGTGAAAATTCCAATATTATTTCACTTCCAAGACCAAATCAACCAGCTTTTATGACAAGAAAATGTGGTAATAATGACACATCTTCTTCTTCTAATGCTAAGTCTGATGGTGTCTCCAACAATCAGCTTACAGTCACCATTGAAATGGGTCGATAACGCACTACTAGAAATCAGTCAATTTTTCATCATGAACATGTTCATTGAAAAATCTATCGAAAATATTTCTGATAGAACGTCTGTTGAAAATTGACTGATTTAAGACTGAGATGCCAACAATTTTGACATAGCAATATTGTAGTTTTATCTTTACTTATTCAATGTATATATTCAACTTAAGTATGTACTTTAGAAACATCTTTAGTAATTAAATTATGTATTTTT

>PGSC0003DMG400031860 G-LecRK

CTTTGATAGTACAAACAATTTCTGGTTAGACCAAGAAATATAGTGAGCCATTGTGTATATGAAATCATTTGTTACTTCCTATTGTCTAAACCAAGGCCTTCTGATTTCATACAAGATGAGAATATCTCTTAATCTTCTTGCATTCCTCCTCCTAGTCCTTCATCATCAGTCAAATGGCCAAAATAATCCAAATATTAGCATAGGTTCTACTTTCACTGCTGGAGAAAATGTTTCTTTGTTATCCTTGTCTGGGGACTTTGCCTTCGGATTTTATCCTGTTTCAGATACTCTTTTCCTAGTAGGAATTTGGTTCAACAAAATCCAGGAAACAACACTTGTCTGGTCAGCAAATCGCGACAATCCAGCTGAAATCAGATCAACTATTAGTCTAACAAATGGTGGTCAGCTTGTGCTCAACTATGCTAATGGCACAGTTCAACAGATTTACAATGAGTCTGCAAGGTTGGGAACCATGCAAGATGATGGAAATCTTGTGTTGAGAGACTCAAGTTCAAATAAAGTTTGGGAAAGCTTTAAATCTCCCACTGATACACTTTTGCCAGGACAGACTTTATCATCAGCAGGAAAGCTCTATTCTAATTCTATTTCAGATGGGAATTCCAAGTACTCTAAAGGGAATTTCATGCTAGAAATGCAACTTGATGGAAAATTGGTACTATCTGCTTATCATTTTGGGGATACAGGATATTGGCTGAGTAGTTTAGATGATGAAGGAAACCGTGAAGTGAATTTAGTTTTCGACCAGAGGAATGCATCTTTGTACCTTGTGAAAGATAACAATAACACAATTTTTTCTTTTCCATTGAATGTGCCAACCCCAGTTGAAGACTACTACCACCGTGCAACTATAGACAGTTTTGGCAACTTCCAGCAGTATGCTTATCATAAGACTAATGGTAGCAATTGGATTAGAGTTTTGAAGATCCCTTCAGAGCCATGTATGGTTAATGCTGTTTGTGGGGCTTATGGATTCTGCAGCTCAAATGATAATGAAACTGTAAATTGTGACTGTCTTCCGGGGTATGTCTTCTTGGATCCAAGTAATCCTACCAAAGGTTGTCATCCAGAGACAATGATCAACTTCTGTGCAGGTCATCTATCTAGCGGAAATTTCACAATAGAGGCTATTCAGAATAGTGACATGCCTTACAATGAGGTTGGGGATTATGAAATTTATCGTGATGTTGACGAGGAGGAATGCAAGAAAGTTATCATGGAAGATTGTTATGCCATGGCTGCTTCTCTAATCAATAGGGAATGTCACAAGAGGCGTACTCCTATACTCAACGCCCGTAGAATTTCTATGACCACAGGTAGTGTTTCCTTCATCAAGGTGCCTATTAAGTCAGTTAAAGATGACATATCAACAAAAAAGAAATCCAACACCAGAGCCCATCTCACTGCTGGCCTTATAACTACCAGTTCACTTACTTTCCTATTTGGAGCCTTGGCTTTTTACTATCATCCTGCTCCTTGGAGACTTTTAAAACGAGAGTGGAATCCAAATTCATCAAGAATAGGTATCAATTTCCGCGAGTTCACATACAAAGAACTGCATGAAATTACAAATGGCTTCAGCAAGCAACTTGGAAAAGGAGCTTCTGCAAAAGTTTACTATGGTAATCTAAAGTTAAAGGATGTTCAAGTAGAGATTGCAGTGAAGCTGATGAAAGACGTTGCAGAGCCAAGTGAGAACGTATTCATGACAGAGCTAAAAATCATAGGCAGAACTCATCACAAGAATTTGGTAAAACTATTAGGATTTTGCATTGAGGATAATCATTTCATTTTGGTTTACGAGTTGATGAAGAACGGAGCTTTATCAGATTTTTTGTTCAAGGAAGAAATACTGCCTACTTGGAGTCACAGAACTGAAATGGCATCAGGAATTGCAAGAGGACTGCTTTACTTACATGAAGAATGTGATTCTCCAATCATCCACTGTGACATAAAGCCTCAAAATGTACTTCTTGACAGCAAATACAATGCCAAGATTTCAGATTTTGGGCTATCGAAACTCTTGAAGAAGGATCAAACAAGAACAGATACTTGTGCAAGAGGGACAGTGGGATATTTAGCACCAGAATGGCTAAAGAATGCTCCAATAACACCCAAAGTTGACATTTTTAGTTTTGGTGTAATGTTACTAGAGATCACATGTGGTAGAAGGCATATTGAGCTTAGCAGGGTAGAGGAAGAGAGCGAAGACGATGAAGGTGATGATTTATTGCATGTAAATTGGGTTTCAGGCTGTGTGAGAAGTGGTAGAATTGACAAACTTGCAAGGAGTGATCCTGAAATATTGAATGATATTAAAAGACTTGAACGATTCGTCAAGGCGGGATTGTGGTGTGTCCATCCTGATCCAACTGTTAGACCTTCAATGAAAATAGTGATGCAAATGCTTGAAGGCATAACTGAGGTTGGAGTTCCTCCAATGTTTTATCACTAAATGATTGTTTAAATCATCAACAGGGAGGAGCATTGGAACACGACGAAAAACTGAAAGGAAAAGGCCTCCTTGGCATTGCAAACTGGTAGTGAAAAATCCTGGAGTACTTGTGATATTGTAAAATTAGTCTCTGTCTCAGAATTTGATTGTTAAGTTTTTTCTTTTTTGTTTGAAGTCCTGAATTTAAATTATTACTTTTGCTAGTTTATAATCTGTGCTTCAGTTTCTCCCTTGTCA

>PGSC0003DMG400031861 G-LecRK

GAAAAGAATTCCACACATTCCAGTGACCTCATAATCATAGCCTTCTGCATTATATTATGCAAGTCTTTGAATCATGGCCTTTTGACTAAAAGATTTTAATTCTCCATTTGTATCTAAGTTATAATATCTCAAAAAGAATAAAGTATTTTCCATAAGATTCCAACAATGGTTTCTTGGAAAATCCTGTTATGGTTTGTCTCAATTTATCTTGCATTTATTAGCTCTTTCTGTCAAGAAACTACAAATATTGTCCCTCTTGGTTCCAATATCACAGCTGGAACTGAATCTTCACACTGGCTTTCATCCTCTGGAGATTTTGCTTTTGGTTTCTACCATCTTAACAGTACTGGACTATTCCTTGTTGGAATATGGTTTGATAAAATCCCCGATAACACTCTCGTTTGGTCAGCCAATCGTGATGATCCAGTTCATGCTGGATCAAAGGTTAATCTAACGTTGAGTGGACGCCTCGTGTTGAAAGATACTAATGGGAGAGAGTTTGTTTTGTACAATGGCACAGGCACTAGTCATGATACTATGCAAGATGATGGAAATTTTGTCCTGAAAAATTCATCCTCAGGTGTCTTGTGGGAAAGCTTTGATTTCCCAACAGATACAATTTTGCCTGGTCAATATCTTGATATGGGGCAGGCGCTGTTTTCCAGTGCCAATGGGACAGTAGATTACTCGACGGGGAAGTATAGACTACTGATGCAGACTGATGGGAATGTGGTCTTGTCTGCATATAGAACTGCTGATATTGGCTACTGGAATTCTATAACTGAAAACAATAAGAATGTAAGACTGGTCTTCGACAACACCAGTGATACTTTGCTCATTACAAATGGAAGCTCTATCATATCCAACATGACATTGACAGAGAAGTTGCCTGATTCAGTTCGAGATTATTACCACAGAGCTATGATCACTGATAAAGGAGATTTCCAACAGCTCTTCCACAGAAAAGTAAATGGGAGCGGTTGGAATGTTGCACTGCAAGTTATCGTGAAGCCTTGTATTGTGAATAACATTTGTGGTGTATATGGCTTTTGCCAATCACCAGACAATAAGGAAGCCAATTGTAGTTGCTTGCCAGGCTATTCTCCAAGGGATCAGTACACTCCATCTAAAGGATGTTATCCGAATGAGATGAAAGATTTTTGTGATCCGAATTCTTCCCTTTCGGATGTCTATGTTGAAAGGATCTCTAATGCTGATTTCCCTAATAGGATGTATGCTGAACTAGAAAGGGTATTTGATACTACAGAGGAGCTTTGCAGGCAGGAAGTGTTGAATGATTGTCTTTGTGAGGCAGCTGTGTTTAATGATTCAACATGTTTCAAGAAGAGGATGCCTATACAGAATGCCAGAAGTATTAATCCTGATACAAATAACATGGTTGCATTTCTTAAAATTAGCAACTCTTCGATAACTCAAAACAAAAAGCACTCTATTTCAAAGGGTGTTTTGGTAGCAGGTGTGACAGTATGTTCAGTACTCGCTCCGCTTTTTGCAGCAATTGCTATATATTATCACCCTTTTGTGAAGAAACATAGAGATGCCAAAACCCCTCCAAAAAGAAAGGCAATTGAGATGAATTTAAGGGCATTTTTGTTCCAGCAGCTGCACGAAGCCACCAACGGTTTCAAGAATAAATTAGGCCAAGGAGCTTCTGGTGCTGTTTATAGCGGAATTCTCAAGTTAGAGGATGAAGAAGTTGAAGTTGCTGTGAAGAAATTAGGAAATGGAATTGAACAAGGTGATGATAAAGAATTTTTGGCAGAAGTAAGGGTGATAGGCCTAACACATCACAAAAATCTGGTTCATTTGATAGGCTTCTGCAATGAAAAAAGCAACAGGCTTGTTGTTTATGAATTAATGAAAAATGGGGCCGTGTCAAATATTGTCTTTCGCGATGGACAAAAACCAAGCTGGAAGCTAAGAAGTGACATAGTACTTGACATTGCCAGAGGCCTGTTATACTTGCATGAAGAGTGTGAGAACCAGATCATTCATTGCGACATTAAGCCACAGAATATCCTTCTTGACAAGAACTACTCTGCAAAAATTGCTGATTTTGGTCTTGCAAAGCTTCTTATGAAAGACCAGACGCGAACAAACACGAATTTTAGAGGAACCATGGGTTATATGGCACCAGAATGGCTAAAAAATGTGCCCGTGACAACCAAAGTTGACATCTACAGTTTTGGTGTTTTGCTGCTTGAGATTATTTTCTGCCAAAGGCACATGGACTTGAACCCAATCGGCCAAGAAAATGAAGAACCCGAGTTGATCTTAGTCGATTGGGTTCTTCATTGTGTGAGAAACGAGAAGCTGAGAGCTGTTGCTAGTCTTGATGAAGAAATTATGTTACATTTTCACAATTTTGAGAGAATGACTATGGTTGGCCTATGGTGCTTGTGTCCTGAGCCGAATCTTCGACCATCTGCAGCAAAGCTTGTTCAGATGTTGGAAGGAACTATAGAACTAGGTGTTCCTCCTATGGTTGATACAACTATATGAGCCATTTTCTGTATTATCTTGTAATTTTTCCAACAATTAAGTGTCATTCTGTTTTATTAATCATTAGACCTGGTTAAATTTCATTGTTTTTTCTCATAGTTCTCTGGTGCAAAAGTTAGTGTTT

>PGSC0003DMG400032782 G-LecRK

TCTCTCTCTCTCTCTCTCTCTCTCTCTTATCTCATATCAGCAAAAAGCCACACAAAACCTGCGTTTGCACCAAACAAATGCAATTTTGTTACCTCTTCTGTTCATCACTCTTCTTGCATACGTGCTGTCTTCCTGTGTAGACTGTCTTCCTCTCTGGTTGTTTCATTCAAATGAAATTTTTCAGAGCTTTGACTTGAATGAATTCTTATTGCTTCTTCTGCTCTTGTGGGTCCTTGTTCACTTTCACGTTTTTTGATTCTTGGACCATAATTTCATACCCCATGTGAATGTGATAGACAATGTCATTTATGAGTTCCCTGCCTTCTTTCTTATCATAATTCTCATTCAACTTTTCCCCTTTTGAATCTGAGGTTTTGGTGGATATTTTCCAGATTGTTCTTGTTCCAAATTGGGTAATGGAGTCTTGGATTTCCCTTTATTTAGTAATGATCTTCCTATTTCTGCCTGAGACTTGTATAGCTAGTGTTCAGAATAAAGGCAGACTGAATCTGGGGTTTCAAGGCTCTCAAATGACTTGGATAGACAACGATGGATTAATTCTTGTTTCCAATAGCTCCAAATTTGCATTTGGATTTAACCCTACTAATGATGTTACCTTGTTCCTTCTTGTTGTCATCCATGTAAGTAGCTCTACCATCGTTTGGTCAGCTAACAGAGATTCCCCTGTTCGAAATAATGATAATTTTGTTTTTGATGATACTGGAAATGCCAATTTGCAAAGTGGGAAATCAACAATTTGGTCTACGAATACGGCTGATAAAGGAGTTTCCGCCATGGAATTGAAGGATTCGGGGAATCTAATTTTGGTTGGGAAAGATGGTAGCGTTATTTGGGAAAGCTTCACACATCCTGTTGATACTCTTCTGTCAGGCCAGAATTTCACTCAAGGAATGAAGCTTGTAAGCACTCCTAACAACAATAATTTGAGTTATTCACTTGAATTTAAGTCTGGGGATATGGTTCTTTCTGCAAGTTTTCAGCCTCCACAGCCTTATTGGGCTATGGGGAAAGATGATAGAAGGACTATCAATCAAGTTGGAGGAGGTGTCACTTCAGCAATTCTTGATGGAAATGCTTGGAAAATTTATGGTGAGAAGAGAGTGTTGCTCTGGCAGTTCATTTTCCCTGATGATAAATATCCAAATGCGACTCGGTTGGCTGTTCTTGGAGAAGATGGTTCCATTACTTTTAGCATTCTTCAAGATGAATCGAAGCTTGACTCGGGGACAAGAATTCCACAAGATGAATGCAGCAGACCAGACTCTTGTGATCCATATTTCATTTGTTATAGTGGCAACAAGTGTCAGTGCCCATCAGCCCTTCCATCTTGCAAACCTGAGACTGCTTCGTTTTGCAATAAAGATGTGGAGCTTGTAGATGCTGGGGATAGTCTTGGTTATTTTGCACTTGGTTTTGTTTCTCCATCTGCTAAAACTGATTTAAATGGTTGCAAAGCTTCTTGCGTTGGCAATTGCTCATGTGCTGCTATGTTCTTTGATAGCACTTCAGGGAATTGTTTTATGTTTGATCAAATAGGAAGTTTACAAGGTTCGGTTAATGGAGCTGGTTTCAAGTCATACATTAAAGTATCAGCTAGCCAGGGGAATGGGGATAGCGGTGGTGGAGGTGGAGGTGGGAAGGGACGATTACCTATTGTCTTTGGTATCGTCATTTCTTCGGCTATTGTCATTCTTGGCCTAATTTATGGAGGTATTAGATATCAGAGGAGAAAGAACAACAAGATGCCTGATTCTGCAAAGGGGAGTTCAGAAGAAGACAATTTCTTGGAGGGCCTATCAGGGATGCCGATTCGTTTTAGTTACAAAGAACTTCAGAATGCAACTAATAACTTCTCAATAAAACTTGGGCAAGGAGGGTTTGGTTCAGTTTACCAAGGGGTCCTCCCAGATGGGACTAGACTTGCTGTGAAGAAATTGGAAGGCATTGGCCAGGGAAAGAAAGAATTTCGAGCAGAAGTTAGTATTATAGGCAGCATCCATCATCTTCATTTGGTGAGACTGAGAGGTTTTTGTGCTGAAGGAACACACAGGCTTCTGGCTTATGAGTACATGGGAAATGGATCTTTAGAGAAATGGCTTTTCAAGAAGAACAAGGAATTCTTGCTAGACTGGGATACACGATTTAATATTGCACTTGGCACAGCAAAAGGCCTAGCTTACCTCCACGAGGACTGTGATGTAAAAATTGTGCATTGCGATATCAAACCTGAAAATGTACTTCTTGATGATCATTTCCTCGCCAAGGTCTCAGATTTTGGCCTCGCTAAGCTGATGACCAGAGAGCAGAGCCATGTTTTCACAACCATGAGGGGTACAAGAGGATATCTCGCACCAGAATGGATCACAAATTATGCCATATCAGAGAAAAGTGACGTTTTTAGTTATGGAATGGTTTTGCTTGAGATAATTGGAGGTAGGAAGAATTATGATCCTTCACAAAGTTCAGAGAAATCCCATTTTCCCTCATACGCCTTTAGGATGATGGAAGAGGGGAAATTGGAAGACCTTATCGATAGGAACTTGAAAGTTGAAGAAGAGGATGAAAGGGTCTCCATAGCAATTAAAGTTGCTCTGTGGTGTATACAGGATGACATGTCTCTCAGGCCATCCATGGCTAAGGTTGTGCAAATGCTGGAAGGGATCTCCCATGTCCCTTCACCGCCAACTGCTTCTCAAATGGGGTCACGACTTTTTTCAAGTTATTTGAAATCACTAAGTGGCGAGGGTACTTCATCCGGCACCTCTGCACCATCAGACTGCAACAGCGATGCTTATCTTTCGGCTGTTCGGCTCTCAGGTCCAAGATAGCATTTGCAAGAAGCATAAACATAGTCGTTCATGTAAGGGCTGTTGTAGATGATTGACAAACATGTGCAATGGTTTCATACCTTTTAGTTTTTCTGTATATTAAAATTATCTAGGAAATTTTTTTGCCATATTAGCCATGACAAAGGTTGGAGTTGATTGAGAATGTTTTATTTGTATATATATCAATGCTATAAATGAGTTGAAAATAAACAAAGAAAATGCATTTGGTAACACTCATCCTGGATTTCAAGTACAGCATGTGTG

>PGSC0003DMG400034859 G-LecRK

ATGTACCTGGGATTATGGTACAAAGATGTAGGTCCCGGTACCGTTGTATGGGTTGGCAATAGAAGAAATCCAGTTAATTGTGGTGCATTGCTAGGACTAGACACAGAAGGAGACATTTTCCTCCAAGATATACTGAGAACAACAGTATGGATTCATAAGCCCAACAGTACAGTTCCCTTACTAGTACTGAAGCTCCTTGATTCAGGTAATCTTGTGTTTAGAGATTCTAGCAATCTGACGGCTGGAGAGTACCTGTGGCAAAGCTTTGACCACCCAGTTGACACATTGTTACCAAGAATGAAGCTTGGCTGGGACAAGAAAACTGGCATCGACCGATCCATAAGGTCCTGGAGGACAGAAGGTGATCCAGCTCCAGGTGATTATTTGTTCAGGTTAGACTCAGGTGACTCAGGTCAGTTACCCCAACTAGTTCTTGAGAAAAACCAGCGAATACATTCAAGATGGGGTCCTTGGGATGGACAGAAATTCAGTGGCGGTTATGCCCTCATGGATAACCATGCATTCAGACCAATATTTCATTCCGATACAGATGCGTTTTTTTTTCACATTCGAAGCCAAGAACGATTGGAGTTTGAAATTTTCACTGAATCCAGACGGAAAAATACAGTTCTTGAGGTGGAATAATGCCTACCAATTACTGGGAGGAAGTGAATACACTGAATAAGGACACATGTGACCAGTATAGAACTTGTGGGCCTTATGGAGTTTGTAGTGATGGAGATCTCCCCTGCGGGTGTCCTGATGGTTTCACAGCTGCATCACCAGATGAATGCGGCATGATGAATTTTACGGAGGGATGTAGGAGAAACACCTCACTTAACTACACCAATAAAGACGTCTTTGTGGAGAATACTGGACTGAAGTTGCCTGATAATGCTACTTACTGGGGAATGTTGTACCACCAGGAGTGTGAACAGAAGTGCTTAAATGAAAGATCGTGTATGGCTTATACTAACATCAATATCAACGGAAATGGAAGCAAATGTGTCGTTTGGTTGGGAAATTTACTTGATATAAGACATTCACAGATAGCTGGGAACAACATTTTTATACGGATGGCACCTGAAAAACTAGGTAAAAATGATTGCTTGCTTTAACTGATTTAATTCCTAGATTATATTATTAATATGGTGCAAAACAATGTGTGCAAGAATGACATTTGAAAGGTGAAGTAGTTAATTGTTGTGTAGGTTTGATTATTTACACAGAAACATCAATTAATAGAATTCCCATATTAGTGCTTGATTAAGTGGTTCATCGCAAATTGATGTTTGAAGCATTAGATTGAATGTTGTCACATAATTGCTACACAATCGAGCATAAGATGAAACAACCCTGCATGCATCTAGTTATTCTTATTACAAAATCTTTAACATAATGAGCAAATACCTCTAGAATTCCTATAAAGTTTATGTCAAATCTGAAGCCTAATTTTATGAGTAAGAAACAACTAACACTTTTAACTTTCTTATGGACAGATGCACCTGTTTCCGACAGCAGATGCAAAAATAAATGGACAGTTTTTGCTTCTTTAATCTCAGAGCTGGCACTACTTATTATTTTGCTTATAGCTGTTGTGGTTCTATGTCAAAAAATGAGGACAGAGGTGCTAGGTAAAAAGAATCACCTTGCGCTGCTATTATATTATTTTCATAGTTCTTCATTCCAGACTAAGCTTAGTTTCACTAGAACACTATTATTTAGCAAACACAAGACCTAAGTAAGAAAGCTAGCGTAATACTCATATTAAGGTATGATTCACATCCATTATTTTCTAATTAATTTGCTTTCTCATAGCAGATGAACAAGAAATCAATCAGCCGACAAATTCTCAAGGAGCTTTGCTGCAAGGCACTAATGTCATTGCATATGATTCCAGTGATCTAGTTGCAGCCACTAACAATTTTTCGCTGGCTAATAAGATAGGGCATGGTGGCTTTGGCAATGTTTATAAGGTTATTATTTCTTTTCGACAAGCATGAAAATGCATATTTACAGTAACTCAGACAACAAAATTAATACCTAAATGATACATCTGTAGGGTGTACTAGAGAATGGAGTAGAAATAGCAGTGAAGAAGCAGGAAGTGACTTCATGTCAAGGATTTACTGAGTTCGAGAATGAAGTCAAGTTAATAGCAAAGCTACAGCATCGTAACCTAACCAAATTGTTGGGATACTGCATTAATGGAACAGAAAAATTCCTAGTCTATGAGTTCATGGCGAATAATAGCCTAGACAAGGTCATATTTGGTATGACATCTTAAATAACATTCTTCTTTTTTACTATTGTCTGTGTTGTTTTGATATTGGAAAAGCCACAAAAGACAGAAATTTAGGAGTATCCGAGTGAAATTGAGTAACTGTTTTCTGTACATGTCTAGATCCTGCAAGAAGGGGTACAGTAACATGGCCAATGCGCTTTAACATTATCAAAGGAATCGCAAAAGGATTAGTTTACATGCATCATGACTCAAGGTTGACAATAATCCACCGGGACTTAAAAGCTAGCAATGTCTTGCTAGACAGAGAGATGACACCCAAGATATCTGACTTAGGGCTGTCAAGGGTATTCGAAGATGATGTTGAAGAAAAGACTCAGCATTTAATTGGAACACGGTTAGTATATTATCATTAAGACTACATGTTTATGTATATTTTAGAGTTTTTACAGATGGTTTTATGTTGTGGTTGTAGTGGTAATTATATGCCTCCTGAGTACATGAAAAATGGACATTATTCAACAAAACAGACGTTTTTAGTTTTGGAATCCTGGCATTGGAGATGTGAGTGGCCAAAAGAATTATGATTATCGGCACCCAGTCTATGATATTGGTCTAGTAGACTATGTAAGTACATCATCGATCCCCTGGTTATCGTCATTACTTCACTTAAATTGGTTACATAGAAGTTTCACCTTGTGCTAAAACAATTTACATGATTGATGATCGATCTATTGTTGGTCAGGCTTGGAGAATTTGGAGTAAGGGGAATGCCATAGAACTACTGGATCCAATGATTGAAAAACCAGGTGATGAGAATGAAGTTTTAGGATGCATCCTTGTTGGGCTCTTGTGTTGTCAACGCCGTGCCCAAGACAGAACTTCAATGATTCAGGTGGTTTCCTTACTGGAGGAGAATGAAATGTCAAGGTTAAATTTTGTGCCTCAGGAACCTTAA

>PGSC0003DMG400035841 G-LecRK

GGAATTCTTACTCTTCTAAATGGTTCTGGCAGAGTAATTTGGTCCTCCGATTCCACTAGACATGGGCAAAATCCAGTAGCACAACTCCTGGATTCAGGCATTCTTGTTGTTCGAGATGCAGCCGAGGATAAGACTCAGAACTACATGTGGCAAAGTTTTGACAATCCAGGTGACACAACTTTACCTGGAACGAAGGTTGGGATTAATCTAAAAACTAGTTTTCATCGTTCCATTTCGTCATGGAAGAGCACAAATGATCCTTCCAGAGGTGAGTTTACCTGGACATTCGATACTGGTGGATTCCTGCAAACATTCATTATGAATAGTTCCATTGAACCCTACAGAGCTGGACCATGGAATGTCCGAGTCTTTCCTAATGCACCATCTCGTGACACATCTTGGAATGGTTATAACTATACATACTGA

>PGSC0003DMG400042814 G-LecRK

ATGGAGATCAAAAACAATTATTTTCTCCTCTTTTCTTTGATGTACCTATGCTTCTCTCTCAAAACCCATCTCTCCATTGAAGCTGCTGACACCATTTCTGCAAATCAATCCCTTTCTGGAGACCAAACAATTATCTCTTCAGGTGGAAAATTTAAACTTGGTTTCTTCCAACCAGGTAATTCTTCCAACTACTACATAGGCATGTGGTACGATAAAGTGGTTGAACAAACTGCTGTTTGGGTTGCAAACAGGGAAAAACCAGTTCTTGATAAGAGCACTGCAGAGTTAAAGATCTTAGATGGTAATTTAGTACTGGTGGATGAATCTCAAACTCCAATTTGGTCCACAAATATCAGTTCCAGCAACTCGAGTTCTGTTGTAGCAGTCCTCCGAGATGATGGTAATTTGATCTTGACAGATGGGTCTAATTCAACACCACCACTTTGGCAAAGTTTTAATAACCCCACCAATACTTGGTTGCCTGGTTCTAAACTTTCTTACAACAAAGCCACCAGAACAAAGCAACTTCTTACTTCATGGAAGAGTGCAGACGATCCTACACCTGGGCTGTATTCACTTGAACTCGACCCAAATGAGAAGCAGTATATTATAAGGTTCAATAGAAGTGAGCAATATTGGAATACTGGACCATGGAATAATCGAATTTTCAGAGATGTGCCTGAGATGAGGACTAACTATATTTACAATTTTAGCTATGAGGACAATCAGAATGAAAGCTATTTTACATATTCCCTTTATGATGATTCTATCATATCAAGATTCATTATGGATGGCTCTGGACAGATTAAGCAACTCACATGGTTGAATACTTCGAATCAGTGGAATCTGTTCTGGTCTCAGCCAAGACAACAATGTGAAGTTTATGCATTTTGCGGGCCATTTGCTACCTGTCAGGAAACCAATCCCTTCTGTAATTGTTTGGATGGTTTCAAGCATAGTTCAGAGACTGATTGGAATCAAAATGATTTTTCTGGAGGATGTGAGAGGCAAACGAAGTCGCAATGTGGCAATGGTAAGGGGGAGAAAGATGACTTTTGGATGCATCCCCAGATGAAAGTACCTGAAAATGCTCAAAATATTTCTGCTGGAAGTGCTGAAGAATGCAGATCTACCTGTTTGAATAATTGCACTTGCACTGCTTATACTTATGAAAGTTCATGCTCAATTTGGAATGGTGAACTCTTGAATATGCAGCAACTCCCACAAAATGATGGCAGGGGTGAGTTGATCTATGTCAGGGTAGCTGCATCTGACATCCCAAAATCAAAGAGTAAAAAGGGAATTCCAATTGGTGTTTCTGTGGGGTCTGCTGCTGCCGTACTGATCCTTCTGGGCATTCTTTTTGTTGTATTCCGGAGAAGGCGTAGGCATATTGGAAGTGGGAAAATAGTGGAGGGTTCATTGGTTGCATTTGATTACAAAGACTTGCAACATGCGACTAAAAATTTCTCAGAGAAGTTGGGTGGTGGAGGCTTTGGTTCTGTTTTCAAAGGGAAACTATCTGATTCATCTGTTATTGCTGTTAAAAGGCTCGATAGCATCAGCCAGGGAGAGAAGCAATTTCGATCAGAAGTCAGCACAATCGGGACAATCCAACATGTTAATCTTGTACGCCTTCGTGGATTTTGCTCTGAAGGTAACAAGAAACTGCTGGTTTATGATTACATGGAAAATGGATCCTTGGATTCACATATTTTTACCGAAAAGCAGTCAGATGTTATGGATTGGAAAACGAGGTACCAGGTGGCACTAGGGACAGCTAGAGGCTTGACTTATCTCCATGAGAAGTGCAGGGACTGCATCATACATTGTGACATAAAGCCTGAAAACATCCTTCTTGATGCACAATTATGCCCCAAAGTAGCAGATTTTGGCCTGGCAAAGCTTGTTGGGCGCGATTTTAGCAGAGTGCTTACTACTATGAGAGGAACAAGAGGTTATCTTGCACCAGAATGGATATCAGGGGTGGCCATTACAGCCAAAGCCGATGTTTATAGCTACGGCATGATGCTTTTGGAAATTGTATCAGGGAAGCGAAACTCGGAATACTCTCACGATGGGAAAGTGAAATTCTTTCCTAGTTGGGCTGCACGTGTAGTAGTTGATGAAGGCGATATCCTTAGCCTATTGGACAATAGGCTGGACAGAGCTGCTGATGCTGAGGAGGTGTCAAAAATATGTAAAGTTGCATATTGGTGCATCCAAGATGATGAATTCCAAAGACCCTCAATGGGCCAGGTTGTTCAGATTCTTGAAGGAGTTTTGGATGTGAATTTGCCTCCTCTCCCACGCTCTCTTCAAGTTTATGCAGACAATGAGGAGCACATCGTTTTCTTCACCGAGTCATCCTCGAGCCAGACTAGTTCACAAGCTCAGAGCAAGACTTCAAGTACAACATCTCAATCTAAAAGCATCACAGAGTCTACGAATTCCAGGTCTTGA

>PGSC0003DMG400042886 G-LecRK

ATGGACTTTGTTAACTCAATAACTCCTTTGAAACTAACTCAGTTTTTCATTATTTCCATCTTTTTCTCAATTTTTTTTCAAATTAAAGTTGAATCTGAGACACCCAACCAGTTTTTCAAAGGTTTTAAAGCAAACCCAGATCCCCAAATTTCGAATTTTCAGCCTCTTCTTACTGATTCTACTGGAAATTACTCACTGAGTTTTCTTCGAGTTGAAGAAAATCAACTCACTCTTGCTATTATCCATGTTAAATCTTTCGAGTCGATATGGGTTGCTAATTTGACCCGTTTAGCTAAATGGGCTGACCCGACTGAGCTCTACTTCAATGGCAGCCTTGTTCTGTCTGATTCACGTTCAGGGGTATTTTGGTCAACTCACACTGATGGAGACCGTGTATGGCTTTCAAATACATCAAATTTGCAAGTTCAGAAGGTTGATAATGGAGTGACGTCACCTTCTGTTTTATGGCAAAGTTTTGATTTTCCCTCAAATACCCTTGTGGAAAATCAAAATTTCACTAGTGCCATGACTTTAGTTTCGTCAAATGGGCTTTATTCCATGAGATTGGGCTTTGATTTTTTCGGGTTGTATTTTAAAGGTGAATCGGGTTCGGATTCGGGTCAGATCTACTGGAGGCACAAGGCATTAGAAGCTAAAGCTGATGTGATGGAAGATCAAGGACATATTTATGTTGTATTAAAGTCAAATGGATTTTTGGGAATGTACCAAAATGAGTCAGTTCCAGTTGACATTGAATCATTTAACAGCTTTCAACAACCCGTATCCGGCGTCCGTCGGGTACGGGTCGAACCGGATGGTAACTTAAAAGGGTATTTTTGGGACGGGTCGAGTTGGGTATTGGACTATCAAGCGATAAAGGAGACATGTGAATTGCCTAGCCCCTGTGGGGCGTACGGGCTCTGTCATCCAGGCAAGAGTTGCTCTTGCCTGGATAACAGCACGGACTACAGTATTGGCGGGTGTGCCCCGCCGAATAGTCAGGAGTCCCGCGATTTTTGCGGGGCCTCCTATGATCTTGATCACAAGAAGTACAAGGGGTTATCAAGAAATGGTGTTGAATTGCCTAACACTGAGTTAATGGACTATCAACATATGGTTTCTTTTCAAGAATGTCAAAGTGCATGTGAAGGAAATTGTACATGTTGGGGTGTGGTGTACACTAACACATCTGGATTTTGCTACATACTAAATTACCCCATACAAAGTTTAGTAGGGGTGAGGGATGAATCCAAGATGGGGTTTTTCAAAATGAGGGAGGGTGTAGGAAAAGATAAGATGGGGGTGGGGTTTGGGGTGAGGGTAGGGTTATTATGTGGGGCCATATTGGTATTAGGTGGGGTCATAGGTTTAGGGTATTATAGATATATAAAAAGAAAAAGAGAAAGAGGAGTTAGTGGTTATGTTGATGATGATGGAGTAGTGGTTGGACCATATAAAGATTTGGGAAATGCAAGTTTTAGGTCAATTGAACTAATTGAAAGATGA

>PGSC0003DMG400044943 G-LecRK

ATGGCTTCCATGCTTTCTTTGTTTACTATCTGTCATCTTTTTATTTCCATCGCCCTGTATGTATGTTACGCTGCTGCTACTGACTCCTTGTTTGCATATCAAATCCTCACCAACTTGAACCCTTTAGTTTCATCCAACCATAATTTCGTAATTGGATTTTTTGGTTATTATTCATCGACTCTACCTCTATGCAAGACGTACCTGGGATTGGGGTACAGGAGTGTAGATTCCAGGGCTATAGTATGGGTAGGTAATGAATTGAATCCACTTGGATGTTTTACAGCGCTAGCACTAACCAAAGAAGGATTTTATGTCAAAGATATAGTGCTGGGAACAACGGTATGGATTCATAAGCCCAACAGAACAGTTCCGTCACCAGTTCTGAAGCTCCTTGATTCAGGTAATCTTGTGTTTGGAGATTCTAGCAATCTGACGGCTGGAGAGTATCTGTGGCAAAGCTTTGACCACCCTCTTCACACATTGTTACCAGAAATGAAGCTTGGCTGGGACAAGAAAACTGGCATCGACCGATCCATGAGGTCCTGGAGGACAGAAGGTGATCCAGCTCCAGGTGATTATTTGTTAAGGTTAGACTTAGGTGACTCAGGTCAGTTACCCCAACTAGTTCTTGAGAAAAACCAGCGAATACAATCAAGATGGGGTCCTTGGGATGGACAGAAATTCAGTGGCGGTTATGCCCTCATGGATAACCATGCATTCAGACCAATATTTCATTCCGATACAGATGCGATTTTTTTCACATTCGAAGCCAAGGAGGATTCAAGTTTGATACTTTCACTGAATCCAGATGGGAAACTACAATTCTTGAGATGGAATAATAATGCTACCAATTTCTGGGATGAAGTGAAGACACTCAATATGGCCATATGTGACCAGTATAGCACATGTGGGCCTTATGGAGTTTGTACCGATGGAGATTTCCCCTGTGAATGTCCTGATGGATTCACAGCTGCATCACCAGAAGAATGGGACAAGATGAATTTCACGGAAGGATGTAGGAGAAACACCTCACTTAACTATACAGATAAAAACGTGTTTGTCAAGAACACTGAACTAAAGTTGCCTGATAAAGCTACTTACTGGGGAATGTTGTACCCTCAGGACTGTGAACAGAAGTGCTTAAACAAAAGATCGTGTATGGCTTATACTAACATCAATATCAACGGAAATGGAAGCAAATGTGTCGTTTGGTTGGGCGATTTACTTGATATGAGACGTTCACGGATAGCTGGGAACGACATTTTTATAAGGATGGCAAATGGAAAACCAGGTAAAAATGATTGCTTACTTGTGATATGATTGCTTGCTTATGACACGACAACTAATTCAATGCCTAAATTATTTTTAGTATAGTGCAAAGTTATGTGTGCATGAATGACATTTGAATGGTGAAGTAGTTAATTGTTGGTGCATAAACATCAATAAATAAAATTTCCATATTAGTGCTTGATTAAGTGATTCCTGCAGATTTGTCTTTTTTTTAAAACAAAAAGTATCTATAACTTGATTCTTACATGAACCACTCATCCCTCATCTCTATCAGAGGGAAACAGTTCTATAACTTTTCAAAATGCATATGTATCTCCGTTATATAGAAACGTTAGATGTGTTTCAAGCATTTTACTGAATGTAGTCACATAACTGCTACACAATCTAGCAAAAATTGACACTAACCTGCATGCATCTAGTTGCTCTCATAGCAAAATCTTTAACTTTGTATCAACAGAATATTATGAGCAAGAAAACAACTAACACCTTTTACTTTCTTATGAACAGATGCACCTGTTTCCCATGGGAAAAGAAAATTGAATGAGGGATCGATAGCAGTGATTTCTGTATTAGTTCTGGAAGTACTTATCTCGATTTCATATATTGTGTATACACATCGAGCAAAGAGTACAGAGGTGCTAGGTAAAATGAATCACCTCTCGCTGCTTATATATTATTTTCGTAGTTATTTTTTTCAGCCCAACCTTAGTTTCACTCAAACACCATTCATGAGCAAAACCTCCTTTTAACTCTTGAGACAACAAGAATGAACAGAGACATTGTTCAAAAGCATCTTGTTCTGCCAATACCCTTTAGTTTGTAATGAAAAACAAAATTTACTTAGAGCACACATGACCCTGTAACCTAGCTCGCGGTTCGCTTCCATTATTGTCTGATATATTTGTTTTTGAAAAAGAATATTTTCATCTGAGCAGATGAACCAGAAATCAATCAGCCAGAAAATTCTGTAGGAGCTTTGCTGCAGGGAACAGATGTCATTGCATATGATTCCAGTGATCTAGCCGCAGCCACTGACAATTTCTCGCTGTCTAATAAGATAGGGCATGGTGGCTTTGGCAATGTTTATAAGGTTAGTATTTCTTTTCAACAAGCATGAGCATGCATATTTACAGTAACTCAGACAACACAATTAATACCCAAATGATACATAAATCTGTAGGGTGTACTAGAGAATGGAGTAGAAATAGCAGTGAAGAAGCAAGACGTGGCTTTGCGTCAAGGAGTTGAGGAGTTCGAGAATGAAGTCAAATTAATTGCAAACCTACAACATCGTAATCTAACCAAGTTGTTGGGATACTGCATTCATGGAATAGAAAAATTCGTAGTCTATGAGTTCATGGCGAATAAAAGCCTAGACAAGGTCATATTTGGTAGGTCATCTTAAATAACAACTTTGTTTTACTATTATATGTGTTAATATGATATTGGACAAGCCATAAGAGACAGAAATTTTAGGCCTATCGGAGTGAAATTAATAGGGTAACTGTTTTCTCTACATGTCCAGATGCTGCAAGAAGGGCAACAGTAACATGGCCAACGCGCTTAAACATTATCAAAGGAATCGCAAGAGGACTAGTTTACATGCATCATGACTCGAGGTTGACAGTAATCCACCGGGACCTAAAAGCTAGCAATGTCTTGCTAGACAGCGAGATGACACCCAAGATATCTGACTTTGGGTTGGCAAGGGAATTTGAAGACGATGTTGAAGTCAAGACTCACCGTGTAGCTGGAACATAGTAAGTAGACTAGCACTCATTATCATTAACACTACATGTTTATGCATATTTTACAGATAGTCTTATGTTGTGGCTGTAGTGGCTATATGTCTCCTGAATACATGCAAGCTGGACATTATTCAACAAAATCAGATGTGTTCAGTTTTGGAATTTTGACATTGGAGATTGTGAGTGGCCAAAGGAATTCGATTTATCGGCACCCAACCTATGATATTGGTCTAGTAGGCTATGTAAGTATATCGATCATCCCCTGATTAAATATGGTCACTACTTGTACACCTAAGTTTCACCTTGTGCTAAAACAATTTACATGCATGATTGATCTTTTGTTGGTCAGGCTTGGAAAATTTGGAATGAAGGGAATGCCATAGAACTACTGGATCCACTGATAGATAAACCAGATGATTTGGATGAAGTTTTAGGATGCATCCATGTTGGGATCTTGTGTTGTCAACGGCGATCCCAAGACAGACCTTCAATGGTTCAGGTGGTTTCCTTACTGGAGGAGAATGAAATGTTAAAGTTTAATTGTGTGCCCCGGGAACCTTACTTCTATAAGGAGACGAGTCAGAGTAGCAGGAGTCGCGAAAGTATGAATGGGCTAAGTATTACTGAATTCACCGGAAGATCATGA

>PGSC0003DMG400045997 G-LecRK

ATGACTTCTTTAGTTATGCTTCTTGTTGGTCTCTTATTGTTTCCCCTTTCAGCCATTGCTCAATCTTACCAAAATGTTAGTTTGGGTTCATCACTCACTACAAGTGATGTTACTAGCTTTTGGTCATCCCCATCTGATGAATTCGCTTTGGGCTTCCAAAAAATTGGAAATGAATCAAGGTACTTACTAGCCATATGGTTCAACAAAATCACAGACAAAACCATAGTTTGGTCAGCCAATAGGAATAATCTTGCCTTAGATGGATCCAAAGTTCAGCTCTCCGCGGATGGAAGACTAGTCCTCACTGATCCAAATGGTCAAGAGATGTGGGCTCGTGGCATGGCGAATGCTCAATTGGCCTATGGAGCCATGCTTGATAATGGAAACTTTGTGCTAGCAACTAGTAGTTCAGACACTCTATGGCAGAGTTTTAATGAGCCCACCGATACGATTTTGCCTGGTCAAGTACTTAACCAAGATAACAGTCTTGTTTCAAGTTTCTCTGACACGAATGTATCGAGTGGGAGGTTTGAGTTCATACTTCAAACTGATGGAAATCTAGTGCTTTACACAGTCAATTATCCAGCTGAGGCAACAAATGCTGCATATTGGTCACCTATGTCAGTTGGCAGTGGTTATCAGGTGATCTTCAATCAATCAGGCTTTATTTTTCTTCAAGCAAAGAATGGAACTTTGATAAATTCGATATCTTCTAATGTGGAAAATTCAAGAAGCCAATCAATGTATCATCGAGCGATTCTTGAATATGATGGAGTTTTTAGGCACTATGTCCATCCGAAATCTTCTGGTAGGGAACCGATGGCATGGTCTAGTTTGTATAACATTCCTGATAACATCTGTTTGTCTATCAGGCAAAGTACAGGAGGGGGTGCCTGTGGTTTCAACAGCCTATGCTCAATAGGAACTGATCAGAGGCCGAGATGTGATTGTCCACTCGGATATATCTTGGATGATCCGAATGACAAATTAGGCAGCTGCAGACAAAATTTCTCTGAGCAAAACTGTAACCATGAATCGCGAGAAGTTGAGTCTTTCACATTCCACGAAATGTTGGATACTAACTGGCCTGACTCTGATTATGAGTCTCACCGCGATGTTTCAGAGGATTGGTGCAGACAGAATTGCTTGAATGATTGTTTTTGTGATGTTGCCATTTACAGTGACGACAACGTCTGTTGGAAGAAAAGGTATCCTCTTTCAAATGGGAGAGTAGGTCCAACGATCGGAGGGAAGGCTTTGATAAAAATAAGGAAAGACAACTCCACAGTGGGAACACCAAATGTTGAAATAAGAAAAAAGAAAAATCAGTCTACTCTGATCATTTCTGGTTCTGTGTTGTTGGCCAGTTCAGTTTTCATGAATTTGTTCTTAATTCTTTTGGCCCTATTGTACATTTTTAAGTTTATAGGAAAAAAACGAAAGAGGACTGCACCGTATTCAGCTGTTCCAGGAGTGAACTTAAGAAGTTTTAGCTACAAAGAGCTAGAACAAGCTACAAATGGATTCAAGGAAGAATTAGGAACTGGAGCTTTCTCAACAGTTTACAAAGCGGTTCTTGATGATGAAAATGGGAAAGTCGTTGCTGTCAAAAAACTCCGAAACATGGTAACAGAAGGAGAAGGAGAAGAAGTATTTGAAGCTGAAGTGAATTCAATCAGCAGGACTAATCACAAGAACTTGGTCCAACTCCTTGGATTCTGCAACGAAGGCCAACACCGCCTTTTGGTATATGAACACATGAAAACCGGCTCAATAGCACACTTGCTATTCAAAGATTCTAGGCTTAGTTGGTCCAAAAGGGTACAAGTTGCCATCGACACTGCCAAAGGACTTTGTTATTTGCATGAAGAGTGCAGTACCCAAATTATACATTGTGACATCAAACCCCAAAATGTGCTTTTGGATGAAAATTTGACAGCAAAAATAGCAGATTTTGGTATGGCCAAGCTACTGAAAAAACATCAGACTCAAACAACCACTAGGATACGTGGAACCAAAGGGTACGTAGCCCCAGACTGGTTCAGAAGCATGCCTGTAACTGTGAAGGTTGATGTGTACAGCTTCGGAGTCTTGCTCTTGGAGTTGATTTGCTGCAGAAAAAACTATGAGCAGGATGTGGCAAACGAGAACGAAATGATACTACTGGAGTGGGCTTATGATTGCTATAAAAGAAATAAGTTGCATTTACTTGTAGGCGATGATGAAGAGGCATTAGAAGATATCAAGAGATTCGAGAAGTTTTTGTTGGTTGCAATTTGGTGCATTCAAGAAAATCTAGCATCAAGGCCTAACATGAAGAAAGTCATGTTAATGCTTGAAGGTTCTGTTGAAGTATCAATTCCTCCTGACCCTTTTTCGTTTGTCGATTCTATTTAG

>PGSC0003DMG400046324 G-LecRK

ATGGTTGTTAATTATAGTGCCACTCTTTTCATGCTCAGTCTCCTCATTCTTTCTTCACAGCATCTGATTAGTTGTCAACCTTCTGAAACTTTTATCACCAATTCCTCCATTTCCTGGATCAATAACCGCGCCCGCTCCTATGTAGTGAATTCCACAGATGGAGCCACACTGACACCCATCCTATCGAGAGAAGGGAATGAGGCTCTGTTTGTCTGTGGTTTCTTATGTGATTACAATGGCACTGCTTGCGTTTTTGGCGTCCTTATATTACCAAATACCTATGATTCTTATCTATACTACCCGAGATTAGTATGGTCAGCAAATCGGAACAATCCAGTTAGAACCAATGCAACCTTGCAACTTAGGCAAGATGGAGGCTTGTTGTTGATGGACTCAGATGGCACTTTGATTTGGAATACAAGCACAAGCGGAAAGGTTTCTGTGGGATTCAACTTGACAGTAATGGGAAATCTTGTCCTCTTTGACAAAAGTAACGACATCATTTGGCAGTCTTTTGATCATCCTACAGACACTTTGGTTCCGGGACAAATAATGGTCCCTGGACAAAAGCTGATATCCAGAATATCAGCAACAGACTGGAATGAAGGTATGCTCACACTTGATTTTAAATATTCTGGCCTTTCTGCTTACATTGAGTCAAATCCACGCCAACTCTATGTCTTGAGACTTGGTGTGGAAAAACCTTTCCAGTTTCCTGGACTAGGCTTTTCTTTGAAAGGTTATGGTTCCAGAGGGATGTTAACCATAGGTTTCGAATTTTCAAATGCTCGATTCATGAGATTAGGGGTTGATGGGCATTTAAGGGTGTATCACTGGAGAGGATCTTGGTCAGAGGAGGCTGATCTATTTACATACTACATTGGTGAGTGCGGGTATCCAACCATGTGTGGAAATTACTCTGTATGTGGAAATGGGCAATGTACTTGTCCTCAAACCGCGGTTGGATTTGGTCAGACTAACTTTCTCCAGCAAATCAATTACAGGCAGCCAAACCAGGGGTGTGTCCTTGTCACACCTATCTCTTGTGAACATTCCCAATATCATATTCTTATGGAGCTTGAGGACATAGCTTATATTCCCCTTTATTTTCAGTACGGTACCAACAAAACAGATTTGGAAAATTGCAAAAGGTCCTGCTTAAGTAACTGTTCTTGCAAAGCAGCTCAATTTCAATTTAGTGGTCCTGGTAACACAATAGGTGACTGCGCATTGTTGAATGATGTGTTTTCGCTCACAGACAATTATGGAGCACCCAATAAGACCACTCTTTTCATCAAGATGCAGAACTCTTCTTACCTACAAGCCTCTCCACCACTTATCTCTTCAGAGAAGAAATCAAAGCGTGTCCCAACAATAGTAGGATCCACCATCGGAGCTTCATTTGGTTTGCTTCTCATGGTTTTAACTTGCTTTGCCTATATTTTCAGAAGGAGAAAAGGACTCGAGGAAGACGAGGAGGAGTTCCTTGATCAAATACCAGGAATGCCTACTAGATTTTCCTACGAGGAACTCACTGTGATGACAGAAAATTTCAGCGAGACACTTGGGGAAGGGGGATTTGGTTCTGTATTTGAAGGAACAATGAGTGATGGCACCAAAATAGCGGTGAAGCGTATGCAAGGATTCGGTAATGTGAAGAAATCATTCTTAGCTGAGGTAGCAACAATCGGTAGCATTCAGCATGTCAATTTGGTAAAACTTGTTGGATTTTGTGCTGAGAAATCACACAGACTTCTGGTTTATGAATTCATGGCCAATGTCTCTTTAGATAGGTGGATTTTTCACGGAACAGGGGAAAAATCCCTCACATGGGGTGTCAGGAAAAAAATCATATCAGACGTCGCCAAGGGCCTAGCTTACCTACATGAAGATTGTAACAACAAAATTATTCATTTGGATATTAAACCTCAGAACATTCTTCTTGATCATAATCTCAATGCAAAAGTATCAGATTTTGGGTTATCAAAGCTTGTGGGGAAAGATGAAAGTAATATAGTTACAGCAATGAGAGGAACACCAGGCTATTTAGCACCTGAATGGTTGCATGAGGTCATTACTGAGAAAGTGGATGTTTATAGCTTTGGGGTTGTGATCTTGGAAATCATCTGCGGCCGGAAGAATTTGGATCGTTATCAAGATGAGGATGATATGCATTTGCTTAGTTTGTTCATGAGAAAGGCAGTGGAGAGACAACTCTTGGAAATGGTGGACAAGAAGAGTGAAGACATGCAGCTACATAGAAAAGAGGCTGTGGAAATGATGAAGATTGCAGCCTGGTGTTTACAAAGTGATTATACTAAGAGGCCTTCCATGTCGTTGGTAGTTAAGGTGTTGCAGGGTTTAGTCGCTGCTGAAACCGATTTGGATTACAGCTTCACATTTCCAACAATGACGAGAAAAGTTGCCGGAGCTATTCAAGAAAGGGAAGCTGTGGTTGGTATCAGTTTACCATTTCCATCACAGTTGTCAGGACCTAGGTAA

>PGSC0003DMG401004946 G-LecRK

ATGGGATTTTTTCCCCCGACTAGTTTCACGAATAATTGGTATGTCGGAATATGGTACAGGCACAATGTACCTGATAAGTCTGTGGTATGGGTTGCCAATCGAGCAAATCCATTCAACAACAAATCAGGTGTTAGGTTGAAAATCATCGATACAGGACAACTTGCTCTTCTCACTGCTGACAATAAGAGCATATGGTCTACTAATTCGTCAAGGAGTTTGGCTGTGAAGAATACAGTTGCACAGTTGTTGAACTCGGGGAATCTTGTCGTAAGAGATGTAAATGATGCTAGGCCTGAGAACTTCCTGTGGCAGAGTTTTGATTATCCTACTGATACACTGTTACCAGGCATGAAAATGGGTAAGAACTTTGTGACTGGACAAGAATTTTACCTGTCCTCATGGAAGAAGGAATCTGATCCAGCTCCAGGAGAATATACATACCGTTGTGATCCTACTGGATATCCTCAGAATGTCGTGAGGAAAGGTAAAGTTAAGGTGTTCAATACCGGACCATGGAATGGAGTCTTCTGGAGTGGTGTACCAGGAAGTATACAAAATACTATTTCCACATCTAAGCTGGATTTTGATGAAAAGAAAGCATTTTACAGCTTCACGCTTCTGGCTTCTGTTATGACCAAATTAACCATAAAAAGTAGCGGAGTGCTTCAAATTTCAACGTGGGTTGAGAATAGACAGGAATGGCAAGATTATGTTTCTTCATCGGGCGATACTTGTAACAACTATGGAACATGTGGTGCATATGGTAGTTGCAACAACGTGCACACTCCTGTGTGTAGCTGCTTAGACACGTTTGTGCCTAAAGATCCAAGAAATTGGGCAATGAAAAATTGGTCCAGCGGGTGTGTTAGAAGGAAACCCCTTAATTGCCAGAATGGAGATGGGTTTCTCAAGTACTCTAGTATAAAGCTACCAGATACACAGTATTCATGGTTTGATGTGAGTATGACACTCCACAAGTGCAAGCAAGCTTGCTTGAGGAATTGCTCGTGTATGGCTTACTCAAATCTTGATATACGCAATGATGGGAATGGCTGTTTGCTTTGGTATGGCGACTTAATTGACATTCGAGAATTACCTGGAGGACAGGACATTTACATAAGGGTTGCTACCTCAGAGTTAGGTAACATTTGTGAATTTGATCAAAGTTGGAGTATTAGCTGATTGATTTAAACTTCCCAATTCTCAGATTTAGTTGTATTTTACTTAGATATCTGTTAAAAGTTCACACAGTATATGTTCACTCTTCCATGTCTATTTAGTTTATTTCACCCCTCCATGTTTTGTTCAGGCTCGAAGAAAACAAAACTACTCGTGTTAAGTCTGTTACTGTTGATTGGAGTGACTGTGACTGGCCTGACCATAGGGCTGTATATTTGGAAGAAGAAGAAGAGGAAGATGAATTTGAAAGATGACCTAGACCTCCCACTGTTTACCTTATCAACTCTAACTAAAGCATCCTCTAATTTCTCAGTCGAGAACAAGATTGGAGAAGGTGGATTTGGATCTGTTTACAAGGTAACATCTTAAGTTTATGAATCATTCTTTCAGTAAACAATATCTACTATTATTGACAGTAAACAGATGACACTTACTGAATCACTAAACCAGTTATATGTTGATATTATAGGGAATCCTGGAAGGTGGACACGAAGTAGCTATAAAGAGGCTATCCAAGTCTTCCTCGCAAGGAGTTAACGAGTTCAAGAATGAAGTTATTTGCATTGCCAAACTTCAGCATCGAAATCTTGTGAAGCTTATAGGTTGCTGCATTGCAGGAGGAGAAAAAATGTTGGTCTATGAATACATGTGCAACAGAAGTCTAGATTTCTTCATATTTGGTCTGGTTACTCTCTCTTAACTCGAAGAACCTCTATTAATACAAATAGTAAAGTTGATAGTTTAATCACATTATCAGTGTTCTAACGGAGGACATGTTAACAGATGAAAAGAGGAGCTTGTTACTCGACTGGCCTAAGCGCTTCGACATCATCAATGGAATTGCTAGAGGACTTCTGTATCTCCATCAAGATTCTCGGCTAAGAATTATCCACAGAGACCTCAAAGCTAGTAACATTTTGCTAGATGCTGACATGAACCCCAAGATCTCAGATTTTGGCATAGCTAGAAGCGTTGTAGGGAATGAGACTGGTGCTAATACACACCATGTGGTTGGGACACAGTAAGAAATTTAATACTACTCATTTTTTAGCATTTATATTATGTATATAATATCCTTTACTTAACACTATTTTTGTTTAATGTGTAATGCAGTGGCTACATGTCCCCAGAGTATGTAGTCCATGGTGTTTTCTCAGTGAAATCAGATGTATTTAGCTTTGGTGTGTTAGTACTAGAGATAATCAGTGGTCGGAGAAACAGAGGATTTGCCCGTGAAAGTCACAGTATCAACCTACTCGGACATGTAAGTCTTGTTTGTAATTAAAGACGAGTTTAAATTTTGTGCACTAACAGAGTAAAATAATCCTTAAACAACGAGGTCACTTATAAGGTAGTTAATTGATAACACCTCATAAAAATGATAGTTAAACTGCATTGAACTTAAATCCATCAGGCACATCAACCGCTACATGTTTCTTGATTATGTACTGATTGTGTTATTTATAGGTATGGAAACTTTACAAAGAAGGTAGACCATTGGAGCTAATTGATGGGGATCTAATGGATTCGAGTTATATTTCTGAATTGCTACGCCTTATCCATGTCGCTCTGTTATGTGTACAACAATGTCCAGAAGACAGGCCAGACATGGCTACAGTAATCTTGATGTTGACTAATGATGCTATTTTGCCTCAAGCTAAGGAACCTGGTTTCTTCACAGAAAGCAAAGTCACTTCTGGATGTTCTACAAGCATGAGTTCAACAAACGAAATTACCGTCACACAGTTAGAACCCCGGTAAATTACATTACATGTTCATTCAGATCGTTCAATGTCAACAAGCTTGAATTACAGTGTTTGTAAAGCCATAATGGCCTATGTTATATGGTATCTGTTATTAGCAAGTCTTCAATATGGGTGTGTATCAGTATTTAGTGTGGATACCTTTAGTTTTTATTAATTCTAATGTATTTTAAAGTATTCGAACAAAGTACCTATTCAGCATGTGTACATCAAAATAAATTAAAGATCCATATAAAATAAATCAACACAATGAGAGTTCACTACTTCACAGTAACAGAAGAAGCTATTGTTGTTGGTGTTTAGAAGCTCCAGTCCCATAGTACTCCATTATTTACAGAGTATCAGTTATGTTGTTCCCATTCTCCGAGTTTACACGATCAAACATCAGGCAGTCACTGCACCAACAATTAGAATGATAGCAGAATGGTTTGATTAAAGTAATGAAATTGCCAAGACTAGAGTGATTGGAAGGATTGCAATTCTAATTTCTTACTTTCTTTCTAGGAAATGTACAAAAGATATTATCACTGCAACTGACTTTATAAATGGAAACAACTAAAATTGAAAGACATAAAACGCATTCACTCAATGCGATTTATGAGTAAAATTCTGTTTCTTCTTTTTTCTTGTTGAAATTACTAGATTAACCCTTCTGTTTAATTTTATTTGTACGCTATTTTAAAAATATATTTATTTATTTCATTTATTACTTTTATAATATCAAGAAAAGATAAATATTTTATCTTTTTAAAAAAATGTCATGTCGTTGAAAACTTGAAAGACCCTTCTTAGATACCTAAATAAAAAGAGCTGAGAATGAAAGGCATAAAACACATTCACTGGATTCGATTTATGAGTAAGATTCCGTTTCTTCTTTTTTATTACTGAAATTACCATATTACCCCTTCTGTTTAATTTTATTTGTCCACTATTTTTAAAATATATTTATTTCTTTCATTTATTACTTTTATAATATCAAGAAAAGATAAATATTTTATCTTTAAAAAAAATGTCATGTCGTTGAAAACTTGAAAACTCGTTCTCAGATGCCTAAATAGAAAGAACTGAGAATGAAAGACATAAAACGCATTCAGCGAATGCGATTTATGAGTAAAATTTTGTGTCTTCTTATTTATTGTTGAAATTACTATATTACCTCTTCCGATTAATTTTATTTGTCCACTATTTTTAAAATATATTTATTTATTTCATTTATTACTTTTATAATATCAAGAAAAGATAAATATTTTATCTTTTAAAAAAAATGTCATGTCATTGAACACTTGAAAACCCCTTCTCAGATACCTAAATAAAAAGAGTTGAGAATGAAATGTATAAAATGCATTCAGCGAATGCGATTTATGAGTACAAATCTATTTTTTATTTTTTATTGTTGGAATTACTATATTATCTCTTCCATATTTCATTTATTACTTTTATAATATCAAGAAAATATAAATATTTTTTCTTTTAAAAGAAAAGAAATGTTATGTCGTTGAAAACTTGAAAACCTCTTCTCCAGATACCTAAATAGAAAGAGCTGAGAATGAAAGGTATAAAATGAATTCACTGAATGCGATTTATGAGTAAAATTCTCTTTCTTCTTTTTTATTGTTGAACTTACTATATTACCCTTCCGTTTAATTTTATTTGTAAACAATTTTTAAAATATATTTATTTATTTCATTTATTACTTTTATAATATCAAGAAAAGATAAAAATATTTTATCTTTAAAAAAAGACACGTTGTTGAAAACTTGAAAAACCCTTCTCAGATACCTAAATAGAAAGAGTTGAGAATGAAAGACATAAAACGCATTCAGCGAATGCAATTTATGAATAAAATTATGTGTCTTCTTATTTATTGTTGAAATTACTGTGTTACCCCTTCCGTTTAATTTTATTTGTCCACTATTTTTAAAATATATTTATTTATTACTTTTATTTAGGTAGATGAGAAGGGTTTATTACCTAGTGCAAAAGATATTATCACGGCAACTGACTTTCTAGCAGATGGAAAGACCATTACTTCATCTGATGGAAGCTGTGAGATGGCTAGTTTCACGAATAATTGGTATGTTGATGAAAAGAGAGCATTTTACAGCTTCACGCTTTTATTCCTAACTGTTAGATTAACAGATCCTTGGTTATGACTTATGACTATCACATTAACCAACTTGATAAATAGGAGTTTTTGTCCACTTGAAACCACAAGTGTGAGACCCGGTCTCAATTTGAGACCCGTGCCGACCCCGACAGCCGACGCGAGATCCGACCCTGACCTTGACCCTAACATCTGACCCAAAACCCGACCTCAACCCTAACACTCTGTTGACACCAAATTTTGACCCTCCTTTTTTCTAATTTATTTCGTTGAACTTCTAAATTGGTAACAAGTCGAATACTTTATAATTCACTATTATTTTCCATTTATCACTATTTCTATCGTAATGATCTATATTACTTTAACTTTTATGATATTTATGTTTATCATCTTTTTATTATCATTTTCACTATTTCAAAATTATTATTTTTTATCAATGTTACTATTAGTACTTTATTGTCGTCTTCTTTAAAATAGATTGTCGTTATTTTATTTTAAGTTTCTTACTTACTTAACTATTTTACGTAATATATATTATATTAGTTATTAAATTAAAAGCTCAAATGATTTAATTTAAAGAAGGAAAAATGCCATGAAAGCCTATCCATTTTCAGCCCTTTTTACAATCTTAAAATATGCAACAATTTGGCTATCCAAAATGCAGCCCAATAGGCCCATACGTTTCAGCCAAGTTCACTAAAAAACCCAAATTCGCAGCCCTAATCCAAGACTCACCAGCAGCAGCGCTAAGCATCCCTTTCTCCTCTCTTCCCTCTTCTAAGCAAAGTTCCAATATAGTGCAGCCATTGACAGTTTTCATCCTTACGTTTTGAGCTTAATTTTGTCCTTGTTGAGGCAGCCAACGAATGCTTTCTTGTTTCATGCTTCAAAACGCAGCAGCTGCTTTTCAAATTTCAAACAAAAATCTCACTATTTCGGACGATTATATGCAGCAATATCACTGCCATATTTGACCAATTTCAAACGGATACTTTCCCGCTCAAGCATGCCTTAACCCTAATCTTCACCTATAAATACATCCTTATATATTTACAAAGAAAAAAAAAAGGTTTTCAGTCACCTTCTTCCCCTCTAACTTTACTAGTCTTTACAACTTTAACAAAAAGGTTTTAATCTCTCTTTGCTCCTGTTGGACGACAACAAAGGCATTTCAAAGTTGTTCATTACGAGAGTATAGATTCAAGGCCGATAACAATCCCAGTTCACTGCCCCACAATCGGTAATCTCGACCGCTTTACTTTGTTATATTTTACTCTTTTATCCAGCTCGCCAGCGTGTTTGGTATTAGTTTCTATATGAAAAATGACAAATGATTAAAGGCTGGGCACCGAACCGGAATGGGACCATCGAACCGGTTTGATCGGGTTGTTGATCGGTACCGGGATGAACCGGACAGGAACTACCGGGATGGAGGCTCAGTTCCATCCCGTCCCACTATATACCGGGATAGAACCGGGATGGACCGAAATGGATCGGAACGGGATGGGATAA

>PGSC0003DMG401005047 G-LecRK

TTAACACCAAAAAACACATTTTTCATCTATGTTCTTGCTTCTACTTTCTTCTAATTTCACAAATCCTCGCAATAGCAGCAACAACAACAGATAATACGATAACGATTACAAATCCCCTTACGATATCGAAAACTTTAGTCTCTCAACAAAAAAAATTCGAATTAGGATTTTTCACTCCTGGTGGTCCGAATTCAGACAAATGGTACGTCGGAATATGGTACAAAGAAATCAAAGAAACGACGATAGTTTGGGTTGCGAACAGAGAAAACCCCGTCACCAATTCATCGTCTCCCCCTGTGTTAAAAATAACACAAGATGGACGTCTTGTTATTGATAATAGAGATGGAAATTATACGTTGTCATTAAATTCCGCGAATAATAATACTACATTTATCGCTAAGTTATTGGATTCAGGTAATTTCGTCGTTCTAACAGAGAAAGACGAAATTATGGTGTGGCAGAGTTTTGATTATCCAACGGATACTCTGTTACCCGGGATGAAACTTGGGTGGGACTCGAAAACGGGGTTTAATCGGAATATAACGTCATGGAAATCGCCATTTGATCCTTCTCCGGGTAATTATACGTTTAAACTTGATGTTAATGGTTTGCCTGAAGCTTATTTGACAAACAGAGATACAATTTTCTATAGAAGTGGACCTTGGAATGGTGTTGGATTTAGTGGTGTACCAGAAATGAAGCCAACAGATATTATAGTTTTCGAGTTTCAGATGAATAAGGATGAAGTTTACTACACTTTTGAGGTATGTTCGTCTCTTCACGTATGAGTCTGATTCTTTTTTAGTAATGACAGGTAGCGTTATGTTGAGTCTCATGCTAGTGGTGGGATCGAGGCAATCTTCACTTCATGAACAAGTGTCTATTTCTTATCATGGTATCAGAGTTAGACTTGTCTGTTCAATTCTTCGTTTATATGATGTTAGTGGGGTTAGGGGACTCTTTTGTCTTCTTATACGGTCTTGAGTAATCTTCACCTTATGAGGTAGTTTTTGGGGTTGAGTTATGTCTAACGTCTATTTTTTACATGACAGATTTATCTATGCTCTGATACCATGTTAATTTTGTAAAAACGTGTTTTTACTTACTTAATTATGTTTTTGTTAGGTGCTTGACAAGAAAATATGTTCGAGATTGTTAGTGAAGCATAACGGTTTTTTAGAGAGGTATACTTGGATTCCGACTAGTAACATTTGGAACAAATTCTGGTACGCCCCGAAGGACCAATGTGATTTATACGAAGAGTGTGGTGTTTCGGGAATTTGTAACGCGAATTTATCGCCGGTATGCAAATGTCTAGTAGGGTACAAGCCTAAAAATCAAGTGGCATGGGATTTAAGAGATGGATCAGATGGATGTATTAGATATCATGATTTGGATTGTGAAACTGATGTTTTTAACATATTGAAGAACATGAAATTGCCACAAAGTTCAAGTTCATTTGTTGATACTAAGATGAATTTGGAAGAATGTGAAAAAATGTGTAGGTACAACTGTTCATGCACGGCGTACACCACCGCAAACGTTACGGGTTCGGGTTCGGGTTGTGTCATTTGGACAAAGGAATTAGTCGATATGCGCCAATATTCGGCAGCAGAAGGTGGACAATTTCTCTACGTTAGAGCTGCTTCTTCTGACGCAGGTACATTTTTCATTTCAATCTGATGTTCGATATCTGTTTGAATTGTTATAGAAAATACGTTTTTTGTGGTAAAATAGTACTCCATTTCAAAATAATGAAAAGGTAATTTAATTTGCTTCTATAAAACTTGAATCTGAAACCTGTGGTTATTATGAAAAGAGTGCTTAATATCCTAGCGCTATTCTTAATTTAGTAGGAGAGAGTTGAGTCGAACTCAATTTAATGGAGTGGTTGAATGTTGGTAGAGATATATATGAAATTTTAATGTATTTGTTGGGTTGAATAATAATAATAATAATTGGTCTCATTTCATGAAGTTTGAACTACATATTTTTAAATGATAAAAAAATGGTCGTCTAGGCAATTTTTTACCCCTTTATTTTAGAAATTTGAAGTAATATCATGGATTGGACACGTGGAGATTCGCCACGTCACACTAGGTTGTTAGCACGCCCTTGTTGAGATTTTACAAAACTCTGAAACGATTGTGTAGATGATGAAGTTCAATATTTTATTTTAGGATATTCTGGAACAATTTTGTAGGTGATAAAGTTCGATATTTCTTTTTTAGGAGATTCTAGAACGATTGTATAGATAATGGAGTTCGGTATTTCATTTTTAGGAGATTCTAAAATGATTGTGTAGATGATAGAGTTCGATATTTCGTAATGAAAAAAAAATGCTTGTTTAAAATATCTTGTTAGTATATATTTAATGAGAGTAATTAATAGAAAAGTAGTTGGTGCTATACTTTGAAAAAACAAGAGGTTAGTACATATTATCATTACAATAAGCAAATTATAATTAATTAAAGTCCATTGACTCTTGACAAAGACTTGAAAACAGCCAACTATATATTCGAATATATATTATCTTGCATTATATTTTATTTGATATTGAGAGAGACAAAACATATTCTTTGTGGTAGGCACGTTTTAATTTTCACTTAATTTTCCTTTTTCTTTTTTTGAATTATTATGCATTGAAAAATTAAAGAAAAAAATGTAAACCAAAACATATGAAAAATACCCTCAAAAGTCCTAAATTAGTTAATTATATGGTTTATCTACTATTAGGTGTTTGAATTTTTTTTTCCTTTTTGATAAGATTGGGATGGTTAATTAGTTAGAAAATAATGCACCTACATCTCAAGCTTCTAGAATTAATTATTTTCCATTCAAAAGAAAATTCTACTAATAATACAAAGTTTTTATTCTCCCTTTGGTAAGTTAACTAAAACTGAGTTTCCTTTTTCTTTGTTATCTTATTCTTTCCTATGTATAAGCATGAGATCATATTTCAACTAGAATTTAAGTTAAGTTATATATACAAATAATATGGAAAAAAAATGCTAATAGTAACAGACATATGAATTAAAGGATAAATAAAAACACTCGATAATACACTAATAAAAAAGGTGAGTATAAAAGGTACATTTCTAACTTTATTAATTAGTTTGATGTGCACTTACAGCCTTACATGTTCACGTAAACTTAGTAGTTTTTATCCAGATTCTATAAATATATATATTAATAAATTCACTAAATATTTAGAGGGTCAGTAACCGATTTACATGAATTCAGTAGCTTTTATCCAAACCATCGTTTTCCTTCGGGTGAAGCTATAATAAGTCTGTAGGCAGGTTCGATAGAACTTAGTAGCTTTGACCTATAGTCTATATTTTTGCTTAAAAATCCACTAATCATGTATAAATTATTTATCAATAACTCAGCAAGCTATTTTTTCTAGAATTTATAATTCATAAACTTAAAATTTTAGCTCAGATTCTTATTATAAATATTCATAAACTTCAAATCTTGGATCCGATTCGTTATAAGTTAATGCTTACCTAAAAAAAGAAGGTGAGAAACGGGTTTAAAATTAAAACTTGGTCATAAAATGAACAAAGCGTGTGTAACCTTTTTAAATTTTGTATTTTGTATTGCTAACGCAAATTGTTGCGTGACATAATGAAAAATACATTAATTTTTTGCTGCTTTTATTGTTATATTTTCATGAGGATATCATATCTTTTTTTAATTATTATTTATTTTTTAAAATTTTGACAATAAATGGAAAAACTAAAATTAGTGGACAATTGGTCCTAAAAAGTTAGGTGTAAGTTCACACAAAAGATTCTTTTTGAAATGGATGAGAAGAAATCATTTTTTTGAACCCACTTTATATTCCAAAAATAGTCACTTGGCCGATTATTTTGACTTTGATTAATTTCAAATTTACATATAGTATGAGTTCTACTTTTAATTTTATTTTAACTTTGATTAATTTCAAATTTACATATAGTATGAGTTTTACTTTTAATTTTTCATAACGTTTACTTTCATATTATAAACATAAAAAATAAATACTTTTTGACAAGGACAGTCTTTGGGTCATTTGAATGGTGTACAAGTTACATTAAAATTATAATATTTAAAAATAACTAATGTATAATATTATTTAGTATATATGTATTTGACTTTACATTGGATAAATTAGTATCAACACTTATTTTCAATTTTAACCTTAGATTTTCTAAAGGACCTTGGAAGAAGAGTCTCAAGAAAGGGCATTTTGGTCATTTTTCGTATTTTATCTAGAGATTATTATCTGCATTAAAGGTTGTATAAAAATAATATCACAATTACGGTGTAACGACTAACCCCAAAATCAAACTTTCAATCACACGGAATAATATAATCTCACATTTCATCTCGGGATTGTTATCTATTATTCCAAATATCAAACGACTATAGGAATTACTAAGTTATGTAATGTGCACTAATGTTTTTGGATTCATGACAAAGGAAACTTGACCTTCCGTTCATTTGTATTTTTCGAGCAATTCACACCTAGTTCAAGTTACTCAATCGCGATAAAAAATAATTTTGTGACTTTACTGTTAAAAAAAAATACAACCTTTGATGGAAGACTCATTTTTATGTGTTAATTTTCAGCAAAAAGTGGAAATGTTGGATCTGAAGATGGTTCTGGCAAAACAAAAAGAATTGCTATGGCTACTGGAATTACAGCTGGTGTTGTTCTTGTTCTAATTGGAGTTGTAAGCATTTGCATTTTATCAAAGAGAAAAAAATTATTGGAAGGTCCAATAAGAAACAAAGCAGAACAAAGAGGTATGTTATTATAAGCCTATAATTTCGATTTCAGTATGGTATTCGATATTTACTTAAGGCATAATACATAAAGAATGCACATCTATAGACACCACGACTCCTTTACTGTGTCTTATGGACACCTGACGTTGTTGTGACACATAAATTTTGGAGATGTCTAGATGATCATTTTGTAAGTTGGAATGTACGTTCACTAGACATGATGGTGACGAGTTGAGGTGTCTAACTATACATTACTCAAAATTAGAGGGTTCAGTTGTCAGTTGAGGTCAAGTTAAGGTGTCTATCTATATATTATACAGTAACCATCTCATAATTCATATTTCTTACACTATTTTTGAACCAGGCTCTATTGAAAGAAGTCAAGATCTTCTAGTGAACACAGGCATTATTCCAAGTAAAAGAGAAATTTCTAGTGAAACAGTCGCGGACGAGTTTGAATTGCCATTATTTGATCTCAGTACCTTAGCTATGGCTACTGAAGATTTTTCTGATGCAAACAAGCTAGGCCAGGGCGGTTTTGGTTGTGTTTACAAGGTAAAGCTCATCTTGACGTTAAAAAAGCAAAAACTTATGTTGCTCGGATTGGTCAAAAATGTCATCATATCCATGTCAGATCCTCCAAAAATACATTACTTTGGAAGATGTAACACGGGTGTAGTGACATTTTTCGAGGATCATAGCAATACATCAAAAGTAGTACTTTTTTTCATTTACAAGTTAAAGCTGATCATATATCAGTAAAAAGAATATATGCTGCTCAGACAGGACAAAAATGTCGTTGCATGCACTCGTGTCAGATCCTCCAAAAATGCATCACTTTGGAGGATTGATCTAACAAGGGTGCAATGACATTTTTTGAGTATCTGAGCAATAATGAAAAACTAGTCCTTTTTTTTTTGCATTTACAATGTAAAACTCAACTAGATGTGAAAAAAACCTATGTTGTTCGGATTCTACAAAAATGTCATTGCACCCCTGATCTGACACGGGTGTTACAACATTTTTCAAGGATATGAGCAACATATGAAAAACTAGTTTTTGCCTTTACAAGGTAGACGTGAAAACAACCTATGTTGTTCGTATCCTACAAAAATATCATTACACCCTTGTCGGATCTCCAAGAAAATGCTTCATTTTTGGAGGATCTGACATGAGTGCAGTGACATTTTTCAAGATTCGAGCAACATTGGGAAAGACGGACTCTGTTTTATTGGAAAATGTGAAGACATTCAACAAAAGAACATCTTCTTTTTTTTTTATTATCAGGGAATAATAGATGAAGGTCAAGAAATAGCAGTGAAAAGGCTGTCAAAGAATTCAGGCCAAGGAGTAGAGGAATTCAAGAATGAGCTAAGATTAATTGCTAGACTTCAACACAGAAACCTCGTTCGCCTTCTTGGATGTTGCGTTGAGATGGAAGAGAAAATGTTGATATACGAATACATGGAAAATAAAAGCTTGGATTCAATCTTATTCAGTAAGTCGTTCTTATGATCAGTAACTATGTTTATGTGTGCTATTGGACTTCAAAATTAACAAATATACGCGTCTCAGATAAGCAGAAAAGCTCGCTGCTAGATTGGCAAAGACGATTCAACATTATCTGTGGGATTGCTCGGGGGCTTCTGTATCTTCATCAAGATTCAAGATTTCGGATTATCCACCGAGACCTCAAAGCAAGCAACATTCTCCTTGACAAGGAAATGATCCCGAAAATATCAGATTTTGGCATGGCACGGATTTTTGGAGGCGATGAGACCGAAGGAAATACAAAGAGAGTTGTTGGAACCTAGTAAGTAGAATCACATTATTCCTTATAAATCATTCATAAAAAACACTTCGAAAAGTGTTGGTAAATCGACGTTTTTTCTCTCTTAATGTAGCGGTTACATGTCTCCTGAATACGCGATGGATGGTCTATTCTCTGTTAAATCAGACGTTTTCAGCTTCGGAGTTTTAGTGTTGGAAATAGTAACAGGGAAGAAGAACAGAGGATTTTATTTTCAGAACAATGAAAGAAACCTTCTTGGCCATGTAAGTACACCAAAATTATCACAACTTAGTCATCATTTCTAACTAAGTCGGAACAGAATTTTGAGTTTATGCGTTCTGAATTCTAAAAAAATATAGCTAATTAGTAGGTTCTCGATAAATTATTTATGCATGTTAAGTAGATTTCATAACATAACTACAAGGTTTTTACTAAAGTTGCTACTGAGTTCTGTCGAACCTGTAACTTTCTCTCTAGTTCCGCCCCTAAATGTATAATCCGCTCTGCAGGCGTGGAAACTATGGAGAGAAGGAGGAGCCTCAGAACTACTCGATTCATCAGTTGGTGAATCATTTTCTCCGTGTGAAGTAATCAGATGCATACAGGTCGGGTTATTGTGTGTTCAGGAACAAGCAGAAGATAGACCGAACATGGCAACCGTTGTATTGATGTTAGGAAGTGAAACCGCAACAATGCCTCAGCCTAAACACCCTGGTTTTTGCCTCGGAAGGAGACCTGTTGATGAACACTCGGAAACTATTTACGAAGAAACTTTCACTGTGAATCAAGTTACGATTACCATGCTTGATCCCCGGTAGAGTTAAACGTTCAGACAGATCATAGGCTTGCCCGCTTCTACGTGACAGAATAGAAATAAAGGCAAATTCCACTGATGGGAGTGTGAACATAGAGATTTCACGACGAATGAATGCAATACATGGATGATGGATCCCTGCTGGCAAAAATGTATAGATAGTACATTAGTGTTTTTTCTCCACCCGACCATATCAGGAGGATCAATAGTGTTTTAACTTACCCTCCTATAGTACATTAGTACTACGAGTGGTTAGCAATTTTCTCTTTGTACACTTACCTTGATTTTTTTTACAAAAATAATGTTTTTTTTTGGGTAGTTAATTATAGAA

>PGSC0003DMG401009323 G-LecRK

GCTGCCAAAAATGAAATCTTTAGCCAATTCCATCTTCATCTTAATATTGACCTTATTCTGCAGCCATACAGGTATCATTACCCTTCCATCTGTGGCTGCAATAACCTCAATGAAGCCTGGTGATGAACTCAATCACACTCAGGTTCTTGATTCTGAAGATGGAACATTCAAGTTAGGATTTTTTTCTATCTCACAAACAAATAACTCTTATCTTGGTATATGGTATGCAGGTGATCCACAAGATAAGAAGCTATGGATAGCCAATCCAAATACGCCTTTATTGAACAATTCAGGATTGCTCACTATAGATACTACAGGAACATTGAAAATCACTAGTGGAGGGAAAACAGTTGTGAATATTACCCCTCCTTTATTGACAGGAAGTTTAATAGCTAGGCTTCAAGATTCTGGAAATCTTGTGCTTCAGGATGAAACTGAGAACAGGACTTTATGGCAAAGCTTCGATCATCCTACCGATACGCTTTTTCCAGGTATGAAGCTTGGTTATAACCTTACAACAAAGCAGAATTGGACTTTAACTTCTTGGTTGAGTAGTTATATTCCGGCTTCGGGGGCTTTTACTTTAAGTTTGTAGTCCATTCAAGATGCATTTCAGCTGGTTATACGTCGAAGGGGCGAGGTTTATTGGACTAGTGGCGCTTGGAGGAATCAAAGTTTTCCATTCTTAACTGCATTGCATGATTCTTCTAACAGATATCAGTATAATCTCAACTTGGTGTCTGAAAAAGATGGTGTGTTCTTCCAATTTGATGCGCCAGATGGAAGTTTCCCAAGTCTTGTGTTGACTTTTAATGGGGCTATTCTTGGTGGTGGTGAAGACAGTCGTATATACGCCCTTTATAATGAGTTTTGTTATGGTTATGAAAGTCACGATGGTTGTGTCTCTAATCAGTTGCCTGAGTGCCGAAAGGATGGGGATAAATTTGAGCTAAAGAGTGCATACTTCACTACTGGTAGTGCTACTTCTAATTATGATGATAATGCAAACATTAGTCTTGGTGATTGTATGAAGAATTGCTGGGAGCACTGCAGTTGTGTTGGCTTCATCACCAGCAGCAATGGAATAGGCTGCATAATTTGGAATGGAAATGGGGAGTTTCGAGTTGATGAGAGTGGTAGTGCAGTGCAACAATATGTTCTGGTTTCATCAAAAGGTAAATCCTTATATTAGGAATACACCTTGTTTATGAACTAATCTATTTTGAATAGATCAAGAGTCACCTGTATAACACTCAGCAACCTATGTGTGCAGGAAAACAAAAGAATTGGATATGGATAGTAATTGTTGTAGCTATTGTTGTCCCTATGCTGATATCTGGTTTCATCTGCTACATTATAGTGAGAAGGCGCAAACTACAAGGTAAATCTCTGAAAAAGTGAAAAATGATTAACTAGTAAGCTTTTTTATTAATTATGTTAAGCATGTGTAAATATCTACAAACTTTTCTTGCACAGCTGAGAAAAGAAGGGAGGAAGAGTATATACGTGAGTTGACAGCATCCGATAGCTTCAACGACACCAATATGAAAGAAAAGGATGGAAGGGAAGTCCAAGATTTGAAGATATTTAGCTTTGGATTTGTATTGGCAGCCACAAACAATTTCTCAAGTGAAAACAAGCTTGGAGAGGGTGGTTTTGGACCTGTTTATAAGGTATAATTACATCATATGCCTTCGGAAAACTTCAGTGCCATCATTGCATTAACTGAATTACCTGTAAAATTTTATGACAGGGAAAATTTCCAGATGGGAGAGAGGTGGCAGTCAAGAGACTTTCACGAACATCAGGTCAAGGGCTTGTAGAGTTCAAGAATGAGCTTATACTCATTGCAAAAGTTCAGCACACAAATCTAGTTCGAGTTCTAGGATGCTGTATTCACGAAGATGAGAAAATATTGATATATGAGTATATGCCTAACAAGAGCTTGGACTTCTTCCTATTCGGTTCGTTACCATTTCAAAACAAAGAATTAATAGCTCAAATAAGTTCTGAAAATTTTGACCTGAAATGCTTTCTGTCTTTGACAGATCCTGAAAGAAAGAAGCTACTGGATTGGCAGAAACGTTATGAAATTATCGAAGGAATTGCTCAAGGATTGCTCTATCTTCACAAATACTCAAGAATGAGAGTGATACATAGAGATCTTAAAGCCAGTAATGTGCTATTGGATGAGAACATGAATCCTAAGATAGCTGATTTTGGTATGGCAAGAATTTTCAAACAGAACGAGACGGAGGCAGTGACTGCCAGGGTTGTTGGAACATAGTGAGTAGAACTTATTTCATGATATAAGTTTAATTCATTTAGACATTTTCATATTGACTAAACGGGTAATGTTTCCTGTCCTCTATACAGTGGTTACATGGCTCCCGAGTTCGCAATGGAAGGAGCATTCTCAATCAAGTCTGATGTCTTCAGTTTTGGAATCTTAATGCTGGAAATTGTGAGTGGTAGGAGAACTACTAGCTTACAACAATTCGATCGTCCACTCAATTTTATAGGTTATGTAAGTTTTCCTTGACACCCCTTTTTCCTTTTAAGTACTATAATACATAAAGTTTGAGAATACTGATCATTTTGGTTTAGGCATGGGAACTATGGAAAGAAAAGTGTGCACTGGAATTGAAGGATCCAGCTCTTGGAGATTTATGTGACACTAAACTGCTGTTACGAGTTATTCATGTCGGACTATTATGTGTACAAGAAGGAGCAACTGATAGGCCAACAATGTCTGATGTTATCTCACTGCTAGGTAATGAAAGTATGCCACTGCCTACGCCAAAACAACCAGCATTTTTCACAGGAAGAAATGAAGCAGAATCAAATTCATCGGGTATCAAAGCAGAACAATGTTCGGTTAATGATTGTTCCATCACTGTCATTGAAGCAAGATAGCTTAGAGACATTATCTTGCTGTACTCTGTTGTGCAACACTCAGTAGCTATACATGAAAAGCATGTTTCCTTATGGTATAGATTAGTCAATTTGTACTGTTAGTACTTGTATAGCTGTCTTTAGCTGTGTTTGTAACAATACATAAATTTCTTTCCTACAAAGACCAGAATTATCAATAACTTTTTTTTTTTTTTGCATTTTCTCACTTAAATGTAGTAACATTTATTAGTATAACAACCTTAAGATTGTAAGATTCTCTTTATTTATTAGCTCCTGTATATTTTTAAAATTAGGAGACTCATCAATGATGATATTTTTTTGTGGTCCACATAAATGGATAACTTATCTGAGGATCATAATTCATAAGGATGACTGTAAACTAATCTAGTCCAAACAACTTACTCGACGAGTTAAGGACATTCTTAATCTTGATTCAACAATGCATCAATGAAGTTCACAGTGTATTTCAACAAGTAGATATGATATCTTTTTGAATTATATTAGCAAGCAGCAAAATGAAATAGAGAAATATGAGGAAGAAAAAAGAAATTAGAAAAGTCAAAGAAATAACAGAAACAAAATGCAAATTGATTTCACATTTCAACAAGAAAATTAAAGATTGTCTCTTTCAACAAATTTTCTTACAAGTAAGGTAATATTTCTACTTAGAAAAATCTTCAAAATTTCTGGCTGGAAAATGTTTAATATAATTGTCTATTCTCTAATGCATTATTGGCTTTGAACAGTAAACAACCTAAAAAGGAATAATCTATTATTATCTGCAGCAACATTTCTTTCTTTCTGGCTGCCAAAATGAAATCTTTAGCAAATTACATTTGCTTCCTCATCTTGACATTATTCTGCAGCCAAACATGTATCATTAGCCTTCCATCTGTAGCTGCAATAACCTCAATGAAGCCTGGTGATGTACTAAATCACTCACAGGTACTTGATTCAGAAGGTGGAAGATTCAAGTTGGGATTTTTCTCTATCCCACAAACAAATAAAACTTATCTTGGAATATGGTATGCAGGTGATCCAGTAGAGAAAAAATTATGGATAGCCAATCCAAACACACCTATATTGAACAGCTCAGGATTGCTCACATTAGATTCTGCAGGAACATTGAAAATCACCAGTGGAGGGAAGACAGTTGTGAATATTACCACTCCTTTATTGACAGGAAGTTTAATAGCTAGGCTTCAAGATTCTGGAAATTTGGTGGTTCAGGATGAGACTCGGAACCGGACTTTATGGCAAAGCTTTGATCATCCTACCAGTTGTCTTTTGCCTGGTATGAAGCTTGGTTATAACCTTACCACAAGGCAGAATTGGACTCTAACTTCTTGGTTGGTTAGTAGTACAATTCCGGCCTCAGGGGCTTTTACTTTAAGTTTGGAGGCAATTCAAGATGCATTTCAGTTGGTTGTTAGACGAAGGGGTGAGGTTTACTGGACTAGTGGGGCTTGGAACAATCAAGGTTTTCCATTCTTACCTTCATTCCGTGATTCTGCTACTACTTATCAGTACAATCTCAACTTGGTATCCGGTACTGATGGGATGTTCTTCCAATTTGAGGCTACAAAGGGAAGTTTCCCAAGTCTTGAGTTGTTTTCAGATGGAGCTATTGCTGCTGGAGACGGTAGTATATACACCCGTTATAATAAGTTCTGTTATGGTTATGGAGGTGATGATGGTTGTGTCTCTAGTCAGTTGCCTGAGTGTCGAAAGAACAGTGATAAGTTTGAGCAAAAGAGAGGAGACTTTATAGATTTATCTGGTACTACTACTAGCAATTATGACAATGCAAGCATTAGTCTTGGTGATTGTATGCAGAAGTGCTGGGAGCACTGCAGTTGTGTTGGATTCACCAGTCTTAACAGCAATGGGACAGGCTGCCGAATTTGGAATGGAAAGAGGGATTTTCGAGTCGATGAGAGTGGTAATGCAGTTCAAAGATATGTTCTGATTTCACCAAAATCATCTAAAGGTTAGTCTTTAATTATGTCATAAATACACCTTCTTATGAACTAATTTTTTGGTCGAACAAACACATTTTATTGCAAACACCCAAAAACAGAATACAGTATATCCTAAGTAAAAAGATACAACTGACCACAGCTAACCAAATTTAGAAACATCCCCTACTCTTCCTCCCTTCTATACAGTTTTCTATTTGTAAAGTTCTTGTTGGTGTTTTTTTAAAAATGGGGCAGCACAAAAAGATTGAAAATGTCTCTCAATATTTTATTAATCATAATAGGGCTTTAAATAGCCAACTTGATAACATAATCTGCAGAAAATCTGCATAACAAGAATTTAAAAAGACTAAATAATCTCAACAACTAAAACAACCTAACAAATATTTAAAATTCAAATTCATCCTAAAACTAGGTAGCTTATTATTTTAAAAAATGACTGCAGTAACTAAAAGCAAACTCAACAGTTCTTCTATCCATTCTCCGTCTTTACAGTTTTTCAATTTTCAATATATAGACTGTATTAGTGAAGAATCAACTGTATAATTTTCATCGTGTTACGTGTGCAGGAAAAACTTGGATATGGATAGTATTATCTATAGTTATCACAATGCTAATATGTGGTTTGATTTGCTTGATCAAAACAAGAATACAAAAACTACAACGTAAGTCTCTGAAAAGTGATTTACTAACTAGTATGTTGTTTATCAGTTATTTTAAGCATGTATGTAAATCTACATACTTCTCTTGCATAGGTGAGAAAAGAAAGAAGGAGGAGCATATACGTGAGATGAACGCAGCAGACAGCTTCAACAACACAAATCTGAAAGAAGAAGATGAAAGGGGAGTACAAGATTTGAAGCCTTAAGATCAATGAAGCTCTTGATACCACTGTTGGTGTTTTTTAAAAGTGGGGCAGCACAAAAAGATTGAAAATGTCTCTCAATATTTTATTAATCATAATAGGGCTTTAAATAGCCAACTTGATAACATAATCTGCAGAAAATCTGCATAACAAGAATTTAAAAAGACTAAATAATCTCAACAACTAAAACAACCTAACAAATATTTAAAATTCAAATTCATCCTAAAACTAGGTAGCTTATTATTTTAAAAAATGACTGCAGTAACTAAAAGCAAACTCAACAGTTCTTCTATCCATTCTCCGTCTTTACAGTTTTTCAATTTTCAATATATAGACTGTATTAGTGAAGAATCAACTGTATAATTTTCATCGTGTTACGTGTGCAGGAAAAACTTGGATATGGATAGTATTATCTATAGTTATCACAATGCTAATATGTGGTTTGATTTGCTTGATCAAAACAAGAATACAAAAACTACAACGTAAGTCTCTGAAAAGTGATTTACTAACTAGTATGTTGTTTATCAGTTATTTTAAGCATGTATGTAAATCTACATACTTCTCTTGCATAGGTGAGAAAAGAAAGAAGGAGGAGCATATACGTGAGATGAACGCAGCAGACAGCTTCAACAACACAAATCTGAAAGAAGAAGATGAAAGGGGAGTACAAGATTTGAAGATATTTAGCTTTGGATTGATATTGGCAGCCACAAACAACTTCTCGAGTGATAACAAGCTCGGAGAGGGTGGTTTTGGACCTGTTTATAAGGTACATTTTCAGTATGGTATATCATATATGTACCCCAAAAACTTCTGTAGTACTCTCATTATATGAAATAATAATCTGTAAAAATGTATGACAGGGACAATTTCCTGATGGGAGAGAGGTGGCAATCAAGAGACTTTCAAGAACATCAGGTCAAGGGCTTGCGGAGTTCAAGAATGAGCTTATACTAATTGCCAAAGTTCAGCACAGAAATCTAGTTCGTGTTCTAGGATGTTGTATTCACGGAGATGAGAAAATGTTGATATATGAGTACATGCCTAACAAGAGCTTGGACATCTTCCTTTTCGGTTTGTAACCATTTCAAAATAAGATCTAATCTTTCAAATAACTGTCGAAATGTTTGATGAAATGTTTTCAGTCATTGACAGATCCTGAAACAAAGAAGCTACTGGATTGGCAGAAGCGCTTTGAGATTATAGAAGGAATTGCTCAAGGATTACTCTATCTTCACAAATACTCAAGAATGAGAGTGATACATAGAGATCTAAAGGCTAGTAATGTGCTCTTGGATGAAAACATGAATCCTAAGATAGCTGATTTTGGTCTGGCAAGAATTTTTAAACAGAATGAGACGGAGGCAGTGACTCGAAGGGTAGTTGGAACATAGTGAGTTGAGCTTATTTCTTGATATTGACCTAGAGAACATGTATACATATTCATACTGACATAGGTGGAAAATGTTTTCTATACACAGTGGTTACATGGCTCCCGAGTTCGCCATGGAAGGAGCTTTCTCAATCAAGTCTGATGTCTTCAGCTTTGGAGTCTTAATGCTGGAAATTTTGAGTGGTAGGAGAAATGCTAGCTTACAACAATTCAATCGTCCGCTCAACTTAATAGGTTATGTAAGTGTTTCTTGACATCCTTTTTATATTTGAAGTACTAAATGTACAAAGTTTGAGATAATTGATCATTCTGGTTTAGGCATGGGAACTATGGAAAGAAGGGTGTGGACTAGAACTGAAGGATCCAGCTCTTGAAGATCTTTATGACACTGAACAATTTTTACGAGTAATTCATGTCGGACTACTATGTGTACAAGAAGGAGCAACAGATAGGCCAACAATGTCTGATGTTATCTCAATGCTTGGTAATGGAAGTATGTCATTGCCAATTGCAAAACAACCTGCATTTTTCACAGGAAGAGATGAAGCTGAATCATATTCGTCATCAAACAAAACAGAACAATGTTCAATTAATGATTGTTCCATTACTGTCATTGAAGCAAGATAAGATTAGAGATGTTATTTTGTTGTAATAAAGATTGCAGTAGATCACCAATGTTTTAATGACCTCTAAAGTAACTTGAAGGCTCTTTTTTGTTAAGGTTTAAAAGCTAATTTTTTCTGCTTCATACATATTAGAACTCGG

>PGSC0003DMG401013876 G-LecRK

GAAAAACTTGAGACTTTAAGAAAGGGGACGAGAGGAGTACACATTTTTGTCCATAATTTCCAAAGCAGCTGACGACAACCCTTTTAACCCTTTGTCACGTAATTCCTACTCAAGCAAATCTTTCGTTGGGCTGTCTAATTCTTGTGCCTCTTCGTAGTTTAACCTTAGGCCCAACATCTTAGCCAGCCTCAGATAATTGGTCCATAAGCATTATCTGGGCTATCCATGTTCACAACATTATTAGTACTTATTTTTTAGTGGAATGATTGCACTACTGCAGTTTGATGTCATTTTCATCAATTGGATATTTTCCTTGCGTAGAATTTTCAAATATTTTGACTAAGGTGTTTGATTAGTTGCGTAACTTTTGGTTTGGTAGAATTCTCAAAGACTAATTTTATTTTTTCTGTTTCTCTGAGATGATAACATTGATATACTTATGGATACTTGAATGTTTGTTTGTGTTAATTCGTCTATTCCCTCGACGTGTCAAAACTTTTTAAAAACAGGTATTTGGGAATGAAGATATTGTTGGTTGATTTAGTTCTCTTGCTTTCTTGTTTTCATTTAGTTAGGTGCAATAATACGCGTTCAGACAACTTATTGCCTGGACAAACCCTTTCCTTTTCTGCAGCAGATACTTTAGTGTCCAAACAAGGGAAGTTTGAACTCGGTTTCTTCTGTCCAGGTAATTGTGAAAAGCTTTTTATTGGTATATGGTATACAAACATAAAGCCCAAGACCGTGGTTTGGATAGCAACTAGGGACAACCCCATCCGCCCTCCGTTCAATAGTTCTCATCTAGAGCTGTCTGATGGAAATCTAGGAATCTTTAATGAACGTAAACAAAGAGTTTGGGACTCTAATTTATCCACAGCAGATGTCAATAGGGCAGTACTTCTTGATTCTGGAAATTTCATTTTGACAAATGGCATTGAAATACAGTGGCAGAGCTTTGATTATCCTACAGATACATGGCTGCCTGGTGCGAGGATTGGATTTGATAAGTCCAAGAACACACTCCAGAAACTTACTTCTTGTATGGATGCAAATGATACGGCTTCAGGACACTTCTCTTTGGAATTAGACCCTAATCAAAATGGTGAGTTAGTCATGCTACGTAACCTCGTGGTAATGTGGATCTGGGTGCCTTTGAATCTCGTGGTATCCAACTTCAGTTATAATCCAACTGAACATAGTGCATTTATTACATACAATGTCTTTGGTGAATCTGTCATTTCTCGAATTGTCCTAGATCATAGCAGGTTCATGAGGGAACTCTTCTGGTCTAATGAGCATCAGAATTGGATGGTTGTGCGGTCTGTACCTGTTGATATGTGTGACGCAACGAATATATGTGGTGCTTTTGGAATTTGTGATATAAATACCAGTCCAATCTGTGGATGTTTACAAGGTTATGAACCCAAGTTTCCATTAGGTTGGGATACTAATGATTATTCTGGTGGGTGTGTGAGAATAACTCCGCTGCAGTGTGGCAGCAATAACGATGGTTTTGTTCGTATGCAAAATGTTGAACTTCCTGCATCATCTGAATCAGTGCAAGTGGGGAATGATCAGATCTGTGAATATATATGTTCATCCAATTGCTCATGCAATGCTTATACTTACAGCTATAGTGGAGTGTGCTTATTATGGAATGATGATCTTATCAATCTGAGAAGATTATCTGATAATTCACGTCAAGCAGTGTTAAGCATAAAAATGTTTGAGCTTTCAAACAAAGGTTAGGTGCTTCTAAACTGCTTCTTGTTCTAATTGTGCTTACTTGAAACTGACATTATGTTCATTTTCTCCTCTCCCCATCTATCTTATTGCTCAGGCAATAAAAAGTCATCGTTTGTAGTGCTAGTAGCATCCATCACTTCAGCAATTTTTGTCTGTGGTACATGCTGTTACTTTCTATGGAGACGAAAGCTCAAGAAAAAAGGTACTTCTTTATATTTCCAATAGCTATGCCAGCGCTTTATGAATGTTAATGTGGCCTACATTGGTTGAGGCCATGAGGGTATAGGATCTTGTCTCCTTATATTGTCTTGGCTATCCTCACCTCATGAACTAGCTTTCAATCGAGATTGACATTTTCTTAACAATGAACATATGTGGGATGCCAATAAACGAACCTAGTCCTTCTTGATTCATTATTAAACTGTGCAGTCTTCGTGTGATATTCTAGTTCTTACTAGTAGTAGCAGGTTATCTGACTTAACTGTGTTCTCGCTGTCACACTCACTACAATCATGGATTAGATTATAAGTTAAAATTACCAATGTACAGCTTCATCTATAGACTTCATGTTTCTATTGCCTTTTCAGGGATTTTGAGAAAGATGAAGTTCAGGGAAATGTTACTTTCTGACTCAGCAACTAACATGTCAAAACCTGGAACTTCCATTGGAAAGGGACAAGAAAAGAATGGTGACATTGAACTGAAGTTCTTTGAACTACATGACTTGAAAGCCGCGACAGACAATTTTCCCCCTGATAATAAGCTTGGAGAAGGAGGTTTTGGTCCTGTTTTTAAGGTATTGTTACTACAACCTGAATACCTGCAACACAAATATGCGTTCACACCTAAAGAAAATGGAAAGAAATCCCTCCGGAAATAGCCAGATAGTATGTATCCTGATTTTAGTTAATATCTTTGCTAGCTGTACTTATGTCTTGGAGGTGTGATAAATGTTTTTTTGCAATTTCTAGGGTCAATTGCCAGATGGGCAGCAAATAGCTGTGAAAAGGCTATCAACTCAATCAAGGCAAGGCATAAGTGAATTCAAAACTGAGGCCCTTCTCATTGCAAAACTCCAACACAGAAACCTTGTTAGGTTCCTTGGCTGTTGTGTAGAAGATGAGGAGAAAATGTTGGTATATGAATATATGCCAAATAAAAGTCTAGACTATTTCATATTCGGTATATATGATAATTTACTACCATCTGTTTTTGTGTTAACCCTTTCCTCCTGTCAAATGTGTGCTTACTGGATAAATTTGATATGTACAACTTAGACGAGAGCAGGAGGTCATTGCTTGATTGGAAAAAGCGGCATGAGATCATTATTGGTATAGCTAGAGGGATACTCTATCTGCATCAGGACTCAAGATTAAGAGTAATCCACCGTGATCTGAAAGCCAGCAACATTCTTTTAGATGAAGACATGAACCCCAAGATATCAGATTTTGGTACAGCAAGAATATTTAGCGGAAACCAAGATGAAGCAAATACTCTTCGAATAGTTGGCACATAGTAAGCAAGCAAATACTGAAAGTTCTTTTCCCATACTACTTTGCCTACTCATTATAGCTCCTTTTCTATGCAGTGGCTATATGTCACCTGAATATGCTCTAGCTGGTCTCTTCTCAGTAAAATCTGATGTCTTTAGCTTTGGTGTCATATTATTGGAGATCATTAGTGGAAAAAAGAACAGAATCTCTTATAATAGTGATTCCCCTCCAAATTTGATACGACAGGTAAGTTGTAAAATGCAAATGTTCGAATCTTTCTTTGATTAAGAATCAAATTTATCCATTTCAATGAACTACTGATTCAGGCCTGGGAGCTTTGGAATGATGGCAAGGCCTTCGAGCTGATAGATCCAACGATAGTTGATTCTTGCCCCAGTGAGGAAGCACTACGCTGTATTCAAGTGGGACTATTGTGTTTGGAAGTGAATGCAGGGGACAGACCAACCATGTCATCAGTTCTATTCATGCTAAGTAATGAAGCAACGATTCCCTCCCCAAAGCAGCCATTGATTACCCCTAACTCTGACTCGAGTACAACAGAAACAACACCATCTTCGATCAATGAGGTGACTATTACTGCACTTGATGGTCGTTAGTCAGCTGATGACTGATGATTCAGCCAATCCGTTCAAGTGTGGGAGGGTTATTGACCCATCCATTTACTAGCTCAACGAATTGGGTTGATATGTTGTCCAAATTGATTCGGGAGAGATTGTCAAAATAATTTTCAGAGGACATTTTTTTATTTGATATGTTATATAAATAGTTACAATAAAGAAAACAATAATTTTAGGTATTAAAAATTCATAAAAGAACAAATAAAATAATTAAACCTTACTAAGAATTGAACGAATTGAGTTACAACTCACATTGTAGTCCTGTTTCAGTCAAATTTAATCCACTCAAATTCGACTCAACCCGTCTATTTGAGACCTTTGATCT

>PGSC0003DMG401019239 G-LecRK

ATGAGCGGTAATGGCTGGGGAAGGGGGCAAGTACATCCTTTTGTTCTCAGTTCGTGGGAATTGCTTCCATCACTAACCAGTTCAGCTCGGCTAGTGGGCTTTTGTGTTCGAACCTCGAATCCGGACCTGTTCGACAAGACCAGATCCATCAACAAACCTAGGTCCAATATTTGCGTCTACGAGCTAGCCATCAACAGACGGACATCCTTTTACCGCAGTCTTCCCCTATTGGGCATAGACTTTTCTACGTTAGGCTCCTGTAAGGGCGGGTATGAGAAGGGTCCCCCTCTCCCTTGTCAAAGCTTAGTTATCCGTGAATGAGAACTCGTTTTGCTTGTTGGCTGGCTTATGATGGTTTCCTGTCCCAATTCAACATATTCCTTTTTTTGCTCCTGCTTGGTAGCAGGCTTGTGTGCGAGCTTGGTCCGGTTTCGATTCCTAAAGCCCACGGAGAAGGGTGTGGAGCCTCGCTTTCCATGCTGCTCCTAGTTCGTGGGAATTGCTTCCATCAAATAGCCGTACCATTAACTGATTAAATTAAAAATAGCTACATTGGGGTAAAATGCTTTCTAACAAATGCGACTTCATATCCAAAAAGATTCAAACCCAAGACTTTTAATTAAGAATGAAGGAGTCACATCACCACAACTGTGTCAGTGATTTGATGTTGTTGTTGTAAGCAAATAAGATAGAACATGGCCCCAACATCCATGTGTGGCTGTGATTAAATTCAACTATTCATTATAAACAAATAATCTCAAAACCAAAGAATACGATGAGTATTAATCAAACCAGCCAAGCTAAGGTGTGTTCAAATAGGACTTGGGTCAAGTGTCATCAATATGCTGCATGAACTGGTCATAAGAGTCGGAAGGCAGACTTGCTAAGCACTTCGACATAGAGGAAAGACAACACAACGTCTATTGGATCACAAAGCATGAAGGAATTGCTAAGGGAGAGTTTTCACTTATTCTTGCTCATGTATTTGTATTTTGTTCAACGCAGCATTGGCGCTACAGATACCATAAGTACAACCCACTTTCTGAAAGATGGTGATGCCAATATTACTTCACCTGATGGAACCTTTGAAATGGGGTTTTTCAGTTCAGGTAGTTCCAAAAATCGCTACGTGGGTATGTGGTACAAAAGCTTATCTGTTAGGACAGTGGTGTGGGTTGCCAACAGAGAAGCTCCCTTGACCAGTGGATCAGGTATCCTTAAGGTTATTGAGCCAGGACGTCTCGTCCTTCTCAATGACACTAACAAAGTTGTATGGTCAACTAATACCTCTACATCTGTGCAGAATCCCATTGCACAGTTGCTGGATTCTGGGAATCTTGTCGTGAAACAAGCTGGTGATGATAATTTCCTTTGGCAGAGTTTTGATCACCCAACTGATACACTATTACCAGGCATGAAGCTTGGTTGGAACTTTTTAACTCACAGAGAGGTGTACTTGTCATCGTGGAAGACCCAAGAGGATCCAGCTCCAGGTGATTATACATATCACTGTGATCCTTCTGGATATCCACAGAACATCCTGAAGAAGGGGTCAAATGTTGTCTATCGATCAGGGCCGTGGAATGGTCTTCATTTTAGTGGAACAAGAAACCCAAGAGGCAGTCATTTATACAAATATGAGATATTCTCAAGTAAGACACAGGTTTATTTTGGATATAAGCTCATATCTTCAGTTATCACAAGGTTGATACTTAATCAGAATGGTGCCCTAAAACTTTGGACTTGGGGTGATCATGTATCTGATTGGATCCCTCAACTCTCAATACCCGCAGATAACTGTGATGCTTACAAACTGTGTGGCGCATACGGGACTTGCAACAGTCAAGATTTTACTGTATGTAGATGCTTAGACAGATTTGTGCCCAACAATAGTGAAGCTTGGAAGAAGAGAAACTGGTCAGGTGGCTGTGTTCGGAAAACAGAACTAAATTGCCTTCAAGGACATAGATTTTTGAAGTATTCACATATCAAATTGCCAGACACACGGAATTCCTGGTCCGATGTGACTATGACACTTGAAGAATGCAAGAATACCTGCTCTAAAAATTGCTCTTGTATGGCTTATGCAAATCCTGACATACGCAATGGAGGAAGTGGATGCTTGTTGTGGTTCGAGGATCTGCTTGACATTAAGGTGGTTTCCGATGAAGGTCAAGATATCTACATAAGAATGGCTGCTTCTGAACCAGGTAGGTTTTGTTGTCACTTAAAGATTTTTGATGGATTACCTGAAAATAGTAAAAGAATGTATGGTTGCAGTCACTTCTTTGGAACCAATTGATAGGTATTTAGTTTTCTGTAGAACAAGAAGTTGAGATGCATGCGGCTTGAATCAAAGAAAAAATCTTCTCAAAATTAAAGAGATTTATAGTAACATTCAAAAATTAAAAATGACTTGCATTATATTGAATTCTAGAACCCTTACAACTCAATATCAACAAATTCAAAATGATGTGCAGTACTCCAAAGGCTTACAGAATCTTTTTTAATTCAATATAAAATTATCTCCCTCGTGGATTGGACAGAAAGTGAAATTGATTATTTTTGCGGAATGGATACTTGCAGATAGTCTGGAGAAATCAGATGGAAAGAAAAGAAAAGTACTTTTCTGGATTCTGCCATTATCAGTTGGTCTGATTCTGGTAATTCTTAGCATGTTGATCTTCCATAGAAGAAGGAAGAAGGCTTTACAGCTCAAAAGGAAAGGTAAAATAGGATAAATCTCATTTTTTATTGACATCTTAGCAAAATCAAGGAACATAATTCAAAGAAAAGTGAAGAACTTTAATATTTGGTCTGGATCACAGCTTAAACAAAATTCACTTCGTTTGCAGGCAGGTAGGTCCGGACGCAATGGCAACTACAAGATGGATCATAGTGGTAATTGTGAGGAAGAATTTGAGATACCAATGTTTGACTTATCTACCATAATGAAGGCTACCAATAACTTTTCAATAGACAGAAAGATTGGAGAGGGGGGCTTTGGACCGGTTTACAAGGTATAACTGGTGCACAGGCAAAAACTTCCTGATCCTATAGAGTAAAGTTTCATCTTTAGATTCAGTATAGAAGTTGTATGATGAAATGCATAATGAAATAAAGCTTTGGCTTGTGAAACCGAAGGAGACATACACAATAGCTAAGGAAGAACAAGGATACGGTGGAACACAAACTGATTGTCTCTTTCTAATTGGAACACAATTGACTCTGTGTACTTCCTACATCAAGTAAGGTAATGAAAATATTAACATGCAGAAATTGTGATGTTTAGGGCATACTTGAAGGACGGGAAATAGCTGTCAAACGGCTGTCCAGAACTTCCAAACAAGGAGAAGGTGAGTTCAAGAATGAAGTTTTATATGTTGCAAGACTTCAGCACAGGAATCTGGTGAAGATTCTTGGTTGCTGCAGTGACGGGGAAGAAAAGATGTTGATCTATGAGTACTTGCCGAATGGGAGTCTTGATTCATTTATATTTGGTACTATTCTCCATCCAATATTGTGCAAATTAAATAAATAATTGTGTTTCAGTGATGATGCCTATAGTAATACTTCATTAATCTTTTTTATTTTGAAGTAAAAGTAACTAGTAGTACTTTGGTAATTTTGTTCCAATTATGAAAGCCCCATAAAGCAGGAAAAATATTGATAGACCAAACCTGTTGTGGGAGTATTTAATACTACTTTCTCCTATAACCAATTAGTCAGTTAATGACCCAACAAAAAAAATTGGCAGATGACACACAAAGCAAGGCATTAGACTGGCCTAAGCGATTTCACATTATCAATGGTATAGCTCGGGGACTTATGTATCTGCATCAAGATTCTCAATTGAGAATAATTCACAGAGACCTGAAAGCTAACAACATTCTGCTAGACAAGGACATGAATCCAAAGATATTAGACTTCGGCTTGGCAAAAATATGTGAAGAGGATGACATTGGAGCCATGACAAACCGAGTAGTTGGAACATAGTAAGAACCTTCCACTGGATAGTATTATTTTGATAAGATTTACTGGATAATAATTTGAACCAAAGAAGAATCTTCAGTTAGTAAAGGCATGTTTTCCAGCATTACTAATTATAATGTTGCGCACATTTTAATGCAGTGGGTACCTCTCACCGGAATATGCATTGCACGGAAAATACTCAGTAAAATCAGATGTATTTAGCTTTGGTATTTTAGTATTGGAAATTGTCAGTGGGAAAAGCAACAGAAAATTCTCTCATCCAGATCATAACCTTAATCTACTCGGGCATGTAAGATTTTACTTGTTTTATTTCTTATTTTTTTGATAACCGAGGTGTTGGAGCCAACATACTTGAGCCAAGCGTCTATCAAAAATAGTCTCTCTACCAACGAGGTAGTGGTAAGGCCTACATACACACTACCCTCCCCAAACTCCACTTGTGGGATTATACTAGTTATGTTGTTGTTGTCGTGGTGCCGAGGCCAGCTTGCGCACACCTCAACTTATTCCATGGGATACCTGCAAACTCCCACCACCAATAGGTACCGGGTAACTCTGTCCAACAAGGTTAGGACTGATGGGAAGAAATTGCCTAGTGTTTTTTGTCTCTGCTGGAATTTGAAACTGATCATCGACCACTAGGTCACACCCTTGGGTGCAGTTGTTTAATTTTATTTATTCCTTGGTGTATGTTATATACTAGTAACAAATATCTTGTAAAACGACCCCTGCTAATAAGCTTGCAATATCTACAGGCATGGAAACTTTATATACAAGGTAGGTCAATAGAACTACTTGATGAATGCCTAGGTGATTCTTGTTCAACATCTGAAGTGGTACGATCAATTAGTGTTGGTTTATTATGTGTCCAACAATGTCCTGAAGATCGTCTAAGTATGTCTTCCGCAGTTCTGATGTTAAACAATGAAGGTGTGTTGCCGTTGGCTAAACAACCAGGTTTTTACATAGAAGGAAATGCACCGGGTGGTGATTTCACTCCTAGCCAATCTGCACAGATTACAGTGAGTGACACCCCCATCACAATACTAGCTGCTCGATAGAAGATATTGGTTCACTTGTTTGCAACAAGCTTGCAGTTGCAATGATCTTTTCTTAATTCTCTATAGTAATGTGGTGAAAAAAATGTGTGCATGATTGTGCTTTTTTAGGTGTATCAACTATGTCAAACATATGTAGGAAGCTTGCTGACTGGCTGTTACTTTGATTTGCGCAGTATATCTGAT

>PGSC0003DMG401019242 G-LecRK

ATGTTACTAACTTCTGCAGCACTAGACACAATCACTACAGATAAATCCATTAGAGATGGTGACACAATTGTTTCAGCTGGAGGGGTTTATGAGCTTGGATTTTTCAGCCCTGGAAATTCGAAGAATCATTACGTTGGCATATGGTACAAGAAGATATCACCTACAACTGTTGTGTGGGTTGCAAACAGAGATATTCCACTTAATGACACTTCAGGAATGTTAACACTTAATCCCAATGAAATTCTTGTACTTGTTGATAAATCCAATGTATCAATTTGGTCATCAAACTCATCAAGATCGTTAAAGAATCCAAAAGCATGGCTCCTGGATTCCGCGAACCTTGTTGTTAGTGATGGAAATGATAGAGAACAGGGAATTAATTTGGCGTGGCAGAGTTTTGATTATCCAGGAAATACTTTATTACCTGGAATGAAGGTAGGAATAGATTTGATTACGGGCATGGATAGGTATGTAACGTCATGGAAGAGCACGGATGATCCTACTCCTGGTGAATATGTAGATCGTGTTGATTCACATGGATACCCTCAATTGTTCTTGATGAGAAATTCATCTGTAGTGTTTTGCTCAGGGCCATGGACTGGTGCTGCGTTTTCTAGTAGTCCTAGTAATAAACCAACTTTGTATTATACGTTCGAGTTTGTTGTCAATCAGAAGGAAATTTACTTTAAATACGAGCTTAAGAATGACTCTATGCCCACCAGGGTGGTGCTCAACCGGATGGGGTGATAGAACACTTAATATGGATTGAGCATACTCAGAGCTGGTTTCTCTACTTGACAGCACAACTGGATAATTGTGATCGTTTTGATTTATGCGGACCTTATTCAAGTTGCAACATCAATAACTCCCCTCCATGTGACTGTTTGAAAGGTTTTGAGCCTAGGTTTGTATTTGTTAAACTTTGTGATCATCTCGCGTTAAGCTATTAGAGAGAGTGCACTTTTGAATTTTGATTACTAAGTTATGTTTTCACACACCCCTCATGTGTGTGCTTGATTCTTTTTCATGGACCAAACACGTGTAATTATTTTTTTGCCTTCACAGGTATCCTCAAGATTCTGCAGCAGACTGGTCTAGTGGCTGCATAAGGAGAACTTCTTTAAATTGTACCCATGATGGTTTTCTTAAATTTACGGGTATCAAGGTGCCGGATTCTAGAAACTCCTGGTATGATGAGAGAATGAACCTTGAAGATTGCGAGAAAATGTGCTTGGCTGATTGCAATTGTACAGCCTACTCAGATCTTGATGTTAGAAATGGCGGAAGTGGATGCTTACTATGGTTTGGAGAACTCATAGATATACGAGAGTTCAGCCAAAAAGAGCAAAATCTGTATGTGAGAGTAGCTGCTTCAGAATTAGGTAATGACCCTACTCTGCATAAATGCACTAACTCTTTAGAATTGAAGGAACTCTGCTACCTAATTTACTTACATGATGGACATTAAAGTTGATTAGGTGCTTTAGTAGTTGGTAATATGGCGATTTCATTGTGAAGGATTTCATTTTCGGTGGTACTGGAACGGAAGTAGCTTGCTGCAAAGAGCATTTGAATAATTTCATATCTTACTCTACGACTATATGGATACTAGGACATAACCGGTTAGTAGGATTGCTAAGGGATGAGACACTTCATTCCCTTAAAAGGCTTAAAAACATGCTAGACCTCATGACAGATCAAGGCTGTTGACAGCGGGTGATAATCATATGCATTCTGTCCTTAGTAGTGCCGCTTGTAATCAGTGACACTGTACTTTAATATGTGTCATCTTTCAAAGAAGATCTCCAATTTAACAATGCAGCTTGTAATCAATTAAACATTGACTACTTTTTCAGTAATCTTGTTGATTGAATGCCCTGGTCTACAAGTTACAAAGTCTGCAGGGCTTCATAACAAAATAAGAAGTAATACAATCAATGCAATTAACTACTGATATGGCTAACTCATATGTAAATTGAATACTGATTATGAACTAATATAGAACATGTTCAGACAGGACGCGGAGGAGAAAGAGGTCAGTCCTGATTGGTGTCATTTCAGCAGTGGTAGCAACGCTTATCCTCAGCTTTTTAGCTTGGTTTTACTTCCAAAGAAGGAAAAGAAGAAGAATAGGTAAATATGCCTGAGGTACTCTTGTCTATCTGGCTGACATTGTTCGCCTTGACCAAATTCTCATTGATAGCATGTTTGTGGCAGGCGAATGTATATTGACAGGTTCAAAAGTTGAAAATGAGGACATGGAGCTTCCATTGTTTGATTTAGTTACTGTTACTAGCGCCACTGGAAACTTCTCTTCTGCTAATGTGATTGGGGAAGGCGGATTTGGACCGGTTTACAAGGTAAATCAACTTTGATCTTCAACTTCTAGCTGGAAGTGGCATCTTTAACCCTTGATAGAAAATTGCAGGGTATCCTACCAAATGGACAAGCAATAGCAGTAAAGAGGCTATCGGAGTATTCTGGACAAGGCATTCAAGAGTTAAAAAATGAAATCGTTCTCATTTCCAAGTTGCAACATAGGAATCTTGTCAAGCTATTGGGTTGCTGCCTTGAAGGAGAAGAACGGATGCTAATCTATGAGTTCATGCCCAACGCTAGCTTGGACTATTTCATTTTTGGTCTGCTTTTTCGTGATTTATTGCCTATACTAGTACTTGACAAGCCCTCAATATCTCTGATCACTCAATGATTAGTTCTACATTGTGCAGATCCAAGCAGAAAAGCTTCACTTGGATGGAAGAATCGTTTTGAAATTGCTATGGGAATATCTCGTGGTCTTCTTTACCTTCACCAGGATTCAAGATTAAGAATTATTCACCGAGATCTCAAGACCAGCAACATTTTATTAGATAGTGACATGAATGCCAAAATTTCTGACTTCGGCCTTGCCAAAATTTTTGGTGGAGATCAAGTGGAAGGAGAAACTAAGAGTATAGTAGGGACATTGTAAGATTCTACACATCTTTAAATCGTATAATCTTTATTATATTAGATCAGTACATGAGCATAAACTTATTGAATTGATGTAGTGGATATATGTCCCCGGAGTATATTGTTGATGGGAAGTATTCAGTAAAATCAGATGTATACAGCATCGGCGTAATCATTCTTGAAATAGTTAGTGGCAGAAAGAACAGGAAATTTCGTCATTTGGATCATCATCACAATCTTTTGGGACATGTAAGCACAAAACTTCACTCCCTTAATTCACTCTTAGCTTCCTATTGTGAATACATTTGTATAAATGTGTTGTTCTGTTTGTAATTTAGGCATGGTTACTCTGGAGTGAAGGCAACGCGTTGGAACTGATAGACGAAAGTATCAAAGAATCCTTTTCAGAATCTCAAGTGTTGAGATGCATCCAGGTTGGTTTGTTATGCGTCCAAAAACTCCCCGAGGATAGGCCTACAATGGCATCAGTAGTTTTCTGGTTAGGCAATGAAGGTCTAGTTCTTCCTCAACCAAAGCAACCTGGTTTTTTCATAGAGAGGAATTCAATAGAATCAACAGAGTCATCAACTGATGAAGTATATGTAAGTAGCAGCGTGTCGATAACAGTTCTAGAGCCAAGATAA

>PGSC0003DMG401028902 G-LecRK

TTTTGGCCATTTTGTGTCAATCCTCTAAATAGAAATTCCCCAGTCAGCAAAGATGCAATCTTGGAGTTTGGCACCACAGGAAATCTCTTGTTGATGGATGGAGACACCACAGTGTGGGCATCAAACACCTCGGAAGCAGGCGTTGAGTTGGCTGTCATGTCTGAAAATGGCAACTTCATTCTCTACAGCACCAATCTAAGTACCATTGCATGGCAAAGCTTTTCACATCCATCTGATACTCTCTTGCCTGGTCAATCCTTAACAGTGTCACTAGAATTAACATCATCAAAATTACCTTCTCATGGTGGTTACTACACTTTAAAAATGTTGCAGCAGCCTACTTCACTAAGTCTTGCATTGACCTATAATGTCCCCAACTACTACGATTCATCGCCTGAATTTTACAGCAATTTTTCTTATTGGTCAGGACCTGAAATATCAAATGTGACTGGAGATGTTACTGCAGTATTGGATAAAGCAGGAAGCTTTGGTATAGTTTATGGCTCATCTTCAGATGGAGCAGTTTATGTATATAAAAATGATGGAGATTATGGAGGACTATCATCGGCTGTAAATCAAACGAATTCCAATGTTCCATCTATTCTTCGTCGATTAACCCTTGAGGTTAATGGAAATTTACGTTTGTATCGATGGGATAATGATGTGAATGGATCAAGGCAATGGGTTCCAGAGTGGGCAGCTGTGTCTAATCCATGTGATATTTCTGGAGTTTGTGGCAATGGGATATGTAATTTAGACAGAAGCAAAACAAATGCTTCTTGCACATGTTTGCCAGGGACTTCTACGGAGGGAGATGATGTTTCATGTTCGGGAAATCCTTCAGTGACAGGGAAATGTGGACCTCAGCATGAAAATATGATGTCCCAGTTCAAGATTTCTACGGTTCAGAAAACGAATTATTATTTCTCAGAATCATCTGTTATAGGAAATTATAGCGATAAAGGGTCACTGTCCAAATGTGGTGATGCTTGCTTATCAAACTGTGGTTGTGTTGCATCTGTTTATGGACTTAGTGAGGAAAAGGCTTATTGTTGGTTACTCAGGAGCTTGGAATTTGGTGGATTTGAGGATCCTGTTTCGACCTTATTTGTGAAGGTTGAGGCCAATGCCTCGGTATCTGAAAAACCTGGGAATTCATCAGATGAGTCGAAGAGTAGGCACGATAAGGTTTTGGTACTTCCTATAGTCCTGAGTATGACTGTTCTCATTTTGCTTCTCTGTTGTTTATTGTATATCAATATCCATAGGAGAAGATCTATGAGAAGGGCACTTGAGGGATCTTTGCTGTTGTCTGGAGCTCCAATTAGCTTCAGCTACCGCGACCTGCAATGTTGGACTAACAATTTTTCCGAGTTACTTGGAACAGGTAAAGATATAGAATTTTTATTACTTCTATGATACTAACCTCTCAACTATAGAAATAACAAGCACAGAAAACTGTTCTCTTTCAGATAAGGTAAGAATATGATCAGTAACATTGCTGCTTTGACAGGTGGATTTGGAAGTGTGTATAAAGGAAGTTTAAAGGATGCAACATTGATTGCAGTAAAGAAACTTGATAAAGTTTCACCTCATGGGGAGAAGGAATTTATAACAGAGGTCAAAACCATTGGCTCTATGCATCATCTGAACTTGGTTCGTCTATGCGGTTACTGTTCTGAGGGAACCCAACGGTAAGTTTTAACTTTTTTTCATTGTAATTTGTGATGTATAGATTGGGAAGTACGTAGAATTGTAGCATTACAGTCAAGCACTAGCACAACTTTCACTAAAATTTTCACTCTTTGGCAGGCTACTAGTGTATGAGTACATGAAAAATAGCTCATTAGACAAGTGGATATTTCATTCATTTAGTACAAGAAATAGACTACTGGATTGGGCGTCACGTTTTCGCATAGCAGTTGGCACTGCACAAGGGATAGCCTATTTTCATGAGCAATGTAGGAACAGAATTATACATTGTGATATCAAGCCAGAAAACATACTTCTGGATGAGAATTTCTGCCCTAAAGTATCTGATTTTGGACTAGCTAAGTTAATGGGAAGAGAGCACTCTCATGTTGTCACAATGATCCGAGGAACGAGAGGTTATTTAGCTCCCGAGTGGGTCAGCAATCGACCTATCACTGTAAAAGCTGATGTTTACAGCTATGGCATGCTTCTTCTAGAGATCATTGGTGGTCGGAGAAATCTTGATATGACTTGTGATGCAGATGACTTCTTTTATCCTGGATGGGCTTACAAGGTACACACGCAATCCTCTAATTGTATTCATAGTGTTCCCTCCTTTCTGGTACATGTTCTCTTCACTATCTTTCTCCTTTTTTTCAGGAGATGACTAGTGGAACACCCGTAAAAGTTGTAGATAGGCGACTTAGAGGAGCAGTAGAGGAAAAAGAGGTAACGAGAGCACTGATGGTCGCCTTCTGGTGTATTCAGGATGAAGTGTCAAACAGGCCTTCCATGGGAGAAGTGGTGAAGATGTTGGAAGGATCAGTTGACATGAATATGCCACCAATGCCACAGACAGTTTTGGAGTTAATAGAGGAAGGCTTAGATCAAGTATACAAGGCAATGAAGAGGGAACTCAATCAGTACAGTTCTTTCTCCATTGCTACTCATCCATCATCTAATGCTACATGCAGTCACTCCAGTATGTCACCTAGATAGTTGAAGGTGAAATCGAACACAGTTTTTCCTAGGATTAAGTAATCGTATCATCATTGATTATTCATTCCCCGTCTCCCAAAAGAAGTCCCGGAAAAAGAATAAGAAGGGAACACAACCACTTTTGTATCATGTGTTGCTACAACACCATACACGTTGCAAGAACTCACTGTCTTGTTTTATTGTTTAATTTCACTCAATTCCTCTGGTCTCTAAGATATCTCCACCATTTCCTCTGTATAATTCGTCACTATCTCTTTTTCGGTTTCTTTTATA

>PGSC0003DMG401029444 G-LecRK

GGTAATAGGAAGAGTATTAGAAAACCAAAAAAGATGACCACTAGCAGCAGAAATGTGCATCATTTTGTTCATCTAATTCTTGTTATTCTTCATTGTTTTAATACAGGATTTTGCACAGAAGTAGATAGCATTACCAGCACTCTATCTCTGAGGGATCCTGGGATTTTATCATCTCCAGGAGGCGTCTTGAAGTTGGGATTTTTCAGTCCTCTAAACAGCTCTAACAGGTATGTTGGTATTTGGTACAATTTTTCTGAAACGATTGTCATTTGGGTGGCTAATAGGGACAAACCTTTAAGAGATTCTTCTGGGGTTGTCAAAATATCTGGTGATGGAAATATCTTAGTTATGAACGGGGAGGAGGAGATTCTTTGGTCATCAAATGTTTCAACCTCTCAGGTAAACTCAATTGCCCTTCTCCAAGATTCTGGGAATTTTGTTCTTGTAGATCATCTGAACAATGGGAGCACAATATGGCAGAGTTTTGAACATCCTTCTGATTCAATTGTCCCTAAAATGAGATTAAGTGAAAACACAAGGACAGGGGAGAGGGTAGAAGTAAAATCTTGGAGAAGCCCTTGGGATCCTAATTTCGGAAACTTTTCTTTAGGAATGAATTCTGGATTCATTCCTCAGGTGTATATCTGGAAAGGTAGTCATCCCTATTGGCGAAGTGGTCAATGGAATGGCCAGATCTTTATCGGAGTGCAGGGTATGTATTCTGTGTCGTCTGATGGATTCAATGTAGTGAACGATCGCGAAGGTACTGTATACCTTACTGGCCCTGGTGATTTTGATTTCTTAACGAAATTTGTCTTGGATTGGAAAGGGAATTTGGTTCAATCATATTGGGATGTGAATGAGACAAACTGGAAGATAATATGGTCAGCTCCCAATAACGATTGTGAAGTTTATGGAACATGTGGTCCATTTGGGAGCTGCAATTTGGAGTCGCCGATCTGTTCTTGTCTGAAAGGTTTTGAGCCAAAGCACAGGGAAGAATGGGAAAAGGGGAATTGGACTAGTGGTTGTGTTAGGAGGAAAGCTTTGCAATGTGAAGTAAGGAATAACTCAGGGGATTCAAGTAAAGAAGACGGATTTCTAAAGATAGGGTCAATAAAATTGCCTGATTTTGCAGAAAGGTCATCGACTAGAGAAGACCAATGTAAAAGCCAATGCTTGGGATATTGTTCCTGCATAGCGTATGCATATGACTCAGGTACTGGCTGTATGTCATGGAGTAACAACTTGATTGATATTCAGCAGTTCCAAAGCTCGGGGAAAGATCTCTATATTCGCGTGGCACATTCAGAGCTTGGTATGTATAGTGGTGTTGAATAGAAGACTCATTATCCACCCAGAATAGCTTAATCAGTCATTTGAGATATCATGGACTTATTGAATTCCATATATATACATTAAGTAGCTTACTTCTGTAGAGAACTGATTTTTGTTCACCTTTTGTGATTGCTCTACATTACAGATCATCATAAAGACATAAAGAAAATTGTAATTCCAGTAATCCTTGGTTTTCTTACACTCTGTGTTTGTCTGTTCCTTTGCTGTACATGGATGGCCAGACTTAGAGGTAAAACAAGCTATAAAGTTTTAGTATTATACATTTAGAAGCACTTTGTTTGCTATAGTAGTTAATTTGTATCCCTGCTGATATTAGGAGTAAAAAGGAAGAAGATAAATTTACTTGGTGACAGAAGTGCAGTTCATATGGAAGAGTTACCAGTCTTCAGCCTCGATACGCTTGCAAATGCAACATCCCAATTCCATGAGGATAAGAAGCTTGGTCAGGGCGGTTTTGGTCCAGTTTACATGGTACAATTTTCTTCTCTCTCGATCACTTTTCAAATTTCAGTTGCTGAAATTGCTGATTACTTGTAGGGAAAATTGGAAGACGGGAAAGAAATAGCAGTCAAGAGGCTTTCAAAAGCCTCGGGACAAGGGCTGGAGGAGTTCATGAATGAAGTGTTGGTGATCTCGAAAGTCCAACATAGAAACCTTGTTAGACTTTTGGGATGTTGCGTTGATAAAGAGGAGAAGATGTTGATTTATGAATATATGCCCAAGAAAAGCTTGGATGTGTTTCTCTTTGGTTAGTCCTTGCTTTTAATAAGCATTGTACTTCTCATAGCAGTTCTTTGTTACTCCACATCTATGTTGTTCGGACTCTTCAATAATGTCGATGGATGCTTGTTAGATCCTCCAAAAGTACTGCACTTTTGGAGGATCCGACACGAGAACATCAACAATTTTGGGGAGTCTGAGCAACATTGCTCCACATTTCTTCTGAAGCAAAGTTAATCCCTTCTCCATTCATATTTGACAAACATCTAGATGAAGGCCACCGAGGCATTTTGGATTGGAGAAAACGTTCCACTATAATCGAAGGGGTTGGTCGAGGACTCCTTTATCTTCACAGAGATTCAAGATTGAAGATAATTCATAGAGATCTGAAGCCAAGTAACATTTTGCTCGATAATGACTTCAATCCAAAGATTTCAGATTTTGGCATGGCTAGGATTTTCGGATCTGACCAAGATCAAGCAGACACAAGGAGAGTAGTTGGTACTTAGTAAGTGGACATTGATCTATATCTACCATAGCTTTACACAGTTTTCTCCTGTAAAAAACCATATCATACATACTTACTAACTAATCATTTTTTACACGAATGTGCTGACATTTTTACAGTGGATACATGGCCCCTGAATATGCAATGAAAGGAAGATTCTCTGAGAAATCGGACGTTTTCAGCTTTGGAGTTCTAGTGTTAGAGATCATCAGTGGCCGAAAAAGTACAAGTTCTTGGAATGAGACATCCTCTTTTAGCCTTTTCGGATATGTAAGATTCCCTTTATATGCTACTATTCAAATACTAAAAATGCTCATACTGCAAAAACTATACAGTCAAAACATTTCTACATCGACCTCGTTTGTCCAGATATCTTTTGGTTGCTATAGTGAAATGCTATTATAGAGAACATCCAAAGTAAATTTCACCATTATAGTAACGTGTTGTTATAGTTCGATTGTGTGTACCTTATATTTTTACATTATGATGGTAGAATTACTTTTGCTGTCACTGTGTTTTATGCATCAATAAGTTTGATGATCAATTTTTCACTGTTTTCAGGCATGGATGTTATGGAAAGAACAAGATTTATCAACTTTTATCGATCCATTCATATTGAACACGAGCTCGGAAATGGAGATCAGAAAATGCATACAGATTGGTTTACTATGTGTTCAAGAATTTGCTGAAGACAGACCAAGTATTTCATCTGTTCTTGCTATGCTTACCAGTGAAACTACAAGTCTTCCTGCACCATCACAACCTGCTTTTACTGAAAGACATGATTGTATCTTCAAAATGTGCAATGTAACTAATTGTACTTTGAACAATATCAGTATCACAAACATAACTGGTAGATGACATGAGTATATGAAATTATTATTTTTAAGATATGCAAAATTATCCATTTGTCTCAATTTGTATGATATCTTTGTCATTCATTTTTCGATTACATGTTAAATATTCGTGTTGAATCAAATATCATAATCCACAAGTGTAATATGTGTTTGCCAATCACAGAATATATTATGGGATGTATTTGGAATAAAATGAAAAGGCATTTAGGTTTAATGTCAATTTGTGCAACAATACATTTTTCATTACACTATGTGCATTATTACCACTTATAGTTGACAAGCATGTTGACGTTGAGATGCTAGCTTATTGTCAAGGTCCCCACTCAATTCTCACACTAACGTGGCTTTTGGATTTTCACTCATTTTACTTTTTCGAAATAGTTTAAATATATCAATCATTATATTTCTGGTCTAAGGTTGAATTAAGCTTAGATGTCATATCTTTACATCAATACTCATTTTATTTTATTTTCTTCTTCTTTTTTCAATGAGCATTTAATAATTTAAGGAAAAGAAATGTGTTATTTCAACAAGTTAAAACTTGTCAATTCACAATGTTATTTCAGGAACATACTACACAATGATAGTGATGGGAGAAAAGACTTATATTGTCAGCGTATAGAAGTTAAACTCTACATTTTACATTATTCGTTTTTAAGTGTCAATTTTTGCATTGGCCTAAAAGTCTCGGATAGAGAATGATATGTGGTTTGTGATATTTCTATAGGGTCAACAAATTAAAAATCCTATTTGGCTGCTTGTGATATTAGACTTTGACAACATATTGTGCTTTTGTTCATATAGTGTGGTAATAAGAAAAGTCTTAGAAAACCAAAAAGATGAGCAACAGCAGCAGCAGAAGAAGAAATTTGCAATATTTTGTTCATATAATTCTTGTTATTCTTCGTTTTATCGATACAGGATTATGCAGTGAAGTGGATAACATTACCAGCATTCAATCTCTGAGAGATCCTGGAATTTTATCGTCTCCAGGAGGCGTATTCAAGTTAGGATTTTTCAGTCCTCAAAACAGTACTAATAGGTATGTTGGTATTTGGTACAATTTTTCTGTAACAACTGTCATTTGGGTGGCTAATAGGGACAAACCTTTAAGAGATTCTTCTGGTGTTGTAAAGATATCTCGTGATGGAAATGTCGTTATAACGAATGGAGAGGAGGAGATTCTTTGGTCATCAAATGTTTCAACTTCTCAGGTAAACTCAATTGGCCTTCTCCAAGATTCTGGGAACTTTGTTCTCGTTGATCATCGGGACAATATGAGCACTATATGGCAGAGTTTTGAACATCCTTCTGATTCAACTATTCCTAGAATGAGAATAAGTGAAAATACAAGGACAGGGGAGATGGTAGAAGCAAGATCTTGGAGAAGTCCTTCGGATCCTAATATCGGAGACTTTTCTTTAAGAATGAACTCTGGAGTCATTCCTCAAGTGTATATATGGAAAGGAAACCGTCCCTATTGGCGAACTGGTCAATGGAATGGACAGATCTTTATAGGAGTGCAGAATATGTATGCTGTGGTATCTGATGGATTTAATGTAGTGAATGATCGCGAAGGTACTGTATATTTTACTGGACCTATTCGTGATAATTTTTTAAGGATATTAGTCTTGGATTGGAGAGGGAACTTGGTTCAATCATATTGGAATGTGACTGAGACAAACTGGAAGATAATATGGTCAGCTCCCAGTAACAATTGTGAAGTTTATGGAACGTGTGGTCCATTTGGGAGCTGCAATCATTTGGAGTCGCCGGTCTGTTCTTGTCTGAAAGGTTTCGAGCCAAAGCATAGGGAAGAATGGGAAAAGGGGAATTGGACTAGTGGTTGTGTTAGGAGGAGTGCTTTGCAATGTGAAGTTAAGAATAACACCGCGAATTCAAGTAAAGAAGACGGATTTCTAAAGATGGAGTTGATGAAATTGCCTGATTTTGCTGAAAGGTCATCTACTTCAGAAGACTTATGTAGAAGCCAATGCTTGGGGAATTGTTCCTGCATAGGGTATGCATTTGATTCAGGTATCGGCTGTATGTCGTGGAGTGAAATGATTGACATTCAGCAGTTCCAAAGCTCGGGGAAAGATCTCTATATTCACGTGGCAAATTCAGAGCTTGGTATGTATTCTGGTACTGAATAAAAAACTCATTGTCAGTCATTTCAGATAGGCGTAATACATAAATGTGCCCTTTTACTTGGCCTCAACTCACATCTATGCTCTCATGAACACTCAAAGTTAGATGGCATGGATGTCAGCAAAGGCCAAATCAAAAGACACATTTATGTATTATGCCTTTGAGATATGTTCGACTTAATTGAATTCCATGTATTCTGTAAAGAACTGATATTTGTTCACCTTTAGTGTTTTCTGCAGATCATGGTAAAGACATAAAGAAAATCGTAATTCCAGTAATTGTTGGTTCTCTTACACTCTGTGTTTGTCTGTTCCTTTGCTATACAATGGTGATCAGACGTAGAGGTAAACATGCTAATAAGTTTTTGTATTACACATTAAGAAGCACTTTGTTCGCTATAGTTAACTTGTATTCCTGCTGATATTAGGAGTGAAAAGAGAGGAGGTAGCTTTACTTGGTAACAAAAGTCCAGTTAATATGGAAGAATTACCAGTCTTCAGCCTCGATACGATTGCAAATGCAACATCCCAATTCAATGAGGATAATAAGCTTGGTCAGGGCGGTTTTGGTCCAGTTTACAAGGTACGAGTTTCATCTCTCTCGATCACTTCCCATAATTCAAGTTACTGATATTGGTGATTCCTTGTAGGGGAAATTGGAAGATGGGAAAGAAATAGCTGTCAAGAGGCTCTCAAAAGCCTCGAAACAAGGGCTGGAGGAGTTCATGAATGAAGTGTTGGTGATCTCGAAAGTCCAACATAGAAACCTTGTTAGACTCTGTGGATGCTGTGTCGATAAAGAGGAGAAGATGCTGATTTATGAATATATGCCAAAGAAAAGCTTAGATGTGTTTCTCTTTGGTTAGTCCTTGTTACTTCTCATCTGCATTACTCCACATCTATGTTGCTCGGACTCTTCAATAATGTTGATGGATGCTTGCTAGATCCCCCAAAAGTACTGCATTTTTGGAGGATCCAACATGAGTACATCAGCATTCTGAGCAACATTACTCCACATTTCTTCTGATCCTTTTGTTTAAATTCTCCATTCATATTTGACAAACATCTAGATGAAGCACATCGAGACATTTTGGATTGGACAAAACGTTCCATTATAATCGAAGGAGTTGGTCGAGGACTCCTTTATCTACACAGAGATTCAAGATTGAAGATAATTCATAGAGATCTAAAGCCAAGTAACATCTTGCTCGATAATAACTTCAATCCAAAGATTTCAGATTTTGGCATGGCTAGGATTTTCGGATCTGATCAAGATCAAGCAGACACAATGAGAGTAGTTGGTACTTAGTAAGTGATCATTTATCTATATACAATACCATAACTTTGCATAATTTAACTACTCGAATCGTACTGACATATTTACAGTGGATACATGGCACCCGAATATGCAATGGAAGGAAGATTCTCAGAGAAATCCGATGTTTTCAGCTTTGGAGTTCTAGTCTTAGAGATCATAAGTGGTCGAAAAAGTACAAATTCTTGGACTGAGACATCCTCTTTGAGCCTTTTGGGATATGTAAGACTCCTCTTATAGTTGCTATAGCGAAATGTTTGTTATAAAGAACGTATAATATAACATAACATGAAGAATCAGTTCCAAAAAAGAACTTGGCTGTTATATAATGAAATGTTATTATAGAGGACGACTATTATAAAGAGGTCTGACCGACTATATCATGTACCTTATATTTCAGGCATGGAAGTTATGGAAAGAACAAGATTTATCAACTTTTATCGATCCGTTTATATTGAATCCGAGCTCGGAAATAGAGATCAGAAAATGCATACAGATTGGTTTACTATGTGTACAAGAATTTGCTGAAGACAGGCCAAGTATTTCATCTGTTCTTGTTATGCTTACTAGTGAAACAACAAGTCTTCCAGCACCATCACAACCTGCTTTTACTGAAAGAAGACACTTCAGAATGTGCAATGAAAATAGAGAAACTAAGTTTACTTTGAACAAAATGAGTATCACAAACCTTACTGGTAGATAACATGAGCATATGAAATTACTTTTTAAAGATTTGCACAATTGTTATAGTATGATCCATTCGTCTCAATTTGTATGATACCTTTTGTTTCAAAATGTTTGACAATATTGGGTCCGTGTAAAGCACAGAATATTTGTTATTAGTTACAAA

>PGSC0003DMG402003983 G-LecRK

AAAAAAGAGAAGAAAAGCTAAAATCTATGGCTGCTTTGCTATGGTTATTAATATTATCTGCATTTCATGCAGTAGCACTAGCACAACAAAGGCAGTTCAATATAACTCTTGGCTCTTCTCTAACACCTACTACCAATTCATCATGGTTTTCGCTGTCCAGGCGCTTTGCCTTTGGGTTTTATGAACAAAACAATGGCTATGCTGTTGGAATCTTAATTGTCGGTATGCCTAAGAAGACAGCAGTGTGGACAGCAAATAGGAACAGTCCTGTTGTTCCGAGCAACGCTGTCTTGCTCTTAACTAATGATGGAAGGCTCATTGTACAAGTAGGAGGACAAGAAATCACTGTCGTAAATCTCTCTGGTCAAGTCATTGCTTCAGCTTCCATGCTGGATACTGGCAACTTTGTGCTCTATAATTCTGATCACAGTATCATATGGCAAAGTTTTGACAATCCAACAAACACCCTTTTACCTGGTCAGCATATATCAGCTGGACAAGAGCTGTTCTCGAGTGCCTCGGAAGCAGATGATTCTTTCGGGATTTTCCGTCTCAAAATGCAAGATGATGGGAACCTCGTTCAGTACCCTGTCAATACACCTCATTAGGGATTTTCCGTCTCAAAATGCAAGATGATGGGAACCTCGTTCAGTACCCTGTCAATACACCAGATAGTGCCTCATATTCTTACTACTCAACAGGCACTTATGGAGCAGGGAATAATGTTACACTGAATCTTGATGATGACGGCCTCCTCTACTTACTCAATTCCACTAAAAGTCTAAGAAATCTAACTCAAGGAGGCAACCGTAGAGAAAGGACCATTATCTATATGCTGAAAATTGATGCTGATGGTATACTTAGAATTTATTCAAATTCTCTGAACCAACAGAACAGCTCCGTGATATGGTCATCAACAAATGATAGGTGTAATCCAAAAGGCCTTTGTGGGCTTAATGGTTTTTGCACCAACATAGATGACCAAGCTAAATGTTTGTGTCTTCCGGGATTTGATTTTGTAATGCCAGGAAATTGGAGCGCAGGCTGTGAAAGGAATTTCACCGCAGAAACCTGTCGGTTAAAGGAGAGCACTTCCAAATACTATGCTATGAGAACAGTTGAAAACACAGGATGGGAAGATAGTACTTATGTTGTTTTGGCCGGAACGACCAAGGAAGATTGTGAACAAGCCTGTCTACAGGATTGTAACTGTGAGGCTGCTCTGTTCAAAGACAGAGAGTGTAGAAAGCAAAGGCTGCCTCTGAGGTATGGAAGGAGAGACTTGGGTAATTCTAATTTAGTATTGGTCAAGGTGGGCACTAATTTCATACCCAACGAAGGGGTGCTTTATCAGATGATTGAAGAAACAAAAGGCAAAAAGCTAAGGATAGACATCCTAATTGCCGGCATTACATTGGCTGTTTTTGCTTTGTTGGTGTTAGGCATCTCTGGATTCCTCATTCACAGAAATCATGTCTGGACTTACAGAAAAATCCAGGAGAGTAGAAGTGTTCAGTTATGTGAGGATGTTGCTCCACGAGCATTTTCCTATGCAGAACTAGAGCAGGCAACAAGTGGTTTCAAAGAAGCGTTAGGGAGGGGAGCATTTGGAACAGTATTTAAAGGAATTTTGGCAGAAGACCAGAAAGTGATAGCTGTTAAAAGACTAGACAAAGAATTAGTAGAAGGGGAAACAGAGTTCCAAACTGAAATAAAAATCATTGGAAGGACACATCACCGAAATCTCGTCCACCTCCTTGGTTATTGCCTGGAGGGATCAAGAAGGCTCCTGGTGTATGAGTACATGACCAATGGGTCACTTGCGGACATACTTTTCACAACAGAAAAACAGCCCACGTGGGAGGAAAGGTGTGGCATCACTCGAGATATTGCAAGGGGCCTCCTATATTTGCATGACGAGTGTGACACACAAATCATCCATTGTGATATAAAGCCTCAGAACATACTCATGGATGATCAATACTGTGCAAAAATATCTGACTTTGGAATGGCCAAGCTTTTAAAGAAAGACCAAACAAGAACCTACACAGGGATAAGAGGGACCAGAGGCTATGTTGCACCAGAATGGCACCGGAAACTGCCAGTGACAGTGAAAGCAGACGTTTACAGTTTTGGTGTCCTCCTATTGGAATTGATCTGTAGGCGGAAGTGTGTTGATATGAGCCTTGATGAAAATGAGTCAATTCTTGAATACTGGGTCTACGATTGCTTTGCTGCAGGAGAGCTAGATAAACTTGTCGGGGACGAGGAGGTTGATAGAAGACAATTCGAGAGAATGGTGAAAATAAGCATCTGGTGCATTCAAGACGAACCATCACTTCGCCCTTCAATGAAGAAAGTTCTACTGATGCTGGAAGGGACAGTAGAAATTCCAGTGCCTCCTAGTCCTACTTCTTTTTTAAGTGCCATATAGTTTTGAACTTCTTCACTTTATGGTTTTTGTACTTCAAATTTCAAGTATTCCAACTTCCCCATAGAGAGCTGACCACTTTATCATCCATACTTGAGCTAAATAAAATATAGTTCTATCAAATTTCTACATCATTC

>PGSC0003DMG402004946 G-LecRK

ATTTTAAGGCCTTCAGAACTAGCTAGATTTACACTGAAAAGGTCAGGTTACCTCCAGATATCTAAACCTGCATAGATTAGGTAGCATGTAATAAAGCAGTTCTTAATGGAAAGTTTTGTGCTTATATTGCAGATATTGTTATGTGGTTTTGTATGCTCAATTTCTAGTGCGAAAGATATCATCACTGCAACTGACTTTCTAACAGATGGAAAGACCATTACTTCATCTGATGGAAGTTTTGAGATGGGATTTTTTTCCCCGGCTAGTTTCACGAATAATTGGTATGTTGGAATATGGTATAAGCACGATGTACCTGATAAGTCTGTGGTATGGGTTGCCAATAGAGAAAATCCATTCAACAACACATCAGGTGTTATGTTGAAAATCATCGATACAGGACAACTTGCTCTTCTCACTCCTGCTAATACAATCATGTGGTCTACTAATTCATCGAGGCCTTTGGCTGTGAAGAATACAGTTGCACAGTTGTTGAACTCGGGGAATCTTGTCATAAGAGATGCAAATGATACCAAGCCTGAGAAGTTCCTGTGGCAGAGTTTTGATTATCCTACTGATACACTCTTACCAGGCATGAAGATGGGTAAGAACTTTGTAACTGGACAAGAATTTTACCTGTCCTCATGGAAAAACGAATATGATCCAGCTCCAGGGGAATATACGTATCATTGTGATCCGACTGGATATCCTCAGGATGTCATGAGGAAAGGTAAAGTTAAGGTGTACAGTACTGGACCATGGAATGGACTCCGCTGGAGTGGTGTACCAGGACTGACAAAAAATACTATTTATACATTTAAACTGGATCTTGATGAAAAGAGAGCATTTTACAGCTTCGCGCTTTTAGGTTCTGTTATGACCAAATTAACCATGAACAATAACGGCGTGCTTCAGCGTTCAATGTGGGCTGAGAATAGACAGGAATGGCATGTTTACATTTCTTCACCGGAAGATACTTGTGACAATTATGGAACATGTGGTTCATATGGTAGTTGCAACAACATACTCACTCCTGTGTGTAATTGCTTAGACAAGTTTGTGCCTAAAGATCCAAGAAATTGGGCAATGACAAATTGGTCCGGCGGGTGTATTAGAAGGAAACCGCTTAATTGCCAGAATGGAGATGGATTTCTCAAGTACTCTGGTATAAAGCTACCAGATACACAGTATTCATGGTTTGATAGGAGTATGACACTCCACGAGTGCAAGCACGCTTGCTTGAGGAATTGCTCGTGTTTGGCTTACTCAAATCTTGATATACGCAATGGTGGGAGTGGCTGTTTGCTTTGGTGTGGCGACTTAATTGACATTCGAGAATTACCTGGAGGACAGGACATTTACATAAGGGTTGCTACCTCAGAGTTAGGTAACATTTGTGAATTTGATCAAAGTTGGAGTATTAGCTGAATGATTTAAACTTCCCAATTCTCAGATTTAGTTGTATTCTACTTAGATATCTGTTAAAAGTTCACACAGTATATGTTCACTCTTCCTTGTCTATTTAGTTTATTTCACCCCTCCATGTTTTGTTCAGGCTCGAAGAAAACAAAACTACTCGTGTTGAGTCTGTTACTGTTGATTGGAGTGACTGTGACTGGCCTGACCATTGGGCTGTATATTTGGAAAAAGAAGAAGAAGAGGAAGATGAATTTGAAAGATGACCTAGACCTCCCACTGTTTACCTTATCAACTCTAACTAAAGCATCCTCTAATTTCTCAGTCGAGAACAAGATTGGAGAAGGTGGATTTGGATCTGTTTATAAGGTAACATCTTAAGTTTATGAATCATTCTTTCAGTAAATAATATCTACTATTATTGACAGTAAACAGATGACACTTACTGAATCACTAAGCCAGATATATGTTGATATTATAGGGAATCCTGGAAGGTGGACACGAAGTAGCTATAAAGAGGCTATCCAAGTCTTCCTCGCAAGGAGTTAACGAGTTCAAGAATGAAGTTATTTGCATTGCCAAACTTCAGCATCGAAATCTTGTGAAGCTTATAGGTTGCTGCATTGCAGGAGGAGAAAAAATGTTGGTCTATGAATATATGTGCAACAGAAGTCTAGATCTCTTCATATTTGGTCTGGTTACTCTCTCTTAACTCGAATAACCTCTATTAGTACAAATAGTAAAGTTGATAGTTTAATCACATTATCAGTGTTCTAACAGAGGACCTGTTAACAGATGAAAAGAGGAGCTTGTTACTCAACTGGCCTAAGCGCTTCGGCATCATCAATGGAATTGCTAGAGGACTTCTGTATCTCCATCAAGATTCTCGGCTAAGAATTATCCACAGAGACCTCAAAGCTAGTAACATTTTGCTAGATGCTGACATGAACCCCAAGATCTCAGATTTTGGCATAGCTAGAAGCGTTGTAGGGAATGAGACTGGTGCTAATACACACCATGTGGTTGGGACACAGTAAGAAATTTAATACTACTCACTTGTTAGCATTTATATTATGTATAATAACCTTTATTTTACACTACTTTCTGTTTAATATGTGATGCAGTGGTTACATGTCCCCGGAGTATGTAGTCCATGGTGTTTTCTCAGTAAAATCAGATGTATTTAGCTTTGGCGTATTAGTGCTAGAGATAATCAGTGGTCGAAGAAACAGAGGATTTGCCCGTGAAAGTCACAGTATCAACCTACTCGGACATGTAAGTCTTGTTTGTAATTAAAGACGAGTTTAAATTTTGTGCACTATATGGTAGCTACAGATATTTCTCTATAATAATTAATTGATAACACCTCGTACAAATGATAGTTAAACTGCATTGAACTTAAATCCTTCAGGCACATCAACTGGTGCATGTTTCTTGATTATGTACTGATTGTGTTATTGATAGGTATGGAAACTTTATAAAGAAGGTAGACCATTAGAGCTAATTGATGGCCATCTAATGGATTCGCGTTATATTTCTGAATTGCTACGCTTAATCCATGTCGCTCTGTTATGTGTACAACAATGTCCAGAAGACAGGCCAGATATGCCTACAGTGATCTTGATGTTGACTAATGATGCTATTTTGCCTCAAGCTAAGGAACCTGGTTTCTTCACAGAAAGCAAAGTCACGTCTGAATGTTCTATAAGCATGTGTTCAACAAACGAAATTACCGTCACACAGTTAGAACCCCGGTAAATTACATTACATGTTCATTCAGATCGTTCAATGTCAACAAGCTTGAATTACAGTATTTTGTGCTGTAAAGCCATAATTGCCTATGGTATCTGTTATTAGTAATTATGGGTGTGTTGTAGTTTTTATTAATTCTAATGTATTTTGAAGAATTCGTACAAAGTACCTATTCAACATGAGTACATCAAAATAAACGAAAGAT

>PGSC0003DMG402005047 G-LecRK

CAATGGTTATCACATGAAAAGAACCATAGCAATTGTGACAATCTGCATTGGTGCAATACTCATCGCGCTCTCGTTTGCTTGTTGTTTGGTAATAAGGAAAAGGAGAAGAGGTAAAGGGTACCCCTTAATTTCTTCGTGTGTTTACTACTTTATTGAATCCCCTTAGTGAAATCTTGGCTCCGCGACTTACAATAGTGAGTTACTTCATGTACAGATAAGGAGGACCAATTCACCAGTTTGAACACTTTGACAAGGAATTTAGCATCCTATGAGAACTCTTCAAGAGGTAATGAGATGGATGGAAGTGAACACGTCGATGTGTTAATCTTCGATCTAAGCACTATCATTTCTTCCACTGATGATTTCTCTGATGCTAACAAACTCGGAGAAGGAGGCTTTGGTAGCGTTTACAAGGTACCTCCCGATATTTTACCAATTTCTTACTCATTTGTTGAGCCAATTTTACTTGACAACAAGCTTACATTTTGATTTCAGGGCCAGCTAAACAATGGGCAAGAGATAGCGGTCAAAAGACTTTCGAAAAATTCAGGGCAAGGAGTGGAGGAATTCAAGAATGAAGTAACGTTAATCGCGAGAGTACAACACAGGAATCTTGTGAGGCTTTTAGGATGTTGCATACAAAGAGGAGAGAAAATGTTGATCTATGAATACTTGCCAAACAAAGGCTTGGACAGTTTCATCTTTGGTATATACATTTGCATATTGTGTTAAAATTTTAAAAAATAGGCATGTTACTGATGGATTGCTTATGTATGAATTTCGAATCGGTATTGCAGATAAAACTAAAGGGTCGCAGTTAGATTGGAGAAAACGATTTGAAATCATCGTTGGAATAGCACGAGGATTATCATACCTTCACCACGACTCACGAGTAAGAATCATACACAGGGATCTAAAAGCCAGCAATGTTTTACTAGATGCCTCAATGCACCCGAAAATATCAGATTTCGGAACTGCTAGGATATTTGGTGGTGACCAAATTGAAGCTAACACGAATCGAGTCGTTGGAACATAGTAAGCATCACTTTTATGTATGAACATATGTGTATTTTTCTCAAAGGCACTTAAACTTGTATAAAGTTGAACAAGTAGACACACTCGTCCTACATGATGATTCGCATAAGATACACTCGAACTGATTTGTATTATGCCATGTAAGACGCATATTATGTCATGTAGAACATGTGTGTCTACTTGTTCACACCTCATAGATGTCAGTTGAGGCTAAGTTAAATGACACGACGTTTATGTATTATGTCATATTAAAATTGTGGATGTTGACATAGCTATTGGTTCATTTCAGTGGTTATATGTCACCAGAATATGCAATGGAAGGGCATTTTTCAGTGAAATCTGATGTTTTCAGTTTTGGTGTTCTGTTGTTGGAGATCATTACTGGCAGGAAGAACACTACACATTATCAGGATCATTCATTAAATTTAGTTGGAAATGTAAGACATCTCTTCATCAAGAACTCACCTTTAGTTAAGAAGATGTATCTTGTTTTGCTCTAGCCCTTATACCATTGCTCTAGGGAAGGATATGTGGAAATGTTCATCTCTTCTTTTTTTTCTTATACGGTGGAGTCAGGGCTAGCTTCTGCATACTTTGACTATTACAATGTACCTGCTACCCTCCACCAACATAGGGATGGAAACAATGGCGGGAGCTACATGCATTGGAAGGGTATTCAAATTGAATCCCCTTCTTTAGGAAGCTATGATAGAAACTATTGGGTGTGTCTGAACAAAGTCCCACATGAAAAGTAAAAAAAAACTACTTTAAGATGTCGGATAATCTTAATGTTATGAAGTCTTCTGGAAAAAAACTATGCGGGTTTGGTCCAAGGGAACAATATCATGCCATGTTCAGAGTATCTTATTGGATTATTTTAGCCTAACAACTGATATCAGAGCCAGTGGTTTGGCAGGACAATTATTGAGATGGTAGAGTGATGGAGTGGGGTCCGACTTGGTGCCTTTGACCATCTATGGGCCGATTTACTATCTTTACCTGTAGCCTTAGAGATGCTTACACAAAAAGAAGCTAGTGGCCGTTCGTGACCATAAATACTCTATTGATATGGTGGCACGCAGTGGAATCTCCAGATAAGCCTATAAAGTGATAGGTCATGTGGAACTCAATTTAAGGAGAAGACTGTTGAGTGTACGTAAACAAAGTCTCACATGAAAAAATAAAAAAAATATAACTTATAAGGTATTGGATACTCTTAATATTGTGAGGCCTTTTGAGAAAACCGTTGGGGTTTTGTCCGCAAACTGACAATATCACATCATGTTAAAAGTATCTTTGAGTTATTTTAGCCCAATAGAAACACATTCTTTCGTCAAACATATATGAAATTTTGGACCCCCTTAACCCCATACACGACATTTTTGTATTTTCCTTGAATCCCCTAATTGAAAATTTTGATTCCGCCACTGACCAGAAATGTTAACTTTGCTCTCCCTTTTTTCAGGTATGGGATTCATGGAATGATGACAAAGCTATAGATGTTGTTGATCCCTCATTAGGGGATTGGTATGAAAGTAGTGAAGTTTTGAGATGCATCCAAATTGGACTATTATGTGTACAATCATATGCAAATGAAAGACCAATGATGTCTCAAGTTGTCTTCATGCTTTGCAATGAAACAAAATTGTCTAATCCTGGACAACCTGGTTTTGTGTTCAGATCAAGAAATTCTTCTTCACTTCCTTATTCATCATCAGCTAGTATTGGAAATTCTGTAAATGACATTTCTATTACTGCACAACATGCTCGCTAAGTGTGTGTCAGATCCTTCAAAAGTAGTACCATATATTTTTGTGGATCTGACATGGGTTCATCAACATTTTTGAAGAGCACTGAGCCATATGGTCATTAGGTAATTTATACTTGATCTATTTATTGATTTAGTTTAAGATTTTTTACATTTATCAATGATGTATTTTAACGTGTTATAACAGGTAACGTATCTTTT

>PGSC0003DMG402007101 G-LecRK

CAAGAAGCAAATGTATCATAACAAACAAGCTAAAACAACAAACTAAACATGTCTCCTTCATGGCTTACTACACTTCTTGCCTCTCTATTTCTTTTCTCCCAAATATTTTCTTGCCTTGCCCAAGTTCCACTTGAAAACACCTTCAAATTCGTCAATGAAGGCGAATTAGGACCTTATGTTGTAGAATACAGAGTAGACTATCGCGTTATTAGTATTTTCAATAACCCATTTCAGCTTTGTTTCTACAACACAACTCCTAATGCATGGACATTAGCATTGCGAATGGGCACTGTCCGTTCTGAATCACTCATGCGTTGGGTGTGGGAAGCAAATAGGGGAAACCCCGTTAAAGAAAACGCGACATTCACGTTTGGAACTAACGGAAATCTCGTATTGGCGGATGCTGACGGTCGAATTGCTTGGCAGACCAACACGGCCAATAAGGGGGTGACAGGGTTCAAGTTGTTACCTAATGGTAACATGGTGCTACATGACTCTAAGGGTAAATTCGTCTGGCAGAGTTTCAACTATCCCACTGACACATTACTAGTGGGCCAAACTCTCCGATTGTCCGGTCCGAATAAGCTCGTGAGCCGGGCCTCGGTGAAAAAGAACGTAAACGGGCCTTATAGCTTGGTGGTGCAGCCAAAATTGTTCGCTATCTACAATAGGACTAAGTTAGGCGTGGAATTAGCTTGGTTCGATTTTGGAAACAGTACGTTGGAATCAGTAAAATTGAATAGTGGAAATCAAAGATTGAAATTGGATTATAGATTGGCAAAATCAACAAAGAGAAGTTCACATGTGATGGCATTTACTAAATATAACACAACATTAACATATCTTAGACTTGAAATTGATGGAAATTTAAAGGCTTACACTTTTGTTAGAGATGAAGAAGCAGATGAATTCCGTTGGAAGGTAACTTACCAAATGCTTTGAGAAACTTTTTAAAATAAAAAATAAAATAATATATATTAAAAACGTAACTTTATAAGATAAGGATTTCATTTCCAGTAAAAATAAGTGGAGAAAATGAGTGGGTCCACAGCTATTATTTTCCTTTTTTTTTTTGGTTGAAGGGTGTATTTTATTATTATGATTTCTAGTTTTTTGTTTTCTGTTAAAGGATGTATTTAATTATGTTTTCTACTAGCTATGACAAATGTGAA

>PGSC0003DMG402009323 G-LecRK

ATGAAGCTTGGTTATAACCTCACCACAAAGCAGAATTGGACTTTAACTTCTTGGTTGAGTTTTAGTGTTCCTGCCTCAGGGGCTTTTACTTTAAGTTTGGAGGTTTTTGAAAATGCATTTCAGTTGGTTATTCGTCGAAGGGGTAAGGTTTATTGGACTAGTGGGGCTTGGAACAATCAAGGCTTTCCATTCTTACCTTTATTCCATGATTCTGCTACTATTTATCAGTATAATCTCAATTTGGTATCCGATAAAGATGGGATGTTCCTCCAATTTGATGCAACAGAGGGAAGCTTTCCAAGTCTTGCATTGAATTCAGACGGGGATATTCTTGGTGGATACCGGTTTACATACACCCTTTTTAAAGACTTCTGTTATGGTTATGAAAGCGAGGATGGTTGTTTCTCTAATCAGTTGCCTGAGTGTCGAAAGGATGGGGATAAATTTGAGCAAAAGAGTGGAGACTTTATACATACATCTAGTATTACTACTGAGTTTTATGACAATGCAAGCATTAGTATTGTTGATTGTATGCATAAGTGCTGGGAGCACTGCAGTTGTGTTGGCTTCACCACCGCCAGCAGCAATGGGACAGGCTGCCGTATTTGGAATGGAAATGGGGATTTTCGAGTCTATGAGAGTGGTAATGCAGTGCAAAGATATGTTCTAGTTTCACCAAAATCATCTAAAGGAAAGTCTTTATATCAGGAATACTCATTTTTATGTACTCAATTGTCAACTGTATAACTCTCAGCATGCTATGGGTGCAGGAAAAACTTGGATATGGATAGTAATTAGTGTAGCTGTTTTTATCGCGTTGCTAGTATCTGGTTTCATCTGCTACATTATAATGAGAAGACGCAAACTACAAGGTAAATCACCAAACTGTGGTTTTAGAAGCTAGTAGGTTGTTTAGTCAATTATTTTAAGCATGAATGTATATCTACATACTCTTCTTACATAGATGAGAAAAGAAAGGAGGAAGCATATATACGCGAGTTGACAGCAGTAGACAGCTTCAACACCGCAAATCTAAAAGAAGAAGATGGAAGGGAAGTACAAGATTTGAAGATATTTAGCTTTGGATTCATATTGGAAGCCACAAACAACTTCTCAAGTGAAAAGATGCTCGGAGAGGGTGGTTTTGGGCCTGTTTATAAGGTATATTTGCACTATGTTATATGTAGCTCAGACATTTGCTTACTGCTCTCACTACATAACGAATCAACTATAAAAATGGATGGCAGGGAAAATTTCCAGATGGGCGAGAGGTGGCAGTCAAGAGACTTTCAAGAACCTCGGGACAAGGGCTTGTGGAGTTCAAGAATGAACTCATACTAATTGCCAAAGTTCAGCATACAAATCTAGTTTGAGTTCTAGGATGTTGTATTCACGGAGATGAGAAAATGTTGATATATGAGTACATGCCTAACAAGAGCTTGAACTTCTTCCTCTTTGGTCTGTTGCTATTCCAAAAAAAAAAAAAGACTAATTTTTCAGGTAATTTTTTAAAATTTTGATGTGAAATGCTTTCTCTTTAACAGATCCTGCAAGAAAGGAGCTACTGGATTGGCAGAAACGATTTGCAATTATCGAAGGAATTGCTCAAGGATTGCTCTATCTTCACAAATACTCAAGAATGAGAGTGATACATAGAGATCTAAAAGCCAGTAATATGCTTTTGGATGAAAACATGAAATTTTGGTCTGGCAAGAATTTTCAAACAGAACGAGACGGAGGGAGTGACTGGCAGGGTTGTTGGAACATAGTGAGTTGAGCCTAGTCATTTCTTCGATATAAATCTATGGAACATGTGTACTTATTCATACTTACTGAGTAAAAAATGTTTTCTATCCTCTGCACAGTGGTTACATGGCTCCCGAGTTCGCCATGGAAGGTACTTTCTCAATCAAGTCTGATGTCTTCAGCTTTGGAGTCTTAATACTGGAAATTGTGAGTGGTAGGAGAAATACTAGCTTACAACAAATTGATCGTCCACTTAACTTAATAGGTTATGTAAGTTTTCTTTAAACCCTTCTTTCTTTAATGTACTAAAGTTCACAAAGTTCAACAATACTGATCATTTTGGTTCAGGCATGGGAACTATGGAAAGAAGGGTGTGGACTAGAACTGAAGGATCCAGCTCTTGAAGATCTTTGTGACACTGAACAATTTTTACGAGTTATTCATGTCGGACTATTATGTGTACAAGAAGGTGCAGCAGATAGGCCAACAATGTCTGATGTTATCTCAATGCTTGGTAATGGAAGTATGTCACTGCCAATTGCAAAACAACCTGCATTTTTCACAGGAAGAGATGAAGCTGAATCATATTCGTCATCAAACAAAACAGAACAATGTTCAATTAATGATTGTTCCATTACTGTCATTGAAGCAAGATGA

>PGSC0003DMG402013876 G-LecRK

TTAGTCCTCTATCAGATTTCCCCCAAAGTTTCATTGCCCATCACCTAGGAACTTTCTTTATTCTCCTCTTTCGTTTTTACGTAGTATGAAGATAATATTGGTTCTCTTGCTTTCTTACTTTCATTTAGTTAGCTGCAACAGTACTCGTTCAGATACCTTGTTGCCTGGCCAGTCCTTTTCCTTTTCTGCAGAAGATACTTTAGTATCCAAACAAGGGAAGTTTGAACTCGGTTTCTTCTGTCCAGGTAATTGTGAAAAGCTTTTTATTGGTATATGGTATACAAACATAAAGCCCAAAACTGTGGTTTGGATAGCAAATAGGGACAACCCTATTCGCCCTCCGTTCAATAATTCTCATCTAGAGCTGTCTGATGGAAATCTACAACTCTTGGATGCACGTAAACAAAGAGTTTGGAGTTCTAGCTTATCCACACCTTCAGCATATGCCAATAAAGCAGTACTTCTTGACTCTGGAAATCTTGTTTTGACAAATGGCATTGACAGACAGTGGCAGAGCTTTAATTATCCTACAGATACGTGGCTGCCTGGTGCGATGATTGGATTTAATAAGTCCAAGAACACACTCCAGAAACTTACTTCTTGGAGGGATCTGAATGATCCAGCTTCAGGAAGCTACTCTTTGCAATTAGGCCAGAATCAAAATGGTGAGTTAGTTGTTCAACATAATTTCCAGGAAGAATGGAGGTCTGGGCCTTGGAATGAAGGAGGGTTTGCTATTCTAGCACAACAAAATTATTATAAGGATTTATTCGACTTCAGTTATGATCCAACTGAATATAGTAAATATATTACTTACAATGTCTTTGATGAATCTGACATTTCTCGAATTGTCTTAGAATTTAATGGGTCGATGAAGCAATGGTTCTGGTCTAAAGATCATCAGAGTTGGCAGTTAGTGTGGTCTCGATCTACTGATAATTGTGACCGAATAAATTTCTGTGGTGCTTTTGCAATTTGTGACATGAATGCCGGTCCACCCTGTGAATGCTTAAAAGGTTATGAAAAGAAGTTTAAATTAGATTGGGATAGTAGAAACTATTCAGGTGGGTGTGTGAGAAAAAATCCGCTGCAGTGTGGCAGCAATAACAAAAATTTTGTCCATATGCCAAATGTAAAACTTCCTGTGTCATCTGAATCAATGAAAGTAGGAAATGATCAGATCTGTGAATATATATGTTCATCCAATTGCTCATGCAATGCTTATGCTTACAGCAATAGTGGAGAGTGCTTATTATGGTATAGCGATTTGATTGATCTGTCAAGATTACCTAATAATTCAAGTCAAGTAGAGTTCAACATTAAGCTTTTTGAGCGCTCGAACAAAGGTTAGGTGCTTCCAAGCTGCATCTTATTTCTAATGTGCTCCTTTTCAACCAAAAATATGTTATTTTCCTCCTTTCTTGATCTTCCTTAATGTTTAGGAAAGAAGAAACTACTACTGGTAGCACTAGTAGTGTCCATCACTTCTGCAAGTTTTGTCTGTGGTACATGCTGTTACTTTCTATGGAGAATAAAACTCAAGGAAAAAGGTACATTCTAATTCTTGCTAGTAGCAGTTGAGTAACTTCAGTTTCACTTCTATTTGGCATGACTGAAAGAGGTCAATCAAAATGTAACCTTTGATTTAATTAGTCATGGAAAGGAAATTATATTCTCTGAATCATGCTTAGATAAAAGTTAAAATTGCCCATGTCCAGCTTCTTCTAAACTCATCAATAAACTTCCTGTTTATATTGATTTACATCAGGGATTTTGGGAAGGAAAAAGTTCAGGGAAATGTTACTTTCTAAATCAGCAACTAACTTATCAAAACCTGGAACTTCCAGTAAAATGGGACAAGAAAAGAAACTTTCCATCGAGCTGAAGTTCTTTGAATTACGTGAGTTGAAAGCTGTGACAGACAATTTTTCACCTGACAATAAGCTTGGAGAAGGAGGTTTTGGGCCTGTTTTTAAGGTATTGTTACTACATAAGAAAATATATTGACAACCTGAATAACTGCAACACTAAATTCTACTCAGGCCCAAAGAAAATGGGAAAAAGTCCCTCTTGAGGTGAACAGGTAGCGTAATGGTTTTAACTAAAATCTTTTCCATACTTGTGTTTTAATGGTGTGATTAATTGCATTGCAATTTCTAGGGTCAATTGCCAGATGGGCAGAAAATAGCTGTGAAAAGGCTATCAACTCAATCACAGCAAGGAATTAATGAATTCAAAACTGAGGTCCTTCTCATTGCTAAACTTCAACACAGAAATCTTGTTAGGTTTCTTGGCTGCTGTGAAGAAGACGAGGAGAAAATGTTGATCTACGAATATATGCCAAACAAAAGTCTAGACTATTTCATATTTGGTATGATAACTTCTTATCTGATGTTATGTTAACTCTTTCCTCCTAAGTAGTGCTTACTGGATAAATTTGATATGAACAACTTAGACGAGAGCAGGAAGTCGTTGCTTGATTGGAGAAGGCGGTATGAAATCATTATTGGAATTGCTAGAGGGATACTCTATCTTCATCAGGACTCAAGATTAAGAATAATACACCGTGATCTGAAAGCCAGCAACATTCTTTTAGATGAAGATATGAACCCCAAGATATCAGATTTTGGTACTGCTAGAATATTTAGTGGAAACCAGAATGAAGCTAAGACTCTTCGAGTTGTTGGAACATAGTAAGCACGAAAATACCAAAAGGGATTTCTCTGTCGCATCCTTACCATTTATCAATACTTTGCGCACTCATTATTGCCCTTTCCCTTTGCAGTGCCTATATGTCACCTGAATATGCTCGAGCTGGTCTCTTCTCAGTAAAATCTGATGTCTTTAGCTTTGGTGTTATATTATTGGAGGTCATAAGTGGCAAAAGGAACATGGTCTCTTATAATAGTGATTCCCCTCCTAATTTGATACGACGGGTGAGTTTTAAGATACAAACATTTGAAACTAACTTTTGATTTAATAATCAAACTTATTTGCTAAATGAACTGCTGCTTCAGGCCTGGGAGTTGTGGAATGATGGTAAGGCCTTTGAGCTGGTAGATCCATCAATAGTTGATTCTTGTCCCAGTGAGGAAGCATTACGCTGCATTCTAGTAGGACTATTGTGTTTGCAACTCAACGCAGCAGACAGACCAACCATGTCAACAATTCTGTCCATGCTAAGTATTGAAGCAACGGTTCCCTCCCCGAAGAAGCCCTTCATTACCCGTAACTCAGACTTGATTACTACAGAAACAGCATCATCTTCGATCAACGAGGTGACAATTACCGCACCTTATGTTCGTTAGTTGTCTGACGTGTATATGTATTTCTGTTATCTGAGTATCAAATGTATCTAGTGCTCATTACTTCCCAAAGTATGATATAAGTCTTCAATGATTCAATGCATTTAATGCTAAAAGATTTTCAACAGACTAGTTGATATTACTTTTACAGATACTCATCACAGAAAGATTGAGACAACAGTGCTCTTACTCATGATCATAATATTATTGTTGTATTTGATACATGTCTATCTCATGATCTAATTATGAAACAAAAGTTAAAATCTTACATAAGATAACTTACATGTTTGTATTTTCTTATCCATAACTTTTTGAGGAAAAACATAACCAACTAATAATAGTTAAATATCTCATTTATTGACAAAACTCCACAACCAAATTACTATCCTGCAAATCAATAAAAAAATTGAAAAGTTGTGTGTGGCTAGCAGGATTAGCCTTTGTTGCTTTTAGAGCATTGATGAACACAGAGAAGACTGTTAAAGAAAACATGAAGTGGGGCAATTATCAAAAACAGCTGACAAAGAGCACCATTTAATTCACAGATGAAATAGAAACCCTCTGTGTTTATGCTTCTTTCCTACCAATCCTTTGCCACCCACCTACCTTTAAGTCTGTAACATGGGCCATGGGGAATCAGAAGTCAGTTTTGGTTCCAGTGCTATTAAAGCCCAACCCCACTAGTTTGAATCAAGAATGGACTAGCATGTTAAATGTTGTGTATAACCCACCAATAAATCTCTCTAATACTATGAAAGCAAAGTACAGAAAAAGGAATTCTGTATTTCTTTCAAAATAGTTCTTGAATAGTTAGCATAAGCTCTAGCGACCATGAACTGTTGAAATAGACATATCGTTGACAGATGATGTTGCTATGCTTGCTGATGAAGAGTGTGTAACTGAACTATGATTCTCTGATCTGAATATAAGGCTAGGTTGTTTAGGGAAAGGAAGATTTGTTTCATTGCACAGCATGAAGACCACTTCTGACATAGTCGGTCTGTCATCTGTAAACGGTTGTACACACAAAAGGCCAATGTGGATGCATCTCAAAACTTCACAAGCTTCATATGAGCCACCTAACAATGGATCAACAACATCTAAAGCCTTTTCATCCTTCCAAAGATCCCAAACCTGAAATAGAAACACCACAATGCATCACAATCAAAATTTCAGACAATTTCTTCTCTATAAGAGAAGCGACTATAACACCAAAAATTGAAATTCTGAAATACAAAATCTCGATCTATTCAGCACTTTTAACAACCTGAAAGGGAATTGTTTCCAAATTATTATTTGCAGTACTACCATCC

>PGSC0003DMG402014514 G-LecRK

TTCATGCATTTCTCACTTCACTCCATAACTCCTATGCATCCACACTCACCGTTCCCGTCGCCGTCGTCGGAAGCCAAAAATGGTATCACTTCAAGAGCCATTCCTAAACCGCCGCCAACAGTCATTAATTTTACACATTCACCACTGCCGCCTTCGCCGTCATCACCGAACGCGTCGTTAAAGCCAATAATTAAACCTATTGATGATAAAATCGCTTTACATCCACCGCCACGGCGGCCGATATATACTCATCCGTTACCGAAGCTTCTTCCGAAACTTACGTTACCACCGCCTCCGGTAACGGCGTTAATGTTTCCTGTGAATGGTTCTTCTGCACCAGTTTCTAGAGATGAAACTTCGTCGATTAAAGTTCAATCGCCGTTGTCAAGTGCATCAATCGGATCTGTAATTGGAATTTCTGTAGGCGTTACCGGAGCTTTCTTCATTGTTGCAATTATACTCTTCGTGTGTTGCAGGAATAAGCTTAAATGTGGAAAATTACGTGATCGTGAATCTCCACTGCCAAAAGGTAAAGCTTTTTCAGTTTGATACAGTAGTTGAATCAGTAATATATACAGTAACAACTAAAGCTGCTTTTTCAGGGGAAATGTGTTTCACTTAACCAGAAAATATTTTTCTTTGCTCATGAAGCTTTCAATGTTCTGTTTATCTTCAGTAATTGATTACCTTGCTAGCTTGATATTTTGCAAAGAAATGACGTTTCAGGGTTAAGTACTTCACTTCGCCTGAATTTTATTTCGCCAAAGATGATTTCGATACAACAACGTACCCAGTATAATATCCCACAAGTGGGGTTTAGGGAATGCGATGTATATGCAGCCTTACCTTGCTTGTGAAGATAGAGAGACTTTTTCCACTAGATACTCGGTTCGAGTAATCATTTCAAATATCAAGTATTTAAGCAAATACAGTAAAAAATTAGGTGCCCGGGGTGACCCAGTGGTTTGTACTTGGAACTTCCATGTTCGAGGTCTCAAGTTTGAAATCCCTTGCAAGCAAAAGCAAGGGCTTTGTCACACCAGGGTTGCTTAATGCAAGTTACCTCTCCTGTGTGGTTTGCGAGCTATTGCATAGGAGTAAGAGTTTTACCCTGTGAGCGCTCAAAGGGTAGCTTCTGTGAATTTCCCTTGCCTTTAAGAAAAAAAAATTAAGAAGAGAACAAGTAGCTACATCAAATAACACAATAATTGAAGCAAAGGAAATAATAGGTACTGCTAAAGTCGAAGAAAAAAATTAGTAAGAGGGTAATAATATAACAATACTAATAAGGGAATTAGACAACGCTCAACTACTACTAACCTTCTAACTTAATCATTGATCTTCATATCTTTCTATCTAAGATCATGTCTGCGATAAGCTGTAGGTGAGCCATGCCTTCTCAAATCACCTCTCCCAAAGACTTCTTCGGCCTGCCCACCTCTACGGCTCTACCTATGCTTATACCCACTACAACCAACCTCTCACACCCAGTGAGGTACGCAAGATTTTTTGTAAGGCATGTCGGTATTTAAAAATAAATAATTGATATAGCAACATTATATTAATATCTTGTACAGGAGCAGACCCCTTCATTTGCTGTAATTTTTTGCCTTGCATCTTCTTGATGCCTTCTAATGAAATCTCCTTACTTCATAAAAAAAAAAATTATAAAAATTATTTAAAGCATACTAATAAAAATAAATCAAGTAAACTACTACAGCTTATTATTGTGTCCTATTCTATTTTGGACAAGTAATCAAATTTCTATAATAGGATATAAATTGTATTTTAGTTCAATTGGTTTATCTTATCCTGATTTTGAATAGGAATTGAAGGTTGGAGGTTCAAATCCCTATAACACTTACTTTTATCTAAAAAGGCCCCAAAATATAACTCAAATTTGGGCTTGTTGAGGGTCGAACTCGGCCACACTTTAGAGAATGACTACACCATCTGCACGCAGTAAACCAATAAGCCAAGGAAGCTGTTCAAGTAGTGTCATTAATATTTTTTCTACTTATATTTGAGTTTTCAGATTTTTAGATATAATTATATAATACTAAAAATTTTCATCGAAGCAGTGTCGGTTGACACTCCTTGACATAGGGTGGGTCCGCCCTTGCTCGCACCTCATCGCTGCGGCGTACATTTTCTTTTCACATACCCGAATCATCTCAGCCTCGCTCCCTCACTACCATGGAGGCCACTCCTGCCTTTTTCTTGAATAACTTTGTTCTTGATCCTATTTCTCCTATATGCCCACACATCCATCCCAACATCCTCATTTCAGCTACTCTCATTTTTTGGACGTGCGAGTTCTTGATTGGTCAACACTCTGCCCCTTACTACATAACTAGTCTACCACCACTCCATAACACTTACCTTTTAAGGATTGGTGGCACATTCTTATCATACAAGACAAAAGGTGCTAGCTTCCATTTCTTCCACCCCGCTTCACGAGGATGTGTAACTTCATAGTCAATCTCCCCATTTCCTTGAATTATTGACCCTAGATACTTGAAACTTTCTTTCTTGGGGGTGACTGGTGTATTAATCCTTACTTCTACATCTTCCATGTGGATATGTCATGAACTTAAACTCTAAGTATTCAGTCTTGGTCCTACTCAATCCGAACCCCTTAGACTCCTGGGTCTGTCTCCAAACCTTTAATATATTGTTGACTCTGTTTTATGCCTCGTTAATCAAAACTATGTCCTCTGCAAATAGTATACACCATGGCACCTCTCCTTGGATATGCCACGTCAGTTGATCCATCACCAAAGCAAAAAGAAATAGATTAAGAGTTTATCCTGATGCAACCTCATCATGATTGGGAAGAGACCTGAGTTTCCTCCTATTGTCCTGACTCGTGTCTTGGTACTATCATACATGTCCTTACTTTCCCTAATATAAGCCTCAGGTACACCTTTAGCCTCCAAACATCACCAAAGAACCCTCTAATGACGTTGTTGTAAGCTTTTTCTACGTCAATGAACATCATATGTAAGTCCTCTTTCTCTCCCTAAACTGCTCGATTAATCGCCTAACGAGATGTATGGCTTCTGTAGTTGAAAGCCCTACGATAAGTCCAAATTGGTTCTCGCTGATAGATACACCCCTCCTCAACCTCATCTCCACCACTCTCTCCCAAACTTTCATAGTGTGGGTCAACTAACAACTTGATATCCCTATAACTGGTACAATCCTGAATATCACCCTTGTTCTTGTACAACAAGATCAAAGATGATTTAATTACTATTTAAATTTATCATTATTTTGTTTTATGATTTAGTAGGAGTGGTGACTTTATCTCTAAATTCAACTATTGAAAATTAGTGATCCAAATTGCTATTTTTTCCTCGTACAATTTAATAGTAACTTCTTTAAAAATTTACTAGTAATGAAATTAGTGACTTATGTTTCTAAATTCATTGTGACAGACAATTATCATTTCTTAAATTTTATACTTCTTATGTTCATTTTATATGATTGTGTTTGAATGTGTATGGACTACGGAGTTTTAAGAAGAAAAAGGGACTTTTTGAGACATAATCGAGTATATGTCACATACCAAAAGTATCTATAGGATCTTATGCTTTTAAACATGCCATGCCATTCTGGAAATGTGACAGTTTGGACATTAATCTCTTAAATGTGAAACTTTCTCCGTCTGACCAGATATAATGCTTAAATCTTGTTTTAAGAGGCTCCTTGTTCAGAGCTTGTTTTCATTGATTGCAATTTGCATTTAGAATGGGGTCCAAATTGTTGATATTATGAAATTTAGCAGTTTAATCTATCAACATAAGTTCAAAAAAGCTGTTTACTAACCAGTATGAGTTCAACTTGGTATATTCATTTGCTAATTGTTTCGGTCGATTCATCAGATGACTTGTACCATGTTCTGCGTCAAAGTTTTAGACAGCAGAATGTAGCAAATGGTTCTCCTGTTTCCCCAAAGCAAGTGACACCATTCCGCCCACTTCACTTCTTTGACAACAATGCTCATTCCAATTCCCTTAATTCAGAAGCTGGCAGTCATTTTTCACCAAAAGACCACATAGTGGATTTAGCTGTTTCGGGTGGCAATTTCACATATGAAGAGCTGTGGTTAGCAACTAGTGGATTCTCTACATCCAACCTTCTAGGTGAAGGTGGATTTGGGTATGTACACAAAGGAGTCCTCCCTACTGGAAGAGAAATTGCAGTTAAACAGTTGAAGGTTGGTAGCCATCAAGGGGAGCGTGAATTTCAAGCAGAAGTAGAGACTATTAGCCGAGTGCATCATAAACATCTTGTGTCTTTGGTTGGATACTGCATGAATGGAACAAAGAGATTGCTTGTTTACGAATTTGTTTCCAACAGGACCTTAGAGTATCATTTACATGGTATGTTCTTTCTCTTTTTCCTTAATTCTGGTTGTTGGTCGAATCCTGTATCAATAAGATTTTGTGACCAGCTTAAGGCATTTTTTGAATTATGCTAATATTTGTACAGCAGAGAGTTGATAATTAAGGTTAAAATAAATCTACCTTAACGGGGGCTTCCTTAATGCAGCTTAGTTTCACTGTATTGATTTTGAAAAGCATGTAAAATACACCTCACGAGTTATGAGTTCCATGTTAAATGGACAATCTTTAAGGGGGTACTTAACCAAGCCTTTTAAGTAAGATTCCTCGTGGTATATGCCTATACCACAAAAAGGTGGCCTAGAATTTCATCAAACCATGGTTTTTAACATGAGAAGGTAAAAGAAGAGCTTGCAAATACATGCTACTTCTGTAACCCAAGCCCTTTAATCTTGCATGGTAAAGCACTGCTATTTCTATGCTTTTGGAAAGACCAGCATTTAATATCTTCTGACTTCTGCTGTCCCAGGCACTTCATTTGCAATTTTCCACATACTTTTCTATACCCTTATGGTGTATTATTTTAACACACTTCATGTGAGGTACATTTGATGGAGGATCTTTCATAAGTACCACATTTTAGATGTGTGATTTGTTTCATCATCTGTTTTAAGGTTTGTATAAAGAATGTTCCTATCCACCCTGATGCATGTTTTCTTTTGAACTTTTGTTTTACAGAAATGCTCTTGTATCAGTTTATACTTCTCTGATCACTTCTAAATATCCGATTGAATAGAGTTTCCATAAGTTCCTCTGATATAAAGGTCTTTCTACCAAGATGCTATTTAGAACAGATTGTATGATATGCAGGGGAGGCACAATCCACGATGAAATGGGCATCTAGAATGAAAATTGCTATAGGGTCAGCCAAAGGATTGGCTTATCTGCATGAAGACTGTGAGTGCGCTTCATCTCTGTCTTGGCTTCCTATGTTTGTTCTTAATTTGCATGTTCGTCCATTAGTTGACTAGAGGTGTCCAAAGGGGTAAATTGAATATTAAGTGGCCTGGATGCTTTTTAACAATGTTGAACATGTTTCCACAGGGAACTCTTGTTAGTGTGTTTGTAGATTGTGACAAGTTGGGTAACATAATGGTTTTCATCACATATCTATCTTCACTTTTTCTCCTGAAAAGGATGTAAATCTAATTTTCTCCCTCCCAATAGATACCATTGATCCTCACTTCTATAATACCATTTCTGAATCTCCTCGAGATTTGAGAGCTGAATACCTCGGTTCTAGGGTCTGCTAATGGATAGCAGAAGGCCATAATAATTCCTAAATTGTGAAAGATATAACATGAATATGCATCAGGCAAACTGTACTTTCCCCATATAATTTTATATGCCATGTTCTTTCCTGTCAAATATGAAATCAAATAGTTTTAGTATATTTTCTTCCACTACTACCAATTTAATATAGATGTCAACGTATCTTTGCGCTCTGTAGGACAATTTAATTGTGGTTGCTTGAATTAGCTTGAATAGAATAAATATCACTTCTTTTGATGTAGCAACAACAGGTAATCGTTCTCTGAAGATACTAACTTGTGGCTGCAGGTAACCCCACTATCATTCATCGTGATATCAAGGCAGCTAATATTCTTCTTGATTCTAACTTTGAAGCTAAGGTAACTTTGTGTTTCGCTTTTTGGTATTGCCACTATATTTTCTGTTCTTTTTTTTGAGAACTGTTTTTTTATTTTTTTTGAGAACTATTTTCTGTTCTATCTTTACTATCAAAGAAAGGTCGGGCTGTTTTCTATTATCGACAACTGGCTTCTGAGTACACCTAGAATCCACTGTTGATGTAAATAGTTTTTTGTTCTTAACAATAAAATCCATATGATGAGATTAATTTGAACCCAAATGAGCATCTACTCAGGTGCAAATGTTAACTGCATTATTGATTCTGCTTTTTGTTTCTTTAATTTTCATGTGACAACTAAGAACATATTTCCTCGCAAGTTATTTAACATGAATGGGAACTTTGATTTGTATCCAGGTTGCGGACTTTGGACTTGCCAAGTTCTTATCTGATTCAAACCATCATGTCAGTCATATCTCCACTAGAGTGGTGGGAACTTTTGGGTGAGTTTTATTGATCCTGTCTCTGCTACCATGAATTAGATATCAGAAGGAAAAAAATTGACTAGATCAAGCAAAATTACTATATACAGGTCACCATACTGAGTGGACCAGTCTCTTCTACTTTTATCAAGTAAAAAGTGGTGCTTCCCTCGTTCTAGCGTTAATAATATTCATTTTTTTGGGCTAACCTCATACGTAGCATAGGCTATGCCCTTCCCATGTTAACTTTGTTTCATCCAACTTGCAAACTGATATCTATATCAGATACCTATAGAAAATCGAGCTAGAATGGGACTCTCAAATCTGGGGATTTTAAAGACATTGATGACTTGCATGACCTCGATTTTAGGTCAACCTGCTCATAATGCAGTGATTCCTCTCAGAGTTATAACCGAAGTTTTCCTGCTAGTCGGAGGATGAGAAGAGCGCATGGGATGTTACTGCTGTTTTCCCGTGTCTGATAGTGTTGGGTTTCTTAGTCCTTGATTTTCCCTTACCAACACTGATTGGGGAAAGAGGGGGTTTGGGGGGTTAAGGAGAGGGGCTTGGATTTTGGAAACTGTAGAACTTAAGCTACTTCATGATTAGTAATGATTTCTGGATTACACAGCTTCAGGTAGATGGACCACGTAAAAACTAACAGGGTAAAACCTACATTTCCTGATCACAATTTCTTCATGTATTTGATAAAGCTGACGTCTTGCAGGTATCTGGCCCCAGAGTATGCACAGAGTGGAAAGACCTCAGATAAGTCCGACATCTTTTCTTTTGGAGTCATGCTCCTGGAGTTAATTACAGGGCGTCCACCTATAATCTCAACTGAATCCTCCGCATGCAGCAGTTTGGTCATATGGGTATGTTAACTGTTCCACTTTCTTGACCGCTATTCAGCATTCTTAAGTGCCTAAAACTAATGTTCAACTAGATGCATGAGTAAAACGTTTGTGTGCAACCTTTATAACTGGTTAAATAGCATAATGCTTAAATTTACTTTCTGTCCAACTACACTAATCAAATCCTCAATATGGAATAACATTCATCTTACTATAATCTTATCCTTAATTAGCTATGTATGCACTGTATTTAATCTGAGGCAGAACATGCAAGCACAGCTTATGCAATTCTCTGAGGTGGCCTTTCTATTTATAAGAGATCTTGATCCTCCGGAAGGTTTATCATTTGTGGTTTTAGAAGTACTGGGTGGTTTTATCCTTTATTGCTTTTCTTTTTTGCGAGTGATAAATGGTTGAGAAATTTTATCACGGCAGTAATCAATATCACGGAAACAACCTTTCTACCTCCACGAGGTAGTGGTAAAGTCTGCGTATACTCTACCCTCCTCAGACCCCACTTGTGGGACCTCACTGGGTATGTTGTTGTTGTTGTTTGTATTAATCAATCACTTAAAAAAAGAACCATGGAAAAGCATTGGCTTAAAATCAGAATTTGGCAGACCACCGTCCTTGTACAAAAGAAATGTGCAAATCAAAAGGATCTGGTGTTCTTGAAAAGTTCATCTCTGTCAAGCCAAAGGCATAAAAATTCACGTCCATCTTACAAATTTTTAAAAGAAATTTGTCATCCTTGTACCAAATGATTTGGCTTTTGGACACCCGTTTTGATGATGGAACATATCTCAAGTTTTCTGTTGCATCTTTTACAACTATTAGTTTGTCCAAAGTTTCAGTCTTAAAGCATGGAGTTCACCATGAATTTGATATCATACTCTATCAAATTTTCCCAAAATAAGGCTCCCTTTTTAGGAAATCTTGGAGATACGGGTTCCTGGAATCATTTTGTTGATCTCAAAGATTATTACTGTTGGTTCTACTTAAGTCCCACAATTGATCAGAATTAATTACATAAGTTAATTCGAAATTGAGATCAGAAAGACGGATTTCAGAGTTAGCAGCTTCTTGCATGCTGCCTACTCACAGACAGAATTAGTTATGTCACTTTAGTTAATCAGAATTAACTTTACACCTCTTCCGGGGCATTTTTTTTCATAACCTCGTCATCAAATACAATATTTTGTATTAGGAATCATTACCGTAGTTTCTGGTTTTCATATTCTTGGTTTTTGATTAAGTACTTTTTGAGTGTCGTTTATTAACTGGATGTACACAAATTTCTTGATCAACATATTGGTTTCTTGTAGCAAAATAACAAAATATTTTGGGAACTAGAAGCATCTCTAGGCCGATGCTTAAATGCATTTTCAAAGATATGGATATCATTGTTGTCAACCCTCTTCTTATAGTATACCACTTGTACATAGCATAAGAAGACTCTTTCTGATACACAGGCAAGGCCTTTGCTTAGAAGTGCATTGGATGATGGAAAATTGGACGCCCTAGTAGATCCATGTTTGGGACAAAATTATAACAGTGAAGAGATGGCTAACATGATTGCTTGTGCTGCTGCTTGTGTGCGTCATTCATCACGGAGAAGACCGCGGATGAGTCAAGTAATAACATCTTCTCTTAATCTTTTTCTGTTAGTGCTGGCGGATTTCTTTAAAGGGGTTTATAAACCAAAACCTTCCTACTTATTGTTCAGGTAGTTCGTGCTTTAGAAGGCGATGCCTTTATCCTGGACCTTGATGAGGAAAATAGGCCAGGACAGAGTACCATTTGTGACTTTGATTCTGATGGAAGCAACTATTTTGAGAAGTTCAAGAAAATCCAGCGGAGCACTCTGAAATCAAAAGAAGAATGGTAATACCACCGTGCACCGGGGGCTGTCATGGAAGTACACCAACTGAAAATGGAGTTAGATGAACGAAGCTCTTCTCCCCTTTTTATCTTTCTTTTCACTGATGTAGATGATAATTCGAGGGTGAAATATATGATTGCAACCATCAAAGTTGTCTCACTTAGTGAGAGGATTGATAGTTTGTAGTTACTTGAAAGTGCAAAAAGAGAAGAAACAGGAATTGGCATGAGATATTCAGTATATCACCAAGTAAAAATCTGTGTATAAAGATGCATGTTCAGATGTACATATAGATTTCTTCCCCTCTCTCTTTTCTTATTCCTTTTTGC

>PGSC0003DMG402019242 G-LecRK

ATGAAAGGCCTATTTTCATTATGCATTTGCTGCCAATTTCTCTTCATTTTACTAACTTCTGCAGCATTAGACACAATCACTACAGATAAATCCATTAGAGATGGTGACACAATTGTTTCAGCTGGAGGGATTTATGAGCTTGGATTTTTCAGCCCTGGAAATTCGAAGAATCGTTACGTTGGCATATGGTACAAGAAAATATCAACTGGAACTGTTGTTTGGGTTGCCAACAGAAACATTCCACTGAATGACAATTCAGGAGTGTTAACACTCAATCCCAATGGAATTCTTGTACTTGTTGATAAATCGAATGTCTCAATTTGGTCATCAAACTCATCAAGATCGTTAAAGAATCCAAAAGCACAGCTCCTGGATTCCGCAAACCTTGTTGTCAGTGATGGAAATGATAGAGGCCTGGAAAATAATTTCGCGTGGCAGAGTTTTGATTATCCAGGAAATACTTTGTTACCTGGCATGAGGATTGGAAAAGATTTGGTCACGGGAATGGATTGGCATGTAACGTCATGGAAGAGCATGGATGATCCAACTCCTGGTGAATATGTAGATGGTCTTGATTCTCATGGATATCCACAATTGTTTGTGTGGAAAAATTCATCTGTAGTATTTAGCTCAGGGCCATGGAATGGTATTGCATTCAGTGGTAGTCCTAATAATAAACCAAATACATATTACAGTTTCGAGTTTGTTATTAATCAACAGGAAATTTACTACACATATACAATTAGGAATGACTCCATTCCCACCAGGGTGGTGCTCAATCCGTCTGGTGTGCTAGAACACCTAATATGGATAGAGCGCAGTCAGAGCTGGTTTCTCTACTTGACAGCACAATTTGATAATTGTGATCGTTTTAGTTTATGTGGACCTTATTCAAGTTGCAACATCAATAACTCCCCTCCATGTGACTGTTTGAAAGGTTTTGAGCCTAGGTTTGTATTTGTTACACTATCTGATCATCTCACGTTAAGCTGTCAGAGAAAGCACACATTGGACTTTTGATTACTTACTTATGTTTTCACACGCCCCTCATGTGTGTGCTTGATTCTTTTTTTGTTACCTAATTATGTCTTCACAGGTATCCTCAAGATTCTGCAACAGAATGGTCTAGTGGTTGCGTTAGGAGAACTTCTTTGGATTGTACCCATGATGGTTTTCTTAAATTTACGGGTATCAAGATGCCTGATTCTAGAAACTCCTGGTATAATGAGAGCATGAACCTTGAAAATTGCGAGAAAATGTGCTTGGCTGATTGCAATTGTACAGCCTACTCAGATCTTGATGTTAGAAATGGCGGAAGTGGATGCTTACTGTGGTTTGGAGAACTCATTGATATACGAGAGTTCAGCCAAAATGAACAAAACCTGTTTGTGAGAGTAGCTGCTTCAGAATTAGGTAATGTCCCCACTCTGCACCATTCTCTCGTCTTAACAATATAAATACACAACTCTTAGAATTGAAGGAACTCTACTATCTTATTTACTTGATAGACATTAAAATTGTTTATCAGCCATTAAATGGCTGCACATCTACAATAGTTTCCCCTTAAGGAGGGAGCTGATAATCTAGTACGAACGTATCCATCTAGTAGTGCTGTACTTGCATTTGCCTCTCAGATCATCCACTAGTCCCCTTTGTTGCAGACTTTATTTTGTATAGAGTTTGACACTAGCAGCTATATTCAGAATGAATAACCATATAGGAGAGCAAAGATTGTTGCGTAACAGTTTCCTCCAAGGGATTTTAGGAAAGCTGCCTCTCAGCTTCTGATGTGCTAGACCTTATGGAAGATCAAGGATGTTAACAGCGGGGGATGGTCGTACACACTCTATCCTTTAATAGTTACTGATTAGCAAGTCTGCGGGGTTTCATAGAAAAATAAGAAGTAATACAATCAATGCAATTCATTACTGATATGGCCAACTCATATGTAAATTGAACATATTCAGACAGGAAGGGGAGGAGAAAGAGGTCAATCCTGATTGGTGTCATTTCAGCAGCAGTAGCAACATTTATCCTCAGCCTTTTAGCTTGGTTTTACTTCCGAAGAAGGAAAAGAAGAAGAGGTATATATATACTTGAGATACTCTTGTCTTTCTGGCTGACATTGTTGGCCTTTTACCAATTCTCATTGATATCATGTTTGAGGCGGGCGAATGTATATTGACAGGTTTAGAAGTTGAAAATGAGGAAATGGAGCTTCCATTGTTTGATTTAGTAACTGTTACTAGTGCCACTGAGAACTTCTCTTCTGCTAATGTGATTGGAGAGGGAGGTTTTGGACCCGTTTACAAGGTAAATCAACTTTGATCCTCAATGTCTAGCAAGAAGTGGCATCTTTGACTCTTGATAGAAAATTGCAGGGTATCCTACCAAATGGACAAGAGATTGCAGTAAAGAGGCTATCAAAGCATTCTGGACAAGGCTTTCAAGAGTTAAAAAATGAAATCGTTCTCATTTCCAAGCTGCAACATAGAAACCTTGTCAAGCTATTGGGTTGCTGCCTTGAAGGAGAAGAAAGGATGCTAATCTATGAGTTCATGCCCAACGCTAGCTTGGACTATTTCATTTTTGGTCTGATTTTCATGATTCATTAACTAAATACTACTTGATATGTCCTCAATATCTCTGCATCACTCAACGATTAGTTCCACATTGTGCAGATACAAACAGAAAAGCTTCACTAGCATGGAAGAATCGCTTTGAAATTGCTATGGGAATATCTCGAGGTCTTCTTTACCTTCACCAGGATTCAAGATTACGAATTATTCACAGAGATCTCAAGACCAGCAACATTTTATTAGATAGTGACATGAATGCAAAGATTTCTGACTTCGGCCTTGCCAAAATTTTTGGTGGAGATCAAGTGGAAGGAAAAACTAAGAGAGTAATAGGGACATAGTAAGTTTCTACACATCTTCAAATCATAGAATCTTTATTACATTCAATCAGTACATGAGCATAAATATATTGAGTTGATGCAGTGGATATATGTCCCCGGAATATGCTGTTGATGGGAAATATTCAGTAAAATCAGATGTATTCAGCATTGGCGTAATCATTCTTGAAATAGTCAGTGGCAGAAAGAACAGGAAATTTCGTCATTTGGAGCATCATCACAATCTTTTGGGACATGTAAGCACAAAACTTCATTCCCTTAATTCCCTCTCAACTTCCTATTGTAAATACATTTGTATAAATGTGTTGTTCTGTTTGTAATCTAGGCATGGTTACTTTGGAGTGAAGGCAACGCGATGGGACTGATAGATGAATGTTTGAAAGAATCATTTTCAGAATCTCAAGTGTTGAGATGCATCCAAGTTGGTTTGTTATGCGTCCAAAAACTCCCCGAGGATAGGCCTACCATGGCATCAGTAGTTTTCTGGTTAGGCAATGAAGGCCTGGTTCTTCCTCAACCAAAGAAACCTGGTTTTTTTATTGAGAGGAATTCAACGGAATCAGCAGAGTCATCTGATGAAGGATATCTAAGTAACAACGTATCGATAACAGTCCTAGAGCCAAGATAGACATAAATTCCAACAATTGTAACTTTTGAAATGTATCCACTTTGTCCTGTTTTTGTCAACAGTATTAGTGTATATATGGACTATATCATGAGGTGCATTGAGGAAGATAGTCTAGTATTTCTGTTTTTCATTGACATGTGTATTCTTCGATACTATAATAAACGTAACGAATTGGCCTATTCTGAATAACTTGTAAAAATCATGTCTCCGCTGCTGACCATCAAAAT

>PGSC0003DMG402026153 G-LecRK

CAGAGATATTGTTTTGGGAATGCCTATGGAGTACTTGCCTCATAGAGATAGTTTTGATGAAGATTTAATCACTGAAGATGATAAGAAACGGATTGATGTCCCGTTTTTCAGCTTGAATAGCATACTAGTTGCTACTGAAAACTTCTCAAATGCAAGTAAGCTCGGGAGAGGTGGATTTGGACCTGTTTATAAGGTTTTGCAACTTTATTTTTCTTTCCTTCTTCTGTTCTAAGTTGTTTTACCATTCAGATTTTTTTCTTTCGTTTTCATTATATAACAGTCCGGAGATATATTTATGCAGGGTAAGTTTCTCGGAGGTACAGATATGGCAGTTAAAAGGTTGTCAAGTAATTCTGGACAAGGTGTTGAAGAATTTAAAACTGAAGTAATGTTGATTGCCAAACTGCAGCACAGAAATTTAGTTAGGCTTCTAGGCTATTGTGTTGAGAGAAATGAAAAGATTTTGTTATATGAATATATGCCCAACAAAAGCTTGGATACGTTTCTATTCGGTTAGTATACTTCCCTCCCTACCATGAAACTCGTTGTAGTAATAGTTTTATGATTGATTCTCTCTGAAGTTATCTTACAAACCTTGAAATTAGATCATGCATTCTGTCGATTACTGGACTGGAGGATTCGTTTTGAAATTATATTGGGTATTGCTCGAGGGCTACTTTATCTTCACCAAGACTCGAGGCTAAGGATCATTCATAGAGATCTAAAGACGAGCAACATATTGTTAGATGAAGAAATGAATGCCAAAATATCAGATTTTGGCTTGGCAAGGATTGTTGAAGGCAAAAATACAGAAGCCAATACTAACAAAGTCGTTGGAACATAGTAAGTCTCTTAGTTCTGCTTCACCTTTGATTTGGTGGGAAGAAGATAGATATATACACACCTAATGTATTGGTTTACACAAACTAGATGCTAAAATAACCTATTCTGGAACACTAAAAACAGTGGCTATATGTCTCCGGAATATGCATTGGAGGGACTCTTTTCAATCAAGTCTGATGTATTTGCCTTCGGAGTAGTCCTACTCGAGATCGTAAGTGGGAGGAGAAACATGGAATTTTTCGAAGATGTAAACCTCATTGGTCATGTAAGTACCAAAATTTCCCTCATTAATATACATCTCATTCTACTTAAAGCTTTCATTGACTCAAAATTAGCATTTCACAGGTATGGAGACTGTGGATGAAAGACAGTGCATTAGACATGATGGATCAAACCATAGTTGAGTCGTGCGATGAAAGTGAAGTGCTAAAGTGCGTTAATGTGGCACTCCTATGTGTTCAAGAAGATCCAGCTGAGCGTCCTGTAATGTCAAATGTTGTTTTTATGCTTGGAGGTGAAAGCATAACACTACCAAGACCAAATCAACCAGCTTTTATAGCACGAAGAAATAGTGCTGGTGCAAGTACATCTTCTTCGTCTTCTAAGCTATACAACACTTCCAACAATGGGTTGACAATAACTCAACGAGTAGGCCGATGAAATCAATATACATATAAGTTAAATGCATAACGCACACAAGCCAATAATGTGCCGGTATATCTTTACTTCAATTTCAGTTTGAATTTCTGTTAAATTATTTTGGCACGCAAAATAACCGATTGTGTTGTAGTCTCGGA

>PGSC0003DMG402029906 G-LecRK

CAAAGAAGTACAAAACACTCATGAAGTAGAAATAATTGGGGACTAAGACTATTTCCCACCGTGATCAAAGTAACAGTCTTATTTTCAAAATTATCACTTCCACTTAATTATGAAATGTTTTCATCTTTTATGGATAGTGGTGTATAAAAGATAAAACTCACATTTATATACCATAGTTTTTGATATGTGAAAATGACATACTTCATAAAATATCTTCTTGTTCTTGTTCTTGTTCTTCTCATCTTTCCCCTGCATATTTTAGCACAAACTATACCATTAGGCAGATCTCTCATAGCAAACGATGAAACAACACCATGGCTTTCTCAATCTGGAGATTTTGCATTTGGATTCAAGAAAATTCAAAGTCAAAATCAGTTCTTGCTTTGCATATACTATGCCAAGATTAAAGATGCCACCATAGTTTGGTATGCAAATGGTGGTAACCTCGTGCCAGGAGGCTCTATCGCGGAGTTAAATCCTCAAAAGGGACTAATTCTCCGCGATCCTAAAGGGAAAATGATCATTTGGAGCACTGGTCGGATTGGTTCTAATGTTGCTTATGCTGTCATGAATGATACTGGAAACTTTGTTCTTGTTGGAGTTGATTCTTCAGTTTTATGGGAAAGTTTTCGATATCCGACTGATACCCTTATGCCTACTCAAATACTAGAGATTAACAACAAGCTTGTTGCTCGAAAATCGGAGTCTTTCTTTGTTCCTGGTAGATTTTATCTTCGTATGTTGAGTGATGGAAATTTGGTTCTTGTTACTCAATCCAAGCCAACGAATTTCGATTATGATGCTGAGTATTACAACAGTCATACTTCTGATTCAGGAGATGAAGCCAATTCTGGTTATCGATTGGTATTCGATGAGTTTGGTTCAGTTTACATTTTAAAAAGGAACAATCAAAGACTTGTGTTGACACCTCCCAATGTTCCTTCAATTTCAGAAAATTATCACAGGCTAAGTCTTGATTTTGATGGTGTTTTAACTCATTATTATCATCCTAAGAGTACGGGGGATCAGAAGTGGAGTACTCTTTGGTCTTTGCCTGATAACATATGTCTAGCAATTCTTGAAGATGTTGGAAGTGGAGTTTGTGGATTCAATAATGTATGCAATCTAGGTGAAAATCAAAGACCATATTGTGAGTGTCCAAAAGGGTATTCGTTGATCGATCCAAATAATAAGTATGGAAGCTGCAAACCAAAATTTGTTCCAAGTTGTGATGAATTTGGACAGGGAAATCCTGAAGAATTATATGATTTTGATGTGGTGACAGATGTTGATTGGCCGTTATCAGATTTCGAGAGGATTTATCCTTCTGCTGAAGAAGAATGCAAGAAGGCTTGTTTAGAAGATTGTTTCTGTGCTGTTACTATTTATAGAAGCAATAGTTGTTGGAAGAAGAAACTTCCATTGTCAAATGGGAGAGTAGATACCTCTTTGAATGTAAAAGCTTTCCTTAAAATAAGGAAAATTTGAACATAAGAGCTTCTTAAATTAGATATAATGTATGTCTGCTATAGGTTCATGGTTGAATTCGTCGATCCAACTTGCAATTTTACACAATTCAATAGTTTGGATTTGTAATAATTAATGGACCATATTGTATAACAAACTTTGATTAGCAAATAGTGAAATATTTCATACATTTCAAGAAACATATTGCACTGAGATACAGATACAGGTTATTGATGTACTCTTTAGGTCCAAGCAAATGAATAATGGAAAAAAAAAATGGTTGATCTAAATAGAATCGACAAACGAAAAAGGGTCAGGAGGAATTGATACTTCAACAGAACCTTCAAGCATTAACATGACTTTCTTCATGTTAGGCCTTGATGCTAGATTTTCTTGAATGCACCAAATTGCAACCAACAAAAACTTCTCGAATCTCTTGATATCTTCTAATGCCTCTTCATCATCGCCTACAAGTAAATGCAACTTATTTCTTTTATAGCAATCATAAGCCCACTCCAGTAGTATCATTTCGTTCTCGTTTGCCACATCCTGCTCATAGTTTTTTCTGCAGCAAATCAACTCCAAGAGCAAGACTCCGAAGCTGTACACATCAACCTTCACAGTTACAGGCATGCTTCTGAACCAGTCTGGGGCTACGTACCCTTTGGTTCCACGTATCCTAGTGGTTGTTTGAGTCTGATGTTTTTTCAGTAGCTTGGCCATACCAAAATCTGCTATTTTTGCTGTCAAATTTTCATCCAAAAGCACATTTTGGGGTTTGATGTCACAATGTATAATTTGGGTACTGCACTCTTCATGCAAATAACAAAGTCCTTTGGCAGTGTCGATGGCAACTTGTACCCTTTTGGACCAACTAAGCCTAGAATCTTTGAATAGCAAGTGTGCTATTGAGCCGGTTTTCATGTGTTCATATACCAAAAGGCGGTGTTGGCCTTCGTTGCAGAATCCAAGGAGTTGGACCAAGTTCTTGTGATTAGTCCTGCTGATTGAATTCACTTCAGCTTCAAATACTTCTTCTCCTTCTCCTTCTGTTACCATGTTTCGGAGTTTTTTGACAGCAACGACTTTCCCATTTTCATCATCAAGAACCGCTTTGTAAACTGTTGAGAAAGCTCCAGTTCCTAATTCTTCCTTGAATCCATTTGTAGCTTGTTCTAGCTCTTTGTAGCTAAAACTTCTTAAGTTCACTCCTGGAACAGCTGAATACGGTGCAGTCCTCTTTCGTTTTTTTCCTATAAACTTAAAAATGTACAATAGGGCCAAAAGAATTAAGAACAAATTCATGAAAACTGAACTGGCCAACAACACAGAACCAGAAATGATCAGAGTAGACTGATTTTTCTTTTTTCTTATTTCAACATTTGGTGTTCCCACTGTGGAGTTGTCTTTCCTTATTTTTATCAAAGCCTTCCCTCCGATCGTTGGACCTACTCTCCCATTTGAAAGAGGATACCTTTTCTTCCAACAGACGTTGTCGTCACTGTAAATGGCAACATCACAAAAACAATCATTCAAGCAATTCTGTCTGCACCAATCCTCTGAAACATCGCGGTGAGACTCATAATCAGAGTCAGGCCAGTTAGTATCCAACATTTCGTGGAATGTGAAAGACTCAACTTCTCGCGATTCATGGTTACAGTTTTGCTCAGAGAAATTTTGTCTGCAGCTGCCTAATTTGTCATTCGGATCATCCAAGATATATCCGAGTGGACAATCACATCTCGGCCTCTGATCAGTTCCTATTGAGCATAGGCTGTTGAAACCACAGGCACCCCCTCCTGTACTTTGCCTGATAGACAAACAGATGTTATCAGGAATGTTATACAAACTAGACCATGCCATCGGTTCCCTACCAGAAGATTTCGGATGGACATAGTGCCTAAAAACTCCATCATATTCAAGAATCGCTCGATGATACATTGATTGGCTTCTTGAATTTTCCACATTAGAAGATATCGAATTTATCAAAGTTCCATTCTTTGCTTGAAGAAAAATAAAGCCTGATTGATTGAAGATCACCTGATAACCACTGCCAACTGACATAGGTGACCAATATGCAGCATTTGTTGCCTCAGCTGGATAATTGACTGTGTAAAGCACTAGATTTCCATCAGTTTGAAGTATGAACTCAAACCTCCCACTCGATACATTCGTGTCAGAGAAACTTGAAACAAGACTGTTATCTTGGTTAAGTACTTGACCAGGCAAAATCGTATCGGTGGGCTCATTAAAACTCTGCCATAGAGTGTCTGAACTACTAGTTGCTAGCACAAAGTTTCCATTATCAAGCATGGCTCCATAGGCCAATTGAGCATTCGCCATGCCACGAGCCCACATCTCTTGACCATTTGGATCAGTGAGGACTAGTCTTCCATCCGCGGAGAGCTGAACTTTGGATCCATCTAAGGCAAGATTATTCCTATTGGCTGACCAAACTATGGTTTTGTCTGTGATTTTGTTGAACCATATGGCTAGTAAGTACCTTGATTCATTTCCAATTTTTTGGAAGCCCAAAGCGAATTCATCAGATGGGGATGACCAAAAGCTAGTAACATCACTTGTAGTGAGTGATGAACCCAAACTAACATTTTGGTAAGATTGAGCAATGGCTGAAAGGGGAAACAATAAGAGACCAACAAGAAGCATAACTAAAGAAGTCATGTAAGAAAGAGGCAAAGTTATAAATGGATATTATGTACACC

>PGSC0003DMG403026153 G-LecRK

GTTTACTTTTTCATACCATAAAAAATGTTACTGACTCAAGTAGTTGCTGGAAAAGTTTAAGTTATGCAAATGAGTAATTTTCTCTTGTTGTTGTTTGCAGTGTTGGTGATTATATGTGGATGTGATGCAAGGAGCACGATCGATGGAGGGAGTGAGTTAATAGACAACGGGGAAACACTGGTATCTGCTGGAAAAAAATTCGAAATGGGATTCTTTTGTGATGATGCCGGGTTAAACTGGTATGTAGGCATATGGTATAACAAGAAAGGTCCTCGAACTGTTGTATGGGTTGCTAATTGGAATGATTCGATTCAGAATTCAACAAAAAATAAGGAGAATGTTTCCATGGTCGTTGAAGATGGTAAGCTCAAAGTTTTGAATATTAGTGGACACATTTATTTCTCCTCAGAAGTAGATGGTAACTCTGGCGTTAAAGCAGAGTTGTTGGATACAGGGAATTTGGTTTTATTTGATGAATCAGGAACTAAAATGTGGCAGAGTTTTCGGAATCCAACCGATACATTTCTACCAGGCATGAAAATGGATAGTAATTTGAAGTTGATTGATTCAAAAAATGGCAACTACAAATTCCAATTAGATCAAACAAACAACAAAAAATATGTCATAGTTGATGAGAATGGAATGCATAAAATTCACTGGCAAGGGTCTGTAGGAGGGTTTGCGAAATCAACTGCTTATGAATTCTTCAGATTCAATGATATGCCCTCTTTTGTAACCTTCTTTTTATCCAATAATTCAGTTGAAGCGAGAAGAACATTATTCAATAATTCAGTGGAAGCGACAAGAACATTATTCAATAATAATCGTAGTTTTACTCCGATTGATTTTTCAAGGTTAGCCAATCATAGTAGCAAAGATTCTATTTTCGAAGGTACTAGACTCCTGATGAACTCCTCTGGAGAAATACAATTCTACTTATGGGATAACCAGAGTGGATGGTCCTTGTTATGGTCTGCCCCAAAAGATGAATGCGACGAGTTTAGAAAATGTGGGACATTTGAGATTTGTAACAGCAATGGAAGTTCAGTGTGCAGCTGTTTGCCTGGGTTTCAGAGTGATCCTCCAGAAGATAATGGAGACGGGGAATATTCAGGTGGATGTTCGAGAATATCAGATAACTCATGCAAAGAAGATACATTTGTGCATCTGAGCTCAATGAAATTTGGAAATTCAAATTGGACAGATAAAAAGATTAATACCAGTGAGAACTGCATGAGTCATTGTTTAAGCAGCTGCTCATGCAAAGCATATACATATTATAACAATTCTGGATGCTGGTTTTGGGGGTCGAATCTCAATAATCTCCAGGAGAACTACCCTGGTGGCTTCAACCTCTCTGTCCGCGTGCCCCCTACAGAGGCGTATGTATGTGAAGGGGAGGGGGTTCACATGCACCCATCCTCCCCTTCCTAACCCCTATATAATACTACTATTTTTCTAATTATCTTTTTAAATTTGTATTGGTGAACCCAAGGTTAAAAAGAATACTCTAGTATATTGATAAGTAAGTACAAATAGGGGGTGCATAACTTATAGATTTGTGGTTCAAATTTTTCTATCCACATTAACATTTTTAATTTTTAATTAATATAACAAAATGTACCACTTTTCATAGATCTTTTTAATAATTTATAAATAAAAAGAGGTTCAAATACCCATATTTTTAGGTTGCATTAGCTATGTCT

>PGSC0003DMG400000578 L-LecRK

CAATCATTAAGGCATTAAAATTTACATTCCATCTGAATTACTATATTTTGTTTACTGAAACTGAAATAGGACAAAAAATTGCATTATGCAAGTGAGGACTTTAAGTTCTCTTGTGTTTGGAAGAAAGTGAAATGAAGTTATCTGTAGACCACATCCTTATGTACAACCTTGACAATTGATAATTTGCTACTCAGAAAATTCAAATTCCAAAAAAATTTATCACTAGCCAAGAATTAGCTCATCATGGATTACTTCTTCTGGTTTTTCTTCTCGTTTGCCATACTAGTTAATAGTAATTTAGGTTCGTTCGCGGACGGACCTTCAATTAACGTGACGAAGCATATCTCTTTTCGAGATTTCAGTTCTAGTAATCCCAGATTGAAGCAAGATCTTACACTTCTCGGTAGTGTTATCGTCTCGGATGAAAAAAAATCCGTTCAAATTCCTGATCCTGAGCGAGAAGGTGATGATCTAAAGCATCTAGCAGGACGAGCTATATATTCTTCACCTATCCGTTTCTTTGATCCTCGGACTCAAACACCGGCTTCTTTTGAAACGACTTTCTCGTTTCAATTTGAAGTGAAGTCCAATTCAAGCAATGAGACGTCAGATCAGGGGAAATCTGTTGGTGGTAGTGGCCTTACTTTTATAATCGTTCCGGATGAATTGACTGTTGGCCGTGCTGGTCCGTGGCTTGGGATGCTGAACGATTTGTGCGATGAGGATTATAAAACAGTAGCTATTGAGTTTGATACTCGGCAGAACCCTGAATTTGGTGACCCGAATGATAATCACTTGGGCATTAATTTGGGTAGCATAGTATCAACTGCAGCGATTAATGCTTCGGATGCCGGAGTTCAATTGAATGACGGATCAATTCACAGAGTTTGGATATCTTATGATGGGCGAAAGGGTTTTATTGAAATTCGTCTTGCACCTGATGGCAGAGGATATCCTTCTAAACCAGTTTACTCTGGTTTACTTGATCTTTCACCTTACTTGAATGAGTATATGTTTGTTGGATTTTCAGCTGCTACTGGTAACCATACACAAATCCACAACATCTTGTCATGGAATTTCACCTCAATTAGTCAAGCTTCCCTTCGAATTCCTTCGACAGAGACGTGCCAGAACAAAATCATGCTTCAAAATAGCACACAATCTGAAATTGATCATCGGAAAACGCCTAATAGTTTCTTTATTTTCCTTGCTGTTGTCATTCTTCTGGTAATCGTTCTTATTAACCTCTATTTCAGTAGCTACAAGCGAGATGACAATTCCGACGAAGCATTCACTCTGCCCGAGAAAAAGCAGAGACCGCGGCCACCAAACAAGGCACGTCGCTTCACAATAGCTGAGATCTCCATCGCAACAAGGAATTTCAGCGAGTTACAAATATTAGGCAGTGATGAGAAGAGCATTACGTATAAGGCCACGATCCTGAACGGGTGCAACGTTGTTGTAAAACGATTTTTAACTCAATTTTTCAACACACACGGGTTTGAAAAGCGTCAATTTCACAAGGAAATCAAGGCTATCACCAGAATTCGTCACCCGAATTTGGTCCCGATTAGAGGATGGTGCTATGACAACCAAGAAACAATAGTTGTGTACGATTTCATCCCCAATGGAAGCCTCGATAAATGGCTGTTCGGCGTTGGAGTTTTGCCTTGGACGAGACGTTTTAAGGTCGTTAAAGATTTAGCAGATTCTCTTGTCTACCTTCATTCAAAGCAACTCGCTCACAAAAACGTTAAAAGTAGCAGCGTGTTTCTTGACGTGAGCTTCAGAGCAGTAGTGGGTGATTTCGGGTTTGTGCTTACTTCAGCCGGGTCAACCCGGTTTGAGGCCATGGTGAGTCAGACAGCTGATGTGTTCGAATTCGGAGTAGTCGTGCTAGAGATCATTGCGGGTCGGAGCAGAAAGTCCAACCCGGGAGAACGGGATTTGTTGGATCTTGCATGGGCAATGCATGAGGTACAACAGAAGGAAACTCTAGTGGATCGTAGAATGGGCGCGGTAGTGAACCTGGAGCAGGCGATTCGGGCTTTGGATATCGGGTTGCTCTGCACGTTGAACGAGAACAAAGGAAGGCCTACGATGGAAGAAGTTGTGGAGTTCCTAAATATGGAGAAACCGATACCGGAGTTGCCATCGGGTCGACCCGTTTGTTTGTTCCCGTACAGCAGCACCACAGGCTTATGCAGTGGGTATACTTGTACAGCATTCAAATGAAGTTACGTGTTGCTTGACTAATTGGTTCTTCGGCGTTCCACGTAAGCACCAATTATTCTTATCTATTATACTACAAAATAATGACAGGGAAGCACATGTATATGCTATGTGACGTTATTTTCATGACATAGACAACTATTGTCTGGTCTATGGTCGGAGTTTAATATCACAAGAACTCCTAAAAAGATTTGAAATTATTGTAAATTATGTACTGCAATTCAAATGTTCAATATATAAAATTGTATGTAATCTGTTTATTCCTCCA

>PGSC0003DMG400000789 L-LecRK

ATGCGTTTTTGTTTGCAAAATCTCACAAGGGTCACTCTGTTCTTCATCTTCACCATAAATTCACCTTCCCTTTCAGCTCCTTTCTTGCCTATAAACAATGTAACTCTTTATGGCGATGCTTCATTCACCGCAAAATCAATCACCCTCACCCAAGAACGCAACTGTTCATCATCATCATCAACACCTCCCATTTCTGGCATTGGAAGAGCTTTCTACACATACCCAGTTCGTTTTCTTGATTCTTTAACCAATAACACTGCTTCTTTCTTGTGTACTTTCTCTTTTACTATACTTCCAACCCCTTCTTGCCCTTTTGGTGATGGCATGGGGTTTTTGGTCACTTCTGATGTTGATTCTTTGAGCATCTCTGATGGGTATATGGGTCTTCCGAATCCCGACTCGGAAGATTCCTTCTTGGCTGTGGAATTCAATGCCAATGATAACCGTATTGGTGTTGATACTAAAGAAATTAGGTCTTTGGCTTCTACTAATGTTGATTCAGCTGGGATTGATTTGAAAAGTGGGAAAGAAATGGCGGGTCGGATTGAGTATAAAGATTCAGAGAAAATGATCAGAGTTTGGATTGGGTATGAACTGCAAATTAGGCCTCCTAATCCTGTTCTTTCTACCAGAATTGATGTTTCCAATCAGTTGAAGGAGTTTATGAGGATTGGTTTCACTGCTAAAGGGTCTGCAGTTTATAGCATTAGTCGTTGGCGATTTAGAACGTTTGGATTGATTTCGTCTCCAATATCATCGTCTTGGGATCAATCCGATGAAGGAAACTGTTTGATGTGTTTCCCTGAAGAGGAAATTGGTGGGCATATTTCTGATTCTCATCATAGTAGTAGTAGTAGTGGTAGTAAAAGTTTACTGAAATTGACTTATGGAGGGTTAGCTGCAATTGTCACACTTGTTGGTTGTGCTTTGTCTGTTGTGTTTGTTCTTGCGTTGAGAAGGAAAAAACGCGATAGAGTAGGGGAGACCAATGAACGGCAAATGTGTAGATTACAAGGAAATAGAGTGCCTCAAAGATTGTCATTATCTGAAATAAAATCAGCTACAGAGTGTTTTAATCATGAAAGGATAATCGGAGAAGGAGCATCTGCTGTTGTGTATGAAGGGGAAATTCCTTCTAGGGGATCTGTGGCTGTTAAGAGATTTGTCCAAGGGAGTAGATTGGGTCCTTCACATATTCCTTTTAATACTGAATTTGCTTCTATGGTTGGCTGTTTAAGACACAAGAATTTGATTCAGCTTCAAGGGTGGTGTTGTGAGAGGAATGAATTGGTGCTAGTGTATGAATTCATGCCTAATGGTAGCCTTGACAAAATCCTCCACGAGCGATCGCATTTGACTAAGTTTCTGACATGGGAGAGAAGATTGAACATAGTTATTGGTGTGTCATCTGCTCTTATGTATCTTCATGAAGAGTGTGAGAATCATATAATCCACAGAGATGTGAAGAGTTGCAATATAATGCTTGATGCTGAGTTCAATGCCAAGCTTGGAGATTTCGGTTTAGCAGAGGTGTTTGATAATTCTAAGACAAGGGATGCTACTGTACCAGCTGGAACAATGGGATATTTTGCACCTGAGTATGTCTATACTGGTATTCCAACTGTTAAAACAGATGTGTATAGCTTTGGTGTTGTGGTACTGGAAGTGGCATCAGGTAGAAAGCCTATCGACGAAGGTGGTGGTTTGATTACTGATTGGGTGTGGGACTTGTGGGAGAAAGGGAGGATAACTGAGGCTGCTGATCCTAAACTAATGGGGCGGTTTCAGAAGAATGAAATGGATAGAATGCTCATTGTGGGACTTTCTTGTGTGCATCCCGATCATGAGAAGAGACCGCGAATGAGAGATGTTTTCCGTATGCTTAAAGATGAAGCTCCACTTCTCATTTTACCTCCAATGAAGCCCACTGTGAGAATTCAATCTGTTTTACCAGAGAGCTGTGAAGAAATCATGAATTGGGCTGCAAGAATGGAGGATACACCATGGTCCACTCCCAGAACTCATTTTAGCAAGAACTAG

>PGSC0003DMG400001773 L-LecRK

ATGGAGATTTACTCATTACTAGTTGTATTGCTCTGTTTCTTGGCTGCTTTAGCTGCTGCCACTGAGTTTGATTCTGGGACTTTAACTCTTAGTAGTTTGAAGCTTTTAGGTGATGCCCATTTGGGGAATAATACTATAAAGTTGACTCGTGACCTTGCTGTTCCAAATTCCGGTGCCGGAAAAGTTTTGTATTCTAAACCAGTAAAGTTCCGGCAGCCGGGTCTTGATTTTCCGGCAAGTTTCTCTACGTTTTTCTCGTTTTCTGTTAGTAATTTGAATCCTTCATCGATTGGAGGTGGATTGGCTTTTGTTATCACACCGGATGATGAGTCGGTTGGTGATTCCGGTGGGTATTTGGGGATTTTGGATGCAGAAGGAGGGCAAAATGGGAATTTTGGAGTTGAATTTGATACACTTATGGATGTTGAATTTAAAGATATTAATGGAAATCATGTGGGGTTAGATTTAAATTCAATGATTTCAACTCAGGTTGGTGATCTTGATTCTATTGGTGTTGATCTTAAAAGTGGTGATTTGGTTAATTCTTGGATTGATTATTTTGGTTCTACAAAAAAATTGAATCTTTATGTTTCATATTCTAATCTTAAGCCAAAAGAACCATTTTTATCTGTTACAATTGATATTTCTGAATATGTAAATGATTTCATGTTTGTGGGGTTTTCTGGTTCAACACAAGGGAGTACAGAGATTCATAGTATTGAATATTGGAGTTTCACTTCATCATTTGATACAAATCCTAAATCTCCCCCTGAAGCGGTGGCGGAGCAGCCTCCGCCCCCGCCAACTGCTAGTTTGATGAACCCCACGGCGGATGTAACTACATNNNNNNNNNNNNNNNNNNNNNNNNNNNNNNNNNNNNNNNNNNNNNNNNNNNNNNNNNNNNNNNNNNNNNNNNNNNNNNNNNNNNNNNNNNNNNNNNNNNNNNNNNNNNNNNNNNNNNNNNNNNNNNNNNNNNNNNNNNNNNNNNNNNNNNNNNNNNNNNNNNNNNNNNNNNNNNNNNNNNNNNNNNNNNNNNNNNNNNNNNNNNNNNNNNNNNNNNNNNNNNNNNNNNNNNNNNNNNNNNNNNNNNNNNNNNNNNNNNNNNNNNNNNNNNNNNNNNNNNNNNNNNNNNNNNNNNNNNNNNNNNNNNNNNNNNNNNNNNNNNNNNNNNNNNNNNNNNNNNNNNNNNNNNNNNNNNNNNNNNNNNNNNNNNNNNNNNNNNNNNNNNNNNNNNNNNNNNNNNNNNNCCACCGGCTAGTTTGATGACCCCCACGGCGGATGTAACTACATCCGCCACATCGCCCCCTCCGGAGCAACAACAGATTGCTCCGGCGGACGCTAATGGTACTGCTAATGCAGTACAGAGAAACAGCAGCAAATGTCATAATAGTTTTTGTAAGCAGGGTGCTGGTGCTGTTGTTGGGGTTGTTACTGCTGGAGCATTCTTTCTAGCTTTTGCTACATTGGTACTTATTTGGTTGTATTCGAAAAAATTCAAGAATGTGAAGACTTCAGAGACTATGTTGCCTTCTAATATTATCAAAATGCCAAAGGAATTTAGCTATAAGGAGCTTAAAATCGCGACGAAAGGGTTTGATTCGACTAGGATTATAGGGCATGGTGCATTTGGTACTGTCTATAAAGGGATTTTGTCTGAGACTGGTGATATTGTTGCGGTAAAGAGGTGTAGTCATAATGGACAAGGGAAAGCTGAGTTCTTTTCTGAATTGTCGATAATCGGAACACTTAGACATAGAAATCTTGTTAGGCTACAAGGATGGTGTCATGAGAAAGGTGAAATCTTGTTAGTTTATGATTTGATGCCAAATGGTAGTCTTGACAAGGCGTTATTCGAGTCCAGAATGATCCTCTCATGGCAACACAGGCAAAAAATCTTGTTAGGTGTTGCTTCTGCATTAGCATATTTACATCAAGAATGCGAAAACCAGGTGATACATAGGGATATCAAGAGTAGTAACATTATGTTGGATGAAGGGTTCAATGCAAAGTTAGGTGATTTTGGACTAGCACGACAAATCGAACACGACAAGTCCCCCGATGCAACGGTAGCAGCCGGTACAATGGGGTATTTGGCACCAGAATACTTGTTGACAGGTAGAGCAACTGAGAAAACTGATGTGTTTAGTTATGGGGCAGTGGTTCTTGAAGTGGCAAGTGGGAGGAGGCCAATTGAGAAAGAAACAACAAGTGGTGTTGGATTAAACAGCAATTTGGTTGAATGGGTGTGGGGATTACACAAAGATGGGAGTCTACTATCCGCAGCTGATTCAAAACTGAATCTCGAATTTGATGAAAAAGAAATGACAAGAGTGTTGTTAGTTGGTTTGGCTTGTTCACATCCTGACCCTATAGCTAGGCCAACAATGAGGGGGGTGGTACAAATGTTAGTAGGTGAGGCTGAAGTACCAATTGTACCAAGGGCTAAACCATCAATGAGTTTTAGCACATCTCATTTGTTAATGACATTGCAAGATAGTGTTTCTGACTTGAATGGTATGATCACACTTTCAACTTCTTCATCAGAAAACAGTTTCAATGGTGTTGGTGGACATGATGGCATGGACCTAGTTTAG

>PGSC0003DMG400004396 L-LecRK

AATATTTTGGGCAACGAAAAACTTGAGTCTGCACTTGAAAATATAAAAACATTATAGAGTAGTGTAGCAACATACATTTGTTTAACTATGGCTGTTTTTGCATCTGTATCATACTTTGTTTTGATAATCCCCTTTGTCACTTCATTGTCTTTCAATTTTGATAGTTTTAATCCCAATGATCAGAATATAACATATGAGGCAGATGCTTATCCAAACGGTGTAATTCAGCTCACCAAAAACGAGCAGGGTAGTGGTTTAAACGATAGCATAGGCCGAGCTACATATTCTAAAACACTCTATCTTTGGGACAAGGCTTCTGGGAATATCACCGATTTCACTACTCACTTCTCCTTTGGAATCAATTCACTGGGAAAAAACTTATATGCTGATGGTCTTGCCTTCTTCCTCGCTCCTGCAGGTTCAAGAATTCCTGATAAGAAGTCAGCCGCAGGAGGCGGCCTTGGCCTTGCAATTGATAGTCGACAAAATACATCGAAAAATCATCCTTTTGTTGCTGTTGAGTTTGACACCTTTAAGAACTATTATGATCCACCAGGTGATCATGTAGGTGTCAACATCAACTCCATGGTATCTGTTGTTAATGTGACTTGGTTTAGTAGCATTCCAAATGGTAAGAGAACTGATGCCTGGATTACTTATAATTCAACCTCAAAAAATCTTAGTGTTGTCTTCACCGGTTTCCAACCACAAGGGAATACCACAGTCACTTTCCTGCAGACTCTATCTTACAATATTGATCTGAGGGACTACTTGCCAGAATGGGTCACTTTTGGCTTCTCAGGTGCAACAGGATCCCTCTTTGCATTACAAACCATTTATTCTTGGAATTTTACTTCTTCTTTAAAATATAATGACAACATAACAGATCCATTTGTGCCCCTACCAAGCCCTGTGCCACATCCAGATGTATCCCTACCAACCCCAGATCCAGATCCAGATCCAGATCCTTCAAGCAAAAAACTGTCAGGACTAGTAATTGGATTGATTTCTGGCGGTTGTGTTTCGGTAGCACTATCTGTTTTGATATTGTTTGCATTTTGGAGAAAGAGGAAGGTCAGAGAATATGAAGATGAAGATGAAGATGATATTATTGACAGTTCCATGACTAATGAATTTGAAAGAAGCACAGGACCAAAGAAGTTCTTGTACAGTGAGTTGGTTAGATGTACAAATTACTTTTCGCGGGAAGAGATGCTTGGGCAGGGTGGGTTCGGAGGTGTTTATAAAGGATATCTCAGTGAATCCAACTCCTATATTGCTGTTAAGAGGGTTTCAAGGGAGTCAAAGCAAGGAATAAAAGAGTATGCATCAGAAGTCAGGATCATCAGCCGGTTAAGACATAAACATTTGGTGCAACTCATTGGTTGGTGCCATCAAAAAAGAGAACTTCTACTTGTCTATGAGTTTATGCCTAATGGAAGCTTAGATTACGATCTCTTCAAGGGAAGAAGCCATTTGACATGGTCAATAAGATTCAAGATTGCTCAAGGCTTGGCCTCAGCGTTGCTTTATTTACACGATGAATGGGAACAATGTGTGGTGCATAGGGACATAAAGTCGAGCAACATTATGTTGGATTCCAATTTCAACGCTAAACTTGGGGATTTTGGTTTGGCTAGGCTAGTTGACCATGATAAGGGATCCCAAACAACAGTTTTGGCAGGCACGATGGGTTACATGGCTCCTGAATGTCTCACCACTGGCAAAGCTAGCAAGGAAACAGACGTCTATAGCTTTGGTGTTGTCGCATTAGAAATAGCTTGTGGAAGGAAACCTATTGATCCTAAAGCTGAGGAGCATCAAGTAAATATTGTCGATTGGGTTTGGAGACTTTACGGGATGGGAAATATTCGTGAAGCAGTTGATCCTAGACTCTCATCAGAGATCAATGAAGAGGAGGTGGAGCACTTGCTAATTGTTGGTTTATGGTGTGCTCATCCAGATAACAATTGCAGGCCTTCTATTAGGCAAGCAACTCAGGTTCTTAATTTTGAAGCTCCGTTGCCCACACTCCCTCCAAACATGCCTGTACCGACATATTGCAGTCTGCCACAATATGGACCAACTACTTCATTTGCATCACCATACGACTCTAATGGTCCTGGGATATCTGAAATACACTCTTCAGGGACTAGAGACTACACTGGTTCTTCAAATAACACAGCAGCTTCTGCAGCATCTTCGCCTTCCGCATCACTTTTGTACACACGTTGATATTGTCTTATAAATGAACAGGTCTTTTATTTCCCTGCATTCAATCTTTCAATTGCAGTGTCCAACTTTTGGTCTATGTATTCTTTATTATTTCTACTTTTA

>PGSC0003DMG400004397 L-LecRK

ATGGCTATTTTTGCATCTTTATCTGCTCTCACATACCTTGTTTTGATAATCCCCTTTGTCACATCACTGTCCTTCAATTTTGATAGTTTTAATCCCAATGATCAGAATGTAACATATGAGGCAGATGCTTATCCAGCAAATGGTGTAATTCAGCTCACCAAAAACCAGCGCGATAGTGGTGCAAACTATAGCATAGGCCGAGCCACATATTCAAAAACACTCCATCTTTGGGACAAGGCCTCTGGGAATGTCACAGATTTCACTACACACTTCTCCTTTAGCATAAATTCACAGGGCATGAACATATATGCTGATGGAATTGCCTTTTTCCTCGCCCCTGCAGATTCAAGAATTACTGATAACTCAACCGCAGGAGGCGGCCTTGGCCTTGCAAATTATTATAGTCGACAAAATACATCAAGAAATCAATCTTTTGTTGCTGTCGAGTTTGACACCTTTAAGAACATTGATGATCCACCAGGTGATCATGTAGGCATCGACATCAACTACATGGTATCTATTGTTAATGTGACCTGGTTTAGTAATATTCCGAATGGTACGAGAACTGATGCCTGGATTACTTATAATTCAACCTCAAAAAATCTTAGTGTTGTCTTCACTGGTTTTCAACCACAAGAGAATACCACAGTCACTGTCCTGCAGACCCTATCTTACAATCTTGATCTGAGGGAATATTTGCCAGAATGGGTCACTTTTGGCTTCTCAGGTGCAACAGGAACCTACTTTGCAATACAAACCATATATTCTTGGAATTTCACTTCTTCTTTAAACTATAATGAAAACATAACAGATCCAGATGTACCCCAACCAAGCCCTGTGCCAGAAGATACTCCAAGCAAAAATAAGTCAGGACTAGTGATTGGATTGATTTCTGGTGGTTGTGTTTCAGTAGCAGTATCTGTTTTTATATTGTTTGCATTTTGGAGAAAGAGGAAGGTCAGAGAAGATGAAGATGAAGATATAATTGATGGTTCCATGACTAATGAATTTGAAAGAAACACAGGACCAAAGAAGTTTTTGTATGGTGAGTTGGCTAGATGTACAAATAACTTTTCGCGGGAAGAGATGCTTGGGCAGGGTGGGTTTGGAGGTGTTTATAAAGGATATCTCAGGGAATATGACTCCTATATTGCTGTTAAGAGGATTTCAAGGGAGTCAAAGCAAGGAATAAAAGAGTATGCATCAGAAGTCAGGATTATCAGCCGGTTAAGACATAAACATTTGGTGCAACTCGTTGGTTCGTGCCATCAAAAAAGAGAACTTCTACTTGTCTATGAGTTTATGCCTAATGGAAGCTTAGACCACCATCTTTTCAAGGGAAAAAGCCATTTGACATGGCCAATAAGATTCAAGATTGCTCAAGGCTTGGCCTCAGCGTTGTTTTATCTACGCGAAGAATGGGAACAATGTGTGGTGCATAGGGACATAAAGTCGAGCAATATTATGTTGGATTCCAATTTCAATGCTAAACTTGGGGATTTTGGTTTAGCTAGGCTAGTTGACCATGATAAGGGATCCCAAACAACAGTTTTGGCAGGCACAATGGGTTACATGGCTCCTGAATGTGTCACCACTGGCAAAGCTAACAAGGAAACCGACGTCTATAGCTTTGGTATTGTCGTGTTAGAAATAGGTTGTGGAAGGAAAATGGGCTTGGAGACTTTACGGGATGGGAAATCTTAGTGAAGCAGTTGATCCTAGACTCTCATCAGAGTTCAATGAAGAGGAGGTGGAGCACTTGCTAATTGTTGGCTTATGGTGTGCTCATCCAGATAACAATTGCAGGCCTTCCATTAGGCAAGCAATTCAGGTGCTTAATTTTGAAGCTCCATTGCCCACACTCCCTCCAAACATGCCTGTGCCGACATATTGCAGTTATGGATCAACTACTTCATTTGCATCACCATACGACTCCAATGGTCCTGGGATATCTGGAATACAGTCTTCAGGGACTAGAGACTACACTGGTTCTTCAAATAACACAGCAGCTTCTGCAGCATCTTCGCCTTCCGCATCACTTCTGTACACACGTTGA

>PGSC0003DMG400004401 L-LecRK

GTTGTTTTTGCCTCTCTCAAATACTTCTTTCTTATCCCTTTAGTCACTTCACTCTCCTTCAATTTTGATACTTTTACACCCAATGATCACAATGTAACCTATGAGAGAGATGCTTCTCCACAAAATGGTGCAATTCTACTCACCACAAACCTCCTCAATCGCAATATCAATGCTAGTATAGGTCGAGCTATTATTCAAAACCACTCCATCTATGGGACAAGGCCTCTCGAAATCTCACTGATTTCACTACACATTTCTCCTTTAGCATTAATTCACAGGGAAGAGCTGAATATGGTGATGGTTTTGCCTTCTTCCTTGCCCCTGCAGGTTCAACAATTCCTTATAATACAACAAGAGGTGGAAGCCTTGGCCTTACAAGTAATAATGAGCGACTAAATTCATCCAGTAATCATTTTGTTGCTGTGGAGTTTGACACCTATCAGAACCTGTATGACCCAAAGGGTGATCATGTAGGTGTCGATATCAACTCTATGCAATCTGTTGTTAATGTGACTTGGTTTAGTAGCATTCCAAATGGTACGAAAACTGATGCCTGGATTACTTATAATTCTAACTCAAAAAATCTTACTGTTGTCTTCACTGGTTTTCAACTCCAGTCTCCAAGGGAATAATATCACAGTCACTGTCCTGCAGAACCTATCTTACAATCTTGATCTGAGGGAATATTTACCAGAATGGGTCACTTTTGGCTTCACTGGTGGAGGAGGAAATAGCTTTGCATTTCAAAGAATCTATTCTTGGAATTTTACTTCTTCTTTAGATAATATGACAGATACAGGGGTAACCTTATCAAACACCAAGCCAAAGGATGCACCAATAAAAAGTAAGTTAGGACTTGTGGTTGGATTGATTTCTGGTGGTTGTGTTTTGGTTGCAGTATGTGCTTTGGTGTTGTTTGCATTTTGGAGAAAGAGTAAGGTCAGAGAAGATGATCATGTCTTTGATGGTTCCATGACTAATGAATTTCAAAAAAGTGCAGGACCAAAGAAGTTCTGGTACAGTGAGTTGGCTAGATGTACAAATAACTTTTCACAGGAAGAGATGCTTGGGTGGGGGGGATTCGGAGGTGTTTATAAAGGATATCTCAGTGAATCCAACTCTTATATTGCTGTTAAGAGGGTTTCAAGGGAATCACAGCAAGGAATAAAAGAGTATGTATCAGAAGTCAGGATCGTCAGCAGGTTAAGACATAAACATTTGGTGCAACTCATTGGTTGGTGCCATCAAAAAAGAGAACTTCTACTTGTCTATGAGTTTATGCCTAATGGGAGCTTAGACCACCATCTTTTCAAGGGAAAAAGCCATTTGACATGGCCAATAAGATTCAAGATTGCTCAAGGCTTGGCCTCGGCGTTGCTATATCTACATGAAGAATGGGAACAATGTGTGGTGCATAGGGATATAAAGTCTAGCAATATTATGTTGGATTGTAATTTCAATGCCAAGCTTGGGGATTTTGGGTTAGCTAGGCTAGTTGACCATGATAAGGGATCCCAAACAACAGTTTTGGCAGGCACAATGGGTTACATGGCTCCTGAATGTGTCACCACTGGCAAAGCTAACAAGGAAACAGACGTCTATAGCTTTGGTATTGTCGTGTTAGAAATAGGTTGTGGAAGGAAACCTATTGATCCTAAAGCAGAGGAGCATCAAGTAAACATTGTCGATTGGGTTTGGAGACTTTATGGGATGAGAAATATTCTTGAAGCAGTTGATCCGAAACTATCATCAGAGTTCAATGAAGAGGAGATGGAGCATTTGCTAATTGTTGGCTTATGGTGTGCTCATCCAGATAGTAATTGCAGGCCTTCTATTAGGCAAGCAGTTCAAGTGCTTAATTTTGAAGCTCCATTACCCACACTCCCTCGGAACATGCCTGTGCCCACATATTGCAGCTTCTCACACCATTTACCAACTATTTCACTTTCATCAACACATGAAACCAACGGTACAGAGATCGCCCTTGCCCAAATCCAGCATTCAGTAAACGAAAACAACATTGATGCTTCATTAAAAAGAACATTTTCTTCAGCATCTTCATCTTCAACATCAGTTTTGTATCCAAGTTAATATTGCCCTGTTAAGATCCTGCATTTTGTCTTCCATTAGCAGTGTCCATCTTTTCCTTTCTTACTTTTTGTTGTTTCTCTTCTAATGAAAGAGGATTGAATGGATTACATTTTCTGTAATTTTAACTGTTCTATCAAAC

>PGSC0003DMG400008822 L-LecRK

ATGTTCTTCAATTTTTTCATACTAGTAGCTCTCTTTTTAGTTGATTTTTCACCAGCTTCTTGTGAAGTTGATGCATTCATTTTCAATGGATTCCAATCAGCTAATCTTAGCTTGGATGGCATAGCACAGTTCACATCTACTGATCTTTTGTTGTTAACAAATTCAGGAACACAAAATCAGGGCCATGCTTTCTATCCAAATCCAATTCATTTCAAGAACTCACCTAATGGTTCTGTATTTTCTTTCTCCACTACTTTCGTGTTCGCGATAAGGTCTGATTATGGAAATTTGAGTGGTCATGGACTGGTTTTCGTTATCGCGCCACATAAAGGACTTCAGGGGTCTTTGGCAAATCACTATCTTGGTCTTTTTAACTCTAATAATAATGGGAATGGATTAAACCATGTCATTGGAGTTGAGCTCGATACAATCTATAGCGAGGATTTTGGTGATATCAATGACAATCATGTTGGAATTGATATAAATGGATTGAGGTCTGTAGCAATTCACACAGCAGGTTATTTTGATGATACTGGTTTTTTTCATAACTTGACTTTAATTAGTGGCAAAGAAATGCAAGTTTGGATTGATTACGATGGGAGGACTAAGCAAATGAATGTAACAGTAGCTCAGTTACATATGGAAAAACCAGTTAGACCACTTTTGGCTTTGAAACATGATCTTTCGTCGATTCTTGATCAGACTATGTATGTTGGTTTTTCATCCTCAACAGGTTCAGTCCCAACACATCATTATATCTTGGGATGGAGCTTCAAGACAAATGGGAAAGCTCAAGAACTCTCGCAACTTCCTGAGCTTCCTCGTCTTGGGAGTAAAGGGACATCAAGATTTGTAACGATTGGTTTGCCTATAATCTCGTTGGTTTCAGTTGTTGCAGCTGTGTTAGCGGTGGTTTATTATGTAAGAAGGAAGAAGTATGAAGAAATTCATGAAGATTGGGAACGCGAGTATAGACTACAAAGGTTTAAGTATAAAGAATTGTACATTGCTACTAAGGGATTCAGAGAAAAGGAGGTATTGGGAGCTGGAGGATTTGGTAAAGTTTATAAAGGAGTGATGCCTATCACTAAACTTGAGATAGCTGTAAAGAAGATATCTCATGAATCAAGACAAGGTATGAAGGAATTTATTTCGGAGATTGTAAGCATTGGTCGTATGCAACACAGGAATGTAGTACAACTATTGGGTTATTGCAGGCGAAAAGGGGAGTTGATTTTGATATATGAATACATGTCTAATGGAAGTCTAGACAAGTATTTATACGATCAACCAAGATACACCCTCGATTGGAACCAAAGATTCAGAGTCATTAGAGGTGTAGCCTCTGGACTTTTTTTCCTACACGAAGAATGTGATCACGTAGTGATTCATAGAGATGTTAAGGCCAGTAACGTCTTGTTAGATGGTGAACTAAATGGCAAGTTAGGAGATTTTGGACTCGCGAGGCTATATGGTCATGGGACCGATCCTCAGTCTACTCGTGTTGTTGGTACTCTTGGTTACCTTGCACCAGAGCATACTAGAACTGGCAGGGCAACACCTAGTAGTGATGTATTTTCATTTGGGGCATTTCTGCTTGAAGTTGCATGTGGTAGGAGGCCGATACAGCCAAGACAAGACGGTGATGATCTGATTTTGGTCGATTGGGTGTTCTCGTGCTGGAATAGGGGTAATATACTTGATGCTGCTGATCCAAACATAGGCATTGATTTTGTTCCAGGACAAGTGGAGTTGGTCTTAAAGTTAGGCTTGTTCTGTTCTCATTCAGATCCTTCGTGTAGGCCAACCATGCGACAAATCTTGTTGTTCTTGGATGGTGTTGTGGCCTTACCAGAGTTATCAGAACTCGGTGTTTCATCAGCTAACCTAACATTTGAACATCGCGGAGGTTTTGATGATTTTGTCAAGTCGTATCCAACTTCTTTGGGCTATGCATATTCCGGCTCTCCATCTGTAACAGACTCCTTTCTCTCTGGTGGCCGATGA

>PGSC0003DMG400008897 L-LecRK

ACTAGATTTGACTATATTGTGTGATTATTTTTCATATATACATATTGAAGTTTGGCTGGCCTCAAGAAAATGACATTACAGAGTATACATTGTAGACTTGAACATTGGGAGATAAATAGTCATATTTGTGATTGTAGAAAGTTAACCCCGCGAACATTATTATCAGTTCTATCCCTATAAGAGTTTTGAGCAGACATGGAGAAATTGTTTTTAGCCATGATTTTTCAGTTTCTTGTTTTACTTTTTCATGCTCCATTAGGCTATGCAGCTGCAGAAGATCTCGATTTCACCTACAACGGGTTCCACTCAACAAACTTGAGTCGTGATGGAATAGCAGAGTTCACATCAGATGGCTTGTTAAGGCTAACTAATGCTTCACAATTGCAGGAGGGTCATGCTTTTTATCCTAGACCTATCAATTTCATGAATTTAGCATCTAATGGTTCAGATTTCTCTTTCTCAACCACATTTGTCTTTGCCATAGTGCCTGAGTACTCGATTTTAGCTGGTCATGGTATGTCTTTTGTGATTGCACCAGCTAGAAGGCTTCCAGAAGCACTTCCAAGCTCTTATCTTGGCTTGTTTAATGATTCTAGCAATGGAAATGCCACCAGCCATGTTGTTGCGGTGGAGTTTGACACGCTACAGAACCGTGAATTTGATGATATTGACAGCAACCATGTAGGCATTGACATCAATGGCCTAAAATCTGTAATAGCCAAGCCGGCCGGTTATTATACCAGTAATGAGAAATCCTTCAATAACTTAACTCTTGCTAGTGGCCAGCCAATGCAAGTTTGGGTGGAATATGATTCTGTGGATAAGCATATGAATGTCACAATGGCTCCACTGCACGTTGCAAAACCAGACCGTCCTCTCCTGTCCTTAGTCTACGATCTTTCATCTGTCATGGATGAAAACGTGTCTATAGGCTTCTCAGCATCCACCGGTGCAGTTGTTTCAACACATTATATCCTTGGATGGAGCTTTAAAATGAATGGAGTGGCTCAAGGGCTTGATCTCTCTCAACTTCCTAAGCTTCCTCGAGTAGTACCTAAGAAAGAATCCAAGCTTTTATTGCTTGGTTTGCCCGTGATTTCTTCAGTTTCATTTGTGATTGTAGTCCTGGTAATAATTTATTACATAGGAAAGAAGATGAAGTTTGCTGAATTGCAAGAAGAGTGGGAGCTTGATTATGGACCACACAGGTTCAAGTACAAGGATTTGTACACTGCCACAAAGGGCTTTAGCAGCAAGGATTTGTTGGGATCCGGGGGATTTGGCAGAGTCTACAGAGGGGTATTGCCCATATCCAACATTCAGATAGCAGTCAAGAGGGTCTCTCATGAATCAAGACAAGGGATGAGGGAATTTGTTTCAGAAATTGTTAGTATTGGCCGGCTACGCCACAGGAATTTAGTACCACTTTTGGGTTATTGCAGGCGTAAAGGAGAGTTACTTCTGGTTTACGAATGCATGCCTAACGGAAGCCTAGACAAGTATTTGTTTGACAAACCAAGCTGCACTCTTGATTGGAACCAAAGATTTCGAGTCATCAAAGGCGTAGCATCAGGACTAGTCTATCTTCATGAAGAGTGGGAACAAGTAGTGATTCATAGAGATATAAAAGCGAGTAACGTGTTGTTAGATGTTGAATTAAATGGAAGATTAGGAGATTTTGGGCTAGCAAGATTGTATGATCATGGGACAGATCCTCAAACTACGCATATCGTTGGAACTCTCGGATACCTTGCCCCAGAGCATACAAGAACAGGCAAAGCCACAACTAGCACTGATGTGTATTCATTTGGGGCATTTTTACTTGAGGTGGCTTGTGGGAGAAGGCCAATAGATCAAAGGGTATCGGACGAGGACCTCGTTTTGGTTGACTATGTATTCTCTTGTTGGACTAGAGGTGATATTGTTGAAGCCATTGATCCAAACTTGGGCAATAACTATGTAAAAGAAGAGGTTGAGTTGGTACTAAAACTTGGACTATTGTGCTCTCAGACAGAACCAGCAGCTAGGCCAAATATGAGGCAAGTGTTGATGTACTTGGAAAATGCCCTGCCTTTGCCAGAGTTATCACTGATGCAAACTTCAAATATTAGCTTAACTTTTGCAGGCTTTGATCATTTTGCAATGTCATGGTCTTCTTCTGTTACTGATTCCCTTCTCTCCGGTGGTCGATGACTCGGGCGAAATTTTAATATTTTCCTAGTAAAAATAAAACTTGTTATATAAGCTATATTTCTTCCACCTAAGGTTGTATACTTGGTTCATTTAGGCATCTTGGAG

>PGSC0003DMG400010466 L-LecRK

CGTGTTTCCTTTGGATTCTCTGCTTCATTCACACCCCTCTTCTTCACCCCCATAATCCTCACTCTTCTCCCCCTGTTTCCAGCTTTCTTCAAAAAATTTCACTATTTTCTAAGCTAAAAAAAAAGCTAAAAATCAAATCTTTGTGCAATTTGATCTTCATTGTGCAATGGGGTTTTGCTAAAATCTAGCAATGAAGTCAATTCTTGAAGAACTTTCCAGCAACATTCCTGTAAAAGTCATCCTGTTGTAGTGTTGTTCTCTCTTGTCTACTTTCTTGCATTTTCACTGTTTCTAAGCTCAAAAAATAGCTAAAAATCAAATCTTTGTGCAATTTGATTTTCATTGTGTAATGGGGTTTTGCTAAATCAATTCTTGAAGAAATTTCCTTCTGTAAAAGGCATCATGTTGTAGTGTTGTTCTCTCCTTAATCCTCACTCTACTGTGTCAGTTTTCTACTTTTTTGCATTTTCACTGTTTCTAAGCCCAAAAAAATAGCCTACCCTTTTTGTATCTTCACCAAATTCTTGTGATTTGGTGTGAAAAATCTTACCTTTTATACAATTTGATCTTCATTCTCTGATGGGTTTTGCTAGAATCTAGCAATGACATCAAATCTTGAAAAAAAATTCAGGGACTTTCTTGTATGGTTCATCTTATTGTTCTGTTTCTGTAGTATTAGTTCAACATTTGGTAAAACTGATTTCGATTTTGAAAATTTAACACTGAGTAGTTTAAAGCTTCTTGGTGATGCACACTTGAGTAACAACACTGTAAGGTTGACAGGTGATCTTGCTGTACCAAATTCCGGTGCCGGAAAAGTTCTGTATTCAAAACCAGTAAGATTCCGGCAACCGGGGTTTGATTTTCCGGCGAGTTTCTCGACATTCTTTACATTTTCAGTTACTAATTTGAACCCATCGTCAATTGGTGGTGGTCTTGCTTTTGTGCTTACGCCGGGTGATGAGTTAGTAGGTGATGCTGGAGGGTATATGGGAATCTTGGATGCTAAAGGGACACAAATTGGTACAATTGCTGTGGAATTTGACACCCTTATGGATGTTGAATTCAAAGATATTAATGGAAATCATGTTGGTTTGGATCTAAATTCAATGATTTCAACTGAAATTGGTAATTTGGATTCCATTGATATTGATCTCAAGAGTGGTGATTTGATTAATTGTTGGATTGAATATTCTGGTTCTAATGGAGAGATGGATATATTTGTGTCATATACTAATCTCAAGCCAAAAGAATCATTTTTATCAGTTAACATCAATCTTGCTGAGTATGTAAATGATTTCATGTTTGTGGGGTTTTCTGGTTCAACTCAAGGGAGTACTGAGATTCATAATATTGTGTGGTGGAGTTTCAGTTCATCATTTGATGCAACTCCAAACTCGGGGGAGGTGGTGGCTCCACCACCACCGCCAACGACTAGTTTGATGAACCCAACGGCAGATTCTGTCGCGTTGCCACCACCTTCTATGGCTCCATCAGAGTCTAATAGTAGTACACCAGGTGTATTACAAGAAAAGAGAAGTAGGAAATGTCATAGCAATTTTTGTAAACAAGGTCCTGGAGCTGTTGTTGGTGTGGTGACTGCTGGTGCATTTTTTCTTGCATTTGCTACATTGGTACTTATTTGGTTATACTCCAAAAGGTTCAAGAGAGTTAAAAATTCTGAGATTGTGGCATCTGATATCATCAAAATGCCTAAGGAGTTCAGCTATAAAGAGCTTAAATTGGCCACGAAAGGTTTCATTCCAACGAGAATTATAGGCCGTGGTGCATTTGGGACTGTTTACAAGGGCATTTTATCGGATACTGGTGGCATTGTGGCAGTCAAGAGATGTAGTCATAATGGACAAGGGAAAGCAGAGTTCTTATCTGAATTATCTATAATTGGAACCCTTAGGCATAGAAATCTTGTTAGGCTTCAAGGATGGTGCCATGAGAAAGGTGAAATTTTGTTGGTTTACGATCTTATGTCTAATGGGAGTCTTGATAAGGTGTTATTTGAATCAAGAATGGTTCTTTCATGGTCACATAGGCGAAAAATTTTGATAGGTGTTGCTTCTTCCTTGGCATATTTGCATCAAGAATGCGAAAACCAGGTGATTCATAGGGACATTAAGAGTAGTAACATCATGTTGGATGAAGGGTTCAATGCAAGATTAGGAGATTTTGGATTAGCAAGGCAAGTTGAGCATGACAAGTCCCCCGATGCAACGGTAGCAGCTGGGACAATGGGCTACTTGGCTCCAGAGTACTTGTTAACCGGAAGGGCAACCGAAAAAACTGATGTTTTCAGCTATGGAGCAGTGGTTCTTGAAGTGGCTTGTGGAAGGAGGCCAATTGAGAGGGAAACAACTAAAGTTGAGAAAGTTAGAGTGAATAGCAACTTGGTTGAATGGGTATGGGGGGTGCATAGAGAAGGGAATTTACTAAATGCAGCTGATTCAAGACTTTGTGGTGAGTTTGATGAACAAGAAATGAGAAGGGTTCTAATGATTGGGCTAGCTTGTTCACATCCCGACCCTACGGCTAGACCAACAATGAGAGGTGTAGTTCAAATGCTGGAAGGTGAATCTGAAATCCCAATTGTCCCAAGAACTAGACCTTCTATGAGTTTTAGCACATCACATCTTCTAATGAGTTTGCAAGATAGTGTTTCTGACTTAAATGGTTTGATCACACTTTCCCCTTCATCATCCGAAAGTAGCTTCACTGGTGGTCGCAATGGCAATGGCATGGAGTTGGTCTAAATGCAAGACTAAAAAAGACTTGACTTTTCGAGTTAATGCACCCATAAGAGCCCAACAATCCCACATATGGGGCCACCATCAAGGCGGAACAGACACAATAACTACTCTGATTGGGCGTGGACAAGGAAAAAGTGGCATCTTTTTTGTTTTGTTCATAAATTCATTAATTTTATTCATAATATATATGTACCAAATAAGCTTTAAGATTTAGTAATGTAATTTTCTCCTTTTCCTTTCATATACTATAGTCCATTGAGTGCAAGATTAGAGATATTTAGTGGGGAAAATAAA

>PGSC0003DMG400011071 L-LecRK

AAAAAAAAAAAAATTCTCACTATCCATCACCAAATCTTGAAAAAAAATTCTTGTGTATCGTCTTTTTATAAGGCAAATCAACTTCGATTCTTGAATAATCCACATCACTTCTGCTTCTATATATCTTACTTTAAAATGTTCAATTTTCTCACATTCCTCTGCTTTACCATTTCCTTCTCAATTCTTGCTTCATCTCAACTTGATGAATTCATTTTCACAAAATTCAATCAACCAAACAACAACATAACCTTAAGTGGAGTAGCAAAAATAAGCCAAAATGGGTTTATTCAATTAACCAATGACACAAGTAGACTAATAGGCCATGCTTTTTACTCTTCACCTTTCCATTTCAAGCCTACTAATGCCTCTACCTTCTCTTTCTCAACGTGTTTCGCTTTGGCCATAGTCCCCGAGTATCCAAAACTCGGGGGCCATGGCCTAGCGTTCACAATTTCTCCGTCAAAAGACTTTAGTACCGCTCTTCCGAGCCAGTACTTAGGTCTGCTCAATGCTAGTGATATTGGTAACTTCTCTAACCATATTTTTGCTGTTGAATTCGACACGGTACGTGATTTTGAGTTTGGTGATATTGATGATAACCATGTTGGTGTTAATATTAACAGCTTAGAATCAAATAAATCAGCTGCAGCTGGCTATTTTATTAACGATCAAAATGTGAAACAAGATTTGAATCTTAAAAGTGGGAAAGTCATTCTTGTTTGGGTAGAATATGATTCTGTTACAAAATTAGTTAATGTAACTCTTTCACCAAATTCTTTAAAACCAAAAATTCCTCTTTTATCTTATAATTTTGATGTCTCTCCAATTTTTAAAGAAAATATGTATGTTGGCTTCTCTGCTTCTACTGGTTTGCTTGCTAGTTCACATTACATTTTGGGTTGGAGTTTTAAGTTAAATGGAGAAGCAAAATCACTTGACTTGGATTCTCTACCATCACTTCCTGGACCTAAGAAGAAGCATACAAGACTAATTATTGTCATCTCGATTATAGTGTTTGTGTTCGCGTTAATTTCAATCTTGATTGCTTTTTATACAATACGGAGATTCAAGAATGCTGATGTGATTGAGCAGTGGGAGCTTGAGATTGGTCCTCATAGATATTCGTATCATGAACTCAAGAAAGGTACTAGAGGTTTTAAGGAGAGTGAACTACTCGGGCGAGGTGGATTTGGGAAAGTTTACAAAGGTGTTTTGACAAATTCAAAAGCAGAAATTGCTGTGAAACGAATTTCTCATGAATCTAAACAAGGGTTGAGGGAATTTGTGTCTGAGATTGCGAGCATTGGAAGACTCCGTCATAGGAATTTGGTTCAATTGTTAGGATGGTGTAGACGTCGTGGTGATTTGTTGCTTGTTTATGAGTTTATGCCTAATGGAAGTTTGGATAACTTCTTGTTTGATGAGCCAAAAATGGTGTTGAAATGGGAACAAAGGTTCAAGATTATCAAAGGGGTTGCTTCTGGTTTACTTTACTTACATGAAGGCTATGAACAAGTCGTGCTACATCGAGACATTAAGGCTAGTAATGTGTTACTAGATGGTGAATTAAATGGTAAGTTAGGGGATTTTGGACTTGCTAGATTGTATGAGCATGGGTCTAATCCGTCCACGACTAAAGTTGTAGGTACATTAGGGTACCTTGCACCAGAATTACCAAGAACAGGACGAGCCACAACAAAGACTGATGTGTTTGCCTTTGGTGCTTTGTTACTCGAGGTGGTTTGTGGACGTAGGCCAATTGAACACAAAGCAGGACCCGAGGAGTTTGTTCTCGTGGACTTGGTGTGGGATAAATGGAGAGAAGGTAAAATACTTGACATTGTGGACTATAGATTGAAAGGCGAATACAATGAGAGTGAGGTTTTGATGGTGTTAAAATTGGGACTAATGTGCTCAAATAACAAGCCATTGGCGCGACCAAGTATTAGGCAAGTTGTAAGATACTTTGAAGGTGAAATTGGGATGCCTGAGGCTCCAACAACACAAGGTTCTGATGATCTTGAAGGATTTGGATTCGATGAAAGTGAACGTTGGAATTCCTTAGCATCGACTGGTATTATGATCAAGTCAAGTTCATTTACATGTTTGTCTAATGGAGATGAAAATAGTACTAATTATGTTTCCTTTTCTACTTCACCACTTCCACTTCTATATAGTGGTGAATGTACCTAGATACTAATATGAACTAGCATTGCAAACTTTTCTATATTTTCCTGATGAACACTAATGTACATTGTTTTTTCCTTTTCTTTGTGTTACCTCTTTGTAGTAGTTTTTTTCCCACAGAAAACTTAGGAAGCTTCCCGGTATTTCAGTGAGAACCTGTTTTCCACCATGTGTTTTCCCAGTGTGCTGGTTCGATGGGAAATGGGTGGAAAAATCATGTTTTATAATACAAATTTTTCATTGATTCGCGATGGAA

>PGSC0003DMG400018184 L-LecRK

TAGGAATCAAAATAATTAGTCAAAGGATGATCAGAACTTAGTCATCTTCATCATTTTCAGAGAAACAGCACTCATCTCTAATTCCCTTCAAAGTACCATGTAATTAACACATAATTTCCCCGTTTCCTTCACAAAATTCTCTTCTTCTTCTTCATGCAAACCCCAACAATATGCCACCATTATCACTGTTCTTCATTCTCATCAACCTTACAGTCTTCCCATCCATTTCCGCCGTCGAATTTCTCTTCAACTCCTTTACTGCTAACACCACCCCTGCCCTCAACCTCATCGAGGATGCCCGCCTTGAGCCACCAGTTATCCGCCTCACAAATGACTCCAACCAATTTTCCCTTGGCCGAGCTTTTTACCCATCTCAAATTCCTGTAAAATCCGCTTCAAATTCCACTTCAATTTCGTCCTCTTTCTCTACCCAGTTCATCTTCTCTGTACTCCCTGATGATTCCTCTAGTCCCGGATTCGGAATTGCTTTTGTTCTCTCTGCTTCAACTTCCCCACCTAATGCCCTCTCCAGTCAATACTTTGGACTTTTCTCTAACGCCACCGTTCACACGGTGGCCCCACTTCTCGCCGTCGAGTTTGATACGGGTCGTAACCCGGAATTCAATGACCCAGATAGAAATCATATCGGGATTGATCTTAACAGCATTGAATCTATTGTTACCCAAACGGCTGGGTATTATAGTTCTTCTGGTAATGATAGTGATTCTTTTGTGCCTTTAAATTTGCGGTCTGGGCAAAATATACATGCTTGGATTGAGTTTAACGGACCTGAATTCGAGATTAATGTTACTATTGCTCCGGCGGGTATGTCACGGCCTGTTAGAACTTTGTTGAGTTATAGAAACCCCATAATTGCGAATTACACGTCTGCCCAGATGTATATGGGGTTCTCTGCTTCCAAGACTCAATGGGTTGAGGCGCAAAGACTTCTAGCTTGGAGTTTCAGTGATTCTGGAGTTGCAAGGGATATTAATACTACGAATTTGCCGGTTTTTCAGCTGGAAAATTCCGGGTCTTCGCTATCTCCCGGTGCTATTGCTGGGATTGTTATTGGTTCTGTGGTTGCTGTGTTGGGTTGTTTATTTATATTTTATTGGTTTTGGTGGCGTAAGAAGGAAGAAGATGATGTTATCGAAGATTGGGAACTTGAGTACTGGCCTCATAGATTTTCATATGAAGAACTTAGCCAAGCTACAAAAGGGTTTTCTAAAGATGAGCTACTTGGGGCTGGTGGTTTCGGTAAAGTATACAAGGGAACATTTGCTAATAGCACAGAAGTGGCAGTGAAATGTGTGAACCATGACTCAAGACAAGGAATTAGGGAATTCATGGCTGAGATATCTACTATTGGAAGGCTTCAACATAAGAATTTAGTACAAATGAGAGGGTGGTGTAGGAAGGGGAATGAACTTATGATTGTGTATGATTACATGCCTAATGGGAGTTTGAATAAATGGATATTCGATAAGCCGGAGAAGGTTATGAATTGGGTAGACCGGAGGAGGGTTCTAGCTGATGTTGCAGAGGGGTTGAATTATTTGCATCATGGTTGGGAACAAGTAGTTGTACATAGGGATATTAAATCTAGCAATGTTTTGTTAGATAGTGAAATGAGAGGGAGATTGGGTGATTTTGGGCTAGCTAAGTTGTATACTCATGGTGGTGTCCCTAATACTACTAGGGTAGTAGGTACATTAGGGTACTTGGCACCTGAAGTGGTGACAAGGGCTAACCCAACTGCTGCTAGTGATGTTTATAGTTTTGGGGTGGTGGTTTTGGAGGTGGCATGTGGGCGAAGGCCGATTGACTTTGGGATTGTGTTGGAGGAAGAGGAAGTGTTGATTGATTGGGTTAGACAAAAGTACAGGGAAGGGCGATTATGTGAGGCAGCAGCAGATAAAAGGATTAAGGGGCAGTGTTCAGAGGAAGAAATGGAAGCTATGTTGAAACTAGGACTTACATGTTGTCACCCTGATCCTCTCCGTCGACCTACTATGGGAGAGGTTGTTGCTGTATTGCTTGGTGAGAATGTGGATGCAACGCCAAATGAGTTGATCGTTGAATTAGCACCTAGTGAAAGGAACACGGGAGATAGAAGCGACTGGTCAACTGAGGAATCAGAACCACTTTCGGCTGTGTAGATATTGTACTTTTGTTTCCTTTATTGTTGAATCTTTTTCATCCCTTGAAATTATTGTTTAAAGGTTTGTTCTTTGTATTGTTCTTACATGCTTTTGAACTCTAAACTTAGCATTAGTTGCAAATAGTTATGGATATGCAATTACTCAAGATATCAAAACTGCTATGTCTTTGCGCTACCTTAAAGGGTCAACATCCATGGCAATTAGCAAACGTCTATACGCATTAAGATTCGCCTCGGTGGAATAACG

>PGSC0003DMG400018677 L-LecRK

ATTTTACACTACTCCAACATGCTTCAAAATAGCTTCTTTTATTGACCCTTCCTCTTATTAAAAGCCCTTACATCTCTATACTTCTTTTCACTACACAATATGGAATTTTTGACACCAAAAATCTTCATATTTTTCTCTTGTTTTCAACTCATATCACAAGCCAAAATCATAACATTTGACCAACAATATGGTGACCCTTTTGATCATACATATGTCCCCCTTCTTGAAATCAAACATCCTGCACAAATAAGCAATCAAGCACTTCAAATCACCCCTGACACAGCTTCCTCTGCTTACCAAATGTTCAATAATTCAGGCAGAATCCTATTAAAACGGCCTTTCAAACTATGGGTAGACAATGATATTTCAAGGACAGCGTCATTCAACACATCATTCCTAGTAAACATATATAGACCGGATAACCAAACACCAGCTGAAGGTCTAGCTTTCTTGATATGTCCAAATTTTGATTTGCCACTAAATAGCCAGGGCCAGTATTTGGGCCTAACGAATGCTACAACAGATGGTGAATTATCCAACAAAATAATCGCGATTGAGCTAGATACATTCAAACAAGATTTCGATATTGATGATAACCATATTGGGATTGATATAAACAGTGTTAAATCCGTTAAGTCGGAACCTTTAACACCACATGGAATTGAACTAGCTCCAATAGGTGCTAGATTTTACAATATATGGATACACTATGATGGATTCAAGAAAGTACTTGATGTGTACATTATTGAACAAATGACGAAAAATGGGACAACACCAACAAGGCCAAAAGTACCAATACTAACACACAATGTTGATTTGAGAGAATTTGTAAAGCAAGAATCATATTTTGGATTTTCAGCTTCAACAGGACATTTCAATCAGTTGAATTGTGTGTTGAGATGGAATTTAACAGTTGAGTATTTTCAAGAAAAGAATGATCAAGAAAAGGTACTAATAATAAGTCTAAGTGTTGGTGTATCAGTATTGGTTGTTTTATTGATTTTGTCAGGGTATTTTGGGTATTTTTTTTATAAGAAAAAAAGGGATGATAGGTCACAATCTAATATATTAGGGGCATTAAAGAGTTTACCTGGAATGCCTAGGGATTTTGAGTTTAAAGAATTGAAAAAAGCTACTAATAATTTTGATGAAAAGAATAAACTTGGTGAAGGTGGATTTGGAGTTGTGTACAAAGGGTATTTAGTTGGTGAAAATTTGGAAATTGCTGTGAAGTGGTTTTCTAGGGAAAGTATCAAGGGTCAAGATGATTTCTTGGCTGAGTTGACAATTATCAACCGTTTAAGGCATAAACATCTTGTCAAATTACTTGGTAAGTATTTTGAACTCTTTACTTTTTATGCTTTTTTTTGAGTATTTGTTCTTTGTTCAAGCTCAGATAAAGGAGGAATGTTATGATCTTAAGATTTTGATATATATGCTTATTGAGTTTAAGCGTAATTGGTTGATAGATTCGTCATGTCATTGGAATTTTATTCATGTTCTATGTATAGGTAGCAAAATCTACTAAGATTCTACTGAAAAAAGAGACAGGTTTTACACATAATCATTTCTTCGTTTTGAGGACACTTTTTATTTATTGAGTCGTGTTTCTTATCCTTCCTCGACATAGTAACCGCTTTCATATTTTTTATAACTTTCAAGTACAAGTGCATGTTCATCCATCTTCATAGTCATTTCTTTGTTCCTTTAGTCGTTTTGATAGGACTTTTTTTATTGAGTCGTATTTCTTCTCCTTCTTCAACATAGTGATTTGTTTCATGTTTTTTTTTCAAATTTCATAACTTTCGACCACAAGTGCATGTTCATCCATCTTCGTTTTGAACGAAAGCTTAAATTAAATTTTATTAATTATGGTACCAAGAATATTTTCTTGTGTTCTAAGGAGTTCCATTTTGACTCATGTTAGGCCCATCTCCATCTTGAAAATGGTCCCAATAATTATATCCAACTACTACAAGATAAGTGAAAAAAGTTAAAATCTTTGATGCAACTTGCAAGTGTCAAAAATGGTGTGCTCATGTACCTTTTACAATAGTTTGAAAGTGATAAACATATACTTTTTTGTCTCAAAATATTTATTTATATATTTTTATACGCCTTTTAAGAAAACATTAATTAGTAAAGATATTTAATTATTTCCCCCTTATTTATATTTTAAAACATAATCTCTATTTATATGTCATATTTGAGGTATCTGTAAATTTCAAAAATAATTACTAACTACTAAATATAAAATGAGGTAGGGGTGATTAATTTTAACTTGAATTTTTAAAATATAATCCAAAGTTTCTTGGAGGACCTACTAACTAATAATTTTAGTGCCCTCTTGATGAAAAATCCAAAGTAATAAATATTAAATTCTTAGCTGGAAGACAGAAGAGCACTGATTCCATTTCCTTTGTCTTTTTGGTTTTCTATACCACCAATTTCAAACATAGGGTCCCACACTCTTTTGTTCTTTGTTAGCTTCGGTATAAATATCGAGTAATTCTAATCATTTAAATTAATTATATGAAAAAAATTGTCTAGTATTTTTGTTTCGATTAAATTTGAATCATTGTTTGAACAAACTTTTTTTTGTAGGGAAGCTTCCTGTCTTTGAAAATCACTTTCATTCAAATGTCACCTATTATGATAAGTTTGGTACTTTCTAATATTAATATAATATAATTGGACAATTGCTTCAAAATGCACTCAACTCCAATTAAGACTAAGATTGTAACCCTACAACATGTCATCATAATTGAGAGAGGTCCATATATTCCAAAATTAGGTCCTACAAAAAAGTTGATTGGAACACATACAACCCAACTTTTTTTGCCTTGTAAAGATTTGATATCCTCAAAATAGTATTGATAAGATATATTTATAGATTATTATTATTGTAGTAACAATTTAGCTTATATATGCATTAACAATTTTAACTGCATCTGTGTGAGCTCGCTCAAATTGTCTGTCCTTAATCCGAATAAAGAAGGAGGGTTGTTGAGGATCACTCAGAAACACCATTCTCTATCCCATAAGAGTAGGAGTAAGGCTGCATCAATTCATAGATCATTATTATTGTAGTCATAATTTGGCCTATATGTATATCTACTAATAACAATTTCAACTCTGTCATCAGTGTGATCTCGCTCACATTGTCGGTCTCAAGATTGAATAAAGAAGGAGAGTTGTCGAGGGTCAACCGAAAACGGTCTCTCTGCCTTATAAAGATAGGAATAAGGATATTTACATCTTACCTTTGTCAGACGTACCCCACTTGTAGGATTTCACTGAATTTGTTGTTATTGTTGTTGCACTAACAATTTCAAGTGATCTAAAGTAATTTTTGAACCTCCTTTACCTTCTAATGAGAACGTAACCTAAATGTTGCATGTAGTTATCATTTAGGTGACCTGCAGGTAATGGTATAAAAGTTCTTTTACGCTTTCAGTGTATATATATGGTAATCACATGCATATTATTGTGACAGGATGGAGCCATAAGCATGGTAAGCTACTACTTGTATATGAATACATGCCAAATGGTAGCCTGGACAAACATCTCTTCTCATCAGGGCAAGAAAAAGAGCCACTCAGCTGGAGTGTGAGGTACAACATTGTATCAGGAGTCGCGTCAGCTCTGCACTATCTTCACAACGAATACGAGCAGAAGGTAGTCCATCGCGATCTCAAGGCAAACAACATCATGCTCGACTCAAACTTCAATGCACGTTTAGGTGATTTTGGATTAGCACGAGCAATTGACAACGAAAAGACCTCATATGCTGATGAAGCTGAAGGTGTACTTGGCACGATGGGATACATCGCGCCAGAATGCTTCCACACAGGAAAAGCAACTCAACATTCTGATGTCTATGCATTCGGAGCTGTGTTGTTGGAGCTAATATGTTGCCAAAGGCCTGGAACTAAAGTTAATGGCTTTCAACTCTTTCTTGATTGGGTTTGGTTCTTGCATCGCGATGGGAGAATCCTTGAAGCTGTTGACAAGAGGCTCGAGGATCAATACGTGGTTGAAGAAGCAGAGAGATTGTTACTACTTGCTCTGGCTTGCTCCCATCCAATTGCCAATGAAAGGCCTAAAACACAAACTATAGTTCAAGTTATATCAGGATCAGTACCAGCACCACAAGTACCACCATTCAAACCATCATTTGTATGGCCTTCTATGGTTCCAATTGACATAGATTCATCGAGCATCGTCGATACAATATCCATCACAACTCCTCAGTACAGTTCAGAGAACAACAGTATTGAATATCAAAGCAAGTAGAGATAGCAATAGCTTGATTTAAGGCCACTAAGATATTTTTTTTGCTTATTGTGTAATGACAAGAATAGCAGCTCATGTATATATTTCTCAGCCTTTTATGTTAAATTCCATCCTATACATTGTTCCTTCTGCGGTTTGACTGTTTGATTTGTTGTATTAAAAATGACATGTATTGCATAATTTCTAAAAAGAAGTTGTTCG

>PGSC0003DMG400020326 L-LecRK

CAACAATAGAACAAATTAGCATATTTTCTCTCAATCAAGAAAATATTACCAATGTTGTTAAATTTTGTCATACTAGTGTCTCTCTTCTTTGTACCAACTTCTTGTGAAGTTGATGGATTCATTTACAATGGATTCAAATCAAGTAATATTAGCTTAGATGGCATTGCTAATATTACATCCAATGGTCTTTTGTTGTTAACCGATTCCAAAACACAAGATCAAGGCCATGCTTTCTATCCAAATCCAATTCATTTCAAGAATTCACCTAATGGTAATGCATTTTCATTCTCTACAACCTTTGTGTTCGCGATAACGTCTGATTATCCACCATTGGATGGTCATGGACTAGTTTTTGTTATCGCGCCACAGAGAGGAATTCAGAAGGCTTTGGCAAACCACTATCTTGGCCTTTTTAACTCAAGTAATAATGGGGATAAATCAAACCATGTTGTTGGAGTTGAGTTTGATACAATATATAGCGCGGATTTTGGTGATATGAATGACAATCATGTTGGAATTGATATAAATGGATTGAGGTCTGTAGCAAATCACACAGCAGGTTATTTTGATGATACTGGTTTGTTCAACAACTTTACTCTGATTAGTGGCAAGGCAATGCAAGTTTGGGTGGATTATGATGGCAGAACTAAGCTAATCAATGTGACAGTAGCTCCTTTACATATGGAAAAACCAGTTAGTCCACTTTTGGCTTTGAAATATGATCTTTCGCCGATTCTTAATCAGATTATGTATGTTGGTTTTTCATCATCAACTGGTTCAGTCCCAACACATCATTATATCTTGGGATGGAGCTTTAAGGCAAATGGGAAAGCTCAAGAACTTTCTCAACTTCCTAGTCTTCCTCGTGTTGGGCGCAAAGGGACTTCAAGATTTGTAACGATTGGTTTACCGGTGATCTCTTTGGTTTCAATTGTTGTAGCAACCTTAGTGGTGGTTTACTATGTAAGAAGGAAGAAGTATGAAGAAGTTCTTGAAGATTGGGAACGTGAGTATAGACCGCAAAGGTTCAGTTATAAAGATTTGTACACTGCTACTAAAGGATTCAGAGAAAAGGAGCTGTTGGGAGCTGGAGGATTTGGTAAAGTTTATAAAGGAGTGATGCCTGTAACCAAACTTGAGATAGCTGTAAAGAAGATATCTCATGAATCAAGACAAGGGATGAAGGAATTTGTTTCGGAGATTGTAAGTATAGGCCGTATACAACATAGGAATGTAGTACCACTTTTGGGCTATTGTAGGCGAAAAGGGGAATTACTTTTGGTTTATGAATTCATGTCTAATGGAAGTCTAGACAAGTATTTATACGACCAACCAAGACTCACCCTCGATTGGAACCAAAGATTCAGAGTCATTCGAGGTGTAGCATCAGGACTATTTTTCCTACATGAAGAATGTGACCATGTAGTGATTCATAGGGATATTAAGGCTAGTAATGTCTTGTTGGATGGTGAACTAAATGGCAGGCTAGGAGACTTTGGACTCGCGAGGCTATATGGTCATGGGACCGATCCTCAATCAACTCGTGTCGTTGGTACTCTTGGTTATCTTGCACCAGAGCATACTAGAACTGGCAGGGCAACACCTAGTAGTGATGTATTTTCCTTTGGGGCTTTTCTGCTTGAAGTTGCCTGTGGTAGGAGGCCGATAGAACCAAGACAAGCCGGTGATGATCTGATTTTAGTCGATTGGGTGTTTTCGTGCTGGAATAGAGGTAATATTCTTGAGGCTGTTGATCCAAACTTAGGCAATGATTTTGTTCCAGGGCAAGTGGAGTTGGTTTTAAGTCTAGGCTTGTTCTGCTCTCATTCAGAGCCCTCATTTAGGCCAACCATGCGACAAAACGTGTTGTTCTTAGATGGTGTTGTGGCCTTGCCAGAGTTATCAGCCCTCGGTGTTTCATCTGCTGGTCTAACATTTGACCATCGCGGAGGTTTTGATGATTTTATCAAGTCGTATCCAGCTTCTTCTGTTAATGCACATTCCCCCGCTTCATCTGTAGCAGAATCACTCCTTTCTGATCCCCGATGATTCATGGTGAGTTGGACATAACTTCTTTGGTGGAATATTGATATACTGATCCTGGTAAATATAACATGATTTTAGTGATTATGTACTTCTGATAAATTATTGTAATTCATACACCTGAAACATATTGTGACATGTCTTAAGATATCAGCCATGGTATCCTGATCTTATTGTTTACTTCCAATGCTAAATAAATACTAATCAGAGATTCAGCAATGGC

>PGSC0003DMG400020327 L-LecRK

AAAAAATGGATTAAAAAGATCATTTTTTTGTCATATCAAGAAATGTATAAACAGTTGAACAAATTAGCTTTATATGGTCACAAATCACAATATTATCTCTCAATCAAGAAAATATTGGCAATGTTGTTCAATCTTGTCATACTAGTAGCTCTCTTCTTAGTTGGTATTGCACCAGCTTCTTGTGAAGTTGATGCATTCATTTACAATGGATTCCAATCAAGGAATCTTAGCTTGGATGGCATAGCAGAGTTTACGCCCGATGGCCTTTTGCTGTTAACAGATTCTGAAACACAAGATCAGGGCCATGCTTTCTATCCAAATCCAATTCATTTCAAGAATTCAACTAATGGTACTGCAGTTTCATTCTCCACAACATTTGTGTTTGCCATAAGGTCTGATTATGGAATTTTGAGTGGTCATGGACTGGTTTTTGTTATCGCGCCACAGAGAAGAATTCAAGGAGCTTTGGCATACCACTATCTTGGCCTTTTTAACTCAACTAATAATGGGGATAGATCAAACCATGTTGTTGGAGTTGAGCTCGATACAATCTACAGCGAAGAATTTGGTGATATCAATGCCAATCATGTTGGAATTGATATAAATGGATTAAGGTCTGTAGCAGTTGAAACAGCAGGTTATTTTGATAATACTGGCTTGTTTCATAATTTGACTTTGATAAGTGGTCAGCCAATGCAAGTTTGGGTGGATTATGATGGCAGCACTACGCAAATTAATGTAACAGTAGCTCCATTACATGTGGAAAAACCAGTTAATCCACTTTTGTCTTTGAAAACTGATCTTTCGCCGATTCTTGATCAGACCATGTATGTTGGTTTTTCATCATCAATTGGTTCAGTCCCAACACATCATTATATCTTGGGATGGAGCTTCAAGATAAATGGGAAAGCTCAAGAACTCTCTCAACTTCCTAATCTTCCTCGTCTTGGACGCAAAGGGACATCAAGATTTGTAAAGATTGGTTTGCCATTAATCTCTTTGGTTTTACTTGTTGTAGCAACCTCAATGGTGGTTTATTATGTAAGAAGGAAGAAGTATGAAGAACTTCTTGAAGATTGGGAACGCGAGTACAGACCACAAAGGTTCATGTACAAAGATTTGTACACTGCCACTAAAGGATTCAGAGAAAAGGAGCTGTTGGGAGCTGGAGGATTCGGAAAAGTTTACAAAGGAGTGATGCCTATCACCAATCTTGAGATAGCTGTAAAGAAGATATCTCATGAGTCAAGACAAGGGATGAAGGAATTTGTTTCGGAGATTGTGATCATAGGTCGTCTACAACACCGGAATGTAGTACCACTTTTGGGTTATTGCAGGCGGAAAGGAGAGTTGCTTTTGGTTTATGAATACATGTCTGGTGGAAGTCTAGACAAGTATTTGTACAACAAACCAAGATTCACCCTTGATTGGAACCAAAGATTCAGAGTTATTAGAGGTGTAGCATCAGGACTATTTTTCCTACACGAAGAATGTGACCATGTAGTGGTTCATAGGGACATTAAGGCCAGTAATGTCTTGTTGGATGGTGAACTAAATGGTAGGCTAGGAGACTTTGGACTTGCGAGGCTATATAGTCATGGTACCGATCCTCAGTCTACTCGTGTCGTTGGTACTCTTGGTTACCTTGCACCAGAGCATACTAGAACTGGCAGGGCAACGCCTAGTAGTGATGTATTTTCCTTTGGGGCTTTTCTGCTTGAAGTTGCCTGTGGTAGGAGGCCAATAGAGCCAAGACAAGACGATGATGATCTGATTTTAGTCGATTGGGTGTTCTCATGTTGGAATAGGGGTAATATTCTTGAGGCTGTTGATCCAAACATAGGGATTGATTTTATTCCAGGGAAAGTGGAGTTGGTCTTAAAGCTAGGCTTGTTCTGCTCTCATTCAGAGCCCTCGTATAGGCCAACCATGCGACAAATTTTGTTGTTCTTAGATGGTGTTGTGGCCTTACCAGAGTTATCATCACTCGGTATTTCATCAGCTGGCCTAACGTTTGACCATCACGGAGGTTTTGATGATTTTGTCACGTCCTATTCATCTTCGATGAATGCACATTCCCGCTCTCCATCTGTAACAGACTCCTTTCTCTCTGGTGGCCGATGATTCTTACATTAAATATGTTAGTAACTTTAACTTGTAAGTTGTAATCATGGTTAGCACCTCATCTTAGTTTTAATTGAATCTCTATACTGCTGCACCTCAAATAAGATGGAAATGCA

>PGSC0003DMG400020328 L-LecRK

AAAAAAAATTAGCCATGTTTTTCAAGATTGTGTCCTTGATTGTGTCTGCTCTTTTTGTTGTTACAAATTCTGAAGACTTTGGATTCATTTACAATGGGTTCAATAGAGCAAATTTGAGTCTTGATGGCATAGCACAATTTACATCAAATGGCTTATTAGAGTTAACAAATACTTCTAGGCTACAAAAGGGGCATGCTTTTTATCCTACACCAATTAATTTCAAGAACTTACAAAATGGTTCAAATTTCTCTTTTCCTACCACTTTTGTCTTTGCTATTGTGCCTTCTGTTTTACCTGGTCATGGTATGGCTTTTGTAATTGCACCTGTTGGTGGCCTAGTAGATGCACTTCCAAGCCCATTTTTAGGCCTGTTCAATGATAACACTACTGGGAGAGTCAGTAACCATGTTTTCGCGGTCGAGTTTGATACTTTACAGAACAGAGAATTCAATGATATTGATGGTAACCATGTTGGAATTGATATCAATGGATTGAAATCCGTTGAATCAAAGACAGCAGGTTATTATTATGGTGTTAAATTCAATAACATGACTCTTGCTAGTGGTCAGCCAATGCAGGGTTGGGTGGACTATGATGGTGTGGCTAAGCAAATTAATGTAACTCTGGCTCCAATGAATGTAGCAAAACCAAATGTACCACTTTTGTCATTGTTGTATGATCTTTCACCAATCTTGAATGAAACCATGCATATCGGTTTCTCTGGATCGACTGGCTCGGTTGTTTCAACACAATATGTTCTAGGATGGAGCTTTGAAATGAATGGAATAGCTCAAGGTCTTGATCTTGCTAGACTTCCTAAGCTTCCTCGAGTTGGACCGAAGAAACAATCTAAACTTTTGTTGGTTGCTTTGCCAATGATATCAGCAGTTGTTGTAGTAATAGCCTTTTCTGTGTTGATTTACTATGTCGGAAGGAAGAGGAAGTTTGCGGAATTGCTTGAAGATTGGGAGCTTGAATATGGACCTCATCGGTTCAAATACAAAGATCTTTATATCGCGACAAAGGGATTTGCTAACAAAAAGTTGTTAGGTTGTGGCGGTTTTGGCAGAGTTTACAAAGGAGTATTACCAACTTCTTCAATTGAGGTAGCAGTTAAGAAGGTGTCTCATGACTCAAAACAAGGATTGAGGGAATTTGTAGCAGAAATTGTTAGTATTGGTCGTCTACGCCACAGGAATTTAGTGCAACTTTTGGGATATTGCAGGCGTAAAGGAGAGTTGCTTTTGGTTTACGAATACATGCCTAATGGAAGTCTTGATAAGTTCCTATATGACAAACCAATATGTGCCCTCAGTTGGAACCAAAGATTTCGAGTCATCAAAGGCGTAGCATCAGCACTAGTCTATCTACATGAAGAATGGGAACAAGTTGTAATACACAGAGATGTGAAGGCTAGTAATGTGTTGTTAGACAGTGAATTGAACGCGAAATTAGGAGATTTTGGACTAGCAAGATTGTATGATCACGGGACCGATCCACTCACTACACATGTAGTTGGAACTGTAGGTTACCTTGCCCCTGAGCAGACAAGAACAGGCAAAGCCACAACCATTAGTGATGTATATGCTTTTGGTGCATTTTTGCTTGAAGTGGCTTGTGGGAGAAGGCCAATAGATCCACGAGTATCGGATGAGGACATCGTGTTGGTTGATTACGTATTTTCTTGTTGGAGTAGAGGTGATATTCTTCAATCCATTGATCAAAATTTGGGGAATGAATATGTTAAAGAGGAAGTTGAATTGGTACTAAAACTCGGACTTGTTTGCTCTCAGACAGAACCAACAAGGAGGCGAAGTATGAGGCAAGTTTTAATTTACTTGGAAGATATCGTGCCTCCGCCATTGCCAGAGTTGTCATTGTTACAAAGTTCTGCCCATTATTTTTCATTTCCAGGTTTTGATCATCACTTGTCTATGTCTTCATCATTTTCTCCAGAGAAAGTACTTTCATATCCCTCCACCGATTCGAACTCTCTTCCATTCACTGGTCCATGATTAGAACTCTTTCTGTATTGCCTTGATGAATTGTGATTCCAATCCCAACAGCAACAACAACTGTACTTCAATTCCAAACAAGTTCGTGTTAACTATGTGAATTCTTACTGTCTATATCGTTTCAATTAAGCATGTAATGTTTGTTACGATACAAATAATGGACTAATGTACACTTAGAATGATCCGGTTTCTTTTGATCTATAATGTTGTCGACATGTATTGCTA

>PGSC0003DMG400023671 L-LecRK

AAAAAACATAGTTTTTTGTTATATGTTGATATGTATATTCTAGGCAAGATTAGTTATTTAACTAACTTTGTTCATCTTTTGATCATATTCATATTTTTGGTTCTTAGAGTGGATTGTATGAGTTTCAGCTACCAAAAATTCACCAAGCAAAACGAAAAAGATTTCATTACTACTAACGATTCGTACATAGCCGTTGATGCTATCCAAGTAACTCGCGATGTTCGTGGTGCTTCCATCAGTAATTTGTCTGGAAGGATATGGTATTCACATCCTATGAATCTATGGAACAGAAGGAAAAACAGAACAGCCTCGTTTAATTCAACGTTTGTGATCAATATCAAACCTGAGAGTGATCCATGGGGTGAAGGATTGGCTTTCATATTGACTAAAGAAAGTGGTTCTCACTCAATCCCGGATAACAGTTATGGACAATGGCTTGGGATTGTGAATGAAAGTACTAATGGATCTTCCTCAAACAACATATTTGCAGTTGAATTCGATACAAGGAAAAGTTACCTTGATGATCTTGATGATAATCATGTTGGCATTGACATAAACAGCATCAATTCTGTCAAGCAAGTCTCGTTGATTGATCGTGGTGTTAACCTTTCACTAGCTGTTGATGTTATAGCAAGTGTCCGGTATGATGGAGAATCCAAGATGTTGAAGGTTTATACATTCATGAGCAATCAAACCGGAGACAATGAGAGAAATCCCATCATTTCCATGCCTATTGATCTTTCTAGTTATCTCCCCGAGGATGTTTTCGTAGGATTTTCAGCCTCAACTGGGATATATAATCAGCTGAACTGCATAAAGGCATGGAATTTTACAAGCACAGATATTGGGAACAGTAATGGAGTTAGCCTGTTATGGTTGTGGATCTTGATACCAGTAATGTCAGTTCTTGTTATGTTCTTAGGTGGGGTTTTCGTATATTTCAGCTGGTGGAGAAAAAAGAAGAGGAGGCAAGTGTTAGACCAATCTGAGAATATAGAGATTCAGATTCAGAATTCAGCTACTGCACCTCAGAGATTCCATCTGAAGGACTTGAAACGAGCAACGGGGAACTTTGATCCGAAGAACATTCTTGGAAGAGGAGGATGTGGAGTAGTCCTCAAAGGGATGTTAGATCAAAAAGAGGTGGCTGTGAAGAGATTCTTTAATGATTCAAGTCAAGGGGCAAAAGATCTCATAGCAGAAGTCACAACTATAGGCAATCTGCACCATAAAAACCTTGTCAAATTGATAGGATGGTGCTATGAGAGCAATGAACTCCTTGTAGTCTATGAGTTCATGCCAAATGGGAGCTTAGACAAGTTGATATTTTGCGAAGAAAACGGAAAAGGTTTGAGTCTGAATTGGGAAATAAGGTATGGTATCATCTGTGGGACTGCTCAAGCACTTGATTATCTGCACAATGGATGTCAAAAAAGAGTACTTCACAGAGACATAAAAGCCAGCAACATCATGCTTGACTCAGAACTCAATGCTCAGCTAGGGGATTTCGGATTAGCTAGGACAGTTCAAGTGAATGGAAAGACTCACCATTCAACAAAAGAGATTGCTGGTACTATAGGCTATATGGCACCAGAGAGTTTCCATATAGGCCGAGCCACAGTTGAAACAGACGTCTTTGCATTTGGAGTGCTAGTTCTTGAAGTTGCTTGTGGTCGAAAACCTGGAAATCGATATGAGGAGAATAATTATACTAATAAAGTCATTGAATATGTATGGGACTTGTACAAGATTGGAAGGATCATTGATGCCATAGATGTCAGATTGGATAGAGACTTCAATGAAGAACAAGCCGAATGTGTGTTAATATTAGGATTGGCTTGTTGCCATCCAAATCCTTACGAAAGGCCAAGCATGAAAACTGCTTTACAGATTCTTACAGGGGAATCAGTTCTACCAAACATCCCAACTGAGAAACCTGCATTTGTATGGCCAGCCAGAGCTCCATCGTCCAACGAAGCTGCTAGTGGCTCTCTCCTGGATGGCCAACTTACACAGATTACAGTACTCAGTGGCAGATGACTGCAGAGTATGAGGAACCTCAGCAAGGTTAGGTAACTCTTTAGGAGGCAAAAGATACCTGTAAATTCACATATGTAAAGGCATGAATGTCTCTTTATTTTTTTTCTTTCTATGTATGTGGTGGGGAGTGGGAGATATCTTCTTCCGAGTGTTCGTTTTTCTGCATTATAAATAATTTTGTATTTAGTCAATGTCATAATCAACTTTCCAAGTAAAAACTGAAAACAAAAGCAGCAAGCATTGTAACAAAGGCAGCTAGCTACATGAGTTACTCCTACAGAACATAAATGTTGTACCAAAACCACAAAGAAGTTGTGAAAATGAAAATTTCACGATCCATCATCATCTTCATGTTCTATCTCGAAGAAACTCTTTCGTACTCCATCAATAACAGTTTCTCTGGATTGTTCATGTCTGCTATGTCCATCGTT

>PGSC0003DMG400026646 L-LecRK

GCACAAAAAAACTTAACAATTTGTACTTCCAACAACTCCATGGACAATCCCTATAGCCTAGTTATTCTTGCATTGATCTTTTTCCCTATTCTTGTGAGAGCTCAATCTTTTGATTTTGCTTACAATGGATTCAATAACTCAAATATCATTAGAGATGGAGTTGCAATAATCAATCCAAGTGGTGCACTTAAGCTCACTAATAGATCACAAAATGTTATAGGTCATGCGTTCCACCCCAATCCTGTACCAATTTTCAACTCCAGTACTAAAAATGTTACTTCATTTAGTACATACTTTGTCTTCGCCATTGTCCCTCTTGAGAAAACCTCTGGTGGCTTTGGCTTTGCTTTCACCTTATCTCCATCCCCGGGTTTTCCAGGTGCTCAGGGTGATCACTTTCTTGGAGTCCTCAACAACACGAACAATAGAAAGGATACTAACCACATCTTTATGGTTGAATTCGACACTGTGGATGGTCATGATGAAGGTGTGGATACAGATGGAAATCACATTGGGATTAACATCAATAGCATGATATCTATCGCGTCACAACCAGCCAATTACTATGTTAATGGTACTTCTAGAACAGAGGAGGTACATCTACAAAACGGAGAATCAATTCAGGCTTGGATCGATTATGATGGAATGAGCAAAGTGGTAAATGTAACTATATCTCCCAGAACAGTTCCAAAGCCAATTCGACCTCTGATCAGTGAGGTCGTAGATTTATCTCCTGTGTTGAACCAAACTATGTATGCTGGTTTCTCAGCAGCCACGGGAGATAAAGCGAGCTCTCATTACATCTTGGGTTGGAGTTTTCGGTTGAATGGAGCTGCTGATCCTTTAGATCTTGATAACCTACCTGTTGCCCCACCTGAAGTGGTAAGACCCTCCAAGAATTCCCATCTCAAAAGGGCCTTGATAGGAACATTCTGCTCCGTCGTTATCCTGCTCCTTGGGGCATTTTATATCTATAGAAGGATGAGGCAGCATGAAGTTCTAGAGGACTGGGAGCTGGATTGTCCTCACAGGTTCCGATTTAGAGATCTTTACAAGGCAACTAAGGGATTCAAGGAAAGTGAACTAATTGGAGTTGGAGGATTTGGTGCTGTTTACAAGGGTGTTTTGCCTACTAATGGAGCCGAGGTTGCAGTAAAGAAGATATCGAGCAATTCTCTTCAAGGAATGAGAGAATTTGCAGCGGAGATCGAAAGCTTAGGCAGGTTAAGGCACAAACACTTGATCAACCTACAAGGCTGGTGCAAGAACAAGAATGATCTTCTCCTAGTGTATGACTTTGTCCCAAATGGAAGTCTTGATTCATTACTTTACAGGCAGAAAAGGGACATTGTTCTAACATGGGAACAGAGATTTAACATCATCAAAGGGATTGCTGCAGGGCTTCTTTACTTGCATGAAGAATGGGAACAAGTGGTAATACATCGCGATGTGAAGAGCAGTAACGTCCTAATTGATGGTGAAATGAATGGTCGATTAGGGGATTTTGGGCTAGCAAGATTATACGACCATGGCAAGAATTCACACACAACAAATGTTGTTGGCACAATAGGGTACATTGCACCAGAATTGACACGGACAGGGAAGGCCTCAACGAGCACGGATGTGTATGCATATGGTGTACTACTTCTTGAAGTAGCTAGTGGAAGACCTCCTATTGTCTATGAACCAGGGAAAGGAGCTTTAGTACTAGCTGATTGGGTGATAGAATGTTTACAACTAGGTAATGTTCTTGATGCAGTTGATCAAAGGTTGAATTCAGTCTATGTAGATAAAGAGGTTCAAATGATTTTGAGACTTGGGCTTGTTTGTTCACACCCAAGACCAGAAGCTAGGCCAACAATGAGACAAATAATGAAGTATCTCACTGGAGATGAATCACTTCCAGTTTTTGAGCAGCAGTTGAGCTGTGTTGGATCTGATAGGGTTGATGAAATCACAGCCAAGTTCTTGGAAGTGTTTTCTACTGATACAATCAATATATCACGTCGTTCGCTATCCGTCGGACAGATATCATCTAGTTCACTCCATTCTGGTAGATGAAAACTCTGTATAAAGGACATACACTCTTCTTTTGTTTTGTTTAACCACTTTGTTGGGATTGATTAAGTAGCTTCTATTTACCCAAATCCTGTTTTTCTTCCTCATAATTCTGCAATTTAGATAGGTCATTTATTTCTAGGCTCATTCAGCATTCACAAGTTAACATAATCTCACTTCTATGTCTGCAACATCACTTATCACATCCTATATTTTTGTTTATGTACAAAC

>PGSC0003DMG400026647 L-LecRK

TAATTGTCTCTTTAAATTGAATATAAATATTGCACAAAAAACTTAACAATTTGTACTTCCAACGACTCAATGGACAATTCCTATAGCCTAGTTATTCTTGCATTGATCTTTTGCCCTATTCTTGTGAGAGCTCAATCTTTTGATTTTGCTTACAATGGATTCAATAACTCAAATATCATTAGAGATGGAATTGCAATTATCAATCCAAATGGTGCACTTAAGCTCACTAATAGATCACAAAATGTTATAGGTCATGCTTTCCACCCCAATCCTGTACCAATTTTCAACTCCAGTACTAAAAATGTTACTTCATTTAGTACATACTTCGTCTTCGCCATTGTCTCTCTAGAGAAAACCTCTGGTGGCTTTGGATTTGCTTTCACCTTATCTCCATCCCCGGGTTTTCCGGGTGCTAAGGGAGCTCATTTTCTTGGAGTCGTCAACAGTAGTAACGATGGCAATGATACTAACCACATCTTTATGGTTGAATTTGACACTGTGGATGGTCATGACGAAGGTGTGGACACAGGTGGAAATCAAATTGGGATTAACAGAAATGGCATGAGGCATATCACGTCAGAATCAGCTGATTACTATAATAATGCTACTTCTGAAACAAAACCAGTAGATCTACAAAACAGAGAATCAATTCAGGCTTGGATCGATTATGATGGAATGAGCAAAGTGGTAAATGTAACTATATCTCCCATAAAAGTTCCAAAGCCAATTATACCTCTGATCAGTCGGGATGTAGATTTATCTCCTGTGTTGAAAGAGACTACGTATGCTGGTTTCTCAGCAGCCACGGGAGATAAAGCGAGCTCTCATTACATCTTGGGTTGGAGTTTTCGGTTGAATGGAGCTGCTGATCCTTTAGATCTTGAAAAGCTACCAGTTGCCCCACCTGAAGTGGTAACATCCTCCAAGAATTCCCATCTCAAAAGGGCCTTGATCGGAACATTTTGCTCCGTGGTTATCCTGCTCCTTGGAGCATTTTATATCTATAGAAGGATGAGGCAGCATGAAGTTCTAGAGGACTGGGAGCTGGATTGTCCTCACAGGTTCCGATTTAGAGATCTTTACAAGGCAACTAAGGGATTCAAGGAAAGTGAACTAATTGGAGTTGGAGGATTTGGTGCTGTTTACAAGGGTGTTTTGCCTACTAATGGAGCCGAGGTTGCAGTGAAGAAGATATCGAGCAATTCTCTCCAAGGAATAAGAGAATTTGCAGCGGAGATCGAAAGCTTAGGCAGGTTAAGGCACAAACACTTGATCAACCTACAAGGCTGGTGCAAGAACAAGAATGATCTTCTCCTAGTGTATGACTTTGTCCCAAATGGAAGTCTTGATTCATTACTTTACAGGCAGAAAAGGGACATTGTTCTAACATGGGAACAGAGATTTAACATCATCAAAGGGATTGCTGCAGGGCTTCTTTACTTGCATGAAGAATGGGAACAAGTGGTAATACATCGCGATGTGAAGAGCAGTAACGTCCTAATTGATGGTGAAATGAATGGTCGATTAGGGGATTTTGGGCTAGCAAGATTATACGACCATGGCAAGAATTCACACACAACAAATGTTGTTGGCACAATAGGGTACATTGCACCAGAATTGACACGGACAGGGAAGGCCTCAACGAGCACGGATGTGTATGCATATGGTGTACTACTTCTTGAAGTAGCTAGTGGAAGACCTCCTATTGTCTATGAACCAGGGAAAGGAGCTTTAGTACTAGCTGATTGGGTGATAGAATGTTTACAACTAGGTAATATTCTTGATGCAGTTGATCAAAGGTTGAATTCAGTCTATGTAGATAAAGAGGTTCAAATGATTTTGAGACTTGGGCTTGTTTGTTCACACCCAAGACCAGAAGCTAGGCCAACAATGAGACAAGTAATGAAGTACCTCAATGGAGATGAATCACTTCCAGTTTTTGAGCAGCAGTTGAGCTGTATTGGATCTGGTAGGGTTGATGAAATCACAGCCAAGTTCTTGGAAGTATTTTCTACTGATACAATCAATATATCACATCGTTCGCTATCCGTCGGACAGACATCATCTAGTTCACTCCATTCTGGTAGATGAAAACTCAGTAAAAAGGACATACACTCTTCTTTTGTTTTGTTTAACCACTTTGTTGGGATTAATTAAGTAGCTTAGAATAATATATAACTAGGAAAAAGGTAAGTCTCAAGTCAACATTTTCACTTAGCTGGAGTGCAAAAACCATGTAATTTAGAGAAGAAAAGTTATAAAATTAGTCTAGAAAGTAAATTTGTAATTATAAAAAAAAAAAAAATCTATGTAGAATATTCGTTATAAAATCTATTTGTAATAAAAATGGTGACATGATTCAGA

>PGSC0003DMG400027140 L-LecRK

CTTTAAACCCCACTCTTTCACTTTCCCAACTCTTTCTTCATTCCTTTGTAATCACACAGACAAATCACATGTGGCAAATCAAAGTAGAACTCTACAAAAAGAATGTCTAGATTCAGCTATAAATCATCACCCCACACAATTCTTCAACTCTACAAACAAATTTCAATCTAACTATGCCATGGCCAAACAAATTACACCAACCCTTTTCTTGATTCTCTTTATTTTGCTGTCAAGAACAGTTTTTTCTGATGAATTTGTGTATAATGGCTTTAATGGGGTTGAAGCTAGTAATCTTACTGTAAATGGGATTGCTCAGATTTTAAAGAGTGGCGCTTTAAAGCTTACAAATGAAACATCTAGAGTTGTTGGTCATGCTTTTTACAAAAATCCAATTAAATTCAAGAATTCCCAATCTGGTAAAGTTTCTTCCTTTTCTACTGCTTTTGCTTTTGGGATTGTTCCTGAATATGCTAAGTTAGGTGGGCATGGCTTTGCTTTTACAATTTCAAGGACTAAAGAGATGAAAGGGGCTCTTCCTAGTCAGTATTTAGGTCTTTTAAGTTCAAGTAATATTGGGAATTTTTCAAATCATGTATTTGCTGTTGAATTTGATACTGTTCAAGATTTTGAATTTGGTGATATTAGTGATAATCATGTTGGTATTGATGTGAATGATTTGAAGTCTAATGCTTCAGTAAATGCTAGTTACTTCTCTGAAGGAAGTTTTAGTAAACAAAAGCTTTTTCTTCAGAGTGGAAAGACAATTCAGGCTTGGATTGACTATGATTCAAGTAGAAATGTGTTGAATGTTACACTTTCACTTTCTTCAACAAAACCAGCCTTTTCAATTTTGTCTTTCCCTTTGGACCTTTCTCTAGTTTTTGAGGAATTCATGTATGTTGGATTTTCTGCTTCAACTGGTTTGCTAGCTAGCTCACATTATATATTTGGTTGGAGTTTCAAGATGAATGGGAAAGCTCAATCTTTGGATCTTGATTCATTGCCTGCTCTCCCTAAGCCTAAGAAGGATCAAACAGTCTTGATTGTAGCTACCACGATATCGGCTGTTGTGTTTCTTGTATTCGGACTCGTTTTAGCCTTGTATGTCATTTGGAAAATCAAGAACATAGATGTGATTGAACCTTGGGAACTTGAAATAGGTCCACATAGGTTTTCTTACAAAGAGCTAAAGAAGGCTACTAGGGGTTTTAGAGATAAAGAGTTGCTCGGGTTTGGTGGATTTGGTCGAGTTTATAAAGGAACACTGCCTAAGACGAATACCGTTGTTGCTGTGAAGCGAATACATCATGAAGCAAAACAGGGTCTTCAAGAATTTGTGTCTGAAATCGCGACTATTGGTCGCCTTAGACATAGAAATTTGGTTCAGCTCTTAGGATGGTGTCGGCGAAGAGGTGATTTGTTACTTGTTTATGACTTTATGCCTAATGGAAGCTTAGACAAATACATTTATGATGAACCAAGATTTACTTTAACTTGGGATCAGAGGTTCAAGGTTATTAAAGGTGTGGCTTCTGGTCTGTTGTATTTACATGAAGAATGGGAACAAACTGTGATCCATAGAGATATTAAAGCAGGGAATGTGTTGTTAGATTCCGATATGAATGGACGACTTGGAGATTTCGGACTTGCTAAGTTATACGAGCATGGAGCAAATCCTAGTACAACAAGAGTAGTCGGTACGTTGGGTTACTTAGCTCCAGAATTAACCAAGATCGGAAAGCCTACACCAAGTTCAGATGTTTTTGCCTTTGGTGCATTGCTGCTAGAAGTTGCTTGTGGGAGGAGGCCAGTTGAGGCAAAGGCATTGCCTGAGGAGTTAATCCTAGTGGATTGGGTATGGGATAAGTGGAAGGAAGGTGCCATTCTTGAAGTGGTTGACTCGAGATTAAATGGCGAGTATGATGAGATGGAGGCCATAATGGTGCTTAAATTAGGACTCATGTGTTCAAATAATACACCATCTAAGAGACCTAGCACGAGACTAGTGGTGAGATACTTGGAGGGGGAGGTGACACTCCCAGAGACGCTGGCAGCTCCCGATGAATATGATGGAAAGAAGGGTGGTACTAGCGGTATGGAGTTAGAGGACTTAATAGATTCCTATCCGTCTTCGTCTTATTTTGAGAAAGTAAGTACATGGTCATCAGCTTATGACGGTGAAGGCGATATTGATATTGAAGCTAATTCAGTAACGCCATTAACGGATTCTGGCAGAGAGGACAACAGATAGCCATGCAAAATGGAGGGACCATTCTACTTGCCCTCCCAAACTCCCTCCATATTTTAGTACTTTTTATGGTTTGATTTTTGAACTTAAATGTAAATTATGGCTTTTGAAAAGAAAATGATATTAGTATCAACCATTTCTATGGTTTTGAGAATATTGGTGTTAGTAAAATGTGAATATATTTTTGCAATCAAGTCAATTTTGAATCTTGATGAAGCTTG

>PGSC0003DMG400029735 L-LecRK

GAATGATATTTCCCCATTTTCTCTGCTTCCCATAATTCTTTATCCATGTGAATTAGCTCTGCTTCTTTCTTAGAATGAACCAATATTATCCAAAAATTCTTCTTTCTTACCTGTTATTAACCCTATTTTCTTGTAATCAATCAGTTGCAGAAACTGATTTTGTCCTCAATGGCTTTACACAATCAGATGTGTCATTTTTTGGGAATGTTACAATCGAATCTAGAGTTCTTACTCTTTCAAATGACTCAACTTTTTCAATTGGTAGAGCTCTGTACCCTTCAAAGATTGTCACAAAAGAAGCCAATTCATCTAAAGTTCTTCCCTTTTCAACATCTTTCATTTTTGCAATGGCTCCTTACAGAGACAGGCTGCCAGGACATGGTATAGTTTTCTTGTTTGTGCCACACACAGGTATTGATGGTACTACTTCTGCACAGAATTTAGGTTTCTTGAATTTCACAAATAATGGGAATCCTGATAATCATGTTTTTGGGGTTGAGTTTGATGTTTTCAAGAATCAAGAATTCAATGACATAAATGAGAATCATGTTGGAATTGATGTTAATTCTCTTGCATCAGAGTTTGCTCATGAAGCAGGATATTGGCCTGATGAGAAAAATAAGTATAATAGTGATGGTAGCCTAAATGAGGAGTCTTTAGAGACTTTAAAGTTGAATAATGGAAGAAATTATCAAGTTTGGATTGATTATGCTGATTTCAACATTAGTGTAACTATGGCACCAGTTGGTGTGAAAAGGCCTAAGCAGCCTTTGTTGGATTTTCACCTAAATCTTTCCCAGGTTTTTGAGGAAGAGATGTATGTGGGGTTTACAGCAGCCACTGGAGAACTTGCTCAAAGTCACAAGATTTTAGCTTGGAGTTTTAGTAACTCGAATTTTTCGATAGGTGATGGTTTGATCACACAAGGGCTGCCTTCGTTTGAGCTTCCTGAGGATCCAGTTTATCGATCGAAGGGATTCATTGCAGGTATGACAGTGTCACTCTTGTTTCTTGTTGTGGTCATTGCTGTAGCTTCATGGCTTTTGATTAAGAGAAATAGGAGAATGAAAAAGGAAAGAGAGGATATGGAAGATTGGGAATTGGAATATTGGCCACATAGGATTACTTATCAAGAAATTGATGCTGCAACAAAGGGTTTTGCTGATGAAAATGTGATTGGAATTGGAGGTAATGGGAAGGTTTATAAAGGGGTTTTGGCTGGGGGTTCAGAGGTTGCAGTAAAGCGAATTTCTCACCAAAACAGTGAAGGGGCAAGGCAATTCTTGGCTGAGATTTCGAGTCTTGGTAGGCTAAAGCAAAGAAATTTGGTGTCATTAAGAGGGTGGTGCAAGAAAGATAGAGGCAGCCTGATTGTGGTTTATGATTATATGGAAAATGGGAGCTTGGATAAAAGGCTGTTTGAATCTAATGAGAGAAGCATGCTGAGTTTTGAAGACAGAATTAGGATTTTGAAAGACGTGGCATCAGGGGTTCTATACTTGCATGAGGGATGGGAGTCAAAAGTGTTGCATAGGGACATTAAGGCTAGCAATGTTTTACTTGACAGGGACATGAATGCAAGGCTAGGTGATTTTGGTCTAGCTAGAATGCATGATCATAGTCAAGTGGCTAGCACGACTCGAGTTGTTGGCACGGTGGGATACTTGGCGCCAGAGTTTGCCAAGACTGGCCGTGCCTCCACACAAACTGATGTGTTTGGATATGGAGTGCTAATTATGGAGGTGATGTGTGGAAGGAGGCCTATAGAGGAGGAAGGCAAGCCACCTTTAGTGGATTGGCTGTGGGAACTAATGAGACGAGGCGAATTGATTAACGCCTTTGATAGTCGATTAAGGATTAATCAAGATTTCAATGAAGAGGAAGCTTTAAGGGTGTTGCAACTAGGCATGATATGCGCGAGCCTAGACGCCAAAGCTAGGCCAAGCATGAGACAAGTAGTGAAATTCTTTGACAGGAATAGTGATATTGATGAATATGAAACTGAGGATATGGATGCTTACCTTCTAGAGAGTTTGAGATCTAATACCATGTTGTCCAATTTTTCCTTGAGTTTAAGCCATGGTTCACATCCAACATTTGAAGAAATTAGAGAAGGTTTGTCTTCTTCCATGTCAATTTCTTGGACAAATTCTCTGGTGGATGGTAGATGAGTTTAACATTTGGTTGGTTTATGTCCATCACTCTGATTTAACTTATGTTACCTTCCAAGGCCTAGCTTTGATTGAATTTGCATTTGGCAATGCTGCTTCATGTACAAATTTATATGATGGAGTAGGAAAAAGGATTTGGTTATGACTTGTGAGATATATACATGTATTCATTTCTTTTTTTGTACCTTGATGCTAGTTACAGTGGAAGAGTTTGTTCTCTTTTTTTTTTTTTAACTATAGAATTTGTTCTTTATCTTTTTCATTTTACACGAGATAAG

>PGSC0003DMG400030144 L-LecRK

GTATATATAAAATTTATTTTTTAGCAAAGCACTAAACAGAGTGTTTGGTGTTTATTGTATATATATGCAGTTGCATGTCTAACAATGTGATTAACATATTGTCCAACTATGGTTGTTTTTGTCTCTTTATCTTCTCTCAAATACTTGCTTCTTATCATAATTATCCCTTTAGTCACTTCACTCTCCTTCAATTTTGATACTTTTACACCCAATGATCACAATGTAACCTATGAGAGAGATGCTTCTCCAGAAAATGGTGCAATTCTACTCACCAAAAACCTCCTCAATCGCGATATTAATGCTAGTATAGGCCGAGCTATATATTCAAAACCACTGCATCTATGGGACAAGGCCTCTCGAAATCTCACTGATTTCACTACACATTTCTCCTTTAGCATTAATTCACAGGGAATAACTAAATATGGTGATGGTTTTGCCTTCTTCCTTGCCCCTGCAGGTTCAACAATTCCTTATAATACAACAAGAGGTGGAAGTCTTGGCCTTACAAGTAATAATGAGCGACTAAATTCATCGAGCAATCATTTTGTTGCTGTGGAGTTTGACACCTATCAGAACCCCTATGACCCAAAGGGTGATCATGTAGGTGTCGATATTAACTCTATGCAATCTGTTGTTAATGTGACTTGGTTTAGTAGCATTCCAAATGGTACGAAAACTGATGCCTGGATTAGTTATAATTCTAACTCAAAAAATCTTACTGTTGTCTTCACTGGTTTTCAACTCCAAGGGAATAATATCACAGTCACTGTCCTGCAGAACCTATCTTACAATCTTGATCTGAGGGAATATTTGCCAGAATGGGTCACTTTTGGCTTCACTGGTGGAGCGGGAAGGAGATTTGCATTTCAAAGAATCTATTCTTGGAAATTTACTTCTTCTTTAGATAATATGACAGATACAGGGGTAACCTTATCAAACACCTTGCCAAAGGATGCACCAACAAAAACTAAGTTAGGACTTGTGGTTGGATTGATTTCTGGTGGTTGTGTTTTGGTTGCAGTATGTGCTTTGGTGTTATTTGCATTTTGGAGAAAGAGTAAGGTCAGAGAAGATGATTATGTCTTTGATGGTTCCATGACTAATGAATTTCAAAAAAGTGCAGGACCAAAGAAGTTCTTGTACAGTGAGTTGGCTAGATGTACAAATAACTTTTCACAGGAAGAGATGCTTGGGCGGGGTGGATTCGGAGGTGTTTATAAAGGATATCTCAGTGAATCCAACTCTTATATTGCTGTTAAGAGGGTTTCAAGGGAATCACAGCAAGGAATAAAAGAGTATGTATCAGAAGTCAGGATCGTCAGCCGGTTAAGACATAAACATTTGGTGCAACTCATTGGTTGGTGCCATCAAAAAAGAGAACTTCTACTTGTCTATGAGTTTATGCCTAATGGAAGCTTAGATCACCATATTTTTAGTGGAAAAAGCCATTTGGCATGGCCACTAAGATTCAAGATTGCTCAAGGCTTGGCCTCGGCGTTGCTATATCTACACGAAGAATGGGAACAATGTGTGGTGCATAGGGATATAAAGTCTAGCAATATTATGTTGGATTCTAATTTCAATGCCAAGCTTGGGGATTTTGGGTTAGCTAGGCTAGTTGACCATGATAAGGGATCCCAAACAACAGTTTTGGCAGGCACAATGGGTTACATGGCTCCTGAATGTGTCACCACTGGCAAAGCTAACAAGGAAACAGACGTCTATAGCTTTGGTATTGTCGTGTTAGAAATAGGTTGTGGAAGGAAACCTATTGATCCTAAAGCAGAGGAGCATCGAGTAAACATTGTCGATTGGGTTTGGAGACTTTATGGGATGAGAAGTATTCTTGAAGCAGTTGATCCGAAACTATCATCAGAGTTCAATGAAGAGGAGATGGAGCATTTGCTAGTTGTTGGCTTATGGTGTGCTCATCCAGATAATAATTGCAGGCCTTCTATTAGGCAAGCAGTTCAGGTGCTTAATTTTGAAGCTCCATTACCCACGCTCCCTCGGAGCATGCCTGTGCCCACATATTGCAGCTTCTCACGCCATTTATCAACTATTTCAGTTTCATCAACACATGAAACCAGCGGTACAGAGATCGCCCTTGCCCAAATCCAGCATTCAGTAAACGGAAACACCACTGATGCTTCAATAAAGAGAACATCTTCTTCAGCATCTTCATCTTCAACATCAGTTTTGTATCCAAGTTAATATTGCCCTGTTATAACCCTGCATTTTGTCTTCCATTAGCAGTGTCCATCTTTTCCTTTCTTACTTTTTGTTGTTTCTCTTCTAATGAAGAGGATTCTTTGGAT

>PGSC0003DMG400047213 L-LecRK

ATGTTGCTATTTTTAGTAATTTTTGTTATCCCTTCTTTATATTCATTGAACTTCAATCTCACCAATATCAATCCTTCTTATGTTAATCTCTCTATCAATGTCACTGGAGACGCCTACGTAACGAAACAGGGCATACAAGTTACGCCTAATGAACGTAACATTACATTAAGTAGGAAAACAGGACGCGCCATGTACATTGAGACACTGCAGTTATGGGACAAAGCTACAGGGGAAATGACAGATTTTACTACACATTTCTCTTTTGTTATTGATTCAAATGGTAACGATCGTTTTGCTGATGGACTAGCCTTTTTCTTAGCTCCTGTTGGTTCGAGTATTCCAGTTGGTTCGTTTGGCAGTGGCCTTGGTCTTGTTGATGCTGAGACAGAAAACATGTCATCACATGAGTCATTTGTTGCTGTAGAGTTCGATACACATGTGAATAACTGGGACCCCTCAGACATACATGTAGGCATCAATATAAATTCTATGGAATCTGTTGCTACAACAATATGGAAAAATGACATTAAACTTGGGAAGAAAAATGATGCTTGGATTAGTTACAATGCTAATTCCAAGGTTCTTGAAGTTGTTTTCACAGGATTTGAGAAAAAATATAATCGAGACAAACTAAGCTACCCGGTTGATCTGAGGGATTATTTGCCTGAGAATGTTAGTTTTGGCTTCTCAGCATCAACTGGACAATTGTTTCAGAAAAACAATGTCAAGTCTTGGGATTTCAATTCAAGTTTGAGTTTTGATGCTATCAAAGTTCCACAACAGCCTAGTGCAAGTCCTCCTATACAAGCTCAAGATCCAAGAAATCATCCTATTCAAGGTCAAAATCCAAATGATCCTCCCATTACTCATGTTCAAGAACCCAAGGGTCCTCCTCAAGAAGTACTCAGCACCTCGGGAAAAGGAAGCAAGGGACTACTAGTCGGATCAAGCATAGGTTTACCTATTCTGATTCTTGGTTTGATAACTGCCAGTTGCTTTTTGTGGAAGAAAAAGAAGAAAGGAGATAAAAAATTAGACCATGTCTTCATTGATCTAGATATGGATGATGAATTTGAAAAGGGTACCGGTCCTAAGAAGTTCTCGTATGGTGAATTAGCTCGTGCAACAAACAACTTTGCTGAGGGACAGAAGCTTGGAGAAGGGGGATTTGGTGATGTTTATAAAGGTTTATTGAAGGAATGTAACTCATATGTTGCTGTTAAGAGAGTGTCAAAAGGGTCTAAACAAGGGATAATGGAGTATGCATCAGAAGTGAAGATCATCAGTCGATTAAGGCATAGAAATTTGGTTCAACTTATCGGTTGGTGCCACGAGAAAGAGAAGCTTCATCTTGTTTATGAATTGATGCCCAATGAAAGCCTAGATAAACATCTTTTCAAAGATAAGTCATTGTTGGTCTGGGAAATAAGGTGGAAAATTGCTCAAGCAATTGCCTCAGCTTTGCTGTATCTACACGAAGAGTGGGAACAATGCGTAGTCCATAGGGACATAAAAGCAAGTAATGTCATGTTGGATTCAAACTTCACCGCTAAATTAGGTGATTTTGGGTTGGCTAGGCTTATTGACCATGACAAAGGATCTCAAACAACAATGTTAGCAGGGACCGTGGGGTATATGGCACCGGAATGTATCATGAATGGAAAAGCTAGCAAGGAGTCAGATGTATACAGCTTTGGAATAGTAGCATTGGAAATAGCAAGTGGAAGAAGATCAATAAATATCAAAGCACCAGAAGATGAAGTGAGATTGGTTGAATGGGTGTGGAGCCTCTATGGAACGGGAGAGCTAGTCGAAGCAACTGATCCAAGATTGAATAAAATGTTTAATGAGAAAGAAATGGAACGTTTAATGGTCATCGGTCTATGGTGTGCTCATCCAGACAACAAACTCAGGCCATCAATAAGGCAAGCAATCCATGTTCTAAATTCTGAAGCCCAATTACCAAATCTTCCATCAAGAATGCCAGTTGCAACATATTCACCCCCTCCATTGAATATGTTTTCGTCTCCATTTTCAAATACCTATGAAGTTAATAAGCAGGAGCAGACTATGACTTACACTATTTCCTCAAGTGTCTATACTTCGTCAGCTGCTTCATCGACAAAATCACTTTTGTAA

>PGSC0003DMG401009415 L-LecRK

CTCCATTTTTCACTCATTGATCAATTATTTTATTTGCATTTACAAAAAAGTAAGACAAAGCCAACGTGCTTCAAAGTTTCTGCTCATTGATTCTTTCTTCTTCCCCTGTTTTCTTGCCCATCACCAAATCTACTTCCTCTGTAGCCACCACCATCTCTGAAGGGACTCACTGCTGATGAAAAATGGGATTGTTAACACCAAAAACACTCACCATTTTCATCATTCTCTTCTCTTTACTACACCTCAAACTTCAAGCTCAAAAAATGGAACATTACAACCTAGAATATGCATCTTTTGATCAAACTTTCTTTGACATCTTTGAAGTTGAAAGTCCTGCAACAATCAGTAACTTTGCTCTCCAAGTAACCCCTGATTCTGCTTCTTCAGATTTCAATCTCTACAATAACTCAGGAAGAATCCTCTTTAAGCGACCATTTAAGCTCTGGGATGGTGATGTTTTAGACAATAAAACAAGGGTTGCATCTTTCAACTCTTCTTTTCTTGTCAACATTTACAGACCAAAGAATGAAACAGCTGCTGAAGGGTTAACATTCTTGATTTCACCAGATTTGGAGAAGCCCGATAATAGCCAGGGCCAGTATTTAGGGTTAACAAATGCTACTACAGATGGGAAATCTAGTAACAAAGTTGTTGCTGTTGAGCTTGATACTTCTAAACAAAGCTTTGACCCTGATGGTAACCATATAGGTATTGATGTTCATAGTGTTAGATCTGTAAAGGTTGAATCTTTAACACCCCTTGGTATAGAACTTGCTCCTATGGGTGCAAGATTCTATAATGTTTGGGTACAATATGATGGAATCAAGAAAGTTCTTGATGTTTATATTGTTGAACAAGCACAGAAAAATGGGTCTACCCCACCTATACCAAAAACCCCCATTTTAACATATAATCTTGATTTGAAAGAACATGTAAATCAAGAATCATATTTTGGGTTTTCTGCATCAACTGGGACTAACTACCAACTCAACTGTGTATTGAGATGGAACTTAACAGTTGAGTACTTCCCAGAAAAGAAAAGTCCTTGGTTGAAGATTGTTTTAGGTGTTGGGATTCCTGTAGTGGTGCTTTTGATTCTTGCCGCGGCGTATTTAGGGTATTGTTATCATAAGAAAAGGATTGATAGGTCACAGTCTAACATATTGGGTGCACTCAAGAGCTTGCCTGGAACACCACAAGAGTTTCAGTTTAAGGCTTTGAAGAAAGCTACAAACAATTTTGATGAAAAGAATAAGCTTGGGCAAGGAGGATATGGAGTTGTTTACAGAGGATTTTTAGCTGGTGAGGAGAACAAAGATATTGCTGTCAAATGGTTTTCTAGGGAAAGCATCAAAGGGCAAGATGATTTCTTGGCTGAGCTTACTATTATCAATCGTCTTCGACATAAACATCTTGTCAAATTGCTTGGTAAGTGTTCTTAGTTTTTCTTCAAACATAGATTTTGTGCATTTGTTTGAGTAGAGTATTAGAGTTATATTATTGTGACTGTTTTGTTTATGATCTTCAATAACTGTTAGGGAAATGGTACTGGTAACTGGTAAAGTTGCTGCAATGTGATCGAAAGTTCATAGATTCGGGGTGTGGATATAGTCTCTTAGGCTGCGTAAAGTACACCCTTGTGGTCTAATCCTTCTCAGTATAGCTACAAATTTGAGTCTCAAATAAGGCCTTATCTCCAATATTTCAATTGAAAAGTAAACTATACAAATAGGGTAAAGATTAGTTTTTATTTTTGGGTATGTACATAAAATGTGTGAATACCTTCAAGGCAAAGGAAGTTGTAAGTGTAGAGTAAATTTTGCTCAAAATTAGTTGCATTTTCTTGAATTTCCCTTATAGAATTCTTTCCCCACCCCTCCTGCCATAATTAAGCAATGAGACAAGGATAATGGCGAGAGGATCCAAATTTAGCAATCAGAAACAGAAATTGTCCTCTATGTTTTTTGATTCTCAATGAAAGAGAGAATTCATTACCAGGATTCATTGATGTGACTTGAAAGAATGGAACCAAAAACCTTGTAAAAGTTACTGACATAATTTAAAGCCAAAGATTGGTGGGACATCAAATTTAATACTCCAATTTGTTTGTCATTTTGTCTTTCTTTTTTTTGGGTATAAAAAAGAAGGGAAAAGGTTCAAATATGCCATCGAACTTTTAGAAAAGGCTCATTTATGCCATCCGTTAAAAGTTTGACTCATTTATGTCATTTCTGTTAAAGAAAATGCTCATCCATGCCATTTTTTTTAACTCCGGTTTTGCAAAACCATTTTTTACACACCAAACATGGTTTAACACTTTTCTATTGGTGTTTTTGGATCAAATGTACATTTTTTGCTTGTTTTTCTTTAAAAAACACCAAGAACACATGAAGAACATAAACAACTCCAAAATTTTCAACGTCTTCAATTCTTCATGAATCACCTCAAATTTCAAGAGTAGTTTCGCTCCGAGCTAGATACCAAAACTCAATTTCTTTTGAGCTCGAATTCAACCTAACTCAACAAGACCCCAATTGAAATTTCTTCAAAGTTTCAAAACTTGAACATTCTTTAAATGGTAGATTCTCCAAAACTTCAAATCATTTGAAATTTTGAACATAAACTCAAAAAACACTAGTCAACAAAGATCTAATGTTAAAAACAAGAATCAAAGCAAATTTTGAAATTTTGGGACTGATTTTCGTTTTTTGGAATATATATTTTTAAATATGGGTGAATTTTTTGAACTCTACAAATGATTTTAAAATTTTTATTTTTTTAAAAAAAATAATTAATGGTGTAGCCGAAATTTGAGATGATTTCGTGAAGAATTGAAGACGTTGAAAATTTTTGGAGTTGTTTATGTTCTTCATGTGTTCTTAGGCGTTTTTAAAGTAAAAACAAAGCAAAAAATAGTACATTTGATCCAAAACAACCAATAGAAAAGTGTTAAACCATGTTGGTGTGTAAAAAATGGTTTTGCAAAACCGGCGTTAAAAAAAAATGGCATGGATGAGCTTTTCTTTAACAGCAATGGCATAAATGAGCCAAACTTTTAACGGATGGCATAAATGAGCCTTTTCTAAAAGTTCGATGGCATATTTGAGCCTTTTCCCAAAAAAGAATGGCTTTTACTATTTTATTTCAACTCCTTCTTTTAAACTTTACAACTAATATGTTTGAGACTATAAGATTGAAAAATCTTTTGATATACATATGTTTAAATTAAGACCATAAAAATCTAAAATCTTCTTTATTATCATAAACTCTGCAAGTCAAAAACAAGACAAACATATTGAAACGGAAGGAGTTTATGTAATATTATTATGTTGTTCGGACTCTCCAAAAATGTTGTTGCACATGTATTGAATTCTTAAAAAACAATACTACTTTTGAAGGATCCGATCGGTATCCATCACATATTTAGAGAGAGCGAAGCAACACTAATACAATAGACTCATTAACCGCGTGAGCACATGCCCTTGTCTTTTTATGCTTTTCTTTTTCTAGTAACTAGTAATTGATCTTATTCAACTAATTTATGTAATTTTAACAAGGGACTAGTCTTTCTATTCTTACATTTAGTATGCTAGGAATTTTGACATGGGACTAGTTTTTCTAGTCTGACATTTAGCGTGCTAGGAATGTACTAGGTTGGACCATCTTCTGACAAGCAAATTTCCCCTTCTGAAGAATCAAAGTCTTCTTGAACATTTAATAACTTGGTACTTCTTTTTGTCTTCAATCTCCTTTTCCTTTCTTAGAATTCTCAAGTCATCCCAAGGGTTTGTCAAAGTTTCTTTTACCCTGACTAACTAGATTTGCGTAGCATAGGGCCCAACAAAGTGAGAAACGCTATATATCAAGACCATTTTTTAATGCTTGAAGTCGAGATCTCTAGTTAAGGAATCACACTCCCCGTTAGGCGATAGTAAATGTTTTAATAGTTGATTCATACCATTTGGTAAGTAAGAGTTTAAGCCAAAAATATCAAGGAAGTAAGAAAAACTAATGGAATGCCTTAAAGTTTTGACCAAGAACCAACTTGTTACTATGGTCCCTCCAATTTAGAAAAACAATGCACTATCTTGAAGCCCTTCATTTTGACTTGCTTAAAAGAAGATGATCATCCTTTGCTGACCATGGAATAAGCAGCCATTCGTCAAAGTTCAAAACAATTTTTTATGTCAAAATTTTGGGAAGTTTCCTTCTTGTCCTAACATCCCTTTCATTCATTGTCAACGATTAAGACAAGTTTTGGATGTTAGTTGAATGATGGTAAATTAAATCCAATGAAAAGGACATTAATTCTGTATTTTAGTATACATTCATCTTCAAACACCTAATTTTTGTGTCACCTACTCTTGTCATCTGACCCTACAAAGAGATAAGGTTTGTAAAATACAAAAAAAAGTTTACCATTTGGGTATTCTGCAACTTCTGTGTCTAGCTGTTTTTACTGATAAAAAACAAACATATGCATCTTACAAAATTACATGTTATGGTGACAGGATGGTGTCACAAGAATGGAAAGTTACTGCTTGTATATGAATACATGCCTAATGGCAGTCTAGATATGCACCTCTTTGCGGGACCAGATAAAGAGTCGTTGAGCTGGAACGTCCGCTACAAGATTGTTCAAGGTGTTGCCTCAGCATTGCACTATCTGCACAATGAGTATGAGCAGAGGGTGGTCCATCGCGACCTGAAGGCGAGTAACATCATGCTTGACTCAAAGTTCAATGCGCGCCTTGGGGACTTTGGCCTTGCACGGGCACTTGACAACGAAAGGACCTCGTATGCTGAGGCAGAAGGAGTGCTTGGCACAATGGGATACATTGCACCAGAGTGTTTCCACACAGGGAAAGCCACACAGCAATCTGATGTATACGCGTTTGGGGCAGTGTTGTTGGAAGTTGTATGTGGCCAGAGACCTGGAACCAAGATTAATGGCTTTCAATTCTTTGTTGATTGGGTTTGGTACTTGCACCGCGATGGCCGTATCTTGGAAGCTGTTGATCCAAGGCTTGGAGGTGACTATGTTGCTGATGAAGCAAAGAAACTTCTACTGCTTGGTTTGGCTTGTGCACATCCTATAGCTATTGACAGGCCTAAAACACAGGCAATAGTTCAGATTATATCAGGGTCAGCACCAGCACCAGAAGTTCCACCATTCAAGCCAGCATTTGTGTGGCCATCAATGATGCCAATTGACATAGACTCAAGTGTTATGGACACAACATCCATTACTACTTCTCACTTCAATTCAGGATGGAGTCTTGACTATCAAAGCAGGGAGACCCCAACATATGCAGACCACTCTTTGGTGTAGTTACTTTGATGGTAAACAACTATTACCAAAAGGAAAAATGATTTTTTTGGCCTTCTATTTTTTAAAAAATTTCCTGTAATTATGTTTTCTTGGGAGGAGGTTAAAAATTTCCTTCAGTTTTTTATTTTTTGTTTGATGTTTTTTTCCCTTATCAATTTGTTATTGGGGCATGATGAAAAGATCAGAGCAACCCGTGTACATTCATATAACTCAGTAGTGTTACATTATGTCATATTTACTGTTTTCCTGTCTTTGTCAACCAAAATTATGTAAAAGACACTCATGTTGAGCCAAATTGGAGGAGACAGAAAATGAATAGTGGCGAAGTATCTTTGTTGCCTCAATTGTACTACTACATACAACTTAAAGAATTCCAAGATCTCAAATTGTAATGAAAGATAAGAACTGATCAATTCTTATAAAAGCAATAGATAACAGAAGAACACAATACAAAGATGGATCATAGATGCTTTGTTGATATGCATAAATGCTCTGCCATTTTATAGGCTATTGTAGTATTAGATCGTTGACAATATATAGTAGATGAAATCCCACTAAGATTTATCTACTGTCAAGATCATTTCTGTAAATCAATATAACGGAGAACCTGAGATGCTTGCCAAAGTTTTACAAGTTAGTGCGCATAAGATCAAGAGTTTCCCCCGCCTTTAACTTTGTTGATAAACGGAGCTTCCTATAATCATTGAAAGTGAAAGGAACATAGATACGGGGGTGTTCAGCATCCACCAGCTCAGGTGGTGGTTCCACTGCTGTGTCCCTTGGTCCCAAGAGGAATGAAGCAATAGACACCCGTACTTTTGCTTCCTTGCATATCACTCTATGCTTGACAGTGTGTAATCTTCCATTGCTCCATACCTTCACATAGGCCATAGCAAAAAGCAAGTATCATTGTCTAGTTCTTTTTTCGGAATCTTAGAAGGGTTAAGGGAGTAGGAATTATCTAACTTCCTGCTAAACAATCCCCGCCAGAGGCTAGCATAAGATTGTTGGAACACGTCAAATCTTGAAAATGAACTAGGTAGAGTAGCAGCATAGTTACCGTGCCCATGTCTCCAAGATTCACCAGGAGGGTGTTTGGCCAAGGATCAACTGCTACAAATTCACCAGACTTCTTCATGACCTCAAGACCGCCAACACTTTCATCATCCTGTAGAATAGTAAGGAATCCTGAGTCTGTGTGTGTCTGAACCCCAGAGGAACCTACAGCTTCTGGGGTGAAGTGGTACTTGTTAATCCTAAACTGACAAGGCCAATCCTCAAAGGAAACATTTGGCAGTCCCAACCCTTCACTCATCTTTCGCAGAATGTCCATCATTAGTCCATTAGTTGCTGCAGCATACGTCTTGATTGTCTCTCTGTTCACAGTAGGAATCAAACTGCTTTTAAGCATTCTAGTAAAGCAGTCTTTT

>PGSC0003DMG402009415 L-LecRK

TGCAACTTCTGTGTCTAGCTGTTTTTACTGATAAAAAACAAACATATGCATCTTACAAAATTACATGTTATGGTGACAGGATGGTGTCACAAGAATGGAAAGTTACTGCTTGTATATGAATACATGCCTAATGGCAGTCTAGATATGCACCTCTTTGCGGGACCAGATAAAGAGCCGTTGAGCTGGAACCTCCGCTACAAGATTGTGCAAGGTGTTGCCTCAGCATTGCACTATCTGCACAATGAGTATGAGCAGAGGGTGGTCCATCGCGACCTGAAGGCGAGTAACATCATGCTTGACTCAAAGTTCAATGCGCGCCTTGGGGACTTTGGCCTTGCACGGGCACTTGACAACGAAAGGACCTCGTATGCTGAGGCAGAAGGAGTGCTTGGCACAATGGGATACATTGCACCAGAGTGTTTCCACACAGGGAAAGCCACACAGCAATCTGATGTATACGCGTTTGGGGCAGTGTTGTTGGAAGTTGTATGTGGCAAGAGACCTGGAACCAAGATTAATGGCTTTCAATTCTTTGTTGATTGGGTTTGGTACTTGCACCGCGATGGCCGTATCTTGGAAGCTGTTGATCCAAGGCTTGGAGGTGACTATGTTGCTGATGAAGCAAAGAAACTTCTACTGCTTGGTTTGGCTTGTGCACATCCTATAGCTATTGACAGGCCTAAAACACAGGCAATAGTTCAGATTATATCAGGGTCAGCACCAGCACCAGAAGTTCCACCATTCAAGCCAGCATTTGTGTGGCCATCAATGATGCCAATTGACATAGACTCAAGTGTTATGGACACAACATCCATTACTACTTCTCACTTCAATTCAGGATGGAGTCTTGACTATCAAAGCAGGGAGACCCCAACATATGCAGACCACTCTTTGGTGTAGTTACTTTGATGGTAAACAACTATTACCAAAAGGAAAAATGATTTTTTTGGCCTTCTATTT

>PGSC0003DMG400000295 LRR-RLKs

TTTTTACCAAATGTTCATAGAAACTTAGAAAAAATCACCAAAAAAAATGTCCAAAATTATTTTTTCACTTTTCATTTTTCTGCTTTTCATCATTTCTCATGGAAAATCCCAAGAATCTCCAAATCAAGAAAAGGCCATTTTACTCCAACTAAAACAACACTGGTTTACTTCACCAAATGTCACAAAATGGATTTCATCATCAAATCATTGTTCTTGGGATGGAATAATTTGTACCCAAAATTTAGTTTCTGGTATTCAAATCCCTTTTGGAAATATCTCAAAACCAATCCCAAATTTCATTTGTGATCTTAAAAATCTCACCTTTCTTGATTTGAACCATAATTTCATCCCTGGAAACTTCCCTGATATCTACAATTGTTCAAATCTTGAATTCTTAGACCTTTCTTATAACTACATGGCTGGTAATCTCCCTGATGAGATTAACCGTCTTTCGAGTAATCTAAAATACCTAAACTTAACTTCCAACAACTTCAATGGTGATATCCCTAAAGGAATTGGTGGATTGAGTCAGCTCAAAGTACTTGACCTGAGGGGGAATTTGTTTAATGGATCTTTTCCTGAAGAGATTGGTGAATTGTTGAATCTTGAAGCACTTATGTTGAGCTCGAATGGTTTTGCTCCACAGGCTATACCATCACGATTCACCAAGTTGAAAAATTTGAAAATTTTTTGGATGACAGAAGCACATTTGATTGGAAATATCCCGGAATATATTGGTAACATGACAAGTCTTGAATTTTTGGATTTGTCGAACAATGGATTGAGTGGTAGTATCCCTGATGGTTTGTTTCAGCTCAAGAACTTGAGTATTGTGTATTTATTTACTAACAAATTGTCAGGGGAAATACCACAGTCAGTTTCATCAATGAATTTAGATGTTGTTGATTTGTGTAACAACAGTTTGACAGGAAAAATACCAGAAGATTTCGGAAAGTTGACGAAACTGAGTGGATTGGCTTTGTTTTTCAATCAATTATCAGGGGAAATACCTTTGAGTATAGGCAAATTATCATCATTGGTATCTGTTAAGTTGTTTGGGAACAAGTTATCAGGTGAAATTCCACCTGATTTTGGTCGATTTTCGAAACTTTTTGACTTCCAAGTTTCAGAAAATCAACTTGTTGGAAAGCTACCTGAGGGTATTTGCAATAACAAGGCATTAGCTAGGATGGTTGCTTTTGGCAACAATCTAACAGGTGAACTGCCAAGTTCACTTGGAAGTTGTGACAGTCTGAGGTATCTTCGAGTCGAAAATAACCGACTTTCTGGTGAGGTTCCTGATGGATTATGGACAGGTAAGAGTTTGTCAATGGTTTTGATGAAGAATAACTTGTTAACTGGTCAGCTTCCACATAGAGTGGCATCAAATTTGTCACAAGTTGATATCAGCAATAACAAGTTTTCGGGGGAATTACCAGCTGGTATGGGTACTTGGTACAATCTAAGTGAGTTCAAGGCAAGTAATAATCTCTTAAGTGGTCAAATCCCTCAAGAATTGACTCTTCTTCCAGGGATAACTAAACTTTTTCTTGATGGTAATCTACTATCTGGAAATTTTCCATCGAATATATCGTCGTGGAAGACTCTTGTCACATTAAATAGCAGAAAAAATCAGCTTTCAGGTCCTATACCTTCTGCACTTGGCCTTTTACCAAATCTCATTGATTTGGACTTGTCAAGTAACCAATTTTCAGGTGTTATTCCAACTGAATTAGGTAACTTGAGGTTTACTTCACTTAACCTGTCATCCAATCGCCTGTCGGGTGAAATCCCATCTCAGTTGGAAAATGCAGGTTTTGGTAAGAGCTTCTTGGATAATCCTGGTCTTTGTGCAATTAATCCATCAGTAGAAGTCGCGTCTTGCAAGAGGGAAACTAAGTCAGATAAGTTTCCTGTTGGACTTGTTGCTGCTCTTGCTAGTGTAGCAGCAGTTACTTTCCTTGTGGCAGTTCTGTATGGTCTGTTTGTGTTGAGAAGTCATAGAAAAAGAAAACAAGAATCAGTTTTGACATGGAAACAAACATCATTTCACAAATTAGACTTCACGGAATCAGACATTGTATCCAATCTGACCGAAAACAACATAATTGGAAGTGGAGGATCGGGACAGGTCTACCTCGTGCCTTTAAGCCGATCAGGTGACTATGTTGCTGTCAAGAGGATTTGGAGAAACCAAAGGTTGGATCACAAGCATGAGAAACAGTTTCTTGCTGAAGTTCAGATATTAGGCACGATTCGACACACCAATATAGTGAAACTCCTCTGCTGCATCTTCAGTGAAGAGTCGAAACTTCTTGTCTACGAATATATGGAGAACAGGAGCTTGGATATATGGCTTCACTCAAAGAATAGGTTGAACAATGCCTCAAGATCAACACCACATCTGGTCTTGGAATGGCCTAGGAGGCTGCAAATCGCGATAGGAGCTGCCCGTGGTATTTGCTACATGCACCATGACTGCTCACCACCTATTATTCATCGCGATGTGAAGTCAAGCAACATCCTATTGGATTCCCAATTCAATGCAAAAATTGCAGATTTTGGCCTTGCCAGGATGTTGCTCAAGCCTGGAGATAACACAGTGACAGCAGTAGCTGGCTCTTTCGGATACATTGCACCAGGTATAAGGCCATCAACTTATCTATAACATCATTATAAGTTTAGAAGTTGGTTTAATCTCTTACATGGAAAAAAGTTTCCTATGCTACACAATTCAATAACATATCAAGGGGTCGTTTTGTTCACGTACTAGCTATGCATGAGTTGTAACACGGGTTGTCTATGTAGAGATTGTTATGCAAGTAGTTATTCTTAGTACACTAATTCTCGCGTAACTCATATAAATGCTAATATATTCATGATGGTATTTCAGAGTATGCACGAAAAACTAGAGTGACAGAGAAAATTGATGTGTATAGTTTTGGAGTCATACTATTGGAACTGGTGACTGGAAAAGAAGCCAATCTTGGAGATGAAGATTCATGTCTTGCAGACTGGGCATGGCGTCACCTTCAAAAAGGGAAACCGATGGCTGATGCATTAGATGAGGACATCAAAGAAACGCGATACTTGGAAGAAATCTTCATTGTGTTTAAGCTTGGAATCTTTTGCACCAGCACATTTCCTTCATCAAGGCCAACAATGAAGGAAGCTCTGCAAATCTTGATCCAATGTAGCAACAGTTCACCGACATCTGGAGAAAAGAAAAATGAAACAGAACAGGATGTTTTGCCACTCCTCAAGAACTCAAGGAGTGACAGGATTGAAGAAAATGATGATGTTGGTTTAACATCACTTGTTTGATGGCATTACAATATGATTAGAGTGACGGAGAGCCTTGACGTAACTGGTAAAGTTGTTGTGGAAATAGTCTCTTGCAGAAATGCAGGGTAAGGTTATGTACAATAGATTCTTGTGGTCCGGTCCTTCCCAGGACCTTGCTCATAGTGAAACTTCGTGCACCGGGATGCCTTTTTTTAACAATATGATTAGAGTAAGAAATAGCAAAGAATAAGACCAATAGATAATGTATCAGGTAGTGAAAACTAGCATTGTACATACACTGTATTGGATCCATAGTGTTGCTTTGCATTGTTTTGCTACATAAATTTTGCTTTTCCAATGTTTTCTTTACTCATAAAATAACTGTCAAAAAGGTACATTAAAACCATAAACTATTGAATGGAGGATAGATTTCCTAAAAATCCAGAAAAAATAGAAACAAGTTTGAACTACCTCAGTACTTTAACTTTTGGCAACCATACAACAACATATCCGATGTAATCCCATGAGTCCCAAGCTTTTGGCAACAATACAACAACATATCTGATTTAATCCCACAAGTGGCTCTGGGGAGGGTAGAGTGTACGCAAATCCTACCCATATCTCGTAGAGATAGAGAGGTTGTTTCCGATAGTAACTTTTTGGCAACAATAAATCTTGTAAAAGATCACACAAAAATGTGGCTACTAGAAGAACTTTATTTGTATCCTAGCTTTATTTCTGGCTAGTGAGGAAAAAATTTAAGTTTACACTTGGGATAAAGATGATCCTTAGAGGCACCAGGAGTGAAAAGGTCAGCATATTGATGTTTAGCTGAAGAGAGATGTGCTCATTTGATACATTGAACTATGGCTCACTTCAAAGCTCACTCTTGCATGATGCGGTGCACAAGGAAGCAAGAAATATTTCAATCATCTCGGAGAAGAAAGTAAGAAGAGGGTTCTCTTTAGCAAGAAGAGGGGTGATTGTAAACGTACAAACTTTGACTAGGCACTTTTGGTTGTGTCACTCTGACCATGATCATGATTCCTCGAGATCTGTGTATTGGGAATGCGAGGCAGCCAAAAGTGCAGCTCCAACACCTGAACCATCATTAGAAAGCTCAATGACTATGCTGTCCGCAGTGTCTCCCAGTAACTCTTTCATAGTTTCCTCCAAGCAATTTCTGAACTTGGTGTAACGCTCAAACAATGCACCGTCCACAGCTATGACAGACCTCTGGTTCTCTCCGTCCTTCAAAGTGTCTTTTCCCAATTTCTTGAGGATGCCCACGATTCCTGCTGCAGAAAGACGAGCTCCACGAGAGGTGATAATGTCGCATAGCTCCACAACTATTTTCCTCATTTTCAAGGTAGAATTAGGTACCTGTGAAAAAAATGTATATATAGCAAATAGGATGAGAAAGACGTTCTCCACTTAATTAAGCCTAGCCAACTTAGTGATGTTCAGCCAGGAGCTAGGAATTTGAGTTTATGGGTTCCGGATAAATTATTTACACATATTAAGTGGGGTTTTCAGTACAAATACAAAGCTACTGGTTTTTTGTCCTATCATATATAGGACAAAAAAGATGAGAACTTCAAAGACTAGTAATGTTTCACTCTAAGTCAAATGTTTGATCTTAATACCTCTAAGATATCCTTCAGTTTATTGCCAACCACCTTGAGATCAGCCGACTCGTCGTGATGCATAGCAGCCATATCCGGAGTCCTGCATGTGTGGCAGTAAATAACCTTTTTCACAATATTTGACAATTCAAAGCTTACATATGCTTAATAAACGTACACGAATCTAGTCCAACAAGAAGCATAACTTACAGTGTAAGAGATTAGTGCAATTACATAAACCTTAAGTTGCACTAATTGCATGGAGAAGAATAAGGTTAAAATTGATGAATGCAAATAGGGCCTGCCAAAAAATAAATTACTGCCGTAATAAATGTGTTGGTCAAAGCAATACCCTACCTCAATATGAAAGGAATTTTCAGTTTGGATGGGACATAATCACCGAATAATGAAGCTTCTTTAGCCATTCTACACAATACTCTACGCAAAATTTCTCCAAGATACATCCCGGAGATCATCTTTTCATAAATCTGTGCAGTTTCCGAAAGTTCAACAACAGATGTAAGATAGAAAATGTAGAAAGGAAGGGGATTGTCTTAGTTACCTGCTCCCCGGGGTTTAAACTCTCAACATCAAGATTTTGGTCATATTCTGTTACTGGAAGATGTGATGAGAGGAAATTACCCCATTCCATGTTGATAACCTTTAAAGACCAGTCAAAATTACGTTTAAGTAAACAAACGTGTATGAATCTACCAAGGAAAATATTTTTAGGAAATGAATTAGGCACATATCAAGTCATGAAGGTGTTATGAAATGTCATAAGCATCTACCATCTCTCCGGATTTCGGCAACAGACCATGCCATTTGGGAATCGCATGAGCCCGTTCAACATATGCTGCATTGGTTCCAGTACCCAAAATTACTGCAGCAATGACATCAGGATTATTGTACCGACCTCCTGCTAACGTTCCAACAGTATCATTGACCTAAACAGCCAAGTCAACTAAGAAATTCAGGCTCAGGAATTGCTTAAGATTAAAAATCAGAAATATCACACATACAAAAACCTAATACATGAAAATCATCAGCTTACTTATATTTTATCGAATTTGATTGGGAAAAAAACTTACTAAAGCAGCCACACGCACATCAAGGCCGGCCCTTTCCATTGCTTTTGCTAATTCTCCCACCACATCTTGGCCAACCTATGTTAGATTCCATGAGGAGTAGTTCAGAATTGAAATACAATAAAACGTTACAAAAGGAACTCTTTATAACGTACTTTAAATCAAATCGGTGAAAATGCAATGAAACAATGATCAACTTTGCTTGTATAAAGTTGTATGCATTAAATCCTCAACAATGCCAACCTATGTTAGGCCATGAGGAGTAGTTCATCATTGAAATACAACATAACGTTGGAAAAGGAAATCCTACAACGTATTTTAAATCAACTCAGTGAAAAAGCAATGAAATATACGTTGTCGGACTCTCCAATAATATTGCCGCACCTGTGTCAGATCCTTCAAAATGTACTGCTTTTGAAGGATCCTACAAGCACCTGGCCAGTGGCGAAATTTTTGAAGAGTCTGAGCAACATAGATGGTATAATCATCAACGTTGCATGCATAAAGTTGTATGCATTAAATCATCATCAATGCAAAACCATACATTGAAATGAATGGACTAAATCTAAGGTTGATTGTAGGCATGATCAATCGGAAGTGAAAATCAGAAACTCTGTATGACTAACCACGTCTTCTATGGAGAAGCCCTTTGTCCATTTAATAAGAGTTCCTGATGCAATTGACAATTGTTTCACAGGGAAAGAGAAGGTAAAGCCTAATTCCCTTTGTCTACCAGGTGGAAGATGAAAATCATCACCTTCTGTAGCTACAAATTCTGCAAGTGCCGTGGCAATAAAATCAAATAACACCTGCAATTAACAGTTTGAAGTTTAATAATGTGAGTACAACCTTGATTGTATTAAAAAAGTTGCAGGATGCATGGCTGATACAGCTTAAGACTTACTTCAGATGATGATCCAGCCATCACATTCTGTGGAATTGAAACTTCTTTAACTTCATGTTTAACTATACGCTTTTCGTTCCCACCCAACTGCACACGCATCACACGAAAGTTTGTGCCGCCTAGATCCAATGCATAAAACAGACCTTCTTCGTCCCTAAGATGAAGGAATAAAAGTAATACCAGTCGTTAAACTGAAGATACAAACATACATGGAAAACATACCCCACCCCTAATGCACTACAGAGGTCTATTGGCAGCAGCTTTTGAAAATTTGTCGATACATGTGAATTTTTATACCCAATTTTAGTAAGTTTATCACAAGAACATGGAAATGTTACCAAGTCCCTACCTCCGCCTACTACTATAGGTATGCCATTTTTAAACTTTAGAGAATGTCAAACTGATTCATATGTCATATATGCTTTTCTTATGGATCTAACTTCATTATGTTGAACACAGAGATGTCTGTGTAGACAGCACAGTAACAAATAATGAGCAGACAGAAGAAGATGGAACTGAAAAATTTACACACATTATAAGTGTTAATGAATGGAACTAGCAATGTAAAATTCAAAAAAGGTGGTCAAGATACAAAACAGTAGAAGAGAATATAGAATAAACATCAAACTCAGGATCCATAGATAACAAGTGACCTGATCACTTGTCAGTTAGAAACTTCTTAGTGGACTCCAAGTTCAACAAATACTTCAAATAATAGAGTTTCATTTTTGCTTTTTGTGATTTATAACGATGCTCAAGGTAATCACCTTAAGAACATCAAAAGAATCATAATTTAGGACTAAAAAATAATAATTAAAAACAATTTGAAAAGTACTTAATCAATTATCCAAACATTACGAAAGTGAAATCAATCTCCATGATCACAGCTTTATTCAAAAAAGAGAAATTCTTGACACCCAAATTAGCAAAAATCAGTTTGTGCACATATACACACACAAATACTCATCCAATGTGGAGGCTATCAACTTAGCTAATAAGCATCTTAAGTTTGCCAATGAATCGTTAGTAAAACACACAATCTGATATAACTCTCCACCCCATAAGCTAAAATCAAGAGACAACACTACCAAGATCACAGCTTTATTGAAAAAAGAGAAATTCTTGACACCCAAATGAGCAAAAATCAGTTTATACACACAAAAATACTCGCCCAGTCGGATGGTTATCAACTTAGGTAATAACCATCTTGAGTATGTCGATGAATCAATTGTAAAAACACAAAAAAGAACTGACATAACTCCAACCCAGAAGCTAAAATAAAGAGACCACCTCCAACCGAGGTGGAGCTAGAATTTGATGTTTATTGGTTCTGAACTTGTTGAGAAAAAACTATTAGGTTCATCCGAACCTGTATCTAATATGTAGCTCCACCCCTGCCTCCAAAATCACAACTTTACTAAAAAAGAGACAAATTCTTGAACACCCAAATAAGCAAAAGACAGTTTATGCACATTCTTAAACACCCAAATTAGCAAAAAACAGTTTATCCACATTCATAAACACCCAAATAAGCATAAAAACAGTTTATCCACATTCTCAAACACCCATATAAGCCAAAAAAAAAACAGTTTATGCACATTCTTGAACACCCAAATAAGCCAAAAACAGTTTATCCACACTCTTAAACACCCAAATAAGTAAAAAACAGCTTTTCTTAAACACCCAAATAAGGCACTCAAGGGTGTGGCCTAGTAGTCAACAACGTGGATTGAGCACCATGAAGTCTCAGGCTCAATTCCCAGCTCCTATCTATTCTAGCCTTGGTGGACTCAGTGGCGAATTTATGTGTAAAATTCGGGTCACCCGAACCCGTGCTCCTCCGACTAAACTTGATATATTACTTGTATAATTATTAAAGAACATATTAAATAAACACTTGTTGAACCCGTTCTTAAAGTTGCTTGATGGTTTACTGGTTTGAGAGCTAGATTTTAACACCAAGGTCCTGAGTTCAATCCCAGCCAGCCACAATTCACTTCTAATTTTTTTAGTAACTAATAACACTAGTTTTTACAAAAAAAAACGAAATTCTAAAGCAATGGTGCACCCGTTTTCTTGAAATCCTGGATTCGCCTCTGTGGACTGAATACTGATACTTATTGTTGGTGTGAGATGCCAAGATTTAGTTGAGGTGCGCGCAAGTTGATCCGGATATCAAAACAAAGAACACCCAAATAAGCAAAAAAAAAACAGTCTATACACATTCTTGAACACCCAAATAAGCAAAAATCAGTTTCTTCACACACACAAAACACAAATACTCACCCAGTAGGAAGGTTATCAACATAGCTAATAAGCATCTTGAGCTTACTCCCACCTTCAGAAGCAAGTCCAGCATGCATCTCTACAGTCATAGCATCAGCTACCTGCTTAAGTTTCCCCATTGGAGTTGCACACTTCTCCTCAAAATCCTTCAATAAAGCTTCAACTTTTCCCCACTCCCCAGAATTCTTCATCCTGCGCTTCATTAACAAAACTGCCACCCCACATACTACAGCAGCAGTACACACAACTGATACACCTACAACCAATTTTCCCATATTTTCTTGAAAAAAAATCAAACTTTTTGTGTATATTTGAGTGAATATACTGTAAAGATTGAAACTTTTTTGGTTGATTTTAAATGGGGTTTGTTAAAGATTCAGAATTTGATGTGTGGAGTGACAGAGAAAGAAGAAAAATGGGAGTTGCAGAGGATATTTTATT

>PGSC0003DMG400000358 LRR-RLKs

TTTTTACCAAATGTTCATAGAAACTTAGAAAAAATCACCAAAAAAAATGTCCAAAATTATTTTTTCACTTTTCATTTTTCTGCTTTTCATCATTTCTCATGGAAAATCCCAAGAATCTCCAAATCAAGAAAAGGCCATTTTACTCCAACTAAAACAACACTGGTTTACTTCACCAAATGTCACAAAATGGATTTCATCATCAAATCATTGTTCTTGGGATGGAATAATTTGTACCCAAAATTTAGTTTCTGGTATTCAAATCCCTTTTGGAAATATCTCAAAACCAATCCCAAATTTCATTTGTGATCTTAAAAATCTCACCTTTCTTGATTTGAACCATAATTTCATCCCTGGAAACTTCCCTGATATCTACAATTGTTCAAATCTTGAATTCTTAGACCTTTCTTATAACTACATGGCTGGTAATCTCCCTGATGAGATTAACCGTCTTTCGAGTAATCTAAAATACCTAAACTTAACTTCCAACAACTTCAATGGTGATATCCCTAAAGGAATTGGTGGATTGAGTCAGCTCAAAGTACTTGACCTGAGGGGGAATTTGTTTAATGGATCTTTTCCTGAAGAGATTGGTGAATTGTTGAATCTTGAAGCACTTATGTTGAGCTCGAATGGTTTTGCTCCACAGGCTATACCATCACGATTCACCAAGTTGAAAAATTTGAAAATTTTTTGGATGACAGAAGCACATTTGATTGGAAATATCCCGGAATATATTGGTAACATGACAAGTCTTGAATTTTTGGATTTGTCGAACAATGGATTGAGTGGTAGTATCCCTGATGGTTTGTTTCAGCTCAAGAACTTGAGTATTGTGTATTTATTTACTAACAAATTGTCAGGGGAAATACCACAGTCAGTTTCATCAATGAATTTAGATGTTGTTGATTTGTGTAACAACAGTTTGACAGGAAAAATACCAGAAGATTTCGGAAAGTTGACGAAACTGAGTGGATTGGCTTTGTTTTTCAATCAATTATCAGGGGAAATACCTTTGAGTATAGGCAAATTATCATCATTGGTATCTGTTAAGTTGTTTGGGAACAAGTTATCAGGTGAAATTCCACCTGATTTTGGTCGATTTTCGAAACTTTTTGACTTCCAAGTTTCAGAAAATCAACTTGTTGGAAAGCTACCTGAGGGTATTTGCAATAACAAGGCATTAGCTAGGATGGTTGCTTTTGGCAACAATCTAACAGGTGAACTGCCAAGTTCACTTGGAAGTTGTGACAGTCTGAGGTATCTTCGAGTCGAAAATAACCGACTTTCTGGTGAGGTTCCTGATGGATTATGGACAGGTAAGAGTTTGTCAATGGTTTTGATGAAGAATAACTTGTTAACTGGTCAGCTTCCACATAGAGTGGCATCAAATTTGTCACAAGTTGATATCAGCAATAACAAGTTTTCGGGGGAATTACCAGCTGGTATGGGTACTTGGTACAATCTAAGTGAGTTCAAGGCAAGTAATAATCTCTTAAGTGGTCAAATCCCTCAAGAATTGACTCTTCTTCCAGGGATAACTAAACTTTTTCTTGATGGTAATCTACTATCTGGAAATTTTCCATCGAATATATCGTCGTGGAAGACTCTTGTCACATTAAATAGCAGAAAAAATCAGCTTTCAGGTCCTATACCTTCTGCACTTGGCCTTTTACCAAATCTCATTGATTTGGACTTGTCAAGTAACCAATTTTCAGGTGTTATTCCAACTGAATTAGGTAACTTGAGGTTTACTTCACTTAACCTGTCATCCAATCGCCTGTCGGGTGAAATCCCATCTCAGTTGGAAAATGCAGGTTTTGGTAAGAGCTTCTTGGATAATCCTGGTCTTTGTGCAATTAATCCATCAGTAGAAGTCGCGTCTTGCAAGAGGGAAACTAAGTCAGATAAGTTTCCTGTTGGACTTGTTGCTGCTCTTGCTAGTGTAGCAGCAGTTACTTTCCTTGTGGCAGTTCTGTATGGTCTGTTTGTGTTGAGAAGTCATAGAAAAAGAAAACAAGAATCAGTTTTGACATGGAAACAAACATCATTTCACAAATTAGACTTCACGGAATCAGACATTGTATCCAATCTGACCGAAAACAACATAATTGGAAGTGGAGGATCGGGACAGGTCTACCTCGTGCCTTTAAGCCGATCAGGTGACTATGTTGCTGTCAAGAGGATTTGGAGAAACCAAAGGTTGGATCACAAGCATGAGAAACAGTTTCTTGCTGAAGTTCAGATATTAGGCACGATTCGACACACCAATATAGTGAAACTCCTCTGCTGCATCTTCAGTGAAGAGTCGAAACTTCTTGTCTACGAATATATGGAGAACAGGAGCTTGGATATATGGCTTCACTCAAAGAATAGGTTGAACAATGCCTCAAGATCAACACCACATCTGGTCTTGGAATGGCCTAGGAGGCTGCAAATCGCGATAGGAGCTGCCCGTGGTATTTGCTACATGCACCATGACTGCTCACCACCTATTATTCATCGCGATGTGAAGTCAAGCAACATCCTATTGGATTCCCAATTCAATGCAAAAATTGCAGATTTTGGCCTTGCCAGGATGTTGCTCAAGCCTGGAGATAACACAGTGACAGCAGTAGCTGGCTCTTTCGGATACATTGCACCAGGTATAAGGCCATCAACTTATCTATAACATCATTATAAGTTTAGAAGTTGGTTTAATCTCTTACATGGAAAAAAGTTTCCTATGCTACACAATTCAATAACATATCAAGGGGTCGTTTTGTTCACGTACTAGCTATGCATGAGTTGTAACACGGGTTGTCTATGTAGAGATTGTTATGCAAGTAGTTATTCTTAGTACACTAATTCTCGCGTAACTCATATAAATGCTAATATATTCATGATGGTATTTCAGAGTATGCACGAAAAACTAGAGTGACAGAGAAAATTGATGTGTATAGTTTTGGAGTCATACTATTGGAACTGGTGACTGGAAAAGAAGCCAATCTTGGAGATGAAGATTCATGTCTTGCAGACTGGGCATGGCGTCACCTTCAAAAAGGGAAACCGATGGCTGATGCATTAGATGAGGACATCAAAGAAACGCGATACTTGGAAGAAATCTTCATTGTGTTTAAGCTTGGAATCTTTTGCACCAGCACATTTCCTTCATCAAGGCCAACAATGAAGGAAGCTCTGCAAATCTTGATCCAATGTAGCAACAGTTCACCGACATCTGGAGAAAAGAAAAATGAAACAGAACAGGATGTTTTGCCACTCCTCAAGAACTCAAGGAGTGACAGGATTGAAGAAAATGATGATGTTGGTTTAACATCACTTGTTTGATGGCATTACAATATGATTAGAGTGACGGAGAGCCTTGACGTAACTGGTAAAGTTGTTGTGGAAATAGTCTCTTGCAGAAATGCAGGGTAAGGTTATGTACAATAGATTCTTGTGGTCCGGTCCTTCCCAGGACCTTGCTCATAGTGAAACTTCGTGCACCGGGATGCCTTTTTTTAACAATATGATTAGAGTAAGAAATAGCAAAGAATAAGACCAATAGATAATGTATCAGGTAGTGAAAACTAGCATTGTACATACACTGTATTGGATCCATAGTGTTGCTTTGCATTGTTTTGCTACATAAATTTTGCTTTTCCAATGTTTTCTTTACTCATAAAATAACTGTCAAAAAGGTACATTAAAACCATAAACTATTGAATGGAGGATAGATTTCCTAAAAATCCAGAAAAAATAGAAACAAGTTTGAACTACCTCAGTACTTTAACTTTTGGCAACCATACAACAACATATCCGATGTAATCCCATGAGTCCCAAGCTTTTGGCAACAATACAACAACATATCTGATTTAATCCCACAAGTGGCTCTGGGGAGGGTAGAGTGTACGCAAATCCTACCCATATCTCGTAGAGATAGAGAGGTTGTTTCCGATAGTAACTTTTTGGCAACAATAAATCTTGTAAAAGATCACACAAAAATGTGGCTACTAGAAGAACTTTATTTGTATCCTAGCTTTATTTCTGGCTAGTGAGGAAAAAATTTAAGTTTACACTTGGGATAAAGATGATCCTTAGAGGCACCAGGAGTGAAAAGGTCAGCATATTGATGTTTAGCTGAAGAGAGATGTGCTCATTTGATACATTGAACTATGGCTCACTTCAAAGCTCACTCTTGCATGATGCGGTGCACAAGGAAGCAAGAAATATTTCAATCATCTCGGAGAAGAAAGTAAGAAGAGGGTTCTCTTTAGCAAGAAGAGGGGTGATTGTAAACGTACAAACTTTGACTAGGCACTTTTGGTTGTGTCACTCTGACCATGATCATGATTCCTCGAGATCTGTGTATTGGGAATGCGAGGCAGCCAAAAGTGCAGCTCCAACACCTGAACCATCATTAGAAAGCTCAATGACTATGCTGTCCGCAGTGTCTCCCAGTAACTCTTTCATAGTTTCCTCCAAGCAATTTCTGAACTTGGTGTAACGCTCAAACAATGCACCGTCCACAGCTATGACAGACCTCTGGTTCTCTCCGTCCTTCAAAGTGTCTTTTCCCAATTTCTTGAGGATGCCCACGATTCCTGCTGCAGAAAGACGAGCTCCACGAGAGGTGATAATGTCGCATAGCTCCACAACTATTTTCCTCATTTTCAAGGTAGAATTAGGTACCTGTGAAAAAAATGTATATATA

>PGSC0003DMG400000728 LRR-RLKs

CTTCTCTCTTACCTATCAAACTGAAGAAACTGAAAGAAAAAGAGTTGGTAGAGAGAAAACAGTATAGGGATTAGCATTTAGAACTCTTCTTTGCTTTATCAAGAACAAAGAATCAAAGCAAAAGCTTAGTTGATTCACTCAAAAAAGGTAATCCCTTTAATTTTCTGACACAGAATGTATATTCCCTTTTTTATTTCAAGAATCACTATGATATAGAAGGGGTATAACAAATTAGAGATGCCCATTTATATATATTTGAGAAATGATGAGAATTTTCGAAGTTAAATGAAGGTTGAGCAAGAATGGCGAACCAACAGGGCAGAGTGGTTTTGTTTTGTTTTGTGTTTTTCTTGCTGTTTGACTCTACATTTCAGCAGAGGGTTAGTTCTTCAGCTGAGAAATTTGCTTTGCTTCAACTGAGATCTTCTTTAGGGTTAAGAGCCAAAGAATGGCCTATAAAGGGTAACCCTTGTTTCAACTGGGCTGGGATTCGTTGCAAGAATGGTCGAGTTACTGAGATCAACATATCTGGTTTTAAACGTACTCGAGCTGGGGGTCAAACCCCTCAATTCTCAGTTGATGCTCTCCAAAATTTGACTCTTTTGGAATCTTTTAATGCTTCCAATTTTGTACTTCCTGGTTCTATTCCTGAATGGTTTGGTCTTAGGCTTGTGTCTCTTCGGGTACTTGATCTCAGGTCATGTTCCCTCATTGGTCCTATTCTTCCTTCCCTTGGCAATTTAACTAGTTTAGTTACTCTCAATTTGTCCAACAATGGTCTCACTGGACAAGTGCCTCGAAGTTTCGGTCAATTGTCACGCCTATCAAGTCTTGATCTTTCGCATAATAAGCTTCTTGGGGTAATTCCTGATACATTTGTTTCTCTTAAGAACCTTACCTTGCTTGATATGTCTTCAAATTTCTTCAATGGGGCTATTCCTTCAGATATTGGGACTTTGCTTCAGTTGAAATCCTTGAATCTGTCTGACAACAGCTTTTCTACTTCGATACCCACCCAACTCGGTAACCTTTCTAATTTGGTTGATCTTAACCTCAGCTTCAATAGTCTAAGTGGAGTGGTCCCTGAATTAGGAGGATTACGGAACTTGAAGAGTATGGTAGTTGGGAATAATAGGCTGTCTGGATCGTTGCCTAATGCTTTGTGGTCGATGCCTGGGTTGCAATTCCTAGATGTCTCTGCGAATAACCTCACTGGTATTCTGCCAAATGTTAGCTCCGTTGTCAATGCCACTGGTGCAGTGTTTAATCTTTCTCATAATACGTTTTATGGAAATTTGCCGTCTTTAAACAGAAGCTTCAGTTTTCTAGATCTATCAGGAAACTATTTTGAAGGCAAACTCCCAAATTATGCACAACGCAACGCGTCAATTAGTAGCAATTGCCTCCAAAATGTGACAAGTCAAAGGAATAGGAGCGAATGTGCTTCTTTTTATAGCGGACGTGGCTTGTTGTTTGATAATTTTGGAGAGCCAAATGCCACAGAGCCTCTTCCAGCTTCTAAATCCAACAGAAAGAGTCACAGAAATAAAATCATATTGGCGGCTGTTCTGGGTAGCGTTGGATTACTTGCACTTGTCTTTATCTGTATATTGCTGCTGATTTTTTGCACCCGCAAACGGGGTGCCACGAATCAAAGAGCCACTGAGGTTGGGCCCGGTCCTGCCAGTTCAAGTCCACCACCTCCTGCACCTGGAGTATCATTGAATTTTTCAAGTTTAGGAGATGCATTTACCTACCAGCAGATTCTTCAAGCCACAGGTGAGTTCAATGATGCAAATCTTATGAAGCATGGTCATTCAGGTGATCTCTTCCGTGGTACCCTAGAAGGCGGGACCCTTATAGTTGTCAAAAGAATAGATGTGCAGTCTGCAAGGAATGAAGCATACTTGTCAGAATTGGACTTCTTCAGCAAGGTTTCTCACTCAAGATTAGTACCTTTTATGGGGCACTGTTTGGAGAATGAGAATGAGAAGTTCGTGGTGTACAAGTATATGCCAAATGGAGATCTTTCAAGTTCTTTGTTTAGAAAAAACAATTCAGATGATGATAGTTTGCAGTCATTGGATTGGATAACAAGACTGAAAATTGCTATTGGAGCTGCTGAGGGTTTGTCTTATCTACATCATGAATGTAACCCGCCTCTAGTGCACAGGTATCAAGGACTTGTCATATTACATTTATCTTACTTCCATGTGCCACCACTTTTATGGTTAATCTGAGAGAACTTGAATAGCAATGCTGGACTTATGTGATTTACTATTAGATTTCCATGTTCGCATGTGCACTTATATTTCACTATCATGATTACAAGTTTTCATTTAAACATGTTTATCATATATCTTGGAAGAGTTGCAGGCTACCAGCTACTCTTTCTATGTATATTCATTGGTTATACATAATTGGTGACTCTTGGGCTAGACTCAAGGTAATAAAGGATATGACTTGATGACTTGGAGAGCGCCCTTTTTGGTTTGTACACGTTGGAGATACTAGGAAAAATGCTCTGAATGAAAGCTATCAAATATCAGTGTATTTCGTGTCTCTGAATTAGGCGGGACAAGATTCCCATAGAAGGCTCTCAAATTTTCAGCCAAAAATTCAGTGCAGCGCAGACACTCTGCACGTTAAAGGTGCTTGCAACTTTATGGATTGTGCCCATGTCTGGTCGACTGAAACTCATCTAAAGCCTATTTATAAATGATAAATATTAGATGTCTGCTTCGCTCTTAGCTTCTCGAGATTTATAAAAGGAGGCAATTTATGGTTGATGGTTTTCTGATTTGCTATCAGTTGGAGATCTGTAATGTTTATTGCCTCGTGAGACTATTATTATACAGAGTTCATCACATAAGGAATAGCCTCCACAATATCTTGTACCTGGAGCTTGACTCTATCAACTTGAGGGCTTCACTTACTATTTTTATGAGCTACACCTGCATGTAATAAAATTAAGAGCTTAGTAAACCTAAAACCTGTCCTTTTAGGGCTTAGCAATGTAAAGTCAAGAGCCATGCACTTCTAAAATTGCTGTAAGCTCAGCTTCTATAGTTCCATGCATGGCTGCGATATCTCTAATCTAGGCAGTTTATATTGTGTTAGATTCGGTATGAGGGATGTGGCCTTAGAAGATCTTGATGACATTATATTTCTATTGTGCAGAGATATTGTGAATGCAGAAATAGAAATTTTCTCAAACTAAGAACAGTAATGTTTTTTTAAAAATTGTTTTCTTTTCTTATTATTGAACCTTCTCCATACATCAATCCTTCTTTTTTTGATAACCGTGGTGCCCGGGCTAGCTTGTGCACACCTCGACACAATCCCATCAATCTTAATCAACTAAAGGTAGTCCTCATATGATACTCCTTATTGTACCTTTTATCTCCATGGAACCTAAGCCTGTCAGTGAACTATCTTATTTCTCTAAAACTTTTCATCTGCATGTTTTGGGTTCAACCATTATTCTTTCACCGGGCAACCCTTAGGGATTTGGGTTAGTGGCAAGAGCACAACTCATTATGTGTGGGTTAAGATCACGTTATCAGTTTGAATCCTACCATAGACTAAAGCTTGGTATTTAAGTGGAGAAAGGTAGAAGGCTGGGACCCATTATCCACCGAGTTTTGGATCGTGCGCCAATAGAGCTGGGAGGATTTCTCAGTTACAAACAAAAGATCCTTTCATGGGGAATCCAATAGACTTTCTTGAACCGAATATCCATTTCAGAAGGAGTTTCCAAAGGAGATGCTTCTCTAAATGTCCGTATAGTTTTTCTGCTGCAGTTCACCGTGAAATTCTCAATGGTTTAATTGCATGAATTAGTTTCTTGGATTCAATCTTACCTCCACCCCCCACCCCCAAAAAAGAACTTTATTTCTGTCAGAAATTTTTTAGTGCATCATGTAGTGACATCTTCTTTTTCCTTTGCCTACTATTCTCGCAGAGATGTTCAAGCAAGCAGCATACTTCTTGATGATAAGTTTGAAGTACGGCTAGGGAGTCTAAATGAGGCCTGTGCACAAGAAGGAGAGAGCCATCAAAACCGAATTTCCAGGCTGCTCAGATTTCCACAGTAAGTAGTTGTTGTTGGTAAATTCTATGTTCTAACCATCCTTTTCTTTTTGAAGGCTAACTACTATTACTACTAGGTTGTTATGTAATGGTCAATTTTTGTCTCCCATGATGGATCTTAAAATCAGGCAGTTTTATTGTTAATATATCTGTCTCTCCAATTTGATGTTATAGAAACTACTTTGGCTTGGTTTCTAAATTGGATTTGAGTTCCACAGTTTTGCTGCCAGCTCAAGTCTTTCTGGTGTATATTTATTAGCTCTGATATAAATGAGAATCCTAACGTTCATGCAGCAACCTTACTCTTAACCTTGTTATCTGTTCTTATACAAAAGGGCTTTAGCTTTGTTTCTAAATTATGACAACTTGTAATATAGGCCCTGACTAGTTTAGTCCTGATGATAGTTATCATGCAGCTGCATATTATCTTATTTCTTCTCAGTATTCAAGTTACAAGGAACACCGTAAAGAACAGTCAGTCTAGTCTGAAAGACCATCACCCACTTGTTAATTACTTGTTCCATCCCTCAGTAAATTATTCCTTTTCTACGTTATATGAATCTTCTTGCCTTGATTTACCTATGTCTAGCGGGTGTCACATGGGTGTTGTAGTTTAAGTAGAAATTAGCAAAAAATTGTATGTACCCTTACTGTATACTTAAAACTCATGTATGGGATATGCATGCGTAGCTGAATCCATGTAACATAGTCCTTTGATGTACATGTTTTCACTTAAGGAGAATTTGTTGGTTTTGCTATTTGGTAGGGAATGCATAGTTCTCATTCTTTTAGCTTTCCATCCTGTGCTGAAGTAAGTTTAACTATTTCTTACTCTCATTGTCTAGCTTTAACTATTACCTATTCTCGTTATGCCTACACATTGAATGCAGGAAATTATTAAATTTTGTATTGTGGGGTAAATACTAAATACTTCTGGAACGACCTAGATATAATGGCATGTGAAAAATTGGTTTGAAGTCCCTATGTTAGATGCCAAAGATTTGAGGCCTTCTATTCTATTATTTTAATCTTCATGGCTTCTGTTAAAACTTTCCAGGTTCTCTTATTTTAGAAATTGAACTTACGACCACTGATCATGATTATATTCAATCATAATCCAGAAAGCTTTAGAAGTTTTTGTTTTATAGTGAAAACTTTTCCATATGTATTTCCTGTCTTATGAATGGTTATCACTGTTTGTCCAGCTTTTCGTCGAAGCTTGTTGCATGATCTCAAAAGTTTTGTTATATTCTTTCAACCAGTACTGTTTTATAGTTTATACTGCTTGTTGACATAGTTCTCCCTTTCTTATTTGCAGGACGTCGGAACAAGGTGCATCAGGTATGTGCAACTTTATTTAGTTCTACTTTCCTTAAATAGAGTTTACTTGGATATTCTTAATATTTATTAAATTGGTGCAAGTAACTATTTAAAATTATTTGCGTCACATTTCCGGTATTGTTTGTTGGTGGGTAAAATCTCCTTAAATTGATCTAGGCAACAAATCAAAACTGTAGATTCTCATTAGAAGTAAGAATAAGTGAAACTATTAGGTGGAAATTTACGATATAAAGCTGATTCCAAAGAGCGTAAAGGGCTGGAGATTTTGATGGCGATGTTTTGCTTTACTTTTTGTGGGAGTATGCTTTGTTGCTTGAACTATCACTAGGGAATACTTCAAAGTCTACTTATCTCGTCCTAATATTCTTTCTTACCCGAATGGACAGGTACACCTAGTGCCACTTGTGCGTATGATGTCTACTGTTTTGGAAAGGTTTTGCTCGAGCTAGTAACTGGCAAGCTAGGTATCAGTGCATCCAATGACGCCAGCATGAAGGAGTGGTTGGACGGTACGCTTAAGTACATTAGTATATATGATAAAGAACTTGTCACAAACATTGTGGATCCTTCCCTAATCATAGACGAAGATTTATTAGAGGAAGTATGGGCGATGGCTATTGTTGCCAGATCCTGCCTCAATCCCAAACCGTCAAGGAGACCCTTAATGAGATACATCCTTAAAGCTTTGGAAAACCCTTTGAAGGTAGTTAGGGAAGAACACACTAGCTCAGCAAGACTTAGAGCAACTTCATCGAGAAGTTCATGGAACGCTGCACTGTTTGGCAGCTGGCGTAGCTCCTCAGATGTAGCAGCCGTACCTGCTGCAGCTTCAGCACATAAGTTGGAAGGAACAAGTAGTTTGAAGCAATCCGGAACAACAGGCTCTCAAGGAAGTGGTCCTAATGGGGACAACAACGGTCATTCTTCATCAACTAGACGACAGTCCAAAGAGATTTTCCCTGAGCCATTGGAGGAGCACGATGTAGAGAGAGGACCACAGGCGGAGTAGAAGGGAGGGTGATTGTCATTAATTCTTTGTTAATAATGCAGTAGGATCTGCATTAAGGTTGCATTTTTGTTTGGGGACACCTTGTACATTCTCCGTTTCCGGAATGCTTGAGTTTGCAAGCCATTATCTTGCCCTGGTCTTGAAAATGGAGATGCATAGTTTACCTGGGGACAGCAAACATCAAATTGTTTATTGGGGTTTTGGGATTTTAAGTTCTTTTTCTTTTTCTGGGGATAGAGTTGGAAAACAGGATTTAGTAGTGGTGGTGGTTCTTTTTGTTAATTTTGTTTTTGGGGGGTTTATGTTTGCTTCCAAATTCCAATTCTTTTCTTATAATTCTTCTCTTTTACTAGTCTCTCCATGGAATGTAAATTATACACAATCTTTCATTTTCCAATTGGAGGAGAGGAAAAAAGAAAAAAAAAGAGTATCCC

>PGSC0003DMG400000996 LRR-RLKs

GGCTGTTGTCTCTTTTTTGCCTTAAACTTGAACTGCTTCTAAAATTTTCTACTCCACATATTGCTGCAGGCATGAAGTCTTTTAATGTTTTCCTTTGTGTTATGTTTTCTTGCTTATTATGTGTTCTTTCTGTAGATGCATCTTCCAGGCTTGCTGGAAATGAAACGGATCGGTTAGCCCTCCTCAGCATCAAAGCACAGATAACATATGATCCTTCTGGTGTTACGAATTCTTGGAATGATTCCTTCCATCATTGTAGTTGGCAAGGCGTTACATGTAGTGCTCGACATCAAAGGGTGACTATGCTGGATTTAAGCTCTAAGCAGGTTGTAGGAACATTAGTTCCTCAGATAGGAAATATGAGTTTTCTTAGAGAACTAATTCTTCAAAATAACACCTTTAACAGTCAGATACCACGCGAAATCGGAAGGCTATTCAGGTTGAAGAATTTAGTGCTTAAGGACAACTCATTTACTGGTGATATTCCAGTGGAACTATCAAATTGCTCTAGACTCATTTATCTTGATTTGGATGGTAATTCACTGACTGGGAAAATCCCCGTGGAGCTTAGTTTGTCTTTGAGAAATCTTCAGGTACTTTTTCTTAGATCAAACAATTTAACAGGTGAACTACCATATTCTTTGGGTAATCTTTCATCCCTTATTGCCTTAGCAGCAATAGAGAATAGATTAGAAGGAAGTATTCCGTATTCGTTAGGCCAATTGACTAACTTGAGTTATATCTCTCTTGGTGGTAATATGCTTTCTGGTTCAATTCCCCTTTCAGTTTTTAACTTATCCTCTCTGTATCATCTTGCTGCACCAGTTAACCAACTTAAAGGAACTCTTCCCACTGATATTGGATCCACTCTTCCGAGTCTTCGCATAGTTTTTTTGTTCTCAAACTTATTAAGTGGTGTGCTTCCAAGTTCCATTTCTAACTTAACCAATCTTGAAATTCTCTCTTTATCAAGAAACCAGTTGTCCGGGAAAATCCCTTCTCTGGAAAAGCTCCGTAATCTTCAAGGACTAGCTATGCATTTCAATAATCTTGGAACCGGAAGAGAAGATGACATGGATTTCTTTTCTTCATTGGTTAACATTACCAGTTTTAAAGAACTGAGTTTGAGTGTGAATAATATAGCAGGCCAGCTGCCTAAGAATATTGGCAACTTAACCAATTTCCGGTCTATTGGTTTTGCTAGGAACAAACTATTCGGACGAATACCTGATGGATTTGTTGATCTCAGCAACATGGAAGTGGTAAGTTTGGAGTATAACCAATTGACTGAAGAAATTCCAGCTAGTCTTGGTAAACTTCAAAAGCTCAAGTATTTCTATGTTAACGGAAACAAACTCTCGGGGGAAATCCCTTCCTCTATTGGGAACATAACATCTTTATATGGACTTAATCTAGCTCAAAACAACTTAGAAGGGACTATTCCTTCAGTTCTTGGAAACTGTCAGTTGTTGCAAATGTTGTATCTTTCTCGAAACAGACTCAGTGGGACGATACCAAAAGAGGTTCTTAGTATCTCATCCTTGTCTATCCAACTAGACCTGTCTGGAAACCAGTTAAGTGGATCACTTCCCCTTGAAGTAGGAAGTTTAGTCAATCTCGGGTATCTTGATATATCCGAGAACAAGTTATCTGGAAAACTTCCGAGCACTTTAAGCAGTTGCATCAAGTTGGAAAACCTCTATGTTCAGGAAAACATGTTTGAAGGTGTGATCCCATCATCTTTAAGTTCTTTGAGAGGGATGGAATATTTGGACTTGTCGCGTAATAATTTCTCTGGTCTTATTCCAAGGTACTTTGAAACTTTCATATCATTGAAAAGCTTAAACTTGTCATTTAACAATTTTGAGGGTGAAGTACCTCGAGAAGGAGTTTTCAGTAATGCAAGTGCAGCAATAGTCAACGGAAACAGAAATCTTTGTGGAGGTTCTTCCGCGCTAAAACTACCACAATGCAACTTTCCAACATCAAAGAAGGGAAGATTAATGTCATCAACCTTGAAGATAGCTATATCTATTGCTAGTGCATTATTTGGAGTAGCACTTGTTCTGGTGCTCCTAATTTTGTGTTTCTTAAAGAGGAAACGTAGCCCTTCACTCGATTTATCAGATGATTCGTTCTTGAAAATCTCCTACGGGGAACTTCTGAAAGCAACTAATGGATTCTCCTCAGATTATTTAATTGGTAAGGGTGGTTTTGGTTCTGTATACAAGGGAATTCTTGGTCCAGATGAAAAAACTGTCGCGATCAAAGCACTCGATTTGCAGCATAAAGGAGCTTTGAAAAGCTTCATTGCAGAATGTGAAGTCCTAAAGAACCTCAGGCATAGAAATCTTGTCAAGTTAGTAACTGCATGCTCAGGTACTGACTTTCAAGGCAATGATTTTAAGGCCCTCATTTACGAGTTCATGGTTAATGGTAGCCTAGATGATTGGCTACATTCATTTTCTAATGATGGAAGTCTGCACGTTCGATATCTAGATCTTTATCAGAGAGTAAACATTGCATCAGACATTGCTTTCGCGCTAGATTATCTTCATCATGGAACCCAAACACCAGTTGTTCATTGTGATTTGAAGCCAAGCAATATCCTTCTAGACAAGGACATGACTGCCCGTGTTGGCGATTTTGGTTTATCAAGATTCCTCCAAGAAACATCTCAAAGAGAAACTAGCACTATTGGTATCAAAGGATCAGTTGGCTATGCAGCTCCAGGTAAACATCTCTTTCCCTTTTTTAGCAACTGTCACATTCCAACTTTCCGCCTGACATGTTTAAGATCATACTATGTTTAAGATCGTAATATTCAAAAGTCTTACATACTTTCTTAAACTTCGTGGCAAGTCCAAACCAGACAAACATAAAACGGAGTGATTTTGATATGGACTAATGAATTTTTTTCTGCAGAATATGGAATGGTAAGTGAAGTATCAACTTATGGTGACATATACAGCTATGGAATCATCATACTGGAGATGTTAACAGGAAAAAAACCGACAGACGACGCCTTTTCAAATGGATTGAACCTTCACAACTATGCTAAAATGGCTTATTCTACTGGTAGAGTCATGGAAATTGTTGATCAAATGGTTTACCACAACTTGCAAGAAATGAAAACCAAAGATTACATTGAAGAGTGTTCAATTTGTATGTGTAAAATTGGGATTGCTTGCACTATAGATTCACCAAAAGAAAGAATGAGAATTAGTGATGTTATTAAAGAGTTGCAATTAGTCAAAGAAACTTTATCTTAATGTGATAGAACACTATGATATTAGAAGTAAATCATAAAACATTATCTATTTTGTTCTATATATTGTGAGTTGGTAAAATTTC

>PGSC0003DMG400001185 LRR-RLKs

TCAATGGTAGTTCCTCTTGTCTCCTTGTTTTACAACTTTTTCCTCTTTCTCATACTTTCGTCCTCCAACGTAACTGCTCAACCATCTAGCACAGACACAGACTTTTCATGCTCGTTTAACTCTTCCTTTTCTTGTGATACTTTTATTTCATATCGTGCCCGTCCACCAAACTATTTAGATGTCGGAAGTATATCTGATCTTCTTGAAGTGAGTCGTTTAAGTATAGCTACGGCCACAGGTCTAGCCTCAGAGGATACCGAGCTATTCCCTGACCAGCTTTTACTTGCACCCGTTAAATGCTACTGCAATAGTTCTCATTATTTCTCCAATGTCACCTATCAGATCAAGAAAGGCGACAACTTTTATTCAGTTTCAATAGGAGCTTTTGAGAACCTTACTAATTATCACGTGGTGCAAGATATGAATCCAACACTAGATCCAACCAACTTGACAATTGGCGCAGAAGCTGTTTTTTCTTTGTTTTGTAAATGCCCCACACATTCTGATCTCGAAAAGGGACTTCAGTACCTGGTCACTTATGTTTGGCAGCCTTGGGATGATGTGTTGCCTGTAAGCAATATGTTTGGAGCATCTGCAGCTGACATTCTGGCTGCCAACAATTACAGAAATTTCACTGCTGCAATATATTCTCCTGTCTTGATACCAATAAAACTTCCTATTATTCTGCAATCTTACCCCTCTTCTGCAAGCAGTAGAAAATCCAAACATGGATGGATAGTAATTACTGTTTTAAGTATTATGGGGTTGCTTGCAGTTTTTTCATTTTGCTTGATGGTATACATGCGTCATCTAGACGAGAAGAGGAGGGCTAACTTAGCCCATAACTCTTCTACTCTTGAAACTTCTGAGCTCTTTCATACCAAGAAAGCCTCAGAAGGTGAAATTACGGACCATAAGAACATTCAAGATAAGCTTCTTCCTGGAGTTTCTAGCTACATTGGAAAGCCAATAATATATGACCTGAAGATCATCATGGAGGCAACTGTCGACCTCAGTGAGAGATACAGGATAGGAGGATCAGTGTACAAGGTTACAATGAACGACCAAGTTGTAGCAGTCAAGAAAACAAAACAAGCCTCGGAGGAAATGACAATACTTCAAAAACTACATCACGCAAATCTAGTGAAACTAATGGGCGTTTCATCAGACGATCGTGGGAATTCCTTTGTAGTTTATGAATATGCTGAAAATGGCTCACTGGATCAATGGTTGTTTCCCGGATTATCATCATCTTCTTCTGCCTCTGTCTTAGTGGCATCACTTGGTTGGAGTCGAAGATTACACATAGCCTTGGATGTTGCAAATGCTCTGCAATACCTGCATGAACATACTCAACCTAGTATAGTTCACGGGGACATCCGAACGTGTAGCATACTTCTTGATTCAAGGTTTAAAGCCAAAATTGCAGGTTTCTCTACAGCCAGACATGCCACCAACTCAATGATCCTGAAAGTTGATGTGTTTGCTTTTGGGATTGTGCTGTTGGAGTTGCTTTCAGGGAAGAAAGCAATGGAATCAAAAGATTATAATGATGAGACTCTGATCATCTGCAAAGAAATTAGTGGAATCTTGGAAGTTGAAGAGAACAGAAAGGAGAAGTTTAGAAAATGGATGGATCCAAAGTTGAGCTTTTACCCTATTGATGATGCTCTAAACTTGGCTGCCTTGGCGACAGCATGCACATCGAAGCAATCAGCAGAGAGGCCTAAGATGACAGATATTGTCTTCAACCTCTGTTTTCTTGCTCAATCATCTTTTGAAATGTATGGACCATCTTGGACTTCAGGTGAAGCTGAGGAGATTGTCCAGATTGTTAGTCCAGTAATAGCACGTTGAGTTAAAGATGGAGCGTAAATATAGTTCAAGAACTTTTGTCAGTTCGTCTCAAATTTGTAACATATTCTGGATTATAAAGCCACAGGAACAATAATCAAGCTGATCTCCAGAATATATTGCAAATTCGAGTTATCTTGTCCAGCTTTTCCCTGTATTTCTCTTGCTGCTAAGTTAGTTGCTAGAGTATATTCTTGTATTGGATGTTTTTCTCAACTTTGAAATAAGTGGAGTGCAATTTGACCTGTATTTCTTTCGTCCTCTCAAAAGAAAAACAGAAAAAATTAAGTTTCTGAAAAGACACTGCAAACTCGTTCTTATTTATATTTATATATTTAAGATCCAGACGAGTACATGTTTA

>PGSC0003DMG400001301 LRR-RLKs

TAATTTCTTAGTAGCTAGCTTCTAAAGGCTCCACTCATACTTGTAAAATAACAGACTGGGTTTTCATTTGTCAAGTCAAATGCCTCAACAGTCTTCATCTTCCTTCTCCTTGCTATTTCCTACCTATTTCTCTTGCGCATATTATCAAACACCTTGTGCCAATTACCCAAAACTGCTGTTGGATCCCTAAACCCCCCAGCTAACGATGGCTCGTTTCAGATTTCTCCTTTTTCTTCTGTTGATTCTGATTTCTCTTGTTCTTCCGTGCCTTGATGCTACTGTAGTTGAAGATGATCTCAAGTGTTTAGAAGGATTTAAGAAGTCTTTGGAAGATCCAGATGGGAATCTGAATTCATGGAACTTTAAAAATACTACTGTCGGAGCTATCTGTAAGTTTACGGGTGTTAATTGCTGGAATGACAACGAAAATCGCATCATCAGTCTCTCTCTTTCGAATATAAACCTCGGAGGGAAGGTTACAGAGCCTGTACAGTATTGTGCCAGCTTGACAACCCTTGATCTCTCTGGTAATCGCTTTTCTGGTCCAATTCCTTCTCAAATTTGTACATGGCTACCTTTTTTGGTAACCCTAGACTTGTCCGGTAACGATTATTCCGGCCCTATCCCTGCTGATCTTGCTAAATGCATCTTTTTGAATAAATTATCGCTCAGTGATAACAAGCTTACTGGAAATATTCCCCCTGAATTTTCTAGTTTTGGTAGGCTTAAAAGTTTTTCTGTGGCAAATAATCAACTCTCCGGTAGAATCCCGGCAGCCTTCGATTCTTCTAACTTTAATTTTGAAGGAAATAGTCTCTGTGGTGGACCTTTGGGGAAATGTGGAGGACTAAGTAAGAAAAGCTTAGCTATTATTATTGCAGCAGGAGTCTTTGGTGCTGCTGCTTCTATGTTGTTGGCTTTTGGTGCGTGGTATTGGTTTTTTACCAAGTCTGGAAAGAGGAAGAGAGGGTATGGAGTTGGGAGAGATGACTCTGATAGTTGGGCTGATAAGCTGAGGGCTCACAAGCTCACTCAGGTTATGTTGTTCCAGAAACCGCTTGTCAAGGTTAAATTAGCTGATTTGTTGATTGCCACAAATGGTTTCAGGACGGACAATGTTATCAACTCAACTAGAATGGGTACTACTTATAATGCTGTGTTACGTGATGGTTCTGCACTTGCCATTAAGCGGCTTAATACTTGCAAACTGAGCGAGAAGCACTTTCGTGATGAGATGTATAGGCTAGGACAACTTAGGCATCCTAATTTGGTGCCACTGTTGGGGTTTTGTGTTGTGGAAGAGGAGAAGCTTTTGGTTTATAAACACCTGTCAAATGGTACTTTGCACTCATTCTTGAATGGGAATGCAAGTGAACTGGATTGGCCTACTCGGTTTAGAATTGGTTTCGGTGCTGCTAGGGGCCTTGCTTGGTTACATCATGGTGTCCACCCACCTATCTTGCACCAAAACATATGCTCTAATGTTATTTTCCTCGATGAAGACTTTGATGCTAGAGTAATGGATTTTGGGCTGGCAAGGTTGATGACATCAGATGCAAAAGAGTCTAGTTATGTGAATGGGGAGTTGGGTGAATTTGGCTATGTAGCTCCAGAGTACTCTAGCACAATGGTGCCTTCACTGAAAGGGGATGCTTATAGTTTTGGGGTAGTGCTTTTGGAGTTGGCTACTGGGCAAAAACCTCTTGAAGTTACTGCTGGTGAAGAGGGATTCAAGGGTAATTTGGTAGACTGGGTGAATCAGCTCTCTGCTTCAGGTCGCATTAAAGATGCAATTGACCAAAACATGCGTGGGAAGGGGAATGATGAAGAGATTGTGCAATTCCTAAAAGTTGCTTGTAATTCTGTGGTTTCTCGGCCCAATGATAGGTGGTCTATGTATCAGGTTTATGAAGCACTTCAGAGCATGGCTGAAAAACAGGGTTTCTCCGAACAGTATGACGAATTCCCGTTACTATTTGGCAAAGAAGGTGCTACTAGAAGCCCTGTTTGAAGCCGGAATTGCGAAATTATTACTAAATGTTCCACTCCAAATGAACATGATGCATCCTGTTGTAAGTTTTCAGTTCCTATTGGTGTAAGTGGCATGGCACTGTTTTGATTGGCTCAGAAGTACACCCTTTAAATGTTTCTGTATTATATTTGCAAAATGCACAAATCATTCTTAAGTATTTGCTTGCTTGATATTTGTAGTATAGGAAGCAGCTGTTAATTATTGTTCATGTTTGTGATTACCATTGCCTATAGCATTAGATGTTATGTATTCAAAGGATTCTCAACTGAACGTCTCATTTACTGGAAACTGGTCTATATTACTATTTACTGCCTGTCTAGCTCTTGTTTGGAAGCCAATGGGGCCG

>PGSC0003DMG400002012 LRR-RLKs

TGGAAGTAGTTAGTCAAAATTGTGTCTAAATGAAAGGTTTTTGGTGGTTTACTGTTTCTTCATTTTTATAAGGAAAGTTGGTTTGACTAAAATAACAACCCCTATTGACTTGCTTATGGTATAGGGAAATGTAGGGTCTCTTGCTTTTGATTTTGATGTATTGGTTATTGGTGAGTGAATTTAATGGTGGCAATTTGCATTTTGGGACATTATTGGTGCTGCTGTTGGACTTCAATGAATTTAGTGGCTGGTGTTGTTGCTGCTTGTTCTTTTATCTAGGTTTATATACTCTCTTATTTCACTTACATAATGATTAGTGATGATTGTAGTTTGTGTTTAAAAATTTTGCTAATTCATTTTCTGGGATTCTTGAAATGATTGTGCAATCTTTAGTCTTTCTACTTGGTGGGGTTTGTCTACTTTCTTGGCCCATTCTGAATTATTAGCTTGTCAAAACGCTGTCTTTTTTCCTTTTTAGTTCAGTATTATATGTAACTCTCTGTTAGTGTCACACGGGTTTTTCATTGGGAGGAAATGCAATCTACCATTTGTTTAGTGCTGCTATTTTTAGTTGAGTTAGCTAAAGGAAGCTTAGATTTGGATGCTCTTTTGGAGTTTAAGAAGGGAGTTTTGAAAGACCCTTCTGGGAAAGTACTTTCTTCATGGGATTCTAAGTCATTGGGACCAAATGGTTGCCCCCAAAACTGGTTCGGTATCGGTTGCAGTGATGGCCATGTCACTTCAATTGAACTAAATGATGTAGGTCTGATTGGAGTTTTGGATTTTGCAGCTATTAGTGGCCTAAAAATGCTGCAGAATTTGTCTGTTGCAAACAACCAGTTAAGTGGGAAAATTACTGAGGAAGTTGGGTTGATTATGTCATTAGAATTTTTGGATCTTTCCAAGAACATGTTTAGGGGTTCTATACCCTCTAAGCTGACTAGTTTAAAGAACTTAGTATCTCTTAATCTTTCTTTAAATAGTCTCGATGGAATGGTTCCCACTGGGTTTGCTAGTCTTGAGAAATTGAAGTATCTGGATTTGCACTCTAATGCCTTCTCAATTGATATTATGCTTCTTCTGGCCTCATTGGGGGATGTAGAGTATGTAGACCTCAGTAGCAACAAGTTTGTTGGATCACTTGACCTGCAAGTAGGAAATTCGAGCTTTGTCTCGTCAATTCAGTATCTGAACATTAGTCATAATAACCTGGATGGAGAGCTATTTCCTCACGATGGAATGCCATATTTTGACAGTTTAGAGGCGTTTGATGCAAGTGACAATCAGCTTACTGGTACTATTCCGTCCTTCAATTTTGTAGTCTCCCTTCGAATTCTTAGGCTCGGGAACAATCAGTTGTCTGGATCACTCCCCGAAGCTCTTCTAGAGGATAGCTCAATGATTTTATCAGAATTGGACCTAAGCCAGAATCAGCTTGCAGGTGTGTTGATCCTTTCTTCTTTAGTTAATTTGTTTGGTTGAAGTCAATCTAAGTAATCTATGGTTAATTACTACTCCCTTTGTCCCAATTCATGTGACACTTTTTATTTTTTGAGAGTCAAACAATTTAAGTTTGACCGGAAATTTGCGCGTGGAATCTTCAACTTTTTTGAAATGAAATTTATATATTTGTAAACTACGTAAAAAGTACTATAAGTCACAATAATTGGCAATTCAAAATAATTAAAAGATATATGATAAATTTACGGTCAAAGATAGAATTGTTTGAATCTCGAAATCCGAAAAGTGCCACATAAATTGGGACGGAGGGAGTACTATCTATGATCTGAGGTTGTTGTCAGTTAGATAACAAATGAATAATATTTAGATTGCTATCTGCTGGAAAGCATGCTTGTTTTGGTTGCAATCAAATGTAATGTATAGAACACAATTCCTGAGCACTTCGGAATTGTGTTCTAGAGGAATGCTGTACAATTCTCTGCTTATGTGTAATTATCATACACATTGTTCCCAATTCTTCCCTAGGTTGGAGATATTTCATACTTAGACTCAACTCTAAGGCAATTATTGCTGTACTTGCATTTTAATTAGTGGAATGTTCCAGTGTCTTTTTGATTGTTTTTTTTTGCTGCATTATTAATTTCATCTCTCCTCGAGCCGAGGGTCTATTGGAAATAGCCTCCCTACCCTCACAAGGTAGGGGTAAGGTTTGCATACATGCTACCCTCCCCAGACTCCACCTATGGGATTATACTGGATTTGTTGTGGTTGTTGTTGTTTGCTGCGTTATAAACAAGCTGCATAAAGATATTTTTCATGTGGTTGGAGTTCTGGTGTATTTTAAGTCTTTCGTTGTTCTTTTAATTATTATATTTAGTTGTTCTTACCACAGGCCTAATCCAAGCTACTTCCTTTCCAAACTATTTCATGAAGCCATTTGCTATTTTGGCAGGTCCCATTGGAGGTATAAGTGCTGTGAATCTCAAGCTTCTAAATTTATCATATAACCAGCTCTCAGGCCCCTTGCCTTTCAAGGTTGGGCGCTGTGCCATCATAGACCTGAGCAACAATCGGCTAACTGGAAATGTGTCCCGGATCCAAGGTTGGGGAAATTATGTTGAAGTTATTGTGTTAAGCTCAAATGCATTGACAGGAACCTTCCCCAACCAAACTTCCCAGTTCTTGAGGCTTACTTCGTTAAAGATTTCAAACAACTCACTTGAAGGTGTGTTACCAACTACGTTAGGTACATACCTCGAACTTAAAACAATTGATCTTAGCATTAACCAACTTAGTGGTACTCTCCTTCCCAGCCTTTTCAACTCTACCAAATTGACTGATATTAACGTTTCTTTCAACAAATTTACCGGTAGTGTACCTATCATGGCATTTAACTCAGAGAATCTAAGCTTGATTTCTCTAGATGTTTCACATAATGCATTAGCTGGTCCGTTACCTCCGGGGCTGGACAAGTTCCTGGACATGGTAAACCTGGATCTCTCTGATAACAAATTTGAAGGTGGCCTTCCTAATGATCTCTCAGACAAATTGGAATTTTTTAATGTGGCTAATAACAATTTTTCTGGACCTGTGCCACAAAATTTATGGAGGTTTCCTGACTCCTCATTCCATCCGGGAAATCCCTTGCTTGTACTTCCAAAGCAAGCCAAAGCTCCTTCAGAGGGAGATTCAACTTTAAGCTTGAGAAGTCATGGTTCTCGCATGAAATCTACCATCAGGGCTGCCCTCATTGCTGGGCTGATCTGTGGTGTATCTGTTATAGCTTTGTTGACATTAATAATCTACCGTAAGGCCCATCAGCGAGATGGCGGGAAGGACGATATGAAAGGAACCAAGGAAAAGAAAGGTATGGAACTATTCTTCTTCTAATAAGACAGGTGATCAGATCAACTGATACTAATGCCTTTTCATTTCCACAGGTTTGTCTTTATCTGATATAGAATGTGGACATGACACTAGAGAACACAGCGTGCCAGTATCAACAGTTCAAAATGAGCCATTATCTTCACCTATATCTGTCATGTCGTCCGCCAATCTCTCTCCTTCTAAGGTTCAAGATCAATCCAAAAGCCCAAACTCTCTCAGGGTTTCTTCTCCTGATAAACTGGCTGGAGATTTGCATCTACTAGATAACGCCTTGAAGGTCACTGCTGAAGAATTATCGTGTGCTCCTGCCGAAGCTGTGGGCAGAAGTTGTCATGGAACATTGTATAAAGCTACTCTTGGTTCAGACCAAGTACTTGCTGTCAAATGGCTTAAGGAAGGGATAGTGAAAGGCAAGAAGGAATTTGCTAGAGAAGCTAAGAAACTGGGGAGTATTAGGCATCCTAATTTAGTTTCTCTACTGGGTTACTACTGGGGTCCCAAGGAGCATGAGCGGCTTCTCATATCAAATTACACCGATGCACCATGTTTAGCTCTTTATCTTCTTCGTAAAGGTCAGTCTCCCCTTTTCTATCTATAGAGTATATCACCTGCATCTATTTCTGTAGCTTTTCTGATTTTTGTATGGTGCTGAAATTGTGATCCAGTTTTACCAAAGAAATTAGTAGTACTCCTTCAATGAAATCAAACATAATGGACCGTGAATGCGCACAACCTTGAATGTGTATATTGTTATTGTAGTTGTCAAGATTGCTCATACCTGATATGCTGAAGTTTAATCACCATCTTGCCACTTGATTTCCATTCCGGTGCCTCAAGTGTACTTTACTTCCAAAAGATATTCTAGCTTGCTGGTTGAATATGTATCTGAGACTATTAATACACATTTCACATAGCAACTTTCATGAAATTTATAAATGGAGCTATCTTCCAAAATCTGAACTTTTCAATTCATATATATTAGCAATGTTAGAATTACATATAGCAGAAAAGATACACACCTGATTTTTGCTTTCATGGTAATACTAAAAGGTAAAACAGTTCCACCTTCGCTAATTATGCCTTGATGGTGATAGTGGGAGAGTCAAAAGTCGAAAAGAGGATTCCTCTCTTGTTTCTCTCATTCAAAACTGTGCATGAGAATTTTATCTCACGTGGCTCATTTTGTATTGATTACTTTGGTTGGGCTCGATTGATGTCATTTTGATAAGGGCATCCCGGTGCACTAATTAGAGACTGCTACTTTGAATTCCGCTTATTGCATATAGCACTGTACTTTGTTTTCTAGTTTTGAGTGATGATATGGATGCCGGTTTCATTCTTTGATGTATGCAGATGCAGAACCCTGTAAGCTACATCCTTTATCTCTCGATGATCGACTCAAGATCTCTGTAGATGTTGCTCGTTGTCTAAACTACCTTCACCATGAAAGTGCCATACCTCATGGCAACCTGAAATCCACAAATGTACTTATAGACACGTCTACCGTGAATGCCCTACTAACTGACTATAGTCTTCATCGGTTGATGACATCTGCTGGAACAGCTGAACAGGTTCTAAATGCCAGTGTACTCGGTTATCGTCCACCTGAATTCGCGAGTACAAGTAAACCTTGCCCGTCATTGAAAAGTGACGTCTATGCTTTTGGTGTTATCTTGTTGGAACTTTTGACTGGAAGAAGTTCTGCAGAAATTGTTCCTGGGAACTCAGAAGTGCTGGATTTAACCGAATGGGCGAGATTATTGGCCTTTCAAGACCGTTCAATTGAGTGTTTCGATCCATTCTTATTGGGAAAGCAAAGCAACAACGAAGACATGCATACGATTCTTGATAGCATGCTTCAGGTTGCACTAAGGTGCATTTTGCCAGCTGATGAAAGGCCTGATATGAAATTTGTTTTCGAGCAACTTTGCTCGATAGCGCAATAACACAAGTTGAGGGGAGGGGTAAATTTGAATACTATGTAGTTGTAGATTTGTATCTTAAAAGTTGATATGTCATTCATGTTCCATTTTTAATTCAATTGTTTTAGAAACAAATTCATTTGTGAATTAGCACATTAAGTGATGAATGAAGTTGTTGTATTCTTCACATTTATTGACAAGAAATAAAGTTTTCTTGTAT

>PGSC0003DMG400002689 LRR-RLKs

TACACAAATTCAACTCTTTTGTTCCTTAAATTCAACTTTGTGTTTTTTTGTTCTCTTCCTAACTTCCCGCGCTTTATGCCATAAACCCAAAAAAATGAACCGTTTTTTCTCCTTCATTCTCCTCCTCACACTTCTCATCTCCGCCGTGACCGCCGCCGTAACCGCCACCGGCGACGGCAACGACACCGCCGCACTCCTCCTCTTTCAGTCCCAAACTGACATCCACGGCACTCTCCTTCACAATTGGACACTTCCGGTGAACTCCACCACCGCCTGCACAGCTCAATGGCTTGGTGTAAAATGCATAAACAACCGTGTCTCCGCCGTAATTCTCCCGTCGTTCAATCTCCGGGGACCCATTACTGCTCTCTCTTCTTTACCCTTACTCCGCCTTCTCGACCTCCGTAACAATCGCCTTAACGGAACCCTCACACCCATTACCCAATGTACTAATCTTAAGCTCATTTACCTCTCCGGCAACGATTTCTCCGGCGAAATTCCGCCGGAAATTTCTTCTCTCCGTCGTCTCCTCCGCCTTGACGTTTCAAATAACAACCTTGAAGGTTCAATTCCGACCCAGATTGCGAATTTGACCCGATTACTCACACTTCGTTTACAGAACAATGAACTTTCTGGAACAATCCCAAAATCCCTAAATTCTCTTCAAAATCTTAAAGAATTGAACTTTTCGAATAATGAGCTTTATGGGTCTGTTCCAAATGGTTTATATAATCATTTTGGTGAAAATAGCTTTGATGGAAATGAGGGTTTATGTGGAATTGGAAATTTACCTCAATGTTCATATACAGGGGAAATAGTTCCTTCAAACCCTAGCTCTTTGCCTACAACCTCCACTGCAACAATCGAAGAACCCCACGAAAAATCGCGAAAAGGGTTGAGTAGAGGAGGGGTAGTAGCAATAGTGATGGTGAATGTGGTGGCATTGTTAGTTGTAGTGTCATTTATGGTAGCATATTACTGTGGGAAGTATTCAAGAACTCAGAGTTTTTCAATGTCAGGAAGTGAGTGTGGGAAGAGGAGGAGTAGTTATTCGAGCGAAAAACGAGTTTACGCTAATAACAATGGTGGTGGTGGTGGTGATAGTGATGGTACAACAGCTACTGATAGAAGTAAACTCGTGTTTTTCGATAGGAGGAAGCCATTTGAGCTTGAGGAATTGTTAAGAGCATCAGCTGAGATGCTTGGTAAAGGGTGTTTAGGTACTGTTTATAAGGCAGTTCTTGATGATGGGATTACAGTTGCTGTAAAAAGATTGAAAGATGCAAATCCATGTCCAAGAAAGGAGTTTGAACAGTATATGGATGTTATTGGAAAGCTTAAGCATCCAAATATGGTGAAACTTAGAGCTTACTATTATGCCAAGGAAGAAAAATTGCTTGTTTATGACTACTTGCCTAATGGGAGTTTGCATTCACTTCTTCATGGTAAACATTAGTAAATGTTCATGTCAAAGATTAAAGATTGTTTTGTAATTCAAGATTGTCTCTTTTGGGGGTTGTTAAGGTCTATATTGCTCGGACTCTTCAAAATGTCGATGGGGTGTGTCGGATCCTCCAAAAATAGTGCATTTTTGGAGAATTCTGACGTTGTAATTCAAGATTGTCTTTTTTGGGGGTTGGTAAGGATTATATTCCTTGGACTTTTCAAAAATATCGATGGATGTGTGTCAGATTCTCCAAAAGTCTCTAGAAATAGTGCATTTTTGGAGAATCTTACACGGGTGTGACGTGTTTTGTAATTCAAGATTGTCTTTTTTGGGGTTGGTAAGGGTTATATTGCTCGGACTTCTCATAAATGTTGGCAGGTACGTGTCGGATCCTCTAGAAATCTCCAGGGATAGTGTATCTTTGGAGGATCTAACGCGGGTATGGCAACATTTTTGGAGAGTCCGAGCAAGTTGTAGGTAAGATACTAATATTGATGGGATTTTTTTTTCTTGATTCTAATAGGAAACAGAGGACCAGGTAGAATTCCATTGGATTGGACAACAAGGATTAGCTTGGTGCTAGGGGCAGCTAGAGGGCTTGCTCACATTCATGAAGAGTATGCAGATTCAAGAATCCCACATGGGAATGTGAAATCATCTAATGTGTTGCTTGACAAAAATGGTGTGGCTTGTATATCTGATTTTGGGTTGTCATTGCTTTTGAATCCTGTTCATGCCATAGCTAGATTAGGAGGATACAAAGCGCCCGAACAATCTGAAATCAAGAGGCTATCTCAAAAATCTGATGTTTATAGCTTTGGTGTACTGTTATTGGAAGTGCTCACTGGTAAAGCTCCCTCAGAGTACCCTTCACCGACTAGGCCTCACGATGAGGAGGAAGAATTGCCAGTGGATTTGCCTAAATGGGTTCGATCAGTAGTAAGGGACGAGTGGACTGCTGAAGTGTTTGATCAAGAACTGTTGAGGTACAAAAACATTGAGGAGGAATTGGTGTCAATGCTACATGTAGCCATGGCATGTGTAGTGCCACTACCCGAAAAAAGGCCTACTATGGTTGAAGTTGTTAAGTTGATTGAGGAGATTAGAATTGAACAATCTCCATTAGGTGAAGATTATGACGAATCGCGCAATTCTCTTTCCCCCTCCCTAGCCACCACTGAAGATGGACTCCCCGGTTACTAATTTTCGATCATCGTGGCATTGATGTTTCATAGTCCTAGTAGTATATGTTCAAATTTGCAGCATCTTACTGCAAATTGGGAAGTTCAAAAGCTATGGACTTATGTTTAGTAGTTATTTGATTGCTTTGTGTTATTCCAAATTACTGCACTTTTGAAAAATCTGATGATCCAACAAGCATATAAGAAGTGCAGCAATCAGTCCATTATGTTTTTTTTTAATGTAATGTAATGTAATGTTAATTCTTTGTATTTAATTCATTTTCAAGATTTGTGTTCTAAGGCAAAAC

>PGSC0003DMG400002870 LRR-RLKs

TTTTTTTTATAATATTAAAAATCTCATTTTGCATTGTGGGTAGTTCTTTTCTTGGAATTCTTGAATTAGTAAAAGCAAGAAAAATGAAGGCTATTTTGGGTTATTTGTACTTATTCAACTTCTTGGCTTATTATGCTCATGCTATGCTTACTCCAGCTGGTGTCAATTATGAAGGTTAGTTCTTGTGATGAAATATTGGGCTTTTGTGCTTTGATTATTGTCTTATTTTGTTGTTGTTATTGTTAATTTTTTTGTATGTTATGTATTTGTTATCCATTTTGCATTTTTTTCGTTGTTATTACTGGGACGGAGCTAGCAAGGGCTGAGGTGCCTTCCACGGAAAGTTATATCGTTTATACATGGTTAAAATTATTTTTATGTATACATAGTAGATGTTAAATCCCTTTGGTTTCTTTGTCTGGTTACTTTTTATATTTCGAACTCCTTAGTGAAAATTCTGACTACACCAGTCGTTGTTACAATTTTTTTTTTAAAAGCTAAGGGTTTATTGAAAACAACCTCTTTGAGGTAGGATAAGGTCTGTGTACACTCTACTCTACCCTCTTCAAATCATGCTTGTAGAGTCACACTGAATATGTTGTTGCGGATTTTGAAATTAAAAGCTCAAGATTTTTTTTTTCTTTGGCAGTGCAAGCTTTAATGGAGATAAAGAAGAATTTAGATGATCCTCATAATGTTTTAAACTGGGATGGAGATGCTGTTGATCCTTGTAGCTGGAATATGGTGACTTGTTCATATGATAAATTTGTCACTAGCCTGTAAGTAAAACAATTTTCACTTATTCTAAAAATAGTACACCTAACAAGATTTCTTAATATAGTTGAATTATTGATTATGCAGGGAAAGTCCCAGCCAAAATTTGTCTGGCAAAATATCACCATATATCCACAACTTGACACACCTTGAACTTATGTGAGTACTTACTTAACCAAAGTAAATAAAAAACGTAATACATAAACGTGTCCTTTAACTTGGTGTTATCTATCATCTATGCCTCCAATATGGATGTGCAGAAGTAGGCATACACGTCCTATGTGAAAATTCTCATCCTACGTGTATTATGCTACGTAGGATGTGAGTGTGCCTACTTGTGCACCCCCAAAGTTGGAGGATAGATGTCAGGTGACACCAAGTTAAATGACATATTTATGTATTAGGCCTGAATAAGATATCAAGACATTTGAGTTTTGATTCTTATTCTTAGATGTTAATATGAGTGTCCTGTTTTTACAGACTTCTACAGAGCAATAATATATCAGGATCAATTCCTATGGAGCTTGGAATGCTTAAAAAACTCAAGACAATTGATCTTTCTGATAACAAGTTAACCGGCGAAATCCCGTCATCTTTAGCTCAGCTGAAAGGCCTTCAATACTTGTAAGCAAAGAGAATTCTGAGATCTTATACAAACAAGTTTTTTTGGGGTTAAAAACTAATGATGTTTGTTCTTGATTTTTGATGATTATAGGAGATTAAACAACAACAGTCTAACTGGAGCTATTCCTTTGGACTTGGCCAATATGACTCAGCTCTCACTTATGTAATTTTCTAATACCATTTTCATGTTATGTTATGTTATATGTTCTTTATAACAACATTTTGCTATAGCAGCTAAAAAATATCCAGACAAATGACGTACGTAGTTATAGAGAGGTTTGACTGTATTTTTCACTTTGGACATATATGTATTATGGAGTTGTGTGTTTTGCATGTTTAGGGACTTGTCTTTCAACAATCTCAGTGGTCCTGTACCAAGGCTTCTAGCCAAAACATTCAAGTAAGGTTCTTGAAGAATTCTCTGTGTATCACTTTTTCCTTAATATCAGTAAATGAAACCAGTTTTTTTGTATTTATTTGTCTTTTTTTTAAACCTTATTGACACAGTATTTTGGGAAATCCAATGATATGTGCAACTGGAAAAGAGTCAGAGTGCAATGGAACAACACCAATGCCCCTCTCCTTCTCTTCAAGCAATCCACAGAGTAAGAACTAACCTTATTTTAACTGCCAAATGATAGTTTTTTTGTTTTTTTTATGACCGTATTGTCTGAGCTAGCTTTTGTACACCTTAACTAATTCCACGAGATACCTACCACCTCCCACCAGCAACATGTACCAGGTAACTCTCTCCACCAAGGTTAGGATGGAGGGGAGAATCACCTGAGACCTCATGGCTCTTAACCACTTCATTGACCAGTAGGCCACACCCTTGGGTGCACAACCAAATGATAAATAAGCGCCTCTCTACCTTCACAAAGGTAGGGGTAAGGCTACATACACACCACCCTCTCCAGACCCCATATATGGGATTACACTGGGTATGTTGTTGTTGTTATTGTCGGTCGATATAATGATTATGGTGCTTTTACTAGTCATACAGATTTATGAAGTAAATTTGCTTCACTGGTTTAATTTTCTCTGCTGATAGATTCAATATTTATAGTTCATTCCCAGCTAAAAATATTCTGTGTTATCTCAGATGTTCAGCCTTCTGGAAAACCGAAAACTCACAAGGCTGCGTTAGCCTTTGGAACAAGTCTAGGATGCATCTTCCTACTAATCGTTGGGTTTGGATTCTTTCTGTGGTGGAGACAAAAGCACAATAAGCAAATTTTCTTTGACAGTAATGGTGAGTATTGCTTGATCAGAACATTTTGAAGTTACGGTCTAGCTCATCGATCCTCTCTTATAAACGTTACTGTCATAAATTGCAGAACATCATATTGAAGAAGTGTGTTTAGGCAACTTAAGGAGGTTTCAATTCAAGGAACTTCAGTCTGCCACAAACAATTTCAGCAGCAAGAACATACTAGGGAAAGGTGGTTTTGGTAATGTTTATAAAGGTCGTCTTAGTGATGGGATTATTGTAGCTGTGAAAAGGCTCAAAGATGGAAATGCAGTTGGAGGCAATCAACAATTTCAGACTGAAGTCGCGTTGATCAGTCTGGCAGTTCATCGGAATCTCCTCAGGTTGTATGGATTTTGTATGACACCAACTGAAAGGTTGCTGGTGTACCCTTACATGTCTAATGGAAGTGTAGCTTCGCGTCTCAAAGGTAAACATATCTAAAAACCTGTCTCCTTTGGCCTATAATTACATTATCCTTCTTATTTGCTCAGCTCATAACGTATCATTTCTTGCAGCCAAACCAACCTTGGATTGGGGTACAAGAAAAGGAATAGCTTTGGGATCTGCAAGGGGTTTATTATATCTACACGAACAATGTGATCCAAAGATCATTCATAGGGATGTGAAAGCTGCAAATATATTGCTTGATGATTACTGTGAAGCTGTTGTTGGAGATTTCGGATTAGCAAAGCTTTTGGATCATCACGATTCACATGTCACAACCGCGGTTAGAGGAACTGTAGGACATATAGCTCCGGAATATCTCTCCACAGGACAGTCCTCTGATAAGACGGACGTCTTTGGTTTTGGTATTCTCTTGCTAGAGTTAATCACTGGACAAAGAGCTCTGGAATTTGGTAAAGCAGCAAACCAAAAAGGTGCCATGCTTGATTGGGTAAGTACAACAAATCCCTATAATTATAGTTACATTTCTCTTTTTCAGCATGCTTTGACATGACTTCTTGAGCCGAGGGTCTATCACAAACAACCTCTCTACATCCCAAGGTAGAGGTAAGGTCTGCGTACACTCTACCCTCACCAGACCTTATTAGTGGTATTACACTGGATATGTTGTTGTTGTATATCTGTAAGGTGTGCATTAACACTATCTAAAATATGAACTCTATAAGGTATTAAGAGATGCAATGAACGAAGTACGAACCATCTTTAAATCCTGGATCCGTCCTTCTGTGTTTTTTTACCTGCAGGTAAGAAAGATTCAACAAGAGAAGAAGCTAGACATGTTGGTGGACAAAGACATGAAAAACGAGTACGATACGATAGAGTTAGAGGAAATGGTTCAAGTAGCATTATTGTGCACTCAGTATCATCCAAGTCATAGACCAAAAATGTCTGAAGTTGTAAGAATGCTTGAAGGAGATGGACTAGCAGAGAAATGGGAAGCATCTCAAAGAGCAGAACCAACAAGATGCAGCAGAGCTAATGAATTTTCTTCTTCAGAAAGATACTCTGATCTTACTGATGATTCTTCATTACTTGTACAAGCAATGGAACTATCTGGTCCAAGATGATAATTGAATTAAGACACAATGTCACTTTTGCTAATGGGAGGTGACAAGTCCCTCGTGGAATAAGTCGAAGTGCGTGAAAGCTGGCTCAGACATCACATACATAAAAAAAAAACATGTGTCATATTGAGCTTTGTTTATAAGTTTATACAGAAGCTGCAAATAGAATATCATACATTCTGATTGTAAAGGATAATATAATAACTTTGTAAGGCTGAGAGTTGTTTTTCCTTTTTTGGTTCTTGTTTGTTTGAGTGGAAATGTTTATGGGATGTTATAATTTGCTTATGTGTATATATAATTAGAAATAGAAC

>PGSC0003DMG400002902 LRR-RLKs

AGAAAAGAAACAGAGACGCATTTGCAGATTGTTTCTTTAAGGGGAAGGTGGACATGTCTAGTAAACATGGCCGCCGTTCCCCTAATCCTTCATCTCCTTCCTCAAGCATCGTCTCCGAAAAAACACCATCTTTTCCTTATTCATATCTCCATTTTCTTCCTCATTTCTCTATCTAATCCACAATTTTCTTTCTCCATTTCCGAGGACGATGCCTTGATCAAATTCAAAGAATCTTTAAAAAACACAACATCTTTAGATTCATCTTGGCTTAAAGGTACTAGTCCATGTGACAAAAACAAGAAATGGGATCGTATTCAATGCGAAGGGAACGTGGTTGAAGGCCTCCTTTTAGGTGAAGCTGGCTTATCAGGAGAACTAGATATTGATCCTTTGATTGCACTTCCAGGTCTCCGAGTTCTTGAAATCGCGAATAATTCATTTTCAGGCACAATCCCTGAATTCTTCTTGCTTGGTGCACTTAAATCTATATATATAGATGGAAACCAATTCTCAGGTGAGATTCCTAAAGATTTCTTTGCCAAAATGGGGTCTCTTAAGAAGATTTGGTTTGCACGAAACAAATTCTCCGGTTCAATTCCTGAATCTTTAACCAACTTGAAGTACTTGATGGAACTTCGTCTTGAAAGTAATGCATTTTCAGGACGCGTTCCATCTTTCTCACAAGTGAGTTTAACATCTATTGATCTTTCGAATAACAAATTACAAGGCGAAATCCCTCAGGGGATGTCAAAATTTGGTGCTGATCCTTTTAAAGGAAATGCAGACTTATGTGGACAACAATTAGGGAAAGAATGTAATAAGGGAATCAATAAGACTCAAGGTGCACCAATGTCAAAATTGAAATGGATCATTCTTGGTATAGTAGTTAGTCTTTTATTGGTGACCATTCTTTTTCGCGTCAAGCGCAAGGAAGATCATTTCGACAAGCTTGGGAAAGAGAATCTTGATGAGGGGTTGCACGTTGGTAGCTCGATTAGGAAGAGCATGAGCATTCATAGCAAAGGAGGTGATTCTGGTCGTGGATCAACACGAAGGGGAGGGTCACAAGGTGGTAAGGCTATGGGAGATCTCGTGTTGGTGAACAACGAAAATGGTACATTCGGATTGCCTGATTTGATGAAGGCAGCTGCTGAGGTACTTGGAAACGGAGTGTTAGGATCAGCTTACAAGGCAAAAATGGGAAATGGAATGTCTGTTGTGGTTAAGAGATTGAGAGAGATGAATAAAATGAATAGAGATGTATTTGATGCAGAAATGAGAAAACTTGGAAAGTTAAGACACAAAAATTTACTTCAATTATTGGCATATCATTACAGAAAGGAAGAGAAATTGATGGTATCTGAATATGTTCCCAAAGGCAGCCTATTATATCTGTTACATGGTATGACAATTTCTTTCATCTACAGTCAAACCTCTCTATAACAACATCGTTTGTCTAGATAAGTTTTTGGCTGCTATAGCGAAATGTTGTTATCTAGAACATAAGTTTCATATCTCAACTGTTACTCCCTTTGTATTGAAACATACCTCGATGATGAGTTGGCCTACACACACCTGTTGTTGATTGAGAACAGATATTTGACCAACTCAACATCGAGGTGTGTTTTGATACAAATGGAGTGAATAGTTAAGGTAGGTTAAGAGAAAAATGGATAGTTAAAGTATGAAACTCGTGAAAAAAGGATAGTTTAGGTGTGTTTTTGACCATTATGGTAAAAATCTATTCCGAAGAAAACTTGGTCACTATTATAGTGAAATGTTATTATAGACAATGATTGTTATAGAGAGGTATGACTGTATATCAAATCATTAAGGTCTTCACTTCTTCTTGACATTTGTATGAGTAGATTTTATCTGATAATGTGATCCGATCTATACTTACCTGTCCCTTCCAAGACATCCGGTCACAATTGGGACGATACAAAGAAGATTAGCATGATCCCACAAGTTTTAGGATTCACAGTTTACAGAAGGATTGCTTATGACTGTTAGTACATTCATATACATTTTGTTCTTCTATGTCAACTATCCTAATTTTGAAATTTCGTGCATACTTTCAGGTGATCGAGGGATATCACACGGTGAGCTAAACTGGCCTACTCGTCTAAGGATCATCCAGGGAGTTGCTAGTGGAATGTGTTATCTTCATTCCGAGTTTGCATCGTATGCAGTGCCTCATGGAAACCTCAAGTCTAGCAACATTCTACTCAATGAAAAATATGAGCCACTTCTATCAGATTACGCCTTCTATCCACTGATCAACAACACGCAAATTGTTCAATCCCTTTTCGCGTATAAAGCACCAGAAGCCATACAACATCAACATCTCACTCCAAAATGTGACATATATTGTCTTGGAATCATAATACTTGAAATCCTTACAGGGAAATTCCCTTCTCAGTATCTCAACAATCAAAAAGGTGGAACGGACGTTGTTCAATGGGTGCAATCAGCTATAGCTGACCAAAGAGAATCTGAATTGATTGATCCAGAGATAGCGTCTGCAACAGACTCCGTTGAACAAATGGTGAAACTCCTTCATGTAGGAGCTGCTTGTACTATAAGCGATCCGGATAAGAGGATTGATATGAAGGAAGCATTAAGGAGGATAGAAGAGATAAGTTTAATATGATGAACATAAACCTTCCAAAAGGTAATAGGTTAGTAGATTCGTGTGAAGCTTTAGTTTCGTTACCTGCCAGAGCGATCCTTAGCCTCCTTTGATTCCGTTTTATCTCAGTTAAACTCCACAACGTCTCTCGACTCGTTCAAATGAATGTTGATGCAGGAAACAAATGTCATTTTCTTCAAATACTGGAGCTGAGGTTCATTTGCCAAACAAAACAGAGTTTACTGTATATAAAACACCAAAATGAGACTTCTAAATTACCAAAAACATAAAGTTGATTACTAATTCTACATGTTTTGTTTTATTGTATAGAATTTGTTTCTTTTTTTGTGTGTGTGAGTTAAAGAGGGACATGGATCCTCAAGGCAATGATCAATTTAGACATAGATTTTGAAGACAACCTTCCGGCTTGATCTCCTCCTTTCTGCTTATGTTAGCTCACGTTGTCACACAATCTAATCTAACGCGCTCCCATCAAGAGATCGTTTGATGATTTAGCAAACAAGATGATCTAGCTTTGATTTGTCAAAAAACCTATGTTGTTCGGACTCTCCAGAAGGGTTTTCGTGCACTTGTTAGATCCTCCCAAAATGCACTACTTTTGAAAGATTTGACATGCATCCTATGACATTTG

>PGSC0003DMG400003002 LRR-RLKs

ATGTTTGACACAACTGTATGTGATGAAACAGTGACCAGAGATGCATTGCTCAAGTTTCTTGCAGAGGTATCTAACAATAATAGTACAATTCTTGGTACTAATTCGGGCTGGAACTCGACATCTGATCCTTGCAAGGACAAGTGGCTTGGTGTTACATGTAACGCGAAGACTCTCTCCGTGGAGAAAATCATTCTTGATGGTTATGATTTTTCGTCTGGAACGTTTGATGTTACCACGATTTGTAATGTGGAGCCTATCGCGCACACCCTTTTCTTCCTGAGTCTGAAAAATGGCCTACTTCAGGGGAAAAACTTTGACAGTATAGATAAATGTAAGCATCTAAGACATCTTGTTCTTGGTGGAAACCGGTTCTCAGGTAGCATTCCACAATCATTTTCAAGATTGAATAACTTGAAAAGACTTGACATATCCAACAATAGATTATCAGGTGTTCTCCCTGATTTATCTAGGATTTCGGGCTTGATTGAATTCGAGGCTCAGAATAACCAGCTATCAGGAGAGATACCTGAATTTGATTTCTCAAACTTTGTTGTATTTAATGTCTCTTTCAACAATTTCAGTGGATCGATTCCGCCTGAGGGTGATAGTTTCCCTGTTAGTAGATACTTGGAAAATCCTCTGTTGTGTGGCTCCCCGTTGGCTGGAAAATGCTCATCACTGTCCGAAGATTCAGAATCAGCTGATCATGACAAGCCAAAGAGCAGCTTGTCAAGAGATGAAATTCTCATGTTTTCAGGTTATTTCCTCATAGGCCTGGCTCTGGTTCTTCTTATCCTACTATGCGTCTATAGAAAACGGAAGCAGAAGCAGCAGAAGAAGAAAGAATCAAGAATCGAGCTGATGAAAAAGGTTGCTACAGTTGATCATGACAGTACTACTACATCAATGGATCAATCAAAGAAAGGCGCGAGCAATTCAGATATCTCATCAGCTGTCATTTCTTCCGATGACAGCAACACCGCAGTTTCACAGTCACTCGTGGTTCTTACGAGTCCTGAGGCGAATAACGGATTGAGATTTGAGGATTTGCTTAAGGCTCCAGCAGAGTTGCTTGGAAGAGGGAAACATGGTAGTATGTACAAAGTGATGTGTGACAATCCCAAAATGACACTGGCTGTTAAGAGGATAAAAGATTGGTCAATAACTGGTAGTGAATTCAAGAAGAGAATGCAGAAACTAGACAAAATCAGGCATCCAAATGTGTTGCCTGCTGTTGCCTTTTATTCTTCTAGACAAGAGAAGCTTTTGGTCTATGAGTACCAAGACAATGGCAGTCTTCTCCTACTTCTACAAGGTAATTCACTCTACTAGTCTATTCTTCTGTGTATTGAATTGTGCGACCATTTCATCTAGGCATATAAACAGACACTCGAACTTGGCCTCAACTGGCAAGTAAGCACTCAACTTTGAGAGAGTGCACATCTAGACACCTCAAATTGGTCTCAACTGACGAGTTGAGGTGTCTAGATGTGCATACTCAAAATTGGAGTGTTTACTTGCAAGATGATGCCAAAATTTGAGCGTCTGTTGATGTTTTATCAATTATGACTTTCATTTAAGAACGTAAGTTGCATGAAGGTCTGATTCTTTTTCATCAGCTAAGCATGTGAACATTCTTATTCATAATAGGTGCATAAAGGTTAGATATCAATATGTTCTATTACCATATGAAATTGTGAGACGTGCTCATCTAAAGACTTAAAAACTGTTAAAAAGCGCGAACTTTTGCTTACTTAACGATGTTGGATCAATGAGTAGTATTAAGGCTTTTAGGCATGTGAAAGACTTTATTTAAGGTACTTATTGTCCCCTCATGATTTAAACATGTCAAAATTCTCTTTTAGAACCTCAGTTAGAAACCTCTGTTTGCCTTATTATCTTCAACACTGGTGTATGAATTTGCTTTTCAGGAATCCAGACAGGCCAAAGATTCCATTGGAGTAGCAGACTTTCTATAGCAGCAGGTATCGCCGATGCTTTAGCTTTCATGCATCAGGAGCTTCAACATGACAGAATTGCACATGGAAACTTGAAATCATCAAACATTTTCTTGACCAACAACATGGAGCCAATGATCGGCGAATATGGCCTTATGAGCATTGCAGAAATCAGTACTACTACAGCAAGCAATTCCCATTATGTCTTCCCTATAACTGATGAAAATGTAGGAGCAATCTTTAAAGAAGATGTTTATGCATTTGGTGTCATTCTTTTGGAGTTGCTAACAGGGAAAGTGCAAAACAATGGAATAGAACTTGCAAGTTGGGTTGTTTCTGTGCTAAGAGAAGAATGGACAGTTGAAGTGTTTGACAGAACTTTGATACAAGAAGGTGCTAGTGAAGAAAGGATGGTTAATTTGTTACAAGTTGCTGTTAAATGTGTGAACCATTCTCATGAAGCTAGGCCTAGTATAAACCAAGTTGCTCTCATGGTTACCACTATACGAGACGAGGATGACAGATCCGTCGATGTTTCTGCAAGTACTAGTACTATCATCTGA

>PGSC0003DMG400003175 LRR-RLKs

ATGGGTGGTAGATGGAATACTTATGGATTCAAGTTATCATATCTTGCACTTCTGATTATTCTTTATCACGTTCACGAATGTAGCTCACTCAATTCTGAAGGTAATTATATTCTATGTTTGGACTGTCCTACTGGTTTTGGTGTTTATGTCTTAGTTGTATCTAGTTTGTTATGTCCATTTGAATATTGGATTGGTCTATTTTACAAAACAAGGATTGGCATTGTTAGGACTACGATCTAAAGTGGATTGTGATCCATATGGAATTCTTGCAAACTGGAATCTTGATCATTGTGACCCATGTATGTGGTTTGGAGTACAGTGTCTGGCAGGCAAAGTGCAAATGTTGTAAGTCCCTTGTTACATACGATCATATCTGTTAATGAGTATTCAATTCCATAAAACTTACTTTATAATTGTAATGTTTTGTTTCTTTTTTAGTTAAGAACAAATCTGTTAGATTTATTTTAATGACAACAGATGTGCAAAAGATGAACTCATATGTTTTGACTTTTGAGCTATCTCTCATGTTGTTAATTATTTTCTTTTTCTTCTCTGTTTAGGGATCTTCATGGGAATTCTTTGGAAGGCACATTGGCACCAGAGCTTGGGAATCTCACTCACTTAAGATCAATGTAAGTTCTTACCTACGACAAAATGTTTTTTGCTTTTTTAGTTCATAGGTGCAAATTTGTGGGGAATGAAAGAGAGATTGTTGCTTGTCATGGTTGAAAATTGATTAAAACTTAGTATTGGTTGAAAATAGCCTGGCAGCTTAAAAATACCTCCTTCTTTTGATTTCTTTTTTAACATGCAATTTTTTATAAAAAAAATTGATTTTCCTCTTCTTCTTTTGGTTTTGTGCTTAAAAAAACTCCCTAAATAATCTTGTCTTGTTTGGAAGCTCTTTTGAAAGAGCTTGCCTCTGGACAATGATAGGATTTGAAAGGAGTGTTTCTAAGGATCGTTCTTCCGTTACAATATTCCATAAATCATAGGTTTCAAGATATGAATTCATTTTCACTGACCAAATCTGATAGTTTTTGGTAGTGAAAATTTATGGGACAATGAAAGAGAGATTGTTACTTGTCATGGTTGAAAATTGGTTTAAAATAGCCTTGCGCATTGAAAATAAGCATAGGTTGAAAATAACCCCTAATGAGTTCTAAAAAATGGATTGAAAATAGTCCTGGTGGGTTAAAAATAGCCTTGACAAGTTAAAAATGGGTTGACAATAGTTCTTTAGAAGAACTCACAGATCCCATAAGATCAATGGGGCTTTGGATACTACTGTTGCTATTTTTTTAAAGATGGTGCAGCTAAATAATGTTGAAAATGTCTCTCAATTTTTATTAGGCATAAGAGGGGTTTAATAGACAACTTGAAAACATAACTTGCAGAAAATCTGCATAACAAAAAGTAGTAGATTAGTAAATTGGGTGGAATAGACTAAAGATAATTAGTTCCATGTCTAATAATCATAGTTTTCATCATCTTGCATGTTGTATCTTCTTGTGTGTTGTGTATGACTCCTTTTTTAATCTTAACAATTTAAACATTTTTTTTCTCTATGATTGACTCACAGTGTGCTGTCAGAAAACAATTTTTTTGGTGCCATTCCAAAAGAATTTGGATGACTTCGAAGGTTGGAAGTGCTTGACTTGACAGATAATAATTTGAGTGGGAGAATTCCAGCAGAAATAGGAGACCTGCATGCGCTAAGAAGCTTGTGAGTTCATAATTTAATTTGTGTGCATGTGAAGATGTGTGCTGCCTATATTAGTTTATATATGTGAGTATTCATGAGGAAGAACCATTAAAAAGATTCTTTTGAACCGTGTCTTATACCTTTTCACAGGTTGATTCGTAACAACAACTTGGAAGGAAAAATTCCATTTGAAATTGGGAAGCTTCGTTTGCTCTCCAAATTGCAATTTGACGAAGCACTCACCCCTGCTGCTGGGGGAACATGCTGTTTAAATAGAAAATTTGCACACTGGTAAGGTTGTTTTGGTTTTATTGGACCTTTTATATCTGTGTTCTGGTTCTTGTTACATTTATGTAACCAAATATCAACATGCAAAGATGTTTGAAATGTATGAAATTTGTATTTAAGCCATTTCAGGCTGCCATTATTTTCAGAAATGGTCATAACTCCTGTCATAAGGTTTTCTTGGTGGGAGTGAGGGGGTGGGTGAACACAAGTTGATAAAGTTGAGTATTAATGACAACAAGGAATATTGTGGTTTATGTGAATTCAAATAAAGCCTGGGAGATAGGTGATTTCATAGATGTGGCGATGCAATGACTTGTGCTGCTAATGCTTGAAATTTACTGTCTAGTTTGACATCTAATACTAATACTTGACCATACATCAACTCAAACGAAAAAGCTTTGTAAACTACAAGGAGGTGATTTTCCAGTCATTGCATCCATCCAAGGCTGGGGTTGAGGATGAGGGATGCAATATATTGGAGATCTTGGGATTGATCCTGGAATCAAGAGAGCATCAAGTTATCATATTGGTTCTAATTTATTTTAATTTTTTATCAAGATACTCTTCTCAGATAGCTTTAACTCTTACATAGAATACTGACTATCAAACATAATAGCATGGTTACTTTCAATAGCACTTACCTTCTTAATCAGATATTTTCATTTGCCTGCTTCTCTCTGGTAATTTGATGTCTGTTTCTGGCAGCATTTGTCATGGCAGTTTGATACCATTAAAGACCGTAGACTCCTTTATCAGACCACTTAAAGGGACACTTGTGCGTTATTTCACTCTGTAAGTAACTAAATGGTTTTCAATCACCATGTGTTTCTACTTACATTTTATCACTCTAGAAGCGTCTTTGTGTCAATAAAGGTTCTCCAGACAACTTGGAAGTGGCTCATTGGATAATCAGATAGGGTATTCCTCTGACAACCTACCAAGTGAGTGAAAAAGTTTTACCCCACATCATTAGCAGGAATTATGTTGAAAGTGCACAAAAGATTGTTTTCACTCTTCTTTTTTAGCAGGTTCATCTAGGCCTCACATCATCCATACTATTTAGAACCAGGCGAATATCGCACGCCACAAGTTAGTAGAACAATCTAGCAACCTTGCTGCTGTTTCTGCCAATGCTGGGAAACATTTGGGTCCTATTGTTTCAGTGCCAAGCACCAGAAGTAGTGGGTCATTTCATGCTGTGAAAAATACCTAAGGAACACCTCCTACACCCTTACCTGCCTCGCAACGACAGCCTGAGTCTCAATCCAAACCAAATCCAATGGGACAGTCAAGTAGTGTTGTTAAATAGCCAACTAATAGACAAGCTCCACCTGCCCAAAAATCTGGTGATACATGGAAGTATTTAGTTGGAGGTTTTGGGTTTTTTCTGCTCATTGTTGTTGTAGCCGTGTTCTTAGTTTACAGAAGCAGAGCTGCAAAACCAATAGGCCCTTGGAAGACTGGGTTGAGTGGACAGCTGCAAAAAGCATTTGTCATCGGTACTTCAATTGAAATTTGTGACAAGAATTACTGTTAATATTCTGTCCACATTCTGATATTCAATGACAAAATCACTGTTGGTTGATAATAGTCCTAAATCACAGCCATAAAAGAAGGTATAATCCTGATGTTGATTGCAAAAAATTGTTTCTCATTGCAGGGGTCCCTAAGCTTAACAGGTCTGAGCTTGAAACAGCATGTGAGGATTTTAGCAATATAATTAGTAGTCATGATGCATATATAATCTACAAGGGAACTCTTTCTAGTGGAGTAGAGATTGTTGTTGCCTCGACTACTCTAACTTCTCTCGAAGATTGGTCTGAGAGTTCAGAAATAGCCTTTAGGAAGAAGGTTTGTTGTTTCCATTTACATCAAAAATAAAATATTATCTCTCCATATATCTCACAGTCTTACACTTCACATAATGACATTGTTGTTATCTCTTCATCCTTTTCGGTTGTAAAAGACATGACATTTTGTGATTGTGTTATTGAACAGATTGATACACTTTCAAGAGTGAACCACAAAAATTTTGTTAATCTTATTGGATACTGTAAGGAGGATGAACCTTTCACTAGGATGATGGTCTTTGACTATGCTCCAAATGGAACTCTTTCTGTGCATCTACACGGTAAGATTCTTTGTTGTCCACAATTCAGCTTTAATTACTCACAATGATGGCTGGAGTTAGTTCAGAATAACACTTCTTATTATTTTATTTTAAAAAAAAGATGTTTAGAATGACTTAAAATATAAGTTTAATTGTGTCAATTGAAGAAACCGATCCTCTTGAATGGTCTGCAAGGATGAGAGCCATAATGGGGACAGCATATTGTCTTCAGTACATGCATGATCTAAATCCTCCAGTGCCCCATTCCAGCTTAGATTCAAATGCCATATTTTTGACTGACGATTATGCTGCTAAGGTCTGCTCCATCTTCTTCAATATGTATATACAATTTATCTACCATATTGAACTAAAAACCATTTTTCATTTGTATTTCATATTGAAGTAGAATTTTTTATATTTTCACAGATTACAGAGATTGGTTTCTGGTCAGAACTAATAGCCAAGTCTAAGGACTCTAGGGATGATTTAGAAAATTCTGAGCTGTCAGCTCTTGCTGATCTTGAAACAAATATCTATAGTTTTGGGATTCTGTTACTAGAAATAATTTCTGGAAAGTCACCTTATTCAGAAGAACAAGGATCTCTTCTGAACTGGGTAAATCTCCTTCATATGTTTCCCTTATTTTCTTATACTTGATCGTTACAATGCGTTCCCTAGTCTGCATGCATTTTCTTGTGTGAATGTCTTTTCTTTCAGACTTATACCAATGCAAAAGCATATTGATAACTATTATTTTATTTAGGCGGAGCAATACCTTAAAGATAAGATGAATGTGAGCTCCTTGGTTGATCCAACACTCAAGTCCTTCAAAAATAATGAGCTTATGGCTATCTATGTGGTAATTGAAGAATGCCTTCAAGAAAATATAAGGAAGAGACCAACGATAACTGAAGTCATTGCAAAATTAAGGGAAGCGATTGATATATCGCCTGATGCTGCAGTTCCAAGGCTTTCTCCCCTCTGGTGGGCTGAACTCGAGCTCTTATCTTCTGAGGCAGCTTAG

>PGSC0003DMG400003210 LRR-RLKs

TCGCAGTATGGATCTAAAAACGTGCTGTATTATATTAAAAAAAAAAAACTGACAGAGACTGACTCCCTCATTAAGAATTATTGAACAACCAGCCAAAGCAAAGCATTAAATATTCTCCTATTTTGGTTGTTTCCTAAGTCATTCATTACCAAATACCACACTAATTAAGCATATGATTAGCTGAGGACACATATTATGAAACAACATTCTCTGGCGCCTCAGCCCAAAAATTAGTTCTTGAACTAGCAGCTCAGTGTTTTAAGAGGTCAGAGATGTTCTTGTTTCTGGTTAATTGCCTTTTCTTCTTTGCCAATTTATTTGCAGTCTTTTCGACCTCATCGCCTCAAGATTCTGTTCATCTACTTGGGTTTCACAGCAGTCTTCCAGAGCTCTCTCAACAATTGCTCCCTTGGAATCAGTCTGTCTCACACTGCCGATGGAAAGGTGTTACCTGCTATTCAGACATAACTTCTCATGTCAAATCATTGATTCTAACAGACTTATTGCTACCTGGAACCTTGGACAAGGCCTTTCCTAATCTCTGCCGTCTTCCTCGCTTGGTTTCTCTTGATCTCAGTGGCAACCATTTCACTGGTGGCATCCCTGATATGCTTGCTAACTGCAGCCAACTTGACACGATTCTACTCAATGAAAACAGATTTTCAGGATCCATCCCACCAGAGATTTTCAAATCAAGCAAACTCATAAAACTTGATTTGGGTTTAAACCAGCTCACTGGCACTATTCCCTCTGAGGTAAGCCTTTCTACAAACCTTCAACATCTCGGGCTGTGGAACAATTTCCTTAGTGGAAATATTCCAAAAGAGCTATTTGATTTGCCAAACTTGACACATCTCCACCTCTACACAAACGAGCTGACAGGACCTCTACCCGACTTCCCATTCTCGTGCTCCCTTTCCGAGTTCTTTATTTATGAGAACCGGTTCTCAGGTTCTTTGCCAATTACCCTAGGCAATTGTCACAACCTTACTTCATTCTCTGCATCATCTGCTCATCTTGGAGGAGTTATTTCACCAGAGGTTTTTAGGGACCTTTCAAATCTTGAATTTCTCTATTTGGATGAGAACAACTTTGAAGGGGGAATTCCTGAGACCTTATGGGATGGGAGCTTACAAGAGTTAGCTCTTTCCTTGAATAAATTTAATGGTAGTATATCCGAAAAGATTGGTGGTTCTCTTCAGATAAATTACATTGATTTGTCAGTTAACAAATTAACTGGTCAGCTTCCAAGATCAGTTGGACGTCTGAAGAATCTGAACAAGCTTTTTCTGTATGATAACATGCTTAGTGGTTCATTACCAGCAGAACTTGGGAACTGCACTTCTCTTGTTGCGATAAGTCTCGCCACTAACTTCATCGGTGGGGAAATTCCTCAAGAGCTCTGCAACCTTCACTCCCTAATAAAATTTCAAGTGTACGGAAATCAAATTCAGGGCCAAATTCCAGAATGCATTGGCAGAATAAGTGGACTGGAGGAGCTGGATCTTAGTGAGAACCGATTGATCGGCAAGATACCACCTGGAATCACAAATATGACGAAACTTGTGTTACTCTCTTTGGCTCATAATAATCTTACAGGGGAGGTACCACCTAATATTGGAAAAAACAATTTTCCTGGGTTATTTAAGGTTGATTTAGGTTATAATAACTTCAGTGGACCAATTCCTTCTGAACTATGTAATGGCAACAGGCTTGGAGTCCTGGTTCTTGAAAACAACAGTTTTAATGGAAGCTTCCCAACATATATAGCAAAATGTGAATCCCTCTATAGGGTAAAAGTCCCCAACAACAATCTGCAGGGGAGTATTCCTGACTACATTGAAAAGAATGAGAACATCTCTTACTTGAATGTTCGGGGAAATATGCTTGCGGGAAGGATTCCAGCAGCATTTGGTTATTGGACCAATCTTTCAACGATTGATCTTTCAGAGAATATGTTTTCTGGTTCCATACCTGCCCAAATTGGGAAGCTGCAAAATCTTGTAAGACTGAACATTTCTTCAAACAGACTGACAGGAAAAATTCCTCTTCAGTTAAGTTACTCTGCGAAACTGGAAGAGTTGGATCTCAGCAACAACAATCTTTCGGGAAGAATTCCGAAAGAAATTGCATCATCTTTAGTACTGACAAATCTTCTACTCCAGGACAACAAACTTTCTGGTACTCTTCCAGACACTTTCTCCTCCTCACAAAAGCTTGTAAAGTTGCAGCTCGGGGACAACTTGCTTGAAGGCTCAATTCCATGCAGTCTGAGTAAACTTAGGGAACCTAATGTTGCACTGAATCTGAGCATGAATAAGTTTAGTGGTCAAATTCCTAAGTGTCTGAGCAATCTAGACAATTTAGAAATCCTTGATATATCAAGCAACAACTTGTCTGGAGCAATACCATCAGAAATGGACAAGATGAGGTCCCTTTCGTTTCTCAACATATCATTTAATAACTTTTCAGGGAAGGTACCCATTTCATGGGGAAAACTATTAAGTTCACATCCAGGAACTTCCCAGGGAAATCCAGGACTCTGCTTATCAGACACTGAAAGTAGCAGCTGCAAGCATGTGAAAAAATCACAAAGAAACTGGAAGACCTTGGCTGGTGTAATTAGTGGATGTGTGTTTTCCGTGGCAGTCATAGCTGCGGCAATATACTTGCTAGTGACTCGGATTCAGCACCCTTCCCTGCTGAATAAGCATCGGCTCGTCAAGTATCATGCAAAAATTGAAGATCTGCCAGATCGTATAAATTTTGAGGACATTGTACGTGCAACAGAAGGCTGGAGCGAAAAATATGTAATTGGAAGAGGCAAGCATGGAACTGTTTATAAGATGGAATCTGCGAAATCCAAAAAGCTTTGGGCTGTCAAGAAGGTTGACTTAGCTCAGAGAGCATTCAGCGATGAAATGAGAAGCCTAAATTCTGTCAGACACCGCAATTTGGTAAGATTGGGAGGACACTGCACCAGACATGGATATGGCTTCATTCTAACAGAGTTTATCCCTGGAGGAACTCTTCATGATGTCCTTCACCAGAGAAAACCACCTGTAGTCCTGGACTGGGAGCCCCGCCATCGTATTGCTCTTGGTGTTGCTCAAGGTCTGTCTTACCTTCATCATGATTCTGTCCCTCAAATCATTCACAGAGATCTCAAATCAGATAATGTAATGCTGGATACTGAGATGGAGCCAAAGATTGGAGACTTCGGGATTGCTAAAACAGTATCAGACTCTGATGAAAACTCGACAAACTCCAAAATTGTTGGGACACTAGGATACATTGCTCCAGGTTAGCCACACGTTACTATGCTAGTCTTCAGTAATAATTTAGTTGGCAGATTACTTTTGATGATTTAATAAATTTTATTATTTTCTCCCCCGGTCCCCCCTCCTTCTCCTTGTCTGTCCATCAATTCAGAGAATGCGTACTCAGTTCAATTGACAGAGAAAAGTGATGTCTATAGCTATGGAGTTCTTCTACTGGAGCTCTTCTGCAGAAAAATGCCAGTAGACCCCAGCTTTGAAAAAGGATTAGACATTGTTTCCTGGGTGAGGAAAAACTTGCACAGAAGCGACAACAATTTCCTTCACTTGCTTGATGAAGAAATTAGTTTTTGGTACATAGAAGAACAATGGAAGGCCCTGAAAATGGTGTATCTAGCGCTTCAATGTGCTGAACTCGAGGCAAGCACCAGGCCTGCAATGAGGGATGTAGTAAGGTCTCTTGTCGAGTTAAATCATAGATGTAAAAGTGAAGAAATGAAACTTTGAATAGTGAAAGCCATAATCAACTTTTCCATATTCCAACGTCTTTTTTTTTCTTTGCTTACATAAAAATTGAACCATTTCTCGATTTGTGTGTGTCATCCTTGCATTCAAGGGCTAAATTCCAACATAACCTGATCGTGGGTGTTTCTGTTTAACTTTGTGACATCCACTGCAACATTATAGATTAAGAAAATGCATGACGTTTCAAATCAAATTTTGGTAAAAGCGGTTTTCGCATTCTAGCCATAGCTCTTTTCATAATGGATCACTGTCCTTAATCAACTTTTGCTTGTTACTTTCTCCAGCTGCTCCAAAATCTGAACAGTTTTGAG

>PGSC0003DMG400003211 LRR-RLKs

TAAAAATGATGTTTGCGGGATACATAGAGTTGTGCACGGGATACATTAGGTTTAATACATCCCACGTAGCGGGATACATAGCGTTGTGCACGGGATACATAGCGTTGTGCACGGGATACATAGGTAAAATAAGGGATTTTTGAAATTTTTGAAATAAGTAGGGATAAATGGATATTAAGGTAAGTAAAGGAGTGTAGTTAAGTAATTTGTCCTAAAATAATCTCCTATATTCATTGTTCCCTCGGTTTCACACGTATCTTGAAAAAACAGGCTGACCTCAACTTAAAACCTAAATAATAGTCTCAGGTTCCAGCCCAAAAATTAGTTTTTGAACTAGCTCCTCATTGGATGCAGCTTCTTTAAAAGAGTTCAGAAATGTTCTTGTTTGATCTTGTTCACTGTTTTATCTTCTTTGCCAGCATTGTAGCGGTTTTTTCGGAATCATTGCCTCAAGATTCTGTTCATCTACTTGGGTTTCGAAGCAATCTTCCAGATCCCTCTCAACAATTGCTCCCTTGGAATCAATCTGTCTCACACTGTAAATGGAAAGGTGTTACCTGCTACTCAGACAAAACTTCACAAGTCAAATCATTGTATTTCAGAGACTTCCTTCTATCTGGAATCTTGGACAAGGCCTTTCCAAATCTTTGCCATCTTCCTCGTTTGGTCTCCCTTGATCTCAGTGGCAACCATCTCACTGGTGGCATCCCTGCTATGCTTGCTAACTGCAGCCAACTTGACACGATTTTCCTCAACGACAACAGGTTTTCAGGATCCATCCCACCAGAGATTTTCAAATCCAAAAGGCTTGTATACCTTGAGTTGGGTTACAATCAGCTCAATGGCACCATTCCTTCTGAGGTGGGCCTTTCGACAAGCCTTCAATATCTTGGGCTGTGGAACAATTTCCTTAGTGGAAATGTTCCAAAAGAGCTATTTGGTTTGCCAAACTTGACAGATCTCTACATCCACACAAATAATCTAACGGGACCTCTCCCGGACTTCCCATCCTCATGCTCACTTTCCCAGCTTCTTATTCATCAGAATCGCTTCTCAGGTTCTTTGCCAATTTCCCTAGGTAATTGTCACAACCTTTCTGCATTCTATGCAACATCTGCTCATCTTGGAGGAGTTATTTCCCCAGAGGTTTTTAGGGGCCTTTCAAATCTCGAATTACTCTCTTTAGATGACAACAACTTTGAAGGGGAAATTCCTGAGACCTTATGGAATGGAAAATTACAAGAGTTGGTTCTTTCCATAAATAATTTTAATGGAAGTATATCAGAAAAGATTGGTGGTTGCCATCAGATGACTTACATTGATTTGTCAGCTAACAAATTAAGTGGTCAGATTCCCAAGTCAGTTGGACGTCTGAAGAATTTGTACAAGCTTCTTCTGTATGATAACATGTTTAGTGGTTCATTGCCAGCAGAAGTTGGGAACTGCACTTCTCTTGCTGAGATAAGTCTTGTCAGTAACTTCATCAGTGGGGAAATTCCTTCAGAAATTTGCAGCCTTCAAAACCTAGAAACATTTAATGCATTCAAAAATAAAATTCAGGGCCAAATTCCAGAATGCATTGGAAGAATAAGCGGGCTGCAGGAGCTAGCTCTTTATGAGAACCGATTGACCGGCAAGTTACCGCTTGGAATTACAAATATGACGAAACTTTCATTTCTGTCGTTGGCTCATAATAACCTTACGGGGGAGGTACCACCTGATCTTGGAAAAGATAATTTTCCTGGCTTAGTTAAGGTTGATTTAGGTTATAATAACTTCAGTGGACAGATTCCTTCTAAACTGTGTAATGGCAACAGGCTTGCAGTCCTGGCTCTTGAAAACAACAGGTTTAATGGCAGCTTCCCAACCTATCTAGCAAACTGCAAATCTCTCTATAGGGTAAAACTTCCCAACAACAATCTGCAGGGGAGTATTCCAGACGACATTGAAAAGAATGAGAATATTTCTTACTTGGATGTTAGACAGAATATGCTTGTAGGAAGGATTCCAGCTGCATTTGGTTACTGGACCAATCTTTCAATGATTGATCTTTCAGAAAATATGTTTAATGGCTCCATACCCACAGAACTTGGAAAACTTCAGAATCTTGTAAAACTGAGCATTTCTTCAAACAGACTGACCGGACAAATTCCCTTTCAGTTGAGTAACTCTGAAAAACTGGAGGAGTTGGATCTCAGCAACAACAGTCTTTCAGGAAGAATTCCTAAAGAAATTGCATCATCTTCAGTATTGACAAATCTTCTACTCCAGGACAACAAACTTTCTGGTGCTCTTCCAAGCACTTTCTCCTCCTCACAAATGCTTGTAAAGCTGCAGCTGGGAAACAACTTGCTCGAAGGCCCAATTCCATGCAGTCTGAGTAAGCTTATGCAACCTGGTTTTTCACTAAATCTGAGCATGAATAAGTTTACTGGTGAAATTCCTAGGTGCCTAGGCAATCTTGACAAATTGGAGGTCCTTGATATATCAAGCAACAACCTGTCTGGTGCAATACCATCAGAAATGGAAAAGATGAGTTCCCTTTCGTTTCTCAACATATCATTTAATAGCTTGTCAGGGAAGGTACCCAATACATGGGAAAAACTATTAAGTTCACGTCCAGGATCTGCCTTGGGAAATCCAGGACTTTGCTTAATAGACACGGAAGGTAGCAACTGCAAGCATGTGAAAAAATCACAAGTAAAATGGAAGACTTTGGCTGGTGTAATTAGTGGATGTGTGCTTTCCATGGCAATCACAGTTGCTGCAATGTACTTACTAGTGACTCGGATTTGGCACCCATCCCTGCTGAATAAGCATCGGCTCGTCAAGTGTCAATCAGGAATTGAAGATCTACCTGATGGTATAACTTTTGAGGACATTGTACATGCAACAGAAGGCTGGAGTGAAAACTATGTTATTGGAAGAGGCACGCATGGAACTGTTTATAAGATGGAATCTGCCAAATCCAATAAACTTTGGGCTGTCAAGAAGGTAGACTTAGCTCAGAGAGCATTCAATGATGAAATGAGAAGCCTAAATTCTGTCAGACACCGCAATTTGGTAAGATTGGGAGGATATTGCATGAAACATGGATATGGCTTCATTCTAACAGAATTTATACCTGGAGGGACTCTTCATGATGTCCTTCACCAGAGAAAACCACCTGTAGTCTTGGACTGGAAGTCCCGCCATCGTATTGCTCTTGGTATTGCTCAAGGTCTATCATACCTTCACCACGATTCAGTGCCTCAAATCATTCACAGAGATCTCAAATCAGATAATGTAATGCTGGATTCCGAGATGGTGCCAAAGATTGGCGATTTCGGGATTGCTAAAATGGTATCAGATGAAGAAAATTCAACGAACTCCAATATTGTTGGGACTCTAGGATACATTGCTCCAGGTTAGCCACACGTTACTATGCTAATTTTCAGTATAATTCAGTTGGCAGATTACTTTGATGAACTAATAACTTTTACTACTTTCTTCCCTGGCCCCCTTTCTCTTTTTGTCACTCCATCTATTCAGAGAACGCATACTCAGTTCAACTGACAGAGAAAAGTGATGTCTATAGCTACGGAATTCTTCTTTTAGAGCTTTTCTGCAGAAAAATGCCAGTAGATCCCAGCTTTGAAGAAGGATTAGACATTGTTTTCTGGGTGAGGAAAAACTTGCAGAGAAGCAACAATTTCCTTAGTTTTCTTGATGAAGAAATTAGGCTCTGGAATGTAGAAGAACAATGGAAGGCCCTGAAACTAGTGGATTTGGCGCTTCAATGTGCTCAACTCGAGGCAAGCACCAGGCCTGCAATGAGGGATGTAGTAAGGTCTCTTGTCGAGTTATGAAGAGAGGAATGCAAGATGATTTGAAGGCAGGTGCAGTTCAATCCAGAAGATCAAGAGCTCCTGCATTTCTCTTCTTTATTTTTGGCGTAAAGAACAGTTAGTTGATGATGTTGAATCATTACTAGACTCTATGGCCTCTTTGTAAGTTTTGTTTGAGGCTGTATCTGTAGATATGATATACCAGGTCTTTATTAGGGTGCTGATGAATATAATATGTTACCAAATTCAAAAAGAGAATAGATGTAAAGATAAGTAATGAAACTTTTAGTAGTGAAACCTGTGATCAACTTTCCAAATTCCAACATCTCATTTTTTTCTTTTTTTGCTTACATCAAAAAATTGGACAAAGAAAATACATGATGTTTCAGTTTAACAAAATCGTGATCTTGGGTTTTTCAGTTTAACTTTGTGACATCCAATGTAGCAACATAGACTAAGAAAATGCATGATGTTTCAAATCAATTTTTGGTAAAAGCAATTTTCACCTTCTAGCCATAGCTCTTTTCATCATGGATCACTGTCCGTAATCAGTTTTTGCTTATTACTTTCTCCAGCTTCTCCAAAATCTGAACAGTTTTCAGGTATCACAATTAAAATGAGAGTGTGAAGGAACAAGGCTTTGCTACAAGCTAGTCCTTATCTTATGGACTATAACATTCCGGATTATAAAAGAAATCAAAAGAACAACCCCCACCCAAAAAGATAAGCTTTTGTTGAATTTGGTCGTACTAATGATGGTTCCTGCGAAAGAGTGCAGAAGACCTCCGCCTTACCGGTCTTTCTTTGTGTTCCAAACTTGCAGTAGCAGAATTATTTGCAGAAAATATAGATATGATGAATAGAAATGGCTTGTGGACAATCATTGTATTTCAAACAAATACAGATATTCAAATAAGGTAGATATAACTCAGATATGAGTTACCAACAGGACTTCTAACAAGACTTAACAGACGAATGGCAAATACATAGGAATATCAGATCAGGAGAGTAAAATAATAGTCCTTGAAGAATTGATCAGAATTCAGTAGAAAGGAAATATGAAAGAAGTGCAAGAATAAACTGGAAGAAATACTCTGCAGGTGACAAATAGATGAAAGACATACCTCAGTTACGAAAACTGCAACACCCGAGCACAATACGATGCGCACCTAAATCATCCAAGTAGGAGAAGCCCAACTTAAATGTCCCAGATTCCTTTTTCGTATTTTTTATGTAGTTGACTTGGATGTGCATTACAAGACAACTGTCACAAGATTTTGAGAATTTTTCATGTTAAAATATCCACCAGAATTTGCATTGGATATATTCACATGGCACCACATATGCTGACAAATCTCAGACTTCTAATAATAGAGATTCCATTCTGAGACAGATCCACTAAATTCCCAAGTTTCTGCTCTTAGCAAGTGCAGCACCATCACATATGATGATAAGTCTAGGACCTCTAGTAATAAAGCTTCCCTTCTGAGCCAAATCCACTAAATTCCCAAGCTTCTGCTCTTAGCAAGTGCACTTTTGTGCAAGACGATTAGAGACAAACTCCAGGAATTTGGTGCTTGCTCTAATACCAAAGTTACAAGCAGCTCAAAAGAGAAAATATAAAACACTAAACCACATTAAAGAAAAGCTAGGAAGCTAAAACCAAAATTGAAGAAGAAAAAGTTCCACACTGTTTCTCTTAACAGAACAGGTGAAATTTCTCATAAAGAGCTGATCATATTCTATATCCAATCACTTTACTGAAGTTAGAGTACAGATAAATGTTTGAAAACCCTAGTGTCCAAAGATAGGGACAATAGAAATATGAAGCTGTAAAATGCTCCAAAAAAAAATAATTAAATTACTGAGAGGATTTCCAAAATAGCCTCGAGGTAAAATATTTATACAAGGTTAAACTTGTCTGCTTCCTCAGTCCAATAAAGCAGAGTTCAGGAGTTTTACTTTGAAGAGACCGAACAAATTTAGAGAACTACTCCATTACTGGCTATGTAACCTATGGCTAACTAGATTAAATTAGAAGCTCAATGTGCATAGACGTGGTGTTCTTGGGTTATACTTAAGACCAGCTAACTTGAAGAAGAATTCATTAGTATACATGGTTACGGACTGAGGCACAGTGCAACAGGATAATATTTTCAACTTAAGTAAGAAATTACCATTAGAAAGATAGGATGTCCTTGATCACATCATACAGAATTAGATACCTAATGGAGCTTGCATGTCTGCTTAATTAACACTAAACAATGTTAACTATTGCATATCCTCGCCACTTGAATATTTTCATGCAACATATTTTTGTCATTTGGATGTATATATGTTGAATGAAATTAAGGAACCTAAAAGTCAGTTCACCAGACAATCTATAATCAGTTATGTACCACATCTTTTTACCTAATACATTTTTTGTCAATCAACACCACTATTTATACCCTATACTACATATCAAAGGGACAAGATGAAGTTTGCAAGGATCAATGCCTAGTTTATACTCAGCATACAAATAAAGCTCATATAAGATACAAAACCAACTAAACCTCTTTTCATCTCCCAAATATAGTTTGAGCAATAAAATAGAGAGATGGAAAAGTATTATTTGATCGCGAAATTATGAACCAGAATAGATTGTGGCAGCAACACCAGACCTCAGTCACAGTCATACTAAGTCATCTACTTTGTTGTTTTTCACATACTTTTCTGCTTCAATTTTAATCAAAAGCAGAATGCTGATAGCCTGATATGCATTCAACTACAGATGTATGGATCCTTTTTTTTTTTTGATGACAAGGGAAACCCGCAGGTGCGCACAGGGTAAAACTCTCGCTCCTATGCAATAGCTCGCAAACCACATAGGAGAGGTAACCCGTGGGTGCGCACAGGGTAAAACTCTCGCTCTTATGCAATAGCTCGCAAACCACATAAGAGAGATAACCCGCACTAGGCAAGGCTGGTGCGACGAGCTCGACCCAGAGACAAACCCCTTGCTTTCGCTGGCAAGGGGTTTCGAACTTGAGACCTCCAACATGGAAGTGCCAAGCCCAAACCACTGGGCCACCCCGAAAGGTCAGATGTATGGATCCTTTTCACCTAGATATTCAACATCAAGGCATCTCAGAATTCTCAAAATCAATTTACTCCTGGCAACTTCTAATTTTGAAATCCCTAAGATCATAAAGGGGCTGTGCCCAAGCATTAGAAAACTAGCATTGCCTTCGAAGGTCAAATTATCGATTTAATCAATGGTACAGAGGGCTTTTAGAAGCGGAGAAATTGGGAACAGCATTGTACCTGTTAATTGTTGATGGAGTCATCGGAATCAAAAGTAACTCAAATTAACGAAGACAAATAAATAAAAATCTCATTTCTCAACTCAAACTAAATTGGATGGGCATAAATTTGTGGAACCCATTAAGCTTTTTTCTCCAACTTGC

>PGSC0003DMG400003280 LRR-RLKs

GGAAATTCTGTCTCTCTCTGGAGAGCGGCTCATTGTTTCTCCACAAAAAATGGCCGCTGTCCTCTTCCAGCTCCACCACCTTATTTTCTTATTTGTTTTTTTCCTATCTCATCCAACTCCTTGTGTTTCCATTGGAGATGATCAAGTCCTTCTTGAATTTAAAGAATTGTTATTAAATACCTCACTTCTTGATTCTTCTTGGAAAAAAGGAACAAATCCTTGTGATAATAATAACAAATGGTTTGGTGTACAATGTGATAATAAAAATGTTATTCAAGCCCTACTTCTTGGTGGGGTTGGCCTTTCAGGGAATCTTGATGTTGATGTACTAATTAGCCTTCAAGGCCTTAGAGTTGTTAATCTTGCAAACAACTCTTTCTCAGGAACAATTCCTGATTTTTTTAGGCTTGGTGCTCTCAAGTCTTTATTCATTAATGGAAACCAGTTTTCCGGAGACATCCCTGGAGATTTCTTCTCCAAAATGGGATCTCTCAGGAAAATATGGTTTTCGCGGAACAAGTTTTCAGGGAAAATCCCTGATTCTTTAGCAGGTTTAAAATATCTTTTAGAACTCCATTTGGAGAATAATGAATTCACAGGACCTATTCCAATTTTGTCACAGGCTAATCTAGCAACTATAAACTTCTCAAACAATAAATTACAAGGTTTAATTCCGCAAAGTTTGTCTAAGTTTGGTTCGAACCCCTTTCAAGGAAATCCTGATTTGTGTGGAAATCAAATAGGGAGGGAGTGTAAGGCTGTAGCCTCTGGTGAAAAGTCTGAAGGTTCTGGAAGTACAAAATGGATAATTATAGGCTTAGTGGTTGTTTTGCTATTGGTGGCTATATTATTTAAGTCAAAGCGTAAGGATGATCAATTTGAAAAGCTTGAAAAAGAGAGCCTTGATGAGGCTGTGAAGGTGCACCTTAACAAGAGAAGCATGAGCACGCGCACCTCTATGCGTTCATCAAGGAAAGGACGCTCAAGAAGTGGCAGCGATATGGGTGATCTAGTTGTGGTAAATGATGAAAAGGGTATATTTGGGATGCCTGATTTGATGAAGGCAGCAGCTGAAGTCCTTGGTAATGGAGGATTGGGGTCAGCTTACAAGGCGGTATTGGGGAATGGAGTGTCCGTCGTGGTGAAGAGGTTGAGGGAAACCAATAAATTTAACAAGGAAAGTTTTGATGCAGAGATCAGACGACTCGCTAGGATAAGGCACAGGAACATATTGCAACCATTAGCATACCATTACAGGAAAGAGGAGAAGTTGGTGGTGTCTGAATACATTCCCAAAGGCAGCCTGTTATACCTATTACATGGTATCGATATTTCTTCTTTGCCTCTCATGTCTCATCTTCAATGGTAAAAGAGTCCATGAAAACAGCCTATATATTATTGTGTACAACTTAATGCTAGCTTCTATTTCATGTGCATAAGTAGAGACTATCTCATTATGGGATATCATTTACGTTTATCAAATGGTCTGTTAATCATTCCCTTGAAATGACTCTTTTAAAGTTTTGGGATTACATAATTCATAAAAATGATCTTAGATAGCCCGAAACATTTCCATTCTTGTAATTGAGGTAATATTCACATTATGTATTTAGCATGCCAATTTGGGATCTGCTTCTATCAATATTATAAACTCTTTTTATCCAAAATTTTCAGGTGATCGGGGGATAGCACATGCTCAGCTAAATTGGACTATTCGCGTTAAAATCATCCTGGGAGTTGCCAGTGGAATGAAATTTCTTCATTCAGAGTTTGCATCATATGATGTACCTCATGGGAACCTTAAGTCTAGCAACATTCTTCTTTCTGCAAATAATGAACCACTTCTCACAGATTATGCGTTTTATCCACTACTCAACGATTCACAAGCTGTCCAATCTCTTTTTGCTTATAAATCCCCAGAAGCCATACTAAACCAACAAGTCACTCCAAAAAGTGATGTATATTGTCTTGGAATCATAATACTTGAAATCCTAACTGGGAAATTCCCATCACAATATCTCAGCAACCAAAAGTGTGGAACTGACGTTGCACAATGGGCGTGGTCAGCAATTGAAGAGAATAGAGTATCAGAATTGATTGCTCCAGAGATAGAAACAGAAAAGGATTCCCTTGAAATGATGGAGAAGTTCCTTCATATAGGAGCTGCATGCACTGAGAGTGATCATGATAATAGAATCGACATGAAGGAAGCCATAAGGAGGATAGAAGAGATAACAAATTTAATGTAGTAATTGGTTAGATGATCAGGAGGTTTCAATTTATTTTAGGGAGGAGGGAAAAAAAGGTACATAAACCTGTTTGAGTTCTCCAACTAATGATCCACGCACGAGCTCTTCCCAGTAGAATGTGGGCATTTTACGTTCCCTGAGATGAATGCTGTTTTCATTTGATTTGACTTGCCTGTATCGAACTGGTGTGACATGGGCGTGGATACTTTGTGAGGATTCTTCAACATGCACTTAATGTTAGTAGGAAACTTAAAGTACCCATACGAAGAATTACACAAACTTGTATATTGGATACATTAACATTATGCATTTTCAATTTTTTCTCTCTATAGTTCATTTTTTTCAGATAACCAATGCAGATACTCG

>PGSC0003DMG400003429 LRR-RLKs

AAGCAACACGTCCACACTTCCAAACCACAAAAAAATAAAAAATAAAATGTCTTCTAGTCATTTTCTTTCATTCCTTTTCCTTTCTCTTCTAATTTCCGGGATCTTTTCGGACCTTAACGCGGATAGAGCCGCTCTACTCCACCTGAGTGCGGCCATCAAAGGCCGCACTCTTAGGTGGAACACAACCAATTCAACCCCATGTTCATGGGAAGGTGTCGAATGTGATACTACAATCAACCGTGTAATTGAACTCCGTCTCCCCGGATATGGTCTATCCGGTGAGATGCCCTTAAATTCCATCGGAAACTTAACTGAACTCCGTACTCTCAGCCTTCGGAGTAACTCTTTATCCGGTCTTCTTCCTTCTGACATAGGATCATGTACAGAGCTACGTATACTTAATTTGGAAAACAACGATTTCTCTGGTTCAATTCCAACGACTTTTTTCAATCTGAACAATCTTATACGCGTGAGTTTTTCAGGGAACAGGATCTCCGGTGAGATATCGGACGCGTTTAACAATCTGACTAGGATGAGAACTCTGTATTTGGAGAATAACAACTTTTCTGGCTCACTTCCTGATTTGAAAAACCTCTCACAGCTCAATGAATTCAATGTGTCTTTCAATAGATTAACCGGTTCGATCCCTTCGAGTTTAGGTCGATTCTTAGCTTCTTCATTTCTTGGAAATTCACTCTGTGGCTCTGCTTTCTCTCTTAGTCCTTGTCCTGAAAACAATAGCCTCACCAGTCAATCTGATAAACTTTCCGGCGGAGCAATAGCTGGAATAGTAATTGGCTCGATTATCGGATTGTTCATACTACTATTAGTTTTGTTTATGTTAGTGAGATCAATTTATAGGAGTAGAAAAAGCTTCCGTCAAGTGAATGTGTCTCCTACGCCGAATCAAGTACTTTCAAGTCCTCATGATTCAATAGTGACAGAAAATCATGATATCGAAAATGTTTTTTCGGAGAGGAAAGTTTGTGAAGATAGGACAAAAGGGATGGTGTTTTTTGGGGAAAGTTTTGAGTTGTTTAGTTTGGAGGATTTATTGATGGCGTCTGCGGAGGTTTTGGGGAAAGGGGCAACTGGAACTACTTATAAAGCATATTTGGATAGAGATGTTGAAGTTGTTGTGAAGAGGTTGAGAAATGTTTGTGTTTTAGAGGAGGAGTTTAGGGGAAAAGTGGAGCAATTGGGAGGAATTAGTCATGGGAATTTGGTGCCTCTTAGAGCATATTATTATGGAAGGGATGAAAAGCTTGTTGTTTATGATTCTATGCCTACAAGTTTGTATGCTGTTTTGCATGGTAAGTATCTTATATCTAATAAACTTTAGCTTATTGTTGTTTTAATAATAATATTTGCTACTTGCTTCTTTTAGGTAGTCTCTGGTTTTGGTTAGGGAGTAAATCAAGTTAATTAAAATTCTTAGCTTAGATTATATTAGCTTCTAGTTAATTGTTTTTTAGCTTTAGCTATGAAAACTGAAATTATGTGGAAGAGATGGTATGAAAACAGGGGAAAGGAAGAAGATCATTTGGTAGTTCTCTTTTACATTAGTTAATTGTCATCAGATCCACTGTCGCTACTCGCTATAGGCTTAGCAGAATTTACAAAGAGTGTTAAAATGTAAAGTCACTAGTAATTGCCTCTAAAAATATATTTAGCAGCAATTAAGCTATTGTCGTAAATTAATAGTGGCTAAAGATAACTTTTGATGTTGTGTATATTGATGTTCACATCACAACTAATGAAGGACCCATGTAATATTATTAGAGAGAACAAGAACTTTCTGTGTAGAGGGAAAGCCTCCTTAATTAGTATTTAGTTGAATAGGCCTTCTATCAACTCACATTCTATTAAATGTTACGGAAACGAACTATAGAATTTTACTAGTATTAATTATTTCGTATTCTAACCAATTACTATTACTATGAGTTAGTTTTAACCTTTCTGATTTTGTTAGGCGAAGGAAAAGAAGCATTGACATGGGTAATAAGGAGCAGAATTGCATTAGGAGTTGCCAATGGTATCGAATATCTCCATTCACTAGGCCTTAAAGTCACTCATGGCAACATCAAGTCATCTAACATCCTCCTCACACACTACTACGATGCATATCTATCCGAATTTGGCATAACACAGTTGATTTCTTCAACATCAAACTCCAAAATGACTGGATACTACGCGCCTGAGGTGACAGATATTCGTAATGTTTCCCAGAAAGCCGATGTCTATAGCTTTGGCACGATTCTGTTGGAACTCCTAACGGGTAAAAATCCTAGTAGTGTCATCAACGACGAGGGAATTGACCTGCCTAAATGGGTGAAATGTATAGTTGAAGAGAGGGGGACAACACAAGTGTTTGATCCTGAGTTGATTAGATTTCAGAATTGTGATGAAGAGCAAATGGTTTCATTGCTGCATCTTGCTATTTCTTGTACATCCCACCACCCCGAAAGGCGGCCGCCTATGGCTGACACTACCAGGAGGATCAAAGAAATTGTTATGTGAGTTTTGTTAATGTTTGATCTTCCCCCCTCATTCTTGTCCAAATGAAGGGGCGAATTCATACCTTCAAAATTGGGCACGTGAATATGTGATCTCTCTATAAAACTAGATATTTTATGTATATATATTTTGAATTAATATAATATTATTTATTGCCTC

>PGSC0003DMG400003551 LRR-RLKs

GTTAAAAAGTGCTGGTCCTTGTTTTGAAAAGTGCTCATTTGCACATTCTCCACTTCCTCCCTCACTCTTCTTCTCTGTGTCACTCTGTGCCTCTGTGCAAAGTGAGGGGGCCTCTGCTATGGCCATTAACACTCATCTTCTCCACTTTATCATGCACCCATTTTAACCTTCATTATAAACCAAGCCAACTCCTTTTTCTTATGTTCTTGATTTCTTCTTCTTCTTCTTTCATGGCTGCTTATCTACTTCTGTTTCTTGTTTTTCTCTCTACATTATGCTCTGCTCAGAGAAACTCACAAACCCTTTTGGAAGTTCAAGCGCTTACTTCTTTTAAACTTAGTATTCATGATCCACTCGGTGCACTCACTGACTGGGATTCTTCTTCCCCTTTTGCTCCCTGTGATTGGCGAGGTGTTTTCTGCGTCAATGGCCGGGTCGGTGAACTTCGTCTCCCTCATTTGCAACTCAGTGGGCCACTCACTACCCAAATTGGTAACCTGCGCATGCTGCGCAAGTTAAGCCTCCGGTCCAACTTCTTTAATGGTACTGTTCCAGCTTCGTTATCCAAATGTACCCTTTTGCATTCTGTTTTTTTACAGGGTAACGCATTTTCCGGTAATCTTCCGCCGGAAATATTTAACCTTACTGATTTACAAATCTTTAAAGTTGCCGGAAATCAGTTGTCTGGTGAAATCCCCGGAGAGCTTCCTCAGAGTCTACGATACTTTGATCTTTCGTCAAATTTGTTTTCAGGAGATATTCCGAGGAAATTTTCTGACAGGTCTCAGTTACTTCTGATTAATCTTTCTTATAATCGTTTTTCCGGCGAAATTCCGGCGAGTCTCGGCCGGCTTCAACAGCTTCAGTACCTTTGGCTTGCGTACAACAACTTGGTAGGAACTTTGCCCTCTGCAATTGCTAACTGTTCATCGTTGGTTCACTTGAGTGCTGAAGGAAATGCTATAGGGGGTCTTATTCCGGCAGCCATTGTTGCTTTACCGAAGCTTCAGGTGATATCTTTATCACATAATAATCTCTCTGGTTCCTTACCAGCTTCATTGTTCTGCAATGTTTCCATTTATCCTCCCTCCCTTAGGATTGTTCAGTTGGGTTTCAATGCATTCACGAATATTGTCAAGCAAGAGTCTTCTAAATGTTTTAGTTCTTTGCAAATTTTGGATCTTCAACACAACCAAATACATGGTGAATTTCCTTTGATTTTGACGAACATTTCTGCATTGACGTCGCTGGATGTGTCATGGAATTTGTTCTCTGGAAAAATCCCAAGTGCCATTGGGAACTTGTGGAGATTGGAGGAACTGAGAATGACAAATAACTCATTTGAAGGTGCTCTTCCATTTGAGATTACGAACTGCAGTGACTTGAAAGTTCTTGATCTTGAAGGGAATCGAATGACGGGTGACATTCCAATGTTTTTAGGTGATCTTAGAAGCTTGAAGACATTGTCACTGGGAAGAAATCAGTTTTCGGGTTCCATTCCTTCTAGTTTTAGAAATTTGAGTAATCTCGAAAATCTCAACTTGGGAGGAAATGGTCTCAATGGAAGCTTGCCCGAGGAGGTAATGGGTTTGAGCAATTTAAGCACACTGAATCTCAGTGGAAACAAGTTTTCTGGAAGTATGCCAGTTGATATTGGGAATCTTCAGCAGCTCTCGGTTCTGAATCTGAGTAAAAATGGCTTTTCAGGCACTATTCCTTCTAGCATTGGGACTTTGTATAAGCTAACGGTTGTTGATTTGAGTGGACAGAATTTCTCAGGTGAAATACCATTTGATCTTGCTGGTTTGCCTAATCTACAGGTTATAGCTTTGCAAGAAAACAAGTTGTCTGGTAATGTTCCTGAAGGTTTCAGTAGCTTATTGGGTATGCAATATTTGAATCTTTCTTCCAATTCCTTTTCTGGCCATATACCATCAACATTTGGGTTCTTGACTTCATTACTTGTACTTTCTTTGTCTAACAACCACATAAATGGTTCGATTCCTCCCGACTTGGGCAACTGCTCTGCTCTGGAGAATTTGAATCTTCACTCAAATTCATTGAGTGGCCAAATACCTGCTGATCTCGGACGTCTTTCCCACTTGAGTGTGCTAGATTTGGGTAGAAACAACTTAACAGGTGAAGTCCCAGTAGATATTTCCAATTGTTCATCCTTGACATCACTCGTGCTCGACTTGAACCATCTTTCAGGGAACATACCAGAGTCACTATCCAGGTTATCAAACCTAACTGTCCTTGACCTCTCTACTAACAACTTCACTGGAGAAATCCCAGCTAATCTTGCCATGCTCTCTAGCTTGGTGAGTTTCAATGTGTCAAACAATAATTTGGGTGGCCAGATTCCGGAGATGTTGGGCTCCCGTTTTAACAACTCATTGAATTATGCTGGCAACCAAGGTTTGTGTGGGGAGCCATTGGAAAGAAGATGTGATACATCTGGTGATGGTGGGAATAAATTGATTATGTTCATTGCTGTGGCTGCCAGTGGAGCTCTTCTCCTTTTATCATGCTGCTGCTTATATACTTACAACTTCTTGAGGTGGCGCAGGAAGCTCAAAGAGAAAGCAGCTGGAGAAAAGAAGCATAGTCCTGCAAGGGCTAGTTCAAGGACCAGTGGTGGTCGTGGCAGTGGCGAGAATGGTGGACCCAAACTTGTTATGTTCAATAACAAAATCACACTCGCTGAAACAATTGAAGCAACTAGAGAATTTGACGAGGAACATGTGCTAAGTAGAACACATTATGGAGTGGTCTACAAGGCCTGTTACAATGATGGAATGGTGCTCTCAATTCGTAGACTTTCAAATGGATCACTAGGTGAGAACATGTTCAGAAAAGAGGCAGAATCACTTGGCAGGGTGAAGCACAGGAACCTAACAGTTTTGCGTGGGTACTATGCTGGCCCGCCTAACCTCAGACTACTTGTCCATGATTACATGCCTAACGGGAACCTTGCAACACTGCTTCAAGAAGCATCCCACCAAGATGGTCATGTCCTCAACTGGCCAATGCGCCACCTCATTGCACTTGGCATTGCTCGTGGCCTGGCTTTCCTTCACTCATCCCCAATGGTCCACGGAGATGTTAAGCCGCAGAATGTCCTATTCGATGCAGACTTTGAAGCCCATCTTTCAGAATTTGGCCTAGGCAAGCTTGTAGTAGCCACACCAACCGAGCCCTCCACATCAACTTCAGTTGGTACCCTAGGCTATATATCTCCAGAAGTAGCATTAACCGGAGAAACCACAAGAGAATCAGATGCTTATAGCTTTGGCATTGTTTTGCTGGAACTACTAACAGGAAAAAGACCCTTAATGTTCACACAAGATGAGGACATAGTGAAATGGGTGAAGAGACAATTGCAAAGAGGGCAAATTTCCGAACTATTAGAGCCAGGATTGCTCGAATTGGACCCTGAATCATCAGAATGGGAAGAGTTCTTGCTAGGAATTAAAGTAGGCTTGCTATGCACAGCACCTGACCCACTTGATCGTCCCACCATGGCCGACATTGTGTTCATGCTCGAAGGCTGTCGTGTGGGACCTGATATCGCGTCTTCAGCTGATCCCACATGCCAACCTTCACCCGCCTGAATCCCCCGAGGTTCATTCTAATTCTTTTCCCTTTTTTATTATTAGCTCTGTATAAGTTGTGAGAGTAAATTTGCAATGCAATGCAACGCAACAGTAGCTAGCTAGCAACTTAGTCGTTCAATTTCATGCTTTTTTTTTCTTGTAGATTTTTTTAAGGTAAAAGAGAAACAATATTTCTCTTTTATTGATTCTTGTTTTATGCTATTATTCAAACACCTCACTCCCCATGGTGCCGGGATTTTTCTCCTTTGGCGTGGGGACCAAAATGTAGTGCTATCAATAAGTCAAAGAAAGCTAA

>PGSC0003DMG400003714 LRR-RLKs

GCTCAACACATGGCGTTAGTCTTTGTCAGTGCTGCTTCTTCTTCTCCTTCTGAAACTGAAACTGAAACTTTTTTTTTTTTCAGCTCATACGCTCCTCTCTTCCCCACTTCCATAACAAACTCCATTAACACCCACCATTCTTGATTTTTCCCCCATTTGATCTTCTTCTTCTTCTCTCTATCATTACTCACCACCACTCTTTCTCATGTCAACGGCTACTTTTCTTCTCTCTCCGCCATTGGTGGTGGTGGCACTTACATTGCTATTTTCCTGTGTTTTTTCCATGGCCGAAAACGTTACAGAGACGGAAATGGCAGCCTTAATGTCGTTCAAACGGAATTTGGAAGACCCACTTGGGGTATTAGACGGTTGGGATTTTTCTACCTCATTAGCTCCGTGTGATTGGCGTGGGGTTCTGTGTGATGCCGGCAGAGTTCGTGAACTCCGCCTACCTGGGTTGCAGCTGAGTGGACGGCTAACTGACCAGGTTGGTAATTTACGCCAGCTACGGAGGTTAAGCCTACATTCTAACAACTTTAATGGCTCTGTCCCACGCTCTTTAGCTCAATGTGCTTTGCTACGCGCTGTTTACTTCCATTACAACTCTTTTTCCGGTGAACTCCCGCCGGCTATCTCGAACCTCACTAATCTTCAAGTTTTAAACTTTGCTCACAACTTCCTTTCCGGTCATGTCTCTGGTAATGTTCCAGCAAGTCTCCGGTTTCTCGATCTTTCTTCTAATCTCCTCTCCGGTGGCATTCCGGGTAACTTTTCAGTTGGTTCTCAGCTCGAACTGTTAAATTTGTCGTTTAACCGGTTCTCCGGTGAAATTCCGGCGAGTATCGGAGCTTTACAGAAGTTAGAGTATCTCTGGCTTGATTCTAACCAACTTTACGGCACTATACCTTCAGCAATCTCTAACATTTCCACATTAATACACTTGAGCACTAGTGATAATCATCTGCAGGGATTAATCCCGGCAACAATCGGTTCACTTTCCAGTTTACAGGTAATCTCATTGTCGCAAAATCAACTATCTGGTGTGGTCCCTGCATCATTTTTCTGTAATGGGTCAGTAAATGCTCATGCTATTAGGATCATTGAGCTGGGTTTCAATGCATTCACGGGTTTAACTAAACCTGGGAATGCAACATGTTTGAGTGCTTTAGAGGTCTTGACCCTTCACGGGAATCACATTAATGGTGTTTTTCCTGAATGGTTGACGAATTTTTCGTCGTTGAAGGTTCTTGATATCTCCGGGAATGCAATTTCTGGAACTCTGCCAGATGATATAGGGAATTTAAGGTTTCTTGAGGAGCTTAGACTGGGTAACAACACATTAACTGGTGATATTCCTGCTAGTATTGTGAACTTTGGTTCATTGGGGGTACTTGATTTAGGTGGAAATCGGTTTTCTGGATTGATTCCACAGTTCTTGGGTAAATTGACAGGGTTGAGAATGCTGTCTCTTAATGGGAATCGCTTTTCTGGTTCGATTCCGGATAGTTTGGGGAGTCTTTATCAGCTGGAGTTTTTGGATTTGAGCATGAATGATTTGAATGGGAGTTTGCCTCAAAACCTTATGTTGCTTAGCAATTTGACTAGTTTGAATTTGTCCAGCAACTTGTTTTATGGTGAAATTCCAAGAGGAATTGGAAGCTTGCATGGGTTAGAGGTTTTAAATGTTAGCAATTGTGGGTTCTCAGGGAACATTCCGACTAGTATCGGAAGTTTGTTGAGGCTGACAACTCTTGACGTGAGCAAGCAAAACCTATCTGGTGAATTACCCTTTGACATTTTCGGGTTGCCAAGTCTACGAGTTGTGGCTTTACAAGAGAACATGTTAACTGGGGATGGTCTGGAAGGTTTTAGCAGTTTATCTGGTCTGGAATATTTGAATCTGTCATCTAATGCCTTTTATGGACAAGTTCCGAAGACATATGGTTTTCTTACATCCTTGAAAGTTCTTTCTATGTCTAAAAGTGGTATTAATGGCTCAGTTCCAGCAGAGTTGGGTAACTGTTCCGGCCTTCAAGTGCTTGAGCTTCGAGGAAATAAGTTAACAGGTCAAATTCCGAAGGATTTCTCACATTTATCTCATCTAAGGAAGCTTGATTTGGGCCACAATGGATTAACCGGGGAAATCCCTGAAAACATCTCTAATTGTTTATCTTTGGCTACTCTCTTGTTGGATTCTAACCATATCTCAGGTCACATACCAGAGTCGTTGACGAAATTATCAAACTTGGAAATGCTGGATCTCTCTTCAAATAATTTGAATGGATCAATTCCTTCAAGTCTGTCCCTGATTTCCAGTCTGAAGTATTTGAATATTTCCCATAATCATCTCGAGGGTGAGATTCCAGAGGCATTGGGTTCTCGGTTCAAGGACCCTTCTTTGTTTGCTGCGAATAATGCTTTGTGTGGAAAGCCATTGAAGGAGTGCAATGACGGTAAAAGGAAAAGAAAGAAGTTGATCTTATTCATTGTTTTGGCTGCTGTAGGGGCCTTTTTCGTGGCAGTGTGTTGTTGTGGGTACATCTACAGCCTTATACTTTGGCACAAAAAGTTAAGAGGTTCAGCTGAAGGGAAGAAGCGAAGCCCAGGCCGTGCGAGTTCAGGGGGAGAAGGGGGCCGTGGAAGTGGAGAGAATGGAGGTCCAAAGCTTGTTATGTTTAACAACAAGATCACTTATGCAGAGACTTTAGAAGCGACACGACAATTTGATGAGGAAAATGTGTTGAGCCGTGGAAAGTATGGCCTAGTATTCAAAGCCACTTTTGCAGATGGAATGGTCTTGGCTATCCGCCGCCTTCCTGATACATCAATCGAAGTGAACACATTCCGCAAGGAAGCTGAATCCCTCGGCAAGGTGAAACATCGCAATCTGACTGTTGTTCGTGGATACTATGCAGGACCACCACCTGATGTTCGATTTGTTGTATACGATTACATGCCTAATGGAAATCTTGCCACACTTTTACAAGAGGCATCGCATCAGGATGGCCATGTGCTCAATTGGCCAATGCGCCACCTGATTGCTCTTGGCATTGCTCGTGGCTTAACTTACCTTCATTCAGTTTCATTGATCCATGGAGATGTAAAGCCACAAAATGTATTGTTTGATGCAGATTTTGAAGCTCATCTTTCGGATTTTGGTTTAGATAAACTAAGCCTAGTAGCAACTCCAGCAGAGACTTCAACATCTTCCACCTCAGTTGGAACTCTAGGCTACATAGCACCTGAAGTGGCATTAACTGGACAACCCACAAAAGAAGGTGATGTATACAGTTTTGGGATTGTATTGTTAGAGATACTAACAGGAAGAAAGCCAGTAATGTTCAATGGGGACGAGGACATTGTCAAGTGGGTGAAGAGACAACTACAAAGAGGCCAAATTTCAGAATTGCTCGAGCCGGGGTTGCTTGAACTGGATCCTGAATCATCAGAATGGGAAGAGTTCTTGTTAGGAATCAAAGTTGGTTTGCTTTGCACCATGCCTGATCCTCTTGAAAGACCATCGATGACCGATATTGTATTCATGCTCGAAGGCTGCAGAGTTGGTCCTGACATTCCCTCTTCAGCTGATCCTACAACTCTTCCTTCACCAATGTGATATGAAAATTCAATTCTTTGCAATGAAATCAATTTGCAAGAAAATGTCCTTCATTTTTCTCAGTTCAATATTGTTCATGCATTTCTTCAATGTATTGCTACTCCATTCATTATCATTTTGTTTGTACATTGTTCATTCAAGAGTTGGCTGGTTTTGTGCA

>PGSC0003DMG400003790 LRR-RLKs

TTGAAAACCAAATATTACAGATGACAACATTTCATACTACCTTAGTTTACATCTTGTATGTGCTACTTCTTTTTTCACTTCCATTAAGTATCACATCTTCAGCAAGAACAGAAGCTGAATCTCTTGTCAAATGGAAGAGAAATTTACCTTCTACTTCTTTCTTGGACACATGGTCCATTTCGAATCTCGAAAACTTGTGCAATTGGACATATATTGTGTGTAATGATGGTGGAACAATTTCTGAGATCAATCTTTCTGATGCAGCCCTCTCAGGCACACTTGATCACCTTGATTTCACTTCATTTCCGAGTCTCGTGAATTTCAATCTTAATGGAAATAACTTCAGTGGATCGATACCTTCGAGTATTGGCAATGCCTCATTGCTCACATTCTTGGACCTGAGTAACAATATCTTATCAGGTGTCATACCTGAAGAGATTGGAAAATTGAATCAACTTGAGTATCTGAGTTTTTATAACAACAATATCGAAGGTGTGATGCCGTATCAGATTAGCAATCTGCAGAAGGTAATGCACTTGGATCTTGGATCAAATTTCTTAGAAACTCCTGATTGGTTGAAGATGAGAAATATGCCTGTGTTGACGTATCTAAGCTTTGGTTATAATGAACTGAGGTTAGAATTCCCTGAATTCGTACTTCGTTGTCATAATCTAACATATCTTGATTTATCGATAAATCATTTCAATGGTTCGATTCCTGAAACAGTATTTACCAACTTGATCAACCTTGAAAGACTTAATCTTTCATCTAATTCATTTCAAGGTTCATTGTCACCAAATTTTACTAAGTTGTCTAAGTTAAAAGAACTTCAGCTAGGTGTTAACATGTTTTCTGGTTTAATTCCTGATGAAATTGGCTTGATTACTAGTCTTGAAGTTCTTGTACTTTTCAACAATTCATTTGAAGGAAAGATTCCATCTTCTATAGGTAGACTCATAAATCTTCAAAAGTTGGATCTTCGAAAGAACGATTTGAATTCAACTATTCCTTCTGAGCTTGGCTTTTGTACTAAACTCACTCTCTTAGCTCTAGCAGAAAATGACCTGCAAGGGTCATTGCCTTTATCATTCTCCTCTCTTGCAAAGTTATCTGATTTGGGGTTGTCTGATAATAGTCTTTCTGGTGAAATCTCATCGAATTTCATCACCAATTGGACTGAGTTGACATCTTTGCAGCTTCAAAACAATATGTTTACAGGGAAAATCCCACCCGAAACCAGTCAATTGACAAACCTTGTATATCTTTACCTCTATCATAACAATTTCACTGGTTCCATTCCCTATCAGATTGGGAACTTGCAAAACTTGTTGGATTTAGACTTTTCTGATAACCAGCTTTCAGGCATAATACCTCCAACCATTGGAAATCTGACCAATCTAAAGATGTTGCAACTTTTTCGCAACAATCTCAGTGGAACCATTCCTCCTGAGATTGGAAAATTGATATCCCTTGAAACCATCGACATCAATACCAACCGACTAAGTGGTGAGCTGCCGGACAGCATTTCTGACCTCAGTGAGCTGAAGTTTCTTTCTGTATATACTAATGATTTCTCAGGCAGTGTCCCTAAGGACTTTGGAAAGAACAGTCCTCAATTGTCAAGTGCCAGCTTTGCAAATAACAGCTTCACTGGTGAACTACCTGCTGGATTGTGTAGCCCGAATCTTGAGGAGTTGACGATAAATGGGAACAAATTCAGTGGGAAGTTACCAGATTGCTTGAAGAACTGCACACTTCTAAGACGAGTACGGCTTGAAGGCAACAATTTATCTGGTAATCTTGCAGATGCATTTGGAGTGCACCCGAAACTTGTTTTCCTTTCTCTTAGTGACAACCAACTTTCAGGTGAACTCTCACCTGACTGGGGGAAATGTGAAAATCTCACAAGTCTAAGAATGGATGGAAACAAATTTTCTGGTGTAATCCCAAGTGAGCTAGGGAACCTGAGAGCGTTGCGCGTGTTAGCTTTGGAAGGAAATGAATTGACTGGTGAAATTCCCTCTGAACTGGGAAGGTTAGACCTGCTCTATAATCTCAGCTTGAGTAAAAACAATCTTACTGGAGGCATCCCTCAGTCTGTTGGCAATTTAACTAAGCTCCAGTATCTTGATTTGTCGACAAACAAGTTAAGTGGTAACACACCAGTAGATCTTGGGAAGTGTGAGAGCCTGTTGAGCTTGAATCTTGGCAACAACTCATTATCAGGTGGCATTCCATCTGACCTTGGTAATTTGATGCAGTTGAGTATTCTTTTGGACTTGAGTGGCAATTCACTAACGGGAACCATTCCACAAAACTTGGCCAAACTCACCTCATTAATGCATCTCAACCTCTCACATAACAACCTCTCAGGTAGAATTCCTCCAGCATTATCTCAGATGATCAGTCTTCAGGAGATGGATTTTTCCTACAATGAGTTTTCTGGACCAATCCCAACCGATGGAGTATTTCAAAGAGCACCCGCTAGATCTTTTCTTGGAAACTCTGGTTTGTGTGGAAATGTAGAGGGATTATCCTCATGTAATTTGGATACTCCCAATGACAAGTCCAGAAATAATAATCAAAAGATCCTCATTGGGGTACTTGTACCGGTGGTCAGCCTCATACTTTTGGCAATCCTGTTTGTTGCATGTCTCGTGTCTCGAAGGAAGGCTAAGCAATATGATGAGGAGATCAAAGCTAGCCAGATACATGAAAATACAGAATCTCTGATTTGGGAAAGAGAAGGGAAGTTTACATTTGGTGACATAGTGAAAGCTACTGAAGATTTCAGTGAAAAGAACTGCATTGGAAGAGGAGGCTTTGGAAGTGTCTATAAAGCTGTTTTGCCCTCTGGGCAGATTGTTGCAGTCAAAAGACTCAACATGTCAGACTCAAGTGATATTCCATTGACTAATCGTCGAAGCTTTGAGAATGAGATTAGAACTTTGACAGAGGTGAGACACAGGAATATAATTAAGCTCTTTGGTTACTGTTCCAAGAATGGATGCATGTACTTGGTTTATGAGTACATAGAAAGAGGTAGCCTGGGGAAAGTTCTATATGACAATGAGATGGGAATGGAACTTGGATGGGGTACAAGAGTGAAAATTGTGCAAGGAATAGCTCATGCACTCGCTTACTTGCATCATGACTGCTCTCCGCCCATTGTTCACCGTGATGTATCATTGAATAACATTTTGCTTGAATCAGAATTCGAGCCACGACTCTCTGACTTTGGCACAGCAAAGTTGTTAGCTTCAGACTCATCAAATTGGACCACAGTTGCTGGTTCTTATGGCTATATGGCACCAGGTATGTTACATACTCATTACCTAATGCATCCATATTTCTGTTGATTAAGTAGTATATTCATTGGAAGGATACTTCAATATTATTGTGTCTTTCTTCTTGTGCAGAGCTTGCACTTACCATGCGTGTTACAGAAAAGTGTGATGTTTATAGTTTTGGAGTCGTAGCGATGGAGACTATGATGGGAAGGCATCCAGGGGAGCTTTTAACTTCATTATCAGCATCAACAACATTGTTTCCAGAAATACTTTTGAAGGATGTTCTTGACCAAAGACTTCCACCTCCCACAGGCCACTTGGCAGAGGCAGTGGTTTTTGTGATCACGATTGCCTTGGCATGTACACGTACCACTCCTGAATCGCGACCAACCATGCGTTCTGTAGCACAAGAATTGTCTGTTCAGACTTTGCCTTACCTTCCACAGCCATTGGGGACAATAGAAATGAGCAAACTAACAAGTTTTCAGAAATAATGAGAAGATAGAGCCTCATGTACAGGTGTGAAATGTACAGGCTGTAGAGTTTTAGTGTTGTAATGCAACTTCCTCAAGATCTAGAATCTATATAGTATCAAGGATGAAACCTTGTATCTATTCAAATTACTATGTTTTGGTTGCACTTTTCCCTAGAAAGAAAGAAATAATATCATCCTGTTAATCATGAATGTTACAAAGTTGGTATATTAATTTGGACAAGTATGAATGTTACAAAGTTGCACAACCTTATCCTAGTTTTCACTTTATTGCTAGGCAAAACACTAGCAAAAAGATACAATGTGTTCACTTGTAATAAAAATAGCAACCTAACAAAAATTTGGCAAAGAATACATCACACAGAAACCAGGCATGGTACTTAGCATTCCACTCCACACTGCTTGAGTATAGGTCCCTTCTGCTTTGGTAGGAATGGGTCTATTCGTACCCAAACGAGTGAGAAGATTGAAGCAAGAAGGATTGACCAGAGGACCACAATGGTGGGAGTCCTGTTCTGCCTGCCCATCAATCCCTTTAGGAAAGGATAGAGATGCACAATAACCCAAAAGGCAAAGAATAACTTTCCGAAGAGAGGACCCCATGAACCATAACCATTGTTGATGGCATCCGAAACACCAGCTACAACACCAACCATATTCAAGATGATCAGAGTGGTTGGTGGGATGAGAAGTGTGGTCCATTTGAAAAGGTAAAGCTCCCCAAATTCAGCGTCATCAGCTGCTTTTGATGTTACAGTGAAGTTTGTGTCTACACCAGCAAGAACTTTAAGAAGGCCTTGGAAAACTGCAAACAAATGTGCTGAAACACCACCGATGACCCAAAATTGCTCGTTACGCCACCAGGCCTCAATGCTTACTCCACTCCATCGCAGCTCGAGTACACTTGTGGCTATGATGGAGATAAAGAGAGCCATGAACCAAATGCTTGCAAAGTTGTTAAGCTACAAAACAAACATGGATTTTGATTTATAAAATCTAAATCAAATCTTTATATTACTGAACCAAATGACTTCAGTTCACTTACTGTGGGGACGATAAATTTTCCAGTGAGAAGGCATACAGCTGGAAGGGTGCAGTAAGCTAGCAGAGCAATGGAAGTGAATGGGTAAACAATAGTGTTAATGTATGCTAGCCTTTCCAACCACTTGAGTTTTCCACCCCAGGCATACCAAAGGGGACAATGACGACTCATGAAGATTTCGACAGAACCAAGAGCCCATCTCAGGACTTGGTGCAATCTATCTGATAGATTGATTGGTGCTGATCCCTTAAAAGCGGCTCTCTTCGGACAGCAATAAACCGATCTCCATCCTCTGCAATGCATCTTGAATCCAGTCAAAATGTCCTCTGTGACAGAACCATAAATCCATCCAA

>PGSC0003DMG400003918 LRR-RLKs

TCTATCTGGGAATCTTTTAGTAGGAGAAATACCACAAAACCTGTACAACTTGTCAAACCTTGTATATCTTGACTTACATCATAACCAACTGAATGGAAGTATTCCATCAACCATTGGAAATTTGTCTAACTTGCACTTTTTGGATTTATCACAAAATCTGCTATCTGGGTCAATCCCTGTTGCTCTTGGGGATTTGCAAAATTTAACTCATTTTAATGTATCTTACAACCTTTTATCTGGTGCTATTCCCTCAATTGAGAGCATCAAGAAGTTTGGACCCTCTGCCTTTTTTCATAATAGTGGTCTTTGTGGTGATCCTTTAGAAGTTTCATGCTCTGCCAGTGGCACTACTCCTGCAAAAGGGAAACCAAAATTGAGTGTCTCAGCTATTGTTGCTATTGTCGCTGCTTCAATAATCCTTACAGGCGTTTGTCTGATAACCATCATCAATATGAAGGCTCGACGGAGAAGGAGGAGAGAAGATGAAACATTCGTTGTAGAGAGCACGCCACTGGCTTCAACAGATTCAAATGTCATTATAGGAAAGTTGGTTCTTTTCAGCAAGACTTTGCCTTCAAAATATGAGGACTGGGAGGCTGGCACAAAAGCCTTGCTAGACAAGGAGTCTTTGATTGGTGGGGGGACAATTGGATCAGTTTACAGAACTTCCTTTGAAGGTGGTGTATCAATTGCAGTGAAAAAGCTCGAGACATTGGGGAGAATCAGAAATCAAGATGAATTTGAGCATGAAGTTGGCCGGTTAGGGACCCTTCAGCACCCCAATCTTGTTGCTTTACAAGGGTACTACTGGTCCTCTAGCATGCAGTTAATATTGTCGGAGTTTGTTCCAAATGGAAACCTATACGACAATTTACATGGACTTAGTTATCCTGGAACTAGCACTGGTGCTGGTAACCCAGAATTGAATTGGCCAAGAAGGTTTCAGATAGCAGTTGGAACAGCTCGAGCACTCACATATCTTCACCATGATTGCAAACCTCCAGTCCTTCATCTTAATGTCAAGTCCACTAATATACTCCTGGATACAAACTATGAAGCCAAATTGACTGATTATGGCCTGAGTAGGTTCCTTCCCCTGCTGGATAACTATGGACTAACCAAGTTTCACAATGCAGTTGGTTATGTTGCACCAGAGTTGGCGCAAAGTCTGAGACTAAGTGATAAATGTGACGTTTACAGTTTTGGGGTGATTCTGCTGGAGTTGGTCACTGGGAGGAAACCAGTAGAGAGCCCAGTAGCAAATGAAGTGGTGGTCTTGTGTGAATATGTTAGAGGATTGATAGAAAGGGGTGCTGCTTCAGACTGTTTCGATAGAAGTTTGAGGGGCTTTGCAGAAAATGAGCTGATTCAGGTAATGAAGTTAGGCTTGATTTGTACTTCTGAAATATCCTCAAGGAGACCTAGCATGGCTGAGGTAGTGCAGGTTCTTGAGTCCATCAGGAATGGCTTGGGATCATAGTAAAGAATCAGTAGAGCCATAAGACTATTTGAAACAGCAGCTGTTCATAGAATGAGAAGAAAGCAGTCTAGCTGTTGTGCTAGACTCACTTCTCATGAGATTTGATGAGGGAATCAGAAAGGGTCTCAAATAAGAGAGACGCGCAGACCAATTAATAGTATCAGTCATATTCTTGAATTTGGAATCAGAGTGATTAGGAGGTGACAGCTTTTGGTACAAAGAGAAGATTCTTTAGTATTTGATTTGTGATTCTTTTAATCAATTGGTCATTGTAAGTTTACAATAAAGCAATTCTTTTCCTGTAAGATGCAAAGGTATTTTGATGTCTCATATGCAGCTTGGCAACTGCTTTTTATAACCATTCAATATTGCTCTTCATTTACTGTTAAGAATTACCTTTGACATGCATATATCAACAACCTTTGAAGGGATGAAAGAAATCGATTCATTTCTCGAATTACATATCTAGTGCTAACACAAGAATGAATGTGAACAAATAATGAGTCCATACATCTACCTGTAACTATACAACAATGAGATAATAAGAAAATATCAAGATTTAAGACCGACCTTATTGACACAAAGAATTTGGAAATGGATTTCATTTGAAGCCATTTCTACATGCTATATGTTTTTCCTTGCATCAACTTTGACTGCTAATTTAGCTATCTTCAAAATCCATAAACTGGGAGGAGGAATGCATTTGTGAAAATGACAGTTGGAGAGGACCAAGAACACACTTCTGGAATCATCTGCATACAATCTGGCTTTTTTTTGTTTGTTTTCTCTTCATGTAGTTCTCAGTGCTGCTGCATTCTGATCTCATTATCCGAACATTGATAACTTATGGATCCATTAAATCACATAACACGCGACCCTTAAAATCTCTGCATAAGTTTTGGACCTTTGGTTACGGTTTACTGTAAGGGATCATATCATTAGCAACTTAACTCAGAGACTTACGTAACTCCAAAACTTTCTGGATACGTTGCATAACTGTGGAAATGTCCTTCTCACTTAATGTCGAAAAACATAACCTGAACCACCCAGGTTCAACACAGTGGCAGGAAGAACCAGGAGTTGCATTGATCTTAGCTACATTTAGTAGATTATCCCAAAGCTCAATCTCACCCTTCTCATTATACGACCGAATTAACCTGCTCATGTCAGCCCAACAATAGAAACCCCCACTACTTTTGGTGCACTCAATTCCCAGCTGCTTCAATCCACTGACAAATAGAGAAGACATCTTTCTTAGCCTCTCTCTGTTCTTTTTGATAAATTGTTGTACAAACTTTGCATCTGATAGCATCTGGATGATTAAATGCTGTGTTGGGGCTGAAATGGATGAGAATCTTGTCAACTTTTTCGCAGCAGCTAGAACATTTTCATTGCAAGAATATATGACCCCCACCCTGAAACCAGGAAGAGAAAGGTCCTTAGAGAGACCATACACAATGTGTACCCTGCTCCTGTCAAAATCTTCAGAATCAATTATCTCTGCTATGCTGACAAATTCTTCACGTCCATAAGTTGACCCTGCCAAGATTTCATTGGATATTACATGGATGTTCTTCTCAGTTGCAAAGTCTAAGAGGTTATAAAGTGTCTCCCTAGAGAAAATATTTCCCACAGGATTTGAGGGGTTAGAAATGATAATCCCTCGTACTTTAAGGCCACGTTTCTTAGCTTGGTTAAAAGCTCGGTCAAGAGCGTCAATGCTCAAGTTAAAGTTGTCTGCACTGCGGCAAGGAACAGGTATAATCTCCACTCCAGTTCTCCATTTGACATCCCTATCAAGACTGCAAACGCATTGAGCTCATTTTATACACTATATACGTAAAGATGACCAAAAGATTGAAATTTATTTGGGAAGGAGAATAAAACTCACTCAGGATAATATGGTGAGGGAACAAGAAAAGCATTTCCAGGATCAGCTAGGCAGAAGCTCAGTATCTCAAGCGCAGGAGCGGCACCACCAGTTAGAACCATCTGCGAGGGGCTGAAGGACACTGATCTCTCCAGAGCTTGAGACATAAATTCTCCCACAGCCTGCATACAGGTGAGGAAACATGAAAATCAATTCAAAGAGATACCATAAATAATGTAGAATCACACAAAACACCTATTGTTTTTATGAATCGATCTTCATAGTGGCTCAAGGATGATTTCATATATTGCTGTTATACAATTCTCAAACCCACAATTTTAAAAGGTAAATCCATCAGATCATCTCCCACCCTAGGATACCCGATGACAAAGGTAGCCACTCATATTTCAAAGAGGCCAAAATTGCCTTCCATCTTCAACAGATACTGCAAATCTAGCAATAGATCCTTCAACAGGTTACACAATGATGAGTATCGAGGTGAGTATTTGTCAAGGAAATGATAGATCACCTCTCAGAACAACTGAGTCCATTATTCACTATTAACTTGCTTATAACGAAAACCATGCAAAGTAAAGCAGGTTAAACTATTCTCTTATGTGCTCAAACTAGATTTTTTATCATATCACACTGCAAGGGATCCTATTGTTCCGGTCAGGACAAACATAACTCAGAAAATCATCAAATGCTGAAACAGCATAAATGACTGAGCTAGATCAACAATAATTATGAATTTCCAACTCAAGTTTTACCAATGTAACGCTCAACTAGGAGAAGATATTTACCACAAAAAAACACCCCCATCCCTCTGGATACTGTTTCAATGCTTTGGCTGGTCAAACACATTTTCAGGTTTCAAAATTTTCCTATCATTTCTCTTCTTTTGTATTTTAACCTCCTTTATTAGACAATCATGGGCTCCAAAAGCTCAAGAAAAAAATCAAAACATTCAGAACCGTGTTTATTTTTCAAAGTGTTGCTTAGCGTCGATAGCGTAGAAGTTAGAGCATTGACGAACTTTTCAGAAGTTAATTGGCACTGTAAAATGAACGTGAAGCTTATTCAATGCTTTTCCTTATGTTTGTTGCACTTTATAGACTTAGAAAGCTAAAGACGGGACATACTAAAACCTTGTATACAATAGGTAATTTTGGGATTGTCAAATGCCATAACCAATCCCGCATCCAGGGGTTCACATTTCAAATTAAACCAGCTATAAAGCACACTGGACTAGCGGATGGGTGATTGATATGGAGTAACTGTTACACAAGCATATGAACATAACATTTTTGAAATTATATGGACATGACATGAGCCCATCCTAGGTTACAACTATAAACTAATACCACCCAAATAGGTATCTCGTTGATTTAATCCTATAGTCTAAACTCAAAGTAGCAACTTTAAATTATACATACAAACTTTTTCAAGTAAATTTTTTGTTCAATTTCCTTCTCTAGCCTGTAAAAACTATAACCATATGAACAACTACTAGTACTTTATTAGTTCAAGAAGAACATATTATATCCAATACATGTCAACATTATATCAACAAACAAGTCACTAGTAGTGTGCAAGGAAAAATCAAAAAGCTTACCACTTTCAACTCCAACAACCCATCAAATGGCTGATAAGTAGCTATTCCGGTTATACTCGAATCTTGAGTCATCATCCATCTTGACACATTTTCTGCAAGCCATTCTTGAACTAAATCCAATGACAACTATAACCATACAAATAGAGACAAAATTCAAAATCATTGAAACAACAAACAAAAGTAGATTCCCCAACAAGAAATGTACTGCATTTGGTACATTTCTCTAGAAAATGATTCAATATATCCAAACATTCAAAATTATTAGTACCTTGTTTTCTGCTAATCCTAGCTGAATAACACCATCTGGATTACTTGACTCATCATAAGGGTCTTCAGAAGCTCTTTTCAATCCAACATAGTAAGGGCTACTATCAATTTGTTTTGAAATAATCGAATTAGCTCTAGCAGATACCTGAGATTGCCCAGTAGTTCCCTTCGGTGACAGTAAAACACGTGAGTGAACCCTTTGTAACCCAGACCCAGAAGACACTTCCGGCAAATGGGTCGACGACGTTGACCTCGCCGGAGCCGTAGATTCGCCGTTATTGTCACCACCGGAGGAACGATTTCGTTTTAGGTAGAGTTGCCAAAAATAGAAAAGAGCACAGGGTATAACTGAACCTAAGAAAAGACCACCCCGACCTTGAACTACACCTTGTAAGGGTACTATCACCCTCATCGCCGTCGTGGCTCCTCCTCCGTCTCTGCCGCCGGCGCCGCCAGTTGAAATGGTTGTTGTTGTTGTTCTAGTTGGTGAACGGTCTCGTGACCTTGTCATAACTGTGACATAGTGGGGGCAGAGAGAAAGACGTCACCTTGTCGGAGTATGACGTCAGTCAGCTATCTGACCGGCGAGACGGTGGACGGAGAGTACGAAGGGAGGTGAGTGATGGAGAAGAGATTTGGAGAAAATGGTGGCAAATAGTTGGTCGAATTATTATTTATCTTTTTGTCCGCCTAAAGTTGTTTATTTT

>PGSC0003DMG400003945 LRR-RLKs

ATGAGAAAATTAGGTACATTTGTTCATTGTGTTGTTCTTGTTTGGTGTTTTTGTTTGCTTGAATTGGAATTGGTATCTCCCATCACTGAAAAGGAGATCCTGCTTCAATTCAAGGGTAACATTTCTAGTGACCCTTTTAATAGTTTAGCTTCTTGGGATCCTAATGGAACACCTTGCTTAGACTTTAGTGGTGTTTTCTGCAATTCTGTTGGAAATGTGGTGAAGATTGTGTTGTGGAATACTAGTCTTGGTGGGGTTCTCTCCCCTGCCTTGGCAGAACTTAAGTCTTTGCGCATCCTTACCTTGTTTGGGAACCAATTCACTGGCAATATTCCTGCAGAGTATGGGGGGATTGATTCATTGTGGAAGATTAACTTCAGCTCTAATGCCTTTTCTGGTTCAATTCCTGAGTTTCTTGGGGATTTATCAAATATAAGGTTCCTTGATTTGTCAAGAAATGTGTATAGTGGTGAAATTCCTGTGTCTTTGTTTATGAATTGTAACAAAACAAGGTTCATTTCCTTGTCACATAACAATCTTTCAGGTCCAATTCCAGCATCAATTGGAAACTGTCAGAGTCTTGAAGGAATAGACTTGTCTTTTAATGGTCTTAGTGGAAGTTTACCTTCACAAATTTGTGACATTCCAGGATTAGTGTACTTGTCTGTGAGGAGCAATGCACTATCAGGACTTGTTCAAGAACAGGTTTCTCCATGTCAAAGGTTGGAGCTTTTGGATCTTGGTAGCAATGACTTCTCTGGTTTGGCTCCTTTTGGGGTTCTTGAGCTGGTGAATCTTACATATTTCAATGTTTCAAGTAATGGTTTTGAAGGGGAGATTGTTAACTCTGGGACTTGTAGTCAGCGACTGGAAGTGTTGGATGTTTCAAGGAATATTTTTTTTGGTGAAATTCCCTTGAGCATCTCGAAATGCAGCGCGTTGAAGTACTTGGACATGGCATATAACAGAATTAATGGAAGTATCCCTATTGAGCTTGCAGATTTGAATAGCCTTTCCGTTATGCGATTGGGTGATAATTTGTTAGGTGGGACAATCCCAACGGAGCTTGGTAGTATCGAATGGCTTGCAGTTTTGGACCTCCATAATCTCACCCTATTTGGTGAAATCCCAAATGAGATAAGCAACTGCAAGCTTCTTCTTCAACTGTGAGTATATCACCATCTTAGTTTAATTTGTTTTATTTCATCATTGCTCTGATCTGGAATTGAACCGTCTAATATTTGCTGCAGGGATCTATCTGGGAATCTTTTAGTAGGAGAAATACCACAAAACCTGTACAACTTGTCAAACCTTGTATATCTTGACTTACATCATAACCAACTGAATGGAAGTATTCCATCAACCATTGGAAATTTGTCTAACTTGCACTTTTTGGATTTATCACAAAATCTGCTATCTGGGTCAATCCCTGTTGCTCTTGGGGATTTGCAAAATTTAACTCATTTTAATGTATCTTACAACCTTTTATCTGGTGCTATTCCCTCAATTGAGAGCATCAAGAAGTTTGGACCCTCTGCCTTTTTTCATAATAGTGGTCTTTGTGGTGATCCTTTAGAAGTTTCATGCTCTGCCAGTGGCACTACTCCTGCAAAAGGGAAACCAAAATTGAGTGTCTCAGCTATTGTTGCTATTGTCGCTGCTTCAATAATCCTTACAGGCGTTTGTCTGATAACCATCATCAATATGAAGGCTCGACGGAGAAGGAGGAGAGAAGATGAAACATTCGTTGTAGAGAGCACGCCACTGGCTTCAACAGATTCAAATGTCATTATAGGAAAGTTGGTTCTTTTCAGCAAGACTTTGCCTTCAAAATATGAGGACTGGGAGGCTGGCACAAAAGCCTTGCTAGACAAGGAGTCTTTGATTGGTGGGGGGACAATTGGATCAGTTTACAGAACTTCCTTTGAAGGTGGTGTATCAATTGCAGTGAAAAAGCTCGAGACATTGGGGAGAATCAGAAATCAAGATGAATTTGAGCATGAAGTTGGCCGGTTAGGGACCCTTCAGCACCCCAATCTTGTTGCTTTACAAGGGTACTACTGGTCCTCTAGCATGCAGTTAATATTGTCGGAGTTTGTTCCAAATGGAAACCTATACGACAATTTACATGGACTTAGTTATCCTGGAACTAGCACTGGTGCTGGTAACCCAGAATTGAATTGGCCAAGAAGGTTTCAGATAGCAGTTGGAACAGCTCGAGCACTCACATATCTTCACCATGATTGCAAACCTCCAGTCCTTCATCTTAATGTCAAGTCCACTAATATACTCCTGGATACAAACTATGAAGCCAAATTGACTGATTATGGCCTGAGTAGGTTCCTTCCCCTGCTGGATAACTATGGACTAACCAAGTTTCACAATGCAGTTGGTTATGTTGCACCAGAGTTGGCGCAAAGTCTGAGACTAAGTGATAAATGTGACGTTTACAGTTTTGGGGTGATTCTGCTGGAGTTGGTCACTGGGAGGAAACCAGTAGAGAGCCCAGTAGCAAATGAAGTGGTGGTCTTGTGTGAATATGTTAGAGGATTGATAGAAAGGGGTGCTGCTTCAGACTGTTTCGATAGAAGTTTGAGGGGCTTTGCAGAAAATGAGCTGATTCAGGTAATGAAGTTAGGCTTGATTTGTACTTCTGAAATATCCTCAAGGAGACCTAGCATGGCTGAGGTAGTGCAGGTTCTTGAGTCCATCAGGAATGGCTTGGGATCATAG

>PGSC0003DMG400003958 LRR-RLKs

TATTTTCTAACTAACTCTCTTTTCTCCTTTGACAATTCACACCAATAAACTCCCCATTTTTCTTTCTCTCTACTGTGAAGCACCATTGAAATCCAATGCAAAAGCTCACTAGTACTTCATAAAAAACTCAACTTTTCACTCAAATCTGAAAGTTCACAAAGTTCAGAGAAATGGGGGTTAATTATTTCCTCTTAACTTTCATTTTTGCTCTGTTTGTTCTTGAATTTTCTTGCATAGAACCTGTTTGTGGAAATGCTGAGGTGAGAGCTTTGATGGAGATTAAATCAGCTTTGGACCCAGAAAGCAAGAAACTTTCTTCATGGACAAGTGATGGTGATCCATGTAGTGGTTCTTTTCTTGGTGTGATTTGTAATGAACATCATAAAGTAACAAATATTACTCTGACAGATAAAGGACTTACTGGAAAGTTGTCACCAGCAATTGCTGAGCTGAAGTGTTTGTCTGGTCTTTACTTGCATTATAATTCTTTAACAGGGGAAATACCAAAGGAACTTGGGAATTTGACTGAGTTGAATGATCTTTATCTCAATGTGAACAATCTTTCTGGGATTATACCACCAGAACTTAGAAACATGGCAAGTCTACAAGGTTTGTAGCCTTTGTAAACACCTTTCTGTTTAATTTTGAGGGAGAATGTTTCTCTTTTTTTCGTAGCTCCCACCTTTTTGCTGCCTTTGTGACTCGATCTGTCGTGATTTCTAAAAACAGTTGTATTAAATGATTTGTCATTTTCGAAGTTTTAAGATAAAATAATTATCCTTTTCTCATTCTACTTTTAATAGTAATTGTTCTTGAAGATCCTGTTTTGGTGACTCGAACCGACAACCTTAAAGTTGGAGGTAGAGAGTGCTTACAACTTGAGCAACCCCTGCTATCAGGGAAATTGCTTGCTGTTTCATTTTACTGTTTTTTGTGTGTGCTGCTGAATAATGAGTCTACATGAATTTAGGTAAACAGAGTAGACAGGTTGCAAAATTTCTTAGAGATCTATGCAGATTTTTTGGTTCTGTAAAGATAGGACTTTGTGGGGATTGATGGTTTACTCGTTTTTTTGAGTTCCTTCATCCATAAATTAAACAGTGCAAGTTTGAGAGAAAATAAAGTGTTACTAGTATATATATGAATTGTTGAATTTCCTAACATAAGGAAAGCTTTAAGTACAACAGTAGAAGTGGCTCAAAATTGCTTTAAGTAGCAGATTCGAATTCCAAGTATCCCTTGTTAGAATTCCTACTTCCGCCACTGTTGCACCCACACCCCCTTTTCTGGTCTTTATTCTTTCTTGTTAATTATATTTTTGGTCTGAACAGCATACTGAAATTTCTTTTCCAAATAAGGTGATGAAAGAAATGCAAGAACTTCTTTTTTTCCCCTTTCTACAATGATATGTGCTCTGAGATGATTTGTGCATTTTGTGGTTCCACAGTGCTGGACTTGAGTTGTAACCAGTTGACAGGGAGCATACCAACAGAGATTGGGTTCTTGAAGAAGTTGAGTGTTCTTGAATTGCAACGGAACAGGCTGACAGATAAAATTCCAGAGAACTTAGGAGTCCTCGGGACGTTGAAAAAGCTATATTTGGGATTCAATCAGCTATCTGGTCCGATTCCATTAAAATTAGCTGCTGCTCCCCAGTTGGAGGTTCTAGAAGTACAAAATAACACCCTCTCTGGAGTTATTCCTCCAGGTTAGTTGCCCTTTTTGAGTTCTCAACATTTTATAAATCCTTTTTGCCTATCTTGAAGTTACATCTTTTGAATGTGTTTTGATTTTAAACCTCTTTTTCTTCTTTTTATCTGGACTGTTTGTAGCTCTAAGGAGATTAAATGAAAAATTCAATTATGAAAGCAATCCTGGTTTATGTGGTACTGGATTTACTTCATTGAGAGTTTGCACTGCCTGGGATCATGTCAATGTGAATCAAGTTGGTTCTGATGTACCTGATTCCAATAATAACGGTGTCCCTATAGATGTCCCAGGAGCTGCTCATATTAGTCCGTTGCCTTGTAATCAAACTCATTGTTCAAGATCATCAAAATTTCCTCAAGCAATTATCGTTGCTTCTGTGATTACAGTCACTGTAACTTTGGTAGTAGCTGTAGTTTTCGGTATATTTAGACGAAGAAGGCTAAAACAAAGAGTAGGGAACACAAGTGATGCATCTGATGATAGGCTTAGCACTGATCAGGCAAAGGAAATATACAAGAGAAGCCCTTCTCCACTACTTACTGTTGAGTACTCAAATCGTTGGGACCCTATGACCCCTGAAAAGAGTTGCAATAGCATGTTCAATGAGTTCTTAAGTGGATTCAAGTTCAATTTGGAAGAGGTTGAGTCTGCAACTCAGCATTTTACAGAGACAAATCTATTGGGACGGAGCAATTTCTCTGCCGTCTATAAAGGAATTTTAAAAGATGGATCTACGGTAGCTGTAAAAAGCATCAGCGTGACAAGCTGTAAGTCGGAGGAGACGGAGTTCATGGAAGGATTGAGCTTATTAACCTCCCTGAAGCATGAAAACCTTGTCAAGCTAAGAGGTTTCTGTTGTTCAAAGGGAAGAGGCGAGTGCTTTTTAATCTATGATTTGGCCTCTAAAGGGAACCTTTCTCAGTATCTTGATGTTGAAGTAAACAGCAACCATGTTCTTGACTGGTCCAAAAGAGTTTCGATCATCAAGGGGATTGCAAAAGGTTGGTCTTCTTTGCTTTTGTTCATCATTACATCTTCAGCATTCTCTACAAATTTCTCAGAATATTTTAAAGTTTGGAAATATTATGGTTGTATAATATTTTCCCCATAATGTGGAGTCTTCTAAGAGTTCCATTGCATAGGCAGTTTATTACCACTAAAGTGCTTCTCACAGGAGTCAGGAACAGGAAGTGACTGTAGATTCATTTGCATGGTAATTAGTGGCAGAGTCAGGAGGGGTTCAAAGTATGAAGAAGTAAACACACAAAGAAGCCGAAGGGGTTCAACATTTACTATATATACATAAAAAATAATTTTAACCATGTATAAACAACGCAATTTTCTGCCGACCACCTCGGCCTACCTAGCTCCACCCCTGTGGTTAATATATTATCAAATTACTCCTCCATCCCAAAATAAGTGTCGCTTTAGTTCATTTCACGCCCATTAAGAAAAGTAATACATGCAAGGCATAGTTTACTAAGTCACCCCTAGTTATTAGATGAAGGAGGAGTAGTTCTTTAATATAAAGGTATTGTTGGAAACAAGGAGTACTATTTTGTCTTGACTTCTAAATAGACAATTATTTTGGGATAATTTTTTATGCTAAAATGACACTTATTTGGGACGGAGGGAGTAGTATGAATACAGAGAGAGATTGGACTTATTGCTTTAATTGAGAATTAAAGTTCCCTTGCCATCAAGGAAAGCAGTAGGACATGTCTCAAGAACCTCATTAATATGTGCATTTAATAGTTCTCCTATGTTCACTATTTCGAAACCAGTGCCTTTGATTTGATTAAAGATGAAATAGCACAGGTGCACCATCACTGCATTAACTTCAGTTATTCATATTAGAACAACAGGCAACAGATATTTGGGCTAAGTTTTAGTATAGCTAAGGTCTCTAATTAATAAAATCTTTAATACACTATGATGGCGAATGATTTCAACTTACCAACTATATTGTTTTTGTGAGAATCAGGGAAAACTTACCCATGTTTTTTGCGTTCTATCTTAATATGATTCCTTCATATTTCCTTTTGACTGATACCAATCGTTATGCATCTTTCTAACAGGACTAGGATATCTGCACAGCACCGAACCAGACAAACCTTCCATGGTTCATCGCAACATATCTGTGGAAAAAGTCCTCCTTGATCAACAGTTCACTCCACTAATACTGGACTGTGGCCTACTAAAGCTACTTGCTGATGATGTTGTTTATTCAGCACTTAAGGTCAGTGCTGCACTAGGATATATGGCTCCTGAATATATTACAACTGGACGATTCACAGAAAAGAGCGATGTATATGCATTTGGAGTAATTATCCTTCAAGTACTCTCTGGTAAAGGACTTCTGGACTGCTCAATGAGACTTGCTGCTGAATCTTTTAACTTTGAGAACTTCATTGACCCTAATCTCAAGGGAGCATTCTCCATAAGTGAAGCAACTATGCTAACTAAACTTGCAGTAAGTTGCACCCTTGAGGATCCAGATAGCAGGCCAAGTATGGTTTTAGTAAATGAAGAATTGAACAGGTCTAGTTGTGGTTGAAAAAGGAAAAATTGAAGTCCTTTTTGTTCGCGTTGAAATTTGTTGATGATTATTAACCATAGGGAACACGTGCCATAGCGATAGAGCTGCAGTTTTCTATCTATTATGCAATATCACTTGTTTCTGTATATAATGAGGCAGATTGAACAAATTCTTTTGATGGTTTGAAGGTGAAACTGGGACGCCCGTTGGCATCCTTGACTGTTGTAGCTCTCAGAAGAGATAAAGAGGAAGCTAATGAACTTGTACCATTATTTGATCTTTTCACTGTATTTTAATGTCTTTTTCAGGGCCAATATTCTGATAAATGCATTAGTCTTTGTATCGAGTGATCCAAATTCAATACAGTGCCTATGGCATATGGGTACAGACAAA

>PGSC0003DMG400004146 LRR-RLKs

CAAACTATGAAAAAAATTCATCTGTTTTGCTTGTTACTTTTGAGTGTTTTTTTCTTTGTTTCATCAAATAAGGATGAAGTTTCTATATTGTTAGCAATAAAGTCTAGTCTTGTTGATCCAATGAATCATCTCAAAGATTGGAACTTTTCAAACAATGAAAGTTCTATTCATTGTAAATGGAATGGTGTGTATTGCAACTCAAAAAGCTATGTAGAGAAGCTTGATTTGTCGAACATGAATCTTAGTGGCAGAGTTTCAGATCAAATTCAAGGACTTCAGAGTCTATCTTTACTTAATCTTTGTTGCAATGACTTTTCCACTTCATTGCCAAAGTCATTAGCTAATCTGACATCTTTGAAGAGCATTGATGTGAGCCAGAACAACTTTGTTGGGAACTTCCCTGATGGTATTGGAATGTCGAATCCTGGTTTGAAATATGTCAATGTTTCAAGTAATAACTTCGAGGGGGTTCTCCCTGAGGATCTTGGCAATGCTACGTTGCTCGAAGTCATGGATTTTCGAGGGAGTTTCTTTGAAGGTTCAATCCCTGGATGTTACAAGAATTTGAAGAATCTGAAGTTTCTTGGCTTATCAGGGAATAATCTTTCCGGAGATATCCCTCGTGAGCTTGGTGAATTGAAGGCTATGGAGACTATGATTCTTGGATATAATCAGTTTGAAGGTTCAATACCAGCTGAGTTTGGTAATATGAGTAGCCTTAAGTATCTTGACTTGGCTGTTGGCACTTTAAGTGGACAAATTCCAGCTGAGTTAGGTAAGTTGAAGAACCTCACTACAGTGTACTTGTATCAAAACAGTTTTGAAGGTAAGATTCCACCTGAAATTGGAAATATAACATCTTTAGTTTACTTGGATCTTTCTGATAATAAGATCACTGGGGAGATTCCGAATGAATTAGCAGATTTGAAGAATTTGCAGCTGCTAAATCTCATGTGCAATAACTTAACGGGTCCTATTCCTACCAAACTTGGTGAGTTAGAAAATCTTGAGATACTTGAACTATGGAAAAATTCTTTAAATGGCTCTTTGCCAATGAATCTTGGGAAAAAATCTCCCTTGCAATGGTTAGACGTTTCTTCGAATTTCCTGACAGGTGAGATCCCTCCTGGCTTGTGTGATTCTGGCAATCTAACAAAGCTCATTCTGTTTAATAATTCCTTCTCAGGATCAATCCCATTAGGCCTGTCTAACTGTTCTTCACTCGTTCGAGTTAGAATCCAAAATAATCTTCTATCAGGAATGATTCCGGTTGGGTTTGGAACACTTCCTATGCTCCAAAGATTGGAATTGGCTAAAAACAATCTCACTGGTGAAATTCCTGTGGATTTCACTCTCTCCACCACTCTTTCTTTTATCGATGTCTCCTCGAACCACCTTGAATCATCTTTGCCTTCTAGCATTTTGTCCATTCCTAGCCTGCAAACCTTCATAGTTTCCAATAACAATCTTAAAGGCAACATTCCTGACCAATTCCAAGATTGTCCTTCCCTTTCTTTGCTTGATCTTTCAAGCAACCATTTCTCGGGGAAAATTCCTCAGAGTATTGCTTCATGTGAAAAGCTGGTGAAGCTTAATCTAAGAAACAACCAATTCAGTGGTGAAATTCCAACTCACATTGCCACATTGCCAACTTTATCTATTCTTGATCTATCATACAACTCGTTAGTTGGAAAAATACCAATGGACTTTGGTAGCTCTCCAGCTCTAGAAATGCTTAATCTTTCCTACAACAAGCTTGAGGGTCCAGTTCCAAGAAATGGGATACTCATGACTATTAATCCCAATGATCTGATAGGAAATGCTGGTCTTTGTGGTGGCATACTTCCACCGTGTTCTCAAAGTCTCACCATAACTTCAAATGCGAGAAAGAATCGTGTCAATCACATAATCGTGGGATTCATTGTTGGAATCTCAGTTATCTTAGCTGTTGGGATCATGGTTCTAGCAGGGAGATGGATGTACAATAGATGGTACTTGTGCAACAGCTTTTTCAAGGAATTTCGGTTCAACAAGAACAACTCAGAATGGCCTTGGAGGCTTGTAGCATTCCAGAGGCTTAACTTCACAAGTACTGATATTCTGGCTTGCCTAAAGGAGTCAAATGTGATAGGCATTGGTGGCAATGGAATTGTCTACAAGGCTGAGGTTCTGAGGCCACATTCTGTTGTAGCAGTTAAAAAGTTGTGGAGATCAGATGGAGACATTGAAGCAGGAGATGATCTCGTTGCAGAGGTTGATCTTTTAGGGAAGCTTCGTCACAGGAACATAGTTAGGTTACTAGGCTATCTTCACAATGAGACTGATATCATGATGTTATCTGAATACATGCCTAATGGAAATCTTGGAGCTGCTTTACATGGAAAAGAAGATGGAAAAATGCTCGTAGATTGGTTGTCGAGGTACAACGTTGCCCTTGGTATTGCTCATGGACTCGCTTATCTTCACCATGATTGTCATCCACCAGTCATACATCGCGATGTCAAGTCAAACAACATTCTCCTAGATTCAGATTTTGAGGCAAGAATTGCAGATTTTGGCTTGGCAAGGATGATGCTTCACAAGAATGAGACTGTTTCTATGATAGCAGGATCATATGGCTATATAGCACCAGGTTAGTTACATTTCCTGATAATGCCACTCACAAGAAAATTCATTTCATTTTATATAGTCTCATTTTCCTTTCTCTGTTGTTTTTGGCAGAATATGGATACACATTGAAGGTTGATGAGAAGAGTGACATATACAGCTATGGGGTGGTGCTTTTGGAGCTTGTAACAGGAAAAATGCCATTAGACCCTTTGTTCGGAGAGTCGATTGACATTGTAGAATGGGTTAGAAGGAAAGTAAAGAACAAGGCATCAGAAGAAGCATTAGATGCTGATGTAGCTGGCCAATGTAAACATGTCCATGAAGAGATGCTTCTTGTCCTAAAAATTGCACTTCTTTGCACTGCCAAGCTTCCCAAAGAAAGACCATCAATGAGAGACATTATAACCATGCTTGGAGAGGCAAAACCAAGAAGGAAAAGTATTTGCCAGAATTGGGGTTATAGTACTAGTGCTAATAAAGACAAGTTAATCTTTGCTCATTCACCAGTTATAGGCCTTCTATAAGTACCAAAACCTTATTTTTCCACTTCAATATCTAGTATATCTTCATTTTCATTAGTTCAAATGTAAA

>PGSC0003DMG400004667 LRR-RLKs

ATCAAAAGCTGTCACTTAGCTTACTGATGAAAAACTCATAAACCCAAAATTTTCAACAAACAATCCCAAAAAAAAAAAAAAAACTTCAAAATTTTGAAGAATCATAGCAAAAAAAAGGTAATCTTTGCCAAAAAAATTCAAGAACAATAGCAAAAAAAAAAAAAACTCCATTGTTGATTGTTCCTAGATCTCTAAAACCTCCATTAATGGCACCCCACAAGCTTTTTTTTCTTGTTAACTTCTTCTTCACCTTCATTTCATTCACATTTTCCGATGACAGCACCGTCATGTCAAATTTTCTGAAGGGTCTGTCTAAACCACCATCTGGTTGGGATGCTTCAAAATCTTACTGTGATTGGAGTAAAGTTAATTGTGATAAATCGAGTAATACTGTTGTTTCCATTAATCTAGATTCTCAAGGTCTTTCTGGGGTTTTGCCTTCTGACCTTAATCAACTGTCAAATCTCAAAACTCTTTCAGTTCAGAGGAATTCACTTTCTGGGGTTCTTCCATCTTTCGCAAACATGTCGAATTTAGCTGAGATTTACTTGAATGACAATGGATTCACATCTGTTCCTCAAGATTTCCTTTTGGGTCTTACTAGTTTGCAGATTTTTAGTATTAGTGAGAATGGGAAGTTGAGTCCTTGGCAAATTCCTAGTTATTTGATTGATAGTAACAATTTGGTTAATGTTTATGCTAGTCATGCTAGTTTGTTTGGTGTTATACCTGATTTTCTTGGTAAATTCCCTAATCTTCAGGATTTGAGATTGTCTTATAATAATTTTACTGGTTCTTTGCCTGCTAGTTTTGGGGATTCTGAGATTAAGAATTTATGGCTGAATAATCAGGTTAAAGGGTTATCAGGAACTATTGATGTGATTTCAAGTATGTTTCAGTTGTCACAAGTGTGGTTACATGCTAATTCTTTTACTGGTCCTATACCCAATTTGTCGAAATGCGAAAATATATTTGATTTGCAGTTAAGGGATAATGATTTTACTGGTGTTATACCTGATTCAGTGATGAAACTTCCTAACTTGTTGAATATATCTTTGAAAAACAATAAGTTGCAGGGTCCAATGCCAGAGTTCAGAAAGGAGGTTAAGGTGGAGATTGAGGGTACTAATAGTTTTTGTTTGGATACTCCTAGACCTTGTAATTCACAGGTTATGACGCTTTTAGATGTGGCGGGTGGTTTTGGATATCCAGGGTTTTTGGCTGATTCTTGGAAAGGAAATGATGCTTGTAATGGATGGTCACATGTAAGTTGTGATGGATCGAAAAAGAATGTGGATGTTGTAACACTTGGTAAGCTACGTTTATCGGGTTTTATATCTCCTGCTTTTGCTAAATTGACTAGTTTGAGGAATTTGTTCTTGAATGATAACAATCTTACCGGATCTATACCGGAAAGTTTGGCAGCGTTGCCTATGCTTCTAACTCTTGATGTTTCTAATAACAATTTATCTGGTCCTGTACCTGCGTTTCGATCAAATGTGAAACTTATTAATGGTGGTAACGTGTTTCTTGGAAAGAATATAAGTGATGGACGGGGTAGTGGTGGATCACCGGGGTCTGGGTCAAGTAGTGATGGTGGAAATTCATCTGGTGGTTCGAAGGGATCCTCAGTTCAACCTGGAGTGGTTGCTGGTGTTGTTATATCTGTCGTTATATTTATCCTTGTTCTGTTGTATGTGTCTTACAAATGTTACATAAGGAGACGGCATAAGAGATTTGGAAGGGTTCAGAATCCGGAAAGAAGCAATGATATGGTTAAGCCTGGTGTAGTGATAGGTAATGGATATGCAGGAGTCCCTAGTGAATTACAGAGCCAGAGCAGTGGGGATCATAGTGAGATGCCGATATTTGAAGGTGGAAATGTTGCCATTTCCTTTCAGGTGCTTCGACAAGTGACAAACAACTTCAGCGAAGAAAATATCTTGGGTAGAGGAGGATTTGGAGTCGTTTATAAGGGTGAATTACATGATGGGACTAAGATTGCTGTGAAGAGGATGGAATCTGGCGCGATGGGTACTAAAGGAATGAATGAGTTCCAAGCTGAAATCGCGGTCCTTACCAAAGTTAGGCATAGGCATCTGGTCGCTCTTCTTGGTTCCTGTGTTAACGGCAATGAAAGACTTTTGGTGTATGAGTACATGCCACAAGGAACTTTAAGTCAGCATTTGTTTGAATGGCAAGAACACGGGTGCCCAATTTTGACCTGGAAGCAAAGGGTCACAATCGCATTAGATGTTGCAAGGGGAGTCGAGTATCTCCACAGCTTGGCACAGACAAGTTTTATCCATAGGGATTTGAAGCCCTCTAACATACTTCTTGGGGATGACATGAGAGCCAAGGTTGCAGATTTTGGGTTGGTTAAGAATGCCCCCGATGGGAAATATTCTGTTGAGACACGGTTGGCTGGAACTTTTGGCTATCTGGCGCCTGAATATGCTGGTAAGTCTGTCTTTCCTCTGCTGATCTTTAATCCTTTTAGTTGTAAAACTGGTAGAAAACTAAAACAATATCCTATCATTAGCTTTCATGTTCTTTTTAATCTTCACTATATTAGAGATAAATGGAGAGGTGTTAGCTTCATTGAAGTTCATTTCCATGTTCAAATGCTAGTTTGTTAATCTCGATCTGATTCGTGGATTGATACCGCAAACGTTGTCCAGTAAATTAACTAATGTTAGCAAAAAAGAGTACGATTTGAAAGACAAACATAAGGTAGAAATCTAGAAGCCCACTCATCCGGTTTTCAAATTGAAATTCGGATGTTGGTACAGCGTGTGTGACTGATCCCATTGTATCTAAATAGAAATTTGGCTTTCGGAATATCACTCTTCTTCAAGACATTAAGTTCAACATATCTAGATTTAGTGTTGGCTGTCAATAGTTTTCTGCTCGTAACATGTTTAATTTCCTTTAGAGATCTTGTTCTGGCCTATTTCTTGCATGTCTTAGCAAGGTTGTCTGTGATCATTACAAAATTTGGATTAAACGTGACTGCTCCTCTTTATTTACAGCTACTGGACGAGTCACGACCAAAGTAGATGTGTATGCGTTTGGAGTTGTTCTAATGGAGATCATCACTGGTAGAAAAGCATTAGATGAGACCTTGCCAGATGAGAGGTCTCATTTGGTCACCTGGCTTCGCAGGGTCCTTGTCAACAAAGACAACCTCCGAAAGGCTATTGACCCAACTCTTGATCCTGACGAGGAAACATATGAGAGCATTTGTAAGGTTGCCGAGTTAGCAGGCCATTGCACTGCACGAGAACCATTCCAGCGCCCTGACATGGGACATGCAGTCAATGTCCTTGCCCCCCTCGTAGAACTGTGGAAGCCTACAAGAAATGAAGACGAAGACAGTGGCATCGACCTCCAAATGAGCCTTCCTCAAATTCTTCAGAGATGGCAAGCTGATGAAGGAACTTCAAGAATGTTCGATGACATCTCCTTCAGTCAAACACATTCAAGTATACCCTCAAAACCTTCTGGATTTGCAGATACTTTCAATTCAACAGATTGCAGATGACATTAAGGCAACAATCAAGAAAGGCACACAATTCATTTGGTTAATTGGTTTCGTCTTAACAACGCGTTTTACTTCGTCTTTTCACAACCCTTTTGGTGATAGATCATTAGACCAAAGAAGCTAAGGTGGATAGCAGAGTTTATTTGTGGTTTTGTTAGAAGAGAAAAATTGTATTAAATTCTTTTAATCTTACATAGTTGATGTAAATAGATTTGAGTTCTTGTGTTAAACATGCAAAATATACTCTTCTTATTAAGTTTTTTACTACAAGATTTCATTATTTGTGCCACTAAGTGTGTA

>PGSC0003DMG400004963 LRR-RLKs

CTTTTATTTCCTTCCTCTCTAAGAAAAAAACAATTTAATACAAATGACTATTCAAAAAATTATCACCCTTGTGTTATTTTTTCTTACTTTGACGTCCACTTTTGTTCAAGGTGATCAATCTCAGTTCTTTGTACTCATGAAGAAATTTGTCACAGGGAGTTCTTTGTCTAATTGGGATATTGGAAAACCAATTTGTCAATACAAAGGTGTCGGTTGTGATGAAAGGGGAGACGTTATCAAAATTAACATATCGGCCTGGTATTTATCAGGCCAATTTCCAAGTGATGTGTGTAGTTATTTTCCAAGATTGAAAAGTCTTCATATTGGTCATAATAATTTCCAAGGTGGTTTTCCTAAGTACCTAACAAATTGTTCTTTTTTGGAAGAATTGAACATGACTAAAACATCACTTACAGGACAAATTCCTGATTTGTCACCAATACAATCTTTGAAACTACTTGATCTTTCATGTAATCAATTGACAGGGGATTTTCCTTTGTCAATTATTAATCTCACCAACTTAGTTATCCTGAATTTCAATGAAAATCGCCATTTCAATCCGTGGCGATTACCAGAGGACATTTCCAGGCTTATAAATCTCAAGTGGATGATTTTAACAGCATGTAATATGCATGGGACAATTCCAGTGTCAATAAGTAACATGACTTCTCTTGTTGATCTTGAATTGAGCGCGAATCGCCTTGTTGGTAAGGTGCCTAGAGAACTTGGGAAGTTGAAGAATTTGAAACTCCTTGAGCTTTTTTACAACTTACTTGATGGTGAAATCCCCGCGGAGCTAGGGAATTTGACAGAACTTGTGGACTTAGACATGTCCGCGAATAATTTCACAGGGAGAATTCCAGAGTCTATAAGCCGGCTTCCTAAGCTGCAAGTGTTGCAACTTTACCATAACGCGCTATCAGGAGAGTTTCCAGCAGCACTTGCTAATTCAACAACTTTAACTATCCTGTCTCTTTATGACAATCTGTTTACAGGAGAAGTTCCACAACACTTTGGTCTTTCATCAGCTTTGTTGGCATTGGACTTGTCAGAAAATCGATTTTCTGGAAAGCTACCGCCTTTTCTGTGTAGTGGAGGTAAATTGAGTTATATTCTTTTACTTCAAAACATGTTCTCAGGTGAACTGCCTGATGGATATGTGAAATGTCAGTCTGTTCTTCGCTTTCGAGTGAATTACAATCAGTTGGAGGGAAGAATACCACAAGAGCTTTTTACTCTTCCACATGTTTCAATCATTGATTTGAGCTATAATCATTTTAGTGGTCCAATTCCAACAACAATTGGAAGTGCTAGGAATTTATCAGAATTGTTTATGCAAAGTAACAAGCTTTCAGGTTTGCTTCCTTATGAAATATCTACATCTTCTAATCTTGTGAAGCTTGATCTTAGCAATAACCTTTTGTATGGTCCAATTCCTTCTGAAATTGGTGGTTTAAAAAGCCTCAATTTGTTACTCTTGCAAGGTAACAAGTTCAATTCTTCAATCCCCGAGTCACTTTCTTCGCTTAAATATCTCAACTATCTTGATCTGTCTAACAATCTTTTGATAGGAAAGATCCCTGAGAGCCTAGGTGAATTGTTACCAAACTCTATGAACTTGTCAAACAACTTACTTTCCGGTGCTATACCTCTTTTGTTTATAAAAGGAGGTGTTTTGGAAAGTTTTTTAGGCAATCCGGGACTTTGTGTCCCTACCTCTTTGAACTCATCAAACACAAGTTTTCAAACATGTTCACATAGCTATAACCATAAGAAGAGGAACAACATTGTTTGGGTTATTGGGACATCGGTGGGGATAGTCATTGTTGGATTAGTCTTGTTTATCAAACGATGGTTTGGTAACAAAAAGGAGGTGATGGAACAGGATGATCATTCATTGTCATCATCATTTTTCTCCTTTGATGTTAAGAGTTTCCATCGATTAAGTTTTGATCAACGTGAGATCTTTGAAGCTATGGTTGAGAAGAATATTGTTGGATATGGAGGGTCAGGGGCTGTCTATAAGATCGAGTTGAGTAATGGAGGAGTAGTTGCTGCAAAGAAGCTGTGGAGTCACAAACATAAGCATTCTGTTTCTGAGGATAAATTGGTTTTGGATAAGGAACTTAAGACAGAAGTTGAGACTCTTGGTAACATAAGGCACAAAAACATTGTGAAATTGTACTGTTATTTCTCAAGTTTGGATTGCAGCTTGTTGGTTTATGAATACATGCCTAATGGAAACCTTTGGCATGCTCTTCATGGTGGGAAGTTCGTGTTGGATTGGCCTATTCGTCATCAAATTGCACTCGGAATAGCTCAAGGATTGGCCTATCTTCACCATGATCTTATGCCACCGATTATTCATAGAGATATCAAATCCACCAACATCCTCCTGGATATTGATTATCAACCAAAAGTCGCGGATTTTGGTATAGCCAAAGTGTTGCAAGCCAGAGGAGGCAAAGATTCCAGCACCACGGTCATCGCAGGAACTTATGGTTACTTGGCACCAGGTATAATTTTCCCAATTCATAAGTGTTACTTCAATCTTTGAGCTTGATTGAGTAGATTAAATTAAATCCTTTATGTGCAAATATCATCCCATAGTTGAAGTGTTCATATTGGCCTAATGTTACTCTAAATTGGCCATTTAATATAGTGAAAAATTTAGAACATTGCCTAACGTTAAGAGTACATTTGACCCTTTTTCGTTCTTAATTATGGCTTTTTATGTTGTAAAAAACATCACATATGCTCAAGTTTCTTGATTGAACTTTCATTTAATACATGTTTAAGTGTTGAAATTAGGCTAATGTTACTCAACTTTGGTGTAACAGAGTATGCATATTCTTCAAAGGCAACTACAAAGTGTGATGTGTATAGTTTTGGAGTTGTTCTAATGGAATTGATTACTGGAAAGAAGCCAGTGGAGCCAGAATTTGGAGACAACAAGAACATTGTTTATTGGGTATCAACAAAAGTGGAGACTAAAGAAGGTGCATTTGAAGTATTGGATAAAAAAGTCTCAGATTCTTTCAAAGAGGACATGATTAAGGTTCTACGTATAGCTATACGTTGTACGTATAGTACACCAACACTTCGTCCTACCATGAATGAAGTTGTTCAGTTGCTAATCGAAGCGGATCCTTGCAAGTTCAATTGTTGCAACATGTCAAATAAGAAGAAGAATGACACAGAAGAAGTGATCAATAAGCCACCAAAGAACATATGAGTTGTAAAAAATTTAGTGTATCAAACTAGTATACAAGGTATATAGAATATGTGTACTGAGATATTACACCCTCAAAAGAATCAGTCTTTATGTTTTCTAATAAATGGTACTACTTTTTATTAGTATATATATAGATTCTCCAATTAGTGTTACCTTGCTTTTTTATATTTTTGGCCTGTTGTAAGTATTTGGGAATTTTATGAA

>PGSC0003DMG400004966 LRR-RLKs

GCGCTCTTCATACTCGTCTCCTTCACACGTTTTTCTGAAGGCGGAAGAAAAAAACTTCTCTTCGTCAACTTCTTCGTCAGCTATGGCGGCGCTATTCAATTGCCGGCCGGAACTTATCCACTTCTTCTCTATTTTCTCCTTCTTCCTTCTCTTCCCCGTCGCTTTTTCCGACGAGCTTCAAACACTTTTATCCATCAAATCCTCTTTATCAAACCCCACCACCAATGTATTCCAAAATTGGGAACCCAATACTCCTCTCTGTAAGTTCACTGGCATCACATGTAATTCCGATGGGTCAGTCAAAGAAATCGAGCTTTCGAGTAAAAAAATTTCTGGGTTTGTTCCTTTTGATAAAATATGTTCACTTAATTCATTGGAAAAGCTCTCTCTGGGTTACAATTCACTTTCTGGGGAAGTTACGGATGATTTGAACAAGTGTGTTAGCTTGAATTACTTGGATGTTGGTAATAATGAATTTACTGGGTATTTCCCTTATGTATCTTCACTCAGTGAGTTAACCCATTTTTATGCAAATAACAGTGGGTTTACAGGAAAATTTCCATGGAATTCTTTTGCGAATATGAGTAATTTGATTGTGCTTAGCCTTGGTGATAATCTGTTTGATCGGACACCGTTTCCTGAAGTGATTTTGAAGCTTAATAAATTGAATTGGCTCTACTTATCGAGTTGTGAGCTTGAAGGGGAAATTCCTGAAGAAATTGGAAATCTTACAGAGCTGATTGATTTAGAGCTGTCGATGAATCACCTTACTGGTGAAATTCCTTCTGGAATTACGAAGCTGAAGAAATTATGGCAGCTTGAGTTGTATGAAAACCAGCTTACAGGGAAGTTGCCGGTTGGATTTGGGAATTTGACAAGTCTTGAATATTTTGATGCTTCGGCTAATAATCTGTACGGTGATCTGTCGGAAATTCGAAAGTTGAATCAATTGGTAAGTTTGCAGTTATTGCAGAACCAATTTTCAGGGGAAGTACCAGCAGAATTGGGAGAGTTCAAAAAGCTGGTGAACATTTCTTTGTACACTAACAAACTCACTGGTCAGCTTCCGCAGAAGTTGGGTTCTTGGGCGAATTTTGATTTCATTGATGTATCAGAGAATAGTTTCACTGGTCCAATTCCACCGGACATGTGTAAGATGGGGACGATGAGAGGATTATTGATACTTCAAAACAATTTCACTGGCGGAATCCCTGAAAGTTATGCTAATTGTACAACAATGACGCGGATTCGAGTGAGCAAAAACTCACTCTCCGGTGTGATTCCTGCAGGAATTTGGGGGCTACCAAAGCTCGAAATCCTTGATGTTGCAATGAATGAGTTTGAAGGTACTATTACTTCTGATATTGGAAATGCAAAATCTTTAGGTGAAATTGATGCGGCTAATAACAGATTCTCTGGTGAATTGCCTTTTGATATTTCGAATGCTTCTTCCTTAGTGAAAATTGATTTTAGTAACAATCAGTTTTCCGGTGAAATTCCGGGAACAATTGGGGAGCTTAAGAAGATTGGTAATCTTAATTTACAGAACAACAAGTTTTCTGGTTCAATACCAGATTCTTTAGGCTCTTGTGTTTCACTAAGCGACATTAACATGGCTAACAATTTGCTTAGCGGCTCTATTCCTGTTTCTCTAGGATCATTACCGACTTTAACTTCATTAAACTTGTCGGAGAATCAGCTTTCAGGGAAAATTCCGACGAGTTTATCAAATCTGAAGTTAAATCTTCTTGATTTTTCCAACAATCAGTTGACTGGAGCAATACCAAACTCCCTATCAATTGATGCTTATAAGGGTAGCTTTGCTGGGAATAATGGTCTTTGTAGCCAAAACATTAAGAACTTCCGTAGATGTTATGGTGAATCTGGAAAGCCTAGAGAATGGTACACTCTGTTGATATGTTTATTGGTGGCTGTGATTGTGGTGCTCGTTTCATTTGCGGGTTATTTGTTCTTGAAGAAGAAAAGTCATAAGGAACATGAGAGGTCTTTGAAGCAAAATTCTTGGAATACAAAGTCATTCCATATATTGACCTTCACAGAGGATGATATTCTTGATGGAATCAAGCACGACAATTTGATTGGAAAAGGTGGTTCTGGTAGTGTATACAGAGTGCAGCTTTCAGATGGAACAGATTTCGCGGTGAAACATATTTGGACTTCTGATTCAGGAAACAGGAAAATATCAGGAACAACATCACCAATGTTGGGAAAACCAGGGAAGAAATTGAAAGAATTTGAGGCTGAGGTTGAAACACTGAGCTCAATTCGACATGTGAATGTGGTGAAATTGTACTGTAGCATAACGAGTGATGACTCGAGCTTGTTGGTGTATGAATATATGCCAAATGGAAGTTTGTGGGATCGATTGCATACTTGCAAAAAGATGTCACTTGATTGGGAGACGAGGTACGAGATAGCTCTTGGTGCAGCCAAAGGACTGGAGTATTTACACCATGGTTGTGATAAGCCAGTGATTCATAGAGATGTTAAATCAAGTAACATCTTGTTGGATGAGTTTTGCAAGCCAAGAATCGCTGACTTTGGTCTTGCTAGGATTGCTCAAGCTGATTCAACCAAAGATACAACTCATGTCATTGCTGGAACACATGGATATATTGCTCCTGGTAAGAAGCTCATTATATTGTAGTTGATTTTATACGTAGAATTTGAGTATATCTACTTAACACTAACCATGTATTCATTTTTCACCATCGGTAAAATTGACTTACGAAATAGGCAACTCCCCATATCAAGCAATATGGTATCTTGAAAGATATAGTGAAAACTTGCTATAAACAGTTGGATCACTATGTTGCTCGGATCCTTCAAAAATGTCACTGCACCTGTGTTGGTTCCCTCAAAAATAGCGCATTCTTGGAGGATTTGACATGAGTGTGACAACATTTTTGAAGGATCCGAGCAACAATTGAATTGCACTAATAAGAGTGTGAAACAATATTAGTGTTAATTAACTTAGATCCTATCTTGATTATCTATTCATTTGGTCATAGTGAGTGTCTCATTTCTGGCATCCTTTTTGAACTTATTTGAATTGAATTGGTGTTGCAGAATATGGATACACACACAAAGTGAACGAGAAAAGTGATGTATATAGTTTCGGAGTTGTATTGATGGAGCTTATATCTGGGAAAAGGCCAATAGAATCCGAATACGGAGAGAATGGCAACATAGTTACATGGGTGAGCAGCAAATTGAAGAGCAAGGAAAGTGTATTAAGCATAGTGGATTCAAGCATTCTTGAAGCTTTCAAGGAAGATGCAATCAAAGTCTTGAGGATTGCTATTGTATGCACATCTAGGCTTCCCACATTAAGGCCAACAATGAGAAATGTAGTCAAAATGCTGGAAAAAGCAGAACCTTGCAGATTGGTTGGGATAATTGTAAGCAAAGATGATGGTAGTATCAAGACAGAACAATTGAAGGATCACACCAAGATGTAATTAGATCCCCCGTCTTTGCTCGTTTCACTCTTCAACCTGATCTCTGTCCTCCATGGAACGTCGAATCTTGTAGCAGTATGTAAAGTTTTGAAGATGGCCTCTTGGCCTTGTAACATATAGTAGAAGCTAGATCCATACTAGTATGTGGAATTAGATTGAATTTAGAGATGCATGGCATAAGTGCTATAAGATGTAATACTTGTATGTGCTAAATAAAAATGGTTTAGTTATTCAAATCCCTTTTCACCCTGCA

>PGSC0003DMG400004972 LRR-RLKs

GTTCATGCTCTAAATAACTTCAGGGAAGCTATTTATGAAGATCCACATCTGGTGTTCTCAAATTGGAATGCCTTAGATTCAGATCCTTGTGGATGGTCTGGAATTTTTTGCTCAACGGCACGAGATCATGTCATAAAAATGTGAGAGAAGTGCCTTCTGCATATGGCGTTTTCGTTTTCTTTCTATCTGTTTGTTGGATTGATTTTTCTCAATTGGATGCAACTTATTTCATTACTTTTTGATACATGTTAGTAGCACAACTGAGCCTTTACTAACGAGTTTACAACGTCTCCTGTCTATGTAATTTTGCAGTAACATTTCTGGTGCTTCTTTGAAGGGTTTTCTTTCGCCGAACTTGCATTTGATTTCTTATTTGCAAGAACTGTAAGCTGTTCCTTCCATGAAAAACTTATACTTGTAACTCCGGTGCATAAAAGATTTGGTAAAAGCTTAATGCTATGTATTGGTGATATGCAGGATATTGCATGGAAATGTACTGATTGGTACAATACCGAAGGAGATTGGCTTGTTGAAAAATTTAAAGGTCTTGGATTTGGGTTCTAACCAGCTCGCAGGACCAATTCCTCTCGAGCTTGGCAATTTGTCAAACATTATTAAGATGTTAGTGCAATTTGAGTGTTCGTTTTGTATGTGTACTTTCGTTTTTGACTTGCATGAAGCTTCCTTATCTCATCTAGTTTTGTTATGTCTCTGCAGAAACCTTCAATCCAATGGATTAACAGGTAAATTGCCCTCTGAGCTTGGTAAATTGATATACCTTCAGGAACTTCGCCTGGACAGAAATAAGCTCAAAGGACCACTGCCTTCCAGTAATGGCTCAAATTTTACATCTAGTGCGTATGGGATGTAAGTCGAATATGGACAGTGCTATTTCTTCAATTCTATGCGTTTTTAATCTTTGGATTTGCTGTTATGTAGGTTTTACATTATCATGTTGGTAGAAGATTTAATGTTGCAAATGCTTTTGAGACGTGGTTCATTTGATTTTTCAATCTTGTACACTATGAAGTTGCCTTTTTTCGTCTTTTTGGGGGAGGGGGTGGATATCTGAAGCTTGAAATCTTTTATTTAAGCCGTTAACTATACCCTTACTAAATGATTCTCTCTAAAATGTAATGTTTCCAGGACTTACGAGCATTATACATTCAAAACTTTTGGAATATTTGAAGCTTGATTAAATTTATATATCTTTTTAATCTTCTAAGGAGCCCATAGTATCTCAATTAGCATAAGATTTGCTGCAACATTTTTGGCTGCAGTTTGTCTGGACATTTACAGAGGCTTAAAATTCAAAGATTATGGCTAATGTTTAATCTTTTTGTCGTCTTGATTGATAGTGCTGGACTTCTGTTGACATGTATGTCTGTTGTACTTGAACTTAATTTAATACATTTGTTGTTCTCCTAGAAATTTCACACAATTACTGAGTGTAATGAAATCTTAAAGGGTTCAACATGTATATTCAAAAGTCATCACCGACGTTATCGGCAATATTCTGTATAGTTTTCTGGAAAAATCAGTAAAAAGCCTCCTATGCTTGCTTGAACTTGTAATTGCTCTACATAGATCCCTTTAAGTTTTATCTCACAGTTGGTTTCTTAGTAATCGTCTTTTATTTTTGTTTAGGCATGCCTCTGGTGCCAAACCTACTGGTCTTTGTCGTTTATCTCAACTGAGAGTTGCTGATTTCTCCTTCAACTTCTTCTTTGGTAGCATACCGAAGTGTCTGGAGTACCTTTCAAAGTATAGTTCTTAATTCCAATCCACAAGAATTTTTTATACTCTTTATATCAATTTCTGAGTGCCTATTATTTTGTTCTTTCTATAGAACTAGCTTTCAAGGAAATTGCCTTCAAGCAAAAGATCCCAGGCAGCGTTCTACTGCTTTATGTGGTATGTGTAATCTGCTAGACAACTCTTACACTTTAGCATAATCATTAGCCAAGCCTTTGAATCTTTTATGCATACATTAGTGTGCGAAAATGCAAGCTGGTCATGCTTACTTTTGATTGTATATCCTAAAGAGGCTTCTGCGAGTAAGTTCGCCCATTTCACTAACAGTCGCAGTTGAACAGATCAACTTTAAAATAATTGAATTGATTGTTGTAACTTTTGCATTATAGGAGGCACTCCACCTTTCAAAAGCAATCCAGCACAGAGCAATAATAAGCACCGGACTGTTGAAGGAAGAGCCAAACATAAGTCTGCTTCAAAACCTACCTGGCTTTTTGTGTTAGAAGTGGTAACTGGAGTTATTGCGGGTTCTCTCTTCCTTGTGGCTATTTTGGCTGCCATTCAGAAGTGTAAGAACGAATCTTCTATTATCATCCCTTGGAAGAAATCAGCAAGTATGAAAGACCATTTGAAAGACCATATGGCAGTTTATGTAGGTGAGTACCATGCAATTTCAGTTGGGCGATTACAGTTAAGAATTGCCTTTAAACATTTGTATTACTTGATTCTTTATTGACTTAGGGTGTGTTAGGTATGAAGGAAAAATGGTTTCCTTGGAAAATGTTTTCCAGAAAATAAGTGGATTTCTTACTTATTTTCTTGTGTTTGGTATGTAAGTAAAAAACACCACCCTAATAAGATTTGTATATAATCTAGGCAAATACTATGGGAGGTGGGAGTAGGGGTCCCGGGTGGTGGGGATGGGGGTGAGCTGAGGTGTGTTGGAGGGTGGGGAGGACACAATTGATGTGAAATACCACTTGTGGAACTTGTTTTCCCTACCTCCATACCGAACACGCCCTTAATTCTTTCTCATATAGGATCTGATTCTTCAATTTAGTACTACTAATGTTTCTGTTACTTCAGTTATCTTTACTATTATTGTTGTTAATACATAAATATGCCCTTTAAGCCACATAGGACTCATGTGCCTACTTGTTTAACTTTATACAAGTTTAAGTGTCTGCTTGTGCACATCCAAAGTTAAATGACATAAATGTGAAATGAGGCCAAATTAAAAGGCACATTACTGTATTATGCCTTTTTCTTTTCTATGATGTTTTCACAATAGCTTCCTTACTCTTGTAATTTTAAAAACCTGTTTTGGAAAACATTTTTCTTGAACCGAGGGTCTATCGGAAACAACTTCTCTACTTCCACAAAGGCAAGTTTAAGGTCTGCATACACCTCACCCTCCCTAGACCCGACTTGTGAGCTCACACAGTGTATAGGATCACAAGTTCAGAAGCATTAATTATAATTCACATTTCTATTAGTAACAAATGTTTCTTTGTCAGATACTGAGATGCTAAAGGACGTTGTAAGATATAGCAGACAAGAACTTGAGGTAGCTTGTGAAGATTTCAGCAACATTATTGGATCCTCACCAGACAGCTTGGTCTATAAAGGAACTGTGAAAGGCGGACCTGAATTAGCTGTAATTTCCATTTGTGTCAAAGAAGAGCAGTGGACCACCTATCTAGAGCTTTATTTTCAGAAAGAGGTGAAGCCTGTTCTCTCTTTAAAATACATGGCTACTGCTGAGCTCTCGTGCGCAATAAAAATTTCATCTTTGCTGCAATGCTCTAACTTGTATGGCTTACAAACTCAGGTGGCAGAATTGGCCAGAATAAATCACGAGAACACTGCAAAACTTCTAGGTTACTGCAGAGAGAGCAGCCCCTTCACAAGGATGTTGGTGTTTGAGTATGCATCTAACGGGACTTTATATGAACACCTCCATTGTGAGTGGAAAAAACTATTTTACACATTTGCTTACTGTATGTTTTAGAGGCTGAAATCCGAACTCTTTGAACTTATCACAGATGGAGAAGGATGCCAACTTTCTTGGACGCGGCGAATGAAAATAGTTATTGGCATCGCTAAAGGACTGAAATATCTCCATTCAGAACTTGACCCGCCATTCACTATATCAGAACTGAATTCAAATGCTGTATATCTCACAGAAGATTTTTCTCCCAAGGTAAGCAAAAGATTTCTGTATCACTGCTGCAAAACCAATTCGCTGTTAAACTTTCTGGTTAATCTTACTGTCAATTGTGCTCATCCATAAGACAATTTTAGTTTACTCTCCACGGTTTCTGCTCATTTAATCTTTGAACTCGCCGAGTTTCCTTACAAACAATTTGGGCTTACATTTTTCCATTTTTGTTTCCTTCAAGCTCAACCCTTTGTGCTTTTTAACCCGAGTAATAAAAGGCTGATTTTTCATACCCTTTTCTCTCAAAGTTTAATTTATCTGGTGCAGCTTGTTGATTTTGAAAGTTGGAAAACAACATTTGCCAGATCAGAAAAGAATTCCGGTGCTATCAGTAGTGAAGGAGCAATGTGTGTCCTTCCAAATTCTCTACAGAGCCGTCATCTTGACATGCAGGGTAACATCTATGCATTTGGAGTACTACTGCTCGAAATAATCAGCGGAAGACCTCCATATTGCAAAGACAAAGGGTGCTTAGTAGATTGGGTAAGCATTTCTTTTCATGAAGTTTTTGTATAGGCATAGTACATAAATATACACTCAAACTTGGCCTCAGATGGCAAATAAGTACTCCAACTTTGAGAGTGCACGTCTAGACGAGTTGAGTTGTTTAGATGTGCACTCTCAAAGTTGGGAGTGCTTACTTGCTAGCTTAGGCCAAGTTTGAGTGTCTGCTTGTGTATTATGCCTTTTGTATAATGATAACTCTCTCTCTGTGGAACCTGGACTCGTGCTTTATGGAATCTGAAGTTGACTTTGATATGGCAGGCAAAAGAATTCCTTGAATTACCAGAGGTATTGCCTTATGTTGTAGATCCCGAGTTGAAACATTTCAGATACGAAGACCTCAAAGTGATTTCTGAAGTGGTTAATCTCTGTATCCACCCAAGTTCGAGTAGCAGGACTTCGATGAGAGATTTGTGTGCTATGCTGGAGAGCGACATTGACACATCCATAACTGCAGAGCTCAAGGCATCGTCTTTAGCATGGGCTGAGCTTGCACTCTCATCCTAA

>PGSC0003DMG400005103 LRR-RLKs

CTCATGGCCACCCTCTTGAGAAAAAATTCTCCACTTTCTCTCTCGTGGATTTCATGTTCCCACAGAAACCAAACATCAAACATAGCCATATCAAACTTTCTTCTCTATTCTTCTTACTAGCGAAAAAAAAAAAAATGTCAGCCAGTGCCAACTTTTTCCGGCCATGCCCAATTTTATTGCTCTGTTTTTTCATATTCTTGATTTCTCCTTCTCATCAACAAGATGAGTTACAACTCCTTATGCAATTCAAATCCACTCTGAAAACAACACAAATTTCACATCTTTTTGACACGTGGACACCTCAGAATAACATTTGCAACTTCACTGGAGTTTTCTGTGATTCTGACAGTAAATTGGTTAAAGAAATCATTCTTTCTGAACAGAATCTATCTGGGGTTGTCTCTTTTGATTCACTATGCTCTCTGAAATCACTAGAAAAAATTTCTCTTGGTACCAACTACTTGTATGGTAGAGTCAGTGACCATTTGAAAAACTGTACAGAGTTGCAGTATTTGGATTTGGGTAACAATTCTTTTTCAGGGGAAGTTCCAAATTTGTCTTCTTTAAGCCAATTGGAGTTTTTGAATCTCAATAGAAGTGGATTCTCTGGTTCATTTCCATGGAGTTCACTTGGAAATCTTACTAGTCTAACTTTCTTGAGTTTGGGTGACAATTCATTTAATAAAAGTCCATTCCCTCTTGAAATCTTGAATCTTGATAAGTTATATTGGGTTTACCTTACCAATAGTAGTATTGAAGGCCAAATTCCAGAAGGTATTGGAAATCTTACTCTGCTTGAAAATCTTGAGCTTTCATATAATGACTTGTCAGGCAAAATCCCAGATGGAATAATCAAACTTACTAAGCTTAAACAGCTTGAGATTTATTCTAATGGACTAACAGGAAAGTTCCCTGTGGGGTTTGGGAATCTCAGTAGCCTTGTCAATTTTGATGCTTCAAGTAACAATCTTCAAGGTGATCTTTCAGAGCTCAAGTCTTTATCTCTTCTTGAATCTTTGCAGCTGTTTGAAAACCATTTTTCTGGTGAGATTCCTGTTGAATTTGGAAATTTCAAGTTTACAGAACTGTCATTGTACAGAAACATGTTTTCTGGTTCTCTTCCTCAAAATATTGGATCATGGGCAGAACTTCAGTACATTGATGTTTCTGAGAATATGTTCACTGGTTCAATACCACCTGATATGTGCAAGAAAGGGAGTATGACTGATCTTTTGCTTCTCCAGAACAACTTCACTGGTGGGATACCTTCGAACTATGCTACTTGTTTGTCGTTGCAGCGTTTACGGGTTAGCAACAATTCACTTTCAGGAGTAGTCCCTAGTGGAATTTGGAGTTTGCCAGATTTGGAAATCATTGATCTCACATTGAATCTATTTGAAGGCCCAGTGACATCCAATATTGGTGAAGCAAAGTCTTTAGCACAGTTGTTCCTAGCCTACAACCGGTTCAATGGTCAATTACCACAGACAATATCAGAAGTTTCTTCATTAGTAGCCATTAATCTGAGCGCGAATCAGCTCTCTGGTGATATTCCAGCAGCAATAGGTGAACTTAAGAAGCTTAATACTCTTCATTTAGAGTATAATTTGTTTTCTGGAAGTCTTCCAGATTCTATTGGATCATGTGTTTCTCTCTGTGAAATTAATCTTGCTGGTAATTCACTTTCTGGTGCAATTCCCGAAAGTCTTGGCTCTTTGCGTAGCTTGAACTCTCTCAATCTATCTGATAACACGCTCTCAGGTCAAATTCCAGCGACCCTTTCGAGTTTAAGATTAAGCCTTCTTGATTTGTCAAACAATAGGTTGAGTGGTAGCATACCGGATTCTTTATCAATAAAAGCTTTTAGTAATAGCTTTTTGGGAAATCCAGATCTTTGTAGTGAGAATTTTGGTAGCCTCAGGCCATGTTCATCAGATCCTCACACATCTAGAGACCACAGGACAGTCATGTTGTGCTTGATAGCTGGGGTAGTTGTTCTTGTTCTGTCACTTACGTGTTTCGTGTATGTGAAGTTTAAGCACAATAATCAGAACACTCCCGTAAAGAGACTTGATTCTTGGGATATCAAACAGTTTCATGTATTGAGCTTTAGTGAAGATCAAGTCTTGAAGGCATTGAAGCAAGAAAATTTGATTGGAAGAGGTGGTTCAGGGAATGTGTACAGATTAGTCTTGAATTGTGGAAAACAGTTGGCTGTGAAACACATTGTTAAATCCGATTCTGGTGACCAAAAAAGTTACCGGGACAGCTCAGCCATACTGGTGAAGGAGAATAGAAGATCAAAGGAGTATGATGCTGAGGTGACCACGTTAAGTTCCATTAGGCATGTCAATGTTGTCAAATTGTACTGTAGCATCACAAGTGAGGACTCAAATATGCTGGTTTATGAATACTTAACTAATGGAAGCTTATGGGACCGATTGCACACGTCTCAGAAAGTCAAGATGGATTGGTTGGTGAGATATGACATTGCATTAGGTGCTGCTCAAGGGCTTGAGTATCTGCATCACGGATATGACAGCCCCGTGATGCACCGGGATGTCAAGTCTAGCAACATCTTGTTGGATGAACAGATGAAGCCAAAAATTGCTGATTTTGGATTAGCCAAAGTTTTGCATGTTAATGGTACTAAGGACTCTTCTCAGGTTGTAGCTGGGACGCATGGCTACATTGCTCCTGGTAAATATTTTTCTCTTCCCTTTTCCATTTCATAGTCACTTAAAAATTATTTGGCAAATTCTAAATTTTATTCTTTCTTCATTGTTTGTCACAACAGAGTATGCTTACACAACCAAGGTGACCGAGAAGAGTGATGTTTATAGCTTCGGGGTGGTGCTAATGGAATTGGTGACTGGAAAGAAGCCAGTGGATGCAGAGTTCGGGGAGAACAGTGACATAGTCCAATGGGTTTGTAGCAAGATAAGAAACAACACTAGCATGATCGATTTGGTGGATTCAAGCATTTTCGAGGGGTTCAAAGAAGATGCTGTTGAGGTTCTTAAAATTGCAGTTCATTGCACATCAAGGACGCCAGCATTAAGACCTTCCATGAGGATGGTAGTTCATATGCTAGAAGAAGCCGAGCCTTGCAAGCTGACAAATGTAGTCGTAAATTCACCGAATGAGGATGGCAGGAAACAAGGACTTGCTAACTAATGGCAAGTCATGATGTTGAACTATCTCGTTGAGAGCAATGGAATTTTGCAGAATATTGAACTATGTTGTAAGTTCACAAAGAATGCTTAGTGTACATATGACACTTTTGAAGAAATAGTGTATTTCACAGCTGCATTAATAAATCCAACTAGCAAGTAATGTATTTCACATCCAAATTACTTATACAAACTGAGCAGTTGATCTACCCTACAACCAATTTCATTTTTCT

>PGSC0003DMG400005254 LRR-RLKs

AAAGATTTAAAAGGTAATTGGCACCTTGCTTTTTAGTGGCCTACAATGGAAAGAATGACTTTATTGCCAAAAGGACTTTAGGCAAAGTATGCAATAAATCCTCTTCCCACTCACATTGAAAATCTGTCTTCAGTAGTTCAATTAGTATCTGTCATAATTCTTGCCTACTGGATTGATCCATTTTCAATATGAAGAGGTAGTAAATCAAATGAAGTTTCCCTTTCATTGAACAACACAACTCTCAGATCCCTTTCATGCATATCATATAAAAGTAAGTTAACTTGTGTGCCATATATTTTAGAACTAAAAGAGTCAGTAAGGCAAAATCTTGGCTTTGTCGCAATAACCAAATGAATTTTTTCTATTCTGAGATAGAGTTCTTCCTTTTTCGTTACTTCTTTAGCTGATACTATTTCTGGATTGTATAGTTTTCTTGCATTGTGTTAAAGCTTTTTCATTGTATTTGTTAATGGTTTCTTATAGCAATCTGATTTTATCTCTCTTTCACAGTGGCTAATGAGCTCTGATTTTGTCTAATGCTCTTCAAAAATAGTCAAGGAGAGAGGGAAGAAGTGAAGGACTGCTGTAACTAGGATCTTGCTTTTGCATTTTTGTCAGCCAGATCAGTGAAAACATAATGAAGCTTCAGCCTTTACTGGCAGCTATTGCTTTTCTTATTCTGCTGCTATCAGTATTTCTTCATGTTATAGCTGATCTTGATTCCGATAGACAGGCGCTACTCGACTTTGCAGAATCTGTCCCTCATATACGAAAACTCAACTGGAATTTGGCTCTCCCTATTTGCAAATCTTGGGCTGGTATAGCTTGTAACAAAGATGGAACGAGAGTCATTGCAATCCATCTCCCTGCTGTAGGACTTTTTGGTCCTATCCCAGCTAATAGTATAGGCAAGCTAGATGCTCTTAAAGTCCTTAGTCTAAGAGCCAATTATCTTAATGGAAGTGTTCCTTCTGATCTCCTTTCGATACCGTCTCTCCAATCTGTCTACCTTCAGCATAATAGCTTCTCTGGTGATATTCCTGTTTCTTTATCTCCTCGTCTTGGTGTACTAGATCTCTCCTTTAACTCTTTTACTGGAGAAATTCCCTCCACAATCAAGAACTTGACGCGATTATCTGTGTTGAACCTTCAATTCAATTCATTGACAGGAGAAATCCCCAGCCTTGACACCCTCAGGTTAACTAACCTGAATTTGAGCTACAATATGCTAAATGGCTCCGTTCCAGACCCCCTCCAGAAGTTTCCACTTACATCATTCGCAGGAAACTCTCATTTATGTGGTACACCTCTAAATAGCTGTTCCTCGACCCCGTCACCTTCTCCAGCAGCTGATGGCTCTGCAATCCCTGAAAAACAGAAGGCTGTCCACTCTAAGAAACTAAGTACCGGAATCATCATTGCCATTGTGGTAGTGGTTTCTTTAGTAATGTTTCTCCTAGTTTTAGTAATCTCCTTCTGCTGCTTGAAAAAGAAAGTGAGTCATAGCACTAGCATAATTAAAGAGAAGGTGGCTAATGGGGGAAGAAGTGAAAAACCCGAGGACTTTGGAAGTGGTGTGCCAGATGCTGAGAAAAACAAGTTGGTGTTTTTCGAAGGTTGCTCTTACAGCTTCAATCTGGAGGATCTTCTTAGAGCTTCTGCTGAGGTTCTTGGAAAAGGGAGTTATGGTACAGCTTACAAAGCTGTCTTGGATGAGGCAACAATTGTAGTGGTGAAACGGCTGAGAGAAGTAGGGGTTGCCAAAAAGGAGTTTGAACAGCATATGGAGATAGTTGGAAGGGCTGGGAGGCACCCTAATATCGTTCCACTCCGCGCATATTATTATTCCAAGGATGAGAAACTTCTTGTTAATGAATACATGCCTGCTGGTAGCTTGTCAGCGGCCTTGCACGGTATAAATTGCTAACCTCTATCTAACGTATGTTGCTCAGACTTTCCAAAATGTTGCTGCACCCGTGTTGGATCCTCCAAACATGTACCTGTCCACATTTTTGAAGAGTCCGAGCAGCACAACCTCTAAACTTTTTTCTTGCACATTAATTGCTTTGATCATCTGTAATGATTCATATTTTTACGCAATCTTGATAATGAATGACAGGTAATCGAGGTATTGGAAGAACTCCACTAGATTGGGATTCAAGATTGAAGATATCTCAGGGAGCAGCAAAAGGAATTGCCCACATCCACACTGAAGGTGGTGTGAAATTTACTCATGGAAACATCAAGTCCTCCAATGTACTCCTCACAAGAGACCTAGATGGATGCATTTCTGATTTTGGCTTAACACCTCTGATGAACTACATATCATACAAATATAGATGTGCTGGTTACCGCGCACCAGAAGTTATTGAAACACGGAAAGGGACACAAAAATCCGATGTTTATAGCTTTGGAGTGCTGCTCCTTGAGATGCTAACAGGTAAATCCCCTCTCCCATTATCAGGGCAGGATGAGGTAGTTGATTTGCCTAGGTGGGTCCGGTCCGTTGTAAGAGAGGAATGGACAGCTGAAGTTTTCGACGTAGAGCTATTGAAGTACCAGAATATTGAAGAAGAGATGGTGCAGATGCTTCAGATTGGACTAGCTTGTGTAGCAAAGGTGCCTGATATGAGACCTGCTATGGGAGAAGTGGTTAGAATGATTGAAGAAATTCGACAACCAGAAGGTGAAACGAGACCTTCCTCGGAGGACAGCAGGTCCAAGGACTCTAATGCACAAACACCTGAATGAAAAATGCAGTTTTAGTCATTCAGCTTTCAATTCTTGATCATTTTAAGAATGCAATTTATCTCCATTGCCAACTTTTGCATGTTCTTGGACATACAAGTAGTGTTATTTTTATTTTATTTTTTTATCTTGATGTTATATCCTATTCTAGAAACTTATCAGCTCAACATAAATGAGAACCTTTGTTTTTCTCTATTTTTAAATCTTTG

>PGSC0003DMG400005485 LRR-RLKs

AATAAAGATAAAAAATAAACCTAAAACTAACCTACACTACCCCATTAACCCCTTATACTACTACCCACTCCACTCACAACCCCACAGTCCCACATTAATCCATCACTCCCCTCAACCACACACACGTACCACACACTACTAAACATCAACAACACACCCCTTCAACCCATATACACAGAGTCACAAAAAACGAAGAAGGACGACACTCAAGAAAACAAAAAGAAAAACTTTCCATTTTATTTCATTATCCTCTTAAAAATTCACCAACACATTCACATTCCTCTTCCTTTCTCATTATTATTTTCATATAAACACCATACATATACCACCACACACACCTACAAAATTTGTGGGTATTCTAATTGAGCTTCAGTTAAGATTTTGCGGTATTCAACTTTACCTTATCTTTCTATTTATTTATTTATGGAATTTGATGTAATTTTATGACAAGTTCTTTTTATATTTTGTTTCATTCATGAATTTGATTGGGATGATTAATTTGTTAGTTGTTATATCTCTATTTCATGAATAAGGTAGCTTTAATNNNNNNNNNNNNNNNNNNNNNNNNNNNNNNNNNNNNNNNNNNNNNNNNNNNNNNNNNNNNNNNNNNNNNNNNNNNNNNNNNNNNNNNNNNNNNNNNNNNNNNNNNNNNNNNNNNNNNNNNNNNNNNNNNNNNNNNNNNNNNNNNNNNNNNNNNNNNNNNNNNNNNNNNNNNNNNNNNNNNNNNNNNNNNNNNNNNNNNNNNNNNNNNNNNNNNNNNNNNNNNNNNNNNNNNNNNNNNNNNNNNNNNNNNNNNNNNNNNNNNNNNNNNNNNNNNNNNNNNNNNNNNAAATATATGACCCGTTTCAAAGGTCGGTTTCAATTAAGAATGTGGTTAAACATCTTTCTTTGGAACACTTAAAAAAAACTCAAGGATGCAGTGACGCACCTTGTCCAAATTGTTTTAATAATTATTTTTTGTTAATTCAAACCAAGAGGTGTAAATTGAAATTATTAAGTGTTATGAAATCGAGAGGGGAAGACAGTTATGTCTCTAAATGTCAAGATCGAGAATGCAGTTATGCATCCGGTTTAAGACTAATAAAAATTCAACAATTAATAAAATAAGTGAACCATGGCCTTAGACACAAATTATCCAAAAATAGTAAAATAATGTATAAGTCAATAAAAGCGACCGTGCTAGAACCACGGGACTCGAGGGGTGCCTAATACCTTCCCCTCGGTCAACAGAATTCCTTACCCGAATTTCTGGTTCGCAGACTAATATAAAGAGTCATTTCCTTTTGATAAGGGATTAAACAAAAAGGTGACTTGGAACACCATAACTCAATTCCAAGTGGCGACTCTGAAAAAATAAAATAATCCCTAAATCAATAACGTCACTTAAATTGGAAAAAACCCTTCCGCCGCCTTGCGCAAAAAGGGGTGTGACACTAACAACGATAAATCTTAGGAATAATAGCTTCCATGGTGAAGTCCCTATGGAAATTGGCAACCTCTTGCAACTGCAACATCTTAATCTCACTTGGAATTCATTTACTGGAACCATTCCTGCTAATCTGAGTTACTGTAAAGAACTTAGATCACTTGCTCTAGAATATAATAGTCTGGTTGGTAAAGTTTTACCTGATCAGTTCAGTTCACTTTCAAAGTTGAATTATTTAGGCCTTGGAAGCAACAATCTAACAGGAGGTATCCCATCTTGGATAGGAAATTTTTCGACTCTTCGAGGTCTTTCCCTTGCAATCAACAATCTACAAGGACCAATACCTCGAGATATTGGACGTTTGTCAAACTTGCAAATATTCCAAGTTTATGGAAATCAGTTGAATGGTACAATTCCTCAATCTCTTTTCAATATATCCTCTGTTTACTATTTCTCTGTTACTCAAAACTTGTTGTATGGAGAGCTTCCATCAGATATAGGCCTTACTCTTCCGAATCTTGTGGTGTTTGCTGGTGCTGTGAACGATTTCACTGGACCAATTCCTGTTTCATTATCAAATGCAACAAAACTTGGTGTACTTGAATTGTCTCAAAATAAGCTCACCGGAAATGTCCCTACGAGTTTAGGACAATTGCAAGGGTTGTATAGGATGAACTTTGAGATCAATAGTCTTGGAAGAAACACAAGTGGGGATTTGAGGTTTCTTGATTTCCTAGTTAACTGCACAAGTTTGCAAGTTCTTAGCTTTGAGGACAACATTTTGGGAGGTGAATTGCCTAAAACAATCGGTAACCTTTCCACGAGGCTAGAAATATTTGCTCTTGGTTATAATATGATAGTTGGTTCTCTTCCTACAGGACTAGAAAATCTTGTTAATTTGACCCTTCTATCACTGGATAATAACTACTTGAGAGGCAGTGTTCCTGAGTCTTTAGGTAAGCTTCGACGCCTGCAAGGACTGCTGTTGAATGGAAACAAGTTATCAGGAAGGATTCCATCTTCTATTGGTAACTTAACATCTTTGTCTACTTTACACATTGAGGACAATGAATTAGAAGGAAACATACCTCCGGAGCTCGGGCAGTGCATCCGCTTATCAAGGCTAAACCTTACAGGAAATAATCTTGTTGGTTCCATACCAAAGGAGCTTGCAGGTCTTTCTTCTCTTTCGATTTCCTTAGCCTTAGCGAACAATTCGTTGACCGGTTCCTTGCCAGCTGAATTTGGAAAGTTGATAAATCTCAAGGAAATGGATATTTCACATAACAAATTATCAGGTGAAATTCCAAGTACTCTAAGTAGCTGTGTCAGCTTAGAGCGCTTCATAGCGAATAACAATCTGTTTCGAGGAGAAATTCCTGAATCCTTGCAAGGTTTAAGAGGCTTAGAAGAAATTGATTTGTCACACAATAACATCTCAGGAGGAATACCTGAGTTTCTTGGGAAACTTCCATATCTTAGGAAGCTTGATCTTTCATTCAACGAACTTGAAGGCGAAGTACCAACTGAAGGGATCTTTGCAAATGAAACCGCCGTCTCAATCTCAAGGAATGATGAACTATGTGGAGGTCCTCCAAACTACAATTTTCCTACATGCCCTAAACAAAAAGATGCATCATCAAAGAAACACATTAGTTCAAGGATAAAAGTAGCAATTATTATTTCGGTTGCATTTCTATTTTTACTTTTGTGTTCTTTTGCAGCTTGCTATATAGTAATTAGGAAGTCGAGAAAAAGAGATCTTACTGGACGGTCCTCAAGAGAAAGGCAGTCTGATCATTTTGATGATGAAGAACCAACTTTATTTAAGGATCCAATTTTGACAGCAAAGATAACTTACCAAGATATATTCAAGTCAACCAATGGATTTTCTGAGGATAATCTAGTCGGTACAGGAAGTTTTGGTTCTGTTTACAGAGGAAAGTTTCAGGTTTTTGATAAAGTAATGGCAGTGAAAGTATTGAACCTTCAACAAAGAGGTGCTTTGAAAAGCTTTTCGGATGAATGTAGAGCTTTGAAAAGCATAAGGCATCGAAATCTACTTAAGATAATAGCTGTTTGTTCAAGCATCGATTACCAAGGTAATGATTTCAAATGCATAGTCTTTGAGTTCATGGAAAATGGAAGCTTAGATGACTGGCTGCACTCGAAAGGTGATGAGAAATACCTCAATATCATCCAAAGACTAAACATCGCAATAGATGCTGCTTCAGCACTTGATTATCTCCACAACAATTGCCAAGTATCAATTGTTCACTGTGATTTAAAACCGAGCAACATACTCCTTGATGAAGAGATGACTGCCCATGTTGGTGACTTTGGATTGGCAAAATTTCTCTTCAAATCATCATGGAACAAACATACATCTATTGCACTAAAGGGTTCTATTGGTTATATCCCACCAGGTATGTCATTAATCCATGTTGATTTCTCTTGCATATCTATTTCCCTTATCAGGGGTGCTAACATGTAAGATGTTTGTGTACCTCAATCTCATATACTCCCTCCGTTTCAATTTGTTTGTGTGGTTTTGACTTGACACAAAGTTCAAGAAAAATAAAGAATACTTTCGAATCTTGTCGTCTTAAACTAAAGATACATAGAATGTACGAACATGCCCTTTAATCTATTGAAATTTAAAGAGTTGTCAAAAAGCAAAGAGACACTCTTTTATAAACGGACTATAAAGGAAAGTAAGACAAATGAATTGAAACGAAGGGAGCTTATCTTTCTGATGCTTAAGGTGTCGAATGAACAAATCAATAACTAAGCATTCATATGAGGTTGTCATTTTCATGTGAGTGAACATATGTTCAGTGTCCCAACATAATTCATCATTATCTTGCAGAATATGGATCAGGTGTAAATGTGTCCACTCTAGGAGATGTTTATAGCTTTGGTATCATGTTGCTAGAGTTGTTTACCGGTAGAAGGCCAACCGATGAGATCTTCAAAGATGGCCTGAATATTCACCAATATGTCAAATCCCATCTGCCTAGACGTGTCACGGAGATTGCTGATGCATCATTGCTTTTAGCATATGAAGAACACAATATATATGAGGACAATGCAAGTGATTTGGAAGAAAAGGCAATACTTCAAGATGATGAGTATATATCCAAGCTAAACACCAGCACTATTATACAAGAATGCTTGATCTCAATCATGAAAATAGGACTTTTATGTAGTAGCTCATCGCCAAGAGATCGAATGCCAATAAGCATAGCTTTGAAAGAGATTCACACAATCAAGAACTTATTTCTAGAATCTAAGAGGATAAATAATAGTATAGATAGATCAGATGGTAATAAGCTAATCAAATTATCATCTTCAGAATTTTATTGATGAAGGCAAATATACATAGAATTACAACTTTGAAATAGATATTATTTACAATTAGTACTATAGATAGCCTCACAAAATGTGAATTGTAACCAAGAAATTAAGGGATATGATTGAGTTTTTTTTTTTTTTTTGAGTAATGGATGGAGTTCCTTTTTCCATCCTATCCCATTCACC

>PGSC0003DMG400005896 LRR-RLKs

GAGAAATCAAAGGACAGAAGGAGACAGAGAAAAAAAGGTGTCATGAATTTGTACTTGTCTTTAGGGTTTGATCCACAACTCTTTGCTAAACTCATCTCTCTTTCTTACAAAATTCACTCACATTATTAAAAAAAGGTACCATTTTTCCAACAATTTATAGTTATTATGTACTTGCATCTTTATATGTTACTGTTATGTACTTTGTTCATTTGGGGTTCACAATTCTTGAGAAAAAAGTACTATGTAATTTTATGATAATCTTGAATTTTCATCATGTGTGTATATAAGTATTGTGATTTCAGAGGTGTTGTTATATACTTTTGTACATTAGGTGTATTTCTTGTAAATGTTGTTTCTGGTGTAACAAATTTGAATGATTTCAAGATTTTGAGTGATTTTAGAAATGGATTAGAGAATCCTGAACTATTGAAATGGCCTTCCAAGGGGAATGATCCTTGTGGTCCTCCTGCATGGCCTCATGTGTTCTGTTCTAGTGACAGAGTTACCCAAATTCAGGTTCAGAGTTTGGGTTTAAAGGGTCCTTTACCTCAGAACTTGAATCAACTTGATAAGCTTCAAAATCTTGGCCTTCAGGGGAATGGTTTTAGTGGGAAATTACCTACTTTCAATGGATTATCTGATTTGAAATATGCATACTTGGACAACAATGAATTCGATACGATTCCTGGTGATTTCTTTAATGGTCTTAGCAGTGTTCAAGTTTTGGCTTTGGATTATAATCCGTTTAATGAGAGTGGATGGTCTATTCCGATTGAGTTGCAAGATTCAGCTCAGTTGACGAATTTTTCTTGTGTGCAATGCAATATAGTTGGACCTGTGCCTGATTTTTTTGGGAAATTGCCTTCACTAAGTGCGTTGAAGCTGTCGTATAATCGTTTGACCGGTGAGATACCAGACAGTTTTCGTGATTCAATGCTTCGGATTTTGTGGTTGAATAATCAAGATAGTCCTGGAATGACGGGTCCAATCGACTTAATTGGTACTATGGATCAGCTTATGTTGTTATGGCTTCAAGGCAACAGTTTTAGTGGACCAATTCCTGATACAATTGATGATTTAAATGACTTGAAGGAGCTGAATTTTAACGGAAATCAACTCGTTGGTCTGATTCCGCAAGGTTTGGCAAATCTAGACTTGAGTGTGTTAGACTTGAACAATAATAAGCTCATGGGTCCTATCCCAAAGTTTAGAGCTGCGAATGCTACATATTCCTCAAATTCTTTCTGTCAGCCAACTCCTGGTGTTCCTTGTGTTCCTCAAGTTAATGCACTTTTAGATCTTCTTTGCGGTTGGAATTATCCAGCCAACCTTGCTCCTGAATGGAGTGGCAATGATCCATGTACAGGTCCTTGGTTGGGAATCAGTTGCAATCCTAAAGGCCAGATTACGATTGTTAATCTACAGAACAAGAACCTTACGGGTACGCTTAGCCCTTCACTTGCAAACTTGGATTCACTGCTTGAAGTTCACTTAAAAAGAAATAGTTTGCATGGTCGAGTTCCTGCAAACTTAACAGAGCTCAGATCCTTGAGATTGTTGGACTTAAGTGGGAACAATTTTGATCCACCATTACCGAAATTCCGGGATAGTGTGAAAGTCATTACAGACGGAAATGCTCATCTTGTTGCTAATGTAACTGCGGCAGCTCCTCCTCTGAGTATTAGCCCATTTCCTCCATTGAGCCACAGTCCTAAATCAAGCAAGGAGTTACCATCGAAAAGCCCATTTCCCGGTGATAATCAGCCAACATTACCAGATACACCGCCCTCCCCAGAGAAAAGTTCAGGTTCAGACTCAAAGTCTCCCAGTGGAACCAAAGGGCAGACCACATCAGAGAACCATGATAAGACCATGATCATTGTAGTTGTGAGTGCAGCTTCTTTCGTTTTCACTCTTCTTGCCGTTGTTTTGTATTTCAAGAGCAAGAGAAAACGAGAGAAGGATTCAGGTACTATTGTTATCCACCCCAAAGAGCCATTCGATCAAGATAACATTGTTAAGATCACTATCTTAGAAGATCCCATGATGTACAGCTTACAGAGTGGTACTACAACAACTACTACAACTAGTGGTGGAACAAAAGGTAGTCGAGTGATTGAGATTGGAAACCTTGTCATTTCAACCCAGGACCTTCGGAGGGTAACCGATAATTTTGCACCAGAGAATGAGCTTGGTCGTGGTGGCTTTGGAGTTGTTTACAAAGGCGTAATTGAAGATGGAATTCAAATTGCAGTGAAGAGAATGGAATCTGCCATTATCAACAGCAAGGCATTGGATGAATTTCAAGCAGAAATAGCTGTGCTTTCTAAGGTTCGACATCGCCATTTAGTTTCACTTTTGGGTTATTCCATTGAAGGAAATGAGAGGCTTTTGGTATATGAATATATGTCAAAGGGCGCTTTGAGTAGACATCTTTTTCGCTGGAAGATCCTGAACTTAGAGCCTTTATCGTGGACAAAGAGACTCAATATTGCTCTAGATGTCGCTCGAGGGATGGAGTACCTTCATAATCTGGCACACCAGAGCTTCATACATCGCGATCTCAAATCATCAAACATTCTTCTAGATGATGATTTTAGAGCAAAAGTTTCTGATTTTGGATTAGTGAAACTCGCTCCTGACAAAGAAAGGTCCGTTGCAACAAGGCTTGCAGGAACATTTGGATACCTTGCTCCCGAGTATGCTGGTATGTGTGCATTTTTTTCCATTGGTATCTCTTTCTTAGACACTCGTTTCGACTGTTCTGGTAAATATTATCATTTAAGCATCAATTGTATTTTTTACGCTCGGGTAAGCTGTTACATGAACACATCTCTACCTGAACGAAACCAATACTCTTTTAGTCCCTCTATTTTAAGAAATGCATCAGTTCTATAATGAGAATCCAGACTGCTAAATTGCGCTTCTGACCTTCCAATCCCTTCCTTCTCCAGCCAAAATAGAACAGAGTAACCAATGGAGCATTTACGTTTTTCCTTTTTGTTGTCGGTGTGCGCTAGCTACTAAATTGATATACCTCGGACATTGATGTATCAATCTCGCATCATTCAGTGATATTCTCATTCTGTTCACTATCTTTGTTAAACAATTCAAGTAGAAGGTTTCACATCCTCTCTCCATATGATCATTCAGAGCATAGTTATCATCTTGACTCCACAAAAAGAAACAGAAACTCTATCCCTTTAAAGATACTTTTCTCTTGAGGCATGGTACTGAAATATGTTTGCGTAAAGCTGTTCAACAAGCATGTCATACCTGTTAATTTTGGTGTAGAAACTATTGCTTACTTTATCTTTATGTTACCCGTATATTTTAGTTCTCAGTCTTGTTGAACGAATTACCTAGTCACCTTTCATATATCCCAACTCACATTTTACCTCTCCAGTGACCGGAAAAGTCACAACTAAAATAGATGTCTTCAGCTTTGGGGTGGTGTTGATGGAACTTGTGACTGGACTAACTGCCCTTGATGAGCATCGGTCCGAGGAAACTCGATATCTAGTTGAGTGGTTCTGGCAGATAAAATCGAATAAAGAAAACCTCCTTGCTTCCGTTGATCCAGCTCTTGACGTGAAGGAAGATATCCACAAGAGCATTTGCACTATGGCTGAATTAGCTGGACATTGCACTGCGAGAGACCCCAATCACAGGCCTGACATGTCACACGTTGTGAATGTGCTTGGCCAACTAGTTGAGTCTTGGAAACCCGTAGAAGAGTCGGATGACTATTCAGGAATTGACTACAGTCTACCTCTTCCAGAAATGTTGAAGGACTGGCAGGATGAAGACACTGGAGACTTCACCAGTACTAGTCAAGATAGCAGAGGAAGTATTCCTGCTAAGCCAAATGGATTTGCGGATTCTTTTACATCCAACGATGCTCGATAGGTGAAGTTGGTTTTGGCTACCAAAGTCCATTGCTTCCTTACTATTAGTTTTGCTAGCATTTTCGACATGAATTTTGCTTTTGGCTCGTGAAAATAATCTATGGAGGCGTTGCGTCCAGTCTATATATGGTGAATATACATATACCGGGTTGTCATGAATGATTGATCAAGAGATCAAATTGGTGTCGTCCAGATACAGAGCAAAAAACAGAAAAAACTTTTCAATTTAATGTTGCTTACACAGTTATTTGTTTTGAATGTTGCTTGTAAAAGTTATATTTATAATAGGTTGGTCCAATTTGCTGTATAAGTTATTTACATGACACTCTTGAGAATCTTATGCATGAATTTGAGAGTGATGCTCAAATTAATAATGCAAAAAAAG

>PGSC0003DMG400006065 LRR-RLKs

ATGGAGAAGCACATTTTCTTATTGATTCTTCTCTTTCTAGTTCAATTTTCTGTATCATTTGCTAACTCAAATGTGACAGACAAAGAAGCTTTACTAGCTTTCCAAAATCTTATTACAAGTCCTACTCATTTTTTGGCCAACAATTGGACCAACAATACTTCTTTTTGCTCTTGGTTTGGTGTCAGTTGCAGTCCAAAAAGGCAAAGGGTTGTAGCCTTGGATCTTTCTAGTTTGAATCTTCGAGGTACAATTTCCCCGTCTTTGGCTAATTTGTCCTTTCTCAGAGAGCTTAATCTTGGGAACAACAACTTTCATGGCGCCATCCCTTATGGCATTGGCCACTTGCCTCGCTTGCGAGTGATTGATATTCAAAACAATCAGCTCCAAGGAAGTATTCCAACAAGCCTATTTCAACACCAGAAAGTTCAAAAGATTTCATTGGCTTTCAATAAACGCAGTGGTGAAATGTGGAATGGTTCATGGTATGTACCAGAACTCAAGGTCTTGTATCTCACCAACAATAGTCTCACAGGCATAATCCCGCCTTCTATTGGAAATTCCACAAAGTTGAGGAACTTCAGTTTGTACGGTAATAGAATCAATGGCAACATTCCGAAGGAGATTGGTAATCTTAGCCAACTTGCAGAGTTGTCCTTGGGAGATAATCAATTAACAGGTTCCATTCCTGCAACACTGTTTAATATCTCGTCGCTACTTATTGCATCTCTGAAAATCAATAACCTTTCTGGTCCTCTCTTGCTTGATGAAGGGAATATTGTGTCAAATCTGAAGCATCTAAGTATATCTAAGAACCAAATTTCTGGTATCATTCCTTCCAATATTTGCCAACTCAAAGAGCTCAAAGTGTTGTCCATATCTTTCAACAAGATAACTGGAGATATACCCAAAAATATTGGTTGTTTAGCTAAGCTCGAGGTGTTCTATATCGGAGGTAACGCACTAAGTGGGACTTTTCCTGCTTCATTGGGAAATATATCTACTCTTGAATATCTGGATTGTCCAAACAATTGCATAGGGGGGCAAGTTCCTCCAGAATTAGAGAAGCTGTCAAATTTGAGGCAATTAAACCTTGGCCAGAATTATAATATTATTGGTCAAATTCCGAAGGCTATTTTTAACATGTCCTCTTTGGAAATGATTGCTTTCAATCTCAACAACCTCTCGGGTAGAATTCCAGCCACTACAGGTCTTCATCTTCCGAACCTTGAAAGACTTTACTTGTCTAAAAATCATCTCGAAGGGGAAATTCCTCCGCACATCACAAATGCTTCCAAACTTAAGACATTGGAGCTAAATAGTAACTTTTTCAGTGGCAGTATTCCAACTAATTTGGGAAATCTTCGTGAGCTGCAACTGCTGCACCTAGATCATAATCAACTTACCAATGAACCAAGAGAGCATGAGTTGCTATTCTTCAGTTCTTTGGCGAACTGTAGGATGTTGCGAGATCTAGATATGAGTACCAATATGTTGAATGGTACTCTGCTCAATACTATCGGAAATCTTTCATCTACTATTGAATTCTTTGCAATAATAGATGCATACATCAATGGCCCCATCCCCACAGGTATAGGCAACATGAGCGGTCTAATATCTCTAAGCCTTCATGGAAACAACTTGGTGGGAGGTTTCCCTTCTGAGATTGGTAAGCTTAAACAACTACAAGGACTATATCTAAATAACAATAAATTGCAGGGACATATTCCAGAGGTGGTATGTCGTTTATCTAATTTGGTTCAATTAGCTCTAGATGATAATGAGCTCTTTGGGTTAATTCCAGCATGTATAGGAAATTTTAGCATGCTACAATATCTTTCATTGAGTTATAACAATTTTGCATCAAAGCTTCCTTTGAGTCTTTGGAAGATGAGTGGCCTTCTCCACCTAGACGTGTCACAAAATTCTATAGAGGGAGAAGTTCCACTGGATATTGGAGAACTGAAAGCAATTGTAGATCTATATCTTTCTAGTAACCGCCTTTCGGGCATGATACCAAACAGTTTGGGGGAACTCCAAACCCTGGAGTCTCTTGACCTGTCAAACAATTCATTTTCAGGCCAAATTCCATTATCCTTTGCCAACTTGATAAGCTTAGAATTCATGGATTTGTCTTTAAATGCCTTGGCAGGTACTATTCCTAAGTCTTTGGAAAAACTCTTGTACCTTAAAGCCATCAATGTTTCATTTAATGAATTAGAAGGTGTAATACCTAGTGGTGGTGTGTTTGCAAATTCCACTTTGCAATCATTTCTTGGTAACAAAGGCCTATGTGGAATGCACATATTGGATGTTCCTGCTTGTGCTGTCACTAATCTTGGACATCAATCAAAGTTTAAGAAGCTTGTGCTAAAAATTGCTATTCCAGTGGTTACTTCATCCTTTCTAATATCCTTGTTTGCTTCAATTTGGATGATGAAACGACAAAAGAAGGGAAATTCCAAAGACGTGGAAAAGGTACCGGAGGTCAAGACACATCAATTAGTATCTTATCACGAGATTCAAAGAGCTACAAATTATTTTGATGGATCAAACTTAATTGCTGTGGGAGGTTCTGGCTCTGTGTACAAAGGTACATTATCTAGCGGAGTTGTGGTTGCAATAAAGGTTCTAGATTTGCAAAATGAGGAAGTATGCAAGAGGTTTGATACTGAATGTGAAGTGATGAGAAATGTTAGGCACAGAAATCTTGTATCGGTGATTACTACTTGCTCTAGCGAATACATTAGAGCTTTTGTCCTGCAGTATATGCCAAACGGGAGTCTTGAGAATTGGTTGTACAAAGAAGATTGCCACTTGAACCTTCTTCAAAGGGTTTCCATAATGTTTGATGTGGCTGTGGCTGTTGAATATATGCATCACAGACATCACACTCATATTGTTCATTGCGATCTAAAGCCAGCCAACGTTCTTTTGGATGAAGAAATGGTGGCACATGTTGGTGATTTTGGAATTTCCAAAATCTTAGCTGTAAGCAAGTCCATGGCATCTACCGAGACATTGGGCACTCTCGGATATATTGCACCAGGTATACTAGTCTCTTTCTTTTTTACTACACAAATTATAACTAAGCCCTTCCAAAATCTAAACAAAAAGCAAGTGTTTGTTTGTTTTGTTGTTATATTGGATGAATAATTTTTTTTCATCTTTTGTTCTTTTAAGAATATGGCTTGGAAGGAATAGTGTCCACTAGTGGCGATGTCTACAGCTATGGTATCATGCTAATGGAGGTTTTGGCAACAAGAAGACCAACTGATGCAGAGATATTCAACGAAAATCTTGGCTTGAGGGAGTGGATAAGGCGAGCATTTCCAAAAACTATCATAGAAGTTGTGGATGCCAATCTTTTTCCTGAGGAAGAACAAATCACTTCTAAAAGTGAAATCTGCATAATTTCCTTGATAGAATTGGCTTTAGATTGCGCAAAGGAAAGGCCAGAGTCAAGGTTAACCATGAAGGATGTAGTCAAGAGGCTTAACAAAATCAAGAACACATTTTTGGAAATATGA

>PGSC0003DMG400006280 LRR-RLKs

CCAAAGCCAAAAATTACTATCATATTTTATCAGTATCAATGGAATCTTTCAAAATACTTGTTGCATCTTCTTCTTCTTCTTACCTTTGATGTTTTCCGAAGTATTCGAACTTCAATCACTAGTTATCTAAATGGCTACTTCTAACACAAGTCTCTTCTTTTTCGCGTATTTCTTACTCGTGTTCCTCATTACTCCATCTCAATCGCGTAACGTGTCTCTGAGACGACAGGCTAAAACTCTAGTTTCATTGAAACAAGCATTTGTAGTATCCTCTGTTCCTAGTACTCTGTCTACTTGGAACATGTCAAATTATATGTCTATATGTTGTTGGACCGGTATCACCTGTGATGATACCAAATCAGTAACTACCATTGATATATCCAATCTAAACATTTCTGGCTCTTTATCACCTGATATTCATGAGCTCACTAGACTTCGCGTCCTGAATATTTCTAACAACTTGTTAGGTGGGAACTTGAGCTGGGAGTATCGAAAGTTTAACGTACTTCAAGTGTTGGATGCTTATAACAACAATTTCACTGGTCCACTCCCCTTGGGAGTTACTCAACTTCTGCAGCTGAAGTATTTGAATTTCGGGGGTAACTACTTTTCAGGGAAAATTCCTTTGAGTTATGGTAGTTTTAATCAGCTTGAGTTCCTGTCTCTTGCTGGGAACGACTTGCACGGTCCTATACCAAGGGAGCTGGGCAACGTTACGAGCCTCAGGTGGTTACAGTTGGGTTATTATAATCAATTTGATGAGGGGATTCCGCCAGAGTTGGGAAAACTTGTTAATTTGGTTCATCTAGATCTTTCTAGCTGTAACTTGACGGGTTCGATTCCAGCAGAATTGGGCAATCTTAATATGTTGGATACTCTTTTCTTGCAAAAAAATCAACTTACTGGTGTATTTCCTCCTCAGCTAGGGAATTTGACAAGGTTAAAATCTCTTGATATATCCGTCAATGAACTCACAGGAGAGATACCAGTTGACTTGTCAGGACTCAAGGAGCTCACATTGTTGAACCTCTTTATCAACAATTTGCACGGTGAGATTCCTGGATGTATCGCGGAGCTGCCAAAGTTGGAAATGTTGAATCTTTGGAGGAATAATTTCACTGGATCGATTCCTTCTAAGCTTGGGATGAACGGTAAACTAGTTGAAATTGATCTTTCTAGTAATAGACTCACTGGCTTGATACCAAAATCTCTATGCTTTGGGAGGAATTTGAAAATTTTGATTCTGCTTGATAATTTTCTGTTTGGACCTTTACCTGATGATTTTGGGCAGTGTCGAACCTTGTCCAGAGTCAGAATGGGACAGAATTACTTGAGTGGATCAATACCAACAGGGTTTCTTTATTTGCCTGAGTTGTCACTGGTGGAACTGCAGAACAACTACATCAGTGGACAACTCTCGAACGAGAAAACCTCAGCGTCTTCTAAACTTGAAGGGCTGAACCTGTCGAACAATCGCTTGTCTGGTGCACTTCCCAGTGCTATTGGAAATTATTCAGGGCTGAAGAATCTTGTGTTAACTGGAAATGGTTTCTCAGGAGATATCCCTTCTGATATTGGCAGACTAAAAAGCATCTTAAAGCTGGACCTGAGTAGGAACAACTTCTCTGGCACAATCCCCCCTCAGATTGGTAACTGTCTTTCCTTAACTTACTTGGATTTGAGCCAAAATCAACTTTCTGGTCCTATCCCAGTTCAAATTGCTCAAATTCACATCTTAAATTACATCAATATTTCCTGGAATCACTTCAACGATAGCCTTCCCGCGGAGATTGGCTCGATGAAGAGCTTAACTTCAGCAGATTTTTCCCACAATAACTTATCTGGATCAATACCTGAAACAGGACAATATTTATATTTCAACTCAACTTCCTTCATTGGCAACCCTTATCTCTCTGGATCCGACTCGACTCCTAGCAACATTACATCCAACTCACCGTCAAAACTTGGAGACGGAAGTGACAACAGAACTAAGGTTCCAACAATATACAAGTTCATATTTGCATTTGGGCTCTTATTCTGCTCCCTCATTTTTGTTGTCTTAGCAATAATCAAGACAAGAAAGGGGAGTAAGAATTCAAATTTGTGGAAGCTGACAGCATTTCAGAAGCTTGAATTCGGAAGTGAAGACGTCTTGCAGTGCTTGAAAGACAACAACGTCATAGGGAGAGGTGGAGCAGGGATAGTGTATAAAGGAACTATGCCAAATGGTGATCATGTCGCGGTGAAGAAGTTGGGAATAAGCAAAGGCTCACATGATAACGGCCTATCTGCTGAACTTAAAACATTAGGAAAGATCAGGCATAGGTACATTGTGAGACTGCTCGCGTTTTGTTCAAACAAGGAAATCAACTTGCTAGTTTATGAGTACATGCTAAATGGAAGCTTAGGTGAAGTGCTTCATGGGAAGAACGGCGGGCAACTCCAATGGGACACTAGGCTAAAAATTGCCATAGAAGCTGCCAAGGGCCTTTCTTATTTGCACCATGATTGCTCCCCTATGATAATCCACCGCGATGTCAAGTCCAACAATATATTGTTGAACTCTGAACTTGAAGCTCATGTTGCAGATTTTGGTTTAGCCAAGTACTTCCATAATAATGGTACCTCTGAGTGCATGTCTGCAATTGCAGGGTCTTATGGTTACATTGCTCCAGGTAACTAGTTTGTTCATACATTAGTTGATTTAATTTCTTCTCGACTGGCCTATTATGCACTCGGACCAATCTACATCCTACTAATAATGGTCTTTACATGTTATCTTTTTGTATAGAATATGCATACACGCTGAAAATTGATGAGAAAAGCGATGTGTATAGCTTTGGAGTGGTGTTGTTGGAGCTTATAACAGGACGAAGGCCAGTTGGTAATTTTGGAGAAGAAGGAATGGACATTGTACAATGGGCGAAAACGGAGACAAATTGGAGCAAAGAAGAGGTGGTGAAAATCTTGGATGAGAGGCTAAAAAATGTTGCAATTGTAGAAGCTATGCAAGTATTTTTTGTAGCAATGCTTTGTGTTGAAGAGTACAGCATTGAGAGGCCTACAATGAGGGAAGTAGTCCAAATGCTTTCTCAAGCTAAACAACCAAATACTTTCCAAATCCAATGATCTAATTGTGGCTCTACTAATTCTTATAGTATATCTTTTTTTGTTTCTTGTGTGCTTGGGAAGACCCCTTTTGTTAGCTTTGCAAACAAATTGCAATTATGTTAGATCTTAGGGCATGATATCTAACTATCCTCTCAACTTGGAATAGTGTATTGGATGTGTAGAACTAGTATTAGTATCCGCGTGATGTGTGGCGAATATCAAAAGAAATTGATTACATAAAATGAATCTCTCTGTTTC

>PGSC0003DMG400006364 LRR-RLKs

ATGAATCTTGCAAATTATGGGATTAGAGGTAATCTTCACCATCTAAATTTCACTTCTTTTCCTCATCTTCAATTGATTAACTTTTCCAGTAACTCACTCAATGGTACACTTCCAAAAAACATGTTCAATCTTTCAAGATTAAGTTTCCTATATTTGAGTTACAATCAATTTGATGGAGTGATTCCACATGAAATAGGGCTATTGAAGAATCTCAAGCAGTTAGACTTGTCGATAAATCGTTTTATTGGCTTTATCCCTTCATCTATTGGTAACTTGATTAAACTTGAGTTGTTATTTTTATCAGGTAATGAACTCTATGGCCCTGTCCCTTCAAGTTTTGGTAACTTAAAATCCCTTGTGCATTTGTGTTTGCTTAGAAATAAACTCAATGGATCACTTCCAAAAGAGTTTGAAAATCTTACACATATGCAAACTTTTCAAGTAGCAGAGAACAATTTTTCTGGTCATTTGCCACAAAATGTGTGCCTTGGTGGATCACTTGCTAAATTCATAGCATATGACAATAAATTTATTGGTCATGTCCCTAAAACCTTGAAAAATTGCTCTACGCTATCAAGAGTTAGGCTTGATGGTAACCAATTGTCTAGCAATATATCAGAAGCTTTTGGTGTCTATCCAAGTCTAGTTTACATGGATTTGAGTCACAATAAACTATATGGCGAGTTATCGTCACAATGGGGCTTTTCTCATAATCTAACAAGTTTGAAGATCTCCAAGAACAACTTGTTTGGTGCCATACCAGTTGAAATAGGAAATTTGATCAAACTAGAAATGCTTTATCTCTCTTCAAATCACTTGACCGGCGAAATTCCTAAAAGTTTAGAGAGGCTAACTCTCTTGCTGGAACTTGATTTACATGAGAATAGAATTTCAGGAGAAATACCTATAGAAGTGGGGAAGTTGTCTAAACTTACAAAACTTGACTTAGGTGCGAATAATATGAGTGGAACGATTCCAGGAGAAATAGGAGATTGTAGGCAGCTTTGGTACTTGAATTTAAGCAAGAACATGCTGAACACAACTATTCCTTCTAATTTAGGTAACTTGCACTCTTTAGCCTATCTTGATCTCAGTTATAACATGCTCAGTGGTGAGATACCGTGGCACATAGGGAGTTTGAGAAGTTTGGAAAGAATGAATCTTTCGCGTAATAACTTGTCTGGTTCAATCCCGCCTAGTTTCAATGAACGCGTTAGTTTGAGGTCAATTGATATATCTTGCAATCAATTGGTTGGTCCCCTTCCCAAGATCGTGGCATTTCAAAATGCATCGCGTGAAGAATTAAGAGATAACAAGGATTTATGCAGCAATGATCATACTGGTATGAGACCTTGCTCTTCATTGAGGAAAAAAGAGAGAACAAGTAGAAAGCTTATACTGACCTTAACTCTGTCTCTAATTTTGGTTGCCTTGCTTCTACTTGTGATTACTTGTATACTTTTTCGAATGAAAAGGAAAAGAAATAGAAGTATTCAGCTTAGAGAACCATCCAGAAATTCTTTTTCAGTTAGGGGCTTTGATGGGAAAATAGCACATGAAAACATAATTGCAGCAACAGAAAACTTTGATAGCAAATATTGCATTGGGAAGGGAGGACATGGAAGTGTCTATAAGGTTGAGTTACCATGTGGTCAAGTTGTTGCAGTGAAAAAAGTTCATGCTTTAGAAGATGAAGAATCTGATGACAACTTGATCAAAAGCTTTTCAAAAGAGATACAAGCTCTAGTAAACATCCGCCATCGGAATATTGTGAAGCTTTATGGGTTCTGTTCACACGCGCGCCATTCATTCTTGGTTTATGAGTTGTTGGAAGGAGGAAACTTGTCACAAAACCTTAGCAATGAGGGTAGAGCAAGAGATTTAAATTGGCTCAAGAGGGTTGATATTGTCAAGGGGATAGCAAACGCGTTGTGTTACTTGCATCACGCGTGTTCACCTCCTATAGTTCATAGAGACATATCAAGTAACAATGTTCTCTTAGATGATGAAGGCAATCCCCATGTTTCTGATTTTGGTACTGCTAAACTCTTAAGGCCTAACTCCACCAATTGGACATCGTTTGCTGGAACTCTTGGCTATGTAGCTCCAGGTATTTAGTCTTCTTTACTTTCTAAACTTCGTGTTAAGTCAAAATCAGACAAACAAATTGAAACAGTAACTAGGAATTCATTACTTCAAAATCGTTCACTCTTTCACACAATTTTCTAATTTGTTTGTAGAGCTTGCTTACACAATGAAGGTAAATGAGAAGAGTGATGTTTACAGCTTCGGGGTGCTATCACTAGAAGTTATTCTAGGTCATCATCCAGGTGATATTATTCAAGCTATATCATCATTTTCGCCAGCATCAGGAGTGAATGGAACGCTGTTAAAGGAATTGATTGACAAACGGACACTAGCTCCTGGAAAACAAGAAGCAGAAGAATTGATCAAGATCACAAAACTAGCATTCGCGTGTCTTCATCAGAGTCCTCTAGCTCGGCCAAGTATGAAGCAAGTTTGTGCATCTTTATCAAAAGAAAATTGGCCTTCAAAAGGCTTGTTCTCCACTGTTACATTAGGCCAGTTGCTTGAACCAACACTACTAAGATGTTGA

>PGSC0003DMG400006502 LRR-RLKs

CAGCACTTAGCCATGGCCTATTATCTTGACAAGATCTTTCTTCTCCTATTTCTCTTCACACTTAGTTCTTCCTCAAAATCAGATAATATGACAGATATTGAAGCACTCTTTGCTATCAAGAATGAAATACTTGATCCCTTTGAATCTTTGATTTCATGGAATGAGTCTGTTCCTTTATGTCAATGGCAAGGTGTGGTATGTGGAACGCGAAGTCAACGAGTCACTGAACTAAACCTTCTTGACCATAAGCTGGGTGGTGTGTTATCACCATTTGTTGGAAATTTGAGCTTCCTTGTACGACTAGACATCGAAAACAATACCATTTCTGGCAATATACCTCCTGAATTAGGCCGTTTGACAAGATTAAGCCATCTATATTTGCAAAACAATTCATTACATGGTGAAATTCCTGTCAATCTATCATTTTGTAGTAACCTCGTTGAGTTATGGGCACACAAGAACAATCTTGTAGGTGTGCTTCCGAAGGAGCTTGGATTTCTTCCCAAACTCAAATATTTTGATTTCAGTTACAACAAGCTCATTGGTGAAATCCCGAAATCTTATGGTAACTTTTCAGGTCTCTTGGAAATGTACCTTTTGTATAATGATCTTGTGGGTAAAATCCCTGATGAGTTAGGGAAACTGAAAAGCTTGGAAATATTTGACGCAGACTTCAATAGGTTATCAGGTAGAATTCCGAGCTCACTATTCAATCTTTCCTCCTTGAAAGTCATTGATGTGTCAGATAACCAATTCGAAGGTACTCTGCCTCGGGATTTAGGGATCAACCTTCCTAACCTGGAGTGTCTCAGCATTAGTGAGAATCAATTCAGTGGATCGCTTCCTTATTCACTTTTCAACCTCTCAGGACTCGCGTACCTTCTTGTTGGCAAAAATAACTTAGGAGGACGTGTTCCTAGGTTTGATAAGTTGCATAATTTGTATATTTTAAGCATGGAAAATAATCAGTTTGGGAGTGACAAATCTGATAATGAGTTGACCCTCATGTCCTCCTTAACAAATGCAACTAACCTGAACTGGCTGTATCTGCAAAACAACGCGTTTGGAGGGACATTTCATGAGTTTTTCAGCAACCTTTCAAGTAATCTGGTATATGCAGACTTGTCAAGGAATCGGTTTTCAGGACATATCCCGATAGACATTGGGAAATTCATCAACTTGGAAGAGCTTAGGCTGAGCGGAAACCAACTCACAGGTATCATCCCTCCGACTATTGGGAAGCTTTATAAGCTACGATATTTATATCTGTCCCAAAACAAAGTCTCAGGTAGTATACCATCTTCCATTGCAAATCTGAGCCTTTTAACAGAGTTAAATCTTGATCATAACAATCTTAGAGGTAGAATACCTTTAAGTGTTGGAAAATGTCAAAATTTGTTGTATCTTAACCTGAACCGAAACTATTTAATCGGTTCAATACCCAAAGAAATTTGTCTCATCACTTCCTTAGTTGTTCTAAACCTTTCTAGTAACAACTTGACTGGCGCCCTTCCCAAAGAAATCGGTAGCCTGAAAAATCTCTTATCATTAGATGTCTCTTTGAATAATCTCACTGGTGAAATTCCTAGTACACTTGAGAGTTGTGTGGCTATTGTTAATTTAACAATGAAAAGGAATTTATTTCGCGGGATCATTCCTTCTACTTTCAATTCACTAAAGAGTCTTGAAGTTTTAGATCTTTCACAAAATAAGCTATCAGGTATGATTCCAAAGTACTTGGAAGGATTTGCCTTGCATTTCTTAAACTTATCGTTTAATGATTTTGAAGGTGGCCTTCCACAACGAGGCATCTTTGAGAACGCGAGTGTAGATTCTTTCGTTGGAAATCCTAGAATCTGTGGTGGTGTACCTGGCTTGAAACTTTCCAATTGCAATTTCAGTCATTCAAAGAAAATAAACTTTAAGCTTGTTATTTTAGTAATATTAGGAATTTTGGGGCTAGTTGTTATGGTTTTTGCTTTTTTCTTCTATCGATTCAGAAGGCCCAAAAGAACATTTCCTTCATTAGATAACAATCTAAACCAACTCATAGCTATGTCCTATCAAAGTATTCTCAAAGTTACCAACGGGTTTTCTGCGAGCAACTTGATTGGTGTTGGGAGTCATGGATATGTTTACAAGGGAATATTAGAAACGGATGGTAAACACGTTGCTATAAAAGTGTTGAATCTTTTACAATATGGAGCTATCAAAAGTTTCATAGCAGAATGTGAGGCCTTGAGAAATATTAGACATCGGAATTTAGTTAAGCTACTTACTGCATGTTCTGGTGTTGATTATCGTGGCAACGAATTCAAGGCATTGGTTTACGAGTTTATGGCCAATGGAAGTTTGGAGGACTGGTTGCATCCAGATAATTCAAGACCAAATGTTCAGCCAAGAAGGTTGGGATTTCTACAAAGGCTGAATATTGCTATTGATGTTGCTAGTGCAATTCATTATCTTCATAATGATTGTCAAATATCAATAGTTCATTGTGATCTCAAACCAAGTAATATACTCCTTGACAATGAATTGGTTGCCCATGTTGGGGATTTTGGATTGGCAAGATTCCTATATCTTACTGATGAAACCACTTGTAGAATTCAGACAAACTCTTCAACTTTTAAAGGATCTATTGGCTACATAGCTCCAGGTAAATTTCACTCTATATAAATGCCTCTAAAGACTTTAGTGACTCTGGATGCAATGACATTAACTCAGTTGCCGCTATATTTTCATTGCAGAGTATGGTATGGGAAGCGAGGCATCAACGCAAGGTGATGTGTATAGTTTTGGTATCGTCTTACTTGAAATGTTAACCGGGAAAAGGCCAACGGATGACATGTTTGGCGGTGATTTAAGCCTTCATGATTTTGGTAGGACTGTCATGCCTGATGGAGCACTTGAAATAGTAGATCCACTTCTTAATTTGGAAGAGGAAGAGATAAGCAGAGAGAGGTCTCAAATTCCAAGATTCATGAGAAGGCAAAAAATGGTAGAGGGTTTAATCTCCTTATTTGGAGTTGGTATTGATTGTTCAATGTATGATTCAAGTAAAAGAAAAAACATGAAGGAGGTTGTTAGAGAGTTGTGTTCTATAAGAGATTCTCTTGTTGGTTGTACAATATGAAACACTTTTCTTGTTTTGTTTTTGTCTTGTAAGGTTCCAATAATATTTAATGATGAATCATTTG

>PGSC0003DMG400006563 LRR-RLKs

AGTAAACATACAGAGCTAAGGAGATCAAGTGTCATTACATTTCTGTATATTTAGGTCTCTGAAATGCATAGCATAGTTATTTAAAATTTATGTTTACTTGTGAGTAAGATTTTGGAACTTGCGTTGTATTTTTAAAGGTGTGAAATTATGAACCAATGCAGGTATTCTTTAAAGCCAATAATTGAAAAGTTGGATAAACATATGTTTTTTGCTCATGGAGCTGTTAATTGTATAACGCCTGGCCATTTCTTTCATGATTACGGTGGTGTCTTGTGAGGACCTTGATTGTTCCAACATATACCTGCATTTACCTCTAAGCATTAGGCATATAAGGAAGAAGATGCCTAACTCTTTTTTTGTCAGCTAGGATTCAAACTCTAGATTTCATGGTTCTATATTCATCGATCACTAGACCACACCCTTGGGTTTGTATCCCTTTTCGACTTTCAGCTTGTGCATTTCTTCATCTGTTGTATTCCCTTTCTCTCTTTTGATAATGTTTGACCCTTTATTTGCCTTGTGGTTTCTCTTTCCTATTCAGTCTATCAGGAAAATGGTGTGAGGGTTCAATAAGTTCACTAAACATCTGAAGACTAAGATGCTAATATTGATAGTGATCAGTTCAATTTCCACCTTTAACTTTTCTAAATTGCATCGATCCTAGCAGAGCTCTTATTCTATTTTTACTTTTCTTTACATGTTAGATGATGATGGAGAGAAAGGGTATGAAGGTATAACAGATATAATGAATTCATGATACATGTTGTGAAAAGACAACTTACCAAAAAGTTATATATGAGAAAGGAATAAACCGAAACATATTTTGGACTCCCACATTATGTTTGAATTTTTGAAAGACAGATTATGGAGTATTTTATCATGATAGAACATGTTGGTAGAAATAAACATGAAGTTAGGTCCTCATCTTGCACACTTATTGAAGATCAAATGCTGCTTCTGGTATTGCAGAAGTGTCAGAAATTGCTCTTTGCAGGGATCCATTCCTGATTTCAGCAGAATTGCAAACCTAAGCTATCTGTAAGTGCGAGTTTGTTATTGTTTTGTTCTTCTTCTTTTGTTAAGCTTTATCTGTATATTTTTACTTCAAGATTGTTCAACTAGTAGATCTCTGAAAATGTCTAGCTTCTGCTTGCTATACAGTGATCTCAGCTGGAATCAACTTTCTGGATCTATACCACAGAATAAGCTTTCCAACAATATGACAACTATGTAAGCCTTTTGATTTGGCCTCGATAGTTTCTTCATATAGTTATATAAATTGACAGATGTTGGCTTTTGTTTCCACAGCATTTGAACTGGATTGCCTGGGTTCCTTTTACTTTTTAGCATATTCTAATGTATATTTCAAATGCATGATGCAGTATTCTATCACATAACCGTCTTAATGGATCTATACCCAAGAACTTTTCTCTCCTTCCTTCCCTGCAGAAACTGTGAGAATCTGGACTTCGTCTTGCAATATCTACTATTTCACTGAAGGAAATGTACCATTAACGAATTGGTGGGACAAATTTGTCACAGGTCTCTTGAGAACAACTTTCTGAATGGTTCCATTTCCACTGACATTTGGCAAAACAAGATCTTTAATACAACTTCCAGACTTGTGATGTAAGATGTGAAGCAGAAAAATAACAGATCTTATTTCAATAGAAGTATTTACTTCCAGAGTCAAACCTTTGAATGAGGATTTCTTTCTAACAACTCTTTCATTGATTTTGCAAACCTAGTGATCTGCGGAATAATTCAGTTTCAAACATTTCGGGTGCTTTTGAACCTCCTGTAAATGTCACACTAAGGTATGATTTCTTCTTTCAAATTCAAAAATAGAACAATTTGAATGTCGGTTTATGTCCTCATGGGCCACATAAATAGGACAAACAAATGGGTAAAGAAACTAGAATAACAGAAGCAATATGCCATACCAAAAAGGGAAAGTACGTCGATTTTCCTCGTGTGTGTGACAACAATTACTTAAATTACTTGAAGAAGCACAAATGCATGGAATCACTTCTGTTATGTTTTATCCACACACACCGATACACATGCTCTTAGCAAATTCATGTATGGTGCTTTATAAAATGTAAACAAAATGGAAAGAACAAGGAGTTGAGAGAACCTTTGTGGTTTTCAGTTTTTTCATTGTTTGAAAAATTTAAGTATGCTTAAAAAGGAGGTATAGATAGAAAAGGGAATTCTGAAATTCTTAGTCACATATTTTCATCTAGTTGCAGTGGAAAAGGATGAAATGATGAAGGGCCCTATAATATTGGTTATGGTCCTGATCAGCGTAAATCTTGAAAAAGTACAAAGTTTTACCCAATAATATAAATGAGAAGTGTGTGTGTGGGGTGGGGGGGGGTGGGGGGTAAAAAGATATAAATAGAGGGAAACTATGAGCATAAAAGGAGTTTTTGTCTTTTGTTGAGTCTCTCATTTATTCGTTGAAAGGAGGCCTGAACAAATGGAGAGGAAGGAAGTTTCTGTAAAATTTGAGTCCACTACATGCCTTATGACACATCTAATCTAAGTTGCTCGGACTCTTCACTTTTGGTGCCGCACCCGTGTCGACAGGATATGGGTGTGGGATTCGTGTCCGATTTTGGGTATTTGGACCAAAATTGATGGAGAAAGTCTGGACAGTTTTAATGATTTTTATAATCAATACAAAAGCTAAGGTGAAATTGAAGAAAATGGAATACCTTTTATATAGAAATTTCTATGTCACTCATTTTCCTTTGATCTCCTTCCAAGATTCTCTTCAAGTTCTCCACATAATATCTCATAATATAGGTTTTATAACTCTTTTAGATATTTCAATTATATTTAGGCGAATCCCTGCACCCGTATCCATTCATGGATCCATATCCCTGAATCTTAAAAGTTATATCATATCATGAAGGATCGGACCTTTAGATCCGCACCCGTATCGGACACCCACACCCTAGTCCAAGCAACTTAGATTCTAATAGTTTGTGAAACTGAGGGAGTTTTTCTGCAGTATATCACATCAAGACAAGTGATATAAACTGGTATCATTAAATTCTTGAAGAAGGAAAACATCACATTTAGGAAAGTGGAGAATGGAAATTGTGCCAAACAAAAAGAGTTATTGGTCTAAAATTACTTGTACATCTTTGTTACTGTCTCTGGAGTTCATGTTTTCAGTTATGCACTCATTCTTGTTGTCCAATTATTCATAAATATCTGATTTTGACAGTGAATTGAGGTCATCTTTCAAAAATTACTTACTGATCTCTTTCTATCTCATGAGGTCAATTCATCTTGAATTCTTAATTGAAAATCTGCTCAATGTTTCTCAAAAGAAACATGTCAATAAAAATGTTTTAGATTTTGTACTATGTTAAAATTCATTTATTATTTTCATAGTCTTCCCCTAACTTTAGCAGACGATTTCCAGGTTCCAAAGCAATCCCATCTGCAGCAGTACAAGTATCAGAAACATTGGTAAATACTGTGGCCCTGACTTTGGAGCAGATGACGACGAACCTAGTAATTCAACAAATCCAAGTGGTGTTTGTCCTATTCATGCTTGTCCAACTGACAACTATTATGAATATGTTCCTGCATCACCTAAACCATGCTTTTGTGCCTCACCTCTTAGAATTGGGTACAGGTTGAAAAGTCCTAGTATTTCTTATTTTGATCCGTACAAGCAACTATTCGAGTCTTATGTGACTAGTTCCCTTCAGTTGGATCTTTACCAATTGTGGATCGATTCATATTTTTGGGAAAAAGGACCTCGTCTGAGGATGTATTTGAAGCTTTTCCCGGTGGTTGGGGACAATACATTTAATGAAAGTGAGATTCTTCGTATCAGTGAGATTTTTGCCTCTTGGGAATTCCAAGGAAGTCACCTATTTGGACCATATGAGCTCCTGAACTTCACACTCCTCGGACCTTATTCACACTGTAAGTATTCACAGCTTAAATAGATTCTAATTCAACTTGTAACAACTTAGGCTGTGTTACTACAATTGCATGACAATTGAATCATTGACATGCAGCATGTGGATAAGATTGAAATATTTGACTATCATTGCCACTTCTAGCCAACAAGTATAAGGATCTTAATTCATATAATCAAATAGTTTAGATAAGCAACAAAAGACTATGATCCATGATCTTCTAAGGACTTCATATATATTGGGTTGAAAAAGACAATGAAGTCTTTGTTTATATCCAAAGTTTTTTTAAGACTTCAGATATTTTACGTTGAGGAACCTTGAAGGATCTCACTAACTTGATTAAAAGGTTGAAACTTCAAGGGGTTCAAACTTAGATAGTTTGGTCCAGTCTCTCTTCAAGAGCTTCAATTCTGGAGGTCTCAGCCCTTACAGTTTTTGTTGGGATGAAAGTGGATAAAAAAAAATGGACCTTAGGCCTCCCAAAAAGCTAGCTCATGAGGAGGGGATTGTCCAAAATCATATAAGGAGAATAACCCTTGTCACTCCAACACCTTGCTGCACTCCCAAGAATTGACATTGGAGCGTGGACAATATAATAAGGGGGGGTTCAACACCGGAAACAATAAATTGGGATCAGTGTGGATCTGAGACCATGTGAGAAATAAAGAAGAAAAATATTACTCCCTCTATCCAGTTTATATGACTTACTTTTCTTTTTAGTCAGTCCCAAAAAGAATAACACATTTCTATATCAAGTAATAATTTATTAAAATGTCAATTTTACCTTAATGAAATGATTTATAGCCACACAAATATCTATCACTTTTTTTTAGACCACAAGTTTTAAAAGTCTTCCTTTCTTTCTTAAACTCCGTGCCAAGTCAAACTAACTCATATAAAATGGGATGGAGGGAGTATTAAAATTATGTATTCTTTTATTACATTGAGCCCTATTTATAGATTCTACATTACAATTCATTTCCAAGTAGGACAATATACACAATCCTATTCGAACACTCCCCCCTTAAGCTGGTGCATACAAGTCAGATGTACCGAGTTTGTTAAGATGTAATTAATATGAGGATCAGTGAGGGACTTGGTGAAAATATTTGCAAACTCATCACTCGACTTCACAAATTTTGTAACAATATCTCTTGGGATCATCTTTTCTCTGACAAAGTACCAGTCAATCTCAATGTACTTAGTCTTCTCATGAAATACCACCGATTTGATGTGATATCAAGGGCTGCTTGGTTATCACAGACAAGATAATGGCTCAATTGTTTGATTTCCTTTGTACAAGCTTTGATTTCTTATATTATCTGCAGTGAATCCTGAAATGAAAGGTAAAAGACAAAGCAAAGGCGTTGTGATAGCTTGCATTGTAGCAGCAGGTGCTTTTGCTGCCTTTGTTTCCTCAATTGCCACTATATTAATCACAAGACGACGTGCTAAGTATCAAAAGATCTTGTCAAGAAAACGTTTGTGTAAGTCCTCTTATTAGATTAGCTATAATTTTACTTTATCAAGTCTGCTATTTTTAATTCTTTCTGCAAAATTGATGACTATGGGTTTTTGTTTGCAGCTTCAAGCCTCTCTATAAAAGTAGATGGCGTTAAAAGTTTCACCTTCAGAGAAATGACTTCGGCCACTAACAACTTTGATACCTCAACTCAAGTTGGTGAAGGAGGATATGGTTCCGTCTTTAGGGGAATTTTAGCTGACAAAACAGTTGTTGCTATCAAGCGAGCAAAAGTAGGATCCCTGCAGGGGCAAAAGGAATTCTTGACCGAAATAGAACTATTATCAAGGTTACATCACCGTAATCTGGTTGTGTTGCTTGGATACTGTGACGAAGAAGGAGAGCAGGTATGACAAAAGCTGAAAACCTTTTCAATTCCAACTGTTTTAGTTACTAATAATTCACACCATTCATCATTTTTAATCATAGTTAGTCATATTTTTTTAGTGTAGCATCATAAACCTTCTTGTATTCTTTCTCACTTCAGTTGCTAATGATCAGACTGAATGTGTAGCAGACTAAATGCTTTCTTGAGCTACTGAAGTCTTCTTGTTTAGCACAAAGAGTGATATAGTCGAGCAATTATGAGTCTTCTGTATACATTTGCAGATGCTGATTTATGAGTTCATGCCCAATGGAACTTTGCGAGACTGGCTTTCTGGTAAGAACTGTTTCCTGGTTGTTTCTAGCTTTCATATGATTAATCCTCCTCAATTTCTTTCTTTTAGTTTAGTAGTGAAGATCCAATGCGGAGTTGCAATGTTTATAGATGTTCCTTGCACATGTTGTGTCATTAGTAGTACTATGGTGAATTGATAATACAGGCTTCAGTATTTTATTGTGTTTGTGGAGAAACCCTTGTCAAGGTGGTGATCGATCTAAAAGTTGCATAATGGAAAAGATATTGCAGGAAATCTAATAGTCAGAGAAACATAGATTCCATCTTCTTTTTCCTTATCAGAACAAGAAAAGTTGTTAGTTTCTTTATGCGGCTGTGATAAACTTAAAATTGACATCGGTCAAGAAACTTTTGTATTGCTATCTAAGTAGGTGATTTCCATCATTTGACAGCCAAATGTAAAAAAAATTTGAAGTTTGGAGCAAGGTTGGGGATCGCACTGGGGGCATCTAAGGGCATCCTTTACCTCCATACTGAAGCTGATCCACCCATATTCCATAGGGATATTAAAGCCAGCAACATTCTTTTAGACTCTAAACTCACTGCCAAAGTTGCTGATTTTGGATTGTCTCGGCTTGCACCTGTCCAGGATGATGAAGGTCTTCTGCCTAATCATGTGTCAACAATGGTGAAGGGGACACCTGTAAGTCTGTCTATGCTCTAGATTTAAGGAAATAATTGTGCAATGATCTTACTGATTTTCTTATGTATATGCACAGGGTTACCTTGATCCCGAATATTTCTTGACCCGTAAAATGACAGATAAAAGTGATGTCTACAGCCTTGGGGTTGTATTTCTGGAGATCTTAACTGGCATGCATCCTATCTCACATGGAAAAAACATTGTCCGTGAGGTATTTCTGTCGGCTGAACACCTATTGGAATACAAAAACAGTGAATGTTTGTCTAGCATTTTAGCAATAAAAGCATAGGTGTATGTATATAACCTTTAACTGTCCATGACAGGTAAAAATGGCTCATAAATCGGGAGTAATGTTCTCCATTATGGATAAGAACATGGGATCTTACCCCTCCGAATGTGCGGAGAAGATGGTGGAGTTGGCCCTGAAGTGTTGCCAGGACAAACCAGAAGACAGACCCTCAATGCTGGAGGTGGTTAGGACGCTAGAAACCATACTCCAAATGATGCCATATACCGACACCGATCCTTTGGATAATAAAGCCAGTTTCAGTGAAACTACATCATCAGCTTCATTTTCTAATACTAGAAGTGGAGACCTATTTATATCATCATCCAATGTATCAGGTGGTGATCTTATTAGTGGTGTTACTCTTAATATTACACCTCGTTAACATTAGCTTGTCTTCCACTGCATAAACATTGACTTGTACACTATGACTAGTAGGAAGTCTATACGACTTGTCTTTTATGTTGTATACGAGTGTACAATAATACCATCTTACTGCTTCTAGATTTATGTATATTCCTCAAGAATCTGTTGTGACAGACAACCCTTGTATATGTATAGTACATTATACATGAAGTGCCTTATCACTGTCCATAGAAATGCTACATGTAGTTGGTTATCATCCTTGCA

>PGSC0003DMG400006923 LRR-RLKs

TTTTGAGGACTGTGAAGTAAAAAATATTGATGAGAAGCCCATCTCTTTCTTCTGCAAGAACCGAACCCAACGGAAAGGTCACCATTAATGGCGGAAACCTCATGAAACTCGCGACACTACCTATCTTCTTCTCATTCAATGCATGTAGCATTAAATTCCTTCTTCTGTCTATAGCAACCCAACTTCTCCATTTTTCTACATTATATATACTACTCTACTTTACTCTTCTCCACTTTTGTGCTTCTTTTAATTTCACCTCTTAAGCCGTACAAGACCCATTTATTAGTGCTTCATTATAGAGTTCTCTGTTCTTTTTGCTTCTTTCTCTTGACCCCCAGAGTGGGTTTTTGTTCTATTTGTTTTCTCTTCTTCCAAAATATCTCTGCCAGGAAAAAGCAAAACATTCCTTCAATGTGCAGTTTAGTTTGTTTGTTTGTTTTTTCTTGATGGGTCTTTGTCAAGATTTGTGTTTGGTTCAGAATTAGTTTAATTTTCACCCAAAAATATTCTATCCATTTGGAGTTTTCCGCAGATTTAGACATAAAAAAATGCTTGACGTCTGCAAGAATTGAAGCTCAAATTGTTCCCTTCAGTTTATTCATAAATTTCTGTTGAGTATTTAGAGGACTGTGTTAAATGGGGTGCCTAAAGATATTTGTCATAGTTGTATTCATGTGGGCTTTATTGATCCCAAATTCTCATCAATTAGGGTCCTATGAAACTCAAATTCTTCTTCAGTTACGGAGGCACTTGGAATATCCAGTGCAGTTGGATGTTTTGGAGAATTACAATGGGGATTTTTGCAGTTTGGGTTCAACATTGAATATGAGCATTATTTGCGAGAATAACTCTGTTACTGAGCTCAAAATCAAGGGAGATAAGCTAGTAAAGGTCAATGAATTCCATGGAGTTGCAGTTCCAAACAATACCTTATCTGAACGTTTCTCCATTGATTCATTTGTTACCACTTTGACAAGGTTAAGCAGTTTAAAAGTTCTCACTTTGGTTTCTTTGGGCATTTGGGGACCTCTCCCTGTTAAAATTCATCGGTTGTCTTCTCTAGAAGTTCTGGATATGAGTTCAAATTTTCTTTTTGGTAAGGTTCCATCTGAGATGTCTACAATGGTGAAGCTTCATACTTTGACATTTGATGGCAATTTTTTCAATGAGAGTCTCCCTGAGTGGTTTGATTTGTTACCTAATATCACTATTTTAAGCATGAAGAATAATAGGTTGAAGGGTAAATTTCCTCATTCAATTTCTAAAATCACAGGCCTAACTGATATTATTCTGTCACATAATGCTCTTTCTGGTGAATTACCTGATTTAAGTGCTTTGTCCAATTTGCATTTGTTGGATTTAAGAGAAAATCATTTGGATTCTGAACTGCCACTGTTGCCACAAGGATTGACCACTATTCTTCTTAGTAGTAATGCTTTCTCTGGTGAGGTCCCAGAAGAATTTGGCAAACTAAATCTACTTCAACACCTTGATCTGTCAAATAATGCTCTTACCGGAACGCCGCCTGCTGATTTGTTCTCTTTGCCAAGTATTAGTTACTTGAATTTAGCGTTTAACGTTCTGAGTGGTTCACTTCCTGAACATCTCAACTGTGGGAGTGAACTTGGTTTTGTTGATATTTCTGATAATAGACTGCTTGGTATGCTTCCTTCTTGCTTGAACGCCAGTTCGGATAAGCGAATTGTAAAGGTCAGTGGAAACTGCTTGTTGGATACACAATATCAGCACTCTGAGTCCTACTGCAAACAAGCTAGTTTGGCAAAGAAACAAACCACTGGAAAAGAGATAGCAATATTGGTAGGTGTTGTTGGGGGAATTGTAATACTTGTGGTGTTTTTGGCTGTTGTTCTTCTCATCTTTTGTAGAAGACAACATGCACGTCACGATGTGGACCGGTACATGTTTCCCAAAGTTGTGCAAGATAATGGGCAACCCAATATTTCTGCTGAGCTCCTGGCAAATGCTAGTAAGTTGTAGTTTGATTTCATTTTCAGTGTCAACTGTAAATTTTTTTCCGGGATTTGATAAGAGTTTCTTTAACTTATTTTACCTCAGGAATCATTTCTCAAACAACAGCACTAGGATCACAAGGTGCCCCATCATATCGGGTGTTCTCCATGGAAGAGTTGGAAGAAGCGACAGAAATTTTTGATAAATCAGCATTACTCGGTGAAGGCTCCATTGGAAAAGTACTGCTCTGAAACTATTTATGACTCCCAAATAATTGTAGGAAGAGGACTTAGCGTATTGTTTCCTTGCATATATCCATTCAAAATTCATTTTATATTTTGTCGGAAGAATAATATTTTTGTTGCTTATTTAGCTTTTGTCCATATGATGCCATATTATTCCAGAGGTCTTGATCTCCGTTATCAATTGAAGCAAATCAAATTAGTTTTGTAGGTAGGTTTCATTGAATAACTCGTATCAGACAATTAAACAGATGCACAAGTCGTCAATATAGGCAGTTGATTAAACAGTTTTTTGCCTTTATGATTGATTGTCTGTGTTCATGAATGCTATATGCATCTTGACTTTGCAATTTAATACCATATAATGGTGGTCACAGTTAAAAGTAATGTTCGTTTTGATTTGGTATCTACTGAACTTTGACAATTAATATCAATAAAGTCAGTGCGCTTGATTCGCTTATTTAAAGTGCATATGCAGTGAGTTGATATCTCTTCAAGCCTTCTGTCCCAGGATTTTTTCTTTCTTGGGATCAAGTTGGACAATTTTTATTTAACTAACACCCTTCTATGTCATTCTCTCTATCTCAGATTTACAAGGGAAAATTAGAGAATGGAACCTATGTTGCTGTAAGGGAATTGACTGTTCATAGACGATGCACTAGTTGGAATTTCAAACTTCGAATGGATTTGCTGTCAAAGTTTCGCCATCCCCATTTAGTTAGCCTTCTGGGTCACTGCATTGATGATGGGGTGCAAGATGACTCAACAGTCCACAGACTTTTTCTGGTTTATGAATTTATTCCTTGTGGAAACTTCCGTGCTCGTCTTTCAGGTCAGTAGCGTAAATATTTTAGTACCAACAGTTGCATAGACATATACTTGATTGAATATGCAAATGCTGCAGGTATATTTCTGGTTATGCTGATTGCTAATTTTCAGATAACTATTTAGTATGATAGTTTTGGTGAATCTGTCTTTTTTTGAATATTGTGCGGGGGAAGGGGGTTAAGGATGAGAAACACATGAATTTATTGCGTCTTTTGTTATTTATTCTCTAGTAAGTACAAATACAGAGTATTGTTATCTTCTGGATAGTTTCTTTCCGTCTATTTGTCCAAGCTTGCGATCTGAAATTTTACAACAAAGTGAAAGACAATTATTGTAGTTTGATGAAGCAATCTGTGTTAGCGTTTCATTGATTTGTCGAGATATACTTGATCGTATGTTCTTTGCATGGTCTGTTAGCCGACACTTGCACTGCAAAATGTTTTGACCATCTTTTTCTTTGTGTTGTTCATTTTATTTATCCCGATATATCCAATGAACCATAGGGAATACATGTTGACCTTGTGTAAATCATCATTTTCACTTGCAGAAACTACTCCAGGAAAAGTCCTCAACTGGTCAGACAGGTTGGCGGTTCTGATTGGTGTTGCCAAGGCTGTGCACTTTCTCCACACGGGTGTAATCCCTCCTTCATTCGGCAATAGCTTGAAGACAGATAGTATATTGCTTGATGAGCATCAGATTGCAAAACTTAGTGATTATGGAATGTCAATCCTTATAGAAGAAAGTGAAAAAGTTGAGGTATGTTGAAATGATATTCATATGATGAAGGGTAGGGGTAAGGTCTGCGTACACCCCACACCCTCCCCTGACTGTTATTACACTGGGTTGTTGATGTTTTAGGGTAATATCTGATACTGATGATATTCTTTTTCAGGCAAAAGGAGATGGCCACAACGCCTGGTATGTAACTCTTTTGCCCAAATAGAGTCATACATCAGTCTGTTTGCAACTTCTATTCCTAGAAAAACTGATAGATTTTTTCTTAGTGCCTCTTCTGACTATGGCATCCCTTAGAGATAGATAAAAATGCTTGGCAATGACAGTTCAAGAATGTTGAGCAGAGACATATTCTCAACATTAAAATTTTGTGCATTTCTTCTCATGCAGGAACACGAGGAAGAAAGAGGATGATGTGTATAACTTTGGTTTCATATTACTGGAGTCGCTGGTGGGTCCTTTTTTTAGTGGAAAAGGAGAAACATTTTTGCTCAATGAAATGGTATGGAACTACTAATTTTGATTTTTGATATGCAACTCCTTTCTCATTGGCAGTAGCACAGAATGCTCTTCAGCATTGTTTCTACTGTGGATCACAAACTTTCGTCCCGTTTGTAGTTAAGGAGTTTTGAAATGAATGTCTGTGACACTTACTATGACTTGGAAGGTAAGCAGTAAAGTCTTTTTTTTTCCGATGACAGACATCCTTTGGTAGCCAAGATGGTAGGCGTAAAATTGTGGATCCTGCTGTGCTAACCACTAGCTCCCAAGAATCTTTGTCAATTGTAATATCACTCACCAACAAATGCATCTCTCCCGAGTCACAGAGTCGCCCCTCATTTGAGGATGTCCTCTGGAACCTACAATATGCAGCTCAAGTCCAGGCTACAGCCGATACAGATCAAAGATCTGATGCAACTTCATTGTCATGAGGTATTCAAGAAATTCAACTGCATCTCATCAGCAATACTTGCTCATGCCGTTGTCTCAATTGGGATGCAGGCTGAATGCATGATGTTTGTGAAGCACAAAATGGTTGCAAACCTCAGCCTCATGTTCAATCTTCTAGTCCTCGTTATCATTTTCTTTTGTATAAAAATGTAAATTTGTCGATTCTTTGGATTTTAGAATTTAGAAGTGGGTGATGTTGCAAAACAGAGACTAGTCTTCACATGAATTTGTATTACCCTCCTTCAGAAAGGATAGAAATGAAAGTGACAGAAGGCTTAGTAGCCGTGCAACCTCAAAAATTGCCCTTCATGCTTATGTGTAAGATAAGCATACATGTATATCAACATCAGACTAATAACTGATCTTCATTGCCTTGGTACTTGCATTGATCCAGATATGAGAAAATCAAAGAGACTAGAGTCCCTCCAATTCTTCCTAAAC

>PGSC0003DMG400006942 LRR-RLKs

GGTGCTTACTCAGGGGAGTCCACTCCATTAACAGGACTAACACTAAATTAAACTTTACAAATTCTCTAAATCCATAATTCCAAGGTTCCACTTCATCATTCAAAGTCAAATGTCATCTTCCATTCTTTTGCTTTTTTCATACACTTATATTTCTCATGTAAACAATTCATCAAACACCTTCTATGCATCCCTACTCCCTAAAGCTCAAAGATGAAACTTTTCAGATTTTTCAATCTTCCCCTTATAGTATTCATCTTATTGATTCTTGTATTTCAACCCTTTCTTCATTGTACTGCTATTGCTGGTGCTGTAGCTGAAGATGATATAAAGTGTTTAAAAGGAGTCAAGAACTCCTTAACAGATCCAAAAGGGAACTTGAATTCATGGAATTTTGCTAATTCTACAGTGGGGTTTATATGTAAATTTGTTGGTGCTTCTTGCTGGAATGACCGTGAAAATCGTCTCATAAATCTTGAACTTCGGGATATGAATCTTGGAGGTAATGTTCCAGATTCGTTAAAGTATTGTCGAAGCTTGCAAACTCTTGATCTTTCTGGTAATCGGATTTCTGGTTCAATTCCTTCTGATATTTGTACTTGGCTACCTTTCTTGGTAACCCTTGATTTGTCGAACAATGAATTTACCGGTTCTATTCCGTCTGATCTTGTTAGTTGTTCCTATTTGAATAAATTGATGCTTAATGATAATAAGCTTTCTGGGAATATACCACCTCAGTTTTCTAGTTTGGGGAGGCTTAAGATATTTTCTGTGGCAAACAATGATCTTTCTGGTAGGATTCCGGAAGCTTTTGATTCGGTTGATTCGTTTGATTTTGGAGGAAATGATGGGCTTTGTGGTGGTCCGTTGGGGAAATGTAGGAGGCTTAGTAAGAAAAGTTTAGCTATTATTATTGCTGCTGGTGTGTTTGGTGCTGCTGCTTCTCTGTTGTTGGGTTTTGGGGCTTGGTATTGGTATTTTACGAAGGCAGGGAAAAGGAGGAAGATGGGGTATGGATTAGGGAGAGTTGACTCGGAAAGGTGGGCGGATAAGTTGAGGGCTCATAGGCTTACTCAGGTTACATTGTTTAAGAAACCGCTTGTGAAGGTTAAGTTGGCAGATTTGTTGGCTGCCACGAACAATTTTAGTACGTCGAGTGTGATAAACTCGACTAGGACAGGGACTACATTTAGAGCTGTTCTACGTGATGGTTCTGCGCTTTCTATTAAACGGCTTAAAGCTTGCAAGCTGAGTGAGAAGTTGTTTCGAATGGAGATGAATGGACTAGGACAAGTTAGGCATCCTAATTTGGTGCCACTTTTGGGCTTTTGTGTTGTTGAGGAGGAAAAACTTTTGGTTTATAAGCACCTGTCAAATGGTACTTTGTATTCATTATTGAAAGGGAGTGCAAGTGTGTTGGATTGGCCTACTAGGTTTAGGATTGGCTTGGGTGCTGCAAGGGGCCTTGCTTGGCTTCATCATGGTTGTCAGCCACCAATATTGCACCAAAACATATGTTCTAACGTTATTTTCCTTGATGAGGATTTTGATTCTAGAATAATGGATTTTGGGTTGGCAAGGCTAGTGACACCTCCAGATGCAAAAGAGACTAGTTTTGTGAATGGGGAGTTGGGTGAATTTGGCTATGTTGCTCCGGAGTACTCAAGCACAATGGTGGCTTCACTGAAAGGGGATGCTTACAGCTTTGGGGTTGTGCTTTTGGAGTTGGCTACTGGGCAAAGACCTCTAGAAATCACTGCTGCTGATGAAGGTTTTAAGGGAAATTTGGTGGACTGGGTAAATCAGCTCTCTGTTTCTGGTCGGATTAAAGATGCAATTGATAAGCATATTTGTAGAAAGGGCCATGATGAAGAGATTGTAAAATTCCTCAAAATTGCTTGCAATTGTTTAATTTCTCGGCCCAAGGAGAGGTGGTCTATGTATCAGGTATATGAAGCACTGAAGAGCATGGCTGAGAAACATGGTTTCTCTGAACATTATGATGAATTTCCCTTACTATTTAACAAACAAGAGACCAGCAGTCCCATTTGATACATGAATGGGAAAGCACTATGAGATGTTTCACATCTGGTGAACACAAATGCATTATGGTATGTTCTTCACTTCCCTTAGGTGCAAGTGGAATCAGGCACGATACATATGGTTCTGTGAAGTACTACCACAGATATTGCTGTATTATACTGTGAAGTGTCAAAACATTCTGAGGTGTTCTGCTCGAAATTTATGTTACAGATAGCTAATATTATTGGTATCTTAGGTTTATGTTTCTCTTCTGCTCATACTTATGCATTAAACAAGACAGTAAGCAAAAGCTGTCACAGTTTATTTGAAAGCTATAATACCTACTAATTTAACTGTTTATAACTGACCTGGGCATTGGCCTTTTTAGGGTCGAGATATATTCAGTATGTCCCGTACAGTCATTTATGGTTTGTAAGATCAAATCACCCTAGGTTTCCCTAATGGATTGGTTCGGTAGAATATCTGCCTCCACACTGATATTAATTCATCTAATCGGAATGCTATCAAGCAGAGTTATTAGTAATTACCCCTTCTGATTAATTTTGGATGATGTTGATCAGTAATTGTAGGTCCTTTTGCAAAAATTGTGTCTGATTGGGAATAATTTTCACTAAAATGGAAGTTGAATGGGCTTTCTGAAAGTTATGTGCCCTCTTAAATTTGATTTGGCAGCTGAAATTTTCATTCACTTGATTCTAATCTGCTTCTTTACAATCTCAGAAACATAATGCACAATTATAGTTTCCAAGTAGGAAAAAAAAATTCTTGTTGACAGTTGAAACTCTTTGCAATAGAAGTTATCAAGCCTCTTACTTTTTCCCTCAAGCTCAAGCTCGGATGCGTGTTTCCATATGCTCTCCTGCTTCCCTGAAAAGTCATTTTGTAATATTTCAAAACAACTACTTGTAGTCTAATAATAGTAAATGAAGGGAAAAGGTCTGTTAGCAAGTTATCTCATATCTGTC

>PGSC0003DMG400007234 LRR-RLKs

CTCATCTTTACATACCTTATTTTCCTCAACCAATTTTTTCTTACAATCTCTTAACCATGAACCATTTCTTTCTCACTACAATTTTCTTCTTTCTCATACAATTTTCACTTGTTTTCACTACTTCAAATTTTCCTCTTCAACTCATTTCACTTTTATCCTTAAAATCTTCATTTCAAGATCCAAACAATACATTTCAAGATTGGAATCCTACATCAACTTTCTCAAATTTTGGTTCACAGCCCTTTTGGTGTTCATGGTCAGGCATCAAATGTGACACAAAAACAAGTCACATTACATCACTTAATCTCTCAAAGAGGAACCTTTCTGGTAAAATACCTCAAAATATTGCATCCTTGATACACTTACATCACTTAAATTTAAGTGGAAATTCATTTGATGGACCTTTACCATCCATCATTTTTGAATTTCCATTTCTTAGAACACTTGATATTAGTCATAATTATTTCAATTCAGCATTTCCTCTTGACGGGCTAACGAATCTCAAGTCCCTAGTACACTTGAATGCCTATAGCAACAGCTTTACAGGCGTGTTGCCTGTAGAGCTTGTTGGGCTACAAAATCTTGAGTATTTGAATCTTGGTGGCAGTTACTTCACTGGTGAAATTCCGGCAAGTTATGGGAATTTCAACAAGTTGAAATTCCTACATTTAGCCGGAAATTCACTGAGTGGGAAAATTCCAGAGGAATTGAGATTGTTGAACCAACTCGAACATCTTGAAATTGGATACAACAATTACACTGGAAATGTTCCTCCTGGATTTTCCTCGTTATCGAATTTAGCTTATCTTGACATCTCACAAGCTAATCTTTCAGGTGAAATTCCAATTCAGCTTGGAAATTTAACAAAGATTGAATCTCTATTTCTTTTCAAGAACCATTTTATCGGTACAATTCCCTCGAGTTTTTCCAAACTCACATTGCTGAAATCACTGGATTTATCGGATAATCGTCTCTCAGGCAACATTCCTTTTGGATTTTCAGAGTTGAAAGAGCTTATCATGTTGTATTTGATGAACAATAACTTATCAGGTGAAATCCCTGAAGGGATTGGTGAGTTACCAAATCTCGAATTGTTAGCTCTTTGGAACAATTCACTCACTGGAATTTTACCGCGAATGTTAGGATCAAATGCTAAGTTACAAAAACTTGATGTGTCTTCAAATAATTTGACTGGTCCAATTCCCCAAAATCTCTGCCTCAGCAACAACTTAGTTAAACTCATTCTGTTTTCGAACGAGTTTACTGGTGAGATTCCTTCATCCTTAACTAACTGCAACGCTTTATCCAGGTTGAGAATCCAAGACAACAAACTCAATGGCACGATTCCATCAGGATTCGGTTTTTTGCCTGATTTCACTTACATGGACATCAGCAAGAACATATTTTCGGGGCAAATTCCAAAAGATTTTGGAAATGCCCCGAAAATGATGTACTTGAACATTTCTGAGAATTTTTTCGGTAGTAACTTGCCTGATAATATCTGGAACGCCCCGAATTTACAGATATTCTCAGCAAGTAACAGTGGCCTTGTTGGAAATCTCCCTGATTTTAAAGGATGTCAAAATCTCTATAAAATCGAACTCGAAGGAAACAATCTCAATGGTAGTATTCCATGGGACATTGAACATTGTGAAAAGCTTATTTCTCTAAATCTTCGACGAAATTCATTTACAGGAATCATTCCTTGGGAAATATCTGGTATTCCTTCGATAACAGACGTTGATTTGTCACATAATTTTCTCACAGGAACAATCCCTTCCAATTTCGAAAAGTCTAGCACGTTGGAACATTTCAACGTGTCGTATAATCAGCTCACTGGTCCATTGCCTTCTTCAGGGTCTATGTTCTCAACGTTCCATCCTTCGTCTTTCGTAGGAAACGAAGGTCTTTGTGGTACTATCATACATAATCCGTGCAGGACGGATGAACTCAAAGATGGAAAGACAGATTTTTCGAACCATTCAAAGAAAACAGATGGTGCATTGATAGTATGGATTGCGTCTGCAGCATTTGGAGTAGTAATTATCATTCTTATTGTGGCAATTCGATGTTTTCATTCAAATTACAATCGAAGATTTCCATGTGACAAGGAAATTGGACCGTGGAAGTTAACAGCATTCAAAAGGTTGAATTTCACAGCAGATGATGTACTCGAAAGCATAGCAATCACAGATAAGATCATCGGTATGGGGTCTGCAGGTACTTAACGGATCTATATTGTTAGTTATAATGAGTTCAGTTGAACCCGCAATTTTTCATTAAGTATATGTGGTGGAAAAATCATACAAAATTGGTGTACATATATATTTTTTTTAAAATGCTAACATCACCTAGACAAGTTGGCAATGTGGACTTTTATCAATATAACTCTCAAATGGAAAAGTATAAGATAGACTAGATCTTACCTTAAAATTGTTACATAAATATTTAGTTTTGAGTGCACTTTCAAATAACTTCGATCCTTAAAGCCATTAAGATTATATTCTAGATCTGCCTCTATCTATAGATATAATGTACAATTTCCACCTTGAACTTAGAATGGGGTCCGCGCGTAGTGGATGATCTGTAGTGTCAGCAAAGTCGAAATGTACAAAGTCTCTTAAGCAATAGTAGAATAAATATTACATTTCAAAATCATGTTTTCAAATAAGTTGGAATCTTGAATACTAAGATTATATTCTAGATTCGCCTCTGTCTGTGGGTATAATGTACCATTTCTAGGTGAAACTCAAAATATGGTCCGCAACTAGTGGATGATTTATAGTGTTAGTGAAGTCGAACATATATAGAGAAATGTGCAAAATCTCTTTGACATTAGTAATTTTAAAAGAAGCTTGAATCTTGAATACTAAGATTATATTCTAGATTTGTCTTTGTCTATAGATATAATGTACAATTTCAAGTGAAACTCAAAATGGGGTCCGCAACTAGTAGATGATCTATAGTGTTAGTGAAGTCGAACATAGAGAAATCATGTGCAAAAATCTCTTAAAAGTACATGAGTAACTTCCAAAATAGTTTGAAAGCTTGAATACTAAGGTTATATTCTGAATCCACCTTTGTGTGTGCGTGTCTATATATAGGTACAGTGTACAAGGCGGAAATGCCAGGTGGCGATATCATAGCGGTGAAGAAGCTATGGGCGGGGAAGCAACATAAGGAGACGACGAGGAAACGACGTGGCGTGTTGGCAGAGGTGGATGTGTTAGGCAATGTAAGACATAGGAACATTGTGAGACTACTAGGTTGTTGTAGTAACAATGAATGTACTATGTTGTTGTATGAGTACATGCCTAATGGTAGCCTTGATGACTTGTTACATGACAAGAACAAAGATGCAAATTTGGTTGGTGATTGGTTTACTAGGTACAAAATTGCACTTGGTGTTGCACAAGGGATTTGTTATCTTCATCATGATTGTGACCCTGTCATTGTTCATCGTGACCTAAAACCTAGCAACATTCTTCTTGATGGTGAAATGGAAGCTAGGGTTGCTGATTTTGGGGTTGCTAAATTGATTGAATGCGACGAATCGATGTCCGTCATTGCTGGATCTTATGGCTACATTGCACCCGGTAAGTAATTTCGAATCTCCCTTAATTTTTTAAACTATTTTCTAAGGTTGATTTAGCTGTCAAGAGGTTACAATTTAACTGACAACGTGCACGAATTTGAAATCTAATAGTTTCATGTTGTGTTATATGTATGTTTGTAGTCTTTATAGGGGTCCATACTCCGTATTAGTGGCGTTTCTATAAAACCTGTCACGAATTATTATAAATCGTGATGATTGAGCTAGTTGTCCGTAACATTATTACTTACGATGGGTAGTGGGAACCCCTAAAATATTACTTTTAGTGACAACTTTTAAGGTGGCCATGAAAAAAACATCACCGAAAATCACATTTTTCATAGTCGATTTTTTAAAAAAATTATCTTAATTTTAATTTCATATTGTTTTTCCAACTAACACTTTAAACATTATTATTATCTTTAACACTCAAAAAGTTCATCAAAATGTTTCCGATTTGCCAATATTAGTAACAATAATGATTATATACCCAGTTAAATTGTTTTCTATAAATTTATCAATATTAAGCTTGATATTTATTAAAAAGATAGACATATGAATCTTTATTATATAACTTTTTGAAAGATATTTTTCATTAACCAACCAAATATTAAAACACAATTTCATAAAAATATTTCCCCACAAAAGACTTGTGTGAACAAAAAATCGGATTTATTCAAGCTTATTATTTTAATAAATAAATAAAATAATTAAGGTTTGTTTTGTGCTTTTTTTATTTTTTTTACGTTGTGTTATGTTTATCCAAAAATATTTCGTCATCGATCGTGTCGGTTTAGTCCGTTGTGTAGGGTGACGAAAAGGGTCATGACTTGCGTGCTTAACGTGGTATCATGTCATATTGTATTTTTGGTGAACTTCATCATTTTGTTTGGTCCTTTTCAATGTGTCCACTTTCTTTGTGAATCCTCTTCCTTGGACGGACTTCAGAAATTCAGTAATTTTTGTACGGTTATATATTTGTATTGAGAAATTCAGTTTAAAGTGTGTTGTTGGTTCAGTGATGATAGAAAAGGCCTAATTTGAGCTCTTTCAAGGGGGGGGGGGGAATATTTCCGTTGATCTTGTATAAGGGTGACATGGCACCTAGAGGCGAACTCACATGTATAGGTGGGGTGCACGTGCACCAGTTAACTTTGGAAAAAATCATGAACATATGTATATTTATTTTGAGAAACTGACAACAACTAGATATAATTAACAGTGTTCCCATGAAAAGTATAGAATGACACTTGGAACAGGCCAATGCATCCGCCCTCGCGACCTAGGTTCGAGCCTAGTCAAAACCACATTTTATAGTTTTTATTATAAGTGATTTCTAGATAAGGGATTCAAGTTTAGAATCATTCAGTAGAACAAACCCATGATGCACCCAAAGTCTTCAAATCTTAGATTCGCCTCTGATGGCACCACCATTACCATTTTTATTTTTGCTCTTTTTCTCCGTGATTTACTACTTATAATTGTTTTAACCTTTTAGGGACCATGTCCACTATTGCGCGTGAATTATTCCCTTGGCTCTTGAGGCTTTTGATTTTCACTTTATGTTATTTCTTTGGGCCCACAAAGATTTTCCCAAAGTATAATATATGAATCTTCATGATTATAAAACCCTTTAATTTATTAGTCCTAGTACTATATTCTTTATTAAACATATAAATTCAAAAGTCAAATACTTTGGGCTATAAAAGCCAAAATGTTATGATAACTCGATTTCTTGTGTGCAAGATAGTTGTGATCTCACTTAATTATCTTACTTGCTAGTGCTACTTGTAAAATAAAATCGAATCAAATATAATCCTATCAAATATGTTTAGGATTTAATCTTTTTTACACTATATATAATAGTAAATAATATAAGATTCAGATCACTAATCTAACTTTGTACATTTATATATTTTTATGATGCAGAATATGCGTATACGTTGCAAGTGGACGAAAAGAGTGACATTTATAGCTATGGAGTGGTGCTAATGGAAATTTTGTCCGGAAAAAGATCGGTAGATTCGGAATTTGGCGATGGAAATAGCATTGTGGATTGGGTAAGGTCCAAGATGAAGATTAAAAATGGTATAAATGATGTTTTAGATAAAAATATTGGTGCTTCGTGTCCTCGAGTGAGGGAGGAAATGATGTTGATGCTTAGGGTGGCGTTGCTATGTACTAGCCGGAACCCGGCTGACCGGCCTTCCATGAGGGATGTTGTGTCCATGTTACAAGAGGCCAAGCCCAAGAGGAAGTTGCCCGGAAGTGGTGGTGACAATTCAATTGCTCTTGTTTGTTTGGCACAAAAGGCCAATGTGGAGTGTTAATATAATTGGAGATTTGGGGTTTCTTTTCTTTAGTCTTTTTTTTTTTTCCAATTGTGGGGGATTGGTATAATCGTATAGGATAAGTACTCAATGTTTCCTTTTAAAACTTTAATTAGTGGAATTTTAGGTATTATGATTTCTGAG

>PGSC0003DMG400007517 LRR-RLKs

GGCATGTTGTTTTTGCTCTCACTTTTTTACCACCAAAAACCTCTGTTTTTTTATTTGTTTTTTTGAAAGAGTCTGACATAAAGCTGTTTGAAACTTTGAGTGAGTGAGTGAGTGAGTGAGTGCAAGTTTTCTGTAAAATGAAGCAGCAAAGCATGAGGAATTGTACCACTTTCCAATGATTTGTGTTTTTATTTGTGAAAAAACCAAGTCCCAAATTCTTCTTTTATAAAAAAAGATCTCATTTTTACTCTTTGTTTAGCTGTTTTCTTGTTTAAGACAAATGGAAACAAGAAGTGTTTTTGGTTTTGTGGTGTTGTTGAGTTTCTTGGTTTACTCTAATGCTATGCTTACTCCTGGTGGTGTTAATTATGAAGGTTAGTTCTTGTTATTTTTAATAAAGATTTAAACTTTGAGACATTATTGGAATGAGATTGATCATATTAGAAAATATCAAGAATTTTAAAGTATGATTTACTCCTGTTGGTGTTAATTATGAAGGTTGTTTCTTGTGGTTTTTCTCATAAAGATTGAAACTTCATGACTTTTTTGTTGAAAAAATCATATGGGGTGTTTGAAATTTTTGAAAGGAATGAGATTTTGATCAAATTTTAGAAATATCAAGAATTTTGAAATATGCTTACCCTTGTTGGAGTTGATTATGAAGGTTAGTTCTTGTGGTTTTTCTCATGAAGATTGAAACTTTATGACTTTTTATTTGAAAAAATCATATGGGGTGATTGAATTTTTTGAAAGGAATGAGATTTTGATCAAATTTTAGAAATATCAAGAATTTTGAAATATGCTTACCCTTGTTGGAGTTGATTATGAAAGGTTTAGTTCCTTGTGGTTTTTTCTTCATTGAAGATTGAAACTTTATGACTTTTTATTTGAAAAAATCATATGGGGTGATTGAATTTTTTGAAAGGAATGAGATTTTGATCAAATTTTAGAAATATTAAGAATTTTGAAATATGTTTACCCCTGTTGGAGTTGATTATGAAGGCTAGTTCTCATAAAGATTGAAACTTTGATACCTTTTTGTGAAAAATTCATAAGGGGTTTCTGATTTTATTTTTTTGAAAGGAATGAGATTTGAACAAATTTAAGATATTTCAAGAATTTCATCTATTTTAACTTTCTTGTTATGTTGATTTTGGACAGTGCAAGCTTTAATGGAGATCAAGAACTCATTACATGATCCACTTGGTGTTTTGAATTGGGATGAAAGTTCTGTTGATCCATGTAGCTGGAATATGATCACTTGTTCAAATGATAAATTTGTCACTGGCTTGTAAGTATACCCGATTGTCCGGTATCTGCATTGGAGCCCGACTGACAAAACCTTTCTATTCCTAATACTTGAACCTGAGACCTCTAGTTAAGGATGAAAGTGCTTACAAGTTTTCTTGAATTGTGCAGGGCTTGTTCAAGCCAAAGTTTGTCTGGGAAAATATCACCATACATCCATAACTTGACTCACTTGGATCTACTGTGAGTACTTTATATGAAGAAAAAATAATGAAATTTCAAATGATTGATTATTTGAATCAAGAATATAGAAAATAGTAGTACTAATATTACTTTTTAACTTTTTTTTTTAAACAGGCTTCTGCAGAGCAATAATATAACAGGACCAATTCCAACTGAACTTGGAATGCTTCAGAAACTTAATACAATTGATCTTTCTGATAATCAACTCACTGGGAAAATCCCTTCATCTTTAGCTCAATTAAAAAGCCTTCAATACTTGTAAGCAAAGAAAATTTTAAAGATTTTGATATAAAAATGAGTTCTTGAATTATAAAAAGTTAATGATTATGGTTTTTTTTTCTTTTTCAATTGTGTTATAGGAGACTAAACAACAATAGTCTTAGTGGAGCTGTTCCTTTGGAATTGTCTAATATGACTCAACTTACTCTTCTGTAAGTTCCTTAGCCTCATTTAGTTTTGTTTATTTTCTCATACTTGAAGGGGAGCCTCGGAGCAACAGTAAGCTTGTCTCCTTGTTATCTATATAGGTCACGCGTTCGAGTCAGGGGTAGGCTGCCTATAGTATACCCTTTGGGGTGCGGCCTTTCCCCAGACCTTGTGTGAACGTGGAGTGCTTTGTGCACCTTTTTCCTAATTGCTTTCTGTTTTCATGAGTATATTTTGATGGGATTTTTGTTGTTTTTTTCTAGAGACTTGTCTTTTAATAATTTGAGTGGTCCTGTGCCAAGGCTTCTAGCTAAAACATTCAAGTAAGTTTCTTGAACAATTCTTTGAATTTCATTTTTTTTCCCTTTGCTAAAGATTAGAGAAAATAAATCATTTTTTCTCTCATTGATGCAGCCTTTTGGGAAATCCTATTATATGTGCAACTGGAAAAGAACCAGAATGCAATGGAACAACAACATCCATGCCTTTGCCCTTCACTTTGAACAACTCACAAAGTAAGAATCTTGAATCATCCTCCTAAGTTTTTTTTCGAGCCAGCTTGTGTGCCCCTCGACTATTCCACAAATCTTGGTTGTTGTCAAGTCTCCTGCTTTCAACGTCGTTATGATTAACTACAACTCTATTATAAGGATTCATCATGAAGTCTTGGATAGTTTCTGTGTTGTACTATTCACAACTGAGAAGTCTCTTGTTATATCTCAGATGCTCAGCCTTTTGGAAGATCTAAAACTCACAAAATTGCCTTAGCCTTTGGGACAAGTCTTGGATGTATCTGCCTACTAATTATTGGGTTTGGAATATTTCTATGGTGGAGACAAAAGCACAACAAGCAAATTTTCTTTGACATCAATGGTGAGTATGATATGCAGCTAAAACCTATATTGCTCGAATCTCGAGATACTAGTAATGTCTTGCTTTAACCAAGAGGATCATGCTTGTTGATAATTTCAGAACAACATCATGAACAAGTGTGTTTGGGAAACTTGAGGAGGTTCCAATTTAGGGAACTTCAAGTTGCCACAAACAACTTCAGCAACAAGAAGATACTGGGAAAAGGCGGTTTTGGAAACGTGTACAAAGGTTGTCTTAGTGATGGCACGATTGTTGCTGTAAAAAGGCTCAAAGATGGAAATGCAATCGGAGGAGACAAACAATTCCAAACAGAAGTTGAAATGATCAGCTTGGCAGTGCATCGCAATCTCCTACGTTTGTATGGGTTCTCCATGACACCAACTGAAAGGCTTCTGGTTTACCCTTACATGTCTAATGGAAGTGTTGCTTCTCGTCTCAAAGGTAAAAAGAAACAACCTTAGAGAACTAAAGATTCTGCAAGTATTCTTAAATATGGTGGGATGGTTGAGATACCGTCATCCTTAACCATAGTAGGCTTATCTGTGTGAACCGGAATAGTCGAACTAGTGGACTTCAGATACCTGATGGTAGATTTCTCTATTTCTGTTGTTTCTTGCAGCCAAGCCACCATTGGACTGGACTACAAGAAAAAGAATTGCATTAGGAGCTGCAAGAGGTTTACTATATCTACATGAACAGTGTGATCCAAAGATTATCCACAGGGATGTGAAGGCAGCAAACATATTGCTCGACGATGACTGTGAGGCTGTTGTGGGAGATTTTGGATTGGCAAAGCTATTGGATCATCGTGATTCACATGTCACAACTGCAGTTAGGGGAACTGTAGGACATATAGCTCCTGAGTATCTCTCTACAGGACAGTCCTCTGATAAAACAGATGTTTTCGGGTTTGGAATTCTCCTGCTAGAGTTGATATCTGGCCAGAGAGCTCTAGAATTTGGTAAAGCAGCAAATCAGAAAGGCATTATGCTTGATTGGGTAAGTATTACACATTCCAAGATGTGACAAAACCCTCCGTTTCAATTTTTCTTACTTTTCGTTTTAGTCTGTTTAGAAAAAAAATGCCTACGACCATTCTATATCTACTAAGTATGTTTTTTTATTTCAACATTCTATTTTACCTTCAAAGGGTATTTGGTGACAAGATTCAAAAGTCGTATTAACATACTGAAACTTTGTGCTCGATCAAACTAAGACAAACAAAATGAAACGGAGAGAGTATCTCGATTCTTGTTGGGGTTGATAATATCAATGTGCATATTTGTTGCAGGTAAAGAAGATTCACCTGGAGAAAAAGCTTGATTTGCTGGTGAGCAAGGACTTAAAGGACAAATATGATCGGATCGAGCTTGAGGAAATGGTACAAGTAGCTCTAATATGCACTCAGTATCACCCTAGTCATAGACCAAAAATATCAGAAGTGGTAAGAATGCTCGAAGGAGACGGACTTGCAGAGAAATGGGAAGCTTCTCAAAGAGCAGAATCTACGCGCTGCAGAGCCAATGAGTTCTCCTCTTCAGAAAGATATTCTGATCTTACTGATGATTCTTCATTGCTTGTTCAAGCAATGGAATTATCAGGACCAAGGTGATGTAGATTATACAAGATAGTCAATTAGATGAAACAAAAAAGGGCATCTGAGATAGAATGCTGAAGTAATATATGTATGTTATTGAGCTTGTTTTTTATTTCTTCTTTAGATAACTGAGAAGTCCTCGAGGGTCAATAGTGCACGGTTCATAACTCAGTGGATCATAGACACCCCCCTTTACTCTTCTCCACTTATATACCAGACGTGGTAGGGTTCGATTGAGCTTGTATATATGATCATATGGAAGCTGCATAAAGGAATGGGATATGTTCAGACTGTCTAGGAAAAAGATTTATGAGCAACTGACCTCCCCAACCACACCCACAACCCCTACTTTTTTTTCCTCTTCCTGTTTTGAGTGATAATTTTTCTGGGATGTGGAATTGTAACTTTGAGCTTGTAAGAATAATATTGTTGAAATTCAATTTCAGCTTATAAATATTGTATCATAATTCTCATTGAAAAATATTGTTCAAAGGGATTATCTTTTCCATTCAAAA

>PGSC0003DMG400007634 LRR-RLKs

TATATATTCATTTTTATCAATTTGGCATCTTCTTCACACCTCCAATGTTGACCAAACTTTCCAAAATCAACTCATAATTCTAACCCCCACCTTTTTTTTTCATGGAGTTTTCCCATTTTCCGACAGGCATCTTCGCCGGAAAACAGTCCCTTTATCTCATTTTACTACTAAACTTCATACCCATTTTCGTAACTTCACAATCTCCGGCCACTACTGAACGTGACACTCTACTGAAGATTAAGCGTCAATGGGGTAACCCATTAGCACTTGACTCATGGAACTCTACTTCTTCTCCTTGTAGCTGGCCGGAGATTGAGTGTGATGATGGGAAGGTTACCGGAATAATAATTCAAGAAAAAGATATTACTGTTGAAATTCCAAGTTCTATTTGTGAACTTAAAAATCTTACTTTTCTTAATCTCCGGTTGAATTATCTTCCCGGGAAATTCCCAACTTTTCTTTACAAATGTTCCAATTTGCAGCATTTGGATCTATCTCAGAATTACTTTGTGGGTTCCATTCCGGAGGATATTTACCGGCTTGGGAAACTCAAGTATCTTAATCTGGGTGGTAATAATTTCACCGGAGATATCCCGCCGTCAGTCGGGAATTTGACGGAGCTGGAGACCTTGTGTATGAATTTGAACTTGTTTGATGGGAGTTTCCCGGCGGAAATCGGCAACTTAGCCAATCTTGAAAGTTTGGGGTTGGAGTTTAATGAGTTTTCTCCGATGGCACTACCGCCGGAGTTTGGGAAGCTGAAGAAGATAAAGTATATTTGGATGAGAGATACGAAATTGATCGGTGAAATCCCTGAAAGTTTCGGGGATTTTGAAAATCTTGAGCTTATTGATTTTGCTTACAACAATCTGGAAGGAAAAATCCCTTCTGGGTTGTTTCTGTTGAAGAATTTGACGATGATGTACTTGTACGGTAATCGGCTTTCAGGTCGAATCCCTGAAACTTTTGATTCATCAAAGTTGATGGAACTTGATGTTTCCAACAACTATTTAACTGGAACAATCCCAGAAAGTTTTGGAGGGTTTAAACATTTGGAAATTATGAATCTGTTTGGTAACCAATTGTATGGAGCAATTCCAGAGAGCATAGCCAAGATTCCATCTTTGAAGGTATTCAAAGTGTTTAGGAATAAACTGAATGGAAGCTTACCATCTGAAATGGGACTTCATTCGAAACTCGAATCTTTTGAAGTTTCACTGAATTCCTTCACTGGAAACTTGCCAGAGCATTTGTGTGCTGGAGGAACTCTGTTTGGTGCTGTAGCTTATGCTAACAATTTGTCTGGTGAAATACCAAAATCACTCGGAACTTGTTCTAGTCTACGTTCGATTCAGCTCTACAAGAATCAGCTTTCGGGTGAGATTCCATCTGGGGTTTGGACTTTGGTTGACATGACAAGTTTGTTGTTGAGTGATAACTCATTTTCCGGGGAGCTACCAAGTAAAATTGCATTCAATTTTACGCGGTTAGAGATTAGTAACAACAGGTTTTCTGGTGAAATTCCTGTGGGGATATCTTCTTGGAGGAGTTTAGTGGTGCTTTTGGCAAGTAACAATTCATTTTCAGGTCGGATTCCAGTGGAATTGACTAGTCTTTCTCAGATAACTCAGCTAGAGCTTGATGGTAATTCGTTATCTGGAGAACTTCCAGCCGATATAATATCATGGAAGTCGTTGACAATATTGGATCTTGCCAGGAACAAACTCTCTGGCAAGATTCCAGCAGTTATAGGTTTAATCCCTGATCTTGTTGCTCTAGATTTGTCTCAGAATCAATTTTCAGGTCCAATTCCACCTCAACTAGGTGTAAAAAGGATCACTAGCCTTAACGTATCATCGAATCAACTTACCGGGAATATCCCTGATGCATTTGCTAACCTAGCCTTTGAAAACAGTTTCTTGAACAATCCCAGTCTTTGTACCACCAACTCACTTCCATATCTTCCAAGTTGTAATAATGCGAAAGTCGCCAATTCCAAAAGATTGTCCCATAGAGTCCTAGCCCTAATTCTAGTCCTTGCATTCGCTGTTTTCCTATTTAGTGTTGTATCGACCTTGTTCATGGTTAGAGACTACAGGAGGAAGAAGCATAAGCGCGATGTTGCAAGCTGGAAGTTGACTTCATTCCAGAGGTTGGATTTCACAGAGGCGAACATTTTGTCAAGTTTGACGGAAAATAACATGATTGGAAGTGGTGGATCGGGTAAGGTGTACAGAATTTCCATTGGCAGACCAAATGAGTATGTTGCTGTGAAGATGATTTGGAGTGATAGGAAAGTAGACTACATTCTGGAAAGGGAATTCTTGGCTGAGGTTCAGATATTGGGATCAATTAGACATTCCAATATAGTGAAGTTATTGTGTTGTATATCAAGTGAGGATTCAAAGTTGCTCGTTTACGAGTACATGGTAAATCACAGTCTTGATGGATGGCTTCATGGGAAGAAGAGAGTTTCCTTGAGTAATAAAGTCATGGATTGGCCAAAAAGATTGGAGGTTGCCATTGGAGCTGCTCAAGGACTTTGTTATATGCACCATGACTGCACTCCTCCGATCATTCATCGCGATGTAAAATCAAGCAATATCTTGTTGGATTCTGATTTCACAGCCAAAATAGCTGATTTCGGGTTAGCCAAGATTTTAGAGAAGAAAGGAGAGCTTAACACCATGTCTGCTGTTGCAGGTTCCTTTGGTTACATAGCTCCAGGTAAAAGCAAAATCCAATCTTGAACTTCGTATTACCTCAATTTCGTCAAAAGAATGACTAACTAATAATCTACTGTTTGGTTGCAGAGTATGCTTATACAACGAAAGTGAACGAGAAGATTGATATCTATAGCTTCGGGGTGGTGCTATTGGAGCTAGTGACAGGAAGACAACCAAATTTTGGGGACGAGCATACAAGCCTGGCAGAATGGGCATGGAAGCAGCACGGAGAAGGGAACACTGCCATCGATAATATGTTAGATACAGATATCAAAGAAACATGTTACTTGGAAGAAATGAAGACTGTGTTCAGACTCGGGCTTATATGCACGAGCAACTTACCTGCAAACAGGCCTTCCATGAAAGAGATTCTGCAGATACTTCATCGATGCAAGTCCTTTAGGTATTCTGGAGGAAAATCACCAGATACAGAGTATGATGTTGCTCCTCTCCTTAGCGGGAACAACAGCGAAAAATACATTGCTAGTTACAAACGTATTAATTCCAATAAGGTAATAGATGATAGTAGTGATGATGGTCTAATAATTTCCACTGTGTAATATTGGAGATGCCAAAGTAAACTGAGTTGTTGTAAAATGAGTGCCACTTAAGTAACGTACAAGTTTTGAGGATGTTGTTAGAATAGAAAACCCCTATAAGAACGTTGCTTTTCTAGTGTACTCTTGAATTTGTTCACTGCTTGTAGATTGAGTGAAATGCATATTTAACTTTTGCTAGTTATGTAGATTGGAAAAG

>PGSC0003DMG400007709 LRR-RLKs

CATCAATCCAATGCAATCCCTTCAGTCCAATCACTCAATAGAAAGTTAACTTTGTAAAACCCAATTGCTTACTTCGTTTCTTTTCACATACATAATAAACAAAGGGATTCTCTTTTAAAAAAACTGACCACATCTCTGTATCTGTGAGTTTTTGGCCTATGTTTTGGCTTTTTCTGTTGTGTTTGTATCTTTGTGGTATATATTACTTAGTTAATAATAGAGGGAGAGGATGTTGATCTTGAATGGATATGGTGGAGAGTTTCATTGAAGGAGAAAAATAGCTCAAACATAGAATTTGTTGGGATTAGTTCATATTCCAATATTATTAGATCTCAAGGTTACCACCTTAGTCTTTTTTCTTTATCAAGAAGATTCCCTTTTTATTCATTCATTCTAATTATAGATGGATGCCCATTTCTCATAATTGTGTAAAGATAGCAACTTTGGTGTTTTGAAGAAATATTGGAAGCTGAAAAGAGCAGGATTTGTGGTTGCTATTGAGTGAAAACAGCATCTATCTTTCTCTCTCTTTTTTTTTTAGCAAATTGGTTCTGTTCTGATTCTGAGAGTGACCCTTTATGGATGAAGAAGTGGTTTTGATGTAAAGAATGTATTCTTGTTGTCTTAATGTTTTATGGGGAGGTTTTGTTAGGCAAGTAATCCATTTTCTGACAAGGTAAATTGTAGAAAAGTAGAGAATGGACAGAAAGTGGTGTATGTTCTTGTATGTGGTTTTAACATTGATGGAGAGTTGTTATGCTTCTCTTTCCCCTACTGGTATCAACTATGAAGGTAAACTTCTTTCCTACATTTCCTTTCTCTCTTGGGGGTGAAAATCATCTTCATCTCACATAATTATATGCAGTTGTTGCCTTGATTGAAATTAAGAAGGCCTTACATGATCCTTACAATGTTCTTGAGAATTGGGATGTCACTTCTGTTGACCCTTGCAGTTGGAGAATGGTTACATGTTCAAATGATGGATATGTTTCTTCTCTGTAAGTAGTTGAGCTTCAGTAATTTTCACTCTACATAATCATAACACCTAAAGTAAACTCTTTTTTCTTCCCTTCATCAGTGGACTACCTAGTCAAAGTTTATCTGGAACCTTGTCACCTGGAATTGGAAACCTCACTAAATTGGAATCAATGTAAGTTCTTCACTGTATTTTTCTCTTAGTACTTCATTTGATTATATGTTAGCTAAATAGTAATACTATTCCCAATATCGTGTAGATTGCTGCAAAACAATGCTATTTATGGTCCTATTCCTGATGTGGTTGGTAATTTAGAAAAGCTTCAGACACTTGATCTCTCCAACAACAAGTTTGATGGCGAAATACCTGCTTCTTTCGGAGATCTCAATAATTTGAATTATTTGTGAGTTCAGGAAATTTTTCTTATTTTTCACTGTTTCTTATCAAACTACACTTGAATCTTATTGTGTATCTTCGCTGTTGAGCAAGACAGACGATTAAACAACAATAGCCTCACAGGAACCATTCCTCAATCTCTCTCAAACATTGGAGGCCTCGCTCTCGTGTACGTCTCCATACCCTCCTTTACCATCGAAATTTGTTATGCTTATCCTTGAAAGACCGGTCTAAATTACTCTAATGTTCTGTTTCTTCCTCGTACATTTTACTGATTTAGCTGATCTTGATTGTAGGGATGTTTCTTTTAATAATCTAAGTGGTCCATTGCCCAAAATATCTGCAAGAGCTTTCAAGTGAGTCACCTTTTCTTCTTTCAATACATAATTGCTGCAGTTGAAGTAGTAATTCTCAATATTAATGAAATTGACTGGTTCATGACTCTGTATATATACTTCTCATTAACAATAATTATGTAGTGCTTCTAGGATAATGGAAAGTATTTATTTTTGATAGATTAGTGAACCTTATTATATTGAATTTAAGTTTCTCATCATGATTTTCAGAGTTATTGGAAATCCTTTGATTTGTGGCCAAAGTTCGGGGAACAATTGTTCAGCAGTCTATCCAGAACCGCTGTCCTTCCCGCCAGATAGTTTAGGAGGTCTGCATAACATCACTCACTTTCATTCACCTTCCGTTCCTTTCATTTAAAAATGGAAAATAGTAACAGACACAGGAGGTTTTGCTATCTCTATTGTTGAAATTGTATTGTCTGTTTTCTTCAATTTCATAGATCAAAGAGCAGGAAGTAAAAACCATCATGTGGCTGTTGCTTTTGGAGCAAGTTTTGGTGCTGCTTTCTTGGTTTTAGTAGTCATAGCACTGGTTCTTTGGTGGCGCTACCGACATAATCAGCAGATCTTCTTTGATGTCAATGGTTAGCTGATTTTCTTTTGCTTTTTATCCATTTAAGATAATTCAATTTCTTTGATTTTTTGTTTTTTCTATTTCTGATTCATTTGGTTATGTACATATTTTCTCGAATGTATTAAACTCCGTAGATTTGTCATTATTTGATTATTTTTTTTCAAGTAAGGGTGTTTCAGAGTTTGGTGGCTTGTGGTTTAACTGGTTATCTTAGCTTCTCAATATACATCTTTCATTTAATCTGACATAGAAAAGGTTTACATTGACATTTGATTGTAAATCTAGTAGTCTAGTATTATGAAGGACCCAAATTTGCAAGTAGTAGATATCTAACAGCAGTTCTACATTTACATCGAGAATATGTTGAATAAAATGCTATAATCCACATCTTGGACTAGATAACTTCCCTGCTTTAGACCAGCTTAATTCATATCCTGACCAACCTTGTCAGGTGAATCAATTGTTATTTGCTCCCCCTCCCATTGAGAAGAAGTTTTCCCTGACAAACTTATAGGGTGGTTGCATATTGTGGACAAATTTTGGGGACTTGTCACTACTCATTCTTTCTCATCCCTTTCCTTGAGCATTAGTATTTTTCGTCCGTCTGTCACCTGCTATTTCTACAGCTGTATCTGAGCAATGATTTCTCAAATGCAGAACAATATGATCCAGAGGTATGCTTGGGTCACTTGAAAAGGTATGTCTTTAAAGAACTGCGAACTGCAACTGATCATTTCAGCTCAAAAAATATTTTGGGTAGCGGAGGATTTGGAGTTGTGTACAAGGGACGCTTAAATAATGGAACTGTCGTGGCTGTCAAAAGATTAAAGGACTACAATGCTGTTGGTGGTGAAATCCAATTTCAGACAGAAGTTGAGTTGATTAGTTTGGCTGTCCATCGAAATCTTCTCCGTCTTTGGGGTTTTTGTTCAACTGAAAGTGAACGGCTTCTTGTTTACCCCTATATGCCAAATGGAAGTGTAGCAGCACGATTAAAAGGTCAGCCTCTTGGCCCGCATGTTTTGTTAGTAATAATTGCTATGCCTCCTAGCTTCAGATCATTCCTCCAACATATATGTTCTGTGAAGTTTGACCTTTAAGGAAGGTGTTTGTTAACTAATTTCAACTGTGGTTACCTTTCTTTTACATTTTTAGGTGCCTGGTCACACCTTTCTAACTCATTATGGGAAGTTAACTAGAAAATACTCTCTTAAAGCTGCACAATTTTCTATTACAGAAAGTGAGGCTTAACAGACTAACGTATTATCTTACAAGAACAAATCTTGTGAAGAAATGGACATTGGATCTAATTCAAACTTAAAAGTTAGCTCATGAGGTAAGTATTGCTCAAAGACCACAACCCATTCCCTCAACCTATGTGGGACTTCTTAAAGGTAGATGAGCTTAGCTGATCTTTAATTCCATACGCTGTAAAGTGTAAATCAGTTTCTTTTCTCCGATATGAGAGTGTCACTAGGGCTTGAATACCAACAAACAAAAATTGGATTCTTTTCCCTTAAGCTGGGACTTGATAATATTGCTTGTTTAAGGACTCCCCTCTTTGCCTTCTCTCTATCCAAGTTAAAACTTGTTAATTTTAGAGAGGTTATCCTCTTCTCATTCTCTCAGTTTTATGCAATTTGGAATAATGAAGGAAGAAAGTGAATTAAAATTAAAATTTCACCATTGCACATGATAAAGCTTTTGAAAAGAGAAAGAAGTTTATTCATCTTATACAACAATTTGCTTCTAAAAAGCATTATGAGACTTGTCATAATTTATGGCTTAAGGAGATTAAAGTGGGTTTGAACCATTCACGTAGTCCATATTTTACCCATATTTATGTAGATGGATTCCAGTTTATCTTCCTTTAACCCTGAAAAGAGCATATGCATCCCGAGCTTTTTACGTTACGCTTCAATAGAACGAATTATCTTTACTTCAATAATAGATATTAGGCCTTTCAGCAGGTTAGTTCACGAATTCTCCTATTTCAGTAAGAGACGTGTTTGGGCCTGCTTTGAAGACAGCCAGTAACTACTAGCACAAGATCCAGTATTTGAAAAGGTTCCATACCCATAAGGGTTCCAAGCTTCAACCCACTGCTCGAGCAACTTGCTATCTATTTTAACTCAATCCACCCAAATCCGGTCCAATGCACCCATTTGCCACCACTATAACTGATACTATCTTTTATTTTTTAGTACAAGCTTTTAATGTTTGACCCATCCTATTTTGTGTCCCTTATCATATTTCTTCTTCTTGAAGTATATGGTAGGCCATCATCAGCGACCACAGTGATGCAGTTTATTCTGGCGAACATTGCCTTCCATTAACCATGTCTTTTGTGGTTCCTAAATTTGCAGATCACATCCATGGCAGGCCAGTTTTGGACTGGTCAAGGCGGAAAGGAATAGCAGTAGGTACAGCAAGAGGGCTAGTATATTTGCATGAACAGTGTGACCCCAAAATTATCCATCGTGATGTCAAAGCAGCGAACATTCTGTTGGATGAGGAATTTGAAGCAGTTGTTGGAGATTTTGGGTTAGCAAAGCTCTTGGATCACCGGGATTCTCATGTAACGACTGCTGTCAGGGGCACAGTTGGTCACATTGCTCCAGAATATTTGTCAACGGGTCAATCGTCTGAAAAGACTGATGTGTTTGGGTTTGGAATCTTGCTGCTTGAGCTAATTACAGGTCAGAAGGCTGTGGACTTTGGACGAGGGGTGAACCAGAAAGGTGTTATGCTTGACTGGGTAAAGAAACTTCATGCAGAGAAAAAGCTGAACCTTATGGTGGACAAAGATTTGAAGAACAACTTTGACCGGATTGAACTTGAAGAGATGGTTCAAGTTGCTCTTCTGTGTACCCAATTCATTCCAACTTATCGGCCAAAGATGTCGGAGGTATTAAGGATGTTGGAAGGAGATGGATTAGCTGAAAAATGGGAGGCTTCACAAAAAGTTGAGACACCTAGGTACAGAACATCTGAAAATACACCGAAGAGATATTCTGATTATATAGAAGAATCATCACTAGTAGTCGAAGCAATGGAGCTTTCAGGACCTAGATGACAACATCTGATTCTATCCTTGAGTTGAAATATGAGAAATAGCTGTTGGAATGCCTGTATTATTAATCTGTAAAAAAGAACACAAAGCTCATGTATAATTCGATCAACACCTAACTTCATACTTCTCTCCCTCTCAATTTTGATAAAATGTGCCATTCATTTTTTGAAGATGGGGGCACAAGTCAGCCATCCTCATGTACAGTGGTCTGATTGTCATGGTGGATGAATCATGAATAAGGTTTTCGGTTTTCTTCTTTTGTTCATGTACTGAAATGTTTAGTCTCGTCATAGTGAAGGGAAAAATGAAAATATGAAGAAAGGTAGTAGTAGGTAGGGGTGTCAAAAATGAGCCGTTTAATCGGATCATAATTTTATGGGTTGAGCTCAACAT

>PGSC0003DMG400007862 LRR-RLKs

TTTTGTTATTTGTGGGTGATTGAGTCTTTTTCTTGCATTTTTATGCTTGAGGAAAACACTGACATTGATGTTTTGTTTTATTTGGTATTTGCAGTTTTGGGAACTTTTTTGTGTTGGCTATGCAGAATTGTTTTCCAAAGAAAGATTGAATTGAATTGAAGAGAAAAGATTGTATTTTTGGGGCAATGGATGGGAAGAGATTTGTTGTATGCTGTTTGAATTTGGAGATTTTGGTGTGGATTCTTTTAGCTTTTGCTGTTCAGCTTTCAGATGAAGCTAACCCTGGAGATGGTAAGGAACGACAGTCGCCAAGTCTTCGTAGGCGTTTGGACATGAATTGAGTGATATACGAAAAAAAAAAGATTAGTATTTGATGTTCAGTTGAAAAATAGTAGTATTTAGAAACTGAAGTTGTGTTTCGACATGCATTTTAGTTTTTAAAAAGAGTTGTAAAACAATGTTTTGAAAAGAGGGATAACAAATTCATGTCCAAACGGGTAAGTAATTTCCTGTGTAAATGCTCATAAGGGAATTCCAATTTCATGTTTTGTGCAATGCTGATTATGTGAGAATATGTAATTATCTAAGCGAGTTGTTGATTTGAGCTCAAAAGTGAATGCATTTGCTTATTGTCCTTGTATATTTGAACTAGTAGGTGGTCTTCTTGCTTCTTATAAAGTTTGTACCTTTTGTTTAAAAAAAATTGAGGATAGGGTAAATGGTCGAGGGATTACAAGGTGGAGAATCGAACTCTCATCAATAAGGTAAAAGTTCTAGTAGCCAACCAGTTCAGCTACTAAGATTTCCCTATAAAGTTTGTACCTTCACTTGCATTGTCCTGACATTTTATTGGGTTTGTAAAGTTGAGGACTAACAGCTTTTGCGCCTATTCTGGTTAATTGCCTCTCAGGGAAGTTCGTAGTTAGCCTTTGTTAATTGATTGGGTTCTAAATTTCCACATGGTTTGTTTCTGTAATTGATTGGATTCTAAATATTTATATGGTTTCATGGATCAGTTTCTGCACTTTCTTCTTTATGTTTGGAATTTCTTAACAATTCTCCTTTTGTATTGGTTAAGTTGCTGCTATTAATAACTTATATGCTTCTCTGGGATCGCCATCCCTTCCGGGCTGGGGTGCTGTTGGTGGAGACCCATGTAATGGACTTTGGCAAGGGGTCGAATGTGACAATGGAAACATACTGACAATGTAAGGACTTACCATTTACTACATTGCGTAGATTTTTGTGGAACTGAAATGCTAATCCTACTTATCATGTTGTAGTCGACTCAATGCTGCTAACTTGGGAGGTGAACTTGGGGATAACTTAGGCTCATTTTCTTCTCTCAAAATAATGTAAGCTTCTCTCATATCTCCTAACTTAAAAGTTTGCTTGTTTGTATGCATGTGTTTTTCATAGACATAATACGTGAACACACTCAAACTTGGCCTCAGTTCGCAAGTAAACACTCCAACTTAGAGAGTGGACATCTAGACACCTCAACTGGAAACTAAACACTTCAACTGGTCCCCACTGCATCTCGTGGTTTGGAGGTGTTTAGATGATCATTTTGTTAGTCGGAGTGTTGAACTGACACGAGTTGAGGTGTCTGGTGCATACTCAAAGTTGGAGTACTTACTTGCCAGTTGAGGCCAAGTTTACGTGTCTGTTATGTATTATGCCTTTTTCCTATTATTAAATGTTTCCATTTAGTGGAGTCCACTGTAATGATATGTGAATACTAATTGTGGTCATCTCCTGTGGCAGAGATCTGAGCAACAACCATATTGGCGGAAGCATTCCGTCTAATCTGCCAGTTACCTTGCGGAATATGTAAGTTCTACCATTTGACTTATGAAGGATCCCCAGAAACACGAGAAAAAGAAAGAGGACATATTGATAAACCTAGCTTTACTTATTGAGTAAAACAGTTGATATTCTTTACCTTGGACAGTTTTCTTTCAGCTAACGCATTCACTGGAAGTATCCCTACTTCTTTGTCTTCCCTGAGTCAACTTTCAGCTATGTAAGTATCACTTTTAGTGTAGATAAAAAGTTGCTTACTTCATAAAGAATCAAACTCAGGTATTTCGCTGTGTGTCTGCAGGTCTCTGAATGCCAACCATTTGACTGGAGAAATTCCTGATTCCTTTCAAGGCCTTACCGCTTTGGTCAATCTGTATGAAAGCAAACAAAAGCTTTTACATACTTCCATTTAGTTTTTTTCATATGAATAACTGGTTTGGGTGTTTATCTTTCAACAGTGATTTGTCAAGTAACAATATAAGTGGATCATTGCCACCATCTGTAAGAAATTTGTCTTCCTTGACCACTCTGTGAGTAATTCAACATTATACAGTTTGTTTATAACTTCATAGTTTGATCAATATGATAAAACTAACTTCTTGATTTCCAGATATTTGCAGAATAATCAGCTTTCTGGAACTCTTGATGTTTTGCAGGATCTTCCCCTTGCAAATTTGTACTCTTCCAAACTAGATTTTACTCGAATATAATTCAGTTCACTAGAATGATAGAATCTTACTAACATCTCTTCACACTCCCTGTGTCCTTTATAGGAATGTGGAAAACAACTTGTTTTCTGGGCCCATACCTCAAAAGTTGTTCTCTATTCCCAATTTCAAGTAAGTACATTCATTTTATGCTTTCATATACTCGTTTCATTTCTGTTTATGTTGAGTCAACATCTTGTATATTAAAATGTAATCTTGGTCACATTTAATATCAATGTATCCCTCGTCGTTTGTTGGAGATATCATTTGACATGTAACATATCCCACAAAAAAGATATATAATCATAGAAGACAATAGACATGTATGTCGAAACCGCGGAAACAAATATCTTTTCAAGTTCAATGTTACTTAATACTGCTCCTTATTTTCCAATAATGCCTCACTTTTGTACTATGTACCAGTGGTTATAACCAAACATATAACTTTTCTAGCTCCCAATATCCCCCTTTAAAGCTTGAGAACTTGTATCGTCTGTGCTATGCTTCATACACACAAGTAAGCCTTTCAGAAACTGCTTGTTTACAAGTTCCTGAAATCTTACTTAGTAAAAAATGGATGTGCTTGTATGCTATGTTGCTCGGACTCGTATTGGATCCTCCCAAAGTAATCCGACACTGGTGCAACAACATATTTGGAAAGTCCGAGCAACATAGCTTGTATGCTGCAAGGTCATTTTGTACAAAATGACCAGAAATATAGGCTTAATGTCTGACATGGATATGAATCCTAAACTTGCCCTTAGCATAAATGAAAGCTCAATGAGTAAAAGTAAGAATAAAACAAATAAGAAATTGACCTGAAAAAAGAAAAAAAGAATCCTGACAAGCATGTCATTTAAATACATGAATTCTATCATATGGCCAGTCCCTCAAAACTTGGAAGTAAACAACTCATATAGACACAATAAATAGACAACAGATATAGGTATAGATTTACAAACCGTAGTACAATATGCATATGAAAACATACAAACAGTTAAGTTATGATATTCATGACCTGTAAAATACACAAAATTGCAAGTTATTTAGTTTAGCAAGCTGGCTCTTGGTCTAAACTTGATCATAATTTAAGTTCACATTCCATGGTGGCTATACTCTAGTTAGGTCACCCACCTTAATCATATATCGAGCCCCTAACTTTCAGGAGGCAAAAATAACATGGAAGTCAGGCAGTCTTATTTATTAGCTGACTTAAATGGTACAAGTTGGTAGATATATTTGATATATGCAAAGATCAATGTATCAGTTGTTGGATCATTTTGTTACCATCTGATACATGCCTTGTCGATTATGATCAGTGATGGTTTGTACTGTCTTAGTTACCGGTTTTGTTTTTCGTGCAGAAAAGATGGAAATCTTTTCAACAGTAGTATTGCTCCTTTACCTCCACCCACATCATCAACAGCACCTCCCATTTCTGGGTCTCCAACTTCAGCACAGACGCCATCTTCTCAAAGCAAACCTAAAACACAGGCCGATGGACCATCAGCATCTGCAGAATCAAGTTCAACAGGAAAAAAAAAGTCTTTATCTGCTAAAAGGGTGGTCTGGATATCAATTGCAGCAGTTTTGTCATTTATCATATTGGTTTTAGCAATCCTTCTCTGTCTGCCAAAGTGCCTCAGGAAATGGCACGAGACTCAAAGAACGCCCAAACACCATGAAATAGCCCCTTATATGGGTGCTAGAGAGAATCTTGGAGATAGTGGTTCATTAGTCCAACCTGGCCATGATATAGAAAAAGGTGCTACAAAATACTCAATATCTATGTTATTTTATGATATCTTTTATCACTGTTTCTTAAAGTTCAACTTTTCCCCCTTAGGGTTCGTTTGGTAGGATGTATTAGAGAAATTAATACATGTATTAGTGATTGTATTATTTAATCCTTTGTTTGATAGGATTTTCAGCATATGTATAACTAATACTATTTAATATTATAGGGTGTATAACTTATACATAACCAACCATGGTATTAGTTAAACAATGGTTATTAATACATGGATAAGTATGGTTAAAGACAGAACTGTCCTTAATATATCAAACTAAACCGTGGATAAGAAACTATATCTAGCATAACTAACCCCAACATAATTAATCCCAACATTACTCATCTCGGCACTACTAATACACCCTATTTCAATACTATTATACGTGCTACCAAACGACCCCTTAGTTACTTGAAAAAAAAAAGATACTGGGTCTGTTGTGTTGATGCACTTGAGCCAAGGGTCTTTCGGAAATAGCCTCTCTACCATCACGAGGTAATGGTAAGGTCTGTATACACTCTACCCTCCCCAGAACCTACTTAGTGGGATTTGACTAGGTATGTTGTTGTATTGGAAGTCTGTTGTGCAGCATTTAGAACTGAAAAGTTAATGAATCTCAAAGTTTAACCTTCTACTCTTTAGTCTTTAGTGATAAGTGGTTTTCTTATGTGTTAAACAGCTCCCCTAGTTGTGACGCCAAAAGAAAAGCAACCAAGAAGACCGGTTCCCGTTCCGAAGCCACAAGATGAGCAGCAGGTAAATGTGCAAACCATGAGTGCAGTGCCAAAGAAGGATGATAGTGAGATAAACCTTGGCAGATTGGATATTGATTTGATGCCACCACCTCCGCCTCCTCCTCCGCCGACTCTTCCGCAAGAGAGAGTTATTGTCAAGCCGATTGCGCCTGCTGAAGATACAGCAACGAGGCTTCCAACCAGGCCACTTCCACTAACTTCTGTGAAATCTTACACAATTGCATCACTTCAGCAATACACTAATAGCTTCTCTCAAGATAATCTGTTGGGATTAGGAATGCTGGGTAGTGTATACAGGGTGGAGCTTCCTAATGGAAAGGTACAACTTTCAATCGGCATGAATTTATTATGATGTACCAATTTATTAACTGTAGTTTCCTACTTCCACTAGATTGTCGAGAGTCTTGACTTACTGCTTTTTCTCTCCTTTTATTTCTCCATCATGTTTGGTATTCTGGAATGCTAAATTATCAGCAAGTAGCATCAAAATTGCCTCTTTATCTCTAGTATAAATTGTTAGGTTGCTTAGATTTATCAAGAATGGTTGTGATACAAGCACAATTATATGCAAAAAGGCCTTTTTTAATGGAAAATTCACCTTCATATATCCTTAATTAGAGCACTCCTTAGCAAATCTATTAAATGAATTCTACCTATTTTATCTGATATAATGTTTCTTGAATTACTTTAGCTAGTTGAGACCTGCTTAAGGACAAAAGTTGTAATTGCAATATTTTCCCTCTAGGTTTCTTCTAGTTCTTTTCTTTTCCTCGTTCCAAATTAATCTTCAACTAGTATTTTTTTTCCAGTAACCAACATCATCGATGAAATCTTGTCCTAGAAGTGTATTTCAAATTTGTTAAAATGAAAAACAGCCTTTTATACTATTTTCTGCCTTGACTTGTCTCACTTTATAAGAGGAACCGCCAGATACTGCTGCATCTTTTTTCTGGTCATGATTTTAACTATTAAAGGTAAGTATATGCATGCTGTTTCATTTATATAAATTACTTGGGGATGTATATGCATTTCTCTGGCATTCAAATCTTACCGCTCTTCCTTCACTTGCACTATGCACCACATCATTCAAGTGGTTGCAACATTTCACTTCTGACACATGCTCCTTTTGGATGCTTCTGCAGTTGCTTGCTGTCAAAAAGCTGGACAAAAGAGTTTGCGATCAACAAAAGGATGATGAGTTTCTTGACCTAGTAAACAATATTGACAGAATTCGTCATGCTAATGTTGTTGAGCTGATGGGCTATTGTGCAGAGCATGGTCAAAGGCTTCTGGTCTATGAGTATTGCTGTAACGGGACACTACAAGATGCACTGCATTGCGACGATGAATTCAAGAGACAACTTTCATGGAATACTCGCATGAGGATGGCTCTTGGAGCTGCAAGAGCGTTAGAGTACGTCAAATAACATTACATACATTTAATTATGCACATCATGTTTGTGCACACTCGCGTTCATGCGTGTGTGTATAAGTGTTTTGGGACCTATAATAAGTGTTTCTGCACACACTATATGATAAATCAGTGCTATTAAATACTGAACAATAAAATGGTTCTGGTATAGTGATAAATTCAATAACAAGCTCATTTTAGTCCTCAACTTTCTTGTGTTGTGTATATTACATGATAAAAAGCCACACAAACGACCTGGAATGGTCTTAACTATATGCAAAATGTTAGATGTAGCTGTCTTTTGCTTCCTCTCCTTTCTTTTCTTCTGCATGTCCAAGAAAAAGATGAACAAAATTAAATTAACTGGTTTGCACTGCTATGCTGTAAAAGGGAAGCCCGATGCACAAGGCATCCCGCATTAGCAGTCTGGGGAAGGTCCATACACCAAGGGGTGTGATGTAGACAGCCTATTCCAATGCAAGCAGTAGTGGCTGCTTCACGGCTCGAACCGGTGACCTTATAGGTCACACAGAGACAATTTCACAATTGCTAAGCAGTCTGATTCTCATTTTTTAATTAGATTTGCATGAAAAAGTGAGGTAGACTGCTAGAGTATGTTTAAATTAATAATCATTCTTGTTTGATGAAATGCAGGTATCTCCATGAAGTCTGCGAGCCACCTATTATTCATAGAAACTTCAAGTCTGCCAATCTTCTGCTTGATGAGGAGTTGGCTGTACATGTTTCAGATTGTGGTTTGGCTCCACTTATATCATCTGGTGCTGTGAGTCAGGTAACTTGTGACCCCAATGTTATGGAGTAGAAAGAATTGAAATGCTTCCTTGGATTATTAGTTTCCTCTATGAGCTGCAAGTGGAAAGAAAAGTGAATTTCTAGTGTTTGATAGAAGTGGACATGGGAGCTGAATATATAATTTAAAAAAGGAACAGTAGTGATGGTCAAAGCGTTTTCAACAAGATGTTGGAATAAGGAGCAGTAAGAATGAATTCCCGTAAATGTGTAAAGTATATTCTTTGGGAAATGGTTGGTTCACACCCTGTAGGATAGGGTGAAGTGTTGTCGAAGGCGCGCTTAAAGCACGCTTGAGCCCTGAAGCGAGGCTCAAAATGTGTTTAGCATTTCACATCGCTTAATGTGAGATTAGGGGTCTATAGGAAACAACCTTTCTACAGTTGAGGTAGGGGTAAGGTCTGCGTACACTCTACCCTTCCCAGACTCCACTGCGTATGTTATTGTAAGTAAACACAAATGGTGAAGGTGCTCTTCTCATGATGTGTTTTATATGAACAGAAAATTGCTCTAGGATGCATGTGGATCATCTATCTTGCTGTTATTTGTTGTTCTTTTGATAAAGTAGTCTTGCTTAGTCTTGGGTTTGATTACTGTTACGTTCAATTAAGGTATGTTAAAAACTATATCAAGTTGACATTACACTGATTTTTCAGTTGTCGGGACAACTTCTCACAACATATGGCTACGGTGCTCCCGAATTTGAGTCAGGGATTTACACTTCCCAAAGTGATGTTTACAGTTTTGGTGTGGTGATGCTGGAACTAATAACAGGCAGAATGTCATATGACCGGTAAAAAAGCCTTCTGCATCCTTTTGAGCTAATATAGGTGTGCCTCATTTCTAAGTTAACACACCTGCATATCATATTCAGGACACGGAGTCGAGGAGAGCAGTTCTTGGTCAGGTGGGCAATCCCTCAGCTACACGATATTGATCTCTTGACAAGGATGGTTGATCCTGCTCTCGATGGAAAATACCCCATTAAATCACTATCACACTTTGCTGACATAATTTCTCGCTGTGTTCAGGTGAGTTATTCTGTTGTTCCTTCATGTGAAACACATAAATGCTTACTAATATACATTGTCTAATCGCTGAGTGGACCGAATAGTCTTAGATTGTTCGTCATCTCAATCGGTTGGAAGGATATTTCTAGGATCTAGTTGTGGAATAGGTTTGCCTTCAAATGTCACTTAACTTTTAAACTCTTATACAGCATGAGCGAGAATTCAGGCCACCAATGTCAGAAGTTGTTCAGGACCTCATACAAATGATAAGAAGAGAGTCTCACAAAAGATCTGATGGTGAATGATTTGAGTTAGACGCGCATACTGGGAATTCAGCCGGTGAGAAACTAAGATTATTTCTTCTTCGAAGGAGCTGAGATACTCGGAAAAAGAAAGGCGGTTTGCAGGCCAAGCACCGTGCTAAAGACAACTTCAAACTGAAGCAAAATGCACCTCAAAGAAGCGCGATTCATGAAAATACCAACACATGCAGGCGACTTAATGACCTGTAAAAACAGCTTGCTCATCTGATTCTTGTCAAGCTGCATCATTTTAATGTTCATTTAGGCTTACATGTGAGCACCAATGCATATAATTCATTGATCTTGACAAAATGGTATATTTTCTTGAAGTTGTATATCTTTACTTGCTGCAACATGTACAAACTTCATTTGGTTGTACAAGGATACCTTCATTCTTTCTCATGTACTTGTGGCACATGCCAAAGTTATTTACTACTAGTCATTACCTAAGGAGTTGGAGACCTACAACCACTCAACCATGTAACAATATGTAAA

>PGSC0003DMG400007948 LRR-RLKs

TTAATAATTAATCCTAAAATAACTTATTTTCGACCGTGAAACATTATGGCCATGGCTGACACAGAAACAGAGAGAAAAATAGCCAGACTAAATATTGGAAGTCCCCACTTTTCTTTTTTCTTTCTCAAAAATCTTCACGTGAATCTGACTCTGCCTTTTGCCATATCTTCTTAGACAGCTGCTTCTCTCTTCACTCTTCCACCTCTACACATATTCCACAACAATTCAATCTGGAAAAAATTTCTTAAACAAGATGAAAATACAACAATTCTTACTACTTACCATTCTTCTGTTTTTATCTGTTTCTTTTCATTCCACTGTTTCGTTGATTCCTTCTGATGCATCAGCATTGTTAGCATTCAAATATAAAGCTGATTTGGACAACAAACTTGCATTTTCTGCTAATACAAGTTCCAGATTCTGCAAATGGAAAGGGATACAATGCTCGGAGAAGAAAGTGATTCGTATTGTTATCGAAAGTTTTAGCTTACGAGGTACGTTTCCAGCTAATACATTGTCTATGCTTGATCAACTCAGAGTATTGAGTTTACAGAACAACTCACTTACCGGTCCGATTCCGGACCTTTCTCCACTTATTAATCTCAAAGTTCTTTTCCTGGACCATAATTTGTTCACCGGTTCAATACCAGCTTCCATTTTCACTCTTCATAGGCTTAAAACCCTTGATCTCTCCTATAATAATCTCACCGGTTCAATACCCGTTGCAATCAACGGGTTGAACCGGTTGTACTATCTCCGGCTTGACTCGAACCGGATAAACGGATCAATCCCACCGCTGAACCAATCTACCCTCCATATCTTCAACATTTCTCACAATGCCCTTTCTGGTCCTATTCCGGTAACGAAAACGTTATCCCGGTTTAAAACGGCGTCGTTTTCAGACAATAAAGGACTTTGTGGTGAGATCGTACACAAAGAATGCCGTCCAATACAACCGTTTTTCAGTCCATCCACCGCTGCCTCCACTAAAATTACGCCGCCGCCGTCTAAAACTCCGGCAGAACTCGGCCAAAACGAAGAGTTACGACAGGGTAGCCCGTTAAACCGTAAAGAAAACAAATCACATAAAAGATCCTTACTCATTATTGGAGTTTCAACAGCTTGTTTAGTCCTTTTATGTTCAGTCATACTATTAGCATTAGCGTCGAAGAAGCACAGGAACTCGAAGAAATTGGGAGAAACTAAGAAATCCGTATTCGATCCAAGTGTTAGCGGCAATGCAGAAGCAGTAATAAGAATTGAAGAAGACAACAATGAACTGGAAGAGAAGGTGAAGAGAGTACAACAAGGAATGCAGCAAGTGATGGGTAAAAGTGGGAGCTTAGTATTCTGTGCAGGTGAGGTGCATGTGTATACCCTGGAGCAGCTGATGAGAGCTTCTGCTGAGCTTTTAGGTAGAGGAACAATGGGGACTACTTACAAAGCAGTGCTTGATAACCGTCTGATTGTATGTGTTAAGAGATTGGACGGTGGGAGATTGGCGGGTACAAGTCAAGAAGAGTTTGAGCAGCATATGGAATCAGTGGGCGGGCTTAGGCACCCGAATTTGGTTCCTTTTCGGGCTTATTTTCAGGCCCGACAAGAAAGGCTTTTGGTTTATGATTATCAACCCAATGGCAGCCTGTCTTCTCTTATTCATGGTACGTTCTTCTCCGTATATCTTGTACTTATTATTTATTAAGATGCGCATAAACTATACGAAAAATATTATTAATGCAATTATTTTGACGAGAAGTAAAATATAATAAAATGAACGGAGTATTTTGAAGGTGTTGTCCAATCAAATAGGGTCCATACACCATACCCTATTGTTACAGTTTCTCAACCCCAATGTTCTCCCAAACACAGAGAATCTTGCAGCTGACCCTGATGACATTTTTGGACACATATTTCTGTCATCGTAAAATACTTTTATATTTGTTTAGTGTCCTAATCATAGATTATAGTATTATTATTGAAGCTAGTTCTGTCTGCATTAAGAATGTTATTGTTTCACGTGTGGTTGCGTAACGTGAAGTTAATAAGAGTACGGTGGTTAGCATTAATAATTCATAAAGTAATATAAATAATAGCTTTATTGCATATACTGAATTTTTTGGTCAATCTGACGTGGCATATGAAATAAAGTTTGACTGGTCAACCTGCCACGTGTACAGGTTCCAAGTCATCAAGAGCAAAGCCACTTCACTGGACTTCATGTCTGAAAATAGCAGAGGATGTAACTCAAGGACTTTCCTACATCCACCAAGCATGGAGGCTTGTCCATGGCAATCTCAAGTCATCTAACGTCCTCCTCGGCTCCGATTTCGAGGCGTGCATAACCGACTACTGTCTCTCCATCCTAGCTGTCCCCTCAGACGATGACAATCCTGATTCTGTAGCTTATCAGGCTCCAGAAATTCGGAAGCTGAATCACAACAACCACCACCACCACCGCCAAGCTAGTGCCAAGGCTGATGTCTATTCGTTTGGTGTTCTTTTGCTTGAACTTCTAACGGGTAAGCATCCTTCAGAACATCCATATCTAATGCCAGATGATATGTTACATTGGGTGAAGTCTACTAGAGAGGATCATGATGGTAGCATTGGAGAAGATAGTAAGTTGGAGATGCTTCTTGAAGTTGCTATGGCTTGTAGAGTGAGCTCACCAGAGCAAAGACCAACAATGTGGCAAGTGTTAAAGATGATACAAGAGATTAAAGAAGCTGTTGTAATGGAAGATAGTAATGAAATGGACCTTCTCACTGGCCCTCCTAAATCCTAATTCTTATTTAAATCTTAGTTATTATAAAATTGAATTTGTATTATATTCACTACATGAGACTTTTCAATATTAGTAATTGCAAAAGAAAGGAACAAAAGGGGAAAGACTTATACTTTTGTTACATGACCCTCATTCACCCTATTATATTGGATAACTATGTCTCACTTCCAAACAAATCGGTATGTTTTATGCTAGTTGTCAGCCTACTACAACCCTCCTCCTTTATCTGGTCTCGGGATCAACATGTTTGTTGTTGGCAGTGTTTTTTCTCAATAACATATTTCAATTTGAC

>PGSC0003DMG400008146 LRR-RLKs

AGTTGTTCAAGTTTGTATCTTTCTTCTATTTGTACTTAAAACTGCAATTTTTATCACTTCTGCAACTTATTTGTTGTTTTTGGTACATGGGGTTCATTCTTGAGTAGTAAAGATTGCATCTTTATTGTTCTGTTTCATTTTCAAGATTCTTGAAATGGGTCATTTCCTTTAAGTTGTTCAAGTTTGTATCTTTTGTTTTCTAAACCCTTCTGTTTCTACTCTAGTTGTCTCTACAAGGTTTCTTGGTTTTTGGGTTCTAGCATTTTGTTTCTTTATGGTTTCTTGATTGTTCCATTTGGGCTTTTTTCTTTTTGGGGAAAAAAGGGAAGAAATTGAAGATTTTCTTGAAAAAAACAACTTAAAAATGGTAACTTTGAGGTTGTTCACTGTTTTGCTGGTGGGGTTGTTGTTAAGGTCTCAATTTGGGATTGTAAAGGCTCTTCATGAACAAGCCATTTTGGAAGCAATTGGCAAAGAGCTAGCAATACCTGGTTGGGATCTGAATAGCACTGATTTCTGTTCTTGGCATAGCATTAGTTGCAGCAGCTCAAACAGTTCAATGGTGGAAAGGCTGAATCTTTCGGGTTTTCGATTACAAGGTAATGTGACTCTAATATCTGAGCTCAAAGGATTGAAATGGCTGGACCTTTCTAATAACAATTTCCAAGGTTCAATCCCACAAGCATTTGGGAATTTGTCTGAGCTTCAGTTTCTTGATTTGTCTTTTAACATGTTTAGAAACTCTATTCCTGGTGAATTAGGTAAGCTCAAGAACCTTAGAGCATTGAATCTTTCAAACAATATGCTCACTGGTTCAATACCTGATGAGCTTGAAGGGATGGAGAATTTGCTTTATTTCCAATTATTTACCAATAAATTGAGTGGTTTTATTCCAATGTGGATTGGGAATTTGACTAATCTTAGAGTGTTTGCAGCCTATGAGAATGAGTTTAGTGGTGATATTCCAGTTAACTTAGGCTTACATTCTGAGCTTTTGTTGTTAAATCTTCATTCAAATCAGCTTGAAGGGACAATTCCTGAGAGTATTTTTACTATGGAAAAGCTTGAATTTTTAGTTCTGACTAATAATAAGTTGACTGGCACTATTCCTGATTCAATTGTGAATTGCAAAGGCCTTTCAAGCATTAGAATTGGGAATAACAAGCTGATAGGTGGTATCCCTAAAGGAATTGGAAATATAAGTAGTCTTGCTTATTTTGAAGCAGATAATAATACACTTTCCGGGGAAATCGTCTCAGGATTTGCGAAATGCTCTAATCTCACTCTTCTTAATTTAGCTTCGAATGGATTTTCCGGAACTATCACTCCAGAGTTTGGTGAGCTCGATAATCTACAGGAGTTGATTGTTCCGGGGAATAATCTCTATGGAGAGATTCCAACTTCAGTTCTGAGGTGCAAGAATCTCAACAAGCTTGATTTAAGTAACAACAAGTTCAATGGAACCATTCCAGGAGATATATGCAACACAACCAAATTGCAGTTCTTGTTGTTGGGACAGAATTCCTTGAAAGGGGAAATACCCCGTGAGATAGGGAACTGCGTAAAACTGCTTGAGTTGCAAATGGGGAGTAACTATCTTACTGGAAGCATCCCTTCTGAGATTGGTCATATGAAGAACTTACAGATTTCATTGAATTTGAGTCATAATCATCTCCATGGACAGTTGCCTAAGGATCTTGGAAAACTTGACAAGTTGGTTTCTTTAGATGTATCTAATAATCAGCTGTCGGGGAATATTCCGTTGGAATTGAAAGGCATGTTGAGCTTGATAGAGGTTAACTTTTCGAGCAATCAATTCACAGGACCAATACCTGCTTTTGCTCCATTTGAGAAGAGCTTAAATTCAAGCTTTCTTGGAAACAAAGGGCTCTGTGGTGAGCCTTTGAGTAGTGATTGTGGATATGATTTTGAACACAACGGTAATCATCACCGAGTTTCCTACCGGCTCATCTTGGCTGTTGTTGGTTCTGGTTTGGCAATTTTTACAGTTGTCACAGTGGTTGTTTTACTATACATGATGAGGGAGAAACAAGAGAAAACCACCATGGAAGATGGAAATACCACTGATGAAACTTGTAGCAAACCAGTGATCATAGCAGGGAATGTTTTCGATGAAAATCTCAAGCAAGCAATAGATTTTGATGCAGTTGTAAAGGCAGTCAGGAAAGATACAAATAAGATTAGTACTGGAACTTTCAGTGATGTGTATAGAGCAGACATGCCTTCCGGGATGATTTTGTATGTTAAGAGCCTAAAGTCGATGGACAAGACTATAGTTCATCACCAGAGCAAGATGATCAGAGAGCTTGAAAAGCTAAGTAAACTCTGTCATGATAATCTGACGAGGCCTCTCGGGTTTGCAATCTATGAGGAAGTTGTTCTACTCTTACATCAATACTATCCCAATGGGACATTGGCTCAGTTTCTTCATGAGTTTAGCCAGAAACCTGAGTACGAACCTGACTGGCCTACAAGACTTTCCATTGCCATTGGCGTTGCAGAAGGGTTAGCGTTCCTTCATCATGTGGCCATAATCCATCTTGATGTTTCTTCAGGGAATGTGTTTCTTGATTCTAAATTCACCCCTTTGGTTGCTGAAGTTGAAATATCCCGACTTCTTGATCCATCTAGAGGGACTGCAAGTATTAGTGCTGTTGCTGGCTCATTTGGATACATTCCCCCAGGTATATTCTATGGCTTGAAAGACTTGGCTAATCATTTTACTACCTCGTGTTTCAATTTGTTTGTCTTACTTTCCTTTTTAGTCGGTTTAAAAAAAATGTCTTTCCTTTCTTTGTAACTCTTTAATTTTAACTTTACACCTGACATTATTAAGACCATGGACATTTTGTATATTCTATCTGCCAGACAAACAAATTGAAACGGAGGGAGCATTTTCTTTTTGTGGAAAATATCTGGTGTCTGACTAGTTCCTTTTTCTTCAACAGAGTATGCATATACAATGCAAGTAACAGCTCCCGGAAATGTTTATAGCTATGGCGTTGTTTTGCTCGAGATCCTTACCACCCGACTACCCGTTGATGAAGCTTTTGGTGAAGGAGTTGATCTGGTCAGGTGGGTACATGGTGCTTCTGCGAGAGGAGAAACACCAGAGCAGATACTTGATGCAAGACTTAGCACCATTTCTTTTGCTTGGAGGAAAGAAATGCTTGCAACTCTAAAAGTCGCATTGATGTGCACCGATACCATTCCAGCAAAACGACCAAGAATGAAGAAGGTGATAGAAATGCTTCAAGAGGTTACACAGAGTTGAATCAAAACTCGTATCATCCAAAAGTCTCTGCTTAATCTCAGACCTGCAGATCATGAAAACCCCTGTAGAAATACAAAACGGTATGAATTCACTGTTCTTAGCGTAGACTAGTGGTCTAAAAATAGTTAAACTATGTATATTGTCTGAGGTTGCTGTTCTTCTCTGATGATCAGCTCCGAGGTTGTAATTTTGATTTTCATATTGGAGCGAGTAAGGAGGTTATATGGTAAGGGTTAAAACCAGAAACAGTTCTTGATATGTACCATTCTGTACAAGGCCTTCAATCATATATACTTCTTTTTTCTTGTCTATGTGTAAATGGAGCTTTGGCATTTCATATGCAACAGTAAAATCTTTATGTATATATAATATACAGTAGATGTTGAACCCTCTTGGTG

>PGSC0003DMG400008296 LRR-RLKs

TTATGTACTCCTAGGTTTCGAGTAGGAGACGAAATTCTTGTTGTCTTTTCTTGTGTGGGGGTACATACAACTTGTTCTTTAACACGGAGCATCCTAGATTAGAGGCATGCTTGGTATTTTGTAACAAATTCATAGAATGCAAGTCATTTTCCTTGGTGTGAATGGATGACTAGTTGACTGCCACTGAAAGCAGTGATTCACTGTACGATTCACTAAGAAATGTGTATAAAGGCTTCATTGTCAGCATAAGTTTTTTCAGTGAGTGAAGCTTCTTTCTTTTGAAACAATGATGTCAAAGACAGTTGTATATGCATTAGCCATATTTTCTATTACCTTTCTCATACCTCTGTCATCTGGGCAACATCCAAGCTTTGAAGTTGAAGTTGCTGCTTTGAAAGCTTTCAAGAGCTCAATCTCTGACGACCCTTTCGGTGCACTTGTGGATTGGACGGATGCAAATCACCATTGCAACTGGTCTGGAATCATATGTGATCCTTCTTCAAGCCATGTCATCAACATTTCGCTTATTGAGACGCAGCTCAAAGGCGAAATTTCTCCGTTTCTAGGAAACCTCTCCAAACTCCAGGTTCTTGATCTAACTTTGAATTCATTTACTGGAAATATCCCACCCCAGCTGGGGTATTGCACAGACCTGGTTGAGCTCGTTTTTTACCAAAACTCTCTTTTTGGTGAAATTCCTGCCGAGCTGGGAAACCTAAAAAAACTGCAGTATATAGATTTTGGAAATAACTTTCTAAATGGGAGTATACCTGATAGTATATGCAACTGCACTGAATTGTCGTTAGTAGGCTTCATCAACAACAATCTCACAGGCAAATTACCGTCTGAAATTGGTAAGTTGGCCAATCTTCAGTTGTTTGTAGCTTATACAAACAATTTGGTTGGTTCTATTCCTACCTCCATTGGAATGCTTACAGCTTTGCAAACCCTTGATCTGAGCGAAAACCAGTTATCTGGACCTATACCATCAGCAATTGGCAACTTGTCAAGTTTAGGAATTCTTCAGTTACATCTCAACTCTCTTTCTGGGAAAATCCCATCTGAACTTGGCCTCTGTATCAATCTTTTTACCTTGAACATGTATACTAATCAATTTACTGGAAGCATACCTCCTGAGCTTGGAAATTTAGAAAACTTGCAAGTGCTCAGATTGCATAACAATAAGTTGAACTCCAGTATACCTGCCTCAATATTCCACCTGAAGTCATTAACTCATTTAGGACTCTCACAGAATGAACTAACTGGTAATATTCCTCCCGAGTTGGGATCTTTAACGTCACTAGAAGTGCTTACCCTTCACTCAAATAGGTTATCGGGGGAGATCCCATCAACTATAACAAACCTGGCAAACTTGACATATTTGTCTTTGGGCTTTAATTTATTGACTGGATCACTTCCATCGGAATTTGGGTTACTCTATAATCTGAAAAATCTAACTGCAAATGATAACCTCCTGGAAGGATCTATTCCATCGAGCATAACAAATTGTTCTCATCTTCTTGTTTTAACCCTCGCTTATAATAGAATAGCGGGGAAAATACCAAATGGGTTGGGGCAATTGTCCAATCTTACATTTTTGTCTTTGGGATCAAACAAAATGATGGGGGAGATTCCTGATGATCTCTTCAACTGTTCAATGCTTGAAATTCTAGATCTGAGTGATAACAATTTCAGCGGGAAACTCAAATCAATGATTGGCAGACTCTCTAAACTTCGAGTTTTAAGAGCTCGTACTAATTCATTTCTTGGGCCAATCCCACCAGAGATTGGTAAACTGAGTCAACTCGTGGATTTAGTGCTTCACGAAAACAGTTTCTCAGGTGCGATACCACCAGAAATTTCAATGCTTTCAAACCTTCAGGGTCTTTTGCTATCTGACAACAAGCTGGAAGGTGAACTTCCTGTGCAGCTTTTTGAGCTCAAACAGCTCAATGAACTCCGGCTGAAGAACAATAATTTCTTTGGTCCACTACCGCAACATATATCCAAACTAGAATCACTGTCGCTCATGGATCTGAGTGGAAATAAGCTTAATGGCACAATCCCAGAGAGCATGGCGAGCCTCCGCAGATTGATGACCGTAGATCTTTCACACAACCTTCTTACCGGAACTATTCCCAGAGCAGTACTTGCCAGCATGAGAAGTATGCAACTTTACTTGAACGTTTCCAGCAATTTGTTGCACGGAGAGATCCCAGATGAGATAGGCGTGTTAGAAATGGTTCAAGAGATTGACATGTCAAATAATAATCTGTCAGGCAGCATTCCCAGATCCCTAGAACGCTGCAAAAACTTATTTTCACTGGACCTATCAGGAAACATGCTCTCTGGTCCTGCTCCAGGTGAAATTTTCACCAAATTAAGAGAGCTTGTGTTCTTGAACCTCTCAAGGAACAGATTAGAAGGTAGTCTTCCTGAAATAGCAGGATTATCACATCTTCGCTCTCTTGATGTTTCACAAAACAAGTTCAATGGAATCATTCCCGAGAGATTTGCCAATATGACTGCATTGAAATATCTCAACCTTTCTTTCAACCAACTTGAAGGTCACATTCCAAAGGGGGGTGTATTCAACAATATAAGGTCGGAGGATTTACTGGGAAATCCATCTCTATGTGGAACAAAGTTTCTCAGTCCATGCAACATAAAAAGAAACCGAACAAGTTCTCATGGTTTTTCCAAGAAAACCTGGATCATACTTGCAGCACTTGGATCTGTTTTCAGTCTCATTCTTCTTGTCCTGGGAATAATTTTATTTCATCGGTACATGAAGAAGAAGAAAAAAGTGAATGACGCGGAGGATACAAATCCAAAGTATACCGCAGCATTGAGCCTCCAGAGATTTTATCAAAAGGATTTGGAACATGCTACCAATAATTTCAGTCCGGAAAACATTATTGGAGCCAGCAGTTTAAGTACTGTGTACAAAGGTACACTGGAAGATGGGAAGATTGTAGCAGTTAAGAAGCTGAATCACCAGTTCTCAGCAGAATCTGGTAAATGCTTTGATAGGGAAGTCAAGACTCTGAGCCAACTCAGACACAGGAACCTAGTTAAGGTGCTAGGTTACGCTTGGGAAAGCAAGAAGCTAAGGGCTTTAGTTTTAGAATACATGGAGAATGGGAACTTGGACAACATGATTTATGGTCAAGTGGAGGATGACTGGACGTTGTCCAATAGGATTGATATTTTAGTTTCAGTTGCAAGTGGACTGTCATACCTGCATTCAGGCTATGATTTTCCAATAGTGCACTGCGACATGAAGCCTTCAAATATTCTTCTGGATAAAAATATGGAAGCACATGTGAGTGACTTTGGGACAGCTAGGATGTTGGGTATTCATCTCCAGGATGGGAGCAGCATATCATCAGCATCCGCATTTGAAGGAACTATTGGTTACATGGCGCCAGGTAATAATTCTTACAACTTCTTGACCCACACTTTACTAACTAGATTATGTGTTACAATAGCATGAATACAAGATGTCTACAATGTAAACGAGCTTATATGGGTACCACAAATAATATTTTGTCAATGGTGAATGTTTCAAATACATTCCATTTTAGCATCATTCAGTAGTAACATCACTTCTGTCAATCACAATATGCAGCACAGCAGATTTTTTTTTCCTTTCAATCACAAAATACTGATTTAACTTTTATGTGGAAGAATAGACAAGCAAGTAAATGATGCAATGTTTAGAACATATAATCTGTAAAGTATTATTTTAAATGATAATGTCAGTTTTAATCTATCCAAAAATCTAGCGCCAGCTTTTACATGCTCCTAATTTAATTTGAGACCTCACTTACATGAAATGTTATCTATGCAACAGAGTTTGCATATATGAGGAAAGTGACCACAAAAGTAGATGTATTCAGCTTTGGTGTCATTGTGATGGAGATCATTACAAAAAGAAGGCCAACAAGTCTTACCGGAGCAGATGAATTACCAATGACTTTGCATCAAATTGTTCAGAATGCCCTTGCGAATGGCATAAACAAGCTAGTTCAGATTGTGGATCCTAATCTAGCTTCATATGTCTCCAAGAAACAGGATGTACTAGAGGGACTTCTTAACTTGGCTTTGTCCTGCACCTCTGCTGATCCTGAAGATAGACCTGACATGGAACAAGTTCTGTCTTCTCTTTCAAAGCTAAGTAAGATGGACTGCATGCCTTCACATTTGGTAAAAGATTAAATTTGATGTAGAAGAAGCTGGAATATCATCACTCTTAAAGTGAACCACCACCAAGTTTCTCATGGGGCTAATGGAACTAGTTCATAACTTAAAATGTATTTACTATTTAAATAAATTTTCCGTATTTCAAGCCATAAATGATCTCTCGAGTATTCCTGCTTTACTAGAGAGATATAATGGAACTGTCCCGTCAGATTATTTTGAAGTTGACATGCATCATGAAAATGAACAA

>PGSC0003DMG400008403 LRR-RLKs

AGTGACCATATGTATGAAAAATCAAATTAGTTATTTGCTGATCTGAAGCAAATTTTTGAACAATGGCAGCCACTCGGCTGTTACTCTGCTTTATCATCTTCTCGTCTGGCATCGATTTCATATATACTGTTACAGATCCCAGAGATGGTAAGAACTTTGTATAAAATTGTCGGATCCAGTATTTAAACTTTATGAGTTTTACTATTTGTTGCAGTTTTTGTGAACTTTTACTTATAAATTTATGTTCTGTGTTGAAAGTACTGAGTTCAGATAATAATGTTGTTATGAATAAGTGTGAACTTGGTTGATTGCAGTAACGATACTTCGATCACTGAAAGATCAGTGGGAAAATACGCCGCCTAGTTGGCAGAAATCGGATGATCCTTGTGGAACATCTTGGGAAGGTGTAACCTGCAACAACTCAAGGGTAACTGCATTGTGAGTATGCTTTTTCTCTTCCAATATCTAGGGATCGTTTGGTTTACGTTCTAGTTATACGATGACTAATAACACATGAATTGTTATGCATGATTGTATAATTTAAGTGCATTTCCGCCTTTAGATGCTCTTATCCACGTATTACTCGTACATGTATTGTTGTTCCACATTCTATCTGTGAAAAATAATATATAGATTCTCGTCTAAGTTATACAAGGTTTTAGTTTCCAATATTAAACTTCGCATGACTAGTGCAGAGTTTAATATCGGAAACAAAACATCATGTTAGTTATATGGAAACTATATTATCTTACAAGTTTTAAGATTGAAAACCAAATGTAGTATAAGTAATACGAAAATTTATACATAAAGTTAATTTTTTAGGAAAAAATACATAATTAGACATCCTTCGCTATCATATATATATCATTACCTATACTTTCAAAACGTATCTTACCACTAATTAAGAGTTTCCTAACATATATTGAGGAAGATGCACTTAGATACATGTATTTTTGCTACCAGATACAGTAAAATAGGGAGAGAGGAGAGCGAAATGGGGAGGGAGGCGAGCGAGATATGAGAGTGGTGGGTGAGATGGGAGAGACGCGAGAGACAAACGAGATCTGTTAAGTATCCTAGATACATGTGAATTCACTTGGATACAATATATCTAGAACAATTACACCTAATTTTTACCCTATGTATCCCGAGATGCATGTTCTGGATGTATGTGGACGCAACAAATGGTAAGATACATAATATTGCAAACTAGAGTGTATCAAAGTAATTAGCTCCTAAATTAGTGGGATTTATGTAAGTTTTTCACCTTTTTATACTTCAAACCAAGCCACATCTTAGTTTTTGAAATTGTTGTTTCACCAACGAAGCAGTTGTTAACAATTGATTGTGTTTATTGTTTCTTTCAGGGGATTATCAACAATGGGACTGAGAGGAAAATTAAGCGGTGACATTGGAGGGCTCACTGAATTAATTTCCTTGTGAGTTTGGTTTTTAAATCCAATAAACTTCTGTTGATTATTAGCAATAAGGTTGTATTTGTCACCATAATTTGCTCACTTCAGTTGCTAAACTTGTATTGCAGAGACCTGTCGTTCAACCGAGGACTTACAGGTTCTCTTTCTCCGCGGATAGGAGATCTGCAGAAACTAAACATATTGTAAGACTACGAAAACTTGATCATCTTGATTGTATAGTGTATACCTGCTATGTACATAAAATTGACATTTATAATATTTTCGTCATAGAATGATCATTTGATCAGCTTTCTTCCTCGTAATTTGTATCATGATTCAATGACATACATAACTTACATGTACGTGAATATATCTATTTACTTTTCAGAATCCTTGCGGGCTGTAGCTTCAGTGGTAGTATACCGATGGAGTTGGGAAGACTTGCTGAGTTATCATTCTTGTAAGAATTTCTCAGCCATATTAACTTGTGATCATTTTAGGTGTATCGTCAACTCATTTAGTTCCATGTCAATTTCTTTGACAGGGCACTTAACTCAAATAATTTCACCGGTGAGATTCCACAAACTCTTGGCAATCTGTCAAAACTGTACTGGCTTGATCTAGCAGATAACCAGTTAACAGGACCAATTCCTGTTTCAACCTCCTCAAGTTCGGGGTTAGACCTTCTAAAAAAAGCAAAACACTTGTGAGTAAACTTTATATGTCCATATTACATGCTGTTCTTAAGTTGGGATATATACCTCTTAAGGTCTGACTTCTACGTTATGTCATTATCGCAGCCATTTTAACAAAAACCAACTTTCAGGCTCCATTCCGGACATTCTTTTTAGTGCTGACATGGTGCTAATACATGTGTAAGTCAAGAAACTAAAGGGTGATTCACGCTATTCTTCCTGCCCTCCTTCTCACTTTGTGAACTTCTGATATGGCAGACTGTTTGATGGAAATCAACTTAGCGGGAGCATTCCATTTACACTAGGACTTGTTCAGACACTTGAGGTTCTGTGAGTAGAACTGACCGTCTTATTTATCCTATCTTTCATGCTACTTCACCTTCTGTTTTGGATTATATGTTTGTACTCTTTTATCTGTTGCTGGTATTCATGTCACCTAACATATACAGTCGGCTCGATAGAAATGCCTTAAATGGTAGCGTCCCATCAAACCTCAACAATCTCACAAGCGTTGTAGAGTTGTAAGTATACTTTTTCCCATCATTGCTCACTCAATTTTCTGCTTCAAATCAATAACTTGTAGATTGAAGTTGTACTTTTGTGCACTTATTTCTGTTTTCAGGAACTTAGCTCACAACGAACTGTCCGGTCCACTACCAGATTTAACTGGAATGAATTCCCTTAATTATCTGTAAGGAAAATATCCAGAATCCTGATTTTCCCTAGTTGTTTCTCAATGTTGAAGTGATACATCATATTTTGTTTCTCTTCCAGGGACTTGAGCAATAATTCCTTCCATAAATCGGAGGCGCCTATATGGTTCTCGACTTTAGAGTCTCTCACCACTCTGTGAGTATTAATCAGTTATGAATGCACCTATTAGTTTCCCAAAAAGTTTGATGAATCTAAGTAGGAAGAACTGATGTAATGCTCAGAGTCATTGAGTATGGCTCACTCCATGGATCAGTGCCTCAGAAACTTTTCGCTTTACCTCAATTACAACAAGTGTGAGTATCGCGATCCTAACTGAATATTCCATGTGTATTACCTGCAATGAAGAGGTTGTAATGATCGTAATAACTTATGCAGGAAACTGAGGAACAACGCATTGAATGATACGTTGAACATGGGTGGCATCAGTGGCAGGCAACTCACACTTGTTGATTTACAAAACAACGAGATCTCCTCAATAACACTTGGCTCTGGTTACAAAAATACCTTGATGTGAGATGGCTCATTTCTCCTTTTTCGTCAATTAGATTATGTTTTGTCTCTTTCTTTCTCCTTGATTGCTTTGTGGTTGTCGTCAGACTGATTGGAAACCCTGTATGTGACACGGCCCTTGGAAACACAAACTACTGCCAGCTTCAGCAGCAATCTGCAAAGCCTTACTCAACCAGCCTAGCTAACTGTGGGAGCAAATCATGTCCTGCTGATCAAAAAGTTAGCCCACAAAGTTGTGATTGTGCTTATCCGTACCAGGGAACATTCTACTTTAGAGGACCCTCTTTTAGGGAGTTGTCCAATGATAATACGTTCCATTCACTCGAAATGAGCCTGTGGGTGAAGTTGGATCTCACGCCCGGTTCAGTTTCTCTGCAGAACCCCTTCTTCAACATTGATGATTATCTTCAGGTGCAGTTAGAATTATTTCCACCAACAGGAAAGTATTTTAATAGATCAGAGGTTGAGAGGATTGGATTCTCCTTGAGTAACCAAACTTATAAGCCGCCTCACGAGTTTGGACCGTACTATTTTATAGCATCTCCTTACACTTTCCAAGGTATAATTACTGAGTTTCTTTACAACTGTAACTTCTAGACCAATATACTGACTTAATACTAAAACTGTTTCAGCCGAACGTGGAGAAACTTCTATTAGCTCAAGACAAGTTATAGGAATAGCTACCGGGTGTACCATCCTGGTCCTCTTGCTTGTTGCTCTTGCAATATATGCTATTCAACAAAAGAAACTTGCTGAAAGAGCTATTGGATTGAGTAGACCATTTGGTAATGTCATCTCTTATAGAAAAAAACAGATGAGTTATAACTATATATACATTCAAGCCTCTCTATAACAGTCATCCTATATGATAACATTTCACTATAACGACCAAGTTTTCTTTAGAACTGATTTTTCATGTTATGTTGAATTATATGTTCTCTTTAACAATATTTCGCTATAGCAGCCAAAAAAATATCAGAACAAACGACGCCTTTATAAAGAGGTTTGACTGTATGTCCAGAATGCAAGTAGGATTGAACTCAAATGTCTGATTACTTTGACCTTCCTTGTCCAGCATCATGGGCTCCAAGTGGCAACGACAGTGAAGGTGCACCACAACTAAAGGGAGCAAGATGGTTTTCTTATGATGAACTTAAGAAATGCACGGGTAACTTCTCGGAGAGAAATGAGATTGGTTCCGGTGGCTATGGGAAGGTTTGCTATGAAAATCCATTGTTTTAGAATAAATTAAAGAGGATAAACTCTTCAGCTGCTTTTCTTATTCGCGTGTGTTCTGTTTCCTAGGTCTACAGAGGAACCCTCGCGAATGGGCAAGTAATTGCTATCAAAAGGGCTCAACATGGTTCCATGCAAGGTGGACAAGAGTTCAAGACGGAAATTGAGTTGCTTTCACGGGTTCATCACAAGAACCTAGTTGGTCTGGTTGGATTCTGTTTTGAGCAAGGCGAACAAGTACTGGTATACGAGTATATGCCAAATGGTTCACTCAGGGAGAATCTGTCAGGTAAGTTACATCCTCCTATACATTAAAGGCTACTTCCTATTTATAGTAATTTACACTAAAAAGTTCCTAATATTTCCATTTCTCTGTAGAAGTTAATTTCCACTATGCTCTACAGGGAAAACGAGCATTTATCTAGATTGGAAGAGGAGACTTCGTGTTGCGCTTGGTTCAGCCAGAGGATTAGCTTATCTGCATGAGCTTGCCAATCCTCCTATAATTCACAGAGATGTGAAATCAACTAACATTTTGTTGGATCAGAACTTAACAGCAAAGGTCGGGGATTTTGGCTTGTCCAAACTTGTATCAGATAGCTCAAAGGGTCATGTTTCTACTCAAGTTAAAGGCACATTGGTCAGTACCACATCCTGCACATTTTCGCTTTCTATTTCAAGAAAAAGATTGAATTTTGTTGAACTCGTGAAGCATGAAATTGAATTTAAGACTATTGACTACATTGCAGGGTTATCTAGATCCTGAGTACTATATGACTCAACAACTAACCGAGAAGAGCGATGTGTATAGCTTTGGTGTTGTAATGCTAGAACTCATAACTGCAAAACAACCAATAGAGAAGGGGAAATACGTTGTTCGAGAGATGAGAACAGCAATCAATAAGAACGACGAGGAGCATTATGGCTTGAGCAATATGATCGATCCAGTCATTAGAAACATGCCAAATCTAATTGGTTTTACAAGGTTTGTGGACGTGGCGATGCAATGTGTTGAAGAGGCTGCTGCTGATCGTCCAACGATGAGTGAAGTGGTAAAAATGCTTGAAAGTATTCTGCAAAATGATGGTCTGGAAACAAATTCTACGTCTGCATCATCATCGATAACAGATTTTGGGACAGCAATAGCTGCTTCTAGACATCCTTATAATAAAGAGGCTCTACAGAGAAAGGAGATAAATGATACTCATGCTTTTGATTACAGTGGTGGATACACTCTTCCAACAAATGTGGAACCAAAATAGTGACAACCAAAAAAAAGGACTTGTCTCTACTGTCTCTCATTCTTTAATTTATTCTCTATTAAAAAAATTGGAAAATTGGTGAAAGGGATAAGAGAATTTAGTTGTTGTATTGTCCCTATATTGGCTACTTTTCGCGAAAACAATCACGTATATTATCAAGATCGAAGTTCAGATTTTCTATTGCCAAGGCTTACAGTCTAGTAGTATCAGTGAGAGTCTCCTATACACTAAGTGTGAGTTCGATTTTTACTGTCCCCTATCCTCAAATGTAATAGAAAAGAGTTATAAGGAAAAAAAAGATTTTGTACATCAAGGCAAAGATTTTGTATATCAAGGCTTACAGTCTAGTGGTACACGTAAGAGGTCTTCCATACACTAAGTGTGAGTTCGATTTTTTACTATCTCCTTTCCCGTTCCATGATTCTACAATGTAATAGAAAAAAATAAAAAATTTAAAAGTTCAATTTTC

>PGSC0003DMG400008581 LRR-RLKs

ATGAATTTATTAACCATTTTCATACTCTTCTTGTTCTACTATTACAGTTTTTTTGCTAACGGGGTGTTTGCGAAAGTTGATTTGAATGATGAGGTGTCTATTCTACTTTCTATTAAAGAAAGTCTCGTTGATCCATTGGATCATCTTCGTGATTGGACAGTGCCCAATAATGCAGCTGCAGGGAACAATAGAAGTATTATTGTTCCCTGCAGCTGGACGGGAGTTGAATGCAACTCCCATGGGGCTGTCGAGAAGCTTGATCTGTCCCACATGAATCTCACTGGCACAGTGTCTAATGATATCCAAAAGTTGAAGAGTTTGACTTCTCTCAATCTATGCTGCAATGAGTTTTCATCGCCATTGCCGAAAAGTTTGTCAAATCTAACTGCATTGAGAAGCATTGATGTGAGCCAGAACTATTTTGTCTATGACTTCCCTGTAGGTCTTGGAATGTCAGAAGCGCTTATGTATTTGAATGCTTCGAGTAATAACTTCTCCGGTTATCTTCCTGAGGATATTGGAAATGCTACATTGCTCGAGACATTGGATTTTCGAGGGAATTTCTTCGAAGGTTCAATTCCTAAATCTTATAGGAATTTGGGGAAATTGAAGTTTTTAGGTCTTTCAGGGAACAATTTGACTGGCTACATTCCGGGAGAGCTGGGGCAGCTTTCGTCGCTTGAGACTGTAGTACTGGGATACAATCTTTTTGAAGGTGGAATTCCAGCTGAATTTGGAAATCTGACAAATCTCAAGTATCTTGATCTTGCTATTGGAAATCTTGGAGGTTCAATTCCTTCTGAGCTAGGAAAGCTGAAGCTGCTTGATACGATTTTCTTGTACAAGAACAAGTTTGAAGGCAAGATTCCGCCAGAGATTGGCAATATGACTTCATTGCAGTTGCTTGATCTGTCTGATAACATGCTTACGGGTGAAATTCCAGCTGAGATTGCTGAACTTAAGAATTTGCAGCTTTTGAATATAATGTCGAACAAATTATCAGGTTCAGTTCCCCCTGGGATTGGAGGTTTGACTCAGCTGGAAGTAGTTGAGCTGTGGAATAACAGCTTATCAGGTCCTTTGCCAAGTGATCTTGGAAGAAATTCGCCATTGCAATGGGTGGATATTTCATCCAACTCATTTACCGGTCCAATTCCAGCAGGATTATGCGCCAAAGGTAACCTCACTAAGCTTATCATGTTTAACAATGCTTTCTCTGGTCCAATTCCAACTGGTTTATCAACCTGCACGTCCCTCGTTCGTGTAAGAATGCAGAACAATCTTCTTTCGGGGACGATCCCAGCAGGTTTTGGTAAACTTGGGAAACTACAGAGGCTGGAACTGGCAAACAATAGTCTGACAGGACAAATCCCAAGTGATCTTGCTTCTTCTACCTCTCTTTCTTTTATTGATTTCTCTAGAAACCACATTCAGTCTTCTATTCCTTCATTCATTCTTGCAATTCCAACTCTTCAAAATTTCATTGCCTCCGACAATAAGATGACAGGCGAAATCCCGGATCAATTCCAGGATTGTCCTTCTCTTACTGTTCTGGATCTATCCACAAACCATTTCACTGGAGATCTTCCAGCTAGCATTGCTTCCTGTGAGAAGTTAGTAACTTTGAACCTTCGAAACAACCAATTAAACGGTCCAATTCCAAGAGCTATTTCCATGATGCCCACATTAGCTATTCTAGATCTATCCAATAATTCTTTAACTGGTGGGATACCTGAGAACTTTGGCAATTCACCAGCACTAGAAATGCTGAATGTTTCTCATAACAAACTGGAGGGTCCTGTTCCGGAAAATGGCATGTTGAGAACAATCAATCCAGATGACCTGATTGGAAATGCCGGTCTATGTGGTGGAGTGCTTCCTCCATGTTCTCATAATGCAGCATACACATCAAAGCAGAAGAGCTTGCACGCGAAGCACATCATCACAGGATGGCTAACTGGTGTAGCAGCTCTTCTGCTTTTTTTGACAGCAGGTCTTGTAGCTAGATCTCTATACAAGAGATGGCATGAAAATGGGAGCTGCTTTGAACCAAGCTTTGAGATGAGCCGTGGTGAATGGCCATGGAGGTTGATGGCATTCCAGAGGCTAGGATTCACTAGTAATGACATCTTGGCTTGCCTTAAAGAGTCAAATGTCATTGGAATGGGAGCAACAGGGGTTGTATACAAAGCTGAAATGCAACGCGAGAATATGGTTGTTGCAGTAAAGAAGCTGTGGAAATCTGGAACTGATATTGAGATGGGGCACAGTGATGATCTCGTAGGTGAGGTGAATGTTCTTGGAAAATTGAGGCATCGAAATATTGTCAGGCTTCTGGGGTTCCTTCACAACAAACGTGATGCAATGATACTATACGAGTATATGCAAAATGGTAGCCTTGGAGAGGTTTTACATGGTAAACAAGCAGCAGGGAGACTGCTTGTTGATTGGGTCACTCGATATAACATAGCCCTTGGAGTGGCACAAGGCCTTGCTTATCTCCATCATTACTGCCATCCACCAGTTATTCACCGGGATGTCAAGTCAAATAACATACTTCTTGATGCAAATCTAGAGGCAAGAATTGCAGATTTTGGTTTGGCAAGAACGATGCTCAAGAAGAATGAAACAGTTTCCATGGTTGCTGGATCTTATGGATACATAGCCCCCGGTAAGTCCTTCAAATTTTTCATCAATGTTTACTCTAGAGTGATTTTTGTTACTAATATTTAACAATCATTTATCTGGATTTTGACTTTGCAGAGTACGGATACACGTTGAAAGTGGATGAGAAGAGTGATATTTACAGCTATGGAGTTGTTCTCATGGAGCTCCTAACGGGTAAACGACCTCTAGATCCTGAATTTGGGGAATCTGTTGACATTGTAGAGTGGTTTCGGATGAAGATCCGAGACAACAAGTCTTTAGAAGAAGCACTTGACCCACATGTAGGAGCAACACAACATGTTCAAGAAGAAATGTTGTTAGTTTTAAGGATAGCAATTCTATGTATAGCCAAACTCCCCAAGGACAGACCATCCATGAGGGATGTTTTGACGATGCTCGAAGAGGCAAAGCCTCGGAGAAAAAGCAGTAGCAACAGTGGAGATAGCCATGATGCTACTACTAACAAAGATAAGCCAATATTCAGTACATCACCTGTAAATGGTCTTTTGTAA

>PGSC0003DMG400009068 LRR-RLKs

CGTGATTAAGAGTGGTTAATTTCCTTTCCCTTTTTCGTAATAGTAATGTGTGTCTTCCTTTAGTGAAGAGCTTCAAAAAAATGGCGGAAATGGTGGAAAATGCTCTTCTGGTGCTCCCTGCCTTACTGGCACCCACTTCCCCTCTTCTTCTAAATCCAAGCAAATTGTTAAAGAATTGATTTTGTGTATTACTGTTAACTTAACTTCTTCATATCATAGTAGTAGTAGTTCCACTTTTGTCTCCACGTGCTTTTCCCACTCTTTTCGCTTCGTTGTTAACCAAATACATACATATACCCCATTTCTTTATTTGCTTTCTTCTTCTTCAAATTAACTGCCTTTCAATCATTTATTGTCTCTCACCCACTCCCCCCCTTCTTCTCATTTAATCATTGTCTCTCTCTAGCTACTTACTTAATTCTGCATTTACTTCCCATCATTTTCCTCTTCATCTACTCCTTCTTTTCAGTTTACTCTACCAAGTGTTTGATTTTTTGCCTCACTGGTACTTGCTAGTATACTATACAGTTTCTGAATTTTGTGGGGTTTTCCGTTTTCTCCATTTTGATGTGATTTGGCTATCTGGGTATTGCTATTCAGGTTCATTTCAATGTTCTTTTTGTTATTGCCGACAGTTTTCAAATTATGCATACTTTGCTTTAAGGTCAAGAAAAGTGTTAAAAATTCAATCTTTTTATGTGTATGTGCTTGGAAATAGGTGAAGAACGTGAAATGCCTGTTTGTTCCTTTTTCTTTTTCTTTTTTTTTTCATTTTTTGGCAGCCCCAAAAGTTGACGGAAGAAATTTGTAGCTGGTTCTTGTTTTTACTTTAGCTATTCTGTTTATTGAAACCTAACTTTTACTTAACTTGCGTGGTTGACCCAAAAACAAAATGTGTTCTGAGTGTCTACGTAACTTTAGCTGCCCTATAAATTGTATATTTATGGTTAAGAAACGGTTCTCTAGTTTAAATGGTAATTATTTATAGTGAATAATGAACATCATACCGTTTCAATGTCTGATTCCATTTTTATTTTTTTTTGAAATTCTTGTAGAGATGAAGGGTGCTGTTACTAGAAGCTGATGCCTTTGTAGTTTAAGCTCAGATGAGGAGAAGGATGGCTACAAAGTTCCTGTTTTTTTCAGTTTTATTCTGCACAGCTCTGTTCTGGCTAGCAAGTAGTGAACCATATGAAGACAAAGAAGCGCTTCTTGATTTCCTTAATAATGTTAATCATACCCGAAACCTGAATTGGGATGAGAGAACTTCTGCTTGCAGTTCTTGGACTGGAGTCACCTGTAATCATGACAAATCAAGAATTATAGCAATTAGGTTGCCTGGAGTTGGATTCCGTGGTTCGATTCCTGGAAACACTCTTAGCCGCTTGTCTGATCTTCAGATCTTGAGTTTAAGGTCTAATAGTTTTAGTGGTAGTTTACCTACTGATTTTGCTAAACTTGGAAATCTTACTTCTATTTATCTTCAATCCAATAACTTTCAAGGTCCATTGCCTGCTGATTTTTCTGCTTGGAAAAGCCTTTCGGTTTTGAATTTGTCAAATAATGATTTTAGTGGGAGTATACCTTCTTCAATATCGAACTTGACTCATTTGACTGCTCTGGTTCTTGCTAATAACTCGCTTTCTGGCAGTATTCCTGATCTCAATCTCCCTAGTTTGCAAATATTAGATCTATCCAACAATAACTTCACTGGAAGCATCCCCAATTCTCTTCAAAGATTTCCTGGTTCAGCTTTTGCTGGTAACCAGCTTTCTCCTGCAAATTTTTCACCTTCATTTCCTCCAGTTCCTCCTCCAAGTGTCCCACCAAAAAAGAAATCGTTTAAACTCCGTGAACCTGCAATACTTGGGATTGTAATTGGAGGGTGTGTTTTAGGTTTCTTGGTAGTAGCTGCAGTGCTGATTATGTGCTTTTCGAAGAAAGAGGGTAAAAGTGGGGCCACAGAGAAATCAATCAAGAAAGAAGATATAGTCAGGAAGGGAGTTTCCAGCAGTCAGCATGGAGTAGGGAATCTCGCATTCTTTGAGGGTTGTAATCTTGCATTTGACCTTGAAGATCTGTTGAGAGCTTCAGCGGAAGTGCTTGGAAAGGGAACATTTGGCACTACATATAAGGCAGCTTTGGAGGATTCTACCACAGTTGTGGTGAAGAGATTGAAGGAGAGTGTGGGAAGAAAAGATTTTGAGCAACAGATGGAGGTTGTTGGCAATATCAGGCACGAGAATGTGGCTCCACTGAGGGCATACTACTATTCCAAGGATGAGAAGCTCATGGTATATGATTTCTACAGCCAAGGGAGTGCATCTTTGATGCTGCATGGTATGTTGAGCACCCAACTTCTACTACTCTTGCAGTTTTATTTTTGAACCGAACCATAGAGAACTGATTATCAGGACCAAAAAGATAATGCAGATCATTTTTAGCTATGATGTGTCTTGTGCAGTAGTATTTGAAATTGCTTGTAGGCTAATTCCAACGTAACCAATTACCTTTGAATTAATAGAGTATCTGCTGTTGTTGTCCTCTTGATAGTTTTTTTAATCACCAATAGTTTATAGGACAGCATAAGTGTCTCGAAGCCATAACTACATGCCACTTTTAGGCCATGTCTTAGTGCAACTTCTGTTTTAATTTGTCAAAATTCAAGAACATCATGTTGCTCGCGAACTTTGCTTTGGTGTTACAGTCAACTTTTATCATTATTAGGTGGATAGTCTCTTAACTGCACCTTCAAGTATTCCTATCATTATATCTGATGCACTCTGTAATTGCAGCAAAGAGGTCTGCTGATCGAGTTCCTCTGGACTGGGAAACCCGACTACGCATTGCCATTGGTGCAGCTAGAGGCATTGCTCAAATACATGGACAGTCTGGGGGAAGGCTTGTCCACGGGAACATAAAATCCTCAAACATATTCCTCAACTCCCAAGGTTTCGGTTGTATTTCTGATCTTGGCTTGGCAACTATAATGGGTCCAATTGCTACACCAATCGTGAGAGCTGCTGGTTATCAACCTCCAGAAGTTACAGATTCAAGAAAAGTATCTCAAACCACTGATGTCTACAGCTTTGGCGTCCTAATACTTGAACTTCTCACTGGAAAGTCACCTACACATGCCACAGGTACTAGCGACATTGTCCACTTAGTCAGGTGGGTACATTCTGTCGTTCGTGAGGAGTGGACTGCTGAAGTATTCGATGTGGAGCTTTTGAGGTATCCAAATATAGAAGAAGAAATGGTAGAGATGTTACAAATAGGTTTGACTTGTGTATCCAGAATGCCAGAGCAGAGACCAAAAATGACAGAAGTAGTAAAGATGGTAGAGGGTGTTAGGAGAGTGAATACGGGGACTCGAACATCCACTGAAGCTTCTACACCGAACCTGACACCACCGATGACTGAAATAGGGTCGTCTTCCATTACACACTGATATTTCTTTGTTATTTAGTGATTAGTTACTGTTCTATATGCTCAATTGTTTCATTGTTATAAGTTTTTTCCTCAGACTTCTGTCAAAACATTATTTCAGCTACAAAATTGAATAGAAATATTTACTTGATC

>PGSC0003DMG400009180 LRR-RLKs

CCAATATTAATCAAAATAAAAATAGGAAAGTTCTGACAAAGCCTTTCTTCTGTTCCCGAGCTGAAGCTTGGGGTTTTGTGTTTTGTGGTGCCCGGCACTGGTGGTGAAAGAAAACTCCGACGGCTGTGACTGAGGTGGAAGAGGAGAAAACGACGCCGTGAAGATGAGGTTGCTGCTGCTGCTGCCGATACTCATCGGCATTTTTCTGTATTGTGCTAATTATGCCATTGCCGACACAGATCCAAATGACGGTCAGCTATTGCTTTTAATCATTCTCATTTATTGTTCCTATCTGTTTGTCTGTATGCTTACTGATCATTAATGGAGGTTGATTGATTTGGAAAGTCAATGAGTAATCTACCTGACAATGAACTCGGCAACCGTCGCAAATTTGTTTGATATTTTCCTTACCATATATATGTAAAAATGCGACAAGTTCCTTTCTGATCAGGGCATGAATCATATAGAGGGAGCTCTCTTGGATTCTTATATATTATGTATGTATTGGCCATGTCGGTGGACTTTTAGTACTGTTGAATCTTTGCATTATTACTTCCATTCATTCTGAGAACTGACATATTGAAACTTCGTCGGACATATTTGACTGAGTTACCATTTTCAGCCAAGTTGCTCCAGAAGATTTTGGCTTTTCTTCTTTCTGCTGCTTGGAAGCGGGACAGGGTAGTGGAGGTGGGCCGCCAAAACAAGTGAGGTTAAAAATGATCAAGGGCATTATAACCTAAAAGACTCTTCTCCCAGTGATAATACACATATAGGTTCATGGCTTGATCAACGCACTTGTTGCGTTTCACCATGTTTATTGATGTCTTGTAGATATACCACTCGTTAACAGTTTGATTACTCTTCAAGCTATTATTAATGAAAAGAAACTTTTTGTTTGAACAGTATAAGAATATGTTGACCTGTAAGTATGCCTAGCTGATTTTTTTGGTGGCGCTTTGATTTTCAGCATCTGCTCTGAGAGTTCTGTATAGCAGCCTAAATTCACCTGGACAGCTAACAAAATGGAATGCCAATGGTGGTGATCCTTGTGGAGAGTCCTGGACAGGTATTACTTGCTCGGGCAATAGAGTTACTGAAATGTGAGTTCCTCTCTTCCTTACTGTTGTAAGTAATTATTGTTGATGTTTTCCTTCTTTATCTTCTTCATCATGTTTCAATTCTTGGAATTGTTGACTTGGTGATTTCTGATGCTTTCTTCCCTTTTCATTTTCCTCCAGCAAGATATCAGGTCTTGGACTCTCTGGTTCACTGGGCTACCAATTAGCAAGTCTTACATCGGTGACTAATTTGTAAGATAAATGGCCTTCAATAATGTCCTCATAAATGTTTCTTCGATCTATATCTATATCAAATTCTTACACACTTCTAATTTTGCAGTGACATAAGTAATAACAATTTGGGAAACCAGTTACCTTATCAGCTTCCTCCAAATGTGCAAAGATTGTATGTCTCGTAGTTAGTATATTTTCATCATTCCAGATGCTATTTTTCCACCTATCTTTCTTAAACCTTGCCTTGCAATGCAGAAACCTTGCTGCTAATGGTTTTAATGGTGCCCTTCCATATTCCGTTTCGCAGATGACTTCCCTCCGATACTTGTATGTAAAATCTTTGAATTTGACTGCAATTTGCTTTCCAGCAATAAGCTTTTGGTTTAAGGTTGAGCTTTTCAATGACTTTAATATTTTTATCTGGCTGTAGAAATGTCAGTCATAATCAAATTCAAGGACAAGTGACTGTCGCGTTCGACTCTCTGTCTTCTCTCAATACATTGTAAGCCGTATCTTCTCTCATGCCATTGCTTGTGTGTTTTAATATTACAAAGTTGTGGTCATAACTCATATATGAGGTGTTGTAGGTAAAACATGTCATTCTTACTTCATTAGTATAAACTTTTCTTCTCCCATCCGAGTTGTTGACTTGTTGCTTTGGTAAAGAAAGTCAATATCATGAAAGACAGTATCTGATCCTTGTTGTAATCTGTAAACTTATGTGGTTGTCCCAGTTAAGTCAACTAAGAAGCTGTATACATTTCGAGATATTCTTAAGTCAAGTCTTAAAAACCTCTAAATGACTAGTTATCATTCTCTGTCTCATAGTTTTTGATTGATTGATCTGTATGCTTCAGGGATTTCTCATTCAATGCTATGACGGGTGATCTTCCTCAAAGCTTCAAGGCACTGACTAGTATGAATAAAATGTGAGCTAATAAATTTTCTGCTTCGTCCTTGTAGACAATTGGTTGAGTCCTTCAGTGGTGAAGTTTACATGTTGAAGTATGCAGGTATTTGCAGAACAACCAGTTTACAGGAACTATTGATATCTTAGCCAATCTTCCTCTTGATGATTTGTGAGTTTTCAGTCTATGCTTCTTTTTAGATGTTATCCCTGATGTTTTATCTTATTTATATAGACTCGATTGAAAGCATGGTTGAAAGTCATGTGTATTCCTTTATGCTTTAGGAATGTTGAGAACAACCGTTTTACTGGCTGGGTTCCTGATCACTTGAAAGGGATCACGAAGTAAGTATATCATGTTAGCGTAATAAATTTTGAAACTTATGTCAGAAGTATTTTGAATTTATTCAGGGTTTGTGACATTTCTGTTACAGATCAAATGGTAACTCATGGAACTCAGGATCTGCACCCCCTCCTCCACCTGGCACACCCCCTGCAAGCAGGCCGCACCACAAATCTGGAGGCAATAATAGCCCTACTGATGGTGCAAGCAGTGGTGATGGTGGTGGTAAATCGGGGATAAGGGGTGGAGCCATTGCAGGCATAGTGATATCTGTTCTAGTTGTTGGGGCAATTGTAGCATTCTTTGTTATCAAGAAGAGATTGAAGCGGTCATCGACGGACATTGAGAGACATGATAATCAGCCATTTGCTCCTCTTGCTCCCCCCCAGGAAGTACATGGTACTTCTTTTCATAATTTCTTGGTTTTATCAGTTAAGATGCCTTTTAGACCCTTTCATGGCCTCTGTGATTGTTATTTAACTGTGGAACATATCAATCAACCACACTATCATCAACCTCATTCCTGTATCCTGCTTAAAAAGGTGTAAAATAGACCTGCTGCATAAAATGGAAATTTTATGAAAGTGAATTTGGAGAATCAATCTTAACCTTTGTGCTTTATCACTGCCTGTGGTGTATAGCTAATAGTGGATCAACACAGCTACTGTCTTCGCTATAAAAGAAAAACACCAAAGTAAAAACAAACTTTATATCGAAATGTAAAACCGCTAAGGCTGATAGGTCCTATCTTCCATGGAAGCAAAACTTCTTGCTTACTGAAAAGTTCCAAGAACTTGATTGTAGAAATCTGAACTGATGTTTAGACTCTGTATTGCCTGTTGAAGCTGATAGTGGGATAAAGCGAACTTCAGCTTTTTTGATAATGAAGAAAACTTCAGCTTTTTGATCTCTTTCAATGGATCTGGTTCTATGCTTGTTATAAGTTAGCTCCCCTTGATGGTTTTCCTTCCTATAAAACTACTATTCATAAATCTTATTGACCTTTATAGAACAACTATTGCTTCAAAATGCAACTCTCAGCTAAGTGAAAGTTTCTCCCATTCTAAACCATTTGTGCTCACCACCAAATTCACCTCTTTGCAAATTCTAATTCTTTGCTTTTCTCAACTCCATCACTCGTGGTGCATAGTTATGCCTTTTAAGCCTCATCTCACTTCCGTATCTCATTAATCCGAACCTGAGTATGCATTTGCCTGTTTGTAACACTAACAGTCTTGTGCTTCTTCTCTGTCTGGATTTTCTGTTCATTCATTGTCCTTGTTTTTTACTGCAGAATTGAAAACTAATGAAGCTTCCTCTGCGCCAATAGTGAAAGTATTTGAAGCTCCTGCAGTAGTAAATCTAAGACCTCCACCTATTGAACGCCACAAGTCATTTGACGAAGTTGACATAGCAGCAAAACCAATTGTCCCACCCAAGAAAGTTAATACAGCTAAAATAGATGCAAGACAATATTCAATCGCAGACCTACAGATGGCGACTGATAGCTTCAATGTTGATAACCTTATTGGCGAGGGATCATTTGGGCGTGTTTATAGGGCTCAGTTTGATGATGGAAAGGTATGATTTCTCTCTTATGACTAACTGACTATCTATGTGAAGAAAATGGCTCTCTCTAATTGGATAAAAAATTCATATCATGAATTTTTTATGTGCTGCGAATGAATGTTTAAGCAAGATAGTTGAAATGACAGATAAATGTTCAATATGAAAAAGTACCTATCTAAAGAAAATGAAAGAAAGAAGTTTGATCATTAAAAATTAGTTGAAAAATTATGTTTGTTCAAAACTCAGTTCGGAGCTACTATCATCTTGTCCTGTGTAACTGAACCATCCTCGTAAGTTCCTTTCTAATTTTTGAAATGCCATAGGTTCTTGCAGTCAAGAAAATCAACTCTTCTGCACTCCAGAATCCTGAAGATTTTCTCGATATTGTTTCTGAGATATCACGGTTGCACCATCCAAATGTTACTGAGCTGGTTGGATATTGTTCAGAGCACGGGCAGCATCTGCTGGTTTATGAATTCCACAAAAATGGTTCTCTGCATGATTTCCTGCATCTGTTAGATGAGGAAAGCAAGCCTCTAACATGGAATAGCCGAGTTAAGATTGCACTAGGCACAGCAAGAGCACTAGAGTGAGTTTCCATGATCTACAAAGATATTTGATGCCATTTTATAATGTTCACGGAGCTCATAACAAGGTTTTATATTTTCCTTGTACCATTGGTTGAAATACTAGCTCTGTCATATATTTTCTGGTATGTGTAGTCAAACAATAATGAAGTTTCTTTTCCTGGGGGGGGGGGGGGGGGAAAAAAAGAGTTTTGGAANNNNNNNNNNNNNNNNNNNNNNNNNNNNNNNNNNNNNNNNNNNNNNNNNNNNNNNNNNNNNNNNNNNNNNNNNNNNNNNNNNNNNNNNNNNNNNNNNNNNNNNNNNNNNNNNNNNNNNNNNNNNNNNNNNNNNNNNNNNNNNNNNNNNNNNNNNNNNNNNNNNNNNNNNNNNNNNNNNNNNNNNNNNNNNNNNNNNNNNNNNNNNNNNNNNNNNNNNNNNNNNNNNNNNNNNNNNNNNNNNNNNNNNNNNNNNNNNNNNNNNNNNNNNNNNNNNNNNNNNNNNNNNNNNNNNNNNNNNNNNNNNNNNNNNNNNNNNNNNNNNNNNNNNNNNNNNNNNNNNNNNNNNNNNNNNNNNNNNNNNNNNNNNNNNNNNNNNNNNNNNNNNNNNNNNNNNNNNNNNNNNNNNNNNNNNNNNNNNNNNNNNNNNNNNNNNNNNNNNNNNNNNNNNNNNNNNNNNNNNNNNNNNNNNNNNNNNNNNNNNNNNNNNNNNNNNNNNNNNNNNNNNNNNNNNNNNNNNNNNNNNNNNNNNNNNNNNNNNNNNNNNNNNNNNNNNNNNNNNNNNNNNNNNNAGGTTTTTTTTCGGGGGGGGGGGGGGGATAGAATGAAGCTTATGGAATAGTTACTTTCTTCCTGAAGAGGAAGTGGAGAGTTATATATTGTGAACTCCAATACTAAGCACTTAGTAATTAAAGCTGTGTTCCTATGTTTAAATCAGACTTAATCATTTATTATGAAGGCCATTAAATAATTCAGATTTCTGGAGCATTGATGATCAGTAACATGCAAGAAGATGATACATTATCTGAACATTAAAAGAAAAGAAAATATGTGTTGCGAGCCATTGTTTATCCACCCTGTTTAATATACTGATCTTTCCATCTTCTTCTTCTCAAGTTTCTAGAGTAGTAGCTAAAATAAGCATCCTTTGCAGGTATCTGCACGAAGTTTGTTCACCATCTTTGGTTCACAAGAATATCAAGTCTGCAAATATTGTACTTGATGCAGAACTCAATCCTCATCTATCAGACTGTGGATTGGCAAGTCTAATTGCTGATGCAGATCAGGTAAGCTATAGCTCCTAGCTTTAGGATTTCCTTTGCTAAGTCTCATTTGTTGGATGTTGCTATATCTTCTAATGTATGAATGTGGGTTCTGATTGACTAATGCATATGCATAAGGGTGCTCATGCACATGCCTATATGCCTGTACTTGCATCCTCTTTGCCTATTCATCTGTCCTTGTAGAAAAGAAACCAGAAATTTGATTCAGATCTTCGATATAAGTCATCCACAGTATCATAACTATAGAAGGCGTATACTCTTTACCATGAACTATCAAATATCAAAAAAGGAATTTTGTTGCCATGTTAAAAATTAGATTGACTCAGAAGCTGAGAAGTATCTACTAGTTTTTCGCAAGGGGTCTTATTTCCTCTTTTTCAGCTAGTATAGATTTTCATCATAAAGCCAGTTTCCTGAGATATTCACATGACTATCAATCTCTTGTCGTAGGCATTGAACCATAATACAGGATCGGGATATGGTGCCCCCGAAGTTGCTATGTCCGGTCTCTGTACCATAAAAAGTGATGTGTACAGCTTTGGAGTGGTTATGTTGGAACTTCTTACCGGACGAAAACCTTTTGATAGGTAATGTTATTTAGCATCCCATGATATTATGATGTAAGTTTCTTAAAGAAACAGTACAAATTTGAAGTTTCAGAAAATTTTCTGATGATATTCTCTGTTCTCTTAATGCAGCGCAAGAACAAGATCTGAACAATCTCTAGTTAGATGGGCAACACCTCAGCTGCACGATATTGATGCCCTAGCTAAGATGGTTGATCCAGCACTCGAGGGGCTCTATCCTGTTAAATCTCTATCACGCTTTGCTGATGTAATAGCTCTCTGTGTCCAGGTAATATTTCGAACTGGAAGTGAATGATTAGCTTTCAGAAATCTGCTACTTAGTTTTCATTAATCTAGTTCCTTAATTATCTCAATGGACCTCTCATTTTTTTTTCTTTGTGGCAGCCTGAGCCCGAGTTCAGGCCACCTATGTCGGAAGTGGTTGAAGCATTAGTTCGGCTAGTGCAACGAGCAAACATGAGCAAGAGGACATTTGGTGCTGATCAGGGTACTTCAAGGGGTGAAACAGATGGCCCGGACTATGAACCATAGTTTCCATGTGACGTTGAGTACTCAACATGTTGCCTATTCTAACACGTGGAGGTGTTTTTTTTTTTGTTTCTATTCATGAGAAATGACTTCGAGCCTGTTGTTTTGCAGACTTGGAGTGCATTGTACCAAAACGAGACGCTAAAATCTGTTCATAGTGCATAATTTATTTTCAGTTCTTTTCCTCTTCCTTCTAATTTGTTCTTACCAGCGGGATTAAAGAAATAATATTATTTCATATTTTCATATGGTGTCACATTACAAACTATCATCTGAACAAACATGTATGTACTAGTGCCTTATTCCAACA

>PGSC0003DMG400009181 LRR-RLKs

GGAGGAATTGAGGAGAGGAGAATAGTATGGTAAAGTCTCTCTCTCTCTCTCTCTCCCACGTATATTACACGCTTTCAAATATCCATAGAATAAACAAAGTCCACCAGCCATCTATCTATCTAGCCGTCCCAAAAAGTAAACGCTTTATTTCAAAAACTCATACCTCACCTTTTTGCTACATAAAAATTGCACCTTGTCAACATCAACTACACTCTCTCTCTTCTCTTCATGGGGTTAGACTAAAGTCTGCACGATCAACAGGACAAAGAATTGCCAAAAAAATACCCACATGGTTGATTGGAGTATCTTGGCATCTTCAACTGTCTTTCTTCTTCATCTTTAGTTGTCTAAAAATAATAATGATCATGGATAATAAAGACAGCCGCCACCAATCTGGTCTGACATATCATCAATTTCGAGCAGCCATGCTTCTTACTTCCTTTCATTCTTCTTCTTCTTTACCCCACCATAAATCTTTCTATCCAATAATATTGATAGTTGTTCTACTTCTTTCTTCTATTGCTACCATCTGTCATGCTTCCTGCAATCAGCTTGATCGTGACTCTTTGTTATCATTCTCTGTTGGTATCTCTTCACCTTCTCCTTTGAATTGGTCCTCTTCCGAGGATTGCTGCACCTTGTGGGAAGGTGTTGGTTGTGATGATAATGGCCGAGTAACCGCTCTCTGGCTTCCCTCAAGAAGCCTTTTTGGAAACATAACCCCTGCTATTGCAAATTTGAGCAAACTCTCTCAACTCAGCCTGTCAAATAATCGATTTTTTGGTCCCCTCCCAGATGGATTTTTCAAGTCATTTAGTAGCTTGCAGATCATTGACTTGAGTTATAACCGTTTATCAGGACGATTGCCGCTATCAGATAGATTGCCATCACCCATCAAGACAGTAAATCTCTCTAGCAACCACTTCAATGGGACAATTCTATCATCATTTCTCGAGCCAGCAATTAATTTAGAGAGTTTCGATATTAGCAACAATAGCTTCTCTGGTCCAATACCCTCCTTTATTTGCAGCTACTCAGCCGCTGTCAGAGTTCTTGATTTCACTTCCAATGACTTCAGGGGCCAGATGCCCCAAGGGTTTGGGAGTTGCTCCAGTTTGGTTACTTTGAGAGCAGGATTCAACCATCTTTCGGGATTCATTCCTGATGATATTTATAGTGTGTCAACACTGCAAGAAATCTCTTTACCTGGCAATAAATTTTCTGGACCCATCCCAGAAAGCATTGTCAACCTTGTCAACCTCAGAATCCTCGCACTCTATGGCAATGAATTGACAGGTTTGATCCCTCAAGATATTGGAAGGCTCTCCAGATTGGAACAGTTGCTCCTCCATATAAACAATCTAAATGGCACTGTGCCGCCATCGCTGATGACTTGCACCCGTCTCACTGTGCTCAATTTAAGGGTCAACTTCTTGGAAGGTGAACTCTCAGCTCTTGATTTCTCCAACCTTAGTCGACTTGGCATAATCGACCTTGGAAATAACTTCTTCACTGGAAGCATTCCACAAAGCCTTTTTTCATGCAGGTCATTAACTGCAATCCGTCTGGCTACTAACTATCTGACAGGAGATATCTTGCCTGGCGTAACGTCTTTACAAGCTCTGTCCTTCCTCTCGGTTTCCAATAACAGCCTAACCAATTTTGCAGGGGCAATAGAAGTCCTTAAGGGTTGTAAGAATCTTACCACACTAATCCTCACCAAAAACTTTTATAATGAAACGTTGCCTGATAATGGGAACTTGATTGGGTCTGAAGATTTTCAAAATCTCCAAATTCTGGGTTTAGGTGGTTGTAATTTCACCGGACAGATACCCACATGGCTGGTTAAACTTGGGAGGGTAGAGGTTCTAGACCTTTCTATGAATCAAATCACAGGCAAAATTCCAGGTTGGTTGGGGACTTTACAGAATCTATTTTACCTGGATTTGTCTCAGAATTTTCTATATGGAGGCTTCCCAGTTGAACTTACTCAACTGCAAAGACTCGCATCACAAGAGGCTGCTGATCAGGTCGACAGAGGTGCTTTAGAATTGCCTGTGTTTGTTCAGCCAAACAATGCCTCAAATCAGCAATATAATCTACTGTCAAACCTGCCACCAGCTATTTACCTTGGAAACAACAACCTTGATGGCAATATTCCAACTGAGATTGGCCAGCTGAAGTACATTCATGTTCTTGATCTGAGCAAGAACAACTTTACAGGAAACATTCCAGAAACAATATCCAACCTCACTAATCTGGAGAAACTAGACCTCTCTGCCAACAACCTCTCAGGTGAAATTCCTTCTTCACTCAAGGGTCTTCATTTTTTGTCTTCTTTCAGTGTTGCTCACAACAATCTTGAGGGACCTATTCCAACAGGAGGTCAGTTTGATACATTTCCTATCACTAGTTTCTTAGGTAATCCAGGACTGTGTGGTCAAATCCTGCAGCACCCTTGCCCTGATCGATCAGGAACTACACAACCTTCTGCAGTTAGAAAAACCGCAAAAAGGAAAATCTTAATTGGACTCATCCTAGGAATCTCCTTCGGCATTGCCTTCACAGTCATCATCATAGCATTCTGGATATTCTCCAAGAGGAGGATCCTTCCAAGAGGTGACGCCGAGAAAAATGACTTGGAAATTGTGTCCTACAACTCCACTTCTGGATTGTCTGCTGAGATTGGAAAGGATAACAGCATGCTTGTTATGTTTCCAACCAATAAGGATCAGATTAATGATCTCACAATTTTTGACATTCTGAGAGCCACGAATAACTTCAACCAAGCAAACATAGTAGGTTGTGGAGGCTTTGGTCTGGTCTATAAGGCAACTTTAGCAGATGGAACTATGTTGGCTGTTAAGAAACTTTCAGGAGATACGGGTTTAATTGAAAGAGAATTTAAAGCAGAGGTGGAAGTTTTATCCACTGCCCAACATGAGAATTTGGTCTCTCTTCAAGGTTACTGTGTGCATGATGGCTGTAGGTTGCTGATCTATTCCTATATGCAAAACGGAAGTCTGGACTACTGGTTGCACGAGAAAACTGATGGAGCATCCCTACTTGATTGGCCAACTCGACTGAAGATTGCACAGGGGGCAAGTTGTGGGCTGGCTTACATGCATCAGATATGTGAACCACATATTGTTCATCGTGACATAAAGTCCAGTAACATCCTCCTTGATGAAAAGTTTAAAGCGCATGTGGCAGATTTCGGATTGTCCAGAATGATTCTTCCTTACCAAACTCATGTCACCACTGAACTGGTTGGTACCCTCGGCTACATCCCACCAGAGTATAGTCAATCATGGATAGCCACTTTGAGAGGAGATGTCTATAGTTTTGGAGTTGTGATGTTAGAACTCCTGGCTGGCAGAAGACCTGTGGACATGAGCAAACCAAAGATGTCAAGGGAATTGGTTGTATGGGTGCACCTAATGAGAAATGAGGGTAAACAAGAAGAAATATTTGATCCTATACTGAGAGACAAGGGTTTCGAAGAAGATATGCTTCAAGTGCTTGATGTTGCCTGTATGTGTGTCAGCCAGAATCCTTTCAAAAGGCCAACTATAGCAGAAGTGGTTGAGTGGCTCAACAGAGTAGTATCCAACGAGGGAGCACCTAAATAAGGTGGTGGTACACATAAATTTGATGTAGATGTGAGTGTCTGTATAGGTGAACTTTTGCTTTTTACACGTAGGAAGTTAACATATAGGAATAAATGTATAAAGTGAACAATTTCCTCAATAAATTACATTTACGATTTTGCGTCATGGCTAACATTGGATGACAGGTAATG

>PGSC0003DMG400009196 LRR-RLKs

GGTTATGCCTCAAATTGATGGTGGTTTGATGTGGTTAGGCAAAGGAAGGATGATGACGGAGGTGGAGGATACCGGCCGGTCACCAGTTACGTTTGCATGTTCTTCATCATCATCATCATCATCAAGAAGAAGATCATACAATGCATTGTGTCTCATTTTCCTTGTTGTTGTTGTATCCTTGATCACATTTGTATCGGCTCAACCGGACGATGACGACACCGGCCCAGATGATCATCATTATCTTCCCGATGCAAAGAGCTCTTCGGAGGCCCTCTTCAACTTTAAAAGCTCACTATCTACATCTTCAGGTAAAGGAAAAGATGTTCTCGGGTCTTGGGTTCCATCGACAAGTCCCTGTACCGGGAACAATGCCAATTGGTTGGGAGTAATTTGCTTCGAAGGGGATGTATGGGGTTTGCAACTAGAAAACTTGGACCTCTCAGGTGCAATTGATATAGATTCATTGCTTCCTTTGCATTTCCTCCGGACCCTAAGCTTTATGAACAACAACTTCAAGGGGGGAATGCCTGATTGGAACAAACTTGGTGCCCTCAAGTCATTGTACTTGTCCAATAATAACTTCTCTGGTCAGATCCCCGATGATGCCTTCAAGGGTATGACTTATCTCAAGAAACTTTATTTGGCCAACAACCAATTCACTGGTAACATTCCCACCTCCCTCGCTACCTCCTGCCCCAGGCTCTTCGAGTTGACCCTCGAAAACAACAAATTCACTGGCTCTATACCTGATTTCCGCCGTGGCCTTCTCAAACTCCTCAATGTATCCAACAACCAACTCGAGGGTCCAATCCCTCCTTCTCTTTCTTTAATGGATCCAACAACATTCGCAGGTATCATTGGTTAATTTAATTATGTACCGATCTAGCCTTTTTAATTTATTATTATTATACATATACATGACAGGGAACAAAGGCATGTGTGGGAAGCCTTTGGAATCAGTTTGCAACTCCCCAATTCCAGAAGCAAATACTACTACTCCGCCTAACAGTCTCAACTCTACCATCAGCACCCAATTATCTGGTGATATAAATAAGAAGTCTCCCTCTCTCCTGAGTAGAGTGATGTTGATCGTTGCAGTTTGTTTGGTGGTCTTATGTCTCGTGATTGTGTTAATTCTTATCATTCGCCGCCGCAGCAGCAGCCAAAATAATTCACAACTCACCAGCCGGGCTGTTGAGTCAAATAATATTGATGGGGATCAAAACATGACCTTTTCATCATCAGCTCTTCCTGATATGACGATGAGCGGCAATCCAACTTATGCAAGACACGATAACAATAACAAGGCAGTAGAGGCACCACCAGCAGCAGCAGCACCAGGCGCAACGGTGGTGGGTAAGTTGTCCTTCGTTAGAGATGACAGGACAAGATTTGATCTACAGGACTTGCTAAGAGCATCCGCGGAGGTGCTTGGCAGTGGGAACCTGGGGTCCTCTTACAAGGCACTACTTATGGATGGCCAAGCCGTTGTGGTGAAGAGGTTTAAGCAAATGAACCATGTCGCTAAAGAAGACTTCCACGAGCACATGAGGAGGTTGGGCAGACTAAGCCACCCGAACCTACTCCCCCTTGTCGCCTATTATTATAGGAAAGAAGAGAAGCTGCTTGTTTATGACTATGCCTCCAATGGTAGCTTGGCCAGTCACCTCCATGGTATGTATATGTATATTAATTAATTAGAGTATATACGTGAGTTTAAATTATTAATTACTCGGCTCTGTGTGTGTATATATATGTAGGCAATCACTCTAGGCTGGACTGGTCAAGCCGCTTGAAAATCATAAAAGGAGTTGCAAAGGCGTTGGCCTACCTTCACAATGAGCTGCCAAGCCTAGCACTTCCGCATGGTCACCTCAAGTCATCAAATGTACTTCTGGACAAATATTCTAATCCGTTGTTGATGGATTACACATTGGTGCCCCTGGTGAATCTTTCCCAAGTCCAACATCTTCTAGTGGCTTACAAAGCTCCCGAGTATGCACAACAAGGCCGCATCACAAGGAAGACTGATGTTTGGAGCCTTGGGATTCTAATACTAGAGACCCTCACCGGTAAACTCCCAACAAACTACCTTGCTCTCAGCACTGGCTATGGCACCGAATTGGCTACCTGGGTCGACACCATCATTAAAGATAATGAATCGGCTTTTGACAAAGAGATGGACCAGCTCACTCATCAAGGACAGATCCAAAAGCTCTTCAACATTGGAGTGGCTTGCTGTCAAGAAGATTTGGATACAAGGTGGGATCTCAAGGAAGCTATTGAAAGTATACAAGCCTTAAATGACAAGGTCGACGATCAAGTTGATGTTGGATTTTAATTTTAACATCGATCGTACGTACAACAATACATATACTCAACCCCATTAGTTAATTCCCCTCTCTATATTTATATGTGCATTACATATCATGTTTGTACACATTTTCATTGTATAGTACTTCTCTAAATTACTCAAGGCGTAACTAGACTTGCATGCCAATCGATCGGTCTCTCACTCCATAAGTATTCGACCCTACCATTTGCAAGTATATATGTACATCTAGAACTTTGTTTAGCTCTAAAATTT

>PGSC0003DMG400009238 LRR-RLKs

ATGAAACACAAAACATTTGTTCCTTTGTTCCCTCCCCTTATTTCACCTGAGCTCAGAGAGTATATGCAAATCTTGTACATAGATTTTCTTGAAACCCATTATCATTTTCTTGAAGTTTTCAGCTTTCTGACTGTCAAAGCCAAAAACAAAAAAAAAATGAACTTAAGGTATGCTCCCCATATAGTTATTACTATATTCTTGTCTTTTTCTCTTCAAATTGTGTTTTCTCAATTGCCCCCAAATCAAATTAATGTCATGAGAAGTGTCTATGATCTGTTTCAGAATGATACTGGTGCTTCTTTTGTATGGAATGGAACTGATAAAGCTTCAACCCCATGTTCTTGGAAAGGGGTTTCTTGCAATTCTAATAATTCTTCCCTTACCAAAGTCACCTTTTCATTGTTCTCTATTTCTAGCTCTGAATTCTTGCCTTTTATTTGTCAAATTGATACTTTGGAGTCTCTTGATGTTTCCCAAAACTTTTTGAGCTCTATCCCAAATGAGTTCATCACTGTTTGTGGGGGAATTAGTGGGTTGAAATTGTTGAACTTTAGTGGGAATAAATTGGGGGGTTTTTTGCCTACTTTCACTGGTTTTGGAAAGTTGGAGTCTTTGGACTTTTCTTACAATAACATGAATGGGAAAGTTGACTTGCAGTTGGATGGATTAGACTCACTCAAGAGTTTGAACCTTAGCTTTAACATGTTTAATGGTCCAGTTCCTACCAGTCTTGGAAAATTTATTCTTCTGGAGGAGCTTCATCTTTCTGCAAATTCTTTTGAAGGTGAATTCCCCACTCAAATTGTGAACTTTGGCAACTTAACTTTGATTGACCTATCTCTAAACAGCCTATCTGGTGTAATTCCTGATAGATTAGGTGAACTTTCCAAATTGCAAGTTTTGATTTTGTCAGCCAATAATTTGAGTGGCACAATCCCACAATCCCTTAGGAATATCACAACGCTGACGCGTTTTGCTGCAAATCAGAATAATTTTGTTGGAAATATACCCTTTGGTATAACCACATACCTGAGGAATTTGGACCTCAGTTTTAACAGATTAAATGGTACTATTCCTCAGGACCTCTTATTCCCAATGAATTTGCAGTTTGTTGATCTCACTTCCAATAAGTTAGAGGGACCTGTTCCTTCAAACATGTCGATAAATTTGATCAGGTTGAGATTGGGGCAAAATGCTTTGAATGGGTCATTTCCATCTGCTTCTTTTGAAAGCCTTCAAAGTTTGACCTACTTGGAACTGGACAACAACCAGTTAACTGGACCAATTCCTTCTGAATTGGGAAAGTGCCAAAAGCTGGCCCTTTTGAACTTAGCCCAGAATAAGCTGAGTGGTGTTATTCCTGTTGAGTTGGGTGATATTTCTAATCTTCAGGTACTGAGTCTTCAGTCCAATAATCTAGTTGGAGAAATTCCAAGTAACATTTCACAGTTGAACAGATTGCAGAGGCTCAATTTCAGCTCGAATTCATTGACTGGTTCCATTCCAAGTTCGTTATCAAGTTTGAGAAGTCTCACAAACTTGAATTTGCAGGGGAATAATCTAAGTGGTCGAATTCCGGTTGACATTAGTAACTTAAATGTGCTGTTAGAACTCCAATTTGGAGGGAACCAACTCAGTGGGCCTATTCCTGATATGCCTTTGAGTTTACAGATTTCTTTGAATCTGAGTCACAATCTTTTTCAAGGACCTATACCAAGTAGTTTTTCTAGATTGACTTCATTGGAAGTTTTGGATCTCTCTTACAACAGGTTCTCGGGTCAGATTCCAGACTACCTGACGGGAATGGGAGGCTTGACTAGGTTGGTGCTTTCAAACAATCAACTGTCTGGAGTTGTTCCAAAGTTTGGAAGCTTTGTCAGTGTTGAGACAGATGGAAATGGGGGTCTGATCTATCCTTCCCCAGTTGTGCCACCACAAGCTGCTGCAAAAAAGAGAAAATCAATAGTTGTTGCAGTTGTTGTCCCAATTGCATGTGTAGCAACAATTGCCGTTTTCGTGGTGATTGCTATTTCAATCTCTAGAAGATATTACAGGATCAATGATGAACATTTTCACTCAGGATTAGAAATTTCTCAATCCCCTGTCGTTCAGGGAAAGGTTTTAACTGCAAACAGCATTCACAAGTCAAATATAGACTTCACAAAGGCAATGGTAGCAGTGTCTGATCCCTCAAATGTTGTGTTCAAGACAAGATTTTCAACCTACTATAAAGCTGTTATGCCATCAGGTACAACCTATTTTGTCAAGAAGCTTAATTGGAGTGACAAAATATTTCAGTTGGGAAGCCATGAACTATTTGGGGAAGAGCTAAAAAATATTGGAAAGCTGAATAATTCAAATGTCATGATTCCACTAGGCTATCTTTTGGCTGCTGATAGTGCTTATCTCTTCTACGAATTTGCCCCCATTGGCTCATTATATGATGTTCTTCGTGGTAGCTTGGGCTACTCGTTAGATTGGGCAAGTCGCTACAGCATAGCTATTGGAGTAGCTCAGGGTTTAGCTTTTCTTCATGGATGTGAAAAGGGTCCAATCCTCCTCCTCGACCTTTCCAGCAAGAGCATTCTCTTGAAGTCCCAGAATGAGGCTCAGATAGGAGACATTGAACTCTACAAGGTGATGGATCCTTCGAAGAGCACAGGCAGCTTTTCTGCTGTTGCTGGTTCTGTCGGTTATATTCCTCCAGGTAATGAACTATTACTTGGTTAATGGTTAAAGTTTAAGATCCTTCGACATTGATTTATACTCCTTAAGGGTAATGTGACGTTTTTTGTCCTAACTGTCGTACCATGCTTTAAACTCATCTTTTTGTGTTTAATCTTCATATAGTTATTGTTGTTTTCGAACTTAAACACTTCATTATATCCTCCCACCATGAAACGGTGTTGAACTGTCTTTCGATATCTAAAGTGAAACAGTGTGTCAGTATAGCCTAGAGAGGGAGGAGCGGGTTAAAATGTAACATGAGAAGCCTCAAAGTTGAGCATATATTAAGTGATGCATTTCCTTTTTGTGTTACAGAATATGCATACACAATGAGAGTCACAATGGCTGGAAATGTATATAGCTTTGGGGTTGTCCTGCTGGAATTGTTGACAGGAAGGCCTGCAGTTAGTCAGGGAACCGAGTTAGCCAAGTCAGTGCTGAGCAACTCTGAGAAGCACAGCAAGTGGGATCACATTCTGGATTCTAGCATTTCTAAGACATCGCTCAACATTAGGAGCCAGATGTTAGCAGTCCTCAAATTGGCTCTAGCTTGTGTCAGTGTATCACCAGAAGGCCGGCCAAAGATGAAGAGTGTGCTTCGAGTGCTTCTCCATGCAAGGTAATCGGATGTCAAAGGAAAGTTAAGGAGAAAACAGCCAAGATGGTTTTCTTCTTTTTCTTTTGGGGAGGAGCTGCAAGTGGTGAAGGTGTTGTGAGCTATGCAGAATATGCCAGTTTTGCCAGTTTATACATATTATATTAATGTAAACATATAGATATATGATGATAGTAGTTACATAACTTTGAATATGATGTTAGTTCTCAGTGTAACATAGTCCATTTTTCTTGTTCATGTGTAATATATATATATATTATATATAAAAGGTCTTTGGGAGTGAATCCAATTT

>PGSC0003DMG400009380 LRR-RLKs

AAAGCTTACACTATTTTAACAAGCTCAGCAAACACCCATTTCACCATTATTACAGAAAAATCTCTGTATCTTCTTCCTGTGCATGAAAAACCATGCAAATGAAACTATTACTCTTCTTTCTGAGTACATTCCCTTTGATTTTTGCTTTAAATCAAGATGGGTTGTATCTACAAAAACTGAAACACTCTTTGTCAAGCTCAGATCAAGGGGTATTTTCTACTTGGTATGAAAATGATCCTACCCCATGTAACTGGACAGGTGTCACCTGTAATGACGCCGGAGATTCTCCTTCCGTCGTCGCTGTTAACCTCTCCGGTGCTTCTCTCGCCGGAACCTTTCCGGTCTTCCTCTGCCATCTTACTTCGCTTTCGTCACTCTCGCTTTCGAATAATCTTATTAATTCTACTCTTCCGGTTTCTATTTCTGAATGCCGTAGCCTCACGTACCTTGACCTTTCTCAGAATCTCATCGGTGGAACTATCCCCGACACTATTTCCGATCTTCCTTACCTCAGGTAACTCAGTTTGAACCTCAATTTTGCATTTCCCCTGTTTTTTTTCCTTTCAATTTTTCGAAAAATATCAACTAATTGGGACTTACATTTAGCCATATATGCGTATTTGGGTCTGTTTTAAGGGTGAATATAAGATCTATAGTTAGTGTATATAACATGGTATGTGTAGAAAAAGACAGTCCAGTGCACAAAACATCTCACGTTAGCAAGCTGCGGGGAAGGGGCTGTAAGCAATGATTTAGCTGAACCAACATAACCCGTATGGATTTGCTTGTTGGTGTTATAGTAACTCTAAATTATTGATTGTTTGAATAAAAATGCAGATGGAACGAAATGAATATTATGATCTTTTATTGCAGATTTTGTTATTCTTGTTGTTTCATTAGTATATTATGATCTTTTATTGCAGGTACCTTGATCTTAGTGGATGCTATTTTTCGGGGAATATTCCGGCAAGTTTCGGAAGATTCAGGCAACTGGAGACTCTTATTCTGACTGAGAACATTCTTACTGGTGAAGTTCCTCCTGCATTAGGTAATGTAACGAGTCTCAAGACACTTGAACTTGCTTACAACCCTTTTGCGCCGAGTCAGTTTCCTCCTGAACTCGGTAACTTGACGAATCTTGAGACATTATGGCTAAGTATGTGTAATCTTGTTGGTTCAATTCCAAAAAGTATTGAGAAATTGAGTCGATTGACTAATTTTGATGTGTCCAATAATGGACTGGTTGGGTCAATACCAAGTGCAATTTTCCAACTTAATAGTATTGTTCAAATTGAGCTCTACAATAATTCCCTCACCGGAGAATTGCCTTCGGGATGGTCTAACTTGACCAGGTTGAGAAGATTCGATGTGTCGACTAACAAGTTAAATGGGACTATTCCTGATGAGTTGTGTGAGTTGCCACTTGAGTCACTCAATTTATTTGAGAATCAATTTGAGGGGTTTCTTCCAGAAAGTATAGCTAACTCTCCGAATCTGTATGAGCTTAAGTTATTCTCTAACAGATTTTCTGGTTCATTGCCTAGTGAACTGGGGAAGAACTCGGCTTTACAGTATCTTGATGTTTCATACAATACATTTTCTGGTAAAATTCCTGAAAGTTTGTGTGAGATGGGAGCTTTAGAGGATCTTATAGTGATATATAATTCGTTCTCTGGGAGTATTCCGGCTAGTCTTGGCAACTGCCGGAGTTTACTTAGGGTCAGGTTTCGGGATAATAAGCTATTCGGGGAAGTCCCAACTGAGTTTTGGAGTTTGCCTCAGGTTTATCTCTTAGACCTTTTTGGCAATGCATTTTCAGGAAATATATCACACATGATTTCTGGTGCCAAAAATTTGTCTAACCTACAAATATCAAGAAACAAATTCTCAGGGGTTATACCTAGTGAAGTAGGAAAGTTGAAGAACTTAGTTGAGTTTTCTGCGAGTCATAATGAGCTAACGGGAGAACTTCCAGACACATTAGTGCAGCTAGGGCAGTTAGGTACCCTTGATCTTAGTTTCAATGAGCTATCAGGGAAAATCCCCTTCGGAATTCACACAATGAAGCAACTCAGTGAGCTTGACTTGGCAAACAATGGATTTTCTGGGGAAATTCCGGAGGAAATTGGGACTTTGCCAGTGCTTAATTATCTTGATCTTTCTGGGAATTACTTCTCAGGGGAAATCCCACTCAGTCTGCAAAGCTTGAAGCTTAATAAGCTAAATTTGTCTAATAATCAGCTGTCAGGGATGATTCCTGCAGTTTTTGATAAGGGTGTTTATAGAGACAGCTTTCGAGGTAATCCAGGTTTGTGTCAAGGTGTTGCTGGTCTTTGTCCTACCAAAGGTAGAGGACAGCATGAAGGATACTTATGGACTTTGAGAGCTATCTATACAGTTGCTGGCTTCGTTTTTCTTGTCGGGATTGCTATGTTCATTTGGAAGTATCAGAAATTCAAGAAGATTAAGAAAGGAAACACTATGACAAAGTGGACATCATTCCATAAGCTTGGATTTAGTGAATTTGAAATACCCGTTGGCCTAGATGAAGCTAATGTAATTGGCAATGGAGCTTCAGGAAGAGTGTACAAAGCTGTCCTAAGCAATGGTGAGGCAGTAGCTGTCAAGAAGCTATGGGAGAGAACAGTTAAAGATGAAACCCCTTATGGTGCTCTTGAGTCTGATAAAGACGAGTTTGAAATTGAAGTTGAAACTCTGGGTAAAATTAGGCACAAGAATATTGTGAGATTGTGGTGCTGTTGCGTTACTGGGGATAGCAAGCTCTTGGTATATGAGTACATGCCAAATGGAAGTTTGGGCGATTTGCTGCACAGTTGCAAGGCCAAATTGTTGGATTGGCCGTTGAGGTTCAAGATCGCTTTAGATGCTGCTGAGGGGCTTTCTTATTTGCACCATGGTTGTGTTCCTCCAATTGTTCACCGTGATGTTAAGTCAAACAACATATTGCTGGATGATGAGTTTCGGGCCAAAATTTCAGATTTTGGTGTGGCAAAAATTGTTAAAGCAGACAGCAAAGGTGACGTTGAATCCATGTCTGTAATTGCTGGTTCCTGTGGTTACATTGCACCAGGTATTGATTTGTCCAAACTATATCACTGGCTTTAGTCATTTTTACCGATATATTGGCACATCTAGCTTATATTTTTGGAAAGATTGCAATTTCATTTTAAAGTAGCCAAAACTAACATAAGTGAAATTCATTGATTGCAGAGTATGCATATACTCTTCATGTGAATGAAAAAAGTGACATATATAGCTTTGGAGTGGTCATTTTGGAGCTGGTGACAGGCAAAAGACCAGTCAGTCCAGAATTCGGGGAGAAAGATCTAGCTACTTGGGTACACACGACGTTGAACGAGAAAGGAGTTGATCAGTTGCTCGACCCAAATCTAAACTCCAGCTTCAAAGAACATATATGCAAGGTTCTTGATGTTGGTCTACGTTGTCTTAACCAAACTCCAGCTAATCGCCCCTCAATGCACAGAGTGGTGAAAATGCTCCAAGAATCAGCTCCTTATAATGTGCCAGAAATGGAAAACAAGAATGGTAAACTTTCCCCTCACTTTCCAAAGTCAGTCTAGTTAATATATTAGATCAAGTAATAGAGCAGGAGTGTTTCTTGTCAAGAATTGGCAAGGAAGAATCTGCATAATGAAAAATTAGTAGTCTTTTTCAAGAATTAAGCATATTGTACTCTTTGAATTCTACAACCCTTTATGAATCTATATTACGAGGATTCATCATGAATTAACAAATTTTACGCTAGCTGTCAGCCTACCGCAACACAGATGGGGTGGGTACTGAAAGTTGTAAGTATTCTTAACTAATGCCTTTGATTAAGTTTGTTAAACTTAGGTCCTACTTTGCTGTCCATCATTCACTGAGCTGGGGCCTTTAGAAGTAAA

>PGSC0003DMG400009561 LRR-RLKs

AAAAATAATATTCTTTCTCCCACCTTTCTTTTGGCTTCAAAACTACATAAAAAGGTAGCATTCTTTCATTTGTAGGAGCCCTTTGAGAGGGCGCGCACCTGCAAGTGCGCGCACCAGAAAAAAAAAAACAAAAAAGTTTTAAAAAATAAAGAAGACTGTCTCTCTTCTCTATGCAATCATGGCATCACAAAAAAACTACAAAAACAAAAACAAAAACAAACATGTCCTCCTCCTTGTTATCATCATGTGTTCATTAGCATTTGTGACAGAGGGAAAATTATCAGAGCCTGAGGTTCTTCTCAAGTTTCGTCAATCGTTAAAGTTCGATGGAGATCCCTTCTCGACGTGGGATATTAATGTACCACCTTGCATCAAAGAGAACAATAAGCCTAAATGGAACAATCTATTTTGCGAAAGTGGAAAGGTTTATGGTTTGAATCTTGAAAATCTAGGGTTAAGTGGTACCATTGATTTGGACATACTTAAAGACTTGCCAAATTTGAGAACCATAAGTGTTTTCAAAAATAAATTTGAAGGTTCCTTACCAATATTAAATAAATTGCCTGCATTAAAGACTGCTTATTTCTCAAATAACAAGTTTTCTGGTCCAATTGATCAAAAGATATTTGAAGGGATGAATTCATTAAAGAAACTTCACCTAGCAAATAATGAATTTACAGGACAACTCCCTCCTATTTTTGGTGATATGCCAAATCTAAGGGAGTTGAATATTCAAAACAATAAGTTTGAGGGTCCAATTCCTCCTTCCTATTCACATCTATATTTTCCTGCTTATGATGGTAATTCCTACCTCTTTTCCTTTTTTATATTTCATTAATTATTGTGTCCAATGAATTAACACTTTTTTTGTTTATCATCATTATGCCATGTTTTTCTTCCAATCCTAGATCATTAAAAAGATCATATCATTGGCTACATAATAAACAAAAGATCATTAGATTCGGGTCATCCATGTGTGTCAACTCGGATCCATGTAAATAATAAGGGAAGAGACACAACTATTCAGATTATTTCTGAAAAGTAGTAGTACTCCATCTCTTTAGAATTATAAAAGAATTGTCTTAATAGAAACCAACAACACTTTTTTTTTTTTTTGAGAAAAAAAATAACTTGATAGACAATCCTCTCTTCTTCTTTCAAGTTCTTGTGAAAGGATTTCATAGAACAAACTTCTGTAACAGGTACGCGATTTTAGACTATGGCTTAATTAATATGATGTATGTAGTTCGTCTTTATATGTGATTACATCTAGTGAAACCAAATTCAGTATTGATATATTATCTAACTAAACATAAGTATTTTTCTTCTTCTTCTTCTTATAAAATAGGCAATGATGGTCTTTGTGGTCCTCCTTTAGCAAAATCATGCAATAAGGAAGATGAACAAAAAAAGCAAGAATCATCATCATCGTCATCGTCATCAGGTTGGAAGATTGCCCTTATTGTGGTTATAGTAGTTGTAGTGATAGGCATTGTTGTGTTTGCACTAATCAATCGTCGCAAGAAGAATCACCAACAAGAAGTAGTACTTGGAGGATCATCATTAAGTTCTTCCTCACCAACAAGTCAAGACCAAAAACTAATCCCTCAATCTCATGATCATCTAAACAAAATGGAACAAGGCCAATCATCAGCTGCATCTACTCCTGATCGCGCGTCTAATGATGGCGGTAAAAGAGCTGAAGTTGCTGGACAAAAACTTCTATTCTTGAAAGATGACATTGAGAAATTTGACTTACCTGATTTGTTGAAAGCCTCTGCTGAAGTATTGGGAAGTGGAGTGTTTGGTTCAACTTATAAAGCTGCACTTAGTACTGGTCCTGTCATGGTTGTTAAGAGGTTTAGACATATGAATAAAGTTGGTAAAGAAGATTTCCATGAACATATGAGAAGGCTTGGAAGATTGAGTCATAAGAACTTGCTTCCTGTTATTGCTTTCTATTATAGGAAAGAGGAGAAGCTTCTTGTCTTTGAATATGTCAACAATGTCAGCCTTGCTGTTTATCTTCATGGTACCTATCTCTACTCTCTTTTGTTTCAATTTGTTTGTCTCGTTTTAACTTGACATACAGTTTAAAAAATAAAAAAAAACCTTTGAGACTTAAAGATACTTAAAATGTAGTAGAATATCCTTTAATTTAATTTTGTGGTATGAAAAATTTCATCAGTAAAGTTGGAATTAAAAAGTTCCCTAAAAGAAGAAGAAGAAAGAAACATTCTTTTTGAAATCAACTAAAAAGATAAAATAGAGTATTAATTATCCTTTCTGTCCCAATTTAAATGTCTTACTTTCACTCTTGTTCTGTTCTAAAAAGTGTGTCGGTTTCTATATTTAGTAAGTTTTCAAATTGTGACATTCTACTTGCTTAGTTTAAGATCACAAAATTAAAAAAAAAAAATCTTTATTTTTTTTAAAACCTATTTACTCACTTAGATTAAGACACTTAAATTGAAAGGAAGGATGTACTCCCTGCATCTATCTTTACTTGTCCAGTATATGGGATGTCCAATAATACTTGTCCAGTTTATAAAACCAATGAATAATTTTTCATTTTATGTCTATTTTACTTTTGCTATTAAATACTATTCATTTTTCAATGTTTAAATGACTTATATTTAAATAAGAGTGATTTAATAAAACTACCCCTATCATTTACTGCTTCATAATCCATGTGCCAAGTCAGTAGTGTACGACGAACAGAGGGAGAGGGAGTAATATTTTGTTCATAGTTTTGACCACCTTTTCTTGATTTTTCAGGTAATAGCAAATCACGTGGTAACCAAAGTCTTGATTGGCCAACTCGTTTGAAAATTGTGAAAGGAGTAAGCAAAGGAATTTTATACCTTTACAATGAGCTACCAAGCTTAACATCACCTCATGGTCACCTCAAGTCTTCAAATGTTCTTCTAACTGAAAACTTTGAGGCAGTTCTCACAGACTATGCCTTACTACCAGTAGTAAATGCTGAACATGCACATGAACACATGATCTCATACAAAGCACCAGAGTTAAAACAATCAGGCAAAATCAATAGAAAAACTGATGTTTGGACACTTGGTATGTTAATCCTTGAGATCTTGACTGGAAAATTCCCTTCAAATCTTTTAGGCAAAGGAACACAAGATAGCGATGATTTAGCGACATGGGTGAATACTACTCTTGGAGATGAATCATCAGAGAAAGAAGTGTTTGATAAAGAAATGAAAGGAACAAAAGATTGTGAGAGTGAAATGATGAAGTTATTGAAGATTGGACTAAGTTGTTGTGAGGTTGATGTGGAGAAGAGATGGGATATAAAAGAGGCAGTTGAAAGGATTGATGAAGTAAAAGAGAAAGGTGATTTCAGCTCTAATGTTGCAACTAATGAAGTTGACAATATGCATACTTCAAGATAAAAAAATATTTTTGATTGATAAGCTCATGATCATAATATGGAAATTTTGATTATATTGGATGCATATTGCATGACTATATATACTTTGGGTATATGATTTTGTTTTGTCATGGTGGAGGATAAATTGAACCATGATGTGGGGTTACTACAAAAGAAAAAAAGAAAAAAAAAGAAAAAGAAAAATAAGTGATAGAAAAATGAAGTTACTTAATTTTTGTTTGCATATTGTAAATTTGTCCTTTATGAATCTATAGTCTAGGTGTGTTTCTTTTATTTATTTTTATTATTCATTCATCAAGCTTATTTGGGACTTGCTTTACAAAATTGGC

>PGSC0003DMG400009843 LRR-RLKs

TTAGAGACTATTTTGACCCTACGGCCAAAATTAATGGACCATATTTGTCATTTTCTTTCTTCTTCACCCTACCCCTCGTTTCCCACTTCACTCTTCTCCATAGTTCCTCACTGTCTCCGAAAATCCAGTCCATTTTTCAGACCTCCTTTCACGCACAAAATGACAACAACAAACACTAAACTTGGCCACTAGTCAACTAAACCAAATCACTCGACATTTTTAGTTGCATAACTTGCAATGTCTCCATTGATGCTTCTGCTCTGTTTTCTTCTTCAACTAATCTACAGTTTCTACTTTTGTTCTTCTTCTTCTGCTGCTGCTGCTACCTCTACCCCTTCTGCTCCATTCAACTCTTTGCTTCCTTCCGACGCTGTTTCGTTATTGTCATTCAAGTCAAAAGCTGACCTTGACAACAAGCTTCATTACACACTTAACGAGCGTTTTGATTACTGTCAATGGCAAGGAGTGAAATGTGTACAGGGTCGTGTTGTGCGTTTAGTTCTTCAAGGGTTTAGTCTCCGGGGTACTTTTCCGGCGAATAGTTTAACTCATCTTGACCAACTCCGAATACTCAATCTTAGGAATAACTCACTTTCCGGTCCTATTCCTGATCTTTCGGGTCTTCCTAACCTGAAAACTCTGTTTCTTGATCATAATTTTTTCTCCGGAACTTTCCCCTTTTCTGTTCTTTCTATTCACCGTCTTGTTATTCTTGATCTTTCTCGTAACAATCTCACTGGTTCACTCCCTGTTAGGCTTACTGTTCTTGACCGGTTGAACTATCTCCGGCTTGATTCCAACTGGTTTAACGGTTCGATTCCGCCGTTGAACCAGACCCAACTCCAAATTTTCAATGTGTCTAATAATAACCTCACTGGTCCAGTACCTGTTACTCCAACTCTGAAGAAATTTAACATACGCTCGTTCTTACGGAACCCTAGTCTTTGTGGTGAGGTTGTTGATAAGCCGTGCCGTTCGGCACCGTTCTTTGATTCGCCTTCCTCTGCTGCCTCTCCGCCCACACCGTTATACCAAAACGCACAGTCGCAGGGTATTCTTATTAGTCCTCCACCTCAGCACAAACACAAAAAAGTTGGTGTCGTTTTGGGTTTTGTAGTTGGAACATTGATTTTGATTGCAGCTGTTCTCTGTCTTTTTGCTTTTGTGAAAAAGAGAAGAGAAGAAACTGAAACTGAATCAAAAGCAACAAAATGTACCATTGAGACCATCACTAATAGTGCGGCGAACGCAACGGTTTCTGAGCCAGATGATAGTAGCCAAGAGATTAAATTAGAAAAAGAAATGAAAGTGCTTCAAGCCCCCAAACAACAGATGAAGAGTGGAAATCTGATATTTTGTTCAGGGGAAACAGAGTTGTATTCTTTAGAACAACTAATGAGAGCATCAGCAGAGTTGCTTGGAAGAGGCACAATTGGAACAACGTATAAAGCTCTGATGGCTAGTCAGTTAATCGTTTCAGTAAAGCGATTGGATGCTGGTAAAACTTCGATTACGAGTGCAGAAGCGTTTGAGCAACACATGGAGTCAGTTGGGATGTTGAGGCACCCAAATTTGGTTGCAGTCAGAGCATACTTTCAGGCTAAACAAGAAAGGCTTGTTATCTATGATTATCAGCCTAATGGCAGTCTCTTCAATCTAATTCATGGTAAAGTTTCTAACTTTTTAATATATCATTACTAGTTCTACAAATTATTCATCTGAAATGGATCAATGTTTATTTTTCATTTTTGGTAATTCTGCTAATTGGTCAATGTGAAGTCAATTTGGTGGACCTGTGATGGAGAAAAAAGTGGAAGAGTGAGTTGTCTGTGGCTACTGGCTAATCACTATCTAGCTTCTTTTTTTTTTCTTTACAGAAATTAGTTGTAGTGTTTTGGATCTTTCAGTGGTTGCTATGGTTGTAGTAGCAGGGGACTTTTTTAGCTTTTAGTGTGGATGCCTAACATGGCTCCCACTGAGTTCACTACCGACAATTGCAAAGTTTGTGTTTACAGGGCAGCCAAATAGTAATGTGTTTTAGTTTCAGGCTGTCACTGACTGACCGGTATGGGTTGATAGCATTATATTCCTCAATTTTTTTTATCATGCATTGAGGCAAATGGTCCTTGGCTCTTGCGTCTGTCTTTTTGGTATTTATGAGTATGTTTGCTTCTTCTTTAAGTTGAATAGGATGTTGACAGTAACTTTTTGGTGTTTTCCTTTTACTTGTCCAAATGTTCTCTGTTTCTTAAATATTAAATTTACTCAAGTGAACGTTAACGTTGTTAGATAGAAGGATTAAGGGAAAGAACTAGAACATCCTTGGGAATTCTTAATGTAGTGACAGAAATGGTATGAATTTGCTTAATTTACATTATTTTGAATCCTAGGTATTGCATTTTTTGTATCCCCTTGTGATACGCTATCAAGAAATTGGAATTTCACTTTCATAGATTACTCTGCTCAAGAAGATTGATTCCTCTATATAATTCTGTGTGTTTAAGTGGTTGGTCTTTTCACCTAAATCCTGATAATCTGTTGTTCGTTTACTTATATATTGTGTTCAATTTTCTATTGTTCGTGGCAGGTTCAAGATCAACAAGGGCAAGGCCTCTTCACTGGACATCGTGTCTAAAAATAGCAGAAGATGTGGCTCAGGGCCTTGCTTACATCCATCAAGCGTCAAAGCTCACTCATGGCAACTTGAAATCCTCCAACGTCCTTCTAGGGTCTGATTTTGAGGCCTGTCTAACAGACTACTCCATCATCGCGCTTGCAGATATTTCTTTGGAGGATGATCCTGATTCAGCATGCTATAAAGCACCAGAAGTGCGCAAATCAGCACGAAGAGCCACGCCTGGGTCTGACGTATATGCCTATGGCATTCTGTTATTAGAGCTTTTAACGGGCAAACCTCCATCTCAGCATCCACATCTCTCTCCCCCTGATGTACCGGATTGGGTTAGAGCTATGAGGGAGGATGATAATGAGGAGGATAGATGGCTTGCAATGCTAATTGACCTTGCTAGTATATGTAGCTTGACATCACCAGAACAGAGACCAACAATGAGGCAAATCTTGAAAATGATACAAGATATAAAGGATAGCGCAATGGTGGAGAATAACAAAAGAGATGCACATAATGGATACTCATAGCTTCTTCTGAATTGCAGTTACACCATTAATCAGCGAGGGTACATGGTCAATTGTAAGTAAGTGTCATCTCCCCCTTGGCAAAGTGCCATTGATCGACAACATATCTGTAATTGAGTTTTTTCCAGATAGACCAAATGACTTATCAACTATAATGGTGCTTGTACCTTCACTACTCAGCAGTAAACACCTAAAACTGACAGGTTAAGGTCTATCTGCTAGAACATGGTCAACTTAGTCACTGAAGATTATGACACATTTTGTTCCCCTCGTCGAAATCCCTCTTTTTTGGTCACACAAACATATGGTGCCTATATTTTTAGGGCATATTGTAGTAGGTGGTTTTGGTTGATTCTTTTTCTAAAGAACTTCAAAATGTATTCGCTTTACCTTAATGGCAGAAGAATATTGTATAATCTTGAATGGTAAAGCTAACCCAAATTCTGAACTTTCCCTTCTCTGCATGGTTGGTTAGTATCAACTTAGCAGGAGAAGAGTTGTTTAATG

>PGSC0003DMG400009941 LRR-RLKs

CTTCAAATACCCTTGAACCTATCCTAAAATTTGTACAAACCAGACTAAACTAACAGTGTAATAATGTCACTCCCCAAAAAAATATCCCTTTTCCTCCAAATTTTCATTTTTTTTGTTTTCTCCATTAATGCAAACTCTGATCTTGAAACCCTTTTGAAGCTCAAAGAATCCATGGTTGCTCCTGGAACTTCTGCACTTCTTGATTGGAACAACAACACAAATTACCCTTTTTCCCATTGTTCTTTTTCTGGTGTTACGTGTAACAATAACTCTCATGTTATATCTATAAACATCACTAATGTTCCTCTATTTGGTACTATTCCACCAGAAATTGGTCTTTTACTAAATCTTGAAAATCTTATTATTTTTGGTGATAATATTACTGGTACACTCCCTTTAGAAATGTCACAACTTTCTTCTATTAAACATGTTAATCTTTCTTACAACAATTTTTCTGGTCCTTTTCCTAGAGAAATCTTGTTGGGGTTAATAAAGCTTGAATCTTTTGACATTTATAACAACAATTTCACTGGTGAACTTCCTACTGAGTTTGTAAAGTTGAAAAAGTTGGAAACTTTACATCTTGGTGGAAACTATTTTCATGGTGAAATACCAGAAGTTTATTCTCATATTGTAAGTTTAAAGTGGTTGGGTTTGGAGGGAAATTCACTAACTGGGAAAATACCAAAGAGTTTGGCTTCGTTACCAAATCTTGAAGAACTTAGATTGGGCTATTATAATAGTTATGAAGGGGGTATTCCATCTGAGTTTGGTAATATTAGTACACTTAAACTTCTTGATCTTGGAAATTGTAATCTTGATGGTGAAGTTCCTCCAAGTCTTGGAAATTTGAAGAAGTTGCATACTTTGTTTCTACAAGTGAACAGACTTACAGGTATTCATGAACTCACTATTTGTTAATTAGTTATACAATCAAATCTATCTTTAATAACGTCGTTTGTCTGAATATATTCTAGTTGCCATGGTTGAATATTGTTGTAGGAAACATATTTGTCACAAATCATTTTCCAAGATGTAAGACTGAAAATCAAATTGTGTTGAAATACTCAATAGTAATAGTTGTTTCTTGAGTGACGTGCAGAGTGCCAGTATATAAGTAAAAAGTGAATGGAGTGAGAATATCATAAAAAGTTTCTTTGAGAAAAACTGTATTTATAGTGAAATGATGTTAGAGTATGTGTTAATAGAGAGGTTTTTGTATAACATAGATGTATATGTGCAAAATATTATGTATATAAAGTTTTGAAGTCATATATCTGATAAATTTTAGAATCTTGAATTCGTAAAACTTAAATCTTGGATTCGCTTATGATTTACAGGTCGCATACCTTCTGAACTATCTGGTTTAGAGAGTTTGATGTCGTTTGATTTGTCTTTTAATCAACTGACCGGAGAAATACCAGAGAGTTTTGTGAAGTTGCAGAATTTGACATTGATTAACTTGTTTAGAAACAACTTGCATGGTCCAATTCCCCCTTTTATTGGTGACCTTCCAAATCTTGAAGTGTTGCAGATTTGGGGAAACAATTTTACTCTTGAATTGCCCGAAAATCTTGGGCGTAACGGGAGGTTTTTGTTTCTTGATATTTCTATTAATCATTTTACTGGAAGGATACCACCTGATTTGTGTAAAGGAGGGAAGTTAAAGACACTGATTCTAATGGAAAATTACTTCTTTGGTCCAATTCCTGAACAACTTGGTGAGTGCAAATCGCTTGCTCGAATTCGCGTTAGGAAGAATTACTTAAATGGTACTATTCCAGCTGGTTTTTTCAAGTTACCTGCATTGGATATGCTTGAACTTGACAACAACTATTTCACTGGTGAGCTGCCAACGGAGATAAACGCGAATAATCTCACTAAACTTGTACTTTCCAACAACTGGATCACGGGGAACATTCCTCCATCATTAGGGAACTTGAAGAATCTAGTCACTCTATCACTTGATATGAACAGATTATCTGGTGAAATTCCTCAAGAAATTGCGAGTTTGAATAAACTCGTGACCATCAACTTGAGTGGTAACAATTTAACAGGTGAAATCCCAAGTTCAATTGCGCTTTGTTCAGAGCTAACATTGGTTGACTTGAGCAGAAACCAACTGGTTGGTGAAGTGCCAAAAGAAATCACCAAGTTAAATAGCTTGAACGCTCTGAACTTGTCAAGAAACCAACTGAGTGGCGCCATTCCTGGAGAAGTCGGAGTGATGAATGGCTTGACAGTTTTGGATCTTTCTTACAATGATCTTTCTGGAAGGAGACCGACCAACGGACAACTAAAGTTCTTCAATGACACTTATTTTGTAGGAAATCCAAAACTCTGTTCGCCTCATGCTACTTTTTGCCCGTCAGCTTCCAATTCACCACAAAACGCGCTCAAAATCCATTCTGGGAAGTTCACAACTATCCAATTGGTGATTACAATAATCATCTTAGTCACTGTTGCATTGCTGTTGGCAGTTACCGTGTTGTTCATCAAGAAGGAAAAGTTCAAGAATTCGAAACTTTGGAAGTTAACAGCATTCCAGAAACTTGATTTCAGAGCTGAGGATGTTTTGGAGTGTTTAAAAGAGGAGAACATAATTGGGAAAGGTGGAGCTGGCGTTGTGTACCGAGGGTCTATGTCAAATGGCATCGACGTTGCAATTAAGAAACTTGTAGGCCGAGGAACTGGACACCATGATCACGGATTCTCAGCTGAAATCCAAACACTAGGAAGGATCAGGCACAGAAACATCGTACGATTACTAGGATATGTCTCAAACAAAGACACAAACTTGTTGTTGTACGAATACATGTCGAATGGGAGCTTAGGTGAAATGTTACATGGTGCCAAAGGAGCACATTTGAAATGGGAGACGAGGTACCGTATTGCTGTGGAAGCTGCAAAGGGATTGTGTTATTTGCACCATGATTGTTCGCCTTCGATTATTCATAGAGATGTCAAGTCCAATAATATTCTGCTGGATTCCGATTACGAGGCTCATGTTGCTGATTTTGGCCTAGCCAAATTCTTGCAGGATGCTGGTGCATCAGAGTGCATGTCCTCTATTGCTGGCTCATATGGTTACATTGCTCCAGGTAAACTAATACGGTTACAATATTTATCATATTTATACAATTATGAATTTCCGTATTAGTTTTCTTTTATTACTGGCTCATATGGTCATCATATTTATACAATTTGGTGGCATATAATGTAGTTACAGGTAAAAATTAATGATAATCTTAAAATGTGATACTAATAACTTGCTATAATAGATTAAATTATGATGATACTATGAAGTTTGTATGCTATTGTGCATATAACCTTGAGCTATGTTGCTCAAACTCTTAAAAAATGTCAGTGGGGTACATGTCGGATCCTCCAAAAGTAATACTTTTTTGGAGGATTCGACACGGATGCAACATCATTTTTTGAAGAATACAAGCAAAATAGATCCTTAAGTCCTTTTATATACCTGTGTTATTGGTCAATATTTTCCTATTTCATGAAAGATACACTTCACTAACTTTTAGTACCAATTACCAAGACCTATGAACTTTTAGAATTTACTGAACTGGGGTTAAAAATAGTAGGTATTTAAAAGCTATGCTCCAGGACAAAATCTATTTAAAACATTTGTCCAAAACATTAAATTGTGCAATTAATGTTTTTCTGAGGTGCATTTTTGTTTGGATATGAAACAGAGTATGCATACACATTGAAAGTTGACCAAAAGAGTGATGTATACAGTTTTGGAGTTGTACTGTTGGAACTTATCACAGGTCACAAGCCAGTTGGTGAATTCGGGGACGGTGTAGATATAGTCAGATGGGTAAATAAAACAATGTCCGAATTATCTCAGCCGTCTGATGCAGCCTCAGTTTTAGCAGTCGTTGACTCGAGGCTACATAGTTACCCTCTTGCAAGTGTTATCAATTTGTTCAAGATTGCTATAATGTGTGTTGAAGAAGAGAGTTGTGCTAGGCCTACCATGAGGGAAGTTGTTCACATGCTTACAAATCTTCCTCAGTCTACTACTACTACTCTCCTTGCCCTTTGAAATTGCGCGGATATCAAGTGTCTGGTTGAAAACTCGTGGAGTTTGAGGCCGGGAACACGAGTCTCATGAGTCTATTTGGGTACGGGGAACAAAAAAATTGTGCAAGTAGAGATATGTATTTGAAATGTATTGCTTGTTTTTATATTTTTGAAGTTGTAGGCAAGTTGTTGTATAAAAAGGGGCATGAAAAAAAGAAGGAAAAAAACTCCTTTAATTACTTATTATGGGTATTGAGTATGAATATGTGGCCTTAGGAAATAAATTTCAAGATGGCAATTAGTCAATACTTGCTTTTAATAACATGGCCTATGTAATGTTCTTAAGAGTTTGCAAAGTTAAAACTTG

>PGSC0003DMG400010055 LRR-RLKs

GACTGACAAGACAGTAATAAAATGATTTTCTGGCATACAAAAGCTATAGTTTTGTTCTCAATCTAAAAGATTATTAGAAGAAAGAAAATCCAATAGAGGGGTTTGTCTCTCTGGTATTCATTTGAGTTTCTGTTTCTTTCTTGAAATTTAATGGAAAAGAAAAGAGAGATAAAGGTTGCATTCTTGTATGTGGCTACATTCTTGGTTCTTTGGAGTTCTGCAGCAGGATTGCTTTCTCCTAAAGGTGTCAACTTTGAAGGTAATTAGTATTATTTTGATAAAATTCTGCATAATGTATACTGAAAATACTAAGTTTTCATCTGAGATGTTCCTCTTTTTCTTTTTTTTTTCTCAGTGCAAGCTTTAATGGCCATAAAAGTTGCTTTGAAGGATCCTCACGGGGTTCTTGATAATTGGGATAGTACTTCTGTTGATCCATGTAGTTGGGCTATGGTTACTTGTTCTTCTGAGAGTTTGGTTATTGGCTTGTAAGTATTTCTTGCAAAGATGAAATCTTTTTCTTTGTTTGTGTCTTTGCAATTCAAAAGTTGAATTATTTTTTTTACCTTTCTGTATTATGCAGAGGGGCACCAAGCCAGAATCTGTCTGGTACTTTTTCTCCTAGCATTGGCAATTTGACAAATCTTCAGATTATGTGAGTTACTAAAATCAGTGATTTGAACCTTTCTTGATTGTTTCTGTTACTTTTTTGAATAACCATTTTTTTTTTCTTTTGAAACAGATTGCTGCAGAATAACAATATAACAGGACCAATCCCAAAAGAAATTGGAAGGCTCTCAAAACTTCAGACTCTTGATCTTTCTGATAACTTCTTCACTGGTGATATTCCTGTTTCTTTGGGACACTTGAGTGACCTCAAGTACATGTAAGAATCTTAACAGAGAGCCAGTTCTCATACTAAACTTCATTTTTCTATCTTCCAACAGTTTCAATATCAGTTTCCTTATATTAATTTGTTCAGGAGGCTCAACAATAACAGTCTATCAGGAGAAATTCCAGTCTCATTAGCCAACATGTCACAGCTTACTCTAGTGTAAATCCTTATTGCTTTACTTATCTTTTTCCTTTTATCCTTTTCAAAACTTGAGATTGATTTTGGTATTTAACCTTTTGATTCAGGGACTTATCCTTCAATCACTTGAGTGGACCGGTACCTAGGTTCCCTGCTAAGAAATTCAAGTAAGCTAATTAACAGTTTATTCATGTCTTGTTAATAGAAAGCTGGAATGTTAAGAATAATTTATGTTGTTCTGTGATTGTAGCATTGTTGGAAATCCATTGATCTGTGAGACGGGGTCTGAACCAGACTGCTATGGAATGCAATTGCTGCCAATGTCAATGACATTGAATGGTTCAGAATGTATGCATTATAGAATGCCATTTTCCAAATCTAACCATTTTTTCTTACCTGTAATTTTTGATTTTTCATTTCTTCTGTTTCACCTCACTTTATTGCTCCGTGTTTGTTATCCAGCTTCGCCTTCTGGGAAGCAAAAAGGTCATAAAATCGCGCTTGTATTTGGCTCGACTCTTGGTTGCATTTTTTTGCTCGTTCTTGGAATTGGACTCTTTCTATGGTCAAGGCATAGACACAATCAACAAGCCTTTTTTGATGTTAAAGGTTGGTCATCTGTTCTAAACTTAAGCTCAATGTTCTCATGAAAGTTATGATCATTGCTTCATACTCATTTGTATAATTTGCTATTATTAGATCGGCATCATGAAGAAGTTTCCCTTGGAAATTTAAGAAGATTTCAATTCAAGGATCTCCAAATTGCAACCAACAACTTCAGCAACAAAAACATTTTAGGAAAAGGTGGTTTTGGTAATGTTTATAAAGGCCATCTCCCAGACGGTACTCCTGTGGCTGTAAAGAGGCTAAATGATGGCAATGCAATTGGTGGTGAGAAACAATTCCAAACAGAAGTCGAAATGATAAGCTTAGCAGTGCATCGTAACCTCCTCAGGCTCTATGGATTTTGTATGACACAGTCGGAGAAGCTTTTGGTTTACCCTTTTATGTCCAATGGTAGTGTAGCTTCGCGTCTCAGAGGTAAAAAAAACATATGAAATTAGCAAAGTGGCATTTCAGATATCTTATTTGTCTTTTGCTGAAATATAGTCCATGTTACTTTCTATAGTAAAACCTGTGTTGGATTGGGGAACAAGGAAGCGAATAGCTATAGGAGCTGCTCGAGGACTGTTATATCTTCACGAACAATGTGATCCAAAGATAATTCATAGAGATGTTAAAGCAGCAAATATATTGCTCGATGACTTTTGTGAGGCGGTTGTGGGAGATTTTGGGCTGGCAAAGCTTTTGGATCACCAGGATTCACATGTCACAACTGCTGTGCGCGGTACAGTAGGGCATATTGCCCCTGAATACCTGTCAACAGGACAATCATCTGAGAAAACAGATGTGTTTGGATTTGGGATCCTTCTACTTGAGCTGATCACAGGAATGAGAGCTATAGAATTTGGAAAAGCAGCTAACCAAAAGGGAGTAATGCTTGATTGGGTAAGTATCAATTGACATGCCTCAAATAAGTTCAACACTTTCTATATTTGAAAACTATTTAACTTTAGACTTCTCATTTTACCCTTAACGACATGCTCTTATAGTCAGAGAAATGTTATGGCATGTTTCGCGACATTGTATATATCCTGTTAAGTGTAACATTTGGTTTACATTATATCAGGTTAGGAAAATTCATCAAGAGAAGAAACTTGATGTCCTTGTTGATAAAGATTTGAGAATCAACTATGATCGAATCGAGCTAGAGGAAATGGTTCAAGTTGCTCTCCTCAGCACACAATATCTTCCAGGACACAGACCAAAAATGTCTGAAATCGTTCGAATGCTTGAAGGTGATGGACTTGCAGAGAGATGGGAAGCATCCCAAAAATTTGATGGTAGTAACAAGTACAAAACTAAAGAGCTATCCTCATCCGAGAGATTTTCAGATCTCACCGATGATTCATTGTTACTTGTACAAGCCATGGAGTTGTCTGGTCCTAGGTGAGCAAGTCATTTACGTATATTTATAAAAAAAAGTAGAATTCACACTTAGAGAGGTAATTTATTAAACTATGTGGTAATGTATAGTAATGAACCATCAAAGATTCAAAGAACTTTGATGTGTTCTGTTATTTTTTTTGTTTCTTTATTGTATATGTTCATCTTTAAGTTAAATGATTTCAATCGAGAGTTCGATTTTTTTTCCTGGTACAACCTGTATTTG

>PGSC0003DMG400010230 LRR-RLKs

AAAAAAGAGAAAAAATCTTGTAGAGAGAGAAACAAGAGAGAGGAAAAAACTTTTTTTTTCCCACCATTTCCCCATATTTTCCCTCTTGTTCCTCCTCAACCATATTTCTGAGAAAAAAAATAGTTGAAAAGAAGCCCCTGTTTTGAAATCACCCTGCAAACTGCAGTCAAGAAAAAAAAGAAAAGATATTACACTATTTTTCACAAAAAAAAAGATTGATTTTTTCTTGTGTGATTTCAATTTTCTTCTTCAATCTCTGTTATTTCTTTAATTTTTTTCAAAAAAGAAAAAAATAAATTTTGTCCCACCACCTGCATTTTCCTATTGCAGTTATTTGATTTTTGGCTTTTTTTCAAGAAATGTGACGAATTGTCATTTCCAGTTCTGCAATAAAAAGCAGAAAAAGATTTCATTCTTGTCTGGTAAGTTTTACTCTTTACATTTTACCATCTTTTGGGTTTTTGGATTTTTATTTTTCTTTTGATTGCTTGCAAATTCAGTTATTCCCTCATCATCTTTTCTTTTTCTTTTTCTGTTTCTCTCTTTTTTTGATGGGGCTTGTGAAGTGTTCTTAGGTTTTTCACCAACCCCTTTTCAAGAAACATATTTTTACTAATTTTTTTCTTTTGTGTTTATTCATTTTCCAGATAAAAATTGAATCTTGGAGATTAATCAATCGAGTTTGTGGAAGTGTTTGGTGAGGGAGATGTTGTTATAGTCTAAATTTTAGAAACTGGGAAAAATGAGAGACCCCAATTTTGTCCATTGTATCAATAGGGTTAATAGTTTTTCTTTTAACTTGTAATATAGTTTTACATTGTAAAAATTGAATGATAAGCAAACAGTGGTGTATTTGCTCTCCTTTTTTTCAACTTGTTTATATTTACGAGTGAATTCAATTGTTTGGAGAGTAATTTTGTACAAGAAAGCTGGCGATTTTGTATAGAGTCCTCGCTGTTGACAGATATTGTTACTGCCTCAACAACTGATCCATATGGTTGTGCAAGCTATGGGTGGTAGATGGAACACTTATGGAATCAAATTGTCGTATCTCGCTCTTTTGATTATCATTTTGGATATTCGCGGTTGTTGTTCTCTTAATTCGGAAGGTAATTCTTATACTTGAGTACAGTTCTATTGGTTTTGTAGTTCTGTCTTAGGCTAGTTTTAAGTGTTTGGTATGTTTTCTTGTTTGCTAATTATTGTTGGTGTGAGAAACAGGATTGGCACTATTGAGATTCCGCGTGAAAGTGGATTCTGATCCGTATGGCATTCTTGAAAACTGGAATTCAGATCATTGTGACCCGTGCTTGTGGTCTGGTGTCCAGTGTATGGACGGGAGAGTGCAAATGTTGTAAGTCTCTTGTTAAATATACCCAAAAATTTCCTAGTGTGAATCCTTTTCTTATAAATTTATATGTGTTGCTTTGCTATCGACAAGCCTGTTGAAATGATAATGACAGACAAAGTGCAAACACTGAGTTGTGAGTTTTTCTGTTTGTGGTTCTTCGGATATTGTTAACTCATTTGTCTGTTCTCTATTTAGGGACCTTCATGGATATTCGTTGAAAGGAAAACTCGCAGCAGAACTTGGAAATCTGACTCACTTAAAATCACTGTAAGCTTCAGCTATACTTATATGCTTCTCATTTGTTTCTTTAGCTCATATTCCTGACCTCATCATCTGAAAAAAGATTGTTCCAATCATTGATGGTTTTTTCTACTTACGAAATGCTTTCATGATTCTGGTCTTAGTCTAGAAATAATATTTAGTGTGCTATGTCATGGAACTAATACTTCCCTATCTTGGTAGACACTATCTTATCACTATACAATTTGAAATTGCACTAAATTTACATGTCTGGTCAGTACAGGTTTCCCTTTTTCGTTCCCATTTGTTACTTAGGAGATGTCTCAAAATATAAAGAGCGTCATATACATTGGGATATTTAGCATATTTGGTCATGCAAATTGCACCAAAGAAAAACTTTACCTTTTTTTTTTCCCAAAGAAAAATGTATAAAGTGAATTGGGTAGATTTCTGGTAATCAATGTTTCACCTTCTGTTCTGCATCTATGTACTATGTATTAATAAACTTTCTTCCTCTTTGAATGCTTTTCTTCCACAGTGTGCTTTCTGAAAACCGTCTTTTTGGTGCTATTCCTAAAGAATTTGGACGCCTCAGAATGCTGGAAGTGCTTGACATTAAGAACAACAATTTGACTGGAAGAATTCCAGCAGTAATAGGAGATTTGCAATCACTTAGAAGCTTGTAAGTTGATAATTTGTTGTGTTTCCGTCTTATAATATATTTGTATGTATTTATGCTACAACATTGCTAACAAAATGTTTTTGAATTGTGCTTGATACTTCTTTTCCAGGTTGATTAGGGACAATAACTTCGAAGGGAAGATCCCTTTGGAGATCAGAAAGCTTCATCTGCTCTCGGAATTGCAATTTGATGATTACCTCACCTCTGGTGTTGCTGCCGGAACTGGTTGCATAAATAGAAAATTTGGGCGTTGGTAAGACTTCCGTAGTTGTTTGGTCCTTTTATCTTCATGTTTCAGCTGTGCTGCATTTTAAAGTCGCAGACTTGTTTCTATGTTCAATTATATTGAAAATGCATAAAATCCATCATTTGAGGAACATATACCAATATACTACTTTCATACTAACTACCTCCTGCTATAAGTTTGGTATTAACATTAAATTAAGTGCAAGCAATTATCCCATGGAGTTGCACTGAAGATCTTTGAGACATTTTCTATGCATAGTGTGTCCCCACTAATCTCTCATACTTTCCTTACAATAGTGAAGCAACTATGATGGTGATTGTCGAAAGCAATGCCATGGGGCACTTGATACTTTCCTGTCTCCAACCCTAATATTCATAGGGGATTAGAGTTTAGTGTTTCTAAAGTTGTGATTCATCTTCTTATTCATCAAGATCTGTTTATGTGGAAATTTGACTGATCTATTGCATAATATTTGTTTATTTTGTCGACTTTTCTGTACATTAACCACTAAAATTGCTGGAAATGTGGTTACTCTTTCCCATGAGTAGAAGAACACTTGGATATTATCATATTCCTGTTTCGTAGCATGAAACTTCAGCCAGCTATATGAATAAATTTATTCTATTCATCCCGTTGACTCTGTTTGAATTCTGGATCGAGATACTCATCTTGGACAAGTACCTTCAGCTTTACCCTTGTAACTGATGTCAGATTACAGACTACCTCTTAATATGCTAATGTTTTATTTCTCAATCACGCACTTTTAAGTTCCATACTCCGCTCTGAAAATTTGGTGCCTTTTTCTTGCAGCATCTGGCATGGCAGTTTGTGGCCATATAAGACTATAGGCTCCTTTCTCAGACCAATTAAAGGGATGCTAGCATGGTATTTTAGTTTTTTCACACTGTAAGTAACTAAATAGGTTTTTTCCACCTTATGCTTTCTAAATAAGTTTAATGGTTCAGAAGCAACTTTTTTGTTTCCTAAAAGGTTCCCGCGTTTTTTGGATGCTCAGGCAGACTTCTGCTCTGACAACCTACCAAGTGAGTGAGAATATTTTGTGACATATCTTTTATTGGATTTCAAAGAATTGTACACAAAAAAAAAAAAAGTATTTTATTCTCTTTTGTTTTTGCAGGTTCAACTAGGCCTCACAATATCCACGATGTAGAGAATCAAGCAAACATTGAACGCCGTAAGTTAGCTGAGCAATCCAGTAACTTTGCTGCTGCTCCTGCTAACGGAGGGAAACCATTGGGACCTGTTGATCCTGTTCCAAGCAGCAGAAGTAGTGGTTCATTTCGTGCTGTGCCAAGTACTGAAGGAGCTCCTCCTCCTCCTCTGACTACACCATCACGGACCACACCTGATCATCATCCGCCTCCAAATCCTGGGGGACACTCTAATGATGCTGTTAAACAGCCAATTGCTAGTCAAGCCCCACCTGGTGGAAAGTCTGGAAGTACATGGAAGTATATAGGAATTGGCATTGGGGCTTTTCTGGTTGCTGTTTTTGTGTGCCTGATAATCATCTGCAAGAGCAAAGCTGCGCAAACAATAGGCCCTTGGAAGACTGGATTGAGTGGACAGCTGCAGAAAGCATTTATTACAGGTACTTTTGATTTAATATTGTGTCAAAGCTGATGAATCCATATGTTCTAGTTCAACTGTTGGATTCATTTTTCTCTGGTGGATATAATCCGAATCTTCAACGCCTACAATAGTTTTTGTTTCATACTCCAAGAAGATGTTAAAAGCATGTATTAATACTCCTGTGCTATATTGTTTCTCATGTCTTATGCATGAAATCAATTTTGCAGGGGTCCCAAAACTAAACAGGCCGGAGCTTGAAAATGCATGCGAGGATTTTAGCAACATAGTTTGCCATCAAGATGCATTTACAGTCTACAAGGGAACTCTATCGAGTGGAGTAGAGATTGCTGTTGTGTCAACTGCTATAAATTCTTTGAAAGATTGGTCCAAGCGTTCTGAATTGGCCTTCAGGAAGAAGGTTAGTAGTTGCCATCTCCTTCACAAATAAAAAACTGCTTCATAACTTATTACCATCTCACTCTTTCTGTTGAAATATCCTAGTCGTCTCGGTTTTTTTACTTATAAGATTTCTACTTATTGTGCGCTTGGGTAGATCGATTCACTCTCAAGGATAAACCACAAGAATTTTGTTAATCTAATTGGATATTGCGAGGAAGATGAGCCTTTCACTAGGATGATGGTCTTCGAGTATGCAACAAATGGAACACTTTTCGAGCATCTGCATGGTAAACTAGTTAATAGTTCAAGTTTCAAGCTTCAAGCTAATCATTTTTGTGATTTGGTTTTGATTAATGCAGTCTACATGTTTTGTTGTTTCAGATGATGAACTTGAACCTCTTGACTGGCCTGCAAGGGTACGAGCTATAATGGGAACAGCATATTGTCTCGAGTACATGCACAATTTGAATCCTCCTCTGTCGCATTCTGATGTGAATTCACACTCCATATTTTTTACAGACGATTATGCTGCTAAGGTATGCTTTATTTCGTATACCACATTGAATTTAGAAACGTTTTACTTCGTGTAATGTCTTGAATTGAACGGATCTTTTTTTATTTACACAGATTACAGAGCTTGCTTTCTGGTCAGATATAATGGTGAAGTCAAAGTCCTCTAGTGGTGATCTAGAGAACTCTGAGCTGCGACCACTTTCTGATCCCGAAACAAACGTCTATAACTTTGGGATTTTGTTGCTGGAAGTAATTTCTGGAAAGTCACCTTATACGGAACGAGATTCTCTTATGAGCTGGGTATCTATCTAACTCTATTTTATACGAAAATCCCTTATTTTCTTCGGTGTTTAATATAGACTAGTGAAACGTTCTTCTCTGCTTTTGTTTAGGCGGAGCAATGCCTTAACGATAGACAGTTGGTTGATCCAAAATTGAAGTCGTTCAAGAATAATGAGCTTATGGTTATGTGTGAGGTGATGAGAGAATGTCTTGGAGAAGATTCAAGGAAGAGACCAACAATAAAAGAAGTGATTAAAAAGCTAAGGGAAGCGATTGATATCTCCCCTGATGCTGCAGTTCCAAGGCTATCTCCCCTGTGGTGGGCTGAACTTGAGATCCTATCCTCTGAGGCAGCCTAATATACTGACTATTGTGTCTACTTTTGTATGTAAGTAGAGAGTTGCTTTGCTATGCATAAATGTTCTGAATTATACTTCACTTAATGTCAACTGCAAGTCCTGTGTACCACATTTTTCTGTTCTAAATATATTAGTTCATTCTTTT

>PGSC0003DMG400010348 LRR-RLKs

ATGAACACCAACATTATTCCTAGTACCTCAATACAAAATTCTGAGGATATTTCAGAATTACCTATGACCACCACCAACATCTCTGGTTATTCATCAACGGATAAGAATTCTTCGAAAATAACTTATCCAATTACCGTTGATGACTATTCGTTATCTTATATATATTTCATGGGATGCAAAGATAATACTAGTGAGAAATTACAAAATCACTCCATCAAGTCTAAGTTGATGGTTGGTCAGCAAAATGGAATTTTGAGTTCCTTGGATAGTGAATTCCCACAAAATACAAGGTTTATTTTCCTTTCCCTATTTTAATTTCAAAAATACATATGTTATTGATGATATTGCCAATTATTATTTTTTTATTACATATTTTCAAACTTTTATTTAATCAATTCTTAGATGTAAGTAAATTTTGTTTTTGTCATTTTGGTTTAGCAGTAGGAATGGCGGTCCTATAAGAAGGTCACCTAATAGGACAAGCTCAAGATACCAACCATTTGGCAAAGAAATTGATCCAAACAATCAAGGATTAGATAGGCATGTTTTCTCTCTTTTATTCTCCAAATTATCATCAATATATTAATTAGTAATATTGTATCTCCCATTTTCAATCACATTAGTTTACATATTATATGAGTGTGTTCGGTACGGAGGAAACATATGGTTTTCAATTATTTTATGTTTGATTAGTGGGAATATTTTCTCATGTTCTTTGAAAATGAGAAAAATGATAAATAGTTGCATCTATCATATATATATATAATAAAAGATCTCTTTAGACAATGACAACGCTCAATATAATTCATCTCATCTTGTTTATATTTTTCAGGATTCGTCAAAACAGTTCAGAGGCGATCTTCAACACACCACACAATTTCAATGAACATACTCAAGGGTATGGCTATTCCAATGAAGCGATTACATTGAGAAGTTTGAAGAGTAGGAGAGAAAATGAAGATGAAGCTCCTGCTCAGTTACAGTCTTCTTCAATACCCCAACCAACTTGCCTCTATATTGAATCTGCCTAGTTCGCAAATAAGGCTCAAAATACTTGTACAATGTGTCTTGGACGTTCAAGATATATCATTTTTAGAACAATGTTTTATTGCCTCAAACAGTATATATATATATGGAACAAAAAATATGGAAAATAGTGTTCAGCAAATAAATATTAATCCTTTTATGTTCGATTCCTCTGTGTCTTCATTTCTTCGTCATTAAATGCAACGTCGCCACAACTAACCAGAAGTGATCATAGGACATTTAGGTTGACATTTTTGGCAAACATCTCATAAAGGCAATATTAGCACTGACACTGACGAAGCAAGAAATGTGATAACTTCTCGTTGTTGGGTTAGGATGTTGCATAATTTGTCACCTCTATATATATTGATTCTACAGTTACTTCTTCGAACATTGCAAGGTGAATACTACTGGATATATGGATCGTAAGACAATTGTGTTTTACGGTTTAGTGTCGAGGTGAGTGTCATGATGTCAGATTTTGTCAGTGGCTTACACTTACATACAAGTAAATGATGTGCTAAACCTATTAATTATTACCAATCTCAAATAGCATTGATGAAAAAATTCTCTCAAGTTTCATAAACAAACATACAATGTTTTTGCACTGTATTTTACTGACAATAAGAAGATACTCCTATGTGAAACCGTCATCAGCAATGAAGGCTCTGTTGTTAAGAGAATTTGTGCTTTCCCGATCTAAGAAAAACTTGGACCAATATGTTGATTTTGCCACATCAAACCATAATTTATACTGTAAACACCCAAATAAAGCCACAGGAACACCAACACAAGCTGCAATGAGCTTTGGTTCCCAACCCATATGATTACTATAGACCAAAAAGAATGTTGCTGCAAAAGCCAAGAGCATGCTAACTATCGAGACAAACAGCATTGTGAGTCCAAACAATAACTTGGCTGGCAACGACACAAGAAAATCATCTTCAGCATAGTGGGATGTCACAATGGACAAGAACATAATGATGGAAACTATTGAGGTGAATAGTGCCACTGCATCTGAATTGACAAAGATCCTAAATCCATACAATTTTAGCAATATGGGAATGCAATTATCACTGTTATTACCGCCTGGCACAGTTAATCCAGCAGCAAAGACCATGGTGGCAATCAAAGTAGCGACAATCATACAAGAATTTGCAGTGTCTTTCATCCACCTTTTCCCTTCTTTCAGTAATAACTTATGCTCTTCTGTGAATATTTCTCTAGGGGTTTTCCCATCTTTGTTTTGTTAGTGCAAAGTTTTAATGATAAACAACTTATTCTTGCATCAAAAAGGAAAGAATCAAGAATGTGTACATCCTTGACTCTGTAAAAAAACAATGGAGGACATATATGCCTAGCTTCAATGACTGATGATCCCTGGATTTGGCACAAGAATTTGGCCATGCAAGCATGCGACTAATAGAGAAATTAGCTAAAAATGAGCTTGTAATTGGACACCACCACCACTTTTGTCTTTTTAGTTATTCCAAAACTATCCTTGAAAATAAAAATAATTCAAAGTAAAAATATAAATATGAACAATTAACTCTCTTGAACTTATCACTTAGAAAAGGTAAATGAAAGACAAAAATGGAAGAAATTTTTAACAACTCTCTTGAACATTGAATAATTGAAAAAATGTCAAAAGTTCAAATAGTATGGACTAGATGGAGTAAAATATATAGTTTATATTTTATTAGTCCATATATCATAAAACAATTTTTTTTTTTGTTTCTAGTCAATAGTAATATGAATTTTACACATCATATTAGGCCTCTTATTGTAATTTGAAGTAATTCCGTTGACCCATCCCTGTGTAACACACTTGTGGAGATGTGTAACACATTATGAAGGGTAGAGAGGACTATTGATAGACCCAAAAATCACGCCAAAACTAATGGCAGTATTCAAAATGCAAGGCCACAAGTAGTGATCAGAAAGACCAAGATTCTTGCACATTTCATTGAAGGCAAAAAACATTTTCCCTGTAGGTCACTAATGCCATTTTCTAGTCTTCACTAAAAGAAGTTGTTGAACTCCAAATTGTCACTCTAAACCAAAAAATACTCCAGTCATCAATTCTTGTTTTAGGAAAAAGAGAAACCTGTCAGTTATAGTTATGGGAAGAAGTTGCGATCTTCTCTTTGCTCTTTCGGTTTTCATTCTGCTCCATACTTCACTATCTATTAGTACTGATGAAGCTGCTCTTCTTGCCTTCAAATCTCACATTTCTTCTTCTAATCCTAACAACATCTTAGCAACCAACTGGTCTTCTTCCACCCTTGTTTGCACCTGGATTGGAATCACTTGCACCTCCCCACACCATCGAGTCGCAGCTTTAGACATTTCTAGTATGCAACTTCATGGTACCATTTCTCCACACCTTGGAAACCTCTCATTTCTTGTTTCCCTCAACATTAGTAACAACTCTTTCCGTGGAGATTTGCCCAAAGAGTTGGCTCATTTGCAGAGGTTGAAATTGATTGATGTCTCAAGCAATAACTTCACTGGTGCCATTCCATCATTTTTAAGTTTGCTAGTAGACCTTTGCATTTTGTGCCTGTCGAGCAACCAATTTTCAGGAAAAATCCCATCTTCACTTTCCAATCTAACAAAGCTGGAGGTGTTGAAAATAAAGGGAAATCTTCTTGAAGGAGAGATCCCTCGAGAACTTGGTGATCTTCATTACATGACTGCTTTAAACCTGGGAAGTAACCACCTTACTGGCTCTATACCACCATCAATTTATAACATTACAACCATGCGAATCATTGCTCTTACCAACAATAATCTTAGTGGTGAGCTTCCAACAACTATATGTGACCATCTTCCAAACTTGGAAGGGCTTTGCCTCTTAGAAAACATCCTAGGTGGCGTTATTCCACCAAACTTGGAAAAATGCAAAAACCTCAAAATTTTGTCATTGTCTGTAAACGAGTTCACTTGGAACCATACCAAGAGAGTTAGGCAACATAACAGATCTTACAGATATAGCTCTTGGAATACAGCACTTGGGAGGTAGTATTGCATCTCCTAGCAAGAAATTTTGAATTTTCTTCTTTTTCACTCTTCGACCAATTTGGTACTCATATATATTTATATTTCCCCACTTTACAGGAAAGATACCAGTGGAGCTAGGTAATCTTAAGAAACTACAGATTCTGTCATTAAACGACTGTAAGCTTACTGGTTCCATCCCCGCAAGCATTTTCAACATGTCAGCACTGCAGCTCCTAGGAATATCAGAAAACAGGCTTTCAGGTACTCTACCTTCAGATTTAGGTCGTGGAATGCCCAACCTAGAAGGATTCTCTTGTTTTCAGAATAGTTTGAGTGGTTTTCTCCCTGCTTCTATCTCAAAGTCTTCTAGACTCAGTGTACTTGAACTCTCCTATAACAGTTTTACAGGCCCAATTCCTGAATCAGTTAGTGACTTAGAATACATTGAGATTTTGAACTTTGGGGGGAATAATTTTTTCAGCAATTCAGCATTGAGCTTCTTGACATCTTTGACAAACTGTAGGAAACTGAAAGAAATCACTTTTGCCGAGAATCCCTTGGACGGTTTTTTGCCTGCCTCCATTGGGAATTTCTCTGATTCTCTGCAAATTTTCCAAGGATGGTATTGTAAACTGAAAGGCTTCATTCCTGGAGAAATTGGCAATCTTACCGGTGTGATAAAGATGGATTTGTCACAAAATGAGTTGACTGGATATATTCCAAAAACCATCCAAGGCTTGTCAAACCTTCAAGAACTTTACTTAGGTGGCAACATGATAAAAGGAACCATACCAGATGTTATGTGCAATTTAAATAAACTTGGAGCATTAGATTTGTCAAAAAATCAAGCTTCTGGTTCCATACCACCATGCTTAGGGAACGTTACCAGTTTGAGGTATCTTTATCTAGCTAACAACAGGCTGAACTGGACCTTACCTTCAAGCTTGGGGAGCCTTCAAGATCTCCTAGAATTCAATGTTTCGTCCAATTTATTAACTGGGGAAGTTCCTATAGAGATTGGAAATTTAAAGGTTGCAACAATAGTTGATCTGTCAAAAAATGATTTTAATGGTAAGATCCCTAGCACACTAGGGGGTCTGGATAGATTGATGAAACTTTCTCTAGCACATAATAAATTAGATGGGCCTATTCCAGATTCATTTGGCAAAATGCTGGCCTTGGAATTCTTGGATTTGAGCATTAACAATGTTAGTGGTGAAATTCCAAAGTCATTAGAAGCTCTCGTGTATCTCAAATACTTGAACTTCTCATTTAATGAACTCAGTGGAGAGATTCCCACGGGTGGTCCTTTTGCAAATGCCACAAGTCAATCTTTCTTGAAAAATTATGCCCTTTGTGGCGACTCTAAGTTTCATGTTTCACCATGCGTCATCAAATCTCCCAAGAGGTCAAAGAAAAAAAAGGCAATATTGGTTTTGGGAGTTGGTATGCTATTTCTTGCATTAGCCCTCACATATGTATTTTTGAGATTGAGAAAGGAAAAAAAGAATGCAGGTCAAGCTGATGTGTCTCTGATAAAATGTCATGAAAGAATTTCCTATTATGAACTTGAACAAGCAACCGAAAGATTCAATGAAAGCAACTTGCTTGGTAACGGGAGTTTCTGCAAGGTCTACAAAGGGAGACTAAAGGATGGTACTCTTTTGGCAGCAAAGGTATTCAATGTGCAATTGGAGGGTGCATTCAAAAGTTTTGATACAGAATGTGAGATGTTGCGCAATCTCCGCCACCGAAATCTGACCAAAGTCATCACCAGTTGCTCCAACCTTGATTTCAAAGCCTTAGTGTTGGAATACATGTCCAATGGGACACTTGATAAATGGCTATACTCTCACAACTTGTTCTTGGATTTATTACAGAGATTAGCTATAATGATAGATGTTGCATCTGCAATGGTCTATCTCCACAGTGGTTGTTCAAATCCGGTGGTGCATTGTGACTTAAAGCCAAGCAATGTCTTGCTAGATCAAGAAATGGTTGGCCATGTCAGTGATTTTGGCATTGCAAAATTGTTAGGTGCAGGGAAGACTTTTGTTCAAACAAGGACAATAGCAACCATTGGATATATTGCTCCAGGTATTAAAACTTTTACATCTTTCTCATGTTTTCAGAATACCCAAAACATTAATTCTTTTCCCCGATATATATGATCAGTTTTAGCCTATTACTCGATAGTTAGGAAGCATTTTTGATCATTTTCAACCCATGATTGCAGAGTATGGACAAGATGGAATGGTATCCACGAGCTGTGATGTTTATAGTTTCGGCATCCTGATGATGGAGACATTTACAAGAACCAAACCAGGTGATGAAATATTTACTGGAGACTTGAGCATACGACGTTGGGTAAGTGATTCTTTTCCATATGAGATTCATAAGGTGGTGGATGCTAATTTGGTAGAGCTAGAGGATGAACGGATTGACGCAAAGATGCAGTGTTTGTTGTCTATCATAGAGTTAGCTTTGAGCTGCACTTTAGTGTCACCTGATGCAAGAATTAGTATGGAAGATTCTCTTTCAACACTTCAAAAGATCATGCTCCAGTTGGTCAATAGTCACCGCTAGGTGAAATTAGACCATCCTCTTATGATTGATGCTTGATTACCTATGTCAATTCTGCTATTTAAGATGAATTCAGTTTTGTACTTCATATTATGTACTTTTTCACTTTCTAAACATCACAGATTTCTTTCAACCATATGCTAGCTGGAATGGTGATTTTCCAGCTACTGGAGTTCACAATTAGTCTTTCTAATCCATAATTAATTGTCTCCTGGATATCTTTCAGAACAAGTTTTATGACGTTGCTGATAAGAGCTTATGCCTTTTGGTGTTTTGATGATCTCCTCACAAATGCAGGGACCTGGTCCTCTGCAGAGTATGCTCGTCCAGTGTCGAAGTGCGGTACAGCTGTAAAAGCTGTCTGCGCAAAGTGCGGTACAGCTGTAAAAGTTGTCTGCGTCAGAGAGTTGGTGAAAGCTGCAGAGTACTTCACCAATCAGAAGCAAGGAGGTTGCTTGTACGTTTGGTAA

>PGSC0003DMG400010350 LRR-RLKs

ATGCAACTTCATGGTACCATTCCTCCATACCTTGGAAACCTCTCATTTCTGGTTTCCATTATAATCGACAACAACACTTTCCATGGAGAGTTGCCAAAAGAGTTGGCTTATTTACAGAGGTTGAAACTGATCTCGGTAAGAAGTAATAACTTTACTGGTGCCATTCCAACATTTCTAAGTCTGTTACCAAACCTTCGCATTGTGCATCTTTCGAGCAACCAATTTTTCGGGGAAATCCCATCTTCCCTTTCCAATATAACACAGCTGCAAGTGTTGGATATGTCCAAAAATTTTCTCGATGGGGAGATCCCTCAAGAACTCGGTGAACTTCATCACATGACTTTGTTCAATCTGGAAAATAACCAGCTTACTGACTCTATACCACCATCAATCTTTAACATTACAACAATGAAAAAGATTGGTCTCACCTACAATAATCTTACGGGTAAGCTTCCAGCAACTATATGTGACCATCTTCCAAACTTGGAAGAGCTTCACCTCTCAGCCAATCACATACATGGCGTTATTCCACCAAACATAGGAAAATGCAGAAAGCTCCAAATCTTGTCATTATCTAGAAATGAGCTCACTGGAACTGTACCAAGAGAGATAGGCAATTTAACGGCACTTACAAGCTTATACCTTGGAACTTTGCATCTGGAAGGTATTATGATACTTACTTTCATTTTTATCTTCTGCCAAATTGGTACTAATCATGTATTTTCCTCACTTGACAGGAGAGATTCCAGCAAGTATTTCTAACATGTCAGAACTGCAGAACCTAGGATTTGCAAGAAACAGGCTTTCAGGTAATTGTTGTGTAACAAAGAATCTGAAGTGTTTATGTAAATATATGACACGTGAGCCTAACTCAACCCTAAAAACTAGCTCATGAGGGGAGGATTCCCCAATTCCATATAATCTGACCACCTGTCTATTCCCCAAACAATGTGGGACTTTTAGATACTCTAACACCCCTTTCACACCCAGGTCCAACCGGAGCGTGGACCGGCAGCCCAAAGGGGATGGCCTGACCCAGATACCATGTAAAGATACGACATCCTCTCCTCATGAACTAGCTTTTGGAGGAGAGTTAGGCCCATGTGTCATATCATACATGGTATTAGAGCCACGTCCATTCTCGTTTGGACTCCCAATATTCACTCCAGTTAGGCCTAGGCAAGAAGGGGTTGTTAGAGAGGTTAAAAGTCCCACATTTGTTGGGAATGAACAGGTGGTCTGCTTATATGAACTTGGGCAATTCTCGCCCATGAGCTAGCTTTTGGGATTGAGTTAAGCCCAGGTGCCATATCTTTACGTTCTCAAGTTCTAACATGTCAACTCATATATTTCCCCCTCTTGACAGGAGAGATACCAATGGAACTAGGTTATCTTCAGAAACTACTGTTTCTGTCATTAGACACAAATGAGTTGACTGGTTCTATCCCTGCAAGCATTTTCAACATGTCAGCACTGCAGATCTTAGGAATTGCAGAAAACAGGCTTTCAGGTACTCTACCTTCAGATTTAGGTCGTGGAATGCCCAACCTAGACGGATTCTATTGTTATCAGAATAGTTTGAGTGGTTTTCTCCCTGCTTCTATCTCAAATTTTTCAAGACTCAGAGTACTTGAACTCTCTTATAACAGTTTCACAGGCCCAATTCCTGAATCAATTAGTGACTTAGAATACATTGAGGTTTTGAACTTAGGGGCGAATAATTTTGTCAGCAATTCAGCATTGAGCTTCTTGACATCTTTGGCAAACTGTAGGAAACTGAAAGAAATCACTTTTGCCGAGAATCCCTTGGACGGTTTTTTGCCTGCCTCCATTGGAAATTTCTCTGATTCTCTGCAAATTTTCCAAGGATGGTATTGTAAACTGAAAGGCTTCATTCCCGGAGAAATTGGCAATCTTACCGGTGTGATAAAGATGGATTTGTCACAAAATGAGTTGACTGAATATATTCCAAAATCCATCCAAGGCTTGTCGAACCTTCAAGAACTTAACTTAGGTGGCAACATGATAAAAGGAATCATACCAGATGTTATCTGCAATTTATATAATCTTGGAGCATTAGATTTGTCAGGAAATCAAGTTTCTGGTTCCATACCACCATGCTTAGGGAACATTACCAGTTTGAGGTATGTTTATCTAGCTAACAACGGGCTGAATTTGAGCTTACCTTCAAGCTTGTGGACTCTTCAAGATCTCATAGAATTCAATATTTCGTCCAATTTATTAAGTGGGAAAGTTTCTATGGAGATTGGAAATTTAAAGGTTGCAACACTAGTTGATCTGTCAAAAAATGATTTTAGTGGTAAGATCCCTAGCACGCTAGGGGCTCTGGATAGATTGATGAAATTTTCTCTAGCACATAATAAATTAGATGGCCTATTCCAGATTCATTTGGCAAAATGCTGGCCTTAGAATTCTTGGATTTGAGCAATAACAATCTTAGCGGTGAAATTCCAAAGTCATTAGAAGCTCTCGTGTATCTCAAATACTTGAACTTCTCATTTAATGAACTCAGTGGAAAGATTCCCACAGGTGTTCCTTTTGCAAATGCCACAGGTCAATCCTTCTTGTCCAATTATGCACTCTGTGGTGATTCTAAGTTTCATGTTTCACCATGCGTCATCAAATCTCCGAAGAGGTCAAAGAGAAAAAAGATAATCTTGGTTTTGTACATCCTTTTAGGAGTGGGTATACTATTTCTCTCATTAGCCCTCGCATATGTATTTTTAAGATGGAGAAAGATAAAGAAGAATGCAGATCAAGCAGATGTCTTTCCGGTAAAAGGGAAATATGAAAGAATTTCTTACTATGAACTTGAACAAGCAACAGAAGGATTCGACGAAAGCAACCTGCTTGGTAGTGGGAGTTTCAGCAAGGTTTTCAAAGGGATACTTAAGGATGGTACTCTTTTGGCAGCAAAGGTATTCAATGTGCAATTGGAGGGTGCATTCAAAAGTTTTGATACTGAATGTGAGATGTTGCGCAACCTCCGCCATCAAAATCTTACAAAAGTCATCACCAGCTGCTCCAACCCTCACTTCAAAGCCTTGGTATTAGAATATATGCCAAATGGGACACTTGAAAAATGGCTATACAATCACAACTTTTTCTTAGACATGTTGCAGAGATTAAGTATACTGATAGATGTTGCATCTGCAATAGACTATCTCCACAATGGCTATTCAACACCTGTGGTGCATTGTGACTTGAAACCAAGCAACATCTTGCTAGATCATGAAATGGTTGGTCATGTCAGCGATTTTGGCATTGCAAAATTGTTAGGTGCAGGGGAAGATTTTGTTCAAACAAGGACAATAGCAACCATTGGATATATTGCTCTAGGTATCGTAATTATAGATTAGTACTCAGTTTTAGCATGTTCCTCGGTAACTAGGAAGTTTTTTTGTGATTATCTTTTACCATGATTGCAGAATATGGACAAGATGGAATAGTCTCCACCAGCTGTGATGTTTATAGTTTAGGCATTGTGATGATGGAGACATTCACAAGAAGGAGACCAAGTGATGAAATATTTACCGGAGGAATGAATATACGATGTTGGATTAATGATTCTTTTCCGAGTGGTATTCATAAGGTGGTGGATTCTAATTTGATACAACCATGGGATGAACAAATCGATGCAAAGATGCATTGTTTGTCATCTATCATGAAATTAGCTTTGAGTTGCACTGTAGTGACACCTGATGCAAGAATTAGCATGACAGATGCTCTTTCAACACTCGAAAAGATCAGACTCCAGTTTGTCAATAGTCTCCGCTAG

>PGSC0003DMG400010351 LRR-RLKs

GATTAGGCAAAACACAAAACTATTCATATTGACTAATGTGCCCAGAGTTTATGTAATCAAAGTACTAAATACTCGTTTAGTCTCATATTACTTGGCCTTGTTAACACTAGTAAAAGTTGACACTCTTTAAGTCTTAACCTAAAGTATACTCCAAAAACTAATGGCAGTACTCAAAATGCAAGGCCACAAGTACTTATCAGAAAGACCAAGACTTAAATTTGAGGTTCAAGTTTCTTGCACATCTCATTAAACATAAAATACATTTTCCCTGTATTGTCGCCATTTTCTATTCTTCACTAAAAGAACTTGTCGAACTAACTCGATATTGACACTCTAAACCAAAATATACTCCATAGACTAATGTCAGTATATTGGTATTCGTGTGAATTTGATATCAATCATGACTATATGAAAAGTCCACATATGGTGAATAATGCCAAAGAAATATGACTTGAGAATTTCCATGGACTGGTTGAGGGGAGGAGACTGGACTGTTGACTCAGACTCACTAATACAACTTGTAGTACTAAAAAAATTTACACTCCAATGAATTTGATATCATTAGATATACACATAATCTAAGATCATTTATTTGCCTCATCGATCAATTCTTGTTTTAGTAAAAAGAGAGATATCAATCTGCTATTAGTTATGGGAAGAAGTTGCAATCTTCTCTTTGCACTTGTGGTTTTCATTCTGCTCCATACTTCATTCTCTACTGTTCCAAATATTAGTACTGATGAAGCTGCTCTTCTTGCATTAAAATCTCACATTTCTTTTAGTCCTAACAACATCTTAGCAACCAACTGGTCTTCTTCCACCCCTGTTTGCACCTGGATTGGAATCACTTGCACCTCCCGACACCATCGAGTCACAGCTTTAGACATTTCTAGTATGCAACTTCATGGTACCATTCCTCCACACCTTGGAAACCTCTCATTTCTTGTTTCCCTCAATATTAATAACAACAGATTCCATGGGAATTTACCCCAAGAGTTGGCTCATTTGCAGAGGTTGAAATTGATTGATGTCACAAGTAATAACTTAACTGGTGCCATTCCAACATCTTTAGGTTTGCTAGTAGACCTTCGCATTTTGCGCCTGTCTAGCAATCAATTTTCCGGAAAAATCCCATCTTCCCTTTCCAATCTAACAAAGCTGGAAGTGTTGAGAATAACGGGAAATTTTCTCGAAGGAGAGATCCCTCGAGAACTTGGTGATCTTCATTACATGACTGCTTTAAATCTGGAAAGTAACCATCTTACTGGCTCTATACCACCATCAATTTATAACATTACAACCATGCGAATCATTGGTCTTACCAACAATAATCTTACTGGTGAGCTTCCAACAACTATATGTGACTATCTTCCAAACTTGGAAGGGCTTCACCTCTCAACAAACATCCTAGGTGGCATTATTCCACCAAACTTGGAAAAATGCAAAAAGCTCAAAGTCTTGTCATTGTCTGTCAATGAGTTCACTGGAACTATACCGAGGGAGTTAGGCAACATAACAAATCTTGCAGAATTATACCTTGGATTGCTGCACTTCGAAGGTAGTATTACATCTCCTAGCAAGAATATTTAATTTTTTTCTTTTTCACTTTTCCACCAATTTAGTACTCATATATATATTTCCCCACTTAACAGGAGAGATACCAGTGGAGCTAGGTAATCTTAAGAAACTACAGATTCTGTCATTATCCGGCTGTAAGCTTACTGGTTCCATCCCCACAAGCATTTTCAACATGTCAGCGCTGCGGGCAGTCGTACTTGACGAAAACATGCTTTCAGGTAATCTACCAGCAGATTTAGGCAGTGGGATCCCCAGCCTAGAAACATTGTTTTGTGCAAACAACTATCTGAGTGGATTTATATCTGCTACTATCTCAAATGCTTCAAGACTCAGTGAGATTGATCTTTCAATCAATAGTTTCACAGGTCCAATTCCTCAATCACTTGGTAACTTAGAAAACCTTGAGGTTTTGGACCTGCAGGTGAACAATTTTGTCAGCGATTCAGCATTGAGCTTCCTGACATCTTTGACACACTGTAGGAAACTGAGAGAACTCACCTTTGCTAAGAATCCCTTGGAAGGTTTTTTCCCTGCATCAATTGGAAATTTCTCTGATTTTCTGCAATTTTTCGAAGGATGGACTTGTAAACTGAAAGGCTCCATTCCTGGAGAAATTGGTAATCTTACCGGTGTGATAAGGATAAATTTTTCACAAAATGATTTGACTGGACATATTCCAAAAACTATCCAAGGCATGCTGAACCTTCAAGAATTTTACCTAGAGAGCAACAAGATAGAAGGAGTCATACCAGATTCTATCTGCAATTTAAAGAATCTTGGAGCATTATTCTTGTCAGGTAATCAATTTTCTGGTCGCGTGCCACCTTGCTTAGGAAAAGTTACCAGTATGAGATATCTTTCTCTAGCTGATAACATGCTTAATTCGAGTTTACCCGAAAGCTTGGGGAGCCTTCCTGATCTCATAGAGTTCAATTTTTCATCCAATTTATTGCATGGCAAAATTCCTATTGAGATTGGAAATTTAAAGTCTGCAACACTCATTGATCTGTCAAAAAATGATTTTTTTGGTATGATCCCTAGCACTCTAGAGGGTCTGGATAGATTGATTAGTCTTTCTCTAGCACATAATAAATTAGATGGGCCTATTCCAGATTCATTTAGAAAAATACTGGCCTTGGAATACTTGGATTTGTCCAGTAACAATCTTAGTGGTGAAATCCCAAAGTCATTAGAAGCTCTTGTGTATCTCAAATACTTGAACTTCTCATTTAATGAATTCAGTGGAGAAATTCCCACGGGTGGTCCTTTTGCAAATGCCACAGATCAATCTTTCTTGTCCAATTATGGGCTCTGTGGTGATTCTAAGTTTCATGTTTCACCATGCGTCATCAAATCTCCCAAGAGGTCAAAGAAAAAAAAGGCAATTTTGGTTTTGTACATCCTTTTGGGAGTAGGTATGCTATTTCTTGCATTAGCCGTCACATATGTATTTTTGAGATTGAGAAAGAAAAAAAAGAATGCAGGTCAAGCTGATGTGTCTCTGATAAAATGTCATGAAAGAATTTCCTATTATGAACTTGAACAAGCAACCGAAGGATTCAATGAAAGCAACTTGCTTGGTAACGGGAGTTTCAGCAAGGTCTACAAAGGGATACTTAAGGATGGTACTCTTTTGGCTTCAAAGGTATTCAATGTGCAATTGGAGGGTGCATTCAAAAGTTTTGATACAGAATGTGAGATGTTGCGCAATCTCCGCCACCGAAATCTGACCAAAGTCATCACCAGTTGCTCCAACCTTGATTTCAAAGCCTTAGTGTTGGAATACATGTCCAATGGGACACTTGATAAATGGCTATACTCTCACAACTTGTTCTTGGATTTATTTCACAGATTAGATATAATGATAGATGTTGCATCTGCAATGGTCTATCTCCACAGTGGTTGTTCAAATCCTGTGGTGCATTGTGACTTAAAGCCAAGCAATGTCTTGCTAGATCAAGAAATGGTTGGCCATGTCAGTGATTTTGGCATTGCAAAATTGTTAGGTGCAGGGGAGACTTTTGTTCAAACAAGGACAATAGCAACCATTGGATATATTGCTCCAGGTTTTTAAACTTTTACAGATTCCTCATGTTTTTAGAATACCTGAAAAATATTAATTATTTCCCCATAAATATGATCAGTTTTCACATATTCCTCGATAATTAGGAAGCTTTTTTTTATATCATTTTCAACCATGATTGCAGAGTATGGACAAGATGGAATAGTATCCACGAGCTGTGATGTTTATAGTTTCGGCATCCTGATGATGGAGACATTTACAAGAATCAGACCAGGTGATGAAAGATTTACTGGTGACTTGAGCATACGACGTTGGGTAAGCGATTCTTTTCCAGATGAGATTCATAAGGTGGTGGATGCTAATTTGGTACAGCTAGAGGATGAACGAATTGACGCAAAGATGCAGTGTCTGTTGTCTATCATAGAGTTAGCTTTGAGCTGCACTTTAGTGACACCTGATGCAAGAATTAGTATGGAAGATTCTCTTTCAACACTTCAAAAGATCAGGCTCCTGTTTGTCAATAGTCGCCGCTAGGTGGAATTAGACCATCCCCTTATGATTGATGCTTGATTACCTATGTCAATTCCGCTATTTAAGATGAATTCAGTTTTGTACTTCATATTATGTACTCTTTCACTTTCAAAACATCACAGATTTCTTTCAACCATATGGTAGCTGGAATGGTGATTTTCCATCTACTGGAGTTTACAATTAGTCTTTCTAAATATTGACACTCTTAAGTCTTAACCAAAAAACACTCTGAAGACTAAACAACACTATAGGGCCTCCAGGTTGCCTGTAGCATAGCCAATACCATTTTCCAGTCCATAATAATGCAGTAACCTGTTCCTATCATCTACATATCCTTTCATA

>PGSC0003DMG400010400 LRR-RLKs

GGCATTTAAATCGAGCCTTCGATTGGAACCTCACCACATTTTGTCCAGAAATTGGAGCTCAGACACACACTTCTGCAGCTGGTATGGAGTTTCATGCTTGCACCAGAGAGTTGTAGCTTTGCAACTTCCTAATTTGGCCATTCAAGGCAGAGTTTCTTCAGAAATTGCCAACTTGACTCAACTTGCTCTTCTTGACCTTAGTAGCAATGATTTGAATGGCAATTTACCAAGTGAGTTGGGCTTCTTGCAACAACTGAGACTTTTTAATGTGACTGGGAATTCTCTTGATGGAACTATTCCTCTGAATATAAGCCGTTGTCGTGTACTGCAAGAACTTCATTTATCAGATAACATGATCAAAGGAAGCATACCTCAAGAACTTGGCCTGTTAAGTCAACTTAGAATTTTGCGTCTCAACGACAATAATCTGACAGGAAAGATTCTACCTTTATTAGGAAACTTATCAAAGCTGGAATATTTTTATCTGCATGAAAATGATTTGGAAGGTGAAATACCTGATGAAATTGGTGATCTTACCAATTTGAGGCTCCTTTCCTTTCGTGGCAATGATTTCACTGGAGCGATACCTGCTTCAATTTTCAATATTTCAAGATTGCAAATTGTGGATCTCTCGATAAATGCCTTATCAGGGGAACTTCCAAGTGATTTGGGTAATTGTTTTCCTATCCTTGAACATCTTTTTCTGGACAGCAACAGGATCACGGGGAAAATCCCGGTTTCTTTATCAAATGCATCAAAGATTACTCATCTCTTCTTCACTGAAAATGATGTGCAAGGAAATATTCCAAATGAGTTTGGTAAGTTGCATGAACTAGTCTGGTTTGAATTTGAATATAATCGAATTTCTGGTGCCATTCCATCCAGCTTATTCAATATTTCGTCTCTTGAAATCCTCAAAGCCAGGCATAATTACTTAAATGGGCATCTTCCATATGATCTTGGTACTTGGCTACCTAACTTGCAGGAGATTTTCCTTTGGAGATCTACCCTCTGCAATATGCAATGCTTCAAAGCTTTGAAAATCTTGAAGTTGCAAACAACAGTTTTACTGGACCGATTCCCATGATGTTAGGTAACCTGGTGAACCTTATAACTCTAAATCTGCAAAATAATCTGTTGGAAAACAAGCCTGGAGCCACACAACTTGACTTCCTGAATTCATTGGTAACCAGCAGGAATCTGGTGTACCTGATTCTTGAATCTAATCCTTTGAATGGAATTTTTCCAGAATCAATAGGAAATTTATCAAGCAATATCAGGGTCCTTGATGCAGCAAACTGTGGAATTAGAGGCAGTATGCCAACAAACATTGGAAACTTAAGTGGCTTGATTTTCCTGGGATTAGCAAGTAATAACATTGTTGGCAATGTGCCCCCCTCGTTTGTTGGCCTGCAAAACCTGGAAAGGCTGTATCTTACTAGAAATAAATTAGAAGGGACATTTCCAGCTGAACTGTGCAGCATAGGAAGATTAGGTCTACTGCATTTGGGAGAAAACAGACTGTCAGGAGAAATACCCTCGTGTATCGGAAATCTGACTGAATTAAGAGTAATGTCAATTGCTGCCAACAATTTCAGCTCAAAGTTACCTTCAAGTTTCTGGAGATTAGTAAAGCTTGATGGTCTGAACATATCGAGAAACCTGCTGCATGGTTTCTTCCCTTCTGATGTGGGAACTCTGAAAGCAATGAGCATCATGGACCTCTCTTTCAATAGGTTTTTTGGTGAAATACCCAATTCCATAGGGTCCCTTCAATATCTAGTTTCACTAGATATGTCAAGAAATGCATTCCAAGGTCCCATACCGGACACATTTAGCGATCTAATAGTTCTTGAGGGTCTGGATTTGTCTACTAATGCTCTATCAGGTGTGATTCCCAAATCTCTTGAGCTATTGCGAGATCTCAAGTATCTCAATGTTTCTTTCAACAACTTGCAAGGAGAAGTTCCAAAGAAAGGGGTGTTTGCAAATGTTAATTACCGGTTCTTGATGGGAAATCCAAGACTTTGTGGAGCAATTGACTTATATATTCCTATATGTCCAGCTCAAGGTAGAAAGACAACAAGGAAAAAGTCCATTATTCTTAGAACAGCAGTACCTGTTGCTGTAACATTTCTGATATTGGTCGCATTGTTTTTCACATGGATGATTTGGTCAAGAAAAAAACATGTGATTGGAAACAATGAATCGGATTACCCTCCTAGAATTGCACATCAGAAAATCACTTATTATGAGCTTCTCCAAGCTACAGAAAATTTCAGTCAGTCCAAGCTAGTAGGAAGTGGAAGTTCTGGTACAGTCTACCAAGGTACATTTTCTGATGGTGCTGTCTTTGCAATCAAAGTTTTTGATATGCAATGGCAACGGGATTTAAGAAATTTTGATTCAGAATGTGAAATTTTGAGCAACGTAAGACACAGAAACTTGGTCAAGATAGTGTCGACTTGTTCAAATTTGGATTTCACAGCAATTGTGCTGGAGTACATGCCTAATGGAAGCTTAGACAAAAGATTATACTCTGATGAAAATTGCTTGAGCCTGGTTGAAAGACTCAATATAATGATCGATGTAGCACTTGCCATGGAGTATCTTCATCATGATTATACAGTGCCCATTGTGCACTGTGATCTGAAGCCAGGTAATGTTCTTCTTGATCAGGATCTGACTGCTCATGTAGCAGATTTTGGAATTGCAAAAATGTTGGCACAAGAAGGAAACATGGCTCAAACCAAGACTCTCGGAACTATTGGTTACATTGCTCCAGGTATGATAATCATGTCTTGAAATGAAGGTGTCCTATGTTGGTTACTTTTTTCCTTTATGTCTACCATAACTTTATACTGTAACATGGATAACTACACATTTTAACTGGGAATTTGTTGAAACAGAATATGGTTTAGACGGGCAAATTTCCACTAGCAGTGATGTTTATAGCTTTGGGATTTTGTTGCTTGAGACATTCACAAGAAAGAAGCCTACGGATGACTTGTTTAGCGGAGATTTAAGCTTGCATAAATGGACATCTCTCTCCTTTCCTGATGCTGTACTGGACATTCTGGATGCTGACCTTGTTAGCGACATTGGCTTTACAACAGATGAAAACCAAAGCGAAATCAAACAATTGCTGGTTTTGATAATAAATGTTGCATTCCTGTGTTTGAAAGAGTTGCCAGAGGAAAGGATAAACATGAGAGAGGTTGTGGTGCAGCTAAACAAGATAAGAGCTGAATTAACCAAGCTTTAACTCAATCCCATTCAAAATTTCAGAAAGAGGATTGATGCAGCAACAAGCAAATTCAGTTTTCCACCTTCTATGACCTGTCAAAACATCTGTTATTTCTTAATATTTTCTTTCTGTTTAGTGTAGGGGCATTTTAATATTATAAGAATGTTACATTTTAGCATTGATCGTTTGCCCCTAAGTCAGTCATTACTCCCTCTATTCATTATTGCATTAGAATATGCATTGGTTGACTTTCATTTGTTGTAGTATGGTGGGAGTTAATGGAAATTCTTTTGCTGCCGTTTGGACCATTAAGTTGGTGCATTACTCCATAGAGTAATTGCATATGTTACTAATTCCTTGGTGCAATTGCAGCAGGTTCAGCCGTTGGTGGCTGTAAGCTTTTTTATCCCATTACTTCTTACTCTTTACTCCTCAATTATTC

>PGSC0003DMG400011160 LRR-RLKs

AAAAAATCTCTCTTCCCTCTCTCTGAGTCCTCTCACTTTCACTGATTTTCTTCATATCATCTCCACACCTGATTCTCTTACACCAAAAACACAGTACAAAAAAATTAGTAAAAAAAAAATATTTGAAAAAAAAACACAAACAAACTCAGATAAAGCAAAAGGAGAAACAGAGTTGGTACAAATACCAACAAAATAGACAGACTGATAAAGAACCTGTTTCTTTTCCTTTACACTCCCTCTCACTTTTTTTTTTAGTTTTTGACTGAAAAAAGACACAATTTTTAATCATTTATATGAACCAAAATGTTAAATAAATTCATGGGGTTTGCTTAATATATGTGTTGAATAATTTGATTTCACTTTCTTGGGTTTTTTTTTTTTATCGTCAATGGCTTCCATTTTCTTGATTTTTGTTCTCTCTTTGCTTCTGTTTTACCCTTTTCTTTGTTTTTCATATGAAGCTAGGAATCATGAAGGTAAACAAAAAAAAAGAATCCCATTTGCTTATTTTTCAGTTTTTTGTTTTCAAAATTTGATCTTGTTGTTTTGATGTATAGTGGAAGCTTTGATTGCTATAAGGAGAAGTTTGGATGACCCAAAAAGAGCTTTAAGTAATTGGGATGAAGATTCTGTTGATCCTTGTAGTTGGGCTATGATCAGTTGTTCTTCTGAAAATCTTGTTATTGCACTGTATGTTTAATAGTATTTTTTTTTTTTGGAATTTTATGGTACTATATGATTATATTGGTTGTTAAAGATTGAAACTTTTACTGTTTTATCAACTAAAAAAGTGTTTTTTTTTGTTTGTAGAGGAGCTCCAAGTCAAGGTTTATCTGGGGTTTTGTCTGGGATGATATCAAATCTTACAAACCTTAAACAAGTGTAAGAGACAATTTTTTAATGTTTTTGTTGTATGTAATTTCAAATTTTGATGACCACATTTGGTTAATTTTCTTGTATGTTGAAAATTGTGTAGTTTATTGCAAAACAACAACATTTCTGGGCATATTCCAAGAGAGCTAGGGAAACTTCCTAATCTTGAAACATTGGATCTGTCTAATAATCATTTCTCTGGTCATGTTCCTGATTCTTTGGGTCTGTTAAACAGTCTTCAGTATCTGTAAGTGTATTAAACTTTCAAAATTTTCATTTTTGTGTGATTATGATTATGTTTTTAGAGTTGAATGGTCTAAAACACACATGCAGTATCACGTTTTTGCGAGTTTGCTACGAATTATCACCTAAAACATATCTCAACGCCATCTCATAGTTGTGTTTTGATAAATAGTTGGTGATGGTTTAAGAAGGAAAATTGAACACATGATAGTTTAGGTAGAAAACGTGCGAAATCTTGATACTTCAAAGTGTGGTTTTTACCATTATCTCTTGTTTTTCCGAGTACCCTTTTTCCTTTAATTGTTGTTTTCTCATGTAGACAATGTTTAAGTGCTATCTTATTCTTTGACTATTATGTTTTTGTTTTTTGTTGATGTCTTGTTCTGGGGTTTGTCTTGTAATGGAGTAGAAGGTTGAACAACAACAGCTTGTCTGGTGCTATTCCTCTTTCTTTAGCCAGGGTCCCTCAGCTTGCTTTCTTGTGAGTATTGCTTCTTTGAATCTATGACTTTTGTTTGATTTTGTTTTATTTTTTTGTCTGATTGTCCTGAGAATTGGAATTATCTAGTCACTTTCTTGAATCAGCCTTATGTTAAAGCTTTTTTTGTTTAAAGGAAATTGAATAGTTTGTTCTGTTTAGGGGTATTATTATATTTTAACTTATTATTCACTTTTATTTTATTTAGTATGACGATGATATGCAATGAGCTTAAATGAAAGTGGGGATTCATATTGTTGACTCCAACTTGTTTGGGACTGAGGCATTGTTGTCGTCGTCGGATAGTGTTTTTTTTTTTTTTTGTATAAAGAAATGATTAGTGTGTTCTGATTAGGGGTATTATTATATTTTAACTTATTTTTTACTTTTATTTTATTTAGTATGACGATGATATGCAATGAACTTAAATGAATGTGGGGATTCATTATGCGGACCCCAACTTCTTTGGAACTGAGGCGTTGTTGTTGTCAAATAGTGTGTCTGTTTTGTGTAAAGGAAATGAATAGTGTGTTCTGTTCTGGGTTATTATTATATTCCTACTTATATTTCACTTTTATTTAATCTAGTGTGACGATGATATGCAGTGAGCTTAAATGAAAGTGGTTATTCATATTGTCGACGCCAACTTGTTTGGGACTGAGGCGTTGTTGTCATTGTTGCCGTTGAATAGTGTGTTCTGTTTAGGGGTCTTTTTTATCTTCTAACTTATTTTTCACTTTTATTTAATCTAGTATGACAATGATATGCAATGAGCTTAAATGAAAGCGGGGATTCATATCGTGGACTCTAACTTGTTTGGGACTGAGGCGTTGTTGTTTTTGTTTTCATTGTTGAATAGTGTGTTCTGATTTTATGAACCTCTTATTCTTTCAGGGACTTGTCTTTCAATAATCTGAGTGGACCTATTCCTAAATTTCCTACGAGAACTTTCAAGTATGGATTCTCTGCCCAACTCTTATTATGTTACTTATAATTGTGTTCTTTTGTACATACAAATTGTCTTGTTTAAGTTCTAATTGTGATTTTTCTTTATCCTTGATGTTTTCGTAGTGTCGTGGGTAACCCTTTGATATGTGGAAATCACTCTGCTGAGACTTGCTTCGGATCGGTTAACCTGATGCCCCTTTCGTTCGACATTGATTCCACCGGTAAAGTTTAGAAAAGTTTCTCATGATTCAACAATGTTTAGACATAAATACAATTTAATTATCTTAAACTTCTTTCTGTTGTAACCAGGTAAGCGAAACTCGAAAAGACTAGCCATTGCTATAGGACTTAGTGTCAGCTTCATCTCTCTTTTCCTCTTGGGATTCGGATTCATAATTTGGAAAAGGAACCAAAACAGGAAACAGTCTATTCTGAACATTAATGGTGAGTCTTAGTAGCATTTTGTGTATTTGATGACGGTTCAAAGTTATTTACGTTTCGTATTTTTGTTCCAGATATGCAAGAGGAGGATCTAGTTAGACTAGGGAATCTCAGAAGCTTCACATTTAAAGAACTTCAACGTGCGACAAACAGTTTCAGCTCTAAGAACATTCTTGGTGCTGGAGGATTTGGCAATGTTTACCGTGGAAAGTTGGGGGATGGCTCGTTGGTCGCGGTGAAACGATTGAAAGATATTAGTGGAACTGCTGGAGAGTCACAGTTCCGGACGGAATTGGAGCTTATAAGCTTGGCAGTACATAGAAATTTGCTCAGATTAATTGGTTATTGTGCTACCCCAAACGAGAGACTTCTCGTGTATCCTTTCATGTCTAATGGCAGTGTGGCATCAAGACTTAGAGGTAAACACCTTGTGTACTCTTTTCAGCCAATCGCATGCCTTACACATTGTACTCCTTCGTTTCTATTCGTTTGTCTGGTTTTGATTTGACACGGAGTTTAAGAAAGTAAAGAATGATTTTTGAATCTTGAAGTCTTAAACTAAAGATATGGAGAGTGTACCAATATGTCTTTTAATCTTGTGGTCTTAAACATGCCATGTGGTCTTAAAAAAATTGATGGAATGTACACATTCTTAATCCATTTTTCGCGATGACAGGAAAACCAGCACTAGATTGGAATACTAGGAAGAGAATTGCAATTGGAACTGCAAGAGGTCTTCTTTACCTCCACGAGCAATGTGATCCAAAGATAATCCACAGAGATGTGAAGGCGGCAAATGTCCTATTAGATGACTTCTGTGAAGCTGTTGTCGGGGACTTTGGCCTTGCAAAGCTCCTCGATCATGCTGACTCTCATGTTACCACAGCTGTTCGTGGAACAGTTGGCCATATAGCGCCAGAGTACCTTTCCACCGGTCAATCATCCGAAAAAACGGATGTGTTTGGATTTGGTATCCTCTTATTAGAGCTTATAACTGGGATGAGAGCTCTTGAATTGGGAAAAACAGTAAACCAGAAAGGAGCAGTACTTGAATGGGTATGTATGTTGCTTTATTCATCTACCAGTTATTGAATTCTATGTTGCTCGAACTCTCCAAAAATGCACTACTTTTGAAGGCTTCGATACGCATGTGTTTTCATTTTTGAAGAGTCCGAGCAACCAATATTTCCTTTAAATAAAAGTGAAAAATGGAGCTCCAGTAGCATCACTTACTTAGTGATTTTTTAACCTTTGTTAGTCGGACTCTTCAAAATGTAGACTGGTGCATATCGAATCCTCGAAAAAACTAGTACATTTTTTGATAATCTGATACGGATGCAACAACATTTTTGTAGAGTTCGAGCAAAGTAGGTTTTAACAAAAAGAGTCTTGCTTAGATTTTCTCATGAACTGCTGCTTAACAACAAGTGTGGCTTGATTGCAGGTTAAGAAAATGCAGCATGAGAAGAACGTCGAAGCTCTCATCGACAGAGAGCTTGGGAGCAACTACGACAGGATCGACGTAGGGGAGATGTTACAAGTGGCTATACTCTGTACTCAATACCTACCGTCCCACCGTCCCAAGATGTCTGAGGTGGTCCGAATGCTCGAAGGTGATGGCCTTGCTGAAAAATGGGCTGCATCTCATAACTATGATGGCTTTTCTCTAACCAAAAACAGTAACAAAACTCAGACTAAACATTCTATGGTGCCTCTTGGATATGATGACAATGATCATGATCATTTGACCGTGTTTGGGATGATGGATGATGACTATGACACACATGCCATGGAGCTTTCCGGTCCAAGATAATAGCGAAATATCATCGAGGTCTGGAGAGGGTACGATGTACACAAACCTTACCCCTATTTCTATGGGATAGAGATGCTATTTCCGATGACCCTCGGCTCGTAATAGCAAAATATCCAAAAGGAAAAGAAATTGAACATAGGTACCTAATTCTTTTCAAATTTTTATAAGTTAGTACCACAATTAACTCTATCAAAAAGCATGTTGGAGAGGAAAAAGAAAGTAGTTCTTTTTTAAAATTTTTTTTTGTTGGTTTTCTTTAGATGGTGTGAAAATAGGAGTTTGCTATGTTTGATAAAAGTACACTTTTAAGGAGTTTATGGGACTAATGTAAATAACCCTCTTCATCAAAAGTATCAATCTTTA

>PGSC0003DMG400011205 LRR-RLKs

ATGTCTATGATTGTTTCTGATCCATTTCATGTATTAGAAAGTTGGAATTCTTCCAACATTCATGTATGTAATTGGACAGGAATTGTATGTGACAAGAAAATTCATAGAGTTGTTGAGCTTGATCTTAGTCATCATTCGTTACGTGGAACGATTTCTCCTTCTCTCTCTGGGCTCTCTTTCTTGCAGATTCTTGATTTGTCTGGTAACTTATTAGAAGGACAAATTCCAGCTGAATTAGGCAATCTTGTTAAGCTAAATCAGCTTAGTTTGTCTTCTAATCTCCTCGAAGGAGAAATCCCAAACGAGTTAGGATTCCTTCATGAATTGAAGTATCTTGATTTGGGAACAAACAATCTTTCTGGTGCAATTCCTCTGTCTTTGTTCTGTAATTGCTCTGCTTCATTACAGTATATGGACCTTTCTAACAATTCGCTTCGCGGTGAGATCCCTATGAATGATCATTGTGAGTTGAGTGGTTTGAAGTTTCTTCTTCTATGGTCTAATGGACTTGTTGGTGAAGTTCCTATTGCTCTTTCGAAATCTACAAAGCTAGAATGGCTTGATCTTGAATCGAATTCGTTGAGTGGTGAGCTTCCTTCTGATATTGTAAGTAAAATGCCAAAGTTACAGTTTTTGTATTTGTCTTACAACAACTTTGATAGTCATAGTGGTAATACTGACCTGACGCCTTTTTTCGCGTCTTTAGTTAATTCTACTAACTTGCAAGAACTTGAGTTAGCTGGGAATAATCTTGGTGGGGAGTTGCCTCCTATTATCGGTAACATCTCCAAGAACCTTGCACAGATACATCTCGACGATAATCTTATCTATGGTCCTATCCCAACACAAATTTCGAGTCTTGTAAATATTACTCTATTGAACTTGTCTAGTAATCATCTGAATGGTACAATTCCTCCTGAGTTGTGTCGAATGGGGAAACTTGAGAGGCTTTATCTGTCGAATAACTCACTCTCTGGTGTCATTCCTTCTGCTTTTGGAAATGTTTCTCATTTAGGCCTTCTTGATCTGTCGAAAAACAACCTCTCTGGATTAATCCCAAATACCTTTGCTAATCTTCCGCAATTAAGGAGGCTTTTGTTGCATGATAATCACCTCTCTGGAACGATTCCCCCGAGCTTAGGTGATTGTGTTAACTTGGAAATACTTGATCTTTCTCACAACAGAATCACGGGGACGATTCCAAGTGCAGTTGCTGGATTGAGTAGTTTGAAGCTCTATCTGAACTTGTCTAGTAATTACTTACATGGTCCAATTCCTTTAGAGCTCAGCAAAATGGATATGGTGTTGGCAATTGACTTGTCTTCGAATAATCTTTCGTCTAATGTCCCTTCACAGCTTGGCAGCTGCATTGCTCTCGAATACTTGAATCTATCGCGTAATTCCTTAGAAGGACCTCTTCCATCTTCGATAGGAAGATTGCCTTACCTTAAGGAAATCGACGTTTCATTCAATGTGTTGAGTGGGGAAATACCACAGACATTTCAAGCATCGTCGACTTTGCAGAACCTCAACTTCTCCTATAACCATTTCTCGGGGAACGTGACAGATACTGGTGCATTTTCATCACTAACCATGAGCTCATTCATGGGAAATTTTAAGCTTTGTGGCTCAGTACAAGGGATGAAAAGTTGCCATAAAAAGAGGGGTAGACATTTCATTATCACAATCCTCCTTTCATTGCTTATTACTCCTATATTTTGCATAGTCGGGTACCCTCTCCTCCTAAGATCAAGATTCAGGAACCAATTCGCGAGTTTTAATAAAAGCAAGGTAATTGAAGATGTGGAGCAAGGAGGTGGGAGGAAAGAGCAAAAGTACCCGAGGATATCTCGTATGCAGCTGATTGAAGCAACGGGAGGTTTTAGCAGCTCGAGTTTAATAGGAGCAGGACGATATGGGCGTGTTTATAAAGGAGTGCTGAAGAACAACATGAGCATAGCTGTTAAAGTGATTGACACAAAAGGAGGTGGAGAGATATCAGGGAGTTTCAAAAGGGAATGTCAAATCTTGAAAAGGACTAGACATAGGAACTTGATAAGGATCATAACGACATGTAGCAGACCGGACTTCAAAGCTCTCGTGTTCCCTCTGATGTCAAACGGAAGCCTAGAGAACCATCTGTACCCGAGCCATGGGCTGAGACATCAACTTGACTTAGCTCAATTGGTTAACATTTGCAGTGATGTAGCAGAAGGAGTGTCGTATCTCCACCACTATTCGCCTGTTAAAGTCGTGCATTGTGATCTAAAACCAAGCAATATCCTTCTTGATTACAACATGACAGCTTTGGTGACTGATTTTGGGATTTCAACATTAGTAAAAGCTGTTGAGGAAAATACTAATCCTATCGACGAATCAGTTAATTACAACTCAACAGATGGATTACTTTGTGGATCAATTGGTTACATTGCTCCTGGTTAGTAACAAATAATAGTTCTATTCTATGATTCTATCTAATTCATTTCAATTTATATGACTAAAGTTTCTGTTTCTTTGATTGTAGAATATGGAATGGGAAGGAGGGCTTCAACAAAAGGAGATGTATTTAGCTTTGGAGTTTTGTTACTCGAGATTGTGACAGGTAAACGTCCCACAGACGTTCTATTTCAACAAGGCTCGAGCTTACACGAATGGGTGAAGAGTCAGTATTCACATAAACTCGAGGCTATGATTGATGAAGCATTACAAAGATGTACAACAGTTCAACGCGATAGCAATTTATGGCACGATATCGTGTTAGAGTTGATTGAACTTGGCTTAATGTGTACTCAGTACAATCCATCAACTAGGCCAACAATGTTGGATGTTGCACTTGAAATTGGAAGGTTAAAACAGTATGTTTCAAGTTCTTCAAACAGTGTTACTGAAGATGTCACAACAAAAAGTTGAACAGTTTTTTTTTTAATATCTAACAATGTACCTTTTAACTTTATTCATTGTACTTCTATGTTCTTTTTTTTTTACCTTTTTAATTGTTCCTTCTTGGGGTGCGGACCTTCTCTGGATTCTACGTGAACGTGGGATGGTTCGTGCACCAAGACTACCCTTC

>PGSC0003DMG400011509 LRR-RLKs

ATGAAAACATGGAATGCATCCACACACTTTTGCCATTGGTCTGGTGTAACTTGTGGACGTAAACATGTTAGAGTCATTAAACTAAACGTCGAGAACCAAAAACTAGATGGCCCATTGTCATCCTTCATTGGGAACATGAGTTTCCTCAGGTCTCTATACCTGTCGAACAATAGCTTCCGTGGTGAAATCCCTTCAGAGATTGGTCGTTTGAGAAGACTACACCGATTGTATCTTGGTAACAACTCATTTCATGGAGAGATCCCCTCAAATTTATCAAGGTGCTTGAATCTTGTATCCCTTGTCCTTGAAGGTAACAAGTTAGTAGGAAGCCTTCGTCCGGAGCTAGGTTCCTTGTCCAAACTCGAGTACTTGTTACTCACTCGAAACAACTTGACAGGTGAAATCCCTTCTTCTTTTGGAAACTTAACATCATTGATAGGATTCTATGCACCATTGAATAATCTTCAGGGGAAAATCCCTGACTCATTTGGGCAGCTGAAAAACTTAGAAATTATCGGAGTAGCTGCAAACCAATTGTCAGGTACTATCCCTTCTGAAATCTTCAACATATCTTCCATCACAACATTTGATGTTGGAATGAACCAAATCCAAGGTACACTTCCTTCAAGTTTGGGAATCACCCTCCCGAATCTTGAATTATTTATCATCGGTGGGAACAACGTTTCTGGATCGATCCCATCTACATTGTCCAATTCATCAAAGCTAGTGTATTTTCTTGCTGGAAGAAATCAACTTACTGGAAGTGTACCAAGTCTTGAGAACCTGAATGAGCTTCAACAGTTAACAATTCCCGGGAATTATCTTGGAACTGGAGAATCAGATGATTTAAGTTTCATCGCCTCTCTGACTAACGCATCTCGTTTCAGAATTTTGGAGATTCAGTTCAATAGTTTTGGAGGGGTTCTGCCTGCATCCTTCAGAAATCTTTCAACTGAGCTTCAGGTTGTGCAGTTGTCATACAACAGAATACGTGGTAACATACCGTCTGAGATTGGAAAGTTCGTCAACGTGGAAGAATTTCAGGTGAGGGAAAACTTACTTACTGGTACTATTCCAATTAATTTTGGTAAAGTTAAGAAACTGCAGATTTTGGATTTGTCTCAGAATAGATTTTCAGGGAATATTCCATCATCTCTAGGAAATCTTAGTGTAGTATCCATCCTTCTTCTGCACGATAATAATCTCACCGGTGAGATCCCAGCAAGTTTAGGAAATTGCAATTACATGATTGAGATATATGTTGCTAAAAATAACCTGTTAGGACAGATACCTAAGGACCTTTTTGCTCTCTCTTCCTTAGTTGCTGTAGACATTTCTGAGAACCATTTAGATGGTTTCATTCCTCTTGAAGTTGGAAATATGATAAATCTTGAATATCTTAATGTTTCCGTAAATAACTTGACTGGTAAAATCCCAAGTACTATTGGTAGCTGTGTGACACTAGAAGCCTTAGACATGAAAGGGAACTTCTTTCAAGGGATTATTCTTCCTTCTTTCAGCTCTTTACGAGGTCTCCATGTTTTAGACCTTTCTAGAAATAATTTGTCTGGCCAAGTACCAAAATACTTGGAAGATTTCAAGTTTCAACTCTTGAATTTATCATTCAATGATTTCGAAGGAGTGTTACCAAATGAAGGCATCTTTAAGAATGCAACTGCAATATCAGTAATTGGGAATCCAAAACTTTGTGGGGGTGTGCCTGACATACATCTTCCGGAATGTGACATCAAAAGATCAAAAAAAATCGGTTCGAGGTTTATCCTCAAGATTGTCATCTCTGTTGTCTTTGGTATACTAGGACTTGGTATGCTTGTCACTTTACTATTTTGCTTCTTGTTAAAGAAACCGAAAAGGGTGCCTGTTTCAAGTTCATTGGGTGAGTCTCTCATAAATGTGTCTTACAGAAGTTTGCTTCAAGCTACTAATGGATTCTCTGAAGACAACTTGATTGGTGCTGGGAGCTATGGATCTGTTTATAAAGGAACTCTCGATGGAGGAATGGTTGTCGCAGTTAAAGTGCTAAACCTCTCGCGACATGGAGCTTCCAAAAGCTTTATGGCTGAGTGTGAAGTCTTGAGAAACATTAGACATCGGAATTTGGTTAAGGTTCTCACAGCATGTTCAGGAGTTGACTATCGAGGGAACGATTTCAAGGCTTTGGTTTACGAGTTTATGGTTAATGGATGCCTCGAAGACTGGTTGCATCCTAGTCCAAGTGAAGATACAAGTCAGGCAGCTGAAACAAAAAAGTTGAATATTCTTCAAAGGCTCAACATTGCTATAGATGTTGCTTCTGCAATCGATTATCTTCATCTCCACTGCGAGACTCCAATAGTTCACTGTGATCTCAAGCCAAGCAACATTCTTCTAGACAACCAATTGGTCGGGCACATTGGTGATTTTGGTTTGGCAAAGTTCCTTCAACCAACTGCACAAAATTCTTCTATTTCGGAAGGAAGCTCTGGTCTTGTTAGAGGAACAATTGGTTACACTGCTCCAGGTAACAATTAATACATAAACAAACACTCGAACGCCCTTTTTTTCTCTCATTTCTTCTATTTTTTCTTGAAAATGATAAATTCATAGCAAAACCACACTGCTTAACCATAACTTTCTTCGACAGAGTTACTTTATCAATATCATAAACCATCTCTTAATTGTTACACACATACTGCCATTACTTTTATAAAATCAAGTCATACTAAGTTATGTCCCTATTAGGAAAGTTATTGTTCCTTGTCTTTTTTTCTTTTGTGAATTTTAGAATATGGGATGGGAAGCGAGCTATCGACTTGTGGTGATGTGTACAGCTTCGGGATCCTTTTACTAGAAATGTTCACCGGAAAAAGGCCTACTGATGGAATGTTTAGGGATGGTTTAGACCTTCCAAGCTTCGCTAAACATGCATTGCTTAATGGAGCTATGGAAGTCATAGATCCATCCCTTATTTACGGAAGTGAAGAAGACGAGAAAGGTAAAAGCACAAATATCTACCAGAATAAGGTGTATCTGGTTTCTGTTCTTCGAGTCGGGGTTGCTTGTTCTGCATATTCTGGAGCTGAAAGAATGAACATCACTGAAACTGTTTCCCAATTGTACTCTATCAAGGAAGCTCTTCTGCAATCAGAAAAAAATTCAGGAGCACGTTGA

>PGSC0003DMG400011580 LRR-RLKs

AATAAAGGTTGAATCTTTCTCATTTTTATCCAACCCTTTTTCAAAATCTTGCATGTATATAGGAAGATCCCCACCATTATTGGAAAAATGTTGGAGGTTCTTGGTTTTAGATATAGTATAATCTTTGATTTTATCTTTTCCTCAATTCATGAGTTTTATGGTCTTTGCTCTGTCATCATCTAAAGGAATTACTTGAGAGTTCTTTGCTTTTTCTGGGTTTTTGGTGGAATTCAGCTCTTTTTGACAAGGTGATGGGCTCAATAGCCTAGAGGTATGCTAAAGCTTAGATCTTTAATGAGTTTCTTATTTATTCCTGAATTACTCCTTTTATTTAGTTGAGTTTCTGGGCATTTCTGAGTTGATTGGATTCTTTGAAGTTAGTTGATTAATTGGTATGCATAGAACTTTTTCTGGTTTGTCTAGTTTATCTAGTTAGTATTGTCTCTTCTTGATGATTGACCTTAAGGGTGTGGAGGGTAGGACTTGGATTAGGGTAGTAGGTGGCAATGCGTTGTCTTGTTGTCCGTTCATATTAGTAGTCATACTATTGCTCATTCTGTTACTATCTATTTGATTCCTTATACCTCGATTGTGTACTGTTTTGTTGTAGTTTTTGTTCTGCTATTCTCTATTGTTTTGCTATTATCTGTTGATTCTTGTACTTCCATTACTTCATTTTCTGGACTGTTTTTTCTTGAGCCGAGGGGCTACCGGAAACAACCTCCCTACCTCTGAGATAGGGGTAAGGTTTGAGTACACTATAGTCTATACCCTCCCGAGTCCCCACTTTGTGTGACTACACTGGTTATGTTGTTGTTGTTGTTGTATTTTATAAGTATTGACACATAGTTGGATTAACAGGTCAAACTGAGACTGTCTTATTGGAAGTTTGTTATGATGTCTGGAGATTTTGAATCAAGAAGTGGTTTAGTTCTTCTAATTTGGATAAGTGCTTTGGTGGCTGCTGTACTATTGCTGGTTTCTCCTGCCGAAGGATTGAATCAGGAAGGAATGTACCTTCTTGAATTGAAGAAGAATTTCCAGGATTCGTTTAACCATTTGGGGAATTGGAATCCAAATGATGAAACTCCTTGTGGCTGGGTAGGTGTGAATTGCACTTCTGATTATAATCCAGTTGTGCAATCTCTTTATCTGAGCTATATGAATCTTTCAGGCACGTTGAGTTCTAGCATTGGTGGTCTTGAATATTTAGCTTATCTTAACCTGTCTTACAATCAATTCACCGGGAACATTCCCAAAGAGATTGGAAATTGCTCAAAATTGCAGTCTCTCCAGCTCCATTTTAATACTTTTTACGGTCCGATTCCTGCTGAACTGTATAACCTGTCTAATTTGAAGGATGTGAACATGAGCAGCAACATGATCTCTGGTCCTATTGCTGAGGAGTTTGGGAAACTCTCGTCTCTAGTTACTTTCGTTGCATACACCAATAATCTTACCGGTCCAGTGCCTCGATCTATTGGAAGTTTGAAAAATTTGACAATATTTCGAGTGGGGCAGAATGCACTTTCTGGAAGTTTACCTGCAGAGATAGGTGGTTGTGAGAGCTTGGAATCTCTTGGTCTAACACAGAATTGCCTGGAAGGAAACATACCAAAAGAACTTGGCATGCTTAGTAAGTTGAAAGAGCTTGTGCTTTGGGGAAATCAGTTCTCTGGTTATATTCCAAAGGAACTTGGAAATTTAACACAAATTCAATTGCTTGCTTTATATCAGAATAACCTGATTGGAGATATTCCTGCTGAAATAGGTAAACTCAAAACTTTGACGAAGTTATACCTATACCGGAATGGATTAAATGGTAGCATTCCAAGGGAGATTGGGAATCTTTCTATGGCAACAGAAATTGATTTCTCAGAAAACTTTCTGAAAGGAGAAATTCCAGTTGAGTTTGGCCAAATAAAAAGTCTGAAATTGTTGTTTCTTTTCCAGAATCAGCTCGAAGGCGTTATACCAGATGAACTTACTACATTAAAGAACTTGATTAGTCTTGATTTGTCAATTAATCACCTAACAGGTCCTATTCCATTTGGTTTTCAGTATCAAAAGGAATTGGTTCAGTTACAGCTTTTTGAGAATTCTTTAACTGGTACTATTCCTCAACGTCTTGGCATTTATAGTCGACTTTGGGTGCTTGATTTAAATAACAATCAACTCACAGGAAGAATTCCTCCATTCGTTTGTCAGAATTCCAATTTGATTTTGCTGAATCTTGCATCAAATAAATTGCATGGATATATTCCATCTGGTGTTCTTAAATGTGATTCATTGGTGCAACTTCGTCTGAACGATAACAGGCTAACAGGGACATTTCCTTCTGAGTTGTGCAAATTGATAAACCTGTCTGCTGTTGAACTGGGGCAGAACAAGTTCACTGGTCCAATACCTCCTGATATTAAATATTGCCAGAAGTTGCAGAGGCTTGATTTCTCAGGCAATAGTTTCAACCAGTTGCCAAGGGAGATAGGAAATCTTACCAGGCTTGTGACATTCAACGTCTCAGCAAATTCGCTCACTGGACCGATACCTCCAGAAATTCGGAACTGCAAGGCACTACAAAGACTGGATCTCAGCAAGAACAGATTTACTGATGTTATACCTGATGACATTGGTAGCTTGTCGCAGCTAGAACGTCTTTTGCTTTCAGAGAATAAGTTATCTGGAAAGATACCAGCAGCATTGGGGAGCCTCTCTCATTTGACTGAGCTGCAGATGGGCAGTAATCTATTGTCAGGTGAAATACCATCAGAGTTGGGTAATCTTAGTGGCTTACAAATTGCAATGGATCTTAGCAACAACAATCTCTCTGGCAGTATACCACCCAACCTTGGTAATCTTATCCTATTAGAGTACCTATATCTCAACAACAATCATTTGAGTGGTGAAATACCAAGCACATTTGGAAACCTGACAAGTCTCTTGGGCATTGACTTCTCATACAATGACCTGACTGGACCACTGCCGGATATACCACTCTTTCGGAACATGGATATCAGCAGCTTCATTGGAAACAAAGGTCTTTGCGGTGGGCCTCTAGGGGAATGTAATGCATCTCCAGCTTATGATGCCAATAATCCTCCTAGAGTCGAGAGTGCAGATTCTCCTCGAGCAAAGATTATTACTGCAGTTGCCGGTGTAATTGGTGGAGTTTCTCTTGTTTTGATTGTGGTAGTTTTGTACTATATGAAGCAGCATCCAGTTGAGATGGTGGTAACCCAGGATAAAGACATGTCATCTTCAGATCCAGACATTTATTTCCGTCCCAAAGAGGGTTTCACTTTCCAGGACTTGGTTGAGGCTACAAACAATTTTCAGGATTGTTATGTTCTTGGAAGAGGAGCCGTCGGAACAGTTTACAAAGCAGTCATGCAGTCTGGACAAACGATTGCAGTCAAGAAGCTTGCTTCTAACAGAGAGGGTAATAACATCGACAATAGCTTCCGTGCAGAGATCTTAACTCTAGGAAAGATTAGGCATCGTAACATCGTAAAGCTATATGGATTTTGCTATCATCAGGGTTCTAATTTACTTCTCTATGAGTATATGGCCAGAGGTAGCTTGGGTGAACTGCTTCACAGTACATCTTGCAGATTGGATTGGCCAACTCGCTTTATGGTGGCTGTTGGAGCTGCTCAAGGACTTTCTTACTTACATCATGATTGCAAGCCCCGGATCATCCACCGTGATATAAAGTCCAATAATATTCTGATTGATGAGAAGTTTGAAGCTCATGTTGGTGATTTTGGTTTGGCAAAAGTTGTTGATATGCCTCAATCTAAGTCGATGTCTGCAGTTGCTGGCTCATATGGCTACATAGCCCCTGGTATGTCTTTCTGTTTTCTCCCGGACAACTTCTTATGTTCCCTTGTTATCTATAGAAATATGTCGAGAAATGTACTGCCATTTGATCCTTATTCCCACATATTTATCTGTATCTTCACTCTGAAAGTGTTACCTGTTTTTGTTGATATGAATAGGATTTTCTTGTTGAGATTAGATAGTTAGAATTGTATGCAACGTCTATTTCTTCGATTATCTGAAGACAACAATGAAATGCTTTTAACTTTTATCTGTGTATACTTCTCATTTTATATGTGTTCTATGCCAGGAGAAGCCATACAAACTGCTCTATGTATCTTAGGCGAACTTTGGGGACTCTTCCTGATACATATGGAGCATTCTTGTTCCCACATATTTATGTTCAGTTTAGATATTCGACCATGTGTTGTTCACTTTGAAAATCTAATTCTTCTCTTTGTGATATGAAATGCATATCTTGTTGAAGTTTGATAGTTAAAATTGTTTACGAACAAGTTAATTTCTTTGATGAAATGACAGAATACAAGAAAAAATGCTTTTAACTTTTCTCTGTCTATGTAGCTTTTAATATCCCTCTCGTAATTTTCATTCTGTGTTATAAAACCATTTTTTCCTTTTCAATTGATGTTTTCTACCTTCTCTGTTGCAGAATATGCTTACACCATGAAAGTTACGGAAAAATGTGATATCTATAGTTATGGGGTAGTCCTTTTGGAGTTGCTAACTGGAAAAGCACCCGTACAGCCCCTAGAACAAGGCGGTGATCTTGTCTCATGGGTGAAGCATTATGTTAGGAACCATTCATTGACACCAGGGGTACTCGATAGCCGATTGGATTTGGAAGATGTAATTACTGTTAGTCATATGCTTACAGTTCTTAAAATTGCTCTAATGTGCACTAGCATGTCCCCTTATGATCGTCCTTCAATGCGCGAGGTTGTTCTGATGCTGATTGAGTCTGATGAGCAAGAAGGGAACTTTCTCTCATCTCCAGTTTATGATCTCCCTCTGAAAGATAATTCCTTCTAGATGCATGTAAGAAGCACATCTCAATTAACTTTGATAATTTTTTAGTACTAATTCTCCTTGTATTTTTTTTCTTAATGGCTAAATTATAGATATTCCCATGTATATTTCTCATAGTGAAAATTTCATCAATCACTTTCTAGAACTAGTTTGGAGGTGATTGATGTTATTTCAATAGTTCTCCATCTGATGGTTATGGAAAATAATATGCTTTATGCTTTTCTCACTATATATTGAGGGCTACCTAGAT

>PGSC0003DMG400011833 LRR-RLKs

CTCTTCTTCTTGTTTCCCTTACTTTTCACTACTACTACTACTGATGCAAGTCCCAATGGGGAAAAAACTTTTTCAGCCTTTGTACATCACTACTACTTTCTTGATTTTTCTATCTTTGATTCTTGTTTTCCCTGTTTTAGCTGTTGATCCTTTTACACAAGGACTCTTGAGCTTAAAAACTGAAATTCTTGATAATTCCAACAGTTTGAAAGATTGGATTTTGCCTTCTGGTTCAACTGATAAAATTCATGCATGTTCTTGGTCTGGTGTGAAATGCAATGAAAATTCTTCTTTGATTATTGGATTGGATCTTTCTGTGAAAAATCTTGGTGGGGTTTTGTCAGAGAATCAATTTAGTGTTTTTTCAGACCTTGTTGAGCTGAATTTGAGTCACAACTCATTTTCTGAGAAACTCCCTGTTGGGATTTTCAAGTTGAGAAATTTGAGAAGCTTGGATATTAGCAGGAACAATTTTTCTGGTCATTTCCCAAGTGGGATTTCAAATCTTGATTCTCTTGTCATTCTTGATGCTTTTAGTAACAGCTTCTCTGGTCCTTTGCCTAAAGATGCTTCAGAAATTGAGTCACTCAAGGTACTGTTAGGGTTTGCTTTGTTTTTTTGAGAATTATGGACAACTTTTCTTCCACATCTGATCACATAACAACACACTAGCATTCACAAGTGTAGTCGTTAAAAGAGTCTTGTTTATAGATGGAGAGATAGTTCTCTCCATAGTTTCTATACTTTCTTTTAGAGTTTTTAATGGTCAAAAACAAGTTTGAATTATGTTTTTTCACGAGTTTCACACTCTAACTATCAATTTTAATACATAGATGGTAATAGTTCAGGTAAAAAAAAGTGAACAACTAATAGTTGGATTGTGAAATTTCCAAAAATAGATAGTTTAGCCGCATCTTTCTTTTATACTATTAGTTTTATGCCTGTTTTAGTATAGTTCTGTTTGTCTATTCATTTATCGTCGTTCAAGATTTGCTTTGTTGGATTCTGCATGACGCGTTGTTATTTCGATCCCAACAGGTACTGAATTTTGCAGGGAGTTATTTCAGTGGTCCAATTCCTTCTGAGTATGGCTCATTCAAGAATCTTGATTTCATTCACTTAGCTGGAAATTCTTTGAGTGGTAAAATACCACCAGAATTAGGGATGCTGAAAACAGTAATTCATATGGAGATTGGTTATAATTTTTATGAAGGAACCATACCTTGGGAACTTGGGAATATGAGTAAGCTTCAGTATCTTGATATTGCTAGTGCAAATTTATCTGGTTCAATACCAAAAGAGCTGACTAATTTGACCAATCTTGAGTCACTTTTTTTGTTTAGAAATCAGCTTAGTGGTAAAATCCCATGGGAGTTTGGAAAAATCATATCTTTATCAAGTTTGGATCTTTCTGATAATTATCTTTCAGGCCCAATCCCCGAGAGTTTCTCGGAGTTGAAAAATCTAAACTTGCTTAGTGTTATGTACAATGACTTAAGTGGAACTGTTCCTGAAGGAATTGCAAAGCTTCCACAGCTTGATACACTTCTTTTATGGGACAATTGGTTTAAAGGTTCACTCCCAAAAGACTTGGGCAAATATTCAAAGCTGAAATATGTTGATGTTTCAACAAACTATTTAGTTGGTAGCATTCCACCAAGTATATGTTCAGGTGGGATGTTACAAAGGTTGATCCTTTTTTCAAATAATTTCACTGGTGGACTCTCTCCATCACTTTCCAATTGTTCCTCTCTTGTTCGGATCCGTATCGAAGACAATTTATTCTCAGGTGATATCTCATTGAACTTTGGCAAGTTTCCTGATTTGTCATATGTTGATATGTCTAGAAATAGGTTCACTGGAGGGATTCCTACTGATATTGCCTTAGCTTCCAATCTTGAATACTTCAATGTGTCTAATAATCCAAATCTTGGAGGCGTAATCTCGGAGAAGACATTGTCTTTGTATGCACTACAGAATTTCTCAGCTACAAATTGTAGCATTTCAGGAGATTTTCCTCCTTTTGGACCTTGCAAATCCTTACTGGTTCTTGAATTGAGTACGAACAATGTTTCCGGGATTCTCCCTCAAAGCATAAGCAACTGCCAAAATCTTGTGAGCTTAGACTTGGCTAACAACAATTTAAGTGGTCAGATACCTGTTGGACTTGCTAGCCTTCCTGATATTAGTGTTGTAGACCTGTCACACAATAGTTTTAGCGGTTCTATTCCGGCCAAGTTCGGAAGTTCTTCCAGCTTGCAACTTCTTAATGTGTCATTCAACGATCTCTCAGGTTTGATTCCTTTTGATAAGAGTTTTAAGGTGATGGATAGTAGTGCCTTCTGGGGTAATCCTAAGCTTTGTGGGACACAGTTGAGGCCTTGTCGTGGACCGAATGGACTGGAGTTAGGAAGCAGAAAGACACAGAAGCTGGCATGGGTTCTCATAACCTGTGGAATCATAGTGTTGGCTATTACAGCTGCATTTTTCGGAGTATTTTACTTCAGAAGAAGAGGTCAAGGGCAGTGGAAAATGGTTTCTTTTAGTGGATTTCCTCGGTTTACAGCAAATGATGTTTTGAGGAGCTTCAATTCTATTGAAGAAGCCACAGATATGGTGCCACCCTTGGCAGGTTCTGATTGCAAAGCTGTTCTGCCAACAGGAATTACCGTCTTGGTGAAGAAGATCGAATGGAGACCGGAGAGAATGAATGCTATGATGGACTTGATATCAAGAATGGGCAATGCAAGGCACAAGAATATGACCAGATTGTTGGGATTTTGCTACAACAAGTGTATGGCTTACCTGTTGTGTGACTACTTGCCTAATGGAAATCTGGCTGAAAGGATCAGGACGAAGAGAGATTGGGCGACCAAGCATAAAATCATTGTGGCAGTTGCTAAGGGGCTGTGTTACCTGCATCATGATTGCTACCCCGCGATACCTCATGGGGATTTGAAGACAAATAACATTGTGTTTGATGAAAACATGGAACCTCATTTGACTGAATTCGGTGTTAAATTCTTGATTCAATTAAATAATGGTCCGTCGGTAGCAAGAGTTGGAAATGAAGCAGGTATGCACCTATACTTTGATGTAGTCTGCATTAATATTTGTTAATTAGTTAGCAATCTTAATATAAGATTAGTTATTAGACAGAATTCGGTTGCTCCAAGGAGACGCGACTTGACTGTTGCTCCAAGGCTCCGCTTCAATTCTAATTGAAGTGTTTAATGATCTTAATTTGAATGACATAGCGAGAATTCCCTATTACTGGTTGTGAATCTTGTTTAATTGTGTCAAACAGGTGAAATCGAACGAGCTATACAGGAAGAACTTTACAGGGACATATACAACTTTGGTGAGCTCATTCTGGAAATCTTAACCAACGGGAAGCTGTCAAATGCAGCGACGAGCATTCAGAACACCTCCAAGGAGGTTCTTTTGAGAGAGGTTTTGGATGAGAATGATGTTGCTCCATCGAGCTCGGTCCAAGAGGAAATAAAACTGGTTCTTGAAGTTGCTTCACTTTGCACCAGAGTTAGACCCTCAGATAGGCCATCAATGGAAGATGCACTAAAGCTTGTTACTGGTTTGAAGAAACAAAGCTAATGATGTTCTTGAACTAGGATGCCAATATTTAGACATGATTAGAAATTTTGAAGCACAAAATATGTTGTAAAAGTCTTTTATTTCAGTTGTTTTTTGTTACTTCATCTGGTCAAGAAGTTGATGCCTTTTCTTTTGTCAGGAATAGAATGTTTTGATGTAAAGTAGAGGAATGTAAGTAAGTTCTATCCTCAGCAATATAAGGTCTGTTATTCTATTTTTTTTTTTTTTTTTTTAGTTGTTTAAAAACAAAATATGTTGTAAAAGTCTTTTATTTCAGTTGTTTTTTGTTACTTCATCTGGTCAAGAAGTTGATGCCTTTTCTTTTGTCAGGAATAGAATGTTTTGATGTAAAGTAGAGGAATGTAAGTAAGTTCTATCCTCAGCAATATAAGGTCTGTTATTC

>PGSC0003DMG400011932 LRR-RLKs

ATGTCACTGCTAGCATTCAAGAATATGATCATTGATGACCCTTTCAAGATCATGGACTCCTGGAATGAAGCTATACATTTCTGTGACTGGCCTGGGGTCTCATGTGGTCACCGTCATCGCCGAGTTACAGTTTTAAATCTTACTTCTCTGAAATTGAGGGGTTCTCTGTCACCAAGCATTGGTAATCTGAGCTTTCTTTATATCCTTAAGCTCCAAAATAATAGTTTTTCAGGTGAAATCCCATCAGAGATTGGCTACTTACATAAGTTACACGTTTTACGCCTTGATAATAACTCATTCACTGGTCATATTCCTTCAAACATTTCTGGTTGCTTCAATCTCGTTTCTGTTGGTCTTTCGTATAATATGATGGTAGGCAAAATTCCAGCAGAATTAGGCACATTGTTGAAACTCAAACAACTTTCTCTTGTTTCTAACTATTTTACAGGAGGAATTCCGCCTTCATTTGGTAATCTTTCTTTACTTGATACCTTTTCCGCAAGCAAGAATAATTTGTTAGGGAAAATACCTGATGAATTATGTCAATTGCTCAACTTAAAGTACTTTGTAGTGAATGAGAACAACTTGAGTGGTACCCTGCCTCCTTGTCTTTTCAATCTTTCCTCTATCGTGGCCATAGATGTTGGAACAAACCATTTAGAAGGAAAATTGCCTCCATTGCTTGGTATTACTCTTCCTAGTTTAGAGTTCCTTAGCATTTATAGAAACAATGTAACTGGAAATATTCCGGTAACATTGTCAAATGCTACGAATCTTCAGTCTCTTATTGCTAGTAGAAATGGACTCACAGGAAAAGTACCACCCCTTGGAAATTTGCTGAAAATGCGGAGATTTTTAGTTGCTTTCAATTATCTTGGGAAGGGGGAGGATGATGACCTTAGCTTTCTCACCACATTGGTGAATGCCACCAGCTTGGAGCTTTTGGAGCTCAATACAAATAATTTTGGTGGAGTCTTACCTGCATCTGTTAGTAATCTTTCGACTGAGATTATAGAGCTTTCGCTGTCTTATAATCAGATTTCTGGAGAAATCCCAAGAGGGATATCCAATCTCAAGAAGCTTCAGGCTTTCTTTGTAGCTTACAACAGATTTCACGGTGAAATTCCGTCTGAAATTGGTGATCTGATATATTTGCAAGAGCTGGCTTTACTTGGAAATCAGTTCTCTGGCAAAATTCCAATCTCATTGGGAAATTTAGCTTCTTTAACCAAACTCAATTTAAGAGAAAACAATCTCCAGGGGAGAGTTCCTTCCAGTTTAGGCAAATGCCACAATTTGGAACTGTTGGATCTTGGTTCAAATAACCTTAGTGGGTTCATACCATCAGAAATTCTTGAACTCTCATCCTTGTCGGAGGGCGTGGACCTATCTCAGAACCAATTAACTGGGTTCCTTCCGATGGAAATTGGGAAGTTGAGAAATCTAGGCTACTTAAACCTGTCTGATAACAAGTTGCAGGGCCAGATTCCTACCAGTATAGGTACTTGTGTGAAACTTGAAGCACTAGACTTGAACAACAACAACTTTCAGGGTAGTATTCCTTCAACCATGAATAGTTTGCGAGGCCTGGAACTTTTAGTCCTTTTCTCACAACAATCTGTCAGGTGGAATCCCGGGATTCCTGAAGGACTTTAAATTTCTGCAGATTTTAAATCTGTCTAGCAATACTTTGGAAGGTGCTGTCCCAACAGGAGGCATCTTTAGTAATGCTACTGTGGTTTCCATCATAGGAAATAGAAATCTCTGTGGGGGTGTACCAGAGTTGGACCTTCCGGCTTGCATTGTAGAGGTAAAGAAAGAAAGGAAATCAGGTTTTCCCTTGAAAATAGTAATTCCTGTAGTTAGTGGGCTTATAGGATTGACATTCATAGTTTGTTTTCTTGGCATACGACAGTTTAGTAGATCGAGAAAACCAACACCTACAGATATACCTGAAAACTCAACCTTAAGAGTATCTTATCGGTGTCTACTTAGGGAAACTGATAGATTTTCTGCATCAAATTTGCTTGGAGTGGGTGCTTTTGGGTCTGTATATAAAGGAATTTCTGAACACGATGGTACGTTTTTTGCTGTTAAGGTACTGGACCTTTCACACCATGCAGCTTCCAGGAGTTTTTTAGCTGAATGTGAGGTTCTAAAGAACATTAGACATCGTAATCTTGTAAAGGTTCTAAGTGCATGCTCAGGTATTGATTATGAAGGTAATGAGTTCAAGGCCATAGTTTATGAATACATGGATAAGGGAAGTCTACAGGACTGGTTACATTTCACTACTCAGGAGAAATCTGAGTCACAGGAAGAGCACAAGAAGCTCCGGTTTATTCAAAGATTAAACATTGCCATTGATGTTGCCTGTGCTCTGGATTATCTCCATAATGATTGCCAACCCCCAATAATTCACCGTGATCTGAAGCCAAGCAATATTCTTCTCGATGAGAACATGACTGCACATGTTGGTGATTTTGGTTTGGCAAGATTTGTTCCACCAGCAATTCCAAACTCATCAGCAAATTCAAAAAGCTCTACAGGTGTAGGGGGAACTATTGGTTATACACCTCCAGGTAATAATAAATCAAGGTTTCTGCTGGCTGTCATATGTATGTCTTTTGATAAGAATATCTCTAATTCTGTAGTATTTCTTGTTGACAGAATTAGGTATGGGAAGCGACGCCTCAATCTATGGAGATGTATACAGCTTTGGCATTCTGCTCTTGGAAATGTTTACCGGGAGGAAACCAACTGATGAAATGTTCAAAGACAATTTAAACCTTCATAACTATGCCAATGCTGCTTTGCCAGACAGAGTGATGCATATTGCAGATCCCATTCTTCTCCAGGAAAGAGATGAACTTGGAATGAAATACAAAGTTGATGACAATACAAGCAGTGCTGGTGACATATTTCTGTCATTCTTAGTTAAGGTGATTCAAATCGGAGTTTCCTGCTCCGTTGAATCTCCAAAAGAAAGAAAGCGGATCAGTGATGTCGTTGGAGAACTCAATTCATTAAGGAAGTTGTTCTTGGAGCAAGCATACCCGAAAGAAAAGGTTATAAGCTTCAACCTGCATTGA

>PGSC0003DMG400011989 LRR-RLKs

CACAACTTCATCACAAAACCAAAAACAATCAATCATTTAGCCTTGATGATGGTTCCCACAATATTTTGTATCCTTCCCTTCTTATATCTTTTCATAGTTATAACTTTTGCTTCCACTGAGGAAGCAACTGCTCTTCTTAAATGGAAAGCAACATTCCTAAACCAGAACAATTCCTTACAGGCTTCATGGACACTAAGTAGTCCTGCTGGTACCAAGAATTCTTCCAGAATTGAAGCGACGAGTTCTGATGCATGCAGGGGCTGGTATGGAGTTGTATGCGTTAATGGTCGGGTTAACAGGTTGAATATTACAAATGCTAGTGTCATTGGCACACTGTATGATTTTCCATTTTCATCACTCCCTTTTCTTGAATATGTTGATCTTAGTATGAACCAGCTCTCTGGCACCATCCCACCTGAAATTGGTAAGCTCGCTAATCTTGTCTACCTTGACTTGTCTGTCAATCATATTTCGGGCACAATCCCACCACAAATCAGTTCACTAACAAAGCTTGAGACCCTCCACATCTTTGGCAACCAATTAAATGGTTCCATTCCGAGGGAAATAGGTCGCCTAAGGTCTCTTACTGATCTAGCTTTGAGTTCTAACTTTCTTAGTGGTTCAATTCCTGCTTCATTGGGGAATTTGAACAATTTGTCTTTTTTGTATCTTTATAATAATAAGCTTTCTGGCTCCATTCCTTCAGAAATAGGAAAACTTGTCAATCTTGTTGAAGCTTATATTAGTAGAAATCAATTAACAGGTCATATTCCTCCAGAAATAGGTAACTTGATCAATGCAAAATTGTTCTATGCTTTCTACAATAACTTGTCTGGTCCTATTCCTGATGAAATAGGGAAAATGAAGTCACTTGAGTACTTAAGCTTCCAGACAAACAATCTCTCCGGTCCAATTCCCAAAACAACAGGTGACTTAACTGAGCTCAAACTCTTGCACCTTTATTCCAACGAACTTTCTGGTCCCATTCCTAGTGAGTTGGGGAAACTGAAAAAATTGAATGATCTGCAATTATCCAATAATCAACTTACCGGTCCTATTCCTGGTTCATTTGGCAATTTGAGAAACTTGCAATTTCTGTATCTCCGTGCCAACAAACTTTCTGGTTCTATTCCAAAAGAACTTGCATATTTGGATAACTTGGTTATTATGCAAATGGATGAAAATCAGTTTTCAGGCCATTTGCCTGAGCATCTTTGCCAAGGTGGGAAACTTGAGAACTTCACGGTGAATAGTAACAAACTGACAGGGCCAATACCAAGAAGCTTTAGCAAGTGCTCGAGTTTCAAAAGAGTTCGCTTGGATAACAACAGTTTTACTGGGAATTTATCTGAAGCTTTTGGCATCCATCCACACCTTTACTTCATTAATTTGAGTGAAAATGATTTTCATGGTGAACTCAGCAGCAACTGGGGGAAATGCAAGAGTTTGACTGATTTGCGGGTAGCCAGAAATAACATTAGTGGTAGCATACCACCAGAGATTGGAAACCTCAAAGGGCTTCAAGGACTTGATCTTTCATCCAATCATTTGGTTGGACAGATACCTAGGGAATTAGGAAAATTGACCTCTCTGGTTAAGCTTTCTGTACAAAACAACAATATTTCTGGCAATATTCCTATAGAACTTGGGTCACTGACAAAGTTAGAGTCCCTAGATCTGTCAAACAATAAATTGAATGGGTCGATCCCAACGTTTATAGGAGATTACCAGCACCTATTTCACTTGAACCTGAGTTGTAACAAGTTTGGGCAGAAAATTCCAAAGGAGATAGGGAGGATAACTCATCTTAGTGTACTTGATTTGAGCCATAATCTTCTTGATGGAGAAATACCTGCTCAGTTAGCTAGTTTGCTGGACTTGGCGAACTTCAATCTTTCCCACAACAGCCTTTCTGGGCGCATTCCTGAAGAATTTGAAAGTTTGACCGGTTTGCAGGATGTTATCTTGTCATACAATGAGTTGGAAGGTCCAATCCCTAATAATAAAGCTTTCATGAATGCCTCATTAGAAGGTAATAAAGGTCTTTGTGGTAATGTGACTGGATTCCAACCTTGCAAAAGGCCATCTTCTATTGTGAAGAAGCACTCAATGGCGAAGGGACTTAAACTCATCCTCATCACTGTACTTCCTGTTATGGGAGCACTAGTACTACTCTGTGCTTTCATTGGTGTTCTCTTTATGTGTAATAAAAGAAGGAGAGTTAGAGACGTTGAAAGAAGGGATTCCAGTGATGATGTTGATGGTTTGATTTCCATATCCACATTAAATGGAAATGCATTGTACTGGGATATCATAAAAGCCACAAAGGAGTTTGATGCAATGTTTTGCATCGGAAAAGGAGGATCTGGAAGCGTTTACAAGGTAAAGCTTCCAACGCTTGAGAATGTAGCTGTGAAGAGACTTCATTCTTCATTTGAGGTTACCCATCGCAAAAGCTTCATGAATGAGATAAGGTCATTGACAAGGATCAAGCATCGCAACATTGTGAAACTCTATGCCTTCTGTTCAAATGCACAACACTCATTCTTGGTTTACGAGTACATGGAGAAGGGTAGCTTGTCTAGTATTTTGAGCAGTGAAGTAGAGTCCAAGAAATTGGATTGGCTTACAAGGGTGAATGCCATCAAAGGTGTTGCTTATGCTTTATCTTACATGCACCATGATTACTCACCCGCGATTGTTCATCGAGACATATCAAGCAGTAATGTTTTGCTTGATTCCGAGTATGAAGCACGTGTTTCAGATTTTGGCATAGCTAAGATTCTTAAGCCAGACTCATCCAATTGTACTGCACTTGCAGGCACATATGGCTATGTAGCACCAGGTAAGATCTAACTTCTTATCCTTTTTTTAGTTCATGAGTTATTTCTGCACAGTCTATGTTGTTTCTATCTCATTTGGAACTTTATTGTTTCAGAGCTTGCCTATACAATGAAGGTGACGGAAATGTGTGACGTCTATAGCTTTGGAGTATTAGCATTGGGGGTAATCAAAGGAAAGCATCTTGGGGATTATATTACTGTGCTAGCAAATTCATCGTCTAGAGATCATGTGCAGCTTAGCGATTTGCTAGATGAACGCCTTCCATATCCTGAAGATAGCGTAAAGAGTTGGTTTTATCATCAAGCTAGCAAGCTCATGTTTGCTTGAAACTCCAAAATCAAGGCCAACAATGCACTTCATCTCCAGTATGTTGTCAATGGATCCACCTGTTTCTCAAATAACCTCTATGAAGGCGAAAAGCGTCCCCGAGATGTAAGGTGTCTATGTATGGGGAAAAAAAGAAGCAAGTTTTGACACCAAGTGGTTGTGAAGTAAGAATAAGAAGCTATTATTACTGGGACTAGTATTTGTCTATTTTATTGTTAGTTGTGCTTATCTAGTTTAGTGTTTCAACCTGAAGGATTCTGAGCTCTTGTACTAGAGGTAATACAATATCAAATAAAATGTAATATTTCAATTTTCTTGTGCCAATTAGTGCTAATAAAAAT

>PGSC0003DMG400012218 LRR-RLKs

TAGCTATTGTGTATTTATGAGAAAATATTCGAAGTTTGATCAATCATAGCGACGCTGGTAAAGGCCCACACTTCCTTTATTCCGCCGCTCCTTCTATTATCTCCTTCTCCATTTTCTGCTTGATATTTTGTTCCCTTATAATTTTAACATATAGTTTTTAACCCCAAAATTCTTCTTCTTCAGCACATAAATTCTCTGTTTCTAGACACCAAATCTCTCTTTGAACCCAAGTTTTATTTTTTATAACTAAGTTGGTGACACTTAATAGTAGATTCCACTCAAAGGGGTTGTTACTGATTTAGATTCTTGAATTCACTCTTAAGCTTTGTTAAGTTTTCTTTAGTGTTTATCTTTCCCCCCTCACTTGTAAAAAAGACAACTTTATCTTTTTCCAGCTTTTTTTTTTGTTTGTTTTCATCTACTGTCTTGCGTCTTTTCTTGTTTTCTCTTTCAATTTTGTTTGGATTATTGCTTTTTGAATCTTTTGCTTTCTGGGTTTTTCTTCTTTTTGTTGATTGCTTAAGTTCTTGGAGACTGTTTCTTTAGATGGGTTCCTTCAGTTCTGGGGTTTTGTAGAAACTTAGTGTGTTCTTGTGGTGAGGAATTATTTGGGGGAAAAGATGAGACTTTGTGTTGTATTTCAAGTTTTAGTCACTGTTGGACTAGTTCTTGGGATCTGTGATGCTTTTCCATCAAATGAAGGTGTGTATTATCAACTCAAGTTCAGATCTTGATGTAGTTACTGCTGCAAAGATTCAATTTTTGTCTTTTAATTGAGCATTTAGTGTTGTAATGCAGTGTTGTTTCGGAAAGATCTTGTATTTTCTTGTTTTGAGATACTTGCTGTATTTTGATGTCATATATTCTTGCTACCATGGTTCAAATTGTCTATTGGAAATAGTCTCTCTACCCTGCAAAGGTAGGGGTAAGATCTGTGTAGTGCGTACTATGTTATTTGTTGTTGTTGTAGATAGTTGCTGTATTTTATGAGATTGTAATTCAAAGAGTAGGTAGTTTATTGTGAGTGAATCTGATTGGGGATTCTTTGTTGATACAGTTTATGCTCTCAATACCTTCAAGGAAGCTATATATGAAGACCCTCTTCTGGTTTTGTCAAATTGGAATGGTTTAGATTCAGATCCTTGTGGTTGGGCTGGAGTCTTTTGTTCTATGGCTCAAGATCATGTCATAAAAATGTGAGTTAATTGTCTTCAATTTTATGTGCTATACATGGATAGGTTGTTTGGTTAATGATTTTTCTCAACTAAAATGCAATCTAGTGAATCTTTTATGGACTCAAGTTTAGTGCAAATGAGTCTCTAATACGCATTTCCACTGGATCGTGTGACAGAAACATTTCTGACGCTTCGTTGAAGGGATTTCTTTCACCCAACTTGCATTTACTCTCCTCTTTGCAAGAATTGTAAGTTATTTCCGAATAGATTTATAAGAATTACGTAGTTCTGGTTGTTCTTTTATGTGGTTTTGTTAAAGGTTATGCTATGCACTGGCAATATGCAGAATTTTGCATGGAAATGTGCTGATTGGCACAATTCCTAAGGAGATTGGCCTATTGAAAAACCTGAAGGCGTTGGATTTGGGTGCTAACCAGCTAACAGGACCAATTCCTTCCGAGCTTGGCAATTTGACAAACATTATGAAAATGTTAGTCTAACTTGAATTTTTTAGGTTCAGTTGTACTTGCCTTTACCTATCCTTGTTGAAGCTCCCTTACTTTTCTAGCTTGCATGTGTCTCTGCAGAAACCTACAATCTAATGGATTAACAGGGAAATTGCCTGCTGAGCTTGGTAACTTGAAATACCTTGAGGAACTTCGGTTGGATAGGAATAAACTCCAAGGACTAGTGCCTGCTAATACTGGATCAGATTTTACATCTGGTGTACGTGGAATGTGAGTTGATTAGCTACATCTTGTACTTTATCGTTTTTCTTTTGAATTTTCTGGTATGTGGATAGGGATTAATCCATGATTCTCTGTTTACTTACAAGATATTTTTGGTTGTCCTTAGAGCGTTTTCCGTCTTGGCCCTCTAGGATTCTGTCAGATATTTGAAGGATTGAGCGACCCAAGAATCTAATTTTCTGATCTTTATGACAGTCATTGTCGGGGATGATGATGACAGAGTTCAGAAAAGAGATGCTTATGGGATGATGTCTGTTCTTTCTGTCAGCATTGTTTGATATGTACTGCCTGCAGATTGACACTGATGGCTGTTTTACTTGAATCTAGTGTAGTACATTCAGTGTCAGCATAGAAAGTGGGGGTTAAACAGTGTTGTTTGATGGTGAATTGGGGTACCCCAATGGAGTGTTTTTTTGCTATAGCCTTTAATATTTGACCAAATGAGAGCAATGTTACTGAGTTCAGAATGTATGTTTCAAAAGTTTAACGATGATGTTCTTGGAAATGATCTGTATTTCTATAAAGAAACAATGAGTAAACTGTTTGATGTAGCCTTATGTCCATCACATGTATCTTTATCTGACTGTTTTTTTCCTTTTCTGTTTTACCATTTTAGGTATGCCTCTGGTGCCAGCACTACTGGTTTTTGTCGCACATCCCAGCTCAAAGTGGCTGATTTCTCATTCAACTTTCTTATTGGAAGCATACCAAAGTGTTTGGGTTACCTTCCAAAGTCAGTTCTTAATTCTACGCCATAAATCTTGTTTTATCTTATCTTCTCTATTCCTGAATGCCTGTTAGTTTTGTTGCTGTAAATCTATATCCAAAGTAGCACTCTGATGCTCTACTAGTGTTGTTTCTCTTACCTTTTCGGTTTCTTAATGCCTGTTATTTTGTGTGTCTATAGATCTAGCTTTCAAGGAAATTGCCTTCAGCACAAAGACCCGAAGCAACGGTCTGCTGCTTTATGTGGTAAGTAAACAATCATAGTTACAATTTTCAACACTACGCATTGTTTGTGCAGCTCAGTCTAGGAGAATTGATATCTCAACCTTTTTTCTTTAATATTTGCAGTCAAATAGATTTATTTATTTTAGTAAGAACTGAACTGGTTGTTTTGACATTGCATTATAGGTAGCACTCCGCCTCCCACTAGCCATGCATCGACACTCAGCAATAAGCACCAGCCTGTTGAAGAACCGAGACATCAGACTTCTTCCAAGCCCACTTGGCTTTTGGCTTTAGAAGTGGTAACAGGAGTTATGGTGGGTTCTCTCTTCATTGTAGGTCTTGTGACTGGCCTTCAGAGGCTTAAGAACAAATCGTCAATCATACTTTGGAAGAAATCAGGAAGTGAGAAAGATCATATGACAATTTATGTAGGTTAGTACTATCATGCAATTGAACTTGAGCACTTAGTACTTTCAAAAGCATTGAGCCTAATCTTCGTTTCTGTTTACCGCTGATGTTTCTTTGTAGATACTGGTATGCTGAAAGATGTTGTAATATACAGCAGACAAGAACTTGAGGTAGCTTGTGAAGATTTCAGCAACATCATTGGATCCTCACCAGATAGCATAGTTTACAAAGGAACCATGAAAGGTGGACCTGAAATTGCTGTAATATCCCTTTGCGTCAAAGAAGAGCACTGGACAGCCTATCTAGAACTTTATTTTCAGAAAGAGGCGAGACTCAATTCTTTTTTCATGATCGCTATTTATCTCTTCCAAAAGTTTGCATCTTAAGTTCCATGCTCTAACTTCTGTTAATTTACATTTAGGTGGCAGAATTGGCCAGAATAAATCACGAGAACGCTGGGAAACTTCTAGGCTATTGCAGAGAGAGTAGTCCCTTTACGAGGATGTTGGTGTTTGAATATGCATCTAATGGGACATTATATGAACACCTTCATTGTGAGTAAAGTGAAATAGTTTTCGCATGCACTTCATTTTATGTTTTCCAAAGCGAACTCCTAACTCTTTGAATGTATCATAGATGGAGAAGGATGCCAACTATCTTGGACACGCCGGATGAAAATAGTTATTGGCATCGCTAAGGGACTGACATATTTCCATTCAGAACTTGACCCTCCATTTACTATATCAGAACTGAATTCAAATTCTGTATATCTCACCGAAGATTTTTCTCCCAAGGTAAACGAGAGACCATTATTTATCATTATCGTATAAAACAGTCCTAAACAACTCATCCCATTTTTATTCACCTTAAGCTTGTGTACCCACCCCATTATGCTCTTAAGCAGTCTACAAAAGGCACTGGCATTTTATAATTGTCTTCTCAAAGTTTATTTCATTTGGTGCAGCTTGTTGATTTTGAAAGCTGGAAGTCGATAATTTCCAGATCAGAAAAGAATTCAGGTGCTATCAGTAGTGAAGGAGCAATATGTGTCCTTCCAAATACTCTCGAGAGCCGTCATCTCGATGTCCAGGGTAACATTTATGCATTTGGAGTACTTCTACTGGAAATAATCAGTGGGAGACCTCCATATTGCAAAGACAAAGGGTGCTTAGTAGATTGGGTAAGCATTTCTTTTCATGTGAAATTTTTCCTACTGTAAAAAGACTTCCCTGGTAGAACCAGCTTCTAAACGCGTTGGCAGGACCTAAAGTCGACCTTTTATGTGCAGGCTAAGGAATTTCTTGAAGTGCCGGAGGTTTTATCCTACGTTGTGGATCCCGAGTTGAAACATTTCAAACAGGAAGACCTCAAAGTGATATGTGAGGTGATCAACCTTTGCATCCACCCGACTACGAGTAGGAGGACATCCATGAAGGACCTGTGTGCTATTCTGGAGAGCAACATTGACACTTCCATCACTGCAGAGCTCAAGGCGTCTTCGTTGGCATGGGCCGAGCTTGCACTTTCATCCTAACTGACAGAAGGCTAACACAGTATATATATCTGAAATATGTAATTGTCTAGCGCTTTCTCCTTTTCGTTTTTTGACCTCTCGATTTCGTTCCCAATGTGTTCTGGAGTGATGGTTGAGTTCATGTAAATTTTGAGTCTGTAGGATAAAATTAGAGTTGAAGAGAAGCTACTGTTGATTCTTTGTAAACAAAGTGGTTCTTGTTAAAATGAATGAGTGTTCATAGTGAATGAGGCACCAACCATATTTTTTGAGTTTCAAAGAACTCGCTCGAGTTCTCGAGTCGCAACTCAAAATATAGGTTTTGGACTTAACTCAACTCCAAAAGAAAAGTACAATACAGACTTTCTT

>PGSC0003DMG400012307 LRR-RLKs

TTTTTCCATGAACAAGAAGAACACAAGTGAAAAAATCAAGAAAATGTGAGAATTTTTCTACAAAAAAAATTCCCAGTGTCTTGGTTTTGGCATTCAAATGGAAAGTTAAAGCGCAAAGAACAAAAAGAAAAGCTTCTTCTACTACAGATTCTGACTAGCAGCTACTTTCATTTTAAAATTACACCCTTTTGACCCTAACTTCCCCTATTTCAAACCTTGTGAGTGACCTCCAATTTTTTTAATCACATTTTTTCAGAACCCCACATTGCTTTTTGCAGACACCATTCTTGGGAATTTGACTCTTTTTTCTCACACCTATATACATATATATACACATGCATCTTCATATTGTTGTTGTCAGAGTGCATTTCTCTTTTTTGTTTGTTTCTTTCTTCTTCTTTTTTCACTTTAAGCTTGTGTTAAGCTGCTGCTACTTCCTTTTATGAGGTTTTGAATTAAAGTTGTGAGCTTTAGATTCTTGTGGGTTGAGGGTGTTGTTGGAGAGTTTTTGAAATAGTTGAAAAAGTTTTGAACTTTAGAATCTTGATGCTTGCTTACTTCATCTTATGTGACTTATAAAAGCTAAGTTCTTGAGGGGGGTTTTTTGAATTAAAGTTTTGAGCTTTAGATTTCCTTTTGTGGATTGAGGGTGTTTGTAGAATGTTTGAAATAGTTGAAAAGTTTTGAACTTTAGATTCTTGATGCTCTTTTCTTTTTCTTATGTGTCTACAAAAGCTAAGTTCTTGAGGGTTTTTTTTTGAATTAAAGTTTTGAGCTTTAGATTTCCTTTTGTGGATTGAGGATGTTTGCTGAGTTTTTGAAAAGTTTTGAACTTTAGATTCTTGATGCTCTTTTTGTTCTTCTTATGTGACTACAAAAGCTAAGTTTTTGAGGGGTTGGGGGCATTCAGAGTCTTTGAAATGGTTAAAGTTTTGAGCTTTAGGGAGTGGTGGGGGATTGTTTCTTTGGGTTTCTTGGTGGGATTGTTTCTGTTGTCACAAGGCACTGTTGCTCTTCTTGAAAATGATAAGCAAGCATTGTTGGATTTTGTTAATCAGTTGCCTCATTTGCACCCTTTGAACTGGGATGCAAATTCTTCAGTTTGCAAGAACTGGACTGGAGTGGGGTGTAATGAAGATGGTTCTAGAGTGATAGCACTGAGGTTACCTGGGGTTGGGTTCAATGGTCCAATTCCTAATAATACACTGAGTCGTTTGACTGCATTGCAGATCTTGAGCCTTAGATCTAATGGTATCAATGGCACTTTCCCTATGGACTTTGATAACCTCAAAAATTTGTCTTATCTTTACCTTCATTACAACAATTTCTCAGGTCCTCTGCCTTTTGATTTCTCTGTTTGGCAAAATCTCACCAGTTTGAACTTGTCAAACAACAGATTCAATGGCACCATTCCTAGCTCGATATCGGGTTTAAGTCATCTTACTGCCCTTAATCTTGCTAACAACTCACTTTCTGGTTCAATTCCTGATCTGCATTTGCCCAACTTGCAGCTACTGAATTTGTCAAACAATAACCTTATAGGTACTGTGCCTAAATCACTTCAAAAATTCCCAAAGAATGTGTTCATTGGTAATAATATGTCTTTACTTGACTATCCTGTTTCGAATTCTTCTATTGTTTCCCTGCCTCAGCAACCAAATCCAAAATTCAAGAATGATGGTAAATTGAGTGAGAGGGCTTTGCTTGGGATTATAGTAGCTAGTAGTGTAATTGGGATTCTTGGATTTGGTTTCTTGATGGTTGTTTGTTGTTTTAGAAGAAAGAAAGATGATGGTTCGTTTCCCAGCAAGATGGAGAAGGGAGATATGTCTCCTGATAAGGCGATTTCAAGGAGTCAAGATGCTAATAATAGATTGGTTTTCTTTGAGGGTTGTAACTATGCATTTGATTTAGAAGATTTGCTTAGAGCTTCCGCGGAGGTATTGGGGAAGGGCACGTTTGGTATGGCGTATAAAGCAATATTAGAAGATGCTACCACAGTGGTAGTGAAGAGGTTGAAAGATGTAGGTGCTGGGAAGAAGGAGTTTGAGCAGCAAATGGAGGTTGTGGGAAGTATAAAACATGAAAATGTAGTCGAGCTTCGAGCATATTACTATTCAAAAGACGAAAAGCTTACTGTCTCTGACTATTTCAGTGAGGGTAGTGTTGCTGCAATGCTCCATGGTATGATCACTTTATGCACTCTCTAAATTTTATAGTTTTGAACTTTTCAACTTTGTCACATTTTGTTAGCATATTCATACTGCTCCGTTATTTGCTTTGAATCAGTAGATTAGTTAGTATTTTTGAGTATGATTGTTAAAACAAGTTATTTGTTATCCCAAACTATGTTGACTCTAACTAAGTTGCTACGACTCTTCAAAAAATGCCGCGGGGTGCATGTCGGATCCTCCAAATAGTGCACTTTTAGAGTATCCGACAAGGGTGTGACAACATTCTTGGAGAGTCCGAGTTACATAGTCCCTAAGTGAGTTCTTACAATCAGACATGTTAGTTTAGTGAATTGAAAAACATAAAGCAATTAAAAACAACATAGAATCTTCCAAACTGAAGGCCAACTAATATAATTTGTTGTGTTATGGGTAGGGCCAAGATGGGAATCCACATATATTATATGCTCCATATATTCTCTTTCCAGCTTAATAGTTAGTTGTACCCACAGGATGATAATATATATATCCGTATCAAAAAAAAAAAATGTGGATGATCCTAACATGTGGTTTACGCATCTCTGTGGGGCAGTGACTCCTTTCTGTTAACAAAATAAGCAAAATCCTATCTCTATTTTAAAATCCATACAAATCATTGTCACTTCTCCTTTGTCACTAGAGTCTTTCAGCTATTACACTATTTGACTGAATTATTTCATAGCATGTGGATAATCATTTCTGCTTTTGTTATCTGTTAGTCTTTTGTGTTTCAGTTACCGCATTATTTCGTGTCTTTGTTGTTACTGTTCTTTTTCTTGATGTCCTGTACTGCTTTCCGTATTAATTGACATAATTTCTCCACTGCTGTTCTTTCTTTTTTCATAACTACTTTGATTTGTTGCAATTGAACCGAGAGTCTTTTGGAAACAGCCTCTCTACCTCCACGAGGTAGGAGTAAGGTCTGCGTACACTCTACCTTCCTCAGATCCCACTTGTGGGAGTAAGCCGGTTATGTTGTTGTGGATAATCATTTCTGCTTTTGTTATCTGTTAGTCTTTTTGTGTTTCAGTTACCGCATTATTTCGTGTCTTTGTTGTTACTGTTCTTTTTCTTGATGTCCTGTACTGCTTTCCGTATTAATTGACAGAATTTCTCCACTGCTGTTCTTTCTCTTTTCATAACTACTTTGATTTGTTGCAATTGAACCGAGAGTCTTTTGGAAACAGCCTCTCTACCTCCACGAGGTAGGAGTAAGGTCTGCGTACACTCTACCTTCCTCAGATCCCACTTGTGGGAGTAAGCCGGTTATGTTGTTGTGGATAATCATTTCTGCTTTTTATCTACTTTCTATAGCAAGTCAAAATAGTTCATGATTTTGTCGAGGTCCCAGCGTCCTTTTTCCCAATTCTTTGTCCACCAGGCAAATGCACATGATGTCAAAGTGCATGAATGGCTAAGCTTTTCTGATAATTTATTTGGAGTTGGTTGGCATCTTTCATTACAATTTTCACTGGTGGGACCTGATTTAATTAACTAGGAGTTTAGGTTAAGGAATCCTAAGCTAATGTCAACCTTAATATGCTTAGAATCTATTGAGAACATTACAACAAAATTCTTTTTGGTTATTTGCTTATTTACTCCCCTAATTTTGCTGTTAGAGCTAGGACTTCTGTTTGTTCTTCTCAAACGAAATGAAGTGAAATGGAAACTTGTAGAAAGTGTTATTTCCTTTTGTATCTCAGTTATGCTAACCTAATATTTCATCCGCCGTAACTTTTGTGAACAAAAGGAAAAGAAAAGAAGATCATTTGTGTAATGAGTAGCTTCCCACTTAGATGATAAGTCCTGCAAGGGATATTCTAAACTTTTAAGCACTTCGCGATTTGCTTATTCTAAGTTAAAGGATCGACCAGGAGCTACAAGGATACTGATTACTCTAATATTTGGCATGTTATAGAATGATAATTAAAATATGAACAAATGAGAGACAAAGGATTGAACGCGCGATCCTTTGCTTTTGTTAAAATTGGGATGGTTACCTGTGGTATCAGAGGTCGTTTGGTAGGCAGGATAAGGTGAGATATTCTAGGATTGACTGTGCTACCAGTTATTCTCCTTTTCAATTCTCTACCTTTTGTTAGGCAGTTCATATCATTGTGATCTCTGTGTTACCTGTTATCAAATATTGCTTTTGCTATTCGATCGTGTTGACACCTATTAAAATGCAACTTATAGGTAAACGGGGAGAAAACAGGATCCCTTTAGACTGGGAAACTCGACTTCGGATAGCAACTGGAGCAGCGAGAGGTATTGCTAGGATACATGCTGAGAATGGTGGCAAGCTAGTCCACGGAAACGTCAAATCCTCAAATATCTTTCTGAACTCTAAACAATACGGTTGTGTATCTGATGTTGGTCTGTCAACAATAATGAGTTCGTTAGCTCATCCAGTTGCACGTGCTGCTGGTTTTCGCGCTCCAGAAGTAACAGACACACGTAAAGCCACTCAACCTTCTGATGTGTACAGCTTTGGTGTGTTGTTATTAGAGCTCTTGACAGGCAAGTCCCCCATCCATACAACAAACGGAGATGAGGTCATACACTTAGTCCGATGGGTTCATTCAGTTGTCCGAGAGGAGTGGACTGCTGAAGTATTCGACTTGCAACTATTGAGGTACCCGAACATAGAGGAAGAGATGGTGGAAATGTTACAGATAGCAATGTCTTGTGTGGTAAGGATGTCAGATCAAAGACCAAAAATGTTTGAGGTTGTTAAAATGATCGAGAACGTTCGTCCCACGAGCTTAGAGAATGAGCATTCATCTGAAGGAAAAGCAGAAACTTCAACACCAAGAGCAGAAGCTACACCTGTGCCTGATACACAATCTGATTCTCAATGAGTTGGCCATTTTTATCATAGTTTACCATCTTAATTCATGTTGAAAATGCTTTACTTCACTCTCTCAGTGAAATATTGAATACTTTGTAACTTAATCCAATAGGCACTTTATTTTAAGGTATTCTTGAAATAGTACTTTGATTACAAATCTAAGTTCTCTTTCTCACTCTCAAAGTACCATCTCTTCCATGTTCTTGATGCAATGATCACAAGAACATGGTTGATGTGGGACGAATTCAAGATTTAGAGTCAGTAAGTTCGGAGTAGGTGTGCATTATTTTGTATGCATTTGTATCTTCTTCTTTTTTTCGATTGTAGTAGTACTTGTTTTCTTTCCTTCAACTATTGAGTTATTTGTAATGCTACTTGTTCTCTTCTTAATTGTTGTAAACATGGTTTTTTTTCTTCG

>PGSC0003DMG400012443 LRR-RLKs

CTTTGTCTAATCTTAATAATAACTGTTCCTTTGATTTGAAATCTCAATCTCTATCAGCTTAATTAAAAACTGTACAATCTTTTTTTTTCTTTATTAGATTCTGTCCCTCTTTCTTGAAATTAACTAAAAGAGTTTGCTTGAGTTGAAAGTTGGGTTCTTGCTTCTCTTACAAGTGATGGGGTGTTGAAAAAATCTCATCTTTTTCATTTTTTCTTCCTAGTTTTAGGCTTTGAGTTTTGTAATCTAGTGTATTTATTTATGATTTGGAGTTAAGGGTTTGTTAATTCATACAAGTAGTTAGAGGCTTGCTTTAATTACAAGTAGTTGGTAGCCTTTTTTCTTTGAGATTTTTGAGGTGTTTTCTTGGGATTTGTGTTAATTGAAGTTTCTGCAACTGGGGTTCTCTTCTTCCTTTACTATAATAATAAGGGTGGTGGTGCCCTGCTGCTACCGTTAAGTGCCTTGATTTTTTCAGTCTTGGGATCTGATTTGATTTTTGGTACTAGGGGATCTGAGGAAGGTTCTTTCTTGGTGCTAATTTGCATCAAAGGTGGAAACTTTTCTTCATTTGGAACTGTGAATTTGGAATACATGGTGGTGGAATCTTGATTTTGAGCTGTTTATATAAAGATTTGTGTGTTTTCATCAATGGTGATTTGGGAGTTTCTGCCAATGAGTTTTCTGTGTTGGGTGTTTTTGGCTTATCTTTTTTGTACATCTTTGAGTCTTGAAACCCCAGTTCAAACCTGTCATCCATATGATTTGTTGGCATTGAAAGAAATTGCTGGCAATCTAACAGATGGGGTTATTCTATCAGCTTGGTCTAATGAACCTAATTGCTGTAAATGGGATGGGGTTGTCTGTGGTAATGTTTCTGCTCAAAGTAGAGTGATCAGGCTAAATTTGTCAAGAAAAGGTTTGAGAGGTGTGGTTTCACAGTCCTTGGAGAAATTGGATCAGTTGAAATTGCTCGATCTTTCGCACAATCACTTGGAAGGTGGATTGCCTTTGGACTTGTCCAAATGGAAGCAGTTGGAAGTTCTTGATTTGAGTCATAATGTGTTGCTTGGACCAGTGTTGAGGGTGTTTGATGGATTGGAATCAATCCATTCTCTCAATATATCAAGCAATTTGTTCACTGGAAATTTCAGTGAGTTTGGTGAATTTCCTAACCTTGTCGCATTTAACATAAGCAACAATTCGTTTACTGGTAGTTTCAAGTTTGAAATTTGCAGTTTTTCCAAGAAGCTTAAGGTTCTGGATATATCACTTAATCATCTCACTGGTGATCTTGGAGGACTGGATAATTGCAGTTCATTGCTCCAGCAGCTACATGTGGATTCTAATGATCTCGGGGGTCACCTTCCGGACTCGTTGTATTCAATGACATCTTTGGAGCAACTTTCACTGTCTGCCAATAATTTCTCAGGCCAGCTAAGCCCACAGCTTAGTAAGCTTTCCAAACTGAAATCCTTAGTTTTATCAGGAAATCGCTTTCATGGTTTGCTTCCTAATGTGTTTGGTAATTTGACATTGTTAGAACAGCTAGCTGCACATTCTAATAGATTTTCAGGACCATTGCCCTCTACGATTTCGTATCTTTCTGTGCTTAGGGTGCTTGATCTTAGGAATAATTCTTTGTCTGGTCCTGTTGATCTTGATTTTACTAAATTGACAAGTCTGTGCACTCTTGACCTTGCAACTAACCATTTCAAAGGTAACCTTCCTGTATCGCTCTCTAGTCGGGAGTTGAAAATCATGAGTCTTGCCAAAAATGAATTCACCGGGCCAATTCCTGAGAACTATGCCAACCTTTCATCCCTTGTGTTCCTCTCATTGTCCAATAATTCCCTTTCGAATTTGTCTGGAGCTTTATCTGTTCTGCAGCACTGCAGAAATCTTTCAACTCTTATTCTCACCAGGAACTTCCGTGGTGAAGAGATTCCGAAAAATGTGAGTGGATTTGAGAACCTGATGATATTTGCATTGGGGAACTGTGGTTTGGATGGGCGAATTCCGATATGGTTATACAATTGCAGTAAATTGCAAGTGCTTGACCTGTCATGGAATCATTTGGATGGCGAAATTCCTCCTTGGATTGGTGAAATGGAAAAATTGTTCTACTTGGATTTCTCGAATAATTCACTCACAGGTGAAATCCCAAAAAATTTAACTGATCTAAAGAGCCTCATTTCCCCACACAACTACGCCTCTAGTCTGAATTCTCCCACTGGTATACCGTTGTTTGTCAAGAGAAATCAGAGCGGTAGTGGTTTGCAGTACAATCAGGCTTCAAGCTTCCCTCCGTCTATCTTATTGAGTAATAACAGACTAAACGGGACAATCTGGCCCGAAATTGGTCGTCTGAAACAACTTCATGTCTTGGATCTCAGTAAGAACAACATTACGGGGACTATTCCTAGCTCGATTTCAAATATGGGGAACCTGGAAGTTTTGGATCTTTCATGTAATGATCTCAATGGATCAATTCCTGCTTCATTCAATAAGCTCACATTCCTTTCAAAGTTCAATGTAGCTAATAATCACTTGCAGGGAGCGATTCCAACAGGGGGCCAGTTCTTGAGCTTTCCCAACTCGAGCTTTGAGGGTAATCCTGGACTTTGTGGAAAAATCATTTCTCCTTGTGCTGCCAGCAATTTGGACCTCCGACCGGCTAGTCCTCTTGCTTCTAGTAGCAGTAAGCTTGGCCGAGGCGGAATTATTGGAATTACAATCAGCATAGGGGTAGGAATTGCACTTCTTCTTGCAATAGTGCTGCTTAGAGTGTCTAGAAGAGATGCTGGTCACCAGATTGGGGACTTCGAGGAAGATTTCAGCAGACCACCTCGATCGTCTGATACTTTTGTTCCTTCTAAGTTGGTACTTTTTCAGAATTCTGATTGCAAGGAACTGACTGTTGCAGACTTGCTTAAATCTACAAACAACTTCAACCAGTCGAACATTGTTGGATGTGGAGGATTTGGTCTTGTTTACAAGGCAGAACTTCCTAATGGCATAAAGACTGCGATCAAGAGGCTTTCTGGGGATTGTGGTCAGATGGAGCGTGAATTCCAAGCTGAAGTGGAAGCCCTCTCGAGAGCTCAGCACAAAAACCTTGTATCCCTTCAAGGTTACTGTCAGCACGGGAGTGATAGATTACTGATATATTCTTACATGGAAAATGGAAGCTTGGACTATTGGCTACATGAAAGAGTTGACGGGAGCTCATTAACATGGGACATTAGGTTAAAGATTGCACAAGGAGCAGCTCACGGATTAGCATATTTGCATAAGGAACCAAATATAGTTCATCGCGACATTAAAACCAGCAACATTCTTTTGAATGAGAGATTTGAAGCTCATCTAGCTGATTTTGGACTATCAAGGCTGTTGCATCCCTATGATACTCATGTGACGACAGATCTCGTTGGAACCTTAGGATACATTCCTCCTGAATATAGTCAAACATTAACTGCTACTTTTCGGGGGGATGTTTACAGCTTTGGTGTTGTTCTACTTGAGCTTCTGACAGGCAAGCGTCCCGTGGAAGTATGCAGGGGGAAAAACTGCAGGGACTTGGTATCATGGGTTTTTCAACTGAAATCTGAGAACAGAGTGGAGGAAATATTCGATACATCGATATGGGATACAAGTTACGAGAGGCAGCTTTTGGAGGTGTTAAGTATAGCTTGTCAATGCATAGTGCAAGATCCACGCCAGAGGCCCTCGATCGATCAAGTTGTCTTATGGCTCGAGGCAATCGGAAGTGTAAAAGAAAGATGAGATTCAACTAGTTTTTCCTTGTAGACTTGGCTTCTTTTGCTTACAAAGTGAAAAATATCAGTGTAAAGCAACAGAAAATTTCAGTTTTATAGAGTTAGTTATAAAACTTATGAAATGCTTTGTTTGGTTCACTTTGTTTCATACACAAAAAATAGTGATAGTTGTGTTTGTATACATATGCTTAACTTGGGGACAAGTCCTTTACCAATATACAACTAATTACTCCCA

>PGSC0003DMG400012514 LRR-RLKs

AAGAAATTGGTAGCTTGCAAAATCTGATATACCTTTCTTTGAGAGACAATAAATTGCAAGGATCAATACCTAGCTCAATGAGCAGCGTGACATCTTTGGAATTTCTAGACCTTTCTCATAACAATGTTTCAGGATTAATTCCCACGTCTTTGGAGAAGCTTCAAAATCTCAAGTATTTCAATAAGTTAGTTGGTGAAATCCCCTCGAGCGGTCCATTCAAGAACCTCTCCGGTCAGTCTTTCATATCCAATGAAGCATTGTGTGGTTCTCCAAGATTCCTGTTCCCCCATGCCACACTAGTACTTCAAAGTATCTATCCAAGATGAAAAAACTACTTGCTCTAATTCTACCAGCTGGAGTTGCACTTGTATTAGTCTCCATCGCCTTTGTGATTATATTGTATGGATAAGGTGTATAAGAGGTAAAAGAAAGCAAGCAGTCGATTCATCGTATGTCGTCTCAACAAGAGGAAGAATTTCATACTATGAATTGCTCCGAGCAACTGATTCACTCAGTGAGAGTAATTTGATTGGTTCTGGGAGTTTCGGCTCTGTTTACAAAGGAATCCTCAGAGAAGGGACTTCTATTGCAGCTAAAGTTTTCAATCTGCAATTACAAGCGGCATTTAGGAGTTTTGATACAAAATGCCAATTTTTGCGCAACCTCCGTCATAGGAATCTCACCAAAGTCATCACTAGTTGTTCAAACCTTGAATTTAAGGCTTTGGTACTAGAGTACATGTCCAATGGGAGCCTCGATAAGTGGTTATATTCGCACAACTACTCCTTATACATCAACCATCATGATATTTCGCTTCAAATAGGCTAGTTTTGTGTCTTTAATTCATAACATAATAGCTTCAAGTGGTACTCTTTTAGTTTCAATAACTTGTCCTATGTCCAGAATTTCATCCAAGTGGTACTCTTTTGGTTTCTTCATCTCTCAAGTCAATTTCACTATTATTCGATTGCTTTCTGAATTGCTCTTAACAATTGTTGTCCAGGTCTCAACTTTCATTAGTGATTTCATCTTGAATCTTCCTCTACTACTCTTTTAGGTTTTGTCTTTGCTGGTTTCTTTACTCAAATTTCCCAAGTTGTTGAATTCTTTCATGTTCGTTTCGTTTTGGGCTTAATAAGCAAAGGTGGCTTTTTGTTATTTCTGAACTTAAAACTTATAAATGTTAAGAACTAAGCTACATTTATTTAGAGTAATGTGCCTGACAATGTTCGAAAATTAGGAAACTTGTATTGTTTTCAGCTTGAGTAGGGGGCTGCACACCGCGCAAGTACAGAGTGTAGACAAGGAGGGACTTTTTCTTTTGTTGAAAGGTATTGGCCCAAATCTTTTAATAAATCTCATATTGTTAGGATCTTGATTAGATTCAACTGATGTGTCGAATCTCCTGAATGAAGGATTCAGATCTTCACCATGGATGATAGAACTCCTCCCGGAAGTGAAGGCAGGAGTAAAGCCTATTGATTCACCAGTTTGCTAATGTACTTGGTAAAGCCTATTGATTCACCAGTTTGCTAATGTACTTGGTAAAGCCTATTGATTCACCAGTTTGCTAACTTACTTGTTGGCTACCCGCACTTTGTAACCAGAATAGGGCTCTGGGAGGGAGTTGTCAGTGTATTGGGAGTTGCTTTGCTATGCGTTATTCATGGTGCTAGAGAAGACTTTATTTGAAGACAGTAATGGTGCATATACATTCCGTACTTCTAACCCAACTCAAGCCGAAGAAACTCATATATCTTTTGACTCTATTTATGTAGGCACCTGATATTTATGAGGTTTCACTCACCCCAGTTACCTATTTGAGGTCTAGGTCAAGATTTTAATAATCATGACGGTCCTTGATCCAACAGGATATTGGCTAGATGGAAAGCTTGGACCTCGAGAATCTGTGTACCGCCATTTTTCCTTGGACTGTGAAATCACGTGTTTCCACTCTTATTATCCTACATGGATATATTTTTGTCTTAATTATTAAAATTTGCTCTCAATTTAATTCTTCTACTCACTCAATATTTAATACAAATATGCAGAAACCCCAAATAGATGAGCAAATGTGTTGTGTTTGAAGTACTGATAGCATATGACATACATTGGCACGCGTTGATTAAACATCACATATGCAATAAAGTGAATAATATTATATATACAAGTGACAAGTGAAACTAACAGCCACCTCTTCTTGCACTGTAATTACACTGCTCAAATCTGGAATTTGTTCCTTAGCATCTCAAATCTGAATTGGACAATGCCAGAACGTACATCAAACGTGTTGAAGAGCTGGGTAAGGAGAGGAGGCACCAAGAGCCAGAAGAGATGGTGGAGGTTAATACCATCATGTATATGGTGGTCAGTTTGGAAGGAAAGGAACAGTAGATGTTTCGACAATAGATCCAATTCCATACACAAGGTCAAATGGAATTGTATAGTATCTCTACTTTTTGGTGTAAACAGTTGTGTATAGTAGATACAGACCAGATTGTAGATTTGATAGGAAGTTTGTAATTGACACTTTTTGGTGGTAGCCAGCATGCCCTTAATGCTGAAGAATACAACCTTACCAGTTTCAAAAAAAAAAAAATATTATATATACACGCTTGTATACAAGGTTGATACGGTATTTACACTAAAGTGAATAATATTATATACTGGGGTATAGAATAATATTATACTGTTGTGGAAATGATTTAAGGTGTAAACTGAAAATATGCTTAGAGGGTGCAACGTGTTATGTTTGGAAGGTCACATTATTCCAGAAACGAAATACACATGCATTTGGAGCCAATGTGTTTTACGAGTGCCTTATCTTTTTGACATATGAATTTGTATTGTTGTTTATTCAAATGTATAATGAGACAATCAATGAATGTGTTTGAAAGGAAAATGTTTTTTTCAGAAAACAAGTTGATTTCTGACTTATTTTCTCATGTTTGGTTAGTAAGTGAAAAATATTTTTTAGTATTTGGTTGGTAAATGAAAAATATTTTTCAAAAAATATCTTTTATTATTGGTTAGAGCTTAGAGACTAGAAAATACTTTTTAGGAGATTAATTTCTAACCCAAAGATCCGAACCTGACAACCTGAACCAACACACGACCTCAATGACCGATTCGGGATCCGAACCCAATACCCAACATGTGACCTGACCTCAACCCCAACACCCGAATTAAAACACGACCCCGATGATCAATCCGGGGATTGGACCTCAAGACCCAACCCAAGACTTGGCCCAGACACCCGACGCATGACCCAACCCCAACACCCAACTTCCAAATCGAGACCCGACCTCGACTCTTGATGCGGATCCGACTTTTGAACTAGGACCCAACCTCGACACCAGATCTAGGACCCAGGACCCAACTTCGACACTCGAACCAAGACCCAACCTCGACCTTGACTCCGGACCCAGGATACGACTTCCAAATCGAGACCCAATTTAGAATCCCGACCTAGGACCCGATTCCTGATCATGTACCCGACCTTGACACCCAACTCGAGATAGGAGTTCCAAATCGGGATCTGACCCCGACACAATAAATGATCCAGGATTCGACCTCGACTACCGAATCTGACTTCCAAATAACAACTCGACCCCAACACCCAACCCGGAATTCGACTTTCAAATTGGGACCCGCCTTTCGAATCAGGATTTGACCCCTACATCCAACCCCAACTCCCGACTACGAAGATCAATCTGGGACCTAAAACCCAACACCTAACTTGGGACCCGACCTTGACTCTCAATTCGAAACACAATTTCAAAACCCAATCCCAAAATCTAACCTCGAATCTTGACCCAGGACCCGACACCCAAACTAGGATTCGAAACCAACCCAACCCAACCCAAAAGTTGGGTCTCAAATTGAGGTTGGGAGTCGGGTCTCGAATCGAGAATCAGGAGTTGGATCTAGCATTAGTGATGAGAATCGGTACCCGAGAGTTGAGTTAGGGTGGAAAATTGAAGTGTGCTGAAAAAACTCTTCGACGCATCCTAATTCTCATTGAATTCAGGGCCCAAATCATTAATAAGGACTCATCCTCGACCATTTGGTTTCATATTAGATAAAAGGAAACTAAAAATGGGGCACTTGTAAAGCAATTGAAAAGAGACCAGAACTACGTGAAAAGATGTTTCATATATAAGCAAACATTATTATTGTGGACCAAGGGAACTTATATACATAAAAGAAATGGCCTAAACCCCTTTCACTCATTTAGCTCCACCCCTGCGTCCGATACTCAACACACATAACTCATTCTACAGTTCTTTGTAAAATTGTGTATAATTAGCCTTGTAAACATTCAAAATCCTTAATATACCCTTTTCTTTTCAAAATCCCTCTTCGTTATGAAACTAAATATCTATGGACTATATTTAGAATATTTACTTCAAAATCATCACAAAAAAATGTAAAAGATGATTAATACTACTAACTACTATTCCAGGCTGTTATTTGCTGAAGAAAATTTTGATTTGATCAACAAAACGTTACTAACATTAGTCAAGTCACACGGTCCCCTTAGAGGAAAATGGAGAAAAAGACAATGAAGACCAAAATGGAGATCAACTGATCACTGCTATTTAAAGTCTCATAGCCCCATTGTAACTAGCCATGCCAAGTTCTTTCTAATTTGTAAAGTACTAAAATGGAGAAAGCCACCAACACTTTTTTCTTCTTGTTTCACTTGGTTGTGGCTAGTATAGCCATGACTCACACCAATATTACCAATGATCAATTAGCTCTTCTTTCTTTAAAATCCCAAATCATTTCTGACCCCTTTCACTTGTTGGATGAAAGTTGGTCTCCAACTACTTATGTCTGCTATTGGGTTGGCATTACTTGTTCTTCCCACCATCGAGTGACGATGCTGAATATTTCCAACATGGCTCTTACAGGCAAAATTCCTCCTGATTTCGGGAATCTCACATTTCTTGTTTCTCTTGACTTGAGTGGCAACAATTTCTATGGAAATTTGCCTCAAGAAATGGCACGTCTACGTCGGTTAAGGTTTGTTATGCTTGGTTTTAACAACTTCAATGGGGAGGTTCCTTCATGGTTCGGGGTCTTACACCAACTTCAAGTTCTGAATATAAGGAATAATAGTTTTAGTGGTTCCATTCCCTCTTCAATTTCTAATATCTCTACGCTTCAAACTTTGGATTTGACATACACCTCCTTAGGGGGCCAAATGCCAAAAGAAATTGGAAATCTTCAAAATCTAAGGGTTTTACGAATATCTGGTAACAATCTTATAGGTTCTATCCCTCTGTCTCTCTCAAATGCCACAAGGTTGATGACTTTAGAACTTTCTGTCAATTACCTTGAAGGAAACATCCCCAAAGAGATTGGAGAACTTCAAAACCTAAAAGTTTTGGCAATACAAGCTAATCAACTTATAGGTTCTATACCATTCTCGATCTTCAACATTTTCGGAATTAAAGTCATTGGATTTTCATATAATACCTTATCAGGAGATCTTCCTACTGATATGTGTGATCGACTTCCAATGCTGAAAGTGCTTTATCTAGGTTTCAACGAGCTACACGGTCATATGCCTCTAAGCTTGGCAAACTGTTCAAAACTTCAAACCTTTGTTTTATCTGATAACAAGTTTGATGGACCCATACATAGTGAAATTGGACATTTAAGTAATTTGCAGGATTTGTCTCTCGAAAATAACCATTTTGAAGGTTTGGCATTGAATACTAGCTTGTGGTTATTTTTCTTCTAACTAACTACTTCCTGAAAAATTTCAATCACATAATTTTTCTTTTGCAGGGAGAATTCCACAAGAAATCGGGAATCTTGATAATTTGGTGTCCATAAATCTTATGGCAAACCAGATTACCGATTGTATCCCAATCTCCATATTCAATATCTCGTCACTACAGTTTCTCTCATTGGATGACAACAATCTCCTTGGGCCCTTGCCAGGGGAGGTTGGCAACTTGACCAAATTGCAGTATCTACTTCTTAATGAAAATATGCTTACAGGTAAGTTATTGCAGCTATCTTGCTACAAGTGAACTAACAAAGTTTAAAATCATTACACGTTGTTGCTTAATTATATCTCCATACATACATTGACAGGTGAAATACCGAAAGAGGTAAGCAATCTCATTGACTTGGCTGAAATGGGGTTGGGTTTCAACAAGTTTACTGGTTCACTTCCAATAGAGATCTTCAACATATCAGGGATAGAAACAATTCAACTTACTTCCAATAATCTGACAGGAACCCTACCATTAAACATGGGTTCTACGTTACCCAACATTGAAGTTCTTTATCTAGGTCGTTTAAATCTTTTTGGGACTATACCTCATTCTCTCTTCAATTGTTCCAAACTCACTCATCTAGACCTTGCTGTAAACAGACTCACCGGTATGATTCCAACCTCTCTTGAATATTTGACTCAACTCAAGTACCTATTCTTGGGGGGAAACTATTTAATAAGTGACACTAAGTTAAGCTTCTTGACTTCCTTATCAAATTGTAGAGATTTAGAATTTCTTTCTTTATCTTTGAACCCTCTGAATGGAGTGATTCCAGTTTCTATAGGAAACCTTTCCAGTGCTTCTCCTCTATGGTTTAGAGCTTTTGAATGTGGAATTAAAGGGGAAATTCCTAAAGGAATTGGGAACTTAAGCAGCTTAGTAGACCTCGACCTTTCTGGAAATGGCATAACTGGATCGATTCCCGCAACAATTAGAAATTTGAGATTCCTTCAGCGCTTTAAGTTGAGTCAAAACAAACTTTCTGGATCTATTGGAGAAGATCTATGTAAATTGCAGAACTTAGGCTATTTACACTTGACTCAAAATAAACTTTCAGGATCTATTCCTAATTGTTTGGGGAACTTAACTTTCCTTCAAGAGATATTTCTAGGTTCCAACAAATTGTATTCGAACATACCAGCAGGTTTAGGGAACCTCAAAAATCTCTTGAGGCTAGACTTATCCTCAAACAACTTGGGTGGCTCATTACCTCTGGAGATTGGAAACCTAAAGGCTGTCATATATATGGATCTATCAATGAATGCACTCTCAAATGGCATACCCGGAGAAATTGGTGGCTTGCAGAATCTAATACACCTTTCTTTGAGAGACAACAAGTTGCAAGGATCAATACCTGGCTCAATGACCAGCATGTCAGCTTTGGAATTTCTAGACCTTTCTCAGAACAATGTCTCAGGATTAATTCCCAAGTCTATGGAGAAGCTACAAAATCTCAAGTATTTCAACGTTTCATTCAACAAGTTGGTTGGTGAAATCCCCACTGGCGGTCCTTTCAAGAACCTCTCCAGTCGGTCATTCATGTCCAATGAAGCATTGTGTGGTTCTATAAGATTTCGTGTTCCGCATTGCAACAATAGTACTTCAAAACATAGATCCAAGAGGAAAAAGGTACTTCTATTTCTACTAGCTGGAATACTAGTTGTATTAATTCCTATCGTCTTTCTGCTTGTATGGATAAGGTATATAAAAGGTAAAAGAAAGCAAGCAGTTGACTCATCCTTTGTCGTCTCAACCAGAGAAAGAATTTCATACTATGAATTGCTCCGAGCAACTGATTCACTCAGTGAGAGTAATTTGGTTGGTTCTGGGAGTTTCGGCTCTGTTTACAAAGGAATCCTCAGAGATGGGACTCCTATTGCAGCTAAAGTTTTCAATCTGCAATTGCAAGCGGCATTTAAGAGCTTTGATACAGAATGTCAAGTTTTGCGCAACCTCCGTCATAGGAATCTCACCAAAGTCATAACTAGTTGTTCCAACCTTGAATTTAAGGCTTTGGTACTTGAGTACATGTCCAATGGGAGCCTCGATAAGTGGTTATATTCGCACAACTACTCCTTAAACATTAACCATCGACTAAGTATAGTGATAGATGTGGCATGTGCGTTGGAGTATCTCCACCATGGCTGCTCATTGCCCGTGATTCACTGTGATTTGAAGCCTAGTAACATCTTGTTGGATGAGGATATGGTTGCCCACATAAGTGATTTTGGCATTTCAAAACTGCTTTCTGATGATGAGAGTGCTTCATACACTAAAACTCTAGCAACATTGGGCTATATTGCACCAGGTATGTACTCTCTTAACTAATTTCTTTTGAATACTTCTCTCAAATGCTGACTTTAAAAATTGGCTGATTGTAGAGTATGGATTGGAGGGATTGGTGTCAACAAAATGTGATGTTTATAGTTATGGCATCATGTTGATGGAAATATTTACACGCACAAAGCCTAGTGATGAAATGTTTGATGGAGATTTTAGCTTGAAGCAATGGGTGAGCAATTCACTCCCACAGGCAATAATGGAAGTTATCGATGCCAACTTAATAACGCCACATGATAATCACTTAATGAGAAAGATAGATTGTGTGGTATCCATCATGAAAGTGGCAATGGATTGTTGTGTTGAATCTCCTAAAGGAAGAATTGACATGAAAGATGTTGGGAGGTTAAAGAAGATCAAGATTCAACTTCTTGCATGTTGA

>PGSC0003DMG400012588 LRR-RLKs

ATGGGCTCATCTAGTCTCTTCATCAAGTACTATGTTGTCTTGCTTTTCATGATGAAATTATGTACCCTGTCCATAGCCGAAGATGGTTCTACGGTAGGCTATCATGGCACAGAAAGGGAGGCTCTTTGGGCGATTCGAGCTGAATTCAACAATCCTAAATTGCTCCAAAAAAGTTGGAACAGCCTCATGTGTTACATGAATAACACTCCAAATTGGTATGGTATTAGCTGTATCAACGGTCGAGTCACTGGACTTGTGCTAGAAAATCTTGGCCTTACTGGGAAACTCGAGGCAAATGCACTTGTCAATCTCACTCAATTACAAACACTTAGCTTCAAGAACAATTCCATATCAGGGAACTTGATGAACTTCTCCAACAACCAAAATTTGAACAACATTGATTTATCCGGTAACAGATTTGACGGAGAAATTTCACCTTCTCTTCTAACACTTAATTCGTTAGAGTCATTGCAGGTACAGAATAACAAATTGAACGGTCCAATTCCCAGTTTTAATCAGTCAACTTTGAAAGTTTTCAATGTCTCAAACAACAATCTTTCTGGTGAAATTCCAAGCACAATAGCTCTTCAAAAATTTGGTCCGTCCTCGTACTTAAGTAACCATGATTTATGTGGTCCTCCTTTTACTAACACAACGTGCATCACGATCAAGGATAATGAGTCATCTACACCAACTCCAAGTGATAGTTCACATAATGAATCTTCAAATAGTTCAAAATTATCCATGTTAACTCCAATCTTGGTAGTAATTAACGTTGTGGGCATTGTTGTTCTTTTATATCTAGTCATTTACTACTTTAAAAAAACAAAAAAGCTTAAGAAAATGCTGATGAATAGTAATACAAGATTAACGGAGGTTGATAAAATGGAGACTAGCAAAGCTGAGATCACAACAACGCCAACAGAAACCAGGAGTGTGGAATCAAGAAGTGTGGCATCAGAGGTTGAATTAGAGAAAGGCAAATTGATATTCTTAGGTAGTGAAATAAATTTTGAATTAGATGACTTACTGAGAGCTTCAGCAGAAGGTTTGGGAAATGGAAATTTTGGAAATTGTTACAAGGCAATGTTGGTAGATGGACCAACTGTTGTGGTGAAGCGTTTGAGGGTTTTGAAACCTTTAACAAATGACGAATTTGTGAGACAAGTAAGAGCTATTGGTGATATAAAGCATCCAAATTTGTTGCCTATTCTTGGTTATTACTATACAAGGGATGAGAAGTTGCTTCTCTTGAAATCTGCACCTAACGGAAGCTTATACAACCGCATTCACGGTATCTTACTGTGATCCACTTTTCTTTGTAGTCTATCATATAACATTTATAATTACTCAAATACATAACTGATTTTTTTTTTCCGTGGTTGGTTGAACAATAGGAGGAAAAGGGACTAGAGATAGGATACCATTTCGATGGAGCTCAAGATTATCCGTTGCTCGTGGTGTTGCTCGAGCACTAGAGCATCTACACCTGAACGTCAGTTCCTCCCAATTCGTTGTTCCCCATGGCAACCTAAAATCCACCAATGTTTTACTTGACGAAAATGATGATGTTCGTGTTGCTGACTTTGGTCTCACTTCCCTTATTGCACTTCCCATCGCTACTCAACGAATGGTTTCCTATCGATCACCCGAATATCTGGCTTCCAAGAAAGTGTCCAAGAAATCTGATATTTGGAGCTTTGGTTGCCTTCTCCTGGAATTATTAACAGGTATTTGTTTTTCTTGATTGATCAGGCAGAATCAAATATTTGCGTATATATACTACATTATTAGTTATCTAGCGATGTATATATTTAGAGCATACAATACATATATTTTGATGGTATATTATATTGATCACATTGTTGTATACCATTGATATATATAAGTAAATAGAAAATGTACCATCGGGCTGATACTCAAAACCACTTATAAATCAAATTATAATAAGAGACTTTGACACAAGTGTGTGCGCTATAATTGTAACAGGTAGAATTTCTTCACATTCTGCTCCACCGGGGGTGACCGGTGCGGATCTGTGCAGTTGGGTTCATAGGGCGGTCCGAGAAGAATGGACGGCTGAGATTTTCGACACGGAAATTGCTGTTCAGAGAAGTGCCAACTCAGGAATGTTGAAATTGTTGCAGACTGCTATAAGCTGTTGTGATAAGTCGCCGGAAAAACGACCGGAGATTAGTGAGCTGTTGAGGGAAATGGAAAGTATTAACGGGGCGGTACCCGAATCGGAAGACGAAGATGATCTCTCCTTCTCAATGGATCAATCTATGACAGATGACTCCATAGCAACTGCAACTCCGTCTCGATAA

>PGSC0003DMG400012594 LRR-RLKs

ATGCTGTAAGCAGTAACGAAATACTATTTCCATGGACAGAATCCCTCTGCCGCAGATGTTCCCATTGACAATTTAAACACATTTTGAACCTCCTTTTCTTTTCTCTCAAAAGGTGTAGCTTGTGGATGCACTTATTAATTTTTAATGTGCTACATAGCTATGTGATGACTAATGTATAGCCCCTTAATTTTTACAGATTGTGTGGTCTGTAAGTTTTAGAACTGATTTGCTTACGACCTCTTTTTCCTAATTGCAGGGAACTTTATAGTAATAATATAAGTGGAAGAATTCCTTATGAACTGGGGAATTTGACGGAGTTGGTGAGCTTGGATCTTTACCTGAACAAGTTAGTTGGTCCAATCCCCGACACTTTGGGCAAGCTTCAGAAACTACGTTTCCTGTATGGGTTACACATCCTTTTTCCTCTTCTATATGTTTGTTTTTCTTTTTCTTTTGCTTGGTTGAATAGTTTCTTTAGCCATGAGCTTTTATTTTCTGAACATTTTGCTTAGTCAAGAAAGGCCACCTTCTATTTTAGAACATGTTAGTGGCTCTGTACTCTGTAGTACTATTTCTCTTGTTTAGTTATATGGAAATAGGGATAGACTTTGCTCCTTATTGGCCAAGTGGCAGACATGAAGATAAAAACATTTCAAACACCTGGAGTATCATTGCCAATGAGACAGCATCATAATTTTCTCTGTGGCTTAAATTATTCATAAAGAGAGAGAGAGGAAGAGGAGGCGGAAGAAGAAAACCTGAAGCAGCTCATAGAAGTACCCAAAGTGGGTGTCTTAGGGTGTCCAAAGTGCAGGACATGCATGTCTGAAAATTCAAAAAATATACAGGAAGTTAGACACACACATGCTCAATGTGTGTCCAACACCAGTACTTGACAATGAAATTTGTCCATCATCCTTGGCAGTCATCGTTCCTATTTCCCTTATATAGGCTTGCAAAAGGCTAGGTGGATTTGAGTGGTTGATAAAATTGTGGTTGCTGCTCTCTATAAAGGCTCATTGGTGTGCTGCTGAACAGGATAAAAAATTTACATGCTATTCTTGTTATTTCCCGAAGCTCTGAAAACTAGATACTTCTGATGATCATATATATGAACATAAATCTCAACCTGCTAACAGTGTCAAGAAGCTTTCTTAGATATTGGGCATGAATTTTGTAGCATCTTCTAATACTACTATTTCTTCTTCATTAGTCAAGTATTTACTACAATGTACACATGGACTGATTAATTAGCCGAGTTCTGTCTGTTTAAGCTTGATCAAACTTTTTTTTTCTTATTGTTATTGTTTTCTTTTTGCTAAATTGAAGGAGGCTTAACAATAACAGTTTGACTGGGCAAATTCCGATACTTTTAACCACAGTCACTTCACTTCAAGTGCTGTAAGTATAAAGCTGGCACTTTTTTATCATGTGGTTTTTTTCGCCATCAGTATTTTCTAATTCTTAGGTACTAATGTTCATTCCTCGCAATGCAGTGATTTGTCAAACAATAATTTAACAGGACCTATTCCAGTCAATGGTTCCTTCTCGCTTTTTACTCCTATCAGGTCTGTGAAAAGCACTAGTTTATATTTTAAGTTCTGTCACTGGAATCATGATAATGAAATTCTTACATCTGAATGTCTATATTTCAGTTTTGCTAATAATCCATTGGATACTCCTCCAGCTTCTCCACCTCCACCTATTTCTCCAACACCCACTTCTCCAGGTTTGTTAAAATAATTTATGTTTTATGGGTTAAGGCACTGCGCCCCCCCCCCAAAGGAAGAAGGCAAAAGGGCTGACTGTTTTAGGGGAAGAGATTTGCACATTATAATGAGTTTTAGTAGTGTATGTGAGATAAATTTTACAGGTGATTTTGGGGTTATCTACTGTTTGATAGGTTTGAATAAATAGATTAAACTTACAAGTGATATTGAAGAGTGGATAGAAGTGTCCTACATCAGGAAATGCAACTGTATGCATTGGATATTGTGACTATAAAAGGACCCAAACTTCAGAATTTATTGTCATACATTCTCTCAATTCTTTCCCCATGTGTCTTATTTCTCACATGGTATCGGAGCTTTGGTGAAAAATTAGTTTCAGTGCATTTACCAGCAACACGCGGGAGAAAAATCTCCTTGCTGTGCAATTCTCATTGACCATGACTGCTGTCAAAATGTATCCTTGCCGTCTCTGACTACTGTACATAAATCAACACCACCAAAAGGCCGAATAACAAACAAGTACCTGTTGTTATGAATAAACTGAAGGATCTTTACCTATTTCTGCATCTTTTCTATTTTGTTAGCACTTTAGCTTTACAACATAGTCCTCGGTGCGAGTGTAAATGGTGAAACTACCATATCCCTTTTCTCAATTTTTTTTGAACTACTTGTGTTAAGTTTTTGTCATCCATCAAAAAAAGGGAATTAAAACTTAAATTTTGAAATTCCTAGGTGTGGGAAACAGTGCAACCGGGGCGATTGCTGGCGGTGTTGCTGCAGGGGCTGCTCTTCTGTTTGCAGCTCCTGCGATTTTGCTTGCCTGGTGGCGCCGAAGGAAACCGGAAGACCACTTCTTTGATGTTCCAGGTGGGAAGCCTCAATCAAACACCTGAATTAGTTTTTTGTTTTCAAATTTAATTTTAATTTAATGAGTTCTTGCAATTTTACGTTCTTTTGCAGCTGAGGAGGATCCAGAAGTTCATCTTGGACAACTCAAGAGGTTCTCCTTGCGCGAATTACAAGTTGCAACTGACAGTTTTAGCAACAAAAATATACTTGGAAGAGGAGGATTTGGCAAGGTTTACAAGGGCAGGTTGGCTGATGGCACTTTAGTTGCTGTTAAAAGATTAAAAGAGGAGCGTACTCAAGGCGGGGAGTTACAGTTTCAGACAGAAGTTGAAATGATTAGTATGGCCGTGCATCGAAATCTACTTCGTTTATGGGGCTTTTGCATGACTGCAACTGAACGGTTGCTTGTTTATCCATACATGGCGAATGGAAGTGTTGCTTCACGCTTAAGAGGTATCTCTTAGTCTTGAATTTTCCATCCTTCTTTTTCTTTGCTCCATTGAGTCGAAATACATCAGATCAAGCACTTGAAAAAGCTGAAAAATGATAACAACGTTTTTTTTGAACTTTTAAGTTATGGTCTGGTAATATAAGTTATCTGGAGAAGTTATTGCAGATGTGTACATCATTGTGTAAACTTGGCATTATTGGATATTGTCGTGGTTGTATTAGGAGTTAGTTTGTTATAGTTAGAATATTCAATCCCGTAGTTAGAAGTTGTTGTTCTTTGTGATTTTACCCGTGGTCTATCAAGCAATTTCAAGAGCATTGCTATCCTTGATGTCCTACCAAATATTAAGATTTCCGTTCAAGTTTTATGTTCTGAATTCATAGTGTCAGATATGGAAAGTAATCTTCATTTATGCATGAACCGCATATATCAATTAATTAAATTGCGTTCAACCTAAAATGGGAACTTGAACCAGTTTACTTGACATCTTGAACTTGTCCTTACAGTAAAATTTTTTTCTTGTAGAGCGGCCAGAGTCAGATCCACCACTTGGCTGGCCAATAAGGAAGTGTATAGCACTTGGATCTGCAAGAGGCCTTGCTTATTTGCATGATCATTGCGACCCTAAGATCATCCACCGTGATGTCAAAGCTGCAAATATATTGTTGGATGAGGAGTATGAAGCAGTTGTTGGGGATTTTGGGTTAGCCAAACTCATGGACTACAAGGATACTCATGTTACTACTGCTGTACGTGGTACAATTGGGCATATCGCCCCGGAGTATTTATCCACGGGTAAATCTTCAGAGAAAACTGATGTGTTTGGCTACGGAGTTATGCTTCTAGAGCTCATAACTGGGCAAAGGGCGTTTGATCTTGCTCGACTTGCAAATGATGATGATGTCATGTTGCTAGATTGGGTAAGGATTTACTAACCTACATGCTCTAGTTATTCTACTTCTAGCTTATGTCTTTTTCTCTTTTCTTTATACGTTGAAACTTTATACTACTTACAATTGCTAATTTTTATCGTGCAACTGACTTCAAACATGGAAGTGAGCACGGGTGAGAAGATCTCACTTTCCTGCATGCATCTCCCTATAAGCAGAACTCTATCATTGCTGCACTCAACATGATGAAAAGCTGATTTGTAGACCCACTACAAATTAGGACTGGCGAATAAGACACTGTCATGTCCATTCTCTTCCCCTGCGTTTTCCTTTAATATCCCCAATCCCGTTCCTGTTCTGTTATCCCTTCCCTACCCCCGTATGTAACCCACATCCCATAATCTCGTTTTATATCCTCATCCATCAATGAGAAATCTCAGCTTCATCTCATCTCCCTCACTCTTTTGCACATATTACACACACTCATAGCTCTCTTTGTTCTTTACTCAAGTCTCAGGGCTGCAGTTCTGTTTAAAGTAAAAAGTTTTATATTGACCCCCCCTCCCCCCAAAAAAAATAAATTAATAATAAAAAAAAAATTGTTAGCTCTAGTAGGTTTTGCATTGCAGTTTGGCATGTATGATTTCTGTAATCTTTGATACTACATCATATGAAACATCCAAATTTCCAAAATAATTTGATTCCTGGGACTTGTATTTGTGCACATTCTTCGTGTAATCTTTTTGTTTCTCTTTCTAGAACTTAACCATCTTGCGAAAACATTTGTTGCATTAGCTGTATGGTCAGGGTGGATCACTTTCATTATCGTGTAGTAGCAGATTGTTCTGGAATTGCTTTTTAGTGTTAAAATCTTTTTCCCTTTGGTTAGCAAAAGCACTCGTATAGCATATTGCTATATCAGAAGATTTAGATACTTTTTCCATGCCATATTGACTGTCGCCTTCTGTGCAGATAAAACTACATCTCTTTTTTTTTCCGTTGTATGGAATTGCTAGGCTATTTCTCTGTATTTGCATCAGAGAGTAGGTGAGATAGAGAATACACTTTACGAGAGGGGACTTCTTTGGGAGGATCCTCTAGAAAGGGAACCCACCCTTTAGGACTTGGGAGTGCAGAAGATGGTCAATTCTTTCACTATGGTTGAACCGTTGAAGTATAGAAAATTGAAAGACAAGTGGCTATCTGAACTTGGAAAGTAAATAAAGTGGCTATCTGAACTTGGAAATTCAATATGGTTCAAATATGAGCACAGAGAATCAGGAGCCTCATGCTCTCTGATTTTCTGTCTTTTCAGTCTCCCTATAGGGCTGCTTATCAAAAATCTCAACAACTTCTATGGTGAATGCTTTGTTATTTTAGTGATGAAATTAATTATTTGCTGACATGATGAGGAAAGTCCGCATGCACTGTGATTTCTTTTTAAAAAATGGCACTCATGTGAAGAAAATGGTTATGTACCTTATTTGACCTCAAAAATTGTGTTAAAATTTCAGGTAAAGGGACTTCTTAAGGACGAGAAGTATGAAACATTAGTAGATGCAGATCTTCAGGGTAATTACAATGAAGAAGAGGTGAAACAGCTTATTCAGGTAGCTCTACTCTGCACACAGAGCTCTCCTATGGAACGTCCAAAGATGTCGGAAGTGGTCAGAATGCTTGAAGGTGATGGCCTTGCTGAGAGGTGGGAAGAATGGCAGAAGGAAGAGATGTTCCGTCAAGACTTCAACCATGCTCACCATCCTCACACTGATTGGATTATTGCAGACTCCACTTATAATCTACGACCCGATGAGTTATCAGGGCCAAGATGA

>PGSC0003DMG400012817 LRR-RLKs

GGCTTAATCACTTTTTCCAGAAAAAAATCCACTAATCCATTCATCTCCTTCAACAATGCATCTTCAAATCTTGCTTATACTTCTATTACCTACATTGATTCTCTCAATAAACCAAGAATCCCTTTATTTACACACCATAAAGCTTGGATTTGATGACCCAAATGGTGTTTTTTCAAACTGGAATCTCCATGATAACTCCTCACCCTGTAACTGGTTTGGAGTAAAATGCGACTCTTTAACTCGTTCTGTTACATCTATTGACCTCTCCAACACCAATATCGCCGGCCCATTTCCGGCTTCTCTTCTTTGCCGGCTCAAGTATATCAAGTACATTTCATTCTACAATAACTCCATTAACTCGACACTTCCGGTGGAGGAGTTATCCGCTTGTAAATCTCTTGTCCATCTCGATTTAGCTCAAAATTTGTTAGTGGGTAGTCTTCCATCGAGTTTGGCGGAGCTTCCTGAGCTGAAATACCTTGATTTGACCGGAAATAACTTTACCGGCGAAATTCCGGCGCGTTTTGGGGCTTTCCGGCGACTTGAAGTTCTGGGTCTGGTTGAAAATTTGTTAACTGGGACTATCCCGCTGGAGATTGGAAACATTTCGAGTTTGAAGCAGCTGAATTTGTCGTACAACCCGTTTTCTCCGGGTCGGATCCCGCCGGAGATTGGGAACCTCACGAATCTCGAGGTGCTATGGTTAACTGACTGTGGGTTAATCGGTGAGGTTCCGGGTACATTAAGGGGATTAAATAAGCTTGTTAACTTGGACCTTGCGTTAAACAACTTGTACGGTCCGATTCCGAGCTGGCTCACTGAGTTAACTAGTGTTGAGCAAATTGAGCTGTACAATAACTCGTTTTCCGGCGAGTTTCCGGTGAATGGGTGGTCAGATATGACCTCGTTGAGGCGGGTCGACCTGTCGATGAATCGGGTTACCGGGTCGATCCCGAGCGGGTTGTGTGAGTTGCCACTTGATTCACTCAATCTTTATGAGAATCAATTGTATGGTGAGTTACCTATAGCCATTGCAAATTCACCCAATTTATATGAATTAAAGCTCTTTGGTAATAGGCTAAATGGAACTTTACCTAAAGATCTTGGTAAATTTTCGCCATTGGTATGGATTGATGTTTCAAACAATGAGTTTTCAGGTGAAATTCCGATGAATTTGTGCGGAAATGGGGTTTTAGAGGAGGTTTTGATGATAGATAACTCGTTTTCCGGTGGAATTCCGGTGAGTTTAAGCCAATGCCGGAGCTTATTACGTGTGAGATTAGCTCATAATAAGTTCTCAGGTGATGTCCCTGTGGAATTTTGGGGGCTGCCACGCCTTTTGCTGCTTGAGTTAACGGACAATTCATTTTCTGGTGTAATCGCGAAAACTATAGCTGGTGCATCGAATTTATCAGCTTTGATTTTGTCAAAGAACGAATTTTCGGGTAATATTCCTGAAGAGATTGGCTTTTTGGAAAGTTTGGTTGATTTTGTGGGAAATGATAACAAGTTTTCAGGGTCGTTACCGGTTAGTATAGTGAATCTTGAGCAATTGGGAAGAATGGATTTTCACAACAATGAATTAAGTGGTAAGTTTCCTAGTGGGGTTCATTCTTTGAAGAAATTGAATGAATTGAACTTCGCAAACAATGATCTTTCTGGAGAAATTCCCCGGGAAATCGGGAGCTTGTCTGTTTTGAACTATCTTGACCTATCAGGAAACAAGTTTTCGGGGGAAATTCCAGTTGCGTTGCAGAATTTGAAGCTCAATCAGCTGAATTTATCGAATAATGGCCTTTCGGGTGGTATTCCTCCTTCATATGCAAAGGGAATGTACAAGAATAGCTTTCTGGGGAATCCTGGTTTATGTGGAGATATTGGAGGTTTATGTGATGGAAAAGATGAAGGTAAAACTGCTGGTTATGTATGGTTACTGAGATTGCTTTTCATACTTGCTGTTTTGGTGTTTGTAGTTGGGGTAGTTTCGTTCTATTGGAAGTATAGGAATTACAAGAAAGCAAAAAGGTTGGATAGATCGAAATGGACCTTGACGTCGTTTCATAAGTTAGGTTTCGATGAGTATGAAGTACTGGAAGCTCTAGATGAAGACAACTTGATTGGTAGTGGTTCTTCCGGGAAGGTTTACAAGGTCGTTTTGAGTAATGGTGAGGCTGCTGCTGTGAAAAAACTTTCAAGAAGTTTGAAAAAAACAGATGAGAGTTGTGACATTGAGAAAGGTAACTATCAGGATGATGGATTTGAAGCGGAGGTTGAGACATTGGGCAAGATTCGACACAAGAACATCGTTAGGCTATGGTGTTGTTGTACAACAAGGGGTTGCAAACTTTTGGTTTATGAGTACATGCCTAATGGAAGCTTGGGCGATTTGCTACACAGCAGCAAAAGTGGGTTGTTGGATTGGCCTAAGAGATTTAAGATAGCTATGGATGCTGCAGAGGGACTCTCATATTTACATCATGATTGTGCTCCTCCGATTGTTCACAGAGACTTGAAGTCGAACAATATCTTGTTGGACGGGGAGTTTGGAGCTCGAGTTGCTGATTTTGGTGTGGCAAAGGCGATTGATGTCGATGACAAGGGAACCACGTCTATGTCAGTCATTGCAGGGTCTTGCGGTTATATTGCTCCAGGTTAGAGGATTAATCCTTCACTTTGGGAGTTCTACTAACATGATTTTGTTGAATTTGGTTTATGCCTTCCATTGCAAATTGTAGTTAAGATGTATGCCTAAATATAACATATTTCATATCTTGTTATATTATAATTAAGCGTCGAATAAGGTCTGCTGTACACTCTACCCTCCTCAAGCCCCACTTGTGGGATTACACGGGGTATATTGTCATTGATGATTGTTAAACGTCCAATATCTATAATTAGGCAGACAGCTATATATGAGTAATTTTGGTCCATTCTTGAGCTATGTTGCTCAAATCCTTCAAAAAAATGTGGCATGTCTTTGTCGGATTCTCCCAAAGGTATTCAATTTTGGAGGATCCAATACAGGTGTAACAACATTTTTGGAGGATCCGAGCAACATAGTACTTGAGTGCCCTTATACATTAACACTCAGGTTTGCTCCAAGAACAGGTCATTCTTTGTTTACATTTGCTTTCTGGTTCTCGTAGAGCTCCTCCGGTTGAACTAAATTCTGTTCGTCTTGCAGAATATGCCTACACACTTCAGGTGAACGAGAAGAGTGATATATACAGTTTTGGCGTGGTAATCCTCGAGCTAGTGACAGGGAAACTCCCTGTAGGTCCTGAATACGGGGAAAAGGATTTGGTGAAGTGGGTTTGCGCTACTCTAGACCAGAAGGGTATAGATCATGTTATTGACCCGAAACTCGACTCTTGTTTCAAGGAGGACATAAGCAAAGTCCTAAAAATTGGCCTCCTCTGCACTAGCCCCCTCCCAATTAACCGACCCTCGATGAGAAAAGTCGTAAAAATGCTGCAGGAAGTTGGTGGCGGAGACCAGCTCAAGACAGCGTTAACAGATGGCAAGTTGACCCCTTATTACCACGAAGACGCATCAGATCAAGGAAATGTAGCTTAAAGATAGTCAACATATCTTTGCTTAGATGTAGCGACATGGACAGTGATGATTCATATGGTCGAGACTCGAGACCAACTTGTCTGGCATTGAGGCATAGTCGTTGTATCGTGAAAGTGAAGATGTGTGAATTTTTTTATTTGCTAATTGCTTAATACTTTGCTATCAATACTTGATTTTTGGTTTAATTTTGGAGAGAAGGGGAAAACAGAGTTGAGATGAAGTGTAAAAGGATG

>PGSC0003DMG400012929 LRR-RLKs

GCTGCCTGCAGTAACACCTTGAAAACCGTACCTCATAGAAGCAAAAAACAAGAGTGTGGTATTGTTGCTGGTTAAATGATGAATGTTGTTGTAGTTGTGATCCTTAGTTTTTTCACATTTTCAATCTCTGTTATATTTTCATTGAATTCAGATGGTGTTTCTCTGTTAGCTTTGAAATCTGCAATTTCTAATGACCCATCTGAGTTTCTTTCTTCTTGGTCGGAGTTAGACTCATCAGCATGCCATTGGAATGGAATCACATGCGATGAAAATCAGAAGGTGACTTCTATTTCTCTGTCGTCGAAGAATTTGAGTGGTTACATTCCTTCTGAAATCGGAGCACTTTCTTCTCTATCAATTATTTCTCTTTCCTACAACAACTTTTCAAAACCAATCCCTATTCATCTCTTCAATGCTACTTCCCTTCACTCCCTTGACCTCTCTAACAATGCTTTCACAGGATTCTTACCTCAACAAATCACATCCCTTATAAACCTTAAACACCTTGATCTTTCTGCTAATTTTCTCAATGGCTCACTTCCCAAAGAGCTAACTCATCTTACACTTCTCACTGGAACTTTAAATCTTTCATACAACAGATTTACCGGCCAACTTCCGGCGAGTTATGGTAAGTTTCCTGTAACATTGAGTTTAGACCTTCGGCATAACAATCTGACCGGAAAGATACCTTCAATCGGGTCGTTGTTGAATCAAGGGCCTACTGCATTTTCCGGTAACCCTTTTCTTTGTGGGTTTCCATTGGAAACTCCGTGTGCAGAACCAGAAGCTCAAAAACTACCACCGAACCCGAACCCTAATGAGGAGGAGGTGGGGTTTGTAGGAAAACGGAAATCGGGAAATGGGTCAATGGTGGTTTCTTTGCTCTCCGGCGTATCGGTTGTGATTGGGGTGATGTTTGTTTCAGTGTTGGTGTTGAGGAGAAAATGGAAGTTGGATGAGGATAAAATAGGAAAAGAAAAGTTGGAGAAGCCAGAGGAGGGGCAAAAGGGTAAATATGTAGTGTTAGACGAGGGGTTTGGGATGGAACTAGAGGATTTGTTAAGGGCTTCAGCTTATGTAGTAGGGAAGAGTAGGAGTGGGATAGTTTACAAGGTGGTGGCTGGCGCTACGGTTGTTGCTGTTCGACGATTAAGCGAGGGGGATGTCACGTGGAAGTTCAAGGAGTTTGAGACGGAGGTGGAAGCCATTGGGAGAGTACAACATCCAAATGTGGTGAGGCTTAGGGCTTATTACTATGCAAGTGATGAGAAATTGCTGGTAACTGATTTCATCCGCAATGGCACCTTGCACAATGCTTTACATGGTAATCTCACTTCATTCTTACTTGGAATTTTCTTTACTGTTAAAAGAAAATTCACTCTAAACATACTACTTACGGAGTATTATTCTAAATCACTATAAGGATAAACTTTTAGTAGTGCTGGAGTAAGTTATCCTGCCTATGCGAGTGAGAAAGGGTGTACTTTTTTGGAGAAAATTAGATCAAATCCTCATAAGCAGTATTATCTCTGTGTCCATGGTACTTCAGGGGTTTTATTAATATATCAGACAACCTTGAAGGTGGCATTGTATTTGATATTTATGCAAAAAAAGAAGTTTAATGTCTACTCCATTGAAGTAAGTGTTTACAACCTAAGATGTCCTTAAAGCTGCTCATGATTATTCGTTGAAGTATTCTCTAGGCACATTTAACAATTTCTGTCATGGTTTCAAGCATTCTTTGATAAATATTAATGATCCCAAATTATATGCTTTTAAGTCGTTAGCTGCTTACTGTCCTTATCTTCTTTTGATCAAGAATAGGCAACCAAATGTAAATTATGATGAACAAGAACTAGTTTGGGGTAGTGGGGTGGGGAGGGGAATAATACGTGCATGGTTCATTTGTTTTTATCTGAGCAATTGGCATTTGTGTTTTCTGCTTCCTGGAAAACATCTATGCAAGTGTTAAGCTATTGAGACCTTTTGGGCTCCACAATTCAACATCTGTTGGTTTATTGTAATTCACTAGTTTCTGGTCCCCATTATGGCATTTTCCCCTTGTTTTCTGCTTCCTAGGGAGTACATTTTATGTGGCCTAAGATACATTGAGAGAGGCTTTAAAGTTGAAGTAAACATTCGGAAGTTACACTAAGGAAAAAAACATTAGATCTATCAATTGGATGTTAAGCTATTCGTTCGATTCACATTCATTAGGAATTGAACTAAGCATGGTTACACATAAAACGTGAATCCAAAGTCTGAAATTTTTGCAGAACAAAATATCGACTAATCATAAAATGTTGTAGTTCCTCCTTGTAACATCTATCATTGTCAAGTTCACTTTAGGATGTTGGACTTTATGTAAAAGAGGCATTCAGCTGACACCGAGAAGAAAACTATTACACCACAGGTTATTTTTGTCTTCAAATTCTTTTTTAACTAAGTGGAGAAGTCCACGACGTTAAGCTTTCTGATAGGTAAATATAAAACAAAAAAGATAATAAGTAAGACACGAAATTTCATCAGATTCCAAGACTTGGTAAGTCATTCCCTGTTTCCTTTAGTCTGTTTGTCTTGTTATTCAGAGGTGTTGTTTAATGTACCTATAATTTGTATTGTGGTTATAGAATCCCTGAATAGGCAGTTACTGGACTTGTGTTAGCCTTGTATTTTGCTCTCTTTATTTTCAGAACCTTGCACTTCACAAAAACATTAATGTTCATTTGAACTCCAGTCCGTTTTTCTTAACTTCAATCTTTGTGTACAAGTCTTCTCCACATAATCCTTACCTTTTTGCTCGTACTTTGGTATTGTCAATTGTGAACCAATCCTATATAAACACAAGTGGAAATGAAGATTGCACCTACTTTGGGGTGGCTTCTTGAGAATGTGGTTCCTCTAGTTCATATTGATAATTTTTTGTCTCGACAAAATACCAACTGACCTATTTCTTGTTTTCCTTGACTCAGACTTCCATTGTTTCAAGCACATATTTTTGTACCAATTTCAGGATTTGAAGTGAAAAATGTTCTATTCGTTATTCTGAAAAGCTGTCGTCGATTTGGCATGAATCACTTTTTAATTTATCTTCCACAAGTCCAAAAATATTTTTTGGATTTCAAAACTTGCATTTTCCGGCAAAAAATTTCTCATGGAAAAAAACACATACACAACTAATTTCTCTAGAGAACTGGGTGATTTTCTTTTTATGTTGGAACATTATCCTTGTTAATCAATCTTTTTATGTTTATTTTTTACAATTGCAGGAGGACTTGGTAATAACTTGCCACCTATGTCATGGGCAGCTAGACTGAAGATCGCTCAAGGCACAGCTCGGGGTCTAATGCACATTCATGAGTGTAATCCCAGGAAGTATGTTCATGGGAACATAAATTCAACAAAAATCCTTCTTGATGATGACTTGCAAGCCTACATATCTGGTTTTGGATTGACTCGCCTTGTTTCAGGCTGCTCCAAGGCCATAAACAGTAGTACCAAAAAGCTAAGTACAAGTCAAATCATAATAAGTCCTCAAAACTCAACTTCGTCTTGTACTATGTACATGGCACCTGAGGGTCGAGTGCCTGGCTCCAAGTTCACCCAGAAAAGCGACATCTACTCATTTGGTATGGTACTTTTGGAGATTCTGACTGGTCAGTTGACAGATGGAGGATCGGAGGTGGAGGATGATGGGAAGGGGCTTGAGAGTCTTGTTAGAAAGGTTTTTAGACAAGAACGACCCTTGTCTGAGATCATAGACCCTGCTCTCTTGAATGAGGTTCATGCTAAAAAGCAAGTTGTTGCAGCATTTCATATCGCTCTCAACTGCACAGAACTGGATCCTGAGCTTCGGCCAAGGATGAGAACAGTTTCTGATAACCTTGATCGCATCAAACTGCAGTAAAAGAAACTTGATCAATGTAAAGGCTTTCAATGTTTTGCTGACTGCCTAATGTAGAGCTGTTTAGTAATTGTGAATCAGAATTTTCAGTTTGTGTTGTGCCAAATCCTACATTTGGCGTTTCTATTGTAACTCTACTAATTCAATGTTCTATTTGTTGAATCATCAATTGAAAATTGCTTGTGCTAGCGAATCTTACAAAGCTATTTGTGTTTGATGGTACGGACATGATCTGAATTCCAGATAACATGCAAAGGAGCATGGTCAGACCCTGCCTAATCAAATGATGTAGTTCAATAGTTGGAAAGTTCTGTATGAATTCTTGATTGCCCATCACCCTATTCAATCTTTTGAAACTACACTCCTCTATTCTCTCATTCTACCAAGTGAAGTTGCTAC

>PGSC0003DMG400012955 LRR-RLKs

CACCCCATTCCAACAGAGAAATACAGAGAGAAAGAGCAAAAATACAAAAGAGACACCAACTCATTTGCCACTCTACCCCGAAATATAAAAACCCTAATTTCCCTCTGTTTTTTCTCCTTTCCCAACTCTTTCCCCACTCCCCACTCCCCACTCCCTCTCTCACCACCCACCTACCCTAATCTCTCTCTCTGTCTGAAGTCTGAACTCTTTCTTGAATTTCTATGGTGCTCTCTCCATAGTCATCCTTAAGCTTTTCTTCCTTTTACCCTTTTTCAGATTTTCCTTCTTTATCTCAATGCCTCATTTTTAGCACTTAAATCACATTCTCAGCCCAATGACTCTCTCTCAATCTCTCTCTTTACTCCTCCTTGTACTTGTGTTTCGACTCAATGGGTCTTCTGCTGCAGAAGATGAAGTTCGATCACTTTTGGAGTTCAAAAAGGGTATTAAAAACGACCCATTAAGTAAAATCTTCAGTTCATGGAGTCAAACGGGTCTTTCAAATCTATCAGCATGTCCAAAGTCATTTCATGGTGTCGTTTGTGATGAAAACTCCGATTATGTTTTCTCCATTTCTCTTGACGGGCTTGGGCTTGTGGGTGACTTGAAGTTCTCAACTCTTAGTGGGTTGAAGCAGCTCAAAATTCTTAGTCTTTCTGGTAACTCTTTTACGGGTAGGGTTGTTCCTGCTTTGGGTTCTATGTTAACTCTTCAGCATTTGGATCTCTCTGGAAATCAGTTTTATGGACCAATTCCAGCACGAATTAATGAGCTTTGGGGTTTGAATTACCTCAATTTGTCCAATAATAATTTTACATTTGGTTACCCAAGTGGAATTAGTAATCTTCAGCAGTTGAGAGTACTGGATTTGCATAACAATGGGCTTTGGGGTGATATCGGGGAGTTGTTTTTGGAGTTGAAGCGTATTGAGCATCTTGATTTGAGTAACAACTCGTTTTTCGGATCTCTTCCTACGAGCCCGGAAAATGTATCGCTCTCTTCGACAATTCAGGTCATGAATTTGAGTCACAACAAGTTGGGTGGTGGTTTTTTTCCTGGAAAATTGTTGGAGGCATTTGAGAATTTGATGGTTTTGGATTTGGGGAATAATGCTATAATGGGGCAGCTTCCTTCTACCGGGTTTATGCATAATTTGAGGGTTTTAAGGCTTGGGAATAATCAGCTTTATGGATTGATTCCCGATGAGCTCTTGCAGGGAACAGGTCCATTGGAAGAACTGGATCTTAGTGGCAATGGATTTTCAGGTTGGGAAATGGATATCAGTTTTTTGGAATATTTGCATTTCTTAGATCAGTATATGTTGTTGTTGTTGAATGATCATAATCAAATACACCTTTTATTTGTATAGTGTTTTTGTGGATGCCTTGAACTTCATGCCTCTTAATTTTTGGAGGTGGACTAAAATTGGAAACTTTCTTGTGTTTTAGGCATTGAAACTTTTACAGTTATGCAAAGTTGCGGGGACTCTTCACTTTTGATGCCGCACCGTGTCGGATTTTCCAAATATACACTACTTTGTAGAATCCGACACGCACCCGTTGACATTTTTGAAGAGTCCGAGCAACATAACAGTTGTGAATGTAAAATCTCGTTCTTTTTGTTTTCTTGTCAGGTTATAATTAGTTTTCTTGTGTGAGTAGTGGTTGACAATCTATATAACAGAGAATAGAAATCTTCAGCTTACGCCAAATACAGAAAAGTTTGCTTTTTTGTGTAAGTAGGTTTTAAAAAAAATGTTTTAACAAAACCAAATTTTGAACAAACTTTTACCGCCAGCTTTTTTTGTTTGCCTCAACAGAACGTGAGCAGCGTGGACTGCATAGACTAATTTGAATTGCAAAGCATGTTCAATAGAACTTGCCCATAATAAGTAGTCAACAAAGTGTTTCCTAATTGTAGTAAATTTAATAAAAGCCCCTAATGGAATAGTTGCATGAATGATTGCATCATTAGTAGCCATAAATAATCTTTTTTGCAATGGTATAACTGTATGCTCAAAGTTCATGGCAAGCTAGCTGGAACAATATTTAGCCCAGTGTCATCAAAGGCGCGCTTAAGCCCTGAAGCAAGGCTCAAATCGTGTTGAGCGCTTCGCTTTGCTTAATGTGCGCTTCTGTGTCGTTATCAAGGTTCTAAGGCATACTTTTCCTTATGTTTATATATTTGTCATTCATGCTTAGGACTAAACATATATATTTTTATTTTTTCTCCCTTTTCGCCTTTTTCCATTAAAGCCCAATCAAACCTTAGAGCTTTTTTACTCTTTTTGCTTTCGATAAGACTGATTTAGCCTTTATCTTGAAAAGTGAAAATGTTGAATTTTTTCTTCTGCTTCCAATATCCAATGCTATAGCACAATATTTGTACCTACAGTAAAATGAGTTTTATTTCATTAATGAGCTATTTGCTCTTAATATGCTCATCTTTCTATTTTGTTATCACTGCATTACACCGTCCAGGGCTCCCCCCTTCCCTCCAGTGTAGGCCACTCATGATGACGTTGGGCCTTAGAATCAGAAAATAATTGATGACTACATTTCTTTTTTGATAAGGAATTGATGCCAACATTCCTGTCTTCATTATTATCAAATCAGTACTCAAATTAGAGAAAAGAATGGAGTTGGTGAGACTTGATTATTGAAGAAAAGCAGAGCAAATTTACAAGGGTGATAATTTGGGTCTCTGAGCTTAAAACTCCTCGGTGACAATCTGATAGGGAAGTTTGTTCTGTTTCATGAGCTACAGAGAGGGTTCACAATTTCATATTTCTTTAGTTACACTTGGATATCTTTCATCATAGGACCAGAGCGTGGTTCCACTTAGAGCCTAAATGTGGAGATCCTCTGTCCAGAAGTTGAATGGTGCCGTTGAGTGATGACTGATATTTCATTCTTCTCATCTTGCATTTATTCTTTGAGGAAGCGACTCTCAGATCAGCTAGATAGTTTTGTATACTTGTCTCACTTATATACTTTTTTGCCTTCTAATTTTTTTTTGGCTATATAGTATTTCTACCATATTATTTAGCTAGAAATAGTTGCACTTAGACCTGAAAGTTAAATTGGATGTTTAATAACAATGCACATTTTTTTCAATAATATGAGCTCTTTTTCTGTTGGACTTGTAAACTTATTCTAAAGAGATATTTAAAGTTTAATATTTCGTTTCTACGCATGCAGGTTCAATTCCGATTGTGAATTCAACAAAGTTGAGAGTTCTAAATATTTCATCGAACCACCTATTAGGTTCACTGCCATCCTCCATAGGAAATTGTGCAGTTGTGGATCTGAGCAGAAATATGCTCGTGAATGGTATATCAGCTATTGAGAGCTGGGAAGCCAATTTGGAAATTATTGATTTGAGTTCAAATAGGTTAACTGGAAATATTCCCACTATAACCTCTCAGTTTCAACTATTGACTTCACTTAATTTCGGAAACAATTCTCTTGAGGGAACCTTACCTTCAGCACTGGACACCTTGCCAAGACTGGTTAAACTTGATCTCAGCACCAATAAACTTGGTGGACCGATTCCATCTACATTTTTTACTTCGACGACATTGATGAACCTAAATATATCTGGAAATCAGTTGTCAGGATCAATTCCTCTTGAAGGCTCTCATGCTAGCGAATTACTTGTTCAATCACCATACCCTGCACTGGAGTCTCTTGATCTTTCAGAAAACACTTTAACAGGTAATTTGTCTTCTGCCATTGGTAATTTGCGGAGGCTTCAAGTGCTAAATCTTGCTAAGAATCAGTTATCAGGCATGCTGCCTACTGAATTGGGTGACCTTAGAAGTTTGGAATTCCTTGATATATCTAACAACAATTTTAGTGGTATGATCCCCGAAAATCTTTCATCAAATTTGAGGGTGTTTAATGTGTCTAACAATGAATTAAGTGGTGCTATCCCTGATAACTTGAGAAACTTTAATGAAAGTTCATTTCGTCCTGGAAACTCAAATTTGGCAATCCCAAGCAACTGGTTACATGATAATCATGGCGATCCTGATCAAAATAGTCAGCACCATCATAACTCGAAATCCAGTATCAGGGTGGCTATTATTCTTGCCTCAGTTGGAGCTGCTTTAATGATTGGTGTTGTTCTCCTGGCTTACCATCGACAACGGTTCCAAGATTTCCACTTACCAAGTGGATTTAACAGCCAATCTGCTGGAAGGGATGTTAAACTCGGGAGGTTCAGCCGGCCTGGAATTTTCAAGTTCCATGGCTCCTCAGAACCACCACCAACTTCCTTGAGTTTCTCTAATGATCACTTGCTCACAGCGAACTCGAGATCTCTGTCTGGGCAAATTGAGTCTGGTACAGAAATTGTTGAGCATGTTTTTCCAGAGGGAGTAACTGCAGTTTCAGCATCTACGCATCTTGGCACTGTGGGTAATAATCCTGCAACATCTGGCCAAAGATCCTCTCCAGGCTCTCCAATAGCTTCCTCCCCTCGTTTCGTCGACACAGTTGAACAACCGGTGACATTGGATGTGAATTCACCGGATCGATTGGCTGGGGAATTGTTCTTCCTGGATGGTTCACTGTCATTTACTGCTGAGGAGTTATCTCGTGCCCCTGCTGAAGTTCTGGGTAGAAGTAGCCATGGCACATTGTATAAAGCTACTCTGAATAGTGGGCATGTTCTTACTGTCAAGTGGTTGCGGGTTGGGTTGGTAAAAAACAAGAAAGAGTTTGCAAAGGAAGTTAAAAAGATTCGATATATTAGGCACCCAAATGCTGTTCCTTTACGAGCATTTTACTGGGGACCTCGGGAACAAGAAAGGCTTATTTTAGCAGACTACATACCAGGAGATAGTTTGGCGTTGCATCTCTATGGTAAGATTTACTTATGAATAACTTCAACAATGATTTTCCATGTCTTCCATTGTTGTTTGGTTACTGTTATTAGAAAGTTACCTTTTGCTGGTCTGGTGGGTGATTATCTTGTGTTAGGACCAATATCTAGTATCCTTCGCATAAATAGTTTCTATAGCAGCATAGCTTATTTCTTGTGGTGGTTTTCTAATTGCTGAGATCTCGTGTTTTGTGGGATTGTTTGATAAAAATTATGGAGGATAAAGTTGACTTGGCCCTATTATTTATTTTTGGTTGGCCTCTCTTGCAAATGTCCCTCTTTTACATGTCTTACAATATGCAGATAGTGATGAGATCTTATTTGGATCTGTCGTCTCTTATGAATCCAGTGGTAGAGTTGTTGATCAAATGGAAAAGGAAAAAAATACAACTGCAATTGTAGAATCTGAATGTTTTTTACAAAATTATGTTTGTTAGAATGTATGTGCAGATCTTTATGTGTCATCTGGTAACCTTCAAAGGAACTCCAAAAAGGTACAAAAGATCTCTGGTTCACTAATTGCCGTACGCCAACAGCCACACATCAATAGCCCATCAAAATTAGGGAGTAGACCTATTCTACGATCTCGTTAACATGACCTAGAATGTAGTGATAGTTGCTCAACAATCTGCTTTAAATAGCATTTCCATTGTTTCTGAGTCATTCATGATGGTAAAACTTGATAAATATGGAGAGTTGGCTTTGTCATAGTATCCCCGGTGCAGTGAGGACTGATTACTTGTAGACATGTTGGGGGCTGTTGGAATCTTTTACTTTTTGTAATTTTTCGCAGTTTTAAATTTTTAATGAATGTCTACTTTTTACAGAGACAACCCCTCGGAGGTACTCCCCGTTATCCTTCAACCAAAGGTTAAAAGTTGCCATTGAAGTTGCTCGGGGTTTGGCATATCTGCATGAGAGAGGGCTGCCTCATGGAGACTTGAAACCTACAAATATAATCTTGGTTGGTGCTGATTACAGTGTCCGCCTCACTGATTATGGTCTCCACCGCCTGATGACTCCAGCAGGGATTGCAGAGCAGATACTGAACCTAGGGGCTCTAGGATACCGTGCTCCTGAACTTGCTACTGCGACCAAACCTATCCCATCATTTAAGGCTGATGTATATGCTCTTGGTGTGATTCTAATGGAATTGCTGACGAGAAGAAGTGCTGGAGACTTAATCTCAGCACACTCTGCTGCAGTTGATTTAACTGATTGGGTTCGGTTGTGTGATCAAGAAGGTAGGGGAATGGACTGTATCGATAGGGACATCGCAGGAGGAGAAGAACACTGCAAAGCGATGGATGATCTGCTAGCTGTGTCATTAAGGTGTATTCTTTCGATAAACGAAAGGCCAAACATCAGACAAGTTGTTGAAAATCTGGGTTCCATATCTGTGTGAAATTTTTGTACATGTTGTCTTGGAATTACAATTCTTTTTTCTTTTCCTTCACATTTTTTATTATGTCCTATTGGTTGCCTTCTAATGTTGGTCAATCCTTACATATTCCCCATTTTCTTTTCCTTCCCCCCTTCAATTTTTGTGTAAAGGGATGATCAGTCCATTGATTGATTGGTTGACATTTTTCTCTAATTAGAAAAGATTGCAATTATTTATGTGTAATTTATTTTTTGC

>PGSC0003DMG400012992 LRR-RLKs

ATGAAGTCACTTGAGGTTTTACTCTTAGATAGAAACAATCTTTCTGGTCCAATTCCAACAACACTAAGTGACTTAACTGAGCTCAAAACTTTGTACCTTTATTCTAATCAACTTTCTAGTTTTATTCCTAGAGAGTTGGGAAATTTGAAAAGCCTCACTTTTTTGGATTTATCAATTAATCATCTTACTGGTCCTATTCCTAGTGAGTTGGGGAATCTGATAAATCTCACAGAGATAGATTTATCTAAAACTCAACTTACTGGTTCAATTCCGATTACTATAGGTAACTTAACTGAGCTCAAAATTCTATACCTATTTTCTAACCAACTTTCTGGTTTTATTCCTAGAGAGTTGGGAAATTTGAAAAGCCTCACTGATTTGGATTTATCCACTAATCATTTTACTGGTCCAATTCCTGGTGTGTTGGGGAATCTGATAAATCTCACAAAGATAGATTTATCTAAAAGTCAACTTACTGGTTCAATTCCGATTACTATAGGTGACTTAACTGAGCTCAAACTTCTATACCTTTTTTCTAACCAACTTTCTGGTCCCATACCTAGTGAGTTGGGTAATTTGAAGAACCTCAATGGTCTGTCCTTATACGAAAATCAACTTACTGGTCCAATTCCTGCTTCTTTTGGCAATTTGAGAAGCTTGCAATTTTTGTATCTCAATGCCAACAAACTTACTGGTCCAATTCCTGCTTCCTTTGGCAATTTGAGAAACTTGCAATTTCTGTATGTGAGTGTCAACAAACTTTCTGGTTCTATTCCAAAAGAACTTGCATATTTAGATAACTTGGAAGAGCTGATAATAAGTGAAAATCAGTTTTCAGGTCATTTGCCTGAGCATCTTTGCCAAGGTGGGAAACTTGAGAACTTCACGGTGAATAGTAACAAGTTGTCTGGGCCAATACCAAGAAGCATGTCCAAGTGCTCGAGTTTCAAAAGGGTTCGCTTTGATAACAACAGTTTTACTGGAAATTTATCTGAAGCTTTTGGTATCTATCCGGAGCTGCAATTTATTAATTTGAGCAAAAATGATTTTCATGGTGAACTCAGTAGCAACTGGGGGAAATGCAAGAAGTTGACTGATTTTCGAGTAGCCAGAAATAGAATTAGTGGTCACATACCACCAGAGATTGGAAATCTCAAAGGTCTTCAAGGACTAGATCTTTCAGCAAATCATTTGGTTGGGAAGATCCCTAGGGAACTTGGAAAATTGACCTCTCTAGTTAATCTTTTTCTGCAAAACAACCAGATTTCTGGCAATATTCCTATGGAACTTGGATCAATGACAAAGTTAGAATCCCTTGATCTATCAAAAAATATGTTGAATGGGTCGATCCCAACGTTTATAAGAGATTACATACACTTGTTTCAGTTGAACCTGAGCAACAACAAATTTGGCCAAAATATTCCAATGGAGATAGGGAAGATAACTCAGCTTATTGTACTTGATTTGAGCTATAATCTTCTGGTTGGAGATATATCCCCTCAGTTAGCCAATTTGAAGGTCTTGGTAAACTTAAACCTTTCCCACAATGGCCTATCTGGGCGCATTCCTAAAGAAGTTGAAAGTTTGACTGGTTTGCAGGATGTCGTATTGTCATACAATGAGTTGGAAGGTCCAATCCCCAATAATAAAGCTTTTATGAATGCTTCATTAGAAGGTAACAAAAGTCTTTGCGGCAATGTATCAGGACTCCAGCCCTGTGAAAGACCATCTCCTGTGGTGAACAAGCACTCAATGGCGCAAGTCATCCTCATCACTGTACTTCCTGTTATGGGAGTACTACTACTGCTATGTGTTTTCATTGGTGTTCTCTTTATGTGTAATAAAAGAAGGAGAGTTAGAGACGTTGAAAGAAGGGATGGTGATGGTTGGCTTTCAATATCCATGTTAGATGGAAAGGCATTGTACAGGGACATCTTAAATGCCACAGAAGAGTTTGATGCAAAATTTTGCATCGGCCAAGGAGGACACGGAAGCGTTTACAAGGTAAACCTTCCATCATTAGGGAATATAGCTGTGAAGAGACTTCATTCTTCATATCAGAATACACATCCTAAAAGCTTCATGAATGAAGTAAGGGCATTGACTGGGATTAAGCACCGAAACATTGTGAACCTCTATGGCTATTGTTCGAAAGCACAACATTCACTCTTGGTTTACGAGTATGCGGAGAGGGGGAGTTTGTCTAGTACTTTGAGTGATGAAGTTGAGTCCAAGAAATTGGATTGGTTTAAAAGGGTGAATATCATCAAGGGTGTTGCTTTTGCTTTATCTTACATGCACCAGGATTGTTCATCGAGACATATCAAGCAGTAATGTTTTGCTTGATTCAGAGTATGAAGCTCATGTTGCAGACTTTGGCATATCAAAGCTTCTCAATCCAGACTCATCCAATTGCACAACGCTTGCAGGGACATATGGTTATGTTGCACCTGGTAATTAAGTTTAAAATATTTCATGAACTAAAGCCAAACACTCCATTGTTTATATCTCATTTACACGTTTATATTGTTTTGGAAATTGTATTGTTCAGAGCTTGCATATACTATGAAGGTTACGCAAATGTGTGATGTATATAGCTTTGGAGTATTATCTTTGGAGATAATCAAAGGAAAGCATCTTGGGGAATACATTACCGTGCTAGTAAATTCATCGACTATAGATCATGAGCAACTTAGTGATTTGTTGGATGAACGCCTTCCATATCCTGAAGATAGAGTAAAAGAGGTTTTGATTTTTATCATCAAGCTAGCATGCTCTTGTTTGCTTGAAACTCCAAAATCGAGGCCCACAATGCACTTCATCTCTCATAAGTTATCATCGATGGATGTACGTCCATCTACTCATGTAAAGCTAGTTCAATTCTTTAATCCGTGA

>PGSC0003DMG400013198 LRR-RLKs

GATTCTCTCCATTTACACATCCTCACATGCTATGTTTCACTACTCTTTCTTCTACTAGTATTTATCTCTTTCTTTTCCCTTTTCTCTTCATTTAAACTTGAAAAAAGTTGAGGGGAGGAAGAAAAAGAAGAGTTTAGCATTTCTTTGTTTAAGTTCTTCTTCTGATTGGAGATTCTGAACTTGTTTTAAGTTTATGGATAGGTGGGATACTTCACTGAGTGGATTTTACAACTACCCTTTTCGATTCTTGAATTTATATTCAGAAATTAAAGATGGGTCTAGCTGGGTTCATGACAAGAAAAAGGAAAAATGGAAGCTTTCTTCTTCTTGGCATAATACTCTGTTCTTGTTTGTTGTGATTGTGTTTTCTGTTTTTCCTATTATATCAGCTGGGAGGAATTCAGATGGGGTTATTGTGACTCAATCTGATTTTCAAGCTTTAAAAGCTATTAAGCATGAATTGATTGATTTTAGGGGAATATTGAAGAGCTGGAATGACAGTGGTCTTGGAGCTTGTGCTGGTGGTTGGGTAGGAATTAAGTGTGTTAATGGGGAAGTAATAGCTATACAGTTACCCTGGAAGGGATTGGGTGGAAGAATATCTGAAAAAATTGGGCAATTACAAGCTCTTAGAAAGCTTAGTCTTCATGACAATGTTATTGCTGGTCCTGTTCCAACTTCTCTTAGCTTCCTACCAAATCTCAGAGGTGTTTACCTTTTCAATAATCGGCTTTCGGGTTCAATCCCACCATCAATTGGAAGAATCCCTCTTCTACAGACTCTTGATCTTAGCAACAATCAGCTCAGTGGTACTATCTCTCCCAGTCTTGCAAACTCCACCAGGTTATACAGACTCAACTTGAGCTACAATGCACTTTCAGGGTCAATCCCAGTAAGTTTCACTCAATCTCCTTCTCTTACTTTTCTTGCACTTGAACATAACAATCTATCTGGATCAATTCCTGATACTTGGGGCAGTGTGGTTGTGAACAAGTCTTATCAACTTCAGTATCTTACACTTGATCACAATCTTTTATCTGGGAAGATTCCAGTTTCAATTAGTAAGTTAAGCATGCTTGAGGAGATTAATCTTAGTCATAACCTTATTAATGGGACTATTCCTGATGAATTAGGTAGTCTCTTAAGGCTTACTGTTCTTGATTTATCTAATAATACCATAAATGGAACTATTCCTGCTAGTTTCTCAAACCTTTCAGCTCTTTCTACTTTGGATTTAAAGAGTAATCTTCTTGATAGCCAAATCCCAGATACCATGTATAGAATGAAAAACCTTTCAGTGCTGGATTTGAGTAACAACAAATTCATTGGTCATATTCCAGCTACTATTGGAAATATTTCTAGGCTCACCTCACTTGATTTATCTGGAAACAATTTCACTGGTGAAATTCCAAACTCTCTTGTTTCTTTGGCAAATCTGACTTCTTTGGATGTGTCTTATAACAATCTTTCTGGGATTGTCCCATCTCTTCTTTCTAGAAAGTTCAATGCAAGTGCTTTTGTTGGAAATTTAGAGCTTTGTGGATATAGTCCATCAACTCCATGTGCTTCACCACCCCCTCAAACTCTTCCTTCTCCTGTTAGTGGGGTTGTCAAGCCTCACCGCCATCGTAAACTTAGTACCAAAGATATCATTCTCATAGCATCAGGTGCTCTTCTAGTTGTTCTGCTTCTTTTGTGCTGCATGTTACTTTGCTGCTTGATTAGGAAAAAGGCAAATTCGAGAGCGAAAAATGGTAGCAAAGCCGGTGGCCTAGCTACCACTACAGGAAGAGGTGCGAAGTCAGTTCCAGCTGTAGGAGGAGCTGAAGTTGAATCAGGTGAAGCTGGTGGAAAGCTAGTCCATTTCGATGGACCTTTCGTGTTTGCAGCGGACGACTTGTTGTGTGCCACTGCAGAGATCATGGGGAAGAGCACTTACGGGACAGCATATAAGGCAACATTGGAGGATGGTAATCAAGTTGCAGTGAAGAGGCTACGCGAGAAGATCACGAAAGGGCAGAAAGAATTTGAAGCTGAGGTTGCTGAATTAGGAAAGATTCGACACCCAAATATTTTGGCGCTTAGAGCTTATTACTTGGGACCAAAAGGAGAAAAGCTTCTTGTGTATGATTATATGTCTAATGGAAGTCTCTCATCCTTTCTTCATGGTAAAATTGTCTACACTCACTCACTGTTTGTTTGTATATTCCTTTTGGTGTTCCTTATCATGTCGACAATCCTAACATCATACTTGTGTTCTCTATATGCAGCTAGAGGTCCCGAGACAACGATAGACTGGCCAACAAGGATGAGGATTGCTATCGGTATAACAAAAGGAATCTGCTTTCTGCATACCAAAGAAAACATAATTCATGGGAATCTTACATCAAGCAACATTTTACTTGATGAGCAAAACAATCCAAAGATTGCAGATGTTGGACTCTCCAAGCTTATGACCACTGCTGGAAACACCAACGTGATTGCCACTGCAGGCACGCTAGGTTATCGTGCACCAGAGCTTTCTAAAATCAAGAATGTAAGCACCAAGACTGATGTCTATAGTCTTGGAGTGATCATTTTGGAGCTCTTAACTGGAAAATCCCCTAGCGAGGCAACGGATGGTCTCGATTTGCCTCAGTGGGTAGCTTCTATTGTGAAAGAGGAGTGGACTAATGAAGTGTTTGATGTGGAACTTATGAGGGATGCACCTAATATTGGTGATGAGTTGCTTAACACATTGAAACTAGCTTTGCATTGTGTTGATCCCACACCAACTGCTCGACCCGAAGCTCCGCAAGTATTACAGAAATTGGAGGAGATAAAACCAGAGGTGGTGTTAGCAGCCACTAGTTCTGGAGATGATGGCACAACAGTTCAAGAAAAGAGTGATGAATAATAATCTCACAAGGCTGATTGGAAATTTTTGAGCTTTTTACTTGATATCTATTTGTTTATTCTTTCATTTTTTTTTACAGTTAGTAGTGGAAATGCTGTTTTATACCAAGATCTACTAAAAAACTGTATATAGGAGATTGTGATTGCTGCATAGGTTTTACTGATGAAGAAAAACATTTTATTCAATTCTTTCAATAAAATAAACG

>PGSC0003DMG400013489 LRR-RLKs

GTTACTAACACATTCCTAGATCTCACGGTTTCCAGAATTCCAGGCTTTTTTAGTCATTGTTTTGTTTTTTTTCCTTTCTCGAGAAAAATTCTGTCTTTATTTCCTTTTAAGCTCAAAATGTTGCCAGTTTTAGCATAATTGAGTAAAAGAAATAAACTTGGCATTTGGGTGTTTTGAGGTTTTTCCAGTTCTTGAAAGTGCTAGTTGTGTAAAGACTGAAACTTTGTCATTTGGGTGTTTGAGGTTTTTTCAGTTTCTAGAAAGAGCTAGTTGTATAAAGACTGGAACTTTATCATTTGGGTGTTTGAGTTTTTTCCAGTTCTTGAAAAGCTAGTTGTGCAAAGACTGAAACTTTATCATCTGAGTGTTTGAGGTTTTCAATCTTGAAATGATCTAGTTGTGTAAAGACTGAAACTTTGTCATTTGGGTGTTTGAGGTTTTTTCAGTTTCTAGAAAGAGCTAGTTGTATAAAGACTGGAACTTTATCATTTGGGTGTTTGAGTTTTTTCCAGTTCTTGAAAAGCTAGTTGTGCAAAGACTGAAACTTTATCATCTGAGTGTTTGAGGTTTTCAATCTTGAAATGATCTAGTTGTGTAAAGACTGAAACTTTGTCATTTGGGTGTTTGAGATTTTCCAATTCTTGAAAGCTAGTTGTGTAAAGATTGAAACTTTGACATTTGGGTGTTTGAGGTTTGCAATCTTGAAATGAACTAGTTGTGTAAATACAGAGACTTTGTCATTTGGGTGTTGAGCTTTTTCCAGTTTCTTGAAAGAGCTAGTTATGAAAACACTGAAACTTTATCATCTGTGTGTTTGAGATTTTCAATCTTGAAATGAGCAAGTTTATCAGGTGTTTGTGGAGGGGTTGAAAATGGATAAAAAGGGCATTTTTTTTAATGATAAATTACGAGGTTTTGTTAGAAAAATTACTTTGTTGAGTAGGTGAGATTGTAGAAATTTGTACTGGAAAGAATGTTGTATCTGTTTTGGAATGTGGGGCTGCTACTGTTTTGTACATTGATGGAGAGTTCTTCTGCTACACTTTCTCCTGCTGGTGTAAACTATGAAGGTTAGCTTCTTTCTTGAAATTTCTCACTTGTACTGAAAATTTCCTTCTATCAAATTAAGCTTAGAGTAGAAGAACTTGTTAATTCCTCTGGATCAGGGTTAGAGCCACATGGGTGCAAGGGGTTCATCTGAATCCCCTTCGACGGAAAATTAAACTGTATATCTAAGGTTACAAATATTTTTATGTATATATAATAGACATTGAATCCCCTTGTATTCGTCCTGTGTTTACTTCTTCATTATTTGAACCCCTTAGTCCTTTGCAATGTATATGAGTATGTGACTATGTGTTCTATCTTTTTTCTTTATGATTTCCTCTCGTTATATTGAATCTTTGATGCAATGACAAACTGAAAAGTGAAAACAGTGGAAGTAATCTAATAGAGTGACTGTTTTTCCATATAATAATGTGCAGTTGTAGCTTTGACGGAAATTAAGAAGGCCTTACATGACCCCTATAATGTTCTGGACAATTGGGATGTGAATTCTGTAGACCCTTGCAGTTGGAGAATGGTCACTTGCTCCTTGGATGGCTATGTGTCTGCTCTGTAAGTTGTTTGGATGAAAGAGTATTTTAGCAGAAACTTAGTTGGCTCATGAAAAGGCCTGATTTAATCCAGTAAATTTATTGCTGAATATTTTCTCTTGCATCAGAGCACTACCTAGTCAAAGTCTAGCTGGCACCTTATCGCCTGGAATTGGCAATCTTACTAACTTGCAATATGTGTAAGTTCTTGACTATATACATCTCTTAGTTGCTGCTACTTGGTTCCGTAATACCTAAGGTCACTTCTTATATGATTTTCTTAGTGTTATTAAGACATTTTCTCGCAAACTGTTCTTGACTTGTTCATGTAGATTGTTGCAAAACAATGCCATTTCTGGTCATATTCCTGATGTGATTGGGAAGTTACAAAAGCTTCAGACACTTGATCTTTCCAACAATAAATTTGAAGGCGAAATACCCACTTCCTTCGAAGACCTGAAGAATTTGAATTATTTGTGAGTTCATGAATTTTCATTTTCACTTATTATTATTCACTTTCAAGACTCCAACCATATATTTCACTCTGTCAGATTGAATACATCATGTTACCTATTGACTCAAATCTTCTGTTTCGTCCTCATCCCTGTGATTGACTTAGCTGATCATTGCTTGCAGGGATGTCTCTTATAATAATCTAAGTGGTACCCTTCCTAAAATATCAGCGCGAGCTTTCAAGTAAGTCAGCCTTTTCTATTGCCGTAATATACTCAGTTATTTCATTTTCAAGTAGTTATCTTAATATTATTGAAGTTAAACTATTGCTGTACGGCTGTACCTGTAAATCTCGATGCTTAGTTGATAAAAATCCATTATTAAAAATACATTCACAAATCCTTTTGGGTAAACTGGTACTTGAAAAGTATTTCATTAGTATTCCGCACTATAGTGAGATTGACTTCCTCTTCATGGCTTTCAGAGTGCTCGGGAATCCTTTAATTTGTGGTCAAGGTTCTGAGAATAATTGCTCAGCAGTCTATCCAGAACCACTGTCCTTCCCACCTGATAGTCTGGCAGGTACATATAGCATTTTCATGCTCAAGTCCCTTCCCTTGCACCTGAAAATGGACAACTGTTAAAACATATTATACATACTGAGGAAGTTTTGCTATTTGTGTTATTGAAAATGTGTTGCATCTATTATTTTTGATATCACAGATCAATCTGGTGCTGCACGCAAGAGCCGTCGCGTGGTTGTTGCGTTTGTTGCAAGTTTTGGTGCTGCTTTCTTGGTCATAGTAGCCCTGGCTTTGCTTCTTTGGTGGCGCTACAGAAATAATCAGCAGATCTTCTTTGATGTCAATGGTTAGTTGATTTACTTTCCCCCCTCTTCTCTATGATAATTTGAGATTTTATTAGTTCTGCATTTCTGATTCTTTTCCTGCCACAGATATATAAGTTGAGCCAAGTGAGCTTTTCATTATTGTCTGAGTGATTGGTGTTAGGTATCATGTGAATATTTTTTAAGCTAGGATGTTACAGAGTCACCTGCTTGTGGTTTGAATCTTTATTTTAATCTCTAAAAAGATTTTTGTGATTGTTGATTTTAAATTCATTTGTTTTGTTTTAGAGAGGACATGAGTGGTCAAGTAATAGGAATTGGGGATAAATTCTGTGTTTACCATAATCAATAAACCTGTTGAATATGAAGAACTTTCGAACCCCGTCAATTAGAAAATTATGCCCTTTGGCATACTCAGTAGTTCTGGTCAATTTTCCTAGTAAGGACTCAGAAAAGTTGTTTGAATTTCGTTGTTATCGGGACAAGTCCTGATCCACTACAGGCTACCTTAAACATTTAGTTCCAAACATTGACAGAATAATTTGTTAATCACATATCTGAATAATTACACTTGCAGTTTCTTGTGCAAGTTGAAGTAAATATTTCTGTAAAAAGGAAGATGGTGTTATGTTCACTCTCTCGGGCCACCATGCCTGGTGAATTGCCTTGCCACATTTATTGCGTTGAAAGTTACTCTTGATAATCTTGTACTTTGGTTGTCTGATTCATATAAGTGGATCAGTTTTGGGACCTTTTTGCCAACCTCTATTATTTTCTCCCTGTCTCACTTCTCAAAAGATTTTCTCCATCTGTCTCTAACTAGTCCTCAAAGCTGTATCTGAGTAACGACTTCTCAAATGCAGATCATTGTTATCCAGAGGTATGCTTGGGTCACCTGAAAAGGTACACCCTTAAGGAACTGCGGACTGCCACTGACCATTTCAGCTCAAAACGTATTTTGGGAAAGGGAGGATTTGGGGTTGTGTATAAGGCCAGCTTAAACAATGGTACTGTTGTGGCCGTCAAACGATTGAACGACTACAATGCTGTTGGTGGTGAAAATCAATTTCAGACAGAAGTAGAATTGATTAGTTTGGCTGTCCATCGGAATCTTCTCCATCTTCTGGGGTTTTGTTCAACTGAAAGTGAACGGCTTCTTGTTTACCCCTATATGCCAAATGGAAGTGTAGCATCACGATTAAAAGGTTAGCTCTTGCCTCCATGTGTTAATACATAATAATCACTATTGATTTCAGATAATGAATCCAACATTTCCATTTATAAAAGTTAAATATTCTCACATGGTGTTTGTTTGTTACTACTCCCTTTATTCATTTTGTTTGACAGATAATTAATAGGATACAGAGTTTAAAATTTTAATTCCACTCATTTATTACATTGGAGGAGAAGAATTTTTTTAAATAGTGGTTATTATACTTGTTAAAGAGAACATCATATAAATTTTAGAATGTGGATAGTATGAATTTGAGTTCTTAAAGTTGTAAAAGGTGTACGGACAAATATAATACTAGAATAGAGCCAAATTTTGGACCATGTCTAATTTAAAAGAAGTCATGTTAGTGAACAGTTCCTACTTCCCAATGTATTGCCATTAAACATCAGTGAGTGCTCTGAAAACCAGCAACCGCTGTCATTGGTGAATTGAGGCCATAAGATTCTGGTTAAGTACAAGCTTAGATGTTACGACCCACCCTTACCTCAGTCCATCCTTGAGCCCGCGGTGGGCTGCCCACCTTAACATACTCTGTACTGCTGTAGTTATATGATAAGACATTATCAGTTTGCTGCTGGCTGAAGCCATTGTTGTATGTTTCTGCATGTTCGAAAGCCAAGCTCATTTGCTTTTCTGGAACTTCATCAGCTACTTTTTAATAGTTTCCAAGTAACTAAATTTTGTGTCAGTTGAAGCTGATACATATATTGGCTTGATTGAAAATGCATTGTGGTTTCTAAATTTGCAGATAGCATAAATGGTAAACCAGTTTTAGACTGGTCAAGGCGGAAAAAGATAGCACAAGGTACAGCGAGAGGGCTAGTTTACTTGCATGAGCAGTGTGATCCCAAAATTATCCATCGTGATGTCAAAGCAGCAAACATTCTGTTGGATGAAGAATTTGAGGCAGTAGTTGGTGATTTTGGGTTGGCAAAGCTCTTGGATCACCACGATTCCCATGTGACTACTGCTGTTAGGGGCACCATTGGCCACATTGCTCCAGAATATTTGTCTACTGGCCAATCGTCTGAAAAAACTGACGTTTTTGGGTTTGGAATCTTGCTACTTGAGCTGATTACAGGCCAGAAAGCTATGGACTTTGGTCGAGGGGCCAACCAGAAAGGTGTTGTTATGCTTGATTGGGTGCGTCTCTCCTCTTTCTCTTTCTCTGGATGTTGTACTGAGTACTTTTTGAAATATGTACTGTGGGTTGTGAGAGGAATAAGCTATACAGAGGCAGGTTGTGCGTCTCTCCTCTCTCTCTCTCTCTCATAGCTGTTTTCTTTTCAGTTATAAGGAGTTGTAGGTTCAAACCTCAGATATAGCTAGCAAGTATTGTGTACATCTAGTTTACAAGGCTCTTTTTCCATTAGCATACTTCAACGAACTCTCTCGTTTAATTACTGCTAATGATATGCAGGTGAGAAATCTGCATCAAGAGGAGAAATTGAACCTTATGGTAGATAGCAATTTGAAAAATAACTTTGACCAGATTGAGCTAGAAGAGATGGTTAAAGTTGCCCTTCTATGCACCCATTTCAACCCTTCTCATCGCCCAAAAATGTCCGAAGCATTAAGGATGTTAGAAGGCGATGGATTGGCTGAAAAATGGGAGGCATCACAAAAGCTCGAGACTCCAAGAAACCGAGCTTTTGAACATACACCACAAAGATATTCCAACTTCGTCGAAGACTCTTCACTAGTTGAACCAATGGAACTTTCTGGACCTAGATGACATCCATTTTTCGTTCTCGATTCTTATCATCCTTGAGTTGCTAAAGTTGAAGTGAGAAGTATTTCATTGCTGTTGAATGTATGTATTCATTTTGTAAAAGAAAAGGGAATAAGTCATGTGTAAGCTGAGTTCATATATGATATATTCCCTCTTGTTCTTTCAATTTTTGAAATGATATGCTATGTAATTATGATTTCATTTGTTCTTTTTATAAGAAAGATTTTGAGATTTATGAGTAATAAAACGCCATAGGCTTGCCATTCTCATTAAGT

>PGSC0003DMG400013500 LRR-RLKs

ATGGTTGGTTCTTTACCTCTCAAAATCGGAAATTTAAAGGCTGCAACACTGATAAATCTGTCAATGAATCAATTCTCAAATGGAATTCCTATAGAAATTGGAGGATTGCAGAATCTGGTGCACCTTTCTTTGAGGCACAACAAGTTACAAGGATCTATACCTGACTCATTGAGCAACATGGTAGGTTTGGAATTCCTAGACCTTTCTCATAAAAATATATCGGGAACCATTCCTATGTCTTTGGAGAAACTTCAAAACATGATGTATTTCAATGTTTCTGTCAACAAGTTGCATGGTGAAATTCCCTCGGAGGGTCCGTTCAACAACCTCTCGAGTCAGTTTTTCATCAACAATGAAGCATTGTGTGGTTCTTCAAGGTTTAGTGTCCCGCCATGCCCCACATCATCAAAGCATAGATCAAATAGGAAAAAATTGCTTGTTCTATTTCTTATCCTCGGAGTTGCACTTGTATTTGTTCCTATTACCCTTTTGTTTGTATGGATAAGGTATAGAAGAGGTAAGAAATACCCTCAACAAGATGATTCATTGTCTATTGCAACGAGAGGTAGAATTTCATACTATGAATTGCTCCAAGCAACTGATTCTCTCAGCGAGAGTAATCTGATTGGTTCCGGAAGTTTTGGCTCTGTCTACAAAGGCATTCTCAGAAGTGGGACTGCTATTGCAGTTAAAGTGTTTAATCTGCAACTGGATGTGGCATTAAAGAGTTTCGATACAGAATGTGATGGTTTGTGCAACCTTCGCCATAGGAATCTCATAAAAGTCATCACTAGTTGTTCCAACATTGATTTTAAGGCTTTAGTGCTCGAGTATATGCCTAATGGGAGTCTTGAGAAGAATTTGTATTCTCACAACTACTTCTTGGACATCCAGCAAAGACTATGCATAATGATAGATGTGGCATGTGCATTGGAATATCTCCACCATTGTAACATCCGGCAATACGAAATAATTGAAGAGGCTTAG

>PGSC0003DMG400013525 LRR-RLKs

GTAAACTTCCATCCAGCTGCAAGTTGTAGCTCATAATAAACCTCTAAATTATGTCATAAATTTCAATTCTCTTAGCTTAATTCCTTCAAATTCTTCATTAATGGGGTTTTCATCGAAATGGGTTTTTGTATTTACGTTATTTTCATTCATAAAACTTTCAATTTCTGATCCAAGAGCAACAGAAGCAGCTCTAATTTGTACAAACAGAACAGCTGCACAATCCGATCGACAAGTTTATGTAGCAACTTTTCTAGAAGCCATGGATTCCATTACGCCATTAGTGGGGCGAGACAAATTTGGAGCAGTAATTAACGGAACAGGGAATAATACAGTTTATGCGTTTGGTGAATGTATGAAGGACTTATCACAAACAGATTGTAATCTCTGTTTTGCTCAGTGCAAAACTCAAATCCTACGATGCTTACCTTTTCAAAGATTAGTTACTGGTGGGAGATTATATTATGATGGGTGTTTCCTAAGGTACGATTATTATAGATTTTTTAATGAAAGTTTGAGTTCTGTTGATAGGACGATCTGTGGGAACACGAGTTTTTCTGGGAATCAGAGTTTGTTTAGTGAAAATGTTGGGAAATTGGCGAGGGATTTGGAGTTTGGGGGTTTGAGGAATGATGGGTTTTTGACTGCAGTTGTGAGTACTGGGAATTTGACTGTTTATGGGTTAGCTCAATGTTGGGAATTTGTGAATGGAAGTGCTTGTCAAAGGTGTCTTTCTGATGCAGTTTTGAAGATTGGTTCTTGTACTCCTAAGGAAGAAGGAAGGGTGTTGAATACTGGTTGTTATGTTAGGTATTCTACTCAGAGGTTCTTTAATAATTCTGCAAGTGATACTCCTCCTGCTGGAAATGGAGGTATGTTCTTTCTTTCTTGAATGTATGAAATGAATAAATTGTTTTCGTTTAGTTGTGCGATCATTTCACCTAAAAGTGTAAGCTATTAGAGATAGTACTTTTTCCGTGACATATGTCATATCTTTACATGGTATCATTGGGTCTGACCGTGAAGGGGTGTTAGAATGGTTAAAGTTTCACATTGGTTGGGGACTGACTTGAGCAATTCTCTCGTGATGAGCTAGCTTTTGGAGTTGAGTTAGACTCAAATGTCATATATCTTTACGAATATCATGTTTAATTGTGTAATCATCTTATTTAACATATGCTTGCGATGTTTCTTGGTTTCATTAGTTATTTCAATCTTATCCAATATTCAATGATCTTAAATGTTTAATTTTGTTTTTGTAGATTTGCAATGATACTTGCTATAAGCCTAAAACTATTGGATGTAGAGCAAGAAAAAGTTCAACTTGTTCTGCCCAATTATTATTTTCTAAAAAAAAGATACCTACTTTGTCATTTGGAATGATTTGAAAAACGGTGGTTGGAAATCGAGGTGATTTTGACATTTCTTGAAAAGTTGCTCTTGTTATCTGTTTAAAATGTGGTAGTTTGTATTTCTTGGATTTTAGTTCCATTCCAGGATTTAATAACTATCTAGACATTACTTGTGCTGATGTTTCTTTTTGTCATCTGATAAGTTGAAATGGTTGACATATATATATACATGAAATGAATGAAGAGTGAATCAGCATTGAGGCAAATGCAGACAATGAATCTTTTAAAAACTGTGATAAAATAAAATGTTTATACAAAATTATGTCATTCATGTTGTCGTATTATGATTAATTATCAGGAGGATCGAATCGTTTGGCTGTTATTCTTGCAACAACTCTGGGTATTTCTGCTTTCTTGCTATTTGTATCGGCTGTTTCATTCTTTGTGCGGAGGAAGGTTCTAAAACAAAAGAGAGGTACAAATTTCTAAGTCAAATCATTTTCTTTCTTTCCGACAAAAGAAATCTCATTTAAGTCTGATTGATCAAATGGCAATTTGACTTGAATTTGCTTCGATGATATAGAGAAGAAGCAGCTAGGAGCTCTCATAAGAACGGTTAATAAATCAAAACTGAATATTTCCTATGAAACTCTTGAGAAGGCGGCAAATTACTTTAACAACTCGAATAAATTGGGACAAGGAGGTTCTGGTTCTGTTTACAAGGTAATTTTCTTTGAACATTATATTCATTTTTCTTATACTAATGAGATTTGCCTGACAATAGTCGTGTTTATACATGAAATAGGGGATCTTGCCAGACGGGCAGGTTGTTGCTATAAAGAGGCTCTTCTTCAATACGAGGCAGTGGGTTGATCATTTCTTTAATGAGGTCAATTTGATTAGCGGCATCCATCACAAAAACTTGGTAAAGCTATTGGGTTGCAGCATTACAGGGCCGGAAAGTCTTCTAGTGTATGAGTACGTGCCCAATCATAGTCTTGCGGATTACGTCTTTGGTAAGTATCCAATCCACCTAAATTTATGAAATCTTAAATTGGTGTAGTCGTCTCTCGTTATGTAAATGCCTTTGTTTTGCACTGAATCAGACACAAAGAACCTTCAGCCACTTACATGGAGTCAAAGATACAAAATTGTATTGGGTACTGCTGAGGGATTGGCTTATCTTCATGAAGAATCGAAATTAAGGATCATCCACCGTGACATAAAGTTGAGCAATGTTCTTCTAGATGAAGATTTCATGGCAAAAATAGCTGATTTTGGTCTAGCCAGGTTATTTCCTGAAGATAAGAGCCACATTAGCACAGCTATTGCTGGCACTCTGTAAGTTCAATACCTTTTTCCGGTTACTTGTTAATTTTTCTTTAATCAATACACTAAGAAAGTGCTGAATTAATAAGTTTTGGCATTGCAGCGGTTACATGGCTCCAGAGTACGTTGTACGTGGCATACTGACCGAAAAAGCTGATGTCTATAGTTTTGGAGTTCTTGTTGTCGAAGTCGTGTGTGGAAAGAAGAACAATTATGTTTTCAAGAACACTAATTCTCTTCTGCAGCAGGTACTTTCTTGTGATATTTTTTCTTTCCTTAACGGACATTAATATAGTATTTACTCAACTCATGTCATTGTGGTTTCTAGTTGTGGAACTTGTATGGGATGGGGAAATCTTATGAAGCAGTTGACCCTTTATTAGAAGGTAACTTCAATAAAGAGGAGGCATCTAAGCTGCTGCAAGTAGGTCTTCTTTGTGTACAAGCATCAGCAGAACTACGACCCTCCATGTCAAGTGTCGTGAAAATGCTCACCGAGAACCATGAAATTCCTCAGCCAACACAACCACCATTCCTCAATTCTGGTAGTTCAGAATTTAGTCCATTTAATCTGCATGGCAAAAGATTTTTCGGTCAACCAAGCTCCTCTACACAATCTTCTGGGAATAAATTGACGGAGAGTTGGATTGAGCCCAGATAAACTAAGTTGATCATCTGGAATTGGTATTTGATTTTGTTCTCAGCAAAGGCCAACTTCTTGAAGGTGAATTGTTCAGCCAACTCTAGTTTTTTCTGTGATGAACTAGTACTCACTACTCACTCTCCATTTTCTTTACTGATATTTGTTTATAGTCTTATTTAGTCAACTTCCTTTTGTGAATACCCCAGTGAGCTATTTCATTTCTAATGTAAAAATTTTGTATAGTATCTGCAGGAGTTCCAATTAACTATATCTACTGTTTTCGATTTTGGATTTTGGATTTTCGATTTAGTAATTTTCATCAAGGATTATGCTTCACAATTGTCATCATCTTGTCTTAGAAGTGGCTATTCAGAATCCCCTGCCTAAAGTCACAGGCATTGGTGAAACTATTCATTACAATTACATCTACA

>PGSC0003DMG400013682 LRR-RLKs

CTAAGCTTCTCTCTCCCTCACCTCGCTCCCATCTCTCTCCTCTTTGTGCTTCCCCATGCATGCCCACACAATCCCATTTCCCTAGAGATGTTGAAATTCTTTTTGCATGCCCTCTAAATCATAGTATTTTCTCATTTGTAATTGCCCTTTGTAAAGGGCGCGCACCAACAAGTGTGCGCACCCCTACAACTAGTCTATGCAATGTCGGTGGCTTATCGTTATAGCAACCACAATCGCCAGTATCGTCATCACCATCACCATCTCTTAATCCTCGTTGTGTTATTATTACTAGTCGTTGTCCCGATCAAATCAGATAGTTCTGAGGCAGAAATCCTCTTTCGATTTAGCAAGTCCTTGCAAAAAAATGGCGCAACAGCTAATTGGAATACGAAAGTATCGCCTTGCGATAAAAATACAGATAAACCCAATTGGGATAGTGTTATTTGCGAAAACGGATTCGTTTTCGGGTTGCAATTGGAGGGTAAAGGCTTAAGTGGGACAATAGATGTGGATGCTCTTAAAGATTTGCCTAATTTCAGGACCATAAGTGTTATGAACAATAATTTTGAAGGACCTTTACCGAATTTAAGTAAACTTGCGGGGTTAAAAACGGCATATTTCACAAATAACAAGTTTTCTGGCCAAATTGACAATACTCTTTTTGAAGGGATGCATTGGTTAAAAAAACTTCATCTTGCTAATAATCAATTAAGTGGTAAAATCCCATCTGTTTTGGGTCAATTGCCTAAGCTTACGGAATTGAGAGTTGAAAACAACAAATTTGAAGGCCAAATACCTGATTTTGTTCAAGAGAGACTTATGGATATGAATTTTGCTAACAACTCTCTTGAGGGTCCAATTCCTCATGGCCTTACTAGTCTCAAACCTTCAGCATTTGAAGGTAACTTTTTTTTTCATGTGTAGGATAAATTCCATTTTTTGAAATTATTGTAACACTTATCTAGCTACGACGTACACAAATGTCTACATTTTAAAATGTAGTCGTAAAGATATGTCATGTATGCCTAACTCCACCTTAGAAGCTAGCAAGGATTGATCAAGTTCATATCATCAGCAGATCATCAATCCATTTCCCAATCAATGTGAGACTCTTTAACACTTCTTTCACGCCCATCTCCCAAAACGGAGATGAACCTGACTCTGACACCATATAAAGATATGACGTTTGCATCTAACTAAATCTCAAAAGTTAACTCACGAGGAAAAAATTGTCAAAGTCCATATTAGCAGACAATCAACACATTCCCCATATGGCGCTTTAACCCACATTCTAACAATAGTTCTTCGTCATATTGGCATTAGAAGAATTGTAGTTCATAACAATTTTTATATAGTTTTTAAATATCTAAATTTTAACTTCAAAATATTAAGTTAATCTAAACATGCTCTCATTTTTGTTCATTTGGTACAATAATACAGGTAATGATCTTTGTGATGGTCCATTATCGAAATGCACGAGTGAACCAAAAGTAGCCCTCTGGACAATTATATTGGTGGTTATCGCAGTTGCAGCAGCAGTAGCAGCAATTGTTGTTGTTATCATCATTCTTCGTCGGGGCAAGCAGACACCCGAAACAGAAACAAGACCAACGCCTACACCCTCTGGAGCTGCAGGGGGCGCTACTAACCAAACAGGAGCGCCCTCAGCAGCAGATCTAAACAAAATGGAACAGGGCAGCAATCAAGCAATGGCTGCTCCTGCTCCTGACCAGTCTCCAGAGGGGACCGGTGGACTAAATAATAATAAGAGGCCTGAAGTACAAGCAATACAAAAACTTCTATTCTTAAAAGATGATATTGAAAAATTTGATTTGCCAGATCTACTAAAGGCTTCAGCTGAAATATTGGGTAGCGGTGTATTTGGTTCGACTTATAAAGCTGCACTTAGCATGGGACGTGTTATGGTTGTCAAAAGGTTTAGGCAAATGAATAATGTAGGCAAGGAAGATTTTCATGAGCATATGAGAAGGCTAGGTAGGTTGAGTCATAAGAACTTACTTCCTGTTGTAGCTTTTTACTATAGGAAAGAGGAAAAGCTTTTGGTCTCAGAGTATGTCAACAATGTCAGCCTTGCAGTTCATCTTCATGGTACGTATTTCATACTTTTAAGTTTCCGCGCTAAAATTAGATCATAAAATCATGATTTGCGCTACCAGTTTATTATATTAACTCAATCTATTTTGACCTTTTTTTTATTCAATTCATTTCAAAACAGAGTTGAGATATGCAACTCAAATCTTTTGAAGAGGGCTCATTTGTTGTTCTCTCTACCTACTCTACTAGCTAGAGTGATGATCTGATGTGTGATTATCTCATCTAAGATATTACATTTTTACTATGATGAATTCTGTGATCATCTCATTACACATTGTTTATTTTTTTCAGGTAATAAATCGCGGGGGCAACCAAGTCTTGATTGGCCAACTCGATTGAAAATTGTAAAGGGAGTAGCCAAAGGACTATTGTACCTCTACAACGAGCTACCAAGCTTGACAGCACCCCACGGTCACCTCAAGTCCTCGAACGTACTTCTAAACGAATCCTACGAGCCCCTCCTCACGGACTATGCATTACTACCCGTAGTAAACCTAGAACACGCTCAAGAACACATGATTGCTTACAAATCTCCTGAATTCAAACATAGTGGTCGTATCACACGTAAAAATGATGTGTGGACGCTTGGAATATTAATCCTCGAGATCATAACGGGCAAATTCCCGTCTAATTTCTTGCAACAAGGCAAGGGGAGTGACACGGATTTGGCTACGTGGGTAAGGTCCGTGGTAAATGAGGACATGATGGAGGTAGATGTATTCGAGAAGGAAATGAGAGGTACGACGAATTCAGAGGGGGAAATGATGAAGTTATTGAAGATAGCTTTGGGTTGTTGTGATCTTGATATGGAAAAGAGGTTTGATATGAAGGAAGCAATGGAGAGGATTAAGGAAGTAAAAGAGAGAGATGGGGATGATGATTTTTACTCTACAAGAGGATTATCTGATGATTATACGCAAGTTTCGATGAACATTTGAAGAGTGTTGGTTATACATACTCATAGTTCGTGTGTAACACCTTTTTATTAATTTGTTCAAAAATCATCAATGGTAAATATGGGATGGGACTTTTATTTTGTTTGATGCATGGTTGGGGGTCGTATATATTGGTCAATGAAACCACAAAATAAATAAATATGAAAACAAAAAAAAGAGATAGAAAAAATGAAGTTCAGTGTTTTTTTGTTGCGATTCAATTGAAGCAAAGGAACATGTAAATTGTCCTTGATGAATCTAGTTTAGCTAGTTGCCTTCACTTTTTTTCTTTTTTGAAACTTTATCTAGTTTATTATATTTGGGGTTGTAAAAATTAATCCAAAAGCATGCAAGAACAACGAATTATGGATCTTTCAATTGCAG

>PGSC0003DMG400013953 LRR-RLKs

ATGGTTGTGGCCTTGGATCTTCCTAATTTGCAACTTCAAGGTACAATTTCCCCTTCTTTGGCCAATTTGTCCTTCCTCAATGTTCTCAATCTCCAGAACAACAGCTTCCACGGTGGCATCCCTTATGGCATTGGCCATATGCCTCGCTTGCGAGTCATTGACGTTAAAAACAATCAGCTCAACAGAAGTATACCAACAAGTCTATTTCAAAACCGGAGAGTTCAAGTAATTTCATTGGATTTCAATGAACTCAGTGGTGAAATCTGGAGAGGCCCTTGGTATGTACCCGAACTCATAGTCTTAAATTTTAGGAACAATAGCCTCACGGGTATAATCCCTCCTTCTATTGGAAATGCCACAAAAATGATGATATTCAGTTTGTCTGGGAATAGAGTCAGCGGCAACATTCCAAAGGAGGTCGGTAATCTGATCCAGCTTGCATTTTTTTCCTTGTTCAATAATCAATTAACAAGTTCCATTCCCGCAACACTGTTTAATATCTCGTCACTACTTGTCGTAACTCTGGCAAAGAATAGCCTTTCTGGTCCTCTCTTGCTAGATGAAGGCAATATTGTGTTGAATCTGTACTTTTTAAGTATATCTCACAACCAAATTTCTGGTTGCATTCCTTCCAACATATGCCAACTCACGGAGCTCCAAGTTTTGTCCTTATCTTTCAACAACATAATTGGAGACATACCCAGAAATATTAGTTGTTTATCCAAACTCGAGAAGTTTTATATTGTTGATAATCTAATAAAAGGACTATTCCATTCATTGGAAATATTCCACTCTACATACTCTTAACTGTCGAAACAATTGGATAGTGGGGGGAATTCCTCCGGAATTAGGGAAGCTATCAAATTTGATGAAATTATGCTTTGACAATAATTCTAATCTTATTGGTCAAATTCCAGAGGCTATTTTCAACATATCTTCTTTGGAAGTGATTGGTTTCAGTTTCAATAACCTCTCCCCAACCTTGAAGGACTTTACTTGGGAGTCAATCAGTTCAAAGGGAAATTCCATTGTTTATAACAAATGCTTCCAAGCTTCAGATACTGGAGCTAAATAATAACTTTCTGACAGGAACTATTCCTAATAATTTGGGAAATCTTCGTCAGCTGCAAATGCTGCTCCTATATAATAATCAACTTACCAATGAATCAAGAGAGCGTGAGTTGTTACTCTTCAATTCTTTGGCGGACCGTAGGATGTTGAAATATCTACAAGTGGGTAACAATCCCTTGAATGGCGTTCTGCCCAATTCAATTGGGAATCTTTCATCTACTATTGAAAACTTTCATATAGAAGATGCACACATCAATGGCCTCATCCCCAAAAGTATAGGCAACATGACCGGTCTTACATCCCTAAGCCTTGACGATAACAACTTGACAGGGAGTATTCCTTCTGAGATTGGTAAGCTTAAACAACTCCAAGGGCTGTATCTATATAACAATAAATTGCAGGGACATATTCCAGAGGGGGTATGCCTTTTATCTAATTTAGACATATTAGCTCTAGAAGGTAATGAGCTCTCTGGATTAATTCCAGAATGCTTTGGAAATCTTAGCTACAACATCTTTATTTGGATTCTAATAAATTTTCATCAAAATTTCCGTTGAGCCTTTGGAAGATGAATGGTCTTCTCTATCTAGACGTGTCACAGAATTCAATAGAGGGAGAAGTTCCCCAGGATATTGGAGGACTGAAAGCCATTGTAGAACTACATCTTTCCAGTAACCACTTTTCAGGCATGATATCAACCAGATTCGAGGAATTCCAAAATCTGCAGTATCTTGACCTATCGAACAATTCATTTTTTGGCCGAATTCCATTATCATTTGCCAACTTGATAAACTTGGAATTCTTGAATCTGTCTTTAAATGCATTGTCAGGTACTATTCCTAAGTCATTGGAAAAACTCTCATACCTTAAAAGCATTAATGTTTCATTTAATGATTTAGAAGGTGAAATACCAAGTGGTGGTGTGTTTGCAAATTCCACTCTGCAATCATTTCTTNNNNNNNNNNNNNNNNNNNNNNNNNNNNNNNNNNNNNNNNNNNNNNNNNNNNNNNNNNNNNNNNNNNNNNNNNNNNNNNNNGGAATGCACATATTGGAGATTCCTGCTTGTGCTATCACTAATCCTGGAAAACAATCAAAGCTTAAGGAGGTTTTGCTAAAAATTGTTACTCCAGCGGTTACTTCATCCTTTCTGATATTCTTGTTGGTTTCAATTTGGATAATGAAACGACAAAAGAAAGGGAAGTCCAAAGATGTGGAGAAGGTACTGGAGATCGGGACTCATCAATTAGTTTCTTATCACGAGATTCAACGAGCAACAAATAATTTTGATGAATCCAATTTAATTGGTGAGGGAAGCACTGGCTCTGTGTACAAAGGCACATTATTTGGTGGAACTGCAATGGCCATAAAGGTTCTGGATTTGGAAAATGAGCAGGTATGCAAAAGGTTTAATACCGAATGCGAAGTGATGAGAAATGTTAGACACAAAAATCTTGTTCCAGTGATTACTACATGTTCTAGTGACTATATAAGAGCCTTTGTTTTGCGATATATGCCCAATGGAAGTCTTGAGAATTGGCTGTACAAAGAAGATCGCCACTTGAACCTTCATCAAAGAGTCACTGCAATGCTTGATGCAGCTATGGCAGCTGAATATCTACATCATGGTCATGTCACTCCAATAGTTCATTGCGACTTAAAGCCATCCAATGTACTTTTGGATGAAGATATAGTGGCTTATGTTGGTGAATTTGGAATCTCTAAAATTTTATCCATAAGCAAGTCCATTGATGGTGTTCTTCCCGAGCTGCATGTTAATATGATGCTTTTTGCTTACCTGTAG

>PGSC0003DMG400013956 LRR-RLKs

ATAAGAGCCTTTGTTCTGCAATTTATGCCCAATGGAAGTCTTGAGAATTGGCTATACAAAGAAGATCGCCACTTGAACCTTCATCAAAGAGTCACTGTAATGCTTGATGCAGCTATGGCAGTTGAATATCTACATCATGGTCATGTCACTCCAATAGTTCATTGCGACTTAAAGCCAGCAAACGTACTTTTGGATGAAGATATGGTGGCTCATGTTGGTGATTTTGGAATCTCTAAAATTTTAGCCATAAGCAAGTTCATGGCTTATAGCGAGACATTGGGGACTCTTGGATACATTGCACCAGGTATAAAAAATCTACTCTCATTGATTTTCTCTTATCATAATAAAGCCTCTCCAAGTTATAGAAGTAAAAAGAGAATTATTGTTGTATTTCAATTGAGTAACTTTTCTTCAATCCATTTTTAAGAATATGGCTCAGAGGGAATAGTGTCCACTAGTGGTGATGTTTATAGCTACGGAATCATGTTGATGGAGGTTTTGACGAAAAGAAGGCCAACAGATGAAGAGATATGCAATGGAAATCTTGACTTGAGGAAATGGATTACACTATCATTTTCAGGTAGTATGATGGACGTTGTGGATGTCAATCTTTTTTGTGAGGAAGAACAAATCACTAGTAAAAGTGAAATGTGCATAGCGTCCATGATAGAATTGGGTTTAGACTGCACAAAGGAAACGCCAGAATCAAGAGTGACCATGAAAGAAGTAGTGAAGAGGCTTAACAAAATCAAGAACGCATTTCTGGAAACATAGAAGTGATCAGCATCTGTTCTTTACTGAGCTGCAAATCAGTTGAGAAAGTAATTATGTGAGGGCGTTAGGCCAGCGGTAACAAGGAATGCAACTCATGCATACTATTAAACATATCTTTTTCCGCAGTCCCATAGATTAAGAAGTTAAGCCTGCTAATGTATTAGTGAGAACTAAAATGATATGATTAATATGTAATTCACATTAGATTTCTTATCTTTAACACATTTTGTGCACTCTCTGTTTAATAGATCAAAAAGTGCCATGATCCACTTTTTGGCAGTTTTTTCCTCACTTGTTTCTGTGAGGATCAGGAACTCGGTAATTAAACTGCAGTTTATTCTCCTATTATATTTTATCTACAAGTTAGGCATGTGTACAGATTGTTTGCATCAGTAACAGATTGATACCGAATTCAAAGGAAAAATATATGGTGGCTCATTTTGCAGCTGCTATATTCAGGATGGAGCCGGTAATA

>PGSC0003DMG400014281 LRR-RLKs

ATGGCTACTGGGTTTTCTTCTACTACTGCCATTGTTGTCTTTCTTCTCTGTTTCTGTGTTTCTAATGTTGTTTTACAAAGAACAGAGGAGGACCCAGAAAGACAAATCTTGTTTTCTTTCAGGAGTTCTCTTGAAAATCCTTATGTTTTGTCTACATGGACCCCCACAATTTCACATTGCAAATGGGATGGTGTTTTCTGCCAAAATGGTCAAGTTGTTTCCCTTATTCTTTCTTCTTTGTCACTCAAAGGGCCTATTTCACCCCACATTGCTTCATTGAAGAGTTTAAGAGTGTTGGATTTGAGTAATAACCAGTTGTCTGGTGAGTTACCAATTCATCTCAGTGAACTTTCTCTTTTAGAAACAATTAAACTTGGTTCTAACTGTTTTACTGGAGAAATATCGCCGGAGTTTGGACGGTTGACGGAGATGAAGTTACTTGACCTTTCAGGCAATGCACTGACTGGAAAAATTCCAGCCCAACTTGGTCAGCTTACTAAGCTACAAGTTTTGGCACTTGGAAATAACCTTCTTTCTGGTTCTTTATCTGCAACACTTTTTACAAAGCTTCAATCTTTAACTTCTTTTGATGTTTCTAATAACACCCTTTCTGGTATTATTCCACCTGAAATTGGGGGATTGAGAAGCCTTACTGATCTTTACATTGGGGAAAACAGATTCTCTGGCCATTTGCCAGCAGAAATTGGTGAACTTTCAAGACTCGAGATTTTCTTGGCTCCTTCATGTTTACTTGAAGGTCCATTGCCTGAATCCATCTCAAAGTTGAAATCTTTGAAAAGATTTGACCTTTCTTATAACCCATTGAAGTGTTCAATCCCAAAAGCTATAGGTAGTCTTGAGAATTTGACTATACTGAACCTTGCTTACTCTGAGATCAATGGTTCAATACCTTCTGAGCTTGGAAAGTGTAGAAACTTGATGTCAGTAATGCTTTCTTTCAACTCACTCTCTGGGTCTTTGCCTGAAGAACTTGCAGAGTTGCCTGTGCTATCTTTTTCTGCAGAAAACAATCAGCTTTCTGGTGCATTGCCTTATTGGCTTGGAAGATGGACTCAGATGGATGCATTGTTGCTTTCGAGTAACCGGTTTTCTGGGAAAATTCCGGCTGAGATTGGGAATTGTTCTATGTTGAGTCATATTAGTTTGAGCAATAACTTACTGACTGGTCCAATACCTAAGGAGCTTTGCAATGCCGTGGCACTCGCGGATATTGAGCTTGGCAATAACTTTCTCACAGGCACCATTGATGACACATTTGTCAAGTGTGGCAATCTGAGTCAGTTGGGATTAATGGATAATAGTATTGCTGGGATGATTCCGGAGTATCTATCACAGCTTCCTTTGGTGGTTCTTGATTTGGATTCCAACAATTTGACAGGTCCAATTCCGGTGAGTCTCTGGAATTCGACTTATATGTTGGCGTTTTCTGCTGCGAATAACCGGTTATGGGGTACTCTACCGGTGGAGATTGGCAATGCTGTGTCATTGCAGAGTCTTGTTCTTAGTAACAATCAGATAACTGGTGTCATTCCTAAGGAGATTGGTAATTTAACCTCTCTTTCAGTGTTGAACTTGAATTCCAATCTTCTTGAGGGTTACATTCCTGATGAGTTGGGTGATTGTGTCTCTCTTACAACATTGGATCTTGGGAATAACAGGCTGCGTGGGTCGATACCGGAGACACTGGGGCATCTGCCTCAGTTACAGTGTTTAGTCCTTTCACACAATGATCTTAGTGGTGCTATTCCTTCTAAGATTTCCAAGTATTACCGGCAAGTTAGTATCCCTGATTCGAGTTATGTGCAGCATCATGGAGTTTATGATCTGTCTCACAACAAATTGTCTGGCTCAATACCTGAAGAGCTAGGGAGTTGTGTTGTTATAGTGGATCTTTTGCTCAGCAATAACATGCTATCTGGAGAGATACCACGATCACTCGCCCGCCTGGTGAACCTCACTACGTTAGACTTAACCGGGAATTTGTTAACAGGCACCATTCCAAAGGAATTTGGCAACTCACATAAGCTGCAAGGATTTTATCTGGGGAATAACCAGCTTACAGGATCAATCCCTGAAAGCATTGGCCAAGTAAATAGTTTGGTGAAGCTGAATTTGACAGGGAACATGCTTTCAGGTCCAATACCCTCAAGTTTTGGGAAGTTAAATGGGCTGACTCACTTAGATTTGAGCTCAAATATACTTGACGGTGAACTTCCTCAGTCGCTTTCCAGGATGGTAAACCTTGTTGGCCTTTATGTTCAGCAGAACAGGCTTTCGGGCAGTCTAGATAAGCTATTCTCAAACTCTGCAGCATGGAGACTTGAGATTATTAATTTGGGTACTAACTCCTTCACTGGAGACTTGCCACCATCATTGGGCAACTTGTCGTATTTGACGTTTCTTGATCTCCATGCAAACAGTTTAACTGGTGAAATTCCAGTTGAGCTAGGAAATCTTGTGCAACTTGAGTATTTGGATGTCTCAGGCAACAGTCTCTCTGGCAAAATTCCCGAAACAATATGTGCCCTTCCCAATTTAGATATACTAAATTTTACGGATAACAAACTGAAAGGAGCTATACCGAGGAATGGAATTTGCCAGAATCTTTCAAAAGTCTCAGTGGCTGGGAACAAAGATCTCTGTGGGGGAATCGTGGCTCTAAAATGCCCTGCCAATAGTTTTGTTAAAAGATCATTGTTGTTTAATGTCTGGGGAATCTTATCAGTGGTGGCTGGGACTATAATTATCACTCTTACTATTGTCATTGTGATAAGGATATGGGTTAATAGAAGCAGCAGAAAGAGTGATCCCGAGGATAGCAAACTTGACAGTGATGATCAGCACCTTTATTTCTTAGGCAGCAGCAAGTCGAAAGAGCCTCTTAGCATCAATGTGGCCACGTTTGAGCAGCCTCTTCTCAAGCTGACCTTGGTTGATCTTCTTGAAGCCACCAACAACTTTTGCAAGACTAAAATTGTTGGTGATGGAGGGTTTGGAACTGTCTATAAGGCTACTTTACCTAATGCTAAGACAGTTGCAGTTAAGAAGCTAAACCAAGCAAAAACTCAGGGTCACCGCGAATTTTTAGCTGAAATGGAAACTCTAGGCAAAGTGAAACATCGAAATCTTGTTCCTTTGCTTGGGTACTGTTCTTATGGTGAGGACAAAGTGCTTGTTTATGAGTATATGGTTAATGGGAGCTTGGACCACTGGTTGAGAAACCGTACTGGAACACTTGACGTGCTGGATTGGAGCAAACGACTTAAGATTGCAGTAGGAGCTGCTCGTGGTTTGGCATTTCTTCATCATGGATTCACACCACACATTATTCACAGGGACATTAAGCCAAGTAACATCTTGCTCAACGAAGACTTTGAGGCCAAAGTTGCAGATTTTGGGTTGGCAAGGTTGATCAGCGCCTGTGAGACGCATGTAAGCACTGATATTGCTGGTACATTCGGTTACATTCCTCCCGAGTATGGGCAGACATGGCAGTCTACAACAAAAGGAGATGTTTATTCTTTCGGCGTGATCCTTCTTGAATTGCTGACAGGGAAAGAGCCAACAGGACTCGATTTCAAAGATGTCGAAGGTGGAAACTTGGTTGGTTGGGTACTCCAGAAGATCAAGAAAGGTCATTCAGCTGATGTTCTTGATCCAACAATACTTGATGCAGATTCAAAGCAGATGATGCTTCAGACATTACAGATTGCTACAATATGTCTCTCTGATAATCCGGCTAATCGCCCGTCTATGCTCCATGTCTTCAAGTTCTTGAAAGGAATCAATGGGGAATAA

>PGSC0003DMG400014576 LRR-RLKs

CTTGCATTAATTGCCCCACAATTGTCACATTATTTTCAACATTAATTACTATCACCACCTTTCTCTCTTCTTCACCTTTCACTCTGTTTTTTCTCTTCACTCTTCAATGGCTACTACTACTCTACTCATGAACTTCCATTACACTTTTCTCTCATTCTTCTTCCTCTTCACCCTTCTTGAATCTTCCCCAAACCCAGATATTGCACCATTACTAGCTTTCAAATCAGCTTCAGATGAAACAAACTCACTCTATAACTGGAACTCATCAACTCACCCCTGTTCTTGGACTGGTGTATCTTGTCTTAACAATAGAGTTTCTAGACTTGTTCTTGAAGGTCTTAATCTAAAAGGGTCTTTCCAAAATCTTGTATTTCTCAAAGAGTTAAGAGTATTGAGCTTGAAATATAACAATTTTTCCGGTTCTGTTCCGAATCTTTCGAACCTTACTGCTTTGAAGCTGCTGTTTCTTTCCCATAATGAGCTTTCTGGGGAGTTTTCTGAGTCGTTTACCTCGCTTTTTAAGCTTTATAGACTTGATCTGTCGTATAATAAGTTTTCCGGTGAGATTCCGGCGAAGGTGAATCGTTTGACCCATTTGCTTACTCTTCGTCTTGAAGGAAATGGGTTTTCCGGTGAGATTTCTGGGGTTAATCTACCTAATCTTCAAGAGTTTAACGTTTCTGGTAACAAACTCGTTGGAGAAATACCCATTTCGCTATCTCGTTTCCCGGAATCTGCATTTTCGAAGAACCGGGTTCTTTGTGGTTCTCCGTTGCCAAACTGTACGGCGGCGGTTCCTCGTGAACCGTCGTCGTCGGCAGGAGCTTTAGCTTCTCCGGTTAGTCCTAAAACTACCATTGCATCTTCTCCGAGCTCATTGCCGGTGACCTCTGCAACTTTGAGTCCAAAAAATACCCATCATAGTAGTGGTAAAATGAGCTCATTGGCCATAATCGCCATTATACTAGGTGATGTGTTGGTTCTATGTGTTGTGTGTATTTTTCTCTACTGTTTCTTCTGCATTAGGAAAGTTTCTTCTCAGAAAAATGGGTCACACATTCTTGAAGGTGAGAAGATTGTGTACTCGTCTAGCCCGTATCCGAATACGGGCCAGACGAGCGGGTTCGAGAGAGGGAAAATGGTGTTTTTCGAAGGGGCGAAGAGGTTTGAGCTTGAGGATTTGTTGAGAGCTTCAGCTGAAATGTTGGGAAAAGGGGGATTTGGGACTGCTTATAAGGCTGTGTTGGATGATGGAAATGTTGTTGCTGTGAAAAGATTGAAAGAGTTGAATGTTTGTGGGAAAAGAGAATTTGAGCAACAAATGGAAGTTTTAGGAAGACTTAGACACCCAAATTTGGTTAGTTTGAAAGCATACTACTTTGCTAGAGATGAGAAGTTGTTGGTGTATGAGTTCATGACTAATGGCAACTTGTTTTGGCTTCTTCATGGTAAGTTTTGTTCTTCAAGAAATTCAATAGTACTAACATATTTTAAAATGGAGCAACTTTGCTGGATTACACTGAGTATGTTGTTCTTGTTATTGATATGTGATTGTGTTGTTGAACTTGTTGATTTTGAAAAGAAAAATGTGATCTTTAGGATTGTTTTGTTGAAATCGATTGATTTGGGTGATAAAGTTTGGCAACTAGTAGCAAAAGTCACATGTGGTTGGTTTGTGTCTTGTTTTTTTTTTTTGGATTTTAGTTGCTTTTTTTGGCTGCAACTGTCAATAAGGCCTGGTCTCTAATGCATATAAGAATAGAAACCTAAATTTGGATAGTTTGAAAGCATATTACTTTGCTAAAGATTGAGTAGTTGTTGGTGAGTTTGGTGTATGATTTCATCACTTGTTTTGGCTTATTCATTTGTTTTGTGTCTTTTTTTTTTTTTTTGGATTTTAGTTGCTTTTTTTGGCTGCAACTGTCAATAAGGCCTGGTCTCTAATGCATATAAGAATAGAAACCTAAATTTGGATAGTTTGAAAGCATATTACTTTGCTAAAGATTGAGTAGTTGTTGGTGAGTTTGGTGTATGATTTCATCACTTGTTTTGGCTTATTCATGGTAAATAGTATTCAACCTTTTGTTCTTTAAGAAATTCATAGTACTAAAATATTTGTGAATTACTTGTTTCTTAAAGAAAACTGTGATCTTTAGTTTTTTGTTGTAGAATAGATTATTGATTGACCAATAAGGATTATAGTGGAGTTGTAAGTACTTCGTCATCTTTAATCAGACATTTCAAGTTCGAGCCTTGAGTATGGACTTGCCCTTGTTAGGTAACGGTTTGTCCCTATGTGGGACTTCCTAGCGCAAACCTGTATCCAGCTCCAATGCAGTTGCCGTACACCGGATGGGTAAAAAGGAGAAGATTATTGATTGATTTGGTGGTAAAGTTTGGCAAGTAGTATAGCATAATCACATGGGGTTTGAGTTTGGCCTTTTGTCTTTAATTGCTTTTGGTCTGCTACTGTTAACCAGGACTAGCCTTCTGATGCAATAGGAATAAAGATGGGATGCCTCAATACTAGACGGTATTTGCTTTAGTTAATGTTTCTATTTACATAATTGAAACTAGCCATGTGGGGATACTGAAAATCTATGTTGCTCGGACTCTTCAAAAATGTCGATGAGGCATTTTTGGAGGATCCGACATGGGTAAGTCCTACATAGTCTTTTCAAGCTAAAACTATAGAATATTGGAGTTTCTGGTCAAGGTATGTTGGTTCATTAACAATCTTTGTCACATAAGCTATTCTTGATTTTCAAATGTAGGGAACTTAAATTATTTAACAATCTTTATCATCTTATTCTTGAAAAAGCTTTTAAAGGAATCTAATTTACTCATGATGGTCCTAAAGTGACAGAAGATTAGAGCAAAGGACTTGTTGTACCAGATGAATAGAATTATGTGTTCCCTCTGATTTTTGGGCCAAGCAAAAGAAAGAAATTCACACTTTTTAGGCAAGAGATGCCCTACTGTCCCATTGTGCTCTACAGTTTTTTTGCTTGTCTTTTTGACCTCACTTAGTATTCATATCTTGCAACTGTTTTGGTAGGTAGATAGTGAAATATAACCTCTTGTTTTTTTCTGATTCAAATGGTAAAGAATCAACTTCTTGTTGAATTGCACCTGTCATTCATGTAACAGGAGACAGCTGCTTCATATGTTAAATTTTCTCATTTTAGAAATATGAACAATTATGTATTACTATTCCTACCATCAAAGTCTATTGTGTCAACTTCATACTCCTTGTCCTCTCTGCCCATGTTTTTGTTCTTTTGTCTAGTTAAAAGTTGCCCCTTTATCTTTCTATAACAATTCTCCCTCCGTTTCAATTTGTTTGTCTGGTTTTGATTTGGCACGAAGTTTAAGAAAAGTATAGAAGACTTTTGAATCTTGTGGTCTTAGTTCTTAAACATGTCATGTGGCCGAAAACGGGAAGAGGCAAATAAGCGAAAATGTGCAAGTGCTCCATTTTATAGTAGTTTGAGTAAATGTTTTGATGGGCATGAATCTTCTTGCTACAGGTAACAGAGGGCCTGGAAGAACTCCTTTAGACTGGACAACAAGGTTAAAGATTGCAGCCGGAGCAGCTCGAGGTCTAGCCTTCATCCACAACTCATGCAAGTCCTTGAAACTCACTCATGGCAACATCAAATCAACTAACATCCTCATTGACAAGAGTGGCAATGCACGTGTCTCCGACTTTGGTCTAGCCATCTTCGCCACACCATCTTCCGTCCCAAAAACCAATGGCTACCGAGCCCCTGAAGTCGCATTGGATGGTCGTAAAATAACTCAAAAGTCCGACATCTACTCATTCGGGGTCCTATTACTCGAGCTGCTAACCGGTAAATGCCCCTCAGTCGTGGACAATGGTAGTGGCCTCACAACCGGCTACGGAGGGGTCGTGGACTTACCGAGATGGGTGCAATCCGTGGTGAGGGAAGAATGGACTGCGGAAGTGTTCGATTTGGAGTTGATGAGGTATAAGGACATTGAGGAGGAAATGGTGGGGCTATTGCAGATAGCTATGGCATGTACTTCAACATCACCAGATCAAAGGCCTAAAATAAATTATGTTGTGAAAATGATTGAAGAGTTGAGAGGTGTTGAAGTTTCACCTTCTCATGACACTGCTACTGATTCTGTTTCTGACTCTCCTGCTGTTTCTGAAGATAATACATGTGGTGCAGCTAGTCAGTAGTAAGAGTTTTTTTTTCTTCTTATTTTTCTAGTTTGTTATTCTGAGTTATGTAAATTAACCTCTTAGTTTTAATCCTTTTATCTATAAGATATAAGAGAAAATTTAGGAGAAGTTTTATA

>PGSC0003DMG400014917 LRR-RLKs

GTTCATTACTACTATTTTTATTTGTATCTCAATTTTGTCAAAAATAAAAAAATCCCCAACAAATGATGTGGAGGATATCCGTTTTTCTTGTTCTTTTACTCAATTCTCATACCGGATTTTGTCTAACTTCCGACGGACTTTCACTTCTATCTCTAAAATCCGCCATGGATGATGGCGGTGGTGGTACAGTTTTCTCCGATTGGAATGAAAACGATGATACTCCTTGTACTTGGACTGGCATTTCATGTGCTAACATTTCTGGTTCTTCAGAGCCACGTGTCGTTGGGATTACTCTTTCCGGTAAGAATCTCCGGGGATACTTATCGTCTGAGCTGGGAACGTTACTCTATCTCCGGCGACTGAATCTTCATGGAAATAATATTTACGGTTCTATCCCTGACCCTTTATTCAATGCGACGTCGCTTCATAGTATTTATTTGTATGATAATAATATCTCTGGTATACTTCCTCCTTCTGTTTGTAATCTTCCTCGTTTGCAGAATTTGGATATTTCTGATAATTCACTTTCTGGAACTTTCTCTAAGGATCTACGTAATTGTAGGCAGTTACAGAGGCTTATTCTAGCTCGGAATAAATTCTCCGGTGAGATTCCGGTGGGGGTGTTCCCGGAGTTGGCGAATTTGGAACAACTTGATCTTTCGTCGAATTTGTTCAATGGTTCGATTCCTGAAGATATTGGTGAATTGAAGTCACTATCTGGAACTTTGAATCTTTCTTTTAATCATTTTAGTGGAAGAATTCCCAAATCTGTAGGTGATTTACCTTTAACTGTTAGTTTTGATCTAAGGAATAATAATTTATCTGGTGAAATTCCTCAGACGGGTTCTTTTGCTAATCAAGGTCCGACGGCTTTTTTGAATAATCCCTTGTTATGTGGATTTCCTTTACAAAAAAACTGTAAAAATAGTTCAAACAATTCAACACAAGTTCAGGGTTCGTCTGATAACGAGGGAACGAATTCGCGCAAGGGGTTAAAACCTGGATTCATTATATTGATATGTTTGGCTGATGCATTTGGAGTGGCATTTATAGGACTAGTGATTATTTACTTGTATTGGAAAAAGAAAGACTCTGGTGGTTGCAGCTGTACTGGAAAAGGGAAATTTGGTGGAAATGAGAAGAGAATGCTTTGTGGTTTCCCATGTATTAATGGATTTCCAAATAATGATTCGGAAGTTGAGTCTGAGAAGGGAGGTGGCGGTGGGGCTAGTGGTGGTGGTGTTAGTGGTGGGGAGGGTGATCTTGTGGCTATTGATAAAGGTTTTAGCTTTGAGCTTGATGAGCTGTTGAGGGCTTCTGCTTATGTATTGGGAAAGAGTGGGTTGGGGATAGTGTACAAGGTTGTGCTTGGGAACGGTATCCCAGTGGCAGTGAGGAGGTTAGGGGAAGGTGGGGAACAGAGATATAAGGAGTTTGTCGCGGAGATTCAGGCAATCGGAAGGGTTAAGCATCCTAATGTTGTGAAGTTAAGAGCTTATTATTGGGCTCCTGATGAGAAACTTCTCATTAGTGATTTCATTTCCAACGGCAACTTGGCTTCTGCTCTTCACGGTGAGCTTTTCATTATAGTAAAACATCTTCAGTTTTGTTTGTCTAGTAGATATAACCTGTTGAAAGAATCATTAACGCATTCATATTTTAGATATCGGGGTATACTATATTCTTATCATTTAGTAGCATGATACACTGATGGATGTTTCGTACATTCCAAGATCGATATGTGTAAATTGTAATTCTTTTGCTGGATTGGTATTTTACTTATTGATCACTCTTTTGAGGAGTGGAAATACAGTACAAGACTTTTGGTATTAGGTTAGTTAGGAGTTTGGGATGTTCTAGTATTTATTCACTAGTATAACTGTTTAATATATTCCCAATGACAAAATAATGACTTTTAGCTGTTGTGTGTTCCTTGATGATTTTGAATAGCAATTATTCATATGCCATCTAGTGATAATTTTATATTGAATGTGTCCCATGTGCTTGCAATCCTTGAGTGAAGGATGAGTTTTGTTTGCGACTTGATAATGTGTAGGATTTTTGCCTAATTGTATCAGCTGATAGGATAGTCTTTATTATGTATCATAGATGAAAAGTTATCTGCATAAAGGGTAAACCAACAACCTAACATTTTCCATCCATTTCAATGGATGTTAAAGCACATCGTAGTATATTAACTTATAATGATCCAGCCCAAGTCAATTTGCAGACTTTAACGATTTACTGCCGATGCAATAATCATATGTTGACAGCATATCGAACCTGGTAGAAAGGGCATTTAGGCATGTTTTGAAGAATTTAGTCTCTCTTTTTTCATGTGACATATGGTTGACTGCCCCGATAAACCTTGCGGAAAATGATTTATCATCAATTAGAAGCAAACCAATAAGTACAAGGGGCCAAGAACTGAATTATGTCATGTACTAGGTCAATTCCTTAAGTTGTGGAGTTACGTATTCTGTCTCTCAAAAAAGTCTAGATATTACATCGTTTTATTATGTCTTTTAGAGATGTATTACCTGATTTGCATATACTAATATTCAAGAATTTGCGAATCGCTCCGCCTGGTGCAGTTTGACTTTACAGATGTTTAATTTTTCGAAGCAAACTGGAATCTGCAGTATCTGGTCTGGTAACTACCATCTAGCATGAGCGGGACTTATACTCCCTTGAGATGGATTATAGACTTCGAATTTTTAAAAATTAGTGTTCTGCAATTGAATTTTGCACTGTTGCTGTTGGTATAGCTGCTTGAATGTTACATTTTTAAAAAGAATTAGATTTTTTCCCCTGATACAAATTGTTGTTTACATTTATGAATATCTACTTCACTGCCTCATATACATGTCACTATGAATCTATAACACATTCGTTACCCAAGAGTAACTGCTTGCTTTCTGAGACAAATATATTTGATGACCTTTGGTTCTATGGTACCTGATGTTCCCAGTTTCTTAATTTACCTGTTTTTAAATTATGTAGCTGGAATTTGACAGTACAGTAACTTACCTTTTGCATTCTGAAGAAGTTGAGGAGTGTTTTGTTTCTGCATCTCTTTTTCCAAATTGTTTTGCCATGCTAGATCTGTATGTATTGACGTAGTGACAGTAGTTTCATGCTCTAGCTGATTGAGATTAGCCGAATTAGTGTGAACACCGATTTGCCTATTGAGCTGAAAAGTTATTACCTTGTATCTTCTTCGTCATACTGTGGTAAATGTTGTGATGACAATCTCGTTTGGCTTTATACAGGGAGAAATGGGCAACCATCACCTAGTTTAACATGGTCAACGAGACTGAAAATTGCTAAGGGAACAGCGCGAGGTTTAGCCTATCTCCATGAGTGTAGCCCAAGGAAATTTGTCCATGGAGACGTCAAACCTTCCAATATTCTCCTTGATACTGAACTGCAGCCATATATATCTGATTTTGGCCTCAATCGATTAATCAATATCACGGGAAACAACCCCTCCTCTTCTGGTGGATTTATGGGTGGAGCTCTTCCTTACTTGAAGCCTGCCCAACCAGAACGTCCCAATAACTATCGAGCACCAGAAGCTCGAATCACTGGTAACAGACCCACCCAAAAATGGGATGTGTACTCATTCGGGGTGGTCTTGCTTGAACTGCTGACTGGAAAGTCTCCTGACCTCTCAGTTCCTACAACATCCACTTCAACGGAGGTTCCAGATCTTGTAAGATGGGTAAGAAAGGGATTCGAGGAGCAAAATCCACTGTCGGATATGGTAGAACCAATGTTGCTTCAAGAAGTCCATGCTAAGAAGGAAGTACTAGCTGTATTTCATATAGCTCTTGCATGCACCGAGGCAGATCCAGATATTCGACCAAGGATGAAAACTATCTCAGAAAACATTGAGAAAGTTGGAGCTTAAACTTTTCGGTACACTCATGTCAAGGAGTTCCTGAGTTTCATGGACTTCTTTAATCTTTTTTTTGCAGTAATTGCTCCTATTATTTTTTTTCTTTTAGAGGTTAATCAAATCTTTAAACTTAACTACAGTACAAAATACCAGTGTGCTGCAACAAAATTCCTGTCATTTTTGAGAAGTTAATTAGTTGCTTATCTCTTTAATCTATTTCTGTAATTCTGATGTTAAAACAACTTTTCATGGCCAA

>PGSC0003DMG400015081 LRR-RLKs

ACACAACAAGTTGCAAGGATCTATACCTGACTCAATGAGCAACATGATAGGTTTGGAATACCTAGACATTTCTCATAATAATATATCAGGAACCATTCCTATGTCTTTGGAGAAACTTCAAAACTTGAAGTATTTCAATGTTTCCGTCAACAAGTTGCACGGTGAAATACCCTCAGAGGGTCCTTTCAAGAACCTCTCGAGTCAATTTTTCATCAACAATGAAGCATTGTGTGGTTGGTCAAGGTTTAATGTCCGGCCATGCCCCACCTCATCAACGCATAGATCAAATAGGAAAAAATTGCTAGTTCTATTTCTTTTGCTAGGAATAGCACTTTTATGTGCTCCTACTACCCTTGTGTTTGTATGGATAAGGTATAGAAGAGGTAAAAGATCTCCTCAACGAGCTGATTCATTGTCTATCACAACAACGGAAAGAATTTCATACTATGAACTGCTCCAAGCAACTGATTCTCTTAGCGAGAGTAATCTGATTGGTTCTGGAAGTTTTGGCTCTGTCTACAAAGGCGTTCTCAGAAGTGGGGCTGCCATTGCAGTTAAAGTGTTCAATCTGCAACTGGAAGCGGCATTCAAGAGTTTTGATACAGAATGTGAAGTTTTGTGTAGCCTTCGCCATAGGAATCTTGTGAAAGTCATCACCAGTTGTTCCAACCTTGATTTTAAGGCTTTAGTGCTCGAGTATATGCCTAATGGAAGTCTTGATAAGTATTTGTATTCCCACAACTACTTCCTAGACATCAGACAGAGACTAAGCATTATGATAGATGTGGCATGTGCGTTGGAATATCTCCATGATGGTTGCTCCTCTCCGGTGATTCATTGTGATCTGAAGCCTAGTAACGTCTTGCTGGACGAGGATATGGTTGCCCACCTAAGCGACTTTGGTATTTCAAAACTGCTTGGTGAAGATGAGACTGATTTATACACTAAAACCTTAGCAACATTTGGTTATATAGCACCGGGTACGTCTCTTTGTTTTTGCTAATTTGAACATTGATTTTTCTTCTTATTCTTTTTTTTCCAATCAAGAAAGTATATTGCATCTTTTAATTAATTGTTTGACTGTAGAGTATGGACTGGATGGACTGGTGTCAATAAAATGTGATGTCTATAGTTACGGGATTATGTTGCTGGAAACATTTACTAGGAGAAAGCCTAGTGATTTTGAGGGAGATCTTAGCTTGAAGCAATGGGTGAGTTATTCACTCCCAAAGGCAGTACTGAACGTTATGGATGCCAACTTGGTTACACCAATGGATAATCGCTTACAGAAGGAGTTAGATATCGTGGCATCAATCATGAAAGTAGCATTAGATTGTTGTGCTGAATCTCCGACAAGGAGGACAAACATGAAAGATGTTGTAGGGATGCTACAGAAGATCAAGATTCAACTTCTTGCATGCTGAGCATGTTCTCTGAATCTCTTGTTCCGAATTGATCACTTTTTTGTTTTGTTTTAGCACTTGATTTGTGCATGATTCTGTTTTCAGAAATGCT

>PGSC0003DMG400015157 LRR-RLKs

TTCAGAACCACCAAGCAACTAAAAGAGACATCCTACTCCTCAAGCACCACCAGTGTGGTGCCTCATCCAACTCCCAATTCCACCACAATACTGCACCAATCCGATGGCCTTCACAGCCTCACATATACACCTTTCTTTTTTGTCTTTTTTCACTGTGATCCTTCTTGTTCAAGCAAAACTCAACCTTTACTCACCTGATCACAGTGCTCTTTTGCTTGTCCAAAAAGGCTTAGGCATCCCTGCTCACCGCAATGCTCTTGAGAACCCATGCAACTCCGTTGGAATATCATGCGAAAAACGACTCACAAACAATTCATACGTGCTAAGAGTCACAAGGGTTGTCTTCAAATCCTACGGGTTGAAGGGTACTCTGTCTCCTGCCATTGGGAGGCTTTCTGAGCTCAAAGAACTGTCCCTCCAAAACAACAAACTCTTTGACAGAATACCAACTGAAATCGTTGACTCCAGGAAATTGGAAATCTTGAACCTTCAAAACAACCAATTTTCTGGCAAAGTCCCACCTGAATTATCATCTCTAGTCCGCCTTCGTACCCTTGACCTTGGCTCTAATGAGTTATCTGGGAACCTGAACTTCTTGAAATACTTTCCTAACCTTGAAAAACTCTCCCTTGCTGATAACATGTTTACTGGCAGAATACCTCAATCCTTGAAATCTTTCAGAAATCTCCGGCTCCTCAACATTTCAGGCAATAGTTTCCTTGAAGGTACGGTGCCTGCCGTGAGTCAAGTTGAGCACTTATCAGCAGACTTAAATCGAAAAGATTATGTTCCTAAGCGTTACATTCTTGCTGAGAATGAAAGAAGGCCAAGGCACACCCCTGCAATGGCACCAGAGTCCAAATCAGGATATGCCCAAGCTCCAGGACCTAGTACAGTTGTAGTACCAGTAAAAAAAGACAAACACAACAAGATCAGAAGGAAGGTAGGCGCCTGGATTCTTGGATTCTTTGCTGGAGTTTTCGCTGGGGGCTTATCGGCGTTGGTCTTCTCTGTCCTCTTCAAGGTGCTTATGTTTTTAATCAGAGGGATCAGAAATGATTCAGGCTTAACAATTTTCAGTCCATTGATTAAGAAAGCCGAGCACTTGGCCTTTCTGGAGAACGAAGATGGATTGGAATCACTAGAACTCATTGGGCAAGGTGGATGTGGAAAAGTTTATAAAGCTGCGTTACCTGGAAGTGACGGAAAGATTATAGCCGTAAAGAAGATCATACAACCACCAAGGGATGCTGCAGAACTCACTGAGGAAGATAGCAAGGCTATGAGTAAGAAAATGCGCCAGATTAAATCAGAAATCAAAATTGTAGGTCAAATCAGACACCGGAATTTGCTTCCCCTACTGGCACATATGCCAAGACCAGACTGCCATTACTTGGTCTATGAGTACATGAAAAATGGGAGCTTACAGGATACCCTCCAGCAAGTCAGAGAGGGGACACGGGAACTAGATTGGTCTGCACGTCACCGAATTGCAATGGGAATAGCTGCTGGACTCGAGTATCTCCATATAAATCATACTCAGCGTATAATTCACAGAGATCTAAAGCCAGGCAATGTCCTCCTTGATGATGACATGGAAGCTCGAATTGCGGATTTTGGCCTTGCAAAGGCTGTCCCAGATGCTCATACACATATTACAACTTCAAATGTGGCAGGAACTATAGGATACATCGCTCCAGAATATCATCAGACACTGAAGTTCACTGATAAGTGTGATATATACAGCTTCGGGGTGCTGCTAGGCGTGCTAGTTATGGGAAAGCTTCCATCTGATGAGTTCTTCCAGAACACTTCTGAGATGAGTTTAGTGAAATGGATGAGAAATGTCATGACTTCTGAGGATCCAAACAGAGCAATTGATCCAAAGCTGATGGGTAATGGAAATGAGGACCAAATGCTTTTGGTTCTCAAGATTGCCTGTTTTTGTACCCTGGAGAATCCTAAGGAGAGGCCTAATAGTAAGGATGTTAGGTGCATGTTGATGCAGATCAAGCATTAAGTATGTAGTGGATAACTTACAGTAAATAAAAGTTTGTCATGAAACTAAGAGAATATTTCTTGTGTATGACGGTCCTTGTGTATTCTGTTACATCTCCAATATGAGAGACTGTCCTTGTGTGTTTTGTTTCCTCTCCAAAATGAGAAGTTTATATTTAGCTGTATGAAAGTTCTTTGCAGAAGTTTCCTCTGCTATATTATAAAGCATGGTACGAAATGTGAATCGAATTCTTTCCACTAGA

>PGSC0003DMG400015162 LRR-RLKs

TTATGACACCTTGGTCCGCACATTCTACACTCACTCTTTCCTTCCCCTTTAAACAGTCATGCATTTACATAAAGACAGTGACTGTTTTTTCTCTCTCTAAAATTTCATCTTCTTTCATGGCTTCTCGCCGGAAAATCAAGAATTCCCGCCGGGAACTACATTCTTGATTCGCATTCTTCTTTAAAAGGGTCGGAACAAGAAAACAAAGGGTGTTTATAAATGGCGGTGTTTTTAAGGTTTGTTTTTTGTGTTTTATTCTTTGCTTTACTCGGATTGAGTCGTTTCCGGGTTTTTTCAGAGCCTACACAGGATAAGCAAGCTCTCCTTGCTTTTTTCTCTCAGATCCGACATGCGAATCGGGTCCAATGGAATTCTTCTGCTTCTGTTTGTACTTGGTTCGGAGTTGAATGTGACCCGAATAACTCTTTCGTTTATTCACTGAGACTTCCTGCTGTTGGTCTCGTCGGAAAAATTCCTTCTAACAGTTTGGGCCGGCTGAGTCAACTTCGAGTTCTTAGTCTTCACGCCAATCGTCTATCTGGTTCAATTCCTTCGGATTTTTCTAACCTTAAGCTTCTTCGTAGTCTCTATCTTCAAAAAAATGACTTTTCCGGCGAGTTCCCGGAGAGTATTCCCGGGTTGACCCGGTTAAACAGGCTGGATCTTTCGTCGAATAACTTCACTGGTACTATCCCTTTCTCTATCAATAATCTGACCCATTTGACTGGACTTTTATTGCAGAACAATAGCTTTACTGGTACTCTTCCTAGCATTAATCCGCCTGGTTTAGTGGATTTCAGTGTTTCGAATAATCAGCTTAATGGGTCAATTCCTACTGCACTTTCCAAATTCCCAGCTTCCTCTTTTACTGGTAACATTGATTTATGTGGCGGCCCATTACCCCCATGTACCCCATTTTTTCCTTCTCCTTCACCTTCCCCTGAAACAGAACCAAAAACCCCTCCGTCAATTAAGAAATCCAAGAAGCTTTCAACAGCTGCCATTGTTGGAATAATTATAGGTTCAATAATTGGGGTTCTCCTACTCTTGTTACTCCTTTTCTTCTGTTTAAAACGAAGAAAAAACGACACATCAAAGGTACAAAAACCTCCGGTAGCATCAAGAGCTATCGGGGCAGTAACAGGGGCAGCGGCTGAGGCAGGTACATCGTCGTCTAAAGACGATTTAACAGGTGGTTCAGGTGAAGGGGAAAGGAATAAACTTGTGTTCTTTGATGGGGGTGGGTATAGTTTTGATCTTGAAGATTTGTTAAGAGCTTCTGCTGAGGTTCTTGGAAAAGGGAGTGTTGGAACAAGTTATAAAGCTGTTCTTGAAGAAGGTACTACAGTTGTTGTTAAAAGGCTTAAAGATGTCGTTGTGCCCAGAAAAGAATTCGAGCAACAATTGGAAGTAATGGGGAAGATGAAACATGAAAATGTGCTACCTTTGAGAGCATTTTACTATTCTAAAGATGAGAAATTGTTGGTATCTGATTATATGCCTGCTGGAAGTTTATCAGCTCTTCTCCATGGTACGCTGCTCTTATTATTTTTAGTTAAAAGTATGTCAAACTTGTTCGTTTTAGTGTATAATTTGCTCTATTATATGTTACTGCAATTCTGTGTCTGTTGTTTGCTCTGAACACTTAAAAGTTCCATTTTTTCACCAATGTTTTAAAACTTTAATGGCAATAAAGATAGTAGCATTCAATTCTTGCTAGATTTTAGTGACTTTGTTCTGATTCTTGAAGAGTCTTGAAGGGGGTTAGGTGGGTTATAGGTATGAAAAAGATTGAATTTTTATTTTGTTTCTTTTTTTGGAAGAATAGATGGGGCTTCAATCTAATACACGTGCAGATCATGTACACATTGTAGCCATAGTCTTACAGCCCTTCTGAACTGGCATTGCATTATTGTTTCACTCAATTATTGAAGTTTGGTACCACTTAATTTACTTCGTTAGTACATTTATTGTCAATTCAACCGCAAATTGCGCATTTTGTTTCTATAATTCCTTAAGCTACAGAACATTCTTTTTTGAGAGAAATGGAACAATGACTTTTTTATTATGATTTATTCGAATCGAGAGCATATCGGATACAATCTCTCTATCTTAACGGACACACTTGTTAGATTACACTTTTACTCTTTCTTTTTTATGAAACGAACGTTACAATGCAACAGACTAGCATGATTCCCTGCCTTAATAAATGATGAAATGAGTTTAAAAAGTGGAAAAAGAAGGCTTCTTTCTTGTTCAAAGATTTACCTTTTAAATTGCTACATTGCTTGTGTCTATGCCTTAAGGTCTTTGCTCCATAGTTCTCAATTTTTTGCCACAATCAACAAACTACCTCTATTATTGTAGTTATATAATATATATATTTTTTTTATTATACATAGGGGAAATGGGGGAGGGAATAACAAAGTGTTGTGGAGTTTGGTGATATCAAATATATTAATTAAATATTAATCTTTTGTTTTCTAATCCCACAATTTTCAAATCATGATTTAATTAGTAGTAATTGCTGTAAGTCTAAACAATACTACCATACTTAAAAGTTTATATTACTTTGGCTTTAAGCTATTTGGTGGTTATAACTGTCTGATTATTATTCATCTAATCTTGATTCTTTTGCTTTACTTTATTCATCATTGTTTCTTGCTGTCCCCCATTTTTCTTCTTTTGTTGAAATAACGTGCTGCTTTACTCTATTGTCCATTGATTCTTGCTGTCACCCATTTTCTTCTCTTTTTTTTTGGTGAAGTAAAGTGCAAAAGAAAAGCTAATAATACAATGTGTCCTCTTGAGTCTTGAGTAGTTCAACTGCTTTGAATTTGAATCCAACATATTAATTTGCATTGACATTAGTTTGTTAACAACATAACCAGTGTAATCCTACGAGTGAGGTTTGACTTTTGAGTAGGGTAGGATGTACACAGACCTTATCCTTACCTTTGGTTGTCATTGGTTTGGTTGTCTAATTAAAATGTTTTCTTTTGGTTAATTCTTGTAGGTAGTAGAGGATCAGGGCGTACGCCACTTGACTGGGATAGTCGAATGAGAATAGTGTTAGGTGCAGCAAGAGGCATTGCGTACCTCCATATTTCAGGAAAAGTTGTACATGGAAACATCAAGGCTTCGAATGTTCTCCTTAAACAAGATAACCAAGATGCATGTGTATCCGATTACGGATTGAACCCTCTTTTCTCTACCTCAGCACCAGTGAACCACCGTGTTGCAGGATATCGTGCACCTGAGGTTCTTGAAACTCGAAAAGTCACATATAAATCCGATGTGTATAGTTTTGGAGTGTTAATGTTGGAGCTCTTAACGGGGAAAGCTCCAAACCAAGCTTCACTAGGTGAAGAAGGCATTGATTTGCCTAGATGGGTCCAAAGTGTTGTAAGAGAAGAATGGACTGCGGAGGTGTTTGATGTTGAGCTAATGCGATACCATAATGTGGAAGAAGAGATGGTACAACTACTTCAGATAGGTATGGCATGTGTTGCTACCATGCCAGACCAACGACCTGCCATGACCGAGGTGGTTAAGATGATCGAAGAAATGAATCATGGTGATACAGATGATGGTCTTAGACAATCCTCCGACGATCCCTCTAAAGGTTCTGAAGGCCAAACACCTCAGGAGTCTAGAGGTTCACCTCATGGTGTAACACCATAAAATGAAATTAATCATAAAAATGGATGGCTTGATTTGTCAAAAATTCTCAGAAAAGCCTAAAGTAAATTGTATTAGATTCTTAGTGGGGGGAGCCTTTTATGTGTTTTTATTTTATTTCATTTTTTTTTCTTCTTGAAATTATTATTTTGAGAAAATTTGTGGAATGATTTTTACAGATGGTTAGTTGAGAATTTGAATATTTCAGTGAAGTGATTGGGAAATTGAAATTTG

>PGSC0003DMG400015276 LRR-RLKs

GCTGTCTATAACTATTTTTTCAACTTTTTAATGTGTTTGACATTGTTTGAATTATACCCCCAATGTTCAATTTTACTTCACCTACTACATACAACTGGTAAATTCTCAATCTATTAGAGTATTACTTACTTATCTATTATTCTTCTACGTCAAGGTTGCCACGACCAGAAATGCGTAGGACGATCCTATTTTTCGCGTTGTTTCTGGTGGTGGAATCAGCCGAATTTTTCCCAAATGAAAGAGATGCTTTGTTGCAGCTGAGAGATAGTGTAAATTCAAGTGTTCTTGATTTACATTCACATTGGACAGGGCCTCCTTGTTACAACAACCAGAGTAAATGGGCTGGAATTTCGTGTTCTAATGGGCATGTAAATCAACTTGTTTTAGAAGGAATTCAGTTAACTGGCTCACTTCCACCAACATTCTTGTTGCACATAGCATATTTGACAAAAATCAGCTTTAGGAACAATTTATTGTATGGTCCTGTTCCAAATCTCACAGGACTAATCTACCTGGAGTTTGTTTTTCTGTCAGAAAATCGATTTTCGGGTTTGATTCCGTTTGGGTACACTGAGTTAAAGAAACTAACAGAATTGGAGCTCCAGGGGAATGATCTTGAAGGTTCAATTCCACCATTTGATCAACAGACTTTGATTGATTTCAATGTGTCGTCGAATAAACTGGAAGGACCTATACCTGAAACTCCTGTCCTGAAGAGATTTTCGAAGAGTTGTTATGAGAATAACGCGAATTTGTGTGGTGAAGTTGTAGGAATACCTTGTTCAATATCACCTGCAAGTCCACCTAAAGCACCAAGTGAAAAGAAAAAGAATGGCCTTAAAGTATGGAGTATTGCTTTGATTGCAGCAGCAGTAGCAGTACTTGTTCTTATTTCAATCATGTTCATTTTGTTCCGTTACTTTAGAAGATCTAAAAGAATGGAAACACAAGGAGAAGACAAACAAGGTAACAGTTTTATTGTATCATCTATGTTATCCTAGCTAGTGCCAGTTATTCTGAGGGGAAAGTTATTACTACTTAGATTGAATTAGATGATGAGGTTATCTGGTGTAGGGCTGTTTTGGTGATACGGAACGAGACAGGGGAGAGGAAGTTGACTGTATTAGAGGGCTGACTGTATTTTTTTCGAAATAGTTTAACCATTTTTGTTAATCGCTAAAAGAAGGTCATTCAGAACTCGTTGGCTCCAGAATGGGGCAGCTCCAGAGGTGCTAAAGATGGTTAATTCGTGATAGGAAGAAAACCAATACATGTCCATTATGAAGTCGAATTCAATCATTTCTAGCTCAATAAGATGAGAACTAGTATCTCGCCCCTGAATTGCGGCTATACAACAACAATAACATACTAGTATAGTCCCACAAGTGAGGTCTGGGGAGAGTGAAGTGTACGTAGACCTTACCCCTACGTTTGTGGGATAGAGAGACCGTTTCTGATAGATATTGTGGCTATACAACATAGAACTAGATTAGCTAATTCAATTAAACTTAAAAGATCCATGTGATTATGGAGCAGTTAATATTGTTGTTATTACATCTTTTAACATTTCCAATTTCCATTTTGACAGGTAATATTTCAATAGAGAATGTGGTAAAGAGAAGTTATTGGTCCAAGAGTACAGAAGATCCAGAAAAAACATTGGACTTAATATTCTTTGAAAAGGATATGTCAGTCTTTGACATGGATGACTTGTTAAGAGCATCAGCTGAAGTTTTAGGAAAAGGAAAGCTAAGCACAACATATAAAGCAATCCTGGAATCTGGTTCTGTTGTTGCTGTTAAGAGACTCAAAGAAATGAATTCTTTGAGCAAAAAGGAATTCACACAACTGGTGCAGTTGCTTGGGAAATTGAAACATGAAAACCTTGTGGAGATGTTGTCCTTCTACTATTCCAAGGAGGAGAAGCTCATTGTTTATGAATATGTTCCACAAGGAGACTTGTTTGGGCTCTTACATGGTTGGTTCATCAACTTTCTTTTACTTTTGGGAAAATGTCTAAAGTTAACCCCTCCTGAACGAAATACCTTAATTTTATCTTTCATTATACTTTGAGATTATTCATATCCTTATCCTTAGTAAAACGTCTCGTGTTATACTTCGAGGTCATTCATATTCATTTCTGATTCTAAACTATGAACTCATGATCAAATGACTTTTGAGGTGGGCTTTCCCTTCTCTATAGTGCTACTGTGAAAATTGAAATTGCTTAGAGGTAGGTGTAAGGTCTGCGGACACTCAACCCCTTGTGGAATTATACTGAACATGTTGTTATCGTCTTATAGTAGTAATAGAGGAATATCTTTTACCATTTTCCCATCTTTTTTCCCCTTAAGATTCAACCTTGTAAAAACTTATGATTCTTGATTTCAGAAAACAGAGGAATTGGGAGACAACCACTAAACTGGACAGCAAGAATATCCCTCATAAAGGATGTAGCAAAGGGTCTAAATTTCTTGCACCAGTCCTTACAATCACATAAGGTGCCACATGGGAACATCAGATCAAAAAATGTTCTAATCCAGCAAGACCAAAGCAAGAACTACCATTCTAAGCTGACAGATTATGGATTCTTTCCTTTGCTACCATCTAAAGAATCCTTAAGAAAAATAGCAGTAGGAAAATCTCCAGAGTTCTCTCAAGGGAAAAAGTTAACAAGAAAAGCTGATATCTACTGCTTTGGTATTCTTCTGTTGGAAGTGATCACTGGAAAAGTACCAGGTGAATTGTTTTCACCAGAAAATGGAGTTACTAGTTATGTTGATGATGATTTGTCAGAGTGGGTAAGGACAGTGGTGAATAATGATTGGTCAACAGACATTTTGGATATGGAAATACTTGCACAAAAAGAAGGATATGATGAAATGTTGAAGCTTACTGAATTGGCTTTGCAGTGTACAGATGAGGCACCAGAAAAGAGGCCTAAAATGAGTGAAGTGTTGAGAAGAATTGAAGAGATTGAGCTTGAACCAAAGAATTTAGTCACTGATGTGGAAGATGATTCAGTCTCTTAGTTGAAGACTTGGAACTTTCCCTCCCAAACTGTAATCAAGCCGTAGCCGCAACAACAACATATCCAGTATAATTTCATCAGGTGGAGCTTGAACAATATATTCAGTATAATCTCATCAAATGAAATCGGAGTCTGAAGAGAATTTCAAGTAGAGTCTGAACAATATATTCAATATAATCTCATCAAGTGGAGTCTGAAAAAAATAGAGTGAATGCATACATTATCGTAACCCAATCCTATGGGAGTAAAGAAAGGGTAAAATATGCACGTACTTTACTCTGATTCCAACCAATGGGATCTTTGAGAGAATAGAGTATGCGCATACCTTACCCAACAAGGATAGAGAGACTGTTTCCTAAATTGTAATCAAGAAAATAGATTAGAAATTTAGGAACAGGCTTTTAG

>PGSC0003DMG400015391 LRR-RLKs

CAAGAATTTGGTGTCTTCTAAGTGCATGACAGCTATAATCTTTTTTATGCAGGGATATATCATACAACAATTTCACACAAATATCTACTTCAGTATGCGATTCCTCTACCTTGTGAGCATTGTTCCTCTTCATTTGGAACCTACATTCATCTACTATTTCTTTTTTTTTATAAGTGAAAATTTTGGTCCTTTAACTACATTACTTACAAACCTTCCCTCTTTAACAGAAACTTAGCTGCCAGCTATTCAAATACAATAAACACGCTCAACAACTCGTAAGTCCTAAATGTTGTCTTATGCATCAACTGCTAGTTTGTTTGATGCTTTCTGTCTATTGTCTATAATTATGTGGATTAATCTATTTGAAATTGCAGGGATGCCTGGTGTTCTATGAAACCCCTCATTTGTCCCACAGAAACGAAATGTAAGAGTTACAGCCTATAAACCTTTTTTTTATACTTGGAAATTCTTAACTTGACCCTTGAGAGGGCCTGAAGTAGCAGTTTGTCAAAGGACCTCAGGGGTGGCAGCCTGTGCACTTTTAAGTACAAAAGCATAACCTTGGAGTTGAGAGATAATTTCACAACTCTTCCTCATGTTATGCTCTTTATGTCAATTGCAGATACTTCGCTATTCATAAACTGTGGAGGACCTAGAACTTCATTTGAGGGAAATGATTATGAGGGGGACACGACCGACAGAGGTCCATCATATTTTTCTTCACCCTCCGACAGATGGGCTTTCAGCAGTTCAGGAGTGTACGTGGGTCTTCAAGCAGCCAGTTATGTTGCAACAAATACATTTTCACTGAATGTGTCCGGTCCTGACTTTTACAATACAGCACGCCTTGCCCCTAATTCTCTCAAATATTATGGGCTTTGCCTACGAGGAGGTAGCTACAGAGTGCGCCTTCACTTTGCTGAAATAATGTTTTCTAATGACTCAACATATAGCAGCCTTGGAAGGCGGATATTTGATGTAGCAATACAAGTAAGTTTGAGGCCTAACTTCTTTTGTAAATAAGGAGTATGTAATCTAAAATGTTGATTTATTTAGTTCTGATATGATTGGCAACTTTCTGATTTAATATTGAAGGTACCAATTTGTTGGAGATAATGTGGTATATTTTGATTTCTTACACAGGGGAGAGTAGTTTTGAGGGATTTCAACATTATGGAGGAAGCTAACGGTGTTGGAAAAGTCATTACCAAGGACTTTGCTGATGTTACTGTTAGTAGCACTCTGGAGATTCACTTGTATTGGACCGGGAAGGGGACTAACGCTATTCCAGACAGAGGTGTATATGGACCGCTGATTTCAGCTATCACAGTGACACCAAGTCCGTATATCTCTGCCTATAGACTTTACACTTATCTTCCCACAATATTATCTTAATAAAATCCCATTTTCTTCTTGCAGACTTTGATGTTGATACAGGCAGTGGGTTACCTGTCGGAGCTGTTATTGGCATTGTACTCGCTTCAATTGTGGTGGTTCTGCTAGTTTTATTTGCTCTGTGGAAGAAAGGCATTTTTGGAGGGAAAAATAATCAAGAAGAATTAGGTAATACATACTTAAAAGAATTTTTTCACTCTTTCCATTTGATAAAATGGGGATTCTTCTGTTGTGATAAATAGAATTTTTTACGAGATTTCTAGCTGCGACAGACAAATAATTAACTCCCCGTAATTGACCAGCATTGAGATGATTGTTCTGTTGGTGGATCGACATCATGACATATCATAGATAGTTAAACTGTCTTCATCATACAGATGTTCATTCTTTTAAAAATTGTTGTTCATCATACAGACGTTTTAGAAGATCTATATATACTGACGTTTGCAATATTCTTCTCATTTACAGAACTCAGAGCCCTTGATTTACAAACAGGTCATTTTAGACTTAGACAAATTAAAGCTGCTACAAATAACTTCGATCCTGCCAACAAAATTGGTGAAGGAGGGTTTGGACCAGTTTACAAGGTATCAATTCTCTATCCATACTGATTTTGCATAAGGACAGCGGGTTTTTGGTGATATTTCCTACTGGTTATCCTTGATGTCACGTTTATAGGAAAATAGAGTAGTAATAGTTGAAGTAACCCGTATCCTTCTTCTGTCTGTAATATGATCGTTGTGAATTAATCTTGCTATCTGTAGGGAGTACTCGCCGATGGTGCCATCATAGCTGTTAAGCAGCTGTCTTCCAAGTCGAAGCAAGGAAATCGTGAATTTGTCAATGAGATAGGCATGATTTCAGCTCTACAACACCCAAACCTTGTCAAGCTTTACGGTTGTTGCATAGAAGGGAACCAGTTATTAGTGATATACGAATACATGGAAAATAACTGTCTTGCTCGAGCTCTCTTTGGTAATGCTTTGCAACAACATTTTAAGAATCATTAATTAGGCAGTGATAGTTTATGCTTTACATTCCTCTAATTCTCGTTTCTCAGGCCGTGACGACCAGAGGTTGAACATAGACTGGGCAACCAGAAAAAGGATATGCTCAGGAATAGCAAAAGGGTTAGCTTACCTTCATGAAGAATCAAGGTTGAAAATCGTACACCGAGATATAAAGTGTACCAATGTGCTTCTTGATAAGGATCTGAACGCTAAGATTTCAGATTTTGGATTGGCTAAATTAGATGAAGAAGAGAACACTCATATTAGCACCCGAATTGCTGGAACAGTGTAAGCCTTCTTATCATAACTGTCTCTGTACTATCATGTGATGAGAGAGTTACAGTTCTGTGCGTTTAAATTTGTTCATCATCAACATTACTATCGTTACTTTGAAGTATCTTACGTTTTGTTTTCTTTTTTTGAAGAGGTTATATGGCTCCGGAGTATGCAATGAGAGGCTACTTGACAGATAAAGCTGATGTTTATAGCTTTGGAGTTGTTTTATTAGAGATTGTCAGTGGGAAAAGCAACACAAACTACAGGCCAAAAGAGGAATTTGTTTACCTTCTTGACTGGGTAAACTATCTTCTTTACTTTCTTAAACTCTGTATACGTCAAAACTAGACAAACAAATTGAAACGAAGGGAGTATTTGCTTTGATACCTCCAAAGAAAATATTAGAATCAGAGTTGTTTCATTAGTTCTGATTCTTTTATCCTGATATGCGGATGTCAATGTTTGTTTTCTCTTTGTCATGCAGGCTTACGTCCTACAAGAACAGGGAAATCTGTTAGAACTAGTGGATCCACGTCTCGGTACCCATTACTCCAAAAAAGAGGCAATGCGGATGATAAATATTTCTCTTTTGTGCACAAATCCCTCTCCTACTCTCCGACCATCCATGTCTTCAGTAGTGAGTATGCTCGAAGGAAAACTCCCCGTGCAAGCACCAATAATCAAGCGCACTACATCAGACGATGAAATGAGATTTAAATCGTTTGAGAAACTTTCACATGACAGTCAAACAACACAAGTATCAACATACTCTCAGGACAGTCAAGGACAAAGTGTGAATGCACCTTGGAGTGATTCCTCAGTCTCAGTGTCTGTCCCTGGTAAAGATGGAAATGTTACTTCCACAAGTAGGCTTCTTCCAGATCTTTATAATGTAAACTTGGATTGATTCATTGACAACTACATTTTGATTTCCATCATGTAATTATCTAACTCTATCTTCACACAGGTCCCACATAATTTCTTTTCGTGTATACTCCAATTTGTTCAATTGTGGTCCATGATGTTTTTGTAAAATGTATGTTACAATTCTTATTAATTAGTTCAATTGAAAT

>PGSC0003DMG400015458 LRR-RLKs

TAAGAAAAAAAATGAAGACATAAATTTCACAACATATCCATACACCTAATATCCAAAAAAATGTTAAAGATTGCTCTTTTCTTCTTCTTCCTCTTCTTTTACTCAGTTTTATCTAATCAAATCTCCAATTTAGAACACAAAGCTTTACTTGATCTTAGATCATCTTTAGGCATCAGAGCAAAAAACTGGCCAAGAAAATCCAACCCTTGTACCAATTGGACTGGAATCCATTGCAACAATGACAAAGTTGTGGCTATTCAGTTGACAGGACTAAGAAGAATTCACAAAGGGAACATGTCTCCACAGTTTGCTGTTGATTCTTTAGTCAATTTTACTCACTTGACATCATTCAACTCATCAGGTTTTGTTCTTTCAGGGCCAATACCTGATTGGTTTGGCCAAAAGTTAACTCAACTCAAACAACTTGATCTTAGTTCAAGTTCAATTCTTGGTTCACTTCCAACCTCATTAGGGAGTTTGAGTAATCTTGATTTTCTGTCTTTGTTTAACAATTCAATTACAGGAACTGTTCCTTTAGAACTTGGAAAGTTGAGTTCTTTAAGGGTTCTTAATCTTTCTTGGAATTCACTTACAGGTTCAATTCCAAGTTCTGTTTCAGATATTAAGAATCTGTCTGTTCTTGATTTGTCTTCAAATTTCTTGTCTGGACAAATCCCAGTTGAGTTTGGTTCATTTCTTGGACTTGAGTTACTTAATTTGTCAAATAACAGTTTAAGTTCTTATATTCCTTCTGAAATTTCAAATCTTTCTGCATTAGTAGAACTTGATCTTGGCTATAATTACTTGTCAGGGTCATTGCCTGAGGGTTTATTAAGAATGAGAAATTTGAGTAAATTGTTAGTTGGAAATAATGAGCTAGAGGGTGCATTGTCAAGTACTTTGTTTACTTCAAATGATTCTGTGTTGGAGTATATAAATCTTAGTTGGAATCGATTCGAAGGCAAAGTTCCTGATGTTTTGTGGTCATTGTCTGATCATTTAAGGGTTCTTGATATATCAGGGAACAATTTTACTGGTGTGGTTTCAAATAGTACCTATTTTAGTGTTACTGGTGCTTCATATAACCTTTCTAATAATCTTTTCTATGGAAATGTGACGTTTGATGTTCGGAGTTTTCAGTCGATTGATTTGTCTAAGAACTATTTTCAAGGACTAGCACCTGATAAATCTGGTGTTCGAGTTATGGATAATTGCTTTGCGGGGTTATTGAATCAGAGGAGTTTAGATGATTGTAGTAAGTTTTATGCTGATAGGGGTCTAGTTTTTGTTGATGGTGGAATCCAAACTCCGGTGCAAAGAGCCTCGAAGGGCGGGCATAAACGTTTGTTTATCATGGTTGGAGTTTTTGGTGGAGTTGGTGTTCTTATGCTTATGGCACTAGTCATTTTATTGTTTTGGAAGGTATGCTATAGAGGGAATTCAAACGAAAGAGTCATATCGAATGTTAGTCCTGTTGTGGAAGGAGGCAATAATCAGGCACCAGTAAAGTTCTCTCGTGATACATCTGGTTCAATTGAATCATTTACTTATGATCAGATACTAAAAGCCACAAGCAACTTCAATGAAACAGATCTTATCAAGAATGGTCATACTGGAGACATTTTCCGAGGAATATTAGAAGGCGGTGTCACCGTAGTTATCAAGAAAGTTAATGTACACCATATCGAAAAAGAAACATACAAGTTGGAGTTGGATTTGTTAAATTGGACAACACACCATAGGCTTGTCCCTTTTTTGGGACATTGCTTAGATCATGGGAATGAGAAGTTTTTGGTCTATAAATATATGCCAAATGGAGATTTGTTCAATTTGTTGGCCCGGGATACGGATTTAGAAGACGAAAATGGACAGTCTTTGGATTGGATAACAAGATTAAAAATTGCAACAGGAGCTGCAGAAGGCTTAGTTTATTTGCATCATGAGTGTAATCCTCCCCTTGTTCATAGGTATTTTTTTCTTCCACTTTGTTGCATTCTTGTTTGATAATGTTGTACACCAAGCTAATTATATTCTGCATTTTCTGGTTCTTGAAGAGATGTTCAAGCTAGCAGTATCCTTCTCGACGATAAATTTGAAGTAAGGCTGGGAAGTTTGAGCAACGTGTGTGCACAGGAAGGTGATAGCCAGAATTTGATCACAAAGTTTTTCCGCACGACAAAGTAAGTATTACTATGATCATAGTTTGATCCTAATACAAGATTTTTCAGAAATGTAATAGACATTTAGTTAGATTTATGTGTTTACATTTCTGTATATTGCTTTTCGTTCAATGTTCTAAGATATATTATGTTTCTTCTTGACAATAAACAGGAATCCTGAGGGAGAGCCTGCTGGTATGTACTATTCTTTTCGTAAAAGAACGTATAGTTGAATAGAGTTATTTATCTGAATTTCTCATATGCTGCCTTTATACATGGACAGGATCTTCATTTACAAGTTGCAGTACTTATGATGTATACTGTTTCGGGAAGGTCTTGCTCGAGCTTGTAACTGGTAAGGTTGGCGTAAGTCATTTAGATGATGCCTCAACCAAGGAGTGGTTGGACAAACTTCTTCCTTTCATCACTATACGCGAGAAAGAACTACTGTCCAAGATAGTCGATCCATCTCTAATTATAGATGATGACTTACTTGAAGAGATGTGGGCTGTAGCAATAGTAGCCAAAGCATGTCTTCATCCGAGGCCTTCTAGACGTCCTGAAATGAGACGCGTCCTGAAAGCTCTTGAGAACCCGTTCAAGGTTGTTAGGGAAGGAAGTTTCAACTCTATAAGGCTACGAAATACTTCATCGAGGCGATCATGGAGTGCTGCTTTTCTTGGTAGTTGGAATCATAGCTCACAGGATAGTGTTAATGCTAGTGGCCAAATAAGCAAAGAAGCTGGCATGAACCTGAAACAGTCAGGAAGAGTAGGATTATCTCATTCTATTAGCAGTGGGAACGAACATTCATCATCGTGTAAGAGATCATCGAGTGAGATTTTCCCCGAACCTCAGGACACACTAGATATAGAGAGACAAGACATGAGCTAGCCATCCAGCACGAATCTATGCCAAATGCATTTCTGCAGGCAGGGGTTTTGTAAGTATCAGGAGCTTTGATTTTTTATTAGATGAGCGGCAGTACTATGATCATTCTGAAGACATGGGGCCTACACGCTACACACATGTTATGCGGTTGGTAAACTCTTTCCTTCTGCAACAGCCGATCAAATCGGATCAGTATTTTTGAATATGTTGTGTTACATCATGTACATGTATTCTGTTTTCCTCTTTTCTTTTAGTGTGTCACAAATGCTCATGTGCTTTGTAAATTTTCAAAAAGTTTACTCTTGTTCCTGTGATGATAGAACAAGTATATAATTGTGTCAAGAGTGTACACTATCTAATCCAGATTCACATTTCCATGATCTCATATTTTGCAATTGCATGAAAAAGTATGATATAAATGAATAATAAAACCTGCACTTTCTTGAGCCGAGGGTCTATCGGGAACAACCTCTCTATTGTAGGTAGAAGTAAGGTCTGCGTACATCCGGTCATCTCCAGACCTCATTTGTGAGATTACATCAGGTATGTTGTTGTAATTGGATAAATAGGGCATCAAAACTGTTATAGTGTGTCAATCCCTTGGTGGCCTAAATAAGGACAAGTAACTCAATACATGTAGATGCACCTGAAAGTTACAATGAGC

>PGSC0003DMG400015666 LRR-RLKs

CAAATTCCCAACTAGTAGTATTTCTTTATCTTCTCCAGCTCAGACACAGACACACAGACACACAGTAGACACAGTCCATTCTTCTCTCTCAAAATCAACCAACAAGAAGATGAACAACATGTTTACCACTACTTTGAACAAGAATTGAATCAAGAAATCAAATTACCCAGAAAAAAATCCTGTTTGGGGGTTTTGGGTTTACCCAAAATCTTCAAAAAATGAGCTCAAAAATCTTGTTTTTATTCACCATTTCTGTATCAATTTTCACTCTTTTCTTGCCGACAAACTCCGACATTGTTTCCGATCGAGCTACCCTTTTATCTATCCGATCTGCTCTTCGTGGACGTTCACTTTTATGGAATATCACAAGTCCAACTTGTTCATGGCCTGGTGTTATTTGTTCACCTGATAAATCTTCTGTTTTAGAGCTTCATCTACCTGGAATGGGACTTTTAGGCCAAATCCCACCTGGGCTTTTCTCAAATTTGACTAAATTGAACTTTCTCAGTCTCCGGTATAATGCACTTTCCGGTGTAATTCCGGCGGACTTGTTTACTTCACTTCATGATTTGCGTAACTTGTACCTTCAAAATAATCTATTTTCCGGTCCAATTCCTGATTCAATCTTCTCTCTTACCAATCTTGTCCGCCTTAATCTTGCTCATAACAATTTCTCCGGTTCAATACCGGAATCTTTCAACAATTTGACTAGTTTGGGTACACTTTACTTGCAGGGAAATGGGTTTTCGGGTCAGATACCGGATCTGAATTTACCTGGTATGGTTCAGTTCAATGTATCGAATAATCAGCTAAACGGGTCAATTCCTAGTAAACTTGCGGGTCAACCCAAGGATGCTTTTTTGGGTACTTCACTTTGTGGTAAGCCACTTGATTCATGTGATGGTAGTAGTAGTAGTAGTAGTAGTATTGGAGAAGGTAAAAAGAAGAAACTTTCTGGTGGAGCAATTGCTGGTATAGTAATTGGTTGTGTTGTTGGTTTGCTTTTGCTTCTTTGTTTGCTGTTTTTCTGTTGTAGAAAAAGGGGTAAAAAAGAAACAAGATCTGCTGATGTTGGTGCTGTTAGTAAGCAGGTTGAAGTTGAAATGCCTGAAGAAAGAGGGGTTGAGAGTAATGGTGGAAAAGATGGATTCTTGGGTAGTGCAATTGCAGCTATTGGTGTTGGAGGGGGTAATAAGGATAAAGGTAAAGCTGAGGCTGTTGTAAATGATGGAAAAAGTTTGGTGTTTTTTGGGAAAATGGCCAAAAATTTCAATTTGGATGATTTGTTGAAAGCTTCTGCTGAAGTTTTGGGAAAAGGGACATTTGGTACTGCTTATAAAGCTGCTTTGGAATCAGGGATTACATTAGTTGTGAAGAGATTGAGAGATGTTACTGTACCTGAGAAGGAGTTTAGAGAGAAAATTGAAGATGTTGGGAAAATGAATCATGAGAATTTGGTTCCTTTGAGAGCTTATTATTATAGCAGAGATGAGAAGCTTCTTGTTTATGATTACATCTCAATGGGAAGCCTTTCTGCACTTTTACATGGTGAGAAAAATAACTTTTTTTTTCTGTCATTCTTTTTCATGTCATTATATTGCTGTTATGCTGTTCCTTGAATCGAGCGTCTATCGAATCATGTCATTATAGTGCTGTACTGTTGTTTACCATTTACATGATCTGCAAATTTTAGTTGTCTGATTGACACTGATACTGCTGCTTTCTATGTTGCCTAGACTCTTAAAATGTGGTGCTTGTCGGATCCTGGATCCTCCAAAAGTAGTGTATATTTGGAGGATCCAAACACGAGTCCGAATAACATAGACTGTTTTTGCTGATTTTATGAGGCTAGTAAGTTTGGATGACCATTTGTTTTCTTGGATCTTGTTATCTGTTACTGAACTGGTGCTGTTGTGTAAACGTTGCTGCATTTGGCTGATGTGTTGTAGTTGCATTATTGTGCCCAATTGTGTTCAGCCACTAAAATTTATGTTTTTTTTTTTACATTATTGTGCTGTTGTTGCATTGTTGTTCATGTTTAATCAGATTTTATCGTAATCTAATGTGATGAACATTGTGTTTAGTGAATATGATATCTGACATGTTTTATATATAACTCAACCTGGAATGGCTTTACTATGTCCAATTGTTGCTGTTCAAACTCGTGTTCATCATTTAGATTGAATGTATACTTTATCCACAAGTGATCTGCACAATTTTTAGATTCCTAAGCCCGAGAGTCTATCAGAAACTACCTCTCTACCTACAGAAGGTAGAGGTAAGGTTTGCGTGCATCCCACCCTCCACAGACCTCACTTGTGAGATCTCATTCGGTATGTTGTTGGAAGAGTTTAGGAAGAAGTCGTAAAATTGAATATATGTGTAGGACAGGTTCACTGCGTTCAACAGGACGAAACAGGATCATTTAACATTTAATCTGTTTGTATGGATTTAAGCCAAATGTACCTTTGTTAACTTATCGACTATATGTTGTGTAAAAACTATGAATTATTTGGGTGCTTTATTCGGCATATTCCACTGCTACTATATCATGTCACAATAGTTTTGTCATTCAACTTGTCTTTTCTATAGGCAACAAAGGAGCAGGTAGGACACCGTTGAATTGGGAAACAAGGGCTGGCATTGCCCTTGGAGCTGCACACGGGATTGCCTACCTTCATGCACAAGGGCCCTCAGTATCACATGGCAACATTAAGTCATCCAATATCCTCCTCACCAAATCATACGAGGCCCGTGTTTCAGATTTTGGCCTTGCTCAACTCGTCGGTCCATCATCAACTCCCAACCGTGTTGCAGGTTACAGGGCACCAGAGGTAACTGACCCTCGTAAAGTCTCCCAGAAAGCAGATGTCTACAGCTTTGGTGTCTTGTTGTTGGAACTTCTAACAGGGAAGGCTCCCACACATTCGGTACTGAATGAGGAAGGTGTCGACCTACCAAGATGGGTGCAATCAGTGGTACGAGAGGAATGGACAGCCGAAGTCTTTGACCTTGAGCTCCTTAGGTACCAAAACGTCGAGGAGGACATGGTCCAGCTATTGCAAGTTGCAGTTGATTGCACTGCTCAGTACCCGGACAGACGCCCTTCAATGGCCGAGGTCACTACTCGCGTTGAAGAACTCTGTCGTATGGATTCTGGAGGTGGTGACATAATTGACAATGATGCAGAAGTGCAAACTGCATGAAATTGAACAAGAAAAAATAGTTCCATCACATTATTTCAATTTTTTTCTTTTTCGTTTTCGATTGAGCATTTCGATTACTTTGTTCATTCTTGGAGTTCTTCAGTGAAGATAAATTGTTCATTTTATTTTATTTTTTTGAATTTATTTTCCAGTAGCAACATAGTTTTATTAAAGATCTGGTTGGTAGTTGAAATGGAGCTGTTTTTTGTTTTATAGATATTGATGTATTCTTTTTTGAAGGTGTGATGAACTGTATAAATAATTGAGATGACCTGTAGTTACA

>PGSC0003DMG400015806 LRR-RLKs

ATTTATGAGCTTTGTGCTTGAGGGGTAGTCAACAAAGTGTCAGCTTTCCCAAAAAGAAAAACCCCAAGAAACAAGTTCTTCCACTTCCTTCACTCATCACATTTTCCTCTTTTTTCTCTTTCATTCAATGGCACTTCAGAGGGTTCTTCTTCTCTCCTTCTACATTGTCTTTTTCATAGCCTTCACTACTACTTCTGCTGCATCTGATGTTCAGCTTCTGTTACAGAACATTAAACCCTCACTTCAGGGTAACACTGATAATTTGTTGTTATCTTCATGGAATATTTCAGTTCCTCTTTGTCAATGGAGAGGTCTCAAATGGGTTTTCACCAATGGTACTTCTTTGCTCTGTAGTGACCTATCTTCACCTCAATGGACAAGTCTTTCTCTTTACAGAAACCCATCTTTGCATCTTGTTTCTCTACAGTTACCATCAGCTAATCTCTCTGGAACACTCCCTAGGGAGCTTGGTGAGCTTTCTACACTGCAAAGTCTTTATCTTGGTGTGAATGGACTACATGGGTCCATCCCTCTTGAGCTTGGCTACAGCTCTTCTCTTTCTGATATTGAGTTGAGTGGCAATAGTCTTACTGGGACACTTCCAACTTCAATATGGAATCTTTGTGAGAAGATGGTTTCTTTCAAGATTCATGGTAATTCTTTGTCTGGTTCTCTTCCTAACCCTGCATTACCTGATGCTACTTGTAAGAATTTGCAGTTTCTTGATTTTGGGCAAAACAGGTTTTCTGGGGATTTTCCTCTGTTTATAACTAGGTTTAATGGTCTTAAAGAGCTTGATTTAGGGGATAACAATTTTTCTGGTCCTGTTCCTGAGACATTAGCTGCATTAAAGCTGGAAAAATTGAATCTCTCACATAATAATTTTAGTGGGGTGTTGCCAAATTTTGGGGAATCTAAGTTTGGTGTGGATGTTTTTGAGGGAAATAATCCTACTCTTTGTGGACATCCTTTGGGGCCCTGTAGTGGAAGCTCTGGAATGAGTCCTGGTGCAATTGCTGGTATTGTTATTGGTCTAATGGCTGCATTGGTGGTTGTGGCATCACTATTGATTGGGTATTTTCAAGGGAAAAAGAAGAGACATGCTGATGAAGAAGAGGAGGAATTTGAGGAAGTTGAAGATGAGGAAGCTGGTTGTGGTGAGGGGAAGCTCATCTTGTTTCAGGGAGGTGAGCATCTGACTCTGGAGGATGTGCTGAATGCGACGGGGCAAGTCATGGAGAAGACTAGTTATGGGACTATTTATAAGGCAAAATTAGCTGATGGTGGAACTATAGCTTTGAGGTTGTTGAGGGAAGGCAGTTGTAAGGATGGAGGTACATGTTTAACTGTGATTAGACAGCTGGGAAGAGTTCGACACGAGAATTTAATTCCTTTGAGAGCCTTCTATCAAGGGAAGAGAGGAGAGAAGCTTCTCATATATGATTATCTCCCAAACAGGAATCTTCATGAACTCTTACATGGTAAATTCTAGCTTCTTTATGTTCTCATGTTTGCTCTTTCAAGTTTTACTAATGAAAATCAATGCAAGCTCAATTACGTTGACGTTATTGTATTAAGTGTTGCCTCATAGGGAAGTAGCAATAGTTCTTAGGCTCATTTTCAGCAGCTTTGGTTATTTGCTTGAGAGACTAAGATATGTGGGCTTCAAGTTGTTGCCTTTTCACACGGTGGGGTTGTAATTGAAGTATTAGTGCATAAATAGATCATTTACAACAGCCTGAAAAATATAAATGTAGCTGAACTTAGCATCTTAAGAGAAACTGAATTAGTTATTGAATGCACAATATAGAGATATTACGTTACACGAATATAGAGAATGTTCCATTTATTCTGACCGATATCCTTTTCGCTAAGTTAAGTTGATCCAACCTATATCCTCTCCTGTGCATCATCTTCCAATACACGAACTGCCATATGTATACCTGTTTGGTCAACAAAGAAAAAGTATCCCTATCATATATACTTACTCATCCAAGAAAAATGGAAAAAAAAAGAACTAGCACATTATCATTAAACTGCCGATGCAACTATCAGATGCCTTGATCAACCATTCATTTATAATTTCGCTTTAAAGAACTCTGTGGCCCTAGATCAACTGATGAAAGCTCATTTCCAAGCATTTGAGCTTGGTGTAGTAAGCTTCTTTAGTTCCTTTTGCTTGATGCAATCTATTGATGCTCTGGTTGTTTCCTGTTCCTCATTTCTTGTACTGCTAATTTAGCAGTAGTTTGGTCATTGAGAGTAGTTTGCGCCAATAAATGTTGGAACCCCACAAATTTTTATATCCTTGTTCGTTCAACCTTAATAGCTCTAAGGATTAGTGAATTTGATTATGATATTTTCTGATTGTCTACAGAATCAAGAGTTGGGAAGCCAGTGTTGAATTGGGCTAGAAGACACAAAATAGCATTGGGCATAGCCAGAGGACTAGCACACCTTCATGGTCTGGAAACTCCCATTACTCATGGGAATGTTAGATCAAAGAATGTTCTTGTTGATGAATTCTTCGTTGCCAGGCTTACTGAATTCGGACTTGACAAAATAATGATCCCAGCTGTAGCTGATGAAATTGTATCAGTTGCAAAAGCTGAAGGTTACAAGGCACCAGAGCTGCAAAGGATGAAGAAGTGCAACTCAAGAAGTGATGTTTACGCTTTTGGTATCCTACTGTTGGAGATTCTTCTAGGAAAGAAACCTGGTAAGAATGGTAGAAACGGGGACAATGTTGATTTGCCTGCACTAGTTAAAGTTGCAGTTCTGGAAGAAACGACTATGGAGGTATTCGACATGGAGCTTTTGAAGGGAATTAGGAATCCAATGGAAGAAGGTTTGGTACAGGCCTTGAGACTTGCAATGGGATGTTGTGCTCCTGTTGCTACTGTAAGACCGTCGATGGATGAAGTAGTAAGGCAGTTAGAGGAGAATAGACCTAGAAACCGGTCTGCGTTGTACAGCCCTGCTGAAACTAGGAGTGGAAGTGGTACTCCCTTTTAAGCTCATGTTGTGACTTTAGTAATGAATATTGTCGCATATTATGCCTGTTTTTCCTCCTAAAGGAAAGAGGGCTAAATGGCTATAGCTTTCTTACTTACTATTTCAACTATTTACCTTTGGCTTTGCATTGTTGGGGAATATCTTATCATAGACAAACTGACAATAGACAATGTGTCACATTAGGCATCAAATTGGATATTACAGTGTGCCTTTCTTTTTTTCAATTTGTTCCTGCATTAGCCTACCTTCTATAAATATATATATGCAAAAGAAATTCAATG

>PGSC0003DMG400015814 LRR-RLKs

ATGCCAATGTCGAGGCGATTCTTGGACCTCCCAAATCTTGTTGTCTTTCTAACCTTTTTTCTTCTATTACATAACTTTGCTCTTCCTTCAAATGAAGTTGATGTCCTGTTTTCTTGGCTTCATTCTACTAATTCACCAATTCCTCAAACATTCTCAAACTGGAATCCTTCAGATTCCACTCCTTGTAAATGGTCTCATATAGTTTGTTCTTCTAATCTTTTCGTTACACAAATTGATATTCAGTTTATTCAGCTGGCTCTTCCTTTTCCTTCTAATCTTTCGTCTTTACAATCCTTGCAAAAACTGATCATTTCTGGTGCTAATCTCACTGGAACAATCCCACAGGATATTGGAGATTGTGTTTCACTTGTAACACTTGATGTTAGTTCAAATGGTCTTGTTGGTACCATACCGAAAACTATTGGTAACCTTATAAATCTTGAAGATTTGATTTTGAACTCTAATCAACTAACAGGGGAAATTCCAGGAGAGGTTGGCAGTTGCATTAACCTGAAAAATCTTATTATCTTTGACAATATGTTTAGCGGAAATCTTCCTTCCGAACTTGGAAAACTTGGAGTTTTAGAAAATATAAGAGCAGGAGGGAACAAAGATATTAGTGGGAAAATTCCGGATGAGCTTGGGAATTGCAAGAACTTGATAGTGTTGGGACTTGCTGATACGAAAATTTCAGGTTCGCTTCCTCCATCGTTGGGTAACTTGGGAAAGCTTCAGGTATTATCTATTTATACTACAATGCTTTCTGGCAAAATACCTTCAGAAATAGGCAACTGCTCAGAGCTAGTTGACTTGTATTTATACCAAAATAGTCTGTCAGGTTCACTGCCTGCTGAGTTGGGAAAGCTTCAGAAAGTAGAAAAGATGTTGTTTTGGCAAAATAATCTTGATGGGCTAATTCCTGATGAAATTGGGAATTGCAAAAGTTTGGTTATTCTTGATCTTTCTTTGAATTTTTTAAGCGGGAGCATTCCGTGGTCATTTGGGAACCTTACTAATCTCCAAGAGTTGATGATTAGTAACAACAACATTTCTGGTTCAATCCCATCTGTTCTTTCTAATGCGACAAACTTGTTACAGTTTCAGATGGATACGAATCAGATTTCAGGCTCAATTCCTCTGGAAATGGGGCAGTTGAAGGAGCTAAATATATTCTTTGCTTGGCAGAACAAGCTTGAAGGAAGCATTCCTCCTGCATTGGGTGGTTGCAGAAGCCTACAAGCTCTGGATTTATCACATAATTCTCTTACTGGTAGCTTACCTCCTGACCTTTTTCAGTTAACTAATTTAACGAAGCTTCTGTTGATTTCCAATGATATTTCTGGTTTTATCCCACCTGAGATAGGTAATTGTAGCTCTCTTATTCGTATACGACTTATTGGCAATAAACTTAGCGGACAAATTCCAAGAGAGATAGGATTTCTAGACAACCTAAGTTTTCTTGATCTCTCTGAGAACCACCTTACAGGATCCGTTCCTGAGGAGATTGGCAATTGCAAGACACTGCAAATGCTCAATCTATCTAATAACACTCTAAGTGGGAATTTACCTAGCTCTCTTTCTTCTCTCAGTAGGCTAGAGATTTTGGATGTTTCCTTGAATCAGTTTAACGGACAGATTCCAGCAAGCTATGGTCAACTTGCTAACCTTAACCGGCTTGTTCTTAGTAAGAATGCCTTCTCTGGATCGATTCCCCCAACTCTCGGCAATTGTTCAAGCCTTCAATTGCTTGATCTCAGCAGCAATGAGTTCTCAGGGAACATGCCAGTGGAATTGTTTGACATTCAGACACTTGACATTGCCTTGAATTTAAGCTGGAACATCTTGAGTGGGGTAGTCCCACCGCAGATATCTGCCTTAAACAAACTTTCAGTACTAGATCTTTCCCACAACAAGCTTGAAGGAGACCTATTGTCTCTTTCGGGGCTTGAAAATCTGGTTTCCTTGAATGTTTCCTACAACAATTTTACCGGCTACCTCCCTGATAATAAGTTATTCAGGCAATTATCATCAGCAGAGATGGCTGGCAACAAAGGTTTGTGCTCATTGGGACATGATTCTTGTTTCTTGAGCAATGTTGAAGGCGGGGGAATGATGAGTAATAGCAATGTAAGACGGTCATGGAGGCTGAAATTGGCAATTGCACTCCTTTCTGTGGTGACTATAGCTTTGGCACTCCTTGGAATGCTGGCAGTGTACAGAGTGAGAAAAATGAGCAGAGAAGATAATGATTCTGAATTGGGAGGAGGGGATTCATCAGCTTGGAAGTTTACTCCATTCCAGAAGTTGAATTTTTCGGTTGAACAAATTTTGAGATGCCTAGTGGAATCCAATGTTATTGGAAAGGGATGCTCCGGCGTAGTTTATCGTGCAGAACTAGAAAATGGTGAAGCAATTGCAGTCAAGAAGCTATGGCCAACCACATTGGCAACTGGATACAACTGCCAAAATTCAAAGTCAGGAATTAGTGGATGTGTTCGCGATTCTTTCTCAACCGAGATAAAAACCCTTGGCTCCATCCGTCACAAGAACATCGTTAAGTTCTTAGGTTGCTGCTGGAATCAAAACACTAGATTGCTCATGTATGACTACATGCCTAATGGAAGCCTAGGTAGCCTTCTACACGAACGAAGTGATGGATGTTTGGAGTGGGAATTGAGATACAAGATTGTCCTAGGTGCAGCTCAAGGGCTTGCTTATTTACACCACGATTGTACACCTCCAATTGTTCATAGGGACATCAAGGCCAACAACATTCTCATTGGCCTTGACTTTGAGCCATACATTGCAGATTTTGGTATAGCCAAACTCGTCGATGATGGAGATTTTGCTCGATCTTCCAATACAGTAGCTGGCTCCTACGGATACATAGCACCAGGTTCGGCACAAGTCGTAGTACCACTTTACAATATTACACTATAATTATATACATTATATCTTTAATACACATGTATATATTATCTTGTGCAGAATATGGATACATGATGAAGATAACAGAGAAAAGTGATGTTTATAGCTTTGGAGTAGTTGTGTTAGAGGTGTTAACAGGAAAGCAGCCAATTGATCCTACAATACCGGATGGAGTACACATTGTGGATTGGGTGAGGCAGAAAAGAGGCAGTGATGAAGTTTTAGATGTGAGCTTATGTGCTCGTCCCGAATCAGAAGTTGATGAGATGATGCAGACTATAGGAGTAGCTATGCTGTGTGTTAATCCATCACCAGATGATAGACCAACGATGAAAGATGTCGCAGCAATGCTGAAAGAGATAAGACTTGAAAGAGAAGAGTACCAAAAAGTTGATATGCTCCTTAAAGATGGAGGAGACAATAAGAGTAGTAGTGATGGAGGGCCATCTTCAAAGATGCATAGCTTGTACTTGCAAAGCAATAATACAAGCTTTTCTGCATCTTCATTGTTACATTCTTCTTCCTCCAACACCAAGAATATTGGTTTCAAATAGACTTTTGTTTCTTTCTTATAGTTAGATTTGTTTTCTCATTTTTCTTCCCGGG

>PGSC0003DMG400016015 LRR-RLKs

TTTTTTTCCAATATATATCCTTTTTCCATTTCTCTTCATAGTAGTATGTTTCAAGTGCATAGTAACATCATGGAGAAACACATTTTCTTATTGATACTTGCTATCTTAGTTCAATTTTACTTTGTTTCTTCTATATCAGCTACTATTTTCTCAAATGAGACTGATCAAGAAGCTCTATTAGCTTTTCGAAATCTTGTTACGAGTGATTCTAGTCAATTTTTAGCCAATAATTGGACCAAAAATACTTCATTTTGCTCTTGGTTTGGTGTCACTTGTAGTCCAAAAAGGCAAAGGGTTGTAGCCTTGACTCTTCCTAATTTGCAACTTCAAGGCACAATTTCGCCTTCTTTGGCCAATCTATCCTTTCTCATAGAGCTAAATCTCACGAACAACAACTTCCATGGTAACATCCCTTATGGCATTGGCCACTTGCCTCGTTTACGAGTGATTGATATTCAGAACAACCAGCTCCAAGGTAGTATTCCAACAAGTCTATTTCAACACCGGAGTGTTCAAATCATTTCATTGGCTTTCAATAAACTCGGTGGTGAAATGTGGAACGGTACATGGTATGTACCGGAACTCAGAGTCTTAAATCTCAGGAACAATACCATCACAGGTAGAATCCCTCCTTCTATTGGAAATGCCACAAAGTTGATGAACATCAGTTTGAATTGGAATAGAATCAACGGCAACATTCCAATGGAGATCGGTAATCTAAGCCAACTTGTAGAGTTGTCGTTGTCTCGTAATCAATTAACAGGTTCCATTCCTTCAACATTGTTTAATATCTCCTCCCTTCTCGTCGTGTCTCTGGCATACAATAGCCTTTCAGGTCCTCTGTTTCTTGATGATCGACGTAATGTTCTTTCATCAAACCTCGAGCATATAGGTGTATCATACAATCAAATCACTGGTCACATTTCTTCCAACATCTGCCAATTCAAAGCTCTCAAAGTCTTGTCCATATCATACAACAACATAACTGGAGAAATACCGAGAAATATTGGTTGTTTAGCCAAGCTCGAAGAGCTTTATATCGGTTATAATGCAATAGATGGAACAATTCCTACTTCATTAGGCAATATTTCCACTCTTCAAAAACTTCATTGTGGAAACAATCACATGGAGGGAGAACTTCCTCCGGAATTAGGAAAGCTATCAAACTTAAGACAAATCAATTTCGAAGAAAATTATAATCTTATAGGTGAAATTCCAAATGCTATTTTCAACATATCTTCTTTGGAATTCATTGCTTTCACTTTCAACTACCTCTCAGGTAGAATTCCAAATCTTCTTCATCTTCCAAACCTTATACAACTTCTCTTAGCAAACAATCAGCTCGAAGGTGAAATTCCTCGGTACATCACAAATGCTACCAATCTTGAGCTATTGGAACTATCAGATAACCTTCTCACAGGCAGTATTCCTTATGATTTAGGAAATCTTCGCGAGCTGCAAGAACTTTTCCTACATCATAATCAACTTACTGAGTTGGGATTCTTTGATTCTTTGGTGAAATGTAGGATGTTGAGATATGTACAAGTGGGATCGAATCCGTTGAATGGTGTTCTGCCAAGTAGTATTGGCAATCTTTCATCTACTGTTGAATACTTTCATATTGGAGATGCACAAATCAATGGATTCATTCCCACTAGTACAGGCAACATGAGTGGTCTTACAACGCTAGTTCTTCAAGATAACAATTTGACAGGAAACATTCCTCGTGAGATCGGTAAGCTTAAACAACTCCAAGGTTTATTTCTAGTTAACAATGAACTGCAGGGGGATATAGCAGAGGTAGTATGTGATTTATCGAATTTGGTTCGATTAGCTCTGTCTGAAAATGAGCTCTCGGGGGTGATTCCGGAATGTCTAGGAAGTCTTACCATGCTACAACACCTTTTTTTAGGTTCTAACAAGTTTGAATCAAAGCTTCCTTTAAGCTTTTGGAAGATGAGTAGTCTTCTCTATGTAAACATGTCGCGTAATTCTATAGAGGGAGAAGTTCCATCAGATATCGGAGAACTTAAAGCTATTGTAGCAATTGAAATCTCTGGTAACCACTTTTCGGGGATGATACCAAGCAATTTGGGGGAACTTCAAAACTTGAAGTTACTTTCCTTATCGAACAATTCGTTTTCAGGTCCAATTCCATTATCCTTTTCAAACTTGAAAAGCTTGGAATTCTTGGATTTGTCTTTGAATAACTTGTCAGGTACTATTCCTAAGTCTTTCGAAAAGCTTTTGTACCTTACAAGCATCAACGTCTCGTTTAATGTTTTAGAAGGTGAAATACCTAGTGGTGGTGTGTTTGCAAACTCCACCCTGCAATCATTTCGCGGGAACAAAGGTCTATGTGGAAGGCAAATATTGGAGGTTCCTGCTTGTGCTGTCACTACTCCTGAACAACAACAACCAAAATCGAAGAGGCTTGTGCTAAAAATTGTCACTCCGGTGGTTATTTCATTCTTTCTGATATTCTTGTTGGTTGTCTCAATTTGGATAATGAAACGAAAGAAGAAAGGAAAGTCCAAAGATATTGAAAAGGTTCCGGAGATGAGGACTTATCAATTGATTTCTTATCATGAGATTCAACGAGCAACAAACAATTTTGATGAATCCAATTTGATTGGCGTGGGAGGTTCTGGCTCTGTGTACAAAGCCACATTACCTAGTGGAATTGTGGTTGCAATAAAGGTACTGGATTTGGAAAATGAGGAAGTATGCAAAAGGTTTGATACTGAATGTGAAGTGGTGAGAAATGTTAGACACAGAAATCTTGTTTCGGTGATCACTACGTGTTCTAGTGATCACATAAGAGCCTTCGTTCTGCAATATATGCCCAACGGAAGTCTTGACAATTGGTTGTACAAAGAAGATCGCCACTTAAACCTTCGTCAAAGAGTCACCATAATGCTTGATGTAGCTATGGCAATTGAATATCTACATCATGGTAATGACACCCCTATAGTTCATTGTGACCTCAAGCCAGCCAACGTTCTTTTGGATGAAGATATGGTGGCGCGTGTTGGTGATTTTGGCATCTCAAAGATTTTAGCTGTAAGCAAGTCTATGGCACATACAAAGACATTAGGCACTCTTGGATATATTGCACCAGGTATACTACTATACTAGTCTCTTTCCATTTCACTTATGTATCGAAGACCTTTCATTATTGTTACATCGGATGAGTAATGCTTTTTTCATTTTCATTCTTTTAAGAATATGGCTCGGAGGGAATAGTGTCCACTCGTGGTGATGTTTACAGTTATGGCATCATGTTGATGGAGGTTTTGGCAAAAAGAAGGCCAACAGGTGAAGAGATATTCAACGAAAATCTTGGTTTGAGGGAGTGGATAACGCGAGCATTTCCAAGAACTATGATGGAAGTTGTGGACGCGGATATTTTTCATGATGGAGAAAAAATCACTTCCAAAAGTGAACTCTGCATACTTTCCATGATAGAACTGGCTTTAGATTGCACAAAGGCAACACCAGAATCAAGGATAACCATGAAAGATGTAGTCAAGAGGCTTAACAAAATTAAGAACACATTTTTGGAAACGTAGAAGTTAGTTAGAATCTCTTTTAATGTTGTTTTTTCTTACAAGTTGTAAGTTGATACTCAATCTGTTTGTCATACTTTTTAGTCCGTTTTAAAAAGAATGCCTCTTTTCTTTTTTGGGAACTCTTAAATTCTAACTTTCTACATGTGTATGACATGTTTAAGACCACGAGATTAAA

>PGSC0003DMG400016164 LRR-RLKs

GGGGTAACTAAGAAATACTGCTGTGAGTGACAGGAAGAGAAGACTTAAACGTAGTAATAATTGCAGTATGCAAAAGAGAGAACCTTTCTTGAACTTCTTCTCCATTGACTACTACTTTTTTACAGCACAACAAAGAACAGCTTTTATTTGTGTTTTTTTCTTCAAGATTTGCTTTGTAAAGCTTATTTTTTTGCTTAATTCAAAGTAAAGTTTCAATCTTTGGTTCAAACTCTGCTCTTTCTTTCACAGCCTAAAAAGATTGGTGCTTGCTTAAAGGTACTTAACTTACTCAGCTCTTGTTTTTGTTTTGTTCTGTTTTTGTTTTCAGATTCTGTCATTTACTGTTATTTGTTTGGTGTATTGAGAATGGTGTTTTGACTTTAGATATGGTGGGTTTGTCTTTTCTGATTTGACTTGACATATGGAAGCAAAAAAGTGAACTTTTTTCTTGGAAAAATTTGTTCTTGATGGATTTTGTGAATTTGTGGATGTGGGGTTCTTTTCTTTTTGGGTGGAGAGGGGAGTTGCTAATTGTTTGTTTTTTGTGATACATATGTGAAATAGAAGGAGGTCAAGTTTAAATTGTTAACCATAGGCAAGGGGCATAGTTGATTTCGGATATGGTCTCTCGACTAATAGTTATTATGTCAGAAAGTTCCTTTCTTTGTCGAAGAAAATTGGAGTCAAGAGGAAAGACTTTATGAGATAACAACGGTTAGTTGAGTGACCATGTGAGAAATCAAACCAAAAGGGACAGTGGAAAAAAGAATGAACTTGGATTTTTGTGAGTCTGGTTAAGTTTTAGGAATTACAATGGTTGAGAAGGAGATTTGAATATGTACACAGTCAGTGTTTTAGCTACAACAACCTTTGTTGTTTCATTTGTTGGAGTTTCTTTTGAAGAACTTGTGTCTTTCTCCTTAATAAGTTACTGAAAGTTTTTTGTAATAGTTCATTACTCACTTGATGTATTTTGAGAAAATGGGGGAATTATATATCAGTTTTGTTATATCTTCAAAGAATTTCTTGTGGTTATCTGATAGACATAACCAATAGACATTCAAAGAGTTGCTGGCGAAAACATCTGGTTTTTGTTAAAGAAAAGGATAGGTTTGTCGGTTGTTTGAAAGTTGTGGATCAATTCTTCCTTTGGATGGTTGACATTATTGGTAGTAGCTGAAATATTGTTTGGATGGAGAAGACGAGTGTTTAGTTGTTCTTTCAACTCCTAATAGCTTTTACGATTCTGCAATACTGCTACAGTTTGCTTATATGGGGTTTGCAGTTCGAAGTACGTCGTTGCATGCTTTACATTATTAATAGAGGGCATTATGTAATGCTGTATTATGCTTTTCATGTGATTGATCTATATCATCAACCACTACCATCTTCTTCTTGTTCTTTGTACTTCAATTATCGCACTTTCTTTGGTGTAGTTACTGGTCCTTTTCTTAGAATGCTTTGTCATACTTTCTATAGTATTTTGCTATGATCCCTTCACTTCAGTTATTTCTTTTTCGATCTGCTTTGATATACTTTTCCTTGAGCCGAGGGTCTTTTGGAGACTGCCTCTCTACCTCTATGAGGTAGGGCTAAGGTCTGCATACACTCTTGACTCTACCCTCTCCTCATTTGTGGTATTACACTGGGTATGTTGTTGTTGTTCTTGCATCAAATATCTCTTTTACATATTTAATTAGGCTTAGCTGTTCTCCTTGAGGTTATACTAATTTTGATCGTAGTAACCTTTCTAATATATTATGCAGAACTGCCATTTGTCATTACCAAGGAAGATAGTGCTTTATGTTACAGTGATGAAGGGGAACAGGTTTACTTGATCTATAGAATTGAACAGCAACTGCAATATGGAACTGTTCCTTGCAATTCTGATATTAGCATGTCTCTACTCATCTGTCTCACCTGATCAGCAGGGTACACATTTCTCCCATTTTCTACTTTAGCAAACTAACAAAGAGATGCCCTTAGCTGTCTTTAGTCTTTTATCTATTCAAAGGTATAAATAAAAATTTAGAGGGGTTAGATAATTTTGCTTGTGCTGACTTAATGTTGTCTGTATTCTGCTCGTAAAGTAGTGTGACATTCGTTATTTTGTAATCTCTTGATCAGGAGATGCACTGTTTGCATTGAAGAGTTCACTAAATGCTTCAAATGATCAACTTTCTGATTGGAATCAGAATCAAGTTAATCCATGCACTTGGTCAAAGATAACTTGTGACGATAACAGCAATGTCTTGATGGTGTAAGTTGAAGTTACTTTGCGAGCCATATTGATCACCATGTACATGATAAAATTATGGTCCCAAACACTTGAAAAGTTGGTTAACTACCATTCTGAAATCCTATATTTAATGTTGTCTGTGCTTTATTTGCAGGTCGTTGTCAAATATGGGGTTCACTGGCACTTTAACTCCAAGATTAGGTGTTCTAGTGCATCTTAATACTCTGTACGTATAGAGAATTTGTGGAATTATTTTCAATTGAATGCTAATGTGTATCTATGATGTACTTATAGAGATCAGGTGTGCACTTTAATCATAACGTCAGTAGATAAATCACTCCATTTACGAAAACTAGGTAAGTGTGATCAACTGAAACAAACAAAAACATACCAGTGTAATCCAACAAGTGGGATCTGGGGAGGGTAGATTTTGCTTGTGCTGACTTTAGGTTGTCTCTGTACTGTTCGTAAAGTAGTGTGTCATTCGTTATTTTGTAATCTCTGACCTTTGATTTGGATGATAATTTTGAGTTTCTTGTTCTTAGGGGAGGAAATAACATTGAACTTTCAATTTGAAACCATTGTGGAAAATGTGATGGTGAAATAAAAATATGGTTTGTTAGGTTATGTGCTGTCAATTTTCTGCTGAATGCCTCTCAAAGTGAGAGAGTCTTTCCATTATATAGAGAGCATAAATATTACACATCCCTATAGATCCGTCATTCCTATCCCTAAAGATCAGCTATTCTATCTCTACAAATCTGCTATTCAATCCCTCTAAATTTGCTTTGCTATCCCTCCAACTCAACTATTCTATCACAATATTTATTCCTCTGATTCTACAACAAATCTATATGATAATACTAATTCATATCACTATATAATTACATTAATATATACAATTATCCATCATGCTAACATGTGCTACTCTAGAAGTCTATATTTATCGATTTCCTTCATCAAGCTGATATTAATCTACTCATGATATTGGCACATTACTAACTGCTCTTGTTTGGGGACTGGTGATTGCTTGTATCTTTGCCCGTGCATGGTTTGGGATCTTAGCCTTTTCCGATTGTACTACTTTCCTACCCACACAAGCATGTGTAGAAATCTACTCGCAATTTTGGTTCTCAATATTCCAACTAAAGAGTAGTATTCTTTTAAGTTAATAATTAGGAAACGACATTCTTGAGTGTTTGGTCATGCTTTTGCCAGAGTGTTGGATATTTCATCTTCATTTCTTGTCCGTTATCACATTATTCTCGTAGTGCCTCTGATAAGCATTCTAAGGTCATAACTGGTAACATTTTCTAAGATCTCCTAAAAATAAGTTTTCATCCTTGTCTGTAGGAAGTTCTACTTTAGTGAGATCTGTGAACTCATCAATTTTCTTGTCCTTAGTTGTGTTATGTATTGAACTGACTGGGCGGTCAATATCCTGGCAGTTCTTTGCAAGGAAATGGAATAACTGGTAAAATTCCTGAAGCATTAGGAAATTTAACCAGCTTGACCATGTTGGATCTGGAAAAAAACAGTTTGTCAGGAGAAATACCAGCTTCCCTAGGCAATCTAAAAAAGCTTCAATTTCTGTAAGTAATATTATTTCATTGATTATTGTTTTTTTTAAAACTGAGTTTTACATTTGGTGTTTGACTTGGGAGTCTATTATAATGTTTCTTTGAATAGGTTTTTGAGTCAAAACAATCTGTCTGGGACTATCCCTGAGTCACTTTCAGGCCTTCCCAACTTGATTAACCTGTAAGTGATTCCTTTAGCCTGATCTTTAATGTGCTTACAACGTGTGCCAAGTGATATTGACCTGACCCGGTGATATCTCTTTTCCAGTCAGCTTGCATTAAATGATCTTACTGGAAAAGTTCCTGACAGTTTGTTCCAAGTCCCGAAATACAAGTATGCAGATCTCAATTCTTGCATTGTTTAATTTGATGATAGATTGAAGCTGCTTAATAATTTGAACATTTTAAAAAGGTATAATACATAAATAGACACTCAAACTTAGTCTCAACTGGCAAGTAAGCGCTCCCAACTTTGAGCGAGGACATCTAGACACCTCAACTTGGTCTCAACTGACAACTACACACTCTAACTCGTCCCCACTGTGTCTAGTGGACACCTGATGCTGACGTGGCATCTAAATTTTGGTGGTGTCTTGATGATCTTTTTGTAAGTTGGAGTGTTCAACTGACACAATGAAGACGAGTTGAGGTACCTAGATATGCATACTCAGTTGGAGTGTTTACTTTCAAGCTGAAGCCAAGTTTGAATGTATGTGTATGTATTATGCCCTTTTTATAATAGCCCCTGCCTCTTTTGCATGGGTCAGAACTGTTGATATTAGTTTAACTCAGTTTGCCTACTAGTTTTTCAGAGAACCATTTGAATTGTGGCTTAAACTTCAGTCGTCGTTGTGTATCTGACAGTGAAGGTAATTAATTGAAAGAGCAAATACCACATTTTTGAACTACAAGTATTTACCTTAGAAGTTAGACGTTGCCGGATATGTTTGGCAGTTGGCTGGTTTATGCACCCTGGTACAATTCTAAATTCTAAAGTTGATATTCTGGCAGGTTCTCCCAGTAAATCAAAGACTGGTCTCGTAATTGGTATCGTTGGTGGATTTCTTGGAATTATCCTTCTTGGGGGATTAATGTTATTTTTCTGGCGGGGGAGGCACAAAGGCTACAAACGTGAAATCTATGTTGATGTTCCAGGTCCTTATCACCTGTTTTACACAGTATTAACTTCAATATGTGTTTGTTAGCTCATCAAAGAAATACTCCCTCCATTTCAATTTGTTTGTCTTGCTTGCTTTTTTAGCTCGTTTAAAAAGAATGCCCCTTTCCCTTTTTGGCTACTCTTCAATTTCAACTTTTCACGGGGCATGTTTAGGACCACAAGATTCAAAAGTCTAATTTACTTTTTTAAACTCCATGCCAAGTCAAAACAAAACAAACAAATTGAAACGGTGGGAGTATGTGTTAGCTCTCCTGAAAAAAAAAGAAAGAAGAAGGCTGAAGTAGTTACCTATTGCTACATTTCTTTGTTCCATCTTACATTGGAGACTAGATTTTTTATGAGCATTACTTTAGTTCACTTGCAGTGTAAGATGGTATAGGTCCATTGAGTATGTTGTGACGTTTTTCTTGTGTCTGTTCGGTGTGCCCAGGTGAGGTTGACAGACGAATTCCATTTGGTCAATTGAGGAGATTCGCATGGAGGGAATTGCAACTCGCTACTGATAACTTTAGTGAAAAGAATGTTCTTGGACAGGGAGGTTTTGGCAAGGTTTATAAAGGAGTGCTCAGTGACAGCACTAAAGTTGCTGTGAAACGCTTAACTGACTATGAGAGTCCTGGGGGAGATGCTGCATTTCAGCGTGAAGTTGAGATGATTAGTGTTGCTGTTCATAGAAATCTTTTGCGGCTTATAGGATTTTGCACCACACCGACTGAACGCTTGCTAGTGTACCCGTATATGCAGAACCTAAGTGTTGCCTATCGTCTACGAGGTACTTATTTTTCCACAATAGTGCTTAGATCTTGGTTTGCAGTTTTCCCAAGGCTTTCCTTGTTCATCTCAATATCAAATTTTCCTGGGATCACTTATTTCTTTTGCTCCAAACTCAGCTGAATTTCTATAGAAGTCTGGAATTTGATGGTCAAATGCTTATGTCAATAGTCTAACTGTCTAATAGCCTATTTGGCCAAGCTTCTAAAATCAGCTTATTTTGATTTTAGGCTATTAGACTCCTATCAAAGGTCTAATCACCTACTTTTTCCTAGTGATCTTTAGTAGAGTAGACAAGTTGAACAGATAATAAAGGAGGGTATTTGCGTAGGTACTCGTATTCCTAATATTCTTAGAAGTGTGTGAGTTCTTGAAAATTTGGTTCTCTTCAGTCTTCAAGAGCCATAGTTGGATTGTACCTCGTCACGAGGGAACTTCCATCACGTATGAATTGACCGATATCCATAATGAGAGCTTAAATGAGTTCATGAATTCATTCTTGCGCATTCATCATTCCCCCCGAGGTGACTTTTTTGAAAACCTTGCATGAATTGAAGATGACACATGCAGTAGTCGGTCAAACTTTAAAATTCAGCATTTATCTCCCAGAGATGATGAGATTTTGGTGGCTTCTTATTTGTAGTATTTTGTTAAATGCAGAACTTAAACCTGGGGAGTCTGTTTTAATTTGGCCAACCAGAAAGCGTGTGGCACTGGGCACTGCACGTGGACTTGAATACCTGCACGAACACTGTAATCCAAAAATTATTCACCGTGATGTTAAAGCAGCCAATGTTTTACTGGATGAAGATTTCGAGGCTGTAGTTGGTGATTTTGGCCTGGCAAAGTTAGTTGATGTTAGGAAAACCAACGTGACAACGCAAGTTCGGGGTACTATGGGCCATATAGCTCCTGAATACTTGTCCACTGGGAAATCATCAGAAAAAACTGATGTTTTTGGATATGGAATCACGCTTTTGGAGATTGTAACTGGTCAACGCGCTATAGACTTCTCACGCCTAGAAGAAGAAGATGATGTCTTGTTACTTGACCATGTGAGTGTACATAACTCAACGACTGATATTAACTTGTGATTCTATTGGTCTGCATATTACTTTTAACGTGTCCTCTAAAATTATTCAGTTTCTTCTAAGTTTATTTTTTTATCTTATACTTTCACTCTTCTCGAAGCTGCATATTGGTGGATGGCTCATTGTCCATCACAAGACATTCTCATGAATTATATACGTAGTTCTTCTTTTTTATGTTGCTTTCTTGGTATATGTTGTAATTTTATATTCAACAGGAAGGAGGGACAAAAAAGAAGTTTCATTTTCCATATAGCACTCTTACGCAAATTCTTTTGCATGTCAAACATAGGTCTTAATACTCAGCCTCTTTTGTCGGATACCCTTTTTAGGTGTAAAGTAAATTAGTGAAAAAAGGTGAAATAACATTGAGAGGACGCAAAATTTCTTAATTAGGGTAGAAGGATAGTAGATAGTGCATTTTCTCGCTATCCGTCCATTCTAGTAGTTGTAGTAGTGCTCCTTTAGTTTCTTGTCCTATGTTTTTTGTTACCATCTGTTGTTTCTTGTATTTCGCTTGTCGTATTATTTTGATGTAGTTATTGTTCAGCATGCTTTGCTATGCTTTCACATCCATTACTTTTTCTGGACTGCTTTTGTTTGTTTTTCCTTGAGCCGAGGGTCTATCGGAGACAACCTCTCTATTCTGAGGTAGGGGTAAGGTCTCTATGTAGACTCTATCCTCCCAGATTCCACTTTGTGGGAGTACATTGGGTATGTTGTTGGTTACTGAATGGCTGCATGTGTCATCTTGAAGGTCAAAAAACTGCAAAGGGAGAAAAGGTTGGATGCTATTGTCGACCGCAACCTACATAAGAACTATAACATGGATGAAGTAGAGATGATGATTCAAGTTGCGTTGCTGTGTACTCAAGGATCACCAGAGGATCGTCCAGCAATGTCAGAGGTAGTACGAATGCTTGAAGGAGAAGGACTTGCTGAGCGGTGGGAAGAGTGGCAGCATGTCGAAGTTACACGTAGGCAAGAATACGAGAGATTACAACGACGATTTGATTGGGGCGAAGATTCAATCTATAATCAAGATGCAGTTGAACTGTCTGGTGGAAGATAACAAAATCTGTATCTCCAAACAAGTACATAATTTCCTGTAATAGTCTCTCCTTGATACATATAACTTGTCGCTAAGTGGCAATCATTCCACGTTTTCATATTGCTTGTGTCATATCTTGATTGTATTGTTTTTCGTATGCTTCTCTTATTTAATCGATTTTTTCCCTGTATGATTGGCCTGTGCAGCGACATAATATATATCGACTTTCTTTTTGTTTCAATTGTAACATCAATCTTTGTTATAGTGATTTGAGTTTTGC

>PGSC0003DMG400016654 LRR-RLKs

CTCCCACTCACACCAGAGAGAGAGAGCGCTTTAATAAATGTTTAACTGACTCGCGGTCGCCGGAAAACTCATTCTCCCTCCGGTGGATTTTTGTCTTAATTTCATTTTTGGATAAGGACAAGTATCCGGCGCCGGTGGTTTCTTTCCCGGAAAAAATCTCAGCTCACATTTTGTGCTGCTTTTAGTTTGGAGGTTGAAACTTCTCTTTCTCGCTAAATTAAACTCCAACCGAGCTGTTTTTTTCAGTGAATTCGTAGGTTTATGCGTTTTTTATTTGTCTGTTCTTCGTTTGGGTCTTTTATAGTTTACTCTTTTTTAGCAGATTTGCACAAACTTTTTTCCAGTTTCAGCTGTATTTTAGAATTCTAGGGTTTTGGGTTGAATTTTTCACCACTGGACCCATTGCGTGCTTATTTAGTTGTAACTGGTCATTTCAACTCCTATTTTGGAAGTTGATTTTTTTTTGTCTTGAAAAACAAGTAGGGCCTCAACATTGTCTTCAGCTGACCTTGTCTATGGAATTATAAGCAACATTGGTTTGGAAATGAGAAAGGTGTTTGTTACTGTGTTTTTGTTTCTGTTGGTGGAGGCTGTGTTTGTGGAAGCTTTGAACCCATCTTTGAATGATGATGTACTGGGATTAATTGTGTTCAAATCGGACGTTCAAGATCCATATGGAAAGCTGAAATCTTGGAATGAAGAGGATGATAGTCCTTGTAACTGGGACGGAATCAACTGTAACCCCAGATCCAATAGGGTGACTGAAGTTGTTCTTGACGGGTTTGGTTTATCTGGAAGGATAAGCAGAGGCCTTTTACGCTTGCAGTTTCTTCACAGACTTTCACTAGCAAAGAACAATCTTACAGGTAGCATTAGTGTTAGTCTTGCTCAACTTGCTTATCTCAAGTTTCTTGACTTGAGTGAAAATAACTTATCTGGTAACATTCCTGGCGATTACTTTCAACAATGTGGGCCGCTTAGATCAATTTCTTTGGCTAACAACAAAATTTCAGGGCAGATCCCAGAAAGTTTGAGCTCCTGTGTTACACTTGCGTCCCTTAATGTATCGTCAAATCAGTTTTCAGGGTTGGTGCCTTCTGGGATTTGGTCTTTGAATGGACTAAGGTCTCTTGATTTGTCAAACAATTTATTGGACGGCGAAATTCCAGTGGACATTCAGGGTCTGAGTAATTTGAGAGCATTAAACTTGGGGAGAAATAATTTCAAGGGTGAAATTCCGGATGAAATTGGAGGTTGTTTGCTTTTGAGGTCAATTGATTTGAGTGAAAATTCTCTCTCAGGAGAGCTTCCAAGCACAATGCAGAAGCTTAGCTTGTGTAGTGAATTCATTTTGAGACGTAATGCTTTTGTGGGCATTGTTCCTAAATGGATTGGAGAAATGAAAAGCCTTCAGATTCTTGACGTTTCTGAGAACAACTTCTCTGGTATCATTCCAACTTCAATAGGGGAACTTGAGTCACTGAAAGTATTAAAGGTGTCACGGAACGGGATTTCTGGAAGCTTGCCTGAGTCCTTGAGCAGTTGTGTTAAGCTTCTGGAATTGGATGTTAGTCATAACTCTTTGGCGGGTAATCTTTCTACTTGGGAATTTAAATTGCAATTAATTCAGGTTATGTTTTCCGTGAATAAAATTAACTGCCAATCTAAATAATGCATTGCTACACAACTCTTATCACAATCTTGTAGCAGTGAGAGTTATCTGGTGAAATTTAATTAACTCTCTGAAACTTCTCTTGCAGGCACCATTCCTAAGACTGTCGGCCAATTGAAATCCTTGAATATTCTTGATTTGAGTGAAAATCGCTTATATGGTATTGTCCCTGTGGAAATTGGAGGAGCAAAATCTCTCATGGAACTTAGTCTGGAGAAAAATTCCTTGACTGGAGAAATTCCTAGTTCAATTGGCTATTGTTCTTCATTAGTATCCTTGTAAGTTCTCGAACCTCCTTATGTTTCATTAATTTGTTTTTGGACATTGCCTTTCTCTGCTTGAAGATTCTTAGAGGATAGTATAGTGGTATCTGGTGTTTAAGTTCTCTATAGTTTTCAGGCCAGGGAATATTGTGTTAGAGGCAAGTTCAAATACGAAACCATGTTTGATATTCTCAATCATTTTCTGCTGAAATTTAGTGGCTCTAGTAATCATGTAGAGTTTATAATTTGCAATGACAATAAAATGGGATTATCAGCTTCTATTATAAAAGCTAATTGCAGAGTAGTATTAGTGGTAATGTGCCTAACTGTATCTGAATATAATTTGAGTGGTTATGACAGTAATATCTTACATGAGGCATGAAAGTAACAACAATAATATATTAATAGGCTCTCAATGATCTATCTATTTATATCCATTATGTTTTGGCTGGTGTTACCTGAATACGAATTTTTTGCTACAATTAGTTAATGTTTTACTTCAATCAACTTGGCAGGTCTTTATCACATAACGGTCTAACTGGTCCTGTACCTGAAGCCCTAGCAAAACTGACTTATCTTCAAAGTGTTGACTTATCCTTTAACAAATTAACGGGAGTGTTGCCAAAGCAGCTTGGAGATCTTGGCCATCTCTCCTTATTCAACATTTCACACAACCAGCTACAGGGCGAACTACCTTCTAATGGTTTTTTTAATACTATTTCTCCTTATTCCGTGTCAGCAAACCCATCTCTATGTGGGGCTGCTGTCAATAGGTCTTGTTCTACAGTCATGCCCAAGCCAATTGTTCTGAATCCCAACTCTACTGATTCTTCTCCCAACACTGTCCCTCAAAGTATTCGCCATGAGAAGAAAATCCTAAGCATCTCTGCCCTCATTGCTATTGGTGCAGCTGCTGTTATTTTTGTTGGTGTGATTGCCATTACTGTGCTCAATATTCGTGTCCGGTCAGCGGCACCTCGCTCTGCTGCTGCAGCCCTTACATTTTCTGGTGGAGATGATTTTAGCCATTCTCCATCCACGGATGCGAATTCTGGTAAGCTTGTCATGTTTTCTGGTGATCCCGACTTCAGCACAGGGGCACATGCTCTGCTTAACAAAGATTGTGAGCTTGGGCGAGGTGGTTTTGGAGCAGTGTATCGCACTGTCCTTGGGGATGGGCATCCAGTGGCCATTAAGAAGCTTACCGTTTCCAGTCTTGTTAAGTCTCAGGAAGATTTTGAGAGGGAGGTTAGGAAATTGGGAAAAGTACGTCACCATAATCTGGTCACCCTTGAAGGTTATTATTGGACACCATCTTTACAGTTACTTATATATGAATTTGTGGCTGGTGGAAATTTGTACAAGCATCTCCATGAAGGATCTGGTGGGTGTTTCCTCTCATGGAATGAGCGGTTTAACATCATTCTCGGGACAGCAAAAAGCTTGGCTCACTTGCACCAAATGAATGTAATCCACTACAACCTAAAGTCCAGCAATGTCCTAATTGATAGCTCAGGTGAGCCTAAAGTTGCAGATTATGGCTTAGCAAGACTATTACCCATGCTTGATCGATATGTCTTGAGTAGCAAAATTCAGAGTGCACTCGGTTACATGGCACCTGAATTTGCATGCAAAACTGTGAAGATAACCGAGAAGTGTGATGTGTATGGATTTGGAGTTCTTGTTCTGGAGGTAGTCACTGGAAAGAGGCCAGTTGAGTATATGGAAGACGACGTGGTAGTACTGTGTGACATGGTACGTGGAGCATTAGAAGAAGGCAGGGTAGAAGACTGTATAGATGCAAGGTTGCAAGGAAAATTTCCAGCAGATGAGGTGATTCCTGTGATGAAGTTAGGATTAATCTGCACATCACAAGTCCCATCAAACCGGCCTGATATGGGAGAAGTGGTTAACATTTTGGAGCTGATTCGATGTCCTTCAGAAGGCCAGGATGAACTTGTATGATGTTCACTAGATAAACACAGATTGTAAGATAAATGTCTCATCATGGCAAAGAAGAATGCGGTTCATTTATTCAGAGAAGTTGGAATTTGTTGTTGTAGAATATTTTGCTCTTAATTGCTATAAAAATCCCTCCTTTGAATTACATCATTTGTTATTTGCTTCTCTATGCCACTCATGGATTCCTTTTCACCTGGTTCCAATGTTGGACTCACATGCTTATTATTACTCTTTTCTTTCTTCTCAGGGTTTAAGTCATCTGCCGCTCCTTAAAATTGTGCTCATGTTTCACTTAGACACA

>PGSC0003DMG400016685 LRR-RLKs

GAACATTTCTTCTTCTTCTCCACCATTTCTGCTCTGCTCACTCTTTCCCCTCCACCATTGAAGAAACATGCGTTTTCTTCTCCTCTTCTTCCTTTCCCTTATTCTCCATTTCCATCTCCTCCACTTCACCACCGCAAAACCACCTTACGTGCCGGAATACCGGGCATTACTCTCCCTGAAAACTTCCATTACCGATGACCCACAATCTGCTCTTCTTTCATGGAATATCTCAACAAGTCATTGTACATGGAGAGGTGTCACGTGCGACCGGTATCGTCACGTGACTTCTCTCGACATCTCTGGTTTTAATCTCACCGGTACTCTCACGCCGGAAGTTGGTCATCTCCGTTTTTTGCTCAATCTTTCTGTAGCTGTTAACCAGTTCTCTGGACCCATTCCTATAGAGCTCTCGTTTATACCAAATCTGAGTTACCTTAACCTCTCTAACAACATTTTCAATTTGAGTTTCCCTCCCCAGCTTACCCATCTCCGGTACTTGAAAGTTCTCGATATTTATAATAACAATATGACCGGTGACCTTCCGGTTGGGGTTTACAATTTGACTAATCTTCGACATCTTCATTTGGGTGGCAATTTTTTTAGTGGCAGTATTCCACCGGAGTATGGTAGATTCCCATTCCTAGAATACCTTGCAGTTTCTGGAAATGCGCTCGTCGGTATGATACCACCGGAGATCGGAAATATTACCACACTTCGTGAGCTTTACATTGGATACTACAACACGTTTTCCGGTGGGTTACCGGCGGAAATAGGGAACTTGTCGGAGCTCATTCGGTTAGATGCTGCAAACTGTGGACTTTCCGGTGAGATTCCGCCGGAGATAGGGAAGCTTCAGAAATTAGATACACTGTTCTTGCAAGTGAATGGTCTTTCTGGGTCTGTTACACCGGAATTGGGGAATTTAAAAAGCTTGAAATCTTTAGATCTATCAAACAATATGCTCTCCGGTGAAATACCGTTCACATTCACAGAGCTGAAGAATCTAACCCTGCTAAATCTTTTCCGTAACAAGCTTTACGGGTCGATACCGGAGTTCATAGAAGATTTGCCAAAACTGGAAGTATTGCAGCTTTGGGAAAACAACTTTACCGGAAGTATTCCACAAGGTTTAGGCAAAAACAGTAAGTTAACAAACGTTGACATCAGTACCAACAAATTAACCGGAAATTTGCCCCCAAACATGTGTTCCGGCAACAAGTTACAGACGTTGATCACTCTCGGAAACTTCTTGTTTGGCCCAATTCCAGAATCTTTAGGTGAGTGTCAATCACTTAATAGGATTAGAATGGGAGAAAATTTTCTAAATGGGTCTATTCCAAAAGGGCTATTCAGTTTGCCGAAGCTTTCACAAGTAGAACTTCAAGATAATCTTCTCACTGGTACATTTCCAGTGACTGGTTCTGTTTCATCAAGTCTTGGACAGATTTGTCTGTCGAATAATCGTTTCACGGGGCCTTTGCCATCGAGCATTGGAAATTTTACTGGTGTTCAAAAGTTGCTTCTTGATGGGAACAAGTTTTCTGGTCAAATTCCAGCTGAATTAGGGAAATTGCAGCAGCTGTCGAAAATGGATTTTAGTGGTAACAGTTTTTCAGGCCTGATTCCACCGGAGATAAGCCGGTGCAAGGCTTTAACTTATGTTGATCTTAGTAGGAATAAGCTATCTGGTGAAGTTCCTACTGAGATCACTGGTATGAGGATACTGAATTACTTGAATGTATCGCGGAATCAGTTAGTTGGGAGTATTCCTGCACCTATTGCAGCAATGCAGAGTTTAACCTCGGTTGATTTTTCGTATAACAACTTATCTGGATTGGTTCCGGGTACTGGTCAGTTCAGTTACTTCAATTACACATCATTTATTGGTAATCCAGATCTTTGCGGACCCTATTTGGGTCCTTGCAAAGAAGGTATTGTTGATGGGGTTAGTCGACCTCATGAGAGAGGTGCATTTTCGCCTTCTATGAAGCTTTTACTTGTTATCGGGTTGCTTGTTTGCTCGATTGTGTTTGCTATTGCTGCAATTATTAAGGCTAGATCGTTAAAGAAGGCGAGTCAGGCTCGTGCCTGGAAGCTTACTGCTTTCCAACGCCTGGATTTCACTTGTGATGATGTATTGGAATGTTTGAAAGAGGATAACATTATTGGTAAAGGAGGTGCTGGAATAGTATACAAGGGGGTAATGCCAAATGGTGAACTTGTTGCTGTTAAAAGGTTGCCGGTTATGAGCCGTGGTTCTTCCCATGATCACGGGTTTAATGCCGAGATACAGACACTTGGGAGTATTCGACATAGACATATTGTTAGATTATTGGGATTTTGCTCAAATCATGAAACAAATCTTTTGGTTTATGAGTACATGCCTAATGGGAGCCTTGGTGAAATGCTTCATGGAAAGAAAGGTGGTCACTTGCATTGGGATACCAGGTATAAGATAGCATTGGAGGCTGCAAAGGGACTTTGTTATCTTCATCACGATTGCTCGCCTTTGATCCTCCATCGTGATGTAAAATCAAACAACATTCTTCTGGATTCCAGCTTCGAAGCTCACGTTGCTGATTTTGGGCTTGCCAAGTTTTTGCAAGACTCGGGAACATCAGAATGCATGTCTGCTATTGCTGGTTCTTATGGCTACATTGCACCAGGTATGCTCTCAAATCTTTACTTCTTTGTAGTCTTGGTTTTGATGTGTGTTGTTCCTTTCAAAAGAAGTTTATTATTGTTGTAAGTTGACAACTAAAATTGTGATTTATCAGTCTCTTCAAGTGTTTATTATGGTACTTCTTGATCCTTTAAAGCTGGTAATGATCAGTCTCCTTGTTAAAACATGGAACAACATGACACGAATATCAGTCCTTGATTGGTAGTTAACAAGAACAACAACAACAATGATAACTACGTGTCGAAAATGAATGCCTCTGTTCATAGATGAGCTGCTAACTGGTGCTTTGATGAATTCAAATTTGGTTACTTGCTGTTTGGCGTTAAGTTGTTTCATTTTGACTTACAAAGTTTGCTTATTGCTTGAAACACAATCTCTCTTTTTTTGTATGTGCCTAATTTCGTTTTCATTTGAACTGTGCAGAATATGCATACACACTCAAGGTAGATGAGAAGAGTGATGTATACAGCTTTGGTGTGGTTCTGTTAGAATTGGTGAGCGGGAAAAAGCCAGTTGGGGAATTTGGTGATGGCGTTGACATAGTCCAATGGGTAAGGAGGATGACCGATGGGAAAAAAGAAGGAGTTCTAAAGATCCTTGATCCAAGACTCTCAACAGTTCCCCTTCATGAGGTGATGCATGTGTTCTATGTTGCAATGCTGTGTGTCGAAGAGCAAGCTGTTGAACGCCCCAAAATGCGTGAGGTTGTGCAAATGCTAACTGAGCTTCCCAAGCCATCTGGTCCAAAAACAGAAGATTCAACAATCACCGAGTCGCCCCCATCATCAGGTCCTGCATTAGAGTCTCCCACTTCGACTCCCGGAGACACGAAAGACCAGTACCACCATCAGCCATCACCTCAATCTCCTCCACCTGACCTACTCAGCATATGACCTACAATGTTCCCTTCTAATAGAGGATGTAGTTCTCTTCCTTTAGTTTTATTATCAGTTTTATTGGGGGTTGGTTTATTTATTAGGCTATAAGCTTTTCCTAGGAAGTGAGTTTCATTGTACAGTAAGATTGGCTAAGGTATCTGAGTAACTTTATCCTTCAAATTTTCCCATTTGCTGTTTAGCTTTTTGATATTGAAAGAATC

>PGSC0003DMG400016709 LRR-RLKs

CGACACCTATAGATACACACTGCCAGTGGAGTCTTCTTGTATAAATTTTTCTCTCTCTCTTCATTAAAAGGAATGCTTGGAATCGGAGAGTCTCTTAACGATGGTGACACCTACCGCTACTGACGATGTTACCTTCTTGTCATTTCCCTGTCTTTCTTTCGCCATGCTTCTTCTTCTTCCTCTTACCTCTGTCTCTGCACAACCCCTTTAATTCTTGACTAACAAGAACCAAAACAGAAAATGAATAGAAATAGCTGTAAGCAAGACACCCTTTTCACCCTTTTGCCAGTAGTGGTGATTGTGGTAGTGGTTTTTTACTCATCTTTTGTTGTGGAAGGAAACACTGATCCGAATGATTGTGAGTTTCTTCACTGGTTGAAAATCTTGGATCCGTTTTTAGAGCTTCATTAGTGATTTTGAGCTTAGTGGTTTGGGTGTTTTTTCTGTTTTCTTGATTGTGTTGAAGGAGGAGGGTTTTTTTTGTTTTTTTGTTTGTCACTCTGTGAATGTTGTGATATAAGAATGCCTTTCTTGTGATTGTTTGAATGCCTGATTTGGTTAATGGGGGTAATGTATTTGTTTGCCCTTTTTCTGGGATCTTTTTCTTCCCCTTTTTGTTGGGTTTCCTAGTCAATGCAAGTCAAAGATTGATGCGTTAGTACAAAAAATATGGATTCTTGATCTTTATACTAAAAATTATATGACTGGGATTTTCTGTGATTATAATGTGAGGACGCATGCAATTGAATTTGAATGCTAAAACTGGTTGATTTGGTAAGATAATATGTTTTCTTGCCCTTTCTTGGTGTCTTTCTTGTATTTAATTAGTTTCAGAGCCAACACTAGCTTAAGATTGAGGCTGTACTTGTTGAATCTTGATAAGATATATATTGAATCTTGGTTTTCTTGTTGGAGAGATGTTCTGGCTTTGGCTTTGTCGTTGCCTGACTCTTGTAATCTGCAGTGCAAGCTCTTCAGGTCTTGTATGCTTCACTGAATAGTCCTTCACAGCTCTCTAATTGGAAGTCGAATGGTGGCGATCCCTGTGGTGAGTCTTGGAAAGGAATCACTTGTCAGGCCTCTTCTATTGTTTCAATGTAAGTTTGCATAGCATTAACTTTGTGCGCATCATTATACATTTTAGTTATGATTTGATTGATATAATGGTTGGTTCTTGCTGAAACAGTCAAATTCCTGATTTGGGACTCAATGGAACGTTGGGATACATGCTAAATGGTCTTACCTCGTTAAAGAACTTGTAAGAGCTTGTCCTTTTAGTGGTAAAGGGTAATATAGAGGACCTACTATCACAATAAAATTGAACTTCTGCAATCTGTTGTCTTCAGGGATATGAGCGGCAACAACATTCATGATTCTATTCCATATCAGCTTCCACCCAATCTCACTAGCCTGTAATGTTGACTCCTTGTTATGAGTTTGTTTTGTTTGAAAACTTCCCGGTTTATGACTGTCTCACTGGAAATATAATCATTGTCAATGCAGGAATCTTGCAAACAACAATATAAGTGGCAACATTCCGTATTCCTTTGCTGGCATGGTTACACTTAGCTATCTGTGAGTGGCCAGACTCACTAGGCACTTTCCAAAACAAAAATAACATTTGCTTGGTCTTCTCTTTATTCTATTGAATCATTTCAGGCAACTATTTAAAATAGCTTACCCTACGATTCATTTTTTCAGGAATCTTAGCCATAACTCACTCTCTCAAACAATTGGAGAATTGTTCACCAATCACTCCGATCTGGGAGCTTTGTAAGTGTTCTACTTAATCTTTAATTGATTGATCCTTTCTGGTAATTTTTTTTCCTTTTCTTCTTTTTTTCTAATAGAGTTCATTCATTGCCACACTTGAAGGGCCTCTTTTCCTACACCAGTCCAATTAGTTACTAGCTTGAATTCATAACACCATGGTGTCTCCGACCCACAGTAGCTGGCCCTTCATTTGTTTTGACGTGCCTGCAATGTACAGACATGACTGCTTCCAAATATGTTCTCAAAACAAATATAGGATATAATCATATACAAATGCCTTTTTAGAATCATTATCTGCTTAATTTTAATGAACAATGAAGTCTGGAGGCTTGTAAAGTCCAACATTTTATTTTCCTTCGTAGCCTTTTTATGTGGAAATTCGTTTTGATTGTATTAGGATAGGTGCCCTGCTCCTCTTTGAAGCTTTCAGTTGCCTTTTGTTTTCATTTATTTGTTCTTTCAGAATCCCATTTAAGTAAACCTACATTAGAGCTCAGTACTCCTGCTACAAAGAGCTTTCTGGCGATATTGATTCCTCATGTCTTTTGAACATCTGTTTCACTAGTTAATAGTTATAAATTTTCACTGATTTGCTGCTACTGCAGGGATCTGTCCTCGAATAATTTTTCTGGAGATCTCCCCTCTTCATTCAGTTCGTTGACAAATCTTTCCATACTGTAAGCATAATAACTTTCTACTAATAGGATTTGCTGTTCCGTTACTTGACTTACCCATGAAATTGGCATCTCTTGTAGTCATCTGGAAAACAATCAGCTAAGTGGCCCTCTCAATATCCTGGTTGGTCTACCGTTGACTAATTTGTAAGTTTCAATACCTCTTCTTGAATTAATGGGCTTAGCTCGTTCGATTCAATGCCAGTAGAGTGTTTTATGGGTAAAACAAAACCTTTTTGCAGAAATGTTGCAAAGAATCATTTCAATGGATGGATCCCGAATGAGCTTCTATCTATCCCTAACTTCATGTAAGTAATTTCACATCCTATATATTCTTACATGGTTTAGAATGTTACAGTTTCAAGGGGTGTGACATAACATCTAATATGACAGATACGATGGGAATTCCTTTGCAAATGGTCCAGCTCCTCCCCCACCACCATACACTGCACCTCCTCCTGGAAGATCTCGCAACAATCGTAGCCATTCTCCTCCTAGTTCACGTACAAGACCAGGTTCTGATGGCCAGCCATCTCCAGAAAATGGTGATAAAAAAAAGGGATTGACAACTGGAGCTATCATTGGCATAGCTCTTGGATCTTCTCTTGCAATTCTTTTTGCTCTGGTAGCACTTGTATTTTGCTTCCGGAGGGGCAAGGGCAAGGAAATTGCTACAAGACCCTCTACAGGAAGTCTTCCAGTTGGCACAGATAAAGGTGTGTTCCCTTGAACTTTTTTCCTTCCTGACTCAATATTGTCATTGGTGCTAATACCCTCAGATGACTAACAGTTGTGGTGTGATGTGCCATCTTAATTTGGAAATTGGTACAAACTTGGTTGATTTCTCATGTATGCTGTTTAATCTTAGTGAATATGGAGGTACAAGAGCAAAGGGCGAATCCTGTGGCTGCTGTTGCTGACTTGAAGCCCCCACCTACCGAAAAGGTGACTGTTGAAATGATACAAGGGAAGAATGGATCTGTCAGGAGAGTGAAGTCTCCAATAACTGCATCTGCTTATACTGTTGCTGCATTGCAAACAGCAACAAACAGCTTCAGTCAAGAAAATCTCATTGGTGAAGGATCTCTTGGGCGTGTTTATAGGGCAGAGTTTCCTAATGGGAAGGTAATTTCAGATGTTTAACGTTGTAACTTCTTACCTTTCCTACTTGTGATGTGCATTCAAAACTATTTGAAGCTGAACAATATTGAATAAATTCAATCTGTCTCTCATCGACATCTTTCATTTTTGATGCTTCGGAGAATCACATACTTGTGAGTTGTATAACGTTCATCAGATACATAAGGACATCAAAAGGGGTATATCAAGATGTTGGCCACTCCACCTATCACCCTATTGCTAAGATGCTTTAACATAGTATCAAGCCGTTACACGTGTGGGACAATGTTGCGTTACCTAATGTCCAACCTCGGGTAAATGAGCCTGTAAAGATATGGTAACTCAACCCCAAAAGCTAACTCATGAGGGGAGGATTGCCCAAATCCATATAAGGAGACCACCAGTCCATTCCCTAACCAATGTGAGACTTTTACCCACTCTAACACCCCACTTCACGCCCAGGCCCAGGCCCGCTGGAGCATGGACCGGGAGCCCAAATGGAGATGAACCTCTGATACCATGTAAAGATATGGCACCTGGGCTTAACTCAACCCCAAAAGCTTAGCTCATGAGGGGAGGATTGCCCAAGTCCATATAAGGAGACCCCAGTCCATTCCCCACCCAATGTGAGACTTTTACCCACTCTAACAAAGGCATCCAAAGTAGTATATAAGGTTCTTGTCCTACTATATCCATTGCCTTAACGTTATGGTTGAACCCTTAGATCCTTTCACAATATTTGTCAAACCGGGCAAACACTAAAAGGCTACATACTTGATCTATGTGTTACTAAACCTGAGTAGAAGTTAGTAGACTTCTTGATACTCTGCTCGCATGGGAGTTAAAGTATTGGGTCCTCTTGCTAGTACATAATCTGTTTTGAGTGATTCTCTTAACTGAGATTAAGTCCTTGATACTCTTTGATTCAACTTGCTCTTTTAGATCTTTCACAAAGTTGATTGCCCACAGTAGGCTCATATTCTGATGAACACATGTCAATTGTTCACTCCTCTGGGTGGTCTGGCCACATCTTTGCTTTTATCTGTCTTGTGATGTTTGATGGGTGTAATTATATGGTCACTTTGATGGTCATCTCACTTCTCTTCTCAGCCTGAATTAACTATGATACTATAACCTTGTCCTTTGGTTACATGATGCAGTCCTAAGCAAAATTATTCTTAGCAGTTTCAAAGTTTGACATTATATATCTTCTACTTTGTGGTTTATTATTGGATGTGCTACTGGTTATTTCTTTTTGAATGAACCAATTGTGTTGACTATTTCTCTGCTCCAAACTTGATGCCAACTGTGTTCCAATAAATGATTTAAGAATTTCATGTGTCAACAACATTTTTGATGAGGAAACTTTTAGTTTTTCATTCATGTAGTTTCTACGGAAGGTGCCAACTGAAAATCTCAATTGATCTTTTGAACTGTCCTTACCAGATAATGGTCATCAAAAAGATCGACAGTGCTGCACTGTCGCTGCAGGAGGAAGATAACTTTCTTGAAGCTGTCTCAAATATGTCAAGGCTGAGACACCCAAACATTGTTCAATTGATAGGATATTGTGCTGAGCATGGACAGCGTCTTCTGGTTTATGACCATGTTGCAAATGGTAGCTTGCATGATATGCTGCACTTTGCTGATGAGAGGAGCGAATTGCTTACTTGGAACGCCCGGGTTAGAGTAGCACTTGGCACTGCTCGAGCTTTAGAGTATGCTATAGCTGTATAGTATTTTACCTCTCTTGCCCTTTTCTTGTTGGGTTAGCTTAAAACTTGACCGTCTTCTTTTCAGGTACTTGCATGAAGTCTGTCTACCTTCTGTTGTACATAGAAACTTCAAATCCGCAAATATTTTACTAGATGAAGAGCTCAATCCCCATTTATCAGATTGTGGGTTAGCAGCTTTGACTCCAAACACAGAACGCCAGGTGAGTAAAGAAGATTTAATCTACGTCGTCGTTTTAACAGAAGGTGTTGATAATATTGGGTGGTACGTGATGCATATAATGTTTGTGAGGTGATGTCAATCAACTTACCTAGTACTGTAGTAGGAGCAATATTAATAAGCAACTATTTCTAGGTAGATCTTTACAGACTTAGTCATGAAATTGGAGATTCTGATTGTGGCAGCTTAGTTTTGTTTTCTAATTCTTCTGGAGCAATCTTGTTAGATGCTTAAACACTTGGCTTACCATAGTGCACAAACTTTAAGGAAGACCCCTTCCCCCTGCTTGAAATTCCATTTAGTTCTGTGAATTATTTCTGTGAAAAGAGAACCTTCTTGTACAGAAGTTCGGGTGAAGGAAGCTAATGTGGGAAGGCTAATTGACTCTGTTATGGGCATTGCCTAATATTGTAATTACTTCTGAATGTTGACAGGTGTCATCCACACAAATGATTGGTTCATTCGGTTATAGCGCTCCTGAATTTGCTTTATCTGGAATATACACGGTAAAAAGTGATGTATATAGTTTTGGTGTGGTGATGTTGGAATTGTTGACAGGAAGGAAGCCACTTGATAGGTAAACTAGATCTTTTTACATTGCTAGTCAACTTTTACACTATTGAATTCTTTTAATGAATTCATTCTTGTTAACAATCATGTAGTTCAAGGGTGAGATCGGAACAGTCCCTTGTGAGATGGGCTACGCCTCAACTCCATGATATAGATGCCTTGTCAAAAATGGTTGATCCTGCACTAAATGGCATGTATCCTGCAAAGTCTTTGTCCCGTTTCGCTGATATTATTGCACTCTGTGTTCAGGTGAGCTTTACTTGCATGTTCTTTCATTTGGAACATAACAGTTGTTAACATGTGAAGCTTAAAAGATGAAACCACTTCTAAAATATCTTTGATGGAGGCGTAAGTCAATGATCATAGGGAATTTACCCACTTAAGGACCAAACTTGACATTCTAACCCGTCAAATAAATGATACACTACCAGTTTTTGCATTCTATATAAATCCAAATTCAGGGGCGTGGATAAATTGGGACCACCTGCACTCTGAGTTTGCTCATGGCTCAGTGCTGAAGTGAATCGATGTGCTTCAGCAAATAGATGACCTTTCTTACACATACCCGGTGATTCTTGGAATGGCGAGAGCACCCATCTTGGTTAAGTTAGTTTGGGAACTTCCAAGGCGTAATCCTTGCATACCATGGTCCATAGTGGCTTTGGGTGCTTAGTCTTTAAATGAGAAATTCTTAATTAAGCTTTTCTTGATCTAAGGTGACTTCAAGACTCACGAAGAAAAATCTGATGGACTCGACAAAATTATAATGAATACTTGCTCATTGACATATCTGAAGTGAATTACTTTACTTGCAATAATTTGAATGCGACTGTTTTCTCCTTGGTAATAACTTTTGACATCTATTTCCTGTACAGCCTGAACCAGAGTTTCGACCCCCAATGTCTGAAGTTGTGCAAGCACTGGTTCGGTTAATGCAAAGGGCGAGTGTGGTCAAAAGGCGATCCAGTGATGAATCAGCTTTCACATACAAGACTCCGGATCATGAGACGGTTGACATGCCATAGATAAACATACACGACGAAGCAGCCTTTGCAATTTTGTAAGTAACTTCTGTGAAGGTCAAGAGCAACTATGGCAGATAAAACTCCGAGGTGTCAATACCAGGGACATTAAAAATGTAATTATTTTTCACATCTTCCTCGTTAAAAAAAGAAGAAGCTTTTTTGCATCCCAATTTGAGGTACATAATTCACTGACAATGTCCTCCAGAGTATATTTTCAAACCATCTTGGGCTACTGGATATAATTTATTGTCAATAAGAACATTGCTACACATTATCATTCTGTGTTTGTTTATATTACAGTTTTTCCAATAACTTTTGGCTAGCCTAAGAGCTAACTGTAGTGTGCATCGGTCGGTTCGGTT

>PGSC0003DMG400016872 LRR-RLKs

ATGTCTGGAGTATATGACTCAAGAACTGGTCTAATTCTGATTTGGATAGTTATCTTGTTGGCTTCCGCGATGATGGTTTGTCCTGCAGAAGGATTGAATGCTGAAGGAATGTACCTGCTTGAGTTGAAAAAGAATTTAAACGATGAGTTTAACAATCTTGAAAATTGGAATCCTAGTGATGAGACGCCATGTAGATGGAAAGGTGTAAATTGCACTTCTGATTATAATCCAGTTGTGCAATCTCTTGATTTGAGTTTTATGAATCTTTCAGGCACTTTGAGTTCTAGCATTGGTGGTCTCGTAAGTTTAACCATACTTGACTTGTCTTTCAATGGATTCACGGGAGAAATTCCGAAAGAAATAGGGAACTGCTCGAAAATGCAGAGTCTTCAGCTTCATGATAATGAATTTTATGGGCAGATTCCTGATGAATTGTATAACCTTTCTCATTTGAAAGACTTGAATTTGTTCAATAACATGATATCTGGTCCTATATCAGAGGAGTTTGGGAGACTTTCTTCTTTAGTTTCTTTTGTTGCATACACCAATAATCTCACTGGTTCACTCCCTCGATCACTCGGGAAACTAAAAAAGTTGGAAACATTTCGAGTGGGGCAGAATCCGCTTTCTGGAACCTTACCTGCTGAGATTGGTGATTGCAAGAGCTTACAAGTTCTTGGTCTTGCACAAAACAATATAGGAGGAAATATACCCAAAGAGATCGGTATGCTTAGAAGATTGAAACAACTTGTACTTTGGGACAATCAACTCTCGGGCTATATCCCGAAGGAACTTGGAAACTGTACAAAACTTGAACTGCTTGCTTTATACCAGAACAACCTGGTTGGAGAAATTCCTGCTGCAATAGGTAAGCTCAAATCTCTCAAGAGGTTATACCTATACCGGAATGGATTAAACGGTACAATTCCAAGGGTCATTGGGAATCTTTCGTCAGCGATAGAGATTGACTTCTCAGAGAACTATCTAATAGGAGACATACCAATTGAGTTCAGTCAAATAAAGGGACTGATATTGTTGTATCTGTTTCATAACCAGCTCAATGGTGTGATACCTCGGGAACTTTCTAGTTTAAGAAAGTTGGAACGTCTTGATCTGTCGATTAACGACTTATATGGGTCTATTCCATTTTCATTTCAGTATCTCACTGAATTGGTTCAGTTACAGCTCTTTCAGAATTCTTTAAGTGGAACTATTCCCCAAGGTCTCGGGAACTATAGCCGACTTTGGGTTGTTGATTTTTCTAACAACTACCTCACAGGAGGAATTCCTCCGAATATTTGTAGGAATTCTAATTTGATTTGGCTTAATCTTGGTTCAAATAACTTGCATGGAGACATTCCATCTGGTGTTATTAAGTGTGACTCATTGGTACAACTTCGTCTTGACGGTAACTGGCTACAAGGGAGTTTTCCTTCTGACTTGTGCAAACTGAGTAATCTATCTGCTCTTGAATTAGGACAGAACACGTTCGGTGGTCTAATTCCTCCTGAGATTGGAAACTGTCAAAAGTTGCAAAGACTTGATCTTTCAGGCAATTATTTCACACATGAATTGCCTCAAGAGATAGGGAATCTTGAAACACTTGTGACGTTTAATGTCTCGTCTAATTTGCTGACAGGTCAAGTACCACAAGAAATTCTGAAATGCAAGGCGTTGCAACGACTTGATCTCAGTAGGAACAGCTTTAGTGGTGCTATACCTGCTGAAATTGGAAAACTAGCACAGCTTGAACGTCTTTTGGTTTCGGATAATAAGTTCTCTGGAAAGATACCAGTAGCATTGGGGAGGCTTTCTCGTTTGAACGAGCTGCAGATGGGTGGAAATTCATTTTCAGGTGAAATACCATCAGAATTGGGTGATCTTACTGGTCTACAGATCGCGATGGATCTTAGTGACAACAATCTCTCTGGTTCAATACCACCTAAGCTTGGAAATCTTATCCTACTAGAGTGTCTCAATCTCAACAATAATCATCTGAGTGGTGAAATACCAATCACATTCGGAAACTTGACGAGTCTAATGAGCTGTAACTTTTCATACAATAACCTCACTGGACCATTGCCGGATATACCACTCTTTCAGAATATGGATGTCAGCAGCTTTATTGGAAACAACGGGCTTTGTGGTGGTCGTTTAGGTGGATGTAACGAATATCCTCCGTTCAATTCTGATCCTCCAATAAAAAATGCAGGTGCTCCAAGAGGAAAGATTGTTATTGTTGTTGTGGCTGTTGGTAGTGGTGTTTCTCTTGTTTTGATTATGGTAATTTTGTATGTGATGAAGCGGAAACCAGTTGATCAGATGGTTGCATCAGTAAAAGATAAGAACGCGTCATTTCCAGCTTCGGACATATATTTCCCTCCCGAAGAGGAGTTCACTTTCCAAGACTTAGTTGAGGCTACAAACAGTTTTCATGACAGTTATGTTGTTGGAAGAGGAGCTGTTGGGACAGTATACAAAGCAGTCATGCAATCTGGACGAAAAATCGCTGTTAAAAAGCTGGCTTCTAACAGAGAGGATAATAACATAGAAAAGAGTTTCCGAGCAGAGATTTCAACTCTCGGAAAGATTAGGCATCGTAACATTGTAAAACTGTATGGCTTTTGCTATCATCAAGGTTCTAATCTGCTTCTTTACGAGTACATGGACAAAGGTAGCTTGGGTGAATTGCTTCACGGTGCATCTTGTAGCTTGGATTGGCCTCAACGCTTCATGATAGCTCTTGGGGCAGCTGAAGGACTTTCTTACTTACATCATGATTGCAAACCACAGATCATCCACCGCGATATAAAGTCCAATAATATTTTGCTCGATGAGAAGTTGGAAGCTCATGTTGGTGATTTTGGTCTTGCTAAAGTTATCGACATGCCTCAAACTAAGTCGATGTCTGCAATTGCTGGATCATATGGCTACATAGCCCCTGGTATGTACATTTCATTATTTTTTCATTAGAACTTGTGTAGTATAGCCCGTCTACTGTCTGCGAATAATGTCAAGAAACATATGCCTGTCTAGATTGCTCTAACAAGACTAATATATTTTAGGTAAGCAATGATTATAACTCTTCATCGAAAAATCTCACGTATAGTTCCTCCCATGAGCTGACTAGTGTATGTTAGTTTTCCTCACACTTTGACTAGCAAATAGTACTAGTGTTTCCTTGGTGAACGTTTTTGCTTCCTTTGTTTATACTCAACAACAACATATCCAGTGAAACCCTATAAAGTGCGATCTGGAGAGAGTAGAGTGAACACAGACCTTATACTCATACCTCGTAGAGGTAGAAAGACTGTTTCCAAAAGACCCTCGACTCAAGTGAGGTGAAGAGAACAATACAACTAGTAGAAAAAAGTGCAGAGCATACATGCAAGGAATGATATTTGAAGTAAGGGTAATACTATTTGATGAACTATTTGAAGAGAGTAAGCAGACGCGTCTACTTTGTTTAACGGTTCTGCTTTTCCCATTGCAGAATATGCTTACACCATGAAGGTAACAGAAAAATGTGATATCTATAGTTACGGAGTAGTCCTTTTGGAGTTGCTAACAGGAAGAACACCGGTGCAGCCACTCGATCAAGGAGGTGATCTTGTCACTTGTGTGAGGCATTATATTAGAGACAATTCGTTGACACCAGGGGTACTCGATATCCGATTGGATTTGACAGATAAAACTACTGTTAGTCACATGCTTACAGTCTTGAAAATTGGTCTAGTATGTACTTGTTTGTCCCCGGCTGATCGCCCTTCAATGCGCGAAGTTGTGTCAATGTTGATGGAGTCTGATGAGCAAGAAGGTAACTTCATCTTGTCTCAAAGTTGA

>PGSC0003DMG400017291 LRR-RLKs

TTTCTCTTCTCTATTATTATTACTATACTATAACAAAATGCCAATAAGGAAGTGACTGAAGAGCTGAAACTCTTTGGATTTTTGTACTGTTTTTTGTACAATCTAATAGTAGCTGGAAATGAGCATAATGAAGATTTTGTTTCTTCAATCTCTGTTATTTTCTCTTTTTCTTGTTTTCTTCACAGTTTTAGCAGCAACTTCAAAATCTAAGCTCCCACAAGAAGAAGGTAACCCCATAGTAATCTTTTTCAGTTTCTACATGTTCCCATTTTGTTTTCTTGGAAAATGTCAGAAAAAGTTTCAAACTTTTTGTGTTTTGAAGTTTGCTGCTTAGTTTTTTCAATTACCCATATGTTCAGTTTGTTGGAAAATTGAAAAAGTTTGGAACTTTCTGTTCAGTGAAGTCTGTTGTTTAGTTTTTTCAATTACCCATGTTTCATTTTCTTGGAAAATTTCAGAAAAAGTTTGAAACTTTCTGTTCATTGAAGTCTGCTGCTTAATTTTTAGTTTTTACTTTCTCAATTTCAATGTATAAAAGTAACTCTCTATTACTTTTTCAGGAACTTTTTAATTTCTAACTTTCCAGTTTACACGTGACATGTTTAAGAGCATAAAATTAAAAAACAATTTGGTACATTTGACAAATCTTTTAATTGAAGGCCATTTAAAAATTGTTCTTTGCTTTTTTTAAACTTCGTGTCGAGTTAAAATAGGACAAGCATATTGAAATGGAGAGAGTACAAATTACTCATATGTTTCATTTTCTTGGAAAATATCAGAAAAAGTTTGGAGCTTTCTGTTCATTGAAGAGTGCTGCTTAGTTTTTTCAATTACCCATATGTTCATTTTCTTGATGTGCAGTGATAGCATTGAAAGTGATTGCAAAAAAGTTTGGGAAAAAAGATTGGGATTTTAACAAAGATCCTTGTAGTGGAGAAGGAAATTGGAGTACTGCTATTACTGTTAAAGGATTTGAGAGCTATGTTGCTTGTGATTGCTCTTTTAACAACAATTCCACCTGCCATATCACCTCCATGTATGTTTTAGTCTGTTCCCTATTAAATTGAAATTATGTTGTTCGGACTCTCCGAAAATGTAGCAGCACCTGTTCAACTCCTCCAAAAATACACTTCTTTTGGAGGATCTGTCACATACTTGACTATATTTTTGAAGAGTCCGAGTAGCATAGAGGTGAAGATTTGATTTTTTTTAAAGCAGCAACAATTAAATCTGCAGTTTTAAGTAGCTTCTTTGCAAGAGAAAAGATTAAAACTTTTCTTTTTACAAGTTTGAAAGGTTGAGACTTTTTACAAGTTTGAACTGATTAGTGTGTGAATAGGAAATGAAATGTGTGGACTTTAAGTACCACTTTTCTATGAGTTGACTGAAAATTGAGTCAACTTCTTATATGGCAGCCCAGCTGTAATGAGGTGTCTGGAGTGTGGACATTATTTTTGTAAAGTATTTCAAAACTCAAAAGACATTTCACTTTCAAAAAAATTCTACAAGATAATTTTTAAAAAATACTTGTTGTAACAAATTACATGTCTTATAATTCAGGTTACTAATTGAATTTTGTTATGTATAATTAATTACCTGTAATTAGTTTCTAGTTATCTTGAAAAGTACTAATTATCTTTTGAAACAGTTAGTGTAATGAAGTTAAAGCTCTTTCTTCAAATTACCAAACAAATCAACATTGTATACTTCCTGTTTATGTTAAACTTGCATGTAATGCAATGAAGTAGTCAAAAACTAACACTAAATATGCCACAGAGCTTTGAAAGCACAGAATATTTCAGCAAATATTCCACCAGAATTCGCGCAACTTCGCCATCTTAAGCACTTGTAAGTCCTTGTTACAGAATTATCACTCAATAGACTGAAGCACCTGATATTACTATGTTGAAAACTTAGGAGCTGCACTAATCCAACTACTAAATGATGATTATGCAGGGACCTCAGTCGCAATTATCTCAATGGTTCAATCCCATTTCAATGGGCGTCACTGCGCTTGCTTGACCTGTACATACTATTTCTCACCTAAACTATTCTTCATTAGTGCTTAAATTTTTCTCTTCTGGTTTAGAATAAGCTATTTGACATGAAAATAATTGTTATTTGAATGTGCAGTTCTTTCATGGGAAACCACTTGTCTGGTCCATTTCCAAAAGTTCTCACCAGAATTACGACGCTTAGGAACCTGTAAGATCAAATTTCTCTTCTGATGCCTACTTGTCATTTTCAGTGTTGTATACTTGCACTTAATCTGTTTTACTATTGAATTTAAGTTTTATTTATGACTGCATAAAGTATTTACACAATCGAGTTTAAGTTCTATTCGCTGACTATATAAGTAATTACAAGTAAAATCTCTATGATAAACACTAATTGGTAACCTGATAAAAATGTTAACTTAAGTCATTTACCATTTCAATATGGCGATTCATTGATTCAACATATATATTATACTGTAATTTGGTACTGAAGTGAGAATGCGGAAAAAAGATCTCTATATGTCATGTCTGTCTGCAACAACAACATACCTAGTGAAATCTCGTAGGTGGTGTTTGGGGAAGCTAGAGTGTACGCAGAACTTATTTCTTCCTCGTAGAGGTAGAGAGGCTGTTTGTGAAAGACTTCGGCTCAAGTAAAACATATCAAAACAGATTCTCTTTTCCTAAGGCATTTTATTTCAAAAGTTAACTGCTGCTCTGACAAGAATTTTTGATCACCTGTATTTGATTCCATGAGAAATAAAATGCCTCTTCTATAACTTTCAATACAGGAGCATTGAAGGAAATAAATTTTCAGGAACAATACCTCCAGAAATCGGCAACTTGGTTCATATGGAAAAACTGTAATTCTCTTTTTCCTGTTTTAATTTTGTTCTGGAAATCTTTATTCGTTTCACGTTGTATTAATGCATCTGTGCCTTAATTCTCAGCGTGCTATCGTCAAATAAATTAACCGGAGCTTTGCCAGCCACACTAGCCAAGCTAACCAACTTAACTGATATGTGAGTTCACAGTTTTAAACTCTTTCTCATTTGGTCATTCCTTCTGTCCCTTGTGAATTGTGATGGATGATCACATAATTAACTTTGCCGATATATTAGGAGGATAAACGACAACAATTTTACTGGAAAGATACCAAAATTTATCAGCAGCTGGACAAAAATTGAGAAACTGTAAGCTTTCTTTTTTCTTTACTTTTCAGACTTCAGTAGATGAAGTCTAGTTTCTAGCTTTATAACAAAAGGGCTTCCCGGTGCACTAAGCTCCCTCTATGCGCGGGTCTAGGGAAGAGTTGGACCACAGGATCTATTGTACCAGCCTTAGCCTGTATTTCTATGACAGGCTGTTTCCACGGCTTGAACTCATGGAAACAGCCTCTTTCTGGAATATTAGAAGTTTATACCCCAATTTTTCCTGCATATTTACAGTTATTGCTGTGATCTCATGAACCTCCAATTATACTATAATTTTTGCAGGCATATACAAGGCTGTTCTCTTGAGGGTCCCATTCCTTCCTCTATATCTTTCCTGACAAGTTTAATTGATTTGTAAGCCAATGTGAAACTCCTTTTTGATGTCATTTGTGGTTGCTTTGAGAAGTAAAATAATTTGTTATTTTTTTCTTTCATGAATTTCAGAAGGATCAGTGATCTAAAAAGTGGAAAATCTGGTTTTCCACCATTAGATAACCTAGAATCCATAAAAATACTGTGAGTTTCTTTACTTTTTCTTCTTTTAATGGACTGCTTGAAGGACATTTCATTTCGATTTCGGAATCAATTTCTGATCAAAGATTCTGAATTCCTCTTGTGCAATGCAGAGTACTAAGAAATTGCTTAATTCATGGAGAAATGCCTGAATATATTGGGGAAATGAAGAAATTGAAGACACTGTAAGTTTGAAAAAATTCAATCACTTTTTTTTGGTTCAGTTTTCCTGCATGTTGATACCTTTTCTGTTTTCATGCTATATAGTATATACACATTCTTCAAGATTCTGAATCTTTTTTAGCTGAAATTTGTTTTTCCATGGAACTTACTATTTTCTGCTTTGTGATTTTAGGGACCTTAGTTTTAACAGCCTGTCTGGTGAGATTCCATCAACTTTTGTACACCTATCCAAAGTAGATTTTATGTAAGTGGATATTCTTCAGATATCTTTATTTCTTTGTAGCCAGATGCTTATCAATATTAGGCTTCTGTTTTTTTCTTTTCTCTAATTGTTCTGGTACTTTGCAGATATTTGACAGCGAACAAACTTACTGGACTTGTTCCTGGATGGATTTTAACCAGAAACAAGAACATGTGTGTGTTTTCGCAACTTAATTCGTACTTCCCTTTTTAGTCCGTCCCAAAAAAGAATAACACAATTCCATATTTAGTAACAATTTTAACTTTAAAATGTTCACAAACATCGATGACTTGTTTTAGACCACAAGTTTTAAAAGTATTTCTTTCTTTCTTAAACTCCATGCCAACTCAATCAACATCACATAAATTGAGATGGATGGAGTATAAATGTGCAAAAAACTTGATTGCTCTTTAAAGTTTTCACTTACTGACCTACATGCTTTTATGCAAGTAGAGGCTTTAGAATGTATTCAATATTGAGTAATCTATTTTTGCCAAAAAAAGAATGGTGAAAAGGAACTAAAAAAATGTAAAAGATCATCTCTAAGCTTCAAAAAGTTATTAGTATTATGGACTTGCGAAACTTCTAGTCTAATTTTGTTGTCTATTTTGCTTCACAGAGATGTTTCAACTAACAATTTTACATGGGAAACCTCACCTTTTGAATGTCCCCGTGGAAATCAGTAAGTTGTTTGATAAAATTGTTTTTCTTCATTTGACTTATTTGTTTGTGATAATTTTCTAACATCAAGACGATTTATTCAGAAACTTGGTGGAGAGCTATTCTGCATTGGGACAAGAATCGTAAGACTAATGTGCCTATAATAAATTTATCTGTTGCACTGATCTTACAGAGGAAACTATTGTTCCTTGAGTTCACCTATAACTAAGAGTGAACTCTTACCTACTATAGAAAACTTTTGTAGCACTCTTAACAACTTTTTCTTTTGTTTTCTGTGTTGCAGAAACAAAATTCATCCTTGCCTGAAGCAGAATTTCCCTTGTTCTGAGCCAATAGATCAGCGTAAGTAAACTTTTATGTGTCCATTTCTGTAATTGATAGCACAACAAAATCTGTCTCCTGTCACATAAATAGGTGTACTTGTAGTGTGAGGTCAGCTAGTGGCAGAAAATATAAACAACAACAACATAGCCAGAGTAATCCCACAAGTGGGGTATGGGGAGGGCGGTGTGTACGTAGTCTCACCCCTGTTAGAGTGGGTTAGAGTCCTACATTGGTTGGGGAATGGATTGGTAGTCTGCTTATATTGACTTGGGCAATCCTCCCCTCATGAGCTAGCTTTTGGGGTTGAGTTAGGCCCAGGTGTCATATCTGACTCCTAGTAACAACAAAAAATACAATTTAAGAGTTTGGGACTGACTGCTGTCTTCTGATCTGTCCATTGTGATTTTCTGCTGAATTCTTGTTAAGAGTGCCCTTGTCTACTGGTTTGCAGAACAATATTCTATCCACATTAATTGTGGAGGCAAAGAGGTAACCATCAAGAATGGTACAAAATACACAAACTATGAAGCTGATTTAGAACCCAGAGGTGCTTCTATGTACTATTCAAGACATAACTGGGCGTTTAGCAGCACAGGAAATTTTATGGACAACGACGTTGAGTCTGATGTCTATATCAATACCAATGTCTCTGCCCTGCAAAATGTCAAGGCCCCTGAACTTGAACTATACACTACTGCACGTATATCTCCTCTCTCCCTCACATATTATGGACAATGTCTAATGAATGGAAACTACACTGTCAAGCTCCATTTTGCGGAGATAATATTCACAAATGATACTTCATTCAACAGCCTTGGAGAGCGTATATTTGATGTCTACTTGCAGGTAAATTTCATGCTCTGGGGGAAGGTGAAATGTGTGTGAATTTCATGTTTGCTGCCACCAGTGTATTTATTCAATCTTCTTGAGACAGGAAAAGCTGGTGTTGAAGGATTTTAACATTGCAAAAGAGGCTGGTGGGCCTGGTAAGCCCATTGTGAAGACGTTCACTGTGAATGTGACAAGCCACACGCTGAAGATACATTTTTACTGGGCAGGAAGGGGGACTACAGGAATTCCATTTAGAGGGGTTTATGGTCCTCTAATTTCAGCTATATCAGTTGTTAATAGTATGTTCCTTTACTCTCTTCAAGTTTATATCTGTCTTGCTTACAGTTGTATGCTCAACTCTTCTTTAATGAATGCCATAGGGATCATAGGACATTATTCTAATGTACTAGTCCTTAAAAAAACCTTTCCGAGACCAAATTACTCTACATACCAAGGCTTAATAAGTTTTGTTAACCATCTTGATGTTATAAATTCTATTCCTACTGATTCATTATGCATCTAAAACCGCTGTCAACATATTCTTTCAGGAAGTTTATGTGTATCAGAACCTAAGAAACTTCATATACCCTTAAATTTCATGATGCATTTCGAATACAGTGGGGAGACCACTTATAGTGTCCCCAGTTCAGTTCCTTAATCTTGTCTGGTAGTGGAAGCATCAAGACTTATACTAAGACTTCAAAGGAAAATGTTTCAAATTGCATATGCAAGTCAATTAAAGCTTGCTAAAATCATCACAAGTTTTGAGATGTCATATTATTTTTTCCTTTCCGTGCAGAGTTTCCCCCTCCTCTGCCTGCTCGTCTGCCTGCTCATCTGCCTGCAGATCAAAAGAGTAAAATTAATGTTGGCATACTAGCCGGCATAATAGCTGGATCAGTATTTCTTGTCCTTCTGATTATTGGTATCTTGTACAAGGGAGGCTGTCTAGGCGAAAATGTATCTACTGATAGAGGTAATAACTACAAGTATTTTACTTTTGAAAGTTTAGTATCCATTTATTTGACCTATGAGTCAGCAATAATGTTCTCAAGTTAAATGCTAAACTACATAATTACTTTCTAGCCAGTATTTCAGTTCATACCAAATTATATCAGCTCAAATACTTGAAGTTATGCTTTACGTTCTATTGACTTGCTTGTTCATCTCATCAATCATTTATTCATTTTAGGCTTTCAAAAATCATTAATGGTCTGAAGCACTTGACCAAAGTTTATATGAACCTTGAAGAAACATAGTGTTATAGCCATTAGCTTCCACTGAGATACTTTTCCTCTTGAACAGAACTTAAAGGACTGGATCTACAAGCTGGACTATTTACTCTCAGACAGATCAAAGCCGCGACAAAGAATTTTGACCCAGCAAACAAGATAGGGGAAGGTGGTTTTGGTTCAGTTTATAAGGTAATCATCTTTTCCTTGTCTATTTTAAATCAAACACATAACTATAAATGTCATTATATTAGCATGAGAAGGATCATAATGACATGGTGCAGATGCCTCTTAAGAGAGAAAATGAAAGACTAAAATCTTAAATTTCATAAACCTTCAAAGGAAGTGTGAATATTCGCCTGTAATGAGGAGACTGTACAGGCAATGCTCTATTACCATATGTGTTTCTATAGCCCTGAGAAGTTGGAGTAATTTACTGATCTTTTTCCATCAACTAAGATGCAGAGTTCTTATCTTCGCGACATAGGATTCAATCTATTATAATCTTACTTGCTGATCATAAGCTGCAGAAGTCTGAATAGAGTTTTGAATCATGACTTTTGTACACAGGGTCTATTATCTGATGGAACAGTAATTGCGGTGAAGCAACTTTCTGCAAAGTCAAAACAAGGAACTCGAGAATTTTTGAATGAAATAGGGATGATCTCTGCTGTACAACATCCAAATCTTGTAAAGCTTTATGGATGTTGCATTCAAGGAAACCAATTATTGCTAGTCTATGAGTACATGGAAAATAACTGTGTATCTCGTGTTCTTTTTGGTAAATACTTCCTAACTCTGATCAATCAAAATATGTCTGCTGCTTATGCACTTGACATTGTTTATCTAACTGAGCTGGTATATACAACTTTTTCTTTTTGTTTGTCTCTCTCTTACAGGCAAAGGTCCAATTGGCAAAATGAAATTAGACTGGTCTACTCGGAAAAAAATTTGCCTAGGAATTGCAAGAGGTTTAGCATATCTCCATGAGGAGTCGAGCCTGAAGATAGTCCACAGAGACATCAAGACCAGTAATATTTTGCTTGACAAGGACTTCAATCCAAAAATATCTGATTTTGGCTTGGCAAAACTACATGAGGATGATACTACCCACATTAGTACCAGGATTGCAGGAACTGTGTAAGTATTATGAAACTAGTTCCTCTTGTTCCGATCCCATTAGCAACAACATTTGGTTGATATGTTCCATAGAAAATGGCTCAACATTTAATCCACTCATTTATCTTGAAACACTCGTGGTACTGTTCTCATCCGAAAGCAACTTTGCAGGTATAATCTGCCAAAAGTAAAGCAAGCCAAGAACTTATTACCTGCCTAAAGAAACTTAACTTTTCTAGATATTAAACAGCCTTTGTAGAACAGTAACATAGAATTTTATTCGTCAATGGTAAAACCACCTCTCTTTCCTTACCATATTTGTAACTCATTGGTTTGCTAATTATTAGCACCTCAACGTCTCACTTCAGTCATAATTTTACATTTTGCAATTCAACATAGCTATAGGTGAACTTTCAAGAAAATATTTTCTTGCTACTTCAAATTTTAAGAAATGTGAAGATTGTTTGTATCAGAATTCTCTGTAATTAGCAAGTAACTGTATTAATGATGAGGAAACAAAGATCTAATAAGTTGGAACATCTTTGGTATTTGGTGGTACAGGGGTTACATGGCTCCAGAATATGCGATGCGAGGCTACTTAACTAGCAAAGCAGATATTTACAGCTATGGAGTTGTTGCATTGGAGATTATTAGTGGGAAAAGCAATACAAATTATAGGCCAACGGAAGATTGTGTTTATCTTCTTGACTGGGTAATTTTAATATTTCTCTCTACAACAATAAACCCAATGTAGTCCCAAAAGCGGGGAATTTCACTACTTCTCTCTATTCGACCAAATATCTGGGCTTCCATCATCCTTAAATCACTTCCCCTTCTAGTTTCATCTCAACCAGAGAACGTTTTATCAACATTCCGAGTAATGCTAATAGCATCTTATTAATCTTCCCCTGCATCAACTTGTACATTTAGAGCCAAAGTACTTTAAAATTGGCTTTGGCAAATGAACATGTCATACTTTTTTAGTACATTAACTGTTCTTCATTTTCTGTTTTGATTAGGCTTATGTTTTGCAAGAGCGCGGGAGTATATTGGAATTGGTTGATCCAGACTTGGGTTCTGACTACTCGTCACAAGAAGCAATTGTGTTACTGAATGTTGCCCTTTTATGCACTAATGCAGCTCCAACACTTAGACCAATAATGTCTCAAGTGGTAAGCATGCTCCAAGGCCAAACTCTTGTTCAGGATGTCCTTTCTGATCCAGGGATGTCAACAAGCGGTTCAGGATTCAGATCAACGAGAAGTCACTTCTGGCAAACTCAAAGCCTAACTGATGGTACGCTTACTGATTCCACTCTCTCAACTGGAAATGCGGAAGAAATTGGCATTCTTCAAAGAGTAGAGCCTAATGTATCTAACAGTTAAACAAAATGTCTGTTAACACAAGCAAGTGTATATATGAACATCAAAATGTTTAATGAAGATTAAGTTTAGTCTTTACAAAACTGCTAACTGGTCCTCCTTTTATAGTACTTCATTTGTTTATTTTCTAAGCCAACCTCATTGTTGCATAGTTCTATCAATTCCTTCTTTATTTTTC

>PGSC0003DMG400017645 LRR-RLKs

TGTCACAAAAATTGTACAACTAGACTAGCTATAGACAAACACATCTGTACTCCATTATTTAATCTCATCATTTAAAAATCCCTTTTCTCTATGTATCATTACTCACTCTATTATAGATTCTTGAATTATTTCATTCCTCACTTCTCTTTAATCACTTCAAAAAAACAAAAAAAGTGCTCACTCTTTTGTGAGCTCAAAAGAGTTTTTTGGGATTTTGAGTCTTTTAGCTATGAAGGGTTTCTTGTTTCTTGGTTTGTTTTTGTTTTTTCTAGTTCCATTTTCATCATCTCAGTTAGTTCCTGCAGAAAGTAGAATCCTTTTTCAAATTCAACAGTTTCTTGAAAATCCACCTGTTCTTCAAGAATGGAATAAATGGACAAATTTTTGTTTTCTCCCACAATCTCCTTCTCTTGTTATCACTTGTTCTGGTAATCATATAACTGAGTTAACAATTGTTGGTAACAAAAAATCCCCTTTTGAGTCTTTAAAATCCTCTTCTCCACAAGCCCTTTCTGGGAAATTTTCAATTGATTCATTTTTCACTGTTGTTACTAAGCTTTCTAGTTTAAAAAAGTTGTCTTTAGTTTCTCTTGGTTTGTGGGGTTCATTGCCTGCTAAGATTAGTAGGCTACATTCACTTGAGGTACTTAATATAACTTCAAATTTCATTCTTGGTGAGATTCCATCATCTATAGTGAATTTCAAGAACCTGAAATCTCTTGTTTTGGCTAGAAATTTGCTTAATGGAAGTGTACCAGATCTGAAAGGGTTAAAAATACTTGAAGAGTTGGATTTGAGTGGTAATAATTTAGGACCAAAGTTTCCATCTTTAGGTGATAACAACAACAATCTTGTTTCACTGAATTTGAGTAACAATATCTTTAGATCAGAAATACCTAATGGACTCAACAAGTTTACTCATTTGCAAAATCTTGATTTGTCTTCTAACAAACTTGTTGGTCCAATGCCTGCTTTCCTGTTTTCACTTCCAGCAATTCAGTCTATTAGTATTGCTAAGAATCAGCTTAGTGGTGCTCTTCCAGGAAGTGTGTCTTGTAGCAACAATCTAAAATTTGTCGATTTTTCGAGTAATTTATTGATGGGAAAGTTGCCTGCTTGTCTTGGATCAAGTTCAAGTTCAAGAAACAGGACAGTGATTAATGTTTGGAATTGTTTGTCAAGTACTAGTACTAAGTCTCAACATCCACATACATTTTGTGAGAAACAAGCTATAGCTGTTAAGCCCCCTCCAAGAACTAGTGATGAGAAGGAACAATCCACAGTTAAGCTTGGTGTTGTCTTAGGACTCATTGCTGGAATTGTTGTAGTTGTGGGTGCAGTTGGTTTCCTGATCTTTTTCATTGTCAAGAAAGTTGTGAGAAATAGAGATCAAGGATATAAAAATGACAGCTTTGCTTTTGAGAAGAATTCAACCCTTTCCAAAACTGCTGATGGAGGTAAATTTCCTTAATGATTAGGGTCTATCGGAAACAGTTTCTCTACGTGTCAAGGTAGGGGTAAGGTCTGTGTACACTCCCCCACTAGACCCCACACTGTGTATGTTGAATCTTGAATAGTGATAGGGGTCTATTGGAAACAACCTCTCTACCTCCCGAGGTAGGGGTAAGGTCTGTGTACACTCTATCCTCTCCAAACCCCACTTGTGGAATTACGCGGGGTTTGTTGTTGAATCATGAATAGTGATAGTGCATCATTATGACAATTTTATTGATAACTTGATTTAATCAGGGAATGCAAGGAGGACAATGAGGATGGTATCATTAGGACTTCCACCATATCATGTTTTCACTTTAGAGGAAATGGAAGAAGCAACAAACAGTTTTGATCCAACTAATCTGGTTGGAGAAGGTTCCCAGGGTCAGGTAAAAATGTTGATTCATTATAAATAGCAAAAAAAATGATATTTATTACTATATCCTGTCCATATTATTCGTCATTTTGGACGCCGAACAATTCCATTCTAGTCAAATACATAATCTACATGAGTATACAAGTCCGTAGATACTCATATTGCTATACTTCTAGTCTAGACCCAAAAAGTCACCCTAAAAGTCAACCTGAACAACACTTGACATTTTAGAAAAACTAAAGATAGTAATATGGATATGCTTTTCTTAAATCTTGGTGAATACAAACTTGTGTTTTGAACATTGGGATTGATGTTATTATTAATGACAGCTATATAGAGGTTGGCTTAGAGATGGATCAGTGGTCCTAGTGAAATGTCTGAAATTGAAGCAGAAGCATTCACCTCAAATTCTGCAGCAGCACATGGAGATGATATCAAAGCTAAGGCATCGACATTTAGTAAGTGTACTTGGACACTGTGTTGTTACTTACCAAGATCATCCAAATACAGCTAGCACTGTATTTATTGTGCTTGAAAACGTCGTCAATGGATCACTAAAGGATCATCTTAGTGGTAAGCTATTACGAGTTGATTTCTCTCATGGCCACTCAACTAATAGTTATTACCTCAAAAGTCACTTTCGAAAAGGTGACTTTTGAGATAATGACTATTAGTTGAGTTACTATCTAATAGTCATTACTTTCTTCAGTCGGAACGATGAAGGACTGAATTTTATATTCCGGGATTCGTTCTTTGGCAGATTGGAGGAAAAGGGATGTATTGAAATGGCCACAAAGAATGGGAATAACGATGGGCATTGCAAAAGGAATTCAGTATTTGCACACTGGAGGAGTCACTGGAAATGATATAAAGCTTGAGAATGTTTTGTTGGATGAAACTCTTACTGCTAGAATAAGCAGCTACAATATATCGTTGCCACCTAAGGTAATGCGATTAAATCACTATACGTAAATCTTGCAGAAACTGTATTTTTGACCATCTAAATTTTTACAGGTTGGTTCGGAAAGTCCTCTTGCTGGACCCGACCATTTTACCAGGTATTATTCTGCAGCTACATTCACCTTACTTTCTCTATTTCGTGCTATCATTTCTGAATTGTCTGCTAATATAAACTTCATATTCAGCGCGAAAGAAGCAGAAAAGGAAGACATTTATCAGTTGGGAGTTATTCTACTAGAAGTTATTATTGGTAGACCAATCAACTCTCGAAGCGAAGCAGAGGACCTTAAGCTTCAGGTATCAATAACAATATCCTAATAACTCTCAAACGATAGAAATATATCATATCGTCACTCAACTAATAGTAATTATTTCAGAAAGTCACCTTTCTTCTTGATTCAGATGTTCTTCTGAAATAATAACGATTTGTTGAGTGACCATATGAGAAATCAACTCGAGAAATAAGAGATATGCGATAAAGAAATTAGAAGAATGCACACGAATACAAATGTCTTACATGATATGAAGCAATACACATGTTTACATATTGCATAAGAGTCTAGTCTATTTACACAATCTATCTAACAACCTTGAACCAAATTGCCTTTTATGACATTCAGGTGGAGACTGCCTTAGCGGAATCACCATCAAAGCTACGAGATTTAACAGATCCTTGTATACGAGGTACATTTGCATATGACTCATTGAAAACGACAGTCCAGATAGCAATAAACTGCCTTGAAAAAGAACCAAGCAGGCGGCCATCAGTCGAAGATGTTCTCTGGCATATGCAATACTCAATTCAGGTTCAAGAAGGAACAACCAACAGTGGAAACCTCAGTGGGAACCAAAGTGGGAAACTTAGTGGAAAAATAAGTGGAAATCTTAATAATAAGTTGTACTGATAAGAGAGAACTGTCATTATATTGTTATATCTTGTTGTTTAGTCATTTGTATGCATCTTGTATAGCATAGTTCAGTAACTTACTGAATAAATGTTTTTGCGTTCTACATCACAAAAAAAATGTACCCAAGTTTACAATACAAAGCAAATTTTCATGGAAGTTGAGAATCTCGGGGATCTCGCGAAAAACTTATCAACACCCGGTGTTTACATCAACTCAATGAAGACATAAAATGTATCTTCATGTGCTTTGTTATGTCTAAGTCCTTCAAAATAAGTAACTAGCGGAAGATATAACTAAAGTAAATAACAATGCAGTCTCTACAGTCATGATATGATTCAATATCGTATCAGAGCAGGCAAAAATCATCGGTTCATATCTCACCACCAGAAATTTTCATGTGCGCTTGGCACATTATAAGACCTTTACACGTGAGGGGGCGTGTTGAGGATATAATAAAGTAAACACTAAAAGTATGTCTCTCCAACAATTGAAACTTCTAGATGAAATGCTCACAACATACCTTTGTTGACTCAAAGGGTACTTCTTGAAGTATAACTTCGCCTTTCCTATACCATTTTCAGCAGCCACGCCTGCCACGGTGGCTGCATGCAACAGCAGAATCTGCTGTTCATCGTTGGTGCTCCAAGAGCACCGACATCAATTACCAATTCCAGCCACAACAGCTGAATTGGCTGATCAGGCTGCTGCTTCTATCTCTGCATTCTCCAGCACCCAAAGAACAATAGCAGAGTAGTCAAGTATTCCCAAGTATCTTTCTCTCCAGCCATTACTTTGTCGACACTACTAAAAAATCACTATTTTCTCCAAGAAATTTCTCACTGATATTTTGATTAAATCGGTGAGAAAAGAGTAATGTTTTCGTACAAAAAAACTAGAAAGTTTACCACTATTTCATTGAGAATCTGTTTCCCACCATGCATATTCCCAGTGAACTGATTTGGTGGAAATGAGTGAAAAACATGTTCTATAACACAAATTTCTCACTAATTCACCGTGGAAAAAATCTATTACTACTACTCATCATTAGTACTTCATCCGTTTCATCTTATGTTGAACTTGATTGATTAGGCGCGGCTAAGAGCAACATGGGAAGTCTTAAAGTTCAATTTTTTCCAAATATAGAAAAAGACCCTTTTTATAACGTAC

>PGSC0003DMG400017690 LRR-RLKs

CACATGTTAGTTGGCATGTCTCATCTTAAGCGGCATGGTGGGCAAATTCTCAGAATAACTTGCTAATGATCTTTTTTTTTAATTTCAGGGAACTTTATAGTAATAACATAAGCGGAAGAATTCCAAATGAACTGGGGAACTTGACAGAGTTGGTTAGTTTGGATCTTTACCTGAACAACTTAAATGGTCCTATCCCTCCCTCATTGGGCAAGCTTCAGAAACTACGCTTCCTGTACGAGTTATACATTCTATTTCCTTCCGGTTTTTTTCTTCTCTTTTTGTTTCTTTTTCAGTTTGGTTATTTTTTTTCAATGTTTCAAATCTTTAGCATATATATCTGAAGTTTAGTTAAATCAGAAATGATCACCTTCTATTTTTATTTTTATCAAGAAAGCAGCGTCTATGTTAAGGATCCGTTTGGTCATAGATTTCCCAAGTAAAATTTGGGAGAAAATTTGGCATATAGTGTTTGTCCATACAATTTGCCATTATTTGGCAAATAACCCAAATTCCCAATACTAGAAAAAACTAGTATTTGGGCCAAATCTCATTATTTGGGATCTTTTAAAAATTTAAATTCTACACCAAACTTTTATGTTTTACAAAAACACCCTCTATAGTAGTTGTTTGCGTTGTATTACATAATTTTTTACGTAAACACCAAACACCAAAGTAGTGATGAAATATTCAGTGAATATGAAGTTGATCATATGGTTATTGATGAAAATGATGAACAATTGGCTCAGGGTAACAAAGTCATGTGCTTTGTCTACTTGACGATGTATGGAATGATGCTTGTTGCACTCACTCTGAACTACCATATTGCTCCAATGTCATGGACACTGCTTGTTATTTGTTGCAACGAAGATCCAATTGACTTGTAATATAAACTTATGGTTAGTTTTGATAGTTTTTAAAACTTTTATAAATCATATTTCCTAAAAAAGTGAAATATGTTTCCCAAATACTATGGCCAAACACATAGTGAAATTTCACCCAAATTTTAACACAAATAATATTTGCCAAAAATATTTGGAAATCTATGGTCAAACGCTAGCTAAATCATGCAAATGGGGTTGTAGTTTCTAATTCTCTAAGCAAATCTGTATGATGTTCTGTAACTTTTCTGTTTATTATGTTTACATTGATGGGTTCTGTCCAAAAAAAATTGATGTGTGGAAGCATTATGAAAATCACTACTCCTGTCAGCGAACAAACCTTTGTTGTTTGAGAGGTCAACAGGGAATATATCTGTATGATTCATTCTGTCTTTGCTCTGTAGTGGAAAGTTAGTTGATATGCTTGGCAAACCTTGTTCTTGTTTCCCAAATACTGTATGGTTGCTGAGTGAAGACCAGGTGACTGGACTGGACTCTTTTTTTGTACGACAAACAGAGATAAATAAGTAATAATATAAAATGTTAAGAATAATATGAATCAACTTGAATAAAAATATATTCTAAAAGTTTCAATTATTATCTGATTAACAAAGTTGTGCTTGCTTGGTCTCTCTCGATATAGACTCACAGCCGTGCAGCTCAACTGAAAGTGACAATTCTGAGATACTTCTCATCTTATCAGGAAAATGGCCTGTGGATTTAGTTCATTGCTAAAATACTTTATCTATGAACTGTGATTCTCAACCTGCAAACACTATCAGCAAGTTTTTTCTAGCATATTGGAATTAATCTTTCTGGAGTATCTAGTCCTATATATATAAATGTATTTGTCTTTGTCTACCTTTAGCACATGTACGCTACATTTAGCACATGTACTGAGTAATGAGCTCTAGACCAGTATTCGCTTTGACGAACTTCTTTTCCTGTCTTGCTTACTCTTGATCAACCTTCCTTTTATCCTTTTTTTTTCTTTGCTGAATTGAAGGAGGCTCAATAATAACAGTTTGAATGAAGGTATTCCCGTGTCTCTAACCACCATTGTTGCACTTCAAGTACTGTAAGTATAAAGTTGTTCCTTTATTTATCAATGCTGTATGTTGTCTGTGTTGTCTAGTTCTTCTGGGTTACTTATTGTCATTGCCTTGCAATGCAGTGATCTCTCAAACAACCATTTGACAGGACCAGTTCCAGTCAACGGTTCCTTTTCACTTTTTACTCCTATAAGGTGTGTTGGAAGCATAAATTTGTAGTTCCTTCTGTTCCTGAAGTTACAATAGTGAATATCTAAAGATTGAGTACCTATATTTCAGTTTTGCTAATAATCAGTTGGAAGTTCCTCCAGTTTCTCCACCTCCTCCCCTTCCTCCTACGCCCTCATCGTCATCTTCAGGTTTGTTGACATAATTTGTGTTTATGCTGTTATGAAATTGTGCTACCTCTCCTTTTTCCCTATTCCAATTTAGTTTTCATGAAAATCCCTTAAGTTATTCACAATAAGGGGAGATGGCAATGGGAAGAAAGGAAAGGCCTGGTTGCCTAGGGAAAGCTGAAGAGATGAGCCCCAGACTGGATTCCCTTATGAAAGCACTTTCTTTATGAAAAAAACTACCATTACGTTAGGCACTATAAATTAGTTTATTTAGATCAATTATACAGGTATTGGGGTCTTGTGGTTATCCAGCATTCATGACTTTCCAAAATAAAAAATTCAAATAAGCTTGGAGATACTATCAACATCTTTTTCTTTGTTGGAAATTAGATTTACTGTAAACTCTGAAGTTCGAGTGATACGTGTGTCAAAAGTACCATATATATTTTAGAATTGCCTGTATTTTAAAATTTAGGTCATACAAATAAAAAGTTAACTCAATCATCTTTTCAGTGGGCAACAGCGCAACTGGAGCTATCGCTGGAGGAGTTGCTGCAGGCGCTGCCCTTCTATTTGCAGCTCCTGCAATTTTTATTGCTTGGTGGCGTCGGAGGAAACCGCAAGACCACTTCTTTGATGTTCCTGGTTTGGCAAAACACATTGAATCTGAGTTAATTGTACTTTCAACTTGGATGTGTTCTCACTCTTTATAACTACTTTTGCAGCTGAGGAGGATCCAGAAGTTCATCTGGGACAACTCAAAAGGTTTTCCTTGCGTGAACTACAAGTTGCGTCGGATAATTTTAGCAACAGAAATATACTCGGTAGAGGTGGATTTGGTAAGGTTTATAAAGGCCGGTTAGCTGATGGCTCTTTAGTTGCAGTGAAAAGACTAAAAGAGGAACGTACTCAAGGTGGAGAGTTACAGTTTCAGACAGAAGTAGAAATGATCAGCATGGCTGTACACCGAAACCTACTTCGTTTACGGGGATTTTGCATGACACCCACTGAGCGGGTGCTTGTTTATCCGTACATGGAGAATGGAAGTGTTGCATCACGTTTAAGAGGTATCTCTTTTTTAATTTTCATGTGCTTGCCTCTACTTTTTGCTCACCTTTAGGCAGGCTCACAACAACCACAACAACAACATACCCACAAAGTGGCGTCTGGGGAGGGTCGAGCATATGCTGACCTTTCCTCTACCTCAGATGTCAGGTAAAGAGGTTGTTTCCAATAGATTCTTGGCTCAAGACAAAAATAGTCTAGAAAAAGACATAATTGAAGTACAAGAGAAAATAGATAGCAACAAAAACAACAGATAGTATCAAAACAGTACTGCACAAAGTAATACTGTAATAGAACTACATAAAACATCATATATCAACAAAAATCGAAGAACAAGAAACTACAAGCATAGTACTACAACTACTAGTAAGGGCAACAATATAACGCACTCCTACCTACTAGTATGGATGTACTTTTACCTACTAATCTTCTACCCTAATTTGTGTCCTCTACAACTTTCTATCTAAAGGTATGTCCTTGGTAAGCTGCAATTACGCCATGCTGTGTCTAATCATCGCTCCCCAATACTTCTTCGGTCTACCTCTACCTTTCCTAAATCCATCCATAGCCAACCTCTCTCACCTCCGCCCTTGGCATCCGTGCATCTTTTCATCACATGTCCAAACAATCTCAACCTCACTTTCCGCATCTTGTCTTCCACCAATGTCACTTCCACCCTGTTTTGGATATCCAAATCCTTAATCCTATCTCTCCTAGTAAACCCACACATCCATCGCAACATCCTTATCTCTGCCACTTTCATCTTCTAAATGCTGGCCAACACTCTGACCTTACAACATAGCTGGTCTAACCACAGAAAAAAAGCCAATAGCTAATGTTAATGACAAGCCTTTAGTCTTTTAACTTTTCTTTCATAAATTTGTAACAAATTTGAAAAAGTTACTGCAGAAGTGTAAGCCAGTGGTGGAATCTTCACATCAATTTCATTGTGCTATTAGTTATTATAGTGATTGAATTTTTTATGAGTTCTCATGATGGGTATGTATATTTAACAAGGTGTCAATGTTGTTTATGATTTTGCTTGTGTTCTATGGACCAATTTCACAAGTGGTGCTATCCTTGATATCTTACTGAATATAGAGACTGCAGTTCATGATAAAATTTATTTAATGAGCACATCTGAAAGTCAAAAAGAATTAAATGTCTTGAAAGAGAATACATATTTTTATTATTGTATGAAACTGGATAAGTAATGCCTCATGACTTGGTGTTTCATGATTTAGAGTGATATTGACCCAAAAAGATCAGTTTTCTTTTTATAACTTATTCAAGCTTAAATAGGAACCTGTTTCCAGTTTGCTGAATAGCTTGAATTTATGCTCATGGGATAATTTACATTTTTCTAGAGAGGCCTGAATCAGAGCCCCCACTTGACTGGCCAAAAAGGAAGCGTATTGCACTTGGATCTGCAAGAGGCCTTGCTTACTTGCATGATCATTGTGATCCTAAAATTATTCATCGTGACGTCAAAGCCGCAAATATCTTGTTGGATGAGGAGTTTGAAGCAGTTGTTGGGGATTTTGGGTTAGCTAAACTCATGGACTACAAGGATACTCATGTTACCACTGCTGTACGTGGTACAATTGGGCATATTGCCCCTGAATATTTATCTACTGGTAAATCTTCTGAGAAAACTGATGTGTTTGGCTATGGGGTTATGCTTCTAGAGCTCATAACTGGGCAAAGGGCTTTTGATCTTGCTCGACTTGCGAATGATGATGATGTCATGCTGCTAGATTGGGTAAGCATTTATACTGTCCTGTATGCTTAAGTTATTTTGCTTCTGCTATCAATCAGCAATTTGGATTTGCTTAAAGATCAAGTCACTTCTGCTATCAGTGAACAGTCTTGGAATGTTCTGAAAAGATCAAGTCAACTCTCTCTTCTCTTCTATAAGTCAAAGTTTGATGTCTTTTTCTACTGTAGTGTTAGTTGTTGAACACTGATGTGCTCATCCTGAAGAAATGGTGTGGTACGAGACCTGCATCGTTGCTTATTATTTATTGTGGTTTTGTGGTGTGAAAGTAACATTCGTTGCAGTTCCCTGTGAGTAATTTTTTTTAAAAAAGGCTTAGATGTTTCAGGAGGATTTAACCATTCATTGCTAAAATTGCACGAGCATGTAGGTTATAATTAGAGTGGATTACTTTCATTATCTTATTCTTGCAGGTGCTTCAACAGTTAGGTTGCCAAATTCGTCCCCTCAATTTGTGACTAATCTAGAAAATGTGCATCTCCACTTCTCATTTCTGATGCTATCTGAAGATTAGAAAAATATACTTTTTTCATTTTCATTCAAATAAGCTAAGGTGTATCTGCAACTGCCATATTAATGTCACTTCAGTCATGTTGTTTTTGGGGGTAGATATAGTATACTTTAGGACAGGTACTCATTCAGGAGCATCCTTGAGAAAGGCTAGTCTTATGTGGGATCGCCATCCTGGAGAAGGAAAAGGAGATGGTCAAACTGTTAGGAAGTGACTATTAAAATTAAAAGCTTTTCTGAACCTCTTGCGTCTCTGATGGATACAAAAACTGTAATAGCAACATGAACACTAATAGACAGCAATGCCACTAATGTCAATCAGAAACACTTGGCTGTGTTTGAGCCTCATGCACTGATGTGCTGTTTTCTTAGTCTCCCTTCAGATCTTACTAGCTTTTCTAGTGACTGTGCTCTTAGAGATGGCACATATGGTTGCATGCATGGCTATTATTTGACTTATAACATTCCACTGGATATTCAGGTGAAGGGACTCCTGAAGGACAAGAAATATGAAACATTAGTTGATGCAGATCTTCAAGGTAATTACAATGAAGAAGAAGTGGAACAGCTTATTCAGGTAGCTCTACTTTGCACGCAGAGTACGCCTACGGAACGTCCAAAGATGTCAGAAGTTGTAAGAATGCTTGAAGGTGATGGCCTTGCTGAGAGGTGGGAGGAATGGCAAAAGGAGGAGATGTTCCGGCAAGATTTCAACCATGTCCACCACCACCATACTGATTGGATAATAGCTGACTCCACTTCAAATATCCGACCGGATGAGTTGTCAGGGCCAAGATGATCTTTCATTTATTCTGCCATCAGCACATATAGGACCGACTCTTTTGAGGGAAAATCCTCTTTGTATCCTTGTATCTGTAATCGGTATATGTCATCTTCTTTTTTTTGGGTTAATTGGTATATGTCATCTTTGTGCATACCATTTTCCTTCTTTTTTTGTTTCAGTCAGTTATATTGTATTTGATTATGTCCAAATGTGTATAGGTGATTGGGTCATTACAACAGGTCTGAACATGAAAATCAACTTATAGCAGGAATGGCTGTAGAAAGCATCTTTGCACCTTCTCTTTTTCTAATGTATCAGATAGAGTGGCGTTTGTTAAAGAATATACACTTTGTAAACTTAAAGTAATCATAGTTTCTTTCATGTTCAGTGACAGTTTGTCTCCTCCTCTTGATGGCTGAATTTTGTAGTATTATAG

>PGSC0003DMG400017713 LRR-RLKs

CAAAAACAATCTTTTAATTATTAGCAATGATGATAGTTCCCACAATTTTTAATTTCCTTCAGTTTATTACTATCTTATATTTTTTCACAGTTACATTTGCTACCACTGAGGAAGCAACTGCTCTTCTAAAATGGAAAGCAACTTTCCAAAACCAGAACAATTCCTTATTGACTTCGTGGACGCTAAGCGGTCCTGCTGGTACCAACAGAATTGGAGCGGCGAGTTCTAATGCGTGTACGGACTGGTATGGAGTTACATGTTCTAATGGTAGGGTAAACATGTTGAATATGACAAATGCTGGTATCATTGGCACACTCTATGATTTTCCATTTTCTTCACTCCCTTTTCTTGATTATCTTGATCTTAGCATGAACCAACTCTCTGGCACTATCCCACCTGAAATAGGAAAGCTCACTAATCTTTTCTATCTTGACTTGTCCATCAATCAGATTTCGGGCACAATTCCACCACAAATCAGCTCATTAACAAAGCTTGGGACTCTCCACATCTTTGTCAACCAATTAAATGGTTCCATTCCGGAGGAAATAGGTCACCTAAGGTCTCTTACTGAGCTAGATTTGAGTTCTAACATTCTTAATGGTTCTATTCCTGTTTCTTTGGGGAATTTGAACAACTTGTCTGTTTTGAGTCTGTATGAGAACCATCTTTCTGGTTCCATTCCAGCAGAAATAGGAAAACTTGTCAATCTTGTTGAACTTTTACTTGATTCAAACAACTTAACTGGTCATATTCCTCCAGAAATCGGAGAACTTGTCAATCTCGATAGAGTTTATCTTGATTCAAACCAATTAACTGGTCATATTCCTGCCGAAATAGGGAAGATGAAGTCACTTGAGGAGTTGAGCATAAACACCAACAATTTTTCTGGTCCAATTCCCAAAACTATAGGAGAATTAACTGAGCTCCAACTCTTGTACCTTCATACCAACCAACTTTCTGGTCCCATTCCTAGTGAATTGGGGAATCTGAAAAAACTGAATTATCTGCAGTTATCCACTAATCAACTTTCTGGTCCGATTCCCGATTCTTTCGGCAATTTGAGAAACTTGAAAATTTTGTTTCTCCGTAACAACAAACTTTCTGGTTCTATTCCAAAAGAACTTGCATATTTGGACAACTTGGTTGTGATGGAAATGGATGAAAATCAGTTTTCAGGCCATTTGCCTGAGAATCTTTGCCAAGGTGGGAAACTTGTGAACTTCACGGTGAATAGTAACAAGCTAACAGGGCCAATACCAAGAAGCTTGAGCAACTGCTCAAGTTTCGAAAGGGTTCGCTTTGATAACAACAGTTTTACTGGGAATTTATCTGAAGCTTTTGGTATCTATCCGGAGCTGCAGTTCATTAATTTGAGCGAAAATGATTTTCATGGCGAACTAAGCAGCAACTGGGGGAAATGCAAGAATTTGACTGATCTTCGGATAGCCAGAAATAGAATTGGTGGTCGCATACCACCAGAGATTGGAAACCTCAAAGGGCTTCAAGGACTAGATCTTTCATCCAATCATTTGGTTGGGAAGATTCCTAGGGAATTTGGAAAATTGACCTCTCTGGTTAATCTTCTTTTGCAAAACAACCAAATTTCTGGCAATATTCCTATGGAACTTGGGTCATTGACAAAGTTAGATTACCTTGATCTTTCAGACAATAGATTGAATGGTTCGATCCCAACGTTTATAGGAGATTACCATCACCTGTTTCACTTGAACCTGAGCAACAACAAATTTGGCCAAAATATTCCGAAGGAGATAGGGGGTATAACTCAACTTAATGTACTTGATTTGAGCCATAATGTTCTGGTTGGAGAAATACCCCCTCAGTTAACCAATTTGAAGGTCTTGGAAAGCTTAAATATTTCCCACAACGGCCTTTCTGGCCACATTCCTGAAGAATTTGAAAGTTTGACTGGTTTGCAGGATGTTGTCTTGTCATACAATGAGTTGGAAGGTCCAATACCTAATAATAAAGCTTTCACAAATGCCTCATTGCAAGGTAATAAAGCTCTTTGTGGCAATGTAACAGGTTTCGTGCCATGCAACATACCATCTTCTATGGTGAAGAAGCACTCAATGGCGAAGGGACTTAAACTCATCCTCATCACTGTACTTCCTATTCTGGGAGCATTAGTGCTTCTTTGTGCTTTCGCTGGTGCTCTCTTTATGCGTGACCAAAGAAGAAGAGTTGGAGACATTGAAAGACGGGATTCCATAGACAAAGATGATGGTCTGCTTTCAATATCCTCATTACATGGAAATTCATTATACTGGGATATCTTAAAAGCCACAGAGGAGTTTGATGCAACATTTTGCATTGGGAAAGGAGGGTTCGGAAGTGTTTACAAGGTAAATCTTCCATCATTGGGAAATGTAGCTGTGAAGCGACTTCATTCTTCATTTGAGATTAAGCATCGCAAGAGCTTCATGAATGAGGTAAGGGCATTGACAGGGATTAAGCATCGGAACATTGTGAGACTCTATGGCTTTTGTTCGAATGCACAACACTCGTTCTTGGTTTACGAGTATGTGGAGAGGGGGAGTTTGTCTAGTATTTTGAGCAACGAAGTAGAGTCTAAGAAATTGGATTGGCTTACAAGGGTGAATATCATCAAAGGTATTGCTTATGCTTTATCCTACATGCACCACGATTGCTCACCCCCGATTGTTCATCGGGACATATCAAGCAGTAATGTTTTGCTTGACTCCGAGTTTGAAGCTCGTGTTTCTGATTTTGGCATAGCTAAGATTCTCAAGCCAGACTCATCCAATTGTACTGCACTTGCAGGCACATATGGCTATGTAGCACCAGGTAAGATCTAACTTCTTATCCTTATTTTAGTTCATGAACTATATCTGCACACTCTATATTGTTTCTATCTCATTTGCAACTTTATTGTTTCAGAGCTTGCCTATACATTGAAGGTGACGGAAATGTGTGACGTCTACAGCTTTGGAGTATTAGCATTGGAGGTAATCAAAGGAAAGCATCTTGGGGAATACCTTACTCTTCTAGCAAATCCATCAACAAGAGATGTGCAGCTAAGTGATTTGCTAGATGAACGCCTTCCACATCCTGAGGATGAAGTAAAAGAGTTTTTGGTTTTCATCGTCAAGCTAGCAATCTCGTGTTTGCTTGAAAATCCAAAATCAAGGCCAACGATGCACTTCATCTCTCATATGTTATCAATGGATCCACCCAATCATCAGATTCCATGAAAGTGAGATTCCTCGAAATGTAAGGTGCCCTGGTATGGAAAAGATTATGCTCAGCACCAAGTGGTTGTGAAGTAACAAGTATGTTGATCAGAACTTTCACAGCAAAATATTATTAAGATTAGCCTTTCTGTATTGTTGAGTTGCTTATATAGATTACTTATTTCAATTTGAAGGTTTCAGAGCCTTTGTACTAGAATATATACAAATTACAGTAATATACTGATGTTAAGTAATAAAAGCA

>PGSC0003DMG400017864 LRR-RLKs

ATGAATATGAAGAAGTTGGAAAGAAGAAATACTATTGTTATTGTTATGTTAGTTGTTGCATTTGATCACTTTACTGCTTCTTTGGCTAACATTGCCACTGATGAAGCTGCTCTTCTTGCCTTCAAATCTCACATTTCTTCTTCTTCTGATCCCAACAACAACAACGTAGTCGTCTTGCAAACCAACTGGTCTTCTTCAATTCCATCATCAGTTTGCACCTGGATAGGGATCACTTGCAACGGGCGCCATAGAGTCACTGCTTTAGACATTTCTAGCATGCAACTTCATGGAACCATTCCTCCACACCTTGGAAATCTCTCATTTCTCTTATCCCTTGACATTAGTAACAACACTTTCCATGGAGATCTCCCCGAAGAGTTGTCTCATTTGCGGAGGTTGAAATTGATTGATGTTACAAGAAACAATTTTAGTGGTACCATTCCATCATTCTTAAGTTTGTTACCTAACCTTCGATTCGTTTACCTTTCAAACAACCAATATTCAGGGGAAATTCCATCTTCCCTTTCCAATCTAACAAATCTTCAAGAGTTAAGAATACAGAGGAATTTTCTTCAAGGAAAAATCCCTCCTGAAATTGGCAATCTTCGTTACTTGACTTTTTTAGACCTGCAAGGTAATAGACTTACTGGCTCTATACCGTCGTCAATCTTCAACATGACTTCCTTGACAACACTTGCTATAATACACAATCGTCTTGTCGGTAAACTTCCTGTAGATATTTGTGACAATCTACCAAATCTTGAAGTACTTTTACTCTCAACAAACAACCTGGATGGCCTAATACCACCAAATTTACAGAAATGCTCAAAGCTACAACTGCTGACATTGTCTGGCAATGAGTTCACTGGACCTATACCAAGAGAACTTGGGAACTTAACTATGCTCACAGTCTTGCATCTTGGAGAAAACCATTTAGAAGGTACGTCATATGAATTACTCCCTCTGTTTCAATTTGTTTATCTAGTTTTGACTTGGCACTGGTTTAAGGAAGTAAAAAGGACTTTTGAATCTTATTGTCTGTAAAATGTACCAAAATACCCAATAATCTTAGAGTCTTAAACAGGTCGGGTGAAAAGTTGAAATTAAGAAGAAAAGACGAACTGACTAAAAAGGGAAGTAAGACAAACAAATTGAAACGGAGGAAGTATATCTTTATGCTTTTGTGTTTATTTTAAGATTTCTCAAAAGCTAAGCAATATCAATATGTACTAATGTTTATGCATTTTACTCTAATTGACAGGAGAGTTACCGGTCGAGATAGGTAACCTACAAAACCTCCAAGTACTTGGGTTAAGAAATAATAAGTTAAATGGTACTATTCCTGCAGAAATTTTCAACATTTTAGCATTGCAGATATTGACAATGTATGGAAATCAGTTATCAGGTAGTTTACCTTCAGATTTAGGCGTTGGAACGCCTAGTCTTGAAGAAGTTTACCTTGGATCAAATGAACTCAGTGGACGAATTGCTCCTACCATCTCGAATTCTTCAAAGCTAACACTTCTAGACCTTGCAGACAACAAGTTTACAGGTCCAATTCCTGATTCACTTGGTTCATTAGAGTTTCTTGAAGTCCTATTCTTGGGAGGTAACAATTTCATTAATGAGCCATCTTCTTCTGAGTTAAGATTCGTCTCGTCCCTGACTAATTGTAGATATTTAAGAGAAGTGGTCATTGAAGATAATCCTTTGAATGGTTTTCTTCCATCCTCTATTGGAAACTTTTCTGATTCTTTTCGGATGTTTGTTGCACGTAGAACCAAACTGAAGGGCACAATCCCGGAAGAAATTGGAAACCTGAGTAGCTTAGGTGTGCTAGCTCTGTCTCATAATGACTTGACTGGTTCGATACCTGAGCAGTTGAGAAGCATGAAGAATCTTCAAGAATTTTACTTAGAAAATAACAGCCTAAGTGGAACAATACCTGATGATATCTGCAGTTTACGAAATCTTGGAGCTTTAAAATTGACTGGAAATCAAATATCTGGTTCAATTCCAGCATGTTTAGGAAATGTTTCTACTTTAAGATATCTTCATTTAGCTTTCAACAGATTGACTTCCACCTTACCTGAAACCCTGTGGAGCCTTCAAGATCTCTTGGAACTCAATGCATCAGCAAATCTGTTTAGCGGGCACATACCTCCTGAGGTTGGTAATCTAAAGGCTGTGTCACTAATTGATCTGTCAAGAAATGATTTTTCTGGTAATATTCCTAGTACTATAGGGGCTCTTGAAAAGTTGATTAGTCTTTCCATGGCACACAATAAATTGGAAGGGCCTATTCCGAGTTCATTCGGGAAAATGGTGGGCTTGGAATTCTTGGATTTTTCTTACAACAATCTTACTGGTGAAATTCCAAAGTCACTTGAAACACTCTCACATCTCAATTACTTTAACATCTCTTTTAACAAATTAAGAGGAGAAATTCCTAGTAGTGGTCCATTTGCAAACTTCACCAGTCAATCTTTCATTTCCAATTCTGCACTCTGTGGTGCTCCTCGCTTTAATGTGTCGCCATGCCTCATCAAATCCACCAAGAAATCAAGGAGACACAGAGTTCTAACAAGTCTGTATATCGTATTAGGTATTGGATCAATGATTCTCACATTGGTTCTAGGATATGTTTTGTTAAGGTGGCAAAAGAGAAGGAAGAATTCAGGTCAAACAGATGCATCTTTGGTTAAAAGACATGAAAGAATCTCTTATTATGAACTTCAGCAAGCAACGGAAGGATTCAGCGAGAGCAATTTGCTTGGTACCGGAAGTTTCAGTATGGTATACAAAGGGATACTTAAAGATGGGAATCTTTTGGCAGCTAAGGTATTCAATGTGCAATTGGAGGGAGCATTCAAGAGCTTTGAAACAGAATGTGAGATACTACGCAACCTTCGCCATAGAAATCTAACAAGAGTGATCACTAGTTGTTCCAACCCAGACTTCAAAGCCTTAGTATTGGAATACATGCCCAATGGAACACTTGATAAGTGGCTACACTCTCATGACTTGTTCTTAGACATGTTGAAGAGGTTGGACATTATGATAGATGTGGCATCGGCATTAGATTATCTCCACAACGGGTATCCAACACCCGTGGTGCATTGTGACTTGAAGCCAAGCAACGTCTTGCTCGATCAAGACATGGTTGGTCATGTGAGTGATTTTGGCATTTCAAAATTGCTAGGTGATGGGGAGACTTTTGTACAAACAAGGACTATTGCAACCATCGGATACATTGCTCCAGGTAATAGAAATTTCAAACTTGTCATCCTTTAGATTTTCACATTTCAAACTTTATCTAACAAGATTTCTCCCATGATTGCAGAGTATGGACAGGACGGAATAGTATCCAAATGTTGTGATGTTTATAGTTTCGGTATCATGATGATGGAAACATTTACAGGTATGAGACCAAGTGATGAAATGTTTACGGGAGATTTGAGCCTAAGATGTTGGATTAATGATTCTTTTCCTAATGGAGTTGTGGATGGCAATTTGTTAAGGCCAGAGGAAGAACATATTAAAGAGAAAATGCAATGTGTATTGTCTATCATGGAATTGGCTTTGAGTTGCACTTTAGTGTCACCTGATGCAAGAGTTAACATTGAAGATGCTCTTTCAGCCCTCCAGAAAATAAGATATCAGTTTGTCAATAGTTGTGCAAGGACTTAA

>PGSC0003DMG400017943 LRR-RLKs

AAGAATTAATAGTGGGAATTTCCACTTCGGAGGGTTGCCAATTGTGTGTGCCTTATGAATGACAGCCAATGAGTAGTTGCAGAAGGTGAATGTTATAGTTAAGCTAGCATAAATGGTTGAAAATGATAAGCAAGAAGTAGTTGGAGAGGTAACCTCGAGATCTGTGCTAGGATAGCTTAGATTCCGTTTGCTTCACGAAGAAGCTCAAACATTATTCTTTATTTCTTTTGAAAAGAAAAGTTTTTTCCTTCCTACTGCTTTGTTTTTTCCTACCGGTACCACCAGCTGCTGCTGCAGAGGGAAGTAAAAAGAATAATCTGATTTTGCAAAGTAAAAAAAATGCCGTACGTTGGAGTAGCAAGTTTAGAGGGGAATTGTAAGGCACAAGACTTCGACCTTTTTTCTCAATTCTCAACCGTTATATATTAAGCTCTCACCCATTTTCTTCATTGCAGTAGAGTTCACCATTTTTCAAGGTATACTCTTTCCCCTATTATTTCAACTCTCTACTACCTTTGCTTACATTAGTGTTGATACTCCATTTTTCAAGGCACACCCCTCTGGCTCTGGCCATAGATTGTGTGTGGTAAGTGTTGGTTCATCAAATTTACTCATTATTTCAACTTCATTCAAACATGTTTTCACTCATGTCCTAACACAACTAAACTTTCAAATTTTGGATATGACAACCAATTTCTCAACTTAAGACATTTAAACTACTTTCTTTGTCTAATGCTATGATGGAATGTTGAAATTAGGAAATAGAGGGTGGTAATGCAGAAAAATCAAATCTTTTTCATACATGTCCTGTCTGCTCGGTATGGGATGCATCTAATAAGTTCATAGGGCTTCAGAATTTAGGATGTTTATCTCTAAACAAGGTTTTTACTCTTATTATGAATCCAACTGCTAAGCTTCTTTTCGCCTTGGTTGACTACATATGAGAACGCCACAAAGTGTCCTTTGCTCTACGGGGAATTAATTGTCTCTTCATTTCATACTGTGATGGGGATATTTTCCATATATCTAACTTCTCTGCCACATTTTTGTTTACCATTGTTACTAGGTGATGCAAATGGATGACATACAAAGTTGATATCTTATGGCAACTTGAGTTGACTTGGCAAGTCCAGGAGTAAATAGATTTAAATGATGAAGCTACACCCAATAATAGATCTACTTCCTCTTTCTCTCTTCCTATTTTTGCTCCTCTTCTCAAATGTTATTGCTGACCTTAGCTCAGATAGACAAGCACTACTTGACTTTGCTTCTGCAGTACCTCATCTTCGAAACTTCAAGTGGAACACTAACTCTTCCATTTGCACATGGCATGGCGTAAGTTGCAGCTCAGATGGCACTCGTGTAGTTGCACTTCGGCTTCCTGGTATTGGACTTTATGGTCCTATTCCAGACAATACCATAGGAAGACTGGATGCACTAACAACCCTCAGCCTTCATTCCAATGCCCTCACTGGAAATCTTCCTTCAGACATCACCTCTCTTCCGTCCCTCCGTTTCATATTTATACAACAAAACAAATTTTCTGGTGAAATACCTTCTTCTCTATCTCTACAGCTTAACTTCATTGATCTCTCTTTCAACTCCTTCTCAGGGGAAATTCCAACAACAATTCAAAATCTGACACATCTTACTGGTTTAAACCTACAAAACAACTCCCTCACAGGATCCATTCCTAATGTAAACCTACCAAGGCTTACGCAATTGAATATGAGTAATAATCAACTCAATGGTTCAATTCCACCATCCCTTGCAAAGTTTTCTGCTTCTTCATTTCAAGGAAATTCTCTATTATGTGGACAACCCTTGACTCAATGCCCTTCTTTTGCGCCTTCACCGTCTCCATTCCCTTCAATTCCACCTTCTCCTTTGAGTCTAACACCACCATCTCGTTCTCCGAGCGTCCTGCCAGCCTCACCAACAATTCCTGAAAACCATAAAGGCAAGAAAAGCTTGAGTACACGGGTTATCATTGGCATTGTTGCAGGAGGTGTTGGAGGGATCCTATGTCTAGCTGTGTTGATTTTCTTGTGTTGTATGAAGCGATATTATACTAAGAGAGGTGTACAGCAAAGAAAAGACTTTAATGGAGGAGGAAGTCCGAAGCAAACAGAGGACTTCAGTAGTGGAGTACAAGCAGCTGAAAAGAACAAATTGGTTTTCTTTGAGGGTTGTTCTTTCAATTTTGACCTTGAAGATTTGTTGAGAGCCTCAGCTGAGGTTTTGGGTAAAGGGAGCTATGGAACAACCTACAAGGCCATCTTGGAGGAGGGAACGACTGTTGTTGTGAAACGGCTGAAGGAAGTTGTTGTTGGGAAACGAGAGTTTGACCAACAGATGGAGACAATTGGCACTGTGGATCAGCATCGAAATGTTGTTGCTCTTCGTGCTTATTACTTTTCCAAAGATGAAAAACTTCTTGTCTATGATCATGTACCGGCAGGCAGTTTATCCACTCGAATGCATGGTAACATTTCTTCCTTCTTTTGTCCTCCCATTCTTTTAGCTGTCATACCCATCCAGGTGCTTGGGAATATCCTTAGTTACTAAGACTGGCAAGAAAATCAGCTGTAAAAGTTGCACGAGATCATAGATTCCGAATGTGAAATGAAGTACCTATATTCTTTAGAAAAGCCCCTAGACAAGCATTATAAATAAACACTCCTGAAATAGGCATGCAAGTTTGTCTTGAAGACTTTATAACATGTGATATTGGTTTCATGCACTCATAGTTCTCCATATTATTTTGAAGCAAGATGCAACTTGCAAGTTGTTGAATGTCAGATTAGTTGTTCGGTGCTATACATAATTTTCAATCTAACTGTTTATACAAGAAAGTTCTTTTCTTTCATTGCTCCTTGTTTATTGATTATAGGCAACAAGGACTTGGGAAGAACACTGGATTGGGAATCTAGATTGAGGATTGCCCATGGAGCTGCAAGTGGTATTGCCCATATCCATGCTGTTTCTGGTGGCAAATTAATTCATGGAAATATCAAGTCATCCAATGTGCTTCTCACCCACGATAATAATGGATGCATCTCAGATGTTGGTCTTACACCTTTAATGGGTTTTCCTACTATCCCATCAAAGAGTGCAGGATATAGAGCACCCGAGGTGATAGAGACCAAGAAATGTACTCAGAAATCCGATGTTTACAGCTTTGGTGTTCTGCTTCTTGAGCTCCTTACTGGGAAAGCACCAGTGCAGCCACCTGGCCATGATGAGGTGGTAGACTTACCAAGATGGGTGCAGTCTGTTGTGAGGGAGGAATGGACTGCTGAGGTATTTGATGTTGAGCTCATCAAGTTTCAAAATATCGAAGACGAAATGGTGCAGATGCTGCAGATTGCAATGACTTGTGTGGCAAACGTGCCTGAAACAAGGCCTGACATGAGTCAAGTTGTCCAGATGATCGAAGATATCCAACAAATTGATTCTGGAAATAGGCCATCATCTGAAGATAACAAGTCTAGGAGCCCAACTAGTCCAACGCCATGATAACTCATGCATTTCTTACTACTCAATTCTATTACCAAATGTGTATTTGAACGTCTTTTCTTGTTTTCAAGTATAATTCAGACCTTTTAGATACTATCATTCTTCATCACTGCCTATATCTAAAAATTCAATCCTATTATGTCTAGCTCCACTTTTATTTT

>PGSC0003DMG400018024 LRR-RLKs

GGTTTCACTACGGACGAACAATCTCAAGGGATCCTTACCACGCGAGATTGGCAACTTAACCAAGATGCAAATTTTAATTCTAGACGAGAATAGGTTTACTGGTACGTATACATTCAATTTAAGTGATAGCAATCTGAACAGTTTAAGAGGTAAACTCATTCTTTCTTCAGGTGAAATACCCAAAGAGATAAGAAATCTCGTTGAGTTGGAGAAGCTTGGTCTTGGATCTAATAGTTTTAGTGGTTCACTTCCAATGGAGATCTTCAACATATCACGATTGAGACATATTGCTCTTACAAACAATAATCTATCAGGAACTCTACCACCAAACATAGGTTCTACCTTACCCAACATTGAAGGACTTTATCTGCATGGCTTAACCAATCTTGTTGGGACTATTCCTCATTCTATCTCCAATTGTTCAAAGCTTACTATTCTAGAGCTTTCACAAAACAAAACTCACAGGATTGATTCCCAATTCTCTTGGATATTTGACTCATCTAAACATCTTAAACTTGCGGAGAAAACAATTTAACCAGTGATTCATTTTTAAGCTTCCTCACTTCCTTAACCAATTGCACAAGATTAACAGTTGTTTCTCTATCTTTCAACCCTCTAAATGCCATGCTTCCGGTCTCCGTAGGGAACTTCTCCAAATCTCTTCTAACCTTTTATGCCGATGCTTGCAACATTAACGGCAAGATTCCAAATGAAGTTGGAAACTTAAGCAACTTATTAAACCTTGATCTTTCTGATAATAACTTGATTGGATCAATTCCTACATCAATTCACAACTTGAGAAGCCTTCAGCGCTTCAACTTGAGTTACAACAAGCTTACGGGATTTATTGGAGATAATTTATGTAAATTGCAACATTTGGGAGAAATTTACTTGGGACAAAATCAACTTTCAGGATCTATTCCTAATTGTTTAGGGAATGTTACTTCCCTTAGATTGATATATCTTACTTCCAATAAATTGAGTTCCAATATACCAACAAGCTTAGGAAATCTTAAAGATCTAGTGGTTCTTGACTTATCGTCAAACAACATGGTTGGTTCTTTACCTCTGGAAATTGGAAATCTAAAGGCTGCAATATATATGGATCTGTCAATGAATCAATTCACAAATGGAATTCCTAGAGAAATTGGAGGATTGCAAAATCTGGTATATCTTTATTTGGAACACAACAAGTTGCAAGGATCTATACCTGACTCAATCAGCAACATGGTAAGTTTGGAATTCCTAGACATTTCACATAATAATATATCGGGAACTATTCCCATGTCTTTGGAGAAACTACAATACCTCAAGTATTTCAATGTTTCTGACAACAAATTGTATGGTGAAATACCCTTGGGGGGTCCTTTCAAGAACCTCTCGAGTCAGTTTTTCAGCGACAATGAAGCATTGTGTGGTTCTTCAAGATTTAGTGTCCCGCGATGCCCCACTTCATCAAAGCACAGATCAAATAGGAAACAAATGTTTGCTCTATTTCTTATGCTGGGAATTGCACTTGTATTTGTTCCTATAGCCTTTCTGTTCATATGGATAAGGTATAGAAGAGGTAAAAGAGCTCCTCAACAAGCTGATTCATTGTGTATTGTAACAACAGAAAGAATTTCATACTATGAACTGCTCCAAGCAACTGATACGCTTGGCGAGAGTAATCTGATTGGTTCTGGAAGTTTTGGCTCTGTCTACAAAGGCGTTCTCAGAAGTGGAACTGCCATTGCAGTTAAAGTGTTCAATCTGCAACTGGAAGCAGCATTCAAGAGTTTCGATACAGAATGTGAAGTTTTGCGTAGCCTTCGCCATAGGAATCTCGTGAAAGTCATCACTAGTTGTTCCAACCTTGATTTTAAGGCTTTAGTGCTCGAGTATATGCCTAATGGAAGTCTTGACAAGTATTGTATTCGCACAACTACTTCCTAGACATCAGGCAGAGACTAAGCATTATGATAGATGT

>PGSC0003DMG400018171 LRR-RLKs

TAAACAAATCCGAAAAAAACACCCATCTAAAATTCCTCCCCACTCTCTCTTACATAATGGCCTAGCTGTTCTTGGATCCGGAGATTCTGGCAACTTTTTCAAATTTTTATATGTAACCTACATAAAATCCATTAGCCCACAAGACATGATTTGGTTTTCCTCTCTTTGTCTCTCCTCCCTAGGCTTAAACCTCTACACCAAAACCAAGAACAAGGCTTCCTTTCTCTGCTTAATCTTGAAGTTTGTAGCAGCAGTCACATATTTGTTCTAATCTAATGCCTGTTTATTCATGGACCCTCTTTTTCTTTTTCTCCTTCTTTCTCTTTACTACCTCCTCTGCTCTCAACCCTCAAGGTCAGGCTCTTCTTTCATGGAAGACAAGCTTAAATGGATCATTAGATGTGTTGAGTAATTGGGATCCTACTGATGAAACTCCATGTGGTTGGTTTGGTCTTAGTTGCAACTTCAATAAGGAAGTTGTTGAATTGGAATTGAAGTATGTGGATTTGCTTGGGATTGTTCCTTCTAATTTTAGCTCATTGGTGTCATTGAACAAGCTTGTTTTGTCAGGGACTAATCTTACTGGTGTAATTCCAAAAGAGATTGGGATGCTTCAGGGACTTAAGTTCTTGGACTTGAGTGACAATGCATTGACAGGTGAAATTCCAAGTGAAATTTTTCACTTGCCAAAGTTGGAGCAACTTCATATCAATTCAAACCGGCTGGTCGGATCGATCCCAGAGGACATTGGTAACCTCACAAGCTTGGTGTGGCTTATATTCTATGATAATCAGCTGAGTGGTGGAATTCCAAGTAGTATTGGGAATTTGAAAAGGCTGGAGATTATTAGAGGAGGTGGAAACAAGAATCTTGAAGGTCCATTGCCTCAAGAAATTGGCAACTGTAGTAATTTGGTTATGTTAGGATTGGCAGAAACCAGCATTTCTGGTTTTCTACCTTCTTCTCTTGGCCAGCTGAAAAGACTCGAAACGCTTGCTGTTTACACCTCCTTGCTCTCTGGACAAATCCCACCGGAGCTCGGGGATTGCTCCAAGCTCCAAAACATTTATCTTTATGAAAATTCACTCACTGGTTCAATCCCAGCTCGGCTTGGTAACCTCAAGAACCTTCAAAACCTCCTTTTGTGGCAGAATAATTTGGTTGGAACAATCCCTCCCGAGCTTGGGAACTGTCAACAATTACAAGTCATTGACATTTCAATGAATTCATTAACCGGAAGCATTCCGGAATCATTTGGGAGATTGAATTCAATGCAAGAGTTACAGCTGAGTGTTAATCAGATTTCAGGTCGGATTCCAGCACAAATTGGCAACTGCACTGGTCTTACTCACATTGAGCTCGACAACAATGAGATAACCGGTTCCATACCATCGGAATTCGGAAACTTGTCGAACTTAACTTTGCTGTTCTTGTGGCAAAATCGTCTTGAAGGGAAAATCCCTTCTTCAATTTCCTCGTGCCATAATCTCGAGGCTATTGACTTGTCGCAGAATGCATTGACTGGTTCGATTCCTAAAGAGATATTCGATCTTCAGAAGCTGAACAAGCTACTCCTACTGTCTAATAATCTCTCTGGTCCCATTCCACCAGAGATAGGGAACTGTTCATCGCTGATTCGACTTCGGGCAAATGATAACAAGCTGACCGGATCACTGCCACCGGAGATTGGGAAATTGAAGAACTTGAATTTCCTTGATGTCGGATCCAATCACCTCACCGGGATCATTCCACCGGAATTATCTGGATGCCGGAATCTGACATTTCTAGACTTGCATTCCAACTCAATCAGTGGCAATTTACCGGAGAATTTGAATCAGCTTGGAATCCTCCAGTTCATTGATGTTTCTGATAATTTAATTGAAGGCACTTTGAGTCCGAGTTTCGGCTCACTGACTTCACTCACCAAACTTGTTCTGGGTAAGAACAGGTTTTCTGGTCCTATTCCAACTCAACTCGGTTCCTGCATGAAGCTTCAGTTGATTGACTTGAGTGGTAACCAACTCTCCGGCGAGATTCCGGCGAGTGTTGGTAAAATCCCTGGTCTAGAAATTGCACTGAACCTCAGCTGGAATCAGCTTTCCGGTGAAATTCCGGCAGAGTTCGCCGCATTGGACAAGCTTGGAGTGCTTGATCTTTCTCATAATCAGCTCTCGGGGGACCTCCATTTTCTCGCGGACCTACAAAATCTCGTGGTTCTGAATGTCTCGCACAATAATTTATCGGGGCACGTGCCTGACACCTCATTCTTCTCTAAGCTCCCACTCAGTGTACTTGCTGGCAATCCAGATCTATGCTTTCCCGGCAACCAGTGCTCCGCCGACAAGGGCGGCGGTGTCCGTCGTACCAAAGCGGCGAGGGTGGCGATGGTGGTGTTGCTTTGCGCAGCTTGCGCTCTCCTTGTGGCAGCTCTCTACATTATCCTCAGCGGCAAAATACGGAACCGCAAGGCCCACGATTACGATCTAGATGGTGACAATGACGTTGAACTTGGCCCACCATGGGAGGTCACAGTATACCAAAAACTCGACTTATCTATAACTGACGTCGCAAAATGTTTAACCGTTGGTAACGTCCTCGGCCGTGGTCGGTCCGGAGTTGTGTATAAAGTGAATATTCCGTCAGGATTGACGATCGCTGTGAAAAGATTCCGAGCATCGGACAAGCACTCGATGTCGGCATTCTCGTCGGAGATTGCCACGCTGGCACGAATTCGACATCGGAACATTGTTAGATTACTTGGATGGGCAGCAAATAGAAAGACTAAGCTATTGTTCTATGATTATTTACCAAATGGTACATTAGGTTCATTTTTACATGAAGGTTTTGGAGGGTTAATTGAGTGGGAAACCAGATTCAAGATTGCATTAGGGGTGGCTGAGGGATTAGCTTACTTGCACCACGACTGCGTTCCACCCATCCTCCACCGCGATGTCAAGGCTCAGAACATTTTGTTAGGTGATCGTTACGAGCCGTGTTTAGCTGATTTTGGACTTGCTAGACTCATGGAGGAGGAAAATTCTTCAGTTACAGCAAATCCTCAGTTTGCTGGATCTTATGGCTACTTTGCCCCAGGTAAATATTTCTCTTCTTCCTAGTTCAAGTTCATTTCTTTCGAAAATCGACTAGTTATGATTCATTTATTACTTTAGTTATTAGGGGAATTCTCCTCTAATGCAATTATGTGAGGTTTCATTTATCACTTAGTTCAAGTCCATGGTGGAGCTAGAGGGTGGAAGGGAGTTCATTTGAACCCTCTTCGTCACATAATCACACACATATATAAGAACTATTTTTTCAATATATATATATATGGAAATTGGAAGGAAGTAATCTCAAAATACGCTTGATATCAATACAGACTTAGGTAAAGATAGTACAAATGGAAGAAAAAGATCCATGTAGGTGATAGCTTAATGTTAAGGATATTTTAAAGACTCGTATGAAACTTTAATCATGATTATTATTAGTATTTTATTTTTTTACTGATGATATGATGTGAGCTTAAATGAAATTAGGATTCATATAGCCAACCCCAAGTTTGCCTTTGGATTTTTTCCGTTTTTTCCTTTATAGAAAAAAGAAGAACTGAATGCGAGGTAGCAGAAGAGAACAGATATAAAGCACTGCAAACAGAGAGCAAGAAGGAGAATTCAGTTTAAGTTGCTTCCATACGGATGGAATTGTGTCACTATTTTGACAATATATGCATGTGAAGCCATGTGCTAGCTTTAACTATGTGCAAAACCAAGTTTGTCATGACATACTTGGCAGCTACAGTGCTATTTCCCCATCTTTTTGCCTGTGTATGGCCCCCCACTAAAGATACTTAGGATGGGGAATACTCTAAAAAAACCATAACCAAGTTAGTGGTCACACTTTCACAGCCAAGCATGGACACATTGCTGCTGTCTCCATACCCCTTTGGCCATTGCTTGTTGTCCGATTGCGATATGTGAAAAATAATCGCCCAACAAATGCCTTTGTTATGTTTCAAAAAGCATCAAGATCAAATAGGTGATAAATATCTGGCATGAAAAGGACTAAACTCTGTTCTATTGTTTTGTCTTTTGCAGAGTACGCTTGTATGCTAAAGATTACAGAGAAGAGCGATGTTTTTAGCTTTGGAGTTGTGTTGCTAGAAATCATTACTGGGAAAAAGCCAGCCGACCCATCCTTCCCTGATGGCCAACATGTTATACAATGGGTTCGCGACCACCTAAAGAGCAAGAAAGATCCAGTTGACGTAATTGATCCAAGGTTACAAGGACATCCTGATACACAAATTCAGGAAATGCTTCAAGCCCTTGGAATAGCCTTGTTGTGTACTAGTAACCGGGCTGAAGATCGTCCAACAATGAAAGATGTGGTGGCATTACTCAAGGAGATTATTCATGAGCATGCAACAGGAAATGAAGCTAAAAAGACATTGAACAACTCCTCAAAAATGTCCGATACTTCATCATTTTCCTACTCGTCTTCATCTGTCACTCCAGCTCAGTTGCTACGCCTCCAAGGGTCTTTTAATTGCTCTCTTACTCACTCATCCTCCTCCGTTAGCTATAGCACCAGCAACCAATAATAATGTAGCGATAGTCATGATAGATCTGTTATTCTTTTGAACTAGATAGTATTAGCAACCGTAAAGAGCTAGGTAGAGGGAACCTTATTGTCAATTTAATTGTTAAGCAAATGAAAATGAAATCCATTGTACAGCAAAGTCAAAATTTTCTTTATCTCCC

>PGSC0003DMG400018253 LRR-RLKs

TCATTTCTGGAGCTCTTTTATTCACTCTGTTTGGTTTTTCATCATTAATGGGCAGGGTTTCTTGAATATTTGGAGATTATAGGATGGAGGGGTGTTCTTAAATAGTTGAAAGAATTGATTTTTATGCAGTTGGTTGGTGCTGGTGTTTGTTTTCAGTAAGATTAGCTACAAAAGATCCTAAAGAATTGATTTTTATGCAGTTGGTTGTGCTTTTTTTCAGTATGAGGAGTCGTAAAAGATGTCAAGAATATTTTTCACTACTGGGGTGATGAAATTTTTTGTCTTTATCATAATACTTCTGAGTTGTGGTTTTTTTGTGGTGGTGTCAAATGCAAGAACACTGCCTGAAAATGATCAAGTAGAGAGGTTATTAGACTTCAAGAAGTCCTCTGTTGAATCTGATCCAAATGGATTCTTGAATGAATGGACTTCATCTTCTTCAAGTCCTTGCACTTGGAATGGGATTTCATGTTCAAATGGTCAAGTTGTTGAGCTTAATCTGTCTAGTGTTGGGCTTGGTGGTCCACTTCACCTCACTGATCTCATGGCTTTGCCTACTTTGCTAAGAGTTAAGTTTAATGGCAACCATTTCTATGGCGATCTTTCCTCAATAGCTAGTTCTTGTGGCTTTGAGTTTCTTGATTTATCAGCTAATAGTTTTAGAGCCTTTGTTGAAGTCTTGTGATAAAATTAAGTACCTAAATGTCTCTGGGAATTCAATTCAAGGGGTTGTTCTAAAGTTTGGACCTTCACTGTTGCAGCTTGACTTATCAAGTAACACCATTTCAGATTTTGGCATTTTAAGTTATGCACTGTCCAATTGCCAGAATTTGAATCTGCTTAATTTTTCCTCAAATAACTTAATTGGAGAACTCAATGATCTTGATTTTGGGACTTGTCAAAATCTCACTGTCCTAAATTTGTCTTTCAACAACCTTACCTCAACTGAGTTTCCAGCCTCTTTGGCAAATTGCCAGAGCCTCAATACACTGAACATTGGCCATAATTCGATCCGAATGGAGATTCCTAGCAAGTTATTGGTGAAGTTGAAGAGCTTGAAGCAATTGGTGCTAGCCCACAATCAGTTTTTTGATAAAATTCCATCAGAATTGGGGCAGAGTTGCAGCACACTTGAGGAGGTTGATCTTTCTGGAAACCGGCTTACTGGTGAACTTCCTTCAACTTTTAAATTGTGCTCCTCACTTTTCAGTCTCAATCTTGGTAACAATGAGCTATCGGGTGATTTCTTACATACTGTTATCAGTTCATTGACAAATCTTAGGTATCTTTACTTGCCATTCAATAACATAACTGGTCATGTACCAAGGTCTTTGGTTAACTGTACAAAACTTCAGGTGCTCGACCTCAGTTCCAACGCATTGATTGGGAATGTACCTTTTGAGTTGTGTTTAGCAGCCTCGGGGTTTCCACTGGAGACGATGTTGTTAGCCAGTAATTATCTTACTGGAACTGTGCCTAAACAGCTTGAACATTGTAGGAACCTCAGGAAGACTGATCTCAGCTTCAATTACCTGACTGGTTCGATCCCATTGGAGATTTGGACCCTTCCAAATCTTTCGGAATTGGTAATGTGGGCTAATAATCTCACAGGTGAAATTCCTGAGGGCATTTGCATCAATGGAGGAAACCTTCAAACACTCATTCTCAATAACAATTTCATCTCAGGGGCACTTCCACAGTCCATTTCCAATTGTACCAACTTGGTTTGGGTTTCGTTGTCCAGCAATCAACTTTCTGGAGAGATACCACAAGGAATTGGGAATCTTGCAAATCTTGCTATCCTTCAGTTGGGTAATAACTTACTTACTGGACCGATCCCTCGGGGATTAGGTAAGTGCAGGAACTTGATATGGCTAGATTTGAATAGCAATGCTCTAACAGGTTTGATCCCTCTTGAACTTGCTGATCAAGCTGGCCATGTTAATCCGGGAAAGGTTTCTGAAAAACAGTTTGCTTTTGTGAGAAATGAGGGTGGAACTGAATGCAAGAGTGCTGGTGGACTGATTGAGTTTGAGGGTATCCATGAAGAGAGGCTTGAGATTTTGTCTATGGTTCACTTGTGTCCATCAACTAGGATCTATTCTGGCAGGACGATGTACACCTTCACGAGCAATGGGAGCATGATTTACCTTGATCTTTCCTACAATTCTTTATCAGGAACTATTCCTGATGAACTTCCTTCAAGTTCTAAATTTGGGACACAACAATTTAACTGGAACCATTCCATTCATCTTTGGAGGTCTAAAGATAGTTGGAATTGTTGATCTGTCACATAACAACCTACTGGGATTCATTTCACCCTCGTTAGCAAGCCTTTCATTTCTTAGTGATCTTGATGTTTCAAACAACAACCTATCGGGAATGATTCCTTCTGGTGGACAGTTGGCTACATTTTCAGCTTCAAGATCCGAAAACAATTTGGGCCTTTGTGGCGTCTCCCTTCCTCCTTGTGGCCCAGGCAATGAACACCATTCATCCAGTATTTACCATAACAAGCCTACAACTATAGGAATGGTGGTTCGTATTATGGTATCGTTTGTATCTATTACTATTATTATCCCACTTGTTATCTGAAATATAAGAATAACTGCACCACAAAGTCTAATTGAATTACCCATTTCGATAAGATAATATATCTTCCAAAGACCCCTAAATAAA

>PGSC0003DMG400018263 LRR-RLKs

GAGATATTATCCTCTTGCTGTTTAGAGAACAGTTTAATCCTCAAAACAAACCAAACACTTGTGGTTTCTCTTTCTTCCCATTACTCTTCTTTTCTCTTTCTTTTTAGTACACATATTGGCCCCTAAGAAGAAACTAAGGTTGCATGGTGGTGGATTTAAACAATGGTCCAAACAAGAACAATCCAACTGATCATCTTTTGTCTGCTTATTATTTCTATGCAAGTTTTGGTGCAGCTGGGAGCTTCAACAAATTCTGGTGACTGTAAGTCCTACTTATCACATCTCTTTTGTTGTTTTTTTAACTGTATAATTATGAATTTAACTTATATACACCATATCGCGAATAAAAAATAAAGTTATACTAGTACCTAGCACAGTTTAACATGTTATATCGTAAGTTACTTATTGTAAATAGTTCTTGTTTTAGATGATCTGAGTTATGAAAGCAAATTGTTTTCATTTCTTTTCTTTTTTTTTTAGTAGGGAATATATTAGTGTCCACTATTTTAACTCCAATTTATTTTGATCCCTTTTTCGAGTTCATTACTTAAAAGGTCATTTCAATTATTGAACATAGTTATGTTTTTCGTTGTTTTGTATCAGAAAAGTCAAATATGATTTTCTTCATCAATTTCATTCCGTATGTTCTGAATTTTTTTAAAAATAATTAGTGTATGGTATTTTTGTTCAAAAATAAAATGATTGGTGCAGATGTTGTGTTAGAGCCACTGAAGGAAGAGTGGGAAAACATACCTCCAAGTTGGGATGGGTCTGATCCTTGTAACGATCCATGGGAAGGCATCGACTGCAACAATTCACGTGTTATTTCTATGTAAGCTTTTTTTTTTTGTGATTAAATTAAAATTGTTAAAAGAGAATCATATTAATCGGAAATAATAACAATATTACCACTCTTTAAATTATTCCTAAATTTATGAATAAAATGTTGAAATTAAGAAATAAAAACACAAGTGCACTCTGTCAAAAATTCCATTTTTCCTTAGCTCGACAACGTCACATGTTTAATTCAATGTTCACCCACCTCATTTATGCCAGCTAAATAATATCTTCGATTTTTAAAAAATAAAATAAAATTTAAACACTAGAATAATTAAGGAACTATTTCCAAAAAAACAGCAAAATATTTTTTTTCTTAAAAAATCTTAATTAGTAAGAAAGTACAAAAATATGTAACTGTTACTGTATTTACTCTTCCATTTTGGTCAGTTCATGTGTATATATAATAATAAAACTAAAATGCTTATTATCTAATGAAACTAGGTTAAAATTAAATCATATTAATTTTGTTAAATGACATTGCTAATTGTAATTTGAGGTACTAATTAATGATTGTGAATTTCAGAAAATTATCAAGCATTAATCTGAAAGGGGAGCTATCTGGAGATATCGAAGGCTTGTCTGAATTGCAGATATTGTATGTCTTCTACTCATCATAATTGAAAGTGTTAAAATTTGGAGAAAAAAAATTGTCTTAATTATGTTCGCTCTGTTTCAATTTATTTGTCTGATTTGAATTTGACACATAATTAATGAAAGTAAATAAGATTTTTGAATCTTGTGTGGAATATGACCATTTTTGTTTTTGTGCTCGCCAAAGTGTTCCTAAAGGATTCATTACAAAAAATAATAGTTTAATGATTTTATTTGGTGCGATAAAATGAAAGTTGAGTGATCATCTGTGAAATTAATCCTGTTCTTGAAGCACCATTTTGCTTGTGATCATGCATTGCATGCACACAAATATTTTTGACTGTTGAAAAAGTTTTAAATTTCTGTTCTGATTATCTGTTTCCTTAATAGGGATCTATCATACAACAAAGGCTTGACAGGTTCACTCCCTCAATCAATTGGAAATTTGAAGAGTTTATCAATTCTGTAAGGGTATTATTACATGATAATGATTAAATGTTGTTAAATTCACCAGCTGATTTCATTAGAGTAATTGCTCCGAGCTGCCTCGCAGGATCCTTGTTGGTTGTGGGTTTTCTGGACTGATACCAAACACAATCGGATCCCTTAGCCAGCTGGAATATCTGTAAGGATATCAATTTCTTCTGTTTGCAACTTTTTATGCTTAACGAATTCAAGATATTGAGTATTTGAAGAACTTTTAGTCCTTTATGTTTTGTATAAAATTGGAAATTTGGCATGAGATGAGTTGGATGGAGTAACTTGGTCAGTGAGGATTTATATAGCCGACCCCAACTTGTTTGGGGACTAACGCTTAGTTATTTTTGTTGTTGTAGTAATTCTTGTATCCCTTTACGTTGTCAAACTGTGAAGTCTTAGAGCTTCAGGTCTCTAGATATGGTATACTAGATTGAAATATTTTTCCTGTTTCTTCTACTGGGCTACATTTCTAAAATACTATCTGAATCATGCAGATCTTTGAATTTAAACAACTTTATCGGACCGATACCAGCTTCTGTTGGTAATTTATCAAAGTTACATTGGCTGGACTTAGCTGATAATAAGCTCAGTGGACCTCTTCCAGTTTCACCTGGTCTAGACATGCTTGTTCACACAAAGCATTTGTGAGTAATACTGATTACTGTCAGTTTCTTAAGTAACACCTTCGTCTCTCATTTTCATCTGATATACCACTTACCTCGCAGTCACCTGGGGGGAAATCAGTTCTCAGGTGAGATTCCAGATCAGCTATTCAGCTCAAATATGACTCTCAAACATTTGTAAGTTCCACTGAACTGGAAGTGTATATATTTTTGAGAATTATTTGATGGATCAACTTCATTATTGTCCCACTCCAACGCTATGGTGCTAGCTCTGACACTTGTTGCAAACTACACCCGTTTGGCTAATGGTTTCTTCTGTTTGGGTCTTGTAATTGCACAGGCTGCTTGACCAAAACCAGCTTACTGGAAAAATACCCCCTTCATTAGGACTCGTGCAGACTCTGGAGGTGGTGTAAGTATTTCCAAAATCCATCATGCTCTTAAAGAATTCTAATAAGTGTCCATCTTTCTTGCTTTTGATTACCTGGTGAAGTTTTATTAACTAGATACCCATGTCCACTGTAGTCGTCTTGACAGGAATTCATTAGACGGATCGATTCCATCAACCCTCAAAAATCTCGCACTTATGAGTGAACTGTGAGTGGACAAGCATTATGCTAATATAAAAAATCCTTTTTATGAGTGTATCAGACCATCTATAAATTTTCCGTATCACTTGCAGCTTCTTATCAAACAATAAATTCACTGGCCCCCTGCCCAATCTTGCTGGCATGAATGCCCTCTACTACTTGTAAGTTCCTTGAAACTCTTTTCTGCTAAGAAAATGCAGCAATCTTGATATTGTGATAATTACAGATTATGTCTTTCGGTTACTTTAATTTGATTTTGTTGCCTATGGTCATGTTTTGCAGAGATATGAGCAATAATACATTCAGCTCAGCTGATTTTCCGCAGTGGTTTTCAACTTTGCAGTCATTGACGACTTTGTATCTATCTCTCTCTCGCTTTCTCTTTTTCTCTTAACATGCATGTGTGTATCTGTCCCTGAGTTCACGTGCCTCAGCAGAACTCACATGCGTCTGAGTGTATATGTTATCCTAATGAAGTGGTTCATAGAATTTGTTATGCACCTCATTTGTATTTGTTCTGCTCATAGGGTAATGGAGAACACTCAACTTCAAGGAGAAATTCCGCCAACCCTCTTCAGTCTTTTTCAGTTGCAGACTGTGTGAGTGTTCTTTTACTTTTAGTACATGGTTCATGTCATGCAACTATTCTCATTTTCGCTGTGTGCTAGTCTTGTACAAAGTTTGAAAGGTAGTACAAAGTGACCTAGTATATAATTCTATGTTTTGCATTTCGATTTTACGACAATGAGGTGGTTATTTATGACAGCAACTTGAGGGGAAACAAGATCAATGGAACCCTGAATATTGCATCCAACTATAGCAGTCAATTAAAACTGATTGATTTGCAGAACAATTCCATCGATGCATATACAGTAAGACCTGGATTTCCTTTTCAAATAATGTAAGTCCTCAAGCCACTTTTTCTCCACTTTAGCAATGAGAAAGAAATAATGGATCATTCATGTTAGGACATTTGATTCGTCTGAATACAATTTAGTCGTTTAACAATCTCAGTTTTCTACTTGGTCACTTGGCAGACTTATGCATAATCCAGTTTGTTATGAAGAAAGATCAGAAGACTATTGTGGCATTTCCCAAACGAACTTTGTATATTCAACTCCACAAGAAAATTGCCTTAAAACACAATGCAGTTCTGATCAAATTCTTAGCCCTACCTGTAAATGTGCTTATCCGTACACAGGCGACTTATTCTTCAGAGCTCCTTCCTTTTCTGACTTGACAAACACAAGCATTTATGAGTCACTTCAGAAGTCTATGCTATCTTCTTTTAGCCCAAATCAGGTGCCAGTAGATTCAGTTTCCTTGAGTAATCCAAAAAAGAACTCAGAGTACTACCTTGTGTTGCACCTGCAAGTTTTTCCATTTGGCCAAGATCATTTCAACCGTACAGGGATTGCCACAATTGGATTTGCACTTAGCAACCAGACTTACAAACCCCCACCAAAGTTTGGACCATTCTTTTTCAGTGGTGAAAGCTACAAATACTTCGAAGGTGATTGGTCATTTCATTTCATTGATATGTAAATGTATTTCCTATGCACAAGTTTCATCTGATAGCCGTACTATTAGTCGTGGTATTACTCTTGTAGTTTATTGCCCTTTGAGTTAGGTTATCGTCTATAGTTTTTTTACACTTCGATATATTGTTTTCAGTTGGATCAAGAGGAACTCACAAGTCAATCACTAAAGGAATTATAATTGGAGCAGCTGCTGGTGGTGCTGTCCTTTTAATTTTATCAGTAATAATAGGAGTTTTAGTTTGCCAAAAGAAAAGAGATCAAGAAGCTATTAGAAAGAGTAACCCTTTCGGTGTGTTCTGTTACTGTTCTTTAGTGTTGCTTCTAACTTTTTATTTCGCTGATGGTTATCGTCTTTTTTATCTGTCCAGCTACATGGGATTCTATAAAAGACAGCGGTGGTGTTCCACAGTTAAAAGGTGTCAAATGTTTCACATTTGAAGAGTTGAAAAAATACACCAATAATTTTTCAGAAAGCAATTACATTGGATCTGGTGGTTATGGGAAGGTTGGTTTCTATTGACTTTTGATAATACAATGTTATATTATGATGACTGGTTAAAAAGCTGTTGCTTTAATTCTATCCATACTATCTGCACCATCTTTCGGGCAAGCTAAAGTTAGTTGATCACTTAAAACTTCTCGTGAACTTACAGGTATACAAAGGCACACTTCCAGATGGACAACTTATTGCAATTAAAAGAGCAGAACAAGCATCTAAGCAAGGCGCCCTCGAGTTTAAAACTGAGATTGAGATTCTTTCAAGGTTTCATCACAAAAATGTTGTCAGCCTTGTAGGATTTTGTTTCAGGCAAGGTGAACAGATGCTGGTCTATGAGTTTATACCAAATGGATCCTTGAAAGAGAGTCTTTCAGGTATTTTGAACTCTCATCGACTGTTGTTTTGAAAATCAGCTGTAGGTTGGAATGTCTGGGCATTGATAAGTTAGTGTTAACTTCTGCAGGAAAGTCCGGGATTAAGTTAGATTGGAAGAGGAGACTTCGGATAGCCCTTGGAGCAGCCAGAGGTTTGCAATATCTTCATGATCATGTTGACCCTCCTATCATCCACAGAGATATCAAATCGAATAACATCTTGCTGGATGAGCGCTTGACTGCAAAAGTTGCTGATTTTGGTCTTTCCAAGACGATGTATGAGCCAGACAAGGGTTACGTCAGTACTCAGGTTAAAGGAACAATGGTAAGTCCAGACTTTACAAATGTTGTCATATGGTAGCATTTCTCAGTTTCTATATTTCTTAAGTCACTGGAAACTCCTGTCCTTTAATAGTTATAGACTGGTGATCGTCTTGGTTCAATTCTTAAAAGTTAACACTGAAGTTGCGTCCTGATCAATTACACATTGTGCTGAACTTATGTATGTAAGTTCATACAACGAAAGGAGTTCATTGGTTAAACTGCTTAAAAAGACTAAAACAACAGTTGATTTGTAATTTTCGCAGAGTAGAAAACTTGAAACCTGAACACTTGATATCAACATTGTTTGAAGAATCTGTAACAGAACATTTAATTAATACGAGCTGCTTTGTACATTTATCGGCTTAGGGCTACATGGATCCAGAGTACTACACAACACAACAGTTGACTGAGAAGAGTGATGTCTATAGCTTCGGAGTTCTGCTGCTAGAGCTTATAACAGCACGGAGTCCTATTGTGAGAGGAAAATACATTGTGAAGGAATTGAGGCAAGCGATAGACAAGTCTAAAGACATGTACAACATTGATGAGTTTGTTGATCCTGCTATCCCCTCAAACATGACACCTATAAGCTTCAGGAAGTTTGTGGATCTAGCTTTCACATGCCTCGAAGATGCAGGAGCTAATAGGCCAACAATGGCTGAGGTAGTGAAAGAGATTGAGAATATCATGGAAATAGATGGATTGAACAGTTGTGCTGAATCCAGATCAACTTCTGTGAGCCACGAAGGAACCAGCGCAACCTTTGATCATCCTTACAGTGAGGAAAGCCTAAATCGTTACAATGGGGTCACCAATTCGCGATAAACACCAACTCTTCTCCTGGAACCAGGTACATTGTCATAGAAATGGAAAGTGATTCAAGCAACTGTTGTGCTTTTGTGTTAGCTGTGTACATTCTGTGAACCTCTGTGTGCGGCGTAAACAGCCTTGTATGTTTCAACTGTTTGCTTCTGATAATCCAGGAGCTAGACTCCGTGGATTCAAAATGAGTGATTTTATTGTACGTATACATTTTAATTTTGATTTATTTTTACCTATGTATACATAATACGAATCAAAAAGTAATGCTTCAATTGAACCTACACAGAACCACCCTAGATCAGGAA

>PGSC0003DMG400018264 LRR-RLKs

ATGGTAGTTTCTATGGCAATGATGCTGCGAATTTTCGTCAGTTTTCTGGTTGTTTTGATCCAAGTTCTGTCAGTAGCAGCACGAACTAATCCTGTTGATTGTAAGTTGCTTTTGTTCTTCTCTATCAAATGTTTCTTAACCAGTGTACAGAGATATTTGATTTATCTGTGAGTACTGGTGTTCAAATCTTAGTTCCTTTTTGTAATCATATGAAGAACTTGTCTCTTAATTGCAATAATCTTGAGGGAACATTAAATATTTCCTGTTCTCTTCTAGATATGGTCAATTAAGTGTTCTAGTGACGAACTGATTCAATATTTTATACAGCTGCTGCTTTAAAGTCTCTCAAGGATTCTTGGTATGGTCCTTCCAATTGGACAGGAGAAGATCCCTGTGGGAGTAGTTGGGAAGGAGTTGGATGCAGAAATTCACGAGTGATTTCTATGTAAGAATTCGGTGTGTATCCTTTTGTCAACTGCTTAAAATTACAGAAATCATATGGTTGATGCCATCTTTCTCAATTTCAGAACACTGTCAAGTATGGATTTGAACGGTCAGCTATCTGGAGACGTCCAAGGATTATCTGAATTGGAGACTTTGTAGGTCTTCCTTTCACATTTCGTGTTTCTACTTCACTCCCTTTTTTCTTTTGACGGAGCAATACTTTGTCAGTTCTAATTATGGACTGCTCCTTATAGGGATCTATCATACAACAAAGATTTGACCGGATATCTTTCTCAGTCTATTGGAAAATTGACAAAGTTATCAAATTTGTAAGTGAATGTGAATAATCATCACTCGAGTATTCACTTACCAGCTAATCTCGTGATAATATGCTGATGTTTTCTTGATTTCATGACAGGATCCTTGTTGGTTGTGGATTTTCTGGACCAATACCAGACTCCATAGGATCTCTGACCCAGCTGTTTTTTCTGTAAGCATTATGTGGCACTCTTTGTACTTTGAATGTGCGACCAGTACGTGACATATTTTACCAGAATGCAAGGGTCATACACTATCTGCTTAAAATTGTTAACGTAATGTGTGCTTTTGGTGCTAGCTGTACTCAAATTTTGGTCCATCTCTGGGACAGGCAATGCACTTCTAAACAGAGAAATCAGTTCATGAGGAGTTCAATCCATACTGCTATTCTTGTGTTTTATACGTTATCAAAAGTTGAACTGTTGTGTTTACCATATCTATCTGTATATGATTTCTTTCAACCCTTCTTCTGGTATATGCTCTAAAACAGCCCCATTCTGTATTATGCAGATCTTTGAACTCTAACAAATTTACTGGAGGAATACCTGCTACCATTGGCTATCTGACAGAGCTTTATTGGCTCGATTTATCTGACAATCAGATAGAAGGAAACATTCCAGTCTCTAGTGGGAGCAACCCTGGTCTTGATATGCTTGTTAACACTAAACACTTGTACGCATCTCCTTTTTCTTGATGTTGTTTCGTCTCTCAACTGCATTTTCTAAGATTATTTACATTTCTTCAGTCACTTTGGGAAGAATCAGCTATCCGGTGAAATTCCAGCTCGTCTTTTCCATTCTAAAATGACTCTGATACATCTGTAAGTTCCATCGAATTGGAGAATGGAGTGAATGAATAATTTTGTCCGTAAATATTGGGGACCTCACTTGTAGTTGAGCTTGCCTTTTGATATTCTTGTTAGTGCATGTTTTTACTTCTGAGAGTAGTCATGTGAAAGAAGCTTGATAATTATATCACAAACAACTTTCCTTTACGGTTTCCTCATGCTTGTATATTCCAACTTGTAGGCTAGTTGAGAACAACAGGTTTACCGGGGTAATCCCAGATACATTAGGATTTGTCCAGACGATGGAAGTGCTGTAAGTATTGCTATCTTGTGCTTTCCCTTTATCATCTAGTAAAGTTGCTTTCTGAATTGTTTCTGGTATTGCAATAGTCGTCTTGACAGGAATTCATTCAGTGGATCTGTTCCACAAAACCTCAACAATCTCACACATGTCAATGAACTGTAAGTAGAAAATGAAACAATTTTCTGAACCTACTGTAGGTCCTCTTGATTTTTTGAAGTAGTATATTTTGACCATGTATTACAGCGACATATCAAACAACAATTTGAATGGCCTTTTGCCGAATCTCACTGGCATGAATGTCCTCAACTACTTGTAAGTTGCTTATCCTAAGTTTTGTGTCTTTATCCCTTTGAGTTGACAATCCATTATTCTCAACATAGTACTATAAATATTTCTCCCCAATCACGATTCACGGGCACTTTTTTTTTGCAGGGACATGAGTAGTAATACATTTAATGCATCAGATATTCCATCATGGATCCTAAGCCTGCAATCATTAACAACATTGTATACTCTCTCTCTCTCTCTTTTTTACTAGATTTGGGAACGTGCGTTGCACGTTTATCCCAAAATACTTATATAAATTTTGTATTGCAAATGACATATTTTATCCCCTTCAAATCTTTGGTATTTACAAAGAAAAATCTATCATTGTTTCTATAGGAACTACATATATCTTCCTAATTAAAAAAAATATACCTTTAACTATAAAATTGACTAAAAGTTAATAAAGTAAAAATAACTAACTATTGCACAAATTCAATATTCAAATGAGGATACTAAAACACCACATGCTAGGCTAATGTATACAAAACAACAAACCATGATAAATTCTCAATAATAACAACACAACAATAGAAAGTTGTACAATAACAACAAACAAGAAAAATAAATGTTAAACTAAACAAATAAAAAGAGACAAGAGAAGAGAGCATAGAGATTTTCTCATATTTTTTTTCAATTGGTTTAAGTGTGTAAATTATTGTGTCAAATGTCATTATTTATAGGATAGCATTGAGGAACACAAAGAGTTGTCATAAATATGATATTAATCCATTTGAGATTATTGATGAGAGTAAAAACATAATGAATATAATTGGTGGGAATTACATACATATAATGAAGAGTTAGTGGTAATTTATGTATACTAATATGATGATTTACTATTTACTACTATTTAATTTATTAAACAATTTATAACATTTAAATATAATGTAGAAATAAAACCGTTATCTAAGGATAAAATATGTAAAAGAATTCAAATATATTAAATGTGTAGTGAAAAATCATTATGACTAATTATTAAAAAATTCATAGTAACTATGATTATGAAAAATCATTACTATTACTACATGGAAGATGTAAAGATGGTTGTTTTTTTCTTGCTAATAATAACTTACAATTTAATAATGAGTATTGAATTGTAAGTCTAGCTGGAAATAGGAGTGTTAACTACCATAATACGAACAATCAAAATTCATTGAATCCCAAAATAAATATTAAATGAAAAAATAATATTTTAAATAAATTCAAGAAAATAAATTAATAGTTAAAGGGTATTTTCGTAATTCAACTTTTGACTTAGATTCTTCCCACTTTTAATATAACTAGTTCTCGACACGTGCGTTGCACGTGCATCCCATATTAAGTAATATAATTCATAATTAAATTGGAGGTTAATGAGCAACAAATTATATGGATATGCTCCATGTATGAAATTCAAAAGTCCATGTTATATGGATATGCTCCATGAAATGAGGAAAAGAAGTTGCATGTTATCAGAAGTCCAAAAGTCCAGGTAAGATGGATATACTCCATGTATGAAATTAATTAAATGAATTTAATCATTAATAACTAACGAAAATTATAATGCCATAACTACAGAATCAAAATGCAATTGCTAGACTTCTAATTCAGTAAATAAAATATGATTATTTCATTAAATTTACTGGTTAATTAAATATTTAATTATATTTTAATTTAGTGTGGGGTAAATTCGTAATTCAACTTTTAAGTTATGAGCTTCCCACTGTTAATATAAAATATAAAATAATAATATGATTTATTGGTTAATTATATTTTAATTTAGTGTGGGGGTAAAATCGTAATTCAACTTTTAAGTTATGAGCTTCCACTTTTAGTATTACTAGTATCTGGACACGTGCCATGTTATTTTGTTCAAACTTATTCATTAAGATGGAGGGAATATATAATAGTTAAATATATTGGTTTTAAAAAATGAAAGCTTATACTTACTAAAATAAAAAAATTACAACACATATCAATTCTCTCTTTACATATAAAGTAGAAACAACTGCTAAAAGTATAACAAGAATATTTGAATCCTAAAATTAAAATATTACATGTTGGAATAATAATACATAAACATTGTGCATTACAATAAATCTGTATTGAACTTTGTTTAGTGAAGTTGATGTTACTGGCTAATCGAACTCTAGACGCCAATAGCTTTAGATACTTTCCATTCTCTCAAATTGTTGCTTTCTCTTCAAGAACAAAAAAGAAGATGAAATTAATTAATAAATTAAAATTATGCAAGAGCCTATAAAAATGGTGAGAAAGAATTCAAGAATCCTTTCTTGCCTTTTATAGCTATTTTTTGAGTAAGAATTCAAGAATCCTTTCTTGTTTTTTTATAGCTATATTTCAATTTTCATTCCTTGAACCTTTTCTTTCATTAGTCTACATTTAATTATTCAATGCAACTCTTCCTATTCTCCAAACTATTTAGCAATGAATAACGCACACTTAAATTTGATTAATGGGTAAATCATTAAGGGCTACATTTAAATTGGACTTTTGAATATCTACAGATTCACAAATCTCATGTAAATTTAAATACAATTACTACTATTTCAATTAGATTAGGCATTTTATTTTAATTAAATATAAGAGAAAAATTGTAATCTAACTTTAAACACTTGTTTTATTTTTCTCCATTTTTATTCATGTTTAGTACATTCACCTCAGACTAATATTCTACTTTGAACAGAAATAACACAAAGCAATAAATATATAGAGATACATATTCCAAGCCATAATAAATGTCATTATTTTTTATTATGGATACATTCACCTCAGTTCTCACTCCATGAATTTCCGTAACATTTCATTAATTTGACTGGAAACATAATAAGTATCATTATCTTTTAGTTGGAATATGAATGTCAAAGAGTGCCCTCTCTTTTTTTGTGTGGATAATTTAGTGATTTTATATTTATGTATATTAATTTTGAAGAGATAATACATTTTTATTATTATTTATTTAGATTAGATGTTGTATCTATTACATTTTAATTAAGTATAGGGGTAAAATTGTAATTTAACTTTGAAGATAGGTGCTTCCCACTTTTAGTAATTAGTAAAAATTACTCCCTCTATCCCTAATTACTTGTTCACTTTTGAATTGACACACCTATTAAGAAAACAATTATTGACATAGTGAGTGTACCATTTTACCCCTATTATTAATTATAAAGTGGATGGATTAAAAATTTAAGATTTTCAAGAAATTTTACCTTTTTCAAAGTATTTAATTGAGGGTATAGTAGGTAAAATTATTTTGTCCTTTCTTGATTTGTCAAAATGAACAAGTAATTAGGGATAATTATAAAAGGAAAAGTGGACAAGTAGTTAGGGACAGAGGGAGTAGTAAATTAGTAATATATATAATATATATATAATATAATATATGATATGATGTGTGTGTGTATGCACGACCATGAGCAGGTGCATGTGTGTATGGGGCTCGTGTATATGCCTGCATCCATTTGAATATGGACTGATTTTATTTTTTCTTGTTTTTATGGATTCAATTCCTAGATGTTCTTTTTAACCGTTTGACTGACCTTGTATTGTCAGGGTGTTGGAAGACACGGGACTTCAGGGATCAGTTCCAGCTAACCTTTTCAGCCTACACCAGTTGCAGACAGTGCAAGTGTCCCTTTTACTTTGTGTCGTTTTTGTATGAAGTATAGATTGAGTATCTGAAGTTGAATTTATCTTTGCTGAAATTTCCGAACTCTGTATTATGCTGATAGTTGTTGACAATGTGCTCGTTGCTTACAGCATCTTGAGGAATAACAAACTCAACGGCTCTCTTGAAACTGGAACCACATATAGCAACCAGTTGAAGCTCATTGATGTGCAAAATAATCTTATCGAATCATTTACTCAAAGACCAGGGTATCCCTTTCAGATAAAGTACGCACTATTCCATGATTCTTCAAGCCATAAAATCTCCAAGTTGATAAGAAACACCATACGCGAGACAGTTTTTTGCCTAGCTTGTTGTGTCGTGTGATTATTCTATGCTCAACTTGACTATATACTCGTTTATTTACTTTTTCCTATGCAGGCTTGCAGGCAATCCTATCTGCAATGAAAGAGGAGAAGGGAGAAAAACTTACTGTATGGAAACTCAACAAACTGAGACATACTCAACCCCACCAGAGAATTGCTTGCCAACTGAGTGCAGCTCTAATCAAGTTTCTAGTCCTACCTGTAAATGTGCCTTTCCATATACAGGAAACCTAGTCTTCAGAGCTCCTTCCTTTTCAAACTTGGGAAATAGAACTACTTATGAGACTCTCCAAAAGTCTTTGTTGCTTTTCTTCCAAAAACGTCAACTGCCTGTGGAATCAGTTTCCCTCCGTAACCCAATAAGAGATTTAAATGACCACCTTGTAATACAGCTGCAAGTTTTTCCATCTAGCCAAGATCGTTTCACTTGGACTGGTGTTTCTGGAATAGCTTCTGTGCTTAGTAATCAGACTTTCAAGACCCCATCGTCATTTGGACCTTTCTTCTTCATTGGTGAAACCTATAAATACTTTGTCGGTAATAGAACTATTTCCTTTTCCTATTTATCACTCATTTCTCATCTCCTACATATATGTGCAGCCTAATCTATAATTACAGAAAGTTGGTGGCTGAAGAAACAATATATATTTGAAATCTTGATGATTAAGTTTGTATGTACTTTCGTTTGACAGAAGAATTTACTGTATTTTTAAGACTTTGAACGTGAATAGCAACTATTACAGATCTACTGTTTTGATAACTATATCAATCCTTTCTGAAGTATTCAATTTAGAAGTACAAGACCCAAAAAAAATTGTCTAGTCATTGAATTTGAGACAAAGATACGATAAAAAAACTAAAAATTTAGGCTACGTGCATGTGATGGATCAACTTTCGTTCATTGTTACTAGTGATTTTTCTGTTAGAGTTTGCATAAACTTCAATGGATTTATTAATAACGTTTTAGTATAGACGCAGGTTATTCTGATCGGTCTCCTTAGAAAAACTGGTTCATTAATTGACAGCTGAAAATTCAAATTATTGTTTGGAACATTATGGAGCTACATCTAAACGTTTCATGTTACTGAAGTCTGAGGCTGTGCTTGCTGTTTTCTAGGGGAATCCACAGGATCAAAAAAATCAGTTAGTACAGGCATTATTATTGGAGCTACAGTTGGTGGTTCTGTCCTTGCAATTTTAGCACTAATCATTGGTATTTATGCTTTCCGGCAAAAGAAAAGAGCTGAAGATGCTGCAAAGAGGAGCGATCCTTTTGGTAAGAGTTGGAACTAACTATTAGGCCATTTACCTATTACTTCATCTTACGGTTGTTGTTGCTATCTTCCTTCTTAATGCAGCATCCTGGGACTCAAATAAACATAGTGGTGCTGTTCCACAGCTGACAGGAGCTAGATTTTTCTCATTTGACGAACTAAAAAAATGGACTAATAATTTTTCAGAAACCAATGTTATTGGTTGTGGTGGTTATGGAAAGGTCAGCTTTTATGCTCTGATACACTAAGCATGTGTTCTGTTAGCAGTACTGCAGCCTTCATTCGTTCTGATCTTAATCATCTTAGACTTAGTAGTTAGTACTACCAGTTTCTTTGTCAAAGTGTCAGGCTATTTACAGACGATAATGAACTGATTGTATATTTGGATGGAGAAGTCTTGATTCTTCCACCCAAGATAATTCACATATTTTGGTTTCGGATGTAATACAATAGCTCTAATCTATTTAATTAATAAGATCCGGTAAGTATGGATATTTTTTATCTTGTTTGTACCAGCTTCTCTTTCCACTGAAAAGAAAATTTCTCGACTACAATTGAAACTGTATTAAGTTTTTAAACTGTAGAAGTGTGGCTCTACAGGTTTACAGAGGAAGTCTTCTGAACAGAGAATTGGTTGCAATTAAAAGAGCTCTACAAGGGTCTATGCAGGGCACACATGAGTTCAAAACTGAGATAGAGCTTCTCTCAAGGGTTCATCACAAGAATGTCGTTGGCCTTGCTGGCTTTTGTTTCGATCAAGCTGAACAGATGCTTGTGTACGAGTACATTCCTAATGGCACCTTAAAAGACGGTCTTTCAGGTATATGTTGATCCCTTCTTGTTGATTAATTGAAACTAGCTCCTTTTAACTTAACCAAAATGCAAATGGAATGAAATTTGTACTTGTAGGTAAGACTGGGGTCAGGTTAGATTGGATGAGGAGATTGAAGATAGCCATTGGAGCAGCCAGAGGTTTGCAGTATTTGCATGACCATGTCAATCCCCCCATTGTACACAGGGACATCAAGTCCAACAATATTTTGCTAGCTGATCACCTAAACGCAAAAGTTGCTGATTTTGGCCTGTCCAAGCTTCTAGGTGAAAGGGGTCATATTACCACTCAAGTCAAAGGAACAATGGTAAGTAAAGATTCTATAAACACCTGCCTGAACTTGTTTCGTTTTAGAATAATGTATGCTCCCTCAAATTTCTGATGTCCAGAAAGTTCAGAAATTAAATGTGTCGATAATTTCTAGTCCTGTAATAACCATTACAAGCACTATTTCATTGTCTAGTAAGAACACCTGAAGCTCCTCCTTTCTCATCGACCACCCTTCCCTCCACTAAAAATGAGTAAATAAATAAGTAGGAACAGGAAAATACGAAAACGTTTTATTTATTTCTTGATATTCTCTCAAGGTAAGGGACTTGTAGGTAATGATGCTTAAAAACACCAAATTTAAGGAGAAATACTACCTTGATTAGGAAACAATGAGTTATTGCCCATAGCAGATAGAAGCTTTTTAACGTTCATATAATGTGATCTTGCCATGTATGAAAGGAAACAAGTTGAGATGTTGCAGAAAAATGTGTCAGTATTTACCTTATAATCTATGTCCTTTTGATGGTTGCCTTTTATTTATTAGGGATACATGGATCCTGAATATTACATGACAAACCAGTTGACAGAGAAGAGTGATGTTTTTAGCTTTGGAGTGGTGCTGCTTGAGATAGTCACAGGAAAAGTTCCTATTGATAAAGGAAGATATATTGTCAAGGAAGTGAAGGATGCAATGGACATGACAAAAGATTTGTATAACCTTCATGAAATTTTGGATCCTGCAGTTTGTTCGGGTGCAACTCCTAGAAGCTTAGAGAAGTTTGTGGATCTAGCACTCAAGTGTGTTGAAGAAAAAGGAGTCAATAGACCTAAAATGAGTGAAGTGGTGAAACAGATTGAATATATTATGGAGATCGAAGGCCTGAACCCCGATGATGAATCTGCATCTACTTCAGCAACCTATGAAGGAGCAAACAAAGGCTTGAACCATCCTTATACTGAAAGTCTCTTTGTCTATAGTGAAGCATAA

>PGSC0003DMG400018367 LRR-RLKs

CTCTGTTAAAACAATGCTTTGCTTGCATTTTTAGTTTTTGTTCTTTTGTTTTTCAGCTCAATATCATGATGTTTTTTCTCAGCTTCTTCCTCATTATTTCTTCATAATTCACTTCATATTTGGTGGGTTTTTCCTTCTTGTTACTGTACTTCAATTTTTTACTTCTCCTTCTTCAACTGTCCAGAGTTAGTTTTTTTTTTTTGTTTCTAGATTTGTTCTATCCTTTTCTCAATTTTTGAATACCCATTTTGTTATCAGTTGGGTGTCATTGATTTGCTTCTTTTTGGTTGTGTTATGATGAATTTTGGGTTTGATTGAAGTACTGATTTTTATTTTTGGGCTCTTTCAGATTGGAGGTTTATGTATTGGTTATGCAGAGTGTGTTTCCAAAGAAAGAGTAGATTGAAGAAAAGGGTTACAGTTTGGATCAATGGGTGGGAAGAGATCTGTGTTAGGGTGCTTGAATTTGGAGATCTTGGTGGGGATTTTGTTAATTTTTGCTGTTCAGCTTTCAGATGAAACTGACCCCGGAGATGGTAAGGAACAACAAATATTAGTATTATATTTCGAGTTGTTTTGCGTTGAAGGAGAGGAATGCTAATTTTCATGTTTTGGGGAATGCTGGTTATGAGGGACTGAATTATGAGCTGAGCTAGTTGTTAACTTGAACTTTATTTTGGGTAAACGTTTTCAAGTCGGGGGCGAGCTGGAGTTATCGAAATTGGTCGGTTAAATGAATAGTCTTTATCCATTTCCGGAATTAAGTTGGGGCATGACGTTAGATATGAGGTTGCATAGGAAATGGATTAGGTAGAAGATTAGTAGGTAATGGAGTGTTGTCTTGCTTATCCTTTGATGCTGGTAGTCGTAGTATTTCCCTTGGTAGTTTCTCGTCCTTGGATTTGTGTTACTATTATTGCACTGTTTTGTTGCGGTTACCGTTTGTTTTTTTCTTCTCAGAATGCTTTGCCATGCTTTCTATAGTATTTCGCCACTATTTCTTCACTTCTGTTATTTCTTTTTTCGATATGCCTTTTCTTGAGCTGAGGGTCTATTAGAAACAACCTCTCTACCTCTTAAGCTAGGGGTAAGGTCTGCGTACTCTACCCTCCCCAAACCCCACTTGTGGGATTACATTGAGTATGTTATTGTTGTTTGAATTAATTTACAAACAACTCCATCATACACAAACAAAAAACCTTAAAAATAACCTAATCGTTTCACTTACATTCTCAAAAAACTTGCATGCCGCAGTTACCAATTCTACTAACAGTGGACAAGCTGTCATTATTTCAGAAAAAAAAAAGAAAAAGAGACAGTCCCAGGAAATTGAAGGGGGAGAAGAGGAAATGAAGTACTGACTCCTAGGTTGCTAGTCAAGTGTAGGGGGCTATGGCCTCGTCAAGTGATATGTAAATTCAACACATATTATTGTCCTAAATATCTACTAAATTGCCATCTTGATCGTGTAAACTTTGCTTGGTTTGAAGTTAGCATGCTTTAAATGTTTGGAATCAAATTTTCTAGTTCATTCATGATCAATATCAACAATATTCTTATATGGTTTGAAATTTCTAACTATGGCATTTTCCTTTCTGCTGGTCTAGTTGCTGCAATTAATGCCTTACATGCTTCCCTGGGATCACCTCCCCTTCCTGGCTGGGGTGTCGGTGCAGACCCATGTGATGGACAATGGCAAGGGGTCGTATGCCAGGAGACAAATATAATATCCATGTAACTACCTCTTATTCGTTGCAGCATTTCATGGATATTTGGAAGTCGTAATGTTAATCCTTCTAATCATCTTCTTTGCAGTCAATTAAATGCTGCTAACTTGGCAGGTGAACTAGGGGATAACTTAGCATCATTTTCTTCTCTCAAAACGATGTACGTGTACCCCACAGATATATCTCCTTCGACATATATTTGGTTGTGTTTTCTGTGCATGTGCTGCATATATTATGTGGATGCTTCCTTTTCTAGAGGTTTCTTTTATTGTGTGACATGTGAACAATAATTGTGGAAACTTCTTGTGGCAGAGATCTGAGTAACAATCACATCGGCGGAACCTTCCCATCCAGTCTGCCAGTTACATTGCAGAATATGTATGTATTATCATTATCCTAGAAGGGGAAACAGAGAATGAAGCTTGAACTATCACAGCTTACCTTTTGAGCAAACCATTTGATAGTCTTTACCTTTTGGCAGTTTTCTTTCAGATAATGATTTAGCCGGAAGTATCCCCAGTTCTTTATCTTCCCTAAGTCAACTGTCAGCAATGTGAGTTTCAGCTTATATTTTAGATTGGAGATTTCCCGTTACCGTAATCATGGTTCCAGTCAGCTATACTTCTCTTTTGTTTCAGGTCTCTCAATGGAAACCAACTGACTGGAGAACTTCCTGATTCCTTTCAAGGCCTTACAGCATTGGTTAATCTGTATGACACCAAGACAAAATCATTTCAATATTTCCATAGAGGTTTTTAATTGTCCATAACTGGTGGATTAATTGTATACTTGCAGTGATTTGTCAAGTAACAGTTTCAGCGGAGCTTTGCCGTCATCAGTAGGAAACTTGTCCTCCCTGACCACTCTGTGAGTGATTCTACATATTGTTGAATTGGTTTGTAATTTATGACCAATAGAAAAGCAGCTTATTTCTTTCTCTCCAGACATGTGCAGAATAACCAGCTTTCTGGAACTCTCGATGTTTTACAAGATCTTCCCCTCGCAGATTTGTACTCTTCGTCACCATTTTTAACTCAAATATTAATTCCAGGTCCATAGAATGATATAATATCGTACTAATATCTCCCATCTTTTGTGTCCCTCCAGGAATGTGGAGAATAACCTGTTTTCTGGGCCCATACCTCAGAAGTTGTTATCTATTCCCAATTTCAAGTAAGTACATTTCACTTTCTGCTTCTAATATTGTTGTTTATCTTCCAGCATGTAATCTACAATATATTCTTGGCCACCTATTACATTCTTGTGCAGTTGTGCCTCTTTGTTCGACACAATCTGCTAGACAGATATTCAGAAACTATGGAGACAGATTCCTTGTTGAAGTTCGATTATTACATGCCAAACTGCTCTGTACTTTTTGTTAATACTTAATACCCTCTTTCTTAAATCTGAGTGACAACCTTGATCAGCTTGTTATTCATGGACAATTAAGGCAGATAAGATCTGAAGGGCTACAATCGTTAACATAAACAATAATTGTACTCCACCCTCCTAAAGCTGCAGCACATGTATCATCTGTACTCAGCTTCATATTCCCTACCCACTTCAGACCATACTAAACTTTTAGTTAACATACTGGTGAATATCAAACTTCACTAAAGGGGAGAGTTTCTGTGCAAAATGACAAGACTTGAATAGGCTTCATCATCTCATTAAGTATAGATCAGCGGAAGAGTTCAGAGTGGCTTCTTTATTACACCCAAGTTTGTTGGTTGGCAAAAAAATGCAATTGCTATTGGCAAACTTCTCAATTTTCTGAACCTATACGTTTTAGACATTTTATGTGCGTAGAAATTTGTGTGTGAGTGTGTGTGTGTGTGTGTGTGTGTCTGTTTTTATGTAAGTTTGTGAATGCAAGACCTCGTTAAATTCCTATAAGCTAGAAAAAACCTTGATCTGAACTACATTACTCTTTCTCATAAGCAGGAAAAATATTATGAAATGTGTGTCGTGCTAGTCTGTATGTTAGCTGCACCATCTTTAACAATAGTATTTTAGAAAATTGTTGAGTCCCACATCGTGTAGCATGAGGAAGTTGACTCCTTTATATGGTCTTGGGCAATCTTCATCTCTTGAGCTAACTTTTGGGGTTGAGTTAGGCCCAAAGTCTTTTTTATACTATCGGAGATCTGATATGTTGAGGAGTATATTTATGTTGCAAAACCAGATTATGCTTACAGGTTTATGGCACTGACTTAGATTAGAATTGCTAGTGGGCACACTGCATATCAACAGCTTGAGTCCTAAATGTTAGAATAATTTCAAATTACATTAGGTAAAGTAGCCAAATTTTGATATCATCAGCACAATTACCTAAAAAAAGAATAATCCTAACACTATGCTGCTCAATTTAACAATTGGATAAAAAAACATCTAGCCTACACAATAAATTGGAAAAAATGTGACGTATACAAGCTAAAGGATTATAGTAGCACAAAACCATATGACACCCATATATGATGTATATGTCACGCTGGACCATGGACTGCCAAGCTGGATCATGGACTGCATTTTGGTCATATTTAAGTTCACTAGCGGGCTAACATTTATTTAAGTTTTGAACACAGGCGTGATTTTCATATCATGTTGTTCAATAAGATTTGTTTGCCGGGAAATTTGATTTATTGAACAATATTGGTAAGGGTACATATTGATCCATGCTAGGAAGATGTATCAGTTGTTTCATTATTTTTTTCAAGTGTCTTATATAGGTTTTGACTCTTGATTGACTTTAATTATGAGAATTGATGGCTAATGCTTTCATGTACTTACTAATTTTGTTCTTGACGTGCAGAAACACTGGAAATCCTTTCAACAGTGTCTCTCCTTTGTCTCCACCCAACAGTAGTATTGCCCCTTTGTCTCCACCTACTTCATTTACGAGACCCCCTCCAGCACCACCAGGACCACCCTTTTTTAAGCCTCCAACTTCAACACAGACACCACCGACTTCAGACAGAAAACCTGGACAAAAAGCTGATGGACCATCAGCAACTGCAGAATCTAGTTCAAAAGGAAGCAAAAAGTCTGTTAAAAGGGTAGTCTGGATATCCATTGCATCAGTTTTGTCATTTATAATATTGGTAATAGCAATCCTGCTCTTTCTGCCAAGATGCTTTAGGGAAAGGCAAGAAACTAATTGGTCCAGACGGCACGGAATAGCTCCTTATGTGGGCTCCAGGGAGAATCGTAGGGATAATGGTTTATTGGTCCTACCTGGACATGATGTAGAGAAAGGTACTGCAAATTATTAAACCTTTCAACTTTTATATCGTGTGTATTGTTGCTAGTATTTGTTAGTCTGGCTTTTTTCCTTAGTTCTTACAGGGAAAACTAGTACTCTTTAATTTAGGCTCTATTTGAATCTGCAGTCTTGACATCTAAGTATGCAGCTAAATATATAGTGGTTATCCATGTGCCCAGGGGCGGACCCACCCTTACCCAAGGCCTTCGTCAGAAAATTACATGGTATATAGAGGTCAAATTTTGATTTAATAAAATATATACAAGTATTGACACCTCTTGACACAAGTTAAAGGTTCAGTCTAGTGGTTAAGACATTGCTTAATGACTACAACCATTGACACCCCTTAATGACAATGGCCGCTACAGTATTGTCAAATTCTCTATTACAATCATAAACGCCCTACAAAAGATTCGTGAATTTGATATCTGTTTTTGCAACATTCAGAACTAATAAGCTATGACTCTTAACCTCTTCTACTCTTTGATGATAAGTGGACTTCATATGTTTTGAACAGCTCCTCCTCTGGTGAGGCCAAAGGAAGAGCAACAACCAAGGAGACCGGCCCGCACTCCCATGCCACAGCAGGAGCAGGAGTTAAATGTGCAAAACCTGAGGGCAGTACCAAAGAAGGATACTAGTGAGATAAACCTAAGCAGAATTGATATTGATTCGATGTTGCCACCACCTCCACCCCCACCACCACCTCCGCCACCGCCACCACCCTTTTCTCAAGAGAGAGTTATTGTGAAGCCAATTCCACCTGCCGACAATACTTCAATGAAGTTTCCCCACAGGCCACTTCCACTAACTTCTGTGAAATCTTACACAATCGCATCTCTTCAGCAATACACCAACAGCTTTTCTCAAGATAATCTGATAGGATCAGGAATGCTGGGAACTGTGTACAGGGCAGAGCTTCCAAAAGGGAAGGTATAAGTTATGAATGATATATAATGTACTGTAATTTGATACTCGAAATTACCAACTGTTTTCCTACCCTGCTGTATTCTCTTGAACCTAGATTGTCAGCTTTTCTCTTCTTATAATTTCCTTTTTTGGGGGTCTTTACGGTGTTCTGTGAGCACTAAAGATGCCTCTTATCTGTTGTTGCATTACGTATCTTGCTCTTGCATTTCTATGCTTAATTGCTTAAGAATGATTCACGGACTGATCGAATCATATGTAAATATGGTTTTGTTGGTGGAAATCCAAAGTCATATATATTAGGGTAAACTGTATTTCATAAATACAGATAAAAGCAAAATGACTTCCAATTGCTTCATAAGATATTTCTTGTAACTTAAATTTTCTTTAGGACGTTTTTTGGTTCTTATCCTATATAATCTTTAAGTGGCATTTTCTTCCAAATAAACAACACCATTGATCAACTGTTGTCATAGAACAATCAACATTCAAGTTGCATGAGCTTATCATTCTGATTTGTACCTTAGTTTTGTCATTTGCTGTATAGCAGGATCTGCCAAATACTGATGCATCTTTTCTAATGAGAAACTATTTTATAATCAGAATATACAATGTCCCTGTTTTATTTTCAGTGTACATTTATTTATCTGGCATGCAATCTTAACTCTCATCCCATTGCTAACATTATTTGAATCATTTGGTGGTCACATTGTGTTACTTCTGACACTTGACCCTTTGAATGCTTATGCAGTTACTTGCTGTCAAGAAGCTGGACAGAAGAATTTCCAATCAACAGAAGGATGATGAGTTTCTTGACCTAGTAAACAATATTGACAGAATTCGTCATGCTAACGTTGTCGAGTTGATGGGCTATTGTGCAGAGCATGGTCAAAGGCTTCTGGTTTATGAGTATTGCAGTAGTGGGACACTACAAGATGCGCTGCACTCTGATGATGAATTCAAGCAACAACTTTCATGGGATACACGGATCCGGATGGCTCTTGGAGCTGCAAGAGGCTTGGAGTAAGTAAAATAACCAGTCACATATACATAGAAACCACATGCAAATATACGAGTATATACATATGAGCTATGGAAAATCTGATGTTGAAGAGTTATTTGGCCGTGATCTATAAGAAATTTCTGTAAGCAGTACTAATAATAATTTAGCACTGGAAGGTAGCAAAGAACTCAAATCACAAACTAGTCCTGAACTTTCATGCCATGTACATACTACATGACATAAATGATAAAATACTGTTGGGAGATGGTCTTTATAAGTATATGTTAACATTATATTTGGTGCTGTATTTCGGTTTATACTACTTTCCATTCCTCCTCCACATGAGTTCATACAAGGGATGCTGAACACGATTGTTGAAAGAAAATATCCTCATAGTCATCCCACCCGTTAATTATTATTAGTGGGATTGCTAGGTGATCTAATTCTTAGATTTGCATGACAAGCGAGTTAAAGCAAATTTTAAATCTCATGTATTAATTATATTTGTTCTTTTATGCAATGAAGGTATCTCCATGAGGTCTGCGAGCCACCTATTATCCACAGAAACTTCAAATCTGTCAATCTTCTCCTTGATGAAGAGTTGGCTGTGCATATCTCTGACTGTGGTTTAGCTCCACTGATATCATCAGGTGCCGTGAGCCAGGTAACTTACCATCCAAATATTATGGATCAGACATTTACCTCTTGGGGTGGAATGGTCATTTTCTGCTTGCATTTCCAGTTGAAAGAAAAGCAAGTTTCAGGTTTATGATGGAAGTGAAAATGTGGACTAAGTAGTAGTTGTTAAGAAAGGAGTGTAATGATGATCAAAATGCTTCCAAGTAACACACCGGAGTAGAGCAGTAATTAACTACAAATGCAGTTTCTCGTAAATGGATAACTTATTGTTTAGAAATAGTTGATTTTGTAGTAGTGCCAGAAGTTATACGGTCAATTTCAGAAGGGTCAAGTCTTTTTCCCTGTAAATCTGGAAATCAGGAAACAAATCTAGCTAGTCTGAGACTACCTGGTTGCCCAGCTAGTCTGAAGTATTTGTCATGTTTTAGGCTTTTATAACATTTCAATGCTTCACGAGATCTAATAGACACTAGTGGAATTTGTAAGCAGGAGAGTAGATCCTTTCCAACATATGCTGGACTTGCCTTCTCATCTTCACCCCTTGACAAAGAGATGAAATTTAGTGATCCTTGAAACTGTTCCCAACATTCTTCAGAAGTAGTTTTGCCGCTGCACATTTAATATGATGGTGACAAGCTCCTGTATTTTTTTCCCTGTTGGAGAATTTTGACTTCTAAAGATGTGGGAAGAACAGTTTCACTATGTTAGTTTGACCTAGTCATACATAAAGACACTCTACAGGTGCTTTTGTTAGATGGGTTTTCAGAAACTTAAATTCTATTTTATTTTTATGAAGTCCACATTATCTCCTCTCATTAATAAGAAACAGTAAAACACAAGATTGAACTAAACAAGACATCACGGATATGACTGGTTTTTTCACTTGTACTTCGTATGATTATAGATCTGCTACTCTAGGAGCATGTGGATCATCAATCTTTGCTGAAACCTTACCTTTTTTGTTTTTCTTATGCACTGTGCCGAAGAATTTGAAATTGAATAAGTGAAGTTGTACTTGATTTTCAGTTGTCAGGACAACTTCTTACAACCTATGGCTATGGTGCCCCAGAATTTGAGTCAGGGATTTACACTTCTCAGAGTGATGTTTACAGTTTTGGCGTGGTGATGCTGGAACTCTTAACGGGCAGAATGTCATATGACCGGTAAGGCCATTTGCATCTCATTGAATAAATTTTGTTGTCATGTTTTCTCCCAGCTTTATCATTACCTATATATCATATTCAGAACTCGAAGTCGAGGAGAGCAATTCTTGGTCAGATGGGCAATCCCTCAGCTACATGATATTGATGCCTTGACAAGAATGGTTGATCCTTCTCTCAAAGGAAAGTACCCACTTAAGTCATTATCACATTTTGCTGACATAATTTCCCGCTGCGTTTTGGTGAGTTTTTTAATCCTTTCTTTTATTACAAGTGAGTAATTTGTCCAAATCTCCATTTTCTTCAATCCTTAATCAAACTTTGGGTACAAATGAGGAAAAAGGTAACATGGAAAATGATTTATTTCTGTCACAGAAAATAAAATAAAAAGTAAAAGGAAGGAGCTGTACTGTGTCAAAATAGGAACTCTACACATGGTGTCTTTTGTTTTTTATAGCCACTAGATTATGAATTATTTACCAGTTTAAGGACATCTGTGAATCTGTAGTATGAGATGTACCAGGTTTATAGGCATAGTTAAAAGTTACAGGGAGTGGAAACATCCCATGTCGATGTTCCGATATCGTTTTCTGCTCAGGTGTCTGAATGCCTTTTCATGACTCGTTGAAGAGTGTGTGTATGTTGTAGTTGAAGCAAACAACACTAGGAAAGTCTGGATCAATGAACAATTTACCAATAATACCCTATTCACCAAATTAAATCTTGTGGCAGAATACGCATGGTTAACTAGCACACAATCACACACCAAACAAATAAATAATAAAAATAACACAAGGAATTTACTAGGTTTGTCAAAACCCATTTCCTCGGGGAAGAAGCGGAGAGAATTTTCCACTATAAAAGATAAGAAAACAGTACCAAGAGATCCCCTATTACAACCCTTTTAAATACCCAAGTGTTTGCCCAATCCAACTGCCCCAGTCTAACAATATAAGCAAATAATAGTGGTTCGAAGCTTTTCGGGGTTGGGTCACATTTCGGATCTGCCAGAATCCAGACACAAAGCTGTAGAAGCAACAGATATATTATATTTGCTTTCCTTCTTCTGCTTATTGTTCTCCTGTCTTATTGTTTATTCGGTGGGTGTTGGATTTTTCACTAGGTAATATGCAATTTTTTGTACAACCAAATGCTTGTGTATGTGGGGGAGAAGAAGAATGAGGCACCACTCTATTGATTGGATATTTGATTGCAACTGATATATCCATTGTATTTATATAATGGTCAAATTTTCAATGTTTCAATGAACAGAAATCGATACTGATAAGTTAAAAGGAACCAAGTAGTTGGTGTAACTCCAATGTGAAGCCTGTTCACAGTGTCCTCCCCATTATCATCTTTCAGTTACCCTTCTAGCAATGACAGAGAAGAGATTAAATGCATCATATGATTTCTTAGGGACTATTTCCATTTGGCTCCACCTTCCATATCCTATTCCCGCTATCTTATAGATTGATCGCACTCACAATTGTGTGCAGAACAACTAGTACAGAGTACTATATTACTGTTAATCAAGGAAGAGTGAAGTAGTTTTCGTGTGGCTGTTTTCTGAAGCCCATCTCATATCTCTTCCTAGCATGACTAAGATTTAGCTGTTGTATAGGATTTGCCTTTGCATTTTGTTGTAACTATCTTTAATATTTCCTTGCAGCCTGAGCCTGAATACAGGCCACAAATGTCAGAAGTGGTTCAAGACCTCATACAAATGATAAGAAGAGAATCTCCCAGTAGATCCGATGAAGAATGATTTGAACCAGACATGCATGCTATAGCAGATTCAGGCATGGAGTAACCAAGACTTCCAAGATTTTTTTACTTTTGCAAAGAACTTAACTCGAACTGAGATACTTGGAACCAAAGATTTTGAGGCTGTAGATGCAGTGCCCTGCCAGCAAAGCTTCATACCTCCAAGACAAAGGGCTGGCTAGAAACTGCACGATTTCCACCAACACAGCTCGGGTTATCCTGATTCTTAGTACAATTTTTTGCAATCATTCTAATGTTCATTTATAGTCCAAACAGAAACCACAATGCATAAAATGCACTAACCCCATTGTTTTGAGAGCTTATTTATGGTTGTTAATTGGTATATCTGAAGTTGTAAATCTTTATTTGTTCTGCTGTACATTTTCAATTGTTTGTACAAGAATATTCTTCACACTTTCAATATGGACTTTCTGTAGCACATTTTTCAGCCTTCAGTATCATCATGGATATATTTAATAATCACTTGAAG

>PGSC0003DMG400018752 LRR-RLKs

TTGAGAAGTTGTTCTGTCTGGTTCAAATCACGAATTTCTTGAGAAAAAAAAAATCCTTGACAAGATGGAGAAAAGCTATGTTCTTTTCCTTATCTTATTAGTATTTCAAGAATCAATTTTTTCCCTATATGCAAATTCCACAAATGAAACAGACATGGAAACTTTGTTATCTTTTAAACAACTCATTACAAATAACTCTTCTTTCCTTTCCAAGAATTGGACATCAAATACTTCATTTTGCTCTTGGTTTGGTATCACTTGTAATCCACAAAACCAAAGAGTCATAGCTTTAAATCTCCCAAACATGAATCTTCAAGGCAAAATTTCACCATCAATAGCTAACTTATCCTTTCTTACTAAACTTAACTTTAGTAACAACATTTTTCAAGGCACTCTTCCTTATGAACTTGGAAATTTGCCTCTTTTGGAAGTCATTGATGTTCATAACAATCAACTAGAAGGAACTCTTCATCCTTTTGTTGGTAACATTACAAAGTTAGAGAGATTGAGGTTAGATGGAAATAGATTTAGTGGTAAAATTCCTACTGAGATTGGGAATCTAAGCCAACTTGTTGAGTTAGATTTGTCACATAATCAGTTTTCAGGTTCAATCCCTGGATTGATCTTCAGCATGTCGTCGTTACGTGCTGTGTATTTGGTTAACAATAGTCTTTCTGGTTCATTCTTGGTTGATGAAATGAAGGGTGTGATGAATCTTGAAGTTATAGACCTTAGTTATAATCGAATCATCGGTGAGATTCCTTCAAGACTTTGTCAATTCAGTAAGCTAAGAACTCTTGTTCTGTCTTACAACAATTTAACAGGTCAAATTCCGAGAAATATTGGTTGTTTATCAAGGCTTGAGAGTTTTTATATAACTCAGAATGCAATAAGTGGTACAATTCCACTTTCATTGAGTAATATTTCCACATTGCAATTTCTTGGTTGTGTGAACAACCATATAAGTGGTACAATTCCTAAAGAATTAGGAAATTTACCGAACTTGAAGATGTTGGGATTTGATTTCAATAATCTTACTGGTGTAATACCTGAATCTATCTTCAATATATCTTCTTTGGAGTACATTGCTTTCAGTGATAACGATCTCTCAGGGCGAATTCCAACCACTCTTGGACTTAAACTTCCAAACCTTAAAGGAATTTTCTTGCCTGATAATCAACTTGAAGGAGAAATCCCAATGTATATCACAAATGCTTCTAACCTTATTGAGCTAGAGTTGTCCTATAACTTGTTCACAGGCATAGTTCCAAGTGATTTGGGAAATCTACGCCAGCTCGAGTTCTTGAATCTTGGTGGTAATCAGCTTACAAATGAACCAGGACAACAAGAGCTTGGATTTTTGAACTCTTTGGTGGATTGTAGGAGGCTGCAGTTCCTGATATTGGCTAACAACCCTTTGAATGGCGTCTTGCCTGATTCGGTTAGTAATCTTTCGTCTACTATTGAAATGTTTAACATAGAGAATGGTCAGATAAATGGACAAATTCCAAGAGGAGTAGGCAATATAAGCAGCATGTTATCGCTAGTCTTGAACGGTAACCAATTGACAGGAACCATTCCACCAGAGATTGGTGAGCTAAAGCAACTTCAGAGGCTTTATCTAAGCAAAAACAAGCTACAAGGTTCAATTCCAGAGGAAATATGTGACTTGGTAAACTTGGGAGATACATTTCTACACGAAAACGAGCTCTCTGGGGCTATTCCAAGTTGCATTGGAAAACTCACTAGGCTGCAAAGGCTTTCATTTGGCTTCAATAAGTTCACCTCAAGCCTACCTTCAAGTCTATGGGAAATGGATAGCCTGATTTTTCTAAACGTGACGCGAAATTCAATACAAGGGGAACTTCCTATTGACATTGGAAAATTGAAATCTATTGAAGGTATAGATTTTTCAAGCAACCAACTTTCCGGTGTGATTCCAAGTACATTTGGAGACCTTATAGGCCTGAGGTATCTTTCACTCTCAAACAACTCGTTACGGAGTGCAATTCCTTCATCATTTGGCAGTTTGTTAAGTTTGAAGTTCTTGGATTTATCTTCAAATGAGTTATCAGGAAACATCCCAAAGTCCTTAGAAAAACTTCAGTTCCTTAATGAAATCAATCTTTCCTACAACCATTTAGAAGGTGAAATACCTTCTAGTGGTGTGTTTGCAAACTCTTCTTCTCAATCTTTTGTTGGAAATAGAGGTTTATGTGGAAAACCTATATCAGAAGTTTCTCAATGTGCTACGAATAGTGCAACGAAACGATCAAAGTCTAGAAAACATGTGCTAGTAGTTGTCATTCCTGTGATTGCATCAATTCTCCTCATATTTGTTGTTCTGTTCGTATGGATTAAGCGAAGATCAAGAAGAACAAAACTTCAAGACCATGATCAAGAGTTAACAGAAATCACAACACACCAGTTGATCACATACCGTGAACTTCAACAAGCAACGGATAGTTTCAGTGGCTCTAACATGATTGGTTCAGGAGGTTCTGGTTCTGTTTACAAGGGCATATTGGCTAACGGAACAATGGTCGCGATAAAGGTTCTAAATATGCAAAATGAGGAAGGATGCAAGAGGTTTGATACCGAATGTGAAGTGATGAGAAGCATTAAACATAGAAATCTTGTCAAGGTGATCACTACATGTTCCAATCAATACGTTAGAGCCATCATTCTCGAGTATATGCCAAATGGAAGCCTTGAAAGTTGGTTGTATGATAAGGAACATCAAGTCTTAGACATGTTTCAAAGAGTTAGCATAATGCTTGATGTTGCAATGGCATTGGAATATCTTCACTATTGTTATGATACACCTATAGTCCATTGTGACTTGAAGCCTCAAAATGTTTTATTAGATGCTGACTTGGTGGCACATGTTGGTGATTTTGGCATTTCAAAGATTTTAGGTGAAAACAACTCCATGGCACAAACTAACACATTGGGCACTATTGGCTACATTGCACCAGGTAAACAATTCTTCTCTGACTACTCTACCTTATATAAGGCATATTTTTGGTCATTCTTCTCTCTGAAAAATAGAATATTGGTCAATTAAGTTGATTTCGGTGCTACTTACTATGCCTGAATCTAACGCTAGGCTAAGTCTTGAACCTTTGAGCTGACCGTTCACTTTCTTAGTAGCAAGAGGGCTTCTCTTTGCCCCTTTATTGGTTGTATCAACTAAACTGCCCTTCCATATAGATCGTAGCAACATGAAACTATTATTAATAGATTCCTCGCTTCCTATTAATCTTGAAAAGTTCTAGCTTAGTTCTTTTTTTTTTGTTAGAATTATCTTTTTAATTAGACCGGACCCACTTTTATACTTATCCTACAAGTGCTATTGGGGACCCACTCAAAAAAAAAAAACTTGAACTTAGAAAAGTCCACCCCTTAATAGATTAGGGTAGAATCAACTGGATTTAACTTATATCTACTGATAGTATAGAATATATGTTTAATTCGTTATAACATATTAATTTCCTTTCCTTTTTGTGTTTTTAGAATATGGCTCAGAGGGAATAGTGTCCACGAGTGGTGATGTTTATAGTTATGGTATAATGTTAATAGAGATATTGACAAGAAGAAGACCAACAAATGAGTTTTTCAATGAAAACATGAACTTGAGGCAATGGGTCAGTGAATCATTTCCAAGTTCTTTGAAGACAGTTGTTGATGAAAATATTTTATTTGGAGAAAATGAGATTTTCATATTTAGTATGATGGAGTTGGCTTTGGAATGCACAAAAGAAAGACAAGAAGAGAGGGTAAATATGAAAGATGTAGTGAATAGACTTGGTAAAATTAAAGAGGACCTCTTCTTGGAAACAAAGAAGAAAAAATCTTAGGAGTAAAATTTTGTTTATTTTTAGTTTATGGTTTAGTTATTCTTT

>PGSC0003DMG400018811 LRR-RLKs

ATAAAAGGGGCACTAGTAATTTTAGTAATTGAAAAGCAAGTGTATAGAGAGAGATAATGAGAGAGAAAGAGTAAGTACACTACTACTGCTACTATCCCATATAGCTGTAATGTTGCAGGTCTGATTTTTGCAGTTGCAGACCCCCTTCTTGGCACAAGCTCTTTTAACTTTTATCTTCTCAAATAATTCTCTCTCTCTCTCTCTCTCTTTTTTCTCTTTTTACATTGTGAGGAAACTGAATACCCATTGTATGTATTAGTGTGAGGCCTATCTGCCACAAGGATGTGATGGAACACTATGCTTCCTCTGCTAAAACCCCACAACCCCAAAACTCTTTTTCACTTCACATTTAATCACAATTCCTCAGTGAAATTATTCTGTTGCTCTCTCTAATTTCAATTTCAATGTCGGTAAGTCCAAGAACTGGTTTTTCAATTCAAAGGAGCTGAGTTAGTGCAAACACTTGAGGTTTTGAGTTTTGACAGAGACTTGAGTCTCAGAGAAACTACCATGACATCATTTTTACTTCAAAGATGTAATCTTTTCTTTGAGGTTCTTCTTCTTTTGGGGTTCTTGAATTTCTTCAGCTTTGGTTCTGTGGTGTCTGATGATGGTGAGTGAGTAGAGTAGAGTAGTATAACATTCTGCTTCTTATGTTCTAGTTTAATGTTTTGTTTAAATGTTGAAAAGACAAAGTGTCCTTTTTTTAATCATTTTTTAAATGGTGTTTTTGATTAATCCCACGTTTTGTAGTTGTTATTTGTTAAAGGTTTATTTTTTTGTCTCATTATTATAATAATAATTGGAAAATAGGTTCTGCATTGTTGGAGATTAAGAAGTCAATTAGGGACGTGGAGAATGTGTTATATGACTGGACTGATTCTCCTTCATCTGATTACTGTGCCTGGAGAGGTGTTACCTGTGATAATGTCACCTTCAATGTTGTTCAACTGTAAGCCATAACTCAAAAAACACTCTTGTTACATTATTATCATTTGGGATTCTTTAGTATAAAGTTGTAATCTTTTGACATTATCTTGTAGTAATCTTTCGAGTTTAAATCTTGATGGGGAGTTGTCTCCTGCAATTGGACAGCTCAAAGGCCTTATATCTATGTAATATCTTCTCCCATTATCCCACATTTTCCCTTTTTGTTTGATCTTTTGACTTAGTGTACATTTAGATTATGTTTGTTAATTTATTTATTTTTCGTAGTGATGTAAGGGGAAATCGCCTTTCTGGCCAGATACCAGATGAGATTGGTGACTGTTCAGCACTGAAAAACTTGTAAGTATAAAATGCTTCTGATTCTTGTGTTATTATTTGGAAAATAAGTAACCATTTTTTTCCCTTAGGGACCTATCCTTCAATGAGCTTTATGGTGATATTCCCTTTTCCATATCAAAACTTAAGCAACTGGAATATCTGTAAGTTTTGATACTCTTCTTCTTCTAGGTGTTGTATCATTTGCTTTCCAAGATTGTCAATTGATCATGCTTGTCTTCTTCAACTTAGGATTTTAAAGAATAATCAGTTGATTGGACCAATTCCATCTACATTGTCACAGATCCCCAACTTGAAGGTCTTGTAAGTATATTCTCTTCTGCTTTGTCATGATATTGGTAGATTATGAATAATTTCAATTTAATCCAAGAACTTCCTCCAGGGACCTAGCTCAAAATAGGTTAAGTGGAGAAATTCCTAGGCTGATATACTGGAATGAAGTCCTGCAGTATTTGTGAGTGTTTTAATTCAGTGCTCCTCTTCTCTTGGTTTCTAGAATACCTTAGGAGACTACCATTTTGTATATGGATGATCATATCTGTTGTATGTTTTTCATTGTATAGGGGACTGCGTGGCAACAACTTGGGTGGATCACTTTCTCCTGATATGTGTCAGCTCACCGGCCTGTGGTACTTGTAAGTTTGTAATCTTTATGCTCTTCAGATCTTACTATAGCTCCTTTAGCTGATGACATTAACCATCGTTCATTGTTTTGTGCAGTGATGTTCGGAACAATAGTTTGACTGGTTCCATTCCTCAAAATATTGGCAACTGTACTGCTTTCCAGGTTCTGTAAGTATCTATAACAAATGAATGAAGTTGGACTATTTTCTAAATGTTTGGTTGGCATAACACCTTGTTTTGTGCTATCAGAGATTTGTCTTATAATGACTTGACTGGAGAGATTCCTTTCAACATTGGTTTCCTGCAAGTAGCGACCTTGTAAGTTTATGCTGCTTCTCTTTATCACAAAGCCATTCATTATATGGTTATAAGTTTGAAGTGTACTTTCATCATTCCAGGTCTTTGCAAGGTAATCGCCTTTCAGGGCAGATCCCTTCTGTCATTGGATTGATGCAAGCTCTTGCAGTTTTGTGAGTTTTTGTGTCTTGTTATCTCAATCCAATGCTACTGAATCATGAACGAGATTTCTTGGAAACCAAAAAATGCATCTGTTATTTAAGTTTTTCTGACCCTTTTACTGACAGGGACTTGAGCTGCAATATGTTGAGTGGAACAATTCCTTCAATTCTTGGGAATTTGACTTACACAGAAAAATTGTTAGTACTATAATAATTTAATTTAAGAGCAATTTGGATCATTTTGTGCTTCCTAAATTGTATACTGGATCAATTACTGTAAGTTCGCATTGTATTGCAGGTATCTACATGGGAACAAGCTATCTGGTTCCATTCCTCCAGAGCTGGGAAATATGACAAAGCTTCACTACTTGTATGAATGCTTTCTATTAATCATTTTTTGTTGGCTTTGTTTTGTTCTTCCTGTTCAAACCCTTTTAAATGAATGCAAGCATTTGTTTGATTAGTGAGCTTTTAGGCAACCACTGATTTGATTGATACAAAGCTGTTTTGAGAATTTTTTATAAGAGACTTTCCTCAATCTCAAGGCTCAAAGATGGTAATTTGCAGAGAATTGAATGATAACCAACTTACTGGACGCATACCACCAGAACTTGGAAAGCTGACGGAGTTGTTTGACTTGTAAATCCCGTTTCTCTTCATCTTCTACTTTGGACTTGTTAACATCATTATTTATTTACTTATGTGGTATGTTTCAGAAATGTTGCAAACAACCACCTTGATGGGCCCATACCTTCCAATCTTAGCTCATGTACCAATTTGAATAGTCTGTGAGTGTTTTTCATATCCGATGTGTTTCAATTATGTATGATCATGCTTGTTTGGGAGTTATTGACACCTAATTTGGTTGCAGCAACGTTCATGGAAACAAATTGAATGGTACAATTCCACCTGCTTTTCAAAAGCTGGAAAGTATGACCTATCTGTAAGTTCTTACTTTTTGAATTTTTCTTTTAGAGAAATATGTGCAAGGTTATCTAGTTATCATTCTACCCATTCGCGCTGTTAAGGAAGATTGTAAACTTACTGTGCCAATTTTATATTTAAATTTTACCAACGTTGTATTATTTTGCACAGAATTCATTTTATTTATCTTGCTCTCAAATCGTTACTGGCTTATCCTTGCATGAAGAATCATTTCTATTGTCGAGAGCTTTCAGCTATGTGTGTCATATAATTTTTTTAATTATTAACTAGTTTATTTAGAGCCTCTGCATCATCTAGTTATGCGAATTTCAATTGCTTATTCTGCATACTCTGAGCATTAACTGCATCTTTTTTTTTGTTAATTGCTCAAGTAATCTCTCCTCCAACAACCTCAAAGGCCCAATTCCAATTGAGCTTTCTCGTATTGGGAATGTAGATACACTGTAAGTGCAAACTTTCTCATCCACTTTCATTTCTCTCATTGCAATTATGGTTGCGGGAAAAGCACGTTTTGACAGTTTAAGAACTCTTCAACATTTTTTTGGCTTAGGGACTTATCAAACAACAGGATCAGTGGTCGTATACCTTTGTCCCTCGGTGATTTGGAACATCTTCTTAAACTGTGAGCATAACCGTCAGGTTGTTATGTTATCATCATATATCTGTTGTACTTACATTCCTTTTGTCAATGCTGTAGGAACTTAAGCAAGAACGAAATAAATGGAAACTTGCCAGCTGAATTTGGCAATTTAAGGAGCATCATGGAGATGTATGGAACCTTGCTAAATTTAGTTACTTCAAATTTATGGTTTGCTTGATTTTCAGCTTTTTGACCGCACTCTTAATTGTAGTGATCTGTCAAGCAATCACCTCTCTGGTCCCTTGCCTCAGGAACTTGGTCAGCTTCCAAATCTGTACTTGCTGTAAGTACTTTAGATTTACTTCGAGACTCTCATCCTCTTAGCCATTGGTAATAATCTGTACAGTGAATAAGTATGAACTTCTAAACTGGGAAAGTTAGATTTTAAATTATTTTGGATGCCATTTTCAAGAAAGTAGAGATGAAGTGGTGTTGTTGCGATTGGTTTATATTATCTGGCTACATATGTTAATTACTTTGGTGTTGTCTCAGTTTTTCTTTATATTGCATTATTGTACGGGGCTCATAGGGCATCATTTCTCTATCTTCTTCTTTTCCTTGAGGCCTTATTCTACAAAATAATGAACAGGAAACTGGAAAACAACAATTTATCAGGCGATGTGATGTCCTTAGCCAGTTGTCTCAGTCTAAATGTCCTGTGAGTTTTCAAGCTAGTAGTATTACACCAGTACAAATGTTTTGTAAATCTAATCAACCGTTTTTTGGCAGAAATGTCTCATACAATAATCTGGGAGGGAATATTCCAACAGGCAATAATTTCTCTAGATTTTCACCAGACAGGTAAGTGGAGCGATGAAGATTCCATTCATTGATCTTCCTCGTCATATGCCAAGTTCACACAAGTCACAAGCATTTATTGGTTTGTTTTTAATTCTTTGCTTCTAATTTTTTCCTTTTGCTATGTTTCCAATAAAGCTTCATAGGAAATCCAGATCTGTGTGGGTATTGGCTCACTTCTCCCTGTCATGCATCTCATCCAGCAGAGCGAGGTATGATCAAACTGTAGCCATCATTTGACCTTTATTCTATTGCATTCTTGAAGTTTCATTTCACTTTAGACACTTGCAACATTTATTAAGCGCGATGGACAGATATTGATTAATGAGGAATTACCCCTTGGTTGAGCAAATTTAATTCTGTGTTAGTCTGGTAGTAGGGTGCACGATCACGGTTTCCTCTGTTCACAATCCTTGATTGTAACACTTGGATGTGTGCAGTAACACTTGGATGTGTGCACATATTAAATTAACATGAAATAATATTCATTTGCTGGAGTTACATTGACGTAAAGATGCCTTAGCTGTCAAATGAAACTGCATTAGTTTTATCCCCATCATCAGTACATTAAGGGGTCATTTGGTAGAGTGTATTAGTAAAAATAATGCATGCATTAATTACATGTGTTAGTAATCCCTTGTTTGATGCATTTTTTCATGCTATGTATAACTAATGCTATACACTCTATTGTGTATTGAGGAGTGTATTACTAATACCATGGATTTCTAGGTATTAGTAATGCAAAGGATTTAATACATGCATTAATATGGTTAAAGACCCAATTACCCCTCAAAAGTACTTCTACATCTTTTTCACCATAATTGTGAAGGATACTTTTGTAAATAAATATTTTATACAATGCATACTATTTTCATACACCAAACCAAACAATGCATAAGAAATAATATATGCATAACTAATACAAGTATAATTAATACATGCATTACTAATGCTTGCATTACTAATACACTCTATTGGGTATTATTTTTATACACTCTACCAAATGACCCCTAAGTGCATAAATATTTTAACAGTTGTCTAATGAAAAAAGATGAATTCATTGGACAATTGCAGTTTCAATTTCTAAAGCTGCTATACTTGGTATTGCTCTGGGTGGCTTGGTGATTCTTCTGATGATACTAGTAGCAGCATGCCGGCCACAGAATCCTGCACCTTTCATGGAAGGATCTATTGATAAACCAGGTACAATATTTTCTGGACGGTTGGATAGTGTTTGGAGACGTTCATGTCTTAAGGACAGTCATCAGAGTTAATTGAAGTTGCCATGTATTGATTATTTAACGTTTTTGATGAACAGTTTATTACTCATCTCCAAAGCTTGTGATCCTTCATATGAACATGGCACTTCATGTTTACGAGGACATTATGAGGATGACTGAGAACTTGAGTGAGAAGTATATAATTGGTTGTGGAGCATCAAGTACTGTATATAAATGTGTTTTGAAAAATTGCAAACCTGTAGCTATCAAGAAATTGTACTCTCACAACCCGCAATACTTGAAGGAGTTTGAGACTGAACTTGAGACAGTTGGGAGTATTAAGCATCGTAATCTTGTCTGCCTCCAAGGATATTCTCTTTCTCCATCTGGCCATCTTCTTTTCTATGACTACATGGAAAATGGTAGCCTTTGGGATTTGCTTCATGGTTAGTAAATCCAAAATGGTTTAGGTGATTTGGTTCATTGAACTTGTGTTTTAAGCATTTCGTAATAATCCCTCTTATATCTTTTTTGCAGGTCCTACAACAAAGAAGAAAAAACTTGATTGGGTTACTCGCCTTCGAATTGCATTGGGATCAGCTCAAGGGCTTGCTTATCTTCACCATGATTGTAGCCCTCGAATTATCCACCGTGATGTTAAATCATCAAATATCTTGTTGGACAAAGACTTTGAGGCTCATCTGACTGATTTTGGCATTGCCAAAAGCTTATGCATATCAAAGACCTATACGTCCACATACATTATGGGAACCATTGGTTACATTGATCCAGAGTATGCTCGCACTTCTCGCTTGACAGAGAAGTCTGATGTTTACAGCTATGGAATTGTTCTATTGGAATTGCTCACTGGAAGGAAAGCTGTAGATAATGAATCTAATCTACACCATATGGTATGCTCTTGTAATTTAGTTAATATGAACTTGTGCTATGATGTATCTTCATATAATTATATAAAGATTCAATTGATCTCAACAGTTTTGCATATGTTACAGATTCTAACTAAGGCAGCAAACAATGCTGTAATGGAAACAGTGGATCCTGAGATCACAGGCACATGCAAAGATCTTGCAGATGTGAAGAAGGTTTTTCAGCTTGCCCTTCTATGTTCCAAAAGACAGCCTGCTGAGAGACCAACAATGCATGAAGTGGCAAGAGTACTTGAAAGCCTAATACCCGTCGCTGAAACGAAACAGCCAAATCCAACCCCCTCACTTGCATTACTTCCATCTGCTAAGGTACCTTGTTACATGGATGAATATGTCAACCTCAAGACACCCCATCTAGTGAATTGTTCATCCATGAGCACTTCAGATGCCCAACTTTTCCTCAAGTTTGGAGAGGTCATATCCCAGAATAGTGGCTGAAAATAACATGAGAGTAGATTTCTTGAGATTTTGTGTAAAAATGTAGTGCCATTATCATATTATTATTGTAGGTAGTTGTTGTAAGATGATGCATGCAATAGTGGTCCAGTCTACTTTTTCCACTACATAGGTCTAGTGTGTGTAAAAATATTTCACTTTTTACCATGATGAAATTGGAAGAGGTAGCACTTAGTAGAGTATTGTAATATTGGTTTTTGGGACCACTGATGCTGAGTATGGACTATACTGTCTGTAAAATTTTTGGCACACACTTTGAGGTGGCCTTAGCACTGCCCCTCTGTGTATAGCAATAATGGATTGCATTGTGGACCTAAGTTTGACAGTCTGCCGCTCTCTCTTGCAGCTGCAGTACATTAGTGCATTTATTTGGTGGACTACTCAGAAACACGAAAAATAATTTTGACATTTTAATTATTACTCTACTTCCTCAGATTATTAAAT

>PGSC0003DMG400018849 LRR-RLKs

ACTTAACCCACTAAAATCATCATGAAACACACTCTATTTTTCCAGCACTCCTAAAAAGCATTTTTCTCTCAAATTTACCTCTCTTTACATACCCCATTTCTCAACCATTTCCACCATAAACAACATCACCATGAGACACTTGTTTCCTTTCTTTTTTACTACAGTTTTCTTCTTCCTAACAAGAATTTCACTTGTTTTCTCCATTACCAATCTACCTCTACAACTTATATCTCTTTTGACTTTAAAATCTTCTTTGCATGATCACCACAATACCTTCAATGACTGGGATCCCACCCTCGCCTTCGCTAGACCGGGTTCTCACATTTGGTGTTCCTGGTCTGGTATCAAATGTGACAAGAAAACTAACCAAATCACCTCTCTTGATCTGTCGAAGCGCAATCTTTCTGGTACAATTCCAGAAGATATCAGAAACTTGGTACACCTTCATCACTTGAATTTGAGTGGAAATGCTTTGGAGGGTCCTCTTCAAACAGTCATTTTTGAATTGCCTTTTCTCAAGACACTTGATATTAGTCACAACTTGTTTAATTCAACTTTTCCCTCTGGTGTATCAAGGCTCAAGTCACTAACATACCTAAATGCTTATAGTAATGACTTCGTAGGCCCTTTGCCTGAAGAAGTTGCTCAAATCCCAAATCTTGAGTACCTTAACTTTGGGGGAAACTATTTCAAGGGTGTGATTCCAAAAAGTTATGGTGGCTTAGCAAAACTGAAGTTTTTGCACTTGGCTGGAAATTTATTGAATGGTCCTGTTTTGCCTGAGTTGGGTTTTTTGAAGCAGCTTGAACATGTAGAAATTGGTTATCAGAATTTCACTGGAGTTGTCCCTGCTGAATTTTCTTCTTTGTCAAATTTGAACTATCTTGATATCTCATTAGCCAATCTTTCTGGTAATCTTCCTGTTGGACTAGGCAACTTGACAAACCTTGAAACTCTGCTTCTTTTCAAGAACCATTTTTATGGTACAATTCCATTGTTTTTTGCCAGACTGACATCCTTGAAATCATTGGATTTGTCTGACAACCATCTCTCCGGAACAATCCCCGAAGGGTTTTCAGGATTCAAGGAGCTTACGGTCTTGAATTTGATGAACAATAACTTAACAGGTGAAATCCCACAAGGCATTGGTGAGCTTCCTAATCTTGAATTATTGGCTCTATGGAACAACTCACTCACTGGGATTTTACCACAGAAGTTGGGTTCAAATGCAAAGCTACAAAAACTTGATGTCTCTTCAAATTATCTCTCAGGTCCCATTCCACCAAATCTTTGTCTCAGCAATAGCTTAGTGAAACTTATCCTGTTTTCAAATCAGTTTATTGGTGAGATCCCTTCTTCCTTAGCAAATTGCACTGCATTATTCAGGTTTAGAATTCAAAACAACAGACTCAACGGGTCAATACCATTAGGCTTCGGTATTTTGCCCAATTTAGCTTACTGGGACTTGAGCAAAAACAATTTCACTGGTCCAATTCCAGAAGATTTAGGGAATACTGCGACATTGGCGTATTTGAACATATCGGAGAATCCATTCAACAGTGAATTACCGGAGAGTATCTGGAGTTCACCATCTCTACAGATCTTTTCAGCTAGTTACAGTGGCCTTGTCGGGAAAATCCCCAATTTCAAAGGGTGTAAAGCTTTCTACAGAATCGAACTGGAAGGAAACAATCTCACAGGAAGCATTCCATGGGATATTGAACATTGTGAGAAGCTCATTTGCATGAATTTTCGACGAAATTCGCTAACGGGAATCATTCCTTGGGAAATTTCTGCAATTCCTTCCTTAACAGAGGTTGATTTGTCTCACAATTTTCTCACTGGAACAATTCCTTCGAATTTCGCCAATTCCACTACCATAGAAAACTTCAACGTATCCTACAATCAGCTCACTGGTCCAGTTCCTTCTTCAGGCTCAATTTTCTCCAGCCTACATTCCTCTTCCTTCATCGGAAACGAAGGCCTCTGTGGTACTGTCATTCAAAAGCCTTGCGGGACTGACGGACTCGCAGCCGGAGCAGCGGAAATCAAGCCACAGCCGAAAAAAACAGCTGGCGCTATTGTCTGGATCATGGCAGCAGCATTCGGTATTGGTTTATTCGTCCTCATCGCCGGTAGCCGCTGTTTCCACGCGAAATACAGTCAGCGATTTTCAGTAGAGAGAGAAGTTGGGCCATGGAAATTAACCGCATTTCAAAGGTTAAACTTCACGGCGGATGATGTACTTGAAAGTTTGACAATGACAGATAAGATCCTGGGAATGGGTTCAACAGGGACAGTGTACAAGGCGGAAATGCCAGGCGGAGAGACCATAGCAGTAAAGAAACTATGGGGAAAACAGAAGGAAACGATCAGGAAAAGGCGAGGCGTTTTAGCTGAGGTGGATGTGTTAGGCAATGTGAGGCACAGGAACATCGTGAGGTTGTTCGGCTGTTGCAGCAACAACGAGTGCACCATGTTGCTGTATGAGTACATGCCTAATGGTAGCCTGGATGATCTGTTGCATGGTAAGAATAAGGATGCTAATTTGGTGGCTGATTGGCTCACTAGGTACAAAATTGCTCTTGGGGTCGCTCATGGAATTTGTTATCTTCATCATGATTGTGATCCCGTCATTGTTCATCGTGATCTTAAGCCTAGTAATATTCTTCTGGACGGTGATCTGGAGGCTAGAGTGGCTGATTTTGGTGTTGCTAAGTTGATTCAGTGTGATGAATCTATGTCCGTGATTGCTGGATCTTATGGCTACATCGCACCAGGTAAGTTAGTACTCCCTCCGTTTCAACTTGACATGAAATTTAACAAAGTAATATTCAACTTTTTATCGTGTTCTTTAATTTAATTATTAATGGCAGGAATATTTGTCTGTTTGTTTGATGGAATTACATGATTGTTTTGGAGTTCAAGTTTTTTTCTTGTGCGGCTTATTGCTATCCAAAATTATTTTGTTCAGATCAGTTTAGTTCGTTCTGTAGGGTGACGAAAAAGAGTTCACTTGCGTGCTAAAACGTGGTGTCATGTCAATAGCATGATGTATATTGTTTTTTTTTTCGTCCACTATCGTCATTAAATCTTCTTTGCCCACACAATTGTCAAATTCCTCAACTGCTTCATACTAGTGGAATCTTCTTTTAATCATAACCAGATCTTGGAGGCACTAATTCATTTTTTGAATTTGAGCGCCACAATATAATATAAATATCTAAATGTTGAAACTTTTATTAGGAAGTTAAAATAATATTCTTATGAAGATTCTAATATTAGTGTAGTGAAACAGGTAATATAATTTTTTTTCTTAATAAAGGGTATCAAGTATTTCTTTTCTCATACTTCTTTATGACAATTGGTGCAAATATTTATTCTCTTTGTTTTTGTCCATGATTTTGTGGTGCTTAAAGAGGGCATGGCGCACTAGTTGCGCGTGAATAACTCCATTAGCCTTTAAGCTTTTATTTTACACTTTTTGTTATTCTTTTGGACCCACCAAGATTTTGTCTTAGTTACTTATCTGTGATTAACCCTTTGATTTATTAGTATATAATATAGTACTAGTACTACATTCCTTGACTTTAATGCTTAAAGTCAACCATTTTTGGGCACTAGTAGACAGAGTGTTATGATAACAAATTATTTTTATATAGTTATTGTACTAAATATGTTTGTTGATACAGAATATGCATACACATTGCAAGTTGATGAGAAGAGTGACATCTACAGCTACGGGGTTGTGTTACTTGAAATTTTATCAGGGAAAAGATCGGTGGAACCAGAATTTGGCGATGGAAATAGCATTGTGGATTGGGTTAAGTCGAAGATCAAGACTAAAAATGGCATAAATGATGTGTTGGATAAAAATGCTGGCGCTTCATGTCTTTCAGTTAGGGAGGAAATGATGTTGTTGCTGAGGGTGGCATTGCTGTGTACTAGTCGTAACCCGGCTGACAGGCCATCGATGAGGGATGTTATATCTATGTTGCAAGAGGCTAAGCCCAAGAGGAAGTTGCCTGAAACTGGTGATAATGCAATTGCTGCTATTCCTTTGGCTCAAAAAGCTAATGTGGAGTGTTGATATTATAGGAGGAGATTTGGGGTTGTTTTGATTTTACAACTTACTGATGAATTGTGGGGGAAATTGGTATAGGAGAAGTTTAAATAGTTTTATTTGCTTCTTAAATTTTTGCATAGTAGGAAATTCCCGAGTGTTGTCATTCCAATGTTGTGATGAAATGGGTATCTTTTCCAACAAATTAATCTTCGATAGCATACTTGTCTAGAC

>PGSC0003DMG400019051 LRR-RLKs

ATGATCGGTCTTACATCCCTAAGCCTTGACAGAAACAACTTGACAGGGAGTATTTCTTCTGAGATTGGTAAGCTTAAACAACTCCAAGGGTTGTATCTAGATAACAATAAATTGCAGGGACTTATTACAGAGGCCGTATGCCATTTATCTAATTTGGTTCAATTATATCTGAATGATAATGACCTCTTTGGATCAATTCTAGAATGCTTAGGAAATCTAAGCATTCTACTACATCTTTATTTGGATTCTAATAAATTTTCATCAAAATTTCCGCTGAGCCTTGGGAAGATGAGTCGTCTTATCTATCTAGACGTGTCACAGAATTCAACAGAGGGAGAACTTCCATCAGATATTGGAGGACTAAAAGCCATTGTAGAACTATATCTTTACAATAACCACTTTTCAGGTGTGATACCAACCAGATTCGGGGAACTCCAAAACCTGCAGTATCTTGACCTATCCAACAATTCATTTTTTGGCCAAATTCCATTATCCTTTGCCAACTTGATAAGCTTGGAATTCTTGAATTTGTCTTTAAATGCATTGTCAGGTAGTATTCCTATGTCTTTGGAAAAACACTCTTAAAAGCATTAATGTTTCATTTAATAATTTAGAAGGTAAAATACCGAGTGGTGGCGTGTTTGCAAATTCCACTTTGCAATCATTTCTAGGGAACAAAGGTCTTTGCGGAATGCACATATTGGAGATTCCTTCTTGTGCTATCACTAATCCTGGAAAACAATCAAAGTATAAGGAGGTTGTGCTCAAATTTGTTACTCCAGTGGTTATTGCATCCTTTCTTATATTCTTGTTGGTTTCAATTTGGATAATGAAACGACAGAAGAAAGGAAAGTCTAAAGATGTTGAAAAGGTACCAGAGATCGGGACTCATCAATTAGTTTCTTATCACGAGATTCAACGAGCAACAAATAATTTTGATGAATCCAATTTAATTGGAGAGGGAAGCTCTGGCTCTGTGTACAAAGGCACATTGTTTGGTGGAACTGCAGTGGCCATTAAGGTTCTATATTTGGAAAATGAGCAAGTATGCAAGAGGTTTGATACTGAATGTGAAGTGATGAGAAATGTTAGACACAGAAATCATGTTCCAGTGATTACTACATGTTCTAGTGACTTTATAAGAGCCTTTGTTCTGCAATTTATGCCCAATGGAAGTCTAGAAAATTGGCTGTACAAAGAAGATCGCCACTTGAACCTTCATCAAAGGGTCACTGTAATGCTTGATGCAGCTATGGCAGTTGAATATCTACATCATGGTCATGTCACTCCAATAGTTCATTGCGACCTAAAGCTAGCCAACATTCTTTTGGATGAAGATATGGTGGCTCATGTTGGTGATTTTGGAATCTCTAAAATTTTAGCCATAAGCAAGTTCATGGCTTATACCGAGACATTGGGCACTCTTGGATACATTGCACCAGATATAAAAAAATCTACTCTCTTTGATTTTCTCTTATATCATAATTAA

>PGSC0003DMG400019415 LRR-RLKs

ATGGCCTATTATCTTGACAAAATCTTTCTTCTCTTATTTCTCTTCACACTTAGTACTCCCTCAAAATCAGCTAATATGACAGATATTGAAGCACTCTTTGCTATCAAGAATGAAATAATTGATCCCTTTGAATCATTAAGTTCATGGAATGAGTCTGTTCCTTTATGTCAATGGCGTGGCGTTGTATGTGGAACGCGAAATCAACGAGTCATTGAACTAAACCTTCTTGACCATAAGTTAGCCGGTGTGTTATCACCCTTTGTTGGAAATTTGAGCTTCCTTATACGACTAGACGTCGGAAACAACACCATTTCTGGCAATATCCCTCCTGAATTAGGCCGTTTGACAAGATTACGCCATCTATATTTGCAAAACAATTCATTTTATGGTGAAATTCCTGTCAATCTATCGTTTTGTAGTAACCTCGTTGAGTTATGGGCGTACAAGAACAATCTTGTAGGTGTTCTTCCAAGGGAGCTTGGTTTACTTACCAAACTCAAATATTTTGATTTCAGTTACAACAAGCTCATTGGTGAAATCCCGAAATCTTATGGTAATTTTTCAAGTCTATTGGAAATGTACCTTTTGGATAATGATCTCGAGGGTAAAATCCCCGATGAGTTTGGGAAACTGAAGAGTTTGGAAATATTTGACGCGGACTTTAATAGGTTATCTGGTAGAATTCCGAGCTCATTCTACAATCTTACCTCCTTGAAAGTCATTGATGTGTCATTTAACCAACTCGAAGGTACTCTGCCTCGGGATTTAGGGATCAACCTTCCTAACCTTGAGTGTCTCAGCATTAGTGAGAATCAATTCACTGGATCGCTTCCTTATTCACTTTTCAACCTCACAGGACTCGCGTACCTTTATGTTGGAAAAAATGACTTAGGAGGACGTGTTCCCAGGTTCGATAAGTTGCATAATTTGTATATTTTAAGCATGGAAAATAACCAATTTGATAATGACTTGACCCTCATGTCCTCCTTAACAAATGCAACTAACCTCAACCGTCTGTATCTGCAAAACAACGCGTTTGGAGGGACATTTCATGAGTTTTTCAGCAACCTTTCAAGTAATCTGGCGTTTGTAGACTAGTCAAGCAATCAGTTTTCAGGAAATATCCCGATAGACATTGGGAAATTCAACAACTTGGAAGAGCTTAGGCTGAGCGGAAACCAACTCACAGGTATCATCCCTTCGACTATTGGGAAGCTTTATAAGCTACGATATTTACACCTGTCCCAAAACAGAATTTCGGGTAGTATACCCTTTTCCATTGCAAATCTGAGCCTTTTAACTGAGTTAAATCTTGATCATAACAATCTTAGAGGTAGAATACCTTCAAGTGTTGCAAAATGTCAAAGTTTGTTGTATCTTAACCTGAACCAAAACAATTTAATCGGTTCGATACCTAAAGAAATTTGTCTCATATCTTCCTTAGTTGTTCTAAACCTTTCTAGTAACAACTTGACTGGCGCCCTTCCCAAAGAAATCGAAAGCCTGAAAAATCTCTTATCATTAGATGTCTCTTTGAATAATCTCAATGGTGAAATTCCTAGTACACTTGGGAGTTGTGTGACAATTGTTAACTTAACAATGAAAAGGAATTTATTCCGCGGGATTATTCCTTCCACTTTCAGTTCACTAAAGAGTCTTCAAGTTTTAGATCTTTCTCAGAATAAGCTATCAGGTATGGTTCCAAAGTACTTGGAAGGATTGGCCTTACATTTCTTAAACTTATCGTTTAATGATTTTGAAGGTGGCCTTCCACAACGAGGCATCTTTGAGAACGCGAGTGTAGATTCTTTCGTTGGAAATTCTAGAATCTGTGGTGGTGTACCTGGCTTGAAACTTCCCAATTGCAACTTCAGTCATTCAAAGAAAATGAACTTTAAGCTTGTTACTTTAGTAATATTAGGAATTTTGGGGCTAGTTGTTATGGTTTGTGCTTTTTTCTTCTATCGATTCGGAAGGTCCAAAAGAACAATTCCTTCATTAGATAGCAATCTAAACCAACTCATAGCTATGTCCTATCAAAGCATTCTCAAAGTTACCAACGGGTTTTCTACGAGCAACTTGATTGGTGTTGGGAGTTATGGATATGTTTATAAGGGAATTTTAGAAACGGATGGTAAGCACGTTGCTATAAAAGTATTGAATCTTTTACAATATGGAGCTATCAAAAGTTTCATAGCAGAATGTGAGGCCTTGAGAAATGTTAGACATCGGAATTTAGTTAAGCTACTTACTGCATGTTCTTGTGTTGATTATCGTGGCAACGAATTCAAGGCATTGGTTTACGAGTTTATGGCCAATGGAAGTTTGGAGGACTGGTTGCATCCAGATAATTCAAGACCAAATGTTCAACCAAGAAGATTAGGATTTCTACAAAGGATGATTATTGCTATTGATGTTGCTAGTGCAATTCATTATCTTCATAATGATTGTCAAATATCAATAGTTCATTGTGATCTCAAACCAAGCAATATTCTCCTTGACAATGAATTGGTTGCCCATGTTGGGGATTTTGGATTGGCAAGATTCCTACATCTTACTGATGAAGCCACTTGTAGAATTCAGACAAGCTCTTCAACTTTTAAAGGATCTATTGGCTACATAGCTCCAGGTAAATTTCACTCTATATAAATGTCTCTAAAGACTTTAGTGACTCTGGATGCAATGACATTAACTCACTTGCCGCTATATTTTCATTGCAGAGTATGGTATGGGAAGCGAGGCATCAATGCAAGGTGATGTGTATAGTTTTGGTATCGTCTTACTTGAAATGTTAACCGGGAAAAGACCAACAGATGACATGTTTGGCGGCGATTTAAGCCTTCATGATTTTGTTAGGAATGCCATGCCTGATGGAGCACTTGAGATAGTAGATCCACTTCTTATTTTGGAAGAGGAAGAGATAAGCAGAGAGAGGTCTCAAATTCCAAGATTCATCAGAAGGCAAAAAATGGTGGAGGGTATAATCTCCTTATTTGGAGTTGGAATTGTTTGTTCAATGTATGATTCAAGTAAAAGAAAAAACATGAAGGAGGTTGTTAGAGAGTTGTGTTCTATAAGAGATTCTCTAGTTGGTTGTACAATATGA

>PGSC0003DMG400019698 LRR-RLKs

TCTTCACATTCTACTACTCCTTTCCCTTTCTCTATTTGTCTGTTTAAAACTTCTACTAATCCTATCTTTCTTTTCAGAAAAATATGGAACCCCTTCAAGAACTCAAGCTATAGATTCAAGAAAATCTCCATTTAAGCTATAAAGTTTCAATCTTTGAAGATGAAAGCTCACAAAACTGTGTTTTACCAACACCCTTTAAGCTTAAACAAGCTTTTCTTTGTTCTTCTTCTTATCTTCTTTCTTCCTCCAGCTTCACCAGCTTCTGTTAATGGTCTTTTTAAAGACTCCCAACAGCTTCTTTCCTTTAAAGCTGCACTTCCACCAACCCCTACTCTGCTTCAGAACTGGTTGCCATCTACTGACCCTTGTAGTTTCACTGGTGTTTCATGCAAGAATTCTAGAGTTTCTTCTATAGATCTCAGTAACACTTTTTTAAGTGTGGATTTCAGTTTGGTCACTTCTTATTTGCTTCCTCTTTCTAATTTGGAGTCTTTAGTGTTAAAGAATGCTAATCTTAGTGGTTCTTTAACTTCTGCTGCTAAATCCCAATGTGGGGTTTCTTTAGACTCCATAGATCTAGCTGAAAACACAATTTCAGGACCTATTTCTGATATCTCTAGCTTTGGTGTTTGTTCAAACCTTAAGTCTCTTAATCTTTCTAAGAATTTCTTGGACCCTCCTGGTAAAGAAATACTTAAAGGTGCAACCTTTAGCCTCCAAGTTCTTGATCTTTCTTACAATAATATCTCAGGTTTCAACTTGTTTCCATGGGTTTCATCTATGGGGTTTGGTGAACTTGAGTTCTTTTCTCTCAAGGGTAACAAACTAGCTGGAAGTATTCCTGAATTAGACTTCAAGAATTTGTCACATTTGGATCTTTCTGCAAATAATTTCTCAACTGTTTTTCCTTCATTCAAAGATTGCTCCAATTTGCAGCACTTGGATTTGTCATCCAACAAGTTTTATGGTGATATTGGTTCTTCACTTTCTTCATGTGGGAAGCTCAGTTTTCTCAACCTTACCAATAACCAGTTTGTAGGTTTGGTCCCTAAACTACAAAGTGAAAGCCTACAGTATTTGTACTTAAGAGGGAATGATTTTCAGGGTGTGTACCCAAACCAGCTTGCTGATTTGTGCAAAACTGTGGTGGAATTGGACTTGTCATACAATAATTTCTCAGGTATGGTTCCTGAGAGCCTTGGTGAATGTTCAAGTTTGGAACTTGTTGATATTTCCAACAATAATTTCTCTGGTAAGTTGCCTGTTGATACTCTCTTGAAGTTGAGTAATATGAAGACTATGGTCTTATCATTCAACAAATTTGTTGGTGTTTTGCCTGATTCTTTCTCTAATTTGCTGAAATTGGAGACTTTGGATGTGAGTTCTAATAATCTCACAGGGGTTATTCCATCTGGGATTTGCAAAGATCCTATGAATAACTTAAAAGTGTTGTACCTTCAGAATAACTTGTTTGAAGGCCCTATACCTGACAGTCTAAGCAACTGTTCACAGCTGGTGTCACTTGATCTTAGCTTTAATTACTTGACTAGGAGAATACCATCTAGTTTGGGGTCATTGTCAAAGCTGAAAGATCTCATCCTTTGGTTGAATCAGCTTTCAGGGGAAATCCCACAGGAGTTGATGTACTTGCAGGCTTTAGAAAATTTGATTCTTGATTTTAATGACTTAACTGGACCAATACCTGCAAGTCTTAGCAACTGTACCAAGTTGAATTGGATTTCATTGTCAAATAACCAATTGAGTGGTGAGATACCGGCTTCTCTTGGTCGTTTGTCGAATCTAGCCATTCTTAAGCTTGGAAACAACTCAATCTCAGGGAATATACCTGCTGAGTTGGGTAATTGCCAGAGCTTGATATGGTTGGATCTCAATACTAATTTCCTGAGTGGCTCCATTCCACCGCCTTTGTTCAAGCAATCTGGAAATATTGCAGTGGCATTGCTGACCGGAAAACGATATGTGTATATTAAGAATGATGGGAGTAAGGAGTGCCATGGAGCAGGGAATTTGCTGGAGTTTGGAGGGATTAGACAAGAACAGCTGGGTAGAATCTCAACAAGGCATCCTTGCAATTTCACAAGAGTTTATAGAGGTATCACTCAGCCAACATTTAACCACAATGGCTCTATGATATTTCTTGATTTATCTTATAATAAGTTGGAAGGTAGTATCCCAAAGGAATTAGGGACCATGTACTATCTGTCTATATTGAATTTGGGGCATAATGATCTGTCTGGTATGATTCCTCAAGATCTTGGAGGCTTGAAGAATGTTGCAATTCTTGATCTGTCATATAATAGGTTCAATGGCCCGATCCCGAATTCCCTCACCAGTCTTACATTGCTTGGAGAGATTGACCTGTCAAACAATAATCTCAGTGGAATGATTCCTGAATCTGCACCATTTGATACATTCCCTGATTATAGGTTTGCAAATAATTCTCTCTGTGGGTATCCTCTCCCCCTACCTTGTAGCTCGGGGCCGAAATCGGATGCAAATCAGCATCAGAAATCTCACCGCAGACAAGCATCGTTGGCAGGGAGTGTGGCGATGGGTTTGTTATTTTCCCTCTTTTGTATCTTTGGTTTGATTATTGTTGCCATAGAGACGAAGAAGAGGAGGAAGAAGAAGGAGGCTGCCCTTGAAGCTTATATGGATGGTCATTCACATTCTGCAACTGCAAACAGTGCCTGGAAGTTTACGAGTGCTCGTGAGGCGTTAAGCATTAACCTCGCAGCATTTGAGAAGCCTCTCAGGAAGCTCACATTTGCTGATCTTCTCGAAGCCACCAATGGTTTCCACAACGACAGTCTTGTAGGCTCTGGTGGTTTTGGTGATGTCTACAAAGCTCAGTTGAAGGATGGCAGTGTTGTAGCCATTAAGAAGTTGATACATGTCAGTGGACAGGGTGATCGAGAATTCACTGCTGAAATGGAAACCATAGGGAAGATCAAGCACAGGAACCTTGTCCCTCTTTTGGGCTACTGCAAAGTAGGGGAAGAAAGACTACTGGTTTACGAATACATGAAGTATGGAAGTCTTGAAGATGTCCTGCACGATCGGAAGAAAATCGGGATCAAGCTGAATTGGCCTGCAAGAAGGAAAATCGCCATTGGAGCTGCGAGAGGGTTGGCTTTCCTACACCATAACTGCATTCCACACATCATTCACCGGGACATGAAATCAAGTAATGTCTTGCTTGATGAAAATTTGGAAGCCAGAGTATCTGATTTCGGAATGGCAAGGTTAATGAGTGCTATGGACACTCATTTGAGTGTCAGCACTCTTGCAGGCACTCCCGGATACGTTCCTCCTGAATATTACCAAAGCTTTAGATGTTCAACAAAAGGAGACGTTTATAGTTATGGTGTCGTATTACTTGAGCTTCTAACCGGCAAACAGCCAACAGATTCAGCTGATTTTGGTGACAACAATCTTGTCGGATGGGTAAAGTTGCACGCTAAAGGAAAAATAACTGATGTCTTCGATCGAGAGCTATTGAAAGAGGATCCAAGCATTGAGATTGAACTTCTACAACACTTAAAGGTAGCTTGTGCTTGCTTAGATGATCGACATTGGAAACGTCCCACGATGATACAAGTTATGGCTATGTTTAAGGAGATTCAAGCAGGGTCAGGCATGGATTCGACATCCACAATCGGAGCTGATGATGTCAATTTTAGTGCAGTTGAAGGAGGGATAGAAATGGGGATAAATGAAAGTATAAAAGAAGGCAATGAGCTAAGCAAACACCTTTGAATGCAGCAGCACTAAATCAAGAGTTTGTTGAAAGCTCACAAATTTTCCAAAATCATATGCAAAGAATTGCCAGTGTAATTTTTAGCCAATTATTGTATGTACCACTAGTTCCCATCCATAAAATCTTGTGTTATTATACATAAAAGGAAGGTTGTTTTTCTTTTTTAAAACATGTAACAGTGTTGTATATAAATAGTGTATGAGTTTGTTTTTGC

>PGSC0003DMG400019801 LRR-RLKs

ATGGTTTCCATGACTATTATTATAGAATTGCTAGTTTTGTTAACGTACCTCTTGGATACTAATGCATCGAACATTTCAACTGATGAGGCTGCTCTTTTAGCTTTCAAAGCTCATATAACTTCAGATCCAAATGGAATGTTGTCGAAAAACTGGACCAAAGGGACTCACGTTTGCAATTGGATAGGCATATCTTGCAGTATAAGGCATCAAAGAGTGACATCACTAGACCTCAACGGCCTCAGACTAAGAGGTTCAATTGCCACAGAAATTGGGAACCTCTCATTTCTTAGTTTCCTGGACATTGGCAACAACAGCATCCACAGCAAAATCCCTGATGAACTCGGCTATTTGAGGAGGCTAAAGTACTTGTCCTTGTCAATGAATAATCTCACAGGTCAAATTCCAGCAAGTCTAGGATTACTGAGAAGGCTCGAATCACTTGATCTTTCTGAAAATGATCTCCTTGGATCTGTTCCTTCATCTATCTTCAATGTCTCTTCTTTGAAGGTCATCAATTTGATTTATAATGCACTGTCTGGGACTTTGCCGAATGACATTTGCCATAATCTTCCGCGGCTGGAGGGATTACAAATAGCAAGAAATCAACTCTCTGGCCAACTACCAAGCAACTTAAACAAATGCAAAGAGCTTACATACTTGTCCTTGTCATATAACCAGTTCACAGGTAAAAATACTGAACCAATTTCTTTTTCTGCTACTTTTGGTTCATATTGACAATAAAGATTGCAACAATTTTTAGTGAAAACAATGATAAGCTGTTGAGAGGTATAACTTATTGTAACGTCTATTACAGGAAACTTGCCTAGAGACATGTGGAACTTGACAAAACTTCAACATCTATTTCTTGGATGGAATAACATAACAGGTACTTATAATGTTCTAGTTTAACTGAATCTGCCTACAACTGCATTTATTGCTGGACTTCTCACCTCGTAGCACTAGAGATCTTTGACATTTCCTAACTTGCTCTTCAAGTCATTTATATGAAATTAGTATTTGCATTTCAAAGGGTAGTAGCAGAAAATGTTAGTATAGCTCTTTGTTCTTCATTTTCAAACAACAACAGCAACTACTCCATAGGTTCAATGCTAGTTTGTGATGTGAATCCTCATTATCCATGTTGATCCATTTAGTCCCATCTTCAAGAGGACGTTAAAACAGATTGCGAGGATTGCTGAACTATTTAAAAGGCATACAACTAACAGTTTTCTGTTATTTATAGTAACTTATTGACTGTAAAGCTGTTATACAGTATTGGACATTTCAGTGTAGTAACCACACAATTCGTCATATATTTTACAGGACATATACCAAGTGAAATTGATAACCTATCAGCAATTCGAAGATTAAGTCTACGACGGAACAATTTGGTGGGAAGTCTTCCACCATCTATTGGAAATTTGTCAAATCTAGAGATGATAGACCTAGGTGACAACAGTCTATATGGTGGCATTCCTCAGGAGTTTGAACATCTTGTAAATTTGAAAGAAGTGTATCTTGGTCCAAACAGATTATCGGGTGAAGTTCCAAGGCGTATGTACAACATTTCTGGACTAAAAAAGATTTCATTTGTAGCAAATACGCTTTCAGGAACACTTCCCCCAAACATAGGCCATACTCTTCCAAATCTTGAAGGCCTTTACCTTGGGGAAAACCAGTTCACCGGGCTGATTCCTACCTCCATTGTGAATTCCACCAAGCTTATCCAGCTTGATTTTGGTCGCAATCTGTTTAGTGGACCTGTACCTATGAATCTTGGAAAGCTACAACAGCTTCAATTCATCAACTTGCAAGTAAACCAATTGACGAATGATCAATCTACAGGCGAGTTAAGTTTCCTGACTTCATTATCTAATTGCAAGTATTTGAAAACCGTTCAAATAGGAGGTAATCAATTTAATGGAAGTCTTCCAAAATCTCTAGGTTCAGGAAACTGGTCCTTTTCCCTTGAATATTTCATTGCCATTCAAAGTGGTATCACAGGAAAAATTCCAACTAACATCAGTAACTTTAGGAACTTGGAGTGGTTAACATTGGGAGACAATAAGCTTATAGGATCTATTCCTCCAGATTTGGGGAACTTGAGGAATTTGAAAAGATTTAGCCTGGAGAAAAACAATTTAGATGGTATTATACCAACTAGCTTGTGTAACATGGAAAACTTGTACCAAATTATCTTGGGGAAAAATCAACTTTCTGGTGAATTACCAAGTTGTTTTGGGAATCTTTCTTCTTTAAGAGAATTCTACTTAGATTCCAATACACTAATTTCCCATATTCCATCAACTTTTTGGAGGAACAGAGATATTTCAATTCTCACTTTGTCCTCCAATTTGCTGAATGGGTCTCTTGCAGTAGAAATGGGGAATTTAAGGAGTTTGCGGATATTACATTTACATGGTAATCAGTTCTCTGGTCAGATTCCTAGCACAATTGGCCAACTACAAAATTTGGTTAGCCTTTCATTGTCAAAGAATATGTTAGATGGTTCCATACCTGAACTATTTGAAGATTTGGTTTCATTGGAACACTTGGATCTATCTAGCAACAATCTATCAGGCATGATCCCCAAGTCCTTGAGGAATCTTGAACATCTCATTTATTTCAATGTCTCGTTCAATGGGCTCATGGGTGAAATTCCAGATGAGGCCATTCGTAAATTTTACAGCTGAATCATTCATCGGTAACCCTGCATTATGTGGATCATCACGCTTCCATGTGATGCAATGCAGAGCCACAAGTCTTGAAAGAAAAAGAAAGAGTAGAGTCTTAACTTCTGTTCTTGCATCAGTTTCCTCAGGAGTTGTAGTCACAACCATTTTCATCATTTGGTTTCTGAAATGCCGAAAAAGGGGTACGGAACTTCCTCTAGTTGATACATTTGGTCAGGTACATAAGAGGATTTCGTACTATGATATTTCTCAAGGGACAAACAACTTTGATGAAGCAAACTTGATTGGAAGGGGTAGCCTTGGTTTGGTGTACAAAGGGACACTAGCAGATGGAATGGTTGTGGCAATAAAGGTATTCAATACAGAACTGCAACACGCATTTAGGAGTTTTGAAGTGGAGTGTCAAGTTTTACGTAGCATTCGACACAGAAACCTTGTTAAGGTGATAAGTAGTTGTGCCAATTTTGATTATAAAGTGTTGGTGCTAGAGTACATGCCCAATGAAAACCTCGAATGTTGGCTTCACTCTACTGACAAATTATTGGATTTAACTCAAAGACTAAAGGTGATGATTGATGTGGCTTCTGCTGTGGAGTATTTACATGGAGGTCATTTGTTTGTAGCCGTCCATTGTGATTTGAAACCAAGTAACATACTTTTGGATGGAGATATGGTGGCAAAAGTGAGTGACTTTGGGATATCCAAACTTCTAACAGCTGAGACGCTAATAGCACATACCAAGACTTTGGGTACTATCGGCTACATGGCACCAGGTAAATGTTATTATCATGTTCATTTGAAGATTTTAAATGACGTTAATTATTTTCACTAATGATTGCGTGATTTTTTTGCAGAGTATGGATCAGAAGGCAAAGTATCAACCATGGGAGATGTTTATAGCTTTGGTATTTTGTTAATAGAGACTTTTACAAGAAAGAGCCCCGTGGATGATCTATTCGTTGGAGACTTCACCTTGAAAAGATGGATATGCCAATCATTACCAAACCGAGTGGTGGATGTAGTGGACATTAATCTATTTTCACTGGACGAAGAAAATCTTACTTCTAAAGAGAGATGCTTCAAATCAATCATGGAATTAGCTCTTGAATGCACAAACGATTTGCCTGAAGAGAGAATTTGCATGGAAGATATTACTCTGCGACTCAAGAAGATCCTAACTCAATTTCTGCAAAATGTTGTAACAAACTAA

>PGSC0003DMG400020594 LRR-RLKs

CCCCAACACCCCCACCCCACATCTCCATTCTTCCTTATTTATGCACCCACCACCCTAACATCTCCACCACAGCCCCTTACAATCAATGGCTTTCCACCACCACCACTACCTCCTCCTCCACCTTCTCATTTTCTTCACTTCCCTTTCTTTCACCTTCTCAGATGATGCTGATGTTATGTCCAAGCTTCTTGCAGCTCTTTCTCCAACTCCTTCTGGTTGGTCAACTTCTAAGCCTTTCTGTTCTTGGACTAATGTCATTTGTGACAAGTCTTCAGCTACTGTAACTTCAATTAATCTTGATTCTCAATCTGTTTCTGGTTCTTTACCTTCTGAAATTACTCAACTTTCCAATCTTAAAACCCTTTCTCTTCAAAAAAACAAACTTTCTGGCCCCTTACCTTCTTTTGCCAACATGTCAAAATTAGCTGAACTTTTCTTGGACAATAACCAATTCACTTCTGTTCCTCAAGATTTCCTTTTGGGGGTTCCCAATTTGGTAACTTTAAGCATTAGTGAAAATGGGAGACTCTCTCCTTGGCAAATCCCTATTTATTTAACTGAAAGTACAAATTTGGGATCTTTATATGCTAGTAATGCTAGTATTGTTGGTGTCATTCCTGATTTCTTTGATGTTTTTCCTAATTTACAGAATTTAAGATTGTCTTATAATAATCTTACTGGTTCTTTGCCTGCTTCTTTTGGTGGTTCTGAAATTATGAATCTTTGGTTGAATAATCAAGTTAAAGGGTTATCAGGTAGTATTGATGTGATTGGAAGTATGACACAATTGTCTCAAGTTTGGTTACATGCTAATTCTTTTACTGGTAGTATTCCTGATTTGTCAAAATGTGAGAATATATTTGATTTGCAGTTGAGGGACAATCAGTTTACTGGTATTGTTCCTGAGTCTGTGATGTCTCTTCCCAAATTGTTGAATATTACTTTACAGAACAACAGGTTGCAGGGTCCAATACCTCAGTTTAAAGATGGGGTTGAGGTTAAACTTGAAACTACTAATAGTTTCTGTAAGGACACTCCGGGACCGTGTGATCCACAGGTCACTACACTTCTTGATGTGGCTGGTGGTTTTGGATATCCATTGTCTTTGGCTGAGTCTTGGAAAGGGAATGATGCTTGCAATAGTTGGAGTTTTATAAGTTGTGATACAACAGGGAAGAATGTAGCTGTTGTTACCTTAGGTAAGAGGGGATTTTCGGGTACTATTTCACCTGCTTTTGCGAATTTGACTTCATTGAGGAGCTTGTTTTTGAATGATAATAATCTTACTGGTACGATCCCGGAGAGGTTGACAACTTTGCCTAGTTTACAAGTTCTTGATGTGTCTAATAATAATTTGTCTGGACCTATACCATTGTTTCCACCTAGGGTGAAGTTTACTCATACTGGTAACTTGTTTCTCGGGACAAATATAACTACTGGTGGTGATGGTGGGGGAAGTGGTTCGGATGGTTCTGGATCGAATTCCAGTGGTCGAGGTGGAAGTTCATCTGGTGGTTCGAAAGGGCCTTCAGTTGGAATGATTGTTGGTGTTGTTATAGCAGTTGTTCTTTTTGTTTTAGTAGTGTTGTTTGTGTCTTACAAATGTTACATGAAGAGACACCATAAGAGATTTGGAAGGGTTGAGACTCCAGAGAAAAGCAATGAAATGGTTAAGCCTAGTCTCCCAAGTGTCGTGGGTGGTTCGAATGGATATACTGGTGGCACCAGTGAGTTACAGAGTCAGAGCAGTGGCGATCATAGTGAGATACCAGTTTTTGAAAATGGAAACGTGGCCATTTCCATCCAGGTCCTTCGACAAGTGACGAACAACTTCAGTGAAGAAAATATCTTGGGGAGAGGAGGGTTTGGAGTTGTTTATAAGGGTGAACTACACGATGGGACTAAAATCGCGGTGAAGAGGATGGAATCTGGGGCAATGGGTACCAAAGGAATGAACGAGTTCCAAGCAGAAATTGCGGTCCTTACCAAAGTCAGACATAGGCATTTGGTTGCTCTACTTGGTTCATGTGTTAATGGAAATGAGAGACTTTTGGTGTACGAGTATATGCCGCAAGGAACTTTGAGTCAGCATTTGTTTGAATGGCAAGAATTAGGCTACAAACCTCTTACTTGGAAGCAAAGGGTAACGATAGCATTGGATGTGGCAAGAGGGGTCGAATACCTTCATAGCTTGGCACAACAAAGTTTCATTCATAGAGATTTAAAACCGTTAAACATCCTTCTCGGAGACGACATGAGAGCCAAGGTTGCAGATTTTGGTTTGGTTAGAAATGCCCCTGATGGAAAATACTCTGTGGAGACACGTTTGGCTGGAACTTTTGGCTATCTTGCACCTGAATACGCGGGTAAGCATTTCTACTCCTTGTATCATTACGTCTTTCTTTAACTGAGAACTGATTTGTATGCCCCTAGCTGTAAACTTCAGATTTTAGCAAATTTTGAAATTATTCGATCATCTGCATTGCTAGGCATTGTTGTCTTCATTACATTTAAAGACATTTGGCATATAATGTAGAGGCAATCTCCATGCGTAAATGCTGCTTTGCAATTTTTATCGAACTCATGAACTATATACTACTATTATAAAAGATAAGTTTCCATTTCGTGTGGATCAGCATCTGAATAATTATCCTATTACATCTCCTCTTTGTTACAGCTACTGGACGAGTCACAACCAAAGTAGATGTTTACGCCTTTGGCGTTGTTTTGATGGAGATCATTACTGGTAGAAAAGCGTTAGATGAAACAATGCCAGATGAGAGGTCTCATTTGGTTACATGGTTCCGCAGAGTCCTTATCAACAAAGAAAACCTCCGAAAGGCTATTGACTCGACACTTGATCCTGATGATGAAACATATGAAAGCATCTCCAAGGTTTCCGAGTTAGCTGGCCATTGCACTGCTCGGGAACCTTTTCAGCGACCTGATATGGGACATGCTGTTAATGTCCTTGGTCCCCTTGTAGAGCAGTGGAAGCCTACAACAAGTGAAGATGATGACGGCTATGGCATTATTGACCTCGACATGAGCCTTCCCCAAGCCCTTCAAAGGTGGCAAGCTGATGAAGGAACTTCAAGAATGTTCGATGACTTCTCAATCAGTCACTCGCAGTCGAGCATACCCTCAAAACCTTCCGGATTCGCAGATACTTTCAGTTCAACAGATTGCAGATGATATGCAAAAAATTTGACAACAGTTCAAGCCATGCCATGCCACACCTCGCGCGGTTTTAATCCTTTGTGGATCAACATAGAGTATGCCAATATTCATTTGATTAATTGGTTTCTTATTGTCAATGTCATTACTTTTTATGTGATTACATATTAAGTTAATATAGTCTTGATGTAAGGTGTGTGATAACAGTTGTAAAGTCAGGTGAATGTCACATTTTTCTTTTGCCTTTTTGGAGACAACAGAAAATTGTAACAAATTCTTTCATATTACATTGTTAATGTAAATCAAATTTCGAGCTTCCCCCTCC

>PGSC0003DMG400020697 LRR-RLKs

ATAATGGAGAAAGTCTTCACATATTTTTACTTAATAGCACTCATGTTGCTTTACTTGGGTGTGGCTAGTTTAGCCATGACTCAAACCAACATTACAACTGATCAATCCGCTCTTCTTTCCTTGAAATTCCAAATCATTTCGGATCCCTTTCACTTGTTGGATGAAAGTTGGTCTCCAGCTATTTCTGTTTGTCATTGGGTTGGAGTCACTTGTGGTTCTCACCACAAAAGAGTGACGTCATTGAATATTTCCAACTTGGATCTTATAGGCAAAATTCCTTCTGATTTTGGGAATCTCACATCACTTGTTTCTCTTGACTTGAGTAGCAATAATTTCTACGGAAAATTGCCTCAAGAAATGGCACATCTTCATAGGTTAAGGTTTGTTGAACTTAGTTTTAACAACTTTAGCGGAGAGGTTCCTTCCTGGTTCGGGGTCTTACACCAACTTCAAGTTCTGACTCTAAGGAATAACAGTTTCAATGGTCCCATCACCCTCTCTTTATTTTCTAATATTTCCACACTTCAAACTTTGGATTTGACATACAATTCCTTAGAGGGCCAAATACCAAAAGAAAAAGGAAATCTTGAAAACCTGAGGGTTTTAAGATTATCTGGCAACAAGTTTATAGGTTCTATCCCTCCGTCTCTCTCGAATGCCTCAATGTTGATGACATTAGAACTATCTCACAATTTCCTTGGAGGAAACATTCCAGAAGAGATTGGAAAACTTCAAAACCTAAAACTTTTGTCAATAGAATCCAATCAACTTACAGGTTCTATACCATTCTCGATCTTCAACATTTCTGGAATTGAGATCATTGCATTTACAAATAATACCTTATCAGGAAATCTTCCTTATGATATGTGTAATTGTCTTCCAATGCTCGAAATGCTCTATCTAAGTGACAACAAGCTATATGGTCATATGCCTGTAAGCTTGCCAAATTGTTCGAACCTTCAAATATTGTCTTTATCTAATAACAAGTTTGACGGACCCATACATAGTGAAATCGGGATTCTTAATAATTTGGTGATTTTAGCACTGGATACTAACCATATTTCTGGTCATTTCCCAATCTCCACATTCAATATCTCCTCATTACAGGTTTTGGTACTGAATCACAACAATCTGATTGGATCCTTGCCAAGGGAGGTTAGCAATTTGACAGAATTGCAAATTCTAGATCTTGCCGATAATAATCTTACAGGTATATATGTATTCAAATCATTACTAGCTAGCAACATAATTTCTTAAATCTTAGTAATTACTAACTATACATATACATACATTGGCAGGTGAGATACCGAAAAAGTTTAGCAATCACATTGAGCGGTTTGATCTTAGTTTCAACAAGTTTACTGGTTCACTACCAATGGAGATTTTCAACATATCGGGTATGAGAGAGATTCAAGTTATTTTCAATAATGTGACAGGAACTCTCCCATTAAACATAGGTTCAACGTTACCCAACATTGAAGTTATTCATCTGGGCAACTTAAATCTTTATGGGACTATACCTCATTCTCTCTCCAATTGTTCCAAACTCACTAGTCTAGACCTTTCCCTCAATAGACTAACCGGTATGATTCCCAACTCTCTCGGAAGTTTGACTCATCTACAGTCCCTAAGCTTGGCAATAAACAATTTAATGGGCGATAGTAGGTTAAGCTTCTTGACTTCCTTGTCAAATTGTAGAGATTTAAAAATTCTTTTCCTCTCTTCGAACCCTCTATATGGTGTGCTTCCAGAATCCATAGGGAACCTTTCCAGTACTTCTCTTCTAAGGTTTAGAGCTATTGATTGCAAAATTAAGGGGGAAATTCCAAAAGGAATTGGGAACTTAAGCAGCTTATTAGAGCTCGATCTTTCTAGAAACGGTTTCACTGGACCAATTCCCACAACAATCAGCAACTTGAGATCCCTTCAGAGCTTGAAGTTGAGTGTCAACAAACTTTCTGGATATATTGGAGATGATCTATGCAAATTACAGAACTTGGGTTATTTAAACTTGACTCTGAATCAACTTTCAGGATCTCTTCCGAATTGTTTGGGAAACTTAACTTCCCTTCGGGAGATGATTCTAGGTTCCAACAAATTGCGTTCAATCATTCCAAAAAGCTTAGGAAACCTCATAAACCTCTTGAAGCTAGACTTATCCTCAAACAACTTGGGTGGCTCTATACCTCTGGAAATTGGAAATCTAAAGGCTGCGATATATATGGATCTATCAATAAATACACTCTCAAATAGAATTCCTAAAGAAATTGGTAGCTTGCAAAATCTTATATACCTTTCTTTGAGAGACAACAAATTGCAAGGATCAATTCCTGGCTCGATGAGCAGCATTTCAGCTTTGGAATTTCTAGACCTTTCTCATAACAATGTTTCAGGATTGATTCCCAAGTCTTTGGAGAAGCTTCTATATCTCAAGTATTTCAACATCTCATTCAACAAGTTGATTGGTGAAATCCCCTCGAATGGTCCTTTCGAGAACCTCTCCGGTGAGTCTTTCATGTCTAATGAAGCATTGTGTGGATCTCCAAGATTTTGTGTTCCCCCATGCCACAGTAGCACATCAAAGCATAAATCCAAGAGGAAAAAAGTGATTGTTCTTGTTCTACCGGGTGCACTTGTATTAGTCTCCATCGCCTTTGCCCTTTTATGGATAAGGAATACAAGAGGTAAAAGAACCGATCCTCAGCAAGCAGTTGATTCATCCTTTGTCGTCTCAACAAAAGGAAGAATTTCATACTATGAACTGCTTCGAGCAACTGATTCACTCAGTGAGATCAATTTAATTGGTTCTGGGAGTTTCGGCTCTGTTTACAAAGGCATCCTTGGAGATGGGACTTTCATTGCAGCTAAAGTGTTCAACCCACAATTGCAAGTGGCATTTAGGAGTTTTGATACAGAATGTCAAGTTTTGCGCAACCTTCGTCATAGGAATCTCACCAAAGTCATCACCAGTTGTTCCAACCTTGAATTTAAGGCTTTAATACTCGAGTACATGCCCAATGGGAGCCTCGATAAGTGGTTGTATTCACACAACTACTTCTTAGACATCAATCATAGGCTAAGTATAATGATAGATGTAGCATGTGCATTGGAATATCTCCATCATGGTTGCCCATTGCCCGTGATTCATTGTGATTTGAAGCCTAGTAACGTCTTGTTGGATGAGGACATGGTTGCCCACCTAAGTGACTTTGGAATTTCAAAGCTGCTTTCTGAAGATGAGAGTTCTTTATACACTAAAACATTAGCAACGTTTGGTTATATTGCACCAGGTACGTGATCTCTTAATCAATTGCTTTTGATTATTTCTCTTTCCTTTTTTCCACTAGTAAATGTTGAATTTCAAAATTGATTGATTACAGAGTATGGACTGGAGGGATTGTTGTCAATAAAATGTGATGTTTATAGTTACGGGATCATGTTGATGGAAACATTTACTAGGAAAAATCCTAATGATGAAATGTTCGATGGAGATTTTAGTTTGAAACAATGGGTGAGAAATTCACTCCCACAAGCAGCAATGGAAATTGTCGATGCCAACTTAATATCGCCACATGATAATCACTTAATGAAAAAGCTAGATTGTGTGGTATCCATCATGAAAGTGGCAATAGATTGTTGTGTTGAATCTCCTAAAGGAAGGATTGACATGAAAGACGTTGTGGGAAGGTTAAAGATGATCAAGATTCAACTTCTTGCGTGTTGAGGAAGATCTTATTAGAAACTTTTCTTCCAAATCATCTCATGTGTAACAAACATTTGCAATAATAGTTAGCTTCAAATAAGTTATAGCGTTATGTCTTTCATAACATAGAGAATTGATGGGCATGTATACCTCTTTGGTCTTGAATTTGTACAGTGGTTTTGGCCTAATACCTTTACTTTTTTCCATCTTTTTAACTTTAACAATGTAAATAGAACTCTCAATATTGAATAACGGAATTTCG

>PGSC0003DMG400020744 LRR-RLKs

ATGTCACTACTAGCATTCAAGAATATGATCATTGATGACCCTTTCAAGATCATGGACTCCTGGAATGAAACTATACATTTTTGTGACTGGCCTGGGGTTTCATGTGGTCGACGTCATCCCCGAGTTACAGTTTTAAATCTTACTTCTCTGAAATTGAGGGGTTCTCTGTCACCAAGCATTGGTAATCTGAGCTTTCTTTGTGTCCTTAAGCTCCAAAATAATAGTTTTTCAGGTGAAATTCCATCAGAGATTGGCTACTTACATAAGTTAAAGTTTTTACTCCTTGATAATAACTCGTTCACTGGTCATATTCCATCAAACATTTCAGGATGCTTCAATCTTGTTTCTGTTGTTCTTTCATATAATATGATGGTGGGCAAAATTCCAGCAGAATTAGGCACATTGTTGAAACTCAAACAACTTTCTCTTGTTTCTAACTATTTTACAGGAGGAATTCCGCCTTCATTTGGTAATCTTTCTTTACTTGATACCTTTTCCGCAAGCAAGAATAATTTGTTAGGGAAAATACCTGATGAATTATGTCAATTGCTCAACTTAAAGTACTTTGTAGTGAATGAGAACAACTTGAGTGGTACCCTGCCTCCTTGTCTTTTCAATCTTTCCTCTATCGTGGCCATAGATGTTGGAACAAACCATTTAGAAGGAAAATTGCCTCCATTGCTTGGTATTACTCTTCCTAGTTTAGAGTTCCTTAGCATTTATAGAAACAATGTAACTGGAAATATTCCGGTAACATTGTCAAATGCTACGAATCTTCAGTCTCTTATTGCTAGTAGAAATGGACTCACAGGAAAAGTACCACCCCTTGGAAATTTGCTGAAAATGCGGAGATTTTTAGTTGCTTTCAATTATCTTGGGAAGGGGGAGGATGATGACCTTAGCTTTCTCACCACATTGGTGAATGCCACCAGCTTGGAGCTTTTGGAGCTCAATACAAATAATTTTGGTGGAGTCTTACCTGCATCTGTTAGTAATCTTTCGACTGAGCTTATAGAGCTTTCGTTGTCTTATAATCAGATTTCTGGAAAAATCCCAAGAGGGATATCCAATCTCAAGAAGCTTCAGGCTTTCTTTGTAGCTTACAACAGATTTCACGGTGAAATTCCGTCTGAAATTGGTGATCTGATATATTTGCAAGAGCTGGCTTTACTTGGAAATCAGTTCTCTGGCCAAATTCCAATCTCATTGGGAAATTTAGCTTATTTAGCCAAACTCAGTTTAAGAGAAAACAATATCCAGGGGAGAATTCCTTCCAGTTTAGGCAAATGCCACAATTTGGAACTGTTGGATCTTGGTTCAAATAACCTTAGTGGTTTCATACCATCAGAAATTCTTGAACTCTCATCCTTGTCGGAGGGCATGGACCTATCTCAGAACCACTTAACTGGTTTCCTTTCGATGGAAATTGGGAAGTTGAGAAATCTAGGCTACTTAAACCTGTCTGATAACAAGTTGCAGGGCCAGATTCCTACCACTATAGGTACTTGTGTGAAACTTGAAGCACTAGACATAAGCAACAACAACTTTCAGGGTAGTATTCCTTCAACCATGAATAGTTTGNNNNNNNNNNNNNNNNNNNNNNNNNNNNNNNNNNNNNNNNNNNNNNNNNNNNNNNNNNNNNNNNNNNNNNNNNNNNNNNNNNNNNNNNNNNNNNNNNNNNNNNNNNNNNNNNNNNNNNNNNNNNNNNNNNNNNNNNNNNNNNNNNNNNNNNNNNNNNNNNNNNNNNNNNNNNNNNNNNNNNNNNNNNNNNNNNNNNNNNNNNNNNNNNNNNNNNNNNNNNNNNNNNNNNNNNNNNNNNNNNNNNNNNNNNNNNNNNNNNNNNNNNNNNNNNNNNNNNNNNNNNNNNNNNNNNNNNNNNNNNNNNNNNNNNNNNNNNNNNNNNNNNNNNNNNNNNNNNNNNNNNNNNNNNNNNNNNNNNNNNNNNNNNNNNNNNNNNNNNNNNNNNNNNNNNNNNNNNNNNNNNNNNNNNNNGGGCTTATAGGATTGACATTCATAGTTTGTTTCTTGGCATACGGCAATTTAGTAGTATCTTATCAGTGTCTACTTAGGGAAACTGATAGATTTTCTGCATCAAATTTGCTTGGAGTGGGTGCTTTTGGGTCTGTATATAAAGGAATTTCTGAACACGATGGTACGTTTTTTGCTGTTAAGGTACTGGACCTTTCACACCATGCAGCTTCCAGGAGTTTTTTAGCTGAATGTGAGGTTCTAAAGAACATTAGACATCGTAATCTTGTAAAGGTTCTAAGTGCATGCTCAGGTATTGATTATGAAGGTAATGAGTTCAAGGCCATAGTTTATGAATACATGGATAAGGGAAGTCTACAGGACTGGTTACATTTCACTCCTCAGGAGAAATCTGAGTCACAGGAAGAGCACAAGAAGCTCGGGTTTATTCAAAGATTAAACATTGCCATTGATGTTGCCTGTGCTCTAGATTATCTCCATAATGATTGCCAACCCCCAATAATTCACCGTGATCTGAAGCCAAGCAATATTCTTCTCGATGAGAACATGACTGCACATGTTGGTGATTTTGGTTTGGCAAGATTTGTTCCACCAGCAATTCCGAACTCATCAGCAAATTCAAAAAGTTCTACAGGCGTAGGGGGAACTATTGGTTATACACCTCCAGGTAATAATAAATCAAGGTTTCTGCTGGCTGTCATAAATATGTCATTTGATAAGAATATCTCTAATTCTGTAGTATTTCTTGTTGACAGAATTAGGTATGGGAAGCGACGCCTCAATCTATGGAGATGTATACAGCTTTGGCATTCTGCTCTTGGAAATGTTTACCGGGAGGAAACCAACTGATGAAATGTTCAAAGACAATTTAAACCTTCATAACTATGCCAATGCTGCTTTGCCTGACAGAGTGATGCATATTGCAGATCCCATTCTTCTCCAGGAAAGAGATGAACTTGGAATGAAATACAAAGTTGATGACAATACAAGCAGTGCTGTCGACATATTTCTGTCATTCTTAGTTAAGGTGATTCAAGTCGGAGTTTCCTGCTCCGTTGAATCTCCAAAAGAAAGAAAGCAGATCAGTGATGTTGTTGGAGAACTCAATTCATTAAGGAAGTTGTTCTTGGAGCAAGCATACCCGAAAGAAAAGGTTATAAGCTTCAACCTGCATTGA

>PGSC0003DMG400020832 LRR-RLKs

ATGGACACAAGTTATACTCTCCTCTTTGCTCAACTTTCAATTTTCATTCTGCTTCTTAACCATACTTCACTTGCTAATATTAGCACTGATGGAGCTGCTCTTCTTGCACTTAAATCACACATTTCTTCTCATCCTAACAACATCTTAGCAAGCAACTGGTCTTCTTCAACCTCGGTTTGCAGCTGGATTGGAATCACTTGCAGCTCCCGCCACCATCGAGTCACTGCTTTAGACATTTCTAGCATGCAACTTCATGGTACCATTACTCCACACCTTGGAAACCTCTCATTTCTTGTTTCCCTCAACATCAGTAATAACGCTTTTCATGGAGATTTGCCACAAGAGTTGGCTCGTTTGCAGAGGTTGAAAGTGATTGATGTCACAAGCAATAACTTCACCGGAGAATTTCCGTCATTTTTAAGTTTGTTACCAAATTTACACATTATGTACCTCTGGAGAAACCAATTTTCAGGTAAAATTCCATCCTCCCTTTCCAATCTGACAAAACTGCAAGTGTTGTCAATACAGAACAATTTTCTCCAAGGAGAGATCCCTCGGGAACTCGGTCATCTTCGTTACCTGAGTATCCTAGACCTGCAATATAATGGACTTACAGGCTCTATACCATCATCAATCTTTAACATTACTACAATGCAAGTAATTGCTCTTACAGGCAACAATCTTACTGGAAAGCTTCCAAAAACTATATGTGACCATCTTCCAAACTTGGAAGGGCTTCACCTCACAAGCAACTCCCTAGATGGAGTTATTCCACCAAACCTTGAGAAATGCCGAAAACTTCAAATATTGTCATTGTCTTACAATGAGTTTATTGGAATTGTACCAAGAGAGCTGGCCAACTTAACAGCTCTTACAGAATTATATCTCGGAGATCTGCATTTGGAAGGTAGTATGATATTTCCTTACAAGAAAAATGAAAATTTTCTTCCTTTTCTCTCTTCCGCTAAACTGGTAACTCATATACTTCCTGAAATTGACAGGAGAGATCCCTCGAGAAATCAGTGATCTTCGTTACTTGACTATCCTAGATCTGCAATTTAATCAACTTAGTGGCTCTATACCACCATCAATCTTCAACATTACCACAATGCAAGTAATTGCTCTTACGGGCAACAATCTTACTGGAAAGCTTCCAAAAACTATATGTGACCATCTTCCAAACTTGCAAAGGCTTTACCTCTCAAAGAACTCCTTAGATGGAGTTATTCCACCAAGCCTTGAGAAATGCCGAAAACTTCAAATATTGTCATTGTCTTACAATGAGTTTATTGGAATTGTACCAAGAGAGCTGGCCAACTTAACAGCTCTTACAGAATTATATCTCAGAGCCCTGCATTTGGAAGGTAGTATGATATTTCCTTACAAGATAAATGAAAATTTTCTTCCTTTTCTCTCTTCCGCTAAACTGGTAACTCATATACTTCCTGAAATTGACAGGAGAGATCCCTCGAGAAATCAGTGATCTTCGTTACTTGACTATCCTAGATCTGCAATTTAATCAACTTAGTGGCTCTATACCACCATCAATCTTCAACATTACCACAATGCAAATCATTGATTTTACTGGCAATAATCTTACTGGTAAGCTTCCACCAAGCCTCGAGAAATGCAGAAAGCTTCAAGTCTTGGCATTGTCGCTCAATGAGTTTATAGGAACTATACCAAGAGAGATAGCCAACTTAACAGTTCTTACAGAATTATATCTCAAAGCTCTGCATTTGGAAGGTAGTATGATATTTCCTTACAAGAAAAATGAAAATTTTCTTCCTTTTCTCTCTTCCGCTAAACTGGTAACTCATATACTTCCTGAAATTGACAGGAGAGATCCCTCGAGAAATCAGTGATCTTCGTTACTTGACTATCCTAGATCTGCAATTTAATCAACTTAGTGGCTCTATACCACCATCAATCTTCAACATTACCACAATGCAAATCATTGATTTTACTGGCAATAATCTTACTGGTAAGCTTCCACCAAGCCTCGAGAAATGCAGAAAGCTTCAAGTCTTGGCATTGTCGCTCAATGAGTTTATAGGAACTATACCAAGAGAGATAGCCAACTTAACAGTTCTTACAGAATTATATCTCAAAGCTCTGCATTTGGAAGGTAGTATGATATTTCCTTACAAGAAAAATGAAAATTTTCTTCCTTTTCTCTCTTCCGCTAAACTGGTAACTCATATACTTCCTGAAATTGACAGGAGAGATCCCTCGAGAAATCAGTGATCTTCGTTACTTGACTATCCTAGATCTGCAATTTAATCAACTTAGTGGCTCTATACCACCATCAATCTTCAACATTACCACAATGCAAATCATTGATTTTACTGGCAATAATCTTACTGGTAAGCTTCCACCAAGCCTCGAGAAATGCAGAAAGCTTCAAGTCTTGGCATTGTCGCTCAATGAGTTTATAGGAACTATACCAAGAGAGATAGCCAACTTAACAGTTCTTACAGAATTATATCTCAAAGCTCTGCATTTGGAAGGTAGTATGATATTTCCTTACAAGAAAAATGAAATTTTTCTTCCTTTTCTCTCTTCCGCTAAACTGGTAACTCATATACTTCCTGAAATTGACAGGAGAGATCCCTCGAGAAATCAGTGATCTTCGTTACTTGACTATCCTAGATCTGCAATTTAATCAACTTAGTGGCTCTATACCACCATCAATCTTCAACATTACCACAATGCAAATCATTGATTTTACTGGCAATAATCTTACTGGTAAGCTTCCACCAAGCCTCGAGAAATGCAGAAAGCTTCAAGTCTTGGCATTGTCGCTCAATGAGTTTATAGGAACTATACCAAGAGAGATAGCCAACTTAACAGTTCTTACAGAATTATATCTCAAAGCTCTGCATTTGGAAGGTAGTATGATATTTCCTTACAAGAAAAATGAAATTTTTCTTCCTTTTCTCTCTTCCGCTAAACTGGTAACTCATATACTTCCTGAAATTGACAGGAGAGATCCCTCGAGAAATCAGTGATCTTCGTTACTTGACTATCCTAGATCTGCAATTTAATCAACTTAGTGGCTCTATACCACCATCAATCTTCAACATTACCACAATGCAAATCATTGATTTTACTGGCAATAATCTTACTGGTAAGCTTCCACCAAGCCTCGAGAAATGCAGAAAGCTTCAAGTCTTGGCATTGTCGCTCAATGAGTTTATAGGAACTATACCAAGAGAGATAGCCAACTTAACAGTTCTTACAGAATTATATCTCAAAGCTCTGCATTTGGAAGGTAGTATGATATTTCCTTACAAGAAAAATGAAATTTTTCTTCCTTTTCTCTCTTCCGCTAAACTGGTAACTCATATACTTCCTGAAATTGACAGGAGAGATCCCTCGAGAAATCAGTGATCTTCGTTACTTGACTATCCTAGATCTGCAATTTAATCAACTTAGTGGCTCTATACCACCATCAATCTTCAACATTACCACAATGCAAATCATTGATTTTACTGGCAATAATCTTACTGGTAAGCTTCCACCAAGCCTCGAGAAATGCAGAAAGCTTCAAGTCTTGGCATTGTCGCTCAATGAGTTTATAGGAACTATACCAAGAGAGATAGCCAACTTAACAGTTCTTACAGAATTATATCTCAAAGCTCTGCATTTGGAAGGTAGTATGATATTTCCTTACAAGAAAAATGAAATTTTTCTTCCTTTTCTCTCTTCCGCTAAACTGGTAACTCATATACTTCCTGAAATTGACAGGAGAGATCCCTCGAGAAATCAGTGATCTTCGTTACTTGACTATCCTAGATCTGCAATTTAATCAACTTAGTGGCTCTATACCACCATCAATCTTCAACATTACCACAATGCAAATCATTGATTTTACTGGCAATAATCTTACTGGTAAGCTTCCAACAACTATATGTGACCATCTTCCAAACTTGGAAGGGCTTCACCTCACAAGCAACTCCCTAGATGGAGTTATTCCACCAAGCCTCGAGAAATGCAGAAAGCTTCAAGTCTTGGCATTGTCGCTCAATGAGTTTATAGGAACTATACCAAGAGAGATAGCCAACTTAACAGTTCTTACAGAATTATATCTCAAAGCTCTGCATTTGGAAGGTAGTATGATATTTCCTTACAAGAAAAATGAAATTTTTCTTCCTTTTCTCTCTTCCGCTAAACTGGTAACTCATATACTTCCTGAAATTGACAGGAGAGATCCCTCGAGAAATCAGTGATCTTCGTTACTTGACTATCCTAGATCTGCAATTTAATCAACTTAGTGGCTCTATACCACCATCAATCTTCAACATTACCACAATGCAAATCATTACTCTTACTGACAATAATCTTACTGGTAAGCTTCCACCAAGCCTCGAGAAATGCAGAAAGCTTCAAGTCTTGGCATTGTCGCTCAATGAGTTTATAGGAACTATACCAAGAGAGATAGCCAACTTAACAGCTCTTACAGAATTATATCTCGAAACTCTGCATTTGGAAGGTAGTATGATATTTCTTTACAAGAAAAATGAAAATTTTCTTCCTTTCGGAGCTCTGCATAAGGTAAGTGTAACTCATATATTTCCCAAAATTGACAGGAGAGATACCAATGGAGCTAGGTAATCTTAAGAAACTTCAGGTACTGCAGTTAGCAGAAAATGAGCTTACTGGCTCTGTTCCTGACAGCATTTTCAACATGTCAGCACTGAAGATTATAGATTTTGGACAAAACAAGCTTTCAGGTACTCTACCTTCAGATTTAGGTCGTAGAATTCCCAACCTAGAAGTATTTCTTTGTGGAGGAAATAATCTGAGTGGTTTTATCTCTGATTCAATCACAAATTCTTCAAGACTCAGAATGTTTGACATCTCATTCAACAGTTTCACAGGTCCAATTCCTAAATCATTTGGTAACTTAGAATACCTTGAGATTCTTAACTTAGAGATGAATAATTTTATCAGTGATTCATCATTGAGCTTCCTTACATCATTAACAAACTGTAGGAAACTAAGAGCACTCCGATTTAATGAGAATGCATTGGATGGGGCATTACCAGCGTCTGTTGGTAATTTCTCAAACTCGATGCAAAATTTTCAAGGAAATGGCTGTAAGCTAAAGGGCGTCATTCCTCGAGAAATCGGTAATCTTACTGGAGTGATACATATGAGTCTGTTTAACAATAAGTTGGCTGGACATATTCCAAATACTGTCCAAGACATGTTGAACCTCCAAGAATTTTATCTACATAGCAATGAGATTAAAGGTACCATACCAAATGTTATATGCAGTTTAAAGAATCTTGGTGCATTAGACTTGTCAGGAAATCATTTTTCTGGTTCAGTGCCCTCATGCTTAGGGAATGTAACTAGTTTGAGGTATCTTAATCTAGCTTACAACAGGCTGAATTCAAGATTACCTGCAAACTTGGGGAGCCTTCAAGATCTCATTGAATTAAGTGTTTCATCCAATTCATTAAGTGGTCATATTCCTCTGGAGTTGGGAAATTTAAAGGCAGTAACACTAATTGATTTGTCAAAAAATGATTTTTCTGGTAAAATTCCTAGCACTTTAGGAGGTCTAGCTGAATTGATTAATCTTCCTCTAGCACATAATAGATTAGAGGGGCCTATTCCAGATTCATTTGGCAAATTGTTGGCATTGGAATTCTTGGATTTGTCTTATAACAATCTTAGTGGTGAAATTCCAAAGTCATTAGAAGCTCTAGTGTATCTCAAATACATGAATTTCTCATTCAATAAACTCAGTGGAGAAATTCCCATTGGTGGTCCTTTTGTAAATGTCACTAGTCAATCCTTCTTGTCCAATGATGCACTTTGTGGTGACTCCCGGTTTAACATAAAACCATGTCAAACTAAATCTACAAAGAAGTCAAGAAGAAAAAGAGTGCTTATAGGTTTATATACGCTGTTAGGGATTGGGTCGCTCTTTGTGTTGGTAGTTGGATATGTGGTGTTAAGATTGAGAAAGACAAAGAAGAATGCAAGTCAAGCAGATGTGTCTCTGGTAAAAGAACATGAAAGAATTTCCTATTATGAACTTGAACATGCAACAGAAGGATTCGACGAAAGCAACTTGCTTGGTACAGGGAGTTTTAGTGTGGTCTACAAAGGGATACTTAAGGATGGTACTCTTTTGGCAGCAAAGGTATTCAATGTGCAATTGGAGGGTGCATTTAAAAGTTTTGATACGGAATGTGAGATACTCCGGAACCTTCGCCACAGAAATCTGACCAAAGTTATTACAAGCTGTGCCAACCTTGATTTCAAGGCCCTGGTGTTGGAATACATGCCCAATGGGACACTTGATAAATGGTTATACTCCCACAACTTGTTCTTGAACTTATTACAGAGATTGGATATAATGATAGATGTTGCATCTGCAATTGACTATCTCCACAATGGCTATACAACGCCTGTGGTTCATTGTGACTTGAAGCCAAGTAATGTCTTGCTTGATCAGGAAATGGTTGGTCATGTCAGTGATTTTGGCATTGCGAAATTGCTAGGTGAAGGGGAGGCTTCTGTTCAAACAAGGACAATTTCAACCATTGGATATATTGCTCCAGGTATATTAAAACTTCTTAAAGTTTTTCTTGTTTCTCCTTTCAGTTTTGGCATATTCCATTTTAACTAGGAAGCTTTTATGATAATTTTTTAAAAATAATCGTAGAGTATGGACAAGATGGCATAGTATCCACGAGTTGTGATGTTTATAGTTTTGGCATCCTGATGATGGAGGCATTCACAAGAAGAAGACCAAGTGATGAAACATTTACCGGTGAATTGAGCATACAGCGTTGGGTTAGTGATTCCTTTCCGGGTGAAATTCATAAGGTGGTGGATTGCAATTTGGTACAGCCAGGGGATGAACAAATTGATGCAAAGATGCGATGTTTGTTATCTATCATGGAATTATCTTTGAACTGCACTTTAGTGAAACCTGATGCAAGAATTAGCATGAAAGATGCTCTTTCAACACTCAAAAAGATCAGGCTACAGCTTGTCAATAATCGGCATTAG

>PGSC0003DMG400020848 LRR-RLKs

ATGGGAAGAAGTTGCAATCTTCTCTTTGCTCTTGCAGTTCTCATTCTACTCCATCACCATACTTTACTTGCTACTCTTCCTAATATTAGCACTGACGAAGCTGCTCTTCTTGCACTTAAATCACACATTTCTTCTTATCCTAACATCATCTTAGCAAGCAACTGGTCTTCTTCCAGCCCGGTTTGCAGCTGGATTGGAATCACCTGCAGCTCTCGCTACCATCGAGTCACTGCTTTAGACATTTCTAGCATGCAACTTCATGGTACCATTCCTCCACACCTTGGAAACCTCTCATTTCTCGTCTCCCTTGATATCAGTAACAACACTTTCCATGGAGATTTGCCAGTAGAGTTGGCTCATTTGCAGAGGTTGAAACTGATTAATGTCAGAAGTAATAACTTCACCGGAACCATTCCATCATTTTTAAGTTTGTTACCAAACCTACGCTTTGCGTACCTCTCGAGCAACCAATTTTCGGGGAAAATTCCATCATCTATTTCCAATCTGACAAAGCTGGAATTGTTGACAATACACACAAATTTTCTCGAAGGAGAGATTCCTAAAGAAATCGGTGATCTTCGTTACTTGATTGTCCTAAACATGCAAGATAATCAGCTTACTGGCTCTATACCACCCTCAATCTTTAACATTACTACAATGCAAGTAATTGCTCTTACTGACAACAATCTGACCGGAAATCTTCCACGAACTATATGTGACCGTCTTCCAAACTTGGAAGGGCTTCACCTCTCAAGCAACTACCTAGGTGGAGTTATTCCACCAAACCTGGAGAAATGCCGAAAACTTCAAATACTGTCTCTGTCTTACAATGAATTTACTGGAACTGTACCAAGAGAGCTATCCAACTTAACAGCTCTTACAGAATTATATCTTGGAATACAACACTTGGAAGGTAGTATGCTATTTCCTCACAAGATATTTGAAATTTTCTTCCTTTCCTCTCTTCACCAAATTTGTACTCATATATTTCCCAAAATTGACAGGAGAGATACCAGCAGAGCTAGGTAATCTTAAGAAACTACAGCTGCTGATGTTAGACCAGAATGAGTTTACTGGTTCTATCCCTGAAAGCATTTTCAATATATCAGCAATGCAGATCCTAGATTTTTCAATGAACAAGCTTTCAGGTACTCTACCTTCGGATTTAGGTCGTGGAATGCCCAATTTAGAAGAACTTTATTGTGGAGGAAATAATCTGAGTGGTTTTATATCTGATTCAATCTCAAATTCTTCAAGACTCAGAATGGCTGATCTCTCAGACAATAGTTTCACAGGTGTAATTCCCGAATCACTAGGTAACTTAGAGTACCTTGAGGTTCTGAACTTGGAGCTGAATAATTTTATCAGCGATTCATCATTGAGCTTCCTTACATCATTAACAAACTGTAGGAAACTAAGAGCACTCCGGTTTAATGATAATGCATTGGATGGGGCATTACCAGCGTCTGTTGGTAATTTCTCAAACTCGCTGCAAAATTTTCAAGGAAATGGCTGTAAGCTAAAGGGCGTCATTCCTCGAGAAATTGGTAATCTTACTGGAGTGATATATATGAGTTTGTATAAAAATGAGTTGACTGGACATATTCCAAATACTGTCCAAGACATGTTGAACCTTCAAGAATTTTATCTACAAAGCAACGAGATTGAAGGAACCATACCAAATGTTTTATGCAGTTTAAAGGATCTTGGTGCATTAGACTTGTCAGGAAATCATATTTCTGGTTTGGTGCCATGCTTAGGGAATGTTACTAGTTTGAGGAAACTTAATCTAGCTTACAACAGGCTGAATTCTAGATTACCTGCAAACTTGGGGAACCTTCAAGACCTCATAGAATTCAATGTTTCATCCAATTCGTTAAGTGGCCATATTCCTCTGGAGTTGGGAAATTTAAAGGCAGTGACACTCATTGATTTATCAAAAAATGATTTTTCTGGTAAAATTCCTAGCACTTTAGGGGGTCTAGCTGAATTGACGGATCTTTTTCTAACACATAATAGATTAGATGGGGCTATTCCAGATTCATTTGGCAAAATGTTGTCACTGGAATACTTGGATTTGTCCTATAACAATATTAGTGGTGAAATTCCAAAGTCATTAGAAGCTCTTGTGTATCTCAAATACATGAATTTCTCATTCAATAAACTCAGTGGAGAAATTCCCACTGGCGGTCCTTTTAAAAATGTCACCAGTCAATCTTTCCTATCCAATGATGCACTCTGTGGTGACTCCTGGATTAATGTAAAACCATGTCAAAGCAAGTCAACAGAGAAGCCAAGAGGAAAGAGAGTGCTTATCAGTTTATATACATTGTTAGGGATTGGATCACTCCTTGTGTTGGCCGTTGGATATGTGGTGTTAAGATTGAGAAAGACAAAGAACAATGAAAGTCAAGCAGATGTGTCTTTAGTAAAAGAGCATGAAAGGATTTCCTATTATGAACTTGAACATGCAACAGAAAGATTCGATGAAAGCAACTTGCTTGGTACCGGGAGTTTTAGCATGGTCTACAAAGGGATACTTAAGGATGGTACTCTTTTGGCAGTAAAGGTATTCAATGTGCAATTGGATGATGCATTCAAAAGTTTTGACACGGAGTGTGGGATACTCCGGAACCTTCGCCACAGAAATCTGACCAAAGTCATCACCAGCTGCTCCAATCTTGATTTCAAGGCCCTGGTGTTGGAATACATGCCCAACGGGACACTTGATAAATGGTTATACTCTCACAACTTGTTCTTGAACTTATTGCAGAGATTAGATATAATGATAGATGTTGCATCTGCAATGTACTATCTCCACAATGGCTATTCAACGCCTGTTGTGCATTGTGACTTGAAGCCAAGCAATGTCTTGCTAGATCAAGAAATGGTTGGTCATGTCAGTGATTTTGGCATTGCAAAATTGTTAGATGCAGGGGAGGCTTTCGTTCAAACAAGAACAATTTCAACCATTGGATATATTGCTCCAGGTATATTAGTTAAGCAATTCTTTCCCCTAGATATGGATTGATGCTTTCAGTTTTGGCATATTCCATTTTAACTAGGAGGCTTTTATGATCATTTTTTACCATGATTACAGAGTATGGACAAGATGGAATAGTATCCACAAGTTGTGATGTTTATAGTTTTGGCATTCTGATGATGGAGACATTCACAAGAAGAAGACCAAGTGATGAAATATTTACCGGAGAATTGAGCATACAACGTTGGATTAGTGATTCCTTCCCAAGTGGAATTCATAAGGTGGTGGATTACTCTTTGGTACAGCCAGGGGATGAACATATTGATGCAAAGATGCAATGTTTGTTATCAATTATAGAAGTAGCTTTGAGTTGCACTTTAGTGACACCTAATGCAAGAACTAGCATGAAAGATGCACTTTCAACACTCCAAAAGATTAGGCTCCAGCTTGTCAATAGTCAACACTAG

>PGSC0003DMG400021008 LRR-RLKs

TTCCTTTCAATATGCTATCTGTCATGGTATTTGTAGTGGTTGTAATCAAGGTGTACTGGTATTGATTTCTTGTTCTAAGTGAACCCCATTTTACCTTGTGGTGTTAGAATGAAGTTTATCAAGTTCTTGATGATTTTTGTTCTTTATTTTGGCTCTGTCATTGGGCAGCTTCCCTCTCCGGACATATTGGCTTTGCTTGAGTTTAAGAAAGGTGTCGAACATGATCCAACGGGGTTTCTTCTTGAATCTTGGAATGAGGAGTCTATTGATTTCAATGGATGCCCTTCATCTTGGAATGGGATAATGTGTAATGGTGGTAATGTTGCAGGAGTTGTTCTTGACAACTTGGGTTTAGTTGCTCATGTTGATTTGAGTGTATTTGCTAATCTTACAAAGCTTGTTAAGCTTTCAATGGCTAACAATTCAATTGCTGGGAAAATGCCTAATAACATAGGTAGATTTAAGAGCCTTCAGTATCTAGATCTCTCAGGTAATCTATTCAACTCTTCTTTACCCCCAGAGGTTGGACAGTTAGGGAGGTTAAAGAATTTGTCATTAGCTGGTAATAACTTCTCCGGTATGATACCGGAGACAATTTCAGGGCTTGTGTCTGTGCAATCTTTGGACTTGAGTCGCAATTCTTTTTCTGGGGGTCTACCTTCATCTTTGATGAAGTTGGCTGGTTTGGTATACCTGAATCTATCTGTTAATGGATTCACAAAGGAAATTCCTAAAGGATTTGAGCTGATGGAAAATCTTGAAGTGCTTGATTTGCACGGAAATATGTTTGATGGCAATTTGGATGCAGAAATGTTGCTGCTTACCACTGCAACTCATGTTGACTTGAGTGGAAATTTGCTGGTGAACACTGCTTCTCAGCAGCAGAAATTTTTGCCAGGTTTATCGGAGTCGGTGAAGTATTTAAATCTTAGCCATAATCAGCTAAAGGGTTCACTTGTAAGTGGAAATGAAGCTCAGATATTTGGCAACTTGAAGGTTTTGGATTTGAGCTACAATCAGCTCTCAGGGGAATTGCCTAGTTTCAATTTTGTTTATGATCTACAGGTGCTTAAACTTGCCAACAACTTATTTTCCGGATTTATTCCGAATGATCTTTTGAAAGGAGATGCATTGCTTCTCACTGAGTTGGATTTAAGTGGAAACAACCTCACTGGTAAATACATGAAACCCTTTGGTAAATAAAGTTATCTGTTTGCTAACAACTCTTTTCGCTGTTTTTCTCAATCTCTTCATTTTGTAGGTTCAATAAGCATGATAACATCCACAACATTGCGAATGCTTAACTTATCCTCCAATGCTCTTTCTGGTGAACTTCCCTTTGTGACTGGAACTTCTGCGGTGCTTGATCTCTCGAAAAATCAGTTCAAAGGTAATCTAACTAGAATGCTGAAATGGGGGAACATTGAATTTCTTGACCTTAGCCAAAACCGTTTGACTGGAAACATCCCTGAGGTTACGGCTCAGTTTCTGCGTTTATATCACCTCAACCTTTCGAGCAATACCTTGACAGGATCTCTCCCGAAAGTTATTTCACAGTTTCCTAAGATCACTGTCCTTGATCTCAGTTTCAACCAGTTGGATGGGCCTCTTCTTACTTCTCTACTAACTTTGCCCACTATTGAAGAACTTCACCTACAAAACAATGCACTTGTTGGAAGCATTGATTTCTCTCTTCCGGCTAGTACTCCTAAGCTTCGTGTTCTAGATCTTTCTCATAATCAGCTAGCTGGTTACTTTCCTGATGGATTTGGGTCATTGATTGCACTTCAAGTACTCGATATATCAGGAAATAATTTCTCTGGATCTTTGCCTACTTCCATGGGTAATGTTAGCTCTCTTACTTCTTTAGATGTTTCAGAGAATCATTTCTCGGGTGAACTGCCAAAAAACCTGCCTAACAGCCTCCAAAGCTTTAATGCATCACTCAATGACTTCTCTGGTGTCGTGCCTGAGAATTTGAGAAAGTTTCCTTTATCATCTTTCTACCCGGGAAACTCTCAACTTCAATTTCCAAATCCTCCCTCAGGATCTGGTCAAGCTTCAACAGAAAAGCATAAGAGCAGGCAACTTAAAACTATTATCAAAGTGGTGATCATTGTTGCTTGTGTGGTTGTTCTTGTTGTTTTAATCATGCTAGCCATTTTTGTATTTTACTTGCGGGCATCGAGGAACTCTCAGGCACATGTTACTGACAAAGACATCCGCCGTCAAGCCCAATCAAATCCTTCTGGTTTTAGCAGTAGAGAAGGTGCAGTAGTTTCAGCACAAGATGTTACAGCTGCTTCACGGAAAGCATCATCATCAGAAGTAATTAGTGCTGATGAGAAAACGACAGGTTTCTCCCCTTCGAAAACAAGTCATTTCACTTGGTCACCAGAGTCAGGGGAATCATATACTGCAGAAAGCCTTGCTGATAATTTGGCTGGAGAGCTGTTCTTCCTTGATGATACAATCTCTTTTACAGCTGAGGAACTATCTAGGGCCCCAGCTGAAGTCTTAGGCAGAAGCAGCCATGGGACATCTTACAGGGCAACACTAGATAACGGGTTGCTCTTGACTGTGAAATGGTTGAGAGAAGGAGTGGCCAAACAGAGAAAGGATTTCACAAAGGAGGCTAAAAAGTTTGCCAACATTAGACATCCAAATGTGGTGGGATTAAGAGGTTACTACTGGGGTCCCACACAACATGAGAAGCTTATTCTTTCAGATTACATATCTCCCGGAAGTCTATCAAATTTTCTATATGGTAAACTTTATCTTCTTGCATCAACCTTGCAGTTCCTCTTATGCATATCGTTTGCTATAACACTTTGAACTCATAAAACACTTATAATCTTTCCTTCTTTTGATATTTTTTAGTATTTCATGGCAAACAATTTGTCAGTTTGATATCTTGACAGTTCTTGTGGTGCATGTATTTTCTCAGTCTGTGATACACGAATTAAGGATAAGAAGCCATTTCCATTTTTAGTTAGCATAGTGAGTCCAGCTAATTCATTCATTCATTATACAGCTTGTAATCAGTTCATTCACAATCATATATGAAACTACACTCTTTCTCTTCTGAATCTATTCTCTCAATATTTCTCCTTCTTTAGCTGTTCTTCTTGAGGGAACTCCTGAAACAGTCTCTTGATCCTCGAATTCCTGCAATCAAGTAACAGTTTGTATTATCATGAAATATAGTATCACTTTCACTGTGGAATTCTTCAACTTAAACTGCTGAAGTCTAGCTTCTGATTTATTGTTGGAAAACTTTAAACTTGTCCAAGGCCCTATACACTACGTCCGACCGGACGTGTAAGCAGGCCTTTACTTTCTTAATATCTCTGCACTGGCAGTAAGGTTCAGCAATTGTGGTACTTTGCTCATAAAAAGGTCACAATGAACTGCAAATACTTCAATGCTCAAATACCTTATTTTCGATTGGTGATTCTACCTGCTCAACTTCTCTACTATTTTGTATGCTTTTGTGTAGTTTGTATAGTGTAAGGTTTCTGTATATACATTTAACGTGCGGCACCTTTGCAGATCGACCTGGAAGAAAAGGTCCACCACTAACCTGGCCCCAAAGACTCAAAATATCAGTTGATATTGCGCGTGGCCTGAATTATCTCCATTTTGACCGTGAGGTTCCTCATGGAAACCTTAAAGCAACCAACATCTTGTTGGATGGTCCTGATCTTAATGCACGAGTTGGTGATTACTGCCTTCACCGTCTTATGACTCAAGCCGGTACAATTGAACAGATTCTTGATGCTGGAGTGTTAGGTTATCGTGCACCAGAGTTGGCTGCATCGAAGAAACCAGTGCCTTCCTTCAAATCAGATGTATACGCTTTTGGAGTCATTTTGTTGGAGGTCCTAACTGGGAAATGTGCAGGTGATGTTGTTTCTGGTGAAGATGGTGGTGTAGACTTGACTGATTGGGTAAGGTTGAAAGTTGCAGAAGGTGGAGGATTCGATTGTTTTGATAATGCATTGTCATCTGAGATGGGAAATCAAACAACAGAGAAGCAAATGAAGGAGGTTCTTGCAATAGCTCTAAGATGCATACGTTCGGTATCTGAGAGGCCAGGTATCAAGACTGTATATGAGGACCTTTCATCTATTTAGTTTTCCTCATTTTGGCTTTTTGCTTGGGAATCAAATAGCTCATTGGCTTCCCTTGAGTTGTAGAATTAGCCAGTTTAGTGTGTGAATGGGCTAGTTTTATTAACCATTGTTTGTTGTAAATTGGGGCATATTTATAATGTATACAACTTTCACCCCCCACCCACCCCCC

>PGSC0003DMG400021017 LRR-RLKs

ATGGAGGAGTACATTTTCTTATTGATTCTTCTCTTTCTACTTCAATTTTCTGTATCATTTGCAACCTCAAATGAGACAGACCAACAAGCTCTACTAGCTTTCCAAAATCTTATTACAAGTCCTAGTCATTTTTTGGCCAACAATTGGACCAACAATACTTATTTTTGCTCTTGGTTTGGTGTCACTTGCACTTCAAAAACCCAAAGGTTTGGGGCCTTGACTCTTCCTAATTTGCAACTTCAAGGCACAATATCCCCGTTTTTGGCCAATTTGTCCTTCCTCGATGTTCTCAATCTTGGGAACAACAACTTCCATGGTGGGATCCCTTATGGACTTGGCCACTTGCCTTGCTTGCGAGTGATTGATGTTCAAAACAATCAGCTAGAAGGAAGTATTCCAACAAGTCTATTTCAACACCAGAAAGTTCAAGTTATTTCATTGGGCTTCAATAAACTCGGTGGTGAAATGTGGAAAGGGCCATGGTGTGTACCGAAACTCAGAGTCTTAAATCTCAAGAACAATAGCCTCACGGGTATAATCCCTCCTTCTATTTGAAATGCCACAAAGTTGCTTAACTTCAGTTTGTATGGAAATAGAGTTGGTGGCAACATTCCAAAGGAAATTGGTAATCTGAGACAACTTGAATTTTTGTCCTTGGTCGATAATCAATTAACAGGTTTCGTAACTGCAACATTGTTTAATATCTCATCACTTCGTGTCCTATCTCTTGGAATCAAAAGACTTTCTGGTCCTCTCTTGCTTGATGAAAGGAATATTGTGTCAAATATAGAGTTTCTAAGTATATCTAACAACCAAATTTCTGGCTGCATTCCTTCCAACATATGCCAACTCACAAAGCTCAAAGTCTTGTCCAGAGCTTTCAACAACATAACTGGAGAAATACCCAGAAATATTGATTGTTTATCCAAACTTGAACAGTTTCTTATTGGTGATAATCCAATAGAAGGGACTATCCCCACTTCATTGGGAAATATATCCACTCTTCAATATATTTATTGTGGAAACAATCGCATAGTGGGGAAAATTCCCCAAGAATTAGGGAAGCTATCAAATTTGAGGCAATTAAACTTTGCCTCTAATTATAATCTTATTGGTCAAATTCCAGAGGCTATTTTTAACATATCTTCTTTGGAAATCATTGATTTCAATTTCAATAGCCTCTCAGGTAGAATTCCAGTCACTACAGGTCTTCATCTTCCAAACCTTAAAGTACTTGAGTTGGGAGCCAATCAGATCAACGGGGAAATTCCATTGTTCATAACAAATGCTTCCAAGATTGAGAAACTGGTACTAGCACGGAACTTTCTCACGGGCGCTATTCCTAATAATTTAGGAAATCTTCGTGAGTTGCGATTACTGTTCTTGCATACTAATCAACTTACCAATGAACCAAGAGAGCATGAATTGTGATTTTTCAATTCTTTGGCGGATTGTAGAATGTTGAAATATCTAGAAGTAGGTTCCAATCCGTTGATTGGCATTCTTCCCAATTCTATTGGGAATCTTTCATCTACTGTTGAATACTTTGAAATAGGATATGCACACATCAACGGCCTCATCCCCATAAGTATAGGCAACATTAGCGGCCTTACAGCCTTGTACTTTGAAGACAACAACTTGGCAGGGAATATTCCTTCTGAGATTGGTAAGCTTAAACAACTCCAAGGGTTGTACCTATCTAAAAATAAATTGGAGGAACATAAGAGGCGGTATGCCATTTATCTAATTTGGTTGAATTAGGTTTGGTTGATAATGAGCTACCTAGATTAATTCCAGAATGCTTAGGTAATCTTAGCATGCTTCAAAAGCTTTATTTGGGTTCTAATAAATTTTTATCAAAAATTCCCTTGAGCCTTTGGAAGATGAGTAGCCTTCTCTATCTTGACGTGTCACATAATTCAATAGAGGGAAAAGTTCTCGAGGATATTGGAGGACTGAAGGCCATTGTAGAACTACATCTTTCCGGTAACCACTTTTCAGGCATGATACCAAGCAGATTCGGGGAACTCCAAAACCTGCAGTCTCTTGACCTGTCGAACAATTCATTTTTTGGACAAATTCCATCTTCCTTTTCCAACTTGATAAGCTTGGAATTCTTGAATTTGTCTTTAAATGCATTCTCAAGTACTATTTCTATGTCTTTGGAAAAACTTTCATCCCTTAAAAGCATTAATGTTTCATTTAATGATTTAAAAGGTGAAATACCTAGTGGCGGTGTGTTTGCAAATTCCACACTACAATCATTTCTCGGGAACAAAGGTCTATGTGGAGTGCACATATTGGAGATTCCTGCTTGTGCTATCACTAATCCTGGACAACAATCAAAGCTTAAGGAGGTTGTGCTAAAAATTGTTTCTCCGATGGTTATTTCATCCTTTCTCATATTCTTGTTGGTTTCAATTTGGATAATGAAACGACAAAAGAAAGGAAAGTCCAAAGATGTGGAGAAGGCTTTAGAGATCAAAACTCATCAATTGATTACTTATCACAAGATTCAACAAGCAACAAATAATTTTGATGAATCCAATTTAATTGGTGTGGGAAGTTTTGGCTCTGTGTACAAAGGCACATTATCTGGGGGGAAGTGTAGTGGCAATAAAGGTTCTAGACTTGGAAAATGAGCAAGTATGCAAGAGGTTTGATACTGAATGCGAAGTGATGAGAAATGTTAGACATAGAAATCTTGTTCCAATGATTACTACATTTTCTAGTGACTATATAAGAGCCTTTGTTCTGCAATTTATGCCCAATGAAAGTCTTGAGAATTGGTTGTACAAAGAAGATTGTCACTTGAATCTTCATCAAAGAGTCACTATAATGCTTGATGTAGCTATGGCAATTGAATATCTACATTATAGTCATGTCAATTAGATAGTTCATTGTGACCTAAAGCCAGCCAACGTTCTTTTGGATGAAGATATGGTGGCTCATATTGGTGATTTTGGAATCTCTAAAATTTTAGCCATAAGCAAGTTCATGGCTTATACCGAGACATTAGGCACTCTTGGATACATTGCACCAGGTATAAAGAATCTACTCCTTTGATTTTCTCTTATTATAATTATGACTCTCCAAATTCTAATGGTAAAAAAGGTAAGTCTTTATTTATGCAGAATTATTGTTTTATTTCAATTGAGTAACTTTTCTTCAATCCTTTTTTTAAGAATATGGCTTGGAGGGAATAGTGTCCGCTAATGGTGATGTTTATAGTTATGGCATCATGTTGATGGAGGTTTTGACCAAAAGAAGGCCAATAGATGAAGAGATATGCAATGAAAATCTTGACTTGAGAAAATGGATAACACAATCATTTTCAGGGACTATGATGGATGTTTTGGATGCTAATTTATTTTCTAAGGAAGAACAAATCACTTCTAAAAGTGAAAGTTGCATCGCCTGCACGATGGAACTAGCTTTTGATTGCACAAAGGAAATGCCAGAATCAAGGATAACCATGAAAGATGTAGTCAAGAGGCTTAACAAAATCAAGAACACGTTTCTAGAATCATAG

>PGSC0003DMG400021018 LRR-RLKs

ATGAGTGGTCTTCTCTATCTAGACGTGTCACAGAATTCAATAGAGGGAGAAGTTCCATCAGATATTGCTGGACTGAAAGCCATTGTAGAACTACATCTTTCCTATAACCACTTTTCAGGCATGATACCAAGCCAATTGGGGGACCTCCAAAACTTGAAGTCTCTTGACTTGTCGAACAATTCGTTTTCAGGCTCAATTCCATTATCCTTTGCCAACTTGATAAGCTTGGAATTCTTGAATTTGTCTTTAAATGACTTGTCAGGTACTATCCCTATGTCTTTGGAAAAACTCTCATCCCTTAAAAGCATTAATGTTTCATTTAATGGTTTGGAAGGTGTAATACCCATTGGTGGTGTGTTTTCAAATTCCACTCTACAATCATTTCTCGGGAACAAAGGTCTATGTGGAATGCACATATTGGAGATTCCTGCTTGCACTATCACTAATCCAGGAAAACAATCAAAGCTTAAGGAGGTTGTGCTAAAAATTGTTACTCCAGTGGTTATTTCATCCTTTCTAATATTCTTGTTGGTTTCAATTTGGATAATGAAACGAAAGAAGAAAGGAAAGTCCAAGGAAGTGGAAAAGGTACCGGAGATCAAGACTCATCAATTAGTTTCTTATCACGAGATTCAACGAGCAACAAATAACTTTGATAGGTCAAATTTAATTGGTGAGGGCAGCTCTGGCTCTGTGTACAAAGGCACATTATCTAGTGGAACTGTGGTGGCAATAAAGGTTCTGGATTTGGAAAATGAGCAAGTATGCAAGAGGTTTGATACTGAATGTGAAGTGATGAGAAATGTTAGACACAGAAATCTTGTCCCTGTGATTACTACTTGTTCTAGTGACTATATAAGAGCCTTTCTTCTGAAATATATGTCCAACGGGAGCCTTGAGAATTGGTTGTACAGAGAAGAGTCCCACTTGAACCTACTCCAAAGAATCACTATAACGCTTGATGTGGCTGTGGCAATCGAATATCTACACCATGGTCATAACACTCTGATAGTTCATTGCGACCTAAAGCCAGCCAACGTTCTTTTGGATGAAGATATGGTGGCTCATGTTGGTGATTTTGGAATCTCTAAAATTTTAGTCGTAAGCAAGTCCATGGCACATACCGAGACATTGGGCACTCTTGGATATATTGCACCAGGTAAACGAGTCTCTTTCTATTTTCAGAAAAATCACAATTGAGCCCTTCCAAATTCTAGAACTAAATAAAAGCAAGTGTCTGTTAATACATAATTATTGTTATATTCAAATGAGTAATTCTTTTTCATTTTTCTTCTTTTAAGAATATGGCTTAGAAGGAATAGTGTCTGCTAGTGGTGATGTTTACAGTTACGGCATCATGTTGATGGAGGTTTTGACCAAAAGAAGGCCAACAGATGAAGTGATATGCAATGAAAATCTTGACATGAGGAAGTGGATCACACAATCATTTTCAGGGACTATGATGGACGTTGTGGATGCCAATATTTTCTCTGAGGAAGAACAAGTCACTTCTAAAAGTGAAATCTGCATAGCCTCCATGATAGAATTGGCTTTAGACTGCACAACGGAAATGCCAGAATCAAGAATTACCATGAAAGATGTAGTCAAGAGGCTTACCAAAATCAAGAACACATGGGCCACGAGGGGTTAG

>PGSC0003DMG400021184 LRR-RLKs

TTTTTTTTTAGAGAGAGAAGCTGAGAGTTGTAAGAGAGAGAAAATATGTCTGCAAGAAGACAAATGTTGAGTGTCACTGCCATTTCTGGGAGCTGTGAGGTTTCTCTTTCATTCATATACAAAACAAGACTAGGGTTTTACCATTGTGGTTTGGTTTTTTACTGAGCTCATAGAAATCTTGAACTTCATTTATCAAGATTTCTAAAGGGCTAAATACTTTGCTGGCCCATTTCATTTTCTTGTACTTTAATGCAGATTTTTTTTGAGGCCACCAGTTGAGTGCATTAAAGAACTTGAGGTTCCAAGATAACCTGAAGCTGTAATCATTGCAAGAAGTTGAATTTCTTCAGTAAATCTTTGAAAAAGACTCTTCTTTTTTTGTTTTTCTTCTAGTTGAGCTGATGTTCTTGGTAAGGGAATGCTGAACTTGTTTTACTATCTTGAATCTTTGAAAGTTCAGTCTGTACTTTGTTTTTGTCACTCATTTCAGCTTGTATACCTAGAAAAGTTTCTATTTTGAAAAAGACTGGTGATTTTGGGTTGAAGTTGTGTACTTTTCCCCCCTGTTGTTTTACTGCTTTGAACTGTGATGGTTGTTGTGAAAGATGGTTTTCATATGTTGAGGTTTTACTTAAAGTGATTCTCATACTTAAAGCTGCTCCTTTTGGATTTCTTGGAAGGTTGAAGATTTAGTTGTATTTTTGGGAGTGAGTTTGTTGTCACTTGGGCTAGTAAAGCTTTTTCTGGTGTTGCAATGAACCTATTTAGGTTTTTGATGCTTGTGCTTTGTTTTGGTTCTGCAATGGGGCAGCTTCCTTCCCAGGACATTTTGGCCCTGCTTGAATTTAGGAAGGGTATCAACCATGATCCAACTGGTTATGTACTGCAGTCATGGAATGAGGAGTCCATTGATTTCAATGGTTGCCCTTCATCTTGGAATGGTATAATGTGCAATGGTGGTAATGTTGCAGCAGTTGTCCTTGACAACATGGGTTTATCTGCTGTTGCAGATTTGAGTGTGTTTGCTAACCTCACAATGCTTGTGAAACTCTCTATGGCTAACAATTCAATTGCTGGACAAATGCCAAAGAAAATTGGTGATTTTAAGAGTCTTGAATATCTGGATATTTCAAACAATCTATTCAACTCCTCTTTACCACCAGAGATTGGGAAAATAGGAAGCTTGAAGAATCTTTCACTTGCTGGTAACAACTTCTCTGGCCCAATCCCTGATACTATTTCAGAGCTCATGTCAATTCAATCTTTGGATTTGAGTCACAATTCTCTTTCTGGGCTACTTCCTTCTTCTTTAACCAAGTTAAATAATTTAGTATACCTCAATCTTTCTCTTAATGGATTTACAAAGAAAATCCCTAAAGGTTTTGAGCTAATGGCTAACCTTGAAGTTCTTGACTTGCATGGAAATATGCTTGATGGTACTTTGGATCCAGAGTTCTTACTGCTTACTACTGCAACTTATGTTGACCTAAGTGGAAATTTACTCGCGAGTGCCGCTTCTCAACATGAGAAATTTTTACCAGGCATATCATCGTCAGTCAAGTACTTGAGTCTTAGCCATAATCAGCTTACGGGGTCACTAGTAAGTGGCGGTGAAGCTCAGGCATTTGGAAACTTGAAGGTTCTGGATTTGAGCTATAATCAGTTGTCCGGGGAATTACCTGGCTTCAATTTTGTTTATGATCTTCAGGTCCTGAAACTCAGCAACAACAGATTTTCTGGTTTCGTTCCAAATGATCTTTTGAAAGGAGATACTTTAGTTCTCGCAGAATTGGATTTAAGTGGAAACAACCTCACAGGTATATTCCAAACTGCTCTCTGAAATTCATTTTGCAAATTAAATAGTATTTCGTTGTGTTGAACTTGAGTTACAACAAAGTAGTTAATTTCTGGTCATTTTGTTAGTGTGCTTTACTCAGTCCCTTCATTTTACAGGGTCGATAAGCATGATAACATCAACAACTCTGCGCGTTCTTAACTTATCCTCCAATGCTCTTTCTGGTGAACTTCCACTGGTGACAGGTAGTACTGCAGTTCTTGATCTCTCGAAAAACCAATTAGAGGGGAATCTAACTAGAATTCAGAAATGGGGAAATGTTGAATTTCTTGACCTCAGCCAGAACCAATTGACTGGAAACATCCCTGAGGTTACAGCTCAGTTTCTGCGATTAAATCGTCTAAACCTTTCTCACAATGCTCTTACTGGATCTCTCCCAAAAGTTATCACACAGTTTCCCAAGATCACAGTCCTTGATCTTAGTTTCAACCAGCTGAATGGACCTCTTCTCACTTCTCTGCTAACATTACCCACTATTGAAGAACTTCACCTTCAAAACAATGCACTTGTTGGAAATATTGATTTCGCTGCCCCCTCTGCTACACCCAATCTCCGCGTTCTTGATCTTTCTCATAATCAGCTTGCTGGTTCTTTTCCTGAAGGGTTTGGTTTGTTGACTGCGCTTCAAGTACTTGATATAGCTGGAAATAATTTCACTGGGTCTCTGCCTACATTGATTGGTCAAGTTGGTTCGCTCACTTCTCTAGACATTTCACAGAATCATTTTACTGGTCCTCTGCCGATGAACCTGACTGATGGCCTCCAAAGCTTCAATGCATCACTCAATGACTTATCTGGTGTTGTCCCTGATAATTTGAGAAAGTTTCCTTTATCATCTTTCTACCCTGGAAACTCTGAACTTCAATTTCCAAATCCTCCCTCAGGATCTGGCCAAGCTTCACCAGAAAACCAGAAAAGTCGATCACTTAAAACTATAATCAAGGTGGTAATAATAGTTTCTTGTGTAATTGCTCTTATTATTTTGGTCTTGCTAGCCATTTTCTTTTATTATATACGAGCTTCAAGGAAGCGTCATCCTCGTGTTGCTGAAAAAGTTGTTCATCGCCAAGCCACATCAAATCCTTCTGGCTTTAGCAGTAGGGAGGGTGCCGGTGGTGCAGTTGTTTCAGCTGAAGATCTTATGACTTCACGGAAAGGATCATCAGAAATAATTAGTCCAGATGAGAAAATGGCTGCTATAACTGGTTTCTCCCCTTCAAAAGGTAGCCATTTCTCTTGGTCACCAGAGTCTGGAGATTCGTATACTGCAGAAACTTTTGCAAGATTGGATGTGAAGTCTCCGGATCGTCTGGCTGGAGAGCTGTACTTCCTTGATGATACAATTTCTTTTACACCTGAGGAACTTTCCAGGGCTCCGGCTGAAGTCTTAGGTAGAAGCAGCCATGGTACATCATACAGGGCAACACTGGAGAATGGGTTGCTCCTGACTGTGAAATGGTTGAGAGAAGGAGTAGCGAAGCAGAGGAAGGATTTCGCAAAGGAGGCTAAAAAGTTTGCCAATATTAGGCATCCTAATGTAGTAGGATTAAGAGGTTACTACTGGGGTCCCACACAGCATGAGAAGCTCATTCTTTCGGATTATATATCTCCTGGAAGTCTTGCAAGTTTCCTCTATGGTAAGCTTCATGTATTCTAAACTCTCATATTATGTTTTGCTAATAAAGGTAAAGGCCTTGTGTACCTGTTCTTTAATCTAGTCATTTTCCTCTGACTGTGTGATATATCTTACAGTTCATATGGTTTTCCCATGCACATTATATATGAGCACTTTAATCATTGCTGTGTGATCCTGTAGTGGGAATTTGTGTGAACGACAGTATTTAGTCCTTGGTTTGATGTGGTAATACTTTAGTACATCTAGAGCCCAATATATTATGCATGTCTTTTGTTTATGTAACTTTCACACCAAGTAGTAATACAACAATCACCATGTGCTCTTCGTGATGGTCACTACATAGGCTGTGAGGGATTTGCAATATGGTACTTATTATTTCCAGTAGCTAAAATTCTGTATGATTGAGTTCTTAATGGATCTGAAAATTGTGATGTTTAGCTTTTGTTTCATGATTAACTTTGTTAGAAATCAGCTAGAATATGATACATTTTCCTTCTTTTATGAATCCTTCAAATGGACTTGCTTTCTATTCATTTGACTAACTAAATGTTTTCATTCAGTTGATAATTATATATGTGGTCATTCTCAATATACTTCTCTAATGTGCTGCACGTTTGTAGATCGGCCTGGTAGAAAAGGTCCACCACTAACCTGGCCCCAAAGACTCAAAATATCAGTTGATGTTGCACGTGGCTTGAACTACCTCCATTTTGATCGTGAGGTTCCACACGGAAACCTTAAAGCAACCAACATCTTGTTGGATGGGCCTGACCTTAATGCAAGGGTTGCAGATTACTGCCTTCACCGCCTCATGACTCAAGCCGGCACAATAGAACAGATTCTTGATGCAGGAGTGTTAGGTTATCGTGCCCCTGAGTTGGCTGCGTCGAAGAAACCACTGCCTTCCTTCAAATCAGATGTATATGCTTTTGGAGTTATTTTGTTGGAGCTGCTAAGTGGGAAGTGTGCAGGTGATGTTGTCTCTGGTGAAGATGGTGGAGTAGACTTGACTGACTGGGTAAGGTTGAAGGTGGCAGAAGGTCGTAGTGCTGATTGTTTTGATAACGTGTTGTCACCTGAGTTGGGAAATCCAGCAATGGAGAAGCAAATGAAGGAGGTTCTTGGTATAGCCGTGCGATGCATTCGTTCTATATCTGAGAGACCTGGTATCAAGACTATATACGAGGACCTTTCTTCTATATAGTTTGTACCAAACTCTACTTTGGGAATCAAATAGCTCATTGACTTCCCTTTTGAGTAGTAGAATTAGCACAGTTAGTGTGTGAATGGGGCTAGTTTTATTGACCATTGTTTGTTGTAAATTGGGGCATATTTAATGTGTAAACAAACTTTTCTCCCCTTTTGGCCTATGAGCTATCAATGTTTGTCAGTCTTTCGGAGTAAATACTTGTGACTTTATTTTCCAAGTTTTGCAGCACAGTTTCACAGAACATTCTCAGATGGTCAATCAAACACAATGCTGACATCAACATTGCAGTGCAAACTGACCGAAAAAGGTGTTCAAGAGAAAGTGCCAGGTTATGAACAAATGCTACTGGTTTTTCTTGTCCACTATTCAATAAAAGTTCAAATGCAACATCATTATAAGACACCTAAGGTGGAAAAAGGAAAAGTTACAAGAACCTAGTTATGCCACTAGTTACTATGCAAGTGAGGAGAGGGGTGGGATAAAGGTATGAATTTAAACAACAGGAACTCTAAATTACATGAAAATACATCATAAGATGCAACATATTTTACTGTTGCAACTTTAACAATGTATGTTCGTCAAACCTGAATCCAGCCATCTCTGCCTCCTTGCAGATATCCTTGCACTTAGCTACCCTTCCAGATGCCATATAAACTTTGATCAAGCTGCAGTATATGTCAAAAGTTGGACTAAAACCTGCTTCCTTAAGTTTAGAGAAATAGCGTACAGCCCTTGAGAGATCGTTCAAAGCAACAAAAAGCTTAATAACTACACGATAAGCTGGAAGATCAAAAAGGCAATTTGAAGCCTCCATTTCCCATAGCATGTCTATTGCCTCTCTATAGTTGTTCGCTGAATAACGACAGTGAATCAGAGTAGTATACATAACTACACTTGGATTACCACCAGATTTCCTGAACCAGTCATATAGTCCCTCCACGATCTCAAATCTTCCCAGCCTAATGCAAATCTTCATGATAGCTGTACAGTCCTGTGGTGTCAAATTCAGATCATCTCTCACAGCAAGCTCCTCTAATAATACCAAGACAAGTGTGCTTTTATCAGGATCCTTACCAAGCTCTAAGATTAGCTTAGCATACACCCCAGTATCAAGCACTCTGTTAGATTCCTTAGCCATTGAAAGTAGCTTCAAAGCAAGCTTCAGGCTTCCTCCTTTAATATAACCCCTTAACATTGCTTCATACACACCCCGACTTGCCAAGCCAGTGAACTTGTCCAAATCAAATGGCACTTTCAACTCATTTGACCCTGCCAAAACTTCAATGGTAGAACCAAGAATATGATCATCTGGGAAAAGATGGGGTTGTTTCTTCACCCAACAGAATGTTTCAAGAGCTCTCTCCGGAAGCCCCAAGTAACCCAACTCTCGGATTGTCAACGACAATGATCCTTTCCTAAGAAATGGACTCCATTTACTGAGAACCTTTGATACATTTTCCTCAGTTGAAAGGCCTTGTATCTCTTTAGCCAAACCAATCAAGAAAGTAGGATTCTTGTACACTTGCGTTGATATGGATTGCCTTGAACGATTTGTGAATCCCTTTTGTGTAGGAAGTCCTAAAGGTCGAATCTTGTAAGGAAGGGGCAGAGGTAGAGGCCTCTCTCTATCCAATTTTCCAGGCTTCTGAGGGATCCTCCCTTTAAAAAGAGATGAAATTGCTTCCATCTCATCCGACTCCCATATAATTCCCCCATCATCATCACACATCCTTCCTTTTTCTACTTCCAGCTCCAATTCTTCATCCATTCCATCAGAACAAGCCTCAGTTGCAACTCCCTCATCCTCCTCGTACAAGACCCTATTAAAACTATAGTCCATATCTGGTGGTAGCTTGGGTCGTCGAGGATTTCTAAGGTTCTTTGGAAGTTTTCTATGATTAGCTCTGTTATTAGATACAGCCATGAAAACAACTCTTCCCTTTCTTCTTCCAAAACTTAAAGGTGTTCCAGAAGCAGCTGTACTGGCACTACTAGACTGCGAGATAGAATATGTGGGTGAAGACCAAGTAGAGGACTCATAAGTGGCCGAGAACATGCAATAACTCCTGCTGAGCATTTGGTCGAGCAGGTGGTCGGCACAAATTTATAAACAAAGGAATATGAGAGACCTTATAGTGTGACGGAGTATAAAACCAACTGTTAAATGAAAGTTGAGCAGTATTTTTGACATATAAGTTACTAT

>PGSC0003DMG400021353 LRR-RLKs

AGCATTCCTCATATGAAAAATCCACTCTTTATGGTCATTTCTTGCTACTAAATCTTCACTTATAAAAAAAAAGATTAAATTTTTTGGATTACTCTGTAAAAACTACTGTGTTGTCAGCAAGAACAACACCATAGTAGTAGCAACTAGCAAGCATTGTGCTTTTTCTCACTCCAATTATCACTCTCTTTCTCCTTTTTTGTTTTGGGGTTTAGTGATCTGAACCCCATGTTTTCCACCTCTTATTGTCTCAAGATTTGACATGGGGGTCTTCAAGAATCAACCTCATTAGGTGGCAAAGATTGTAGCTTTCTTGATTTTTGTTGTAGTGGGGTTTTTCTTTGTTGAATGTTAATGCAAATGCTCAAATGTTGTTGCTTTGTGGATCATCTATGGAGGTTTTTTTTCACCACATATGGTTTGTTGTAGCTCTGTTTCTTGCTTTTGCTTCACTTGTTTTCACTGTCACTGACCCTAATGACTTGTCTGTTATCAATGAGTTTAGAAAAGGGTTGGAAAATCCTGAGGTCTTGAAATGGCCTGAAAATGGTGGAGACCCTTGTGGTTCTCCAGTTTGGCCACATATTGTGTGTAGTGGTAGTAGAATTCAACAGATTCAAGTTATGGGGTTAGGGTTAAAAGGCCCTTTACCTCAAAACTTGAATAAGTTGTCTAGGTTGACACATTTGGGGTTGCAAAAGAATCAATTTAGTGGAAAGTTGCCATCTTTTAGTGGTTTGTCTGAATTGAGCTTTGCTTACTTGGATTTCAATCAGTTTGACACAATCCCATTAGATTTCTTTGATGGACTTGTGAATCTACAAGTTTTGGCTTTGGATGAAAATCCTCTAAATGCCACTAGTGGTTGGTCATTGCCAAATGGGCTTCAAGATTCAGCTCAGTTGATCAATTTAACAATGATAAACTGCAATTTAGCTGGTCCTTTGCCTGAATTTCTTGGAACTATGTCTTCTTTAGAGGTTTTGTTGTTGTCTACAAATAGGCTTTCAGGGCCTATTCCAGGTACTTTCAAGGATGCAGTGCTGAAGATGCTTTGGTTGAACGATCAGTCGGGTGATGGTATGAGTGGTTCAATAGATGTTGTTGCAACAATGGTGTCACTTACACATCTTTGGCTTCATGGGAACCAATTTTCAGGTAAAATCCCAGTAGAGATTGGTAATCTAACAAATCTGAAGGATCTCAATGTGAATACAAATAACCTTGTTGGATTAATCCCTGAAAGTTTAGCAAATATGCCATTAGACAATCTTGATTTGAATAATAATCATTTTATGGGACCAGTTCCTAAGTTCAAGGCTACTATTGTTAGTTTTATGTCCAATTCTTTTTGTCAAACCAAACAAGGAGCAGTATGTGCCCCTGAGGTTATGGCACTTTTAGAGTTTCTTGATGGTGTGAATTATCCTTCTAGGCTTGTTGAATCATGGTCTGGAAACAACCCTTGCGATGGACGTTGGTGGGGAATAAGCTGTGACGATAACCAAAAAGTTAGTGTCATAAACTTGCCTAAGTCTAATCTTTCCGGGACCTTGAGTCCTTCAATCGCGAACCTTGAATCCGTTACTCGCATTTATCTTGAATCAAATAATCTTTCTGGTTTTGTTCCATCTAGTTGGACTAGTTTGAAATCTCTGTCTATTCTTGATTTGAGTAATAACAATATTTCTCCACCTCTACCTAAATTTACCACCCCTTTGAAACTTGTTCTAAATGGAAATCCAAAGCTGACTTCTAGTCCTCCTGGAGCAAATCCTTCACCAAACAACAACACAACGCCTGCAGCATCACCTGCGTCGTCTGTACCATCTTCACGACCCAACAGTTCAAGCTCTGTGATCTTTAAACCCGGTGAACAGTCACCCGAAAAAAAGGACTCAAAGTCAAAGATAGCCATAGTTGTGGTTCCTATTGCTGGTTTTCTACTTTTGATTTTTCTTGCTATTCCACTGTACATTTATGTCTGTAAGAGGAGTAAAGATAACCATCAAGCTCCAACTGCTCTTGTGGTTCATCCTAGAGATCCATCTGATTCGGATAATGTGGTCAAGATTGCGATCGCCAATCAGACTAATAGAAGTCTTTCCACAGTAAATGCAAGTGGTTCTGCTAGCATACACAGTGGTGAATCCCATATGATTGAAGCTGGGAATTTGCTCATATCGGTTCAAGTACTTCGGAATGTGACTAAGAATTTTTCTCCGGAAAATGAACTTGGGCGTGGTGGTTTTGGTGTGGTTTATAAGGGAGAATTAGATGATGGGACACAAATAGCTGTCAAAAGAATGGAGGCTGGTATTGTTAGCAACAAAGCTTTGGATGAATTTCGATCTGAAATTGATGTTCTCTCAAAAGTACGGCATCGACATTTAGTGTCTCTACTGGGATACTCTGTTGAAGGTAGTGAAAGAATTCTGGTTTATGAGTACATGCCACAAGGTGCACTTAGTAGACATCTTTTCCGCTGGAAAAAGTTCAAGTTGGAGCCTCTCTCTTGGAAGAAAAGGCTTAATATTGCCTTAGATGTTGCTAGAGGTGTGGAGTATCTACACACACTAGCACATCAGAGTTTCATTCATAGAGATCTCAAATCCTCAAATATCTTGCTGGGTGATGATTTCCGAGCAAAAGTATCCGACTTTGGACTCGTGAAACTTGCTCCTGATAAAGAGAAATCTGTGGTCACCAGGCTGGCCGGAACTTTTGGATATCTGGCACCTGAATATGCTGGTAAGCAATTTTTTGCTTACTTTGTCAAAGAGAAAAGATGATTTTTTAAAATGCATGTTTATATGACAATTTTTACTTTTGCAGACACCTAGAGTTTGCTCAACTCAATTCTGATGTATAGCATTTTATTTTTTTGCATACCAATGTACTGTTTTCCGAACTAACATGAGATGTTTATTTGATCCAGTTACGGGTAAAATCACCACGAAAGCTGATGTCTTTAGCTTTGGTGTTGTGCTAATGGAGTTGTTAACTGGGATGATGGCACTAGATGAGGATAGACCTGAGGAAAGCCAGTACTTGGTTTCGTGGTTCTGGAATGCCAAATCCTCTAAAGAGAAGCTTATGACAGTCATTGATCCAGCTCTAGACGTGAAAGATGAGATAACCAAGAGCATTTCCACCTTAGCTGAACTTGCTGGTCATTGCACTGCAAGAGAACCTGGTCAACGGCCAGATATGGGCCACGCTGTAAACGTGCTCTCCCCACTTGTTGAGAAATGGAAGCCTCTTGAGGATGATCCAGAGGACTATTGTGGTATCGACTACAGTCTTCCCCTCAATCAAATGGTCAAGGGCTGGCAAGAATCGGAAGGAAAAGACTTAAGTTACGTGGACCTCGAGGACAGTAAGGGTAGTATCCCTGCAAGACCAACTGGATTTGCAGATTCATTTACATCAGCTGATGGTAGATAATGGAGGTACTTCTATGTAGTAGATGTAGATATCAATTTTCTTTGTATTGTGTGAGATTTTGATCGTATTTTCCACGTGCCTTCGTTCATTTCTCCCCCTTCAATGTGAATGTATTAGATATTCAAACTATGTGTAGCCTCAGTTCCTTCTGTAGATACAAAATAGTGATGAAGGAGAACTATACATCGGTTAAACTCCTCGCTCAGTCACTATTTTCATCTTCTATATACTATTGTGAGATTTAAGAGCATTTCTTACAACTTTGCTTATTTTCGTCGTCTAGTTTGCTGCTTCAGAAGTTCTGTGTTGAGAAACATACTGATGTTGAGATAGCCTTGTAATAGAATATGTCGCTTGATTCGAT

>PGSC0003DMG400021394 LRR-RLKs

AATTTGGCTGTAAAGTGCCAAACACTTGCAACAAGAACACTAGTAAATTACATTTATTATTCATTTCTCTTTTCTCTTGTTCATTCCTCCAATCTCTAACAACGTAGATGCAATTCACAGGCTCAAGTAGTTTTCCTTTTTATCTAAAATTGATCATCGTTTTGTTGGGGACATTTTGTGATGCCGATCCTCGGATTTCAGAAGCAGGGCAGGTGTGTGGCACTAACATGACATTGCCGTCCATTATCATCCCACAGTATACTAAAGAAATGCAAGTGATTACCCAGCTTGTGAGTGAACATGGATGGGGCAGCAGCTATGCTGTTAATTCAACAGATATTTCGATATACGCTTTAGCAAATTGTTACCAAGATCTTCCACCTAACGACTGTGTTCAATGCTTCCTTATAAGTTGTTCTACACTCCCTACTTGTCTCCCTGCTGTCTCTGGTCGTGTATATCTTGATGGTTGTTTCATCCGTTATGATTATTACAACTTCTTCGGGGAAACTACTGATTCTTTCAAAGATAAAGTGAACTGTAGCAGTTCGTTTGGAAGGGCTGTTACTGATTCAGATCAATTGGCTTTGGCTGCTGCAGCTGGAAATTTGATTGAAAACGTGACGAAAAAAGCAATTAATGAGAATGATGGATATGCAAAGGGGAAATTGAATGGAGTTTATGGATTGGCACAGTGTTGGAGAACAGTGAGTAAACAAGGTTGCAGAGAATGTTTGGACAAGGCAAGTGAAGCTATAAAGGGATGTTTGCCTAATAGGGATGCTAGAGCTTTAATGGCTGGTTGTTATTTGAGGTATTCCACTCACAATTTCCTCAGTCATCATTCTCAGGCTACTAATAAAGGTATCTATATCACTTCTCTTTTCACCATACCAACTCTATTCTTCGGTTAATCCATAGACTTTTGAAAATTTACTTCCTCATCAACATATCTCTTTTTATTTCTTCTTTATTAAGGGCTCGTTTTGTAGGTGGATAAGAAATAAGTTATTTATGTTATTAATTTCTGCATAACTTATAATACTACAATTGGCAAGTATATAGGAAATAAGGTATTCATTTATAAAATTAATAGGACATTTGAATGACAATTTAGAAACTCGCATAAATAATATATGTATAAGTTATGAGATAATTTTTATATATATTCTACAGAATAGAATGTGGAATAAGTATTAGCACATGTATAACTAATATCTGCATAAAGTAATACATAAATTCACTCATAACTAATCCTACATTACTAATACCTCTATAATTCTCACCAGCAACCAAACAACCTAAGGGTGTGTTTGTTATGGAAGAAAACATTTTCGCTACAAAAAACATTTTCAAGCTGTTAAGGATTATAATTTGTAAGTAGCCAACAATGACGAGAATTGCTAAGAAAAGAAAACACTAAAAAATTTCTTCAAGGGATTCAATAATACAGAGAAAGAAAGATGAAAGAATGATAAACAACTTCAGATTAAGTTGTGGTACCTTTTTTTTTGTTTTTTGTTTGAGTATTTTAGATTGGCTTAATGAAAATAAAGAAGTATGAAAGGGAAGTGATGGATACGTACGTTCGGTGCTTTGCTGTTAAACATTATTTTTCAATATAGATCTAGCAGATCATAAATCTTATTTTTAGCCAAAGAAATTTCCTATTCTTTGGGTTAATACTTCATTAAATGGTGTCTCAACATGTAATGTTTTGAAAATAATCCCCGCATCAACTCCTAGTTTTCTATTTTGGAACACCTCTAGTTATGGGAATTACATGTTTAATGATTGATTGGAAAAACCTTCATTGAAAAATTGATTTCAATCTCAAAAAATTCATCTTCAGAAACTCATTGCATCTCATGAAAATGTCTTGAACTTATTATTTTCAATACAAGGAAAAATGTCCAAACAAGATCTTAGAACGATATATATCCTGTTTGCTAGCTTTACTAAAGGCATTTTATGAACGTGTTTCCCCAAAAGAATTTGATATTCTCCTACCAAACACTCAAAATTATTTTTTTAAAGTATTTATTTCAAAATATAAAATCATATATAATTTCCGTCCTGTTTGATTATCTCAAGGAAAATAAAAAATGATATTATAGGAATATCTTGACTATGTGGGATGGTCAAACTAACGCCATATTTGCTTTAAAAAACAAGAATGGTCCAAAGTCCGAACTTATATGAGTTCTCTAACATCCTTCCTTTTGTCTTATTCCCCTAATCCAACTATAATTCACATTGTACAAAAAAAAAACAGCATAAATCATTATTTTAAACGGATAGCAAAATTAAATAATTCATAACACATTAGCACCTAAGATTAATTAAATTGTTAAAAAAAGCTCAACTGTGGAAATAATTCTTATTTGATTTACTAAACAACGCAAGTTAGACTCTATGTTTTTTTCAAGTACTAAAGCCAAATATATGCAAATTGGAAGGCAGAAAATTTCAAACGTATCGATAATGACATCCATATGACCTACTCCTACAAAGATGTATTATAAATATTTCTGTTGGGTCAAGTCTATACGGATCGACAGAGACATTTTAATGGTATGTAAAATAATTATTTTTCTTTGTTGTTATATTTTGATTTTGGTGAAAAAGATTGGAAGTATATATTTATATATATACAACTAGCTGAATTAATAAATGTAGAAAGTATAAAATATAAAACATCAATTGTGCTATTGTGGAATTGACTTGTCTTATAAGTTGAAGTAGAACCAGCTCCTCTCCTCTCGGACTCAGGTGTAGTTCAGTTCCAATACTAAACATTTTCTTAAAAGAAAAATAGAAAATAAATATTCAAGAAAATAATACTCCCTCAGTCCCTTTTTATTTGTACAATTTGGACTTCTTACTTGTTAGTTTTGACAAATCAAGAAAAAATAATATATATTTTTTTACGTATTATACTCTCAATTTATTTACAGAGTTGATATTTAAAAAAAAGATAAAACCTCTTTAATAAGTAAATTGATTTAGAATAGTAAACTCATTATGTTAATTATTATTTTTCTTAATAGATGTATCAAATAAAAAGAGTATTATTGAACAAACAAAGTTTTTACTGATATACTAGACTATAAGGGATTGGGTTGTGAGAGTCTATAAAAACAATCTCAGAACTGACAAATTAAAATCTGCTACCTTATACTCTCAAGACCCCATTTTGTCAGATTACATCAATCAAATATTCTGTTGTTATTGTTGTAGTATACTATAAAACTTTATCTGGCTCCTCTTTGACTGCTAAACTTGAAAGCTTGTTCTTTAGAGACTTTCAACTCTGTTTTTTTTCTTTTCTCTCAAGGACAATAATATACTTGCTTTTGACTAATTTGTAAATAAATGCTCTTGGTTTCAGGCTTAAGCAAAGGTGTGATAATAGTTATCGTTTTTGCAATGATAGCTTTCTCCATGCTGTTTCTCTTTGTTGCGTACGTAGCTCGTAAAAAATCGTTAAAGCGAAAACAAGGTATCATAAAAGAAACTAACCTTAATTTGTCAATTTAGACAATCGATCAGTCGTATATGTTATAAAATAATTACATAAATGTTTGCAATTTTTCTTGGCCACAGAATTCATTAATCTTGGCAAGATATCGAATTCATACAAGAGGTCCAGCTTGAACTTTAAATATGAATATCTTGAAAAGGCAACACACTACTTTGATCCATCAACAATAGTAGGCCAAGGAGGAAATGGTTCTGTATACAAGGGGACTCTTCCTAATGGGAAGGTTGTTGCAGTTAAGAGACTGTTTTTCAATACAAGGCACTGGGTTGATGAGTTCTTCAATGAGGTTAACCTTATCCATGGGATTGAGCACAAAAATCTTGTGAAGTTGTTGGGTTGCAGCATTGAAGGCCCCGAGAGCCTCCTTGTCTATGAGTTTGTGCCTAATAAGAGCCTAGATCAATACCTTAATGGTACGTTAATATCGATGAATTTATATCTATAAATGCAGTGACAATTAGGGACGACTTCTAATTGAGTTGTTCAAATTTTTGTTGTGTAGATAAGAATAAGATAAAAGTTCTAAGTTGGAAAGAACGATTACACATCATAGTCGGAACAGCAGAAGGCCTTGCGTTCCTCCATGGAGGTTCAGATATGAAAATCATCCACAGGGACATCAAGAGCTCCAATGTACTTTTGGATGAAAATTTTGAGGCGAAAATTGCTGATTTTGGACTAGCTCGGTGTCTTGCAGCTGATAAAACTCATCTTAGCACTGGAATTGCTGGGACATTGTAAGAATGATAAAACAGCATTCACAGCCTTCTTTTAACTACTTAATTATAACTAGTGATTTTATTTGTTGCGTAGAGGATATATGGCGCCTGAATATCTCGTAAAAGGACAACTCACAGAGATGGCTGATGTCTATAGCTATGGAGTGCTTGTTCTTGAAATTGTTTGTGGAAGAAAAAACATTTCTTTGGCAGAGGATGACTCTGGATATCTACTGCAAACTGTAAGAATTAATGTTGTACATTTCTCTTAAAAGTTTGTTGGTAACAAATTGAACCTGTCATTTTTTATTTCAGGTGTGGAAACTGTATACAACAAATCAAGTGACTAAAGCATTAGACCCCATCTTGAAAGATGATTTTTCACCAGAAGAGGTATCAAAAGTCCTCAAAGTAGGGCTTTTGTGCACTCAAGCTTCTGTTACATTAAGACCATCAATGTCTGATGTTGTCCAAATGCTAACAACATTTGGACAACCGATTCCGGAACCATGCCAACCACCATTCTTACGTTCAAGCAGTAGCATAAAAAGATTCGTCTCCGATACACTATCCAAACTTGATGAATCTTGTATCTCCTCTGCCATGCATAGCTCATCAGATGGACCACACATTAACTGATCAAGAATTCTTTTCAAAACCGTCTCTACTGGAAACATATATACAAATTAAAAGAAGTCAAATTTCATGATGTATGTAAGTGAAAATGAAAATTTTGCACAACATTTTTGTACCGTTCGAACTAGTAGTGATTACATGTTGCATTTTTACGATTTGATTTTACTTTAACTCTTCG

>PGSC0003DMG400021609 LRR-RLKs

CACAAAACACACAAGCCCACTACACTACCAATCTTAGTCATTATTTCTCTATCTACTATCACTCTTGTGAATGCAGTTACTTCATTTTCCTTTCCACCTTGTACACATGAATATCTAACAATAATCCCAAACTCAACATTCATTCAATGAAACTTGTACTCCATTACCTAAAATTCACACCTTTTTCCCTTTTTCATTTCCATTCCCTTCTCTAACTACTCTGAAAAAAACATTTGTTTGCTTCTCTTATTGTAGGAACTAAAAACACCATCATAAAAATGAGATCTTTTAGTCCCTTTTTAGTTCTAATACTTTATTTTGGCTTATTGTATGGATCTTTATTAGCTGATGATACTACTATGCAGCTTAATGATGATGTTTTAGGCCTCATTGTGTTCAAATCAGCTCTTCTTGATCCTTACTCAAAACTCTTGTCATGGAGTGAAGATGATAACTCTCCTTGTGCATGGGAATACATCAAATGCAATCCAATGAATGGTAGAGTCACTGAGCTCAACTTAAATGGCTTAAGTTTATCAGGAAAGATTGGTAGGGGGCTCGAGAAGTTGCAATCTTTAAAGGTACTATCTTTGTCAAACAACAACTTTACTGGTGCAATTAGCCCTGAGTTAGCCTTGTTGACAAATCTTGAAAATCTCAACTTTAGTCATAATGGCTTATCAGGGAATATCCCTGGATCTTTTTCAAATATGACATCTTTACAGTTTCTTGATCTGTCTGAGAATGCTTTATCTGGACCTGTCTCTGATACCATGTTTGATAACTGTGGTGATTCGCTACGTTATCTTTCTCTATCCGGGAATTTTCTTGAAGGTTCATTTCCTAAAACAGTTTCAAAATGCACTAGTTTGAATCATCTCAATCTTTCAAGAAACCATTTTTCTGGTGACCCTGGATTTTCCGGGGGTATCTGGGGATTGACAAGGCTTAGAACATTAGATCTTTCACATAATGAACTCTCTGGATTAGTACCAATTGGTGTTTCAGTTTTACATCAGTTGAAAGAGTGTTTGTTACAAGGGAATCATTTCAGTGGAGATTTACCAGCTGATATTGGTTTCTGTCCACATTTGAATAGATTAGATTTGAGTAATAATCAATTCACAGGACAAATTCCAATGTCACTTCAAAGGGTGAATGTTCTTTCTTTTCTCAGTTTATCAAACAATATGATAAATGGAGATTTCCCTCAATGGATAAGTAACATGAGCAGCTTGGAGTACTTGGATCTTTCAGGCAATAGTTTAGAAGGAGCATTGCCTGATTCAATAGGAGATTTGAAAATGTTGAAGTACTTGAGTTTATCTGGTAATAAATTGAGTGGAAACATTCCAAAGTCTATGGTGTATTGTACTAGTTTATCGACTATTCGGATGAAAGAAAATGCATTGACTGGTAGCATTCCTGAAGGATTGTTTGGTATAGGATTGGAAGAAGCAGATTTTTCAAGAAATGAGTTAACTGGTTCAATCCCTCCTGGTTCTGGTAAATTCTTTGAATCACTTCAAGTTCTTGATCTATCAGGAAACAATCTTACTGGGAATATCCCAGCTGAAGTGGGACTTTTCTCCAAGTTGAGATACTTGAATCTTTCTTGGAACAATTTTCAATCTAGATTGCCTCCTGAAGTTGGATACTTTCAGAATCTAACGGTGTTGGATCTTCGACACAGTGCTTTAGTTGGATCAATTCCTGGTGATATATGTGATTCTGGTAGCTTGGGAATTCTTCAGCTTGATGGAAATTCATTCACTGGACCTATTCCTGATGAGATTGGAAATTGTTCATCCCTCTACTTGTTGTAAGTATATTTATTCTGATTTATCTCATTATAGTATGGTACAGCCTCTCTATAACAGTCATTTCACTATGAGAACCATGTTTTTTGTGGAACCAATCTCTCATGTTATGTTATCTATTACAACATGTTGCTATAGCAGCAAAAATAATATCGGAACAAACAATGTTGTTATAGAGAGGTAGCAAGGGATATGTTTATTTGAAAATGAGGTTCTGTATCACTTTTGCTTTGTCTATTTTCCCTGTAGTTCTTGAATTGAATGTCAAATTTGTTAAATGCAGGAGTTTGTCACATAATAATTTAAGTGGCTCAATACCAAGGTCCCTTTCAATGTTGAAGAAGCTCAAGATTTTGAAGCTAGAGTATAACCAATTGAGTGGTGAAATACCACAAGAACTTGGCAAGTTAGAAAATCTTCTGGCTGTTAATATATCCTACAACAGGCTCGTTGGACGCCTTCCAATGAGTAGTATATTCCAGAATTTAGACCAGAGTTCATTGGAAGGGAATTTGGGAATTTGTTCACCCTTGTTGAAAGGTCCCTGTAAGATGAATGTGCCAAAGCCTTTGGTTCTTGATCCTTATGCATATGGAAACCAGAGGGGAGGTCAAAACCGGGACGATGGAACTTCAAGAAGCAATAACAAAAGTTTCAAGAACCATAGATTCCTCAGTGTTTCATCCATTGTTGCAATCTCTGCTGCAGCTGTGATCGCGGTTGGAGTCATGATAATAACCTTACTCAATGCTTCTGTTCGAAGGAGGATCACATTCGTTGACAATGCCTTGGAAAGTATGTGCTCAAGTTCTTCTAAGTCAGGGAGCCTTGCCACTGGAAAGCTTGTGTTATTGGACACCAAATTGTCCCCGGATTGGACTAATAGCAGTCTTGAATCAATCCTAAACAAGGCATCTCAAATCGGTGAAGGTGTCTTCGGGACTGTTTACAAGGCTCCATTGGGAGGGGAAGGAAGAATAGTAGCTATCAAGAAGCTTGTAACATCAAAAATCCTCCAATATCCTGAGGACTTTGATAGAGAAGTTAGAGCTTTAGCAAAAGCAAGGCATCCAAATCTAATATCCTTAAAAGGGTATTACTGGACTCCTCAACTTCAGCTTCTAGTATCAGATTATGCACCAGAAGGAAGTTTACAAACCATACTGCACGAAAGGCCTTCGTCTTCAACTTCCCTTCCCCTGTCTTGGTCCACTCGATTCAACATTGTGCTCGGAACAGCCAAGGGACTAGCACATTTGCACCATGCGTTTAGGCCAGCAATCATACACTACAACATAAAGCCTAGCAACATCCTCCTTGATGAAAACTTCAACCCTAGAATATCAGATTTCGGGCTGGCAAGGCTAGTGACAAAACTCGATAAGCACATGATAAGCAACAGGTTCCAGAGTGCACTAGGCTATGTAGCACCTGAATTGGCATGCCAGAGCTTAAGGGTGAACGAAAAGTGCGACGTGTATGGTTTTGGAATGTTGATTCTTGAAATTGTCACAGGTAGAAGGCCAGTTGAGTATGGAGAGGACAATGTGTTGATACTCAATGATCATGTGAGAGTTTTGCTTGAGCAAGGCAATGTGTTGGAATGTGTTGATCCAACTATGAAAACTTATCCTGAAGAGGAAGTTTTGCCTGTTTTGAAGTTGGCATTGGTGTGCACTTCTCAAATACCATCAAGTAGGCCTTCAATGGCTGAAGTGGTTCAAATCTTGCAGGTCATCAAAACACCTGTTCCTCAAAGAATGTCAGCATACTAAACATGTCACAAACTTGATGAATTTCCCTTTCTCTTTGGCTTTCTATCATCCTATTGTTGTTATGTTAACCTCCTAATTTGATGTTTTTGGACTAGATAGTGATAGAGAGAGTTTTGTACTTCTCTGTCATTTGATGAAATATATGAGAGGTTGGTTCAATGACTTCAATCATGTTTGTTG

>PGSC0003DMG400021789 LRR-RLKs

CTTTAATGTTTTGTAGACATATTTTCATGGAAGATGGTGTCTTATGTCACAGTGATAAATTGCAACAAGGATACTTGACATAAAGTGCTATATCCGAAAGATGGTACTGTTCTTTGCAATTGTGATATTGGCATGTCTCCGCTCATTTGTCTTACCTGATGGGCAAGGTACCTGGTTCTTCCAGTTTTACATCATAAAAACCAACTTTGTTTTTATTTATCTACTCAAATGGATAAAGAATAATTTAGATGTGACCACTATATCTTGTTTACATTTTGCTTCTGCCGGTGAAATGTTGTCTTTGTTCTTCTAATAAAGTAGGGTGACATTTGGTTTTTTATACTCTGGCCCATTTTAGGAGATGCACTGTACACATTGAAGAACTCACTAAATGCTTCAAATGGTCAACTTGCTGATTGGAACCAGAATCAAGTTAGTCCTTGTACTTGGTCCAAGATCACTTGTGATGATAAAGGCAATGTCATTATGGTGTAAGTACTATTGTTAGAGCCATTAGAATCACTGTGTGCATATACTATGATGTTACTGTTTTTATGCACTAGAAAGGTTAATCTTTGGTTGGTCAACTTCCTTTACTGGATGTAATTTTGATAATGTCTGAATCTGTATTTGCAGGTCATTATCAAGTTTGGGATTCTCGGGCACTTTAACTCCCAGAATAGGTGTTCTAAAGAATGTGAATACACTGTACGTTTAAAGTACTCGGGCTGAGTTATTATTGATAGAATGATGTCTTGAATGCTGCCCACATATAATTTTCTAGATGCTATTTATTGCTATTTTAAAATGTTTATTTAAATTTTGTTTCTGAACATACAAGTTTGATGTCATGGCATTGCACTAGAACAAAAATCCCTTCAAATCATCAGATGTTAACTAAAAGTTGAGAACACGATGATTAATATTTTCAAGGGAGCTTCACTTAGCGGAAGGGGGAAAAAGAAAAAGGAAAGTACCTACATATTGTGGTGTTATAAATGAACAAGGAGAAATATTTGGCCATTTCACAAAGGCAAACAGCTTTAAGAGAGATGGAAGCTGTATGAACTGATCTAGCAAATGGGAGTCTCTGTTCGGATGGCTGTCTTTTTTCTTGTGGTAACTGTGTGGTGTCTCTACACTTGTAACTGCTGGTTCCGTTGGACTAAGAGCAACCTAAGCCATTTCACGAAGACTAGAAATGTGAACTATACAAGATTAATCCTCAGTATCAGTGACTTTATCTTTTTTCCTGAGTGGTCATTCCGTCACATTGGTATTTGCACTTTGTCTACTTCTTAGGGTCCTATTTTAGTATTTGACTTATATATATTTCAAATCAATAATTTGGATGCATAACCATACGTAAGGGTACATAGTTCTTTGATGTAGCTCTTGATGAACTTTGATTTGTTAGTATGTTGTTGAGCTCAAAATTGTAAGGTCTATGACTAAAATTTTTGGGATGATTTCTTGTCTTCTTAGATCCATGACTAAAATTTTTGGGATGATTTCTTGTTTTCTGCTTCTAACCTAGCTGACTATGTCGCATACACTCCCTGCAGGTCCTTGCAAGGAAATCGTATAACTGGTGAAATTCCCAAAGAGATTGGAAATTTAACAAGCTTGACAATGTTGGATTTAGAAAATAACCGTCTTTCCGGAGAAATACCAGCTTCGTTGGGCAATCTTAAAAAGCTTCAGTCTCTGTAAGTCATATTATTTCATTTAATCATTTGAGATATAACTTAGTTTACTTGGTGACTGGAATAGGTGTGAACTCTATTATTCCATTGAGCAGGTTTTTGAGTCGAAACAATATCACGGGGACTATCCCTCAGTCACTTTTGGGTCTTCCAAACTTGATGAACCTGTAAGTTATTTATCTTGCCTTTTCCTTATTCCAAGTATAAATTACTTTTCTGAATAGGAGAATAATACATTTTCTTTTGGGATGTCAACTTCAGACTTCAGTGATCCTTAACCTGTTACCTGTATCATTCGATATTTGACTTGATTCTCTGATTCCGCTTTGCAGTCAGCTTGGTTCCAACGGTCTTACTGGCCAAATTCCTGAACAGTTGTTCCAAGTTCCAAAATACAAGTACGATTACAGTTCTTACTTTAGACAGTAGCTCAAGGCTTCTTAGTTTTTGCCTTCTCTACTAATGCATAATCTTTAATGTTAGTATAACTTCATTTTATTCTAGTTTTACAGGGAATCATTTGAATTGTGGCTTAAATCTCACTCATCATTGTGAATCTGCTAATGGAGGTAAGTTCTACCAGCAAATATGTGAATTTTTGAACTGTATATGCTTCATGCTTCTCTGTATTCTGGAAGAATGAAAGTAATGTAGCTAAATTTTTTGAATTTTACTGCCATTATCTTGATTTTATTTGTTATTATCTTTTAGCTAGAAAGGAGGCTTTATCTTTTAGTTTACTTTATTAGAGAAAAAACTTTAGTATTTTCAAGATGTCTAGCTTGTTAGGGGTTTCATATTCTATATCTAGTATAAGTGTGTACTTTGTTTGGAACGACACTTGATGAATAGACTCTATTAATTTCCTTAATACAACAACAACAACATACCCAGTGAAATCCCACTAGGTGGGGTATTAATTTCCTTAATATGTCATCTATTTTTTCTTGTTTCTTTCCGTTCCCATTCTTACTTAGTCCCTTCTCCCTTTTCTTGGTGCTATTTCCTCCTTGTACTCAAGACATGGAACAAGATTAAAGTTCTTAGTGTTATTTAGAGTTATACTTGCCATTGGCTGTGGGATATGTTTGTTGTGTGGTATACTGGTATCTAATTTAATGGATGGAAGAATCAAGCACCTCATGGTGCTGCCTGCACTAAAAAGTATGAGTTTCTTTCCATTAGTACATTTAATCTTGGAGAGGTGTAACTTGTAACACTGACTTACTTGTAGGTGCCAAGCATTCTTAACGGAGGGATAATGATGTAGCTGAAAAATATAGCATTATTTACTACCTTTACTGTTGATCTATTGTCGTCATCTTTTAGTTAAAGTTTGAGGTTTGATATATTTTTTTTAGAAATACTATCATATTATAGTAGGAGAGAATTTTAATAATTTTAGGAGTTGTAATGCTTAAAAAAAAGAATTTAGTAATTCTAAGAGTTGTAATGATTCTAAAAGAAAAATTATTTTAACTCTAAGAGTTGTAATGATTAAAAAATAGAATTTTAACAATTCTAAGAGTTATAGTTATCTAGGTTCCCTAATCTATAATTAGAGTCCATAATTCCTTTGCTTGGAACAACATTTGATGAGTAAATATTGTTGATCTCTTATTATATCTCTCGTCCCCTTTTCTTCTCGTTCTCTCCCTTAATCATCATGCTCATTCTCTCTCCTTTTGGTGCTACATCTCAGAAAGTTTCTGAAGTTGATTACCTATCCGATGTATTACTATTCGTTTAGCAAGGAAATCGTCATTTAACATATGCACCTTGTTGAACTTTCTGGACAAACCAAATTCTAAAGTTGAATTTCTTGCAGGTTCTCCTAGTAAACCAAAGACTGGTCTCATAGTTGGTATTGCTGTTGGATTTCTTGGACTTCTCATTCTTGGAGGATCTGTGTTATTGTTCTGTAGGAGACGGCACAAAGGCTATAGTCGTGAAGTCTTTGTTGACGTTGTAGGTTTGTGTCACATATTTTTGACAATAATTTCATCAGTTTAATGTACTTGTTATTCTTCGTAAGGGAAGAAGATATTCTGGCCATGCATTGTTGTATCTTGTAATTCCATCTTAAGTTAGTGACCTCATTCTTTATACACCAGTACTTAGTTTTCATTTACCTTCAAGATGGTATGATTCCTTTGAGTATATTATGATATTATATTTTATGTTTTATTTGGCGCACTCAGGTGAGGTTGATCGACGAATTCCATTTGGTCAATTGAGGAGATTTTCATGGAGGGAATTGCAGCTTGCTACTGATAACTTCAGTGAAAAGAATGTTCTTGGACAGGGGGGTTTCGGCAAGGTCTATAAAGGAGTGTTCAATGACGGCATTAAAGTTGCCGTGAAACGGCTAACTGATTATGAGAGTCCTGGGGGAGATGCTGCATTTCAGCGAGAAGTTGAGATGATAAGTGTGGCTGTTCATAGGAATCTCTTACGGCTTATAGGATTTTGCACCACACCGACTGAACGCCTGCTAGTATACCCATATATGCAGAATCTAAGTGTTGCTTATCGTCTACGAGGTATTGTTTCTTCATAGCATTGCTTTTATTCATTGTAGTTTCTTTTACATCCTGAGTATGCTATTTTCCCTTACTGTGCCTGTTTCATCTAATTTATTTTATCCCAATCAATAGATCCTTCATTTCATTAGTCTTACATTTCCTGTGGCGCATATTTTGATTTTGTCAGCTTGGTTGTAAACGATCACGAGTCATTTCCCCGTTCGATCTGCGAAGAAAGCGTGCATTTGTCATTGTACGGTCTAAATTGACTATTAAAGTACTGTAGAACTGCTGATCCGGTAAAAGAACTTGTACATGGCATCAAGCTCGAGGGGCATGTTTATAGAGCTAATGTGTGTAGCTTAATTGGCAGAAGCATAAACCCTTTAATTTGTGTTTTCATGCTTCAGAATCTTATTCTTTTTCCTTCTCAAAAAAGAATCTTATTCTTTCAACTTTGAATGAAGTGCTAGAAGTTTAATGCCTTAGGGTTGCATCACCATGAGTACAATTTGTATGCTTCCTTTTTTCTCCCAGAATATGGTAGGGAAAGACAAGTTCTGAGCAGATTTAAGGGTTCTGTAGAAGTATTTTCTGATATTCATTGAAGGGGTTGAGTGTTCGGAGATTGCAATTTTCTTTGGTCAAATCAAGAGCAATACTCAAATCATACCTCATTAATCATTAATAGATGTATTCAGTTTTTTTTTAAAATCATTAAAAGATGTATGTGCGTTAAGTTGGTATCACAGATCGTGATATGAAAGAGCTATGTGACTTCATAAATTGATTCTTGTGATCCTGGCTTGAACTAGAAGGTGTTACATGAAGTAGTTAGTGCAGCTTAAGATTTCAGCAGTTTTCTGCTAGAGTTGATGTGTCTGTTATTAGTTTTCAGTGGTGTCTAATTAAATTTTTAATCACTTACAGAACTTAAACCTGGGGAGCCTGTTTTGAATTGGCCAACTAGGAAGCGTGTGGCAGTGGGTACTGCACGTGGACTTGAATACCTACATGAACACTGTAACCCAAAGATTATTCACCGTGATGTTAAAGCAGCTAATGTGTTACTAGATGAAGATTTTGAAGCTGTAGTTGGTGATTTTGGCCTGGCAAAGCTAGTTGACGTTAAGAAAACCAATGTGACAACTCAAGTTCGTGGTACAATGGGCCATATAGCTCCTGAATACTTATCCACTGGCAAATCATCAGAAAAAACTGATGTTTTTGGCTATGGAATCATGCTTTTGGAAATTGTAACCGGCCAACGTGCAATAGACTTCTCACGCCTAGAAGATGAAGATGATGTCTTGTTGCTTGACCATGTGAGTGTACTTAACTCAATGATCTTCATATATTTTCAAATGGGCATCAAATTATTGCATCAAACTTTTCATATATCTTATCTCTACTGACCAATCTTTGTTTCTCTCCATATTCTATTGGGACGACTCATTGCATGAGAGCCTCTTGGATGGATATACATTCGTTTATCTTTCTCCTATTGGTGAAACTGCTGCAATTTTTCTATGATAATTGAAATGGTGTAAACAATTATTATTTTTCGCGTGGCACCAGGGTACAAATGCTTTGAATATGAGCTAGGTCTTAATATTGAGCCTTTTAGGCACAAGAAATATGATGACTATATATATTTGATGCTAATTTTTTCTCTCTTTCAAAAGTGTGGATATGTGGACCCTGAGCCTCTCTTATAACTTTACATGATTATTCTTTAAGCTTCATAGGACCTTTATGTAGTAATAGTTGCTAAAAAGAGATCGTCTTGAAGGTCAGAAAACTGCAAAGGGATAAAAGACTGGATGCAATTGTAGACCGTAACCTGAATAAGAACTATGAGCTGGATGAAGTGGAGATGATGATTCAGGTGGCATTGCTGTGCACTCAGGCATCACCAGAGGACCGTCCAGCAATGTCGGACGTGGTTAGGATGCTGGAAGGAGAAGGGCTTGCTGAGAGATGGGAAGAGTGGCAACACGTTGAAGTCAACCGTAGACAAGAATACGAGAGACTCCAGAGACGATTTGATTGGGGAGAAGATACAATGTTTAATCATGATGCTGTTGAGTTGTCTGGTGGAAGATGACACATGAGTAATAACTTTGAACCACAAAAATGTTTCTTGATATATAGATGCCTGGTTGCTAAATGTTTTTTTTAAAATTATTTATGGAAAACCCCTTGTTCTAACTAGGCAGATCAACAGTGGACCTGTATTTATGTAAGTTTTAGTCATTATTTTCATGTTCCATTAACTACTCCTGGGAGTGGGAATTTGATTCCTCAAATTGTTACATATTTTTCCTCACTAAACTATTTTTCATTTCATA

>PGSC0003DMG400022000 LRR-RLKs

ATGGTTTCCAGGACTATAATTATAGCATTCCTAGTTTTGTTAATATACCTCTCAGTTACAAATGCATCGAACATTTCAACTGATGAGGCTTCTCTTTTAGCTTTCAAAGCACAGATAACTTCAGACCCCAATGGAATATTGTCAAAAAACTGGACAAAAGGAACTCACATTTGCAATTGGATAGGCATATCTTGCAGTAAAAAGCATCAAAGAGTGACATCATTAGTCCTCACAAGCTTTGGATTTAGAGGTTCAATAGCAATAGAAATTGGGAACCTCTCCTTCCTTAACAAGTTGGATGTTGGAAACAACAGTTTCCACGGCCAAATCCCCGATGAAATCGGGAGTCTGAGGCGTTTAAACTTCTTGTCTTTGCAAATGAATAATCTCACTGGTCAAATCCCAGAAAGCCTTGGATTTCTCACAAGGCTTCAACTTCTTGATCTTTCTGAAAATGATCTATTTGGAAATGTTCCATTTTCCATTTCCAATGTGTCTTCCTTAAAGATCATTGATTTGGGATTTAATCGGATTAGTGGGAATCTCCCAAGGGGATTTTGTACTAGGCTTCCAAATCTGCAAGGTTTATTTCTCTCAAAAAATCAATTAGCAGGCCAAATCCCAAGTGGGCTGAATCAATGCACACAACTTATTTACCTTTCCTTGTCATATAATCAGCTCACAGGTAATTAGTAATGTTAATTCAATTCTTGGACTATATATACTTGATTCAGAATCATTTAATCCTTTTACACTGTTATGAGGCTATCTGATTGTGTTCATATATGTATTAACAGGAAGTTTGCCAAGAGACATGTGGAATTTGACAAAGCTTCGAGAATTATATCTTGGATGGAATAACATAACAGGTATCTTGGTTCTAAGTCTAATAATTTCTAGTTAATTAGATATTGTAGTAGTAGCAAAAAATGTTCATATAGCTCCTTGTTATTCATTTTCAAACAACAATGACTACTACTACATCGGCTCAATGCTAGTTTGTGTCAACTTTATGAATCCTCATTATCCGCGTTAATCCATTTAGTCCCCATCTCAGTTCAAGGTTCAATAATTTAAGTTTTCTAACATTAAAATTTCTCTATATTTTCTGTTAACATACAAATATCTAACAATATTAAAAGCATCCTAGCGAACATTGAGTAAGTTGGAGGTACCAACATGATTAAGCTTTAATATCAACTAGTTGGTATCACCTTTATGGATCTTACCATTCCATTTTGCTCTACTTTTCACTGAGTCAAAATGGATTCTAACTTATAGTTGGTCTTTCTTTCAACTTCCTCCCATGTGATTTTAGATCTACCTTGTCCTTTTTAACAAATCCAATCACCATATATAGTTTCATACCTGCAGACCAGTGCATCTAGAGATCCATGTAAGACTGAACTAACCATGGTAGTTTCTAATGTCTCATCTCAAGTGACTCACATATCCATCTTAATATAGATAATTGACTGACGATCAAATAGTTTAAGCTTCCTTTCTTCAGTTACCATATAAGAAGAGAGGAAAAGAGAATAGCAGAAGTAAATGAATAGTTCTCATGTTGCCAATATTTAAAATTTAAATGCCCTAATTAACAGAAAACCTTTCCTAGGCTTGTTATTGTTCGTCTTCAAGAGGACGTTGAAACAGAATACAAGGATTGCCGAACTATCCGTGATTCTAATTCTCTTACACAATATTGTACGTTTTAGGGTTGTAACCACACAATTTAATATATATTTTACAGGACATATCCCAAGTGAAATTGATAACCTATCAGCAATTCGAAGATTAAGTCTACCACGGAACAATTTGGTGGGAATTCTTCCACCATGTATGGGAAATTTGTCAAATGTAGAGGTGATAGACCTAGGTCAGAACAGTCTACATGGTGGCATTCCCCTAGAATTTAAACATCTTGTAAATTTGAAAGGATTGTTTCTTGGTCAAAACAGATTATCGGGTGAAGTTCCAGGGCCTATGTACAACATTTCTGGATTAGCAATGATTTCATTTGTAGGAAATGGGCTCTCAGGAACACTTCCCTCAAACATAGGCCATACCCTTCCAAATCTTGTCGGCCTCTATCTTGGGAACAACCAGTTTACAGGGCTGATTCCTACCTCCATTGTGAATTCCACCAAGCTTATCCAGCTTGATTTTGGTCGCAATCTGTTTAGTGGACCTGTACCTATGAATCTTGAAAAGCTACAACAGCTTCAATTCATCAGCTTGCAATTTAATCAATTGACGAATGATCCATCTACGGACAAGTTAAGTTTTCTTACATCATTATCTAATTGCAAGTATTTGAAGACAGTTCAAATAGGAGGTAACCAATTTAATGGAAGTCTTCCAAAGTCCTTGGGATCAGGCAACTGGTCCTTTTCCCTTGAATATTTCATTGCCACCAATAGTGGTATCACAGGAAAAATACCAACTAACATCAGTAACTTTAGGAACCTGGAGTGGTTAAGCTTGGGAGAGAATAAGCTTATAGGTTCTATTCCTCCAGATTTGGGGAACTTGAAGAATTTGCAAAAGTTTTACCTGCAGATAAACAATTTAGATGGAATTATACCAACTAGCTTGTGTAACATGGAAAGCTTGTACCAAGTTATCTTGGGCAAAAATCAACTTTCTGGGGAATTACCAAGTTGCTTTGGAAATATTTCTTCTTTAAGAGAACTCTACTTAGATTCCAATGCACTAGTTTCCCATATTCCATCAACATTTTGGAGAAACAAAGATCTTTCAGTTCTCGATTTGTCCTCCAATTTGCTGAATGGATCTCTTGCAGTAGAAATGGGAAACACAAGGAGTTTGCGGATGTTAGATTTATCTGGCAATCAGTTCTCAGGTCAGATTCCTAGCACGATTGGCCAACTACAAAGCTTGGTTAGCCTTTCATTGTCAAAGAATATGTTAGATGGTCCTATACCTGAACTATTTGAAGATTTGGTTTCATTGGAATATTTGGATCTATCTAGCAACAATCTAACAGGCATGATCCCCAAGTCCTTGAGGAATCTTGAACATCTCATGTATTTCAATGTCTCGTTCAATGGGCTCATGGGTGAAATTCCAGATGGAGGGCCATTCATAAATTTTACAGCTGAATCATTCATGGGTAACCCTGCATTATGTGGATCATCACGCTTCCATGTGATGCAATGCAGAGTCACAAGTCTTGAAAGCAAAAGAAAGAGTAGAGTCTTAACTTCTGTCCTTGCATCAGTTTCCTCAGGAGTTGTAGTCACAACCATTTTCATCATTTGGTTTCTGAAATGCCGAAAAAGGAGTACGGAACTTCCTCTAGTTGATACATTTGGTCAGGTACATAAGAGGATTTCGTACTATGATATTTCTCAAGGGACAAACAACTTTGATGAAGCAAACTTGATTGGAAGGGGGAGCCTTGGTTTGGTGTACAAAGGGACACTAGCAGATGGAATGGTTGTCGCAATAAAGGTATTCAATACAGAACTGCAAAATGCATTTAGGAGTTTTGAAGTGGAATGTCAAGTTTTACGTAGCATTCGACACAGAAACCTTGTTAAGGTGATAAGTAGTTGTGCCAATTTTGATTATAAAGTGTTGGTGCTAGAGTACATGCCCAATGAAAATCTCGAATGTTGGCTTCACTCTACTGACAAATTTTTGGATATAACTCAAAGACTAAAGGTGATGATTGATGTGGCTTCTGCTGTGGAGTATTTACATGGAGGTCATTTTTTTGTAGTCATCCATTGTGATTTGAAACCAAGTAACATACTTTTGGATGGAGATATGGTGGCAAAAGTGAGTGACTTTGGGATATCCAAACTTCTAACAGCTGAGACGCTAATAGCACATACCAAGACTTTGGGTACTATTGGCTACATGGCACCAGGTAAATGTTATTATCAAGTTCATTTGAAGATTTTAAATGACGTTAATTATTTTCACTAATGATTGCGTGATTTTTTTTGCAGAGTATGGATCAGAAGGCAAAGTATCAACGAAGGGAGATGTTTATAGCTTTGGTATTTTGTTAATGGAGACTTTTACAAGAAAGAGCCCCGTGGATGATCTATTCGTTGGAGACTTCACCTTGAAAAGATGGATATGCCAATCATTACCAAACCGATTGGTGGATGTAGTGGACATTAATCTATTTTCACTGGACGAAGAAAATTTTACTTCTAAAGAGAGATGCTTCAAATCAATCATGGAATTAGCTCTTGAATGCACAAATGATCTGCCGGAAGAGAGAATCTGCATGGAAGATATTACTCTGCGACTCAAGAAGATCCTAACTCAATTTCTGCAGAATGCTGTGACAAACTAA

>PGSC0003DMG400022139 LRR-RLKs

GTTTTCTTGAAGATTTGGAGATTGTAGGATGGAGGAGTGTTCTTAAATAGTTGAAAGAATTGATTTTTATGCAGTTGGTTGGTGGTGGACTGGTGGTGTTTGTTTTCAGCAAGATGAGCTATTAAAGATTCTAAAGAATTGATTTTTATGCAGTTAGTGGTGTTTTTTACAGCAATATGAGTCATAAAAGATGCCAAGAATATTTTTCACTAAATGGGGTGTTGAAAAATGTTGTCTTTATGATAATACTTCTGAGTTGTAGTTTTTTTGTGGTGGTGTCAAATGCAAGAAAACTAGCTGAAAATGATCAAGTAGGGAGTTTATTAGCCTTCAAGAAGTCCTCTGTTGAATCTGATCCAAATGGATTCTTGAATGAATGGACTTCATCTTCTTCAAGTCCTTGCACTTGGAATGGGATTTCATGTTCAAATGGTCAAGTTGTTGAGCTTAATCTGTCTAGTGTTGGGCTCAGTGGTCTACTTCACCTCACTGATCTCATGGCCTTGCCTTCTTTGCTCAGAGTTAATTTCAATGGCAACCATTTCTATGGTAATCTTTCCTCAATAGCTAGTTCCTGTAGCTTTGAGTTTCTTGATTTATCAGCTAATAATTTCTCTGAGGTTCTTGTTTTAGAGCCTTTGTTGAAGTCTTGTGATAAAATTAAGTACCTAAATGGCTCTGGGAATTCAATTAAAGGGGTTGTTCTAAAGTTTGGACCTTCACTATTGCAGCTTGACTTATCAAGTAACACCATTTCAGATTTTGGCATTTTAAGTTATGCACTGTCCAATTGCCAGAATTTGAATCTCCTTAATTTCTCCTCAAATAAGCTTGCTGGAAAGCTAAAAAGCTCCATTTCTTCTTGCAAGAGTCTATCTGTTCTTGATCTTTCGCGTAATAACTTAACTGGAGAACTCAATGATCTTGATTTTGGGACTTGTCAAAATCTCACTGTCCTAAATTTGTCTTTCAACAACCTTACCTCAGTTGAGTTTCCACCCTCTTTGGCAAATTGCCAGAGCCTCAATACACTGAACATTGCCCATAATTCGATCCGAATGGAGATTCCTAGCGAGTTATTGGTGAAGTTGAAGAGCTTGAAGCGATTGGTGCTAGCCCACAATCAGTTTTTTGATAAAATTCCATCAGAATTGGGGCAGAGTTGCAGCACACTTGAGGAGGTTGATCTTTCTGGAAACCGGCTTACTGGTGAACTTCCTTCAACTTTTAAATTGTGCTCCTCACTTTTCAGTCTCAATCTTGGTAACAATGAGCTATCGGGTGATTTCTTACATACTGTTATCAGTTCATTGACAAATCTTAGGTATCTTTACTTGCCATTCAATAACATAACTGGTCATGTACCAAGGTCTTTGGTGAACTGTACAAAACTTCAGGTACTCGACCTTAGTTCCAACGCGTTTATTGGGAATGTACCTTTTGAGTTGTGTTTAGCAGCCTCAGGCTTTCCACTGGAGATGATGTTGTTAGCCAGTAATTATCTTACTGGAACTGTGCCTAAACAGATTGGACATTGTAGGAACCTCAGGAAGATTGATCTCAGCTTCAATTACCTGACTGGTTCGATCCCATTGGAGATTTGGACCCTTCCAAATCTTTCGGAATTGGTAATGTGGGCTAATAATCTCACAGGTGAAATTCCTGAGGGCATTTGCATCAATGGAGGAAACCTTCAAACACTCATTCTCAATAACAATTTCATCTCAGGGGCACTTCCACAGTCCATTTCCAATTGTACCAACTTGGTTTGGGTTTCGTTGTCCAGCAATCGACTGTCTGGAGAGATGCCACAAGGAATTGGGAATCTTGCAAATCTTGCTATCCTTCAGTTAGGTAATAACTCACTTACTGGACCGATCCCTCGGGAATTAGGTTCGTGCAGGAACTTGATATGGCTAGATTTGAATAGCAATGCTCTAACAGGTTCAATCCCTCTTGAGCTTGCTGATCAAGCTGGCCATGTTAATCCGGGAATGGCTTCTGGAAAACAGTTTGCTTTTGTGAGAAATGAGGGTGGAACTGAATGCAGGGGTGCTGGTGGACTGGTTGAGTTTGAGGGTATCCGTGAAGAGAGGCTTGCGATTTTACCTATGGTTCACTTCTGTCCATCAACTAGGATCTACTCTGGCAGGACGATGTACACCTTCACGAGCAATGGAAGCATGATTTACCTTGATCTTTCCTACAATTCTTTTTCAGGAACCATTCCTGATAATTTAGGTTCATTGAGCTTCCTTCAAGTTCTGAATTTGGGACACAACAATTTTACCGGAACCATTCCATTCAACTTTGGAGGTCTAAAGATAGTTGGAGTTCTTGATCTGTCACATAACAGCCTACAGGGATTCATTCCACCCTCGTTAGGAGGCCTTTCATTTCTTAGTGATCTTGATGTTTCAAACAACAACTTATCGGGAACGATTCCTTCTGGTGGACAGCTGACTACATTTCCAGCTTCAAGATACGAAAACAATTCAGGCCTTTGTGGTGTCCCCCTTCCTCCTTGTGGCTCGGGCAATGGACACCATTCATCGAGTATTTACCATCATGGGAATAAGAAGCCTACAACTATAGGAATGGTGGTTGGTATTATGGTATCTTTTGTATGTATTATACTACTTGTTATTGCCCTTTATAAGATTAAAATGACTCAGAATGAGGAGGAGAAGAGAGATAAATACATTGATAGCCTTCCAACTTCTGGCAGTAGCAGCTGGAAACTATCAACCGTTCCTGAGCCTCTAAGCATTAATGTGGCAACATTTGAAAAACCATTGAGGAAGCTGACCTTTGGTCATCTTATAGAAGCGACCAACGGGTTCAGCTCTGAAAGCATGATTGGCTCTGGAGGTTTTGGTGAAGTCTACAAGGCTCAGCTGAGAGATGGCAGTACTGTTGCAATCAAGAAGCTCGTGCACGTAACAGGTCAAGGAGACAGAGAATTCATGGCTGAAATGGAGACTATTGGAAAAATCAAACATAGAAACCTTGTCCCTTTGTTGGGATATTGCAAGATTGGGGAGGAAAGGCTTCTCGTATACGAATACATGAAATGGGGAAGTCTTGAATCTGTTCTTCATGATGGGGGGAAAGCAGGGATGTTTCTTGATTGGCCAGCTAGAAAGAAAATCGTGATAGGATCAGCAAGAGGATTAGCATTTCTTCACCACAGCTGCATGCCACACATAATTCATCGCGATATGAAGTCCAGTAATGTACTTTTAGACGAAAACTTTGAGGCACGAGTTTCTGATTTTGGAATGGCAAGGCTGGTAAATGCTCTGGATACTCATTTAAGTGTGAGTACACTTGCTGGTACTCCAGGTTATGTTCCTCCAGAGTACTACCAAAGTTTCAGGTGCACCGCAAAAGGGGACGTGTACAGTTATGGTGTCATACTGTTGGAGCTTCTTTCAGGGAAACGGCCAATAGATCCCCGTGTATTCGGTGATGATAACAATCTTGTTGGATGGGCAAAGCAACTTCATAACGAGAAACGAAGTCATGAGATTCTTGATCCTGAGCTGATAACAAACCTATCTGGTGATGCTGAACTCTATCATTACTTGAAAGTTGCATTTGAATGCCTTGATGAGAAATCTTACAAAAGGCCAACAATGATACAAGTGATGACAAAGTTTAAAGAACTACAAACTGATTCAGAAAGCGATATTCTTGATGGCATTTCAGTGAAAGGTTCTATACTCGAGGAATCGCAAGAAAGGGAACCGTAAATCAATGATATTTGATCTTTTTGTGGTTAGAATTCTGTTCTTTTTTTCTCTTTTTTTGGATTGCCTCATGGATAGTAGTAAAAACTTTTATCTTTAATGGTTCTTCACCATTGAGATTGATAAGAGTTTATACTTCAGATTTTGCCTACTGTGTAAATATTAGAACACGACGATGAATGGGAGATGTGAA

>PGSC0003DMG400022189 LRR-RLKs

ATGGACAATCTTCTCAAGTTCATTGTAAGTGTTTTACTTTCTCTTCTTTTTATCAGATTCTGTAATGGAGGCGAGTTTATCGAATCCAAATCATTTATCAGTTTTCTTGTTGCTATTGATCCAAACAATGTACTAAACGTTGATCGAAATGCTTTGTTTTCTTACCCTTGTTACTACAAATTACATGGTGTTAAGTGTGATTTACGTTCCGAAGAAATCATAGAAATAAGACTTGAGAACATGAATCTTAGTGGTGTAATTGATGCAGAGTCACTCTGTAAACTCTCAAACTTGAGAGTTCTTAGCTTAGCTAGGAATAAGATTGAGGGAACTGTTCCTGAATCGATATCATCGTGTAAAAGTTTAACTGTTCTTGATTTAAGCAGCAACTTTCTTAGTGGAAACTCATTAGTACTAATGCAGTCATTGACAAGGATCAAGAATCTTAAAAAACTTAACATTTCTAAGAACAATTTCACTTCCTCACAACGTATGATGAGTCTACACGCGTTGAGACGTGTAACTATTAAAGAATCTTCATCTTCAACTACAGTAAGTGACGATAAGAAGACAAATAATCATCATGTGAGTCTGATGGTTTGGGTACTCATATCTCTTGGAACTGTGATTGTGTTAGTAGTTTTAGTATTCTTGTATATGAGATGTGTTGGGACAGTTAAGGATAAGGAGGTTATAAAGGAAGTAGTTGATTATCCGTCTCCTCGAAAAACTCCCTCAGATGAAGTTGTGGATGGAGTGTGTAACACTGAAGAGAAAAGCTCAGAACTTTACTTCTTCATGGAAGATGTTGAGAAGTTTACGATGGACGATCTTCTTGAAGCAACTGCTAACTTAACAAAACAAGGCATTTGCAGCAGCCTTTACAAGGTTCATATCAGTAGAAGTGGTGTGTTTGCTCTCAAGAGACTGAAGAAACTACAAGTGGGGTTCAAGGAGTTCTCGCGAACGATGAAAAAGATAGGGAGCTTGAAGCATCAAAATGTACTTCCACTTGTTGCTTATTACTCTAGCAATGATGAAAAAATGCTCATCTACAAATATCAGAACAGTGGAAGTCTACTAACACTTTTCGAAAGTAAGTAGTCAAGAACTTCAAATTTCTCATCTTGTAATTCAGAATTTTATATCATATTGTTGTTGTTTATTGTTGCAGATTATGCAGAGGGGAAATGGAACTTCCCGTGGAACTTAAGGCTGTCGATTGCAGTTGGCATAGCAAGAGGTTTCGCGTACATATACAGAAGTTCAAAGAAAGGCAATGTCATTCCTCATGGCAACATTAAGCCATCAAACATTCTATTAAGCGATAACGGGGAGCCATTGATAAGTGAGTATGGCTATTACAAATTCTTAGACCCTGACAAAAGTTGCCTCTATAAAGACAATGGTTATACAGCCCCTGAGAAAACGTTGACAGAAGAAACTGATGTCTATAGCTTCGGGGTGATTCTGCTGGAACTGCTAACAGGTAAAGTTGTGGAGAAGACAGGATTAGACCTTGTCAAATGGGTGAAATCTATAGTGAGGGAAGAATGGACTGGAGAAGTGTTTGATAGTGAAGTTGCCAACTATGAAATGTATGCTTTCCCTCTACTAAACGTTGCACTCAAGTGTGTGGATCGTTTGCCTGATGAGCGACCTACTATGACAGAGGTCTTGGAGATAATCGAAGAAATTGTGAATGATCAAGAAGACATTTCTCCTTCTTCTATGACTTCTTTTGAATCCACCCCTGGTTCAATGAAGCATGTGTATGCCCTTTAG

>PGSC0003DMG400022264 LRR-RLKs

GTTCAATAATAATAATAGAAAAATTCAAAGTGCTCTTGGATCCAGATCTTCTTCTGTTTTTCTTCTATAAACAAAACATGGGGTTCCCCCATTTACAATCCCAAAGTCCTCAACTTTCTCTCTTCCCTCTTAGCAGCAGCAAACAATGCCTGCTCCTTCAAGAAATCCCTTTCTTCTTCTTCTTTTACTTTTCCTCTCTTCTTGTTATTCACTTGATGTACAAACTCAAACACTTCTTGCATGGAAAAAAACATTAAACATAACTTCCAATGATGTTTTAACATCATGGGATTCTTTAGATAAAAGTCCTTGTAATTGGTTTGGCATAAACTGCAATTCCAATGGTCATGTTGTTAGTATAAGTCTTAAATCAGTTGATTTACAAGGTCCATTGCCTTCAAATTTTCAGTCACTTAAATTCTTGAACACTCTTGTTCTTTCATCTGTTAATCTTAGTGGTCCAATCCCAAAAGAGTTTGGAGATTATCTTGAACTTAAGTTCATTGATATTAGTGATAATTCAATCACTGGTGTCATTCCTCAAGAACTTTGTAAGCTTATAAAGTTACAAACTTTATCACTTAGTTCAAATTTTCTTGAAGGTGATATCCCTTCAGACATTGGGAACTTGTCTGATCTTAAGATGTTCTTGATTTATGACAATCAACTTAGTGGTGAAATTCCAAAGGGTATTGGAAAGTTGAAGAATCTTGAAGAATTTAGAGCTGGAGGGAATCAAAATCTTAAAGGAGAACTCCCTTTTGAGATTGGAAACTGTTTGAACTTGGTTTTTTTAGGCCTTGCTGAAACAAGCATTTCAGGGAATTTGCCAGTGTCAATTGGAAACCTGAAAAAGATTCAGACAATTGCAATTTATACAGCTCTTTTGTCTGGACCAATTCCAGAAGAGATTGGGAATTGTAGTGAACTACAGAACTTGTATTTGTATCAGAATTCGATCTCTGGTTCGATTCCAAGGAGTATAGGGGAACTCAGGAAGCTTCAAAGTTTGTTGCTTTGGCAGAATAGTATAGTTGGTGTGATTCCAAATGAGCTTGGGAATTGTAAGGCTATTACAGTTATTGACTTGTCTGAGAATCTCTTAACAGGTAGTATTCCTACAAGCTTTGGTGAACTTTCAAGTCTTGAAGAGTTGCAGTTGAGTGTCAATAAGTTATCAGGTACAATACCTACTGAAATATCAAATTGTACTAAATTGTCTCATTTGGAAGTTGACAATAATGGCATATCAGGTGAAATTCCTAATGAAATAGGTAAGTTAAAGAGCTTGACTTTATTCTTTGCTTGGCAAAATAATTTAACAGGTGAAATTCCTGTTACATTATCTAACTGTGAGAATCTTCAGGCTATAGACCTTTCTTATAACAATCTTTTTGGTTCCATACCAAAAGAAATCTTTAGTTTGAAAAATCTAACAAAGTTGTTGCTTCTTTCCAATGATTTGTCTGGTTTTATACCAACTGATGTTGGAAACTGCACAAATTTGTATAGACTTAGAGTGAATAGGAATAGGCTAGGGGGTACTGTTCCATCAGAAATAGGAAAGTTGAAGAATTTGAATTTTCTTGACATGAGTGGCAATCACTTTATGGGAGAAATTCCTTCAGCAATATCAGGATGTCAAAATCTTGAGTTTCTTGATCTCCATTCGAACGCGTTTACTGGTTCTTTACCAGAGAAACTGCCAGGGAGTCTACAGTATGTGGATATTTCAGACAACAGGCTTTCGGGTTCGTTAAGTCCAAGTGTTGGTTCTTTAACAGAATTAACAAAATTGAATCTTGGGAAGAATCAACTTTGTGGTAGAATTCCAGCTGAGATAGTTTCTTGTAGTAAGCTTCAGCTGTTAGATCTCGGGTACAATGGCTTATCAGGCGATATACCTAAAGAGTTGGGTCGAATACCATCCCTTGAAATCTCTCTTAACCTTAGTTGTAACCAGTTTACAGGAGTGATTCCAAGTGAGTTTTCAGGTCTTAGTAAATTAGGAAACCTTGATCTTTCCCACAACAAACTCACCGGGAATTTGGACGTTCTCACGAACCTTCAAAACCTTGTTTCACTCAATATATCATTCAATGACTTCTATGGAAAACTTCCCAATGCTCCATTCTTTCATAATCTTCCTTTGAGTGATCTTACTGGAAACCAAGCTCTATACATTTCTGGAGGGGACGTGATCCAAACGGGGCCTGCTGGACATGCTAAAACCACCATGAAGCTTGCTATGTCAATCCTAGTCAGTATAAGTGCTGTGCTGGTGCTGCTAGCCATCTACACATTGATCAGGATGCAAATGGCGGCAAAGTATGGACCAGAAGTTGATACTTGGGAAATGACTCTTTACCAGAAGTTGGACTTTTCAATAGATGACATTGTTCATAATCTAACATCAGCTAATGTCATTGGCACTGGGAGCTCTGGGGTTGTCTACAGGATAATGACCGAAAATGGGGTGACTTTAGCAGTCAAGAAAATGTGGTCATCAGAGGGATCAGGAGCATTTGGTTCAGAAATTCAAACACTTGGTTCAATCCGGCACAAGAACATAGTACGCCTTCTCGGCTGGGCTTCAAACCAGAACATGAAACTTCTGTTCTATGACTACCTTCCCAATGGGAGTTTGAGCTCATTGCTCCATGGTGTTGGCAAAGGAGCAGCAGAATGGGAGAACAGATTTGATGTGGTCCTTGGAGTAGCTCATGCACTTGCATACTTGCATCATGACTGTGTACCCCCTATAATGCATGGAGATGTCAAAGCAATGAATGTATTGTTAGGCTCTAGAATGGAGCCTTACCTGGCCGATTTCGGGTTGGCAAGGATTGTCAACACTGATGTTGATGCAGACTTGTTGAAGGAGAGCCAAAGGCCTCATCTTGCTGGTTCCTATGGATATATGGCTCCAGGTAATTATTTTTACTGCATTTTTACTCATTATCTTAGATTGATGACGGAGAACGTAAAACTAGTCAATTTATGATGAATCCTCACAATACAAAAGATATTGTTGGTGTTGTACTCATTAGTCCTTATATCAGCTTGATTAGTTGCATCAACTTTTCTTGAATTGACAAAGACTCTTGCTTATTATGTGTCCACAGAACATGCTTCGATGCAACGGATCACAGAAAAGAGTGATGTATACAGCTTTGGTGTGGTCCTCTTGGAGGTACTAACAGGAAGGCATCCACTGGACCCAACGTTGCCCGGGGGTGCACACTTAGTACAATGGATACGCGATCACTTACAAAGCAAGCGCGATCCAAATGACATTCTTGATCCAAAGCTTAGAGGCAGAGCTGACCCTGAGATGCATGAAATGCTGCAAACTTTAGCTGTTTCATTCCTGTGTGTGAGCACGAAAGCTGATGATCGTCCGATGATGAGGGATGTTGTAGCCATGCTCAAGGAAATTCGGAATGTTGATCCTGTTGTGTCCGAATCCGACTTGTTGAAGAAAAAAGCAAGTGTAACACCTCTTCCAAAATCACCTGGCACCAAGAATGTGGATTCTCAAATGTCTTGTAGCTGCTCTTTTGTATTTTCTGATAACTCTATCTCCAAATAAGTCATCAACACAGTGAACAAATTTGTGAATATAGCATGTTTTATTAGCAGTATGAGTTAGAGCAGATTATAGATTTTAAAAGGAATATCATGTATCCAATTAATTTAGCATAAGCATTATAGAGTGAAA

>PGSC0003DMG400022425 LRR-RLKs

CTTTTGGTTGTTGGAGATTTGTCTGTTGTTTATGTTAAAGTGAAGATCTTAGTGTGTTTTGCAGAGTGTTTTCTTTTTGCTGGAACTGTGAAGCAAAGATTTGAAGGAAAAAGGACAGAGGAAGAGGAAGCAGGAAATTTGTGTTCTTGGAGTGTCCCTTGGTGATGATTGGTCACAAGATGTGTGGACTATATATGAAGATTTTGTTTGTATTTGCAGTTTTTGCTGTTCAAAATTGCCAGGGGTTTACTGATCCTCGTGATGGTATTAATAAATCTCTCTGTGGTTTGAAATTTGTGTATTATACAGTATAGAATGTCTCAATAGTTTGAATTTGTTATCTTTTCTATTAGCTTTATTGAAATTACAGTTTGATGTTTTGTGTAAAAGAAAGGTGAAGTTTATGACTTTGTGTTCTTTTCTTGCTCTCCAGTATTTGCCATAAATAGTTTATATACTAGTCTGGGCTACCCCTTGCTTCCTGGATGGGTGCCATATGGATTAGACCCTTGTGGTGATCTATGGCATGGTGTACTCTGTGTCAATTCCAATGTAACTGGAATGTAAGATTTCCTTCTTTTTTTTTTTGCTTGTGTTTCTCACATGGTTTGTATACTTTATAATAGATTTCTTTTCTTTTTTTGGCAGAGTACTAAATGATTCAAATTTGGGAGGTGAATTAAGTGAGGGCATAGGAAGCTTTGCTTCAATCATACAAATGTTGGTACCATCTTTATCTTCATCATTGCATTGATTTAACTTATGTTGCTGAAATTTGCCTTGTCATTTGGTTCTGTGCACTTTGTCCTTCCCTGCATCATTTGGCACCACTATTACATTGTTTAATTCTATGGCTTTAGTTAACTGCTACGTATTAAATGTCTACGCTTTGTGTTATGATCAAACCAGAGATCTCAGCAACAACCATATTGGAGGCAGTATACCATCCAATTTGCCTGTCACCCTCAAAACCTTGTAAGTTAATAATCTTTGTCCTTACTGAGCTTCTTCACCACTTTATAAGTTATTTGCTCCACATACATCCACATTTTTAATCCTATTATGGTATTTGTGTATATACAGGTCTCTCTCTGGTAACCAGCTTACTGGGAGCATTCCAGATAACATATCCTCGTTAGGGCAATTAACAGACTTGTGAGTTATATTTGACTTATCTTGATATACTTTTTCTATGACCATGCTTTTGTTGCCATTACATTTAATTCCTTCCTTGTAGGTCGTTGAACAACAATCATCTCAATGGAGTCATTCCAGATGCCTTTCTGCAACTTAAGACTTTGATTAACATGTACTCATACTTCTTTTAGCTTTTTACCTTTTCGCAGTGATGAAACTACTTTTCTTGGTGTTGTTTGTATTTATGGCTTCATGCTCTTATTTTGCAGGGACTTATCTGGGAACAGTTTAAGTGGTCAGCTGCCTCCTTCAGTGGGGACTTTGTCATCACTTAGTACCTTGTATGTATTTCTCCTTGTTGGTATGATAATTAGGGAAATATTAGGAAATGAAACTACTTTTCTGGCTATGAGGAATGAGTGCTTTCCTCTCTCTAGACACTGAATAGAAGTTCAAATAAGTCCTTAATTCCTTCAGAAATTTAACATAAGTTCTGCTCATACATAATTTAAACTGATAAATTTTGTAAAGATAAATATAGTTTTTGACATTTCATTGCATATGTAACTATTGTATGGTTGTGCTGTATGTTGGTTTGTCAAGATGTGTTTTCTGTGAACAAGCTAGAGGAATGATATTTGAGGATAATACTTCACAAAGTTGATTAATTTTGTCTAGAATCTCAATTTGCAACCATTCAACTTAATATACTTGGATGGTAAAAGGATGTTAACATAATTTCTCAACTGAACTTGTGAATATACTTGTATTTATGGTGAATTGTGATACTCTAGTTTTGATTTTTGAATATATACTTTTGGCTCTATGCAGACACTTACAGAACAATCAGCTTTCTGGGATCCTTGACGTTTTGCAAGATCTTGCTCTAACGGATTTGTAATCTCCTGGACTAATAAATTTTCCATCATTATAGATAATGTCATCAACTATGTCATTTTCTGCTCAATGTGTGCTTAAATTTGGTCACTTTCTTGTTCTGCAGAAACATAGAGAATAATTTGTTCTCTGGGCCTATACCACCAAAATTGTTGAGCATCCCGACCTTCAGGTAAAAACCTAGGAATTTTTTTGCCCTCTTTAGTTTGAGTTTTCTTTTAAGCTGATGAAATCCATTTTATTCTATCTTCAACTTCTGTAATATGAATTGGAGTATCTTATTTCTCTTTGTTTTAATTTGGCTTCATCTCTCTGTGTACTTCCATCAATTTAAAATTGGATAAGCATTGGTCACTGTGGTAATGGGACAATGTGACTTAAAAAATTTAAAACAAAAGGGAGTTGTCTCTTGGGCTAGAGATTGCACTATTTAGATCTAATTTAATTGCCAATTGTGCTTGCAATGAATATTTATGGGAAGCCAGTATAACTGTTGTTTCTAATAAACTTGACACCATTCCATGAGCTGACCATGAAAATGTTGGTTTTACAAGTTTTAGTTTCTATAACTTTGTCATGCATCGATCTGATAAATAAATCAGTACACTGGCTGATAAATGGGTAAAGTAGCAAACTTGAAGATATGGCATGTAATGATATTACTAAATAGTATGGTCTTTTTGATGAAGACTGCATAGTACACCTTATGAACCAATTCTTTACTCCATCAATCAACCATCCTCAATCCCAAATTAATTGTGGATGGCTATATGAATCCTCTTTATCCGTTCTACCTTATGCATCATCCTACATGTCACTCTTAACTTTATCTAAAATTCTATTTTGTGCAAGATTGATAATAGATATTTAGTTGGTGTTTGTTGTGAAAGCTCTTTTGGAGTTCTTTTAAAGACCTTATTAAAACCATGTTTGCTTTAGTTCTTATAAGACATTTTTCATTTCATCAATGAAGAAGGGGGTTTGAGGGATTTTCATTAAAAAACAATTCTTCTTCCACTTATAAAAGTGCTCACAAATCTTCAAAAGATCAACTGTGTAAACACATTTGTTTTGCACATCTACAAAAATATTTCTCCAGAACCATTTGATTGGTTAAAATGAAATTACAATGGATAACCAATCAACATGATGCATTTTTGTGATAGACTAGCCTTAGTCATGGCCTCAATTGTTTAGTTGATACTCTTTGGGGGAAGCGAACTATGGTAGTCTTTTCTTCTTCATGAAAGGTTCATTTGTATACTTATAAATTTTGTTGCCTTACATGGTGAAATCTATGTTTTTTATCAGACGTGCAGGGAATCCTTTTAATACTACTATCATCCCTTCACCACCCATGATATCGCCTTCTCCAGCACCATTTGCGCTTTCTCCAGATTTAGCCCCTTCACTGCCATATATTGTACCGCCAGGACAGGAATTACAGCCCTCTGGAAGGCAAAAGAGTAGTAATTCAACCAAACATGTCAAATTGATTGCTATAGCTGGATTAGTATCCCTTGTCATACTTGGATTGGGTGTTTGCCTATTGATGTCACGATTCTGCAAAAGAAGAAGAGAGACCCAAAAAGAAACCAAGAGACATGAAACATATGATCATAGTTTGCCCAAGGCCAAGGGTAACCCTAAGCATGATCAATCCCTGCAAAAACCATATTATGATGCAGAAAAAGGTAAATAGTTCATATGTGTACTTTCTTCTTATAGATTTATGTTTATCGTCAGTTACCTGTTGTTTTTAATGGCATTATGATGACAAAATGGATATGTCCAAAAGAAGCAGGTTTGAGGCCAGTGGCTGGAAATGGGAAAGCTCAGGGTAGGAACATGTCAATGAATACTTCAGAGGCTGTGCAGCAGAAAGACGTGAAGAGCACTATATCATATACAGACGATGAGCTGGGCTCGGAGATGGAGTCAATGATATCGGATGTTCTTCCACCACCGCCACCACCTTTTCTTCAAACTCTTTCTGCTGAGAGGATGGTTGTAAATCCAATTGTTCCCCCAATTACCTCAGGCAAGCATGCTATGAAGAATGTAAACTCTGCAGAGTTCTTCACCATTGCCTCTCTTCAGCAGTATACAAATAGCTTTTCTCAAGATAATCTTATTGGAGGAGGCATGCTTGGAACAGTCTACAGAGCTGAGTCTCCCAAAACAGTAAGCATCTTTATTTTCACGAGCTTCAAATCAGTAAATACTGTGTGCTTGCATGCATTGTGACTGCATCAGTACCACATCTTGATCTTCATTTCACTTTTTTAAAGGTGGAAATTACTCGTTATTTTTTTATTAAAAGCATGGTTTTCTGTGACAGACATTATATCTATTTTACATTGAACTTGTAGATGTTTTGTGGGTCTGTAATGTTTGACTCGAGTTTTGCATAATAGAAGTAGTCCAACCTTGTAGATGGCAACAGTTTCCATTAGTAGACTGTGTAGATATACCTGACATGCCTCCTGATTCATTAAATTAGTTTCCTGTTTTCTCAATAACCACTTGTACTTGTCATGTTTGTATATAACCTATTTAGTTGCCGTACAAGGCTAGAAGATGCTGAGACTTTGCTATTTCTTGATGCCTTTTGTTTTGTTAGCTATTGGCGGTCAAGAAATTAGACACTGCTATGACCAAGCGTCAAAGTGATCAAGAATTTGTTGAAATGGTTTCCAATATCTCTAAACTTCAGCATGAAAACATTGTCAAACTTGTGGGTTACTGTTCAGAGCATGGACAACGGTTGCTTGTCTACGAGTACTGTAGAAATGGTACCCTCCACGAAGCATTACACTTAGATGATCAGATCCACAGAAAACTTTCCTGGAGTACACGTGTACGCATTGCACTTCAAGCAGCAAGAGCATTAGAGTAAGTACGAAAAAAAAGCTAAATGATATTAGTTTATAAGTGCTTGTAACTTGTTTTAATGAATTGTGATGGTTAAATAAATTATAAATTGGCTGTTAAAGTCAGCTCTGCATGTTTCCCGGGCTTAATGTTTGCTACACCACAATTAATTTTGTAGTTTTATCCTTAGCTTGTAACCGTTTAGAGATTGAGTTGCTTCCTGTTTCACTGAAAGGTACTTGCATGAAGTTTGTCAACCACCTATTGTCCACCTGAACTTCAAGTCCACTAATGTTCTCCTTGATGATGAACTTGCTGTACATATGTCAGATTGTGGTTTGGCACCTCTGATGTCATCTGATTCCATAAGACAGGTAATCCCTTCAATCCTCAAACTAGAAACAAGAATAACAGGAGATTTGACTTGCAAACTGTAGTTATTCTGAATTCGTAATGGGAAAATTTATCCTTTTGGACTGTGATCTTCATTGAACTTGTAAACACTTCCTTCTCAACTTCTAGAAACTGATTCATTATTCATTAGTATGAAAACTAGGTGCCAATTGGCTGCTAAAAATTGGTGCATTCTGTTTAGAAATGGTTGTATGTTTCTATTTCGAGGGGATAAAAATGAAAGGACAAGACATGGTATTTCTGCTAATTTTGGTTTGCATCAAAAAGCTGTCCATCTCATATTGCATTTATCGATGAATAAAGTGTTGATAACTTCAGTATAACTTTCAGTTGCAAGGATCCGGCTATGGTGCTCCTGAACTTGAATTGGGAAACTATACTTGTCAGAGTGATGTTTATAGCTTTGGAGTTGTTATGCTAGAACTTCTCACAGGAAGACAATCTTACGACAGGTTAATTTCTTTTCTGACAGATATATTTTAACACCTTGCTAGTTGTTATTGTTGCAACTACATATTTTTCTAAAATTATACGACTCCTGACTTTGAGCCCATTAAAATATGATTTCCAGGTCCCGGCCTAGAGGGGAGCAGTTGCTGGTTCGATGGGCTATTCCACGGTTGCATGATATAGATGCATTATCAAGAATGGTTGATCCTTCCCTAAATGGTTGTTATCCATCCAAGTCTTTGTCTCGGTTTGCTGACATAATTTCCTTGTGTATTCAGGTTAGTCTCTTCCTTTACAGGTTCCTCATAGCTCTTTAATGAAGTTTAAAGTCCCACTTACAGTTTTGAGTTACCGCGTGTGAGAAACTTTACTGGTTATTCTTCATAAAATGAGACTTTAATCCAGTTAAATTCCAGGGCTTTGGTAAATGTCTGTTTTGGATAAACCTACTTTACAGGCTGCATATAACTATGTCCAACAAGAAAACAAACTCACATTTGAGCTATTTAGGCTTAAGTAAATTTTAGAATCAACTCAGTCTAGGGAGAGAAGAGGTTCTTTGTTAGTAAATTCGCCATTCTGCTATAGCTAAAAAACACACACATAATATTTGAACTGATGTTTTTCGAGCTGTCGCCTTTCATACTTTTCTCATGTCTTCTTACTTTTTTGTTGGGCAGTCGGAACCTGAATTCAGGCCACCAATGTCCGAAATCGTGCAGAATCTTCTGCAGATGATCTAAAGATATCATCAAGATTGAGTCTTTTAGCATAGATGGCAAGGATTCAAGGAGAAATCTTTACACAACATTTTCATTAAGCTCAAATATTGTGTGCTAACATGAGTAATGTAATGTAAATAGCATTATTATGTTTTGCAACATTTTCATGTGTGATTCTTTTTGACATATCTATGAGAAAGTAGAAAAAGACTCTCATTCTTTTAGTGTAATCGAGGAGTTTGTAATATATTACCAGCCAATTCTTTATGCAAGTTTTGTGATCATTCGAAG

>PGSC0003DMG400022565 LRR-RLKs

CCAAAACTAGCTGCAACCCTGCCAATAACACTCATCTTTCATTGTCTCACTCTCACAATGATACCCTTAATCTCCATGGCTACTTCTTCATCTTCTGTAAATTTCTTCTTCTTCTTCCTCTTTCTCTTCACCCATTCATTCTTTTGCTTTTCTCAACAAAACCCAGAAATTCAAGCACTTCAATCCTTCAAACTTAGTATTCATGACCCACTTGGAGCACTCACAAGTTGGGATTCTACATCTCCTTCTGCTCCATGCGATTGGCGAGGAATTTTTTGTTCAAATGATTCCTATGTGAGTGAAATTCGTCTTCCACATTTACAGCTTAGTGGATCTCTCACTACCCAGATTTCTGATTTGCGTATGTTGCGTAAGATTAGTCTTCGTTCAAATTTCTTTAATGGAACTATACCTGCTTCAATCTCCAAATGCAAACTTCTGGATACTGTTTTCTTACAGTATAACTCTTTTTCCGGTGAAATCCCGCCGGAGATTATGAATTTGACTGAGCTTGAAACGTTTAATGTGGCTGGAAATCAGATGTATGGTGAAATACCCAGTGAGTTGCCGGTGAATCTGAGGTATTTTGATGTCTCGGAGAATTTGTTTTCCGGTGGTATACCGGAGAAAATATCGGACCTTTCTCAGGTTATTCTGCTTAATCTGTCGTACAATCGATTCTCCGGTGATATTCGGGCGAGTTTTGGTAGGCTTCAACAACTTCAGTATCTTATGCTTGACTACAATGAGTTAGAAGGAACAGTTCCTTCTGCAATTTCGAACTGTTCGTCGTTGGTGCATTTGAGTGCTGAAGGGAATGGTATTACTGGTGTTATACCGGCGGGGATTGCTGCTCTTCCGAAGATTCAGGTGATAAATTTATCGCATAATAAGCTTTCTGGATACTTAGCAACTTCATTTTTCTGCAATGGTTCTGTGTATCCTCCTTCCCTTCAGATTGTTCAGTTGGGTTTCAATGCCTTCGCTGAGATTCTTCACCCACAATCGTCTAGATGTTTTAGTTCATTGCGAGTTTTGGATATTCAACACAATCAGATACATGGAGATTTCCCTTTTTTCTTGACAGACAATTCTGCATTGTCGTCCTTGGACTTGTCAGGGAATTTGTTTTCGGGTACAATCCCGAGTTCCATTGGGAATTTGTTGAGGTTGGAGCAACTGAGAATTGGAAATAATTCATTCGAAGGTGATATACCGGTTGGGATAACGAAATGTAGTAGTTTGAATGTTCTTGATCTTGAAGGGAACAGATTCATAGGGGAAATTCCAGCTTTTATAGGTGATCTTAAAAGCTTGAAGATTTTGTTCATGGGAAGAAATCAGTTTTCGGGTTCAATCCCTTCTAGTTTTAGCAATATCACTAGTCTTGAAAGTTTGAATTTGGAAGGAAACCGTCTTACGGGAAGCTTGCCTGAGGAGTTGATGTTTTTGAGCAATTTAAGCACATTGAATCTTAGTGGGAACAAGTTTTCAGGTAGTATTCCCGTTGGTATTGAGAACCTTCAGCAGTTATCGGTTCTAAATCTGAGTAAGAATGGCTTCTCAGGGACTATCCCGTCTAGCATTGGAACTTTGTATAAGCTGGTAGCTCTTGATTTGAGTCGACAGAATTTATCAGGCGAATTGCCATCTGTTCTTGGTGGTTTGCCTAGTTTACAAGTGATTGCTTTGCAAGAAAACAACTTGTCTGGAAATGTTCCTGAAGGTTTTAGTAGCTTAATGGGTTTGCAGTTCTTGAATCTATCTTCCAATTCATTTTCAGGTCATATACCGTCTACGTTTGGGTTCTTGACATCGTTAGTTGTCCTCTCGTTGTCTAAGAATCACATATCTGGTTCAATACCTCCTGACTTGGGTAATAACACTGCTCTGAAGATTTTAAATCTGAGGTCAAATTCATTGAGTGGCCAAATTCCTTCTGATCTTGCACGTCTCTCCCGTTTGAGTGTGTTGGATTTGGGTAGAAATACCCTGACAGGTGAAATTCCAGAAGTTATTTCCAATTGTTCATCCCTCACATCAGTTGTGCTGGACACGAACCATCTTTCGGGAAACATTCCAGCATCACTCTCCAGCTTATCAAGCTTAATTACCATTGACCTCTCCGGAAACAACTTGAGTGGAAAAATCCCAGAAAATCTTACAGTGCTCCCAAAGTTGGTGAACTTCAATGTATCGAACAATAAGTTGGAAGGCCAGATACCAGTGAAGCTCGGTTCTCATTTTAACGATCCATCGGATTTCAGTGGCAACCAGGGTTTGTGTGGGGAGCCATTGAATAGAAAATGTGAGCGAACTGGTAATGGTAAGAATAGACTGATTATGTTTATTGCAGTGTCTGCTAGTGGAGGTCTTCTCCTTGCATCATGCTGCTGTTTTTACATATACGCCTTGCTGAGATGGCGTAGGAAGCTCAAAGAAAAAACAGCTGGAGAGAAGAAGCATAGCCCTGCAAGGGTTAGTTCGAGGACCAGTGGTTCTCGTGGCAGTGGCAACAATGCTGGGCCAAAACTTGTTATGTTCAATAACAGAATTACAGTCGCTGAAACAATCGAAGCAACTAGGGAATTTGATGAGGAAAATGTTCTTAGCAGAACACGCCACGGAGTCCTATTCAAGGCATGTTACAGTGATGGAATGTTGCTCTCGATCTGTAGACTACCAGATGGATCACTAGATGAAAACACATTCAGAAAAGAGGCTGAATCGCTTGGCAGAGTGAAGCACAGAAACCTAACTGTTCTACGCGGGTACTATGCTGGCCCTCCAGACCTTAGACTACTTGCATATGATTACATGCCAAACGGGAACCTTGCAACACTCCTTCAAGAAGCATCTCATCAAGATGGTGGTCATGTTCTCAATTGGCCAATGCGACACCTCGTTGCACTTGGCATTGCCCGTGGGCTTGCTTTCCTCCACGCAGCCTCTATTATTCACGGAGATGTTAAACCACAAAATGTACTGTTTGATGCAGACTTTGAAGCCCATCTTTCAGACTTCGGCTTAGACAAGCTTACTGTAGCCACACCAGTCGAACCTTCCTCTTCAACATCAGTTGGGACTATAGGCTATGTGGCGCCAGAAGCAACATTAACAGGAGAAGCCACAAGGCAATCAGATATTTATAGCTTTGGCATTGTATTACTGGAACTGCTAACAGGGAAAAAGTCGTTAATGTTCTCGCAGGATGAGGACATTGTGAAATGGGTAAAGAGGCAATTACAGAGAGGGCAAATTTCAGAATTGTTAGAGCCAGGATTGCTTGAACTGGACCCTGAATCATCAGAATGGGAAGAGTTTTTATTAGGAGTAAAAGTAGGCTTGCTTTGCACAGCACCGGACCCCCTTGATCGTCCCACCATGACCGACACTGTGTTCATGCTCGAAGGTTGTCGTGTTGATCAAATTTCACTCTCCTGAAGAACCACGACCGGCAATGGTGACACGTAAAACCCAAGGTCTCATCCTAATTCTTTGCCCTTCCTATTGACTCTTTGATTTTTTGAGGAGTTGCATTGCAATGCAACAATTGCTAGCACTAGAACTTAAGTTTTTTGTTTAAGGAAAAGAACAAAAAGAAAATATTCTCAATTTTTTTTAGCTATAGCTATTTTATTGATTTTGGTTTTCTATTACAAAAGTGCCTCTTGAGTTGAAAACAATGAAATGAATTCTTTTTAAGTAATTCAAG

>PGSC0003DMG400022632 LRR-RLKs

CAAACCTCAGTCAAATGACAACAAGCACAGCCTCTTCATTCTTGTTTTTGCTCTTCCCCCTCTGTTCTTCTTCTTCTTCCTAACCCCCTTCTCACTTCACTTCTCCACACTCAACCCACCCACCATGACAGCAAAATCAACTAACCCCATCTTCTTCTTCACTCTTTTCTTCCTCATTTTCACATTAACCCCATCTTTTTCCATTGAAAATGATGTTAAATGTCTTGAAGGTATCAAATCTGCACTTTCTGACCCTTTAAACAAGCTCTCATCTTGGTCATTCTCCAACACTTCTGTAGCTTCAATTTGTAAACTTGTTGGTGTTTCTTGCTGGAATGAGAAAGAAAACCGCCTTCTTTCTCTTCAACTTCCTTCAATGTCTTTATCCGGTTCACTCCCACCTGCCCTTCAGTTCTGTACTTCTCTTCAATCTCTTGATCTCTCTGGTAACTCTTTTTCCGGTCCAATCCCGGTTCAAATCTGTTCTTGGTTACCTTATCTCGTCAATCTCGATCTCTCCAGCAACTCTTTCTCCGGTTCCATACCCCCTGAGTTTATTAATTGCAAATTCTTGAACACCCTTGTTCTAAATGACAACAAACTTACTGGGTCGATCCCATTTGAGATCGGCCGGCTCGACCGGTTGAAACGGTTCAGTGTGTCCAACAATGGCCTTTCCGGTTCAATTCCTGATGATTTAGACCGGTTTTCGAAAGATGATTTCGATGGGAATGATGGGCTTTGTGGGAATCCAATTGGATCTAAATGTAGTAATTTGAGTAACAAAAATCTTGTTATTATTATAGCTGCTGGTGTTTTTGGTGCTGCTGCATCACTGATTCTTGGATTTGGGATTTGGAGATGGTTTTTAGTTCAGCCTAGTAAGAAAGATAGAGAATTCGGTGACGGTAAAGGTGGTGGTGGTATCAGTGATGATTGGGTCGATAAATTGAGAGCTTACAAGCTTGTTCAGGTGACATTGTTTCAAAAACCTATTAATAAGATTAAGCTGAATGATTTACTAGTAGCTACAAATTCATTTGCTAGTGAAAATATTGTTATTTCAACTAGAACTGGTATTTCTTACAGAGCTATGTTGATTGATGGATCAGCATTGGCTATTAAAAGGTTGAGTAGTTGTAAGTTAAGTGAAAAACAGTTTAGGTCTGAGATGAATCGATTAGGACAGCTTAGACACCCGAATTTAGTCCCGTTATTGGGATTTTGTATTGTTGATACTGAAAGGCTTTTGGTTTATAAGCATATGCAGAATGGAAGTTTGTATTCGCTTTTACATGGTAATCTTAGTACTGGTATTAGAAATGATAGTTCTGAGCTTGGTTGGTTAGCAAGGGTTAGGGTTGCTGCTGGTGCAGCTAGAGGACTAGCTTGGCTTCATCATGGGTGTCAACCGCCATACGTGCATCAATACCTTAGCTCGAATGTGATTCTTGTAGACGATGATTTTGATGCTAGGATCACGGATTTTGGGCTTGCAAGGCTTATTGGTTCTGCTGACTCGAATGATAGTTCGTTTGTGAATGGGGATTTGGGAGAGTTTGGTTATGTAGCTCCTGAGTATTCGAGCACGTTGGTTGCTTCAATGAAAGGGGATGTGTATAGTTTTGGAGTAGTGCTGCTTGAGTTGGTGACAGGGCGGAAGCCTCTTGGTGCGGGTAATGCAGAAGAGGGATTCAAAGGTAGTTTAGTCGATTGGGTTAATCAGCTTTCGAGCTCAGGTCATAGTAAGGACGCGATTGACAAAGCTTTTGCTGGAAGTGGTCAAGATGATGAAATTTTGCGAGTTCTTCAAATTGCATGTTCGTGTGTGGTTTCAAGGCCGAAAGATAGACCTTCGATGTACACAGTGTATCAGTCCTTGAAGAGCATGGTGAAAGAACATTGTTTCTCTGAACACTTTGATGAGTTTCCAATCAACTTGACCAAAGAAAATCATGATCATAAAGATTAGGTTATAACATTGAAATGATTAGACAAATGTTTATGAATTTTGAATATTCTAGTTATGGCATTTGTATTTCTTGATAACTCTGTATAGTTTGAGAAGTAAATGCATGATCTTTGTACTTTCTTGACACATTTT

>PGSC0003DMG400022900 LRR-RLKs

AAAAATTATACTAATTCTACCTAACAAAAAAACAAGTAAATTTTTCTTGAGATAACAAATGAAAAATCCAGAAATTTACTTCATAACGGGGACGAATATAGTAATTTTGAAGAAATAACATGGACATTTTTCGTGTTATGATCAAAACATGCCTTCTTTGCATCGTCTTCTTGATCAATTACCCCGTGAAAATAATTTCAGTTTCCGATACTGATGCTTTGTTGAAGCTGAAACAATCATTTACAAATGCAGCGTCATTGGAATCCTGGAAACCTGGAACGGATCCGTGTGACAAAAATATACGTTGGTTGGGTGTTTTTTGTCAGAAAAAAATGGTTACTGGACTTCTTCTTGCAGAAACGAATTTATCTGGAATTATAGACGTTGAGGCATTATCACAAATGCCTGCTCTGCGTACACTTAGTTTTCAAAGCAATTTATTTTCTGGACCAATTCCGGAATTCAATCGAATGGGTGCTTTAAAAGGTCTTTATTTGTCGGAAAATCAATTTTCTGGTGAAATTCCATCAAATTACTTCGAAAAAATGTTGTCGTTGAAGAAGTTATGGCTCTCCGACAACAAATTTTCGGGTGAAATTCCGTCTTCATTAATGGAGCTGCAATATCTCATTGAATTGCACCTGGAGAACAATCAGTTCACAGGTCCAATACCGCCAATCTCGAAGCAGTCAACATTAGAATCAATCGATCTTTCTAATAATAACTTGAAAGGGGAAATCCCTGTTTCGTTATCAAGATTCAATGAGAGTTCGTTTAAGGGGAATTCCGAGCTTTGTGGTGAAAAATTAGGCAAAGCTTGTAATCCAACCATAAATAGCAGCAACGACAACAACACTAATAATAACGCAAATAATGGCACTCAACAAAGTCCGGATAGTAATTCGAATAAAAGCATCTCAATGTGGATTATGATATCATCAGTTGTTCTTTTATTGATCATGATCATAGTAATATATCTCATATGTCGCTACCAACAAAATCGCCGTGCTTCAATTGAAAGTTTTGATGAGCCATCACTTGGGAGGCGAATTTCGAGTGATAGCAAGAAGTCATTTGAACTAAGTAGACGTGGCAGTTCGATTAGAAAAGGATCGATAATGGGAAAACGTGTAGGAGATTTGACAATGGTGAATGATGATAAAGGTGAGTTTGGTTTAGCAGATTTGATGAAAGCTGCTGCAGAGGTGTTAGGGAATGGGCCATTGGGTTCTTCTTATAAGGCTATGATGTCAAATGGATTGACAGTTGTGGTAAAGAGGATCAAAGAAATGAACAAGATCGGAAAAGATGGTTTCGATGCAGAGGTTAAACGTTTAGGAAGATTGAGACATAAAAATATTCTAACACTTTTGGCTTACCATCATCGCAAGGAAGAGAAGTTGTTTGTGTATGAGTACATTCCCAAAGGAAGCTTGCTGTACATATTGCACGGTAAAGTACCTACCTCTAATTTTATCAAGTCATACTAGAAATGCATGGTAAGACTGTGTACAATAAACTCATTTTTGTTAATTTTGTGCATGTGCTTAGAATAGGAGACCAAAAGCCAAAAACTTCTACAAGTAAGTAAGTTTACATTGTATTAGTAGTCTAGTTGTCAAGTGTTTCATATTAGTTGAGGGAACGGAAAGTAGTCTCCTTGTATGATCTTGGCAAACTATACGTCTATATCCATGAACTAGCCTTGAAGTTGAGTGAGGTCTAAAGGTCCATTTGATTCACAAGTTATCAAAGCCAGACTCGTATGTACTCTTGATATCCCGATGTAGAGTTCCCATCTTACATTGGGATTATTCGCACCTTGTGGTTTTTTTCAAGACCACTTTTTGAATTATGTTCCCGTTAGCCCAGTGGGTTAAGGGGAACATAAGTTAAATAGAGACCTCAAGAAAGGTCATTTCTACAGTTGTCCTGTTATTTTCTCCATGTTAATTAGTTGAGGAGTTGAGGGACGGATTTGAGCTATGCTTAACTTATGAACTAGTTTTCGAGGTTTAATTAGGCATAATATTCATTTTCTTCGCGCTAATTAGTCTGTGCATGATTACTTATAAAGATTGCTGCTGCTTTATATGAATGATCACAGGTGAAAGAGGGCTGCCCCATGCTGAGCTCACCTGGCCAGTGCGTCTAAATATTATACAAGGAGTAGCTGAAGGACTAAATTATCTTCACACTGAACTTGCTTCTTCTGATTTGCCCCATGGCGATTTGAAATCGAGCAATATACTCATAAGCACAAACCACGAGCCAATACTCACAGGTTACGGATTCTGCACCTTAATGAACAACGCGCATGCAGTTCAAGCACTAATCGCGTTCAAGTCACCAGAAGCAGTACAAAATAACCAAGTTACACCCAAATGCGATGTTTATTGCCTTGGAATTGTGATTCTTGAGATAATAACAGGGAAGTACCCGTCTATATATCTTAATACCGGTAAAGGAGGGATAGACATAGCACAATGGGCGAAATCCGCGATTGCTGAGGGAAGAGAAACTGAATTATGTGATCCAGATATAATAGCAAGCGCGAAAGATTCAATGAGTTCAATTAAGGAACTAATTCACATTAGTGCAGCATGTGTAGAGAGTAATCCACAACAGAGGATAAATATTAGAGAAGTCATTAGAAGAATTGAAGAAATACAACAACAAAACGGAGTAGGCCAAACAGGACAAACAACAACAACAACGCCTCAGTCTCAAACGGGACAAGCAATTGAGATAATGCAGTCTCCAAGAGATGGAGATGCTGAAATACAAGTGTCGGAATCCTAACAAGGAACAGCAAATCATTAGGAATTTTAGCAGACTCCGGTAATTCAAGGCGTGGTTAATATTGCAGCTTCTCCTATGGTCATCAGACTAATGATAGTGTTTTTTTTTCTTCATTTTTTAAGGTTCGTTAACTAGGGATAGTACATATATTACTGACTAAGTTTTAGTTTCATTTTGTAGTCTGATGACCATTATCTCCCCTTGTAGAACAAGAAATGAGAGGAGAAGCTGTATCCTGTGTAGAGATGAAAGAGCTTACTGCTTGCTTTAGAATTGTATATGCAATACAATTTCCTGTTTCTACTACGTTGAGTACCAACTATTGTAGACTAAAAATTCGTAGCGTACTTTGACCAACTCCAAGGGATACTGTCATTTTCAACCAGTAACATGTATCAGGTAACTCTGCTTCCAAGGATAGAACACATGAGAAAGAAATTACCTAGTGTTTTTATCCATATTGATATTTGAACCTGAAACCTCATGATTCTCGACCTACTTCATTGATCACACTCTTGCGTACCTGTATGTTAATTTTTTTCACACAATAATAAACGGAGAAGTATGGAAGGCCCAATCAGATAACTATAAC

>PGSC0003DMG400023004 LRR-RLKs

CTACTGCCCAAAAGAAAAAAAGAAACCAACTGAGTCAGTATAGTACTTCATCTCTGAAACAAAGCAAGAACCTCACTCTCTCACCATTTCCAGTCAAAGGTAGGTAACAACTAACATGGGTTTTCCTCCATTATTCCTCTGTTTTCTCTTCATACTCTTTTCACTCTCTGTAAGTGGAAATGCTGAGTTACAACCTCTAATGGACTTAAAAGCCAGTCTTGACCCTGAAAATGCTTTACTCACTTCATGGACTGTTTCTGGTAATCCATGTGATGGTTCTTTTGAAGGTATTGCTTGTAATGAAAAGGGTCAAGTTGCTAACATTTCTCTACAAGGTAAAGGCCTTAAAGGGAAGTTGTCACCAGCCATTGCTGGGCTAACTCACTTGACTGGACTCTACTTGCATTATAACTCATTGTATGGAGAGATACCAAGAGAAATTTCTAATTTGGTTGAACTCACTGATTTGTATTTGAATGTTAATGATCTCTCTGGTGAGATCCCTCAAGAACTTGGAACCATGTCAAAATTGCAAGGTCAGTGACTAAGAAACTAACTTCTCTGTTTTGTGCGTTATGCTTTGCTTGGTTTTTGGGCTATTTTAGATTTTGAAGTTACAAACTTGAAACTTGAAAATTTGAATTTTTGTTACTATGATAAAAATAAGATATTTTTACATTTTCTTTGTTCTTTCTGAATATTTTGTCCAAATGTGTTGAGCTCTAGTTGGAAAAAGTAGGGGCTTCTGCTGTTAATCATTTATGTCTCACTGAAAAAATTGAAGTCACAAACTGGAACTTGTTCCATTTGGGAAATCATAAAGATTGTTTTTTTCCCCCAAATTGTTTTAGAGTTCAATTTCAGTCTAGTAGTTCCATCCCTTGTATTCCTTTCATACTACTAGTTTCTTGTTCTATGATTTTGTATTTTGGTTATCGTATGATTTTGTTGTAGTTACTGCATAGAATGATTTGTCAGGCTTTGTATAGTATTTTGTCCTGACTTCTTCTCTTCTATTATTTCCTTTTCTGTACTACTTTGAACTTTTTTTTCTTGAGCCAAGGGTTTATTGGAAGCAAACTCTCTACCTCCAAGGTAGGGTAAGGGTAAGGTCTGTGTATACTTCACCCTTCCCGGAGCCTACTTTGTGGGATTACATTGGGTATGTTGTCGTTGTTGTAGTTCATCCCCGTATGTTCTAAAGGTGGCTTTAAGTTTTGTTTTTGGGTGTTTATTTACAACCTCATTTTGATAATGGAGTCTGTCTCACTTTAGTTTATTGTCTTTTTCCGTATTGTACAGTTTTGCAACTATGCTATAACCATTTTACTGGTAGCATACCCACTCAACTAGGGACTCTAACAAAGCTCAATGTCCTGGCTCTGCAATCCAATCAGTTAACTGGTGCGATCCCCGCTAGCTTGGGTGACTTGGGAGTGTTGATGAGGGTGGACTTGAGTTATAATAGTCTTTTCGGTTCAATTCCTTCAAAACTTGCGGATGCTCCCCTACTTGAAGTCTTGGATATCCGAAACAACAAGATTTCTGGCAATGTGCCTCTTGGTAATGAATTTTCTCGCCCTACATTGTACTCTCTCCGTTGGAGTTTCAAGAGTCAATCAAGTTTTTTTTTAGTGGCATTTCCATATATGCCTTTTAGATAATTTTAGTTGTTAATTATTGTGTTTCATAGTACTTATTACGTAGTTTTCAAATATGAGTTTTTTTCAAAAAACTTGAAGCTTCTATGTCCGATTCATGGTCAAAGTTTAGACGTTTGAATCTTGAAATTCCAACCGTGCCACATAAGTTGAGGCAGAAGGAGTTATGTATATGTAGCCCCATAAGAGTCGTACGTATAGTTTCTTGGTAAATTATATCTTATCTAGTCTTTTGTTTTCTGTCTTGTGAAGCTTTGAAGAGATTAGTTGGAGGTTTCCAGTATGAGAACAATCCAGGACTATGCGGTGTGGGTTTCCCATCCTTGCGAGTTTGTACATCTTTGGATCGCTCGAATCCTAATAGACCAGAACCCTATGGAGTTGGCTCAAGTGGTTTATCTACTAGAGATATTCCAGAAACAGCTAATCTCAATCTAAATTGCAGCGGAAATGCCTGCACAACGTCGTCTAAAACCTCACAGGCATCTGTTGTTGTAGGGGTTATTGTGGTTACTATTATTGCATCTGCTATAGGAATTCTTTCCTTCACACACTATAGAAGACGAAAGCAGAAACTTGGAGGTGGACTGGAAATGTGTGATAGCCGTCTCAGTATCGATCAGGCCAAGGAAGTTAATAGGAAGAATGGTTCTCCCCTAGTCAGTCTTGAGTACTCGATTGGTTGGGATCCGTTGGCTGAGGGCCGGCGTTATGGTGGAGTTTCCCAAGAGATTCTGCAGAGATACAGATTCAATTTGGAAGAGGTGGAGTCAGCTACACAGTACTTTGCTGATAAGAATCTATTGGCTAAGAGCAACTTTTCAACTACATATAGAGGGACTCTAAGAGATGGATCACTTGTTGCTGTTACGAGAATTGCCAAAATTAGCTGCAAGTCAGAAGAAGCTGAGTTTTTAAGAGGACTAAACATTTTGACGTCTCTGAGACATGATAATTTAGTTAGGCTGAGAGGTTTCTGTTGCTCTAAAAGCCGGGGAGAATGTTTTCTCGTTTATGATTTTGTGCCAAGAGGAAACTTGTTGCACTACCTAGATTTGAAGGAGGACGAAGCTCATGCTCTTGAATGGTCCACGAGGGTATCTATCATTAGTGGCACTGCAAAAGGTCAGTTTTTGAATCAACTCTATCTTATAAGATCTAAAGTACAGTAGTTTGACATTATATTACTTTTTTTGAGCTTACTGTCAACAATTCAATAATGCCTTTTTATCTATCTTGAGCAGGCGTTGAGTATTTGCACGGGTGCAAGGTGAATAAGCCAGCTCTAGTGCATCAGAATATCTGTGCTGAAAACGTGCTCCTTGACCAACGATTCAAACCATTGCTTTCAAATTCAGGCCTGCATAAGCTCCTCACAAATGATACCGTCTTTTCAACACTCAAGGCAAGTGCTGCCATGGGATATTTAGCTCCAGAGTACTCCACTACAGGCCGATTTACTGAGAGGAGTGATATTTATGCCTTTGGAGTGCTCATTTTCCAAATTTTATCTGGCAAGCGAAAATTCACCAGTTCAATGCGTGCTGCTGCTGAATCGTCCAAATTCCACGACTTGATGGACGTGAATCTCCGTGGAAGGTTCTCTGAGTCTGAAGCAACAAAACTAGCGAAAATCGCTTTGTTATGTACACATGAATGTCCAGAGGAGAGACCAACCATGGAAACAATCGTACGAGAAATTGGTAATTTAGCTAGTCCCTCCTAATGCTAATGATCTTCTTCCATCTACTATTGCCTGTGGCAAATTTTCAATCCTCCTACTTATTAGCTGGAGATAGATTCGTACTAACCTCTCACGAGTATTGGAGCTTCAAATATTTAGTCAGTCTGAGAACTTCTTCGACTGAGGACTTGACCATTTTGTGAATGTATCTTGTATGCCATTTAACCTTAGGATTTCTCATGTTGGAGGATTTTCCTCCGTTTATCTCATGGATTCTAATTAATCTGTTTGTAAAAAGATGAATGTTCTTCTATTTACCTACGATCATTGCTTCGTTTTTGTTTTGTGCCTA

>PGSC0003DMG400023057 LRR-RLKs

ATGTCGATGTTTAACAAATTATTGTTGGTTTGGTTTTTTGTTGTAACACTGGTTCTTGGATATGTGTTTTCTGAAACAGAGCTTGAGGTTCTGTTAGAGATAAAGAAGTCATTTCTTGATGACCCTGAAAATGTTTTGTCAAATTGGTCTGATAAAAATCAAAATTTCTGTCAATGGAGTGGTGTTTCATGTGAAGAAGATACATTGAAAGTTGTGAGATTAAATCTTTCTGATTGTTCAATTAGTGGTTCAATTTCACCTTCTTTTGGCTTTTTGCATGACCTACTCCATCTTGATCTTTCTTCTAATCTATTATCAGGTCCCATTCCACCTACTCTGTCTAATCTTTCTTCTTTACAATCTCTTCTTCTTTATTCTAACCAACTCACTGGCCCCATTCCCAATGAAATTGGCTTGTTGAAGAATCTCCAAGTTCTTAGAATTGGTGATAATGTTGGCTTAACAGGGCTAATTCCAAGTAGTATTGGTGATCTTGAAAATTTGGTTACTCTTGGTTTGGCTTCATGTAGTTTGAGTGGTATGATTCCTCCTGAGTTGGGAAAACTCGGGCGGGTTGAGAATATGAATCTGCAAGAAAATAATCTTGAAAATGGAATTCCTGCTGAGATTGGTAACTGCTCTAGTCTTGTTGCGTTCAGCGTTGCAGTTAACAATCTCAATGGATCAATCCCGGAGGAGTTATCTATGCTTAAAAACCTTCAAGTTATGAATTTAGCGAATAATAGTCTTTCAGGACAGATTCCTACTCAGCTTGGAGAAATGAATGAGCTGCAGTATCTCAATTTGCTTGGTAATCAGTTGGAAGGTTCGATACCAAAGTCGTTGGCAAAGTTGAGTAATGTTCGGAATTTAGACTTGTCTGGTAATAGACTTACTGGTGAAATTCCTGGAGAATTTGGGAATATGGATCAGCTACAGTTCTTGGTTCTGACTAGTAATAATCTTTCGGGTAGCATACCGAAAACAATATGTAGTTCTAATCGAAACAGTAGTTTGGAACACATGATGCTATCTGAAAACCAACTTTCTGGTGAGATTCCTGTGGAGTTGAAAGAATGCATTTCACTAAAGCAGCTTGATTTGTCTAACAACACGCTTAATGGTTCGATACCAGTAGAGCTGTATGAGTTGGTTGAGTTAACTGATCTCCTACTTAACAACAACACATTGGTTGGTTCAGTTTCTCCGTTGATTGCAAACCTAACCAATCTGCAAACATTGGCATTGTCGCATAATAATTTTCATGGCAATATACCGAAAGAGATTGGGATGGTCGAAAATCTTGAGATTCTGTTTCTATATGAGAATCAGTTCTCTGGAGAAATACCAATGGAGATAGGGAATTGCTCCAGATTGCAAATGATCGATTTTTATGGGAATGCATTCTCAGGACGCATTCCCATTACAATTGGAGGGCTGAAGGAGCTGAATTTCATTGATTTTAGACAGAATGATCTCTCTGGTGAGATTCCTGCCAGTTTGGGAAACTGTCATCAGCTGAAGATCCTTGATTTGGCGGATAATCGTCTATCTGGTAGCGTTCCTGCTACGTTTGGTTATCTTCGAGCATTGGAGCAGCTTATGCTTTATAACAACTCACTTGAAGGTAATCTTCCTGATGAACTGATCAATCTTTCAAACTTGACAAGAATAAATTTTTCTCATAACAAACTGAATGGTAGTATAGTTTCCCTGTGTAGTTCGACGTCTTTTCTTTCGTTTGATGTTACAAATAATGCATTTGATCATGAGGTTCCACCCCATTTAGGATATTCACCATTTCTTGAAAGGTTAAGGCTAGGAAACAATCGTTTCACCGGAGAAATCCCTTGGACATTGGGATTAATCCGTGAGTTATCACTGCTCGATCTCTCTGGAAATGAATTGATTGGTTTAATACCTCCACAGCTTTCTCTGTGTAGAAAGCTCACACATCTCGATCTCAATAACAACCGGCTTTATGGATCAATTCCATCTTGGCTCGGAAACTTGCCACTCTTAGGTGAGCTTAAGCTGTCCTCAAATAAGTTCTCGGGACCTCTTCCTCGAGAATTGTTTAACTGCTCAAAGCTTTTAGTGCTATCTCTAGAAGAGAATTCTCTCAATGGAACTCTGCCCCTTGAAATTGGTGAACTCAAGTCCCTAAATATCTTAAATTTCAACAAAAACCAACTCTCTGGTCCCATTCCATCTACCATTGGCAACTTAAGCAAGCTATATATACTCCGTCTCTCGGGAAACAACTTAACCGGTGAAATACCAAGTGAGCTTGGAGAACTCAAGAATCTTCAAAGCATACTGGATTTGAGTTTCAACAACATCATTGGTCAAATTCCTCCTTCTGTTGGGACACTTACTAAGCTTGAAACACTTGATCTATCTCATAACCATCTCACCGGAGAAGTTCCTCCTCAAGTCGGTGAAATGAGCAGTTTGGGAAAGCTTAACCTCTCCTACAACAACCTTCAAGGCAAATTGGACAAACAGTATGCACATTGGCCAGCTGATGCATTCACGGGGAATCCACATCTTTGTGGAAGTCCTCTACAAAATTGTGAAGTCTCTAAGTCCAACAATCGGGGTTCAGGACTAAGTAACTCGACAGTTGTGATCATTTCAGTGATATCGACTACTGTGGCTATCATACTTATGTTGCTTGGAGCTGCCCTCTTTTTCAAGCAAAGGCGAGAAGCCTTTAGAAGAGGAAGCGAAGTGAATAGTGCTTACTCATCCAGCTCTTCTCAAGGACAAAAGAGACCACTTTTCGCAAGTGTAGCTGCTAAGCGTGATATCAGGTGGGATGACATCATGGAAGCAACAAATAATCTAAGCAACGACTTCATAATCGGATCTGGTGGTTCAGGAACTGTCTATAAAGCTGAGCTGTTTAACGGAGAGATCGTGGCAATCAAGAGAATACCAAGCAAGGATGATCTCTTGCTAGACAAAAGCTTTGCAAGAGAAATCAAGACACTTTGGAGGATAAGGCACAGACACTTGGTGAGGCTACTTGGATACTGTAACAACAGCGGAGAAGGTTCCAACGTATTGATTTATGAGTACATGGAGAACGGAAGTGTGTGGGATTGGCTTCACAAGCAGCCAGCTAATAATAACAAGAGGAAAACATGCCTCGATTGGGAGGCAAGGCTGAAGATAGCGGTGGGATTAGCACAAGGAGTTGAGTACCTTCATCATGATTGTGTACCCAAGATCATTCATAGGGACATCAAATCAAGCAACATTCTACTAGACTCAAACATGGAGGCACATTTGGGAGATTTCGGGCTGGCTAAAGCCGTTCATGATAACTATAATTCCTACAACACTGAATCAAACTTGTGGCTTGCTGGTTCATTTGGTTATATTGCTCCAGGTATATATCATTCCTGAATTCTCGAGTATTAGCATTCGAAAATAGTTTTTTATTTTTTTTATACTAAGTTGCTTATTTGTTCGCAGAGTATGCTTATTCTTCGAAGGCAACAGAGAAGAGTGATGTTTATAGCATGGGGATTGTACTGATGGAGCTAGTGAGTGGAAGGATGCCAACTGATGGGAGTTTCGGAGAAGACATGGACATGGTGAGGTGGGTTGAATCGTGCATCGAAATGTCTGGGACAGTTAGGGAAGAGCTCATTGATCCTGTTTTGAAACCATTGTTACCTAATGAAGAAAGTGCTGCACTTCAGGTGCTGGAAATAGCACTTGAATGCACCAAAACTGCACCTGCTGAAAGGCCATCTTCGCGCAAAGTTTGTGATCTGTTGCTACATGCATTCAACGACAAGGTGGTTCATTCTGATAAGATGAGTCCAGACAATTACGTCTAG

>PGSC0003DMG400023192 LRR-RLKs

ATGACAACTTTGTCATATTTTTTCATTATATTCTTCTTCTTGTTAAGTTGTTTGAGTCCAAGAAGAGTGAACTCTCTGTCTTCTGATGGAAAAGCTTTGCTCTCTCTTCTCAAAGCAACTTATGACCCTTATGCTAAATCATCATCATCTTTTGTACTTTCTTCTTGGAATGCTTCAACTTCAACTCCATGTTCTTGGCAAGGCATTTCTTGTTCTCCTCAACAAAGAGTTATTTCTGTTTCTATTCCAAATACATTCCTCAATCTTTCTTCTTTTCCTTTTGAGCTTTTTTCACTTTCATCTCTTCAGCTTCTCAATCTTTCTTCTACTAATATATCTGGCTCAATACCTTCATCTTTTGGTTTGTTTACACATCTTAGGCTTCTTGATCTTTCTTCTAATTCTCTTTCAGGACATGTTCCTTCAGAACTTGGTGGACTTACCTCACTTCAGTTCCTTTTCTTGAATTCAAACAGATTGAGTGGTAGAATCCCATATCAACTTGCTAATCTTTCTTCATTAGAAATACTCTGTCTCCAAGATAATCTCCTTAACGGGTCGATTCCGAAATATTTAGGCTCATTGGTATCACTCCAACAGTTCAGGATTGGAGGGAATCTAGAGTTGAGTGGTGAAATACCTGCAGAGCTAGGAATGCTCACCAATCTTACGACGTTTGGTGTTGCGGCTACTGGACTTTCAGGGGTTATTCCACATACATTTGGTAATTTGATCAGTCTACAAACTCTGGCAGTATATGATACTGAAGTGTTTGGTTCAATACCTCCTGAACTTGGGATGTGTTCAGAGCTCAGGAACTTGTATTTGCACATGAATAAGCTTACTGGTCCAATCCCTCGTCAGTTGGGTAAGTTGCAAAAGATTACTAGCTTGCTTTTATGGGGAAATTCGCTTACTGGCCCTGTCCCGGCTGAGCTGTCGAATTGTTCATCCCTTGTGGTTCTTGATGTCTCTGCAAATGACTTGTCTGGCGAAATTCCGGGTGATTTAGGGAAGCTTGAAGTTCTTGAACAGCTTCACTTATCTGATAATGCACTTAGCGGGGCTATTCCAATGCAGTTGAGTAACTGTAGCAGCTTAACAGCTCTTCAGTTGGATAAGAACCTATTATCCGGCACGATTCCTGAGCAAGTTGGTGAGTTGAAGTACTTGCAGATTTTCTTGTTGTGGGGGAATTCGGTTTCAGGAACTATCCCGGCTGCTTTCGGAAACTGTACTGAACTGTATTCACTTGACCTCTCAAGAAACAATCTCACTGGTTCAATACCTGAAGAGATATTCAGTTTGAAGAAGCTGAGTAGGTTGTTGCTTCTTGGGAATTCGTTAACGGGAAGATTGTCTCCGAGTGTGGCAAAATGCCAGTCTCTGGTCAGGTTGAGGATTGGCGAAAACCAGTTTTCTGGACCAATACCTGATGAGATAGGACAGCTGCAGAATCTTGTGTTTCTTGATTTATACATGAACCATTTCTCTGGTGAATTGCCTTCTGAAATTGCCAACATTACGGTTCTTGAGCTGTTGGATGTGCATAACAATTACCTAACCGGGGAAATACCATCATCGCTGGGAGAGCTCGTGAATTTAGAGCAGCTTGATCTCAGCAAGAACAGCTTCACTGGTGAGATTCCGTGGAGTTTTGGTAATCTCAGTTACTTGAACAAACTTATTCTTAGCAATAATCTGCTTACTGGTCCAATTCCAAAGTCATTTAGCAACTTGCAGAAGTTAACTCTACTTGATTTGAGCTCCAATAGTTTATCTGGTGCAATTTCACCTGAGATTGGTTACATGACAAGCTTAACAATCAGTTTAGATTTGAGCTCAAACCGATTCACAGGTGAACTCCCTGACACGTTATCCGGATTGACACTGTTGCAATCTCTTGATATTTCTCATAATATGTTGAGTGGAAGAATCACAACTCTAAGCCTCTTGACCAGTCTCGCTACTCTAAACGTATCGTATAATAATTTCTCAGGACCTATTCCTGTAACACCCTCCTTTAGAACTCTAACATCAAACTCTTTTCTTGAGAATTCACTTCTTTGTGAATCAACTGATGGCTTCACTTGTTCTGCACATATAACTAGAAGGAACGGATTGAAGTCTGCAAAAACCATAGCTTTGGCTGCTGTGATTGTGACTTCTGCATCCATAACAGTCGTGGCAACGTGGTATCTTGTGACAAGGAAGCATAGATATGAATTTGAGAAATCACCTGGCATGTCGGTCTCTGCAATAGGGACAGAAGATTTTTCCTATCCGTGGACTTTCATCCCTTTCCAGAAGCTCAATTGCACCGTTGACAACATCTTGGATTGCTTGAAAGATGAAAACATTATAGGGAAAGGCTGTTCGGGAGTTGTTTATAGAGCAGAAATGCCAAATGGCGAGTTGATTGCAGTGAAGAAGCTCTGGAAAACAAAGAAAGATGAGGAACCAATAGATTCTTTCGCTGCTGAAATTCAGATTCTTGGTCACATTAGGCATCGAAACATAGTGAAGCTCCTCGGATATTGTTCAAACAGGAGTGTTAAGCTTCTTCTCTACAACTACATTTCAAATGGTAATCTTCAACAGCTCTTGCAAAGTAACAGGAACTTGGATTGGGAAATCAGGTACAAGATTGCAGTTGGATCAGCTCAAGGCCTTGCTTATCTTCACCATGACTGTGTGCCAGCAATTCTTCATAGAGATATCAAGTGCAACAACATACTCCTCGACTCTAAATTCGAGGCTTATTTAGCAGATTTTGGACTTGCAAAGCTGATGAATTCTCCAAATTATCATCAAGCTATGTCCAGAGTAGCAGGCTCTTATGGCTATATAGCTCCAGGCAAGTATCTTTCAACTTTTATTTCTCATCTTAACCTAAATGTTGTGCAGCTATATGATGCTTTAAATCCTTGTGTTGAATCGTGTGATCATCGCATCTCTAAAAGCTTAAGTTATTAGAAAGGATACACTTTTCTCTACTAAATCATATCTTCAACACATGCGGCTCTGATTCTTGCCAAGCACGTGGATAATGGGTGACGCTTAGACTTCAACCTAGGATACCATGTTGAAATGTGTGACTCTCTCATTTAGTTTAAGTTATTAGAGATGGTACACATTTATTTACTTCATTATATCGTCAACAGTTCCCAGATATAGGTCAAAACGAGAGTCACAACGTTATGAGCTATAGTCAAATTTGGAAGGATAAACTAAACAAGTTATGCTTCATAGATGTTAACACTATTTTTTACCACTATGGTTGCAGAATATGGATACACAGTAAATATAACAGAAAAGAGCGATGTCTACAGTTATGGAGTGGTGCTGCTGGAAATATTGAGTGGACGTAGTGCGATTGAGCCTCAGATTGGTGATGGACAACACATTGTCGAGTGGGTGAAGAAGAAGATGGGAAGCTTTGAACCAGCTGTTACAATTCTTGATTTGAAGCTACAGAGTTTACCGGATCAAATGGTGCAAGAAATGCTTCAAACACTTGGAATAGCTATGTTCTGTGTGAATTCATCACCCACTGAAAGGCCAACTATGAAGGAAGTAGTGACACTTCTAATGGAAGTGAAGAGCCCAACTGAAGAATTTGGAAAAACTTCTCAGCCTTTAATCAATCAGTCAACTCAAAGTTGA

>PGSC0003DMG400023282 LRR-RLKs

AATGAACCATCCCTCTTTCTTCATTTTCCTTTCACTCCACAATAACACTCTCTTCCTTTTAACAAAATTTTTTTCTTCTCATTAGATTTCTGAATTCTGCTGCTCAAACATGAGACTAATTAAAGCACACATTGTTACTATTATATTTTAGGTAAGTAACTCTGTTTCCTCTTAACTCTTTCTTCATTATTTTTGTAATATTATGTATGCAACAAATTCTTGAGTTATAGTATAACATGTTCTTTAATAGCAAAAGATGCATTCTACTTTTTGTTGTACTTAATGTCTCTGTTTCTGTTAAATTTAAGAGTTAAATGATTTTAGGCACAAAAATATATGATTTTGGTTGTTAGGACTCAATTATGAAAGTTATATGTTTTTTTTTGTCTGTCACCACTCACCACTTTCTCTTTTTCTCTTCCTGTATGTATATGCCATACTCATTTCCTCTTCTTTTTTTTTTTTTGATGATTTGTCATATTAGAATCTCCCAATTTGTTCATTCAATAATAAGTGTGTGGGTCAATTTAAGCGCATCTTGATTAATTTTATGACATATTTGCCATCTGTTATTCATGCATTCAGATTAGCAAATCAGAAACAATTAAGATGAAAAGAAAAAGAATTTTCTATTCACACCTTTAAAACCCAACATCATCTCACATCAATAATAAATAGCATGTAACTCTATTTACCAAGGTTAGGACAGATGAAAAGTAATAACCTAGTGTTTTTTGTTTCTACTGGGATTTGAATTTATCTCATAGTTTTCACCTACTTCATTGATTAATAGGCTGCACTGGTGGTGTAATAAGTTTAGAACTATTCGGTTCAAGATTTTAATCTTTTCTGGTGCTATGTTCTATCCCACTCTAGTCAGATCCTTCCCCGGATATGAAGAATTTGATTGTAGGAAAGATGAAAAAACAAAGGCCGCTCTTTAGTATCTTTCCTTATAACAAAAGACTATAAAGTCAAGCGAAACATTATATACTTGAGTCACTTCAAAGAAAGATGTAAATTCATTTGCAATATAGTAAACCCACAACTAAATACAAATTGACATGTTAAACACATCAGCCAATCACCCAACATCCTTTGTCTGTGTATATCTGTTTAACATCTTTCATATGTGCTTTACTCAATTCATAAGCTATCTTATTCTTTTTAAGTGATATTTTTAATTTCTTATTATACAACAAACATTTGATTAGGTATACATTGCTTTTAAAGTAAAAGGGGTTTGTTATATGTTTATTCTTGTTTCTATACTTTATTGTTTGGTTACTGATTGCTACTTCTCTTCTTTCAGTTACTCATTAAGAATGGGCATCCATCATATCTGTTATGGAGTTTGTCATCCCCTTGTGAAATGACTCTTTAGCCCTTCCCATATTCAATCATGTCTAGAATTCAAACTCTCCATTCCCTCCAAATCCTTGTTATTTTCTTGAAGTTTCTTGTTTTCTCTAATTCATTGTCTGTTGAGACAGAAGCACTTCTTGAATTTAAAAAGCATCTTGTTGACCCTTTGAATGTCTTGGAATCTTGGAAATATTCAGATTCCCCGTGTAAATTCTATGGTATTCAATGTGACAAACACACTGGCTTGGTCACTGAAATTTCACTTGACAATAAGTCCCTTTATGGAATTATTTCCCCTTCAATATCTGTCCTCCAAAGCCTTACTTCTCTAGTGCTTCCTTCAAATTACCTGTCTGGAAATCTTCCAAGTGAATTGGCGGATTGCACCAATCTCAAAGTATTGAATGTTACTGACAATAACATGAACGGTACGATACCTGATTTATCTAGCTTGGCTAAGTTGGAGGTGTTAGACTTGTCAGATAACTGTTTTTCTGGAAAGTTTCCAGCTTGGTTTGGAAAGTTGACTAGTTTGGTTGCATTAGGCCTTGGTGGAAATGAGTATGATGAAGGTAAACTTCCTGATTTATTTGGGAAGTTAAAGAAAGTTTATTGGCTTTTCTTGGCAGGGTCAAATTTGACAGGACAAATCCCGGAATCTATTTTCGAAATGAAGGCACTAGGTACATTAGATATTTCGAAAAACCAGATATCCGGGAATTTTCCAAAGTCGATAAACAAGCTGCGAAATCTCTTCAAGATTGAACTTTATCAAAACAATTTGACCGGGGAATTGCCAGTAGAACTTGTAGACCTTATACATCTGCAGGAAATTGATGTTTCTAGAAACCAGCTGCATGGGACTTTGCCTAAGGGGATTGACAACCTGAAGAATATAACGGTTTTTCAAATATTCAAGAACAATTTCTCTGGGGAAATCCCTCCAGGCTTTGGAGATTTGCAGCATCTTAATGGCTTTGCTGTGTACAACAACAGTTTTACTGGAGAAATTCCAGCTAATCTTGGCCGGTTTTCGCCACTGAATAGCATAGACATATCTGAGAACAAGTTTAGTGGTGCATTTCCAAAATATTTATGTCAAAACAACAACTTGCAAAATTTGCTTGCTGTTGAAAACAGTTTCACAGGGGAGTTTCCAGGTAATTATGCTTCTTGCAAGACTTTAATGAGATTGAGGGTGAGTCAGAATCAACTTTCTGGGAGAATTGCTGAGGGTTTATGGGGGCTTCCAGAAGTCACAATGATAGATTTCAGTGATAACAACTTCACTGGAACTGTATCTCCTGGAATCGGCGCTGCGACTAAGTTGAATCAGTTGGTATTATCAAACAACAGATTTGCAGGTGAGCTTCCAAAAGAACTTGGAAAGCTCACACAGTTGGAAAGGCTATATTTGGATAACAATGAATTCTCTGGCATAATACCATCTGAACTTGGCTCATTGAAGCAAATTTCATCTTTGTATTTAGAGAAAAACTCACTTTCCGGGTCGATACCATCGGAGTTGGGTGAGTTTCCTAGGTTGGCAAACTTGAATCTTGCTTCAAATCTTCTCACAGGCAACATACCGAATTCGTTGTCAATGATGGCCTCGTTGAACTCTCTGAATCTTTCGAGTAACAAACTCAGTGGCTCAATTCCAACAAGCTTGGATAACCTGAAGTTATCTTCATTGGACCTCTCTAACAATCAGCTAACTGGAAGAGTTCCTACAGATCTTTTGACGGTGGGAGGAGAAACGGCATTTATAGGAAACAAGGGGCTTTGTGTTGATCAAAGCATCAGGAATGTCCGAAGGAACTCAAGCATCGGCGCTTGTAGTGGAAAGGCTGCTCAGGAAGTGTTCATGAAGAGCAAACTAGTTGTGTTTTGCATTGTATTACTTTCTTTGGCTGTTCTAATGGGTGTTTTCATGCTTGTGAGTTACTGGAAGTACAAGTGCAATGCTGAAGCTGATTCAGAAAAGTGTCTGGGACATTCAAATGGAATGAATCCAAAATGGAAGCTTGAAAGCTTCCAACATGTGGAACTTGATATAGATGAAATATGTGATGTTGGTGAGGACAAGCTGGTAGGAAGTGGGGGCACAGGAAAAGTTTATCGGTTAGATTTGAAGAAAGGTTGTGGCACAGTGGCAGTAAAGCAGCTTTGGAAAGGAAATGAGGTGAAAGTTTTGACAAGGGAAATGGACATTCTAGGAAAGATCAGGCATCGAAATATAGTTAAACTGTATGCCAGTCTAATGAGAGAAGGTTCGAATATGCTGGTTTTCGAGTACTTGCCAAATGGTAATTTATTCGAGGCACTACACCGGGAAGTCAAGGCTGGGAAGACAGAATTAGATTGGTACCAGAGGTATAAAATTGCAGTTGGAACTGCAAAGGGAATCGCTTATTTGCACCATGATTGTGTCCCGCCTATTATTCATAGAGATATCAAGTCAACAAACATTCTACTTGATGAGGAGTATGAAGCAAAAGTTTCTGATTTCGGGGTTGCAAAAGTTTCAGAAATTTCCTCTAGAGTCTCCGAGTTCAGCTGCTTTGCAGGCACTCACGGCTATTTAGCTCCTGGTATGTTTCAAAGAGGATAAACCAAATACTATGTTTCCACTTCTTTCTATTTTCACTCTTGCTAGTTTCCTAACATCACATTTAAATTTGTTTTATTGACAGAGATAGCATATACTTCCAGAGTGACAGAAAAGAGCGATGTATACAGTTTTGGAGTTGTTCTATTAGAATTAGTAACCGGAAGGAAACCAATAGAGGAAACCTACGGAGAAGGCAAAGACCTGGTTTATTGGGCATCAACTCATCTTAATGACAAAGGAAGTGTTCTGAATATTCTTGATCAAAAGGTGGTTTCCGAGCTTATCCAAGATGACATGATCAAAGTTCTGAGGATTTCTGCCCTATGTACGACTAAGCTTCCTAATTTACGCCCCAGCATGAAAGAAGTAGTAAATATGCTTGTTGATGCTGAACCTTGAACATTTAGTCTTCAAGCAAATGCGAGAAAAATGAGAAGGATCCTTTGTAGTTTTCTCCTTTTCCCTCTTACCATATAGTTTAAATTCCAAACTTCAGTATAGTTTTAGTTTTCTCATTATGGATATGCCATGTAAAATTGTATCCAACTTTGTCTGTACAAGTTCATTTTGTTAAATGGAATGCATATTTCTGGTTGAGCTTATAA

>PGSC0003DMG400023283 LRR-RLKs

ATGATTGTCTTGAACTCTAGTGAATTGTTTGGTCCATTGTCACCAGCAGTTGGTAATCTTAGTTTCCTCAGGGTACTTTGGCTCAGCAGAAACAGTTTCACTGGCCAAATCCCTGGAGAGATAGGAAAGCTATCGAGATTACGGAGGTTGAACTTAGCAAACAATTCATTTTCTGGTGAAATACCAAGAAACATATCGCGGTGCTCCAATCTTAATTATATTCATTTAGGTAATAACAATCTGAGTGGAAGCATTCCAACAGAGTTGAGATCATTGAATAAACTTGAGAATTTACTACTTCAGTTGAATAAATTGACTGGAGAAGTTCCAGCTTATCTAGGTAATCATTCTTCACTCTTGGCAATTGCATTAGGAGATAATCAATTACATGGAAAGATACCTGATATTTTTGGTGGACTCAAACATTTGGTGTTTCTTGATTTTGCTTTGAATTATCTGTCTGGAGAGATTCCTATGTCAATTTTCAACTTGTCATCATTGCATACACTTCAGATAACTTTTAATAAACTTGAAGGGGCTCTTCCATCAGATATGGGGTTCAATCTTCCAAGTTTAAAGTTCTTTCTCCTTGTGGGAAATCAACTGACAGGAAAGGTTCCTACTTCAGTACTGAATTCTACATCACTTATGGAATTTGGTATCGACGGGAATAATTTCACTGGACAAGTACCTGCCTTTGGAAACCAAAAGGATTTGTACTGGCTTGGACTTGCTGAAAATCATTTTGGAAATGGAATGTTGGATGATTTGAAGTTTATGTACTCCCTACAAAATTGTACTAGTTTAGAGCAGCTAATATTGGAAAACAACAATCTTGGAGGAGTACTGCCAAGCTATATAGGCAACATGTCCAATCTTTTACGGCTTTCTGTAGGGGGGAATCTCATCCAAGGAAACATTCCCACAGAGATTTTACAGCTTGAAAACCTGCAAGTACTTGGTCTCGAGAAAAATAATTTTACTGGTATGATTCCTGAGTCTATTGGACAACTAAGCCAACTAAGTAGGTTATACATTCACCACAACAGATTTTCAGGTGAGATTCCACATTCTCTTGGAAATTTGACGAGGTTGATTGAACTCGATTTGGGATCAAACAACTTGCAAGGTACTGTGCCTTCAAGTCTTGGAAGCTGCAAGTTCTTATCACTCTTGTATCTGAATGGAAATCAACTTAGTGGTCTTATACCGAAAGAACTGTTTGAGCTCTCCCTTATAGAATTTGACCTCTCAAACAACCACTTGACGGGGTATTTTCCTGTAGGGATTGGCTCTGGAAACTTGACTGGATTGATAAACTTGATCTACATGAACCATTCATACAACAATTTATCAGGTGAAATACCAAGCAGCTTTGGCACTTTGACTAGTTTGAGGGAGCTATACCTTGGTAACAATGCCCTTCAGGGAGTTATTCCTGCATCCTTGAGTTCTTTGAGAAGTCTTGAATATATGGATCTTTCTCATAACCACTTTGTAGGGAGGATCCCAAAGTTTCTGGATGAACTTGTCTCCTTGAAGTTCTTAAATTTGTCTTATAATGATTTGGAGGGTGAGGTGCCTTTGAAAGGAGCTTTCAGAAACATGAGTGCTGTTTCACTTGTTGGGAACAGTGAGCTATGTGGTGGTATTCCAGAGTTCAAAATGCCAAAGTGCAGCAACAAAGTAGCGTCAAGAAGACGAAGGTTATCTCATAGATTGATAATAGTAATGCTGGTTATTGGAGGATTATTGGCAGCAACCACAGTGGCCCTTCTTATTTTCTTGTGTGCTAGAAGAAAAAAGAAGTCCACTTCATCGGAGAATTCATCGTTGGACGTGATCCCAAGAGTGACTTACAATAGTCTTTACAAAGAAACTAATGGCTTCTCTACGTCAAACATGATTGGTTCGGGTGCATTTAGCTTCGTGTACAGAGGCATTCTAGAGGAAAATGGAAAATTTGTTGCCATAAAGGTGCTCAAACTCCAAGTTAGGGGAGCTTCTAAGAGCTTTCTTACTGAATGTGAAGCACTAAGGCATATCAAGCATAGAAACCTGGTAAAACTCTTAACATCATGTTCAAGCATTGATTACCAAGGTAATGACTTCAAGGCTCTCATTTATGAGTACATGGCCAATGGTAATTTGGCAAATTGGTTGCACAATAGAAGTACAGATGGTGAGGAAAATCATGAGCCTAAAACTTTGAACATGCTACAGAGACTCAATGTAATAATTGATGTGGCATCTGCATTAGATTATCTTCATCATCAATCTGGAACGCCATTGACACATTGCGATATCAAACCAAATAATGTGCTTTTGGATGAAGATTTTGTGGCTCATTTAGGTGACTTTGGACTTGCAAGGTTTCTGCCTGATGCTGCAAATCTGCTCTCCTTAAGCCAATCAGCCAGCTCTCTCAATATAAGAGGAACTATTGGATATGTTCCACCTGGTAATTAACTCTTCTCCCTCTCTAAACATCATTTTATATTTATGAAGGTCCTCTTACTTATCAAGTTTGTCAATTCATGTATGTGCAGAATATGCTACATGCAATACGTTTTCGACCTATGGAGATGTCTATAGTTATGGAATTCTACTGTTGGAAACATTTACTGGACGAAGTCCAACTGATGAGATTTCCAAGGATGGTCTAAACCTTCATGATTTTGTTAAGAGAGCCATACCTGAGCAAGTGAAAGATGTTTCTGATCCTAAACTTGTCTATGATGAAATGGGCAGACTGATCAGCAACAATAAAACAATCGAGTGCCTCACATTGATTTTTCAAGTTGGGATTGCTTGTTCAGTGGAATCTGCAAAAGATCGCATGGACATTGCCAATGTTGTTAATGAACTGAATGTAATCAAAGATGCCTTCCTGAGGAACTAGCAATATTGGCTATGAAGTTTTTGAGTTTCATTATAAAGATACTACTTATATAAGTTATATACTCTTTATCCACTTTTCCCTTCCCCTTTTCCTGCTTCTCCTTATGTAATGGAAATT

>PGSC0003DMG400023329 LRR-RLKs

CAAAATTGTTCTATTCCTACTTTCTTGAATGCTTTGTGTGTCCCACTTTACACATTACACTAAACCCCCAATTGCCATTTGAATCTCACCTATTCATCATGTCATCAATTATCTACTATTACTCTTCCTTGTATAACTTAATCATTCATTCAACTCTTCAAAATCCATGGCTATTAGTCCAGGTCACTTTTCTTACTAACTTGTATATTTACATATTACTATTTGCATATATACACGTTGTGAACCTCATATAGTCGACCTCAACTTGTTAGGGACTGAGGCGTAATAGTTGTTATACATGTTGTGAAGCTTATATAGTTGACCTCAACTTGTCAGGGACTGAGGCGTAGTAGTTGTTATACATGTTGTAAAGCTTATATAGTCGACCTCAACCTGTCAGGGACTAGTAGTTGTTATACATGTTGTGAAGCGTATATAGTCGACCTCAACTTGTTATGGATTGAGGCATGGTAGTTGTTATACATGTTGTGAAGCTTATATAGTCGACCTCTAACTTGTTAGGGACTGAAGCATGGTAGTTGTTATACATGTTGTGAAGCTTATATAGTCGACCTCAACTTGTTAGGGACTGAGGCGTAATAGTTGTTATACATGCTATTATATAATTTTCTTGATATTTTTTTGGGTATATTGAACTAAGTTAATTTTGCCTTAGCTTTTAAATGAACCTATGTTAAGGGGATTCAACAATCTAATATATATATGTGCCTATCATTTAACATACGTATGTCATTTAGTTTTTGTACCTTTGTCATCCCTCAACCCCTCTCCAATCTTTTCTATCTATCATTAGAAGTAGGACGGGTAATATTCCGGTGAAGTGGATAAAAATTAGTTTAAAGGCTTTATGTTTTGATGATCATTTGAGAAATTAACTCTTAGGAAGACTAGTCGCTCAGAAATAGTCATTTTGATTCTTTTATTCTTACTCAGTATGAGCATTAATTCTTATTGGTAGCACAAAAAGAAGTGGATACTTCTTGAACTAATTGGTAATATTTTCATTGTGTTGCAGCTACAAGGCCTATTTCTTTGACATCATATTCTCCGACCTAGACTTGTTTGAGATTGAGGCGTAATTATTGTTGTTGTTGTTATATAGAGTATTGGAAACAAGGAGAAGGATCAATTAGAAGCTCTCCTCTGGTTGATAGAGAAAATTCAATGCCTACAGCCTTTTCGTTCCGGAATTTTCACTTGAACTTGCTGCTATTCCTACTTTTATGCAGCAATGGTCTCTCCTCAAATGATTATGATGTTGAAGGTATGGAGTTATAGGTTTTGGTTACTGTACCTGTGAAAATTTCATATGCTTGCAAAATAATGTGCTAATAAAATCGCGATAACTGTTATTCTCTATTGAAATCATATATCATATATACATCATTTCAGGTCATGCTTTAATTGAGTTGCTGAGGGCTCTGAATGATTCACACAATAGAATCAAAGATTGGAACATTGATTTTGTGACTCCATGTTTCAGTTGGTCTCATGTTACATGTAAAAATGGAAATGTCATATCCCTGTAAGACTAATTCTTTCCTTTCTGTTATAAGTAAATCAAGTAGATTCATATAGTCGACCTCAACTTGTTTAGGACTGAGTCGTAACTATTGTTGTAATCAAACAGGAACTTAGCTTCTAATGGATTCTCAGGAACACTTTCACCGTCGATAACCAAATTGAAATTTTTGGTTAGCTTGTAAGTATACTGCACTTTGTATTAAAATTAGAGTTACTGGTCAAAAACACACCTAAAGTATCCCTTTTTTCGAGTTCCCTATCACCAACTATTTATCAAAAACACACCTCAACTATTAGTTGCTCCCTTTTTCCTACCTGAGATATCACCATCGTTCAGGCAGGAAAAATGAAATGTTGCTAATACTTGATGTCTCGCGAACCTAGGAAAGTTTTGTGAAGTGTTGCTAAAATGTTAATGGAACTGTGCAGGGATTTACAGAACAATGATCTATCTGGTGCCTTACCTGATTACCTTAGCAGCATGTCAAATCTACAAAACTTGAATCTTGCAAACAATAATTTCAATGGCCCTATTCCACCAGCTTGGGGTCAGCTCTACCAACTTAAATATTTGTAAGTACTTGTTTTTCTTTGTTTCTTTGTTCTCTCCGTTTTCCTAAAATATCAAGGAATGTCACTCGGACTCTTCAAAAATGATGACAGGTTCGTGTCAGTTCCTCCAAAAGCAGTGCATTTTTGGAGGATCCAACACGAGTGTGACAACAGTTTTGGAGAGTCCTCTCTGACCTTAACATAGCTCTTATTAAATAGTTCTTGTAACAAAGTCTTTCTATGATATGTCCTTTTGTCCTGCATCAGGGTCATGAGGGGAAATCACATAACTGGAACTGTTCCAAATTCACTTACAAATATCACAGGCTTAAAAGAACTGTAAGAACTATGTTTACCTTACGGGCATAATTTACTCTTTGTTTGAAATTTTCGGTTGGATTGAGGCCTATGTTGCTCAGACTCTTCAAAACTACTATAGTATGTGTGTCGAACCCTTCAAAAGGTACCATTTTGGAAAGTCCGAGCAACATAGAATGAGTTGTTATTTGATGTTTTCATTTATTTCTGGTTGTAGTTCTTACTTGTTTTAATTCTTCACAGGGACCTGTCCTCCAATGATCTGACTGGAGGTATCCCACTGCAGTTGTTTTCAATACCAACATACAGGTAATGTCATCATTTATTCTTCTTTCTTGCCAACTTGTTTAGCCTTCCCATTTTGATGTTGAGCTCTCAAAAAGGGCTACTTGATGCACAAAACAGCCATGGGGGTGTGACATAGGCAGTCTACCCTGGTGCTAGCATCAGTAGCTTATTCCACAGCTCGACCTCATGACCTATAGGTCATATGGAGACAACTTTACGTTGCTCCAAGGCTCCCTTCAACTGATAAAAAAATGAAAATTTGTTACTCATGACTGCATTATTCAACTTGAAGCTATTATTATTGACTTAGTTGCTTAACTATGTTGTTTTAACAATCATTCTATACGTCGTAATCAGTTTTTCAGGAACTCATCTTGCTTGTGGCAATAGATTTCAGAAACCCTGTGTTTCTAGTTCCTCTGCTCCAGGTTAATAGTCTTGTCACCAATGCATCTTTTAGTTCTTAAATCTTTGCCAACCAAATGATTATTGCTCAAGTGATATTTTTTTCAACAGCTTCTAGCAGAAGACCAAAACTTGAGGTTGCTATTACTGGTGCAAGTTGTGGTGCATTCCTTCTTCTCTTAGTTGGGGCTATCGTCACATATCGTTGTAATTATAAACGTAAACTCAAACAAGATCAATTTTTTGATGTGGAAGGTATATGCTTTACTCCTATTTCTTCAACAGAAAGAGTATATGGTTTTTCTGTACTGAACAAATTTCCTTAATAACTTCAAATGCCATCTACTTGTATGGCATTTTAGGTGACGTTGAGCGCAACATTTCATTAGGCCAACTAAGAAGGTTCTCATGGCGCGAACTTCAGATTGCAACGGACAATTTCAGTGAAAGCAACGTTATAGGACAAGGAGGTTTTGGTAAAGTTTACAAGGGCTATCTTTCAGACAACACAAAAGTGGCAGTGAAAAGACTCACTGATTACCACAATCCTGGTGGAGAGGCTGCATTTCTAAGGGAAGTGCAGCTGATTAGTGTTGCAGTTCATCGAAATCTCCTACGTTTGATCGGGTTCTGTACAACCTCTTCTGAGAGAATTCTTGTTTATCCATTTATGCAGAATCTTAGTGTTGCATATCGGTTAAGAGGTATGCCCCTTATCACTCAACCACTTTTTTTGACATTCTTGATACTATTCAGTCTGTTAAGTCTGTGTTTGTGTTTTTTCCTCTCTTTTCGACAGATTTAAAACCTGGAGAGAAAGCTCTAGATTGGCCAACGAGGAGACGGATAGCATTTGGAGCAGCACATGGTCTAGAGTACCTACATGAGCATTGTGATCCTAAGATCATTCATAGAGATCTAAAAGCAGCAAACATCCTCCTAGATGATAACTTTGAACCTGTTCTTGGAGATTTTGGACTAGCAAAGCTAGTTGATACAAAACTGACTCATATTACAACTCAAGTTCGCGGAACAATGGGCCACATTGCTCCAGAATACTTGTCTACTGGAAAATCATCTGAGAAAACAGATGTATTCGGATATGGTATTACACTTTTAGAACTTGTAACAGGGCAACGTGCAATTGACTTTTCGCGTCTTGAAGAAGAGGAAGATGTGTTGCTACTGGATCATGTAAGTCCCTACTTCCACACACTTTTCAATTTTTGAAGAGTTAGAGCAACATAATGGATACCTTTTTACTTCTACAGATCAAGAAACTACTAAGAGAGAAAAGGCTAGGTGACATTGTGGATGGAAACATGAAAACTTATGAACCCAAAGAAGTAGAGACAATCCTTCAAGTTGCATTGCTATGCACCCAAAGTTCACCTGAGGAACGTCCGAAAATGGCACAAGTAATCACCATGCTAAAGGGTGTAGGTTTAGCTGAAAAATGGGCAGAGTGGGAGCAACTTGAAGAAGTAAGAATTCAACAATTTTCGCTCATGTCGCGACAGTTTATGTGGGCTGAAGACTCAACACATGATCAAGAAGCTATACAATTGTCTCAAGCAAGATAAACAAAAGGAAGATGCAGAGAAATGTATGTATAAGTACTTCTGCAATAAAATTTAAGAGTAGTTTTAGCTTCTGCTGCAGCTAGCTAGTTATGTAACATGTAAGAGTAATTTTATCATACATCTAGCTAGTTATGTATCATGTAAGAGTAAATTTATCATACGTATATAAGGCGCTAAGGAGTCTTTTTTTAATTTTACGATGTATATATATAGAGAGAGAGACCATCTATTACTGAATTCACTCAATTCCTAGCTCGTTGTACTAAAGTTCCTGCTCGATACTAATACTAAAGTTCCAT

>PGSC0003DMG400023550 LRR-RLKs

ATGTTGAGATATCTAGAAGTGGGTAACAATCCGTTGAGCGGCGTTCTACCAAATTCTATTGGGAATCTCTCATCTACTATTGAAAAAAATCATTTAGTAGATGCACACATCAATGGCCTCATCCCCACAAGTATAGGCAACATGAGCGGTCTTACATCCATAGACATTCAAGACAACAACTTGGCAGGGAATATTCCTTCTGAGATTGGTAAGCTTAAACAACTCCAAGGGTTGTATCTAAGTAACAATAAATTTCAGGGACATATTCCAGAAGCGGTATGCCATTTATCTAATTTGGTTAAATTAACTCTGCAAGTTAATGAGCTCTCTGGATTAATTCCAGAATACTTTGGAAATCTTAGCATGCTACAAAAGCTTTATTTGGCTTCTAATAAATTTTCATCAAAATTTCCGTTGAGCCTTTGGAAGATTTAGTGGTCTTCTCTATCTAAGTGTGTCACAAAATTCAATAGAGGGAGAACTTCCATCAGATATTGGAGAACTGAAAGCCATTGTAGGACTATATCTTTACAGTAACCACTTTTCAGGTGTGATACCAACCAGATTCGGGGAACTTCAAAACCTGCAATATCTTGACCTATCGAACAATTCATTTTTTGGCAAAAGTCCATTATCCTTTGCCAACTTGATAAGCTTGGTATTCTTGAATTTGTCTTTTAATGCCTTGTCTGGTACTATTCCTATGTCATTGGAAAAGCTCTCCTACCTTAAAAGCATTAATGTTTCATTTAATGATTTAGAAGGTGAAATACCGAGTGGTGGCGTGTTTGCAAATTCCACTTTGCAATCATTTCGCGGGAACAAAGGTCTTTGCGGAGTGCACATATTGGATATTCCTGCTTGTGATATCACTAATCCTGGAAAACAATCAAAGCTTAAGGGGGTTGTGCTAAAAATTGTTACTCCACTGGTTATTTCATCCTTTCTGATATTCTTGTTGGTTTCAATTTGGATAATGAAACGACACAAGAAAGGAAAGTCCAAAGATGTTGAAAAGGTATCAGAGATTAGGACTCACCAATTAGTTTCTTATCATGAGATTCAACGAGCAACAAATAATTTTGATGAATCCAATTTAATTGGTGAGGGAAGCTCTGGCTCTGTGTACAAAGGCATTTTATCTAGTGGAACTGTAGTGGCCATAAAGGTTCTGGATTTGGAAAATGAGCAAGTTTGCAAGAGGTTTGATACTGAATGCGAAGTGATGAGAAATGTTAGACACAGAAATCTTGTGTAACATCCGACAATTTGAAATAGCTTTGAAGAGGCTTAGGATTGGAAATTTTCATTTTTGGAAAGGATTTAAAAATCTGGAAGAATTTGGCTAAGTGTAGAAATCAGTGATTTTTTGGCCAACTTTGAGAAGTCATAACTTCCAGCTCAGGATGAGTTAGGAGTAGTTCCAGTTATGGTTGTGAAGCTTGTGGAATGATCTTTCCAACGCCGCCGAGTTTGCTCGATGACGAGTTCGTATGAGTGAGATATGCCCTTTGAAAGTTGGGAAGTTGACAAGGAAATCTGTCCGGGAATCATTTAGGGCATTTTAGTCTTTTCCCTAGCCAATTCTTTTGGGATTATATTGTTGTATAGGGATGATCTTTGGATCATTTTGTCCCATTTTAAAGAGAGAAAGTTAGGGTTCTTGAGAGTGAAGAAGAGAAGAANNNNNNNNNNNNNNNNNNNNNNNNNNNNNNNNNNNNNNNNNNNNNNNNNNNNNNNNNNNNNNNNNNNNNNNNNNNNNNNNNNNNNNNNNNNNNNNNNNNNNNNNNNNNNNNNNNNNNNNNNNNNNNNNNNNNNNNNNNNNNNNNNNNNNNNNNNNNNNNNNNNNNNNNNNNNNNNNNNNNNNNNNNNNNNNNNNNNNNNNNNNNNNNNNNNNNNNNNNNNNNNNNNNNNNNNNNNNNNNNNNNNNNNNNNNNNNNNNNNNNNNNNNNNNNNNNNNNNNNNNNNNNNNNNNNNNNNNNNNNNNNNNNNNNNNNNNNNNNNNNNNNNNNNNNNNNNNNNNNNNNNNNNNNNNNNNNNNNNNNNNNNNNNNNNNNNNNNNNNNNNNNNNNNNNNNNNNNNNNNNNNNNNNNNNNNNNNNNNNNNNNNNNNNNNNNNNNNNNNNNNNNNNNNNNNNNNNNNNNNNNNNNNNNNNNNNNNNNNNNNNNNNNNNNNNNNNNNNNNNNNNNNNNNNNNNNNNNNNNNNNNNNNNNNNNNNNNNNNNNNNNNNNNNNNNNNNNNNNNNNNNNNNNNNNNNNNNNNNNNNNNNNNNNNNNNNNNNNNNNNNNNNNNNNNNNNNNNNNNNNNNNNNNNNNNNNNNNNNNNNNNNNNNNNNNNNNNNNNNNNNNNNNNNNNNNNNNNNNNNNNNNNNNNNNNNNNNNNNNNNNNNNNNNNNNNNNNNNNNNNNNNNNNNNNNNNNNNNNNNNNNNNNNNNNNNNNNNNNNNNNNNNNNNNNNNNNNNNNNNNNNNNNNNNNNNNNNNNNNNNNNNNNNNNNNNNNNNNNNNNNNNNNNNNNNNNNNNNNNNNNNNNNNNNNNNNNNNNNNNNNNNNNNNNNNNNNNNNNNNNNNNNNNNNNNNNNNNNNNNNNNNNNNNNNNNNNNNNNNNNNNNNNNNNNNNNNNNNNNNNNNNNNNNNNNNNNNNNNNNNNNNNNNNNNNNNNNNNNNNNNNNNNNNNNNNNNNNNNNNNNNNNNNNNNNNNNNNNNNNNNNNNNNNNNNNNNNNNNNNNNNNNNNNNNNNNNNNNNNNNNNNNNNNNNNNNNNNNNNNNNNNNNNNNNNNNNNNNNNNNNNNNNNNNNNNNNNNNNNNNNNNNNNNNNNNNNNNNNNNNNNNNNNNNNNNNNNNNNNNNNNNNNNNNNNNNNNNNNNNNNNNNNNNNNNNNNNNNNNNNNNNNNNNNNNNNNNNNNNNNNNNNNNNNNNNNNNNNNNNNNNNNNNNNNNNNNNNNNNNNNNNNNNNNNNNNNNNNNNNNNNNNNNNNNNNNNNNNNNNNNNNNNNNNNNNNNNNNNNNNNNNNNNNNNNNNNNNNNNNNNNNNNNNNNNNNNNNNNNNNNNNNNNNNNNNNNNNNNNNNNNNNNNNNNNNNNNNNNNNNNNNNNNNNNNNNNNNNNNNNNNNNNNNNNNNNNNNNNNNNNNNNNNNNNNNNNNNNNNNNNNNNNNNNNNNNNNNNNNNNNNNNNNNNNNNNNNNNNNNNNNNNNNNNNNNNNNNNNNNNNNNNNNNNNNNNNNNNNNNNNNNNNNNNNNNNTTGTTACTCCAGTGGTTATTGCATCCTTTCTGATATTCTTGTTCGTTTCAATTTGGATAATGAAACGACAGAAGAAAGGGAAGTCCAAAGAGGTGGAGAAGATACCGGAGATGGGGACTCATCAATTAGTTTCCTATCACGAGATTCAACGAGCAACAAATAATTTTGATGAATCCAATTTAATTGGTGAGGGCAGCTCTGGCACTGTGTACAAAGGCACATTGTTTGGTGGAACTGCAGTGGCCATTAAGGTTCTGGATTTGGAAAATGAGCAAGTATGCAAGAGGTTTGATACCGAATGTGAAGTGATGAGAAATGTTAGACACAGAAATCTTGTTCAAGTGATTACTGCATGTTCTAGTGACTATATAAGAGCCTTTATTCTGCAATTTATGCCCAATGGAAAGTCTTGAGAATTGTCTGTACAAAGAAGATCGCCACTTGAACCTTCATCAAAGAGTTTCTGTAATGCTTGATGCAGCTATGGCAGTTGAATATCTACATCATGGTCATGTCACTCCAATAGTTCATTGCGACTTAAAGCCAGCCAACGTTCTTTTGGATGAAGATAATGGTGGCTCATGTTGGTGATTTTGGCATCTCTAAAATTTTAGCCATAAGCAAGTCCATAGCTTATACCGAAACATTGGGCACTCTTGGATACATTGCACCAGGTATAAAAAATAATCTCTCTTTGATTTTCTCTTATCATAGTTAAGCATTCTAAAAGTAAAAAAAGCAAGTCTTCATTTATGCAGAATTATTGTTGTATTTCAATTGAGTAACTTTTCTTCAATCCTTTTCTAAGAATATGGCTCAGAGGGAATAGTGTCTTCTAGTGGTGATGTTTATAGCTATGGCATCATGTTGATGGAAGTCTTGACCAAAAGAAGGCCAACAGATGAAGAGATATGCAATGAAAATCTTGACCTGAGGAAATGGATAATACAATCATTTTCAGGGAGTATGATGGACGTTGTCGATGCCAATCTTTTTTACGAGGAAGAACAAATCACTAGTAAAAGTGAAATCTGCATAGCGTCCATGATAGAATTGGGTTTAGATTGCACAAAGGAAATGCCAGAATCAAGAGTGACCATGAAAGATGTAGTGAAGAGACTTAACAAAATCAAAAACGCATTTCTGGAAACATAG

>PGSC0003DMG400023554 LRR-RLKs

ATGTGGAAAGGGCCATGGTATGTACCCGAACTCAGAGTCTTGGATCTCACCAACAATAACCTCACGGGTATGATCCCTCCTTCTGTTGGAAATGCCACAAAAATGATGAACTTCAGTTTGTTTGGGAATAGAGTCAGCGGCAACATTCCAAAGGAGGTCGGTAATATGAGCCAACTTGCAGACTTGTCCTTGACTGATAATCAATTAACAGGTTCCATCAGTGGCGAATTTAGAGCCAAAAAAAAGGGCACCCGCACCCATGGTCTTTTCGTAAAATTAGTATATTTTATATGTATATTTTTTAAGATTTTGTATAATATTATATTCTTGCACCCGTACTACAAGAAGTCTCATGGTTAAATGTTGAGCTTTTGACCTTGAGGATTAAGGATCAATTTTCACTTCAATACATTTTACTTGGTGCACCCATGTTCGAAAAATCCTAGATTCGCCTCTGGGTTTCATTCCCCAGAACTGTTTAATATCTCGTCGCTACTTGCCGTATCTCTGGCAAACAATAGCCTTTCTGGTCCTCTCTTGCTAGATGAAGGCAATATTGTGACGAATCTGTACTTTTTAAGTATATCTGACAACCAAATTTCTTGTTGCATTCCTTCCAACATATGCCAACTCACAGAGCTCAAAGTTTTGTCCGTATCTTTCAACAATATAACTGGAGACATACCCAAAAATATTGGTTGTTTATCCGAGCTCCATGAGTTTTATATTGGTGATAATCTAATAAAAGGGACTATTCGCACTTCATTGGGAAATATTTCCACTATGCATAATCTTAATTGTGGAAACTATCGCATAGTGGGGCAAATTCCTCTGGAATTAGGGAAGCTATCAAATTTGAGGAAATTAAAGTTTGCCCATAATTCTAATCTAATTGGTGAAATTCCAGAGGCTATTTTCAACATCTCTTCTTTGGAAGGGATCGATTTCAGTTTCAACAACCTCTCAGGGAGAATTCCAACCACTACAGGTCTTCATCTCCCCAACCTTGAAGGACTTTACTTGGGAGTAAATCAGTTCAAAGGAGAAATTCCATTGTTTATAACAAATGCTTCCAAGCTTCAGCTACTGGACCTAAGTTATAACTTTCTGACAGGAACTATTCCTACTAATTTGGGAAATCTTCGTGAGCTGCAAATACTGCTCCTATATAATAATCAACTTACCAATGAATCAAGAGAGCGTGAGTTGTTATTCTTCAATTCTTTGGCGGACTGTAGGATGTTGAGAGATCTAGAAGTGGGTTCCAATCCGTTAAATGGCGTTCTGCCCAATTCAATTGGGAATCTTTCATCTACTATTCATAACTTTAATATAGAAGATGCACACATCAATGGCCTCATCCCCAAAAGTATAGGCAACATGACCGGTCTTACAATCCTAAGCCTTAGAGGAAACAACTTAGCGGGAAGCATTCCTTCTGAGATTGGTAAGCTTAAACAACTCCAAGGGCTGTATCTAAGTAACAATAAATTGCAGGGACATATTCCAGAGGCGGTATATCATTTATCTAATTTGGTTGAATTAACTCTGCAAGTTAATGAGCTCTCTGGATTAATTCCAGAATGCTTAGGAAATCTAAGCATGCTACAACATCTTTTTTTGGATTTTAATAAATTTTCTTCAAAATTTCCGTTGAGCCTTTGGAAGATGAGTGGTCTTCTCTATCTAAGTGTGTCACGAAATTCAATAGAGGGAGAAGTTCCATCAGATATTGGAGGACTGAAAGCCATTGTAGAACTATATCTTTACAATAACCACTTTTCAGGAAGGATACCAAGCACATTCGGGGAACTCCAAAACCTGCAGTATATTGACTTGTCGAACAATTCATTTTTTGGCAAAATTCCATTATCCTTTGCCAACTTGATAAGCTTAGAATGCTTGGATTTGTCTTCAAATGCGTTGTCAGGTACTATTCCTAAGTCTTTGGAAAAACTCTCATACCTTAAAAGAATTAATGTTTCATTTAATGATTTAGAAGGGGAAATACCAAGTGGTGGCGTGTTTGCAAATTCCACTTTGCAATCATTTCTTGGGAACAAAGGTCTTTGCGGAATGCACATACTGGAGATTCCTGCTTGTGCTATCACTAATCCTGGAAAACAATCAAAGCTTAAGGAGATTCTGCTAAAAATTGTTACTCCAGTGATTATTGCATCCTTTCTGATATTCTTGTTGGTTTCAATTTGGATAATGAAACGACATAAGAAAGGGAAGTCCAAAGATGTAGAGAAGGTACCAGAGATCGGGACTCATCAATTTGTTTCTTATCACGAGATTCGACTAGCAACAAATAATTTTGATGAATCCAATTTAATTGGTGAGGGAAGCTCTGGCTCTGTGTACAAAGGCACATTGTTTGGTGGAACTGCAGTGGCCATAAAGGTTCTCGATTTGGAAAATGAGCAAGTATCCAAAAGGTTTGATACCGAATGCGAAGTGATGAGAAATGTTAGACACAGAAATNNNNNNNNNNNNNNNNNNNNNNNNNNNNNNNNNNNNNNNNNNNNNNNNNNNNNNNNNNNNNNNNNNNNNNNNNNNNNNNNNNNNNNNNNNNNNNNNNNNNNNNNNNNNNNNNNNNNNNNNNNNNNNNNNNNNNNNNNNNNNNNNNNNNNNNNNNNNNNNNNNNNNNNNNNNNNNNNNNNNNNNNNNNNNNNNNNNNNNNNNNNNNNNNNNNNNNNNNNNNNNNNNNNNNNNNNNNNNNNNNNNNNNNNNNNNNNNNNNCTAAAGCCAGCCAACATTCTTTTGGATGAAGATATGGTGGCTCATGTTGGTGATTTTGGAATCTCTAAGATTTTAGCTGTCAGCAAGTCCGTGGCACATACAGAGACATTGGGCACTCTTGGATACATTGCACCAGGTATAAAAAATCTACTCTCATTGATTTTCTCTTATCATAATTAAGCCTCTCCAAGTTGTAGAAGTAAAAAGAGAACTATTGTTATATTTCAATTGAGCAACTTTTGGTCAATCATTTTCTAAGAATATGGCTCAGAGGGAATAGTGTCTGCTAGTGGTGATGTTTATAGCTACGGAATCATGTTGATGGAGGTTTTGACCAAAAGAAGGCCAACAGATGAAGAGATATGTAATCAAAATCTTGACCTGAGGAAATGGATAATACAATCATTTTCAGGGAGTATGACCGACGTCGTGGATGCCAATATTTTTTCTGAGGAAGAACAAATTACTTGTAAAAGTAAAATGTGCATAGCCTCCATGATAGAATTGGCTTTAGACTGCACAAAGGAAATGCCAGAATCAAGAATAACCAATGAAGGATGTAGTCAAGAGGCTTAA

>PGSC0003DMG400023587 LRR-RLKs

ATGGAGAAGCACATTTTCTTACTGATTCTTCTACTTGTAATTCAATATTACTCTGTTGCTTCCTCAAATGAAACTGATCAACAAGCTCTACTAGCTTTCAAAAATCTTATTACAAGTCCTAATCATTTTTTGGCCAATAATTGGACCAAAATACTCTTTTGCTCTTGGTTTGGTGTCACTTGTAGTTCAAAAAGGCAAAGGGTTGTGGCCTTGGCTCTTCCTAATTTGCAAATTCAAGGCACAATTTCCCCGTCTTTGGCCAATCTGTCCTTCCTCAGTGTTCTCAATCTCGAGAACAACAGCTTCCGCGGTGGCATCCCTTATGGACTTGGCCACTTGCCTCGCTTGCGAGTGATTGATATTCAAAACAATCAGCTCAACGAAAGTATACCAACAAGTCTATTTCAAAACCGGAGAGTTCAAGTAATTTCATTGGCTTTCAATGAACTCAGTGGTGAAATGTGGAGAGGGCCATGGTATGTACCCCAACTCAGAGTCTTAAATCTCAGGAACAATAGTCTCATCGGTATAATCCCTCCTTCTGTTGGAAATGCCACAAAAATGATGAACTTCAGTTTGTCTGGGAATAGATTCAGCGGCAACATTCCTAAGGAGGTCGGTAATCTGAGCCAACTTGCAGACATGTCCTTGACTGATAATCAGTTAACAGGTTCCATTCCCGTAGAACTGTTTAATATCTCGTCACTACTGTCCATAAATCTGCGACACAATAGCCTTTCCGGCCCTCTCTTGCTTGGTCAAGGGAATATTGTATCAAATCTGGAGCATTTAATTATATCTTTCAACCAAATTTCTGGTTGCATTCTTTCCAACATATGCCAACTCACAGAGCTCAAAGTATTGTCCATAGTTTACAACAACATAACTGGAGAAATACCCAGAAATATTGATTGTTTATCCAAACTCGAGATGTTTCTTATTGGTGGTAATCCAATTAAAGGGACTATCCCCACTTCATTGGGAAATATTTCCACTCTACAATATCTTTATTGTGGAGAAAATCACATAGTAGGGAAAATTCCTCAAGAATTAGGGAAGCTATCAAAGTTGAGGGTATTAGTCATTGAGGACAATTCTAATCTTATTGGTCAAATTCCAGAGGCTATTTTCAACATATCTTCTTTGGAAGTGATTGCTTTCAGTTTCAATAACCTCTTGGGGAGAATTCCAACCACTACAGGTCTTCATCTTCCGAACCTTAAAGAACTCTACTTGGGAGACAATCAGCTTGAAGGGGAAATTCCATTGTTCATAACAAATGCTTCCAAGCTTGAGAAATTGGAGCTAAATGATAACTTTCTGTCAGGAACTATTCCTACTAATTTGGGAAATCTTCATGAGCTGCGATATCTCTTCCTACATCATAATCAACTAACCAATGAACCAAGAGAGTGTGAGTTGTTATTCTTCAATTCATTGGCGGACTGTAAGATGTTGAAATATCTACAAGTGGGTTTCAATCTGTTGAATGGCGCTCTGCCAAATTCTATTGGGAATCTTTCATCTACTATTCAAAACTTTCATATAGGACATGCACACATCAACGGCCTCATCCCCACAGGTATAAGCAACATGAGCGGTCTAATGACCCTATACTTTCAAGGAAACAACTTGACAGGGAGTATTCCTCCTGAGATTGGTAAGCTTAAACAACTCCAAGGTCTGTATCTACGTTACAATAAATTGCAGGGACATATTTCTGAGGCAGTATGCAATTTATCTAATTTGGTTCAATTATATCTGGAACATAATGAGCTCACTGGAGTGATCCCAGAATGTATAGGAGATCTAAGCTTGCTACAACATCTTTATTTGGATTCTAATAAATTTTCATCAAAATTTCCGTTGAGTCTTTGGAAGATGAGTGGTCTTCTCTATCTAAGTGTGTCACGAAATTCTTTAGAGGGAGAAGTTCCATCAGATATTGGAGAACTGAAAGCCATTGTAGGACTATATCTTTGCAGTAACCACTTTTCAGGCGTGATACCAACCAGATTCGGGGAACTCCAAAACCTGCAGTATCTTGACCTATCCAACAATTCATTTTTTGGCCAAATTCCATTATCCTTTGCCAACTTGATAAGCTTGGAATTCTTGAATTTGTCTTCAAATGCATTGTCAGGTACTATTCCCAAGTCTTTGGAAAAACTCTCATACCTTAAAAGCATTAATGTTTCATTTAATTATTTAGAAGGTGAAATACCGAGTGGTGGCGTGTTTGCAAATTCCACTCTTCAATCATTTCTCAGGAACAAAGGTCTATGTGGAATGCACATATTGGAGATTCCTGCTTGCGCTATCAATAATCCTGGAAAACAATCAAAGCTTAAGGAGGTTCTGCTGAAAATTGTTACTCCAGTGGTTATATCATCCTTGTTGATATTCTTGTTGGTTTTAATTTGGATAATGAAACAACAAAATAAAGGAAAGTTCAAAGATGTTGAAAAGGTACGGGAGATTGGGACTCATCAGTTAGTTTCTTATCATGAGATTCAACTAGCAACAAATAATTTTGATGAATCCAATTTAATTGGTGAGGGAAGCTCTGGTTCTGTTTACAAAGGCACATTATCTAGTGGAACTGTAGTGGCCATTAAGGTTCTGGATTTGGAAAATGAGCAAGTATGCAAGAGGTTTGATACCGAATGCGAAGTGATGAGAAATGTTAGACACAGAAATATTGTTCCAGTTATTACTACTTGTTCTAGTGACTATATAAGAGCCTTTGTTCTGCGATATATGCCCAACGGGAGTCTTGAGAATTGGTTGTACAGAGAAGATTGCCACTTGAATCTTCATCAAAGAGTCACCATAATGCTTGACGTGGCAATGGCAATTGAATATCTACATCATGGTCATGTCACTCCAATAGTTCATTGCGACTTAAAGCCAGCCAACGTTCTTTTGGATGAAGATATGGTGGCTCAAGTTGGTGATTTTGGCATCTCTAAAATTTTAGCCATAAGTAAGTCCATGGCACATACCGAGACATTGGGAACTCTTGGATACATTGCACCAGGTAGACTACTTTCTTTCTTTCTCACTTATATAATAACCTTTCTATTTGATTTTTTTCTTATCTAAATCAAGCCCTTGCAAATACTAGAACTAAATAAAAAGCATGTCTTTGTTAGTACGCGCACTATTGTTATATTCAAATGAGTAATTCTTTTTCATTTTCATTTGTTTAAGAATATGGCTCAGAGGGAATAGTGTCCACTAGTGGTGATGTTTATAGTTATGGCATCATGTTGATGGAGGTTTTGACCAAAAGAAGGCCAACAGATGAAGAGATATTCCATGGAGATCTTGACTTGAGGAAATGGATAACACAATTATTTTCAGGGACTATGATGGACGTTGTGGATGCCAATATTTTTTCCGAGGAAAAACAGATCACTTCCAAAAGTGAAATCTGCATCGTGTCCATGATAGAATTGGCTTTAGACTGCACAAAGGAAATGCCAGAATCAAGAATAACCATGAAAGAAGTAGTCAAAAGGCTTAACAAAATCAACAACACATTTCTGGAAACATAG

>PGSC0003DMG400023612 LRR-RLKs

ATGGAGGGGCAAATTCCTCCGGAATTAGGGAAGCTATCAAATTTGAGGAAATTAAGCTTTTCAGAGAATTATAATTTTATTGGTCAAATTCCAGAGGCTATTTTCAACATATCTTCTTTGGAAATGATTGATTTCAGTAACAATAACCTCTCGGGTAGAATTCCAACCACTACAGGTCTTCATCTTCCTAACCTTAAAGTACTTGAGTTGGGAGTCAATCAGCTGGAAGGGGAAATTCCATTGTTCATAACAAATTCTTCCAAGCTTCAGATACTATCGCTAATTGATAACTTTCTCACAGGAACTATTCCTACTAATTTAGGAAATCTTCGTGAGCTGCAAGAACTGTTCCTACATACTAATCAACTTACCAATGAACCAAGAGAGTATGAGTTGCGATTTTTCAATTCTTTGGCGGACTGTAGGATGTTGAAATATCTACAAGTAGGTTCCAATCTGTTGTATGGCATTCTGCCCAATTCTATTGGCAGTCTTTCATCTACTATTCAAAAATTTGAAATAGGAGATGCACACATCAATGGAATCATCCCCACAAGTATAGGCAACATGACCGGTCTTACATCCTTAAGCCTTGGCGGAAACAACTTGACAGGGAATATTCCTTCTGAGATTGGTAAGCTTAAACAACTCCAAGGGTTGTATCTAGATAACAATAAATTGCAGGGACATATTCCAGAGGCAGTATGCCATTTATCTAATTTGGTTCATTCTTATCTAGATAATAATGAGCTCTCTGGACTAATTCCAGAATGCTTAGGTAATCTTAGCATGCTACAAAAGCTTTTTTTGAGTTCCAATAAATTATCAAAAATTCCAGGGAACCTTTGGAAAATGAGTGGTCTTCTCTATCTAAGTGTGTCACAAAATTCTATAGAGGGAAAAATTCCATTAGATATTGGAGAACTGAAAGCCATTGTAGAAATACATCTTTCTGGTAACCACTTTTCAGGCATGATACCAAGCACATTCGGGGCACTCCAAAACCTGCAGTCTCTTGACCTATCAAACAATTCATTCTTTGGCCAAATTCCATTATCCCTTGCCAACTTGATAAGCTTGGAATTCTTGAATTTGTCTTTAAATGCATTGTCAGGTAGTATTCCTATGTCTTTGGAAAAACTCTCATACCTTAAAAGCATTAATGTTTCATTTAATGATTTAGAAGGTGTAATACCAAGTGGTGGTGTGTTTGCACATTCCACTGTGCAATCATTTCTGGGGAACAAAGGTCTTTGCGGAATGCACATATTGGAGATTCCTGCTTGTGCTATCACTAATCCTGGAAAACAATCAAAGCTTAAGGAGGTTCTACTAAAAATTGTTACTCCTGTGGTTATTTCATCCTTGCTGATATTCTTGTTGGTTTCAATTTGGATAATGAAACGACAGAAGAAAGAAAAGTGCAAAGATGTTGAAAAGGTACTAGAGATCAGGACTCATCAATTAGTTTCTTATTACGACATTCAACAAGCAACCAATAATTTTGATGAATTCAATTTAATTGGTGTGGGAAGTTCTAGCACTGTGTTCAAAGGCACATTATCTAGTGGAACTGCAGTGGCCATTAAGGTTCTGGATTTGGAAAATGAGCAAGTATGCAAGAGGTTTGATACCGAATGTGAAGTCATGAGAAATGTTAGACACAGAAATCTTGTTCCAGTGATTACTACTTGTTCTAGTGACTATATAAGAGCCTTTGTTCTGAAATTTATGCCCAATGGAAGTCTTGAGAATTGGTTGTACAAAGAAGATCGCCACTTGAACCTTCATCAAAGAGTTACTGTAATGCTTGATACAGCTATGGCAGTTGAATATCTACATCATGGTCATGTAACTCCAATAGTTCATTGCGACTTAAAGCCAGCCAACGTTCTTTTGGATGAAGATATGGTGGCTCATGTTGGTGATTTTGGTATCTCTAAAATTTTAGCCATAAGCAAGTCCATGGCTTATACAGAGACATTGGGTACTCTTGGATACATTGCACCAGGTATAAAAAATCTACTCACTTTGATTTTCTTTTATCATAATAAAGCCTCTCCAAATTCTACAACTATAAAAACAAGCTTTTATTTATGCAGAATTATTGTTGTATTTCAATTGAATTAACTTTTTTCAATCCTTTTTAAGAATATGGCTCGGATGGAATAGTGTCTGCTAGTGGCGATGTTTATAGTTACGGCATCATGTTGATGGAGGTTTTGACGAAAAGAAGGCCAACAGATGAAGAGATATGCAATGAAAATCTTGACTTGAGGAAATGGATCACACAATCATTTTCAGGGAGTATGATGGACGTTGTGGATGCCAATCTTTTCTCCGAGGAAGAACAGATCACTTCAGAAAGTGAAATCTGCATAGCGTCCATGATAGAATTGGGTTTAGACTGCACAAAGGAAATGCCAGAATCAAGAATAACCATGAAAGATGTAGTCACGAGGCTTAACAAAATCAACAACACATTTCTGGAAACATAGAAGTCA

>PGSC0003DMG400023614 LRR-RLKs

ATGCCTAGAAATATTGGTTGTTTATCCAAGCTCGAGAAGTTCTTGATTGGCAATAACCCAATAAAAGGGACTATTCCCACTTCATTGGGAAATATTTCCACTCTGCAATATATTTATTGTGGAAGCAGTCGCATAGTGGGGCGAATTCCTCCGGAATTAGGGAAGCTATCAAATTTGAGGCAATTAAGCTTTGACAACAATTCTAATCTTATTGGTCAAATTCCAGAGGCTATTTTCAACATTTCTTCTTTGGAACTGATTGGATTTGCTTTCAACAACCTCTCCGGGAGAATTCAAGCTACTACAGCTATTCATCTTCCGAACCTTAAAGAACTTTTCTTGGGAGTCAATCAGCTGGAAGGGGAAATTCCATTGTTCATAACAAATGCTTCCAAGCTTGAGACACTGTCGCTAGCACGGAGCTTTCTGACAGGAACTATTCCCACTAATTTGGGAAATCTTCGTGAGCTACGAAATCTGTTCCTATATGATAATCAACTTACCAATGAACCAACAGAGCGTGAGTTGAGATTCTTCAATTCTTTGGCGGACTGTTGGATGTTGCGATATATATCAGTGGGTTCCAATCCGTTGAATAGCGTTCTGCCCAATTCTATTGGGAATCTTTCATCTACTATTGAATACATTTATATAGGAGATGCACACATCAACGGCCCCATCCCCACAAGTATAGGCAACATGAGCGGTCTTACATCCCTAGTCTTTAATGAAAACAACTTGATGGGGAGTATTCCTCCTGAGATTGGTAAGCTTAAACTACTCCAAGGGCTGTATCTATTAAACAATAAATTGCAGGGACATATTCCAGAGACGGTATGCCATTTATCTAATTTGGTTCAATTATATCTGGACGGTAATGAACTCTCTGGATTAATTCCAGAATGCTTTGGAAATCTTAGCATGCTACAACATCTTTATTTGGATTCTAATAAATTTTCATCAAAATTTTCGTTGAGCCTTTGGCACGAAATTCTTTAGAGGGAGAAGTTCCATCAGATATTGGAGGACTGAAAGCCATTGTAGAACTACATCTTTCCAATAACCACTTTTCAGGCATGATATCAACCAGATTCGGGGAACTCCAAAATCTGCAGTATCTTGACCTATCGAACAATTCATTTTTTGGCCAAATTCCATTATCCTTTGCCAACTTGATAAACTTGGAATTCTTGAATTTGTCTTTAAATGCATTGTCAGGTAGTATTCCTAAGTCTTTGGAAAAACTCTACTACCTTAAAAGCATTAATGTTTCATTTAATGATTTAGAAGGTGAAATACCGAGTGGTGGCGTGTTTGCAAATTCCACTCTGCAATCATTTCTTGGGAACAAAGGTCTATGTGGAATGCACATATTGGAGATTCCTGCTTGCGCTATCAATAATCCTGGAAAACAATCAAAGCTAAAGGAGGTTCTGCTAAAAATTGTTACTCCGGTGGTTATATCATCCTTTCTGATATTCTTGTTGGTTTCAATTTGGATAATGAAACGACAAAAGAAAGGGAGGTCCAAAGATGTGGAGAAGGTACTGGAGATCGGGACTCATCAATTAGTTTCTTATCACGAGATTCAACGAGCAACAAATAATTTTGATGAATCCAATTTAATTGGTGAGGGAAGCTCTGGCTCTGTGTACAAAGGCACATTGTTTGGTGGAACTGCAGTGGCCATAAAGGTTCTCGATTTGGAAAATGAGCAAGTATCCAAAAGGTTTGATACCGAATGCGAAGTGATGAGAAATGTTAGACACAGAAATCTTGTTTCAGTGATTACTACGTGTTCTAGTGACTATATAAGAGCCTTTGTTCTGCGATATATGCCCAATGGAAGTCTTGAGAATTGGCTGTACGAAGAAGATCGCCACTTGAACCTTCATCAGAGAGTCACTGTAATGATTGATGCAGCTATGGCAGTTGAATATCTACATCATGGTCATGTCACTTCAATAGTTCATTGCGACTTAAAGCCAGCCAACATTCTTTTGGATGAAGCTATGGTGGTTCATGTTGGTGATTTTGGAATCTCTAAAATTTTAGCCATAAGCAAGTCCATAGCTTATACGGAGACATTGGGCACTCTTGGATACATAGCACCAGGTATAAGAAAACTACTCTCTTTGATATTCTATTATCATAATTAAGCCTCTCCAAATTCTAGAAGTAAGAAAAAAAGCAAGTCTTTATTTATGCAGAATTATTGTTGTATTTCAATTAACTTTTGTTGAATCGGCCTTTTCTAAGAATATGGCTCAGAGGGAATAGTGTCTACTTGTGGTGATGTTTATAGCTATGGCATCATGTTGATGGAGGTTTTGACCAAAAGAAGGCCAACAGATGAAGAGATATGCAATGAAAATCTTGACTTGAGGAAATGGATCACACAATCATTTTCAGGGACTATGATGGACGTTGTGGATGCCAATATGTTTTCTGAGGAAGAACAAATCACTTGTAAAAGTGAAATATGCATAGCCTCCATGATAGAATTGGCTTTGGACTGCACTAAGGAAACGCCAGAATCAAGAGTGACCATGAAAGAAGTAGTGAAGAGGCTTAACAAAATCAACAACACATTTCAAGAATCATAG

>PGSC0003DMG400023638 LRR-RLKs

ATGAAGAAGCACATTTTTATATTGACTCTTCTCTTTCTAGTTCAATTTTCTATATCAATTGCCTCCTCAAATGAGACGGACCAAGAAGCTCTACTAGCTTTCCAAAATCTTATTACAAGTCCGAGTCATTTTTTGGCCAATAATTGGACCAAAAATACTTCTTTTTGCTATTGGTTTGGTGTCACTTGCACTCGAAAAAGGCAAAGGGTTGTGGCCTTGACTCTTCCTGATTTGCAACTTCAAGGCACAATTTCCCCGTCTTTGGCCAATTTGTCCTTTCTCAGCGTGCTCAATCTCCGGAACAACAACTTCCATGGCGGCATCCCTTACGGACTTGGCCACTTGCCTCGCTTGCGAGTGATTGATTTTCAAAACAATCAGCTCCAAGAAAGTATTCCAACAAGTCTATTTCAACACCAGAGAGTTCAAATCATTTCATTGGCTTTCAATAAACTCGGTGGTGAAATGTGGAAAGGGCCATGGTATGTACCCGAACTCACAGTTTTAAATCTCAGGAACAATAGCCTCACAGGTATAATACCTCCTTCTATTGGAAATGCCACAAAGTTGCTGAACTTCAGTTTGTATGGGAATAGAGTCAGCGGGAACATTCCAAAGGAAATCGGTAATCTCAGCCAACTTGAATTTTTGTCCTTGTTCGATAATCAATTAACAGGTTCTATTCCTGCAGCACTATTTAATATCTCCTCGCTTCTTGTCGCATCTCTAGCATTCAATAGCCTTTCCGGCCCTTTCTTGCTTGATGAAGGGAACGTTGTGTCAAATCTAGAGTCATTAACTATATCTAACAACCAAATTTCTGGTTACATTCCTTCCAACATTTGCCAACTCACAGAACTCAAAGCGTTGTCCATATCTTTCAACAAGATAACTGGAGGAATACCCAAAAATATTGGTTGTTTATCCAATCTCGAGGAGCTTTATATTGGTGACAATCCAATAAAAGGGACTATTCCCGCATCATTGGGAAATATTTCCACTCTGCAAAATCTTTATTGTGGAAGCAATCACTTGGAGGGGCCAATTCCTCCAGAATTGGGGAAGCTATCAAATTTGAGGCAAATTAGCTTTGTCCAAAATAATAATCTTATAGGTCATATTCCAAAGGCTATATTTAATATATCTTCTTTGGAAATAATCGATTTCAGTTACAATAACCTCTCGGGGAGAATTCCAACCACTACAGGTCTTCATCTTCCAAACCTTAAAGAACTTTTCTTGGGAGTCAATAAGATCCAAGGGGAAATTCCATTGTTCATATCAAACGCTTCCATGCTTGAGATACTGGGGCTAAACAGGAACTTCCTCACAGGCAATATTCCTACCAATTTAGGAAATCTTCGTGACCTGCGACGCTTATACTTAGAGGGTAATCAACTTACCAATGAACCAAATAATCATGAGTTGCAATTCTTTAATTCTTTAGTGGACTGTAGGAAGCTGCAATATCTAACAGTGGGTAACAATCCATTAAATGGTATTTTGCCCGATACTATTGGAAATCTTTCATCTACTATTGAATCAATTCACATGGGAAATGCACAAATCAGTGGCCTCATTCCCACAGGTATAGGAAACTTAAGCGGTCTAATGTCGCTAGCTTTTGTGGAAAACAACTTGATGGGAACCATTCCGTCTGAAGTCGGTAAGCTTGAACATCTCCAAGGTCTGTATCTATATAGCAACAAATTGCAAGGGAATATTCCAGAGGTAGTTTGTCATTTATCTTATTTGGTTACATTATCTTTGCATGTTAATGAGCTCTCTGGGGTGATCCCAAAATGTTTAGAAAATCTTACCACGCTACGAGTTCTTTCATTGAGTTCTAACAAATTTTCATCAAAGCTTCCTTTGAGTCTTTGGAAGATGAGTGGACTTCTCTATCTTTTCATGTCACGAAATTCTATAGAGGGAGAAGTTCCACAGGACATTGGAGGATTGAAGGCCATTGTAAAACTAGATCTTTCTGGTAACCACTTTTCCGGCATGATACCAAGCCAATTGGGGGACCTCCAAAACATGAAAGTTCTTGACCTGTCAAACAATTCATTTTCAGGCTCAATTCCCTTATCCTTTGCCAATTTGATAAGTCTAGAATACTTGAATTTGTCTTTCAACGCATTGTCAGGTACTATTCCTAAGTCTTTGGAAAAGCTTTCATACCTTAAAAGCATCAATGTCTCATTTAATGATTTAGATGGTGAGATACCTAGTGGAGGTGTGTTTTCAAATTCCACTTTGCAATCATTTCTTGGGAACAAAGGTCTATGTGGAGTACACATATTGGAGATTCCTGCTTGTGCTATCACTAATCCTAGAAAACAATCAAAATCTAAGGAGCTTGTGCTGAAAATTGTTATTCCAGTGGTTACTTCATCCTTTCTGATATTATTGGTTGTGTCAGCTTGGATAATAATGAAACAAAAGATGAAAGGAAAGTCCAAAGACTTGGAAAAGGTACCAGAGATCAGGACTTATCAATTGGTTTCTTATCATGAAATTCAACGAGCAACAAATAATTTTGATGGATCCAATTTAATTGGTACGGGAGGTTCTGGCTCTGTGTACAAAGGTACATTATCTAGTGGAACTGTGGTGGCAATAAAGGTTCTCGATTTGCAAAATGAGGAAGTATGCAAGAGGTTTGATACTGAATGTGAAGTGATGAGAAATGTTAGGCACAGAAACCTTATTTCAGTGATTACTACTTGTTCTAGTGAATACATAAGAGCCTTTGTTCTACAGTATATGCCCAATGGAAGTCTTGATAATTGGTTGTACAAAGAAGATTGCCACTTGAACCTTCTTCAAAGAGTCAACATAATGCTTGATGTGGCTGTGGCAATTGAATATCTACACCACGGTCATGATACTCCAATAGTTCATTGCGACCTAAAGCCAGCCAACATTCTTTTGGATGAAGAAATGGTGGCACATGTTGGTGATTTTGGCATCTCTAAAATATTAGCTGTAAGCAAGTCCATGGTACATACTGAGACATTAGGCACTTTTGGTTACATTGCACCAGGTACGTACACTAGTTTCGTTCTTTTTCACTTATCATTAACCCTTTCTCTTATCAGAACTAAGCCCTTCCGAATTCTGGAACTAAACAGAAACCAAGTCTTTCATTGTACAAAACAATTATTATCTTCCGATTGTAATTTATTTTTCATTTTCATTCTCCTAAAGAATATGGCTTGGAAGGAGTAGTGTCCATTAGTGGTGATGTTTACAGCTATGGCATCATGATGATGGAGGTATTGACAAAAAGAAGACCAACTGAAGACGAGATATTCAACGAAAATCTTGGCTTGAGGCAGTGGATAAGACGAGCATTTCCCGGGACAATTATGGAAGTCGTAGATGCCAATCTTTTTCATGAGGAAGAATCTGTCAATTTCAAAAGTGAAATATGCATAGCCTCCATGATGGAATTGGCTTTGGATTGCACAAAGGAAATGTCAGAGACAAGGATAACTATGAGAGACGTAGTCAAGAGGCTTAGCAAAATCAAAAACACATTTTTAGGAACATAG

>PGSC0003DMG400023639 LRR-RLKs

ATGGAGAAGCACATTTTTTTATTGACTCTTCTCTTTCTAGTTCCATTTTCTATATCAATTGCTTCCTCAAATGATACGGACCAAGAAGCTCTACTAGCTTTTCAAAATCTTATTACAAATCCGAATCATTTTTTGGCCAATAATTGGACCAAAAATACTTCTCTTTGCTCTTGGTTTGGTGTCACTTGCAGTTCAAAAACGCAAAGGGTTGTGGCCTTGGCTCTTCCTAATTTGCAACTTCAAGGCACAATATCCCCGTCTTTGGCCAATTTGTCCTTTCTCAGAGAGCTCAATCTCGAGAATAACTTATTCCATGGTGGCGTACCATACAGACTTGGCCACTTGCCTCGCTTGCGAGTGATTATTGTTCGAAACAATCAGCTAGAAGGAAGTATTCCGACAAGTCTATTTCAACACCAGAGAGTTCAAATCATTTCATTGGCTTACAATAAACTCAGTGGTGAAATGTGGAAAGGGCCATGGTATGTACCCGAACTCAGAGTCCTAAGTCTCAGGAATAATAGCCTCACAGGTATAATCCCTTCTTCTGTTGGAAATGCCACAAAGTTGCTGAACTTCAGTTTGTCTGGGAATAGAATCAATGGTATCATTCCAACTGAGATTGGTAATCTTAGCCAACTTATAGAGTTGCATTTGTTCAATAATCAATTAGCAGGTTCCATTCCCGCAACACTGTTTAATATTTCGTCGCTAATTAGAGCATCTCTAGCAAGCAATAGCCTTTCTGGTCCTCTCTTACTTGATGAAGGGAATATTGTGTCAAATCTGAAGTATCTAAGTATATCTAAGAACCAAATTTCTGGTTGCATTCCTTCTAACATATGCCAACTCACAGAGCTCAAAATTTTGTCCATATCTTATAACAATATGATTGGAAAAATACCCAGAAATATTGGTTGTTTATCCAAAATCGAGGAGTTTTATATTGGAAATAATCCAATAACAGGGACTATTCCCACTTCATTGGGTAATATTTCCACTCTGCGGAATCTTTATTGTGGAAACAGTCGCATAGTAGGGCAAATTCCAAAGGCTATTTTTAACCTATCTTCTTCGGAAATGATTGATTGCAGTTACAGTAACCTCTCGGGTAGAATTCCAACCACATCAGGTCTTCATGTTCAGAACCTTAAAGAACTTTTCTTGGGACACAATCGGCTCGAAGGGGAAATTCCATTGTTCATAGCAAATGCTTCCAAGCTTGAGATATTGGGGCTAGAAAATAACTTTCTCACTGGCACTATTCCTACTAATTTGGGGAATCTCCGTGAGCTGCAAGAACTGTTCCTACATGATAATCAACTTACCAATGAACCAAGAGAGCATGAGTTGCAATTCTTCAATTCTTTGGCGGACTGTAGGATGTTGCGATATCTACAAGTGGGTTCCAATCCGTTGAATGGCATTCTGCCCAATTCTATTGGGAATCTTTCATCTACTATTGAAAACTTTCATATAGAAGATGCACACATCAATGGCCCCATCCCCAGGGGCTTACTCAACATGAGCGGTCTAATAGCCCTAAACCTTGGAGAAAACAACTTGGCGGGAAGTATTCCTTCTGATGTTGTTAAGCTTGAACAACTCCAAGGTCTGTATCTAAATAACAATAAATTGCAGGGACATATTCCAGAGGCAGTATGCCATTTATCTAATTTGGTTCAATTAAGTCTGGGTGGTAATGAGCTCTTTGGATTAATTCCAGAATGTTTAGGAAATCTTAGCATGCTACAAGCCATTATGTTGAGTTCTAACAAATTTTCATCCAAGATTCCTTTGAGCATTTGGAAGATGAGTAGTCTTCTCTATCTAATCATGTCACAAAATTCTATAGAGGGAGAAGTTCCACAGGATATTGGAGGACTGAATGCCATTGTAGGACTAGATCTTTCTGGTAACCACTTTTCAGGCATGATACCAAGCCAATTGGGGGACCTCCAAAACATGAATACTCTTGACCTGTCAAACAATTCATTTTCAGGCTCAATTCCATTATCCTTTGCCAACTTGATAAGCTTAGAATACTTGGATTTGTCTTTAAATGTGTTGTCAGGTACTATTCCCAAGTCTTTGGAAAAGCTATTATACCTTAAAAGCATTAATGTTTCATTTAATGATTTAGAAGGTGTAATACCCAGTGGTGGTGTGTTTGCGAATTCCACTCTGCAATCATTTATTGGGAACAAAGGTCTATGTGGAATGCACATAATGGAGATTCCTGCTTGTGCTATCACTACAACTGGACAACAATCAAAGTCTAAGAAGCTTGTGCTTAAAATTGTTATTCCAGTGGTTGCTGCTTCTTTTCTGATATTTTTGTTTGCTATAGTTTGGATAATGAAACGACAGAAGAAAGCAAACTCCAAAGATGTGGAAAAGGTACCGGAGATCAGGACTTATCAATTGGTTTCTTATCACGAGATTCAACAAGCAACAAATAATTTTGATGGATCCAATTTAATTGGTGTGGGAGGTTCGGGCTCTGTGTACAAAGGTATATTATCTAGTGGAACTGTGGTGGCAATAAAGGTTCTCGATTTGCAAAATGAGGAAGTATGCAAGAGGTTTGATACTGAATGTGAAGTGATGAGAAATGTTAGGCACAGAAATCTTATTCCAGTGATTACTACTTGTTCTAGTGAATACATAAGAGCCTTTCTTCTACAGTATATGCCCAATGGAAGTCTTGAGCGTTGGTTGTACAGAGAAGATCGCCACTTGAACCTTCTTCAAAGAGTCACCATAATGCTTGATGTGGCTCAGGCAATTGAATATCTACACCATGGTCATGAAACTTTGATAGTTCATTGCGACCTAAAGCCAGCCAATGTTCTTTTGGACGAAGAAATGGTGGCACATGTCGGTGATTTTGGGATCTCAAAAATATTAGCTGCAAGCAAGTCCATGGCACATACTGAGACATTAGGCACTCTTGGTTACATTGCACCAGGTACGTACACTAGTCTTATCCCTTTTCACTTATCATTAACCTTTTCTCTAATCAAAATTAAGCTCTTTCGATTTCTAGAAATATATAAAAATCAAGTCTTTGATTGTAAAAAACAATTATTATCTTTCGACTATAATTTATATTTTTTTCATTCTTCTAAAGAATATGGCTTGGAAGGAAGAGTGTCCAGTAGTGGTGATGTTTACAGCTATGGCATCATGATGATAGAGGTATTGACAAAAAGAAGACCAACTGATGACGAGATATTCAATGAAAATCTTGGCTTGAGACAGTGGATAAGACAATCATTTCCCAAGACAATTATGGAAGTTGTGGATGTCAATTTTATTCATGAGGAAGAACATTTCAATTCAAAAAGTGAAATATGCATAGGCTCCATGATGGAATTGGCTTTGGATTGCACAAAGGAAATGCCAGAGTCAAGGATAACAATGAGAGACGTAGTCAAGAGGCTTGACAAAATTAAGAACACATTTTTGGGAACATAGAAATTAGCATCTTTTAAAGTGGTGTTCTTTTTTAGTTGAAGTTAATATGTTGATCAGCTTTTTCTGTTTAAATGGTTTATATAAGGAATGAAGTTCGAGGATTAAAATGGAGCAAGGTTGGGATGGAGTACCAAAATGAAATTTTGGTGCAAGTTAGAGGGGCTATCTATGTATTTGACCTTGTGTGAATGTACG

>PGSC0003DMG400023643 LRR-RLKs

ATGGAGAAGCACATTTTCTTATTGATACTTCTCTTTCTACTTCAATTTCCTATATCAATTGCTTCCTCAAATGATACTGACCAACAAGCCCTACTAGCTTTCCAAAATCTTATTACAAGTCCCAATCAATTTTTGGCTAATAATTGGACCAAGAATGCATCTTTTTGCTCTTGGTTCGGTGTCACGTGCAGTGCAAGAAGGCTAAGGCTTGTGGCCTTGGCTCTTCCTAATTTGCAACTTCAAGGCACAATTACCGGGATTAGGGCATTGCTTGTCAGAGAGCTCAATCTCGAGAACAACTTATTCAATGGGGGCATCCCTTACGGACTTGGCAAGTTGCCTCGCTTGAGAGTCATTGATTTTCAAAACAATCAGCTCCAAGGAAGTATTCCAACAAGTCTATTTCAACACCAGAGAGTTCAAATCATTTCATTGGCTTTCAATGAACTCAGTGGTGAAATGTGGAAAGGGCCATGGTATGTACCAGAACTCAAAGTCTTAGATCTCGCGAACAATAGCCTCACGGGTATAATCCCTCCTTATGTTGGTAATGCCACAAAATTGATGAACTTCGATTTGTCTGGGAGTAGAATCAATGGTACCATTCCAATGGAAATTAGTAATTTGAGCCAACTTGCATCTTTGTCCTTGGTTGATAATCAGTTAACAGGTTTTATTCCTGGATCACTGTTTAACATTTCATCACTACTTGGCGTAACTCTGGGATTGAATCACCTTTCTGGTCCTCTCTTGCTTGATGAAGGGAATATTGAGTCAAATCTGAAGATACTAAGTATATCTTTCAATCAAATTTCTGGTCGCATTCCTTCTAACATATGCCAACTCACACAGCTCAAAATTTTGTCCATATCTTTCAATAACATAACTGGAGACATACCCAAAAATATTGATTGTCTAGCCAAGCTCGAGATGTTCTTAATTAGTGAGAATGCAATAAGTGGAACTATTCCTACTTCATTGGGCAATATTTCCACTCTGCAACATCTTGATTCGGGGGACAATCGGTTGGAGGGACAAGTTCCTCAAGAATTAGGGAAGCTATCAAATTTGAGGGTGTTAAACTTTCACAAGAATTTCAATCTTATTGGTCAAATTCCAAAGGCTATTTTCAATATATCTTCTTTGGAAGCGATTGATTTCAGTTTCAATAACCTCTCAGGTAGAATTCCAGCCACTACCGATCTTCATCTTCCGAACCTTAAAGAGTTTATGTTGGCATTCAATCAGCTCGAAGGAGAAATTCCTCAATACATAGCAAATGCTTCCAAGCTTGAGATACTGGCGCTAAATTATAACTTTCTGACAGTTGCTATTCCTACTAATTTAGGAAATCTTCGCGAGCTGCAATCACTGTTCCTAGATCATAATCAACTTACCAATGAACCAAGAGAGCCTAAGTTGTTATTCTTGAATTCTTTGGTGAACTGTAGGATGCTGAAATATGTAGATGTGAGTTTCAATCCGTTGAATGGCGTTCTTCCCAATTCTATTGGGAATCTTTCATCTACTATTGAAGTATTTGATATAAGAAATGCACACATCTATGGCCTCATTCCCACAAGTATAGGCAACATGAGCGGTCTTATATCCCTAGTCTTTAATGAAAACAACTTGACAGGATATATTCCTCCAGAGATTGGTAAGCTTAAACAACTCCAAGGTCTGTACCTATTTAACAACAAATTGCAGGGACATATTCCAGAGGCGGTATGCCATTTATCTAATTTGGTCGAATTAGAAATGGATGGTAATGAACTCTCTGGATCAATTCCAGAATGTTTAGGAAATCTTGGGATGCTACTATATCTTTCGTTGGGTTCAAATGAATTTTCACCAAAATTTCCCTCGAGTCTTTGGAAGATGAGTGGACTTCTCTACCTAAACATGTCACATAATTCTATAGAGGGAGAAGTTCCACCAACTATTGGAGAACTGAAAGCCATTGAAGAACTACATCTTTCCGGTAATCACTTTTCAGGCCAGGTACCAAGCAGATTGGGAGACCTCCAAAGATTGAAGTCTCTTGACCTATCAAACAATTCATTTTCAGGCTCAATTCCATTATCCATTACCAACTTGATGAGCTTAGAAGACTTGGATTTGTCTTTAAATGCATTGTCAGGTACTATTCCTAAGTCTTTGGAAAAACTCTCATACCTTACAAGCATCAATGTTTCATTTAATGATTTAGAAGGTGAATTACCCAGTGGTGGTGTGTTTGTAAATTCCACTCTGCTATCATTTCTCGGGAACAAAGGTCTATGTGGAATGCACATATTGGAGCTTCCCGCTTGTCCTATCACTAATCCTGGACAACAATCAGTGTCTAAGAAACTTGTGCTAAAAATTGTTACTCCAGTGGTTATTTCATTCTTTCTGATATTCTTGTTGGTTTCAATTTGGATAATGAAACGAAAGAAGAAAGGGAAGTCCAAAGATGTTGAAAAGGTTCTGGAGATCAAAACTTATCAATTGATTTCTTATCATGAGATTCAACGAGCAACAAATAATTTTGATGGATCCAATTTAATTGGTGAGGGAAGCTCTGGCTCTGTGTACAAAGGCACATTCTCAAGTGGAACTATGGTGGCCATAAAGGTTCTGGATTTGGAAAATGAGCAAGTATGCAAGAGGTTTGATACAGAATGCGAAGTGATGAGAAATGTCAGACATAGAAATCTTGTTCCAGTGGTTACTACTTGTTCTAGTGACTATATAAGAGCCTTTGTTCTGAAATATATGTCAAATGGGAGTCTTGAGAATTGGTTGTACAGAGAAGCTCGCCACTTGAACCTTCTTCAAAGAGTCACTGTAATGCTTGATGCGGCTATGGCAATTGAATATCTACATCATGGCAATGACACTGCGATAGTTCATTGTGACATAAAGCCAGCCAACGTTCTTTTGGATGAAGATATGGTGGCGCATGTTGGTGATTTTGGAATCTCTAAGATTTTAGCCGTAAGCAAGTCCATGACACAAACCGAGACATTGGGCACTCTTGGATACATTGCACCAGGTATACTAAAATTATAACCTTTCTATTTAATTTTTCTCTTATCAAAATCAAGCCCTTGAAAATTCTAGGACTAAATAAAAAGCAAGTCTTTGTTAGTATGAGCATTATTACTATATCCAAATGAGTTAGTTCTTTTTCATTTTCGTTCTTTTAAGAGTACGGCTCAGAAGGAATAGTGTCGGCTAGTGGTGATGTTTACAGTTACGGCATCATGTTGATGGAGGTTTTGATGAAAAGAAGGCCAACAGATGAAGAGATATGCAATGAAAATCTTGACTTGAGGAAATGGATAACACAATCATTTTCAGGGAGTATGATGGACGTTGTGGATGCCAATCTTTTTTCCGAGGAAGAACAAATCACTTGTAAAAGTGAAATCTGCATAGGGTCCATGATAGAATTGGGTTTAGACTGCACAAAGGAAATGGCAGAATCAAGAATAAGCATGAAAGAAGTAGTGAAGAGGCTTAACAAAATCAACAACACATTTCTCGAAACATAG

>PGSC0003DMG400023953 LRR-RLKs

TTAATTCTATTCTTTAAGATTGGAAATGAGAGATCAAGATCGAGTTTTTACTAGTTTTGAACCATACCCTTTTCGAGTTGTTAACAAGTTTTTAACTCGAAAAGGAGGAGCTGGTAGATGTATAATTTCTCATCTTGCAAAAAGATTCTTTCTTTTTGCTCAACTTTTCTTGTTATTTCAGTTACTGATATTTGGGATTCAGCCAACTTCAGGTGAAGATTGGGATGGGATAATTATTACTGCAGCAGATTTTCAAGCACTTCAAGCTTTTAAACAAGAATTGGTTGATCCAAAAGGGTTCTTGAAAAGCTGGAATGATAGTGGATTTGGAGCTTGTTCTGGTGGATGGCTTGGTATTAAATGTGCTCAAGGACAAGTTATTGTTATTCAGCTTCCATGGAGAGGTTTAGGTGGAAGAATTACTGAAAGAATTGGACAATTTCAGTCTCTTAGAAAACTCAGCTTACATGATAATGTCATTGGTGGTTCAATTCCTTCTACTTTGGGACTCATACCTAATCTTAGAGGACTTCAACTTTTTAACAATAGGTTATCTGGTTCTATTCCTGCTTCTCTTGGTTTATGCCCTCTTCTTCAAACACTTGATCTCAGCAATAACTCATTCTCTGGTGCAATTCCAGATAGTCTTGTTAATTCAACTAAGCTTTATAGGCTTAATCTTAGCCATAATTCTTTGTCTGGTTCAATCCCCACAAGTCTCACACAATCTCCCTCAATCATCTTTCTTGATCTCAAGTATAACAATCTTTCAGGCTCAATACCAGATACTTGGGATGGAAATGGAAATGGAAAAAGACTCTTCCAACTTCAGTCTCTTACACTTGATCACAACTTCTTTTCTGGAAGTATGCCTGCTTCCTTAGGCAAGTTGAATGAACTTGTTGAGCTTTCACTTAGTCATAATCAATTGACTGGAGTTATCCCAAGTCATATTGGGGGACTCTCTAGGCTTACAACACTTGATTTGTCTTATAATGCCATTAATGGAAGCTTGTCTGATAGCTTCTTGAATCTATCCTCACTAGTGGTCTTGAACTTGGAAAGCAACCAACTTGATAACCAAATCCCAGCAGCCATAATAAAATTGCAGAAACTCTCAGTTCTCAACTTGAGGAGTAACCACTTTAGTGGTGATATTCCAGTCACTATAGGAAACATCTCTACTCTTAGACAACTAGATTTAGCTCATAATAATATAAGTGGAGAAATCCCAGCTTCTTTAGATACTTTACCAAATCTTAGTGCCTTCAATGTCTCATATAATAATCTGTCTGGTCCTGTTCCAACTCATCTCGCACGGGAATTCAACTCGAGTGCCTTTGTGGGTAATCTTCAGCTATGTGGTTACAGTGCATCAACCCCATGTCCAATCTCTCCAGTTAGCCCATCCCCCGAAACACCAAAAAAGCAACGTCGTAAGTTAAGTACTAAGGACATAATTCTCATAGCAGGAGGGGCACTTCTCATAATCTTGGCTATGCTATGTTGCATTCTCCTATGCTGCTTGATCAGGAAAAGATCTGCAGCTGAAGCAGGGAAGGATGGTCAAAGCACGAGTAGGGCGGCTGCAGCAGCCAGAGGTGAAAAGGGTGTTCCACCAACTGCTGGAGAAGTTGAAGCAGCAGGAGGAGGAGATACAGGAGGAAAGCTTGTCCATTTTGATGGACCTATTGTGTTCACAGCAGATGATCTTCTTTGTGCAACCGCCGAAATAATGGGAAAGAGCACTTACGGTACGGTGTACAAGGCCACGTTAGAGGATGGCGATCAAGTTGCTGTGAAAAGATTGAGAGAGAAGATCACAAGAGGTCAAAGGGAATTTGAGAGTGAAGTCAATATTCTTGGCAAGATCAGGCATCCAAATCTTTTGGCCCTTAGAGCATATTACATGGGACCTAAAGGAGAGAAACTTCTTGTTTTTGACTACATGCCTAAGGGAAGTTTGGCAACTTTTCTTCATGGTAAGCACAAAATCAGTTTTCCATTTCACTTTGTATTTGTGAGAACTTCAGATTATAAAAATCTTACTGTTTTTTGCAGCTCGCAGTCCTGATACACCAATTGATTGGGCAACAAGGATGAGAATAGCAAAGGGTACAGCAAGGGGATTGCTCTTCCTTCATACAAATGCCAACATCATTCATGGGAATCTTACATCGAGCAATGTTTTACTTGATGACAACACAAATGCAAAGATTGCAGATTATGGTCTTTCACGCCTAATGACTGCTGCTGCAAATGCAAATGTTATTGCAACTGCTGGAGCACTTGGGTATCGGGCACCTGAACTTTCAAAACTCAAGAAAGCAAACACAAAGACAGATGTATACAGCCTTGGTGTTATTATACTGGAACTTCTAACTGGGAAGTCACCAGGAGAAGCAATGAATGGTGTGGATTTGCCTCAATGGGTTGCATCAATTGTGAAAGAGGAGTGGACTAATGAGGTATTTGATTTAGAACTAATGAGGGATGCATCCGTTATCGGTGATGAATTATTGAATACCTTGAAATTGGCTTTGCATTGTGTTGATCCCTCACCATCAGCTCGACCAGAAGTTCAGCAACTTCTCCATCAACTAGAAGAAATTAGACCTGAAACAGCTACCAGTTCTGGGGACGACGGTGCTGCAGCTCCCTCAGCAAGCGATGATTAGATATTAGATTTCTCATCTAAGTCATTTTTAGGCAGGTGTGTTAGGAGTTAGGTGCCACAATTCTTTCATTCTTGAATTAGGTTGATTAAGAGAGTGATTTTTGTTCTTCTACTGTCTGTAGATACCCACCCAAGTGTGCTATTTTGTTCTGTTATTTTGAGCAGCAATGGAATCCTCTTAATTTCTGGCCCTAGAGACAGAAAATACACTAAAAACTCCACACACATGTCTATTTTCAGGTCATTCTTCTATACAAC

>PGSC0003DMG400024485 LRR-RLKs

ATGTTGCTAGTACTTGCAGGTTCAGAAACTCTGGAAGTGAGACGAGCATTAGTTAAGTTTATGAACAAGATATCACTTGGTAACATTTCAAAAGAAGTAAACTTTGGATGGGATTTATCATCTGATCCATGTACCAACAAATGGGAAGGGATAACTTGTGATTCAAGAAACCAACATGTAAAAAATATTGTTCTTGATCAGAAGAATCTCACTGGAATTCTTGATGCTTCTTTTGTCTGTGAGGCTACTTCCCTTGCTGTACTGAGCTTGAACGAAAACGAAATTGTTGGAACATTACCTCAAGAAATATCAAACTGCAGACGTCTCACACATTTTTATCTGCGTGGCAACAAATTATCAAGCAACTTACCAAGCTCTCTTTCGCGGTTGAGCAATTTGAAAAGGTTTGTAATTTCAGACAATGCCTTCTCGGGGCAGATACCGGATATGTCAAGAATCTCAGGGCTGATAACTTTTTTGGCTGAAAGAAATCAGCTAACAGGGCAGATACCAGAATTTGATTTTTCCAATCTTGTAGAATTCAATGTCTCATACAATAATTTAACTGGTCCAATACCTGATGTCAAGGACCAGTTTAGTTCCAGCAGTTTTTTGGGTAATCCTGGATTATGTGGAGTGCCATTGCCAAGCATATGTCCACCCTCGCCACCGCCACCTCCCCATCCAGTTGATAAGGAGAAAAAAGTTTCATACTTTATTTATCTTGGCTATGCTATTCTTGGGCTGATTATCATACTTCTACTTGTATGGAAGCTTTTTAAGTGCATAAGGAAGAAAAAGAGCAATAGTACTCCAATGGATCACAACAAGACTATTAGCTCTTCTGGTGTTACAAAGACTCCTAAAAATAGGTCAGAATATTCAATAACATCTAGTCCCGAAAACTCTATGCTCTCAGCATCATTTGAAATCCTTTCAAGTCCTCTGGCTAATAAATTGAGATTCGAGGACTTGCTTCGAGCTCCAGCTGAATTGATTGAGAAAGGAAAACATGGAAGTGTTTTCAAGGTCAATGTTGATGGAGTGACATTGGTAGTAAAGAGGATCAGTGGTTGGAATATTTCGAAATACGATTTCAAGAAGAGGATGCAGAGAATACATAGAATGAAGCATCCTCATGTGTTGCCCCTTGTTGCCTTCTATAGTTCCAAACAAGAGAAGCTAACAGTTTACAAGTACCAACAGAATGGCAGTCTATTCAAGCATCTCCATAGTAAGTTTTATTAGTGCAAACTTAATTATTCTCACTTGAATTTCAAGCCATAGAATGATACATGAGTAATAGGCCTTCCTGTAAGAACTATCAAGAATTGATATCAGTAAAGGAATATGTTAGATAGAACTATAGTTCAATTTCCAGATCCAGGGTCTAGCTAAGTCAAAATCTTTTTAAGTATTTTGTCGGTCTACGTTGCTTTGACTCCAAAATGTTGCCACACCTGTGTCAGATCCTCCAATAATGCATTACTTTTGGAGGATCCGACACACACCTGTATGACATTTTTGAAAAGTCCGAGCAACATTGTTGTCGGTTGATTAAGTAAGAAAGAAACTTTACTAATTAGTATTATTAGTTTGGGCAGCAAGATATAGTCATGTTTTCATTGGAATCTAACTTCTGATTCTGAACCTAGTTCAAAAGATAATGTCAACATATCAGCTTATAGGATTTTAGGTATTTACATTGGCAATACAAAGCTAACCTATAATAGCAAGTCAATTACCTATAATTAGTTACCATACAAGTGATCTAATCGTATAAGTATCTTTTACGCAGTCAGTGAATAGACCTTAAACTCTTAAAAATAATCAAGTAGTAACTAGTTTGTCTATTATTCTTTGAGAATGAACATGGAAAGGTAAAGTTACTGCATAACTGTTCATTTCAGCATCTCAGGGCAGTAAAGTTTTCGAGTGGGCGAGCAGATTAGCTATTGCAGCCAGTGTCGCAGAGGCCTTAGCATTCATGCATGAAGGCCTTCAGAATGACAATATTCCTCATGGAAACATGAAATCCACCAACATTATGCTGAACGATGACATGGAAGCCTGTATCAGCGAGTATGGTCTAATGCCCAACAATCAAGACCAGTCATTTGTTGCTCAAAGTGATCATAGCATCAGAGAAGATGATTCAGTTGCAATTACTACACGTAACACCTTCAAGATGGATGTTTATAGTTTTGGAGTGATTCTTCTTGAATTGCTCACAGGGAAGCCTGTTCAGGCTAGTGGATATGATTTGTCTAGGTGGGTAAATTCAGTAGTTAGAGCGGAATGGACTGGTGAAGTTTTCGACAAGTCCCTTATTACAGATGGTACAAATGAAGAAAGAATGATCAATCTGTTGCATGTAGCTCTCAAATGCATAAATACATCACCAGATGCAAGGCCTAACATGAAGGAAGTTGCGTTCATCATCAATTCTATTAAGGAAGATGAAGAGAAATCCGTGTCTGCGTTCAGCTAA

>PGSC0003DMG400024661 LRR-RLKs

TTTGTATTAAAAAGTTTGGTTTGGTCCATTTTGCAGGCCCTCCCTTTCTGAAATTCAACTCCATTTTCCCATGTTTATTCCTTGCACAGAGATATTGTTTCTGAACTACCAAAACCCCCTTTCTTAAACATAGTAAAATACCATTTCTTTCTTGTTTTGCACAAATTTCTATAACAACAACAACTCATCTGTCACACAACAAAAAATAAATAAAAAAATAGAACAAATAGTACTGGAAGAAATCAAGATTCAAGATTGAGTTATGTCAGAAGAGGAATCTGATATTCTTCTTCCTGTTGTATTATTCCACTTCTTGCTTCTAATCACTACTGGTAAGTCTGAGTTTTTTTTTTTCGATTTTTCAAGATTGTACTCAGAATTCTGATTGTTGTTTTATGAGGTTTTTAGCTGATTTGTAATACTAAAGTTTTGATTTTTATTTATAGTTTGTGGAGATACCCTGGAGAGTGACAGGCAAGTGTTACTGAGTTACAAGTATTTTCTTGAACATCAAAATCCGGTTAATAAAGGATACAGGCATACAGAATGGAATGCTTCTGATTCGTCTCCGTGTAATTGGCGTGGGGTTTTTTGTGATGGTGGAGTTGATCGTGTTACTCGAATTGATCTCTCTGGTGATAGTTTGGCTGGGAATATGTTTTACAACTTCTCAGCTATGACAGAGTTGACATATATTGATTTGTCTATGAATACAATTGGAGGGTCTATTCCTGCAGACTTAGGCCAATGTAAAAACCTGAGATTCTTGAATTTATCGCATAATATTATTGATGGGGAGCTTAATTTGACTGGTTTGAACAATTTGCAAGTTCTTGATTTGACCATGAACAGGATTCATGGTGAGATCAGTTTAACTTTCCCTGGAATTTGTGATAGTTTGGTGGTTGCTAACATTTCGAATAACAATTTTACTGGTGAGATTGGAAGTACTTTTGATCAGTGCTGGAATCTTAGATATCTTGATTTGAGCTACAATAACTTGACTGGAGGATTGTCATTTGGTTTTGATAAGCTTAAGGAGTTTTCGGTGTCTAAAAACAAGTGTAATGGCTCTCTGCCTTCGTCATTTTTCACTCAAAACTGCACCTTGCAGGTGTTGGATTTATCAGAAAATGGATTCGTTGGAGGAGTGCCTAAAGAGATATCGAATTGTAAGAACTTGGAGGACTTGAATTTGTCTGGAAATAACTTTTCAGGGCCAATTCCTGAGGAAATTGGATCAGTTATGAGTCTTCAAGCACTTTACTTGGGAAGCAACAATTTTTCAAGGGATATTCCAGAGAGTCTATTAAGTTTAAGTAACTTAGTGTTTCTTGATCTTAGTAGAAACAACTTCAGAGGAGAAATACAAGAAATTTTCAGGCAATTTACACAGGTAAAGTTTCTTCTGTTGCATGGTAATTCTTATACTGGAGGTATAGTTACCTCTGGAATTCCCAACTTAGTGAACCTTTCTCGATTGGACTTGAGCGATAACCAATTCTCCGGTCCATTGCCAGTTGAACTTTCCAAGATGAAAGGGTTGAAGTTTCTGATTCTTGCATATAACCACTTTAATGGAAGTATACCATCAGTATACGGGGATATTCCGACACTTCAGGCCCTTGATCTTTCATCTAATAAGTTAACTGGCTCAATACCACCAAGTTTAGGGAAGCTAAGCTCACTTTTGTGGTTGATGCTTGCTAACAATTCACTGACCGGTGGAATCCCACCCGAGTTGGGAAATTGCAGTAGCTTATTGTGGTTGAATCTTGCAAACAATCAGCTTTCTGGTTCAATTCCACCTCAGTTAGCAAGAATTGGCTCAAATCCTATGCCTACTTTCTTGTCGAATAGGGCTAAGGATAAGGTGACTGCTGGCTCGGGGGAGTGCTTTGCTATGAAGAGATGGATACCAGCTGATTACCCTCCATTTAGCTTTGTATATCCTCTCCTTACCAGGAAGAATTGTAGAAGCCTATGGGATAAATTGCTCAAAGGGTATGGTTTATTTCCAGTGTGTGAACCGGGTAGTAATGTTCGTTCAAATCAGATATCAGGCTATCTTCAACTTAGTATGAACAAATTTTCTGGTGGGATCCCTCCTGAAATTGGCAGCATGCAGAATTTCAGTATGCTTCATTTGGGTGTAAATGAATTCGGTGGCACGCTCCCTTCAGAGATTGGAAAAATGCAGCTAGTAGTCCTTAATATTTCACAAAACAGAATATCTGGTGAGATTCCAAGCCAGATTGGCAACATTAAGTGCTTACTGAATCTTGATCTGTCCAGCAACAATTTCTCTGGTCTGTTCCCAGCTAGTTTTAGTAACTTGACTGATCTAAGCAAGTTCAATATCTCTTACAACGCGTACATCTATGGTACTATACCAGAAAGTGGGCAATTAGCCACATTCGAGAAGTCATCATACCTTGGTGTTCCATTGCTGCACCTTCCACCTTTCATTGATAACACTAGGAATAATACTATAAACAAGGGTGGAAGTTTCAAAAGGCCTACAAAGGTTGGTACAGTTTTGGTATTCATGGCTTTGCTACTGGCTTTGCTAGTTTGTGGACTTATGACACTTGTTATCTGCCTTGTTCTAAAATCTCCGATAGATACACCAGGATACCTACTGGAGGATTCAAAGGGCCGACACGATCTTGCATCGAGTTCAGGTGCATCCTCACCATGGTTGTCTAATGATGTTAAGGTTATCCGCTTGGACAAAACAAGCTTCACACATTCTGACATATTGAAGGCCACAGGCAGATTCTCGAATGACAGAATTATAGGGAAGGGAGGATTTGGGACAGTTTATCGCGGAGTCTTGCCTGATGGAAGGCAAGTGGCGGTGAAAAAGCTACAAAGAGAGGGAATTGAAGGGGAAAGAGAGTTTAGAGCTGAAATGGAAGTACTAAGTGGAAATGACTTTGGTTGGCATCCGAATCTTGTAACACTTTACGGTTGGTGCCTTAATGGATCAGAGAAATTGTTAGTCTATGAATACATGGGAGGTGGAAGCTTAGATGAGATCATCACAGATAGAACCAAATTCACATGGAAGAAACGACTCAACGTGGCAATTGATGTTGCACGTGCTTTAGTCTTCTTACACCACGAGTGTTACCCTTGTATCGTCCACAGAGATGTCAAGGCTAGCAACGTGCTGCTAGACAAAGACGGAAGGGCAAGAGTCACAGATTTTGGCCTAGCTAGGGTTATGGATGCTGGAGATAGTCATGTTAGCACAATGGTTGCTGGTACAGTCGGGTACGTTGCACCAGAATATGGACAGACATGGCAGGCCACAACAAAAGGCGATGTCTACAGTTATGGAGTGCTGGCAATGGAGTTAGCCACAGGGAGACGCGCTGTAGATGGAGGCGAGGAATGTCTAGTTGAATGGGCGAGACGTGTGATGGGAGACGGAAGGCATGGATTCACCAGAGCCATAATACCAGTTTCTCTTTTGGTATCAGGCCTAGCAGAGGGAGCAGAGGAATTGTGTGAGTTGCTTAGAATAGGAATAAGGTGCATTGCTGATAGTCCTCATGCTAGGCCTAACATGAAGGAAGTATTGGATTTGTTAATCGCGATTTTGCGCAGCCAAAGATCAGGATCCAGTCGTAGCACTTCTCCTTCATTTTGATCAATTTTTGTTTCAAATTACACAGCAAAGTGTCAATAGTTTTTTTTTCTTCACATTTGTAACATACACACATTACATTCTTCTTGTAAACAGTTCTTTTTAGATTCTTTGTTCAGGCACTCTTGTTTGTTAGGATCATAAGAG

>PGSC0003DMG400024795 LRR-RLKs

CCTATGGCCAACCCTGTCCTACAACACTGTGTCAAGTGTCCATCTTGACAATGAATCTAATTGAAGACAGATATATTTGGAGGAAAATCAAGATTTCTGAAAATTCCATTTGTTTCTTTCTTTTACATAAAATTCTGCAAGAATTTAACTGTTGTGCATTTCCAACACCAGCAAACCATGATAAAGAAATCATTGCAGATAACCTCTTATGACTACATTTGGTGCAAAAAAGAAAACTGAAATAGAAGTACAGTAGATCAGAGAGTACATACACACCCTCATTGTCATCATAATCATAATCATGTCAGGCAAACTAACAGTCATCTTATTCTTCTTGTTTCTCAACATATCTTTTTTATTCCCAACATCCATTTCTGGTTTAAATCAAGAAGGGGTTTCTTTAATGTCATGGCTTTCCACTTTTAACTCTTCTGCTTCTGTTGCACCTTTCTCTTCATGGAATCCAAGTCATGAAAATCCATGCAAATGGGATTATATCCAATGCACCAGTAATGGATTTGTTTCAGATATCAAGATCAGGTCGATCAATCTTCCCGCGATATTCCCAAGTCAAGTTCTTTCTTTTCACTTTCTGGAAGTGCTAGTTTTATCAAATTGTAACCTCACTGGTGAAATTCCTGCTTTTATTGGAAACTTGTCATCGTCTCTGAGGATTTTGGATCTCAGTTTCAACGCTCTAACAGGAAGTATTCCGCCTGAAATAGGAAGATTATCACAGCTAAAGCAACTTTTGTTGAGTTCTAATTTCTTTGAAGGTCAGATACCAAAGGAGATAGGAAGATGTTCTGAATTGCAGCAGCTTGAGCTCTTTGACAATCAGTTTTCTGGAAAGATACCTGAAGAGATCGGTCAGTTAACGAGTCTTGAAATCTTTCGTGCTGGTGGGAACGTGGGAATTCAAGGAGAAATCCCAATGCAGATATCAAGCTGCAAAGAGTTGGTCATTTTGGGTCTTGCAGATACCGGTGTTACGGGTCAGATTCCACATAGTATAGGTGAACTCAAGAAGCTGCAGACTCTTGCTGTTTACACAGCTAATTTGACTGGTGAAATCCCTCCAGAAATTGGGAATTGCACTTCTTTAGAAGAATTGTTTGTCTATGAGAATCAAATTACAGGAGAAATTCCAAGTGAACTCGGTCTCTTGAAGAATCTTAAGAAGGTTTTGCTTTGGAAGAACAATCTGACAGGGGAAATTCCAGGAAATCTTGGAAACTGTTCAAGTTTGAAAGTGATTGATTTCTCTGTGAACTATCTATATGGTGAAATCCCTCCGTCTTTCGAGAATTTAGGCACATTGGAGGAGCTTCTATTATCAGAAAACAGCATTTCTGGAGAAATCCCATACTATATTGGCAACTTTTCCAGCTTGAAGCAGCTTGAATTGGACAACAACAACATTTCAGGTGTGATTCCACCTACCATTGGGAAACTAAAGGAGCTAAATCTGTTTTTCGCTTGGCAGAATCAACTTCACGGGAGCATACCGACTGAGCTAGCAGATTGTCGGAAACTTCAGTCTTTGGATCTTTCTCACAATTTCCTTACAGGGTCCATCCCAAACACTTTATTCAATCTCAGGAACTTGACTAAGCTGCTGCTGATATCAAATGTTCTGTCCGGTGGAATTCCACCTGATATTGGAAATTGCACCAGCTTATCCAGATTACGCCTTGGATCAAACAGGTTAGATGGTCCAATTCCTCCAGAAATAGGGCGTCTACCGAGTTTGAGTTATCTTGAACTGTCGGAAAATCAATTCACTGGATCAATCCCTCCAGCTATTGGCAACTGTCCGCAGCTAGAAATGGTTGATCTTCATGGAAACAATCTTCAAGGAACAGTTCCTTCCTCTTTTGTGTCCCTCACTGGTCTTAATATATTAGACCTATCCATGAACAGAATATCAGGTAACATTCCAGAAGATATAGGGAAACTCATATTACTGAACAAACTCATACTGAATGGAAACAACATAGACGGAACGGTTCCTAAATCACTTGAACTCTGCCAGGATTTGCAGTTGTTGGATTTAAGCAGCAACAGACTCGTGGGCTTAATCCCTGAAGAGATTGGTAACCTCCAAGGACTAGATATTCTGTTCAATGTAAGTCGTAATTTTCTGACCGGACAGATTCCTGAAAGCTTCTCTAACCTCTCCAAGCTAGCCAACATGGATATTTCTCATAACATGTTGACTGGAAGTCTTAGAGTACTCAGTAACCTTGATAACCTTGTTTCTCTGAATGTTTCATACAATAACTTTTCTGGTGTTCTTCCTAACACAAAATTCTTTCAAAGTCTCCCTCCTAGCGCTTTTATTGGTAATCAAGAGCTCTGCACTGACAGAGCTGCATGCCACTTAAGTGGTGATCACCATGGATTAAAATCCATAAAGAAGATCACCATTGCTATTGTACTCAGCATTTTCATGACCATGCTAATTGTGACAGCTTCTATTGCCATCTTTATCCGAACACAGGGAGAGATATGTCAGAAGGACGACGAAGAAAATGGTTTGCAGTGGGAATTCACCCCATTCCGAAAGCTTAGCTTCTCTGTAACTGATGTAGTGCCAAGACTTAGTGAATCTAACATTGTTGGAAAGGGCTGCTCAAGCTTTGTTTATCGCGTTGAGACTCCATCAGGACAGGTGATCGCGGTGAAGAAACTATTGGCTAAGAAAATCGGTGAGGTTCCTCAGAGGGACTTTTTTTCTGCAGAAGTTAGAACACTGGGATCAATCAGGCATAAAAACATAGTGAGGCTTCTTGGATGTTGTAATAATGGCAAAACAAGATTGTTGTTGTTTGATTACATTAGTAACGGGAGTTTATCGGGACTTCTCCATGAGAAGAGGGTATTTCTGGATTGGGATGCTAGGTTTAATATCATATTAGGAGCAGCTCAGGGCTTAGCTTATCTTCACCATGACTGTAGTCCTCCTATAGTCCATCGTGATATCAAAACCAATAACATTTTGGTAGGTCCACAGTTTGAAGCTTTTCTTGCAGACTTTGGACTTGCAAAGCTATTAAACACATCATCAGATACTTCAAGAGCCTCCACCATAATAGCTGGTTCCTATGGGTATATAGCTCCGGGTGAGTACATTTCCTCAAATTTCCGCCCTATATATTCAAATATTTATTAAGTGCTCAACATTTTGATAACATGATTATGCTAGTACTATTGAATATTAGCTTTAAAGAAACACTTGAAGTTCACTATATTAGTCTTGTACCCCGTGAAATTCGCTGTGCAAGACATATAGTAGGTATATATACTATGTTTGGACAGCATTCTAAATGTTTTGGTCCTACATTGCAGAATATGGATATAGTCTAAGAATAACTGAGAAGAGTGATGTCTATAGCTATGGCATTGTGCTTCTCGAGGTCCTAACGGGGATGGAACCAACTGATTCCCGGATACCAGAAGGCAGCCACATTGTCACTTGGGTCAATCAGGAGCTAAGAGTAAAACATAAGGAATTCACAACAATTCTTGATCAGCAATTACTTTTAAGGTCTGGTACACAAACTCAGGAGATGCTTCAAGTTCTTGGGGTGGCACTACTTTGTGTTAACCCATGCGCGAATGAAAGACCAACAATGAAGGATGTAGCAGCAATGCTAATGGAAATCAGGCACGAAAATGAAGATCTTGAGAAGCCTAATCGAGGAGTGGTACCGAATCCTAAAGAAGCAATTAGTTGTCCTAGTTTCTCTGGTTCATCTCAACCTCTCAATAGATCCCCTCCTCAGTAACTCTAGTTAACTTAGCCTCTCTCTAGTTCTAACTAGTTGATTCTTTCAAGTAACTAATAAGGATGTATAGATTATCAATATTCATTTCACAAAATAAGAACATTACTCTTTCTAGTCCCGACTATAAGTCGTTTTGACA

>PGSC0003DMG400024890 LRR-RLKs

ACCATTTAATTTAGGCTAAAAACTAACTTTGATCTTCTATGTAATGTGTTGTTCTTCAGTGAGAGATAAAAGGAAAAAAAGCAAAGAGAGGAAAAGGGGAGGCTTAGCTTCTTCCTTGATCTTGAAGACTCACTTTCTTTCTTTTGAAGATGTTTGATTCTTTAAATCTATCTATACCTGAAGTTTCTCAGATTCCCTTAACTTTTTTCTCCTTTTACATTGCCTCAAAGTCTCATAATAACTCACTTTGTTTCTTCTTCAAGTCCAAACAAATTCACTTTCCTTGAAAAAACTTCCAAGAAAATATGTTGCCACTTATTCTTGTCACACTCTTTACTCTTGTATGCACTTCCATTTCCTCCTCTTCCTCTTCTATTAGTACTGATTTTCATGCTTTGTTGAGTCTAAAAAAAGGATTTGACTTCTCAAATTCTGTTTTGAGTTCATGGGATAGTTCAAATCCTAGCTCAGTTTGTTCTTGGATGGGAATCAAATGCTTACAAGACCGAGTTGTATCGATAAACTTATCCAATATGGAGCTTTATGGTTCTGTTTCCCCTGCTATTTCAAGGTTAGATAAGCTTGTTGAGCTTTCAATTGATGGAAACAACTTCACTGGTGAAATCAAGATTGAGAACATGAGGAGTCTTGAGTCACTCAACATTTCAAACAACATGTTTAGTGGGAGTCTTGATTGGAACTACACAAGTTTGGCTAATTTGGAGGTTCTTGATGCTTACAACAATAACTTCTCATCTTTTCTTCCTGTTGGAGTTGTTAGCTTGGAGAAACTCAAGTACTTGGACTTAGGTGGCAATTATTTCTATGGACGAATACCGGAGAGTTATGGGGATTTGATTGGGTTGGAATACTTACAACTTGCTGGAAATGACCTGCATGGAAGAATCCCTAGAGCTTTGGGGAATCTTACCAACTTGAAAGAGATTTATTTGGGATATTTCAATGTTTTTGTAGGTGGGATTCCTAAGGAATTTGGAAAGTTGGAGAATCTTGTTCACATGGATATTTCTAACTGTGAATTAGATGGTCCAATTCCTCCTGAATTAGGGAATTTGAAATTGTTAAACACACTTTTTCTACATATAAATCTTCTTTCAGGTCCAATTCCAAAGGAACTAGGCAACCTTACTGGCTTAGTCAATCTTGATCTTTCAGCCAATGCACTTACTGGTGAAATCCCATTTGAGCTCATTAACCTTCAGCAGCTACGCCTTTTCAACTTGTTCATGAACAAATTACATGGTTCTATACCGGATTTCATCGCTGATTATCCTGACCTGAAGATTCTTGGTCTCTGGATGAACAACTTCACCGGTATAATCCCGGAGAAGTTAGGCCAAAATGAGAAGCTACAAGAGCTGGATTTGTCTTCAAATAAGCTCACTGGTACAATTCCTAAACACTTGTGTGCTTCTAAGCAACTCATAATCTTGATTTTACTGAAGAATTTCCTCTTTGGTTCAATCCCAGAGGATTTAGGCACATGTTTAAGTCTTGTTAGGGTGAGATTGGGACAGAACTACTTGAATGGCAGTATTCCAAATGGATTCATTTACATGCCTGACCTAAACTTAGTTGAGCTTCAGAATAACTACCTGTCTGGTACCTTATCAGAAAATGGTGACACTTCTTCAAAGCCAGCCAAGTTAGGTCAATTGAATTTATCAAACAATCAACTTTCAGGTTCTTTGCCATTTTCACTTTCTAATTTCTCTTCCCTTCAAATCCTTTCACTTGGAGGGAACCAATTCTCTGGTCCTATTCCTACTTCTATAGGTCAACTCTCACAAGCTTTAAAAATTGACCTCAGTCAAAATTTTCTTTCTGGTGAAATCCCCCCTGAAATAGGCAATTGTGTCCACTTGACTTATCTAGACTTGAGCCAGAACAACTTCTCTGGCTCAATTCCACCTCGAGTATCCGAAATTCGAATCTTGAACTACTTGAACTTGTCAAGAAACCACTTGAATGAGACCATACCAAAGTCAATTGGTACCATGAGGAGTCTCACTACTGCTGATTTCTCATTCAATGATCTATCAGGCAAGCTGCCAGAATCAGGCCAATTTGCTTACTTCAATGCTACTTCTTTCGCGGGCAATCCTCAACTCTGTGGTTCTTTATTGAACAACCCTTGCAATTTCACACTAATCACTGATCCACCAGGAAAGTCTCATGGAGATTTCAAGCTGATCTTCGCCCTCGGATTGTTAATCTGTTCACTAGTCTTTGCAGCTGCAGCTATCATAAAGGCCAAGTCTTTCAAGAAAACAGGCGCGGACTCTTGGAAAATGACAGCATTTCAGAAGGTTGAATTCAGTGTTGTTAATGTTCTTGAATGTGTAAAAGATGGCAATGTGATAGGCAGAGGAGGAGCCGGGATTGTGTACCATGGGAAAATGCCAAATGGAGTTGAAATTGCAGTGAAAAAACTGCTTGGCTTTGGGAACAATAGTCATGATCATGGTTTCAAGGCGGAGATTCGGACACTAGGCAACATTCGACATAGAAACATTGTAAGACTAGTTGCTTTTTGTTCCAACAAGGAAACGAATCTCCTTGTGTATGAGTACATGAGGAATGGTAGTTTAGGAGAGGCGTTACACGGGAAAAAAGGCGGATTCCTTAGCTGGAATTTGAGGTATAAAATTGCTATAGAAGCTGCTAAAGGATTGTGCTATCTTCACCACGACTGCTCGCCATTGATCGTTCACAGAGACGTCAAATCCAACAACATTTTGCTGAATTCAAACTTTGAAGCTCATGTTGCTGATTTTGGACTAGCAAAATTCTTGGTAGATGGTGGTGCCTCAGAATGCATGTCTGCTGTTGCTGGCTCCTATGGTTACATTGCCCCTGGTACATTACTTTTAACTTTTACTCACACATTCTCAATAATCAATCGTTCATATTAAGTGAAATTAATAACCTCGAAAGATAAAATACAACTTGGTACAACATGTCAAAATACACATTGTATAGAGGGGACAATTATTGTGTAAAAATCATATATGTACGTTGGTTAGTTAATTTAGTTAGGTATGTAGACATTACCTGACAAAGTGATTTTTGCCTGGTCGTACATACAACAACAATTAGTCTTATCAATTGGTCATAAAATCAAACTTTTAAGATAAAATATTATATTGCATTTTGTTTCTTTTAGTTTGGAGTTTGTTGAATTCGATAGAAATTTTGGGCTAAAGTGAAGAGGCCCAAGATCACCCGTGATTGAAGGCATAGAGTTTAGGGCTTTGTTGGAATTTTCTGGGTAATCGTAAATCGTATCATGGGCCCGTTTTTCGAGAGAGCACTTAATTTCTGTGCCATAGAGAGAGTTCTTAGTGGGGGCCCAATAGAAGATCACTCTGCTAAAGTAGGTCAAAAAACTGACTTGACTGACACACAAAGATAATTGGGCCCCCTCGATTGTCGTGCTTGCCAACTCAACATGCACTTGTTCAAAAGGAATTCAACTAAATGGAATAGAGTATATAATATATAGTCACAATATTACATATTTTACTGTATAACTCTTTAAAGTAGTCAACTTTTTTAAAATTAAAGGAGGGAGAGAAAGCAATAGTGTGAATAGCGATGAAGTCAAGATATTCTATTTGTCACATATATTTGAGGGGTGTTAACTGATCTCAAATATTGTTTAATGCATTTATATTAAATTTGTATATACATTGTCATAAATGTTAACCTATTTTTTTAAAAAAATTGTTGCAGAATATGCATATACTTTGAGAGTAGATGAAAAGAGCGATGTATATAGTTTTGGTGTGGTTCTATTGGAGCTGATCACCGGGCGTCGTCCGGTGGGTGAATTCGGAGACGGAGTGGACATAGTACAATGGAGCAAGAAAGTGACAAATTGTAAAAGGGAACAAGTAACACACATTGTTGATCCAAGGTTAACCACTGTGCCTGAAGATGAAGCCATGCATTTGTTCTTTATTTCTATGCTATGTATTCAAGAAAATAGCGTAGAACGTCCAACAATGAGGGAAGTCATTCAAATGTTGTCCGAATTTCCACGACAATCATCAGAATATAATCAACCTTCGTCTTCAAAAGTTGTTTTACAGAAGCTAAAAAGCCTTGAGAATGACCAAACTACTTGTCCGAAGATTCGGAAAGAACTTTTGGTCTAACCAATTTTCATTTTTATATAATATGGTCCTCTTGGCAAGTTGTTTTAATTTTAACTTTGTATTATTCTTTATGCTTATTTAAGGTTAGGGGTTTTGTGACAAAGAGAAGTAGGAATATAATTAGTACTATGTATTACTAAGTAGTTAATTAGTTGCTTTTGATTAGTTGTTAGGGTACTTGGGGGTGTTTTAAGTTCTTGGCAAAGGCAGACCAGGCCCAATTAAGTTGGTTCTGAATGCTTTTTGTATCATATGGTTATATACAGGTATATATTCTTCCTTTAACGGGTGAATTGATTCTCGATCCAATAATGTTTAATTGTTTATTGTTGGTCATGTTGTTATGGATGAATTGGTTCTCG

>PGSC0003DMG400024954 LRR-RLKs

GGAAGGTTGGTTTTCGATGAGCTGTGTCTTCTTCTTCTCTCAATATTTTATAAACTTCCCTTCACATGGTCCCCTTTTTATCATAAATCTACATTTATTCTTCTAACTTTTTTTTCTATTCAATTCCTAAAAAAAAACAGAGATGTGGGAACATTTAGAGACACCATGAAATTCCAAACCTTTGATCTTGTTCTTCTTCTTTTTCTACTACTTGAATATCAAGCTTCTGGGCTTAACAATGATGGACTTCTGTTACTGACATTCAAGTACAATATTCTGAGTGACCCTTTAAGTGTTCTGCAGAACTGGAATTCTTGGGATGATACACCATGTTCATGGAAAGGGGTAGGTTGTGGAAATCCAAATGCTTCAGACCCAAACCTTCGAGTGACAAATTTGTCTCTTCCTAATTCTCTGCTTCTTGGTTCTATTCCTTCTAGTCTTGGTATGATTCAATATCTTACAAATCTTGATCTTTCTAATAATTCCATTAATGGCTCTATTCCTCTTACTCTCTTTAGTGCCCCTGAGCTTCAACGCCTTGATTTTTCTAACAACAGAATATCCGGCCAGTTGCCTGAGCTTGTCGGACATTTAAATAATCTTCAGTTTCTTAACCTCTCCGGTAATGCTTTGACGGGAAGATTACCGGCAAATCTTGGTAATCTAACTGTTGTTTCTTTGAAAGATAATTATTTTTTTGGTGCTGTTCCTTTTGGGTTTGATTCAGTTCAAGTTTTAGATCTGTCTTCAAATTTGATTAATGGGTCATTGCCTCCTAATTTTGGGGGCAGTAATCTCCGTTACTTTAATGCTTCTTTTAATAAACTTTCTGGAGATATCCCACCGGAATTTGGCAGCAAAATCCCTCCAAATGCAACTATAGATCTCTCTTACAACAATTTCAGCGGTGCAATTCCAGAGTCAATTGTTTTTATCAACCAAAATAGAAAAGCTTTTTCTGGTAATCCAGAATTATGTGGTGCGCCGCTGAAAAATTTATGCCCAATTCCGTCAACAGTAACAACTCTGCCAAATTCCCCTGAACCAACTTCTTCCCCAGCAATTGCAGCCATTCCGAAACCCACAGACTCAAACCCAGCCGCCGAATCTCCAAAAGAGGGCAAAAATGGGTTGAGAACTGGCACAATTATAGGCATAATAGCAGGGGATATTGCAGCAGTAGGAGTTTTAGCACTGATTTTCATGTACGTATACCGAGCCAAAAAGAAGAAAAGGAACATAGAAAATAGCATTAAAAAAGAAGCTGAAACTGCAAAAGATTTTGACTGGGCATCGTCAGCATCATCAGAAGAATACAACTGGTTAAGGTCATGGACTTGCTTGAAAAAACAAAGGCACGGAGATGACGACGAATTATCAGAAGCTTCCCATTCAGAAAGTGAAAATTCCCAATTGGGTCATCCTCAGCAGAATCACGTGCATACAGAGCAGAAAACAGGTGAATTAGTAACGGTTGATGGAGAAAGGGAACTCGAATTAGAAACACTACTAAAAGCATCTGCATACATTTTGGGAGCTAGTGGGTCGAGTATAATGTACAAGGCTGTGCTAGAAGACGGTACTACACTGGCGGTGCGACGAATTGGTGAAAGTGGTGTGGAACGGTTCAAAGATTTCGAGAATCAAGTAAAGGTAATTGCTAAATTGGTGCACACAAATTTGGTTAAAATCCGTGGCTTCTATTGGGGCGCTGAAGAGAAATTGGTCATCTACGATTTCGTTCCTAATGGTAGCCTTGCCAATGCACGTAACAGTAAGTCCACTTTTTCCCATATACATTATGTTTTACATCAAATAACGTTACAATTATTGAGTAGAATCGCGTCTTTTGTTATTTGTTTTACTATTTTGCTCCTGTTTTACAGTGGAGAATAAATTGAAACAAAATTGACTAGTCATTATGTGTGTACATTGTACCTAATCATCTCTAATTCTTAATTTGTTTCAATAAGTATCTTTTGAATAATTGCGAACATTTTGGAATATAAAAGTTCGGAGTAGGTTTGTACTTTTAAGACATTTGGTTATGCTTAATATTCAAAAGAAAGACGTAACAAGTCACTCAAATTAAGTTCACACATGTATAAAATTGTGCGCAAATTAGAAAAATATTATGAATTTCTGTTAAAATGCGCGGGAAGTAAAGGAGATAGTGGACCTTGGGGTGGAGTAGTTGAAACTTTAGTGTTTGTTTTTAATGTGGAAAGTAGCAGTAGTTCCCTCACTGCAAGTCAACCGTGTCTGTTTCTAAATGGGCGCCCCACAAATTCATAAATACCGCCAAACGCCCCCTCTTCTACTGTCACTTACATGGAAAATGACTACCTTTTAAAAAAAAAATATTACTCTGTAGGCCATAAGTATGCTCTTTTGTTATGTCAAATTCATGCATTTCTTTTGCCCACAACACCAATAATGTTAAAAGAATCTTCACCTTATTTACCATTTATAGTATGTTGTTTATTTTATAGATATTATTGTGAGAATCTTAGCGCTCAGTTGATTGACTACCTGATGTAATTATAACGTGTGCATTTTCATGGTTTCAGGAAAGGCTGGTTCTTCGCCATGTCACGTACCCTGGGAAATCCGGCTGAAGGTTGCTAAAGGTGTGGCTCGTGGGCTCACTTACATTCATGAAAAGAAGCATGTCCACGGCAACTTGAAGCCCAGTAACATTTTACTAGGGGAAGATATGGAGCCCAAGATTGGTGATTTTGGAATTGAAAGGCTTGTAACAGGAGACAGTAGTCATAAGACCTATGGATCTGCCCGTAATTTTGGAAGCAAAAGGTCCACAGCATCCCGTGAGAGCTTCCAGGACTTTACATCTGGGCCTACTCCAACCCCAAGTCCAAGTGCATTGGGCATATCTCCTTACCATGCACCCGAATCGCTTCGTAGCCTCAAGCCCAACCCGAAATGGGATGTATTCTCATTCGGGGTAGTCTTGCTGGAGTTGCTAACTGGAAAGGTGATTGTCTCGGATGAAATGGGACCCGCATCGGTCATCGGTGCTACCACATCAGCAGAGGAAGAGAAGAGTAAAGTGTTACGGTTGGCTGATGTGGCGATACGTGCTGATGTGGAAGGGAAAGAAGATGCTTTGTTGGCACTAATGAAAGTAGGATATAGTTGTATATCACCAACACCTCAGAAGAGACCAGGCATGAGAGAGGTTGTTCAAGCATTGGAAAAATTTCCAACTAATTCAACCAGTTCACCATACTATTATGGACCTTAATTAGCACTGATTAAAAGATGGACACAATTGTAGAGACAAGAGACTTTGTATGTAATGAGACTGACATCTTTGTTATTTTATTTTTTTTCAATTTTTTTGGTTTGTGTCTTAATATAGTATTAGCCATTGGTTTTCTTTTTTAACAGTGATTAGTGTAATTGTCCGTGCTTTGTTGTGTTGTAGTTTTCCTAAACCTTATGGTGACTTTGGATTACCAATATATATGACAACTTATTATGTG

>PGSC0003DMG400025076 LRR-RLKs

TAATCTCTTTGCTATTTAAATGTATAGTTTCATTTGTTTGCTCTTTAGTTTCTCAAGAACTGTCACTCTCACTCAAAATCCTCTCCGCCACCGTCAACCGCCGTCAACCACCGTCCGCCGCCGTCATGAAAAAATGCAAAATGACAAATGTGTTTGGTTCATGCTAGTGACCATTTATCTTCACCACCATCATTAGTAAGCTACTAGCTAGCGCCGCCGTGTACGGAGGAGCAAGGAGCAAGGAGGAGCTGTTTTCTCCAGTAAGAGAACTTTAATAGTAGCAGTTTAGTAAATGTTTCATTAAAGATAAACAAGAACTTTTTAGTTCACATGCATGGTAAGTAAATTTTTTGAGTTAGTGTGAGTTTGAAGCAAACCACCATTAACACACCACATGTATGTTGTCTTGTTAGAGTGAACTTTCAAACACTATTATCATTTTTTTCAACAACCCCATTTCTTGACTTTGATTTTCACTCAATAATGGAGAAGAAAAGCTACCATTTTTACACTTTCCCTTCATGTTATTTCTTGTTTTTGTTGATTTCTTGGTTAAGTTTGAGTCCAAAGATTGTATTTTTAGTGAGCTCTGTTTCTTCTGATGGACAAGCGTTGCTTTCTCTGCTTAAAGCAGCTGAACCTTATACTAAATCATCATCTTCTGTGCTTTCTTCTTGGAACCCTTCAAGTTTAACTCCATGTTCTTGGCAAGGGATTACTTGTTCTCCTCAAGAAAGGGTTATTTCACTTTCTATCCCAAATACTTTCCTAAATCTCTCGTATTTGCCTTCTGAACTCTCTTCCCTTTCATCTTTGCAGCTTCTTAATCTTTCTTCTACTAATATTTCTGGAACAATCCCTCCATCTTTTGGTTCTTTTTCTCATCTTAGGCTTTTGGACCTTTCTTCTAATTCACTTTTAGGTTCTATTCCATCAGAACTTGGTAGGCTTAGTTCACTTCAGTTCTTGTTCTTGAATTCAAATAGATTGACTGGTAAGATTCCACCAGAGTTGGCTAATCTTTCTTCACTAGAAATCTTTTGTCTTCAAGATAATCTCCTCAATGGGTCAATTCCATCACAGTTAGGGTCCTTAGTGTCACTCCAGCAGTTTAGGATTGGTGGAAATCCATATTTGAGTGGTGAAATTCCTGCACAGTTAGGCCTGCTCACCAATCTTACCATGTTTGGTGTCGCGGCGACTGGTCTTTCGGGTGTTATTCCACCTACGTTTGGGAACTTGATCAATCTTCAAACACTGGCAATTTATGATACTGAAGTATTTGGTTCGATACCGCCTGAACTCGGGATGATTTCGGAGCTTAGGTACTTGTATCTGCATATGAATAAGCTCACTGGTTCGATACCTCCTCAGTTGGGTAAGTTGCAAAAGCTTACTAGCTTGCTTTTATGGGGAAATTCATTAACTGGACCTATACCAGCTGAGGTTTCGAATTGTTCGTCCCTTGTGATTCTTGATGTTTCCGCGAATGAATTGTCCGGTGAAATCCCTGGTGATTTGGGGAAGCTATTGGTTCTTGAACAGCTTCATTTGTCTGATAATGCACTTACCGGTTCTATACCGTGGCAGTTAAGTAATTGCACTAGCCTGACAGCTCTTCAGCTAGATAAGAACCAATTATCAGGGCAAATTCCCTGGCAAGTTGGTAAGTTGAAGTACTTGCAGAGTTTTTTCTTGTGGGGGAATTCGGTTTCAGGAACCATTCCAGCTGCTTTTGGAAACTGTACTGAGCTATATGCACTTGATCTTTCGAGGAACAAGCTCACTGGATCGATCCCTGAGGAGATATTTGATTTGAAGCAGCTGAGTAAATTGTTGCTCCTTGGGAACTCGTTAACTGGGCGATTGCCTCGAAGTGTTGCGAGATGCCAGTCTCTGGTGAGGTTGAGGCTTGGTGAGAATCAGCTTTCAGGACAGATTCCTAAAGAGATTGGTCAATTGCAGAATCTCGTGTTTCTTGATTTGTACATGAACCATTTCTCTGGAGGCTTGCCTTCTGAAATTGCCAACATTACAGTTCTTGAGCTGTTGGATGTGCACAACAATTACCTAACTGGGGAAATACCACACCAGATGGGGGAGCTTGTGAATTTAGAGCAACTTGATCTTAGCCGGAACAGCTTTACCGGTGAGATCCCTTTGAGTTTTGGCAATCTCAGTTACTTGAACAAATTTATTCTCAGTAACAATCTGCTTACTGGTTCAATTCCAAAGTCTTTTAAGAACTTGCAGAAGTTAACTCTACTTGATTTGAGCTCGAATAGTCTTTCTGGCGAGATTCCATCTGAGCTTGGCTATGTCACAAGCTTAACAATCGGTTTGGATTTGAGCTTGAATCGTTTCACGGGTGAACTTCCTGAAACATTGTCCGGTTTGTCACAGTTGCAATCTCTTGATATTTCTCATAATTTGTTGAGTGGAAGGATCACAATTCTAAGCTCCCTGACCAGCCTTACTTCCCTGAATGTTTCAGACAACAATTTCTCGGGGCCTATTCCTGTCACACCATTCTTTAGAACTCTTACTTCAGATTCTTTTCTGGAGAATTCACTCTGTCAATCAGTTGATGGTTATAGTTGTTCTTCACATATAATGGGAAGGAATGGGTTGAAGTCACCGAAAACTATAGCTCTGGTTGCGGTGATTCTGACATCTGTAGCCATAGCAGTCGTGGCTACATGGATTCTTGTGACGCGAAACCACAGATATGTGTTTCAGAAGTCCCAAGGCATGTCAGCCTCTTCGGTAGGAGCAGAAGATTTTTCCTACCCTTGGACTTTCATCCCCTTTCAAAAGTTCAATTTCACCATAGACAACATCTTGGATTGCTTGAAAGATGAAAACATCATTGGGAAAGGATGTTCCGGTGTTGTATACAAAGCAGAAATGCCTAACGGTGAGGTGATTGCAGTGAAGAAGCTTTGGAAAACAAAGAAAGACGAGGAGCCAGTAGATTCTTTTGCAGCAGAAATTCAAATTCTCGGACATATTAGGCATCGAAACATTCTGAAGTTATTGGGATATTGTTCAAACAAAAGCGTTAAGCTTCTTCTCTACAACTACATTTCAAATGGCAATCTCCATCAGCTCTTGCAAAGTAACAGAAATTTGGACTGGGAAATTAGGTACAAAATTGCAATTGGATCAGCTCAAGGCCTTGCTTATCTTCACCATGACTGTGTTCCAGCCATTCTTCATAGAGATGTCAAGTGCAATAATATACTGATCGACTCCAAGTTTGACGCTTATATTGCAGACTTTGGACTTGCAAAGCTAATGAATTCGCCAAACTATCACCACGCTATGTCCAGTGTAGCAGGATCTTATGGATATATTGCTCCAGGTATGCATGCACTTAGAATATTTTCAATTACTAATATTTCAACATAAAAGCCAAATCGATCATAACATCTTTTGAATATTTACTTTATGAGTCATAGTTCGTTGAGTGTGACACAAATATTTGTCAACGGTTGCAGAATATGGATATACAGCGAATATAACAGAAAAGAGCGATGTCTATAGTTATGGAGTGGTGCTGCTGGAAATACTTAGTGGGCGTAGCGCTGTTGATTCTCAGATAGGTGATGGGCTTCACATCGTTGAGTGGGTAAAGAAGAAAATGGGAAGCTTTGAACCAGCTGTTACAGTTCTCGACACTAAGCTACAAGGTTTACCCGATCAAGTGGTCCAAGAAATGCTGCAAACGCTTGGAATAGCGATGTTCTGTGTCAACTCATCGCCGGTTGAAAGGCCAACAATGAAGGAAGTTGTGGCACTTCTAATGGAAGTAAAGTGTTCACCTGATCAAGAATTTGGGAAAACTTCTCAGCCTCTAATAAAACAGTCCTCAACTCAAAGTTGATTAATTTTGTTATGTTTTTTTTTTTTTTTTGCTTTTGTCTAAGAGGAGTAGGAATTAGTTTTCTTTAGTTCCCAAGATATGATCAACTCATTGTTCCAAGCATTGGATAATGTCTTGGGGTCCATTGTGTACATATTTCAGAAACATTTATGTAGTTTATTTGCTTCTTCATTCTTCCTATCCATCTCTTCCATCTCTGTGCTGAGATTTTAGCTTTCAAAATGCTTGCAGTTGCACCATATAATATTTTAGTTTACCATAGTTTCAAAAAAAGAGAAATATAGTTTGACCTATTCAGAAAAATTGTATTGTTGGTGTATGAATGTGGTTTCAAACCTTCCGGTGATGCTAAAAGTAGTTTCAGAAAAGAGAAATATAGAAGAAGAAATCAAAGAAGGGGAATCTTGGAGCAACGGTAAAATTATCTCATGGGTTCAAACTGTGGAAGCAGGCATTAATGCATTAGGTTAAATTGTCTACATCATATCCTTTGAAGTGTGGCCTTTCTCTGAGCCTTGCTAACACATGATGTTTTGTTCATTGAGTTGTTTGTAGAAAGAAGAAAACAAACAACTCTTAAAACCAAATATAATACTCATGTAATGTCATTAGAGACTATTGGTCTAGTGGTAAGAGTGTAGCGCGTAATGCGTGGGTAGTTGTGTC

>PGSC0003DMG400025330 LRR-RLKs

TTTTGAATTCTCCACTGGTGTCTGGACAACGGTTCTTCTCTCTTTTTTTTTTCTTTCTTTCTATTATACACAACTTTCTAGACTACTTTCAAAGTAAAGTCTTCTCTTTTTCAAAATACACACACCCCCCTCCCCTCTTCCTCTTTTTCTCACTTCGTGCTTGAAATTTCATTATTATGGATTTTCCATCTGTAAGAACTGTAATCATACCCATTGTTTCTATCAAAACCCCACCATTTTAGCTTTACTCTTTTTTTTTTCAAGATCTGTATTGATTAGTGAACTTGCTACTACTCCACATTGCACTTGATATTTATAGTCCAAAGTGAAGAAGTAAAGCTCAAATCTGCATTTTTCATCAAGAAATCGAAATGGGTGTCTCGGATTTGATCTTTTTGGGACTATTTACGATGACCATTTTCAGTAGAAGCTGTTTTTCACTAACCCCAGATGGTAAAGTTTACTTCTTTGTGGGTTTTATCTAGTTTTTTCAGCATTGTATGTTGTTCTTGAGCTAAAAAGATTGTACTTTTTTAGGTTCTGCTTTGTTGGAAATGAGAACCCTTTTGAATGATACAAAAAATGTTTTGAGCAACTGGAATGATTCGGATGAGTCGCCTTGTAGGTGGACAGGCATTTCTTGCTATCCTAATGACCAAAGAGTTCAATCTATGTATGTTCAATTCACTTTTTTTTGTTTTCACTATCAGTGCTTTATTCACTTTGTTTAGTTCTGTTTGCTTGGATTCATTTTCTGCATTTTGTTTCTGAATTTTCAGCAACTTACCTTATATGGAACTTGGAGGAATTATATCTCCTAGCGTTGGTAAACTCACCAGATTGCAGAGATTGTAAGTCCCATAGTCATTGAGGCAATGCAACAAAATTGTATTAGGGCTAAAGTTTCTGTCTTGCTACTTGCCTTGATAAACTATGTGTTCTTTCCTAAAACCAGGGCACTTCACGAAAACAGTCTGCACGGTGTGATTCCTAATGAGATTGGCAATTGCCCTGAACTCAGAGCTATGTAAGTAGGAGTTTCATATATAACTGAGTGATAACATGTTTTCTTGGAGATTCCACATGTTTCTTGATGAATTTGTCACAAGCTAGGTGTTATTTATGCGTACAGATACCTCAGGGCAAATTTTCTTCAAGGAGGCATACCATCAGATATTGGAAATCTTTCAATACTGACCATATTGTGAGATATATATATTTTAATTCACCATCTTATTATATAGTTAATGAGGTTGTTGTTTTAATGATTGGTACCTTTTAATCCTTTTTAGAGATTTATCAAGCAATTCACTTAGGGGTGCTATACCTTCTTCTCTTGGTCGTCTCAGACATCTTCGTCTTCTGTAAGTCCAATTTCTGTTCATTGTAAATTTTTAATTTGATAACTTAATGCATTACACACTGATGTGTTAAACATGAAAATCAATTCAGGAACTTGTCAGCCAATTTCTTCTCGGGTGAGATCCCAGATGTTGGAATACTGAGCACTTTTGGTAACGATTCGTAAGTTCTCCTCCATTGGAATATAATCCCACATTTGATTTTGCCCTCTGTGTTCATCATTTTCTATATACTCTAAATAAATTTAGACAGATGGTTCAAAACTCAGAAAATACAATTTATCGTGAATAATGCTTCACATTATTATATGTATAGAAGTGCAGCAAAATATTCACATTCCATTTGGCACTACAATCACACTTATAGGAAATACAATACAAGATAAATCAGGTCTTTGCACACCAGTTTTCTGGTGATTAACTGCTGTTTGCTGCATAGGCTAGGTTTGTGGCAGTGTTACAATGACACCCAGCAAATACAAACTAAGCAGAATACCCATCAAAAGATTGATGTATTAACATAACAAGTGCAGTTGAGATTTTTTACAACTGAACAAACCGAATCTATCAATTGATAGTCCTTTAATCATGATCCTAGAAGTTAAATACCATGTGATAATTGAGATGGGAACTACACTAGCAAATTGATGGTGCTTCAATTTTAAAGATGGTGATTAATTTGTAAGTATACCTGACTTAAATTTATGGTACTCCAATCATGCAGTATGTGTTGCATATGTTAAGATAGACTTTTTGGTGTTAAATTGCCTAGTGGGTGATTGGCTGCTACATAGTTGAAACTAGGAATCTAGTCCTGTATCATGATCAGTTTTACTTGTGTACCTTGCAAAAATTATTCTTCCCCATCATCTCAAAAGGGAAGCTTGATGCTCTCAAAGTCTCAAATAGTTAAGCTTATAAAGTATTTTCAAATGTCACAAGGGCAAGAATGTATGTGACGGGTCAATTCACGTAGGTTTCTTTCTGGAGTAAATTCTTAGTCATCCATTGGTAAGGGTGTTGCATCACCAACACTAGCTAGAAACAACTTGTTTAACTGGAGTCTCGTGAACTGACCTGCCGTTTACCTTTCTTCTTCATAGTCTTGGGAGATAATATTTCCAGTGTTAAATTTTTCTATTAATGGCTATCTTATAGCAGACTATTCTGATGAGGATAATGCTTATGCTTAGGTTTATTGGCAATTTAGATCTCTGTGGTCAACAAGTCCGCAAGCCTTGCAGAACATCTATGGGATTCCCTGTAGTATTGCCTCATGCTGAAAGCGATGAAGCTGCAGGTTGTTATTTGCTACTGCTTCATATCTATTTCCTTTTAGAAGTTTACTGGCTTTTGGAACTTCCCTTTCCTCCACCAACTGAATCTATTTTTTTTCTCCTTTACTTCTATGTAGAAAGCTTGATTTGTTGCTAAATTAATGTGATTTCTGCAGTCCCTACGAAGCGATCCTCTCACTATATTAGAGCAGCAGTTATTGGTGCAATATCTACATTGGGTTTTGTACTCATTGTGTTCTTTATTTTCCTCTGGGTTTGGTTGTTATCTAAGAAGGAAAGGACAGCAAAAAAATACACAGAAGTCAAGAAACAAGTCCACAAAGAGCCAAGTAAGTGGGTGAATAGTACAATACCGTTTCTTTTCCTTCTCCCAATTTTTGTCTTAATTTGTGGAGGCTTCAGGTGCCAAGCTTATTACTTTCCATGGAGATCTTCCTTATCCTTCATGCGAGCTAATAGAGAAGATTGAATCCCTTGATGAGGAAGATGTTGTTGGGGCAGGAGGGTTTGGTACTGTATATCGGATGGTTATGAATGATTGTGGAACATTTGCTGTTAAAAGAATAGACCGGAATCGTGAAGGCTCTGATCAAGTCTTTGAGAGAGAGTTAGAAATCTTGGGTAGCATCAAACACATTAATCTGGTCAATCTACGAGGTTACTGCAGGCTTCCTACTGCAAGGCTTCTCATCTATGATTATTTGGTCATGGGAAGCTTGGACAATTTCCTACATGGTATGTTTCCAGCTTTTCAAAATTTGTCCCCAGATCCTTCCTCCTCTCCACATCTCCTCTTAAGCTTATACAAGACATCAAATATGGAAAGTGTGTATATTTTTGCCATACATTTTTTATTTGGTTTATCCCGAATACAACCATTATTTTCAGTCATCTGTTTTTAAACTGGAGAGGGGGAAAAGCACTATTGCTTTCATTTAAACAATCTGTTTTTCTTGGCTATTCATTGCATGCATCTTTATTGTATTTTTGCAGAACGTGTGGATGATCGGTTGTTAAACTGGAATGCACGTCTGAAAGTAGCTCTTGGTTCTGCAAGAGGATTGGCCTACCTACATCATGACTGTTGTCCTAAAATAGTTCATCGAGATATCAAATCAAGCAACATACTCCTTGATGAGAATCTAGAGCCTCGTGTTTCAGACTTTGGCCTTGCCAAGCTTTTGGTGGATGAGGAAGCTCATGTCACGACTGTGGTTGCGGGTACTTTTGGTTATTTAGCACCAGGTATGCATATTCACTTAAATCAGCAAGGCATAAAGATAGATTCACACATATGTTGCCTTTTAATTGCAAGAATGGTTTATTTTCTTATTTTTTCGCTTGGATAAACCTTAACCTATTCTTCTTTTTATAATCATCTATATGTAGGTTTGACTTGCTTAACCTCTATAAATTGAAATTCCAAACTCACCTCATCTTCTAATGAAATGTCTGACCAACGATTTCTTGTTTGTATATTGTAATATAACCAGAGTACTTGCAAAGTGGAAGAGCCACAGAGAAATCTGATGTGTACAGCTTTGGGGTTCTCCTTTTAGAGCTTGTGACTGGAAAAAGGCCAACAGATCCATCCTTCGTAAACCGAGGCTTAAATGTGGTTGGCTGGGTATGCATGCTTCCCTTGAAACTCACCAAATTTTTGAACTTCATGAAGACTCATTAATTTTTTAAACTTCCCTTTTCTGTATTTAGCTGTACAATATCCACATATTGATAGTGTCACAATTTGAAAAATGAACAGATGAATACCTTGCTGAAGGAGAATAGACTGGAAGACATCCTTGATGCGAGTTGCAGAGAAGCCGATGCAGAGACGGTAGAAGCAATTATAGAAATAGCTGCAAGATGCACTGATGCAAATCCAGATGATAGACCTACAATGCAACAGGTGCTGCAATATTTGGAGCAAGAGGTCATGTCACCATGCCCGAGTGATTTCTATGACGAGTCTCATTCAGATTATTCTTGATGTATGGAATAATTTCCTGGGACCAAATATTTAGAAACAATTTGCAGCCTGATGCTGAAATGTGTAATCTGCCATTGCAGATACTCTGTATGCTCCTTGCTTTTGGAGCCCTGCTGCTTTTCTTCCTAATGTCTTTGTGCATCACTTGTGCCTTTTTATGTAGATACCTGCAGTTAGTTTTAAAGCTACAATTATGAGTCCTAACTAGCCATGGCATTATGTAGTATTTATAATCTATTTATCACTAGTGAATTTTTCCTTCACCTATAATAGAGTTTAGGTTCCTCCTAAGAAAGTTCTTTAAGAAAATTTGATGAAATATGTGAGGACACTCAACTACTGGTGGATGTCATATCTGCAAAGCATATTTGTATGCCACTCAATTATCA

>PGSC0003DMG400025635 LRR-RLKs

ATGCGTCTTATATCACGTATTGGTGCTCCAATACCAACAACGACATTTCTGAGAGGTATCATCTTTTGCTTCTTGACATTTTCCAGAATGGAAATTTTCACAGCGAGTTTTAACTTCCACACCTTTCTGGCTATACATGCTGTGTTTCTTGTGTTCCTATTCTCATTTTCTCTCCAACATATTGTATCAGCTGCTTTTCTTGGCAATGAAACTGACAAACTTGCCTTACTTGGATTCAAGTCCCAAATAACTGAGGATCCAGCAAGAGTTTTCGCCTCTTGGAACGATTCTGTTCATTTCTGTCAGTGGACTGGAGTAAAATGTGGCCTGAGACACGGAAGAGTCATCCGTTTGAATCTCAAAGGGCGAGGCTGGCAGGTATGATCTCAGGTAATCTTGGAAACCTCTCTTTTCTTAATTCTCTTGACCTTGCAGAAAATGCCTTCCATAACAAAATTGAGGCTTCTTGGATTGTCAGTAAATAGCTTATCCGGGGAGTTCCCTCCTCCGTTGTACAATTTGTCTTCCCTGGAACTAATATCACTTTCCTTTAACAACTTTTCTGGTAATCTCAGATCCGGTTTAGGCAACTACTTCCCTAATCTCCAAATACTTTACTTGGCGAAATGTCAATTCATTGGTTCCATACCGTCCTCTTTGGCCAATGGTTCAAAATTGCTAGAGCTTGATTTTCCTGTAAACAATTTCACTTGAAATATACCCAAGGGTTTTGGTAACTTGCAGATTTTCTTGTGGCTCAACGTTTGGAGTAACCAACTTGGATACAGTAAGCATGATGACCTTGACTTTGTAATTTTTCTTACCAACTACCAGCAGTCTACAAATGCTCCATTTTGGAGACAACCAGTTTGTAGGTACATTACCTCATTCAACAGTCAACCTTTCTAGTCAGTTACAACGCCTACTCTACTAGGAAACAGAATTACTGGAAGCATACCCAGAGATATCTCAAACCTGGTGAATATGCATCTACTCGAGATGAGTAACAACAATCTTACAGGTAGCACTCCAGGTTCTATTGGAAGACTGACAAACTTGGGAGGTCTCAACTTGGGTAACAGTCTCTTGACAGGGGTCACCCCTTCTTCAATAGGAAACCTCACTGAACTCATCTATCTTTATTTGGGACTAAACAGGTTAGAGGGTAACATACCTTCAACATTGGGAAACTGTAGCCAACTGCTAAAATTGGGTATTTCTGAAAACAACCTAACAGGCACTATACCACAACAACTTTTTGCTCTCTCATCCCTCACAAATATTTATGCATCTTATAACTCTTTGACAGGTCCATTACCAGTCTATATTAGTAACTGAAGTTATCTTACTTATCTTGATTATTCTCACAACAATTTTTCGGGGATGATTCCACGAACTCTTGGAAAATGCTTGGCCTTGGGGGATCTATTTGAAGGGAAACTCCCTCCAAGGGACCATTCCCAATTTAGAAGATCTGCCGGATCTCCAATCCTTGGATCTTTCCCAAAATAATCTATCAGGGCCAATTCCTCACTTCATTGCAAGTTTTACTTCCTTACTCTACTTGAACTTATCTTTTAACAATCTAGACAGTGAGGTTCATGTCACCGGAATATTTTCAAATTTGAGTGCAGATGTATTAAACTGGAACTCCGAGCATTTTGGTGGGATTCAAGAGCTGCATTTACGACCTTGTGTTTATCAAAAGACACAGAAGAAGCATGTACTTCCCTCAAGCATGATACTTCACAGAATGAATATCATAATATGTGGCCTCTGCACTTCATTATCTTCACCATGAGTGCCAAACACCCATGATCCACTGTGATATAAACCACAGAACATTCTTCTTGATGAAGATCTCACAGCTCATTTGGATGATTTTGGTTTGGTGAGACTCATTCTTGATGAAGATCTCACAGCTCAGTTTAGCTCACTTGGAGTTATGGGAACCATTGGCTATGCAGCTCCAGGTGGTGTTCCTAGGTTAACAAAAACTGTGATATGAAATTTCTTCAAACAATAGTTGTTTATTTATAAGCTAAATTTGCTTGTTGCAGAATACGGCATGGTAAATAAGGTCTCCGCTTTAGGAGATATGTACAGTTTTGGGATTCTCATATTGGAGATATTCACTGGAAGAAGACCCACAGACACTTTGTTCCAAGCAAGCTCTAGTCCTCACCACTTTGTCGAAACAGCGTTGCCAGAGAAAGTTATGGAGATTCTAGACAAAACAACTTTTCATGGTGAGATGAGTAAAGCAACCAACGTCAAAGAACACTGGGGCAGTATCAAGAAAGAAGAGATGGAATGCTTGGTTAGCATACTTGAGATTGGTGTCGCATGTTCAGCTGAATCCCCAAGAGACAGATTGACCGTGACACAAGTTTACAGTAAGTTGACACTAATGAGAGAACAATTTCTACAAAGGGGAGGTTGAGGCCACAGGCTACAAGTGAAAGCATTGGATTTCATCTAAAATGGTTTAAATTCTGTCTACTTGGTAAGAATGCTAACATTCTGTCCAGCATGCAGAAGAACTAAGGTAGTGTCAGATTTCTTGTTACTTAATGGATTTAGTCTAGAAACATCAGATATTAGTTCTTAATGTCTACATCAGATAATACGTTTCATTCATGAATGTAAAGAATCAGATGTTTTAGGAGATCATAAGGCAAGATCAAATGGTCATGGTGAGAATGTTCCTAAAAGGTTGTGATTCTAGACACAGATGAGGTATTCTTGTCCCAAAGTTGGAGCATCATGATCAATACTTGGAGGGTTAGTGAGAACACTGTTGATCTTTGCGTTGCTTTCTTAGCTTTGCTTGTCGTTTGTCCTTGTCCTTTATTTGAATACCAATGCCATGCCCACATGACATTGTTTAGCCTTCATCTTGTCAAGTCCACACCAAGTTGCCTTGCCCTCAAGTCCTTGGTCCTCACTTACCACTTTGGAATCAAGCCTTGGTGCTATTGTGCACACCTTAGCTTTGTCAACCGTCCGTCGCCAACTTAG

>PGSC0003DMG400025771 LRR-RLKs

CAACACTTTCTTTTTTTTGCCTTAACAATTCTCCATCTCTTGCTGCCTTTTTGGTCACCAAAACAACTTCTCCAGTAGTTGGTTTACCTCTTCCTCTTTCTTCTTATGTATATAGAAAGTAATCATGAATAATCACTTCAAGAAATGATGACTATCTCAGAGCATTACTATTCTTTCTGGTCATTTGGGATCTTTTTCTTTCTTCTTTTATGTCCTTCATTTGGTCTCAACATAGATGGAACTCTCTTGTTGTCTTTCAAATACTCTGTTCTTGATGACCCTTTATCAGTTCTTGATAATTGGGACTATAATGATGCAACCCCATGTTTATGGAATGGGGTAACTTGTGCACCAGACATGTTTAGAGTCATTAGTTTACTACTCCCAAATTCTAAGCTTATTGGTTCTATTCCAGAAGAATTAGGATTTATTCAACATCTTCATACTATTGATCTTTCAAATAATTTCTTGAATGGAACACTTCCTCTTTCACTTCTCAATGCTTCTGAACTTCAAGTTCTTTCACTGTCAAACAATGCTATTTCTGGTGAGCTTACTGAGTCTATTGGAGTGTTAAAGAGCTTAAAGGTTCTAAATCTTTCTGTCAATGCATTTGTAGGAAATATCCCACAAAAGCTAAGTTCTTTGCAGAATCTAAAAGTTGTTTCTTTGAGTAAAAACCTATTTTCTGGTGCAATACCAAGTGGATTTCAGTTTGTTGAAGTGTTGGATCTTTCTTCTAACTTGTTGAATGGAACATTGCCAGATGATTTTGGTGGTGATAGTTTGAAATACTTGAACTTTTCATCAAACAAGTTATCTGGTTTAGTGTCACCACAATTTGCCAAGAAAATACCAACAAATGCTACAATTGATCTTTCTTTCAACAGCTTTACAGGTGAAATCCCAGAGTCATTAGCTTTATCAAACCAGAAAACAGAATTCTTTGCAGGAAATACAGATCTTTGTGGCAAACCACTCAAGAAACTCTGTACAATCCCTTCAACACTTTCATCTCCACCTAATATAACCACTAATCCTCCAGCAATTGCAGCCATTCCAAAAGAAATTAACTCAACCCCATTTCAAGATTCTGATGGAACAACTCAAACTGCAGCTCAGAATCAACAACAACATGGGCTAAAGCCAGGTACCATAATGGGAATCATAGTTGGTGACTTAGCTGGTGTTGGTGTTCTTGCTATTGTCTTTCTTTATGTCTACAAATTGAAGAAGAAGAATAAAGCAAGTGAGGCAACCATAGAAAGCTCAATTGACAAAGATCAAAAATACAACAAAACACCAGAACCATCAATACTTGTTGTAAAAGAAAAAGACACCATTTTTCCTAGTTGGCCATGCTTGACCATAAATCAAGAAAGTTCAGATACTGATGACAGCAAAAACCAAGAAAGTGAAGACCAAATTGAGTATGAAACAGAACAAAAGAATGAGAAAAACAAGAAAACAGAAAGGTCATTTGTAATGGTGGATGGTGAAACAGAATTAGAGCTAGAGACTTTGTTGAAAGCTTCAGCTTATATTTTAGGGTCAAGTGGAGCTAGCATAGTTTACAAAGCTGTGTTAGAAGATGGAACTGCATTTGCTGTTAGGAGAATTGGAGAGAGTGGTGTTGATAAATTGAAGGATTTTGAGCAACAAATTAAAGGAATTAACAAGCTAAGGCATCCAAATCTTGTTCGTGTTAGAGGATTCTATTGGGGAGATGATGAGAAATTAGTCATCTATGATTATGTTACTAATGGAAGCTTAGCCAATATTGGTTACAGTAAGTTACTTTTCTTTCATCTTCATTATCTTATCATAGTTATAGTTTTGATAAATCTTTTCTGTCCTTATCATAATTACCCCCAAATATTCTGATTAGTCAAGTGTACTTGACATTTAATGTCACTGGTATATTTGCAGCAACATTCCTTGATTTGTCACTTAACAAAAACATTAAAGCCTATAGTTTAGCGGTAGTAATAAATATTGTACTAAGGGGTCGTTTGATAGCTGGTTAGAGTTATGCATGTATTAATCATGTAGGAATTGGTATTATTTAATTAAGTGATTAGTTACTCCACATTCTATGTTGTATAAAGTAATACATAGCTTTCCTCGTAACTTATATATGTATTAGTTATAAGGGTTTTTAAATTACAAACCAGACAACTCCCAAAGATAATACAACTTACGTCTAAACCAACTACCAAACGAGTACTTTTTTAAGTACACTGTCTGTCATTAGAAAAACTAGTTTAGTTTTAGTGTACATATACTACTCATAGGCATGGATTTTCGTTACTAAATTTTGCAGATACTATACTTCATATGTGGTTGTGTTTGAGGCTCCCAATGGTCCTCCATGTGACATTGTCATAATCCGTGAAAGTCCCCAAACTATACTACAATACATATAATACGTCAAAATACATGTGCACCCCTTCTCCTCCATATTGTAGTACCACAAACGTTCCCGAGACTTGATTTTTTCAAACTCCACGTTTGATCAGATTATACTGAGTATAATATTATTGTTTAAATTTTTCAGAAAAAGTTGGTTCCTCTCCTTACCATTTATCATATGAAGTCCGGCTCAAGATAGCAAAAGGAATTGCAAGAGGATTGACCTACATCCACGAAAAGAGACAAGTACATGGCAATATTAAACCTAGTAACATTCTCTTAACACCAGATATGGAGCCCATAATAAGTGACATTGGGCTTCATGGGCTAATGCATGGTAAAAACACCTCTAAACCGGATAATTCGGGCCGGCATTTCGGTAGCAAGCGGTCCACATCCTCCTCCCGGGACGGTTTAAATGATCAACCGGTTCATGGAAGTCCTTACATTGCCCCAGCCGGGTTCGTGGTAGGGTGCACGTCCCCCTACCACGCCCCGGAGTCGTTGGAGAGTCTCAAACCTAGTCCTAAATGGGATGTTTACTCATTTGGGATTGTGTTGCTTGAGCTTTTAACCGGGAAAGTATTCTCGGACCGGGAGTTGAGCCAGTGGACCACCGGTTCAGTATCGGATGACGTGGACCTGGTTTTAAGGATGGCTGATGTGGCGATTAGAGCTGACGTGGAGAGCAGGGAAGAGACCACGGTGTCATTGTTTAAGTTAGGTTTTAGTTGTGCATCATTGAATCCACAAAAAAGGCCTACAATGAAAGATGCCCTTCATGTGCTTGATAAAGTACCAGGTTATTCACATTACTAATCATTATGTTGGGACCAAGAAAAGCTATAAAAAGCTGTTTAGTTCCCTTTAATTTGTTCTTCTTTTGACTTGGAAAGATTTTTGCTCCATAAAGATGAGAAGAACCGTTGTGTTGTAATTGGTGAAAGTGTTTGTAAGTTTGACAAATTGATCAACAGTGTATTTATTTGAGTTTTTTAATAATAGTAGTTTTTTTTTTCACCCGC

>PGSC0003DMG400026004 LRR-RLKs

TTCTTCACAAAAGCCAACACTCCTTTCTTGAAATTTGTCACTAAAACCAACTTTCCCTACAAACTTTTCCTTCCATTTTTCTTGCAAAAACCAAATATGGGTTTTCTCTCTGTTCTCTCTTCCATTTTCCTTTTCATCCTAATATTAACATTCCCACAACAAAGGGTTTCTTCTGAAGCCATTAATGATAGTACTAAAACTTCTTTATCTTTGCTTATGACTATCAAAGCTTCTTTGGACCCAAAAAGCCTCATTCTTTCATCATGGTCACCTACTTCTACTGACCCATGCAATGATTCTTTTGAAGGTATTGCTTGTAATGAGCTTGGTCAAGTTGTTAACATTTCTTTGCAAGGCAAAGATCTTACTGGAAAAATCCCACCTGAGATTGGTCAGCTTCAAAGCTTGTCTGGATTGTACTTGCACTTTAATAAGCTACATGGGGTTGTACCTAAAGAAATTGCTAACTTAACTCAGCTCTCAGATTTGTATCTTAATGTGAATAATCTCTCTGGTGTTATTCCTCCTGAGGTTGGCAACATGTCTAGTCTTCAAGGTAGGTTCTTTTTTGCTTTGTTGTCTTGACTAGTTATGGTAAAAAGCCTCAGTCTTGATGCTTAGTTTTGACTTAATTTGAATGAATGTTTGAACTTTGCCAAATAGATCTATACCTCTATTAATGAAATGGGTAGCTCAATAAAGAGTCCAATAAGTGATAATGCTGTTAGTTATTGAGTAATTTCACAATGCCTATGGCTTTTCTTGAAAAAAGATTGGATTTTTAGTACTCATTAGTCATTGAAAGTCTATATTAAAAAGGTTCTTGAAGTATCTTTTATGAGGTGATTTTTGTTCCTATGTAGCTAACAGCTAGGTGCTTAGAACATGAAACACTTGAAAAAGCCTCCATCTTTGGTTTTATGTTAAGCTTTTCTCTGTCATTTGTCTCTTTCCCCATTGATTTGAGCTTCAATATGGGCTAATGCTTGAATAAAGTTTCAACTTTCCTGGCAAACTTGCAAAATTCCTAGTCTCTTGTTTGGGTGGCTTTTTCTGCATTTTTGGTGGAATAAATCTCAAATAAAAGTAACCGAGCTCCATTTGTTTTCTGTTTTTACACAGTACTGCAACTATGTTATAACCAGTTGACTGGAAGTATACCTAATCAACTTGGAGCTGTGAAGAAACTGAGTGTTCTTGCTTTGCAAGTCAATCAGTTAACTGGAGCAATTCCTGCAAGCTTGGGGGATTTGGAGATGTTAACCAGGCTTGACTTGAGCTTCAATAATCTTTTCGGGTCTATTCCAGTGAAACTCGCGGATGCTCTAAAGCTAGAAGTTCTTGATATCAGAAATAATACCCTTTCTGGCAATGTCCCTCAAGGTAGTTCTATTTGATTTTCTTGATAGTACCTTTAATCCTATTTCATTGCTTAATATGCTTAAGAAATAAAGAAATCTTAGGTTCTTTGATTCCCCCTAATATCAATCCAAATGTTTACTAATATTAGACATTATAGTAGAACTAACTCTCTTTTTGGTCTTTTTGGTTTGGGCAATGGATGAAATGAAGCCTTGAAGAGACTGAATGAAGGATTTCAGTATGCAAACAACCCAGACTTGTGTGGAATTGAATTTTCCTCCTTGAAACTCTGCACTGTATCTAGTCTAAACCAAAACAGACCACAACCATTTGAACCTGGTTCGAATCGTCTTCCCACAAAAGACATTCCAGAGTCTGCAAATGTACAAACAAAACAAATGAGTCAGTCAAGAAAATCACAAACTGCTGTGGTTGTTGGTGTTATTGCACTTTTTGTTGCAGTAGCAGTAACTGGACTTTTCACATTTTCGTTGTACCGTCGAAGGAAACAGAAAATTGGAGGTACTCTTGATACCTCCGATAGGCGACTCAGTACTGATGAGGTGAAGGAGATCTCTAGAAGAAGCGCGTCGCCCCTCATCAGCCTGGAATATTCCAATGGCTGGGATCCTCTAGGCAAAGGCCGAGGTGGAAGTGCTTTCTCTCAGGAAGTATTTGAGAGCTTTATGTTCAATTTGGATGAAGTTGAGTCAGCTACACAGTACTTTTCAGAAGCAAACTTGTTAGGCAAGAGTAATTTCACAGCTGTTTATAAAGGGACATTGAGGGATGGGTCTGTTGTTGCTATTAAGTGCATTTCCAAGACTAGCTGCAAGTCAGATGAAACTGAATTTCTCAAGGGACTAAAACTCTTAACCTCATTGAAACATGAAAATCTTCTGAGGTTGAGAGGCTTTTGTTGTTCAAAAGGTAGGGGGGAATGCTTTCTGATCTACGATTTCGTTCCAAATGGAAATTTGTTGCAGTATCTTGACGTGAAGGATGGAAAAGGAAAGATCCTTGATTGGTCTACCAGATTTTCTATAATCAAAGGCATTGCCAGAGGTGAGCTACTGTTTGATTAGTCCCATGTTTGATGTTTTCTTTATAGCTCTTTGACGACCAGATTTGAATAACAACATTTGTTATCTATTGTTTAACAGGTATTCATTACTTGCATAGAAACAAGGGAGGCAAACCTGCTCTAGTCCATCGAAATATATCAGCTGAAAAAGTACTTGTCGATCAACACTACAACCCATTGCTCTCGGATTCTGGTCTGCATAAACTACTAGCAGATGACATTGTCTTCTCAACACTCAAGGAAAGTGCTGCAATGGGATACCTAGCTCCTGAGTATACAACCACTGGACGGTTCACCGAAAAGAGTGATCTTTATGCATTTGGCATGATCATATTTCAGATTCTCTCCGGAAGACGTAAAATCACCCCTTCTAACCACCAAGGGGCAGAGTTGTGCAGATTTGAAGACTTCATTGATCCGAATCTTTGGGGCAACTTTGTTGAAGCGGAGGCAGTTCAGATGGCAAAGGTAGCTCTACTTTGCACGCACGAGTCACCAAATCAGAGGCCAGACATTGAAATCGTCATGCAAGAACTGAATGATTTAACCTCCAGCAAGTCGTAAAAACTATTGCAAACAACATCCAAGTTCTCCAATGACCATTAGGAGCCTGATATTAACATCTTACTGATGCCAGGAGTAGGCACCATTTTCCCATCCTTTTGATCCTGCATGACTTGTGCCTGAGGACGACACATGAATGCCAGGTGCCTGTTCCACATAGTTTATTAGTACTAATGCACCTCGACGTAGGTAGGCCTCAAGTGATCAGTTCTCTTGTATTATGTAACTTAACCATAATTCTAAGCAAGTCTTCTCCACTTGCAAACTAGTGTACCAGAAGTGCAATTTCTATTTAGTATCATATGAGTTATGACCTCTTGC

>PGSC0003DMG400026072 LRR-RLKs

CTAAGACCTCATTTTTCTATCATCACTAAATGGACTAAAAACTAACAAACTCTGCATTCAGAAACTTCTTTTTCTTTTATCTATTTCTGGTGACTGACACTGCAAATTCCCATTTTCCCTTTAGAGGGGTCTCATTCAAACTTTTGAATTTTTTGGGTATTCAAGAAAACAAAAAGTTTGAAGCTTACTGCCTCTTTATGTGTGAATAAATAAGAGAAACAGAAAATCATCTTACTATTTCAGGTATAACTCATGTTTTCTTTAAGCCTTATTCATTTGTTAGTCTTTTAATTACATAAAAATCCAAACTTTATGTGTTATATGAGTTATCAGATGATTCTTAACAACTAAGCTATCATTTAGCCATAGATTTTAGATGATATTTTGAAAATTTATATTCCTATCTGTTTGGCCATGAAATTTGATATTTTTGGAGTTTTGGGACTTTCCCTCTTAAAACTTCAAAATCTATAGCCAAACAGGACCTAAAGGATGCATGTTACTTATGATTTCTTAAGAAAGTTTACTTTTTTGTTTCTATGGATATCTCTGTTTCTGTAAAATTATGGAAGCTGTTTGTTTTCTGTCAATTAGTTGATTTCCATGTTCCTTGTTTGCCTATTGTAGTTTGCTACAACTTTCTGGGAAAAAAAGTTGTTTTGTTATTGTTACAAATATGGAAATCCTAGATTTCAATCTTCTTGCTTATATTCCTATAATTCTTGCAATTATGTTTCGAGGGGTCTATTGAAAAAAGTCTCTACCTACCCCAGATTCCACACTTGTGTTTTTGGGGTAAATGAGTATCTTCTTTCATCTGGACAGTCTGTAAAATAAGGTGAAACTATTATATTTTGAGTCACTTTAAAGGGAGATAATTTAATTTTTATATAGTGTAATTCACAACTAAATGAGAAATTGACATCTTAACATAAGTTTATTTCCTTCCCAACCCCACCCAACAATTCCATCTTTTATAATCTAGACTTCCTGTATCTGTAAGTTTTCTTTTTCTTGAACTACAATTTTTATTTCATGTTGGAAAGTTATATGAAGTATATGACAAACATTTTGTTAGGTATACATACAACACTTTAAAAAAAGACAGCTTTTTTTGAATTGGATATCAGTATTATCTTGTTTTTTTTAATAGACCACTTTATCTTAGTGTTTGATATCTGAGTTTCCACTGTTTTTCTTGCAGCTGCTTCTTGGATGGACACCCATCATGGCAATAATGGAGTTCTTGATTGGATTTGTGTCCCCTTTATGAAATGACCCTTTTGCCCTTCTCAAATTTCAAGAATGGCCAGAATCCAAAAACTTCAAAACTCTCTCTCAATTCTTGCTGTTTTCTTGTTTTTGAATTTTTTTGTCCAGACATGTAAGTCCTTAACCTCAGAAAGTGAGGCTCTTTTGCATTTCAAAGAGCAGCTTAATGACCCTTTGAATTACTTGGATTCTTGGAAGGACTCAGAATCTCCTTGCAAGTTCTATGGAATTACATGTGATAAAAATACTGGTCTTGTTATCGAAATTTCCCTCGATAATAAGTCGCTTTCCGGGGTGATTTCACCTTCAATTTTTTCACTCCAAAGTCTTACTTCTCTTGTGCTACCTTCAAATGCACTATCAGGGAAACTTCCTAGTGAAGTTACCAACTGCACCAGTCTCAAAGTCTTGAATGTCACTGGAAATAACATGAACGGAACGATACCTGATTTGTCTAAGTTGACTAACTTAGAAGTGTTGGACTTGTCGATTAACTATTTTTCCGGGGAATTCCCATCTTGGGTTGGAAATATGACTGGTTTGGTTGCGCTAGGGCTTGGAGATAATGACTTTGTTGAAGGTAAAATTCCTGAGACTCTTGGGAATTTAAAGAAAGTGTATTGGCTTTACCTGGCAGGTTCAAATTTGACAGGAGAAATCCCGGAATCCATTTTTGAAATGGGGGCACTAGGTACATTGGATATTTCAAGAAATCAGATTATTGGGAATTTCTCTAAGTCAGTAAACAAGCTGAAAAATTTATGGAAAATTGAGCTGTTTCAGAATAAGTTGACAGGGGAACTTCCTGTAGAACTGGCAGAGCTCTCTCTTTTGCAGGAATTTGATATCTCTAGCAACCATATGTATGGGAAGCTGCCTCCAGAAATCGGGAACCTGAAGAAGTTAACAGTTTTTCAAGTATTCATGAATAACTTCTCTGGTGAAATTCCTCCTGGTTTTGGAGATATGCAGCATCTTAATGCCTTTTCTGTGTACAGGAACAATTTTTCTGGGGTATTTCCAGCAAATCTTGGTCGGTTTTCCCCCCTGAATAGCATAGACATATCTGAAAACAAGTTTACTGGTGGATTCCCAAAGTACTTGTGCCAAAATGGGAACCTGCAGTTTTTGCTAGCTATTGAGAACAGTTTCTCAGGGGAGTTTCCGAGTACTTATTCATCGTGCAAGCCTTTACAGAGATTGAGGGTCAGTAAGAATCAACTTTCTGGGCAAATTCCTAGTGGTGTATGGGGACTTCCAAATGTATTTATGATGGATTTCAGTGATAACAAATTCAGTGGAACTATGTCCCCGGAAATTGGGGCTGCTACTAGCCTGAATCAGTTGGTATTATCAAATAACAGATTTTCAGGTGAGCTTCCAAAAGAACTTGGGAAACTCACACAATTGGAAAGGCTTTACTTGGATAACAATAACTTCTCGGGCGCAATACCTTCTGAACTTGGTAAACTAAAGCAAATTTCATCTTTGCATTTGGAGAAAAACTCATTCTCGGGCACTATCCCATCAGAATTGGGTGAATTTCCTAGGCTTGCAGACCTGAATCTTGCTTCAAATCTTCTTACAGGTAGCATTCCCAATTCCTTATCAACAATGACCTCTTTGAACTCTCTGAATCTTTCACACAACAGACTCACTGGTACAATACCAACTAGCTTGGACAATCTGAAGCTGTCTTCATTGGATCTCTCAAACAACCAGCTATCAGGAGAGGTATCATTAGATCTTTTAACACTGGGAGGAGACAAGGCATTAGCTGGTAACAAAGGACTTTGTATCGACCAAAGCATCAGATTCTCGATCAACTCAGGCTTGGGCAGTTGTGGTGGAAAGGCTGCTAAACATAAGTTAAACAAACTAGTTGTGTCTTGCATTGTGTTGCTTTCCTTAGCGGTTCTAATGGGTGGGTTATTGCTCGTCAGTTACCTAAACTACAAGCATAGTCATGAAGTAGATCATGAAGAAAAACTGGAGGAAGCCAAGGGAACAAATGCAAAATGGAAGCTTGAGAGTTTCCACCCTGTGGAATTTGATGCAGATGAAGTTTGTGATTTCGATGAGGACAATTTGATCGGAAGTGGAGGTACAGGAAAAGTTTATCGGTTAGATTTGAAGAAAGGTTGTGGAACAGTAGCAGTGAAGCAGCTGTGGAAAGGGATTGGTGTGAAAGTTTTGACAAGGGAAATGGAAATCTTGGGAAAAATCAGGCACAGAAATATAGTTAAGTTGTATGCCAGTCTAATGAAAGAAGGTTCAAATATCTTGGTTTTCGAGTACATGCCAAATGGTAATCTTTTCGAGGCACTACACCGAGAGATCAAGGCTGGCAAGCCAGAACTAGATTGGTATCAGAGATATAAAATAGCCCTTGGAGCTGCAAAAGGAATTGCTTATCTGCATCATGATTGTTACCCTCCTATTATTCACAGAGATATCAAATCAACCAACATTCTACTTGATGAGTATTATGAAGCAAAAGTTTCAGATTTCGGGGTTGCAAAAGTTTCAGAAATTTCTTCAAGGGGATCCGAGTTCAGTTGTTTTGCAGGCACTCATGGTTACATGGCTCCTGGTAAGTTTCAAGAAAACCTTAAATTGAGTGACCATTTTGTCTCTCCTTGTTACTTTCCATTCTTGAATTTGCATTTACTTTCCTATATGTTTGCTTGACAGAAATGGCATACACTCTCAGAGTGACCGAAAAGAACGACATTTACAGCTTTGGTGTCGTGCTATTGGAGCTAGTAACTGGAAGAAAACCAATAGAAGAAGCCTACGGAGAAGGAAAAGACCTGATCTACTGGACCTCAACTCATCTAAATGACAAGGAAAGTATTAACAAAGTTCTTGATCAAAAGGTGGTATCTGAACTTGTTCAGGATGAAATGATCAAAGTGTTGAGAATTGCCACACTTTGTACTACTAAGCTTCCTAATTTGCGTCCCAGCATGAAAGAGGTTGTCAATATGCTCGTTGATGCTGAACCTTTGACGTTTAGGTCTTCAAGCAAATCTGAGAAAAAAGGGAACAGTTTTTCTGAGGTCTAGTTTCTCTTCTTTTTTTTTATTGGGTTTTCCCTGTTGATAGTTGCTCTGTAGTATGATTCTCAAGTTAGTTTTAGGTATCTGATATACATAAATTTATACCATGTGATGTGATATTCCACTTTCTTTATGTAAAAGTTCGAATTGTGTAAGAAAAATGTACTTTTTTCTGCTGTGATTGGAAACTTCAGAAGTGGGATTAATCAGAAATGTGGGTAGTTAATGTCCATGATTAAGAGTCCCACATCGGCAAAAATACATTAGATATGTTGAGTATATAAATAAGCAGGCCTTACACTCAAATAACATGTCTTAAAGCCGCATAGGTGTAGACTCAAAGCTAACAATATTGCCTTACACAAAAACTTGTTTTGTTGTAAACTTTGAATCTGATTCAATGAATATCCACATAGATAGTTGGAAGCAAAACTGATATACATAAACATTTCATACATTTGTATAAGAGGGATGGCAACCAACAATGTTACTCTTTTATCTTTTGTGATCCTCCTCAGTATTACATGGTACATTCTGCAGATTTTTCCTGCTCTCTATGACATTACGGTTCCAAATCTTCACTCCTTTGCTTCAAGAAGTTGAGTTAATGCAGCATCAATTTGAGCAAATACCTCCTCTTTGGAAACGCTCCCGTCCACCTGAAATTAAAATAGGTCAGAGAAGATAGAAGTTTTAACACTGAGAAATGATAAGAACACGTTCATGCACGCGTATTGAAAGAAAGAACATACAGTATCCTTGTTCAGTGAAGAAAAAGAGAAGGGTAAGCAAAAGGGGAGTCCTTGATTAAGTATATTTCCCCCCAGATTGTTTGGTATGAATGTAAAACTCAAAGGATTTGCTCTTCCTTTATCTAATTGAGGAGAGATGAATAATTTTAAGATAAAAGTGGATGTATAGCCCTTTTGCTTTGGCTTCCCTTGTTGTTTTAACAACTGTGTGAGTATGGATCAGTCTGGATGAACTGGAACATAATTTAAATTGAACATTTTGTAACCTACATGTCAAACAACATAATCTTATAACCATATCATTATTCTTTCGCCAATTAGAAAACCAAACTACTTTTTCTCCAAACCATTTCCAACCAACCCCCTCCACATTGTCTTTTTTATGGACTATATTCCTGTTAACCAAGACCTTGGAAACCATCAAACAACCCAACAACTATGAATCAAGCATATAAGAATGATTCCACGTTTTGAAGGTGAGTGTTCAATCTGCATTCCCGAGTGAAGCAACTGCACCACATCTGTGGCAAAAAGAATACCTGGAATATAGTATCTTTGTACATTGAGAGAACTGATTCCACATTTTGATGGTGAGTGTGCAGACGCAGCTTCACCTGTAAATGGTAAACTGAAAAAGAAATAAGACAAAAATGAAGCTGATCACAACATAAATTTATGGTGTTTTAGAAACCTATACTTAAGTCAATTATCTACAAGGGGACTTGAGTAAGCAAGGGAAAAGAACATCAAGAATGATAAAG

>PGSC0003DMG400026749 LRR-RLKs

CTCAAAACCACAGGTTACTAGTACTAATATTTGCACTAATGGAGAAAGCCTTCACTATTTTTCTCTTAACAATACTCTTGTTGCTTCACTATGTTATGACCCAAACCAACATTACTACTGATCAATTAGCTCTTCTTTCTTTAAAATCTCAAATCATTTCGGATCCCTTTCACTTCTTGAATGAAAGCTGGTCTCCCGCTATTTCTGTTTGTCATTGGGTAGGAGTCACTTGTGGTTCTCATCACCAGCGAGTGAAGTCCTTGAATCTTTCCAACATGGCTCTTACAGGCAGGATTCCCCGTGATTTTGGAAACCTCACATTTCTTGGTTCTCTTGACTTGGGAAGCAACAATTTCCAGGGATATATGCCTCAAGAAATGGCATATTTGCATCGGCTTAAGTTTCTTGATTTAAGCTTCAACAACTTTAGAGGGGAAAATCCTTCTTGGTTTGGGTTTTTACACCAACTTCAAGATCTAAATCTTGGAAATAATAGTTTCATTGGTTCCATCCCTTCTTCATTTTCTAATATTTCCACACTTGAAACTTTGAATCTGAATTTCAATTCCATAGAGGGTGAAATCCCAGAAGTGATTGGAAGTCTTATAAACCTTAGAGTTTTAAGCTTGTATGGAAATAATCTCATAGGTTCTATTCCTCCGTCACTCTCGAATGCCTCAAGGTTGGAGGCATTAGATTTATCTCGTAATTTACTTCAAGGGAACATTCCAGAAGGGATTGGCAATCTACACAAGATGAAATTGTTGTCCATACAACATAATAAACTTACGGGTTCTATACCGTTCACAATTTTCAATATTTCTAGAATCAAATTCATTGCATTTACAGACAATAGCTTATCAGGAAATCTTCCCAATGGTTTATGTAATGGTCTCCCAATACTCAAAGGGCTTTATCTATCCACAAACAAACTTGGTGGTCATATGCCTACAAGCTTGTCAAATTGTTCACAACTTCAAATATTGAATTTATCAGGTAATGAGTTTGACGGACGAATACATAGTGAAATTGGAAGATTGAGTAACTTGCAGATATTGTATCTCGGATCTAACCATTTCACAGGTACGATCCTAGCTACAACATTATTAATTCTTTTATGAATACTTGTAATATTGAACTTGAACTGCAGGAATAATTCCACAAGAGTTAGGAAATCTTGCTAATTTGGCGGATTTAGGCATGGAGGATAACCAGATTAGTGGCTCTATCCCGATCAACATATTCAATATCTCCTCACTGCAACGTTTTGGACTATGGCGGAACAATCTCAAGGGATCCTTACCACGAGAGATTGGCAACTTAACCAAGATACAGATTCTAGATCTTAGAGAAAATACATTTATTGGTACGTATTTGAAGTGATAGCAAAATTATAAACTTGTACTAGCTTGAACTAACTCATTCTTAATTTCTTTAGGTGAAATACCCAAAGAGATGAGCAATATGATGGAGTTGGAGGTACTTAGTCTCGGGTTAAATAGTTTCAGTGGTTCACTTCAAATAGAGATGTTCAACAGTACTTCACGCTTGAGGATAATTTCTCTTACAAACAATAATCTATCAGGAACTCTACCATCAAACATAGATTCTGTCTTACCCAACATTGAAGAGCTTTATTTGGGTGCCTTAACAAATCTTGTTGGGACTATTCCTCATTCTATTTCCAATTGTTCAAAACTTACTAATCTAGAGCTTTCAAATAATAAACTCACTGGCTTGATTCCCAATTCTCTTGGATATTTGGCTAATCTACAGTTCCTAAACTTGGCAGCAAACAATTTAACAAGTGACTCATCGTTTAGCTTCCTCACTTCCTTAACCAATTGTAGAAATTTAACAATTCTTTTTCTATCTTTGAACCCTCTAAATGGCATGCTTCCAGTATCTGCAGGGAACCTTTCCACTTCTCTTACAATGTTTTACGCCAGTTCTTGCAATATTAAAGGGCGAATTCCAAATGAAGTTGGGAACTTAAGCAACTTGTTAGAGCTTGATCTTTCTGGAAACAACCTGGTTGGATCAATTCCCACATCAATTGGCAACTTGAGAAACCTTCAACGCTTCAACTTGAGTTACAACAAACTTACAGGATTTATTGGAGATTTAATTTGTAAATTGCAGCATTTGGGTGCTATTTACTTGGGACAAAATCAACTTTCAGGATCTCTTCCTAATTGTTTAGGGAATGTTACTTCCCTTAGATTGATACATCTTGGTTCCAATAAATTGATTTCCAATATACCACCAAGTTTAGGGAATCTTAAGGATTTAATGGAGCTTGACTTATCCTCAAACAACATGGTTGGTTCTTTACCACCAGAAATTGGAAATCTAAAAGCTGTGACACATATGGATCTGTCGATGAATCAATTCTCAAATGGAATTCCTAGGGAAATTGGAGGATTGCAAAATCTGGTGCATCTTTCTTTGAGACACAACAAGTTGCAAGGATCTATACCTGACTCAGTGAGCAACATGGTAGGTTTGGAATTCCTAGACATTTCTCATAATAATATATCAGGAACCATTCCTATGTCTTTGGAGAAACTTCAAAACTTGAAGTATTTCAATGTTTCCGTCAACAAGTTGCACGGTGAAATACCCTCAGAGGGTCCTTTCAAGAACCTCTCGAGTAAGTTTTTCATCGACAATGAAGCATTGTGTGGTTCTTCAAGTTTTAGTGTCCCACCATGCGCCACATCATCAAAGCATAGATCAAATAGGAAAAAAATGCTTGTTCTATTTCTTGTGCTGGGAATTGCACTTCTTATGCTGGGAATTGCACTTGTATTTGTTCCTATTACCTTTGTGTTTGTATGGATAAAGTATAGAAGAGGTAAAGGTGATCCTCAACAAGCTGACTCATTGTCTACCGTAACAAGAGAAAGAATTTCATACTATGAACTGCTCCAAGCAACTGAGTCGCTTAGCGAGAGTAATCTGATTGGTTCTGGAAGTTTTGGCTCTGTTTACAAAGGTGTTCTCAGAAGTGGGACTGCCATTGCAGCTAAAGTATTCAATCTGCAACTGGAAGCGGCATTCAAGAGTTTTGATACAGAATGTGAAGTTTTGCGCAGCCTTCGCCATAGGAATCTCGTGAAAGTCATCACTAGTTGCTCCAACCTTGATTTTAAGGCTTTAGTGCTCGAGTATATGCCTAATGGAAGTCTTGATAAGTATTTGTATTCGCACAACAACTTCCTAGATATCAGCCAGAGACTAAGCATTATGATAGATGTGGCATGTGCGTTGGAATATCTCCATCATGGATGCTCCTCACCCGTGATTCATTGTGATATAAAGCCTAGCAACGTCTTGCTGGATGAGGATATGGTTGCCCACCTAAGCGACTTTGGTATTTCAAAACTGCTTGGTGAAGATGAGAGTGATTTATACACAAAAACTTTAGCAACATTTGGTTATATAGCACCAGGTACGTCTCTTTGTTTTTGCTAATTTGAACATTGATTTTTCCTCTTATTCTTTTTTTTCCAATCAAGAAAGTATATTGCATCTTTTAACTAATTGTTTGACTGTAGAGTATGGGCTGGATGGACTGGTGTCAATAAAATGTGATGTCTATAGTTACGGGATCATGTTGCTGGAAACATTTACTAGGAGAAAGCCTAGTGATTTTGAGGGAGATCTTAGCTTGAAGCAATGGGTGAGTTATTCACTCCCAGAGGCAGTAATGGACGTTATGGATGCCAACTTGGTTACACCAATGGATAATCGCTTACAGAAGGAGTTAGATATCGTGGCATCAATCCTGAAAGTAGCATTAGATTGTTGTGCTGAAACTCCGACAAGAAGGACAAACATGAAAGATGTTGTAGGGATGCTACAGAAGATCAAGATTCAACTTCTTGCATGCTGAGCATGTTCTCTGAATCTCTTGTTCCGAATCGATCACTTTTTTTGTTTTGTTTTAGCACTTGATTTGTGCATGATTCTGTTTTCAGAAATGCTAGTTGGTTGCAATAAATGTCACTGTTTCCTCTTAAA

>PGSC0003DMG400026760 LRR-RLKs

CCAAATTTCAAGAAAAATAGAAAAAATAAATAAAAATTTTGTTTTTGTTTTTAGAATTGTTTGGTTGGTGGTGGGTATTGGATCCAATGTCACCAAGTAAATGTTTTTAGTTATCCAATGTGTTTTTCTTTTTAAAATTCAGCAAAAAAAAAAGTGTATTTTATTAGACACACAAATCATTGTTTTTAGAAGCATTTTTGGGTGTCTCTGCAATCTCTGAGGTCTGTCATGTGAAGTGTAGGGAAAAATAATTCAAGAAACCAAAACCCCAAGTTTTGAATTTCTTTATTTTATTATATGGTATTTTTTTTGTAATTTTTTTGGGTTTTGGGGTTTTATGAAAAGTGAAAGTAGTAGTAAAGAGTGAAAAAAAGGGTAGTGATGGTGATGAGGAAGTTCAATAAATTGAGTTTGTTTTGGAGTTGTTGATTTTATAGTTTTGTGGTCAAAGTTTTTGTTTGTCAATTGATGATAATGACACTAATTGGTAGATCTGAAGATTCTTGTGTTGATAGTGATCTTGTTAAAACTTTTATAGACAGAAAGGAAAATGTCTTTTGGATCTTTTACATTTCAAGATTTAATCATCAAATAAGAAAATACTCTACTTTTGTCATATGAAAATCCAAAAAAAAAAAAAAACTTCATTTTTAATGAACTTTCTTGTTTCCATTCATCAATGGTGACACCCCAACCAAATCAAGAATCCATTTTTTTTCTAGGTTTCTAGTAGATCCCTTTAGTGGCAAGACTCCATTTTTAATTCTGCATATTAAACTTTGTAGTTTGGTGACACCCAACCAAATCAAGACTCCATTTTTAGCTCTGCATTTTGAACTTTCTAGTTTCCCATTCATCAATGGTGACACCCAACCAAATCAAGAATCCATTTTTGCAGACCCCTTTAGTCTCAAAACTCCATTTTTAACTCTGCACTTTGGACTTTCTTGCTTCCCATTCATCCATGGTGACACCCAACCAAATCAAGAATCCATTTTTGCAGAGACCCCTTTAGTCTCAAGACTCCATTTTTAGCTCTGCATTAGATCCCTTTAGTGTTAAGACTCCATTTTTAGCTCTGCAGTAGATCCCTTTAGAGTCAAGACTCCATTTTTATCTCTGTAAGTTGAGTATTTTAGTGTCAAGACTCCATTTTTAGCTCTGTATTGAGCATTTCAGTGCTCAAAAATTGTCAAGTTCAATGAAAATCAAAAACCCCTTTTTCCTCTAGAATTGAACAACTTAATGCTCAATCATTCAACAATGGAAATGTCAATTTCAATGGAAATCAAGAACCCAGTTTTGTTCTTAAGGTACTGTTCTAGTCACAAATTCTTGTTTTTCATTTTCTCAATATTTTCCATAGTTTTCCCACTTGTAAGATCAGATGATGCTGAAGCTCTTTTAGCATTGAAATCCACAATAGACCCCTCAAATTTTCTTGATTGGAAAAAAGGAACTGATTTTTGCCAATGGCAAGGTGTTAAAGAATGCATGAAAGGTAGAGTTACTAAGCTTGTTGTTGAAAATCTCAATTTGAGTGGTTTATTGGACCAAAATGCTCTCAATCAACTAGATCAATTACGAGTTTTAAGCCTCAAAGAAAACTCCATTTCCGGCCAAATCCCAAACCTTGCCGAACTTTCCAATCTGAAATCCCTTTTCCTCAACAACAACAACTTTTCCGGTGAGTTTCCGGCTACCATTACCCTCCTCCACCGTATCAAAGTTGTGGTTCTCGCCGGAAACAAACTCTCCGGTGACATTCCTCCGGCCTTAACAAATCTATCCCGGTTGTATGTTCTTTATCTAGAAGACAATCACTTCACCGGAGAAATCCCACCGTTGAATCAGCTCAGTTTAAGGTTCTTTAATGTGTCAAAGAATCAACTTTCTGGTGAAATCCCGAATACTCCGGCGTTATCTCGGTTCAATGAATCATCTTTCGCCGGAAATGTCAAATTGTGCGGAGTACAACTGCATAATAACTGTCCTTCAACTAGTCCATCGTATCCCATCATTCCAGAAGATGTATCTCATCATAAAAAAAGCAAAAAGCTTATCATCATCATTGTCCCAGTGGTCGCTGGAATCTTGATCCTCTGTGTTTTGATGGCGATTTTTATAGCATGTGTTCGAAAACGAAGAAGAAAAACTCAAGATGGAAAGAGTAAAGGAGTTGAAGCCGGTGGAGGCGGGGGAGGGGGAGGGGAAGAAGAGGGGACTAGTCGCGGTGGCAATGGCAGTGGTGGTGGTGGTGGCAGGGATGGTAATAATGGTGGAAAAGAAGGGGCATTTTCATGGGATCAAGGAAGTGGAGGACTAGGGACATTGGTGTTTTGTGGACCAGGTGATCAACAAATGAATTACACCTTAGAGGATTTGTTGAAAGCATCAGCCGAGACATTAGGGAGGGGTATAATAGGAAGTACATATAAAGCCGTGATGGAATCCGGCTACATTGTGACGGTGAAGCGGTTGAGGGATTCGAGGTTTCCGAGGTTAGAGGAATTCCGGAGGCATGTGGAGATTATTGGTAGGCTAAGACATCCTAACTTGGTCCCACTCCGAGCATATTTTCAAGCTAAGGAAGAACGTTTGCTCGTTTATGATTATTTCCCTAATGGCAGCCTCTTCTCCCTCGTTCATGGTATGTATTCCGTCATTCATTTATTTATTTATAGTATATTTTGCTTTTATCCGTTTTTGTGTACTGAACTCTCGTGAAGGAGGAAAAAACAGGTATAGTTGAACTCATGAATAGTGAAAGGAAGTTAAAAAGGGATAGATTTGGCGGAGTGAACTTGCACTTGAGTGGTATAAGGTGGAATAAGATAAGGGTGAGGTAACTTATTTACGTCTAGTATATGCATCGATTCAATAGATGCATTACATGTGTTTTTTTACTAAATTAGTTATCGAATAGTTTATCCTCTCTTGAGGTTAAACTAGTTTAGTTTTGTGCAAATACCAAGTGGGGACAAATTCTTTTTCATGAGATCACTAGTTTATAACTCAATCTCTCGAAGATTCGGCTCCTTTATTGACATGAGTCATTTTCTTATTTTGAATTCAAATTTGTTACCTCTTTATGCTTGGCTGTTTTGTGGTGGAGCATTGGAGGACTTTAGTGTCACTTTATTTTCCTATTTTTTTGTGTTTATATTTGTGTACTTTGCACATTTTCCTTGAATTGTTTATACTTTTTAATATGTTTCTTTATCTTCTTCAGTATTCTGTCCCTTCTTATTTTGTTTTTTTGTTATTTTTGAATAATTTCCATATAATTGGTTTGGTAGTGTATCACTTTCTGGTCTCTCCTGTGGGTTCTGTTTGGTGATGGTGGGGAAGAGAGGGCCAACAAAGGAAGCTTGTAGGAGCCAACAAAGAATTGAGATAGCTGGCCAAGTAGGACAAATATGTAAATTTGTAAAGGAAGTTCCAAACTTGCTTTGGTATACTGATGTGAAGATTATAGAAAAATATTGTTTAAATAATTAAGTTAAAAAAAATGAATAATCAAATAAGTAAACTCTTCTCATCTTGTTCTCTACGACTACGGATGCTATACGAAAAGAAGATAAGGAACAAAAGGGGAGTATAGTTATATATTGGCCTTTTGGAATCTTACAAAGGATGATATTTCGTTCAAATGAATTGTGAAAAAAGATGTTTAGTGTGTTTTTGCATTATTTTGTTGTTGTTACTGTTCCTTTTATGAATGCTTTGTGCTTTCCTTATTATTTGCCATATTTCTTCAATGTCATTTTCCCTTTCATAACTACTTTGTTTAGCTGCACAAGAGCCTAGGATCCTTCGGAAACAACCTTTCTATCTCAACGAGGTAAGGTCGCGTATACTCTACCCTCTCTAGATCCCACTTGTGGGATTTCACTTTGTTGTTGTTGGATTGTGAAAAAGATGGCCCTCTGGGAGGAGGAAACTCAAATTAGGAATCAAAACAATGATGTCACCGCTACACTCGCATATTTTGGCCTTATCATTTATTGCATAGAGCCAAAAAAGTTGTCCACCATATATGGAGATAAGGCATGATACTTTGAAATTAGAAAGACCCTGCATATTGCTCAAATTTGCATACGTAAGAGCTTCTTTGCCTTGTGCTGATTTCAGTACTTCTCTGGTAAGGTGTTAATCGTTAAATTGACCAGTGAGCTTCCAAAGATACCTGGAAAAATCAACATTTTCCTTCTGATTCAAAAAGAACAACAACAACAACGTACCTAGTGTAGTCCCACAACTACGGTCTGGGGAGGGTGGTGTGTACGCAGAAAAACAGAATTTTTCTGACACTAGTCACTAGCGATTAGAAGTTCCATTGCTATTTATAAAGTTTCATAATTGTCTGTACTATTTAGCTTTTAACATCTGCTCAACATCTCTTGTTCTCTCCACATTATATCTTGTGATATAGTTTGAGATGTTCTAAATTTGATTTGTTCCTAGTAAACTGAAGTTGCACCTACGTTGTATTCCAGAGTGTGTTCAATATGATATTTACTTTTCTTCTATTTAGGGAAACTAGAATGCAATACACCATTGACAAGAACAAAAAAGTTCAATATCATATAAAACTATTGTCAACCGACATGTTGAACTCCGAGATCCCTACATGTAAGTATAGCAAATTGGGTAATAATCAAGAATAAGGTAGTCTAGATGCTAGTCTTTTCTTCCAAGTAAAGTTGTGGTGATGCAGTGAAAGGCATTGAAAATGAGTTGCTTTGTTAGGCCATTAGACCATTTCAGGCACTAAATCACCTATGGCGGAATCACAGAGCAAAAGGCACACCGAAGTTTATTTTTGGAATAAGGACAACGTGGAAGGTGCTATGCAACAACAACAACATACTCGGTGTAATCACACAAAGTGGGGTAAAGAGGTGGTTACAATAGACCCTCGGTTCGAGGAAAAAAAATTCAAAGCAGTTCGGATGGAAGGTGATTTATGCGTCTAGCAATTGTATACGCCGTATGCAAGTACCACCTCAAATAGTGTGTTACCTTTCATGTGGTCTTATCTTGTTACTAGTCATTGTAAGACCACATCTGATGTTTAATACATGGGGGTGCCAATCGAATATTTGAAGAGAACATATTCAAAGTTAGCCGGGACTCTCTTATTTTCATGTATCTGGACAACCAATATGTTTTGTTATTCAAAAAATATTCCAGATGATAACTAAGCTATATTTTTACAATTTGCTAACTTTTTCAGGGACAAGGGCTAATAGTGGTTCAAAGCCTCTTCACTGGACTTCCTGTCTCAAAATTGCAGAGGATGTGGCTACTGGATTGCTTCATATCCATCAGAACCCTGGTCTGACTCATGGAAACTTGAAATCATCAAATGTACTCTTGGGGGCCGACTTTGAGTCGTGTCTTACAGATTATGGCCTCATGCCATTTAGGAATCTAGATTCTCCTGAGGAATCAGGTGCTTCTTCCCTCTTCTATAGAGCTCCTGAGTGTCGTGACATTCGAAGGCCACTTACCCATCAGGCCGATGTCTATAGCTTTGGTGTCCTCCTATTGGAGCTCCTCACGGGAAAGACTCCATTTCAAGATCTTGTTCAAGAACACGGATCAGACATACCGAGGTGGGTGAAATCTGTTCGAGAAGAAGAAACTGAATCAGGTGATGAACCTGTTTCTAGTAATGAAGCCTCGGAAGAGAAACTCGGTGCTCTCTTAAACGTAGCTATGGCTTGTGTTTCACTTGTACCCGAGAATCGACCCACAATGAAGGATGTTTTGAGGATGATCCGGGATGCTAGAGCAGAAGCTCAAGGGTCCTCCAATAGCAGTGACCATTCACCAGGAAGATGGTCAGATACTGTCCAAAGCTTGCCAAGAGATGAACATTTGAGCATATGAGTTTGTACTATACTTATGGAGGCCCCGAATAACGTGTCTTGATTCTTTTGGAAGGCAGAAGTTACTTCGTAAGTTAAGCATCATCGATGTATCTGAGTTCTCAGAGATTTGTTGATTCAGTTCTTGTAAAGTCTCAAGGGCTTTCTACTTTTCTTGTTTTTCTAGGTTTAGTTGTTTGTATGTAGGAGATTTGCTTTTATACCACTTGGATTGACATTCACTAGTTGATGATACATTCCTGATTCAAAAGTTGTCATTGTGTTTTGTGAGAGATGTAATTGTAATTGAGGTTCCAACACTTCTTTAACATAGTTTCTTATTCTAGTGGTAAGAGTGTATCACGTGATGTGTGGATTAGGAGTACGACAACCATTCAAACCTTGCCCTAGACAAAAATATGATTGCTACTTTCTTGACTTGGGGCATTACATTTCGG

>PGSC0003DMG400026792 LRR-RLKs

CGGCGTTACACTCTCTCTTCACTCCAAAACTTGTCTAATTTTGACTATACAAAATCATTTGAATCCACCTCCTCAACAAAGAACCAGCCTTGATGAACTTTTCTCTCTCACTACTCACTGTCTTTCGTCACTCTCTTCTTCTAAAACTGTCCTCCCCCTTTTAAGTTTCTCTCTTTCTTGATTCCAACTTGTCTGCTTAAACACCCTTTTTCTTCCTCTCCATCACACACACATTCTTGCAGAATACCCTTTTTCTTCTACTCTATCTTCAACCACTGACAATGTTTCTTTTATTCACTCAATCTATTCTTTAACCTTTATGTTTTGTTTTCTTATATATGCAATTTACATATCTCTTCACCATATATTTCATTTTTGTTGGAGACAAAATGTTGTGGCATATAATAACTGTTTTATCTTTAGCTCCATTTCTGGTACACTCTTTAGACCCAGCATTCAATGATGATGTAATGGGTTTAATTGTTTTGAAAGCTGGATTCACTGACCCTAAATCTAAACTTGCATCTTGGACAGAAGATGATCCTACTCCTTGTAATTGGGTTGGTATAAATTGTTATCCTCAATCCAATAGAGTTTCTGAGATTTTACTTGATAATTTCTCTCTTTCTGGACATATTGGGAGAAGTCTCTTGAGGTTACAATTCTTAAAGGTTTTGTCTTTGTCTAACAATAACTTTACTGGCAATATCAATCCCATTTTATCCCAAATACCTAGCTTGAGGGTTATTGATTTGAGTCACAACACTTTGTCAGGTTCAATCCCTGATGAATTTTTTCAACAATGTAGCTCTTTACAATCTGTTTCTTTTGCAAACAACAATCTTACTGGTGAAATCCCTCAATCTTTAACATCTTGTTCAACGCTCCAAAGGGTTAACTTTTCATCCAACCGCCTTTCGGGTCCGTTGCCTTCAGCTTTATGGTCCTTAACCTCTCTTCAATCTCTTGATGTTTCTGATAATTTAATGGAGGGTGAGATTCCTAAAGCAATTGAAGGTTTATATTCTTTGAGGTCAATTAATTTACATAAAAACAAGTTTACTGGTTGGCTGCCTGAAAATATTGGAAACTGTGTGCAGCTCAAGTCAATTGATCTCAGTGACAATTTGTTGAATGGGGAACTTCCTGAGTCTATGAGAAGATTGGGTTTTTGTACCAATATGGAGTTAAGATCAAATTTGCTTAATGGGCAGATTCCAGATTGGATAGCAGAGATGAAAAGCCTCAAGGTTTTGGATCTTTCTGCGAATAATCTTTCTGGTAGGATTCCAACCTCTATGGGAGATGTTTCATTGCTAAAGGAACTCAATTTGTCAAACAATTACTTCGTTGGAAGCTTGCCTCGGTCCTTGATGAAATGCAATGACCTTGTAATCTTGGACATAGGCAACAACTTCTTGACTGGTAACCTTCCTTCTTGGACCTTTGAATTGGGTGTAGAGAGTATTTCTCTTTCGGGGAATAGGTTTACCGGGCATATAGATTATCCGTCCATATCCATGGATGCTTCTTATCGAAGCCTTCAAGTTTTGGATTTGTCTTCCAATGCATTATCTGGTGAAATACCAGCTGCCATTTGGAATATCAGTAGCTTGCAGGTCTTGAATATTTCCAGGAACTTTTTGTCTGGTACTATTCCAGAAGCTGTAGGTAAACTAAACGCAACTCGAATTCTTGATTTAAGTCACAATCAACTAAACGGAAGTATTCCCAGTGAAATTGGGAGTGCTGTTTCATTGCTGGAACTGAAGCTGAGAGAAAATCATTTGAGTGGTACAATTCCTGCAGATATAGCGAATTGCTCTGCTCTAAGTTCATTGTAAGTTTCCTGCCACTTCCATCTCATTTTTATTCAATGTGTTGAAAACTTTTGTATTTAGCTTTTCGATCATGTTATCTTCGTTTAAATTGCAGTTATGTGCATCATGATGTTTGCATTTGTTATGTCTGTGTGATTCTAAGAGTCATTTGTTCTGCATTTTCGATTGTGTAGGGACTTGTCACACAACAACCTCACTGGCCCTATCCCTCCAGAGATTGCTAAGTTGACCATTCTTGAAGTGGTAGATTTCTCCTTTAACCAATTTTCTGGAAGCTTACCTAAAGAACTAACCAATCTTTCGCACCTTGCCACATTTAATGTCTCCCACAACCATCTCAAAGGTGAACTTCCAGTAGGTGGCTTTTTCAACACCATTTCTCCATCATCTGTTGTTGGTAATCCATCCCTATGTGGTTCTGTTCTCAACCACTCCTGCCCTGCTGTCCATCCAAAGCCTCTTGTACTAAACCCCAATTCATCTGACCCGAATCATGCTTCTGTTACTTCGCTTGGTCATAAGAGAATCATGCTCAGTATATCATCTCTCATTGCCATTGGAGCAGCAGTTTTCATAGCACTTGGAGTAGTGGTCGTTTCTATTCTCAATTTGCATGTGCGCTCTTCTATGGCACTCTCTGCTGCAACTTTCACCTTATCTGGTGGCGATGACTTTAGCCATTCACATGGTACAGAAGCCAATCTAGGCAAGCTTGTTATGTTTTCTGGTGATGCAGATTTTGTTGTTGGAACTCAAGCGCTGCTCAACAAGGATAATGAACTTGGACGTGGTGGTTTTGGATCTGTATACAAGACAGAACTCGGAGATGGACGTTCTGTTGCCATTAAGAAGCTCAACATTACAAGTTTGATCAAGTCCCAAGAAGATTTTGAGAGGGAAATGAAAAGTCTCGGAAGCATCAGGCACGAGAACCTTGTGGCACTTGAAGGTTATTATTGGACACCTTCTCTCCAGCTCCTGATTAATGAATATGTATCTGGAGGCAGCTTGTACAAATTACTCCATGAGGAAAGTTCCAAAAGGAGTCTCTCTTGGCAGCAAAGGTTCAACATTATTCTTGATACTGCGAAAGGCTTAGCCTATCTGCACCAGCTGAACATAATTCATTACAATATGAAGTCAACCAACGTCCTCATAGATGATGGTTCTGCCAGCACAAAAGTTGGAGATTTTGGTTTGGCTCGTTTATTACCAGTCTTGGATCGTTACATATTGAGCAGCAAGATTCAGAGTGCACTTGGATACATGGCTCCTGAATTTGCATGCCAGACGGTGAAGATAACTGAGAAATGTGATGTCTATGGATTTGGCATCTTGATTCTTGAGGTGGTGACAGGGAAAAGACCCATGGAGTACATGGAAGATGACGTGATTGTATTATGTGACATGGTGCGGGGAGCACTAGAAGAAGGCAGGATAGAAGAATGCATTGACGGACGGCTCCAAGGTAACTTCCCTGTCGAAGAGGCAATTCCAGTTGTGAAACTTGGTTTAATATGTGCATCTCAAGTTCCATCCAATCGACCAGACATGGAAGAGGTGATCAAAATTTTGGAGCTAATTCGATGTCCGTCAGAAAGTCCAGAGGAAATAGAGTAATGTTATAAGGTGTTTGTGGAAAACCAGATTATAGAGAATGAGTCATGTGGTGAACAGGAATTCAGTTTTGTTTGTTACATTTGGAGAAGGTCACTATCCTCTTGTTATTTTGTCATATCCATACATGTTAAATCTTTATATGTTATTTAAATAACTGGATTCCTCTTCACCAGCTCCAGTTTGGGACCCTTGTTAGATTTGGATAGACTGGCATTGTCTTTATATAGTATTCAGACATCCAACTTGTTGTCTTGTGCTTGTTTGATGTTGTTGTCCATACAAAGCTGAAAAAAATACATCAGTTGTATTCTTATAGAAGAAGACCCACATGGATTCCTAATGAAATGAAAACATCAAAGTAGCCCACCACTCA

>PGSC0003DMG400027071 LRR-RLKs

CAACCATTTTTGATTCATATAAACCACCAATACACACTTATAAAAAAATCCATCAACCACCACAGCCTGGTTGCCTCATTCACCTCTCTCTTCCCTCCGTCCGGCCTACCGGAAAAACAAGAATCTCAATGGCTTCAAATTTCCACTTTTTTCTCTTATACCTTGTGACCCTTTTCCTTTTTGCTCAAGCAAGACTGAATCTTTATCCACAAGATCATGCTGCACTTTTGCTTGTTCAAAAAGACTTAGGCATCATTGCTCTTAACAACCCCTGTACGCTCGCAGGAATATCCTGCGAGCGTAGACCGGGTAACAGAACACAAGTTCTGAGAGTTACCCGTATTGTATTCAGATCCAATGGATTGAAGGGAACTTTGTCTTCTGCTATTGGCAAACTCTCTGAGCTCAAAGAGCTTTCTCTTTCCGACAATCAACTATCTGAACAAATCCCAATTCAGATTCTTGATTGTCGGAAATTAGAGATTCTTGAACTTCAAAGAAACAGATTTTCTGGAAAGATTCCGTATGAATTGTCATCTTTAAACCGTTTAAGGGTAGTTGACTTTTCATCGAATGAGTTTTCTGGGAATCTTGATTTCTTGAAATACTTTCCTAATTTGGAAAAACTGTCTCTGGCTGATAATATGTTCACTGGAAAAATACCCTTTTCATTGAAATCTTTCAGGAATCTTCGTTTCCTCAACATTTCAGGGAATAGTTTCCTTGAAGGTCCAGTACCTGTCATGAGTCAAATTGAGCATTTATCAGCAGATTTGAATCGAAACAATGCCGTTCCCAAACGTTACATTCTTGCTGAGAATTCAACAAGGTCAAATCAGATATCTGCAATGGTGCCTGCTCCAGCTCCAGCACCGGTCAATCGTGTTGTACCGGTGGTACATAAACGTAACAAGAAGAAAAGGAAGTTAAGATCTTGGTTTCTTGGTTTCCTAGCTGGAACTTTTGCTGGGGGTATATCTGCTGTGATCTTTTCGTTGCTATTCAAGCTCGTAATGTTCTTCGTAAGAAGGGGAAAGAACGATAGTTCAAGTTTAACGATATTTAGTCCGTTGATCAAGAAAGCGGAGGACTTGGCTTTCTTAGAGATGGAAGATGGAGTGGCATCACTTGAAATGATTGGAAAAGGTGGATGTGGAGAAGTTTATAGAGCCGAGTTACCGGGGAGTAATGGGAAGATTATAGCTATAAAGAAGATTATACAATCCCCAATGGATGCTGCAGAGATCACGGAGGAAGATACTAAGGCGTTGAACAAGAAAATGCGTCAAGTTAAATCAGAAATTCAAATTGTAGGTCAAATCAGACACCGGAATCTGCTTCCATTACTGGCGCATATGCCAAGGCCAGACTGTCATTACTTGGTGTATGAATATATGAAAAATGGGAGTTTACAGGATATCCTGCAGCAAGTCACAGAAGGCACAAGAGAATTAGATTGGTTGGGACGACACAGAATTGCAGCGGGGGTAGCTGCTGGTCTTGAGTATCTCCATATAAACCATACTCAACGCATAATTCACAGAGATCTAAAGCCAGCAAATATCCTACTTGATGATGACATGGAAGCTCGAATAGCTGATTTTGGGCTTGCAAAGGCAGTTCCAGATGCTCATACACATATTACGACTTCAAATGTTGCAGGAACTATGGGATATATTGCACCAGAATATTATCAGACACTGAAGTTTACAGACAAGTGTGATATATACAGCTTCGGTGTGGTGCTAGCTGTGTTGGTTATCGGAAAGGGTCCATCGGATGAATATTTCCAACATACTTCTGAGATGAGTTTAGTTAAGTGGCTGAGAAATGTAATGACTTCTGATGATCCTAAAATAGCAATTGATCCTAAGCTGAGAGGGAATGGATATGAGGAGCAAATGCTTTTGGTTCTCAAGATAGCTTGCTTTTGTACTCTCGACAATCCAAAGGAGAGGCCTAACAGTAAGGATGTTAGGTGCATGTTAACTCAGATCAAGCATTAGCTAAGTCATACGGGGTATACGTTGGTTATGTAACTAAGAATGTGTTCTGTATTTCAGTTTTTGTTGTAATTGTGTTATACATGTCAAATCTTGTATGTTTACTGCAGTGTTAATAATATTATTTCCACTTCGCTACTTAAATTACTAAAACTCTA

>PGSC0003DMG400027115 LRR-RLKs

TTCTTGTGGTGTGTGTAAAAAAAACACTGTCCCTATCCCATTTTTATACTTATCAAAAAGCAAAAAAAACAGGGATGTCAACTGAAAAATCCCATTTTTTCTAACAAAAAAATGGAAAATCAAGACCAAGTTCAAGACTTTACCTGTTTTTACCAGCACCCATTAGGGGTTTTCATGTTTTTAATCCAAGAAAGAAATGTGGGGTGTGTTAAAAAGAAGGAAAAATGGATACCATTTATCCAAATTTTGTTGTTTTTGCAGCTGATGATTTGTGGAGCTCAGTTAGCTTTAGGTCAAGTTTGGGATGGTGTAGTTATCACTGAGGCAAATTATCAAGCACTTGAGGCGCTTAAACATGAGTTAGTTGATCCCAGAGGGTATTTAAGTAGCTGGAATGATAGTGGTTATGGAGCTTGTTCTGGTACTTGGGTTGGCATAAAATGTGCTCAAGGACAAGTTATTGTTATTCAGCTTCCGTTAAAAGGATTTGGGGGAAGAATCACTGAAAGTATTGGTCAATTTCAAGCACTCAGAAAACTTAGTTTTCATGATAATGAAATTGGTGGTTCAGTGCCTTCATCTTTGGGTTTTTTACCCAATCTAAGAGGTGTTCAATTGTACAATAACAGGTTTTCTGGTTCTATTCCTGGTTCACTTGGTTTATGTCCAGCTTTACAAACACTTGAACTTAGTAACAATTCTTTATCTGGTGCAATCCCTGATAGTCTTGTGAATTCATCTAAGCTTTATAGACTTAATCTTAGTTACAATTTATTGTCTGGTTCCATTCCTGTTAGTATAACTCAATCGCGCTCTCTAGTCTTTATTGACTTGAAATATAACAACTTGACAGGGTCTATTCCGGATTCTTGGGGTGGAAATGGAGATAGACAGTTTAAATTAGAGTCGTTGACGCTTGATCATAACTCTTTGTCTGGTGGAATTCCTGTTTCTTTTGGGAAGTTGAGTGAGCTTCTTGAGATTTCGTTGAGTCATAATCGTATCGTTGGTGTCATCCCAAATGATATTGGAAGGCTTAGTATGGTTAGGAATCTTGATTTTTCTTATAATGAGATTAATGGTAGTTTGCCAGAGAGTATAACCAATCTGTCCTCCCTCATGGTTCTCAACTTGGAAAGTAACAATCTTGATGGTGAAATCCCGTTGGATGTAAAGAAGTTGCAAAAGTTGTCGTTTCTTAACCTGAGGAATAACCGGTTTAGAGGTGATATTCCAGCTGCTATCGGGGACATTTCTGGTCTAGTCATGATAGATTTGTCTCGTAACAATCTAACCGGAGAAATCCCAGAGTCTGTTAGTGAGTTACCTAATCTTAGTTCCTTTAATGTTTCATATAACTCTCTGTCTGGTCCTGTTCCTACTTATCTTGCTAAGAAATTCAATTCAAGTGTGTTTGTGGGTAATGTTCAGTTATGTGGCTATAATACTTTGAATCCATGTCCTGTTACGCCTTCTCCAGTTACACGAGGTAGTAAACGAAGTATTAAAGACATTATTCTCATAGTAGTAGGGTGTCTTATAATAGTCTTGTTTCTACTCTGTTGCATTCTTCTCTGCTGTTTGATCAAGAAAAGAGGTGTGGCCAAAAAAGCAAAAGATGTAAAGGATGTTCCTCCAACTGCCGGAGAAGTTGAGGCGGCTGGGGATAACGGAGGAAAACTTGTCCATTTTGGTGGAGGTATGATGTTCAGTGCAGATGATCTTCTATGTGCTACTGCAGATATATTGGGAAAGAGTACTTACGGTACGGTGTATAGGGCTACATTAGAAGATGGAAGTCAAGTTGCAGTGAAAAGAATGAGAGAAAAGATCACGAAAGTTCAGAGGGAATTCGAGAAAGAAGTCAACGTTCTTGGAAAGATAAGGCATCCGAATCTTCTGGCTATTAGGTCTTATTACTTAGGTACGAAAGGCGAGAAGCTTCTTATTTTCGACTACATGCCTAAAGGAAGTCTCTCAAATTTTCTTCATGGTAAGCAGAAAACAGATTTCCATTTGACATTCTTTTGCTTCTGTTTTTTTGAGAATCAGATACTTTCTCTGTCTCATTTTAACGTAATGAAAATTGGATTATGTATCATATCGGAAACAACTTCTCTAACTCACACAACGTAGGTAAGGTCTACGTACACTCTACACTCCCCACTCCCCAGACCTCACTTGTGAGATCGCATGGGGAGGAAATTGTTATGGAGGGGGTAATATAGCTTTATTGTTTTTGTTGCAGCTCGTGGACCGGATACACCAATTGATTGGCCAACAAGGATGAGAATAGCAAAAGGGACGACGAGGGGATTGCTGTACCTTCATACTGATGCCAACATCATTCATGGGAATCTTACATCAAGCAATGTTCTACTTGATGAAAACACAGATGCAATAATTGCAGATTATGGCCTTTCACGCCTTGTAGCTGCTGCTGCAAATTCAAATGTTATGGCCACAGCAGGGGCACTTGGTTATCGTGCACCTGAGCTTAATAAGCTCAAGAAAGCCAACGTGAAGACGGATGTCTACAGCCTCGGAGTCATCATACTTGAACTCCTAACTGGAAAGTCTCCAGGTGAGGCAGTGAATGGTGTAGATTTGCCTAGAAAGGTGGCGTCAATTGTGAAAGAGGAGGGAACAAAAGAGGTATTCGATTTGGAACTAATGAGGGATGCATCCATAATTGGTGATGAATTGTTAGTTACACTAAAATTGGCTTTGCACTGTGTTGATCCCTCGCCATCAGCTCGACCAGAACTTCAGCAAATTCTCCAGCAACTGGAAGACATTAGGCGTGAAACGCCAACTGCAGGTCCCTCGACAAGTTACTAGATACTCAGTTTTTGATTTTATTAACATGATTCTTTCATTCTTGAATTTCATTGCTAGAGTACTAGATCTGTTTCAAGTGTCTTTAAATAAAAGTGTGCAACATTGTGAAAAGTGTTTGAGCTTAATGGATTCCTAAATACCATCCCCTTTTGGCCTAAAGAACATAACTTGAGATTGAG

>PGSC0003DMG400027467 LRR-RLKs

ATGGGACTAATAGGTGAGTTTCCGCGAGGCATACAAAATTGCACTTCTTTAACTTCTCTGAATCTTTCAAACAACGAGCTATATGGAACCATCCCTTCCGATATATCAAAACTAATCGTATATGCTGTAATCGTTGATCTCTCGAGCAACCTATTCTCAGGCGCGATCCCAAGTGACATTTCAAATTTCTCTTTTCTCAATTCCCTCAAATTAGAAAACAATAAACTAGAAGGACTGATCCCACCTGAAATTCGCTTGTTACAGCGACTCAAGACGTTTAATGTTGCAAATAATATGTTGACAGGGCCAGTGCCAAATTTTGTCAATGCTACTTTTCCAGATGAAAGTTATGCAAATAATCCAGGACTTTGTGGTAGTCCTCTGGAATTATGTGTTACAGAACGCGTAGTAAGCCGTCATATTCTGTTTGCTTGTGGTTTTGTGACGGGTTGGTCGCTTTCTACATTACTTGGTGTATATCTTTTTTTCTTTGGTGTAAAGAAGATGCTTCTATTGATCAACAAGAGAACAAAGGTAATGGTGATTGATGGAAATGAAGTAAACAATGACCCAAAGGTAATTATGTCTACCTGCTCAATTATATACAAGTTTAAGTATCTATTTGTGCACATTCAAAGTCGGAGGGCATAGATTTTAGATGAGGCCAAGTTAAAGGGAATGATTATATTATGCCAATTTGGAAACAATTTCACTTAGTGACCAGCTTTTATAGCCAGACAGTGTTATGGCACGTTTAAACTACAAGTTTAAAAAAACTACCTTTTTTTTTTTGAAAATTTGGGCCCTGTCAAACTATGTTAGATAAATTGCGACGAAAGGGGAACTACTTAATAAAAAAACTGTTGTTGTAGAGGAGAAATAAAATGTGGTAATGGTAAATATGTTTTTTTCCTCAGATTTTAAAGCTGGAGAAGATTGTTACAAGAATGAGTTTTATGGAACTATCAAAAGCGACTTTGAATTTCAGCCAAGACAATGAAATCGGGAACGGAATGCTGGGGAAATTGTACAAAGCACTAGTCCCAAATGGATGGACTGTTGCCATAAAGAGGCTCTAAAGTTCCGAGGATTTGGAGGAAGAGTTTGTTTCGGAGATCACAACTCTAGGTATTTTGAGGCATCCGAATTTAGTGCCCCTAATCGGATTTTGTTATGAGAGGGACAAGCGACTTCTGGTTTACAAATACATGCCTAATGGAAACTTACATGAGTGGCTGCATTCAACTGACGACAAGGCAAGACTCTTGGACTTCCCTCTAAGGGTTAAAATTGTTGTTGGTGTCGCGAAAGCCCTGGCTTGGCTTCATGATGGTGGAAATTTCGATCCCAAGTTATCCAATTTTTGGGAGGCAACATTTGTAGAACCAAATGACATAGGTTCAAACTTGAGTCTATCGCCAATGGTCGAGTTTTTAGACTTTACAACATACAAAAAAGATGTCTATAAATTTGGGGTTGTGCTTCTCGAGATTTTAACGGGGAAGGAATCTTACGAATTGAGCTGCTCGAGTCTAAATTTGTCGAGCAGCTCTTTTGCCAGTCCTCTTGATGTGGATAAACTACTGTTAGGCCAAGGATTTGATAATATGGTAATGCAACTACTTGAACTTGCAAGTAACTGCATGAAATTCATCCCGGATCAAAGACCAACGATGCAACAAGTGTATCAGATTGTTGCTGCGATTGCTCGAGTCCATGACCAAACAGGGGACTCTGAAATACAATTGCACTTGGACTAG

>PGSC0003DMG400027468 LRR-RLKs

TTTCAACAATGTGTAACTGCAACATTCTTTACTTAGTACTACCATTTTTCCTACTATTTTGCAGCTTTTCTCTAAGATGCAACGCGGTGCAGAGTGATATCGATTGCTTGAAATCGATTAAAGATTCGTTGATAGACCCTTTAAACATCTTGAATTCTACATGGAAGTTCGATAATCAGACAGAAGGTTTTATATGCAAATTTACAGGAATACAATGCTGGAATGCCGATGAGAACATAGTTGTATCCATTAGTCTTCCAGACATGGGACTAATAGGTGAGTTTCCGCGAGGCATACAAAATTGCACTTCTTTAACTTCTCTGAATCTTTCAAACAACGAGCTATATGGAACCATCCCTTCCGATATATCAAAACTAATCGCATTTGCTGTAGACATTGATCTCTCGAGCAACCTATTCTCAGGCGCTATCCCAAGTGACATTTCAAATTTCTCTTTTCTCAATTCCCTCAAATTAGAAAACAATAAACTAGAAGGACTGATCCCACCTGAAATTTGCTGGTTACAGCGACTCAGGATGTTTAATGTTGCAGATAATATGTTGACAGGGCCAGTGCCAAATTTTGTCAATGCTGCTTTTCCAGCTGAAAGTTATGCAAATAATCCAGGACTTTGTGGTAGTCCTCTGGAATTATGTGTTACAGAACGCGTAGTAAGCCGTCATATTCTGTTTGCTTGTGGTTTTGTGACGGGTTGGTCGCTTTCTACATTACTTGGTGTATATTTTTTTTTTGGTGTAAAGAAGATGTTTCTCTTGATCAACAAGAGAACAAAGGTAATGGTGATTGATGGAAATGAAGTAAACAGTGACCCAAAGGTAATTATGTCTATCTGTTCAATTATATACAAGTTTAAGTATCTATTTGTGCACATTTAAAGTCGGAGGGCATAGATTTCAGATGAGGCCAAGTCAAGTTAAAGGGCATGATTATATACTATGCTAATTTGGAAACAATTTCACTTAGTGACAAGCTTTTATAGCCAGACAAGTGTTATGGCACGTTTAAACTACAAGTTTAAAAAACTACTTTTTTTTTCTGAAAATTTGTGCCCTGTCAAACTATGTTAGATAAATTGCGACGAAAGGGGAACTACTTAATCAAAAATTTTTATTGTAGAGGAGAAATAAAATGTGGTAATGGTAGATATGTTATTTTCCTCAGATTTTAAAGCTGGAGAAGATTGTTACAAGAATGAGTTTTATGGAACTATCAAAAGCGACTTTGAATTTCAGCCAAGACAATGAAATCGGGAACGGAATGCTGGGGAAATTGTACAAAGCACTGGTCCCAAATGGATGGACTGTTGCCATAAAGAGGCTCCAAAGTTCCGAAGATTTGGAGGAAGAGTTTGTTTCGGAGATCACAACTCTAGGTATCTTGAGGCATCCGAATTTAGTGCCCCTAGTCGGATTTTGTTATGAGAGGGACGAGCGACTTCTGGTTTACAAATACATGCCTAATGGAAACTTACATGAGTGGCTGCATTCAACTGACGACAAGGCACAATTCTTGGACTTCCCTCTAAGGGTAAAAATTGTTGTTAGTGTCGCGAAAGCCCTGGCTTGGCTTCATGATAGTGGAAATTTCCATGTCGTGCATAGTAACATTAGCACACAATGTATCTTGCTCGATGAAATCTTACATATTGAGTTGCTCGAGTCTAAATTTGTCGAGCAGCTCTTTTGCCAGTCCTCTTGATGTGGATAAACTACTGTTAGGCCAAGGATTTGATAATATGGTAATGCAACTACTTGAACTTGCAAGTAACTGCATGAAATTCATTCCTGATCAAAGGCCAACGATGCAACAAGTGTATCAGACTGTTGCTGCGATTGCTCGAGTCCATGACCAAACAGAGGACTCTGAAATACAATTGCACGTGGACTAGAAGCAGGGGCGGAGCCACCAAAGGGACGTGTAGCTTTGGCTCAAACCATGTATTTGTGATGAAGAACCATTTCATATGTATAAATAATCCATCTAGACTCATGTGCTACAAATTCTGGCTCCGCCTCATACTAGAATCACCAGCTTTTGGAACCGTGTTTCCTATGGTTTCTTTGTTTAGCAAGAATGTATGTCTTCCTGAATTGTTGTTTTCAATATTGCAATAACTATGCTTCAGAATGATCATCATTA

>PGSC0003DMG400027482 LRR-RLKs

ATTTGGTGCCTTTAATCGGCTTTTGTTCTGAGAGGGACGAGCGAATTCTGGTTTACAAATACATGCCTAATGGAAACTTACATGAGTGGTTGCATTCAACCGACGACAAGGCAAGACTCTTGGACTTTCCTCTAAGGGTTAAAATTGTTTTAGGTGTCGCGAAAGCCCTGACTTGGCTTCATGATGGTGGAAATTTCCATGTTGTGCATGGTAACATAAGCACACAATGTATCTTGCTCGATGAAAATTTCGATCCTAAGTTATCCAATTTTTGGGAGGCAACACTTGCAAAAACAAATGACATAGATTCAAACTTGAGTCTATTTCCAATAGTCGAGTGTTTAGACTTTACAACATATAAAAAAGATGTTTATCGATTTGGGGTTGTGCTTCTCGAGCTTTTAACGAGGAAAGAATCTTACATATTGAGCTGCTCGAGTCTAAATTTGTCGAGCAGCTCTTTTGCCAGTCCTCTTGATGTTGATAAAGTACTGTTAGGGCAAGGATTTGATAATATGGTAATGCAACTACTTGAACTTGCAAGTGATTGCATGAAATTTATTCCGGATCAAAGGCCAACGATGCAACAAGTGTATCAGACTGTTGCTGCGATTGCTCGAGTCCATGACCAAACAGGGGGTTCTGAAATACAATTGCACTTGGACTACAAGCAGGGGCGGAGCCACCAAAGGGACGTGTAGCTTTGGCTCAAACCAATGTATTTG

>PGSC0003DMG400027499 LRR-RLKs

GTGACTGTATTCTTTAAAAATGCATGCAGCTTCTTCAACCCCATCATTTTCTTGAAGTTTACAGCTTTAATTCCTCAAAGAAATCAGCAAAAAAATTGCATTTGCCATGGGAATTGCAAAAACAGTGCAGGGCTCATGTTGGCTTCTTGTATTCATGTTGTTTTTTCTGTTTAATTTATCAGATGAGTTCTTAGAACCTTCACAATCTGAAGCTATAGTGAAAATCAAGCAGCTTTTGAACTTCCCTCAAGATTTGAGTAGTTGGAGTGACAATACAGACTTTTGCAATAGTGAGCCAAACACTGCCTTGACACTCATGTGTTATGAAGATAACATAACACAGCTTCATATTTCTGGCTATAACTGGTTTCCTAATTTGCCTCAGGGGTTTTCTACAGATACCCTTTTCTCTAATCTTGCTCTTTTGCCTAATTTAAAGGTACTTTCTTTGGTTTCTTTAGGTTTGAGAGGAACTTTACCTAAAAAGATTGGATTTTTATCTTCTTTGGAGATTATTAATATCAGTTCAAACTTCTTTTATGGTGAGATTCCAGGAGAGATTTCATATTTGAAGAGTCTTCAGACACTTATATTGGATGATAATAAGTTTACAGGTCAAGTTCCTGAAGGGGTTGGTTTACTTCATTCTTTGTCTGTTTTGAGTTTCAAGAACAATTCATTTTCAGGTTCATTGCCTAATTCTTTGTCAAATTTACAAACCCTAAGAATCCTTTCTATTTCTGGTAATAATTTTTCTGGGGTTGTTCCAAACCTTCATAATCTGTCAAATCTACAAGTTCTTGATTTGGAAAGTAACAATCTTGGACCTAATTTCCCTAATATTCCAACCAAGTTGGTATCTTTGGTACTTAGGAAGAACAAGTTCAGTTTAGGGGTACCAAAAGAACTGAGTTCATGTTATCAGTTAAAGAAACTGGATATTTCTTCTAATGAGCTTGTTGGACCCTTTTCACCAACAATTTTATCATTGCCATCACTAAGCTATTTGGATATTTCTGGAAATAAACTAACTGGGAAGCTTCTAAAGAATGTGACATGTAGTCAAGACCTTTCTTTTGTGAATTTATCATCAAATTACTTGACAGGGGAGTTGCCTGATTGCTTGCACCCTAGTTCTGGTTCCAAGATTGTATTGTTTTCTGGGAATTGTCTGTCGAATAAAGAGCAATGGCAGCATCCTTATTCATTTTGCCATAATGAAGCTTTGGCTGTTAGTATTGAGCCTCATAAGGGAAAGGTTAAAGGGGGAAATGGTAAAGCTGTTCTTGCATCAAGTATGGTAGGAGGTTTTGTTGGAGTAGTAGCGATTGTCGGTTTGGCTTTCGTGGTTGTTAGAAGAGAGTATGCCAAGCAGAAAGCTAGCAAAGCCCCTCAAACAAGATTGATACTTGAAAAGGTCTCTCCTGCTCACACTCTTAAGCTGCTTAATGATGCAAGTAAGCTCTCTCTTTCTCTCACACAAACACATTTTGTTAATGATTTCAACTTGTGAAAGTTTTTAGATCCTGATAATTGTAAATCTAATGTTCTGTAAGGTCTATGTACACTCTACCTTCCCCTAATCCCGCCGGTGGGATTACACTGAGTATGTTGTTGTTGTTATTGTTGTCATTGTACTTGAAAAGGCTATAAACATTTGAATTTAAGCATTATTCCACAAAAGGAGTTTCTTTATATATAGATTGAAAATATCAATTTGCTTCAGGAAATTGTCCAATACTGTGTTATTGGAATAAACTCTGTTCTTGCCTAAATTTCAAGATGTATCTTGATCTTTGATTCTTGATAGGGTATATTTCTGAAACGAGGAAGTTGGGATTGCTTGGGGCCCCTCCTTATCGAACGTTTGTCTTGGATGAGCTTCGAGAGGCCACTAATAACTTTGATATATCAAATCTAATTGGCGCAAGCTCCAGTGGACAGGTATGGATTTCCTTACTAAATACTGTAAGACTGTTTGAACAATGTTAGTCCACATGGACTGTTTTGTAGTTCATTCCTCAATTCTCTTGCATCTATGGTGAATGCTGATTGTGCTTTTCTATACTCTGAACAATAGATTTACAAAGGGCGTCTCACAGATGGCACTGTGGTTGCTATAAAAAGCATAAAAATGAGGAAGAGGCATAGTGTCCAGTCCTACACTCACCAACTCGGGCGTATTTCAAAGATCAGATACTGCCACTTAGTTAGTACTATTGGACACTGCTTTGAGTGCTACCAAGATGACTCAAGTGTTAGCCGAATATGTTTAGTCTTTGAATTTGTGCCAAATGTGACGCTGCGGGGGGTCATATCAGGTATATAGTCTAGATAACTTCATCTGCTTCGCGAGTTATCATTTGGCACAACAACTGCAACAGTTTATAATTGAGCATTTAGTTTACTTATTTGACTATTTCTACAATATTACATAAATTATACTAAGTCTAAGTAAATGACTTCTTGGTATGTAGACGTTATAGAAAACAAGCATCTTTGCAACAATCTGTTCCATGTTGTTTTGATATATATTACTAGATTGTCCTCAATCACGACAAGTACCTACTACATTGTACTTCTTGTCAAAAGTGCTGATGAATTTTAAGGAGACATGAAATGCTTTTAATTACTTTCTCCTAGATTATGCAACGCGTTGTTTCTTCTTTGCCCGATGCTTAAATGTACAGGTGTTTATTGTCTTTAAGACAGTTGTTCTGAATTCTTTGCAAATGCTCGGTTTTGTAGGAGCAAATTCAGCTCAAAAGTTTACATGGACGCAACGGATGTCAGCTGCAATAGGGATAGCAAAGGGAATTCAGTTCCTTCATACTGGAATAGTACCAGGGATATTCTCCAATCAATTGAAGATTACAGATGTATTACTGGATCAGAATTTCCACGTAAAGATTAGCAAATACAATCTGCCTTTGTTAATTGAAAATAAGAAAATGGTATGCAAACCTCTCTTCTCTACTATATTCTCTATTAATCACTCATTTCATTCTTGAACATTCACATTTTGTTATTGCAACTGTTCTTAGGATACTGGACCGTCTTCTAGTGGATCAAAGGGAAATGATGGGCAAAGGTAAATGATCTCATCATTCTCAGCTGCTACTGCTGCATTTTCGCTCTTCATTACTCCACTTTTATGGTTCGATTTCAGAAGCACCTATTGGAAACTAATCTTGTTATACTTCTAAATGCAAATTGGTCTAAAAAGGTTTTAGAGAATCTTAAGTGTATCTCGACCAATTCTAGACAACACCTCTCCAGCCACTGCTATTTATTGGTTATGATTCTTGAAGACAATATATAGAGCCATAGACTTGGACCGTCTTAGGAGATTCCAAGATAGTGCTCTACCGTTTAGTAGACTACTAATGAATTCCTTCTTTTGTTAATATCTTCTTGAATGCAGGTTAAAATACGAGGAGAAAGATGATGTGTATGACTTTGGAGTAATCTTACTTGAAATCATATCGGGAAGGACAATCGATACAAAAAATGATATAGATGTTTCTAAAGACATTGTAAGAGTCACTTATGACATTTATGAGAAATTCTTTCATGTCATCAAGTTGAAGAACTATTTTGTTAAAACCTTTTCTTGTTGTTCTTATTGCAGCTAATAGTTAGCTTAACAGCTGACGAAATAGCTCGGAGGAACATCATTGATCCAGCTGTTCGGAAGGAATGCTCAGATAGTTCTCTTAGAACATTAATGGAACTATGCATAAAATGTCTTTCAGATGAACCATCGCAACGTCCTTCTGTAGAAGACTTGATATGGAACTTGCAGTTTGCAGCTCAGGTCCAAGACCCTTGGAATAGAGACACTTATGGCAACCAAGAGTCCCCTGGTCATGTGTAAGAACTTCCAATGATCAATGGACCTTTTCTTGTACAAAATGTAACATTATACCACTAGGATAGTAGTTCCAATTATTTAGTAGTCTCTATTGCAACACTTTTGGCTATATGAGGTTTCAGTACTCTTTGGAGGTTAAACTCTACATTTGAGCCAAAATTTGTGAGCAAATGGCTATTTCTTGCACTTTTTTTTTAACGAACCTAAAATTTAAAAGGTGGGCCTTTCACCGACCATCCAATGGACCATGGACTAAACGGCACCCAAAAATGTATGGGAAAATTTTCAAAATGTCAATATTTTAGCATTTAATGGGACCTCTTATCAACATTTTCAATATTTATT

>PGSC0003DMG400027536 LRR-RLKs

CTCTGCGCACACACAACTCAGTAGAGAGAGAAAGAGAAGCAGTACAAAAATGTGAAAAGTTGCAGCAGATAAAAAAAAAAAAGGGGGTTCATCAGAAGAAGGAGAAGAACTAAACTGAACTGAGAAATACAAAAAAGGGTTGTTCAGTGAAGAGTAAAAGGGGTAGCAAAATCTTGGTAAACAAGTGTAAAAATTTGATCTTTTTTTGTTGGGTTATGAGATGAGATCGATTCTTGGGTTTTGAGCTTTGTGGGGGTGTTTATCTTGGTTCATGGTAATGAGGGGTAGGGTTTTCTTGAATTTTTGGGTAAATGCTGGCTTTTTTGAGTTTGCTTTACTTGGATCTACGAGATTGTTTTGTCAGCTGGCTGCTTGATTTGTAAAAAAAAAAACAGTATTTTGATGGTGAAGGTGATGGAGAAGGATGCTGTGGTGGTGTCACTGGTGGTATGGCTAATCTTGGTTGTACATCATCTTAAGCTCATTTATGCTAATATGGAAGGTTTGTGGGACCATCATCCTCATTTAGTTAATTTTTAGGTCTATATTTGTTGTTCTATTTTGTGAGATGAGTTTCAAATGTCGAAACTTTTCGTACTAGAAATTAAATATTTGCTGAAAGTGAAATGAGTATGTGCAAAATTGGATTTTGAATTTGACTAATGTGTGAGTTGTTGAATTTTCCCAAAAGGGTAAGCAAGATGTGCAGTGAGCTCCCTGCATTGTTACGGTGGCAAGCAGTTTAGTTTTAATGTCAGTTATATGTTAAAGTTTTTAGTTATCATTGCCATTCAACCTTAGTGCATTATGTTTCCTATACAAGCTTGAATAATTTGATGTGATTTAATTTGGTTGTTGCCAAAATCTTGATAAACTGTTTTATTCAGTGAATAGATTAATTTTTTATGTGCATTTTGGTGGATTAAATATCGTTTATGTTTTCCTAAATGGATACACGAACAGGTGATGCATTGCATAGTCTACGCGTCAATTTACAAGATCCTAACAATGTGCTACAGAGCTGGGACCCTACCCTTGTTAATCCTTGCACATGGTTTCATGTTACCTGCAACAATGACAACAGTGTTATAAGAGTGTAAGAATCTGTTGTCTGGTTTACCTCCGTTTTAATGGATTTGCAATTCCACTCTCTTGTGGTGGTGAGCAAATCTAATTGCAGTTTGTGCTTCCATAACACTTGTTATATGATCAACCTTTCCAGTGATTTAGGAAATGCAGCTTTATCTGGTCTGTTAGTCCCACAGCTTGGCCTTTTGAAGAATTTGCAGTACTTGTAAGTCTCACTTCATGAACTATGTTTGGAATTATTTTACGAATTTAGACTTGGAAAACTGGCATTAAGTGTATTGGTTTTTCCAGATGTTGAAGATTTTTCATATTACCTCTAGAATTTGCTGCTGGAAATACCATAGATGGCTTGCATATTAACTTTCGCATAAAACAAAGCTGTTGTAAGTGGCAAAAGCGAGCAGAGAACACTTTCTCCTCTCCATCTCTTGTTTCACTTGGTGTTGTAGTCAATTGCATAGTTGTTAATGTGTTCCCTGAACAACACAAATTTGAATGTATACTTCATAAAAAAAGATTTAAATTTGTGTTTCACAATACATACTAGTAAATTAAGAAACTCGGATACTTGGATGAGACTAGCAGTGAATTAACTTATGTGGCAAAGTAGTGTCTGTAATTGCTTGTGAAAACATTGTAGTTGAACTTAAATTTCTGTCATACACTCCCCTATCCAAAGTAAGCATAGATTCTAAATACCATTACTAATGCCAGTACCCTAAGCACTCTTTATTCTTTTGGTTTCCCACCCGGTGTCCGGTACCCACATTGGAGCCTGACTAAATCCGGATTCGCGCTGGGAAGTCCCACATTGGGGGTAAAGCGCTCCCTAACAAAGGTGACTCCGTACCCAAGAGGGCTCGAACCCAAGACCTCTTGGTTAAGGATGAAGGAGTACTTACCACTCCACCACAACCCTGCTTGGTACCCTAAGCACTCTTAAACAAACTTCCAACTCATATCAAATCCTTTATTCCCTCTTTCCTTGTCCCAGGTAGCGTCTAATCATGATTTAGGATGATGGAATAGGATAAAGCAAACAAGAGAAACAAGTCCATTGTAAAAGTAAAGCAAAATCTAGACTTAGAAAGAGGAAATATGGCTTGAGGGTAGGTTGATATCTTGGAGGAGAAGATCTGAAGTTTTGTGGTCATATAACAGATGCTATTCTGGTTTTCACCACGAGTTTGAATCATGATTCAATTAAGCATAGATTTTGTACCTGTGTTAGCTTAGTTTGGAGTTTGATAATATACCGTTCACTTAAATTGTCAACGTGCTATAGGCATATTCTGTCTCAAACTGGCTTTCAAGTTCTTATTGTAACCCATGCTCTTTTGAAGATGCAACTGTATGGAAGTTCACAGAGACAGTGAGTGCATGTGCACCTGGTGAAGCTGGGCATTACATGATGTATACTGATGCATCAGACAATGTTTTAATGATGGGCATAATTATTACCTAGTGACTGCTTCGTTCTGCTCACACATTCATATATTCCTATGGATGCCTAGATTTGCTGGAGGCTTGTGTAGTATGTGTGCTAAGTATTTTCCAAGTGTATGCAAGTGACAATCACATCAAGAATGAAACAATATAAAAAGAACATCAAGAATGCTGATGGATTTTAACGATCACTGGGCATACACACTGTTTAGATGAAGTTCGAAATAAACATAATGACAGACGTTAATTTTTTATTGATGAGCTATAAAGATTGTATAAAATGTTTGACGAGCTGATCCTTTGCAGCACATGGGATGACAGGTTTTTATCTAGCAAAATGCTTGTAACCAAATATAAGAAAATTCCTCCATGACATATACATTGGTAACTAGTTATAATTGTTAGCTTGGAACTCCCAGACCTTTCTCATTTGTTGACATTTAACTGTAGCTTGATGCCAGTTTTGATGAATGTGTGAACTTTGCTTCATTATCTTGGTTGTCCCATTATCCTAGGGATGTCATTGGTATGATCTGTGGAATTAATTCATTCTCTGGTTTTCTTGTCACAGGGAGCTTTACAGTAATAACATAAGTGGTCTAATACCGAGTGATCTTGGGAATTTGACTAATTTGGTCAGCTTGGATCTCTACTTGAACAACTTCGTCGGTCCCATCCCAGATTCCTTGGGCAAGCTGTCGAAATTGAGATTCCTGTATGTATTTTTATTCTATATTAGATTTATGTGGATATTGTGATGCCAAATGTGTATGTCATCAACCAAGATGGACCTGCTTTTGCCGGCATGTTGAGAAAGGAAAGGAAACCTGAATTCTTTTCTAGGTTTCACATAAGATTACCTCTATTAGTATTAGGTTTCACACTAAGTTTTGATGGTAGAAGGTATTTGAGGACTTTATGGGACTGATATCATTAATAAAACAGATCTTACATATAAGTCTTTTTATTTTCTCCCTTTGTTCTAATCCTTATATAAAGGAAAAGAGTAGAAGCCTATGCTTTGTCTGATATTTGGGCCTGATATTGTAAAATTAAGCAGAGTTTAGTAATTGGTGTTTTATTTTTCTCCCATTGTTGGGATCCTTTTCTAGAAGAGAAAAGAAGTTTAGAATCCCATGCTTCATCTTACATTTTTCACTGCTGAAGTTTCTCTTGCTTAATGCTCCACAAACTTCTTTCTTTTGGCTTGACTGTCTAAGAAGTGTACCAAATAAGTTCTATTATGCAGACTCAGCATGAAGCCTCATTAACTGCATCTTGGGACTTACTATAGAACTATTGCAGTACTGCCAATGTTGACTGACATGATTTAGTCCAAAAAGTGTTTCAAGTTTCTGTTTGTCAGGGGGGAAATGTTTTGAGGCTGTTTATTGTTGTTCTGAACTGACAACTTCATTTCCAGCTGAAGTGCCGGTTCTGGCTTGAAGTTCGAATATGTTTTACATCCACATATAAATAATTTATTAAGATGATTCTACTTGTCAATAAATAAATAAATTAAGATGATTCAATTAGTAGCCTTTTCTTCTGTTTTATTCTCCAATTATTACTTATACCAAAAGTGAAGCTCATTCCTGTACACCTTTGTGATTTTTTTGCAAGGATAGTCGGCTCAACAATAATAGCTTAACTGGTAACATCCCTATGTCACTGACTAATATCTCATCACTACAAGTGTTGTAAGTACACTAGTTATTTTGTGACTTAACTTAGATAATTCTTTCTGTCTTCCATATCTTTTCATTCATTTCCTTTTCCTTCTAGTGCATATACAGATATACTCATGCTCAATTCTTGTCTCATCTGTATATAGGGATCTGTCAAACAACCGTCTCTCAGGTGTTGTTCCAGATAATGGTTCCTTTTCTCTATTCACGCCTATCAGGTATATTTCATTTAAGGTGGTGCACCATCAAACTGGCTGGTGGTTTTAGTATGCTGCTATTTGCTAATATATTTTGCTTGAATTTACAGTTTTGCGAATAATTTAGATCTTTGCGGGCCTGTAACTGGACGCCCTTGTCCGGGATCTCCTCCATTCTCCCCTCCGCCTCCATTTGTTCCACCACCACCAATTTCTGCTCCAGGTGGTTCTCTAAATTGGTGTGGATAGGTTGTTTTCCCTTTTTTTTTTTTTGCTTGTTTGAGTTTGGTGATTTTTGTTGGTCCATCCATCTTGCCTCTCTTAGTTTTTCCTAACCTAAGTGAGAATTTGTGCTATAGAAGTACTGACAAGAAAGGGGCAGAAGAGGAAAGCCCATCTTAGCTAGTCAGGCATTTTGGTTAAGTAGTTCTTATGGGAAACTGGTATGCACGAGACTACATTTAGTCTCTAAGCATTCTGGTCTTTATACAAATTTAATTCAGCATTGTGGAGATCTTTTCTTTGGTCCCCTTGTAAATTATTATCTGTGGTATTTGAAGTCGAGTATGTCTGAATGAAATTAATCATATTTAAGCCAGAACTTGTGAAGATTTTCTTTGCTTTGTAAAATCTATCCTGGAAATTAGATCTGAATGATAGATACATATTTGTGTTCTTTCTTTTTGAAGTGATTTAAATAATGAAGGGCCAAAATTGTGTGTTAAATATAGTCTGGATCATATGAAGGAAATTAATCAAATTTATGGGAACTTCAGGAGGAAATGGTGCAACTGGAGCAATTGCTGGAGGTGTAGCTGCTGGTGCTGCTCTACTATTTGCTGCTCCTGCCATTGCATTTGCCTGGTGGCGCCGTAGAAAGCCACAAGAATATTTCTTTGATGTACCAGGTTAGCAGTATTCACAATACCCAACCATAAGTCCATAACTCCTACTTGCTCCCTCGCGTGTTTGTGGTTTCTCTTGCATGTTTATTTTTTGGCTCCATAATTACCGTTTTTGCTTAAATTTATTGCAGCTGAAGAAGATCCTGAAGTTCACTTAGGTCAACTGAAAAGGTTCTCCCTCCGAGAGCTACAAGTTGCAACCGACAGTTTTAGCAATAAAAATATTCTGGGTCGAGGTGGATTTGGTAAGGTATACAAAGGACGCTTAGCAGATGGATCATTGGTGGCTGTTAAGCGGCTTAAGGAGGAGCGTACTCCTGGAGGGGAGTTGCAATTTCAAACAGAAGTTGAGATGATTAGCATGGCAGTGCATAGGAATCTTCTACGATTGCGTGGTTTCTGTATGACACCAACTGAAAGACTGCTTGTCTACCCCTACATGGCGAATGGAAGTGTTGCATCATGCCTGAGAGGTGACGCCTTCTGAAATCTATCACTCCGTAACATATTTGGATGTTCTCACCTTTAATTTAGAGGGTATTATTACATAATGCAAGAACATCTTTCGCTGGTTAACGTTCTATCTGGCATAACATACTCTTTATAATAAAACATATTCTTGTATGTTATTCCTGTAACTTGGTAAAGGTAGAAATATAATTTATTTTGTATTCTCTTGACAAAATAATTTATTTTATCAGAACGACCGCCTTCTGAACCGCCACTTGATTGGCCAACGCGAAAGCGCATTGCTTTGGGGTCTGCCAGGGGATTATCGTATTTGCATGATCATTGTGACCCTAAGATTATCCATCGTGATGTGAAGGCTGCAAATATATTGCTAGATGAAGAATTTGAGGCTGTTGTTGGAGACTTTGGTTTGGCTAAACTTATGGACTACAAGGATACACATGTTACAACTGCTGTGCGTGGTACAATTGGGCATATAGCTCCAGAATACCTGTCCACAGGGAAGTCTTCAGAAAAGACTGATGTTTTTGGGTATGGGATCATGCTTCTGGAGCTAATCACCGGCCAACGTGCTTTTGATCTTGCTCGGCTTGCAAATGATGACGATGTCATGTTGCTTGACTGGGTATGTTGTCATGCCTGCTTTACATGTAACATGACACGAGTACCATAATGTGCTCATTTTTTAATCTGTACAATCATAACACTAGCTGATTAATAAGTGTTTCTGCCTTTAGCAGGAATCTTTACATCTATGTCTGAACTTGTTGAGGTTCTTGTTTCAGGTGAAAGGACTCCTCAAAGAGAAGAAATTGGAAATGCTGGTTGACCCTGATCTTCAGAACAAGTATGTGGAGGCTGAGGTGGAGCAACTGATCCAAGTAGCATTGCTTTGTACACAAAGCAACCCAATGGATCGGCCTAAGATGTCGGAAGTGGTGAGAATGCTTGAAGGTGATGGTTTGGCTGAAAGATGGGATGAGTGGCAGAAGGTAGAAGTTCTCCGGCAGGAAGTGGAACTTGCACCACATCCTGGTTCTGATTGGCTTGTTGACTCAACAGAGAATTTACACGCAGTGGAATTATCGGGTCCAAGGTGACCGACCTCGTACTACCCCAAGTAAAGAAAATCAAATCATTAGGAACTTTTTATTTTTGTTGACCTTGTGAAAGTTTTTCTTTGTGAAATTTAATGAAAATCTCCAGTTGTAAGCCCATGAAGTGTGTCTGTTACATTGTGCATACAAATTTGTACTGACAGGAAATTCCAAATCCATGCCCTGACAAGTTGGCACATAGGCGTAGTAGTTAAGTGCTTTAAAATTGTCCAACACAGTACACTACTGTCAAGCAGACTCTTGTTTTCAGTGACATTGAAAATGCTATCAATTTGACAAGGTTATTTACAAGATTTTGGATTTTACAATCAAAGAATAGTGCTTCACATTCTCTTACGGTTTTTATCATAATCTCGTGCTACAAGATAATGGCAGATATTTCATCAATAGGGCACTTGCAGAGTTTGTTTTGCCACTCTATGCCCAAATTTTGGGCTCTTTCTGACAATTCAGATTAGTACCAGACGCTCTGATATAAGTACAAGATCTGCTCAGTGATACTTTTCGATCATTTCTGTCATTATTTTTTACTCCAAAGAAAATTTCTTACATGCAAATTTTTTGAACTGCATACTCTTCAACATACAAGGATCCATCGAATGTACTACAGACCAAAGGGCAATAGAATAAACAGAAGAATCTCAAAACTCACAATCAACAAACAAACCAGAGCCTGCAGAAGGTAGAGAAATACACTTCACTTATCATGGCTGGACCTGATATTATCTTTGAATGACTGTGAACGAGAACCAGGATTAGAGCCATTTGTTGCATTAAACAGGTTTGGATATGAATAATATGTTTCATCGTCCATGGAAGGCAATCTCTGTGGTTGGTGAAGACCCTGACCAGGATGATGAATATGGGAGATGGGAGACTCAACACCCACTGTTGAATCATCGTATGAAGGAATATTCGACACCTGATGTTCCAATGGACTGCCATGAAGTGGGCTGGGGATACCATCAGAGTTTGACGATTGGCGTTGGAGGTCTAAGTGAGGCTTTTCTAGGATGTGAGATTCTTGTTTCTCAGCATAAAAGGGGTTCTCACTGGGGCTTACAGGCACCTCTTCGCTCCTTTTCCTGGTAAGCTCGCTGATTCTAACCTGAGCAAGGCTAGCTGCTGAGCGAGCAGCAGCAGCTGCACGTTCGGCTGAATCAGCAGCTGCTTGGGCAGCAGCCAATACATCTTGCAAATCTACATCAGTCTCAACTTTGGTCTTTGTAGTAGAGGGTGGTTGAGTACTTTGGGTTGTGGTTCTTTCTGTTGAGGAGAATGGAGGTGATTTTGGCGGGGGAACCATGAATGGCAAAAACTGTTTCTCTTCTTCCGGTGCAGATAAACTATCAGCTGATCCAAAATCTTTTGAAACTACCTTCTCTTGCAAAAGTTCATCTCGCTCCACATCTTGTTTGTGAGATTTGAGCTCACCGTCTTCAGAAGACTTAGCTACCTCCTCATCGGAGTCAGATAGTGCAGAGGCTGGGAAAGGAAGCATTTCAGGGGCAGAAACTCCACCGGTGCTAGGTCGTAGTGGCTGCTTAGGGACTTCTGGAAAATCTAAGTCAAAATCAGCCTCAGATTCCGTTTGTTTATCAAAGACTTGATCTGATGCAGAGTGCTGCACCTCATCCAACCTTTCTTTGGGAAGTGGCACCTTAGCCCCACTGACAAATTGTGTGGGCCCATTCTGCAAAGCAAAGGGAAGAAATCATGATGATTGAAAAGAGGACAAAATCAAAGCCAGTACAGACTGAAATATACCAGCAAGTCCTCATGTGACTTCAACAGCTCAGTTTCAGAAGCACTTGGATCCCAATCTAGCTGATGCTCCTCCGCAATTTCCTTTAGCAGCTTCATTTTCACATCAGGAGCAGGAGCTCGAATAGACAGCAACTCTATCAACTGTCAGAGACAAGAAAAAAGAGAGAAAAATCAGAAACAAGAAACAGTTCCTCTTTCTCGTTTAGCTATGGCATAAAGGAGAACTAAGGTAACCTGGCGATTGACTCCGCATTCTGGCATGAGCTCTGTAGCAGCAGCAATGAATTCCTTCCCATATTTACCAGCAAACATCATCTGAACCTGCAGCAATTCCGGAAGATCTGCACATCTTGGTGCAGCAAAACAAACACTTGAAATAGCTTCTTTCAGATCTAGAGGGCATTCCCTGGACATTATAGAGTTTGGGGGTTCAGCTCTTTCCAAAAATTCAAGTTAAGAAAATTATCAACAAAATTCAACTAGAAGTCTAGAATATTAAGACTTGGAAACAAAACCTCCACATTGCAGGACAGTCAAGTTCAAGAGTATCAGTTTGCTTTAAAACAAATATCGAAGATGAAGCGCTTTGGGTCCATCTTGGAACTAGAGATCCACATTTATGGTCTCATGTAGATGATCAAATTATTTTAAGGTTTCACACTCCCATAACACAAGCAAGAATGTATTGTTCATTCTATGCCCTAGATACTTCTCGGTCATATAAGCAGAAAGTTAACGTAGTTAATCCTTGTTTGGCTAAAATTGACTTTATGCAAAAGGATCAAACATGCTGCTGCTCGGATACATGCACATACTTCCATTTAATTGACCTACAACCTAGACATTTCAAAGTTGGTAACTTTTTTTCAGAATGCATGAAATTTAGCAAATTAAAGGACTGTGATTCATTAATTGAACTAGAGTCGCTCCATGAAAAAGTATCCATATCTAGGGCAGATAATTCTCGCAAACAACAGCTAGGGTGGAGTATAGTAGATCTGAAAGTTTGGTATTGAAGGAGATATGGAACTAAATGTTAACAGCAGCATAGACAGCAAAGGCAAGAGGCAAGCAAGGAAGACAGAGAGGAACATTTGGGACCCGTTTGGCCATAGATTTCCCAAGTAATATTTGGGGGTAAATTTGGCAAATAGTGTTTGTCCATACAATTTGCCATTATTTGGCAAATATCCCAAATTCCCAAATACTAGTTTTTTCTAGTATTTGAGCCAAATCTCATTATTTGGGATCTTTTAAAAATTGAAATTTTACACCAAACTTTTATCTTTTACAAAAACTCCCTCTATAGTAGTTGTTTGCGTTGTATTACATAATTTTTTTACGTGAACCAAAGTAGTGATGAAATATGAAAGTGATGATATGGTTGTTGATGAAAATGATGAACAAACGGTCCAGGGTACCAAAGTCATGTGCTTTGTCTACTTCACGATGTATAGAATGATGCTTGTTGCACTCACTCCACACACCACATTGCTCCAGTGTCATGGACACTACTTGTTATTCGTTGCAACGAAGATCCAACTGACTTATAATACAAACTTATGGTTAGTTTTGATAGTTTTTAAAACTTATGGGTATAAATCATATTTTTTCTAAAAAAGTGAAATATGTTTCCAAAAGATTATGGCCAAACACATGGTGAAATTTCACCCAAATAATATTTGCCAAAAATATTTGGAAATCTAT

>PGSC0003DMG400027555 LRR-RLKs

CAGCTGAAGAAGATCCTGAAGTTCACTTAGGTCAACTGAAAAGGTTCTCCCTCCGAGAGCTACAAGTTGCAACCGACAGTTTTAGCAATAAAAATATTCTGGGTCGAGGTGGATTTGGTAAGGTATACAAAGGACGCTTAGCAGATGGATCATTGGTGGCTGTTAAGCGGCTTAAGGAGGAGCGTACTCCTGGAGGGGAGTTGCAATTTCAAACAGAAGTTGAGATGATTAGCATGGCAGTGCATAGGAATCTTCTACGATTGCGTGGTTTCTGTATGACACCAACTGAAAGACTGCTTGTCTACCCCTACATGGCGAATGGAAGTGTTGCATCATGCCTGAGAGGTGACGCCTTCTGAAATCTATCACTCCGTAACATATTTGGATGTTCTCACCTTTAATTTAGAGGGTATTATTACATAATGCAAGAACATCTTTCGCTGGTTAACGTTCTATCTGGCATAACATACTCTTTATAATAAAACATATTCTTGTATGTTATTCCTGTAACTTGGTAAAGGTAGAAATATAATTTATTTTGTATTCTCTTGACAAAATAATTTATTTTATCAGAACGACCGCCTTCTGAACCGCCACTTGATTGGCCAACGCGAAAGCGCATTGCTTTGGGGTCTGCCAGGGGATTATCGTATTTGCATGATCATTGTGACCCTAAGATTATCCATCGTGATGTGAAGGCTGCAAATATATTGCTAGATGAAGAATTTGAGGCTGTTGTTGGAGACTTTGGTTTGGCTAAACTTATGGACTACAAGGATACACATGTTACAACTGCTGTGCGTGGTACAATTGGGCATATAGCTCCAGAATACCTGTCCACAGGGAAGTCTTCAGAAAAGACTGATGTTTTTGGGTATGGGATCATGCTTCTGGAGCTAATCACCGGCCAACGTGCTTTTGATCTTGCTCGGCTTGCAAATGATGACGATGTCATGTTGCTTGACTGGGTATGTTGTCATGCCTGCTTTACATGTAACATGACACGAGTACCATAATGTGCTCATTTTTTAATCTGTACAATCATAACACTAGCTGATTAATAAGTGTTTCTGCCTTTAGCAGGAATCTTTACATCTATGTCTGAACTTGTTGAGGTTCTTGTTTCAGGTGAAAGGACTCCTCAAAGAGAAGAAATTGGAAATGCTGGTTGACCCTGATCTTCAGAACAAGTATGTGGAGGCTGAGGTGGAGCAACTGATCCAAGTAGCATTGCTTTGTACACAAAGCAACCCAATGGATCGGCCTAAGATGTCGGAAGTGGTGAGAATGCTTGAAGGTGATGGTTTGGCTGAAAGATGGGATGAGTGGCAGAAGGTAGAAGTTCTCCGGCAGGAAGTGGAACTTGCACCACATCCTGGTTCTGATTGGCTTGTTGACTCAACAGAGAATTTACACGCAGTGGAATTATCGGGTCCAAGGTGACCGACCTCGTACTACCCCAAGTAAAGAAAATCAAATCATTAGGAACTTTTTATTTTTGTTGACCTTGTGAAAGTTTTTCTTTGTGAAATTTAATGAAAATCTCCAGTTGTAAGCCCATGAAGTGTGTCTGTTACATTGTGCATACAAATTTGTACTGACAGGAAATTCCAAATCCATGCCCTGACAAGTTGGCACATAGGCGTAGTAGTTAAGTGCTTTAAAATTGTCCAACACAGTACACTACTGTCAAGCAGACTCTTGTTTTCAGTGACATTGAAAATGCTATCAATTTGACAAGGTTATTTACAAGATTTTGGATTTTACAATCAAAGAATAGTGCTTCACATTCTCTTACGGTTTTTATCATAATCTCGTGCTACAAGATAATGGCAGATATTTCATCAATAGGGCACTTGCAGAGTTTGTTTTGCCACTCTATGCCCAAATTTTGGGCTCTTTCTGACAATTCAGATTAGTACCAGACGCTCTGATATAAGTACAAGATCTGCTCAGTGATACTTTTCGATCATTTCTGTCATTATTTTTTACTCCAAAGAAAATTTCTTACATGCAAATTTTTTGAACTGCATACTCTTCAACATACAAGGATCCATCGAATGTACTACAGACCAAAGGGCAATAGAATAAACAGAAGAATCTCAAAACTCACAATCAACAAACAAACCAGAGCCTGCAGAAGGTAGAGAAATACACTTCACTTATCATGGCTGGACCTGATATTATCTTTGAATGACTGTGAACGAGAACCAGGATTAGAGCCATTTGTTGCATTAAACAGGTTTGGATATGAATAATATGTTTCATCGTCCATGGAAGGCAATCTCTGTGGTTGGTGAAGACCCTGACCAGGATGATGAATATGGGAGATGGGAGACTCAACACCCACTGTTGAATCATCGTATGAAGGAATATTCGACACCTGATGTTCCAATGGACTGCCATGAAGTGGGCTGGGGATACCATCAGAGTTTGACGATTGGCGTTGGAGGTCTAAGTGAGGCTTTTCTAGGATGTGAGATTCTTGTTTCTCAGCATAAAAGGGGTTCTCACTGGGGCTTACAGGCACCTCTTCGCTCCTTTTCCTGGTAAGCTCGCTGATTCTAACCTGAGCAAGGCTAGCTGCTGAGCGAGCAGCAGCAGCTGCACGTTCGGCTGAATCAGCAGCTGCTTGGGCAGCAGCCAATACATCTTGCAAATCTACATCAGTCTCAACTTTGGTCTTTGTAGTAGAGGGTGGTTGAGTACTTTGGGTTGTGGTTCTTTCTGTTGAGGAGAATGGAGGTGATTTTGGCGGGGGAACCATGAATGGCAAAAACTGTTTCTCTTCTTCCGGTGCAGATAAACTATCAGCTGATCCAAAATCTTTTGAAACTACCTTCTCTTGCAAAAGTTCATCTCGCTCCACATCTTGTTTGTGAGATTTGAGCTCACCGTCTTCAGAAGACTTAGCTACCTCCTCATCGGAGTCAGATAGTGCAGAGGCTGGGAAAGGAAGCATTTCAGGGGCAGAAACTCCACCGGTGCTAGGTCGTAGTGGCTGCTTAGGGACTTCTGGAAAATCTAAGTCAAAATCAGCCTCAGATTCCGTTTGTTTATCAAAGACTTGATCTGATGCAGAGTGCTGCACCTCATCCAACCTTTCTTTGGGAAGTGGCACCTTAGCCCCACTGACAAATTGTGTGGGCCCATTCTGCAAAGCAAAGGGAAGAAATCATGATGATTGAAAAGAGGACAAAATCAAAGCCAGTACAGACTGAAATATACCAGCAAGTCCTCATGTGACTTCAACAGCTCAGTTTCAGAAGCACTTGGATCCCAATCTAGCTGATGCTCCTCCGCAATTTCCTTTAGCAGCTTCATTTTCACATCAGGAGCAGGAGCTCGAATAGACAGCAACTCTATCAACTGTCAGAGACAAGAAAAAAGAGAGAAAAATCAGAAACAAGAAACAGTTCCTCTTTCTCGTTTAGCTATGGCATAAAGGAGAACTAAGGTAACCTGGCGATTGACTCCGCATTCTGGCATGAGCTCTGTAGCAGCAGCAATGAATTCCTTCCCATATTTACCAGCAAACATCATCTGAACCTGCAGCAATTCCGGAAGATCTGCACATCTTGGTGCAGCAAAACAAACACTTGAAATAGCTTCTTTCAGATCTAGAGGGCATTCCCTGGACATTATAGAGTTTGGGGGTTCAGCTCTTTCCAAAAATTCAAGTTAAGAAAATTATCAACAAAATTCAACTAGAAGTCTAGAATATTAAGACTTGGAAACAAAACCTCCACATTGCAGGACAGTCAAGTTCAAGAGTATCAGTTTGCTTTAAAACAAATATCGAAGATGAAGCGCTTTGGGTCCATCTTGGAACTAGAGATCCACATTTATGGTCTCATGTAGATGATCAAATTATTTTAAGGTTTCACACTCCCATAACACAAGCAAGAATGTATTGTTCATTCTATGCCCTAGATACTTCTCGGTCATATAAGCAGAAAGTTAACGTAGTTAATCCTTGTTTGGCTAAAATTGACTTTATGCAAAAGGATCAAACATGCTGCTGCTCGGATACATGCACATACTTCCATTTAATTGACCTACAACCTAGACATTTCAAAGTTGGTAACTTTTTTTCAGAATGCATGAAATTTAGCAAATTAAAGGACTGTGATTCATTAATTGAACTAGAGTCGCTCCATGAAAAAGTATCCATATCTAGGGCAGATAATTCTCGCAAACAACAGCTAGGGTGGAGTATAGTAGATCTGAAAGTTTGGTATTGAAGGAGATATGGAACTAAATGTTAACAGCAGCATAGACAGCAAAGGCAAGAGGCAAGCAAGGAAGACAGAGAGGAACATTTGGGACCCGTTTGGCCATAGATTTCCCAAGTAATATTTGGGGGTAAATTTGGCAAATAGTGTTTGTCCATACAATTTGCCATTATTTGGCAAATATCCCAAATTCCCAAATACTAGTTTTTTCTAGTATTTGAGCCAAATCTCATTATTTGGGATCTTTTAAAAATTGAAATTTTACACCAAACTTTTATCTTTTACAAAAACTCCCTCTATAGTAGTTGTTTGCGTTGTATTACATAATTTTTTTACGTGAACCAAAGTAGTGATGAAATATGAAAGTGATGATATGGTTGTTGATGAAAATGATGAACAAACGGTCCAGGGTACCAAAGTCATGTGCTTTGTCTACTTCACGATGTATAGAATGATGCTTGTTGCACTCACTCCACACACCACATTGCTCCAGTGTCATGGACACTACTTGTTATTCGTTGCAACGAAGATCCAACTGACTTATAATACAAACTTATGGTTAGTTTTGATAGTTTTTAAAACTTATGGGTATAAATCATATTTTTTCTAAAAAAGTGAAATATGTTTCCAAAAGATTATGGCCAAACACATGGTGAAATTTCACCCAAATAATATTTGCCAAAAATATTTGGAAATCTATAGCCAAACCCTAGCTTGGTATCCTATACTATTTTTATTAGGAAAAAAAATGCCAAAGAACCCCCCCCCCACCCCACCCCACCCCCGACTTGTAGCTTTTTAGTTAGTGAGCACCTAAACTATCACTCGTATTAAATACCCCTGGAATTTGTTTTTTCAACCGTAGCGTACACCTACGACGTTTAGATCCCTCAACCGTGTATACACGCACGTGACACGGTGAAAATATGACAAATGTGGATGTCCATGTAGATAAATAATGCCCCCACCCTCTCTAATATTTACCCTTTTTCAAAAGCTTTTTATATAATCCCCCAATTCACACAAATCAACCCTAATTACCCGATCTTCTCTTTCACACAAATACTCTCTAAATAATCAAATCCTAAGAAGTTTAACAGTCGGAGTTAAGGGTAGATTTTGAACATTTCTGGCTCTGCTATTTGAATGTTTTCTCCTCAAATAGTACAGTCCATTTGGGTGAAGCTTTTTATTTGAGTGGATTTTTCTCCGATCTTGTCTGAAATTCTTATCAAGTAAGTGATTATCCATTCTTAAAATGTATTTTTGTTTGTTTTTCCTCGAATAGCCTTGGTCAGTTACAATCCAATTACGTCCGGGGTAGTCATATGACCATGAAATTCTCACAATCGGAGGACACCCACATGCACAAACAAGCATCCTATTATAATTCATGGATTCATCAATTGCAACTTTAGAAAGACGGAGGAGAGATAAATTATAGGCAAAGAAAGAAAATGATGGCAAAGATAGGTATTTCAGAAAATTGAAAGTGATGGAGAAGAAGGGTTTAGGGAGAATAAATAGGAAGAGAATGAAGAGAGATGCTGTGCAAAAAAGGAAAAAAATAAAGAGAAACAAGAATTTTTAATGTTTTTCACCATTGAAGTAATATGATGCGGCAAAAAATTTACTCCCTCACGCACCTAGCTGAGCTTGTTGTCAGTTTCTATGCCAGGTGGACGTTGTAGGTGTATGCGACGGTTGAAAAAACAAGTTCCGGGGGTATTTCATATGAGTGATAGTTTAGGTATTCACTGAATAAAAAGCTACAAGGTCGGGGGGTCTCTAGCATTTCTGCCTTTTTATTATTATTAAACTGGTGTATGCGACGGTTAAAAAACAAGTTCCAGGGATATTTCATATGAGTGATAGTTTAGGTATTCACTAAATATAAAGCTACAAGATCGGGAGGTCTCTAGCATTTCTGCCTTTTTATAATTATTAAACCTCATTCAGCTGACTATCCTAGTCTATATCAGGTGTAACCCAGTTTATTGACTTATTGCAGCTAGAGTAAAGTTAAATGGAGCTATTGTTTTAGAGCACAATACAGCTATAAAGATACATATGACCAGCATAACTACACAAGATGACAATATACTTTACTTATGAAAACGGAAGAAGAAAAATAAGCTAAAAGGCAATTCTACAAGTAGGCCAACCTTACAGCTGGGATTTACATCTGCCAACGGTGTCAAAAAAATCATATAACCACGAAAACCAAAAACTTGCTGTTCTATAGGCACTTCGCCAAAACATCATAGGTGATCATCTAATGCATCATAAAGGAACCCTTTTATATTCTTTGCCATTCGCCCCATACATTCTTTATCCTAAGAGCAACTAAAAGCATATTCAGTTGCTCATCAATAAACTGAAGAACCATCTTAGGATACTGTAAATGTTCACACCATATAGAATATCAAGAACATTAACACAACAGATAACTCATTACCTTTGTGCTTCAATAATTGGAAGACGAACCGATATAAGCTCGCAGAATAGCTCAACAATCTCCTGCGCTGCCATCATTTTCTCTTCTCTTATGATATGCTCTACCTAAACAAGCAAACATCACAGTCAAGCTCATTACTCTGTAAAACCATCTATCAAAATACATAATAACCTTTCAGTATTCAAATGCGAAGAACACACAGCCTAGCACAAATTATATCAGCATGAATCGATGAATTTAAGCAAAATAAGAGTCAGTTTATACCCGAATTCGAGCAGTAGCTTCCTGACCAGTCTCTAGTAGCTTAGCTATTTCTTTCCTCATCTGCTTCAGCTGAAACTCCCTACGATTCCTCAGTAGTTTGATCCGAGGAATTGTCAATTTCAACATGGTTTTACTGCAAATTGATAGGTACCATCAGCAAATTCCCATATAAAAAAGACAACATTTTTCTCAAAAATTTGAAAAATTCGAAACCCCATGATTTCAACAATACAAAATTTCAAATCTGCATCATAAGAACCCTTAACTAAAAAGATCCAAATCATAAAATCTGACAAGAACCTTCAATAAAAGACAGGAAAGTGGGAAATTGAGGATTACCATTTGGCAGCTTTGAAACCTTTGCTGAAGAAAGAATCCATTATAGACTTGATTCAGATCTTCAATATCCGTGTACCCTTTAAAATTCTTGCAGTAAAACCTTATGAATTATCACTTACCTAAAACCACAAAATAATTAATCAAACTTGGTAGTATTTCAGATTAAAGCAAAGAATACCTTTTTATTTTTCAAAAAAAGAAAAGAAAAAGAGATCAAAGATCAAAACTGATGACGAAGTAGTTAAGAAAATATGTGTTTTTGTGGAAAGCACAGAAAGTGAACCTGCTCCGCTATGCATATAAAGACAAATGGAACGCGGACAAAGCTTTCAATGTCTACGACAACAAGCAGCTTCCTTAAGCTCTAAGCATTATTGGATTTCAACTTTCAACGAGTATTATTTTTGG

>PGSC0003DMG400027581 LRR-RLKs

ATGAAACTACAAGATTTACGTGCAACTATCGTCTTTCTTCTATCCTTGCTAGCAATTCTTCCTCACATTATTGCTAATCTTGATTCAGACAAACATGCTCTACTTCAATTTGCTGCATCTGTCCCACATCTTAGAAAACTCAACTGGAACTCTGCTTTGTCAATCTGCAATTCTTGGATTGGGATCACTTGCAACAAAGATGGAACGCGAGTAGTTGCAATTCATCTGCCTGGTGTTGGACTTACTGGCCATATCCCAGCAAACAGCATTGGCAAACTAGATGCTCTCCAAGTCCTTAGCCTAAGAGCAAACAATCTAAATGGAAATCTTCCTTCTGATATTCTTTCAATCCCTTCTCTTTACTCTATCTACCTTCAACATAATAACTTCTCTGGTGATATTCCTGTTTCCTTTTCATCAACACTTGGTGTAGTAGATCTCTCGTTTAACTCTTTTACCGGAGAAATTCCCCCCAAGATCAAGAATTTGCCTCGACTATCCATGTTGAACCTAAAGTTCAATTCACTATCTGGATCAATTCCCAATCTTGATGTGTCAAGGTTGAGTTTTCTGAATTTGAGTTACAACATGTTAAATGGCTCAGTTCCATATTCCCTCAGGAAGTTCCCACTTTCTTCTTTTGTGGGGAACTCTAATTTATGTGGAACACCTTTGAGTAGCTGTTCTTCGAGATCTCCATCACGTAAAGGTGACAACTTTAAGAAACTTAGTAATGAAATTATCATTGCAATTGCAGTAGGTGGTCCTTCTGTTATAATTCTCCTTGTTTTATTCATCTACTTCTGTTATTTTAACAAGAAGGTTGATAACAACACAAGCATGGTTGAAGAGAAGAATGAGAAGTTAGAAGACTTTGGAAATGGGGTTCAAGATTCTGAAAAGAACGAGTTAACTTTCTTTAAAGGGTGCTCTTATAACTTTGATCTTGAGGATCTGTTAAGTGCTTCTGCTGATTTTCTTGGTAAAGGGAGTTATGGAACTGCTTACAGAGTTTCTTTGGATGAGGTATCGATGATAGTTGTGAAAAGGCTGAAGGAAGTAAGGGTTGTTAAAAAGGAGTTTGAACAGCATATGGATATTGTTGGAAAGATTACAAGGCACCCAAATATAGTGCCATTTCTTGCTTGTTATTACTCAAAGGATGAGAAACTTCTTGTTTGTGAATACATCCCTTATGGTAGCTTGTCATCCGCCTTATATGGTATAAGCTCCTTGACTCTTCTTATATAAAACTACTTCTTAGAACAGATCATGCTTGCTGCACATTATGCTGTTAATAGCTATTCCAATTCCGGCTTTATGCTTACATGGTCTTAGCTTAAATAGAGCGACATGATTATTGGTGGTGATTCATATAGCCGAGTAGTAGTTGTTGTTTCGTACTGTTACATGATCTTGTTTATAACTAGCTATATGCCTACTGTTAGCTTGTCAACAGCCTTGCATGGTATACAAAATTACACTGTTTATACATGATTAAAATTACTATTTATGTATATATAGTAGATGTTGAACCCTTATGGCTTTTGCGTCTTTTGAACCTCCTAAGTGAAAATCTTAGCTCTGTCACTGACGGTATATGTTGCTCAACTCTGATTGAAACTGCAGTGTCATCGATCACTTGCTACAGATTATTAATGATTCCTGTTTTGGGCTTTATTTTTACATGATATTGGTTATAGCAGGTAATGGAAGAACTCGACTAGACTGGGATACCAGATTGAAGGTCTCTCTTGGAGCAGCAAAAGGAATAGCACATATCCATTCTGAAGGTGGAGTGAAATTTACTCATGGAAACATCAAAGCATCCAACATACTTCTTACAAGGGATCTAGATGGTTGCATTTCTGATTTTGGCTTAAGCCCTTTAATGAATTACACAGCCATCAAGAATAAGGCTGCTGGTTACCATGCTCCAGAAGTGATTGAAACGCGAAAAGGCACACAAAAATCAGATGTTTATAGCTTTGGAGTGTTGGTACTTGAGTTGCTCACTGGCAAATCTCCTCTGCCATTGCCAAGGCACGAGGACGTAGTCAATTTGCCTAGATGGGTACGAGCTGTTGTTAAAGAGGAATGGACAGCTGAAGTTTTTGATGCAGAGCTGATGAAGTACCATAATATTCAAGAAGAGATGGTGCATATGCTTCAAATTGCACTTTTATGTGTAGCAAAGGTGCCTGACATGAGGCCTTCAATGGATGAAGTGATTAAAATGATTGAGCAAATTAAACATCTATAA

>PGSC0003DMG400027585 LRR-RLKs

GAGAGAGAAAGATAAAGAGAACGTTAATAAATGCTCCCACTGACCCGCCGTCCCCGGTAACACTTTACTTTTATCTACTTTCTCCGGCTTATTCCTCCCTACCTCATCAACAACCTTCAACTTCACCGGCGCTTTACATTTTCCGGCGAAACCCCAAATTCACTCTCATTGTCTCCTTCTTCTTCCCTTCAGATTTCCCTTTTTCTGCTGCCAAAATTCAACTGTTTCTTCCACTCATTCATCTACATTTTCTTCTTCTCCACTTTGTTTCATTAAAAATTCACTCTTTCCGGATTTCTAGGGTTTTTTAAGAACTGGGTGTTTTTAGAAAAAAATTCTTACCCATTTTATTGGCGGTTACGTTCTTCTTGTGCTAAAGATTGTATCTTTTGCTTGTTCTTGTATATAATTATGCATACCCATTTGGGGTTGCTGATGGGATTATGAGTAACTTATATATTGCTTTGTGATTATTGAGCTGGGAGAATGGAAATGAGGAAGGTTTTTGGTATAGTAATATTGTGTATTTTAGTATCTCCTATTTTTGTTAAAGCCTTGAACTTATCTTTTAATGATGATATACTGGGGTTGATGGTATTCAAGGCTGATGTTCAAGATCCACAAGGGAAGTTGGTGTCTTGGAATGAAGAAGATGATAGCCCCTGCGGCTGGAATGGTATCCATTGCAACCCCAGATCCAATAGGGTCTCTCAGATTGTTCTTGATGGGTTTGGACTTTCAGGTAAGATTAGTAGAGGTCTTATGAGGTTGCAATTTCTTCGAAAGCTTTCTCTTGCGAAGAATAATTTTACAGGGAGTATAAGTAGCAGTGTTGTTCAACTTGCTTATCTAAGGATTCTAGACTTGAGTGAGAACAACTTATTTGGGACGATTCCGGGTGATTTTTTTGAACAATGTGGGCCATTGAGATCAATTTCTTTGGCCAAGAATAAGTTTTCAGGGAAAGTTCCAGAAAGTTTGAACTCATGTGTGGCATTGGGGTCTTTAAACTTGTCCTCAAATCAGTTTTCAGGATTGTTACCTTCTGGAATTTGGTCTTTGAATGGGCTAAGGTCACTTGATCTGTCAGATAATTTGCTGGATGGTGAAATTCCAGTAGGTATTGAAGGTATGTATAATTTGAGAGCCATAAATTTGAGGAAAAATCATCTCAAAGGTGAAGTTCCTGATGGAATTGGAAGTTGTTTGCTGTTGAGGTCAATTGATTTGAGTGAAAATTCTTTCTCTGGAGAACTTCCTAAAACAATGCAGATGCTTAGTTTGTGCAATGAGTTGATTATGAAACACAATGCATTAGTTGGTAGTGTACCTGAATGGATTGGAGAAATGAAAAGCCTTGAGATGCTTGATCTTTCTGGGAACAACTTCTCTGGTCAGTTCCCGAATTCAGTTGGGAAGCTTCAATCACTGAAATTACTGAACGTGTCGCGAAATGCTATTTCTGGGGACTTTCCCAAGTCCATGAGCAGTTGTGTTAATCTTATGACACTGGATGTTAGTCACAACTCATTGACGGGTGATCTTCCTCCTTGGGTATTCAAATTAGGATTGCGACATGTTTTGTTTTCTGAGAATAAATTAAGTAGGGGCTTGAAGAATGCTATTGCTTCTTCACTGGAAAACTCTCGACAAAAGCTTCTAGTTCTCGATATCTCTTGCAATGAGTTAGCTGGTGAAATTCCATTTGCTATTGGTGATTTTAACAGCTTGCAGTCATTGAATTTATCTAGAAATTCACTTGTAGGGAAAATCCCTGAGACTGTTGGCCATCTAAAGTCACTGGATGTTCTTGACTTAAGCGAGAATCAGTTAAATGGTAGTATCCCTCTGGAGCTTGGAGGAGCCTATGCTCTCAGGGAACTGAAACTGGAGAAGAATGCCTTGACAGGAGAAATTCCTACTTCTATTGGAAACTGTTCTGCTCTTTTGTCCTTGTAAGTTCTTGTTTCTCTTTATCGTTCATTTATAACTTTCATTTGAGATAATCTCTCTGCTTGGAGGTTTTCATAAGGACAATGTAGTGGTAAGTTATTGTACAGAAGAACCTGATTGATATATAGTTTGATAATTAAGGAACTAGCATTTGCATTCTTTTTCGCAGTTTCCTCTCAGACAACGTTGTTTTGGATTGTCAAAATTCTAGTGACTAAATCAAGCACGAATCCTTATATTAGAGATTCTTTATCATGTTCTACTAATTAGTTAGTTCTAAGCTTCTAGCAATCATTTAGAGTTTCCATATTGAGTTAATCAAGTAGCTTCTAGTGCATGATTTTAGATAAAATCCTTTTTTTTTTCTAGATAAGGCAATTTCCTATACTTAATATTTTACATAAAGTATAATGTAACAACATCACATTACCTGGTTTTCGATGTTTTTATGTGACGTAGCCTGTAAATTTATTTATGAAGTTGTCTTTGAATATCAGCTTTCCTCAAAGCTTGGTTAATCGGTTTTGTTTCCATCAACTTGGCAGGTCTCTGTCACATAATGGTTTAACTGGCCCTGTACCTGCAACCCTTGCTAAACTATCTAACCTACAAAATGTTGACCTTTCCTTCAACAAATTAACCGGAATCTTACCAAAGCAGCTGGTAAATCTTGGTCACCTTGAGTTGTTCAACATCTCACACAACCAGTTAAAGGGCGAACTGCCTTCTGGTGGTTTTTTTAACACTATTTCTCCCTATTCAGTGTCAGCCAATCCATCTCTTTGTGGAGCTGCTGCTAATAGGTCTTGTCCAACTGTCCTTCCCAAGCCGATTGTTCTGAATCCCAACTCCACAGAATCTATTCCTGGTACTATCCCTCTGACTGTGGGCCACGAAAAGAAAATCCTAAGCATCTCTGCCCTCATTGCCATTAGTGCAGCTGCTATTATTGTTGTTGGTGTTATTGCCATTACTGTGCTAAATCTTCGCGTACGGTCTGCAACATCTCACTCTGCTGCTGCCCTTACATTTTCTGGTGGAGATGACTATAGCCCCTCACACAGTACAGATGCAAATTCTGGTAAGCTTGTCATGTTTTCAGGCGAACTTGACTTCAGCACAGGGTCACATGCTCTGCTTAACAAGGATTGTGAGCTCGGACGTGGTGGTTTTGGAGCCGTTTATCGCACTGTCCTCGGAGATGGGATGCCTGTGGCCATTAAGAAGCTTACTGTCTCAGGTCTTGTTAAGTCTCAGGTAGACTTTGAAAAGGAGGTTAAGAAATTGGGAAAAGTTCATCACCCTAATGTTGTGGCTCTTCAAGGTTATTATTGGACTCCGTCACTACAGCTACTTATATATGAATTCATAACTGGTGGAAATTTGTATGATCATATCCATGAAGGTTCTAGTAAAAACATGCTCTCATGGAATGAGCGGTTCAACGTTATTCTTGGGACAGCAAAAGGCATGGCTAACTTACATCAAATGAACATAATTCACTATAACCTCAAGTCCAGCAATATTTTGATTGACAGCTCTGGTGATCCTAAAGTTGCAGATTATGGGTTAGCAAGGCTGTTACCGATGTTAGATCGATATGTTTTGAGCAGCAAAATTCAAAGTGCACTCGGTTACATGGCACCTGAATTTGCATGCAAGACTGTGAAGATAACTGATAAGTGTGATGTGTATGGGTTTGGAGTTCTTGTTCTGGAGATAGTCACTGGAAAGAAACCGGTTGAGTACATGGAGGATGATGTTATTGTATTGTGTGACATGGTACGGGGTGCATTGGAAGAAGGCAAGGTGGAAGAGTGTGTTGACAAGAGGCTGCATGGAAAATTTCCAGCAGAGGAAGCGATTCCTGTGATGAAGTTGGGTTTAATCTGCACTTCACAAGTCCCTTCAAACAGGCCTGATATGGCAGAAGTGGTTAACATATTGGAAATGATCAGATGTCCTTCAGAAGGCCAGGAGGAACTAGTATGATATCCCACTAAACTGATGGAAGTTTTTTAACATATGCTATTAGTCCCGGAGAAGAAGAATCCGCTTGATCTATTCATATTATGGTCATTTAAAGTTTCAATTTTTTTTAATTATGTTATTGGTTCTCTGTTGTTCCGGGTTACCGGATTCCTTTTCACCGGGTTTTAGTATGTGAGGATCCTTTTTTCACGGTTCTTTTATGTAACAATTGATTCTAACATGTAATTCCAGTAACCTCTTGCGTTTGTTGTTCAAATGAAGGAAGGTCCAAATGTTGATAGTGGATGTGCCAATAAGTATATTTCATGACAAAATGATCAAACAAGC

>PGSC0003DMG400027586 LRR-RLKs

GCTCAACAAGGTATATAATTAAAAGAAACCTTGTTGACATATATAGTTATGGGCAGAAGTTGCATTGTCCTCTTTGGTCTTGCAGCTGTTTTCATTCTGCTTCATAACCAAACTTCACTTGCTACTGTTCCTAATATTGGCACTGATGAAGCTGCTCTTTTTGCACTGAAATCACACATTTCTTCTCGTCCTAACAACATCTTAGAAAGCAACTGGTCTTCGTCCAGCCCGGTTTGCAGCTGGATTGGAATCACTTGCAGCTCCCGCCACCATCGAGTCACTGCTTTAGACATTTCTAGCATGCAACTTCATGGTACCATTCCTCCACACATAGGAAACCTCTCATTTCTTGTTTCCCTCAGGATTAGTTACAACGCTTTCCATGGAAATTTGCCAGCAGAGTTGGCTCATTTGAAGAGGTTGAAATTGATTAATGTTAGAAGCAATAACTTCACTGGAGCCATTCCATCATTTTTGAGTTTGTTACCAAACCTACGCACTGTGTCCCTCTGGAGCAACCAATTTTCGGGGAAAATTCCATCTTCCCTTTTCAATCTAACAAAGCTGCAAGTGTTGAGTTTGCAGAGTAATTTTCTTGTAGGAGAGATCCCTCGAGAAATTGGTGATCTTCGTTACTTGACTGTCCTAGACCTGCAATACAATCAGCTTACTGGCCCTATACCACCATCAATATTTAACATTACGACAATGCGAGTGATTGGTCTTACCAACAACAATCTTACTGGAAAGCTTCCAACAACTATATGTGACCATCTTCCAAACTTGGAAGGGCTTTATCTGTCAAGCAACTCCTTAGATGGAGTTATTCCATCAAACCTTGAGAAATGCAGAAAGCTTCAAAGATTGTCATTGTCGTCCAATGAGTTTATTGGAACTGTACCAAGAGAGCTAGCAAACTTAACGGCTCTTATAACATTATTTCTTGGACAACAAGTACACTTGGAAGGTATATAGTATGATATTTCCTTACAACATTATATCTAAAAATTTCGATCTCTCTTCCGCCTAACTGGTAATATTTCAAACAATTGACAGGAGAGATACCTGTGGAGCTAGGTAATCTTAAGAAACTACAGATGCTGGGATTAGCCGAGAATGAGTTTACTGGTTCGGTCCCTACAAGCATTTTCAACATATCAGCACTGCAGTTCCTAACACTTTCACTAAACAAGCTTTCAGGTACTCTACCTTCAGATTTAGGTCCGAACCTAGAAGAATTTTACTGTGGAGGAAATAATCTGAGTGGTTTTATCTCTGATTCAATCTCAAATTCTTCGAGACTCAGAATGCTTGATCTCTCATTCAACAGTTTCACAGGTCCAATTCCTAAATCACTCGGTAAATTAGAATACCTTGAGGCTCTAAACTTGGGAAACAATAGTTTTATCAGCGATTCAACATTAAGCTTCCTTGCATCATTGAATAACTGTAGGAATCTAAGAAATCTCTGGTTCTATAATAATCCTTTGGATGCCATTTTTCCGGCATTTGTTGGTAATTTCTCCGACTCCTTGCACGGTTTTGAAGCAGATGGTTGTAAACTTAAGGGCATGATTCCCCAAGAAATTGGTAAACTTACTGGTTTGACAAAGATGAGTCTATCTAACAATGAGTTCACCGGATTTATTCCAAATACCATCCAAGGCATGTTGAGCCTTCAAGAACTTTACCTAGATAAAAACAAGATAGAAGGACCCATACCAGATGATATGTGCAGTTTACAGAATCTTGGTGCATTGGACTTGTCACAAAATAAATTTTCTGGTTCCGTGCCACCATGCTTAGGAAAATTTACCAGTTTGAGGACACTTCATCTAGATTACAACATGCTGGATTCTAGATTACCAGAAAGCTTGGGGAGCCTTAAAGATCTTTTAGAATTTAGTGTTTCATCCAATTTATTAAGTGGGCAAATTCCATTTGAAATTGGAAATTTACAGGCGGCAACAATCATTGATTTGTCAAAAAATGATTTTTCTGGTAAGATTCCTAGCACTCTAGGGGGTCTAGATAAATTGATTAATCTTTCTCTAGAACATAATAGACTAGATGGGCCTATTCCAGATTCATTTGGCAAAATGTTGGCATTGGAATTCTTAGATTTGTCCTATAACAATCTTGTTGGTGAAATTCCAAAGTCATTAGAAGCTCTTGTGTCTCTCAAATACCTCAACTTTTCATTCAATAAACTCATTGGAGAGATTCCCACTGGTGGTCCTTTTTCAAACGCCACAGGTCAATCCTTCTTGTCTAATGATGCACTATGTGGTGATGCCAAGTTTAATGTGACAACATGTGTCATCCAAACACCCAAGAGGAAAAAGGCTAACTTAGTTTTGTACATCAGTTTGGGAGTGGGTATGTTACTTCTTGTATTAGCCCTTGCATATGTATTTGTAAGATTCCGAAACACAAAAAAGAATACAGGTCAAACAAATGTGTCTCCATTAAAAGAGCATGAAAGAATTTCCTATTATGAAATTGAACAGGCAACAGAAGGATTCAACGAAACCAACTTGCTTGGTAATGGGAGTTTCAGCATGGTCTACAAAGGGATACTTAAAGATGGTACCCTTTTTGCAGCAAAGGTATTCAATGTGCAATTGGAGCGTGCATTCAGAAGTTTTGACATAGAATGTGAGATACTCCGAAACCTTCGCCACAGAAATTTGACCAAAGTCATCACCAGCTGCTCCAACCTTGATTTCAAAGCCCTAGTGTTGGAATACATGCCCAATGGGACACTTGATAAATGGTTATATTCTCATAACTTATTCTTGAACTTGTTGCAGAGATTAGATATAATGATAGATGTTGCATCTGCAATGGACTATCTCCACAATGGTCATTCAACAGCTGTGGTGCATTGTGACTTGAAGCCAAGCAATGTCCTGCTAGATCAAGAAATGGTTGGTCATGTCAGTGATTTTGGCATTTCAAAATTGTTAGGTGTAGGGGAGGCTTTTGTTCAAACAAGGACAATTGCAACCATTGGATATATTGCTCCAGGTATAGTACAACTTTTCAAAGTTTTCTCATATCGTTTAAATACTCAAAACAATTCTTCCTCCCCTAGATATAAATGGATTTTCTTTGTTTTTTGCATATTCCATTTTAACTAGGGAACTTTTATAATTGTGATTTCAGAGTATGGACAAAATGGAATTGTATCCACAAGCTGTGATGTTTATAGTTTTGGCATCATGATGATGGAGATGTTCACAAGAACAAGACCAAGTGATGAAATATTTATCGGAGGCTTGACCATACAACGTTGGGTTAGTGATTCCTTTCCAGGTGAAATTCATAAGGTGGTGGATTCTAATCTGGTACAGCCAGGGGATGAGCAAATTGATGCAAAGATGCAGTGCTTGTTGTCTATCATGGAATTAGCTTTGAGCTGCACTTTCGTGACACCCGATGCAAGAATTAGTATCGAAAATGCTCTTTCAACACTGAAAAAGATTAGGCTCCGGCTTGTCAGTAGTTGGCACTAGCTGGTGGAATTGGACCGGCTACTCTTGTATGCTATTTGATTAACTATGCCAGTAACATTAAACTTTCAATTTTGC

>PGSC0003DMG400027683 LRR-RLKs

GTCGTCGCCTGTCCCCGTTTAACACTCTTTTTCTTTTTTTCAATTTCTTTGCAAGTTTTCAGTTTTAGTTCTCACTGTTAGCCTTCTTTCTTTGACCAAAGAAAAACTGTATCATCAAAAAAATACTCTCTTTTTTTGCCCCTTTTCTTCTTTGTAATTGACCATCAAGATCCAGCATTTGATATCCTCAACTGGGCTGTAAGTATCTTGCATCTATAATTTCTTGGTCTTCTTTTGGTATTTTTTATGCTCAATTTGTTTATTGGTAGAATTTCTTATTAAAAATTGTTTCTTTATTATGTTGTTGCTTGCCTTTGACTAAGGACTTGGTAATTTGTTGAATTTTGAAAGTTGTGTAGAAAGGGTCTGGACTTAATTTATGGAAATTGAAGTTTTTTTGGTATTTTTGCCTAGTTGCTATAAAAAGATTGATTTTTTTTTAATTGTGTTGGTAGCTGCTTTGAATATGGACTTTGTAAATTTGTTTAATTTTGAAAGTTGCATAGAAAAGCTCTGGACTTGTATTTCTGGAAATTGAAGTTGAATTTGGCCTTTTGTTGTTTGATCTTAAGAACCCTTTTTGGGTTTTGTGGATTGTGTGAGATGGGGTTTGTGTCTTATATCAAGTAGAGGTCTTTTTTCTTTTTTGAACTTTCCTTTTGAAGGTCTGAACTTTTGTGCTTGTGGATGTGGAAGATCAAGTAAATTCTGTATCTCGGAAACAATCTCTCTACCATCTCAAGGTAGGGGTAAGGTTTGTGCAGTACCCTCCCTAGACCCGACCTGTGGGATTACACTAGGTTTGTTGTTGCCGTTGTTGTTGTTGTTGTTGACCTTAGTGATTAACTAATTATTACTAACCATTTCTTTAATACATTGGCTGCCTTCTTGAGTACTTTGAACAAGGACAGTGGTTTATTATTTGTTGAAAATTTGAAGGTCATGGTCAAAAAGCTCTCAACTTCAATTAATTTTTTTTGGGATAGGAATTTGCTCTCTTGTTGTTTGATTTTCAGAACCTTTTCTGGGTTCTATTAGACTTGCTTGTTGGGATTCCCTGTATATCAAGTAAGGGGGCTTTTATCAGCTTCCTTCTTGGAGGTGTGAAGTTTGTGCTTGTGGCTTTGTAAGTTCAGATAAAACCAACTTCCATCGTGTGTTTTGTTTATATCAAGTAGGGGTGTTCTTTTTTTTCAACTTCCACGTAAAAGGCGTGAACTTTTATGGTTGTCAATGTAGATGATAAGGTGAAATCATCTTTTGAATGAAGTTTTACTTTTGTTGACGATAGTTTTATGGAAGTGCATCTCTTGGATTGCTGTTATTGTGCTTTTTTCTCTTCCCAAGTCTCTTTTGCTTCCTTGTTTGCCGTTTTTCCCATTAAAATCTGTGAGGAACAGTTTTCTGCTTGTATCTGACTTGTCCTTCAATCTTTATTTTGGTTGTCCTGTAACTGTGGGATTTGAGTACCTCTACTTCTGGAGAATCACTAGTTCGCTTAAGAAGAAAAGTTATCTAATGTTGATATGAAATGAGTCCGAATAGAGCGGATGAAACGTAGATTCATATAGTTTTTTTTTTACTAGTTTGGAAGTGAGGCAGCATTGACTGATTGATAAAACTGATTCATCATCTTATTTTTTTCTTGCAGCCTTACATTTTGCTTCTTAACTCGGGAGAAATTGGATTTGTTCTTGCAATGGATTTTTCGGGGTTGCAAACTAGTCTATAGAGATCTTAAATGTTTCAGTCAACTACTTAATTAGGACCTTCGATTGTTGCGGTTATGACTGGGCCTCTCACCAAGATGGCTTTGGATAACAGAGCTCTTACTACCTTTGCTGCTATTTTGATCTACTTGGTCTTGAGCTGTGCTGTTTGCTCTGCTCTTCAAAGCGACATTGACTGTCTGAAATCTATAAAAGATTCATTTGAAGATCCATTGAAGTTCTTGAACACAACATGGAAGTTCGATAATCAGACCGAAGGTTTCATCTGCAAGTTTGCTGGGATTCAGTGCTGGCATCCTGATGAAACCAGGGTTCTGAGTATCAGTCTTCCTGACATGGGTCTCAAGGGCAAGTTTCCACGTGGTCTTAAGAATTGCACCTCTATTACATCATTAGATCTTTCGAGCAACAAACTCCATGGAAGTATTCCAAATGATATCTCCAAAATTATAGGTTTTATGGTAACGCTGGATCTCTCATCCAACAATCTATCAGGGGATATCCCAGTTAATCTTGCAAACTGCTCTTTTTTGAATAGCCTTAGACTTGACAACAACCAGTTTACGGGTCAAATTCCTCCAGAAATTGCCCTGCTTGGTCGGCTCAAGAATTTTAATGTAGCAAACAATCGATTGACTGGACCAGTTCCACGTTTCATAAATTCAACTTTTCCCCCAGAGAGCTATGCCAATAATGCCGGACTGTGTGGACCACCATTAGCTCTTTGTGAGGGTTATGCTAAGAAATCCCCTACTGGAATCATTGTCGGTGCTGCTGTTGGTGGGGTGACTTTAGGCGCTGTATTATTGACCATCGGTATGTTCTTCTATATGAGAAAGATATCCAGGAAGAGGAAAAAGGAAGATGATCCTGAAGGGAATAAATGGGCGAAGAGTATCAAGGGTTCCAAGGCAATCCAGGTGAGAATTCTGTTCGTGCTTTGTTGATAGGTCATAAATCAATTCTAGTGGGTTGTGGTTTGTGTTTACGGGTTCTACTGTGGCATTGCGATCATATTTAGAACCATAAATTCGTGTTTTAATATAAGCATTTAACTAATGAATTTTTCACCATAAGCTCGTGTGAGTTTTCTCTTGCTCAAGCATTCTTAATTGACTTAAAGTGCCTTATTTTCAGCTTTCAATGTTTGAGAAATCTACTTCAAAAATGAGACTAAGTGATCTCATGAAGGCCACCAACAACTTCAGCAAGAACAATATTGTTGGGTCAGGAAGAACCGGGACATTCTACAAAGCAGTACTTGATGATGGCACTTCACTTATGGTTAAGAGGTTGCAGAATACTCAACACTCAGAGAAAGAGTTTACATCCGAGATGGCTACGTTGGGAAATGTAAAGCACCGTAACCTGGTGCCTCTTTTAGGTTTCTGCATGGCGAAAAAAGAAAGGCTGTTGGTCTACAAAGATATGCCAAATGGCACCTTGCATGATAGGTTGCATTCGGTGAGTGAAGGGGAAAAAACTCTTGAGTGGCCTATGAGAATGAAAATCGCTATTGGAGCAGCCAAAGGATTTGCATGGCTTCACCACAACTGCAATCCTCGTATCATCCACAGAAATATTAGTTCTAAATGCATCTTGCTGGATGTGGAATATGAACCTAAAATATCGGATTTTGGACTTGCTAGGCTCATGAATCCAGTTGACACTCATTTGAGTACCTTTGTAAATGGGGAATTCGGGGACTTTGGTTATGTTGCACCTGAGTACGCGCGAACTCTCATGGCTACACCAAAAGGTGATGTGTACAGTTTCGGTGTTGTACTTCTCGAGTTAGTTACAGGTGAGACACCAACTTCTGTCGCCAAAGCTCCCGAGACCTTTAAGGGGAATTTGGTGGAATGGATCACACAACTCTCTGGTGAATCTAAACTTCAAGATGCGATTGACCATTCATTGTCTAGTAAAGGTTATGACAGTGAGATCTTCCAGGTCCTTAAAGTTGCTTGTCGATGTGTGTTGGCTGCTGCTCCTAAGGAGAGGCCAACAATGTTTGAACTGTACCAGCTTCTGAGAGCCATCGGAGAACGATATCATTTCACAACTGATGATGACATTATGATGCCTGAGAGTGATAGCGGATTCCAAATGGACGAGCTCATTGTTGCTCAATAAGTAAATGAAGCCCTTCGATAATGGTCGAGTAAGATACAATAATTACTAGATCATTTTCTATCCTGTACAAATACCACGGTTCACTATCTATATAATATCATTCTATTTCTACTGTAAGTCCCTTATTCATAGCTACCCAACTATGTAATTTGTGCTTCTTTTTCTGTATTTTTAATGGTGAAGAGATTTCTACAGATTGGTGTTAGTCAGTATTTTTCCTCTTTTCAGATTTACTGCTTGAACAGTGTAGTCTCATCTTCCAGTAATTATAAAATGCAAAGTTGGAAAAAATCCATCACTCAAAGCTGAC

>PGSC0003DMG400027703 LRR-RLKs

TGAATCCTGATATTCAAGCTAAACATAAAAACAAAGGTGCTATCATAGCTATCGTTGTATCAGGTGTTTTTGCTGCATTTGTTTCAGCAATTTTAACTGTGTTAATAAAAAAACGACATGCCAAATACCAAAGTCTCCTATCAAGGAAACGGCTATGTAAGTTATCTCATTTTGGTTATAAGCTACCTTTACTTCATAAGAATATGTTTCTTTCAATTCTTCCCGCAAAATTGACTAGCATAAAATGTTTAATTTCATAATGCCAGCCGCAAAGCTCTCTATAAAATTGGATGGCCTTAGAAGCTTCACTTTCAGAGAAATGACCTTGGCCACTAACAACTTTGATCACTCAAATCAAGTTGGTGAAGGAGGATATGGGGCTGTCTTTAAGGGAATTTTAGCTGACAAAACAGTTGTTGCTATCAAACGAGCAAAAGAAGGATCAGTACAAGGGCAAAAAGAATTTTTGACTGAGATATCTTTGTTATCAAGGGTACATCACCGGAATCTAGTTTCATTGCTAGGGTATTGTGATGAAGAAGGAGAACAGGTATGATAAACCTCAAAACTTCAAAATTCCAAATCTTTTTGAATCCACTCATTAACATGGAATTCATCATGTTGATCCTTTCCTCGGCTATATGAGTAGTTCAGGCTCACATTGTATGCTATAGCTTCTTTTTTTGCTCCGTAGCAATCTCTTGTCATTCGTTCTGACATCTCTTTGGTATATCATCTATGTGAAGCAATTAGTGTGTTTCCTGTTTATGTATTTGCAGATGTTGGTTTATGAATTCATGTGCAATGGAACTTTGCGAAACTGGCTTTCTGGTAAAGAAACTGTTCCCTGGTTTATTCTGATCCATCATCTTCCTCTGCTTTATTTTTGGTTAGGGAAGCTATTGAGCCAGGAGAGGAATTTGATATTTATCATTATTCTGATAATTGTGTTGTGTAAGCTGGCTTGCTTGCGCTTTGACTAATCTATTAAATACGTGCTACCTCCCACTAGCACAAATATCAGATAACTCCATCCTTCAAGGTTTAGGAAGATGGATCACCTCTGTTGGCGTTCCAACTTTAATTACCAAGTTTATTACTCCAAATCTACCACCACTAGGCCACCTTTCTATGTTCCTTAAGCAAATAGTAGCAGTATCTTTGAACTGGTACAGTAGACTGGTAATATATGCTCTTAAACATTGTTTTTTAAGTCTATAGATGGAGATCAGATGTCAAGAAGCAAACTTTGATCTAATAACTGTATGTCTATATGGATATCACCTAATCTAGATTATCTTGTTGTAGTTACTGCTCCGTTACTATCTGCTGTTTTTGTTACTATGTGTTGTTTCTTGTACTTCTGTTACTTCTTTTTCTGGACTGCTTTGGACTATTTTTCCCTTGAGCCGAGGATCTATCAGAAACAGCCTCTCTACCTCTAAGGTAGGGGCAAGGTGTACACTCTACCCTCCCCAGACACCACTTTGTGGGACTACACTAGGTATGTTTTTGTAGTTGTAACACTTGATATCATTTTTTGACAGCTAATTGTAAAGGAACCTTGTCGTTTGCTACAAGATTGGAGATTGTACTAGGTGCAGCAAAGGGCATTCTTTACCTCCATACAGAAGCTCATCCACCAATATTCCATAGGGACATCAAAGCCAGCAATATTCTTCTTGACTCAAAAATGGCAGCTAAAGTTGCTGATTTCGGATTATCTAGGCTTGCACCTGTTCAGGATGATGAAGGTGTACTACCTAACCATGTTTCAACTATTGTGAAGGGAACACCTGTAAGTCCTTCACTACATCATTATGTTCTATTTGACCTCACTCGTATCATTTAAATTTGATGCATTGAGACTCTACTTGTATATGCAGGGTTATCTAGATCCCGAATACTTCTTGACCCGTAAATTGACAGATAAAAGTGATGTTTATAGCCTTGGGGTTGTGTTTCTAGAAATCTTAACCGGAATGCGTCCCATCTCACATGGAAAAAACATTGTCCGCGAGGTATTACTCTCATTGTTAACTTCCCGATTTCCATGTCTTATCGTGTGCATATGTTTGATAGATGATTTATCCGCACACAAAGTGAAATACTGCTGACTGATGCAAGGGTAGGAATAGAGGCAGATTCATGAATTAAACTTGTAATGCTCTTAGTTCTTACCATTGAACACACTACATGTCGCACAAAATGACGCTTTTAGAATTTATATTTTATGGATACATTTTAGTGATTTTACATATGTATGTTATTTTTATGGTGCATCCTCCTCTGGGCTGAGAAGGGCTAGGGGAAATGTGAACATAATTTTTACGCAATCTTTCCTTGCATTTCTGCTATGAAGCTTCACACCTCGAATCTAGCTCCACCGTGTTTAATAGTCCCTTGTACCATCAATGGCAGGTAAATTTGGCTCATGGCTCTGAGAAGATGTTCTCAGTTATGGACAGTACAATGGGATCTTATCCTTCTGAATGTGTGGAAAAGTTTGTAGCTTTGGCCCTCAAGTGTTGCGAAGACAAACCAGAAGACAGACCTTCAATGTTGGATGTGGTCAGGGAGCTGGAAACTATTCAGTCTATCCTCAATCTGATGCCAGATATTGACACTGATTCAGTAGATTCTAAAGCAAAATTTAACGAACCAAAGACATCGTCTTCATTTTCTGATTGCACTAGCAAGGACGCGTTTTTATCGTCCTCCAATGTATCGGGAGCTTACAGTATCAGTGGTGTCAGTCTTACTATGCCTCGCTAACAGAATAGACAATAGTTGCTCATGTTTTGTACTAGAAAAAATAGCATGTAGAATAGAGAGTTGTACCATTTAGCTTCTGTTTTATATATGTATATTTCAAGGATTTGTTAACAAAAACCTCTTTATCATAGCCTTCTGAGTTCTGACAGTATGTGTATGCTATCAAAATCCAAAACTTTGCTACAAAATGGACTCTGAGATAGGTACTGCTATTTGTTTGCAAGGAGTTGCACCAACCAATTCCTCAAAACAACTGACAACAAACAGACAATGGCAGAGCTAAAGGGGCGGAAGGGGGTTCATCGCCAGAAAATCACACTGTATATATAAAAGGTCGATCTTTTTATCTATATAAATGATGAACCCTCTTGTAATCCAACATCATTACCAGAAATAGAAATATTATCTAAGCTATGGCTACAATGAGAGTCGTACAACAAAACAGGCAAAACTCGTAGTTTTGAAGTAATGTTAGCTTTAGATCACATAACATGTCATCGTAAACCACACAAAGGAACAGAAAACAAATCCTCAAATCAATATGAGGAAACTCATGTTTTGTTCTATGGAACCAAAGTTAGACATGGTTGACACACTCCAAGCTCCTACAAGAAGTCAGAAGACTATTTCTTGGACTAAAAAGACAGCAAAGTTCATTTTTATTTATAGATCATAGTAAGAGAGCAGCGGGTAAAATGGCTCTACCGGGAGGGAGGGAGGGAGCCGGTTTTGGATTGGTGTGTATTTGAAACAAATATTGATAATCACTTTGAAGCAGCATCAGCAGTATGCATACTGTCATTCTGGTCCAGCAACCTCTCTTTCTCTCCAACTAGTGTGTCTTCAACACGAACAACCTTTCCTTCATCTGGTTTATCTGCTTTCTGTGATCCTGTGAACTTTGCTAACATGACGAATGTCATTCCCCCAAGAATGTTATTTGCGCATCTCAAAGCAATAACCGAGGTCTTAAATACCAAAGGAGGAAGACCTTTAGCTAGCACAAACTCAACACCGTTAAGTGTTTGATATCTTAAATTACTACTGAGACCCATGTGTATTGCCCAAGTTGCTGCATTCAAAAGTGTAGGAGGTGGCTTGTTTGGTGTTTCGAAATTCGGATCCATCTTCTTCCTCAACTTGATTAAACCATTAGATAATGCAGTCCCAACAAGCCCTGCTGCAAATCCAACTGCAGCAAACTGAGTTCCTTTGTACACAAATGTACCAAGCCTACTCAATAAACCGTAAGAACCAGACTGAAACATGTGGCTCGGTGGACAACTGGCGAACATCAAAGGGAGAGTTTGGATCGAAGCAGATGCCGTTGGTGCCAACAGATACATCAACACAAAATTCATAATTGAACCAACAACAAGTGTCGAAAACACAAAATCTAATTCATTAAGCCCAAAATTAGGCCTTGATGCCATATCTCCAAGAACATTAGCACTAACACCAACCAATTCCTCCATAAGAACCTTAAAAGGAAACTGTGGATCTGCAGCCACCCTTGATCTCCAGCCATTAACAAAAGCACCAATTGGTCCAATACTATCCCAAGAAGACTTTGAATCATCTGAATCTCCACCTTTACTCCAATCACCACTACCCCCACCCCCACCACCCCCACTACCCCTTCCAATCCCAATACTTCCTCCATCACCTCCACCAACACAAGGAACTTGATACCTCAGGTGATCCACTTGTTTAGACACAATATACCCAGTTTTCAAAGGGTTAAAACAAACCCTATTATCACAAAATCTGGCTGAAACACCAGAAAGATCACGACTTTGATTATGATTCTGATAGTAACCCGTAAGAGGAGAACAACGAAGCTGAGACATCGCCACCATTTTCTACCAACCAACCTAATAAAATGAGAAAATAGAAAGTAATGGCTATATTATTCATCCCATCAACAAACAAAGATTCAGATTCACTAAAAGATCAAGAAAGACTCTTTCTTTGACACAAATAATATATATAAATACATACACACACAACAAAAAAGATGCAACAGAGTTAACAATAAATCAAGAAAAGGAAAAGGGGTTACCTTTTATGAAGAGATGTGTTCAGGGGCTGCACATCAAGTGATCTGAGCAGAGCAAAAAAAGATGGCTGCAGCGGCTAAAGTGACTAAAATGGGGATCTGCATGAAAGTGGGAATCTTTATAACCTAAATAAAAATACAGGCGGCAGGGCCGAAGTTAAAGAAAAAC

>PGSC0003DMG400027710 LRR-RLKs

ATGAAGTACCTGGAAGATTGGGAGAAGGGTGATCCATGTACATCTAACTGGACAGGAATTCATTGTTTTAACAAAGTTGGAGCCAATGGCTACTTGCATATCAAGGAACTGTATGTCATTATATTCTTCTCTCTTATTAGTTGTTATGTTTTCTTCTTCTTTATACCTGGAATTGCTGCACTTGAGTAGAGGGTCTTTCGAAAACAGTTTCTCTACCTCCACAAGGTAGTGGTAAGACCTGCACACACTCTACACTCTCCAAGCCCCGGGATTTCACTAAGTATGTTCTTGTTGTTGTAATTAGGTAGATTCAAAGCCAATTTACTCTAGCAAAACTTGGGTTGGTGTTTGAAAAACCCGAACCAACCTTGCCTATGTACACCTCTACTTTGGGCCATAAAAATACCCTTGATGTTAGAAAATCTTTGTATCTTTAATCGAGTTCTAGTGCGGTGGATTTCACGTGTAAGAATATATCAGGAAAAAGGTCTAAATTTACCCCTTTGCTTTTGGTTGTGGTTAAGAAATTACCTTTCATTATGCTTCAAGTTGGAGTTCCTCTAGCATTAACAGCAAAAAAAAGAGACTTGGTTCTTTCGATATCTGCATAATACTCCTGCAATCCAGTTAGCAGCATAACTAGTATCTCCAAGGAGGGACCTGGTTCTTTTGTAGTTGTTTGTTGATCATCAAAATCAATTTAAGCTCATCTCAACAGTTTGCTAACGACAAGGGCATGAATAACTAAGTTAACTCATAAATTAGAGCACTTAGTTGTAAACTCTAGCAAAAGATGCGTCTTTACTACTTTGAGTTATGACTATCTTGTATTCTTGTGTGAGGGTGATGCATGACGCATTAAGCATATTATATTTCCCCTTTGTGCAGCCGTATGATGGCTATGAATCTTTCTGGAAGTTTAACACCTGAGCTTGGACAACTATCTCATCTTCAGTTTCTGTGAGTTGTTACACATATTACTACGTTGTACAAATATTTTGATTTTAAGATATAGTAGGGTTTATTAATTTTTCTATCTGTTTTACTTATTTTTATCTTTATAATTCACTTTCTGAAAGGAATTTCATGTGGAATGATTTGACTGGGAGTATACCGGAAGAAATTGGAAACATCAAGTCTTTGAAACTTTTGTAAGTTGAACGAAATTTGATCCGTTTTTTTTTGGTCCCATCTTCTTGATTTTGCAGTCACCCTACATGTTTCCCCGAATGCTTTGAACCCTGAACTATCTCATCTAGGAGATCTAGATAATCATTTGGCTATGATGGTGGTATACTTTATGTTGGATAGAACAGGTGGATTTATGATTTTAAGTTTGATGGTTTCAATCTTTATGTTCTTATCACGGAGTCCATTACATTTTCAAAACTAGGGGTTCAAAGTTCAATATTTTTAAATTTATAGTGATTTGTCATGTATGTATCTGTGCTTCCCGTCGAAAATGTTGAGTTCAGTTGAACCTATTAATTAGATCCTTCATCCGCCACTGGATAGAATTGAGGAACATTTTTTTTCTTGAATTTCTTATGAAAATTGTGCTTAATGTAAGATTGTTTCAAGAAGAAGTAAAACCTAATATGCTTGGAAAGATCATGACTATGTCGATTGGTTTAATCCCTTCGGAATTATGTCCATGCAGGCTCTTGACTGGAAACAAACTTTCAGGATCTTTATCTGATCAGCTTGGGTATCTTCCAAACCTGAGAATATTCCAGATAGATGAGAACCAGATTTCCGGAAGAATTCCTAAATCCTTCTCCAATTTAAACAGTGTCCAACATATGTAAGTGCAACTCATTTGATTGTTTTAAACAGTGGAACAAGTCTAAGGTTCTACCTTTGTAATCTTATGTGATATATCATGTGTCTACAGCCACTTCAACAACAACACACTGATTGGCCAAATCCCTCCCGAGCTTTCCAATTTGTCCACTCTGCTTCACATGTAAGTTTCTTAGTTTCTCTTCTTCAATAGACATGCATACTCTCAGTAGAAGTTAGAGCGAAGCAAGTTTTTGTAACTTCTAAAATGACCTTTATCTCTGCTTATGTCTGTAAACTAACGGAGTAATGTTTTTGTGATGATACTTTCTTTTTGTCAACAGGCTTATGGACAGCAATAATCTTTCAGGATACCTTCCTCCTGAGTTTTCAATATTACCACGTCTTCGAATAATGTATGTCCCTACTCACACACACATATATGATATGCATATATTGAACCTCATATGCAACTTTCATTAGCTGTATTTTGCTCAAGAGGTATAACTAACTGTTACAAAAAAACAATGGTTATTCAGAGGCAACTTTAAACTGAACTTTCTGAAGTAACTAGTTATAAATTTGAAGGAATGCTGATTCCTTACTACCTATGAGTGAGACTGTCTATGTTTGTCTTAGTTGAAGTGGCGTTATTGGCCAGATGATCGAACAACTGAAGCTGAGAACATTTCGTTACAGATCTATCTCAGAACCCTCTAATTATCATCTTCAGATCTTTCTGTTGATTATAAAATTCATCGATAATCTTTGCTTGTAATATCCAATGATAAAAAAATGATTTTATATAAGACAAATATAGTAATCCTTGTCATACAATAACTGCTATGCCATTTGCAGTTCCACTGAGAAGTGCATGGGTTGAAATTTATGATGAAGTTCTGAATTTTCTTATGCAGACAACTTGATAACAATAATTTCAGCCGTTCCGAGATTCCAGCTTCGTATGGGAACATGTCATCATTAGTGAAACTGTAAGTGCTCCTTGAAAATGTTCCTCTCTTTTATTCTTCCCCTATTCTCTATGACTCAGCATTATAAGTTATACATTTGAAGGACAACAATAGTAAATAATATGTACAAACATAAGCATATATTCATTGTGCACAATGTATATGAAAGAGAGAATGTTCAGGACATCTTATTATGACGGTTCGGGTAGATAGAATAAGATGTAATAGTTGAATCGTACTGTCACCTAATCAAATGGAGATTCAATGCTTCTGATAATATTGCAGTAGTCTCAGAAATTGCACGTTGCGAGGATCCATTCCTGACCTTAGCAGAGTACCGAGTCTACACTATCTGTAAGTTCAAGTTTTGAAATTGTTTCTTCTGTTCCCGTTGATTTGTGACCTTTTTACTTCATGCAACATGATTATAAGATTTTCTAAAATGTTAACACTTCTTGCTTACTGTTTGTGCAGTGATCTCAGCTGGAATCAACTGTCTGGATCTCTACCAAATAAGCTTTCCAACAATATGACTTCTATGTATGCTTTCTATTTCATGCTACAGATTTCTTCATATATCTAGCATCTTAGTTCAGCCATATTTTGTGTTAGTTCATTATCTCATGTTGTGAAGAGTTCTACATCAGCCCTTTAAGAAACATTACTAAAGAGCTAGCCTTTGGGGTTGAGTTAGGCCCTACATCCATAACACAAGTTAAACATAATTTTTATCATGATTGCACATTTCTCTGTGTTTTTCCTGTCATTTTCCCTAATCTAATGTGTTGTTGTACATTCTCATGCAGTATTCTGTCACACAATCGTCTTGACGGATCTATTCCGGAGAGTTTTTCATCTCTTCCTTTGTTGCAGAAACTGTGAGGATCTTCACTGTGTCTTGTAATACATTCTCTTTAGTACTGTTTCTTTAACATTTCAAAGCTCAATTTCGTATATAACAAATTGATTTGACGGACTTGTCACAGGACAATGGATAACAACCTTTTGAATGGTTCCTTTTCTACCGACATTTGGCAAAACAAGTTTACTTCAACGTCGTCGCTGTTGATGTAAGACCTGAAGCTAAATTGTAACATCAAATATCTGATATGAAATGTACTATCATTTACACAGAAGTCTGGGAGTCTTTTTAACCCTTCTATCTTCTTTGCAAAACTAGTGATCTTCGGAATAATTCACTTTCGGACATTTCCGGTACTCTTGAACCTCCTCTGAATGTCACTCTAAGGTAAGGTTTTTCTTTCCATATTCAACGACAAAACCTAAGAGAAGCCATTGAATTACTAATTTTATGATCTGACTATCACTTCATATCCACATTTTCAATAATAAAAGACATTATTTTCAGGTTGCAGGGCAATCCTGTCTGCAGAAATGAGAATGTAAGAAACATCGTCAAGTTCTGCGGCTCTGAAGCCGGAGCAGAACATAAGACGAATAACTCAGTGATTGTAACTGGTGCTTGTCCTATTGTTGCATGCCCAAAGGATAACTATTATGAATATGTTCCAAACTCACCTGTTCCCTGCACTTGTGCCTCACCTCTCATAGTTGGCTGGCGGCTGAAAAGTCCAAGTTTTTCTTACTTTGATCCATATGTGCATCAATTTGAGCAGTATATGACTAGAGATCTTCATTTGGACCTTTACCAAATGTTGATTGAGTCAAATTCTTGGGAAGAAGGACCTCGTCTACGAATGCAATTGAAGCTTTTCCCAGTAGTTGGGGTCAGTACATTTAATAAAAGTGAGGTCATTCGTATCAGCGACATTCTGCAATATTGGGAAATATCTGTAGTTGACCTTTTTGGACCTTATGAGCTCCTTAGTTTCACTCTGGAGGGACCTTATTCCTACTGTAAGTTCTCAGCGCTTCATTGTGCTTCATGGTTGATCTTTGTTATCATGATTAGTATCTTCCTTCCTCAGAATTTTGGTTTATCATATGATGCATGTTTTTCTGCAGTGAATCCTGATATTCAAGCTAAACATAAAAACAAAGGTGCTATCATAGCTATCGTTGTATCAGGTGTTTTTGCTGCATTTGTTTCAGCAATTTTAACTGTGTTAATAAAAAAACGACATGCCAAATACCAAAGTCTCCTATCAAGGAAACGGCTATGTAAGTTATCTCATTTTGGTTATAAGCTACCTTTACTTCATAAGAATATGTTTCTTTCAATTCTTCCCGCAAAATTGACTAGCATAAAATGTTTAATTTCATAATGCCAGCCGCAAAGCTCTCTATAAAATTGGATGGCCTTAGAAGCTTCACTTTCAGAGAAATGACCTTGGCCACTAACAACTTTGATCACTCAAATCAAGTTGGTGAAGGAGGATATGGGGCTGTCTTTAAGGGAATTTTAGCTGACAAAACAGTTGTTGCTATCAAACGAGCAAAAGAAGGATCAGTACAAGGGCAAAAAGAATTTTTGACTGAGATATCTTTGTTATCAAGGGTACATCACCGGAATCTAGTTTCATTGCTAGGGTATTGTGATGAAGAAGGAGAACAGGTATGATAAACCTCAAAACTTCAAAATTCCAAATCTTTTTGAATCCACTCATTAACATGGAATTCATCATGTTGATCCTTTCCTCGGCTATATGAGTAGTTCAGGCTCACATTGTATGCTATAGCTTCTTTTTTTGCTCCGTAGCAATCTCTTGTCATTCGTTCTGACATCTCTTTGGTATATCATCTATGTGAAGCAATTAGTGTGTTTCCTGTTTATGTATTTGCAGATGTTGGTTTATGAATTCATGTGCAATGGAACTTTGCGAAACTGGCTTTCTGGTAAAGAAACTGTTCCCTGGTTTATTCTGATCCATCATCTTCCTCTGCTTTATTTTTGGTTAGGGAAGCTATTGAGCCAGGAGAGGAATTTGATATTTATCATTATTCTGATAATTGTGTTGTGTAAGCTGGCTTGCTTGCGCTTTGACTAATCTATTAAATACGTGCTACCTCCCACTAGCACAAATATCAGATAACTCCATCCTTCAAGGTTTAGGAAGATGGATCACCTCTGTTGGCGTTCCAACTTTAATTACCAAGTTTATTACTCCAAATCTACCACCACTAGGCCACCTTTCTATGTTCCTTAAGCAAATAGTAGCAGTATCTTTGAACTGGTACAGTAGACTGGTAATATATGCTCTTAAACATTGTTTTTTAAGTCTATAGATGGAGATCAGATGTCAAGAAGCAAACTTTGATCTAATAACTGTATGTCTATATGGATATCACCTAATCTAGATTATCTTGTTGTAGTTACTGCTCCGTTACTATCTGCTGTTTTTGTTACTATGTGTTGTTTCTTGTACTTCTGTTACTTCTTTTTCTGGACTGCTTTGGACTATTTTTCCCTTGAGCCGAGGATCTATCAGAAACAGCCTCTCTACCTCTAAGGTAGGGGCAAGGTGTACACTCTACCCTCCCCAGACACCACTTTGTGGGACTACACTAGGTATGTTTTTGTAGTTGTAACACTTGATATCATTTTTTGACAGCTAATTGTAAAGGAACCTTGTCGTTTGCTACAAGATTGGAGATTGTACTAGGTGCAGCAAAGGGCATTCTTTACCTCCATACAGAAGCTCATCCACCAATATTCCATAGGGACATCAAAGCCAGCAATATTCTTCTTGACTCAAAAATGGCAGCTAAAGTTGCTGATTTCGGATTATCTAGGCTTGCACCTGTTCAGGATGATGAAGGTGTACTACCTAACCATGTTTCAACTATTGTGAAGGGAACACCTGTAAGTCCTTCACTACATCATTATGTTCTATTTGACCTCACTCGTATCATTTAAATTTGATGCATTGAGACTCTACTTGTATATGCAGGGTTATCTAGATCCCGAATACTTCTTGACCCGTAAATTGACAGATAAAAGTGATGTTTATAGCCTTGGGGTTGTGTTTCTAGAAATCTTAACCGGAATGCGTCCCATCTCACATGGAAAAAACATTGTCCGCGAGGTATTACTCTCATTGTTAACTTCCCGATTTCCATGTCTTATCGTGTGCATATGTTTGATAGATGATTTATCCGCACACAAAGTGAAATACTGCTGACTGATGCAAGGGTAGGAATAGAGGCAGATTCATGAATTAAACTTGTAATGCTCTTAGTTCTTACCATTGAACACACTACATGTCGCACAAAATGACGCTTTTAGAATTTATATTTTATGGATACATTTTAGTGATTTTACATATGTATGTTATTTTTATGGTGCATCCTCCTCTGGGCTGAGAAGGGCTAGGGGAAATGTGAACATAATTTTTACGCAATCTTTCCTTGCATTTCTGCTATGAAGCTTCACACCTCGAATCTAGCTCCACCGTGTTTAATAGTCCCTTGTACCATCAATGGCAGGTAAATTTGGCTCATGGCTCTGAGAAGATGTTCTCAGTTATGGACAGTACAATGGGATCTTATCCTTCTGAATGTGTGGAAAAGTTTGTAGCTTTGGCCCTCAAGTGTTGCGAAGACAAACCAGAAGACAGACCTTCAATGTTGGATGTGGTCAGGGAGCTGGAAACTATTCAGTCTATCCTCAATCTGATGCCAGATATTGACACTGATTCAGTAGATTCTAAAGCAAAATTTAACGAACCAAAGACATCGTCTTCATTTTCTGATTGCACTAGCAAGGACGCGTTTTTATCGTCCTCCAATGTATCGGGAGCTTACAGTATCAGTGGTGTCAGTCTTACTATGCCTCGCTAA

>PGSC0003DMG400027967 LRR-RLKs

ATGGATCCCAAAATACTTTATTTCCTTCCTCTGGTATATCTCTTCACAATTACTTTTGCTTCCACAGAGGAAGCAACAACTCTTCTTAAATGGAAATCAACTTTCAAAAACCAGAATAATTCCTTATTGGCTTCTTGGACACTACGTCATCCTGCTGCTGCCAAGAATTTTTCGAGCCATGATCCATGCAGGGACTGGTATGGAGTTAAATGCTTTAATGGTCGAATAAACAGGTTGAATATTACAAATTCTGGCGTCATTGGTACACTCCATGATTTTCCATTTTCATCTCTACCTTTTCTTGAATATATTGATTTTAGCATGAACAACCTTTCTGGTCCCATTCCTGCAGAAATAGGGAAACTTACCAACCTTGTCGAAATTGATCTTGATACAAACCAGTTAACAGGTCATATTCCTCCGGAAATAGGCAACTTGATCAATGCAAAATTGTTCTATGCTTACTCCAATGAATTGTCTGGTTCCATTCCTATTGAAATAGGGAAGATGAAGTCACTTGAAAGTTTAAGCTTACAGAAAAACAATCTTTCTGGTCCAATTCCAAAATCTTTAGATGACTTAACTGAGCTCAAAATTTTGTACCTTTATTCTAACCAACTTTCTGGTCTCATTCCTAGTGAGTTGGGGAATCTGAAAAACCTCAATGAGCTAGATTTATCCGATAATAAGCTTAGTGGTTCAATTCCAATTACTTTAGGTGACTTAACTGAGCTTGAAACTCTATACCTTCATTCTAACCAACTTTCTGGTCCCATTCCTCGAGAATTGGGGAATTTGAAAAACCTTAACGATCTGGATATATCCGATAATAAGCTTAATGGCTCAATACCAATTACTCTAGGTCACTTAGCTGAGCTCAAACTCTTGCATCTTTATTCCAACCAACTTTCTGGTCCCATTCCTAGTGAGTTGGGGAATCTGAAAAAATTGAATGATCTGCAATTATCCAATAATCAACTTACCGGTCCTATTCCTGGTTCATTTGGCAATTTGAGAAACTTGCAATTTTTGTATCTCCACACCAACAAACTTTCTGGTCCTATTCCGAAAGAACTTGCATTTTTGGATAACTTGGTAGAGCTGATAATAAGTGAAAATCAGCTTTCAGGCCATTTGCCTGAGCATCTTTGCCAAGGTGGGAAACTTGAGAACTTCACGGTGAATAGTAACAAGCTTACAGGGCCAATACCAAGAAGCTTTAGCAAGTGCTCGAGTTTCAAAAGAGTTCGGTTGGATAACAACAGTTTTACTGGGAATTTATCTGATGCTTTTGGCATCCATCCACACCTTTACTTCATTAATTTGAGCGAAAATGATTTTTATGGTGAACTCAGCAGCAACTGGGGGAAATGCAAGAGTTTGACTGATTTGCGGGTAGCCAGAAATAACATTAGTGGTAGCATACCACCGGAGATTGGAAACCTCAAAGGGCTTCAAGGACTTATTCTTTCATCCAATCATTTGGTTGGACAGATACCTAGGGAATTAGGAAAATTAACCTCTCTGGTTAATCGTCTTTTGCGAAACAACCAAATTTCTGGCAATATTCCTATGGAACTTGGATTATTGACAAAGTTAGATTACCTTGATCTATCAGACAATAGATTGAATGGTTCGATCCCACCGTTTATAGGAGATTACCAGCACCTGTTTCACTTGAACCTAAGCAACAACAAATTTGGCCAAAAAATTCCAATAGAGATTGGGAAGATAACTCATCTTAATGTACTGGATTTGAGCCGTAATCATCTTGTTGGAGAGATCCCCCCTCAGTTAGCCAATTTAAAGGTGTTGGTAAACTTAAATCTTTCACACAATGGCCTATCTGGGCGCATTCCTCAAGAATTTGAAAGTTCAACTGGTTTGCAGGATGTCGTATTGTCATACAATCTGTTGGAAGGTCCAATCCCTAATAATAAAGCTTTTATCAATGCCTCATTGGAGGGTAATAAAGGTCTTTGCGGCAATGTAGTAGGAATCCATCCCTGCGGAATGCCGTCTTCTGTGGTGAAGAAGCACTCAATGGCGAAGGGACGTAAACTCATCCTCATCATTGTACTTCCTGTTATGGGAGCACTTGTTTTACTCTGTGTTTTCATTGGTGTTCTCTTTATGTGTGATAAAAGAAGAGCTGGAGATGTTGAAAGAAGGGATGGTGATGGTTGGCTTTCGATATCCATGTTAGATGGGAAGGNNNNNNNNNNNNNNNNNNNNNNNNNNNNNNNNNNNNNNNNNNNNNNNNNNNNNNNNNNNNNNNNNNNNNNNNNNNNNNNNNNNNNNNNNNNNNNNNNNNNNNNNNNNNNNNNNNNNNNNNNNNNNNNNNNNNNNNNNNNNNNNNNNNNNNNNNNNNNNNNNNNNNNNNNNNNNNNNNNNNNNNNNNNNNNNNNNNNNNNNNNNNNNNNNNNNNAAGGGCATTGACTGGGATTAAGCACCGGAACATTGTGAACCTCTACGGCTATTGTTCGAAAGCACAACACTTGCTCTTGGTTTATGAGTATGTGGAGAGGGGGAGTTTGTCTAGCATTTTGAGCAACAAGGTTGAGTCCAAGAAATTGGATTGGTTTAAAAGAGTGAATATCATCAAGGGTGTTGCTTTTGCTTTATCTTACATGCACCAGGACTGTTCACCCCCGATTGTCCATCGAGACATAAGTAGCAGTAATGTTTTGCTTGATTCTGAGTATGAAGCTCGTGTTGCAGACTTTGGCATAGCGAAGCTACTCAAGCCAGATTCATCGAATTGCACTGCACTTGCAGGCACATATGGCTATGTTGCACCTGGTAAGTTTTAAATATTTCATGAAAGCCAAACACTATTGTTTTTATCTTATTTACAAGTTTAAATTGATAATATTGTTCTGGAAACTGTATTGTTCAGAGCTTGCATATACTATGAAGGTTACACAAATGTGTGATGTCTTTAGCTTTGGAGTACTATCATTGGAGATAATAAAAGGAAAACATCTTGGGGAATACATTACCGTGCTAGCAGATTCATCGACTATAGATCCTGAGCAGCTTAGCGATTTTCTGGATGAACGGCTTCCATATCCTGAAGATAGAGTAAAAGAGGCTTTGGTTTTTATCATCAAGCTAGCATGCTCTTGTTTGCTTGAAAACTCCAAAAATTCAAGGCCAACAATGCACTTCATCTCTCATAAGTTATCATCAATGGATGCACGTCCACCTATTCATCAGAGAAACCCCCATGTAAGGCGAGCAATATAA

>PGSC0003DMG400027971 LRR-RLKs

ATGGATCCCAAAATACTTTATTTCCTTCCTCTGGTATATCTCTTCACAATTACTTTTGCTTCCACAGAGGAAGCAACAACTCTTCTTAAATGGAAATCAACTTTCAAAAACCAGAATAATTCCTTATTGGCTTCTTGGACACTAAGTCATCCTGCTGCTGCCAAGAATTTTTCGAGCCATGATCCATGCAGGGACTGGTATGGAGTTAAATGCTTTAATGGTCGAATAAACAGGTTGAATATTACAAATTCTGGTGTCATTGGTACACTCCATGATTTTCCATTTTCATCTCTACCTTTTCTTGAATATATTGATTTTAGCATGAACAACCTTTCTGGTCCCATTCCTGCAGAAATAGGGAAACTTACCAACCTTGTCGAAATTGATCTTGATACAAACCAGTTAACAGGTCATATTCCTCCGGAAATAGGCAACTTGATCAATGCAAAATTGTTCTATGCTTACTCCAATGAATTGTCTGGTTCCATTCCTATTGAAATAGGGAAGATGAAGTCACTTGAAAGTTTAAGCTTACAGAAAAACAATCTTTCTGGTCCAATTCCAAAATCTTTAGATGACCTAACTGAGCTCAAAATTTTGTACCTTTATTCTAACCAACTTTCTGGTCTCATTCCTAGTGAGTTGGGGAATCTGAAAAACCTCAATGAGCTAGATTTATCCGATAATAAGCTTAGTGGTTCAATTCCAATTACTTTAGGTGACTTAACTGGGCTCAAAATTTTATACCTTTATTCTAACCAACTTTCTGGTCTCATTCCTAGTGAGTTGGGGAATCTGAAAAACCTCAATGAGCTAGATTTATCCGATAATAAGCTTAGTGGTTCAATTCCAATTACTTTAGGTGACTTAACTGGGCTCAAAATTTTATACCTTTATTCTAACCAACTTTCTGGTCTCATTCCTAGTGAGTTGGGGAATCTGAAAAACCTCAATGAGCTAGATTTATCCGATAATAAGCTTAGTGGTTCAATTCCAATTACTTTAGGTGACTTAACTGGGCTCAAAATTTTATACCTTTATTCTAACCAACTTTCTGGTCTCATTCCTAGTGAGTTGGGGACAGGGCCAATACCAAGAAGCTTTAGCAAGTGCTCGAGTTTCAAAAGAGTTCGGTTGGATAACAACAGTTTTACTGGAAATTTATCTGATGCTTTTGGCATCCATCCACACCTTTACTTCATTAATTTGAGCGAAAATGATTTTTATGGTGAACTCAGCAGCAACTGGGGGAAATGCAAGAGTTTGACTGATTTGCGGGTAGCCAGAAATAACATTAGTGGTAGCATACCACCGGAGATTGGAAACCTCAAAGGGCTTCAAGGACTTATTCTTTCATCCAATCATTTGGTTGGACAGATACCTAGGGAATTAGGAAAATTAACCTCTCTGGTTAATCTTCTTTTGCGAAACAACCAAATTTCTGGCAATATTCCTATGGAACTTGGGTTATTGACAAAGTTAGATTACCTTGATCTATCAGACAATAGATTGAATGGTTCGATCCCACCGTTTATAGGAGATTACCAACACCTGTTTCACTTGAACCTGAGCAACAACAAATTTGGCCAAAAAATTCCAATAGAGATTGGGAAGATAACTCAACTTAATGTACTGGATTTGAGCCGTAATCATCTTGTTGGAGAAATCCCCCCTCAGTTAGCCAATTTGAAGGTGTTGGTAAACTTAAATCTTTCACACAATGGCCTATCTGGGCGCATTCCTCAAGAATTTGAAAGTTCAACTGGTTTGCAGGATGTCGTATTGTCATACAATCTGTTGGAAGGTCCAATCCCTAATAATAAAGCTTTTATCAATGCCTCATTGGAGGGTAATAAAGGTCTTTGCGGCAATGTAGTAGGAATCCATCCCTGCGGAATGCCGTCTTCTGTGGTGAAGAAGCACTCAATGGCGAAGGGACGTAAACTCATCCTCATCATTGTACTTCCTGTTATGGGAGCACTTGTCATCCTCATCACTGTACTTCCTGTTATGGGAGCACTTGTTCTACTCTGTGTTTTCATTGGTGTTCTGTTTATGTGTGACAAAAGAAGTAGAGTTGGAGATGTTGAAAGACGGGATAGTGATGGTTGGCTTTCGATATCCATGTTAGATGGGAAGGCATTGTACAGGGACATCTTAAACGCCACAGAAGAGTTTGATGCAAAATTTTGCATCGGGCAAGGAGGACACGGAAGCGTTTACAAGGTAAACCTTCCATCATTAGGGAATATAGCTGTGAAGAGACTTCATTCTTCATTTGAGAATACACCTCCCAAAAGTTTCTTAAATGAAGTAAGGGCATTGACTGGGATTAAGCACCGGAACATTGTGAACCTCTATGGCTATTGTTCGAATGCACAACACTCGCTCTTGGTTTATGAGTATGTGGAGAGGGGGAGTTTGTCTAGCATTTTGAGCAACAAGGTTGAGTCCAAGAAATTGGATTGGTTTAAAAGAGTGAATATCATCAAGGGTGTTGCTTTTGCTTTATCTTACATGCACCAGGACTGTTCACCCCCGATTGTCCATCGAGACATAAGTAGCAGTAATGTTTTGCTTGATTCTGAGTATGAAGCTCGTGTTGCAGACTTTGGCATAGCGAAGCTACTCAAGCCAGATTCATCGAATTGCACTGCACTTGCAGGCACATATGGCTATGTTGCACCTGGTAAGTTTTAAATATTTCATGAAAGCCAAACACTATTGTTTTTATCTTATTTACAAGTTTAAATTGATAATATTGTTCTGGAAACTGTATTGTTCAGAGCTTGCATATACAATGAAGGTTACACAAATGTGTGATGTCTTTAGCTTTGGAGTGCTATCATTGGAGATAATAAAAGGAAAACATCTTGGGGAATACATTACCGTGCTAGCAGATTCATCGACTATAGATCCTGAGCAGCTTAGCAATTTTCTGGATGAACGCCTTCCATATCCTGAAGATAGAGTAAACGAGGCGTTGGTTTTTATCATCAAGCTAGCATGCTCTTGTTTGCTTGAAACTCCAAAATCAAGGCCAACAATGCACTTCATCTCTCATAAGTTATCATCAATGGATGCACGTCCACCTATTCATCAGAGAAACCCCCATGTAAGGCGAGCAATATAA

>PGSC0003DMG400028123 LRR-RLKs

CAGTTACAAGAGAGCATAATGTCTAATGCAATGTCAAGAAAGAATATGGTTGATCCAGCAATTAGAAATTCGTGCTCGGATGAATCATTGAAGACGATGATAGAGATATGTTGCAGGTGTTTGGAACAAGATACAGAAGATATGCCCTCAGTTGAAGATGTAATATGGAATTTGCAGTTTGCTGCTCAAGTTGAGGATTCATGGAGAAAAGACTCTTCAAGTAGTGATGCCTCTCCAATTTCACATCTCTACAGCTTCAAACCCAACTTGTCGCGCAATAGCAATAGCAATAACAATAGCAATGGCACGTCTGTGTGACAAGCTATTTCTAACGTGGATAACTAATCTCGATACTATTATTCCATCCTGTGACAAGAATAACTAAACTAACAATTTATGTAATAATTTTCTTTGCATAGCTTATACAAAAAAGTAAGTTTTCATACAATCTTTTTCCTAACAAGCAGATGCCATAGCATTTTCCTTGTGTATATACCAACCCAACCTTAATAGTACCTAGATATTACAAGAGCAACATATAGTGCAGAAACCTATAATAGACATGGATGATAAAAAACAAAAACAAAAGAATGCAAAGAGGGACTAGTTAATGGAAAAGGGAAGCATTACAAATCGATTTTTTAGACAGTAGAAACAGAATATCTATTCGTTTGCAGGAGGAGTTTCCATTTTCTGAACTTGAGGGGCTGTCACTGCCACTAAGCTAAGATCAGATGAGTTTGAATCTCCAGATCTTTCATGGATCTCTCGGTACTCCTGGCATATGGCACATAAGTGGCAGAAAAAGTGTGTGGCGAAGTCTCCACATGGAGCTTCCTGCATATATATCAGGACACACAAAATTGTTAAATGCAGAAATACTGATGTAAGCAGTTTGCATCTTTCTATTCAGGCATGGAACCATAGTATTTCGTAGAACAGAAAAGGGGGGCGGGAGCTAATGGGAAGAGAGGAAGGTGAAGAATTCAGGGTAACTGGTTATAGATGCTGCAGACTCATCTGATAATAAACTCAAAACTAAGATTTCTAACTCGGACAACTTCTCAATGTAAACCAAACAGAAAAGGAAAGTACAGTCAGTTCTCTTTCTACATGGAGTCCTGATTTTCCTGATTTTCAAGTTGTTGATAGTTTATCGAACTGAAGTCCACCACTACTAATTCAAATGAAGTATAATCATCTCTGCCGTGGAGTCGATTACACACTGTAACTCTTTAATCTTTTGTAGCTTCTACGGATAACATCCAATTCAAAAGAATGTATGTGGAATTTTAAGACATTTCTGATTAATTTTTCTTTCTAATGAAAAGGTTAACTTCAAACTATCATCATAGAAGTCTTAATATAGTCATTAATATTAATATATATCACACCATAATGGGGAGTTTATGCATAGACGTAAAGAAACATCCTCAAAAGGTGTGCACATCAGACTATCCTCCTATGCATAGCCCATGTTATAGGAGATACAACTATGAATATGGAAGGTTAGATTGAAGGTAAATTTTCAAGCAATCTAACTTCACTCATTTTTTCAAATGTCTTTCCTGTTGAATCAATAGAGCTACTGATAATATGCTAAACCACATTCTCATCCATGTGCACTATAAACAGCAAAACAACAGAGATGGCTACCTGAAGATTGTACTTTGATCTCAATGTCTTTCGATAGCCACAGGCATAACATGCAACAAAGCACCCAGGCAAACCCAGAAGAATGCCTTCAGTTAAGAAGCAGAAAATTGTGTTGAAAAAAGCCCACAAAATAAAGTGAGTTGTACATGATCCCATCAAGGTTCCTGACCCCAAGAAGTCCGCATTTTTTCCAAACAGAAAGCAAGGGCAAAGAAGTCCAATAAAACCTGTGCTCAAAATTTTGCAATTAGTGCTAACTCGAGGTTGTCAACCAACATCTAGTGTCATTCCTCTGGGATGACAGAAGAGCCATTGCTTAGGAACAGGTGTTGGAATTTTAAACATAAACACACATAATTAACCCTCAAAAGAGAAGTTCCACAATCACACTTTCATCTTATTATATGTTCAGAAAGGAATGATTGATCTAGTCCCATCGGCACTGCTGAAAACCAGTTTCGGGAAATTCCGGCAGTTGTATTATTTATAACTTCATGAATATGACAAACAGTGAAACCACAAGGGGGAGCAAATAGCAAATAATCTGGAAACAGAATGGAAGCAGTGTTGACTGTTGAGAAAAGAAATGCATGTGTGTTATCCAGAACTCTATGGAATTACACTACACTGGGTATGTTGTTGTTCATAAGAATATATACATAAATCCTGTGGGATTACACTACACTGGGTATTATTTTGTTGTTGTTCATAAGAATATATACATATATCCTTATGGGTATACTTTTGTCCCAGCAAAAGAGCACCAAGAAGGTTCTGTATATGTACAGTATACAAAAACTCAGATACCCCTTTATAAATGCAGGGCACTGAGGAACTCTAGCAAGGCACCTAAATCATACACACTTAAGCATGTTACACCATAAATAAAAAAAGTACAGGCACTTCTAAGTTCCTAAGTCTGGATAGGACTAGTTTTGCAAATTTGGTAAGAAAGTGTATTTTCAAAGGCATACCAACTTCAAGCTAAATTTTTATCTAAGAAGCATATAAGCAGAGAAAAATACTTCATTCCATTATAGAGCCCGAGGAAAACTCAATTAAATGTATGGCTTATAATAGCTTCCTTAGTGGTTCACTCAGAGATTTATCATTGTTTTCCATCTAATAAGATAAGAGGGTAATTAATCATGAATCTTATTTTACTACCCAAAAATGCACCTTTCATTCACTTTTTTCCCTTCTCATAAAGCAAGATCACAGGGACGAGAGTAGCGTACAGCTCTGTGGATCATCAAAACAAGCACAGATTCCAGAACTCCACTGAGCATGGTCATCTTTTTTGGTCTGTTGAATTTCTTCCGCAGTAGAAACTTGTGGTACTTCTGCATCCGACTGATCCAGAGGAATATAAGGTGGAGGTGAATAATTCTTGTGCCCTACCATTATTAGTCAGGTGCAAACAGCAACCTGCAGCACAAATTCAAGAGAAAGCAAAAAATATCAATTAAGAACAAGCGAGTTACACATAAACATCCAATCAGTTCTCTGATTTCATGAAGGACGGTTACTTTTTGACTAAGTTGCTCGGACTCTCCAAAAATGTTGCCGCACCCGTGTTGGATCCTTCAAAAATACATTATTTTTGGAAGATCCGACACGCACTTGTCGACATTTTTGTAGAGTCCGAGCAACATAGCTTGTGTGGGGGGGGGGGGGGGGGGGGGGATTTTAAAAATCCTTAGGGGTACATGCCCCCCTCCCCCCTTAAATTCATCTCTGAATGTCTAACACTTACTAGATTCCCTTTGCTAAACTCTACAGAGTTCTTGTTGGACATTGGAACTTAGATAGTTTGTCTATACTGTACGGGTTCAACAAATATATTTCAATATCTTTGTCAAATTACTCCAAATTGATGTAGTTTAATAAATATAAACGGCATCTATAATATTCTCTATTGGTAATTTTATAAATACATCGTAGATTTATTCTTTTTAGTTCTTCTATATGTGTATACTCCAAATTAACACTTTCCTCTAACCGTGCTTATAAATGTCTGGTACTCATTTTCACAACTATAGTGTCTTATCACAGGTGAAATCTCGAATACACCCGTGTTAGTTCTGCATTAAATTGTAAAACCTTAACGTATATAAGAGAGTTATCCTTCGTATTACTTATCAACTAAGTTGCTCGGACTCTCCAAAAATGTTGCCGCACCCGTGTTGGATCCTTCAAAAATACATTATTTTTGGAAGATCCGACACGCACTTGTCGACATTTTTGTAGAGTCCGAGCAACATAGCTTTTTGATGAAGTTATAAATCATCTGCCATCGTTTCAAAAGAAAAGAAACACACACACAGATATTGACAGGAAAGAGAAAAAGAAAGATGCTTAACATTACAAAATCGATTCTAAGTTGTCAAACTGAAAAATGTTTACCAACAAAAAATTAGCACTGTGTAGATTGACCACATTGCAGATTGGCTCCACTAGGACTGACGATTATTTTATCATGTAATGCTTACGATTAAGTATTTCCAACTTTCAATTATGGGTGTTCCTTAGAAAGGATTGCACCCAATTTGTGCACTTCCTTTATTTTTTGACCTTGTAATCACAAAAGGAAGCTTTCCTAGTGCTTGCCTGTTTTACAATGTGAAAATTTCGGGCCAGCTTGCTCGCACCTCAACTAATTTAACGGGACACCTACTACCTCCCATCAACACAAGTACAAGATAACTCTATCCACTAATGTTTGGACATATGGGAGGAAATCATCTAATATTATTGCACTTGGAGCCTCATGGTTCTCAAACCAATTCATTGACCATTAGCCCACAGCCTCACAACCTTGGTGCAACTATTCAGAATTTTTTATCAATAACTATTAACCATAGCCATTTCTGATTTAAAAAATTAGACTAGATGCACCTGATACCTGTTGCTACTTGTTGCTGGAGGGAGGTAACACGTATCCCGTGAAATTAGTCAAGGTGAGTGTAAGCTGACTCAAACACGACACGATTATCAAAAGAAATTGTAATGTGGTGTAACGGTAATTAGAAGGATTTGACAAATGATTAGCAATTCATGACAGTATGTTTCGATTCATCTATCGGCCTTAATTTTTTTTCATTTTCTTCAAACAATTGATAAGCAATACTTCAATGCAATTATTACTCAGATCTCAACTATAGAGCCATGACAAATAACGACATAGTATCGAAATTCAAGCTTTATTCATGCAGTGTAAACGAATTTAAAGCATTGAATCCAAATATCAAAAACAATATCGTCTATTTTACCGCTAAGAATCATTTGGGATGAGGTTTTCGTTTTTTTTTTGCTCGTAAGAGAAAATAAAAAGGCAAAATCTGGAGAAGAAAGAGACAGACCGGAACTTACAGCTCGATGTCGCCGGAGAGAAGGATATTGCTGAGGATTTCGCCGGAGAAAGAGATCGCCGGGCAATGAACTTCGCCGAAAAACAAAACAAGGAGTAGTTTCCAGAGGACAAAAAGTGGCGTTTTGTT

>PGSC0003DMG400028147 LRR-RLKs

TCTCACTCCAATTATAACACTGTATATTACTATCTCTCCATTATTCTTGGCTTTGTAGTGATCTGAATCAACCCATGTTTCCCACCTCTCCTTGTCTAAATTCTAACATGGGTTTGTACAAAAAACAGCCATTAGTTGGCACAAAGAAATCAACTTTATTGATTATGTGATTCTTGTCAATACATGCTGTCTAGTGGTTCTTGGTGAAATCATCATCAATGGAGGCTGTTCAGCTAATGGGGTTTGCTCTTACACTCTTCATTTCACTTTTTTCAGTTGTTTACAGCGTTACAGACCCTAATGACCTTGCCATTATCAATGAGTTCAAGAAAGGGTTAGAAAATTCTGAGCTTTTGGAATGGCCAGTTAATGGTGATGACCCTTGTGGTCCTCCAGCTTGGCCTCATATAATTTGTACTGGAAACAAAATTCAACAGATTCAAGTTATGGGGTTGGGGTTAAAAGGCCCTTTGCCTCAGAACTTTAATAAGTTGTCTAAATTGACTAATTTGGGGTTGCAAAAGAACAAATTTAGTGGAAAGTTACCATCTTTTGGTGGTTTGTCTGAGTTGCGCTATGCTTTCTTGGATTTCAACATGTTTGATAGTATTCCATTAGATTTCTTTAATGGGCTTGTGAGTTTAGAGGTGTTGGCATTGGATGATAATCCTTTGAATGCAACTACTGGTTGGGGTTTGCCTAATGAGTTGCAAAGTTCAGCTCAATTGACTAACTTGACTTTGATGAACTGTAATTTGGCTGGTTCTTTGCCTGAGTTTCTTGGAAATATGTCTTCTTTAGATGTATTGTTGTTGTCCAAGAATCGACTTTCGGGGACTATACCAGGTACTTTTAAGGATTCTGAGCTCAAAATGCTATGGTTGAATGATCAGTCTGGTGATGGCATGAGTGGTTCAATTGATGTGGTTTCAACAATGAGATCACTTACTAGTCTTTGGCTTCATGGGAACCATTTTTCAGGTAAAATTCCAAAGGAGATTGGTAATTTGACATATCTGCAGGATCTTAATGTTAATAGCAATGATCTTGTTGGTTTAATTCCTGAATCTTTAGCAAATATGTCATTAGGCCATCTTGATTTGAATAATAATCATTTTATGGGGCCAATACCAAATTTCAAAGCTATAAATGTTAGTTATCAATCTAACTCTTTTTGTCAAGCCAAAATTTGTGCACCTGAGGTTATGGCACTTTTAGAATTCCTTGATGAGTTGAATTATCCATCTAAACTTGTTGAATCATGGTCAGGAGATAATCCTTGTGACGGGCCGTGGTGGGGATTAAGTTGTGACATTAACCAAAAAGTTATTGTGATAAACTTGCCTAAGTCCAATCTTTCTGGTACCTTGAGTCCTTCAATTGCAAAATTAGATTCTCTTACTCATATTTATCTTGGATCTAACAATATTTCTGGTTCTATTCCATCTAGTTGGACTAGTTTGAAACATTTGGTTCTGCTTGATTTGAGTAATAACCACATTTCCCTTCCTCTGCCTGAATTTACTCCCCCCTTGAAACTCGTCCTGAGTGGAAATTCGTTACTGAACTCTAGTCCTCTTATAGCAAGCCCTTTGCAAAAGAATAGTACATCCACTTCTGTTTCACCCTCTTTGCCGACCAATAAGTCAAGCTCTTCCAAGTCTAACTTAGTCATTTTTGTGGTTCCTATTGCAAGTTTTGCACTTTTAGTTTCTTTTGCTATGCTACTATATGTTTACGTTCGAAAGAGGAGTATGGATAGGCACAAAGGTCCAACTTCTCTAGTGGTTCATCCTAGAGATCCTTCTGATTTGGATCGGATGGTCAAGATTGCAATTTCCGATGAAACTAAAGGAAGTCTTTCGATACTAACTGGAAGGGGTTCTTCTAGTATACACAGTGGTAAATACCCTGTGATGGAAGCTGGCAATTTGGTCATATCAGTTCAAGTACTTAGGGATGTAACCAAGAACTTTGCTCCAGAAAATGAACTCGGGCGTGGTGGTTTTGGAGTGGTTTATAAAGGAGAATTAGATGATGGAACAAAAATAGCTGTCAAAAGAATGGAGTCTGGAGTAATTAGCAGCAAAGCGTTGGATGAATTTCAATCTGAAATTTCTGTTCTTTCCAAAGTCAGACATAGGAATTTAGTGTCTCTGTTGGGATATTCTGTTGAAGGCAATGAACGGATTCTGGTTTATGAACACATGCCACAAGGTGCTCTTAGCACGCATCTTTTCAACTGGAAAAGCTTGAACTTAGAGCCTCTTTCGTGGAAGAGGAGGCTGAATATTGCATTAGATGTTGCTAGAGGAATGGAATATCTACATACATTGGCTCATCAGTGCTTCATACACAGAGATCTCAAGCCCTCAAATATCTTGCTGACTGATGATTTCCGAGCAAAAGTATCAGACTTTGGACTCGTGAAACCCGCCCCTAATGGAGAGAAGGGCTCTGTGGTCACCAAGCTAGCTGGAACTTTCGGATATTTAGCACCTGAATATGCGGGTAAACATTTCATTTCCTGTTCTTTTATTCCTAACATGACATTTTATTTTGTAAAACAGGAAACAGATGTTCTATTTGCCATTTGATGTTCTCTTTGGTATGAGTACAGTTACATAAAACTCAAATAGTTTAGTTACTGTTGTCTGTTTTGGGGTGGATAAGTTAAGGACTATAATAAATCTGTTCACCGATAAGATGATATTGAATATCAACTTTAGTGACTAATTCTTGTAGACACCTAGGATTTTCTTTAGATGATTCTGTCTTTTCCGTCTGCTGGTTTCCCAACTCATTTTGCCTGCTGTTTTAATACAGTTACTGGTAAAATCACCACTAAAGCCGATGTCTTTAGTTTCGGTGTTGTCCTTATGGAGTTGTTAACTGGATGGATGGCACTTGATGACGACAGACCAAATGAAAGCCAATACTTAGTTGCATGGTTCTGGAACATCAAATCCTCTAAAGAGAAACTTATTGCAGCAATCGATCCAGCTCTGGATGTGAAACAAGAGAGTACATTTGAGAGCATCTACACCGTTGCAGAACTTGCTGGTCACTGCACAGCAAGAGAGCCTGGCCAACGCCCTGACATGTCCCATGCTGTGAATGTGCTCTCCCCGCTCGTTGAGAAATGGAAGCCTCTTGAGGAAGATTCAGACGACGATTGTGGTATCGATTACAGTCTTCCACTCAATCAAATGGTCAAGGGCTGGCAAGAATCAGAAGGAAAAGACTTGAGTTGCGTCGACCTTGAAGACACTAAGGGTAGTATCCCCTCAAGGCCTACTGGATTTGCTGAGTCCTTTACATCAGTTGATGGTAGATGAGAAAAAAGGTACTGCACTACTTCTAGTGTAGTTGATCTGATATTCTTATTCAGTAAATGTGGATAGCATTTTTGTTTGTTTTTGTGTGATATTTTGATCCTGTTTTTTATCTATGCTTTCTTTCACTTTTATTTGTAGAAAATTTTACTCCAATGTATAGTCTTTTGAAGTTATGTGGGATGTAGATATGGTATAGTGATGATTGATGAATGAAAACTGTTTCCA

>PGSC0003DMG400028429 LRR-RLKs

TGAATTCTAAAAAGGACTTTTTTCTGAAAGCTCTGCACCAAAATCTTTCTTTTGGGTAATTTAGCTTTTTGGGTCCAAATGGTAACTTTGCTACTCATTTTCTCATTTTTTCTTCATTGGGTGGCCACCACTCCAGCTTCCATGGGTTTTCTTTCACCTAAAGGTGTCAACTATGAAGGTAAAGATTAAAACTTTACTATAATCCAATCATTTCTTGTTTTTTTTTCCAGTGAAAAATCCAAACTTTATTGCTTAGCACTTGTTTATTTTTGTTTTCATAGTTTAGAGTGGTAAAGATGAAAACTTTACTACAATCCAAACATTTGTTGTTGTTTTTTTTCCAGTGAAAAATCCAAACTTTGTTGTTTAGCACTGTTTATAAGCATTTATTTGTTTATGTTTTCATAGTTTAGAGTGGTAACTTTACTTAAAGATTTGTTGAATTTATTGTGGGGTAAGCAGTGGCAGCATTGATGTCTATGAAGAATAGAATGAGAGATGAATACCATGTGTTAGATGGATGGGATGTAAATTCTGTTGATCCTTGTACTTGGTACATGGTTGGTTGTTCTTCAGAAGGCTTTGTTATTTCTCTGTGAGTTTTCTCCCTTTCTCTTACATCTTGTCCATTTCTTTTCCCTTTCTGTGAAAATAATTTATTATTATTCTTTGGTTATATATTCTTGGCAGTGAAATGGCAAGTATGGGATTATCTGGCACACTTTCGCCTAGTATTGGGAATTTAACTCATCTTCGGACGATGTAAGACTAATTTGACATGAATTTCCCACTTTATTCTTATTTTGATGTGATGCTGCTCATATATATCTTGGAGATACTTGAAGTTGAAATCACATAGTACTTGTTAATAATTACTAATATTAGTAAGAGTAACATTAAGTGCTTTAATTGCAAGATATGTTGAAAAATATCTTTTAATATGATCTGTTAAAGTCATTTTTGCATTGGTTTTTGTTGTAGGTTGTTGCAGAATAATCATTTATCGGGTCCAATTCCTGCTGAAATTGGACAGCTCACTGAACTGCTGACCCTTGATTTGTCTGGTAATCAGTTTGTTGGAGGAATTCCTAGGGCGTTGGGTCGCCTGATTTATTTAAATTACCTGTGAGTGTAGGCTAAACTCTTGATTTTGTTCTCTCTTTATTATTCATATGTTACAGTTCCCCTTGTTGGCTGTGTATGTAATCCTCTGGTGTGTCAAATTCTCAGGAGGCTTAGCAGGAACAGATTGTCTGGACAAATTCCTAAACCTGTGGCATACCTTTCAGGTCTTTCATTCTTGTATGTCTTCCATCCATGACATTTATATGTCCAATACAAGGGCTATATGAGAGTTAAATTAACTGTGGATGGTGTTGTGTAGGGACTTATCATTCAATAACCTTAGTGGTCCAACTCCAAAAATTTTGGCCAAAGACTACAGGTAAGATCCAAGGAAATATTATTATATCTGTCTCAATGTTTGGTCCTCAATAGTCGGTGAACTGGAAGAATCCCTCGGGTCTCTAGTTTGATGAATCAAGGAATTCAGGGTTTACACTTTTACTATGAGGAGGATGGTATATGTCACAGTACTGTGCCAGAATCTTATTCATTAATGGAGTTTAAGATAAACACCTTGATCTTAAATTCAACTGCAAAATAGAAGCTTGATTCAATATTGTTGCATCTGCGACATGCCATATGGCGCAGTGCCACACATAATGTTTCTCTCGTAATAAAGTTATGCTTTCTATAGTTTTAGTAACTGTTATCTTGCATTCCAGTTTGTTGTTGTGGTATGTCCCTTCACTGGAAACCAGCAGTCAGTTCTTGCAGACCTGAAAGTAATATCTCAAATCTCATTATCAAAAAGAAGAAGAAAAGAAGAAGTAATATCTCAAATCTCCATATCAAAAAAAGAAGAAGTAATATCTCAAATCTTATCATTGAAGTATTATCTTCTTACCTGGTCTAATGGAACTTCCATGATGTTTGTTTGATATAATTCCTATATTTCTAATCTGCTAATGTCAAGCAGAATATTTGCTTGAGATAATGTCTGATGTTATAGAGGTTTTCTATCCTACTTGCAATGCTTGTTGATTATCTTCCCCCTGGCATCATACTAGCAACCCTTCGGGTTCTTTTTTATTCTATTTCAGCTTATATTTTGGTTGTTGCCGCCTTAGTATTGCAGGAAACAGATTTCTTTGCTCTGCATTGACCACACAAATTTGTGGGGGTGTGCCAAAACCAGTAAATGGTATTGTTGAAATTTTACCAGAACTTGTCATTATATTGAGGCTGATGTTTTTAATCAAACTCTATTTTTACCATGCAGAAACATCTTCAGACAAGAGGATTAGCAATCATCATCGATGGGTTGTTTCTGTTAGCATTGGGGTCAGTTGCACATTCATAGTTGCAGTCACGCTTCTTGTTTGTTGGGTGCATTGGTACAGATCTCGTGTCCTCACAGGATATGGTAATTACAAAATTAGAAACACATCTCCGATTTCTTGCACATTAAGACTAGATACATTATGAGTCTCTGATCATTTGATCTGCAGTGCAACACGATTGTGAGTTCGCCACTGGCCACCTGAAGAGATTTTCATTTCGTGAACTGCAAATTGCCACCGGAAATTTCAGCTCTAAGAACATAATAGGACAAGGTGGATTTGGAGTGGTTTACAAAGGATACCTTCCAAATAGGACAGTAGTGGCTGTCAAAAGATTAAGAGATCCAAGTTTCACTGGAGAAGTGCAGTTTCAAACAGAAGTTGAGATGATAGGCTTGGCTGTGCATCGGAACCTTCTACGTCTGTATGGATTCTGTACGACTCCCGAGGAAAGATTACTTATTTACCCTTATATGCCAAATGGGAGCGTTGCCGATTGCTTGAGAGGTTTGCTTTGAAGTATATTTGCTTGGTGCTTTTGCTGAAGCATTCTTTTTTCTTGTCACAATGGATTTCGTAAACTGATCAAGTTTAACATATGCATTTCAATTATTCCTAGTGCTCTCAATGTTAATCTGACAAAAACTCTTGAAAGGTCATGGTTTATGGAAATCGAGGACTTTGTGCTATATTTTTACTGTACATCTGAAAAAACCAGGGCCCTATGATGAAAACAACTTTTGATTGATTATCTTAGATAGTTAAATTGGATTTACATTGTCAATGTTTCTTGTGAAAGTGGAATGAAATCTATTATCAACATATCAATCAACAATGCATCAAACCCAAACTAGTTGGGGTTGACTATATGAATATTTTGTATCCATTGCAATTAAGACTCCTGATGTAGTGCATTACCTAAGACTCTAGCAATCTATTGCGACCTTACGTGCTATTAAGCTTAATTTTTATGTGACAAAGTTTCTATCCATCTTCTGAAACTCTAATGCCATTTATGGTATCGACTTACATTATCCATGAATAAAATAAATTGTTATTTGGCAGTAAGGTAACTGTCGTGTTGTCATTAATTTTTAGCATGACCAATTCAAGTCAATTTTTAGGTATATGAACACAAAATCAAGTTGGAATTATTGGAATGTTAACGCATTAGCTTGATGTAGTGACTGTGTCATCTTTTGCTCTGTTCTCACATAATCTTTTGTTATCTACCAGACAATGGCCGGGATAAACCATTTTTGGATTGGAGAAAGCGCATGCATATAACTCTTGGAGCTGCCAGAGGTCTTCTGTATTTGCATGAACAGTGTAATCCTAAAATAATTCACAGAGATGTTAAGGCTGCTAATATTCTGCTTGATGAAAGTTTTGAAGCTGTTGTTGGTGATTTTGGTCTTGCCAAGCTTTTAGACTGTAGAGATTCTCATGTTACAACTGCAGTTCGTGGTACTATAGGTCATATAGCACCAGAGTATTTATCAACTGGTCAATCATCTGAGAAGACAGATGTCTTTGGATATGGGATACTACTTTTGGAACTCATTACAGGACACAAGGCATTAGATGCAGGAATTGGTCAGGGTCAGAAAGGAACTATTCTTGACCGGGTGTGTAGCTTCTAACCCTCCTACTGTTTGAAGCTTTATATAGTTGTGCATATAATTTCTAGACTTTATTTGATTAACCAGTTAATAGTTAATCTTCTTGCAGGATTTTGTTTGAGTCATAATCAATATCAATATTTGACTTGAAATTTTTCACTAAAATTGACGCAATAACTAATGTCTAACGCAGAAGTTGCTCCACATCTTATGAGAGAAGTACAGTTCCAAAATTCTGAATCATTGCTCGAAAACTTGTTCTTTTTACTGACTGCATAATCTGTAGGAATTATGGACAACATATTCAAGGTTTTTGAATCACTTACTGATCAGTGCAATAAAATTATTGCTGTGTATTACCTTACTGTTGCATACTTAAGCCCTTAAATATAAATTATTTGGTTGATTTTTTATCACATATAATGTTACTCTTGCATACTTTAAGCCCTTGATTAACTTATTTGGTTGATAATTATTACATTATCATGCAAGTCTGAACAACCATGTATTATGCTAGAATAGTTGTGTGAAAAAAGGTACCCGTTCTATGCTTCTTCAATTATGATTTTTCTTCCCTCTCAGGCTGTTTTATGTGGAGTAAAGATTTTCAGAACATGATAGAGCTCAGAGTTTGTTATGTCTATTATTCGGGCCTATTCATTCTGACAACCTATCCAATTTACAGGTCCGGAATTTATTTGAGGAAAAGAAAGTGGAAATGCTTGCGGATAGGGATCTAAGAGGATGTTTTAATGCAGAGGAGCTAGAGAAAACAGTTGAAGTGGCTCTACAGTGTACTCAATCTAACCCCAATAATCGGCCTAAGATGTCGGAAGTTTTGAGAATCCTGGAAGGCGTTACAGAACAAATGGGGCATGTGGACGAATCACAAGGTGGAAGCAACACATGTCAAACAAGGGCTTTCAGCTTCTCTAGGAATTTTAGTGATATTCATGAAGAATCCTCATTTACCTTTGAACCAATTGAGCTTTCAGGACCCAGATAACAAAAGTAGCCTGCACTGGCCTTGTCATAATAAGTGTGCAATCTTGATATATCTAGTTCAGCCATACATACGAGTATGTTGTTGCTTTAAGTCAATGTCCTAGTACTCTGTAAGTTACTGTAATTTGTAAATTTTGTTGCAAAAGCATTCATTATACTAGTTCTTGGTGGTTCACAACCTGTGTTAGAACTAGCTAGACTAACTTCACTTTGGAACTTCATTCTTTGGAAATGACAAACTTC

>PGSC0003DMG400028435 LRR-RLKs

ATGTTCCAACTCAAGCCGAATCTTTTCCTGCAACAACACCAGCCAGGATTTCATATATACATTGAAATTGGAAGGGCAACTCAAGGGTTCGACCAATGCAACTTGCTAGGCTCTGGAGGTTTTGGCTCTGTTTACAAGGGCATGTTTGAAAATGGGATGACTTTGGCAATCAAAGTGTTCAATTTACGGATTGAAGGTGCATTCAAGAGTTTTGATATTGAATGTGAAGTTCTACACAACCTTCGCCATAGAAATCTTGCCAAAGTTATCAGCAGTTGTACTAACATGGATTTTAAAGCATTACTTCTAGAGTACATGCCCAATGGGAGCCTGGAGAAATGGTTACATTCTGATGATCACTTCTTGGACATAATCCAAAGATTAGACATTAATGGTTGATGTTGCATCAGCTTTGGAGTATCTCCATCATGGTTATTCAACAGTCGTTGTACATAGTGACTTGAAGCCTAGTAATGTCTTGCTAGACGAAAGATTGGTTGGACACGTGAGTGACTTTGGCCTGGCCAAGTTATTCGGAGAAGGGGAATCAGTTGCTCATACTAAAACACTTGCTACAATGGGGTACATTGCACCAGGTAAATTTCTTTTAGTCCTTTTGTTACCACATATAAAGTGTTATTTTACATCACATTAGAACTTGGTGAAAGCATAATCAAAATGGCCAAGTTAGCTCATAGATTGAATCTGTATATTATTACCTTTAATCTGCCAGAAGTCATATTGTTTCAATTGATTACCTTACTCTTGTTTTATTCCCATTATACCTTTAATTTCTTATTTGGCTCGTCAAGAAACTAATTACTCTGGTAACATTGGATCAGAGTATGGATCAGTAGGATTAGTTTCTACAAGATGTGATGTGTACAGCTACGGCATTATGCTCATGGAAACTTTCACAAGAAAAAGGCCATATGATGAAATGTTTCAAGAAAATTTGAACATGAGGAGTTGGGTCTGTAATTCACTACCTGTCGCACCAGATGATATTATTGATGACACCTAATTTCTGTTAGGATTTTGTCCTGATTTTCTTACCTTCTTTGAAGTTTATTTTTTCTAAAAGAAAAGACAAAACATTACCATAAATGTTTTTACTTTTTCTTGAAAGGAAAATATAATATTCTTTTCATATTTGGTTTATTCTTTCCTTTTAGGAAAATGTTTTTGGAGTAGCTAGTTCTTTTCTAGTAGGAAAGGTTTTAGGACTCTAAAAATATAGGTTTGTTTCTTCTAACTCAATAACAATAACATCCACAATGTAGTCGTATAAGAGTCTTGTTTATGGGGAAATTTAATCTCTCAACAGTTATAATATTTTTTTCTTTTAAATTAGTTTTCATATTAATAATATTATGTTTTAGTATCGCTTTATTTGTCATCTAATTTATCACCGTCTTGATTGCAATTGTAAGCTTCCGCATGACGCCCAACTCACCTTCATAACCCAACAATTTCAACCTGAAGAGATTGATTTTGAGAAAAAGTTGTGTTTTGTGTCCTCTATTTTGGACTTGGCATTGAATTGCACAGTTGAATCTCCTAATGAGAGGCCGAACATGAAAGATGTCCTGGCAAATATCAATAAGATCAAGCATGAATTTCTTCACAGATGATGCACCAAGGTTTATGCAAACGGTAGGCATCCAACATAGCTCCTAGATTTAAACTTTTCTTGTTTCTTGTGCTTTCCTCTCGTGTATATTTCAAGTTTGAGTGAGTGTGTCAAATAACTTAAGTACATAACATGTTTACTACAATTGGAAAGGAAGAATGCCCTGTATAATATGATTAATGAGGTTATTATGTTAATTTCAACTACAATAGTACACAAAGAGTGGTGTATAATGTTGAACTAAAATTGAACTTTTCTTTGTTTATTGCTTTCCTCTTGTGTATGTACCAATTTGAGTGTTTCTCTCAAATATAAGTGTAATATAGGACAAGTATATATAAGATGTGTACTAAAGTTGTTATAGAAGAATGCCCTGTACAATATTAATGATCGTTATTGTTTTGATGTATCTACTTTGCAGATGTGAATCCCTCCGCTTAAGCTTTTTAGATGATATAGTCAGACACTTCAATACTCCTTTATGTTACACCACGCTGCTCCTTTCTCAAATAACTTGATAGAACATTCAAAAAAGACGTAATCTAACACCCTTTTTTGGGACAGACTTTAGTAAAGTCAAGTCGACCAACTGTGGAAAATTTATAACTAGGGGTGTACAAAATCGAACCAAACCGCAAATTAAACCAAAAAAAAAACCCGACTAGTGGTTTGGTTTGACTTGGTTTGGTATTGGAAAAAAAACCCGACTATATTTGGGTTGGTTTGGTTTTAACTAAAAAAATCAACCCGAGGCCAAACCAACCCGACGTTATATATATAATTTTAAAATATTATTTTATACATAAAATAGATATAATTTTTAAATATTTCTTATACTTTTTCATAGTTTTTGTCTTTTAATATATTATTTCAAGTTTGTAACTTAGAATTATGAATGTTCCAATAAAGATTATAGTCCACATATGTTGGTAATTATAATAAAGCTCAAATCAAAATCAAATTAATGCTAATGCAAAAAGAAAATCAATTCAACACTAATAATGACAATAATATTGAATATTTGTTCTTTAGTTTTACATTGGTTTAGACAATTAAAATACATAATCTAATTTTAATTTCATTTAATATTTAGTCATGTAACTAATACTTATTGAACTTATTTTAGCATGATTTAGTACTTTTAAATTATGATCATTTTCATTATGAATTGTTAATTTGCAATTTTTGTTTTACGCGATTTCATTATTATTATTTTTGTTGGATATTCTAGTGTCATTAATTATATCACATTTTGTGTTATTTTTTTAAGAAACAACTTAGATAGTTGTATTTTGATAGGACTAAAGAAATATTTGAAATACAAGTAAATTATATGTTTGTATGAATACTTTACCAGAAAAACCCGAAAACCCGAGGTTGAAAAACCCGAGTTTTTTTGGTTTGGTTTGGTTTATAAATTTTAAAATCCGACATAAATGGTTTGGTTTAATATTTGAAAAACCCAAACCAACCCGACCATGTACACCCTTATTTACAACCCCTTATCTACGGCATTGCTTGGCCCTCTTTCCTTGATAATAATTAAAAATCAAGGAAAAAGGTTTGCATAACATTAGAAAGGTATGCTGTACTGTCCATATGGTTTAAACAGGAAATAATAAGAAAACCTTTCATCATAATAAGAGCTCAGGTAAAGATCGGATATTACCGTTGACAATATAATTAAATAATATAAAAGTTTGTTCTCTCTAATCGCTTAAGTTTTTAGATGAGATGGTCACACAATTCAACAATTATTAGTGGAAACTAATTTTTCTAGTACTGTTCGAACTAGATAAGTATTAGGTAAATTATATAAATTTGTATTTTCTGCTTATTTAAAATACAAGAAAATATACACTAAAAAAATATTTTTCTTCCGACCCTACTCAATCACTGAATTTAATGCCTATTTGGAAGGAGAAAAAAAACTCTCTAAGAAGACCTAGTAAATCCAACTAATTTTCCATATACTGCTCTTTGATATAGCACATGAAGTCAAGTCACTCTGGAAAATTTATAGCCCCTTATCTACGACATTGCTTGGCCCTCTTTTATCCCTATTTAATGACTAGCTTGGTTTGATTTTTCATTTAAAAATAACTAAATGAATTATATTGATGTATTAAATCTATAAATCAACCCAAATTAACAAAAAGTGTTTTTCAATTTTTTGGTTTTTTTTATAATTATACAATAATTGAAAAAGAGATCAAATATATTTTCACTTACATATGTGATAGGCTAAAACCTAAAGAGATCAATTTTACTTTATATTACGAAAGAATTTTAGAAAGAAAAAAAAAAGACAAAAAAGAGAGAATTTTGCTTTAAAACAATTGACAAATAATAAGAAGAGAGAAAATAGGAACAATAAAATAATGAACTTTTTATTTTCAAAATATATATGATAGATTAAATATAGTCAAGAACAATGATAAAAAAATGATTAATATCATCGTGTACACAGGTTGAATGTAGATGCGAAGGAGAAAATTTAGGCAAAAAAAAGAATATATGCATGTATGAAAAAATCTTAAAGAGATAAACAAAAAAGTAAAAAAAAAAAAAGTTGAAAAAGTTGTTATTACCAAAGAACTGTTTAAAATAAGAAGAGAGATGAAGAAATGATAAAAAATAATTATACTTTTTCAATAAATAAAGGAAGGAAAGAAAAAACATTTCTTGAAAAGGCAAAGAAAAAAAAACGAGTAAAAGAGAACATGGCTAATAAATGATAAATAGATTTTAAGAAATCTTTTTTTTGCTAAAAAGTTATAAGTGGGCTATTTATTGCCAATTTTAATTATAAAACATGTCATGATGTGCCATTTTTTCTTTTTAATATAGTGTACAATTATTATTGGATGAAGGGATTACCATAGTAAAGAAAAGTTCTAAAGAAATTTGGATGTCTGATGAAATAATATATTATCTCAAAACAAAGACTTGGGGACAAAATTTTAATAAAGTCAAGCCAACCACTGTGGAAAATGTACAAATTACAACCCCTTTATCTACGGCATTGCTTTTGTTCCTTTTTTTTATACAAATAAGGGACCCAGTCCTTAGCAACCTTTTGTCCCTATTTAGCTCAACACTTGTTTCCTTATCTTATGTGCAATCACATGGAGAAAGCATACTCTTTCTTTCTTTCACTACTACTCTTGCACCTATACTTGTTAGAGACATGCATTAGCAAAAACATAGTAATTAACACAGATCAATCAACTCTACTTTCATTGAAAGCTAGTTTTACTCTAAACTCTTCTCATCCCTTAACTCAAAATTGGTCTTCTCAAGCTTCAGTTTGTGATTGGATTGGAGTCACTTGCGGCTCTCGCCACCATAGGGTGAGAGCACTAAATGTATCCAACTTGAACATTCTTGGAAACATTCCACCACAATTGGGAAACCTCTCATTTCTCGTTTCGCTTGATCTTAGTAGTAATAATTTCCAAGGAGACCTCCCTCAAGATTTGTCTCGTTTACGTAGATTGAAGGTGATGAATCTAGCAAACAATAACTTCACTGGCCAGATTCCCAAAGATTTTGGTAACCTTATGAGGTTAGAGAAGTTGGTTCTAGTCAGTAATAAACTTAGTGGCTCAATTCCATTCTCCATTTTCAACATTTCCTCGTTGGAAATTTTAGCTCTCACATATAATCAATTATCAGGGAGTCTACCTGTAGATATATGTCGTCGTCTTCAAAAAATAAAGAGTATTTCAATCATTTCCAACCACTTGAGTGGTCAAATTCCAACAGGTTTGTCTAACTGCACAGAATTGTATGAATTGTCATTGTCATACAATAACTTCAGTGGAACCATTCCACCAGAAATTGTAAACTTGGAGAGGCTTCAGTTCTTAAACCTTGGGGGAAACAACTTGCAAGGTACATTCTGCATACTTTAATTTCTCAATGTGAATATATAACATCCTACTTTGCATATATATGTAAGCCCACAATATTTAGGGTGTGTTTGGTACCAAGGAAATTTTCCTACTTTCCTATTTTTTTATGTTTGGTGCTTAGCATGAGGCTAAGCACCAAAGTTTGTTAATTGAGAATTAACACCTTGAGCAACATCATAAAAATTTGCAAATTTGTTTGCCGTTTGTTGAAATTTGCAGGCACAATTCCAGCAAAGATTGGTAATTTAGGAAAGATGCAGCAATTACAATTGGAGAACAATCGCATCGTGGGTTCAATACCACGCACCATATCGAACTTATCATCACTATGGATGCTTAATTTGAACACAAATAATTTATCAGGTACCTTGGTCTTGTTTGTCATGGAATTAATTGTTGGTTTTTATCAACTCTTATATATACTGATGGGCACAATCAAATAAAAGTAATTTAGATATATTCTTTCGTGTGGCAGGTATTATACCCAGAGAGATTGGAAATCTTCACAAACTGGAGATACTTTATTTGCAAATTAATAAGTTAAGTGGTTCCATACCGGAAGGGCTCTTCAATATCTCTACACTAACAGAGGTGTCAGTTTTCTTCAATAACCTTTCGGGTAGTCTTCCATCTGCTTCAGGCTACTGGCAAACAAATCTACAGCTTTTGAATCTTGGTCAAAACAACATTGGTGGAGTAATACCCACTTCAATCTCCAATTCATCAAATCTAGAGATTATAGATCTTAGTATAAACAAATTCAGTGGCCAAATTCCTAACTCGTTGGGGGATTTGAGGGATCTTGAATATCTGGACTTGTTCGTTAATAACTTATCGTCTCCGCACCTAAGTATCTTAACTTCATTGGTGAACTGCAGATATTTGAAAGAAGTAATATTAGGGGAGAATCCTCTGTATGGTGTTCTTCCAGATTCCATTGGCAATCTCTCCATTTCTATCATGGAACTTGAGTTATATAGTTCTGAAATTAGAGGCCAGATACCATTAGGAATTGGGAATTTAAGTAACTTAAACACTTTGTCCCTATCCGGCAATGACTTGACTGGATCAGTGCCAACAACCTTATGTGATTTACACATTCTTCAAAGGCTAGGACTTGGTCTTAATAGGTTGAGTGGATCCTTGCCAGAGTGCCTTTGCAAAATGTCATCGTTGGGCTTGCTTTCTTTGTACTATAATCGGATTTCTGGTCCGATACCATCTTGCATTGGTAATGCAACCTCTTTGAGAAACGCTTATCTAAATTCAAATAGGCTCACTAACATACCCACAAGTCTATGGAGCCTCAAATATCTTTTGGTGCTTAACTTGTCAAATAATTCTTTGATTGGTTCTATACCTCCAGAATTAGGGAACTTGAAAGCCATAACATCCATAAATCTGTCCAGGAATCACCTTTCAGGAAGTATTCCCTCCACAATTGGAGACTTGCAGAGCCTAGTTTATCTTTCCTTGGCTTATAATGAGTTACAAGGTTCTATTCCAGAGTCACTTGGGAAAATGATAAGTTTAGAATCGGCGAATCTATCCAATAACATTCTTTCAGGTACCATTCCAAAATCATTAGAGTTGCTTCGGTATCTGAAGAATTTCAATGTATCATTCAATAGATTAGAAGGTGAAATCCCAAGTAAAGGACCTTTTCTCAATTTCAACTCTCAATCTTTTATGGGAAATGGAGGGTTATGCGGCGGTTTAGTTTTCCAACCTTGTATGACTAGGTCTTTTCATCATTCAAGGAGAGGCAAACTGCTTCTGATTATACTTGTCTCACTGGGAGCTGCGGTAATGGTACTTGGCTCAATCGTTGTGTTTATGTTAAGGAGACGTTGGAATAGAAATATTCCAACTGAAGCTGAATCCTTTGCTGCAACAACACTAGCCAGGATTTCATACATTGAAATTGAAAGGGCAACTCAAGGGTTTGACCAATGCAACTTGCTAGGCTCTGGAGGTTTTGGCTCTGTTTACAAGGGCATGTTTGCAAACGGGATGACTTTGGCAATCAAAGTGTTTAATTTACAGGTTGAAGGAGCGTTAAAGAGCTTTGATGCTGAATGTGAAGTTTTACGCAACCTTCGTCATAGAAATCTTACCAAAGTTGTCAGCAGTTGTACTAACTTGGATTTTAAAGCATTACTTCTTGAGTATATGCCTAATGGAAGCCTAGAGCTGTGGTTACATTCTGATGATCGCTTCTTGAATATGATCCAAAGATTAGACATAATGATTGATGTTGCATTTGCTTTGGAATATCTCCATCATGGTTATGAAACAGTTGTTGTACATAGTGATTTGAAGCCTAGCAATGTCTTGCTAGACGAAAAGTTGGTTGGACATGTGAGTGACTTTGGCCTGACCAAGTTATTAGGAGAAGGGGAATCAATTGCTCATACTAATACACTTGCTACAATGGGCTACATTGCACCAGGTAACTTTCTGTAAGTCCTTTTATCATCGCCTATGATGTGTTATTTTACATCAAATAATTAGTTGACTGGTTCTAATATAATGCATCACATTAGAACTGGGTGAACGCCTAAGTGGATGGCTTATTTTAATTGGAAGTTATCACCTAGTGTTTTTTGTCACTTTGTTGGAATTTAATCCACTTCATTGACCATTAAGCCACAATTAGCCTTGGGTATAAATATAGATAGTATTTTGATTTCTTTGGCTCGTCAAGAAACTAATTACTCTGGTAACATTGGATCAGAGTATGGATCAGTAGGATTAGTTTCTACAAGATGTGATGTGTACAGCTACGGCATTATGCTCATGGAAACTTTCACAAGAAATAGGCCATATGATGAAATGTTTGACGGAAATTTAAGCATGAGGAGTTGGGTCTGCAATTCAGTACCTGCAATGCCAGAGGATATTATTGATGTCACGTTAATGGAACCAGAAAAGACTCACTTTCAGGAAAAGTTGTGTTGTGTGTCTTCTATTTTGGAGGTGGCGTTGCATTGCACAACTGAATCTCCCAATGAAAGGTTGAACATGATAGAAGTCCTTGCAAATATCAAGAAGATAAAGCAGGAATTTCTTCGCAAATGATGTACGTAGCAGTGGTAAATATCCAACTTTTTAAAAGTATTCTGATGTCATCTAGATATATTATTTCCTTATTCAAGAATAGAAATGATTATTGTCTTGTGAGCTTGTCTATATCGTACAAATATATTGAATTGGATATTCTGGTTTTCCATACAAATATATCCCCTTGTATAAGAAAAAACAAGAACTAATTGAAGCATAACTTAATTCATGATTATAAGTAGAAGAACCCAACATTTTTGCCTATGGAAATTTGAATTGTGATGAATGTTGAAAACTGAAGTAAACAACTCAAGTCTGTGTTACTATTAGTTTATTTTTGTATGTGCTGTGCCAGCTCAAGTTTAATCCTAATTGAAAATTTGTGCAGGTTGTACGTGCTGCCTTTTCTCCTGGTTTAGCAAATTTGGGTGATACCAATTGAGATGAAAAATCAACATTTGGTCCGAGATTTGAACTTTTCTTGTTTCTGTTCTTGTGAATATTTCGACATTTGGTAAAATAAGTCTAATGTATAAGATGTGTACTACAGTTGGAATAGAAGAATGCCCTGTATAATATTGGATTGTTAATGTATAATATTGAACTAAAA

>PGSC0003DMG400028436 LRR-RLKs

CCTAAATATTTATTTCTTGAAAATTATATATACCTCTCACTTTCTTATTTGTCCCTACCTGAAAACTTAATTTGTTCCCCAATCACTTGGAGAGATCATACTCTTTCTTGCTTTCACTACAATTATTGACACTTTACTTGTTCGAGGCCATCGACATAAAAACAGATCAATCAGCTCTACTTGCCTTAAAAGTTAGCTTTACTCTAAACTCTTCTCATCCCTCAACTCAAAATTGGTCTTCTCAATCCTCCGTTTGTGATTGGATTGGAGTCACTTGCAGATCTCGTCACCATAGGGTGAGTGCACTAAATATATCCAACATGAACATTCTTGGAAACATTCCTTCAAACTTGGGAAACCTCTCCTTCCTCGTTTCTCTTGATGTGAGCCAAAACAATTTACATGGAGACCTCCCTCAAGATTTGTCTCGTTTGCGTAGACTGAAGGTGATGGATCTTGCGTTCAATAATTTCAGCGGAGAGATCCCAGTCATGTGGTTCGGTTTCATTTCAAAGTTAGAAACTTTGAGTGTGAGAAACAATCGTCTTCAAGGCAATATCCCTAAGGAGATTGGGAATCTTGAGTATCTAAAGGAGTTGATTTTATTTGAAAACCAATTTACTGGCTCTATCCCGCTCTCCATTTTCAATATGTCGTTGTTGGAAATTTTAGGTCTATCACATAATCAATGAACTGGGAGTCTTCCTGTAGATATATGTCGTCGTCTTGCAAAAATAAAGAGCATTTCAATCATTTCCAACCACTTGAGTGGTCAAATTCCAACAGGTTTGTCTAACTGCACAGAATTGCATGATTTTTCATTGTCATACAATAACTTCAGTGGAACCATTCCACCAGAAATTGTAAACTTGGAGAGGCATGAGCTCTTAAATCTCGGGGGAAACAACTTGCACGGTACTCCTTTCGTCCACTTTTACTTGTTCAATTTAGAAAATAATTGTATTTATATTGCTGCTTAAGGATGGCCATCCCATTTGGCAAAATACATCTAACTGGTCTTTAGTTTTCATTTTGATACTTCATCTCAACCTTGTACGTTCCATTTTAACCCTCCCAAAGCTATCATTTTATTTTCACTTTATAAAATAAGTTTATCCATATTCAAAAGCCTTCTAGCATGAGGCTAATTTCGAAGTTTGTTAATTGAGCAACAGGATAAAAGGTTTACAAGTTTGTTTGCCGCTTGTCAAAATAATTTGCAGGCACAATTCCAGCAAAGATTGGTAATTTACGAAAGTTGCAGCAAATACAAGTGGAGAACAATCAAATTGTGGGTTCAATACCGCGCACCATATCCAACCTATCATCACTATGGGTGATTAATTTGAACACAAACTATTTATCAGGTAGCATACGTGGTCTTGTTTGGTTATCCGGAGGCAATAAAATAAATTAATCGAGATATATGTACTCTTTCATGTGTCAGGTGTTATACCCAGAGAGATTGGGAATCTTCACAAATTGGAGACACTTTATTTGCAATTTAATCAATTAAGTGGTAATATACCAGAAGAGCTCTTCAATATCTCTACACTTACAAGGGTTTCTCTCCTCTCTAACCACCTTTCGGGTAGTCTTCAATATGCTTCAAACTACTGGCCAACAAATCTAGAGATTTTGAATCTTGGTGTAAACAACATTGGTGGAGTTTTACCCACCTCAATCTCCAATTCCTCATATCTAATGAGATTATATCTTAATGGTAACAAATTCAGTGGTCAGATTCCTAACTCATTGGGGGGTTTGAGGCATCTTGAACATCTGGACTTGTCCAATAATAACTTATTATCTCTGCACCTAAGTATCTTAACTTCATTGACAAACTGCAGATCATTGAAAGAAGTTTACTTAGACCATAATCTTCTCGATGGTGTTATTCCAGATTCCATTGGCAATCTCTCCACTACCATTATGGAATTTTATTTAGACAGTTGTGAAATAAGGAGTCAGATACCATTAGGAATTGGGAATTTAAGTAACTTAAACACTTTGTCCCTAACCCACAATGACTTGACTGGATCAGTGCCAACAGCATTATGTAATTTACATATTCTTCAAAGGGTAGCAATTGAGGATAATAGGTTAAGTGGACCGTTGCCCCAGTGCCTTTGCAAATTATCATCCTTGGGCATGGTTGATTTTTCTAATAATAGGATTTCGGGTCCGGTACCGTCTTGCATTGGTAATGCAACCTCTTTGAGAAATGTTTATCTAAATTCAAATAGGATCACTAACATACCCATGAGTCTATGGAGCCTCAAAGATCTTTTGGATCTTAACTTGTCAAATAATTCCTTGGTTGGTTCTATACCTCCAGAGTTAGGAAAGTTGAAAGCCATAGCATCCATAGATCTTTCCAGGAATCACCTTTCAGGAAGTATTCCCTCCACAATTGGAGACTTGCAGAATCTATTTTATCTTTCGTTGGCTTATAATGAGTTACAAGGATCTATTCCGGAGTCACTCGAGAATACGATAAGTTTGGAATCAGCAAATCCATCCAATAATTTTCTTTCAGGTATGATTCCAAAATCATTAGAGTCACTTCGACATCTGAAGGATCTCAATGTATCATTCAATAGATTAGAAGGTGAAATCCCAAGTAAAGGACCTTTTCTCAATTTCACCTCTCAATCTTTTATGGGAAATGAAGGGTTATGCGGCGGTTTAGTTTTCCAACCTTGTATGACTAGGTCTTTTCATCATTCAAGGAAAAGCAAACTGCTTTTGATTATACTTGTCTTCCTGGGAGCTGCAGTAGTGGTACTTGGCTCAATCGTTGTGTTTATGTTAAGGAGACGTTGGAATAGAAATATTCCAACTCAAGCTGAATCCTTTACTGCAACAACACTAGCCAGGATTTCATACATTGAAATTGAAAGGGCAACTCAAGGGTTCGACCAATGCAACTTGCTAGGCACTGGAGGTTTTAGCTCTGTTTATAAGGGCATGTTTGCAAATGGGATGACTATGGCAATCAAAGTGTTTAATTTACAGATTGAAGGTGCGTTAAAGAGTTTCGATGCTGAATGTGAAGTTTTACGCAACCTTCGTCATAGAAATCTCACCAAAGTTATCAGCAGTTGTACTAACTTGGATTTTAAAGCATTACTTCTTGAGTATATGCCCAATGGAAGCCTAGAGCTGTGGCTACATTCTGATGATCGCTTCTTGAATATGATCCAAAGACTAGACATAATGATCGATGTTGCATTTGCTTTGGAATATCTCCATCACGGTTATGAAACAGTTGTTGTACATAGTGACCTGAAGCCTAGCAACGTCTTGCTAGATGAAAAGTTGGTTATGTGAGTGACTTTGGCCTGACCAAGTTATTAGGAGAAGGGGAATCAATTGCTCATACTAAAACACTTGCCACAATGGGCTACATTGCACCAGGTAACTTTCTTTTAGTCCTTGAATATCGCATATGATGTGTTATTTACATAATAATTAGTTTACTATTACATCACATTAAAACTCGGTGAAAGCCTGAGTGGATAGCTTAATTAAAGAAATGTAATATATCACAGATTAAGCGTGAACAAAAATGGTATAGAAAACATTGAAATATGACTGTCAAACTCTATATGCATGCATCTACGAATTTCTTTTTATCAAAAGTTCCCTCAAAGGATATTAAATAGATGAGCCAAAGGGGCATAGAGGGGGACTACGCCTGACCTCATCAATACAAATTTACGATACTAGGAGTGAGGAAAAGAAGAAATGTGGGGGGGGGGGTTAAGGGTTGCCCCACATCAATTAAGCATACAAAAAGATAAATTCTAGGATTTTAGAGCAAGCAGAGAAGACTCAGCCAGGATCAAAAAGAAAGCTGTTCTCTTAACTTAGATTCTAATGTTAGGCTTGCCAACTACATCATTCTTAAACGGAAGATTTATAGGCCTAGTCTCAGTGAAGAAGGTAGTCATTTTGCACTATTCAGCCAAGTTAGCAAGCATGTGAGCAGGAGTGTTACCTTCTCTTAAGGTATATGTGAAAGTGAAACGACTAAGTGAACTCATAGATTGGATCTGTCTAATATCATCTTTTATCTGCTAGAAGTCCATATCGTTTCGATTGACTGCCTTATACTTGTTTCATTCCAATTATACTTTCAGTTTTTAAAGTCCTTATCTACTCAAGGTAAATTCAACTATTTTCAATATCTAGTAAATATATATATATATATATATACACATACAGACACACACACATATATATGTATAATATTTTCATTTCTTTATCGGGCTCATCAAGAAACTAATTACTCCAAATATTGGATCTCAGAGTATGGATCAGTTGTATTAGTTTCTAGAAGATGTGATGTGTACAGCTATGGCATTATGCTCATGGAAACTTTCACAAGGAAAAGGCCATATGATGAAATGTTTCAAGAAAATTTGAACATGAGGAGTTGGGTCTGTAATTCACTACCTGTAGCACCAGATGATATTATTGATGACACCTTATTGCAACCTGAAGAGATTGATTTCAACAAAAAGTTGTGTTGTGTGACCTCTATTTTGGAGTTGGCATTGAATTGCACAGCTGAATCTCCTAATGAGAGGCTCAACATGAAAGATGTCCTGGCAAATATCAAGAAGATCAAGCATGAATTTCTATGCAGATGATGTAACAAGGTTTATGCAATGGTAGGTATCCAACTTTTTAAAATATTCTAATGTCATATGGAGAAATTATTTCCCATTTCAGGAATAGAAAGATTACTGTCTTGTCTATACCATGTAGAAATGGGTTTTTTCCATACAAATAATAATAATAAAAATAAAAAATAAAATAATAATAATAATAATATAATAATACTTTTTTTTTTATTAATATAATAATACTTAAATCAAGGAGAACTTCATTAGTTTTGAAATTGGTCACATAATCAAGGAGAACTTCATTAGTGAGTAAAGTAAATCAAGAATTCATGAGGACCAACATTTGGAAATTATGATGAATGTTGAAAAGGGAAGCAAACAACTCATGTATCCGTACAATTAGTGTATTTTTGTGTGTGCTATGAGTTTTTATCTGCAATGACTGCCAACTCAAGTTTGATGCCATTTAAATTTTTGTACAGGTAGTGCCGCCTTTTATGCCTGTCGTTGGCAAATTTGGGTGACACCAATTGAGTTGCCAACAAAGATTTGGTCTGAGATTCGGACTTCTCTTATTTCTTGTGCCTTCCTCTTGTGCATGTTTCAAGTTTGAGTATGTTTATGTAACAAATCTAATATATAAGATGTGTACTAAAGTATAATACCGGTGATCATGAATTATTATTGAGATAGCAGCTTGAATAGTCTATGAAGA

>PGSC0003DMG400028437 LRR-RLKs

AATTTTCAAATGGTATTCCAAGAGAAATTGGAGACATGCAAAATCTGATACACCTTTCTTTGGCACAAAACAAGCTACAAGGATCTATACCTGACTCAATTGGTAGCATACCGAGTTTGGAATTTCTAGACCTCTCGAATAACAATCTATCTGGATCAATTCCGATGTCCTTAGAGAAGCTTCGGTATCTCAACTATTTCAATGTTTCCTTTAACAGCTTGCAGGGTGAAATTCCCTTTAGTGGTCCTTTCAAGAACCTTTCCAGCCAGTCATTCATGTTCAATAAAGCCTTGTGTGGTGCTCCAAGATTTCGTGTCCTCCCGTGCCCCACTTCTTCAAACCATAGATCAAAGAAGAAGAAGTTACTTCTGATAGTCTTTCCTCTACTGGGAGCAGCTGTGATAATTGTTTTTGTAACATTGGCGTTTGTGTGGATGAGGTACAGAAGAGAAGGAAATGTTCCAATTGAAGCTGATTTGTTAGCTACAATGGAAAGAATTTCATACTATGAAATCATACAAGCCACTAATGATTTTAGTGAGAGTAATTTCATTGGTTCGGGGAGTTTTGGTTCTGTTTACAAAGGAATTCTCAGAGATGGGACTACCATTGCAGTTAAAGTGTTCAACCTACAAGCTGAAGGTGCATTCAAGAGTTTTGAAACAGAATGTGAAGTTCTTCGCAACCTTCGCCATAGGAATCTCACCAAAGTCATTAGCAGTTGCTCCAACCTTGATTTCAAGGCTTTAGTGCTCGAGTACATGCCTAATGGGAGCCTTGAAAAGTGGTTGTATTCACACAACTACTTCTTAGACATCCTGCATAGACTAAGCATAATGATAGATGTGGCATGTGCATTGGAATATCTCCACAATGGTTGCTCAGCACCTGTGATTCATTGTGATCTGAAGCCTAGCAATGTCTTGCTAGATGAAAACATGGTCGCTCATGTAAGTGACTTTGGCATTTCTAAACTACTCAGTGAAGATGAGAGTGATTTACACACTAAAACCTTAGCAACATTCGGCTACATTGCACCAGGTAAATCAAATCTCTTATACTTCTTTTTTCAGTATTCTATCTTTGTTTGCCTTTTATTTCGATTATATGATATTATGATAAACCGTAAGTGTGCTTTTATTGTAGAGTATGGAAGGGAAGGATTGGTGTCGCCAAAATGTGATGTTTATAGTTATGGGATCATGCTGATGGAAACATTTACGAGGAGAAGGCCTAACGATGAAATATTTGATGGAGATCTTAGTTTAAAGAAATGGGTGAGTGATTCACTCCCAGAGGCAACAATCAAAGTTGTAGATGCGAATTTGCTAACGCCAGAAGATGAAAAATTCATGGACAAGATAGATTGTGTAGCATCCATCATGAAAGTGGCACTGGATTGCTCTGCAGAATCTCCTGAAGAAAGGATCAACATGAAAGATGTTGTAGGAATACTACAAAAGATCAAGATTCAACTTCTTTCATGTTCTGCAAGTACATAAAAACATTTGATTCATGTCACTTGAGGGTATAGAAAGTGTTTGTTAACTTTTTAGCTGCATACCTAGACTGTTTTCCTTTCTTGCTGCATAAGATTCTTTTCTGTAACACTAATGTAATATCGG

>PGSC0003DMG400028438 LRR-RLKs

ATGAAGAGAAAAATCAATTTCCTTTTCCCTCTTGTAGTTTCAGTAATATGGCATCATTTGCTGATATTATCAGCCACTGCAGAAGTCAATCTCAAAACTGATGAAGCTGCTCTACTTGCTTCGAAATCCTATGTTACTTCTGATACTTACAATATCTTGACAAGCAACTGGACCTCAACTACTTCCGTTTGCAATTGGATTGGAGTCACCTGTGGCTCTCGCCATCAAAGGATCATAACCTTGGATATTTCCAATATGGGATTAATAGGTACCATCCCTCCTCACTTGGGAAATCTGTCCTTTTTGGTTTCTCTCGATATCAGTAGCAACGGCTTCCAAGGAATTCTGCCACGAGAGTTGGCTAATTTGCATAGATTGGAGTTTATAAATGTCACTTCTAATAAGTTCACAGGAGATATTCCCTCATGGTTCTCTCTTTTGCCAGAACTCCAGCACTTGCATCTAGCTTTTAACAGTTTCACAGGTATTATTCCACCAGATATTTGTAACGCTTCCAAACTTGAGAGTTTAGTCTTGGGATTTAATCAGTTACAAGGGGAAATTCCAAATGAAATTGGTAATCTTCAGAACTTGACTTGGTTGAGCTTGGGATCAAATCAGCTAACAGGCTCTGTACCACTTAGCCTGTACAACATTTCATCGTTACAACGTTTGGTTTTGACAAAAAATAGTTTGTCTGGAAATCTTCCTGTCGATATATGCTTGAACCTTCCAGAACTTATGGTGCTTGCACTTTCTGACAATGAATTTGATGGTCAAATTCCTTTGGGGATAGATAAGTGTTCAAAGCTTCAGATTTTGTCATTGTCATTCAACAAGTTCAGTGGATTGATACCTAAACAGATTGGGAACTTGAATGTGCTGTCTATTTTATATCTTGGTTGTAATGACTTGAAAGGTGAGTTATACATATTTGCTTTTAGGTTCCTTCTTTGTGCATACATGAAACATATTTTTTGAGTCATGTATGCACTCCTTAATTGATATACTGCAGAAAGTAAACTTTCTATTTATGATTTAATTTGCAGGTGAAATTCCAGAAGAGATTGGAAATCTTCGTAATCTAGAGATATTGGATGCACAGAACTGTAGCCTAAGTGGTCCTTTACCATCTTCTATAAGTAACATAACTTCACTTCAATCACTCAATCTTTATGGCAATAATCTCTCTGGCACATTGCCCCGGTATATATGTTTGAACATGCCTGATCTCAGAGCATTTGATCTTGGGAATAATTTATTCAGTGGAAACATTCCAAAAGAATTTGGAAACTGCACTTCACTTACTGACCTCTTCTTAAGAGAAAACAACTTAACAGGTATCATCTTGTCATTTTATTTTTCCAAGATTTGGTGTATGTTCCTATGCTTTTCATATCCTAATGAGTCGCTCAAATGTCATCAGGTGAGTTACCAAGGGAGATTGGTAATCTTTTCAATTTGGGAAGATTAGATCTACATTACAATTTCTTAACAGGTCCTATTCCATCTACAATATTCAACATGTCAAACATAAGGGGCATTTCATTTTTAGGGAACTTCTTTACTGGAAGTCTACCATCAGATATAGGACTTGGTCTTCCTAATCTTGAAGAACTTTATCTTGGTTACAATAACCTTACTGGAGCCATCCCTAATTCTCTCTCAAATGCTTCCAATATTTTCCGGCTTGGAATTGGATATAATGACTTTTCTGGTCATTTTCCGAGGTCGTTTGGTAATCTAAGATGTTTAGAGTACCTCAACGTGAACGGCAATCATTTCACAAGAGAGCCTTCATCTCCAGGATTGATTTTCTTTGATTCCTTGACAAATTGCAGACATTTGAGAAAACTGTGGATAGGTTATAATCCACTCAATGGAAATCTTCCGGCTTCTATTGGAAATCTCTCAAGTTCTCTCGACTACATTTATGCTGTTAATAGTGAAATCAGAGGCTACATTCCCAGTGAAATAGGAAATTTAAGTGGTTTGTCTTTCTTGTTACTTCAAGGCAATTACTTGAGTGGGTTCATTCCAAGAACAATAGGGAATTTAAAGAATCTTCAAGCCTTGAATTTGTATGACAACAAAATGATAAGTGGACCTATCCCTGAGGAACTCTGCAATTTAAAGAAACTAGGATTCTTGAGCTTGGGAAATAATGAGCTTTGCTGTTCTATACCAGCATGTTTAGGGAACATTACATCTCTTAGATACATTTACCTAGGTTCCAACAAATTGACTTTTAGCATACCTCCCAGCCTGTGGAACCTCAATGATCTCTTGCATCTTGATGTATCCTCAAATTCCTTGAAAAGCTCTCTACCTCCAGAAATTGGAAATCTGAAAGTCGCAACGTTGCTGAATATATCGAAGAATCAAATCTCAGGAAGTATACCGAGCACAATTGGAGGCATGCAAAATATGGCTGAACTTTCTTTTGCAGAAAATAGACTAGAAGGACCAATTCCAGAGTCAATGGGGAACATGATCGCCTTGGAATCGTTAGACTTGTCTCATAACAAGCTCTCTGGTGGCATTCCCAAATCATTGGTAGCTCTTTCACATCTCAACTATCTTAATGTATCTAACAACAGATTGAGTGGTGAAATTCCAATTGGAGGTCCTTTCGTGAACTTCTCTTATGATTCATTCCTTTCAAATGAGGCATTATGTGGTGCTGCTAGGTTGCAGATACCAGCATGTAGGTCTAATTCTCCTAGTAGGAAAAGAAAAAAGATGGTGCTTCTTATTGTGTTGATCTTGTTGGTGGCTTCTTCAGTCATAATGCTCTCAGTCATGCTCACCATCTTTTTGGTGATTAGAAGTCGAAAGAGGAAGACAAACATTGCTACTCATCAGGCGGATGCCTCTCCAGCCACGGTACATGGAAGAGTTTCATACCATGACCTTCAACAAGCAACCGACAGATTCAGTACAAGCAACTTGCTCGGTTCTGGAAGTTATGGCTCTGTTTACAAAGCAACATTTGGGAGTACTATAGTGGCCGTTAAGGTATTCAACTTGCAAACAGAAGGCGCATTCAAGAGTTTTGACACAGAATGTGAAGTCTTACGCAACCTTCGTCATAGAAATCTGACCAAAGTCATCAATAGTTGCTCCAGCATTGATTTCAAAGCCTTGGTATTAGAGTACATGCCTAAGGGAAGCCTGGATGATTGGCTTCATTCTGAAACTTGTTCCTTAGATATAATGAAGAGAGTAGACATAATGATTGATGTGGGATCAGCATTAGACTATCTTCATCAAGGCTATTTTGTGCCCGTAGTTCACTGTGATCTAAAGCCTAGCAATGTCTTGCTCAACGAAGAAATGGTTGCACATGTGAGTGACTTTGGCCTGGCAAAGCTATTAGGGGTAGGAGAAAGTATTGTACAAACCAAAACACTTGCAACCATTGGATACATTGCACCAGGTAATAATCATGTCAACAAGCTTCATTTCTCCTTCAAATTTCAACCATTTTAAAGATACCATATTCTCTTGATAAATTGTTTTATTTCAATGTAGAGTTTGGATTGGAAGGACTCGTATCTACAAGGTGCGACATTTACAGCTACGGTGTCATGTTGATGGAAACCTTTACGAGAAAGAAACCAACGGATGGAATGTTTGCTGAAAATTTAAGCCTGAGGGAGTGGATAAGACAATCATGGCCACACGGAATGGACGAGATAATAGACCCCGAGCTTATGATACCAGGGGAGAAGAACAAAACAGGAAAGGTGCAGTTCTTATCATCCATCATGGAATTAGCTCTGAGGTGCACTTCAGACTTGCCAGAGGAAAGAATGAATATCAAACATGTTCTTGCTCAACTGAAGAACGTCAGAACTATGTTGGAAAATGTAATAAATCACTAA

>PGSC0003DMG400028473 LRR-RLKs

ATGAATATTAGCACTGATCAATCTTCTCTTCTTGCCTTGAAATCCCACATCACTTCAGATCCTTTTCATGTTCTTTCGACGAACTGGTCTTCTTCTACCTCTGTTTGTGATTGGATTGAAATCACTTGTAGCTCTCGCCATCAACGAGTGATTGCACTGAATATTTCAAACATGGGTTTATCGGGTACTATCCCGCCACAACTTGGCAACCTCTCCTTTCTTGTTTCACTTGATCTAAGCAAAAATGATTTCTGTGGTGAACTTCCATCAGAATTTTCTCATTTACGGAGATTGAGAGTCATTAAACTTAGTTACATCAACTTCACCGGAGAAATTTCAATTGGAATAGCCCCTCTTCCGAGTTTGATACGGTTTTCCATGGGATACAATAAATTACTCAATGGCTCCAATGTGCTCTCTAAATTCAACATCTCAACCCTTGAATATTTGGATCTCAGAAATTCTGGTTTAACTGGTGATTTCCCTTCTGATTTATGTCATCGTCTTCCTAGATTGCAAAAGCTTGCACTTGGCAACAACATGTTAAGCGGAGAGGTACCAAGAAACATATCTGAATGCTCAGAGCTTCAACTCCTAATGTTGTTTCAAAATAACTTTGTTGGAACAATTCCAAGAGATATAGGTAACTTGCATCAGCTGCAACATTTAGATCTTGCAGATAACACGTAAAAGGTATGGCTTTACCATTTTTCTCGGGACATTAGATTTAACTTATTTAAAAGAAAATGTATTTTAGAGAGTTGAAGTGTTTGGACAATCTTTAAAAAAATGTGAGTATTTTTTTAATAGTAGCATAAGAAGATTTGTTTGTTTTCACTAATTGCTCCGTCACTTCTCCTCTCGCTTTGTATACTTGTAGCCGGTTTGACCAATCTTCCGGCATACCCATTTGGAAGTTAAGCTCTTGTTAAATATATGATTTTTATGTGATATTTCTGTTTAACAATATGACAGGCACAATTCCTGATGAGATCGGCCATCTTTATAACTTAAAGAACATATTCATGGATAAAAATTATTTAACAGGCTCAATCCCTTTAACCTTATTCAACATTTCATCACTTGAAATGCTATATATGAATGATAACAAGCTTGAAGGACCTTTACCAAGACAGGTTGGAAATTTTACTATGCTTGCCTGGTTTGATCTTTCCAATAATTACCTGGCAGGTACGGGGAAAAAATCTAATTCAAGTCATGTTAGAATTTTTTGTTTTTTGGGATACGTCTTCTGATCATAATTATACTGAATTTTTCAGGTGTAATTCCACATGAAGTTGGTAACCTTCAAGAGTTGACAGACCTTCCATTGTTTTACAATGAATTTAGTGGGTCGATCCCTATTGGTATCTTTAATATCTCATCTCTTGTAACTATTGGACTCACAGAAAACCACATTTCAGGTAATCTTCCATCCACTATAGGCAATGGGTTACCTAATGTTGAAAAAATATTTCTAGGTGGAAACAACATTAATGGTGTTTTACCTGGATCCATCTCAAATTTGTCTAAGCTCACCTTTCTCGAACTCTTTAGTAATGAATTTACTGGTTCAATTCCTGACTCTCTCGGAAATTTAAGACAACTTGAACTTCTCAACTTGTACGGTAACTCCTTTACAAGTGAATTGAGCTTCATTACTCCTTTGGCCAATTGTAAACACTTGAGAATATTGATATTGTCTTTCAATCCTCTAAATTCAATGCTCCCGAGATCCATTGGCAACATCTCTTCTCTTCAATCATTTATTGCTGTAGGCTGTAACCTCAAAGGCAATGTTCCAAGTGAGATCAGAAATATGAGAAATTTATCTACTTTGTCTCTGGGAGGTAATAACCTTACTGGAATTGCCCCGACGACATTAAGTTCTTTGAAAAACCTGCAACGGCTTTTTCTTGATGCAAACAGAATAACTGGTCCTTTCCCAATTGTTTTATGTGAGCTACCCCACTTGGGGATACTAAGCCTTTCACAAAATAAAATTTGGGGAGATATTCCTAGTTGCTTGGGGAACATGACATCCCTAAGGGAGGTTTATCTTGATTCCAATAACCTCACTGCTAGCATACCTTCAAGTCTATGGAACCTCAAAGACATTTTGAAGATGAACTTGTCTTCTAATTCCTTCATTGGATCTCTACCACTAGAAGTTTGAAACCTCAAAGCTGCACTACTTCTGGATCTTTCCAGGAACCAAATATCAGGCAACCTTCCAAGTACATTGGGAGGTCTACAGAAATTGATTCAACTATCTTTGGCTCATAACGGAATTGAAGGATCTATTCCTGAGACATTTGGGGAACTCCTAAGTTTGGAAACATTAGATCTTTCATATAACAAAATGTCTGGTGTGATTCCAAAGTTATTAGAGGCACTTAAGCAGCTAAACTCCTTTAATGTCTCATTCAATAGGTTAGATGGGGAAATTCCGAGTGGGGGACCTTTTCTTAATCTCCCTTACCAGTCTTTCATGTCGAATAAAGGATTGTGTGGTAACCCTCAAAAGCATGTCCCAGCTTGTCCTTCTTATTCAAAGAATTAGTCTAAGTCGAAGAAAAGAAATCTGATATGGATTATAGTTGCCTCTTCAGTTATCTCTGTAATAGGGCTTGCTTCAGCAATAGTTTTCATGTTGATGAGACGCGGGGGTAAAACGATCAATGTTGAAGTTGAGTGGTCTCCTGAGGTAGCACCACAAAGATTTTCTTACTATGAACTTCAAAGAGCAACTCAGAGTTTTAATGAAAATAACTTGCTTGGTAGTGGAAGTTTTGGTTCTGTTTATAAAGGGATGTTGGCTGATGGGATGATTATAGCTGTTAAAGTTTTCAATGTGCAGATGGAAGGAACATTTCAAACCTTTGATAGAGAGTGTGAAATCTTGCGGAATCTTCGTCACAGAAATCTCACAAAGATCATTAGCAGCTGTTTTAACTTGGATTGCTCCAGGTGAATTCTTTATTTAACTAGTAACAATGTTATGGTTTAATGCATTAAAAATGCTGACTTATTTCCTTTTTTCTTTTTAGAGTATGGTCTGGAAGGCCTTATATCCAAGAGGTCTGACGTTTATAGTTTTGGCATCATGCTGTTGGAAACTTTTACCAAGAAGAAACCCAGTGATGAAAAGTTCGCAGGAGATTTGAATTTGAGATGCTGGGTGCATAATTCACTTCCTGATGAGTTGGATCAAATCATAGATGCTGACTTACTAACAGTCAATGAACAAAACTTAAGTCAAAAGCTGCAATGTGTGTCATCCATTATGGAGTTAGCCATGAATTGCACAACTAGTATTCCAGTTGAAAGGATGAACATGACTGATGTTGTAGAAACATTGAACAAGATCAAACAAAAGCTTTCTTACTGTTATTGAACTACCTAACAACATGACATCAGGTGCTTTTCATATCCCTTCTGAAAAGAACGAGGTTATAAGTTCAGTAGCATTTGACCATCAGTAAGTGTTCCTTGTTATTGCAGGAGATTATGTGGCTATTGGAAGTTAAAAGGATTGAAGTTCATATACTTCTTACGTCAAGAAGTTGCTATGATTTTGGGACTACGTACTATTATAAAAAGCCGATGATAGCTTGTGTACTTCTTCCTTTTTCTTGTGTAAACCAAGAGGGCCTTTCCATAACCTGATGTTCTAAGCACCCTTCCAATTTGATGATACAAGGAATGTAAGCCATTTAAATAAACTATGTTGAATCCATTGTATATATCTCAAGTTCTCCCAAGTATTCAACTGACTTAAAAAAATTTCTCCAAAGAATAGAGCAGCAAGATATGCATATTCCTTGATAGTTTTCACTAGTTACTTTGACATCGGAGTTCCTTTCTGTTACTGATCTATCCATATATATATAGTCAACTAGTCAAGGTTTTACCTTGTTATGGCATTGAACATGCATGAAGGAGAACTTTCATACAGCCACAGATAAATGAAGGGTAAACAAGAAAATATCTTAGGCAGAAAGGGAGTTGAATTTGAAGTTCTTGATTTGTACAGTCAAGGGTGTATGAACTGGGTGCCAAAGAGGGCAACTAGTGGGGCAAATAGTGAGTCGAAAAAGGGTTGTAAATTTTCCAGAGTCAACTCAAATAGTTTATTTATTAATGACTGCATGAAGCCTGGAATAAATTCATTAATTCAAATAGTTTAGGGTATTGATGCAAAGATCCAACCTAGGAAAGTTAACGACCTAGCTAGAGAGTCAACTCTTATTCCACTCTTTTGTATGTCTGTATACTGATAACCACAAACAAATTTTCTATGCCAAAATCTAGATAGCTGTGTTTAGCAACATCTCACCATTAGCAAAGTTTCCTTCTCTTTAAGTAACAGATTAATTTAGTTCAATATATGGTGCAAAGTCAACTTCCTTAGTGGAGTAATATATTACTTAGGCCATAAACTTATATTTTAATGTATTAGCATTTATTAGCTAGCTACCAGATCTTGAATAAATTTACTTGCACAATTAACGGACTAAATAGTTGCTAGAAGCATTTTACAATTCCAAGTATTATGTTGGTTTCAATAATATAGAACTATGTACTAGCATTATATTTTCGATTATAAATCTAATCAGTTTCAATAGGTATAGATCATTATATTGCTGCAATGATCAAACCAGAAAACTCATCAATGGAGAAAACATCCTCCTCCTCCTTTCCGCTTCTAGAAATTTTATTAATATCTTTTCATGTTATAACGACATGCTTAGCCATGAATATCACTATATCAGCACAGATCAATCTTCTCTTCTAGCCTTGAGAGAACACATCACTTCAGATACTAATCACATTCTCTTGACAAACTGGTCTTTTTCTAACTCAGTTTGTAACTGGATTGGAATCACTTGTAGCTCTCGTCATCACAGAGTTATTGAACTGAATATATCAGGTATGGATATTACTGGTACCATCCCACCGGAACTTGGCAACCTCTCTTTTCTTGTTTCACTTGATCTAAGCAAAAACAATTTCCATGGTGAACTTCCACCAGAGTTGCTTCGCGGAAGTTGAGAGACATTGATCTTAGTTACAACAACTTCACCGGAGAAATACCAAAAGGAATAGCCACTCTTCCAAATTTGAAATGGTTCATCTTGAGATGCAACAAATTACTCAATATCTCATCTCTTGTAGTTCTGGGAATCTTAGGAAACCACATTTCGGGTAGTATTCCTTCCACTATAGGCAATGGCTTACCTAATATCGAAGGAATTTATCTAGTTGGAAATAACATCAATGGTGCTTTACCTAGTTCCATCTCAAATTTGTCTAAGCTTGAATTTCTCGAACTAGGTGGAAATGAACTTACGGGTTCAATTCCCGAATCTCTAGGAAATTTAAGACTACTTAGAAATCTCAACTTGTACGGTAACTCCTTCACAAGTGAATTGAGCTTCATTACTCCTTTGGCCAATTGTGAAAACATGACAAGATTGTTATTGCCTTTCAATCCCCTAAATGCAATGCTTCCAAAATCTGTTGGCAATCTTTCTTCTCTTAATTATTTTTCGGCAATAAGCTGTAACCTCAAGGGACAAATTCCAGATGAAATTGGAAATATGAGAAAGTTATCTACTTTGGAGTTGGATGACAATGACTTAACTGGAATTATCCCGACAACGATAAATTCTTTGAAAAACCTTCAACGGCTTTCACTTGGTGCAAACAGAATAAGTGGTCCTTTCCAAATTGTTTTATGTGAGCTACCCAACTTGGGCATGCTAAACCTTTCACAAAATCAAATGTGGGGAAATATTCCTAGTTGCTTAGGGGATGTGACTTCTCTAAGGGAGATTTATCTTGATTCCAATAACTTCTCTGCTAGCATACCTTCAAGTCTATGGAACCTCAAAGACATTTTAAAGTTGAACTTGTCATCTAATTTCTTCAATGGTTCTCTTCCACTAGAAGTTGGAAACTTCGAGGCTGCAACTCTTCTGGATCTTTCGAGAAACCATATCTCAGGCAACATTCCAAGTACATTGGGAGATCTACAGAAATTGACTCAACTATCTTTGGCTCATAACAGAATTGAAGGATCTATTCCTGACACATTTGGGGAACTCATAAATTTGGAAGCGATGGATCTCTCGTATAACAAAATGTCTGGTGTGATTCCAAAGTCATTAGAAGAACTTAAGCAACTACGCTCCCTTAATGTCTCATTCAACAGGTTACATGGAGAAATTCCGAGTGGAGGGCCATTTGTTAATCTCACTTACCAATCTTTCATGTCGAATGAAGGATTGTGTGGTAACTCTCAAGAGCATTTCCCAGCTTGTCCTTCTAATTCAAAGAATCATTCTGAGTCAAAGAAAAGAAGAATGATATGGATTGTAGTTGCCTTTTCAGTTATCTCTGTGATAGTGCTAACTTCAGCAATAGTTTTCGTGTTGATGAGACATCGGGGCATAACAGTCACTGCAGAAGACGAGGGATTGCCTGAGGTAGTACCACAAAGAATATCTTACTATGAACTTCAAAGAGTGACTCAGGGCTTTGATGAAATTAACTTGCTAGGTAGTGGAAGTTTTGGTTCTGTTTACAAAGGGACATTGGCAGATGGGATGATTGTAGCTGTTAAAGTTTTCAATGTGCAGATGGAAGGTACATTTCAAACCTTTGATAGAGAATGTGAACTCTTAAGGAATCTTCGTCACAGAAATCTCACAAAGATCGTTAGCAGCTGTAGTAACTTGGATTTCAAGGCATTGATACTTGAGTACATGCCAAATGAGAGCTTAGACAAGTTGCTATATTCTAGAGATTTCTGTTCAAGTATGATGCAAAGATTGAATATCATGGTTGATATTGCATCTGCTCTGGAATATCTCCATCATGGTTACTCAGTAGGAGTTATTCATTGTGATCTGAAGCCTAGCAACGTGTTACTTGACAACGACATGGTGGGACACCTGACTGACTTTGGCATTGCGAAACTTTTAACCAAAGAAGAATCTATTGCTCACACTAGTTACATTGATCCAGGTGAATGCTTTATTTAACTAGTATAACAGTGTTATGGTTTAATGTGTTAAAAATGCTGACATAATTTATTTTTTATTTTCTAGAGTATGGTCTGGAAGGCCTTATATCCAAGAGGTCTGATGTTTATAGTTTTGGTATCATGCTGTTGGAAACTTTTACCAAGAAGAAACCCAGTGATGAAATGTTCGCGGGAGATTTGAATTTGAGAAGCTAG

>PGSC0003DMG400028474 LRR-RLKs

AACACATTTTCTCCCATTCACTCAGAAAAAAATGAGGATTTCATTAATGGAGAAAAATCCCTCTTTTCAGCTTCTAACAATTTTGTTCGTTCTTCTTCATGCTCTAGCAACTTGCTTAGCCATGAATTTTAGAACTGATCAAACTTCTCTTCTCGCGTTGAAATCCCAAATCACTTCAGACCCTTATCAAATTCTTTCAACAAACTGGTCTTCTTCTGCCTCTGTTTGTAACTGGATTGGTGTCACTTGTGGCTCTCGTCATCAACGAGTTACTTTACTCAATATTTCAGACATGGGATTTTCGGGCACCATACCGTCACAACTTGGCAACCTCTCTTTTCTTGTTTCACTTGATCTAAGCTACAACTATTTCCATGGTGAACTTCCACTTGAGTTTTCTCGTTTGCGAAAATTGAGAGCTATTAATCTTAGTTTCAACAACTTCACCGGAGAAATCCCTAAATTCCTCGGTGATTTTCAAGACCTTCAAATATTGTCTCTTGAAAACAATAGTTTTAGTGGGTTCATTCCTTCTTCTATCTCAAATATGAAGAATCTTGGATTCCTGAATCTGAGGTACAACAATTTAGAAGGAAATATTCCAGCTGGAATAGCCGCTCTTCGGAGTTTGAAATGGTTGAGCTTTGGATTTAATAAACTCAATGGTTCCAATGTGCTCTCTATGTTCAACATCTCTACACTAGAATATTTGGATCTCAGAAATGCTGGTTTAACTGGTGATTTTCCATCTGATTTGTGTGGTCGTCTTCCAAGATTGCAAAAACTTGGACTTAATTTCAATATGTTAAGTGGAGAGATCCCAAGACGTATATCAGAATGCTCGCAACTTCAAGTCTTACTGTTGATGGAAAATAACTTGATTGGAACAATTCCAGGAGAACTTGGGAAATTACAACTGCTGCAACAATTAGCTCTCGGAAATAACAAGTTAGAAGGTATGCAGCTTCCAATCTTAACAATATCTACATATATAGTAATAAATCGAAGATATTGTTTTGTGCTGAGCACATGCATTCACTTTAATTTTCAGTTTCGACAAGAGGTTAAGTGTTCAATTTTCCTCTCTTAAACAAGCATATGTATGGTTTTGACCTGGTGTGAGAATTTGTTACATCACCTTTGCTTATACCTGTCTGTTAAATTATTTAATGAGGTATTGAGATTGGTTCAACTTAATTCCTTAAGACTTTCATTCGATAATTTTGCATAACAATTTGACAGGCACAATTCCCAATGAAATTGGTCATCTTCATAACTTGAAGCAATTGGGATTGGAACAAAATGCATTAACAGGCTCAATCCCATTAAGCATATTCAATATTTCATCACTTCAAGTTTTATCAATGTGGGACAACAAGCTTGAAGGGCCTCTACCAAGAGAGGTTGGAAATTTGACTATGGTTAACGTACTTGATCTTGGAATGAATAGCTTGACGGGTATGGAATAATATTTTACCACAGGATTTCAGAATGCTAGTTCTTTCAATACTTCTTCTAATGGTAATCTTACTTCATTCTGCAGGCGTACTTCCAGATGAAATCGGTAACCTTCAAGAGTTGCTGATGCTGAAGTTGGATTTCAATAATTTTAGTGGGTCGATCCCTATTGGTATATTTAATATCTCAACTCTTGTATCTATTACACTCACGCAAAACCGCATTTCAGGTAATCTTCCTAACACTATAGGCAGCGGGTCACCCAATCTTGAACGAATTTTTCTAGGTGCAAATAACATTGATGGTCTTTTACCTAGTTCCATCTCAAATTTGTCTAAGCTTACTGTTCTCGAACTCAGTGCGAATGAACTCACGGGTTCAATTCCTGACTTTCTAGGAAACTTAAGACTTATTGAAATCCTCAACTTGCAAGGTAACTCCTTCACTAGTGACTCTTCAATGTTGAGTTTTATTACTCCTTTGGCCAATTGTAAATACTTAAGAGAACTGATATTGTCTATCAATCCTCTAAATGCAATACTTCCAAAATCCATAGGGAATCTTTCTTCTCTTCAAACGTTTGAGGCAATAGGTTGTAATCTCAAAGGCCATATTCCAAATGAAATTGGAAATTTGAGAAATTTATCTTATTTGAAGCTGGATGAAAATGACTTTACTGGAATTGTCCCTTCGACAATAAGTTCTTTGGAAAAACTTCAACAGTTTTCACTTAGTGCAAACAGAATAAGTGGTCCTTTCCCAATTGTTTTATGTGAGCTACCCAACTTGGGCATGCTAAACCTTTCACAAAACCAAATGTGGGGAAGTATTCCTAGTTGCTTAGGAGATGTGACTTCTCTAAGGGAGATTTATCTTGATTCCAATAACTTTACTGCTAGCATACCATCAAGTCTATGGAACCTCAAAGACATTTTGAAGCTGAACTTGTCTTCTAATTTCTTCAATGGCTCTCTACCATTAGAAGTTGGAAATCTCAAGGCTACAATACTTCTGGATCTTTCCAGGAACCAAATCTCAGGCAACATTCCTGGTACATTGGGAGGTTTACAGAAATTGATCCAGCTATCTTTGGCTCATAACAGAATTGAAGGATCTATTCCTGAGACATTTGGGGAACTCATAAGTTTGGAAGCATTGGATCTTTCAAATAACAATATTTCCGGTGTAATTCCAAAGTCATTAGAGGCACTCAAGCAACTACAGTCCTTTAATGTCTCATTCAACAGGTTACACGGGGAAATTCCGAGTGGTGGACCTTTTCTTAATCTCCCTTACCAGTCTTTCTTGTCGAATGAAGGATTGTGTGGTAATCCTCAAAAGCATGTCCCAGCTTGTCATTCAAATTCAAAGAATCATTCTAATTCAAAGAAAAGAAGAATGATATGGATTGTAGTTGTCTCTTCAGTTATCTCTATAATAGGGCTTGCTTCAGCAATAATTTTCGTGTTGATGCGACATCGGGGTAAAGTAATCAAAGGTGAAGATGAGTGGTCGCCTGAGGTAACACCACAGAGATTTTCTTATTATGAACTTCAAAGAGCAACTCAGGGGTTTGATGGAAATAACTTGCTAGGTAGTGGAAGTTTTGGTTCTGTTTTTAAAGGGACATTGGCAGATGGGATGATACTAGCTGTCAAAGTTTTTAATGTGCAGATGGAAGGTACATTTCAAACCTTTGATAGAGAATGTGAAATCTTGAGGAATCTTCGTCACAGAAATCTCACAAAGATCATTAGCAGCTGCTGTAACTTGGATTTTAAAGCATTGGTACTTGAGTACATGCCAAATGGGAGCTTAGACAAGTTGCTATATTCTCAAGATTATTCTCTAAATATAATGCAGCGATTGAATATCATGGTCGATGTTGCATCTGCTCTGGAATATCTCCATCATGGTTACTCTGTACCTGTTATTCATTGTGATCTGAAGCCTAGCAATGTCTTACTTGACAAAGACATGGTGGGACACCTAACTGACTTTGGCATTGCAAAACTTCTAACAAAAGAGGAATCTATTGCTCAAACTACAACCTTTGCCACAATTGGTTACATTGCTCCAGGTGAATCATTTATTTAACTTGTAACAAAGTTTTTGATTAACAGTTAATCAGATTTACGCCTTTTAAGTTCAGTTGCTGACGTATTTTAATTTTTCTTTTCTAGAATATGGATTGGAAGGCCTTATATCCAAGAGGTCTGATGTTTATAGTTATGGTATCATGTTGCTGGAAACTTTTACAAAGAAGAAACCTAATGATGAAATGTTTACGGGCGATTTGGATTTGAAAAGCTGGGTACATAGTTCACTTCCTGATAAGCTGGATGAAATCATAGATGCCGACTTACTAACAGTAGATGAACAAAAGCTAAATGAAAAGTTACAGTATGTGTCATCTATCATGGAGTTAGCCATGAATTGCACAGCTAAATCTCCGGCTGAAAGGATGAACATGACTGATGTTGTAGCTGCATTGGAAAAGATCAAGCAGCAGCTTTCTTCATGTTACTGATCGAATGGCTTAATGACATGACATCAGGTACTTTTTCCATATCACTTTTGGGAAGTAAAAAGGAAATATGTAAATTTCATTAGCATTTTAATCAGTAATTGCTCTTCAAATGCTTTTGCAAGAGACTATGTGGCTATTGTAACTGTCCTTTGTTGAGATGAGTCTTCTGTACAATGTATTGCTAATTAAGGATCAATAAAATGCAGTGCTTCTTAATTTTAAGTGAAGTGATCCCCATCTGGCCTTATAAATATCATCCAAAAGAGTCATTTGCCCTTGCTTTAATCTTGTATCCAGTGAAAATGATAAGTGCTAACACTCCTAGATGTTTTGATGATCTCCTCACAAGTGCAGGGACCTGGTCCTCTGCAGAGTATGAATACTCGTCCAGTGGTGAAAGTGTTGAACAGCTGTAAAAGTTGTTCGTGTCAGAAAGTTGGTGAAGTTGCAGAGTACTTCACCAATCAGAATGGGCGTAGTTGTTTGTACATTGGACAATGGCTTTTCATGGTTGATATATTCAACACTCAAACAACAAATTCATTATTCATTCAAAAAGAGATTTTTCAAGCTTCAAAAACAGCAGGGACCTGATCGATTGCTTAAAGTGCTTCCTGTATCAGTTGTTACTTTGTTTGCTTGTAATAGTTCTTATATGTATGATTCGTTTCACTTGTGAAAGAAACGTTTCAAACTTGTGTAAATCATTGTAAGAACTAGGTTAAGGGTAACTTAGTTTCTTACTCTAAAGTCTATATTAATCAGTTAGGATTAGTATAGTGGGCAGTGTTGTATTTGCTCTGGCTCAGCTAAATGATAGGAGATATTGCTTAGTGTGGAGATTAGCAGGCTAATCTTCATTTTGTAAACTGGCTTTTACTTTTGCTTGGAGAAGATTAGTGGAAGCAGTTTGAAAAGTCCTATTGAGAGCAGGTCGTGGTTTTACTCCCTTGAGCAAGGAGGTTTCCACGTAAAGTTGTTTGTGCAATCTTTACTTTCAGCATTTATTTTACTGTTTGTTATTGTAATTGTGCTGAGGACTTGGTCCCATCAAAATCGGTG

>PGSC0003DMG400028478 LRR-RLKs

CAATTATATTGGACTTACTTACCTCCACTTGAAATCCCCTTAATACATCTACTCTTATCAACTCTATATGGCATTTACATCAACTACTTCCAACTGAAAGAAACAATCACAAGTCCCACAAACAAAAAATGAAAATCTTGTTCACACTCTTCATCTTTCTTTCTCTTTTCACATTCTCAAAATCAGAATCCCCTTGCAAAAACAACACAGATTTAGATTTATATCTTGTTTCCAAGGCTTTCAGTTCTGTTTATGGTTTCAACATCTCTTGGTTCAACTCAAACTGTTCTTCTCCAATAACTGAAATAAACTTATCTTCAAGAAACCTCACTGGCATAGTTTCATGGAAGTACTTAACAAATTTGTCTCATCTTCATACTATAGATCTATCAAATAATTCCTTAAAGGGTTCTGTTCATCCTCTGTTTTGGTCCATTTCATCTTTGGTTCAAGTCAATTTGTCAAAGAATAAGCTAGGTGGGGCTGTTGCTGTAGCAAGATCATCATCATCTATTCAAAGACTTGATCTTTCTTTCAACAGGTTTACCAACTTAGGCTCTGTATTTTATGGTTTTCCTAATCTTACTTCTCTTGACCTTTCACATAATGATATCAAGTCTTTACCTTTTTGGTTTACCAACCTAACTAAGCTTGAAAATTTGAGCATTTCTAGCTGTAACATTTATGGTAATCCAGAACCCCTTTCACATATCAAGTCACTGAAGCATTTAGATGTTTCTGTCAACCATATGGATGGTAAATTTCCTAATGATTTTCCTCCTCTGTCTAGTCTCAATTTCTTGAATATCTCATTCAATAACTTCACTGGGGAAATACCCAAAGACCAGTTTGCTAAATTTGGGAACTCTTCTTTCATTCATGCTGGGCACTTGCAAATCAAGAATCTCTTACCAAATCCCAAGAATTCCTCACAGTTACACATTAAAAATCATAACTTTACTACCCCAACACACAAAAAAATGTTGCCTTCGAGACACAAACCTATCAAACCAAACACACACAAGAAAAGGCCCAAATCAAGAAAGAAAGTTTTAACCATAGCTATTTCCTCTGCTTCAGCATTTCTAATCTTGGCCATGGGGTTAGTCATTTTGTGTCTATACAAGAGGAGAAAAACATTGGCTAGGAAGAACAAATGGCTAATTTCAAAGCCAATACAAATACCATTTAGAATGGACAAATCAGGGCCATTTTCATTTGAAACAGAGTCAGGGAATTCATGGGTTGCTGACATAAAAGAACCAAGTTCAGCAGCTGTAGTTATGTTTGAGAAGCCATTGATGAACCTAACATTCAAAGACCTCATAGCAGCCACAACTCACTTTGGAAAAGAGTCATTGTTAGCTGAAGGAAGGTGTGGGCCCGTTTACAGAGCCGTACTTCCAGGTGACCTCCACGTGGCAATCAAGGTCTTGGAACATGCTAGGGAACTAGCCCATGATGATGCTATTATCTTGTTTGAACAACTCTCCAGGCTTAAGCATCCTAATCTGTTGCCTATTTCTGGTTACTGCATTGCAGGTACTAATCTTCTTTCACTTATTACAGCATCTTCTGTTAATAAGTACAATTTATGTGATACAATTCGAATTCTAGCAGTAAAAAATATCTAGAGGTGAGCATATTGAATGCATTCACGTGGGAGGATGGGATGAGAAATAGGACAGACAGTAGTAAATGAGACAGAAATCTTTGCAACCATAGAAACGTAGGCAGTAGTTATGATTACATACAACAAGATGAAGCAGTAAATAGGAATGCGTGACTTATTTGGATGTTAGTACTATGACTACATCATTCAAATTATATGTGCTTTTAATGCTCTTTGTAAATTGTGTCCATTTTTTACCCTACAAAGTCAAACTTTCAAATCTTATGCCAGATTGGGATTTGAAAAATACTGAAACTGCAGTATGGTTTGGTTCGCAGAATTACGATAACCAATGATATTCTATGAATTGAGATATTTCACGACATTTATTTATCGTGAAATTAAAATGGTAGAATAAAAAAAAATTGTGATTAATTAATATCTCAAACTAAACATGGAAAAAATTAATTTTTAATTTTATTTCGAGATTATTATTTGTATCACATATATCACCACGACATTTATGATGATGCTCTTGGTAAATTGTGTCCATTTTGCACGATAAGGTCAAATCTTTAAAAATCTTACCAGATTGGGAGTTCTACTGTTCAGCCTTGATTTATGATTTTTGGAAGAATGTTGCACAAATATTATCCATAAAGTATTATTTTATGGAGGCGTTATTGAATTAAAAAGATAAGAGGATATTTTAAAAGCATTAATGGTTCAGAAGAATGTCATTTTCAATTGAAAATGCTTAAAGTTGGAAAATTATTTCATGACATGCATTGATTGATCAATTGTAGTATCAAAGTCCTTGTCAAATATTTTTGGAAGACTTTTCTAGCTAGGAACCAAATTGCTCCATTTGATAAGAACAGAATCCAATTATATTTTCTTTCAATTAAGTCATATTTAAAGCTAATTTAATACTTCCCTTTGATGCAGGGAAAGAGAAGCTAGTCTTATACGAATTCATATCCAATGGTGACCTCCACCGATGGCTACACGAGCTTCCAACTGCTACCACCAACGTGGAAGATTGGACAACCGACACGTGGGAGTTACAAAACGGTTCTCAAATTACGTCACCCGGGAAAATGGAATGGCACACGCGCCACCGTATTGCCGTTGGCGTAGCGCGTGGACTAGCCTACCTCCACCACGCGCAGTCGAAACCTGTGGTCCACGGTCATTTGGTGCTATCTAATGTCTTACTAGCCGATGACTTTGAGCCTCGGATAGCCGATTTTGGATTGAGCCAAAACCAAGCCGAGGGATCTAGTGAGACGGATGTGTACGATTTTGGAGTCGTATTAGTAGAGCTATTAACAGGGAAAATCGGCTCGGATGAGACGATTAAATGGGTGAGGAGGCTAGTGAAGGATGGACTTGGAGCTGACGCGCTTGATTCGAGACTAAGACTCGGTGGTGACTCAGTGAGTGAGATGGTGGAGTGCCTCCGAGTTGGGTACTTGTGTACGGCGGAGACCCCAAATAAACGGCCTAGAATGCAACAAGTTTTAGGTTTGCTAAAAGACATACAGCCCCACTACCAACTCGGGAAAACGCAAAATTTATAATCTTTAGGTAAACCACTCTCACTCCAGCGCGTGATTTACACGACGAGGAAGATTGATTGGCAGCATTGCATTATTGTACTCCCATGTGTTTGGTTGACCATTGAAAAGTCAAAAGTACAAGGCATCTATTCCACTTCATCAATGGTATTTAGTCACGTGCAGATCACGTGATGAAGTGCAGTGAGAGGAACATGTTTGCTTGCTTGTCTCATTTTATTCTCTCTTTTTTTTTTTTTTTTTTGTAGTTTTGTATAGTATTAAATTAGAAAAATGAAAAAAAATTGTGGGTAAATCTTTTTTTTTTATATATAAA

>PGSC0003DMG400028688 LRR-RLKs

TTTTGCTTCTCTATACAAACTCTCCTCATTCTTAGCTTTCCAGTTTTCTGTGGGCCCTTTTTCTCATGTGCTTCCTTCTTCTTCCTACTCTCAACTCTGCAAAAAACCCTACCACATTGCACAATGTTTCTCTTCTAATCTTGAAAAAAGATTGAATTTTTGCTATAAAGAACAATTGGTTTCTTGTTTTTCTTCATTGTCTTGTCAAAAAGTTTCAATCTTTGCAAAAGAAATGAACACCCTTCTTGTGTTAAAGCTTGTATCTTTACTTGTTCTATAAAGAACACTTGGGTTCTTGTTTTGTCAAAAAGTTTCAATCTTTACAAAAGAAATGAGCACCCTTCTTGTGTTAAAGCTTGTATCTTTACTTGTTCTGTAAAGAACACTTGGGTTCTTGTTTTTTATTTCATTGTGTTGTCAAAAAGTTTCAATCTTTACAAAAGAAAACACCCATCTTGGTTTAAAGCTTGTAGCTTTACTTGTTCTATAAAGAACACTTGGGTTCTTGTGTTTTTTTCATTGTCTTATCAAAAAGTTTCAATCTTTACAAAAGAAATGAACACCCTTCTTGGTTTAAAGCTTGTAGCTTTACTTGTTCTATAAAGAATACTTGGGTTCGTGTGTTTTTTTCATTGTCTTGTCAAAAAGTTTCAATCTTTACAAAAGAAAACACCCATCTTGGTTTAAAGCTTGTAGCTTTACTTGTTCTATAAAGAATACTTGGGTTCGTGTGTTTTTTTCATTGTCTTGTCAAAAAGTTTCAATCTTTACAAAAGAAATGAACACCCTTCTTGGTTTAAAGCTTGTATCTTTACTTGTTCTTGGTATTTCTGCATTTTTTTTGGGTGTTGAGTCTCAAGATGATGATGTTTCAGTGATGTTAGCTTTGAAAAAGAGCTTAAATCCACCTCAAGAAGTTGGATGGTCAGACTCGGATCCATGCAAATGGAACCATGTGGGTTGTTCAGATAAACGGGTCACCCGGATCCAAATTGGGCGCCAGAACATTCAAGGTACTCTCCCTCCAGAAATCTCTAAACTTACTGAACTTGAACGTTTAGAGCTTCAAGGGAACAATATTTCTGGGCCTTTGCCTAGTTTGAAAGGTTTGAGTTCATTGCAAGTGTTACTGCTTGGTGAAAATCAGTTTAGTTCAATTCCTGCTGATTTCTTTACTGATATGTCTTCATTACTATCTGTTGATATGGATAAGAACCCTTTTGTTGGTTGGGAGATCCCTGAAAGCCTTAGAAATGCTTCTTCACTTAAAAACTTTTCAGCAAATTCAGCAAATGTAATTGGGAGAATTCCTAACTTTTTTAGTCCTGATGAGTTTCCTGGATTGGTAAATTTACATTTGGCTGGCAATAACTTGGAAGGTGAATTGCCCTCCAGTTTTTCTGGCTTGCTGCTTGAGTCTTTATGGTTAAATGGACAGAAGCTTAATGGAGGAATTGATGTTCTAACAAACATGACATTCCTGAAGGAGGTTTGGTTGCATTCCAATAACTTTTCGGGGCCGTTGCCGGACTTTTCAGGATTAAAGGCTTTGGAGACATTGAGTCTTAGAGATAATGCCTTTACTGGTCCCGTGCCAAGTTCTTTGATGAATCTTGAGTCTTTGAAGCTCGTTAATTTGACGAATAATTTCTTTCAAGGACCAATGCCGGTGTTCAAAGGTTCTGTTGTTGTTGATTCGGCAAAAGGTACAAATAGCTTTTGCTCGTCACAACCAGGTGATTGTGATCCTAGAGTAAATACATTGCTTTCAATTGCAAAAGCAATGGATTATCCAACAAATTTTGCGAAGAACTGGAAGGGAAATGATCCCTGTGCTGATTGGTTTGGTCTTACTTGCAGTAATGGTAACATTACTGTCATAAATTTCCAGAAAATGGGGCTTTCAGGAACAATTTCTCCTGAATTTGCTTCACTGAAGTCGCTACAAAAAATAGTTCTTGCAGATAATAATCTGACTGGTACAATTCCAGAAGAGCTAACCACATTGACAGGCCTTACAGAGTTAGATGTGTCAAACAATCAAATTTATGGAAAGGTACCAGCTTTTAGGAAAAATTTGATTCTGAAATTTAGTGGTAACCCTGATATTGGAAAGGATAAGAGTGATGCACCTTCCCAAGGCAGTTCTCCAGGTGGTTCAACAGGCTCTGATGATGGAAATGCCCAAGCAGCTCGTAAGAAGTCCAATCGCCGGGTTGGAATTGTGGTGTTTTCTGTAATTGGAGGTGTATTTATGCTTTGTTTGATTGGAGTGGCAGCTTTCTGTCTGTACAAAAGCAAACAAAAGCGTTTTAGCCGAGTGCAAAGTCCAAACACCATGGTGTTGCATCCACGTCATTCTGGTTCTGACAATGATAGTGTGAAAATCACAGTTGCAGGATCTAGTGTCAGTGTTGGTGCAGTTACTGAAACTCATACTGTTTCTGCTAGTGAAGCCGGTGATGTTCAAATGGTTGAAGCAGGAAATATGGTGATTTCCATCCAAGTACTGAAGAATGTGACCAACAATTTTAGTGAAGACAACATACTAGGCCGGGGAGGCTTTGGCACTGTCTACAAAGGTGAATTGCATGATGGGACTAAAATTGCAGTCAAAAGAATGGAAAATGGAATCATCACTGGGAAGGGACTAGCTGAATTCAAATCCGAGATTGCTGTTTTGACCAAGGTCAGACATAGGCATCTTGTTGGACTACTTGGGTACTGTCTTGATGGAAATGAGAAACTTCTAGTCTATGAATACATGCCTCAAGGAACATTAAGCAGTCATCTCTTCAACTGGGCAGAAGAAGGGCTTAAACCCATGGAATGGACAAAACGATTGACCATTGCATTAGATGTTGCTAGGGGTGTAGAGTACCTCCATAGTTTAGCTCACCAGAGTTTCATTCACAGGGATCTGAAGCCTTCAAATATTCTTCTAGGGGATGATATGAGGGCTAAAGTTGCAGATTTTGGCCTTGTCAGACTTGCCCCAGAAGGGAAAGGTTCTATAGAGACAAGGATTGCTGGGACATTTGGATATCTGGCACCGGAATATGCAGGTATGCTCTTTGCTTACTTATATGTGTATGCTAAATTGCTGATAAAATACTTGTATGCAATTTGTGCCTAGATAAACTTGGTCTTAAGTCCAATACCTCTAAAGTATGAATTATGTGAACCTTAGGGGTTTGATCTCAGGATGCTTCCAGAGAATTAGCATTCATTTGACTGATGAGAGTCAGATGATACTAACTGTACTCCAAGAAACTTAGCCTGTTTTACATTAATCAACTTTTGCCCTATAATAGGTCTGTAACAGTATATATTAATGTTTTGTCTTTTTTCTCAGTCAATTATACTTGTGCTCATGGTTAAGAACCTGCCATTCATACATTGTCATATATCAAATATCCTTATTGATTCTGTCATCCCCTTTAATGCTATGATTACTTAGTTTCTAAAAGTGGATACCCATCCTACTGGAAACACTTTTTCCTGATGCATTTGCTTAATGCCACCGTTTTCTACTTTGGCCTCATCACACACTTTAAACATGTTATTATTCCTATGACAGAAGATCCTCTTGTTTTCTTTTCCCCTTAGTGGAATTTAAGGATAGCCAAATTTGTTATTTGCTCACTTTCCTGATTGTGATTTGTGCATAGGAGTTCTTGGAATGTAGATACATCATTATCTAAACTTGTATTTTATCTTGTTTCGTCTGCCAGTAACTGGGAGAGTGACAACCAAGGTGGATGTGTTCAGTTTTGGTGTGATTTTGATGGAGCTCATCACAGGAAGAAAGGCTCTTGATGAAAGCCAGCCTGAGGAGAGTATGCATCTAGTAACATGGTTCCGAAGAATGCATCTGAATAAGGATACATTCAGAAAGGCAATTGACCCCGCAATTGACCTTAGTGAGGAAACACTCACCAGTGTCAGCACTGTAGCTGAGCTAGCTGGTCACTGTTCTGCAAGGGAGCCATATCAAAGGCCTGACATGGGCCATGCTGTCAATGTTCTTTCATCTCTTGTAGAGCTCTGGAAACCATCCGACGAGTGCTCAGAGGACATATACGGCATTGATCTCGATATGTCATTACCCCAAGCGCTTAAAAAGTGGCAGGCTTATGAAGGTACAAGTCACATGGATTCATCCTCCTCTTCTTACCTTCCAAGCTTGGATAACACTCAAACTAGCATACCTACTCGCCCCTACGGATTTGCCGAGTCATTTACATCTTCTGATGGTAGGTGAGAAGTTGACACTCTTTGATCTAGGTTTGGTTGTTCATTCTCTCTGTTTTCTTCCTCATCGCGAGTCACACATCATTTTTTTCTTCTTTTTTTTTGCTTTTAACTCTGATGCTTTTACCTTAACTCATTGTTAGTTTTGGCACTGTGGTGAATAGGGATTCTTTTTAAGATTTTTGATGTCTTTTGACTCTTATTGTATGATGAATCTCTCTCTCTTTTCCATGTGAATACATTGTGGTTTGGTTGGAAATGAAAGGGATGCCATCAATTACCAGTAGATCATGGTACTTTTGTATTCTTGAATTGTTTTTTTGTTAGAAAAATTTGTAGGTGCAAGCTGTATTGAAATGTAAAGTGGCTTTGAAGTGTGCAAATCATAAAGTTAGCTTTTTGCTTATGATGCTTTTTCAAGCAAAATGTAAAGTTTTTTGTTATGATTACAGATCAACTGCTG

>PGSC0003DMG400028801 LRR-RLKs

AACACATCCTAGAACTACTAAGTACACACTTTTCATCACTCCTACAAAGAACACCATTCTTTCTTCTTACTTCTGCTACTACATTTCTCTCTGCTATGAATTTCTAGCAGTTCTTGGTTCTTGCGCCATATTCAGCTAAAACAAGTAGGCACTGCCCATTTCAGTTGGCTAGTTCAATTTCAGTTTCACTGTCAAATTTGACAATGGTGTTGTTAATCATCTTCATAATTCATGTTTTCATGTTTGTAACAGCTTCGTCGGATCTTAGCTCCGACCGTGCTGCACTTTTAGCTTTTCGCTCCGCCGTGGGTGGCCGTACATTTCTCTGGAACACCACCAGTACCTCCCCATGCAACTGGGCTGGTGTACAATGTGAAAACAACCGTGTTACAGTTCTCCGGCTACCGGCGTCAGCTCTCTCCGGTACCCTTCCGGTGAACACAATTTCCAACCTCACTCGTCTTCGTACACTTAGTCTTCGTCTCAATCGCCTTTCGGGTCCTCTCCCTTCTGATCTCTCAAATTGTGTTGAGCTCCGTAACATTTATCTCCAGGGTAACTTTTTTACCGGAGCTGTTTCTTCTTCCTTCTCTGGGCTTCACTCTTTAGTTCGGTTAAATCTCGCCGAAAACAATTTTTCTGGAGAAATTCCATCTGGGTTTAACAGTTTGATCCGATTGAGAACATTTCTACTTGAGAAAAACCAGTTTTCTGGGTTTATGCCTGAGCTAAAGTTTTTCCCAAATCTCGAACAGTTTAATGTGTCTTTCAATAGATTAAATGGGTCTATTCCGAAAAGTTTGGAGGTAATGCCAGTGAGTTCTTTTACTGGGAATTCTCTATGTGGAAAGCCTATTAATGTTTGCCCAGGAAGTAAGACTCAGCCTGCTATAGCTACTGATGGAATCGAAATTGGGAATTCAAATAACAAGAAGAAGAAGCTATCTGGTGGAGCTATTTCCGGTATTGTCATTGGGTCTGTTGCTGGATTTTTCATTCTTTTGTTGATTTTGTTTGTGTTAGGTAGGATGAAAACCGGTGATAAAACAAGGTCTCTTGATGTTGAGACAATTAAGTCACCGGAAACAGAAGTTCCCGGGGAGAAACAAATTGAGAAACCCGATAATGGAGGTGTAAATAATGGGAATTCAGTTGCTGTAGCAGCACCTGCTGCTGCAGTATTGAATAGTGGTGAAGAGAATTGGGGAGAAAATGGGGTTAGGAAAAAGCTTGTGTTTTTTGGAGATTATTATAAAGCATTTGAATTGGAGGATTTGTTGAGAGCATCTGCTGAGGTTCTTGGAAAGGGGACATTTGGAACAGCTTATAAGGCAGTTTTGGAGATTGGTACAATTGTTGCGGTTAAGAGATTGAAAGATGTGGCAATATCTGAAAGGGAGTGCAAGGAGAAAATTGAAGCAGTTGGAGCAATGAATCATGAGAATTTGGTGCCTCTCAGAGCTTATTATTTCAGCAGGGAGGAAAAACTTCTTGTCTTTGACTATATGCCAATGGGAAGCTTGTCTGCTCTTCTTCATGGTAAGTTTTAGTACACTCTTTTACTGTCAATTCATGTTGTTCGGACTCTCTAAAAATGTTACCACATCTGTGCGAAATCATCAAAAAATGCACTACTTTGGAGGATTCGACACACACTCAGTGGTATGGATCCTCTAACAATACACTACCTTTTTGGAGGATCTCACATGCACTTAGCAGCATGGGTCCTCAAAATATGCACTACCTTTGGAGGATCCAACACTCAGTGGCATGGATTCTCAAAAAAAAATTCACTATTTTTGGAGGGTCCGACATGTATTGGTCGATATTTTTAATGAGTTCAAGCAACATAGCTACCAATATCTGGTGTTTTACTTGCTTTTCCCTTATTCAGTAGATGCACTATATTTGTTCCAGATTTTGACATTTTGTTAGAGTAATTAGTTTGACTTGTTGCTAACAGTGTAACATGCAATGTCTGTAGTATTTGGCCCCTCAATCCAATGAAAATGTGATCCTAGAATTAAGCATTAATTTCTTATCATTACCTTAGTCTTCATTATGTAAAGCATGCTTCTAATATAAGAAAGTATGTTAAATCTAAAGTTGTTTGTAATTAAGAGCAATAAGTAGTACTCCATTTCATTTAGTGCTTTCATTATAGCAAAAGGGTCCTTTGTTCTTTATAGATACCCTGTCCATATGCAGGGGAAAGGTGAACAAGCACACATATAATCTTTCACTTGGCCCTACAGTAGTAGGGTCTATCATTATATGTTCTTTTTCTATCTAGAGCATCTGTATAGGACTAGAATTTTGAATCTCCGAAGCTAATTATTCAACTGGTGATTAGTTTCCTGGCTTGTGAATCTTGCATGAACATGTACATTTTTCGCCTTTGGTATACTTGAAACTTGTTAAAGTGACTAAGAAGCTGTAATGAGCTGTATGCTGCAACTCTACTTTCTATTCACTACAATTTCTTTAATTTACTAGGAAGCAAGGGGGCTGGCAGAACACCATTGAATTGGGAAATCAGGTCCAATATTGCCCTCGGAATTGCACGTGGCATCGAATACCTTCACTCTCAAGGTCCTGATGTCTCCCATGGGAATATCAAATCATCCAATGTTCTTCTCACCAAATCATACGAAGCACGTGTCTCAGATTTTGGCCTGGCTAACTTGGTTGGATCGCCATCCTCCCCTACTCGAGTTGTTGGATATCGTGCTCCAGAGGTAACTGATCCTCGCAAGGTTTCCCAAAAGGCTGATGTCTATAACTTTGGAGTATTACTCTTGGAGCTTCTTACTGGAAAAGCACCTAGTCATGCCCTCTTAAATGAAGAAGGTGTTGATTTGCCAAGGTGGGTTCAATCTGTCGTTCGTGAAGAATGGCCTTCTGAAGTTTTTGATATTGAGCTCCTCAGGTACCAGACTGCTGAAGAAGAGATGGTCCAACTCTTGCAGCTAGCAATAAATTGTACAGCTCAATATCCTGATAAGCGACCTTCAATGGCTGAGATCTCTAAACAGATTGAAGAGCTTCGACGCCCAACCTCACATGATAAACAAGAACTCTCTGATCTTGCCAATGCAACAGAGTGAAGTGGTCATTATTAGTCCCCCTCGAGATGTGCAGTAATTAAGCTAAACTCAGATTCATCACCAATCTTTTTTAAGCATATTAAACTTGTTGTTTGTTGGGGGAGGGGTGTCTCATCTTCTTTCAGTTTGAATTAACTGATCATGGCCCCTTTTTCTCTATGTTTCTGTAGTAGACTAGTGGTTACCCCTTTCATTTTCCTTTAGATGAGGGGCCAAAGCCAATATCTGTACCATTTATTCTTGTTGGATGTATAATGCTTTCTGTAATTATTACTGCTTCTGCTTTTACTTTTGCTTTTTCAGTTGCAACTGTCGATGGCTATTCTGTGTGTATATGAGGCATGCATGCATGTGGTTCTTGATATGCTGAAGCCAGCAGGGAGATTGCATTGAGAGTTTTCATAGATGCTGAGTTGGCCAAATATTG

>PGSC0003DMG400028845 LRR-RLKs

TGTGTGGCTTTGGATTCTTGTAGCTGAATAGTGGTTTTTCTTTAGAATGGCAAAAGCATCTGACCCTTTTCTTCTTTTAGTGTCTTTTTCTCACTCTAAAAAGAGTCTTTCTGTGTTGTTGGTGTGGGAGAGATGAGGTTGTTGTTGCTACTGGCAGTAGTACTGTTGGTGGCCATTTTGGCGAATTTTTCAAGCTTTGTTAATGGTGATACAGATCCATCTGATGGTTAGCCCCTTTTGATTTATTCATCATTCTCTTTATTTATTTCTTTTTTAGTATGTGAAGTAGTAAGTACAAGTTCCAGTTTTCTTGTTCTCAAGGAAAAATTTAGCTGAGAGTGGGCTTGAGTCTTGACAACTCATGAAATGAACAAACCCAACTGAAAAAACTTGTTTCTTTGAAGCCCCATGATATTGAAAAGAAAATAGCATTTGAAAACACTAGGTCTAGGCTTGGTAGATAGAGTTATGTGGTACATGTTGTTGGTATATATCCTTGGAATTAGTTAGTTGCACACAAGCTGGCTCGGACACTGTGGTTATCATAAAAAAGAAATCTTTGAAAATGATGGATATAATAAGTCTTGCTTGGGCTGCATCAATGGTAGTCTTCACTTTCGAGTATGGGAAATACTCATTACTTATGGCTGAACAACCCCTTAGTCTTGTCGACTAGATTGAGTCATAGTTATTGTAATTGTTGAATTTACTGTTGCCGATGCACTTAATAGTTACAACAACATATCCAATGAAATCTTGTAGCTGGGGTTTGGGAGGGTAGAGTGTATGCAGACCTTACCCTACCTCGTGGAGATAGAGGGGAGACTCTTTCGGGAAGATGCACTTAATAGCTAATAGAAAACGTCAACCTTCTCACAATAATTTTCTTTCATTAAAAATTGAACAAGTCTGGGTAAAAGGTAAATCCTTACAGAAACTTGTGGTTTGTGAATGTGATTGGTTTTAGTTACTATTGCCAGCCTTATATAAGCAATTACTGGATAATGTTTTCTACTTTATGTGGTAAGGGGTGACAAACACCAATGAGTTTGAAGTTGATCAAAACCTTAATAAAGCTATATAATTTACTTTTGGACTCCTGATGATCCAAAGAGTTTCACAGTAATGGTTTGGTTAAGTTGCTCGCTACTTTGTACCATGCTGGTCAAGTTGCCGTAGCTGCACATTCTTTTAACAGATTGTCCTATGTAAAATCTGTCTCTTGAGCCGTTCTCTGTCCATATATAACTATTTCTTTATCATTAAAATTCCTTACACAATCACATATGTCTTGATTTTAAATTAATCTAGTTGACTTCTGTTTGGCTTTCAGCATCTGTTCTGAGAGTCCTGTATACCACCCTAAACTCACCAGGGCAGCTGACAAAATGGAATGCCAATGGCGGTGATCCTTGTGGAGAGTCCTGGACAGGCATTACCTGCTCGGGCAATAAAGTTACTGAAATGTGGGGTTCGGTCTTCCTTTTTGTCTTCATCTCCATGTTGTTTCAACTTTTTGGAATTATTGACTAGGTAAACTGATGATTAGTTTCTTTTCATTTTCCGTCAGCAAGATGTCAGGTCTTGGGCTCTCAGGTTCATTGGGGTCTCAGCTAGATAGTCTAAAATCTGTGACCAATTTGTAAGATAAATGGCCTTTGTTTGTTCCTTATTAATTGGATTTATGTTGAATTCCTGAATGTGGTCATCTTTGTGAATGTTGCAGTGACATAAGCAATAACAATATGGGGAGCCAGTTACCTTACCAGCTTCCTCCAAATGTGCAAAGATTGTATGTTTGTTTATTTGCATATCTTCCTCCTTGCAGGTTCTATTTTCTTCCTCTTATCTTTCTTAGAACTTGCCTTGCAGAAACCTCGCTTCTAGTGGTTTTAGTGGTGGCCTCCCATATTCCATTTCACAGATGACCTCCCTCCAGTACTTGTAAGTGAATGCTTAAATTTAATATTTGCTTAAATACTTGGTCCTCTTCATTCTCTTTGGGGCTCCTTATGATTCTTTGCAATGTTTATGCCTGGTTGTAGAAATGTCAGTCATAATCAAATTCAAGGATCAGTGGTTGTCTCGTTTGACTCGCTCTCTGCACTCAACACATTGTAAGGCCGATTCATCTCCTTCCATTACACCAGAGTTTTCATATTGCTTAATTATGATCTGAATGAGAATAGTTTGCTGTTCTCAAATGTTATGATGAAAGTACACTTATTCTATTATTTAAGTTGTCTCGATTTGAGAAAGAGGTTAAACACACTTGTTCCATTTCCCACATTACTGTGCAGAAGTGAGTCATGTCTTTGGTCTGCTGCCAAAGTATTGAATTATGCCAATTGAGCTTAGTTGCTACTTAGATTGTACTTATTCTTAAGCAAGTATTAAAAGCCTCAAAGGTGCGCTTATGCCCTAAAACGAGGTTTAAAACGTGTTGAGCGCTTTGCCTTGCTTAATGTGTGCTTTAGTGTCATCATCAAGGTCCTAAGGCATACTTTCCCTTGACTAAACAAGTGATATTTCACTTTATCGTAAAAAAATCCAATTTCTTTGTTCATATATTTGTCATGCATGCTTATAATTATTAGTCTTGGACTAAACATATATATACTTGTATTTTTTTCTCCCTTTGCACCTTTTTTTCATTAAAGCTTACACTTTATTTGCGCTTTAAGCCCCAATGGACCTTAGAACTTTTTTGTGCTTTTTGCTTTTGATAACGCTGGTACCTTATCATATTTTCTTGTTTAAAACATAATAGTTTGAAATTTATCATTCTCGGTCACAATTTTGATTGATCTTTTTAATGCTTCAGGGATTTCTCTTTCAATTCTATGACGGGTAATCTTCCTCAAAGCTTCCAGTCACTAACTAGTATGAACAAAATGTAAGAAGAGTTTTCATACTTCATCTATCTATTTGCAAACTTCCATCAGATATAGAAGTTTTGTTGTTATAATATGCAGGTATTTGCAGAACAACCAGTTTAGTGGAACTATTGACGTCCTGGCCAATCTTCCCCTCGGAAATCTGTGAGTTGTATGCCACATGTGTAGATATTATGTCTTATGTCTACATCTCCTTCAGTTTCAGTTAAATCCATTGGTTGAAAACCATGTTCATTCTTCTATTATTTAGGAATGTTGAAAACAACCAATTTACTGGCTGGATTCCAGATCAGCTTAAAGGGATAGTGCAGTGAGTGTCTTAGATTGCAATAATTTTCATTCTTCTTTGCAAATTCTTTGGGGAGTTCTTTCTAATTGTTTTCTGGATTTATGAATCTTTCATAACAGATCAAATGGTAATTCATGGATCTCTGGACCTGCACCCCCACCTCCACCTGGCACACCTCCTGCAAACAATCCCAATGGAGATCACAAATCCAGAGACAACAGCAGCAATTCGGGCAGTAGGAAATCAGGTATAGGGGGTGGAGGTGTTGCAGGAATAGTGATATCTATTCTAGTTGTTGGAGCAATAGTAGCATTCTTTATTATCAAGAAAAGATCAAGAAGGTCGTCCACAGACATAGAAAAACTTGATAAGCAGCCACTTGCTCCTCTTACTCCACAGAAAGTAGAAGGTATTTGGCTTTCATAATTCTTGATATTGTTAGAAATGTCTTGAATTCTCTCTTAACTCTTTGGATATTTCTTAACGTATAATTATAATCAATTGCGGTGTTACTATTTACTCAGTTGCTTGGGAATATGGAAAATGATGTCTAAGCCTCTATCCTTACCTTGCTAATGATTGGTCATAAACCATGCAAGTAGTATCAAAATTTTGCAAGGGGTTGTCATTGATACAGAAGGAAGCAGCAAAGCTGGAAAATATTTATCTTCGAAGTGGAAAAACTGTATGGATTTGAAAGATCCTAAGTTCTGAACAAACAAAAGTTTCATACTTTATTGGTAGTCCCGTAGGTATAATCTTTAGAAATTCAAACTGATCTTTTGGAGGAGGTCCTAAATTACGTGGTGCTTTTAAAGCTTATGATTATTTTTTGACAAATTAACATCAGCTACCCTTGATGGTTTAACTTCCTGGGAGACAAAAACTCTTGAGAATATAGTTGACCCGAAGAACTCTTGCTTCAGACTGTAACTGTCAGCACAGTGAAGGAGTCATCTTCAGGTTCATATTTGGGAAGTTGCTTTCTCCTTTTTACTCCATCAAAAGCCCACAAAACAAACCGTCTCTTACAATACGAGTTCCTTCTCATTTGCAACTTCATCAGTAATTTTCCAGTTCATGCTTATAACTTTCATGCCTCATTTCATTATTGACTTCAATGATGTCATGGAATCCGAGTTGGAGCTTGTATTTGCTTATGTCTAACTTTTTCATTGCTGTGTTGCTGTTTTTTTAAAAAGGTAACGTCACTGTTATATTTTGTTTGTTCCTGGAATTCCTACTCATTGTCTGTTTTTGGTTGCAGAGTTGAAACCTTATCAAACCTTCTCCACAGTAAGCATGGGCACATTTGAAACTCCCACTGCTGTAACTCTAAGACCTCCACCTATTAATCGCCTTAAGTCCTTTGATGAGGTTGACATGTCACCGAAACCCATTGTTCCTCCCAAGAAAATTAATACAGCTCAAATCAGGGCTACACAGTTTTCAATTGCAGACCTACAGATTGCTACTGATAGCTTCAATGTTGAAAACCTTATTGGCGAGGGATCAATTGGGCGTGTTTATTGCGCTCAGTTTGATGATGGCAAAGTAATACTTCTCTCTATGAGTACCCATTAAAAAAACTTTTTTTCTTTCTTATGACTATATAAGGTGATTTAGTGATGCTAACACATGTTTAAGCAAAAATGTATTATCAGAGAAACTACATACTTATTGTGTTCCTTTCTGTTATTTGAAATGTTGTAGGTTCTTGCTGTCAAGAAAATCAATTCTTCTGCCCTCCAAAATCATGAAGATTTTCTCTATATAGTTTCTGAGATATCACAGTTGCACCATCCAAATATAATTGAGCTTGTTGGTTATTGTTCAGAGCATGGACAGTACTTGCTAGTTCATGAATTCCATAAAAATGGTTCTCTGCATGATTTCCTGCACCTGTCAGATGAAGACAGCAAGCCACTGGCATGGAATAGCCGAGTCAAGATTGCACTTGGTGCAGCACGAGCACTAGAGTGAGTTTTTGCAATCTCAGAATTATTTTATCATATTTTATATTCAATTGGTTACAGTTATGCGGGACTATTTCCTTATTGATTCATTCCAGAAAGGATGACACGTTGCTGTATTTGGAAACAATTCACTTTATATTTCCCATTTTTACCATTAGTGAGAAGTTTTATAGCAACACAAATGTTATGGTATGTTTAAGGCCACAAGTTTCAAAAGTTTTATAGTCAGACAAATGCTATGCCATGTTTAAGATCGATCACAAGTTTCAATTTTTTTTTATCTCTTTCTTAGATTCAATGCCGATTGGAACTATGTCACATAAATTGGAAGGGGGAAGTAAAATTCATGGAGCATGCAGCAATGCTATGTTTTTCTCTTTATTTATTGATGGCTAATTACTAGTAGTTTACATGAATCTTCTGCTTATCTATATCGGAGATTCATAGAATAGATCTTCTGGGTAGAAGAAAAGGGGATTCTGTGAAAGTCAATACCAAGCACTTAGTCGTGACTCGTGATGAACTTTACCAGAGGATCCATAGTAATATTGCATTTAGAGGTCATACTTGTTGGACTAAAAAGACCACTATAGAGTTTAGTCTTTTTTAAACATTGATGACTAGTAAGTTAACATAAAGAAGAGTTCATTTATTCCTTTTAGTTAGAGCCTAGTATCCTAAATATGCCGGTCTTGCCAATATAAAGCTAATCAAAACTTCAATGCTCCCATCATTTTGTTTTATATGGTGGTGTTTGAGTGGGGACAGAGTTCAAAAAGAAAGAAAGACTTCTGGAACTTGTGGTCTTAAACTTGCCATGTCATTTGTGTGGCTATAAAAGCATGTCATTAGGGTAACAATGAATATTTGAAGTTAAAATTGTTTCCAAATATAGGAAGGCGTCAATCTTCTTGAAAAGGCTTAAAAGGAAATCCGCCATATAAAATGATACAAAGGGTGTAACAAATTTCAATTTATGTGGTTTTCATATCTGTTCATTTCCTGTTTTCCATACACAAATTTCTTCACTAGCTAATATAGGCATCCTTTGCAGGTATCTGCATGAAGTTCATTCACCATCATTGGTTCATAAAAACATTAAATCGGCAAATATTTTACTTGATCTGGAACTCAATCCTCATCTATCAGATTCTGGATTAGCAAGCCTTATTAATGATGCAGATCAGGTGGCTTTAAGCTCCTTACTTTATGCATTGTTGCAAAGTGTTTCGGTGGATTCGATATAGAAGCATGCATGCACATAAGTACACATGTATTATATAGGCATGTTTTGACATACTATTGGCTGTGTATGTGTCCTTTTCAAGGAGATAAGAAATATCAAAACACCACATGGTTGAAACGTCATTTGTTTCAGGAGAAGCTTTGTTTTGTGCACAAACCTTTGAAACATGAAATCATTTACAACAGTTTCTGGGGTTCATAAATTGGATTCCAAGAGAAGTTGTCAAGTAAAAAACAGAAAATTTTATCCAAATCTAGGGCATGGAATTATCCAGTATATTCTATAGAAGTAACACTACCATATAACTCATTTTAATGAACTAGTCTTTCTATGGTCCCTGTTGCTTTATCTCCTGCTCTTCTCTACTTACTCAATTGGTAAAATTGTAGCTGTTTCTTCTTCTTAAAAGATTCACTTCATTATCATCTTCTCAAGATATTTGGAAGTGATCATCTTTGCAGGCACTAAACCATAATACAGGGTCTGGATATGGTGCACCTGAGGTAGCTATGTATGGGCAATGCACCATAAAAAGTGATGTCTACAGCTTTGGAGTTGTTATGTTGGAACTTCTTACCGGACGGACACCTTTTGATAGGTAAAGTTGCTTAGGCATCATGATATTGTGTCAAATTACTTAGAGACAAACAGGAAGAAATTCAGAAAGTCTTAACTTTCTTGATCAATATGTATGCTCTTGATGCAGTGAGAGACCAAGATCTGAACAATCTTTAGTTAGATGGGCAACGCCACAACTCCATGATATTGATGCCTTAGCTAAGATGGTTGATCCAGCACTCAAAGGGCTCTATCCTGTTAAATCCATTTCCCGCTTTGCTGATGTAATTGCTCTTTGTGTCCAGGTAATATTTTGTATAGACGGTGTGCCTAATTCATAAATCTGCTGTGTTTATCAAGAACTTGATTCTGTTAGTTCAGTCAACACTATCTCATTGTCATGTATACTATTGTGGCAGTCTGAGCCTGAGTTCCGGCCACCTATGTCAGAAGTGGTTGAAGCCTTAGTTCGGCTAGTTCAACGAGCAAATATGAGCAAGAGGACATATAGTATTGATCGGGGTTCAAGGGGTGAAACAGATGCTCTAGACAACCAACCTTAGTTCCGATGTAATGTAGCCTTTTCTACCAGAATGTCTCTCTTTATAGTTTGTTGTTTATGGGATCTATCAGCTTATCTGTTCAGCAAGATCTTGTTGGTTGCATTGGTTGTACACACTGCTAGATACTAAAATCTGTTCATAGTGCATATTCAATTTTCATTTTTCCCTGTATCTGCT

>PGSC0003DMG400029146 LRR-RLKs

GGAAAATGAAGGATCATTCCTATTAGAAAAAAGTTACTCAAGCTTTATTTAGTTCCTCTTTTTCCATGCAACTTATGATATAGAGCTTTCATATCTAGCTAGAGAGAAAAAATGGCCAGCATAGGATCAAAATCGATCATGTGTTTGATCGTGTTCGTGCTATTTTACATGCAGCAAGTCAAAAGTAGCTCTGAGCTCGAGCTACTTTTATCGATGAAAACATCGATGAAGGATCCATTAGGATCTCTTCATGATTGGATTCCGCGGTCTCAATCTTTTTGCCATTGGAATGGTGTTGTTTGTGATGACTTGTCACATGTTGCAAAAATTGAGCTATCAGGGAAGAACTTATCTGGAAAATTATCCGAGACTATTTTCAACTTTCCATACGTTGAATCGATTGATCTCTCGAATAATCAACTCTATGGAGAAATTCCAAGCAATATCTCAACTTGTTTGGCGCTGAGATTTCTCAATCTGAGTAACAATAACTTCACAAGTCTACTTCCTCAAGGCTCGAGAATCCCACTTCTCGAGACATTGGATCTCTCAAACAACATGATTTCGGGGAAAATCCCCGAAAATATTGGTTTATTCTCAATGCTCAAAGTTCTTGATTTTGGTGGAAATGTGTTGGTTGGAAGCATTCCAAAGTCTATTGCAAATATTTCTAATTTGGAGTTCTTGACTCTTGCTTCAAACCAACTAATCGGAGAAATCCCTCGAGAACTAGGTCTTTTGAAGAACTTGAAGCTTATTTATTTGGGATACAACAATTTCTCTGGTGGAATACCGGAGGAGATTGGTGAATTGTCTTCTTTGTATCATCTCGATCTCGTGTACAACAATCTCACAGGTGAAATCCCTTCATCTTTAGGTAATCTCACCAATCTTGAATACCTTTTTCTTTACATAAACAAGCTCACTGGCCCGATTCCAAGATCTCTATTTAATCTCAAGAAAATCATCTCGCTTGATTTAAGTGATAATTTCTTGTCTGGTGAAATTCCTGAGCTTATTTCCCAATTACAAAATTTGGAAGTTCTACAATTGTTCGCGAACAATTTCACTGGTAGAATTCCAAATACTTTATCTTCCTTACCTCGACTTCAAGTTCTTCAATTATGGTCTAATAAATTATCTGGTGAGATTCCCAAGGACCTTGGGAAACACAACAATCTCACAATTCTAGACCTTTCCACCAATAACCTCACGGGTAAAATACCCGAAACCATATGTTACTTCAACCATCTTTTCAAACTCATACTCTTTTCAAATTCACTTCATGGCGAAATCCCTGTAAGCTTGAGTCATTGCAAAAGCTTACAAAGGGTCCGCCTTCAAAACAACCACCTCACCGGTGAATTGTCCCCAGAATTCACAAAACTCCCACTTGTTTACTTCCTTGATATCTCGGGCAACAATCTATTTGGCTCCATTAGCGAAAGACGATGGGATATGCCATCACTTCAAATGCTAAATTTGGCTAAGAACAAGTTTTTTGGTACCTTGCCTGATTCTTTTGGTAGCAAGAAACTCGAAAACTTGGACTTATCCGAAAATGATTTTAACGGTACTATTCCTAAGAATTTCGGAGAGTTATCAGAGTTAATGGAGCTCAAACTACGAAGCAACAAGCTTTCAGGCGAAATCCCCAATGAATTGTCTTCATGCAAGAAGATTGTGAGTCTTGATCTGAGCCATAATCGATTTTCAGGTCAAATTCCAACTAGTCTCTCCGAAATGCAAGTTCTTAGCCTTCTTGATTTGTCAATGAATGAATTATCAGGTGAAATCCCGCCTAATTTGGGGAAAGTTGAATCACTTGTCTTGGTTAACATCTCTCACAACCATTTCAGTGGATATTTACCGTCCACCGGAGCTTTCCTAGCCATAAATTCAAGTGCGGTGGTTGGAAACCAACTATGTGCCCGTGGGGACGACATTACAAGCGGCTTAACGCCTTGTAAATCACTCAAGAAAAGCTCAATTTGGTGGTTCTTCTTGACATTTCTTCTTGGAATACTAGTTTTGCTAGTATTCTCGGCTTTAGTTATTGTATTTATTCAAAGAAGAAGAGAATTGAAGCTAAAGAAAGTGGAGAGTACTACTCAAGATGGTAATAATTGGGAAATTCAATTCTTTGATTCAAAAGCCTCAAAATCCATCACCCTAGATGACATCTTAGGGATTGGAGTTTCGTACAAAGGGTTTTATTCTGAAATAAGTAACATGCAAGTATTCGTGAAGAAGTTGAATGTGAATATTCCAACAAGTTTTTGGACAAACATCCAAGAACTAGGAAATATTCGACATCCAAATGTTGTGAAGATATTGGCTGCATGCAAATCGGAGAAAGGAGGTATTTTGGTTTATGAGTATGTAGAAGGGAAAGATTTGAGTGAAGTGATTCGAGTCATGAGTTGGGAACGTAGACAAAAAGTTGCTATTGGCATATCTAGAGCATTGAAATATTTGCATTGTTCTTGTTCACAAAGCATTTTCATCGGTGACCTTTCTACCCGGAAAGTTATCATTGACGGAAAAGATGAACCTCGCCTGAGATTGAGTCTTCCCACTACGTCCTACGTTGGTCCAGGTAAATTTACCATTTTACCCTTGTCATCTACGGTTACTTCTCTTGTTTTGACATTTTTTAAGAGTTTAAGTAATATACAATCGTAAGCTAAAACATCATATATTCTATCAGGTAACATGTCTTATTTGCAAGATTATAAGCTTACTTACATACATCTTTCAACTGACTAGATCATGTAAATTAATTTATACTGATAATGCATATAACTTAATTAACTCGTTAGTTTAATAGTTAACAACCTTTGTTATAACTTAAACAAACCACATATAGTTTCTAAAGAGTGTAACGTAGCTATGATGACTTTAATGAAAATTGATTTTTTCAGCTTTAGCATTGATGTGATAGGAATGTCATGTTATGTTGAAAATTATTTGATCTAACTAAAAGAAATTTTGTATTTTTCTAATTATTAGAATAGAAATAATTCTTAGAATTTACGTACTTGGTCTTTATTTGGTTTCTATTTTAACTTTAGTTCTCATAGTCTCTAGGAGTAATGATAAGGTCAAGGAATTAAGCAAACAAATGTCATGGAGGCAATCATGAATGTTTTTTTCTCTTAAAATAAATGTAGAGACATAAATCTTATCATTCCTTTCTTTACGTAGAAAACAATAAATCTAGTCTAGTGTAACAAAAGTCATGACCTTTTATGCCATGCACGTACAATACTGTTACTTAATTACGAGTTACAACTTTTCTCATAAATTGTAGAATACAATGGAATATCGGAGAGGAGCGATATATATGGATTTGGGCTTGTTTTGATCGAGTTATTGACTGGAAAAAATCGCGGTGATGCTGAATTTGGGAAGCGTGAGAGTATAGTTGATTGGGCGCGATATTGTTACTCAGAATGTCATTTGGATACATGGATTGAACCACTATTGAAGAGTGATGCAGTGAATAATCAGAATAAAATGGTGGAGATGATGAATGTGGCTCTTCAATGTACTGCTAGTGAGCCAGCAGCAAGGCCTTGTGCTAGTGATGTTGCAAAGACATTGGACTCTTTTGTTAGATCAAATTCTTGTGGTTTAGGGTTGAAAAGGTGTGGTAGTGTTTAGATTTTGGGATTTCTAACTTCAAGTCCAAGATTTTTGTTTTTATGTTAATGTTACTTTCTTTTTTTTGGTTTACTTTGTATTGGGCATAAATTAATGAGTGTTGCTCAATGTTTTGGATGTTATTTGAGTGTAAAAATGTAGATATGAAACTAAATGCTAATATATAT

>PGSC0003DMG400029442 LRR-RLKs

TTTCAACACTCACTAAAAGCTTACTCTCTAAGCTCTCTTCTTTCATTTTTTTTTCTATCTATCTATTTTCTCTATTCATCATGGACAACATGAACACCAACTCTGCTATAACATTGCTCATTGTCTTGTTTTTTTCCTTGATTCTTGCATCCGTAAACGCGGTATCTTCCTCCATCAAAACTGATGCTGAATCACTCTTGCTATTCAAGAACATGATACAAAAAGACCCTTCTGGTGTTTTATCAGGATGGCAACTTAAAAACAATCCATGTTCATGGAATGGTGTAACTTGCAATAGTCTTGGAAGAGTAACAAATCTTGATCTTCAACAATCTGAACTTGTTGGAGAGGTTTCTTTTTCTCCTTTCAATTCTCTAGATATGTTAACTGTCCTTAATCTATCATCGAATTCGTTTTATGTAAATGCATCAACCTCTTTAACTCAGTTACCTTATAGTTTGAAACAACTTGAACTTTCATTCACTGGCTTAGCAGGCTATGTACCTGAGAATTTCTTTGCAAAATGTCCAAATCTTGAATATGTTAGTCTTTCTTTCAATAACATAACAGGTTCTTTGCCTCAGAATTTCTTGTTGCATACTGATAAGTTACAGTATCTTGCAATGGACTATAATAATCTTACAGGATCAATTTCTGATATCAAGATTGAGACTTGTAATTCATTGTTACGTCTCGATTTGTCAGGGAATCAGATAATGGATTCAATTCCTTCTGCTTTATCTAATTGTACAACACTTCAAGAATTGGTTTTGGCTGAGAATTTCTTCTCTGGTTCAATTCCAACTTCTTTTGGTGAGCTTATAAGTTTGCAAAGATTGGATCTTTCTAAGAACCATTTATCCGGTTGGATCCCATCTGAATTAGGAAACTCATGTAGTTCACTTGTGGAACTCAAGTTTTCTAATAACAATATCACTGGTTCAATCCCGAATTCATTCTCATCATGTTCTTCTCTTCAAAATCTTGATTTGTCGAATAATAATCTAACCGGTCCATTCCCTGATTCCATCCTACAGAATTTAGCTTCTTTAGAGTCTTTGCAAATGAGTAGTAACAAGATTTCTGGTTCATTTCCTGCTTCTCTTTCATATTGCAAGAAACTCCGCGTTGTAGACTTTAGCTCCAACATGATCAATGGGATGATTCCACCAGATTTATGCTCTGGAGCTTCATCATTGGAAGAGCTGAGAGCACCTGATAACTCACTTTATGGTCCAATTCCATCTCAATTGTCTCAATGTTCACAGCTCAAGACTATTGATTTTAGCTTGAACTATTTGAACGGTTCGATTCCATCAGAGCTAGGAAAACTTGAGAAACTTGAGCAGCTTATTGCTTGGTACAATAGTTTGGAAGGGAGCATACCAGCAGAGTTGGGAAAATGCAGCAACTTGAAGAATCTTATACTTAATAACAACTACTTGAGTGGGAAAATTCCAGTTGAATTGTTTAATTGTGGTAATCTTGAGTGGATTGCTCTCACAAGTAATGGACTAAGTGGTGAGATTCCGAAAGAATTTGGCCATCTTTCAAGGCTGGCTGTTTTGCAACTAGCAAATAATAGCTTGAGTGGTCAGATTCCAAGTGAATTAGTCAATTGTAGTAGCTTGGTTTGGTTGGATTTGAGTAGCAATAGGTTAACCGGGGAGATCCCACCGCGACTTGGTAGGCAGCAAGGGGCAAAAGCATTGAGTGGGATTCTATCAGGTAACACTTTGGTGTTTGTTCGAAATGTAGGAAATTCGTGCAGAGGCGTTGGAGGATTGCTTGAGTTCTATGGGATACATCCTGAAAGGCTTTTACAGGTTCCATCATTAAAGAGTTGTGATTTCACTAGGCTGTATTCTGGTCCAGTTCTTAGTGCTTTTACTCGGTATCAGACTATCGAGTACCTTGATCTTTCGTACAATGAGCTTCGTGGGAAAATCCCTGATGAATTTGGTGATATGATAGCTTTGCAAGTTCTTGTTATATCACATAACCATTTGTCAGGAGAGATTCCTTCATCACTTGGAGGGCTAAAGAATTTGGGAGTGTTTGATGCATCACATAACAGATTGCAGGGTCAAATTCCTGATTCATTTTCGCTTCTTTCTTTCTTGGTGCAGATTGACTTGTCTAATAATGAATTGACAGGACAAATTCCACAAAGGGGTCAGCTCAGTACACTTCCTGCAAGCCAATATGCAAACAATCCAGGACTCTGTGGGGTTCCGTTGTCTGAATGCCAGTATAACTCACCTGCCACGAATACTGGAGACGGAGGAGGAGGAAAAAGAAGTTCAGCTGCATCATTGGCTAATAGCATTGTTCTTGGAGTTCTCATTTCCATTGCCTCAGTATGCATTTTGATTGTTTGGGGTATTGCTATGCGTGCAAGGCGTAGAGAGGCTGAAGGAGTGAAAATGCTTAGTAGTTTGAGTACTAACTATGCAGCTTCATCATGGAAGATTGACAAAGAGAAGGAGCCATTAAGCATCAATGTAGCAACTTTCCAAAGACAACTTAGGAAGCTGAAGTTCTCACAACTTATCGAGGCCACGAATGGATTCTCAGCTGCAAGCCTCATCGGTTCTGGAGGATTCGGAGAAGTCTTCAAGGCAACATTGAAGGATGGATCAAGTGTTGCAATCAAGAAACTCATCAGGTTAAGTTGCCAAGGAGACCGAGAGTTCATGGCTGAAATGGAGACATTAGGAAAGATCAAGCACAAGAACCTTGTACCCCTCTTAGGATACTGCAAGGTTGGTGAGGAAAGACTTCTAGTTTATGAGTTCATGGAATATGGCAGCCTAGAAGAAATGCTTCACGGGAAGACAAGAATGCCTGATAGAAGGATCCTAACATGGGAGGAAAGGAAAAAAATAGCCAGAGGAGCAGCCAAAGGACTCTGCTTCCTTCACCACAACTGCATCCCACACATCATACACCGCGACATGAAGTCAAGCAACGTGCTATTAGACAACGAAATGGACGCCAGGGTGTCTGATTTTGGGATGGCAAGGCTGATAAGCGCGCTGGACACACACCTCAGTGTGTCCACACTAGCAGGAACACCAGGTTATGTGCCACCTGAGTACTATCAGAGTTTTAGGTGCACAGCAAAGGGAGACGTGTACTCGTTTGGAGTTGTGCTTCTTGAACTCTTGACAGGGAAAAGGCCAACAGATAAGGAAGATTTTGGTGACACAAACTTGGTAGGATGGGTGAAAATGAAAGTAAGAGAAGGAAAATCAATGGAAGTAATTGATCAAGAATTGTTATCAGTGACTAAAGGGAATGATGAAGCAGAAGTAGTAGAAGTTAAAGAGATGGTAAGGTACTTAGAGATAACAATGCAATGTGTAGAGGATTTTGCATCAAAAAGGCCTAATATGTTACAAGTTGTGGCTATGTTGAGAGAACTTATGCCTGGAAGTAGTTCAAGCAACAGTGGCTAAACAGCAAACCAGAAACAAAAAAAAGGAAAAAAAAAAACCAAAAGAATGTAAAATTCTTTCTTGTTTTGTTTGTGCAAGTATTCATCTGCATTTGTATTTAGTCAACAACTTCTCTTATTTTTTTGCAATATTATTCCCATGTTTTAAGAATCTTGTGTATGGATATTGATAACAT

>PGSC0003DMG400029644 LRR-RLKs

AAAAAACAGAGCAAAACCCAAGAAACCAAAAAGAAGGGATTTTTACTTAGTTTTTTGATATGCAATTGCAAACCCAAGAAGAAGTTTTTGGAATTTGAGGCAAAGATGTCTTCTTCTTCTTCAAATCCCAACAAAAAATACCATTTTTTTCTTTTCTTGGTTACCCTTTTCTTCACTTCTTCTTCTTCTTCTGCTGCTGTTACTGATGTTGTTGCACCATTTTCAAATTGGGACATTCTTGATGATACCCCATGTAAATGGTCTTTCATAAAATGCAACTTTCAAGGTTTTATTACAGAAATCAATATTCAGTCTATTCATCTTGAACTTCCTTTGCCTACAAATCTTTCTTCATATAAGTACCTGAAGAAGCTTGTTATTTCTGATGCTAACATCACTGGTACAATACCCTTTAACATTGGTGACTGTTCTTCACTTGTTACTATTGATTTGAGCTCAAATGGTCTTGTGGGTACCATTCCTTTAAGCATTGGAACCCTTGTAAACCTTCAAGATTTGATTTTGAACTCAAATCAGCTTACTGGGAGAATCCCAGTTGAGATTGGGAACTGTAGAAACTTGAAAAATCTTGTTCTTTTTGATAACAGGCTTAGTGGTGGTTTACCAAGTGAAATAGGGTTGTTGTCAAATCTTGAAGTGTTAAGAGCTGGTGGGAATAAAGATGTTACTGGTAAAATCCCAAATGAGTTTGGGGATTGTGGGAATTTGACTGTTTTAGGCTTAGCTGATACAAGAATATCTGGTTCTTTGCCTGTTTCTTTGGGTAAGCTTAAGAATCTTGAAACTTTGTCTATTTATACTACAATGTTGTCTGGTGAGATACCTTCTGATTTAGGTAACTGTACTGAGCTTGTGAACTTGTATTTGTATGAAAATAGTCTTTCTGGTTCAATCCCATCTGAGCTAGGGAACCTTAGAAAGTTAGAAAAGTTGCTTCTTTGGCAGAATAATCTTGTTGGTGTTATTCCAGAAGAGATTGGGAATTGTACTAAGTTGACTATGATTGATTTGTCTTTGAACTATTTGTCTGGGAGTATACCATTATCTTTTGGTGGTCTTGTTGTGTTACAAGAACTTATGCTTAGTAACAACAATGTTTCTGGTTCAATCCCTTCTGTTCTTTCACAATGTACAAGTCTTGTGCAGTTGCAGTTTGACACAAATCAGATATCAGGTTTGATTCCTTCTGAGTTAGGGAATTTGACTAGTCTTGTTGTGTTTTTTGCTTGGGATAATCAACTTGAAGGAAGTGTGCCTTTAACTTTGGGTAGTTGTAGTAACCTTCAGGCATTGGACTTGTCACATAACTCACTTACTGGTAGCATTCCTCCAGGCTTGTTTCAGTTGAAAAACCTTACTAAGCTACTGTTGATTTCTAACGATATTTCGGGTACTATACCTCGAGAAATTGGATATTGTAGCTCATTAGTGAGGCTAAGACTTGGAAACAACAGGATTGCTGGTGGAATTCCGAAAGAAATTGGAGGTCTTAAGAGCTTAAACTTTCTTGATTTGTCCGGTAATCGCCTTTCTGGACCAGTTCCTGATGAGATAAATAGCTGTACTGAGCTACAAATGGTAGACCTCAGCAGCAATACACTGGAAGGTCCCCTGCCTAACACTCTGTCTTCACTATCAGGGATTCAAGTCTTGGACGTTTCGAATAACCGGTTTGGAGGGCCTATATCGGCTAGTTTTGGGCGTCTTGTGTCCTTGAACAAGCTGATTCTCAGTAAGAACTCGTTCTCAGGGTCGATACCTCCATCTATCGGCCTGTGTTCGAGTCTTCAATTGCTTGATCTTAGCAGCAATGAGCTCTCGGGAGGCATACCGATGCAGTTAGGAAAGATTGAGTCCCTTGAAATCACTCTCAACCTTAGTTTCAATGAACTAACAGGTCCAATTCCTGCTGAAATTTCCTCACTAAGCAAGCTGTCGATACTTGACCTTTCGCACAACAAGCTTGAAGGGAACTTGAATCCACTAGCGAGGCTCGATAATCTTGTCTCACTGAATGTCTCATACAACAACTTCACTGGTTATCTTCCGGACAATAAGCTCTTTCGACAGTTACCATCATCAGACCTAGATGGAAATGAAGGTCTATGTTCATTCGGAAGGCCTTCGTGTTTTCTCAGCAATATCGATGGAGTAGGAGTAGCTAAAAATGGAAATGATGAGGGAAGGTCAAAGAAGCTCAAATTAGCCATTGCATTGCTAGTCATCATGACAATAGCAATGGTGATCATGGGAACTATTGCTATCATACGAGCACGAAGGGCGATGAGAAGGGATGATGATTCAGAGATGGGAGATTCTTGGGCTTGGCAGTTTACTCCATTCCAGAAGCTCAATTTTTCAGTGGACGAAATACTCAGATGCCTTGTGGATACTAATGTCATTGGAAAAGGGTGCTCGGGGATGGTCTATCGTGCAGATATGAACAATGGTGATGTCATAGCAGTGAAGAAGCTTTGGCCAATAACCATGACTACAACTAATGGAGGCAATGATGAGAAGTGTGGGGTACGTGATTCGTTCTCTGCAGAAGTTAAGACGCTTGGTTCAATTCGACACAAGAACATTGTTCGATTCTTGGGGTGTTGTTGGAACAGAAGCACAAGATTGCTCATGTATGATTACATGCCAAATGGGAGCTTAGGAAGTCTTCTCCATGAGAGAGGTGGTAACCCTTTGGAATGGGAATTGAGATACCAAATATTGCTCGGGGCAGCGCAAGGGCTCGCCTACTTGCACCATGACTGTGCTCCTCCAATTGTTCACAGAGACATCAAGGCCAACAACATTCTCATTGGTCTTGAGTTTGAGCCTTACATTGCAGACTTTGGCCTCGCAAAACTAGTCGATGATGGTGATTTTGGTCGTTCCTCCAATACAGTTGCTGGTTCTTATGGCTACATTGCTCCAGGTAAACATGTTTTTTTATTTCATGTCAAGTTACAATTCGTCATCATTTTCTAATTCCTAGCATATTATTCCGAGTATACATTTTGCTTAGTTATAGTATAAGCATTGATTGCACATTGCAACAACAACAATATACCCAGCGTAGTCCCATGCAATGCAGTCTATGAAGGGTAGAGTGTACGCAAACCTTATGCCTACCTCAGAGGTAGAAAGGTTGTTTCCAATAGACCCTCGCTTCATGGAAAAACAAAAACAAGGCAATCTAGAAAAAGAAGTAACAGATGCGACAACGATTACACATTGCATTTATCACAATTTTTTCAAAGAATATCAGTAGTTCTTGATGATCATTGATCCAAGTGCTCCAATGTTGTTCATTTTTCATTACAGAATATGGCTATATGATGAAGATAACTGCAAAGAGCGATGTCTACAGCTACGGAGTGGTTGTATTAGAAGTCTTGACAGGTAAGCAGCCAATCGATCCAACAATTCCCGAAGGAGTGCATCTAGTTGATTGGGTAAGACGGAAAAGAGGTGGAATTGAAGTCCTCGATCCAAGTCTACACTCGAGACCAGAATCAGAAATCGAGGAAATGTTACAAGCATTAGGAGTAGCCCTGTTATGTGTCAACTCCACTCCTGATGAAAGGCCTACAATGAAAGATGTAGCAGCAATGCTAAAGGAAATCAAACATGAAAGGGAAGAATACGCCAAAGTCGATGTTTTACTCAAAGGCTCTCCAACAACAACAACTACTACTGATCATAATCAAGAAAACAACAACTCATCAAAGGGTGTTTTAGCAACATCTTCATCAGGACAGAAAGCAAGAAGTATGTATACAATGAGCAATAACACAAGCTTCTCTGCTTCATCATTGCTTTATTATTCATCATCATCATCAAATGCCAAAAGTCGATTCTTGGGGTGTTGTTGGAACAGAAGCACAAGATTGCTCATGTATGATTACATGCCAAATGGGAGTTTAGGAAGTCTTCTTCATGAGAGGGAAGAATACGCCAAAGTCGATGTTTTACTCAAAGGCTCTCCAACAACAACAACTACTACTGATCATAATCAAGAAAACAACAACTCATCAAAGGGTGTTTTAGCAACATCTTCATCAGGACAGAAAGCAAGAAGTATGTATACAATGAGCAATAACACAAGCTTCTCTGCTTCATCATTGCTTTATTATTCATCATCATCATCAAATGCCAAAAGTGGTGTTTAAGTGATTCAACAAAACTTATTACATTTGCATTGAATAGAAGTTTGTCATATTAGTTACTATTAGTTTTAAGTTTCAATAATCCTTTGTTTGTTTGTTTGGAATTTTCTGTAAAGAGAATTAGAAAGGTAGATATGTGTGACAAGTTGAAACAAAACAAGCCAACTTATGCTAATTTGGCTTTTTGAGCATTTTTAAGCTTTCATGTAGCTACTAAAGTCAACTTTATTCCTCCTGGTTTTAAGGTTCATTTTTGAAGCTTTCAAACAAATATTTG

>PGSC0003DMG400030035 LRR-RLKs

GGCTAGGGTACCAAATGAAGTTTCCAAGTAAAGTCCTTGAATAAACAAGTGAAGATTATAACAAAGGGCTTTGTCATATTATGATGATCTGTCGTTAGCTTTTACTCAGTCATCAAACTTTGTGAATCTTTCAATCTCTCTCTCAAGTCTCAACCATTTTTCTACGTCATAGTTTTTTTGAATAGAAGAAGTAGGCAACTTGTATTGAAAATTCTATCTGAAACTCAAAAAAGAAAAATTCACGTTTTTCACAGTAAACCTTTCTTCTCCTTGTTTTTCTATTGGCAAAGAAGACAATACCATTTTTGCCTAAAGTGAATAAAGTCAAAAAAGAGCTTCCAGTTTTTGTTTGCTTGGTTTTGTTTGTTTTTTACGCGTTTTTCGAACAAACTCAGTCTCCTATTTGTTTTTCCCTCTTTCTTCTACTATTTTATTTACTGGGTTTATTCAATTTCTGGGTTTTGGAGCGGTGGACTTTTAATTAATTGTGAATTTTAGCGATTGTTTTTCAGTTTTGGGTAAAAGATATACTAGTGGGTTTGTTTATGTTTTGTGTGAAATGAATTGGCAATGGAAATTCAGTGAGAATAGCTTGTTGTTGGTTATTGTGCTGTTCTTGTGTTTGGTGCATCAGGAATTGTGTTTCTGTTGGTCTCTCAATGATGAGGGTATTACTCTTCTTCTTGTTATTATTTGCTATTTAAGACTTACTTTGTTAGAAATGGCAATAAAAAATTTGTTATTTTGCTTAAAGTTTTAGTCTTTATTGCTGCAGGTTTGGCACTGTTAAGATTCAGGGAGAGAGTGGAGAGTGATCCATTTGGGGCTTTGTTGAATTGGAAAGATGGTGATGGAGTGGAAAATCCATGTACTTGGTTAGGAGTTGGGTGCGTTAATGGGAATGTTGTGTCTCTGTAAGTACTAAAAATTGCTCTTTTCTCTTGTTATTTGGTTCTGCAGAAGTCTTCTTTTTCAATTTTGATCTGGTTGGCTTATATAGATGATTCTAAATTTGGTCTCTGATTGAATTCTGAAACTTTTGTTCCATGAGTTGGCGGTGATTCCAGAAGGGGTATGATCAAAAGACAGATAAATGAGAAGCTTTTGGCACCAATTCCTGTTGAAGCTTTAGTATAATGAACATCTTTTGGTGTTTTGTACCTGGAGTGTAGCTAATGTGTTCTTGTAAAGAGGGAGCTATTTTAAAATTTGATTTTTATTACTCAATATTTTGAGAGGCTAAGCACAATCTTATGATATGTTGGTGATCAAGATAATTAAACATAGCCTCGAGAATGGATGTAAATAAGTTGATGTCTAAATTCAGAATGATGAAGCATCTGACCGTGGGTCTAATGGAAGGGAGAGATTTGGAAAACCCGCCCTCCTCTAGAAAGGGCAAACATGGCGTTAGATATAGTAACAAAGATATAAATATTATCAGAGATCAAATAAAAGGAGGCAAATTTCACATACTACCTTCGATGTCAACTTTCTCCAGTTTATTTATTTTTCTCAAATAAATGTAAACTAGATAATTAGAGCTACATGTATAACATGATCTCTTTGTTTCTTGACCTGATACATGATTCTATATTTCTATGAATTAACTGGCAGTCGAATTTTGTGAAACGGGTAATTATATCATAGATAGAATATATTCATCCCTAGCTAACTTTAGAAACGCATTTTACTGTAGTCATTTTCGTAAATGCAATAATTTTCATACACAATTATCTCCATTTATTATCATGAATATAATTTTCAAGTCGGTAAGTAGCATAGTGTCTAAGTTGAATTGGCTATTGTAAAAATTATCCATGATACTGATGTTCTAATCTGTGTTTTTTATCTACTCACAGGGACTTGAAAGATCTCCATCTATGTGGTACCCTAGCTCCTGATCTCGCGCACTTGGTTCACTTGAAGTTTATGTATGTGCACATTTAAACTAGATGTCTTCATTTTCCTAGTCGCTGTCATTACAAACTCTTAATTCAAGTCTTTTTCTTTAAGATAATTCGCAACTCACTGGAGACTTCTTTACAGAATTTTACATAACAATTCCTTTACTGGGTTCATACCTGATGAGATTGCAAAGCTAAAAGAATTGGAGATTCTGGACCTGGGATCTAATAACTTTAGCCAGCCGCTGCCTTCCGAGCTTTGCCATAATCCCTCTCTGGAAATAATGTGAGTTCAAACAGAGGTGGTGTCATTTTCTAATTTTACCACCTTGTTCCCTTACTTACAGAGTTACACTAATGCAGCTTATCAGACAACAATGAGCTGTTTAGCAGCATATTTCCTGGAACCGATAGACTTGAAATGGTCCCAAATGTCCAAGAAAATAACAACCACGCTACGAGAGCCCAACCTTTGTCACTTCCCAATGGGAGATTGATATTAAGGTAAACATCTTTCAATCAAATGAAATGTATATTTTAAGAGCAATCATGTGTATTTGTAACTTTAAATGTCCAACATAGTAATAGTAAAATGACTTAAATCATTAGATTCAGTTATTGGAATACATATTCCTGGAATAAGTATGCATCTGTAGCAATTCATTGCTTGTCCTTTTGACCCCCCTGATGCTTGCGAAGACGTTTAGAAGTCCCATCAATTTTCCCTGTCTTCTAGTTGTCTTTCAGTTTTCTTATGGTTTGCCCTTCACATTGAACATCTCTTGTTTCCCTTAATGGCCTGATAGGTGATTGTTAAGCACATTTTGATTCATGTTTGCACAAACAATGTTCAGTTAATTTTCTGATATTTCTGAAAAGACACTACTAATCTTTTCTTTTCATCTTGTGATAATTTTCCTCATTTTCAATCAAGCTTTGCACTAGCTTCTTTTATGGGTATATACTTCTATGGTAACAACTTACATAGTTATCATGAAATTTAAATTATTTGTAGGACTACTGAAGCACTTGGAAATGTAGCTGGAAGAAAATTACTAAGAGTATCATCTTTTTCTGTCCTGAATAACTTCCAGAAGCAATTTCTGCCAGTCTTCCGACTTCTAGACCGTCGTGAAGTTGCTGTTGAGAACAGCGCACCTTCATCATCACCATCTCCATCACCGAGTCCTTCTCTTTCCCCGTCTCCTACACCGGCAATTACTGCAGAACCTCATATGGCTCATTCACCAAGATCCTCTCCTTCTCCATCTTCACTTCCCAGTCCTGCAGAGAACAGCAGTAGAGTTGAAGATGCCAGACGAAGTAATCATCGAGTTTTAATATTGTCTGCAGCTATAGGAGGTTCCGTTCTACTAATAATATTGATAACTGGCACTATCGTCTTTCGAAGCAACAAAATGGCTGTTGTAAAGCCTTGGGCCACTGGATTGAGTGGACAGTTGCAGAGAGCATTCGTTACTGGTAATTTTCTATCTTTCGATGAATGAATAGTATTTCTATTTTTAGAACCAAAATAGTTGTCTTAAAAATCAGATCCATAAAATAGAAGCTTATATGAACATGTGTAGCTGTAAAACCAAAATTTTTCCTTAAAAGTATTACCCAAATGGGAGAAGCTTAAATGGACCTTTTCGCTCCCATTATGAGCTTGTTACAAGCTTTCTTTTCTAATTGTTAATGATCTTTTCCCTTTTCCCTTTTTTCTTCTTGATATCTCGAAACTTTTGTTTATCAATATGAGCAAATAGTCTTAGATAGAGAGCTGAGTTCTATCATTCTTACTGTGAACAGGTGTGCCAAAGCTTAAGAGATCAGAACTTGAGTCAGCCTGTGAAGATTTCAGCAATGTGATCGGTTCCTCATCTGTTTGCACATTATATAAGGGCACATTATCCAGTGGAGTTGAGATAGCTGTGATATCTCGTACAGTGGCATCTACTAAGGATTGGTCAAAAGATATGGAAGCCCATTATAGGAAAAAGGTGATTTAGATTTTCCAGTTGGAATTTTACATCTCATTTTTTCCAATATGCGAGTTTCTAAATGCTTTTCTCCTCCTTTTACCAGATCGATACATTGTCAAAGGTCAATCACAAGAACTTTGTCAGCCTCCTTGGGTTTTGTGAAGAAGAGGATCCATTTACCAGAATGATGGTTTTTGAGTATGCTCCTAATGGGACCCTCTTTGAACATCTTCACAGTAAGTTTCTGATTCTAATGGTGTATCTGTTGTGATTTACTCAATATTGTTACCCAACTCGTCTACATCTGCTTGATTGAACTGCACTAGGATATACTTTTATATAGCAGGTAAAAACATACCTGCATTTGGTGTTTACACAAACCATATTAATTTACCATTGTTAAGTGATAAATTGACGTGAGAATGTGTATTAATTAACTTTGGTTCAATGATACTAGTGGGTGTGAAAACACATCTTATGTATTCATTGGGTGGTGTAAACACCAAGTGTGAATAAGTTTTTTCTCTACCTCGAGTATGATGTATACGGACGTGTCTTTGGTCTCGTGTCGATAATTTGTCTGTAATTTGAGAAGTAACCATTTTCACTTTAGCCTCATAAGATGTGTTCACACCTGTACAAGACATCTTTGGTATTCACTTTACATGCTCTTTCAACAATTTATGTTTAGCTTTCCCTGTGGTAAACAACTATTCTTATGTAATCTATTATGACCGAAAAGATCTCTAATTCTAGTGTTTTGTAGTAAGGGAAGCTGAACACTTGGACTGGGGGATGCGAATGAGGATAACAATGGGAATGGCATACTGCCTTGAGCACATGCACAATTTGTCGCCATCGTTACCTCATAAAAACTTGACCTCCTCTGCCGTATACCTCACAGAAGACTCTGCAGCCAAAATATCGGATTTTGGGTTCTGGAATGAGACAGATGCAGCTGAAATGGTGTCGAGTCCAGAAAGTAATGTTTACAGCTTTGGTGTCATTCTGTTTGAGATGATGACTGGTAGGCTCCCTTACTCAGCAAACAGTAGCTCTCTTGACGACTGGGCATCAGACTATCTTAGGGGAGGAGAATCCCTGAGAGAAATGGTTGATCCAACTTTAAATTCTTTCCAAGTAGAGCAGCTAGAGAGAATAGCTGTTGTGATTAAAATGTGTGTACATCCCGAGCCAAGACGAAGACCGACAATGAGAGAGGTTTCTTCCAGATTGAGAGAGATAACAGATATTGGACCAGATGGGGCAACCCCAAGACTCTCTCCTCTATGGTGGGCTGAGCTTGAGATTATATCAACTGAGGCTTGTTAATGCATAGGTACCTCAGTTTTGCCTTTAGATGGTAATTAATATATAGGCTGTATATACAAAGATTTTTCTTTTGTGTGGATAACAAGAGCATATCGACTGTGGGGGCTAAAGAGAAGTTGTTTGGGTACAAAAATGGAAAATTTAGTAAGTAAAAAAAATTTCTTTTTTTATTTTATATTTTAGTATGTAAGTAATTTCTAAATCGCCTTAATTTGCTTTCTTGGCGGCTTGGGATCAATTCTATTGCTTATGAAAACTATAAGGACCCTAGACAAAAGGGATAAAACCTAAGGGGTCGTTTGGCTTTATCGCATTAACAAAAAC

>PGSC0003DMG400030113 LRR-RLKs

CTTCAGTCTTCATTACCATATGAAAAGAAACACAAGGGAAGCATCATTTTTAATATGCTCCAATCAAGATCTAACCGTTTTTTGCTTAATTTGCAAATCTTCTTCCTAATTAGCCTGTTTTTCCAAGAAACCTTACAATTTTCGGTCAGGGAACAGTCCATATTACTAAGCTTGAAACGACACTGGAGGGATTCCGAGTTCCTCCAGTCATGGGACTTGAATTCTTCAGCATGCACTTGGTCAGGAGTTTTGTGCATCGATGACCGGGTAGTGACTCAACTAAATCTTGGAGAAAAGAACATTACTGGAAAAATTCCATCAACAATAATATGTCAGCTCAAGAACCTTAACTTCATTGATCTCTCCAACAACAACATTTCAGGAACTATTCCAGTAGGCCTGAAAGATTGCTCAATGTTACAACATTTGGACTTGTCCAATAATTCTTTGATCGACCGGATTCCAGGTGAGTTGTTTGGGATGAAACAGTTACTCAGTTTGTATCTTAATGGTAACATGTTGTCTGGTGAGATGCCAAAACAAATATCATCTCAACTGGAAAATCTTGATCTTTCTGAGAATCACTTGAACGGTTCGATACCAGAAGATATTGGAAACTTCAAAAATCTTTTGAAATTGGATCTGTCACATAATTCTCTCAGTGGTTCAATTACAAGCAAGTTGTTTCAGTTGCATGATTTGCGTCACCTGTCGCTAAGTTCCAACTACTTGTCCGGTGTGATACCTGATGAAATGGACTTGTTTAGTTTATACGATATGGATCTTTCACATAACCAATTGACTGGTTCTATACCAAAGGGATTTCAGTATTTACCTGGACTGCATGCCCTGGATTTGTCTTATAATCAGTTATCAGGAGACATTTCAAAAAGTATAGAATATCTTAGGCCTAGAAATACTTTAAGGCTTTGTTCCAACAAATTCTCAGGAAGAATTTCTTCTGAATTTGTTAAGCTAACATATGAAGTGAATTGCTTTGATGAGTCCAATCTTTGTTCTACATCCAACAACCGAAAACTTTTGAGATCACAACACTTGATTATTCCTATTGTAGGAGTGGGGTTGGCTATACAGTTATCATGGATCTTTTACAGAACAAGAAAGCATTGGTGGAAAACAAAGAAGCAGAATGTCAAAGATGACATGAAGTTCATTTCATTTCAGAAGTTGAAGGTGACTACAGAAGACATTTTGTCTAGCTTGAAAGACAAAAACATAATAGGAAATGGAGGATCAGGGAAGGTCTATCGAGTTGTGATAGACCAAACAGGCAATACTTATGCTGTTAAAAGCATAGGGCATGGAGAAAAATCAGGTGGAAGACCCCAAAAGGAGTTCCTGGTAGAAGTTAGAACCCTTGGTAGCATTCGACACAACAACATTGTCAAGCTCATCTGTTGCATCTCAAGTTTAGACAGAAAGCTTCTAGTCTATGAATACTTTGAGAAACAAAGCCTCGACAAATGGCTTCACAGAAAGAAAAGTGTAGTATCATCGGGTCAAAGTAGTACCCCGGCCCTGGATTGGCAAAAGAGATTAAAAATAGCCACTGGTGCAGCGCAAGGACTCTGCTATATGCACCACGACTGTACTCGATCCATCATTCATAGAGACATTAAGTCCAGCAATATCCTTCTGGACTCAGAATTCAATGCAAAAATAGCAGATTTTGGACTAGCAAAAATACTATCCAGGAGGGATGATAATCCTGAGACAGTTTCTGCTATTGCTGGAACATTTGGTTACATTGCACCAGGTATATTACAGTTTAAATATCTATTGTACCTCGAGAATTTCACAAAAGAATTATACTCCAATATTAAATTTTAATGTTTGGTCCTCACAGAGTATGCCTCGACATTTAGAGTGAATATAAAGACAGATATCTATAGCTTCGGAGTGGTGCTATTGGAATTAACAACAGGGAGGCAACCCATTCTCAGAGAGGACCAAATGAATCTAGCACAATGGGCTCAACAGCGTTACAGAGATGGGAATTTCATTGTTGCCCTTGATGAAGAAATCATGGAAACAAGTAATGTGGAACAAATGAGAGGTGTTTTTAAACTAGGACTAATGTGTACCGGAGCATCACCATCTAGTAGGCCGTCAATGAAGGAGGTCTGCAACATTCTTCAAAGTCTCAGAGACCCCATATTTTGAGAGAATAGAATGTTTCGCCTTACATAGATAAATGTAGTTGTATTTTTTTTAACAGAATGAGAAAGTCGTGTTGATAACGATTTCTTCCAGCAATGTATACTTTAGTAATACTGTCAGAGGACATTTTTTATCCTCAGACAGATCAATAGCCTCATCTGGTTTAAGTGGTGTCAAACACAAGGGCTTGTAAGCTTTACCTTAGAATGTGATTCCTTGTTCATACTTGAGATGTCGCGAGGCAAGAGCTAAGTGTCGTGGCAATTACAGTGTATATATATATATATATCTTTTTTTTAATTCTTCAGAACTCTCCAAGGAAGCTTTTGGAACATACAGGTTTGATGAACTGCAAATGCCTTCTTTCAAACATAAATCCTTCTCATGATAATCTTGTAGGAAGTCTAGTTCTTGTGGCAGAGGTTGTAGCGGCTGGGGATTATCCTGGGGTTCCCAAAAAATAAAAAATAAAAATTCTAACAGTTTTACACAATTGATAAAAGCTAAAACCATTGATTAGATATTCATTAATGGTCTAGGCTAACTTATATGATTGGAAAAATGAAATGTTCCTATAAGCGATCAAGTTTTTCTACTCGTTAATTCTTTTGGTACTTGTTTTAATTAGTTTGTTAGAATTGTTCTGACCCTCAGTGATACTCCTGATGGAGCAGGGTCAAGTAGGACTAACAGACTTCTAGGTTCACAAGCTTCAGAAGATGAGGTGAACATGACATCTTAG

>PGSC0003DMG400030117 LRR-RLKs

ATGTTCTTCCGCATTTTTTTCGTCCTAATTAGTCCGTCATTGCAAGAAAGTTTACAATTGGAACAGTCCGTGTTGTTAAAATTGAAACAACACTGGAGCGATTCGGAGTTCCTTCAGTCATGGGATTTGAATTCTTCAGAATGCACTTGGTCAGGAGTTTCGTGCATCGATGGCAGGGTAGTGACAGAACTACATCTTGGAGGAAAAGACATTACTGGAACAATTTCATCGACAATATGTGAGCTCAAGAACCTTACCTTCATTGATCTCTCCAACAACAATATTTCAGGAATCATACCAGTAAGCCTGAACGATTGCTCAATGTTACAACATTTGGACTTGTCCAATAATTCTTTGAATGATCGGATTCCAGGTGAGTTGTTTGGGATGAAACAAATACTCAATTTATATCTTAATGGTAATATGTTGTCTGGTGAGATGCCAAAAGAAATATCATCATCCCAGCTGAAAAATCTTAATCTCTCTGAGAACTACCTGAACGGTTCGATACCAGAAGATATTGGTAACTTGAAAAATATTGTGAAGTTGGACATGTCACATAATTCTTTAAGTGGTTCAATTACAAATCAGTTGTTTCAGTTGCATCATTTGCGTCACCTGTCGCTAAGTTCCAACTATTTGTCCAGTGTGATACCTGATGAAATGGACTTGTTTAGTTTGTATGCTATGGATCTTTCTCATAATCAATTGACTGGTTCTATACCAAAGGGATTTCGGTATTTACCTGGATTACATGCTCTGGATTTGTCGTATAATCAGTTATCAGGAGATATTTCTCAAAGTATAGAACATCTTAGGCCTAGAAATACTTTAAGGCTTTGTTCCAACAAATTCTCAGGAAGAATTTCTGCTGAATTTGTGAAGTTAACATACGAAGAGAATTGCTTTGACGAATCAAATCTTTGTTCTACATCCAAGAACTTATCCGTCCCTGTCCTACCAAGCTGTTCCTCTGATGATGAGGTTCGAAAATCTTCGAGACCAAAACACTTGATTATCATTATTTCTGTTGTAGGAGTTGGGGTAGCTATACAGTTAACATGGATACTTTACATGGTCAGAAAGCATTGGTGGAAAACAAAGAAGCAGAATGTCAAAGATGACATGAAATTCATTTCATTTCAAAAGTTGAAGGTGACTACAGAAGACATTTTGTCTAGCTTGAAAGACGAAAACATAATAGGAAAAGGAGGATCAGGGAAGGTCTATCGAGTTGTGATCGACCAAACAGGCAATACTTATGCTGTTAAAAGCATAGGGCATGGAGAAAATTCAGGTGGAAGACACCAAAAGGAGTTCCTGGCAGAAGTTAGAACACTTGGTAGCATTCGACACAACAACATTGTCAAGCTCATGTGTTGCATCTCAAGTTTAGACAGAAAGCTTCTAGTCTATGAATACTTTGAGAAACAAAGCCTCGACAAGTGGCTTCACGGAAAGAAAAGAGCAGCATCACTAGGTCAAAGTAGTACCCCGGCCCTGGATTGGCGAAAGAGACTAAATATAGCCATTGGTGCAGCTCAAGGACTATGCTATATGCACCATCATTGCACTCGACCAATCATTCATAGAGACATTAAGTCCAGCAATATCCTTCTGGACTCAGAATTCAATGCAAAAATAGCAGATTTTGGACTAGCAAAAATACTATCCAGGCGGGATGATGATCCTGAGACAGCTTCAGCTATTGCTGGAACATTTGGTTACATTGCACCAGGTACATAACAGTTCAAACATGTATGAATATTCAATCTTTCAAAATCCAATTCTACTATATCAAATTTTAATATTTGGTCCTGACAGAGTATGCCTCGACATTCAAAGTGAACGTAAAGACTGATATCTATAGCTATGGAGTGGTGCTATTAGAATTGACAACAGGGAGAGAACCCATTCTCAGGGATGAGCAAATGAATCTAGCACAATGGGCGCTACAGCGTTACAGAGAGGGGAATTCCATTCTCGAGGCCCTGGATGAAGAAGTGATGGAAACAAGTAATTTGGAACAAATGAGAAGTGTTTTTAAACTAGGAGTGATGTGTACAGGAGCATCACCATCTGGTAGGCCATCAATGAAGGAGGTTTGCTATGTTCTCCAAAGCTGCAGAGAGTCTAATATTTGA

>PGSC0003DMG400030119 LRR-RLKs

CTTTGCTGTTTTTTCTTGGCCACAGTCCACAGACGTCAGTGAAGTCAATTTTTGTTTCTTATCTTTTTAAAATTTAACAAGTCAAACTTTGTTAGCATTATACAAATTTTCGTCCATGTTGACTGACTTGTGTTGTATAATACACGGTTTCTTCAAAGCTAATTCATGATCCACAAAATAAAGAGGACAATCAAACCAAAGGAGCCATGAAAATCAACCATATTCATGTTGTAAAATCTACATTCTCAATTCCCAGGAAACAAGTTTACCTTTTCATTGTCTCCTTACTTGCCTTTATGCATTTCCAGGGCGGTTTACAGAGTGACGTCTACGAAAAGGAGAAGAACATTTTGTTGGAGCTGAAGCAACTATGGGGATTGCATCAATTATGGGATTCCGATTCCGATTCCTCACACTGCAGTTGGTATGGTATAGGCTGCAGAAATGGTTCTGTTACAAGCATCTCGATTCATGAAGGGAATCTACTTTCCGGAACAATTTCACTAATCATATGTGAACTCAAAAGCCTGCAATTGATAGACCTGAGTAGCAACTATATCAGAGGAGAATTTCCTACCTCCTTGTACAATTGCTCAAAGCTGGAATATCTTGATATTTCTTGGAATCAATTCCATGGACCACTGCCTAGTGACCTTCATCGTTTGTCGAGGCTAATCCATTTAGATATTGCTGGAAATAACTTCAGTAATATCCCAGAAGCCATAGGTCAGCTTTCTGAATTGCAATATCTATCTCTTAAGTTCAATAATTTCAACACTTGGATATCCCGAGAAATTGGAAATTTATCAAAGCTTGAGACCTTGGATATATCCTATATTGACAGCTTTAAGCAGGCAACAATCCCAGAGGAATTAAGAAGACTGAAGAAGTTGACAGCCCTTTTCATTGTACGGTCAAATTTGAAAGGGGAGATTCCAGAAACATTTTCCGCTCTTTCAGGTCTCGAAGTCTTGGATCTCTCAATGAATTACTTAAAGGGGTCTATCCCAAGTTACTTGTTTGAATGGAAGAAGTTGACGAATCTTCATTTGGAGTTTAACCAATTTTCCGGTAGGTTACCAATGTTGGATGCAAATATGGAATTGATGGGGAGAACTCAAGAAATGTTTTCTAATCTTTCTGGTCTCCAAGTCTTAGATCTCTCAGGAAATCACCTAAATGGTTCAATTCCAAGCTACTTATTTCATTGGAAGAATTTAACATCTCTTCTTTTGGGGGATAACCAATTTTCTGGGAATTTACCTAGTATAACTGGAAATTTGAGATTAAGAACATTGGATCTTTCATCGAATCAGCTTTCTGGACATATTCCACAAGAGTATGACAAGAATGAATACTACTTTAGAAATAATCTGAATCTATGTAGCAAATATACCAACCATTACATCCAAGTTCCACGTTGAAGTGGAAAAACAAGAGTTGTAATTGCCGTTGTGGCCCCTGTTGCATTTTTAGTCATGATAACATTATTATGTTACACATTTAAAAAGAATTGGACTTTCTCCACAGACCAACTCATGTGGAGGAAGAGGCATGAGAACCAAGATCCTGAGTGGATGTTCATTTCTTTCCAGCAGTTAGGATTCACAGAATTTGAAATTCAGGTGAATATGACAGAAGAAAATTTGATTTGAAGTGGAGGATATGGGAAGGTCTATCGAGTTGGTGTCAATCCTAATGGGAACTTTGTGGCAGTTAAGAGAATATGGAACAAGAGGAATTTAGACCACGGGCTTGAGAAACAGTTCCTTGCAGAAGTTGAAGTATTGGGTAGCATTCGACATTCTAATATTGTGAAGTTGTTATGTTGCATTTCTAGAGGAAACTCAAAGGTTTTGGTGTACGAGTACATGGAAAATCAAAGCTTAGACAAGTGGTTGCATCACAAGAGAAGAAGTGAAAAAGATATTACTGCCTCAGCACATTGTGTATTGGAATGGCATACTAGACTGCAAATTGCAATTGGTGCTGCCCGAGGTCTATGTTACATGCATCACGAGTGCTCTACGCCAATCATTCATCGTGACATTAAGTGCAGTAATATCCTTTTAGACCATGAACTGAATGCCAAAATTGCAGATTTTGGACTAGCTAAGATTTTGGCCAAGTGGGGAGAGACAGAGACTGCTTCTGCTATCGCTGGCACGTTTGGCTACCTAGCTCCAGGTATATTTCACTGTTCTCATAATGAAATAAACAAAAACACACCAGAGTGTACGTATGTATCGTGTTTCTCATTGATTTTATCCTAATACCTTGTTATTCTAATTGGAATTTCAGAGTATGCATACACGTCGAAAGTGAATGCCAAGATTGATGTCTACAGTTTTGGTGTGGTGTTATTGGAGCTGGTTACCGGGAGAGAACCAATTAACGGAGACGAGCATATAAACCTTGCACAAAGGGCATGGAACCACCATGAAGAAGGCAATCCAGTGGTTGATGCTATTGATGAAGAAATCAAGGAAGCATGTTTCTTGAACGAAATGAGTTCCATGTTCAAGTTGGGGATTATTTGCACCAGTACTGTACCTTCTGCTAGACCTTCCATGAAGGAAGTTCTGCAAATTCTATTACTTAGCCTTAATACTTCTTCCTAGCAATTGGCTGCGAATCCAGGATTATCGTTGCTTCTTTATTTGCACTGAAAAATTAAAAGTCCAAA

>PGSC0003DMG400030169 LRR-RLKs

CTTTTCTTCTTCAAGAATTTAGGGTTTCTCTTCTCCACCGACGGGAGGCGTTATTTTCCTCCTCTTCTTCACAATTCTTCTCGAATTTCTTCTATATATAAGTGGGGTTCTCGTTCTTGATCCTTTTGATTTCTTATTCAAGCTGCTCGTATGGGTGAAAACGATTCAGAAAGCTCTGAAAGTGCCAAACCCAAGAATGTATCTGAAAATTCCGACGGAGGTGATGAATTGACGGGTTTAGAGGAAGATTCGAGCAAGATGGTTTCTGGGTTTGATTCAAGTTTTGATATTTCGGGGAAAAGTTTGGATTTTCCATTGTTGGAAGGTGTAGAAGGTGGGGTTGAAGGTTTGTATATGTATAAGAATGTATTTAATTTGATTCCTAAGGCAATTGGTGCACTTGGGAAGGTTAAGATACTTAAATTTTTTGGGAATGAGGTTAATTTGTTTCCAACTGGAGAGTTAAGGAATTTGGTTGAATTAGAGAGTTTGCAAGTGAAGGTATCATTTCCTGGGATGAGTGGATTGGATTTGCAGAAGTTAAAGAATTTGAAAGAATTGGAGCTATGTAAAGTTCCCTCTAGACCTTCAGCTTTCCCACTTTTAAGAGATATTGCTGGTCTTAAGCGCCTGACTAAGCTCTCTGTCTGTCATTTTTCTATCAGGTTAGATCTGTTTCTTCTTGCAATAATTCTATCTAAGTTTATTTACGCTAAACTCATGTTTACTATAGACAAATTACCTGTAGTTGGCTGTCAGTTAACTTGATAACATAAAATTTCTGACTTTGTCAATGTATAAATATTAAACCAATTCGATGATGTTTCTCTGTGAACTTGTCTATATTGTTGCACTAAATTCATGTGCAGGTTGCAGCTCAATGTGCTTTTGGTGTTATTGGAAGATTAGGTTGCATCTGCACTTTTTCGCTTTTGCTGCATTCCTTAAAGTAATTTGAATTTAATAGAACAAGTTTATTGCTAAACTATTGTTTTTACAAGTGCTAGCTGCTGCAACTATAAGCCCCAAGTTAAATAGTATTAAGCAGGGTACATGATCACACAAACTTCCATTAATACAGGGGAAACTAGGTAGGTAGAGCATCCTACCAGGATGTTTTTTGTGGTCTCTTTAAGGAGGTAAAGACAGGGAAAAAGTATTTGCAGGTTTAAATGGTTAGTTTCAGTTTTCGGGTACAAAATTTTCCTGTTAGTTCAAACTTGTTTTGTAGATTGACATGACCAAGCAAAACAATTTTATGCATGATTAGTAAAAGTTTTACAAAAACAACAGATCTTCTCAAAGAAATTTTGGATGTGGTAACACAGCTGGAGCATAAAAGAAAATAAAGACTTGTTTTTGCTGCTTAGATGCTTCATGAGATTGAAGTTTGTTAATTAGGACCAAAAGATATGCACAAAATTTTGTAAGTTGTTTATCGGGAGTTTAGACTTTAGCATTCTTCAAAGGGTATCAGGTTCTACTCTAGGACCATACTAAGTTCAACCAGGATATGATACGATATTAGGATTAGTCGATAGTACATAGCATAACTATCCCTTGGTTAAGGGTCGTAAAGAGTGTTGTTTTTTAACTATTTATTTGTGTTTCCTTTAATTGAAATTTTTATTTTTTGTTTATGGGATTTCTTATCCTATAGTGCATCTTAGTTAGGTTAAAATCATCTTATTATTTTACCTCATGTAATTTGACAGCTGAAGAGGTTATATGGTTTGCCTTTATAATGGATAAATTTATTATTTTGAGGTTGGCATTATGCAGCCACCAGAAGACTTTTCCCTTCATCTCCGTCTTGTTGTTTCTTGCAACTACAACGAGTTAAAGATTTTGAGACATTAAAAATCTGGCTGGTTAATCAATTTGCAGCATATTGGTTGCTGTTGTCTCCTGGCAATTATACCCTCTCAAGGTCCACATTTTGTTTGAGAATTATCTTTTGGTCCCTGAAGACATCATTCTACTTTTGATTAGCAGGTGTTATGAAAAGCCTATGCACGAGTACATCACATAATTTTAAAGTTATAGACGTCTTAAATCTCACAGGTCTCCAATAGGAAAAAAAGAAAAGGGAAGTATATGAAGTGACTTTGTTCGCTTGGAGTATGGAATATAGAATATGCTGCCTCAAGGAGTTTGAGGTGCAAAAGGTGTCCACACACTTCACACTTCAGAATTGCGAGCCTGTTGCTGTCACATCTTTCCATCACAGTAGTATGTCCTATAGCTAATAGCAGATTAAGTGAACTTTTGCTTAGCATGTGTAATTTATCAGTTGTATATCATTGCTTAATGTTGGTAGGACTGGAAATTAGTCGTTTTTGCAGTTGGAAGGAGGATTCAAATTGGAAATAGGTGTTAGGATTTGTATTTTATAGTGTTCCTTTTATTGGGTGCAGTTACGCCAGAATATTGACTCTTCAGATAAAAGTCATGTTTAAGTTGTTTGACACTTGATACATTTTGTTAGATGAGGTCTGATATCTAAAAATATCTGTACATCACATTTTATTGTTTTGATATCTGAAAGGCTAAAACTTCATTCTGCTACTCTTTTGGCAAGCAGTGCTTTACTTTTGGTTGCTAGAGGTTGAAAGTTGTGAACCATAAACTTGTAGATTCAGTTCATATCTATAAACCATTTTATCTTTTTTTTATCATTGTACTCCATTATAATGCCATTGTTCCTTGCTGTTCTTGCAGATATTTACCTCCTGAAATAGCCTGCCTCTCTAAATTGGAGTGCCTTGATCTTTCATTCAATAAGATGAAGAATTTACCTGTTGAGATAACTCATCTTAACTCCTTACTGTCACTGAAGGTTGCTAATAATAAACTGATTGAAGTACCTCCGGGGTTGTCTTCTCTGCAAAGATTAGAGAGTTTGGACTTTTCTAACAATAGATTAACATCCTTGGAGAATCTTGATCTCCTTTCAATGTATAATCTGCAGAGTCTCAATCTTCAGGTATGTGTTACTGAAATTACAGTTTCTTTCTAGTCTTGTTGCTTTTAGGCTTTTTCTCTAGTTTCTTTTCCTTCTCTGATAAATGTTTATAACATGAGAGGCATCTACTCTTAATGTGGATGAAATTCTTGCATCCCACTGGAAAAAAACCCGAAATGTATTTGTCTCAATTGTCTAGTCAAGTAACTTGATATATTATAATTACCTTATCCTGACCCATATATCTTCATTGAAATTTGCTGAAGCACAATAAGCTTCGTAGGTGGTGTTCGATTCCTTCATGGGTATGCTGCAATTTGGAGGGGAATTTCATAGATTTGTCCAAGGATGACACCACCAGTTCGTCTTCAGAAATGGACGTGCTTGAAAGTTATGAACAGGAAACTTCTGAGAATACCCAAAATGGTGAGTTCCTCGGAATCTCTTGTCTCCCCTCCTTATATCTATTGCTATTTCTGCACATTCTGATTCCTTTGAATCTTGTTTGTTGCCCTTCTAAGAAGTGTTGTTCGGAAGGGTAGTACCTTTGAGAAGTCTGATGTGTGAACTTTTAATTGTGTAGGAGTTTCTATCAAATTGTCAGGTCATTTATGTGGCTCATCACCCAGTCATCGATGTTTTCGGCCTCGAAAATCCAAGAAATGGAAGCGGCAGTATTATATGCAACAGAGGGCTCGTCAAGAGCGATTAAACAACAGCAGGAAATGTGTTGCCTGCAAACATTCTAAGCTAATTGATGATTCTCTGGTGGAGGCCTCATCAAGCATAGTGGATGATGATACACATGATAAAGAACTGATTCCTGAAGAAGCTGAGTGCAAAGGCTCTTTAGCTAGTGGTATTGATGAACATATCAGATTAAAGGAGGACAATTATATAGGAAGACCTTCATGTGTTGCTTCTGACTCCATTGAAACATGCATAGATATTCAGAATTGTAAAACATGTGATGCTTCAGTAGGTTCTGTTTCTGATGCTGCTGATGTGGCTGAGGAAAGTTTGTCTTCAGAGGTGTCTAATAGTCCTCCAAAATCGAAAAGGCACCTGGATGGGGTCATTGATAATCCAAAACCATGCAAAACCCGCAGACCAACAGATCATTCAGAGGTATCCTGCAAATACAGCATGATGTCTTTCTGTGGTATTGATGATTATTTACCAGATGGCTTTTATGATGCTGGGCGAGATCGGCCATTTATGTCGCTGAGGAGTTATGAGCAGAATTTGCATCTTGATTCACGGGAAGTCATTTTAGTTGATAGGTAACTTCCATGTTTGCATGCCATTAATAGACTAATAGTTATGGACCTCAACTACTAACTTCAGGGAATGCTTTGCCTCTCTACTCATTTTGATTACTTATTTAATCTGTTTAGGCAAAGAGATGAAATGCTGGATGCCATTGCTCTACGTGCTCAAGCTCTAATCTTTCATTTTAATCAGATAGATGGTCTGTTTAAAGATAGAGAACATGTTGCTGTAGACAACTTGCAAATTGCATCACTGCTTGCCCTTTTAGTGTCGGATCATTTTGGTGGAAGTGACAAAAGTAATATTGTACAGAAGGCAAGGAAAGATGTATCTGGGTCGAACTATAGCAAGCCTTTTGTCTGCACATGCCCTACTGGAAATGATGACACTACCAGCATGGTCACAAAAGAGAGTCCTAGCATCTCAGAAGATATACTTTTCCTTAATCTTTGTGAGAAAGCACTTCACTCAATAAAATCAAGGCAAAATTCTATTGTTGTTCCTATTGGGAGTTTGCAGTTTGGTGTATGTAGGCATAGGGCCTTGCTTATGAAGGTATTTTGTCATCAGACATGTTTCTAATTAGATATGGTGGTTCTGAATTTAGTTTATTATTATGAACTGATAAATTTCATTTGTCTTCTGCAGTACCTGTGTGACCGAATAGAACCTCGAATATCTTGTGAGCTTGTTAGGGGTTATTTGGATTTTTCACCGCATGCGTGGAATGTGATCGTGGTAAAGAGGGGTGAATCATGGGTTCGAATGATAGTTGATGCGTGTCATCCCCTTGATATCAGAGAAGAGACAGATCCAGAATACTTTTGCAGGTCAATGATTTTCCAATTCTGTGTTACTTTCTTCATGACTAATTTCTGGTACTACAAGCATTTATATCAATACACGTGTATGACGATATTGCATTTTCAATTTAGCCAGTGTGATTATTGGTTATTTTGGGGTGAAATTGCCAGCTTCTTGAGTAGTTCCTTAGTTACTAAAAATCTCTAGTTTACCTGAGTTATGTCGGAATGAAAATGGCCAGATCACTGAGCTTGCTGTTTCTTTCAGGTATATTCCACTGAACCGGATAAATGTTCCTGTTGTGCCAGATGCTAGTCCTGGTCAAGTTAGTTCTTTTCCTTCTCTAACCGGTGCTGATAAAATTCATAAAGCTCCCTCCTCAACCCTCGTCCCGTGTAAATTGGGATCCCTTGAGACTCTGGCAAAGGTGTGGTTCCTGATTGCCTTTAAGTTCTCCAGTTATTATTATGCAAATATACTTAAGTTCTTATTCCAAACTTCAATGACTTGAATTAAATATGTTTTATTCTTAGAGTGACTGAACTACATGTTTATGTGTTAACCCACAACAATTAGAAAATGATTTTTCTGGCTTTATTGTACTTTCCATTATTGACTGGAATTTACCAGTCTATCTGGTCTTGTGCTTTTGATTTGTTTTGGAAATTTGAAATAGATAAGAAGCAAAGCAATCCTTTTTGACTTATTCTGCTATTCAGCTGGAATCCATTATATTGCAGGCTTATGCTTTTCATTTTGCAAATTTTTCATTTATCTGCTGATAAAACTTGATTAATTAGTGACACGTATTATACCACTAGAATGTGGGAATTTTTCTCGGCTAGCATCTTTATTGACGAGGCATAGTTGATTTTTTTTTAAATTTATTCTGCTCAATTAACCTGAATTGAGCGGAATGAATATAAAAGAGTCATATGACTGATACCGACTAGTTCAGGATTGAGGTTTAGATGATTGAATGAACATTTTTGCCACTGCAGGTGCGAACTCTAGAGATGTCCAAGTCCACAGCGGATGAGATTAAGAATTTTGAGTTTAACTGTATTGGAGAAGTTAGGGTGTTGGGTGTCTTAAACAGTTCGTGCATTGTCAAATATTATGGTCATCAAATATCTTCAAGATGGGTTCCTTCATCAGATGGCAGTTCAGAAAGTCGTACTTTGCAGTCTGCCATTTTGATGGAGCATATTAAAGGGGGTTCACTAAAGGTATATGTTTTCCTAGAACTCTTCTCGTACCTGACTTTCCTCTCCTTTATTTCCTTTCCCTTTTGAGGTTGGAAGGGTATAAATAACACCTGATTGCCTTTTGTATTCAGCTCATTTTACTTTCTTGTCCTTAACTGGCAGAAACATGTAGATAAGCTATCTAATGCTGGTGAGAAGCGTCTTCCCGTTGAGCTGTCGGTATTTATTGCCCGAGATGTAGCAAGTGCGTTGACAGAGCTGCATTCTAGGCATATCATACATCGTGATATAAAAAGTGAAAACATCCTGATAGATCTGGATAAGAAGAGAGTTGATGGTACCCCTACAGTAAAGCTATGCGACTTCGATATGGCAATTCCCCTTCGTTCATACTTACATACTTGCTGCATTGCCCATGCTGGAATTCCTCCACCAGATGTTTGTGTTGGTACACCTCGTTGGATGGCTCCTGAGGTCTTTCAGGCGATGAATAAGCGCAATATCTATGGGTTGGTAAGTCACCAATCCTTCTGACCAAAAAAAAAGTCACCAATCCTTGAAGTTAGAATCCTAACCATTGAATTCTCTACTAACAAGTACCAATTAAATGCCCCACCGGGTGCTCTGATCTGCTAAAACAATGTGGTTCTTTCTGATTTGCCAATTTGTCACCCTGGGGTAAAATCAATTCAACACAATCAAGAACTTACCACCTTCAAAAGACACACATTTTCTACTAAAAACTGATATATCACTCATTCCTCTATTAAAGTTCAGCTTTATCTTGTCATTATTAACTTCAAATGTTTTAAGCTAGAAAAGAAAGCATATTTTGTCTGCTCTATTGCTGCACTGCTACTGATTAAAGAACAATAATATGTACTCATTGACAAATTCAAATTTTCACCTTACTTTCTTGGTAATTTGTTTGTTTAAATTTCCTCATTTTTGATGCTTGAATCTATTTCGGAGATGAGGATTTTTGTTTTTGAGAGACTATTTTTATATGACGGCTAATCTCAGAGGATGTATCCTGTGGCAAAAGGGGTTGGGGAATATTATCTCTTACTTCTTTGAAGTTGAAAAGGTCTCTAGGTTATAGATGAGTTGCTTCAAGGTAAAAAGGTTTTTCTTGCTTTGACTTTGCAGGTTCCCCGTTTCAATAGGTAGATGAAAGCTAAATAGGACCTAAAATAAGTCCACTATGGCCAAAAAATATCAGGAGGGGCGGGTGGATTATTTCACCCTTAACCCAAATTCAATAGATGGGTAAATTGGCTAGAATGGGTTATTGAACAACAAAAAAGTATTTATAGGTGCTAATTTCCTGTATTATCTTTTTCCTTTTGAAGATGAACAAATGAGTTTAATTGCTTTGAGTTGGTGACTTGAAAGATAAAGTGAGTTTTTGTCATGTCTAAACTGCAATTCTAGCTTAATGTATTCAAAGGTTTTCCGAACTAGGTTTTTACTTGTCTTACTTTGGATGTAGGGTGCTGATATTTGGTCGTTTGGGTGCGTCCTTCTGGAATTGTTAACACTGCAACTTCCTTACTCAGAGTCATCCGAGTTAGATATCCACCATTCTCTTCAGGTATTTCTTGATCTCTAACCTTCGATTATTAATGCTCTAAATACCATGATGAGTACCTAGTGTGCTTGAAATATCATTGTGTGATGAGGACTTCAGAAGTTCCAAAGTCCCCTGTTGGACAATCTTTATTATGGCTGCCCTTGGTTGCTCAAAAGGCTAATGGTTCTAGCATCTTGGTTGTTGGATATTTCGGAGCTCTCATGGTTTTCTTCCTTATATTGCTTTCGTTCATAAGAACTGCCTGCAGAAGTACACTTGTCGCATGCACATTTAAAAAAAAACAAGTTTTTGTTTGAATTGCTTTAAATGGTAGAAAACAATGTAATTCATGTTTAACACATGCAGGCGGGTAAACGGCCACAATTAACCGAAAAATTAGAGGCAATGGCTGCGTCCAAAGCAGAGCTAGAGGATCTAGCAAAGTCTTGTTCTAGCTCGGACCTTGACAAGAAACAATCTGAATCTCGTATCCTGAAACTTCTCGTTTCTATTTATCGTTGGTGTACTGAGAAGGATCCAAACGACCGTCCCACAGCAGAGAACCTCTACAACCTCTTACTTACCTGTGCTAATTCCCTCCCTTCACAACAAAGTCAAGAAGATTCTTGATCTTCCTCAAATTTTGGTAGCCTTAGGATGTTGTAGATTAGCATTAGATTATTCAGCAGGAGTTACAGACCCAATACTTGGGTCTGTTGACCCCTCATTGAGCAATGTAACATGTTCTTAGTCTTATTTGAAATGGAAAACCATACGGACGTATTCGCGTCCATAGGACATTTATGATGGTCTTGTTGTGTGTGTGTGTTGTGTATCAGTTTCTTTAGTCCTTTTCCAGCTTGTAGTTGTATACATTTGTACCCATCTAATGTATTCATCTTCACTAATTCCAATTGATGTTAATTATGAATGTATTCTGAGTTTCTAAGACTTGTTGGATTGTGC

>PGSC0003DMG400030298 LRR-RLKs

ATGTGGAGAGGGCCATGGTATGTACCCCAACTGAGCGTCTTAAATCTCAGGAATAATAGCCTCACGGGTATAATTCCTCCTTCTGTTGGAAATGCCACACAAATGATGAACTTCAGTTTGTATGGGAATAGAGTCAGCGGCAACATTCCAAAGGAGGTCGGTAATCTGAGCCAGNNNNNNNNNNNNNNNNNNNNNNNNNNNNNNNNNNNNNNNNNNNNNNNNNNNNNNNNNNNNNNNNNNNNNNNNNNNNNNNNNNNNNNNNNNNNNNNNNNNNNNNNNNNNNNNNNNNNNNNNNNNNNNNNNNNNNNNNNNNNNNNNNNNNNNNNNNNNNNNNNNNNNNNNNNNNNNNNNNNNNNNNNNNNNNNNNNNNNNNNNNNNNNNNNNNNNNNNNNNNNNNNNNNNNNNNNNNNNNNNNNNNNNNNNNNNNNNNNNNNNNNNNNNNNNNNNNNNNNNNNNNNNNNNNNNNNNNNNNNNNNNNNNNNNNNNNNNNNNNNNNNNNNNNNNNNNNNNNNNNNNNNNNNNNNNNNNNNNNNNNNNNNNNNNNNNNNNNNNNNNNNNNNNNNNNNNNNNNNNNNNNNNNNNNNNNNNNNNNNNNNNNNNNNNNNNNNNNNNNNNNNNNNNNNNNNNNNNNNNNNNNNNNNNNNNNNNNNNNNNNNNNNNNNNNNNNNNNNNNNNNNNNNNNNNNNNNNNNNNNNNNNNNNNNNNNNNNNNNNNNNNNNNNNNNNNNNNNNNNNNNNNNNNNNNNNNNNNNNNNNNNNNNNNNNNNNNNNNNNNNNNNNNNNNNNNNNNNNNNNNNNNNNNNNNNNNNNNNNNNNNNNNNNNNNNNNNNNNNNNNNNNNNNNNNNNNNNNNNNNNNNNNNNNNNNNNNNNNNNNNNNNNNNNNNNNNNNNNNNNNNNNNNNNNNNNNNNNNNNNNNNNNNNNNNNNNNNNNNNNNNNNNNNNNNNNNNNNNNNNNNNNNNNNNNNNNNNNNNNNNNNNNNNNNNNNNNNNNNNNNNNNNNNNNNNNNNNNNNNNNNNNNNNNNNNNNNNNNNNNNNNNNNNNNNNNNNNNNNNNNNNNNNNNNNNNNNNNNNNNNNNNNNNNNNNNNNNNNNNNNNNNNNNNNNNNNNNNNNNNNNNNNNNNNNNNNNNNNNNNNNNNNNNNNNNNNNNNNNNNNNNNNNNNNNNNNNNNNNNNNNNNNNNNNNNNNNNNNNNNNNNNNNNNNNNNNNNNNNNNNNNNNNNNNNNNNNNNNNNNNNNNNNNNNNNNNNNNNNNNNNNNNNNNNNNNNNNNNNNNNNNNNNNNNNNNNNNNNNNNNNNNNNNNNNNNNNNNNNNNNNNNNNNNNNNNNNNNNNNNNNNNNNNNNNNNNNNNNNNNNNNNNNNNNNNNNNNNNNNNNNNNNNNNNNNNNNNNNNNNNNNNNNNNNNNNNNNNNNNNNNNNNNNNNNNNNNNNNNNNNNNNNNNNNNNNNNNNNNNNNNNNNNNNNNNNNNNNNNNNNNNNNNNNNNNNNNNNNNNNNNNNNNNNNNNNNNNNNNNNNNNNNNNNNNNNNNNNNNNNNNNNNNNNNNNNNNNNNNNNNNNNNNNNNNNNNNNNNNNNNNNNNNNNNNNNNNNNNNNNNNNNNNNNNNNNNNNNNNNNNNNNNNNNNNNNNNNNNNNNNNNNNNNNNNNNNNNNNNNNNNNNNNNNNNNNNNNNNNNNNNNNNNNNNNNNNNNNNNNNNNNNNNNNNNNNNNNNNNNNNNNNNNNNNNNNNNNNNNNNNNNNNNNNNNNNNNNNNNNNNNNNNNNNNNNNNNNNNNNNNNNNNNNNNNNNNNNNNNNNNNNNNNNNNNNNNNNNNNNNNNNNNNNNNNNNNNNNNNNNNNNNNNNNNNNNNNNNNNNNNNNNNNNNNNNNNNNNNNNNTCTTGAATTTGTCTTTAAATGCATTGTCAGGTACAATTCCTAAGTCTTTGGAAAAACTCTCATACCTTAAAGGCATTAATGTTTCATTTAATGATTTAGAAGGTGAAATACCGAGTGGTGGCGTGTTTGCAAATTCCACTTTGCAATCATTTCTCGGGAACAAAGGTCTTTGCGGAATGCACATATTGGAGATTCCTGCTTGTGCTATCACTAATCTTGGAAAACAATCAAAGCCTAAGGAGGTTGTGCTAAAAATTGTTACTCCAGTGGCTATTGCATCCTTTATGATATTCTTGTTCGTTTCAATTTGGATAATGAAACAACACAAGAAAGGAAAGTCTAAAGATGTTGAAAAGGTACCGGAGATCGGGACTCATCAATTAGTTTCTTATCACGAGATTCAACGAGCAACAAATAATTTTGATGAATCCAATTTAATTGGTGAGGGAAGCACTGGCTCTGTGTACAAAGGCACATTATTTGGTGGAACTGCAATGGCCATAAAGGTTCTGGATTTGGAAAATGAGCAAGTATGCAAGAGGTTTGACACCGAATGTGAAGTGATGAGAAATGTTAGACACTGAAATCTTGTTTCAGTGATTACTACATGTTCTAGTGACTATATAAGAGCCTTTGTTCTGCAATTTATGCCCAATGGTAGTCTTGAGAATTGGCTGTACAAAGAAGATCGCCACTTGAACCTTCGTCAAAGAGTCACTGTAATGCTTGATACAGCTATGGCAGTTGAATATCTACATCATGGTAATGTCACTCCAATAGTTCATTGCGACCTAAAGCCAGGTAACGTTCTTTTGGATGAAGATATGGTGGCTTATGTTGGTGATTTTGGAATCTCTAAAATTTTAGCCATAAGCATGTCCATGGCTTATACCGAGACATTGGGCACTCTTGGATACATTGCACCAAGTATAAAAGAATCTATTCTCTTTGATTTTCTCTTATCATAATTAAGCCTCTCCAAATTCTACAAGTAAAAAGAGCAAGTCTTTATTTTTGCAGAATGATTGTTGTATTTCAATTGAGTAACTTTTCTTCAATCCTTTTCTAAGAATATGGCTCGGAGGGAATAGTGTCTACTAGTGGTGATGTTTATAGCTACGGCATCATGTTGATGGAGGTTTTGACCAAAAGAAGGCCAACAGATGAAGAAATATGCAATGAAAATCTTGACTTGAGGAAATGGATTACACTATCATTTGCAGGGAGTAGGGGCGGCTCAACATAATTTGTGGCCTAAAGCGAAATTTTTATTAGAGGCCTAAAATCTATATAAATTTTATCTATATCTATTTATTTTTCTAAATTTCTAGACATAAATTACTACTTAATATCTTTTATATAAATGATTTTTCAAACACTTTTTTTTTAATTATTAGTTTTAAGTAAGATTTTATCAATTCAACATTGAAAAACATCCTTTTATTGAGAAAATTATTATATGAATTATTAGCACAATTTTATAAGTAATAAGTTGATAAACTTTAAAGATAAGAGTTAGAATTATAGAATTTGATTCAAAAAGTTGATAACTTGGTATACCTACTTTAATATGAAAATTAACAAATAAATAAAAAAAAGAATTTGTAATGCACTTTGTTCAACACTTTTCGACTATTGATTCATATTTTTGAGAGTGTTTACTCTAACTATCAAAGATTTGAAGAAATAGAGTGCAAAAAGAGTTACAAATTAAAGGATAAATAATGTGAGTGTTAACGAAGAAAATGTAAGAATCCAAAACAATGTTTCCTTAATTTCTTCTATTTGAATGGGGTTAAGATACATAATTTTTTATTTATAAAAATTTGAGGCCCCCAAAAGGCCAATGCCTAAAATTTTATAGGCAAGAGCCGGCCCTGGCAGGGAGTATGATGGACGTTGTGGATGTTAGTCTTTTTTCTGAGGATGAACAAATCACCAATAAAAGTGAAATCTGCATAGCGTCCATGATACAATTGGGTTTAGACTGCACAGAGGAAACACCAGAATCAAGAGTGACCATGAAAGAAGTAGTGAAGAGGCTTAATAAAATCAAGAACGCATTTCTGGAAACATAGAAGTTGAGCAGCATCTCTTTTGATATGATGCTTTTTGTTTCTTCAATAAAGTTGTTGTATAGTACTACTTGAAGTCATGTATTGTTTGGTAATGTAACATCCGACAATTTGAAGTGGCTTTGAGGAAGCTTGAAATTTGGAAATAGTCATTTTTGGAAAAAAAATTTGAAATCTGGAAATTTGGCTAAGTGTGGAAATCAGTGAGTTTTTTGGCCAACTTTGAGCAGTCATAACTCTTAGCTCAGGGTGAGTTAGGAGTAGTTCCAGTTATGGTTGCGAAGCTCGTGGAATGACCTTTTCAACGCCACCAAATTTTCTCGATTCCGAGTTCATATGAAGGGAAATCCGTCCGGAAATTTAAGGGCATTTTGGTCTTTTCACAAATCATTTCTTTTGAGCTTATATTGTTGTATTAGGCTGATCTTTGGATCATTTTTGTACCCATTTTGAAAGAGAAAGAGTTAGGGTTCTTGAGAGTGAAGAAGAGAAGAAGAAGAGAAGAAGAGGAGAAAGGAGATCAACAAGTTTCCGTCGTGGATTATCGTCGGGGGTGATCCCTATCAAGGTATGTGAGTTTTCGTGTGGGTTGATCCTTTCCCTCACACACACCAAGCTTATTCTATGATTTGAGAAAAGATTGGAATTTGTTGTTGAAGTTATTGTTCGTTGTGAGACATGATTGGGTTGTGTATTTAAGTTGAAACCTATTGTATGTTGAGGGAATTATGATTCTAAGAGTTATGACCCTTTTGAAGTTGATTCATCATCAATTCTCAGCAAATTTTGGCTATCGTTCCCCATCTACGGGCGTGACCTACGGACCGTAGATCGATTGACGGTCCGTACTGGTCAACCGTCAATCGGGACAGAAGTTGGGTTTTTGGGGCATCGATCTACGGGCATGACCTACGGTCCGTAACCTGGACCTACGGACCGTAAGTCCGGACCGTAGGTCAACACTTAGTCATTTTAAAAAAAATGAATTCTGGGGAAGTTCTGTCACGATCTACGGACCTGCAGGACGGACCGTAAGTCCATCTACGGTCCGTCGATGGCGTCCGTAAGTGCCATCTATGTCACCAGAAATCTAAAGTCTCAGCCAAGTTTCCCCATTTCTTTTCCCACTTTCTCATGCATGTTCCAAAGTATTATACCTATATTTTACAGTTGTTTCCGACACTCCAAAGTACGTCTAAGAGTTCAAGAATCCCTCCATAACATATGATCGAACACCTTGAATTCATAATTCCGGTTCAAGGAAAGATAGTTCCAAGTCAAGTGAAGTTAGAAGCCAAGTAAGAGTCTCAAGTTTCAAGTAGAGTAAGAGTCTCAGTTTCAAGTTAAGTCATGAGTTTAAAGTTGAGTTCGTTTCTCAAAAGTTATTAGGGAACTATGTATTCCCAAAGAAGTTTTAAAAAAATGTCTTTACATTTAAATAAGGTTGGAAACTGAGATTTCCAAAGGGGCCTTCGGGCTAGTTTTTGAGTAATTATCTCAACTAAAGAAAAAATGTGTTTAAAGACAATTATGAGCCAAAGTATTTCTGGGAGTAGTATTGAGCACCGAATTGGGGACACGAGTTCTGATTTACTCAACTCTCCATAAGAACCATGTGCCAACATGGGATTTCTCGGATCATACTTTTTAGATGATCACGAAAGTTAAGCTAGTGGATCCACTAAGTAAAGTTAGCTTCCTATATCGCGGCAAGGTATAGGATGGTCCTTGGGCAACGCGAGGTAAATCGTTGTATCATCACTAGGGCTCATAGTGGTGGTTGTCGGTTAAAGAACCTCCCACATAAGTTATATTACTTTTATATAAGTACAGTTGAGTTGTTATTGTTTTATCATTAACAAGCTAAGTTGTTTATACTGTTTTACAAGCTTTATATATTGCATGCGTCTTATTGCTTTAAATATTGAGTTTAGTTAGTCATGAGTCGTACAGAGCCGAGGTAAATGTTCTCTTGCATTTCCTTTCAAGCTTAAGTTGTGTTTTGGCTTTCCCGCTCACATACTCGTACATTCCATGTACTGATGCCAGTTGGCCTGCATCGTTTGATGATGCAGATTCAGGTACCCCGGATCAGCATCCTGCACGTCGTTGATCCAGCTGAGCACTCCAGAGTCAGTGGTGAGCCTCCTTGCATTCCAGAGGACTCAATTATTTTGGTGTTCTTAGTTTTTGTCTTATTAGGATGTTGCGGGGTCTGTCCCAACATCCATCTCAGTATTTTAGAGGCTTCATAGACAGACAGTCAGTCAGCTAGTTTCGAGTCTCTTATCTATATATATATGTAAATATTCTATTTTGAGATTTGGGTTGCCTTATGGCCATATTTTATAAGTTAAGTTGTTTTTGTAATATTGCATTGCATTGAGTTATTCTGTTGAGTTAGGTTTCCGCTGAGTTAAGAAAGCCAGGCCGAGGGTTCGCTTTGGGCCAGAAATGGTCTCCGGGTGCCGGTCCCGCCCAGGGTGTAGGCTCGGGGTGTGACAGGTAAACTGTGTGGGAATAGACATGACACAGCCACGTTGGCGCTTTTGTCAATCAACAACAACGTCTCAATGTCAAACAAGTTTGGACATATTTTTGTTGATAGTTTACTTAAAATATTGACTACTATTTCATGACGAATTAAAATAAAACCTCTTAACAAGAATTCTTGTATCTATTATGAGTTAGTCGACTTTTATTCCATTTTGTGAAACACAACCGAAGTAGTGTCAACTGTCAAGGAATAAATGGATAAATTCCTTTTCTTATTTTCCTCATCCATATTAATCTGTTTTCACATTTTGGAAATTTGAGACTATTTATTTAAATTAATGAAGGTATTTAGCTGCAACTGTTTTGCTTTTTTTTTTGCAAACTCGGAAATCTCTCTTTTCGCCTCTCTCCTTCATTTGTATTTGCCTCAGAAGTTTTACACAGTCAATGTGTAGAGCCTAAACTCTAATGTGATTAAAAACATGAATAGATTAAGCACCTTATCCATTTTTTTTAATCTCTTCCGGCCCGACTTGGTTTAAAAAAAGTTTTCATATGACAATTTCCATTCATAAAAATAATTTTGTTATTTTGTTACTTTGCTTATTATTGACGAGTGTAATTCATGTATTAGAGTATGTTAATAGAACCAATTTAAATCAGGATAAGATGTTTAGATCGTCATAAGCAAAAGTTAAGGTTCACAATACGATATAGTATTATGAAACAATACGTAACAACAATAGTGAAATCAAGGTCTATTCTAGATTTTTTAAAAAAAAAAGAGGAAGAGAAAATACAGGTAAAAATAAGATTTCGTGTACTCAAAATTGATTTTGAGATACATAATTCTCTACATAAGATACTTGTGTTTAAGAGACATTATATTAAGTTAATGTATTTAGATGATCACTGGATACATAATTTTAGTATCAAACGGATAATCAAGTTAAAGTTTAAGTATCTATCGATGTATTTTAGCAAAAATAATGGCATTTAAACAATGAATGTTGCCCTTTTCATTTGAATTTGATTGAATATATTATATTTTGGAATAAAAAGATGAAAAATTTAAATAACATTTGGTACAAGATTTGTGTATCTTTGATTGTTGTAGCAAAATGAAATTCACGAGCTGAATGATTTGCTCCAGATTTTTCTTTGTATCACTAGGAAACAAAGATAGTGCTTCACAAACTAAATGCCCTTTTTGTTGGGTGTCTTTAATCTAAGAAGAAGATGTTACCCTTTTCTTTTGAGTTTTTTTATTTTTGTACTTCTGTCTTTGTTAAGAGCGTAAATCATCTATTTAAAAAAGGGATTGTTTACGTTGGAATAGTTACAAAAATTGTTCCCTCCCTTATTTTAAATTAGCATTTATCCCTTTTACAGATAGCATTGCACCAGGTATAAAAGATCTTCTCTCTTTTATTTTCTCTTATCATAGTAAAGCATTCTAGAAGTAAAAAAAGCAAGTCTTCATTTATGCAGAATTATTGTTGTATTTCAATTGAATAACTTTTCTTCAATCCTTTTCTAAGAATATGGCTCGGAGGGAAAGTGTCTTCTAGTGGTGATGTTTATAGCTATGACATCATGTTGATGAAAGTCTTGACGAAAAGAAGGCCAATAGATGAAGATATATGCAATGAAAATCTTGACTTGAGGAAATGGATAATACAATCATTTTCAGGGAGTATGATGGACGATGTTGTGGATGCCAATCTTTTTTCCAAGGAAGAACAAATCACTAGTAAAAGTGAAATCTGCATGGCCTCCATGATAGAATTGGGTTTAGATTTTACAAAGGAAATGCCATAA

>PGSC0003DMG400030443 LRR-RLKs

ATTTGATCCATCCCTAAATTTATATTCTTTCTTTCTCTCTTTATAAACTTTGTGTGAAGTTAATTAAATTTGGGCGAACAAGAGAGTACTAATTCAAATCGGCGAGCTTTGAAACCAGAACGCACACTACAACATGCTATCCCTGGGATTCTCACTTCCTCCTCCTTCCAATAATAAACTAATTGTTACCAACAAAAACCAATATAATTTTCGTACCAATTTAGCCAAGGTTTGTTCTAATAATGTTAATGCAGCTGGGGATCAATTAGTGACACTGCCTGAGGGCCTCAAGCATGTTGCAGTGATCATGGATGGTCATAGAAGGTGGGCAAAAAACAAAGGCTTGACCGTAAAGCAAGGTCACCGCGCTGGCGGAGAGAAAATCCAAGTCCTCACCCGCCTTTGTAGCCAGTGGGGAGTCAAAGTTCTCACCATTTTTGCATTCTCTACAGAAAATTGGGTTCGCCTCAAAGTAAGTCATTGTATTCCTTCCATCCCAATTTACATGACACAGTTTGAATTTCGAGAGTAAAATAGTATAATTTTGATTGTGATTCCGGGGAATCTCTAACTTTTGAAATAAAATTTACATATTTGAAAACTAGGTTAAAAATACTATAAGTCACCATAATTCATAATTCAAGATGTTTACATGAAAAAATCCGGTCAAATAAAGATTTATTTGAATCTCGAAATTTGAAATTAGGTGCACCATAAATTGGGGAATGAGGGAATAACTTTAATGACGATCTATCAAGCATGTTAAAATAAATCATCAAGAAAATAAATTAAATAAAGAATAACAATAACATATTCAGTGTAATTTTATAGGTGAGGATCTGAAAATGATATAATGTACGCTGAACTTACCCCCACCTCGAAAAGATAATAAAAGTTGTTTTTAAAGATCATTGACCCAAGTAACCTGTACAAAATCAACGAATATATATATATCAGAGAAAGTGACTTTCTTTTCTAACAATGAAATTAATTATACAAGAAATGAATCATATGTCTGATTATATTTAACTATATATAATTATAATTGCATATGATTTGTTTATTTAATTTCTAATACACCAATGTTTTGTTTTATTTGTTAACAACGCAACTCTAAACTTGAGTGCCGCAGGAGGAAGTTGATTTTCTTATGAAGTTGTTTTTAGAGCTGATAGGCTCACAAGAAAACTTGGATGAGTGGACAAGGTATTCTATATATGTGATAAATAAAAATATCTTGTGTGATCAGCTATCCGACTATCTATATAAGAGATCGTTAATTCCACCATTTTAATGCAAAATGGTTCCCGAAATTAATTCCCCATGGTTTAGTAAAAAAAAAGGTGGTATTTGATATTCATTTTGAAATTCAACTTATTCGAATTAAATTATATTATGTGATAGGGATGGTAGACGAGTATCATTTATTGGGGATAAATCAATCCTTTCCAAGTCTTTACAGAAAGCTCTTACTGTAATGGAGGAGAGAACAAAATGTAATTCTGGACTACATGTTATAATGGCAATCAATTACAGTGGACGACATGATATATTACAAGCAACCAAAAGCATTGCCAGTAAAGTGAAAAATGGACATTTAACTGTAAAAGACATTGATCAGAGCCTATTTGAACAAGAACTGGAAACTCATTGCGCCGAATTTCCTGAACCAGATTTGTTAATTAGAACAAGTGGAGAGAAAAGAGTTAGCAATTTCATGTTATGGCAATTAGCTTATACTGAATTCTATTTCGCGAATAAGTTATTTCCTGACATGGAAGAAGCTGATTTTATGGAGGCTTTAATGTCGTTTAAGTCGAGACAGAGACGTTATGGTGGACAAAATATCTAGTAATATTTTCAAGAACTTCCATTACTGAGAATTAAACCTAGCTCGAAATATGAAGAATTATTGATTAGATCTCTTGAATTTCCTCTTGCTATTAGTACTATTAAATAATTTTATGGTGGTCGATTGTATCTATACCAGCACTATATAAGCATGAAATATCATCATGCTAGTTATGTGGTTTGTGCTGAGTTTTTCTTTTCTTTTCCCATCGCCCCCATTAAATTTGTATCACACACCGAAGGACTCGTGGCACTCCAACAGGATATTCATATTTTCATAAAAAAAGAAGAAAAAAAAAGGAGGGGGACGGGGTGGGGACAACGCCGGAAAATGAGACTATAAAAAGTACTCAAAGTCATATGCTTCCAAGATGTCAATCACATCACCATCATTCATCATTTTGTTGGTTCTAAGTGATGAGTGGATATAAAATTTTCATTTTTTACATCACATTTGATTTTTGCTAGACTATAATATAGTAAATTAAATTATAAAATTGAGGCGAAAAATATAGTATATTAATTAACTATCAATAATAATATCATATATTAATTGAATTAGTATTAAATCGTAACTTGTCTTTTTAGCAAATCCTTAAATTTATTTTAGATGAGGGAAGTTCACTTTCTTAAGTCAAATGTCAACTCACTAGTCATTAATCATCTTATTATTCAAAGATCCAGTGAATTTCATTACAATTCCATGGAAAATAAAAAGATTTGACAAACAAATAAGAAATTATTTGGTGTTCTCCTTTCTCTTAAAGGGAGAGCCGTTTACCTATACAAGTCTATATTTGTCGCACTCTTAAAGTCATTGGCCATACTTTGACTCAAGGGGTGGTATCCTGTTTGATTGGTAAATCTAGAACCTTTTTATTATTATTATAGTCGTGGTATTCAGATCACCTTATAAGCACCTTGTTAATTTCACAAGATATGTTACCTTCCATCAACACAAGTACCGAGTAAGATTTGCTAAGAAAGTATCCTTTCCCTTCACATGATCTTCTTTTTTCAGTACAAGTTATATGCACTAATTGAGTCAAAATAACTACACAACCATGTCGCTTGTAAAATAATTCATGTAAATATGACAGTATATATTATTGATAACCCGAGAAAAATAATATCTAATCTGATATAATAGGTTAAATTACACTGAAAATGGTTGTAAAAAATTTGTACACTACCATTGTGTATAACTTTAAGCCCTTTTTCTCAAACACTTTGGTACCCCACATAAGGAATAATGAACCCATTATAATTAGCAGAATCAATTGTGACTCAGTATTATCCTCTTTGTCCACATATCACCTAACCTTCCCAATCCCCAAATGGAGCTTCAAACTACTTTTGTTCATTCAAGCTCTCTTTTTTTTCTGTCATTCCATGCTATTCTCTTCATGTTCCTATACTTATCTTTTCCAGAGTGTGCTTCAGCAGGCATACTTGGGAATCAAACAGACAAACTTGCATTACTTGATCTTAAGTCCCAAATAACTGAAGATCCACAAGGACTCATGGATTCTTGGAATGCCACTCTCAATGTCTGCCAATGGCCTGGTGTTACATGTGGCCATAAACATCAAAGGGTCATTTCTTTAGATCTCAAAGACCATAGATTGGCTGGCACTATATCACCTAGCATTGGCAATCTTTCATTTCTTCGAATTCTTGACATCTCGGATAACTCCTTTCATGGTGTAATCCCTCCAGAGCTTGGCCAATTGATCAGACTTCAAACCATGAACTTAAGCTTCAACTTCTTGAGAGGTGAAATCCCGCTTACCCTATCTCGTTGCGTTAACGTTGTTAACCTTATTCTAGACCATAATATCCTTGAAGGACACATTCCTACTGAACTTGGTTCATTGACAAAGCTGGAAATGTTGTACCTCAAGAACAACAATCTCACAGGAAATGTTCCGAATTTTGTTGGAAATCTTACATCTCTAAGAGAGCTTTACATTTCCTACAATGACTTGGAAGGTGAATTACCTGAAACAATGGCCAATATGCGAAGCTTGATTGAGTTAGGTGTTTCAGTGAACTCTCTGTCCGGAGAATTCCCCCCTGCACTCTACAATCTTTCATCTCTTACACTTTTAAGCTTGTCATTTAACAAGTTCAGAGGCAGACTTAGACCAGATATTGGCCTTGCCTTCCCAAATCTTCAGAGACTGTATTTGGCCAACAATTACTTCACAGGTTCAATACCAGCTTCATTGTCCAATTGTTCAGATCTTCTGCGGCTTGATATTCCTCTAAACAATTTCACAGGAAATATACCATTGAGTTTTGGAAACCTCAAGAATCTCTTATGGCTTAATGTTAACGATAACCAGCTTGGAGTTGGTGCACCTGATGATCTAAATTTCATAAATTCTTTAACCAATTGCAAAATGCTTGAGTTCCTGGACATTGCAAACAACAAATTTGGAGGTATGTTGCCTTACTCTATAACTAATCTCTCGACAACTTTGACAAAGCTACTGATTGGATACAACAGGATAAGTGGAACCATACCTAGAGAGATCTCCAACCTTGTCAACTTGGATATGCTTAGCATCCAAGGAACTCTTATCAATGGTAGCATTCCTGATTCTATTGGAATGTTATCAAACTTGAAAAATCTGCACATGGAATCGAATCAGTTGACAGGAAATATCCCCTCTTCCTTAGGAAACATTAGAGGATTGCTGTACATTTATTTGCAAGATAATAGCTTGGAGGGAACCATACCTTCAAGTTTAGGAAATTGTACATCTCTCCAAACCTTAGACATCGCTCAGAATAAACTCAGCGGTAGTATACCCAAACAAGTCATTGCACTCTCCTCCCTTTCGGTACTCCTTAATATGTCTTACAACTCTTTGAGTGGTCCCTTGCCTGTTGAAATAGGTAACCTGACCAATCTTGCTGCACTTGACATATCGAACAACAAACTGTCTGGTGAAATTCCACACTCACTGGAGAGTTGTTCATCCCTTGAAATACTCTACCTGCAAGAAAACATCTTTGAAGGAACGATTCCTCCTCTGGATGATTTGAAAAACATTCAGTATCTTGATCTTTCACGTAATACCTTGTCAGGCAACATTCCTAGGAGTATCACCAAGCATGTTTCATTACAGAACCTCAATTTGTCATTTAACCACCTTGACGGAGAGGTGCCAGTTCAAGGTGTTTTTTCTGATGCAAGTAGAATTCAAGTGATGGGAAATATGAATCTTTGTGGGGGAATTGAAGAGCTGCATTTGCATCCATGTCTCAAGCATGCAAATAAAAGACCCCAGAAGCACATTGCCCTCATTTTGGTGCTGGCACTGGGTACTAGTGCTGCATGTTTAACTTTGTTGCTGCTAGTTTCGTATTGCTGCGTGAAAAAGGGGAAGCATAGGCCTAGCACAGCATCTTCATTCAGGAAAGGATACACCCAAGTTTCCTATGAAGAACTCTTAAATGCAACTGGTGGGTTCTCTTCAAATAACTTGATAGGATCAGGAAGTTTTGGCTCCGTTTACAGAGGAAATCTTTCTCCAGAAGGAACAATTGCAGTAAAGGTACTAAAACTTGAAAAAAAGGGTGCTTCGAAAAGCTTTTTGGCTGAATGTGAAGCCTTGAGGAACATAAGGCACCGGAACCTGGTGAAGATCTCAACTGTTTGCTCAAGTGTTGATTTTGATGGCAATGATTTCAAAGCTCTCATTTATCCATTTATGGAAAATGGGAGCCTAGACGAGTGGTTGCACCCGAAGGAGGGACAGATGCTGCAGAAGAGATTGAGTATCTTACACAGATTAAACATCACAATAGATGTGGCTTCAGCACTGCATTATCTTCACAGTCAATGCCATACATCTATAGTTCACTGTGATCTTAAGCCAAGCAATGTTCTACTGGACAATGATCTAACTGCTCTTGTTAGCGACTTTGGTTTGGCAAAGTTCCTGTCTGACTCGGGTCAAAATGCCGATGTAAATCAGTTCAGCACAAGTGGGATTAAAGGCACAGTAGGTTATGCTGCTCCAGGTAAATATTATAGTGGGGATCTTAGTCTAAATATTGTCTGAACTCTGAAATCCATTTTTTTTCTCCATAAGACCATAACATAATAACAATTTAAGCAATTACATACTTCCTTGAGAGACTCCGGATGCTCCTTGTCCTAAGCCCCAGGCATCCGTTACCTTTGCAGATCACTACCTCATGATTTTGTGCTATTGATTCTTATAACCAGAGGCAGAGGCAGCTTTAGTTTAATGGATTCAACCAAATCTAGTATTTTCGACACTGAATATGGATATATAAGTTTAAAAAACTACTTAAATTACAATAAATATCATATTTTGAACCCATAATTTCAGAAGTACAATGAGTTCAGTTCTAAGAATCTTAGACGTTGAACCCATCAAATTTAAATCCTGGATGTCTCTTCTCTCAAATATAATTCCTGACTAGTCATAGTTTTTCTAAGGACCTTAATAAGAGGTTTCAGCTTTCTAGGGCATTTTACATAACATTCTCTATTGGAAAATCGATACAAGTAATCTGTGTTGCTTAGGCATTTTTGACAAATGTTTAGGATCCCTAATATTACTATGAAACACCTTCTCTGATACATCAAATTTTTCAGAATATGGCATGGGTGGTCAGGTATCATGTCAAGGAGATGTCTATAGCTTTGGGATTCTTCTATTAGAGATTTTCACTGGACGAAGACCAACCAGTGAACTATTTGAGGACAATGAAACTCTTCACAGTTTTGTTAAGCAGGCATTGCCCGGTCAAGTGATGGATGTTGTGGATCAATCTGCTTTGTACGAGACAGAACCAGGGGATCTTATGGATATACTCAGCTGCAGAAGTGACTTCTCGGATGAATTTGTTGAATGCTTAGTTTCTATTCTTACTGCTGGAGTTTCTTGTTCAGAGGAAACTCCACACGCCAGAATAAGCATGGGTCAAGTTATTTTGGATCTCATTTCCATCAGAAACAAATTGAGCAGAATTTTAGTTCATTCAGAGAAAGTGAAAATCTCAAGAAAAGGTGAATCATAA

>PGSC0003DMG400030474 LRR-RLKs

AAAAAACAGATTCATTACAAAAAAGCTGAAATTCCTTTTACCATTTCTCCACCTCTCACTCTTCACTTTGTACTTGGCACTAAACACTCCTCCTTCTTCTCCATTAACACAGATCCAAATCTCTTACTACTTCAGCTCAAGCTAAACGGGTCGGGTCGGGTCTTATTCTGCGCAAAATGTGGTTACTGCAACTCGATGAATTCGATTTGATGTTCATGTTTACTGTTAATTTCACCTCAGAACCTCAGTATTACTCCTAGAACTTCAAGTGCTTTCTCTATTTTATAGGTAATTACACTGCGATTTCATATTGTTTGTTGAAATTTATAATTTTGTATTTGTTTAGCTTGATTTCTTTTGTGTTTTCTGCTAATTTTGGTGGCAGATCTGTTTGCTTATTTGTTTCCTATTTGTTTGGCATTGTTAGATTGAACAGTCAATTTTTATTTATTTTGAAATATGATTTTTGAAGGTTTTCTGATAGTATTTTTTTTTTTTGTAGCTTTAAATTTGATTTTTAAAAATTAATTAGGAAATCCGTGATGAATTCATGCAGAAAATTAGACTCATTGGAGGGAGGAGTATTAAGTAGCGGCGGAGCTAGGATTTTCACTAAAGTGATTGAAAAATAAAGAAAGAAATATAATGATAAACTATGTATGATTTTTTTCTATTTCTTTTTTTTTGAACTTGTATATATGGTATGATTTTCCAGCTTTCACCCCTGGCTCCGGCGGTGGACTATTAAGTAGCAAGACCATGTTAGGTTCTGTATGTTGTTCAGTCTTCAGTGTTTGCTTCTTTGATAATGTGGAAGTACAATGAATTGAGAAATTGAAAATGTTGTTAGTCATTATGAACTATCATTGGATTATATTTGTGTAAAGGAAGCTTAGGATTCCCTATTACTGGGACCTCATATGAGTTGGGGTTGTTCAATAGTCTTAATTAACCCAACTCTCTTTGACATTTACTTGAAGAAGAATTATGAATGTGGTAATAGGTAATCCCCAACCCCCAAAATAGTAATAATAATAATAATAAAAATCCCATGATCTGGTAGATTTTTTGATGTAGTTCTTCTTCATCAACTGTTTCAAGCTCCAACCTTGAATAAGAATTATCATGACATCAGGATTGAAAAACTTGTAATTAAATGGGTTAGTCTAGTCCCGGGTTGACTTGATGGAGTGGTTATTTTCCTTCCCGTTGCTTATTAGATACTATCATGTTAGAATACTTAAGTGCTAGCTATTTGTATGTTTTTTTTTTTTTCTCAAAACATCTCTTGTTTATTAATAATATAATTACAAAAGAGGGTGACCTAAGGCCTAACACCATTTCTCACACGTAACACCAAAAAAAAAGAGTTTCCATCCCGGAAGGTTTCTTCTTCAGTTAGAGTTTTTACCAAAACCGAGGGTGGCTCATAATGCCAACCAGAATGGTTTTCTTTTTTTGAACCTAAAGGAGGCAAGGTTAAGTTTATCCATCCTCAGAAGGCTTATTTGTATGTATTAGCTCTTGCTCTTCTACTTCAAGACTCTTTTGTTCATGAAATTGAGAATGGAATATTTGTGAACCAAAGTTTTTTCTGTTAAATGTTTCTTCTTCTACTATATTCTTTTCTAATCTCGCTAATTCTTCTGTTTTGAGCTGTTTAATATTGTACATGGTTTGGTATATCTTAGCATCTTTTTATTTTATTTTGGTGTAATCATTCATCCATCTGTAACATTTTCGCCCACTTCATTGTTAAGGTTTTCGTTATTTTTGGCTTAGAAAGTGATATGGCAAGGTTTCTGGTTTTTGTTTCCAGCATCTCTTTTGCATTTTGTTTATTTCTTGGCATCAGTATCTATGAGATTGAATAGCTTTTGAAGTAATTATAAGTTGTTAGTATGGTGGATATGATTTTTAACCAAGTCAGCAGAGCTTTCATCATACTATGTTATTTGACTGACTGTTTTTTTCTTTTTTTTATTCTTCTTTTTATTCCATGAGAACATGATTAGTTTTATTAATCTTCTTCTGTAGATGGTGAAAAATTAGTTAAATGTAGAACAATTACTATTTGGCATTCATAAGAAATTGTATAAGTCTAGTTCATTGCTAAGTGCTTTTGGAACGAATGTGCAAAGAGAAACATTAGTTGGACACATAATAATTGTTAATTGGAATCCATAAAAAGTTATCCAGTTTTAGTTCATGAAAATTTAGTGCAGATTCCAAACATATTTCTTGAGGATATTCTTATATCTTTGACTGCATAGATGCTTGCTATGTCAAAATGTCTTTGTTTATCCTTCCTCTCTCACCTTTGGTGTTTGTTTGGTTCTCAAAATGTTTATCTACTGAATTATTCCAGGACGACCACAATACTGGCAGCATAGTGGCTTTCAATCTGTTGAAGGTTTCCTCATGAGAATCTACCACACCAATCAACTGCTACATTCTCTTTTTGTTGTAGCAACTCTTCTTTCGCTGTGCCGTGGTGATCTAAACTCAGACAAACAAGCCCTTCTTGACTTTGCTGCAGCAGTCCCACATGGCTGGAAGATCAACTGGGATCCTGCCACTCCTATTTGCTCATCTTGGGTAGGCGTTACTTGTAGTAATGATGGAGCCCATGTGGTGGAAGTACGACTTCCAGGAGTTGGACTCATTGGTTCACTTCCAAAAAATACTTTTGGGAAGATGGATAGCTTAAGGATTATCAGCCTTCGATCCAATCGTCTCAGTGGAAACCTTCCTTCCGCTATCACTTCACTTCCTTCTCTCCAATTCCTGTTCTTACAGCAAAATAACTTATCTGGACCTATTCCTTCTTCTCTTTCAAATAAACTGAATGTTTTGGACCTATCTCATAATTTCTTCACTGGAAAGATACCTGAAACAATTCAGAATTTGTCTCAGCTAACAGGTTTAAGCCTCCAGAATAACTTGCTTTCTGGACCCATACCCAATATTACTCTTCCAAGGCTAAAACATATTAATATAAGCCACAACAATCTTAATGGTTCTATTCCATTATCCCTCCAAAAGTTTCCAAATTCCTCCTTTGTGGGGAACTCACTCTTGTGTGGACTACCCTTAAATCCTTGTTCACCTGTTCTACCACCATCCCCTCCTTCCCCTTCCCCTTCCCCACCTGCACCCTCCCCAAACACTCAAGAAAAAAGTTCAAAAAAGAAACTTAAACTAGGAGTTATTCTTGCTATTGCTGCTGGAGGGGTTGTGCTGGTCTTGCTGCTAGTTGTGATCCTCGTTTTGTGCTGCCTGAAAAGAAAAAGGGGTGATGGCAGAGGTGTGCGGAAAGGGAAATCTCCGGGTGGAGGAAGGAGTGAAAAACCGAGGGAAGAGTTCGGTAGTGGGGTGCAAGAACCTGACAAAAATAAGCTTGTTTTCTTTGAAGGCAGCTCTTACAATTTCGATCTCGAGGACCTATTGAGGGCCTCAGCTGAAGTGCTAGGAAAGGGGAGTTTTGGAACTGCTTATAAGGCTATTTTGGAGGAGTCATCAACAGTGGTGGTTAAAAGACTAAAAGAAGTCATTGTGGGGAAGAAGGACTTTGAACAGCAGATGGAAATTATTGGTAGAGTTGGACAGCATCCAAATGTGGTTCCTTTACGTGCTTATTATTACTCCAAGGATGAGAAGTTACTGGTGTATGGTTATTTCTCAATGGGCAACTTGTCAATGCTCTTGCATGGTAAGGCATCATATGCTCTCATCGTCATTTCCTTTTTCTATTTCTAGCGTTTGGTGGAGTAACTTGATTCACTCTTTGTGATTGAATGCCTGATAGTAAGCTTCTAACACAATACAAAATGCACTCAAGTAATGTCCTAAACGAATCTAAACAATGTGAGTTTGAACTTTTAAACTTGTGAACTGGTGTGAAAAGAAGCATTCTTTCTTGTAGCTTATTGTAAAGTTAGAGAAGAAAAAGTCTTCTTTAGAATATATGCATCAGTTTGATCTTTATAGGTCTATTATATGAGTATAAACAGCTCTTATGTTATTCTGTATCTATTGTTTCTGTAGCAACATATTGACACTTGCTGGCTTTCTTTTTCTTATCACTGGTAGTTTATTGCTCGCACTATCTTGCTGTTTGTCTAACTGGAAAGCTGCTAGTGCTGAAGAATTTCATGTATGATATACCTATTATCTATCCTGGAAGACTACACCAGAAGTGAATTCATCTTCATTCACTAACCTGCATAAACTTATCAATGTTAAGTATACTGCCAATGATGTTACGTACTGCTGATATTTCTGTTCTCTAACCCTATCAAACCCCACTATCATTTAGGAAACAGGACTCCGGGAAGGACTCCATTGGACTGGGAGTCGAGAGTTAAAATATCTCTTGCAGCAGCAAAGGGTATTGCCCACATCCATTCAATGGGTGGTCCAAAATTTACTCATGGAAATATAAAGTCCTCAAACGTCCTACTCAAGCAAGATATGGAAGCTTGCATCTCAGATTTTGGTTTAGCCCCTATAATGAACTTCCCTGCAGCCCCATCTCGATATCCAGGGTACCGTGCTCCTGAGGTGATAGAAAGCCGAAAGCATAGTCATAAATCAGATGTCTACAGCTTCGGTGTCTTACTACTCGAGATGCTTACTGGTAAGCAGCCTATGCAATCACCAGGGCGTGATGACATGGTTGACCTTCCTAGGTGGGTCCAATCAGTTGTCAGGGAGGAATGGACGTCAGAGGTTTTTGACGTCGATTTGATGAGATTTCAAAACATTGAAGAAGAGATGGTACAAATGCTGCAAATTGCTATGGCATGTGTTGTAAAGGTCCCGGACATGCGACCCAACATGGAGGAAGTTGTTAGAATGATTGAAGACGTCAGGCAATCTAACTCCGAAAACAGGCATTCTTCAGAAAGAGAATAGATCCTATGATTCAAATGTGCATATTCCTTGGATTTCTGACTTGCAATTTGTAAGTCCTGTGAAATTCATTCGATTCGATTACCCAGTGCAACAACATACATGTTTTCTTTCCCTAAATTTTCTGATTCAATCCTATTGACAAGTGTATCTAGGAAAGTGTGGATTTGTTTTCCTTAGGAGATATAGTAGACAAGTTAATCATCTTCCTTTTTCTTTTCGTTTAAGAACTCTTGTCATTTGTTGTAAGATTCAATCTGCACTGCACTTATTAACATCTGATTTATGCAACATGTCGAAGCAATTCATTTGCTGTGTGTTTGGATTTTGGTGTGAAATCATTTTTATGGCATTCAAACACCAAGTAATT

>PGSC0003DMG400030572 LRR-RLKs

CTATTTTCCCAACGTCTCTTTATTTTCTTAGCTGAAATCAACTTTACCTTTTCTCACTCTCAAAAGTCTCACCTTTCACACTCTTTTTTCTCTCTTAAGCTCCATTCTTGGTTATCTGGTGTACAAAAGGTAAGGTCTTTGAGACACCCTTTTGCTCATTTTTCATTTTTCAGCTGTCACTGTTCTTGATTTCATCATATTTGTAGATAAAAATGCGATCTTTTTTGTTTTTTTGCATTTTCTTGATTCTGTGTAGTTGTGTTTTTGTTACTTCTTTGAATGATGAAGGGATAGCTCTTTGGTCATTCAAAAAAGGTATTGGACAAGACCCTGAAGGGTCACTTAAAAATTGGAACTTTTCTGATGAAACCCCATGTTCTTGGAATGGTGTTACATGCAAAGACCTAAAAGTTGTATCTGTTAGTATACCTAAAAAGAAACTCACTGGGTTTCTTTCTTCTAGTCTTGGATCTCTTACTGGGCTTAGACATGTTAATTTGAGAAGTAATTTGTTTTCTGGTAGTTTGCCTGTTGAGCTTTTTGAGGTTCAAGGTTTGCAAAGTTTGGTTCTTTATGGGAATTCCTTTTCTGGGGTTATACCTATTGAGGTTGGGAAGCTAAATTATCTTCAAACTTTGGATTTATCACAAAATTTCTTGAATGGGTCTGTGCCTATAACATTACTTCAGTGTAAGAGATTGAAGTTTCTTTATTTAAGTCATAATAACTTTACTGGTGTTGTACCAGAGGGGTTTGGGGGTAATTTGTCTGCATTGGAAGAACTTGATCTTGGATTCAACAAATTTGATGGTAAAATTCCAAGTGATTTGGGAAATTTGTCTAATTTGCAAGGAACTGTTGATTTGTCTCATAATATGTTTAATGGTTCGATTCCGGCTAGTCTTGGTAATCTTCCTGAGAAGGTGTATATTGATCTGACTTATAACAATTTGAGTGGTCCAATTCCACAAAATGGTGCTCTGATTAACAGAGGACCTACTGCTTTTATTGGAAATCCTGGGCTTTGTGGACCTCCGTTGAAAAACCAGTGTTCTGCACAAAGTGATGCAAGTTCACCGTCCTCGGAACCTTTCTTGCCTAATAACGTACCTCCCTTAGATGGTGCTGGAGGGGATGGGAATGGTAGAGGTCTGAGTAGAGGTGCTGTAATTGCGATAATCGTGGGGGATGTGGTTGGAATTTGTGTTATTGGGTTGTTATTCTCGTATTGTTATTCGAGAATCTGTGGTTGTGGGAGGAAAAAGGATGAATCCGGATTTGGTTTTCAGAAGGGGGGAGGGAAAGGAAGGAAAGAGTGTTTATGTTTTAGGAAGGATGAGTCGGAGACGTTATCGGAAAATGTTGAACAATATGATCTAGTGGCACTAGATAATCAAGTGGCATTTGATCTTGATGAACTTCTCAAGGCATCTGCTTTTGTCCTTGGTAAAAGTGGTATCGGCATTGTGTACAAAGTTGTGCTTGAAGACGGACTTAACTTGGCTGTGAGGAGATTAGGCGAGGGTGGTTCACAGAGATTTAAGGAATTCCAGACGGAGGTCGAAGCAATTGGAAAACTGAGACATCAAAATATTGTGACTCTTCGTGCCTATTATTGGTCTGTCGACGAGAAGCTACTGATATATGACTTCATTCCAAATGGAAACCTTACCACCGCAATTCACGGTATGCTTCAAACTTAGGCTGTTGCATTCTCGTTTATCGTTTTTTCTTTGATTTCCACAACTGTTTTAAGCTTTCGTGATTATTAAGTTATTAACTTGTCTTTTTGCTTGTGTAATCAACTCTTAGGAAAACCTGGTATGGTGTCATTTACACCGCTTTCGTGGTCCATTCGACTGAAAATAATGAAGGGAACTGCAAAAGGTTTGGTTTATTTACATGAGTACAGTCCCAAGAAATATGTTCATGGTGATCTAAAGCCATCTAATATATTACTTGGGCACGACATGGAGCCCAAAATCTCCGACTTTGGACTTGGACGTCTTGCTAATATAGCCGGAGCATCTCCTACGTTGCAATCCAACCACATGACCTCTGACAAACCACAACAAAGTAAGCAAGGCAGTGCACTATCAGAGTCCGGGACAGTTACATCAACTACAACATCTGGTTCTTGCTACCAAGCTCCCGAGGCATTGAAAGTGGTGAAACCATCTCAAAAATGGGACATTTACTCGTATGGAGTGATCTTGCTTGAAATGATCACGGGAAGAACCCCGATTATCCAAGTAGGTTCTACAGAAATGGATCTAGTAAACTGGATTCACTGGTGCATTGAAGAGAAGAAGCCACTTTCGGATGTATTGGACTCATATTTAGCTCAAGATGCTGATAAAGAAGAGGAAATGATTGCGGTTTTGAAGATTGCAATGGCGTGTGTTCACAGCAGCCCGGAGAGAAGACCCTCAATGAGATACATATCTGATGCATTAGAAAGACTGCAAGCTTCCTCTGAATGAACGATCGAAACAGAACCTGAAGGCTACACATTGCAGAGTAAGGATCACAGCATCTAACCGATAACTATGGAGCACGATCACTGCTTGGCGCGCGATCATATTGACTAACAGTAGTAATCTCCACTTGTCTGTTTGAATATTGGCCTTCTTGCTCTCTAATTGCAAGGTTTTTGTTGTGTCTATTTGTTGTTTCCTTCTATTGATGAACATAAGTTATTTGTTCTCTTTATTACTACTACTATTCATCTTTGTGTGTCACAAATGAGTTTATGTAAATAAAAATGATTGTCAAAATAATAAATAAGATAGGTGTAAGCAATAACTATGTCTGTTGTGAGTCTGCAAG

>PGSC0003DMG400030625 LRR-RLKs

ATTTACTCAAAACAAAAGAATATACACCAAACGGATTACCCACCCCCTCCTCACTTTTGCCTTTCTCACTCTCACTGAGTGAAACCGCAAACCAAACAGTTGGTGGGCATTAGATTAAGGAAGGAAAAATGCGTCTTCTTTTTCTTCTTCTTCTTGTTATGCATTTTACCGACTTTTCTGCCGGTAAACAACCACGGTTACCGGAATACCAGGCTTTGCTTGCCCTGAAAACTGCCATTACCGATGACCCACAGTTAACACTTGCCTCATGGAACATCTCCACCAGTCACTGTACGTGGAATGGTGTCACGTGCGACACGCATCGTCACGTGACCTCTCTTGATATTTCTGGGTTTAATCTTACCGGTACTCTTCCGCCGGAAGTTGGGAATCTTCGTTTCTTACAAAATCTGTCTGTTGCTGTTAACCAGTTTACTGGACCCATTCCTGTTGAAATCTCCTTTATTCCAAATCTCGGTTACCTTAATCTTTCTAATAACATATTCGGGATGGAATTCCCTCCGCAGTTAACCCGTCTGCGTAACCTCCAAGTCCTTGACCTTTACAACAACAATATGACCGGTGAACTTCCCCTTGAGGTGTATCAGATGACTAACCTTCGACATCTACACCTCGGCGGGAACTTTTTCGGTGGCCGCATTCCTCCGGAGTATGGAAGGTTCCCGTCTCTAGAGTACCTCGCAGTTTCAGGCAATGCACTGGTAGGAGAGATACCACCGGAGATTGGAAACATCACTACACTTCAGCAGTTGTATGTAGGATACTACAATACCTTCACCGGTGGGATTCCCCCGGCAATAGGGAACTTATCGCAGCTCCTCCGGTTTGATGCTGCTAACTGTGGACTTTCGGGGGAGATTCCACCGGAGATTGGGAAGCTTCAGAACCTTGACACTCTCTTCCTGCAAGTGAATTCTCTGTCTGGGTCATTAACTCCGGAGATAGGTTATCTGAAGAGCTTGAAATCTTTGGATCTGTCGAATAACATGTTCTCTGGCGAGATACCGCCAACATTTGCGGAGCTTAAGAATATCACTCTTGTTAATCTTTTTCGGAATAAGCTTTATGGGTCAATACCAGAGTTCATAGAGGACTTGCCGGAGCTAGAGGTGTTGCAACTTTGGGAAAATAACTTTACGGGAAGCATTCCACAGGGGTTAGGCACAAAGAGCAAGCTCAAAAATGTTGATCTCAGTTCCAATAAATTGACTGGAAATTTACCCCCAAACATGTGTTCCGGTAACAATCTGCAGACAATTATCACTCTAGGGAACTTCTTGTTTGGCCCAATTCCTGAATCTTTGGGTAGGTGTGAATCACTTAATCGGATTAGAATGGGAGAGAATTATCTGAATGGGTCAATTCCAAAGGGGCTCTTAAGCTTGCCACGTCTGTCACAAGTTGAACTTCAGAATAATATTCTCACTGGTACATTTCCTGATATTTCTTCCAAATCTAATAGTCTTGGGCAGATTATCCTTTCAAATAATCGCCTAACTGGACCTTTGCCGCCAAGCATTGGAAACTTTGCTGTAGCCCAAAAATTGCTTCTTGATGGGAACAAATTTTCGGGACGAATTCCAGCAGAAATAGGAAAGCTTCAACAGCTATCCAAAATTGATTTCAGTCACAACAACTTTTCTGGACCCATGGCTCCGGAGATTAGCCAGTGCAAGTTGCTGACTTATGTTGATCTCAGCAGGAACCAACTTTCGGGTGAGATTCCTTCTGAGATCACAGGTATGAGGATACTCAACTACTTGAACTTATCGAGAAACCACTTAGTTGGGAGTATTCCTTCCCCTATTTCTAGTATGCAGAGTTTAACTTCTGTTGATTTCTCATATAACAACTTTTCTGGTTTAGTTCCTGGAACCGGGCAATTTAGTTATTTCAACTACACCTCATTTCTGGGCAATCCAGATCTTTGCGGACCCTATTTGGGCCCTTGCAAAGAGGGTGTTGTTGATGGGGTTAGTCAACCTCATCAACGAGGAGCCTTATCGCCTTCGATGAAGCTTTTACTTGTTATTGGTTTGCTTGTCTGTTCTATTGTGTTTGCTGTTGCTGCAATTATAAAGGCCCGATCTTTAAAGAAGGCAAGTGAAGCTCGTGCCTGGAAGCTCACTGCTTTTCAGCGCCTAGATTTTACTTGTGATGATATTTTGGACAGCTTGAAGGAGGATAACATTATTGGAAAAGGAGGTGCTGGTATAGTCTACAAGGGGGTAATGCCGAGCGGGGAACATGTAGCAGTTAAGAGGTTGCCAGCTATGAGCAGGGGTTCCTCTCATGATCATGGGTTCAATGCAGAGATACAGACTCTTGGGAGGATCCGACACAGGCACATTGTTAGATTATTAGGATTTTGCTCGAATCATGAGACAAATCTTTTGGTTTATGAGTACATGCCTAATGGAAGTCTTGGGGAAATGCTTCATGGCAAGAAAGGCGGTCATCTACATTGGGATACCAGGTATAAGATAGCCGTGGAGTCTGCAAAGGGTCTTTGCTATCTCCATCACGATTGCTCTCCTTTGATCCTCCATCGTGATGTGAAATCAAACAACATTCTGCTAGACTCCAGCTTTGAAGCTCATGTTGCTGATTTTGGACTTGCTAAATTCTTGCAAGATTCAGGGACATCAGAATGCATGTCTGCTATTGCTGGTTCTTATGGGTACATTGCTCCAGGTATTTTGTAAAAGCCTTCGTATCTTGAAATTCTTAGTTTCTAACTGTGTTCATATTTATTTTTCTTGTCACATCATGCTATGCGAGTTTAGTGCCGAAGTTACAAACTTACTGATGAACTAAAATTTATCAGCTGAGACTAATTATCAATCTGTTAAAATGAAGCTTAGGTGTTCTAAACATCATAAATGTGCTATCCTGTAGTCTTGTATTTCCACTACTAATAATGTGACGATGAAGGACAGCATTAAGACATGGATAGTGGGGGTCCTTGACTGGTAGTAAACTAGAGGATTGGGCCTTCTTACTTTTGAAAAGGATCTCTAACTACTTCGGTGATGGTTAGGTTGGAATTTATAAATGCTTATGTCACTTGATAAGTAAGGCAACTGTGTTGCATGCTCCTGAAGCAAAGCAACATTAAATAAGTGGTACTGGTACTGATGAACTATTTACACTGTTGGAACGGGTAAGGGGATGTAAAACTCTGGTTTCTTGTTTAGTTATCTAACAAACCTACTGATTGCTAGCAATTATTTGTTCTGTTCTATCGTCTACTTCTCATTAGCATCTTATTTTCGTTTTTGCAGAATATGCTTACACGCTTAAGGTTGATGAGAAAAGTGATGTATATAGCTTCGGTGTGGTGCTATTAGAACTGGTAAGTGGCAAAAAGCCAGTTGGAGAATTTGGTGATGGTGTTGACATAGTCCAATGGGTTAGGAAAATGACTGATGGGAAAAAGGATGGAGTTCTCAAGATCCTTGACCCAAGACTCTCAACGGTTCCCCTTAATGAGGTGATGCATGTCTTCTATGTCGCATTGTTGTGTGTTGAAGAGCAGGCTGTGGAACGCCCCACCATGCGAGAGGTAGTGCAAATACTAACTGAACTTCCCAAGTCACCAGGCGCAAAATCAGATGACTCAACCGTCACTGATCAGTCACCCCCACCATCAGACTCTGCATTAGAGTCCCCAACCTCAATTCCCGAGGAAACAAAAGACCACCATCAACCAACACCTCAATCACCTCCACCTGACCTACTCAGTATCTGATTTGCAATGTTCTTGAAATAGGAGTGTTTTATTTAGTTTGATTCTCTAGTTCTATTATGATGAATTGTGCTAGCTTTGTTCTTTTTTTAAAAAAAATCGGGTCTTTTTAGGCTCAGGGGTTTATTCTAACTCTAAGATGGGTTTAATGCTCAGAAGTTTTCCTCTTGTACAGTAAGATTGGTAGGGTTTTCTCAAGTGTATTAATGGAAAAATTGCCCTTCATTTGCTA

>PGSC0003DMG400030626 LRR-RLKs

TTTTATGCCATATAATATTTTGAGTCCAAAAACCTTATCATTACTCTCTGCTTCAAGTTACTGCAACTCTTGTCTTGTTCTTCGCCATTAGAGTCCAATACTAGCTTTTTTTTCTTCTAATGTGTGAACACTCTTCACGCTTTCAGTACTATTTCTTCTCCTTAAACTCTGCTTCCGTTACACTTGTAATCCATGCAAAAATTCCCCCCACCTCTTTCTAACTCTTACCTTCCCAACATTCAACCTCCATTTACAGATGTTTATGATTAATCGGTTCTTGCCTTTTTAAGGACTTGGAGGTAGGGGAGATTCATTCAGTCATCAATGGATTACATGAAGCTTCAATTTCTGATACTCGTTTGTTTTTTTCTCTTCATTGTTCCGGCGAGTTCATCGCCTCGGGATATCGCTATTTTACTCCGGGTTAAGTCCGGCCAACTCGATGACCCGAATGGGTTGCTTGATGATTGGAACGGGTCTGCTCCAAATGCGCCTTGCAGCTGGAACGGGATTAAGTGTGATCGGAAAACCGGTCAGGTTGTGTCCATTGATTTTGCAAGTTTTGGAATCGCTGGTCGTTTTCCTGCTGATTTCTGCCGGATTTCGACTTTGCAGAAACTCAATCTGGGTGATAACAGTTTTGGTGAGTCCATTTCCTCTGACTCATGGTCACTCTGTTCGCATCTACACTTTTTGAATATTTCTTTAAATTTCTTTGTTGGCCGGCTGCCGGAGTTTATTACCAAGTTCGATAACTTGACCATCCTTGATGTTAATTCAAACAATTTCTCCGGTGAAGTTCCGGCGAGTTTAGGCCGTTTACCAAAATTACAAGTGCTGAATATAGCTAACAATCTCCTCAATGGTTCAATTCCTGAGTTCTTGACGAATCTTACCGAGTTGACTCGATTGGAAATTGCTGCAAATCCGTTTCAGCCAGGTCCATTGCCTTCCTCAATCGGCCGACTCGGTAAGCTTCGAATTTTCTATGCTCGGTATGCGAACCTTATTGGAAATTTTCCAGATTCTATCAAAGACCTGAAATCTATTCAGAATTTTGATGTGGCAAACAACAATCTCTCCGGAAAAATTCCAGAAAGCTTCGGAGAACTCAAAACCATACAACAAATAGAGCTCTTTGGAAACCATTTTTCAGGCGAATTGCCGGACATGTTTTCCGGTCTTGGTTCTCTTTCCAGGTTTGACGCTTCTGAGAACAATCTCACCGGGAAAATACCTGAAACCCTTGCCCATTTGCCATTGGAATCTCTAAATCTCAATGATAACCAATTAGAAGGCGAAATTTCAGAAAATTTAGCTCTTAACCCAAATCTTAGTCAGTTCAAGCTTTTTAACAACAGATTTTCAGGTACTTTACCTCAAAATTTTGGTTTAAGTTCAGATTTAGATGAGTTTGATGTCTCCGGCAACAATCTAAAAGGTTCTTTACCTCCCAACCTATGTTCTAGAAAGAAACTTAGGATTTTGAACCTGTTTGATAATAAGTTCAATGGGCCAATCCCAGAATCCTATGGGCAGTGTTATTCACTATCATATGTGCGTATCTATAACAATCAATTCTCTGGTGAATTACCAACTGGTTTCTGGGGATTTGATGGATACACATTTCTTGAACTGCGAAACAACAACTTTCAAGGTTCAATTCCAGCTTCAATCTCCAATGCTCGAGGCCTGACACAAATTCTTATCTCCGGCAACAAGTTCTCCGGGGAATTGCCAGCAGAACTATGCAATTTGGAAGAGGTTGTGATCATGGACATTAGCAAGAATCAATTATCAGGGGAGTTGCCTTCGTGTATCACAAGATTGAAAACATTACAGAAGCTTGATCTTTCACAAAATAGGATCAAGGGTCAAATTCCCAAATCAGTTAGTTCTTGGAATGACCTGACTGAATTGAATTTAGCTAACAATCAATTGACAGGTGAAATTCCTGGTGAGCTTGGGACGTTACCGGTCTTAACCTACTTAGACCTTGCCACAAACTTGCTTTCTGGCGAAATTCCATCTGAGCTGAGCAAACTCAAGCTCAACAAATTCAATGTATCAAATAACAGGCTGGAAGGGAAAGTTCCACTTGGGTTTGATAACGATTTTTTCGTCTCAGGTTTACTGGGCAATCCGGATCTTTGTAGTCCGGATCTTAAGCCTCTGCCCCAATGCCGAAGACCCAAAAGTGTAAGCTTGTACTTGGTGTGTATTTTATCAGCTTTTGCCTTCATACTTGTCGGGTCACTTGTTTGTGTCTTACTCAAGGCCAGTAAGCTGCTACCAATCCGTAGCAAGCGTAAAAGTGTGTGGAGAATTACTGCATTCCAACGCGTCGGTTTCACAGAGAGAGATGTGTTAGATGCACTGATAGAGGAAAATCTCATTGGAGCTGGTGGGTCGGGTCGTGTGTACCGGGTCAAATTGAAAAACGGACAGATGGTAGCAGTGAAGAAACTTTGGGCGGCTAAACGGGAAAGAGAATCCGAGGAGGTGTTCAGGTCTGAGGTGGAGACATTAGGGAGAGTCCGGCATGGAAACATAGTCAAATTACTATACACTGGCATTGGTGATGACTTTAGGATATTGGTGTACGAATACATGGAAAATGGAAGCTTAGGAGATGTATTACATGGGGAAAAAGGTGGCTTGTTATTGGATTGGCCGAGGAGATTTGCCATAGCAGTTGGAGCAGCTCACGGATTGGCCTATTTGCACCATGATTCTGTGCCAGCAATAGTGCACAGAGATGTTAAGTCTAATAACATTTTGTTGGACGAGGATTTCAGGCCCAAAGTGGCTGATTTTGGGCTAGCTAAGGCAATGCAACGGGATGCTGAGGAGAGTGAACAAGCCATGTCCCACATTGCTGGTTCCTACGGCTACATTGCACCTGGTAACCTTAAATTCCTTATTGTTTTTCACCATTTATTTTAGGTCCAATGACTGTTAGAAATTTTCTAGTCTTGTTAGAAATTTATTTGCCAATAAAAAATGTTGAAAAAATCTAGTGCTAAGGAGACTATTTAAAAGAAAAAAAATATTGAACCAAGATAGGCATAAATGAAGTGGTATGGACAGCAGGAATTCATAATGCTAACCTCTACTTGTTTGGGATCAGGGCCTAATTATTGTTGTCGTTCAACTTTTTTGTTAACTTGTTGTGAAAAGATATTTAATATCTTGCGTTTGGCATACCAAGGACATCCAAAAATAGTTTTGTGGAAGTTTTGGTTAACTTCACATACTATCTTAAAATTTTGATTTGATTTGATTTGAAATTCTTTTATATGAAGCCAGAGAAGATTTTAGGAGATGATTCAAACTTAGAAGTAGATTGGAAACTTTAATAATATCCATTTAATTGTAGCATCTAGTTTCATTCCTCTAACTGGACGTTGTTTCTGAAGTCTGCATATTCTTGTTTGGCTTGCTAATGCTTGATTATATGTTTAACAAGGGGATTTGTCAAAGTAAAATCAAAGAATGATTGGGCATCCCATTTTTGAAGTCTGCATGACTATTTAGTTTTAACTACCTGTAATATTTAATCCTGAGTTTCGTTTGGGATTTGTCAAAGTAAAATCAAAGCATAGTTGGGCGCTCCTTTTTCTGAAGTCGGCATGACCATTTATTCTACACCGGTAGTAATATTTAATTCTGAGATCACATCAAATTATTGAATGATTTTGTCACCCAATGTTTGTTAACACAGTCAAGATTTGTTTGAAAGAATAAAAAATAGGAGGGTCATGTGTGTCCTGTCACCTAGCATTTGTTAACATATTCAAGATTTGATCTAAAAGAATAGGATTGAAAGAGACATGTGCCTAAGTATTGTTACTTATTTATCTAATTTATCGATATCTTAAATTCTATTTACTCCCATAAGTCCTCTTTGATTGGACATACAGTTTCATCTTTTTTGTAAAGCTGACACTATCCGGTCTGTGGCTCTGCCCATGTGCCCTTCCTTTGCAAAACTGAGAGAATACAGAAACCAATATCGTTGATGGATATTTTTTGATGGCCTTTTGATACTAGAAAACATTCCGGTTGCAATGACAAACAGTTTGTTAAGTTGATCTTGGCTACTTAACGATTAGGAAGATGCTAAAGTTTGCTAAATATCTCATGGGATGGCACATGTGTTATCCAAGGTTTTCAGCTGCACGTTTAAACTTAACCTCGATTCCTTTGTTTCAATTGCAGAATATGCATACACTCTGAAGATCACTGAGAAGAGTGATGTTTATAGCTTTGGTGTGGTACTGTTGGAACTAATAACTGGTAAAAGGCCTAATGACTCCTCTTTCGGAGAGAACAAGGACATTGTCAAGTGGGTGTTAGAGGTTGCAACATCGTCTAAGAAAGATGAAGGAACTGGCCATATTGTTACGTGCGCAAGTGGTATTCTTGATTTGAACCAGCTAGTTGACCAGAGAATGAATCCATCTGCAAGCAATTATTCAGAGATTAAAAATGTTTTCGATGTGGCTTTGCTTTGCACTTCAGCATTGCCTATTAATAGGCCCTCCATGAGAAGAGTTGTTGAGTTGCTGAAGGTTATCCCCTCTGCTCGCTCTAAAACAACCCATTAGCTTTTCACAGTTTCAACATGTCCTTGCTCAAGTAACTTGATACTATTGTAAACTGTAAAAGAATGTGTGAAATTAGGTGTGTTTAGGTCTGCTTCATAGGTCGATTTAGTTTGCTTGCTGTGGTTTAACAAGTACCTTGTTAGATACTCCAGTACTACACAGAGTAGCTTAACTAAAAGTTTGCTGACCCTGTAAAACTCATGTTGTAACGACTAAACCTTAAGAGTTTTTGTTGCAATATATGAACTTTACTATCATATACAAGTATTCTATTCTGTATCCATG

>PGSC0003DMG400031028 LRR-RLKs

ATGGAGGTGAGCGTGAAGATGAAATTCCCCTCACAAGCACTACTGTTGGCTCTATTGCTTGTTTTACCGATCGTTTTAGCTCTCACCGAAGAAGGTAACTATTTTTTTGAACACCTAGAAGCTGTTTTGGTGTTTTCTAGCTCCTCTGTTTTTATAGCAGAAGGTAGTGCTATGTGTGCTATCTAGGAAATACGAAAGTTATTGTTCGAATTTAGTTTTTATTTTCAGTTTCTGGAGCAGCATTCCATGCTATTTAACTCTGATAACGAAAAATCCGTGTTATTTAAGATATATTTAGGCTCCAGTTTATGGCATAACTACTGGAATACTAGTTGATGAAAACTGAAAATGGTTATGACTGCTTGCGCAGGCAAAGCATTAATGTCGATCAAGGCATCGTTTAGCAACGTGGCAAACGTGTTGCTAGATTGGGATGATGTCCACGACGAGGATTTTTGCTCATGGCGAGGCGTGTTGTGTGGAAATTTCTCCATTTCCGTCGTTGCCCTGTAGGTTTTCCATCCTTGTCTCCTAACTTTTACTGATACACCAGAACAAAGAAGTAGCTGGACTCTGATGACCTGCTAGCTATTGTATAGCACTTTGTTATTTTAGCTAAAACTTATACGTCTTTTATTTAAATTTGCCTTCTCTGCTTGCGAAGGAATCTGTCTAATCTTAACTTGGGCGGGGAAATTTCACCAGCCATTGGAGATTTGAAGAATCTAGAGACATTGTATGGTCCATGTTCTCTTTTACTGTTCTTGGTCCATTGACTGTTATTTTTCCGTGCTAATATTATATTCCAATTTTAATGACAGAGACCTTCAGGGAAATAAATTAACTGGTCAAGTCCCAGATGAAATTGGCAACTGCATTTCACTGATCTATCTGTAAACTGTTTTTAACCTATGCTTTTAATATCTATTTCTTGCGTGTCAGTAAATTTAAAGTGTTCACACTAGTTGATTTGCTATTTTCTGCAGTGATTTGTCTGATAACTTGCTCTATGGAGATATACCTTTCTCAATTTCTAAGCTCAAGCAGCTAGAGTTATTGTGAGTTATTTAATCACGTTAAATTGATGTCTTCTCATATCTAGTTGATATGGTTATGATGAATTAATTCATATGTGGTACAGGAACCTGAAAAACAACCAGTTGTCTGGCCCAATCCCATCCACATTAACTCAAATCCCTAATCTAAAGACGCTGTGAGTTTCATGACTTTAGTTTATCTCACTCAAAATTTAGTCCAATATACATGGTTACTAATGGTTATTTGATGGTGAAGTGATCTGGCTCGAAACCAGCTCATTGGTGAGATACCAAGGTTGATCTATTGGAATGAAGTTCTGCAATACCTGTGAGTGCATGTTCCTGGTGTTTTGGAGGTTTTCCTTTTTGTTTGTGAAATTTACAATGTTATAATTACCTTCTGTTTTGCAGTGGCTTAAGAGGCAACATGTTGACAGGAACATTGTCCCCTGATATGTGCCAGTTGACTGGTTTGTGGTATTTGTATGTGCTCTGCTACAAGTTGTCTATACGGAATGCTGTGTTGTCTGTTTGGTGTTATGTTTATTCTAACAATAGAAGTTTCATATTATTTCAGTGATGTGCGGGGCAATAACCTCAGTGGAATAGTTCCAGATAATATTGGGAATTGTACAAGTTTTGAGATACTGTCAGTGTCCTCTCCTCGCTCTTGTAACATTAAGCTACGGTTTTTACTCCTACTGCTGCCCAATTATAACAAAGTCTTTTTTTTCGTGATTTCAGGGATATCTCATACAATCAGATAACTGGAGAAATTCCCTACAATATTGGATTTTTACAAGTGGCTACCTTGTTAGTGAATTACTTTTAAATGATTTTTAGCTGTTGTCATTTTTACCTTGTCTGTTCTGTTATGGGGTTCATTTCTGTGAATTGTAAATGGTGATTGTGGTCCTTTCAACAGGTCTTTGCAAGGAAATAGGCTAACTGGGAAGATCCCAGAAGTGATTGGTCTAATGCAAGCTCTTGCTGTTCTGTGAGTATTCATACTATTACATTTTTTTTTTAACAACTCCATTTCTTACTATTTACTACTTGCAAGTTAGAAAGTTTCATGATGTTTGTTTCCTCTGTAGGGACTTGAGTGAAAATGAGTTGGTGGGACCAATTCCTCCAATCTTTGGCAATTTATCCTACACTGGGAAACTGTAAGTTTTCACCACTATTTTAATGCATATACTTTTCTACGTGTGGCTCGTTTGTCTGATTCACTTGTACATTCAACAGGTACCTGCACGGCAACAAACTTACAGGGCCAATACCACCGGAGCTAGGAAATATGTCTAAACTTAGTTACTTGTAAGAGGGTTAATTCACAATTTCAGATGCATTATGCGGTCACCTTTTTTTTTTGCTTCATGCATTGAGCTATCTTATTTACAGGCAATTAAATGACAATCAGCTAATGGGGCGAATTCCCTCCGAACTTGGCAAACTGGACCAGTTATTTGAATTGTGAGCCTTTTATTCTTGTAGTATATAAATATGGTTCCATTTCTCTTAGAATATTGTGTTCTTAGCTTTCCTTTACCAGGAATCTTGCAAATAACAAGTTGGAGGGACCAATTCCTGAAAATATCAGCTCCTGCTCGGCATTGAATCAACTGTAAGATTGTTGCGTATCCTTTTGGAACTTCTTTTTTTTTTCTTTCTCTTGCTATTTCCATTCATTCTGAGTCTAAACATATTTGTTTCTTTCTTTCAGTAATGTTCATGGCAACAACTTAAACGGGTCCATTCCTTCAGGGTTTAAGAATCTTGAGAGCCTGACATATCTGTATGTTATATGCTTCTGGTTTGGCTAGTGATCATGAAAATGATTTTGAATTTCTTTCATATGGTACAAACTCATTGGTGATATTTTGTTGATATATACCTGGCAGAAATCTTTCAGCAAATAAATTTAAAGGTCACATACCCTCTCAACTAGGGCGAATCATCAACCTTGATACATTGTGAGTTCTTGACTGTCAAGGTCAATAAAATTTATTAGTTCAAGTGATTTCTTGTACTAAAACTTGTATTCTTTAGGGATCTCTCTGGCAATGAATTTTCTGGGTCTATCCCTGGTTCTATTGGAGATTTGGAGCATCTCCTCACACTGTAAGATTTTAAACATTTTGATGTACAAGGAGTTTGTATCATATTGTGACTTGAACATAATCACAATATCACTGCATATTGTAGGAATCTGAGCAGCAATCATCTTGATGGACAAATTCCTGTAGAATTTGGCAATCTGAAAAGTATACAGACCATGTGAGTTTCTGCTGAGAGCTTTGGTTACACTGCCTTCAAAATGTGTACTTTTTTTTAGTATACTGATTTTCTTGTTCTGAGATTGGTTTCTATTCTTTCAGTGATATGTCATGCAACAAGATTTCTGGTGCCATCCCAAAAGAGCTGGGACAGCTGCAGACCATGATAACTCTGTAAGTACATTTAGTTTTTGTCAAGTAAATTGATGTAGTGTTTAATCAGTTGTCCTTAATATCAGTCAATTCTAATGAATTTTTGATTGGCTTTGTTTCTTTATGAAGTACTTTGACAGCTAACGATCTTAGTGGAGCAATCCCTGACCAATTGACCAATTGTTTCAGCCTAACTAGTTTGTGAGTTCATCTATCTTTGCCTTAACATCATAGACATTCTAATTCTTCCTACAGTTACTGATCTTATGCCATTTTCCTTTAGGAATATATCCTACAACAATTTTAGTGGTGTTGTTCCTCTTTCACGGAATTTCTCGCGGTTTGCACCTGACAGGTATTTTCCGTATAGAATGAAGCTTCATCATTATATATGAGCATTAGTACAAGTTAATGATCATCTTACTATGGTCATCAGCTTTTTGGGGAACCCATTTCTTTGTGGCAACTGGAAAGGCTCAATATGTGACCCCTATGCACCAAGGTCTAACGGTATGACTTCATTTTTGTGCATTGGTTAGTGAATCTTCTGGTATGCTAGAGTCATGTGCATTAAAATGACTTGTTACTTTTGCAGCCTTGTTCTCTAGAACAGCTGTTGTTTGCACAGCATTGGGTTTCATAGCACTCTTATCCATGGTTATAGTGGCAGTGTACAAGTCCAACCAACCACACCAGTTTCTGAAGGGGCCTAAGACCAATCAAGGTAAAAATTAGTACATGTCCATTCTGTTCTTTTGCTTTTCAGTACTTTCAGGTATTTATGTTAGCTTTTATCTTGTTTCCCTCTAATTCCCTAGGTTCCCCCAAACTTGTGGTTCTTCACATGGATATGGCCATCCATACATATGATGACATTATGAGGATTACTGAGAACTTCAATGAGAAATTCATCATAGGATATGGTGCTTCCAGCACTGTATATAAATGTGTTTTGAAAGATTCCCGACCGATTGCCGTTAAGCGACTTTACACTACACATCCGCACAGCTTGCGAGAGTTTGAGACTGAACTGGAGACCATTGGAAGCATCAGGCATAGAAACCTTGTTAGCTTGCATGGTTACTCCCTTTCCCCTCATGGGAATCTCCTTTGTTACGACTACTTGGAGAATGGTTCACTCTGGGATCTACTTCATGGTTAGTAACCCTCTTTTCCTTGTAATCTTTTATAAAGCATCTTCATGTAAGACAGTGTTGACTATTAGTTGATATTTGTTACTGTTTTCCCTGTCGGAGAATGGTGCTATAAGCCAGGATTGATTTGAAAATGGCTAATTATCAACTGAATACATGTCATAGGGCCTTCCAAAAAGGTGAAGCTTGACTGGGAAACACGTCTGAGGATTGCTGTTGGTGCTGCTCAGGGTCTTGCTTATCTTCACCACGATTGCAACCCAAGAATCATCCACAGAGATGTGAAATCTTCAAACATTCTTGTTGATGAAAATTTTGAGGCTCATCTTTCTGATTTTGGGGTTGCAAAATGCATCCCTTCTGCAAAAACTCATGCATCAACTTTGGTGTTGGGCACCATAGGTTACATTGACCCTGAGTATGCCAGGACTTCCAGGTTAACTGAAAAATCAGACGTCTACAGCTTTGGCATTGTTCTCCTAGAGCTTTTGACAGGAAAGAAACCGGTTGATAATGACTTGAACCTGCATCAGCTGGTATTATTCTCCTCTTGTAGTCTACGTTGTTGCTTGTAAAAAAGATTTAACTCAGACTGGATATAGAAAAGAACAACTTATCTCAAATTTTCCTACCTTCCTCCAGCATTTGCGATGTTATCATTCCATCAAATGTCTTTTGTCTTTTAACTCCTGTATTATATTTGTTGTCTTTATAGAGTACTTTGATTTGTGTTACAGATAATGTCAAAGGCGGATGATAACACCGTGATGGATGCTGTTGATCCTGAGGTATCTGTTACATGTATGGACTTAACACATGTGAGGAAAACTTTTCAGCTTGCGTTGCTGTGCACAAAAAGATTTCCATGTGAGAGGCCAACGATGCATGAGGTTGCTAGGGTACTTGTTTCCTTGCTTCCTCCCCCGCCAACCAAACCTTGTTTAGACCCACCTCCCAAATCCATTGATTATACAAAGTTTGTGATTGGGAAAGGACTACCGCAAGTTCAGCAGGGTGATGATTCCTCCGAAGCACAGTGGCTTGTTAGATTTCAAGAAGCTATATCCAAAAACTCCCTTTGACTTGGGTTCATAGACACACTTCATATCAAATCAGAAGCGAATATGGAACATATATATGTTGGCCAAAGGTTTTTTGATGTTAGAGATTTCTCCTATAAGATGGTTAGGTACACGGTAATAGCTAGTGTATTTCACATTTGTAGATGGAATGTTTTATGGTGGATGAGATCCACAATAGGTTAGATCTACAACCAGTGCAGTTCAACTTAAATCTTTGTTGA

>PGSC0003DMG400031103 LRR-RLKs

CAATTTTATCATCACCATATAGTTGTATTCATTCACATATCACACTCACATGTTCTAGTACTCCTTCCATTCTGTAAAGTTCTAGTTGACTCTCTCCCTTTGTGGTCCTTTTCCATTTTCTTGCCTTATCAATCAATCAACTTTTTTAGCCAATTTTTTTCTCTGTTTTCGCATCCCTTTTTTATCCTCTTATTTAATTTTTAGCTAAACCCCATTTCTTGATTTTTAGTCTTTTGCATGTCCTGAGTAGCATTATTTTTCAAGATTTCATATTATAGCTAAACCCCATTTCTTGATTTGTATTATCTTGTCTTTTGCATGTCTTGAGTAGCATACTATCCAAGATTTCATATTATAGCTAAACCCCATTTCTTGATTTTGTAAAACCTTTGGTGATTTGCATGTCTTGAGTGGCATGTTTTTGTGGTCATTAAGGTTAAAACAATTCAAGATTTCATTTTTTCCTTTGCTTCTTGAAATTGGGTTGAGTTGTATACTGAGAGTTATGGGCCCTGGAATCTCAAAACTTGCTTAATGGTATCTGTTTCCTGGTCTGCCATTAGCTCTGTTACAACTTTTGGGCATTGCCATACTCATAAGTTTCCTAATTTTCATCCATAAGAAAAATATTGACATGACTTTAGTGTAAACTAAACAGCATGTCATTCATGCTCTTTTATCCTTGGAAATGGCAAGTTATAGTAACTCTGCAATCCAACAGCATTTGTTTTTTGTGCTGTTGATTCTTCTTTTCTTTACTGGTTTTGCTGAATCTTTAAATGAAGAGGGACTTATTCTTTTGGAGTTCAAAGAATCCCTTAATGATCCTGATAACAATCTTGAAAGTTGGAATTCTTCAAACTTGAATCCTTGCAAATGGGATGGTGTTAAGTGTAGCAAAAATGATCAGGTGATTTCTCTCAATATCGATAATCGTAACTTGTCTGGTAGTTTTTCTTCAAGAATTTGTGAACTTCCTTACTTGACAGTATTGAATGTGTCATCAAATTTCATTTCTGGTCAAATTCCTGATGATTTTGCGTCGTGTCATAGTTTGGAGAAATTGAACCTTTGTACCAATAGGTTCCATGGAGAGTTTCCTCTCCAATTATGCAATATTACCTCACTTAGACAGCTTTACTTATGTGAGAATTACATTTCTGGTGAAATCCCTCAGGATATCGGAAACTTGTCACTTCTTGAGGAGCTTGTTGTTTATAGTAACAATCTTACTGGAAGAATACCTGTTTCAATAGGTAAATTGAAAAAACTTAGGATCATCAGGGCTGGGCGAAACTATCTGTCTGGCCCTATTCCTGCTGAAGTTAGTGAATGTGATAGTTTACAAGTGCTTGGTGTAGCGGAAAATAGGCTAGAAGGGTCTTTTCCAGTAGAGCTACAAAGGCTTAAGAACCTCATCAACTTAATTCTTTGGGCTAACTCATTCTCTGGTGCAATTCCTCCTGAGGTTGGGAACTTCAGTAAGTTGGAATTACTCGCGTTGCATGAGAATTCATTCAGTGGACAAATTCCTAAAGAGATTGGGAAGTTAACTAATTTGAGAAGGTTGTACATTTACACTAATCAGTTGAATGGAACAATTCCATGGCAAATGGGGAATTGTTTGAGTGCAGTTGAGATTGATTTATCCGAAAATCAGTTGAGAGGAAGCATTCCGAAAAGTTTGGGGCAACTTTCTAATCTGCGGTTGCTTCACCTTTTCGAGAACAGATTACATGGGAAGATTCCAAAGGAGCTTGGAGAGTTGAAGCTGCTTAAGAATTTTGACTTGTCTATAAACAATTTGACAGGTAGAATTCCAGCAGTGTTTCAGCATCTTGCATTCTTGGAGAATTTACAACTTTTTGACAATCACCTTGAAGGTCCTATTCCGCGATTTATTGGCCTCAAAAGCAACCTCACTGTAGTGGACCTGTCCAAGAATAATCTTAAAGGAAGAATACCTTCAAATCTTTGTCAGTTTCAGAAGTTAACATTCCTGAGCCTCGGGTCTAACAAGTTGTCTGGCAATATCCCTTATGGCCTAAAGACTTGCAAGTCCCTCGAGCAGCTAATGCTAGGAGATAACCTGCTCACTGGGAGTTTTTCTTTTGATCTTAGCAAGCTGGAGAACCTTTCAGCTCTTGAGCTTTTTCACAATAGATTTTCTGGATTGCTACCACCAGAGGTGGGCAACCTTAGAAGATTAGAGAGGTTGCTCTTGTCAAATAACAACTTTTTTGGGCAAATTCCTCCTGATATTGGAAAACTTGTGAAGCTTGTTGCTTTTAATGTTTCCTCCAACCGGCTCTCGGGTGATATTCCTCATGAATTAGGGAATTGTTTAAGTCTCCAAAGGCTTGATCTTAGCAAGAACTCGTTCGCTGGAAATCTCCCCGATGAACTTGGTAGGCTAGTCAACTTGGAACTGCTTAAGCTATCTGATAATAAGTTTAATGGACAGATACCTGGTGGCTTAGGTGGACTAGCACGACTAACGGATTTGGAAATGGGAGGCAATTTCTTCTCTGGTAGTATTCCTATTGAGCTTGGCTACCTTGGAACTCTTCAGATTTCTCTGAATCTGAGTCACAATGCTCTCAATGGATCGATTCCTAGCGCCCTCGGGAACTTGCAGATGCTTGAAACATTGTACTTAAATGACAACCAGCTTATTGGTGAGATTCCAACTTCAATAGGGCAGTTGATGAGTCTGATAGTATGCAATCTTTCTAACAATAACCTTGTGGGATCAGTACCAAACACGCCTGCCTTTAAGAGGATGGACTCGAGCAACTTTGCTGGAAATGTTGGGTTATGCACGTCCGATTCCATCCACTGTGATCCTCCTCCGGCCCCTTGGATTGCTCCAAAGTCAAACTGGTTGAAACACGGTTCTTCTAGGCAAAAAATAATTACTGCTGTTTCTGCGACTGTTGGGATGATCTCTTTGGTATTGATTTTAGTTATATGTAGGATCATTAGAGGCCACAAAGCAGCTTTTGTGTCCGTGGAAAATCAAGTAAAGCCGGATGACTTGAATGATCATTACTTTCCACGAAAAGGGTTCACGTATCAGGACCTTGTCGATGCTACTGGAAATTTCTCAGACAGCGCGATCATAGGAAGGGGAGCTTGTGGCACTGTCTACAGAGCTCATATGGCTGATGGTGAGTTTGTTGCAGTTAAAAAGCTGAAGCCACAAGGAGAAACCGCGAGTGTTGACAGCAGCTTCCAAGCTGAATTATCTACACTTGGTAAGATAAATCATAGAAATATTGTCAAGCTATATGGTTTCTGCTACCATCAAGATTGCAATCTTCTCTTGTATGAGTACATGGGAAACGGAAGCCTCGGGGAAGTACTTCATGGGAATAAGACAACTAGTTTACTAAATTGGAATAGCAGGTACAAAATTGCCTTAGGAGCTGCTGAGGGACTATGCTATTTGCACCATGACTGTAAGCCACATATCATTCACAGGGATATAAAATCGAACAACATTTTGTTGGACGAAATGCTTGAGGCACATGTAGGAGACTTTGGCTTGGCAAAGCTAATTGACTTCCCTTACTCAAAATCCATGTCCGCGGTTGCTGGTTCATACGGCTACATTGCCCCTGGTAAGACCGGTGTTTAGGCACTTAAAACGATACTCTTTATTAGCATTACTTACATACAGAAGCTAACTAAGTTCTAGTTTTTTCTTTTTTTCAGAATATGCATACACAATGAAAGTGACAGAGAAATGCGACATCTATAGTTATGGCGTTGTTTTGTTGGAATTGATTACTGGTAGATCTCCAGTTCAACCCCTTGATCAGGGAGGCGATTTGGTGACTTGGGTAAGGAGATCAATTCATGAAGGAGTGGCACTAACTGAGCTTTTTGACAAAAGACTGGATGTTAGTGTTGCAAGAACAAGAGAGGAGATGTCTCTAGTTCTCAAGATTGCTATGTTCTGCACCAATACATCCCCTGCCAATAGGCCTACAATGCGAGAAGTCATCGCCATGCTGATTGAAGCCCGGGAATTTGTGTGCACTTCACCACCATCACCATCCTCCGAAACTCCTCTGAGTGAAGCTGATGCTAATAAAGGTGCATAAAAACAACTTCCTCATCTTTCAATCTTTTCACTACAGTCAAATATCTCTATAACAGTCATCCTCTATAACAACATTTCACTATAACAAACGGGTTCTCTATGGAACTAATTTTCATGTTATGTTACATTACATGTTCTCTATAACAACATTTCACTATAACAAACGGGTTCTCTATGGAACTAATTTTCATGTTATGTTACATTACATGTTCTCTATAACAACATTTCACTATAACAAACGGGTTCTCTATGGAACTAATTTTCATGTTATGTTACATTACATGTTCTCTATAACAACATTTCACTATAACAAACGGGTTCTCTATGGAACTAATTTTCATGTTATGTTACATTACATGTTCTCTATAACAACATTTCACTATAACAAACGGGTTCTCTATGGAACTAATTTTCATGTTATGTTACATTACATGTTCTCTATAACAACATTTCACTATAACAAACGGGTTCTCTATGGAACTAATTTTCATGTTATGTTACATTACATGTTCTCTATAACAACATTTCACTATAACAAACGGGTTCTCTATGGAACTAATTTTCATGTTATGTTACATTACATGTTCTCTATAACAACATTTCACTATAACAAACGGGTTCTCTATGGAACTAATTTTCATGTTATGTTACATTACATGTTCTCTATAACAACATTTCACTATAACAAACGGGTTCTCTATGGAACTAATTTTCATGTTATGTTACATTACATGTTCTCTATAACAACATTTCGCTATAGGAGCCAAATATTGGAACAAGTAAAACTGTTATAGAGATGTTTGATTGTATATCATTTAAACACATATCGAATCAATACCTGTGTTTATAGCAGCCCCAAAAGAAAGGAATTCTCATTTTTCATGTCCATAATTCTAAGTTTAGTTAATAATAAAAGGAATTATTCAATACTGAATGGATGTTCACAAAGAAAAATCAGCTAAATCACATGCATGTCTGTTTACTAACAGCAACCAGTTTCTACATTTTTTTAGTAGCATTTGTGTATTCATGCTATACATGTTTGATAATTTTGATTCAGAATGTTGCGCATACTCTCCAAAAATATCGTCAGACACTACTTTTGGAGGATCCGACATGCATCCAGCTGATATTTTTTTGGAGAGTTCGAGCAACATAGATTCAGATTATATCATCCACAAAAAACTAAAATGCATCCACTTTATGTTTCTTGCAGGTTCTAATTAAGCAACCATAGACCAAAAAACAGAAAATTGAAGTTGTGGCTCATCTCAAGCTCTGCCAAATTTCTAATAGAGTTAGAAGCATCTTATTATTCATGTACACACAACTACCTTTGATACCATACTTCCCCTCTAATAATTATTAGTTGGCATTTCTTATCTATTTATCTTCTAAATTATATCTACTTCTATGATCTGTCTCCACATTAACTGCC

>PGSC0003DMG400031228 LRR-RLKs

CAAACCTGCAGGAGTCTTAGGAACTGCAACTTGCAAGGAACAGTTCCTGATCTAAGCACAATACCACACCTTCTTTACTTGTAAGCATACTTCATCGAGAAATTCATGCGTCCAGTTAAACCACATGTTGAGGCTTATGTTCTTGTTGTTCTAACCTCTTGGTAATGTTGATATCTGCTTAATAATTCTTATATGTCATCTGCTTAATAATTCTTATATGCCAGATGTAGTTGAACTCATCTGCCACTCAATTTGTGCTGAATATAGAAAGTTAAATTTACAGCCAGTGGCAATTCATCTGATATAAATTTCTCATGTCACAGAGACCTTAGCAGAAACCAACTCACGGGAAATATACCTTCCAATAAGCTGTCTGATAATATCACAACCATGTAAGTTTCTGGAAGTGTTTAACTGCTGATATCTTTCCTGGTTTGATTCACCCATCTTTAATCACACTTCCTTCCCCCCCCCCCCCAACTCTCTATCGATATCTGTGAATAGGTGGTTGAATGGACATTAGTGTTTTAACATGCACTCTTTTCTTCTTGATGTTTGAGTAACTTGGTCTTGGTATCAGACCCTCAATAGTGAACTAACTACTTTGATGATGATTGCAGCATTCTGTCAGGAAACATGCTTAATGGTTCTATCCCTTCGAACTTTTCTGTCCTTCCAAATCTGCAGAGGCTGTAAGGAACTTCATCTGCTCAATCCTTGTTTCTTGATCCTCTGTCTGTTTCAAATTTCATGTCGATACTCTAATAAATGTAGTATTGACACTACCTTTCCTTAAATTATTTTTCTTGTAGGTCACTCAACAATAATAGGTTAAGTGGTTTTGTGCCAACTACCATTTGGGAGAATAAGACATTTTCCCCAGGCGCAAAACTTAGATTGTAAGTGTATCATTTGTACATTTTCCTCTTATCAGTAATTTGATTTTCATGTTTCTGTTGTCTGCTGCCCCACAATCTATTGGCCAGTTAAAGTTTTTAACTGTAAGATACAGACAAAAGCTTTTTGTAGAGAATCAAGCCAAATTTGTATTCCTATTTCATTTTGGTAGCTAGTCAGTCCACATGTAGTTTTGGATATACAAAGAAAGTATCCCGGAAAATAATAGTACAATCAAGGAATAGTGTGTCCATTGTCAACAAGGGTTTCCTATCTCTATACAGTTAATATATAACAAGGAATAATATGTCTGGTGTTTTGTTTCGTTCGAGGTAATGTAGCATGTATTCTTTCCTCTTACTAAACCAGGCCATGTGACTAGAAAAGGCCCCTCAGATGTTAACATATCAGATAGCCAGGCTGATCCAGTTGTTCTCCTTTGTGCAGGAACTTTCAACGTAATTTTCTCTCAGATATTTCGGGTATTCTTGATCCTCCTCCAAATGTCGATATTATGTAAGACCTTGTTTCCGGTAGTCTTGTACATACATATTTGCGGACTTACTTTCAGTTCTCGGCTTTTATAATTATATGTGGCGAATCAAGCTCCCCTAATTCAATTGCTATCTAACTTTCATTATCCTATTTTTTATAAGCTTCAAATTACAAAGTAATTCTGTGGAGTCTTAGTTTCTATTTCAGATTTCTGTGATTCAACAGAACTACTATCATGAAGAAGGCAGAAAAACTTGCTATAGCATATCCCCATGACAACATTTGGGATAGTTGCATAACACTGCCTTCATTTGCTGTTTTCCTGCTAAACTCTTTCTATCAGTATAGAGATTTACTTCGAATATCCATTGATGAAGATAGAGTAGTGAACCAACACAGTTTCAACTTCTTTTATTTTGTCATCGTTCAAAACTTCATTTCTTGTCAGAAGATGGGTAGTTTTCTTGCTAGCAGTGCACTTTATATGGAGCTCGAAAACAATGGGGTCAAATTCATATATGTGAATTTCTTTCCATATGATATGGTAATATGACAAAGCTAGGGTTATGGGGGAGAAGTTGGTGGCTATGGGAGACTGGGGAAGCAGTGGGGATGCGACCAATATGTGGGATAGGACCGCTAGTTGCATAAGGGAAGCGGCTAGAGAGGTCTTGGGGGTTTCAAAAGGTAACTCTAGTGGACACCGAGGAGATTTCTGGTGGAATGACAAAGTCCACGTAAGGTAGAGGCAAAGAAGGTGGCTTATGTGAAGTTGCTGGAGAGCAAAAGGATGAGGAGGATAAGAGGGCGACGAGAGAAGTATATAAAGTGGCAAGGAAGGAGGCGAAGTTGGCAGTTATGGAGGCAAAAATGGCAGCTTTTGAACGCCTGTATGAGGAACTAGGGGACGAAGGCGGAGATAAGAGGTTGTATAGGCTAGCCAAGGTGAGAGAGAGGAAGACTCGAAACCTGGACCAAATAAAGTGCATTAAAGATGAGAAAGGCAGAGTATTGGTGGATGAGAGGCAAAGTTGGCAGTTAAGGAGGCAAAAATGGCAGCTTTCGAATGCTTGTATGATGATCCCTGCATTATAATCTTTTTATTTTTATTACAATTCTGGCATAACTTGCCTCTAAACCAGACAACTAACTGGTTACATACCGAAGTAGTAACAGGCTTTCCAATTTGATGGAGAGATCACGCCTCAGCAATACCTTTCAAGCTGTATATATTTAATTCTAAAAACAAATTGTTCAACTTCTATCATGGTTAAATAAATTCTGGTTCACCTACTAACTTTATTTTTCTGAAGAGCAGCTTCACCTACTAACCTTATACATGTTTGTGTGCAGTTTTGTGTGCCATATTTGCTGATTTCATGAATAACATAAATAATGAGTTTGAGTTTTGTTTTTAACCCATTTACTGGTACTTTAACGTGCAAATATTTTCAATGGCATAGAATCCCTACACACATTTTGCTTTATCACTCCTTTATATTTCTGATTAAGTATTCATTAGGTGTGACAAAGAACTGTAACTTTCAGGCTTTATGGAAATCCTGTTTGTGGAAATGCAAATGAGCGCCAGATTACCCAATTTTGTAAATCAAGAGACGGAGGTGAAGAGTATGGAGGCTTGAACAACTCTATTCCAAGTTGTGCAGCCCAGCTGCCTTGCGATGTGGATTTTGAACATGTCCCAGCATTAATGGATGGCTGCTTTTGTGCAGCACCTTTTGGAGTTGGTTTGCGTTTAAGAAGCCCTAGCATTTCAGACTTTCCACCACATTATAGTGACTTTGAACAGTGGATAACAAAAAGCGTTAATTTGAATGATTATCAACTGCAAATTGATTCGGTTTCATGGCAAAGCGGTCCTAGGCTTAGGTTCTTTTTGAAATTCTTCCCTCCACGTGCAAATGACTCTGGAACCAATGATTTTGGCAAATTTAATGACAGCGAGATAGTACGGATCGCAAACAAGTTTGCCTTTTTTAATCTTACTGGAAGTGATATCTTTGGCCCATATGACCTGCTCAATTTCACTGCCATGGCATATAATTCTGGTATGTCTCCAAATCCATTATGTTCTTGCATAAGTAGGATATAATAATACATTAACATGTGGACATTTTCAGTTGTAGTTCTGTAATGATTTCCTGAGGAATTTAAAGTAAAACCAATGTGAACAATAATAGAAACAAACGTCAATCTCGTTTGTTTTTTACAACTTAGATCATGCACGGAGAGTGGCAAAAACTCAATTTTTTTAGTTTAAATTTCTCTAGGGAGGCACAAGAACCCAAGGTTGAGATTAGCATGCAAGCCGAGAGAAACACCTTTATTGTAAAACTTGTTTCTTCAGTTCTTATTTTGTGTTTAATGCCAAATATTGGATACTTTTCAACTGGACTGGAAGTCTGGAACTCTTGTGCACCCGCATGACTAATTTTTCTTTTGTTCTTGCTGAATCTTGAATATAATTATTATGATTATTATATTTCTTTTATCTTTACTGGTCATGCTCGGATAAAAAATTCCCTTTCCCATTGTGCTCCATTTCGTTTCTGAAAATGTCTATTTATATCGTTTACCCTGTGATATAGAACATAGTTTTTTCTTCTTGTCATAGAAGTATGCGAGTATGAAAATTCAATGAAATGGTTTAGATATCATGCATGCTTTGAAGCTTAACATCCTCATGAACATGGTAACCTGACAGAGATGCTAGTAATCTCTTAGATACTCAGTAAGGGTCAAGAGACTAAATTGACTACCTTAGTACAAGTTCTTGATTAAGTAAAATCTTAGGTGAACCATATGGTACTGTCAAAACATGTGAATATCCCCATTCAGAAACTCAAGATGAATTGGCTACTTTTGTTTTATTTTTGGTGTATGAAGAATTTGTTAATGACGCAGAATCTGTGTGAAGAAGCCTTATAGAATCTTTGTAAGCAGAACAAGGATTTACTTTTCTTTTTTTTTCTATTTGAGATGTCTCGCAACACAGCCTTTGTCCTGAATATAAATATATCATTGTTAAATATATGCTATCATATAAGAGAGCCACTCTGTTCTTGTCTTTCTTTCTGCTTTCTCAAGATCAAATAACGAATGTTTTTCCTTCTTGCAGTCCTTTTTCCTCCGTTGGGAGGAGAGAGAAGCAGAAAAAACAGAGGCACTGTGGTTGGAATTGTCTTAGGGTCCATATTTGCTGCAGCTGTGCTACTTATGGCCTTAATATTCGTGTTGTTTAAAATGCGGCGGAGGTTACACGTTTCAAAAGATCAACCATGTGAGTTAATCTTCAGTAATTTTATGTGACTTTTCCTCAGCTATGGGGAGTTTCTTCTCCATTTCTTTCATTCACCCCATTAGGACCTGTTGGTTAAGTAGTCAATACGTTTAACTTTTCTTGGGTATTCTGTTGAATCGCCAAAAAAGAGGACATGAAACCACATTCTTTGAGTGCTTCATGCATATAATGCAAACTTTGCACCTGATAATTTCCAATTTCCATCTTCTGAAAAACAAAAAGATGCCAAAAACCATAAAATCTGTCTTCCTTACTGTTATTTTCTTGTTTAAGTTCTCTATTTAATCAAGAAATAACACAGAATTGCCAGTGTCAACCATCTTTTTCAATGTGGCAAATTATTTTTCTTAGTATGTATACTTGTAATTTAGCTATGATACTGCTTGATGTTAGTTTTTTTAAGGAGAGAGTATGGTGCTTTAATTCCTCACATGGACAGATAACTTTACGGTTTTATGGGAAGTTTGTGCGTATTTATGATGAGTAAACTCTCCGGCAATGCCCCATCACCCTTAGAACAAAAAGTGGAATAGAAATAAATAGGAAGTGTGTGCTGGACAATTGGCATGTATATCTAAGTAATACTCTTGTTGTAGTGCTTCTGTTATTTGTACATCAATTAGACGATTATTTATGTTCTTTTCTTAGTCAGATTAATTGATATTCTTAAAGTGAACAAAGTTAGCTGAAGCCAACTATTGTATCAAAGGTTCATTTGCATTAAAGTTATGCTACCATCACCTAAGCTTTTTCGTTTTCATTCTTTTTATGTCAACTTTTTATGCTTAAGTTTTCAATGAATATCTGATTTTTAATAAATTGAATCTTGTGGCAATGATGTATTTCTTGCTTAAACAAACTTACCCAGGACCACATGGCACTCATAGTGATATTCCTTTCATTTCTTTTGTAGTACCAAAATTTCCAATGCGAATAGAAGGTGTTAAGGCAATGGGTTTTAAGGAATTAGAGGCAGCAACTAACAATTTTAGCAGTACTACTGAAATTGGCCAAGGAGGCTATGGAAAGGTCTACAAAGGCACTTTGGCAGAGGGAACAATAGTGGCAATTAAACGTGCACAGCAAGGCTCATTGCAGGGTGAAAGAGAGTTCTATACTGAGATAGAATTACTTTCACGTGTGCATCATCGAAATCTGGTTTCCTTGGTGGGATACTGTAATGAAGGAAGTGAACAAGTAAGATTCTCCAAACTGGAAATGTAATAGACACACACACACACACACACAGAGCCAAGTCCATGTTGCTATTCCTGCATCCAGATGGGAGAGGGGAGTATGTTCGGTGGAATAAGTTCTTTGATGTATCAGTTCAGCCACTTGTTGAATTCTGATTGCCCAGTTCTGGTTGAATACCTAGGTCAAAATTGAATTTTTTTGCATATCCTTGATCATTTTGGATGCCTCATTGCCGTTGTGCAGAACCAATCAACCATAATATCAATTACCTTCTACAACTTTGAGAAATCCATTTGATATCACTCTAGATGCAAATGAAATTACAAGAATCACAAATGATAGTTAAATTGCAGGGCAAAGTTCAGACTTAGCTGATTCATCTGTTAAAGTAAACTCGAGCAAAAGTCTTCTGAGATAGTATTTTCATCATACTTCTTTTTTTGTCACCAATTATTATCACACGTTAAGTGATTCAATTTGTGAGCATATCCAGTGTACAGTACTTAGATCACCTAATATTAAGTTCAACCAAATTTCCATCTATGAGTAGCTCATCAGTTACATCCCTCTTATACTTGCGTAGTCTGGGCACATTAGTGAAATGCATGTCTTCTGAAAAAGTAATCAAATAATCAAGCAGATGGAAGGATAAGAACGGATATTATGTTTTATGTTTTTTTATTCTACTTTTGTAATAGTTGGTTACATGTGCATGCAGATGTTGGTGTATGAGTTCATGCCAAATGGTTCCTTGCATGATCTTCTAAAAGGTATGTGATGTATCCTTCCTTTGCCTATGTTGTCATTTGATTTTCTTTTGTTAGAATCAAGCACATCTATTCTCCTCTTATCACTTATTTTTTCCTGGTGTAGTACTTCTTATACGAAAAAGAGGGTGAAAGATATGTCCCTCTAGTAAAGAATATTGTATAGATGAAAGTAAATCAAAATTATCTAGTCACTCAGATTTTCTTTTCTGAAGTTTTTACTTCATTTTGACTCCTGAAAGTCACTGTTAGTTTTCCTACTTTCTTCACTTGGGTGGTCATCAACAAGTTTATCACATGGAACTTTATATTCTTAATTTTAGTCATGTGTCAGTTCCAATATGCATATCAAACAGGTATCTCCAAAGTACAATTATTAAAGATTTTTTTACTTATTTTTACAGCTAGATATGGGGAATGTCTGAGTCTTGGAACAAGGTTATACATTGCTTTAGGTGCTGCCAGGGGCATCCTCTACCTCCATACTGAAGCTAATCCTCCAATAATCCATCGTGATGTCAAAGCAAATAACATACTGTTGGACTCTAAGTTCACCGCAAAAGTTTCTGATTTTGGAATCTCAAAGCTTGCACCATTACCTGATGCTGAAACGAGTGGGCATGTATCAACTGTTGTGAAAGGAACTCCTGTAAGGCTTTACTCGTCTTATATGTTGAAGCCTGCAATAGAAAAGCTAGAAAATCGCAATGAGATAAGATCTGCAACTTGAGTAATCTAGGTAGCATTGCTACAGACCAGGAAAACTTCATTCTCT

>PGSC0003DMG400031752 LRR-RLKs

GTCAGCTTTTACTACTCCCTTTTCCCAAAAAGAAAATTGCAGTCTTTCTTCTTCTTCTTCTTTGTTCTTTCTTGATCCCCAGAAAACCAAAAAATGAAATCTTCAATTTCAATAATGTTTCTTCAAATCTTGGTTACCCTTTTGCTCCCAACTTTGATTTTCTCACTTAACCAAGAGGGTCTTTATTTACATAATGTGAAGCTTGGATTTGATGACCCTGATAATGTTCTTTCAAACTGGAATGAATATGATGATACACCATGTAACTGGTTTGGTGTTTCATGTGACCAATTGACTCGCACTGTTACATCATTGGACCTTTCTAATGCTAATGTTGCTGGTCCTTTTCCGACTCTGCTTTGTCGGTTGAAGAAGCTTCGTTACATTTCGTTGTATAACAACTCTGTGAACAGTACTCTTCTTGACGATTTATCTGGGTGTGAAGCAGTGGAGCATCTTGATTTGGCGCAGAATTTCTTGGTGGGTACACTTCCGGCGAGTTTATCTGAGCTCCCAAACTTGAAATATCTTGACTTGTCGGGTAATAACTTTACCGGAGATATTCCGGCGAGTTTTGGTTCTTTTCAGCAGCTTGAAGTTCTTGGGTTAGTTGGAAACTTGCTTGACGGGAGTATACCGGCGTTTCTCGGGAACGTTACGACGTTGAAGCAGCTGAATCTGTCGTACAACCCGTTTACTACGGGTCGGATCCCGCCGGAGCTGGGGAATCTGACGAATCTTGAGGTTTTGTGGCTTTCCGACTGTAATTTGATTGGGGAAGTTCCTGACACATTGGGGAGCTTGAAGAAGATAGTGGATTTGGACCTAGCTGTTAACTACTTGGATGGGCCGATTCCGAGTTGGCTCACTGAGCTGACTAGTGCTGAACAAATTGAGCTGTATAACAACTCGTTTACCGGGGAGTTTCCGGTGAATGGGTGGTCGAAAATGACGGCGTTGAGGCGAATCGACGTTTCGATGAACCGGGTAACTGGTACGATTCCGAGGGAGTTGTGTGAGCTGCCACTTGAGTCACTCAATCTTTATGAGAACCAGATGTTTGGTGAATTGCCACAAGGCATTGCAACTTCACCAAACTTGTATGAGTTGCGCCTTTTTCACAACCGTTTTAATGGGAGTTTACCTAAACATCTTGGAAAAAATTCACCTTTGTTGTGGATTGATGTGTCAGAAAACAATTTTTCTGGTGAAATTCCGGAAAATTTGTGTGGGAAAGGGTTGTTGTTGGAGCTTTTGATGATAAATAACTTACTTTCCGGTGAAATTCCGGCGAGTTTGAGTGAATGCCGGAGCTTACTGCGCGTGAGATTGGCACACAACCAGTTATCCGGTGATGTTCCGGAGGGGTTCTGGGGTCTGCCACACCTTTCTCTGCTTGAGCTCATGGACAATTCACTCTCCGGTGATATCGCGAAAACTATAGCCGGTGCTTCAAATTTATCAGCTTTGATTTTGTCTAAGAACAAATTTTCAGGTTCCATCCCAGAGGAGATTGGTTCTCTGGAAAATCTTCTTGATTTTGTGGGCAATGATAACCAGTTTTCTGGGCCTTTACCTGCAAGCCTGGTGATTCTTGGACAATTGGGAAGGCTGGATCTTCACAACAATGAGTTAACTGGTAAGCTTCCAAGTGGGATTCATTCTTTGAAGAAATTGAATGAATTGAACTTGGCAAACAATGATCTTTCTGGAGATATCCCCAAGGAGATTGGGAGCTTGTCTGTTTTGAATTATCTTGATCTATCAGGGAACCAGTTTTCAGGGAAAATCCCAGTGGAGTTGCAGAATTTGAAGCTCAATCAGCTGAACTTGTCGAATAATGACCTTTCGGGTGATATTCCCCCTGTTTATGCAAAGGAAATGTATAAGAGTAGCTTTTTGGGGAATGCTGGTTTATGTGGAGACATTGAGGGCTTGTGTGAAGGAACAGCTGAAGGTAAAACTGCTGGTTATGTTTGGTTATTGAGGTTACTCTTCACCCTTGCTGGATTGGTGTTTGTTATTGGGGTTGCTTGGTTCTACTGGAAGTACAAGAATTTTAAGGAAGCTAAAAGGGCTATTGATAAGTCTAAATGGACTTTGATGTCGTTTCATAAATTGGGTTTCAACGAGTATGAAATCTTGGATGCTCTTGATGAGGACAACTTAATTGGCAGTGGCTCTTCTGGGAAGGTTTACAAGGTGGTTCTGAGCAAGGGTGACACTGTTGCGGTGAAGAAGATTTTGAGAAGTGTGAAAATAGTAGATGAGAGTAGTGATATCGAGAAAGGTAGCTTTCAAGAAGATGGATTTGAAGCAGAGGTTGAGACGTTGGGGAAGATTCGACACAAGAACATTGTTAAGCTATGGTGTTGTTGTACAACTAGGGATTGCAAACTTCTGGTCTATGAGTATATGCCTAATGGAAGTTTGGGTGATTTGCTACACAGCAGCAAAAGTGGCCTTCTAGACTGGCCTATGAGATCTAAGATAGCCATGGATGCAGCTGAGGGACTCTCTTACTTGCATCATGACTGTGCTCCACCTATCGTTCACAGAGATGTTAAGTCAAATAACATCTTGCTGGATGGTGAGTTTGGAGCTCGTGTTGCTGACTTCGGTGTTGCAAAGGCGGTCGATGCCAATGCTAAGGCAATCAAGTCCATGTCTGTTATTGCAGGGTCTTGTGGTTACATTGCTCCAGGTTAGTGGATTAACTTTTTCACTTTTCAGCTTCAGTTTCATGAGTTTGTGTTGAATTTGTTCATTTTGAGTTTTACTAGCAAATTGTTATGCCTGTAAGTTTATGGATTACTGTAGCCTTTCAAAAATGAGATCTAGTATTACTGTAGCTATGTGGCACCTATTGTGTAGAAGGCTAGATCTATATACTACGATATTTCCTAGATATTCCACATTTGTTAAGTAATTGGCTCATATCTGGAGTACCCCCTAATACAGTAACAGTGTACCATTGCTCTAGATCATAGTTTGTGAACAGTTAAGTAGGCTCCACGAGTTTCCGGCTCTTGAGCACAAAAGACTGTATTAGGACTGTAGAAAAAGATATACTTAGCTTTCCACTTATGATGTTTGAAATCTTAACAGCTAGCTATATTGTGGGCGGCTGCTAAGTTTAATAACCACCCACCCATATTTTTGGTTTCATGTGGTCATTCTTTGTTGTGTTGTTTCCTTGCAACAACTTAGTTGACTGAATTCTTTTTTGCATTCCAGAATATGCATACACACTGCGGGTGAATGAGAAGAGTGATATATACAGCTTCGGTGTGGTCATCCTAGAGCTTGTAACGGGGAAACGCCCTGTGGATCCTGAGTTCGGGGAAAAGGATCTGGTGAAATGGGTGTGCAGTACATTGGACCAAAAGGGCATAGATCATGTAATTGACCCTAAACTTGATACTTGTTTCAAGGAGGAGATATGCAAGGCCCTAAACATTGGCCTACTCTGCACTAGCCCTCTCCCAATTAACCGACCCTCGATGAGACGAGTCGTTAAAATGTTGCAAGAAGTGGGTGGTGGGAACTTGCCCAAGGCTGCCTCAAAGGATGGCAAGTTGACACCCTATTACTATGAAGAAGCATCAGATCAAGGAAGTGTAGCTTAAGAATGTCTCAACAAATTTTGTCCTTTAAAAGTGAGACTCAATTAACTTCTTTGTTCATCTGCTAACTACTCAATTCTTTCCCACTAGATGTTTGGGTTGAAAGAATTGAAGTTAAATGAAAAAAAGGTGAGAACAAAAGAATATTAGGAAAAAAAAATGGAGTTGGAAGAATCTTGTTTTTCATTGTAAAAATCCCATCCAACTACCAAAAGTTATTGCAGTAAGTATTTTACTATCTGGCACTAAAAGATCTTAAATGTGCTACCTATACTCTATTTCCTTG

>PGSC0003DMG400033049 LRR-RLKs

GAGGAAAACAAGATTGTAATTAAGTGACACACTAATTATCATGTCTGAAGTGGAATCTCATGTTCTTCTTCTCATTGTATTACTCAATTTCTTCATTCTCATAACAGGTATGTATATATCCTCATGTCATATTTGATAGCTTTAATTTGATCAAGTTTAGTATTAGAGTCTCATTTTGTACATAATTAAAAAGAATCAATTTTTTAAACAACATTTATTGCAGCAAGACTAATGTTTTTTTTTACTTAGCAAAATCATAGGACCAGTCTTGTTTGTGTTACTCCAACACCAAGCATTAAATGTTGTTCTGTTTTGTTTTTGGGAGAATTGACTTGTATTTATTGTAGCATGATTAGTTTATGCAAAATATTATATAGGTAGGCTTGTTTGTGGAGACTCAATTGAGACAGACAAGCAGATGCTGCTGAACTTGAAGTCATTTCTTAAGGAACAAAATCCTGTTGACAAAGGATTCAAATATAACCATTGGAATCCTACAGATTTGTCACCATGTAGATGGCCTGGAATATCATGCAGTACCTCTATCAATCGTGTCACTGGAATCGATCTCTCGGAAAGTAATCTGGCTGGGAAATTGTTTAATAACTTCTCAGCCATGACAGAATTGAACTCTCTTGACCTGTCCAAGAACACATTTTCAGAGTCCATTCCATCAGACTTAGGCTGGTGTAGAAATCTGAAGTTCTTGAACTTGTCACACAATATCATTGTTGGTGAGCTCAACTTGACAGGCCTGAACAAGCTGGAAGTTCTTGATTTGACGATGAACAGGATCCACGGGCTAATGATCCCTGAGATTTGTGACAACTTAGCTGTTGCAAATATTTCTAACAACAATTTCACTAGTGGGAGTGGATTTGAATTTTCTCACTGCAAGAAACTGAAGTATCTTGATCTGAGCTACAATTATTTGACAGGGAATCTGTCGTTCGGGCTTGATATGCTGAATATGTTTTCAGCATCTCATAACAACTTAAGTGGCTCTCTGCCTTCCTGGATTTTCACTCAAAACTGCTCTTTGCAAGGTTTGGACTTATCAGAAAATATGTTCTTTGGTGAATTGCCTACGTCTATTGCGAACTGTAAACGATTAGTAGAGTTGAATTTGTGGGGAAATAGCTTTTCAGGGTCAATCCCTAGAGAGATTGGATCAGTACAGAGTCTTAAAGAACTTTGCTTGGGAAGTAACAACTTTTCAAGTGATATTCCAGACACTCTATCAGGTCTAAACAAATTGGTATTTCTGGACCTAAGTAGAAACAACTTTGGAGGAGAAATACAAGAAATTTTCGGGAAATTGACACAGGTAAGATTTCTCATGCTGCACGGAAACTCATATATTGGAGGCATAGTGTCATCAGGAATTCCAAACTTGGTGAACCTTTCGCGGTTGGACTTGAGTGATAACCACTTCTCTGGTCCATTACCAGTTGAAATTTCTGAGATGAAAGGTTTGGAGTTTTTGATTCTTGCCCACAATCAGTTTAGTGGCAATATACCTTTAGAATATGGAGATCTTACTGCACTTCAGGCTGTTGATCTTTCCTCTAATAGGATCAATGGTTCAATACCACCAAGTTTCGGGAAGCTAAGGCCACTACTGTGGTTGATGCTTGCAAACAATTCATTGAGTGGTGAAATCCCATCAGAGTTGGGGAACTGCAGCAGCTTGTTATGGTTGAATCTCGCGAATAATCAACTTACTGGTCCAATTCCTCCTCAATTAGCAAGTATTGGTGCAGATCCAATGCCTACTTTTTTGTTGAATAGGGGAAAGGAGAAGCTCACTGCTAGCCCAGGGGATTGTTTTGCCATGAGGAGGTGGATACCAGCTGACTACCCTCCATTTAGCTTTATATATCCTCTATTGACAGGGAAGAGTTGTAGAATCCTGGGGGATAAGTTGTTTATGGGAGATGGTTTAATGCCACTCTGTGAACCTGGTAGTAATGTCCGAAAGAATCAGGTACCGGGCTATATTCAACTTAGTGATAACAAATTGTCTGGTGAGATCCCTCCTGAGATTGGCAACATGAAGAAAATGAGTATGATGCATTTGGGTGCAAATGAATTTTCTGGAAGGCTCCCTTCAGAGATTGGACAAGCGCACCTAGTAGTCCTTAATGTTTCACAGAATAAATTTTCTGGTGAAATCCCAAAGCAGATTGGCCATCTTAAGTGCTTGCTAAACCTTGACCTTTCATTTAACAATTTTTCTGGTCCATTCCCAGTTAGCTTTAGTAACTTGCATGATTTGAGCAAGTTCAACATCTCTTACAACCCATACATCTATGGAGCTGTACCAGAAATCGGGCAACTGCTTACATTTGATAAGTCATCATTTCTTGGTAATCCGTTGTTGCGTCTTCCATCCTTCATGCATAACTCTAACAACAACACAGAACGAAACAAGAATGACAATCACAAAAAGCCCAAAAAGGTGGGTGCACTTTTGGTAATTGTGGTTCTGGTACTAGCTTTCCTAGTCTGTGGAGTCATGTCACTCCTTGTCTGCCTCCTTATCAAAGTGCCAAGGGGTTCCCCAGGAATCTTACTGGAGGATACAGAGGGCAGACATGATTCTCCATCAAGTACTAATGCATCCTCCTCACGGTTGTGTGCATCCTCATCACGGGGTTCTGATGATGTTAAGGTTATCCGTTTGGACAGAACAAGCTTCACACATTCTGATATACTGAAGGCCACATGGAACTTCTCGAATGATAGAATTATCGGGAGGGGAGGATTTGGGATAGTCTATCGTGGAGTCTTGCCTGATGGAAGGGAAGTAGCAGTGAAGAAGCTACAGAGGGAGGGAATCGAAGGAGAAAGAGAGTTCAGAGCTGAAATGGAGGGGCTCAGTGGGAATGGCTCCGGTTGGCCTCATCCCAACCTTGTAACTCTTTATGGGTGGTGCCTTGATGGATCAGAGAAACTTCTAGTCTATGAGTACATGGAAGGTGGTACCTTAGAGGACTTCATTACAGATAGAACAAGGTTTGCATGGAAGAGCAGAATTCAAGCAGCAATCGATGTAGCACGTGCTTTAGTCTACTTGCACCATGATTGCTATCCTTGCATTGTCCACAGAGATGTCAAAGCTAGCAACGTGCTTCTTGACAAGGACGGAAGAGCAAAAGTCACCGACTTTGGCCTTTCTAGGGTCATGATTTCTGAACATACTCATGTTAGCACAATGGTGGCAGGGACTATTGGTTATGTTGCACCAGAATATGGGCAGATAATGCAGGCCACTACAAAAGGTGATGTCTACAGCTATGGGGTGCTAGCAATGGAGCTAGCAACTGGGAGACATGCTATAGATGGGGGTGAAGAATGTCTAGTTGAATGGGCGACAAGGGTCATGGGAGACAGAAGGAAAGGGTTCACCAGAGCCATTATACCAGTTGCTCTATTGGTACCTGGCCTAGTAGAGGGAGCAGAGGAAATGTATGAGTTGCTTAGGATTGGGATAAGGTGCACAGCTGAGACGCCTCATGACCGGCCTAACATGAAGCAGGTATTAGATATGTTGATTAGTATTCCTAGCAGCCAAAAGGGATCCAGCCGTAGCTTTGGATCAAGTCGTAGCACTTCTCCATTATTATGATAGACTTTCTTGGAACCCAGAGATAAAAACGTCAATAGTTTCTTTACAAGATGTGTAACATACCTAGCAGATTCTTCTTGTAAAACAGTTTTTCTTTTTTCACATTTGTTAGTCTAGGAAAATGTAAGTCAATTTTGTTTGTTTGGATCACAAAATGTTCTCAAGGATATCAATTTTTGCTATGAAATTGAAGCACACAAATATTTGACCCACCTACAAAGTTACACTACAGTAGGTGGCAAGATTGTTCCCCTTATCAGGTACATTATTTGTCTGGTGCAATTCAAAGTCTCTATTTCAAACAAGATTCCTCTACCAGGGACTGTGCAATAGCAACTACAAACAGGAGTGTCTCCAAATTTCAACTTCTGGACCACATCTGATGGAAAATTTTAGTATACTTCCACTACATCAACACACCACCAATTGTCAACGCCACAATACTCATTTTCATATCAGGCTAGCTTCATCATCCGTGGGAATAATGTGGATATGCACTCAAGCATCATACTGATTTCAAGTTAAGGCTCAAAATTTTAAGTCACTCTCTGATGTATAAACACTGCTAATAGAAGAGTACAAATTCTTGTCTATCTGCTTAAGAAAAAGAAACTACAGATAACGTTTAGTGAAACTCTTAATGGTGTGAGACCTTTTAAGAAAAACATTGCGAGGTTGGATGAAAACTACCAGTATCCTACCATGTTAAGAATGTCTTTGGGCTAAACATAACCATGTATACCCCCATTAATCTAGAGGTCACCTATG

>PGSC0003DMG400033667 LRR-RLKs

CAATTTGGCATTTTCTTCACACCTCCAATGTTGACCAAACTTTCCAAAATCAACTCATAATTCTAACCCCACCTTTTTTTTTTCATGGAGTTTTCCCATTTTCCGACCGGCATCTTCGCGGGAAAACAGTTCCTCTATCTCATTATTCTACTAAACTTCATACCCATTTTCGTAACTTCACAATCTCCGGCCACTACTGAACGTGACACTCTACTGAAGATTAAGCGTCAATGGGGAAACCCAGTAGCACTTGACTCATGGAACTCTACTTCTTCTCCTTGTAGCTGACCGGAAATTGAGTGTGATGATGGGAAGGTTACCGGAATAATAATTCAAGAAAAAGATATTACTGTTGAAATTCCAAGTTCTATTTGTGAACTTAAAAATCTTACTTTTCTTAATCTCCGGTTGAATTATCTTCCCGGAAAATTCCCAACTTTTCTTTACAAATGTTCCAATTTGCAGCATTTGGATCTTTCTCAGAATTACTTTGTGGGTTCCATTCCGGAGGATATTCACCGGCTTGGGAAACTCAAGTATCTTAATCTGGGTGGTAATAATTTCACCGGAGATATCCCGTCGTCAGTCGGGAATTTGATGGAGCTGGAGACTTTGTGCATGCATTCGAACTTGTTTGATGGGAGTTTTCCGGCTGAAATCGGAAACTTAGCCAATCTTGAAAGTTTGGGGTTGGAGTTTAATGGGTTTTCTCCAATGGCACTACCGCCGGAGTTTGGGAAGCTGAAGAAGATAAAGTATATTTGGATGAGAGAGACGAAATTAATCGGTGAAATCCCTGAAAGTTTTGGGGAATTTCAAAATCTTGAGCTTATTGATTTTGCTTACAACAATCTGGAAGGAGAAATCCCTTCTGGGTTGTTTCTGTTGAAGAATTTGACGATGATGTACTTGTTCAGTAATCGGCTTTCGGGTCGAATCCCTGAAACTTTTGATTCATCAAAGTTGATGGAACTTGATGTTTCCAACAACAATAATTTAACTGGAACAATCCCAGAAAGTTTTGGAGGGTTTAAACATTTGGAAATTATGAATCTGTTTGGTAACCAATTGTATGGAGCAATTCCAGAGAGCATAGCCAAGATTCCATCTTTGAAGGTATTCAAAGTGTTTAGGAATAAACTGAATGGAAGCTTACCATCTGAAATGGGACTTCATTCGAAACTCGAATCTTTTGAAGTTTCACTGAATTCCTTCACTGGAAACTTGCCAGAGCATTTGTGTGCTGGAGGAACTCTTTTTGGTGCTGTAGCTTATGTTAACAATTTGTCTGGTGAAATACCAAAATCACTCGGAACTTGTTCTAGCTTACGTTCGATTCAGCTCTACAAGAATCAGTTTTCGGGTGAAATTCCATCTGGGGTTTGGACTTTGGTTGACATGACAAGTTTGTTGTTGAATGATAACTCATTTTCCGGGGAGCTACCGAGCAAAATTGCATTCAATTTTACGCGGTTAGAGATCAGTAACAACAAGTTTTCTGGTGAAATTCCTGTGGGGATATCTTCTTGGAGGAGTTTAGTGGTGCTTTTGGCAAGTAACAATTCATTTTCAGGTCGGATTCCAGTGGAATTGACTAGTCTTTCTCAGATAACTCAGCTAGAGCTTGATGGTAATTCGTTATCTGGAGAACTTCCAGCCGATATAATATCGTGGAAGTCGTTGACAATATTGGATCTTGCCAGGAACAAACTCTCTGGCAAGATTCCAGCAACTATAGGTTTAATCCCTGATCTTGTTGCTCTAGATTTGTCTCAAAATCAATTTTCAGGTCCAATTCCACCTCAACTAGGTGTAAAAAGGATCACTAGCCTTAATGTATCGTCCAATCAACTTATCGGGAATATCCCTGATGCATTTGCTAACCTAGCCTTTGAAAACAGTTTCTTGAACAATCCCAGTCTTTGTACCACCAACTCACTTCCATATCTTCCAAGTTGTAATAATGCGAAAGTCGCCAATTCCAAAAGATTGTCCCATAGAGTCCTAGCCCTAATTCTAGTCCTTGCATTCGCTGTTTTCCTATTTAGTGTTGTATCGACCTTGTTCATGGTTAGAGACTACAGGAGGAAGAAGCATAAGCGCGATGTTGCATGCTGGAAGTTGACTTCATTCCAGAGGTTGGATTTCACAGAGGCGAGCATTTTGCCAAGTTTGACGGAAAATAACATGATTGGAAGTGGTGGATCGGGTAAGGTGTACAGAATTTCCGTTGGCAGACCAAATGAGTATGTTGCTGTGAAGAGGATTTGGAGTGATAGGAAAGTAGACTACATTCTGGAGAGGGAATTCTTGGCTGAGGTTCAGATATTGGGATCAATTAGACATTCCAATATAGTGAAGTTATTGTGTTGTATATCAAGTGAGGATTCAAAGTTGCTCGTTTACGAGTACATGGTAAATCACAGTCTTGATGGATGGCTTCATGGGAAGAAGAGAGTTTCCTTGAGTAATAAAGTCATGGATTGGCCAAAAAGATTGGAGGTTGCCATTGGAGCTGCTCAAGGACTTTGTTATATGCACCATGACTGCACTCCACCGATCATTCATCGCGATGTAAAAACAAGCAATATCTTGTTGGATTCTGATTTCACAGCCAAAATAGCTGATTTTGGGTTGGCCAAGATTTTAGAGAAGAAAGGAGAGCTTAACACCATGTCTGCTGTTGCGGGTTCCTTTGGTTACATAGCTCCAGGTAAAAGCAAAATCCAAGCTTGAACTTCGTATTACCTCAATTTCGTTAACATTATATCAAAAGTATGACTAACTAATAATTTACTGTTTGGTTACAGAGTATGCTTATACAACGAAAGTGAACGAGAAGATTGATATCTATAGCTTCGGGGTGGTGCTATTGGAGCTAGTGACAGGAAGACAACCAAATTTTGGGGACGAGCATACAAGCCTGGCAGAATGGGCATGGAAGCAGCACGGAGAAGGGAACACTGCCATCGATAATATGTTAGATACAGATATCAAAGAAACATGTTACTTGGAAGAAATGAAGACTGTGTTCAGACTTGGGCTTATATGCACGAGCAATTTACCTGCAAGCAGGCCTTCCATGAAAGAGATTCTGCAGATACTTCATCGATGCAAGTCCTTTAGGTATTCTGGAGGAAAATCACCAGGTACAGAGTATGATGTTGCTCCTCTCCTTAGCGGAAACAACAGCGAAAAATACATTGCTAGTTACAAACGTATTAATTCCAGTAAGGTAATAGATGATAGTAGTGATGATGGTCTAATAATTTCCAGTGTGTAATATTGGAGATGCCAAAGTAAACTGAGTTGTTGTAAAATGATCATAGAATAGTGCCACTTAAGTAAAGTACAAGTTTGAATTTCTAGGTACTCTTGAATTTGTTCACTGCTTGTAGACTGAGTGAAATGCATCATTTAACTTTTGCTAGTTATGTAGATTGTAAGGAAGTGTGAAATTGGAAAATGAAAATGCCAATTTATATAATCAAAGAGAAATGACAGTAGAGTGTGAAATTGGAAAATAAAAATGCCAATTTCCAGGTCAACACGTAGTTCATTAAGAAGATTTGGTTTTGACTGAAAATGACTCAATAGGAGGGTATGGAGAGGCGAT

>PGSC0003DMG400033914 LRR-RLKs

CTGGCAAGTGAGGCCAAAGTGACCACAATACATGTCCATGTCAGTGGCCACTACCTTCTAACCTGCCCGGAGAAATTCAACTCTGCGAACTGACACACGCATAGGCATCGGTGTTTCTTTTGTTTGGCACAACTGGGGGAAAACACCGGGCAAACCACCCCCTTTTAATTAAATCTTCTGATTACAGAAGTGGTTTAGGGTTGGGATGGGAATCGTGTGCAGTTCTCAGCTCCACACCCCCGTGTGACAACTTAGCTTATGCTTTTGCTTTTAAATCACTATGTTTCCTTTTGAATATCCGATGTGGGACAAAGAGGCCATTAAAGTGATTTTGTATTGCATTATGAAGTGAAGGCTGTATTTTGTTCTCTTTGACTGCCTCTGTTTTTATCCAACCTCCATCGAAGTGGGAGCTTACACACTGTATAAAAGTAGAAAAATGGAGAGGTTCGCGCTTAACTACTTCTCCCTCCCCTTTTCACTATTTTTAAGTATTTCCTCTGTCACCCATCAACTGAAAGAAAAGAATAACCCAACTATTATAGCACTTGTATACTTTTCAAGAGGAAGAATCAAACCCCAGACCTTCTGATTCAGTACAACGCTGAATGATAAGCAGTGGGGTTTTGCCTCACTTGTATTCATATTTGTATACCCCAAACCTTCACAAGTTTCAATGAAGGTTCAGCAAAAATGAGGCACCAACTGGGTTACCTTTCTTTCCTCCTTTTCTTGCTTTTTCATTCTACATTCCAGCAGCACGAAGATAGATTGGCTTTACTTCAATTGAGATCGTCTTTAGGATTGAGAGCCAGAGAGTGGCCTATTAAGGGTACCCCTTGTTCAAATTGGAGAGGTATTTCTTGCAACAATGGTAGAGTTACTGGGATTAACATTTCTGGTTTTAAGCGAACACGAACTGGGAATCAAAATCCTCAATTCTTGGTTGATTCCCTTCAAAATTTGACTCTGTTGGAGTCATTTAATGCATCTAATTTTGCACTTCTGGGTTCTATTCCTGATTGGTTTGGCATTGGACTTGCATCTCTACAGGTTCTTGATCTTAGCTTCTGTTCAATAACTGGTGCTATTCCTTCAACTATTGGTAATTTGAGAAGTTTAGCTGTTTTATACATATCTAATAATGGTCTTAGTGGAATGGTGCCTTCAAGTTTGAGTCAATTGTTAAGTCTTTCAGTTCTTGATGTTTCACACAATGCACTTGGTGGGTCAATTCCTGCATCATTTGGGTCTCTTGGGAATCTTACTTTGCTTGATATGTCTTCAAATGTTTTGTCTGGGGCTATTCCTCCTGTAATTGGGACTTTATCTCAGTTGAAATTCTTGAATCTTTCTCGGAATATGTTGTATGGGAATCTGCCGCCTGTAATTGGGAGATTCAGTTTTCTTGATATATCAAACAACTATTTTCAAGGCAAAGTTCCTCAGTACTGGCAAAGCAATGTGACCCTTAATAGAAATTGCCTACAAAGATTGGAAAGTCAAAGGGATACAAGAGAATGTGCATCTTTCTATGATGAGCGGGGCTTGATATTTGACAATTTTGGAATGCCTAATGCTATGGAGCTTCCGGTTCTTATATCTGAAAAGAAGAGTCCCAAAAATGTGATCATATCAGCTGCTGTTCTAGGGGTTGTCGGATTTATTGCTACTGTGATGCTCTTTGTAGTGCTGCTGATTTGTTGCTGCAAGAGGGGCGTGACGAATCAAAGGGTCACTGGTGTAGGGCCTGTTCCTACTACTGCAAGTCCTCCCCCACCTCCAGGACCATCATTGAAATTTTCAAGTTTAGGAGTAGCATTTAACTACCAGCAGATCCTTCAAACAGCCGGTCATTTCAGTGATGTAAATCTTATTAAGCATGGACATTCAGGTGATATTTTCCGTGGAATCCTGAAAGGTGGAGTTCATGTAGTAGTTAAAAGAATTGATTCCAGCTCAGTTAAAAAGGAAGCATACTTGTTAGAATTGGACTTCTTCAGCAAAGTTTCTCATTCTAGAGTAGTCCCTTTTCTGGGACATTGTTTAGAGAGTGAAAATGAAAAGTTTCTGGTATACAAATATATGCCAAATGGAGATCTCTCTAGTTCTTTGTTAAGAAAAAACAATGCAGATGATGATAGTTTGCAGTCACTTGATTGGATAACGAGACTGAAAATCGCTATTGGAGTTGCTGAGGGTTTGTCTTATCTGCATCATGAATGTAATCCACCTCTTGTGCACAGGTATTTTAGAGCTTGTTGTGATGCATTACTCTTACTTCCACTTATCACGCTTAATCTGACAACTTTTAAATTGCAGTAACAAACCAAATCTATTATCTTTATACTGCCTATCTGAATCATAAGTTATCATTGTGATGGGACACTTGATCATGAATCTTGGAAGAGTAGTTGCATATTCCTCTACATATGTTTCTTTGGTTGTAGACTTTTGGCGAATTAGCATAATAAATGATATGCATGTTTTATTAATACATGATATTTCAGAAAATTAAAATCATCCATTATTCATGAAAGATGAAGTGCTTAAGTTTCCTCCATACTAATTGGCGTCACTGAAAAAGCCTCTTCTTACTGATAAAATTTGGCCAAGTTAATGCTGGCACCATTTTGTACTTCATGAATTTCAACTGTCTATCAGTGCCAATTAGGTCAATACTATAAGAGACATGAGTGAAAGATATAAGCTAGAGCTGGATGGTTTGGCAATTAAATATACTCTGAGGTGTTCTTTTTTTGGAACTAAGTGGCCACATGATACAAACCGGAGCTGGATATTTAGCAATTATATACAATCCAAAACTGTTCTTAGCAGTGATCTAAAAGCTTGCTAGGACACATGAATTGGGCAAATCCAATCATAAACTATTTACAGAAAATATGAAACCAAACTTTGTCTTGAGATACCCCTGTTACAACTTACAAGAGTAAGGGTGAATAGAATAAGTCCTTTACTCTTTTGGCCGTGTATTACTTCACAATCTTGGGCTAACTACTCCAGACGTACTGAGTGAAACCTAATGACAAAAAAACTCCTATTGGCAACTCGGTCTAAATTTTGACAAAACTCCTATCACTGCCATCTTTCCTGAACAACTTGGATCACAGCCCAAATGTTAACCCTTCTTCGGAATTGTCCTTTTTAGAAAAGATTTCTATTTAATTTTCTCATATTGTTTCATCAGAAGCTTGTAGAGAATGATAACTCAATGAACAACTTTGTGGATTTAATAACATGTCTGGTTGTTTGAAATTGATCCAGTATGTAGATAAATAGCAAAATATCAGATGACAGCTGCGACAATTAGCTTCATGAAATTTTACAAAGACAAAGAAAAGGAATAGACAAGACTTCATCCTTGTTTTGCTCTTCATTATATGTTCTCTATGTGTTATTCACGTGCAATAAATTACAAATTTTCCATAACAAGACGATTTGGCCAATGAGGGGCCATGGTTATGTACACTTGTACTAGCTATTCTCTCTAGTTGCATGCATGTCTATTTCCACTTTAGCAAGTTCTGGTTTCTTCTAGATTCTATATCTAGTGACATGGCTTAAGGTTCACTGTTACTAGATCTCTTACTGCTCTAGCATATTAAAATTTATTTCTGAAACTACGGCGAATTGAGGTAGCTTTGCTTATTAATTTTTTGAAAACTTTCTAACATCGAGGTCTTCTTGTGTTTGACATGGGTGCCGCTTTTGATACTGAGATTTTATGAGATCGCTAAACTTGTTTAGTTGAGTACTAGCTGGTAAATATGTTAGGTAGGTCATTTAGAAATCTCATAATGTAGAAGTGACCTTTTCTTATTTTTTCATCAAAGGATTTCTAATAGGATAGCACTTTGACTGACTTTTTTCATTTTAAAACATCAGCATAGCTCCTTGACATATCACATACTCGTGATAAACTATGTACAGATTGATTGAGCTGGTATAACCGTGACTATCAAATTTATTTCTTGGATTTCCTTTGTACTTCAGTAGACCCATCGAGCTTCTTTCTGAAGTTAGTTTGAATATTTACTAAACTTGCAAAACAGGTTTAGATATCACCATGATTTATGTCCATTTCTTCCTTCCAATTTTAATCATCTTCCAATTTTAATCATGAATGTACTCTATTTCACTATAGGTTACTACACTTTCCTCAAGCTGTTCACCACATTTTTTCAGTTAAGTATTATTTTTTTTATGTGGCAACTAGAATTTCTTAAGCCAGCATGCATAAATAATTATATTCTTATCTTCTCTCTGTTTCTTTTCCACCATACCACGTGGTAGACAAAAGATGGTCTAAGCTGGAACTTCCCAAACGCCTCATAATAGACCATACTTAAGGTAGTGCATGAAAAGGAGAAACCAATTGCACAAATTTCTGCATTTTCCCTGAGCTATATCAGTGAACTGTAAAAAAGACGGATTTTATACTATTCTTAATGGTTACATTGTCTTTCTGCTGCATTGCAGATGAAAATTCACATAATCTCGGTTATATAATTTATGCCTTGGGGTACTATGGTGAAACACCTTTTTTGAACCTTGTCTTGCTATATTTACAGAGATGTCCAAGCAAGCAGCATACTTCTTGATGATAAATATGAAGTACGGTTAGGGAGTTTGAGTGAGGTCTGTGCACAAGAAGGAGACACCCATCAAAACAAGATTTCCAGGATGTTGCGGCTGCCACAGTGAGTAGTTGTTGCTTTTCTTTTCATTTTATTTTATATGACAATATCTTTATTTGGGATTGAGGCATAAGT

>PGSC0003DMG400038191 LRR-RLKs

ATGGGTTCCCTGCCAATGGAGATTGGAGATATAACAAACCTTGCAAGTTTTGACCTTTCTTACAACAATTTCTCCGGCATGATTCCAAGCACCATAGGGAAATGTTTGGCCTTGGAATTACTTTACATGCAACACAATTCATTTGAAGGGTCCATCCCGTACATAGCTGACTTACAAAACCTACGGGAGCTAGACCTTTCAAGAAACAACTTATCAGGAGAAATTCCACCTTGGACTATGAATCTTTCCTCCCTACTGTACCTGAATCTTCCAGACAACAATTTAGAGGGTGAGGTTCCCGTCCAAGGAGTATTCTCAAATTCAACCGCATTACAAGTTGAGGGGAATCCCAAGCTTTGTGGTGGAATCCAAGAGTTTGATTTACCACCTTGCCCGAAGCAAGAAAACAATAGAACAAAGAGGAAGCGTCTTACACCCGTTCTGACTACAATTATCAGTATTGTTTTATTTGCTGCAATAGCATGTTTTCTGCTTTCTATGTTTTTTATTAAAAGATCCAGAAAACGAAGAGATGCAACATCTACATCTGGGCATCTCATCTTCCCCATGATTACATATAAGGAGCTTCATGATGTAACTAATGGTTTTTCTTCAGATAGGTTGTTAGGATCAGGCAGTTTCGGCACTGTGTACAAAGGAATTCTTCACTCATCGAATGAAAATCCAGTTGCAGTCAAGGTACTCAACCTCCAAAGTCACGGTGCATCTAAAAGCTTCATGACTGAATGCCACGGCTTGAGAAACACCCGGCATAGAAACCTGGTGAAAGTCAGAACTGCATGCTCCAGCATTGACATCAAAGGAAATGACTTTAAGGCCATAATGTATCAGTTCATGGAAAAGGGAAGCTTGGAGGAGTGGTTACATCAAGAATACTGCAATGGTTGCTGA

>PGSC0003DMG400039680 LRR-RLKs

ATGGGAATAGGCATTCTTGGTTCTTTTCTGCTTTTGACTGTGTTCTTTAGGCCTTTGCTTTGCCAACAGCCAAACACTGATGGATTTTTTGTGTCTGATTTTCTTCAAAAGATGGGGGTTGCTAAAGTTCACAACTTTTCAGCTCACTTTTGTTCTTGGAAAGGGGTGGGATGTGATTCCAAGGATGAAAATGTTGTTAATTTGACAGCTAAAAGTTTTGGTTTATCTGGTGTGATTCCTGATAACACCATTGGTAAACTCACAAAGCTGAAGTATTTGGATCTGAGCAACAATAAACTCACTGGTTTGCCTTCTGATATATGGAGTTTAGGTTCACTAAAGTACCTTAACCTCTCACATAACCATATCTCTGGTGACCTTTCAAGTAACATAGGCAATTTTGGTGGTCTTGAAATCATGGACTTTTCTGTCAACAATTTCTCTGGGAAAATCCCAGATGCCATTAGCTCCCTTTCAAGTTTGCATTCTCTTAATCTTAGTAAGAACTTGTTTGATTCAGAAATCCCTTCTGGAATTTTGGGCTGTCATTCTTTGGAATCCCTTGATCTTTCAGAAAACAGACTCACTGGTTTTCCTAATGATTTTGGTTCTACATTTCACAAGCTCAAGTTCTTGAATCTTGCAGAAAATGAGATTCTTGGTAAAGATTCAGATTTTTCAAGAATGGGTTCAATCACTCATATCAATATTTCAGGCAATCTGTTTAAGGGTTCTGTTGTTGGTCTTTTTGAGGGGCCATTGGAAGTGATTGATTTGAGTAGAAACCAGTTTGATGGTCACATTTCTCAGGTAAACTTCAGTTCCAGCTTCAATTGGTCTCATTTGGTTTATCTTGATTTATCTGAGAATCACCTTAGTGGAGGGATCATTAGAGAATTAAAGAATGCTCGAAATCTTCTATACCTTAATCTTGCAGATAATAGATTTTCACATCAAGAATTTCCACAAACTGATATGTTATCTAGCTTAGAGTATCTGAATTTGTCTGGAACTAGTTTGATTGGCCAAATTCCTCAAGAGCTCTCGTCGTTGAGTCGCTTGAAAATCCTTGATATATCCAAAAACCATCTTAGTAAACGCATTCCTCCGCTGAGCAATAGAAATCTCCTGGTTCTTGATGTTTCATACAACAATTTGACTGGTGATATCCCATTGCCACTTGTAGAGATACTTCCAAGGATGGAAAGGTTCAATTTTTCTTACAACAATCTAACTCTTTGTGCTACTGAATTCTCCCCTAAAACACTCCTTTCGGCGTTTATCGGGTCATCAAATGGATGTCCTATTGCTGCCAACCCAGCTCTCTTCCACAAGAAAGCTCCGAAACACAGAGGACTTAAACTTGCTTTGGCATTAACTTTCTCTATGGTTTTTCTGCTTCTTGGTTTACTGTTCTTGGCATTTGGTTGTCGAAGGAAAACCACGATATGGGCGGTTAAACAGAACTCTTACAAGGAAGAACAGACTATTTCTGGTCCTTTTTCATTCCAGACTGATTCAACAACTTGGGTTGCTGATGTTAAGCAAGCAAACTCAGTGCCTGTTGTGATTTTCGAGAAGCCACTGTTGAATTTCACATTTGCAGATCTCTTATCTGCAACTTCTAATTTCGATCGAGGCACATTGTTAGCTGAAGGGAGGTTTGGTCCTGTTTATAGAGGCTTTTTACCAGGTGGCATTCATGTGGCAGTGAAAGTTCTTGTTCATGGTTCCACTATGACAGACCATGAAGCTGCAAGAGAGCTCGAGTATCTTGGCCGGATAAAACATCCTAATCTTGTTCCATTGACAGGATACTGCTTGGCTGGTGAGCAAAGAATTGCCATTTATGATTACATGGAGAACGGGAATTTGCAGAACTTGCTTCATGACTTGCCACTCGGGGTTCGAACTACTACTGAAGACTGGAGCACGGACACGTGGGAAGAGGAGGATGATAATAACAGTATTCAGAATGTTGGTTCTGAAGGATTGTTAACCACTTGGAGATTTAGGCACAAAATCGCGCTTGGCACTGCTAGAGCACTTGCATTTCTACACCATGGTTGTTCACCTCCCATTATACATAGAGATGTCAAAGCCAGCAGTGTTTATCTTGATATGAACTTGGAGCCTAGACTATCCGATTTCGGATTGGCCAAAATCTTTGGCACGGGACCCGAGGATGAGATCACTCGCGGTTCACCTGGATACATTCCCCCAGAATTTCTTCATCCAGAGAGCAGCTCACCTAAATATCCTACACCAAAGTCAGATGTGTATGGATTTGGAGTCATCCTTTTCGAGCTAATTACCGGGAAAAAACCAGTTGAAGATGATTATCCACAACACAATGATGCGAATTTGGTTGGTTGGGTTAGAGGATTAGTAAGGAACAACGAGGGATCAAGAGTTATCGATCCAAAAATTCGTGGAACTGCATCACAGACACAAATACTAGAAGCCTTGAAAATCGGTTATCTATGCACAGCTGAAGTTCCCGCGAAACGACCAAGTATGCATCAAGTAGTTGGACTGCTTAAGGATATTGAGCCTACACAACAATGA

>PGSC0003DMG401000515 LRR-RLKs

TTTTATCATATAACTTAACTCAGTATATTGTCATTTTGGTTTCAGGTGTGGAAGCTGTACACAACAAATCAGGTAACTGAAGCATTAGATCCTCTATTGAAAGGCGATTTTCCGCCAGAAGAGGCATCAAAGGTACTAAAAGTGGGGTTGTTGTGCACTCAAGCTTCTGTCGCTTTAAGACCATCAATGAGTGAAGTTGTCCAAATGCTAACGTGTGAGGGCTATCAAATTCCAGAACCATGCCAACCACCATTCTTAAATTCGAACTTATTAGTTGGCGGTTCAGTCAAATCCAGTATAAGAAGTTTAGTATCAAGTGCACTCTATAAACTTGATGAATCATCATCTTATGCCACCACCACTGGGTCCTCAAGTATGCAAAGCTCATCAACTGGACCACACAAAAGTGATGAATTCCTTTTAAAAGAGTCTGAAAATAGGAAATAATTGCAAAGACAAATATAGAAATGGAAATTTTGAGAGATTTTTTATCAGTACATTTTTTGTACAGTAATAGAATTGCTGTAGTTGTGCATTTTTATGATTTTATTTTTGCATTTTTGGTGAGTGATGATACAAGAAAGTGTAGGGGTGGCAATTTCAGCCCATATAAGTGAAATGAAACCATTTTGCCCATATTAAATTAAATGGATCACTCATTTTC

>PGSC0003DMG401002925 LRR-RLKs

TGGACTTGTAGACTTGAATCAAAACCTGTCGCAAAATTTTTATCATAAGCTTGGTGAGGGGTCAAACATGTCTATTGAAGGTTATGGAATGAGCAATGCTGGTGGTTCTGGTGCAATGTCGGTGGATAACAGCAGTGTTGGATCAAATGATTCACACACTCATATTCTAAACCAGCAGGGTAATCACATCCACAATAACGATTCTGTTGCTGCTAGCGTAGTCAGGGGGAGAGTTACTAGGTTGAGCGATGATGCCCTTGCTCAAGCCTTGGTTGATCCTCAATTCCCCACTATTGGGCTTGCGATTTATCATGAGTGGACAATTGACTTGAGGAGGCTTACCATTGGACAAGCATTTGCTCAAGGATCTTTTGGGAAACTCTATAAGGGAACTTATAATGGTGAAGATGTTGCCATCAAGCTTCTAGAGAGGCCAGAACATGATCTTGAGAAGGCTCACTTTATGGAGCTGCAATTTCATCATGAGGTCAAAATGTTGGCAAATTTGAAGCATCCAAATGTAGTACGATTTATTGGTGCATGCCGTAAACCCATGGTGTGGTGTATTGTGACTGAATATGCCAGAGGGGGATCAGTGCGTCAGTTCCTCACGAGGCGGCAAAACCGAGCTGTGCCATTGAGGTTAGCAGTTATGCAGGCCTTGGATGTGGCAAGGGGGATGGAGTATGTGCATGGCCGGAATCTGATTCACCGAGATCTGAAATCCGACAATCTACTGATTGCTGCTGACAAGTCAATCAAGATTGCAGACTTTGGTGTTGCTCGAATTGAGGTGCTGACTGAAGGAATGACACCAGAAACTGGCACTTACCGCTGGATGGCTCCGTAAGCATCTAGACTCTTTAGCAGCCTTGTCAAATCTGGTGTCTTGATCCAGTTTGAGTTTATTGCCAAACTGTATAATAGAGCTCTTAAGCTTTTTATCCTATTCTTGTGGGCTATTTTTTCTCTGTCTGCATTACTTTCTGTAATTTTTTTTCAAAATTGCTTTGATATGTGTTTCTTTAGCCAAGCATCTTACGGAAACAACCTCTCTTCTTCCACGAGGGCAGGCTAAGGTCTGCTTTCTCTCTACCCACCCCAGACCCCACTGAGTGGGATTTCACAGGGTATGTTGTTGTTGTAGCATGCTGAATTCCCTTTATGGAATAAATTTTATTAGAACTGAAGTGAAGAGCATGGTGGTTGATGGGTTCAAGTCCTGTATGTCCTTTTGATGCCACCTCACTGATTGTGATTGTATTTTCTATTGTATACCTAGTGTTTCTTCCAGTTAACTTCTTAATTTAGTTCTGTACCTGTTAATGAAGGATAAAGAGTTGATTGACTTTACTTTCTCTTTCAATTCCTTTCCCTCCTTTTCCTTTTCCTTCTTGTCCATGACTTGTAACTTGATTCAAGTTAGACACACTCAAGGGTGTCACTTAGTTGTCAATGAAGTAGGAGCAAACCTTTGGAGATTAGGGTTCAATCATCCCAGAGACAGAAAACACCAAGTGTTCTCTTCCCATATGCCTAAACCTTAAACCTTTGTGGGTATAGTTACCCGAAATCGAGATGCATGCAAGTTGGATTGGATACCATTTTCAGAAAGAGTAAAATTGCATATATCTGAAAATGGGTCTAGGATATTAACTACCACACTGGGTTCACTTTCAGGATGTGTCTGAGAGATATGAACCACAATTCTATTCTTATTGAATATGTAGATGAAAGTTCTATGGGAAGTATATTATCTCTTTCTCTCCTTTGTGAGTGTCGACCTTCCAGCATCATCTTTAGCATCACCTCTTTCTTCAAAGCAGTGAGGCTATTTTGGAAATAATGTATGGGTAAGAAAGAGAATGGGGAGGTGCGATTCACCAGAGATCTCTCTCTTTATAGGATGACATACATCCACAAGAGTCGCATTGTTTGTTGCTGCTGGAGAAACTTAAAAAGAGCAAGGGAATGACATGTCTACCAGAAACATTTCTTGTTTAGAGTGGAAGTTGGTTGCCGAATGAAAGTTTTTAAGTTGTTAGTAGTTGAACAGCATTCCTGTTTGCTTGGCTCGTGCTAACTTTCTCAAAGCACTGAAATGAATCAGTTAAAGAAGGATTGTTTAGGACATGTTATTTGGACATACACTGGTGTCTCTGGAATCTGCAGGGCTCTTGTTAACTTTCTCAAAGCACTGAAATGAATCAGTTAAAGGAGGCCTGTTTAGGACAATTTAATTGGACATACACTAGTGTCTCTGGAATCTGCAGGTAGATTTTAAGTTGTGCAGGTGGAAGTCATTGCTCTGTTGGAGGATGGCAAAACAAAACATCGATTTTTTTTTATGACAAAGGAAACCCGCAGCTGCTATCCATTGGGTGTGTACACGGTAAAACCCCCGCTCCTATGCAATAACTCACAAAATTCGGAAGAGAGGTAACCTGCACAAGGCAAGCTCGGTGCGACGAGCTTGACCCAGAAGACAAACCCCTTGCTTTTGCTGGTAAGGGGTTTTGAACTTGAGACCTCCAAAATGGAAGTCCCAAGCTCAAACCACTAGGCCACCCCAAAGGGTAACTGACATATCTATTCTGATGTTTTTGGAAGGAATTCTAGGAGGGCCTTACAGAAACTAGACTCTACAACGATAATATGATTCTCCGCTGAGTTACTACATAGACTTTTTTCATCATTAGACAATCTCTACAGAAAGGTAGAAGTGGAATATTTGGTTCAATCTTCAATGAATCTAAATCATTTATTGGAACCTTCACTATTCTTTAGGAAGAGGCCTCTTTAAAAGCATAAAAAGCGACAAGGTGATGTTTCTATAATCTACAACGATTATGTGGATTCTGCATTGAGTTGCTACATGGACTTGTTCCGTCCATAAGACTATTTCAATCTCTATGGAAAGGTAGAAGTGGAATACATGGCTTAATCTTCTCATATTAAAAGCAAAAAACGATCCATGGAGTAAAAAGCTTGGAGTGAAGCATACAAGTTATGGAAAAGTATAATGAAGTACTATCCACAAAGAAATTTAGTAATCTTCTCATGAACTTGAAACATTTGTTGAAACCTCTAATTATTGAGAGTACTGTTATTAGAAGCGAGAAACACAAAAAGCGACAAAGTCCATGGAGTTTGAAGCAAATAGCGAGAAGTGAAGCACAATGGCTATAAGAAGTAAGTAATGCCGAACATAAAAGGATTTAGTACAATAAAGATGCAAATAAAATAAATGGTTTTTGCATCTAGTTTACTTAATATTGATATTGATCTTGCTCTTCAAAGTCGAGATTCAAAAAAGTAGATCCTTTCTCGTTAGGATCATTTCGCACGTCATTGTTTAAAGCAGTGCAAGGCTTCAAGAAGTAATCCCATTCTTCAAATTGCTTTAAGCGATGGCTTTGAATAATAATGTGGGAGAGATGTTGTGACACATGATGAAATCAATGCCCAGTAAGAAGAATGATAAGAATGCCTTATTTGGTATTAAGTCTTACAACACGTGTTGCTAGACCTGATTTGACAAATGCCTTATATAAGTTGAAGTAGATCTTTTAGAAACAAAAAGAGCATGAAAACCTCATAATGATCAATCAATTAGTATAACTTGTTAAATTACGGAGATCCTACGTCAGGCCGATTATGACTTAGTATTTCTGACATCTAACTTTTAGTGCCTCACATACTATGTTGTATCGGTCCTTTGCATTACATACATGTTAAAATGGGTGTGTCATGGTTGCGCGTACTCTTCAAAATACACAAATAATTTACAAAAAAATGCACGTACCTATAATGATACTTACATCTGATAAAATTCATACTCAAGGACGAGTCCAGACAAGAAGATTTAGGATTTAAAGTTAATCGGTTCCCAATGATATCAAGTTCAAAAAGTAAATTATTATATGTGCCCAAGGGGAATCGAGCCTTCAATGAGGAGGGCAATAAAGCTCAAGCAAGTTCCCATTTATGCCGCTGGACCAAGAGCTTACTTTGTTAGTAGATCCCAACAAATAGTTTTATATATTTAATTGACTGGATTCCACAAAACCTACACATGATACTTTAGGTCCGCCCTTGTTCAATAGCAGCAACAACATACTCGGTGCAATCCCACATGTGGGGTTTGGGGAGAGTAGAGTGTATGCAAATGTTACCCTTAACTTGTTGAGATAAAAAGGTTTGTTCTGAAAGACCCTCAACTTTAAATAAAACTGCCAAACAGTTTGCAAAAGGGAATACAAATATTCGGAGCAATCACAATGACACATTTATGTAAAATGGCAAGTGTTACATAACAAACAAGTGAAATAATGCAATAATCGACACACAAAAAACTTCAAGAGTAGTGCTAATACTACTGATAAGAATACCAGTGAAAGCCTATCTCGCAAAGAAGCAAGAGAATGCTCAATTACCAACTAACCATTTACCCTAATACACGACCTCTATACTCTCCTATCAAATGCCATGTCCTCGGTAAGATCTAGATGCACTCCGTCTAATCACTTCCTCCCAATACTTCTTCGGCTCGACTTTACCTCTCTTGACGTCCTCCCTTGCCAACAACTCACACCTCCTCACTGAGGCGTCTATACTCCTCCTCCTCCTCACGTGTCCAAATCATCTTAGTTTCACTTCCCTCATCTTGTCCAACATGGAGGCCACTCCTACCTTGTGTCGAATATCTCCATTCTTAATCTTGCCACTCTTAGTATGCCCACACACCCACATCAATATCCTCATCTGAACGATTTTCTTCTTCTGAACTCGAGATTTCTTAATAGGCCAACACTCCGCCTCATAAAACGTGGTCGGTCTAACCACTACTCTGTAGAACTTACCCATCTATACTCCATATAATATGGATTTCATGTTATTTTAGTTGTGGAAGAATAAGTATATTTGAAACATTTTGGATTTCTCCACTAAAACTCATCACCTCTAAGAATGATAGAAAAAACGAAAGCATCAAATCTATTTAGGTGGCACGGGTGTCCACATGTGCTTGATTTGCCAAGTTTGTTAATATGTACTATGAAACGCCAAATTAATTTTAGAAGTATGCATTAGGTATGTAAGAATATATTTTACTATTTGTTTTATGTATTACGGTGCATGTATAGGCTGACTGTTGTATTGATTCATGCATTAGCGCCTGATTGTTAAACTGTTGAAAATTTATTGACATCTATTGGAAGATCTTCTTGGTTGGCACAATTTGACCTAGAGGCTGCAAAAATTATGACAACTAAGAAAATTACTTCATTTTGCTCACTTTATTGTTCTACTGATACATGGGGTTTTCCTGTATTTAGGGGTCGTTTGGTGTGAGGGATAAAAATAAATAGTCCTGGGATAAAAAAACATGGGATTAAATTTTGGTGTCTTGTTTGGTTGCCATGTTTGGGATAACTTATCCCACTATTTATACCATAGTGATGGGATAAGTTATCCCATATACATGGTGGAATAAGTAATCTCAGGATAACTAATCTCAGGATAATTAATCATGGGATAACTTGTTCCCAACCAAATGACCCCTTAGAGTCTTATTTAAGAGTTGGTATATTTAGTGACTCTTTTGAGAAAAAAATTTCAAATCACGATTTTCCTTAGGATTGTGCTCTTAGAAGTTCAGTTATTTGTGTCTCAGGCATATATGGCAGTTAGAATTTAAACCTTGAATTAGATCATCAATTCAGGTTGGTTTAGAGAATATGGTATAGAGTTAAATAAATCCCTTTCAATCTGGTGGAGATTGATTGCTATATCCAGTTGAGTGTAGATTACTGATGATAGAACTGGCATTTTTTTATTTCTTTTGACCTGGACAGAAGTATCACCATCAGTTTCCGTTTGGGTTTGTACCTTACACAATGTTAGAGAGGAAAGCATATTCCCTATAAGCTAAGTGAGTTGGAGATTAAGCTTTTGTTTTCTTGAGATAATGTGATAACTGGAGAAGTTTCGATTTCTAATATGACATTTCTTTCAGACTTGATCAAAGATATGTGTTCATGTCTGCCTGAAATATTTTCGAAATGGTTGCGATCAGAAAGGATTTAGGAGGTTGACCTGGTGAGTGGTGGTAATTTTATAGTCTTCTCTGTGTGATAATAGGCTAGTATGCATGAATGACTTTCCTTGGGGAAATAGACAAAAAATAATATGCTTGCTTTTGTTAATTTGATGTGATAATGAAAGATCTTGCCAAAACAGGTGTGGATCGAATCTCAATATCTGGTAAAACTCGAATACGTGCAGCCAGTCTGACTCTTTGTATAATTCTAGTATGCTCAAATTTTTATTCTGATTAACTGAGAAATGAAGTTACTTTTCAAGTTTACTCTTGAGAAATTAGTTATTCACATTCTGCAGATCCTATCCCGTTTAACTTTGTGAATTATTGGCAAGTGTGTCTGGTTTCTTCTCTTTTCAGAAAAAAGAAAAATGGATGTTCAAAGTTGTTAGCAATAGCATCTGTGTTTCGTTGTGGATCGCTAAGCGTTTGACTTTGTTTTGTAGGGAGATGATCCAGCACCGGCCATACACCCAGAAAGTTGATGTTTATAGTTTTGGCATTGTTCTGTGGGAGCTCATAACAGGGTTGCTTCCTTACCAGAGCATGACTGCTGTGCAGGCTGCTTTTGCTGTTGTCAACAAAGGCGTCCGTCCAACAATCCCCATTGATTGCTTGCCTGTCTTATCTGATATCATGACCTGCTGCTGGGATTCTGACCCCGATAATCGGCCAACTTTCTCTCAGGTGGTCAAGATGCTTGAGGCGGCTGAGACAGAAATCATGACTAATATCAGAAGGGCACGTTTCAGGTGCTCCCCTCATCCTACGGCTACAGGGTAAGAGGAAACTGGAGGCAACTGATTTTGAATGGCAAAACTGATTTTGAGTCTGCCATGTGTATCTATCAGGCACAATTTCTTTTCAGACAGAAAAAAGGGTTATAGTTGTGAGAAAAAACTAAACATTTCTCTGCTGTATTTTGTTTCTTTTCTTTAATCCGTGTATTTAAGATGGTGACTAGTTGGTAACACAGGATATCTGATGTGCATCTATGCATGTCAGATTATGGCTGTATTTGCTGAATTTGTTCAAAGTTATATCAGGTTCTGAATTA

>PGSC0003DMG401006401 LRR-RLKs

GTTAGCCTTCGACATGGTTCCCAGAGTATTTTATCTCCTGCCTCTCTTATATCACTTCACAATTACTTTTGCTTCCATTGAGGAAGCAGCTGCTCTTCTAAAATGGAAAGCAACTTTCCAAAACCAGAATAATTCCTTATTCGCTTCTTGGACACTAAGTCATCCTGCTGCTGCCAAGAATTCTTCCAGCCATGATCCATGCAGGGACTGGTATGGAGCTAATTGCTTTAATGGTCGGATAAACAAGTTGAATATTTCAAATTCTGGTGTCATTGGTACACTCCATGATTTTCCATTTTCATCTCTACTTTTTCTTGAATATGTTGATCTTAGCATGAACAACCTTTCTGGTCCCATTCCGGCAGAAATAGGGAAACTTGTCAATCTTGTTGAAGTTGCTCTTGATACGAACCAGTTAACAGGTCATATTCCTCCGGAAATAGGCAACTTGATTAATGCAAAATTGTTCTATGCTTTCTCCAATGAATTGGCTGGCTCCATTCCTATTGAAATAGGGAAGATGAAGTCACTTGAGAGTTTAAGCTTACAGAGAAACAATCTTTCTGGTTCAATTCCAAAACATTTAGGTGAATTAACCAAGCTCAAAATTTTGTACCTTTATTCTAACCAACTTTCTGGCCCCATTCCAACAGAAATAGGGAAACTTGTCAATCTTGTTGAAGTTGATCTTGATACAAACAAGTTAACAGGTCATATTCCTCCGGAAATAGGCAACTTGATTAATGCAAAAATATTCTATGCTTTTTCCAATGAATTGTCTGGTCCCATTCCTGCAGAAATAGGGAAGATGAGGTCACTTGTGGATTTACTGTTCCAGAGAAACAATCTTTCTGGTCCCATTCCTAGTGAGTTGGGGAATCTGAATAACCTCAATGATCTGGTTTTATCCGAAAATCAACTTACTGGTTCAGTTCCTACTACATTTGGCAATTTGAGAAACTTGCAGACACTGCATCTCCATACCAACAAACTATCTGGTTCCATTCCAAAAGAACTTGCATATTTGGATAACTTGCGAGTTCTAACAATGACTAAAAATCAGCTTTCAGGCCATTTGCCTGAGCATCTTTGTCAAGGTGGGAAACTTGAGAGGTTCACGGTGACTAGCAACAAGCTGACAGGGCCAATACCAAGAAGCTTGAGCAACTGCTCGAGTTTTAAATGGGTTCGCTTTAATAACAACAGTTTCATTGGAAACTTATCCGAAGCTTTTGGAATCTATCCAGAGCTTCAGTTCATTGATTTGAGCGACAATAATTTTCATGGTGAGCTCAGCAGCAACTGGGGGAAATGCGAAAATTTGATTGATCTGCGTGTAGCCAGAAATAACATTAGTGGTAGCATACCACCAGAGATTGGAAACGTCCAAGGGCTTCTTGGACTTGATCTTTCATCGAATCATTTGCTTGGGCAGATTCCAAAGGAATTTGGAAGATTAACTTCTCTGGTAAGGCTTTCTGTGCAAAACAACAATATTTCTGGCAATATATATGAGGAACTTGGGTCACTGACAAAGTTAGAGTCCCTTGATTTATCAGACAACAGATTGAATGGATCAATCCCAACGTGGATAGGAGATTTTGTGCGGTTGTTTCAGTTGAATCTGAGCAACAACAAGTTTGGACAGAAGATTCCAAAGGAGATAGGGAGGATAACTCATCTTACTTTACTTGATTTGAGCCATAATCTTCTTGATGGAGAAATACCTGCTGAGCTAGCCAGTTTGCTGGACTTGTCAAACTTGAATCTTTCCCACAATGGTCTATCTGGGCGCATTCCTAAAGAACTTGAAAGTTTGACTGGTTTGCTGGATGTTGTATTGTCATACAATGATTTGGAAGGTCCAATCCCTAATAATAAAGCTTTTATGAATGCTTCGTTAGAAGGTAATAAAGGTCTTTGCGGCAATGTAACAGGATTCCAGCCCTGCAAAAAGCCATCTTCTGTGGTGAAAAAGCACTCAATAGCAAAGGGACGTAAACTCATCCTCATCACTGTACTTCCTGTTATGGGAGCACTAGTACTGCTCTGTGTTTTCATTGGTGTTCTCTTTATGTTTAATAAAAGAAGGAGAGTTAAAGACGTTGAAAGAAGGGATGGTGATGGTTGGCTTTCGATATCCATGTTAGATGGAAAGGCATTGTACAGGGATATCTTAAATGCCACTGAGGAGTTTGATGCAAAATTTTGCATTGGGCAAGGAGGACATGGAAGTGTTTACAAGGTAAACCTTCCATTATTAGGAGATATAGCTGTGAAGAGACTTCATTCTTCATTTCAGAATACACATCCCAAAAGCTTCATAAATGAAGTAAGGGCATTGACTGGGATTAAGCACCGGAACATTGTGAACCTCTACGGCTATTGTTCAAAAGCACAACACTCGCTCTTGGTTTACGAGTATGTGGAGAGGGGGAGTTTGTCGAGTATTCTGAGCAATGAAGTTGAGTCCAAGAAATTGGATTGGCTTAAAAGGGTGAATATCATCAAAGGTGTTGCTTTTGCTTTATCTTACATGCACCACGATTGCTCACCGCCAGTTGTTCATCGAGACATATCAAGCAGTAATGTTTTGGTTGATTCTGAGTATGAAGCTCGTGTTTCAGATTTTGGCATAGCGAAGCTTCTCAAACCAAACTCATCCAATTGCACTGCACTTGCAGGAACATATGGCTATGTTGCACCTGGTAAGGTCTAAATTTAATTTCTATTTTATTCAGACTTCGTGAACTATAGCCAAACACTCTATACTGTTTTTATCTCATTTACACATCTAAATTGTTATACCATTTTTGGAAATTGTATTGTTTCAGAGCTTGCCTATACAATGAAGGTTACGGAAATGTGTGATGTCTACAGCTTTGGAGTATTAGCATTGGAGATAATCAAAGGAAAGCATCTAGGGGAATATATTACTGTGCTAGCAAATTCATCGACTATAGATCATGTGCAGCTTAGTGATTTGCTAGATGAACGCCTTCTATATCCTGAAGATGAAGTAAAGGATGTTTTGGTTTTAATCATCAAGCTAGCAAGCTCTTGTTTGGTTGAAATTCCAAAATCAAGGCCAACAATGCACTTCATCTGTCATAGGTTATCATCAATGGATGCACGTCCACCTACTCATGTAAAGTGATCTTTG

>PGSC0003DMG401009161 LRR-RLKs

ATCATTTTTCATAGTAGCACAAATTATGAAGATAGCTGTTAAAAATTTCATCTTTTTTTACTGCTGCTGTTTCTCTCTGTGGGGTTATGCTGTATATGCTCTCACTTCTGATGGCACCGCTTTACTTTCCCTTTCTAGTCACTGGATTGGTGTACCCTCTTCTTGGAATGCTTCAGATTCCACTCCATGTTCTTGGGTTGGTGTTGAATGTGATAACAACCACTTTGTCACCTCTTTGAACCTCTCTGGCTATGATATCTCTGGCCAATTGGGACCTGAAATTGCTTATTTGAAGCACTTGCTTACCATTGATTTGAGTTATAATGCTTTCTCTGCCTCCATTCCTTCTCAACTCACTAATTGCACTCTTCTTCATTACTTGGATCTATCCTACAACACCTTTACTGGACATATTCCTTCAAACATTGGAAATTTACACAAGTTAACCTACATAAGCCTCCTCTCTAATTCATTGACTGGTAATATTCCTGATTCCTTGTTTTCTATCCCACATTTGAAAACTATCTACTTTAACCAAAACAGGTTAAATGGTTCCATCCCATCCAGCATTGGTAACTTGACTCACCTTTTATCTCTCTATCTTTATCAGAATGACTTGTCCGGTCCAATTCCGTCTTCCATAGGAAACTGTACTAACTTGCAAGAATTGTATTTGAATGATAATCATTTAGTTGGTAGTTTGCCTGAGAATTTTGATAAGCTACAACACCTTGTTTATTTGGACTTGAGCAATAATAGTCTACAAGGAAGTATCCCTTTCACTTTAGGTAACTGCAAACATCTTGACACTCTGGTCTTGTCATCCAACACTTTCAATGGGGAACTTCCTCCAAGTTTGATGAATTGTACTAACCTAAAAGTTCTTGCTGCTTTTTCTTCTGGTTTAAGTGGCCCTGTTCCTGCTTCTCTAGGCCAGCTCACAAAGTTGGAGAAGCTTTACCTTACTGACAACAATTTCTCTGGAAAAATACCACCTGAGCTGGGGAAGTGCCAGGCCTTGCAGGAATTACTTCTACCGGAAAACCAACTGGAAGGTGAAATTCCTAGTGAACTAGGATCACTCAGCCAGTTGCAGTATCTCTCTCTATATTCCAACAAATTGAGTGGTGAGATTCCCCGCACTATTTGGAAGATTCAAAGTCTTCAACATCTTCTTGTCTATCGGAACAACCTGACTGGGGAGTTACCTCTTGAAATGACTGAGTTAAAACAATTAAAGAACATATCCTTATTTGAAAACCAGTTTACCGGAGTTATACCTCAAGGTTTGGGGATTAACAGCAGTCTAACGCTGCTGGATTTCACAAACAACACCTTCACAGGTCCTGTTCCACCTAATCTTTGTTTTGGAAAGAAACTGGAGAAGCTTATTTTAGGTTATAATCATCTCGAAGGTGGCATTCCTTCTCAATTAGGACAATGTTATACTTTGACGAGAGTAATTCTCAAAAAAAATAATCTGTCAGGTGCCATTCCAGATTTTGTTAAGAATATAAATCCTATTTTCTTGGATCTCAGTGAGAATGGCTTCAGTGGGAAAATATCCCCAAGTTTGGCAAATCTTGGAAATGCTACATCAATTGACTTATCAGTGAATAAGCTTTCAGGGTTTGTACCACCAGAGCTGGCAAACCTTGTCAACCTCCAGGGTTTGAATCTTTCGTACAATGGTTTGGAAGGTGTACTGCCATCTCAGCTTTCAAACTGGCAGAGACTGTTGAAGTTTGATGCAAGTCATAATTTGTTGAGTGGATCGGTCCCATCCATTTTTGGAAGCTTGGAAGAGCTATCCATTTTGAGTCTGAGTGAGAATAATCTTTCAGGAGGTATTCCAACCTCCTTGTTTGCACTCAAGAAGCTCTCCAAGCTGCAGCTTGGTGGAAATGCACTTGGTGGGGAAATTCATTCAGCAATTGCGACTGCATCAAGAGAAACACTGAGGTGTTTAAATCTGAGTAGTAACAGGTTGACAGGTGAACTTCCTGCGGAGCTAGGAAAATTTACTTTCCTGGAGGAGTTAGATATAGCTGGCAATAACATTTCTGGAACTTTAAGAGTTCTTGATGGAATGCGTTCATTGATATTCATCAATGTATCATACAACCTCTTTTCTGGTCCAGTACCTGCACATCTGATGAAGTTCCTCAACTTAACTCCTACCTCTTTCTCGGGAAACTCGGGCCTTTGCGTGCACTGTGATCCAGAAGAAGGCTCAAATTGCCCTGAGAATATCACGTTAAGGCGCTGTGATCTTCAATCAAACAATGGAAGGCACCTGAGTGTAGCAGAAACGGCAATGATTGCACTTGGGGCACTGATATTTACCATTTCCTTACTCCTAGTAATTGCTTACATGCTTCTGTGGCGCAAAAGTTCTGGGAAAGGAGTTGCAATCTCTGCTCAAGAAGGTGCATCGTCCCTGCTTAATAAAGTATTGGAAGCTACTGGGAACCTAAACGATAAGTATGTCATTGGGAGGGGAGCTCATGGAGTTGTATATAAGGCCATCTTGGGTCCAGGGAAAGTGTATGCTGTGAAAAAGCTGGTGTTTGTTGGTATAAAGGATGGAAGCAGAAGTATGGTTAGAGAAATTCAGACAATTGGAAAGGTTAGGCACCGCAATCTTGTCAAATTAGAAGACTTCTGGCTGAGAAAGGATTATGGGCTGATTCTTTATAATTACATGGAGAACGGGAGCCTTCATGATATTCTCCATGAGACTAAGCCACCTGTAACATTAGAATGGAGCGTTCGGTACCGAATTGCTATTGGAGTTGCTCAAGGGTTGTCATATCTCCATTTCGACTGTGATCCTGCCATTGTGCATCGAGACATCAAGCCCATGAATATCTTGTTGGACTCTGATCTGGAGCCTCACATATCAGATTTTGGTATTGCCAAGCTTCTGGATCAGTCTGCAGCAACTTCCGCCTCCAATGCCCTTCAGGGCACTGTTGGATACATGGCTCCAGGTATGTAATTTTCATTTTTTTCTCACACTTCCTTTCAGTTAGCTGTCTCTTTAGAGATCTAGTCATCTGTAGAAAAATAATATTGTCTGTGTCTTGTCTGTTTTTGTTGACAGAAACTGCATTTGCAGCTACAAAGAGCAAGGAGTCAGATGTTTATAGTTATGGTATTGTCCTGTTAGAACTTATAACTCGAAAGAAGGTGTTGGACCGTTCACTCTATGGGGATACAGATATTATGTGTTGGGTTAGGTCTGTTTGGACTGAAACTGAAGAAATTGAGAAGATTGTGGATCCAAGACTTTTGGACGAATTCATAGACTCAAGTGTCATGGAACAAGTAATTGAAGTGCTTTCACTAGCTCTTAGATGTACAGAGAAGAAAGTCAGCAAAAGACCCTCAATGAAAGAGGTAGTCAAGCTATTAACAAGGTCAAGTTCGAGTATACGAAGCAAGTACTAGCTAGCAGGGACATGTAGTATCTAGTGTATCAGTCAGAGGGGGTTTCATATAAACGCCTGATGTGTTCTAGGTACTCTATTTTCCAGATTAAAGTAGTCCTAGCTTCCAGCTTTCTGAATCCATGTTACAAAATGCTCCCTGCAGGTTCAGTATCTTTTTATTGGTAGAGGGTGATAAAAATATATTTACTTCTGTAGTTTAGAAAGAGAAGGCGTGTGCCTTTAGCTGTAAATGTTCTTTTCCATCACATGAACATGGATGAAAACTAAAGATAACTTACTGATTTTCATCTTAGTTAATCATATTCTCAGCAAAAAGCAGTTCCTTTTGCCATTCCTAGCAGGATTTGTTATGGTAAAGCCATATTGTTTTCTCCCTAGCTATTTGCAACTGTACAATTACTCCAATAAAATAAGCTTTC

>PGSC0003DMG401013731 LRR-RLKs

GCTTTGCTGTCTGGTGATGCTTATCCTGCCTGCTTCTGATAAGAATTATTTACTTGCAGGGACCTGAGCAACAACATGTTGAATGGTTTAATCCCAGATACATTTAGTCAGTTGCACTTTGATAACATGTAGGTACCTTGAAGTGAAACCTTTACAAGTTGAAGCATAAATGTTGTAATGTTTTTTTTTTTCCTCAAAAGTTTTCTTTCAGTAACAAGAAGCAAAAGTTTTATCTGTGATGCAGGTTTCTTGGTAACAACGCATTAAGCGGAGCAATTCCCAGCTGGGTGTTTAGTAGTGATAAGAACATGTTGGTATTTGCTCAATTTGTTCCGTTTTCCTCATCTAAGTAGTAAACTTTGGTTATTATCATGCCAGCACACTTCAAAGTTAACAGTTTTTCCCCAAAACATTTTGATTGCTGAAAGGACGCATTTGCAAGAATTTGGTGTCTTCTAAGTGCATGGCAACTATAATTTTTTTTTTGCAGTGATATATCATACAACAATTTCACCCTTATATCTACTCGAGGATGCCAGTCCTCTATCATGTGAGCATTGTACTTCTTTGTTTGGGAACCTACATCCGTGTATTATTTCTATTTTTATTAGTAAGAATTTCGGTCATTTAACTACATAACTTACAAACCTTCTTTCTTCAAACAGAAACTTAGCCGCCAGCTATTCAGATACAATGAACACGTTCAACAACACGTAAGTCCTAAATGTTGTCTTATGCATCAGGTGCTAGTTTGTTTGATGCTTTCTGTGTATTGTCTATAATTATGTGGCTTAATCTATTCGAAATTGCAGGGATTCCTGGTGTTCCATGAAACCCCTCATTTGTCCCACAGAAGCGAAATGTAAGAGTTACAGCCTATAAAAGGACCTCAGGGGTTATGTCTGGATCATTTACCACTGAACCAACTCTTGGAGTGGCAGCCTGTGGACTTTTTAAGTACAAAAGCATAACTTTGGAGTTGAGAGATAATTTCACAACCCTTACTCATGTTACGTTCTTTATGTCAATTGCAGATCATTCACTATTCATAAACTGTGGAGGAGGTAGAATTACATATGAGGGAAATGATTATGAGGAGGACTTGACCACTAGGGGTCCATCATATTTTTTTTCGTCCCCTGATAGATGGGCATTCAGCAGTTCGGGAGTGTACGTAGGTCTTGCAAATGTTAGTTATATTGCAACAAATAGATTTTTACTGAACATATCCGGTGCCGACTTTTACAACACAGCACGCCTTGCCCCTAATTCTCTCAAATATTATGGGCTTTGCCTACGAGGAGGTAGCTACAGAGTGCGACTTCACTTTGCTGAAATAATGTTTTCTAATGACTCAACATATAGCAGCCTTGGAAGGCGGATATTTGATGTAGCAATACAAGTAAGTTTGAGGCCCAACTTCTTTTGTAAATAATAAATATGTAATGTTGATATACTGAGTTCTGATATGATTGGCAACTTTCTGATTTAAGATTGAAGGTACCAATTTGTTGGAGATAACGTGGTATATTTTGATTTCTTACACAGGGGAGAGTAGTTTTGAAGGATTTCAATATCATGGAGGAAGCTAACGGTGTTGGAGAAGTCATTATCAAGGACTTTGATGATATTACTGTTAGTAGCACTCTGGAGATTCACTTGTATTGGACCGGGAAGGGGACTAACGCTATTCCAGACAGAGGTGTATATGGACCGCTGATTTCAGCTATCTCAGTGACACCAAGTCCGTATATCTCTGCCTATAGACTTTACACTTATCTTCCCACAATCTTATCTTAAGAAAATCCCATTTTCTTCTTGCAGACTTCAAAATAGGTCGCAGGTTACCTGTCGGAGCTGTTATTGGCATTGTACTCGCTTCAATTGTGGTGGTTCTGATAGTTTTATTTTCTCTGTGGAAGAAAGGCATTTTTGGAGGGAAAATTAATCAAGAAGAATTAGGTAACACATACCTCAAAGAATTTATTCACTCTCTACATTTGATATGGCGATACTTCTGTTGTGATAATCAGAATTATGTACGAGATTTCTAGCTGCGACATACAAATAATTATCATAGATGATTGTTCTGTTGGCAGATAGACATCATGACATATCATAGATAGTCAGCACACAACAAAATGTTCATAACATATCATACAGATGTTTATTCTTTTAAAAATTGTTGTTCATCATACAGATGTTTTAGAAGATCTATATATACTGACGTTGATATTATTCTCGTTTACAGAACTCAGAGCCCTTGATTTACAAACAGGTCATTTTAGACTTAGACAAATTAAAGCTGCTACAAATAACTTTGATCCTGCCAATAAAATTGGTGAAGGAGGGTTTGGACCAGTTTACAAGGTATCAATTCTCTATTTATATCTATCCGTACTGATTTTGCGTAAGGACAGCTTGAGGTTCTGTGGTGATGTTTCCTACTGGTTATGAAGGTCTGTAATATGATCGTTGTAAATTAATCCTGCTGTCTGTAGGGTGTACTCTCCGATGGGGCAGTCATAGCTGTTAAGCAGCTCTCTTCCAAGTCAAAGCAAGGAAATCGTGAATTTGTTAATGAGATAGGCATGATTTCAGCTCTACAACACCCAAACCTTGTCAAGCTTTACGGTTGTTGCATAGAAGGGAACCAGTTACTGGTGATATACGAATACATGGAAAATAACTGTCTTGCACGAGCTCTCTTTGGTAATGCTTTGCAACAACATTTTGAAAAGCATTTATCATGCAGTGATAATTTTATGCTTAATACTCCTCTAATTTTCCTTTCTTAGGCCGTGACGACCAGAGGTTGAACCTTGACTGGGCAACCAGAAAAAGGATATGCTCAGGAATAGCAAAAGGGTTAGCTTACCTTCATGAAGAATCAAGGTTGAAAATCGTCCACAGAGATATTAAGAGTACCAATGTGCTTCTTGATAAGGATCTGAATGCTAAGATTTCAGATTTTGGATTGGCTAAATTAGATGAAGAAGAGAACACTCATATTAGCACCCGAATTGCTGGAACAGTGTGAGCCTTCTTATCATAACTGTCTCTGTACTATCATGTTTTGATGAGAGAGTAACAGTTCTACCCATTTGAATTTGTTCGTCATACATTAACATGGTCAGTTTGAAGTATCTTACATTTCTTTTTCCTCTTTTGAAGAGGTTATATGGCTCCGGAGTATGCAACGAGAGGCTACTTGACAGATAAAAGCTGATGTTTATAGCTTCGGAGTTGTTGCTTTGGAGATTGTCAGTGGGAAAAGTAACACAAAACTACAGGCCAAAAGAGGAATTTGTTTACCTTCTTGACTGGGTAAACTATCTTTTTACTCAACTGTCATTATTCCCAAATACTAAAATCTTTAAAAACTGCTGCAACATCATGTATTATAAGTCTTGCATGTACATGTTTAGTTTAATTCCACTCTGAATTATAAGCCAGAAAAAACTTCTCTTGTTAGTGTTTTGTGCAAATATATACTCCATCGGTTTCAATTTGTTTGTCTTACTTTTATTCATAGTGAGTTTCAAAAAGGTCTCTTTTCTTTTTTGCCAGCTCTTTATTCTAACTTTCCACGTGACATATTTAAAACCACATGATTTAAGGGTATTTTGGTACATTCTACATATCTTTAGTTTAAATTAAAACCAGACAAACAAAGTGAAACGAAGGGAGTATTTGCTATGATACCTCCAAAGAAAATATCAGAATCAGAGTTGTTTCATTAGTTCTGCTCACATTCTTATAGATGCATCTTTGCGTATTTAGTTTTTCTTTTCTCCAATTTCATCTTGCTTTGTATGTCAATTTTGTTTTAGGCTTACGTCCTACAAGAACAGGGAAATCTGTTAGAACTAGTGGATCCACGT

>PGSC0003DMG401017953 LRR-RLKs

GGTCTCCGGAAAATTTTCATTTATCATAAGAGTTGTTGAGCCAGTATTGTGGTGATGGGAAAAGAAATGGGAGTTGAAAAGCTTCTGGTTTTGTTTTTGGTACTAATCTCAATCAGATATTCAGAGCAATTACAATCTTCTCAAGTGCAAACACTTTTGAGAATTCAACATCTTTTGAACTATCCAACAGCTTTGAGTAGTTGGAGTAATGAGACAGATTTCTGTAAGATTGAGCCAAGTTCAACTGTTACTGTTGTGTGCTACGACGAAAACATAACTCAGTTGCATATAATAGGGAGAAAAGGGACTTCCCCATTACCTAGAAAGTTCTCCATTCGTTCATTTCTCAACGCGCTTGCCAAGCTTCCTAGCTTGAAAGTCCTTAGATTAGTTTCTCTTGGTTTATGGGGGCCGTTGCCTGACAAACTTTCAAGACTTTCAGCTCTCGAGATACTTGACTTGAGTTCAAATTACTTTCACAGTGGCATACCTGAAGCAGTTTCATCCTTAACGGACCTCCAAACTTTAGTGTTAGATGGAAATAGGTTGACTGGCGGAATTCCAGATGGCCTTGGTTCACTTTCAGTTTTGGCAGTTTTGAGTTTGAAGAGCAATTCACTAGATGGACCTTTGCCAGATACACTGCATGATTTGAATAATCTTCGTGTACTTTCACTTTCGAAGAATAATTTTAGTGGTGATGTCCCTGATCTTAGCAGTTTACAAAATCTTCAAGTTTTAGAATTAGAGGACAACTCATTTGGTCCAAAGTTTCCTCAAACTGGAAGTAAAATTGAAAGTATTGCTCTAAGGAATAACAAATTCACTGCAAACATACCAGAGAAAGTTCAGTCCTATTATCAGCTTGAACATTTTGATATTTCTTCCAACAACTTTATGGGGCCATTTCCTCCGTCGTTGCTATCTTTGTCATCCATCACTTATCTGAATGTTGCAGGAAACAAGTTAACAGGAATGCTCTTTGAAGATAATCCATGTAACACAGCATTAGATTTTGTTGATCTGTCTGGTAATCTGTTGACAGGATCATTACCTAATTGCCTTCTGTCTAGTTCTAGGAACCGGATTGTGCATTTTTCCAATAACTGCTTAGCAACCGGAGATGGTACTCAACATCCATTTTCATTTTGCCGGAATGAGGCTTTGGCTGTTGGTGTCTTACCTCAGCATCAAAGACAAAAACATGCTTCTAAAGTGATTCTTGCCTTGATCATATGTGGTAGCATCACCGGTGGAGTCATACTCGTATGCCTGACCATCTTGATTGTCAAATCTTTTCTTGCAAAGAAGGCTTCTGCCCAAAATACCCCACCCAGATTTATTTCAGAAGATGCATCATCTTCATATACCTCCAAGTTATTCACAGATCCAAGTAAGGTTCCTCTCAATAACCAAATTACTGTTTAAAATAGAAGCTTAGGGCAAATATATCTGCTCTCTGCGCCCTAGATGCCTGAAGGCGAGGGATTTCCTTCAATTCCCAAGTGCCAAGTATACTAACTTGATACCTTTGTATTTCTTGTATAAGGATACATTACTCAAGCAATGAAGTTAGGAGCACTTAGTCTTCCATCTTATAGAATATTCTCTCTAGAAGAACTTGAGGCGGCAACGAACAACTTTGATACAACAACTTTTATGGGTGAAATTCCTAATGGCCAGGTATGGTATATGAATTCTATTATTCTTTCCTGCTATAGCATTGAGGTCTTTGGGTGTTATACTTATGGCCATATTAGTAATGTAGCTCACGTGGCAGATGTGGACAAGTCATTCCCAAGAAAGGGAGTTTCAAGTATTTGGGGTAATAATCCATGGAAATGGAGAGATTGATGATGATGTTACACATCGTATTGGAATGAGGTGTATGAAATGGAAGCTTGCATCTAGTGTCTTGTGTGATAATAATGTGCCACCATGTTTTAAAGGCACGTTTTATAAAGTGGTGGTCAGACCAATCATGTTGTATAGGGCAGAGTGTTGGCCGTTCAAAAACTCGCATGTCAAGAAGATGAGAGTAGCGGAAATGAGAATGGTGAGATGAATGTGTGGACACGCTAGGAGAGATATGATAAGGAATGAAGCTATTCGGTACAACATGGGAGTGGCCAAGATGAGGGAGGCAATACTGAGATGGCTCGGGCAAGTGAAGAGAAGATGCACAGATGTTCCATTGAGGAGGTGTGAGAGGTTGGCTATAGTGGTATTAAGAGAGGTAGAGGTATGCCAAAGAAATTTTGGGGAGAGGTGATTAGACATGACATGACTTACGTGTAGCTCACCGAGGACATGACCCTAGATAGGAAGATATGGAGGTCGAGGATTAGGGTAGAAGGTTAGTAGGTCAAGAGTGGTGTCTGGTTTCCCTTTTTAGTATTGTTATTACTACCCTTCTATGATCCTATTCTTCGATATTAGTTTTACTGGTTTTTTCCTTTGCTTCAGTTATCGTATTATTTTGTTGTTGTTGCTACTAATCTTTTCTTCATTGTTTTCAGCATGAATTCTTCGACTATGTTTGCTTTACACACTTTTTTTCTATATGTTTTACTTGAGCCGAGTATCTTTCAAAACAACCTCTTTACCTTCACAAGGAAGGGGTAAGGCTGCGTACACCCCACCCCACTCTCCCCAGATCCCACTTGGGATTATACAGAGTATGTATCAGTATGGGTGTTATATTAGAGAGGTGGTGTGGACATGTAGTAGTAATGCCATCTGTTTGTTCTCTATCATTTTGGCACAAAAGGATGAGTTTGACTCTTGAAGAGTAAACCTATTAATATTACCAGCTGTTCTCGAGGACTAAGGCTCACAAACGTTTTCTGTTTTGGATTCAGATGTACAGAGGGCAGATGAGAGATGGTTCCTACATTACGATTAGATGCCAGATAATGAAAAGAAGCAACACTAGGCAAAATTTTATGCATCACATAGAGTTGATATCCAAGCTCAGACATCAGCATTTGGTCAGTGCTCTTGGACACTGCTTTCAGTTTTACTCGGATGATTCAAGTGTCAGCAGAATTTTTCTCGTCTTCGAGTATGTACCTAATGGGACCCTAAGGAGATGGATTTCTGGTAATTACTTGACAAGCAGATATATACCCCTTCCCTATTGTTGGATGTAAATGTGCTTCAGTAATTTGATGTTCTCTGTTGCTTAATTGCAGATAAACATGCTAGACGGAAACTAACCTGGACACAACGTATAGCAGCTGCTACAGGAGTAGCGAAGGGACTACAATTCTTGCATAATGGAATTATCCCTGGTCTCTTTTCGAATAATTTGAAGATAACAGATATTTTGCTGGATCAGACCCTTGTCGCCAAAATTAGTAGCTACAATCTGCCTATTCTGTCGGAGAATGTCGGGAAGGTGTGAC

>PGSC0003DMG401021514 LRR-RLKs

CAATAATTCACTACATAGCTCGCTATAAGTTCAACAGTATCCAAAACAAGCAACGTACTGAATGATGATGGTTCATAGAATATTGCTCTTTTTTTTATTTAGAGTTTCTTTTGCTGCTTCCACTGAGGAAGCAATAGGTCTCCTCAAATGGAAAGCAACTATCGAAAACCAGAATACTTCCTTATTTGTTTCTTGGACAATAAGTGTCAAGAATTCTTCCAGCCATGGAGAAGCAAATCCTGATGCATGCAGGGGCTGGTATGGAGTTAAATGTTTTAACGGTCGGGTAAACATGTTGAATATTAATAATGCTGGTGTCATTGGTACACTCCGTGATTTTCCATTTTCATCTCTCTCTTATCTTGAATATGTTGATCTTAGTATCAACCAGCTCTCTGGCATCATCCCTGCTGCAATAGGAAAGCTCACTAATCTTGTCTATCTTGACTTGGGGTCTAATCAGTTTTCAGGCACAATCCCACCACAAATCGGCTCACTAACAAAGCTTGAGACCCTCCGCATCGTTAACAATCATTTACATGGTTCCATTCCAGGGGAGTTGGGGAATTTGCAAAACCTCACTAGACTTGCTTTGTACGATAATCAACTTACTGGTTCTATTCCTGCTTCATTTGGCAATTTGAGAAACTTGCATATTCTGAATATACGTGCCAACAAACTTTCTGGTTCCATCCCTAAAGAGCTTGCATATTTGGATAACTTGGTAGTGATGATAATGAGTAAAAATGAATTTTCAGGTCATTTGCCAGAGAAGCTTTGCCAAGGTGGTAAACTTGAGAATTTTACAGTGAGCAGTAACAAACTTTCAGGGCCAATACCAAGAAGCTTCAGCAAGTGCTCGAGTTTTAAAAGGGTTCGCTTTGATAACAACAGCTTCACTGGGAATTTATCTGAAGCTTTTGGTATCTACACGGAGCTTCAGTTCATCGATTTGAGCGACAATGATTTCCATGGTGAACTCAGCAGCAACTGGGGGAAATGTATGAATTTAACTAATCTGCAGGTGGCTAGAAATAACATTAGTGGTAGCATACCACCAGAGATTGGAAACATCAAAGGGCTTCAAGGACTTGATCTTTCATCCAATCATTTGGTTGGGCAGATACCTATGGAATTTGGAAAATTGACCTCTCTTGTTAATCTTTTTTTGCAAAACAACCACATTTCTGGAAATATACCTAGGGAACTTGGGTCACTCACAAAATTACTGTCCCTTGATCTATCAAACAATAGATTGAACGGGTCGATCCCAACATTTATAGGAGATTACATGAACATGTTTCTCTTGAACCTAAGCAACAACAAGTTTAGCCAAAAAATTCCAAAGGAGATAGGGAGGATAGTTCATCTTACTGCACTTGATTTGAGTCATAATCTTCTTGATGGAGAAATACCCCCTCAGTTAGCCAGTTTGCTGGACTTGGCAAACTTGAATCTTTCCCACAATAGCCTCTCTGGCCGCATTCCTGAAGAATTTGAAAGTTTGACTGGTCTGCAGGATGTTGTATTGTCATACAATGAGTTGGAAGGTCCAATACCAAATAATAAAGCTTTTACAAATGCCTCATTGGAAGGTAATAAAGGTCTTTGCGGCAATGTAACAGGATTTGTGCAATGCAAAATGCCCTCTTCTATGATGAAGAAGCAATCAATGGCAAAGGGACGTAAACTCATTCTCATCACTCTACTTCCTGTATTGGGAGCACTGTTACTATGTGCTTTCGTTAGCATTCTCCTTATGCGTGTTAAAAGAAAGAGAGTACTTAGAGACGTTGATAGAGGAGATGATGGTTTGCTTTCGATATCAATGTTAGAAGGAAAGGCATTGTATAGGGATATCATAAACGCCACAGAGGAGTTTGATGCAACATTTTGCATCGGGAAAGGAGGGCATGGAAGCGTTTACAAGGCAAATCTTCCATCATTAGGGAATATAGCTATAAAGAGACTTCATTCTTCATTTGAGAATTCATATCACAATAGCTTCATGAATGAAATAAGGGCATTGATAGGGATTAAGCACCGGAACATTGTGAACCTCTACGGCTATTGTTCGAATGCACAACACTCGTTCTTGGTTTACGAGTATGCAGAGAGGGGAAGTTTGTCTAGTATATTGAGCAACGAAGTTGAGTCCAGAAAATTGGATTGGCTAAAGAGGGTGAATATTATCAAAGGAGTTGCTTTTGCTTTATCTTACATGCACCATGATTGCTCCCCACCGATTGTCCATAGAGACATATCAAGCAGTAATGTTTTGCTTGATTCCAAGTATGAAGCTCGTGTCTCTGATTTTGGCATTGCTAAGATTCTCAAGCCAGACTCATCCAATTGCACTGCGTTGGCAGGAACATATGGCTATGTTGCACCAGGTTAGGTCTAACTTTACATTTCCTCTTTTTTTTCAAACTTTATGAACTATAGCCAAAGACTCCATTCTGTTTTTATTTCATTTACACATTTGAATTGTTACCTCTTTAGTTCAATTTGTTTGTCTTACTTTCATTTTTAGTATGTTTAAAAAAAGTTGTCTCTTTCCTAATGCAACTCTTGAGTTTCAACATGTCATATAGCATGTTTAAGACCGTGAGATTAAAGAACATTTGGTTGCAATACATATCTTTAGTTTACAATCACAAGATTTAAAAGTCTTCTTTTACTTCTTAAATTTTGTGGCAAGTCAAACTAAGACATACAAATTGAACTTAGGGAGTAATACTTTTTCGTAAATTGTGTTGCTTCAGAGCTTGCATATACAATGAAGGTTACACAAATGTGTGATGTCTATAGTTTTGGAGTATTAGCATTGGAGATAATCAAAAGGAAAGCATCTTGGGGAATACATTACTGTGCTAGCAAATTCATCGACTAGAGATAACGTACTGCAGTTTAGCGATTTGCTAGATGAACGCCTTCCATATCCTGAAGAGAGAGTAAAAGAGGTTTTGCTTTTTATCATCAAGCTAGCAAGCTCTTGTTTGCTTGAAACTCCAAAATCAAGGCCAACAATGTACTTCATCTCTCATAGGTTATCATCAATAGATACGCATCCACCTATTTATGGAAGGCGATATGAATAACCTTTGACACCAAGTGGTTGTAAAGTTCCACAAAATGTTATTACTGGGACTAGTATTGTTGGTTTAAATCTTAGGGTGCTTAACTGGTTTATTGTACTAGAGTGTATGCAGTGTAATGTTGAAGGTAATTATCTAATTTCCATAGTATTTTGC

>PGSC0003DMG401023552 LRR-RLKs

ATGTTGACATATCTAGAAGTGGGTAGCAATCCGTTGAGTGGCTTTCTGCCAAATTCTATTGGGAATCTCTCATCTACTATTGAAAAAATTAATTTAGCAGATGCACACATCAATGGCCTCATCCCCACAAGTATAGGCAACATGAGCGGTCCTATATCCCTATACATTGAAGAAAACAACTTGATAGGGAGTATTCCTTCTGATGTTGGTAAGCTTAAATGACTCCAAGGACTCTATCTATCTAACAATAAATTGCAGGGACGTATTCCAGAGGCGGTATGCCATCTATCTAATTTGGTTCAATTATATCTGTATGGTAATGAGCTCTCTGGGTTAATTCCAGAATGCTTAGGAAATCTTAGCATGCTTCAACAGCTTTCTTTGAATTCTAACAAATTATCTTCAAACTTTCCGTTGAGCCTTTGGAAGATGAGTAGTCTTCTCTATCTAAGTGTGTCACAGAATTCAATAGAGGGAGAAGTTCCATCAGCTATTGGAGGACTGCAAGCCATTGTAGAACTACATCTTTTCAGTAACCACTTTTCAGGCGTGATACCAACCAGATTAGGGGAACTCCAAAACCTGCAGTATATTGACCTATCAAACAATTCATTTTTTGGCCAAATTCCATTATCCTTTGCCAACTTGATAAACTTAGAATTCTTAGATCTGTCTTTAAATGCGTTGTCGGGTACTATTCCTAAGTCATTGGAAAAACTCTCCTATCTTATAAGCATTAATGTTTCATTTAATGATTTAGAAGGTGAAATACCGAGTGGTGGCGTGTTTGCAAATTCCACTCTGCAATCATTTCTCGGGAACAAAGGTCTATGCAAAATGCACATATTGGAGATTCCTGATTGTGTTATCACTAATCCTGGAAAACAATCAAAGCTTAAGGAGGTTGTGCTAAAAATTGTTACTCCAGTGGTTACTTCATCCTTTCTGATATTCTTATTCGTTTCAATTTGGACAATGAAACGAAAAAAGAAAGGGAAGTCCAAAGATGTGGAGAAGGTACCGGAGATCGGGACTCATCAATTAGTTTCTTATCATGAGATTCGAGGAGCAACAAATAATTTTGATGAATCCAATTTAATTGGTGAGGGAAGCTCTGGCTCTGTGTACAAAGGCGCATTGTTTGGTGGAACTGCAGTGGCCATTAAGGTTCTGGATTTGGAAAATGAGCAAGTATGCAAGAGGTTTGATACCGAATGCGAAGTGATGAGAAATGTTAGACACGGAAATCTTGTTCCAGTGATTACTACATGTTCTATTGACTATATAAGAGCCTTTGTTCTGCAATTTATGCCCAATGGAAGTCTTGACAATTGGTTGTACAATGAAGATCGCCACTTGAACCTTCATCAAAGAGTTTCTGTAATGCTTGATGCAGCTATGGCAGTTGAATATCTACATCATGGTCATGTCACTCCAAAAGTTCATTGCGACCTAAAGCCAGACAATGTTCTTTTGGATGAAGCTATGGTGGCTCATGTTGGTGATTTTGGAATCTCTAAAATTTTAGCCGTAAGCAAGTCCATGGCTTATATGGAGACATTGGGCACTCTTGGATACATTGCACCAGGTATACTAGTCTCTTTCTATATATTTTCACAAAAATCAGAATTTAGCCCTTCCAGAATCATAAAAGCAAGTGTCTGTTAGTACATAATTATGCTATATTCAAATGAGTAATTCTTTTTCATTTTCATTCTTTTTTAAGAATATGGCTCGGAGGGAATAGTGTCTTCTAGTGGTGATGTTTATAGCTATGGCATCATGGTGATGGAGGTCTTGACGAAACGAAGGCCAACAGATGAAGAGATATGCAATGAAAACCTTGACTTGAGGAAATGGATCACACTAATCATTTTCAGGGAGTATGATGGACATCGTGGATGCCAATCTTTTTCGTGA

>PGSC0003DMG401027271 LRR-RLKs

GTTTTCTTCATTTCTTGATATGGCTTCTTCTTAGATAGTACTCACTACTGCTGTCTATTTCCTAAATTTAAGGGCTTCCTAGCTTTTTGGTTATTTTACATTTTCAAGATTCCTCAAATGGAATACTCCCTCACATTTCTTGAGCATTTCAAGACTCAATCTTTTTCACTCTAAACTCCCTTTTGTCTTCTACACCCCACCCCCACAATCTCCAATTCTGCAGAAAGTTTCATGTGATGGATTCTTGATGGGAAAAGGAGGAAGAAGTTGAAGAAAAGGTGTTTTCTTTGTAAACAACCTCATAACCATTTCAAAGATTCAATCTTTTTCACTGTAAACCCCTTTTTTCACCCCCCTTAATCTCATCAATGGATTCTTGATTGTTCTATCTAGTTGTTTTGGAGGGAAAAGGAGGAAGTAATTGAAGAAAAGGTGTTTTCTTGAAAAGAAAAAAACTCAAGAATGACACTTTTGAGGTTTCTGTTAACTGTGTTTTTTCTAGTTGGTTTGTTATCAAGATTCCAACTTGTGAATTCCCAGTTGGTTTTTCATGATCAAGATGTAGTGGAAGCAATTGGGAAAGAGCTATCAGTACCTGGATGGGGTGTGATGAACACAACTGATTTCTGCTCTTGGCATGGAGTTATTTGCAGCTCAAACAGTTCAATGGTGGAAAGACTTGATCTTTCTGGTTTTAGATTACAAGGTAATGCAACTCTAATATCTGAGCTCAAAGGTTTAAAGTGGCTAGACCTTTCTAATAATAACTTCCAAGGTTCTATTCCATTTGCATTTGGGAATTTATCTGAGCTTCAATATCTTGATTTGTCTTTTAACAAATTTGGAAACTCAATACCTAGTGAATTAGGTAAGCTCAGAAGCCTTAAGGCATTGAACCTTTCGAAAAACTGGCTCACAGGAGCTATACCTGATGAGCTTGAGGGGTTAGAGAATTTGCAAGATTTTCAAATATTTACTAACAACTTGAATGGTTCTATCCCAATGTGGATTGGTAATTTGACCAATCTTAAGGTTTTTGCTGCTTATGAGAATGAATTTAGTGGTGGTGTTCCGGTTAACTTAGGACTGTACTCTGATCTTTCTTTGTTGAACCTTCACTCAAACCAGCTTGAAGGTACTATTCCTGAGAGCATTTGTGCTAATGGGAATCTTGAATTTCTTGTTCTAACTCAGAATAAGTTGACCGGAACGATTCCTGATTCGATTGGGAATTGTAAAGGACTTTCTAGTATTAGGATTGGGAATAACAAGTTGATTGGAGGTATCCCTAAATCAATTGGGAATATCAGTAGTCTTACTTATTTTGAAGCAGATAGTAATAGTTTGTCTGGTGAAATTGTCACAGAGTTCGCGAAGTGCTCGAATCTCACTCTCCTTAATTTAGCTTCAAATGAGTTTAATGGAACTATTCCTCCTGAGTTTGGTCAGCTTAACAGTTTGCAAGAATTGATTGTTTCTGGTAATAACCTTTATGGAGAGATTCCAACATCAGTTCTTAAGTGCAAGAATCTCAATAAGCTTGACTTGAGCAACAACAAGTTCAATGGTACCATACCGGCAGATATCTGCAATACGTCCAGATTGCAGTATTTGCTGTTGGGGGAAAACTCGATTCGAGGGGAGATACCTCATGAGATAGGGAATTGTGTAAAGTTGCTTGAGTTGGAAATGGGGAGTAATGAACTTACTGGAAATATCCCTCCTGAGCTTGGTCATATGAAGAACTTGCAGATTTCGTTGAATTTGAGTCATAATCGTCTCCACGGAAAATTGCCTCAGGATTTAGGAAAACTTGACAAGTTGGTTTCTTTAGATGTATCTGATAATCAGCTCTCTGGGAATATTCCACCGGCATTGAAGGGCATGCAGAGTTTGATAGAGGTTGATTTTTCGAGCAACCAATTCACTGGTCCAATACCTACTTTTGCACCATTTCAGAAAAGTCCTAATTCAAGTTTTCTTGGAAACAAAGATCTCTGTGGTGATCCTTTGAGTGGTGGTTGTGGGGATTTAAATGGTTATGACCACAGCGTTTACGGTCACAAGGTCTCTTACCGAATCGTTTTGGCTGTTATCGGTTCTGGTTTGGCAGTTTTTGTATGTGTCACAGTGGTTGTTTTGCTGTTCATGATGAGGGAGAAACAAGAGAAAGCTGCCAAGGAAGCAGCTGGAAATACCATTGATGAAATTTCTAGCAAACCAGTGATACTGGCGGGAAACGTCTTTGTTGACAATCTAAAGCAAGCAATTGATTTTGATGCAGTTGTAAAGGCAGTCATGAAAGACTCGAATAAGATTTGTGTTGGTACTTTCAGCACTGTCTATAAAGCAGACATGCCTTCCGGGATGATTTTGTCGGTGAAGAAGCTAAAGTCAATGGACAAAACTATAATTCATCACCAGAGTAAGATGATCAGAGAGCTTGAAAAGCTTAGTAAGCTCGTTCATGATAATCTTGCTCGTCCTATCGGATTTGGAATCTATGAAGATGTCGTTCTTCTGTTGCATGAATATTATGCCAACGGAACGTTGACTCAATATCTTCATAACTCTAGCCAGAAACCTGAATATAAGCCCGACTGGCCAACAAGACTTGCCATTGCCACTGGAGTTGCAGAAGGATTGGCATTCCTTCACCATGTTGCTATCATCCATCTTGACATTTCTTCCGGGAATGTGCTTCTTGATTCTAATTTCAGGCCTTTGGTTAGTGAAGTTGAAATCTCTAGGCTTCTCGATCCATCTAGAGGGACTGCAAGTATTAGTGCTGTTGCTGGTTCATTCGGATATATTCCTCCAGGTACATATCATTTGAACACACATACTCCCTCTGTTTCAATTTGTTTGTCTGATTTTAACTTGACACAAAGTTTAAGAAAGTAAAGAAAACTTTTGAATCTTAGTGTCTTAAACTAAAGATATGTAGAATGTACCAAAATGCCCTTTAATCTTGTGGTCTTAAACATGTCACGTGAAAAGTTTGAATTTAAAAGTTGCCAAAAAAGGAAAAAGGCATTCTTTTTTAAGTATGACAAACAAATTGAAACTATTATCTCTCTTTTCGCATTTGTAGAGATTATAAATCTGTTATTGACTTGTTTTTTTTTTCTAAACAGAGTATGCATATACGATGCAAGTAACAGCTCCTGGAAATGTTTACAGCTATGGGGTTGTTTTGCTTGAGATCCTAACCACCCGACTACCTGTTGATGAAGCTTTTGGTGAAGGAATAGACTTGGTTAAGTGGGTACATGGTGCACCTACAAGGGGGGAAACACCGGAGCAGATACTCGATGCAAAGCTCAGCACCATTTCATTTTCTTGGAGAAAGGAAATGTTAGCAGCACTAAAGGTTGCGTTGCTATGCACTGACATGACACCAGCAAAGCGACCAAAGATGAAGAAGGTGGTAGAAATGCTGCAAGAAATCACGGAAAGCTAAACAAAGTTCGTATCATCAAATCAAAGTATGTCTTGATCTCTGTCTCTTCATATGTGAATGCTCCTGTAAAAATACAAGTGTACGACATTCTTAGTCGAGAGGACAAAATGGTCGAACTCTTAAGGGTATGATGTGTCTTACTTCAATCATATGATCACCAGTTCTTTTTCTCCAAAACATGTAGATGGAGATGTAGATTAATTTCTCACAAGGAGAATATGTTTACAAATAGCATTATCTTTCATTTTCCATTTCCCCTTTCTGGTACATTGCTTCCAAATCAAGTATAGATTAAATCATAGTGAAAGATTAAGGGAATGCTACTATTCAAC

>PGSC0003DMG401029358 LRR-RLKs

CTCTTGTTTTATAATCCCCAAGTGTGCCATCCATTCTTATTTGTCTATTCACTTATTCCAACTTCCATATGTTTGAAAATTCTGTGATGAGAACATCAAACCAAGAATTCAAGAATCATCGCTAAATGAAGACAAAAATTGAAACTTTGTCATTGCTACAATATGTGTGAACCATGGAGAAAACCAATTCCAGAACAAGACCTTTCTTTTTCATCTGCGCATTTCTTGGCTTTCTATTGATATTACTTAGCAGAATTTGTGTCTGCAGTGCTGCAGAAAGTGATTTGTATTGCTTGAAATCAATAAAAGATTCATTACAAGATCCTTGTAATTCCTTGGGCTCTTGGGATTTCAGCAATGTTACCGAAGGCTTCATCTGCCAATTTAGAGGAATTGAATGCTGGCATGATTATGAGAATAAGGTAATGAACATTAGACTTTCAGACTCAGGACTAATAGGTGAGTTTCCACAAGGCATTCGAAATTGTACAAGTTTGATTGGTTTAGATCTTTCAGACAACAACTTGTATGGAACTATCCCTTCAGATATTTCCGTAATAGTTGAACATGTTACAACACTTGATCTCTCAAGTAACAAGTTTTCTGGCAATATACCACCTGACATAGCTAATTGTGAATACCTTAACTTTCTAATGTTGAACAACAATAATCTAGGAGGTGAAATTCCAAGTAGAATTGGTTCTCTATCTCGCCTTAATACATTCAGCGTAGCCAACAATAACTTGACAGGGCCAGTGCCATCATTTGTTAACATACAAATCACAGCTCACAGTTTTGCAAACAATTCAGAACTTTGTGGGAAGCCCTTGAAAGAATGTGAGTATTATGAGGATTCTTGGATTTGGAAACATGTAGATCGTGCTTCATTTATCGAGGCATTTGTGTGTGGTTGGGTACTTTTCTTTACATTAGTTTTAGTCCTTTGCTTCTTTAAATTTCCTGTCAAGGTTATCAACAAGATAGTCTCATTAAACATATGGAAGCTGAGGAAAGAGGACTTAACTTCAGGAAGTGAAGAGGATTTAAGCAGCCACCAGAAGGTAAAGATTTTGCTAACTTTCCGTCTTTTCTCCTGCATTGTTAAGATACATATTTTTGCTGTAGCTCTAGTTTAGTTATTAGTCCTTACAAATCCATCATGGTATATATTCTTGACTAAAGAGATGAATCGAGGTATAAATTCTTTATTCGATGATCGAATGTGATTGAAGTAAGCTTCTCAAATTTCTGTTCTGTTCTGTGTATGTTCCAATGATTCAACTTGCTGTTTTACAGATATTAATGTTGGAGAAGTTTGTTACAAGAATGAGCTTCACCGAGCTGGAAAATGCAACTTCTGATTTTAGTGAAGATTATTTGGTAGGTAATGGAATGTTGGGCAAAGTGTACAAAGCAATTCTACCAAATGACTGGACACTTGCAATCAAGAAGCTGAATGACTCAGAGAACTTGGAGGACGAGTTTGTCTCGGAGATTACAACTCTTGGTGGTCTGAGGCATCGGAACCTATTGCCTCTTATAGGTTTCTGTGTTGAAAGGGAAAAAAAACTTCTCGTTTACAAATACATGTTTAATGGAAATCTTCATGAATGGCTGCACTCGAACGAAGACAAGGCCAAGATTTTAGACTTCCGTCTTAGAGTTAAAATTGCACTTGGGATAGCAAAAGGTGTAGCTTGGCTTCATCATGGTTACAAATTGCACGTTGCACACGGAAGCATAAGCACACGATGTATCTTGCTAGATCAAAATTTTGAGCCCAAAATATCCAACTTTTGGGAAGCAAAAATTTGGAGTAAGAATGATACAGCATTAAGTTGGAGTCTTTTCCCAGTAGCTGAATATTCATGTTTAGGTTCTTACAAGAAAGACATCTACTGCTTTGGCGTCCTGCTTCTCGAGCTGGTTACAGGGAAGGAACCGCATGAATTGACAAGCTCGAGAAATCTGTTTGATCACTCGCCTTGTCTACTTGATGCAGATAAAGATTTGCTTGGGAAAGGAATCAACGATTTGATCCTCGAATTTCTAGAGTTATCTTGTGACTGTGTGAAGTTCTTTCCTAATGAAAGGCCAACAATGCTGAAAGTTTATGACAAACTCAAGACAATTTCTCAAGGTCGAAACGACGACTAGGTATAGAAAACTCCCTTACATTCAACTGGCATTTGAAGTCTATATATAGGACTAGTACTATTGTTTATAGTTTACACTATTGGAAAGTTATATTGATTGCATTTGTGGTTTTTAAAAGAATCTACTGATGCTGCATTGACTAGTTAACTTCAACCCCTGTACCTTTAACTAGAGGTCTCGAGTTTGAGTCCCTAGGGGCTAGGAATGGAAAAAATCACGTTGAGAGCGCTGCTCCTAGAAATGGACCCTTCAATGCCGCGAATTTGACTTTAATGGGGCCCTTGATTCCCCTGCTACTTGATCAGTAATATGTATTAGATGCTAACTACATTATTGAGTCTACTCAGATTGAGGCACCTAGATTGAGGCCACTATATGTGTGTT

>PGSC0003DMG401030114 LRR-RLKs

AAATATTTTGTTAAAGGAGAGTATCAAGTCATGGTGGATAAGTTAAAAGGGGGAGGGAGTAAATTAGTAGCTTGACTTGGTCAAAAGGAGACAACTAGCCAATTATTATGGTTTGATTCTTCAAAGCCAAATAGTTGGCTCTAAATGACTATTTTATTGTTGGAATAAGAGTTGACTTGACAGACTTCATTATCAAATGAAAAGAAACACAAGGAAAGCATCAAATTAACATGATCCAAGCAAGCTCTAAACTTTTTTGGCTCAATATGCAAATCTTCTTCCTAATTAGCCTGTCTTTCCAAGAAACCTTACAATTTTCAGGCACGGAACGGTCCACATTACTAAGCTTGAAACAACACTGGAGGGATTCGGAGTTTTTCCAGTCGTGGAATTTGAATTCTTCAGAATGCAATTGGTTAGGAGTTTCATGCATCAATGACATGGTAGAGACACAATTAAATCTTGGAGAAAAGAACATTACTGGAACAATTCCATCAACAATAATATGTCAGCTCAACAACCTTACCTTCATTGATCTCTCCAACAACAACATATCAGGAACTATACCACTAAGCCTGAAAGATTGCTCAATGTTACAACATTTGGACTTGTTCAATAATTCTTTGAGCGGCCGGATTCCAGGTGTTGTTGTTTGGGATGAAAACACTAGTCAATTTGTATCTTAATGGTAACATGTTGTCCGGTGAAATGCCAAAGCAAATATCAGCATCCCAACTGAAAAATCTTGATCTTTCTGAGAACCACTTGAACGGTTCGATTCCAGAAGCTATTGGTAACTTGAAAAATCTTGTGAAATTGGATCTGTCACATAATTCTCTAAGTGGTTCTATTACAAGCAAGTTGTTTCAATTGAACCAATTGAGTCACTTGTCGCTAAGTTACAACTACTTATCTGGTGTATTACCTAATGCAATGAACTTGTTTAGTTTGTATGATATGGATCTTTCTCATAACCAATTGACTGGTTCTATTCCAAAGGGACTTGCAAGTTCAACTGGACTGGATGCTTTGGATTTGTCGTATAATCAGTTATCAGGAAACATTTCAGAAAGTATAAAACATCTTAGGCCTAGAATTAGTTTAAGGCTTTGTTCCAACAAATTCTCAGGAAGAATTTCTTCTGAATTTGTTAAGCTAACATATGAAGAGAATTGCTTTGATGAGTCCAATCTTTCTTCTACATCCAATAGCTTATCTATCCCTGGGCTACCAAGCTTTCTCTCTGGTGATGAGGTTCAAAAAATTTCGAGATCACAACACTTGATTATTCCTATTGTAGGAGTTGGGGTAGCTATACTGTTATTATGGATCTTTCACAGAACCAGAAAGCATTGGTGGAAAACAAAGATGCAGAATGTCAAAGATAACATAAAGTTCATTTCATTTCAAAAGTTGAAGGTGACTACAGAAGACATATTATCTAGCTTGAAAGACGAAAACATAATAGGAAATGGAGGATCAGGGAAGGTCTATCGAGTTGTGATAGACCAAACAGGCAATACTTATGCTGTTAAAAGCATAGGGCATGGAGAAAAATCAGGTGGAAGACCCCAAAAGGAGTTCCTGGCAGAAGTTAGAACACTTGGTAGCATTCGACACAACAACATTGTCAAGCTCATGTGTTGCATCTCAAGTTTAGACAGAAAGCTTCTAGTCTATGAATACTTTGAGAAACAAAGCCTAGACAAGTGGCTTCACAGAAAGAAAAGAGCAGCGTCACCCGGTCAAAGTAGTACCCCGACCCTGGATTGGCAAAAGAGATTAAAAATAGCCACTGGTGCAGCTCAAGGACTCAGCTATATGCACCACGACTGTACTCGACCCATCATTCATAGAGACATTAAGTCCAGCAATATACTTCTGGACTCAGAATTCAATGCAAAAATAGCAGATTTTGGACTAGCAAAAATACTATCCCGGAGGGATGATGATCCTGAGACAGCTTCTGCTATTGCTGGAACATTTGGTTACATTGCACCAGGTATATTACAGTTCAAATATATATCGTACCTCGTGAATTCTCAAAAGAATTTATCCAACTCTAATATTTTTTAATGTCGGGTCCTCACAGAGTATGCCTCGACATTCAGAGTGAATATAAAGACAGATATATATAGCTTCGGAGTGGTTCTATTAGAATTAACAACAGGGAGACAACCCATTCTCAGAGAAGAGCAAATGAATCTAGCACAATGGGCTCAACAGCGTTACAGAGATGGGAATTTCATTGTTGAGGCCCTTGATGAAGAAATCATGGAAACAAGTAATGTGGAACAAATGAGAGGTGTTTTTAAACTAGGACTGATGTGTACTGGAGCAACACCATACGGTAGGCCGTCAATGAAGGAGGTCTGCAACATTCTTCAAAGCTTCAGAGACACCATATTTTGAGAAAACAGAATGTTTAGCCTTACATGGATAAATGTAGTTGTATTTTTTTAACAGAATGAGAAAGTCTCAATAGCCTCAGCTGGTTTAAGTGGTGTCAAACACAAGGGCTTGTAAGCTTTACCTTAGAATGTTACTCCTTGTTCATAATTGAGATGTAGCGAGGCAAGAGCTAAGTGTCTTGGCAATTACAGTGTTTGTATATATCATTCGGGAGATTAGACAGATTTTAAGCACACTGAGCATTAAGGTCTAACAATGCTTTAGGAAGGATAATCATGATGCAGATGCTTTAGCTAAATGAGTAATGGAGGTTAGACCCATATCTTTTTTTAATTCTTCAGACAACCTGCCTATTCCCAACCAATGTGGGACTTTTACCCACTCTAACACGGGAAGTAGATTTCTTTCATGAATAGAGATGCTGGTAACAGAACCATTTCTGCAGCCTATAGCATACCAACTGCAGTGTGAGGAACTGGAATCCAATAATTGATGAAATCCCCATAGTTGCTTCAGCTCCAACAAAATGCTCTTCTCATTTTCGTAGACGTCACTCTGTAAACCACCCTGGAAATGCATAAAGGCAAGTAAGGAAACAATGAAATGGTAAACTTGTTTCCTGGGAATTGAAAATGTAGATTTTACAACATGAACATGGTTAGTTTTCATGGTTCCTTTGGTTCGATTGTACTAGCTACCTCATTCTCTACAATCTCTTGTTTTATTTTTTTCAAGTGTTTTGATTGGATAGTTAATTAGCTTTGAAGAAACCGTGTATCATACAACACAAGTCAGTCAACATGGACAAAAATTGTTGCTGAC

>PGSC0003DMG402002925 LRR-RLKs

GTTCACTTCTCTTTAGTTTTTACTTCGATTTTTCTCCGGAGCTCCGGTGGTCGGAAAGTTGATCGGCGTCGGAAAACAAGTAGGATTATAAATTTATAATCACAATCGAGGTAAATTTGCTTTTTTCTCCTTAATTTTGCCCGTTGATCATTCATTTTTTCTGAATTTTCTGTGTTTTAATTAAAGTTTAATTTTTCCTGTGATTCTGTAGTTAAAGTTTGAAGCTTTGGGAAGTAGTTTGAGTTAGAGATGAAAATGGTGAACTTTTTTTTTACTTGGAATTGAAACCCCACCCCCCACCCCACCCCATAATTGGAAGAAAAATGTAGGGGTGAAAGTTGAATTTGGAGTTTTTTGTGATTTTGGAGTTGGTTTGAATGTGTATCTTTCTGGTTTTTATGTTTAAAGTTTGGAGCTTTTTTGCTTAGGGAAGTAGTTTGAGCTGGAGATGAAAACATGTTTTGTTTAATTTGGAATTGAAACAGAAAAATTTGTCTTGTTTGTTTGATTTTTGATAAGTCTGAAACTAATAAAATGAAAGAGGTTAGAATGATGACATTTTGTTCTATCATTATTGGAAGAAAGATGTAGATGTAGGGGTGAAAGTTGAATTTGGAGTTATTCATCTCTTTTATTTTATTAGTTTTATTTTTTTTTTTGATTTTTTTTTCTGTGGTTTTGGGAAGTTCAGCATATTGGTTTATGCTTTAAGTTTGGATTCTTTTGGCTTATGGTGATGTTTTGTTTTACTTGGAAATAAAAACCCCATATTTGGAAGAAAAATGTAGAGGGTCAAAGTTGAATTTGGAGCTTTTTTTCCTTCTTTTGTCTTGTTAGTTTTATTTTTCGACTAGTTCTTTTTTTTCTGTGATGGTGGTGTTGGTTGAAGGTTTATCTTGATGATTTATGTTTAGTGTTTGGAGCTTTTTTCTCCCAAATTTTAAACATAAACCAAGAAGATAAACTTTCAACCAACTCCAAAATTACAGAGAAAAAAAAGAAGCTAATCAAAAATAGAAATAATAAGAAAAATGAGAAAATAAAACTCCAATTTCAACTTTCACCACTACATTTTTTCTTCCAATTATAGGGGGTTCAATTTCAAGAAGAACAAAAGGTCACCGTTCTTATCTTTGTGCTGGAGATAAGAATAGTAACCTTTTGTTTTACTTGAAATTGAACCCCCTATAATTGGAAGAAAAATGTAGGGGTGACATCCATGAGGTCTCAGGTTGAATACCCCAAAGACAAAAAACACTAGGTGATTTCTTTCCATCTGTCCTAGCATTGTTAGATAGAGTTGTTTGGTACTTGTTGCTAGTGAAAGTGACAGGTATCTCGTGGAATTAGTTGAGGTGCATGCAAGCTGGTCTGGACACCAAGGTTATATAAAAAGAAGAGTTGAAAGTAGCTCACCCTGTGAGGCTACTTAGAAGAATTGATTTTTGATGTACCTAACCTTTTACATTTTTCATGTTTTGCAGTGTCTGTGGAGGTGTGGTTTGATTCTAGAAATCCTCGTGCAAAAATAATAAGTCGGATTGGAACTTTTCTGCTGTAGCTGCTTTGTCAGGAAAAATGATGGAGGCTCCAAAATCTTTTGGACTTACAGGCTTGAATCAAAACCTGTCGCAGAGCTTTTACCATAAGCTTGGCGAGGGGTCAAACATGTCTACAGAGAGTTATGGGATGAGCAATGCTGGAGGTTCTGTTGCAATGTCGGTGGATAACAGCAGTGTTGGATCAAATGATTCTCACACTCATATTTTAAACCACCAGGACAACCACATCCACAATAACTATTCTGTTGCTGCTAGCATAGTCAGGCGGAGAGTTTCTAGATTGAGCGATGATGCCCTCGCTCAAGCCTTGGTTGATCCTCAGTTTCCCACCATTGGGCTTGAAATTTATGATGAATGGACAATTGACTTGAGGAGGCTTACCATTGGACAAGCTTTTGCTCAAGGATCTTTCGGGAAACTGTATAAGGGAACTTATAATGGTGAAGATGTTGCCATCAAGCTTCTGGAGAGGCCAGAAAATGATCTTGATAGGGCTCATCTTATGGAGCAACAATTTCAGCAGGAGGTCATGATGTTGGCAAATTTGAAGCATCAAAATATAGTACGATTTATTGGTGCATGCCGTAAACCCATGGTGTGGTGTATTGTGACTGAATATGCAAGAGGGGGATCAGTGCGTCAGTTCCTCACAAGGCGGCAAAACCGAGCTGTGCCATTGAAGTTAGCAGTTAAGCAGGCCTTGCATGTGGCAAGGGGGATGGAGTATGTGCATGGCCTGAATCTGATTCACCGGGATCTGAAATCCGACAATCTGCTGATTGCTGCTGACAAGTCAATCAAGATTGCAGACTTTGGTGTTGCTCGAATTGAGGTGCTGACTGAAGGAATGACACCAGAAACTGGCACTTACCGCTGGATGGCTCCGTAAGCATCTAGACTCTTTAGCAGCCTTGTTAAATCTGGTGTCTTGATCCAGTTTGGGCTTATTGCCAAACCCTATAATAGAGCTCTCTCTGTCTGCATTACTTTCTGTATTTTATTTCTTTTTTAAAACTCCTCTGATATGTGTTACGTGAGTCGAGCATCTTTCAAAAAAAACCTCTCTACCTCCACGTGGGAAGGGTAAGGTCTGCTCTCAGTCTACCCACCCCAGACCCCACTGAGTGGGATTTCACTGGGTATGTTGTTGTTGTTGTAGCATGCTGGGTTCCCCTTTATGGAACAAAATTTTTTAGAACTAAAGTGAAAAGCATGGAGGTTGATGGGTTCAAGTCCTGTATGTCCTTTTGATGGCACCTCACTGACTCACTGATTGTGATTATATTTTCTATTGTATACGTGCTGTTGCTAGGAGGGAGAAATTATTTGTTTCAGTTAGCTTTTTAACATAGTTATGTGCCTGTTAATGAAGGTTAAAGAGTTGAGTGACTTTATTTTCTCTTTCAATTTTTTTCCCTCCTTTTCCTCTTCCTTCTTGTCCCTGATTTGTAACTTGATTCAAGTTAGTCACATCCAAGGGTGTGGCTTAGTTGTCAATGAAGTAGGTGCAAACCGTGGAGATTAGGGTTCAATCACACCAGAAGTGCTTTCTTCCCATATGCCTAAACCTTAAACCTTGGTGGGTATCTATTCATGTCTGTTTATTGATCCAAATAAATGGTAAAGTGCCTTAGTTGATAACTCCCAGCACCAACCACGTCATATTCCATGAACTTATCATCCCAAAGTCATTATTAACAGGAAAAGAACTAATCTTAAAAAAAAAATAGTTCTTTTTCTGTCAATAATCACTTTGAGCTGATAAGGTTCATGGAATATGACGTGGTTGGTGCTGGGAGTTATCAACTAAGGCACTTTACCATTTATTTGGATGCATTCTCCCCGATGAAGATGATAAGTTCTACATTGTGGTGGAAATTTTGGCCTTTTAACAGGCATCTTAGAGAAGCATGGGAGGCTTTACTTCAAATCTTAGAAGAGGCACAAAAACACTAGGTGATTTCACCTGATACTTGTGTTAGTGGGGTATTACAAGTACCTCATAAAATCATTGTCGGCATGCACTAGTGATCTTCGAGGATTTCTCGATTATCAAAAAGAAAAATCTAAATAAGTTCTATTCTTAGACATAATTAAGTTTCAACACTTTTATGCCTCCCAACTTATTTATGCAGTTGTTATAATTGATATATTTAAGCGACTAATTAATGGATTCAAAAACAAAAACTGGTTACACCATAACTCAAGTATGGCTCTTTTAGTTTCTTTGCTAGCTCTGAGTTGGAAAGAAGTTCGTTCTTCTTTGGCTGTTTCCGAAAGGAATTCCAGGAGAGGCCTTACAAAAACTAGACTCTACAACGATAATGTGATTCTCCACTGAGTTACTACATAGACTTTTTCCATCATAAGATCATCTCAATCTCTACGGAAAGGTAGAAGTGGAATATTTGGCTCAATCATCCATGAATCTAAATCATTTGTTGGAACCTTCGGTCATTCTTTACGGAGAGACCTCTTTTATAAAGCATAAAAAGCAACTAGGTGATGTTTCTAGAATCTACAACGATTATGTGGATTCTCCATTGAGTTGCTACGTAGACTTGTTCCATCCATAAGATTATCTCAATCTCTACGGAAACGTAGAAGTAAAATACTTGGCTCAATCTTCTCACGACCCTAAATCATTCGTTGTAACCTCCGAGCATTCTTTCTGGAGAGAAGTCTTATTAAAAGCAAAAGGCGATTCATGGAGTTTGAAGTGAAAAGCTAGGAGTGAAGCATGCAAGTTTTGGGGAAGTAAAAAGAAGTACCAAACACATACAAATTTAGTCATCTTCTCATGAACCTGAATCGTTTGTTGAAACCTCCAATTACCGAGAGTAGTGTTTTTCGAATCGAGAAATGCAAAAAATGGCAAGGTCCATGATTTTGAAGCAAAAAGAGAGAAGTGAAGCGCACTAGCTATAAGAAGTAAGTAATGCCAGACATAGAAGGATTTACTACTATAAAGATGCAATTAGAATAACTGATCTTTTCATCCAATTTACGGTAATATTGATATTAGTCTAGCTCCTCAAAGTCGAGATTCAAAAAAGTAGACCCTTTCATTAGGATCGTTTTACATGTCATTGTTTAAAGCCCACACTTCTCTGAAATGCATGGCTTCAAGTGGTAACCCTTACTTCAAATTGCTTAAAGCGATGGCTTTGAATAACACTGCAGGAGAGATGTGATACATGAAGAAATCAATCTCCGGTAAAAAGAATGATAAGAATGCCTTATTTGGTATTTAGTCTTACAACACGTGTTGCTAGACCTGATTTAACGAATGCCTTATATAAGTTGAAGTAAGTCTTTTAGAAACAAAGGGAGCATTAAAACCCCTTAGTAATCAATCAACATGAACAACTTGTTAAGTCAGGGAGAGCCTACGTCAGGCCGGTTATGACTTAGTATTCCCGACGTCTTACTTTTTGTGCCTCACATACTATGTTATACCGGTCCTTTGCATTACACACCCGTTGAAATGGGTGTGTCGTGGCTGTGTGTACTCTTTAAAATACACAAAAAAAATTAACAAAAAAAGAAAAATGCACGTACTTATAATGATACTCTCGTACAACTGATAAAATTCATACTCACAAATGAGGCCATACTTAGAAGCGGGTCTAGAATATGAAGTTTATGGGCTCCCAACGATCTCAAATTAAAAATAAAAAATTATTATAAGTGGTCAAAGGGAATCAGACCCTCAACGAGGAGGGCAACAAAGCATAAGCAAGTTCCATTATGCCGCTGGAGCAAGAGCTTACTTTGTTGGTAGATCCCAACAAATAGTTTTTATATATTTAATAGATTTTTTCAATAAAAATACAAAAGTTAGTGGATTCCCCAAAACCTACACCCGATACTCTAGGCCCACCCATGTCCAACAACAACAACAATGTACCCAGTGCAATCCTACAAGTGGGATTTGAGGAGGGTAGAATGTACGCAAATCTTACTTTTACCTTGTAGAGATAGAGAGGCTTATGAAAGACCCTCGTCTTTAAATAAACATGCCAAAACAGTTCAAAAGTGGGATAAAAATATGGGAGCAATCAGAACGACACGTAATGCAAAATGGGAAGTGTTACATGACAAACAAGAGAAAGAACGGTAACGAAAACAAAATAATGCATAATCAATGCACAAGAAACTACACGAGTAGTGCTAATACTACTGGTAAGAACACCATTGAAAGCCTATCTCGCAGGGAAGTGAGAGAATGCTTAATTACCAACTAACCTTCTACCCTAATATGCGACCTCCATACTCTCATATCTAACACCATGTCCTCAGTAAGATCCAGATGCACTCTATCTAATCACTTACTCCTAATACTTCATCGGCCTGCCTTTACCTCTCCTGGCATCTTCCACTACCAATTGCTCACACCTCCTCACTGGAGCGTCTACACTCCTCCTCCTCACATGTTCACTTAGTCTCACTTCCCTCATTTTGTCCAACATGGAGGCCACTCCTATCTTGTATCAGATATCTCAATTCTTAATCTTGTCTCTCTTAGTATGCCCGCACATCTACCTCAACATCCTCATCTCAACGACTTTCTTCTTCTGAACATGAGATTTCTTAATAGAACAACACTCAGCCCCGTAAAACATGGTCGGTCTAACGTCTACTCTGTAGAACTTTCCCGTCCATACTCCATATAATATGGATCACATAGTACCTTTAGTTGCTGAAGAAGTAATATTTGAAACATTTCAAAATTTTGGATTTCTCTACTAGAATTCATTACCTCTAAGAATGATAGGAAAAACGAATGTATCAAATCTATTTAGGGGGAATGGGTGTCCACATGGGCTTGATTTGCCAAGTCTGTTCATATGTATTATGGAACACCAAATTAATTTTAGAAGTAATGCATAATTAGTACGTAAGAATATATTTTACAATTTGTTTTATGTATTGCGGTGCATGTATAGGCTGACTATTGTATCAATTCATGCATTTGCGCTTGATTGTTTAACTGTTGAAAATCTTTGTTTTATTGATATCTATCAGAAGATCTCCTTGGTTGGCTCAATCTGACATAGAAGGTGCAAAATAATGACAACGAAGATAATTATACTTCAATTTGCTTAGTTTATTGTTCCATTGATACATGGGGTTTTCCTGTATTTAGAGTCTTATTTAAGAATGGGTATATTTAGTGAATCTTTCGAGAAATTTTTTCTTATCATGATTTTCCTGGAGTCTTTAGGATTCTGCTCTTGGAAGTTCAGTTATTTGTTTCTCAGGCATATATGGCAGTTAGAATTTAAACCTTGAATTAGATCACCATTTCAGGATGAGAGAGAGAATATGGTATAGATTGATTGCTTTATCCAGTTTATGAGTATGGATTACTGATGATAGAACTGGCATTTCTTGATTTCCTTTGTCCCGGACAGAAGTATCACCATTAGTTTCTGTTTAGTTTTGCTCCTTACATAATGTTGGAGACGAAAGTATATTCTTTATAAGCTAAGAGAGTTGGAGATTAAGCTTTTGATTTCTTGAGATAATGTGACAAACTGAGAAGTTTCCATTTCTATTAGGACATTTCTTTCAGACTTGAATCAAAGTAAGTTTTCATGTCTGCCTGAAATATTGTTGAAATGCTTGCGATCAGAAAGGATTTAGGAGGTTGACCTTGTGAGTGGTGATAATTTTGGTGACCTAGTGCAGTCGTGAAACACCACATCCCTGCGATGGTGGATGATGTAGCAGTGCAATATAGTTCTTCTCTGTGTGATAATAGGCTAGTATGCATGCTAGACTTTCCTTGGCGTAATAGACAGAAGAATCTGCTTGCATTTGTTAATTTGATGTGATAATGAATGATCTTGCCAAAAAGAAAATGATATTGAGCGATCTTTGCCAAAACAGGTGGGGATCGGATCTCAATATCTGGTAAAACTCGTATACGTGCAGCCAGTCTCTGACATTTGATTCTGATTGACTGAGAAATGAAGTTACTTTTCAAGTTCACTCTTGAGAAATTAGTTATTCACATTCTGCAGATCCTATCTTGTTTAACATTGTGAATTTTTTGGCTAGTGTGTCTGGTTTCTTCTCTTTTCAGTAAAAGAAAAACGATGTTCAAAGTTGTTAGCATCTGTGTTTCGTTGTGGACCGCTAAGTGTTTGACTTTGTTTTGTAGGGAGATGATCCAGCACCGGTCATACACCCAGAAAGTTGATGTTTATAGTTTTGGCATTGTTCTGTGGGAGCTCATAACTGGGTTGCTTCCTTTCCAGAACATGACTGCTGTGCAGGCTGCTTTTGCTGTTGTCAACAAAGGCGTCCGTCCAACAATCCCCATTGATTGTTTGCCTGTCTTATCTGATATCATGACCTGCTGCTGGGATTCTGACCCCGATAATAGGCCAACTTTCTCTCAGGTGGTCAAGATGCTTGAGGCAGCTGAGACAGAAATCATGACTAATATCAGAAAGGCACGTTTCAGGTGCTGCATTCATCCTACGGCTACAGGTTGATTCGGGAAAGAGGAAACTGGAGGCAAAAAAATGTTGAATGGCAAAACTGATTTTGAGTCTGCCATGTGTATCTATCAGGCACAATTTCTTTTCAGATAGAAAAAGGGTTATAGTTATGAGAAAAAACTAAACATTTCTCTGTTGTATTTTGTTTCTTTTCTTTAAACTCTGGATTTAAGACAACGACTAGTTGTTAATACAGGATATCTGATGTGCATCTATGCATGTCAGATTTTGGCTCTGTGTTTGCTGAATTTGTTCAAAGTTATATCAGG

>PGSC0003DMG402004902 LRR-RLKs

TCCTATTAACTATTCCGATCCGCCATTAATCTAATTCTTCAGTCAGCTTTACTGACCATATATTTATAGATCATCCATCGGTATCATTGATCATATACAAGAAGATTTGCAGATCGTCCTTTTCTTTCTCCTATTAATTTGTCAAGAGGATTACTCTCTCTTTTTTCTGATAAAGCCTTGATTTTTTTTCGAGTTCAGAGACATAGACGAATTTAGAATTTAAATTTGATGGATCAACCTTTTAAGTTATGACTATTGAATCCATCTGATCACATTATGAGTTCGAAATTTAATGTTACAGTACTATTTGATAAAATTCCGTATTAAAATATTGGGTTCAATTGAACGCGAAGAAGGTGGGGGCAGGATGAGCTTGGTGGTAAAGGTGGCCCTACTCTTCTTCATTTTGCATGGCTGCAGCTTCATCAATGCAGACACAGATCCATCCGACGGTCAGCTTAATTCTACTGCTACGTAGCTTAATTATAAAGAAATTAATATATTCACGGCACTTATGATTAATTTGGAAGTTACTAACAAAGTTTCTCTGTAAATAAGAAAGAATTTCGTTGTCAACTTATAATGTGAGCGTGGATCACTTGGTGAACAACTTTTTTTTGTCTGGTGTTCGATATCTTGCTTTATTGGCTTGCACTAAATCGAATTTGTCTATCCTAAAACTCGTTAGAGTCCTATCTGTAATTTTTCATTCTCAAAACTCAAACCTGAGACTTTTTATAGCTAAGCTAGGTTGCTAATTTTCACTAAAAAGCCGGACGATAAGCCAAAGAGTTCACAACATGCAAAAAAAAAAGAGTTATAAAATTTAGTAACAAAATAGTGGAATTCGATTTATATATTTTTGGAGCTGTTCTTAGTCTCTGTGTGTATATTTATATGTTCTCAATAATTTCGTTGGTGGCGTGCAACTTTGGTTATCAGGTTCTGCCCTGAGAGTGATGTATAGCTCCCTAAATTCACCGGGGCAGCTTACAAAATGGAGCTCATCAGACGGTGGTGATCCTTGTGGAGAATCCTGGAAAGGCATTACTTGCTCGGGCAATAGAGTAACTGAAATGTAATTATGTTCTCTTCGCCATCCTCCTATAGTTTTTCGATATACACTTTATTTTTGGTAACTTCTCGTTAGTTAATTGATTTGGCAATTGCTGATTTTCAGTTTTCGTTTCATTTTCCTTCAGCCAGATATCCGGTTTAGGACTATCCGGATCAATGGGATACCAACTAATGAGTTTAACATCCGTAACCAACTTGTAAGATGATCAATGGTTTCAATTACTATTTCCTTACTTCGTTCTGTTATTTGTATTTACATCGTAAAATGTTGTAATTTGCAGTGACATAAGCAACAATAATCTCGGAAACCAGATACCTTACCAGCTTCCTCCAAATGTAAAGAGACTGTATGTGTTTTTATGCCTAATTTCTTCTTTCTTTGCTCAATTCCAGGCTCTAGTGTTTTTTACAAAGTTGCTTACAAAATCTTTCTGTCATTTGCCTCGTATTGCAGTAACCTTGCTGGTAATGGTTTTACCGGTGTCCTACCTTATTCCGTTTCACAAATAACTTCCCTTCAATACTTGTAAGTGAATTTTTAAAATTAAATTTACATGTATAGTATTATTACACTCGTTGTTCTGATTTTCTCTCCGTTGTATGTGCCTGATTGTAGAAATGTCAGTCATAATCAAATTCAAGGAGAACTAAATGACATGTTTGGATCACTTTCTTCTCTCAACACATTGTAAGGCCTTGTCTCGATTTCTTCCTTTTCTTATACAGATTGATATTACCCTGTGATGATTGAGATTTTCAATGTATCTTGTGCTATTCTCGATAGATAAATATAGTATCTCTTCTGTTTATCATGTGAACTGTTCTCACGGAGTGAAAGTATCTTTTCTTCTTCTTGGTTGATTATATTATGTTATATCCCGTGTTTCTTACTTTGATTGTTTACATTTATGCTTCAGAGATATCTCTTTCAATTCAATGACCGGTAAACTTCCTCAAAGCTTTAAGTCACTGACTAGCATGAAGAAAATGTAAGACGCGTATCTTCTGATTCATTATTGTACTGTATTGGTCCAGTGTCATTGAACTTTCTAGTATTGTTATGTTGTGCAGCTACCTGCAGAACAACCAGTTCACTGGAAATATTGATGTCCTCGCCAATCTTCCTCTTGACGATCTGTGAGTTATGCAACATCTTTGTGTGTGTGTGAATGTTATCGATCTCTGAATGTTCCATTTCAAGGCTTTCGCAGTTACTATATTGTTGATTCCTTTATGCTTTAGGAATATTGAAAACAATCAGTTTACTGGTGGGATTCCAGAGAAGTTTAAAGGGATAATGCAGTGAGTGTTTCAGTCTGCAAATTATTTTTCATCCTTGAATATTTGTAGAAAATAACTTTCTAACTGAATTCGCGATTTGTGATACAATTGTAGAAACTCTAACAGTAACACAGGGAGCTCAGGGCCTGCAGCCCCTGCGAACAAGTCCACTCAAAGTCACAAATCAGGAGGCAATGGCAATTCTTCAAGCGGTGGAGGTAAAAGTGATAGTGGTGAGAAATCTGGTATCGGTGGTGGAGGCGTTGCAGGCATAGTTATATCAGTTTCAGTGGTTGGAGCAGTTGCAGTAATCTGTATTATCAAGAAAAGACGCAGGAAGTCGTCAACAGACATAGAAAAACTTGACGTTCAGCCATTTGTTCCTGAGTCACATGAAGTACAAGGTATATAATTTTATGACTTCTTTTTTGGGTCAATTGAGAACATAGCTGGTTAGTTTGACTGATACTTCAATATCTCAACATTTATACACTCTTATGGTGGATAGGATCCTTTTCTTGCCATGTTATCTAATTTGAATATAGAGTTGAATTTATTGTGGTACACAAAGGGGAGCCCTGGCGTAACTAGTAAAGTTACTTCCATGTGACTAGGTTCGAGCTGTGGAAACATCCTCTTGCAGAAATGCAGGGTAAGGTTGCATACAATAGACCATTATGGTCCGACCCTTCCTCGTATCCCACATATAGCGGGAGCTGCACCAGGCTGTCCTATTGCAAATGTAGTTGTCTTTATATAAACACACTTTTAACTCTGGATGCAAGTTCCAGCAAAGATACTGAATCACCTTAGGTGAATTTTGGAAAATTAGTTTTCCTTTTGAACTCGTCGTCCTTCATTTCATGTTCAAGTGAGCATGTCTATAATGATGAATTATCTTTCCCTTTTGCAGCTTCATTTGTAACCTGGTATTTCATACATCTTAAGCCTTACCTCAATTGTTTTAAGTTCTGTAGCATCATTCCATCAGAGTTCCAGCTAATATTTTCCCTTTTCTGGTTACTAGTATTATTGATTTTTAATCATGTGTTAGTTTTGGAACCTATATTTTTCTATTCATCACCTATTTTGGCTGCAGATACGAAACTCAGTCAAGCTCCGTCCACAATAAGCATGGAAGCTACTGAAACTCCCACTATAGTATCTCTAAGACCTCCACCTATTTATACTCATAAATCCATTGGTGGAGATGACATTTCAGAAAAACTCATTGTTCCACAGAAGAAAACTCGTACAGCTGAAAGAAAAGTTGTGCAATTTTCAATTGAAGACGTACAGATAGCTACTGACAGCTTCAGTGTAGAAAACCTTATTGGTGACGGATCTATTGGAAGTGTATATCGAGCTCACTTTGACAATGGCAAAGTAATATCTATTAGTTCTGTTTGTCGTTTAACTGTTTCTTGTTGCAGCTAACATTACCTATGCTGATGTAAGTGATTCTGATCGTTTAAGTTCCTATTCCATAGGTTCTCGCTGTCAAGAAACTTTATTCATCTGAGCTCCGGAATCCTGAAGATTTTCTAAAGATGATTCCTGAAATATCCCAGCTGGATCATCCAAATGTAACAGACCTAATTGGTTATTGCTCAGAAGATGAGCAGCACCTGCTAGTTTATGAATTCCACAGAAATGGTTCTCTGCATGATTTCCTGCATCGGATGGATGAAGACAACTCGCCACTAGCGTGGGATAGCAGAGTTCAGATTGCACTTGGCACAGCAAGAGCACTAGAGTGAGTTTCATAATTTACAAAATTACTTTCATTAGGCATCACATAAGATTAATGCAGCTTGGAGAAAACTTTTGTTTGCTTGGTTTCAGATATTTTTTTATGGTAAACTACCCTATCTATCATAAATCTTCTAGTTTTCCTTAAAACATTGATGATAGTAAGCATGTAGAAGTGTTTCAATAATATTTCGAATAGAAAAGCAATTATATTGTTTTAAGTTGTAGAACACCGCGTATGCTGGCTTTCCACTAGATCTAATCAGAATAAATTTTCTATTTTGGCCGCGACCTCTCCAATTCTTATGAGTAGGTTTATTCTAGTAGCTAAATCTAAGCTTCCTTTAATTTGCAGGTATCTACATGAAGCTTGTTCTCCATCTTTAGTTCATAAAAATATAAAATCTGAAAATATTTTACTTGATGCTGATCTCAATCCCCATCTATCAGACAGTGGGTTAGCAAACCTTATGGCTGATGCAGATCAGGTAACACTTTATAAATTTGTTGTTTTGCCTTACTCTAAAGTATAATTTCCTAAAACTCCACATCAGTATCTTCTTAATGGCTTCAACTTGGACCCCTTATTTTTGCTCCGTGTAGGGATTGAACCATAATATAGAGTCTGGATATGGCGCACCGGAGGCTGCTATATCTGGGAAGTGTACCACCAAGAGTGATGTATACAGCTTTGGAGTTGTTATGTTGGAACTTTGTACTGGTCGAAAACCTTTCGATAGGTAAATGTTAATTTAATGATATTGAACCCAAAATCTGTTTAAGACAAATAATAATATACGTGTGTTCTTCTCTTCTCCACTCAGCTCAAGATCAAGATCTGAACAGTCTTTAGTTAGATGGGCAACCCCTCAGCTTCATGATATTGATGCCCTGGAAAAGATGGTTGATCCAGCTCTCTCTGGATTATATTCTGTCAAATCTCTTTCCCGGTTTGCTGATGTTATTGCTCTCTGTGTTCAGGTAATACTCTCGTTGTTTCAATTCATGTGACTAGACACAAAGTTTAAGATAGAAATGCAAACATTTGAAACGTGTGGTCTTAAACATGACATAACATCTCATTAGAGTAATATGAGAAGTAGTTCGTTCATTTTGTACTGAAAGCTGCTACATGCCATGTGTTCAATTTTCAATAGAGTACTCCCCTAAATCGCGCTCACTATCACAATGGTTTGTGTATGTTGTGTCAGCTTGAGCCTGAGTTTCGGCCACCTATGTCTGAAGTGGTTCAAGCATTAGTCCGGCTAGTGCAACGAGCAAATATGAGCAAGAGGTTACACAATAATGATCAGGACTGAAGAGATTTGCTCAAGACCTGCTAATTGTTACACCAAATGGTTCTCTTAAACTGTTTGCTGCTATGGGAGATCATCTGATGCTGATCAGCCCAGCAATATCTTCTTCGGTGCAGTTGTATTTGATTAAATACCATATGCTAAAATATTTCTCCTACATAAGCTCCTCCACTGTTGTAGCTGATTCCCCATTTGTTATGTTGAGCTAA

>PGSC0003DMG402005482 LRR-RLKs

AAATGCCAATTGAATAAATATAGTACAAAGGATAAAAAGAAGAAAAGAAAGGATTTTTCACACCAAAAAGCACAGAATAAACCAGCTAGCTACCTCATTCTTGAATACATTGTCTGTAAGAAGTATGGTTTTTTCTCTAGACTTCAAGATTCCAATGCCATTCTGTTCTTTTGTTTTCTATCTAATTACCCTCCTTTCATTCATGTTCAATATCTCACATGCCTTGAAGTTACAGAATGGTAATGAAACTGATATCCAAGCCTTACTTGCCATCAAAGAAAAGGTAACTCAAGATCCTCATGGTATTTTTACTTCATGGAATAATTCAGTCCATTTCTGCAGCTGGGAAGGTGTCACTTGTGGCCATCTTCATCAAAGAGTCACTAAACTGCACTTAACATCACTTGACTTAGTAGGTACTTTGTCACCTTTCATAGGAAATCTTACTTTTCTTACTAGTCTCAAACTTGAGCTCAACAATTTCCATGGAAAAATCCCACCACAGGTTGGTGGTTTGTTTAGGCTTCAGCATCTTAGTTTGACTAATAACTCATTTTCTGGAGAGATACCTGTAAATCTTTCTAGTTGTTTGAATCTTGTTATACTTGGTGTTGGTTGGAATCAGTTGTCTGGAAAGATTCCATTTGAACTTGGTTCATTGCAAAAGCTTGAGAGATTGCAAGTTCATAACAACAACCTCAATGGACCAATCCCGGAAACCTTAGGTAATCTTTCCGCAATCAAATCACTTTCCTTATCGGTTAACAATTTGGAAGGAACTATTCCAAGTTCTCTTAGTCAAGTGAAGACATTGAATGTTCTTGGTTTGGGTATAAATAAGTTATCTGGTATTGTACCTGCTGAAATTTTTAATCTTTCATCTCTTGAGATATTCACAGTTTGTTATAATCAGTTGTATGGAACTTTGCCATCAGACTTTGGTTTAAGTCTTCTTAAGTTGAAAGTTCTTAAGATTGGTCATAACTGGTTTACTGGACCCCTTCCAAAGTCATTATCTAATGCTTCAAATCTAGTTGAACTTGATGCATATGGTAGCAATTTCACTGGCAAAGTGTCAATTGACTTCGGGGGGTTATCGGATCTTTGGTGGCTGATTCTTGCATCAAATTCCATTGGAACAGGAGAAGTTGATGACTTGAGTTTTTTCAATTCTTTGAGTAGATGCAGAAACCTGAAAGTTCTGGATTTGAGTGATTGCAAATTTGGAGGTGAGCTACCTGATTCTATTGCAAATTTATCGACGACTCTTCTGTCACTAAGATTGAGAGGTAATCAACTATTTGGTTGTATTCATTCAGGAATAGGGAATCTTGTTAACTTGACTGAATTGCAACTGCAAAAGAATGATTTTTCTGGTAGCATTCCTGAAGTCGTTGGCAATCTTCGTAGGCTTCAGTTACTGGATTTGTCCGAAAACAAGTTTTCAGGAAGTATTCCACCTTCAATGTCTAACATGACTCGGTTATATTCACTTCATTTGGAAAAGAATGAGTTAACTGGTAATATTCCCCTTACTTTTGGCAATTTCCGGTACTTGCAAGATTTAGACCTTTCTCAGAATCATCTAAGTGGTACCATACCAGATGGATTTATGAGTTTGTCTTCCTTAACAAATTCTCTTAATCTTGCTGACAATCAATTGTCTGGTCCGCTGTCAGTTGAAATTGGAGCTCTCAATAATCTTGGAAGATTGGATATTTCTAATAACGTGTTGTCTGGGAAAATTCCTAGTAGTATAGGGAGGTGTGTAGCTTTGGAAAGCCTTGTTTTAGCTGGCAACTTCTTTGAAGGTATAATTCCTTCATCCATTAGTTCTTTGAAAGGTCTCGAAGAATTGGATCTTTCGCGCAACAATTTATCTGGGCAGATTCCAACATCTCTTCAGCTCATTTCCTTAAAAAAAGTGAACTTGTCTTTCAACCAGTTTGAAGGTCAGTTGCCAACTGAAGGTGTTTTCAGGAACGCAACTGCAATCTCCGTATCAGGGAATAGGAAACTTTGTGGGGGTATACCGGAACTGGAGTTGCCAATATGTCCGAACGCTGACCCTGATGGTAGTGATAAGTCCAGAAGCATTAAGCTAATGATCCCTCTTCTTTCTGGACTTGTTGCACTGGTTTTCATCATGTCTTTAGTCATAATCATTCGGTTAAGAAAGGCAAGGGGAGAGCCTTCTTTAACATCTTCACCGGTTACTTATGAGAGCCTATATAGAGCGACAAATGGATTCTCTTCTGCCAATTTGATTGGAAATGGTAGCTTTAGTTCTGTTTACAAAGGGGTACTTGATCCAGGTCAATGTATGGTTGCGGTGAAGGTGATAAACATTGACCAACAAGGGGATACTTCTAAGAGCTTCATGGCTGAGTGTGAGGCCTTGAGAAACATAAGGCATCAAAATCTTGTTAAGATCTATAATGCCTGTTCAACTAGTGATTTTGAGGGTAATCCGTTTATAGCTTTAGTTTACGAATACATGCCTAATGGGAGCTTAGAAAGTTGGTTGCATCCTATTCCGGGAGCTGATGCCTCAACTAATGAGGTGAGGATACTAGGTCTAGTTGAAAGACTAAGCATTTCTATTGATGTGGCTTGTGCACTGGAATATCTTCATAACCATTGCCATAATCCAATTGTGTTCTGTGATTTGAAGCCAGACAACATTCTTCTCGACAATGACATGACTGCCCATGTAGCTGATTTTGGATTAACAATGTTTTTCTCAGAAACCATGAGCAAATATAGTTCAATTGGATATGCTGCACCAGGTAACGTAGTTGATAGTCCCTTGAATTTTAGACTATCCTTGTTCAGAGCACTAACAGTCCTCCATGTTTACAGTATTGCTGCAGTTTTAGTCCGCCCTCAAACTTGTCTCTTTTTTTATTGGGATTGAGAGGTTCATTGTGTTAGATTTCTCGTGCAACAGTACAAAATTTTCTTTCCTTGCAACGTTATTTGTCTACAATAAACTGATATAGTTCGAAAAACAAAGGTAGATTCTTGGTTTAGCTGCAATGATGCTTGGACTAGTTGTACAGAACGAAGAGCACATACGCTTCGTCTCACCTTCTCTCTCATGGTGTTATTGTTTCTACTAAACATAAAGGATGATGACTGAACATTTTATACACTAAATTATGGACAAAACCAATGTATTAAATAGACGGACATTTGAGGGAGGACCAAAAGTGCATGCACCAATATTGCGAACATAGGGACTACAAGTACTCCGTGTGAAAAAAACAAAGATGACTTTTAAGTCTTAGTCCAAAGATAAGGGACTATTTTGGGTCTTTTCTCGTGTCTTCTTGAGAAAGTTTTCATATTTGATTTTACATGTTTTAAGTTGATTACATGTTTTCTTTCCTCTGTGGGCAGAGTACAGCATAGGAGGTAAGGCTTCCGAATTTGGTGACGTTTACAGCTATGGGATCCTTCTACTTGAGATGTTTACTGGAAAGAGACCTACTGACAGCATGTTCGAAAATGGACGGAGTCTTCACAGTTTTGCTAAGACGGCTTTGCTTGATGAAATAGTAGACCCCATGCTTCTACCTAGCAACAGCAGAGAAAGACAAGAGGCTGAGGAAGAAGGTGTACTTATCAATCAAGATGATACTAGTATAAAACAAGCGCAGGAATGCTTGATCTCGATAATACAGATTGGAGTTGCATGTTCAGCTGAATCCCCGAGAGAAAGGATGGATATCGGGGATGTTGTCAAGGAACTTCAACTAATTAGAGACATTCTCCTTGCTTCTCATGCAATTCACAGCTCCACCAGCGGGTAAATTCGCTCTACTAGCTTTGTTTTCAATCGTGATATTGAGTTCCTCTGCTAGATATTGAGAAACTAGTTTCTAAACACATGTGTTGCACGTGTATTGTCCTGTGTAAATTCTTGTAGATCCATTAGAACGATGAAAGTTCTATGAAGATATGCATGTGTTTACCATGTAAATTTTCTTCTTCTTGTTAATCGTTGTCTTTCTTTCCTTGGTTTCAGGAGCTTGAGGTTCGAAGGGTCTTCCAGTCGTTCTGTTACTAGTAACTGGCAAAATTTCACATCATTTCGTTTGCCTTGATGTTTTTAGCTCTTATATAACAAATTACTTTAGCAGATTGAAGATGTTAATCATCAGTTCAATCATTTGTTCAGTTGGCCTAATGCTGGTTAGTTTAGTTGAGCTTAAGCAAATGGAATTGGAATTGTCTATGTATATGATGTTATGACTTGGCCAAACACATACACGACCCCGTTAGCTCGTCTCGAAATCTC

>PGSC0003DMG402008032 LRR-RLKs

AAAAAAAGTTCTCTTTCAATGCATTGCTTCATTAAAATCCCATTACTCATTTCTGGGTAGTCTTTTTTTTTTTTCTCTCTGTATTTGTTTGCCATGAAAAATGCCGTCTTTATGGAAAGCATTTTCACTATCTTCAAGTTCTCAACACTCTTCTTCTTCTACTACTACTGCTAGTACTACTTCTTCTCCGGCCGATTCACCGAATACCCGGCGGATTTACGGCGGCGGAAGGAAGCTTACTAGGCAGAGGAAGTTGAGACATGTTAGTGATGATGATCTTGGTTTGCGGAGACCTAATATTCAGGCACTGATTATTGATGAACGTTCAAAGTCTTTGCCGGGTTCTCCTGATTCTTACGCTGATTTAGGGTCGGGTTCTAGGTCGTCTCACCACCACCTCCGTCACTGTTCTAACTCCTCTGCTGTGCCACTGCCGTTGCCGCTCCCTGAGCTCAATTCTCTGCCTAAACAGAACTCGGTTGACTCTAATTTGCCGGCTAGAGTTGACAGGGAGCTTTTAAGTCCTCCTCTTGCCAGGTTAGTCTTGGAAATTGAACCCATTGTTCAGAATTTATACTCAATTTCAAATTTTAGTCTTACTTTTCGTTTTAGTCTGTTTCGAAATGAATGAGTTTTCCATTTTTGGCAACTCTTTAATTGTAACTTTCCACCTTACATATTTAAGAGATTAAATGATATTTTGATATATTTTATGTATTTAGTTTAAGATCACGAGATTCAAAAGTGGTTTTTACTTTGTTAAACTTCGTGTCAAGTCAAAACCAGTCAAACAAATTGACATGCTGGGTATGTTTATCAAACTTTTGAGTTTTTTCTTACATGTATGTTTTATGCTTTAAGTAAAAAAAATGCTACCTTTCTGAGTGAAGTATTGTTTAGCCGATTCATACTGGTTTGGGATTGAGGGGTAGTTAATTTCGGTTTGATGATGAAAGCTTTTCACCATAGGTAACTCTGTATGTAAAGGGATTTAGTAGTAGTAGGTAGTTGGTTCGAATTTGAAGTAGTTTTTGGGGTGGATTGATTGCAGTAACACTTTATAGGTCATTACCATAAAAGAACCATTTTTTCCTTTTTTGGGTTGTGGATTAACTGCAGTAACCTTTATAAGGTTCACCATAAAGGTTGGTTTGGAATTGTGAAAGAGCATGTGGTTTGTAGTTGTTTCAAATGATGATTTGTAGTGTGCTACTTTGTTTTTGGGGTAAAATGTTCGATTAGCTATGATGATTTTGTAAGAACAAAAACCGAACGCGGAAATGTTAACCTGTTGCGAGTTGCAGGGAAACTTTTGGTCGAACTCCAGCTGAGGTGAAAAGTTCTCAGCATCCAAGATCATCTACTCCAACTTATCGGCGTAGGGGGTTCCCTCAGGATCTGAATGCTGAGGGCGTTGAATTCAGGTTGAATGTTCCTGCTAGAAGTGCCCCGAGTAGTGGTTTTACAAGCCCCGTCCGAAGCCCAAAAAGGTTTAGTACACAGGACCTTTTTCATCATCCCTTGCATCAGGCCTCATCATCACCTTCAGAGGCTTACTCTTTCCAACTTTCACCCACAAGGATTATAAATAGCGCTGATCATTCTCCACTGAGTAGCCCAATTTTGCCAAGTTCAGCTAGCAGAATCAGGAACAGCAGAAGTGGTGCTGTGCATTCACATCATAAATCTCTTCCGGAAAGTTCTCTAGGTTGGAATGAAGCTAACAACAACAATGTGCATCCACTCCCCCTTCCACCAGGAGTTCCACGACAACCAGAATCATGTACCATGCACAGCAATATGGATAAACCATGTGTATCACCAGCAAAGGGTCAATGGCTAAAGGGGAAGCTTTTAGGTCGTGGTACATTTGGAAGTGTATATGAAGCTACTAACTGGTAAGGGATAGAGCTTGCTTAGTAAGTAACTCGGTAGGAATGTAACATTTGCTTCTTTTGTTTTTCTTCCTTTTACAGTGAAACCGGTGCTTTATGTGCAATGAAAGAAGTTGATCTTACTCCTGATGATCCTAAATCTGCTGAATGTATAAAGCAGTTAGAACAGGTGACGTCCTCATGTCTTCTTAGGCTACATAATCAAAATTTGTATATAAACTGCAATTGTATGAGATACACGGTGAGAGAAAACATATTTTAAAGGCCAAATCTTTGTTAAGTGAAGTGAACTTTAAAACAAGTACTTACTAAGCATGTTATATAATATGTCCTGGTTGTCTAACTTGCATTAGCAAATTAGTCAACTTAAGGTTATGTCCAACTTTGAGCTAACTATCATGTGCCTGAAGATGCAAAAATCAATATCGTATCCATGGAGCTACTTTTATTGATCATATTCTTTGACTTCATAAGCCTCCTTCACGTTTTGCATCATTGAGAAAATATTTTCGGAATGTTGGTAACTGACTTGGGAAAAAAGTTTCATATTGTGGAGTTTTTTATTCTACTTGCTTATACATAGTTTGTGATGGATTGCAAAGACTATGACATGTTTTTCACTTGTTTATGCTATTCTAAATCGATTCAGTTTCTTCCTTACTTCTGCTGTAGGAAATAAGAGTTCTCCAGCAGTTAAAGCACCAAAATATTGTTCAATATTATGGCAGTGAAATAGTAAGAACATTTATCTTTCTCTTGCCATGTTCAGGTATTTCTTTAGAATATTGAAAGATAAATGTTGTTGTTGTTGTTCAGATGGAAGACCGCTTTTGCATATATTTGGAATATGTGCACCCCGGTTCAATTAATAAGTATGTTCGGGAACATTGCGGAGCTATGACAGAGTCAATAGTTCGAAACTTTACTAGACATATTGTATCAGGGTTGGCTTACTTGCATAGCACTAAGACAATACACAGGTAGATACTGGGTTACATCGATGTGGTGTATGTCACTTGATTTAAAGATTCATTAACGGAACACTAAAGCGTCTGTGTTGCAATTCCTTGTTATAAGCAAGGAAGAAACTATTTTCTTTTGCCATTTTTTTTTAAAAAACTTTTCGTTGACAGTGATGCTCAGTCTAATGGTTGGTTTCAATGATTTTTTCTGAGTAGGGACATTAAGGGAGCAAATTTGCTTGTAGATGCATCTGGTGTTGTCAAGCTTGCAGATTTTGGGTTAGCAAAACATGTAAGCTTTCAGAACTCTAATGCCAAACTGAAAATGCTACTTTCTATGTTCGTTCTAATCTAGTATTTTTCTTGTAAACCCTGATCTGTGGTGAAAATGAATGAGCTTGATCTTTCATCGATTATTTTTCATGCATATCTTACCGCTGTAATTTATAACGTTTCTCTTTTTCCCAGCTTTCAAGCTGTGCAACTGATCTTTCTCTGAAAGGGAGCCCTCACTGGTTGGCTCCAGAGGTGGGCATTGTTAAGCTATTTTGCGGCTACTGCCTCTGCCTGGCTTCAGTTTTCTTTCCCCTTTATTAATTGGCTTGTACTTATGTCTGGACTTAACTTTATATGCAGGTAATGCAGGCTGTGTTGCGCAAAGATGCCAATCCAGAGCTTGCCTTAGCTGTTGATATATGGAGCCTGGGTTGTACTGTTATTGAGATGTTTACTGGACAGCCCCCTTGGGGAGAACTTAGTTGGGTAAGTGTTTAAAGCATCTCATTTTGAGAACAATGTAATCGCATTACAAGCTTAAATATATTTGTGAGCTTTTAGGTTGTCAATATAGCATCATAGCTTGTTAATATACGAGCAAGGCTGACTGATTGGCTTAATCTCAATAAAGCCAGAGCTATGTTCTTACTTTTCATGCTTTCATGAATTGGTTTCTGTAAGTGTCTTAATGTTCATGCTTTGGCTTTATTAGGTGCAAGCAATGTTCGGTGTCTTGAACAAATCACCACCTATACCAGAAAAATTATCACCAGAAGGAAAAGATTTCCTTCAGTGCTGCTTTCGGAGGAAACCTGCAGATAGGCCATCAGCTATGATGCTACTTGAACATGCTTTTTTACGTAGTACTAGTTCCCTTGAGCATAGTATCAATGTTGCTGGTTGCTCAGAAGATTATCCGGGGAAGAAATTCCATGTGAGTGGTACATGCTGCTAGCCTGCTACTGATGTGACATTAAGGGTTAACGTCACTGTATTATTTGTGAGATTCTCTAAAAAAAAATTCCAAATGATTTTAAACTCTGAATTCATTACATTTCAGGATACCCTGAGTCCTAAGAACCCAATCAACCATAAAAAGGAACAAAAACCACTGTTACCAGGAACATCTGGCAGGCATGCAAAATCACCATGTAGCAGGTAATAACTGACTTGTGTTTGTCATGTTATATACCAAACACTTTATTCTGTGAGATTATTTTTTTCGCGTATTAGTATCTACAATTGCAGCCAGATACTATATCTTGTGTTGTGCCATTGACTGTCACGTCCATCTTATGCAAGTTAAACTGTGTCTCTTCCAGTGAAACTTGCCGGCAAACTCAACCTGAAACCTGTGACTATGGAGCAGCTTCTCACCATTCTCCACGTTCTGCACTTGAAGTCTTTCCCTGTATTTCTTCAATGGAGCTGAACTCGAGTTCACGTGCAGCCAGCCCATCAAGCATCCCTAGTAGCTTTCGCCTGGGACCTGAAAACAGAAGTCCTTATAGAATCATATCATAGGAAAGGAAATTCCAAACCTCTGTATAAGATCATAAATGAATGGAGAGGGGGAAAACTCATGTACAGATCAGGAGAATTAGATACAGAGAACTTCAGTTGGATGATTAGTCAAAATGTTGCCATATGTGGGAGAGATCTCAATTCCCGTATGCATCCACATCTTCTGTGTAGTATTGATGTTTGTATTTGAAGGCAAACATATCTTTGTAACAGATTCATGAAGTACTGAAGATGATTCTGAAGTTAATAGTGTATTTTAGATTGAAAACATGATTTAGTGTACACAATTAAATGATTTATGCTAAATTCAATGAGTATCTTTTTCCCTCATTTTTACAACCAA

>PGSC0003DMG402014186 LRR-RLKs

GCAGCAGCAGCTGCTGCCATGACGGCTACTGGAAAGGGAGGAGAGATCGGTGGAAATGGTATTAAAAAGCTGATCTTTTTCGGGTCTGATAGATCGTTTGATTTAGAGGATTTGTTGAGAGCTTCAGCTGAGGTTTTAGGGAAGGGGACGTTTGGAACAGCTTACAAGGCAGTGTTGGAGATGGGCACAGTTGTAGCTGTTAAGAGATTGAAGGATGTAACCATATCTGATATGGAGTTCCGGGAGAAAATCGATAAAGTTGGACAAATGAATCATGAGAATTTGGTTCCTCTTAGGGCATATTATTACAGCAGGGAGGAAAAGCTTCTTGTCTATGACTACATGCCAATGGGAAGCTTGTCTGCACTTCTTCATGGTGAGTCAAAGTTATACTTTTTCCTTATAAATTGATGATTTTTTGTTGTTGTAAATACCTGAGGGCAATCTCTATTGCTTTTCTTGCTTTAGTCTGTGGATTGGTTAGTTGGACTTGTTAGCTTATTGTAGTACTACATGCAATGCATCTTTTTGTGGGAGAACATCTAACTGAAGTGTTGCATTTCGTATGAATTTGAACATTTCTTTTTGATGAATTTTAAAAGCACGAGTTCAATTATTGATAATGTCCTCTCATTTCTCTAGATTTGGTAGAGTTGTATCTGTGAAATGTTGCTTTTATAGCCTGTGAAACTCGGAAACTTATTAAATTGCTAATAGATTGTAAGATGTTTGCTATAATTTTATTCGTGCTATTCACTACTAAGCTCTGTAATTTTCCAGGAAATAAGGGGGGTGGCAAGACTCCACTGGATTGGCAAGTCAGGTCAGGCATTGCCCTTGGCACTGCTCGGGGCATTGAATACCTGCACTCTCAAGGTTCAAACGTCCATGGGAACATTAAATCATCCAATGTTCTTCTCACCAAATCATATGATGCACGAGTGTCAGATTTTGGCCTAGCCCAGCTGGTTGGACCTCCAACTTCCCCAACTCGAGTTGCTGGATATCGTGCTCCAGAGGTAACGGACCCACGCAGAGTTTCCCAAAAGGCAGATGTATATAGCTTTGGTGTATTGCTTCTGGAGCTTCTTACTGGGAAGGCACCCACTCATGCCCTCCTGAATGAGGAAGGTGTTGACTTGCCAAGATGGGTGCAGTCCATCGTTCAAGACCAATGGACTTCACAAGTTTTTGATATTGAGCTCCTCAGGTACCAGAGCGTTGAGGAAGAAATGGTGCAACTCTTGCAGCTCGCAATAGATTGCTCGACTCAATATCCCGACCACAGACCTTCTATGTCTGCTGTCGTTGAGCGAATTCAAGAGCTCTGTCTCTCAAGCTTGCGGGTTACTCAAGAACAGTCTGACTCTGTCAATGAATCTGATTGAGGTGGTCCTTGGTCCTTTTACATGTGCAGTACTTGAACTAAAGAAGCACCATCACCAATTTACAGAACCCCCTTTAAGCATATTAAACTTTAATGCTTTCTGTTGGTTAGTAGGGCCTATCTTCTTTCATTTAAATTTTAACAATTTATGCCCCTTTCTTTTTTCTCAGTTCTCCATTGTAGTTGAGCTAGAGCCCTTTTCATTTTCCTTTGTGTGGGGGGCCATTCTCTGTACCATTTGTTCTTGTTGGGTGGATAATGGTTGCTGTAATTATTACTTTCAGCCTTTACTTTGCTTTCACTTTAACCAACTTTGTTGAGAACTTAATACCTACCTTTACTTTGTAGTACTTTCATTTAGGAGGTTTGTTGTTTGGTAGTAGTAAATTAGTGAAGTTTTTTTTTTTGGTTTCCCACCTGGTGTCCCGTAACTACATTGGAACTCGACTATTCAGGGGAAGCGCTCTCTACCAAGGATTGCCATGCTTTGTTTGTTGATCAGGACAAAGTTTCTTTGGTTTCACGGATTGTGTACCCTCTTTTACTTCGTTTTTGA

>PGSC0003DMG402015976 LRR-RLKs

ATGGCAACAACGGAAAGAATTTCATACTATGCACTGCTCCAAGCAACTGATTCGCTTAGCGAGAGTAATCTGATTGGTTATAGAAGTTTTGGCTCTGTTTACAAAGGCGTTCTCAGGAGTGGGACTGCCATTGCAGTTAAAGTGTTCAATCTGCAACTGGAAGCAGCATTCAAGAGTTTCGATACAGAATGTGAAGTTTTGCGTAGCCTTCGCCATAGGAATCTCGTGAAAGTCATCACTAGTTGTTCCAACCTTGATTTTAAGGCTTTAGTGCTCGAGTATATGCCTAATGGAAGTCTTGACAAGTATTTGTATTCGCACAACTACTTCCTAGACATCAGGCAGAGACTAAGCATTATGATAGATGTTGCATGTGCTTTGGAATATCTTCATCATGGATGCTCCTCTCCGGTGATTCACTGTGACCTGAAGCCTAGTAACGTCTTGCTGGATGAGGATATGGTTGCCCACCTAAGCGACTTTGGTATTTCAAAACTGCTTGGTGAAGATGAGAGTGATTTATACACAAAAACCTTAGCAACATTTGGTTATATAGCACCGGGTACGTCTCTTTGTTTTTGCTAATTTGAACATTGATTTTTCCTCTTATTCTTTTTTTTCCAATCAAGAAAGTATATTGCATCTTTTAATTAATTGTTTGACTGTAGAGTATGGACTGGATGGACTGGTGTCAATAAAATGTGATGTCTATAGTTACGGGATTGTGTTGCTGGAAACATTTACTAGGAGAAAGCCTAGTGAGTTTGCGGGAGATCTTAGCTTGAAGAAATGGGTGAGTCATTCACTCCCAGAGGCAGTAATGGACGTTATGGATGCCAATCTGGTTACACCAATGGATAATCGCTTACAGAAGGAGCTAGATGTTGTGGCATCAATCATGAAAGTGGCATTAGATTGCTGTGCTGAATCTCCGGCAAGAAGGACAAACATAGAAAGATGTTGTAGGGATGCTACAGAAGATCAAGATTCAACTTCTTGCACGCTGAGCATGTTACCGGAATCTCCTTTTCCAAATTGA

>PGSC0003DMG402020263 LRR-RLKs

CTTTAGTGGCCTAGGTTAAGACGGCCCTGGTGGCACAGTGCTTTTCTGAGACTCTGTGAAAAGAAAACTGTCACCATTAAAGCCAGAAACTTGAATCGGTCCCCTGTTTTTTCTCCTTGACTTGCTCTGCATGCAACACTAAATGCTTTCTCTTTTGTCAACTACCCTCAATTTTGCTCTCTGTTTTTTTATTTTCAATATTGTTCTTCTCCCAAGAAAAAAAGAACATTCCTTTAATTTGCAATTATAGGTCAATTTATTCGCATTTTTAATGGGGTTTTATCAAGATTGTAGCAGCATTAACTGTTTGGTAGCTCAATCTTGACTTGTCTGCACATTTAGACTTTAATGCTTTTCATTTTCACTGAATTAGAAAGGGTCCACCCATGTAATTTTCTTATTCATGATTAGCCAAGAACGTAAAATGGGTTGCTTCAAGATTGTTGGAAACTTGGCTCCATTTTGAAGTCACTGTTAAATGGGTTTCTTCAAGCTTTTTGTGATAGTGTTGGTTATTTGCGCTTTCTTTATAGCAAATACTTATCAGTTACAGTCCTATGAAATTCAAGCTCTTCTTCAGTTGAGGAAGCACTTGGAATATCCAGTTCAGTTGGATGTTTGGGAGAATTACCATGGAGATTTTTGCAGCTTGACTTCAACACTGCATATGAGCATTACCTGTCAAGATAACTCTGTTACTGAACTCAAAATCAAAGGAGATAAGTTAGTTAAGTTGAATGAATTTCATGGATTTGCAATTCCAAACCAAACATTATCTGAGGGTTTCTCCATTGATTCTTTTGTTACCACATTGACAAGGTTAAGTAGCTTAAAGGTTCTTACTTTAGTTTCTCTTGGTATTTGGGGACCCCTTCCTGATAAAATTCATAGGTTAGCTTCACTTGAACTTTTGGATATGAGTTCAAATTTTCTATTTGGTTCTGTTCCTTTTCAGATGTCAAGATTGATTAAGCTTCATACTTTGACATTAGATGCCAATTTTTTCAATGAAACTTTCCCTGATTGGTTTGATTCCTTACACAATCTTACTATATTAAGCTTGAAGAACAACAGGTTGAAGGGTCAGTTTCCTTCTTCAATGTCTAAAATCACAACCCTCACTGACATTGTTCTGTCACACAATGTGCTTTCTGGTAAATTGCCCGATTTAACGGCCTTGTCTAAGTTGCTTTTATTGGATTTAAGGGAAAATCATTTGGATTCTGAATTGCCACCCTTGCCAAAAGGACTTACCACAGTTCTTCTTAGCAGTAACGTTTTTACGGGTGAAATCCCAGTAGAATTTGGCACACTAAATCAACTTCAACACCTTGATCTGTCGAATAATTCTCTCAGTGGAATGCTGCCTGCTGGCTTATTCTCTTTGTCTAGCATTAGCTACTTGAATTTAGCGTCTAATGTTCTTAGTGGTTCACTTCCTAGCCATCTCAATTGTGGGGGTGAACTTGGTTTTGTTGATATTTCTGATAATAGATTAGTTGGTGGGCTTCCTTCTTGCTTGAACACTATTTCAGATAAGCGAATTGTTAAGGTTGGTGGAAATTGCTTGTCTCTTGACACTCAATATCAGCATTCAGAGGGCTATTGTAAACAAGCAAATTCGGATAAGAAACAGATCACAGGAAAACAAATAGCATTGTTGGCGGGTGTTATTGGGGGAATTGTAATTGTTGTGGTGTTTCTGTTAGTTGTCCTTTTCATCTTCCGTAGAAAACGACATGCGCGTAACATGGTGGATAAGCACATTCCTCCGAAAGTTGTGCAAAATACACAGCCCGGGGTTCCCTCTGAACTCCTAGCAAATGCCAGTGAGTTGATGTTTTTCTTATTTCATTTTTAATCTGTTTAATTAATATGAAGGAAATGTTCTCCAGAAAAGCTTAATCTCAATCTTCGAATTCTCAGGAGTCATTTCTCAAGTGGCAAACACAGGATCACAAGGTTCCCCATCATATCGAGTGTTCTCCATGGAAGAATTGCTAGAAGCAACAGAAAATTTTGATCAGTCAGCATTACTTGGTGAAGGGTCAGTTGGAAAAGTATGGCTTCAAAACAATACTTATGGTTCTCATTGATAATATATATCTTGATGTTTTCCATTCAATCAATGGTCACGCTGAATTTTGGCGCAAAGTGATATACTTGCTGTAAAACTTGAATAAACGAAGCAGGACGTAGATTTTGTTGCATAGACCATGTCCAACAATGATAATGCAATGAATGCACTAGTTCACCAAAGTACAAAATGGATTTAGGTGTTTTTTTTTTTTTTTAAAGTGCTTGCCTATCATATTCGCATTCTACATGTGCATCTGTGACTTCTTTTGATGATAAAAACTGGTGTCACCAATAATGCTTTTGTGCATTTGATTTCATATCTAATGAACACATGACAGTTCAACTAAATATATCGAATGCCCTTAAAGGCAAAATTAGATAGTCTCACACGTTTGAGTGCCTATGCAGTGAGTTGACATCTGTGCTATTTTGATTGACATAAACAGGTGTTAGCCTTATTATGTCTTGCTTGAATAAGACAGTTATTTTAGTTATCAGCAATTCCCTCAATATTTATCTCCATTTCAGATTTACAAGGGAAGATTAGAGAATGGAGCTTACATAGCTGTAAGGTCACTAAATGTGTGCATAGTTCACAAAAGTTAAAAATTGTTTTAGGTTTTTTTTTTTTTTTTAAAGTGCTTGCCTATCATATTCGCATTCTACATGTGCATCTGTGACTTCTTTTGATGATAAAAACTGGTGTCACCAATAATGCTTTTGTGCATTTGATTTCATATCTAATGAACACATGACAGTTCAACTAAATATATCGAATGCCCTTAAAGGCAAAATTAGATAGTCTCACACGTTTGAGTGCCTATGCAGTGAGTTGACATCTGTGCTATTTTGATTGACATAAACAGGTGTTAGCCTTATTATGTCTTGCTTGAATAAGACAGTTATTTTAGTTATCAGCAATTCCCTCAATATTTATCTCCATTTCAGATTTACAAGGGAAGATTAGAGAATGGAGCTTACATAGCTGTAAGGTCACTAAATGTGTACAAAAGATACTCTAACCGGAACCTCAAACTACGATTGGACTTACTGTCAAAGTTTCGTCACCCCCATTTAGTTAGCCTTCTGGGTCACTGCATTGATGGTGGAGCACCAGATGATTCAACTGTCCCCAGAATTTTTCTCATTTATGAATATATCTCTAATGGAAACTTCCGTGCTCATCTATCAGGTCAGCTAAGTAAATCCTTTAGCATGAGCATTGCATGTGTTTTGATGCTTGGTTGAATATGGAAATATTATATGATGCACATATGGAGTGCTGGTTGGATTCTTTTCTCCCAGAAATGATTGGGATACCCTTGTTGATGAATCTCTATCTAGCATTGCAAGTATTTTTGCTACTTATGAGCTTGAGGCATCTTATTCATCATATTAATTTCACACATAGTGGTATTCAGTATGCAATTACTGTCAGTTTGATTGAGCAACAATGTAATGATTCACTTGCACGAAGTTCTGTTGGTTGAATTTCATACTGAGTTGGACCTTTATTTTTGTTGTTATATACTTTCATTTAACCCTGCATGTCCAATAAGTACAGAAAATACAAGAGACACGCCATAACGTATTCAGCTTATCTCAAACACTTGTACTTGCAGAAAATAGTTCAAGAAAGATCCTCAAGTGGTCAGACAGGTTGTCGGTGCTGATTGGTGTTGCCAAGGCAGTGCACTTTCTTCACACAGGTGTAATTCCTTCTTCATTTAGCAACCGCTTGAAGACGAGTAACATATTGCTTGATGAGCATAACATGGCAAAATTAAGTGATTTTGGGATGTCCATTCTTATGGAAGAAAGTGAAAAGGTGGGTATTTTAATATTTTGTGTGAAAGGTGAAAATTCAAAATTTAGGATAGATGGTAATTTCTCATACTCAATATGTCTTTTTTCAGGCAAAAGGAGATGACGTCACCTCCTGGTATGAAACTCTTCACTCATGTATTCCGTAACGTGCCTTACCAACTTCCTTTTACTAAACCAACTGGGATTCTATTTCAAGAGTGTCAACTAGAGGCATATATATCATCTACTATAGCTGTTTGCAGGATATAAATCAGTGATATTTTATTCTTGTGTAGGCATATGACAAAGAAAGAGGACGATGTTTATAACTTTGGTTTCATATTACTGGAGTCGCTGGTTGGCCCTTCTGTCAGTGGAAAAGGAGAAGCATTTTTGATGAATGAAATGGTATGAAACTGCTATATTATTATGTATGCAACTCCTTTGTCATTGACAGTAACGTAAAGTATTCTTCAGCAATTTGTTCCCTTTAGGATCCTAAACGTTGTTGCAATGTTGGATAAGTAACAAAATTTTCGTTCGCTCTGCAACCTAACTTTACAATCAGTACTAATATAGATTAACAGAACTACACACATTATTCAAGTTTTATTAGGTAAACCTTCTGATATGCAAGTACCATAGGGAGAAAAGTTGAGCCTGATATACAAGAGTCAAAAGTCTTGATTTTTTCACTGGTGTGTTACATAAATATCTCTGGATTTTCTCTTTGAATGACAGGCATCCTTTGGTAGCCAAGATGGTCGACGCAGAATTGTGGATCCAATAGTGCTAGCCACTAGCTCCAATGAGTCATTGTCAATTGTAATATCCATCACCAACAAATGCATATCTACAGAGTCATCAACTCGCCCCTCGTTTGAGGATGTTCTCTGGAACCTACAATATGCTGCTCAAGTTCAAGCTACAGCTGATGCAGATCAGAAATCTGATGCTACTTCACCGTCATAAGCTATTTGAGAAGTACAACTGCATCGAATCAACGATATTTAATTTGGTCAGATCATAGGAGAATGTTAGCTGATTAATTTCATGATGGTTGTGGGAGTACCAACCGGTAGGCTACTAGCCTTAAAGCTCAAGTTTACGCTACAGGTCCTCCCATATTTTCTTTATTTTGTATCAGAATTGTAAATTTTGTTGATTCTAGTAATAGGTGATGTTTTCATATAGTATAGACCCTCGTTTTAAATTCTATCATTGCAAGTAGGCAATTGAACTCGTAGAAGTCTAGATTCAGACAAGACAATACTTTAGTCATCATATATATGTTTGGTGCAACGGAAAAAAGATATGGATACCGCAAATGCAAG

>PGSC0003DMG402026747 LRR-RLKs

GACTTATCATCAAACAACATGGTTGGTTCTTTACCTCCGGAAATTGGAAATCTAAAATCTGTGATACAGATGGATTTGTAGATGAATCAATTCTCAAATGGAATTCCAAGAGAAATTGGAGGATTGCAAAATCTGGTGCATCTTTCTTTGAGACACAACAAGTTGCAAGGATCTATACCTGACTCAATGAGCAACATGGTAGGTTTGGAATTCCTAGACCTTTCTCATACTAATATATCAGGAATCATTCCTATGTCTTTGGAGAAACTTCAATACCTCAAGTATTTCAATGTTTCTGACAACAAATTGCACGGCGAAATACCCTCGGCGGGTCCTTTCAAGAACGTCTCGAGTCTTTTTTTCATCAACAATGAAGCATTGTGTGGTTCTTCAAGATTTAGCGTCCCTCCATGCCCCACTTCATCAAAGCATAGGTCAAATAGGAAAAAAAATCTTATGCTGGGAATTGCACTTGTATTTGTTCCTATTACCCTTGTGTTTGTATGGATAAGGTATAGAAGAGGTAAAAGATCTCCTCAGCAAACTGATTCATTGTCTATGGCAACAACGGAAAGAATTTCATACTATGAACTGCTCCAAGCAACTGATTCGCTTAGCGAGAGTAATCTGATTGGTTCTGGAAGTTTTGGCTCTGTTTAAAAAGGCGTTCTCAGGAGTGGGACTGCCATTGCAGTTAAAGTGTTCAATCTGCAACTGGAAGCGGCATTCAAGAGTTTTGATACAGAATGTGAAGTTTTACGCAGCCTTCGCCACAGGAATCTCGTGAAAGTCATCACTAGTTGCTCCAACCTTGATTTTAAGGCTTTAGTGCTCGAGTATATGCCTAATGGAAGTCTTGACAAGTATTTGTATTCGCACAACTACTTCCTAGACATCAGACAGAGACTAAGCATTATGATAGATGTGGCATGTGCGTTGGAATATTTTCATCATGGATGCTCCTCTCCCGTGATTCACTGTGATCTGAAGCCTAGTAACGTCTTGCTGGATGTGGATATGGTTGCCCACCTAAGCGACTTTGGTATTTCAAAACTGCTTGGTCAAGATGAGAGTGATTTATACACAAAAACCTTAGCAACATTTGGTTATATAGCACGG

>PGSC0003DMG402029358 LRR-RLKs

ATTACAAGAAAGTTGAGTTCTTAGCCTTACTAGTTTGACTGACCCCATTGCTACAAATAACTCAATAAGTTTAAAAAAGTGATATTTTCCTTCTTGATTTGCTAGGAAGTCTCTCAAGAATATCAAAATATGGAGAAAACCAATTTCAGAACATTTCTTGGTTTTGTATTAGTATTACTACTTAGCAGAATTTGTGTCTGTTGTGCTTCAGAGAGTGATGTTTATTGCTTGAAATCAATAAAAGATTCATTACATGACCCTTTCAATTCTTTGGGCTTTTGGGATTTCAGCAATGCTACTGAAGGTTTCATATGTAAATTTTCAGAAGTTCTTTGCTGGCATCCTGATGAGAATAAGGTACTGAGCCTCTCACTTTCAGGCTATGGATTAATAGGTGAGTTTCCTCGTGGCATTCAAAATTGTACAAGCTTGACAAGTTTAGATCTTTCAGGGAACAGCTTATATGGAACTATCCCTTCTGATATTTCAGCAATAGTTGAACATGTTACAATACTTGATCTCTCAAATAACACGTTTTCGGGTGATATACCACCTGATATAGCTAATTGTCAATACCTTAATGGTCTAAAGTTGGATAATAACTATCTAGAAGGTGAAATTCCAAGCAGAATAGGCTATTTGCCTCGCCTTAAGACGTTCAGTGTAGCCAACAATTACTTGACTGGGCCAGTGCCATTGTTTTATAGAGAAAGTATCACAGTTGAGAGTTTTGAAAACAATTCAGAGCTTTGTGGGAAGCCCTTGAAAGGATGTGAGGATTCTTCGATATGGAAACATGTAGATCATGCTTCGTTCATCAAAGCGTTTGTGATTGGTTGGGTACTTTTGTTTACATTGGTTTTAGTCCTTTGCTTATTTAAATTTCCTACCAAGGCTATCAACAAGATAGTCTCATTAAACATATGGAGGCTAGGGAAAAAGGAACATTTAACTTCAGGAAGTGAAGAGGAATTAAGCAGCCAACATAAGGTAAAGATCTTACTAAATTTGCTGCACTGCTTGTAGTCTGTAAAGTTACCTTTTCTTCTGTAGTTCTAATTTTGTGATTAGTCCTTGCTAGTCCATTATGTTATAAATTCTTTATTCGATCGATGATCAAACATGATAGCAATAAGGTTCTCAAGTTTCTGTTCTGTATATGTTCAAATGATTCAACTTGTTGTTTTAATTACAGATATTAAAGCTGGAGAAGTTTGTTACGAGAATGAACTTCACGGAGCTAGAAAATGCGACTTCTGGTTTTAGTGAAGACTATTTAGTAGGAAATGGAATGTTGGGCAAAGTGTACAAGGCAATTCTTCCAAATGGCTGGATACTTGCCATCAAGAAGCTCAATGACTGGGAGAATTTGGAAGACGAGTTTGTCTCGGAGATTACAACTCTTGGTGGTCTGAGACATCGGAACCTATTGCCTCTTATAGGTTTCTGTGCTGAAAAGCAAGAAAGACTTCTTGTTTACAAATACATGGCTAATGGAAGTCTTGATGAATGGCTGCACTCGAATGAAGTTAGCGCCAAAATTTTGGATTTCCCTCTGAGAGCTAAAATTGCACTTGGGATAGCGAAAGGCCTGGCTTGGCTTCATCATGGTTACGAATTGCACGTTACACATGGAAGCATAAGCACGCGATGTATCTTGCTAGATCAAAATCTTGAACCAAAAATATCCAACTTTTGGGAAGCAAAATTTTGGAGTAAGAATGATTCTGCATTAAGTTGGAGTCTTTTCCCAGTAGCTGAATATTCTGGTTTAGGCTCTTACAAGCAAGACATCTACTGTTTTGGTGTCGTGCTTCTCGAGCTGGTAACAGGGAAGGAACCACATGAATTGACTAGCTCAAGAAATCTATTCGATCACTCGCCTTGTCTACTTGATGCAGATAGAAATTTTCTTGGGAAAGGAGCTGATGATTTAATCCTCCAATTTCTTGAATTGGCTTGTGACTGTGTGAAGTTCTTTCCTAATGAAAGGCCAACAATGCTGGAAGTGTATGACAGACTCAAGAATATTTCTCAAGGTCGAAGGGACTGGATACAAAAGATACCATTATCAACCGATATTTCAAGTAAATATGACTAGTGCTTATGTTTTTATGGTTACTGAGAAAAACAAATTAGGAATACACTACATTAAAGCATGAAATCCGTGCTTCGAGTCTGTATGCTGATAGCTTCTGTTGCAACTTTAATATATATATATGAATACGTATATAATCTACAACATTGGAAAGCCTTACTGACTGCAGTTTTCAAGTCTGGCATTAACATCTTGCAGTTTCTTTAGTTAGATAGTTTTTATTTGCATATTGTAAAGAAGAGAAGTTGACTATGAATCTGCATATACTAATAGAACAAATCAATGGTATTTAATAGTTGACTTCATTGTGCTATATTGTGTAAGGTTGACAGAACAATATATAAATCATATTATTGAAAGCTTT

>PGSC0003DMG402030114 LRR-RLKs

CCAGAAAACGATTATCAATAACATAATCAACATGTTTTTGTTTGTAATCTAAAGAACTGCCAATTATTTATTTGAATTATTAAAATGGGAAAACTCAATTAGCTCAAATTCAACATAATATTAATAAGAAATAGCTTTTGAACCTAACTCAACTGTAAAAACAAGCTAAGAATTGTCCGAAACTATATAAAGAGACCAAATCCATCATCCCACTACAATTTTTGTGCAAGCACAAGAAAGTGCACCTACAAATTCATTTTCACAGTCTCTTTTTCTTTCTCACATGACTTCACATATTTCATAATGAGGGTATCAAGTCATCGTGGGTAAGTACAAAGGAAGGGAGGGAGTAAATTACTAGCTTGACTTAGTCAGAAGGAGACAATTATTATGGCTTGATTTTTCAAAGCCAAATACTTGGCTCTGGATGACTATTTAATTGTTGAAATATGATTTCTTTATTAAATCCACACTGACTTCACAGACTTCATTATCAAATGAAAAGAAGCACAAGGGAAGCATCAATTAACATGTTCCCAGCAAGCTCTAACCATTTTTTGCTTAATATGCAAATTTTCTTCCTAATTAGCCTGTTTTTTCAAGAAACCTTACAATTTTCGGGCACGGAACAGTCCATATTACTAAGCTTGAAACAACACTGGAGGGATTCGGAGTTCCTCCAGTCATGGAATTTGAATTCTTCAGTATGCACTTGGTCAGGAGTTTCATGCCTCGATGACCGGGTAGTGACACAACTAAATCTTGGAGAAAAGAACATTACTGGAACAGTTCCATCAACAATAATATGTGAGCTTAACAACCTTACCTTCATTGATCTCTCCAACAACAACATTTCAGGAACTATACCACTAAGCATGAAAGATTGCTCAATGTTACAACATTTGGACTTGTCCAAGAATTCTTTGAGCGGACAGATTCCAGGCGAGTTGTTTGGGATGAAACAATTACTCAATTTGTATCTTAATGGTAACATGTTGTCTGGTGAGATGCCAAAACAAATATCATCTCAACTGGAAAATCTTGATCTTTCTGAGAATCACTTGAACGGTTCGATACCAGAAGATATTGGAAACTTCAAAAATCTTTTGAAATTGGATCTGTCACATAATTCTCTCAGTGGTTCAATTACAAGCAAGTTGTTTCAGTTGCATGATTTGCGTCACCTGTCGCTAAGTTCCAACTACTTGTCCAGTGTGATACCTGATGAAATGGACTTGTTTAGTTTGTATGATATGGATCTTTCACACAACCAATTGACTGGTTCTATACCAAAGCGATTTAGGGATTTACCTGGACTGGATTCTATGGATTTGTCTTTTAATCAGTTATCAGGAGACATTTCAAAAAGTATAGAACGTCTTAGGCCTAGAAATACTTTAAGGCTTTGTTCCAACAAATTCTCAGGAAGAATTTCTACTGAATTTGTAAAGTTGACATACGAAGAGAATTGCTTTGACGAGTCCAATCTTTGTTCTACATCCAACAACTTATACATCCCCGGCCTACCAAGCTGCTCCTCTGGTGATGAGGTTCGAAAATCTTCGAAACCAAAACACTTGATTATTCCTATTGTAGGAGTTGGGGTAGCTACACAGTTATCATGGATCTTTTACAGAACAAGAAAGCATTGGTGGAAAACAAAGAAGCAGAATGTCAAAGATGACATGAAGTTCATTTCATTTCAAAAGTTGAAGGTGACTACAGAAGACATTTTATCTAGCTTGAAAGACGAAAACATAATAGGAAATGGAGGATCAGGGAAGGTCTATCGAGTTGTGATCGACCAAACAGGCAATACTTACGCTGTTAAAAGCATAGGGCATGGAAAAAAATCAGGTGAAAGACCCCAAAAGGAGTTCCTGGCAGAAGTTAGAACACTTGGTAGCATTCGACACAACAACATTGTCAAGCTCATGTGTTGCATCTCAAGTGTAGACAGAAAGCTTCTAGTCTATGAATACTTTGAGAAACAAAGCCTCGACAAGTGGCTTCACAGAAAGAAAAGAGAAGCGTCACCCGGTCAAAGTAGTACCCCATCCCTGGATTGGCAAAAGAGATTAAAAATAGCCACTGGTGCAGCTCAAGGACTCAGCTATATGCACCACGACTGTACTCGACCCATCATTCATAGAGACATTAAGTCCAGCAATATACTTCTGGACTCAGAATTCAATGCAAAAATAGCAGATTTTGGACTAGCAAAAATACTATCCAGGCGGGATGATAATCCTGAGACAGCTTCTGCTATTGCTGGAACATTTGGTTACATTGCACCAGGTCTATTACCGTTCAAATATATATCGTACCTCGTGAATTCTCAAAAGAATTTATCCAACTCTAATATTAAATTTTAATGTTTGGTCCTCACAGAGTATGCCTCGACATTCAGAGTGAATATAAAGACAGATATATATAGCTTCGGAGTGGTGCTATTAGAATTAACAACAGGGAGACAACCCATTCTCAGGGAGGAGCAAATGAATCTGGCACAATGGGCTCAACAGCGTTACAGAGATGGAAATTTCATTGTTGAGGCCCTTGATGAAGAAATCATGGAAACAAGTAATGTGGAACAAATGAGAGGTGTTTTTAAACTAGGACTGATGTGTACTGGAGCAACACCATACGGTAGGCCGTCAATGAAGGAGGTCTGCAACATTCTTCAAAGCTTCAGAGACACCATATTTTGAGAAAACAGAATGTTTAGCCTTACATGGATAAATGTAGTTGTATTTTTTTAACAGAATGAGAAAGTCTCAATAGCCTCAGCTGGTTTAAGTGGTGTCAAACACAAGGGCTTGTAAGCTTTACCTTAGAATGTAATTCCTTGTTCATAATTGAGATGTCGCGAGGCAAGAGCTAAGTGTCTTGGCAATTACAGTTTATGTATATATCATTCGGGAGATTAGACAGATTTTAAGCACACTGAGCATTAAGGTCTAACAATGCTTTAGGAAAGTAATCATGATATCCGCATGGAGGTTAGAATCATATCTTCTTCTTTTTTTAACTCTTCAGAACTCCCTAAGGAAGCTTTTGGAACTTACAGGTTTGATGAACTGCAAATGCCTTCTTTCAGATATAAATCCTTCTCATGATACATGGGGATAATCTTGTAGGAAGTCTAGTTCTTTTAGTTAAGTTGAATGGGATCGTCCATTTTCGAAAAGGGGGCACAACTTGAAGGCATAATTGATTGGAAAAAAAATAATCGAAAGATATAGGAGAGTGTTTTTTAAATGAATAAGTCCTTATTGTAGTTGAGGAGATGATAATCAACTTCAAGAAGAACTTTCATCTCAAAATACTAAAAAAAAGTTTTTCATACTATCAATGTTATTGAATTGCCTCTGCATATGAATATACTTCATAATAAATGTGGGACTTCTCACCGCGAATCTGAATTTAGTTAAACCTCAATGCATGTACCGGACATTGGGTGAAAACAAAAAAGGATAGTATGAAAAACTATCCCAAGTGTTGCAAACCTATTAGCTGGAACTGTCCTCGCACTGAATGTAGAAATGCAATATTGATGGCTCCTCTAATGAAAATCCAGACCCCAGATCGTGGACCTTTTTGTATCAGAGATATTATGGGAGAGTTAATTTATGCTACTACTAATGTCATGAATCATGATTAACACTACTAGCCTATGTGCTGAGGCTAAAGCAATAGAAAAGGAGTTGGAATATTCTGTGAGCCATCAATACAACCTCTTATAATAGAGACTGATTCAACCAAGTTGCGGATTTTTTAGCTAATTATGTGTTCATTATCAATATCTACTATATGAAGCAAAGAAATTATTGAACATTGATAAGCAACAAGTTCCCGGGTGATATTTTAAAAAAAAGGCAGGAACTAGCAACAAGAACAACAAGCAGGCTACTCAATCAACTATCACAATAGACTATCCATACAAAATATACAAATCTTCATGAGTACAACCAGAGTAATAACAAAATACAGATGGTGAAATTAACTTGACAAATCCTCCAAATATTTTCTTATCAAAAGTGGTATCAATCTCAAACAAGGTTCAATCTTTTGGGGAGGAAATATTAGAGGACTATCTCGGGGGTTTTTGACACTTTACTCGCTTCTTGGAAACTATTTTGAGGATGGTAACATGTTACACCCCAACTTAGGGATGCAACTAGCACCTGGTGCGACACTTGACCCTGAGTCGATCCTTGCTGGACAACTACGCTACCAACATAACACCAGCCACAAAATCAAAAAAACAAACAAGTATATTTCAACTTAAAAATAAGGCCTCTATCGGCAAGGACCCTGTCAACATAATCTTCAATAAACTCAACTACTCCCAGCAATTCATATAGAATAGAAACTAAATCATTTATTGCCCGTCAAGGTCACTGTTATATCTAGAAACATATAAACTGAACACATATAGTCATATATGATTAAGGCCAACAAGCCACTAAGCTAAAAGGGCAAAACAACAAGGCCAGCATAGCCAACAACACACATCTATTCAATCGAAAGTTTGCAAGCCTCTAAGAGTAATATACTTGGTCGGGACCCTTATACTTCCTGTCCCGACCTACCTATTAGTTGAATATGACAAAACACTAGCAAACCTTTAAATTTGGCAACTCTAGAAGAAATAGAGCTGACCAATCTTCTGGAACTCAGCTACACTATTGAGGAGGCTTGTTTAGCTGTCTGTCAGGATCTGCAGGCAAAAATGCAGCACCCCTGCCCCAGGTATGAGGCGTCATTATGAATTGATATATCGAGTATAAAAGGCAATAGAATAACTGAGCAGGTATCATAATGTACCAAAAGTAATCAGGTAAGGAAGTCAAGAATTAGATGAGTGTACTGACCTGTAACACCTTAGAATTTGGCAAGTGTGGTTTGGAAAGTGTCATGGTTTTAACGGGCAGGTTCATGGACAGTGAAGTGGTCCACGACCTGCCAACCCATCCGTGAACCCTACTCCCCAAGGCTCAGGAGAACTTGATCCCTCACGGACGCCTGAACGGTCTGTGGAAGGGTAAATGTCCAGTTGATAGGGCCGTGGTGATTGGCAATAAAATCCCATTTTTTCTCTCTTGTTCACAATCACCTTCATGGACCGTGAAGTGATAACCGGTCCGTGAAGGCCTCCGTACAGGGGTCCCGTTAGCTTTTAAGTCAGGGCTATTCTAGTATTTTCCTATGCGTCTAATACCTAAGCTATGTTGTTTTGACCCCTATTCTAAGACCTATATAAGTATTTTAATTCCTAAAACTACCCATTCTAAAATCATTCTCCTAAAATCTCAAATTCCCTTCAAGTTCTTCCCCCAAATTCCTCTCTCTCAAGCAAAGAAAGAAAAAGCTAGGGTTTCACTTCAAGGTCTCCAATTCACCATAAAATTTGGGCCTTGATCATCAAGATATGTTACTATTCACCTATGGAGTCTTTTCCTCCATTGGGTTCCCAAAGGTCCCAATTCTACAAGCTAAATTTTCAATTCAAAGTTAGGATTTTATTCAAATTCCATGAATTCTTCTCAATTATGATTCAATTAGTTATTTCCTAGCTTCTAATGCATGAATTGAAGTATTTACATGATTTTGAGTTGAATTCATATGAACTCATGCATCCATGATTTTTACTATGAACACATGATGAAACTTATGAACTATGCAATATGAATATATGAAGATATGCATGTTTTTGTGAAATTAATGTATGACTTGAGGATGCTTTAGATGAAAGTATCAATAAGATTGTGAAAAGTTTTCTCACATGCTACTATGATCATGATTGTAAAAGGTTTTCTTACATAAGGTTGAAAGGTTTTCTCACCTAACTTGAACTAAGATTTAAGGAGCTATTCTAAATGACTTTAGCTTGGATTGGTAGCTTAAACGTGCCTTTCCATGGATGATGAATATGATTAATGACTATTCTGTGGGATTTTACTTAGCACCGAGAGGATATTGAGATGGAGGCTCTCCAATTAGTAGAGGTCGGGTCTCTAGTAGCAATCTCCTTGTCCCTAACTACGTGCCCCCGTAGGATTTCTTAGCTAGTGGATCCACGTAAACTAGAAGTTTATGGTCCTTACTTAGCAAGTAGGACATCTCGAAATAGACACCGGATTTCATGTTTTAGAGCACATGTTCCATATCGGTTAATGGAAAACATCCCACAATAATGAACTAGATTATTTTCAAAGTATCTCACCAAGTGTTTTAAATATGTTAACCTTGCATTGACCATGATTATGACTTGTGTTTCTAACACCGTATCTTATGCTTTATCTTGGCTATTGCATATCATGTAAAGTCTTTTTAGCATGATTTCATATTGTTTGTGCATTGTTATCATACTTAGTATATTCCATGTACTAATGTATACTTGTGCCTATATTGTTTCCCTAATGTAGGGTCCGATGCTCCTACCTCGCATACTTGTGGCTAGTTGATCTCTCTTGTCGATTTGAAGATTGGTGAGTCATCATGTTCCGAGGACCAAGACACTTTTATAGCTTTTCTTTATTTCAGTTTTCAGACATTGTACGTGGCGTATGGGTTATATCCCAACCACTTCTTTTGATGTATTAGATGGCTTGTTGAAACTATGTTAGACTTCCGCTTTGTCAAAAAAATTTATGATATGAGAATATTTCTTTGAAATCTTTATTTCTATTATCTAGTATGATACATGGTAAGTGACTAGCATAGGGCCTCTCGGGGTTCTATACGCTGTATTACACCTAGGGTATACTTTGGGTCGTGACAAACTTGGTATTCAGAGCACAAGGTTTAGAAAGGTCCTAGGATGGCTCACAAGCCACGTCAAGTAGAGTTTGTTCATGAGTGTGAAGCGCACCATATTTATGAATAGGAGGTAATAGCCTATAAGATGTTTAGGAAATTCCACTTCTTTCTTTATTCTAAAGTCGGGCGATAGAGTATAACTCTGTATGTCTCTCTCATAATCCTTACCCGATGCTTTTCGGGATATGGCTCCTCGAAGAGCCTACGTTAGAAGGAATGCTGGAGAGAATGTGGAATAAGAAGCTCATCAATTTCTGGCCGACCCTTTTGCCGAACAAATGACTAATGCGGAGTTCTGATTCGTTTTTCATGTTTTGCCCTCAAGTTCATAGGTATTGCAATCACATAAGTTTTATGTCAAACTAATTGTAATGATGTGAATATCAACAAGGAGTGATATGCAGCTAATTACAAAGCCATTAAGGCATCAACAACGTAATGAGCTATCAATACCCCATCTAAGCAATGACAATTTATCCAAGAAGATTCTTACGTCGCTTTTTAACATGTATCACCAACTAAACAAGAATAATCCCAATAAGGAAAGCTCACATGTATACATATATATAACTCTAGCAACCAACTCCATCAACAACTATTATGACTATGAGTACTATGGTGATAAAGAACAAGTCGCATTACCTATACGACAAAGAAACTTGCCGAAAAAAGTTAAGCTTTAACATACCTTTTTTCATCGAATTTACTCTGTCATATTACTTCTGCCCGCACCTTATGATACTCGCTGCAGACACCCAAAGAACTTTCAAGAACTCAAAACACAGCCAGATCCCTACTGACACAACTTCCACAGAGCTACACCGCACAATCCACAGAGAAAAAATATACCAAAACAAAGAGCGCGAGACGTAGAGCAATTTCCGTTAAAGACCCAAGACAAGATCATAAATATATATATGGATGATTTAAAGGTTGAATATCTCGC

>PGSC0003DMG400000386 LRR-RLPs

ATGGTGTTTGAGAATGATAGAATCAAGAATGCTTATATTGCTTTACAAGCATGGAAACAAGCTATCATTTCAGATCCCTTTAATATCACTCTCAATTGGGTTGGTCCTAATGTATGCAGCTACACCGGAATCTTCTGTGCTCCGGCTTTGGATAACCCCAAAATCCGTACAGTTGCCGGCATTGACCTTAACCATGGTGACATTGCAGGGTACCTCCCGGAAGAGCTTGGGCTTCTTACAGATCTTGGGATTTTCCATATCAATTCAAACCGTTTCTGTGGTACAATACCAAGAAAATTCAGCAAGTTGAAGATACTGTTCGAGCTGGATCTGAGTAATAACAGGTTTGCTGGGAAGTTTCCTTATGTTGTTTTGAGTTTACCCAAGTTGATATTTTTGGATATTAGGTTCAATGAATTTGAAGGTAATGTACCTTCACAGCTTTTTGATAAGCCATTAGACGCCATTTTTATTAATCATAATCGATTTGCGTTTGAGTTGCCGGAGAATTTTGGGAATTCGCCGGTTTCTGTGATAGTTCTGGCGAGTAACAGTTTCCATGGGTGTCTTCCGGCGAGTATTGGGAATATGAGTAATCTTAATGAGGCGATTTTAATGAACAATGGGTTGCGTTCTTGTTTGCCGGCGGAGATTGGGGTGTTGAAAAATTTGACTGTATTTGATGTGAGCTTTAATCAGTTGATGGGTCCGTTGCCGGAGAATTTTGGTGATTTGGTGAATTTGGAGCAATTGAATGTGGCACATAATATGCTTTCTGGGACAATTCCGAAAAGTATTTGTCAGCTTCCTAAGCTTGAGAATTTTACGTATTCTTATAATTTCTTCACTGGTGAACCGCCGGTGTGTTTGGGGTTGCCGGAGTTTCATGATGAACGGAATTGTTTGCCCAACAGACCGGTGCAACGCTCTCCGGGACAGTGTAAGGCCTTTTTGTCTAAAAAAATTCATTGTAGTGCTTTCAGGTGTCATAAATTTGTTCCTGTTTTGCCACCTCCTCCGCCACTTTCACCGCCCTTGCCTGCCCCTCCGCCACCTGTTTATTCGCCTCCGCCACCAGTTTATACTCCTCCTTCTCCTTCCCCACCACCTCCTCCACCTCCAGTTTATTCACCACCGCCACCACCACCCTCTCCACCTCCTCCATCACCTCCCCCGCCTCCACCACCAGTCTATTCACCTCCACCCCCACCTCCATCTCCTCCTCCACCATCACCTCCACCACCACCCCCGCCATCTCCACCTCCACCACCAGTTTATTCATCACCACCACCGCCACCATCTCCCCCTCCACCATCACCTCCACCTCCTCCCCCACCCGTTTATTCGCCACCACCATCTCCACCTCCTCCATCACCATTGCCTTATTGTATACGCTCACCTCCACCACCAAATTCCCCACCTCCTAATTCTCCACCTCCACCGCTTGCTCACTCGCCTCCACCCCCTTCACCTTACTATTACAATTCACCAGCACCTCCGCCCCCAAATTCGCCGCCTCCACCACCAAATTCGCCTCCTCCTCCACCAGTATGCATATACTCATCACCTCCGCCACCAAATTCACCTCCTCCACCACCAACATACGTATATTCTTCACCTCCTCCACCTCCTTCCCCATTGCCTTGTATAGAACCCCCTCCACCACCTCCTCCTTGCATTGAACCACCCCCACCTCCCTCCCCTTCACCATCACCTCCTCCACCACCAGTTTATCATTACAATTCTCCGCCCCCACCATCTCCATCTCCTCCACCACCAGTTTATTATTACAATTCTCCACCCCCACCATCACCACCACCTACTCCAGTATATGAAGGGCCATTGCCACCAATAATAGGAGTTCAATATGCTTCACCACCTCCACCACCCTTTTATTAA

>PGSC0003DMG400000819 LRR-RLPs

CTCTTATTCTACCTTAGCTTTCTATAAACATTTTTAACAAGCACCTTCCTTCTTTACTTCTTTGTTCCCTCCATTTCCCTGTTTGATTGGATACTTTTACCATTTTCCACCACTGCCCCAATAAAACAAAAAAGTACACATTTTTTTTTATATTTATGAAATGAAAGCATTTTAAGATACCCCCTCTTGGAGTTTCTTCAATGTTATAAGTTCTAGTTTGGCTTTAAACACTTTTCTTGCATAGTTGACAAAGTTTATTACTTTCAAGATTTTGTCAACCCAACTCACTTTCAAGCATTTTTTTGGGTTTTAAAGAGTGATGGGGGTCTTCAAGAACTTGTTTATGTGATAAAGTTGTTATCTTTGATCACTTTGTGTACTCATTTTCAAGAAATTTTAGCTTCAAAAGAGTGATGGGGGTCTTCAAGAACTTGTTTGTGTGACAAAGTTGTTTTCTTTTAACACTTTGTGAACTCATTGTGTGATAAAGTTGTTATCTTTGAACACTTTGTGTACTCATTTTCAAGAAAATTTAGCTAAGATAGGGGTCTTCAAGAATTCTAACTAGTTTGTCTATGTGACAAAGTTCTAAACTTTCAAGAATTTGTCAACTCATTTTCAAGAAACTTTTTTGTCTACAAAAAGTGATGAAGGGTTTCAAGAATTCAAGATTGTGTGTGTTAGTTATCTTCTTTCTCCTCCATTTTGGTGTTTGGTGCAATGGTGATACTGAAAACATGGGAGCTCAAATGGAGAAAAATGAGATAAATTCTCTTTACTCAACCATTCAAGGCTTTGTTGGTAAATGGTGGAATGGTTCAGATCTTTATCCAGATCCTTGTGGTTGGACTCCAATACAGGTAACACAAAAAAAATTACATAATTTTTTTCACTTTTTAGTTACTTTTCTGCCTTAAGGATTCTGTTAAGTAAAAAGGTAAAAAAAAAAAAAAAAAAAAAACTATCTTTTTTGTGCATATGGTTCTAATTGTCCTTTTATTATTTTGTGGTATACATGACAAGATATGACTTCTCATTGTATATAGATTGTTTTAAAGCTTTAAGTGGCAATTTTTTACTATGACATATAATCAAAGAAATTGATAGTATATTTGGAATGATAGCTAATTAGCCATAGAAGTTTGATGAACCAAACTTCAATAATGAATATCAGTACATTCTTGGCTTGTTTTGTTTAGCAGTTTTGACAGACCAACATAGACAGAATATGTTTTAAAACACAATGATAATACAGGACAGGACAGAGCTTATACTAAATATACTCCGTTCGTTTCAATTTGTTTGTCTGGTTTTGATTTGACACGTAGTTAAAGAATGTACCATAGTTTTCTTTAATCTTGTGGTTTTAAACATGTTATGTAGAAAGTACAAAAGATGCATTCTTTTTGGAACATACTAAAAACGGAAGTAAGACTAACAAATTGAAACGGAGAGAGTGAGAGTGATATAGTTGATGCATTATACTCATATTTCTGATTTGTTTTGTTTCTATTTCCTCTGTCCAGGGTGTATCTTGTGACCTGTTTGATGGTTTTTGGTATGTTACTGACTTAAACATTGGACCTATACATGATAATTCTCTAAGTTGTGCACCAAATGTGGAATTCAGCTCCAATCTGTTCACATTAAAGCACCTAAAATCTCTCACATTTTTCAATTGTTTCGTGTCGCCTCGCCATCATCCAACTTCAATCCCAACAGAGAGGTGGGAGTTTCTTGCTAGTAGTTTGGAGTCACTCGAGTTTCGATCAAATCCCGGTCTCATTGGCCAAATCCCAAGTACATTTGGTAGACTCATAAAGCTCCAATCTTTAGTACTAGTAGAAAATGGACTCTCTGGTGAAATGCCACAAAGTCTTGGGAATTTAGTGAACTTGAAAAGACTAGTTCTTGCTGGAAACAAACTCAACGGCGAAATCCCGGATACTTTTGGAAGGTTCACTCAACTTTTGATATGTGATTTAAGCAGAAACAAACTATCTGGTGTCTTGCCTAAACCATTATTTGGAGGTTTGATCTCACTTTTGAAACTTGATTTGAGTTACAACAACTTACAAGGCAAAATCCCACAAGAGATCTCACAATTGAAGAATTTAACACTTTTAGACCTTAGTAACAACAAGTTATCAAGTGGATTACCTAAGTCAATTCAAGAAATGACATGTTTGGAGGAGTTAGTCTTGTCAAACAACCCAATGGGTGGATTCTTGGATGTTCTTGATTGGTATAGCATGATAAAGTTAACAACTTTGGATTTGTCAAACATGAATTTGACAGGTGAGATACCTAAGTCTATAGTAGAGTTAAAAAAGTTAAGATTTTTGGGACTTAATGACAATAAACTCATTGGTAAAATCCCAAAGAATCTTGAAAATTTGCCTAATGTAAGTGCTATTTACCTATATGGTAACAATCTGACAGGTGAACTTCAATTCTCTCAATGGTTTTATAGGAAAATGGGAAGGAGATTTGGGGCATGGGGGAATCAAAATTTATGTTACACTTTTGGTTTGGTGTCAAGTGATGTTCCATTTGGGGTAAAACAGTGTCAACAAGAACTCAAGTTTGTTATTGATGTTGAGATGAACTCAAGTTTTGTGGATTTTCAATCTTCAATGGGATATTTGAATGATTGTGGGATTTGGAGATGTTTTGTAGTTGAACTATTCATGGTTATTCTACTATTGTTGAAATGAAAGGAATTTGACAATAGTGATTTACCCTTTTTTTTTTTTGATATTTTGTTGTTGTTTTACTTGAAATGTACCACACATTTTGGTTTGTCTTTTAACATATGAATGAATGAATGTACATATTTTACC

>PGSC0003DMG400000888 LRR-RLPs

ATGGAAATCAATGATGTTCGGATTGTGGGGATCTGGGGAATGGGGGGAGTTGGTAAAACGACAATAGCAAGGGCTATTTTTGATACAAACTCAAATCGATTTGGTGGTGCTTGTTTTCTTGCGGATATTAAAGAAAACAAATGTGGAATGCATTCTTTGCAAAATATCCTTCTCTCAGAACTATTGAGGAAAAGAGATAATTACGTGAATAATAAGGAGGACGGAAAGCATATGATAGCTCGTAGACTTCGGTTTAATAATAAGGAGGACGGAAAGCATATGATATTTCGTATGTATTGAAAAAAACCAGTGCATTTGTATGTATGTCTAGTCGAAAAGCTTTTCCTGCTGGGATATAACGAAAAGCTTTGCAAACGACAATAGCATGGGCTATAATTAACAGAAGCTGTAATTATGGTTTTCAGTACAACCCATGTACTGGTTTTTGTCAATACATACGAATGTTACCTTTCTCAAAAAAAAAAAGTAGCTTTTCGACTCGACTCTAAAACCTGAGAAATTTGAGACTATTGTGTATAAGCTTTGGCATTGTATTAATTTATTGAGTGATAACATACGCAGGGAGTGTATTTATACACATACAAGAGTTAGACTTAACCTAAGACTAGTTCTTCTAATACAATAGGGAGACTAACTAGTTGGATTCACTACTCTTACCTTGTCTCTGAAGAATATGGTGGTCCTTTGCTAGAGTCTTTGTATGAATGTCAACTAGTTGATCTCTTGATCTGACAAACTGAGGATTAGTTACAAAGTGATAATCCAATTCTACATGTTTAGTTCTAGCATGTAGAACTGGATTTGTTGTCATACATAGTGCACTCATGTTATCACAAAATAGCGTAGGAGCAGACTTGATGTGTATCCCTTTATCCATGTGAGCTCAACAGCTGTAGCTGCTAAGGCTCTATACTCAACCTCTGTACTTGACCTTGGCTAATTCTTTTAAAATTTGGATAATCATAATGAAATTATAGCCAATTGTCAAGTTTAAGGGGTTTAGATATTCTACCTTTTATTTATGGTAGTAACTCTTTTATTTAGTTTCTTTCTGTGTTTTGAATAGGAGCTTTTGCCGTCTCTACGAAAGCTAGATCTCAGCTACTCCAAAAGCCTGATGCAAGCACCAGATATCACAGGGATGCCAAATTTGGAGTATTTGAATTTGACGGATTGTATGAGTCTTGAAGAGGTTCACCATTCCCTAGGATATTGTGAAAAACTCATTGAGTTAAGTTTGACTTGGTGTTCGAGGCTTGAAAGTTTTCCATGAGTTAACGTGGAATCTCTGCATCTAGTGGCTTGCTCTAGTTTAGACAAATTTCCAGAAATCCTCGGAAGAGTGAAGCCTGAGTTAAAAATTTTCATGCATTTCTCTGGGATAAGGGAACTACCATCATCTATTGTTCAGCACAAAGCTTGTCTTCCAGAGCTAGATTTTAGTTCTAAGGAAAATAAAGCAGCAGTTCTAAGTTGAAAGGTTTGGTGTGTCGCGTTGCTCAGAACTTGAAAGCTTGCCAGAAGAGATAGGCTATTTAGAATACTTGGAGGTGCTTGGGCTCACAGTCTCCGCCTCCGCCTTCAATCTGTAATTTGTTGTTTCAGAATATGTCATCATTGCTGCACGACATCTCTGCTTCAAATTCTGTCACTAAGAGTTCTTACTACCAGTGGGAATATCCCAAGTTGGTTCCACCGTCGGGGAATTGGTAGAATTGTATCGGTCAATTTGCCTGAAAATTGGTACGTTAGTTATAACTTCTTGGGATGACCCAACAATTTGCCTCATCCAACCATTCAGATTTTGATACCAAACCTACTATTAATATTTTCTTGGTATCTTTTGCTGGCTTATGGAAAAACACCAAATGACTATGGACTCCTTGTGTTATGTTTATCTGGAGAAGTGAAGGAGTTTGGACTTCGTTTCTTGTATAAAGTTGAACAAGAGCTTGATGCCTTGTTACAAATGAGGGAAAATAATAATGAACAGATATGAGCACCATGATGAAGCCAGTTTCTCCTCTTCGAAGAAACAAATATCTTATTTTAATATAGGTGGGAGCTGTCTTTGAGAATCTGCAGCAACAAGTAGAGGCACCAGTTTATTCAGAAACTTTCCAGGTATTTCCCGCAAGCCCAAAATTTTGGCATAAAAATAGGTATATACCTCATGATTTGTTTTTCCTTACTAGTTGGTAGTGAAATATAGTTTACGTATGTATTTATGTCAGTTAATCTAAGAAACTCCCTAAATGAAAGAATTGAATTATGAAAGATGTTTGAAGTTCTAAGGAGTCCAGAGAAAAGGCAAAGGGTTGTTATTACTTATTCCCTAAGGTATCCACTGATCCATGGGTTTAAAATTTATTAAAATATCATTTTGATGCAAGGAAATCTCTGTTTTAAATACAAATATCTCTCAGGTTATGGCAGGGAGTTTGGGAGCACAATGTGAGACAGTGGAGGAAGCTGGAGTACACGGTGAGGTTAGTAAATCACCTCACATCTTATGATTTGCATCATATCGTGGTGCTCCATCTGTGCAAGCCATTATTCTGCTTTTCTTAGTACCACGTTTCAGATTTTGATGCTAAGGTTGTACTTCAGTTTTTAACTATACCTGTTCCCTTGATCATGTCTTTCTGCAGATGTATACCCTACCAATATAAATATTTCACCGTGTCATCGTTGGTAGATGAGGATGCCTTATAAGTTCTTCTAACAATCCACAGAGAAAAGGCAAAGTGTTGTTTCCAAAGGTAGCCACTGATCCATGACTTCAAAACTAATTACTAGTTCAATTTGATGCCCTCTTGTTATGGAAGGGAGTTTGGGAAAAAGATAAACTCATTGAAGGTCCAGCACAGTGTGAAACAATGGAGGAAGCTGGAGTACGAGGTTAATTTGAGGTTCGTTTCCCTCGTGATCTCTTTCACCTCTTAGTTTTGTTTTAAATTTTCTATTACATCAGATAAAATATGTTGCCATTATCGAAAACATCAGAATACTAGCTGTGATAGAGTTGGCTTATGACCTTCCATCTCTCTCTCTCTCTCTCTTTTTCGTAAATATTGAATGCTAGATTAGTAAGCCAAGACTGAATGCACCAATTCATGATTAGATGTTGTAATCGTTGTTGATTTAGGAATATGCGACCTTTTTATCTTACTGTCTTCTGTCTCGCTTGTTGGCTTTGATCTTCACAAAAGTAATAACAAGTAGCATCTATGTTTTGATTACTTATTTCAAGTGGTTTTAACAAGTTTCACTCTTTTGGTTTTGCAGTGCCTGTGAAGGCACATTGTTTTCTGCGCTCGTAACAGAACAACCAGACCCTTTGCCTGAGAAAGAGATTCACGGAACGACAAAAGTTATCCGTTCTCTTCAAATCTTTATATATCAATTTACTGGTCTTATTAGTAAAGGTGACACAACCCCTCTCTTGGATCAGAATACCACGTTTCATTTTGAAATAAAATTATGTCTTCACATTGCTGATGAGTAGCCTGTCCAAAATCATATAGCAGATGAATGTGCGCGGGAAGCAAGAAAGCTATGCTAAAAAGGGTCAATGAAAGAAACATTGGAACTACTATATCTACTACTCTGCATATATATAGTTTCATAATCAATTATTGGTAATGGTAAGGAAGATTCAGAAAAGCTGTCGCAATTGTAGACACTAATATTTGTTTTAATTAATTTGTCAACTCTTCTCTATTGCCCTTTTCCAAAGTTAGCTGTTATCTGGTGACACAAAAATTGTTTAGATTATTTTTATTGATTCAGTCTGTTGAAGACTGAGCAATACTTTGCTGGATTAAAGACTTTTAGCTGTCCATTGTGTACATATTACGATGAAGTTTTCGTGTAGGGGACTGGTCTTTGAAGGAATTTCCTAATACCACAATATTTCTAATCAACTGGAGTAGACATTAAGAAGCTTCACAGAAATTTCCTATTACTTCTATAGTTCTATTAATATTCAATGAAAATGTACCACATATAATTTGACTAGATCCAACAATAATAATTTATTTAACTGGAGTAAACCGAATGCTCATAGAAGCTTTTTGTTTCTTCTTTCCTGTATATAACTTTGTCCAAAATATCATCAATTGATCTCATCCATACAGTCTATTTTTCTTTTCAGACAATTAATCAAATATATGGCATCATCTTCTTCTTCTGTGAGTAATTCACAGTACTGTACTCGATGGAAGTACAATGTCTTTCTAAGTTTTAGAGGCAAAGATACCCGGAGAAATTTTACGAGTCACTTGTACGAAGGTTTGGACAATAGGGGAATATTCACCTTTCTAGACAATAAAAGGCTAGAGAATGGCGATTCCATTCCACAAGAGCTCTTGAAAGCTATCGAAGAGTCTCAAGTTGCACTCATCATTTTCTCAAAGAATTATGCTACATCCAGGTGGTGCTTGAATGAACTAGTGAAGATCATGGAATGCAAGGAGAAGGAAAATAATGGAGAAACAGTCATACCGATCTTCTATGATGTGGATCCATCACATGTTCGAAACCAAAATGAGAGCTTTGCAATAGCATTTGCCGAACATGAATTAAAGTATAAGGATGATGTTGAGGGGATGCAGAAGGTGCAAGGATGGAGGAATGCCCTAAGTGCTGCCGCAGATCTAAAAGGATATGATATCCGTCACGGGTGAGTTAAAAACACATAAATTGTTTTTATGAAAAAGGAAATGGAATAATTTCCATCCAAATTTAGATGCTTCTACGAAACCTAGACTTATTCTACAATACAACAGGAATCTATACCTTCAAAATGAGTTAATTTAGTTAGAGAGAATACACAATAATCTTCAAAATGTGCAAGTTTTATACTTCAACTGTAGAGTGTTGCTGTTTCCCTGAACTACAATGTTTCCTATTTGATTAACAATATGAAATACTTACTACTACTTGACTTTCACATTTTCCACAATCAATCTTCTTATTTATGCAGGATCGAATCAGAGAATATTCACAAGATCGTCGATCACATCTCTTCCAAATTATGCAAGACTTCTTATTATTTATCTTCTTTGCAAGATGTTGTGGGAATAAATGCTCATTTAGAGAAACTAAAATCCCTACTTCAGATAGAAATAAATGATGTTCGGATTGTAGGGATTTGGGGAATAGGTGGAGTTGGTAAAACGACAATAGCAAAAGCCATCTTTGATACTATATCTTATCAATTTAAAGCTTCTTGTTTTCTTGCGGATGTTAAAGAAAATGCAAATTATAATCAATTGCATTCTTTACAAAATACCCTTCTCTCTGAACTATTAAGAAAAAAAGATGATTATGTCTATAATAAGTATGATGGAAAGTGCATGATTCAAAGTAGGCTTTCTTCTAAGAAGGTGCTAATTGTGCTTGATGACGTAGATCATGTTGATCATTTGGAGTATTTAGCAGGTGATGTTGGTTGGTTTGGTAATGGCAGCAGAGTTGTTGTAACAACTAGAAATAAACATTTGATAGAGAAGGATGATGCAATATATGAAGTATCTACACTACCTGATCATGAAGCTATGCAATTGTTCAATCGACATGCTTTCAGAAAAGAAATTCCAAATGAATGTTTTATGAAGTTCTCATTGGAGGTAGTAAATCATGCTAAAGGCCTTCCTTTAGCCCTCAAGGTGTGGGGTTCTTTGTTGCATAATAAAGGCCTAACTCAGTGGACAAGCACTGTAGACCAAATAAAGAAAAATTCTAGTTTAGAAATTGTTAAAAAGCTCAAAGTAAGTTATGATGGGTTAGAGCTTGAAGAGCAAACAATATTTTTAGATATAGCATGTCTTTTACGAGGAAAAGAAAAAAAACTAGTCATGCAAATCCTTGAGAGCTGTGATTTTGGAGCAGAACACACATTGGATGTCCTAATTGATAAATCTCTTGTGTTCATCTCAAAAAATGATAAGATTGAAATGCATGACTTAATTCAAGATATGGGTAGATATGTGGTAAAAATGCAAAAAAAATCTGGAGAACAGAGTAGACTCTGGAATGTTGAAGATTTTGAAGAAGTGATGGTCAACAATATAGTAAGTACTAAGTAGGCTAACAATGCAATGATATTTATTAGTCTTCATGTTTTACTTTTTATTTAAAAGAAAAAAATTGTTGGTACTGTCTCAATTACTACTTCATTTTATCATAACATGATTTTTTTTTTTTCAATTTGACTCTTTAGAACTTGTCAAAAATTGTAATACTAGAAATTAGAAATATAGAATTAATTGCATAAAGTATTTCTCTTTAGCAACATTTTTCTTAAGAAAGTCCTAACTAATATTTTACCTCTTTTTTTTTTTTTTTTTTTAGTGTTGCTAGCTTTTCAAGCGAAATAATAAATAATGCAAAATATGATGAAATTTGGCATTGCAAGGAGTTATAAAGTAGTCATGAGATTTTAAACAAACATTTGAAGTTGTGCTTTGTAGACTAATGGATCTTTTATCAGGGGACCAACGCAATGGAAGCAATTTGGATACATGGTGTTCAAAAACTATGCTTTAGTGAAAAAATCATGAAAAAGCTTAGATTATTATCCATCATCAGCTTTCATACACATGATGACTCCATTGAGTATCTGCCCAACAACTTGCGTTGGTTTGTTTGTCATGATTATCCTTGGGAGTCATTATTAGAAAATTTTGAACCTAAAAAGCTTGTTTATCTTGATCTCCAGCACAGTTTGCTGCTTCAATTATGGACTGGAACAAAGGTACAATTACTGAATACAGTTTTATTTATGCTTCTTTTTGTCAAATACATCTTGACCATTCTAATTGAAAAACTTGACATAAAAATAAAGTAAAACAATTTCTAATTTCACACAAAATATTGTGTTGACTCTCATAATCTGAATGTCCTTACTCATTTTTCGTGTAAGGGAGTAAAATTATACGATTGCCCACTAGCCTAATTCGCATTCTTTCTATGTTCTGAATTGCAGCATTTGCCGTCTCTGCGAAAGCTAAATCTTAACCACTCCAAAAGGCTGTTGCGAACACCAGATTTCACGGGGATGCCAAATTTGGAATATTTGAATTTGTGTAACTGTAGTAATCTTGAAGAGGTTCACCGTTCCTTGGGATGTTCCAGAAAACTCATTTGGTTAAAGTTGTATAATTGTGAATGTCTTAAAAGGTTTCCATGCGTTAATGTGGAATCTCTTAGATCTCTGGATCTACGATATTGCTCTAGTTTAGAAAAATTTCCAGGAATCCTCGGAAGAATGAAGCCAGAGTTAGAGATCAAGATGAGACGCTCTGGGGTAAGGGAAATACCATCATCTTTTATTCAGCAACACGCATGTCGTCTTACAGAGTTAGATTTGAGTTTTATGAAAAACCTTGTATCTCTACCAAGCAACATTTGTCAGTTGAAAGGTTTGGTGAAGCTAACTGTATCCTGCTGCTCAAAACTTGAAAGCTTGCCAGAAGAGATAGGTGATTTAGAAAACTTGGTGACGCTTAATGCCGAATGCACTCTTATTTCACGAGCCCCGTCTTCCATCACCTGGTTGAACAAGCTTAAATACTTGAGTTTTGCATATCAAGAATCAAATTTTGGGGTGTCCTTTGTGTTCCCTGAGGTGAATGAAGGGTTACGCTCATTGGAATCTCTGAATCTCAGTGACTGCAATTTAATAGATGGAGGACTTCCAGAAGACATTGGATGCTTATCCTCGTTGAAAACAATGTATCTCAATAGAAATAAATTTGAGCATTTGCCTCGAAGCATTGCCCAACTTGGTGCTCTTCAATCCTTGTGCTTATCAAATTGCATGGGGCTTAAAGAGTTGCCAAGTTTCATGGGGATGCCAAACTTGGAGACTCTGAATCTCAGTAACTGCAATATGATAGATGGAAAACTTCCGGAAGATATTGGATGTTTATCGTCTTTGAAAGAGTTGAATCTCAGTGGAAACAGTTTTGAGTATTTGCCTCGAAGCATAGCCCAGCTTGGTGCTCTTCGATCCTTGGACTTATCAGATAGCAAGAGGCTTACACAGCTGCCAGAATTTCCACAACAATTAGATACAATAGGTGCAGATTGGACTAATGATTCGATCTGTAATTCGTTGTTTCAGAATATCTCATCATTGCAGCATGACATCTCTGCTTCAGATTCCTTGCCACTAAGAGTGTTCACCACTAGAGGGAATATCCCAAGATGGTTCCACCATCGGGAAAATGGTAGAAGCATATCAGTCAAATTGCCTGAAAATTGGTATATAAGTGATAACTTCTTGGGATTCGCTGTATGTTACTCTGGCAAATTAATTAACGCAGAAGCTCACTTGATTTGTTATGATGGGAGGCCAGTGACGTGTATCACCCAGAAACTTTCCTTATCCAACCATTCAAAATGTCCTGCAGAATCTGCTATTCATTTTTTCTTGGTACCTCTTGCTGGCTTATGGGACACATCTAAGGCAAATGGAAAAACACCAAATGACTATGGGCGCGTTAAGTTATTTCCCTCTGGAGAAATGAAGGAGTTTGGACTTCGTTTGTTGTATAAAGATGAACCTAAGATTGAGGTCAGTTTCTCCTCTTCTAAGAAACAATGA

>PGSC0003DMG400001016 LRR-RLPs

ATGTTCCTAGTAATTTATGCAATTATGAAAAAAGCCAATTGCTCTTTCCTCTTGACACTCTTGTTGCTTCACTATGTCATGGCTACTTCAGCCATGAACCCAACAAACTTTTCTACCGATCAATCTGCTCTCCTTTCCTTAAAATCTCAAATCATTTTGGACCCTTTTCACTTGTTGGATGAAAGTTGGTCTCCAACTACTTCTGTTTGCCATTGGGTCGGAGTCACTTGTGGTTCTCGCCACCAGAGAGTGAGATCACTCAATATTTCCAATATGGCATTAATCGGAAAAATTCCTTCTCATATCGGGAATCTCACATTTCTTGTTTCCCTCGACTTGAGTCACAACAACTTGTATGGCAATTTGCCTCAAGAAATAGCACATCTACGTCGATTAAGGTTTCTTAATTTGGGTGTTAATATGCTTAGCGGGGAGGTTCCTTCATGGTTTGGGAACTTACACAAACTTCAAGTTCTGAATCTTAGGAATAATAGTTTCAATGGTTTCATCCCCTCTCTATTATCTAATATTTCCGGGCTTCAAACTTTAAATTTGAGATTCAATTCTTTAGGTGGTCAGATACCAGAAAAGATTGGCAAACTTCAAAACTTAAAGGAATTAAATATAGAAGCTAATCGACTTACAGGATCTATACCATTCTCGATCTTCAACATTTCTGGAATAAAGGTCATTGCATTGACAGATAACACTTTATCAGGAAATCTTCCTCCTGATATGTGTGATCGACAATCAATGTTGGAAGAGCTTTATCTAAGTTTCAACGAGCTACGTGGTCATATGCCTCTAAGCTTGCCAAACTGTTTGAAACTTCAACTATTGTCTTTATCAGTGAACAAGTTTGATGGACCTATACATAGTGAAATTGGACATTTAAGTGACCTACGTATTTTGTATCTCGGACAGAACCAATTTGAAGGTTTGTTGTTGAGTGCTAGATTTTCTTCTAACTAACTATTTCCGAAAAGAAACCATCACACATAAATTTATATGTATGTTATTGTTTTGACAGGGACAATTCCACAAGATATTGGGATTCTTGATAATTTGGAGGTTTTGGGGCTAGAGAGAAACCATCTAACTGGGAGCATCCCAATCTCCATATTCAATATCTCCTCGTTGCAAACGATGTCATTGGAAATGAACAATCTCATGGGATCCTTACCGAGGGAGATAGGCAACTTGAACAAATTGCGGATTCTATATCTTGGTTACAATATGCTAACAGGTATGTAAGTAGTCATTGGGTTATTGAAAACATAAATTCAAGCAACTTGAACATTAGTCACGGAAGTGGTCAAACAACTAACGTAAAAGAATTGGCAGGGACAATACCACAGGAAATCGGGATTCTTGATAATTTGGTGGAATTAGGACTAGATAGCAACCAGATTACGGGTCATATCCCAATCTCCATATTCAATATCTCCTCGCTGGAGACGTTGTATTTGGAAAGGAACCACCTCATTGGATCCTTGCCAGGGGAGGTTGGCAACTTGACCAAATTGCAGAGTCTACGTCTTACTGAAAACAAGCTAACAGGTATGTACATATATAGTAAAAAAAGGTACTCCTTTCATCCCAATTTATGTGATGTAGTTTGACTTAATACAAAATTTAAGAAAGAATGTCTTTTGAAACTTGTGGATATTTGTATGGGTACAAATAATTTTTAAAGTTAAATTGTTACTACATATAGAAATATATTATTCTTTTTTGACGGACTAAAAAGAAAGTAAGTCGTATAAATTAAGGCAGATGGAAACTAAAGTATCATTACTAGTTGTACCTTTACTATTTTAACAATCATACATTTGCAGGTAAAATACCAGACGTAAGCAATCTCGTTGAATTGGTTGAAATTTACTTGGGTCTCAACAAGTTTACCGGTGCACTTCCAACGGAACTCTTCAACATGTCAGGGATGAGAGTGATCGAACTTGTGCGAAATAATCTGACAGGAAGCGTCCTCCCATTGAGCATAGGTTCTATGTTACCCAACATTGAAGGTCTTTATCTATCTAGTTTAAATCTTTTTGGAACTATACCTCATTCTCTCTTCAATTGTTCCAAACTCATTTATCTTGACCTTTCTTATAACAAACTAACCGGTTTGATCCCAAACTCTTTCGGATATATGACTCATCTAAAGTACCTAGGCTTGGGGGGGAACAATTTAATGAGTGACACTAGGTCGAGCTTCTTTTCTTCCTTATCAAATTGTAGAGAATTACAATTTCTTGATGTATCGTTGAACCCGTTGAATGTTGTGCTTCCAGTTTCAGTTGGAAATGTCTCTCTTGTAAAGTTAATGGTTGTTGAATGTGGACTTAAAGGGGAAATTCCGAAAGGAATTGGGAACTTAAGTAGCTTAATAGACCTCGATCTCTCTGGAAATGGTTTGGTTGGATCAATTCCCACAACAATTAGCAACTTGAGACTCCTTCAGAGTCTTAAGTTGAGTGGCAATAAACTTTCTGGATTTATCGGAGACAATCTATGTAAATTGCAGAACCTGGGGTATATACATTTGGATCAAAATCAACTTTCAGGATCTCTTCCTAATTGTTTCGGGAACTTAACTTCCCTTCGGGAGATATTTTTAGGTACCAACAAGTTGCATTCAAACATTCCAGCTAATTTAGGAAACCTCAAAAATCTCCTAAACCTTGATTTATCCTCGAACAACTTGACTGGCTCTTTACCTCGAGAAATCGGAAGTCTAAAGGCCATGATACAGATGGATCTTTCGATGAATAAACTCTCGAATGGCATACCTGAAGAAATTGGTAGCTTGCAAAATCTGATACACCTTTCATTGAGAGACAACAAATTGCAAGGTTCAATTCCTAGCTCAATGAGCAGCATGTCAGCTTTGGAATATCTAGACCTTTCTCATAACAATGTCTCGGGATTAATTCCCAAGTCTTTGGAGAAGCTTCTAAATCTCAAGTATTTCAACGTTTCATTCAACAATTTGGTTGGTGAAATCCCCTCGAGCGGTCCTTTCAAGAACCTATCCAGTCAGTCTTTCATATCCAATGAAGCATTGTGTGGTTCTTCAAGATTTCGTGTCCCCCCATGCCATAGTGGAACTTCAAAGCATAGATCCAAGAGAAAAAAAGTACTTGTTTTGATTCTACCGGTTGGAATTGCACTTGTATTAGTGTCCATCGCCTTTGCGTTTGTATGGATAAAGTATATAAGAGGAAAATCAACTGATCCTCAACTAGGAGTTGATCCGTCCTCTTTTGTCTCAACAAGAGGAAGAATTTCATACTATGAATTGCTCCAAGCAACTGATTCACTCAACGAGAGCAATTTGATTGGTTCAGGGAGTTTCGGCTCTGTTTACAAAGGCATCCTAAGAGATGGGACTTTCATTGCAGCTAAAGTGTTCAATCTACAATTGCAAGCGGCATTTAGGAGTTTCGATACAGAATGTGAAGTTTTGCGCAATCTTCGCCATAGGGATCTCACCAAAGTCATCACTAGTTGTTCCAACCTTGATTTTAAGGCGTTGGTACTCGAATACATGTCCAATGGGAGCCTCGATAAGTGGTTGTATTCACACAACTACTTCTTAGACATCAAGCATAGGCTGAGTATAATGATTGATGTAGCATGTGCATTAGAATATCTCCACCATGGCTGCTCATTGCCCGTGATACACTGTGATTTGAAGCCTAGTAACATCTTGTTGGACGAGGACATGCTTGCACACCTAAGTGACTTTGGCATTTCAAAAATGATTTCTGAAGATGAGAGTGCTTTATACACTAAAACTTTAGCAACATTGGGTTATATCGCACCAGGTAAGTTATCTTAATTAATTGTTTTTTTTTTTGAATATTTTCTTTTTCCTTTTTCTGCCAAGAAATGTTGAATATAAAATTCTCATTGTAGAGTATGGATTGGAAGGAATGGTGTCAACCAAATGCGATGTTTATAGTTATGGGATAATGTTGATGGAAACATTTACAAGGACAAAGCCTAGTAATGAGATGTTTGATGGAGATCTTAGCTTAAAGCAATGGGTGAGCAATTCACTCCCACATGCAGTAATGGAAGTTATCGATCCCAACTTAATAACACCACAAGATAATCACTCAGTGAAAAAAATTGATTGTGTGGTATCCATCATGAAAATAGCAATAAATTGTTGTGTTGAATCTCCTAAAGGAAGGATTGACATGAAAGATGTTGCCGCGAGGTTAAAAACCATCAAGATTCAACTTCTTGAATGTTGAACAAGTTCTCAGGTTTAACAAATGTTTGCTAGAACATTGATCCTTTAAATATGCTATAGATTTACATCTTTTCATCACCAACATAGAGAACAAATGTTATGTATACCTCTTCCATCTTGAATTTTTGAAGTGGTTTTAGTCTAATCCCCCATTACTTTATTTTATATTTCCTTTTTATCGTTGATAGAACTCAATACATAACTTCTAATGGCTTTCATTAAAATATGTTTCATATGTCACTACAACAAAAAACAAATTTTAGTCGCGATAAATATACACATTAACAAAGAGTGCTAATTTTTTTACCAGCATTAGTTAATTGTCATTGGATTCAATGTCGCTATAGGCTTTAGGGACATATGTCGTTTGAGTCTCAAAATAGAATGACACCAAGTAAATTGAGATAGATGGAAACATTGTTCGTGACAAGACAAAAAAACTCGTTAGGTATGGGCAAATAACAATAATTATACAACCATCAACCTTCATCTATATACCTACTATATCCCAATACTGGCCATATAACATGTATGCAAGACAAAATTAAACGCCACTACTTTGGTTCTCTGTTACGTATATATACGAAGAAATAAAAACTCATTTCCAGAAGATCACTATTATATTATAACAATGGAGTCCATAATTAGATGGAGAAAACCGAGTCTGAAACCAATTGAAACTAAGGATAAAGCCATTGACGCATCTGAAGATCGAAGAGCCATTGAGTCTGTCATCGAGGCTATTTCATGACCCCAACTTCAACGTGCACGTTCTAGCTATTTTGGGCATCAAAACAAGAGCCAGCACTGAGCAAGCTCCCAGAGAGCATGTGGCTAATACTTTGCTTAAACATCCTGTTTCTCCAGCCTGCCGTTACCAGTATAACACTTTGTTTGTCTTACTTTCCATTTTAGTAGGTTAAAAATGAATTTTCTTTTCGTGACAACTGACAACCCTTTAATTCTAACTTTCTCATTACATGTTTAATTAGACGACAAAATAAACAACACTTTGGTACATTCTATGTGTATTTAGTTTAAGACCGTAAGATTTAAAAGTCTTCTTTACTTTCTTAAACTCGGTGTCAAGTCACAAATAAACAACTTTAAACAGAAAGAACACTATTTTCTGATTAAAATCACCAAGATGAATAGATGTAAAAGACAATGAATTGATAACCTGCTGCTATCAAATAAAAACATACATTACACTTGATGGTAGCCAATAAAAATTGAAACTACAAGTACTTAAGAACAAGTCTAGTGTACTTCTCCTCAGTAAGTTCAATACATAAATATTGTAAACTAATAAAAGGGAGAATTGTCAAAAAGCATAAAACGCTCGTATTCAAAGATCTTGTTTTACTTTCATAGCCCTCATGGTTAATTCACTGCAGGTTATTGATGAGGGAAAAGGCGGAGAAATGAAATGGGTGCGGACAAAGGTGGACTTAGATCAGCACATTGTGGTGCCACAAATTAATGAACAGAACCTGCAGGAATCACCATACAAATTTGTGGAGAACTTCTCTTATTTCCCTTCTCTTGGAGTGTACACACCAAACTGCTGATCCAGATAAAGTTCCACCTATTCATGGAAGTAAAAAGAGGCTTATTGATTCTTCAAAGCATTCCAATAAGGGATTGCGGCGACATGTTACAAAAATTTGGTCTTTTATGATATTGTTTGGGTACACAAAAGTGGATGTGTTGATGTTTATGGCTACAACTATGTTCTTGAAGGACACAGCTACACCGATTAAGAGTAGACCTGGTTATGAGTCTAATCCTAGGTGATTTGTTTACCGGACAATAAGTCTTGACGATATGAAATTGGTGAAAAATGCATAAAATATGTTAAGCCCATACAAATTAATTCTATATTGTAGCAACATCCTTCTCTTCAATTTTGAAATGTAGATGTTTATAGCGTGTTGTTTGATGTTGTTGTCGGACTTTGTCATTTCTTGCACAGACAGTAACGGATGTCGCTTTGGGAGTAACACAAGATGGTTTGTCTATCTATCTTAATAGAAGATATGGTAAGTTTCTGTTTCATGTATTTGGACTTATTAAGCTTAGTATTAAGAAGTCCCACATCGGTAAAGGGATGGGAAATTGGTCTCCTTATATGGACTTGGACAAACCTACCCTCATGAGCTAGGTTTTGGGTTGAGTTAGGCCCAGGTGCATATCTTTACATGGTATCAGAACTAGACCCATCCCAAGTTGTTGTTCACCGATGTTGGCCCCCATTTAGAAGTGTTCACGCTCCAGTTGAGTTCTGGGCGTGCGGGGAGTGTTAAGAAGTCTCACATCGATAGAGGGATGAGAAATTGGTCTCCTTTATGGACTTAGACAAATCCCCTTCATGAACTAACTTTTGAGGTTGTGTTAGGTCCATATCTTTACACTTAGTTGATCGAAAATCTTACGGAAGAAATCTTTAATGCAAAATTTTGCAGGTGAGGTAAGAAAGACAGGAGCAACAGAGAAAAAAATAATCTTCCGATGAACATTAGACTTAGATCAACTATACTTATGAACCTAAGACCGGCAGTCGGAATTCAGGTGATGGAATCCATTCAAATTTTAATCAGCACTTAAGTTTTCTTAGATTATATACTTAAAGTTATTAACATTTTTTTGGGGCTTTCTTTCATGTTTAATTAATTAGCAAATAGGCTTTTGCTGATATGATGGAGAATGACACTGAGGCAAAATGGGGGAATTGGATTGGTTTTGTTCTTTTCCCCTTAAAAGTTGCATTACGAGATGATCCATTAGATTATATCAGAGAAGCTAAGGCAACTATTGGTCGAAAGAAGAATTCACTTGAAACTTTCTACACATTTTCTATTTCAGAATTAGCACTCAAGATTTTTGGGATTAAGGTAACTAATTTTCATTCTCAGTTCTTAATCCTACACATGAGATATCTTCATTGTAAGATGTGACTCTTTTTTCAGACATCAAGTTCGATATCGCACAGAACTATCACCAATACAACAATGTGCTTTTCTTGTCTGCCTGGTCCTCAAGAAGAAATTTGCTTCTGTGGGCATCCTTTGGCTTACATTGCACCCAGTTCTTATGGCTCATGTAGGTGTTTTACTCTTACTAATTTCAAGTTTTTGCTTCCATTCTCAGCTAGATTTTGGGGTGTGAATTAG

>PGSC0003DMG400001072 LRR-RLPs

TATCTCTTCTTCTCCACAAAACTTCATCTCTCTCATCTCACATTTTGCTTAAAGATTCCATTTTAAAACCAAAAATAAAAAAATTCCCCAAACCCCATTTCTCAAATCCATTTTTCTTCATAGATCTTCACTTTTTTCATCTCACATTTGGTTAAAGATTCAATCTTTGAACAGAAAAACGCATAAAAAAACAAGCAGTCTTGATTATCAGATGACTCATGTGAGATGAAGGAGAAAACTCAAACACCCATTTGCTCTCTGTTTTTTCTTGTGCTCTGTTTTTTGTGTTCAGTTTTTACTGAACACACAAATGCTGAGTTTACTGTCTCTGAAAATGGCCCTTTAACTGATTCTGAAGCTCAGTTCATCAAACACAGGCAGCTTTTGTATTACAGAGATGAGTTTGGTGATAGAGGTGAAAATGTGAAAATTGATCCATCTATGGTGTTTGAGAATGATAGAATCAAGAATGCTTATATTGCTTTACAAGCATGGAAACAAGCTATCATTTCAGATCCCTTTAATATCACTCTCAATTGGGTTGGTCCTAATGTATGCAGCTACACCGGAATCTTCTGTGCTCCGGCTTTGGATAACCCCAAAATCCGTACAGTTGCCGGCATTGACCTTAACCATGGTGACATTGCAGGGTACCTCCCGGAAGAGCTTGGGCTTCTTACAGATCTTGGGATTTTCCATATCAATTCAAACCGTTTCTGTGGTACAATACCAAGAAAATTCAGCAAGTTGAAGATACTGTTCGAGCTGGATCTGAGTAATAACAGGTTTGCTGGGAAGTTTCCTTATGTTGTTTTGAGTTTACCCAAGTTGATATTTTTGGATATTAGGTTCAATGAATTTGAAGGTAATGTACCTTCACAGCTTTTTGATAAGCCATTAGACGCCATTTTTATTAATCATAATCGATTTGCGTTTGAGTTGCCGGAGAATTTTGGGAATTCGCCGGTTTCTGTGATAGTTCTGGCGAGTAACAGTTTCCATGGGTGTCTTCCGGCGAGTATTGGGAATATGAGTAATCTTAATGAAGCGATTTTAATGAATAATGGGTTGCGTTCTTGTTTGCCGGCGGAGATTGGGGTGTTGAAAAATTTGACTGTATTTGATGTGAGCTTTAATCAGTTGATGGGTCCGTTGCCGGAGAATTTTGGTGATTTGGTGAATTTGGAGCAATTGAATGTGGCACATAATATGCTTTCTGGGACAATTCCGAAAAGTATTTGTCAGCTTCCTAAGCTTGAGAATTTTACGTATTCTTATAATTTCTTCACTGGTGAACCGCCGGTGTGTTTGGGGTTGCCGGAGTTTCATGATGAACGGAATTGTTTGCCCAACAGACCGGTGCAACGCTCTCCGGGACAGTGTAAGGCCTTTTTGTCTAAAAAAATTCATTGTAGTGCTTTCAGGTGTCATAAATTTGTTCCTGTTTTGCCACCTCCTCCGCCACTTTCACCGCCCTTGCCTGCCCCTCCGCCACCTGTTTATTCGCCTCCGCCACCAGTTTATACTCCTCCTTCTCCTTCCCCACCACCTCCTCCACCTCCAGTTTATTCACCACCGCCACCACCACCCTCTCCACCTCCTCCATCACCTCCCCCGCCTCCACCACCAGTCTATTCACCTCCACCCCCACCTCCATCTCCTCCTCCACCATCACCTCCACCACCACCCCCGCCATCTCCACCTCCACCACCAGTTTATTCATCACCACCACCGCCACCATCTCCCCCTCCACCATCACCTCCACCTCCTCCCCCACCCGTTTATTCGCCACCACCATCTCCACCTCCTCCATCACCATTGCCTTATTGTATACGCTCACCTCCACCACCAAATTCCCCACCTCCTAATTCTCCACCTCCACCGCTTGCTCACTCGCCTCCACCCCCTTCACCTTACTATTACAATTCACCACCACCTCCGCCCCCAAATTCGCCTCCTCCTCCACCAGTATACATATACTCATCACCTCCGCCACCAAATTCACCTCCTCCTCCACCGCCAACATACGTATATTCTTCACCTCCTCCACCTCCTTCCCCATTGCCTTGTATAGAACCCCCTCCACCACCTCCTCCTTGCATTGAACCACCCCCACCCCCCTCCCCTTCACCATCTCCTCCTCCACCACCGGTTTATCATTACAATTCTCCGCCCCCACCATCTCCATCACCTCCACCACCAGTTTATTATCATTACAATTCTCCGCCCCCACCATCTCCATCACCTCCACCACCAGTTTATTATTACAATTCTCCACCACCACCATCACCACCGCCTACTCCAGTATACGAAGGGCCATTACCACCAGTAATAGGAGTTCAATATGCTTCACCACCTCCACCACCCTTTTATTAATTCTTTCAGAAATTTCACTAACTCTTATCCTCAGAGCACAAAATAGTTCCTACAATTATTATTTCCTCAACTCTTCAAGCCATCAACAAAATCGAGCTAATATGTGATCACGAACCTTAAATGGTACGACAGAAAGAAAGATAGAGGAGGTAAAGAGGAGAAAGATTCAATCTCCAATGATTTTATTTTTGAATTGTTATCAAATAGATTGAAGTTGTGGAGAGAATAAATTGGCAAAGTAAAGATTGAAACTTTGCTGGAAAATGTATGTATAGAGAGATCATAATCTTGAATTAGAAACCAGTTTACAGTGCTTGCTGCAACTGGTTTTATTTAATGATCATACTTTGATTTCTTTGTAATTTTTTCTTTTTTTTTTTGGATCTTCCAAACTATAGTATAAGATGAGCTTTTTATTTTCTTTTAAAACTTGTATTGATTCCTCATAATTCAGATGAT

>PGSC0003DMG400001252 LRR-RLPs

ATCAATTTAGAAAATTTTAGAGAGAGAAACAAAAACAACACCCTCAGGATGGGCAGCTTCATGTACATTGTCATCCTGCTTCTTGCCATAATCTCAACCGTTCATTCCTGCTCGCCATCAGATCGGGCAGCATTATTATCTTTCCGAGCTGCCCTAAATGAATCTTACTTAGGAATTTTCAACTCATGGAAAGGCTACGACTGTTGCCATGAATGGTACGGGGTGAGTTGTGATCCCACAACTCACCGCGTAGCAGACATAAACCTCCGCGGTGAATCCGAAGATCCACTCTTCGAAAAAGCTCATCGTACCGGTTACATGACCGGTACGATCTCTCCATCTATATGCAAATTAGAAAGACTATCAAGTCTCACTATAGCTGATTGGAAAGGCATCACTGGACCTATACCATCATGTATTACATCCTTACCCTTCCTACGAATCATCGACTTAGTCGGAAACAAGCTCACCGGCTCAATCCCTTCAACTATAGGTCGATTAAGTCGACTCACAGTGTTAAACGTAGCTGAAAATCAACTTACTGGCTACATACCTCGATCATTAACCAACTTATCATCTTTAATGCATTTAGATCTCCGAAACAACTTCATCTATGGAAACATCCCAACAAATTTCGGAAAATTACGGATGTTGAGTCGAGCTTTACTGAGTCGAAACAAGTTAACAGGTCATATACCGTATTCGATTTCGTATATTTACAGGCTTTCAGATTTGGATCTGTCATTGAACAAACTTTCCGGTAGCATACCTCCGTCTCTAGGTAAAATGCACGTACTTGCAACGTTAAATCTTGATGGTAACAATATTACTGGTACTATACCACCTAGTTTGATTAATTCACGTATCAACATTTTGAATTTGAGCAAAAATGCTATTGAAGGTTATATTCCTGATTCATTTGATGAAAGATCTTATTTTATGGTAATGGATTTATCTTACAACAAATTAAAAGGAAGGATTCCAAAATCGATATCGTCGGCGACGTTTATCGGTCACTTTGACGTTAGTCATAACCATCTTTGTGGCCAGATTCCGGAAGGTTCTCCGTTTGATCATCTTGAAGCTTCATCTTTTACTTACAATGATTGTCTATGTGGGAAGCCTCTTAAACCTTGTCACTAAAGTGTGACTATTATTAGGTAAATTTAATTTTACTTTTCTTTTTTGAGATTTTTATTAACACTACATATTGGTGAATATGTTTGTATTATTATGTGATACTGAAAAGTAAAGTGTTTTTTTTAGCATATATTTGTAATTTTAAATGTATAATTGTGCCTTGAGATATATGAAGGCTTTCTTA

>PGSC0003DMG400001829 LRR-RLPs

ATGGAGTACCAAAAATTGGTAATGGTTCTTTCTATACTTTTTGTCCAACAATCTCAGCTTACGTATTCTGGGAAACATCTCTGTCCCCGCGATGAAGCTTTTTATTTGCTTCAATTTAAGCAAGAACTCACATTTCTTGATACAAGATCTTTTTACTTTCATAATGTCTGTGACGGCGAGACCCAAGCCAAGACTTTTTCCTGGAATGTTACAGGAGATTGTTGCGAATGGGATGGTGTTACTTGTGATGGATTTACGGGTCATGTCATAGGCCTCGATGTTTCTTCGAGCTGTCTCATGGGAACCATCAATGCTAACAGTAGCCTCACAAAACTTGGTCATCTCCAAAGACTCAACCTTGCCTTCAACTACTTGGATGACTTTCCACTTGGAAATAGTATTAGTGAATTCACTAGTTTGACGCATCTCAATCTTTCAGATTCTGGATATATGAAAGGGAAGATGATCCCACAAGGATTATCCAACTTGTCTAAGTTGATTTCACTTGACTTCTCCAGGAATTACATCCAAGTTGGCCGAACAACCTTCACTAGTTTGGTGCACAACTTAGCCAACTTAGAGGTACTGTTGTTTCACAAAGTGGACGCACCATTTGAGTTACCTAAGAGTTTTCCTTCTTCCCTTAGGAAATTAAGTCTTGAGGGCACCAACATGTTTGGGAAAATAACTGATTCTCAGCTTTTTCATCTACCAAACTTGCAAGTACTGAGATTGGGATGGAATCCTTCATTAACCGGCACTTTGCCAAATTTTAATTGGAGTTTCAGTAAAAGTATTTTAGAGTTGGACTTTTCCAATACTGGCATTTTTGGGAAGGTACCTGATTCTATTGGAAACCTCCATTCTCTTTGCTATTTAGATCTCTCTTACAACCATTTATCCGGCTCGATTCCAGAATCCATTGGCAACCTCACTGCAATCACAGAGTTGACACTTTCACGTAACAGCTTCACTGGTAATGTTCTCTCAACTATTGCAAAATTAAACAAACTCGTTTATTTAGAACTCTATTCCAATTATTTTCAAGGCTTGATTCCAGAATCCATTGGCAACTTCACTGCAATCAAAGAGTTGTCACTTTCAGATAACAACTTCACTGGTAATGTTCCCTCAACTATCGGAAAGTTGAACAAACTTAACTACTTATATCTTAATTCCAATAATTTTGAAGGCTCGATTCCAGACATCTTTGCCAACTTTTCAGAGCTATCTTTTTTAGATTTTCACGGTAACAATTTCACTGGCCCATTCCCCTGTTCAATTGCAACCTTGACTCACCTGCAAAGGTTAGAGTTGCAAATCAATTCACTAACTGGTCCACTTCCCTCTAATATAACCGGACTTCAAAATCTAGACATACTTGATTTGTCGTTCAATTATTTCACTGGCACAACACCCTCTTGGTTATTCCATCTTCCGTCCTTGACTTCTTTATCTGTTCAAGCCAATCAATTAACTGGATTGCCAAATGAGCTCAAGAGAACTTCCTCAAAATACTCTAGAATTGATCTTTCGCACAACAAGCTGCATGGCGAAATCCCTGATTGGATGTTATCCATGAGCATGAGAAGTCTGGATCTCTCTCATAACTTTCTCACAGGCTTTGAAAAACAAGTGTGGCGCGCAGAGAACTTGTCCTACCTTAATTTGGAGAACAATTTTCTTCAAGGGCCTTTGCATCAATCCATTTGTGACTTGATTAACCTTGGAATCCTCATTTTGGCTCAAAACAATCTCAGTGGTTCAATCCCAGGATGTTTAGGTAACTCCAGTAGGTTCATTTTTATTTTAGACTTGCGAATGAACAATTTTCATGGAGAGATACCAACATTCTTACATAGAGGGTTAGAGTATCTTGGTTTATATAGCAACCAATTGAGGGGACAAGTCCCCCGATCCTTAGTTAACTGTACAAGATTGGAAGCTCTTGATTTGGGGAACAACATGATAAATGACACGTTCCCCATTTATCTGCAGAAACTACAAAACCTACAAGTTTTGATTCTGAAATCAAATCTCTTTCACGGTCCAATTGGTGACTTGGAGTCCGAATTTCCATTTCCTAAGTTACGGATCTTTGACCTTTCCTTTAATGGGTTCACAGGAACTTTGCCATCTAATCTTTTCAAAAGTTTTAAAGGTATGATGGATGAAGAAAAAACAGCAAGTACAAGCACTCGCACGGATTACCTTTACAATATTAGTTTGGTGATAAAAGGCAATGAGTATGATATGAGAATCACGTCAATTATGACGAGTGTTGATTTATCAAGCAACAGGTTTGAAGGAGATATTCCAAATTCCATTGGAAATCTGAGCTCACTTGTGTTACTCAACCTGTCCCACAACAGCTTCCGTTGTCGTATTCCTGCAGAATTTGCAAAGTTGCAGGCGCTCGAAGCATTAGATCTCTCATGGAACAGACTCATTGGAGAAATTCCAGGTCAATTGTCGAGTTTGACATTTCTTGAGGTCTTAAACCTTTCATACAATCATCTGGCTGGGCGCATTCCTATTGGGAAACAGTTCAATACATTTCCAGATGATTCATATTGTGGAAATCTTGATTTATGTGGATTTCCACTATCAAAGAAATGTGGAAACAATAATATGCGAGATGAGTCAACACTTGAACAAGATGATGATGATTCATTTTTTATGAGCGGGTTCACATGGGAAGCTGTTGTAATAGGTTATGGTTGTGGTATGATATTTGGATTGTTGATGGGAGGCCTTATGTTCCTACTGGAAAAACCAAAATGGTTTTCGAACTTTGCTGAAGATATTGCTCAACAAATTGCTGCTAAGAAGCGAACAAGGCAAAAGAAGAGACGTCAAAGACGTGGTCTAAGAATGAATTAG

>PGSC0003DMG400001831 LRR-RLPs

ATGGAGTACCAACAATTGCTAATAGCTTTCTGCTTCTATTCACTTTTTATAATTTTTTCCCAACAATCTCAGCTTACATATGCTGGGAAACATCTCTGTGCTCGCGATGAAGCTTTTTATTTGCTTCAATTTAAGCAAGAACTCAGAGTTGATTCAAATTATGAGTACTGTGACAACAAGGCCCGAACCAAGAATTTGTCCTGGAATGTTACAGGAGATTGTTGTGAATGGGATGGTGTTACTTGTGATGGATTTACGGGTCATGTCATAGGCCTCGATCTTTCTTCGAGCTGTCTTTGGGGAACCATCCATGCTAACAGCAGCCTAAAAAAACTTGGTCATCTCCAACGACTCAATCTTGCCTTCAACAAGTTGGGCGAGTTTCCACTTGGAAATAACATTAGTGAACTCAGTAGTTTGACGCATCTCAATCTTTCGCATTCTGGAACTTTGATGCACATCCCACAAGTATTATCAAACTTGTCCAAATTGGTTTCACTTGACCTCTCCTGGCATTTGGCTGTACTATTCCCATTTGGTCAAACAACCTTCACAAGTTTGCTTCAAGACTTAACCAATTTGGAGGTACTGTTTCTTGACAACGTATACGCACCATTTGAGTTACCTAAGAACCTTTCTTCTTCCCTTAGGTACTTAAGTTTTCAAGGCACCGGCATTTTTGGGAACCTAAGTGAGTCTCAGATTTTTCATCTACCAAACTTGCAAGTGCTGCTATTGGGAGAGAATCCGTCATTAACAGGCACCTTGCCGAATTTCAATTGGAGTTTCAGTGGAAGTATTTTAGAGTTGGACTTCTCCAATACTGATATATTTGGGAAGCTACCTGATTCAATTGGTAATCTCCATTCTCTTTTACATTTGAACATCTGGTATTGCCATTTATCCGGCTCTATTCCGGAATCCCTTGGCAACCTCACTACAATCACAGAGTTGAGACTTCCAGGTAACAGCTTCACTGGTAATGTTCTCTCAACTATCTCTAAATTAAGCAAACTTGTTCATTTAGATCTCTCTCACAACCATTTTCGAGGCTCAATTCCAGAATCCTTTGGCAACCTCACTGCAATCAGAGAGTTGCGACTTTCAGATAACAGCTTCACCGGAAATATTCCCTCAACTATCCAAAAGCTAAACAAACTTAGCTTCATATCTCTCTCTTCCAATAATTTTGAAGGCTCGATTCCAGACATCTTTGCCAACTTTTCAGAGCTAGATACTTTATATTTTGAAAGTAACAATTTTACTGGCCCATTCCCCGATTCAATTGCAACCTTGACTCACCTTGAAAGTTTAGACTTGCAAAACAATTCACTAACTGGTCCAATTCCCTCTAATATAAGCGGCTTTCAAGAGCTACAAATACTTGATTTGTCGTTCAATTATTTCACTGGCACAACACCCCCTTGGTTATTCCATCTTCCATCCTTGCTTTCTTTATATGTTCAAGACAATCAATTAACCGGGAAATTGCCAAATGAGCTCAAGAGTAATTATCTAGAATACTTTAACATTGATCTTTCATACAACAATCTGCATGGAGATATCCCTTATGGGATGTTATCCAAGAGCATGGGTAGTCTGGATCTCTCTCATAACTTCCTCACAGGCTTTGAAAAAAAAGTATGGCACTCAGACTCCTTACAATACCTTAATTTGGAGAACAATTTTCTTCAAGGGACTTTGCATCAATCCTTTTGTGACATGATTAACCTTGTATTCCTCATTTTGGCTCAGAACAATTTCAGTGGTTCAATCCCAGACTGTTTGGGTAACTCCAATAGCTCCATTTCTATTTTAGACTTGCGAATGAACAGTTTTCATGGAGAGATACCAAGATTCTTACCTACAGGGTTACAGTATCTTGGTTTATATGGCAATCAATTGAGGGGACAAGTCCCGCGATCCTTGGTTAACTGTACAAGCTTGGAAGCTCTTGATTTGGGGAACAACAAGTTAAATGACACGTTCCCCATATGGCTGGAGAAACTTCCGAACCTACAAGTTTTGATTCTGAAATCAAATCTCTTTCACGGTTCAATTGGTGATTTGGAGTCAGAATTTCCATTTCCTGAGTTACGAATCTTTGACCTTTCCTTCAATGGGTTCACAGGAACTTTGTCATCTAATTTTTTCAAAAGTTTCAGAGGTATGATGTATGTGGATGAAGAAAATACAGGAATTACTCGAGCTACTAATAGGACTCACAGAGATTACCTTTACCATGTTAGTTTGGTGATAAAAGGTAATGAGTTTGATATGAAAATCACGTCAATTATGACGAGTGTTGATTTATCAAACAACAGGTTTGAAGGAGATATTCCAAATTCCATTGGAAGTCTGAGCTCACTTGTGTTACTCAACCTGTCTCACAACAGCTTCTGTGGAGTTATTCCTGCAGAAATTGCAAAGTTGCATGAGCTCGAAGCATTAGACCTTTCATGGAACAGATTCATTGGGGAAATTCCAGGTCAATTGTCGAGTTTGACATTTCTTGCGGTCTTAAACCTCTCCTACAATCATCTGGCTGGGCACATTCCTACTGGGAAACAGTTCAATACATTTCCAAATGATTCGTATTGTGGAAATCCTGATTTATGTGGATTTCCACTGTCAAAAGAATGTGGAAACAATAATGAATCACCACTTGAACACGATGATGATGATGATGATGATGATCATTTTTATGAGCGGGTTCACATGGGAAGCAGTGGTAATAGGTTATGGTTGTGGTATGATAATTGGATTGTTGATAGGAGGCCTCATGTTTCTACTGGAAAAACCAAAATGGTATGTGAACTTTGCTGA

>PGSC0003DMG400001832 LRR-RLPs

GTACATATAAAAGAGAAAAAGTTACAACAAAGATGTAATCAACAACTAGCGCAGGAAAATGAGAAATTTCAAAAATATATACTCTAGAAAAATTCCTTCAAAATAATCACAGTAACTAAAAATACTATATTTCTCACAGCTATAGAGAAGCCAACAAATAGTTGACTTCTTTGCTAAACAGGCCTCTTCAAGTGGAAGTGGTACCTTCTATTATTTTTTTCATCAATTTCCTAAAGAGGGGAATAGACTATTTCTACTAGATAAGGGACAACTTCCTAGCATAAGGAGAAGATACAACAAATGTAATTTTTTTTGTTAGTTGAACCATTGTTTTTTGCTTATAGTATGAAAGTCTATGTCACTTTAGCCTTTCTGTAGATATTGCTGCTCAGAAGCAAAGAAGGCAAAACAAGAGACGTCAAAGAAAAATAAGCTTTAGACATAGCTACCTCTGCACTCTGTTTTTGGATGGATGGATGGATGTCTAAACAAAAATAGGCTTTAGACATAAAAACACCATTTTTCATGAAAAGGCCCTCAAAGTTTGAATTCAAAAATGAGGATTTGATGTTGAATTCATGAAGAACTAAATGGGTAATGGAGGATTGGGGTTGAAATCACTTACCCAAACGTTATTTCTTAAAATCGCCTACTCCAAGCTTCCAAGGTCCAAAAATGGAAAAATGGGGAAGAAACCCAAATTGGGAAATTTCATTAAACTGCCAAGCCATTTGGTCTTCGCAATCGCGGAGACCAGACGGAATAACCCTTCACGAACGCAGCCACTCTGACCCTCCACGATCGCGGCCAAGCTCCACCAACGCGAAGGAGGCACCAGAAGACAGCAAACAGCAAATACCAAGGCATAAAAACATTCACCCAATGGACCCTTGGAATCCCGAGCACACAAACTACTACCCAACTAAACTCGACGTTTCGGACTCAATGGAATCATCGAAATATGGATTCGAGGTCTCCGTGACCCGGCACCCATTAAAGGAATTAATATCTCAGGTGATCACTCAACTAATTCTTTTCTCTGAAAGTCACTCAATTTTGATTTTTAACTCAAAAGTCACTTAACTATGAATTATTTTCTCATAAAGTCACTCAACTATAGATTTTTTTCTCAAAAAGTCATTCGACTTTGAATTTTAACTCAAAAGTCACTCAATTATGAGTATTTTACTTACAAAGTCACTCAACCTATTCAAATAATTTTTTTAATTAAAATTTACTTTAATGAAATTTTTATTCCTAAGCAAAAATAAAAAATCAATAAAAAAGTATCCATTTGTACTGCCAATTATATTATGCTTAAAAATATCCAATATCACCGTCAATATACTATACTTAATTATTTTTTTAAAAACTGGAATAATGTTCCTATAAATACGTGTTAAAAGTAGTGTAGAAAATAATAATAAAACTCATGGAATTTAAAGGTTCTTAAAATAATTAATGCTAACTATTATATTTGGGGGGAAAGGTAAAATAAAAGGATCACTATTATATTTTTAGTGGATCGAGTCAAGTAGGTGGGTCATCTTTTTACAATTTTAGTTAGAAATAAAAATTTCATGTAAGTAAATTTTAATTAAAAAAATTAATTAAATAGGTTGAGTGACTTTCTAAAAAAAGAACGCATAGGTGAATGACTTTCTGAGAAAAGAAATCATAGTTGAGTGACTTTTGAGTTAAAAACCAAAGTTGAGTGACTTTCTAAGAAAAGAATCAATAGTTGAGTGACTTTTGAGTTAACAACCAAAGTTAATTGACTTTCTGAGAAAACAATCAATAGTTGAGTGACCATCTGAGATATTACCTACTCGTTAAAGCCACAATTTACCAACTTTCACCCAAAATATTAAAATAATATTAGTGGCAGTAAAGTTTTGAAAAAAATATTAAAATTGTAGTAGTGGTAGTAAAGTTTTTGAAAAAAATATTAAAATTGTAGAGTGGTAGTAAAGTTTTTGAAATTGTATTAGTGGTAGTAAATTTTTTGAAAAAAATATTAAAATAATATTAGTGGAAGTAAAGTTTTTGAAAAAAAAATATTAAACTTGTAGTTGTGGTAGTAAAATTTTTGAAAAAAATATAAAAATTGTATTAGTGGTAGTAAAGTTGTTTGACTAAATAAAGGAGGAAATTGTGTTAATTGAGTTGGGTGAACAACAATAATGCATTATCGATCACACTGTATGTATAAAAACTAGTTCTAAGAATGTGCATGTGTAACACGTTTGACCTCTACTGCTATATAAAGATGGAGTACAAGTTCGTAGCTAGTTAGATATAGAGAATGGAGTACCAACAATTGCTAATAGCTTTCTGTTTGTATGCACATTTTATACTTTTTGTCCATCAATCTCAGCTTACATATGCTGGGAAACATCTTTGTGCCCGTGATGAAGCTTTTTATTTGCTTGAATTTAAGCAAGGCCTCATCGTTGGTCGAAATGATTATTACAATGGTTGTGACAGCAAAGCCCAAGCCAAGACTTTGTCCTGGAATGTTACAGGAGATTGTTGTGAATGGGATGGTGTTACTTGCAATGGATTTACGGGTCATGTCATAGGCCTCGATCTTTCTTGCAACTTTCTTAGTGGAACCATCTATGCTAATAATAGCCTCACAAAACTTGTTCATCTCCAAAGACTCAACCTTACCTTCAACTACTTGAATGACTTTCCACTTGGAAATAGTATTAGTGAACTCAGTAGTTTGACGCATCTTTTGGCAATCTCACTAGAATTAGAGTGTTGAAACTTTCATTTAACAACTTAAATGGTAATGTTCTCTCAACTATCTCAAAATTAAACAAACTCGATCATTTAGATCTCTCTTACAACGATTTTCAAGGCTCGATTCCAGAATCTTTTGGCAATCTCACTAGAATTAGAAGTTTGGATCTTTCAATGAACATCTTCACCGGAAATGTTCCCTCAACTATCGGAAAGCTCAACAAACTTCACTCCTTATCTCTTTCTTCCAATAATTTTGAAGGCTTGATTCCAAACATATTTTCCAACTTCTCAGAGCTATCTTTTTTAGCTTTTGACACTAACAATTTCACTGGTCCACTCCCCTATTCAATTGCAACGTTGACTCGTCTTGAAATATTATACTTGCAAAACAATTTACTAACTGGTCCACTTCCCTCTAATATAAGTGGACTTCAAGAGCTACAACAACTTGATTTGTCCTTCAATTACTTCACTGGCGCAACACCCCCTTGGTTATTCCACCTTCCATCCTTACTTTCTTTATATGTTCAAGACAATCAATTAACCGGGAAATTGCCAAATAAGCTCAAGAGTAATTATCTAGAATACTTTAACATTCGTCTTTCATACAACAAGCTGCAAGGTGAAATTCCCGATTGGATGTTTTCTACAAGCATAGTTAGACTGGATCTCTCTCATAACTTCCTCACAGGCTTTAAAAAACAAGTATGGCACTCAGACTCCTTACAATACCTTAATTTGGAGAACAATTTTCTTCAAGGGACTTTGCATCAATCCTTTTGTGACATGCATAACCTTGACTTCCTCATTTTGGCTCAGAACAATTTCAGTGGTTCAATCCCAGACTGTTTGGGTAACTCCAATAGCTTCATTTCTATTTTAGACTTGCGAATGAACAGTTTTCATGGAGAGATACCAAGAATCTTACCTACAACGTTACAGTATCTTGGTTTATATGGCAATCAATTGAGGGGACAAGTCCCGCGATCCTTGGTTAACTGTACAAGCTTGGAAGCTCTTGATTTGGGGAACAACAAGATAAATGACACGTTCCCCATATGGCTGGAGAAACTTCCAAACCTCCAAGTTTTGATTCTAAAATCAAATCTCTTTCACGGTTCAATTGGTGATTTGGAGTCCGAATTTCCATTTCCTGAGTTACGAATCTTTGACCTTTCCTTCAATGGGTTCACAGGAACTTTGTCATCTAATTTTTTCAAAAGTTTCAGAGGTATGATGGATGTGGATGAAGAAAAAACAGGAATTACTCGAGAAAGAACAAGCAATGACACAGATTACCTTTACCACATTAGTTTGGTGATAAAAGGTAATGAGTTTGATATGAGAATCACGTCAATTATGACGAGTGTTGATTTATCAAGCAACAGGTTTGAAGGAGATATTCCAAAATCCATTGGAAATCTGAGCTCACTTGTGTTACTCAACCTGTCTCACAACAACTTCCGTGGTCATATTCCTGCAGAAATTGCAAAGGTGCAGGCACTCGAAGCATTAGATCTCTCATGGAACAGACTCATTGGAGAAATTCCAGGTCCATTGTCGAGTTCGACATTTCTTGAGGTCTTAAACCTTTCCTACAATCATCTGGCCGGGCACATTCCTATTGGGAAACAGTTCAATACATTTCCAAATGACTCGTATTGTGGAAATCCTGATTTATGTGGATTTCCACTGTCAAAAGAATGTGGAAACAATAATGAATCACCACTTGAACACGATGATGATGATGATTCATTTTTTATGAGCGGATTCACATGGGAAGCAGTGGTAATAGGTTATGGTTGTGGTATGATATTTGGATTGTTGATAGGAGGCCTCATGTTCCTACTGGAAAAACCAAAATGGTATGTGAACTTTGCTGAAGATATACAAATTGATGCTAACAAGCAAACAAGGCAAAAGAAGAGACGTCAAAGACGCTGTTTAAGAATGAATTAGCAATATACCAGGTATGTATCAAAACCTTTTGTTTTGAATCAGGAACAGCACTATAGAGGTTGTTGGATAACTATTTTTCCTAAATATTCAAAAACTCAATGTCTCTGCAGAGGTTACTGGAGAATTGGACAATTTAATTAGCAAGTTAGTGTTGATGTTCGTGCAAATATCAAAGGATGCATTTGTTTCTTCACTTTATGCCCATGGTTGCCAAATACATAATAAGTTACAGTTGATTTCTATAGTTTTGTGGCAAATAATAAATTAGCTTTAGACATAGCAACCTCTACCCTCTGTTTGGATGGATGTCTCATAATGTATCGTATTATATTGTATTGTATTGTTA

>PGSC0003DMG400001870 LRR-RLPs

CTTGCATAAATAAAATTGTCGATCGACTGAGTCTGTGTGTGCTCTTGTTACTATTCTTGCGTTCGCTGAAGTTATTGAAGTTTGAGATACCGCTACTTCTTTAACAGGTTAATCCGTTTTATCTTGGGAGGAATTAATCTACAACCTCGGGTACAGTGAGGGGATTAAATTTCTTAAGGAAACACAGTGGTTTCTGTGGACTCGGATTAAAATTCATTCTGTCTATTTCATCTTTTTTTGTTTCTGTATTTTTGACTAACCTAAATACAGGAGATAACACTTTCCACTTGGAAATAGTATTAGTGAACTCAGTAGTTTGACGCATCTCAATCTTTCGGATTCTGGAATATTTAATGAAAGGGAGATGCAGATCCCATCAGGATTATGCAAGAGTTGTCTAAGCTGATTTCACTTGACCTCTCCTGGAATTACATCCAATTTGGCCGAACAACCTTCACAAGTTTGCCGCACAACTTAACCAACTTAGAGATTTTGTTGTTAGACTTCGTGAACATACCATTCAAGTTACCTAAGAATTTCTCTTCTTCCCTAAGGTACTTAAGTCTTCAAGGCACAAATATGTTTGGGAACATGAGTGATTCTCAAATTTTTCATCTACCAAACTTGCAAGTGCTGAGATTGGGATCAAATCCTTTATTAACAGGCACTTTGCCGAATTTTAATTGGAGTTTCGGTGGAAGTATTTTAGAGTTGGACTTCTCCCGTACTGGTATTTTTGGGAAGATACCTGATTCAGTTGGTAATCTCCATTCTCATTGGCATTTGGACCTCTTGAATTGCCATTTATCTGGCTCGATTCCAGAATCCATTGGCAACCTCACTGCAATCAGAGAGTTGATACTTTCAAGTAACAACTTCACTGGTAATGTTCCCTCAACTATCTCAAAATTAAACAAACTCGTTAATTTAGATCTCTCTTTCAACCATTTTCGAGGCTCGATTTCGGAATCCATTGGCAACCTCACTGCAATTAGAGAGTTGACGCTTTCAATTAACAGCTTCACTGGAAATGTTCCCTCAACTATCGGAAAGATGAACAATCTTAACAACTTATTTCTCTTTTCCAATAATTTTGAAGGCTCGATTCCAGACATCTTCGCCAACTTTTCAGAGCTATACTTTTTAGCTTTTGACACTAACAATTTCACTGGCTCATTCCCCTATTCGGTTGCAACCTTGACTTACCTTCGAATCTTAGAGTTGCAAAACAATTTACTAACTGGTCCACTTCCCTCTAATATAAGCGGACTTCAAGAGCTACAACTACTTGATTTGTCATTTAATTATTTCACTGGTGCAACACCCCCTTGGTTATCTGTTCAACACAATCAATTAACTGGGAAATTACCAAATGAGCTCAAGAGCAATTACAAAGAATTCTTAGGTAACTCAAATGATGCCTCCACCACCTCAAATAAATTATTGCCAAAACAAGTATGGCGTTCAATGAATTTGGTGTCACTTAATTTGGAGAACAATTTTTTTCAAGGGCCTTTGCATCATTCTATTTGTGACTTTATTAACCTAAAAGTTCTCATTTTGGCTCGGAACAATTTCAGTGGTTCAATCCCAGACTGTTTGGGTAACTCCAATAGGTTTATTTCTATTTTAGACTTGCGAATGAACAATTTTCATGGAGAGATACCAAGATTCTTACCTACAAGGTTACAATATCTTGGTGTGTATGGCAATCAATTGGGTGGGAGGGGGGGATCCCACGATCCTTGGTTAACTGTACAAACTTGGTAGCTCTTGATTTGGGGAACAACACAAGATAAATGACATGTTCCCCATTTTGGATTGTTGATGGAGGCCTCATGTTCCTACTGGAAAAACCAAAATGGTTTTCGAACTTTGCTGAAGATATTGCTCAACAAATTGCTGCTAAGAAGCGAACAAGGCAAAAGAAGAG

>PGSC0003DMG400002118 LRR-RLPs

CAATCCTTCAGAGATGTCTGAAGATGAGGAGCGTCCTTGTCCTTCAGATCTCACCGGTGGCGCCACTGCAAAAGCAAGGAACTGTTGCTTTAACGCCGCCGTTACCGGTGGCGGAGGGGGGATTTTTAACTTTTCGCCACACCCAGATCAAGTTCTTGAAAATGTGCTGGAAAATGTTCTTTGCTTCTTAACTGATCGCCGTGACCGTAACGCTGCCTCTCTTGTAAGCAAATCTTGGTATCGGGCTGAGGCTTTAACCAGATCCGAAGTGTTTATTGGCAACTGTTATGCTGTTTCGCCGACCCGGGTTACGACCCGTTTCAAAAGGGTGACCTCTGTGGCCATTAAAGGGAAACCTAGGTTTGCTGATTTCAGTTTGCTTCCTCCAGATTGGGGTGCTCACTTTACTCCTTGGGCTTCTGTTTTGGGTGATTCTTATCGTGGGCTTGAGAAATTGTACCTCAAACGTATGTCCATATCTGATGATGATCTAGGTTTGTTGGCGCGTTGTTTCCCCAGTTTCAAAGAGCTTGTTCTTGTATGCTGTGATGGTTTTGGGACTAGTGGACTTGCTATTGTAGCTCGTGATTGCAGGTTAGTATTGTTTTGTCCGGATATAGATCTGTTTATGTGATAACCCTTTTGGTAAAAAGTTTAGATTTTTAATTTCTAGAAGCTGAATTCTTCTTTCTGTAATGGGAAATTAACAGCTTAAACTAGTGTAGATATATTGTGTTATTTGAATTTGTGCTTCATGTGTGCTGGTGTTTGAATTAGTTATGTGATTGAGTTGTTCTTTAACTGTTTTATTTCTGCATTTGAAACTTAATGATACCTTTTGAGGACACTAGCATTGAAGTTATAATATGTGGTGACTTGTTGTATTGGATTTGTGAGTTTGCAAGCTGTGAAATATGGTTTGGTAGTGTGATTTTCTTGTGCATAATAGAAGGTTTTTACTGCAGGCAAATTAGAGTGCTTGATCTGATTGAGTCTGAGGTATCCGACGATGAAGTGGACTGGGTTTCCTACTTTCCTGAGAACAAAACGTGTTTGGAGTCTTTGACCTTTGATTGTGTTGAATGCCCTATTGATTTTGAGGCATTGGAGAAGCTAGTAATCAGGTCACCTAGTTTGAAGAGACTAAGGTTGAATCGGTTTGTTTCTATTACTCAGCTGTACCGTTTGATGATTCGAGCTCCACAGCTTACCAATCTGGGAACAGGCTCTTTTGGCTCCTCAACGGTCATTGATGAACCAGATCCTGATTATGCTTCAGCATTTGCTGCCTGCAAATCCATTGTCTGTCTCTCTGGTTTCAAGGAAATTGTTCCTGAATATCTGCCTGCAATCTATCCAGTTTGTGGCAATCTGACCTCTCTTAATTTAAGCTATGGTGCCAACATTAATACTGAACAATTCAAGTCTGTCATCAGTCGCTGCCATAAGCTCCAAGTATTGTGGGTATGTGGTTTGAATTTTCGCTAAATGTTCTTTTAAAGAAAATTGTACTCTGTAATAGCGTTATATTGAGTTCAATTTTGATGTTTCAGGTATTTGATTCTGTATGTGACGAAGGACTTGAGGCAGTTGCTGCAACATGTAAGGATCTGCGGGAGATTCGAGTTTTCCCTATCGAAGCTCGGGAAGATGCAGATGCCCCAGTTTCTGAAGCAGGTCTCCTTGCGATTTCTGAGGGTTGCAGTAAACTTAAGTCCATCTTATATTTCTGCCAACGAATGACAAATGCGGCTGTTATAGCTATGTCAAAGAACTGCCCAGACCTTGTGGTATTCCGACTATGCATTATGGGTCGGCACTTGCCTGACCATGTTACTAATGAACCAATGGATGAAGGCTTTGGGGCTATTGTCAAGAACTGTAAGAAGCTTACTAGACTCGCTGTATCTGGTTTACTGACTGATAAGGCTTTCAGTTACATTGGACAATATGGAAAACTAGTCAGAACCTTATCAGTTGCTTTTGCTGGAAACAGTGACTTGGCTCTGAAGTATGTGCTCGAGGGCTGCCCTAAGTTGCAGAAGTTGGAGATCAGGGATTGCCCATTTGGAGATTTGTCTTTGCGTTCTGGTGTACACCACTATTACAATATGAGGTTCCTTTGGCTGTCATCTTGTAGAGTAACTCTACAAGGTTGTCAGGAGATTGCTCGCCAATTGCCCCGTTTAGTAGTGGAAGTGATAAGTGGTGATGAGGAAGGGAGTGAGACTGGTGAGCATGTCAATACCTTGTACATGTACCGATCTCTTGATGGACCGAGGGCTGATGTACCATCATTTGTGCAAATACTGTGACCGGCCTTTTTAGAAGATAGGAATTTTCTGCAGTAATTGATTGGGACCTTTGAGAAATGATTAACTATTTTCTTGGGAAGAGATGCTCTCCGGAAATCTGTTGTGACCCATATGAAGGTGCAAGTTTCCTTCTTGAGAAAACGAATGGTGCACCACTTGCAAAGGCAGTGAGCAATCTATAATGTATTCAATGCTGGTCCCCATGGATATCATTGTATATGGTTCAGTACAGAGATTGTTTTTTTCCAGCAAAGCAATCTAGAAGACAAATGTTCTCTGTATCAGTAATCTTTTCTATTCACTTTCTTCTCTTCATATTCATGGAGCCTTGAATTAGTAAGTGGAGTCATTGGTAAGGCCACAGCAAGGGTAGATGTTCCAAGCTGCACGGTGGCTGATCAAATTGCAGCTCAATGAGATTTCTCAAAGGCTTTCGGCAGAACCAACACTTGTGGAAACACCTGCATGCTCGGTCAGGATACTGGCTCGTCAAGTAGTATCTTTTCATACAATTGAGGCCCTTCTTATTTTTTGTTTCATTGTTAGTTTAGAGTTGAAGTTTTTTATTTTTAGTTGGTGATAGCCTAATGGTCAGAAGGATACCTACCGCAGTTGGATCTCTCCATCTCGCTTTCTAGAAATATTTGTTCTTTCCAGGATGTGTTGTTTCGTAGCACCTGACTGTTGAATTTATTATTGTTATTTTTAGAAGAAATGTGAGATGGGTAATAAATGCCAAATATGCATGGACCAGATGATGCTCTAGTGTGTGTTTGTGTTTGTCTTTGCTTTTTCTTTATGTGGAATGATCAGCTTGATAGTGAAGACCCATAAATGATGTGTGGAAAGCCTTGTACAAGTACAAAAATGCAGGACCCTCCTCTCCTTCACTCACTCTCTTTGTGGGATCCATTTATGAGTCTTATCCTCTATTACCTTTCCAGCTGATGAACGGCCCTTCTCTGTCTCATCTCTTGTATCTGAAAGAGCCAAGAGCAGGGAGTAGACTGTAGTAGTGGAAAGTATAGATGGATACTACAGCACACCCCTTAATTTTTAGCTGATGCCAACTTTATCATTATTCTTTTGGATAACAGTCTTGAGGAGCCTGTTGTGTTATCCTGTTTACTTTTCATATGCTGCTGCTGCCTCTATCATTGACTACAACTCAC

>PGSC0003DMG400002426 LRR-RLPs

CTTCTCAAATTTCAACAAATTAACCAAATAAATGGCATCTGCTTCTTCTTTTGCGAGTGATTCACAATACTGTGCTCGATGGAAGTACGATGTCTTTCTAAGTTTTAGAGGTGCCGACACTCGGGCTACATTTACAAGTCACTTGTACGAAGGTTTGAAAAACAGGGGAATATTCACCTTTCAAGATGATAAAAGGCTAGAGCAAGGTGATTCCATCTCAGAAGAACTCTTAAAAGCTATCGAAGAGTCTCAAGTTGCACTCATCATTTTCTCAAAGAATTATGCTACATCGAGGTGGTGCTTGAATGAACTAGTGAAGATCATGGAATGCAAGGAGGAAGAAAATGGACAAACAGTCATACCGATCTTCTATGATGTGGATCCATCAAATGTTCGATACCAAAGTGAGAGCTTTGCAGAAGCATTTGCCAAACATGAATCGACGTATAAGGATGATGTTGAGGGGATGCAGAAGGTGCAAGGATGGAGGAATGCCCTAACTGCTACTGGAAATCTAAAAGGATATGATATTCGTGGCGGGTGAGTTGAAAACACATAATATGTTTCTATTACCCTGAACTATAATGTTTCCTATTTAATACGCTCTTGAACATTAAAAGTATGAAAATACTTTCTCATACTTGACTTCCTCACATTGTCAACAATCAATCTCTAATTTATGTAGGATTGATCAATCAAAGGAAATTGAGCAGATCGTCGACCACATTTCTTCCAAATTTTGCAAGAGTGCTTGTTTTTTATCTTATTTGCAAGATGTTGTGGGAATAAATGCTCATTTGGAGGAACTAAAATCCCTACTTCAAATAAAAATCGATGATGTTCGAATTGTTGGGATCTGGGGAATAGGCGGAGTCGGTAAAACGACGATAGCAAAAGCCATCTTTGATACTTTATCTGATCAATTTAAAGCTGCTTGTTTTCTTGCCGATGTTAAAGAAAATGCAAGAAAGAATCAACTACATTCCTTACAAAATACCCTTCTCTCTGAATTGTTAAGAAAAAAAGATAATTACGTCAATAATAAGTATGATGGGAAGTGCGTGATTCCAAACAGACTTTGGTCTATGAAGGTGTTGATTGTGCTTGATGACATTGATGAAAGAGATCATTTGGAGTATTTAGCAGGTGGTGTCGGTTGGTTTGGTAATGGCAGCAGAGTTATTGTAACAACTAGAAACAGACAATTGATAGAGAAGGATGCTGCAATATACGAAGTGCCTACGCTACCTGATGATGATGCTATGCAATTGTTCAATCAACATGCTTTCAAAAAAGAAGTTCCAGATGAGTGTTTTAAGAAGTTCTCGTTTGAGGTAGTAAATCACGCTAAAGGGCTTCCTTTAGCCCTCAAGGTGTGGGGTTCTTTGCTGCATAAGAAAGGCCTAACTCAGTGGACAAGAACTGTTGACCAAATAAAGAAAAAGTCTAATTTAGAAATTGTTGAAAAACTCAAAATAAGTTATGATGGGTTGGAGCCTGAAGAGCAAAAGATATTTCTTGATATCGCATGTTTCTTCCGTGGGCATCATAGAAAAGAAATGAAATTTTTTGATAGCTGTGATTTTGGAGCTGAATACGGATTAGATGTTCTAGTTGACAAATCCCTTGTGTTCATCTCTATGTATGGTAGAATTGAAATGCATGACTTGATTGAAGATATGGGCAAATACATTGTGAAAATGCAGAAAGATTCCGGAAAACCTAGCAGAGTATGGAATGTTGAAGATTTCAAAGATGTGATGATGGACAATATGGTAAGTAAGCTAAACAATGCAATAGTATTCATATTTCAAAGATATATATATACATACTCTGTTCCTCTAGCTTCATATACTCTAACTAATATTTGTACCTTAAATTTTGTTTCGTGGTGCTAACTCTTCAAGAACTTCAAGTGTAATAATAAATATAATATAAAACATACATGACAAAAATAGCATTTGGGGGTAGACCGGTAGAGTAAGAAAGTAATGATATTGATAAGTGAATTGGACAAGAGGTGGCTATATGTGAAGTTGTGATTTGTTGACTCATTACCTTTTTATAGGGGACCATGACAGTCGAAGTAATTTGGTTGACAGATTCTGAAAAACTATGCTTTAGCAAAGAGGCAATGCAAAATATGCAAAAGCTTAGGATATTATGCATGTCATATCATCCATGGGTTCCTCAAAGGGCTTTCAAAGATACTCCCAAATCCTCTATTGAGTACCTTTCAAACAACTTGCGTTGGTTTGTGTGGCATTCCTATCCTTGGAAGTTATTGCCAGAAAATTTTAATCCCAGAAGGCTTGTTCATCTTGATCTCCGGTGGAGTTCATTGCATTATTTATGGAATGAAACAAAGGTACCATTCTATTTTAGATACTTTCTATTTTGTCTTGTCCTAAAATTATATTTGATTCTTGTGTTTAAATGAAAATAAGTTATTAATTCCATAAACTCAAACTAATAGTTTCTGAGCATGTAATTTCCATAGAAAGTAGTAAAATAAATACGTTGAGTTATAATATAAACATGTCATTTACTATCATTCTTTCTGTGTTTTGAATTGCAGCAATTTCTGTCTCTACGAAGGATAGATCTAAGTGGCTCTAAAAGCCTGAAGAGAACACCAGATTTCAAGGGGATGCCAAATTTGGAGTATTTGAATCTAGAGTATTGTAGGAGTCTTGAAGAAGTTCATCATTCCCTCAAGTATAGCAAAAAACTCATCCACTTAGATTTGCATTATTGTAGAACACTTGAGAGGTTTCCATATGTTAATGTGGAATCTCTTGAATCTATGAATCTAAAATTCTGCTCTAGTTTAAAGAAATTTCCAGAAATCCTTGGAATAATGAAGCAGGGAACTGCCAGAAAGATTATGATGTCATGCTCTGGGATAAGGGAACTGCCATTATCTTTCTTTGATCACCAACCTCATCTTATAGAGCTACATTTGAATGGCATGAAAAACCTTGTATTTATTCCAAGCAGCATTTGTAAGTCGAAAGGTTTGGTGAAGCTATCTGTGTCGTATTGCTCAAAACTTGAAAGCTTGCCAGAAGAGATAGGTGATTTAGAAAACTTGGAGGAGCTTCATGCCAGCAATACTCTAATCTCACGTCCTCCATCTTCCATCGTCCGGTTGAACAAGCTTATAAGATTGAGTTTTGGACAACATAGATCAGAAGATAGAGTGTACTTTGTGTTCCCTCAGGTGAATGAAGGGTTACTCTCATTGGAATATCTGGATCTCAGTGGCTGCAATATAATAGATGGAAGACTTCCGGAAGACATTGGATGCTTGTCCTCTTTGAAAAAGTTGAATCTCAAGGGAAATAATTTTGAGCATTTGCCTCAAAGCATATCCAAACTTGGTGCTCTTGAATACTTAAACTTATCAGATTGCAAGAGACTTACACAGCTGCCAGAAGACATTGGATGTTTATCCTCTTTGACAAAGTTGAAGCTCAACGGAAATAATTTTGAGCATTTGCCTCGTAGCATATCCCAGCTTGATGCTCTTGAATACTTGGACTTATCAGATTGCAAGAGACTTACTCAGCTGCCAGAAGACATTGGATGTTTATCCTCTTTGACAAAGTTGAAACTCAACAGAAATAATTTTGAGCATTTGCCTCAAAGCATATCCAAACTTGGTGCTCTTGAATACTTAAACTTATCAGATTGCAAGAGACTTACTCAGCTGCCAGAAGACATTGGATGTTTATCCTCTTTGAAAAAGTTGAAGCTCAACAGAAATAATTTTGACCATTTGCCTCAAAGCATATCCAAACTTGGTGCTCTTGAATACTTAAACTTATCAGATTGCAAGAGACTTACAAAGATGCCAGAAGACATTGGATGTTTATCCTCTTTGAAAAAGTTGAAGCTCAACAGAAATAATTTTGACCATTTGCCTCAAAGCATATCCAAACTTGGTGCTCTTGAATACTTAAACTTATCAGATTGCAAGAGACTTACAAAGCTGCCAGAATACATTGGATGTGCATCCTCTTTGAAAGAGTTGTATCTCAAGGAAATAATTTTGAGCATTTGCCTCAAATCATATCCGAACTTGGTGCTCTTCGATCCTTGGACTTATCAGATTGCAAGAGGCTTACTCAGCTGCCAGAATTTCCACAGGAATTAGATACAATATATGCAGATTGGAGCAATGATTTGATCTGTTATTCCTTGTTTCAGAATATGTCATCATTGCAGCATGACATCTGTTCTTCAGATTCCTTGTCACTAAGAGTGTTTAAGAGTTGGGGGGAGGGTATCCCAAGTTGGTTCAAATATCAGGGAATGGGCAAAAGTGTATCAGTCAATTTGCCTAGGAATTGGTATGTAAGTGATAACTTCTTGGGATTTGCTGTATGTTATTCTGGCGAATATGATATTGATTGCATCAGAGCTCATTTGATTCCCTTATCATGTGATGATGGGATGTCGTCGATGACCCAGAAATTTGCCTTATCCAACAATTCACAATATTCTAATGTATGGTATATTAAATTTTTGTTGGTACCTCTTGGTGGCTTATGGGATGCATCTAATGCAAATGAAAAAACACCAAATGACTATGGGTGCATTAAGTTAGATGTTTTTGGAGAAAAGAAGACATTTGGAGTTCGTTTGTTGAATAAAGATGAAGCTGAGCTTCACAATGGGATAAGGAAGAGCAGAGATGAAGAAGAGGCCACTTGCTCCTCTTCTAAGAAACAAAGGCAACTCTTTCCCACAAGCTCGGGATTATAGATTAAATATAGGTATATACCTCATGAAATTGTGTTCTCTGTTTATATATAAATTATTTCTTCTCATTCTTTCTTCTATTGGGAATAGGTGATGTACGTATCAGTAATCAGCAGCGAGTTTGGGGACACTGCCTATGAAGGGACATGTACGTACAATTTTGGGTGGCAAAAATTCATTCTTTTTATATTTCATTTCGAAATATAATCACATTTTCCATTTCCTGACAAATAGATAAATTGCAGAAGCTTTGGAATCCAATGGAGCTCAAATAATTTCTCCCCAACCATCAAGTTCTACTGAAGTTGCATAGTCTCCGTTGATATCTACTGTAAAATGTTACATTCTTATTCTATTTTAATAAGTGCAAGAAACACATTTGATTAATCATCTCTAAAATTAATGTAATCTAATTAGCAAAGAACACTGCTATATAATTGAACCCCTCCCTATCCTCATTTCTTGTTCAATGTATTGCCAATGTCGAGCACGAAACGATATTTCACATCCGATTTCAAAAGACGTTCCAACGTTGTGTTCACATAGTCCATTGGCACAACTTCAATATTTGGTGTTATGTTATGCTTTGCCGCAAAATCCAACATTTCTTGAGTCTTCATCCCTCCTATGCAACTTCCAGCCACTAGCTTCCTTCCTGTTTT

>PGSC0003DMG400002478 LRR-RLPs

CTTCCACTTGATTTGAGACAATGCTTTGAGTATTGTGCGGTATTCCCAAAGGACACCGAAATGGAAAAACAAAACCTTATCTCTCTTTGGATGGCGCATGGTTTTCTTTTATCGAAAGGAAACATGGAGCTAGAGGATGTGGGTAATGAAGTATGGAGAGAATTATACTCGAGGTCTTTTTTCCAAGAGATTGAAGTTAAAGATGGTCAAACTTATTTCAAGATGCATGATCTCATCCATGATTTGGCTACATCTCTGTTTTCAGCAAACACATCAAGCAGAAATATCCGCGAAATAAATATGATGATGTCCATTGGTTTCGCTAAATCGGTGTCTTCTTACTCTCCTTCTCTCTTGCAAAAGTTTGTCTCGTTGAGGGTGCTTAATCTGAGTCGCATAGGAAATGAACAGTTACCATCTTCCATTGGAGATCTAGTACATTTAAGATACTTGAACCTCTCTCGCAATAACATGTGTAGTCTTCCAAAGCAGTTATGCAAGCTTCAAAATATGCAAACTCTTGATTTACAATATTGCACCAGACTTTGTTGTTTGCCAAGAGAAACAAGTAAACTTGGTAGTCTCCGAAATCTTTTACTTCATGGTTGCTATAGATTGACTTGTACACCACCAAGGATAGGATCTTTGACATGCCTTAAGACTTTAGATTGCTTTGTCGTGGGCGAGAGGAAAGGTTATCAACTTGGTGAACTAAGAAACCTAAATCTCTATGGCTCAATTAAAATCTGGCATCTTGAGAGAGTGAAGAATGATAAGGATGCAAAAGAAGCCAATTTATCTGCAAAAGGGAATCTGCATTCTTTAATCATGAGTTGGGACAGCGGACCACATAGATATGAATCAGAAGAAGTTAAAGTGCTTGAAGCCCTCAAACCACACCCCAATCTGAAATATTTAGAAATCAATAGCTTCAGAGGATTCCGTCTCCCAGACTGGATGAATCACTCAGTTTTGAAAAATATTGTCTCTATTGTAATTTACGACTGCAGAAACTGCTCGTGCTTACCACCCTTTGGTGATCTGCCTTGTCTAGAAAGTCTAGAGTTACAGTTGGGACCTGCGGAGTATGTTGAAGAAGTGGATATTGATATTCATTCTGGATTCCCCACAAGAATAAGGTTTCCATCCCTGAGAAAACTTTGCATAAAATATTTTCGTAATCTAAAAGGATTGCTGAAAAGGGGAGGAGAAGAGCAATTTCCTGTGCTTAAAGAGATGGAGATTTGCTATTGCCCTATTCCGACCCTTTCTTCTGATCTTAGGACTCTTACTTCCCTCAATATTAGGGATAACGAAGAAGCTACTTCACTCCCAGAAGAGATGTTCAAAAACCTTCTTAGGACTCTTACTACCCGCAATATTAGGGAAAGCGAAGAAGCTACTTCACTCCCAGAAGAGATGTTCAAAAACCTTGCAAATCTCAAAGACTTGAAAATCTCTAACTTCAAGAATCTCAGAGAGCTGCCTACCAGCCTGGCTAGTCTCAATGCTTTGAAGAGTCTGAAAATTCATATGTGTGACGCACTAGAGAGTCTCCCAGAGGAAGGGGTGAAAGGTTTAACTTCACTCACTGAGTTGTATGTCTGTTACTGTAAGATGCTAAAACGTTTACCTGAGGGATTGCAACACCTAACAACCCTCACAAATTTAGGAATTTGGGGATGTCCAGAAGTGGAAAAGCGGTGCGAGAAGGGAATTGGAGAAGACTGGCACAAAATTGCACACATTCCGTATCTGAAAATATATTCCTAAGTATTATTTGTAATTGTTTCTTTGTTTGTGAGTCTTTTTGGTTCCTGCCAGTGGATGTCATTTTTTCTAGGGTTGTCTGTTTGTGAGTCTCTCTCTTTGGATGTAATTTTCTTTTGGAAACAAATCAACAATCTATTTGTATTATACGCTTTCAGAATCTATTACTTCTTATTTGTAATGGTGAGTATCTCTGTTTCCT

>PGSC0003DMG400002618 LRR-RLPs

TAGTAAAGTAAAATAATGGGGAAAGGAGCAATTTTGAAATTGAAAAAAAAACAAAAACAAAGAGTGGTGCTCTTATTCCGGAAAAGCAAGGTGGGAAATCTTAACAGGACTCGTCTCTTTTTCCTTATCAGAATCAGAAGAAGAAAGAAGAAGATACTTTTTATTCGTCTCTCAGGTATGGTGTTCCTCTTACAATCAATCTATAATGCTTATGTTTGTTTGTTTGAGAGGCCGATCCGCCCACCCTCATGATGGATGGGTCAAACCTGTTTTTTTCCATAAAAACAAACCAGACGCTCAACGTCTATCTTTTCCGTTTTCCCTTTTTGAGTTTTGACCAATTGTTAATTAGATGAAAATCAACAATAGGGTAAGAAAAGGGGGTTTGATTGAATTATGCTGGAATATTCTGTCTTGTTAGCATGTGTATTTGTGAACAAGGTTCACCGCACGCGTTTATTTGACCTTTTCCGTATTATATTGGGTCTTGTCTTCTAATTAATTAATCTCGTGCGTTTTTGTGACAGCACAAAATTTGACAGCTATATAGTCTTGCTTGTGGGGCTGTCATTATAGTTCATCTATATTGGCAGCCCAATTTCTGAGGTGTGTCTTCTTTCTCTTCACCGGCAAAACAAATATATCATCATGGCTTACTTGTCTGGAATCCTCCAAAGACCTATAGTTGCAGCATCTGCAGTTGCTATAGCTTCTGTATCTACTGATCTTCATGAAAAATTCTGGCCATCTAAGTCATTACACAACAACAACAACAATTCCCAAGAAAAGGAAAAGATAACATCTTGGGTCTCTCACATATATCTTTCTAAGCTACCCCATTTCTCCTCCTACCTCACCACAATACCTGTTCCCCTGCCTATTCTAAATAACACCAATTTTACTTCTCTTTCAACCCTTCCTTTTCCTACTTCTTCTCTGTTGCTCAACTCTTATCAATCAGCAGCATTGCCCAACTCTGCTAATCCCTCCACATATACATATACACTTCCTTCTTCACCTTCCGAGGTGTTGTATACATGGCACTTACCTGAGCCAAATGCATCACGGAATTCTGATTCTTCACCACTAAAGTCTACCACTCTGGTAGTATTGTTGGGATGGTTAGGTGCAAAGCAAAAGCATCTAAAGAGATATGCAGAGTGGTATGCATCAAGGGGATATCATGTCATTACATTTACTTTCCCAATGTCTGAGATTCTAAGCTATCAAGTCGGGGGAAAGGCAGAGCAAGATATTGAAATGCTTGTGAATCATCTAGGTGATTGGTTGGAAGAAGAGCATGGAAAGAACTTGGTCTTCCACACTTTCAGCAATACTGGATGGTTAACGTGAGTTCTTGATGATACTCATTAGTCTAGGAAGAGAACAGAAACATAACTATTTACATTTTCCTTAATGCTAGCTTGATAATTATTGATTGACTTCTTTTATTCTCTTGGCCTTACAGTTATGGTGTCATTTTGGAAAAGTTTCAGAAACAAGATCATGCTTTAATGAGAAGGATTAAGGGTTGTATTGTTGATTCTGCCCCTGTCGCAGCTCCCGATCCACAGGTATTGCTGACTCAAAAATTGTGATGATAAGATATCATTTTTATGTGCAAGTGTCAGAAGAGTGTTTCTTGGAGAGTAGTCTTATTTCTGAAAGTTAATGCAGAGCACTTGTTTAGAGCTATACATTTCTATTGAAAACCAACCTTTTCATGCTTTTATTGGACAACTATTACAAGTGTCATATATAGTCAGGTTTTAGCCATTCAGAATTGCACAGTTGTTTGGATTCTAAACAGGACCTGCTGTTCTCGTATTTTTATCCTGTTGCTTTCTTTTATCTTAGGGCCTTAACCATGCAGCCTGCTTTTCGAAAACTATTCTTATGCTTGATAACGTGAAAGAATCTTGAGAGTTTAGCTCTCTCTGACAACTTTTTAACAGGAAGCATTCTATGCAACTTTTGTTTCACGGTTCAAGCTTTAAGCCTACTTATTCTGGCTGAGAATAAGCCATCGAGGAATTTCCCCTTGGAGCTACTAAACTGCACACGTCCCTGCGACTACTGGATCTCTCCAGTAACAACTTTGGAGGAATGCTTCCATGAGGCTTCGACAGGCTAGAAAGTCTCACATCCTTTTCAACAATAACAACTTTACTGGAAATATACTGCCTCAAACAGGAAACTTGACTAACTTGGAAGATTTGTACCTAATCCATAACATGCTCTCCGGAAGAATTCCAGTGGAAATAGGAAAGCTTCAAAGGCTGTGTGAACTACCTATAGCTATGGAAACCAGTTGTCAGTAACTACCCCCAGAGAATTGACAAATTGCTTAGTACGCATTGATTTCTTGGCTAATCAATTTTCAGGCTCTATTTCTGATGCCATTGGTAGGCTTAAGAATCTTGTTATCTTCAGCTAACTCAGAATGAGTTGTCTGGTCCGGTCCCATCAAGCTTAGCCTACTGCAGGAAGCATGAAAAACTGGCCTTTGCTGATAACAAACTTTCAGGATAATTGCCACCAACTTTAAGATTTCTATCAGAGCTGGACCTGACTACTCTTTACAACAATTGTTTTGAAGGTCCACTTCCTGAATATCTTTCCCTTCTGAAAAATCTCAGCAAAGTTAACTTTTCTCACAATAAGTTCAGTGGAAACATTTCTCCGCTAGCTGGTTCAAATTCTCTGACAGTTGACATGACAAACAACAGCTTCTCTGGTCCTATCCCATCTGAGCTAGTGGTGTCCAAAAATCTGACCCATCTTCGCCTTTCTAACAATTTTTTCACTGGAAAAACCCCTTCTGAATTTGGACAACTTGAAGATCTCAGATGTCTTGACTTATTGTTCAAAAATTTGACTGGAGATCTAGCACCTTCCCTTGCTGGCCTGAAAACCTTCGCTGTTTTCTTCTTATATCTAATCAATTTTCGGGGAAAATTTCCACATGGTTTGTGGAGGAATGGAAGGTCTTGGTGAGTTCGATCTTTCATTCAACAACTTCACTGGAACAGTTCCAGCAGAACTTGGAAACTCTCCAAAATAACTCAAACTTTTTCTCAGTTACAACAGATTATTAGGTGCCATACCACCAGAATCAGGAAATCTCACAAGTTTAAATGTCTTGAATCTTCAATGAAACAATCTTTTTTGGCTCCATTCCTTCAACTCTTCAAAAATACCAGAAGCTCTATGAGCTGAGGCTCTCTGAGAATAATATAAGTGGCTCAATCCCTTACGAAATTGGCTCTTTATCCGAATTACAGGTGATAATAGACCTGAGCAAAAATCATTCTATTTTATGTGCAAGTGTCAGAATAGTGTATCTTCAAGCATAGTCTTATTTCCAAAGGTTAATGCAGTGCAATTGTTAAGAGCCATACATTTCTATTGAAATCCAACCGTGTCATGCTTTTATTGGATAAGTATTACAAGTGTCATATATAGTCTGGGCTTAGCCATTTAGGAATTGCTCAGTTGCTTGTACTCTTATCAGGACCTGCCGTTCTCTTATTTCGATCATGTTGCTTTCTTCTATTTAGGCCTTAGACATGCAGCCTGCTTTTCAAAATATATTCTTACACTTGATAATGTGGAAGTTGTTCATTCTTGATTTAGTCTTCAAGGGAAGGTGGATGGCCTGGGGGTCAGCTGTGGACATTAGGAGGAGGGAGGGGAAGCTAGTACTTGTTTAGTTGCATATAATAAATGTTAATCTGTGCTTTGGTAAGGTTAAACTAATTTTGTCAAGTTCCCTTCTTTGTTTGTATCAAGGTCTGGGCTTCTGGATTCTCTGCTGCCTTTTTAAAGAAGAATAGTGTTGCAACAAAACGCATCATGACTTTAAACGACAGAGATGTGACAATAGAAGCCAAAACTTCTGTTGAAACTAAGCCTGCAGTAACTGAAGCAGCGTTGCTATTAGTACTGGAAAAGTTCTTTGAGGTGGTTTTGAACCTTCCCACCGTAAATAGGTAAGGAGATGTCTTCCAGAATCACCTTTTCTGCAAATGTTTATCAGCCTGCAGTATTTTTAAACAGCAGTACCAATATTAAATCTACTTTTTGTCTTTGTACTCTACTCAACGTGCGAAGTAGACCTTGTCGATATCATGTCCATTCACCAATGGATTGTAATTACATTTTCAAGTCATTAGTGTGTATAGCAAGTGCCAAAAATAGTTTCGTGTTTGTACTTACATTGATGGTTGTATGAAATTGCAGGAGGCTTTCTGACGTTCTAGATCTATTGACATCCCGGCAACCAAGTTGCCCTCAATTATACATATACAGCAGTGCCGACAGAGTCATTCCTGCAGTTTCGGTTGAGTCCTTTGTAGAAGAGCAACGAAGAATTGGTCGCGATGTTAGAGCTTGCAACTTCGTTTCTACACCTCACGTTGATCATTTCAGAAATGACCCAGAATTGTATACTTCACAGCTTACCCAATTTCTAGAGGACTACGTCCTAAGCTCTTGCAAACAGCCTTCTTGAATAATTCCTAACCACAACACATTGATTCTTTCTGCCCTTTTTTTCCCTCACTTCAACCAAATACTTGGTAAAGCTTCTGTTTTCCAACTTTAAAGCAGATATACATTGATGTAGATTGTTACGTATACAGAGAATTAAGATTTTTTTTTCTTTTCCATTCTTATTTAATTTAGTTGTATCTGTGGTCTCCCAGCAACAGAATTGTTGCATTGTACCCAAAATATACTTTACTTTACTACATATATATAAAAAAGTTGGGCTTAATT

>PGSC0003DMG400002885 LRR-RLPs

ATGTGGCCTCTCTTTTTCATTTATTCACTTTTCTGTTGTGTATTTCTAAGTCTAAGTTTTTCTTCATCCTTTGATCATCATCTTTGCTCTCCCAGTGAAGCTTCTGCTTTGCTTCAGTTTAAGCATTCCTTTAATATATTGGACTTCTCCCCGTGTGATACGTCTTTCCCAAAAACAGTTTCATGGAATGAGAGTAACGATTGCTGCACTTGGGACGGGGTCACTTGTGACATGTTAAACGGTCATGTTATCGGGTTGGATCTTAGTTGTAGTCAGCTTAATGGAACTATTTATCCCAACAGCAGCCTCATCCAACTTCATCATCTTCAGACACTAAACCTTGCTAAGAACTACTTTTATCCTTCTACGATTCCAAATGACGTTAGCCGGTTGAGGAATTTGAAGCATCTCAACCTTTCTGATGCTTACTTTCAAGGGGAAATCCCAACAGAAATCTCATACCTTTCCAATTTGGTTTCACTTGATCTTTCTGTTCCTTCTTATGTGAACGGATTACAATTTGATCAGAGAGCATTTAAAGCAGTGCTTCAGAACTTGACAAATCTAGAGGTACTTTCTCTCTTTGGTGTCAACGTCTCATCTCCGATACCTGTAAATTTGTCTTCCTCCTTAAGGTACGTCGATCTTAAGGTAACAAATTTGGAAGGTGTTCTCACGGAGAGCGTTTTCCTTCTGCCAAAATTGGAAACGCTCTATTTGAGTAATAACTATCTTCTCAAAGGAGTTTTACCAAAGATCCATCCAAGCAGCACATTGTTGGAGTTGGATATTTCATACACAGGAATCTCCGGTGAGCTGCCTGATTCAATTGGCACCTTGAGTTCCTTGACTCGCTTGTACCTCTATGGATGTCAATTCTCTGGCCGCATTCCTGATTCCATTGGCAATCTTACACAAATTAGGTATTTGAATTTTGGTAATAATCATTTCACTGGCAATATTCCTTCCACAATCTCTAAATTGAAGCACCTCGCAGTTTTAGCTCTTTCATCTAACTCCTTCGGAGGTGAAATTCCCAATTTTTTCTCTAACCTCCAAGAACTACGTTATTTAAGTCTTTCTAATTGCAGCTTCATCGGTCCGTTTCCTTCTCCACTTTTAAGCTTGACACAACTTGAAACCTTAGACTTGTCGAGTAATTCCCTATCTGGACCACTGCCTAGAAATGCAAGCATGCTTCAAAAGCTAGCCGAGCTGGATTTGTCTTACAACTCACTGAATGGTACCATACCATCTTGGGTGTTTAGCCTACCTTTGCTATCTTCAGTGTCGCTCCATCATAACCGGTTAAGTGGTTCTTTTCCTCAATCACCTGTGAATCTCACAAACCTTTCTACCCTTGACCTTTCATCAAATAACATCACCCTTGATGCAGGAATACAAATCACTTTACCTAGCCTAGTAGTTTTACAGTTATCATCTTGTGAACTTAGGGATTTTCCACACTTCTTGAGAAATGTAAAGACACTTATGGTCTTGGATATTTCTAACAATAAGATTCGTGGTCAAATCCCTAACTGGTTTAGCGGCATGAGGTGGGACTCCTTGTTGCACCTAAACCTTTCACATAACTCATTAACAGGCCACCTACAGCCAGTGGCGGAGGCAGCATGTAATGAGGGGGGGCACCCCCCTCGGCGAAAAATATTACTATTTATACATGGTTAAAATTATTTTTTAGGTATGTATAATAGATGTCGAACCCCCTTCCACTAGTTCGTGTGTTTACTTTTCAGATTTTGAACCCCCTTATTAAATTTTCTGGCTCCGCCACTGCCTACAGCAACTTCATTATAATAGTCTAGAGTCTCTTGATCTTAAATTTAACTTCCTTGAGGGGGCACTACCTTCATCCATTTGTAACATGAGCAGTCTTAGCCTTCTGGATTTGTCACACAACTACTTCAGTGACTCGGCTCCGCATTGCTTGGGCAGCATGGTTGGGCTATCTGTGCTAGACTTGAGAAGAAATAGTTTCACTGGGAGTCTTCCACCGTTTTGTGCGCAGAGCAATTCATTGAGAACCATTGTCTTGAATGGTAATCATTTTGAAGGAACTGTCCCGATGTCATTGCTCAAGTGTGATGGTCTAGAAGTCCTTGATGTGGGGAACAATGTTATAAATGACACGTTTCCAGCTTGGCTCGGAACTCTTCAAGAGCTGCAGGTACTTATATTGAAGTCTAACAAGTTCCATGGACCTATAAGTACTAGGCTGAAGTTTGGTTTCCCCGTGTTACGGATTTTTGATCTCTCTCATAATGAGTTTGGTGGTTCACTGCCTGCAGAAGTTTTTGAAAACTTCAAAGGAATGATCAAAACAGATGAGAGAGACAAAGGAGAGATCGAATATATGCAACCACAGTCTTACATCGGATTTAGTAACGTAATGTATGAGGTTTCAGTGAGGTTGGTGATCAAAAGCCAGGAAATTCAGCTAGAAAAAATCACCACAATTATGACAACCATAGATCTCTCAAGCAACCATTTTGAAGGTGTCATTCCGAAAACACTAAAGGATCTCAACTCACTTTGGCTACTCAATTTATCTCATAACAATCTCAAAGGAGATGTACCAATGGAACTGGAGAAATTGAATATGCTTGAAGCGTTAGATCTCTCTTGGAATCGGCTCACTGGAAAGATTCCACAGCAATTGACAAGAATGAACTTTCTGTCATTCTTAAACCTCTCTCAAAATCTCCTTGTTGGACGCATTCCTCAAGGTTCACAATTCAACACATTTGAAAATGATTCATATGGTGGCAACCTTGATTTATGTGGTCCTCCTTTATCAAAGAAATGTGGAACAAGTGATCCATCACATGTTCCTCAACCATTGGAGGAAGAAGAAGAAGATGATGAGTCATATTTTTTTAGTGGATTTACATGGGAATCAGTAGTCATAGGCTACAGTTTTGGACTCGTTGTTGGAACTGTCCTGTGGAGTCTCATGTTTAAATATCGTAAGCCAAAATGGTTTGTGGAATTTTTTGACGGCATCTTTCCCAAGAAAATGAGAAGGCCAAAGAAGAGAGCTCAGAGACGACGGACATAATGAAAGATTGAAAGCAGTTCCTTTAACTGTTTGTATATATGCAGTATAATTGTATGTGTTGTTTTAGCATTTAGCTATTGAAAAGTGTCAATGTCCATTTTTTGGTCATTTTTTACTTCATTATCAAGGGATTGGGCTGAAAATTTGCA

>PGSC0003DMG400002886 LRR-RLPs

ATGTTTCAGAACTTCACAAATCTGGAGATACTTTCTCTCTCTTATGTCAACATCTTATCTCCGATACCTGTGAATATTTCTTCTTCCTTAAGGTATGTGGATCTTGCACATACTAATCTGCGAGGTGTTCTCACAGAGAGCTTTTTCCTTATGCCAAACTTGGAAAGACTCCAATTGAGAGGGAATGATCTTGTCAAAGGAGTTTTTCCAAAGATCCACCGGAGCAACACTCTGTTAATGGAGTTGGATATTTCATCCACAAGCATCTCTGGTGAGCTGCCTGATTCAATTAGCGCCTTCAGTTCCTTGAATATATTGTACCTCGAAGGATGTCAATTCTCTGGTTCCATTCCCGATTCCATAGGCAACCTAACCAGAATTACGGAGTTGGCTTTGTCTTATAATCATTTGGGATAATACATATATTTGACCTTAAACTTGGCTTCAAATTTTAACTTTGACCTCCAACTTTCATAGTGCACAAACAAGCACTTTAACTATCCAATCTTTTAATAAATAAACACGTGATTTTTGGAGCCAAGAGCGTGAAATGCAAACGCTGCCACGTGTATTAGCGAGCCACTCAATGGCTGCCAACTAATTAAACACTTTACACGTCCATTTTAGAACTAAAAAAAAAAATAATTAATTAAGGGTAGAGTTGTCATTTCAGCATTTTTTTCATTGTTGTGTGCCTCAAAAATTACTTCTCACTCGCGCAAAATGCGTGCACATCACGCGTTTTGCCAGGTAAGACACGCGTGTTTATTTATTAAAAGGTTGGATAGTTAAAGTGTCTATTTGTGCACTTTGAAAGTTGGAGGTCAAAGTTAAAATTTGAAGCCAAGTTTAAGGTCNNNNNNNNNNNNNNNNNNNNNNNNNNNNNNGTCCAACTCCTTTTCAGGTGAAATTCCTGATGTTTTCTCTAACCTCCAAGAGTTACGTACGTTACTACTTTCTTATAACAACTTCATTGGTTTGTTTCCCGCTTCACTTTTAAGTTTGACACATCTTGAATACTTAGGATTTACGAGTAATTCCCTATCTGGCCCACTGCCTAATAATGTGAGCATGCTTCAAAAGCTAACCAAACTGAATTTGTCCTACAACTCAATGAATGGTACCATACCATCTTCTGTGTTTAGCCTCCCTTTGCTAGGTCAGTTGTGGCTCGATAACAATAGATTTAGTGGATTACCCGATGAGCTCAAAACAAACCCAACATTAGCACGCCTGGATTTAAGCCATAATCAACTCAGTGGTTCTTTTCTTCAATCACTTTCGAATCTCACAAACCTTTCTATCCTTGACCTTTCATCGAATAACATCACCGTCGATGAGGGAATACAAATCACTTTTCCTAGCCTACAAGATTTGTGGTTATCATCTTGTGAACTGAAGGATTTTCCAGTCAACTAAAGGATTTTCCACACTTCTTGAGAAATGTAAAGACACTTGTGTACTTGGATATTTCTAACAATAAGATTTGTGGTCAAATCCCTAACTGGTTTAGCGGCATGACGTGGGACTCGTTGCAGTTCCTAAACCTTTCTCATAATTCATTAACAGGCCACCTACCACAATTTCGTTACTATAACCTACAGTATCTTGATCTGAAATTTAACTTCCTTCAGGGGTCACTACCTTTATCTATTGGTAACATGAGAAAACTTATCTTATTAGATTTATCACACAACTACTTCAGTGACTCAGTTCCACATTGCTTGGGAAGCATGGCTTCACTAGCGGTGCTGGACTTAAGAAAGAACAATTTCACAGGGAGTCTTCCCCCATTATGTGGACACAACACTTCATTGAGTACCATTGTCGTAAATGGTAATCGATTTGAAGGACCTGTTCCTGTATCGTTGCTCAATTGTAATGGTCTACAAGTCCTTGATGTGGGGAACAATGCTATAAATGACACGTGTCCAGCTTGGCTCGGAAGTCTTCAAGCTCTGCAAGTCCTTATATTAGAGTTGAACAAGTTCCATGGACCTATAAGTACGTGTCAGACTAAGTTTTGCTTTCCCAATTTGCGAATTTTTGATCTTTCTCGTAATGATTTCAGTGGCTCACTTCCTGCAAAAGTTTTTGGAAACTTCAAGGCAATGATCAAATTAGATGACGAAGACACAAGAGAGATCAGGTACATGGAATCTATGTTGAATTTGTCATTTGTCAGATCGTATGAGGATTCAGTGAGTTTGGTAATCAAAGGGAAAGATATTGAGCTACAAAGAATCAGCACAATTACGACAACCATAGATCTCTCAAGCAACCATTTTGAAGGTGTCATTCTGAAAACGTTATAG

>PGSC0003DMG400002887 LRR-RLPs

TTTGGTTTCACTTGATCTTTCTGATAGTTATGAATTACAACTTGATGAGAGAACATTTGAAACAATGCTTCACAACTTTACAAATCTGGAGTTACTAGCTCTCCCTCTTGGTGACATCTCATCACCGATACCTGTAAGTATTCATCCCAATAGCAGCCTCTTCCAGCTTCATCATCTCCACACACTAAACCTTGATAATAATTACTTGAATCCTTCTTCTTCGATCCCACATAACATTGGCCGATTGAGGAATTTGAGGCATCTCAACCTTTCTGATGCTTNNNNNNNNNNNNNNNNNNNNNNNNNNNNNNNNNNNNNNNNNNNNNNNNNNNNNNNNNNNNNNNNNNNNNNNNNNNNNNNNNNNNNNNNNNNNNNNNNNNNNNNNNNNNNNNNTACAAATCTGGAGNNNNNNNNNNNNNNNNNNNNNNNNNNNNNNNNNNNNNNNNNNNNNNNNNNNNNNNNNNNNNNNNNNNNNNNNNNNNNNNNNNNNNNNNNNNNNNNNNNNNNNNNNNNNNNNNNNNNNNNNNNNNNNNNNNNNNNNNNNNNNNNNNNNNNNNCAATCCCAAATGGCATTGGCCGATTGAGGAATTTGAGGCATCTAAAACTCTCTGGCTTTGATGGGAAAATCCCAACAGAAATCTCATACCTTTCCAATTTGGTTTCACTTCATCTTTCTGGTTATGGATTACAACTTGATGAGAGAACATTTGAAGCAATGTTTCAGAACTTGACAAATCTGGAGGTACTTTCTCTCTCTGAAGTCAACATCTCATCTCCGATACCTGTGAATATTTCTTCTTCCTTAAGGTACCTGGATCTTAGTCATACTAATCTGCGAGGCATAATTCCCCATGGTTTCTCTAATCTCCAAGAGCTAGTTGAATTGGATCTTTCTAATAACAACTTCACCGGTCCATTTCCCTCTTCAATTTTAAACTTGACAAGTCTTCAATACTTAGATTTGTCTCACAACTCACTGAATGGCACAATACACTCTTGGGTGTTTAGCCTCCCTTCCTTACTTGATTTGAAGCTCCATCATAACCAATTCAGTAGAGTGGCTGATGAGATCAAAACAAACCCGACATTAGTGACTTTGTATTTAAGTCATAATCAATTCAATGGTCCTTTTCCTCGATCACTTGTGAATCTCACAAGCCTGGCCTTGCTTGACTTTTCATCAAATAACATCACCGGCGATGTGGGAATAAATATCACCTTTCCTAGACTATCAGCTTTGTTATTATCATCTTGTGAACTGAAGGATTTTCCACACTTCTTGAGAAATCTAAAGACACTTCAGTTCTTGGATATTTCTAACAACAAGATCCGTGGTCAAATCCCTAACTGGTTTAGCGGCATGAAGTGGGACTCTTTGGAACAACTAAACCTTTCACGTAATTTTTTGACAGGCCATCTAGGAGAATTTCATTATTATAGTCTTGAGTCTCTTGATCTTAAATTTAACTTCCTTCAGGGTCCACTACCTTCATCCATATGTAACCTAAGAAGCCTTAGAATTCTGGATTTATCACGCAACAACTTCAGTAACTCAATTCCAAATTGCTTGGGAAGCATGGCTAAGCTAACGGTGTTGGACTTGAGAAGGAACAATTTCAGTGGGAGTCTTCCACTGTTATGTACTCAAAGCACTTCATTGATGACCATTGTCTTGAATGGTAATCAATTCGAAGGATCTGTCCCTGCGTCATTACACAACTGTGTTGGTTTAAAAGTTCTTGATTTGGGGAACAACGCTATAAATGACACATTTCCAGCTTGGCTTGGAACTCTTGAAGAGCTGCAAGTCCTTATATTAAAGTCGAACAAGTTCCATGGACCTATAAGTGCTAGGAAGAAGTTTTGCTTTCCCAGGTTGCGGATTTTTGATCTCTCTCATAATGCATTCAATGGTTCACTGCCTGCAGATATTTTTAGAAACTTCAAGGCAATGATAAAAAATGGCACAGACAAAGGTAACATTACATATATGGAAACATCAGTTTTCAGGTCATTGGTTGATTCATCGATCAAAGACTGGACTATGTATAAGCTAGAATTGGCCATTGATGAAGTGTACAAGGATTCAGTGAGGTTGATGATCAAAGGAAACAATATTGAGCTAGAAAGAATCAGCACAATTGTTACAGCCATAGATCTCTCAAGCAACCATTTTGAAGGTGACATTCCGAAATCACTAAAGGATCTCAGCTCACTTTGGCTACTCAATTTATCCCACAATAATCTCAAAGGTGATATTCCAATGGAATTGGGGCAATTGAATTCACTTGAAGCCTTAGATCTCTCTTGGAATCGGCTCACTGGAAAGATTCCACAGGAATTGACAAGAATGAACTTTCTGGCGTTCTTAAACCTCTCTCAAAACCAGCTCGTTGGACGCATTCCTCAAGGTTCACAATTCAGCACATTTGAAAATGACTCGTATGGTGGCAACCTTGATTTATGTGGTCCTCCTTTGTCAAAGCAATGTGGAACGAGTGATCGCTCACATGTTCCTCAATCACTGGAGGATGAAGCAGAAGAAGATGAGTCATATTTCTTCAGTGGATTTACGTGGGAATCAGTAGTCATAGGCTACAGTTTTGGACTAGTTGTTGGAACTGTCATGTGGAGTCTCATGTTTAAATATCGTAAGCCAAAATGGTTTGTGGAATTTTTTGAAGGCATTTATCCCAAGAAAATGAGAAATAGGCTACAGTTTTGGACTAGTTGTTGGAACTGTCATGTGGAGCCTCATGTTTAATTATCGTAAGCCAAAATGGTTTGTCGAATTTTTTGACGGCATCATGCCCCACAAAAGGAGAAGGCCAAAGAAGAGAAGGCTGAGACGGCAGACATAATGGAAGATTGAAAGATTGAACTTTTCTGTTTCTGTTTTTTCTGTTTGTTGTTTGTATATTCTTCATACAAACTTAACTGTATGTGTTTTTTTTTAAATTAATGTATCTTTTTCGGAGACTTGTTTGTGGTGGAAG

>PGSC0003DMG400002906 LRR-RLPs

CGAGTGTCCAACCTTGTCAAGCCCACTATTAAAAATGGAAAAAACAACAAATAGACCAGGCCCTTTCGTAAACCTGATATTCTAAGATAGGATTTTCGCCGCTATCTCCGCTTCTTCGCCTTAGCTCTCTTCACTGTTTCTGTACATGAACAACAATGTTGCCTCCATGTTACTGACCTTTATGCTTTTTTCCAGTTGTGTTCAAAATTTTCTTCCTCTTGCTTTCTTTTCTTCTTCTCTCCCTCTTTTGGTCTCTGATTCCCATCATTCTGCAAGTGATTTTGCGATGGAGTGCAAGGGTAGTTCAATTCCTCCATCTCCAGGTTAAAAGCTCCATCTTGACTCCAGATCCTCCCTTTTGCTGAGATCGCTTTGTTGACTGGAATAGCAATAGCATTTTCCCTGTTTTGTTGCCTTCTAACTTCTATGTAATTTTGTTGTTTGTGGAAAAATATATGTTGTAGCCTTCCCTTGTCCCCTATCCCCTTAAGGATGATACTCTTAAATATGGGCATAGTAATTTTTCATTATTTATGAGTCTCCTCGCAAGTGTCTCCCTGTTAAAATTTGTGAATGTGCAGCTCCCTCGGTCAATAGCAAAGTTTGCCACATGATCTGCTAGCTTGTTTCCTTCCCTTAGAATATGTTGGAAAATAACTTGTTTATCTTGTAGACATGATTGAATCTCTGTCACCATTTCTGCAATTATCCATGGACATTCCCATTTCCTTTCTAATACCTTGTACAAAAGCATTGAGTCAGTCTGTATGATTATTTGATTATGTTGTAACTTCTTGCAATGCTTGGATTCCTCCAGAATAGCTTTAGCTTCCGCGACTCTATTAGTAGTATTTTCTATACTTGCTCCTTCCACGTATTCTACATCTCCTTTAGCATTCCTCAAAACGAAGGCATAGGAACTAAGACCCGGATTTCCTCTAGAAGTTCCATCTGTATTGTATTTTACCCACCCATTTGGTGGGAATTCCCATAAGACCCTTTTCACTTTCATTTTAGGTTTGTAGCTTGTCCATTCTATGAGCATATCAGGCCAGTTAACAGGACATGTCATATAGGGTTTCCTGAGTCTTACCAGCATATACATGTTCCTGGTGATATTGTGAATGACTCTTGCTACTGATGTCTTTTTTCCTTCATGCTTCATCCTGTTTCTCTTTCTCCACAATTCCCATATTATAAAACTAGGAAGAGCTCTGTAAAAAGGTCTCATATTTGTCCCAATTTTTGTCCCCCACCACAACATTATAACTTCTCTTAGTTGGAGACCCTCCATGTTAAGACCTGTAAAAGAACAAAAATAGGACCAAGTCCTGTTAGCTGAATCAGATTTCAGAAACACATGTTGAGGTCTTTCACAACACCAGCATTTGGATGGGCCTTCAAGACCCCATCTTCTCATTATGTCATCAACGGGGATTTTAAATTTCCATAATCTCCACATTATGAAAGCCATTTTAAATGGTACACCCTTACTCCACATCCATTTATAAATTTTGTTTTCCTCCGCCTTATGTCTGATATAATGCCAAGCAGATTTAACTGAGAAACTTCCCCTTGTCTCTAGCATCCAACAAGGTTTATCCATTGCTGTTTCTTCTTTTGGAGGATGAATATTCTGCAGAATATGGTTCACCAATTCCTGTGGGAGTATCTCATGTAGTACCTCTGTATCCCACTCCCCTTGTCTAGTTAAATCCCTAACTGTTTTATATGTATCATCACATTGAAATTCTTCTCCAGTAATAGTGTACAAATCCCCTAGGCCCGTCCAGTTATCATGCCAGACTGATACTGATCCTTTCTCCATTTTCCATAAAATTTGGTTTTCAATGAGATCTCTAGCTTTAAGCATTTTTTCCCATAAAATTTGGCTCGGAACTCTTCAAGAGCTACAGGTCCTTATATTAAAGTCGAACAAGTTCCATGGACCTATAAGTATGTGTTAGACTAAGTTTTGCTTTCCCAAGTTGTGAATTTTTGATCTTTCTCGTAATGATTTCAGTGGCTCACTTCCTGCAAACGTTTTTGGAAACTTCAAGGCAATGATCAAATTAGATGGTGAAGACACAGGAAATATCAAGTACATGGAATCTGTGTTGAATTCGTCATTTGTCACATCGTATGAGGATTCAGTGAGTTTGGTAATCAAAGGGCGGGATATTGAGCTACAAAGAATCAGCACAATAACGACAACCATAGATCTCTCAAGCAACCATTTTGAAGGTGTCATTCCGAAAACACTGAAGGATCTCATATCACTTTGGCTACTCAATTTATCTCATAACAATCTAAGAGGTGATATTCCAATAAAACTAGGACAATTGAGTATGCTTGAAGCTTTAGATCTCTCTTGGAATCGGCTCACTGGAAAGATTCCACAAGAATTGACAAGACTGAACTTTCTGGCCTTCTTGAACGTCTCTCAAAATCATCTCGTCGGACCAATTCCTCACGGTCTACAATTCAACACATTTGAAAATGACTCGTATGGCGGCAACCTTGATTTATGTGGTCCTCCTTTATCAAAACAATGTGGAACGAGTGATTCATCACATGTTCCTCAACCATTGGAGTCTGAAGAAGATGAAAGCGAGTCATATTTTTTAGTGGATTTACATGGGAATCAGTAGTCATAGGCTACAGTTGTGGACTAGTTGTTGGAACTGTCATGTGGAGCCTCATGTTCAAATATCGTAAGCCAAAATGGTTTGTGGAATTTTTGATGGACTCATGCCTCACGAAAGAACAAGGCCGAAGAAGAGAGCTCAGAGACGACGAACTTAATGGAAGATTGAGAGGGGCCAAGTTGTCTTTGTTTGATTGAACTTTTTTGGTTGTGTGTTTTAATTTTCTGTTTGTTGTGTTGTATATGTTTGATAGAAACATAATTATATGCATGTGTGTTATTT

>PGSC0003DMG400002908 LRR-RLPs

ATGAAACTAGGACAATTGAGTATGCTTGAAGCTTTAGATCTCTCTTGGAATCGGCTCACTGGAAAGATTCCACAGGAATTGACAAGAATGAACTTTCTGGCCTTCTTAAACCTCTCTCAAAATCATCTCGTCGGACCAATTCCTCAAGGTCGACAATTCAACACATTTGAAAATGACTCATATGGTGGCAACCTTGATTTATGTGGTCTTCCTTTAACAAAGAAATGTGGAACGAGTGATTCATCGCATGTTCCTCAACAATTGGAGTCCGAAGAAGAAGGCGAGTCATATTTTTTTAGTGGATTTACGTGGGAATCAGTAGTCATAGGCTACAGTTTTGGACTAGTTGTTGGAACTGTCATGTGGAGCCTCATGTTTAAATATCGTAATCCAAAATGGTTTGTGGAATTTTTTGATAGACTCATGCCTCACAAAAGAAGAAGGCCAAAGAAGAGAGCTCAGAGACGACGGATTTAA

>PGSC0003DMG400002910 LRR-RLPs

ATGACTTCAATCCCACATAAAATCATTGGCCGATTGAGGAATTTGAGGCATCTAAAACTCTCTGGCTTTGATGGGAAAATCCCAACAGAAATCTCATACCTTTCTAATTTGGTTTCACTTGATCTTTCTGGTTATGGATTAGAACTTGATGAGAGAACATTTGAAACAATGCTTCACAACTTTACAAATCTGGAGTTACTCTCTCTTGGCGACATCACATCACCGATACCTGTAAGTATTCATCCCAATAGCAGCCTCTTCCAGCTTCATCATCTCCACACACTAAACCTTGCTGGCAATTACTTTTATCCTTCTTCAATCCCACATAACATTGGCTGATTGAGGAATTTGAGGCATCTAAAACTCTCTCACTTTCAAGGGAAAATCCCAACAGAAATCCCATACCTTTCTAATTTGGTTTCACTTGATCTTTCTGGTTATGGATTAGAACTTGATGAGAGAACATTTGAAACAATGCTTCACAACTTCACAAATCTGGAGTTACTTTCTCTCTCTTATGTCAACATCTCATCACCGATACCTGTAAGTATTCATCCCAATAGCAGCCTCTTCCAGCTTCATCATCTCCACACACTAAACCTTGTTAACAATTTCTTTAATCCTTCTTCGATCCCATATGGCATTGGCCGATTGAGGAATTTGAGGAATCTGATACTCGCTGACTTTGATGGGAAAATCCCAACAGAAATTTCATACCTTTCCAATTTGGTTTCACTTCATCTTTCTTCTTCTTATGGTAATGAATTACAACTTGATGAGAGAACATTTGAAGCAATGTTTCAGAACTTCACAAATCTGGAGATACTTTCTCTCTATCAAGTCAACATCTCATCTCCGAATATTTCTTCTTCCTTAACGGAGTTGGATATATCATACACAGGCATCTCTGGTTCCATTCCTGATTCAATTGGCACCTTAAAATCCTTGAATATCTTGATCCTCTATAGATGTCAATTCTCTGGTTCCATTCCTGATTCCATAGGAAACCTAACACAAATTACGAAGTTGATTTTATCTCATAATCATTTCACTGGCCATATTCCTTCCACAATCTCTAAATTGAAGCACCTCACACGTTTATATCTTTCGGACAACTCCTTTTCAGGTGAAATTCCTGATGGTTTCTCTAACCTCAAAGAGCTACGTTATTTATATCTTTCTCACAACTCCTTTTCAGGTGAAATTCCTGATGTTTTCTCTAACCTCAAAGAGCTATATTATTTAGATCTTTCTGAAAACAGCTTCACTGGTCCATTTCCCGATTCAATTTTAAGTTTGACACAACTTCATACCTTAGACTTGTCGAGTAATTCCCTATCTGGCCCACTGCCTAGCAACCAAAGCATGCTTCAAAAGCTAACTTTTGTGGATTTGTCATACAACTCACTGAATGGTACCATACCATCTTGGGTGTTCAGCCTACCTTTGCTACATTCAGTGTCGCTCCAACATAACCGATTCAGAGGACTAGCCGATGAAGTGATCAAAACAAACCCAACATTAAAACAACTGCATTTAAGCAATAATCAACTCAGTGGTTCTTTTCCTCAATCACTTGTGAATCTCACAAACCTTGAAACCCTTGGAATTTCATCAAATAACATCACCATTGATGAGGGAATGAATCTACAGTATCTTGATCTGAAATTTAACTCCCTTCTGGGTCCACTACCTTTATCCATTTGTAACATGAGCTCGCTTATCTTATTAGATTTATCACGCAACAACTTCAGTGACTCAGTTCCACATTGCTTGGGAAGCATGGCTAGTCTAACGGTGCTAGACTTAAGAAGGAACAATTTCACAGGGAGTCTTCCCCCATTATGTGCACACAGCACTTCGTTGAGTACCATTGTCATAAATGGTAATCAATTTGAAGGACCTGTACCTGTGTCGTTGCTCAAGTGTGGTGGTCTAGAAGTACTTGATGTGGGGAACAATGCTATAAATGACACGTTTCCATCTTGGCTTGAAATTCTTCAAGAGCTGCAGGTCCTTATATTAAAGTCGAACAAATTCCATGGACCTATAAGTATATGTCAAACTAAGTTTTGCTTTTCCAAGTTGCGAATTTTTGATCTTTCTCGTAATGATTTCAGTGGCTCACTTCCTGCAAATGTTTTTGGAAACTTCAAGGCAATGATCAAATCAGATGGCGAAGACACAAGAGAGATAAAGTACATGAATTCTGTGTTGTATTCGCCATTTGTCACATCGTATGAGGATTCAGTGAGTTTGGTAATCAAAGGGCACGATACTGAGCTAGAAAGAATCAGCACAATTACGACAACCATAGATCTCTCAAGCAACCATTTTGAAGGTGTCATTCCGAAAACACTAAAGGATCTCAGATCACTTTGGCTACTCAATTTATCTCATAACAATCTCAGAGGTCATATACCAATGGAATTGGGGCAATTGAGTATGCTTGAAGCTTTAGATTTCTCCTGGAATCGGCTCACTGGAAAGATTCCACAGGAATTGACAAGATTGAACTTTCTGGCCTTCTTAAACGTCTCTCAAAATCATCTCGTTGGACCAATTCCTCAAGTTCTACAATTCAACACATTTGAAAATGACTCATATGGTGGCAACCTTGATTTATGTGGTGTTCCTTTAACAAAGAAATGTGGAACGAGTGATTCATCGCATGTTCCTCAACAATTGGAGTCCGAAGAAGAAGACGAGTCATATTTTTTTAGTGGATTTACTTGGGAATCTGTAGTCATAGGCTACAGCTGTGGACTAGTTGTTGGAACTGTCATGTGGAGTCTCATGTTTAAATATCGTTAA

>PGSC0003DMG400002914 LRR-RLPs

AAAAAAGAACCATTCTTTGTTTCTTCTCTCTCTAACCTCAAATCTTTCTCTCTCTATTTTTGCCCTACTTTGTTGGGGTTCTTGAAAAAAAACTGGATTTTTGAGGGTTCTTGAAAAAATTTGGATTTTTTCATCTGGAATGCAAATCTTGGTTAAAGATATAATCTGGGTTGTGTTTTTCTTGGTTCAAAATCTGATCTTTTTTGCTTTACAGTTCGGGATTTCTTCTGAAGAACAGACATGCCTACTCTTGTTAATTACAGTGGTGAGTTTCAAAAAACTTCATTTCTTGTTTTTATAAGTTATGGGGTTGTTCATATTTCAATGAATTTTGAAGTTCAGTTACAATTTTTTGATTTTTCATCATGAATTACTGAGATGGAGATGGATTTGTTATGTTTAGATTCATATAAAGTTGTTTATCATGAATAAAGTTTATAACTTTGTATAGGTAGATTATTGTTGAGATGAATATGAGCAATAATTTATTGAGATGAAGATGAATTCTTTTAAGGTTTAGATCCATGAATATAGTCTATAACTATGTATAGGTGCATGATTGTTGATAACTTGAGATGAATATGGTTTATTAATTTACTGAAATGAAGATGATCTTTTAGGTTTTAGATCCAATCATGTTTTTATTTAAAAAAGGAAGTTGGTTTATGAGATAATAACTAAAAGTTGAGTGACTATCTAAGAAATTAACTCATATAACTTTGTATAGGTGCATGATTGTTGAGGTGAATATGAGCATTAATTTACTGAAATGGATAAAAAAGGGAAAAAAGTTATGATAAGACTAGATCCTTTGGCTTTGAAGATTAATTTCTGTGTTTGATAGATCTTTTTTTGTCTTATGTTCTGAATCTATATTAGAGATTGGTAATATATTGAAGAAACTGTGGATTACTCTTTTGGGTTCTGTATCTAGAATCGTGTATTTATCTGATCTGATCTGAAGATTTATGTACATAATAGGTAGATCTAAAGCTTCAATTCACATGTGATAGTTGTTTCTCTGTTATACTTGAAGTTGAAGTGAAAATTCATAAGAAATGGGCATATGGATGAATACTTGTAAAATTCTTTCATTAGACAAAATTTAGTATGCTTGCTTGTTGATTTAAGGGAAAATATAGCTGACAATGTATTTGCTCCTTGGATTTCAGGTGATGATGAGTTCTATTCTGGTGGATCGTTTTGTTCTGCAGATTTGGGTCTCATGTTGTCTTTGGGTCATGCTGATGTTTACTGTCCTCCTAGGAAGAGGGCACGCATTAGTGGCCCGTTTGTTGTCGAAGACCGGTCAAAGGATCCATCTCTTGAAATTCTTCCTGATGAATGCCTATTTGAGATCCTCAGACGATTGCCCGGTGGCCGAGAAAGGGGTGCAGCAGCTTGTGTTTCTAAGCGCTGGCTTACAGTGTTGAGTAGCGTGAGGAACTCTGAAATCTGCAGGAGCAAGAGCTATAACAATCTAAATGACGCAATAATGATCTCTAAAGATGAAGATCTTGAAGTTGAATGTGATGGATACCTTACTAGGTGTGTGGAAGGGAAGAAAGCTACTGATGTTAGACTTGCTGCAATTGCAGTTGGGACGTCTACCCGTGGTGGACTAGGAAAGCTTTCTATCCGAGGGAGTAACTCAGTTCGTGGTATTACTAATGTCGGTCTATCTGCTATTGCACATGGGTGCCCTTCTCTTAGGGTCCTATCATTGTGGAATGTTCCTTCTATTGGGGATGAAGGTCTCTTGGAGGTTGCAAGGGAATGCCGTTCATTAGAAAAGCTAGATCTGAGCCACTGTCGTTCAATCTCCAACAAGGGTCTTGTTGCAATAGCAGAGAATTGCCCGAGCTTGACTTCATTGACTATTGAATCCTGTCCAAAGATTGGAAACGAAGGCTTGCAAGCTATTGGAAGATGCTGCACCAAACTACAGTCTCTTACGATTAAGGACTGCCCACTTGTTGGAGATCAGGGAGTTGCTAGTCTTCTGTCATCAGGTGCTTCAATGTTGTCAAAAGTGAAACTACATGGTCTAAACATCACAGATTTCTCGCTTGCTGTCATTGGTCACTATGGCAAGCTGATTACCAATCTTAATCTTTGTTCACTTCGTAATGTGAGTCAGAAGGGATTTTGGGTCATGGGTAATGCTCAAGGCCTCCAGTCTCTGGTTTCTTTGACAATCACCTTATGCCAGGGAGCCACTGATGTCGGTCTTGAAGCTGTTGGAAAGGGATGCCCAAATCTCAAACATATGTGCATTCGCAAGTGTTGCTTTGTTTCTGACGGTGGGCTTGTTGCTTTCGCTAAAGAGGCTGGATCTCTTGAGAGTTTGATCTTGGAGGAGTGCAACAGAATTACCCAAGTAGGTATCCTTAATGCAGTTTCAAACTGCAGGAAGTTAAAGTCTCTTTCCCTAGTGAAGTGCATGGGAATCAAGGATCTAGCTCTACAAACTTCCATGCTGTCTCCATGTGAATCTCTTCGGTCCTTGTCTATCCGAAGCTGTCCAGGATTCGGGAGCACTAGCTTAGCTATGGTCGGTAAGCTCTGCCCCAAGCTGCATCAATTAGATCTCAGTGGGCTTTGTGGAATAACCGATGCTGGTCTTCTCCCACTCTTGGAGAACTGTGAAGGACTTGTTAAGGTGAATCTTAGCGACTGCCTGAACTTGACAGACCAAGTGGTGCTTTCATTGGCTACGCGACATGGAGAGACCCTTGAATTGTTGAATCTCGATGGATGCAGGAAGGTTACCGACGCAAGTTTAGTGGCAATTGCAGATTATTGCCCATTACTTAATGATCTTGATGTTTCTAAGTGTGCAATAACTGATTCTGGTGTAGCTGCTTTATCCCGTGGAGTGCAAGTGAATTTGCAGGTCCTTTCTTTATCAGGTTGCTCCATGGTATCAAACAAGAGTGTCCCTTCTCTTAAAAAGTTAGGAGAGAATCTACTCGGCTTGAATCTCCAACATTGCTCTGTTAGTTGCAGCTCAGTCGAGCTTCTTGTGGAGGATTTGTGGAGGTGTGACATCCTCTCCTAAACCAAAAGGGCACAAGGAAATGCATCAAAATGGTCAAAAAGAATAAGCAGAAAGCTATAATGGTTCATTCAGCCTGGCAATTACAGAAGCAAACATGAGTGTTCTTTTTTTAGCCCGTATATTCGTGGTTATCTAATACTTCAGTGTCTCGGTTGTTTTCCAGGAGGCTATGGCATCACTTTTTTCCTCCATTTTTTGCAGGCTTCGTCAAATCGAAGCCTGTTTCATCCATTTTTGGAGACAACTCCATCATGCCATGGTTTCCTGAGAATACAGTCACCTAGTACTGAGATTAAAATTTTCACTCATGTTGCTCCTCGAAAGAATCAGGCCGTTTGAATCATTCAGCTACAACATCGACAGACAATATCTATCGATAGTCTTTTAGTGTTTCATTGTAAGTATTCTACTGTGTGTCTTTTAGACTTTAGTTGCTTTGAGCAAATTTCTGTTGTTGTTGTGCCTGTTATTCTAGGGGCTTGGCCATTACAACCTTACCTAGGTTGTTTTTTTTAACCAGTCCCTTGTTTTTTGATAGGATTATCAATTTCCTATTTTTTACCAGGTTCCACATATTCAGATTTGTACTCTTGCAACTATGTTTTACATATATAATGAAGTACTAGTTTGTTTTGAA

>PGSC0003DMG400002937 LRR-RLPs

GAGTGATTTGTAAGAAGAGAACATCCATCCTTTTTTTGGCTGAATACGAAAAAGAAAGATGGGACATTTCTTTTTCTTTTATTCATTTCTGTGTGTTGTGTTGCTAATAAGTGAATGTTTTTGTTCTTCCTTTGAGCATCATCTTTGCTCTCCCACTGAAGCTTCTGCTTTGCTTCAGTTTAAGCAACACTTTGAGATTATATCCTACCCTAATTCCCTATGTCCTATTCATTTCCCAAGAACAAAGTCTTGGAATGAGAGTAGGGACTGCTGCATTTGGAATGGAGTCACTTGTGATATGTTAACGGGTCGTGTTATCGCGTTGGACCTTAGTTGCAGTCAGCTTTGGGGAAGTATTCATCCCAACAGCAGCCTCTTCCAACTTCATCATCTCCACACACTAAACCTTGCTTACAATAACTTCAATTATTCTTCAATCCCACATAACATCGGCCAATTGACGAATTTGAGGCATCTCAACCTCTCTGAGTCTGGCTTTGATGGGCAAATCCCAACAGAAATATCATACCTTTCGAATTTGGTTTCACTTGATCTTTCTGGGTATGAATTACAACTTGATGAGAGAACATTTGAAACAATGCTTCACAACTTCACAAATCTGGAGCTACTTTCTCTCTCTCAGGTCAACATCACATCTCCGATACCTATGAATATGTCTTCCTCCTTAAGGTACGTGGATCTTGGATTCACTAATCTGCGAGGTGTTGTCCCAGAGAGTCTTTTCCTTCTGCCAAACTTGGAAAACCTCAAATTGGGTGTGAATCTTCTTCTCAAAGGAGTTTTACCAAAGATCCCCCTGAGCAACACTCTGTTGGAGTTAGATATTTCACTCACAGGCATCTCTGGTTCCATTCCTGATTCAATTGGCACCTTAAGTTCCTTGAATATCTTGAACCTCGAACTATGTCAATTATCTGGTTCAATTCCTGATTCCATAGGCAACCTAACACAAATTACGGAGTTGATTTTATCTACTAATCATTTCACTGGCCATATTCCTTCCACAATCTCTAAATTGAAGCACCTCACACGTTTGGATCTTTCATCTAACTCCCTTTCAGGTGAAATTCCCAACGTTTTCTCTAACCTCCAAGAGCTACGTTATTTACATCTTTATGATAACAGGTTCATCGGTTCATTTCCTTCTACAATTTTAAATTTGACACATCTTGAAAGCTTAGCCCTGTCGACAAATTCCCTATCTGGCCCACTTCCAGCTAATAAAGCGAGCATTCTTCAAAAGCTAACCCAACTGGATTTGTCTGACAACTCACTGAATGGTACCATACCGTCTTGGGTGTTTAACCTCCCTTTAGCATCTTCATTGTGGCTCCATAATAATCAATTTAGTGGACTACCCGATGAAATCAAAACAAACCCAACATTAGAATACCTGTATTTAAGCAATAATCAACTCAGTGGTTCTTTACACCAATCACTTGCGAATCTCACAAACCTTTCTACCCTTGACCTTTCAACAAATAACATCACCGGTGATGCAGGAACAAATATCACTTTTCCTCGCCTACAAGTTTTGCAGTTATCATGTTGTGAACTGAAGGATTTTCCATATTTCTTGAGAAATGTAAAGACACTTGAGGTCTTGGATATGTCTAAGAATAAGATTCGTGGTCAAATCCCTAACTGGTTTAGCAGCATGAGGTGGGACTCGTTGGAGTTCCTAAATCTTTCACATAATTCGTTAACAGGCCACCTACCACAATTTCATTACCATAGTCTAGAGTATCTTGATCTTAAATTTAACTCCCTTCAGGGTCCACTACCTTCATCCATTTGTAACATGAGCAGCCTTAACATTCTAGATTTATCACGCAACAACTTCAGTAACTCGATTCCAAGTTGCTTGGGAAACATGGCTAGTCTAACAGTGCTGGACTTAAGAAGGAACAATTTCACAGGGAGTCTTCCCTCATTATGTGCGCAGAGCACTTTATTGAGAACCATTGTCTTGAATGGTAATCGTTTTGAAGGAACTGTCCCGATGTCGTTGCTCAAGTGTGATGGTCTACAAGTCCTTGACGTGGGGAACAATGCTATAAATGACACATTTCCAGCTTGGCTCGGAACTCTTCAAGAGCTGCAGGTCCTTATATTAAAGTCGAACAAGTTCCATGGACCTATAAGTACTAGGAAGAAGTTTTACTTTCCCAGGTTGCGGATATTTGATCTCTCTCAAAATGAGTTCAGTGGCTCACTGCCTGCAGAAGTTTTTCGAAACTTCAAAGCGATGATCAAATTAGATGACAGAGAGGAAGGCGAGATCAAATATATGAGTCAATATTTGAATGGACATTCAATATATGAGGATTCAGTGAGGTTGGTGATCAAAGGCCAGTATATGGATCTAGAAAGAATCAGCACAATTATGACAGTCATAGATCTGTCAAACAACCATTTTGAAGGTGTGATTCCGAAAACACTAAAGGATCTCAGCTCACTTTGGCTACTCAATTTATCCCATAACAATCTCAAAGGTGATATCCCAATGGAATTGGGGCAATTGAATATGCTTGATGCCTTAGATCTCTCCTGGAATCGGCTCACTGGAAAGATTCCGCAGGAATTGACAGAATGAACTTTCTGGCATTCTTAAACCTCTCTCAAAATCATCTTGTCGGACCAATTCCTCAAGGTCCACAATTCAACACATTTGGAAATGACTCGTATGGTGGAAACCTTGATTTATGTGGTCCTCCTTTATCAAAGAAATGTGGAACGAGTGATCCATCACATGCTCCTCAGCCATTGGAGTCTGAAGAAGATGAAGATGAGTCATATTTCTTAAGTGGATTTACGTGGGGATCTGTAGTCATAGGCTACAGCTTTGGACTCATTGTTGGAACTGTCATGTGGAGCCTCATGTTTAAATATTGTAAGCCAAAATGGTTTGTGGAATTTTTTGAAGGCATTTTCCCCAAGAAAATGAGAAGGTCAAAGAAGAGAAGGCCGAGACAGC

>PGSC0003DMG400002954 LRR-RLPs

ATGACTCAAGGTATTAACTTCTCCCTTTTTCCCTTTTAACTCTTTTCTAATAGATAGTAAAGGGTTGCTGCCGTTGAAGATAGAATCAGTGTTCTACCAAGTAACGTTATAGATCATATCCTCGAGCTCTTGATGGTTCATGAGGCTGCAAGAACTAGTATTTTGTCCAAAGATTGGAGATATAGTTGGGCCAAGCTTCCCAATTTGGTGCTTGATAAGCAATTCTCCATGAAATTAGCTAAAAAATCTCACAATGTCTTCAAAGAAACAATAGATATGATTCTCTTACAACATATTGGAGATATTGTGAAGTTTTATCTTGATGTTTCAGGAGTACATTTGTCTACATATGCAGATATTGATAGATGGATGCTTTATGTCACCAGAAATGGTGTCAAGAAGCTAAAGCTTAACATGACAAAAGATGGTACTTACAAACTACCTTCTTATATATTTAATTGTCCAACACTGACTCATTTGAAACTCTTCAACTGCAACTTCAAATTACCAAATTCTTTTCTTGGCTTACAGACTCTTACCGTCCTTCGGTTGGAAAAAATAACCTTTGTGCCTGCAATAGAGTTTTATACTATCAATGTGCCCCTTCTTGTCAAACTAACCATAGATTTATGTGATGGTACTCAATACCTCAACATTGTGTTCTCATCTGAGTTGAAATCCTTGGTTGTTCATGAGAGTCACTATAATCTTGACCTAAGTAACAACTTCTTTAAGAAGTGCAAAAAATTGACAGATTTATACCTTGTGATTGATAATCCAATACCTGCGGATAATAGATTAACTTTGGAAAAGCTTCTTTTTAGCTCGCCTACACTTGAGACACTTTGTTTGGGTAAATTTGAACTTGAGGTAAGAATCTAACGTGCTGTTTTACTTAGATATCTTCTTTGAGTGTGATCCTTCTTATTGTATAATGTTCCGTTAAAGCTTTTGACTGCAGGTGTAGTCCCAAAGAGGCTTCCTTGTATACTCAACTGCTTGTGGCATCTTGATTTAGATGTAAACTTCAATAAATTGGGTCAGACTTCTTACACTCTGGAGTTGATTAAAAGTTTCCCCAATTTTCGTGACCTTGAGATTTGGGTAAGAAATAGCAAATATATGTGGTCATACATTCTATATATCAATGGTCTTGTTCGTTTTTGCATTTCACGTACTTAAGAAGCTAAAATGACCTTGTATACCATCAGGACAAAACTACCGATGACATAGTTGAAGGAGTTATGAAATATCTAGACACACCATCATGTTTGGACAAAACCCTCAACAAGCTTGAAGATGTTTCAATCCATTCCTTTAACTATTCTAAAGTTGTACTATATTTCGTAAAATTATTGTTTGCTCATGCTTCATCTTTGTCAAGGATGAGCATCTATCCGGATAAAGCTTCTAATTCCAAAGAAGAATTGAATGTTGCCACAGAGTTGATGCGTTTCCCCAGAGTATCTCCTAAGGCAGAGCTAATCTATCATCCAAATGAAGAGTTGGATTAA

>PGSC0003DMG400002960 LRR-RLPs

ATACACAAGAGTAGTACATTATTTGATATCTTGAGTTACATCTAAAAAATTTTTTTTGTTGTAATTCCTCTCCATATTGACTTGATGGCAGCTTATACTGCTGTAATTTCTCTTCTTCAAACTCTTGATCAACCAAATATTTCAGAACTCTTTCATGATCACAGTACTGAAATGTTCGATTCTCTTCGTGCTACAGCTGAATATTTTCAAGATGTTCTTGAAAACACTAGCAATGAGAAAATCAAATCTTTGGAGGAAGATATTAGAGTTGTTGTTAGTGAGGCAGAGGATGTTGTTGAAATGAAGATTTCTGAAATCATCAAAGAAGAAAGCTGGACATTTGGAATTTTACAACACCATGATATGCTACCAGTTGTTGAAAAAATGGATACCACAAAGAAACAAGTGATGGAGATTCTTTGTCATGATGCTGATCAAATTCTTGAATTAACCGGGGATTCCTTGATTGGCGCTTCTTCTATGAGTGATCCAATGCTGTCAGATCAGTTGGAAGATGATATTGTGCAGGGACTTGATGGTGACTTGGAGATAATAGATAAGAGATTGAGAGGACCAACGTCGGATCTAGACATTGTCACCATATCAGGTATGGGTGGCATTGGCAAAACAACACTCGCTAAAAAAACTTATGATCATCTTCCAATCAGGTATCACTTTGACATTTTTGTTGGGTTACAATATCTCAAGAATTTCAATATAGAAATGTATTGTTAGAAGCTTTACATTGCATTTCAAAGCAAAGAGTTATTGAGAACACAAAAGATTATGATAAGATGAATGACAGTGAGTTAGCTGATCTGGTGCAAAAGAGTCTAAAGGGTCCTAGATACCTTGTTGTTGTTGATGATATTTGGAGTAGGGATGTTTGGGATAGCATATCACAAATATTTCCTAATCGCAACAATGGGAGTCGAGTCTTATTGACTACTAGGGAAACTGATGTAGCAATATATGCAAATACTAGTAGCCCTCATAAGATGAACCTCTTGGATTTAGATAATAGTTGGAAGTTACTCCGTGATAAGGTGTTTGGACTAGAATACGATCATCCTCCTGAGTTGGAAGAAATTGGAAAAAAAATAGCAGAAAAATGTCAAGGACTTCCCTTAATAATTTCAGTGATTGCAGGGCATCTCTCTAAAACGGCCAAGACATTAAAAAGTTGGGAGTATGTTGCCAAAACCTTAAATGAAATCATTGCTAGCCATCCAAATAAATGCTTAGGAGTGCTCGGTTTGAGTTACCATCACTTGCCTAATCGACTCAAACCATGCTTTCTTTCTATGGGTAATTTCCCAGAGGATTTTCAAGTTGATACTTGGAGATTGATCCAATTATGGATCGCAGAAGGTTTCATAAGGGCACCTGCTGGAAGTCGTAAAAGCTTGGAGGAAGTGGCAAAATGTTATTTGGAGGATCTTATCAGCAGGAACTTGATAATGGCTTGAAAAAGGAGATTTAATGGTGAGGTTAAAGTATGCGGAATACATGATCTGCTGCGTGAGTTCTGTTTCATAGAAGCTGAAATGACTAAGTTTATGCATGTTGTGAGAACTCCATTTTCACCAACACAAAAGCCTAATGTTCGTCGTTTCAGTATTCAAAAGGCATTTTTAGATGAGTATGCTTATAAGCTATTACCTCCTGTTGCCAGATCTATGTACTACTTTTCTCTATTGAATCTACCTTTTGAACCCCGTATTAAGCTTCTCGGGATATTGCCCATCTACCGTCATGATCCTATAATACATGGATTTTTCTCTCGTTTCAACCTTCTCAGGGTATTGGCCATCTTCAATACAAATGTCAGGTTCGAGTCATTTCCAGTTGTGATTACAAAGTTATTTCATTTGAGATATCTCCAAATTCGATGTTTTGGCAATATTCCTGAATCAATCTCAGAGCTTCAGAATTTGCAAACTCTAATTTGTGATGGTCATTCTTTAGATATAACTTTACCTGAGAAGATATGGATGATGAAGAATTTGAGGTATATACGTTTAGGGGGAGCCAGTTATTTACCAAGTCCTAGAATAGATAACAAGATGCCAAATCTAGAGGAATTATTCAGTATTTGTTACTCTAGTTGTACAAATGAAGTCTTTTCTAGCATTCCCAATCTAAAGAGATTGACCATTCGTCTACCTTTGAGTGTCGGAGACAATATTCCCTCTCAGCTCATCGATATGTCCAGCTTGAGAAAACTCGAGTCAATCAAGTTTAACTGGTTACTTCCTTTCAAAAACCCCATCAACATATTTGCTTTTACAACATCACTTAGGACGTTGTCTTTAACTGATTGTGCTAATTTTATTTGGAAAGACATATCATCAACTTTTATCATGTTACCAAATCTTGAAGAGCTCAAACTTAAAAGTTGTCGAGCCGAAGATGATGTATGGAGATTGAGTGATAAAGACATTTTCAAAAGCTTGAAGTTGTTGTTACTGAAATACCCAAATCTTAAGTGTTGGGAAGCTAGCAGTGATAACTTCCCAAATCTAAAACGTCTCGTTCTCATGAATTGCAACAACCTGCAAGAAATTCCAACAGATTTTGGGGAAATTTGTACTTTACAATCGATTGAGTTACATAATTGCAGCACTTCTGCTGATGAGTCTGCCATAAATATTGTACAAGAACAAGAGGAAATGGGAAATAATATCCTTAAGGTCTACATCAGCCACGGTAAGTTTTAAATGATAAATAAACCAAGAATAAGATTCCTTATTAGTTTAGGATTTGTTATCCTCTTGACCAACAAGTTTATAAATGTCAGGTACAATGGTGAACTTTCTGGTGGACAATGTATTGCACCTATTAAGTGAAAATGTGTTGTTAATAAAAGCCGGAGATGTCGAATTCAGTAATTTATTGAAAGAAGTGCGATGTCTCAAACAGTTCGTTGACGTTGCATGGAAATACCACAGCGAAAGCAAGCAATGGGATGAATTGGTGAGAGATATTCAACTAACATTATTTAAAGCGGAAGATGCAATTGATGAATTTGTGATTCAATCGAAGCTGAATCGAGAAAAGAATAAAATTGGAAGAGTTTTTGATAAAGTGAGTCATATCAGTAATGTTAGGGCTCTGGTAGCTGAGATCAAAGAGATTCATTATAAGGCGACAAATTTTCGACAAAACAATTAACAAGCTTTTCAGTTTGTGTGTAAATGTTGTAATTATTTTAAGTAAAAAATCGATGATGGATGAAGAAAAATAATTTAGTTATGGACATCTCAATTAGGAGTGTACAAGTCAAATCGAA

>PGSC0003DMG400002961 LRR-RLPs

ATGACAAAGTTTATGCATGTTGTGAGAACTCCATTTTCACCAGCACAAAAGCCTAATTTTCGTCGCTTCAGTATTCAAAAGGCATTTTCAGATGTGTATGCTTATAAAGCATTACCCCCTGTAGCCAGATCTATGTACTGCTTTTCATACTTGGATCTACCTTTTGAACCCCATATTAAGCGTCTCCCTAGATTGCCCATCTACCGTCATGATCCTATAATACATGGATTTTTCTCTTGTTTCAACCATCTCAGGGTATTGGTCATCTTCCCTGCCTGGTTCGGCCCATTTCCATTTGTGATTACAAAATTATTTCATTTGAGATATCTCCAAGTTCGATATTTTGGAAATATTCCTGAATCAATCTCAGAGCTTCAAAATTTGCAAACTCTAATATATGATTGTTATTCTTCACATATGGCTTTACCCGGGAAGATATGGACGATGAAGAACTTGAGGTATATACATTTAGGGAAAGCTAGTTATACACCAAATCCTAGAATAGATAACAATCTTGTGACGGGGATGCCAAATCTAGAGGAATTTTTCAATATTTGTTACTCTAGTTGTACAAATGAAGTCTTTTCTAGAATTCCCAATCTAAAGAGATTGAGCATTTGTGCATCTTCTATAGTCAGAGAAAATATTCTCTATCGTCTCTTGGATATGTCCAGCTTGAGAAAACTTGAGGCATTCAAGCTTTACTGGAGATATATTTTGGAAAAGCCCATCAAAAGATTTGTTTTTCCAACATCACTTAGGAAGTTGATTTTAGCTTGCTGTAGTCATTTTATTTGGGAAGAGATATCATCAACTTTTATCATGTTGCCAAATCTTGAAGAGCTCAAACTTAAATATTGTCGAGCCGAAGATGATGTATGGAGATTGACTAGCAAAGACATATTCAAAAGCTTGAAGTTGTTACTGAGCGAGCTAAATCTTAAGCATTGGGAAGCTAGCAGTGTTAACTTCCCATGTTTAAAACGTCTTGTTCTGAAGAATTGCCAGGACCTGCAAGAAATTCCAACAGATTTCGGGGAAATTGGTACTTTGGAATCAATTGAGTTACATGATTGCAACACTATTTCTGAGGAATCTGCCAGAAATATCGAACAGGAACAAGAGGAAATGGGAAATAATATCCTTAAGGTCTACATCCATAACAACAACAGTAAGTTATAA

>PGSC0003DMG400002980 LRR-RLPs

ATGGAGGTAGTTGGTGGGGCATTTCTCTCTTCAGCTTTGAATGTTCTCTTTGATAGGCTTGTCTCTCATGGTGATCTGCTCAACATGTTTCGGAAGCATAAGGATCATGTTCAGCTCTTACACAAGCTGCAGGACAATTTGCTTGGTCTTCAGATTGTGCTAAGTGATGCAGAGAAAAAGCAAGCATCAAATCGACATGTGAGCCAATGGTTTAATAAGCTTCAGAGTGCTGTGGACGGTGCTGAAAACTTGATAGAACATGTCAACTATGAAGCTTTGAGGCTTAAGGTAGAAGGCCAGGATCAAAATCTTGCAGAAACAAGCAACAAACAAGTAAGTGATCTCAACCTGTGCTTGAGTGATGATTTTTTTCTTAACATAAAGGAGAAGTTGGAAGAAACCATTGAAACATTGGAGGTGTTGGAAAAGCAAATTGGTCGCCTTGGCTTAAAGGACCATTTTGGTTCGACTAAACAAGAAACTAGAACACCTTCAACTTCTTTGGTTGATGATTCTGATATCTTTGGAAGGCAGAATGATATAGAGGATTTGATTAACCGTTTATTGGCTGAAGATGCAAGTGGAAAAAAGCTGACTGTGGTTCCTATTGTTGGAATGGGCGGTGTGGGTAAGACAACACTTGCTAAAGCCGTTTACAATGATGAGAAGGTGAAGGACCATTTTGGTTTGAAAGCTTGGTTTTGTGTTTCTGAGGCATATGATGCTTTCAGAATAACAAAAGGTTTACTTCAAGAAATTGACTCATTTGACTTGAAGGCTGATGACAATCTTAATCAGCTACAAGTCAAATTGAAGGAAAGCTTAAAGGGAAAGAGGTTTCTAATTGTTTTAGATGATGTGTGGAATGACAACTACAGCGAGTGGGATGACTTGAGAAATGTTTTTGTACAAGGAGATATAGGAAGTAAGATCATTGTGACGACACGTAAAGAGAGTGTTTCCTTGATGATGGGTAGTGGGGCAATCACCGTGGGGACTCTGTCTGATGAAATCTCTTGGGATCTATTCAAACGACATTCACTAGAAAACAGGGATCCTAAAGAACATCCAGAACTTGAAGAGATTGGAAAACAAATTGCAGACAAGTGCAAAGGGTTGCCTTTAGCTCTAAAGGCACTTGCTGGTGTTTTACGCTGCAAATCAGAGGTGGATGAGTGGAGAGACATTTTAAGTAGTGAAATATGGGAGCTTCCAAGTTGTTCGAATGGTATATTACCAGCGCTGATGTTTGAGCTACAATGATCTTCCTGCACATTTGAAGCAATGTTTTGCTTATTGTGCAATATATCCCAAAGATTATCAATTTTGGAAAGACCAAATTATTCACCTGTGGATTGCTAATGGTCTTGTACAGCAGTTGCAATCAGGTAACCAATACTTTCTTGAGTTGAGATCAAGATCATTGTTCGAAAGGGTCCCAGAGTCTGAATGGAGACAGGAGGTATTCTTAATGCATGACCTTGTCAATGATTTGGCCCAAATTGCATCTTCAAATCTTTGTATAAGGTTGGAAGAGAACAAAGGATTGCACATCTTGGAACAATGTCGGCACATGTCCTATTCAATAGGAAAAGATGGTGACTTTGAGAAATTGAAATCACTCTTTAAATCAGAGCAGCTGAGGACATTACTTCCAATGGATCTCCAGCGCCCTGATTATGGCCGAAGTAAGCTTAGCAAGAGGGTGCTGCATAACATACTGCCAAGACTAACATCCTTAAGGGCATTATCATTGTCATGCTATCAGATTGTTGAGTTGCCGGATGCCTTGTTTATCAAATTAAAGCTCCTTAGATTTTTGGACCTTTCTGGGACATTGATTAAAAAGTTGCCAGATTCCATTTGTACATTGTATAACTTGGAGACACTTCTCCTGTCATTTTGTGTTGATCTTGAGGAGCTACCGCTGCAGATGGAGAAGTTGATTAACTTGCGTCATCTTGACATACTCGATACTTGTCGCTTGAAGATGCCGCTACATCTGAGCAAGATGAAAAGTCTCCATGGGCTAGTGGGAGCCAAGTTTCTTGTAGGTGGTCCTGGTGGTTTGAGAATGGAAGATTTGGGTGAAGTACATAACTTATATGAATCTCTATCAGTTGTAGAGTTGCAAAATGTGGTTGATAGAAGGGAAGCTGTGAAGGCAAAGATGTGGGAGAAGAATCATGTTGGGCAGTTATCTTTGGAGTGGAGTGAAAGTAGTAGTGCCAACAATTCACAAACTGAAAGAGACATAGTTGATGAGCTACGCCCACATAAAAACATAAAAGAAGTCGAAATCAGTGGATATAGAGGGCTAACATTTCCAAATTGGCTAGCTGATCTTTTGTTTCTTAAGCTGGTGAAATTGTCTCTTAGCTACTGCAAGGTCTGTTATTCCTTGCCAGCATTAGGACAACTCCCTTGTTTGAAAATCCTTTCCATTAAAGGGATGCATGGAATAACAGAGGTGACGGAAGAATTCTATGGCAGTTTGTCCTCCAAAAAGCCTTTTAACTGTCTTGAGGAAATTAGTTTTGAAGATATGCCGGAGTGGAAGCAATGGCACGTACCAGGAAATGGAGAGTTCCCTATACTTGAGAAGCTTTCCATTATAAATTGCCCTGAACTCAGTTTGGAGACACCCATCCAACTTTCAAGTTTAAAATGGTTTGAAGTTAGTGGTTGTCCAAAGGTTGGAGTTGTTTTTGATGATGCTCAACTACTTAGATCACAACTTGAGGGAATGAAGCAGATTGAGGTATTATATATTGGTGATTGTAAGTCTCTTACCTTCTTTCCTTTTAGCATACTGCCCACTACCTTGAAGAATATATGGATATCTGGTTGCCGGAAATTGAAATTGGAGCAGCCAGTTGGTGAGATGAGTATGTTTCTGGAGAATTTGAGACTGGATGGATGTAATTGTATAGATGATATATCACCCGAGTTGATCCCAAGCGCACATACATTGAATGTATCTAATTGCGACAACCTTACTAGGTTTTTGATTCCTACTGCCACTGAAAGTCTCTTCATTCGGAATTGTGAGAATGTTGAAAAACTTTCGGTGGCATGTGGTGGGACCCAGATGACGTATTTGTATATTGACGACTGTTCAAAGCTGAAGTGGCTGCCAGAACGTATGCAGGAACTCCTTCCATCTCTTAAGGAACTGCAACTGATTAATTGTCCAGAAATAGAGTCCTTTCCTCAAGGAGGATTGCCCTTCAATTTACAACAACTTGCGATCAACAATTGCACGAAACTGGTGAATGGCCGAAAGGAGTGGCATTTACAGAGACTCCCCTGTCTCACACAGTTAATCATCTCTCATGATGGCAGTGACGAAGAGATTGTTGGTGGTGAGAATTGGGAGTTGCCTTCCTCTATTCAAACACTTATCATAAACAATCTGAAAACATTAAGCAGCCAAGTTCTCAAAAGCCTCACCTCTCTTCAATATCTATGTATTAAGGGTAATTTACCTCAAATTCAGTCAATGCTGGAACAAGGCCAGTTTTCCTCGCTTCAAAGTCTACACATCGAGGATATCCCTAATCTCCAATCACTTTCCGAATCAGCACTGCCCTCCTCCCTCTCTCAGCTGACCATCATCAATTGCCCTAATCTCCAATCACTTCCTGAATCAGCACTGCCCTCCTCCCTCTCTCAGCTGGAGATCTCCCATTGCCCTAATCTGCAATCTCTTCCAGTAAAAGGGATGCCATCTTCACTCTCTACACTATCAATTTCCAAATGCCCATTACTCACACCACTACTAGAATTTGTCAAGGGGAAATACTGGCCAGAAATTGCTCATTTCCCCACAATAGAGATCGATGAGGAATGCCTTTAA

>PGSC0003DMG400003195 LRR-RLPs

TGGAGCTTTAAATACTTCTTAATTACAAGTTCTAGAAGGAGTCAGAGAAATTTATGAGCCATCCGGGATTGAATCCTTATCTCTCATGCACTTTATCGACCACTAGACCAAATCACTGAGGTGACGGGTGTCTAGTCATTTATATATAGAGTCGCAACCATCTGAATTTTTCCTGTTTGAATGTCAAAGATACTCATAGCACGATCAAACTGTCAGTCAAACAAAAACAATAGAGTTGACATTTAACATATATGCACACAAAATTGCCTAACTATGATGCAAACTGCATAATGATTTGACAATCAATGAGGCTGGATTTTATTACTTGTTCTAGAAGGAGTCAAGGAGATGTTCATGAGCAACCCAGGATTTGAATCCTGATCTCGCATGCACTTTATCGACCACTAGATCAAATCACTGGGGTGACGGATGTCTAGTCATTTATATATAGAGTCGCAATCATCTGAATTTATCAAGTTTGAATGTCAAAGATACTCCTAGCGCGATCAAACCGTCAGTCAAACAGAAAAAATAGAGTTGACATTTAACATATACACACACAAAGTTGCCTAACTATGAATGCAAGCTGCGTAATGATTTGACGATCAATGATGCTGGATTTTGTTACTTGTTCTAGAAGGAGTCAAGGAAGATATTTCATACCTCAAGTATATAAAATCTTGTGAATTATTCCTTCATTCCTCAAGTACTTATTATTTTTGTGCAGATCATGGGCTGTCTATTTCTTCTTCTTCTTCTTCTTCTTGTGTTGCAATATCCACTTGTTATCTTTTCAAGTCATCTATGTCGACGAGACGAGAGTTTGGTTTTGCAAAACCTGAAGCTTAAACTGTCCATAGATGACTCTGCTTCTGCTGATTGTGAATCCAATGGTTACTTTCCCTATCCAAAGACACTTTCTTGGAATAGAACCACTGATTGTTGTACTTGGGATGGTGTAACATGTGATAAAGTTACAGGTTGAATAACTCGTCTTGATCTTAGTTGTAGCCAACTTCATGGAACAATTGACTCCAACAGCACCCTCTTCCAACTTTCTCATCTTCAAAACCTTAATTTAGCTTACAATGATTTCTCACCTTCCCAAATTTCAAGCAAATTTGGATGGTTTCCGAGTTTGACACATCTTAATCTATCGCGTTCTGGTTTTTCAGGGACAGTACCTTTACAAGTCTCATATCTATCTAAGTTGATATCACTTGATCTTTCTGCAGGTATTGAGGAGTTGAGAGTTGAGCCTCACACTTTCAAAATGCTCCTTCAAAACTTAACTCAACTAAGGGAGCTTTACCTCACCTCAACATATATATCTTCATCACTTCCATCAAACTTGTCCTCTTCCTTACAAGAGCTGAATCTTGTCGGAACGGAATTGTCTGGGAAAATCCCTGATGACATTTTCCATTTACCAAGATTGCAAATGCTTAACTTAGGGAGCAACTTGTATCTCACAGGGCATTTGCCAACAACTCAATGGAACTGCAGCAACTCTTTAAGGGAATTAGATCTTTCTTCCTCAGGCTTTTCGGGGAATATACCAGATTCTATTGGTCATCTCAACTCTTTAAGGAAATTAGATATATCAAGCTGCTACTTCTCTGGTAACATTCCTCCAACTATCGGTGATCTCACCGAGCTCACGAGTTTGCAACTCTTCTCCAACAACTTCAATGGTCAACTACCATCAACAATCTTAAAGCTGGTGCAACTTGTTGAGTTTGATATCTCATCTAACAACCTTACAGGCAACATTCCAAATATTTTCGGTAACTTCACCAAGCTCAAATCTTTATCCTTTTCATATAACCTCTTTATGGGCCTTTTTCCATCCTCGGTTACAAACCTAACAGAGCTAGAGAGTTTAGACCTCTCAAATTGTTCTATAACTGGTGCTATTCCTTCTAGCGTCAGCACCGGATTCCCTAACCTCATCTTGCTCTTCCTACCATATAACTCCCTTAGCGGAGAAATACCATCTTGGATATTCAATCTTCCTTCCTTAAAATACTTAGTTTTGAGAGGCAATCAACTTGCTGGTCAACTTAAGGAAGTTAGATACAACTTGTTAGAAGTTGTTGATGTCAGTGACAATAAGCTGAATGGACCTATTCCAACATCATTCTCTAAACTAGTGAACTTGACAACACTTGATCTTTCTACAAACAACCTTTCTGGTGGTCTAGATATTGGGTTGTTTTCAAAATGCAAACAACTTCGACGTCTTGGCCTTTCTTTTAACAATCTGTCTGTATTTTCCAGTCAAAAAGATGCATCCTTGCCTAGTTCTCTTGGAAGTTTGTATGCATCTTCTTGCAACATAAGGGAGTTAAATTTCCTACGAGCTGCAAAATACATTGGTAACTTAGATCTTTCCAACAACAACATGCATGGAAAAATTCCTGATTGGGCATGGTCTAATTGGCAAGTCTCAGTGTCTTATTTGAATCTTTCTAGCAACTTTCTAACGGCTATTGACCTGCTTCATAACTTTGAGAGTCTTTACAATCTTGATATTGGATCTAACTTGATTCAAGGAGAATTACCTGCTCCACCTCCCTTATTGTTCCTCTTCATAGCCTCGAACAATAACTTCACCGGGAAGTTACCTAATTCACCATTATGCAAAATGAGTTCCCTTTTGATTCTTGATTTGTCCAATAACAGCCTGAGTGGAGTAATTCCAAAATGTTTAGTGAACATGAGCACAAGTCTCTCAGTGTTGGACCTGCACAACAATCATTTTAGTGGGACAATTCCAAATAACTTTGGTAAGGGAAGTAGTCTGAGAAGTCTCAATTTGCGGAAAAATAAACTAGAAGGAATGTTACCAAGAAAGCTTACCAATTGCAGAGGGCTCGAAGTAGTTGATCTCGGAGAAAACTTGTTGAATGATACATTTCCCAAATGGTTAGGATCACTTCCTAGGCTGCAAGTTCTCAGTTTGAGATCCAATAAATTACATGGCCCGATTACCATTTCAAGAAACCAAGTCTTATTCTCAAAGCTGAAAATCCTTGATCTCTCTTATAATGACTTCACTGGAAACCTGTCAGAAAGGTTTTTCAACAACTTGAAATCCATGATAATAGAAGATCAAACGGGAATTCCTTTAATGTATATTGGAGAAGTTCGATACCATGATTCCTTGACATTGTCTATAAAAGGACAGCAGATTGAACTTGTGAGGATTCTTTCTATCTTCACAACCATTGATTTCTCAAATAACAGATTTAAAGGTAATGTACCAAACTCCATTGGAAATCTTGGTTCACTTCGCGGGTTGAATTTGTCACACAACAGCCTTCTTGGACCTATACCTCAGTCTTTTGGAAATTTATCTGTTCTTGAGTCGTTAGACCTTTCGTGGAACCAACTTTCTGGGAACATTCCTCAAGAAGTTGCTTCACTGAAATCTCTTGCTGTCATGAATCTCTCACAGAACCACCTGATGGGAAGCATACCTAGAGGTCCACAGCTTGATACATTTGAAAATGACTCATATTCAGGAAATGCTGGGTTATGCGGATTTCCCTTGTCAAGAAACTGCGGAGATAATGAGATGCCTCCTTCTACTGTGACTGATATTGAAGACGAAGAAGATTCAGGTTTTATGGATTGGAGAGCTGCGATAATAGGATGTGGTTGTGGATCAGTGTTTGGGATGTTTATAGTATATATCATTTTCTTAACAGGGAGACCAAAATGGTTTGTGAGAATTGTCAACCAACAGATCAACAGGATCCTTTTGAAAATTGAGGCAAAGAGAAGAATTGAGGCAAAAAAGAGAAGACGGCAGAAAAGAAGAAGGATGAAGTTTTAGTATATACTTCATGCATTATTGTCTTGCAACATACACAGTTTCAGCTTACACTTTGTTTTGGC

>PGSC0003DMG400003380 LRR-RLPs

GAGGAAGTGAAGAAATAAACTATAATACTGTGTGTTCTTCTTTCCCAGTCCACAACAGTTTAATTCTTGTGTCACAAGGATGTATTTCAACAAAGAATTATCTAATCTGAAAGATCGACTCCTTAGCACGGTGCAAGATCAAAATTACTCGGATGTTACAAGGGATCGAATAAATTTCTTTCTATGGGACTTAAAATTCCTCGATTGTTTTCTCCATTTGAAAAGGTTGCCTTTTGCAAGTGAATGCGGTATGCTAAAAGTCTCACAGAAAATCATAGAAATTTGGAAGTTTCAGTGTAAACCCTATGATTGTCCAGATCTATTTCCGTACAAGACTCTGTATGTTTACGATTCATTTGGCTACTGGAAGGATGTAATTTGGAAGACTAAGCAAGAATTCAGAGCTGAATACTCTTTTCCAAATACATCACTTGCAGCCAACAAGGTGGATGATGATGTAGCCCCAAATTTGTGATGGAAGTCATCGATGTTTTTGTGGAGAACCTCAATGTTCTAGTGAAGATCAATGATCCAATTTCGTGGCTTTTTGTTCCAGAACACAAGGAACAAATAGAACAAGTGTTAAAGGAGTTGAAGTTATTGAGATTTTTTGTCTGCTTTGTTTCAAACAAATGTATAGAGCCTCAATACCGACATACTACTTTTTATATTCATGCTTTAATTGAGGCTAGCCACATTGCAATGGTTGTCTGGTTGCATTTGCCAGTCGTCTATGGAAATCAAGACGTGGCTCCAAGTGAAGTCAGTCGTTTGCTCTCTGATTTCATGGAATTGAAGATTAAGTCCATTCAGCCAGTCATCAGCCGCAACAATATTTATATTGATGTCCTGCAAGCTTTGAAGTCAACCATACCACAAGCACAAAAGAAGCATGCAGCTGAGAGTAGCACTGTGGAGATTCCAACACACAGTCTGACGGTTGGTTTGAGTGATCAAATGGCTAACCTTCAGGAGATGCTCTGCCTTCTAAGAGATAATCTCATTCATCTGCCAATACTAGATCTGGAATTTCATCTTCAAGATATGGATTCTGTTATTATTGATGCTGGACTTCTTATTTACTCATTATATGATATCAAGAGGGAGAAGGAAGACACAGTATTGGATGATATGAACCGAGCACTTGGTTTTGATCTTCCAAGAAACATTGAGCCTATCAAGGCGATGGTCTACCTCATCATGCAAAAGGCATTTCAATGTAACTTGCCAAGGATTCATGGACTAGGCTATGTCGATTTTCTATTGAAAAACCTGAAGGATTTCCAAGGCCGTTATTCAGATTCACTCGCTTTCCTCAAGAATCAAATTCAAGTAATTCAAACGGAATTTGAGAGCTTGCAACCTTTCCTGAAGGTTGTCGTAGAAGAGCCACACAATAAGTTCAAGAGACTGAATGAAGATTGTGCTATACAGATAATTAGGAAAGCACATGAGGTGGAATATGTAGTTGATGCTTGTATAAACAAAGGCATTCCTCATTGGTGCCTCGAGCGTTGGCTCCAGGATATCATAGAGGAGATTACTTGTATCAAAGCAAAGATTCAGGAAAATAACACTGTTGACGATACAATGAAGACTGTCATTGCTCGTACATCATCACAACTGGCAAGGACTCCGAGGATGAATGAAGAAATTGTTGGGTTTGAGGATGTCATAGAAAATTTAAGAAAAAAACTACTGAATGGAACCAAAGGGCAAGATGTCATTTCAATTCATGGCATGCCAGGTTTAGGTAAGACGACTTTAGCCAACAGACTCTATTCTGACAGGTCAGTTGTTTCTCAATTTGATATTTGTGCACAATGTTGTGTGTCTCAAGTATATTCGTATAAGGACTTGTTATTGTCCTTGATACGTGATGCTATTGGTGAGGATTCTGATCAGCATAGAGAACTTCATGACAATGAATTAGCTGATAAGCTTCGCAAAACCTTATTGCGCAGAAGGTACCTTATCCTTGTTGATGATGTGTGGGAAAATAGTGTTTGGGATGATTTAAGAGGTTGTTTTCCAGATGCCAATAACAGAAGCAGAATCATTCTAACGACAAGACATCATGAAGTTGCCAAATTTGCTAGTGTTCATAGTGATCCCCTTCATCTTCGTATGTTTGACGAAGATGAAAGTTGGAAGTTGCTTGAAAAGAAAGTGTTTGGTGAACAAAGCTGCTCCCCTCTCCTAAGAGATGTTGGGCTAAGAATAGCAAAAATGTGTGGACAACTCCCTCTTTCAATTGTTTTGGTGGCTGGTATTCTGTCAGAGATGAAAAAGGAAGTAGAATGTTGGGAGCAAGTGGCCAACAATTTGGGTACCCACATTCACAATGACTCAAGAGCCATTGTAGACCAAAGTTATCATGTTTTACCCTGTCATCTTAAGTCTTGCTTCCTTTATTTTGGAGCATTTTTAGAGGATAGAGTGATTGATATTTCAAGGTTAATAAGGTTATGGATATCAGAATCATTTATAAAAAGTTGTGAAGGCAGGAGGTTGGAGGATATAGCAGAAGGTTACTTGGAGAATCTTATTGAAAGAAATCTAGTAATGGTTACTCAAAGGGCCGATTCAGATGGTAAGGTCACAGCATGTCGCCTTCATGATGTACTACTCGACTTCTGCAAGGAAAGAGCAGCTGAGGAGAATTTTCTACTATGGATAAAACGGTAATATGATAAGTAATGGTACTTTCAATCAATCAAGTATTTCAAGTTATATTTGAAAATTAATGATATGATTTTGCTAATTGATATATTCACAGAGATCAGATTACCAAAGCTGTTTACTCTCACAAGCAGCATGCTCACTTGGCGTTCACTGAAATGGATAATCTTTTAGAATGGAGTACGTCTGGCTCACTTGTTGGCTCTGTACTTTTTAAGAATTATGACCAATACTTTGCCGATCGTCCCTTTTTCTCTCATGCTTTTGAAATTTCACACATTTTACCACATTTCAAGTTTCTAAAAGTACTGGATTTGGAACACCAGGTTGTTATTGATTTTATTCCGACTGAGCTCCCTTACTTGAGGTATTTCTCTGCGCTCATTGATCAGAATTCAATTCCTTCAAGCATATCCAATCTTTGGAACCTTGAAACTCTTATATTAAAGCGTAGATCAGCTGCCAGATATAACAGGGTATTACTACCTAGTACAGTTTGGGATATGGTTAAATTGAGATATCTGTATATTCCTTACTTCAGCCCTGAAATAAAAAGCATTACTCGAGAACTCTCCAAAACTTGATGATTTGGAAACCCTTTCCAATTCATATTTCACTCGTGTTGAGGATGCAGAATTGATGCTGAGAAAAACACCTAATCTTCGAAACCTGACATGTAAAATCGAGTGCTTGAAATACCCCCATCAGTACCATGCATTGAATTTTCCAATACGGCTTGAAATACTAAAGCTTTATGGATCAAAAATCGTCAAAACCATCCCCTTTTACATCTCTGCACCAAATCTCAAATACCTGATACTCTGTGGCTTTTACCTGGATTCTCAGTACTTATCAGAAACTGCTGATCATCTGAAGAACCTTGAGGTACTCAGACTGTCCTTCGTTATATTTGGTGGTCATATGAAATGGAAAGTGAGCAATGGCAAGTTCCCTCAACTCAAAATCTTGAAACTAGAATATTTGCCCATGATGAAATGGATTGTAGCTGATGATGCCTTTCCTAACCTTGAACAATTGGTTTTGCGTCGATGTCATGATCTTATGGAGATCCCTTCTTGTTTCATGAACATCCTTTCTCTGCAGTACATCGAGGTAGAAAACTGCAATGAGTCGGTTGTCAAGTCAGCCATGAATATACAAGAAACACAAGTCGAAGATAATCAAAATACTAATTTCAAGCTCGTTCTCATCGAGGTACACTACTGAAAAAAGCTTTATTCTGCATGATTTTGATGACTCAGAAATAATGGCCTAAATTGTATGAACTGTTTTCTCTGTTATCTTTACCTGGTGGCCTCGTTTTACATTTGGGTTTTTCTCTTGGTTTTCTTGCAGAAAGAGACATACCCAAGGCATTTAAACAATTACTTTTACTTCCAGGTACTACTTATTATTTTTTCTTTCTTTCCTTTTTTTGAGTACACCGAATAGATAGATTCATCTTTTTTCTCCTGTCGATATAAAAGGGATAGAATCAATTTCAACTGATATGAAAGAGAAGAAACTTACTGTGACCAGAGAATGTGGATGCTGATGAAGTTCAATTAGTTGTGGAGAAACTGAGAAAGCGTGGCATGCTCTAACAAAATGCTGAACTCTGGTGAACTCAACCGTGATGGATTTCCAAATAATGACAGGGTTGTAGTCCCAACTTGTCAACACAAATGTGCTATACTCATTTTTCTTACTGTAATACCATTTCATGACACACACACAAACATTAACTGTAGTAAAGTTTTGATGGATCAGCTGATATGTTAGTTCAACCCATTGTAATCCCGTTCAAATTCAACTCAAAAAATGCCCATTGAGTTATTCTTTAACAGGGTATCCAGAGTTTGTAGCTGGAGCAATTTGGAATATCGCATGTAATTTCTTTATGAGTTATTTCGTTTAATAAAGATTCTGTAAAACGTCCAACGGCTGTTGCATTCATTGTAAACTAAATACATCTCACAATGTAACTGTTGAACAAATTTTTCATTCAAGTCCTTAAGGTTTGATGTAAGTCATTAGATTTTAC

>PGSC0003DMG400003543 LRR-RLPs

CTTTCTTTGTTTGTTCCCTTCTTTGGACACACAATATATCCTCATGAATCCATTTAAAAACTAATCTTGAAGGCTCTTCTTCTTCATCAATTCTCATTACGATAAAAAAAAACCACACAGAAAAAATCCAGAAAATGGGTTGTTTTTCAATTTCTACTGTTCTGTTCTGTATAGTTCTCTATGCTCTGTTCTTGCATGCGCATTCAACTACTCACTGGCAAGACATTGAAGTCTTGAAGCAGCTGAAAAATAGCGTCGACCCCAATTCAGTGTCACCTGGTTCATGCCTGAATTCATGGGATTTCTCTGTAGACCCTTGTGATAATCTCGGGGGTGAAAAATTCACTTGTGGTCTCCGTTGCGACCTTGTCGTTTCCTCTGTTAGCCGAGTCACTGAACTCGCTCTAGACCAGTGGAATTACTCCGGTTCACTCACTTCTGTCTCTTGGAATTTGCCTTATCTCCAAAACTTAGACATCACTAATAATGCCTTCTCCGGTTTAGTACCTGAGTCACTCTCGAATCTAACTCGGGTTCAACGAGTCGGTTTATCAGGTAACTCGCTATCCGGGTCAATTCCGAGTTCACTGGGTTCACTTTCAAACCTTGAAGAGCTTTACCTTGACAATAATTTCCTTGAGGGAGAAATTCCCCAAAGTTTCAACGGTCTCAAGAATTTGAAACGGCTCGAATTTCAAGGCAATCAGATTACTGGCGAGTTTCCCGAGTTGGATCAGTTAATTAATTTGAATTTCATCGATGCTAGTGATAATGCCATCTCCGGCGAACTCCCGGCGAGTTATCCGGCTTCTCTCATCGAGCTTGCGATGAGAAACAACAGCATAGAAGGTAACATTCCGGCGAGCCTCATCGGACTGAATTTCGTACAAGTTATCGATCTGAGTCACAACAAACTCAGTGGGTCAGTGCCAGCGAGTGTCTTCACTCACCCATCCCTCGAACAGCTCACCCTATCATACAATCAATTCGGGTCGGTTCAGGAACCCGGGATCTCGTTTCAGAATAGTCAGCTTATTGCAGCAGACTTAAGCAACAATGAGATCCGTGGGTTATTACCCGGGTTCCTGGGTTTAATGCCCAGATTATCATCACTTTCACTAGAGAACAACAAGTTATCGGGTATGATACCGACCCAATATGCATTGAAAATGGTGTTTCCGGGTGAAGGGGTGTCTCAATTTGAGAGGCTACTGTTAGGAGGAAACTATTTATTCGGACCCATACCGAGTCCATTGTTGGATCTGAAAGCGGGTTCATTCACAGTGAGATTAGGAGACAACTGTTTGTACCGGTGCCCGTTGAGATTATTTTTATGTGAAGGTGGGGAACAGAAATCATTATCGGAGTGTCAGGCTTTTGGTCCCATCATTCCTTGAGTAATGTGATAATGTCTTTACTTTGAGGTCCTCGTTTTCATTTTGTACTAGTAATTATTGTATCGTTTCAATTTTATCGGAAGGGACCAGTAAGTATTGTATAAATATATTTTTTCCTGCAACCTTACCCCATGTATGGGGTTTTAGATATGATCTATGGTAGAGTTGTTATTGTTGTGAAGAGTGAGACAGAAATATAAGATTTCTGACTTGATGTGTTGAAGTAGTAATTGTGGGACAGTTGAGAAAATTTTATTAGCGGAT

>PGSC0003DMG400003544 LRR-RLPs

CAAATATGGGAATTTTGGGTAGGAAAAAATCTTTCTTGTTCATCTTATTTTACATAATTTTTTCTGTTTTTGTTCCAAGGCCATTAGAATGTTGTCATTTGGTTGATAAAACATCTTTGTTAGACTTTAAACAAAAAGTCACTTCTGATCCTTCAAACTTATTGCAAACATGGACACTTTCAACTGATTGTTGCAAATCTTGGGAAGGTATTGCTTGTAATTCCAATGGAAGGGTTGTTAACGTCTCTCGCCCTGGCCTTTCTTCAGGGGACGATTTCATTATCGATACCTCAATTTCTGGAACTCTTTCTCCTTCTCTTGCCAATCTTTCTTTTCTTGAATTGCTTGATCTTAGTAATCTTAAGGACTTAACAGGGCCAATTCCACCTGAATTTGGCAAGTTATCGCGCTTGAGTTACCTCTTTCTTGATACTAATAAACTCTCAGCATCTATACCTGTCACATTTAAATATCTATACCAACTTAAGAAGCTTTATCTCAGTGATAATACTCTATCTGGTACTATCCCTTCGAGTATTTTCGAACGCTTCCTTTCACTCTCAGAACTTGGTTTATCAGCAAATCAACTTTCAGGTCCAATACCATCTTCAATTGGGAATTTAGTATCATTGACCAAGCTTGATATGAGTCGAAATAAGTTCTTTGGAAGCATTCCAGAGAGTATAGGAAGATTAAAAAATCTTGCATATGTTGATTTGTCTGAGAATCAGTTATCTGGAAAGATTCCAAACTCTATTGGAAACCTTTCCCAAGTATCTCTAATGTATCTGAACCAAAACCAATTGAGAGGAAAAATTCCTTCATCAATATCTGGTCTTAGATCATTGGTGTTCTGCCGTTTATCCGAAAACCAGCTGAGTGGCAGTATTCCACCTTCTATTGGCAATCTTCCAAAGATCGAAAGGCTGATATTTGAGAACAACAAGTTAAGTGGAAAACTGCCTGCAACACTTGGACATCTTGTAACTCTCACTGATATGTACTTTTCCAATAATCTATTTACAGGGAAAATCCCATCAAGTTTTGGTAATCTAAAGAATCTACAGATACTAGATTTGTCGAAAAATAAGCTCAGTGGTGAAATCCCTCATCAGCTTGTGAAGTTAGTGAGTTTACAGACATTGGATCTGTCATTTAATCCTTTGGGGCTTTCAAGAATACCAAACTGGTTCAAGAAACTAAGACTGTTTCGTCTCCTTTTGGCGAAGACTGGGATCAGAGGGAACCTCCCAAGTTGGTTGGCTTCATCGTCATTATCAACACTTGATTTATCAAACAATGGTTTGACAGGAAAATTACCTACATGGATTGGTAACATGACCAACCTTTCACTCTTGAATTTATCAAACAACGCCTTCTATTCATCAATCCCTGATGAGTTCAAGAACCTGACACTTTTGATGGATCTTGATCTTCACTCAAACAGATTTTCAGGCAACTTGATAGCAATTTTCTCCAAGAATTTTCAAGACCCCCTTGGTCACTACAATTCTATTGATCTTGCATACAACATGTTCACTGGACCATTAGACGAAAACAGCGGAAATGAACCTGTCATGGATTCTATTTTGTCACTAAGTTTGTCCCACAATCCATTGACAGGACACATACCAAAATCACTGGGAAATTTGACAACTTTGCAGGAAATTAAACTAGCTGAGAATGGGCTAACAGGTGAAATTCCAACAGAACTTGGGAATGCCAAGGAACTGAAAACAATTTTACTTTCAAATAACAATTTGGAAGGTGCAATACCAGCACAAGTTCTGAATTTGAAAGACCTTCGAGAATTTGATGTATCGGGAAATCGACTAAGCGGTAGTATCCCTCCTCATAAAGCTAATATCCCAAAGTCTGCATTTTTGGGAAATTATGGCTTGTGTGGAGCTCCACTTCCTCCTTGTAAGCACACATAGTGGAAACCTTCAAACACAATATTTCCCTTTTCTTTATTTATTTTTCATTTTCGACATGATTGTTCGTCATAATGTATTGTATTTCAGTTGTAATCACTCAAATGTTATTAGTAAATACTGAACAAAGTTGATTTTTCAAAAGCAACCTTGCAGAATTCTACATCAATATCCTCAATGGTTTAAAC

>PGSC0003DMG400003710 LRR-RLPs

ATGAAAATGCAGTTTTTTCTTCATTTTTTTTCATTTTTCATCTTCTTTGTATTATTAGCACAATGCAAAACTCATCCTATTGATATACAAGGACTTCAAAATCTCAAAAATGGAATAAACCCAAAATCCATTAAACCCGGTTCATGCTTAAACTCATGGAACTTCACTATAGACCCATGTGATCACATTTTCACCGACCGGTTCACCTGCGGTTTCCGCTGTGATCTCATCGTCTCCGGTTTTTATCGAGTCACTGAAATCAGCCTAGACCAAGCCGGTTACTCCGGTTCATTGCCTAATTCATTCAACCTCCCTCATCTCGAGATTTTAGATATGTCGTATAATTCCCTCTCCGGTTCAATCCCGAATTCATTTTCAAATCTAACTCGTCTCCGGCGGCTCAGTCTTTCGAAAAACTCGTTCACCGGTCCAGTTACTTCTTCTATCGGTTCTCTATCCCGTTTACAGGAACTATTTCTCGATAACAACAAATTCACCGGTTCAATCCCGGAAAGTTTTAATGGTTTAATCAATCTGAATCGATTCGAGCTTCAACAAAACAATATCTCCGGTGAGTTACCTAATTTGAATCAGCTCGGAAATCTGATCTTCTTAGATTTAAGCAATAACAGACTCGCCGGAAATTTCTTCACGACAAATTTACCTGATTCGATGATTGAACTCTCAATACGCAACAATTATTTACACGGCGAGTTTCCGTTGAACTTCGAAGAGTTGAAGTTCCTACAGGTGTTAGATCTGAGCCATAATTTGTTTTCCGGGATGATTCCGGCGGTGTTATTTCATCACTCATCTCTTCAGCAGCTTACACTTTCTTACAATAACTTTACGTTTTTGCAAGTGCCGGAAGATTTAGTGCTTCGAAGTAAGCTTATAGCTATTGATCTGAGTTACAACAATTTGCATGGCTTATTGCCGGCGTTTATGGCATCGATGCCGGAGCTGTCGGCTCTGAATTTGGAGCATAACAAGTTTACCGGCGTGATACCGATACAGTTTGCAGTAAAAGTTGTGGCTCCGAGGAATCATACGGCGTCGTTTGAGAGGTTATTGCTGGGTGGAAATTATTTGTTTGGGCCGATTCCTGGGCCTTTAATGGGCCTAAAGCCTGGTTCTGGAAATATTAGTTTGGTGGATAACTGTTTGTATATGTGTCCAGATTCGGTATACATTTGTCATGGTGGGTATCAGAAGTCTTTATTGGATTGTAAAAAGTTTGGGCCTAGGATTCCTTGA

>PGSC0003DMG400003802 LRR-RLPs

ATGTTTGAATTCGCCCGGAAATTGCAGAAGCGAGGCGGAGAGTTGATTAGCGAGTTTTGCAGTGAAATTGGAGTCGAAAAATGGGGTTTTAAGGTACGGTTGCACAATAGGACACCTTCAATTGATATATTGTCGGAATGTCTCATTCATAAAATACTTTGTTGTCTTAGTTTTAAAGAAGCAACGCGGATGAGCATTCTTTCCAAAACATGGCTGCAAGCCTGGTCGACTCTTCCAAACTTGGATTTCACAGTTGATTTTTCTAAAGGCAACGATATGAAAATAGTGGATGATATTATGGGGAGATATAGGGACGGAAAAATCCCTATAGAAAAGTTTGAATTACTAGAGTTCGATGGTGATTCTGAAAATCTCTTCCGTATGATTGATAAGTGGCTTGACATTGTACTTACAAATGGTGTAAAAGACCTTACCATATCATCATATTCATACCCCTTGTTTATTTTTGCAATCTTCACGGTAAAATCTGTGAGAAAATTGGTTCTGGAATCTTGTACTCTTTTGCCTAGTGTTGTTGTGAATTGCAATTCTTTGAGAAAACTTTCTCTATCTTATGTAAGATTAGATGAAAACATGCTTCAGGCTCTACTTAATAGCTGTCCGTTGATTGAATCTTTCATATTTGAGTATTGCAAGGGGTTGGAAACAATGAATCCTCAAAAGATTAAGTCGGTTTCCCTGAAGTTCTTGAAGATTCGGTGTTGTGGGGGAATATGGGAGATTGATGCTCCAAATTTGGTATCACTTAAGTACACCTGGGAAGACATTCCTCAACTTAAAATTGTGAGAGAGTCAAGACAACTGAAGTACTCAAGAATCGTTCTTCAATGCTTAGATAGTGACAATGTAAATGCAGCATGGTTTTGTAAGTTGAGGAAGTTTCTATCAAATTCGATCTCTTGGTCTGAAGTTTGCATTGAATCTTATAGATGCAATGACATCAACATTCAAAATTTGCAACTGGACCATAAAGATTCTAATCCCCACGTGGACGTTTTAAATGTCATCATTATATGGAAGAATGGGGAGAGCTCAACATTTGTGGATGCTTTGGTATGGAGTTGTCAGCCTAGGAGACTCAACGTGGATTCAACAAGAGAAAAGATCACATTTTTCATTGATCATTTGATGTATATGAAGAATTCGAGTCAGTCTACTTCTCATGGAAGCACGCCTCTTCTTAGTCAATTAAAAGGAGTAAAAGCCTACAAATTTGATAGGAAGAAGGAGAGTTGGCATCCTGTGGAACACAAAAGTGGTGAGCTGGTAACAACAAACTCTGAAATGTGGGATTGGTGTTATTTTTTATTAGATTGGTGA

>PGSC0003DMG400003814 LRR-RLPs

TTTTTCATGCTGTAGCCATTAAAGCTTCCACCTATCTCCACACGAGAACTCGTTTGCTTCTCACGGAAAAGCAATACCCAAAAGTCTCTACTTTTTTCCCCTTATATAAACCCCCATTTTAATTTTACTCTCTCTTCACCTCTGCAACAATTAATTAATCATGAGGTTTTTGTTGTTGATGTTTCTGTTTCATGTTATTGTTATTCAACAATGTGGTGCAATGCTTCACCCCATTGATTTCTTGGCTTTACAAGCGATTCGTAAAACCCTCAATGATTTACCCGGTTCGAATTTTTTCTCTTCTTGGGATTTTACTTCCGACCCATGTAGCTTCGCCGGAGTTTATTGTGCTGGTGATAAAGTGATCGCGTTGAATCTGGGTGACCCGAGAGCCGGGTCGCCGGGTCTAACCGGAAGATTAAACCCGGCTATTGGGAAACTCTCTGCTTTAGCTGAGTTCACTGTCGTCCCCGGTAAGGTATTCGGAGTATTGCCGGAAAGTTTTTCTCAGTTGAACAACTTAAGATTTCTCGGAGTTAGTCGAAATTTTCTATCTGGACGGATTCCGGCGGGTTTAGGTAAACTCCGGCGACTTCAAACTCTTGATCTTAGTTTCAATCAGCTCACCGGAAACATTCCATGGGCCATCGGAACCCTCCCGGCGCTCAACAATGTCATTCTCTGTCACAACCATTTATCCGGTTCAGTACCGCCTTTCGTATCCCTTAGATTAACCCGGCTCGATCTCAAACACAATGCTCTTTCTGGTTCGTTATTGCCAATGTCACTTCCTTCTTCACTTCAATACCTCTCAATGTCATCGAACCAGTTAACCGGTCCGGTCGACCATTTACTTACTCGGTTAAACCGGTTAAACTATCTCGACCTTAGTATAAACCGGTTCACCGGTTGTATCCCGGGGAACTTATTTAGATTTCCGATAGCAAATCTTCAGCTACAAAGAAACCAATTCACCAGTCGAATTTACCCGGCCGGTTTAGTAACAATTCCGGTTGTGGACCTTAGCTTCAACCGGTTTTACGGCGAAATATCGCCATTGTTCTCCGGTGTACAGATTTTATATATGAATAACAACCGGTTCATGGGTCAGGTTCCGGGGATCATGGTAGACCGGTTGTTATCGGCTAGTATACATGTTTTGTATTTGCAACATAATTTTCTAACCGGAATAGCTATCAATCCGGCCGCAGAAATTCCGGTTCGGAGTTCATTGTGTTTGCAGTATAATTGCATGGTTTTGCCGGTTCAGACCCCTTGTCCATTGAAAGCCGGGAAGCAGAAGTCCCGGCCCACGGCTCAGTGTGGGCAATGGAAGGGGTAAAATAGACAAATCACAAGTATGTTTTTTTTAATTATGGTTTTGGTGGATTCATTGATTAATTTTGAAATTAAAGGGTAAAAAAAAATGTGGATAAAGAAGAAAAAATTAGAGTGAAAAAGGGTAAGAGTTTTAAGTTTAGTTATTTGTATGTTTTTTTTTGGTATTTCTTGTAATTATAGATATATACTGATTTACCTTATTAAATATGTTGCAAATTTCTTGTTTTTTTTTTTTTGGTTCTTATTTTGTAAAATGTTGCTTTATGGGGTTTTCTTTTCTTTTTTATTCCTGTTTACAACAGCCTTATTGTTCTGATGATAAAAGTGAAGTCTTTAGGGATATAGAGTATCTTTTTCCTGTTTTTAGGAAATTATATGTGATATACAGAGATTGTATACAGCCAAAATACAGCTGCACCAGCATATCTCTGTTCTTTGTGGTACTAAAC

>PGSC0003DMG400003822 LRR-RLPs

TTGAAAACCAAATATTACAGATGACAACATTTCATACTACCTTAGTTTACATCTTGTATGTGCTACTTCTTTTTTCACTTCCATTAAGTATCACATCTTCAGCAAGAACAGAAGCTGAATCTCTTGTCAAATGGAAGAGAAATTTACCTTCTACTTCTTTCTTGGACACATGGTCCATTTCGAATCTCGAAAACTTGTGCAATTGGACATATATTGTGTGTAATGATGGTGGAACAATTTCTGAGATCAATCTTTCTGATGCAGCCCTCTCAGGCACACTTGATCACCTTGATTTCACTTCATTTCCGAGTCTCGTGAATTTCAATCTTAATGGAAATAACTTCAGTGGATCGATACCTTCGAGTATTGGCAATGCCTCATTGCTCACATTCTTGGACCTGAGTAACAATATCTTATCAGGTGTCATACCTGAAGAGATTGGAAAATTGAATCAACTTGAGTATCTGAGTTTTTATAACAACAATATCGAAGGTGTGATGCCGTATCAGATTAGCAATCTGCAGAAGGTAATGCACTTGGATCTTGGATCAAATTTCTTAGAAACTCCTGATTGGTTGAAGATGAGAAATATGCCTGTGTTGACGTATCTAAGCTTTGGTTATAATGAACTGAGGTTAGAATTCCCTGAATTCGTACTTCGTTGTCATAATCTAACATATCTTGATTTATCGATAAATCATTTCAATGGTTCGATTCCTGAAACAGTATTTACCAACTTGATCAACCTTGAAAGACTTAATCTTTCATCTAATTCATTTCAAGGTTCATTGTCACCAAATTTTACTAAGTTGTCTAAGTTAAAAGAACTTCAGCTAGGTGTTAACATGTTTTCTGGTTTAATTCCTGATGAAATTGGCTTGATTACTAGTCTTGAAGTTCTTGTACTTTTCAACAATTCATTTGAAGGAAAGATTCCATCTTCTATAGGTAGACTCATAAATCTTCAAAAGTTGGATCTTCGAAAGAACGATTTGAATTCAACTATTCCTTCTGAGCTTGGCTTTTGTACTAAACTCACTCTCTTAGCTCTAGCAGAAAATGACCTGCAAGGGTCATTGCCTTTATCATTCTCCTCTCTTGCAAAGTTATCTGATTTGGGGTTGTCTGATAATAGTCTTTCTGGTGAAATCTCATCGAATTTCATCACCAATTGGACTGAGTTGACATCTTTGCAGCTTCAAAACAATATGTTTACAGGGAAAATCCCACCCGAAACCAGTCAATTGACAAACCTTGTATATCTTTACCTCTATCATAACAATTTCACTGGTTCCATTCCCTATCAGATTGGGAACTTGCAAAACTTGTTGGATTTAGACTTTTCTGATAACCAGCTTTCAGGCATAATACCTCCAACCATTGGAAATCTGACCAATCTAAAGATGTTGCAACTTTTTCGCAACAATCTCAGTGGAACCATTCCTCCTGAGATTGGAAAATTGATATCCCTTGAAACCATCGACATCAATACCAACCGACTAAGTGGTGAGCTGCCGGACAGCATTTCTGACCTCAGTGAGCTGAAGTTTCTTTCTGTATATACTAATGATTTCTCAGGCAGTGTCCCTAAGGACTTTGGAAAGAACAGTCCTCAATTGTCAAGTGCCAGCTTTGCAAATAACAGCTTCACTGGTGAACTACCTGCTGGATTGTGTAGCCCGAATCTTGAGGAGTTGACGATAAATGGGAACAAATTCAGTGGGAAGTTACCAGATTGCTTGAAGAACTGCACACTTCTAAGACGAGTACGGCTTGAAGGCAACAATTTATCTGGTAATCTTGCAGATGCATTTGGAGTGCACCCGAAACTTGTTTTCCTTTCTCTTAGTGACAACCAACTTTCAGGTGAACTCTCACCTGACTGGGGGAAATGTGAAAATCTCACAAGTCTAAGAATGGATGGAAACAAATTTTCTGGTGTAATCCCAAGTGAGCTAGGGAACCTGAGAGCGTTGCGCGTGTTAGCTTTGGAAGGAAATGAATTGACTGGTGAAATTCCCTCTGAACTGGGAAGGTTAGACCTGCTCTATAATCTCAGCTTGAGTAAAAACAATCTTACTGGAGGCATCCCTCAGTCTGTTGGCAATTTAACTAAGCTCCAGTATCTTGATTTGTCGACAAACAAGTTAAGTGGTAACACACCAGTAGATCTTGGGAAGTGTGAGAGCCTGTTGAGCTTGAATCTTGGCAACAACTCATTATCAGGTGGCATTCCATCTGACCTTGGTAATTTGATGCAGTTGAGTATTCTTTTGGACTTGAGTGGCAATTCACTAACGGGAACCATTCCACAAAACTTGGCCAAACTCACCTCATTAATGCATCTCAACCTCTCACATAACAACCTCTCAGGTAGAATTCCTCCAGCATTATCTCAGATGATCAGTCTTCAGGAGATGGATTTTTCCTACAATGAGTTTTCTGGACCAATCCCAACCGATGGAGTATTTCAAAGAGCACCCGCTAGATCTTTTCTTGGAAACTCTGGTTTGTGTGGAAATGTAGAGGGATTATCCTCATGTAATTTGGATACTCCCAATGACAAGTCCAGAAATAATAATCAAAAGATCCTCATTGGGGTACTTGTACCGGTGGTCAGCCTCATACTTTTGGCAATCCTGTTTGTTGCATGTCTCGTGTCTCGAAGGAAGGCTAAGCAATATGATGAGGAGATCAAAGCTAGCCAGATACATGAAAATACAGAATCTCTGATTTGGGAAAGAGAAGGGAAGTTTACATTTGGTGACATAGTGAAAGCTACTGAAGATTTCAGTGAAAAGAACTGCATTGGAAGAGGAGGCTTTGGAAGTGTCTATAAAGCTGTTTTGCCCTCTGGGCAGATTGTTGCAGTCAAAAGACTCAACATGTCAGACTCAAGTGATATTCCATTGACTAATCGTCGAAGCTTTGAGAATGAGATTAGAACTTTGACAGAGGTGAGACACAGGAATATAATTAAGCTCTTTGGTTACTGTTCCAAGAATGGATGCATGTACTTGGTTTATGAGTACATAGAAAGAGGTAGCCTGGGGAAAGTTCTATATGACAATGAGATGGGAATGGAACTTGGATGGGGTACAAGAGTGAAAATTGTGCAAGGAATAGCTCATGCACTCGCTTACTTGCATCATGACTGCTCTCCGCCCATTGTTCACCGTGATGTATCATTGAATAACATTTTGCTTGAATCAGAATTCGAGCCACGACTCTCTGACTTTGGCACAGCAAAGTTGTTAGCTTCAGACTCATCAAATTGGACCACAGTTGCTGGTTCTTATGGCTATATGGCACCAGGTATGTTACATACTCATTACCTAATGCATCCATATTTCTGTTGATTAAGTAGTATATTCATTGGAAGGATACTTCAATATTATTGTGTCTTTCTTCTTGTGCAGAGCTTGCACTTACCATGCGTGTTACAGAAAAGTGTGATGTTTATAGTTTTGGAGTCGTAGCGATGGAGACTATGATGGGAAGGCATCCAGGGGAGCTTTTAACTTCATTATCAGCATCAACAACATTGTTTCCAGAAATACTTTTGAAGGATGTTCTTGACCAAAGACTTCCACCTCCCACAGGCCACTTGGCAGAGGCAGTGGTTTTTGTGATCACGATTGCCTTGGCATGTACACGTACCACTCCTGAATCGCGACCAACCATGCGTTCTGTAGCACAAGAATTGTCTGTTCAGACTTTGCCTTACCTTCCACAGCCATTGGGGACAATAGAAATGAGCAAACTAACAAGTTTTCAGAAATAATGAGAAGATAGAGCCTCATGTACAGGTGTGAAATGTACAGGCTGTAGAGTTTTAGTGTTGTAATGCAACTTCCTCAAGATCTAGAATCTATATAGTATCAAGGATGAAACCTTGTATCTATTCAAATTACTATGTTTTGGTTGCACTTTTCCCTAGAAAGAAAGAAATAATATCATCCTGTTAATCATGAATGTTACAAAGTTGGTATATTAATTTGGACAAGTATGAATGTTACAAAGTTGCACAACCTTATCCTAGTTTTCACTTTATTGCTAGGCAAAACACTAGCAAAAAGATACAATGTGTTCACTTGTAATAAAAATAGCAACCTAACAAAAATTTGGCAAAGAATACATCACACAGAAACCAGGCATGGTACTTAGCATTCCACTCCACACTGCTTGAGTATAGGTCCCTTCTGCTTTGGTAGGAATGGGTCTATTCGTACCCAAACGAGTGAGAAGATTGAAGCAAGAAGGATTGACCAGAGGACCACAATGGTGGGAGTCCTGTTCTGCCTGCCCATCAATCCCTTTAGGAAAGGATAGAGATGCACAATAACCCAAAAGGCAAAGAATAACTTTCCGAAGAGAGGACCCCATGAACCATAACCATTGTTGATGGCATCCGAAACACCAGCTACAACACCAACCATATTCAAGATGATCAGAGTGGTTGGTGGGATGAGAAGTGTGGTCCATTTGAAAAGGTAAAGCTCCCCAAATTCAGCGTCATCAGCTGCTTTTGATGTTACAGTGAAGTTTGTGTCTACACCAGCAAGAACTTTAAGAAGGCCTTGGAAAACTGCAAACAAATGTGCTGAAACACCACCGATGACCCAAAATTGCTCGTTACGCCACCAGGCCTCAATGCTTACTCCACTCCATCGCAGCTCGAGTACACTTGTGGCTATGATGGAGATAAAGAGAGCCATGAACCAAATGCTTGCAAAGTTGTTAAGCTACAAAACAAACATGGATTTTGATTTATAAAATCTAAATCAAATCTTTATATTACTGAACCAAATGACTTCAGTTCACTTACTGTGGGGACGATAAATTTTCCAGTGAGAAGGCATACAGCTGGAAGGGTGCAGTAAGCTAGCAGAGCAATGGAAGTGAATGGGTAAACAATAGTGTTAATGTATGCTAGCCTTTCCAACCACTTGAGTTTTCCACCCCAGGCATACCAAAGGGGACAATGACGACTCATGAAGATTTCGACAGAACCAAGAGCCCATCTCAGGACTTGGTGCAATCTATCTGATAGATTGATTGGTGCTGATCCCTTAAAAGCGGCTCTCTTCGGACAGCAATAAACCGATCTCCATCCTCTGCAATGCATCTTGAATCCAGTCAAAATGTCCTCTGTGACAGAACCATAAATCCATCCAATCTGTCAAAAAAAGAAAAATGAAAACCAATTTATTAGAAGTTAGCTAAGCGAACACCCATAACTCATAATTTATTCATGAAATATAATGCTAACCTCTTTGCCCCAGTCAGTTTTCTCTTCATACCCACAGCTAATAACATGAATAGCTTCTTTTATGAGTGTTGTCGGGTTTGTCCCTTCAGGAAGCCCACCATCTTCCTTAAGTGTGGAAGCTATGAAAACAGGGGACTGCCCGAATCGTTTCTCAAAGTTCTTTTGGGACATGAGAGACGACTTCTCGAGCTCATCATATCCTTCAAGGCCTTCTTCAATTTCCTCAAGATCAAAGACAGGTCCAGATGGCTTCCTAGTGTAGTTCTTTCCATTCATTGTTTTCTTTTTCTTGCTGTATAGTCCTCCAAGTCCAAGTAAGGCCTTGATGCTCTTCTTCTTGGACTTGGACTTGGACTTTGACTTTCTTGAACCACCACAACAGCAACTGGGCCAGCAATCACATGTCATCTTTTGACGCTTCTCTGAAACAGGTGGATCATAGCCGTATAACGCCGGCCTGTTAAAGACAGTACCTGTTCCAACATAAACTGGTCCTTGAATTCCATCCAGGCCTTTCATGTTGATCTGCCACCATGAAGGGAAAAAAGCATATTAATACCGGGAATAAATCCTTTGTCAACACATTGCTGTATATATAGATCAACTTACATCAAAAAAGACAACATTACGATTGGCATATCGATCGTGGCGATCAATACCATCAAATCTCTGGGGAAATTGGACATAGCATAGCTTCTTTCCAATTTGTGGGTCCATTAAAAAGCACATGGCTTCTCTAACTGCCTTGCTGTTGTTAAGGTAGTGATCACAATCCAAGTTCAACATAAATGGTGCATTCGTCAGGACTGCAGCGACTCGAACCTATTTAGAAAAACAGCACAAATTACATAACATTAAGAAAAGGACTTAATAGGATGTCAAGGAAGCAAGCCAGAAACTAATATACTAATTACTAACCAAAGCATTCATGGCACCAGCTTTCTTGTGGTGTTGATAGCCCGGTCGTTTCTCACGGGAAACATAGACAAGGCGAGGTAGCTCCTTACCTTCAATGTCAAGTGCACCTTCACTGCCAAGATAAACTTGAATCATACCAGGATGGTCTCTGGTGTTGTTACCGGGCCACGGAGTTCCATCTTGCATTACCCAGCCTTCTTCTGGTTTTTTCTGAGCTTTTGCTACTAATGCATTAATTCTTACTTTGAATTCTTCATATTCTCTCTGCCACAAGAGAGTGAGTTTTCAATTTGCAGAAAAAATCTTATTTTAAGCCAAAGAAACTTAAGCCCCCGATAAATGATACTCCTTCTATCCCAACTTATATGACATTCTTTACTTTGGGCACGTCCTAAAATGTTCGACACGATTCCTTCTACAACTTCTTTTTAAACCTTGATATGATTCAGTCAACATTGTAGGCACTGCATCTCCTCTGATATAGTCGTATGAAATGAGACGAAGGAAGTAATAGATACATACCTTCATGGCCCTGCGATCCTTGACAAAGGTAGGCTGAACTTTATCTTTCAAGTAATCAATCTTTTCGTTGAAGTAGATATCAGGTGCTCTAGGCTCTACACTATATTTCTTACAAAATGGCACCCATCTCCTGGCAAATTCAGCTGTTTCTGCTAATGAATCAAACAACAGCATGGAAGCACCATCATCTGATACATAACAGCTAACCTTCTCGACAGGATAATCAACAGCCAAGATTGACAGAACAGTGTTTGCAGTGATAATAGGAGGCTCCTTAAGAGGATCAACAGTACTCACAAACACATCCACTGGGGCTAATCGATTAGGCTCGCCTTCACGTTCAAACCTCAATGTCAATCGATCTAAATAAGTTTCACGGTTTATTGGTGACCATTTGGGGAACTGATCGAGTATCCATGATAATGCAAACCATATTTCACATATAACAGAGATAATCCATAATGGATAGGCATCATAGGCTGGTGTTAGGATCCGGAAATGGAAAAAGAATCCAAGAATTACAAGCCTGAGGATAATGACTATCCGATATGGACTGATTAAACTTGATGGAAGTGGAACTTTTCGCCAGAGTGGTTGCCGAGCTTCAGCCAAACTGCTTGGAAATCAGTTAAGTCGTTATAATATTGAAATATACTGCAAGTTTTATATGCTTAAACTTGATCTGTTTCAATTCATACACACTCATTTCATGTTGAAACATTGTAGTATCTGATTCATTTTGTTAATAATATTATATGTAAAAGTGTTCAAATTATTTAACTATACCAGTCAGTCACTGAGTGATACATAACTACAAGTGACACAGACGATATATAACCGAAAATCCAATATTTAAAAGTTGTAAGGAGAATGAGGAGAAGGTGGAAACAAGACCAGATATAATTTGATCGCTATGAACAAAACCAAATACCAGATTGACTGATTTGTTATAAATTTAAAGTATTTCTATAGCATGTTTTCCTTTTCCTTCTATGTTGTTTTCTCAATGATTCCTCACATAAAATCTTCTCATATTAACGACGATATAACTTCAACACTATCAAAGAGAGGCGGCAATTAGCTTGAATAATCAAACAAACTTATCAATATTTTCTTCCACTACTAAGAATATATGTTAAATTCCCCTTCTAAGTCAAAGGTCTATCTAAAAAGCAAATCTAGTAGGGGTAAGGTGTGGTATATCTTACCCTCCCAAACTTCACTTGTGGGATTACACTGGGTGTATATATCATGTTGAATTCCCCTTCTAAACCGAGCTGTTAATTTCCACTTCTAAACCGAGGGTCTATCAGAAACAGTAAAACTACCCTGGCCCGCAAAGGTAGTGTTAAGGTGTGGTATATCTTACCCTCCCAAACATCACTCGTGGGATTACACCGAGGGTAAACAAGATGTAAAGTGAAAAGAAAGAGTACATACAGAAAGTCATCATCTTCATCAGGTTGATCATTTCCACCATTATTAACAAGACCTCTCTTTTCTTGACGATTCTTCCATTTCTCAACCCTTTCCTTCCATTCTGCATCACCATAACTTTCTTTCTGCTCATCCTCCAATTCCTTCCCAGCAACTGTAAGTGATTTTCCATTGAAATAAAACAAAATCATGACACATATATATAAAGTGCAACAATTACTCATTTCGAAATTAAGTAATAGAGATCGATATAGCTTTACCGCTTCCAACAATTGATTGATTAAAAGCAGCAGCACCATTGGTGTTGTGATTATGCTCTCCATTCTCCTGCCAAACATACAACAACTTGCTTTCAGTTCAAATGAG

>PGSC0003DMG400003834 LRR-RLPs

ATGAGCATTCTCTCCAAAACATGGCTACAAGAATGGTCGAATCTTCCCAACTTGGAATTCACTATTAATTGTTGGGAAGGCAATATAAATACAGCGAACACAGCCATGGAGAGATATAGGAAGGGGAAAATCCCTATACAAAAGTTTGAATTATTAGAGACCTTTGCTAATTCTCATGAAGATTTCCCTCTCATTGATAAGTGGCTCGATATTGCACTTCAGAATGGTGTAAAACATCTTTTTCTTAACTTTAAATCATACCCCATGCCTATTTTAAGAATCTTGGCAGCAAAATCTTTAAGAGAATTGGATGTGCAGGGTAGTATGCCTGATTCGTTATCTAGTGGTGTTGTGAATTGCAAATCATTGAGAAAGCTTTCTCTATCTCATCTAAGATTAGACGAAAACATGGTCCAGACTATACTTAATAGTTGTCCCTTTATTGTCTCTTTAATCCTTGAGTATTGTACTGGGAAGCTTCAAAAGATCAAGTCGGATTCCTTGAAGGTCTTGAAGATACATCATCGCGGGATAGAGGAGATCGATGCTCCAAATTTGGTATCACTTGACTATATGGGGAGTCAAATTCCCGAACTTAATAAAATTGCAAGAGACTCAAAAATCAGGCTTGGATGCATCAACAATTTAAATGCAGCATGGTTTTGTAAGTTGAGGAAGTTTCTATCAAATTCGTCCTCGTGGTCTCATGTTTCCCTTAAATTTATGAATTGCTGTGAGATCAAGATGAAAGATTTGCATATGGACCACATAGGTTCTACCCCTCGGGTGGACGTTTTAAATGTCGATATTCAATGGATGAATCAGAATTTTGTGGATGCGTTGCTATGGAGTTGTCATCCAAGGAGACTCAACCTACTCTCAAATATTACTCCAACTATTACACATTTCATTGGTCGTTTAGTGTTTTTGACAAATTCCAGTCATTCTACTTCTCATAAAAGCTTACCTTGGCATAGTCAATTAAAAGAAATAAAAGCTTTTGATGGAAAAAACCAGTCGCTACAACTCATAAATGGGGAGCTTGCAAAAAGGACCCTTATGGAGGGGGAGAAAGTTTATTTTTTATTAGATTGGTGATGTAGTTAATTTGTTTCTATTTCAAATTGAATGCACTTATTCTTTTCTGCATTGGTGTACAGAGGTATTAGATATTATTAAAACAGAACTTAGAGCATAATTTTAAACTTGGATCTTATAGGCTTGCAAAAGCAAAAACAATTTTTTTAAGAAAGTAATGAGTATGGCAAATTTCTTAAAAGGAAAGAGACTTAATTTCTACATAATTACTAGTGTTTCAACGTAACTACAAATGCCTTCGATGCCATACTTTCTAGGTAGCCCGTTGATTTCTAATTTTCTCAAACTCTGTCTCGTGACATCATAAGAGAAACATAAAGTATCCGACATTGAGTCCAGGATGAATACAATCTCTTCATCACTGGACGTGCATGATGATATAACTAAGGGTTTGTCCTTCCACCACATCGAAGGGAAGTCAATGATTTGAATTTTCCATTCTTCTTCTTTCTGAGTTTGTTCTAAAATCCACAAGTTGAATTATCCATGTTGCCAATATTCATAATCCATGATCGCTAGTTTACCTTTCACCTCTAACAACTCGTAATACCACCACGAGTTAGATGCATTCAATAAATCAATATATTTTGAATTTTCGATTTTAACATCAAATGCATCTATAGTAAGTCTGTCCTGGAAAACAAATTTATAGATAACTCCATTGATGCAAACCCCGGGCTTGCACAGAATAGGAGGGGTTATGCTTTGAGTCTCTCTCCATGATTTGTCTACTCCTAAAGTGAAAATCCAATGTTTTATGTACCCTTTTATAATATGCTTTGTTGTGGAAAGAACTTTGTACTTCTTTTCTTCTGGGTCAAAACCGAATGAACAGTACCACATGGAAGAACCATCATTTAGGTCGGGAAGAAATCTTACTTCTCTTGTACTAGGATTGAAAATTGCAGCAAAAATAATGGTTCCCAAATACAAAATAAACCAGGTCTTGGGTCAGATCTCAGGATCGGATTTTGGAACGGTCGTTGGGGTTGTGTCCTAGTTTGGGTGTTAGGGTCGGGTCCTACATCGATTATCGGGTTGATTTTCGAGTTAAAAATTATTCTCCTAAAGAGAATTTTCTAGTCTCAAACAAAAAATAAAAGATTTTTTCTGGAAAATATTTTTTATTCACCAACCAAACATTAAAAAATATTTTTCAGAAGATATTATCTACTCATCAACCAATAAACATAAGAAAATAAGTTAGAAATCCATTTGTTTTTTCCAAGAAAACATTTTCCTTCGTACCAAACACACCCTAAAGAGTTATAAAAAAGAAAAAAGGACAATCTTTTTTAAATGGATTTAATTGAAAAGTAAGGCAAATATGTTAATTGAAACATATGAAGTATTTTTTTTCGACCTTCTGGAATTTTTTTTAAAAAACAAACAAAGAAGAAAGTCTAGGGGATAATTTTATATAGTTTCTTTTTTACTCTTTTTGCTGACAAAGAACTTTAATATTTTGACCTTATGTAGAAGAAAATATTTTAATTTTTAAGTAGCGCATGATGTTTTCTTCTTTTTTTAAAAAAAATCAAGCATGATGTATTAATTTTTTTGTACATTTATTTTTATTTTTATTAATGAACCATGATACACGTGCAACACACACAATATTAAAAGCATGATTTTTTTTAGAATATAGATTGAACTAACTTTTTATCCTTCATTCAAAATTTTCACAATAGGCAAAACAATCATAATATTATTCGATTTTTGCACACCTCTACTTTTCGATGCGAATCTAAATTTAATCGGAATCTAATGTATAAAAATAATGATTCATTCAAATATTATTTGACCGTGTGACGAGGGAGGGAGATCTTTTAGTAATATAGAGCTCCTTATTTTTTTTTTTTTTTTTTTTTTNNCAGCAAACTCCGGCGAAGGTAAGAGCATTTTTCAGCTTCTCCGGCGAACTCATCAGCAAACTCCGGCGAAGGTAAGAGCATTTTTCAGCTTCTCCGGCGAACTCATCAGCAAACTCCGGCGAAGGTAAGAGCATTTTTCAGCTTCTCCGGCGAACTCATCAGCAAACTCCGGCGAAGGTAAGAGCATTTTTCAGCTTCTCCGGCGAACTCATCAGCAAACTCCGGCGAAGGTAAGAGCATTTTTCAGCTTCTCCGGCGAACTCATCAGCAAACTCCGGCGAAGGTAAGAGCATTTTTCAGCTTCTCCGGCGAACTCATCTATCTTCTCCGGCGACCAGCTCAGAAGAGAGACGGCAGTGCTACTGTTTGCCGCCGGAGATCAACCAGATCTGGCCGGAAATTACCCATTGATCACCGAAACCCTCCATCTTCACTGACCAGGTTCTATCTATCCTTTTCTCTCCCTTTTTCGCTGAACCTAGTTGCTGCTGGGCTCTGTCTTGCAGACTCCTTATCCCGGAACAGTATCCCCAGTTTTTAA

>PGSC0003DMG400003835 LRR-RLPs

ATGGCTGACATATTGCCTGAATGTTTCATTCAAAAAATATTGTCTTCTCTTAGTTTTGATGAAGCAGCGCGGATGAGCATTCTCTCCAAAACATGGCTGCAAGCCTGGTCGACTCTTCCCGACTTGGTATTTAATGTTAAGTTTTGCAAAGGCGATATAAAGATATTGGACACAATTATGGAGAGATATGGAAACGGAAAAATCCCTATAGAAAAGTTTGATTTATCAGAGCTCTTAGGCAATTCTCATGAAGTTTTCCCTCTGATTGATAAGTGGCTTAACATTGCACTTCAAAATGGCGTAAACAATCTTGTCCTTAACTATAAATCATACCCCGCGCCTATTTTAACAATCTTGGCAGCAAAATCTTTAAGAAAATTGGTTCTGAAGAGTTGTACTCTTTTGCCTATTTCGTTATCCAGTGGTGTTGTGAATCGCAATTCTTTGATAAAGCTTTCTCTATCTTATGTGAAATTAGACGAAAATATGCTTCAGACTCTACTTAATAGTTGTCCCTTGATTGTCTCTTTCATCTTTGAGTATTGTTCGGACTTGGAAACAATCGAGCTAGTGAATCTTCAAAAGATCAAGTCGGTTTCCCTGAAGGTCTTGAAGATTCGGTTTAGTGGGAGCATATGGGAGATTGATGCTCCAAATTTGGTGTCATTTGAGTACATGGGGAATCAAATTCCTGAACTTCAAATTGTGAGAGAGTCAAGACAACTGAAGCACTCAAAAATCATTCTTCACCGCTTAGACAATATAAATGCTGACTGGTTTTGTAAGTTGAGGAAGTTTCTATCAAATTCAATCTCTTGGTCTCAAGTTTCCCTTTATTTTTATGAATGCGCGGAGATTAAAATGAAAGATCTGCAACAACACCACAGAGTTGCTACGCCCCAGGTGGACATTTTAGATGTCAACATTCTGTGGCAGAATCGCGAATACCCTTCGTTTGTGGATGCTTTGGTATGGAGTTGTCAGCCTAGGAGACTCAACTTGCAATTAACAAGAGAAATGGTTACAGTTTTCATTGATCGTTTGATGCATATGAAGAATTCGATTCAGTCTACTTCTCATGGAAGCAACCCTTGGTATAGTCAATTAAAAGAAGTAAAAGTCCACAAATTTGATAGGAAGAAGGAGATTTGGTATCCTGTGGAACACAAACTTGGGGAGCGGGCAACAATGAACCTTGATAGATATTATATTTTATTAGATTGGTAA

>PGSC0003DMG400004056 LRR-RLPs

ATGTACAGAGGTGTAGCAGATGCTTCTATATCTCCAATGGAGAAACATGAGAAAGAGGCATTGTACTCTGCTATTCAAGGTTTTGTTGGAAATGAGTGGAATGGTTCATTTCTTTATCCTGATCCTTGTGGCTGGACATCAATTGAGGTACTAATCTTTTGCTCTAGATGTTCCTTTGCTAATTTTCAAAGCTAAATTAGATTACATTAATTTGATATTTTAAACTAAACATTTAGATATTCAAAAACTATATAAAAAATACTATAAATTGTAATTTTTTGCATATTAATATGATGAAGTTAGTCAAAGTTACTCTAAAGAAAACAGGAAAACATGACAACTAAAAAGGAACGAAGAGTATATGATTTCATAATTTGCTAGTTGATTCTAGTCTATGTTTTTTTACCATTATTGACTTTTCTTGCTGTAGTTCAAAATCAAGAAAACAGTAATGTCACAAAGCCATGCATGGATGTCCTCAATTTTTATTTTTATTTTTTAATATAAAGAAATGGATCTTGGGCCTTACTCAACCGCATAAACTACATCATAGGAATTAGGATGAGTATTGTTCAAGACATTTGAGATCAATTACCCATCCCATATTCGACTCTTGTGGGCAATATAATATGGAAGCACAACATAGGTGAACAAAGAATTCGGATGAGCGGACTTTGGTAACATGATAAAAAAAATGGATCTTGGCCTAACTCAAGACAAAAAAATATCTCATAGGGTGAGGCTTATTCAAGACCATATAGGAGATCAATTACCCATCTCATATCCGATGTGGGACTCTTAATAGATCAAACTGTCCTCTCAATGGTTGTTAATTAAAGTGAAAAAGACTTTCTCAAAGTCTTACTTTTTCTGTTCTGGATTTGCAAACAGTGTTATATGAGTGTGTGTAAGAGAGACATCACTAGTTAGGAGCTTTCCAATTTCTAACACACCTTTTCTGGTAGTACTAGTAAATATATATGCTAGTATTCTATTGGTAATTTGCAATATTTATTCCTAAAACCAAGTAGTTATATGTATTGAAATTATCGTTCTTGTATGTTAACTTGACATAACAGTGTAAAAATTCTTTACACCGTAACATAAACTCTTGTCTTAACACTTTAATCTACTCTGTGAACTCTTATTTTTACTATAATTCACAGGGAGTATCATGTGATTTCTCCAATGGCTTCTGGCACGTAACGGATTTAACTATTGGAGACCTATATGACAATTCCCTTCGTTGCTCCCAAGCTGCAGAGTTCACAGAACACTTGTTTAACCTCAAGCATCTAAAAAGACTTTCCTTCTCAAATTGCTTCCTCTCAGATCAGCACAAATCAATTCCCATTTCAAATTGGGATAGTCTCGCGAACAGCCTTGAATCGTTAGAGTTCCGATCAAATCCTGGTCTTATTGGGACAATTCCCACAAGTTTTGGCTACCTTAAGAATCTTCAATCTTTAGTACTGCTGGAAAATGGACTGAAAGGAGAAATACCAGAAGCTTTAGGCAATTTAACTAAACTAAAGCGCCTCGTTTTGGCAGGCAACAATTTTATTGGTCCTATTCCAACTAGTCTAGGAAGATTGACAAATCTTTTGATACTTGACACAAGCAGGAATTTTCTATCTGGTGCATTACCTGTGACTATTGGTGATTTATCTTCGCTCATAAAGGTCGACTTGAGCAACAATCGATTAGATGGAAAATTGCCTCGAGAAATTGGAGGACTAAAGAGTCTAACACTACTTGATCTCAGCCACAACAATTTCTCTGATGGTTTGCCAGAGACATTTCAAGAAATGGATTCCTTAGAAGAACTTGTGTTATCTGACAATCCGATAGGCGATTATGTTACAAGAATTCAATGGAGGAACATGAAAAACTTGGAAATGTTGGATCTTTCAAACATGGGATTGAAAGGGAATATACCAAATTCCATGACAGAAATGAAGAAACTGAGATTCCTCAGCCTCAGTAATAACATTCTTTCAGGAATTATTCCGTCAAAATTCGAGAAAATATCAACTATTAATGCTCTATACTTGGATGGAAATGATTTCACAGGGAAACTTGAATTTTCACAAAGGTTTTATACAAAATTAAGGAGTCGTTTTCGAGCTTCAGGTAATGTGAAACTCTGCTTGTCCCTCGAGTTAACGTCAACAAGTCATGTTCCAGAAGGAGTAAAACAATGTGAACAAGAAAGTACAGTAGATAGTAAAGAATCAGACTCCAATTTAAAAAATGAAGATTTGAGTCAGAATTCCCACCATATTATAGCTTCTTTGGGGCATTCAGGACATGTTGTTAGTCATTTTACATTTTGTTATGTCGCAAGAATAATACTTATGTCTCTTCCATGGATCGTATTTTTATAA

>PGSC0003DMG400004190 LRR-RLPs

AATCCAACGACGCGCAGTGCAAGCTAAATAGCGCGTCGTGTAAAAAAACCAATTTTGAGGAAAATTACCAAAGTGAGGATTCATTTCACACACAGATCTTCTTCCTTCACTTCCATCATCTCTCTCAATCTCGACTATAGTAATGTACGTATAGCTTTGCGTTTGGATTCCGGTTGATTCAGTAATTCTACTCCCTGAAGGTGATTAACCCTATTCAAATGGCTCGCCATGGGCATCTACCTTCAGTATTTCTCCTGATCTTCTCGTTATCCTTGATCCAAAGCGCTCTCGTTCACTCTAAAACCCTCAAACGTGACGGTACACATTTCGATCCATTTACTTTTTAAACCTAATTTATTAGCAATTTAGGTGAGAAAATCAGTTGAATTTGATTGATAATAGTTTGGTTTTGTTATTTTGTAGAGTATGTATGCAGCTTCAAGTGAAATTAATTTTGTAGTTACAGAGAAAATGTAACGATGTTGAACTTTGTAGATCTTATGTATTCTAGGTGCAAGGTTATAGTTTGAGTGGAGTTTAATTGAATTTTAAACGTTCAAGTGAAAATAGTTAGATTTTGTAGATTATATGCAGTTTGTGGTGAAATTTTTGTAGTTTCAGCGAAAACAATATTGAACGTTGTACATCTTATGTAATTTCAAGGTGTAAGCTTATAGTGTGAGTGGAAGTAATTGAATTTTGTAGTTCAAGTTGTAGTAATTGGATATTATCCAGTTTCTAGGTGAAAGCTTATAGTTTGAGTGGAATGAACCTGGATTTTATAGATTATGTGCAGTTTGTGGTGGATTCATGGTTTCTGCGGAAACAATTGAAAGTAAGTGGATTTTATGTAATTTCTAAGAGAGAGCTTATAGTTTGAGTGAAAGTAAGTGGATTTTATGTAGTCCTAGAGCAACTTTATAGTTTGAGTTGTAGAAATTGGATTTTATAGATTTTTTCCCAGCTTCTAAGTGCAAATTTTTAGTTTGAGTTGAAATAATTGGATTTATTCTATAAATTATATGTGGTTTCGAAGTGCACATGTATAGTTGGAAATGATTGGAATTTATAGATTAGATGAAGTTGTAAGTGCAAGTTTATAGTTTGAGCCAAAATAATTGGATTTTATGCATTAGCTGCTTTCTAAGTGTATGTTTATGGCTTTCCTGGAAGTAATTGGACTTTTTAGATTTTATGCAGCTTCTAAGCTCACGTTTATATATTTTGAGTGGAAGATTTTATGCTGTTTCTAAGTTTTACTTATTGTGTTTCATCTGTTCTTACTATAATTTCCTTTTTCCTGAATGTCTGTTTTCTTGGTTAAGTAATTCAAGCTTTGCAATTTGTTGATTTTGTTGAGGAATAGCTATTCGTTTGATGTATAATCTTGATCTGTTCAATGCATTATCTATTATCTGCAATCAATAACTAAATCAGTGCCTTGGAGTGTCATTAACTATAAATGTTTTCCCCTTTTTTTGCTCGGAGAATTCTATTTTCTAGTTGTTACAGGTAAATATAGCTATCCTTGGAAAGTTAAACCGATTTATCATTCAATCAATGCCAATTTAATATCTCTCCATGATTACATTGTATTAGTTAATTAAATGGTAGTATAAGTAAGTATATAATCGATCACGAGTATTTTTATGGTTGTCAGAAATAAGAATGTCCACTGTAGGTCTTGAGCCCATGATCATAAGGTTAATAGCTTTGCTCTCTACACTTAGTGTTGTTGGAAATGTTGTGGAGTTCCAATTATATCCACTCTTTCTCTGTTAACATTTCATTAAAATGATTTACACAGTTTAAGGCCATTGATTGCATTTATGAGAAAACCGAGGAGTTATCGTAGTACCTGAATGTATATGTTTACATGGACTAATTTGAACTTCTGGAATATGACTGTAGTGAAAGCTCTGAATGAAATTAAGGCATCTCTTGGATGGAGAGTGGTGTATGCATGGGTTGGGGATGATCCTTGTGGTGATGGTGATTTACCTCCATGGTCTGGTGTTACATGTTCAACGCAGGGTGATTATAGAGTAGTTACTGAATTGTAAGTTTGCTTCACCAGTTCCAAGTAGCTTTTCTCTAACTAAGCTTCAAATCTGTATGAAGGTACTCACAGAATTGAAATATCTTTACTTGTCACTTTTTCACAATGTTCTTTTCTGTTTTACCACTTGCAGGGAAGTTTATGCTGTTTCAATTGTTGGCCCATTTCCTACTGCTGTCACTAATTTATTAGATCTTACTAGACTGTAAGATTCTTCTTCTCTTTGTCTTTTTACTGGAATCTAGTTTTTACAAAACTTCAGCTACTTCTTACAGATTTTTTAACCTTGTACAAATTGAGAAGTTTCCCTAAGTTCCAATATGGATGTTTTTACACCTTCCAGTCTTAATTGTTCATTAACCAACTTAAAGAATTGTTCTGCTTGGGTGTTGACTGTTACCTCAAAAAATGTGGGCGTTATCTATAATGTGCATAGTCATTTTTTTCAAATTTGATACTACTATTTAAGGCAATAGTTAGTGGACAATTGTGCCTGTACTGTTGGATCTTTTGATTTTCAGTCTGAACTTCCTAATCCATAATATTCTTTCATATGACAGGGATCTGCATAATAACAAGCTGACCGGTCCTCTTCCTTCTCAAATTGGGCGTTTGAAGCGTCTTAAAATATTGTATAATCCTTTACTTGTATACTCCTATTGAAGTATCTATATTCAGTAGCAATTATGAAGTGATATTTATGTTTAGGGCTTATGCTCTGTCTCCGAATAGCCTGTTTCTCCAGTTTCAGGATTAACATATGTTTATCCATTAACACCTAGCTAGATGGTGTTAATATGATTCAACTGTGTTGTACAAAGTGATTAAGTACATGGCTTTTCATCGAGATGAAATGTGTTAATAACAAGGAAAAGCTTACCTGATTAATGATTAATCCCAAAGAAGGGAGTGAGAGGGAGTCTGCCTGATATAATGTGCACCGAAGAAGAAAGATAAATTTTCAACTGAGAGAATATATACTCGTAGGCTTATATTCTAAAAAATAAAAACAGAGTAGGTTTATTATCAAGGAAATGTCACTTAACTAAGAAGGCCTTCTATACTCTTTTTAGGATTGTATTAGATGTATATCTCAACATCTACTTTCTGTTGCCTGTATCCTACCTTTAGGTTTTCCTTTGTCCATAGGAACCTCCACGTTGTCCCAGAATGTTAACTTTAAAAATTACTTTTATTGCTTTGTTACCCAACATACTAATATGAAATTATATCCTCTAGTTTCCTAGTGTGTTGTAATGTTGCATTATTGATTAGTTCCTTCTCCTGATTCTTTTTACCAGAAACGTGAGGTGGAATAAGCTGCAAGATGTCATTCCACCTGAAATAGGTGAACTAAAGCAGTTAACACATCTGTGAGTGCTTTTTGAGCTTCTCTAGCTGTGCTCTTTATCCATAGATCTATTTTGCTGTTATAGACTAATAACTGATATGTATATTTTTCAGCTACCTAAGTTTTAATAATTTCAAAGGCGAAATACCTAAGGAGCTTGCTAATCTTCCTGAACTTCGTTATCTCCATCTGCATGAAAATCATTTTATTGGACGAATTCCACCTGAACTGGGTTCCTTACAGTATCTTCGACACCTGTATGATACCATCTTAACTTCTTCTCTTGTTTAACATATGGAACCTGTGTCACACATGGATCTCACTGTGATCTCATGCAATTCCAGGGATGTTGGTAACAATCGTTTGGTTGGAACTATAAGAGAACTCATACGTATTGAAGGATGTTTTCCGGTTCTCCGCAACCTGTAAGTTGATTATTCTTGTTTAATTGTCCTGTTGCTAGTGTTATGAATTATATGCTAATTCAAGGATAACACGTGCTGCAACTTGCAGATACTTAAACAATAATTATCTTACTGGAGGCATTCCAGCGCAACTTGCAAATCTGACTAACTTGGAGATCTTGTAAGTTTCGGGAAGAAAGTTTCTCCGCGATTAGTGCTTGTCTTTTGGACAAATAACAACTAATTTCTCCTGTACTGGTTGATAAATATTATTATGCTTAATTTGATTGCTCTGGTTATTACAGGTATCTATCCTACAACAAGATGACAGGAGTTATACCATATAGCATTGCTCATATTCCTAAGTTAACTTACTTGTAAGTCCTCCTCTATGCTCTTAGCTACTCATTTATTACACTGTATGTGGACAGTTTGGACACAGAAAATTCAGAATTTTTGAAAGCAAATACAGAAACATTCTTAGCCTTAAGTTGAGATCTCTTTCTTTGATAGATTTTTGGGGCAACCTGGTCTTTGTGCATGATTTAGACGCCACGTTAGATACGATCAGGGACCTGTGTGTTATCATGCAGTAGTTTAGTGCTGTTTTCTGTTATCAATAGATTGCTTTATTAACCTATAAAAATATTAGCATATTGGATGTGATGTCCCAATGATTGTCTAGTGAGTGCAGTCTTTCAAGGATCGTTCTTTCTTAAAAAGATGAATTTTGAGGATTGTTCAACTTAAGGATGAACAAAAACAACATACTTACTACTTAGTGTAATCCTACAAGTGGGGTCTGGGGAGTAGTGTGTACCCAGACCGTACCCCTACCTTAAAAAGGTTGAGATGTTATTTCCAATAGACCCTCGACTCAAAACAGATGAACTCGGGACTAGCTAATGTGGTCTTGCAAGGGTGTTTCTTTCTTAAGGAGATAAATTTTGAGGATTGTGCAATTTATGGATGCTGTGACAAAAACATTAGTAGATGGACTTGATGCTTGTTGGCTGACCTTATTGGACCTTATCCGTAATGATTTCCTCCGACAAGTAGTAATGTATAGAAATTGCCTGGAGAAATTAGATGCAAGAGTGTTCCCATCCTAGTAGGTTCTAATTTTAGCACAAAAAACACAATCCATTTCAATAGTAAAACATCTCATGTACTCAAGATTTCAGTTGAGCTATGTAGCAACAGCACAGTGATACAGGGACGCGTGAATGACATGTTCAACTTTAATTATAGAAGATTGTCTGTTAGAACATAATAGTTTTATTGGTATAATGTGTTAAAAAAACTGTATGTATAATTCAAAAAAATATGAACACTGATGCATGCAAATCTTTAAGCGATTAAACAAAGCCGATTTTGCACCTGGTTGTCAATATCCATTGCATTTCCTACCCATTTTCTTCTTTCCTCCAAAATCCATGGTCTTTCAACACTTACATTCCCCTCCATCCCCCAGATTAGAAGTCATTATTTGTACAAGTGAGAGGCTTGGATAATGCAGAAGGACCTGCATTTTGCTGGTCAAACAACCAAAGAATGTCAAGAGACAAGTCTACTTGTGTCAGATATATATCCCTGCTCAAGATATGCTGTGTTGGAAATGGGAACCTTGTTTTGAAGCTTAACCCACATACTTCCTTTGACAAAACCTTTTATCACAATATTTTCCTTTAAAGTGTTGGTGTTGGAGCTAAACCACACAAGCTGAAACTAGGGCCTTCTCCATTTAAGTGGTTGTTAGTGTTGTCAAAGTGATGCTTGAGCCATGACTTGAGGCTCTGCACAAGTTTGCTCCAATGAAGGTGGTAACTGGTAAGGGATATTTTCCCAAAACCAGCCAAGCAACTTCTTTTCTGAGAATATTAGATTAATTTGTGAGGTCCATAGGTTTCTTCTTGGATAGGGATTAAGCAACAAATATTCAATAGATAGCGAGCAAACCGATATATTAGTTGTTAGCGAGACTGTACTAATGATACATTATTGATTAGGAGTAAGAAGCTGTAAATTTAATGCAAGAAAGCTAGCGACTAAGTCTGAGTTGAAATTTTAAAGGTAGATGTTCCTTAAAGCTTGCCACTCAGATTCAAAATGCAAACTTGATGACCAATGATCCTTGTAAGTTGCAATATCCTAGATGTTCCCTTAAAGCTCAAATGGTCCTTGCAGTTGCATATGTGAGTGTTTTGTTCCATTTATTTATTTATTTATTTTTACGTTCTTCGCCCTTGATAAATCAGGTGTTACGCATCATTCCTTCACTCTTCTTGATTGTATACTGGTAGATTTTGCACCAAAAAGAAAAATAGTTCTGCATGCATTTGATAGTTGATGTCTGAGGCTGAATCTGATTTCAGGTATCTCGATCATAATCAATTTTCTGGGAGGATTCCAGATGCCTTCTACAAACACCCTTTCTTGAAGGAAATGTGAGTGTTCATTTTTTGCTACTGACCATTCTTTGGCTTTGTGCAAATCCTAATAATATGACAAATTATGTCATAGGTACATTGAAGAGAACACGTTCCGGCCTGGTGTAAAACCAATTGGCCTGCACAAAGTGCTAGAACTTTCGGACGCAGATTTCCTTGTTTAGAAAAAGCCAACTCCTGTTATTTGTTCATGAGCAAAGTGTACCTAGGTTGTCGAAAGAAGCAGCCAGTCCCTGTTCTGTCAAGTATCTATTCTTTAGTTGCCAGATGAGAAAAAAGGTTCCATCTTTTTTGTGCAAAGAAGTAATCAACCAAACTTAGTTGCGGTTGCTGCAGTTAAATAGTACCACGTTATGAATCATTATTAAGTAAGTGCGACATACGTCTCAATCAGTTCTCACTTGCCTTGAATTCTTGCATTTGGTTTTTGATTATACATGATGCGATTTCAGAGTTAATATGAAAGGGTAGAGAATATGCTCTGATATGACAGGTTTTGGATAGGTTTTCCATTTTTTCGTTAAATGATACAAATACATACCTAAATTATCGTATTATTTTTTTGAGTTTCACACCCTAATTATAGTTATTATCTATGTGCAAAAACACTTCGCTGAGTAGCTGATGGTATGCATGCTTTATTAGGTGCCAAAGTGGCAGTCCATGTGGATAAATATTTTCACTGGAA

>PGSC0003DMG400004195 LRR-RLPs

CTAAAACTCTTAACATAAATTGTTAAATATTTCTAACCTAAGTAGGTATTACCGAAAATCTTTAACACAGGGTGACATCGAGCTATGGAACATAAACAATTGGTAATTGCTTCACGCTATCTATTGCTCCTTTATTCATGAATTCATGTATATACAATGTAATTTTCTGTTAGAGGGGTTCGAACCTTCTGAGCCCATATGGTTCTGCCCCTGACATCGCTTATCTAGTTAAGCTCAATAATTTGAATATGTTCTTAAAAATATAATTTTTAGAGGCTTCATTGGTCAATTTGTTAATAAAACAAGAATATTATAGAATGGGAACAGTGATATCCTCCATAAATGAAACTCAACTACACTATTTGGTTAAGACTATGTTTGTGACTATAATAGTGGAGAAATAATAGTAGAAAGGCTTTCCATATTATTTGTCATGACATTTTTACTATTTTTTATTGCTTTAATTGTTATGAATTATTTTTCTTGAGCCGAGGGTCTACCAGAAACAATCATTTATCTACACAGATAGGGATAATGTATGTGTACATACATTATCCTCCTCAGATCCCACTTTATGAGATTAAACTTCAGTAGTAATATTCTAAGCAATTACAAAAATATTTCCACAAAAACGAAAAATATACTAAGGGGAGACATAATATTGTGTTTTATAAAATGATGTTGTAGTAATTTGTCTATTTGTTTACTTTCTTTAAAATCAACATGGCTTGTGCAGAAACTTGACGACATGAGATGCTTAATGTATCGAGCTACCGAATCTATTATTGACATAATCATCCCTTTTTTGGCAGGCCAAAATGGCATCTCATCAATTTATTTGTTTCCTAGCAATGATATCTCTTATGATTATCAATATTAGTGCAAACAAGGCATGCCACCCTGATGATCTCAAGGGTCTCAATGACTTCAAAGCTAGCATACATTCGGATACTTCTGGCAGACTAAGCAAATGGATAGGCCAAGATTGTTGCAATTGGCCAGGCATTTCTTGCAACTCTACAACTAATAGAGTTGTAGAAATCTATCTACCTGGTCACTATGTATCTGGTGACGATGAATCTCCAAACTTCGTGTCAAGCACAATGAGTGGTTCAATCTCTCCTTCAATTTCACTTCTCACTTCACTACAAGTTATTGATCTTAACAAACTAGTCGGACTAACAGGGCAAATTCCTGAGTCCATTGGGGTTCTCAAGAATCTCAAAGAACTCAACTTACAAACCAATCAAATCTCTAGTACAATTCCAGAGTCAGTATTCACATTGACGTCTCTAACAACCTTGAACCTTGAAAATAACCACTTGACAGGGGAAATATCGGAGAATATTGGCAACTTACAAGCACTTCAAAAGCTTTTCTTGTCAAACAATTCGTTCACTGGAAAAATCCCAGTTTCAATAACAAAACTACATTCCATTTCCACAATCCACTTAGAAAAAAATCAACTTGTTGGAGAAATACAATTGCCTTTAACTCCTTCCCAATGGCCTTTAATTAAAACTTTGGCTCTTGACAACAATGCACTTACTGGAGCTATACCTGATTCAATTGGTTATTTGACAACACTTTCATCACTCTCTATATCAAATAATCAATTTACAGGACCTATTCCTTCTAGTTTGGGAAACATAAAGCAATTGCAAATATTGAAGGTCAATAACAACAATTTGGCTGTTGAGATGTTGCCAACGACGATTTGTGGTCTTACTGAGCTAAGTGTTTTGTATGTTTCTCAAAACAAGATTCATGGACCATTACCTGGATGTCTCTATTCATTCAAGTACTTACTAGATGTCGATGTACCGTTCAAACGTACTGCCACTCTTCCTATGGGAGTACACAAATAATAAGTAATAGTGATGGAGCCAGAAATTCTGAAGGAAGTCCAAAAATGCCTCAATTTGAAATGTTATGTATGTCTACAACAACCAAAAAAAAAACTTGTGTTTATTTATTATCTCTATGTTATATGAGACTCCAACTAGCCAGAATCACTGGCTAGTATGAAAAAATTCGAATACATAAAGTACTTTCTTTGTTTTATTTTTG

>PGSC0003DMG400004284 LRR-RLPs

AACTTTTTTTCTTTCATTACCTACCATATTTCTTGCAAAATGAGTCTCTTGTTTCTACTTGCATCTTTGTTTCTTTGTGAACTTGTTTTGTCCTCATTTGCAATATGCCCCAGAGATCAAAGTCTTGCCCTTGTAGAATTCAACCAGTCTTTTGTTGTTGATGCATCTGCATCTTTTATTTGTGACAAGCAGTCTTATCCTAAAACGAGTTCATGGAACATGAACAAAGATTGTTGTTCGTGGGATGGAATAATATGTGATGAGATGACCGGTCATGTGATTGAGCTTGACCTCAGTTGCAGCCAACTTGTTGGGAAGATTGATTCCAATAGCAGCCTATTCCAACTCTCTCATCTCCAAAGGCTCAATCTTTCTAATAATGACTTTGACGGTTCTCATATCTCGCCTAAATTTGGTAGGTTAGCAAGCTTGACACATCTTGATCTTTCCCAGGCAGACTTCTCAGGTCAAATTCCTTCTGAAATCTCTCATCTCTCCAAGTTACAGTCTGTTTTTTTATCGTCTAACAGTGAGCTTAGACTCGTAGCTTACGATTTCAAGATGTCCCTTCAAAATTTGACTCAATTAAGAGAGCTTCATCTTAGTGGTGTGAACATCTCTTCCACCATTCCTCTGAATTTCTCTTCTCATTTAACGACGCTGGGGCTGACAAGTACAGGAGTGTATGGGATAATACCTGAGAGTATCTTTAACCTGCCCAACCTGGAAACACTTGACTTAAGTTCCAACGATCAGCTCAATGGTTATTTTCCAAATACCAAATGGAATAGCAGTGCATCTCTCATGGAGCTGAATCTCGCTGGGGTAAATTATTCTGGTAATTTCCTGCCTGAATGTCTTGGCTATCTAACTTCACTGCAGAGATTGGTACTTAGTTCCTGCAATTTATCGGGGCAGGTTCCGAAATCTCTTTGGAACCTCACCCATCTGGAGTATATGGACCTTGAAGATAACCGTCTTGAAGGACCGATATTTCCTCAGTTTACAAGTGGACTGCAGGATCTGAACACACTTAAGCTATCAAATAACTCCCTGAATGGAGAAATACCGTCCTGGATATTCTCCCTTCCATTACTGAGTGAACTAGACTTGAGTAATAACCACTTTTCTGGTCAACTTAAGGAATTCAGTAACACTTCAGTACTAGTCGGTGTTGATATAAGCGAGAATGAGTTGCAAGGATGTCTTCCCAAATCACTTCAGAACCTTGTGAACCTAGTCTGGATTGATTTAGCAAATAACCAGTTGCAATGTCCTCTTCCCAAGTCACTCCAGAACCTCAGAAATCTAAAATTACTTGATCTTTTGTCTAACAATTTTAGTGGGAGTGTAGATGTCAGTGTCTTTTCGAACCTCAAACAACTTTGGTATCTTGATCTGTCATACAATAGTATTTCTCTGATCAATGAGAACAAAGTCAAGTCTACATTGCCCCAATCTCTTGAATACTTATATTTGGCTGCTTGTCAAGTAAAAGAACTGGATTTTTTGAGATCAGCAAATAACTTTTATACCTTGGATCTTTCATATAACAACATTCAAGGAACAATTCCTGATGTGGTATTGTCTAATTGGATGCATTCCATAAAAAATCTTAATCTAGCCCACAACATGTTGACTAGTATTGACCACATTTCATCATTTTCCCAACTGATCTCTATTAACTTATTGTCCAATTCTCTTCAAGGGACACTACCAATTCCACCACCCTCTATAGAATTCTTCTTCATGTCAAATAATAATGTCAGTGGTAAGATCCCTTCTTCTATTTGCAATTTGACAACATTAAAAATCCTAGATTTGTCAAATAACAATTTGAAGGGAGAAATTCCACAATGTTTGGGTAACATGAGTGATCAACTCGAGGTTTTGGATATGCGACGCAATAGTCTTTCTGGGAGTCTGCAAACAACTTTTAGCTTGGGAAGTAAACTCAAAAGTTTCAACTTGCATGGCAATAATCTAGAGGGAAAACTCCCAAGATCCTTGGCTAATTGCAAAGAGTTGGAAGTTCTTGACTTAGGAAACAATCAACTCAATGACACATTCCCTATGTGGTTGGGGACTCTTCCAAATCTACAAGTTTTAAGTTTGAGACTCAATAATTTGCATGGACCAATTAGAACTTCAACTAGTTCGAAACTGTTTCCTCAGCTTCGGATGTTAGATCTCTCTCGCAATGCCTTTACAGCAGAACTGCCAACAATTCTTTTCAGAAATCTGAAAGCAATGAGGAGAATTGCCATTGATAAAACAATGAAGGCACTAGGTGATGAAGAAAAATCATATTACCAAGACTCGGTAACTGTTGTAACAAAGGGAACGGAGTTTGAAATTGTGAGAATCTTGTCTTTGTACACCACTATGGATCTCTCAAACAACAAATTTGAAGGACATATTCCTAGTATGATGGGAGACCTCATTGCACTACGCGTGTTGAATTTATCTCACAATGAGTTGCAGGGTCATATACCACCATCATTTGGAAAGTTATCTGTAGTCGAATCATTGGACCTTTCATCCAACAGGCTTTCAGGAGAGATACCAAAACAACTTGTTTCTCTTACATCTCTTGCAGTTTTTAATCTCTCTCACAATCATCTTGAAGGATGCATTCCAAAAGGGAATCAATTTGATACATTTGAGAACAATTCATATGAAGGAAATGATGGATTGCGTGGATTCCCAGTTTCAGGAGGTTGTGGAAGTGATCGGATACCGGATACAAACAACACAACATTTGTGCCAGATGAAGAAAATGATTCAACATTTCTAAGTGAGCTCAGTTGGAAAGTGGTTCTTATGGGATATGGTACTGGATTGATTATTGGATTCTCCATAGCATATTTGATGCTTTCTTCTCAAAATCCAAATTGGCTTTCTGGGATAGCTGAAGATTTAGAAAACAGAATCATTATGAGAAGGCGAAAGAAGCAGCGAGGTCAAATGCATTACAGAAGAAGAAGAAGAAATAATGGCATCTAGATGTAAGTTCAAAATTTATGAGAGATGTAGTCATGTTAAGCTCTATACTGATTCTTTTTGTTTACTGTAACTGGTATTAAAAGAATGAGATTAGTGTTCTGTATCTGACTCTTTCTTTTGTAACTTTTATGTAGGCAAAGGGTGTATCCAGTTGAAATATCAAGAAGATGATGCCTGCAACAACAAATATATCTTCTTGGAGGAAATTACTTGAGCAAGAGATGCTTTTGATATTTCCTGGAGGTTGCTACTCCTTTTTCTTCATAAAAATAAAATAAATTCTAGGATTGAAGTATGTTGCTTTACTATAAACAGTTGTCACAATGAAGTGTTTTTGTTTTTCCATCCAAGTGTTAGTAAAGAGAAAAAAGATATTTGTCTGCAAAATAATAGCTTTCTTTTGACATGTGTATTGACTTGGAATATTACATTTACCTGGAATACAGTTT

>PGSC0003DMG400004384 LRR-RLPs

CAACCCTCTTTCTCAGGGATGGTCGAAGAGAGGTTCCTCTTCTGAGATCATCCTGAACACAGCAGTCAACAAAAAATATTCAAAATAAAATAAAATATTTTTTTTGGGATTTTAATTCTACATTTCATAAAAAATCCCACATTTTCCATTGGGTTAAATTTCTTGAATTTTTTTTTCAAGAATTTGATCTCTGTTTGCTTTTTCTGGTAAATCTTTCATCTTTACCCCCACTTTTTTAATTTTTGTTTGATTTTGCTTGAGCATATGGTTATCTTATGATGGCCTTTTTGTTTTTCCCAAAAAAGATTCAATTTTTATTAATTCTTGTTGTTTTTATGTCTGCATTTTCAGCTAACAATTGAATCTTGGAATTTTTGAAGACTTGGGGTTTCTGCAATGATGGAGAAAACAGAGGTAATATTGCCTATATTATAGAAATGTTAAGAAAATGAGAGACCCCATGAAGGGGTGGAGCCAGGTAGGGCCGAGGGGTTCAAAATTACGTTGTTTATACATAGTTAAAAGTTTTATATATATGTAGTACATGTTGAACCCCTTTGCCTTGTAGAGGTGTTTACTTCTTTATATTTTGAACGTTGTTAGTGAAAATCCTGGCTCCGCAGTTGACCTCATTGTGCCTAATATAGATAGATTTTGTTTGTAATCTTTTTTATTTTTTGTGTCTAGTGTAGAGTTGTGCAGATAAGCAACTGTGTGTTCGTGTGTCCGAGTGATTTTTGTTACTCGGGGGAGAGTCATTTCATATCAAAAAGCTGGCGATTTCGTGTGAGAATTTGTTCAGAATCCTGGCTGTTGGCAAGAAATGAGGCGTAGATGGAACATTTATGGAATCAAGTTATTGCTTCTCACCATTTTAATTCTTCTTTTGGAGGTTCATGAATGTTGCACTCTGAATTCTGAAGGTAGTTACCATTCTCTAGTTTCTTTCTTTTCTTACTTCTGGAGGACCATAGTGTTCGCAATTCGGGCCAGCTTGCATTAATTCGACTGCTCCACTGTATATCCTCTACCTCCAACCAACATAGGCATCAGTTACTATGCCCATCGAGGCTGAGGCATATGATGTGTTAGGCGAACCAAAGTTCTACATTGGTAGCTAAAAAGATTGGAAATCTACAATTCTACATAAGGTGTAGGATCTCTTAACATGGTGTGAGGCCTTTTGTGAAAACCGTGCGGGCGTGCCCCCTAAGCGGACAATATTGCACCATGTTAAGAGTATTTTTGGGTTGACTTAGCCCAACAAGATGAAGTTACTTACTCCCTACATCCCATTTTATGTGGTCATGTTTTGTTGGACACAAAGTTTAAGAAAGAAAGAATTTTGTAGGTTGTAGTCTAAAACAAGTTATAGACATTTAAGTATGGGTGTAATTTGAATATTAAGATTAAATTGTTTCTAAAGTCTAAACATGATGTTATTCTTTTTGTGACAAACTAAAAGGAAAATGTGCTGCGAAAATTGAGACAGAGGGAGTCGAGTTTTTGGTCTTTGCTGGGATATGAACCTTGATCTTTCATGGTTTTTTACTCACTTAGGGCTGTGTCTTCTGTCTCCCTGGGCCGTGGGCTATTCTCTAGTTTCTTTAGTCTATTCATGTTATTGGCTTATTCTTTGTTTCGCTGATTACTGTTGATTTGAGAAACAGGATTGTCGTTGTTTGAATTCAAATTGAATGTGTATTCTGATCCAAATGGAGTTCTTGAAAACTGGAATGCTGATCACTGTGACCCATGCATGTGGTCTGGTGTCGAGTGCATGGATGGTAAAGTGCACATGTTGTAAGTCTCTTGTTATAGATACTCAAGTTTTCTAGTTAAGATAGTGTCAATAAACTTTCTCTAATGGTTAGTGTTTAGTGATTTGATTCTGGAACCAGTTTGTTGAACTAATAATGATAAACCAAATGCAACAGGGTGAAATTGTTTGTTTCGACTCTTGTGATTCTTCTTATGTCATTAAATTAGTTGACCTATCTTCTTTTTTTCATTCTGTTCAGAAATCTCCATGGATTTTCTTTGGAAGGAACACTAGCATCAGAGCTTGGAAATCTCACTTACTTGAGATCACTGTAAGCTTTAGCTATACTAATATGCCTTTTCTTTAGGCTCCTGCTCATTAAGTCTTAAATGGTATTTTTTACCTGAACCGTCATCCGTAGAAGGTTGTCATAATCTTTGAATAGTCTTCTATCTTCTTATGTAAATTCGAAAGAACGCAAGGGATCGTTTGGTGTATGATGGACAAGCATAGATAGTCCGGGATAAAAAATTAGTAAGGGCTTATCCCTTGTTTGGTTATTAATCCTGGTATAAGTTATATCAGGATTAGTAATTATTATCGGTATAAGTTATCCTCATCAGAGGGTGGAATAATAGTACTGGATTAGTTATCCTAGGATAAAATAGTGAAAATGACAAATATCCATGAAGTCCCTCAAACTCTTTTTCTACTAAATAAGGTGGATGGTGTTTTTGTAAAAGAACTTCTAAGAAATTTATGCAATGCATGTTATTTTTAATACAACAAACCAAACAATCACTAAGCAATAATACATTACCAACATAACTAATCCCAACATAACTTGTCTTCAAACTAAACGACCCCTAAGAGTTATGTTGGTCAAGTAAATTGGGAGGTAGAGACAAAGGTGGATAAGCTTTGGATATCTACTAATCAGCTCTCATCATCTTGTATTCTGCATTTACATTGTTCTCCGCATAAGTTGTGTTTGAATCTTAACAGTATTTCTTCTTGCTGTCTGGTTGCTTTTACAGAATGCTCTCAAGGAATCATTTTTTCGGTGCTATTCCTAAAGAATTCGGAAGACTCGATATGCTTGAAGTGCTTGACTTGAGGGATAATAATTTGAGTGGAACAATTCCAGCAGAAATAGGAGACCTGCAATCACTAAGAATCTTGTGAGTTAATGATTTGTTGTGTGCACGATAAATGGTTTGTTGTGTGGATGATAAACGGTTTGTTGTGTGCAGATGTATATATACATATGCATACACATTCATTATGCGAAGTGATTTCAAAATGCTCATCAATCGTGTTTATTTCGTTTTTCTAGGTTGATTCATAACAATAACTTTGAGGAGAGTGTTCCTTTGGAAATTGGGAAGCTTCATTTAGTCACAGAATTACAATTCGACGAAAATCTCACTTCTGCTTTTGCTTCTGGAACTCTCTGTATAAGAAAATTTCGTCATTGGTAGGATTCTTTTATTTGTTTTAGTACCCTATCTTCATATTCAGCTGGTTTTTTGCATATCAGATGCTGGGTCAGATGCTGGATAGTATATCTTTGTGGCAACACACTATCTAATTGATCTAGCTCATTTCTCAATTTGATATTTTGCATGTCTTACAGCATCTGGCAAGGCAGCATGAAGCCATTCAAGACGATAACCTCCATTATTGAACCAATAAAATGTACACTTTTGCATTACCTCAGTTTCTTCCAACTGTAAGTAATTAAATAGTTTTTCACATCACGAGTCATCTAGTTAAGTTTTATTATTGAGAGACATTTTTTCTATTTCTTTAGGCCATCTCCTGGAACGGGCTTTATGGATGACCGGAAAGACTGCTCTGACAACCTTCCAAGTGAGTAAACACGTCATATTTTTTAAAATAACTCCGAAGAACCGTTTACAAATAATAATTCTTCATTGTTTGCAGGGTTATCCAGGCCTCACATCATCCACACTATACAGCACCAAGAGGAGATTGCACGCCGTAAGTTGGCTGAACAATCCAGTAACCTTGCTGCTGCTCCTGCCAATATCAATGGACCGTTGGGTCCTGTCATTCAAATGCCAAGAAGCAGTGGGTCGTTTCGTGCGGTGCCAAATTCCGATGGAACACATCCTACAAACTTACCATCACCACCTCCACAGCATGAATCTCAAGACAAATTACATCCTGTGGGACAGTCTGCAAGTGCTGTAAAACAGTCGCCTGCTGTGAAAAAGCCAACTAATAGCCAAGGCCCCCCTGCTGGAAAATCGGAAAGTACTTGGAAGTACGTAGTTGGGATTTGCATTGGAGTAATTTTGCTCATTCTTGCTGCTTCTGTGTTCTTAATCTGCCGAATCAGAGCGGCTAGAACAATTGGCCCTTGGAAGACTGGATTGAGTGGACAGCTGCAAAAAGCATTTGTTACAGGTACTCACATCATATTGTGCCGAAAAGTGATACTGCTAATATTCTGCTCCCCACCCTGATAGTAGTTGCATTGCTAACAATATTCTTAGCATAGCGTTACCTTATCAAAAAAAATATTCTTAGCGTAGTGTTGTTATAACAATTTAACTGGAGTAAAGGGAGGTACTGATAATCTTCGTCTTATGAGGAATTAGAGGACTAATATTCTCAGTGTAGCATTGTTATATACCCTTTTTCACTATATGTTTGCTAGGACGAGAAGGTTCAAATGGAACTATTTCCTTCTTTCTCATGAGAAGTGAATTAGCTGAGACGTCAATACTAACGCTTGAAGGCTGGAATCCTCCCCTTATACTTCTAAAATGTCGCTGATCTTTCCTCATACATTTGCTTTTGTCTAGCTTATATCCCCCCCTACAAAGTTGCTTTTCAACGACAACCATAGGAATGTGCAATCCTTTATGTTAATCTATGATATTTTTCGTCTATTTTAATTCTTTTGCCCGTGAAAACATTGTTATAGGGGTTCCAAAACTGAACAGATCCGAGCTCGAAACAGCATGCGAGGATTTTAGCAACATTATTACCAGCGGCGATTCATATATAGTATACAAGGGAACACTGTCTAGCGGAGTGGAGATTGCCGTTATCTCAACTACTATAAGTTCTCTGGAAGATTGGTCCATGCACTCGGAAACAGCCTTCCGGAAGAAGGTTTATCATTTCTTTGCTTTCACAAATTAACCACCTCCA

>PGSC0003DMG400004529 LRR-RLPs

TTTTTGGCCTACGTGGCTGAGTCCGTGACTTCACCTATGTTAGGCGCGTGGGTGCAATTTTTAGACAACACTGACCAATAAAAATTGACCACGTGTTAAAATAAACTTTATTGTTATTTATTGTTCTTCATCTTTAACTAAGTTTTCTTCTTCTTTATTGTTCTTCATCTTTAGAAAAAAAAACCTCCATTATCATTATCATTCAAGTTAAATTGCTCCTTACTTCGAAAAGGTAAAGTAAAATTTTAGTATATTCTTTATATTTACACCTTTTTATTTCGATTGTTAGTTCTGATTTTAAATTTCTTTCACAATTTTTATGTATAAATGTTAGGGTTATGACAAAATCCGAGTTGAATAAGTTGTGTATGGAGCTAGGTGAAGGATGTCCCTTGCTGAGGGATATTGTGTTGTCTCACTGTCGTCAAATAACAGATGTAGGGTTAAGTTATATTGCTAATAGGTGTACCTTGTTGGAAACTAGCCACATGGTCTATTGCCCTGGTATCACCGCTGCTGGAGTAGCCACGGTGATTACAAGCTGCACAAATATAAAGAAAGTTCTTGTAAAGAAGTGGAAAGTGAGTCCAAGAACCTAAAGGAGAGCAGGATCCATCATCTCCTATCTCTGTGTTGATCTGTAGATCT

>PGSC0003DMG400004561 LRR-RLPs

CGGAAATAGGAAAAGAAAATGAAGGAGAAATAGAGAAATTAGCAAGCAACTCATTGGTATGTTACTCTGTCTGTTGTTTCTTGTACCTCGATTAGTATATTGTTTTGTTGTAATTTATGTTCTATTACTATGTGCTATTTTTGTTATTATTGTCTGTTGTTTCTTGTATATTCATTAGTTCCTTTTTGGGATTGCATGCTTAAGATTGTTTTTGTTAAGCCGAGGATCTATCAGTAACAATTTATGTACCTTTGAGGTAGGGGTAGGTTTGATGTAAATTGTTTCTCTTGAGAAGTTGAAAGGAAGGATTTTTTTAAATTCTTTTTAACTACTTTGGCTTGATTAGTGAGAATGGAAAGAGGAAAAGAAAATGAAGGGCAAACAAAAGGGGAAGATAACAAAATCATTGGTATGTCATACAAAATTTAATTATTTTTTTCATTCACATAATTACATATTCCTGTAAGTATGTCTGTTTCTTTGTGTAACTTAGATGGTGAATATAATCTTTATAACCTATAAGACTCAATGATTTAATCTTTCTGATATAAGTTAAACTCTATGTACTAAATCATCTGCTATAGTTGATAAATGTAAGGGATTAAGAACTACGAAAGTAAGAGCTTTTGTTTAGAGAGATAGCGCGTAAACAATGAGAGAGCTTAGTGCACCAGGCCGTCATTTTTTGGTTTAAGATTTGAGGTGTAACAGGACAAACATTTTGGCTTCTCTATTAGGCTATTCTGGAGAAAGTCCTCTGCACTTGTGACAGTTGCTTGTTGTAAAATTTAGAGCTATCACATTTTCGCGAGTTTCACACCTCAACTATATTAGTCGTTCCCTTTTTCTATCTAAATTATCACCATCTATGTATTAACATAATTTGAAACAACATTTGCTCTTGTGGCAGGTGTCATCCGTTGTACTTCGCGAGGACATTGTCAATCTTCTAGATTTCATAGAGAGGTTAAATAGTTGGCGTGTTCGAATTTCTCTTGACTTGTATCAATATAAAGAGTTGACATTGGAGCTGACGTTTGTGTCCGCATTTTGTCATCTTTATTACTCTTTCTTTTTGGAAAGCTGTAATGACGAAATGTCCTGCATATCAAATAAGATTCATGATTTGGTTCAGTCACTGTTTCATCCAAGTGGAGAAGACATGCTGGTTAATATAAAGAATCATGACCTCTTGGAGAATATCAAAAGTTATATTAACTCACATTGCTATGCTGAACCAAGTCGTGTAACCATGACCGAGGATCGTTTGGTTGAACTTTTAGATGCCATCCTCGTGTATCTCCAATGTCTACTCAAGTGTTGTTCTGAATCTATTTTCCCATCAATGACTCAATGTGAGCTTCTTCAGAATGTATTTGGCAATTTAAGAGATTTCCATAGGTTGAAAGTAAATGGTTGCGTTGAGTATGAAACAATCGAATATGTCTTACCTCATCTTCAACTTATGGCTCTGAGAGTAGTAAACATCTGTTTTACTCTTTTGAATTGTCAACTGGACAAATTCGATGAATCAGATGTCTCTCTAGTTAACTCCAAGCTAGCTAATCTACTTGTGGAGGTTATTCCTGTTGAACTGGAGGTTATGCACATATGTTGTACAAATTTGAAAGGTTCAAAGTCGGAAGAAGTTGAACGTTTCATCAAGCAGCTCGTAGAAGCCTCTCCGGACATTCTTAGAGTATCTCTGATTCATCTACAAGAGCACATGGTTAATGCTGGTACTCCTAGAGCTTCTACGTGTAACATTCATGTCATGATAGAGTTCTTATTGATTATTCTTAGTGATGTGCTCAAGGACGTTATTCGTCATGACAAATTGTTTGTTTTCTTGGCACGTGTTAGAGAACTGACCAAGGAGGTATTCGTTCTTGTTCGCAACTTAGAAGAGAATATGAATGAAACAAGTGATGCAAATCTAAACTTGCTGGAAAGTATTGAACTCCTCAAGGAAGATCTCAAGAATGATTTCTTGAAAGCCCGTGCAGACTCATCTCGGCTCTGCTTCCCCATGAGTGATGGACCGCTCTTCATGACTCTTCTGCTCACAAACTTGAAAGACTTGGTCAATTCCAATGCTTCTTCAGTTTCTATGATAAAAGAAGAAATTAAGCAGGTGAAACAAGACCTGGAAATCATAAGATCGTTCTTTGGGTATGGTGAGCAAGAGTTACATAGAGATCTTTGGACTCACGTTCTAGATTTGTCATATGAGGCGGAACATGCCATTAACTCAATTCTTGCCCGAGATCATGGTCTATTACAGCTTATCTTCATACTTCCTGATACCGTAGAAAAGATCAAGCTTGTCAAAAAAGAGGTACAAGAGAAGATCCCCAAGAGCAGCGGTATTATTGTTGCGAACGCTCCCAACAAGCCGGTTGAAAGAAACTCATCGAGTACAGTTGGTAAAATAATTGTAGGTTTTGAGGAGGAGACAGAGTGGATAATTAGGAAGCTCACCAGCGGACCAGCTGAGATAGATGTCATTTCCATAGTCGGCATGCCAGGGCTTGGAAAAACTACTTTGGCATACAGAGTGTATAATGATAAGTCCATTGTTGATCATTTCGACGTTTGTGCTTGGTGCACAATCGACCAAGAACGTAATGAGAAAAAGTTGTTGCAGAAAATTTTCAATCAAGTTATTGGTTTGAAAGAAAGATTCAGTGAGAAGGACATAGATGATGACGTTGCTGATAAGCTGCGGAAACAACTATTTGGAAAACGGTACCTTATTGTCTTGGATGACATGTGGGATACTGNNNNNNNNNGATCAGACGAAAGTTTGGAGTTATTAGAGAAAAAGGTATTCGGAGAAGAACGTTGCCCTGATGAACTAAAGGATGTTGGAGAAAAGATAGCTCGAAAGTGTGATGGGCTTCCTTTAGTACTTGATCTAATCGGTGGAGTCATTTCAAGGAAGGAAAAGAAAGAGGCTTTGTGGCTTGAAGTTCTCGATAATTTGAGTTCCTTTATTTTTAAGGATGAAGAGGAAGTGATGAAGGTTATACAATTAAGTTATGACCATTTGTCAGATCACGTAAAGCCGTGTTTGGTTTACCTTGCAAGTTATCCAAAGGACAAAGATATAATGATCTCCGAGTTGAAAGATTTATGGATTGCTCAAGGACTTGAGATGAAAAATGCAGAAGAAGTAGTGGATGTGTTAATTTCCAGTAGTTTGGTAATACCTTTCAATAATTCTATTTGCAAAATTCATGATCTTGTGCATGACTTTTGTTATATAAAATCTAGAAAGGAAAAGTTGTTTCACTTCATAGGAGGTTCAAACGCTCCGTCTTCTTCTTCAAATCTGATTCCGCGTGGAATTACCATTCATTATAATGAAGGTCTCTTTCATATGGATAGAAATTTTGTGGTGTTTAATCCAGAAAAGAAGAATCCTTATGTTAAACACCTCCTCTCTCTGAAGGTGTGTGATGGGGTTAAGAATCGTCTTTCCTACAAGAGTCACCTAAAACAGTTGAGGCTTCTCAAAAGTTTGGATTTGAAGGGCATAACATTGACAGATTCTTTACTGAATGAAATCGGTATGCTCGTTCATTTGAAGTACTTAATCATTCAGACGAAGGCTAAAGCTCTCCCTCCGTCATTTTCAAACCTCTGCAATCTAGAAACTCTGGTGGTGGATAACCTGGAGGGATCATGCATGATACTATCGCCTTGCTTTTGGAGTCTTGCAAAGCTACGAGATGTGCGGATGAAGTGTTGTGCTATCTTTGATCCAACTGTGTTAGACGAGGAGTCAAGGTTAGAAAATTTTAGAACATTATGTGGTCTCTACCTTCCCGGTTTAGAAGAGACGGAGGATATTTTCAAAAGGTTTCCTAATCTTGAAGCCGTTAGATTCAGGATCAAACAATTACCAGATCGGTCAGGAGAGAAGATTTGTATTCCAAGATTGGATGTCCTTAACGAACTTGAACAACTCCATTTAACTGCTTCTAGGGATTCATTGAGTGAATATACACATGGCTTCCCTTTGAGCTTGAAGAAGGTGATGTTGAAAGGGCTTGCTCTGACATCCGATTCATTGTCAAGAATTGGTAGACTACCCAACCTTCAAAAACTAAGTCTAAAAGACGCAATCATCGAGGAGGGGAAAGAATGGAACATGGAAGATGTCACCTTCAAGAATCTCAAATCTCTAAAATTGATTAATGTGTCATTTTCTGAATGGCAGGTTGATGCAGAGAAATCCTTTCCCGTGCTTGAGAAACTAGATATAACTATATGTGATATGCTTATGAACATACCAGACAGTTTTGGAGATATTGCGTCATTAAAGTTTATCCAAGTATGGTTCAGCCCTCAGCTTGAAGAGTCGATCTTCAAAATTAAGGAATATGTTGAAGAAATGACTGGAGAAGACAAGCTTGAGGTAGAATTCTATGCTTGAGGTATGTTGAAGAATATGTTGAACTTTTACAAATGTTATGTGTTTTCCAATATATTACTTTTAACTTATATGTTCTTTGCGATCAGATCGAGCTAGCTAGCGCATACAAATGATGAACCGATAGATTCTCAAGACAACTTTCTCCATAGATTTGTGCCTAATTAAGAGCTCCCCTTGCTGCAGCACCTCCCATTATGAACACATTTTTGGGGCTTGCTTTCAAAATGGAGACATGACGTGGTGGATTGGTAAAATCCTGTTTAGCCCATTATTATGTTCAACTAAAGTAAACTAAATTCATGTCATTATTTTTCTGTTTCTCCAATTTGCTCTCCTTGTGAAAATCTTCTGGTGTGCTTGTCCTCTCGCCATGACTGACAAACACATTTGCTTTCTTAAGTTCCAGTTATGCTAAGCAGAAATGTCGTCTCGTTGTGAGTTGTCAATCCGACTTTGAAGTAGAGATGGTTAAAGGTATAGAGGAAGCAGAAGAGGATTATGTTTGTATAGAAACAACAAGGAGGCCCCTGAATAAAGCTTAGTGAAGAAGAGTTCATTGCATGCCTTGTATTAGTTTTCGACATTTCTGAAAAAATTGCTTCAGACATGGCCTCATTTGCTTTGTCTCTTGTTGTCATATTATTCTTGCAATTCTGCTTCGGAAACATTAAAAATAACCCCATAAAGAAGGTAGTAAGGTTAGCGTTCATCCTACCTTCCCCAGATCCCACCTGTGGGATTAGAGAGTAAAGTTCTCG

>PGSC0003DMG400004722 LRR-RLPs

ATGCATTCAGAGGAAATGACTAAGATGGATGTTGATGGCACTACGGGAATTCACTTTGAAGAAGATATAATCACGGAAATCCTAAGTAGGTTACCTGTCAAGTCTCTTTTTCGATTCAAATGTGTTATAAGGTCTTGGATGACATTGATCTCTAAACCTTACTTTGTGCAAACGCATCACAATCATTCCAAGAATAACCAAAATTCCCAAAAAAATTGTCTTCAAGAAAATGCTTACTGTAGATACCTCATTAGTTTGGTGTAGTTACTCTTACCACACACGAGTTCATGGAACATGAGCAGAGATTGCTGCTCATGGGATGGAGTCGTATGCAATGACATGACTGGCCATGTCACTGAACTTGACGTCAGTCTATTCAGGGACGAGAGCTTGAACTTGTCAAAATTTTGACCATTCACACCACTATTGTTCTTTCAAGTAACAGATTTGAAGGACTTATTCCAAGTATCATGAGAGAACTCATTGCTCTTCGGGTGCTGAATTTGTCTCATAATGAATTGCAAGGTAATATACCATCATCACTTGGAAGTTTATGTGTGGTCGAATCATTGGACCTTTCATCCAACCATCTTTCAGGAGAGATACCACAACAACTGGCTTCTCTAACGACACTTGAATTCTTAAATCTCTCCCACAATCATCTCGATGGATTCATCCCTCAAGGACCTCAATTCGCTACATTAAAGAAATATTCATATGAAGGCAATAATGGATTACATGGATTTATGCATTGCCAAATTCATCATTGCATATGTTGGAGGAATATCAGGGGTCATAAAGTTTTATGAGCTTAAGTTGGCTCGAACTTCATGATTATTACCCGTTGACATACTAATGACAAGGTGATATATAGAGTTGAAACGAGTTAGATATTTTTTTCTTTTTGTTTTTCAGTAATCACCTCTTATTTGTTTTACTGTCATAAATTCAACTTTCTGATGCAATATAGTTGAAAGTAGTACCTATTATTTTGAAAGTACATAAATTCTGCTTATATGGATGCAGCTAAAGTATTTATGTACAACTATACTTCCTTTCATGATTCATAAATGTACACAACAACCTCATGACCCACTGCAATAGATTACGAAGTACCTTCACTTCAAATTGAATGAGTTGTAGCTTTCATTTGCAAGACGCGTGACTTTTCTGTTACATAACAAATAAGGTGGCTTTCTAAGATAATTATCAATAATTGAGTGACAATATGAGAAATCAACTCTTGATGTCAGCTATATGAAATCTCAGTAGTCATATAGCTTTACCTAAACTCATCTCGATCTAGTACAAAGTATATTAGAGATATAATGTAGTCTAGAACTATTGTTCTATGACATAATTGTTGGATTGGTCCACTAATTTAAAGGTGAGATTGAGAACCAAACATGTCCGCATCATGGCTGATAACACAACAAAACTTTTTGGCCACTTGGTTAAGTCCATAGCTGAATTACATATGTAACCCGTGGATAAAAATAAAAAGCAAAAAGACAAGAACAGAACAACCAAAATTTGCATTGAGAATGAGAACACTTTCTCTGATAAATATGAACAGACAAAGAAGTGTGAATAAGAGAGACAGAAGCCGAATTGGCAACATTTGGTGGCAGTCAAACATGTTTCGCTTTGATCTTTAATATTATGCACTATATAAAATTCTACGAACTCTCTACAATTAGTTCATTGAAACGCAAAATATGATGATGGTTCCCAGCAAAATATCCTCTCTACTTCAGTTTTTCACTCTTTTATATCTCTTTACAGTTACTTTTGCTTCCACTGAGGAAACAACTGCCCTCTTTAAATGGAAAGCGACTTTCAAGAACAAAAATAATTCCTTATTGGCATCATGGACACGAAGTTCTGATGCATGCAGTGATTGGTATGGAGTTATATGCATTAATGGTAGGGTAAACATGTTGAATATTACAAATGCTAGTATCTTTGGTACACTCTATGCTTTTCCGTTTTCATCCCTCCCTTTTCTCGAGTATCTTAATCTTAGCAGGAACAATTTATTTGGCACCATCCCACCTGAGATTGGTTATCTAACTAATCTTGTCTATCTTGACTTGCATATCAATCACATTTCAGGCTCAATCCCACCACAAATTGGCTCACTAGCCAAACTTCAGATCATCCGTTTATTTGACAATCATTTAACTGGTCCTATTCCAGAAACAATAGGTTACCTAAGGTCTCTTACTAAGCTATCGTTGAGTATTAACTCTCTTACTGGTTCTATTCCTGCTTCATTGGGAAACTTGACCAACTTGTCTTTTTTGTTTCTTCATGTTAATCACCTTTCTGGATCCATTCCTGAAGAAATAGGTTACCTAAGGTCTCTTACTCGTCTAGATTTGAGGAATAACTTTCTTATTGGTTCTATTCCTACTTCCCTTGGCAACTTGAAAAATTTGCAAGTTCTGCTTCTCAATGACAACAATCTAATTGAGGAAATTCCTCCATCTATTTGCAATTTGACATCACTGGCAATCCTGTATTTGTCGAGAAACAACTTGAAAGGAAAAATTCTGCAATGCTTAGGTAATATCAGTACCCTCCAGTTTGTGATGATGTCACATAATAATCTCAGTGGAGAACTCCCTTTGTCTATTTGCAATCTAACATCACTACAAGTTCTAAATTTGGGTAGAAACAATCTGAATGGAGCAATTCCACAATGTTTTGGCAACATGAGTGGTCATCTTGAGATTCTTGATATGCAGCACAACAATCTTTCTGGTACTTTTCAAAGCACTTTTAGCATTGGAAGAGCACTCAGAAGCTTCAATTTGCATGGCAATAAGCTAGAGGGGAAAATTCCACGATCCTTGGCAAATTGCCAACGATTGGAAGTTCTTGATCTAGGAGCTAATCTCCTCAATGACACATTCCCGATGTGGTTGGGGACTCTGCCAGAGCTACAAGTGTTAAGCTTGAGATCCAACAAATTGCATGGACCCATCAGAACTTCAAAGTTCATGAAGAACTTGTTTCCTAAGCTTCGAATCATAGACCTCGCTAGCAATGCATTTACAGCAGAGTTACCTGCAAATCTTTTTCAAAATTTCGAAGCCATGAAGAGAGTTGATCAAACTGTGGAGGATTCAAGTAATAAAGAAGGCGGTTATTACCAAGACTCCGTAACTGTCGTAACAAAGGGACTGGAGCTTGAAGTTGATTCAATCTTGTCTTTGTACATCACAATTGATTTTTCAAATAACAGATTTGAAGGACGCATTCCAAGTATTATGGGAAATCTCATTGCACTTCACATGTTGAATTTATCTCATAATGGATTGCAAGGTCATATACCGCCATCACTCGGAAAATTATCTTCAGTTGAATCATTGGACCTATCAGCTAATCATCTTGAAGGAGAGATACCTAAACAACTTACATCTCTTACATTTCTTGAATTCTTGAATCTCTCTCACAATCAGCTCCAAGGATGCATCCCAACAGGACGCCAATTTGATACATTCGAGAAGAACTCGTATGAAGGTAATGATAGGTTACGTGGATTTCCTAATTTGGAAGGTTGTGGAAACAGCAGGTTTCCCAAGACAAACAACACGACACATGAGCTGGAACAAGAAAGTGATTCTACATTTCTGAGTGAACTCAACTGGAAAGTGGTTCTTATGGGATATGTGTGTGGACTTATTATTGGATTCTCCATAACATATTTAATGCTTTCAGCTAAAAATCCCGATTGGCTTTCTAGGATTGCTGATGAACTAGAATACAGAATCCTTATGAGAAGGCGAAAGTAG

>PGSC0003DMG400004725 LRR-RLPs

ATGGGGTCTGGCCAATTCTCATTTCTGCTCTTTCTCTGTTTGCTTTCTTTTTCTTCATCCATACCTCATCCATGCCGCAAAGATCAAAGTATTTCCCTTCTAAAATTCAAGAAAACGTTTACTGTAAATTCCACAGCCTCTGATTCTAATGTCTGTGAAATATTTTATGGTCAGAAGCCATATCCAAAAACGAGTTCATGGAACATGAGTATAGATTGTTGTTCATGGGATGGAGTCATATGTGATGAGATGACTGGCCATGTCATTGAACTTGACCTCGGCTGCAGCTGTCTCGTAGGAAAGCTTGATTCCAATAGCAGCCTTTTCCAACTCTCTCATCTCCAAAGGCTCAACCTTTCTGTTAATAACTTCTCTCGTGTGGATTCAATTCCTCTTCAGTCTCTAGATACTATTGATTTGCGGTCCAATTTGCTTCAAGGTTCACTACCTATTCCACCAATTTCCACGAGATTCTTCCTCATAGCCCATAATAATCTCACTGAAGAAATTCCTTCACATATTTGCAATTTGAAATCATTGGTAATGCTAGATTTGGCGAGAAACAACTTGAAGGGAGCAATTCCGCAATGTTTGGGTAATATCAGAAGCCTCAAAATTCTAGATATGCACAACAACAAACTTTCTGGGACTCTTCCAACAACTTTCAGCAATGGAAGTTCACTTAGAAAACTCAACTTGCGTGGAAATAAACTAGAGGGGAAAATTCCCCGATCTTTGGCCAATTGCAAAGTGTTGCAAGTTCTTGATCTAGGAGATAATCATATCATTGACACATTCCCTATGTGGGTTGGAACTCTTCCAAAGCTTCATGTTTTAAGCTTGAGATTGAATAAGTTGCATGGACCCATTAGAACTTTAAAAAATGAAAACATGTTTCCTGATCTTCGAATTATAGATCTCTCTTACAATGCCTTCACAGGAAATTTACCTACGAGTCTGTTTCGACATCTGAAAACCATGAGAACAATTGATCCATCAAAGAAGAAACTAACATACCTTGAAGACTATTATTACCAAGATTCTATTATAGTCATAAGCAAGGGATATGAGATTGAACTTATCAAAATTTTGACTGTTTACGCCTCTATTGATCTTTCAAGTAACAAATTTGAAGGACATATTCCAAGTATCATGGGAGAACTTACTGCACTTAAGGCGCTGAATTTCTCTCATAATAGATTGCAAGGACGTATACCATCATCACTTGGAAATTTATCTTTGGTTGGATCATTGGACTTATCAGTTAACCAGCTTTCGGGAGAGATACCAAAACAACTTGCTTCTCTTAAGTATCTTGAATACTTCTTCACATCAGAAAATAATCTCACTGAAGAAATTCCTTCATCTATTTGCAATTTGACATCACTGGTAATACTAGATTTGGCGAGAAACAATTTGAAGGGAGTAATTCCTCAATGTTTGGGTAATATTACTGGCCTCGTGGTTCTTGATATGCACAACAACAATCTTTCTGGGACTCTTCCAACAATTTTCAGCAATGGGAGATCACTGAGGAGCCTCAACTTGCATGGCAATAAACTAGAGGGGAATATTCCCCGATCTTTGTCCAATTGCAAAGCGTTGCAGGTTCTTGATTTAGGACATAATCATCTCAACGACACATTCCCCATGTGGTTGGGAACTCTTCCAGATCTACAAGTTCTAAGCTTGAGATCCAATAAATTGTATGGGTCCATTCCACCGTCAAGTGTTGGAAGTATGTTTCCTAAGCTTCGAATCATAGATCTCTCTTACAATGCCTTCTCAGGAAACCTACCTACGAGTTTGTTTCAACATCTGAAAACCATGAGGACAATTGATCCATCCAAGACGGCACTAATTGATGGTTATTATTACCAAGACTCAGTAGTTGTAGTTACAAAGGGATTGGAGCTTGAAGTTGAGAGAATCTTGTTTTTGTACACCACTATGGATCTTTCAAACAACAATTTTGAAGGACATATTCCTGATGTTATAGGAGATCTCATTGCGCTTCGGATGCTTAATTTTTCTCATAACAGATTGCAAGGTCATATTCCATCATCACTTGGAAATTTATTTGTAGTCGAATCATTGAACCTTTCATTTAACCAGCTTTCGGGAGAGATACCACAACAACTCGCTTCTCTTACATCTCTTGCGGTCTTAAATCTCTCCCACAATCATCTACAAGGATGCATCCCAAAAGGACCTCAATTCAATACATTTGAGATCAATTCATATGAAGGTAACGATGGATTATGTGGATACCCTTTTCCTCAAGGTTGTGGAAGTAGCAGGATGCCAGAGACAGAAAACACAGCATATGTTCTGGATGAAGAAAGCAATTCTACATTTCTGAGTGAATTTTGGAATGGGATTCTCATGGGATATGGAAGTGGACTTATTATTGGATTCTCCATAGCATCTTTCATGCTTTCGAGTCGAAATTCCAATTGGCTTTCTAGGATTGCTGAAGAACTTGAATACAGAATCGTTATGAGAAGGCGAAAGAAGAAGCAAGGCCATCACAGAAGATGA

>PGSC0003DMG400004727 LRR-RLPs

CCAAGTTACACTCTCTTCGTATCTCCACATTTTTTAACTTTTACACTAAGGAGCTAAGACTCACATCCCATGATTTTAAATCGCTTCTTCAGAACTTGACCCAATTAAGAGAGCTTGATCTTACTCAAGTAAACATCTCTTCCACCATTCCTCTGAATTTTTCTTCTCATTTCACGACTCTGAGGCTGTCAGAAACAGGATTGTATGGGATAATACCTGAGAGTATATTTCACCTGCCCAACTTGGAAATACTTGACTTAAGTTACAATGTTCATCTCAATGGTTATTTTCCAAAGGCCAAATGGAACAGCAGTGCATCTCTCATAGAGTTACATCTCAGTATGGTGAATTTTTCTGATAATTTTCCAGAGTCTATTGGCTATCATCTAACTTCGCTGCGTTATTTGTCTCTTGTAAGTTGCAACCTTAGAGGGCCTATTCCTGAATCTCTTTCAAATCTCACCCGCATTGAGTATTTGGTCCTTTCATATAACTCCCTGAATGGAACAATACCATCAGGGATGTTATCCCTTCCATCACTAAATACAATAGTCTTGAGTAATAATCACTTATCTGGTCCCCTTGAAGATTTAAAGTCCAATTCACTAATCTCGATATATTTAGAGGGCAACCAGCTGCAGGGTCATCTTCCCCACTCAATTCAAAACCTTGTGAACCTAACAGGCCTTATCCTTTCTTTCAATAATTTTAGTGGTCGTGTGGATGTCAGCTTCTTTTCAAACCTTAAACAGCTTCGGTATCTTAGTCTTTCATATAATAATATCTCATTGACCAATGACAACAACGTTACTTTGCCCGAATCTCTTGATACGTTAGAATTGGCCGCATGTGAAGTGAAAGAGTTGGAGTTTCTAAGATCCGCAAAGCTTCTTCGGCATTTGGATCTTTCTTTCAATAATTTTAGTGGTCGCGTGGATGTCAGCCTCTTTTCAAACCTTAAACAGCTTGTGGAACTTCGTCTTTCATATAATAATATCTCATTGACCAATGACAACAACGTTACTTTGCCCGAATCTCTTGATACGTTAGAATTGGCCGCATGTGAAGTGAAAGAGTTGGAGTTTCTAAGATCCGCAAAGAAGCTTTGGCAGTTGGATCTTTCAAATATAAGATTCAAGGAAGAATTCCTGATTGGGCATGGTCTAACTGGATGTTTCATTGCAAGATCTTAATATATCCCACAATATGCTGACGAGTGTGGATTCAATTCCTCTTCAGTTTGCAGATACTATTGATTTGCGGTCCAATTTGCTTCAAGGATCACTACCCATTCCATCAAATTCCACTACATTCTTCTTCATATCTAATAATAGTCTCACTGGAGAAATTCCTTCATCTATTTGCAATTTGACAACACTGGTAATGCTTGATTTGGCGACAAACAATTTGAAGGGAGCAATTCCGCAATGTTTGGGTAATATTACTGCCCTCCAGGTTTTGGATATGCGCCACAACAATCTTTCTGGGAATATTCCAACAACTTTCAGCAATGGAAGTTCACTGAGCAGCCTCAACTTGCATGGCAACAAACTAGAGGGGAAAATCCCACAATCCTTGACCAATTGCAAACAACTTGAAGTTCTTGATTTAGGAGACAATCACCTCAACGACACATTCCCCGTGTGGTTGGGGACTCTACCAAAGCTGAAAGTTCTAAGCGTGCGATTCAACAAATTGCATGGTCCCATCAGAACATCAAGGATTGAGAACATGTTTCCAGAGCTTCGAATCATAGATCTCTCTTACAATGCCTTCTCAGGAAACTTACCTTCGAGTTTGTTTCAACATCTGAAAGCCATGAGGACAATTGATCTATCATTGGAAGCATCAAGATATGGTGCAGATCGATATTACCAAGACTCGATAGCTGTCGTATCCAAGGGATATGAGCGTGAAGTTGTGAGAATCTTGTTTTTGTACACCACTATCGATCTTTCAAAGAACAAATTTGAAGGACATATTCCAACTATTATGGGAGATCTCATTGCACTCCGTGTGCTGAATATATCTCATAATGGATTGGAAGGTCATATACCGTCATCACTTGGAAGTTTATCTTTGGTTGAATCACTGGACCTATCAAGTAACCATCTTGTAGGAGAGATACCAGCACAATTTGCTTCTCTTACTTCTCTTGCAGTCTTAAATCTTTCCTACAATCATCTCGAAGGGTGCATTCCTCATGGAAATCAATTTCAGACCTTTGAGAACAATTCATATGAAGGCAATGATGGATTACGTGGTTTTCCAGTTTCAATAAGTTGTGGCAATGATCATGTATCAGGGAAAAATTATGCCGTAGATGATGAAGAAAGCAATTCTGAATTTCTGAGTGATTTTCGGAAAGCTGCTCTTATGGGGTATGGAAGTGGACTATGTATTGGAATATCCATAGTATATTTCATGATGTCAACTGGAAAACCGATATGGTTTGCAAGAATCATTGTAGAGCTGGAGCACAAAATAATGATGGGAAGGAGAAAGAAGCAGCGAAGTCAAAGGAATAACAGAAGAAGAAATAATCACTAATCATATTACTGTTATTTCTTTAAGGAAAAGATGTAATAAGTTGTAGACGCATTATCGTTAAATCCTGTTTGCATAAAGTGTGTAGCATCATTTGGCCATGAAAATTGAGGGTGAG

>PGSC0003DMG400005171 LRR-RLPs

ATGAGAAAGCTAAGGCAAGTGGATATATACAATTTGTCCTTGGTATGGGAAGAGAATGATCGAGCAATTTTTGATGAATCTCCAACAACTATGCTAGTGAACTTGAAGACATTCCGCACTTATGTTATACACGTGGATAATAGTACCCCAAGGTTCTGGTGGAGGTTTCCAAATATTGAAGAACTCAGCTGCCTCCGTATTGAAGCTTTACCTAGTTGTCCTTTGTTTCCCTCAGTGGAAGTTCATATTCGGCTTCAATCTCTTAATCTGGTTATAAGCCCTCGGGGATTATTGGATTCAGTTGGTTGGGAGAGCTACTTTGTCTTCCCTTCAAATCTTAGGCATTTGTGCATTGGTGGTTGTTTCCTAACGAAAGAAATGACTTTAAACATTGCAAGATTGAAGAAGCTTGAGAGTCTCACATTAAAGGGAGGATTTCCTTGGGGGGAAATAAAGTACCATTGCTGGGATGTTACAAATGTGGAGTTCCCTGCACTCAAATACTTGGCATTACACTTTATGAGAATGGAAGAATGGAAAGCTTCAGAGGAATCCTTTCCTGTGCTTGAGAAGTTAGTTATAAGAGAATGTCACTACTTCAAGGAGATCCCTATTAGCTTTGCCGATATTCCAACACTGCGACTTATTGTGCTGTTTGAGTGTACTGATTCACTTGTGGCTTCAGCTGTGAACATAAAAAATGAAATAGAAGAAAACACTGGATGTGATAGTCTCCAACTCGGGGTAACAAATTCGTATAAATAA

>PGSC0003DMG400005172 LRR-RLPs

ATGGAAGTTGTTGAGACTTGTGGAAAAGGAACTCTTTCTGCTGGAGAGCCTTCCACCAAACGTAGTATTACATCAATTTATGATGATGAAGAAGTAGTGGGCTTTGAGAAAGATGCAGAAATTATAATGAAGAAATTGATACGAGGAACAAAGGAACGGGATGTTATCTCCATCTATGGAATGCCTGGACTTGGTAAAACAACTTTGGCAAGGAAGGTGTACAACAACCCCACTACTGTTAATTACTTTGATGTTAAAGCATGGTGTGCTGTGTCGCAAACATATAATAGGAGGGGATTATTGGTTGAGATTTTCAAACAAGCAACAAGTAATGAGAATGAAATCAAAGAGGATGTTGACATAGCTGACATGTTGCGCAAGAGTCTAATAGGCAGAAGATACCTCATTGTATTAGATGATATTTGGGATGTCGAGGCATGGGAAGATTTGGGAATATGTTTCCCTGAAGGTGAATATGGAAGCAGAGTAATGGTCACAACACGAATTGAACAAGTGGCTAAGCATCTTCAGGTCCACAGTGTTCCTTATTCTCTTAGCTTTCTAACGTCCGAAGAGAGTTGGGAATTATTGGAGAAGAAGGTATTTCGAGGAGAGAGTTGTCCCCCGGATCTACTGGAAGCAGGGTTACAAGTTGCCCTACACTGTAAGGGATTGCCTCTTGTGGTTGTNNNNNNNNNNNNNNNNNNNNNNNNNNNNNNNNNNNNNNNNNNNNNNNNNNNNNNNNNNNNNNNNNNNNNNNNNNNNNNNNNNNNNNNNNNNNNNNNNNNNNNNNNNNNNNNNNNNNNNNNNNNNNNNNNNNNNNNNNNNNNNNNNNNNNNNNNNNNNNNNNNNNNNNNNNNNNNNNNNNNNNNNNNNNNNNNNNNNNNNNNNNNNNNNNNNNNNNNNNNNNNNNNNNNNNNNNNNNNNNNNNNNNNNNNNNNNNNNNNNNNNNNNNNNNNNNNNNNNNNNNNNNNNNNNNNNNNNNNNNNNNNNNNNNNNNNNNNNNNNNNNNNNNNNNNNNNNNNNNNNNNNNNNNNNNNNNNNNNNNNNNNNNNNNNNNNNNNNNNNNNNNNNNNNNNNNNNNNNNNNNNNNNNNNNNNNNNNNNNNNNNNNNNNNNNNNNNNNNNNNNNNNNNNNNNNNNNNNNNNNNNNNNNNNNNNNNNNNNNNNNNNNNNNNNNNNNNNNNNNNNNNNNNNNNNNNNNNNNNNNNNNNNNNNNNNNNNNNNNNNNNNNNNNNNNNNNNNNNNNNNNNNNNNNNNNNNNNNNNNNNNNNNNNNNNNNNNNNNNNNNNNNNNNNNNNNNNNNNNNNNNNNNNNNNNNNNNNNNNNNNNNNNNNNNNNNNNNNNNNNNNNNNNNNNNNNNNNNNNNNNNNNNNNNNNNNNNNNNNNNNNNNNNNNNNNNNNNNNNNNNNNNNNNNNNNNNNNNNNNNNNNNNNNNNNNNNNNNNNNNNNNNNNNNNNNNNNNNNNNNNNNNNNNNNNNNNNNNNNNNNNNNNNNNNNNNNNNNNNNNNNNNNNNNNNNNNGATATACGCTTGGATAATATGAATGCAAGGTTGTGGTGGAGGTTTCCGAATCTTGAAGAACTCGGCCTCCACGTTAAATATAAACCTAAGTTTCCTTTGTTTCCCATACCAGAAGTTCATACTCGCCTTCAGTCTGTTTCTCTGACGTTCTCATTAAATAGAGTCTTCAAGTCAGTTGGATGGGAGAGGAATTTTGTGTTCCCTTCAAATCTTAGGCATTTGAACCTTCGCGGCTGTTTCTTGACGGAAGAAATGGTTTTAAACATTGCAAGATTGAAGAAGCTTGAGAGTCTCAAATTACACATGGGATTTCCTTTTATGAGATTACAGAGGCTTCGGAACTGTTGCTGGAATGTCACAAATGTGGAGTTTCCTGCACTCAAATACTTGGGATTATTCTTTGTGCAGATAGAAGAATGGAAAGCTTCAGAGGAATCCTTTCCTGTGCTTGAGAAGCTAGTTATAAGTGGATGTTCCGATTTCAAGGAGATCCCCCCAAGCTTTGCGGATATTCCAACACTAAGACGTATTGAACTGACTAACTGCATCGACTCTCTAGGGGTTTCAGCTATGAATATCAGAAGAGACATTGAAGAAAACACAGGATGTGACAGTCTCCATGTTCTAATTGTAAACCGAATATAAATGTATTTAGAAGCAATTCAAAGTTGCTCTATTTCCACTTTTGTATTGTCACATTATCTGTTTTTTTTTTATTAGGAAACTTTTTAACTTGGAAGCTGAGATTGCAATTCAACTTTGCA

>PGSC0003DMG400005226 LRR-RLPs

CTTGATTTCTTCTCTTCCAACATAATTTTAATACTTTTAGTACAAAAATGGGTTATCAAACACTTGCATTGTTTCTTTTATGTTCCTTTCTATGTCAACTTGTTTTCTCCTCATTCTTACCTCATTTGTGCCCGAGTGATCAAGCCCTTGCTCTCCTACAATTCAAGCATATGTTTGCCATTAATCCTAGTGCTTCTCATGATTGTTTCGACATAACAGGCGAATGGATGCAGTCATACTCGAAAACATTGTCATGGAACAAGAGCACAGATTGTTGCTCATGGGATGGAGTTCATTGTGACGAGACAACGGGAAAGGTGATTGAGCTTAACCTCACTTGTAGCAATCTTCAAGGCAAATTTGATTCCAACACTAGTCTCTTTCAACTCTCCAATCTCAAAAGGCTTGATTTGTCTTCGAATAATTTTTCAGGATCGCTCATTTCACCTAAATTTGGTGAGTTTTCTAATTTGACGCATCTTGATTTGTCACATTCAGGTTTTAGATTCGGACCTCACAATTTAGAACTTCTCCTTAAGAACTTAACCCAATTAAGAGAGCTTGACCTTTACGGTATGAGCATCTCTTCCACCTTTCCTCTGAATTTCTCTTCTTATTTAACAACTCTACGACTTAGTGACACACTATTATGGGGGATACTGCCCGAAAGTGTTTTTCACCTTTCAAACCTGGAAACACTTGATTTATCATACAATCCCCAACTCACAATTAGGTTTCCCACGATCAAATGGAATAGCAGTACATCTCTCATGTACTTATCTCTACCTGAAGTGAATTTTATTGGTAAGATACCCGAATCATTTAGCTATCTAACTTCATTGCATTACTTGGATATGAGATCTTGTAACCTCTCAGGTCCTATTATTAAACCTCTTTGGAATCTCTCCCACATACAAAGTTTGTATCTCAGAAATAACCATCTTGAAGGACCAATTTCCCAATTCTTCAAATTTGGAAACCTCAAGTTGTTATCACTTGGAATTAACAACTTTTATGGCCAACTTGATTTTTTATCCTCCAACAGAAGTTGGACTCAACTTGAAGTGTTCGAAGTTTCTTCCAATTCCTTAGTTGGTCCAATTCCTTCCGATGTAAGTGGTCTGCAAAGCCTACAATCATTGTTCTTGTCATCAAACCACTTGAATGGGACTATACCATCATGGATATTCTCCCTTCCTTCACTAGATGAATTATACTTGAATGATAACTATTTCAGTGGAGAAATTCAGGAGTTCAAGTCCAAAACATTGTGGGACGTTACTCTAAAACAAAATCAGTTGCAAGGTCCTATTCCGATGTCACTCCTAAACCAACAGAGCCTATCTTATCTTCTCCTTTCGAAAAATAATCTTAGTGGACAGATTGCTTCAGCTGTCTGCAGTCTAAAAGCCTTGACAGTGCTAGATCTTGGAAGTAATAATTTGAAGGGAACAATCCCACGATGTTTGGGTGAGATGAGTGAACTTTGGGTTGTTGATCTAAGCAACAACAGTCTTAGTGGGGCAATTAATACAACATTTAGTATAGGAAACACGCTCAGAGTCATTAAATTGCACGGGAATAAGCTAGAGGGGGAAGTCCCACGATCTTTGGTCAATTGCAAGAGTTTGGAATGTCTAGATTTAGGTAACAATGAGTTGAGTGACACATTTCCAAAATGGTTAGGAACCCTACCGAATTTGAAGATTTTAAGCTTGAGATCAAATAAGTTCCACGGGCCCATTAAAACTTCAAGGGCTGGAAACTTGTTTGTACATCTTCGAATAATTGATCTATCATCCAATGGATTTAGTGGGAATTTACCTATGAGCCTTTTTGAGGATTTGAAAGCTATGAAGATAAATGATGAGATCACGAAAATCCCACAGTATGTGGGAGAAGACTTGTATTATCATTTTTCTGAAACAATTACAACAAAGGGGTTGGATCTTGAATTTCGTGGAGTTTTGACTACAAACATAGTTATCAATCTCTCGAAGAATAGATTTGAAGGTTGTATTCCAAGTTCCATTGGAGATCTCATTGCACTTCGTACTTTGAATTTATCTCATAATGGCTTGGAGGGTGTCACACCAACTTCGCTGCAATACTTATCTGTACTTGAATCATTAGATCTCTCATTCAACAAAATAAGCGGAGAAATTCCACAACAACTTGCATCCCTCACATTTCTTGAAGTCTTAAATCTCTCTCACAATCATCTTGTTGGATGCATCCCTAGAGGAAAACAATTTGATACGTTTGAGAATAGTTCATACCAAGGGAATGATGGGTTACGTGGATTGCCACTGTCAAAAGATTGTGGTGGTGATGAAGGGGTACCACAAACAACAACTCCATTTGGGCTTGATCAAGGAGAAGAAGGAGACTCAACAATGATCAGTTGGCAGGCAGTTCTCATGGGGTATGGTTGCGGTCTAATTATTGGATTATCCGTAATATACATAATGTTATCAACTCAAAATCCAGCATGGTTTTCGAGGATGGTTGAAGAATTGGACCACAAGATTATTACGAGAATGAAAAAATACAAGAAAAGATATTAGTGTGCAACCTAGCCAGGTATTCAACTTGATCTTTATCATTCATAAGATTATTTTGTATATCTCAATGTATTATATATCTACCTCCTTCATCCTTAAAATTCTTAACTTCAATTCTTCATTTTTTAAAATTGCAGGATTCGAGCCTAATATGTTGTTGGTGATGCAAATATAAGCTTTTATTTTTCACATTTTGTGATGTATCTAAGTATTTATCGCCCTATGAATAAAGACATGAGCTTTCTTCCTTTGAGCTTCGGCCAGAATTTTTAGAACGTTTTTAAATAGAGATTGTTAGCTTGAGTTCAGCCTAATAATAAGTTGAGTAATTAGGCTCAACCGCAAAGTCA

>PGSC0003DMG400005395 LRR-RLPs

ATGGGCTGTTGTGTGAAAGCTAATGCCTCCAAGGAGATTAGAACCCGCAATCATAGACATGTAGGACTTGAAAAGATCTTTCGCCAAGTGCACGAACCTCTACTTTGCATTCTGGTGCCTTTGTGTATCTATTTCTAAGATATCAAGATAGTTACCTATATAAACCAAGTTTCCATCGAAGTTTCCGGGTGCAGGTGTATCCTCAACCTACTATAAACGGGCTTGCTAAATCCAGATTGAGGGAAATTCATGATTAAATGTGTCTTTTAGTAAATATTAACTGTGTTACTCTGTTGGAAATGCAAGGTTAAAGCAGTAACAATTTGTCCAAAATATAATGGCAAACACAAAATTAAAGATATGCTGACAGCTTAACTTCAAAGGGAGATTTTAAGCTATCGTAGGATAAGTCTTAGGAGGAAATAGCCCTGTGAAGGAGCTGTAATGAAAGTCAATTTTTTTGTCAACGAGGTAATATGTGGTGGGTAAAAAGCCCAAGTGTCACACCCGAAATCCACGCTTATTGCTCAAAGTACTTCCATGCTTGTTCTAGTACGTTTAATCACTTTTGGACTTTAGTATCCTTGCTGGTCCCCACACTTCCAACCTTGATCTGATTTTTTCAAGGTTTGAATTTTCGGGGTCTTACACCAAGTATACTAGTGGTGTACAAGAAGTGGACCTACACCAACACATATTTCTGGTACTCCAAAAATCTATTTAAAAAAACAAAAAATCAGTTGTAATCTGTACAAGAGTCCTCAAGACTCTACTGGGATGAAAATCAGTTCTACCTGGGAAAACTGAAAAGTGTCTGCTTCTCATTTATTCTTAAGGGACTCTTCATCTGGGTTTTCACGCTGAATACTATTGATATTGACATATTGTAGGGTGTATTCTCTTAAAAAGAAAATTTTATAAAGAGTTTGTTATATCTGATTCTATCCAAAAAATTGAGTCTGATTATTTCTTTTCAGGTTTTGGCTGATAACCTTATTGAGAATCTCCCGGTGAACCTGGGATTGCTTAAGTCTTTGAAAGTTGCAACACTTGATGGGAATCGAATCAGAGCATTGCCTGATGAATGTAATTTCCCACCGTCCTAGATCATATGTGAAGTTAAGCAGGATTCTCTCTGCTATAGAGTATCACACTCTCTATTCTAACAACATAATGTTCTCATATTCAGTGGGCCAGTTGGTGAAGCTTGAGCGTTTGTCTGTCTCAGCAAACTTGTTAATGAGCTTGCCTGAGACTATTGGGAGATTGCAGAATGTGAGTGATCCATTCTGATGGCTAATACGGTGCCATCTTCCGCAATTCTAAAAAGTGTGCCAGTTAATATATTATTTCTTAGCTTGCTTTACTGCTTTGTTATTTTTTTTTTCCGAGTATAAAGGTGCTAAGGTGTCTAATGAAATGGTGTCAATCTTGTAGAAAATGTTCAGCATATTGATATCCATATTAATGGATCATTTTGCTTACAACTTTCTGTTCTTACAACTAAAAGCCCTACTTTTTTTTCTCTATTTCTATAGTGATGCCGCTTCTTCATGAATTTCTATTATCTTCCTGGTAGTTGATGATGTCTCTTTGCTCTCTTATGTTAGTATTTTGGATTTGGAATAGGGTCCTGCAGCTCCTGGGGGAGGGTGGTTGAATTTGGAGAATAAGTGGCTCTATTGCAGGTTGTCTTGAGGTCCACCACCCTGAGTGCCGTCTGTTCTTCCTTTTTATTTCACTCTCTCCTTATATTGAATATTTTTCTTCTCAAGTTTTTTTCTTCAATGTCATTGATGAACCAAGTGTGAGAAAGAAAGACTCGGAAAATACTTTCAATATACGTAAACTACATTATTACACAAAGCATTGTTTATAGACAATACACTACAATCCTTTTCCATGTATGGCACCATGTACAATCCTATTTGAGCACTCCCCCTCAAGCTGGTTCATGTAAGTCATATGTACCAAGCTTGGTACCTCCACTTGGATATATTGTGTCACTTCATTTATCACTTGGGTAAATAAAAAATGGGCTAAGAATCGATCCATGGTGTAACCCCATCTCTACTAGGAAGTGTTCTAAGTCTCCTCCCATTGTTCTCACTGGGTTTTGACTCCATCGTACATGTTATTAATCCCTCTAATGTATGCCACATGTACACCTCTGGGCTCCAAGCATCTCTATAAAATCTTCCTAGGGACTGTCATATGCCTTTTTTAAGTCAATGAACATCATTTGTAGGTTTCTCTTCCTATTTCTATACTCATCCAACAATCTCATTATAAAATGAATGACTTCCATAGTTGATCGCCCTGACATGAATCTGAATTGGTTCTCCAAAATAGACATATCCCTCTTCCTTCTCATCGTCATCTCTACCACCCTCTCCCAATCCTTCATAGTGTGATTTAGTTGTTTAATACCACTCTATAGTTGTTGTAATTTTGAATATCATCCTTTTTCTTGTACAGCGGGATCATTGTCCTCCACCTCCATTCTTTGGGCAGTTAATTCATCCTAAAATGATATTAAACAGCCTAGTTAGCCATTCAATGCCTACTTCCCCTGCGCTCTTCCAAAAATCCATAGGCATCTCATCTAGTCTGTTCGCTCTTCCTGTTTTCATTTTACGAATTACCCCCTTGACCTCCTCAAACTTTTTACACCTACAATACCCAAAATCACGATGTCTCTTGGAGTGTTATAAATCACTCAACACAATAGTTCTAATTCATGCTCCCCTCTGCATTCAAGAGTACAAGTTCCATTTGACTGTTTTTTTCTAAATTTTAATTCTCTGCGCATTGACAAACAAAGACTCAGAAAATATTATTAAGATACGTAAATATCACACTGAGGTCTACTTATAGATAATACACTACAATCCTTTTCCATGTAGGACACCATGTACAACCTATTGTAACACCAAGAAGTATAGTGTTCACTTTTAGGTGTATATCTGATTCTCATGGTTGCTGACATGGGATTATGAATTAAAGTTTTTCAATCAAGTTAACTAGCTCTATCCTCTAGTAGTGTTCATATCATTCTAAACACTTGCATATACTTTTACTCCAAATATGTTTTGATATTTGTGAGTTCAGTTCAATATGATGATGCCGTTTTTTGTCTGTCAGAGATAGTCGTTCAGCCTATACTTAAAAGTATTCTTAATTCATCTTCTTTGGAAAATGAGGCCTTTGGAATTGACAGTAATGGTTTTTCACTGCAGTTGGTGCTGTTGGACATTTCAAACAATAAGTTAAAGTTTTTTCCTGAATCAATTGGAAGTTGCTTCTCTTTGGAAGAATTGCAAGCAAATGGTACCTTCAAATTTTAGTGTCACTCATCCATTCTTGCTATTTCTCTATTTCCTTTTATTTTCCTGGCACTGGAACTCTGTATCAAATATCCATCGAGGAAAAAAAAACATGTTACCATCAACATAAAATTAAGTGAAATTTATGGCCGATGATAAAATTGGAGTGTTGGGTGACCTAGAAAGCTAAAGGAAAATCTACAAATTTTCTCTGTTAAGCTTATAAGATAATATTCTTGGCACCATCCATTTGAACTGGTACATGATTTTCACGTTCTGTTTCTTATTCCTCAGTGTGGCATAATGAGGTTTACTGTGTTTTTTTGGCAATCTTGCCTCGTGATAACAATATTTCCTTTCAGATAATTCCATTGAAGAGCTTCCTGCATCAGTTTGCAATCTTGTTCAACTAAAATCACTCTGCCTGAACAATAACAATCTCAATCAGGTTAGTGTCTAATTTTACCTGTGTGTATGTGTGTGTGACCCTTGTTATTTTGAGGTCATATCTAGTATTCAGAGTTTACCTCGATTTGGTACTGCTCTCGTGGCCTGCACTGATACAGTGCTTTACCTCTACATGTGCAGTCAACTGCTGTGCATGTGTGTGTGTGTCTGTCTCTATCATCGTCACACATAAAAAGTGCAAAGAATAGTAAACCACGTGGATCAAAAGCTTACCTGTTTGTCACACCGAAAAGCTGAACTTATTGTTGTATTGGTGATACTTGTATGTGCTCTCTCTCTCACTTTCTATCTTAATAATACTATTAAGCAGAGTTAGGAACTCTCTTCGCATGATCTGTAAGCCTAGTTCACTTCCCCGTGTATGAGTGTTAAAAGGAAACTCATACATTAACCTTAAAACGTGCATCTAAAAGCTTTTCCTTTTAATAAGACCTCATGAAGAAACGAAGATAATCAAACAACCAGTAGCATCTACATTGCAAACCGGCAATAAAGCCAAATACGCATAATGATCCACATTTGAGGTCATTGGATTGAACCTCACGTATCTTCCATTTAAAGCATCATCATCCAAGTTTTTGTACTTGTGCCTATCAAGTATATGTTTTTACTAGATAGTCATGCTTCCTTTTGGTAACTGGATTGTTCAGATGCCTCTAAATTTGTTAAGAGAATGCAAGAGCTTGCAGACCATATCCCTGCATGGAAATCCGATTTCCATGGACCAATTTCAACAGGTTACTCACTAATATTCTACTATTTAGTTCTGCAAGTACTAAAAAAATCATTTATGAAGAATTCTATTTGCCTTTTCTTTTTCCAGATGGATGGCTTCAAGGACTTTGAAGCCCGCCGTAAACAGAAGTTTGATAAACAAATTGATTCAAATGTGATAATCAGTTCTAAAGGGCTTGATGAGGGTGTTGACTTGTGATTCCTCTCCGCAGGCGTTTTCTATAAGGCTCCGCGCAATCTTGTTAATCTCAGGTTTACTATTTTTTTGCTTCAATTCCCTGATGGAAAATTTCCCTTTATTTTATGTATGACCATGTAGTAGCCACACTACACAGTTGTGCTCCTTCAGTTGCTGTCAAGCTTTTAGTATTTCTAGTGTCTGTAATGCCTAATTTGGACAATTCATTCTTGTTTACTTCATGCATGTCTCCGTTGCTCTTAGCTCCCAGCCTGGCGTGCCTGCATAGTTGCCTCTTCTCATTTATGTTCCAGTCTTCTAGTTTGTATTACTCTCCATTATACCCGTGCCCCTAAGTGTATATAATTGTGGTAATTATTAGTAACACCAATTTCAAAAGATTGAACACGAAAAAGCTTCTCATTTATGTTCCATGGAAGCAGGACCGTTTTGTCCATTTAGTTTCTGAATTTGATAGGTAGTAGCTAATCCAGTGGCCAAATTAGTTCAAAAAGAAATTGCACTCGTTTTTCTAAACTTCCCAGTCAACTGGTGCTATTGATGATACAATTTTTTTTGAATAAATCATTATATCTTATTTACTGCAGGTTAAATTTGTTCAGGTTGGATGGAATTGCGACAGTTTTGACTTTGATGAAAGGTATTGCTTCTGCTTATTTGGCATGAGATATTTCACACCTTTCTATAGTTTGTCATTGATTGTAAGTTGTGACTTGTGCGTAATTGTATCTATGCTGACTTGCTGAGTATGTTTGTGATTGTAAATGACTCAACTCTAACAAATTCTGTACATTTAGCGAGCTGTATATCACTTTAAAATCAAACTAACGATTTTTGTACCTAAAGATATTTGATTTTAATATGTTTGGATTGTAACGATCCTTTCAACGTATCCAAGAGTAGGGCCTGGTGGCCAATAAAGTGGCTGAAAATTGTGGCCACTTTTTATCTCCGTTCGTCGTTTTAAAATTAATTGAAACTAAGGGCGTGTTTATGATATTTTCATGTTTGGTTGGTTAAATTTTTTGGAAAATATTTGCTCTAGTAAGTATTTCATACGCGAGCCATCATCTAAAATCAATTCATCGATATCAAGTCAGCAACACAAGCATTTATCTAGTGCTATATACTAGGAGTCAAGCTCACATAATCACATTAATCAATTCAGGAAGAAAATCTCCAATTTTTCATATTCATGGCATTGGTACTCAAAGCATTTCATATTCATGGCATTGGTACTCATAGAAATGCTTGTTATCTCTTTTATACCGGACCTCCATTATCCATTAACCTTTTATTCAATTAATAAATACACTTTAACATTTCATAAAAAGTTTCAACTTGCCTTAATTGGAATTCAAAAAGTTAATGTGGATACCTAGGCACCCAAAGACAATGAAGGGTGTAGTAATCGACAAAATGCTTTAGGGAGTTGAAAATAAGCATAGATTTGGAGATTCTCGAATAAGGTAACCTTTCAAAATGCTATTGAATCCACGGGTAGGCAAGATACAACTAACGATTTGCTCTTCTTTTATGCCTTTGAACTATTTCAATTTGGTCATAAATGTAATTTAGATGAGTATACAAGTTCGTAGACACACAAATTGGCACATAATTGGCTCGAAAATTGAGTGAAAAAGTCAACCTTGAGCCTTACTGTCAAATTTGAAAATTTGTTTACCCATAGGTGGGTCAAAAATCCGCTAAAATGGATTGTTAATCGGTCCCCAAATCGAAAGCTACATAAATCATTCTTTGGAGAATTAAAAACCAACCAAATTCCCCAATTCAAAAACCCCCAAATTAAATATCTATGAGCTTGTATAGCAAGGTGAGACCCTCCTCTTATTGGATTCTAATTACTTTAAGTCACAAATAGTATTTTAGGATGAAAATTCATATCTTTCGTCGGATTGTCATAGTACTTCTTGTTATTGATTGTTGAGTGAACTCATTGTTATAGTAATGTGGGGCTGTGGGAAAAAGTTATTGGATGGCAAAGAGTCAATGGAAAAAGGTATGATTAGGTTACTCTCTTCTTTTAGCATGTTTTCTTAAAAGGGTAAAGCTATATGTCAAGCGTACTTTGGATGAGCTACTCATAGCCATAAGTTTAGTGTGTTGGCTTCCTAGAGTTGTTATGTTCGTCCCTTGTTGGGACTCCTATTATAATGATAGCGACCTCGTGCTTAGATACAACTTAGAGATTACTCATATAGGTCGCTTATAGTTGGTTTGCTTGCCTTAGGGCCACTTGATTGATGTTGATGCCTTTGGGCCATATGTTGGTTGTTACTTCTTTCGGGGGGGGGGGGGGGGTTTTGTGTNNNNNNNNNNNNNNNNNNNNNNNNNNNNNNNNNNNNNNNNNNNNNNNNNNNNNNNNNNNNNNNNNNNNNNNNNNNNNNNNNNNNNNNNNNNNNNNNNNNNNNNNNNNNNNNNNNNNNNNNNNNNNNNNNNNNNNNNNNNNNNNNNNNNNNNNNNNNNNNNNNNNNNNNNNNNNNNNNNNNNNNNNNNNNNNNNNNNNNNNNNNNNNNNNNNNNNNNNNNNNNNNNNNNNNNNNNNNNNNNNNNNNNNNNNNNNNNNNNNNNNNNNNNNNNNNNNNNNNNNNNNNNNNNNNNNNNNNNNNNNNNNNNNNNNNNNNNNNNNNNNNNNNNNNNNNNNNNNNNNNNNNNNNNNNNNNNNNNNNNNNNNNNNNNNNNNNNNNNNNNNNNNNNNNNNNNNNNNNNNNTTTGGGATGGGTGGGGGGGGGGGGGGGGGGGATTGTGGTTACCTTCAGAGTTATGGTTCATGTTTTCAAGGTTATATGTAGTTTGCCCAAGGGTAATATAGAGCCTTGTGAGCTCGGATAGATAGTCTTATGTGCTCAGACATATAGAGCCTTTTGTGCTCGATTGTATAGTTATACAACAAGGGTAGTATAGAGCCGAAAGAGTCCTCAACTATACAGAGCTGACGAGATCCTCAACTATACGAGCTAGTCAATTACACAAAGCTTATGGGGCAAAATTGTTGCCAATAGGGGTATAGAGATTGATTTGTACATGTCGAGCTTATGGGGGATAGGGCAAGTGTATTAATTGTTATTTGTACTTGTCGATTTTATAGGGACTTGGTAAGTTGTTGTTTAATTACTGTTGTATGATCCATGTATGAATTGTTGGATTTTGCTGTGACTATTCTTACTCCCTTGCAAGATTCCCTTTAGTATTGTATTTGTATGACATTAGTTGGCCTAGTTAGGCCGAATTGATTGTTGAGCTGAATATGGTTTCTGAGGTTTAAGCCAAAAACATGTAAGTACGTTCTGACTATTTTTAGACTCCCTATTTGATTACTTATAGTACTTTTATCATATCATTATTACTTATTCAATTTTCTTACATACTCGGTATTTATTTCGTATTAATGCTCTATTGTCTACAGGCTCTGCATTCATGCTTGCAGGTACTAACAAACTAGTAGCCTGACTTCTTTAGTAATGTTGCCGAGTTTAGCTAATCAATAAGTTCCACTTCATTCAGAGTTGCGAGGTTTAGAGTTTGAGAGTCTGTGATTATATATAGTTGATGGGTAGGTCGGGGCCCACTCCCGACTAGATACTATATTCTTGAGGCTTGCAGAGCTGGGTCATGTCATAGTACATCTCAGTCGAACGTATTGGCATTACTACTGTGTTATTATTAAGCATGTCATGTTTGTCGTCTATATCATTTTGCTATTGTAGATATTATAATGGTCTTGTCAGCCCGCATAGCAATTGATATGCTATTATTATGAACTGTTGGGTTAGCTAAAATTGTTGTTGCTGCGGGATTCTATGATTATAGCTTTCCCAACCTTGACTTTTA

>PGSC0003DMG400005538 LRR-RLPs

TATTCATCTATTCCACCATTCCTCTGAATTTTTCTTCTCATTTAACAACTCTGAGGCTGTCAGAAACAGGATTGTATGGGATAATACCTGAGAGTATATTCCACCTGCCCAACCTGGAAATACTTTTCTTAGGTTTCAATTATAAGCTAAGTGGTTATTTTCCAAAGACCAAATGGAACAGCAGTGCATCTCTCGTAGAGTTATATCTCAGGGGCGTGAATTTTTCTGATAATTTGCCTGAGTCTCTTGGCTATCATCTAACTTCGGTGCGTTATTTGTCTCTTAAAAGTTGCAACCTTAGAGGGCCTATTCCTGAATCTCTTTCAAATCTCACCCGCATTGAGTATTTGGGCCTTCGCAATAAACTCCCTGAATGGAACAATACCACCAGGGATGTTATCCCTTCCATCACTAAATGCAATAGACTTGAGTTTTAATCACTTATCTGGTCCGCTTGAAGATTTCAAGTCCAATTCACTAATCTTGATAGAGTTAGAGGGCAATCAGTTGCAGGGTCATCTTCCCCACTCAATTCAAAACCTTGTGAACCTACAATATCTCTATCTTTCTTTCAATAATTTTAGTGGTTTTGTGGATGTCAGCTTCTTTTCAAACTTCAAACAGCTTTCGTATCTTAGTCTTTCATATAATAATATCTCATTGACCAATGACAACAACGTTAATGCTACTTTGCCCGAATCTCTTATTGGGTTAGAATTGGCCGCATGTGAATTGAAAGAATTCGAGTTTCTAAGATCCGCAAAGAAGCTTCAGCAGTTGGATCTTTCAAATAATAAGATTCAAGGAAGAATTCCTAATTGGGCATGGTCGAACTGGATCTTTTCATTGCACGATCTTAATATATCCCACAATATGCTGACGAGTGTGGATTCAATTCCTCTTCAATATCTATATACTATTGATTTGCGGTCCAATTTGCTTCAAGGTTCACTACCTATTCCATCAAATTCCACTACATTCTTCTTCATATCCCATAATAATCTCACTGGAGAAATTCCTTCATCTATTTGCAACTTGACATCACTGATAATGCTGGATTTGGCGACAAACAATTTGTGTGGAGAAATTCTGCAGTGTTTTGGAAATATCACTTCCCTCCAGGTTTTGGATATGCGCCACAACAATCTTTCTGGGAATATTCCAACAACTTTAAGCAATGGAAGTTTACTGAGAAGCCTCAACTTGCATAGCAATAAACTAGAGGGGAAAATTCCCCGATCTTTGGCCAATTGCAAAGAGTTGCAGGTTCTTGATTTAGGAGATAATCATCTCATTGGCACATTCCCGATGTGGTTGGGAACTCTACCAAAGCTGCAAGTTTTAAGCTTGCGGTTCAATGAATTGCATGGTTCCATCAGAACATCAAGGATTGAGAACATGTTTCCAGAGCTTCGAATCATAGATCTCTCCTCCAATGGCTTCTCGGGAAACTTACCCTTGAGTTTGTTTCTACATCTGAAAGCCATGAGGACAATTGATCCATCAATGGAAGCACCAAGTTATAGAGGAGATATATATTACCAAGATTCTATTACAGTTGCAACTAAGGGATTTGATCGTGAAATTGTGAGAATCCTGTATTTGTACACCGTTATCGATCTTTCAAGTAACAGATTTGAAGGACGCATTCCAAGTATCATGGGGGATCTCATTGCAGTTCACATATTGAATTTATCTCATAATGGATTGCAAGGTCATATACCGCAATCATTCGGAGATTTATCTTCAGTTGAATCATTGGACCTATCAGGAAACCAGCTTTCGGGAGAGATACCACAACAAATGGCTTCTCTTACGTCTCTTTCATTCTTAAATCTCTCCCATAATCATCTCCAAGGATGCATTCCTCAAGGACCTCAATCTCTCACTTTTGGGAGCAATTCATATGAAGGAAATGATGGATTACGTGGATTCCCTGTTTCAAGAAGTTGCACATATGATCGGGTATTAGATACAAATGATACCGTATCTGTACTAGACGATGAAGAAAGCGATTCTGAATTTCAGAGTGATTTTCGGAAAGGTGCTCTTATGGGATATGGAAGTGGACTATGTATCGGATTATCCATAATATATTTCATGTTGTCAACTGGAAAACCGATATGGCTTGCAAGAATCATTCTAGAGATGGAGCACAAAATTATGAGGGGAAGGAGAAAGAAGCAGCGAAGGCAAAGGAATTACAGAAGAAGAAATAATCATTTCTAGACAAGTTATAAGTTTTGACTTCAGGTAACTAATCGGTGCTATATACTTTTCATTTTTTTCATTTCCAATAATTTCAAAAGGGTCAGCATCAGTTGATGTTTAACTAAAATATTTTGATTACAGCGAGGAGCTAGTGTCTATGGAAGTGGAGAAAGGGAAGAAGAAGTTATGGCAATGAATGGAAACGAATGCCAATCAGTTTTTAAGCAATAATAATTGTCTAACTGAATATTTTCGTTGAACAGTCTTAGATGTGGCTTCAATTCTTACTGTGTAGTGTAAATTACAAATTTATGTAGTGTGTTTCTGTTGATATATAAAATCTATGAAATTTCTCAGTCTTACTGCTTGTCCTAAGATGAGGCAACGCTGGGGCACCTAAATCTCATTCCTAGTTACACTGTTAAGGCACTCATTGTAATTATGGATAATAAGATTGTTACGGCACTTATCAATTATGTGGAAATGCAAATGATAATGCAAGTGAACCTCTGATTCATGCCTAGGTTAAAGAAAACTCTGCTGCTACACTTTTTAGCTACTTCAGAAACAGTAAATTTGTTGTATTAAACTTCATATTATTATGAGACGAAGAAAGTATTATTTTTGACATTTTCCTGGAAAGACTTTGTGCATTGATTTGAGAATGACAACATTTTGTGAACTTGGAATTGATTGATGATTTGATATGGAGGCAGCAGTTCATACTTGACTTTTCCCCTTTCCTCTCTCTTTTTGATAATAAAATTCGCAAGTAATTAATTTCAATATGAAAATTCAAAGTTAAAGGGATTTGTCTGATAAGTAACTTTCATTGAAATTTCCGCAGCAAGACTCATTGCAATTACTTCATAATTCAAAGAACTGCATTAATACTTCTAGGTGCAAGAAAAAAGATAATTGGAGGGACACTCTGTGCAAAAACTTGGTGTTATTTCCCTCTTTGATAGGCCTTACACGCCGAGAATCCAGATTATTCGGGATCCTAATCTGGATATCGAAAATCGAATAGAAAATCAATTTATTTTTAGAAAAAGCCATTTCTCATTATAAATATGAGGTAAACAAGTCAATATCCTGTAATTATACCAAAAAGACACTATGGCATATGACTTAGTCGTGTCTAATATTTGAGAATGACATTTCTCTACTTTCTTTTGAAATTTTCATAGGAAGACTTGTGCATTGACTTAAGAATAACAAACATTAATTTCCCTATTTCTTTTCAAATTTACTATTGGTGAAGGTGGAATTTGGAAATAATTTCCCTATGCAATATGATTCTGGATTTGTTTTTAATTTTTTTTTGTTTGTTTAAGAAAGCCTTTTCTTCTTCCTTTTGGAGTAGCTAGGAATTTGCTTTGCCATTTAGAGCCACTTTTATAACGAGGATATATGATATATATATATATAAGCTCTAAATGAAAAAGTTAGGGGGTATATTAGACCCTTTTTCCTGAATAAGAAAGTAAAACAACGTACAATTATATTTTAAGAAAAAAAGATGATTCTCTACTCCTTTTTTAAGCTTGTTTCTTTAGAGAAATAAAACAACCCAAACTTGACTTGTGTTGAACAATATAAAATCCTAAGTTAGTTCATTGAAACCCAAAATAATAAAATAAAAATATGATGATGGTTCCCAGTAAAATATTTTCTCTACTTCAGTTTTTCACTCTTTTATATCTCCTTACAGTTACTTTTGCTTCGATCGAGGAAGCAACTTCCCTACTGAAAATGGAAAGCAACTTTCAAGAACCAGAACAACTCCTTATTCGCTTCATGGACACCAAGTTCTAACTCATGCAGGGACTGGTACGAAGTTATATGCTTAAATGGTAGGGTAAACACGTTAGAATATTACAAATTCTAGTGTCATTGATACCCTCTATGCTTTTCCGTTTTCATCTCTCCCTTTTCTTGAATATTTTGAACTTAGCATGAACAAATTTTTCAGGAATCATTCCCCCTGAGATCGGTAATCTCACTAATCTTGTCTATCTTGACTTGTCCATGAATCAGATTTCAGGCATAATTCCGCCACAAATCGGCTCACTAGCCAAGCTTGAGACCCTCCACATCTTTGACAACCATTTAAGTGGTTCCATTCCACAAGAAATAGGTTACCTAAGGTCTCTAACTGAGCTAGCTTTGTATAACAATTTTCTTAATGGTTCTATTCCTGCTTCATTGGGGAATTTGAACAACTTGTCCTATATGTATCTTTATGATAATCACATTTCTGGCTCTATTCCTAAAGAAATAGGAAAACTTGTCAATCTTTTTGGAGTTTATCTTGAAACAAACCAATTAACAGGTCATATCCCTCCAGAAATAGGCAACTTGACCAATGCAAAATTGTTCTATGCTCCCTTCAATGAATTATCTGGTCCCATTCCCGCAGAAATAGGGAAGATGAAGTCACTTCAGGAATTAAGCTTACATACAAACAATCTTTCTGGTCTAATTCCAAGAACTACAGGTGACTTAATTGAGCTCAAATATCTACAACTTAATGATAATCAACTCGCTGGCCTAATTCCTAGAGAGATGGGGAATCTAGAAAACCTCAATGGTCTGGAATTATTCAATAATCAACTATTTGGCCTAATCCCTAGTGAGTTGGGGAATCTTAGTATCCTCTATCATCTGACATTATCCAATAATCAACTCACTGGTTCAATTCCTTCTTCCTTTGGTAATTTGAGAAACTTGCGAGTTCTGTATCTCCATGACAACAATCTGATTGAGGAAATTCCTTCATCTATATGCAATTTGACATTCCTCATGTTCTTGCATTTGTCGAGAAACAACTTGAAAGGAAAAATTCTGCAATGTTTGGGTAATATCAGTACTCTCCAGTTGGTGACACTGTCACATAATAATCTCAGTGGAGAGCTCCCTTTGTCTATTCGCAATTTAACATCACTACAGATTCTAAATTTGGGCAAAAACAATATAAAGGGAGTAATTCCGCAATGTTTTGGCAACATGAGTGGTCATCTTGAGGTTCTTGATATGCAGCCAACAATCTTTCTGTGACTCTTCCAATGACTTTTAGAACTGGAACACTTAGAAGCTTCAACTTACGTGGCAATAAACTAAAGGGGAAAATTCCTCAATCTTTGGCCAATTGCAAAGAGTTGCAAGTTCTTGATTTAGGAGATAACCACCTGAATGACATCTTTCCGATGTGGTTGGGAACTCTTCCTAAGTTGAGAGTTTTAAGCTTGAGATTGAATAAGTTGCATGGACCCATTAGAACTTTAGGAAGTGAAAACATGTTTCTTGAGCTTCAAATATTGGATATCTCTTCCAGTGCCTTCACGGGAAACTTACCTACGGGCCTGTTTAAACATTTGAAAGCCATGAGGACAATTGATCCATCAAAGAAGGCACCAAGTGATGGATATTACCAAGACACGGTAGCTGTCGTAACAAAGGGATTTGAGCGTGAAATTGCGAGTATCTTGTATTTGTACACTGTTATCGACCTTTCAAGTAATAAATTTGGAGGGGAAATTCCAAGTATCATGGGGGATCTCATTGCAGTTCATACATTGAATTTATCACATAATGGATTGCAAGGTCATATACCACCATCATTTGGAGATTTATCTTCAGTTGAATCATTGGACCTATCAGGAAACCAGCTTTCGGGAGAGATACCACAACAACTGGCTTCTCTTACGTCTCTTTCATTCTTAAATCTCTCCCACAATCATCTCCAAGGATGCATCCCTCAAGGACCTCAATTCCACACTTTTGAGAGCAGTTCATATGAAGGCAATGATGGATTACGTGGATTCCCTGTTTCAAAAAGTTGCAGCGATGCTCGGGTGTTAGATACAAATGATACCGTATCTGGACTAGACGATGAAGAAAGCAATTCTGAATTTCAGAGTGATTTTTGGAAAGGTGCTCTTATGGGCTATGGAAGTGGACTATGTATCGGATTATCCATATTATATTTCATGTTGTCAACTGGAAAACCGATATGGCTTGCTAGAATCATTGTAGAGATGGAGCACAAAATAATGAGGGGAAGGAGAAAGAAGCAGCAAAGGCAAAGGAATTACAGAAGAAGAAACAATCATTTCTAGACAAGTAATAAGTTTTGACTTCAGGTAACTAATAGGTGCGCTATACTATTCATTTTTTTCATTTCCAATAATTTCAAAAGGCTCAGCATCAGTTGATGTTTAAATAAGATATTTTGATTACAGCGAGGAGCTAGTGTCTATGGAAGTGGAGAAAGGGAAGAAGAAGTTATGGCAATGAATGGAAACGAATGCCAATCAGTTTTTAAGCAATAATAATTGTCTAACTGAATATTTTCGCTGAACAGTTTTAGATGTGGC

>PGSC0003DMG400005539 LRR-RLPs

ACAATTTATGACTATAAGAAGCTAAGACTCACAACCCATGATTTTAAATCGCTTCTTCAGAACTTGACCCAATTAAGAGAGCTTCATCTTAGTGGCATAAACATCTCTTCCACCATTCCTCTGAATTTTTCTTCTCATTTAACAACTCTGAGGCTGACATATACAGGATTGTTTGGGATAATACCTGAGAGTATATTCCACCTGCCCAACCTGGAAATACTTTTCTTAGGTTTCAATTATAAGCTAAGCGGTTATTTTCCAAAGACCAAATGGAACAGCAGTGCATCTCTCGTAGAGTTATATCTCAGGGGCGTGAATTTTTCTGATAATTTGCCTGAGTCTCTTGGCTATCATCTAACTTCGGTGCGTTATTTGTCTCTTAAAAGTTGCAACCTTAGAGGGCCTATTCCTGAATCTCTTTCAAATCTCACCCGCATTTGAGTTATTTTGGGCCTTCGCAATAACTCCCTGAATGGAACAATACCACCAGGGATGTTATCCCTTCCATCACTAAATGCAATAGACTTGAGTTTTAATCACTTATCTGGTCCCTTTGAAGATTTCAAGTCCAATTCACTAATCTGGATATATTTAGAGGGCAACCAGCTGCAGGGTCATCTTCCCCACTCAATTCAAAACCTTGTGAACCTAACAGAACTTTCCCTTTCTTTCAATAATTTTAGTGGTCGTGTGGATGTCAGCTTCTTTTCAAACTTCAAACAGCTTTCATATCTTAGTCTTTCATATAATAATATCTCATTGACCAATGACAACAACGTTACTTTGTCCGAATCTTTTATTGAGTTAGAATTGGCCGCATGTGAAGTGAAAGAGTTGGAGTTTCTAAGATCCGCAAAGCAGCTTTGGAAGTTGGATCTTTCAAATAATAAGATTCAAGGAAGAATTCCTGATTGGGCATGGTCTAACTGGATGTTTTCATTGCAAGATCTTAATATATCCCACAATATGCTGACGAGTGTGGATTCAATTCCTCTTCAGTCTGTAAATACTATTGATTTGCGGTCCAATTTACTTCAAGGTTCACTACCCATTCCATCAAATTCCACAAGATTCATCTTCATATCCCATAATAATCTCACTGGAGAAATTCCTTCATCTATTTGCAACTTGACATCACTGATAATGCTGGATTTGGCGACAAACAATTTGAGTGGAGAAATTCCGCAGTGTTTCGGAAATATCACTTCCCTCCAGGTTTTGGATATGCGCCACAACAATCTTTCTGGGAATATTCCAACAACTTTCAGCAATGGAAGTTCACTGAGCAGCCTCAACTTGCATGGCAATAAACTAGAGGGGAAAATCCCACAATCCTTGACCAATTGCAAACAACTTGAAGTTCTTGATTTAGGAGACAATCACCTCAATGACACATTCCCCGTGTGGTTGGGGACTCTGCCAAAGCTGAAAGTTCTAAGCTTGCGATTCAACAAATTGCATGGTCCCATCAGAACATCAAGGATTGAGAACATGTTTCCAGAGCTTCGAATCATAGATCTCTCTTACAATGCCTTCTCAGGAAACTTACCTTCGAATTTGTTTCAACATCTGAAAGCCATGAGGACAACTGATCCATCAATGGAAGCATCAAAATATGGTGGAGATTCATATTACCAAGACTCGATAGCTGTCGTATCCAAGGGATATGAGCGTGAAGTTGTGAGAATCTTGTTTTTGTACATCACTATCGATCTTTCAAACAACACATTCGAAGGACATATTCCAACTATTATGGGAGATCTCATTGCACTCCGTGTGCTGAATATATCTCATAATAGATTGGAAGGTCATATACCGCCATCACTTGGAAGATTATCTTTGGTTGAATCATTGGACCTGTCAAGTAACCATCTTGTAGGAGAGATACCAGCACAATTTGCTTCTCTTACATCTCTTGAAGTCTTCAATCTTTCTTACAATCATCTCGAAAGGGTGCATTCCTCAAGGAAATCAATTTCATACCTTTGAGAACAATTCATATGAAGGCAATGATGGATTACGTGGTTTTCCTGTTTCAAAAAGTTGTGGGACAACTTATGCCATATCTGGACAGCTAGACGGTGAAGAAAGCAATTCTGAATTTCAGAGTGATTTTCGGAAAGCTGCTCTTATGGGGTATGGAAGTGGACTATGTATTGGAATATCCATAGTATATTTCATGATGTCAAGTGGAAAACCGGTATGGTTTGCAAGAATCATTGTAAAGCTGGAGCACAAAATAATGATGAGAAGGAGAAAGAAGCAGCGAAGGCAAAGGAATTGCAGAAGAAGAAATAATCACTAATAATATTACTGTTATTTCTTTAAGGAAAAGATGTAATAAGTTGTAGACTTACTATCGTTAAATCCTGTTTGCTTAAAGTGTGTAGCATCATTAGGGTGAGATAATCTTTAATGATAAGTTATATGCAATATAAAATCTTATGACC

>PGSC0003DMG400005542 LRR-RLPs

ATGGGGTATGCAGTTATTATGCTCTTAATTGTGTTTTTCTGTCATGAAGTTTCTTCAATATCCATATTACCTCACAAACATCAAACCATTTCACTTCTAAAATTCAAGAACACCTTTACTATAGATCCGTCAGCCTCTGGCGTTTTTCACAGTAGAATCTTATTATTGTCCGAAGCCTTATCCAAAAACGAGTTCATGGAACACGAGTAGAGATTGCTGCTCATGGGATGGAGTCGTATGTGATGACATCACTGGGCATGTCATTGAACTTGACCTCAGTTGCAGTAATCTCACTGGAACCATCGATTCCAATAGCAGCCTATTCCAACTCTCGCATCTCCAAAGGCTTGATCTTTCTTGGAATAACTTCTCCAATTCTCATATCTCCCCTGAATTCAGCAAGTTTTCGAGCTTGATGCATCTTGATCTTTCTGGGTCCTATTTCTCAGGTCAAATTCCTTCTGAAATCTCTCATCTTTCCAAGTTACACTCTCTTCGTATCTCCACATTTTATTACTTTTACAGTAACAACCTTTACAATAAAAAGCTAAGACTCACAACCCATGGTTTTAAATCGCTTCTTCAAAACTTGACCCAATTAAGAGAGCTTGATCTTAGTGGCATAAACATCTCTTCCACCATTCCTCTTAATTTTTCTTCTCATTTCACGACTCTGAGCCTGTCAGAAACAGGATTGTAAGGGATAATACCTGAGAGTATATTTCACCTGCCCAACTTGGAAATACTTGACTTAGGGATCAATCATGAGCTCAATGGTAATTTTCCGAAGACCAAATGGAACAGCAGTGCATCTCTCATTGAGTTAGATCTCCGTGGCGTGAATTTTTCTGATAATTTGCCCGAGTCTCTTGGCTATCATCTAACTTCGCTGCTTTATTTGTCTCTTGAAAGTTGCAACCTTAGAGGGCCTATTCCTGAATCTCTTTCAAATCTCACCCGCATTGAGTATTTGCACCTTAATCGTAACTCCCTGAATGGAACAATACCACCAGGGATGTTATCCTTTCCATCACTACATAGAATAGACTTGAGTAATAATCACTTATCTGGTCCCTTTGAAGATTTCAAGTCCAATTTGCTTCAAGGTTCACTACCCATTCTATCAAATTCCACAGTACTCTTCTTCATATCCCATAATAATCTCACTGGAGAAATTCCTTCATCTATTTGCAACTTAACATCACTGATAATGCTGGATTTGGCGACAAACAATTTGCGTGGAGAAATTCCGCAGTGTTTCGGAAATATCACTTCCCTCCAGGTTTTGGATATGCGCCACAACAATCTTTCTGGGAATATTCCAACAACTTTCAGTAATGGAAGTTCACTGAGGAGCCTCTACTTGCATGGCAATAAACTAGAGGGGAAAATCCCACAATCCTTGACCAATTGCAAACAACTTGAAGTTCTTGATTTAGGAGACAATCACCTCAACGACACATTCCCCGTGTGGTTGGGGACTCTGCCAAAGCTGAAAGTTCTAAGCTTGCGATTCAACAAATTGCATGGTCCCATCAGAACATCAAGGATTGAGAACATATTTTCAGAGCTCCGAATCATTGATCTCTCTTACAATGCCTTCTCAGGAAACTTACCTTCGAGTTTGTTTCAACATCTGAAAGCCATGAGGACAGCTGATCCATCAAAGGAACCATCAAAATATGGTGGAGATTCATATTACCAAGACTCGATAGCTTTCGTATTCAAGGGATATGAGCGTGAAGTTGTGAGAATCTTGTTTTTGTATATCACTATCGATCTTTCAAACAACAAATTCGAAGGACATATTCCAACTATTATGGGAGATCTCATTGCACTCCGTGTGCTGAATATATCTCATAATGGATTGGAAGGTCCTATACCGTCATCACTTGGAAGTTTATCTTTGGTTGAATCACTGGACCTGTCAAGTAACCATCTTGTAGGAAAGATACCAGCACAATTTGCTTCTCTTACATCTCTTGAAGTCTTCAATCTTTCCTACAATCATCTCGAAGGGTGCATTCCTCAGGGAAATCAACTTCACACCTTTGAGAACAATTCATATGAAGGCAATGATGGATTACGTGGATTTCCTGTTACAAAAAGTTGTGGGACAACTTATGCCGTATCTGGACAGCAAGACGGTGAAGAAAGCAATTCTGAATTTCTGAGTGATTTTCGGAAAGCTGCTCTTATGGGGTATGGAAGTGGACTATGCATTGGAATATCCATAGTATATTTCATGATGTCAACTGGAAAAACCGGTATGGTTTGCAAGAATCATTGTAAAGCTGGAGCACAAAATAATGATGAGAAGGAGAAAGAAGCAGCGAAGGCAAAGGAATTGCAGAAGAAGAAATAA

>PGSC0003DMG400005590 LRR-RLPs

ATTGATTTGAGCAATTTGGAGTATCTGGATCTCTCGTATAACAATATAGAGAGCTTTGTGACTACCAAAGGTACACGAATCAATGCAGTTGCTCTAATGTTGAATGTTTTATTTTAACCTTAACACATCCTACAGAATAAAATACATCTCTACGGACTTTAGATTGAAAATTGTGGGATACAAATTAGATTTTAATAAGATCCAACTATTCATTCTTATTTAAAAATGTATATGATTTATTTATTCACCATTTAACAGGTAACAAAAGAATAAGTTCTCTGCGGAATTTACAATTGGGAAGTTCCTACTCCAACTCAAGCAGAGTCATAGAATCCCTAAAATCATTCTCTTCTCTTAAATCACTTTCTTATGAATTTAGTGATCTCACTTCTCCAACCATAATTTACGGTAAGCTCAAATTCATATATACCTTAGCTTACTTCTATCGCCACTCTTTTCTTTTGATATGTACATTTAGTTTTCTTTTCATTTTTTGGACTCTATGATATAAGCATTAAGAAATCTGAGCACGGTGGAGCACCTATATTTGGAGGGATGTTCTTTAAATGACAACTTTCTCCCAAACATTGGACAAATGCCTTCTCTCTTGGTGTTCAGTATAGCATTTGGTGACAATGATGGCACCCTCCCTAATCAAGGTACCCTATTTTTCTCTTTTTTCCCTAATGCCCGCATTTTCATGTTGGAGGTCTGATTTTAGCATGTTCAGTTGATCAAACATTATAATAATTTTTGTTTATTACCAAACACAAGAAAATAAGTAACACAAGCCACCTAATTTTCCTAGAAAATATTTTCCTTCGTATCGAACACATCCATAGAGACATAAAAAAAATATAATAAGATGTCACTTGTTATAGAAATTTGTTTTAGCTAAATGTTTCTTAATATATATATATAGGTTGGTGTGAACTCAAACATATTCAAGAGCTGGATTTTTCAGCCAATAGTTTTCGGGGAACACTGCCTTCATGTTTTGGAAACTTGACATCACTTCGACGATTGAGTTTTGGGGGGAATTACTTCAGAGGAAATATAGCGTCAGATTCTATTTGGAGGAGACTCACATCACTTGAGTACATCGATATTTCATATAACCAATTTGATGTTCCTCTGTCATTCAGCCAATTTTCCAACCATACAAAATTGATCTACTTGGATGTTAGTTCTAATACAATAATTACAGATACCAAATTCCAAAATCAGATCCCAAATTTCCAGTTGGAGTTTTTTGCTATCAAAGGATGTAAACACCTTAAAAAATTGCCTTATTTCCTTCACTACCAATATGACTTGAGGATTCTTGCTATTGGAGGAAATCAATTGCCCGGGATCTTTCCAACATGGTTGTTAGAAAATAACACCAGACTTGCAGGGATTTATAGTAGAGATAATGCTTTCAGCGGATCATTCAAGTTGCCATCGAGTTGTCACCTCCATCTAGAGACAATTGATGTTTCTAATAACAAATTAAATGGGCATATTCCAGATAAAATGAATTCAGCATTCCCAAAGCTTACTTATTTGAACATGTCACAAAATTATCTTGAAGGTCCTATACCTTCCAAGATAAGTAGCATTCATTTACATAATCTAGACCTATCTCACAATTTCTTATTTGGAGGAGTTCCTAGTGATCTGGCAATTGGTTCTCCAGTATTAGCTTACCTTCGATTGTCACACAACAAGTTAAAAGGTCAAATTTTTCCAAAGGATGTCAAACCAAAGATGTTATTCTTTTTGTTCTTGAATGGAAATAACTTTGAGGGACCACTACCTTGCAACATGTTTATCACATCCCTTATTGTATTGGACGCTAGCAACAATAATTTCTCTGGAGAGATTCCGAGATGCATTAGGGATAATACAAGATTGTTACAACTTGATTTATCCAAGAATCATCTCAAAGGTTCAATTCCAGTTGAGATTTGCAATTTAAAGCTCATTCAGGTCTTAGCTATATCAGAAAACAGGCTTTCAGGCTTTATTCCTTCTTGTGTGAATTCTTTACCTCTTGAACACATCCATCTACAGAAAAATCAATTGGGTGGTGAATTGGAACATGCACTTTTCAACTTATCTTCTCTGATAACTTTGGATCTTGGGTACAACAATTTTACAGGAAGTATCCCACACACCATAGGCTCACTTAATACTCTCAACTACCTTCTCCTTAGCAATAATCAATTCGAAGGAGAGATTCCAACTGAGATTTGTATGCTGAACAAGCTATCTATCGTGGATTTATCTTTCAATAAGCTTTATGGTTCACTCCTTCCTTGTTTGAGCTACTTAACACAAGCCAAAAAGGATACAGAGATAAGTAAGACCAATTATCCTACGACTTCTAGGCGTTCATGGATAACTTTTTTGGCCTGGATGCGTTTGAAAAGGCACTACCACGATAGTCATGGATTTCTAAGTGACATATTTCTGATGGATGTGGAAACTCAGGTCCAGTTTTCAACAAAAAGAAACTCATACACTTATAAGGGAAGCATTCTAAAATATATGTCAGGTATGGATCTCTCAAGTAACAGATTAACTGGTGAAATTCCCGTTGAGCTAGGGAAGATGAGTAATATACCTGCATTGAATCTATCGCACAACCATCTCATCGGAAGAATACCAAATACCTTCTCCAATCTACAGAAAATTGAGAGTTTAGATCTTTCCTACAACAACTTGAATGGGAGCATTCCTGTTGGTCTACTTGAGCTAAATTCTTTGGCAGTATTCAGTGTTGCATACAACAACTTATCGGGTGCAGTGCCTGATTTTAAAGCTCAATTTGGAACTTTCAACAAAAGCAGCTACGAGGGAAATCCTTTTCTTTGCGGTTATCCATTAGACAACACGTGTGGCATTCCTAAATTGTCATATACCTCCAAGATTAATGGAGGTGAGGAATCATCAGACTTGGAGGATATTCAGTGTTTTTACATTGGCTTTGTTGTGTCTTATGGAGCGATCTTGTTGGGATTAGCTGCGGCACTCTGCTTCAATCGTTATTGGAGAAAAGCATGGTTTAGGATGATAGAGGCATTGATGTTTTATGGTTACTATTTTCTCTTGGACAACGTTGTTACACCCATCAAGAGTATATGGTACAGGAATGTTGGTTAATTCTAGGTAGTTGTTTTACAATATTGTTGTATGTGTATCACCATTATTAATTATATTGTCATTCTGTTTTATCAATATTCGACAAATAGGATCTGTTTGGAGTAGTTACAACTTCATAATTTAATACCCCCTGCATAGCTTCATTAGTGTTGCTATTTATGTAGCCG

>PGSC0003DMG400005600 LRR-RLPs

CCCTGAGACTTAGTTACCATCACCTTCCATTTGATTTGAGACAATGCTTTGCATATTGTGCAGTATTCCCAAAGGATACCGAAATGGAAAAGCAAAATCTAATCTCTCTCTGGATGGCACACGGTTTTCTTTTATCGAAAGGAAACTTGGAGCTAGAGGATGTAGGTAATGAAGTATGGAATGAATTATACTTGAGGTCTTTCTTCCAAGAGATTGAAGTTAAATCTGATCAAACTTATTTCAAGATGCATGATCTCATTCATGATTTGGCAACATCTCTATTTTCGGCAAGCACATCAAGCAGCAATATCCGTGAAATAAATGTAAAAGAATACCCACATATGATGTCGATTGGTTTCGCAAAAGTGGTGTCTTCGTACTCTCCTTCTCACTTGCAAAAGTTTGTCTCGTTGAGGGTGCTTAATCTAAATAATTTAAGACTTAAGCATTTACCGTCTTCCATTGGAGATCTAGTACATTTAAGATACCTAAACTTGTCTGGCAATCGTAGAATTCGTAGTCTTCCAAAGCAGTTATGCAAGCTTCAAAATCTGCAGACTCTTGATCTACATAATTGCTACTCTCTTTCTTGTTTGCCAAAAGAAACAAGTAAACTTGGTAGTCTCCGAAATCTTTTACTTCATGGTTGCTATGAATTGAATTCTATGCCACCAAGGATAGCATCTTTGACATGCCTTAAGACTCTTGGTTGCTTTGCAGTGGGAAGGAAGAAAAGTTCTCAAATTGGTGAATTACGAAATCTGAATCTCTATGGCTCAATTGAAATCACGCATCTTGAGAGAGTGAAGAATGATATGGATGCAAAAGAAGCCAATTTATCTGCAAAAGAAAATCTGCATTCTTTAAGCATGAGATGGGATGACGTTGAACGTGCACATAGATATGAATCAGAAGAAGTTGAAGTGATTGAAGCCCTCAAACCACACTCCAATCTGACTTGTTTAACAATCAATGGCTTTAGAGGAATCCGTCTCCCAGACTGGATGAATCACTCAGTTTTGAAAAATGTTGTCTCTATTGAAATCAGGGGTTGCCAAAACTGCTCATGCTTACCACCCTTTGGTGAGCTGCCTTGTCTAGAAAGTCTAGAATTATGGAGTGTGTCTGCGGAATTGGAGTATGTTGATTCTGGATTCCCTACAAGAAGAAGGTTTCCATCTCTGAGAAAACTTATTATAGGTTATTTTGATAATCTGAAAGGATTGCTGAAAAAGGAAGGAGAAGAGCAATTCCCTGTGCTTGAAGAGATGGAGATTTACAGGTGCCCTATGTTTGTTATTCCACCCTTTCTTCTATCAAGAAATTGGTAGCTTATGGGGAGAAGTCAGATGCAATAGGTTTCAGTTCGCTATCTAATCTCAGGACTCTTACTTCCCTCCGCATTGGCCATAACGAAGAAGATACTTCGCTCAAAGAAGACATGTTCAAAGGCCTTGCAAATCTCAAGTACTTGAAAATCTCTTTCTTTAAGAATCTCAAAGAGCTGCCTACCAGCCTGGCTAGTCTCAATGCTTTGAAGCATCTGGATATTTATCGGTGTGACGCACTAGAGAGTCTCCCCGAGGAAGGGGTGAAAGGTTTAACTTCACTCACACAGTTATCCATAACATACTGTCAGATGCTACAATGTTTACCGGAGGGATTGCAGCACCTAACAGCCCTCACAAAATTATTAGTTACGTATTGTCCAACACTGGCCAAGCGATGTGAGAAGGGAATAGGAGAAGACTGGTACAAA

>PGSC0003DMG400005972 LRR-RLPs

AAAAATTGTATTGTTATTGAAACCAAATACTACATTAGACATGACAACTTCTCATTGTACATTCATTTACATTCTTCTACTTCTTTTTTCACTTCCATTCAAAGTAATCACATCATCAGCAAAAACTGAAGCTGAGGCTCTTATCAAATGGAAGAGTACTAACTTATCTTCTACTTCCTTATTGGATTCATGGTCCATTTCCAATCTCGAAAACTTGTGTAATTGGACATTCATTGCCTGCAATGCTGATGGAACAATTTCGGAGATCAATCTTTCTAATGCAACACTCTCAGGCTCTCTTGATCACCTTGACTTCACTTCATTTCCGAATCTCACGAGTTTCATTCTCAATGGCAACAACTTTAGTGGATCGATACCATCGAATATTGGCAATGCCTCTAGTCTTGAAGTTGTTGTACTTCTCAACAATTCATTTCAAGGGAAGATTCCATCTTCAATAGGCAAACTTAGAAATCTTCAAACACTTGATCTTCGAAAGAATCGTTTGAACTCAACCATTCCTTCTGAGTTAGGCCTTTGTACAAACCTCATTATCTTGGCTCTAGCAGAGAATGCTCTGCAAGGAACATTGCCTATATCTTTGTCCTCTTTGATCAAGTTATCCCAGTTGGCGTTGTCCGATAATTTTCTTTCTGGTGAGGTCTCATCAGATCTCATTTTACCAATTGGACTGAGTTGATAGCTTTGCAGCTTCAAAACATTTACATTACTGATTTAGGAGATAATCACTTCAACGACACATTCCTAGTGTGGTTGGGAAATCTGTCAGAGCTGAAAGTTTTAAGCTTGAGATCAAACAAATTCCATGGACTCATTAGAACGTCAAGGATTGAGAACATGTTTCCCGAGCTTCGAATCATAGATCTATCTTATAATGCCTTCTCGGGAAACTTACCTACGAGTATGTTTCAACATCTAAAAGCCATGATGAGAGTTGATCAAACAATGAAGGCACCAATATATCTTGGAGTGTACAGTTATTGCCATGATTCTATGACAGTTGCAACCAAGGGACGAGAGCTTGAACTTGTTAGAATTTTGACAACTTACACCACCATTGATCTTTCAAGTAACAGATTTGTAGGGAATATTCCAAGTATTATGGGAGAACTCATTGCACTTCGGGTGTTGAATTTGTCTCATAATCAATTGCAAGATCATATACCACCATCACTTGGAAATTTATATGTACTTGAATCATTGGACCTTTCGTTTAACCATCTTTCAGGAGAGATACCACAACAACTTGCTTCACTTACGTCTCTTGAATTCTTAAATCTCTCCTATAATTATCTCCAAGTATGCATTCCTCAAGGACCACAATTCTCTACATTCCAGAACAACTCATATGAAGGTAATGTTGGATTACATGGATTCCCAATTTCAGATGGTTGTGGCAATGATCGTGTATCTGAAACAAATTACACCGTATCTTCTCTAGAAGATCAAGAAAGCAACTCTGAATTTCCCAATGACTTTTGGAAAGCTGCTCTGATGGGCTATGGCAGTGGACTATGTATCGGAATGTCCATAATATATTTCATGATCTCCACGGGAAATCCAAGATGGCTAGCAAGAATCATCGAAAAACTGGAACACAAATTTATCATGCAAATGAGAAAGAAGCAACAAGGTCAAAGGAATTACAGAATAAGAAATAATCGTTTCTAGACAAGTTATAAGTTTTGCTATTCAATCAAGCTCTACATTGATCTTTAGTTTATTGTAATTTGTAACTATTATGATATCTCATTGAGTGCTTCAGCTTCATGTGATATAATAATACTGTACTATTTCCAGTGTGACTAACTGCAACATGTACTTTAATTTCCTGCCTTTCTTACATTTAGACAGGAAACACAAAGGACCAAATAATAACTGGACAATATATGGGCAGTAGGCAAAAGAGAAAGAAACAAACCATAAAATAGGTTACATAATGAGAAACTAAATACACATTACATATGAGGTATGGTCAAAGAATCTGCCAAATTCATTCCTTGACATGAACCTTCCTCGGGTTCAGGGGACTCGTATACGCACTCCCTACAATATAAACAAGGTACTTGATTACAATTAAGGGTAGAAAACAGAGCAAAGAACACAATTTCAGATAAGTTTGGTCTCACTTGTTAACTAGCATCTATGGATACGAGTGATGAGTGAGACCATGACAAGCGTATTTGGCGCCAAGAATATGGAGCTCGCTACCCAAAGAATGGTCAAAACTAGAGTACAGGCCCTAGCTAGTATCATCATAACGGCTTAAAAATTGGATTATACAAGGATATACATGTTCTGCGGCCTGCAGGCACCAGGAAGAGTATAAGTTGTTATGCAATAATAGAAACGGAAAAATAATAACAATGCACATGCAAAAGATGCCTATTTCACGGAGTTGTGAAAGTAAATAACGTATCATATTTCCTAATCTAGGTATATTTAACTCTCAGGTTTACATAAGGAAACAAAAAAAACATTGAAAGGCAATTCCAATGTGACTAGGCACCTCTCCACTTCCAGCATCTTTCTATTCTTTATCATCCTTTTCTATACTCTGTCTGCTAATCAATCAATCAACTAGACCTCAAATCCTCAATCCTATATCGGTAGGGTTTGGTTTTATAAATTCTCCATATCCAGTCCGGACTATTCAGGTTAATCTCATTCTAATACTAAATAATGTCTTTCAAAGCAAGCAAATTAAAATTCCTCTAAAACTCACATTTATTCCCAAATGAATAACTAGTCTAAATCCAATCAGTTAATGTCACTCATATAGCTCCTTTGTTTCCATTGTATTCTGTTCTTGTATTCCAAAAGAAGTTAAAAATTGTTAAAGTTTACATACCATGCGTCCTCCCACCAAGTCATAATGAGCATAATGCTGACCATTATCATCCCCCAATACTTTGTAGGTGACCAAGTGCTCGGGGATAAGCTTTGCCGTTTCTGCTTGAACCAATGTGTTGCAAGAAAAGAATTGTTAGAAAACTTATCATCCTCTGACAGAAAAATGAGCGGAAGTACTATGAAACATACCGCACACTGCTTCTGGTGGGCAGATTATGTCTTTATCTCCAGCAACAGCCAACACGGGTACATTGGTTTTGTGAAGATTATCCTTGTAGAATATTTTACCACTTCTGTCACGGAGTCCCCCTTCTCGAAAAGCTGTAGTCAACTGCAAAATAAGTTTAGCAGGAATAGTACCTGCCAGAAGATCTATTAGAAAATCTAAGTAAGAGGTACATGTGAAATCTCAAGAGAACATAGAAAGGGCGTGATCCCCAAAGGAATTTTTTAACAAAAGAAAGTGAACTTTCAGGATCACCGTCAGACTACAGCCCTTATAAATCAGTTATGATAAACTATCCTTACTTTGATGGAAACACTCTTCCATTGAAGATCATCTATCAACACCGTTCCACAGTGTCAAGAAAACAACACAGGAGTGAAACTTGTGGGAGCAAAGAGTCAATAGTTCATCAAACAAAGAGTCAAATAAATATATTGGTATATGCAGCTCTTTGTCAGAAGAGAAAATATCTAGCTGGCAAATAAATATACAGCGGATGTTTACTTGATATAATTTTCTTTTTCAATCTGAAGATGTTAATAGTTCTTCGTCGCTCATTCTTTATGCGAACAAATAAGAATTAAGAATCAACTCTCAGTCTCTCACATAGTTTGTTTCCACTGATTACCTTATTGCTCAGAAGCCTATAAGGAAAGACCTTTTTAGTTTTTAGTACCGACTCCTCTTGAGGCCAAGAAGGAGACCAACTAGACGAGACAGACCATTGTTCATACTTCATAGGTTGGTTATAAGCAAGTACAAATCAAAATTCAATTTGGCTTGCCAAAGGACATACTTCAAACTTTGGCTGTTCTTCCTTGTATAACGCTGCATGACTACTGTGAAAGATGAGATATACTAATAATCATAATTAATAAGTACATTTGCCTTCCTAACTGATACCCATGAGGATGTATCTGAAATGTAGAATATTAGTTCAACTCAGACTCAACATATCATCCAGACAGAAAGTGGAGGGGAAAAGAATGTATTGGTGCAAAAACGCTGTAGACAGGAATGTGGGGCATAGCCAGATGGTCATTGAATCATTCTTCAGAACATGTAAGTTTCTCATACTGTTATCTTTAAAACTCATACTAAATATCTATGCCTATACAACTTGAAACCTGCCTGCTCTCCATATAAATAAATTATAAAACAAGAGGACCATAAATTTTTACAGTTCACAATCAATTAATACTCCCTCTGTCCCAATTTATGTGACTTATTTTCCTTTTAAGTCAGTCCCAAAAAGAATGACACATTTCTATATTAAGTAACAATTTAACTATAAAATGTCTATTTTACCCTTAATGAAAATGATTTACAGCCACACAAATTCCTATGATTCATTTTGGACCACAAGTTTTAAAAGTTTTCCTTTCTTTCTTAAACTCCGTGCCGAGTCAAACTACCTCACATAAAATGGGACGGAGGGAGTAAGAAATACTGCTTATTGCCCATTTAAGCACTTACAGAAGTTATTTAATACGAGCTTTTTCAGCAACTCTGGATGCATCATGTCTGCAGCTGATATTAAATCATTAAGCCAAGACAACACATAAGGAGGGCGAGAAGAAAGAGGATAAGCTGCTGCAAGTAATGCTCCTAAAGGGACAACAGGTACATTGAGAGCTTGGGCAGGATCAGCCTGCAGCCAAGGTAAATAAATATAAGAATATGCTGAAAGGGCTTTTTGCTGCACAACGAGAGTTGAAAGACATTTATTTTTAGCTTACAAGGGGTAGGAGCAGTCTGAGTGCAGACTTCGAAGATGTGTAATCCAGTGATGAAGCCAAAGTTACAATAGCAGTTAATCCAGGCTCTCTTCCTTGCAATCCTTTCCAAGATAAAACCAGAAACATATTAAAATTATGTCATATCTGTATGATTCCCGGATGTAGTCAGTCTGAAAATATTTTTTTGAGACTGTGCCCACTGCACATTAACAAGGAATCAGACAATGCTCCAAGTGACTTATCTGTTAGTTGCGAAACCTTTTTACATCTAAAACCAGGTCAAATTCCTTCCAAATCAAATTGTCCCTTACACACTGTTAAAACGTATGGATAAAGTATAAAATCTTGCCTAGCTCCCTATAATCTCAATAAGGTCAAAAGATGGCTCATTTTCAGCTCTGTATGAAAGACGCTAGTAAGCAGATAAGTCCCACTGTTAACCAGATCCTGTTTACCTTGGTGCACCCTATCATGGAATTTTTAAGTTAATAGGAGTACTGCACTTGGTGGCTATTACAACTATTCTCCCAAAGTTGTTGTGAACTATAGCATCTTTCTTATTTTTCAGCAAAAAGCTTGTTGAAGTTGAGTCTGAAGATTTTGCTCCATGAATATTAGGAATTAAAAGAACAATAAGTCAAGAGGAGACCAGGTTAACTCAAAGTTGCACCAAGTGAAGTGGTAGCCTTTGGAATTTAATCCCAAGACTAAATCGTATGATACTCCAGCAGGTTCATGGAATAAATTTCTGGAATTATACTGCCTTCTTTATTTAAATAAAATGTTTAAATATTACTGTTTTCTTATCAACTGCAACTACAGAAAGGATTCTTTGTTACTTTTTCAAAACACTTGCACATTAATTTATTAGTCTCTTAAGTGGCTCATAAGTAGACCCGCCAGATCACGTAAGTGTAACAACATTAATTCCATATAAACAACGAATGAATCTTACCACACTGTGATAGCCAAGCATACAGTAATATTCCTCCCATAGAATGTCCAATGGCAAGAAGTTTACCATCTTTTGGTGCAGTTTGGGCCTTAATGTATTCCATCTAGAATGAAAATAAAATAATTAGACTAGTCAGAATAATAAATAAGAGTGAAAAATCAACAAATTCTGAATTTCACTTGGCTGCGTAATCTCAAGGACAATAATATCTTTATATTCTACAACATGGGAATCTGTCACCTCACCGCAGCAGGAACATCCTCTTCCAGGTAATGATCAAAGTCCCAATCATACTTGACAATCAAGTCAAGCTGTTTCTGGAAATCTTCTATGGTAGTGGTTAGACGTTCATGAAGGTCAACTAACGGAGAAACAGAACGTTGGCCTTCTTCAATGATATTTACTAGCTTTTGGCTAAGATCCTTTACTTGGTCAACAACGGCAGAGTTTTGCCCTGTCTCCAGCAGACTTAACAACTTCCCCCTAGTCTCGTTAAAGCGTTCATAAAGAAAGGAATCCTCTAATAATTTTGATATTTGATCAAATAGTTTAGCAGACATAATCCTTGATTGGCTTTCATTGAGGAAGCCAGAGACTCTTTCTGACAAGCGCATAAATGTTTCCGTAAGCTCGTTTACTACTCTTGACTCATCCCACATTGTTGCAATAGCTGTGGATTCTTCTTTGACCACAGCTGTGTCAGACTCTTCTAAGGTACTCTGGGGATCTGTTGACTGCTGTGCTGCAGAAAGAACTTTGTCAGTAGCATTATCAGCGGCAGCTTCCATCTGTTCTGAGACTGCAATTGCAGTCTTCTCAATGTTTGTGGAATCTGACTCCTGCACACTCAACCCAGCTCCACGAAGTTCAAGAACCCAAGTGTCAAATCCTTCACCAGACATGTACCGGGCGAAAGATGACTGCATAATTATTTTTAAAAGTAAAGACCGTGATTAAATAACTTTTAGAAAGAATTTAAAAAAACCAGCCATACTATGTAAGATTTCTTTGATAAGCAGGATGGACATGGTTGCCTGAAGTGATAAAGAATTCTAAACCTCCACTGAAGAGTAGGGTGCACATACTTGCCTCAAGTGATAAAAATGAACTTAAAAGGATAAGAAGATGCATCAACAGAACTTTAAACAAAAAATAGACAAGAAATAATCCCAGTGAAATCAAGTTGACACTAGTTATTGCATTAACCAGCAGACATTTAGATTTGTCTCACTAAAAAAGAAAAGATAGAAATCATCACTAGCAACTAAGCAAAAAATCGCCATCACAATTCATATTTCAGAAATGTACAATCACAGAGAGTGCTAGCCACATGACTACCATCAACACCTGATTGATGCAGCCATAAAGCCATATAAAAAAAAGGAAGATGAAATAGATAACAAGCCAGTAATAAAAAAATCACACAAGTAAACTTTTATACGCTTGAAATGCATAAAGTATGAGAAAAGGACGAAGTTTTTAACCGACCCACCATTTTGATCACCGCACCACCATGTGTGCAGAAGGAAGTAGATGCAGATGCACTAGGAACTAGTTACCCTAGGACCGTAAACAATCCATTTTGTATTATCTAGGGTCGGGATGTGACCCAAACTACTTACAATGACCCACCCGTTGACACAGGTGGGGAAACTACTAATGTTCTTGATCCCTAATTATATAATATAATATAAGTATCCTTTTGATCAAAAAAAAAAAGTAGTAAGGATAGTACAACATCAAAACCAGGGTTTACTCTAAAAAATTTGAGCATGCCATAGAATTCTACCAGATTCAGAAACAAGAACCACTCCACGAAATAAAGAGTGAAACTTTAGTGGAACCTGACATATCCACTAAGCACTGATAATATACACATTTGACATCTGTCATTATGATCTATTAAATGTGGCTATCAATGAACGATCCATGAAGATCCCCTAGTTCTGTTCACCTATACAGGCATAAATCTGCAAATTAGAGTGTCCCGGCTACTCTTTCACGCAATACTGGACATACAACAAAACAGAAGAATGCACAAAGAATGGAGAGAAGGCAGACGAGTATCATACCCCAGGAGCAAGATCATATCCAATAGCATTAGTCCCCACACCAGACAACAGAAGCAGGGGATGATTTCTCTTTGGAGCCTATCATCATCAAAAATCACACACCCTTAATGCAAAGCTCAAATCCTTGAACAAAAAAATCATGTAACAGAACTGTTAAACCTTCCTAGTATCACTCCCTGTGTTTCATTTTATATAACAGTATTTGATAGAACACAAAATTTAAAATATTCAATAGTTGAATTAGATAAAATGGCGTCAAATATTCTTAATGCCTTCATGTCTTATATATTGGGACCAATTCTATATGACCACATTTACTTCAAGCCGAGTATCTATCAGAAACGGTCTCTCTACCTTCACAAGGTACGAATAAGGTTGCATTCGCTCCACCCTCCCCGGACCTCACCTGTGAGATTACGGGTTGTATTCTCTATGACCACATTTAATTGATCGCAAAATTTAAAATTTGAACATATAGAAATGAAAGTGTCGAATATATTATTTTTTTCAAATTGTCATACAGTTAAGAGTAACAGTTCAAATTAAAAAACATAAAAAATTGAACCTGAGGAGGAGGAATGTAGCGCCAGAGAGCAAGTTTCCAATCGGAATTATTAACAGATACGTAGTGGAGTTCATCGGCAGTGCATAACGATGGCTTATTACAAACCTTCTGGACTGTAGTATCGCCATCATTGGTGGTGAAAGCTCTGAGTTTAAAGGCGGAGTTCACCGTAAAGCAGCAGTTCGGGGAGTTCCGGTATGAAATGACGGCGGGAGGAAGTCGCCGGCGGTGATTTGTGAGGTGGTGGTAATGGTTGTGGTGGCGGAAGATGGCGGCGATTTGAATCGCCGATCGCATATCGGATTGAATTGGATAAACCATATTTTTTGGGATTCGTGACAATTTGGATAAAGAAAATTAGTTTGATAAATCTTTTACTCTTTTGACTTTGGAAGGTTTGAGACGTAAACTGTTTTTTTTTTTTTAATAATTT

>PGSC0003DMG400006027 LRR-RLPs

TAGAGGTGTCTACTTGCTATTTAATTCAAATCAAATTCATTCATTCATTCAACAACACATTTTTACACTGTGAAAAAAAAAACTTCATTTTCTTCTTCATAGCTTGAAAATCTCTATCAAATCCAGCAATGCGGCCTCCTAGCCGCAGCGTATTTATCTCTTTTCTTGTTATCTCCCTCTTGTCGTACCAGATCTCGAATGTGGTAGGACAAGAAGGTGATAACGATATCGATTTAGATGATATTAAGGCTAATGCTAGTAAGTTAAGTTTCGAAAACCCAAAGATTCGCGATGCTTATATCGCCTTGCAATATTGGAAAACAGCAATGTTTTCTGATCCTTTTAACTTCACAGCTAATTGGACAGGACCTGATGTTTGTTCGTATGGTGGAGTGTTCTGTGCTCCTTCGTTAAACGATGATTCGATTAGAGTTGTAGCTGGCATTGATCTTAATCATGCTGACATTGCTGGTTCCTTAGTTGCAGAGCTTGGTCTTCTTACAGATCTGGTTCTTTTCCATCTCAACTCGAATCGTTTTTGTGGAGTTGTACCGAAAACTTTTAGCCATTTGAAGCTTCTTAGGGAGCTTGATTTGAGTAACAATAGGTTTGTTGGTGGGTTCCCTAAGGTAGTTCTTTCATTACCTTCTTTGAAGTTTCTGGATCTAAGGTTTAATGATTTTGAAGGACCAGTTCCTTCCGAATTGTTCGATAAAGATTTGGATGCGTTGTTTCTGAATGACAATAGATTCCGTTTTGGCATCCCAGAGAACTTGGGTAACTCCCCTGTTTCTGTTCTTGTGTTTGCTAATAACGATCTTGGTGGGTGCATTCCCGCGAGTATTGGGAAAATGGGGAAAACGTTGAATGAGTTGATATTGATGAATGATAACCTCACTGGGTGTTTGCCCATGGAGATTGGATTGTTGAAGAAGTTGACAGTGTTTGATGTTAGCTTCAACAAGATTCAAGGCTCATTGCCATTAACTGTGAGCAAAATGAGGAGTGTGGAGGAACTCAATGTGGCTCACAACAAGCTTACTGGGGTTATACCAGCTAGCATTTGCCAACTTCCAAGGTTGCAGAACTTTACGTATTCGTTCAATTACTTCACTGGGGAGGCTCCGGTTTGTGCTGCGACTAGATCTGGTGATGGTCAGGAGAACTGTATAGTTGGAAAGAAGAATCAAAGATCTGCTAAAGAATGCTCTTCGGATGATGCAAAACCGTATGATTGTAAGAAATCGAAGTGTTATAGCCCATATGCTACGTCTCCTTCGACGAAACCAAAGCCAACACCAACACCAAAACCAAGAGGACCACCACCACCTACTTGGAAGTCTTCGGGTTCGCATAACAAGCGATCTCCGCCACCACCTAAATCAACTCCTCTTCCTCCACCAGCACCTTACAAAAAGTCCCCAGCGTATCGACATAGAAGTCCACCACCACCTACTCACAAAATTTCACCTGTAACTCATCATTCACCTCCTCCACCCTCACCCGTATATTACCACCCATCTCCATCGTCCCCACCACCACCAGTTTATTATTCTCCACCACCCCCGGTGTATTACGAGCCACCTACTTACAAGCCTAAATCACCACCACCGCCAACACCGTCATATGAACACCCAAAAACCCCATCACCACCACCACCAACACCATCATACGAACATCCAAAAACCGCATCACCACCAACACCATCATACGAGCATCCTAAAACACCACCACCAACACCATCATACGAACATCCAAAAACACCATCACCACCACCACCAACACCATCATACGAGCATCCAAAAACACCACCACCAACACCATCATACGAGCATCCAAAACCCCAATCACCACCACCACCATACGAACATCCAAAAACCGCATCACCACCAACACCATCATACGAGCATCCAAAACCCCAATCACCACCACCACCAACACCGTCATACGAGCATCCAAAACCCCAATCACCACCACCACCAACACCATCATACGAACACCCAACCCCACCTACCCCACCTTGTAACGAACCACCACCTCCACCACCTAATTCACATTGGGAACCAAAACCATCCCCACCATACACCTACTCATCACCACCACCTCCATCACCATCTCCTCCTCCTCCCACTTACTACTACTCATCACCACCGCCACCACCACCATCACAATCACCTCCCCCACCCACTTACTACTACTCATCACCGCCGCCACCACCACCATTTTACGAAAATATCCCTCTCCCACCGGTAATCGGAGTTTCCTACGCATCTCCACCACCACCAGTCATTCCATATTACTGAACAGGGGTTATCAATTAGCCTTTGATTTTCCATTATCAAAAAAAAAAAAGAATTCATTTTTTTTTATGAATTATGTATACTTTGTTATTTATTTTTTGTAATATTTTTATTTTGAGTTTGATTATATTTATGAAAAATTGGGATTATGGGCGGGAGTTAGTGAGGGGTTTAACTATCAATATTTTAATATTGTTATAT

>PGSC0003DMG400006076 LRR-RLPs

AATTCCCCTTTCAAGCAAAAAACATGGCCGCCACTGCTGACATATTCTCGGAATGTGTGATTCACAAAATACTCTGTTATCTTAGCTATGGCCAAGCATCCCGGATGAGAATTCTATCTAAAACATGGCTACAAGCTTGGTCGACTCTTCCGAACTTGGAATTCTTAGTTGAATATTTCGAAGAAACAAAAATAGTGGACGCAATTATGGAGACATACAGGGAAAACAAAATCCCCATTGACAAGTTTGGATTTTCAAATTGTTCGAAATTCAATCGTCGTCATCAAGTTTTCCCTCTGATTGATAAGTGGCTTGACACAGCTCTTCAAAATGGCGCAAGAGATTTAGTCTATGTAGATCTTCTCTACTCGGTACGATTATATCCTTTTCCTATTTCTAAAGTCTTGGCTGCAAATTCTTTAAGAGAATTGGTTCTGAAAGGTTGTAACCTTATGCATGTTTCGTCATCTAGTGGGTTCGCGAATTGTCATTCTTTGAGAAAGCTTTCTCTATCTTATATAATTTTAAGCAAAAATTTGCTTCAGACTTTACTTAATAGTTGTCCTTTCATCGTCTCTTTCAACCTTGATCATTGTTCGGGGTTGGTAAAGATTGAGCTTATGAATCTTCAAAAGATCAAGTCAGTTTCCGTATCAATAAATAGAAACCAACATGTGAAAATCCAAGCACCAACTCTTGAACACTTGTTTTATAATGGTTATTTATCGGGAAATTTGGTTATCGTTGGATGTCAGAATCTAAAATCTTTAGAGCTATCATATGTGAAGATATGTAATGGATTTCTCCATAACCTTATTTCTAGATCGCAATCCCTAGAAGTATTGAAGATTCAGTATTGTATGGGCATAAGACAGATTGAAACTTCGAATTTGGTATCACTTGAGTATATAGGATATCAAATTCCTGAACTTAAAATTGCGAAAGCTTCAAGTCAATTGAAGCATTCAAAAATCGTTCTTCATGGCAACAACAATTTAGATGCTGCATGGTTTAGTAAGTTGAGGAAGCTTCTATGGAATTCGGCCTCTTGGTCCCAAGTTCTCCTTTGTTTTCCCGAATACAATGAGATCAAGATTAAAGATTTGCAGCAGCACAACGGAGTTGCTTGCCCCCAGATGGACGTTTTAAATGTCAACATTACAAGGTCATATGGGGAGTGTCCAACATTTGTGGATGCTTTGCTATGGAGTTGTAATCCTAGGAGACTCAACCTATCATCAACGGTTGAAATGATTACATGTTTCGTCAGTCGTTTGATGTATATGAAGAATCTGAGTCATTCTTCTTCGGAAAGAAGCAACCCCTGGCATAGTCAATTGAAAGAAATCAAAGTTTTTGATGTGAAAAATCAAGAGCTGCAACTCGTAAGTGGGGAACTGGCGATAAGCACCCTTTCGGAGGACAGGAAGGAAGTTTATTTTTTATTAGATTGGCAATGCAATTAATTTATCACTATTTCAAAACGAATGCACTTGTTCATTCCTAAACTTTATTAATCTGTTTTATGTCTAGCTTTAAATACTTGCACTTTCTCACCATTGTATAAAATTGTGTTGAAGTTGAAGCATAAGATTCTTT

>PGSC0003DMG400006095 LRR-RLPs

ATGGGGACAGCTGACATGTTGCCTGTATGCCTCATTCGCAAAATACTCTGTTATCTTAGTTATGAAGAAGCATCTCGGATTAGAATTCTCTCCAAAACATGGCTACTAGCCTGGTTGGATCATCCCAACTTGGAATTCACAGTCTCGTCGGGCAAAACCATAACAATAGTGGATCATGTCATGGAGAGATATAAGAACGGAAATATTCCTATTGACAAGTTTGAATTACACAACTCTACAAAGTCTGATCATATTTTTCCTCTGATTGATAGGTGGCTCGGTATTGCTCTTCAAAATGGTGTAAGACATCTAGAGTACATAGATGTTGAACTTTCAACATCATACCCCTTCCCTATTTTTAAATTCTTTGAAGCAAATTTTTTACGAGAACTGGTCCTTACGGGTTGTGATCTGATGCATGTTTCGTTATCTAGTGCTAGTCAAGCGGTGACGTGTCATTCTTTGAGAAATCTTTCTCTATCTCTTGTACGCTTAGACAATAACATGCTTCAGACTATACTCAGTAGTTGTCCGTTCATTGTTGATTTAATCATTGGGTATTGTAGTAGGCTGAAAAATATTGAGATACGGAATCTTCCAAATATCAGGTCTCTTGTCATTGGTAATGACCAACCCGTTAAAATCCAAGCACCGTCTCTTGAATACTTGTATTATAGTAGTTCTTGTTTGAATAAATTGAATATTGTTGAACATAAGAATCTGAAAACTTTAGATATATCATGTACAGAGATATCTGATGTATATCTCAACCACCTTATCTCCATACCTTACTGCCTGGAGAAACTGTTATTAGCTAACATTGCTTTTGGAAGATTCGACATTTGCAGGAGTCAATCCCTAAAGGTGTTAAAGATTCATAATTGCAAGATCATAGGAGCAATTGATGCTCCAAATTTGGTTTTGCTTGAGTATAAAGGAAATGATATTCCTCAGCTTAAATTTGCTCAAGAATCAAGACAATTGAGGCAATCACAAATGATTCTTTATCCCTTGTTCAATGTAGATTCAGCATGGTTTTGTCAGTTGAGGAAATCTTTATCAGATTCGACGATCTCTTGGTCTCAAGTTACCATTTACTTTCACAAATGCGAGGAGATCAACATGAGAGATTTGCTACTGCAACCTAGAGTTGCTATCCCCAAAGTGGACATTTTAGATGTAAATTTTTCAAGGCCGAATTGGGAGTTTCCGACATTTGTGGATGCCTTGCTGTGGAGTTGTCATCCCAGGAAACTTAACCTACAATCAACAACTACAATTTTTACATGTTTCATGTATCGTTTCATGCAGATGAAGAATATGCGTCATGGTCAATTAAAAGAAGCAAAAGCCTACAAATTTGATCAGAAAAGTAAGAGTTGGCATCCTGTAGAACACAAAAGAGGAAGAGATTTCCCAATAAGGACCAGTAGCAAGTCGAAGAAATATTTCTTTTTATTAGATTGGTGA

>PGSC0003DMG400006226 LRR-RLPs

GTACAACTCCATAATTCATTGCCCTAACCAAATGGCTACCATTATTATCTCTATTATTTTCATCCTTCAATTATTTTTGCAAAATTGTCTAGTCCTTGCTAGTCCAATTCTTGGCCAAAGTAGACAAGCTTTGGAGATAATCATCGGCGGAGGTGGTGGAGTACCTCCACCGGATAATCAAGATTGTCCACCACCTCCTCCGCCACCAGAACCTCCTTGCCCTCCGCCACCATCGCCACCATTGCTCCTCTTTGAAAGTAAACGTGTCGAGATTGCCTATCATGTCATCCAAAAGTTAAAGTCCAAAATCACATATGACCCTTTACATATTACAAAAACATGGATTGGCTCTGATGTTTGTAACAAGTACAAAGGTTTTCGTTGTGCCACGGTTCCTGACTTAAAGGTAAAAGCACTAGCTGCTGTGGACTTCAATCAGTACCAATTTGATGGCCCTGACCTTACTATTGACGGCTTTCTTGATGAATTACCAGACATAACAATCTTCCACGCAAATACCAACAATTTTAAAGGTACAATCCCCAAGAAAATCGCGAATCTTCGTTATCTCTATGAGCTAGACCTTAGCAACAACAAATATAATGGTGAATTCCCAAAAAACATTGTTTTTGGTGCCAAGAAATTGTATTTCTTGGACCTTAGGTTCAATTCATTTTCCGGGTTGGTTCCCCCTGAAGTTTTCTTGTTAAATCTCGATGTTCTGTTCATCAACAACAACAATTTAATCCAAAAACTTCCTGATAACTTGGGATCAACTCCAGTTCTTTACTTAACTTTAGCCAACAACAAATTCACCGGTCCAATTCCTCGTAGCATTGGTCAAGCTTGTAAATCATTACGCGAAGTTCTGTTCTTGAACAACCAGCTCACTGGATGCTTGCCTTATGAAATTGGGCTTTTAAGTAAAGCCACAGTTTTCGATGCGAGCAAAAATCAGTTAACAGGAAAAATACCTCATTCATTTGGATGTTTAGCTAAGATGGAGATACTTAACTTGGCACAAAATCAGTTCTATGGAGCTGTGCCTGAACTCGTGTGCAAGCTCTGCAATTTGAAAAATTTGACGTTGAGTTACAATTACTTCACTGAAGTTGGACCGGAATGCAAGAAGTTAATTCAGAAAAAGGTTCTTGATGTCAGGATGAATTGCATTCCGGGCTTGCCTATGCAAAGATCAGCAGAGGAATGTGCTGCGTTTTTCTGTAAGCCAAGATCATGTCCTGATGAAAAGTCAATGGGATTTGTTCCTTGTAGCATTGGCAGTTCTGAGCAGAATCAACAGAACGCAAGTGAGACCGCGCCAGCGCCAAGAACTTACAGTGTTTTGAAACCGCATTCGTTGTAAATGCTCTGTTCTTTCCATGTTTGCAGATTGAACTATGGCAGCAGGAAATGTAAATTTGCCAAGTTTAACTTCCTTATCTCTGTTGATGCATATTATGTTTGTTAATTATAATAAAATAATTACGTTTGGATGAGATGATCACATTGTTCAATAGAATTAGAGATGGTAGGTTGTGTAGCACAATGGCTCAACTAATGCATCTATGGTTTTGGATAGCA

>PGSC0003DMG400006255 LRR-RLPs

AAAAAATTTATTTCTCTCTTTCATGAATGAAGCAAGACCAAAGAGGAGGACTTCCACAAGTCAGTCTTCCACAGAATGAACCAAAAGGGATAACATCAAACAAGTAAAAGGAGTTAATTCAATCTGTACAAACGCGTAGTTTTCTTTGACATTTCCCAATAACCCTTCTCCTTATTAAGTTCCTTCTAACTGCGTACTTTAGTCATATTCAAGCTAACAAGTTGCATGAATATGACAACTCTCTACTTTCTATGGCTTTTCTTGATTCCCTTGTTTCAAATCTTATCAGGATATCACATTTTCTTGGTTTCCTCTCAATGTTTTCACGATCAAAAGTCGTTGTTGCTACAGTTGAAGGGCAGCCTTCAGTATGATTCTACTTTGTCAAATGAATTGGCAAAATGGAACGAGAAGACAAGTGAATGTTGCAATTGGGATGGAGTTAAATGTGATCTCTCTGGTCATGTGATTGCCTTGGAACTGGATAATCAGACTATTTCTAGTGGAATTGAGAATTCAAGTGCTCTTTTCAGTCTTCAGTATCTTGAGAGGCTAAATTTGGCTTACAACAGGTTCAGTGTTGGCATACCAGTTGGTATAGCCAACCTCACGAACTTGAAGTACCTGAATTTATCCAATGCTGGTTTTGTGGGGCAAATTCGTATGATGTTATCAAGATTAACAAAGCCAGTTACTCTAGATCTCTCAACTCTTTTCCCTTACTCTGACCAGACACTAAAACTTGAGAATTCCAATTTGAGACATTTCATTGACAACTCAACAGAGCTTAGAGAGCTTTACCTGGATGGGGTTGATCTTTCATCTCAGAGGACTGAGTGGTGTCAATCTTTATCTTCATATTTGCCTTACTTGACCGTCTTGAGTTTGCATGCTTGTCAAATTTCAGGCCCTATTGATGAATCACTTTCTAAGCTTCAGATTCTCTCTATCATCCGTCTTGACCAGAACAATCTCTCTACCACAGTTCCAGGATATTTTGCCAATTTCACAATCTTGACTACCTTGTCCCTCGACTCTTGTAATCTGCAAGGAGCTTTTCCTAAAAAGATCTTTCAGGTACAAGTTTTAGAGATTTTGGACTTTCAAATAACAAGCTGCTTAGTGGTAGTATTCCGAGTTTTCCTCGAAATGGATCTTTGAGGACGATATCACTAAGCAACACCAACTTTTTGGGTTCATTACCAGAGTCCATTTCGAACCTTCAGAACCTGTCCAGGTTAGAGCTCTCCAATTGCAATTTCACTGGACAGTACCTTCAACAATGGCAAACCTTATCAATCTTGTTCATTTGGACTTCTCTTTGAACAACTTCACTGGTTCTATCCCATATTTCCATAAATCCAAGAAACTCGCCCACTTAGACCTATCTCGTAATGGTTTAACTGGTTTCCTCTCCCCGACTCATTTTGAAGGACTCTCAGAGCTTGTCTACAGATATTTAGGGAACAACTTACTCAACGGGATCATTCCTGAATGTATCTTTGAGCTCCCCTCCTTGCAGACGCTTTTACTTAACAGCAATCAATTTGTTGGCCAAGTCAACGATTTTCACAATGCATCCTCCTCTCTGTATGTAATCGATTTGAGCAATAACCACCTGAATGGTTCAATTCCCAAGTCCATGTTTGAAATAGGGATGCTTAAGGTCCTCTCATTGTCTTCCAACTTCTTTAGTGGGACAGTTCCACCTTGACCTCATAGGGAGGCTGAGTAATCTTACAAGACTGGAGCTTTCTTACAATAACTTGACTGTTGATGCAAGTAGCATCAACTCCACCTCTTTCATATTTCCCCAGTTGAGATTATTGAAACTAGCTTCTTGTCGGTTGCAAAAGTTTCCTGATCTTAAAAGTCAGTCAAGGATGATCCACTTAGATCTTTCTGACAACCAAATACGGGGGGCAATACCAAATTGGATTTGGGGAATTGGTAGTGGAAATCTGGCTCACCTGAATCTTTCCTTCAATCAGCTGGAGTACGCAGAACAGCCATACAATGTTTCCAGCTATCTTGTAGTCTTGGACTTTCATTCCAATAAGCTAAAGGGTGACATACCAATTCCACCTTCCTGTGCTGCATATTTGGACTACTCGAGCAATAATTTCAGCAATTCCATCCTACTAGATATTGGCAATTATCTTGGTATATTTCTCTCACTTGCAAATAATCGGCTCGCAGGAAGAATTCCTGAATCTATATGCAATGCCAGCTACCTTCAAGTTCTTGATTTCTCTAACAATGCCTTGAGTGGAACAATCCCACCATGTCTACTGGAATATAGTACAAATCATGGAGTGCTGAATCTAGGTAACAATAAACTCAATGGCGTCATACCAGATTCATTTCCAATTGGTTATGCTCTACAAACTTTAGACGTCAGTGAGAATAAGTTACAAGGTAGGCTGCCAAAATCGCTTGTGAACTGCAAGTTGTTGGAGGTCCTGAATGTTGGAAATAACAAACTTGTTGATCGTTTCCCATGCATGTTGAGGAACTCAAATAGTCTGAGGGTTCTAGTTTTGCGTTCCAATCAATTCAATGGAAGTGTTCAGTGTGATGCAACCAGGAATAACTGGAAAAATCTCCAGATCATAGATATAGCTTCCAACAACTTCACTGGTGTGTTGAATGCAGAATTCTTTTCAAATTTGAAAGGAATGATGGTTGCAGATGATTACGTGGAGACAGGACGCAATCATATCCAGTACAAGTTTCTCCAACTAAGTAACTTATACTATCAGGACACAGTGATAATAACCATCAAAGGGCTAGAGTGGGAGCTTGTGAAAATTCTCAGGGTCTACACAGCTATTGATTTCTCTTCAAATAGATTTCAAGGAGTGATACCAGATACTATTGGGAATCTTAGCTCACTCTATGTTCTGAACCTATCACACAATGCCCTTGAAGGACCAATTCCAAAATTGATTGGGAAGCTACAAATGCTTGAATCACTAGACCTGTCAACAAACCACCTATCCGGGGAGATCCCATCAGAGCTTGCAAGTTTCACATTCTTAGCAGCTTTGAACTTATCGTTCAACAAATTGTTTGGCAAAATCCCATCGACTAATCAGTTTCAAACATTCTCAGCAGATTCCTTTGAAGGAAACAGTGGCCTATGCGGGCTCCCTCTCAACAACAGTTGTCAAAGCAATGGCTCAGAGTCCCTGCCTCCACCAACTTCGCTACCAGACTCAGATGATGAATGGAAGTTCATTTTCGCAGCTGTTGGATACGTAGTAGGGGCAGCAAACACCATTTCACTTCTGTGGTTTTACGAACCAGTGAAGAAATGGTTTGATAAGCATACGGAGAAATGCTTGCTTTGGTTTTCAAGAAAGTGATCCTTAAACCCATAAATGATGAGTTTATTTAATCCTTGCTTTGTTTAAATAAACACCAGGATCCAAATTCAAGTCAGTTTGTTATTTCCCTTGATGAATGTATACAATTTTCTACTATCAGAAACAGCACCCACTTTTGGATTAAATAGCTTCCCACATTGCCTGCAAC

>PGSC0003DMG400006268 LRR-RLPs

AACCAAGCACTTGTCCTATTTTTCTTTTGACTATTTCTTCCCCTCATTTTCATCTTCTTTTCTACCTTCTCCCTTCGCTTCAATCACAAAGTGGAAAAAACAAGTTTGATGGATTATAATTCCTCCAAGACTTGAGCAATGAGATTTTTACACTTTCTATGGATCTTCTTGATACCCTTTTTGCAAATTTTATTAGGTAATGAGATTTTCTTAGTTTCCTCTCAATGTCTTGATGATCAAAAGTCATTGTTGCTGCAGTTGAAGGGAAGCTTCCAATATGATTCTACTTTGTCAAATAATTTGGCAAGATGGAACCAGAACACAAGTGAATGTTGCAATTGGAATGGGGTTACATGTGACCTCTCTGGTCATGTGATTGCCTTGGAACTGGATAATCAGACCATTTCTAGTGGAATTGAGAATGCAAGTGCTCTTTTCAGTCTTCAGTTTCTTGAGAGGCTAAATTTGGCTTACAACAAGTTCAATGTTAGCATACCAGTTGGTATAGGCAACCTCACCAACTTGAAGTACCTGAATTTATCGAATGCCGGTTTTGTTGGCCAAATTCCTATGATGTTATCAAGATTAACAAGGCTAATTACTCTTGATCTCTCAACTCTTTTCCCTGACTTTTACCAGCCACTAAAACTTGAGAATCCCAATTTGAGTCATTTCATTGAGAACTCAACAGAGCTTAGAGAACTTTACCTTGATGGGGTTGATCTTTCATCTCAGAGGACTGAGTGGTGTCAATCTTTATCTTCATATTTGCCTAACTTGACCGTCTTGAGCTTGCGTGATTGTCGAATTTCAGGCCCTATTCATGAATCACTTTCTAAGCTTCACTTTCTCTCTTTCATCCGGCTTGACCAGAACAATCTCTCTACCACAGTTCCTGAATATTTTGCCAATTTCTCGAGCATGACTACCTTGAACCTCGCCTCTTGTAATTTGCAAGGAACATTTCCTGAAAGAATATTTCAGGTGTCAGTTTTAGAGAGTTTGGACTTGTCAACTAACAAGTTGCTTCGTGGTAGTATTCCGATTTTTCTTCGAAATGGATCTTTGAGGAGATTATCACTAAGCTACACCAACTTTTCTGGTTCATTACCAGAGTCCATTTCGAACTTTCAAAACCTGTCCAGGTTAGAGCTTTCCAATTGCAATTTCAATGGATCAATACCTTCCACAATGGCAAACCTTACCAATCTTGTTTATATAGATTTCTCCTTCAACAATTTCACTGGTTCTATTCCATATTTCCAACAGTCCAAGAAACTCACCTACTTAGACCTTTCACGTAATGGTCTAACTGGTCTCTTGTCTAGAGCTCATTTTGAAGGACTCTCAGAGCTTGTTAACATAAATTTAGGGAACAATTCACTCAATGGGACCCTTCCTGCATATATATTTGAGCTCCCCTCCTTGCAGCAGCTTTTTCTTAACAACAATCAATTTGTTGGCCAAGTCCACGAATTCCGCAATGCATCCTCCTCTCCGTTGGATACGGTTGATTTGAGTAACAACCACCTGAATGGATCGATTCCCAAGTCCACATTTGAAATTGGGAGGCTTAAGGTCCTCTCACTTTCTTCCAACTCCTTTAGGGGGATTGTGCCCCTCGACCTCATTGGGAGGCTGAGCAACCTTTCAAGACTGGAGCTTTCTTACAATAACTTGACTGTTGATGCAAGTAGCAGAAATTCAGCCTCTTTCACATTTCCCCAGTTGAACATATTGAAACTAGCTTCTTGTAGGCTGCAAAAGTTCCCTAATCTCAAGAACCAGTCAAGGCTGATCCACTTAGACCTTTCAGACAACCAAATAAGGGGAGCAATACCAAATTGGATTTGGGGAATTGGTAGTGGAAATCTGGCTCACCTGAATCTTTCCTTCAATCAGCTGGAGTACATGGAACAGCCTTACAATGCTTCCAGCAATCTTGTTGTCCTTGACTTGCATTCCAACCGTTTAAAAGGTGACTTACCAATTCCACCTTCCTCTGCTATCTATGTGGACTATTCGAGCAATAATTTAAACAATTCCATCCCACTAGATATTGGAAATTCTCTTGCTCTTGCCTCCTTTTTCTCGGTAGCAAACAATAACATCACTGGAATAATTCCTGAATCCATATGCAATGTCAGCTACCTTCAAGTTCTTGATTTCTCTAACAATGCCTTGAGTGGAACAATACCACGATGTCTACTGAATAATAGTACAACTCTTGGAGTGCTGAATCTAGGGAACAATAGACTCCATGGTGTTATGCCAGATTCATTTCCAATTGGTTGTGCTCTAAAAACTTTAGACCTCAGCAGGAATATCTTTGAAGGGAAGCTACCAAAATCGCTCGTCAACTGCACGTTGTTGGAGGTCCTGAATGTTGGAAATAACAGTCTTTTTGATCATTTCCCATGCATGTTGAGGAACTCAACCAGCCTGAAGGTCCTAGTCTTGCGCTCCAATAAATTCAATGGAAATCTTACGTGTAATATAACCAGAAATAGTTGGAAGAATCTCCAGATCATAGATATAGCTTCCAACAATTTTACTGGTATGTTGAATGCAGAATGCTTTTCAAATTGGAGAGGAATGATGGTTGCAGATGATTACGTGGAGACAGGACGCAATCATATCCAGTATAAGTTCTTCCAACTAAGTAACTTGTACTATCAGGACACAGTGACATTAACCATCAAAGGCATGGAGCTGGAGCTTGTGAAGATTCTCAGGGTCTTCACATCTATTGATTTCTCTTCAAATAGATTTCAAGGAATGATACCAGATACGTTTGGGCACCTTAGCTCACTTTATGTTTTGAACCTGTCACACAACGCCCTCGAGGGACCAATTCCAAAATCAATTGGGAAGCTACAAATGCTTGAATCGTTAGACCTGTCACGAAACCACCTGTCCGGGGAGATCCCCTCAGAGCTTTCAAGTCTCACATTCTTAGCAGCTTTGAATTTATCATTCAACAATTTGTTTGGCAGTATCCCATTGAGTAATCAATTTCAAACATTCTCAGCAGATTCCTATGAAGGAAACAGAGGCCTATGCGGGCTCCCTCTTAACGTCACTTGCAAAAGCGATGCTCCAGAGTTGAAACCAGCACCAAGTTTTCAAGATGACTCTTATGATTGGCAGTTCATATTTACGGGTGTGGGATATGGAGTAGGGGCAGCAATCTCCATTGCACCTCTGTTGTTTTACAAGCAAGGGAGCAAATACTTTGACAAACATTTGGAGAGAATGCTTAAACTGATGTTTCCAAGATACGGGTTCACTTACACCAGATTTGACCCTGGGAAGGTTGTAGCTGTGGAAGAATATGAAGATGAGACCCCAGATGATACCGAAGATGACGATGATGGGGGAAAAGAAGCATCTCTTGGGCATTATTGTGTCTTCTGTAGTAAACTTGATTTTCAGAGAAATGAAGCAATGCATGATCCTAAATGCACTTGTCATATGTCATCATCCCCCAATTCTTTTCCTCCTACACCATCCTCTTCTTCACCTTTATTAGTCATATATCACAAAAAGTTTTGATTCTTAAGCTTCTTCACATCTGCACATTTGTAATCTGTATATAGTTACAGGTAGCACAGGATTGATTCAAAGCAAATTAGTCTCTCT

>PGSC0003DMG400006279 LRR-RLPs

TTTGATCCCCCTCCATCCTGATCCCTGCCCTAATTTCAGGACCTCAAATTTCAAACAACAGGAAGCAAGAACAGAGGAAAATCCATGGAAATTCAGAGTCCGATGAAGCTAAAGAGATTGGTTGTGATCGGAAGGCATAAAAAAGACCGAATCAGTGATTTACCCGATGCCATCCTTCTTCACATCCTCTCTATGGTGCCGGACGGAAAACAAGTTGTGCGAACCAGCGTACTGGGTAAACGATGGAGGTCTCTTTGGATGTCTGTTCCAATGTCAATCGATTTCAACTTCCCCCAAACTGAAAATCTAGTCGTCACACTTGATTACCTGTCGTCCACACTTACAGAACTTCATTACTGTAAGTCGTGCGATAAAATACCCAGATTTACCGTTGGGGGATTTAGGTACGAACACTGTTATGCTAAGTATGTTGATTTATGGGTTAGTTTCGCTACTAAAGTTGCTAAGGTGGAAGAATTTTTGCTTGGATTGTTATCTGTAAATGATCAAACTTATGACTTCCCTTCGTATGCATTTAAGAATTCGTTGCTTAAGGATTTGGAATTGAGGAATTGCCAACTTAACCCTCGTGCTACTGTTGATTGGACTAATCTAAGCTCTCTTAAAATTGGGTATATCGATTTGACAGATGATGCTATGGAAAAGATATTATCTGGTTCCCCGAACTTGAAAAGCTTGGAACTAGATAATTTTTCCGGCCTACAACGTCTCGAAATTAACTCTGTCAAGTTGACACAACTGATGATAGAATCTTATTACAATGACAATGATGAGATTTGGCTTGAAATAGTAGCTCCCCATCTTCAGCATTTAGAAGTTTTGGAATTGTGCAGTGAGATACGCATTGGCCAGAGAAATGTGCCTTCACTTGTCACTGCAATCTTTCGTTTAAATTTTAATTTTGAGAATGAAGAACAAGACTTGGAGATGGAATTTAGCTGTTTGAAGGAACTTCTTCATAGTGTTGCCCATGTCCAGAATCTTGAATTGGGTACCTGGTGCATCGAGGTTTATTCATACACTTCATACTGCAATCTTGAATTTTTATTTACTATTGAATCATCACTATAGTAACTTTAACAGTTCACTACTTATAATCTGATAACTAAGTTGGTCAGTACTAAAGAGTTTGCAATGGATACAACAACATGTAAAATTTGGTTTGATTTACACATATCATACTTTTAGTTATGTTGATAGGTTTTTCCAAGTGTTTAAAGGTCCATCTTTGGATTTGCAAGCGAACCATGAACACGTTGATATGTGTACAGTCATATAGTCCAAGTTGTTTTTATGTTATATCCAACAATGTGCATTTCATTTATTAAGTATGCCATGTTTGTTTGTTTTGACATATTTTGGTTTGTCTGTTTTTGGTTTATCAGTGCGTGTCCATCCTAGAGTTGAAAGGGTGGCGGCCTCCACCATCAAGCCGGAAATTCTTAGAACTGAACCTAGCCGTAATACAATTGGACTTCCCTGGAATTTACAGCTTTCTACAGAGCTCATCGGATCTTGAGACATTGATCATTGGTTGTTGGTACAATCACAATGAAAGAGTAAGTTTCCTTTCGACCTTTATTTATATTACTTCATGTTATTCAAATCAAACAACTATTAGTGGAGGAATAAGCTGATATATATCAACCTATTTCTAATCTTAAGTACTTCAGGAAAGAAAATTTATTTAAGTGTTTTTTTTAATAGCAATCTCCTCAGTCACTTTACCATCCCAACTGTGCTTCCTTCCATGAGTTCTATTTTTGTTTTTTCCAATTCATCGTTTCCTAGGAGATATTGCAGCTTCCCTAGCTTCATTTTTAAAATATTCATCCAGTGTGGCATCCTCCTTGCTTCTCTCAAAAATGTTTGCACCTTGAGAACTTCTTGGAGAAAAATCACTAGGTCTAACATGTAGTTGCTTACTAGTGTTAATAGGAGTAAGAGAATTCACTCTTTCAATATTCGAACTCATCAGCACATCAAGAAGCGCCATGTCTATAGTATTATTAATCTTCTGAACATTAGAACTGCTGCGCAGAATTCTCAGATGATGCATCTGCCAGTTTCGAAATTGTATGACATCCCTGTTTCTGGTGCATCTGTCTAGCACTTGAACTTGCAATTTCTTTTGAACTCAACGCTCCATCTTTCATGATTTCAGGCGATCGCTGCCCCCAAGTGACAACTTCATTCCCCTGAATATTGTCTTCTAATATCTTCTCGTTCCCAGCAGCCTTCTTTAACACCTGATTGACTAAAATCTGTACCTCTGCTTTTGTGCCATGAATGGTCTTACACTCTTACCAACTTCAGAACCTGTAGCATCTGATTTCTCACCCAAATTACGAACACTCTGTGCAGCTCGTCTTGAATTAATAATTTCCCTAGCATTTCCCTGATGCTTAATACCTTGCATATTAATCTGCATACTTCATCTCTATGGCCTCGATGTTTGCATGTACGTCAATATGAAGGTTAATAATCATATGTTATTATTTGGTACACTATCAATTCATTCGACTTTGGCTCTAATGAGCAATCTCACACGTTCAGGCAAAGCTGCCAGTAAGTTGACTTCAACTTTAACTCTTGCTGTGCTTGGCCTTGTTTGATCTGTGATTGCTCTATCAACAACCAAAGGTCTGCCCACTGCTGATGCTAACAAATATAATGCCTCTCGAACGAAAAATTGCACCAGTAAATTCTGAAACGATATCCAAGTCAAAGCAATGGATTCTTCTTCTTTGGGATTAAACCAAGGGGTCCAACGAAAAATTCTAAATTGATAGTCTTCCTCCATGGATTTTCTCCTTCTCACCAATTGATGAAATAGCTATATGAATAGACAGCACCAAAATCCTCATGTGCCGTCAATCGAATTAAAATATGCCGTGAATCTAATGAAAATGTTTCATGCCCCTTTAACCTTTAGTTGTTGTGGAAAATACGCTCTCAACTCTGTCATTGCAAGTTTGCAACAAGATCATATCGCAAATTTTTCTCTTGGAAATTTATACAGAGCACATTGCAAATTTTCCTCCTGAATATCAGATTATTCAAAAACTATAGTAGGTTCTCCATTGATTATCAAAATTGTTGCACGCCCCTCTAGCTTCTCATGCTAGTGAATTAAAGCTGAAAATGTTAACAGTATGTTACCTTGACTTATGATTCAACTCTTTAATCTTTTTTTGTTGTTTGGATGTACCACATTGTAACATTTGCTAATATGTTCCTTGAATTTAAAAATTCTTCACCTTTTTGTTTTATCCACAATCAACTCCTTGCCTTCATGCTTTCTTCTAAGGACAATAATTGGTTAATATAGTGTATTCTTTCCTACAGAATCTGTTGGCAAGGTACACAGATGAGGATGAACTGAGCGAGAGGTTTGAGAAGCATAATGTTAAATGCTCATTTTCACATCTGAAGACCATCACGATCAATAACTTTTCTGGATCATCAAGTGAAAATAAGTTTGTGCTGCAATTGGTAAAATATTTGCTCAAGCATGCAGCCATGCTAAAAGAGTTTGTCATTGCTTCCACATACCAATGGATGTCTCCGGAGTATCGTAAAATGGCACAAAAGCTCTTAAGCTTTCCAATGTCCTCTCCGCACGCTTCAGTTGTCTTTTCTGACATATAACCTTTGTCCCTTACAACATACCTTACTAACAATGTGTTTATGCTCTAAACAGCAAGAAAGTTTGTCGTAAATGTTTTATTGACTTATACTCATTGTTTGTGAGAATAGTTTTTCAAGATTCAGTTATGTGTCCAAGGGCTTTGTTTTTGTTGGCCTATGTTATTTTTAATTGGTGGAATGATTATTTCTCGACTTGGACAGTGAATGACCTTAGGAGGGAAGCAAGGCAGTGGAACCTTGTCCCAGCATGTTTTCTGGGGTATTACAAAAGGAAAGAAATAGCATGTGTATATCTGCAGAGTTCATATTAGATTAGTTTGTATCAGAACTTTTATGTTATTGAAGACTTTATGATAATCTGTATGGTAAATTGTTGAGTCCTGTACTGTTCTC

>PGSC0003DMG400006422 LRR-RLPs

ATGGTCAAAGGAAAAAAGGAAGTGGGAGATGATGATGAGTTAAATTATTCCTACATTTTTGGAGTTTTGAGAGAGAGGGCCATAGTAACGAATAAAGGCAATAAGTACCAGTATGACACCATTTTGGCATTGTTTACAAGTATGGATATGTCTAGCAATAAGCTTTCTGGGGATATTCCTGTAAGTTTAACAGGTCTTGTAGGATTAAGATCGTTTAATTTTTCTAATAATAACCTAACTGGTAAGATTCCAAATGACATTGGTGACATGAAAGTCTTGGAATCTGTTGATCTTTCAGAAAATAAATTTTACAATCAAATCCCACAAAGCTTTTCAAGTTTGTCCACTTTGAGCTTCTTGAATCTATTTGATAATAATTCGTTATGTATGATACCATTGAGCACTCAACCTCAAAGCTTTGATCTCACGAGTTTTCAAGGAAACAAACTCTGCGGACTTCCACTCTTGATGAACTGCAGTTTAGATGACAATATTCCAAATCACGAATATGAAGATGATGAGGGTGACAAAGATGAAGGGGATTGGTTTTACATTTCATTGGTTATAGGATTTGCGCTAAACTTTTGGGGTGTTTGTGGTTCACTGCTTTTCAAGAGATCATGGAGACATGCTTATTTTCGTTTTCTAGATCGTAGCTGGGAAATGTTGCTTGCAAAGATACCAATATGGTGA

>PGSC0003DMG400006458 LRR-RLPs

TTTCTCCTCTCCATTCCCTATAAGTGTTTGTGTCACCACACACAAGTAACGACCCCTCAATTTGTCAATAAACAAATAATAGGATTAGTTAATTAACAAAGGAAGAAGAAGATTATGGAATTTCAATGATGATTTGTGTTAGGATTAAGAGATCATGAAGATGAGGATGGAATTAGTTGAACCACGACGACAACAACAACGACAAGAAGAAAAAGGAGCCTTTGAGAGGAAGAGTAGTAGGGGAAAAAGTACAAGAAGAAAGATGACAATTAGTATTGTTGAAGAAGAAGAAGAAGAGAGCCCAAAGATTGTGGATCTAAGTGGCATGTCATTAGACTCTCTTCCTGTAAATCCTACTATCAATTTGGGAGCTATATCCAAATTACATATTTCCAACAATAATCTTCAGGTTGGTCAGCTTTCGTACACCTCAATTAATTTTTGTGAAATATCTGTTATCTTCTATCAGCAATAAGCAATATGTACTAGATGACTCTGTTCATAGAGGTTAGGATAGATGAATAGATTCACTTAGTGTTTTTTTTTTGTCTTCGCTAGAATTAATTGAACAACCACTTCGTTGATCACTATATGCTATCATGAATTATTTATTTATGAGAATTTTATTCATGATGGAATATTTTTAATCTTTGATTGGAAATTAAGTGAAAAATATCTTTTTTTGAAATAAGTGGTGGCTAAAGTATTTGATATGAAGATAAGATAATTATGAAGAAAAATGTTTTAGTCAAAACCGAGTGAAGTTATTTTCCTCATTTTGACCGGAGGCGAACTAGTTGATTGCTTTGAATTGTATCAGTAGCACAAGGACACTATTAAAAACACTATTGCATTTCATAAATGTACGATTAATTATATATTAGTTTATGTCGATATTAATAGGTGGACGTGCACTACCTATATAGGTGGATCTGCCTCTAATATAATTTTGACCAAAGAATATTTTTCTAAAAATATTTTCTGACAAGCACTCACGACTAGTATTTTGGAAAACATTAATTTTTCAATCATACCAAATACAGTCGTTGCTTCCATTTTTAGTTGCTTTTGGTTCCTTGTACTTTCACAAATATTTAGTTTTGTTTAATTTCTTCCATATTTATGGAAAGAAATATATTTATCATTTACCTTTCTTCTATTTTACGCAACAATAGTTATTATATGTTTCCTAGTTTTCAATTTGTTTATTCAGTGTTGAGTTGATATTGAGTTTCAAAAGGTGGATAATTTTTTTATTGTTATAGTCTTAAACTAAAAATATGTAAAATGTATCAAAATATTTTTTAATTTTGTGGTTTTAAACATGTCTTGTGGAAAGTTAGAATTAAAGAGGTGTCAAAATAAGAATTTAATGTACTACTATATTTTCTTAAAATATTCCCCTTTACTTGATAATGAGTCGTTTATGTGAACTATACCGTATAATATTTTCAGGAATAAAAGGCAGTACTATTTTATTCTGTGTTTAATACAAAACAAACAATATAAATTTTAACGGATCAAATTAAATCCAAATCAACCGTCCTAACGTCATATTTTGTCGTTCATCATTATTTTAAATTGACCCAATAGTAAGGATAAAGTCTATTAGTTTATTTTTTACTTCATTGATACATTTTAAAAAATATATATATAACAACTTGATAGGTTTTCTAATTATTCGTATTAAACGCAATAGTATATTTTTCTGGTTGTTATATGTATGTGTGGATTGTGGATTTACTTAAATATAATTGACAAAATTTCACAAAACACGATAGTACATAACGTAACAACATTTTCGTTAATTATATATCCCGTCTTTGTTTAAAGTCTAATACAATGTATATATCTCTTATTGTGTTTTACAATGTGTATGTGTCTCACTCATCTCTCTTCTTCTATGTATTTGTATTTTATAATGTATATGTGTCTCACAATTTCAATTATAAGTACCATACAAATATCTGATACACAAATACATCGACGGTTTATATTTTTTTTTTATATCAATGTATCTATTTCATACATGTATCGATTTTAAGTAGCATGAATTGTATCATAGATTAACACTTTATCATTTTACTCTCTCTGTCCAACAATAGTAGTCCACTATTGACTTGGCATACCCCTTAAGAAAGAATAAATAATAGGGGTAATATTACTATATTATCTTTTGACTTTATTAAATTTAATGTTTTGAGAAATGTGTTAGATGATAAATAGTATTTAACAGTAAGGGTAAAATAGATCTCTTGATTTTCTTAATTGGACAAGTATTATTGAACAACTATTTTTAGTAGGAAAAATGACAAATATACCCTCGAACTATCGTAAATGGTTTGGAGATACCCTTCGTCATACTTTTGGGACATTGGTGCCCTGTCGTCCAAAAAGTAGAGCATATATACCTTTCATACTAACGGACATACACGTGTCACAATCTTATCCACTGACCCACATTTATTAAATATCGAATCGACAGATAAGATTGCGCCACGTATCCCTATTTAGTCTTCCGTTAGAGTGAAGGGCATATATGCTCTAGTTTTTGGATGAAGGGGCACCAATGTCCCAAAAGTATGACGGATGATATCTGCATACCATTTACGATAGTTCGGGGTATATTTATCCTTTTTCCCTTTTTAGTATAGTGGACAACACTATTGTTGGACAGAGGAAGTACTAGACACAATGATTTTAGGGTAAGTCACATTCTTATTTGGTTGGACGATTTTCAATTATCATGTATCACTTGTGTATATAAATACACAAACACACATCTGTATCTCTTCAAAGTTCCACTCATCTGCCATAGATACATAAATACATTGATACACATATATATCTCGTCAAAGCTAAGCACTCAATTATATTATATCATGGATGCACATTTACATATATACACAAATACACATATATATATCTCTTAAAAATATTCCACTCGTCTTTCATGGATACACAAATACACATTCATAGATACGCAATACACATATATATCTCCTCAAAACTACTCACTCATTTGCAATAGATGCACAAAAATATAGATACACATATACACAAATATACATATGTATCTCTTCAAAGTTGCACTCTAATTTGCCATAGATGCACCAATACATATAGTCCCTCTGTAAAGCGAGAGAGGGGGAGAGGGAGGAGAGTGGCGAATTTGCTACGAAAAAGGGAAATCTCATTGAGTTATGTAATTTTTTAAAACTATAAATAGGTTGAGTAGATATATATACTAGCTATGCTTGATTAATTATGTAATTTTTCATGTATAATTCTCACTTCATTGTTTTTGTTGTATGTTTGACAGAGTATACCAGAGTCTTTAACAGCAAGACTACTTAACTTGGTTGAGTTAGATATGCATTCCAATCAGCTCAACTCCATTCCCAACTCTATAGGTTGCTTATCAAAGCTCAAACTTCTCAACATTTCTGGGAATCTCCTTCTTTCGCTCCCTAAAACTATCGAGAATTGCAGGTTAGTCATCTAAACATTTCTAATTCATAACATTATCACTACAACAAAAGAGATATTTAGTAGCAATTGAATTACTGTCATTGCATTCAGAGTCGCTAAAGCCTTTAGAACATAAGTATAACCTTTTCTTCCTACTTGCAGATCATTGGAAGAGTTGAATGCCAACTTCAATATGCTTACTCATTTACCGGACACCATTGGATTTGAGCTCATAAATCTCAAGAAGTTATGCATAAATTCAAATAAAATAGCATACCTTCCTTACTCCACCTCCCACTTGACCAATCTTCGCGTGTTGGATGCTAGGCTAAACTGCCTTAGATCACTCCCCGATGATCTTGAGAACCTAATCAACCTTGAAATCCTTAACGTTAGTCAAAATTTCCAGTACCTCTCAAAGTTGCCCTATTCAGTAGGATTTCTCATTTCTCTTCATGAACTAGATGTTAGTTACAACAAGATTACTGAATTACCAGATTCCATTGGATGCTTGAAGAAGCTCCAGAAGCTGAGTGTAGAAGGCAACCGCCTTGTTTCTCCCCCACCGGAGGTAGTGGAGCAAGGGATACGTGCAGTTAAGCAATACTTATGTGAAAAGATAAATGGCATGCATGACAAGTCCCCTAAGAAGAAGTCATGGTTTGGAAAACTAGCAAGATGTAGCACATTTAGTGGTGCTAACATACCAAGAGATGATAGAGAGGACTTTGGTGTTCCAAGCAATCGAACTATCGATGTAATTGCCTCACCGCGATTTATGGGAATGCTTTCTCCTCGTCGTCTTCTATCTCCAAAAACCTACTTCTCCAAATAAGGGACATTGCAATTTGCTTGGATATCATATTGTGGTAGCTCTTAGCTTGTTTTCTTTTACGTTTGAAATCGTGCAATGTGTGCTAGATCAAGCTGTACATTATGTTCTGTAGATTTTGTTGTCTTTGAGATTGTACAGATTTTCATAGATGATTAGCCTCATGAGTCATGGGTATGTTTGAGAGGTAGAGGTACAAATTTACCCGAACAGGGTGTTTATATCAAATCATATGAATAATTACATCGAATCAAAATTTGTTATCGTAACAATATCACTTGTACAAAAACTCATTTTGTTCATTAATTATCAAAACTCAAAAAAAAATTATCTTACATAAACAGAACCATTAAAAAAACCCCAAAAAAACAGAACATGACTCTTAGAAAGCAAACAGAACAAATGAAAAACTTGCCAGAATCACAAATTATGAATTAGTTTCTTGCACTTGCGATTCATTTCTTTGCAAATCCGATTCACTTCTCCTGTTAAACGACCATTCTCTAATGGTAGACAAAACTGGCTTCCATGCTCGCGATTTATTTGATCTTTCTCCAAATTTGCACGAAGTTGTGTTAGTTGCACCACTTTTAGCTATGCAACTGAGTTTTCTAGCTATTGGAACTAAATCCATTGGAGAAACACTTGATTGAAATAAGGATTTAGGCAATAAATAGTAAATATGGCCTTGTTCTAACTTAGTATCAGGGTTGAGAGGCTTTGAAGCAGCAGATAAAAGTTGGGCATGAGTGAAGACAAAGTGTTTTTGAGATTTGGCCATGAGTTCATGTACACTAATAGGATGATCAAAATCCTCTACATGACCATTTAAATGTACAACTCTAATAGTCTTTAATATATCACAAGATTTGGGAGAAAAACAACAATATGTTATTTTGATGTACATATTGTTAATTTTTTCCTTATTTCTCTCTAGCTAGTGTGTAAAAGGGCTCCAAAGAAGATGTCCTTTTATACAAGGGGAAAGATGGTGTTGAAACCAGGTTGTGGTGCGGTAGTGGAATTGCTCCACCCTTAATCAAAGATCTCAGTTTCGAGCTCTGGGTATGAAGAAAATCCTATTGGGAGTGCTATCCTCGGATGGGTCTTGCAGTGTGCAAATACAGATTTAATCTAAGCTCCAATGCAGACTCCGAATATTGAGTGGAAAATCAAAAAAATGGTGGTGGGGAAGAGTTGTGGAGATAGATGATGAGAACACGCTATTTAAGTTAGAAATTTTTTTTATTTGCTTCAAGGCTTATGTGACGCTTCCAATAGGATTTTCTTCATATTCAAGGTTCAAAACTGAAACTTCTGATTAAAGATGGAGAAGTCTCATCA

>PGSC0003DMG400006486 LRR-RLPs

ATGGCAAAAAGAGTTTGTGTTGATGGAGAAGACAGAATATCAGAATTGCCTGTGCATATCATTCATCACATCTTATGCCGCACTGACCTCGACGTCGAAGAGGCGGCGATAAGTTGTATGTTGTCCAAGAGATGGTACTATTGCTGGACTTCAAGACCTAATTTGATATTCAATCAATTTCAAGGCTACAAATATAGGCCCCTTGAAAACTATGTCAAGTTGGTTGATCAATCTCTGCGATCCCATGTCGAAAAAAATTTACATCTTGAACAATTCATCCTTACATATAATGGTCAAGAAGTGGATCCTCACGTGGATACTTGGATTGAATTGGCGGTTAAACTCAACGTCACAGAGCTGGGAATTCACCGTTTCGGTTTAACATCATATAACTTACCTGATGTTATTTATGATGCTAAAAAGCTGACAACATTGCAGTTAAGCAGGTGCAATTTTGAATTTGACAAAAGATTTTCTTGTCTTGAGTCTCTTTCTTTGAATCATGTCCATATTTCAGATGCCCAATTACAAAGAGTTATCGACAGATGCCCATCTATCAGGACTCTAAGGACTCGTTTGGTGTAAGGGATAATACCAAATAGTCCCGGGATTAAATTATAGTACCGCTTATTTTGTTGTTTGGTTGGCAAGTTCGGGATAACTTATCCCGGGATTAATAATTAGTACCGGGATAAGTTATCCCTCTCCTTGGGTGGTATAGTAATCCCGGGATAACTTATCCCGGGATAAAATAGGTAAATGACAAAAATGTCTCTTTCAACCCTTTTGTTGTATCACTTTTTACCTTAATGAAGGGCATTTTTGTAAATAAATAAATTGTTCTTAAAATTTATTATTTTGAATATAACAAACCAAACACTCAATAAAAATAATCCAACACAACTTATCCCATCATAACTAATCCCAACATAACTTATCCCATCATAACTCATTTTCAAACCAAACGACCCCTAAGGTTACAGTGTTGTCAGGGTATAAGCAAATGTCATATTTTTGGCCTAGTACATCTCAAGTATTTCGATGCTAGTTATTGCAAACTCCATAGTGTCATAGTCCAAGCTCCATATCTTCAATGCTTTAGATATACAGAACCTACTGAACAGGGCAATCTTCATCCTTGCCAAATTGCTATTTTGGATGGTTACAATACTTTACAAACACTCAAGCTCACTGGTGCCACCATCACAGACCAACAGTTTCGGGATGTATCCTATAAGTTCCCAAACATTTCAGAATTGGATCTAGCAAGATGCTATAAGCTGAAAAATATAGAGATCCAGAGTGAAAAACTCAAGAAATTCACCTTATTAGAAGAGTTGAACAGTCTGGAAAAAATCACGATTCAAGCTCCAAATTTATTGAAATTTGATTTTCATGGTAGTAAAATGCCCTTCACATATATGGATACTTCTTCCCTAGAAAGTGCCCGGCTCAATTTCTTCTTGCCTGGCACAAACTTTGGATCAGTGGACAGCAGTTGGTACACGAGCCTTCAGCACTTTGTGAAAAAGTTCAACTATTCTAAAGATTTCATATTAATAATATTTTGCCACCAGGTAAATCAAAGTAGTTTACATTATTTTTGTGCTTATGTCATTGATTTATGCATTACTTGTTGAATTTGGTTTAAACTATGCCATTGACAAAATTCTTGCAGACCAGAAGCATCCTTATTTATAAAAATCTAGGAGAAATTGTTATTCCACCAAGTCACAAAGTCGAGATATTCATTGCATCCATGCTGCGTGTTGAATCGATCATTGAAATAATAATGTTCGAGTGTCCCTATATCATATCCATACTTCCATGTACAGATAGCAAAGCACTACAGGTAATTTCTTGCTTGTCTCTTGTGCTTGTATAAACTAAGACATTAATTTGAGCTTACAATCAGAAAATTTGTTGTTAAATGTTGATTGCTCGTCTCATAAGCATTATTGTATAATATATCTTATGTTTTTGCTTAAATGCAGGCGATGTCCACACTAAAAGGGTGTATACAAAAGCAAAATTGTGGCAAAGAATGTCCATCCGATACCGTCTGGCATCGTAACTTAAAAGAAGTCATCAGTTGTAGTGGGACATCTGAGGAGGGAATGGCTGCCTCCATGTGGAATCTCTGGTTGAAATCCACGTCTTTATTTGATCAAGTGAATAATTTCGTGTTAGAATGGAAAGATGAAGCCTAG

>PGSC0003DMG400006487 LRR-RLPs

TATAAACCCTAACAGCAAACGGCGGCACCTACCAAAATCAGAAACCCTAATGGAAGAAAGTGTTGATGGAGAAGACAGAATATCAGAATTGCCTGTGCATATCATTCATCACATCTTATGTCGCACTAACCTCGACCTCAGAGAGGCGGCGATAACTTGTGTTTTATCCAAGAAATGGCAGTATTATTGGCTTTCAAGACCTTATTTGCTGTTCCGTCAATGTAAAGGCAACAATTGTATACCCCTGGAAATTTTTGTCAAGTTGATTGATCGACCCCTACGAATCCATGTTGAACAAAACTTACATCTTGCAAAATTCTTCCTTGCATATCGGGATCGAGAATTGGATCCTTATTTGGATCATTGGATTGAATCGGCTATTAAAGTCAATGTTTTGGTGCTGGTGATTCATGCTCCGGATATTGATTCGTCGTATTATAGCTTACCTGATGTTGTTTATGATGCGAAAAAGCTCAACACATTGTGGTTAAGCAGGTGCAAGTTTGAATTTGATATTAGCACCACTCATATAAGATTTTGTTCTCTTGTGGCTCTTTTTTTGTATAATGTTCATATTTCAGATGCCCAAATGCAAAGTGTTATAGATAAATGCCCATCCATCAGGAATATAAGAGTACTGGCTTGCAAAGGTATGAGTAAATTGTATGTTGTTGGACGGTTACATCTACACTCTTTGGCTGTACTATTATGCAAACTCGATAGTGTGATTGTCGAAGCTCCGAAACTTGAATCCTTTAGTTATACAGAATGTATTGATCAGGGAAATTTTCATCCTTGCAAAATTGATATTTTGGATGGTAACAATACTTTACAAACACTCGAGCTCACTGGTTCCAGCATCACAGACCAACAGTTTCGAGATGTATTCCACAAGTTCCCAAACATTTCAGTGTTGCATCTAACAAGATGTTATAAGCTGAAATATATAGAGATTCAGAGTGAAAAACTCAACAAAGTCATCTTATCGAAGTTGGATAGTATCAAAAAAATCATGATTCAAGCCCCAAATTTATCGCAATTTGATTTTGAGGGTGATAAAATGCCCTTCTTATCTATGGATTCTTCTTCCCTGGAAAGAGCCCGACTCAATTTCTTCTTGCCGAGGACAGTAATATCAAACTTTGGAGATGTGGATAGCAGTTGGTTCACAAACCTTAATCACTTTGTTCAAAACTTCAAATATTCTAAAGGTTTGATGTTAGTAATATTTTGCCGCAAGGTACTTTCCATTCTTTTTGTGCCATTGATTTATGCATTACTTGTTGAATTTGGTTTTAAGCTATGTCATTGGCAAAATTCTTGCAGACCAAAAACATCCTTATTTATGAAAATCCAAGAGAAATTCTTATTCCACCAAGTCATAATGTCGAGATACTCATTGTACCCATCCTGAGCATTGAATTGATCATTGGATCATTATTGCTCAACCGTCCCAGCATCATGTCCATACTTCCATGTACAGATAGCAAAGTACTCCAGGTAATTTCTTGCTTTACCTGGAAATCCTTTCAAAAAAACGTCCCTTGTGCTTGTAGACACTAATGCAGTAGAGCTTAAGCATTATTGTATTCTCTATAACTTATGTTTTTGTTTACATACAGGTGTTGCCTGCATTAGAGCGGTGTACACAAAATCAAAATTGTGGTAAAGAATGTCCATTTAGTACCAGGTTTTTCCACAAGCATCGTATATTAGAAGAAATCATCAGTTGTACTGGGACATCTGAGGAGGAAATGGCCTCCATTTGGTATACCTGGTTGAAATCTACGTCTCTAATTGGCAAAGTGAATAATTTCATGTTCAAATGGAAAGAACAAGCTTAGACCGAAGAATCTTGGGGAATTCAGCATATAGTGTTTATTAGTGTGTTCAGAATAAAGTTCAGTTAACCCGCTTGTGGCTCTGTCAAATCCATGAAAGGTTCCTTAACCTTGCTCTAGCTCTAAATAAATTGTGCTCAAATCTGTGGTTTAATTTCATTTGTTCGATTGTACGGAATGAAGTTTAACTGTTAACAAACAGCATAAGAGGTCTTCATTTCATTTTTTTTTTTCTGGCTCTGCCTCTTCTTCACATTCCATTCAATTTCTCTCATGCTTAGACTACTTGGAGGTGATTAAAAAGAAGTGAGCACTTGAGGTAGAGAGATGTAATGATGGATGCATATGCCTCCTACTTGTTCTGAAATCTGGTATATATTCACTGTCATACTTGAAATGTAATATGATTTTTCGTATAACTTTCTTTGTCA

>PGSC0003DMG400006580 LRR-RLPs

TATAGAAACTTTAATTGTGAAGAGTTGTCCTTATGTACTGTTACCAGTGTCTTTTTGGCAAATGGAAAAATTAAGGCATGCTCATGTTGATTGGGCTCTTTTTGGTAATCAGGGACTCTTTGAAGGATCCTCTAAATTGGAAAATTTGAGGATATTAAAGAAAATTAGTGCATTTCCAATTGATAGGGTGGATGTGTTGTCAAGGAGGTGTCCTAATCTTCAACAACTTCATATCGAATTTGATGTACATTCTGCAGATTCTTTTTGTCTCACATTGGAGAATCTTACCCAGCTTCAAATACTTCGCGTTTTCTTTCAGCGGCCCCACATCATTGTATCTGGGTTGCAATTGCCTTCAAATTTAAACAAGTTGGTACTTAGAGGAATTCATATAGAAAGTGCTATTCCCTTCATTGCGGGACTACCAAGCCTGGAGTATCTTCAATTAGAAGAATGGAATTATGCAGAAGGGGAACCAATTTTTCCTCAATCAGAAGAGTGGTGCCTTGGAGATATCACGGTCCATAACCTTAAGTTCTTGAAACTGGTGCGGTTAGGTATCTCAAAGTGGGATGCCTCGGAGGAATCATTTCCCCTGCTTGAAACACTTGTTATAAAAAATTGTAGTAAGCTAGAGGAGATCCCCCTTAGCTTTACCGATATTCCATCATTGAAACAGATTAAGTTGATTGGGTCTTGGAAAGTATCTATGGAGGCTTCAGCTGTGAGAATTAAGGAAGAAGTCGAAGAGATTGAAGGATGTGATCGTATACACCTCGTCAAAGAACTGGCAGATTAGTGAGGGTTTTAATAGCTAATTATTGCAGCCATGCTCATACTGAAGTCTAGCCCATTCTCTTTTATTGTAACACGACTACTATTTGCTATGTTTGTTTTAATTGATAAAACTCTTTTTATGTATGATTGGAGACAAATATATATCAGTTGGTCTTTCACTGATTTCTTGTTTTT

>PGSC0003DMG400006625 LRR-RLPs

ATGAACAAATCTGGAGGAATCAAAGTACCTAAATTCATCGGTCGCCTTAAAGAACTAAGGTATCTCAATCTATCTTCTGCGTCTTTTACTAGTTCAGTATCGTCATTCCTTGGAAACCTCTCAAATTTGTAAGTTCTTGATTTGGGAGGTTTAGATCTTAGCAAAGCTACTAATTATTGGGTTCAAAAAATCAATGATCATATTCCTTCTCTCTTGGAGTTACACTTGCCTCGATGTAACTTTTTAGACTTCCATCTTCTTTTCCGTCTCTACATCTATCTTCCTTTTTGGTACTTGATCTTTCCGACAATGCTTTCAATTCAAGTACATTCCCTCAATGGATATTAAAACTTAATAACCTTGTGCATCTTCACTTAAGCTCCAACAATATCTTTAGTGAACTTCCTGATGAGTTTGCAAAGCTCAATTTATTTGAATATCTTGATCTCTCTTCCATTTATGGTATCAATGGGACACTAAAGAGAAGTTTGGGAAAACTATGCAATTTGAAGACTTTGATCCTCTCCTACAACAACATCTTGAAAACATAATCCATTTAATCGATGCCTTATCTAAATGCCAAAGCAACAGCTTGGACATTGGATTTTATTTTCAATGACTTGACTGGAAATCTTCCTGATGAATTACGCAACTTGTGGAAGCTAAAAGATCTTCAATTGAGGTATAACTCACTGATATGCACAATAGTTGAAACTATAGGGAACTTATCATCCTTAGAGACATTCTACCTCACATCTAACAAAATGAGTGGAAACCTCACCACGAATATTGGGCAGCTCATGTCATTGGTCTCATAGGACATCTTAGAGAATATGTGGGAAGGTATTGTCAAAGAATCCCATTTACTTGATCTTTCCAACTTGCAAGAGTTTTCAATTGGCATGAAACTTGGTACAAATATAACCTTGGCATTCAATATCAGCCTGAATTGGAATCCTTCATTCAATCTGACATTCTTGACCATTCAGACACGCTAATTAGGCACTGAATTCCCACATTGGCTAAAGGATCAAAATGAGCTCAACAATATAATATTCAATACCACAGGGATTTCAAATGTTGTTGCAGATTGGTTTGTAGAGTTGGATTTGAAGCTTGATAATCTAGACATGGCTTACAACAACTTGACAGGGAAAGTTCCCAAAGCTCAAATTTCAAGCCAACGTGGATTTAAGCACCAATCGCTCCGAGGGACCTCGCCCACAATGGTCTTCTAGTGTTACAACATTGTATCTAAGGGATGACTTATTTTCAGGACCTATTCTTGTTCATCCTGTATTCCTAAGATACTTAGAACAAAACTTTGAAGAATTTGGTAAGAAACAACAAAGTTTAAAGAGTATTTTCAAAACTAGAAAATTAAAACTAGTAGAGAAAATAGAATCAGAAACAGATTAGAAACAATATACAAAATATTTTCTCAAAGAAGAAAATCGAGCCCACTGAATGCACAGTGTCCCCTTAAGGAAATTATTCCCCTCAAGTACCCGAGGTTAATGAAATATATCCTCCCAGGATAGAACGATCTTACTCACTGGTGTATCGGTACCTACAATTACGGTGTCAGCGAGCCACTCAACGGCAATAAAGTACACTTAGAATACTAGATTTAGTAATAGTAGAAGAAGTCCAGGAAATTGGTTCTTTCCTTGTTGAAAAATAAGAGGAAACTCTCTATTTATAGACAACAAAGGGTAGTGTGAAAAGGTTCTTATTGTGCCTTACCGAAAGGTCACAACCCTTTGGAAAAGTCACAATCTTTCAAAAAGGTCACAACGTTTCATAAAAGTCACAAGTTTTCATAAAAGTCACAACAATTCATAAAAGTCACAACTTTTCATAAAAGTCACAACTCTTCATAAAAGTCGCAACTCTTCATTTTCCATCCACACCTTTTAAAACCCAACAATCCCCCGCATGAATGGGGAATGACTCCAAGACAAAGAAACGAACAAGTGTGTGTACTTTACAAGTAAGAATTAATTGCATCTGGATAAGTAGGTTTTCCTTTGAACTTTCTGTAGTGAACATATGTCGGATATACTCGGTCAATCGATAGATTTGGTATCTTTGAACCGTCAATCTTTGGTGTATATGTAGACAACCATATGTCACACAATTAACTTTACCGTTTATGGTTCTTACGGTTGTGTTCGTTTTAGCCATGAACACCTCCTGGTTTCATGAGTGTATAGAGGGATGGGCTTTTGACAATCATCATCTTTGAAGCGGCTTACACTTCACACTCACATAGGTGATTTCTAATCGTGTTATCACGTAGGTACACTATTTGGTCAAATCTGCCAAACTTAGCAAATCATTAAAAAACTATAAGCTTTATTAACTCATTAACAAGCCTTAATGTTATATTATTGTTCATAAGCATTATCTTCATCATGAGAATGGATTGAGTTGGTTGACAATGCGAAACTGTCAGTCACAACTTTGTTTTTCTTCTTGAACCTAGCTCTTGGGATCTCCAGTCTACTAGGTAGAGTTACCGCCATCATGACTTGTCCTAAGCCATAAACACATTCCTTTAGATGATCTTTCAACTTTCTCTCTAGTTAGGCCTTTCTGTAAGTGGATCCGACACATATCCTTTGACTTTACATAGTCAATTGTGATAATTCCACTAGAGAGTAGTTTTTTAACGGTATTATGTCTATGTCATATATGACGAAACTTTCCATTATACATCATGCTCCATGCCCCCTACCTATTGCATCTTGACTATTAAAGTGTATGCATACTGATGCCAAAGGTTTAGGCCAAAAAGGAATATTTATTAAGAAATTTTGGAGTCTTTCTACTTATTCACCGGCCTTATCTAGAGCACCAAGCTCAGAGATCATTGTAGAGCTAGCGACACGTCTTTTTGGAAGATTTCCAAGATATTGCTCATCCACCAAGGGTAAATACAAATCCACTCATGGATTTTACTTCATTTGATCCAGGTGATCCAATTTGCATCACTATATCCTTTCAATACTACTGGATATTGTTGTAATGCAAAACATAGTTTTGAGTATGTTGTAAATACCCCAAAACTCTTTTCATTGTCATCTAATGAGTTCGATTGGGATCACTCATGAACCGACTCAGTTCATTAATAACACATGTTATATCTGATCATGCACACTTCATGATGTACATCATACTTCCCAATACTCTAACACAATCCAATTTTGAATCACTTTCACCTTCACTCTTTTGAAATGCAAAGCTCACATTTATTGGAGTCTTGACAATATTGAAATCCAAATATTTGAACTAGTCAAGTACCTTTTCAATGCAATGAGACCGTGACAATGCTAAACATTATGGAGTTCTAAGAATTCTTATACCTAAGATCGCATCAGCAACTCCAAGATCTTTCATATCAAACTTTCTTTTTAGCATACGTTTAGTAGCATTTATGTCATTAGTGTCTCTACTATGATCAACATGTCATCAATATACAAACAAACAATGACCTTGTGATTTTAATGTAAACACATTTATCACTTCCATCATTCTTAAATCTATTTGCCAACATGGTTTGGTCAAGTTTCTTATGTCATTGTTTGGGTGCTTGTTTTAGTCCATAAAGTGACTTAATAAGTTCACACACTTTCTTTTCTTTACCAGGAACCACAAAACCCTCAGGCTGTTCCATGTAAATTTCTTCCTCCAACATATACCGTAACTAATGTAACTAACATCCGAAGGAATGTAATTCTAGTTACTGGTAAGTATGTGTCGACAAGATCAATACCTTGTTTTTGTCTAAAGCCTTTGACAAAAAGTCTTGCTTTATATTTGTCAACAGTTCCATCGACTTTCATTTTCCTTTTGAAGATTCACTTTGAACCCAAAGTTTTATTTCCTGGAGGAAGATCAATTAACTTCCAATGTGGTTGCTCAAGATTGAATCAATCTCATTAAAGACACCATCTTTCCAAAAGGATGAGTCCACAAAAGACATAGGAAATGTTATAAAATCAGATCCAATGGTAGTATATGTCTTTTGACATTTACTACGCGTCCGAATCTATTTATTAAGTACATTCTCTTTTTGTTATTTTCAAGGTCTTTTAGACATTTCGCTAGACTACTCAAGTCTAATTTTTCAGTCCACAGTTCTTAGGTCCTATTTTAATCTTTTTAGGAATAGGAACTTGGACTTTGGCTAGACACCCCCACACTTTGAAATATTTCAAGTTGAATTTCCTTCATTTTCATTTCTCATATGGAATAGATTGTGTCTTGAACAAACAAAACTTTTTATTGCTTTCTTTCTGAAAAGTACAAAACTTTTTTGTTGTTCCCTTTTGCTGTAAGGATAGCTTCTCCACACAAGTTTTATGGTAAACCTGAACTTATAAGTAATGTAGCCATTACTTCCTTAAAAGTTCGGTTTTTGTTTTGATTCTACAATTCCATTAGATTGAGGTGAATAGGGTAGTAATTTGATGGATAAATCCACTTTCCAAACATATTTCTACAAAAGGGGATTTATATTCTCCATCCCTATAACATCTTTTTTTCCCAACTAATTTCAACTTCAATTTTAGTATTATGCAGTTTTTCAAGATTTTCAAGAGTCAAGATCGAAACGGAATTGATGCTACAAAATTTGTTAGGATTCATTTAAATGGCAGGCATTTTTTACCCTTTTTCATAATTGTTATCATTCCAATACTACAGTCTACATTCAGCCCTTGTGGGCCACTTGAATTGCCAAGGAAGGTGGAATCTGCAGCACAGGGGGTTATCTGTTTTCTGGTCCCTGTTACACTATTGATATCCCTTCTTCTAAGGCATGTTCTCACCTTCAGAATGCTTAATTTCCATTTTGTTTATTTTTTATGTCCATTTTCTGTTATCCATGAAGTAGATATTTACTTTGATGCCTTTGCACTTTTTCGATGGTCTCCTATTGTTTTCTGCGTATCAACAATAATCTTGTAGACTAAATCGCCTAACTATTTAGAGGTATTGATGTTATAGACCGTAAATGATTATGCAACTATTCATGCATTTATGTTGGAAGCATGTACATTATCTCCTCTTGCAAGTCGAGATTAGGCCAGTTTGTTATAATTTGGCATGATTTTTCTGCACTGAAATAGTGATTAGTTTGAACTTTGGATGTACTCGATCAAAGATGCCAATTGACCAAATATCAAATAAGGAGAAAGGGAATGTCTATTTCTTGTCTCCATGGCAGGAGATTAGTTAAGTTAGTTATGTTTAGCAATTTTTTTTAAAGTACAACAACAACTATACCGTAGTCCCTGACAAGTTGGGGTCGGCTATTTGAATCCCCACTTCTTCATTTAAGCTCATTTCTTGTATCCAACTGACTTCTCTGCTTTTGCATGTCCAAGGTTCTTGTTGTTCATATGTTGTTTAGGAGTTTGATTGTTTTGGCTGTTGGGATTTGTACTGCAGGCACTTTTACTCTGCCTAGACGTTTCTATTGCTTCATCCCTACCATTCAGCAATAAACATAATTGTATCAAATGCAATTGTCAATAAAAGTTATGAGATATTTTTTCCACCATGAGAAGATGTTGACTTCATATAAAGAATGTAAGCGTAAATCAAATCTAACGGATTAGAATTCCTTTCAACAGATTTATAAGAATGCTTAGCATACTTAGATTCAACACAAACTTGACATTTTTATTTATTGCACTCAAAGGTTTTGTAATTTACATGTCCCAAGTGCTCATGTCACAAACAGTTTGACTCAAGCAAGTAAGAACAAATAAAAATATTGAGTTTGAAAAGCTCTCAGTGAGGTAGCTTTTTCACACAAATATTTGTTCCTATTTCTTACAATTTTGTGTGATACAAACATTGTTTAGAGTCATCACCTTGTCGAATATCCTTTTCGGAAGGACCTTTCCAAAACTTTCAATTTGATTGTTATAGAACTACACATGAACAAAGTTTCATCGGGTCCAACAAAGACAATAGTCCAAATGCTTCTTTTACAAAACATGACAAATGACTATTCACTAATATTCATGTCTATACAAATAGTTATATTTAATAGACATATCTTTTCATGAAAAATCCCAGCATTTTAAGTGACATTTTTTCATAAAAGAACTCAATCATTTTGAACAAGTATACATATAATTTGGTAGTTTCATACTATACATACCGTATACTAGTAAAAGAGAAACTCAGCGGCCACAATATCAAATATACTCCAAGAATTTTGTGGGTCTAACATAATACAAAAGTATCATCGAATTTCATATTTTGTGCTCCAAATTTAAATTAGACACTAAGTCTATTTTTATCTTTGTCATAAGCAAAGAACCCCCCACATTTTAAATATGACCTCAAAAATATTTTTCAAATAGTTGAAAATAGTTTTTCCACATATTTTCCCAGACCATGAATGGCGGCATCAAACCTCTTTTGGAAATATTATTCTACGAAAAAAATATAGTGCTTCAATTCTCTTTGACAATCTTTACATGGTGTCACAGTTCATTTCATAGAAGCACCTATTACAGACACAAAATCACACACTTTAAGTCACTGGTCGTTTTTCTATCACAATTAGACATACCACTTAGTGTTTAGAATACTGATAATAAAGACAAAAACTTTTGTACATTTTTTTCTATTTAACAGTGAAGTTTATAGAAAACAATAAGTACAACACTGACTTATTAGATGAAGTTTTCATATTCTTCAAACCAATTGCATAGTCTTATTTATCCAGAGTAGAAAACTACAGAAGTTTTTACTCAACAAAAATAAGACACAATCAAATTCACACAGAATACAAAACTACAAAAGTTTTTTTTCTCCAAACAGAATAAAACATATTCTTAGATAATCACATAGAAATGCTTTTATTTTTAAATAGTCCTCCAACTATAACATTTTGACATACTATTTTATCAAACATAAAATGTGCTCTCTCATTCTGCAAACTAAAGAATTTTAAAGGAAACATTATTTGCCAAGGAAAATCCATTCCAAAACATATGATTTGCAGCTATATTTCCAAACATACACATGATAAATATCAAAATAATTAACTTTTAGAAACTATTTTCCTCCTTAGATAAATAATAAGAAGGTTACCTGGTGTAATCTTTAACCGGAATACCACCAATTTTTCTAGTACATTCTCAATTCGATCTTGCTAAACAAGGAACCAAATTCTGAAGTAGTAGTGCATTAATCTTCTACTTCCATGTTTTGTTTTCCTTAATCAATAGAATACAAGAAATACCAACCGTATAATAATTTCCTTAAGATTGTTGTTCATCATGTATTCCTAAGATACTTAGAACAAAACTTTGAAGAACATGGTAAGAAACAACAAAGTTTAAAGAGTATTTTCAAAAATTAGAAAATTAAAACTAGTAGAGAAAATAGAATCAGAAACAGATTAGAAACAATATACAAAAATATTTTTTCTCAAAGAAGAAAATCGAGCCCACTGAATGCACAGTGTCCCCTTAAGGAAATTATTTCCCTCAAGTATCCATGGTTAATGGAATATACCCTCCCAATATAGAACGATCTTATGCACCGGTGTATCGGTACCTAAAACCATGGTGTCAGCGAACCACTCAACGGCAGTAAAATATACTTAGAATACTATATTTAGTAGTAGTAGAAGAAGTCCAGGAAATTGAATCTTTCCTTCTTGAAAAATAAGAGGAAACTTTCTATTTATAGACAACAAAGGTTAGTGTGGAAAGGTTTTTATTGTGCCTTACCGGAAAGGTGACAACCCTTTGGAAAAGTCATAATCTTTCGGAAAGGTCACAACCTGTCATAAAAGTCACAATTTTTCATAAAAGTCACAACTCTTCATAAAAGTCACAACTCTTCATAAAAGTAACAACTCTTCGTAAAAGTCGCAACTCTTCATTTTCCATTCACACCTTTTAAAAGTCAACACTTATTCCTCTAAACATTGTGAAGCACTTCCCAGTATAGCAGACTTGGACATCTCTAGGAACAATCAAAATGGCACAATTTCCTTATGCATGGGAGATAGATATGAATCAGTGGACTACCTTAGCCTTCAATAATAACCAACTTATCGGACAAGTTCCTGATTTCTAGGGTAAGCTACCATATCTTTACTAGATAGATATGTCAGAGAATCGCCTATCCATCCAAATTCCAGACTCACTAGGCTCTCTTGGTTATCTGATGTTCTTAAGACTCTCAGGCAATGATTTCAATTGAGAGCTGCCTTCGAGTTTGAGAAATTGCACGAAAATCATTAACAATTATCTTAGTAACAATCAGTTGTCAAGGCTTATCCCAGCTTGGCTAGGGGAAACAATGAGATTACTGTTATCCTTAGTGTAAGGAATAACAGGTTTTCAGGCCACATTTCTTTTAAAGTTTGTAGCCTTTCAGGTCTTCACATACTAAACCTTTCAAGAAACAACTTATTTGGTTCTATCCTCTCTTGTTTTGGAAACTTGAAAGGCTTCAAGGTTGAGCTAACTGATGTAGAAGCTAGACATTATCAAGGGACATTGAAGCTTGAGGCAAAATAAAGAAAAAAAATTATGATAAAATCCTTTACCTAGTAAAAATAGTATCGGCCTCTTGAGCAACAGTTTGTCTAGAGAAATCCCAGTAGAGATAACAAACTTGGAACCTTGAACTTGTCCAGGAATCATTTGAGAGGAAATATACCAACATATATCGGAAAGTTACGTTGGTTTGAAACGTTAGATCTTTCAATAAATCAGTTAACAGGTTTTATCCCACCAAGTATAGCTACTTTAGAGTTTCTGAGTCATTTGAGCTTGTCATAGAATAAGTTGACAGGAAATATTCCTACTAGTACTCAGTTTCAAACCAAAGTTGATCCAACAATTTTCCAAGGAAATGTTGCACTTTGTGGTCCTCCTTTGACTGAATGTGTTGGTGATGGAACTACAACGACATCTGAAAGTGGAAGAAATGATGACGGAGAAACAGATGATGAAGATAAACTTGAGAAAGTATGGTTCTTTGCAGTAGTTGGTTTGGGATATTGTCACGACCAAGTCTACACCCTGGATGTGGCTGGCACTCAAGAACCATTGATGGTCCCCAAGTGA

>PGSC0003DMG400006652 LRR-RLPs

ATGGATCTCTCCTGGAATTGGCTCACTGGAAATATTCCAACTGAATTGACAAGACTAAAATTTCTAGCAGTCTTAAACCTCTCTCAGAATGTTCTTGTCGGTCCAATTCCTCAAGGTCTACAATTCAACACATTTGGAAATGACTCGTATGGTGGCAACCTTGATTTATGTGGTCTTTCTTTATCAAAGCAATGTGGAACGAGTGGTTCATCACATGTTCCTCAACCATTGGAGTCGTATTTCTTTAGTGGGTTTACGTGGGAATCAGTAGTCATAGGCTACAGTTGTGGACTAGTTGTTGGAACTGTTGTGTGGAGCCTCATGTTTAAATATCTTAAGCCAAAATGGTTTGTGGAATTTTTTGATGGACTCATGCCTCACAAAAGAAGGAGGCCAAATTAAGAAGAGAGCTCAGAGACGACGGACTTAATGGAAGATTGAGAGGGCCAACTTGTCTTTGTTTGATTTTATGGACTAATTAGTGAACTTTTCTGGTTCTGTTTTTGTCTTTCTGTTTGTTGTTTGTATATGTTTGAAAGAAACATAATTATATGTATGTTATTTTAATATCAATGGTTATGTTAGTCTGTTTCTGTTTCAAGTTGACAAAAGTGATCTACACAGCCACTCATTAGCTCAGATTTTTCAAAAAGTTATGTATTTCTTAAAGAACCAACACCATCATATTTTAGGAACGCGGGGTTCCTCATATATTAGCTTAGGTACAAATAACGTGAAAGTCAAATATATTCTCAAAAACAAGCTAAAAGAAAGACATATCTGCATTTGAAGTTCGATTTTTGCCTTTTTTATATTCCGACATATTCATTCACTTTGACTTAAAGGGCGTGTTCATCCTTGTAATGAGAATTGAGCATGAATGCTCAAGTGCCTGGACCTCGAGGAGGTGAAAACTTTATAATATGTATTTAACTTGTTTCTTCCATGATTTTAAACATCAAAGAATGAGTTGATTACATCTATAAAAGCAAAAGACAGAAAACAGTGATCCTTCTATACAATTAGTCTCTCAGTTTTCTTTGCCTGTGCATCACTACACCTTTATCTTGAGTCGAGAGTCTATGGAAACAACACAAGAGACAACTAATCGTCTAGAGAAAATGAGTACATAGGGGGTATTATTTTGTTAAGTCTTCTCCGTCCTCTAAAAAGTAATTTGTGGGAAACACTACCAAAAATATGAAGTAACAAGTTGTTAGAAGTGAGTAGTACTCATCCATGGACCACAAGTTGCGTAGTTAGTGATGTCAAGGTTGTATTATTTTCTCTCCATTTATGCCAAATTTGGACAAAATAAATATTGTATAATAGTTACTATCAAACAAGATTTGAAGACTTGAGTCTTTCTGGAAAATGACTCAAAACACTTTGGACAACTTTTTTGCAATCACAAATTAGTTAGTGGTAGATGTGGGGCAAATACCCCATTTTATTTGTCTGGATATGTTTCTTCTCTTTTTCCATTCACACCTTTTATTCATAAAATTATTCAAAACATACTAATTATGCAGGAAATTAAGACACATATCAAACAAAAAATTGAATAAAAAGCAAATAAAAGAATAAAAATAAAATTCCACACGGTATTATTGTTAATTGTTTCCTCACTTGACTTAGTCAAGAAATAGTTTCTCTCATTGGCAATTTCTTAGAAGTTCTGCAGAAATTTTTTCCACATAATGGGACCAATTTCACATCATCTTACCTAACACACAATACGGCGATACTTACTAACCTCGAAATGAACACAACATCTAGAGAAAATGAGGACATAAAGAGTATTATTTTGTTAGGTCTTCTCCGTCCTCCTCTAACAAGTCTTGTGGACACTACAAAATATGAGTAACAAGTTGTTAGAAGTGACTAGTAATCATTCATGGACCACATGTTCTGTAGTTAGTGATGCCAACGTTGTACTATTTTCTCACCATTTATCCCGAAGTTGAAAAAAAACAAATATTGTATAATTGGTACTATCAAACAAGACTTGACTTAAAGACTAGAGTCTTCCTGGAAATTGAGTTGACAACACTTGGGACAACTTCTTTTGCAACCACAAATTAATTAGTACCCCATTTTATACCTCATAGTCCCATGCATAGACAAGTCAACAACGTTACTTTATGAAATAGTGTTTAGTTAACTGTTTACTTTATTTAATGTGGAGGTCGTTGTAATAATTAAGAAATCGAATAGTACTTAATTAATTATAGCATCCTTTCTTAAATATATATGTGGATAATGCTAGTTGATTTGTAAGAAGAGAACAAACATCCTGTTATTTGCTGAAAACAAAAAATGGTCATGAGTCTGTTTGATATTCCACACGACCTAGATTGTTTTACGTTTACCAGACTTTACTTAGGTGAACCAAGTTGCCTAACATGTTTCAAAGACAATGTAAAGACAGGAAAGTAAGAACACAATATTTTACGTGGAAACACCCGGCTCAAAAGGTGTAAAAACCACGACTTGTTCCTCAACAGGATTTACCCCAACTTCACTAAATAACTACAAGTCTCAACCAACAAACAAGATTACAATAGTTTATCTAACCTAAAGATCAACTACTTAATCTCAATTCTACCACACACCTCCAAGGTATGCGTTCTCAGTGTCTTCGAGTTGCCCTCAACTTGAAGACTAAAATCCTAATATCAGTTATAAATTACTGATCTAGAAAATATAAAAATCACAAATACTAAGAAAACTTCCTAGATCTGTTCTTCTATATTCTTTGATCAGCAGCACTTCAGAAATCAATCTCTCAATTTTTTTGCTCTGTGGGTGCGTAATTATTTTGAACGCCTCTTCTCCTATTTATGCTTTTGTTTCCTGAAGTCTTCTACTCCTTCAGTGACTCCTCATTCGTTCAACCACTTTATTTCATTTAGAGTTCTTCTTATACCTTTATTAATTAGAAGTCCTTCTTGAAACAGTTTCTCATCAACTCACTTTGCCAACGTGTTCCTTGCATTTTGTCATGTATCAAAATAATACTTATCCCAACATTTTCTTTTATTCCTTTCTCTGTTTTGTGTTGCTAATAAGTCTATGATTTTCTTCTTCCTTTGATCATCATCTTTGCTCTCCCACTGAAGCTTCCGCTTTGCTTCAGTTTAAGCAACCTTTCAAGTTAAATCCGACTACTCCCAGTGTGATACTTCTTTCCCAAAAACAAAGTCTTTGAATGAGAGTGGAGATTGTTGTACTTGGGACGGAGTTACATGTCACATGTTAACCGGTCCTTAGTTGTAGTCAGATTGTTGGCAATTTTCATCCCAATAGCAGTCTCTTCGAACTTCATCATCTCCAGACACTAAATCTTGCTTACAATGACTTTTATCCTTCTTCGATCCCACATAACATTGGCCGATTGACGAATTTGAGGCATCTTAACCTTTCTCGTTCTTTCTTTGATGCAAAAAATCCCAACAGAAATCTCATATCTTTCCAATTTGGTTTCACTTGATCTTTCTATGAATTGATTACAACAAATTGATGAGAGAACATTTGAAGCAATGCTTCACAACTTGACAAATATGGAGGTACTATCTCTCCTTCTTGTCAACATCTCATATCCCATACTTGTGAATATTTCTTCCTCCTTAAGGTACCTGGATGTTGAAGCTACTAATCTGCGAGGTGTTCTCACAGAGAGCCTTTTCCTTCTGCCAAACTTGGAAACACTCAAATTGAGTGGTAATCATCTTCTCAAAGGAGTTTTTCCAAAGATCCACCCGAGAAACACTCTGTTAATGGAATTGGATATTTCATACACAGGAATCTCCGGTGAGCCGCCTGATTCAATTGGCACCTTGAATTCATTGAATATCTTGAATCTCCAAGGACGTCAATTCTCTGGCCATATTCCTGATTCCATAGGCAACCTAACACAAATTACAGAGTTGATTTTAATCTTATAATCATTTCACTGGCCATATTCCTTCCACAATCTCTAAATTGAAGCACCTCACAAGTTTACTTCTTCGGTCCAACTCCTTTTCAGGTGAAATTCCTGATGTTTTCTCTAACATCCAAGAGCTACGCTTTTTAGATCTTTTTGATAACAGCGGTTCGTTTCCCTCTTCAATTTCAAACTTGACACATCTTCAACGGTTAGACTTGTTGAGTAATTCCTTATCTGGCCCACTTCCTAATAATGCGAGCAATCTTCAAAAGCTAACCCAAGTGGATTTGTATGACAACTTACTGAATGGTACCATACCATCTTGGTTGTTTAGCCTCCATCTGCTAACACTTTTTTTGTCGCTCAGTAGAAATCAATTTAGTGGACTACCCGATGAGCTCAAACCAAATCCAGCATTAACAAAAGTGGATTTAAGCAATCACTTGTGAATCTCACACACCTTTATAACCTTGACCTTTCATCAAATAACATCACCGGTCCATTTCCCGCTTCAATTTTAAGTTTGACACATCTTGAATAGTTAAACTTGTCGAGTTATTCCCTATCTGGCCCACTGCCTAGCAATGCAAGCATGCTTCAAAAGCTAACTTTTGTGGATTTGTCTTACAACTTGCTAAATGGTACCATACCATCTTGGAACCTCCCTTTGCTATCTTTGTTGTGGCTCAATCATAATCAATTTAGTGGACTAGCCTATGAGATCAAAATAAACCCAGCATTAGAATTTTTGCATTTAAATAATAATCAACTCAGTGGTTCTTTTCCTCAATCACTTGTGAATCTCACAAACCTTTTTGCCCTTGACCTTTCATCAAATAACATCACCATTGATGAGGGAATGAATATCATCTTTCCTAGCCTATCATCTTTATTCTTATCATCTTGTGGACTGAAGAATTTTCCACACATCTTGAGAAATGTAAAGACACTTCAGGTCTTGGATAATTCTAACAATAATATTTGTGGTCAAATCCCTAACTGGTTTAGCGGCATGAGGTGGGACTTGTTGCAGTTCCTAAACCTTTCTCATAATTCATTGACAGGCCACCTACCACAATTTCATTACTATAACGTACAGTATCTTGATCTGAAATTTAACTCCCTTCAGGGTCCACTACCTTCATCCATTTGTAACATGAGCAAACTTATCTTATTAGATTTATCACATAACTACTTCAGTGACTCAATTCCAAGTTGCTTGGGAAGCATGGCTAATTTAAGGGTGTTGGACTTAAGAAGGAACAATCTCACAGGGAGTATTCCCATTATGTGCACACAACACTTCATTGAGTGCCATTGTTGTAAATGGTAATCAATTTGAAGGACCTGTACCTGTGTCGTTGCTCAAGTGTAATGGTCTAGAAGTCCTTGATGTGGGGAACAATGCTATAAATGACACGTTTCCAGCTTGGCTCGGAACCCTTCAAGAGCTGCAAGTCCTTATATTAAAGTCAAACAAGTTCCATGGACCTATAAGTACTTGTCAGACTAAGTTTTGCTTTCCCAAGTTGCGAATTTTTGATCCTTCATGTAATGATTTCAGTGGCTCACTTCCTGCAAAAGTTTTTGGAAACTTCAAGGCAATGATCAAATTATATGGCGAAGACACAAGAGAGATCAAGTACATGGAACCATCTGAGGAGCCATCGTACGTATCGTATGAGGATTCAGTGTGTTTGGTGGTAATCAAAGGGCAGAATATTGAGCTACAAAGAATCAGCACAATTACGACAACCATAGATCTCTCAAGCAACCATTTTGAAGGTGTGATTCCGAAAACACTAAAGGATCTCAGCTCACTTTGGCTACTCAATTTATCCCGTAACAATCTCAGAGGCGATATTCCAATGAAACTAGGACAATTGAGTATGCTTGAAGCTTTAGATATCTCTTGGAATCGGCTCACTGGAAAGATTCCACAGGAATTGACAAGAATGAACTTTCTGGCCTTCTTAAACCTCTCTCAAAATCATCTCGTCGGACCAATTCCTCAAGGTCCACAATTCAACACATTTGAAAATGACTCATATGGTGGCAACCTTGATTTATGTGGTCTTCCTTTAACAAAGAAATGTGGAATGAGTGATTCATCGCATGTTCCTCAACAGTTGGAGTCCGAAGAAGAAGGCGAGTCATATTTTTTTAGTGGATTTACGTGGGAATCAGTAGTCATAGGCTACAGTTTTGGACTAGTTGTTGGAACTGTCATGTGGAGTCTCATGTTTAAATATCGTAAGCCAAAATGGTTTGTGGAATTTTTCTATGGACTCATGCCTCACAAAAGAAGAAGGCCAAAGAAGAGAGCTCAGAGATGA

>PGSC0003DMG400006655 LRR-RLPs

ATGTCCAAAGAGTACTTTAGTTATTGGTCTGATGGAGATTCTTTCCCGAAAACAAAGTCTTGGAATGAGAGTAGGGATTGCTGCAGTTGGGATGGAGTCACTTGTGACTTATTAAACGGTCATGTTATCGGGTTAGACCTTAGTAGTAGTCTGCTTTGTGGAAGTATTCATCCCAATAGCAGCCTCTTCCAACTTCATCATCTCCAAACACTAAACCTTTATAACAACCTCTTCAATTATTCTTCAATCCCACATAGCATTGGCCAATTGACGAATTTGAGGCATCTCAACCTTTCTCATTCTCACTTTGGTGGGAAAATCCCAACAGAAAACTCATACCTTTCTAATTTGGTTTTACTTGATCTTTCTGTGTATGGATTACAACTTGATGAGAGAACATTTGAAACAATGCTTCACAACTTCACAAATCTGGAGGTACTAGCTCTCTCTCTTGGCGACATCTCATCACTGATACCTGTAAGTATTCATCCCAATAGCAGCCTCTTCCAGCTTCATCATCTCCACACACTTAACCTTGATAACAATTACTTTTATCCTTCTTCGATCCCAAATGGCATTGGCCGATTGAGGAATTTGAGGCATCTAAAACTCTCTGGCTTTGATGGGAAAATCCCAACAGAAATCTCATACCTTTCCAATTTGGTTTCACTTGATCTTTCTTATAGTTATAAATTCTTACAACTTGATGAGATAACATTTGAAGCAATGCTTCACAACTTCACAAATCTGGAGATACTTTCTCTCTGGTATGTCAACATCTCATCTCCGATACCTGTGAATATTTCTTCTTCCTTAAGATATGTGGATCTTGCACATACTAATCTGCGAGGTGTTCTCACAGAGAGCTTTTTCCTTGTGCCAAACTTGGAAAGCCTCAAATTGAGTGGGAATGATCTTCTCAAAGGAGTTTTTCCAAAGATCCACCGGAGCAACACTCTGTTAATGGAGTTGGATATTTCATCCACAGGCATCTCCGGTGAGCTGCCTGATTCAATTGGCACCTTTAGTTCATTGAATATCTTGAACCTCGGAGGATGTCAATTCTCTGGTTCCATTCCCGATTCCATAGGCAACCTAACCCAAATTACGGAGTTGGCTTTGTCTTATAATCATTTGGGATAATACATATATTTGACCTTAAACTTGGCTTCAAATTTTAACTTTGACCTCCAACTTTCATAGTGCACAAACAAGCACTTTAACTATCCAATCTTTTAATAAATAAACACGTGATTTTTGGAGCCAAGAGCGTGAAATGCAAACGCTGCCACGTGTATTAGAAATTCCTGATGTTTTCTCTAACCTCAAAGAGCTACGTACTTTACATCTTTCTAATAACAGCTTCATCGGTTCGTTTCCCTCTTCAATTTTAAGTTTGACACATCTTGAATACTTAGGATTGACGAATAATTCCCTATCTGGCCCACTGCCTAGCAACCAAAGCATGCTTCAAAAGCTAACCGAACTGAATTTGTCATACAACTCACTGAATGGTACCATACCTTCTTGGGTGTTTAGCCTACCTTTGCTATCTTCAGTATCGCTCCAACATAACCGATTCAGAGGACTAGCCGATGAAGTGATCAAAACAAACCCAACATTAAAAGAACTGCATTTAAGCAATAATCACCTCAGTGGTTCTTTTCCTCAATCACTTGTGAATCTCACAAACCTTGAAACCCTTGGAATTTCATCAAATAACATCACCATTGATGAGGGTATGAATATCACCTTTCTTAGCCTATCATCTTTATTCTTATCATCTTGTCAACTGAAGGATTTTCCACACTTCTTGAGAAATGTAAAGACACTTCAGGTCTTGGATATTTCTAACAATAAGATTTGTGGTCAAATCCCTAACTGGTTTAGCGGCATGACGTGGGACTCATTGCAGTTCCTAAACCTTTCTCATAATTCATTAACAGGCCACCTACCACAATTTCGTTACTATAACCTACAATATCTTGATCTGAAATTTAACTTCCTTCAGGGGTCACTACCTTTATCCATTTGTAACATGAGGAAACTTATCTTATTAGATTTATCACACAACTACTTCAGTGACTCAGTTCCACATTGCTTGGGAATCATGGCTTCACTAACGGTGCTGAACTTAAGAAAGAACAACTTCACAGGTAGTCTTCCTCCATTATGTGCACAGAGCACTTCATTGAGTACCATTGTCGTAAATGGCAATCGATTTGAAGGACCTGTTCCTGTGTCGTTGCTCAAGTGTGATAGACTAGAAGTCCTTGATGTGGGGAACAATGTTATAAATGACACATTTCCAGCTTGGCTCGGAACTCTTCAAGAGCTGCAGGTCCTTATATTAAAGTCGAACAAGTTCCATGGACCTATAAGTACGTGTCAGACTAAGTTTTGCTTTCCCAAGTTGCGAATTTTTGATCTTTCTCATAATGATTTTAGTGGCTCACTTCCTGCAAAAGTTTTTGGAAACTTCAAGGCAATGATCAAATTAGATGGCGAAGATACAGGAAATATCAAGTACATGGAATCTATGTCGAATTTGCCATTTGTCAGATCGTATGAAGATTCAGTGAGTTTGGTAATCAAAGGGCAGGATATTGAGCTAGAAAGAATCAGCACAATTACGACAACCATCGATCTCTCAAACAACCATTTTGAAGGTGTCATTCCGAAAACACTAAAGGATCTCAGCTCACTTTGGCTACTCAATTTATCCCATAACAATCTCATAGGTCATATACCAATGGAATTGGGGCAATTGAATACGCTTGAAGCTTTAGATCTCTCTTGGAATCGGCTCACTGGAAAGATTCCGCAGGAATTGACAAGGATGAACTTTCTGGCAGTTTTTAACCTCTCTCAAAATCATCTCATCGGACCAATTCCTCACGGTCTACAATTCAACACATTTGAAAATGACTCTTATGGCGGCAACCTTGATTTATGTGGTCCTCCTTTATCAAAACAATGTGGAACGAGTGATTCATCACATGTTCCTCAACCATTGGAGTCTGAAGAAGATGAAGGTGAGACATATTTTTTTAGTGGATTTACGTGGGAATCAATAGTCATAGGCTATAGTTTTGGACTAGTTGTTGGAACTGTCATGTGGAGTCTCATGTTTAAATATCGTAAGCCAAAATGGTTTGTGGAATTTTTTGATGGACTCATGCCTCACAAAAGAAGAAGGCCGAAGAAGAGAGCTCAGAGACGACGGACATAA

>PGSC0003DMG400006792 LRR-RLPs

CAACAACAGACTATACGGAAAGTTCCCCTCATGGATATCTGATGGTTTTCAAAATCTTAGAATCCTCAGATTAAGGTCAAATTCATTTTCCGGAGAACTTCCTCTAGGAATGTCAAAACTGAGTTCATTGCAGGTGCTTGATCTTGCAGAAAACAACATAACTGGCGCCATTCCAACAAGCATAGGAGACCTGAACGCCATGGTACACGAGAAAAAAGTGAATGAATATCTGTTATATGGGAAGTATAGAGGGATTTACTATGAAGAAAGCTTGGTAGTGAATTTGAAAAACCAGTTCCAAAAGTACACCAAGACTCTATCACTTCTTACCTCAATAGACTTGTCGAGAAACAACCTCAACGGAGCATTTCCTGTGGAATTGACAAACTTGCATGGCCTCATAGTTCTGAACTTGTCTGGAAACCAAATCAGTGGAAAAATACCAGAAAACATTTCAAGCTTACATCAGTTAGCATCGCTTGATCTCTCGAGTAATACACTGTCTGGTGTCATCCCATCAAGCATGGAATCAATGTCATTCTTGAGCTATATTAACTTCTCAAACAACAACTTATCCGGTATGGTTCCTTATAAGGGGCAGATGACAACTTTTACAGTGTCCGCGTTTGAGGGGAATCCCCATCTTTGTGGAGCTCCACTTGTAGTACTATGTCAAAATGGAAACTCGGACAATAGAACAGTACTAGAGAATGACAGCAGTGATGAATTTCCTGATAAATGGTTTTATCTTAGTGTTGGATTGGGATTTATTACAGGTATTCTTGTTCCTTACTATATCTTGCTCGTTCGAAAGCCATGGAGACGAGCATATTTCAGTTTCTTAGACACAGTCATCGACAGATTTGTCCCAGTTAGAAGCAACAGAACAACCAGCACCAAAAGATCAGATACGGGCACCAGCTAAGGCAAAAGAAGTGTTCAATAGGTACTAGTCCTAGT

>PGSC0003DMG400006880 LRR-RLPs

CACAGAAGAAACACTCTCTGTAAATAGAAAATAGAGAGTAATAGTACTGCATTTTGTAATAGAGAGGAGAAATGGGTCTCACCGGAGTAACATTGTTGGTCACTCTCATGGTGGTGGCCGCCGTCACTTTGCCGGCAAATGCCTGTCTGCCGGCGGACCAAGCAGCATTGATGGATATCAAAGCAGCTCTTAAGGAGCCGTATTTGGGCATTTTCAACACATGGACCGGCAGCAACTGTTGCCAGGGCTGGCATGGTGTTAGCTGTGACCCGACTACTCAACGGGTTGCCGACATTGTTCTTCGGGGAGAATCCGAAGATCCGATTTACGAGAAAGCCGGCCGGTCTGGTTACATGACCGGTTCACTCTCCCCTTCTCTCTGCAAACTCGACAAGCTCACTACACTCATCGTCGCCGACTGGAAAGATATCTCCGGCGAAATTCCGGCATGTGTCACTTCATTACCAGATCTCCGTATCCTTGAGCTCATCGGCAACAAAATCAGCGGTCAAATCCCGGAAAACATCGGTCAGCTGAGTAAACTCACTGTTCTAAACCTCGCCGACAATAAAATCTGCGGTTCAATCCCCGCCTCCATTGTCAACCTCGGAAAACTAAAGCACCTCGAACTCAGCAACAATCAACTCACTGGCGAAATCCCGTCCGATATCGGAAAACTCGGAATGATGAGCCGAGCACTACTCAACAAGAACAAATTCACAGGTTCAATCCCTAACTCAATCACTCAACTTCGCCGGTTAGCCGATCTAGATCTGTCAATGAACCAAATCACCGGTTCACTCCCGGCTCAGCTCGGTTCAATGCCGGTTCTCTCAACTCTAAACCTCGATTCCAACCAAATCTCCAGTTCAATCCCCACTAATCTACTTAGCAGCAGCGGATTAAACATCTTAAACTTAAGCAGAAACTCATTAGAAGGAGAATTACCAGATGTGTTTTGCTCAAAAACGTACTTCACACATCTGGATTTATCATACAATAATCTTCGAGGTTCAGTTCCAAAATCATTATCATCAGCTAAGTACATTGGCCATTTGGATTTAAGCCATAATCATCTTTGTGGCCCGATTCCAAACGGGTCACCCTTCGATCATCTAGAAGCTTCTTCTTTCTCTAACAACGATTGTTTGTGTGGATCGCCATTACGTACTTGTTAATTCTTTTTTGTTTACTTACTAAGTTTGTAATTAATCGAAATTATCGGTTGATTACGCATAAATGGATGTGTAATTGTGTATTATGTTGTTATGATTAAAAAAATATTCTTATGTGTTTAGCTTAATTATTATTATTATTGCTAAGCAATATGATGGATTATGCGGTTCTATCATGCATGCACGTGAAAATGCGTGACAGTTTACAAGAAAAATTATTGAAGAAATTGTGTGCTATTGTTATTAGTAAATTGTAATATAATAAATCCCAC

>PGSC0003DMG400007033 LRR-RLPs

ATGGCCAGATCAGTTAACTCCATTTTCTTAACCTTCTTGTACATAAGTTTAATATACACACTCTCACCTTATGTCAATGCAACAACTTCACAATCTGATATTCAAGTTCTAGTAGCCATCAAGACGTCGATAGACCCTTTTTCAATAACCCCGGATTCCTTCCTTAGTAGTTGGGATTTCAACTTAGACCCTTGTGAAACAACTGGTGCAAGTTTCTTAGGCATTCTTTGCACAGTTCCTGATCAGGTAGACACTAATGCCACGAGCCGGATAATGGAGATTGATCTAGAAGGTGATGGACTAGAGGGGTTCTTGACATCGTCTATCGGGAACTTAACAGAACTTGTTACTCTCAATCTAGGTAGGAACAAGTTCAGGGGACCAGTGCCAGAGAGTATTACTAATTTGAGGAAACTCACAAGTCTTCAACTTTATGAGAATTTTTTCTCAGGTAGTCTTGTTGATGACATTGGTGTCCTTACAAAACTTGAAACTCTTGATGTCTCGAATAACAGGCTATCTGGCTCAATCCCTTCCTCGATTACATCACTTAGAAGTTTAACGCGATTAGACCTGTCTAACAATGAATTCACTGGAAAATTTCCTCAACTCAACGGATTATGGCAACTCTCTTCATTTGACATTTCAAATAACCAAATTTATGGGAATTTACCACAATTCCCATTGAAGATCAAAACACTATCACTAAGCCACAACTTGTTATCAGGACACATTACCCCAGTGCATAAACTTAGACATTTAAGCACATTAGACCTTAGTGATAACAGATTTTCAGGAGGAATAAACAAAGGGATCTTCATGTTACCTGAATTAAGTCACGTAAACGTTTCAGTAAACCGTTTCACAGTGTTTGAGGTGGTAGAATTCTCTGATAAAAAATCACAGCTACACACGTTAGACGTACATGCTAATCGTCTACATGGTCATTTGCCTGTGAACTTGGTTACTTACCCAAACTTAACAGAAATAAACCTTGGACATAACCTGTTTTCGGGTGAGATTCCATCAGAGTATTGGTCAAGACTCGGATTTTCATGGAGAAGTCTCAACTTGGAATATAATAACCTGGAGGGATCGGTGCCTCGAGAGCTCAACAGGACTTTAGAAGGGGTTCAGGGAAGTTTTGAACACAACTGTCTCATTTGTCCAAAGGGATTGCAGCTATGTCATGGACAGAGGACAGCTTCAGAATGTCATGGAGGAGACCATAATTAG

>PGSC0003DMG400007281 LRR-RLPs

ATGCCGGCTCTACGCCGCCAAATTTCTACCCTCTCAACGGTTAACTTGAACAAAACGATCAACAGCTTCATTGGTAACGGAAACCTCCAAGAAGCTCGGAAGCTCTTCGATCAAAGCCCCCACTTGACAAATGTCGTATCATGGAATTCCCTTATCGCTGGTTACTTCAAACACAATCTCATCCAACAAGCAGAGTACCTGTTTGACAAAATGCCTCACAGAGATGTCGTTTCTTGGAACACCATGCTCTCCGGATATCGCAACGCCAACAACCCAGAGAAAGTATATAGATGTTTTTAGATATGAACAGATGTGGAGAAATGAGACCCAATGAGCTCACTTTTGCAGTGTCAATTAGCTCCTTCTTATATTTAGATTACAAGCATTTGATCCCTCAGCTTCAAGGTCTCGTGCTCTGTTTGGGGATAAGTCTCAATGTCTTTGTAGGGTCAGCATTGATGAGGGGTTATGTTGATTTGGATGATTACAGAGGTTTAGCTCGAGTTTTTGATGAGATTTTGGATAAAGATGTTACGCCGTGGAATGTGTTGATTTTGGGTTATATGAAATTTGGGTGCACAAGTGAGGCGCAGAGGGCTTTTGATATGATGCCTATGAGAAATTATTTTACTTGGAGTACTTTGATCAATGGATATATTGAGAACAAGAAGCTTAATGAAGCTCGATTCATTTTTGATAAGATGAGTGAGAAAGATGTGGTTTCTTGGACAGCAATGATAAGAGGGTATGCGCAGTATGGAGAGTTTATGAAGGCTTTGAAATTATTTAAGTTGATGTTGAACTTGGGTTCTCGTCCTAATCATTTTACGTTTTCAACTGTTTTAGATGCTTGCGCAGGCTACTCTGCTGTTCTGGTAGGCAATCAAGTTCATGCGTGCATCTTGAAGTCTGGTTTCCCTCTTGATGTCATCTTGTTGACCTCCCTTCTTGACATGTACGGTAAATGTGGAGACATTGAAGTTGCTTTCTGCATTTTTGAGTCAATCCCGGCAAGGAATTTGGTAGCCTGGAATTCGATCATTGGAGGTTTTGCAAGACATGGGCTTGCAGAGAGGGCAATGCAGGAGTTTGAGAGGATGCTAAAGAGTGGTATCAGGCCTGATGAAATTACTTTTATTAATTTGGTATATGCATGTGGCCATGGTGGGCTAGTCGAGGAAGGTGAGAGAATTTTCAATTCTATGGTAACAGAGTACGGATTGAAAGCAGAGATGGAACACTATGCATGTATGGTGGACTTGTATGGAAGGGCAGGTCAGCTTGAGAAAGCAGAAAAACTAATCGATGGGATGCCTTTTAAGCCTGATGTGGTCGTCTGGGGGGCATTGCTAGGAGCATGTGGCTTGCACTCATGTCTAGAGCTCGGAGAGATTGCAGCAAATGGAATCTACACATTGGAGCACGACCATCCTGCAGTATACTCGGTGCTTTCCAAAATCTATGGAGACAAAGGAGTGTGCAGTGACATAACTGGTTTGGATAAGTTGATGAAGAAATGGNNNNNNNNNNNNNNNNNNNNNNNNNNNNNNNNNNNNNNNNNNNNNNNNNNNNNNNNNNNNNNNNNNNNNNNNNNNNNNNNNNNNNNNNNNNNNNNNNNNNNNNNNNNNNNNNNNNNNNNNNNNNNNNNNNNNNNNNNNNNNNNNNNNNNNNNNNNNNNNNNNNNNNNNNNNNNNNNNNNNNNNNNNNNNNNNNNNNNNNNNNNNNNNNNNNNNNNNNNNNNNNNNNNNNNNNNNNNNNNNNNNNNNNNNNNNNNNNNNNNNNNNNNNNNNNNNNNNNNNNNNNNNNNNNNNNNNNNNNNNNNNNNNNNNNNNNNNNNNNNNNNNNNNNNNNNNNNNNNNNNNNNNNNNNNNNNNNNNNNNNNNNNNNNNNNNNNNNNNNNNNNNNNNNNNNNNNNNNNNNNNNNNNNNNNNNNNNNNNNNNNNNNNNNNNNNNNNNNNNNNNNNNNNNNNNNNNNNNNNNNNNNNNNNNNNNNNNNNNNNNNNNNNNNNNNNNNNNNNNNNNNNNNNNNNNNNNNNNNNNNNNNNNNNNNNNNNNNNNNNNNNNNNNNNNNNNNNNNNNNNNNNNNNNNNNNNNNNNNNNNNNNNNNNNNNNNNNNNNNNNNNNNNNNNNNNNNNNNNNNNNNNNNNNNNNNNNNNNNNNNNNNNNNNNNNNNNNNNNNNNNNNNNNNNNNNNNNNNNNNNNNNNNNNNNNNNNNNNNNNNNNNNNNNNNNNNNNNNNNNNNNNNNNNNNNNNNNNNNNNNNNNNNNNNNNNNNNNNNNNNNNNNNNNNNNNNNNNNNNNNNNNNNNNNNNNNNNNNNNNNNNNNNNNNNNNNNNNNNNNNNNNNNNNNNNNNNNNNNNNNNNNNNNNNNNNNNNNNNNNNNNNNNNNNNNNNNNNNNNNNNNNNNNNNNNNNNNNNACAACCCCCCCCCCCCCCCAATATATGTGAAGGTTAATGCAATTGGTGCTTATTATGAAGAATGCACAACCATTGATTGAGATAAATATTACATGGCATGCCATCTCAATGTCACAATCCATTTTAGCCTCCTCTCCACTTGTTCTTCCGCTAATTAAACTAAGCAAAAATAACTACAATGAACTAAGCTAATTAAACTAAGCAATATTCCATAACCAACAAATCTATAGAGAAGAGATGAGATGAGAGTCAAAATTTAGGAAGTTGTATTCTGCTCTTTGTCTGATGTTTTAAATCTTCTGATTTTCAGGAAATGGTGCTTTCATCCAAGGTTCATAGCCAAGTCCAAAACAATCCGCCCTGTCTTTATCACATCGGCAAAATGGGAATGTATATTTTCCTTCTTGGAAGAGGATTTGTGTCAGTACCGCTGTTGAAAATTTCGAGCAGGAGAAGCAGTCGTCCAATGAAATCCTTCTTGATGAATGCCTCGTTGAGGTATTCAAACGTCTTTCCAGTGGCCAAGAGAGAAGTGCTTGTGGTTGTGTTTCCAAGCGCTGGCTTATGCTTTTAAGCAGCATCCGCAGGGATGAAGCATTTGCCTTCAAGCCCAATCTGCCTTCAAATGCTGAAGAGGAATCTACTTAACGTACCCAAGTTAAAATTGGTGAGATTGCAGAATCTACTGGTGTAGAAGGTGAGGGATATCTTGCCAGGTCCCTTGCTGGCAGGGAAGCCACAGATGTCAGACTTGCTGCTATTGCCATTGGAACTGCAAACCTTGGAGGCTTAGTAAAGCTATCTATCTAGGGAGACAACCTTTGTCGTGGTGTGACTGATACTGGTCTCAACGCTATCGCTCGAGGTTGCCCTACTCTCAAACATCTTTTCCTATGGGATTTGTCTTCTGTTGGTGATGAAGGCTTATCTGAGGTTGCTCATGAGTGTCATCTGTTAGAGAAGCTTGATCTTTTCCAATGCCCAACAATTACTGACAAGTCTTTACTCGAAATTGCGGTGAACTGCCCCAAAATGACCTCTATAACAATGGATTCTTGTTCAAACATTGGGAATGAGTCTCTCAAAGCTGTCGGTCAATATTGCCTGAACCTGAAGCTTGTTGTATTTAGAAATTGCCCACTCTAGAATGAAGCTGTTATCAAGATGTTTGCCATACTTCTCTGAGAATGCACATCCTGGATTTTTTTGGTCCTGGCTCCTTTGCCAGTAGTTTTTAGCTTCTAGCACTGCATTGCTGAGAAAATTGTGCTACAATAAAAACTTTTTTCACCATCTCTTATGTTCCTGTTTAGTTTGGAGTGAATTTATTTGAAGGTTTTTTTTTTGGGTTGGCCCTTCTCCTGTATCTGCAATATGGAGACACTGTATGAA

>PGSC0003DMG400007606 LRR-RLPs

ATGGACCCTACACATGATAATTGGAAGAAGGTTGAGGAAAATCTGAACTCATTCTTTGGTACGGTGTCCGAACGGTGCCAATCAATTCTGTCTCTGAGCTACAATTATTTGCCCCAGTATTTGAGGGCTTGTTTTCTCTATGTTGGAGGTTTTCCTGAAGATATGGAGATTGATGTTTCCAAGTTGATTAGGTTATGGATTTCAGAGCAATTCGTAAAGGCAAGAAGCAATAAAAGATTAGAAGTGGTGGCAGAGGAGTATCTACAAGAGTTAATTGATAGAAGTCTAATTTTGGCTGATAAACAATGGGTTCATGGAATGATGAGAAGTTGCAAAATTCATGATCTTCTTCGCCAACTATGCCTAAGTGAAACTCATACTGAAAATGTTGTGCATGTCATGAATGGGAAAGTTCTCAAAGCCATAGATGGTCAATGTCGAGTTATCCTTCTGTCAAATTATGAAGAGAAGTATGATTATTTTCCGCGACATAGCAGTGGTATTGTCCGTACCTTTATTTCAATGGGTGTAGCTTTTCCAAAAGAGATGTGTTCCATTATTTCAGAGTTCAAGTTGATTAAGGTGTTGGACGTGTTACCAGTCCTGTACGATTTCTCTAGTGTAATACCTGAGCTTGTACATTTGAGATATGTTGCTGCAAGAATTGAGGAAGGTCTTTCACTAGCCAAATTGAGAAATCTACAGACCATAATTCTTCAAAGATATTTCTCCAGACCAACAACGTTGAAGCAGCCAGTAGATATCTGGAGTATGTCAGAGATAAGACATGTGGATATTCAATTTCCACTATATATATCAAATCCTGTTGAAGCAGAACAACCTTTGTGCCTTCTGAATAACTTGCAAACACTTTATCCCTATAACTCTCCTTTTGTTGAGGAAATCATAAGAAGAACTCCCAATCTAAAAAAGCTAAAGATTTTAGATATATCTGAGCATTCTGATTGGGCTGTTATTCTTGATTCTCTCAGTATTTTACAGGATCTGGAGACACTACACATACAAAACATTGATCCGATGATTATCTCTGAAGACATTTCTCTTCCTAATCTTAAGGAGTTGAATTTATCAAATACTTGTATTCCATGGGAAGTTGTGAATTTGCTGGCTAATTTACCCAATCTTGAGAAGCTCTATGGGCATGCTGCATTCGATGGAACAGATTGGAAAGTAGATGAAGATGTTGTATTTCACAAATTAAAATATCTACGTCTATATACGTGCAGAGATCTGGAAAGGTGGGAATTAGCAGCTGGTAGTGATAATTTTCCGATGCTTGAGAATTTAATACTGTTTGAATTGCCTAAACTAGAGGAGATTCCGGAGAGTATTGGAGAAGTAATGACACTAAAATTCATCCGAATAGTTAATTGCGGCTCTGGTGTAAAGACAAGTGCAAAGAAAATTCAGGAAGAGCAAGAAAGCTTGGGAAATTATGAGCTTGATGTTCAAATCTACCCCTAA

>PGSC0003DMG400007684 LRR-RLPs

ATGAAACTCATCAACTTAGCCCTTAGTGGCTTCTTGTTCATCATTTTCTTCTTCTCAAAACCCTCATTTCAAGCTTCAATCAAATATGAACCAAACCCTAGACTTTTCGATGCTTACATAGCTCTACAAGCTTGGAAACATGTCATGACTTCCGATCCAAGAAACTTCACAAAAGATTGGTATGATTACAATGTTTGTAACTACACCGGGGTTTATTGTGCTCCAGCTCCAGATAACCCTAATATCACCACTGTTGCTGGTATTGACCTCAACCATGCAAATATCTCCGGATACTTACCTGAAAAACTTGGTTTATTGACTGATTTAGCCGTCTTTCATATAAATTCCAATAGATTTGAAGGCTCAATCCCGAAATCTTTTTCTAAGCTTCGAATTTTATTCGAACTTGACGTGAGTAACAATCTTTTTTGTGGTGAGTTTCCTTCCGTTGTGTTATCTCTTCCTTCTTTAAAATTTCTTGATATTAGGTATAACCAATTCGAAGGAAAAGTCCCTTCAACCCTTTGGGATCGAACACTCGATGCTCTTTTCATAAACAACAACAAATTTCAATTTTCATGGCCTAAAAACTTAGGTAAATCACCCGTTTCTGCTTTGGTTATGGCAAATATTAATGTAACGGGGTGTATACCATCAAGTATAGCAAAAATGTCGAAAACATTAAATGAGATTATTCTCTCGAATTCGAGTTTAAGTGGATGTTTACCACAAGAATTAGGGTTATTGAAAAATGTTAGGGTTCTTGACGTGAGTTTTAACAAGCTTGTTGGTGAGTTGCCGGAATCAATTGGTGGAATGAAGAAATTGGAACATCTTAACGTGGCACATAACAAGTTTTCCGGTGAGATTCCGATGAGTATTTGCTCATTACCAAAGTTGGAGAATTTCACTTACTCTTATAATTATTTTTGTGGTGAACCAAAGATTTGTCTCAAGTTGAAGGATAGAAATGATAAGAAGAATTGTATACCATATCGACCATCGCAACGTTCAGAAAGTGAATGTAAGACATTTTATTCACGAGGTCCGGTGGATTGTAGTTCATTTGGTTGTAGATCACCACCACCGCCACCACCGCCACCGCCACCACCANNNNNNNNNNNNNNNNNNNNNNNNNNNNNNNNNNNNNNNNNNNNNNNNNNNNNNNNNNNNNNNNNNNNNNNNNNNNNNNNNNNNNNNNNNNNNNNNNNNNNNNNNNNNNNNNNNNNNNNNNNNNNNNNNNNNNNNNNNNNNNNNNNNNNNNNNNNNNNNNNNNNNNNNNNNNNNNNNNNNNNNNNNNNNNNNNNNNNNNNNNNNNNNNNNNNNNNNNNNNNNNNNNNNNNNNNNNNNNNNNNNNNNNNNNNNNNNNNNNNNNNNNNNNNNNNNNNNNNNNNNNNNNNNNNNNNNNNNNNNNNNNNNNNNNNNNNNNNNNNNNNNNNNNNNNNNNNNNNNNNNNNNNNNNNNNNNNNNNNNNNNNNNNNNNNNNNNNNNNNNNNNNNNNNNNNNNNNNNNNNNNNNNNNNNNNNNNNNNNNNNNNNNNNNNNNNNNNNNNNNNNNNNNNNNNNNNNNNNNNNNNNNNNNNNNNNNNNNNNNNNNNNNNNNNNNNNNNNNNNNNNNNNNNNNNNNNNNNNNNNNNNNNNNNNNNNNNNNNNNNNNNNNNNNNNNNNNNNNNNNNNNNNNNNNNNNNNNNNNNNNNNNNNNNNNNNNNNNNNNNNNNNNNNNNNNNNNNNNNNNNNNNNNNNNNNNNNNNNNNNNNNNNNNNNNNNNNNNNNNNNNNNNNNNNNNNNNNNNNNNNNNNNNNNNNNNNNNNNNNNNNNNNNNNNNNNNNNNNNNNNNNNNNNNNNNNNNNNNNNNNNNNNNNNNNNNNNNNNNNNNNNNNNNNNNNNNNNNNNNNNNNNNNNNNNNNNNNNNNNNNNNNNNNNNNNNNNNNNNNNNNNNNNNNNNNNNNNNNCACCACCGCCACCTCCACCACCACCGCCACCACCACCGCCACCACCGCCGCCGCCACCTCCACCACCACCAAAATCACATTATTACCATTATCCTTAA

>PGSC0003DMG400007876 LRR-RLPs

ATGGAAGGTGTAGCTTTAGTACTTTCACTTCTCCTTATTGTTGTTAATTTAGATGGATTTTCTTGTGTTTATTCAGGAAGTCATTGCATTGAGAGTGAAAAAAATGCTCTTATTAAGTTCAGCCAAGGTTTCAAGAATCCGTCACAAAGTATGTTATCTTGGATGTTTGACGAAAACTGCTGCAGCTGGAAAGGTGTTGAATGTGACAACAAAACCGGGCACGTGATCACTCTTGATCTACATAAGCAATTCTTGCAAGGTGAGTTCAGGACTTCTTTACTTGGTTTGCCTCATTTAAGGCACTTAGACATGAGCCACATTGATTTCCTTGGGGAACGGATTCCTGAGTTTATTTCCAGTTTTAAGGACCTGGAGTATCTCAATTTGTCTAAAACGAATTTTAGGGGAATGATTCCTGAACATCTTGGGAATCTGTCGCGGTTGCAGTTTCTTGATCTGAGTGGTGATTTTTCACTCCAGGTTAATAACCTTGAATGGATTCACTCCCTCTTCTCTATGAAGATTCTTGATTTGAGTGGTGTTGACATGAGAAACGCGGAAAACTGGCTACATGACATCAATTTGTTAAGTTCTCTCTCTGAATTACGTTTGTCTGCTTGTCAGCTTACTACATTTCCTGAATTGCTTCCGACTTCTGTGAATTTCACATCCCTTCGGATTCTTGATCTTTCATTAAACTATTTTAATACTAGGATTCCTAGCTGGTTGTTCAACACAAGCCACAATCTTGTTTACCTTAATCTTTCGAGAAGTCAGCTGAACGGTTCACTTCCAAATGCCTTTGGCAACATGCGTTCACTTAGAGTTCTTGATCTTTCTGGGAAGTCTCTCCTAGGCAACAGTCCACATTCGTTTGAGAAAATGAGTTCACTTACGGGCTATTTACCACCAACCCTGCCATCAAAGCTGAAATACTAGGATATTTCTGATAATCAAATGGAAGGTCCTTTAGCAAGAAGCATCACACAACTTAAGCAATTAGTTGTCCTCAATGTTGCTCGGAACTCTTTCAAGGACTCGATTACAGAACATTTCTTGAATATAACTCATTTGAGAGTGTTAGACTTGTCATCTAACTCATTTATTTTCAATGTAAGCGCAACTTGGATGCCTCGTTTTCAACTTGACTCTCTAAGCCTGCAGTTTTGCGGACTTGGTGCACAGTTTCCCCAGTGGATTCAGACACAAAAAGAGTTGTCATTTATTGATATCTCTCGCACCGATATCTCAGGTGAAGTGCCCGATTGGTTCTGGAATTTTTCTGCCAAGGTTTATCACATAGACCTTTCAGGAAACAATTTCAGTGGAGAAATCCCTGAATTTACAGAAAGAGTTTGTCTTGCAAAATTGGACCTGAGTGATAACAATTTTCATGGTCCTTTGCCTCATTTTTCACCCAAGATGATGACGTTGATTTTGGCAAGGAATTCGTTTAATGGTACCATTGCCCCTGTATGTGAATCACTTGTCATGAATAACTCTCTGAGTCTTCTAGACCTATCTTCAAATTTTCTATCAGGACAACTTTCAGATTGTTGGAGATATGGGAAGAATCTGCAAGGATTGAATTTAGGCCATAATGATCTATCAGGGGAGATACCTCATTCGATTGGAGATCTTACTAACCTCTTCTTTTTGCAATTGCAGAACAACAGTTTCTCCAAAAATCTTCCTTCATCATTGAAGAACATAACTGGACTAAAAATTCTTGATGTTTCTGAAAATAGTTTATCTGGGAATATACCATTTTGGTTAGGGGAGAATTTACATGCTCTTGAGATTCTTATATTAAGCAGAAACAACTATGACGGAATTATTCCCAGGGAGATTTGCCAGCTTAAATACCTATGTATTTTGGACCTTAGTTCCAACGCACTATCAGGAGTCATTCCAAGGTGTCTTGATAATCTCCACACCATGTCCGGAGAAGAGGAAGCCCCATCCTTTACTAATGGACCCTATGCAAATTACAGGATTCATGGAATGCTCGTTCGTAATGGATACTTTAAGGAGTTCATTTTTAGCTGGCCCCGGATAGCAATTGATCTAGCAAAGAACCATTTATCAGGGGAAATTCCTGTGGAAATCACCAGGCTTACTGCACTGAGGGGCTTGAATTTATCAAGTAACAATTTCAGAGGAGCAATTCCAGGTGACATTCACAACTTGCAACTTTTGGAGTCTCTAGATCTTTCCAGGAATCAGCTGTCATGCTCCTTTCCTCCCCGTATGGGACATCTTATGTTTCTTGAGGTCGCGAATTTTTCTTTCAATGATTTGACAGGAGAAATTCCAGTTGGAGGTCAATTCGCCACCTTTGAGAGTACTTCTTATGTTGGGAATGCAAACTTGTGTGGATTCCCAGTCTCACGATTCTGCTCGGATCATTTACATGAGGACATGGTTCACTGTAGTAACAATAATCAGGAGGTTCAGGCCATTGAACACGAAGAAAACAACAACTGGCTCGATGAATTTTCGTTCTACATTAGTATGGGAATTGGATTCAATACAGGTTTCTGGTTATTTTGGGTTACTTTAATGTTGAAAGAGTCATGGAGATATGCCTATATGAGGTTCCTGGAAAACATGGGCAATAAGATATATGTCTTTGCTGCCATCAGATTGAAAAATATTCAAGCAGCAAAGAGGCTGACAAATTTCCCAAATTAAGCTGCTTCTCTTTACTATTCTCTGTTTCTTTTTTTTCCTTTCTGTCTCTTCTTATATTTTGGTTTTG

>PGSC0003DMG400007915 LRR-RLPs

ATGAGGTGGGACTCGTTGCAGTTCCTAAACCTTTCTCATAATTCTTTAACAGGCCACCTACCACAATTTCGTTACCATAACCTACGGTATCTTGATCTGAAATTTAACTCCCTTCAGGGTCCACTACCTTCATCCATTTGTAACATGGGTTCGCTTATCTTATTAGATTTATCACGCAACAACTTCAGTGACTCAGTTCCACATTGCTTGGGAAGCATGGCTAGTCTTAAGGTGCTAGACTTAAGAAGGAACAATTTCACAGGGAGTCTTCCCCCATTATGTGCACACAGCACTACATTGAGTACCATTGTCATAAATGGTAATCGATTTGAAGGACATGTTCCTGTGTCGTTGCTCAATTGTAATGGTCTACAAGTCCTTGATGTGGGGAACAATGCTATAAATGACACGTTTCCAGCTTGGTTCGGAACTCTTCAAGAGCTGCAGGTCCTTATATTAAAGTCGAACAAATTCCATGGACCTATAAGTATATGTCAAACTAAGTTTTGCTTTTCCAAGTTGCGAATTTTTGATCTTTCGCATAATGATTTCAGTGGCTCACTTCCTGCAAATGTTTTTGGAAACTTCAAGGCAATGATCAAATTAGATGGCGAAGACACAGGAAATATCAAGTACATGGAATCTATGTTGAATTCGCTATTGGTCACATCGTATGAGGATTCAGTGAGTTTGGTAATCAAAGGGCACGATATTGAGCTAGAAAGAATCAGCACAATTATGACAACCATAGATCTCTCACGCAACCATTTTGAAGGTGTCATTCCGGAAACACTAAAGGATCTCAGCTCACTTTGGTTACTCAATTTATCCAGTAACAATCTCAGAGGCGATATTCCAATGAAACTAGGACTATTGAGTATGCTTGAAGCTTTAGATCTCTCTTGGAATCGGCTCATTGGAAAGATTCCACAGGAATTAACAAGAATGAACTTTCTAGCCTTCTTAAACCTCTCTCAAAATCATCTCGTCGGACCAATTCCTCAAGGTCGACAATTCAACACATTTGAAAATGACTCGTATGGTGGCAACCTTGATTTATGTGGTCGTCCTTTAACAAAGCAATGTGAAACGAGTGATTCATCGCATGTTCCTCAACAATTGGAGTCCGAAGAAGAAGGCGAGTCATATTTTTTTAGTGGATTTACGTGGGAATCAGTAGTCATAGGCTACAGTTTTGGACTAGTTGTTGGAACTGTCATGTGGAGTCTTATGTATAAATATCGTAAGCCAAAATGGTTTGTGGAATTTTTCGATGGACTCATGCCTCACAAAAGAAGAAGGCCAAAGAAGAGAGCTCAGAGACGACAGACTTAA

>PGSC0003DMG400007999 LRR-RLPs

CTCCAGTAGCATAATAACATCACTCAAAACTTCAACTTCTTCAAGCAGAATGTAAAATCAGTTTGAAGTCCCCAAACATGGCGGTGACGGACTTTTTCGTCGGCGAAATCACCACCGAACTCTTAAAAAACCTTCTGCTAATAGTTAAAAAATCCACTTTATGCCGTTCAAGTGCCGAGAATCTCATCGACAGCATCAATGGTCTCCTTCCAATCATCCAAGAAATCAAACAAACCGGTGTTGAACTTCCACAGATACGTCAAACTCAGCTCGACGATTTCTCCAAACTTCTCCGAGATGGTTACGAACTCGCCGGAAAAGTTGTCCACTCCGGCCGTTGGAACATGTACAGGTGTCACCTTCTCATTCGACTCTAATTACAGTGAATTGAACCTCTTCCTGCTCTAATTAGGGTTATTGAATTGTGCGAATCTGTTAGAAATTACTTAATTTATATTATGCCTTGATAACAGGAACCTACAGTTGGCTAGGAAAATGGAGAGGCTGGAGAAAAGAGTAGCGAGGTTCATGCAAGTTACAATGCAAGCTCATGTACTTGCGGATGTTCATCATGTTAGGTTTAATATGGAGCAGAGATTTGACGTGCTTGAGCATAGGCTTAAAGCTATAAAAATCGGAGTTGACGATAAAAGTGGTGGTGGTGGTGGAGGAGGAGGGTGTTTAGGGGAAGCTGTGAAAAGAATGGAAGAAGATGAGAAATGGTTTGAGAATAGTTTTGTAAATTTAGGTGCTGGGATTGAATTGGGGAAGAGGAAAGTGAAGGAGATGCTGATGGGTGAACAGGATAGAGGTGTGTTTGAGATTTGTGGAATTGGGGGTAGTGGCAAAACTACCTTGGCTAAAGAGATTTGTAAAGATGATCAAGTTAAAAGTAAGTAATTAATCCCAGAAAAGTTTCTCATGTAGAGTACTTTAGCTTTTTGCGGGGTTGATTTCTCGAAAGGTTACTCAACTAATAGTTATTATCTATCACTTTTCGTTCTTGATTCTGATGCAAAGAAAGTGACTTTATTAGATATTAACTATTACTTGAGTGGACATCCGTGAAATCAACTCATGTTTTGCGTTGCTGATGATCGAGTTATTAGAAGTTAATAATCACACAAAAGTTTCTCATTTCATTTTGTTTTTACTCTTCTGTTGCGGATGAAACAGTGGATAAAAGTTGAAAACTTTATTGGCTATTATCTCTATTTTATCAGTTGAATTCTAATAGAGATCAATTTTGCATGTAAAGGACTCATCAAAAGTTTTGATTGGAATACAGTTTAAATTGGTCTTAAAAACCTTTTCTTTAAATAATGGTATGGTGTCTAAAACCGCCATGTTGTTCCAGGTTATTTCAAGGACAAGATTTTCTTTTTCACTGTTTCTCAATCTCCAAACGTGGAGCAATTAAGGAAAATGATTTGGGAAAAGATATCAGGGTGCAATCTCCATGGTTATGGACACGGGGAGATGTTACCCCAGTGGAACCTACAGTACCAATGGAATACGAAATCTGCATCCCCAGTACTCTTGATTCTTGATGATGTGTGGTCTGCATCTGTGCTAGAGCCACTAATTTTCAAGATCCCCGGCTGCAAGATCCTAGTTGTATCGCGTATCAAGTTTCCTCCATCAATCATTGACTGTATTTATGATTTAGAGTTGTTAAAGGAAGATGAAGCTATGCCCTTATTTTGCCATTTTGCATTTGGACACAATTCCTTTCCGCGTGGTTTCAGCCAAAAGCTTGTCAAAGAGGTATTAGTGTATGTTTTTGGCAGCATATTAGTTTTTCTTTGCAAAAATTTCTTGTTGTACTTGAGTTAAGCTAACTAAAGCTTCTTGGACTTTTTAGATTGTGGATGAATGTGAAGGGCTTCCTTTGGCTCTTAAGGTCATTGGATCTTCATTAAAGGGAAAACCTGAGATGTTCTGGATAAGTGCAAAAAACAGATTATCACGATGCCAACCTGTCTGCGAGTCTCATGAACTGCAGTTGCTTGAGCGAATGAAATTGAGTATTGACTGTTTGCCTGAGAAGGTGAGAGAGTGTTTCCTGGACTTGGGTGCTTTCCCAGAAGACAAAAGGATTCCTCTTGATGTTCTAATTAACATGTGGGTGGAGCTACATGATATTGATGAGGAGGAGGCTTTTCACATTCTTGTTGAACTTTCAGACAAAAATCTCCTAAATCTAGTCAAAGATGCACGGTAAGCGGTGGACTCAAATCTCCTCAAACAATCTTTTAATTTTTGTTAGAAGGTGATTGTTTATGGAAAAAATCAATATGTTGATAGTTTTCAATGTGGTCTTCATGTGGCAGAGCTGGAGACATGTATACAAGTTACTATGAGATATCGGTGTTTCAGCATGATGTATTACGAGACCTAGCAATTCATATGAGCAACCGTGATGATATAAATCAGAGAAAGCGATTGGTTATGCCACGAAGAGACGTAAACTTTCCAAGAGAATGGGAAAGAAATGTGGACAAACCTTTCCATGCACGAGTTATCTCTGTGCATACAGGTATTTTAGATCCTTGGTTACTGCTGATTGCTTAATAATGAAAAGTTTATCATGTGATATAAACAGAAATATCTCTCTTTAATGCAGATGAAATGAGAGAAATGGACTGGTTCAGAATGGATTGCCCGAAAGCTGAAGTACTGATTCTCAATTTTGCCTCATCTGAGTACTTCTTGCCTCCTTTCCTGGAGAATATGCCAAAGCTAAGGGCATTGATAATCATAAACTATAGTGCTGGCAATGCAGTTCTTCATAACATGTCTGTATTCAGTCATTTAACCAACTTGAGAAGCCTTTGGTTTGAGAAGATATCTGTCACTCACTTATCTGACTCCACAAATCCTCTCAATAACTTGCGGAAGATATCTCTAGTGCTTTGTGACATGAAAAACAGCTTAGATGAGTCAGATGTGGACCTCCCTGGTTTGTTCCCACAGCTTTCTGAGTTCACAATGGATCATTGCATCAACTTCAATAAGCTGCCATCAAGTATTTGCCGGTTGCATAAGCTCAACAGCCTTAGTATCACTAATTGTGATAGTCTTTATGAACTTCCATCTGATTTAGGTGAATTACAAACTCTACAAGTTATAAGGATATATGCCTGCCCACATCTGAAAAGGCTTCCCCCGGGAATTGGTCATCTGGTAAAGCTGAAGTACCTTGACATTTCACAATGTGTTGGTTTGAGATGTCTCCCTGAAGCAATTGGTTGCTGTAGAAACTTGGAGAAGATTGATATGAGGGAGTGCCCTCAAATTGACAGTTTGCCAAGCGCTCTATCCTTTCTTGAATCATTACGTTGTGTTATTTGTGACGAAGAAGTTTTTTGTCAATGGAAAGATGTTGAGAAGGCTGTACCAGGTCTCTGTGTACAGGTTGCCGAGGAGTGCTATACTCTTGATTGGCTATCTCAGTAATCAAGTTTTCAATTCCCTTATCTTGGGTTTCTTTCGTGTAAATGTAATAAGGCTCAGTCAATTTTCCAGTTCTGTATTTTCCCCTTTGCATGGGGTTCATTGTATCTCTTGTAAATAATATATAGCAATATGTAAATATGTAGATGGACTACATCTCAATAGCTGATCTTTCCTTTTCCTTTAAAACCATAATGCTGGAAGCAAAATTACCACAGGTAATCCAAGACTAAGGCGAGTTATCTGCTATAACATGT

>PGSC0003DMG400008036 LRR-RLPs

ATGAATAGTACTATCAAATTTCTAATGTTTTTCTTCAGTGTCACAACAGCACAACCACTAAAGGGACAGAACCAATTGATTTTTGCAGATCAAAGGCTTGCTATAGTATACCCCATTATACAAAAATTCAAGAATATCATCTCTGCTGATCCTCTAGGTATTACAAAAACTTGGGTTGGTCCAAACATATGTAATTATACTGGTTTTTATTGTGAAAGTCCTCCAGACAACAATTCAGCAATTGCTCTTGCTTCTATTGATTTCAATGGCTTTCAACTTAGCGCGTTGACACTTGATGGCTTCCTTGATCAGCTTCCTGATATAGCTATTTTCCATGCTAATTCAAACAATTTTGGTGGGACTCTGTCTCCTAAAATCGCGAATCTTCCTTATCTCTATGAACTTGATATTAGTAACAACCAATTTTTCGGTCCATTTCCTAGTCCAATTTTGGGCATGAATAGCCTAAATATATTGGATATAAGATTCAATTCTTTCACTGGATCAGTCCCACCTCAATTGTTCACTAAAGATCAATTAGACGCGCTCTTCATCAACAACAACAATTTCATGCAAATGCTTCCTGATAACATCGTGACCAGCCATGTCACATATCTTACTTTAGCTAACAACAGATTTTTCGGTCCTATCCCACATAGTATTTCCAAAATCTTGTCGAGTTTGTCTGAGGTTTTGCTGCTTAACAACTTGTTAAGTGGATGTCTTCCTTACGAGATAGGATTCTTAAACGAAGCAATAGTCTTCGATGCTGGCAATAATAGACTCACCGGTCCACTTCCGTTCTCATTAGGGTGTTTGGAAAATGTAGAAATGTTGAATTTTGCTAGAAATCAGATGTATGGGATAGTGCCTGATGTGATTTGTGCCCTGGGGAATTTGGCAAATTTATCGTTGTCCGATAACTATTTCACAGGATTTGGGCCAATTTGCTTGAGATTGATAGAAAATGGAGTGCTTGATTTGAGGAATAATTGTATTCCTGGTTTTCCATTTCAAAGATCAATAGCTGAATGTGTTGCATTTTTTGCTTACCCAAGGTACTGTCCTCATATGGCTACATATACTTATATTCCTTGTTGGCTTTCAAATTTCAAAACACCTACTCTGGATCTCCCTGAATTGGCTCCTTAGTAAGGGTAACACGGTGTACAAAGCATTATGCGTTAGCATGGTTCGGGAAAAGATCATTAGAAAATTCTGGTATTTGATCATAAACATACATGTCCCATTTTCGCGATCTTTTTTTGTTTTAATTAAATTAAGCTTGTAATTTCCATTGAACAGAGTGTGATATCTCATAATAACTTGCGATCTCTGTTTATGGAGTATTGCTCAGTTGTAAATTTACGTTACTTTTATGTTATGG

>PGSC0003DMG400008185 LRR-RLPs

TATATTTGGCTTTTTAAAAATATATAAATAAATGTGTGTAAGCAGTTTTGGCTTGTCCACTTATGACCTAGTGTTTTGTTCCAATAGCTCCAACTCTATCTGCAACAATGTTTGCAGGTTTTTTGTCCCAACCTGGAAGACCTATACATCCATGGCCTTAAAAGCATAAGTGCTCTATGCTCTCACCAACTTCCAACTGACTACTTCACCAAACTTCAAACATTGTTTGTATCATTTTGTGGAAAATTGAGAAACTTGATGTCTCCATCAGTGGCCACAGGTCTTTTGAATCTCCGAATGCTATGGATACAAGACTGTGAATCAATGGAAGAAGTGATCACAAAAGAGGAACAAGAAGGAGCAGGAATCATGACCTTATTTCCACTCTTAGAACAGCTGCATCTTATTCGTCTGCCTAAGTTGGGGCATTTCTTTCTGACAAACCATGCCCTTGAATTTCCATATCTAAGAAATGTGGAAATTGATGACTGCCCTGAAATGAAGACGTTTGTTCAACAGGGAATATCTGTGAGTATACCAATTCTCAAATATGATGATGAGGTGAAATTCGTAGATGATCTCAATAAATGTATACAGCAGGAGTTCAATAACAAGGTATGTCTTATTCCACTAGCTATATAAGTAATTGCTACATGATATTCTAATCAGGCTCTACTCCTCTCTTTCCGTTTTAGATACTTATTTAAGTGCATCAACTTTCTTTTACTTAATAATCTAGTCATCTCTCTCTTCTTTTGCATTCTCTGTTTCCTGCAGGAACAAGAAGCTAGTGATGACTACGAAGCTAGTGATGACGACGAAGCTGAACCTAGTCATGCCAGAAGTTAGGAAGCTGGTAAGAAAAACTCAATGAAATAATGCAAGTAAAACCTTGAACATCACCTGATACCTATGCTCGTGATGGAGGTAATTAACAAGGAGAGGCAGATCTACATTATTGTAACCTATCGGTTCATGTCAACCTACTAGCTATTGGCTATAGATCTAGTATATGTATTAGTAATATCTAAAAAATAGTTGAT

>PGSC0003DMG400008394 LRR-RLPs

CTTCCTTTGAGTACTCTACAAACAACTACCAAAAGAAGACTTACTTTGAGAAGACTTTGGTGCAGAGGAAGAAGACTTTGTGAAGGAGCAAACACAACAATGGCCGATCCTGTAATTGGTGCTACTGTTCAAGTTTTGCTTGAAAAGTTGATTTCTCTCACTATCGAGGAGGTCAACAGCTCAAGGGATTTCAACAAAGATCTCGAAATGTTGACACAAAATGTATCCTTAATCCAAGCTTTCATTCATGATGTTGAAACACCACAAGTTGAGAAACAACAGTCTATCGAACAATGGCTCAACAGGCTTGAGAGAGTTGCTGAAGATGCTGAAAATGTGTTTGATCGATTCAGATATGAATCTCTCAAAACAAAAGTGGTGAGGAGCCCATTGAAAAAGGTCAGTGGTTTCTTTTCTCATACTGCTTTTAAGAGAAAAATGTCTCAAAAAATCAACAACATCAATAAAGAGTTGACGGCTATCAATAAGGTAGCCAAAGACCTCGGTCTACAATCACTCATGGTTCCTTTTCGGAAAATACTACCAATTCGAGAAACAGATTCCTTAGTAGTTGCTTCTGATATTGTTGGTAGAGATTTGGATGTTGCTGAGATAAAGGAGAAGATTTTGAACATGAGAGAGGAGGATATTGTTCTGTCCACCATTCCCATAGTAGGTATGGGAGGTTTAGGGAAAACAACTGTGGCTAAGAGGATTTACAATGATGAACACATCCAGCAAATCTTTGAAAAGAGAATTTGGTTGTGTCTACCTGAAATGTCTGAAACGAAGTGCTTTCTTGAACAAATCCTCGAATCATTGATAGAGAGGAGAATTGAGGTTGAAAGGAGAGATATAATAGTCAAGAAGCTACAAGATGAATTGAGAGGAAAAAAATATTTGCTAGTCCTGGATGATTTGTGGTGTGTTGACTCTACATCATGGCATGAGTTCGTTGACACCTTGAGAGGAATAAATACATCCAGAGGAAACTGCATTCTTGTGACTACTCGTATGAAGCGGGTGGCATCCACAGTAGCAACAGATCTTCATATCTTGGGGAAGTTAACAGAAGATCATTGTTGGTCTATTTTCAAACAAAAAGCATTTGTTGATGGCAGGGTTCCAGAGGAATTAGCGAGCATGGGCAACAAGATTGTTAAAATGTGCCAAGGTCTACCGTTGGCTGCAAGTGCGTTGGGAGGGCTCTTACACAACAAAGAAAAACATGAATGGCAAGCAATTCTTGATGGCAGCCTCCTTGTTGCAGGTGAAGATGATAATGGAGAAAATAGCATAAAGAAAATCCTAAAACTCAGCTATGATTATCTACCATCTCCACATCTGAAAAAATGCTTTGCTTACTTTGCAATGTTTCCAAAAGATTATATGTTTGAAAAGGACCAACTAATCCAACTCTGGATGGCAGAAGGGTTTCTTCGTCCAAGTCAAGAGATCCCTGTGATGGAAGACGTTGGGCACAGGTTTTTTCAAATCTTGTTGCAGAATTCCTTGCTACAAGATGTTGTGTTAGATGAACACAACACTATAACACACTATAAGATGCACGATCTTGTGCATGATTTGGCTGGAGATATCTTAAAATCTAGACTATTTGATCCGAAGGGAGACAATGGAGAAAAACTTTCTCAAGTTCGATACTTTGGATGTGAGTCACCAACGGATCAAATAGATAAGATATATGAGCCAGAACGTTTGTGCACACTGTTTTGGAGAAGCAATTATACGTCTAAAGATATGCTGTTGAACTTTAAGTTCTTGAGAGTTTTAGATTTGTCCAGTTCAGGAATCAAGGAGTTGTCAGCCAAAATGGGGAAGCTGATATACTTGAGATATCTTGATCTCTCGAACACTGAGATCACAGCCTTGCCCAACTCCATTTGCGAGCTCTATAATTTGCAAACATTTAGAGTCATCAACTGCTTTTCACTCCAGAAACTTCCATATGAGATGAGAAATATGATAAGTTTGAGACACATATATTACACTTCTGTTGACGAAACAAGTGGGCATTGGGGAGGATGGTGTCTTCACAATGAACATTTTCAGATTCCACTTAATATGGGGCAATTGACTAGTCTTCAAACCCTCAAGTTTTTCAAGGTAGGTTTAGAGAAAGGTCGTCAAATAGAAGAATTAGGTCATTTGAAAAACCTAAGAGGTGAATTGACAATCAATGGTCTCCAATTGGTCTGTGATAAAGAAGAGGCTCGAACAGCATATTTACACGATAAACCAAATATCTGCAAGCTGGCATATTTATGGTCCCATGATGAATCAGAAGGCTGTGAGATCAATGATGAGCATGTGTTGGATGGTCTTCAACCGCATCCTAACTTGAAAACCTTAGCAGTAGTGGACTATTTAGGGACTAAATTTCCTTCATGGTTCAGTGAAGAGTCTCTACCAAATTTGGTCAAGTTGAAATTAAGTGGTAGCAAAAGGTGCAAAGAAATTCCATCCCTTGGCCAACTGAAATTCCTTCGGCATCTTGAGCTGATAGGATTCCATGAGTTGGAATGCATTGGACCTGCTTTAGATGGTGTTGAGATTAGAAATATTGGATCAAACAGCATTATCCAAGTGTTCCCATCATTGAAAGAACTAGTATTGGAGGATATGCGTAGCCTTATTGAGTGGAAGGGAGATGAAGTTGGAGTAAGAATGTCTCCCAGGCTTGAGAAGTTGCGGATTACAGACTGTCCATTGTTAAAAAGTATTCCGAATCAATTTGAAATCCTCCGTCAATTAGAAATTAGAGGAGTTGACAGTGAAATGCCATTGTTGAACTTGTGCAGCAACTTGACATCTCTCGTAAAGCTTAGTGTCTATGATATGAAAGAGCTCACTTGTCTTCCAGATGAGATACTACGTAACAACGTTTCTCTTCAACAGATAACGATTTTCAACTGCGGAGAGTTTCGTGAATTGCCACAAAGCTTGTACAATCTCCATTCTCTTAAGAGATTAGACATTTACAACTGCACCAATTTCAGTTCTCTTCCTGTTCCCAATGGAGACAACTATTTGACTTCCCTCGAATTCTTTTTCTTATACAATTGTAATGGATTGATCAGTTTACCAATTGGAATGCTAGATCAATGCCGGTCTCTAGAGTTTTTGTGTGTCAGCTGCTGTAACAACTTGGTTTCATTCCCTTTACATGTGTGGGAAATGCCTTCATTTTCATTTTTGGATATAACAGAATGTCCCAAATTGATTAGTGTACCCAAAGTGGGCCTTCACCATCTCACCGGGTTATTGAGATTGGGAATTGGTCCTTTCTCAGAGATGGTGGATTTTGATGCATTCCAATTGATTTTTAATGGCATTCAGCAGCTGTTGTCCCTTCGTGATCTGGAGGTGTACGGACGTGGGCACTGGGATTCTCTGCCCTATCAGATTATGCAACTATCTGACCTAAGAGAGATCACAATAGCTGATTTTGGAATTGAGGCTCTTCCTCCTAGACTTGACAACCTTACTTCTCTTGAAAGTTTGACTCTAGTGAGGTGCAAACAGCTACAACATCTGAACTTCTCAGATGCCATGCCCAAATTACGGCTTCTGTGTATAAGTGATTGTCCATTGTTAGAAGCTCTGTCGGATGGGCTCGGCAACCTTGTTTCTTTGGAAGAATTATATTTACATGACTGCGAAAAACTAGAGCATCTACCGTCCCGAGATGCCATGCGACGCCTCACTAAATTATGGAACATGGGAATTAAAGGCTGCCCAAAGTTAGAAGAAAGTTTCACCAACTACTCCCAGTGGTCCAAAATTTCCCATATTTCAAATATTGAATTAGGTGGGCGGAGAAGGACAACCGTAAGTCTCGGTTTCTCTTTCACTTACTGAGTCGCTCGCATTTTTAATTAGAATATTATTTTCCACGAGTTCCAAAAAAGAAGTTTGGTAAGTAATGTAATAATTTTTCTTGAAATTATTTTGACATAAAAAAAAAAAGTTAGATTAAATCAATTGCATCTGTTGCTATGCTGGATTCTGCTTGTCCACTTGAACTGAATTGCTGTTATTCTAAAAATTATGTTTCTTTTCCTTATAGGTAGCTGGTTCAAAACGTCTCTGTGGTAAATCAAGCCAAATCGTATGAGCAGCAGGCAATTACGTTAGGTAAATTTTCATTTATAATTTATAGGCACTTTGTAATCTACTTGTTTCTATCTTTCTTTTTGTTGGAAACCAATTTTCGGAGGTCTATGCTGATGAATACAACATTACCTTCCTTAAAAAAAAGTTTAATCATGAAAATTTATGATTTCTGCTCATAGGTTACTAGGTACAAGGACATAGTAGAACGTCGTTGTGGTATATCTAGTCAAAATCCATGGGCAGAATCATGTTAGGTATTTTTATTTAGTTGTTAGAGGAATGCTATAACAAGTTTGTAAAGTTTCTTTTTTTTACTTTCCTTTTCTCTCATGAAATAATTTTAATGTTTAACGGGACTTGTCGTGTTTCACAGAGGAACCACCACTGTTATTTTGTATGAAGAACAATCTTCAACTGTAGACTCATTATGGGACACTTTCATGAGTATTTTCGGGGAACTACAGTCTACATATGAAACATCAAAAACTCAAAAGCGTATAGGGTGCTCAATGAGAAGCTACACGGGGTCTATTCTCCATGTAAGTATTGATAACTCTCAAGGAACAATACCAAGTCTCATATATGGTAGAAATCTATATAAGTTTCTTAATAGACTCATAAAAGAATAATCTTTATGTCTTCTGCAGTTCTGGTTTTATTTTCATGCTTAAGACAGTACAACTTTTATGAATATATGAAACACTTCTCGTTACTTTCACCTCACAGGACAATTTCTCCGTTCATTAAAACTTCCGCAAATGAAGGACTTCAACAAAGATAGTTGTCATGAATATTTTTGCAAAGGGTGACCCATTGGCTCTTAAAATGTCCTTAGCAGTATATGTTCATCTTAAACAGGTTTTCCTCTGTCCTTTTTTAGTGGTTACTTGTAATGATCCGCTTCTCCATTCTCCAAGGAGGAAACGTGTGCCAAAACTAACTTTTTACCATTTCCCGTGTTTCAGATTCATGTTTGAAACTCCAGCATGGACCTTCGAAATGCTACTTGTGTTAGAGATGTGTTTGAGTGTCATGATTGGCAAGGGAAATTGCATGTGGCGGAAATTGGAGTGGAAATACTATTTACTTGTATGACTAAGATTTGACAGCAGAGGCGCCTTTCTTTCACAGCATACGTGCCTTTTGGTACAAATAGAGTTCATTTTTCCAACAAACATTTTGAACTTTTATACTTAGCTGCTGCTTCATTGCCTAGTTGTTAGCTTCTGTAATGTACATCTGCTTTTCCTCTTGAAATTTCTCTTGGCTTTTGTTCGCAAGTATAAACAAAAAAGACAGGGCCTACCATTTGCATTGTTTAAATCCAAACAATATGTATTCAAAAGTCACAAGCTACGTGTGTTATTCCAATTTCGTACAGGACTGCATTTTTGTGTCTTTGTATAACATATCGCATTGCTCATTATTCTTTCAGATAGACCAGATTGCAC

>PGSC0003DMG400008450 LRR-RLPs

ATGCATGACTTTTGTTATATAAAATCTAGAAAGGAAAAGGTGTTTGACTTCATAGGAGGTTCAAACGCTCCGTCTTCTTCTTCTGGTGTGATGGCACGAGGAATTACCATTCGTTATGTTAAACACCTCTTCTCTCTGAAGGTGTATGATGGGATTCATGATTGTCTTTCCTACAACAGTCACCTAAAACACTTGAGGCTTCTTAAAAGTTTGGATTTGAAGGGCCTAACATTGACAGATTCTTTGCTGGATGAAATTGGTATGCTCGTTCATTTGAAGTACTTAATCATTCAGACGAAGGCAAAAGCTCTCCCTCCGTTATTTTCAAACCTCTGCAATCTAGAAACTCTATCGGTGTATAACATTGAGAGACCATGCATGATACTATCACCTTGTTTTTGGAGTCTTGCAAAGCTACGAGATGTGCGGATGATGGAATGTGCTCTCTTTGAACCAACTGTGTTAGATGAGGACTCAAGGTTAGAAAATTTGAGAACATTACAGCATCTCTCCCTTCCCGGTTCAGAAGACATGGAGGATATTTTCAAAAGGTTTCCTAATCTTCAAATCCTTAGTGTCTCTATCATAGCAGAGAAGATTTGTTTTCCAAGATTGGATGTCCTTAACGAACTTGAACAACTCAATTTATCTGCTTCTTGGGATTCATTCCATGAATATACACACGACTTCCCTTTGAGCTTGAAGAAAATGAGGTTGCAAGGGCTCGCTCTGACATCCGATTCATTGTCAAGAATTGCTAGACTACCCAACCTTCAAAAACTAAGTCTAAAAAACGCAATCATCGATGGGGGGCCAGGAGAATGGAACATGGAAGATGTCACCTTCAAGAATCTCAAATATCTAAAATTGGATAGTGTGGAATTTTCTGAATGGCAGGTTGATGCAGAGAAATCATTTCCCGTGCTCGAGATGTTAGATATAATTACATGTGAGAAGCTTATGGACATCCCAGATAGTTGTGGAGATATTGCGTCATTAGAGCTTATCAAAGTATGGTTTAGCCCTCAGCTTAAAGAGTCAATTCTTCAATATTAA

>PGSC0003DMG400008492 LRR-RLPs

CTTTTTCTTGTTACTTGCTTCATAAAATATAATGAAGCCCTCCTCCTCCTCTATACTTTCCATTCTTTCTTTTTCTCTCTTGTTTCTTTTTCAACTCTCTTCATCTGAAAGATGCCATCCAAATGACAAAAAAGCCCTCCTAGAATTCAAAAAAGGTCTAGGCAATCCTGAAGACTTGGGTAATTGGGATCCGAAAACTGATTGTTGTACTGATTGGTACGGACCTACTATCAGATGCGATGAAAAAACCAATCGTATTAACGTCATCGACTTCACCAAAATGGATATGGCCGGATACCTCTCCCCCGCCATAGGAGACCTTACATACCTTACAACATTCAGTATCCACAATGTACGTAATCTCACCGGTCCAATTCCATCAACAATTGTTAAACTGACTAATCTCGAATTCTTCAGAATTAGTCAAACAAACATTTCAGGACCTGTCCCTGAAGTTCTTAGTAAGTTGAAAAGCTTAACGTACATTAACTTGTCCTATAATAAGCTTGTTGGAACTATTCCACCTTCGCTATCTCAACTTCCACACTTGGAGTTTTTAGGATTGGATAGGAACAAACTTAGTGGACCAATCCCGGAGTCTTTTAGAAAGTTAGCACCAAATCTAACGTATCTTTACCTTGGACATAACCAACTTACAGGGATTGTACCAACTTCATTTGCTGGCTGGAGTTTTGCGACAATTGACTTGTCAAGGAACATGCTTGAAGGGGATATTTCATTTTTGTTTGGTAAAGATAAGACTACATTTGAAATGCTTTTGGATAGGAATAAGTTTGAATTCGACTTCTCAAAGTTGACGTTTGGGAAGAGATTATGGAGGTTGGATTTAAATCATAACAAGATTTATGGGAGTATTCCAAAAATAATAGCCAAGAATCCGTGGCAAATCTTGAATGTGAGTTATAATAGACTTTGTGGCGAGATTCCAAAGGGTGAGTATATGAAGAAATTTGAGATATATGAGTATCTTCACAACAAATGCTTGTGTGGTGCTCCATTGCCGCCCTGTAAATAATCATCACTAGTATCCTGTCTATGTATTTCAACTTGTTCTCTTTGAATGTTGATGAATAATTAAGCGACGAAACTCAAGGCTTAAAAATGTACTTTAATTAATTTATGAATGAATGTTATCCTTTTTTAGTAACTTCTCGATGGATGATTATA

>PGSC0003DMG400008596 LRR-RLPs

CTTTATTTAGGTAAAGAAGACTTTGGTACAGAGGAAGAACATTTTGTGAAGGAGCAAACACAATAATGGCCGATCCTGTAATTGGTGCTACTGCTCAAGTTTTGCTTGACAAGTTGCTTTCTCCCACTATTGAGGAGGTCAAAAGTTTAAGGAACTGCAAGAAAAATCTCGAAATGCTGACAAAAAATGTATCCTTGATCCAAGCTTTCATTCATGATGCTGAAAGACGACAAGTCGAAGATCAGGCTGTGGAAAAATGGCTCAAGATGCTTGAGAGAGTTGCTGAAAATGCTGATAATGTGTTTGACGAATTCAGGTATGAATCTCTCAAAAGACAAGTGAAGATTCGAAACAATCCAATGAAAAAGGTCAGTGATTTCTTTTCTCATACAGCTTTTAAGAGCAAAATGTCTCAAAAAATCAACAACATCAATGAAGAGTTGACGGCTATCAATAAGTTAGCCAAAGACCTTGGTCTCCAATCATTGATGGTTCCTCCTCGGCAAATACTACTAATTCGAGAAACAGATTCTGTAGTGGTTGCTTTGGATGTTGTTGGTAGAGACAAAGATGTTGCTGAAATAAAGGAGAAGATATTGACCATGAGAGAGGATGTTGTTCTGTCAACCATTCCAATAGTGGGTATGGGAGGTTTAGGGAAAACAACTGTCGCAAAGAGAATTTACAATGATGAACACATCAAGCAAACCTTTGAAAAGAGAGTTTGGCTGTGTCTACCTGAAATGTCGGAGACTAAGAGCTTTCTTGAACTGATCCTCGAATCATTGACAGAGAGGAAAGTTAAGGTCCAGAGCAGAGATATAATTGTCATGAAGCTCCGAGATGAACTGGCAGGAAGGAAGTATTTGCTTGTCCTGGATGATTTCTGGCGTGTTGACTCTACATTGTGGAATGAGTTTGTGGATACCTTGAGAGGAATAAATACTTCCCGGGGAAACTTCATTCTCGTGACTACTCGTATGGAACAGGTGGCGTCCACAGTAGCAGCAGTAGGTCCTCATAAGTTGGAAAAACTAGCAGAAGATCATTGTCGGTCCATTTTCAAACAAAGAGCGTTTGTTGATGGGGAAATTCCAGAGGAATTAGCAAGCATGGGCAACAGGATTGTTAAAATGTGTCAAGGTCTACCGTTGGCTGCAAGTGTATTGGGAGGCCTCTTACGCAGTAAGGAAAAACATGAATGGCAGGCAATTCTTGATGGAAACCCCCTTGTTGCAGGTGAAGATGGTAATGGAGAAAATAGCATAAAGAAAATCCTAAAACTCAGCTATGATTATCTACCATATCCACATCTGAAAAAATGCTTTGCTTACTTTGCAATGTTTCCAAAAGACCTTGAGTTTGAAAAGGACCAACTGATCCAACTCTGGATGGCAGAAGGCTTTCTTCATCCATGTCAAGAGACCACCGTGATGGAGGACATCGGGAACAAGTTCTTTCAACTCTTGTTGCGGAATTCCTTGCTGCAAGATGTTAAGCTAGATGAGCATAACAATATAACACACTGTAAGATGCACGATCTTGTGCATGATTTGGCTGGAGATATCTTAAAATCTAAACTAATTGATCCAAAGGGCGATGGTGGAGAAAATCTTTCTCATGCACGATACTTTGGATGGGACTCACCAAGTGATCAAATAGATAAGATAAATGAGCCAGGACGTTTGTACACACTGTTCTGGAGAAGCAATCATATATCGGAAGATATGCTGTTGAGCTTTAAGTTCTTGAGAGTTTTAAATTTGTCCAGTTCAGGAATCAAGGAGCTGCCAGCCAAAATCGGCAAGCTAATATACTTGAGATATCTTGATCTCTCCAACACTAAGATCACAGCCTTGCCCAACTCCATTTGCAAGCTCTACAATTTGCAAACATTTAGAGTCAATGACTGCTATTCACTCAAGGAACTTCCATATGAGATGGGAAATATGATAAGTTTGAGACACATTTACTGCTATTCTCGATCTCAGTTGCCACTTAACATGGGACAATTGACTTGTCTTCAGACCCTGCAGTATTTCAATGTGGGGTTAGAGAAAGGGCGTCGAATAGGAGAATTAGGTTGTTTGAAGAACCTTAGAGGTGAATTGACGATCAATGAGCTGGAATTGGTCTGTAGTATTGAAGAAGCTCGAACAGCATATTTACAAGAGAAATCGATTATCTACAAGCTGGCATATGTGTGGTTCCACGATGAACCAGAAGGCTGTGAAACCATTGTTGAGCATGTGTTGGATGGTCTTCAACCGCATACTAACTTGAAAACCTTAGAAGTCAAGAACTATTTAGGTACTAGATTTCCTTCATGGTTCAGAGAAGAATCTCTACCAAATTTGGTCAAGTTGAAATTAAGTGGTTGCAAAAGATGCAAAGAAATTCCATGGCTTGGCCAACTGAAATTGCTTCGGCATCTTGAGCTATTAGGGTTACATAAGGTTGAATGCATCGGAACTACTTTTTATGGTATTGATGGTAACAACATTGGATCAAGCAGCAATGTCCAAGTGTTCCCATCATTGACTGAACTAGTATTGAAGGATATGTCTAGCCTTATTGAGTGGAAGGGAGATGAAATTGGAGTAAGAATGTTTCCTCTGCTTGAGAAGTTGAGGATTAGTAGCTGTCCATTGTTAAAAATTACTCCAAGTCAATTTGAAATCCTCCGTGAATTAAGAATTGAAAGAGTTGACAGTGAAATGCCATTGTTGAACTTGTGCAGTAAATTGACATCTCTCGTAAAGCTTAGTGTTTATGATGTGAAAGAGCTCACTTGTTTTCCAGATGAGATGCTACGCAACAAGGTTTCTCTTCAACACCTATTGGTCTCAGACTGTGGAGAGTTTCGTGAATTGCCACAAAGCTTATACAGTCTCCGTTCTCTTAAGAGCTTAAAGATTTACCGCTGCACCAATTTCAGTTCCTTTCCTGTTCCCAGTGGAGAGAATTATTTGACTTCCCTCCAAAGTTTTCAGTTATGGAACTGTGATGGATTGACCAGTTTACCAAATGGGATGCTAGAGCATTGTCGGTCTCTGGAATCTTTGATGGTCTGCTACTGTAACAACTTAGTTTCCTTCCCTATACGTGTGGGGGAAATGCCTTTACTTTTTGATGTGTATCTATCAGAATGTCCCAAATTGATTAGTGTACCCGCAGGGGACCTTTACCGTCTCACTGGGTTACGGCATTTGGAAATTGGTCCTTTCTCAGAGATGGTGGATTTTGAGGCATTCCAATTGATTTTCAATGGCATTCAGCAGCTGTTGTCCCTTCGTTTTTTGTGTGTGGTAGGACATTTGCACTGGGATTCTCTGCCCTATCAGCTTATGCAACTCTCTGACCTAACAGAGATCCAAATATATGGATTCGGAATCGAGGCTCTTCCTCATAAATTTGACAACCTTGCTTCTCTTGAAAGATTAACGTTAGTGAGGTGCAAACGGCTACAACATCTGGACTTCTCAATTGCCATGCTCAAATTACGGCATCTCCAGATACAGGATTGTCCATTGTTAGAAACTCTGTTGTATGGGCTTGGCAACCTTGTTTCTTTGCAGGAGTTAGTTTTACAGAAATGCGAGAAACTAGAGCATCTGCCATCCGGAGATGCCATGCAACGCCTCACCAAATTACGGTACCTGAAAATTGAAGGCTGCCCAAAGTTAGAAGAAAGTTGCACCAATCGGATTGGCTCAAACTCCCAGTGGTCCAACATTTCCCATATTCCAAAAATTAAAGTAGGTGAAAGCATAATTCAGGACTTAAGTAAGTCTCTGTTCCTCTTTCAGTTTCTGAGTCGATTTTTTTGTTCTTTCTTTTCCACGAGTTACTCTCCTTCTTTTTAACAAACTTTTCATTCCCCAAACTAATACATGTTATGGTACTGAAGAATTGCAGATTTTTTTATCACTCACTTGAAACTAACTGTTCTTCTTCTTCTTCTTATCGTCATCAAAAGAGGAATATATACATTCCCGTCCTTTAATAATGTAATATGTTCCTTCTCATAGGTAACTAGATCAAATGTCATGCTGGAACGTTGCTAGCTGTGGTATATCAAGTCAAAATTTATGGTTGGTAATCAGGCTGGCTAAAATTTTCATTTAGACTTGGTAGAGGAATGCTATAACAAATTTGTCGCTTCTTTTCACTGGTCATTATTGTACTGTTTAACTAGGTTGGCTGTGTTTTGTATGCATGCAATGTAAGCTTGGTTGCGTTTCTAGCACCATGAGGGACGTTCATCCATTTGGTTTAAGGATTCTCTACTCTCATTGGACCAAACATCTCCATGTAAGCTTCTTCCCAGTTATTAATCTCCTAGATCCCATAGTTTCTTTTGTTAATCGTGCAGCAAAAGAGTATCTTTTAATCCTCAAATATTATAATGCTCTTCGATATTGGGTATAGTGTGTTAGCATGTTGGACAGATACTAATAGTTGTACATATCTATGTAATAGCTCTAAATATTTGTGTAATTACTGTGTCAATTGCTTTCCACATGTAGGAGATATCAGGGATAAATGTAGCATATTTTTATAGATATTACAGATATTTAGGGGTGTAAGTGGCAACTTCTAATTTAATGGAAGAGCTCTTAATTTGAGAGCATTCATTCAGCAAATTCGACTCTGGATAAAGACAATTTCTCTTTCTCTTGAGGAAAATGAAGTATTTGATTGACAAAACATTGGCTTATCATGGTTGACACAATCAGTTTCTTTTGCATAAGAGAAGGTGAAACATAGAGGGGGGAAAACGACCTTCTGCAAGACAATTTTATTATCGACTATCTGGTGAAATAGTATAGGTAACAAAGCCATTTCATCATAAGCCTTAGATGCAAGACAGATTTCATTTTGGTAAAAAATGTGACTTCAGTTGATAGTAGAGGCTTTTCCAAATTTTGTTGACGGTCTTCACTCTTCAATAGTATTTTCTTTAATTTTCATGGGACTTATCGGAACATTGTTTACATTGGTGCCTTTGGGGACGGTGATTGCACAAGCTGATTAGGTCTTGTTTTCTCCTAAAATTTCTCTTTTTAGTGGTTTTTTTTTTTTTTTTGGTTGAAAACAGAGTAATATATATGTTATGTAATATGCATCGTTACATTGTTACCAGCAAGTGGCAAAACAGGACACCTACTATTGGTGTTCTGAGGAAGGGGCATACTATTACTAGTACTATTACATGAGTTAATTACAGAGGGCATACTATTACTAGTACTATTACATGAGTTAATTACAGAGGGCATACTATTACTAGTACTATTACATGAGTTAATTACAGAGGGCATACTATTACTAGTACTATTACATGAGTTAATTACAGAGGGCATACTATTACTAGTACTATTACATGAGTTAATTACAGAGGGCATACTATTACTAGTACTATTACATGAGTTAATTACAGAGGGCATACTATTACTAGTACTATTACATGAGTTAATTACAGAGGGCATACTATTACTAGTACTATTACATGAGTTAATTACAGAGGGCATACTATTACTAGTACTATTACATGAGTTAATTACAGAGGGCATACTATTACTAGTACTATTACATGAGTTAATTACAGAGGGCATACTATTACTAGTACTATTACATGAGTTAATTACAGAGGGCATACTATTACTAGTACTATTACATGAGTTAATTACAGAGGGCATACTATTACTAGTACTATTACATGAGTTAATTACAGAGGGCATACTATTACTAGTACTATTACATGAGTTAATTACAGAGGGCATACTATTACTAGTACTATTACATGAGTTAATTACAGAGGGCATACTATTACTAGTACTATTACATGAGTTGCTTGCAAAAGATATCTTAGATAGTCTAACAAATGGAGTTCCTTCTTTGTCTGCTTCTAGTTTGCTGGATACAAACACCGGAGGAACTTCCATGCACAAAAAAGAGTTGATTTGATCCATTTTAGCGCCTTCCTTTGCTAGAGCGTCTTTTTAGTGGTTACTTGTAATGACCCGCGTCTCCATTTCCCAAGAAAGAACTGTGTGCCAAAACTGACTTTTTTCCCCTTGTGTTTCAGATGGTTGAACAACGGCATGGAAAAAAGACTATGAAGTTTCATAGTCATGATTGTCAAGGAAAATTGCATTTCGTAAATTCATTTGCATTTGGCTGGAACTGGAATTGAGATACTATTCTGTATGTGGAAAATTCTGGACTAAGATCTGTCAGCTGAGGACTTTCGTGCATTGTGTACATGCTACACTTGCCTTTTCTTGTATATAGAGATTTTGTCTCCAGCAAACAGTTTTTTAATTATTTTTTTAACGTTGTGTGGGGTTAGGTTTGACCCCTACCACGTCTGTAACATCCAAAAAGTTGCAAAAGCAGAAGGGGACATGGACCTAACTTCCAGCAAACAGTTTATATTCGCTTAACTGTTGTATCATTGTCTAGTTGATAGCTTCTTTAATGTATCTCTGTTTTCCTTTCGAAATTTATCTTGGCTTATGTTAGTGCATACTATGATGGATTAAATATAAAACATGTGTTCCAAATCAACATTTCC

>PGSC0003DMG400008925 LRR-RLPs

ATGGATCCTAATGAGGTAATGAGAAGGTTACCTAATCTTCGGAAATTGAGATGCATATCAGTTGACTCGGAGGACTGCTGTCATTTTCCATCACTGGAGTGTGTGAGTCGACTTGAATCACTCAAGATATTACACTATGGTCAGTCATTTTCTTCTTGTAAGTTCAACTTCCCCTCAAATCTCAAAAGACTGACATTGCCAAAGTTTCAGCTACCATGGACTGAAATTTCAAGAATTGGTAGCTTACCACACCTTGAGGTTCTCAAACTACTTGCCCGAGCCTTTGAGGGTGTGCAATGGGACATGAAAGACGGAGAGTTCCTGAAACTAAGATTCTTGAAGTTAGACTCTTTGAACATTGAACAATGGAACATCTCATCTGATTGCTTACCTTCCCTTGAGCATCTTGTTCTGCAAAGTTGTAAGAAACTTAAGGAGTTCCCTTCATGCTTAGGCGATGTTTCAACGCTACACAGTGTTGAGCTACACTGGTGCAGCCGTTCTTCTGTGAACTCGATTATCTCAATTCAAGAAGATGAGGGGTTCAAGGTCCATGTATACCCTCGAGAATTCGATTCCTGA

>PGSC0003DMG400009029 LRR-RLPs

GGTTATCCCTCGAGTAACAAGACAGTCGAATATAAAAACACTGTTCCCACAAAACACACACACATTGTTTACTTCACTTTTTTCAGTTGAGATTTTCTTCACTTGCACTCTCTCCGGCGGCGCGTCGAACTCACACATTCTCCCTTGTTGTTTTCCGGTGAGTTACAACAGACACTCACTCCAACGATAATCAGAAAAATCCCAAAAAGGTTTTTTCTTTTCTCTTGATTTCTAGGGTTTTTGCGCTTATGCTCTGTACGAGTCTACTTTTTTGGTCATGTGATTACACAGATCTGTGCTGGATTTTAGTAATTTTTCTTGTTTAATGTGCTTTCGTTTGTGGTTTTAGGTAGTTTTTTTTTATGTTTATTATTTTGGGGTTTGGTAGAGTAGGTTGCAATTACTGGGAGGAGGACTACAAACTTCACTGTTAGGTATAAAGTTTTTATTTTTCTCGTGAGGTTTGTTTGTTTGTAGGGGTAAAAGAGGGATTTTGTTTTGTATTACAAGTTTCTGATGTTTTAGCGGCTCTGTAAATTGTATATATGCTGCTCTGCGCTGCTCATATTATTCTTCGGTAATGTATTGTGGGGTATTTTTTTTGTATCCAAGAAAGTTTTTATGTGTTATAGTGTGGTGAAATGGAATACATCTGTTGAGTATACCATTTTTGGTTTTGAGTGTTTTTTACATTGTTGGATATGAATGTGGTGCATTGCAAGGTGGTTTATAAAAGAATTAGAAAATTCAATCTCGTTTTCGATGATGGATGATTGTAGAAACTTTGCCATTCTTTGGACACCTGCTTGGTGTTTTGTGGTGAAACTATTGAGTTGTCAAACAATTATATCTGATTTCTATATGAGGAAGAATTTGAGCTCTTCAAATTTGGTTTCGATATATGTTAGTTTTTCTTACGTGGAGTAGGTGTGCACGAACTTGTTCTTGCCTGGTACTTTGTTTTTGGTTGTTATGTTCTTAACTGCCAGAGATGAGTCTTGTAGTGGTTATATTTCTTGAAAGATAATCACTTTTATAGAAACTGAGACATGATTCACTATATTCCTTTGATTCTAGGTTTTCTTCCAATGGATGCCACAACAGCTAACTCTCAGCGCAGACCATTTTCAATTAAACTGTGGCCTCCAAGTGAGAACACAAGAAAAATGCTGGTGGAAAGAATGACGAATAATCTTTCTAGTCCAACCATTTTCACCCGCAAGTACCGCAGTCTATCCAAGGAAGAAGCTGCCAAAAATGCTGAAGAAATTGAAGATGCAGCTTTTACCATTGCTAATCAACATTATGAGAAGGAGCCTGATGGTGATGGAAGTTCTGCTGTGCAACTTTATGCCAGGGAATGCAGCAAGCTTATCCTGGAAATTCTAAAGAAGATACCCAAATCAGAGGACAAGGAAATTTCAATTTCTGAAGTGGTTCCTACTGTTCAGGAGACCTTTTTTGATATCTCAAAAGGTAAAAGGGCTTTTATCGAAGCGGAAGAGGCTCAAGAACTTTTAAAGCCATTAAAAGAGCCTGGAAATTCTTACAGCAAAATTTGTTTCAGCAATAGAAGCTTCGGTATAGATGCAGCACGTATTGCAGGCCCTATCTTGGCAGCCTTGAAGGATCAATTGAAGGAAGTTGACTTGTCAGATTTTGTTGCCGGAAGAAATGAGGCAGAAGCACTTGATGTCATGAATATATTCTCAGAAGCTCTGGAAGGTTCTAACTTGAAGTTTCTGAATCTCTCTGATAATGCTCTGGGTGAGAAGGGAGTTAGGGCATTTGGAAAGCTCTTTCAGTCTCAAACCAACTTGGAAGAACTATTTTTGATGAATGATGGGATTTCACAGGAAGCTGCAAATGCTGTTAGCGAGCTAGTTCCTTCCACCGAGAAGCTTAAGGTTCTTCATTTTCATAATAATATGACAGGTGATGAAGGGGCTGTTGCGATTGCGGAAATTGTGAAGCGTTCTCCTTTATTGGAAGATTTCAGGTGCTCTTCTACCAGGGTAGGCTCTGAAGGAGGGAGTGTCTTGTGTGAAGCACTTGGGATGTGCTCTCATTTGAAGAAGCTTGATTTGCGAGACAATATGTTTGGTCCAGAAGTTGGTCTTGTGTTGTGTAAGGCACTCAGCAAACATGAAAATCTTACGGAAATTTACCTGAGCTACCTTAATTTAGAGGATGAAGGAGCAATTGCAATAGCTAACGCTCTTAAAGATTCAGCCCCTTCACTTGCTGTCTTGGAGATGGCAGGTAATGATATAACTGCTGAAGCTGCTTCAGCAATAGCTTCTTGTATAGCTGCAAAGCAGCTTCTTGCCAAGTTAAGCTTGGGTGAGAATGAACTCAAGGATGAAGGTGCAATTCAGATTGCCAAGGCACTGGAAGGACACAGCCATCTGATTGAAGTTGATATGAGCAGCAATGCACTAAGGAGGGCTGGGGCTAGGGTGCTGGCTCAAACTGTGCTGCACAAAGATGAGTTTAAATTACTAAATGTTAATGGGAATTTCATCTCAGAAGAAGGCGTTGACGAGTTGAAAGAGATCTTCAAGAAATCTCCTGAAATGCTTGCATCCCTGGAAGATAATGACCCGGAAGGAGAAGACGAGGATGATGAGGAGGATGAGGAGAAAGAATCTGGAGATGAGGGCAAAGACGTTGAGGATGAACTAGAATCAAAACTCAAAAACCTGGACGTCAAGCAAGAGGCAAGTCTTGACACACCAGATAGCAATTAGAATCCAGTACTTGACGACTTCTGCATGTTGCAGTTTTAGTTTTACCATTTTCTTTATGCTAAGCTGGCCATATTATGTTCCTCCAGCAAGTGCTCTAACATATAGTTCCATCAGTAATTTAAATTTGGTGGAATTGAACTGTTGCCTTAATTGGTCAAGTTAGGTCTCTGGCTGAATTATGTAGGGTACGAGAGAGAGAACTGCTATTTATGTTGAACCTTCTTTTGTTATGTAGACTTGATAATCTCCTCTTTTTTTCCTTTTCTTTTTTTTTATATTTCCATGATGCATTTGACTTGTTTTATT

>PGSC0003DMG400009230 LRR-RLPs

CTTGACTGGAACTTGCACACCAAAGACCATTTCTTCATTCTTCACGCGTATTCTCTCATGAAAAACTCTATTTAGACTAATCCTCCCACTTTGTCCTTTCATCATTTTTTCACCATGGCTAAATCTACAATATTTTTTCTCCTTTTTCTTTCATTTTCAGTGTTACTTCACAATGTTACTGCATCAGATGATATTAGGTGTTTGCAAGGACTGAAGGACTCATTTAAGGATCCTAATGGAAATTTGAATTCTTGGAACTTCTCAAACTACTCCATGGGGTTTATCTGCAAGTTCATTGGTGTCGGTTGCTGGAACGACCTCGAGAATCGGGTGCTCAGCCTCTCACTTCCAAACATGAATCTCAGTAGACAGCTCCCAGATGCCTTCAAATATTGCTCCTCTTTGACTTCTCTTGATCTCTCTGGTAACAAGTTCTCCGGTCCCATTCCTTCGGAGATTTGTAGCTGGACTCCTTTTTTAGTAAACCTCGACTTGTCCAACAACAGTTTCTCCGGCTCTATACCTGCTGAACTTGGAAACTGCACCTACCTGAACAAGTTGATGCTCAATAACAACAAACTCTCTGGAAATATTCCACCAGAAATCTCTCGATTAACGAGGCTTAAGGTGCTCTCCGTGGCCAACAATAATCTTTCTGGTAAAATACCACCATTTTCAGGATTGTCTGATTTTGAATATGGAGGAAACAGACATCTCTGCGGTGGACCATTAGCCAAATGTGGTTGAAGATGGCAGAAAGTTAGCCAAGTTTTGGCTAATTGTATCTTTTATAGCCTATCTAGATTTCTGTTTCCTGTCATTTCATAATTGTATTGTATGGTGAAAATGTCTACAATTCCATGCCTTTTAAGTTAACAAATGCAACACTGGCTTCTTCCTGTGTCCATTCCACAAGTTATTGTACACAACACAAAGGAACAGAAATGTTAATACTCGAACTATCATTGCCTTGCTCAAGACTGTTCTTCGCGTACAGAACAATTTATCAACAGAATACTAATCTACCATTTTTTTTACCCACTATACTACATCTAGGTTCCCACCAAACCCAGAAATCTACAAACTAGGCTGAAAAATTGAGTCATAGAAATGGCCGAGAATAATCAACAGATATCAGACATCTCACCAAAAGAATAAAAACAGAAAACACTCTCAAGCTGAAATTGATAAAGGGGAGATGACCGGAAAATCCAAAATCTGGCACACCTATTTTTTTTGGACGGAAATGAACAGCCATAGAGAAGTGCCTCCAACCAATCATTTAAGTTCTTTTGCGCTACTTGAAATGATCCTGCCTGATTATTGCAACAAACTCTGAATGCAAACTGTCCTCTTTCTGCTTCCATATTTCTCCGTTGGCATGTATTAACCATGATATTTGAGGGGAAAGTTGTGCGTACAAATCTGCAAGTAATATATGATATTTAAACTAATCAAAGACGTGCCCAAAAGCAACAAATTTCTGGTTGCAAGAACTGCAACAAACAAGAATATATACATTTAGAATGTCTAGACTATGATAAGTCTTCTATTAATGCTCTACAAAGAGAGTTATCAAGTATCACCTAAGTTTATCCACACCTATTACTACTTCATAACAACAAAGAGGGTTATCAAGTATCACCTAAAACTTTTATCACAATAGTCAGTATGTGTAACCTTTTTGCATTGTTGCACAGACTAGAATGATAAGGCACATGGATTTTTCTGATCAAGCTCAGTGTAAATACCTGTACTACTAGGAGGAGGGTCTGGATCTCTTGTACTATGAAGAACCAATGTACCAGAGAGCACAGTTAAAAAGCCACAAAGCGCTGAAACTATGTTGCTAACACTCTGACCAGACCAATCCTGAGAGAAAGACATGACATTAGAAATGAAAATGAGAGAAGGCCTCCTTGCATCCCGAACATAAACTTTCAGAATACCTTAAACATTATGGCACTGGCTAAAATTGTAAGTGATGTGAACATGGCATAATATATTGGAGAAACCACAGCAGTGTTAAATGTGTCCAATGCCTGCAAGAGTCAAATTCAGGCATTTGTCCAATTTACAATACAAGTCTAAGCCAAGATTCAATAACATGTAGTTTCTATAGTAATCCCTGTAAATATTAGCTCAAGCATTTGAACAAATGCCAATGAGTGTCAGTTCATAAGTTTTCCAATTACAATTGTACTTGTAAAAAGAAATGGATCTTGGGTCTTACTCAACCTCAAAAGATAGCTCAAGAGATCACCCATGTGGTGAGAATTGTTCAAGCCATATTAAAAAGATCTCTTCATACACTACCTCACGCCCAAGAATGGAGATCTAGTGTATGGGCAATTTAATAAGAGGAGCGTTGAAAAATGCTCTTTTACTAATTGTTGGTTTTTAGCAGAGAAATTTTGGAAAATAACAGAGCAAGACTGAGAGGAAAATGCTGCAATGCTATCAAAGGGAACTTTTTTATTCAGCTTACTCTACATTTACGGGGAACTTGAAATAAAAAACAGTAAAAATATGATCATTATCCTAGCCTGGAGTATCTTCATACAATCATATCTTGCCTTCACCACTATCCCAGTTAATATGGCATACTTTACTTTTTAGTCAGTCACAAGTCACAACAAAATGACACATTTCCTTATTTGGCAAATGTTTAATGACATTATCAACATTTTACCCATGCTGGGTCACACTTATTTGGCAAAAGGTCCACAAAAGTGTATTCTTCTATTTAAAAATTCATTTCTTTTAAGGGTAGTTTTGGAACATTGCAAAGTCTTCTCATATTGCATAAATGTTGTGCCAAGTCAAACTAATATCACATAAACGGGAGAAACTAATTACTAGAAATTAAGGCTAGCTAGCTAACAAACTTTAACTAAGATTTATTCAAAAACAACACAGCTAAACAAAGTTACTGTAGTCCAAAAGCATAATTCTAAGGGGTCGTTTGGTTTGAATATGAGTTATGATGGGATAAGTTATGTTGGGATAAGTTATGTTGGGATTAGTTATGATGGGATAAGTTGTCCTGGGATTATTTCTTATTGAGTGTTTGGTTTGTTATATTCAAAATAATATATATTGATAAATTTTAAGAACAAGTTGTTTGTTTACAAAATTACCCTTCACTTTATTTAAACTTTTTAAATATTTCTTTGTGAGCTTTAGTTATTCATTTTTTAAAATAAAACTTTTATTTTATTTAACTTAAATATTTTTGTTTATGTATTCCCATGATAAGTACCATGTGTATTTAAAAAAATAACTTTTTTTCTTATTTAGCTTAAAAATAGAACTTCCCTCTTAAGTTTATTAAAGAGAAGAAAATTTATGAGAAATCAAAATAAAATAAACATAAGAATTAGTCAAAATAATTCATAAATAAAAATTATATTTATATAGTATAATTTTTTAATAGAAAAATATGTATAATTATCATAAAAATAAGGAGTAGTAAATATTATTATACATGGTATTAAAAAAAGTGTAAATATACATAAATGTAAAAAATGATGTATAAAAAGGATTTAAAGGGACATATTTGTCATTTACCTATTTTATCTCGGGATAAGTTATTCCGGGATTACTATACCACCCATGGGGAGGGATAACTTATCCCGGTACTAATTATTAATCCCGGGATAAGTTATCCCGGACTTGCCAACCAAACAACATAATTTTTTTTTTATCCCATCATTATTTCCTTATCCCATCACTATTATTTCTAATCCCTCACACCAAATGAGCCCTCTAATAGTACTCTCAGACTTCAAACATCACTGCATACCAGTGTTTTCAAAAACGTTTTTGGGGCGAGTCTCAGGGCAGGGCGTACCAAAAACGCTCCGGAATGTAAATCAAAGACGTAGATCCATAAGGCGTAAGTTCTAAAATTTGAGGCATAAGCCTCGAGCATAAAAACGTAATTCCTGGGCGTAAAAACGTAGGCCCGGGCGTTTTTAATTTTTTTTAAACCTTTTTTGCTTTAAATCATATATTTTATTATTGGTGTAATGATATTTCTCAAATAGTAATTATTAATACTTTTTTTCTTCATTATTGATTATTAATATTTATCTCAAATAAATATAATTTCTATATATTATAGGTTATTTATAAAAAAATTATTTGTAAAAATCATTGAAAGATTTACTTTTGCTTATTTTATTAGTAATAAAACTATAAAATTGAATATACATGAGACTACGCCCCGTAGATTTATGGGACTTACGCCCCATAGATCCATGGGACTTACGCCCCGTGTCTCGGGGCTTACGCCTCGCCCCATACTGTATAAAGCGCCTCGCCTCACACCCCCGCCTTTTAAAACACTGCTACATACTATAAGGAGAAGTAGAAAGTCTACCTTATTTAAGTAATTTAGTTGAGTAATTATACATGTAACAGAAACCATCACAAACACCCAAGTTTGAAAATTTGCAGCTTGGTTTGAACCCTCCAATGTAAGCTTTATAGCTATACCGATAGCTTTGATGCTCATAACCTACAACAGAGCAACAAACAAACTTAAAATGACTGCTACATGCAATTTTTACCAATTGTACTTATGAGAAGGACTAGTTCCTATATTTTCTTTACTTCTCATGAGGAATAATTTTATCATATTTCATTGGCAACAGAAGCACTCTCTCTAAGTCAGTTCGGCAAAGAGAACTGCAATAGTAGCTCAATTAATACAAAAAAATTAAGAGAGGAACCAATTCATAAACTGTTATTTATCGAGTTTTTTAAGCAAAATACATGAGTATTATTCAAAAGATAGTGAATTCGATTGATAAAGAAGTCATAAAATAACTCGACCTCGAACTAACAATGACTAGATAAAAGCTTATTTCAACTAGCAACTTAGACTTATTTTAAATTTATGTAAAAGACAAAAACTAATAAAATAAAGTAATAGCACTGCAATCTTTTATTGAGTTACACACCTACTAGTCAAGGAAAAACAACAGCAACCCTGGATGCCTTATAAATTATAACAGATCAAAGTATAGTCACTTGTACAAGTATAGTGGCAGGAAAAGACATCTACTAGCATGCAAACACGTTACAGCACCCTAAAGCTCTGGAGTAATATTTCAAGTTCCATACATCCATTTATACCTAAATGAGAAGTTTTCTTTTACTTCAGTGACAACTAGCAATGGATAAGGACCCAGGACCAGTTTGACTCTTCACAACTTAAAAAATATGATCCATTTAGTGGTCAAATTATGCTATTCCAACACCAAGGAGAAAGATAGAATTTGGATAACGTTAAGTTTAAAAGATAAGGCGTCTTGCAAACTGTCAATTTAGCTTCAAGGTTAACTCAGTCAGTTCAGAGCTACAAACTAAAGCACGGTTGTGACACATGTTTCTTCCCTTCCTTCCTTCGCTTTCTTAATTCTTGAGTAGAAGTACAGACTCTGTAGGGATATAGATCAATACAATAGAGGCACTAACCGTCAAGGACCCAAAGATGGAGCAAACACCAATATAGACCATAATATTTGTCCGACCATAGCGAGGCTCACAATACAAGACTAGTACCAATGTTATTGCAACTGCTGATGCAGTGTACAAAAGAAAAGCTGCAAAACAAAGAGTGGAACATCAAAAAAATGATATATCAGCAAGGCAAATAAGAATGATCTTCAAGGAAATACAGCACAGTACATTAGCACACCTGGTTGTGTAGCTAATGCCCATATTTCATCCACTGAATTGAGATCATGTTCACGAGGTGCATGAAGAACAATGACAACGGAACCTACTATACATAAAAGACATCCTAATACTCCCAGTTTCTTCAATTTTTCCTTCAGTATGAAATGTGCTAGAACAGCACTGCAGCAAAGCGCCAAATTATATGGAAGAAAACCGAATAAGCCATTGCCCAGAAAATAATAAGGCCGAGTGGAAAATTTACCTCACAATTATACTTAAAGCTCCAAGCGGTGTCACAAGAACTGCTGGCGCATAAATGTAAGCCACGAAATTTGCAAATTCTCCAACAATCACTGCAAGGATTATGGTAAACATTGCACTCTATATTAGAATAACAAATTCAGGAACATATTCTCTAACATTTAAGAAGCAATTCAAGACTTACTAGTTACCATCCCAATCCACCAAAGAGGCTGCCGTAGATAACCATATCCGCCCGAGCCTTCAGCAAGAAACAAGTATCTTATTGTCAAATAGACAGAGCAACAGAGTTCATATAATCAGCTTTTAAACTAAATTCTATTTAATGCTGCTACAGAAAAGCATCTTCTATATATAAACTTTACAAGATGTCCCCAATATATCTACTTTTAATGTGTAAAGTAGCTGATAAATTACAATGTCAAGGAAATTCATTCAGCTTATTACAGTGCAGAATGATTTAAAAAATCCACTACACTGATCATTTTCAGTTCTTAAAGTCAAGATCCATTTATTATGTCTTGAGCTCATTTCTTCTCTGTAATACATGTATTAGTACTACCTAATACAGCAGTCTGTGTTTATTCGGAAATGATAGATCAAATAATTTGGAATTACCAACAACAATCATTTAGTGAAGCAGAAGCATGAGTGCATATATTTAAATTGTTGGCCCATTCTACAAGCGGTAACTATCAAAGGTATTAAGTTTAAACAGGTGCTGCTGGAGTCGCTCTGATTGCTAAAACAATGTCATTAGTCCTAGATAACTGCCACCGAAATCTAGGCAGCTGACTGGTCAACATTCAGAAGAATATGAATTCTTTTGCAAATACTTCTCCAATATTATTCAAGGACTACTTACTTGCATGATAAGTGTGAAGTGTCAATTCCAATATAAGTAACTTGATATTCATTTTCCCAGAAATGTACACCCAATACTTCCCTTTCGGAGAGAAGTGCAAATTCCATTCCTTGAGTTCCAGAACAAGTTTTCTTTATTCCCAAACGGATAACTGCTTTGTAAGATGCAATATCACCTTTTATAAGCAGTCATTGTGTATTAGTAATACTCATTGTGACTCTACATTTGCTGGAAGTGTTACCCCATTGCTCTTATTTTACAGTCGGGCACCTACAACATGGTGTACCAACAGAGCATAACATAATCTTAAACTGTACTTAAGCAGAAAACAAATGAAGTAAGAATTTTCAGGTGTAAAACATGTGTTTCACCCGCAAAACTAAAATGGGGACAGCAGAAAAGCAGGTACCACCATACCAGTACAAAAGCAAACCCAACATTATCAGTCTGTTACCACTTGATATTTCAAAGTAAGAAGTAAAAGAACAATCTATTCATCAAGCATAATCTCCAATTCCATACATCTATAAACAGCATCTGATATATGGTAAATTGGTAATGCAGTAAAATGGTTAGAAATATCAATCTTCAATCGACTAATTTTGACTTGATCATGTAATAGTCGGTGTCAGCTTTATGAGTCCACTATATTCATTCTACATATCTGGGCCCATTTCAATTTCAATACTACAGGAATTTTATATTAAGAAAAACTAGGGGTTTTCTAGAGTTAAGAGATTCTCCAAATGTCACACTTCTTTTCTGTACACTAACATACAAGGTTTTAACCTTATTCCCAATTAGCTAGCGTCATCTACAATAAATCATTTACTTCAACTGTGATTCTTACCTAATTTGCCTTAGTTGAGAGATTGTAGGGGTTTTGTTGTCATTGTTAGGATCGAAATTACCAAGTGTCATGAAGAAGCCAATAAAGCAAACATTGGACTATGATAGTATAACAAAGAGAAATATACCAAATACATAAAAATTTAACATGAAGAATTCACAAAGATAAAAAAAGAGTACAAACATATAAAGAGAATAACCTCACAAAATTCACTCGTAAAGAAGTTTACTTTTGCCCTAAACAAAATTCTCTTGAGACTACATTGTGCATGCTCTAAATTGAGAAGGAAATCCTCAATGAATAGAAATTCAAAACATGTTCTTCCAAGAAAAGGATTAGCCAAATATGGAAGATTCATACTTTCCTGATAAAATCTAGTTATCCTAAATGTTATATCCAAATATGATAAATAAATTAGGGCAAACACCTACAACAACCTGCGTTTGTATAATTTTAAGCAAGGTCAACCTTCCTAACCTTAGCAAGAAGTAATGCATTGCAACACACACATATAAAGAACTGAGCATATTAAACAACTTTAGCAAAACTACAAGGTAGAAGAAAACACACAAACATCGGCAAACTGAATGGGGACACAACTAAACCTCCATGAATCCACCACTAGAAGCATTGACAAAGTTCAAAATCAACAAAAAGCAATAGATTATTGACAATTAGACCTAAAAACACTTTGCTATCAAGAAAATGAATTACTCACTGGCCCGAGTACCAGAGGCGCCTGCCTTCTGAAGACCCTTCTTCTTGATAATGAAACTGGAACCAATAAAAGCGCTAGACGCCACAGCCAATCCAAAACCAGTCAAATTAGCACTGTACATGATCCCAATATTATATCTAAACTATTGGGCAACAATTTTGGAATTTAATTCTCATCTGAACTTTCCCAAGAACAAAACTTTCAACACCCAAAAGGTTCCCCTGGATATACTCAACAAGACAACACTCAAGTAAGACTAAAATAGCAAGAAAAACGAAATGGGCAAGTTTGAAATACTTAAAACCGCTGTAGCTTTACTCAATTACCCTATAATTGTAGAGCTGAGAATGAAAAAACAGCAAGAAAAACGAAATGGGTATATGAGAAAATTGGCGAAAACAGGTTTGAAATGGTGAGCTAACAGTGCAAGCTCGTTATAGACAATAATGGAGAATAGTATTACAATAAGCATAATTTTGTCATGCTTTGATTCGTTGCTTTGGGTTGATTTGGCTATGGTATGTCTTCGACAATTGGGCATAACTATGAATAAATC

>PGSC0003DMG400009324 LRR-RLPs

AGGAGACCTAGTAAAAAAGAAAAGAAAAAAACATGGCCGACGAAGTTCAGCCAGTCAACCAGAGGTACGGAGAATTGTTGGTATATGCAGTTATCATGAAATATATAAAGTCCACACACTGCTCAGCTGAGCAAACCAGACACTTTGAATTGGTCAATATATCTTTACTGGATTCAATACATGGCGGAAGCTTTTCTTCAAGTTCTGCTAGAAAATCTCAATTCTTTCATAAGAGGGAAACTTGTATTGCTTTTCGGTTTCGAAAAGGAATTTGAAAAGCTGTCCAGTGTGTTTTCCACAATTCAAGCTGTACTTCAAGATGCTCAAGAGAAGCAATTGAAGGACAAGGCAATTGAGAATTGGTTGCAGAAACTAAATTCTGCTGCGTATGAAGTTGATGACATATTGGGCGAATGTAAAAATGAGGCAATAAGATTCGAGCAGTCTCTATTAGGGTTTTATCATCCAGGGATTATCAGTTTCCGTCACAAAATTGGGAAAAGGATGAAAGAGATAATGGAGAAACTAGATGCAATTGCTGAGGAAAGAAGGAAGTTTCATTTCCTTGAAAAAATTACAGAGAGACAAGCTGCCGCTGCTACGCGTGAAACAGGTGTGAGTACTGAGTAATAGTAGCTTAGTTTATATTCAATTTGTTACCACATCGTGTGTTCACCGTGATCTCTACAGTAGGATGGCAATGGGGCGGGGCGGGGTTGGAGGTGTGCAGGTGTGGGGCGCAATCCCAACTTTGAGTCTGCACAAGTAGGTACTTAAATTTGTATAGAGTTAAACAAGTACACACGCATCCTACTTGGTGTCCTACATGGCTCTTATGTGTATTATGTCATATAGGATGCATGTGTCTATTTGTTCAATTTTATACGAGTTTAAGTTCTACTTGTGCACACCCAAATGAAAGTTGGAGGGCGTAGATGTCAGTTGATACCAAGTTAAAAATTCATATTAATGAATTATGCCTTTAAATTATGATTCAATTTTATATCAGTCTGTCCAAAAGATGTTCTAGTGAAAGTGTTAAACTTAGCCTGGATCTGCTATTGGAAGTGTGAATTTTTGTGGCACTAAACAATGCAATGGGTCTGGATTCATTTTTGCATTAACATTTGTTTAGACGATTTTCTTTATCGAATTTTACTGTCTAAAAAGGAAAAAGCAAAGAAATAAGACGTATACAGAGGCTGACTTCTTCATAGGATTTCTCATATAAAAAAAAAACATTGATTTCTAGGATATGGGTTATTTTAAATTACAAATTTGTGAGTTAAAACAGTTCTGTTGGGAAGGATTTAGATACACGTGGATAGTATCTAGAAGTTTTTTAAATAAAAAATTAGCAAATTATGTGGGCTGGGGCGGGTTGAAAACAGCAAACTTTGCAAGGCAGGGCGGGTTGAAATCATTGCAAGTTTGTGTGGGTTTGCCCTGCACCACCCCATCTGCCATTCCTGTCTAAATGTTTGTTTTGTCTATATAATTCTTGCTGACTCATTCTAATGAGCTCAATTGTAACAAATTCTTTGTGTCCACACTACTTGGAACAGGTTTTGTGTTAACTGAACCAAAAGTCTACGGAAGGGACAAAGAGGAGGATGAGATAGTGAAAATCTTGATAAACAATGTTAATGTTGCCGAAGAACTTCCAGTCTTCCCTATAATTGGTATGGGGGGACTAGGAAAGACGACACTTGCCCAAATGATCTTCAATGATGAGAGAGTGACTAAGCATTTCAATCCCAAAATATGGGTTTGTGTCTCAGATGATTTTGATGAGAAGAGGTTAATTAAGACAATTATAGGAAATATTGAAAGAAGTTCTCCTCGTGTCGAGGACTTGGCTTCATTTCAGAAGAAGCTCCAAGAGTTATTGAATGGAAAACGATACTTGCTTGTCTTAGATGATGTTTGGAATGATGATCTAGAAAAGTGGGCCAAGTTAAGAGCAGTCTTAAATGTTGGAGCAAGAGGTGCTTCTATTCTAGCTACTACTCGTCTTGAAAAGGTTGGATCAATTATGGGAACGTCGCAGCCATATCATTTGTCAAATTTGTCTCCACATGATAGTTTACTGTTATTTATGCAACGCGCATTTGGGCAACAAAGAGAAGCAAATCCTAATCTAGTGGCCATTGGAAAGGAGATTGTGAAGAAATGCGGTGGTGTGCCTTTAGCGGCCAAGACTCTTGGTGGTCTTTTACGCTTCAAGAGAGAAGAGAGTGAATGGGAACATGTGAGAGATAATGAGATTTGGAGTCTGCCTCAAGATGAAAGTTCTATTTTGCCTGCTCTAAGACTGAGTTATCATCACCTTCCAGTTGATTTGAGACAATGCTTTGCGTATTGTGCAGTATTCCCAAAGGACACCAAAATGATAAAGGAAAATCTCATCACTCTCTGGATGGCCCATGGTTTTCTTTTATCAAAGGGAAACTTGGAGCTAGAGTATGTGGGTAATGAAGTATGGAATGAATTATACTTGAGGTCTTTCTTCCAAGAGATTGAAGTTAAATCCGGTAATACTTATTTCAAGATACATGATCTCATCCATGATTTGGCTACATCTCTGTATTTGGCAAGCACATCAAGCAGCAATATCCGCGAAATAAATGTCAAAGATTATAAGCATATAATGTCCATTGGTTTCGCTGAAGTGGTGTCTTCTTACTCTCCTTCGCTCTTGAAAAAGTTTGTCTCGTTAAGGGTGCTTAATCTAAGTTACTCAAAACTTGAGCAATTACCCTCTTCCATTGGAGATCTATTACATTTAAGATACCTGGACCTGTCTCGCAATAACTTCCGTAGTCTTCCAGAGAGGTTATGCAAGCTTCAAAATCTTCAGACTCTTGATGTGCATAATTGCTACTCACTTAATTGTTTGCCAAAACAAACAAGTAAACTTAGTAGTCTGCGAAATCTTGTTCTTGATGGCTGTCCATTGACTTCTACTCCACCAAGGATAGGGTTGTTGACATGCCTTAAGACTCTAGGTTTCTTTATTGTGGGAAGCAAGAAAGGTTATCAACTTGGTGAACTGAAAAACCTAAATCTCTGCGGCTCAATTTCAATCACACACCTTGAGAGAGTGAAGAACGATACGGATGCAGAAGCCAATTTATCTGCAAAAGCAAATCTGCAATCTTTAAGCATGAGTTGGGATAACGATGGACCAAACAGATATGAATCCGAAGAAGTTAAAGTGCTTGAAGCCCTCAAACCACACCCCAATCTGAAATATTTAGAGATCATTGCCTTCGGAGGATTCCGTTTTCCAAGCTGGATAAATCACTCAGTTTTGGAGAAGGTCATCTCTATTAGAATTAAAAGTTGCAAAAACTGCTTGTGCTTACCACCCTTTGGGGAGCTGCCTTGTCTAGAAAGTCTAGAGTTACAAAACGGGTCTGCGGAGGTGGAGTATGTTGAAGAGGATGATGTCCATTCTAGATTCTCCACAAGAAGAAGGTTTCCATCCCTGAAAAAACTTCGTATATGGTTCTTTCGTAATTTGAAAGGGTTGGTGAAACAGGAAGGAGAAAACAAATTCCCCATGCTCGAAGAGATGGCGATTTTACATTGCCCTCTGTTTGTTTTTCCAATCCTTTCTTCTGTCAAGAAATTAGAAGTTCACGGCAACACAAAAGCAAGAGGTTTGAGCTCCATATCTAATCTTAGCACTCTTACTTCCCTCCGCATTGGTGCTAACTACAGAGCGACTTCACTCCCAGAAGAGATGTTCACAAGTCTTACAAATCTCGAATACTTGAGTTTCTTTGACTTCAAGAATCTCAAAGAGCTGCCTACCAGCCTGACTAGTCTCAATGCTTTGAAGCGTCTCCAAATTGAAAGTTGTGACTCACTAGAGAGTCTCCCTGAACAAGGGCTGGAAGGTTTAACTTCACTCACACAGTTATTCGTTAAATACTGTAAGATGCTAGAATGTTTACCCGAGGGATTGCAGCACCTAACAGCCCTCACAAATTTTGGAGTAACCGGTTGTCCAGAAGTGGAAAAGCGCTGTGATAAGGAAATAGGAGAAGACTGGCACAAAATTGCTCACATTCCTAATTTGAGTATATATTAGTGAGTTATTTGTAATTGTTTCTTGGGTTATTTTTTTGTCAGTCTCTCACAGGTTACAATTCCTGCCATTGGATTATTATAGCAAAGTTTGACTGTATGAGGAAACATTTAAAAAAAAAAAAGAAGGTAAATCGTTTAACATGACTTTTTTTTTACTGTTTAATTTAATATAAAAGGATTGTACTAATTTTCTATTTGAAAAC

>PGSC0003DMG400009532 LRR-RLPs

ATGGATCGAATTTCCGAATTACCAGAGCCAATTTTAGAGTACATCTTGTCTTTTCTTGTTTACAAAAATGTTGCACAATTAAGTATGTTGTCCAAGACTTGGAACAAGGTTTGCAATTCAATCTCCTATTTGAAATTTGATGGTGATTTTTTCTTTACGCCATCAAAATACAAACCAAAGCTGGTGGATGTTGTGCATCAAACCCTAGAAAATCGTCGAAAGCTGAAGATTGCCGTACACAAGTGTTGGCTACCATTAACGATTGGATCTCATGAGGTATACAATTGTTGGATTGATACACTTGTAGTTTGTAATATCAAAGAACTACATCTATCAATTAAATCTCATCTTTACAAATTGCCTGAGATGATCTTTAATGCAAAAGCCCTAAATGTGCTGAGGCTGATTGGATTTAACATTATTGAATTGCCTTGTAAAAGTATAAATTTGCCATCTCTGCGAGAGTTTTATCTTGACAGTGTAATTTTGGACGACAACTTTCTTCGAGCTATATGCACAACCTGTTCTAATTTAGTGGTTTTGTATTTGGTGGGCTGTCATGGATTTAACAGTTTGCAAGTTGGTGAAAATTTACCTAAACTGAAAAAATTGAAATTGGAATATTCTTATCATCAAGTCCAATTCGTTGATATTAGTTCAACTAATCTTGAACAACTTACCATCTACGGCAATTGTGATTACCTAAAGGCAGTTGAAATAACTGCTTCTAAAGCCCTTAAGACTCTGTACCTCCAGCGTGTGCCGATAACCCAAAAAATTCTTTCTTGCCTTGGAAATCTAGACAAGTGTGAGTTATTTTTTTGCCACACATTGAAGACTCTAAAGATTGCAAGTTTCCTGCTCAAACAGCTGACGGTACATCAATGTCCTAACCTAATCGCTGTTGAATTAGACACTCCTAATTTGATAAGGTTTTCGTACAATTGTCATTCATTGCCAACCCTGCAACTGAAAGCTTCAGCTTCAGCTTCATTGGAAGCAAAAATTTGCTTTGTCTCAGGAACCACTGACAGTGATGTCACAAAATTTCTTGGTAACTTCAATCAGTCCATGGTTGTTGAATTAACATGCAAAAGTGACAAGGTAATAATCTTTTTAATTCCCACATATTCTTTTAGTTGTTGTTATATATTATGGATTTCTCACAAGTTGTTTACTTTTGCTGATACAGGCACTAGTTATACCAAAAGACATGAGAGGAAGCTTGCTTCCTCCCCTTTACGGTACTAATCGTGTTCATGTAAATATTCGAAGTCCATTGATAAACTGTTCGATTGTCGACATTCTTGATAGTATGCTTTGGATTTCTCCTCAACTGGACACCCTATCTTATTTCAAACCTGACTTAAAGATCCTAATGGTAAGACATTTATCTTTCAAAGGTACTGACTTTGTAAAAAAACGTTTTTTTAGATGCAAGGTTTCTTGCAATTCTTTTCCCTTTTAACATGCCTATTTCTTTCTTTCTTTAGCAGTATAAGTAATTAAGTCTGCAGTTACTTGAACTAACAAGTCAATTAAGTCTAGTATAAGGATACTGAGATTCTGAAATTCTTTCGCGTCCCCCCAAAAGCTGTTTATTTTAAAAAAATTGCGAGAATATATGTGAAAATTTCGGGTTAATAATTATTGGATTGTGCTTTGTCTCTGCGGCTAGTGGATAAATTTGAAGTTATATGTAGCTGGTTAACTCAAAAAAATATGTTGTTAACAAACAAGACAACGAAATCAGGGTCATAAGATTGTATTGAAAATTGCACTTCTTAACAAGCCGTGCATGCTACATTTGCAGCAATAACTCTACTAGAAAAGAGACTGGTGTTTCTGATATTTGTTTTCATGACTGATTCTTTATGAAGACAGTTTTTATATTCATGCGTTTGTGTATTGTCAATACTTATTGACTTTTCTGGTTTTACAGTTTATATATGAAGATGCATTAGATGAAGATGAGAAGCCTTGTTGTACATCTCAACCTTGGAAATGTTGGAGGCATAAGTTAAAGCAAGTCAAAATACAAAACATTACATGTATGGAACAAAAACAGGAGTTGAGAAACTACTTGTACGCAAACCCAGATATCCCGGAGATAATTGAGGTTCCTGGCCCAGTATTCATCGATCCCACAGTATTCATCCACAGATTCGTAATTGGATAA

>PGSC0003DMG400009755 LRR-RLPs

ATGGACAAACCAAGTTATAAAGGAGATGGATATTACCAAGACTCGGTAGTTGTTGTATCAAAGGGATTGAAGCTTGAAGTTGTGAGAATTTTGTCTTTGTACACAGTTATAGATCTTTCAAACAACAAATTTGAAGGACATATTCCTGGTATTCTGGGAGATCTCATTGCGATTCGGGTACTGAATTTATCTCATAATGGATTGCAAGGTCCTATACCACCATCACTTGGAAGCTTATCTGTAGTGGAATCACTGGACCTTTCGTTTAACCATCTTTCGGGAGAGATACCACAACAACTTGCTTCTCTTACGTCTCTTGAATTCTTAAATCTCTCTCACAATTATCTCCAAGGATGCATCCCTCAAGGACCTCAATTTCGTACCTTTGAGGGCAATTCATATGAAGGTAATGTGGATTACGTGGATTCCCCATTTCAAAAGGTTGTGGCAATGATCCTGTGTCGGAGACAAATTATACAGTATCTGCGCTAGCTGATCAAGAAAGGAATTCTGAATTTCTCAATGATTTTTGGAAAGCTGTTCTGATGGGCCATGGCAGTGGACTATGTATTGGATTATCCATAATATATTTCATGGTCTCAACTGGAACTATGAAATGGCTTGCAAGAATCATTGAAGAACTGGAACACAAAATTATCATGCAAAGGAGAAAGAAGCGGCAAAGTCAAAGGAACTACAGAAGAAGAGATAATCATTTCTAGACAAGTTACCAATAAAGAAGGATTCGATTTCAGAACTTCAGGTATTCAAGCTAAGATCTACACTATCTTTATTAGTTTATTCTAACTACTAATATATTAGTTTACACGTTTTTTGTTAATAACAACAACTAATTAATGCTCAATATGAAGCATATTCGTGTTTCTTTCAACTTCAAGAAGCTAAAGATTTGAAAAGCTTGGCAGTTGTTATAACAGTGGCTACTAAATTCAATCAGTGATCAACTCTCAAGTATTTGACTCACTATCTTATTTTTTCAGATGCCAAGAATAAGAAGATGTTGGCGTAATGGATTTGATTTTAGTTTCTTTTTAGTTTAATAAGTAAGCTTCTTCTTAACTTGGAAAATGTCATATTATGAATTTTTGATGATATATAAAGTTTATTTAAGTTTCTAATGTTATTCAATAAATGTTGTGTGTATTGAATGAAATGTATACATTTCTCTGATCCCTAACTTACAATTCAAAGAATCTAGGAGGGACTTAGTCATGAATAAATTTTTTGAACAAGCTGAAGAATGTGTTATCTTGTCCCTATTTAGCTCGAGAAAGAAGTGAAGTCAGAGTGTTGGGTGTCCCTGGATTGCTCACTGCTGCCTCAACGTGTGCTGAGGTTTTGCAGCAGTCTAAAGAGAGAGCGTTGAGAAGAGCTAAATTAGAGGAAGATATTTATCGTGTGATAGAGGAAAGAGCATTAACAAGGAAGAATAACGAGTTCGTGACAAGTGATTACATTTACTGGCCAAGGGAATTGCTTTAA

>PGSC0003DMG400009899 LRR-RLPs

ATTGTCTAATAAATTTCTCTTTGATTATTGAGCAAAATTTTGTCACCAATGGAAGGTGTAAATTTACTACTAGTCCTTCTTCTTGTAGCTAGTAGTACTATATTAGATGGATTGTTATTATGCAGAGCTATGAATTCAGACATACTTTGTGATGAAAATGAAAAAAATGCTCTTATCAAGTTCAGACAGGGTTTCAGGGATCCATCACAGTCAATGATCATGTCGTCTTGGATGGTCGAGGAAAATTGCTGCAACTGGAAAGGTGTTGAATGTGACAACACAACCGGATACGTGATCACTCTTGACCTGCACCAGCAATTCTTGCAAGGTGAGTTTGGGACTTACTTGCTTGGCTTGTCTCATTTAAGACACTTAGACTTAAGCCAAAATGATTTCCTTGAGGCCTTGTTTCCTGATTTCATTTGCAAATTTAAGTACCTAGAGTATCTCAATTTGCACAAAACCAAGTTTAGGGGATCAATCCCTGAGTGTCTTGGAAATCTGTCGCGGTTGCAGTTACTTGATCTTGGTGATGGTTTATCACTCCGGGTTGATAGTCTTGAATGGATTCAGCATCTTTCCAATATGAGGATTCTTGACCTGAGTGGAGTTGACCTGAGCAATGCCAAAAACTGGCTACATGACATCAATTTCCTAAGGTCTCTCGTGGAATTACGTTTGTCTTCTTGTCAGCTTCCTAAATTGCTTCCTGTTTTTGTAAATTTCACATCCCTTCAGGTTCTTGACCTCTCGTTAAACTATCTGGATGTTCCATTTCCTAGCTGGTTAGTCAATACAAGCTCTCAGAGTCTTGTTTATCTTAATCTCAAAAGAAGTCTGCTGCATGGTTTACTTCCAAATACCGCCTTTGGAAACATGTACTCACTTAGAGTTCTTGATCTTTCTGACAACTCTATCCGAGGAAAGCTTCCGTCTTTTGAGAACATGACTTCAGTTACTTCTCTTGATATCTCAAGGAATTCTCTTGTAGGCAAATAACCGCCAACCCTGCCTTCAAAGTTGAAACATTTGGATCTTTCATCCAATAATCTGGAGGGTCCTTTAGCAAGAAGCATCGCACAACTTAGGCAATTAGTTGTCCTCAATGTTGCTTGGAACTCTTTCAATGACTCAATTACTGAACATTTCTTGAATTTCAGTGATTTGAGAATGTTAGACTTGTCATCCAACTCAATTATCCTCAATATAACTGCAACTTGGATGCCTCCTTTTCAACTTGACATTATTGGCCTGATGTCTTGCCAACTTGGTACACAGTTTCCTCAGTGGCTTCATACACAAAAGGATTACTCATTTATCGATATCTCTCATGGTAGCATCTCAAATGAAGTGCCTGATTGGTTCTGGAGTCTCTCTGCAAAGGTACAGCACATGAACCTTTCTCTCAACTATTTCAGAGGGGAAGTACCTCAAGTTACATCAAGACTTGATAGTCTTGAAAAACTAGACCTGAGCCAAAACAACTTTTCTGGTCCTTTACCTCATTTTTCGTCCAAGATGAAGATGTTGATTTTGTCCCGGAATTCTTTTAATGGCGCCATTTCCCCTGTATGTGAATCACTTGTCATGAATAATTCTCTTATGTTTTTAGACCTATCGTTCAATAATCTATCAGGAACACTTTCAGACTGTTGGAGATATGGGAACAATCTGGCAGTATTAAAGTTAGGCTATAATAATCTGTCTGGTGAAATACCTCATTCTTTTGGATATCTTGAAACTCTCAGGTATCTGCAACTGAAAAACAACAGGCTGTCCAAAAATCTGCCTTCATCACTGAAGAACCTACAAGGACTAAAAATTCTTGATCTTTCCGGAAATTGTTTATCTGGAAAGATACCATCTTTCTTAGGGGAGACTTCACAAAATCTATTGGTTCTTTTGTTAAGAAAGAACAAGTTCGATGGAAACATTCCCCTACAGATTTGCCAGCTCAAAAACTTGATTATTTTGGACCTTAGTTCCAATTCTCTATCAGGCACCATTCCAAAGTGTGTTAAAAATTTCCTAATGATGGCTGGGGTAGAGGAGCTCCCTTCCTTGGTTTATGGTCCCTATGAAGAATACAGGAAGGATGTGGCGTTGATGCTAAAAGAGAGAGGCTATGATTATAGTTTTGCCTTTATGGCGGTGATTGATCTATCAGATAACCATTTATCAGGGGAAATTCCTGAGGAAATCACCACACTTGTTCAATTGCAGGTATTGAATTTATCAAGGAATAATTTAACTGGAGCAATTCCTTGTAAGATTAGTAAAATGCAATCTTTGGAAGCTCTTGATGTTTCCTGGAATCGTCTGTCATCCTTCTTGCCTTCCAGCATAGTGGAGCTTTCCAGGCTCGAGATCGCGAATTTTTCGTTCAATAGTTTGACAGGAAAAATTCCAATAGGTGGTCAATTTTTAACCTTTGAAAATAGTTCTTATATTGGGAATCCAGATTTGTGTGGCATGCCACTTTCCAAAAGCTGCTCGGGACATTTTCTTGAGGACATTACTCACTGTAACTCTCCCAAGAAAAAAGACGTGCAGGCCATTCATCAGCATGAGGAAGACAATTGGCTTGATGAATCATCATTCTATATCAGCATGGTAATTGGTTTCATCACAAGTTTCTGGTTATTTTGGGCTACTCTATTAGTGAAAACATCTTGGAGACATTCCTATATGAGGTATCTTAATAAAATGGGCAACAATATTTATGTGTTTGTAGCCATAAGATTACCAAATAAGCATAAACCATCAAAGATGAAAGATTGAAGTTCTTTCAGGGATTGCTGAATAAGGTAATGAAGGTCTTATTTCTGGTGCAACTTTCTTTTGTTGTTTTGTTTTCTAGGAGCTCTTCTGGTTTTGTTTTTGTAAAATCTTCTAAGTACTATATTGTGTATGCCTAGTTGCATTGTTTCATCTCGTGGGATTGTACTAGGTATGTTGTTGTTGTAGTCATAATTGTATTCTATGATATGTTTTGTGGATATCTAAGTAACTTGAGAATACTATTACTAATGTTTTT

>PGSC0003DMG400010183 LRR-RLPs

ATGTCATTGTCAATGACCATCTTGTATTTCATTTGTGTTGTTTTGACCGAGTGAGTATAAAGGCGCTATATTGCAGCTGCATTTCCTCCTAAAAAGCAGTTGAGTTTAAAATACAAGACAAAGCAAAGCAAAAATAGTTAGTCAACACTTACCTTGCATGCACCAGAGAGTCGTGACTTTTGAAATTCAATTTCAAGAACTGGATAGTTTTTTAATGCATATTTTAAAGAAAAGCTTAGGTGGATAGAGAACCAAACTCAGAATCAATGTGCTAACAAAATATTTTATTGTTGATATATCATTGGCAGCTTATTTTATAATCGTGTGTCTTATACCACGTAGTGACTCTCCATTACCAAAAACAACATTCATCTCTTGCTCCTTGACATTTTCCAGAATGGAAATTTTCTCAATGAGTTTTAATTCCCACACCTTACTGGCAACGCATGCTGTGTTTCTATTCTCATCTACTCTCAAATATGCTGCAGCAGCTACTATATACTTGGAAACGAAACTGACGAACTAGCCTTACTTGGATTCAAGTCCCAAATAACTGAGGATCCATCAAGAGTTTTCGCCTCTTGGAACCAATCCGTTCATTTATGTCAGTGGACTGGAGTAAAATGTGGCCTGACACAGGAAAGGGTGAGTCTGGCAGGTACAATCTCAGATCATCTCGGAAATCTTTCTTTGCTCAATTCTCTTGACCTGGCAGAAAATTCATTCCATGACGGAATTCCTCAAGAACTCAGCAGGCTGTCAAGGCATCAAAACCTGAATTTGAGTTTTAACTATCTAACAGGGGAAATTCCAGTTAATCTATCACATTGTGTTAACCTCAAGAGCCTTGTGCTGGACCACAACACTCTTGTGGGACTGATTCCTTACCAAGTTGGATCTTTAACAAAGTTAGTGAGATTGTATCTCAGAAATAACAACCTGACAGGAATCTTTCCAGTGTCTATTGGAAATCTTACATCTTTAGAAGAGTTGTATTTATCGTACAACAGTCTAGAGGGACAAGTGCCAGCTTCTTTAGCTCGATTGACCAAGTTGAGACTGCTTGGATTGTCAGTAAATAGCTTCTCCGGGGAGTTCCCTCCTTCCTTGTACAATTTGTCATCCCTGGAATTAATAGCACTTTCCTTCAACCACTTTTCTGGTAATCTCAGATCTGATTTAGGCCACTACTTCCCTAATCTCCAAAGACTTCACTTGGGGAACTGTCAATTCCATGGCTCCATACCATCCTCTTTGGCCAATGCTTCAAAATTGCTACAGCTTGATTTCCCTGTAAACAAATTCACTGGAAATATACTAAAGGGTTTTGATAACTTGCAGAATTTGTTGTGGCTCAATGTTGGGAGTAACCATCTTGGATACGGTAAGAATGATGACCTTGACTTTGTAAATTCTCTTACCAACTGCAGCAGTCTACAAATGCTCCATTTTGGAGACAACCAGTTTGTAGGTACATTACCTCATTCAACAGTCAACCTTTCTAGTCAGTTACAACGCCTACTCTTCTTCGGGAACAGAATTGGTGGAAGCATACCCAGAGAGATCTCAAACCTGGTGAATTTGAATCTACTCGACATGAGCAACAATAATCTTACAGGTAGCATTCCAGATTCTATTGGAAGACTTGCAAACTTGGGAAGTCTTGACTTGCGTAACAACCTCTTGACAGGGGTAATCCCTTCTTCAATAGGCAACCTCACTGAACTCGTCTATCTTTATTTAGGATTTAACAGGTTAGAGGGTAACATACCTTTAAGGCTTTAA

>PGSC0003DMG400010325 LRR-RLPs

ATGGTGGAACAAGAAGCAGTGTACAATGTTCTTGAATCCATTAACTCTGATGTCTCTTGGCGTTCCCTTTTCCCTGATGATCTCTGTTCTTCTTCCCCACATGGGGTTATTTGTGATTACTTTGTTTCCACTACAAACTCATTTACTCCTCATGTAACTGAGATTAGTCTTGGTTATGTTTCAGATTATGCTCCTAATCCACCTTGTACTCCTAATAGCACACTTAATGCTTCCCTTTTTACCCCTTTTTCTTATCTAAGAAAGATCTTTGTTTACAAATGTTTTACTGAAAATGAGGTTTCTTTGATTGACTTTGGATCTTTGCCTTCTTCTTTGGAGGAGGTGGTTTTCTTGGAGAATCCAGGTCTTGTTGGTTCTTTAAGTGATAAAGTTGGAAACTTGATGAGTTTGAGGAGGTTGGTTTTGAGTGGGAGTGGTGTTTATGGGGAAATCCCAAATGGAGTTGGTGGTTTGGTCAATTTAGAGCAGCTGACTTTATCAAGAAACAAGATTAGTGGTGAAATTCCAGTGGGGTTGATGACATTGAAGAAACTCAAGATTCTTGATTTGAGTCAAAATGAGCTTGAAAGTAGTGTTCTTGAATCTATAGGGAACTTTACTCAACTTTTGAAACTTGACTTGAGTTACAACAAGCTTACTGGGAAAATCCCTCAAAGTTTGAAGGGTTTGAAGAGTTTGGAGTTTTTGGACTTCAGTTACAATAACTTTTCCAATTCTGGGGTTCCATTGTTTTTGGCTGAGATGTCTAGTTTGAAGGAAGTCTACTTAAGTGGAAACTATCTTGGAGGGGAAATTCCAGAAATATGGGAGAATCTTGAGGGTATTGTGGGGCTAGGATTATCAAAAACAGGACTAGTTGGGAACATTCCTGTTTCTATGGGGGTGTACTTGAAAAACATATGTTATTTGGGGCTTGATAATAACAAGCTGGAAGGGGCATTGCCTGAAGAGTTTGGGGCATTGGAATATGTGAATGAGTTACATTTGGAGAACAATAATTTGAGTGGTAGACTTCCATTCTCTCCAAAATTTGTATCCAAAGTTGGAGAAAAGTTGAAGTTGGGAGGCAATCCTCAACTCTGTGTAGATGAAGGCTTGAGAATATCTAATGTTAGTGTTAGCTTGAGGCTACTAAAAATGTGCAACAATGCTATTAGTGCCAAATTTGTCCCATTCTCTGGGGCATCTTTAGTACTGGTGCCTGTTTCCCATGTTGTCTTTTTTCTTGGAATTTTGTTTCTCCTGTGCTAG

>PGSC0003DMG400010328 LRR-RLPs

TACAAACATTATCTTATGTAGAACCCATGGAGAAAAGACTGCGACTTTCCAACAAATTTGCTTTAAAAAAGGTGTTTTTCTCAACCCCATATCTCTTTTCTCCAGTCTCTTGTCATTCTTGTGGTTTGATCAATTTTTTTGTTTCTAGTGTCAAAAGGAGGTCTGCATTGCACAAGAAGATCGTGTCAGTCAGTTACCTGATGAAATTCTGGTATATATATTGTCTTTCTTTTCTGTTAAGGAAGCTGCAGATACAAGTCTCCTCTCAAGGCGTTGGCTAAACTTATGGACATATATTGATCACCTTGATTTTGATGCTACAAAATACTTGGATCAAGTGTTCGTGCAACCCAAGCTACAAAAAGGGCATATGGACAAGTATCTGACATGGGTCAACTCTGCTTTGCAAAAGTTTAAAGGCCAGAGACTTGACCAATTTCGTGTTTACTATGATTTAAGCAAATTTGCTCAGCACGACATTGACAAATGGCTTGAACTTGCCTTTTCTAAGAAAGTTCAAAGACTAGAGTTAGACTTATTAAGGTGTGGATCATTCACTCGTAATTTTGATTCCTGTTATACTTTTCCGGAGCGGCTCCTTGGTCTTGGAGATTCTTCTTCATGCTCTAATAACCTCCAAATACTCTCACTTATCAACCAGAATTTCAAGTCTCTAAAAGTGCTTATATTGAAATCTGTAAATGTGACGGGCCAAGTTCTTGAGTTCTTTCTACGCAATTGTCCATTTCTTGAAAGAATGCAAGTTCATGGATCAGGAACATTGAAAAATTTGGAAGTTGTTGGTCCGTCCCTCAAGTTGAGACACTTGGAGATATATAGTTGCTTCAATGTAAAATCACTTAAAATTTGTGATACAAATCTTATATCATTAAGGACTTCAGCAGGAAAAGTATTACTGCTTAAGAATGTCCCGATGCTGGTTGATGTAGAGATTTTGGGTATGCCATACACCCATTTATTGGATGACATATCCTCTCGTCTCTTGTGCGTTCTCTCTCAGCTGGAGGTCCTTACAGTAGATGCTTGTATATCATTGGTTAGTGAACAAAGTTTGTGTCTATCTACTCTTGTTTTACTAGCCTAGCACTAACTGGTGCCTTATCTGTAGTTTGATTATCCGCACGAAGATTTGGAGCATTACAATTTTCCTGAACTTACGAAGCTCAAGAAATTTGTTTTTATTACATTTGGAAATTCACTAAGAGTGTCAGGCCAGAGTCTCTTAGGTTGCACGTCTATAATAAGGGCTGCCCCACAATTGAAGGAATTTGAACTACAGGTAAGTATCATGCCTTTTCCTCTTTGGTGTACAAAAGTCTGATTTATTCATATTAGCAATTCTGGTTCATGCAGTTATTGATGCTACTTCTTTAATAAACTTAGTTATTTAAGTCGTTGAGCTTGAGTTGAAACAATTGTGAAGAGCTAAACTTGAATAAATGGTACTCTTGCATTTTAACTCATCTGACTGTGCCAATATAAAAGTCCTGCCCTTACATAGGTGTCATTTTAGATAGGTAGTAGACAGCTCTGTCGTCCCTGAGCTTTGATTTATTATCTGTAACAAAGAGCTTAAAGTCTCATCATTTCATCAAAGAGCTTAGAGCCTCATCATTCCTTCAGTTTCTTGAATATTTTTACTGAGTTACTGCACTTGTTTAGCGTAATGGAATGACCTGAGTGTTGGGATTGTGGTTTCAGTTGTCGATTGTGTTGCTCCAAAATCTTAGGTTTCCACCTGTTTGTTGCCAGTATATTATATGATGGAATTGGGAAGTACTATACAGTGATTATTATGTGTTTAATAAAACTACTTTTTTGAGAAAAGGTAACTTGTATTATAAACACAGGAATACTCATGACGTATTCCAGTAATACTTACAATCAGAAATACAGAAGTGTTGCTGTTTTCCTATAAACTCAAGAACTCTAAGATACTAGGGATATCATCATTTTCATGAGGATATTTTTTTTTTTTTTTTTGATAACCGAGAAATCCGTCAGTGACCCGCCCTTTGGACCAATCATAGCCTTCTAAACTCGGTGGATAATGGGCCCGCCCCTCTACCCTTCTCCACTTAAATACGGGGCTTCATGAGGATATTTATGTTTACACCAAAAACAAGAGCTAAACAATTCAGCTTCATCTTCTCCATAGAGCAGCTCTTGTTCTCAAAACATCTTTGATTCCTTTCCAGCCAAATAGTCCACCAAATGCAGGCTGGGACAATCTTCCATCTCTCCCTGTGCCCTGACTTATTTCCATCCCGATTCCAACATGCCAAAGTTTCCTTTATGTTTCCTGGCATAGACCAACTGATGCCCTTTTTATTGATGAAAATTTGCCGAAGTTTCAATGTCTCCTTAAAGTGTAAAAAGAGATGATTTATTGTCTCTGTCTCACACTCACAAAAAAAACATCTGGAGCACAAATCGAGGCCCTTCTTCATACGGTTATCCTGTGTGAGAACAGCCTGTTTTGCTAAGAGCCAAGTAAAACATGCCACCTTGAAGGGCACCTTAACTTTCCAGATCATTTTCCATGGCCAGTAGCTTGATTGAGTGTGTGTCCTCTGTGAGTTTCTGTAAGCTGTTCCAACTGAGAATCTCTCCTGTGTATCTGGAGCCCATAACAGTTTGTCTTCACTATTACTGAGTTCCTTGTATCGTTCCAGGATATTTAGAAATTCAACCATTCTATTCACCTCCCATTCATTTAAAGGTCTTCTGAACCTTAAATTCCAACCCTGAGCATCTTTTATAAACCTACTTTCTTGTCATAATTATACTGGTTGGTTTCAAATGAGAACAACCAACTAAGAAAATTGAGAAGTAAGTTGTTGCCTAGCTAGGTAGATGGGTGAAGGAGGAATTTAGTGGGGTAGAGGTTTCAGAGATGAGGGATTACAAAATGTTGTTAGAACCGAGGGGAACTTCCAAATTCTAATAGTAGGATTCGAAGGTGGCATCCTGAATCCCAATCAAATGACTCCAGGCTCAAAGAGTCAGAATCGAGCCTGGATGGTATTGTAGAGGAGGCTTAGGGCTTCACATGGTATCTGATGTTCTCTCTCTTGAATATTTATTTTTTCAAAAAATCAAGTAACGATTATATAAGGGAAAAGAGAAAAAATACCCGAACTTGTATCGGATTTCTTGTTACACTTATTCTTTGAAGTTGTTCTATGAATTATTTTGTAGTAGAATAATACCATAGGAAGTGTTGATGTGGCTTTTACTTTGCACCTCCATATAGTGCATGAAAAGACAAATACAAGCTAAATGATAGAGGCTCTAGAGTCCAAAAGATATAATTTTTCTTCTACTTTTTAGGCCCTTTTCTAAATCTCGAGCGGTGATCCACCACTTGTCCCACCAGGCCACCATCCATGCCCCCCACCCCCGTGTCTGAGCCAGGATTTTTTTTTTGTAGTTGAGTAGATACACAAATTATGTTTTTCCTACTCATGAGTGGGTGCACGCACTTGGCTAAAGTTGTGTGCATAATATTTTCAATAAAAATCAAGCAATAAAAGTAAAAAAAATTATCTAGGGTGTGTTTTGTATGGAGAAAAATGTTTATCTATTTTCATGATTGGTTTGTCAATATATTTTGTGAAATATTTTTTCTAGGAAAACAAGCTCCTTAAAAATGAGGAAATTTACTTTCCTAATGGAAGTAGAAAACATAAGTTCCAAAGCAATTAAATCATATTAGCATTTTATAACTATAAAAAACAATCATATAAATCACTATAGACATAGAGAAGTCAATAAAATAATTTTAGTTACTAGTTAAATTAAATAAAAAACAATACATATCACAATAAAAGTAATAAAATAATAATTAAAATAAAAGAATTACCGCGAAACATTATAAAAAAAACTATTGATAATCTCAACCACAAAAAAACTTGAAAGGAAAAACACGAGTACGGAGCCAGAAGGGCAAAAAAATTGACCAAAAATTCCAAAAGGGTTTGAAATCAAAAGGGCAACATTTAAACGGTAGAGGAGATGCTGTTTAAATTAGGGTAGCGCCTTACAAGGCGCAACTCCCTATGCGCCTTACAAGGCGCAAGAAAACAGAGCCACCATCATTGTTTATTTTTTCCATGTCAGTACTTACGGTGGTACTTTTTGCACTACAAAATAGTATAGGGGTCGTAGGGACATCACAAAGAATAAGTGTGTAACTGGAAAACTGAGATAAGTCCATAGGGTTTTTATGCCTTTTCCCACTTTGTTATTTAAAAAAGCACCAAGAGGGCTCCAACCTTTACAAGTAGAAGATTCTGTTGTTCTGTCAACCTCCCAGTCATGCAATCTCCTTCTTAATCTAATATTAAAAGATAGCCTTTCTCTGTATTCTTCAATCACATCTTTTATATGGATAGCTATGAGATGTAATTCTGAATAAGTTATTAAAGCTTCTTGATCCGGCGTACATGAAACTTTATCTTCTTCTTGCAACTAAAAGCTTTCACAAGATTAAAAGAATCAATTCCATAAGTTCTAATGTACTTTCAAACTGTGACATCATCAGTTGTATTGATTAGTTTTGCTTTTGAACATCTCATATCGTTGCCATGCTCTATGTTTAACAGCTTAATCCATAAATTGTTTTGCTCTACTGTGTTTTATCTCCACAAACTAATTCGACAATAGACACTTTTTTTAATACCCAAAGTTTTTATCTATGAACTACTCTTCTTAGAGAATATAGCTCTATTTTTATGCTCTTTGAATCAACTTGCCACATCTTTCTCCTTGCATTATCGAACCGTCTTCGCACCTTAAAAAATGATAGATACAGTTTCTTGCGGGTTGCCAATCTTCTCAACCTTATGTATCACTCTCTGCCATATTATTGCACTTCTATATCTGGCATTAGAGACTTGCTCATGTATGTCTCTGCCTGGATTCTCAATTTAAATGCAGTCAAGTGCTGTCTCAAAGTGTAATAGTAGAGATCTGTGATGCATGATCTTCCGCTCCTCAAGTCCAAATAGAAATAGGGTATAGTCTGCATATCGTGTACATGAAAAATTTCCATATTTTCATTTGTACAACTTTCCCCAACTTTTGACTCCTTTTATCCATACTATATTCATGGCATAATGCAGCATCTTACTGAATGTTTCCATTGCTTAGGAAGAGAATGGGATAACGGGGTTCTCCTATTTAACTGCATATCCTAATTATATTTTTCCATTTGTTTGCAAACCGTGTTGATTCAAAAGCTTCAGTAATATTGCCAGTTTACATGATTGTTCCACCGGGTCAAATCTTTGCTTGATCCTGTGTTGTCCCTAGTCTGAGGTGATAACTTGAGAATGAATCATTCATCGGTGCTCTCATAGTAAAGAAGGCTTCGCTCAAGGAGTGATAGATTTGGACGGAGTCTTGGCTTTGATGCTAAGGAGACGAGATCCATCGATCTGAATAAGTCTAGTACACCTTTCTCAAGTCCAGCTAACAAATAATTCTTGCTACTTTAATTCTATTTAAGGCTCCAAGAATTCATTTTCTAATAACACTGGATCTATAGTTTGTCCTTCCCTTTACAAATACATGCATTTTTTGACCTTGATATCTACTTGCCAATACTTGTTTCATCCTTGTTGTTAGGACCCCACCGGTAGTAATCTCATCCTTTTACCTTGTTCTTAAGAACCAATGCTGTGAATATTGCAGTCCAGCCTCTTGTAAAAATTTCTGTCTCATGAAAGAAGTGCAATGCACTGATAATGTCCCTTTCCAAAAAGTTCCTAGATTTCTGGTAAGAAGACATTATCTGCCCATCTGGCCCTAAAGTTGTTATCACCACTAGACCATCTCCATTACTTCTCCGAGAAAATTTCTTAGTAGCCAACACTCTTCCCGTGATAATAGCTCAAGGTCATTGTAACTGGAAGGCGCCATCATTGCTCGGTTTTACTAAATAACTTCTCATAGCATTACACTCATCTGATGACCTTTGGATTTGGTAGTTCGTCCATCTCTTACAAGTTAGTTTATATGGCTGTACTTTTTATTTGTATTGATCAATTTACAGTAAAACACTTAGTATTTTGTAGGGGTGTCAAAAATGAGCCCACCTATAGGTAACCCGCCCAACCCGCCCATAATTTTAAGGGTTGGGCTTAAGATAATTTGAATTAGGTTTAATCTCAACCCCCATTTTAAGAGAATCCTCAATTGAGCCCAATTTAATTTTCAATTTTCAACCCATTTTAAGACTCTTTATTTGTATTGATGTAATGTATGTTCTTATATTGAAGGTATGAATTACTTTTGTTTTATATTTTTTAGGATTTATCTATCCATTTGTAACTTTTTTTGTAATTTTCTTAAGTTGAAATCTAAAATGTGGTTATAAAACTTAAATAAACAATATATTAAAACTATTGAGATTAAGCGGGTAAAGACCCAACCATTTTTAAGCTCATTTGAGCCCAACCCATTTGAGCCCAAAGTAAACTTGGGCGGGTCAAGACTCAACCCAATTTCTATTTCAACCCATTTTAATATCTCCAATTTCAACCAACCCGCCCATTTGACACCTCTAGTATCTTGTCTCATTCATCTAACCAGAAACACCTCAATTGTTGTTTCCAGGATGTTTTTTAGCATTGGTTATTTGTTCCAGTTCTACCAAGTGCTTCTTTTAGGCTTTTCGTTTTTCTATATCCGTTCACCCTTTTCTGATCATACTTACCAATCCAGCAATCTGATTATGTATAATTTTTCGACATGCATTGTTGTTTGAACTCAAAGTCTGCCTTTTAAGTTCAGTTTCCTTTCTATTAACCATAATCAGATTGCCAACACTATCATAAGATTGCTATTGCTGTTCCGCTAAATCAATGGATCCCTCTATAGCTTTTTTCATTTTCTCGAATTTGCATTAAGATTCGCCTATTCCCAAGTCTGCACTCTGCTGGAATGGGTGTATGATTGGTTATCATCATATGCTTTGAAAACTTGAATATCACATTCCAAAGAAATTGTATACACAAATTTCTGTGTATTCCACCTCATAAGTTTACTGTACATGCTATTGTATTTATACAAAACTAGGTCTAAATTCTGTACCAAAATAAATGAATTAAACAATGATTATTTGTACACTTGGCACCTAATTATTGGACTACTTTGTAACAAACTCTAACAGCCATTTTGCTCTTCATGTGCCTTCCTTGTGCCTAGGTTGAGAATCTTCTAGACCTTGTACAACTGGTTAGGTTGAGAATCTTCTAGACCTTCTGATTAGGTTTCCTAGAACTTTTGTTCTTTTTGGCACAAATCTTTTGGTTCTTGAAAAGAAATGTTGGAGAAGGTTCATTTCGGTTAGCAAGTCCGGCATGCAGCTCGAATATGTTTATATGATAAGTCACTTATGTATTCTTTTGTTTCCTAATTTCTCCAAGTTAAGATTTTTCTCAGAGAGAGTCTAGAACATGGATAATGAGTAGTGGTGATTGAATATTGTTGAATTAGTCAACTTGACTAGGAGGCATCTTACATGATAATGGGAAGTTTTCATTCAGTCGATCAAGGAAGCCATATATCCCGAATCTTAAAATTTAGATCCACAACCGATTTCGAGCAACTTAGTCTAAAATAAGTCGACCCAAACATGTTGAAAGAAGTTGCAATTCTCTCAGAAAATTGAGCACTCCACGTCTGTAATATTCTTGTCCATATTCCCCACTTTCCTTCCTCCGTGTGGTAAATACTTTAGGGAATTTGTAGAAGAACAAAGTAGTAGGAGAAGTCAAATGACATGAGCGATTGATACAATCATGGTTCATCAGTGATTGGTACATAGTTTTATGTTATACTGATTCCCCAAAGTTTAATTCTTTTCATTTAACTTGTGTCCCAAGAAAAAGGCCTGTTCAAAATTTTCTTTTATTAGCTTTACTGACTTATCTTATCATTATTTTTTATTTTAATTTTAATTTGTGTTGTATATATTTACGGCAGGTGATGTCAATCCCCTTCGAGGCAAAGAAAGAGTGTAAGAAAGCAGCCATAAAATGCCCACTCCATCACCTCAAAGTGGTCAAATTTTCGGGGTTTTATACAGGCTTATTTCAACTTGAACTCGTTAGGTACTTTTTGGAAAATGGTGTTGCTCTTGAGAAAATCATTATTGATCCAGGTACTCCATTTTTTTATCGTATCCCTCCGTGCCCTAGCGAAATCAAACACACGCAAGCTTCGAGAAATCTTGCTAAGCTCCAGCTTGAATGTGAAGTGCCTCCGGCTATTGAGTTGGTCATACTTTAATGAAAATAATAATGTTATTAGGGTTAAATTTGTGATACATATCGGAGGTGAATTTTTATTTCTTAGTAGATGTTCCTCATGCTTTGTTTCATTATTGAGTGATATTTTCACTTTTTGCGAGGTTTATATTGCTATATGAAATATGGGGGATTTTCAAAATGAAAATGTCGCAGTTATTCCGCCA

>PGSC0003DMG400010447 LRR-RLPs

CAACTTGCACTTTCCAAGTCTAAAAAGCAGACATCACATTTAACATTTATAATTTCAGAAGCCACCAAGAATTTGTCATCCAAATGTATGTATATATTATGCCATTTTTTGGCACATTCTTTTCTTATAATTTCACAAAAATGAGAGGGCTGTACTACATAGTACTTGTCCTGTTTACTATAAAGCTTTGTTCTTGCAGTGGAATTTGCAAAGAAAATGAGAAAACAGCTTTGCTAAGGTTAAAGAAAGAAGCAAATGATCCCACAAATGTACTATCTTCTTGGGTTGATAAAGAAGATTGTTGCAACTGGGAAGGAGTTTTGTGCCACAATGTGACTGGTTCTGTCATTGAGCTATCAATTAACGGATGGGATTGGTCAGAACTTAAAGGCCTAAAGATTGATGATTTCCAGTGGTTGTCAAGTCTTGAAAATCTTGATAGTTTGGACATGTCTGGTGTGGATTTAAGTGAAGCAACAAACTGGACTGAGGTGATTAGCATGCTTCCTTCTTTGGTTAATCTTCGTTTTTCGAATTGTAGCCTTCATTCAATACCTCCTTTGTTTGATCATAACAGTTCTGTTCTTGAAAATCTTGATCTTTCTTTGAATAACTTTGGCTCTCCGATTCCCGGGTGGGTTTTCAGTTTTGGTAGTCTTGTTTCTCTTGAATTCACTGGAAGTAATTTCACTGGTTCATTTCCTGAAGGTCCTTTTAACTTGACTTCTTTCACAACATTGAGAGCTTCTAGCAACTCCTTTGGTTCTGTTTTGCCTCAATGGTTATTTGATCTTAGTAATCTTGAATATCTTGATTTGAGTTTTAGTGGTCTTGAGGGTCCAATACCAAATGGAGTTGGTGATTTTACCAAGCTTACATATCTTAGTCTTGCTTCCAATAATCTGAACTCCACTATCCCAAATGGCTCTATGGATGCAAGAATCTTGAAACACTTATCATGAAGGATAACAGTCTTGAGGGAACAGTTTCAGACTCGATTTCAAATCTGACATCTCTAACTAGTATCGATATGTCTGAAAATTTTCTTTCTGGAAAGTTACCGAGTGTAATTGGAAAGCTATGGAAATTAGAAGATCTCGATCTCTCAGAAAATCGATTTGAAGGAGAAGTATATGAGGTTTTCAATACTATGAATGATTTTCCTCCAATAGGCTCAGGAAATTGTTCTTCTTTGAGGACCTTAAGACTAAATGACAACAAACTGATTGGAAATCTTCCAAAAAGTTTTGGGCAGCTTCCAAATCTACAATTCTGTTTCATGTCCAATAACAGGTTGGAAGGGGTGTTAACAGAAGAACATTTCACCAACTTGACAAATTTGAAGTTTTTCTCTGCATCTAAGAGTAATTTGACTTTAAGGGTGAGTTCAGATTGGATTCCATCATTCCAAGCTAGTGACATAGTAATTGGCGGTTGGCGCTTAGGCCCTGACTTCCCTTTGTGGATCCAGACTCAACAATCAATCATGAATTTGGACATATCAAATGCTGGTATAGTAGGTGAGGTTCCAACTTGGTTCTGGAACTTATCGTCTCAAATTTGTTTCCTCAACATGTCTCACAACCAGCTCATTGGTGAGATTCCAACTTTCTCAATTTCTGATAGCTCCATGGGAAGTGGAGGTCCTTGGTTGATATACTTGAGTTCCAACAACTTCAGTGGTTTACTCCCACATATTCCAACCATGGTAACAGAGTTAGACCTTTCAGATAACACTTTCTCGGAGGGTTTAACAAGCTTCTTGTGTGATAACAAGAATGAATCTTACATGTTGGAGATTCTACACCTTGGGGGAAATCGTTTCTCCGAAGAAATTCCTGATTGTTGGATGAATTGGCCAGAGTTGAGAGTTGTCAACTTGGGAGAGAATAAGCTTACCGGAGGAATCCCAAGATCCATTGGAGCTTTAAGCAATTTGAAATCTTTGGACTTGAAAAGAAACAAACTTTCTGGTCCATTGCCTTCATCCTTGAACAACTGCACAAGATTGTGGAAAATAGACATGAATGAGAATGAGTTAGATGGAAATTTACCACCTTGGTTTGGTACACTTTCAAAGCTGATAGTCCTTAGTCTTCGGTCTAATAAATTCGATGGTGAACTAGCTCCAGAACTTTGCAAGCTCAATGAACTTCAAATCTTGGATCTTGCCAACAACAATTTGTTTGGAGCTATACCTCGGTGTGTTAATAATTTCACAGCAATGGTGAATGGGAGGAAGGAAATTAGAGATGGGGACGACGAGATGGATTATTCTTATTATGTTGGAGTTTTCAGGGAAAGTGCAAGAGTAGCAACTAAGGGCAACATGTATCAGTATGATACCATTTTGTCATTGTTTACTAGCATGGATCTTTCTAACAATAATCTTTCTGGAAACATTCCTATGAGTCTCACAAGTCTTGTTGGATTGAGATCATTAAATTTCTCCCACAATCAGCTGACTAGTAGTATTCCCAGGGACATTGACAAGATGAAGGTATTAGAATCACTTGATATCTCGAATAATCAACTCTCTGGTGAAATACCACAAAGTATTTCGAGTTTGTCCTCTTTGAGCTACTTGAATGTATCTCACAACAACTTATCAGGGCAGATTCCTGTGAGCACTCAACTTCAAAGCTTTAGCCCTTTTAGTTTTACTGGAAATAAACTGTGTGGGCTTCCACTCGTGAAGCGTTGCAGTAAAAATCCTGAGGTTGAAAATGAAGAAGAAGAAGATGATGATAATGAAGTGGAATGGTTCTACGTTTCCATGGCAATAGGATTTATTGTGGCTTTTTGGGGTGTAAGTGCTCCTTTGCTTTTCATAAGACCATGGAGACATGCCTATTATGGGTTTTTAGATAGAAAATGGAAATCATGGCACACATAATTTGTAAAGATAAGACATTAGGGAGATTTCTATTTATGTATACTGGATGAAATTTGTTTATATCTCATGAAAATCCATCACCATTCGGCGTTCTTTGAATTAAGACATTTTTATATTTGTTGTCTCCATACAAATATGAGAAATTTGACAATTTTGATTTGGCAAGGTTTGAGTTATGGAATGAACAAAGGAAATTAACAGTTCCT

>PGSC0003DMG400010519 LRR-RLPs

TCAGTTCAAGTCCTCGAGACATTTCAAATCAACATGTCTTGGATGACCGTCTGCAGCTAAACCATAACGAAAATCGTATGGTGCTCCAAACAGATTTTTTCCATTTCTATAGCCTACTTCTTCCAAAGCCTTCACAAGTGGCTCCATATATGCTGTAATTTTCCTAAACATGTTAAAAGTAAGTTACTACAAGGAATTTCTGGATGAGTTATTGGCTGAAACAAACTGAATTTCTTGAATCATACGTTACTAGTATGGCGTTGTAAGATAACACCATACTGGCAACATATGACATTTTGATCAATCCAAGGCATATTTTTTTTCTTTTTTTGTAATGGTATCAGGCTAACTTGTATGTGCTTCAACTGGATAACAAGATTGTGCATTTTCAACACTAAGAATAATAAATAATCTAGAAGGTGATAGACTAGAGGGGTTCTTGACATCATCTATCGGGAACTTAACAGAACTTGTCTCTCTGAATCTAGGCAGGAACAAGTTTAGAGGACCAGTGCCAGAGAGTATTACTTATTTAAGGAAACTTACAAGTCTTCAACTTTCTGAGAATTTTTTCTCTAGTAGTCTTGTTAATGACATTGGTGTCCTTACAAAACTTGAAATTCTTGATGTCGCGAATAATAAGCTATCTGGTTCAATCCCTTCATCGACTACATCACTTAGAAGTTTGACGCGATTAGACCTGTCTAACAATGAATTGACCGGGAAAATTCCTCAACTCAACGGTTTATGGAAACTCCATTCATTTGACATTTCCAATAACCAAATTTAAGGGAATTTGCCACAACTTCCATCGAAGATCAAGACATTGTCACTAAGCCACAACTTGCTATCAGGACACATTACCACAGTGCATAAACTTAGACATCTAAGAACATTAGACCTTAGTGAAAACAGATTTTCAGGAGGAATAAACAAAGGGATCTTCATGTTACTTGAATTAAGTCACGTAAGCATTTCAGTAAACCGGTTCACAGTGTTTGAGGTGGTAGAATTCTCTGATAAAGAATCACAGCTTCACACGTTAGACGTAAATGCCAATCGTCTACATGGTCATTTGCCTGTGAACTTGGATACTTACCCAAACTTAACAGAAATAAACCTTGGGCATAACCTGTTTTCGGGTGAGATTCCATCGGAATATTGGTCAAGACTCAGGTTTTCATGGAGAAGTCTCAACTTGGAATATAATAACCTGGAGGGATCGGTTCCTCGAGAGCTCAACAGGACTTTAGAAGGGGTTCAGGGAAGTTTTGAACACAACTGTCTCATTTGTCCAAAGGGCTTTCAGCTATGTCATGGACAGAGGACAGCTTCAGAATGTCATGGAGGAGACCATAATTAGCTGAGAATGTTTTAAAAATTACATTGCATTTTTTTTAATGTTTTCCCAGTTTCCAATATAGTTTATAAGTTGCATTTTTTTTTTATGTTTTCGTAGTTTCCAATATAGTAGAAGTTCCTTAAATTTGTCAGTATATTTTATTTAGATACTTGAACTACAACATGTTTCCTATTAAGCACCTGACATGATGAACTGTGTTAGAGTTTTGCCCTGAATGTATGATATTATCAGTTATAGCAAAGTATATTTGCATAACAATATTGAACAAGAGTATCAAAATTGGTTTAAGTATACGACATGAGCAAGAGTATCAAAATTAACTACTGAAGCAACATGTAAGCCAACGTTTTAAGTGCGAGCTTGTTTTAAGTTCAACTATAAGTGAGCTTGTTTACTTTGTTGGTTCCAAGATTAGATAAATTCTCATAATTTATGGTGAAGCAAGGTTGCTGCAAAGAAAAGTGATTCACTCTCAACTACCTTACTAACATAAAGTCATTCTTTATTTTCTTAAACTCCATGTCAACTCAAAATACATCAAACAAATTTTAAACGGAGGAAGTACCATACTACTAGTAACTGGAAAAATCAAAACAAGCTATAATGAAAGTTGTTTTAATTGCTAAAGAAGCTCATGAAGACAATGACATTACAACTCATTTCTGTTGCCTAGTCTAATAAAAACAATATTTGCCACCTAAAGATATATTTTAGCAGAATTGCATGTCTAAGAACTTAAGTACCTTTTTCAGGGGAAGATGCTGGGAAGAGTCCTTTACAAGCATCTGTATTCCACAAACATGAAATCAAAAGAGCAAGTCAGGCCCGAGACTACAAGCTGCTGTATTTCAAAGCATCTCAATGGATTAGAGAAAAACCTTGACCCACGAGTGTTAAGCTAGTAATATTATAGCTATTGACAGAAGGCTTGAAAATTGTACTACTTATGTAATGATATGTGTAATTCATTGTTATTTCATACTCGTTCATGAGTTTTTGAAGTTGAGAAAGAGATTCTGAAAGTAACTAGTGACTGAAGATTTGAGAATAGCAATTCACACATTTTTTAAGAGCGGGAATTGTGTACAAAAATGTACTATTTCTTCTAAAGTAACGTCTGCATTTTGGAAGAAAATCTCACCTCCTATATGTGTCCGAGTGTTCCTTTGCTGTCAAATGGGGGCATCCCGTACAGAAACTCAAAGCATAGGACACCAAGGCTCTAAATGTCCACACTTGCATCATGCTCCACACTTTCCACTGCATGTTCAACAATTAACATATCAGAAAAAATGTAAAAACAGATTAACAAAATGTTAAAAGCTAAACTCTCATACACGCCCATGTGAAGGTAAAAAGCAATGTATAACATTCACACATAGTCCGCCTTCGATTAAAGGTATGCACTGACCACCCAAAGTCTGCCGTTTTGAGTTCACCCTAAAAGCAGAAATATTAAAGCTGTCATTTTACTGCAATGCATTGTATATACTGTTTAGCAATGCTAAGTTTGTACCTCAATATCTTCTCTGTGTATTGTGTGATTCCCATGACAGTATATTGAAAGTTCGTAAGAATTGAGAAGAAGCAATGCAGAAGAGGCAACGACAAGTTACCGGGTCATAAAAGTAACCATAAGTATTTCAACTTCACGACGAAGCTGATGTTCAACCAGGGACTGTTTTAGCTGGCTCTTGAACAACACTTTTAACGCGACGATATGATTGCTATGTTTCCAAGAAACAACTAACCTTAGTATTGGAATGTGATTGTAGAATGTTAAGTCTATCGAATAGAATTCGAAAAATCTTTTAAGCCATGTCCACAAGATATCATCCCCAACCTCCCACCAATCCATAGCAATTATAACAAATATGTTGAATGAGAGATTCTCACTCGCAGTGCATAAATCCGAGGCATTGAAAAAGTAAGTAGGTTTATCAGCCATTCTTAACATGTAAAGGTTTAAAGGACTTACATACATGACCGAACTTTCCTCATCCAAGAGGCTTCCCAATGTTGAAATCATTAAGTGTCCGCCTCTTTTTATTAGCCTCTGATGAAGTCTGCAGAAGTTGAAAATTCTATAGTCCATTTAAAAGATCAGTTAATTAAAAATAACAATAAATTCTTGTTGTTGTTGTTGTGTACAGTATAACAGATCAGTTAATTAAAAATAACAATAACTTGTTCTTTTAATTTCATGATAACATATGGTTATTTCACATCACAAAGTTGTACGATCCAGCTGAGCAAGAATCAAGATGCTCTTTTGATTCCATTTAGTGCGGGAAGAAGATATGTCAAGGAAGACAATTTGCTTCAAAAGCTCCAAGAATCAGGCTTAAATGCATTGGTAGAAGATTATCTCAATTAACTGCACCTCTGTTCATACAGTCCTCTTAGAGGACTTCATGAACAAGCAATCAACCACAAGTGATCGCAACTAACATACAAGCTTAAACTCAATCAACAATGTGTACAACAAATTTTAACTAACCTCAACATTCTTCCTTTAGCAAGGTGGTACGCTTGTGAAACTATCAAGTACAACTCAACTCAATCTACATACTAATTTATTATAAGCTCAAAAATAACCAAACACAGTGAGGTCAGAGCATGTTACTAGAAATTTTATTTTTCTTTACGACAGAAACAAACATTGTTG

>PGSC0003DMG400010523 LRR-RLPs

GTTCTTTAGTGGTGACTCTCAATGAAGTACCTACTTTCAGCTTACTCTCTCTATACTCACTACTACTGCTACTCAGTACTGATTCCTTTCACACATACTGTGCCTGTAAACCCTGTCCAGGGACCCCCATTTCCCCTTTCCCCTTTCCCCTTTCTCCTTCCTTCCTTAGCTATCTCTCACACAAACACTAATCTTTTTTCACCTCTACCTTACCTCCATGGCAGAATCAGTTCTTGAACCTTGTACAACCTCTTATTCCTTCAAAGTTTCAATCTTTATCCTATTCTTCTTGATTTTCCCTTTCTTGAACCCATTTTCATCTGCATTTCCTCTTTCTTTTGATACTAATGCAACTGAGGCTGTCAATCTTGAAACAGAAGAGGACATGGGTTTGCTTTTGTTCTTCAAGTTACAGTTTCGAGAAACCCCTTTACCAAGCTGGGATGTCAATGTTCCTCTATCAAACTGGACTGGTGTTACCCGGTCTAACCAGACCGGACGGGTCACTGGACTTAACCTCACAAGGTTTAACTTGTCAGGACAGGTTCATCCTTGTTTGTGTAATCTTACTTTTCTTGAAACCCTTGTGTTGTCTCATAATAGCTTTAACAATTCAATACCTTCTTGTTTATGGAAGTTGTGGAGCCTTAAGACCTTAGATCTTAGCTATAATATGCTTACTCTTCTTATTCCTAGTACATTTGCAACAACTATGAGTAAGTTAATTGAGCTTGACCTTAGTCATAACATGTTGAGTGATGAAATCCCAATGTGGATAGGGAATGTCTCAATGTCACTTGAAAAACTTAACTTAGGGTTTAATAGTTTTCATGGGGATATACCTAAGAGCTTGTTGAATTTGATGTCTTTGAAGTATCTTGACTTGTCTCACAATAGTTTGATGGGAAATGTGGGTGATTTTAACCAAGAATTGGTCTCACTTAATCTTGAGTCTAATTTGTTATCGGGTACTTTGCCTTGTTTATATTCGTCAAGGGAATCACTTACACTTCTTAATTTAGCAAACAATTCGATTCTTGGAGGTATACCAACGTGTATCTCGAGTCTTGGGGGTTTGACACAGCTCAACTTGTCACGTAATGAATTACGATATGGTATCTCGCCTAGACTGGTTTTTTCAGAGAGGTTATGTTTGTTGGACTTGAGTTATAATGAGCTATCAGGGAAGATTCCAAGTAGGATTGTTGAGGCATCGGACAAGTCTGGACTTCTACTTCTTGACCTGTCTCACAATCAGTTCTCTGGTAATATTCCTGTAACGATAACAGAATTGAAGAGCTTGCAAGCATTGTTTCTGTCTTATAATCTTCTTGTGGGCGAAATACCAGAAAGGATTGGTAATTTGACCTATCTACAGGTGATTGATCTCTCACATAACTTCCTCACTGGCTCGATTCCTTTGAACATCGTAGGATGTTTCCAACTACTGGTGCTGATACTAAACAGTAATAATCTTTCTGGGGAAATTCAGCCAGTGCTTGATGCGTTGGATAGTCTTAAGATATTTGATATAGGAAACAACAAGATTTCTGGTGAGATCCCACTGACATTGGCAGGCTGCAAGTCGTTGGAAGTTGTTGACTTGAGCTCTAACAATCTCTCAGGGTCTCTAAATGATGCAATAACCAAATGGTCGAACCTCAAATTCCTCTCCCTTGCTCGGAACAAGTTCAGTGGATCTCTGCCAAGTTGGTTGTTTACATTTCAGGCTATTCATACTCTGGATTTTTCTGGAAACAAGTTCTCGGGATATATACCAGATGGTAACTTTAACACTAGTCCAAATTTCTACAACGGCGACATTAGGAAAACCATTCCTGCAGTACCATCAATTTCAGCTCGAAGCCTGGATATCAAACTTTCACTCATTGCTGATGAAACTAGTTTGAGCTTCAACTATAACCTGACAACCACAATTGGAATTGATCTGTCTGACAATTTGCTTCATGGTGAAATTCCAGAGGGTCTGTTCGGATTACATGGTTTGGAGTACCTTAATTTGTCATACAATTTTCTTAATGGTCCAGTTCCAGGGAGTTTAGGGAAGTTGCAGAAGCTAAAAGCACTTGATTTATCACATAATTCTTTATCTGGCCACATCCCTGAAAACATTACTGTCCTCAGAAATTTGACAGTTTTAAATCTGTCTTATAATTGTTTCTCTGGTGTTATTCCGACAAAGCGAGGTTATTGGAAATTTCCTGGAGCATTTGCTGGGAATCCAGACTTATGTATGGAATCATCTGGTAATGTCTGTCAAAGAACTTTGCCTGTAGAGCCAGGGAAGAAATTTGAAGAGGAAATGGAAGAGGGACCATTATCAGTTTGGATTTTCTGTATAAGTGCTTTAGTTAGCTTCTATGTTGGCATTGTTGTTTTATTTTGTTCATCTCGAACAAGAAGCTGTATTCTGCAAACAAAAAGTTTAGCAGGTTGAAAGTGATAAGTGATCACATTGTACAGCAACTTATGAATTGTATCATATAGCAAAACTGCTTCAAGCATATCTTGTTTACTAATCAGCCCTTCATTTTTTTTAACTAGTTAACAATGCTTCACGTTG

>PGSC0003DMG400010700 LRR-RLPs

ATGGGCATTGGAAAACTTGCATTGTTTCTGCTATGCGCGTTTCTCTGTCAACTTGTTTTCTCCTCATCTTTACCTCATTTGTGCCTCAATGATCAAGCCCTTGCTCTTCTTCAATTCAAAAACACGTTTACTATAAATCTTGATGCTTCTTATAGTGGTTATTATCCTAAAACAGTTAAATGGAATAAGAGCGCAGATTGTTGTTCATGGGATGGAGTTAAGTGTGACGAGATGACAGGACAAGTGATTCAGCTTGATCTCAGTCGCAGCCAACTTCGAGGCGAGTTTCATTCCAACAGTAGCCTCTTTAAACTCGTTCATTTCACCTAAATTTGGTGAGCTTTCTAGTTTGACTCATCTTGATTTGTCATATTCAACTTTTACAGGTATAATGCCAGCAGAAATATCTCATCTTTCTAAATTGCACGTTCTTAGTATCGAGAGTGATCCATATGAGCTTAGATTCGAACCTCACAATTTTGAACTGCTCCTTAAGAACTTGACCCGATTAAGAGTGCTCCACCTTAGCTCTGTGAACATCTCTTCAACCATTCCTCTTAAATTCTCTTCTTATTTAACAAATCTAGGACTTTCAGACACGCAGATATACGGGATATTGCCCGAAGGAGTTTTCCACCTTTCCAACTTGGAATCTCTTGATTTATCAGACAATCCCCAGCTCACTGTTAGGTTTCCCACAACCAAATGGAATAGCAGTGCATCACTTATGAAGTTATATCTCAGTGGTGTGAATGCTACTGGTACGATACCAGAATCATTTGGTCATCTAACTTCACTGCATAGATTGAGAATAGCTTCTTGTAATCTCTCAGGGTCTATTCCTAAACCTCTCTGGAATCTCACCAACATAGAGGTTTTGGACCTTGGTTATAACCATCTTGAAGGACCAATTTCTGATTTCTTTAGATTTGGAAAGATCAGGTCGTTATCACTTAGAAATAACAACTTTAATGGCCAACTTGAGTTTTTATCCTTTAACAGATGGACGCAACTTGGATTATTAGATTGTTCAGCCAATTCCCTAACAGGTCCCATTCCTTCTAATGTAAGTGGTATGCAAAACCTATATTCACTCTTCTTGTCATCAAACCTCTTGAATGGGACTATACCTTCCTGGATATTCTCCCTCCCTTCAATAGGGATTTTAGACTTGAGTGATAACCATTTAAGTGAAAATATTCAGGAGTTCTATTCCGAATCATTGTATCGAGTTTCTCTAAAACAAAATCAGCTGCAAGGTCCTATTCCAAAGTCACTCTTAGATCTGCAGGGCCTACAATTTCTTCTCCTTTCACGAAATAATCTCAGTGGACAGATTGCTTCAACCATCTGCAATCTGACAAAACTGTTAGTGCTAGATTTGGGCAGCAATAATTTGGAGGGAACAATCCCACTATGTTTGGGTGAGATGAGTGGACTTTGGATTTTGGATTTAAGCAACAATAGTCTTAGCGGGAAAATTAATACAACTTTTAGTATAGGAAACCAACTTATAGTCATTAAATTTGACGGGAATAAGCTAGAGGGGAAAGTTCCGCAATCTTTGATCAATTGCACATATTTGGAAGTTGTTGATTTAGGTAACAATGAGTTGAGTGACACATTTCCTAAATGGTTGGGAGCCCTACCTTATTTGCAGATTTTGATTTTGAGATCAAATAAGTTCTATGGCCCTATAAAAGATTCAAGGACTGATAACTTGTTTGCTCAAATTCGAGTCATAGATCTCTCAGCCAATGGATTTAGTGGAGATTTACCAGTGAGTCCTTTTTTGAGAATTTTGAAGCCATGAAAATGATTGGATGAGAGCAGTGGAACCCGAGAGTATGTAGGAAATATATATTCTTATTATTACACGAATTCTTTGACAGTGACAACAAAGGGACTGGATCTTGAACTTCCTCGAGTTTTGACTACAGACATGATTATGGATCTCTCAAGGAATAGATTTGAAGGTCATATCCCAAGCATTATTGGAGATCTCGTTGGGCTTCGTACGTTAAACTTATCTCATAATGTCTTGGAAGGTGATATACCAGCATCACTGCACCAGTTATCTGTACTTGAATCATTGGATCTCTCATCCAACAAAATCAGTGGAGAAATTCCACAACAGCTTGCATCCCTCAAATCACTTGAAGTCTTAAATCTCTCTCACAATCATCTTGTTGGATGCATCCCGAAAGGAAACAATTTGATACGTTTGAGAACAGTTCATACCAAGGAAATGATGGGTTACGTGGATTTTCACTCTCAAAGGATTGTGGTGGTGATGATGGGGTAGCACAAACGACGAATCCAGTTGAGCTAGATGAAGAAGGAGGAAGAGATTCATCAATGATCAGTTGGCAGGCGGTTCTCATGGGTTATGGTTGTGGACTTGTTATTGGACTGTCCATAATATACATAATGTTGTCAACTCAATATCCAGCATGGTTTTCGAGAATGGATGTAGAATTGGAACACAAAATTCTTACGAGAATGAAAAGGCATAAGAAAAGATATTAGTGTGTTACCTCCA

>PGSC0003DMG400010940 LRR-RLPs

ATGAACTCAGTGGTGAAATGTGGAGAGGGCCATGGTATGTACCCGAACTCAGTCTTAAATCTCACGAACAATAGCCTCACAGGTATGATCCCTCCTTCTGTTGGAAATGCCACAAAAATGATGAGTTTCAGTTTGTCTGGGAATAGAGTCAGCGGCAACATTCCAAAGGAGGTCAGTAATCTGAGCCAGCTTGCATATTTGCCCTTGTTCGATAATCAATTAACAGGTTCCATTCCCGCAACACTGTTTAATATCTCGTCGCTACTTGCTGTATCTCTGTCAATCAATAGCCTTTCTGGTCCTCTCATGCTAGATGAAGGGAATATTGTGTCCAATCTGTACTTTTTAAGTATATCTTACAACCAAATATCTGGTTGCATTCCTTCCAACATATGCCAACTCACAGAGCTCAAAGTTTTGTCCATATCTTTCAACAATATAACTGGAGACATACCCAGAAATATTGGTTGTTTATCCAAGCTCGATGGGTTTTATATTGGCCATAATCCAATAAAAGGGACTATTCCCACTTCATTGGGAAATATTTCCACTCTGCATGATGTTTACTGTCGAAACAATCTCATAGTGGGGGGAATTCCTCCGGAATTAGGAAAGCTATCAAATTTGAGGCAATTAACCTTTGACAATAATTCTAATCTTATTGGTCAAATTCCAAAGGCTATTTTCAACATATCTTCTTTGGAAGTGATTTCTTTCCATTACAACAACCTCTTGGGGAGAATTCCAACCACTACAGGTCTTCATCTTCCGAACCTTAAAGAACTTGACTTGGCAGTGAATCAGCTTGAAGGGGAAATTCCATTGTTCATAACAAATGCTTCCAAGCTTGAAATGCTGGAGCTAAATAATAACTTTCTGAAAGGAACTATTCCTACTAATTTAAGAAATCTTCGTGAGCTGCGATATCTGCTCCTACATACTAATCAACTTACCAATGAACCAAGAGACTGTGAGTTGTTATTCTTCAATTCTTTGGCGGACGGTAAGATGTTGCGATATTTACAAGTGGGTAACAATCCGTTGAATGGCATTCTGCCCAATTCTATTGGGAATCTTTGATCTACTATTCAAAACATTGATATAGGAGATGCTCACATCAACGGCCCCATCCCCACAAGTATAGGCAACATGAGCGGTCTTACAGCCATAGACATTGAAGGAAACAACTTAACAGGGAGTATTCCTTCTGAGATTGGTAAGCTTAAACAACTCCAAGGTCTGTATCTATATAATAATAAATTGCAGGGACATATTCCAGAGGCCGTATGCCATTTATCTAATTTGGTACAACTATTTCTGAATGGTAATGAGCTCTCTGGATTAATTCCAGAATGCTTAGGAAATCTTAGCATGCTACAAAAGCTTTATTTGAATTCTAACAAATTTTCATCTAAATTTCCGTTGAGCCTTTGGAAGATGAGTGGTCTCCTCTATCTAAGAGTGTCACGAAATTCAATTGAGGGAGAATTTCCATTAGATATTGGAGGACTGAAAGCCATTGTAGAACTACATCTTACCAGTAACCACTTTTCAGGAATGATACCAAGCACATTCGGGGAACTCCAAAACCTGCAGTCTCTTGACCTATCGAACAATTCATTTTTTGGCCAAATTCCATTATCCTTTGCCAACTTGATAAGCTTAGAATACTTGGATTTGTCTTCAAATGCGTTGTCAGGTACTATTCCTAAGTCTTTGGAAAAACTCTCATACCTTAAAAGCATTAATGTTTCATTTAATGATTTAGAAGGTCAAATACCGAGTGGTGGTGTGTTTGCAAATTCCACTTTGCAATCATTTCTCAAGAACAAAGGTCTTTGTGGAATGCACATATTGGAGATTCCTGCTTGTGCTATCACTAATCCTGGAAAACAATCAAAGCTTAAAGAGGTTCTGCTAAAAATTGTTACTCCAGTGGTTACTTCATCCTTTCTGATATTCTTGTTCGTTTCAATTTGGATAATAAAACGACAGAAGAAAGGGAAGTCCAAAGATGTGGAGAAGGTACCGGAGATCGGGACTCATCAATTAGTTTCTTATCACGAGATTCAACGAGCAACAAATAATTTTGATGAATCCAATTTAATTGGTGAGGGAAGCTCCAGCTCTGTGTACAAAGGCACATTGTTTGGTGGAACTGCAGTGGCCATAAAGGTTCTCGATTTGGAAATGAGCAGGTATGCAAAAGGTTTAATACCGAATGCGAAGTGA

>PGSC0003DMG400010950 LRR-RLPs

CATTTTTCTATTAATATTTGCACTAATGGAGAAAGCCTTCACATTTTTCCTCTTGACAATACTCTTGTTGCTTTCACTATGTTATGACCCAAACCAACATTACTACTGATCAATTAGCTCTTCTCTCTTTGAAATCCCAAATTAGTTCAGACCTTTTCATTATTTGGATGAAAGTTGGTCTCCTGCTATTAATGTTTGTCAGTGGGTAGGAGTCACTTGTGACTACCGTCACCAGAGAGTGAAGTCCTTGAATCTATCCAGAATGGCTCTTACCGACAGGACTCCTCGTGATTTTGGAAACCTCACATTTCTTGTTTCTCTTGATTTGCGAAGGAACAATTTCCATGGAAATTTGCCTCAAGAAATGGTACGCTTACGTCGGCTTAAATTTCTTAAGTTAAGTTTCAACAACTTGAGAGGGGAAATTCCTTCTTGGTTTGGGTTTTTACACCAACTTCAAGTTCTAAATCTTGGAAATAATAGTTTCACTGGTTCCATTCCTCCGTCACTCTCAAATGCCTCAAGGTTGGAGACTTTAGAAATATCTGCCAATTTACTTGAAGGAAACATTCCGGAAGAGATTGGCAATCTTCAGAACCTGAAATTCTTGTCCATACAATATAATAAGCTTACGGGTTCTATACCATTCACAATTTTCAATATTTCTAGTATCCAAGTCATTGCATTTACAAACAATAGCTTATCAGGAAATCTTCCCAATGGTTTATGCAATGGTCTCCCGACACTCAAAAAGCTTTATCTATCAACAAACAAACTTCATGGTCATATGCCTACAAGCTTGTCAAATTGTTCACAACTTCAAATATTGTCTCTATCCGGAAATAATTTTGATGGACCAATACATAGTGAAATTGGAAGAGTGAGTAACTTGCATATATTGTATCTCGGATCCAACCATTTCACAGGTATATATGTTTTATCTTTCTAGTTTCTGGACACGTGTGTGGTCATCTAAATTGACATAAATTGTTTCAAATTATCATTGTATTAAAATAGATTTATAATTAAATTACATGAATATCG

>PGSC0003DMG400010952 LRR-RLPs

TTCAAAGTCTCAAAACCACAGGTTACTAGTACTAATATTTGCACTAATGGAGAAAGCCTTCACTATTTTATTCTTAACAATACTCTTGTTGCCTCACTATGTTATGGCCCAAACCAACATTAACACTGATCAATTAGCTCTTCTCTCTTTTAAATCCCAAATTAGTTCCGACCCCTTTCACTTCTTGAATGAAAGTTGGTCTCCTGCTATTTCTGTTTGCCGTTGGGTTGGAGTCATTTGTGGTTCTCGTCACCCGCGAGAGAAGACGTTGAAGCTTTCCAACTTGGCTCTTACAGGCAGAATTCCCCGTGATTTTGGAACCCTCACATTTCTTATTTCTCTTGACTTGGGAAGCAACCATTTCCATGGAAATTTGCCTCAAGAAATGGCACATTTGCGTCGGCTAAAGTTTCTTGATTTAAGTTTAAACAGCTTCAGAGGGGAAATTCCTTCTTGGTTTGGGTTTTTACACCAACTTCAAGTTGTAAAGGAGTAACAGTTATACTGGTTCCATTCCTCCGTCACTCTCGAACGCCTCAAGGTTGGAGACTTTACAAATACCTGCCAATTTACTTCAAGGAAACATCCCCGAAGAGATTGGCAATCTCCACAACCTGAATTGATTATCCATAGAAAATAATCAGCTTACGGGTTCTATACCATTCACAATATTCAATATTTCTAGAATTGAAGTCATTTCATTTTCATATAATAGCCTATCAGGAGATCTTCCCAATGGTTTATGCAGTGATCTCCCAATACTCAAAGAGCTTGATCTATCAATAAACAAGCTTCATGGGTCATATGCCTACAAGCTTGTCAAATTGTTCAAATTAGCAAAAACAAAGAGACGTACCGGGTACAATATAACCAAATGTTGCTAAGGTTTTTGTGTATAAATAAATCACTCTCATCTTCACGGAGCAGTTTTGAAAAACCAAAGTCACTTAGGTGGGTAACCATATCCTCATCCAACAAGACGTTACTAGGCTTTAGATCACAATGAATCACAGGTAAAGAGCACCCATGATGGAAATATTCCAACGCACATGTAACATCTATGATAATGCTTAGTCTCTGGCTGATATCTAGGAAGTAGTTGTGTGAATACAAATACTTTTCAAGACTTCCATTAAGCATATACTCAAGCACTAAAGCCTTAAAATCAAGGTTGGAGCAACTAGTGATGACTTTCACGAGATTCCTATGGCGAAGGCTGCGCAAAACTTCACATTCTGTATCAGAACTCTTGAATGTCGCTTCCAGTTGCAGATTGAACACTTTAGCTGCAATGGCAGTCCCACTTCAGAGAACCCCTTTGTAAACAGAGCCAAAACTTCCAGAACCAATCAGATTACTGATTCACTTAGCGAGAGTACTAATCTGATTGGTTCTGG

>PGSC0003DMG400010964 LRR-RLPs

AGGACAGAATCTATTAAAGGCATAATGGTACCAATTGGCTCAGACCAACATATATGCAAGTGGAGCAAAGCTTTCAGAAACATGCCTTGTCTTAGGTTACTCATTGTCAAAGGGCAGGAGGTCTGGCATCATGACCTTATTTGTGACCCTATTGAGTGTCTCCCCAGTAACTTGAAATGGCTTGATTGGTCATACTACAGTTTTGAATCTTTACCAGCATATTATGAACCAGGAAACCTTGTTGGGCTCCACATGACTTTTAGTTCTCTTGTTGAAGTTTTCAAGGAGCCAAAGGTATTATTTACTCCTTTAGAAACAAATCATATGTGAAAAAAAGAATCATTGCTAAAGCTGAAGTTTCTATTCATTCTCTCTTTATGAACAGGCATTTGATAAGTTGACAGTTCTCAATTTAAGTTTTTCTAGAAATTTAATCCGAACACCCAATTTTTCGGATATTCCAAACCTGCAGAGGATTATACTGAAAAGCTGTGTAAGCTTGGTAGAAGTTCATCCATCCATTGGCCATCTCAGAAAGGTTGTCTTTTTGAATATGGAATACTGCGAAAGTCTCAAATCTTTACCAAGCAGTATTCAAATGGAATCTCTTAAAATCTTCAATCTCTCAGGTTGTGAAAAGTTGGAGAAATTTCCAGAAATTCAGGGAAATATGGAATTACTATCAGAGCTCCTTTTGGCACATACTGCAATATGGGAACTTCCATCATCAGTTGGACTGCTCAGTGGCATCAGCTTGCTCGATTTGCATTCATGTAAATACCTTGTAAGACTCCCAGCCAGTGTCTCTGAGATGAGAAAACTGAAAATTTTGACTTTGAAAGGCTGCTCAAGACTTGCACACTTCCCTGAAAATCTAGGTGATCTAAACCAGTTGGAGGAGCTCTATGCTGGTAACACTGCCATTTGGCAACTACCAGATTCTCTTGGAAACTTAAGCAAACTTAAAGTCCTATCGTTAAGAAGAGGCCGGAAGGTAAAGTATCAAGCTGGTGGCAGTTTAATGCTACCCCCATTCTGGGAGTTTCATGGTTTGAGGGAATTGAAAAGCTTAGATCTCAGTGGATGCAATTTATCTGATAACCAAACTGCTGCTCTTATGAACTTGCCTTCTTTACTAGAACTGAATCTGAGCAGAAACAAGTTTATTTCATTACCTGATATCATCAGTCGACTTTCTCAGCTTCGATATCTCAACATAACACAGTGTCAGGAACTTAAGGAACTTCCCAAACTACCTCAGAGTATAGAGGAATTATATGCAGAAGATTTTTTGGCCAAGCAAAGTATCGCAAAGCTACAAATGTACCCGAGATTGAATTTGGTGTCATTCACAAATTACAGTTTTGATCAACAATCTTATACAAGGGAGAGCAATGGTAGCTCAGTTTTGGATGAGATTCTCGGTTCATTTCTTTCACTCAACATGGATAATGTGGTCCAGTCCTCTCTCAACTCCAATTACAGGGTGACTTGTTCCATCGTCTTTCCTGAATGTGCAATTCCCACGTGGTTTAGGCACCAGAGTGTTAAGGAAAAGATGTTGCTTGAACTTCCTATTAATTGGTATAATGATAAGTTCAAGGGCTTTGCTATATGCTGTGCCACTCTCATGGGAGCAGGTGTGTTGAATCCTGATTCAGGGCTATCCAAGAAGTATGATTATGCTTTCGTCAAAGCTAAATTGATATGCAAAGATGATTTAGAAGACCTTAAAGTGCTAGAGAAAGAATGTAAAGTGGGAACAGCATCTAGAACGTATGGCTTGTGTGTTTGCTTTGTCTACATACCATTGTATGCTTCACTGCAGGTGTCCGGCACAGATGTTGGAAATATTAACCAGTATAGCCTATTTGAGGCATCTATCCATGGGCGCATTGTGAGACAGTGGGGAGTTCATATGATCTATGAGGATGAAAGTACATTTTTTGCAGGAAAATATGGAAGTGTTGGTTTCACACCACTTAAGCTGAAGAGAGGCTAAATTAGCTGTATATGCTGGGATATTGAGGGAATTACCAAGAAACCGAGTTTAAGGTAACTCAATGCCTATTATTATTGAAGTTTAATAAATTGAATTTTGAATACAGTTAGATTAACAATTTCATTTCAATGTGGTTAAGTATTTGGTTAATAATGTTGGTAAAAATTAGGCAAAACTAATGATTTACTTAGCATTAATGAATTAATAAGGCAAGTCAATAATTACGTTAAAATTATTAGTAAATGGCAGTTAACAGGTTTGAAAAATATATGAGTCAATCTGATAGCTTAATAGTTATACCAATATAATAGTTACATTTTTATTTTTTTTTACCTGGATAGCTCTATATGAGGCATGTCTCACCCAAGACAACATAAAGAAAAGAAAAATATAGTTCCCAAACAGATGCTATATGTGCAAAAAAGAGGCTAAAACCCCCAGACACCTTCTTCTTCATTGTGAGGTGGCCTCAGAATTATGGAGCATTTTTTTCGTCTCTCTGGCATCAATTGGACCACACCCCTCACTGTTATGGATGCATATGAGAGTTGGAGTCTATGGAAAGTTGACAAAGCCATCAAGAAGATCTGGATTATGATACCTGCTTGTATTTTTTGGTGTATTTGGTTAGAGAGGAACAAGAGATGTTTTGATGGAGAGTCAACTGCTTTAGGCATTTTGAAGACTAGGTGCATTGAAAATTTGTTTAGTTGGTCCAACTTGTATCCTGTTGTTAATGCAGAACAACTTCAGGATTTTACTAACTCATTAGCTCTGGCTTAGATAGTTTGTATAGTTAGCCAGAGATTTTTTTTGTAAGCTCAAATTCCATATCTTGTAAATTTTGCATCTTCTTGATGTTGTTATTAATACCACTCATCTTATAAAAAAAAAATAGTTACATTTTAAATCACATTATTCTTGATGTTGTATATCCTCGATCATTTTTGTAGCAACCAATTCGAAATAAACTAGTTGATATACTGACCTTCAAGCGGGAACCTAATATTAGGGATGGTTGTGTAACAAATCTTCAAAATCTTAAGTGTAATAGTACCTCTTCCTTGTTAAATTTACTGTATAATATTACAATTAAGAATTAAATTAGCAGTAAGTGTTTTAGAAAACTAAAGACTGTATTAGGCACCACCAAAGGTTGTGATAGGGTGGTAAGTACCCTTTTATCCTAAATCACAGCTCTCGGATTTGAGCCCTGAGATTTGATTGGAGGCACTTTACCCCTCAAAATAGGATTTACCTGCACAAATTCATATTAGTCGGGCCCAGAGGCAGTGTCATTTGTTTGAGCTCAAAGAGGAGACGAACATTGAGCAACCCGTCATATATATTCTGTATGTATCAGGTGCATCCTTTTTCTTTTTCATTATTCTTAATGTTTCTCCTTGTAAATATTAGTTCTATGTTCTTTCCGATCCTTTATATTGGGTTTGATTGCTCAATCAAGCTCTCCGGGGTTTTTGCAATTTAAAATGTCCAAGGAAAAAAAAAGTACTTCATCTGTTTCAATTTGTTTGTCTGATCAAGAGTGTACTTTATATCTTATTCTTAATCTGTGATTTGTAGGATGGTCGGAGCAAAAGTTGATGAGTGCACAAGTAGTGTTGTATCTAGACAGATTTGTGAGCAGGGGAAGCACTTCTCTGTTAGTCATTGATCTGGCTAATCTGGATTCGTGCTGAATAGAGTTACTAAAGGGGTAAAGCGTTTTCTCCAAAAGGTTCTCTTGCATTCACAAGTCTCCAACCCAAGATATCAGGAAAGCGCTACCATATTAATGAGGAGATTTTAAAGATGATGAACTCTCCTCCTCTGCCACAATCGTCATCTCCTCCTCCTCCATCGTCACCACCATCACTTTCGCCGGCTCTTTCTCCGTCTTGTTTGCTTCACTTTCTATCAAAAAATTAAAGTGTCTTTATTCATCTCCTCCAGACTATATTTTAGAATTTTTAGTGTCTTTATTGCTTGTAGCATCATATATGGATTTTCTTATATCGTCTTAATTCTATCCAAAAT

>PGSC0003DMG400011045 LRR-RLPs

ATGGCCGATCTTGTAATTGGTGCTACTGTTAAAGTTGTGCTTGATAAACTGCTTTCTCTCACTATTGAGGAGGCCAAGAGTTTAAGGAACTGCAAGAAAAATCTAAGAATTCTGACAAAATATGTATCCATGATTCAAGCTCTCATTCATGATGCTGAAAGACGACAAGTTGATGATCAGGCTGTTGAACAATGGCTCAAGATGCTTGAGAGAGTTGCTGAAGATGCTGAAAACGTGTTTGACGAATTCAGATATGAATATCTCAAAGCACAAGTCATGAACATCCGAACCAAACTAATGGAAAAGGTCAGCAGCTTCTTTTCTCATACTGCTTTTAAGTACAAAATGTCGCGAAAAATAAACAACATCAATGAAGAGTTGAAGGCTATCAATCAGTTAGCCAATGACCTCGGTCTCAAACCACTGATGGTTCCTTCTCAGAAAATACTTCCAATTCGAGAAACAGATTCCTTAGTTGTTGCTTCGGATGTTGTTGGTAGAGACAAAGATGTTGCTGAAATAAAAGAGAAGATTTTGAACATGAGAAAGGATGTTGTTCTGTGCACCATTCCCATAGTGGGTATGGCGGGTTTAGGGAAAACAACTGTGGCTAAGAGAATTTTCAATGATGAACACATCAAGCAGCAGTTCGAAAAGAGAGTTTGGTTGTGCCTACCTGAAATGTCAGAAACTAAGAGCTTTCTTGAACAAATCCTTGAATCATTGACAGAGAGGAAAGTTGAGGTTCAAAGGAGAGATTTAATAGTTAAGAAGCTACAAGATGAATTGGAAGGAAAAATGTATTTGCTTGTCCTAGATGATATGTGGCGTGTTGACTCTATATCGTGGCATGAGTTCATGGACACCTTGAGAGGAATAAATACATCCAGAGGAAACTGCATTCTTATGACTACTCGTATGAAGCAGGTGGCATCCATAGTAGCAGAAGATCTTCATATGTTGCGAAGATTAGCAAGAGATCATTGTTGGTCCATTTTCAAACAAAGAGCGTTTGTTGATGGGGAAGTTCCGCAGGAAATAATGAGCATGGAGAACAAGATAGTTGAAATGTGTCAAGGTCTACCATTGGCTGCAAGTGTGTTGGGAGGCCTCTTATGCAACAAGGAAAAACATGAATGGCGGGCAGTTCTTAATGGCAATCCCCTTGTTGCGGGGGAAGATGATAACGGGGAAAATAGCATAAAGAAGATCCTAAAACTCAGCTATGATTATCTACCATCTCCACATCTGAAAAAGTGTTTTGCTTACTTTGCCATTTTTCCAAAAGATTTTGAGTTTGAAAAGGACCAGCTTATCCAACTCTGGATGGCAGAAGGTTTTCTCTGTCCATCTCAAGAGACCACTTTGATGGAAGACGTTGGGAACAAGGCTTTTCAACTTTTGTTGCAAAATTCCTTGCTGCAAGATGTTAAGCTAGATGATCTGAACAATATAACACACTGTAAGATGCATGATCTTGTGCATGATTTGGCTGGAGATATTTTAAAGTCTAAACTATTTGATCAAAAGAGTATTGGAGGAGAAAATCTTTCTCAAGTCCGATACTTTGGATGGGACTCACCAAGTGATCAAATAGATAAGATAAACGAGTCAGGACGTTTGTGCACATTGTTCTGGAAAAGAATTATATCTGAAGAACTGTTGTTGAGCTTTCAGTTCTTGAGAGTTTTAAATTTGTCCAGGTCAGGCATCAAGGTATTGTCAGCCAAAATCGGCAAGCTAATATACTTGAGATATCTTGATCTCTCCAGCACTGAGATCAAAGCATTGCCCAACTCCATTTGCAAGCTCTACAATTTGCAAACATTGAGAATCAATAGCTGTTTCTCACTTAGGAAGCTTCCAGAAGAAATGGCAAATATGATAAGTTTGAGACACATATATTGCATCTATCATTACCCAACATATCTTTATTTTGCGGGTCGGAGATCAAGGTTTATAGGTAATTATCAGTTTCAGATGCCACTTAATATGGGGCAATTGACTAGTCTTCAGACGCTACAGTTTTTCTATGTAGGTTTAGCGAAAGGTCGTCGAATAGAAGAATTAGGTCATTTGAAAAACCTTAGAGGTGAATTGACAATCAAACATCTCCAATTAGTCGTAAATAAAGAAGAGGCTCGAACAGCATATTTACAGGAGAAACCAAATATCTACAAGCTGACATATTTATGGACCCATGATGAATCAGAAGGCTGTGAGATCAATGATGAGCATGTGTTGGATGGTCTTCAACCGCATCCGAACTTGAAAACCTTAGCAATAGTGGACTATTTTGGGACTAAATTTCCTTCATGGTTCAGTGAAGGATTGCTACCAAATTTGGTCAAGTTGAAATTAAGTGGCTGCAAAAGGTGCAAAGAAATCCCATCACTTGGCCAACTGAAGTTCCTTCGGCATCTTGAACTGATAGGATTCCTTGAATTGGAATGCATTGGATCTACATTTTATGGCGTTGATGTTAATGATAATGGATCAAGCAGCAATAATGGCAATATCCAAGTGTTCCCATCACTGAAAGAACTAGTATTGCAGAATATGAATAGCCTTATTGAGTGGAAGGGTGATGAAGTTGGAGTTAGAATGTTTTCTGCGCTTGAGAAGTTGAGGATCACAAAGTGTCCACTGTTAAAAGGTACCCCACGTCAATTGGAAATCCTCCGTGAATTAAGCATTGAAGGAGTTGACAGTGAAATGCCATTGTTGAACTTGTGTAGCAACTTGACATCTCTGGTAAAGCTTACTGTGAATGATGTGAAAGAGCTCACTTGTCTTCCTGATGAGATGCTACGCAATAACCTTTCTCTTCAACAACTATTGGTCACACACTGCAGAGAGTTTCGTGAATTGCCCCAAAGCCTGTACGATCTCCATTCTCTTGAGAGCTTACAAATATTCTTCTGCCCCAATTTCAGTTCTCTTCCTGTTCCCAGTGTAGAGAATCGTTTGACTTCCCTCCAAAGTCTTGATTTATCACTGTGTTTTGGATTGACCAGTTTACCAAGTGGAATGCTAGAGCATTGTCGGTCTCTACAGACTTTGAAGGTCAGCTGCTGTGACAACTTAGTTTCCTTGCCTTTACACGTATGGGAAATGCCTTCAATTTCATATTTGGGTTTATCAAATTGTCCCAAGTTGATTAGTGTACCCACAGGGGGCCTTCACCACCTCACAGGGTTAAAGGTATTGGAAATTGGTCCTTTCTCAGAGATGGTGGATTTTGAGGCATTCCAATTGATTTTTAATGGCATTCAACAGCTATCGTCCCTTCATACATTGGAGGTGTATGGATGGACGCGTTGGGATTCTCTGCCTTATCAGCTTATGCAACTCTCTGCCCTAACAAAGATCCATATACATGATTTCGGAATTGAGGCTCTTCCTCATAGACTTGACAACCTTACTTCTCTTGTAACATTACATCTAATGGGGTGCAAAAGGCTACAACATGTGGACTTCTCAGATGTCATGCCCAAATTACGGTATCTGGAGATCCGTTATTGTCTATTGTTAGAAGGTCTGTCGGATGGGCTTGGCAACCTTGTTTCTTTGAAAGAATTAAGTTTACGTAACTGCGAAAAACTACAACATCTGCCATCCAGATATGCCATGCTGCGCCTCACTAAATTATGGAAACTAATAATTAATGGCTGCCCGCAGTTAGAAGAAAGTTGCACCAATCGGAGTAGCCCAAACTCCCAGTGGTCCAACATTTCCCATATTCCAAAAATTGAAGTAGGCGGGAGGATAATTCAGAACTTACGTAAGTCTCTGTTTCTCTTTCACTTTTTGATCTGTAAAATTTTTAAATATTATTTTCCCCGAGTTCCAGAAAAGAAGTTATGCATGTAG

>PGSC0003DMG400011108 LRR-RLPs

ATGTTTCCAAAAGATTTTGAGTTTGAGAAGGACCAACTAATCCAACTCTGGATGGCAGAAGGGTTTCTTCGTCCATGTCAAGAGACCACTGTGATGGAAGACGCCGGGAACAAGTTTTTCCAACTCTTGTTGCGAAATTCCTTGCTGCAAGATGTCAAGCTAGATGAGCACAACAATATAACACACTGTAAGATGCATGATCTTGTGCATGATTTGGCCAGAGATATATTGAAATCTAAATTATTTGATCCAAAGAGAGTTGGAGGAGAAAATCTTTCTCAAGTCCGATACTTTGGATGGGACTCACCAAGTGATCAAATAGAGAGGATAAATGAGCCAGGACGTTTGTGCACATTGTTCTGGAAAAGAAATCTATCTGAAGATATGTTGTTGAGCTTTCAGTTCTTGAGAGTTTTAAATTTGTCCAGGTCAGACATCAAGGAGTTGTCAGCCTCAATCAGCAAGCTAATATACTTAAGATATCTTGATCTCTCCTACAATAAGATCAGAGCCTTACCCAGCTCCATTTGCAAGCTCTACAATTTGCAAACTTTTAGAGCCAATAACTGTTTTTCACTCAGGAAGCTTCCAGTAGAAATGGCAAATATGATAAGTTTGAGACACATATATTGCAAGTATCATTACCAATCATGTACAGGTAAGGATCGGTTTCAGATGCCACTTAATATGGGGCAATTGACTAGTCTTCAGACGCTACAGTTTTTCTATGTAGGTTTAAAGAAAGGTCGTCGAATAGAAGAACTAGGTTGTTTGAAAAAACTTAGAGGTGAATTGACGATCGAACGTCTTCAATTAGTCGGAAATAAAGAAGAGGCTCGAACAACATATTTACAGGAGAAACCAAATATCTACAAGCTGGTGTATTCATGGTCCCGTGATGAATCAGAAGGCTGTGAGATCAATCATGAGCATGTGTTGGATGGTCTTCAACCACATCCGAACTTGAAAACCTTAGCAGTTGTGGACTATTTAGGGACTAAATTTCCTTCATGGTTCAGTGAAGAATTGCTACCAAATTTGGTCATGTTGAAATTAAGTGGTTGCAAAAGGTGCAAAGAAATTCCATCACTTGGCCAACTGAAGTTCCTTCAGCATCTTGAGCTGGTAGGATTCCTTGAATTGGAATGCATCGGACCTACATTTTATGGTGTTGAGATAATTGATAATGGATCAAGCAGCAATAATGGCAATATCCAAGCGTTCCTGTCATTGAAAGAACTAGTATTGAAGAATATGGCTAGCCTTATTGAGTGGAAGGGAGTTGAAGTTGGAGTAAGAATGTTTCCTAGGCTTGAGAAGTTGAGGATTAGTAACTGTCCCCTGTTAGAAAGTACTCCGAATCAATTTGAAATCCTATGTGAATTAGAGATTGTTGGAGTTGACAGTGAAATGCCATTGTTGAACTTGTGCAGCAACTTGACATCTCTTGTGAATCTTGTTGTTAGTAACGTGAAAGAGCTCATTTTTCTTCCCGATGAGATGCTACGCAACAATGTTTCTCTTCAACACCTATTGGTCTCACTCTGCGGAGAGTTTCGTGAATTGCCACAAAGCTTATACAATCTCCGTTCTCTTAAGAGCTTAGAAATTACCAGCTGCACCAATTTCAGTTCTTTTCCTGTTCCCAGTGGAGAGAGCTATTTGACTTCCCTCCAAAGTCTTGAGTTATCCAATTGTAATGGATTGACCAGTTTACCAAGTGGAATGCTAGAGCAATGCCGGTCTCTAGAGATTTTGACGGTCAGTTACTGTAACAACTTGGTTTCCTTGCCTTTACATGTGTGGGAAATGCCTTCACTTTCGTATTTGTGTTTATCAGAATGTCCCAAATTGGTTAGTGTACCCGCAGGGGGACTTCACCACCTCACTGGGTTACAGCATTTGGAAATTGGTCCTTTCTCAGAGACGGTGAATTTTGAGGCATTCCAATTGATTTTTAATGGCATTCAGCAGCTATCTTCCCTTCGTACATTGGAGGTATACGGACGTACGGATTGGGATTCTCTGCCCGATCAGCTTATGCAACTCTCTGCCCTAACGGAGATCGGAATAAATGATTTCGGAATCGAGGCTCTTCCTCATAGACTTGGCAACCTTACTTCTCTTGTAACATTAGATCTGGTGAGATGCAAACGGCTACAACATGTGGACTTCTCAGATGCCATGCCCAAGTTACGGTATCTGGAGATCCGTGATTGTCCATTGTTAGAAGCTCTGTCGGATGGGCTCGGCAACCTTGTTTCTTTGGAAGAGATAAGTTTATGTAACTGTGAAAAATGGAGCATCTGCCATCCAGAGATGCCATGCAACTCCTCACTAAATTATGGAACCTGCGAATTAGTGTCTGCCCACAGTTAG

>PGSC0003DMG400011116 LRR-RLPs

ATGTTTCCTGGGCTTGAGAAGTTGACAATTACAGAGTGTTCACTGTTAAAAAGTGCTCCAACTCAATTGGAAACCCTGCGTGAATTAGAAATTGAAAGAGTTGACAGTGAAATGCCATTGTTGAACTTGTGCAGCAACTTAACATGTCTTGTGGATCTCGATGTCAGTGATGTGAAAGAGCTCACTTGTCTACCTGATGAGATGCTACGCAAGAACGTTTCTCTTCAACGCCTGTGGATCTCAGATTGCAGAAAGTTTTGTGAATTGCCACAGAGCTTGTACAATCTCCATTCCCTTGAGAGCTTACGAATATCCTTCTGCCCCAATTTTAGTTCTTTTCCTGTTCCCAGTGGAGAGAATTATTTAACTTCCCTCCGAAGGCTTCACTTACTTTATTGTGATGGATTGACCAGTTTACCAAATGGAATGCTAGAACAATGCCGGTCTCTAAAGATTTTGAGGGTCAGCAACTGTAAGAACTTGGTTTCCTTCCCTTTACATGTGTGGGAAGTACCTTCACTTTCCATTTTGGATATGTCAAATTGTCCCAAATTGATTAGTCTACCCACAGGAGGCCTTCACCATCTCACTGGGTTATGGGATTTGAAAATTGGTCCTTTCTCAGAGATTGTGGATTTTGAGGCATTCCAATTGATTTTTAATGGCATTCAGCCGCTATTGTCCCTTCATACGTTGTGGGTGTACGGACATGGGCACTGGGATTCTCTTCCCTATCAGCTTATGCAACTCTCTTCCCTAATTGAGATCAGATTAAGTGATTTCGGTATCGATGATCTTCCTCATAGTCTTTGCAATCTTACTTCTCTTGAAACATTACATCTAGTGGGGTGCAAACGGCTACAACATGTGGACTTCAGAGATGCCATGCCCAAATTACATTATCTTGAGATCTGTGGTTGTCCATTGTTAGAAGCTCTGTCGGATGGGCTCAGCAACCCTGTTTGTTTGGAAATGTTAATTTTTTGGAACTGCGAAATACTGGAGCGTCTGCCATCCAGAGATGTCATGCGAAGCCTCACGAAATTACAGTTCCTGAAAATTAAAGACTGCCCACAGTTAGAAGAAAGTTGCACCAATTGGAGTAGCCCAAACTCCCAGTGGTCCAACATTTCCCATATTCCAAAAATTAAAGTAGGTGGGAGGATATTCAGAACTTAA

>PGSC0003DMG400011209 LRR-RLPs

GCGACTACTAGGTGACCATCATACTATGCCTATAAATGACAAGAGATGTTCCTTCCTACTTCATTCACAAACAAGCAAATATGAAGAACAATTCACTTCAACTTATCTTTCTCATATTGTTATCTCAACTTTTCTATTTCACTTGCATAAACCATCACTTTGTTTCCTCTACTAACACCTCTTGCATTGAAACAGAGAAAAATGCACTTCTAAACCTTAAAGAAAATCTCACAGATCCTTCAGACAGGCTATCTTCTTGGATTGATCAACAAGATTGTTGCCAATGGCTTGGTGTAACATGTAACAACAAGACCGGACATGTCGTAAAGATTGATCTTAGGAACAAATACTCAACAGAACATGAATTGAGAGGTGTGATTAATCCATCTTTACTTAAGTTACAACAATTGAGATACTTAGACTTAAGCATGAACAATTTTGAAGGAATCAAAGTACCGGAATTCATCGGTCGCCTTAATGAACTAAGGTATCTCAATCTATCTGGTGCATCTTTTACTGGTTTAGTCTCTTCATTCCTTAAAAACCTCTCAAATTTGCAAGTTCTTGATTTAGGAGGCTTAGATCTTAGTAAAGATACTAATTCTTGGCTTCAAATAATCAATGATCATGTTCCTTCTCTGTTGGAGTTGCGTTTGCCTCGATGTCAACTTTTAAACATTCCTTCTTCTTTTCCATTACTAAATCTTACTTCCCTTTTGGTACTTGATCTTTCCAACAATGCTTTTAGTTCATGTACATTCCCTCAATGGATATTTAAACTTAGTAGCCTTGTGTATCTTGACTTAAGCTCCAACAATATGTGTAGTGAACTTGTTGATGAGTTTGGTAAGCTCACTTTCCTTGAACATCTTGATCTGTCTTCGAACCACGGTATCAATGGGACACTCAAGAGGAGCTTGGGGAAACTATGTAATTTGAAGAGTTTGATCCTGTCTTACAACAACATCAGTGGAGACATAATCGAGTTCATCGATGCGTTATCTGAATGCCAAAGCAATAGCTTAGAGACATTGGAGTTGAGTTTCAATAAATTGACAGGAAATCTTCCTGATGCATTAGGCAACTTGAGGAAACTGAAACATCTTCAGCTGAGGTATAACTCACTGACAGGCACAATACCTGAAAGTATAGGGAACTTATCATCCTTGGAGACATTTTACCTCACATCTAACAAAATGAGTGGGAACATCACCACGAGTATCGGCCAGCTCATGTCGTTGGTCTCGTTGGATATCTCGGAGAATATGTGGGATGGTATTGTCACTGAAGCTCATTTGATGTATCTTTCCAACTTGCAAGAGTTTTCAGTTGGGATGAAACTTAGTAAAAATATTACCTTGGCATTCGATATCAGCCCGAATTGGAATCCTTCATTCAGGCTGACATTCTTGACTATTCAGTCATGCCAATTAGGTCCTGAATTCCCACATTGGCTGAAGGATCAGAATGAGCTCACCAGCCTAATACTCAATACCGCGGGGATTTCTGATGCGGTACCTGATTGGTTTGTAGAGTTGGATTTGAAGCTTGATAATCTAGACATGGCATACAACAACTTGACAGGGAAAGTTCCAAACAAATTCAAGTTCAACTTTCAAGCCAATGTGGATTTAAGCACTAATCGCTTTGAGGGACCTCTCCCACTATGGTCTTCTAATGTTACAACATTGTATCTGAGGGATAACTTGTTTTCAGGACCTATTCCTCTCAACATTTATGAAGCACTTCCCAATATAGGAGACTTGGACATCTCCAGGAACAATCTAAATGGCACAATTCCCTTATGCATAGGTGATATGAATCAGTTAACTACGTTAGCCCTCGATAACAACCAACTTATCGGCCAAGTTCCTGATTTCTGGGGTAAGTTACCATATCTTTACTGGATAGATATGTCAGAGAATCGCCTATCCGGACAAATTCCACACTCACTAGGCTCTCTTGCTAGTCTGATGTTCTTAAGACTCTCAGGCAACAATCTCTCTGGAGAACTGCCTTTGAGTTTGAGAAATTGCACGAAAATGATCAACATTGATCTTAGTAACAATCAGTTGTCAGGGCTAATCCCAGCTTGGTTAGGGGAAACAATGAGTTCACTGTTAATCCTTAGTGTAAGGAACAACAGGTTTTTCGGTCACATTCCTTTAAAAATCTGTAGCCTTTCAGGTCTTCACATACTAGACTTTTCAGGAAACAACTTATCCGGTTCTGTCCCGTCCTGTTTTGGAAACTTGGAAGGGTTCAAGGTTGAGCTAACTGATGTAGAAGCTAAACAATATCAAGGTACATTGAAGCTTGAGGCAAAAGGAAGAACACTTTCTTATGATAGAATCCTTTACCTAGTAAACAGTATCGACCTCTCGAGCAACAGTTTGTCTGGTGAAATCCCGGAAGAGTTAACAAACTTACACAAACTTGGAACCTTGAACTTGTCTATGAACCATTTGACAGGACATATACCAACAGATATCGGAAAGTTACGTTGGGTTGAAACGTTAGATCTTTCAATAAATCAGTTATCAGGTACTATCCCACCAAGCTTAGCTACATTAGACTTTTTGAGTCATTTGAACTTGTCATATAATAAGTTGACAGGAAGAATTCCTACTAGTACTCAGTTTCAAACCAAAGTTGATCCAACAATTTTTCAAGGAAATGTTGCACTTTGTGGTCCTCCTTTGGAACAATGTGTTGGTGATAGGACTACGACGACGACAACATCACAAAGTGGAGGAAATGATGAAGGGGAAACAGATGATGAAGATAAACTTGAGAAAGTATGGTTCTTTGCTGTTGTTGGTTTGGGATACATAGTTGGTTTTTGGGCATTTTTTGGCACTTTGATCTTCAAGAAAAGATGGAGAATTGCTTATTGCAGATTCATTGAGATGTGTATCTTGGAGTTCTATGAAATGTCTTGTAGTTTTGTGAAAAAAATGATAGGAGTCTAATAAGCTATGTTGCTCAGACTCTTCAAAATTGTCGAGGGGTGCATGTGGGATCCTCCAAATATAGTATATTTGTGAAGGATACGACAATATTTTTAGAGAGTCCGAGCAACTTAGCTAATAAGTGGAGCTATAGGAGTTTGTCTCTGCAATTCTGCTATATGCTATGTTTTAGTTTCAATATAGCACATATTACTACTTTTTCTATATGTAAAATAATGCTACCTAAGTATTGGCACTTATAGAAATGCCAGTGTTTTGGGTAGAGTTACGACATGAGTGGTCAGTTGAACACTTTTACGTCGGTCAAAATAATACTTCTTATATATGTATACATGTTCACAAGACTATTGTAGTACATATAATTAGGAAGTTGTTTAAATAGTATCTAAGACTATTCTTATACTTAATTTGCATGCATCTACTTGTATA

>PGSC0003DMG400011353 LRR-RLPs

ATAATAAAACCTTATCTTCTCTGTAGAACACTCACACACACTCTCTCTCTCCTCTTTCTCTCTCTAAAATACAGCAAATCTAAGCTCGTTAATGTCGATTCAGATCTGAGAGAAGAAGAAGAAGGAAGAAAAAGGACTACTGTAAATCTCTCTCTCACTGTGAAATTACACTTTTGCCCTTCCCCCTCCATTACCGTACAACAATGGCGTATGCATTCCCGGAGGAAGTACTAGAACACGTCTTCTCCTTCCTCACCACCGACAAGGACCGTAACGCAGTATCTCTAGTCTGCAAATCATGGTACGAGATCGAACGGTGGTGTAGGAAGAGAATATTCGTTGGAAACTGCTATGCCGTTAGCCCTAGGATCATGATCCGACGGTTTCCGGAAGTCAGATCTGTTGAGCTTAAAGGGAAGCCTCATTTTGCTGACTTTAATCTTGTACCGGAAGGTTGGGGAGCTTATGTTTATCCTTGGGTTTTGGCTATGTCTAGGTCTTATCCATGGCTTGAAGAAATTAAGCTTAAGAGGATGGTAATTACTGATGAGTCGTTGGAGTTGATTTCTAAGTCGTTTAAGAACTTTAAAGTGTTGGTTTTGTCTTCTTGTGATGGGTTTACTACTGATGGACTCGCTGCTATTGCTGCTAACTGCAGGTGAAATTTTTTGTCTTACTTGACATTTCTCTTCTGTTTTGCTCACTTTAAAGTAGTTAGTGGTGAGAGAAGTCTTAATCTGTAAAGTAATTAGATCTAATTTTGGGTATTAGTTTGGACAATGACTTGTATTGTTGTTAAATCTTAAGCTTTTGGATGAATGCAGTAAGCATTTTGGGGGTTTGTTATAATCCCTTGCTAAACACGAGAATGAAGAAGCACTTTTCTTGCAGCTGTTTGTTCTGTTTATAGGATATGTAATGTATTAGTGGAAACGAAGTATCTACTGTAGATGAATTCGTAGTGTGTGAACTTGAAGAACTGACATTTCTTCAAGTGATTATGGTATTATTGGATACGTCTTGCTGCTACATTGCATCAGTCTGGTTCATCTGACTTTAATACTTCAAAGATTAGAGCTTTTTGTTTGTTTCTTTTAGCTTCTTTTTTTTGAGGTGAACTTCTGGGAATTAATTTTTCGGATTTTGATTGAATGAGCTAATGTCAATAATTTCTTGTACTTCCTTCCCCAGAATTCTACATTTGCATTCTATCTTTACAGATCTCTAAGGCCTTATAACAGAAAATGCTAACTTGTGGGTTTTCTTAATTGTTATAGGATTTAATCAGTTGGACTTCCTTTTAGCATTAATTTCTTGTGGTATACACTGTGTAAACATGCCCTTTAGCTTTGCTTTGTTTTGAAGTTGTTGGAGATTTGAATGTTATCTTAATTATCTGGAGATTTCCATATGTTTGTTTCTTGGTTCTCTGGTGGGATTAATTTGTTTTATCAGGAGGGAGTTTTGTAATGGAGTATAGTTGTAGTTCAATGTTGAAATGGTTAAGTGGGTTAGGAATATGAAATGGTGATTTATTGAAATTCCAATAGGAATATGATTCTAAATCTTTCTTATTTATCTTGTTCATATCGTCTGTGATTTATTATCATTTGTTGTTTTCTGTTGAGTCTGTTGATTTGCTTCAATTTATCACAGAATAATTGTGTAGATGGGTATTGTCATCTAACTCCCGTTTTGCTGCAGGAATCTGAGAAAACTAGACTTGGGAGAGAGTGAAGTGGAGGACCTTAGTGGGCATTGGTTAAGCCATTTCCCTGATAACTGCACGTCACTTGTTTCACTTAACATTGCCTGTTTGGCTTCTGAGGTCAGTCTATTAGCTTTGGAGCGCCTGGTTACTCGCTCTCCTAATTTGAGAACTCTTCGGATAAATCGTGCTGTGCCCCTTGAGAGGCTTCCAAACTTGCTCCGCCGCACTTCCCAGTTGGTCGAATTTGGTACAGGTGTTTTCTCTGCTGATGTGCGGTCTGATTTTTTCTCGAACCTGACAGAAGCTTTTTCGAGCTGCAAGCAACTTAAATGCCTGTCTGGGTTTTGGGATGTTGTACCAGCTTACCTTCCAGCTTTATATCCAGTGTGCTCAAGACTCACCTCCTTGAATTTGAGCTATGCTACATGTCAGAACCCTGAGCTTGGCAAGCTCATAAGTCAGTGTCACAATTTGCAGCGTTTGTGGGTATGTTCTAAATGTTTTGAAGTCTTAATTGATGTTCTTGATACTGCAACTGCAAACATGTTTTTTTTTCTTATAAAGAAGAAGAGGCAATGAGATTGTAACATCATACTTGCTGTGATGACATTTAGTATATATTCTTAGTGAAGAATAAGAAAAAACCTAACTGCTACAGATCAACCAATCAACCACAACTTAATCCCAATTTAGTTGGTCTCAGCTTTATGGGTCCTCTTAAACCATATTGCTATATTCAGGTCATTTTCGTTCCGATGCTAAATTATTTGTCTTTACGCAATAATTGCCAAGGATAGGAAAAAAGTAGTTCATTCTGTAGATAAACTTCACCTATGCTGGTTAGGAAAATTCCAGGCTACATATATTTGATATTATATTGCTAAGCAGCTCACTTTCCTTGGAAGTTGAAGCCAATTGTTACATGCTTCATATTGTGATCTGGACGAATGGGTTTCAGTTTTTAATACTTGTGCCTACCCAATGCACGTAAAAATTCATCTAAATGTGATGGTTCTATGCTGAAAAATGTATCTATTCTTCCCTAGTAGCTTCCCCAAACTGGTTCTGTAATATAAACTCCAGTATCAAGTTTAAGTGCTGAATTACTTTTCTTGTAATATATGTGACTGCAGGTGCTGGACTACATTGAAGATACTGGTCTTGAGGAACTTGCTGCCAATTGTAAGGACCTTCAAGAGCTGAGGGTGTTTCCTTCTGACCCATTTGCTGCAGAACCAAATACAACCTTGACGGAGCAAGGCCTTGTAGCTGTCTCAGATGGCTGCCCAAAGCTCCAGTCTGTCTTATACTTCTGCCGCCAAATGACTAATGCAGCTTTAGTTACTATTGCTAGGAATCGACCTAACATGATCCGTTTTCGCTTGTGTATCATTGAGCCTCGAACTCCTGATTATTTAACCCTTGGTTCATTTGATGCTGGTTTTGGGGCCATAGTAGAGAACTGTAAGGAATTGCGGCGACTTTCACTTGCTGGCCTCCTTACCGATCGTGTCTTCGAATACATTGGCGCCCATGCTAAAAAGTTGGAGATGCTTTCCATAGCTTTTGCTGGTGATAGCGATTTGGGACTCCACCATGTTCTCTCGGGTTGTGATAACCTTCGGAAGTTGGAGATCAGAGATTGTCCGTTTGGAGACAAGGCTCTTTTGGCTAATGCTGCAAAGCTCGAGACAATGCGATCCCTTTGGATGTCTTCTTGTTCAGTAAGTTTTGAAGCATGTAAGATGCTAGCTCAGAAGATGCCTAGGCTTAATGTGGAAGTCATAGATGAGAGGGGTCCTCCAGATACAAGGCCAGAAAGCTGTCCTGTGGAGAAACTTTACATATATCGAACTGTGGCTGGACGAAGATTTGACACACCTGGTTATGTTTGGACAATGGATGAAGATGCAGCTGTAAGGTTGACCTGAGCCATACATCAAATGGAAAATTCCAGCAAGACTTCACATGGAAAGATGGGCAGGTATTAGCGCTATGTTTATCCTCCTGCATTTGTTTGTCACGCCTTATGTAGTTGCAGTGCCAGTTGAGGGTTGGAACTTCACTGCTTTGAAGTAGAGACTCCCGAGCTCATACGCCATCAATTTTATTCTTGGCTGAGAGCTACTTTACAGCAGGTAGTTTTTAATAAAGAATTTGAACTCTGAGTTTCTGTGTATCATCTAGAGTAACAATTATTCGTTTTCTTTGGGTGGGATTTTGCTTGGGGGGTTTGGGGGGCATATTTTTGCCATGTATGTGTTATTTCCAGACTTTGCTTCATTATGTAGTTCCCTTTATTTATTCTATTGACTTATTCTCTATGAAATGGAGTGTCT

>PGSC0003DMG400011513 LRR-RLPs

ATAAATAGCAAAACAAACACAAAATGAAAAGATAACTCCTCTTTCTTTCCAATAAGATTAGAACCTTAAACTTCTCCGATGGCCAGCGACGACTCTCATCTGCAAAACGACGTCGTTTCTGTCCACTGTATAAACGACAGTCTCGGCGACGACGAGCTTCGGGCCGTACTGTCGAGGCTCGGAGACGACAAAGACAAGGAAGTTTTCGGGCTGGTATGTAAGCGGTGGCTTCGTATTCAAAGTACTGAAAGGAAGAAGCTTTGTGCTCGGGCCGGGCCTCACATGCTCAGAAAAATCGCGGCCAGATTCACTCGATTACATGAGCTTGATCTATCTCAATCGGTTTCTAGATCTTTCTATCCTGGTGTTACTGATTCTGATCTTTCAGTTATTGCTACTGCTTTCTCTTGCTTACGGATTCTCAATCTTCAGAACTGTAAAGGTTCGTTTTTGTATTTTTAAAAAGTTTTATAGTTTGAAAGTTGTTGAATTTGAATGTTCATTGCTGAAGTGAACCTGAGCTACTGCAGGAGAGGCCAAATGCACCAGTGAGTAGTAGTATGTGTAGTGATATTTCACATTTAGTACTGATTTAAAATATTGAATTTGTCTTTGTGTGGGGTGGTGGAGCAGGGATTTTCACTTAAAGGGTTTGAAATACATAAAACATAATTTAACCACGTATAACCGGTTTGGGTGAACCCCTGGGCCCTTATTAGCTCCGCCCCTGCCTGTGTGGGGTGGTAGGTGTATTGTTCCCAACTTGGACATCAAGGAAATTTTGAGGTTTGACTAAAATCTATTGTTTGAAATGTTAAAGGTTTTGACCCGATATGGCGAAGACTGGCTCTTCCTCTTTTCTATCTTGTAAATAATGGTTTTTATGATAAAAAGGGCGTAATGGGGTTTATGGAGGATTTTAGGTGAATTTATGGTTGGTTTTTTTGAAGATTTGTAGGATTTTGTGTTTCTGTGTAAAGGTTATGTCTTTTTTGCTATGTGGTTTTTGAAGTGGTAGCAAAAGGGAGGATAGTGTAGTTATATCATATGTATTCCAATTTTCAAGAGAAAGTCTAACTTTGTTGCGAATCTAAGGCTCAGATGCTTCCTTGACTTAACTTGCGCGGTTTTCTGGTCCAAATTTACAGGCATAACAGACAATGGATTGGCAGCGATTGGATCAAGTCTCTCTTCTTTACAGTCCTTAGATGTATCCTACTGCAGAAAGATCACAGACAAGGGGCTTTCAGCTGTTGCTGAGGGGTGTCATGACTTGAGAACGTTGCATCTTGCTGGCTGTAGATTTGTCTCAGATTCATTAATGAAAGCTCTCTCCAAGAATTGTCATTATGTAGAAGAGCTAGGTTTGCAAGGATGCACCAACATTACCAACTCAGGTCTCTCTGTTTTGGTTGAAGGTTGCCGGAGGATAAAGTATTTGGACATTAACAAATGCAGCAATATTGGTGATATTGGGATAACTAGCGTGTCTGAAGCTTGTTCATTGACACTTAGGACATTGAAGTTGTTGGATTGCTACAAAGTTGGTGATGACTCCATCTTATCCTTGGCCAACTACTGCAAAAATCTTGAGACACTTGTTATTGGTGGTTGCCGTAACATATCAGATGAGCCTATGAAGTCACTGGCAGCTTCTTGTAGTAATAGTCTGAGGAAACTGAGGATGGATTGGTGTTTGAATATTACTGACTCTTCATTGGACTGCATCATTTCCAAATGCAGAGAGCTTGAGGTTCTCGACATTGGCTGCTGTGAGGAGTTAACTGATGCTGCATTTCAACAATTAGGAAGTGAAAACATTATGTTGGGGATGAAGATTTTGAAGGTCAGTAACTGTCCAAAGATAACTGTTGAAGGAATTAAGAAGCTTATGAAGTCATGTGAATATCTTGAATATCTAGACGTACGGTCTTGTCCCCTAATCACTAAAGCAGGGTGTGAGGAGGCTGGAATTCAGTTTCCTGAATCCTGCAAAATAAACTTCACTGGAAGCTTAGCAGAGCCAGATGTGTTACTTTGAGCAGTACTTGATTTAGCAATGTCTTTGAAGGACTGAAGGGGATCATGCAGTGAAATTATCACATTCTTTGCCTAAGACCTAAAATATGACTTTGGAAGCTCCTGATGTCAGTCCCTTGTTCCCATTAATTTGCCTTTTGTTAAGCATCAAGTGTTTTGTTGTAGATTATTTCCCTGCTTTTCAGTTTTTTTTTCCCCTGCTTTTCAGTTATTTGATATCAAACTCCTGTATATCATCCTTGTAAACTACTAGTAGGCACCATTGTTTCCTGTAGGTGTTTTCAGATTGTGAATGAAATGAGTCCATTGCTGCTAGCTTGAATGTGAACTTATACCAACAATGATGGTTTGATTGATTACATTTCTTGTACGAC

>PGSC0003DMG400011610 LRR-RLPs

ATGGTTAATAGTATATCCCAACAAATTGATCAAACAGCGAAGGTACCGACATATCTTGGTAAATCTGGAGAGAGAGATTACAATGATTCTATAACAATTTCAACAAAGGGAATGGAGTATGAACTGGATAGAATTTTGACAATTCTCAAGACTATTGATCTTTCAAGTAACAAATTTGAAGGACATATTCCAAGTAATCTCGGAGATCTCATTGCGCTCCGGGTGTTAAATTTGTCTCATAATAGATTGCAAGGTAATATACCGTCATCACTTGGAAGTTTATCTTTAGTTGAATCATTGGATCTTTCATTTAACCAGCTTTCGGGAGAGATACCACAACAACTTGTTGCTCTTACATTTCTTTCATTCTTAAATCTCTCTCACAATCATCTCCAAGGATGCATTCCTCAAGGACATCAATTTGATACATTTCAAAATAATTCATATGAAGGTAATGATGGATTACAAGGATTCCCTATTTCGAAAGGTTGTGGAAGTAATTGGACACCGGAGACAAATAATAGGGATTATGAGTCAGATGATGAAGAAAGCAATTCTGAATTTCTAAATGATTTTTGGAAAGCTGCTCTTATGGGATATGGAAGTGGGCTATGCATTGGATTATCCATAATATATTTCATGATCTCAACTGGAAATCTGAAATGGCTTGCAAGAATCATTGAAGAGCTAGAACACAAAATTATTATGCGACAGAGAAAGAAACAGCGAAGTCAAAGGCAAAACAAAAGAAGACATAATCGTGTCTAG

>PGSC0003DMG400011611 LRR-RLPs

ATGAATAATCTCTCTGGAACTATCCCACCTGAAATTGGTAATCTCACTAATCTTGTCTATATTGACTTGCACACCAATCAGATTTCAGGCACAATCCCATCACAAATCGGCTCACTATCCAAGCTCCAGAACATTTACATCTATGACAACCTTTTAAATGGTTCCATTCCTGCCTCATTGAGCAATTTGACCAACTTGTCTATTTTGTCTCTTTATCAGAATAATCTTTCTGGCTCGATTCCTACAGAAATAGGTTATTTAAGGTTTCTTACTTCTCTAGATTTGGGTACTAATTCTCTTAATGGTTCTATTCCTACTTCATTGGGAAATTTGACCAACTTATCTAGTTTGTTTCTTAATGGGAATCACCTTTCTGGCTTTATTCCCGCTTTATTGGGAAATTTGACCAGCCTTTCTATTCTGTATCTTCAGGAGAATGACCTTTCTGGCTCCATTCCTGAAGAAATAGGTTATTTAAGGTCTCTTAGTCAACTAGTTTTGGCTAAGAATTCTCTAAGTAGTGCTATTCCTGCCTCATTGGGAAATTTGACTAGCTTATCTGGTTTGTATCTTAATGAGAATCACCTTATTGGCTCTATTCCCCCTTCATTGGGAAATTTGACCAGTCTGTCTATTCTTTATCTTAATCAGAATAACCTTTCTGGATCCATTCCTGAAGAAATAGGTTACTTAAGGTCTCTTAGTCAATTAGTTTTGGCTAAAAATTCTCTAAGTGGTGCTATTCCTGCTTCATTGGGGAATTTGACCAGCTTGACCTCTATGTATCTTCGTGAGAATCATCTTTCTAGCTCCATTCCTAAAGAAATAGGTTACCTAAAGACTCTTAGTTATTTAGATTTTTCTACTAATTTTCTTAATGGTTCTATTCCTGCTTCTTTGGGGAATTTGAACAACTTGTATCTTTTGTCTCTTTATGCTAACAACCTTTCTGGTTCGATTCCAAGTGAGTTGGGGAATATAGGAAGACTAGTTACTATGTATTTAAATATTAATCAACTCATTGGTTCAATTCCGGATTCCTTTGGCAACTTGAGAAACTTGCAATGGATGTATCTCCACAACAACAATCTCACTGAGAAAATTCCTTCATCTTTCTGCAATTTGATGAAATTGGAAGTGGTGTATTTGGGGAGAAACAACTTTAGGGGAGAAATTCCGCAATGTCTAGTTAATATCAGTGGACTCGAGGTATTGAAAATAGAAGATAATAATCTCAGTGAAGATATCCCTTCGTCTATTTGCAATCTAACATCACTAAGGATTTTAGATTTGGGTAGAAACAATTTGAAGGGAGCAATTCCACAATGTCTTGGCAACATGGGTGGTCATCTTGAGGTTTTGGATATCCACCAAAACAATCTTTATGGGACTCTTCCAGAAACTTTTAGCAATGGAAGTGTACTCAGAAGCTTAGACTTGCATGACAATGAATTAGAAGGAAAAATCCCCCGATCTTTAGTCAATTGTAAAAACTTGGAAATTCTTGATTTAGGAGACAATCACTTCAACGACACATTTCCAATATGGTTTGAAACTTTGCCAAATCTGAAAGTTTTAAGCTTGAGATCGAACAAATTGCATGGACCAATTAGAACTTTAAGTAATGGAAACATGTTTTCTGAGCTTCGAATCATAGATCTTTCTTATAATGCTTTCACAGGAAATTTGTCGACTAGTCTATTTCAAAAATTGAAAGCCATGAGGACAATTGATCAAACAGCAAAGGTACCAACATATCTTGGTAAATCTGGAGAGAGAGATTACAATGATTCTGTAACAGTTTCAACGAAGGGAATGGAGTATGAACTGGATAGAATTTTGACACTTTTCAAGACTATCGATCTTTCAAGTAACAAATTTGAAGGACATATTCCAAGTAGTTTGGGAGATCTCATTGCACTTAAGGTGCTAAATTTGTCTCATAACAGATTGCAAGGTAATATACCATCATCACTTGAAAGTTTATATTTAGTCGAATCATTGGATCTTTCATTCAACCAGCTTTCAGGAGAGATACCACAACAACTTGCTGCTCTTACATTTCTTTCATTCTTAAATCTCTCTCACAATCATCTCCAAGGATGCATTCCTCAAGGACATCAATTTGATACATTTCAAAATAATTCATATGAAGGTAATGATGGATTACGAGGGTTCCCAGTTTCAAAAGGTTGTGGAAGTAATTGGATACCAGAGACAAATAATAGAGATTATGAGCCAGATGATGAAGAAAGCAATTCTGAATTTCTAAATGATTTTTGGAAAGCTGCTCTTATGGGATATGGAAGTGGGCTATGTATTGGAGTATCCATTATATATATCATGATCTCAACTGGAAATCTAAAATGGCTTGCAAATATTTTTGAAAAGTTAGAAGAAAAAATTATTATGAGACAGATAAGGAAGAAGCGAAGTCAAAGGTAA

>PGSC0003DMG400011708 LRR-RLPs

GTGTTACATATGATTTTGTCTTTAAGTCAGTAAAGTTGGTTCTTTGTATATGTGGTTGGGAAAAATTTATGTAGGAGATATTTATCTCAAGTTCACATTTCTATTTTTTTTGCGGGGCCATTGAATGGTAAGTGTTTGGATTGGAATCTCCCTAGTTTGATTTCATCTTCTCTATCATTTGAAAAAACTCAATATGGAGGATGAGACGGAAACTGAAAAATCTCCGATTACTACCCGTGTGATACTTCTTTCCCAAAAACAGTCTTATGGAATACGAGTAGAGATTGTTGTACATGGGATGGATTACAACTTGATTACAGAACATTGGACACAATACTTCATAACTTCACAAAAACCTGTGAATTTCCCAAAAACAGTCTTATGGAATGCGAGTAGAGATTGCTGTACACTTTCTCTCCCTGATGTCAACATCTCATCTCCGATACCTATGAATTTGTCTTCCTCCTTAAGGTACGTGGATCTTGAATATACTAATATACAAATTAGGTATTTGAATTTCACCGGCCATATTTCTTTCACAATCTCTAAATTGAAGCAGCTCACAAGTTTAGATCTTTCATCTAACTACTTTTCAGGTGAAATGCTCGACGTTTTCTCTAACCTCCAAGAGCTTACATCTTTCTGACAACATGAATTATGCGGTTCCTAAACCTTTCGCATAATTCATTAACAAGCCACCTAGCTACCACAAGTTCATTACAACAATCTAGAGTATCTTGATCTTAAATTTAACTTCTTTCAGGGTCCACTACCTTCATCCATTTGTAACATGAGCTAGCTTGAATTTCTAGATTTATCACACAACAACTTCAGTAACTCGATTACAAGTTGCTTGGGAAGTATGGCTAGTTTAAGGGTGTTGGACTTAAGAAGGAACAATTTCAGAGGGAGTCTTCCACCATTATGTGCGCAGAGCACTTCATTGAGTACCATTTTTTTGTTTGATTTTCTTGTTTTGTTTTTAAATTTTCTAGTTGTAGTTTGTATATGTTTGATAGCTACATAATTATGTGTGTGTTATTTTAGTATCAATGGTTATTCAAGTTATGTTAATCTGTTTCTGTTTCAAGTTGACAGAAGTTATCTACACAGCCATGCACTCATTAGCAGGCTCTTCAAAAATGTCAATGGGTATGTGTTGGATCTTCAAAAGTAGTGCATTTTTAAGGAACTAACACAGATGCAACATCATTATATAGAAGAGTCCGAGCAACATAGCTCATTAGTACAGCAAAAAGCAAAAAGCAGTAAGCATATTGAGTTTGGTTCATGATTGAAATCACTTGATTCAATTGTTCATCATTACTCTCCATTTATCGTAATCACTCGTAATTGATGAATCACACTTAAAACGTATGATGATGCTTCACCTACATAAGGTAGAAAAACTAAAAATGTCTACTACTTTGGTTATTTTCCATCTCTTCATGTTTGAACTGTATCATATTTGTCCTTTTCCCCTTGTAGTCTTTGTGGAAATTGGTTTAAAGACTTTGGTGGAATATTGATAAAAGTTTCCCTCGTTACCTTAATGTAAAAAGTCTTATGGAAAATTATAGAATGTACCTAACAAGTTTGAAAACAGCACAAGAGACAACTACTTGGAAATTTCTCTTAAAATTTCTCAACAGGTGGAATGATCTTCACCAAGTTATTCAAAACATATTTGAGATAATTAAAAAATTATTTGAGATAAAGGAATTGAAATAGGTAGTTGAGTCAATAATAAGTTGCTTAATATTTAAAAAATAGAAGTAGAGTGTCAAATGGGGCTACACGTGCAATGCGCGTGCTTCGAGACTTTCCCAAGTTGTGAATTTTTGATCTTTCTTGTAATGAATTCAGTGGATCACTTCCTGGAAAAGTTTTTGGAAACTTCAAGGCAATGATCAAATTAGATGGTGAAGACACAAGAGAGATCAAGTACATGAAACCATCTGAGAAGTCATTTTACATAATGTATGAGGACTCGGTGAGATTGGTGATCAAAGGCCAGGAAATCGAGCTAGAAAGAATCAACACAATTATGACAGCCATAGATATCTCAAGCAACCATTTTGAAGGTGTCATTCCGAAAACACTAAAGAATCTCAACGCTCTTTGGCTACTCAATTTATCCCGTAACAATCTCAAAGGTGATATTCCAATGGAATTAGGGCAATTGAATACGCTTGAAGCTTTAGAACTCTCTTGGAATTGGCTCACTGGAAAGATTCCACAGGAATTGACAAGAATGAACTTTCTGGCATTCTTAAACCTCTCTCAAAATCTTCTTATTGGACCAATTCCTCACGGTCAACAATTCAACACATTTGGAAATGACTCGTATGGTGGCAACCTTGATTTATGTGGTCTCCCTTTATCGAAGAAATGTGGAACAAGTGATTCATCGCATGTTCCTCAACCAATGGAATCCGAAGAAGATGAAGACGAGTCATATTTCTTAAGTGGATTTACTTGGGAATCCGTAGTCATAGGCTACAGTTTTGGACTAGTTGTTGGAACTGTCATGTGGAGTCTCATGTTTAAATATCGTAAACCAAAATGGTTTGTAGAATTTTTTGATGCAATCGTGCCTCACGAAAATAGAAGGCCAAAGAAGAAAGCTCAGAGACAACGGACTTAATGTTGGAATAATCTCATCAACATGCATTTGATATTTTGTTCTTTTACTTTGTCTGAATTTTCTCATCTGTTATTTGCTTCGACTACTGAATTGAAGTTATAAAAGCAGTTTTCTTTGCTTGTGCAGCACTTTATTTT

>PGSC0003DMG400011864 LRR-RLPs

AAGCATTCTTCTTCTTCTTCTTCTTCCTCCTCCTCACTTGGTTTATTCTCAACTAGGCCAGTGTTTTTCTGTTAAACGGTTCCTTCTCAATTTTCCATCTTCTATATGTGATCAAATTCACAAATGGACTACAAATTACTTCTCCACCTGCAAATTTTCCCCTAAACAATCCCACCATTACCAAGAATAACCAAATCTAAGCTGCAATTTAAGCATAAAGCTACAATCTTTGTCCAAAATGATCCCTAAAAGTTCAATCTTGGGAACAATGAAGAAGCAAAAGCCAATTTCAGGAACCCATTTCAACCCCTTTGATCTACTTTCTGAAGAAATCGTCTTTTCAGTTCTTGATTGTCTTAATGACAACCCATTGGACAAGAAATCATTTTCTCTTGTTTGCAAATCTTTTTATGCTATTGAGTCTCATCATAGGAAAATCTTGAAACCCTTTAGGCCAGAACACTTGACAAAGGTTCTTAGCAGATACCCACAAGTTACCCATCTGGATCTTTCCCTTTGTCCTAGAATTACAGACGGTTCACTAACAGTTATAGCTAATTTCTGTAAGGAAATGTTGAGGTCAATCAATCTTTCTAGGTCTAAGTTTTTTACCCATGTGGGGTTGTCGAATTTGGTGATGAATTGTGGGAATTTGGTTGAGATTGATTTGTCAAATGCTACTGAGTTGAAGGATGTGGGTGCTGCTGCATTAGCTGAGGCAAAGAACTTGGAGAAGTTGTGGTTGGTGAGGTGTAAGTCTTTAACTGATATTGGACTTGGGTGTATTGCTGTTGGCTGCAGGAAGCTCAGATTGCTTAGCCTGAGATGGTGTTTGGGTGTTGGTGATCTTGGTGTGGGGTTGATTGCAGTCAAGTGTAAGGAGATTAGGAGTTTGGATCTTTCTTACTTGCCGGTAATGGAGCTTTTCTTCCCTTATAAATTTTGTTTTCGATTATATTTTTGCTCTGTACTTTTGTGTTGGTTATGTTTCTGTTTCTGTTTTTCATGGTAAGTTAGTTTGTTTGTCTAAGTAGTTCATTTGAACATGTTTAGACAGTATGCTCGTCGATTGGTTGGTGTTCTGTTTACTATGATTTTCAGGGTCAAATGATGTGTGTATTCATGTTGTGAATCAATAATCGCTAGATAACCCTGAGGGGTAGCTCGGCTGGTAAAGGCTGAGGGGCAAAGCCCTTTGGTCGCCAGTTCGAGCCCCCTCAGGGCCACTGGGGGCATTACAAACCGCCCGTGTCTCCAGGGGCCCGTGGAACTCACCGGTGTTATCCACCTTGAAACAGTGGGACCCGGAGGGTACAGCTAGTCGGGGAGTGGGTTGGTAAAGGCTGAGGAGCATGCTCTTTGGCGCCAGCTGTAACCCCCGGAAACCACGGTTAAAAAAAAATAAAAAAAATAAAAAAATAATCGCTAGATGTGTTGTGTGATTCTTTTTGAGTACTGCTATTTGATTTCTCTTTGCCATACTCTGCCAAGCTTGTTTGCCAATTGATGGCTTATCAGATTATAGTCTTTCATGAATCCTTGCTTGTGTCATGAACTCGGTTTATTGAATTGTGATGTATCATGAATGCTAAATATGCACTGTATGTTATAGAACAAATGTAGCTTGGTGATGATAATTTATGCGGCTTTGGGAGTGTTAACCAATGATTTAGAAGGTAATGTGCAGATTAAGCTGGCCAAGAAAGTATCATTATAAATAAAATGCTAAGCCGTTTAGGAATTTCTCATTGATATCCTTAACGAAGTCTATGAGAATTGTGACATCCATGACGGAGGGCACACCAACAAGGGTGGGTCAAGACGAGTAATGGCAGGCTTTTAGCTTGGCAAGCTTCCGAAGAAGGGGTGCACATGACACCTTAAGGGAATTCACATCAGAATGGGAAAGAAGAACTTCGTGAGAATATTTTGCACAAGATTATTTTTCACTCTCTATCCTCCGTAAGGGAGGAAAATTTCATGAGTTTGCATTGCTCAACAAGTGCTGACATGTCTTAGGTTCAAAACTAGACGATTTCTTCCCATCTGTATAAGCTTTAGTAGACAAAGTTACCTGGTACGTGTGCTGGTGGGAGGGAGCAGGTCATTGGTGGGTATGCACTAGCTTGCCCAGGCACCAACGTCATAAAAAGGAAAAAAGCTAGTGCTGATGTGTCGTCAAACTCAATGATGATTTTTTTTAAAATTGAACGGAAAATGACATTTCACAGAACATTAAATATTTAATATTATTAAACATTTGCTAAGATTCTTACATTGTCTTAGTTAAATTTATCCCGTGTATTGTGCTTTCTCTACTTGGGAGATATACAAGTTCTGAAAAATGAGTCTGCTTATGCACTTTGCACATCTCTTGCACTTGGCATGCTTCTGACCTTGAGTTTGTTGAAAAGAAAAGATATTGCAGAGTCCAGCTTTTGAATCCTAATGAACACTTCTGCTTGACATTCCGATGCATCTTCAGATGAATAATTGAATGTTCTCTTATCATACTCATGCTGAAATACTCTAAGTTAGTGGAGGTTTTATGCTATAGAACATCTATCCATATTCTCTCGATATCTGTGCACCACTCAAGGATAGCATGACCAAAAAAATTGTGTTTTTAAGCACCTTGGTGCTTTGTCTGATGTCCAGGTCTCAATTACATTTCATGACCATCACAACATGCTCCTTTTTTCATCGTATAATATGGGAAAGTTCTTGTTCATGTGATTCGAAGTCCTTTCTTGACTTATGAAGTTACCACGACTTGGATATGAGCTTAAATAGAGGGACATAGACAATGAAAATTCATATAGCTGGCCCCAACTTGCTTGAGATTAAGGTGTGGTAGTAGTTGTTGGTGCTGTAGTTTCCCACTGCTTTGGTACAGAACTTAAATTCCAGACGTACCAAGTTTCCAGTGACTGACTGTATTTATGAAGTTGTTACATGTAATCTGTTTCTGTGTTCTGTTATTACATTCCATCTCTTTTTCTTTCTCCGTTTTTGTTTTGTTTTGTCTAGGGTTTCTTTATCGATCATACAATAATGTGTTGATCTTCTTATTAAACAGATCACTAATGAATGCTTATCATCGATCTCGAAACTGCAATATCTTGAAGATTTGGTTCTAGAAGGCTGTTATGGAATTGATGATGACAGCCTTGCATCCCTTGAGCAAGGCTGCCAGTCACTGGAGGTATGTACTTTGGAAGACTCTGTTTAACAGACTTTGATTCTCATATTTGCCTGATTGCCTCTCCTTGAGATAAAATGAACAATAAAAAATGAAGCTAGTATTGATGTATGAAATTGCCAAAAGCTCCCAGGACGTAGCTCAATTGGCGAAGGTTGATAAACTTGTGACCTAGGTCATTAGTTCGATCCCCATGTCAAGCGAATCTGATATTTAAGTGGTGAAGGGAGAAGTGAGCCCATCATCCCCGAGTTTCGAAGGCTGCGGTTGGTCCTAAGGGTTGGCTCCTAATGGATTTCTCGGTCATCAAAAAGAAATTGCCCAAAAATTTGTCAAATTGTCCTAGCTTCAAGCACTACTTTTAGATGTACATATATATGACTTTTTTTCCCTAATTGCCTAGTCAATAATTGGAGACCTACTGTGTGAATGCCACTATAAGTACTCTGAAATAGTTAAAGAACAAGGTAACTCAGATTTTTTTTTTTTGATAACCGAGAAATCCGTCTGTGACCCGCCCTTTGGACCAATCACAGCCTTCTAAACTCAGTGGATAATGGGCCCGCCCCTCTACCCTTCTCCACTTAAATACTGGGCTTCACTTTGCATGGTGTGGGGCATGAACCTGGGACCTAAGCCACAAATCCTCTACCTTTTGCCACTTGAGCTAGGCCTTGGGGGCAAGGTAACTCGGATTTTATGCACCCAGAATTCTACCTAATTGTCAGATGTCCCAAAGAAGTGGACTATTGTGCAAGTCAAGTGTTGCCTTGCATGTATCATGCCTTTTGGTCCTTTTTTTCCTCTTAAATAACTTTTTTCTTTAGAAGGTACTATTGACTAATGTCAGTGTATGTATGTTGGCTCTTTTCAGAAACTTGATATGTCGAGCTGTCAGAATGTTAGTCATGTGGGTTTGTCATCTCTTACCAGCTGTGCTGGATGTCTGCGGCAACTTGTATTGTCGTATGGTTCTCCCGTAAGGCTTAACAAACTTGAGCATCTTTCAGTAGTATGAGTTTTAATTTGATTGCTGATTTCAAATTACTGGTATTTCAGGTTACACCTGCTGTCGCAGATAGCTTGCAGAAGCTTTCCAGGTTGCAGTGCGTCAAACTAGATGGTTGCCAAGTTACATGTTCTGGACTGAAGGCTATTGGAAATTGGTGTGTCTCACTTAAAGAACTGAGCTTAAGTAAATGTGTGGGAGTGACAGATGAAGGTCTCTGCTCCCTCGTAACAAAGCACAAAGACTTAGGGAAGCTTGACATTACTTGTTGCCGCAAAATTACTTGCGTATCAATTTCCCATATAACGAATTCTTGTGCCTCCCTTACTTCTCTTAGGATGGAGTCTTGTACTCTAGTCCCAAGAGAAGCTTTTGTTCTAATTGGACAGCGTTGCCAGTACCTTGAAGAGCTTGACCTTACAGACAACGAAATTGATGATGAAGGTCACCCCTTTACCTTAAAGGTCTCAGGCTTCTGTTGTAATTTTGAAAGTTCCTCCTTTATATCTGACGGTGTCCTACATATGCAGGCTTGAAGACCATTTCCAAGTGTGCCAGCCTTTCCAGCTTAAAGCTAGGGATCTGTCTGAACATAACCGATCAGGGTCTTATCCACATTGGCATGTGCTGCTCAAATCTTAAGGAGCTTGACTTATATAGGTATGTATTTCACAGTTCATGGGATATTATTTGAGATAGTGTAACTATATATCTTGTGCAGTTTTTTTTTATATTCTTTTCTTTTTGTGATGGATTGGCTAAACTCACTTTTTAACTATGGAGTACAAGAGATTGAGTGTGCTGATTACTGGGTGATTCTTCTTGACAGGTCTGCTGGAATTTCCGACTTGGGAATATTAGCAATCTCTCGCGGTTGTATTGGCCTTGAAATGATCAATATTGCATATTGCAATCGTATTACTGATGGCTCATTCATATCTATATCAAAGTGTTCAAAGTTGAATACACTTGAAAGTCGAGGTTGTCCTCTTGTCACATCTTTAGGTCTTGCAGCTGTTGCTGTGGGATGTAAGCAGCTTACAACATTAGACATCAAAAATTGCCACAACATTGATGATGCCGGAATGATTCCACTTGCTCACTTCTCGACAAACCTCAAACAGGTAACCACAGTTTCTCTTTCGTCGACATTGACGAACTGGAGCATTGACATCTAATATGAGCTGGCCATCTGTCAATTACACAGATAAATTTATCCTTCACTTCAGTGACGGACGTGGGGCTGTTGTCTCTTTCCAGCATCAGCGGTCTACAGAACATGACAATCCTGCACCTCAAGGGTTTATCTCCTGGTGGGCTAGGAGCTGCACTGTTGGCTTGTGGCGGACTTACTAAAGTTAAGCTGCAGACATCATTTAAATCATTGCTACCACAACCTCTCCTTCAACATCTAGAATCCCGGGGTTGTGTTTTTCAGTGGAGAGAGAAACCATTTCAGGTATCTTTCTCATATTTATCATTACCTACTTTTTCTCTGAAAATATGTGAGGTACTTCTATGGTTGTCGAAATATGCCGTCGTATGAATCAATTAGAATTACATATTTTAGCTTAACATGACATAGTTATTACTTGATCTAGAGGGTCAATGCAGTTTCAGAGGCTTTCTTGTTTGGCATTTAATTGAGACAATAAAATTACAGTTTTGATTATCTTTGTTGTAATAATGAAAATATCCTTTTTGCTTGCTCTAGTCATTAATTATCTTTCTTTTATTGGTTAAACACAGGCTGAAGTAGATCCAATATACTGGAAAATTCAGCTGGACAATTTAGACTGAAGATTTGAGCCATTTATGCGAGTTGTCCTAAGATGGATAGTCGCCCCAGAACCTCCGGAGTCCTTACTTCATCTGTTAGCACTTCATCAGGCAAGGTGCATTGTTCATGGCGACTGTTTTGCATGTTAAGTGTCTGCTACTTGTTCGTCTGTATCCTCTAAACACTCTGTAAATTTCAAAAACCGGGAAAAACTCAATGTCCATTCAGCATTGCCCCGGCGTGAAATCAATTTTGTCAACAATGTTATGAGATGGATTAAACAGATCCAAATCATGTTTATTGTTTAGCAATCAATATCTGGGTTTGTGAATGCTTATTCCTCTTGACTAATGTGATTTCGCGTGAAGTTACTTCAAATTTACTCTCAATTAAATAGTTATTTG

>PGSC0003DMG400011900 LRR-RLPs

TTTTAGGGGATTTGGAGAAAGAGCTATCATCTCTTACATCCATTTTAGAGAAGGAGCTATCATCCATTTTCAGAGATGTCGTGCACCATGAACATAAAATTCCTAAAGATCTTCAGAGACGTACCATCAATTTGGCATATGAGGCTGAGGTTGCCATTGACTCTATTCTTGCTCAGAATAATGCTTTTTTGCATATTTTTTGCTCACTTCCTACAATTTTAAAAGAGATCAAGCAAATTAATGCAGAGGTGACTGAGATGTGGTCAACGGACATTCCTCTTAATCCTCACTACGTGGCTGCTCCATTTAAACATCTGCCAGCTCGACATAGCAATCTTGTGACTGATGAGGAGGTAGTGGGTTTTGAGAATAAAGCAGAAAAACTAATTGCTTATCTGATTAGAGGTACAAATGAGCTAGACGTCATCCCAATTGTAGGCATGGGGGGACAAGGGAAAACGACAATAGCTAGAAAGTTGTACAATAATGACATCATTGTTTCTCGCTTTGATGTTCGAGCATGGTGCATCATTTCTCAAACATATAGCCGAAGAGAGCTATTAAAAGAGATTTTCAGTCAAGTTACGGGCTCCAAGGACAAGGAAGATGAGGTAGGCAAACTTGCTGACAGGTTGAGGAAAAGCCTAATGGGAAAGAGATATCTCATTGTATTGGATGATATGTGGGATTGTATGGCATGGGATGACTTAAGGCTTTCTTTTCCAGATGTTGGAATGAGAAGCAGAATAGTCGTAACAACTCGACTTGAAGAAGTGGGTAAGCAAGTTAAGTACCATACTGATCCTTATTCTCTTCCATTCCTCACAACAGAAGAGAGTTGCCAATTGTTGCAGAAAAAAGTGTTTCAAAAGGAAGATTGCCCGCCTGAACTACAATATGTGAGTCAAGCAGTTGCAGAAAAATGCAAAGGACTCCCCCTAGTGGTTGTCTTGGTAGCTGGAATAATCAAAAAAAGGAAAATGGAAGAATCTTGGTGGAATGAGGTGAAATATGCTTTATTTGACTATCTTGATAGTGAGTGCGAAGAATATAGTCTGGCGACTATGCAGTTGAGTTTTGATAACTTACCCGATTGTTTAAAGCCTTGTCTTCTTTATATGGGGATGTTTTCGGAGGACGCAAGAATTCCAGCATCTAAATTGATAAGTTTATGGATTGCTGAAGGATTCGTGGAGAACACTGAATCTGGGAGATTAATGGAAGAGGAAGCTGAAGGTTACTTGATGGATCTCATTAGCAGTAACGTGGTAATTGTTTCAAAGAAAGGTTATAATGGTAAAATCAAATGCTGCCAGGTTCATGATGTTGTGCATCACTTTTGCTTGGAGAAGGGTAGAGAAGAAAAGTTTATGCTTGCAGTGAAGGGTCAATATATCCAGTCTCAACCCTTGGATTGGAAGGGAAGTCGAGTGAGCTTCAGTTTCAGTGAAGAGCTTTCCAAGTTTGCATCTCTGGTTTCCAAAACACAGAAGCCTTTCCACCAACACTTGAGGTCTCTGATAAGGACAGATGGTGAAAAATCTATTGATATGATTCCCTTCTGTCAGATTAGTGAATTGCGACTTCTTAAAGTCTTGGATTTGAGTTCTTATTATGTGGAGTTTTTGTGGTTAGCTACATTCAAACCACTTATTCATCTGAAGTACCTCGCAGTTCGGGCAGGTGATTTCTATTTTGATCCAGGATCACATCTGCCCCATATAGAAACTTTAATTGTGAAGAATTCTCCTTATGTATGGTTACCAGTGTCTTTTTTGCAAATGGAAAAATTAAGGCATGCTCATTTTGGTTGGGCTGTTTTTGATAAGCAGGGACTCTTTGAAGGATCCTCTAAATTGGAAAATTTGAGGATATTAAAGAAAATTGAGAAATTTCCAATTGATAGGGCGGATGTGTTATCAAGGAGGTGTCCTAATCTTCAACAACTTCATATCGAATTTGATGGACATTCTGCAGATTCTTTTTGTCTCACATTGGAGAATCTTACCCAGCTTCAAATACTTCGTGTTTCCTTTGAGCGGCACCACATCATTGTATCTGGGTTGCAATTGCCTTCAAATTTAAACAAGTTGGTACTTAAAGGAATTCATATAGAAAGTGTTATTCCCTTCATTGCGGGACTACCAAGCCTGGAGTATCTCCAATTACAAGATGTGTGTTTTCCTCAATCAGAAGAGTGGTGCCTTGGAGATATCACTTTCCATAAACTTAAGTTGTTGAAACTGGTGCAGTTAGGTATCTCAAGGTGGGATGCCTCGGAGGAATCATTTCCCTTGCTTGAAACACTTGTTATAAAAATGTGTGGTCACCTAGAGGAGATCCCACTTAGCTTTGCTGATATTCCATCATTGAAACAGATTAAATTGATTGGGTCCTGGAAAGAATCTCTGGAGGATTCAGCTGTGAGAATTAAGGAAGAAATCAAAGAGATTGAAGGATGTGATCGTATACACCTCGTCAAAGAACTGACAGATTGGGAGGCATATAGCAATTTGTCGCAGCTGTGCTTATACTGAAGTCTATTTTGTAGCTAGCCCATTCTCTTATATTGTAACACTACTACTGTTTGTTATGTTTGTTTTGATTGATCAACTCTTTTTATGTATGGTTGGAGACAAATATATATCAGATGATCTTTCAC

>PGSC0003DMG400013644 LRR-RLPs

TTTCCACAGAACCGAACAAGACACATTTTCTGTCTTAGATAACTCATCCTTTCTCTTCCCTTCTTCTTCTTCTCTCTGTTCTCCACATTTCACGTACACTTTCAGAAACATTGTTGATTTACTTTGGACAGAAAAGCCACTAAATAGTGTGAGTCAATTTCTTTCCTTAAATAATTGTTTGGAGTTTTGTTTTAATGGGAAGGTTTTTTTTTCATTCATTTTTATGAGTGTTTTTTCCGTGTTTTTGAAAACAGTTGTTTTACATATAGGAAAATACGGCGAAAAATAAGCTTTACTTTGCATGCCTTCATGTGGGTAAAGGTTTGAATTTGCTGTCATCGTTGATTGGCGCTTTCTAGAAACTTGTAGAAACAAAAAACTTAGCTGCGGTTTTCATTTTTCTTGATCTGCTGATTGGTTTAGATATAGCGGAATTCATCTTTGTTTTTTGCTTTGAATAATACATACACAACGAGGAAAAGGGTAGTCTGAGTCTAATTTTTGTGGCCGGAGTTGTATTTATGTTCAATTAAATTTATCTGGGTGTTTCTTGTTTTCGAGAAATTTTCAGAGGATTATTGAAAATGGGGTCGTCAGGAATGTCGATTGATGAGATTGTGCAAGAAATTATGAGAATTCATGGATCTCTACCGACAAGGCCATCGATTGATGAGGTTGAAGGAGCAAGAGGTTTAATTGGAAATCTGGAGAAAGAAGATCAGTTGAAAATAGAAGCTATCATGAGGCAAAATAAAAGGAAAGGTGTACCAGAAGAGCTGTTCAAAATTTTGCAAGAGATGCAGAGGAACGCTGTTCTTTTTCAGAGTAAAGAACAAAAGAGGGAGGCGTTGAAACTTTTAGATCTCGACAATATTCACAGTGTGTTCGACGATTTGATTCAGAGAGCTACGAAATGTTTGCCTTCTAAATCTCACGATAATGACCCGTCTTCTTCTTTAATTGAACCGAATGATTTATCTTTGGCAAACCAAGTTTCCTTCTCCGGCGGCGGCCTTAATTCTCCGACAACTACAACTGCGGCTGCTTCTTCGTCGAGCTATTATAACTCAACAGAGCCTATTAAAGCGTCTGAACTGTATAGTAGAGATGACAGTTACTTGAATAAGGCAAAAGCCACGTTTCTTGTGGATGGAATTGGTGTTGGGCTTCGATCTGGGGATGCTTTATCAGGGCCTAAGATTGTTGATTCCACCTTAAAGCCGGCGTCAATAAGTGGTATGTTACATCTATTTGCTGCACCGTATTTCTAGTGAATTTGATTATATAATTTGTTGTGTTGACTCAAAGTTCGAATTTTTTGATCAAGAAGACACTGGTTTCTTTGTTTGTAGGTCAAGATGGTGAGAAATTAAGTTTAATTAAGCTTGCTAGTATGATTGAAGTGTCATCCAAAAAAGGAAGTCGAGAACTTATCCTTCGGAATAAGTTGTCTGACCAAGTAGAATGGCTTCCAGATTCTATAGGGAAGCTATCAAGTTTGATCACTTTGGATCTGTCTGAGAATCGAATTACGGTCCTACCCACCACCATAGGGGGGCTTTTATCATTGCAAAAGCTAGATTTGCACTCAAACAAAATTGTTGAACTGCCTGATTGTATTGGAGATCTTCTTAACTTGGTCTATCTTGATCTCAGTGGAAATAACTTGAAGTTGTTGCCCGCGAGTTTTGCTAGGTTAGCTCACCTTCAAGAACTTGATTTGAGCTCTAACATGCTATCAGTACTCCCTGAGACAATTGGCTCCTTAGTAAGTCTTAAGAAATTGATTGTGGAAACTAATGACCTTGAGGAACTTCCTCACACTATTGGTCAATGTACTTCGCTCAAAGAGCTTCGTGTGGACTATAATCATCTTAAAGCTCTTCCTGAAGCAGTGGGACGAATTGAGTCTTTGGAGATTCTGACTGCGCGGTACAATAACATTAGGCTGTTGCCTACAACTATGTCATCCCTTACAAGTTTGAAAGAGCTGAATGTTAGTTTCAATGAAATCGAATCAGTGCCTGAGAGCTTGTGTTTTGCTACTTCACTGGTAAAGCTGAACATAAGCAACAATTTTGCAGACCTGAGATCACTACCAAGGTCCATTGGTAACCTTGAGTTGCTTGAAGAATTGGACATGAGTAACAACCAGATACGCGTTCTTCCAGATTCGTTTAGGATGCTCTCAAGTTTGCGGGTCCTTAAAACCGATGGAAATCCTTTGGAAGTGCCACCAGGAAACATCGTTGAGAAGGGAGCACAGGTTTAATTTTGTTTCGTTTAACTATTATGCTTTCTGAATTAGTGTAGTATTTAATTATCTTAGCTCTTCTCTTGGATATATCAATCACTAATGTTTTATTTCCTCAGGCTGTTGTACAATATATGTCTGATCTGGTTGCCAATAGAGAGGTAAAGGCGCAACCCGTAAAGAAGAAGAAGAAATCATGGACTCAAATATGCTGCTTTTCAAGTTCCAACAAACGTCAGCGCAATGGCAGTATGGACTATGTTAATGCCTGATGTGTTCCACACAAATTTGGTATGATTATATATTTCTCCTTCTTACCAAGTTAGTCATTATGGTACTGAACTTTCCATATTTCATAAACTTCGTGTTCAGTCAAACTACATCCCATAGACTATGGAGGGAGTTTTAAATATATATTGTTAATAAATGGGAATGGTGCATTATACATGGTGCTGATTAGTGTTAATTAGAATAATGAAAATGAAGGGTTGTAAGGGTAACGGCTCACATGATGTCTGTGTTATCCGTTGTAGGGAAATGAGCTGTGATTTGGCTGGACGAAAGCTGCAAAATTTCAGTTGAATTCGAACATTTTTGCCAGCGAGATTCAATCAACTTTCTTGTTGTTCCAGATAAGCTCTAATCCACCACGTGGTTTTTTATTCTCATTTCATGTAATAATTCTTTTAATGCTCTTTCCTGTATTTTCCTCGATTATGATGTAAAATATCAAATGAGATGTAAAATCTTATCTTATTTTTTTACTTTCAATATATTCAATGAATTCCAGAATTTCTG

>PGSC0003DMG400013766 LRR-RLPs

ATGAAGAGTGCTACAATTAGCAATTTGCCCTGTAATGTCTTGGATAAAATTCTTGGGAGCTTGCCTTTGAAAGATGCAGTGAAGACCAGTATCTTGTCAAAAGGTTGGAGGTACAAATGGGTAACGCGTGCAGAACTTGATTTTCGTAGTGAGTTTTTCACATCTTTCAACGACAACCAAGAAGCCAAGAAAATCATTTACCAAGTCCTACGACTGCATCAAGGACCAATACTAAAGTTTACTCTCCAACATCCTAATTTGATATGCTATCGTGATATTTATAATTGGATGCTCTTCTTGTCAAAGAAAAACGTCCAGGAACTCACCCTTCAAATATTCACAGGCAACAAATATCATTTACCTTCTTATCTTTTCACATTTCAGCAACTAAGACATCTGGACCTTGATATGTGCTTCTTCCACCCTCCACCTGATTTCAAAGGGTTTGCAAAGCTTATTAGTATTAATCTTCAACATGTCATTTTTGATCCAACAATATTTAGGAATCTCATAACCAAATGCCCATTGCTTGAAAGTTTGATGTTGACTAGGTGCACCGCATTTGACGTTCTTGAAATTGATGCCCCTAATCTCAAATGCTTCGACTTTCTTGGAACATCGAAGTTTATTTGCTTCAAGAATGTCCCTATGCTTAAAACCGTTGCTGTATGTCTCAATCAAATTTTGATGGATACGTCAACTTTTTGTTCTAATCTTACAAAGTTCTTCCATTTCATGCCTTCCCTGGAGGAACTTGAACTAGGCGGTTCATCATTAGAGGTAATGTACATTTGTGATAATATTACTTATGATAGTTGGTACTTATTATAATCATATGATGTGTCTTCAACCAATTTTTTTACATGCTTCTAGTACTTGAACATGGGAGGTATACCAGAGAATCCCCCGGCTGCTCTAAACAATGTCAAATCCCTATGTATTTCGGACATGTCTTTCAGAAGTGTTGAGGAGGTTTCTAGTGCAGTTTACTTGATTACAAGCTGCCCTAAATTACAAGAACTTACAATTGAATGTGTAAGCACCCTCCTACTTTCTCATAAAACTTATTGATATGGTTTCTTCATCTGAATCCGCCTAACTGTTACCAAATTTTTTGCAGGAAGCAGTAGGCATTGTGGTGGAACCTGTTATACAATTCTTACGAGCCAAAGCAATCTCATGTGGTGCTATGAAGCTGCTTAAAAGTGTGGAAATGCGTTATTTTATTGGTTTTGAGATGGAAATAGAATTTGTGAAGTTTATATTGGCATCTGCACCAGTACTTAAGGAAATCTTCATTTGGAGTAGTGGGCAGTTTCATCGGGGTACACAGATGATGGATGAGATAAAACAATTCCATCGAGAATCACCCAATGTTAGATTCAAATTCGAAGAAATGTGA

>PGSC0003DMG400013898 LRR-RLPs

CCCCTTTGGCTTCAAGGCTTACAAGTCATCGTCCGCTAGTATATATATCTATGAGAACCCTTTTAATGTAAAAGCTCAAAAAAGTTGTATTATTGTTCTTGAAACCAAGTACTCTACCTATTAGACATGACAACTTCTCACAGTACCTTCCTTTACATTCTACTTGTGCTTCTTCTTTTTTCACTTCCATTCACAATTACATCATCAGCAACAACTGAAGCTGAAGCTCTTGTCAAATGGAAGATGAACTTATCTTCTGCTTCTTTCTTGGATTCATGGTCCATTTCGAATCTCGGAAACTTGTGCAATTGGACATCTATTGTCTGCAATGCTGGTGGAACAATTTCCGATATCAATCTTTCTGATGCAGGCCTCTCAGGCACTCTTGATCAGCTTGACTTCACTTCATTTCCGAGTCTCACAAGTTTTTCTCTCAATGGCAACAACTTAAGTGGATCGATACCATCAAATATTGGCAATGCTTCTATGCTCACTTTCTTGGACCTGAGCAACAATATCTTGGAGGGTGTCATACCAGAAGAGATTGAGAAGTTAACACAACTTGAATATCTCAGTTTCTATAACAACAATATCATTGATGTTATTCCCTACCAAATAAGTAATCTTCAGAAGGTATGGCACTTGGATCTTGGATCAAATTTTTTAGAAACTCCTGATTGGTCCAAATTGAGGAATATGCCTATGTTGACATATTTGAGCTTTGGTTATAATGAACTAACATTAGAATTCCCTGAATTTGTACTCCGTTGCCATAATCTGACTTACCTCGATTTGTCGATAAACCATTTGAATGGTTCAATACCAGAAACACTATTCACCAACTTGGATAAGCTTGAACAACTCAATCTTTCATCTAATTCATTTCAAGGTTCATTGTCACCAAATTTTACCAAGTTGACCAAGTTAAAAGAACTTCAGCTTGGTGTCAACATGTTTTCGGGCTTAATTCCTGATGAAATTTCTCTGATCAGCAGTCTTGAAGTTGTTGTACTTTTCAATAATTCACTTCAAGGAAAGATTCCATCTTCTATAGGTAGACTTAGAAATCTTCAACAACTTGATCTTCGAAAGAATCGTTTGAGTTCAACCATTCCTTCTGAGTTAGGTCTTTGTACTAACCTCATTTTCTTGGCTCTAGCAGAGAATATCCTGCAAGGACCATTGCCTATATCCTTGTCCGCGGTGACCAAGTTATCTGACTTGAACTTGTCTGATAATTTTCTTTCTGGTGAGGTTCAATCATATCTCATTACCAATTGGACTGAGCTGACATCTTTGCAGCTTCAAAACAATTCATTTTCACGGAAAATTCCACCTGAAATCAGCCAGTTGAAAAACCTCGTATATCTTTACCTCTATAGTAACAATTTCACTGGTTCCATTCCCTATCAGATTGGTAAGTTGCAAAACTTGCTGGATCTAGATGTCTCTGAGAACCAGCTTTCAGGCACAATACCTCCAACCATTGGAAACCTGACCAATCTAAGGACGTTGCAGCTTTTTCGGAACAACATCAGTGGAACCATTCCTTCTGAGATTGGCAAATTGACATCTCTTCAAACCCTTGACCTCAACACTAACCGGTTTAGTGGTGAGCTGCCCGACAGCATTTCTAACCTCGGCAGTCTGAAGCTTCTTGAGTTCCTTTCTCTTAGTAACAACCAGTTTTCAGGTGTGATCCCTTTGTCTATTTGCAATTCGACATCACTACAAACTCTAGTTTTCGCAAGAAACAATTTCAGGGGAGAGATTCCACAATGTTTGGGTAATATCAGTGGCCTCTCCATTTTGGATATGCGCTGCAACAATCTTTCTGGGATTCTTCCAACAAGTTTTGATATTGGATGTTCACTTAGAACCTTCAACTTTCATGGAAATGAACTACAAGGGAAAATCCCTCGATCTTTAGCCAATTGTAAAGAGTTGCAAGTTCTTGATTTAGGAGACAATCACCTCAATGATACATTCCCCATGTGGTTGGGAACTCTATCAATGTTGCAGGTTTTAAGTTTGAGATCGAATAAATTGCATGGACCCATTAGAACTTCAAGGATTGGAAACATGTTTCCTGGTCTTCGAATATTAGATCTCTCTCGCAATGCCTTCACAGGAAACTTGCCGACAAGTCTATTTCAACATTTGAACGCCATGAAGACAACTAATCAAACAAAGAATGCACCAATTGAAGGAAGCCACTATTACCAAGACTCGGTAGTTGTTGTAATGAAGGGACGCATGTTTGAAGTTGTGAGAATCTTGTATTTATACACGAGTATGGATCTTTCAAGTAACAAATTTGAAGGACTTATTCCAAGTATTATGGGAGATCTCATTGCACTTCGGGTGTTGAATTTGTCTAACAATGGGTTGCAAGGTCATATACCGCCATCACTTGGAAATTTAGCGGTAGTAGAATCATTTGACCTTTCATTCAACCACCTTTCAGGAGAGATACCGGAACAACTTGCTTCTATAACAAGTCTTGCAGTTTTAAATCTCTCCTACAATCATCTTGAAGGATGCATTCCTCAAGGACCTCAGTTTGCTACATTTCAGAACAGTTCATATGAAGGTAATGATGGATTACATGGATTTCCAGTTTCAAAAGGTTGTGGTAATGATCAGGTATCAGAGACAAATGATACCGTATCTGCACCAGACGAACAAGAAAGCAATTCGGAGTTTCTGAATGATTTTTGGAAAGCTGCTCTTATGGGCTATGGAAGTGGACTATGTATTGGATTATCCATAATATATTTCATGATCTCAACTCAAACTCCAAAATGGCTAGCAAGAATCATGGAAGAGTTGGAACACAAAATTATTACGCGAAGTAGAAAGAAGCAACGAGGTAAAAGGCATTATAGAAGAAGAAAATAATCGCTTCTAGACAAGTTACCAACTGAGTAGGATTTGATTTCAGAACTTTGGGTATTCAAGTTAAGGTCTATGCTGATTTTTAGTTTATAGTAGCTATATGTATTGATATATATACACTTTCGTTACAACAACAGTTAATCGATACTTTATACTAGGTATATTTTTCTTTCCAATAATTT

>PGSC0003DMG400013955 LRR-RLPs

ATGAAGCAAGAAACTCTTGAAGATGTTCATACTGGTGATATGGTAGCTTTTATCGAAGAAATCAAAGAAGTAACTCCTGAACATGCAGTGTTCAAAAGTATGTATGCCGATGGTGACCTTCATGAAGAAAGCATTGAGGGACATGTATTGGAGATCGAAGTCTCTAGTCAATCATCTGTCATATCGAGGTCAGTAACTGGCTTAGATTGAACATTGGAAACAAATGGAGAAAATGATGACCAAATAGATGAAGAAGTTGATTTTGAAGTTTCAAAGACAAATGATATGATTTTTGAAAGCTTTGAAGCTTCCAAAAAATCTATCGAGGAGATCATAAGAGAATATGGCAATGGATCCTATGTTGGTGTTGAGACTTCACAAGAAATTGATAGAGTTTCTCTGGATAATATAATTGTCGAGAAGTTAAGAGAAAGGGGGAGTTTGAGAACTCAACAAGGAGATAATTTACATGGATCAACTGTCGGTGAAAAATCTTTATCATTTGAAGAAGGAAGAGGAGTCCGGAAGAAAGTGCTAAGATTTTCACCAAGGCGTGGCTCAAAGCTTCGTTTGAGCTCTTCCATGACAAGTGTGGGGTAGTTTCCAAAAAATCACTTTAAGGGGGAGTGTGAAAAATTAAAGTTGATTTTTTGGAGGTACAAAAAATATCTTAGATATTTAGTAAAACTTTCTAGAATTTTGTAGAGTAAATATCTTTGATATTTATACTTAAGAAATATCTAGATGTTCTAGATAAGTAGATATTTGTAGTAAAGTATAGAAATATCTAGGTTCTTGTTACATAGTTAGAAGATAAGAATATCTAGATTTTTCTTGTAGAAGTATCTAGAAGCATCCCTATACAAGTATAAATAGGGGTGGCATTAGGCATTTGTAAAACAATCTAATAACAACCCAAATAAAAGAGTGTTCTTCCATAAACAAGTTCTCCTTTCTAGCTTCCTCTTCTTTAGTTGCATATTCCAATCTTAGTTAACGATCTTGGGCTAGCAGAAGGTTTCCCAATACTTTTCTTCTGCTATTCTCTACAATTCTATCATTCAGGTCTCCGGCCCAATCGCATCAGTGTACATGTAAAAATTGTTGTCTGAGTTGCGAAGAATTCTAAGACCTGAAGTTGTTGTGGCTTTAAGATACCGTAGTAGTATCCTCTTAACCGCCTTCCAGTGTTCATGGTAGGTTCATGCATAAACTGTGATAGTTTGTTGACAGCAGGCCGTGTGAATGATAAATCCTGTAACTTACCAAGGGTGCGCCTGTAGAGTGTAGCATGAGCTAGCAGGCGGAGAACCATCAGCTGCCCGAGGTAGTTTACCGGAGCACATTGGAGTCAAGACTCCTTTACAATCTTGCATATCCAATTCAGACAAAATATCATGAATGTATTTGGATTGAGAGAGAATAAGACCATGACCATCTGACACTTTTTTTAACTTCAACACCCAAAAAGTAGTCTAAGTAACCAAGATCCTTCAAAGAAAACCCACTTCTATTCCATTTTGTGAAACACAACCCAAGTAGTAGTGTAGTACTGTCAACTGGATAAATTCCTTTTCTTATTTTCCTCATACTTAATAATCTGTTTTCACATTAATCTTTTTGCTACTTATTTTCCTCATCCATATTAATCTGTTTTCGCATTTTGAATATATTATATTTTGGAATTAAAAGAAAAAAAAATAAAAAAAATAAAATAGATTTAAAAAAAAAAAGTGAAATCACACATGACAATGTGTGTAGTAGCGTGTGTCACGTTTGATTTTCTTCAAAAGAAAACATTAAAATTTCTTCCATTTTGTGCTAATTAGTGAACGTATGTTGTGTAACACAGCCTTGTAAGGTTGATTTGAAAGATTGACTCCCATTGACTTTGCTCTTAACAATTATTCTTTCTATATTATGACTTTTATTTCGTTCTGTGAAACACAACCCAATTAGTGTCTTTTTTTTTATTTTGCTATTTATTTTCCTCCTTTTAATCTGTCTTTACATTAATCTGTTTTCACATTTTGGAAATTAGTCCATACTTTTCAGCATTTTATTTGTTTTGGTGCTATATATATAGTGGAGTAACTTTGATCATTGGAAATACCACAAACACAAATTAATTGCCTTGAAATATTCTATATCATATGGAGAAGCACATTTTCTTATTGATTCTTCTCTTTCTAATTCAATTTTCTGTATCATTTGCTTCCTCAAATGAGAGTGACCAACAAGCTCTATTAGCTTTCCAAAATCTTATTACAAGTCCTAATCATTTTTTGGCCACTAATTGGACCAAAAATACTTCTTTTTGCTCTTGGTATGGTGTCACTTGTAGTCCAAAAAGGCAAAGGGTTGTGGCCTTGACTCTTCCTAATTTGCAACTTCAAGGCACAATTTCCCCGTCTTTGGCCAATTTGTCCTTCCTCAGTGAGATCAATCTTGGGAACAATAACTTCTGTGGTGGCATCCCTTATGGACTTGGCCACTTGCCTCGCTTGCGAGTGATTGATATTCAAAACAATCAGTTTAACGGAAGTATACCAACAAGTCTATTTCAAAATCGGAGAGTTCAAATAATTTCATTGGCTTTCAATGAACTCAGTGGTGAAATGTGGAGAGGGGCCATGGTATGTACCCGAACTCAGAGTCTTAAATCTCGGGAACAATAGCCTAACCGGTATAGTCCCTCCTTCTGTTGGAAATGCCACAAGAATGATGAACTTCAATTTGTCTGGGAATAGAGTCAGTGGCAACATTCCAAAGGAGGTCGGTAATCTGAGCCAGCTTGCATTTCTGTCCTTGATCGATAATCAATTAACTGGTTCCATTCCTACATCACTTTTTAATATCTCGTCACTTCTTGTCCTATCTCTGAGAAATAATATCCTTTCTGGTCCTCTCTTGCTAAACGAAGGAAATATTGTTTCAAATCTGAACCTTTTAAGTATATCTTACAACCAAATATCTGGTTGCATTCCTTCCAACATATGCCAACTCACAGAGCTCAAAGTTTTGTCCATATCTTTCAACAATATAGCTGGAGACATACCCAGAAATATTGGTTGTTTATCCAAACTCGAGAAATTTTATATGGGTGATAATCCAATAAAAGGGACTATTCCCACTTCATTGGGAAATATTTCCACTCTGCATGATCTTTACTGTCAAAACAATCGGATAGTGGGGGGAATTCCTCCGGAATTAGGGAAGCTATCAAATTTGAGGCAATTAACCTTTGACAATAATTCTAATCTTATTGGTCAAATTCCAGAGGCTATTTTCAACATATCTTCTTTGGAAGTGATTGGTTTCACTTTAAACAACCTCTCCGGGAGAATTCCAACCACTACAGGTCTTCATCTTCCGAACCTTAAAGTACTTGAGTTGGGAGTCAATCAGCTGGAAGGGGAAATTCCATTGTTCATAGCAAATGCTTCCAAGCTTCAGAAACTGGAGCTAAATAATAACTTTCTGACAGGAACTATTCCTAATAATTTGGGGAATCTTCGTGAGCTGCAAGCACTGCTCCTACATACCAATCAACTTACCAATGAACCAAGAGAACGTGAGTTGTTATTCTTCAATTCTTTGGCGGACTGTAAGATGTTGCGATATCTACAAGTGGGTAAAAATCCGTTGAATGGCATTCTGCCCAATTCAATTGGGAATCTTTCATCTACTATTGAAAACATTTATATAGGATATGCACACATCAATGGCCTCATCCCCACAAATGTAGGCAACATGAGCGGTCTTACAGTCATAGACATTCAAGGAAACAACTTGACAGGGAGTATTCCTTCTGAGATTGGTAAGCTTAAACAACTCCAAGGGCTGTATCTAGATAAAAATAAATTGCAGGGACATATTCCAGAGGCGGTATGCCATTTATCTAATTTGGTTAAACTATTTCTGAATGGTAATGAGCTCTCTGGATTAATTCCAGAATGCTTAGGAAATCTTAGCATGCTACAAAAGCTTTATTTGTATTCTAATAAATTTTCATCAAAATTTCCATTGAGCCTTTGGAAGATGAGTAGTCTTCTCTATCTAAGTGTGTCACTGAATTCAGTAGAGGGAGAAGTTCCATCAGATATTGGAGGAATGAAAGCCATTGTAGAACTACATCTTTCCAGTAACCACTTTTCAGGAATGATACCAAGCACATTCGGGGAACTCCAAAAACCTGCAGTATCTTGACCTGTCGAACAATTCATTTTTTGGCAAAAATTCCATTATCCTTTGCCAACTTGATAAGCTTAGAATACTTGGATTTGTCTTCAAATGTGTTGTCAGGTACTATTCCTAAGTCTTTGGAAAAACTCTCATACCTTAAAAGCATTAATGTTTCCTTTAATGATTTAGAAGGTGAAATACCGAGTGGTGGTGTGTTTGCAAATTCCACTCTGCAATCATTTCTTGGGAACAAAGGTCTATGTGGAATGCACATATTGGAGATTCCTGCTTGTGCTATCACTAATCCTGGAAAACAATCAAAGCTTAGGGAGGTTCTGCTAAAATTTGTTACTCCAGTGGTTATTGCATCCTTTCTGATATTCTTGTTGGTTTCAATTTGGATAATGAAACGACAGAAGAAAGGAAAGTCCAAAGAGGTAGAGAAGGTACTGGAGATCGGGACTCATCAATTAGTTTCTTATCACGAGATTCAACGAGCAACAAATAATTTTGATGAATCCAATTTAATTGGTGAGGGAAGCTCTGGCTCTGTGTACAAAGGCACATTGTTTGGTGGAACTGCAGTGGCCATTAAGGTTCTGCATTTGGACAATGAGCAAGTATGCAAGAGATTTGATACCAAATGCAAAGTGACGAGAAATGTTAGACACAGAAATCTTGTTCCAGTGATTACTACATGTTCTAGTGACTATATAAGAGCCTTTGTTCTGCAATTTATGCCCAATGGAAGTCTTGAGAATTGGNNNNNNNNNNNNNNNNNNNNNNNNNNNNNNNNNNNNNNNNNNNNNNNNNNNNNNNNNNNNNNNNNNNNNNNNNNNNNNNNNNNNNNNNNNNNNNNNNNNNNNNNNNNNNNNNNNNNNNNNNNNNNNNNNNNNNNNNNNNNNNNNNNNNNNNNNNNNNNNNNNNNNNNNNNNNNNNNNNNNNNNNNNNNNNNNNNNNNNNNNNNNNNNNNNNNNNNNNNNNNNNNNNNNNNNNNNNNNNNNNNNNNNNNNNNNNNNNNNNNNNNNNNNNNNNNNNNNNNNNNGAGACATTGGGCACTCTTGGATACATTGCACCAGGTATAATAAATCTTCTCTCTTTGATTTTCTCTTATCATAGTTAAGCATTCTAGAAGTAAAAAAGCAAGTCTTCATTTATGCAGAATTATTGTTGTATTTCAATTGAGTAACTTTTCTTCAATCCGTTTTTAAGAATATGGCTCAGAGGGAATAGTGTCCACTAGTGGTGATGTTTATAGCTATGGCATCATGTTGATGGAGGTTTTGACGAAAAGAAGGCCAACAGATGAAGAGATAGGCAATGAAAATCTTGACTTGAGGAAATGGATCATACAATCATTTTCAGGGAGTATGATGGACGTTGTGGATGTCAATCTTTTTTCTGAGGAAGAAGAAATCACTAGTAAAAGTGAAATGTGCATAGCGTCCATGATAGAATTGGGTTTAGACTGCACAAAGGAAACGCCAGAATCAAGAGTGACCATGAAAGAAGTAGTTAAGAGGCTTAACAAAATCAACAACACATTTCTGGAAACATAG

>PGSC0003DMG400014463 LRR-RLPs

GCTAAAGATGGGGTAATGCAATTGTGCTCACTCTCGTCTTTTTCTGTCTTGAAGTTTCTTCATCCATATTACCTCATCACTCTCACAAACATCAAATTCAAACCATTTCCCTTCTAAAATTCAAGAAGTCCTTTACTATAGATCCCTCAGCCTCTTATTTTTGTCAAGACTTCGGTAAGAAGTTTTATCCAAAAACGAGTTCATGGAACATGAGCAGAGATTGCTGCTCATGGGATGGAGTTGTATGCGATGACATGACTGGACATGTCATTGAGCTTGACCTCGGCTGCAGCGGTCTTGTTGGGACCATTGATTCCAATAGCAGCCTATTCCAACTCTCTCATCTCCAAAGGCTTGATCTTTCTTTAAATAACTTCTCCAATTCTCACATCTCCCCTGAATTTGGCAGGTTTTCGAGCTTGACACATCTTGATCTTTCAGGCTCCTATTTTTCAGGTCAAATTCCTTTTGAAATCTCTTATCTTTCCAAGTTACACTCTTTTCGTCTCTCCACAAATATTGGGTTTGTTGGCCTCCCCTCATATTTTGGGCCCATACAAAGACTCGCGGCCCATGATTTTAAATTGCTCCTTCAGAATTTGACCCAATTAAGAGAACTTGATTTTAGTTTCATAAACATCTCTTCAACCATTCCTCCGAATTTTTCTTCTCATTTCACAACTCTGAGTCTGGGATTAACAGGATTGTATGGGATAATACCTGAGCGTATTTTTCACCTTCCCAACTTGAAAAAACTTGACTTATCCAGCAGTTATGAGCTCAGTGGTTATTTTCCAAAGACCAAATGGAACAACAACACATCTCTTAGGAAATTAGAACTCAGTGGCGTGAATTTTTCTGATAATATACCTGAGTCTCTTGGCTATCATCTAACTTCGCTGAATTATTTTTCTCTTTCGCATTGCAAGCTTTGGGGGCCTATTCCTAAATCTCTTTCCAATCTCACTCACATAGAGTATTTGTACCTTGATGGTAACTTGTTGAATGGAACAATACCATTAGGGATGTTCTCCCTTCTATCACTAATTTATTTAGAATTGAGTCATAACCACTTTTCTGGTCAGCTTGAGGATTTCAAGTCAAATTCACTATTCTATATTGGTTTAAGCAATAATCAGCTGGAAGGCCATCTTCCAAAGTCAATTCAAAACCTTGTGAACCTGACAACTCTTGATCTTTCATCGAATAATTTTAGTGGCAATGTGGATGTCAGCCTCTTTTCAAACCTCAAACAACTTTCGTATCTTATTCTTTCATCGAATAATTTTAGTGGCAATGTGGATTTCAGCCTCTTTTCAAACCTCAAACAGCTTTCATATCTTATTCTTTCTTCCAATAATTTTATTGGAAATGTAGATGTCAGCCTTTTTTCAGACCTCAAACAGCTTTCATATCTTAGTCTTTCACAAAATCATTCCTCATTGACCAATGAAAACAAAGTTAAGGTTACTTTGCCAGGATCTCTTTATACGTTACAATTGGCTGCATGTGAAGTGAAAGAATTGGAGTTTCTAAAATCCACAAATCATCTTGTTTTCTTGGATCTGTCAAATAATAAGCTTCAAGGAAGAATCCCAGATTGGGCATGGTCTAAATGGATGTTTTCATTGCAATATCTAGATCTATCCTACAACATGTTGACAAGTGTTGATTCAATTCCTCTTCAATCACTAGATACTATTGATTTACGGTCCAATTTGCTTCAAGGTTCACTACCTGTTCCACCAAATTCCACAATATTCTTCTTAATATCCCAAAATAATCTCACTGGAGAAATTCCTGCATCTATTTGCAATTTGATGTCACTCCAAATTTTGGATTTGGCGAGAAACAATTGAGAAATTCCACAATGTTTGGGAAATATGACTCCCCTCCGGGTTTTGGATATGCGCCACAACAATCTTTCTGGGAATATTCCACCAACTTTCAGTAATGAAAGTATATTGAGCAGCCTCAACTTGCATGGCAATAATCTAGAGGGGAAAATTCCCCGATCTTTGGTCAGTTGCAAAGACTTGCAGCTTCTTGATTTAGGCAATAATAACCTCAACGACACATTCCCTGTGTGGTTGGGAGCTCTTCCAGATCTGCTCATTTTAAGCTTGAGATCAAACAAATTGCATGGACCAATCAGAACATCAAGTTGGAGTGAGAACATGTTTCCTAAGCTTCGAATCATAGATCTTTCCTACAATGCCTTCTCGGGAAACTTACCTTCGAGTTTGTTTCAACATCTTAAAACCATGAGGACAATTGATCCATCAAGGGAAGCAGCAAGACATGGTGGAAATACATATTACCAAGATTCTATTATAGTTGCAATCAAGGGATCTGAGCGTGAAATTGTCAGAATCTTGTATTTGTACACCGTTATCGATCTTTCAAGTAATAAATTTAGAGGGCAAATTCCAAGTATTATGGGGCATCTCAATGCAGTTCACATATTGAATTTATCTCATAATGCATTGCAAGGTGATATACCGACATTATTCGGAGATTTATCTTCACTTGAATCATTGGACTTGTCAGGAAACAAACTTTCGGGAGAGATACCACAACAACTGGCTTTTCTTACTTCTCTTGCATTCTTAAATCTCTCCCACAATCATCTCCAAGGATGCATTCCTCAAGGACCTCAGTTTTACACTTATGAGAACAATTCATATGAAGGCAATGATGGATTACGTGGATTTCCTATTTCAAAAGGCTGTGGCGATACTCATCGTCTCTAAAGAATCCACGCCCTTTTTTTAGCGTTTAGGGCCGGGCGTCATGAAACAAGAATTCAGAATTTTCTAAGTGTTTCGTGTGTATCAATTCATGAATTTTTGGAAATATGACTGATCGAATATTTTTGCCCAAATATTTATTGAAATCATCTGTAATGACCTAGTGAAC

>PGSC0003DMG400014717 LRR-RLPs

ATGGTGAAAGTGTTTGAAGATCGTGGGGGATGGGCTGGGGCTGAGAGATACAGAAGGTAATGGGATTTGGGGAGGGAGCAACAGAACGAAGTGAGAGAAAAACTTTTTGAATTGATAATTTATCATTAATTAATGAGTTTAAATTTTAAGATAATTATTTAACATCATATATAAGAGATTTGTGTATCTAATTAAATATTTAAAAAATAAAAGTGATATTGTGCGTAATTTTTTTAAACTAGAGGTAAATTAAGTATATATGGCACTAAAGTATGTCTAACTAGGTAGTTTTTCCAATAATGAATTCATGAAAAATGCTTTTTTCTTTTCTAGCTCCAATGGTGAAACAAGGTTTGATCCTTTCCTTTTCAGGAACAAGGTGGGTTAACATTGTATCACATCAAGATGTCGATTCATTTTTTGCCCTCAATGAAACGGAGAAATTTTAACCCCGATTTTTGAGCCTTTCATCTTTCTTAATCAAGACCTAACAACCGCCGAGGAGCACCTCAACATATGATTGTGCACAGTAACTGAGACTCCTATTGCACCTAAGGTCAACTTCAATTCACCAAAACTAAAGTTTATCTTTGTGTAGTCATTGTAGACTCATACAAGTAGGGGAGGAGCCAACATGTCTTTAGGGGTTCATCCGAACCCCCTTCGAAAATATTACTATATATACATGGTTAAAATTATTTTTTAGATGTATATAATAGATGTCGAACCCCCTTCGACTAGTTCGTATGTTTACTCCTTCGAATTTTGAACCCCCTTATTGAAAATCCTGGCTCTGCCACTGCGTACAAGGATATTGAGTAGATAGTTGATGTATCAGACTCGATACTTCAAAAGAATGGAGGAGAAGACGATCTACGGCTGCAAGTGTGCATGAAGACAATATATGAAATAACTTAGCAAACATTCTTTGAAACAAACAGTTCACAATTCATAGATCAGAAAAATAATTTCCCAGCACGTCTAAAACAAGCAAAAATGGAAAAGAAAAACAAAACAGTTCCAAAGGTAGATAAAAAGTTCTTAAGCTTTGCAAACCACACCATCCCTCTGCAATGTCCAACACAAACGAATCAGATCGATACATAACCATCACAACGGTAAAAAGATTATGCAATCACACCTTGATCTTTGCCAACACGAGCTTAAACCTGTCATAATCATTGAGTGAGGCCGTCCTCTTCTGCACGATCGACTTCCTTCCTCAGGAACTGGACTCCCACTTGGTCTTCACATATGCGAGGACATAAAGCAAATGAGAGTTAATCAAGTTCTTACGTAACCCTGTCGTTAGCCAAATGACTCAAATTACCATTTCTGGATGATGAAAACCGTTTTTCCTAATTAAAAATTCAAATTTGAATACTGCAATTCAGCCAACGTACACGTATCTTGATCTCTAGCTCGGCCACTGTTTCCAGTGATATGTATTTTCTGAGGGGACTAATTAACACATAGTCTAGTCACTTTAGTATTTAGTATTGTAACTCTAGTGCTACAAAGAGTTAAGTTTAGTAGTGACGATTCATGAGATTAACCTCTAAATTTTGTGATGACTGAAAATCAGAAAGTAACACAAGATGCTGTTCTATCTTATGAAAAAGATTAGAAAGGTGAAACTCTACTTCTTTTTAAGCTTTGTATTCTAATTGATAGAATTTATTTTCTGTATGATAGTTTTGTTTTACTTGGAAAAAGTGGGAGTAGATGTGCAACACATGGATAAAGACAATAAAAAGACAAAAGCACAACAACCCAAATTTGACTAAAGAATGACAACATTTTCTCTAATAAAATTGAACAGACAAAGTGTGAATAAGAGTTTAGAAAAAAATGGGAGTAGATGTGTAACACATGGATAAAGACAATTAAAAGACAAAAGCACAACAACCCAAATTTGACTAAAGAATGACAATAATATTTGAGTGACTTTTCTCTTACAAAAACAAAGAAAGGTGACTTTCTGAGATAATTATCTATAGTTGAATGACCACGTGAGAAATCAACTCTTGACGTCTACTACATATGAATTAGAGATAATGTAGTTTAGAAAATTATTAATTTAAGACTGAAAATTGTTGAAACAAATATGTCTATATCATATGCTAAGGCAGAGTGGTCATGATCAATGTGTGATAACAAAACTATTTTTGCCACTTCGGTTAAGTCCTTGTCTGAATTAGTGTTGGTCTTTAATTATAATTTACGTACAATATAAAATCCCATGAACCCGAAACAATAAACTCAAAATATGATGATGGTTTCCAGCAAAATACTCTCTTCACTTCAGTGTCTCACTCTTTTATATCTCTTTACAGTTACTTTTGCTTCGACTGAGGAAGCAACTGCCCTCTTGAAATGGAAAGCAACTTTCAAGAACCAGAATAATTCCTTATTGGCTTCATGGACGCCAAGTTCTAATGCATGCAGGGACTGGTATGGAGTTAAATGCTTTAATGGTAGGGTAAACACGTTGAATATTACAAATACTAGTGTCAGTGCTACACTCTATGCTTTTCCATTTTCATCTCTCCCTTTTCTTGAGAATCTTAATCTTAGCATGAACAATTTCTCTGGCATCATTCCACCTGAGATTAGTAATCTCACTAATCTTGTCTATCTTGATTTGAACATCAATCAGATTTCAGGAACAATTCCACCACAAATCGGATCACTAACCAAGCTTCAGATCCTCCGTATATTTGACAACCATTTAAATGACTCTATTCCTAAAGAAGTAGGTTACCTAAGGTCTCTTACTAAGCTATCTTTGGGTAGGAACTTTCTTAGTGGTTCTATTCCTGCTTCATTGAGGAATTTGAACAACTTGTCCTTTTTGTTTCTTTATGAAAATCAACTTTTTGGGCCTCTATTCCTAAGGAAATAGGTTACCTAAGGTCTCTTAGTGAACTAGATTTAAGTAGTAACTCTCTAAATGGCTCTATTCCTGCTTCATTGGGGAATTTGAACAACTTGACTTATTTGTATCTTTCTGATAATCACATTTCGGGCTCTATTCCTGAGGAAATAGGTTACCTAAGGTCTCATAGTGAACTAGATTTAAGTTATAACTCTCTAAATGGTTCTATTCCTGCTTCATTGGGCAATTTGAACAACTTGTCCTTTTTGTCTCTTCGTGAAAATCAACTTTCTGGCTCTATTCCTCAAGAAATAGGTTACCTAAGGTCTCTTACTGATCTTCGTTTGAACAATAACTTTCTTAATGGTTCAATTCCTCTTGAAATAAGTTACCTAAGGTCTCTTAGTTTCATAGATCTGAGTATTAACTCTCTTAATGGTTCTATTCCTGCTTCATTGGGGAATTTGAGAAATGTGCAAAGAATGTTTCTTGATGAAAACAATCTCACTGAGGAAATTCCTTTATCTATGTGCAATTTAAAATCACTTGATATTTTGTTTTTGCGGAGAAACAACTTGAAGGGAAAAGTTCCGCAATGTTTGGGTAATATTAGCGGCCTCCAGGTTATGACGATGTCACGTAATAATCTTAATGGAGAGATCCCTTCATCTATTTCCAATTTAAAATCACTACAAATTCTAGATTTGGGAAGAAATAGTCTGGAGGGAGCAATTCCACAATGTTTTGGCAATATTAATACCCTACAGGTTTTTGATATGCAGAACAACAAATTTTCTGGGACTCTTCCAACAAATTTTAGCATTGGAAGTTCACTTATAAGTCTCAATTTGCATGGCAATGAACTAGAGGGGGAAATCCCTCGATCTTTGGCCAATTGCAAAAAGTTGCAAGTTCTTGATCTAGGAAATAATCATCTCAATGACACATTCCCCATGTGGTTGGGAACTTTGTTAGAGCTGAGAGTTTTAAGATTGACATCGAATAAATTGTATGGACCCATAAGATCTTCAGGGGCTGAAATCATGTTTCCTGATCTTCGAACCATAGATCTCTCTAACAATGCCTTCTCGAAAGACTTACCAACGAGTTTGTTTCAACATTTGAAAGGCATGAGGACAATTGATCAAACAATGATGGTACCAAGTTATGAAGGAGATGGAAGTTACGAAGACTCGATAGTTGTTGCATCGAAGGGATTGAAGCTTGAAGTTTTGAGAATTTTGTCTTTGTACACAGTTATCAATCTTTCAAACAACAAATTTGAAGGACATATTCCTAGTGTTCTAGGAGATCTCATTGCTCTTCGGGTGTTGAATATATCTCACAATAGATTGAAAGGCCATAGACCACCATCACTTGGACGTTTATCTGTAGTGGAATCATTGGACCTTTCATTTAACCGCCTTTCGGGAGAGATACCACAACAACTTGCTTCTCTTACGTCTCTTGAATTCTTAAATCTCTCCCACAATTATCTCCAAGGATGCATCCCTCAAGGACCCCAATTTCATACCTTTGAGAATAGTTCATTTGATGGTAATGATGGATTACGTGGATTCCCCGTTTCAAAAGGTTGTGGCAATGATCCTGTGTCAGAGACAAATTATACAGTATCTGCGCTAGCTGATCAAGAAAGCAATTCTGAATTTCTCAATGATTTTTGGAAAGCTGCTCTGATGGGCTATGGCAGTGGATTATGTATTGGATTATCCATAATGTATTTCATGATCTTAACTAGAAATCCGAAATGGCTTGCTAGAATCATTGAAGAAATGGAACACAAAATTAACATGCGAAGGAGAAAGAAGCAGCGAGGTTAA

>PGSC0003DMG400014718 LRR-RLPs

ATGATTCTCAATGATAACAATCTCATTGGGGAAATTCCTTCATTTGTGTGCAATTTGACATCACTGGAAGTGTTGTGTATGCCGAGAACCAATTTGAAGGGAAAAGTTCCGCAATGTTTGGGTAATATAAGTGGCCTCCAGGTTCTAGTGATGTCACATAATAATCTCAGTGGAAAGATGCCTCCATCTATTTCCAATTTAACGTCACTACAAATTCTAGATTTGGGCAGAAACAATCTGGAGGGAGCAATTCCACAATGTTTTGGCAATATTAGTAATCTCCAGGTTTTTGATATGCAGAATAACAAACTTTCTGGGACTCTTCCAACAAATTTTAGCATTGGAAGTTCACTTATAAGTCTCAACTTGCATGACAATGAACTAGAGGGGGAAATCCCTCGGTCTTTGGCCAATTGCAAAAAGCTGCAAGTTCTTGATCTAGGAGATAATCATCTCATTGACACATTCCCCATGTGGTTGGGAACTTTGACAGAGCTGAGAGTTCTAAGATTGACATCAAATAAACTGCAAGGACCTATAAGATCTTCAGGGGCTGAAATCATGTTTCCTGATCTTCGAATCATAGATCTCTCTAACAATGCCTTTTCAAAAGACTTACCAACGAGTCTGTTTCAACAATTGAAAGGCAGGAGGACTGTTGATCAAACAATGAAGACACCAAGTTATCTAGGAGGTAGATATAACTACCAAGACTCGGTAGTTGTTGTAACAAAGGGATTGAAGCTTGAAGTTGTGAGAATTTTGTCTTTTTACACAGTTATCGATCTTTCAAACAACAAATTTGAAGGACATATTCCTAGTGTTCTAGGAGATCTTATTGCGATTCGGGTACTGAATTTATCTCATAATGGATTGCAAGGCTATATATCACCATCACTTGGAAGTTTATCTGTACTGGAATCATTGGACCTTTCATTTAACCGTCTTTCGGGAGAGATACCACAACAACTTGCTTCTCTTACGTCTCTTGAATTCTTAAATCTCTCTCACAATTATCTCCAAGGATGCATCCCTCAAGGACCCCAATTTCATACCTTTGAGAACAGTTCATTTGATGGTAATGATGGATTACGTGGATTCCCCATTTCAAAAGGTTGTGGCAATAATCCTGTGTCAGAGACGAACTGTACAGTGTCTGCGCTAGAAGATCAAGAAAGAAATTCTGAATTTTTCAATGATTTTTGGAAAGCAGCTCTGATGGGCTATGGCAGTGGAATATGTATTGGATTATCCATAATATATTTCTTGATCTCAACTGGAAATCTAAGATGGCTTGCAAGAATCATTGAAGAACTGGAACACAAAATTATCATGCGAAGGAGAAAGAAACAGCGAGGTCAAAAGAATTACAGAAGAAGAAATAATCGCTTCTAG

>PGSC0003DMG400014719 LRR-RLPs

ATGATGATGGTTTCTAGCAAAACATTCTCTTCGCTTCAGTTTTTCACTCTTTTGTACCTCTTTACAGTTACTTTTGCTTCCACTGATGAAGCAACTGCCCTCTTGAAATGGAAAGCAACTTTCAAGAACCAGAATAATTCCTTATTGACTTCATGGATGCCAAGTTTTAATGCATGTGAGGACTGGTATGGAGTTATATGCTTTAATGGGAGGGTAAACACGTTGAATATTACAAATTCTAGTGTCATTGGTACACTCTATGCTTTTCCGTTTTCATCCCTCCCTTTTCTTGAAAGTCTTGATCTTAGCATGAACAATTTCTCGGGAACCATTCCACCTGAGGTTGGTAATCTCACTAATCTTGTCTATCTTGACTTGAACATTAATCAGATTTCAGGAACAATCCCACCACAAATCGGTTCACTGGCCAAGCTTCAGATCCTCCGTATATTTGACAACCATTTAAATGGCTCTATTCCTGAAGAAATAGGTTACCTAAGGTCTCTTACTAAGCTATCTTTGGGTAGGAACTTTCTTACTGGTTCTATTCCTACTTCATTGGGAAATTTGAACAACTTGTCTTTTTTGCTTCTTTATCAAAATCAACTTTCTGGTTCTATTCCTAAAGAAATAGGTTACTTGAGGTCTCTTACTAATCTATATTTGGGTAATAACTCTCTTAATGGATCTATTCCTGCTTCATCAGTGTGCAATCTGACATCACTGGAAGCGTTGTTTATGTCGAGAAACAACTTGACAGGAAAAGTTCCACAATGTTTGGGTAATATCATTCGCCTCAAGGTTATGACGATGTCACATAATAATCTCAGTGGAGAGATCCCTTCATCTATTTCCAATTTAACATCACTACAAATTCTAGATTTGGGCAGAAACCATCTGGAGGGAGCAATTCCACAATGTTTTAGCAATATTAGTAGCCTCCAGGCTCTTGATATGCAGAATAACAAACTTTCTGGGACTCTTCCAACAAATTTTAGCATTGGAAGTTCACTTATAAGTCTCAACTTTCATTGCAATGAACTCGAGGGGGAAATCCCTCGATCATTGGCCAATTGCAAAAAGCTGCAAGTTCTTGACCTAGGAGATAATCATCTCAATGACACATTCCCCATGTGGTTGGGAACTTTGCCAGAGCTGAGAGTTTTAAGATTGACATCGAATAAATTGCATGGACCCATAAGATCTTCAGGTGCTGAAATCATGTTTCCTGATCTTCGAATCATAGATCTCTCTAACAATGCCTTCTCGAAAGACTTACCAACGAGTCTGTTTCAACATTTGAAAGGCATGAGGACAATTGATCAAACAATGAAGGTACCAAGTTATGAAGGACATGGAGATTACCAAGACTCGACAATTGTTGTAACAAAGGGATTGAAGCTTGAAGTTGTGAAAATTTTGTCGTTGTACACAGTTATCAATCTTTCAAACAACAAATTTGAAGGACATATTCCTTGTGTTCTGGGAGATCTCATTGCGATTCGGGTATTGAATATGTCTCATAATGGATTGAAAGGTCATATACCACTATCACTTGGAAATTTATCTGTAGTGGAATCCTTAGACCTTTCGTTCAACCAGCTTTCGGGAGAGATACCACAACAACTTGCTTCTCTTACGTCTCTTGAATTCTTAAATCTCTCCCACAATTATCTCCAAGGATGCATCCCTCAAGGACCCCAATTTCATACCTTTGAGAACAGTTCATTTGATGGTAATGATGGATTACGTGGATTCCCCGTTTCAAAAGGTTGTGGCAATGATCCTGTGTCAGAGACAAACTTTACAGGGTCTGTGCTAGAAGATCAAGAAAGAAATTCTGAATTGTTCAATGATTTTTGGAAAGCAGCTCTGATGGGCTATGGCAGTGGACTATGTATTGGATTATCCATAATATATTTCTTGATATCAACTGGAAATATGAAATGGCTTGCAAGAATCATTGAAGAACTGGAACACCAAATTAACACACGAAGGAGAAAGAAGCAGAGAGGTCAAAGGAATTACAGAAGAAGAAATAATCGCATCTACACAAGTTACCAGTAA

>PGSC0003DMG400014736 LRR-RLPs

TTTTCACTTGCTAGTCTAAAACATTACACACCATGAAAGTGTTAACAATAACACTATGGGTTCTCTTGTTTTTTATTTTGAAAAATACCAAGTTTGTCATTTGTGTAGGGATTTGTAGAGAAAACGAGCAACGAGCATTGGAGACCTTGAAGAAAGAAATACACGATCCATCGGATATTCCATCTTCTTGGATTGTCGGAAAAGATTGTTGTGAATGGGAAGGGGTCGTGTGCAACAATCTATCTCGTCATGTGATTGAACTACATCTAGGTTATGTAACTCGTCATGAAGAGATTGGGCGAGGTATGTTTTTTGATAGGCTCGAGTTTACTTATCCAAGGATAAATAGCCTTGAGTGGCTGCCAAGTCTTTCGAGTCTTGAGAACCTACAGATGGTAAGTGTTGATCTTAGCAAAGTTAATGAATGGCTACAGGTTATTAACATGCTTCCTTCTCTTGTTGATCTTCGTTTATCCAGTTGTAGTCTTCATCATATAACACCTCTACTTGATCATCACAATTTTTCTTCACTCAAATTTCTTGATCTTTCTTTCAATTACAAATTAAGCTCTTCTGTTCCGAAATGGGTTTTCAACCTCCCAAATCTCATTTTTCTTGACCTGAGTTACTGTAATTTTACTAGCCCATTTCCTGATGGTCCAGTTAACTTGACTTCTCTCACAACCTTCAAGGCTTCTGGAAACTTTTTCAATTGTCGTTTACCCAAATGGTTGTTTGATTTTAATAATCTTGAACATCTTGAGCTTTCTAGTAATATGCTTTCTGGAAAGTTACCAAATGTAATTGGAAAGTTAGGGAAATTAGAATACCTTGATCTCTCATATAATCAATTTGAGGGAGATATTCCTATAAGTGTAACACGTCTTGCAGGATTAAGATCATTTAATCTTTCCAAAAATAACCTAACTGGTAGGATTCCAAATGACATTGGTGACATGAAAGTCTTGAAAACTGTTGATCTTTCAGAAAATCAACTTTATGGTCAAATCCCACAAAGCTTTTCGAGTTTGCCCACTTTGAGCTACTTGAATCTATCTGATAACAATTTGTCAGGTATGATACCAGTGAGCACTCAACTTCAAAGCTTTGATTCCACGAGTTTTCAAGGAAACAAACTCTGCGGACTTCCACTCTTGGTGAACTGCAGTTCGGATGGTAATATTCCAAATAACAAGCATGAAGATGATGAGAGTGACAAAGATGAAGTAGATTGGTTTTACATTTCAATGGCTATAGGATTTGCACTAAGCTTTTGGGGTGTTTGTGGTTCATTGTTTTTAAGAGGTCATGGAGACATGCTTATTTTCGTTTGTATGCTTACATTTCTCATGTTAGCAATCATGTTGTTTCATTGTATATTCAAATGAATGTTTCTTAAGATGTTCAAACAATGTTGTTTCTTTCATAGTTCCAATTTTCTTTTTTCAATCACAGAGTTGTGAG

>PGSC0003DMG400014738 LRR-RLPs

ATGGTGACGGATAAAGGCAACAAGTACCAGTATGACACCATTTTGGTGTTGTTTACAAGTACGGATATGTCTAGAAATAACCTTTCTGGGGATATTCCTGTAAGTTTAATAGGTCTTGCAGGATTAAGATCATTTGATTTTTCTAATAATAACCTAACTGGTAGGATTCCAAATAACATTGGTGAAATGAAAGTCTTGGAATTTGTTGATCTTTCAGAAAATCAATTTTACGATCAAATCCCACAAAGCTTTTCAAGTTTATCCACTTTGAGCTTATTGAATCTATCTGATAATAATTTGTCAGGTGTGATACCATTGAGCACTCAACTTCAAAGCTTTGATCCCACGAGTTTTCAAGGAAACAAACTCTGCGGACTTCCACTCTTGGTGAACTGCAGTTTAGATGACAATATTCTGAATCACGAATATGAAGATGATGAGAGTGATAAAGATGAAGTGGATTGGTTTTACATTTCAATGGCTATAGGATTTGCGCTAAGCTTTTGGGGTGTTTGTGGTTCACTGCTTTTCAAGAGATCATGGAGACATGCTTATTTTCATTTCCTAGATCGTAGCTGA

>PGSC0003DMG400014739 LRR-RLPs

GCTATAAATACATATTACTTGGCCTAGCGAAAAATATAAGCAAACACATACTTTACCCAAATACTTTTCACTTGCAAGTCTAAAACATTACACACCATGAAAGTGTTAGCAATAACACTATGGGTTCTCTTGTTTTTTATTTTGACAAATACCAAGTTTGTTGTTTGTGTAGGGATTTGCAGAGAAAATGAGCAACAAGCGTTGGAGACCTTGAAGAAAGAAGTAGACGATCCCTCAAATATTCTCTCTTCTTGGGTTGTCGGAAAAGATTGTTGTGAATGGGAAGGGGTCGTGTGCAACAATCTCACTCGTCATGTGATTGTGCTAAGCATTTATTTTGATTGGTTTGAGTCGAGATATCTAAGGATCAATAACCTAGAGTGGTTGACAAGTCTTTCGAGTCTTGAGAACTTGGAGATGGAAAGTGTTGATCTTAGCAAAGCTAATAATTGGCCACAGGTTATTAACATGCTTCCTTCTCTTGTTGATCTTCGTTTATTCAATTGTAGTCTTCATCATATAGCACCTCTACTTGATCATCACAATATTTCCTCACTCAAGTCTCTCGATCTTTCTGAGAATCGCAACTTAAATTCTTTTGTTCCCAAATGGGTTTTCTACCTCCCCAATCTTGTTTCTCTTGACCTGAGTGATTGTAATTTTATTGGCCCGTTACCTGACGGTCCAGTTAACTTGACTTCTCTCACAACCTTCAAGGCTTCTGGAAACTCTTTCAATTGTCGTTTACCCAAATGGTTGTTTGATTTGAATAATCTTGAACATCTTGACCTTTCTACTAGTGGCATTGAAGGTGTGATACAGAGTAAGAGCGGAAACATAACAAAACTTAAACATCTTGATCTTTCTAGCAATAACCTCAACTACACTATACCAAATTGGCTCTATCAATGCAAAGATCTAGAATCACTTCAACTTGGATTAAATCGTCTTGAAGGAACAGTTTCAAGTCTGATTTCAAATTTGAGCTCAATAACTATTATTGACCTTTCTAATAATATGCTTTCTGGAAAGTTACCAAATGTAATTGGAAAGTTAGGGAAATTAGGATACCTTGAACTCTCAGGAAATCTGTTTGAGGGAGATATATATGAATTATTCAATGTTAGGAGTAATTTCCTTTCAGTGCCATTGCGAAATACTTCTTCTTTATCTACTTTGAGACTAGGTTGGAATAAACTCACAGGAGCTCTTCCGGAAAGTGTTGGCCAACTTTCAATGCTAGAAGACTTTTCCATCTATAACAATAGGTTGGAAGGCGTTGTAACGGAAAGTCATTTCTCTAAATTAACACATTTACAATACTTTTATGCATCTAGAAATAATCTGACTTTAAAAGTGAGTCGGAATTGGATTCCGTCTTTTCAAGCCACAACAATGTTAATAGGCGGCTGGAATATAGGCCCTTCGTTTCCCATGTGGCTCCGAACTCAGAACCAGATAATGTATGTCGACATATCAGATTGTGGAATACAAGGTGAGGTTCCAACTTGGTTCTGGAACTTGTCTTCTCAAATTCGATTACTCAATCTTTCTCATAACCACTTTGTTGGTGAGGTTCCAATCATCTCAACAGATGAGCAGAGTGGTTCTCCGCTAATGTACTTGGCTTCCAACAATTTTAGTGGACCATTACCATTGATTTCAACCAATGTAAGAGAGCTAGATCTCTCAAGCAATTCTTTTTCCAAAGGTTTATCTAACTTTTTGTGTGAAGCAAAGAATGGATCATACAAATTGAGAATCTTAAACCTCGGGGGAAATGATTTATCAGAAGAAATTCCTGATTGTTGGATGAATTGGCCGGAGTTGAAAGTTTTAATCTTGAGGGACAATAATTTGATTGGAAGCTTACCAAGATCTATGGAGGTTTTGAGTAATTTGCGATCATTAGACTTACGAAGAAATAGACTTAACGGTCCGTTTCCTTCATCCTTGGAAAACTGCACAAAACTGCATAAAATAGATTTAGCTGAGAATGAGTTTATTGGAAAATTACCATCTTGGTTGGGAATGAGGTTTCCAACTTTGATAGTCCTTATCCTTCGGTCCAATAAATTCGATGGTGAATTGCCTCAAGAACTTTGTCACCTCAAAGATCTTCAAATCTTAGACCTTGCAAACAATACATTTGTTGGAATCATACCGAGGTGTATTGGCAATTTGAGTTCGATGGTTAAAAAAAAAAAGGAATTGGAAGATGATGAAGACTTAAATTACTCTTATTATATAGGAGTTCTGATAGAGAGTGCAATAGTGACGACTAAAGGCAACATGTACCAGTATGACACCATTTTGGCGTTGTTTACAAGTATGGATATGTCTAGCAATAATCTTTCTGGAGATATTCCTATAAGTGTAACACGTCTTGCAGGATTAAGATCATTTAATCTTTCCAAAAATAATCTGACTGGTAGGATTCCAAATGACATTGGTGTCATGAAAGTCTTGGAATCTGTTGATCTTTCAGAAAATCAACTTTATGGTCAAATCCCGCAAAGTTTTTCGAGTTTGACCACTTTGAGCTACTTGAATCTATCTGATAACAATTTGTCAGGTATGATACCATTGAGCACTCAACTTCAAAGCTTTGATCCCATGAATTTTCAAGGAAACAAACTCTGCGGACTTCCACTCTTGGTGAACTGCAGTTCGAATGGTAATATTCCAAATCACAAGTATGAAGATGGTGAGAGTGACAAAGATGAAGTGGATTGGTTTTACATTTCAATGGCTATAGGATTTGCACTAAGCTTTTGGGGTGTTTGTGGTTCATTGTTTTTCAAGAGATCATGGAGACATGCTTATTTTCGTTTTCTAGACCGTAGCTGGGAAATGTTACTTGCAAAGTTACCAATATGTTGAGAGATATGGTTTAGGATATTTCTATCTTCAAAAGTTACTGTGCGTGCTTACATTTCTCATGTTAGAAATCATTTTGTATTCAATATGTAGACTTTTATGTCTTAACAACTTCCTCAAATGAATGTTTCTTAAGATGTTCAAACAATGGTGTTTCTTTCATAGTTCCAATTTTC

>PGSC0003DMG400014740 LRR-RLPs

GGCCTAGTGAAAAATATAAGCAAACAACTAATACACACAAAAAATACTTTTCACTTGCAAGTCTAAAACATTGCACACCATGAAAGTGTTAGCAATAACACTATGGGTTCTCTTGTTTTTTATTTTGACAAATACCAAGTTTGTTGTTTGTGTAGGGATTTGCAGAGACAATGAGCAACAAGCGTTGGAGACCTTGAAGAAAGAAGTATATGATCCCTCGAATCGTCTCGCTTCTTGGGTAGTCGGAAAAGATTGTTGTGAATGGGAAGGGGTCGTGTGCAACAATCTCACTCGTCATGTGATTGAGCTAAGCATTTATAGTGATCAGATCGAGTCGAGATATCTAAGGATCAATAGCCTAGAGTGGTTGACAAGTCTTTCGAGTCTTGAGAACCTAGAAATGTATAATGTGGATCTTAGCAAAGCTAATGAATGGCTACAAGTTATTAACATGCTTCCTTCTCTTGTTTATCTTCGTCTATACAAATGTAGTCTTTCAAGTCTTGAGCTCCGTTATGGTGGCATTGAAGGTGCGATACAGAGTAAGAGTGGAAACATAACAAAACTTAAACATCTTGATCTTTCTAGCAATAACCTTAACTCCACTATACCAAATTGGCTCTATCAATGTAAAGATCTAGAATCACTTTTCCTTGGATCAAATCGTCTTGAGGGAACAGTTTCAAGTCTGATTTCAAATTTGAGCTCAATAATTAGTATTGACCTTTTTGATAATATGCTTTCCGGAAAGTTACCAAATGTAATTGGAAAGTTAGGGAAATTAGGATCGCTTGAACTCTCAGTAAATCAATTTGAGGGAGATATATCTGAATTATTCAATTTTATGAGTAATTTCCTTTCCATAGAATTTGGAAATAGTTCTTCTTTAACTGTGTTGAGTCTAGACAATAATAAACTCACAGGAGCTGTTCCGGAAAGTGTTGGCCATCTTTCAATGCTACAAGGCCCTTTGTTTCCCACGTGGCTCAGAACTCAAAAAAAGATATTGAATGTCGACATATCAGATGGTGGAATACAAGGTGAGGTTCCAACTTGGTTTTGGAAATTGTCTCCTCAAATTCGATTACTCAATCTTTCTCACAACCAATTTGTTGGTGAGGTTCCAATCATCTCGACAGATGAGCAAAGTGATTCTTGGCTATTAATGTACTTGGGTTCCAACAATTTTAGTGGACCAATACCATTGATTTCAACAAATGTAAGAGAGCTAGATCTCTCAAACAATTCTTTTTCCAAAGGTTTATCTAACTTTTTGTGTGAAGCAAAGAATGGATCTTACAAATTGAGAATCTTAAACCTCGGGGGAAATGATTTATCAGAAGAAATTCCTGATTGTTGGATGAATTGGCCGGAGTTGAAAGTTTTAATCTTGAGGGACAATAATTTGATTGGAAGCTTACCAAGATCTATGGAGGTTTTGAGTAATTTGCGATCATTAGACTTACGAAGAAATAGACTTAACGGTCCGTTTCCTTCATCCTTGGAAAACTTCACAAAACTGCATAAAATAGATTTAGCTGAGAATGACTTTTTTGGAAAATTACCATCTTGGTTGGGAATGAGGTTTCCAACCTTAATAGTTCTTATCCTTCGGTCCAATAAATTTGATGGTGAATTGCCTAAAGAACTATGTCACCTCAAAGATCTTCAGATCTTAGACCTTGCAAACAATACATTTGTTGGAATCATACCAAGGTGTATTGGCAATTTGAGTGCGATGGTCAAAGTAAAAAAGAAAGTACTGGAAGATGATTTCGAGTTAAATTATTCCTCTCGTTATGGAACTTTGATAGAGAGTGCAATGGTGATGACTAAAGGCAACATGTACCAGTATGATACAATTTTGGTGTTGTTTACAAGTTTGGATATGTCTAGCAATAATCTTTCTGGAGATATTCCTATAAGTGTAACACGTCTTGCAGGATTAAGATCATTTAATCTTTCAAAAAATAATCTGATTGGTAGGATTCCAAATGACATTGGTGATATGAAAGTCTTGGAATCTGTTGATCTTTCAGAAAATCAACTTTATGGTCAAATCCCACAAAGCTTTTCAAGTTTGACCACTTTGGACTGCTTGAATCTATCTGATAACAATTTGTCAGGTATGATACCATTGAGCACTCAACTTCAAAGCTTTGATCCCACGAGTTTTCAAGGAAGTAAACTTTGCGGGCTCCCACTCTTGGTGAACTGCAGTTCGAATGGTAATGTTCCAAATCACGAATATGAAGATGAGGAGAGTGACAAAGATGAAGTGGATTGGTTTTACATTTCAATGGCTATAGGATTTGCACTAAGCTTTTGGGGTGTTTGTGGTTCATTGCTTTTCAAGAGATCATGGAGATATGCTTATTTTCGTTTTCTAGACCATGGCTGGGAAATGTTGCTTGCAAAGTTACCAGTATGTTGAGAGATATGATTAAGGATATTGCTATCATCAAAAGTTAGTTTGTGTGCTTACATTTCTCATGTTAGAAATCATGTTGTTTCATTTTGTATTCAAATATGTAGACTTTTATGTCTTAATAACTTCCTCAAATGAATGTTTC

>PGSC0003DMG400014753 LRR-RLPs

CAAACAACTACCACACACAAAAAATACTTTTCACTTGCTAGTCTAAAACATTGCACACCATGAAAGTGTTAGCAATAACACTATGGGTTCTCTTGTTTTTTATTTTGACAAATACCAAGTTTGTTGTTTCTGTAGGGATTTGTAGAGAAAACGAGCAACGAGCGTTGGAGACCTTGAAGAAAGAAGTATATGATCCCTCGAATATTCTCTCCTCTTGGGTTGTCGGGAAAGATTGTTGTGAATGGGAAGGGGTCGTGTGCAACAATCTCACTCATCATGTGATTGATCTAAGCATTTCTATTGATTGGCCTTATTCGAGATATCTAAGGATCAATAACCTAGAGTGGTTGACAAGTCTTTCGAGTCTTGAGAACCTAGAAATGGAGAGCGTTGATCTTAGCAAAGCTAATGAATGGCTACAGGTTATTAACATGCTTCCTTCTCTTGTTGATCTTCGTTTATACAATTGTAGTCTTCATCATATAACACCTCTACTTGATCATAAAAAAATTTCTTCACTCAAATCTCTCGATCTTTCTGGGAATTACAACTTTAATTCATCTGTTCCAAAATGGGTTTTCAACCTCCCCAATCTTGTTTCTCTTGACCTGAGTGACTGTAATTTTACTGGCCCGTTTCCTGACGGTCCAGTTAACTTGACTTCTCTCACAACCTTCATGGCTTCTAGAAACTCTTTCAACTGTCGTTTACCTAAATGGTTGTTTGATTTGAATAATCTTGAACATGTTGTGCTCTATCGTAGTGGCATTGAAGGTGCGATACAGAGTAAGAGCGGAAACATAACAAAACTTAAATATCTCGAAATTTCTTACAATAACCTCATCTCCACTATACCAAATTGGCTCTATGGATGCAAAGATCTAGAATCACTTGTCCTTGGTGGAAATCATCTTGAGGGAACAGTTTCAAGTCTGATTTCAAATTTGAGCTCAATAATTAATATTGACCTTTCTAATAATATGCTTTCTGGAAAGTTACCAAATGTAATTGGAAAGTTAGGGAAATTAGGATCGCTTGACTTCTCAGGAAATCTGTTTGAGGGAGATATATCTGAATTATTCAATGTTAGGAGTAATTTCCTTTCAGTGGGATTGCGAAATACTTCTTCTTTATATACTTTGAGACTAGGCTGGAATAAACTCACAGGAGCTCTTCCGAAAAGTGTGGGCCAATTTTCAATGCTAGAATACATTTCCATCTCTAACAATAGGTTGGAAGGTGTTGTAACAGAAAGTCATTTCTCTAAATTAACACATTTACAATCCTTCTCTGCATCTAGAAATAATCTGACTTTAAAAGTGAGTCGGACTTGGATTCCACCTTTTCAAGCCTTTGAAATTGAAATAGGCAGCTGGAATATAGGCCTTTCGTTTCCCATGTGGCTCCAAACTCAGAACCAGATAATCTATGTCGACATATCAGATTGTGGAATACAAGGTGAGGTTCCAACTTGGTTCTGGAACTTGTCTTCTCAAATTCGATTACTCAATCTTTCTCATAACCACTTTGTTGGTGAGGTTCCATTCATCTCGACAGATAATGAGCAGAGTGATTATCCGCCATTAATAAACTTGTCTTCCAACAATTTTAGTGGACCAATACCATTGATTTCAACCAATGTAAGAGAGCTAGATCTCTCCAACAATTCTTTTTCCAAAGGTTTATCTAACTTTTTATGTGAAGCAAAGAATGGATCTTACAAATTGAGAATCTTAAACCTCGGGGGAAATGATTTATCAGAAGAGATTCCTGATTGTTGGATGAATTGGCCTGAGTTGAAAGTTTTAATCTTGAGGGACAATAATTTGATTGGAAGCTTACCAAGATCTATGGAGGTTTTGAGTAATTTGCTATCATTAGACTTGCGAAGAAATAGACTTAACGGTCCGTTTCCTTCATCCTTGGAAAACTGCACAAAACTGTATAAAATAGATTTAGCTGAGAATGAGTTTATCGGAAAATTACCATCTTGGTTGGGAATGAAGTTTCCAACCTTGATAGTCCTTATCCTTCGGTCCAATAAATTCAATGGTGAATTGCCTCAAGAACTTTGTCACCTCAAAGATCTTCAGATTTTAGACCTTGCAAACAATACATTTGTTGGAATCATACCAAGGTGTATTGGCAATTTGAGTGCGATGGTCAAAGTACTAGAAGATGATTTCGAGTTAAATTATTCCTCTTATTTTGGAACTTTGATAGAGAGTGCAATAGTGACGACTAAAGGAAACATGTACCAGTATGACACAATTTTGGCGTTGTTTACAAGTATGGATATGTCTAGCAATAATCTTTCTGGAGATATTCCTATAAGTGTAACACGTCTTGCAGGATTAAGATCATTTAATCTTTCCAAAAATAACCTAACTGGTAAGATTCCAAATGACATTGGTGACATGAAAGTCTTGGAATCTGTTGATCTTTCAGAAAATCAACTTTATGGTCAAATCCCACAAAGCTTTTCGAGTTTGACCACTTTGGGCTACTTGAATCTATCAGATAACAATTTGTCAGGTATGATACCATTGAGCACTCAACTTCAAAGCTTCAATTCCACGAGTTTTCAAGGAAACAAACTCTGCGGAGTTCCACTCTTGGTGAACTGCAGTTCAGATGACAATATTCCAAATCACGAATATGAAGTTGATGAGAGTGACAAAGATGAAGTGGATTGGTTTTACATTTCAATGTCTATTGGATTCGCACTAAGCTTTTGGGGTGTTTGTGGTTCATTGCTTTTCAAGAGATCATGGAGACATGCTTATTTTCGTTTTCTAGATCGTAGCTGGGAAATGTTACTTCCAAAGTTACCACTATGTTGAGAGATATGACTTAGGATATTTCTATCATCAAAAGTTAGTGTGTCAGTGTGCTTACATTTCTCATGTTAGCAGTCATGTTGTTTCATTTTGTATTCAAATATGTAGACTTTTATGTTTTAATAACTTCCTCAAATGAATGTTTCTTAAGATGTTCAAACAATGTTGTTTCTTTC

>PGSC0003DMG400014829 LRR-RLPs

CAAATATCGAAATTTTCCTAAACTACTTTGCACGAAAAAACAAAGGGAGTAGAAAGAGGAGGAAGAAATAGTATTCGTAGATTACATCTGAGCTTGACAACAAATTATGCAAGATTTTGAGTGAAACAACCAGACCTTTCATGGTGTTTCTTGTTGGCTGCTAAGAAAACAATGGCGTTTTCACCTGAAAAATCAAATTTCAGGTCACCATTGAAAGGGCCACCATCTCCTAGCTGGAATGATTTGTTGCTCAAGAACAAGAAAGCACTCAATCATGTCCTTTTCACTATGCATCTTCAATCCAGGTCCAATAAGCAAACCCCATCTTCATCCCCTTCTCCACAACTTCGCAAATCCCTAGATTTCACTCTTCCGTCACTTGTATCCGATCCAATTTCTCTTCTTTCTGATCAAATACTTCTTTCTATTCTCTCTAAACTCCCTGATTCACAAAGAAATTGTAATTCTCTTGTATCTAAACGCTGGCTCTATCTCCAAGGTCGTCTGGTACGGTCGATTAAGCTTCTTGATTGGGATTTCCTTGTTTCGGGTCGGTTGTTTATTCGGTTCCCGAATCTTATCCATGTTGATTTGGTTAATGGGTCTCTGATTTCACCGGGGAATGCGGGTGTTTTTTGCTCTAACCGGTTGCTTTCTTCTCACGTGGATTGTAATGCTGATGCGAAAGATTGGTTTCTTAAGGAAAGATTTGTGTTGCCTTCTGATGAGATTGATAAGGGGTTGAGAATACTAGCAAGTGGGTGTCCTAATTTGCGTCGATTATTGGTGGTAAATGCTAGTGAATTGGGATTGTTGGGGGTAGCTGAGGAATGTCCAACTTTGCAAGTGTTGGAATTGCACAGATGTAATGATCAGGTTCTTCGTGGTATCGCGGCCTGTCAGAACTTGCAGATATTGAGATTGATTGGAAATGTTGATGGGTTTTATAAATCTTCGGTTACTGATATTGGATTGACTATTTTAGCTCAGGGGTGTAAGAGGTTGGTGAGGCTGGAGCTGAGTGGTTGTGAAGGGAGCTATGAAGGGATTAAGGCAATAGCACAATGTTGTCAAATGCTCGAAGAGTTGATTCTTCGTGATCATAGGATGGAGGGTGGTTGGTTGTCTGCTCTTTCCTATTGTGAAAATTTGAACACCTTGAGGTTTATGTCTTGTAAGAGTATTGATCATTGTGGATGGTTTGATGAAGATGTAGGATCTTGCCCGACACTTGAAAGATTACATTTCGAGAAATGCCAATTAAGAAATAAGGAGAGTTTGAGAACGTTGTTTCTACTATGTCAAGAAGTTAGAGAGGTTATTTTCCACAACTGTTGGGGACTGGATAATGAAATGTTCAGTCTATCCACTGGTCTAAGGTATTGGTTATATTATTCTGTTCTCGTTTACTTTGCTTGCTGTTGCTGGAAAATTTCGTGCTGGATTAGCATTTTGTTATGCATTTGACATACATGAACTTATTGTGTTTAGGTTTTATGACCTTGAAATGTAGTTCCTTATTTTTGTTAAAAATGACTTTTTAGTGTTGCTGTAACCCCCATCCCCCGCCCCTCGTGTATAGTAGACGACCAAATGAAAGGATAGTTTGCTTGTAATAGTGCATTTTATTTTGGTCATAGTATACCCTTTGATCTTATCTTCTTGGTAAAAGGATAAGTGTGATAGATTTCCTTAGCTATTTGTGTTGGGGATAGATTGTTTTTTTAAGGCCAGTTTGTAGATGCATTTGTAATCATGCACCCTCTTGGTGTTTCAGTTATTAATGTTAAAATAGCCTTACCGTTTCTCAAAGAGTATACGCTTCCATCTTGCGTTAGAGAAAACACATAATGTTCATGGCATGACAATTATGTTTCAATTCATTATTGATGCTTGATTTATCAATATCATGTGTGCTATCTTGGATTATAAAGGGTCCAGTTTTGAGTTTTATACACTGAACTTGCTTCAGTTCTGACTGGCAATTTGGCATCATATTTTGTTTTGCATTCACTTGGAGCAGCAATAAAGTAACTGTCTCCGTGTGACCTATAGGTCACTGATTCGAGCCGTGGAAGTAGCCACTAATGCTTGCATTAGGAGGGTAGGTTGTCTACATCACACCCCTTGTGTGAACAATGGATACTTTGTGCACCAGGCTGCCCTTTTTAGTTCACTTGAATAAAGGGCCTGTAATTAACTAAATAACATTATGTGAGGAATTTGGGTTTCACAGCTTCATTTTGGTGTTATTTTCAACAGGAGAGTGAAGTCTATTTGCCTAGAAAGCTGTTCACTGCTCACAACTGAAGGCCTCGAATCTGCACTCCTTTCGTGGAAGGAGATCCAGAGCCTCAAGGTGATATCATGTGGCAATATAAAGGATAGTGAAATCAGTCCAGCACTTTCTACCTTGTTCTCCACGCTAAAAGATTTTCAGTGGAGACCAGACACAAAATCTCTTCTTTCAGCTGGTCTTGTGGGAACTGGCATGCGAAAAAGAGGCAGTAAATTATTCAAGAAGACGTGCGACTGGAAGTCACTGCCTGGTGCATAGACTGGCTTCCTCATCATGCATTCAGACAATAAATTATTCAAGAAGACATGCGACTGGAAGTCACTGCCTGGTGTATAGACTGGCTTTCTCATCATGCATTCAGACGACTCTACTAGATTTTACCTCCTCATTTATGCATATGAGAACTCTAACACATGAAAGAGGTCAGTGCGACACTTTGCTCCAATACTGTCAGCAATGAGGCCTTGATGTCATTATCCTGTATTTGTAGTCGCTTAAGTATGTAACGTAGGAACTATAGGATCGAAGATCGGCACTTTCTCAGGGTAGAGATGGAGAAAGAGGCCTAAGTTAGTGTATAGAGCTCGATGCTGTAACCTCATGAGCATTCAGCTAAAGCCATCTCGCTGTATCTTTGTCAAGCTTAATGGTATGTATTTCTGTATTGGAACCCCAAAAACTCTTAAAAGGGGTTTTCTTATCTACAATGTTATAAAGAATTTTGGCACCTTTTATGCCATGACTACTGCCTTCCACAATTGTGTGAAAGCATATTTGATATGAACCATACCAGCAAGCATGTCATTTGTGGAGGAGACGAGAACCAGAATAATTTTCTACATCAATGGCATCTATGTC

>PGSC0003DMG400015121 LRR-RLPs

TGGAAATCAAAACCATTTAATGTTTTGCATGATTGATGTGATACGTTTGCACTTTGCAGTGAACACTTGAGAAAAGTGATATCGTAAAGACTTGGGCAAATACCAATTGTTCGGTCAACTCAACAACATAAGTATTGCTATATAGTAATACATTTGAAAAGCAAATCAAGCTTAGAATCAATAGTAACAGCATGAAGAAGAACATTATCTTATTGATTCTTCTCTTTCTAATTCAATTTTCTTTATCACTTTCTTCCTCAAATAAGACTGACCAACAAGCTCTACTAGCTTTCCAAAATCTTATTACAAGTCCTAATCATTTTTTGGCCAATAATTGGACCAAAAATACTTCTTTTTGCTCTTGGTTTGGTGTCACTTGCACTCCAAAAAGGCAAAGGGTTGTGGCTTTGGCTCTTCCTAATTTGCAACTTCAAGGCACAATTCCCGATCTTGGCCAATTTGTCCTTCCTCAGTGTTCTCAATCTCCAGAATAACAGCTTCCGCGGTGGCATCCCTTATGGACTTGGCCACATGCCTCGCTTGCGAGTTATTGATGTTAAAAACAATCAGCTCAACGGAAGTATACCAACAAGTCTATTTCAAAACCGGAGAGTTCAAGTAATTTCATTGGCTTTCAATGAACTCAGTGGTGAAATGTGGAGAGGGCCATGGTATGTACCCGAACTCAGAGTCTTAAATCTCAGGAATAATAGCCTCACCGGTATAATCCCTCCTTCTATTGGAAATGCCACAACATTGATGAACTTCAGTTTGTCTGGGAATAGAGTCAGCGGCAACATTCCAAAGGAGGTCGGTAATCTGAGCCAGCTTGCATTTTTTCTCTTGTCCGATAATCAACTAACAGGTTCCATTCCCGCAACACTGTTTAATATCTCGTCACTACTTGCCGTATCTGTGTTACACAATAGCCTTTCTGGTCCTCTGTTGCTAGATGAAGGCAATATTGTGTCAAATCTGTACTTTTTTAAGTATATCTCACAACCAAATTTCTGGTTGCATTCCTTCCAACATATGCCAACTCACAGAGCTCAAAGTTTTGTCCATATCTTACAACAATATAACTGGAGACATACCCCAGAAATATTGGTTGTTTATTCCAAACTCGAGGAGTTTTATATTGGTGATAACCTAATAAAAGGGACTATTCCCACTTCATTGGGAAATATTTCCACTCTGCATAATCTTCACTGTCGAAACAATCAGATAGTAGGGGGGATTCCTCCGGAATTAGGGAAGCTATCAAATTTGAGGGAATTAAGCTTTGACAATAATTCTAATCTTATTGGTCAAATTCCAGAGGCTATTTTCAACATATCTTCTTTGGAAGTGATTGGTTTCAGTTTCAATAACCTCTCCGGGAGAATTCCAACCACTACAGGTCTTCATCTTCCCAATCTTGAAGGACTTTACTTGGGAGCAAATCAGTTCAAAGGGGAAATTCCATTGTTTATAACAAATGCTTCCAAGCTTCAGATACTGGAGCTAAATAATAACTTTCTGACAGGAACTATTCCTAATAATTTGGGAAATCTGCGTGAGCTGCGAGGACTGCTCCTATATAATAATCAACTTACCAATGAATTAAGAGAGCGTGAGTTGTTATTCTTCAATTCTTTGGCGGACTGTAGGATGTTGCAATATCTACAAGTGGGTTCCAATCCGTTAAATGGCGTTCTTCCCAATTCTATTGGAAATCTTTCATCTACTATTGAAAACTTTTATATAGTAGATGCACACATCAATGGCCTCATCCCCAAAAGTATAGGCAACATGACTGGTCTTACATCCCTAAGCCTTGACGGAAACAACTTGACAGGGAGTATTCCTTCTGAGATTGGTAAGCTTAAACAACTCCAAGGGCTGTATCTAGATAACAATAAATTGAAGGGACATATTCCAGAGGCCGTATGCCATTTATCTAATTTGGTTCAATTATTTCTGAATGTTAATGAGCTCTCTGGATTAATTCCAGAATGCATAGGAAATCTTAGCATGCTACAATATCTTTATTTGGGTTCTAATAAATTTTCATCAAAATTTACGTTGAACCTTTGGAAGATGAGTGGTCTTCTCTATCTAGACGTATCACAGAATTCAATGGAGGGAGAAGTTCCATCAGATATTGGAGCACTGAAAGCCATTGTAGAACTACATCTTTCCAGTAACCACTTTTCAGGAATGATTCCAAGCACATTCGGGGAACTCCAAAACCTACAGTATCTTGACCTGTCGAACAATTCATTTTTTGGCAAAATTCCATTATCCTTTGCCAACTTGATAAGCTTAGAATACTTGGATTTGTCTTCAAATGACTTGTCAGGTACTATTCCTATGTCTTTGGAAAAACTCTCCTACCTTAAAAGCATTAATGTTTCATTTAATGATTTAGAAGGTGAAATACCGAGTGGTGGCGTGTTTGCAAATTCCACTTTGCAATCATTTCTCGGGAACAAAGGTCTTTGTGGAATGCACATATTGGAGATTCCTGCTTGTGCTATCACTAATCCTGGAAAACAATCAAAGCTTAAGGAGGTTGTGCTAACAATTGTTACTCCAGTGGTTATTGCATCCTTTCTGATATTCTTGTTCGTTTCAATTTGGATAATGAAACGACAGAAGAAAGGGATGTCCAAAGATGTTGAAAAGGTATTGGAGATCGGGACTCATCAATTAGTTTCTTATCACGAGATTCAACGAGCAACAAATAATTTTTGATGAATCCAATTTAATTGGTGAGGGAAGCTCTGGCTCTGTGTACAAAGGAACATTGTTTGGTGGAACTGCAGTGGCCATTAAGGTTCTGGATTTGGAAAATGAGCAAGTATGCAAGAGGTTTGATACCGAATGTGAAGTGATGAGAAATGTTAGACACAGAAATCTTGTTTCAGTGATTACTACATGTTCTAGTGACTATATAAGAGCCTTTGTTCTGCAATTTATGCCCAATGGAAGTCTTGAGAATTGGCTGTACAAAGAAGATCGCCACTTGAACCTTCATCAAAGAGTCACTGTAATGCTTGATGCTGCTATGGCAGTCGAATATCTACACCATGGTCATAACACTCCTATAGTTCATTGCGACCTGAAGCCAGCCAACGTTCTTTTGGATGAAGATATGGTGGCTCATGTTGGTGATTTTGGAATCTCTAAAATTTTAGCTGTCAGCAAGTCTGTGGCACATACAGAGACATTGGGCACTCTTGGATACATTGCACCAGGTATAAAAAATCTACTCTCATTGATTTTCTTTTATCATAATTAAGCCTCACCAAGTTGTAGAAGTAAAAATAGAATTATTGTTGTGTTTCAACTGAGTAAATTTTCTTCATTCCTTTTCTAAGAATATGGCTCGGAGGGAATAGTGTCTGCTAGTGGTGATGTTTATAGCTACGGCATCATGTTGATGGAGGTTTTGACCAAAAGAAGGCCAACAGATGAAGAGATATGCAATGGAAATCTTGACTTGAGGAAATGGATTACACTATCATTTTCGGGGAGTATGATGGACGTTGTGGATGCCAATCTTTTTTCTGAGGAAGAACAAATCACTAGTAAAAGTGAAATCTGCATAGCCTCCATGATAGAATTGGGTTTAGACTGCACAAAGGAAACGCCGGAATCAAGAGTGACTATGAAAGAAGTAGTGAAGAGGCTTAACAAAATCAAGAACGCATTTCTGGAAACATAGAAGTCATGGTGTTCTTTGCTGAGCTGCAAATTAATATGATGCTGTTTGTTTCTTTAATAAAGTTGTTGTATAGTACTACTTGAAGTCTTTGAGTAGTTTTGGACCTATTTTACTCTATATATTTGTGATTATGTTTTGAATGCTTGAAAGAAACCAGAAATAAAGGTGTTTAGCTGCACTATTTTGTTTTTTGAATGTGAT

>PGSC0003DMG400015283 LRR-RLPs

ATGCACTCCAATACCAAAACAACATTTACGAGAGGTATCGTCTTTTGCTTCTTGACATTTTCCAAAATGGAAGTTTTCACAACGAGTTTTAATTTCCACACCTTTCTGGCTATCCGTGCTGTGTTTCTTGTGTTTCTATTGTCATTTTCTCTCCAACATGCTGCATCAGCTGCTTTTCTTAGCAATGAAACTGATAATCTTGCCTTACTTGGATTCAAGTCCCAAATAACCGAGGATCCATCAAGAGTTTTCGTCTCTTGGAATGATTCTGTTCATTTCTGTCAGTGGACTGGAGTAAAATGTGGCCTGAGACACGTAAGAGTCATCCGTTTGAATCTCAAAGGGCTGAGGCTGGCAGGTACGATCTCAGATCATCTTGGAAATCTCTCTTTTCTCAATTCTCTTGACCTTGCAGAAAATGCCTTCCATGAAAAAATTCCTCAACAACTCAGCAGGCTGCCAAGGCTTCAATACTTGAATTTGAGTTTTAACTATCTAACAGGAGAAATTCCAGCTAATCTATCACATTGTGTTAAACTCAAGAGCCTTGTCCTAGACCACAACACTCTAGTGGGAAAGATTCCTTACCAAGTTGGATCTTTAACGAAGTTAGTGAACTTGTACCTCAGAAATAACAACCTGACAGGAATCATTCCAGGTTCTATTGGAAATCTTACATCTTTAGAGAAGCTGCATTTATCGTACAACAATCTAGAAGGAGAAGTGCCAGTTTCTTTAGCTCGATTAACCAAGTTGAGGATTCTTGGACTGTCAGTAAATAGCTTATCTGGGGAGTTCCCTCCTCCGTTGTACAATTTGTCATCCCTGGAACTAATATCACTTTCCCTTAACAACTTTTCTGGTAATCTCAGATCTGAGTTAGGCAACTACTTCCCTAATCTCCAAATACTTTACTTGGCGAACTGTCAATTCATTGGTTCCATACCATCCTCCTTGGCCAATGCTTCAAAATTGCTAGAGCTTGATTTTCCTGTAAACAATTTCACTGGAAATATACCTAAGGGTTTTGGTAACTTGCGGAATTTGTTATGGCTCAATGTTTGGAATAACCATCTTGGATATGGTAAGCATGATGACCTTGACTTTGTAAATTCTCTTACCAACTGCAGCAGTCTACAAATGCTCCATTTTGGAGGCAACCAGTTTGGAGGTACATTACCTCATTCAATAGGCAACCTTTCTAGTCAGTTACAACGCCTACTCTTCTTCGAAAATAGAATTGGTGGAAACATACCAAGAGAGATCTCAAACCTGGTGAATTTGAATCTACTCGACATTGGTTCCAACAATTTTATAGGTAGTATTCCAGATTCTATTGGAAGACTTACAAACTTGGGAGCTCTCAACTTGGTTAACAATCTCTTGACAGGGGTCATCCCTTTTTCAATAGGAAACCTCACTGAACTCGTCTATCTTTATTTGGGACTTAACAGGTTAGAGGGTAACATACCTTCAACATTGGGAAACTGTAACCAACTGCTAAAATTGGGATTTTCTGAAAACAACCTGACAGGAACTATACCACAACAACTTTTTGCTCTCTCATCCCTCACAGATATTTTTGCATCTAATAACTCTTTGACTGGTCCATTACCAGTGGATATTGGTAACTGGAATCATCTTACTTATCTTGATTTTTCTTACAACAATCTTTCGGGTATGATTCCACAAACTCTTGGAAAATGCTTGACCTTGGGGGAGATCTATATGAAGGGAAACTCCCTCCAAGGAACCATTCCCAATTTAGAAGATTTGCCGGATCTCCAATCCTTGGATCTTTCCCTAAACAACCTATCAGGGCCAATCCCTCACTTCATTGCAAATCTTACTTCCCTACACTACTTGAACTTATCTTTTAACAATCTAGAGGGTGAGGTTCCTGTCACCGGAATATTTTCAAATTTGAGTGCAGATGTATTAAGCGGAAACTCCAAGCTTTGTGGTGGGATTCAAGAGCTTCATTTACAACCTTGTATTTATCAAAAGACACGGAAGAAGCATGTGCTTGCCCTCAAGTTTATTTTGATAATTGTTTTTGCTGCTTCATTTTCAATCCTGGCCTTGTTAGTAGTTTTCCTTTGTAGGAGAAGAAATTTGAACAATCAACCAGCACCACAAGATAGGTCTAAATCTGCGCATTTCTACCCCAATATTTCGTATGAAGAGCTCCGCACTGCAACAGGTGGATTTTCTTCTGAAAATCTTATTGGCTCTGGCAGTTTTGGAACTGTTTATAAGGGAACTTTTCCCTCTGATGGAACGGTCGTTGCTGTGAAGGTACTCAAACTCCAGCATGAAGGTGCTTCAAAAAGCTTCTTAGCAGAATGCCAAGCATTGAGAAACATAAGGCACCGGAACCTTGTCAAGGTCATCAGTGTATGCTCAAGTTCTGATTTCAAGGGAAATGATTTTAAAGCTTTAGTCTTTCAGTTCATGCCAAAAGGGAACTTGGACGAATGGCTACATCCAGAAAAGGAGATGAATGAGAAGAGTAGCTTGACCACACTTCAGAGAATGAATATCATAATAGATGTGGCCTCTGCACTTCATTATCTTCACCATGAGTGCCAAACACCCATGATTCACTGTGATATAAAACCACAGAACATTCTTCTTGATGAAGATCTCACCGCTCATTTGGGTGATTTTGGTTTGGTGAGACTCGTTCCTGAATTCAGTAATGAATCGGATCTACATCAGTTTAGCTCACTTGGAGTTCTGGGAACCATTGGCTATGCAGCTCCAGGTAATTTTTATATGTAAACAAAAACTATGAATATACCATATGGTTTCTTGTAACAATAGTGGTTTATTTATGAGATGATCAATTTGCTTGTTGCAGAATACGGCATGGGTAGTAAGGTCTCCGTTGTAGGAGATATGTACAGTTTTGGGATCCTAATATTGGAAATATTCACTGGAAGAAGACCCACAGACACTTTGTTCCAAGCAAGTTCTACCCTCCACCACTTTGTAGAAACAGCATTGCCAGAGAAAGTTATGGAGATTCTAGACAAAACAGCTTTTCATGGTGAGATGAGTAAAGCAACCTGTGGCGAAGAATACTGGGGCAATATCAAGAAAGAACAGATGGAATGCTTGGTTGGCATACTTGAGATCGGTGTTGCATGTTCAGCTGAATCCCCAAGAGACAGATTGACCATGACACAAGTTTACAGTAAGTTGACACTAATAAGAGAAAAATTCCTACAAAGGGGAGATGGAGGGCACAGGCTACAAGTGAAAGCATTGGATTTAATCTAA

>PGSC0003DMG400015345 LRR-RLPs

TATTTCTAACAATAAGATTCGTGGCCAAATCCCAAACTGGTTTAGCGGCATGAGGTGGGACAACTTGTTGTACCTAAACCTTTCGCATAATTCTTTAACAGGCCACCTACAACTATTTCATTTCCATAATCTAGAGTATCTTGATCTTAAATTTAACTTCCTTCAGGGTCCACTACCTTCATCCATTTGTAACATGAACAATCTTAAATTGCTAGATTTATCACGCAACAACCTCAGTAACTCGATTCCAAGTTGCTTGGGAAGCATGTCTAGTCTAACAGTGTTGGACTTAAGAAGGAACAATTTCACAGGGAGTCTTCCCCCATTATGTGTGCAGAACACTTCATTGAGAACCATTGTCTTGAATGGCAATCAATTTGAAGGAACTCTCCCTGTGTCGTTGCTCAACTGTAGTGGTTTAGAAGTTCTTGATATGGGGAATAACGCTATAAATGACACATTTCCTGCTTGGATAGGAACACTTGAACAACTGCAGGTCTTAATATTAAAGTCCAATGTGTTCCATGGACCAATTACTACTTGTCAGACTACACTTTGCTTTCCCAAGTTGCGAATTTTTGATCTTTCGCGTAACGAATTCAGTGGCTCAATCCCTGCAAAGGTTTTTGGAAACTTCAAGGCAATGATTAAATTAGATGGTGAAGACACAGGACAGATCGAGTACATGGAACCATCTATGTTTCACACATTATATGATGATTCCGTGAGTTTGGTGATCAAAGGCCAGGATATTGAGCTAGAAAGAATCAGCAAAATTATGACAACCATAGACCTCTCGAGCAACCATTTTGAAGGTGTCATTCCGAAAACACTAAAGGATCTCAGATCACTTTGGCTACTCAATTTATCCCATAACAATCTCATAGGTCATATTCCAATGGAATTGGGGCAATTGAATAAGCTTGAAGCGTTAGATCTCTCTTGGAACCGGCTCACTGGAAAGATTCCGCAGGAATTGACAAGGCTAACATTTCTGGAAATGTTAAACCTTTCTCAGAATGTTCTCGTTGGACCAATTCCTCACGTTCTACAATTCAACACATTTGAAAATGACTCGTATGGCGGTAACCTTGATCTATGTGGTCCTCCTTTATCAGAGCAATGTGGAACGAGTGATCTATCCCATGTTCCTCAACCATTGGAGTCTGAAGACGAGTCATATTTTTTTAGTGGATTTTACGTGGGAATCAGTAGTCATTGGCTACAGCTGTGGACTAGTTGTTGGAACTGTTGTATGGAGCCTCATGTTTAAATATCGTAAACCAAAATGGTTTGTCGAATTTTTTGAAGGCATTTATCCCAAGAAAATGAGAAGGTCAATGAAGAGAGCTCAGAGACGACGGACTTAGTGGAAGATTGCAAGAG

>PGSC0003DMG400015347 LRR-RLPs

GTTGGTAATTAGTGAAAGCTTTTCTTCTTCCTCTGTGATGATGCATCATGTTTGCTCTCCCACTGAAGCTTTTGCTTTGCTTCACTTCAAACAATCCTTTCAAATCTCTGACCAACCTACATGTTGGGATGATATCCCAAAAACAAAGTCTTGGAATGAGAGTAGGGATTGCTGCAGTTGCTGCAGTTGCTGCAGTTGCTGCAGTTAACCGGTTATGTTATTGGATTGGACCTTAGTTGCAGTATGCTTACTGGAACTATTCATCCCAATAGCAGCCTCTTCCAACTTCATCATCTCCGGACACTAAACCTTGCTCACAATGCCTTCTTTTCTTCAATCCCACACAACATTGGCCGATTGACGAATTTGAGGCATCTCAACCTTTCTCATTCATTCTTTGAAGGGGAAATTCCAACAGAAATCTCATACCTTTCCAATTTGGTTTCACTTGAACTTAATTGTTATGGATGTGAACTTGATGAGAGAACATTTGAAACAATTCTTCAAAACTTCACAAATCTGGAGGTAGTTTCTCTCTTTGGTGTCAACATCTCATCTCCGATACCTCTGAATATATCTTCATCCTCCTTAAAGTATGTAGATCTTGGATTTACTAATCTGCGAGGTGTTCTCACAGAGAGCTTTTTCCTTCTGCCAAAATTGGAAATGCTCTATTTGAATTACAATGATCTTCTTAAAGGAGGTTTACCAAAGATCCACCCGAGCAACACTCTGTTGGAGTTGCATATTTCATTCACAGGAATCTCCGGTGAGCTGCCTGATTCAATTGGCACCTTCAGTTCCTTGAATCGCTTGAACATGTATGGATGCCAATTATCTGGCCGCGTTCCTGATTCCATTGGCAATCTTACACAAATTAGGTATTTGGCTTTCAGTTATAATCATTTCACCGGCCATATTCCTTCCACAATATCTAAATTGAAGCACCTCAGCTATCTAAAGCTTTCGTCCAACTCCTTTTCAGGTGAAATTCCCGACATTTTTTCTAACCTCCAAGAGCTACGTTCTTTATATCTTCATAACAACAGTTTCATTGGTTCGTTTCCTTCTACAATTGTAAGCTTGACACATCTTCAATACTTAGACTTGTCAAGTAATTCCCTATCTGGCCCACTGCCTAATAATTTTAGCATGCTTCAAAAGCTAACCCAACTGGATTTGTCGTACAACGCACTGAATGGGACCATACCATCTTCGGTGTTTAGCCTCCCTTTGCTACCTCAATTATGGCTCGATAATAATCGATTTAGTGGACTACCCACTGAGCTCAAAACAATCCCAACATTAGAAAGCCTAGATTTAAGCCATAATCAACTCAGTGGTTCTTTTCCTCAATCACTTGTGAATCTCACAAGCCTTTCTACCCTCGACCTTTCATCAAATAACATCACCATCGATGAGGGAATACAAATCACTTTTCCTAGCCTACAAGATTTGTGGTTATCATCTTGTGAACTGAAGGATATTCCACACTTCTTGACAAATGTAAAGAAACTTCAGGTCTTGGATATTTCTAACAATAAGATTCGTGGCCAAATCCCAAACTGGTTTAGCGGCATGAGGTGGGACAACTTGTCCTACCTAACCCTTTCGCATAACTCTTTAACAGGCCACCTACAACAATTTAATTTCCATAATCTAAGGTATCTTGATCTTAAATTTAACTCCCTTCAAGGTCCAATACCTTCATCCATTTGTAACATGAACGATCTTCAATTTCTAGATTTATCACGCAACGACTTCAGTAACTCGATTCCAAGTTGCTTGGGAAGCATGTCTAGTCTAACAGTGTTGGACTTAAGAAGGAACAATTTCACTGGGAGTCTTCCATCGTTATGTGCGCAGAGCACTTCATTGAGTACCATTTACTTGAATGGTAATCAATTTGAAGGAACTCTCCCTATGTCGTTGCTCAACTGTAGTGATTTAGAAGTTCTTGATATGGGGAATAACGCTATAAATGACACATTTCCAGCTTGGCTAGGAACACTTCAACAAATGCAGGTCTTAATATTAAAGTCCAATCTGTTCCATGGACCAATTAGTACTTGTCAGACTACATTTTGCTTTCCCAAGTTGCGAATTTTTGATCTTTCGCGTAACAAATTCAGTGGCTCACTCCCTGCAAAAGTTTTTGGAAACTTCAAGGCAATGATTAAATTAGACGGTGAAGGTACAGGAGAGATCAAGTACATGGAACCATCTGAGATGTCATCTGATGATTCTGTGAGTTTGGTGATCAAAGGGCAGGATATTGAGCTAGAAAGAATCAGCACAATTATGACAACCATAGACCTCTCGTGTAACCATTTTGAAGGTGTCATTCCGAAAACACTAAAGGATCTCAGCTCACTTTGGCTACTCAATTTATCCCATAACAATCTCATAGGTCATATTCCAATGGAATTGGGGCAATTGAATAAGCTTGAAGCGTTAGATCTCTCTTGGAACCGGCTCACTGGAAAGATTCCGCAGGAATTGACAAGGCTAACATTTCTGGAAAAGTTAAACGTTTCTCAGAATGTTCTCTTCGGACCAATTCCTCTTGGTCCACAATTCAACACATTTGAAAATGACTCGTATGGCGGTAACCTTGATCTATGTGGTCCTCCTTTATCAAAGCAATGTGGAACGAGTGATCCATCCCATGTTCCTCAACCATTGGAGTCTGAAGACGAGTCATATTTTTTTAGTGGATTTACGTGGGAATCAGTAGTCATAGGCTACAGTTGTGGACTAGTTGTTGGAACTGTTGTGTGGAGCCTCATGTTTAAATATCGTAAGCCAAAATGGTTTGTCGAATTTTTTGAAGGCATTTATCCCAAGAAAATGAGAAGGTCAATGAAGAGAGCTCAGAGACGACGGACTTAA

>PGSC0003DMG400015348 LRR-RLPs

ATGTCGTTGCTCAACTGTAGTGGTTTAGAAGTTCTTGATATGGGGAATAACGCTATAAATGACACATTTCCTGCTTGGATAGGAACACTTGAACAACTGCAGGTCTTAATATTAAAGTCTAATATGTTCCATGGACCAATTAGTACTTGTCAGACTACGTTTTGCTTTCCGAAGTTGCGAATTTTTGATCTTTCGCGTAACAAATTCAGTGGCTCACTCCCTGCAAAAGTTTTTGGAAACTTCAAGGCAATGATTAAATTAGATGGTGAAGACACAGGACAGATCAAGTACATGAAAGCATTTGACACATCGTATGAGGATTCTGTGAGTTTGGTGATCAAAGGCCAGGAAATTGCGCTAGAAAGAATCAGCACAATTATGACAACCATAGACCTCTCGAGCAACTATTTTGAAGGTGTCATTCCGAAAACACTAAAGGATCTCAGCTCACTTTGGCTGCTCAATTTATCCCATAACAATCTCATAGGTCATATTCCAATGGAATTGGGGCAATTGAATATGCTTGAAGCGTTAGATCTCTCTTGGAACCGGCTCACTGGAAAGATTCCGCAGGAATTGACAAGGCTAACATTTCTGGAAAAGTTAAACGTTTCTCAGAATGTTCTCTTCGGACCAATTCCTCTTGGTCCACAATTCAACACATTTGAAAATGACTCGTATGGCGGTAACCTTGATCTATGTGGTCCTCCTTTATCAAAGCAATGTAGAACGAGTGATCCATCCCATGCTCCTCAACCATTGGAGTCTGAAGACGAGTCATATTTTTTTAGTGGATTTACGTGGGAATCAGTAGTCATAGGTTACAGCTGTGGACTAGTTGTTGGAACTGTTGTGTGGAGCCTCATGTTTAAATATCGTAAGCCAAAATGGTTTGTGGAATTTTTTGAAGGCATTTATCCCAAGAAAATGAGAAGGTCAATGAAGAGAGCTCAGAGGCGACGGACTTAA

>PGSC0003DMG400015349 LRR-RLPs

ATGGGAATTCTGTTTTTCTTTTATTCACTTGTCTGTTTTTTGTTGGTAATTAGTGAAAGCTTTTCTTCTTCCTCTGTGATGATGCATCATGTTTGCTCTCCCACTGAAGCTTTTGCTTTGCTTCACTTCAAACAATCTTTTCAAATCTCTAACCAACCTAGCTGTTGGGTTGAGATTAATCTGCCAGATATGACCTTTGTTGATATCCCAAAAACAAAGTCTTGGAATGAGAGTAGGGATTGCTGCAGTTGGGACGGGGTAACTTGTGACATGTTAACCGGTCATGTTATCGGATTGGACCTTAGTTGCAGTCTGCTTAATGGAACTATTCATCCCAATAGCAGCCTCTTCCAACTCCATCATCTCCAGACACTAAACCTTGCTCGCAATGACTTCTATTCTTCTTCAATCCCACATAACATTGGCCGATTGACGAATTTGAGGCATCTCAACCTTTCTCAATCTTTCTTTGTAGGGGAAATTCCAACAGAAATATCATACCTTTCCAATTTGGTTTCACTTGAACTTAATTGTTATGAATGTGAACTTGATGAGAGAACATTTGAAACAATTCTTCAAAACTTCACAAATCTGGAGGTAGTTTCTCTCTGTGGTGTCAACATCTCATCTTCAATACCTGTGAATATTTCTTCTTTCTCCTTAAGGTACGTGGATCTTAGACATACTAATCTGCGAGGTTTTATCACAGAGAGCTTTTTCCTTCTGCCAGGATTGGAAAGGCTCTATTTGAGTGACAATGATCTTCTCAAAGGAGTTTTACCAAAGATCCACCCGAGCAACACTCTGTTGGAGCTGGATATTTCATTCACAGGAATCTCCGGTGAGCTGCCTGATTCAATTGGCACCTTGAGTTCCTTGAATATCTTGAATTTTCTTAATAATCATTTCACCGGCTATATTCCTTCCACAATCTCTAAATTGAAGCACCTCACATCCCTAGATCTCTCGTCCAACTCCTTTTCAGGTGAAATTCCCGATATTTTTTCTAACCTCCAAGAGCTACGTTTTTTATATCTTCATAACAACAGCTTCATCGGTTCGTTTCCTTCTACAATTGTAAGCTTGACACATCTTCAAGACTTAGACTTGTCGAGTAATTCCCTATCTGGCCCACTGCCTAATAATGTGAGCATGCCTCAAGAGCTAACCAAACTGGATCTGTCGTACAACTCACTGAATGGTACCATACCATCTTCGGTGTTTAGCCTCCCTTTGCTATCTTCATTGTCTCTCCAACATAATCAATTTAGTGGACTAGCCGATGAGTTCAAAACAAACTCAACATTAGATTACCTAGATTTAAGCCATAATCAACTCAGTGGTTCATTTCCTCAATCACTTGCGAGTCTCACAAACCTTTCTACCCTCACCCTTTCATCAAATAACATCACCTTCGATGAGGGAATACACATCACTTTTCCTAGCCTACAAGATTTGCATTTATCGTCTTGTGAACTGAAGGATTTTCCACACTTCTTGAGAAATGTAAAGACACTTACGGTCTTGGATATTTCTAACAATAAGATTCGTGGCCAAATCCCAAACTGGTTTAGTAGCATAAGGTGGGACTACTTGTACTACCTAAATCTTTCGCATAATTCTTTAACAGGCCACCTACAACAATTTCATTTCCATAATCTAAGGTATCTTGATCTTAAATTTAACTCCCTTCAGGGTCCACTCTCTTCATCCATTTGTAACATGAGCGATCTTAAATTTCTAGATTTATCACTCAACAACTTCAGTAACTTGATTCCAAGTTGCTTGGGAAATCTTCAATTTCTAGATTTATCACGCAACGACTTCAGTAACTCGATTCCAAGTTGCTTGGGAAGCATGTCTAGTCTAACAGTGTTGGACTTAAGAAGGAACAATTTCACAGGGAGTCTTCCCCCATTATGTGCATAG

>PGSC0003DMG400015351 LRR-RLPs

ATGTTAACTGGTCATGTTATCGGATTGGACCTTAGTTGCAGTCTGCTTAATGGAACTATTCATCCCAATAGCAGCCTCTTCCAACTCCATCATCTCCAGACACTAGACCTTGCTTACAATGACTTCTATTCTTCTTCAATCCCGCATAACATTGGCCGATTGAGGAATTTGAGGCATCTCAACCTTTCTTATTCTTTCTCTGGAGGGGAAATCCCAACAGAAATATCATACCTTTCCAATTTGGTTTCACTTGACCTTTTATGTTATGGATGTGGTCTCCAACTTGATGAGAGAACATTTGAAACAATTCTTCAGAACTTCAGAAATCTGGAGGTACTATCTCTCTATGGTGTCAACATCTCATCTCCGATACCTGTGAATATTTCTTCATCCTCCTTAAGGAACATGGATCTTGGACATACTAATCTGCGAGGTGTTCTCCCAGAGAGTCTTTTCTTTCTGCCAAAATTGGAAACGCTCTATTTGAGTGACAATGATCTTCTCAAAGGAGTTTTACCAAAGATCCACCCGAGCAACACTCTGTTGATGTTGGATATTTCCTACACAGGAATCTCTGGTGAGCTGCCTGATTCAGTTGGCACCTTCAGTTCCTTGAATCAATTGAACCTCCGAGGATGTCAATTCTCTGGTTCCATTCCTGATTCCATTGGCAATCTTACACAAATTTGGCGTTTGGATTTCCGTAATAATCATTTCACCGGCAATATTCCTTCCACAATCTCTCAATTGAAGCAGCTCACATGTCTATATCTTTCGTCCAACTCTTTTTCAGGTGAAATTCCCGACATTTTCTCTAACCTCCAAGAGCTAAGTTGTGTATATCTTCGTAACAACAGCTTCATCGGTTCGTTTCCTTCTACAATTGTAAGCTTGCCACATCTTCTAGACTTAGACTTGTCGAGAAATTCCCTATCTGGCCCACTGCCTAATAATTTTAGCATGCTTCAAAACCTAAACGAACTGCATTTGTTGTACAACTCGCTAAATGGTACCATACCATCTTCGGTGTTTAGCCTCCCTTTACTAGTTGAGTTGCGGCTCGGTAATAATCGATTTAGTGGACTACCCAATGAGCTCAAAACAAACCCAAAATTAGAACGCCTGGGTTTAAGCCATAATCAACTCAGTGGTTCTTTTCCTCAATCACTTGCGAATCTCACAAACCTTTCTACCCTTGACCTTTCATCGAATAACATCACCGGTGATGCTGGAATAAATATCACCTTTCCTAGCCTAGAAATTTTGCAGTTATCATCTTGTGAACTGAAGGATTTTCCACACTTCTTGACAAATGCAAAGAAACTTCATGTCTTGGATATTTCTAACAATAAGATTCGTGGCCAAATCCCAAACTGGTTTAGTAGCATAAGGTGGGACTACTTGTACTACCTAAATCTTTCGCATAATTCTTTAACAGGCCACCTACAACAATTTCATTTCCATAATCTAAGGTATCTTGATCTTAAATTTAACTCCCTTCAGGGTCCACTCTCTTCATCCATTTGTAACATGAGCGATCTTAAATTTCTAGATTTATCACTCAACAACTTCAGTAACTTGATTCCAAGTTGCTTGGGAAGCAAGGCTAAACTAAGAGTGTTGGACTTAAGAAGGAACAATTTCATAGGGAGTCTTCCCCCGTTATGTGCGAAGAGCACTTCATTGAGTACCATTGTCTTGAATGACAATCACTTTGAAGGAATTCTCCCTATGTCGTTGCTCAACTGTAGTGGTTTAGAAGTTCTTGATATGGGGAATAATGCTATAAATGACACATTTCCAGCTTGGCTAGGAACACTTCAACAACTGCAGGTCTTAATATTAAAGTCTAATATGTTCCATGGACCAATTAGTACTTGTCAGACTACATTTTGCTTTCCCAAGTTGCGAATTTTTGATCTTTCACGTAATGAATTCAGTGGCTCTCTCCCTGCAAAAGTTTTTGGAAACTTTAAGGCAATGATTAAATTAGACGATGAAGTCTCAGGAGAGATCAAGTACATGAAAACATTCGGGACTGTATATTCCACACCGTATGAGGATTCTGTGAGTTTGGTGATCAAAGGCCAGGATATTGAGCTAGAAAGAATCAGCACAATTATGACAACCGTAGACCTCTCGAGCAATCATTTTGAAGGTGTCATTCCTATAACACTAAAGGATCTCAGCTCACTTTGGCTACTCAATTTATCCCATAACAATCTCATAGGTGATATTCCAATGGAATTGGGGCAATTGAATACGCTTGAAGCGTTAGATCTGTCTTGGAATCGGCTCACTGGAAAGATTCCACAGGAATTGACAAGAATGAATTTTCTTTCCTTCTTAAACCTCTCTCATAATCTTCTTATTGGACCAATTCCTCAAGGTCCACAGTTCAACACATTTGAAGATGACTCGTATGGCGGTAACCTTGATCTATGTGGTCCTCCTTTATCAAACCAATGTGGAACGAGTGATCCATCAGATGCTTCTCAGCCAGTATTGGACTCTGAAGAAGATGAAAACAAGTCATATTTTTTTAGTGGATTTACGTGGGAATCAGTAGTCATAGGCTATAGCTGTGGACTAGTTGTTGGAACTGTTGTGTGGAGCCTCATGTTTAAATATCGTAAGCCAGAATGGTTTGTCGATTTTTTTGAAGGCATTTATCCCAAGAAAATGAGAAGGTCAATGAAGAGAGCTCAGAGACGACGGACTTAA

>PGSC0003DMG400015352 LRR-RLPs

ATGATGGAGACTGAAGAAAACATTGTAACTGTTGGTGCTGGCATAGCTGGTCTTGCTACTTCTTTGGCCCTTCACAGGTAATAATTGCTATTCTAAAATACTTGTTCAATAATTACATCATCTACTTCCAAAAGTTTGATTTAGAAGGGCATATGCATTTGAACGAGCATTTATGCCCTCCACTTTTAGATGTGCATAAGTAGTAATACACTTAGGATGCCAGGTAGATGCAGAACTGTCACGTAGGATGCCACATGGGATGGGTGTGTTTACTTAGCTTGTTCAACTTTTGAATAATTTAACAGTCTACTTGTACCGGCTCAAAGTTGGATGACATAGATGTGGAGCAAAGTTAGATGATATATTTATGTATTATGCCTCTCTAGAATATGATATTTATGATATAATAGCGCTATTAGTTGAGTGACGATTTGAGAAATAACTCGATGGAAGGCGTGTCTTGTATGTAATATGTAGGTTGGGGCTGCGGAGTGTTGTGTTGGAATCAGCAGATTCGTTGCGAGCTACTGGATTTGCATTGGCATTGTGGACTAATGCCTGGAGAGCTTTGGATGCATTGGGCATTGGGGGCTCTCTTAGACAACGTTCCTTATCAATTACTGGGTAAGCTTCCTCTTTATTATTAGTTAGGATTTAGGATTACTAATTGAAGTTCAATTTCGAATTTTTATGAGATTTTGTGGTCATATGTGTTTTCCGGAGTTCTAGCTCTGGAAAAAAAAAGTTACTGATTGTCCAAAATGCATTCGAGTTGATGTGCAGGTTTAAAAGTTTCTCAGCGGATTCAGGTGCACCTATCAAGGAGGTATCATTTGTGGGGAATAACAGGTTAGCATTATACCAATTTTCATATTAGTTGGGGTCGGTTATATAAGTCCTTTCATCTATTCGTGTTTATGTATGTTCATTTTGTTCCAACACTAGATAATCTAGCATGTGAGACAAATTAGGAATTCTCTAAAATTAGAGGTTTTTTGAGTATCGCAATCTATAGAGATGTATAAATTTCTAACCATTGATATAAAGACTCCTCACAAACGTTGAAGTAATGCAAACCTCCGATTTTCACAACATCGATCTTCGTAGAGAACAATCACATTTGTTTACGAAGGACTTAAGTATGAATAGCATATTCAGAGGATGCTACATACCCAATTTTTTGTATTGTTCACAAGTACAGAGTGACGTGATTTTAATGTCCAAATCCCTATGATTTCAACAATGAAATTAAGTTCATTTTGAATAACGTGGTTGAGTATGTCTTCCTTCCGACTATTGGATGGACTGATCATTCGTCAGCCAACATTTTTATTTTTGAAGTCAATAGTCATCTAAAGTCAGTTATATTACGTTGCAGTGTGGAGTATGAAAGTCGTTGTGTGAGGAGAAAAGATCTGTTGGAGACATTAGCTAATGAACTGCCTCAAGGTGCGATTAGATATTCTTCAAAGGTCGATTCCATTGAAGAATCTGGACCATTGAAATTGGTACATCTTGCAGATGGTTCCACTATTAGAACCAAGGTAAATCTTTCTCTGAAATCCGTAACTTGGAATCCCTATTATTGGAAACTTCAACCAACGTGATAATTCAAACGTGTATTTGACCATTATCTCAATTCCTTTCGTTAGTCTAGTACGTGACTAACTTTGATAAGCTTCAATTAAGGCCTTAATTGGGTGTGATGGAGTCAATTCTGTGGTGGCAAACTGGCTTGGNNNNNNNNNNNNNNNNNNNNNNNNNNNNNNNNNNNNNNNNNNNNNNNNNNNNNNNNNNNNNNNNNNNNNNNNNNNNNNNNNNNNNNNNNNNNNNNNNNNNNNNNNNNNNNNNNNNNNNNNNNNNNNNNNNNNNNNNNNNNNNNNNNNNNNNNNNNNNNNNNNNNNNNNNNNNNNNNNNNNNNNNNNNNNNNNNNNNNNNNNNNNNNNNNNNNNNNNNNNNNNNNNNNNNNNNNNNNNNNNNNNNNNNNNNNNNNNNNNNNNNNNNNNNNNNNNNNNNNNNNNNNNNNNNNNNNNNNNNNNNNNNNNNNNNNNNNNNNNNNNNNNNNNNNNNNNNNNNNNNNNNNNNNNNNNNNNNNNNNNNNNNNNNNNNNNNNNNNNNNNNNNNNNNNNNNNNNNNNNNNNNNNNNNNNNNNNNNNNNNNNNNNNNNNNNNNNNNNNNNNNNNNNNNNNNNNNNNNNNNNNNNNNNNNNNNNNNNNNNNNNNNNNNNNNNNNNNNNNNNNNNNNNNNNNNNNNNNNNNNNNNNNNNNNNNNNNNNNNNNNNNNNNNNNNNNNNNNNNNNNNNNNNNNNNNNNNNNNNNNNNNNNNNNNNNNNNNNNNNNNNNNNNNNNNNNNNNNNNNNNNNNNNNNNNNNNNNNNNNNNNNNNNNNNNNNNNNNNNNNNNNNNNNNNNNNNNNNNNNNNNNNNNNNNNNNNNNNNNNNNNNNNNNNNNNNNNNNNNNNNNNNNNNNNNNNTCATCCCATGACTCCTGATCTTGGTCAAGGTGGATGTTCTGCTTTAGAGGACAGTGTTGTTATCGCAAAGTGCCTTGGAGAAGCCTTGATAAAACCAATAACAGAACAAGAAGACGATGAATCTATGAATATTAGGAAAGGACTCGAGAAGTATGCTAAAGAGAGAAGATGGAGAAGTTTTACATTTATCAGTGCTGCTTATTTATCTGGTTTCATACAAGAGAGTGATAACAAGATAATCAGTTTCTTAAGACAACATTTNNNNNNNNNNNNNNNNNNNNNNNNNTTTTGGCTGGAGTCACTATATCAGTTACACTAAGAATAGCTAACTTCGACTGCGGAAGATTAACTGTTTCTTAGTAGCTCAATTGGATCCTTTAGAAGCTATGAATATCCTTTATGGACTTTGTTTAATATCTATCTAGCATACATGTTCTGGTTTAGTTATGTCTAAGAATTGTAAGACCGAATCGGATCCTTATTTTGCAATAATAAAATTGTGTCCTTCATCAAAGATATCCAATAACATCAGTTGTTTAGAAGCAAATATGGTAAAAGCTGCATTTTATTATTATCATGGAAAAACAGTGTAAAAGGTAAATCTCTGAAGTATAGAATAAGTGTTTTCTTATCTGAGATCATTTCGTGCAATTAGATACTACTCCCTCTGTTGCAACTTGTCTGTCTGGTTTTGACTTGACACGAAGTTTAAGAAAGTAAAGAAGACTTACAATTAGAGATATGGACACGTAATATATATCCTGTATAATAAGTAATGTTGGAATAATCTCATGAATATTTTTTGTGGTCTATGTAAGAACATTGTTATTTTTAGCATTTGCTTCGACTACTGATTTAGTTATTTTCCATCTCATCATATTTGTCCTTTTTTTCCCCTGTAGTCTTTGAGGAAATTGGCTTAAAGACTTGGAAGTTTTTTAAACATCTATTTCTACTACAACTACATGGATGTGAAAAGTCTTGTGGAAAATTATAGAATATGACTCACAAGTTTGAAAAATTCCACACGGTATCAAAAATGTTACGTTTCGTCTAAGTAGATGTAGGAGTTAAATCATGTCTTTAATAATTGTTAATTTTTTATTTTTCTCACTCCACTTAGTCAAGCAACAGTATCTCTCATTGGAAATTTGTTACAATTCTGCAGCAAATGTTCTACCGTGTGTCTTACATAATGGGACCAATTCACATTGACTTAGGTGACACACTGCAGAGATAGTCGATAACCTTGAAATGAAACAACATTTAGAGAAAACAAATATGGAATAATTGGTACTAGCCACTAGGGGTGTACATGATCGGGTTGATTTGAATGTTCCAAATATCAAACCAAACTATTTGTGTAAAATTTTAAATTTATAAATCAAACCAAACTAATAAAATTCAGATTTTTTGGGGTTTTTCAACCTCGGGTTGGTTCGCGTTTTTCACTTGTGACTTGTTAACTGGTCATGTTATCGGATTGGACCTTAGTTGCAGTTTGCTTTTGGGTACTATTCATCCCAATAGCAGCATCTTCCAGCTCCATCATCTCCGGACACTAAACCTTGCTTACATGACTTCTATTCTTCTTCAATCCCACATAACATTGGCCGATTGACGAATTTGAGGCATCTCAACCTTTCTGGTTCTTTCTTTGAAGGGGAAATCCCAACAGAAATATCATACCTTTCCTATTTGGTTTCACTTGGACTTAATTGTTATGAATGTGAACTTGATGAGAGAACATTTGAAACAATTCAGAACTTCACAAATCTAGAGGTACTATCTCTCCATGGTGTCAACATCTCATCTTCAATACCTGTGCATATTTCTTCATCCTCCTTAAGGTACATGGATCTTGGACATACTAATCTGCGAGGTGTTCTCACAGAGAGCTTTTTCCTTCTGCCAGGATTGGAAAGGCTCTATTTGAGTTCCAATGAACTTCTCAAAGGAGTTTTACCAAAGATCCACCCGAGCAACACTCTGTTGGAGTTGGATATTTCTAACACAGGAATCTCCGGTGAGCTGCCTGATTCAGTTGGCACCTTGAGTTCCTTGAATATCTTGAACCTCAAAGGATGCCAATTCTCTGGTTCCATTCCTGATTCCATTGGCAATCTTACACAAATTAAGTATTTGGATTTCAGTAATAATCATTTCACCAGCCATATTCCATCCACAATCTCTCAGTTGAAGCAGCTCACATATCTACATCTTTCGTCCAACTCTTTTTCAGGTGAAATTCCCGACATCTTCTCTAACCTCCAAGAGCTAAGTTATTTATATCTTGGTAACAACAGCTTCATCGGTTCGTTTCCTTCTACAATTGTAAGCTTGCCACATCTACAAGACTTAGACTTGTCGAGTAATTCCCTATCTGGCCCACTGCCTAATAATGTGAGCATGCTTCAAAAGCTAACCAAACTGAATTTGTCCTACAACTCAATGAATGGTACCATACCATCTTCTGTGTTTAGCCTCCCTTTGCTAGGTCAGTTGTGGCTCGATAACAATAGATTTAGTGGATTACCCGATGAGCTCAAAACAAACCCAACATTAGCACGCCTGGATTTAAGCCATAATCAACTCAGTGGTTCTTTTCTTCAATCACTTTCGAATCTCACAAACCTTTCTATCCTTGACCTTTCATCGAATAACATCACCGTCGATGAGGGAATACAAATCACTTTTCCTAGCCTACAAGATTTGTGGTTATCATCTTGTGAACTGAAGGATTTTCCACACTTCTTGAGAAATGTGAAGTCACTTTGGGTCTTGGATATTTCTAACAATAAGATTCGTGGCCAAATCCCAAACTGGTTTAGCGGCATGAGGTGGGACTACTTGTGGTACCTAAACCTTTCGCATAATTCTTTAACAGGCCACCTACAACAATTTCATTTCCATAATCTATGGTATCTTGATCTTAATTTAACTCCCTTCAGGGTCCACTCTCTTCATCCATTTGTAACATGATCAAGCTTGAATTTCTGGATTTATCACGCAACAACTTCAGTATCTCGATTCCAAGCTGCTTGGGAAGCATGTCTAGTCTAAAAGTGTTGGACTTAAGAAGGAGCAATTTCACTGGGAATCTTTCTCCATTATGTGCGAAGAGCACTTCATTGAGTACCATTGTCTTGAATGACAATCACTTTGAAGGAACTCTCCCTGTGTCGTTGCTCAACTGTAGCGGTTTAGAAGTTCTTGATATGGGGAATAACGCTATAAATGACCTATTTCCATCTTGGCTAGGAACACTTCAAGAACTGCAGGTTCTAATATTAAAGTCCAATGTGTTCCATGGACCAATTAGGACTTGTCAGACTACGTTTTGCTTTCCCAAGTTGAGAATTTTTGATCTCTCTCATAATGAATTCAGTGGCTCACTTCCTGCAAATTTTTTTGGAAACTTCAAGGCAATGATTAAATTAGAGGGTGAAGACACAGGACAGATTGAGTACATGAAACCACACGAAAATACAATTTACACATCGTATGACGATTCTGTGAGTTTGGTGATCAAAGGCCAGGATGTTGTGCTAGAAAGAATCAGCACAATTATGACAACCATAGACCTCTCGAGCAACCATTTTGAAGGTGTCATTCAGCAAACACTAAAGGATCTCAGCTCACTTTGGCTTCTCAATTTATCCCATAACAATCTCACAGGTGATATTCCAATGGAATTGGGGAAATTGAATACGCTTGAAGCGTTAGATCTCTCTTGGAATAGGCTCACTGGAAAGATTCCGCAGGAATTGACAAGAATGAATTTTCTGGCCTTCTTAAACCTCTCTCATAATCTTCTTATTGGACCAATTCCTCAAGGTCCACAATTCAACACATTTGAAAATGACTCGTATGGCGGTAACCTTGATTTATGTGGTCCTCCTTTATCAAAGCAATGTGGAACGAGTGATCCATCACATGCTCCTCAACCATTGGACTCTGAAGAAGATGAAGACGAGTCATATTTTTTAAGTGGATTTACTTGGGAATCAGTAGTCATAGGCTACAGTTTTGGACTAGTTGTTGGAACTGTTTTGTGGAGCCTCATGTTTAAATATCGTAAGCCAGAATGGTTTGTGGAATTTTTTGAAGGCATTTTCCCCAAGAAAATGAGAAGGTCAAATAAGACAAGGCCGAGACGGCGGATTTAA

>PGSC0003DMG400015353 LRR-RLPs

ATGCGGAAATTGAATTTTGGAAATAATCATTTCACCGGCTATAATCCTTCCACAATCTCTAAATTGAAGCACCTCACATCCCTAGACCTTTCGTCCAACTCCTTTTCAGGTGAAATTCCCGATATTTTTTCTAACCTCCAAGAGCTATGTTATTTATCTCTTTTTGACAACAGCTTCATCGGTTCGTTTCCTTCTACAATTGTAAGATTGACACAACTTCAACACTTAGACTTGTCGCACAACTCACTGAATGGTACCATACCATCTTCGGTCTTTAGCCTCCCTTTGCTATCTTCATTGTCGCTCCAACATAATCAATTTAGTGGACTAGCCGATGAGTTCAAAACAAACCCAACATTAGAACACCTAGATTTAAGCCATAATCAACTCAGTGGTTCTTTTCCTCAGTCACTTGCGAGTCTCACAAACCTTTCTACCCTCATCCTTTCATCAAATAACATCACCGTCAATGAGGGAATACAAATCACTTTTCCTAGCCTAGAAGTTTTACAGTTATCGTCTTGTGAACTGAAGGATTTTCCACACTTCTTGACAAATGTGAAGTCACTTAGGATCTTGGATATTTCTAACAATAAGGTTCGTGGCCAAATCCCAAACTGGTTTAGCGGCATGAGGTGGGACAACTTGTGGTACCTAAACCTTTCGCATAATTCTTTAACAGGCCACCTACAACTATTTCATTTCCATAATCTAGAGTATCTTGATCTTAAATTTAACTCCCTTCAAGGTCCAATACCTTCATCCATTTGTAACATGAACGATCTTCAATTTCTACTTTTATCACACAACGACTTCAGTAACTCGATTCCAAGTTGCTTGGGAAGCATGTCTAGTCTAACAGTGTTGGACTTAAGAAGGAACAATTTCACTGGGAGTCTTCCATCGTTATGTGCGCAGAGCACTTCATTGAGTACCATTGTCTTGAATGGTAATCAATTTAAAGGAACTCTCCCTGTGTCGTTGCTCAACTGCAGTGGTTTAGAAGTTCTTGATTTGGGGAATAACGCTATAAATGACACATTTCCTGCTTGGCTAGGAACACTTCAACAACTGCAGGTCTTAATATTAAAGTCCAATGTGTTCCATGGACCAATTAGTACTTGTCAGAGTAGGTTTTGCTTTCCCAAGTTGCGAATTTTTGATCTTTCGCATAACAAATTCAGTGGCTCTCTACCTGCAAAAGTTTTTGGAAACTTCAAGGCAATGATTAAATTAGACGGTGAAGACACACGAGAGATCATGTACATGATGCCATCTATAATGCCATTTTACTCATCATCATATGAGGATTCTGTGAGTTTGGTGATCAAAGGGCAGGATATTGAGCTAGAAAGAATCAGCACAATTATGACAACCATAGACCTCTCGTGTAACCATTTTGAAGGTGTCATTCCGAAAACACTAAAGGATCTCAGCTCACTTTGGCTACTCAATTTATCCCATAACAATCTCATAGGTCATATTCCAATGGAATTGGGGAAATTGAATAAGCTTGAAGCATTAGATCTCTCTTGGAATCGGCTCACTGGAAAGATTCCGCAGGAATTGACAAGGCTAACATTTCTGGAAAAGTTAAACCTTTCTCAGAATGTTCTCGTTGGACCAATTCCTCACGTTCTACAATTCAACACATTTGTAAAATAG

>PGSC0003DMG400015449 LRR-RLPs

CTCATACAACAACTATATCTCCCACAAAAAAAGTGAAAAAATCAATCTTCATTTCTTATATGATGAACATTTCTTATCTTGAATAATTGTTATTGTGTTTAATATATAGTTCTTTTGTATATCATCTTCCTTAGGGGCAGCCTTGGGGGTTGGTTCTTCCGAGGCAGCTTGTTAGTCGAAAATTCCCTTTGAAAAATGCAGGTGTATCGATGTTTTTTCAGTTTGTTGCTGATCTCTGCTGTTCTGTTTTCTTTCTCTTGTGCACTCTCTGATCATGAAGCATCATTGCTTGCACGACGACAACTCTCAACACTTCTTGAAAATGGTAATTTACCCGATAAATATGAATTCGAGGTGAATGTAGAATACACATTTCCTAATTCTAGGCTTAGACGTGCTTATATTGCGTTAAAGGCATGGAAAGAAGCTATATATTCTGACCCTTCTGAATTCACTAGCAATTGGAAAGGTCCAGATGTTTGTAACTATAAAGGCGTGTTCTGTTCGCCTGCTTTAGATGATCCTAATGTCACTGTTGTTGCTGGGATTGATCTGAATCATGCAGATATAGCAGGTTATTTTCCTGTAGAATTAGGATTATTGACAGATGTAGCTTTGTTTCACTTAAACTCCAATAGATTTTGTGGAATTGTTCCTGAGAGTTTTTCGAAGCTTACGTTGATGCATGAGTTTGATGTTAGTAATAATCGTCTTGTTGGCCCTTTCCCTAAGGTTGTTTTGAACATGTCTAGTCTTAAGTACTTGGATTTAAGGTTTAACAACTTTGAGGGAGAGTTGCCTTCACAGTTGTTTGACAAAGATCTTGATGCTTTGTTCTTGAATGATAATCGATTCGTGTCTACCATCCCTGAGACTCTTGGTAACTCTTCAGCTTCTGTGATTGTGTTTGCTAATAACAAGTTTCATGGTTGTATTCCCAGTAGCATTGGGAAGATGTTGAATTTGGATGAGATTGTGTTTATGAACAATGATCTTGGTGGATGTTTGCCTGTTGAAGTCGGGTTGTTAAAGAATGTAACAATGTTTAATGCTGCTGGGAATTTGCTTTCTGGGATATTGCCAAAGACACTTGATGGTCTTTCACATGTTGAGGAATTGGATATTTCACATAATACCTTAACTGGGTTTGTTCCGGAGAATCTATGTTCATCGTTGAGTTTGAAGAAGTTTGTGTTTTCGTTTAATTACTTTAATGGAGAGGCTAAAGGTTGTGAAGCGCGAACAAGGAAAGATATGAGTTTTGATGATAGGAACAATTGTTTGCCTGGTAGAGCAAAACAAAAGTCTCAAAAAGAATGTCAACCTATTGTGAGTAAGCCTATTGATTGTAGTAAGGCTAAATGTGGAGGGCCTTTAACAAAATCTCCTCCTAGTAAGAAAAATGAAGTGCCACCAAAACCAACACCAACACCAGCACCAAAACCTCGACCTACCCCTAAGTCACCACCCACCCCAAAGAAGGCTTCTCCTCCACCAACACAAGTCGTTTCATCGCCCCCTCTAGTACATTCTCCACCACCACCTGTTCACTCCCCTCCACCACCTCTAGTACATTCTCCACCACCACGTGTCCATTCTCCTCCTGTTTCCTCTCCACCACTACCCGTCCACTCCCCTCCATCACTACCTGTCAACTCCCCTCCACCACCTATAGTACATTCTCCACCACCACGTGTCCATTCTCCTCCCGTGTCCTCTCCACCACCACCTCTTGTCCAATCTCCACCGCCACTACATATCAACTCCCCTCCACTACCTATAGTACATGCTCCACCACCACATGTCCATTCTCCTCCTGTTTCCTCTCCACCACCACCTCATGTCCAATCTCCACCGCCACCTCTAGTACATTCTCCACCACCACCTGTCCACTCTCCTTCTCCACATGTTTCCTCTCCACCACCACCTATCCAATCTCCACCGCCACCTCCAGTACATTCTCCACCACCACCTGTCCACTCTCCACCTCCACCTATTTCCTCTCCACCGCCACCTGTCCATTCCCCTCCACCACCACCCGTCCACTCTCCTCCTCCACCTGTTTCCTCTCCACCACCACCTGTCTACTCCCCTCCACCACCACCCGTTCACTCTCCTCCTCCACCACCACCCGTCCACTCTCCTCCTCCTGTTTCTTCCCCACCACCACCTGTCCACTCTCCTCCACCTCCGGTACACTCACCACCACCACCTTTCGAACATGTTGTCCTTCCACCAAATATTGGTTCAATTTACGCATCGCCACCGCCACCAATCTTCCAAGGATATTAATTAAGCAATGAATGAAGCTTCGTCTCGTTTCCACCTTTTTAGTACGTATAATATTTAGTACATGCAGAATTACATAATCATTTTTCTTAATGTAAGATTATTATCTAATATATGGTGATGTTTTTCTTTCTTTAAACTGATACAATATAATCTTGGAAAAG

>PGSC0003DMG400015603 LRR-RLPs

CAGATCATCATCACTGCAACGCCGCGCTTTATCTCCTCTTCTGGAACTCTCCCTTCGTTCCTCTCACTGTCTAACTCACTCATTGAGTCTCTCCATGGGCTCATCTTCTACATTCTCTCTGTGCTCTCATCCAACGGTACTTTCCTCTACTCTATTAGCTCACTCGAACTTACAATTATGACTAAATTCTCACTTTTTTTCGTAATTTTTAGGTGAATTTTGAATTTAATGGTAGAAAAGTCGGATACGGATTGCAGAAATCGTTGAGCAGTTCATCCAGTTATTGTTTGATTCATATTTCGCCGATTTGTATAAGTAGTAGAAGAAAATGCTTGCAATTGAGACGTGTAATTGTTAGAGCTGCTTCGTCGTCTAGTACTGGTCCGAGACGTTCGACGAATACTCGGAGAGTTTACAAGGAATCACAAGCTCAAACGCCAGCATTGCCTGTTGACAGAATTGTTTCGTTTGTTGTGCCGGCTGGTGCCTTTGTGGTTATCTCTTTTGGTATTGACATTTTGCCTATTTTCATCTCTTTTTCTACTGATAGGCTTATAGCTTTAATTTCTTATGCTGAATTGAGCTTTTGAGGAGGATTTTGCGTTTGAGTACATTCGGTTATCACTACTAGTGTTGGAGCAATAAATAAAATAAGGAGTTAAAAATTACTTCTTTCAATATATGTTACACCTTTTAACTCAGACACAGAAAAAGAAGACTTTTGAATTTATGGTCTAAAGTACTAGAGTACTTGATTATTGGTGCACCTATAAAGTTAAATTGTTTCCAATTATAAAAAAAGTGACATCTTTTGGACAAATTAAAAAAGGAAGTGTGCCACATAAATTAGAACGGATTAAAAAGGGAAGTGTGCAATGTTTGCTTTCCATTATGTATCTGTAATGGTTTTAATTTTCTGTTTTTCTTGTATTGAAATTGTTTTTGCATCCTTTTGCGTCTAGTTTAAGCTCAATTGCAATGCACCTTATAACTGTGGCAAAAAATATGAAATTGCAAATGGTAAATTTGACATTCTTGCTCTTGTGTTTTTCACAATGCCAAGTTAGTGGCAGGCAAGGCTTTAGTATTTGGGGAAAGGGTCTAACTTAAGTGGGTTTCCAGTTTTGGCGGATACTGGAAGAGGTCAGGAATCATGAGGTCTTGTGTGAAAATTCTCAACTTAATAGAAAAGTTTGAAGTTATTGGAGTGCTTGGCAGTATGACAGAGACCTGATATTCAACCATGAATGGAGAACTTAAGGCTCTTTATATGGAACCTTTTAAGCTGTACATCCTGCAACGGATTACCCTCTTTTTGTTGAGTTATTCGTGACTAAGAAAATGAGTTAACACGGTAATGTTTCAGCATTTTAGAGGCTTAGATTTAAAACTTAGAAGTGCAAGGCACCTTACAGCCAAACAGGATTGCCATTTTCATTTTGCAGCCTTTTGATATCATTTTAGATGTTCTGCTGCCTGCACTCGTATGAATGTATTTTGTTTGACATTTTGACTGAGTGACACTGACCTGTTTTTTAGGTTCCCAATTGTTGCAATCATTGTAATCAATTTAATATGCTTTAATTATCTAATATAAATTAGCCATTGAAAATTATATTTCCTCTTTAAGAGCCATTTTCTAACAGCTTTTGAGATATAGGTTCTGCAACTTCACAAATTCATTCTCTTGAACCAACTTTGGTTGTAAAGCGGCTTGCTCAACACAACCATAAATCTGGTTTTCATCTTTTAGTTGCCAAATATAGTTCTATGCTTAAGGGACACCGGACATGTAGTGTGTGGTAGTTTTTTTTTTTTAAAAAAAAATTAGTGAGAAGATGATGACTTGTTCCGCAACTTATGGCATCCTCATTTAACTATCTTTTGAAAATCAAGAGGATTACTAGGAGGAATGCTCTTGTTCTTTTTAGTTGTTTTACTGGTGAAAGTTAATGGAATGTTATTAATATTGTGTAGAAATATTCTTAATGATTATGTTATATTGATTAAGAGCTCGTTTGGATTGACTTCAAAAAAGTAACTTTTTAAAGTGTTGAAAGTTATTTTAGAAATAAGTAGTTATGTGTTTGGATAAAAGTGTTGCAATTGAAATTTTTTTATTGATGTGTCTGGCAAATAAGTGTTGTTAAACACTTTTTTTTACCAAAATGTCCTGAAATACCCTTAAAACTGATAACATTATATAAAAAAAAGGTTTTGAGATGAAGGTCTTCAATGAAGTTGAAGACCGTGCTCATCAACATTCTAAATATATGTTGGATTTTCTCTATGTGAAGATTAGTGATTGAAAGCGAAGGAGGGTTTATTGGTAATAAATGCTTGAGGACACTTCAATGGCATCCTAATCAACAACGGAATATCCTAGAAAGAATAGTTGTGCTGGGGATAGAAATAGAATAGCACTTATATGGATGAGGTCATGAGCCTCCCTTCAGTAACTTACTTCAGTGTCGATCATCCTGTGAAATTTTCATGAAAAAAAAGAGAAAATTGAAATGTTGAAGAGGAACAAGAAAAATGTTGGTGAGAACGTTGGAGAAAACTAGGGCCACAACAACAAAAGAAAGTTGAGGGAAAGAGGAGATCTTAGAGTTTTATTTCACAAGGGCAAATTTGGAATTATATACATGAATTGAGGGTACAAAAGTAATATCCTTGGTTTTAAGTTATTTTTATATTTGCCAAACACTCCTAGAAGTCAAAAACTGACTTAAGTCGATTCAAACAGGCCCTAAGAATCCAAAAGATTAGGAAATAATCCTTATTATTGTAGGCTAAAACTTCTATTTAGGAGTCTTCTTGTATGGTTGTAAGCATCAATTAAAAAGAAGTTGTTATTCCCTTTTCAGATGTTATAGATCTTGCTAATGACTTAAGTAACTTCTGCCTATCAGTGCACGGCATTATGCTACTACTTTTGGGTGGACTTCTCTTTCCTTTTTACGTTTTACTGATCACACTTCTTTTTTCCTTCTGGTGGCACTGCAATTCTTTTTTACTTCTTAAGTTGACTTTACCACCAATCCGAGCTATGTGGATGATGTCCAAGAGATTTAACTTAGCATATACCTTCCGGTGGTTGAATTATAGATTCCTTTACATGTGTCGAAAAACACTTTCTTTCATTATTCAATTACCAGTAACCTTTTAGAAGAAAAAAATTTCAACGCTGCTTTTCTTTTCCATGATCATTCACTACTAATTAGACGATTCAGGTGACTGTTCTTGTTCTAATTACTTTTCTAGTTATTTGACCATCTTGAATATGAAGGAGCATGCTTGACGTTAAGTGAATCATTAAAAATATTGTGTAAATTATTCATAATAATTATGTTTTATTAGTTAGGAGAGCGGGAATTAGTAAACTTTCCTTCTTAATTTAGTTCAGAATTTCTATATGAGTAGCTCCTTGAATGTGGTTAAACCTTTCATTACAAGAGCTCATTTCTCTCTTCTTTTTTTACAATGTCAAATTCTAATAAACAGTTAATCAACCACTGGTATCGTTGCATCTAATTTATCTTAACATTCAAAGTTCTTACATTTCCATCTCCAAAGATGTTGCTCCATACTACTATTCTATATGATGCGATTGAATTCCTCTTTCATTGCCAATTACCTCTACTAATAACTTCTGAGCTCTTTCTGCCCAAAAAATTGAGGAGCTACTTAACTTGCATCTGCAGAAAGAGTAAATGTCTTAGTTTTACCACAGTGTTGCTCTGGTTCAGTCATTAACTATGGGCCCTGGCCATTTCTATTGCTGCACCCTTCCCAATGAACTCGTTGCAAGTTATATCCTTCCAACTTTCGCCTGGCATCCTTTCTCACGGAGGTTGAATCTTGTGAGAATGTGGCTGCTAGACTGAAGATCTTCCTCTTATGAATTGTGACTCGAAATAGAAATCTTAAGTAGAAGGAAGATTCTCTATATCGTACTAATCTATGTGTTCAGTTGACAGTTTTGTGGAAGCTGGTGGAGAAAGTTCTTCTACCCAAACCAGCCAAGACTACATCAGAAGAAAGCAAGTCTACTCAGGGATTGAAGTGGTCCATTGCCGCTGGGACAAACTTGTTACCTGGTTTTGGTGCAAAGATTGAAAGAGAATCCAAGCTGAGGCTTAATGACTTTGCTAAAGAGCTTAGATCTTTCAGCATTGTTGACATGTCAGGTAAAATTGCAGTCAGTTCTGGCTCAGCTGCTTCAGTATTAAAGTTGGTGGTCGTCTATGCTAAATGTTGTCATAATGAATATCTTCCTTCTTTATACCTTTGATAAAAAACCTTCTATCATTTGTAGTTTTCAGAAATGTGATCACGATGTTAAAATATTTGCCTTGACTTGTTAAATTCTTGAGTCCCACCTCTTTAGAAAGTTCAACATATAAAGATCAAAGCTTTACCGTGATTTATGATTAAAAAACATTTTGTGCATACTGATATTGCCTGAAGGAAACTTTCCAATCTTGTTTCATTTATAAGCTTCCCCCACCAAGTGCGCGACACACCAAAACCAAAGTTTTACTGCATCTTCCTTGTATTGTGATTCTACATTCTTTGCATACTGATTAAGGATTAAAATTGCAAGTCACAAGAGGTATTGGATTGAATGTTTTAGGCTTTGGCAGGTCGTAACTTTGGAGATGAGGGGCTCTTTTTCCTAGCAGAGAGTTTAGCATATAATCAGGTAATGTCTTTTGCTCTCTCTACCCCCCTCCCTCTTCCTTTATCTGTGTAGAGTATGTCATTATTTATCATGGTCCATCTTGTTAACCCAAGAGGATAAAGCCATAATCCTGTAATGTAGACAAGTAACAATACAATCCTGGCTTCTTTTTTTCTGGAATAATGTTTTTGGGCAAGTTTGGTTCCTTTACCAATGTACACTGCATTTCCTTTCATGTGATTTCCAGACAGCCGAGGAAGTAAACTTTGCTGCTAATGGAATAACTGCAGATGGATTGAAGGCCTTTGATGGCATATTGCAATCAAATATTGCTCTAAAGACTCTTAACCTATCTGGAAATGCTATTGGGGATGAAGGAGCCAAGGTAATGCACAGATGATATAGGTGAAAATCCAAAGCAGTGAGTTCTTGGAATACAGATGACTAATAAGCAGATCCTAACTTTCCCTTTGCACTTACTTAACTGTTATACCAATTTAACACGTGTATTTATGGTCATGCTTTGTATTGTATTTCTCATCTCCGCCTTGGCACTTCAGGAGCAATCACGTCTATTTGGTGTTAATTTATGTTTGCAGTGTCTATGTGATATACTGGCAAATAACTCTGGTATTCAAAAGCTCCAGCTGAACAGCACTGGCTTAGGAGATGAGGTCTTGAACATGCCTTGAGCATTTAATTGAAATGCTGTTGCTAACTTGCGATGGTATTTTCTGCTTTCTGGTAAATTATTGCTTAAATTTCGCTGCAATGGTTTGTCTGAATTTTGCAGGGAGCCAAAGCCATTGGTGAAATGCTGAAAAAAAATTCAACTTTGCGTGTTCTTGAACTAAACAATAATCTGATTGACTATTCTGTATGTTATTAAGCAGAACTTGTAATTTGTATTTCCTAGTTATTAATTATCTCAATTTCTGAAGGAAAACATGTTTGATTTGGATATGGAAATAGGGATTTTCAGGTCTTGCTGGATCACTTCTTGAGAATAAAACTTTACAATCCTTGCATCTGAAGTAAGCAAAATTTATTCTGCTGTCTTTCTGCGCCCATCCTCATTTGTCGTATGTCAAATGTAGCAGTTAATTATTTTATTTTTCTTTATTGGTTTGGAGGGTGGGGGTGGGGGGAAAAGAAGTGGATTTTAATTTGTGAGGGAAAGCATATCAATTCTTATTTTCCCTTATATCTTTTAATTTATGTGCTTTAATTAACTGTTCAGCGGCAATTATGGTGGTGCTTTAGGGGCTGCAGCACTGGCTAAAGGCTTGGAGAGCAATAAGTCCTTGAGGGTACATTTCTAATCAATTCCTTCCTCCCACTGAGACCCTTCGCCTTTTGCTCTTTTTTTTGTTCTTGTAATTTGACTTGGCAAATAATTGAGCTTTTTTTCAGCCTTGATATCTTGCTTCTGCGTCATTGGACTGTTTCTTAGTATTTAACTTCTGCACTCATGTTTCCCTTCTTTTTTTTTTTGGGAAACAACCTCTCTACCTCCACAAGGTATGAGTAAGGTCTGGGTACACTCTCCCCACCCCAACCCCACTTGTGGGATCACACTGGGTATGTTGTTGTTGTTACTCATGTTTCCATTTGATTGTATAGAAGAGCTCTTCTTTTCAAAAGCTTCAATTTCCTCTGTGACTGCTGACACTGGGATTAAGTGTTAAATGCACTAATGTTATTAGATAAACTTCTTCAGGGGGTTTCTTAATGTTCCTCACCAACAATCCGAGGTTGAAGCTAGATCTGTTCCGCCCAAAATGGAAATGGGCTACTGTTTACATATTACTATATATCATTAACATACACCTGTATATTCAAAGGTGTTTGAAAAGATTCTTTATGTGCAGGAACTTTATATGCAGGGGAACTCTGTTGGTGATGAAGGAGTACGGGCATTAATCTCGGGCCTATCTTTGCGTAAAGGTTTGCTTATTGGATATAATTAACCATTCTCTCTAAACACATATTGGCGTTGTTTCCTGTGCATCTTTTTTGAAAGGTCACCATCATATGTCTGCAGGGAAACTTGTCTTACTTGACCTTGCCAACAATTCAATTACTGCTAGAGGAGCATTTCATGTTGCCGAATACGTCAAAAAAAGCAAAAGCTTACTGTGGTTGAATCTCTATATGAATGACATCAAAGATGAGGTAATTGAACCACTTAGAACTTTTCACAGTTTTTTTCCTTTGTAATGTAGAAGACTCTTCTCAGCCTTGTCCTTCTCACTGCAATCAGTAGACTCTCTTGAACTATTTTATGGTTTTATTTTATGTAACAGAAGCAAAGAATGAGAAAGCTTGAGTTTTTGTCAAGTTGAAACCACTAAAATAAGATAGAAAAGATTTTTGGGTAAGAGGGTGAGGATTGATTGTCGAGAAAAAACAAAAAGAATTTGATTGGTTAGAGCTGAAGGTGATGGCTGAGTCGTTCTTGAAACCAATCATAAGTTCAGATCACATGGTTAGGAAATTTTATAATGGTACTTACCATTAAAGAATATAATTTTAAGACTAGGTTTAATCACTTTGGGGATTCTTGCCAGAAGAAAATTAGAAGGGTTGCATGGTGTTGGATGAACTGAGGTCTTTTTAAATTTTGTCAGCAGTTCAGAAAGATAAAACTCAAATAATCCCAAATCTGTAGGGGTAGCTTTATTTTTTCCCTTCAAATACATATGCATATGGACAGACATATTAACTATGTAAAAATGAATACTTTTATAGTCCTGCACAACTTTAACGTTAATATTTCAACAGTGAGAAACCTAACTGTGCTCCTCTTGGATTATAGTAGAACTGTCTGCATCCACCATAATTAAACTGTGTTCATTGGAATCAAAAGGAAAGCTTTGTGGTCGAAGCAGAAAGATAAAAAATTGATGATTGGAGGAAATTGGTACGGATTCAGTAACAGAAGGTTAAATCTGTTGTGAAGAGTTCAAGGGATTTGGGCTGCAAAGAAGGATTTTTTTCAGAAATTTATGGGGGTGGGTTTGCATTCGACAGGGTAGTTGCTCAGAAGCACAATTAAATTTCTGCTGGAGAAGAGCTCATGTGCTGGCAACATCAGATTGAAACATAAAGTCAAACTTTAACCATATCCTTGGTAGATTAGACCAAACTAGCTAATGTTGATTACCCATTCCGGAAACTTAAGGCTATGGTTTACAATGACAGAGTACAGTGTCTACAAGCAACAAGTCCTGCACAAGAACTTGGAAAGAACTCTCTAAGTTAGCTCGCAAAAAAATTTACTGCCTTCATTTGTGTCAATAAACTTCACATGTAATCAGTAGCCTCAAGTTTCCCACTTCTTCAATTCAACCCAGTTTCTTTTTGTTGGTATGCGATTTCCTTGGTGTTGAAGATTAAAGGCGAAGGCTTTTTGTTTTAAAAGAGTACACAAATCACATGCTTTTCACTTGAGAAACTACCGTTAATATTTGACGTCAAATTTGGGAAGGACCAAAGGTGTCCATTTTTTGCAAAGACGAAGGATTTATTTTATCCACAGTTATATTTGAAGGACAAAGTGTAACCTGTCCCCATAAATAAGGGATCATTTGAGTTTTTACATTGCAGATCCAGACTCGATCTCTTGATATGCATTGACTTTTCCTTCTTTTTTTTTTCCCCCATTTACATCAATATCCATCAGGGAGCTGAAAAGATTGCAGAAGCTTTGAAGGAGAATCGCTCGATAACTAACGTTGATCTTGTAAGAGTTGTTTCCTCTTTTCTATTTTATTCAACATTTCTTGTTTTGTTCATGGAGGAGGTTGGGTTTCTTAATCTTATCAGAGCATAATTTTCTATTTGCTCTGATGGCTGGAATTCTGCGAATTTTACAGGGGGGCAACGACATACATGCTAAGGGTATTAGTGCCATAGCAGAAGTTCTGAAAGATAATTCTGTCATTACCTCTGTAAGTTTCATCAAAATGTTAACTTTTACTGATTTATTGGACAAACTTAAATACATTATACTTCAAATTTCTCACGAACAGTTAGAACTTGGTTACAACCCCATTGGACCAGAGGGGGCAATGGCATTAGCAGAGGTACTCAAGTTTCATGGAAAAGTAAAAGATCTCATGCTTGGGTGGTGTCAGGTAATGCATGGATTGTTTCATATATGCCTTCTTTAACTATCTCTTTTAGCATGCTCACTTAATTTTGTTCCAGATCGGAGCCAAAGGTGCAGAATATTTTGCCGATATGTTGAAGTACAATAGTACCATATCAACTCTAGACTTGCGAGCCAATGGACTTAGAGACGAGGTTTCATTCAAAGTTTATAGTTCTTCTATTGTGCCACAATCTTTAATTTCCCTCAAAGTACATAACCATTTCCTTTTTCAGGGTGCAATCTGCCTTGCTCGGAGTTTAAAAGTGGTCAATGAATCTTTGACGACACTCAATTTAGGGTTCAATGAAATTAGAGTATGCAACTCTTTTCCAGTACAAGTACAACCACCTGTCTTGATAAAGGTATCATCTTGTATTAACTTAGTTTCTGTTGCACACAGGATGAAGGAGCTTTTGCTATCGCTCAAGCACTTAAGGCAAATGAAGATGTTAGACTCACATCTTTAAATCTTGCCAGTAACTTCCTCACCAAATTAGGACAGGTAACATCAAATGATAATATTATTCTTTTATTTTAGCTGTTTTACGCGATTTTTAGTTCCTTACAATATCCGGATGGATTCCAAATTTCAGACAGCTCTCACAGATGCAAGAGACCATGTGTTTGAGATGACTGAGAAGGAGCTTGCTGTTATGTTCTAAAATACTACATTCCTTAGATCAGATATATTTTCCTCCCTACTTGGGGCAAGGTAGATAAGTATTTTTTTTTCTATTCTGGGGGAATAGAATCGCAGCCTACCATGATATTCTTAAGAGTCCCACCTTAGTTCCCTTTTGTGACTGAAATCATTTTGAGCGGGGAACAGTGGAAATTTTGATATCGAGATACACACTGATATTATAAAAAATGTTCCTTTTCTGAAACTTTTTTTTTTGCCTGATAACGTTAGTGGCAAGTACATAGAGATTACTTCTATTCACCATTTCAGATTCATTCTCTGGTTTGAAACTCTTCTGAGTTGTTGTTCTTTCCCTATGGCCTTTTAAGCTTATTTTGGTGTTGCAGGACAG

>PGSC0003DMG400015672 LRR-RLPs

CTTCCTTCCTCTAAAAGCTAAAGGTTCCTCAGTTTTGCTAAAATTTCACACATTGCAGCAAAAAATTCTCTAATTTTCCCAATCAACATGACTCAAGGTATGCAAATTCTCTCTTTTGTTGTTTTTTTTGGATTATGAAAGTGGATTTGAGTCTATAGTTGAAGAAAAATAGCATTTTTCTCTCCAGTTATTTACTAACCTGAAGTGGAAATTTTACTATTCAGATATGAAATTTGAAGTTTTAGAATTTGATAAAGTAACTTAGATTCCTCATTTTATTGAGTTGTGTCTTTCAATTGATCCCAAAAAAGTTTAATTTTACTCCTATTATGTAGTGGAAGAACAAGACACTAAAGAAAGAAAAAAATATGCTTTGAGAATTAGGTTTCAAGGCTTAAGCCCTGAAGTGAGGCTAAAAATGCATACTTTTCCGAAGAGGCGCCACTAAACAAATGACAATTGATATTTCACTTTGTAGTTAAAAAAAATCAATTTATTTGATCATATATGTGGCATTCGTGCTTATAATTATTAGTGTTTGTATTTTTTCTTCTGTTCGCGTGTTTTCATTAAAGCCTATGCTTTATTTGCACGTTATGCTTAAAGCCGTTGTGCTTTTTGGTTTTTGATAACACTTGTAGAAGCACCTTTAGTGGTCGTTTGATAGAGTGTATAAAAATAATGTATAATAGGGTGTATTAGTAATACTTTTATTAGTTATGCTTGCATTAGTTAAACATAGAATATTTTTTATGCATTGTTTGGTTTGGTGTATTAAAAAAAGTAAGTATTGTATAAAAATGTGTATTTACAAAAGCATCCTCCACAATTATGGTGGAAAAGATGTAAAAGTGCTTTTGAGGGGTAATTGAGTCTTTAACCATACTAATGCATGTATTAAGTCCTTTGCATTACTAATACCTAGAAATTTATATACATGGCATAAAAAAATGTACCAAACAAGGTATTACTAATACACAAAGCTAATGCAAGCATTATTTTTGTTAATACACTCTACCAAACGACCCCTTAGTTTGTTCATTTTGTGCACAATAGGATGTGACTTAGTGGTCGATGAAGTATGAGGGAAATCGTGAGGTTAAGAAAATGCTTGGTAATTTCTATTTATCTGCCTAGGATTTGGTGGGCAGAGACTCAGAGTTACATGATTCATGTGTTGGTGGGAGGTACTACATATTTAGTGTAATATTCTATGTGCACGCAAGGTGGCTCGGACATCACTGTCGTTAAAGATAAGACACTACTAAACTTAAGGGACTAATTTGATTTTTTTTTCTCCATATTCTTCTAACAGATGGAACAAGATGTGTCATTGAAGGGGATAGTGAAGATAGAATCAGTGGTTTACCAAGAAATGTTATCGATCGTATCCTTGAGCTCATACCAGTTGAAGATGCAGCAAGAACTAGTATTTTGTCTCGAAACTGGAGATATATTTGGGCAACATTTCCAAATCTGGTGCTGGATGACCGCTTCCTTCATAATTTTATAGGAAAGCTCCAATTCAACTTCACAGAAGCGGTAGATTGTATTCTGTTATTGCATATGGGGAACATTGTGAAGTTTGTTCTTGACACTAGAGGAGTTTTTACTGCAAATTCTTCATATGCAATTATCGATAGATGGATTCTTTATGTCACCAGAAATGGAGTCAAGATGCTAACCCTTCACATTTCAAATGATGATACCTATACTCTGCCTTCTTCTATTTTTATTTGTTCAACCTTGACATATTTGAAACTCTCCAACTGTGTCTTCAGACCACCCAATCCCTTTCTTGGTTTTCAGAATCTTATTACCCTTCGTCTGGAAGGGTCAACCTTTGTGCCAGCCACATCATTTTGTGTTATCAAGGCTCCCCTTCTTGCCAGATTGGATTTGATCTTGTGTCGTGGTACGCAATACCTAAACATTGTTTCACCAGGGCTGAAGTTCTTGTACATTCAGGACAGTCACTCATATCTCGTGCTAGATTGCTTTATGAACTGCAAGAATATAAGACTGTTAAAACTTGAATTTAACAGAGTGGTTGATGATCCAACAAATGATAAAATATCAACTTTGGAGAAGCTTCTTGTTAGCTCTCCTGCACTTGAGGTACTCCGTTTAGATTCATTCTTTGTTGAGGTAAGATATTGACTACTTGTTTCTCTTGATATCTTATAGTGATGGCAATTTCGTATGATTCTAATTATTGTGTTTTCTTCAAAGCGTGCACGCATTGTTCCGAATGGGCAACCTTCTACACTCAACTGCTTGTGGCATCTATATTTAGATGTAGACTTCAGCAAAATGGGTCAGATTTCTAATGTTCTGCAGTTAATTAAGAGTTCCCCCAATTTGAGGAAACTCCGTATTTCGGTAAGAAAAGAAAATATATGCACGATCTGGTTCCTTTTTGCACTTCATAATTGTGAACATGAGAATCTAAAATAGCTTGGTATTTTCTTCATATACATATTAGGTCCATGCTACAAGTGACAATGCTGAAGCAATCTTGAAGTATCTTGATACGCCATCCTGCCTGGAACAACCACTCGACAAGCTAGAGCACATTTTCATCAACTTTTTTAAGAGATCAAAAACTGAACTGCTTTTTGTAAAGCTATTGCTTTCTCGCACTCCATCTCTTTTAAAGATGTGCATTGACCAGGTTGCAGACATAGACGTTGATATTGCCTTAGAATTGATGCGTTTCGCCAGGGCTTCTCCGAGGGCAGAGCTATTCTATTCTCGGATAACTCTGTGATTTGAGTGTTGGAAGGCAGACTGATGTTTATCCGTTGTCAATTAGTTTTGTTAAAACAAGGGATGAATTGGCAAATTTTGTAAAAAATGTGTGGTACCTTTGTGCATATGTTCTTCACTAGATGTTTTTGATGCTCAGCATGCGTCGATGTGTTTGATATGACTAGATGTCTTTTGCTATATTTCTTTGTATTCTACTTCTGATTCAAGTTGCACTGGCTATTGAGAAATAGGGTGTCTGATGGTTTAGCATAAAGTGTTATTTTTTTCACACCATTGTGTATATTGCTTGTTAAG

>PGSC0003DMG400015687 LRR-RLPs

ATGGATGGAGATGTTGTTATTGTTGTTGGAAAGAAGAAAGGACTATGTTCCATTGATCATAAGTGTTAGTAACTATGACTATGTTCCAAAAAGGACTGAAGTAGTTATACATATCAGCAAGAATGTTAAATTCAAAATTTAATCTATTTGCCTCGCGAAAACTTAACTGTTTAGCATGTCTGATTTTAAGTTTCTTCGTGATTTGTTTACAGGTTTCAAGAAACCTGGACAACTAGAAATACTTGATTTGTCATTGAACTACTTGAACCGCAGCATCTTTAAGTCTCTTAGTCAACTTTCATCTCTCAAGTCGTTGTACCTCGCGGAAAACAACATTGAATCAGGTGCTAACATTTGGTTCATCTGTTTTCCCCTTTGCACTATTAGTCTTCCTCTTTTTACAGAACTGTGCTAAAAATAATCCATTTCCTTCTAACATTTCAGGTTCTGAGAGATTATCAGGGTTGGACAAGCTGGAAATCCTGGATTTGAGCTACAATGCACTAAATGATGAAAATGTTCTCTCTGTTCTAGGTAATTAAACTTATAGTTTCTTTTGTGAAAAGCCGTTTTTCCTCAGTAGCTTCCTACATAATTATTCTGAATTCCTCGACAGATCTAAATGTATCTAGAATGACGTTGAAGAAGCTTGATATAAGGTACAACAGATTCCGGAGTTTTATACCAAATGAAGGTATATATACAATTACTGAAGTTTTTCTTCCTTCAATTGATTTTATTCTACCTCCTTGTACATCTTGCTATGTAAATATATCTTATTGGTACCCAATCGAAATCACAAGCTTAAAAGAACTGAAAAGTCACAAAATCATTTACAAATTCCACAGGCTCCATTTGCATTTGATGAGATGTGTATAAAATATAAATACATAGACTTACATGGAAAGATGTTATTTTAATAACTCTCTTTGTCAACATTTTGGTTAAAAATATGACATTCCACCATTTTAGGCGTCATTTGCAAGTCGCTATTGCCATTTTAGTTCAAACATTTACAGTTTAAAGCATACTTTTAATGTTTCTTTTGCATGTAAGAGTTGGGAGCTTTAAGAAATATCGAGTGGTTACCGTTGGATGGAAACACATTGGATGAGAACTTTCTCAGGAGTACTGGTGTAATGTCATCTCTTACAGTGTTATCAGTAGCTAACTGCAACCTCAGTGGAGTCCTACCTCTTCAAGGTAATTTCAGACATACGTAAATAGAACAATGAATAGTCGAGAACAATGAATAATTTCAGAACAATCCATGTTTCTAATATTTTCTGCATAACAACTATATAAGAACATGAATAACAATCATCTTCATTTTATAGGCTTGTGTGATCTAAAATATCTTGAAGAACTAAGTCTCCGCCTAAACTACTTCATTGGAGAAGTCCCTGCATGTCTTGGAAACTTAACACCTCTTCGGGTAATAGATCTCAGTCAAAATTACTTTATTGGAAATATTGCCTTATCTCCACTTCGAAGTCTTGTCCCTTGAATATCTCTTGTTAGCAAACAACAAGTTTGAAATACCAATTTCATTCGAGCCATTTGCTAACCATTCAAAACTTAAGTTTGTCATTGCTTTGTTATTCTAAAAGCTGGACTCCAAATTTCCAATTGGAAGCCTTGTCTTTATCCAATTGCTCTCAAATGCCAAGTTTCCTTCAGTATCAACATCATTTACGGATTCTTTGTCTCTCGAAATGTAACATTGGTGGAAACTTCCCAAATTGGTTGTTAGAAAACAATCCCAGACTAGGAGAAGTTTATTTGGATGGAAATGCATTCAATGGATCTCTTGAATTTCTGTTCCTTCCTAATCTGAAAGCTTTTGATATCTCCAGCAACAAAATTCAAGGACAACTTCCTCCTAACATTGGGTCCATTTATCCGAAACTCATGATATTCAAAATGTCCAACAATATGCTTGAAGGCTTGTTGCCTTCAAGTTTTGGTGGCATGAAAGACCTTCAATGCTTGGATTTGTCAGGCAATAAGCTAACAGGTGATCTACCTATTGGATTGGCTCGTAAAGGATCCAAGTTAAATTTTCTGAGATTATCGAATAACATGTTGAAAGGGGAAATATTTCCAGTTTTGACTAACATCAACAATTTTGCATATTTGTTCTTGGATGGAAATAACTTCTCAGGCCCAATTCCGCAGGCGTTATCCACTGCCTTATTATTTTCACTGGATTTAAGTTACAATAATTTGTCAGGAAACTTACCAGTGTGGCTTGGTGACATATCCTCCTTAGATTCCCTCGCACTGTCCGGAAACCATCTAAGAGGTCATATTCCGCCTGATTTTTGTAGACTTGAAGGACTTCAGGTGTTAGATCTCTCTGAAAACAACCTTGTTGGCACGATTCCATCATGTTTTAGTGCTTTACGAGCCTTAAGACGTGTGTATTTGAGCAAAAACATGTTACAAGGGGAATTTAACGTGTTCTCGAATTCCCATCTACTAATGTTGGATCTTGGAGATAACAACTTCAGTGGTTCAATTCCAAAATGGTTGGACGAAACTTCTGAGTTAACTGCCTTACTCTTGAAAGGAAATCGTCTTCAAGGCACCATTCCTCCACAGTTATGTCATGCAAGTTACTTAAGAATGTTGGATCTTTCTCACAATAATATCTCAGGACCTATACCTCATTGCTTGGGAAACATAATGCAGAATCCATGGGACACAGATGTATACTACAGTCCATCATTTGGATATGGAGACTTCGAGATGTTTGGAGGAAATACCTTGATAGACGTAGATGACTCATTCGAGACAACATCAACAATTTCATTTCCAAATAGTTTTGGCGCGTGGGTTGGAGTAGAGTTCACAACCAAATATAATACTTATTCATATGAAGGTAGCATTCTTAGTTACATGTTTGGTGTTGATTTATCTTGCAACCAGTTAAGTGGTGAAATACCCAATGAACTCGGTAATCTTACTGTCATTCATGCATTAAACTTGTCACATAACCACCTTACTGGAACAATACCATCAGAATTCTCAAATCTCCAGAATATTGAGAGTTTGGACATATCATACAACAACTTGACTGGGAGGCTTCCTACTCAACTTCTAAAGTTGACAGCTCTAGAAGTTTTCAGCGTGGCACACAATAATTTAACAGGTATGACTCCACAGCGTACAGCGCAGTTTGCAACTTTTGATGAAAGCAGTTATGAGGGAAATCCTGTTCTTTGTGGTCCTCCATTGCACATCAGTTGCACAGAGGCCAAAGAGATTCCAACATACCTTCCTGCACCAGATTTCTCCGAAGATGATGCTTCTTTTTTAGATATGGAGTGGTTCTGTATTTCATTTCTTGTAGCATATGCAAATGTGGTGGTTGCAGTGGTTGTAGTACTTTGCGTAAACCCCTCCTGGAGAAATGTCTGGTTTTATTATATCGAGTCTTTCATGTATTCATGTTACGATTACTTTAATTCCAAAAGTTCATGACTTTCATAATTATGTAACATTTCATTTATGTAAATGTATGTGTGAGTTGTTCTTGGAATGTTGGTGCCTATCTAGTCCATTAATACTAGTGAATGTTGTGTTGATTTGCTTGTGCCTTCTTTAAGCATTTGCCAGATTAATTAAATTATTTCGAAATATAAAAATGTGTCATATTTTTTAGAACAGACGAATAGCCAAAATTCTTCAGTGCAATTAATGGTTTGAAAAATATGTATACTGCTCAATGAGCCATTTCTTCTATATGGTCCACTGAAATAAGTGCTAAAGACTTGGTAAACTCCTAAAGTAGCTGAATTTTTTTGACATTAGCACACAAAAATTATAACTAAAAAAAAGAAAGAAGAAGAAAATGGCTAAGGAATTAATTTTTGTTGACTAGTTAAGCACAATAATTATCAAAATTTACTCACCGTTTCAATTTGTTTGTCTTACTTTCCTTTTTAGTCTGTTTCAAAACGAATGTCTCTTTCCTTTTTTGCAACTCTTTACATCAAACTTTCCACGTGACATGTTTAAGACCACATGATTCAAAAATCTTCTTTACTTTCCTAAATTTCGTGTCAAGTCAAAATCAGAAAAACAAATTGAAACGGAGGGAGTATATAGCAATAGAAGTGTTTGCATGAAACTTCCATACAAGTGATACTAATGGGAAATGTTTCCTCATGAGTAAGTGTCTCAAGTCAAGTCAATGATGGTGCTAGTCAACTCAAATTGCAAGCAATGTTTTGATGAAAATTAGAGAAAAATATAGAGATAGAAGAACAACAAGACATATTAAAGAGTAAAATAAAGAAGTTGAAAGGGACATAACAAGAAAAGTAGTTGCACAAAATTCCACTCTAGTCTACATCAATTATTGCACTATTCCTTAAGATGCCTGCTACCTCTCACCAGCACAGATAACGAACAAGCTTCTTTTTCTTATATATAGTTCGGATATATCTTATAGTGTTGTCAACTTGTAGGTAAAGAAGCAAAACATGGCTAATAGCAAGATAGTAGATTGGTGGTTGTTGATTTTGTTCATGGCAAATGGATGGTTGTGTTGTTATTGTTGTTTGAATGAAGAAAAGACTGCTCTTTTGCAACTCAAGGCAAACATAAACTACAGTACAGGCAACGATTACTTGTCATCGTGGGGGGCTAATGAAACCTCGGATTGTTGTCAATGGGAGGGAATTGTTTGCAACAACACCACCAGAAGAGTCGTAGAATTGTCCATTATTGCTAAGCAAATTAGTCAAGAAGAACAACTAAGAAATGTTACAGATAGAGATGATATGTTGAGGAACTGGCTTTTTAATGCATCTCTATTTTTTCCCTTCAAAAATCTCAAAGCTTTGATCATACCTGGACATTCATTAGCAGGCTGGGTGAAGAATGAAGGTATGTTCTCTCTATATACGCTCCCATAATATAGAACGAAAGAGATTATACTACTCCCTCCGTTTTATATTATATGGCGATGTTTGACTAGGCTCAAAAGAACCTTAAAAGCTTGTGGTTGTAAACATGTAATGATAATTCTATTGTGTTGTACTTTTTTTAAACGGACTAAAACAGAAAGAATGTCATATAAAATGAAACAGATAGAGTACTACATATCTGAAAAATGGTTCTAAAGATCCATTTCTAAATACAAAAAGGTGTCATTCTTTTTAAGCGGGATAAAACAGAAATGTTGAAGTTTGCTTTTAGTTACTGCAGTAACTTTTAGCAAAATAAGTTATTTAGTTTGAATAAATTTGAATTTTGAATTTTAGTTAGTTTGTTTATGTTGTTGAGATATTTTAGGTTGTTTTAAATTTCAAATTACGCAGATTATGTGTTTTATGTTGGCTATTTAAAGCCCCTCAATGCTTAATAAAATTATTGAAAGCTATTGAAAGTCATTTCAATCCATTGAGAGACATTTTCAATCTTTTTGTGCTGCCCCATTTTTAAAAAACACCAACAGTGGTATCAAGAGCTTCATTGATCTTAAGGGATCTGTGAAATCTTCCAAAAAACTACCATATATATTTAACCCATAAGGGCTACCCATTTAACCCATCAGGGCTACCCATTTTAACCCTACCTTTTCAACCGGTGCTTTTTTCAATGCACAGGCTTTTCTAAATCATTTTTAACCAAAATAATGGTAAGCAGCAGTCTCTCTTTGAATTCCCCACAAATTTTCACAGACGAAAACTATCAAATTTGGTCTGTGAGAATAAAACCATATTTTGAAACCTATGTGATATGTGAGGTTGTGATGGAAGAAAAACCCTTACAATCTCTCTCTGTGAAGTTCAGATTAAAGTACTTTTCAAAAAAACTTCTACACAAGACAATGTCATCACATGTGTTTTTGAAGAACAAAATCATAAGAAGAAAAAGAAAAAGAAGAAATTTTGCAAGTTCAAAACTAATAGAAAAAAATCCAGCATCAACTACCAAAGCAAAAATTGAGGAGAAGCATTTTTTTCAATTGAAGTAACTAAAGCAAAGGAGGAGTGTTGAAGTTTGCTTTTAGTTACTGCAATAACTTTTGGCAAAATAAGTTATTTAGTTTGAATAAATTTGAATTTTGAATTTTAGTTAGTTTGTTTATGTTGTTGAGATATTTTAGGTTGTTTTAAATTTCAAATTATGCAGATTATGTGTTTTATGCTGGTTATTTAAAGCCCCTCAATGCTTAATAAAATTATTGAAAGCTATTGAAAGTCATTTCAATCCATTGAGAGACATTTTCAGTCTTTTTGTGCTGTCCCATTTTTAAAAAAACACCAACAAGAAAGAATGTCACATAAAATGAAACGGAGGGAGTATGATCGCAATACAAACTTGAATTACAGCTTCCAAAGGTTTAACTATATACTGGTTTAAGAATGTTAAGTTCATTGTACAATCTACTTGACTCTGTCACATAATTTAGCATGTCCATTTTAAGTTTCTTGTTAATTTGGTAACAGGTTTCGAGAAACTGAGACGACTGAGGAAACTAGAAAGACTTGACTTGTCGGGGAATCAGTTCAATCGCAGCATCTTTCAGTCCCTTAGTCAGCTTTCATCTCTCAAGTCACTGAACCTCTCAAGTAACAACAAAATTGGATCAGGTTCTGAGAGATTGTCAGGGTTAGACAAGCTGGAAATCCTAGATTTGACCTATAACAATCTAGAGGATGAAAATGTTCTCTCTGCTCTAGGTAAGTAAACTTATAAAAATGTTCTAAAAAAAGCTCTTCTTTTATCATCCATACCTTCCTAACATTATTATTATGAATTCCTGGACAGAGTTGAATAAATCTAGAACGACATTGAAGAAGCTCGATATAAGGTACAACAGATTCCAGAGTTTTATACCAAATGAAGGTACTTAAAGTTTCTTCTTTCAGTTGATTTTATTGTTAGAAAAACCATAGATTACACGCGGAGATATTATTCTAGGTGTCTCAGTATCAGTGTCTTGGTTAAAAATATGCTGTTCCACCATTTTAGGTGTCATTTGTTAGTTACTATTACTTCACACGTCTACGCTGCACTTTTGTAACACCTATGGCTACATGTTGATGATTCTTTTATGTTTCTTTTGCACGAAAGAGTTGGGAGCTTTAAGAAATATAGAGTACTTATTGTTGGATGGAAACACATTGGATGAGAACTTTCTCAGGAGCAGTACTGGTGTAATGTCATCTCTTAAAGTATTATCAGTAGCTCAATGTAACCTCAATGGAACTCTACCACTTCAAGGTAATTTCAGACATAGCATCGATCCTTTAAATTTTCGTCTTTCTAGATAGTGCTACGTAGGATTGATGTCACGTTAATCTTGTGTAAACGGGACAAAATTTCAGAATTTTGGTTCCATAGTAGTCTATATTTTCTGCATAACAAAATAGAACATGTATAACAATTATCTACTTGTTCATTTTATAGGCTTGTGTGATCTAAAATATCTTGAAGAACTAAGTCTCAGCAGCAACAGCTTCACTGGAAAACTCCCTACATGTCTTGGAAACTTAGCGTTTCTTCGAGTAATTGATCTCACTCATAATCAGTTTACTGGAAACATTGCTTTATCTCCACTTTCAAGTCTCTTGTCCCTTGAATACCTCTTGATAGCAAACAACAACTTTGAAGTACCAATTTCATTCGAGTCATTTGCTAACCATTCAAAACTTAAGTTTGTCATTGCTGACTACAACTCGGTAATTGTACAAACTACGTCTAAAAGTTGGATCCCGGAGTTTCAATTAGAAGCCTTATCTTTATTCAACTGCTCTCAAATGCCAAACTTTCTTCATTCTCAACATCACTTGAGACTTCTTCGTCTCTCGAAATGCAACATTGGTGGAGACTTTCCAAATTGGTTATTAGAAAACAATCCTAAACTTGGAGAAGTTTACTTGGATGGAAATGCATTCACTGGATCTCTTCAATTGCCATTACTTCCTAATCTGAAAACTTTCGATATCTCAAACAACAAGATTCGGGGGGAACTTCCTGCTAACATTGGGTCCATTTTCCCAAATCTCGTGATATCAACAATGTCCAACAATATGCTTGAAGGCTTGTTGCCTTCAAGTTTTGCTGACATGAAAAACCTTAAATGCTTGGATTTGTCATACAATAAGTTAAAAGGAGAGCTACCGACAAGATTGGCTCGTAAAGGATCCAAGTTATATTTACTGAGATTGTCGAATAACATGTTGAAAGGGGAAATATTTCCAGTTTCAGCCAACATCAACAATTTCGAATACTTATACTTGGATGGGAACAACTTCTCCGGCCCAATTCCACAGAAGTTATCCACTGCTCCTTTACGTACATTGGACTTAAGTTACAACAACTTGTCAGGTAACATACCGTCCTGGCTTGGTAATATCTCCTCCTTAACCTCCCTTGCATTGTCCAAAAACCATCTAAATGGTCATATTCCACCTGATTTCTGTAGGCTTGAAGGACTTGAGGTGTTAGATATCTCTGAAAACAACCTTGTTGGTGTGATTCCATCATGTTTTAGTAGTTTCTGGGACTTAAAACATGTGTATTTGAGTAAAAACAAGCTACAAGGGGAATTCAACATGTTCTCAAACTCAGATTTAAGAGTGTTGGATCTCGGAGATAACAACTTCAGTGGTTCCATTCCAAAATGGTTGGGACGTTCTGGTATAACCACCTTACTTTTGAAAGGAAATAATCTTCAAGGCACCATTCCCACAGAATTGTGTCATGCTAGTGACTTAAGAATTATGGATCTTTCTCACAATAATCTCTCGGGACCTATTCCTCGTTGCTTTGGGAACATAATGCAACAAGAATATGTCATAGAAGACAAATACAGTCCAGCATTTGGATATTCAGGATTCCAGACGTTTGGAGGAGACGCTGCGATAGAGGTAGAATTCTCGATCCAGTATTCATCGACAATCATGTTTCTAGATACCTATGTATGGGTTGGAGCAGAATTTATGACCAAACACATTACATATTCCTATGAAGGTAGCATGTTTTATTATATGTCCGGAATTGATCTTTCTTGCAACCAGTTAAGTGGTCACATACCTAAGGAGCTCAGTAATCTTACTCAAATTCGTGCGTTGAACTTGTCACATAACCACATAACTGGAACAATACTATCAGCATTCTCGAAGCTTCTGAATATCGAGAGTTTGGACTTGTCATACAACAACTTGACTGGGAATATTCCCACTCAACTTCTAGATTTGACAACTCTAGCAGTGTTTAGTGTGGCACACAATAATTTAACAGGTATGACTCCACAGCCTATTGCACAGTTTGGAACTTTTAATGAAAGCAGTTATGAAGGAAATCCTTTTCTTTGTGGTCCTCCACTGCTCATCAGTTGCTCGGAGCCTAAAGAGATACCGAAATCACCTCTTGAACCAGAATGTTGTGAAGACGATACTGGTTTTCTAGATATAGAGTCATTCTATATTTCATTTCTTGTGGCATTTGCAAATGTGGTGCTCGTAACGGTTGTAGTCCTCTGGGTGAACCCCTACTGGAGAAATGTTTGGTTTTATTATGTTGAGTATTACATGTATTCTTGCTATTACTTTTTTGCTTCCAAAATGTAA

>PGSC0003DMG400015706 LRR-RLPs

ATGAAGAGTGCTAGGAGATCTGATGTAGATACACTTAGCAACAATTTGCCTCGTGATATTCTGGATGGAATTCTGGGGCACTTGCCTTTGAAAGATGCAGTGAAGAGTAGTATCTTGTCGAAAGATTGGAGGTATAAATGGGCCACACGTCAAGAGCTTGAGTTTGATTATCACTTTTTCGAGTTGTTTGCACATTTTCAAGAAGCTAAAACAATCATTTATCAAGTACTCTTACTCCATAAAGGACCTATACTGAAGTTTAAACTTGGAGGGTCTAATTTGATAAGATTTCGTGATATTGATCATTGGATACTTTTCTTGTCAAAGAAAAATGTCGAGGAATTCACTCTTCGTGTAAGATCAAACAACGATTATCATTTACCTTCTCACCTTTTTACATTCCGGAAACTAAGGCTTTTGGAAGTTCAAAATTGCTTATTCCACCCTCCACCGGATTTCAAAGGGTTTAAGAAGCTTGTTAATCTTGGTCTTCATTGTGTCACTTTTCTTCCGACAGTATTAAACAATCTTATCTCTCAATGCCCGTTTCTTGAAATATTGAAGTTGAATTGGTGCACAAACTTCGACACCCTTGAAATTGATGCTGCTAATCTCAAATGCTTTGAATTTAGAGGAACGTCAAAGTCCATTTGCTTCAAAAATGCCCCTATGCTTAAGGAAGTTACTATATGGCTCAATTCACAAGTTTCGACAGATCTATCAGCTCCTGTTTGTTCTAATCTTATAAACTTCTTCTATTACATGCCTTGCCTAATGGAACTGGATATAAGTGGTGTTTCATTAAAGGTAATAGTCGAGTGAAGTTAGCTACTTATCATAACTAAATTATGTATTTGATTCGTATCTTGAACTAATTTTTGACATGCTTATAGGATTTAACAATGGGAGGTCTGTCAGGGAATTCTCCGACTGTTTTGAACAATGTCAAATCTCTCAGAATTCAGAGAATGTTATTCGAAAACAGTGAGGAGGTTTTAGGTGCTGTTTACTTGATTGCAAGCTGCCCCAAATTACAAGAGCTGACTATTGAATGTGTACGTACTCTCTCCTCCCTAAGGTAATTTGATAATAATTTGTCCTTTGTATCCGGCCTGACAACTTCCTGAATTTTTTTTTAGGAAATAATGAACGATGTTGTGGGACCTGTTGCAGTTGTACAATACTTACAAGACCATCAAAGCTTGTATGATTCTGTGAAGCTGCTTCGGAGAGTGCATATGAATACGTTTAGTGGTTTTGAGATAGAAATGGAATTTGTGAGGCTTATATTAGCATCTGCACCTGCACTTGAGAAAATTTTCTTTTGGAATTTTTCATGTTTCCTTCATCAGCCGGGTAGACAACTGATGGATAAGATGAAGCAATTTCACCAAGCATCACCTAATGTTGAATTCACATTTGAAGAAGTTGTTGCAGAAGATCGTAGACCAATTGAACCCCAAGAAGTCATGGAAGCACCATATGACTTTGATTAG

>PGSC0003DMG400015707 LRR-RLPs

ATGAATATGAGGAGTGGAGGAAGCTCTGGTGCAGATAGGATTAGCAGTTTGCCTTGTGATCTTCTTCAGAAGATATTAGGATGCTTGTCTTTGGATGATGCCGTAAGGACAAGTGTTTTGTCAACAAAATGGAGGTACAAGTGGGTAGTACGTTCCGAACTAGTGTTTGATTATAACATCAACAAAGAACTCCTCCTGAAAACAAGTATTTATAAAGTCCTATTGATGCATCAAGGAGAAGTACATAAATTTAGCCTCTTGTACCCTAGCTCCAACCTTTTTTCTGATATTGATAATTGGATTCTGTTTTTGTCGAACAAAAATGTCCAGGAATTATCACTTGCATTTCTTACATGTTTTCGCCTCCCTTCTCACCTATTCAAATTTCAGGATCTAAAGCACCTGGAACTTAATAGTTGTAAATTCCACCCTCCTCCCAACTTTAAAGGTTTTCGAAGTCTTGTTACCCTTAACTTTCAAGTTGTAACCTTTGAGCCCATGTCATTTCAAGTCTTCCTTTCAAGTAGCCCATTACTTGAGCATGTATTAATGGTTTGCTGCAGTGAGTTTGATAGCTTCATTATTGAGGGCCCCAATCTAAAAATCTTTGAATTCAGCGGCAGAGCAAATTCCCTTTGCTTTAGAAGAACCCCTCTGCTTGAGGAAATTGTATTGGGCATGCCTTTTAACATGTTGGAGGTAATCATCATCATTGAAATATCTGTCTGATGCACTACTAGATTTTCCTCGTAATTTTAGCTAATTTTGTCGCACTTTGTTTTAGTCCTTTACAGTACCAGAGCCTCAGTTAACACTAGACAACATCAAAATTTTGAATTTCAAGAATTTGATCTTTACTATGGTTGACTGGGTTGCATACGTGCTTGACTTGATCAATCGATGCCCCAACATAGAAAGGCTTCTAATTACGGTATGTCTCAGTTTTCATCTTTTGATTGTAAAGAGGTTGATTTTTAGCTTCTTCTTATAATTAATCTTCTTTTTCCCTTCTCAAGTCTGATGCTTTCGAAGCGTGTGCGTGTGTACACATCCCTTGCGATCAAAAGAAATGGCAATGACACATAGTGCAATGAAGCGGCTCAAAAGCGTGGATATGACCCTCCTGTACTAGGTTAATAAAACTGAAACAGAATTCTTGAAGCATGTATTGGCATATGCAACTAAACTTGAGAAAATATACATTGCGACTTCTGCAGATATCCGTCATAGGGGAATGAAAATGATGGAGGAGATGAAGCAATTCCCTTGAGCATCACCTAATGTAGAATTTATCTATCATTCATGAGATTGATCTATCTTCGTAGCAATAACAATATTGTTTATCCGAGATTCTTATTTTCCTTTTTAATTTCATTGTTTGTTTGATCTCTTCATAAAGGAAAATATTTTAATTTTTGGATGGACGATGTTCCTCAGAATTGAGTTTGTCAATGTAATAAAAAGGAACCTAAAGGTTTTTGTATTTAGTTGAAATTATTGAATTATTAGTCATGTAAGGCTATGCATGCGTACTTGTTTAAATATGTATTCATGTAAATTTGAATATTGTCTTTATTCGAAATTCTCCAAAGAATATTGAATACTTTGAAAGTTGATAACCACATTTGCTTAGTACTATTGGTTAGCTTTCCTAATCTCAGTTTTCAGAGGTGCCTACTAACTCTCATTAGTACTCGACCACGAAGAGTACTAGGAGATCCAGTATCGATTCAATTAGCAACTTTCCCCGCAATATTTTGGACGGAATTCTAGGGTGATTGTTGTGTCTTATCAAAAGATTGGAGGAATAAAGGGGTCACACGTCAAGAGCTTGCGTTTGATTATCACTTTTTTCAAGTCGTTTGCACATGGTCAAGAAGCTAAGACAATCATTTACCAAATCTTATTACTCCATAAAGGATCTATACTGAGGTTTAAACTTAGAGGCTCTAACTTGATAAGATCTCCTAATATTGATTGTTGGATACTTTTCTTGTCAAAAAGAAATGTCGTGGAATTCACCCTCCGTGTAAGCTTAGGCAACAAATTTCATTTACATTATTACTTTTTACATTCCAGCCACTAAGGCATTTGGAAGTTCAAAATTGCTTGGTCCACCCATTTCTAAAAATTTGACTCGGAACCTAAGACTCGAGCCTCGAGATCCAATCCCTTACCCGAGGTGAGACTAGATACATGAACTCGAGACTCGGCAACCAAGTGTAGCTCTAGCAATCGAGACCCGGGAGCGACATCTGAGCGTCAATCCTGACTCAGGACCCGATCCAATCTTTAATCGAAACCAGATATTCAAGACTCGAGATTTGAGCCCTAAGGCACAATATGGGCTAGAATTAGATTAATATTGGGCTAAATTGAAGACAACTTTAAAATTTGGGTTATATTTGATTGGACTAAAATCAGCCCAATATAAAATTATCTCATGCCCAAGTCCCAACCTCATAAAATTTTGGATGGATTGGACGGGTTATCAATTTTTGAGCCAGATTCTGCTAACCATTAAGCGAGGAGTTAAAACACCAAACATCACTCTGGCGAAAAATGAGACTAAAAAGTAGAGGGCAGCTTTCGATCTCTCTCCCAGCGGACTCTGCTATCGTTAAGCGTCCAAATTCAACTCTCTAAAGGTGAAATCCCAACATTGACACAAAGTTCGAAACTCGGTAGATAACCGATCTGGCCGTTTTTCTGGTACGAATGCCAATTGATTTCTTCAACAGAATCTAGGTTGAAATTTCTAGCTTATTTGATTTTGATATTCTGTTGTATTTGAAAAGAAAGTAGCAAACAGTCTCGATTAGGGATCAGGTGAAAAGTCATATGTAGTCATGTTTACAGAAATTGTGAATTTTCATTAGTTGGGTAGAGAATAGTACTCTGGCCCTAAGTTGAGATTTCTAGCTTGTTTCATTCGACTATGCTCTTGTATAGGAAATAATTATACCCCTTAACAACAAACCAGCGCGCAACGACTGAAAAGCCGTTGTTGCTTTTGCTTGCAGACTCGAAAATCTCTCTTTCGCCTCTCCTCCCTGTCTCAATTCTGATAGATTAGCTTCTTCCTTCGTACAAGCTACTATTTGTGAGTAACAAACACTTTTCTATAGCAGACCGGGTGAAAAGTCAAATGTAGTACTATCAAGCAGTTTGAAACAGGGGAAATATATTATCAAATGGTGAATTTTCCCTTGTTATCTACCTGTTAACCTTTAGAATGTTATATTCCTTTGTGAAAAAAAGTTGATCTCATGTTTCTATGTTTACCACTATATTTTGTCTAGTGGTAAGTGTGCCACCCCAGTTGCGTGGATTAGACATACATCTTAAAGATTACGACTAATAGTCTTCGCAAACCCTTTGAAGTCAGGTGGAGGGTGGAAGAAGCACATATCAAGTTCCACATGTCTTAGTTGCTGGAATGTAAAAAGATAGGTAAATGATATTTGTTGCCTGAGAATATTTGAAGGGTGAATTCCTAGACATTTTTTTTGACAAGAAGAGCATCTAGTAATAAATATCAGGAAGACATGTCAAATTAGGACATTGAACTGTAAATTTTAGTATTGGTACTTGATGTAGTCGTAGGACTTGGTAAATGATTGTCTTGGCTTCTTGGTTGTCGTTGAAAGATGTGAAAAACTAACTACAAAAATCAAGTTCTGCACGTGTTACCCATTTGTACCGCAAGTCTTTTGACAAGATACTGGTCTTCATTGTATCTTTCAAAAGACAAGCTCCCTAGAATTTTATCCAGGACATTACACGGCAAATTGCTAATTGTAGCACTCTTCATGGTTTAATACACACATTTGCACACAACTCTAATAATTCAAATTTGAAATAAAGAAAATATCCAAATTCTCATAATTACACATTTCAACAATGTATGCATAGTCTTTTATAACTACTAGTTCACATTTTCGATTGCATAGAAGTTTATCATTTTGCCAATATAGAAGGGAGAAAATAATTAATGATGAATGAGAGAGTACCTGAAAAATGTTGGTTGGAGACAATAATGGTGAAAGCATGAATTTAATCTAGCAGATTCAATAATGTGTGTATATATATACATAAAAGGATATTTCACCTCAATACACAATACATTTTTTTGGCGAGGGGGACTAGGAAGCTATTTCCATTTCTAGAAAAAGATCTTGTAGATATCTATTCTTTGCAATTGGAAAACCTTCATTAGATCAATTAGCTCATTCTCATCTTTGGAATGTCAGCTATTTCCATTTCTTGAAAAAGATTTTATAGAAATTTATACTTAACATATGAGATATCTACTCTTGCAAATTTTTTTCACCCAAATCCACCCTGATTACTATCTAAATTATGAATAGATCGGAAGAAAGCATAAATAGGCCCCAAAAATCATTAGGAGTTCCGCCCCTACGACTTTTGTATAGTGGTAAGTGTGCTGCCCATGTTGCATGGATTAGACATATATCTCGGGTTTGGGTTTGCAACCGATGAAACCTACTATTGAAGCCGAGAAGAATACACATCCAGATTATAGTCTTCTCAGCTTAAATAGTAGGTTTCATCAGTTGCAAACCACGATTTATGCCTAATCCACGCAACACAGGCAACACACTTACCACTAGACAAAAACCCTAGGGGCGGAACTCCCAAGGATTTTGGAGCCTATGTCTGCTTACTTCCAATTTGTTTATAATTCTATAATAATCTGGGTGGATTTTGGTAAAAAAAAAAATTGTTAAAGTAGATATCTCATATGTTAAGTATAGATTTCTGTAAAATCTTTTTATAGAAATGGAAATAGTTGTCTGACGTTCCAAAGATAGAGAATGAGCTAATTGCTCTAATGTAGGTTTTGCCAATTGCAAGAATAGATTTCTACAAGATCTTTTTCTTGAAATGGATATAACTTCCTTGGCTCCCCTCGCCAAAAAAAAAAGTGATGTGCATTTAGGTGAAATATCCTTATAGTTGTATATACATATACATTGTTAAATCTTCTAGATTAAATTCATGAAAGCACCATTGTTGTCTCCAATCAACATTTTCCTCATACCCTCGCATTCATCATTATTTTTCCCTCCGATAATGGCGAAATGATAAACCTCCATCAATTGAAAAAGTTGAACTAGCTAGTAGTTATATAAGACTATGCATACATTGTTAAAATGTGTATTTATGAGAATTTGGATATTTTCTTTATTCCAGATCTGAATTATTAGAGTTGTCTACAAACTTGTGTATTAAACCACGAAGAGTGCTACAATTTGCCCTGTAATGTCCTGGATAAAATTCTAAGGAGCTTGTCTTTGAAAGATGCAGTAAAGACTAGTATCTTGTCAAAAGACGGGAGGAACAAATGGGTAACATGTGCACAACTTGATTTTCTTAGTGATTTTTCACTTCTTTCAACAACAATAAAGTAGCCAAGACAGTCATTTACCAAGTCCTACGAATCCATCAAGGACCAATACTAAAGTGTACCGCTCCAAGATTATCCTGATATCCCCCTATTTGACGTGTTATCTTGATATTTATTACTGGATGCTCTTCTTGTCAAAGAAAACTGTCTAGGACTTCGCTCTTCAAATATTCTCTGGCAACAAATATCATTTACCTTCTTATCTTTTTACATTCCAGCAACTAAGACATCTGGAACTTGATATGTGATTCTTCCACCCTCCACCTGATTTCAAAGGGTTAGCAAAGCCTATTAGTCTGAATCTTCAAAGTGTCAGTTTTGATCCAACAATATTAAGGTATCTCACATCCAAATGCCCATTGCTTGAAAGTTTGAGGTTGACTAGGTGCGCCGAATTTGATATTCTTGAAGTTGATGCCGCCAATCTCAAATGCTTCGACTTTCTTGGAACATCAGAGTCCATTTGCTTCAAGAATGCCCCGATACTTAGAAGCGTCATTGTGTGGCTCAACTCAAGTCACGAGTTTTGATGGATCCGTCACCTTTTTGTTCTAATCTTACAAAGTTATTCCATTTACATGTCTTCCATGGAGGAACTGGAACTAGGCGGTTCATCAGTAGAGGTAATGTACATTTGTGATGATATTAATTATAGTCTACAAATGAAAGTTGGTACATATCATAATCAAATGATGTATCTTTTACCGATTTTTGACTTGCTTCTAGTACTTGGCCATGGGAGGTATACCAGAGAATCTGCCGAGTGCTCTAAACAATGTCAAATCTCTCTGTATTTTGGATATGTCTTTCAGAAATATAATGTTGACGAGGTTTCAAGCACAGGTTACTTGATTACAAGCTGCCCTAAATTACAAGAACTTACCATTGAATGTGTAAGCACTCTCCTATTTTCTCATTGAACGTATTAATATGGTTTCTTCATCTGAACCTGCCTAACTGTTATCAAATATATTCTAGGGAGTAGTAGGCATTGCTGTGGAATCGATCTGTTATACAATTCTTACGAGCCAAATCAATCTCGTGTGGTGCTGTGAAGCTGTGCATTATTTTATTGGTTTTGAGATGGAAATGGAATTTGTGAAATTGATATTGGCATCTACACCTGCACTGAAGGAGATCTTTGATGGATGAGATGAAACAATTCCATTGAGAATCACCCAACGTTTTAGATTCAAATTTGAAGAAATATGTACAGGTCTGGAACTTGATTTGTACTTATTGAGGTAGATTTGTCTTCAATATCAGCTTTCCCTTCAACATCAATCGATGAAGTTAATTTCTTATATGATCATCAAGTAATTGAAATAATCTATCAGTGTCACTTTTTTAAATAAGAAAGTCATTCGACTAT

>PGSC0003DMG400015824 LRR-RLPs

ATGGACTATGCAATTGTGCTCTTACTTGTTTTTCTCTGTCATGAAGTATTTACTGTTTCTTCATCCAAATTACCTCATCTCTTTCACAAACATCAAACCAGTTCACTTCTAAAATTCAAGAACACACTTACTGTAGATACCTCAGCCTCTTATCTCTGTCAAGAGCCTTATCCAAAAACGAGTTCATGGAACATGAGCAGAGATTGTTGCTCATGGGATGGTGTCATATGCGATGAGATGACTGGCCATGTAATTGAACTGGACCTCAGCTGCAGCAATCTTGCTGGGACCATCGATTCCAATAGCAGCCTCTTCCAACTCTCTCATCTCCAAAGGCTTGATCTTTCTCTGAATAACTTCTCCAATTCTCACATCTCCCCTGAATTTGGCAAGTTTTCGAGCTTGACGCATCTTGATCTTTCTGACTCCAATTTCTGTGGTCAAATTCCTTCTGAAATCTCTCATCTTTCCAAGTTGCAGTCTCTTCGTCTCTATGGTCGCCTACGACTCATAGCTCACGATTTTAAATTGCTCCTTCAGAACTTGACTCAATTAATAGAGCTAGATCTTACTTCCATAAACATCTCTTCCACCATTCCTCTGAATTTTTCTTCTCATTTAACAACTCTTAGGCTGGGAGATACAGCATTGTATGGGATAATACCTGAGACTATATTTCACCTGCCAAACTTGGAAACACTTCACTTATGGGAAAATTTTCAGCTCAGTGGTTATTTTCCAAAGACCAAATGGAACAGCAGCACATCTCTCCAGAAGTTACAACTCAGTTTTGTGAATTTTCCTCATAATTTGCCCGAGTCTATTGGCTATCTAACTTCACTGCGTTATTTGTATCTTCAAAATTGCAACCTTAGAGGGCCTATTCCTGAATCTCTTTCGAAGCTCACTTGCATAGAGGATTTGTACCTTCAATATAACTCCCTGAATGGAACAATACCATCAGGGATTTTCTCCCTTCCATCACTAAGTCGTTTAGTCTTGAGCAATAACCACTTTTCTGGTCAATTTGAGGATTTCAATTCCAATTCCAATTCACTAATCTTGATTGATTTAAGCAATAATCAGTTGCAAGGCCATCTTCCCAACTCAATTCAAAACCTTGTGAACCTAACAGATCTTGATCTTTCTTTCAATAATTTTCATGTGGATGTCAGCCTCTTTTCAGACCTCAAACAGCTTTTGTATCTTAGTCTTTCATACAATACTATTGATGAAAAGGAAGTTAATTCTACTTTGCCTGAGTCACTTGAGTACTTACTATTGGCCGCATGTGAAGTGAAAGAATTGGCGTTTCTAAGATCTGCAAAACATCTTTCCAACTTGGATCTTTCAAATAATAAGCTTCAAGGAAGAATTCCTGATTGGGCATGGTCTAACTGGATGTCATTGACAACTCTTAATATATCCCACAATATGCTGACGAGTGTGGATACAATTCCTCTTCAGATTGTATATACTATTGATTTACGATCCAATTTGCTTCAAGGTTCACTACCTGTTCCACCAAATTCCACAACATTCTTCTTCATATCCTATAATTATCTCAGTGAAGAAATTCCTTCAGATATTTGCAATTTAACATCACTTGTAATGCTAGATTTGGCAAGAAACAATTTACGTGGAGAAATTCCGCCATGTTTGGGAAAAATCACTGCCCTCCAAGTTTTGGATATGCGCCACAACAATCTTTCTGGGAACATTCCAACGACTTTCAGCAATGGAAGTTCACTGAGCAGCCTTAACTTGCATGGAAATAAACTAGAAGGGAAAATTCCCCAATCTTTGGCCAATTGCAAAAACTTGCAGCTTCTTGATTTAGGCAATAATAACCTCAACGGCACATTTCCTGTGTGGTTGGGAGCTCTGCCAGATCTGCTAGTTTTAAGCTTGAGATCAAACAAATTGCATGGACCCATCAGAACATCAAGGATTGTGAACATGTTTCCTGAGCTTCGAATCATAGATCTCTCTTATAATGCCTTTTCGGGAAGCTTACCTTCGAGTTTGTTTCAACATCTGAAAGCCATGAGGACAATTGATCCATCAATGGGAGCACCAAGATATCATGGAGATACATATTATGAAGATTCTATTACAGTTGCAACCAAGGGATTTTATCGTGAAATTGTGAGAATCTTGTATTTGTACACAGTTATCGATCTTTCAAGTAATAAATTTAGGGGGCAAATTCCAAGTATCATGGGGGATCTCATTGCAGTTCACATAATGAATTTATCTCATAATGGATTGCAAGGTCATATACCGTCATCATTCGGAGATTTATCTTCAGTTGAATCATTGGACCTATCAGGAAACCAGCTTTCAGGAGAGATACCACAACAACTGGCTTCTCTTACGTATCTTTCATTCTTAAATCTCTCCCACAATCATCTCCAAGGATGCATCCCTCAAGGACCTCAATTTCACACTTATGAGAGCAACTCATATGAAGGCAATGATGGATTACGTGGATTTCCTATTTCAAAAAGTTGTGGCGATGCTCGGGTATTAGATACAAATGATACCGTATCTGGACTAGATGATGAAGAAAGCAATTCTGACTTTCAGAGTGATTTTTGGAAAGGTACTCTTATGGGCTATGGAAGTGGACTATGTATTGGATTATCCATAATATATTTCATGATTTCAACTGGAAAACCGATTTGGCTTGCAAGAATCATCATAGAAATGGAGCACAAAATAATTATGGGAAGGAGAAAGAAGCAGCGAAGGCAAAGGAATTACAGAAGAAGAAACATTAATCACTTCTAG

>PGSC0003DMG400015853 LRR-RLPs

CAAAAATATAGATAGAAGACAGCACCACCCACACACAAACCTCATTGTTTTGTACCTCATCCTCCTTCTACCTCTTTTTTCTCTCTTTTTGCCCTATTCAATAGGGTTTCCAAATGATCTCTTGCTACCTGATTTTCTGATGCAATCAGTTGTTCAAGCTCTTTTAATTGTGTTTCTTTTGGTTGATCTGTCAATTTTTGCTCTCCATAGAGGGTTTTTTGGTCCTGGACAGTGCAGATTCTACTTCCCCAGTGTTTTTCAACCATACTCATGTCTAAAGTCTTCAATTTTAGTGGTATGATCTTCATCCTTGTCTGGGTTTTGCTTAAAAATTCAATCTTTATCTTAATTATTGTTTTTTTCTCTAGTGTTTTAGATTTTAGGTTTTTTCTTTTTGAGTTCATCTGTTATTAATTTTATGTAACTTTCTCTTTGAGACTAGTGGGGTTTCTACATCTGGATCCTTGATTTTATCCTAAGTGTGATTTTACTTTGATCTATCTAGATATGATACAATTGAATCTCTCTTGTTTTCCTTTACTAGTGCAAAATTCCATATAAAATGTTAATGATATGCTAATCTAGGAAAAAAATGGGCAATAATGACTGGAAAATTCATTGATGGATATTTATGTTGAAAATCAATTATTTTTTTCCCATGTTTGACTGAAATATTTTTTTCATAGAACTCTTCAAAACTGATTAACCTGGCATATGGAGTATCTTGTATTTCTGTGATATATGCTCTAAATCTGGATCTGATCAGACTTGTATGTGAAGTGGGTTTGGCCTTTCTCAAGATTTGTTTGGTCTCAATTGTGATCTTTGGCTGATGTTCTTTACCATATTCTGGATTTCAGGTGATGATGCTTTTTACCATGGTGGGGCTGTGTACCCAAGTCCCAAGGAATCCAGTCTATTTCTGTCCCTTGGGAATCATGTGGATGTCTATTTTCCTCCTTGCAAGAGGTCTCGTGTTGCGGTCCCTTTTGTTTTCACTGAAAAGAAGCAGAAGTTGTCTTCCATTGATGTCCTACCTGATGAATGCCTTTTTGAGGTATTGAGGCGCCTTTCTGATGGCAAAGAGAGGAGTGCCTCTGCTTGCGTTTCCAAGCGCTGGCTTATGCTTTTAAGCAGCATCCACGGGGATGAAACAGTCATCTCAGATCTCAATCCATCTTTGGAGACCGAGGAAAGATCTATCCAAACCGCCCTTGTTAAGCCCGTGGACTGTGTTAAGAAGGGTGAGGTATTGGACTCTAATGGTGCAGAAGTTGCTGATGCTGAATCTCAAGATATTGAAGGAGAGGGTCATCTTTCCAGGTGCCTTGATGGAAAGAAAGCAACAGATGTCCGACTTGCTGCTATTGCTGTTGGAACTCCAAGCCATGGAGGGTTAGGGAAGCTTTCTATTCGGGGAAGCAACCCAATCCGTGGTGTGACTGATACTGGCCTTAAGGCTATTGCTCGAGGTTGTCCTTCTCTTAGGGCTCTTTCTCTCTGGAATGTATCTTCTGTTAGTGACGAAGGTTTAATTGAGATTGCTCAGGGATGTCATCTCTTAGAGAAGCTTGATCTTTGCCAATGCCCTGCAATTACTGATATGTCTTTGATGGCTATCGCAAAGAATTGTCCTAATTTGACCTCTCTAACGATAGAATCTTGTTCAAAGATTGGAAATGAAACTCTTCAAGCTGTAGGTCGTTTTTGCCCGAAGTTGAAGTTTGTGTCTCTCAAAAACTGCCCACTCATTGGAGATCAAGGAATTGCAAGTCTCTTTTCATCTGCTGGTAATGTTTTGACAAAGGTGAAACTTTACGCATTGAACATCAGTGACATCTCCCTTGCTGTTATTGGACATTATGGCATTGCCGTGACTGACATAGTCCTTATTGGTCTTCAAAACATAAATGAGAGAGGATTTTGGGTCATGGGTAATGGCCAAGGTTTGCAGAAGCTGAGGTCGCTTGCTATAACTGCTTGCCATGGAGTTACTGATTTGGGACTTGAAGCTCTTGGTAAAGGTTGTCCAAACTTGAAGCTGTTCTGCCTGCGAAAATGTACAATCCTGTCAGATAATGGCTTGGTTGCTTTTGCCAAAGGTTCAGTTGCACTAGAGAACCTCCAATTAGAAGAATGCCACAGGATCACCCAGGCTGGGTTTGTTGGTGTTCTTTTGAGCTGTGGTAAGAAGTTAAAGGTTCTGTCCATGGTGAATTGCTTTGGTGTTAAAGAGTTAGCCTGTCGATTTCCATCTGTGCTACCTTGCAACTCACTGCAGTCTTTGTCTATTCGCAACTGCCCTGGAGTTGGTAATGCTACGCTGGCCATAGTGGGTAGGCTGTGCCCCAAACTGACTCATTTGGAGTTGAGTGGCCTTCTTGAAGTAACTGATGAGGGTCTTTTCCCTCTTGTTCAGAGCTGTGAAGCTGGTTTGGTCAAGGTGAATCTAAGCGGATGTGTTAATGTTACAGACAGATCAGTTTCATTCATAACTGAGTTGCATGGGGGAAGTCTCGAGTCTCTTAATGTTGATGAGTGTCCATATGTTACTGATGCGACCTTGTTAGCAATTTCCAACAACTGTTGGTTGCTCAAGGAACTTGATATTTCAAAGTGTGGAATAACTGATTCAGGTATTGCATCTTTGGCCAGTACAGTGCGACTTAATTTGCAGATCCTCTCATTGTCGGGTTGCTCTATGCTTTCGGACAAAAGTGTACCCTTTCTGCAGAAGTTGGGGCAGACACTTGTGGGCTTAAACATCCAGCACTGCAATGGGGTCAGTAGCAGGTGTGTTGATCTTCTCTTGGAACAACTCTGGAGGTGTGATATCCTTTCTTAAGTGCACAGGGTTTAAGGATGAATGCAACTTACAAGTTGAGCTCCTTTGAAGATGGAGCTTAGTAAGGAAGTTTAACTAGTCTTTCTTTGTTAGTGGTAGCATAAAGTTTTTTGGTCCATGATCTTGGGTCTTCTAATAGTCCAGTATCTCGGCTCCTTTTTCCAGGGATGTGGTTGGTTCTCTTTTTTTGCAGCTTCTTCACTAAAGCTGTTTATGGGAATTTTTTTTTCCTGTTTTTCCAACCATACTTCCCTGAGATACAAATGCTGAGTTTTTGGATTAAGAGATTCATGTCTGGTCCTAGTCCTAGTACTTGTTGTCACTAGTTTTTAGCTTCTACCCTTGCATTGATTACCAAATCTTGCAGCGGTAGAAGCTTTTCAATATTGTCTTAGTGTTTTATGTGTGTGCACTTGTAGTTTTACCGAGTCTTTCACGTTGTTGAACCTTTCAACTCGTGATGGTTGGGTTATGTTTGTGTGTTGTTCTTGTGGTTGCTTTTGGGCAGTTGTAGGTATAGATGCCTCTCTTTCATGGGCTTTTGGCCATGGCAGCAAGTTCCTTTTCTGCCATGACAACCCTCTGGGGTTGTTTTTTCTTTTTACCAAGCCTTAGTTTTGCAGGCCTCTCTTGTTCTTTAACTCCCAAATGTAACCTCGTTTTGTACTTGTGGAAATGTTAGTTTGTACTTATGTCTTTTATTCAATAAAAAAAAAGAAAAAAAAGTTAGGTTGTAT

>PGSC0003DMG400015878 LRR-RLPs

ATGGGTGATTTCCCAGAGGATTTTCAAGTTGAAACTTGGAGATTGATCCAATTATGGATCGCAGAAGGTTTCATAAGGACGTCTTCTGGAAGTCATGAAAGCTTGGAAGAAGTAGCTAAACATTATTTGGAGGATCTTATCAGCAGGAACTTGATAATGGCTAGAAAAAAAAGATTCAATGGTGAGGTAAAAGTATGCGGAATACATGATCTGCTGCGTGAGTTCTGTTTGATAGAAGCTGAAATGACAAAGTTTATGCATGTTGTGAGAACTCCATTTTCACCAGGACAAAAGCCTAATTTTCGTCGCTTCAGTATTGAAAAGTATATTGAAGATGCTTATAAGCTCTTACCCCCTGTAGCCAGATCTATGTACTGCTTTTCTAACTTGGATCTACCTTTTGAACCCCGTATTAAGCTTCTCCCTATATTGCCCATCTACCGTCATGATCCTATAATACATGAATTTTTCTCTTGTTTCAACCATCTCAGGGTATTGGCCATCTTTCCTGCCTGGTTTGACTCAATCTCAGAGCTTCTGAATTTGCAAACTCTAATTTGTGATGCTAATATAATGTATATAACTTTACCTCGGAAGATATGGATGATGAAGAACTTGAGGTATATACGTTTAAGGGCAGCCAGTTATTTTCCAAGTCCTAGAATAGATAAGAATCTTGTGACAGGGATGCCAAATCTAGAGGAACTTTTTGGTCTTTGTTACTCTAGTTGTACAAATGAAGTTTTTTCTAGCATCCCCAATTTAAAGAGATTGACCATTCACGCACCTTTAATGGTAGACAATGTTCCATATCATCTCTTGGATATGTCGAGCTTAAGAAAACTTGAGGCATTCAAGCTTTCCTGGAGATATGTTTTTGCAAAGCCCATCAAAAGATTTGTTTTTCCAAAATCACTTAGAAGATTGTCTTTAGCTAGGTGTTATAAGTTTATTTGGGAGGAGATATCATCAACTTTTATCAAGTTGCCACATCTTGAAGAGCTCAAACTTAAATATTGTCGAGCCGATGATGATATATGGAGATTGAATGACAAAGACATATTCAATCTCCAACAGATTTTACTGAGCGAGCTAAATCTTAAGCATTGGGAAGCTAGCAGTGATAACTTCCCTAATCTAAAATGCCTTATTCTGAAGAATTGCGAGTTCCTACAAGAAATTCCAACAGATTTTGGGGAAATTTGTACTTTGGAATCAATTGAATTACATGATTGCACCACTATTTCTGAGGATTCTGCCAGAAATATCGAACAAGAACAAGAGGATATGGGAAATAATATCCTTAAGGTCTACATTCGTAGCTGCCACAGTAAGTTTTAA

>PGSC0003DMG400015957 LRR-RLPs

ATGCTTGGAGTGCTACTGATGCCACCACTTTTAGGTGATTTGGTGAATTTGGAGCAATTGAATGTGGCACATTATATGCTTTCTAGAACAATTTCAAAAAGTATTTGTCAGCTTCCTAAGCTTCAGAATTTTACGTATCCTTATAATTTCTTCACTGGTGAACTGCCGGTGTATTTGGGATTGCCGGAGTTTCATGATGAACGGAATTGTTTGCCCAAAATACCGGTGCAACTGTCTCCGGGACAGTATAAGGCGTTTTTGTCTAAAAAAATTCATTGTAGTGCTTTTAGGTGTCATAAATTTGTTCCTATTTTGCCATCTCCATCACCTCTACCACCAGTCTATTCACCGCCACCACCAACTCCATCTTCTCCTCCACCGCCACCACCACCATCTCCACCTCCTCCACCACCTGTTTATTCACCACCACCGCCACCACCATCACCTCCACCACATGTTTATTCACCACCACCACCACCACCCCCACCATCTCCACCTCCTCCATCACCATTGCCGTATTGTGTACGCTCACCTCCACCTCACTTGCAGGCCTTCGACCACTCCCTTGCAGGCCTCCTCTTCTCCTCCTTCACATTTTTTCTGTCATCTAATGAAACCCACAAATCTAATAGCTTCCTATGCTTTTCTTTTCTATTTTTCCTGTTGGTCCTAAACTCGACGAAGCCCCAATTCATCTGTCTTCGGCCTCTCCTTCTTGATCTCACCAATCTAATCGCTTCCTAA

>PGSC0003DMG400016357 LRR-RLPs

GCTACTGCTTCTCTAAAACAAAATCAGCTGCAAGGTCCTATTCCAAAGTCACTCCTAAACCAGCGTGATCTATATTCTCTTGTCCTTTCGCACAATAATCTCAGTGGACAGATCGCTTCAACCATCTGCAATCTGACAAAACTGTTAGTGCTAGATTTGGGCAGCAATAATTTGGAGGGAACAATCCCACTATGTTTGGGTGAGATGAGTGAACTTATGATTTTAGATTTAAGCAACAATAGTCTTAGAGGGACAATTAATACAACTTTTAGTATAGGAAACCAACTTATAGTCATTAAATTGACGGGAATAAGCTAGAGGGGAAAGTCCCACAATCATTGATCAATTGCACATATTTGGAAGTTCTTGATTTAGGTAACAATGAGTTGAATGACACATTTCCTAAATGGTTGAGAGCCCTACCGAATCTGAAGATTCTAAGACTGAGATCAAATAAGTTCTTTGGCCCTATAAAAGATTCATCGACTTACAAGTTTTGCTCAAATTCGAATCATAGATCTCTCATCCAATGGATTTAGTGGAAATTTACCAGTGAGCCTTTTTGAGAATTTTGAAGCTATGAAAATAAATGGTGAGAAAAGTGGAACCCGAGAGTATGTAGCAGATATGAATAACGTTTATTACGCAAATTCCTTTATAGTTACAACAAAGGGGCTGGATCTAGAACTACCTCGAGTTTTGACTACCAACATAATTATCGATCTCTCAAGGAATAGATTTGAAGGTCATATCCCAGGCATTATTGGAGATCTCATTGGACTTCGTACTTTGAACCTATCTCATAATCGCTTGGAAGGTCACATACCAGCATCACTGCACCAGTTATCTGTGCTTGAATCATTGGATCTCTCATCCAACAAAATCAGCGGAGAAATTCCACAACAACTTGCATCCCTCAAATCACTTGAAGTCTGAAATCTCTCTCACAATCATCTTGTTGGATGCATCCCCAAAGGAAAACAATTTGATACGTTTGAGAATAGTTCATACCAAGGAAATGATGGGTTACGTGGATTTCCACTCTCAAAGGATTGTGGTGGTGATGATGGGGTAGCTCAAACGACGAATCCAGTTGAGCTAGATGAAGAAGGAGGGGGAGATTCATCAATGATCAGTTGGCAGGCGGTTCTCATGGGTTATGGTTGTGGACTTGTTATTGGACTGTCCATAATATACATAATGTTGTCAACTCAATATCCAGCATGGTTTTCGAGGATGGATGTAAAATTGGAACACAAAATTCTTACGAGAATGAGAAGGCACAAGAAAAGACATTAGTGTGTAACCTCGAGGTATCAACTTAATCTCTATCCTTCAGAAGATCACATTTTATATATCCATAAATTTTCTACCTCCTTCATCTTTAAAGCTTTTAACTTTAATTCTTCATTTTTGAAAATTGCAGGATTCAAGTCTACTGCATCGCTGGGATGTAAAAATAGAAGCTTTTACTTTCATAGTTTGTTATTCTGTCGCAACTCAACAAAATTTGATTTCACTTATCTGTAGTATGTAGAAATAAATTCAGCATTTTCAATTGTGAGGAAAAAACAAGTTCATAAAATGCAAAGGGATCAAAAGGGAAACACCAGTAGAGCAAAAAGTTTGAAATAACAGTAGAAAGCACATTCAAATTCCTAAAACAGTTCCAAATTCATCGATATATAATAACGACGACCACAATAGAGTTGGTGTAGCAGTTGAAAAGTTTGAGAAACAACCTTGGATACTAAGGTTTAAATCCTAGCAGAGATATCACATTTAGGAAGAAATCCAGCAGAACTGCTTAATTTGATGAGTCATGACTTCGTATCGACAATTTTACAAACTAAACTCTACTAACTAAAACTGTGGGCTGGAACCTTGTGTATTCCAGCATCTAACATTGTTCTTTTCCACCTCTCTGCGCCAATATAAATTCTTGAGAGTTGCACATAACGCCCAAAATGATTCTGTTGCGACTCAAGCAATAGTTGTGCTACTTCATTTCCCAACTTAATAGCTCCACCAAGTCTACAAGCACCCATCAGAGCCCCCCAAGACAGTAGCATCAGCTTCAAAAGGCATCTTCTTTATGAAGTCGTATGCTTCCTGCAGTAGATCAACCTCACAACCATAACGTTTCATCTTAGCTGCAAGTCCAAATTCATGTGACATTGCTTCAAACAATTAGAACCCGAAATCGACAAGCTTGGCACACAGACAAGCTGATAGAACTGCCACGAAGGCAATCTCATTTGGTTGCAGCCCCTTTTACTCTCATCTTTGTTGAGAATATAATGAAATAATAGCTTAAGCTTTTAGATGAGATGGTCACACACTTCAGCAACCTTGACATACATCGTTAATGCTAGGGAAGAAATCATAGCATTCCATGTGCAAACCTGCTCCATCGCATCAAATACCTGTGTTACAGCAAATGGTTAAAAATACATGCAAGGCCTCGTACTATAAAGAGCACGCAGTTTAAAATACATGCAAGGCCTGCTAACAAGCTTTCATTAAATGATAAGCACGAGCAAAAATTAAAGACCATCGATTTGAGCGACAGAATCAAAGACCACCCCGAAATAGGGACAATCCGTGCAAAAACTTCTATCAATTTAGGACAGAAAATAAGACCTTTAAAATCACGAGAAGTGCAGATCCAAACCAATTTACAGGCGAATTGTTAACAGAGACATCAACTGCACAAAATTCTAAACGAAGAAACAATTAATATAGTGTAATATTGTTGGCTTCGGGCAAAACATGCACAGTTTTCTTCCAAATGTCTCCAATCATATTGCAATCAGTTTCGCTACAGCACCCGCTGCTCTAGCCTAATATGAGTATCACTTCACTTTATACACATCTTCTTTTTCTTAGGAGACAAAATGCATGTGTTATTACAAGTATTGGGACTTTCATTTAAATCTTTCTTAAATTGACTTGAGGATGTTATTTGACTTACAGTGCGATGACTTGAGAAAGATTAAATAGTCTTTACATATTGTGTGACACATTGAATGTGAATGTGTAATTGACAACTCTAAAACTGACTGAAAATGACTAGGTGACCTTCCTGGAATTGATTATTTCAAAGCAAAACTTAAATGTGCTATATAGGATTTGTTATCTGACAATTTCCACCTGCATCAAACAGTCCAGAAGTTGTTGTGGATTTTCTGTTTAAGTACTCCTTCAATTATTATTTTGTTGTCAAGGTTTATACACCAATTATTCATAGTGGTAGGAAGCTCAAAGGAAAGTGTAAAAATATCCCAATAGATAGGGAATGGAACAAGCTTTAAAGGATTTTCGACTGGCATGAAGAGTTTGATTCTTAACTCTAACTCTAAGGTCTTGAATACCTAGAGTTAAGAAATTATGTTCATAACATTCAAGAACTTGAATTATTCTGTGCTCATGGGAACTAAATTAAGACATGGCATACACCTTCTTTTGCTTAAAAAAGTCGACACTACTTAAATAAAAGGACACTAATTATCAACTTGGGACAGAAAGTAAGACCTTAAAATCACAAGAAGTGCAGATCCAAACCAATTAATAAGACGAATTGTGAAAATAGACCTACATCAATTGCACAAAATTCTTAAAATGGTGTGATTTGCAGATTTTAAACTTGAGATCAAATAAGCTGTATGGCCCTATAAATGATTCAAGGACTGAGAACTTGTTTGCTAAAATTCTAGTAATAGATCTCTCATCCAATGGATTCAGTGGAGATTTACCTGTGAGCCTTTTTGAGAATTTTCAAGCCATGAAAATGATTGGTGAGAATAGTGGAACCCCAGAGTATGTAGCAGAAACATATTCTACTTTATACACAAATTCTTTGATAGTGACAACAAAGGGACTGGATCTTGAACTTCCTCAAGTTTTGACTATGGACATAATTATCGATCTCTCAAGGAATAGATTTGATAGTAAGTATTATTAGAGATGTAATTGGACTTCGTACGTTGAACTTCTCTCATAATAACTTGAAAGGTCATATACCAGCATCAATGCAACATTTATCTATACTTGAATCATTGGATCTCTCATCCAACAAAATCGTCGGAGAAATTCCACAACAACTTGTATCCCTCACATCACTTGAAGTCTTAAATCTCTCTCACAATCATCTTGTTGGATGCATCCCCAAAGGAAAACAATTTGATACGTTTGAGAACAGTTCATACCAAGGGAATGAAGGACTACGTGGATTTCCTCTCTCAAAAGATTGTGGTGGTGATGATGGGGTACCACAAGCGACAAGTCAAGTTGAGCTAGATCAAGGAGGAGATTCACCAATGATCAATTGGCAGGAGGTTCTCATGGGTTACGGTTGTGGACTAGTTATTGTATTGTCCGTAATATACATAATGTTAACTCAAAATCCAGTGTGATTTTCGAGGATGGTTGATCAATTGGAACACAGAATCATTACGAGAATGAAGAAACACAAAAAAAGATATGTAACCTCTAGGTATTCAACTTGATCTTTTTCATTCAAAA

>PGSC0003DMG400016424 LRR-RLPs

GTTTTTTGAACATTTTAGTGCAACAATGGGCTACGTAAACCGTATATTTTTTATGCTATTTCCCTTCCTCTGCCATCTTTCTTTCTCCTCATCCTTACCCCATTTGTGCCCCAAAGATCAAGCTCTTGCTCTTCTACAATTCAAGCACATGTTTACTATAGATGCTTTTGATCGTTGTTTCGACATAACAGGCCAACCGTTATATCCAAAAACTCTTTCGTGGAACAAGAGCACAGATTGTTGCTCTTGGGAAGGAGTTTATTGTGACGAGACGACAGGAAAAGTGATTGAGCTTAACCTCGCTTGCAGCAAACTTCAAGGCAAGTTTCACTCCAACAGTAGTGTCTTTCAACTCTCCAATCTCAAAAGGCTTGATTTTTCTAATAATAATTTCTCTGGATCGTACATTTCGCCTAAATTTGGTGAGTTTTCTAGTTTGACGCAACTTGATTTGTCATATTCAGGTTTTACAGGTATAATACCAGCACAAATATCTCGTCTTTCTAAATTACAAGTTCTTCGTATCCAGAGTTATTCAGATGGTCTTAGATTCAAACCTCACAATTTTGAACTGCTCCTTAAGAACTTGACCCAATTAAGAGAGCTTGACTTTGACAATGTGAACATCTCTTCCACCATTCCTCTGAATTTCTCTTCTTATTTGACAACTCTACACCTTCGGAACACACAGTTCTACGGGGTATTGCCCGAAAGAGTTTTCCACCTTTCCAACTTGGAATCTCTTGATTTATCAGACAATCTCCAGCTCACTGTTAAGTTTCCCACAACCAAATGGAATAGCAGTGCATCACTCATGGAGTTAGATCTCTATAGTGTGAATGCTACTGGTAGGATACCTGAATCATTTGGTCATCTAACTTCACTGCGTAGATTGAGAATAGTTTCTTGTAATCTCTCAGGCTCTATTCCTAAACTTCTATTGAATCTCACCAACATAGAGGAATTGTACCTTGGTTATAACCATCTTGAAGGACCCATTTCTGATTTCTTTAGATTTGGAAAGCTCAGGTTGTTATCACTTGCAAATAACATCTTTGACTTCAAATTCCTTCCAATGTAAGTGGTATGCAAAACGTACAATCACTCATATTGTCATCAAACCACTTGAATGGGACTATACCTTCCTGGATATTCTCCCTCCCTTCACTTGACTTGTTAGACTTGAGTGATAACCATTTCAGTGGAA

>PGSC0003DMG400016653 LRR-RLPs

ATGCTTACCCAGGATTTGACATCCAAATCGAAAATTGACCCTGAGCTGATTCAGAACTCGAGAGTTGGGTCTCCCATCAAAGTTGTGAGTTTGGTGGCGAATTGAGAATTATTAGTCGGATTCCAATTAGTATCGGGAATCGATAACCGAAATTTGATTTAGGGTGGAAAGTTTAGGTGTGAGGTTGAAAAAGAGTTTTCGAAAATGTTTTCCTTATTATTTTTTGGGGTAAGTCATTTTCCTTGATTTTGAAGAAATTAAATTTATTTGGAAAACATTTAAGTCAACCAAACATGAAAAATTTGAAATCATTTTCCAGAAAATATTTTCCTTCGAACTAAACACAAAACTTGTGTCTTTATCTTCTTTTTTTTCTCCCCCTTTCAAATCATAGTTTATATAGGTTGGTGATAAGAAAACATGATGGGTATATTATGTATATAAGTTTCTATTTTGATCTTTGTCTGATGTTTTATATCTTCTGATTTTCAGAAAATGGTGCTTTTTTCAAAGCTTCATATCTAAGTTCAAGATAATCATCTTTGCTTTTATCACATGGGCAAAATAGGAATGTAAATTTTTCTCCTAGGAAGAGGATTTGTGTCACTACCGCTTCTAAAAATTTTGAGCAGAGGAATCATCCTTCCATTGAAATCCTTCTTGATAAATGCCTCTTTGAGGTATTCAAACGTCTTCCGAGTGGCCAAAAGAGAAGTGTATGTGCTTGTGTTTCCAAGCGCTGGCTTACACTTTTAAGCAGCATTCACAAGGATGAGATTGCAGAGTTTAATGGTATAGAAGGTGAGGGATCTCTTTTAAGGTCCCTTGTTGGTAGGGAAGCCACAGATGTTAGACTTGCTGCTATTGCTGTTGGAATCGCAAATCGTGGAGGTTTAGCAAAGCTATCTATTCGGGGAGACAATCCGTGTCGTGATGTGACAGATGCTGGTCTCAAGGCTATCGCTCGATGTTGCCCTACTCTGAGAGATCTTTCCTTGTGGAATGTGTCTTTTGTTGGTGATGAAGGCTTATCTAAGATTGCTCATGAATGCCATCTATTATAGAAGCTTGATATTTTCCAATGTCCAACAATTACCGACAATTCTTTGTTGGCCGTTGCAAAGAAATGTAATCACCTGACCTCCTTGATCATAGATTCTTGTTCAAAGATTGGGAATAAGTCTCCCAAAGTTGTGGGTCAATACTGTCTGAACCTGAAGCTCGTCATACTAAAAAATTGTCCACTCATTGGGGATAAAGGAATTGAAGATCTCTTTTATTCAGCCGGTACTGTTTTAACAGAAGTAGAGCTCCAGGCACTACACATTAGCGACATTTCTCTAAAAATTATTTCACATTATGGAACTGTACTGACAAGTCTAGTTGTTGGTGAACTTCGAAGAATAATAGCGAGGGATTTTTGGGTCATGGGCATTGGTCAATGTCTGCATAAGCTGAAGGCCCTTTCAATAAGCTCTTCCAATGGAGTCAATGATTTGGGTCTTCACGTTATCTGTAAAGCTTGCCCAAATTTAAAGTTGTTTTGTGTCCGAAAAAGTGTCATCCTATCAGATAATGGATTGGTTGCTTGTGTCAAAGCTTCAGTCTCACTTCAGAACCTTCAGTTAGAGGAATGCCACTTGATTACCCAGGCTGGATTTTTTGGTATACTTCTGAACTGTGGTAAGAAACTGAAGACTCTTTCCTTGGTGAAGTGCTTGGGTGTGATGGACTTAATTAATGTAGCTCCTTCGAAGGCTCTTTGTTGTAACTCACTGGTATCATTGACTATTCACAAATGTCTTGGAGTAGGCAATGCTACCATTGTTGTAGTAGGTAGTCTATGTCACAAACTAACTCATATAGAGCTGAGTGGGCTTCTTTGTATAACTGACGAGGGGCTCTTCCCTCTAGTTAATAACTGTGCAGCTAATTTGGTGGCGGTGAATCTAAGCGGGTGTGTCAATATTACAGACAAAGCAGTTTCAGCCATAGTTAAATTCAATGGGAGATCTCTGAAGTCTCTACTCGTAGATGGTTGCAGACATGATACTGATGCAACATTAGTAAAAATTTGGAACTCTTGCTGGATGCTTAATGTACTCGATGTTTCCAAGTGTGGAATCACCGATTCAGAGATTACAACTTTGTCCAGTGCTATACAACTTCATTTGCAGATCCTCTCACTGTCTGGTTGCCCTTTGGTGTCAGACAATAGCTTACCCCTCTTGCTCGATTTGGTTCAGAACCTTCAGGGCTTGAACATACAGCAATGTCCCAACATCAGCTTTAGAACTGTTGATGCACTTGTAGAGATAAATCCAGGTTGTAATATACTTTTCTAA

>PGSC0003DMG400016808 LRR-RLPs

GAAGCTTTCATTTTTGAATTGGTACTTGTTTATTTCTTCTTTCCTCTATATAACTAATCTGTTCATCAATTGATCTCATCCATACACTAGACTATTAATCGAATCTATGGCATCATCTTCTTCTTTTGCGAGTAATTCACAGTACTGTCCTCAACGGAAGTACGATGTCTTTCTAAGTTTTAGAGGAGAAGATACTCGAAAAAATTTTACGAGTCACTTGTACCAAGATTTGGAAAATAAGGGAATATTCACCTTTCTAGATGATAAAAGGCTAGAGGATGGAGATTCCATCTCAGAAGAACTTGTGAAAGCTATAGAAGAGTCTCAAGTTGCAGTAATCATTTTCTCAAAGAATTATGCTACGTCAAGGTGGTGCTTGAATGAACTAGTGAAGATAATGGAATGCAAGGAGAAAAAAATTGGACACACAGTCATACCGGTCTTCTGTTATGTGGATCCATCACATGTTCGATACCAAAGTGAGAGCTTTGCAGAAGCATTTGCCAAACACGAATCGAGGTACAAGGATGATGTTGAGGGGATGCAGAAGGTGCAAGGATGGAGGAATGCCCTAGCTGCTGCCGCAAATCTAAAGGGATATGATATCCGTGACTGGTGAGTTGAAAACACATAAACTGAAAAGAAACCATATGTGTATTGCATTATGGGACAAACCTCAAAAAATGAGTCAAAACAGAATCTTAATTATAATTTTGCATATTTCCGAATTATATATTTATATCAGTAAGAAGGAAAACTATTCTGATTGATTTATGTTTATATAACTTATTCACGATTTATGTCTCAAATTGCTCTACTAATTTTTATTCCTGATTCTTTATCTGTATATATTAACAATATTCTTTTTTTATATATAGGATTGAATCAGACTTCATTCAGCATATTGTTAGCCAAGTTTCGTCCAAATTATGCAAGACTTCAGTATCTTATTTGCGAAATGTTGTGGGAATAGATACTCATTTAGAAAAAGTAAAATCCCTATTAGAAATGGAGATCAATGATGTTCGGATTGTTGGGATCTGGGGAATGGGTGGAGTTGGTAAAACGACAATAGCAAGGGCTATTTTTGATACAAACTCAAATCGATTTGATGGTGCTTGTTTCTTGCGGATATTAAGGAAAACAAGTGTGGAATGCATTCTTTGCAAAATATCCTTCTCTCAGAACTATTGAGGATACAAGGTAACCACGTGAATAATAAAGAGGACGGAAAGCACATGATAGCTCATAGACTTCGGTCTAAGAAGGTTTTAGTTGTGCTTGATGACATAGATCACAATGACCATTTGGATAACCTAGCTGGAGATCTTGATTGGTTTGGCAAAGGCAGTAGGATTATTGCAACAACTAGAGATAGACATTTGATGGGAAAGAATGATGTAGTATATAAAGTGACTGCACTACTTAAACATGATGCTATTAAATTGTTCCATCAATATGCTTTCAAAGAAGAAGTTCCAGGTGAATCTTTTGAGAAGCTCTCGTTGGAGGTCATTGATCATGCTAAAGGCCTTCCTTTAGCACTGAAAGTGTGGGGTTCTTTCTTACATAAGAGGGACATAATTGAGTGGAGAAGTGCTATAGAGCAAATGAAAATTAAATCCAAGTCAGAAATTGTTGAAAAACTTAAAATTAGCTACGATAGACTGGAGACCATACAACAAGACATATTTCTAGATATAGCATGCTTCTTACGAGGGGAGGGAAAAAAATACATCATGAAAATTCTTGAGAGTTGTTATTCTGGAGCAAATATTGAATTGAGTGTCCTAATTGACAAATCTCTTGTGTTCATCGACGACTATGATATGATTCAGATGCATGACTTAATACGAGATATGGGTAAATATATAGTGAATATGCAAAAGGATCCAGGTGAACGTAGCAGACTATGGGACATGAAGGATTTGGAAGAAGTGTTTGTCAACAATACAGTAAGTAGACTAAACAACACAATAATATTCAATTTCTAGTTTTCATTTATCAAAATTCATTTATGTTATGTTATCAAGCCACGAATTAGTCTAGTAGCAGCTTAATATGTTTAATCTAACTTTTATTCTTTGGATTAACCTGAAATTTGATCAATACTTCGTACAGAATAACTCAATTAATAATCTTTTTAGCTAATATTACCCAACTATTTATTTTGAGTAACATTTTGTAAATAAAAATTCTCAAATGGAAAAAATACAAGTAAGGAGGGGCAAAAAATACAAGCTAATTGTACAAAGCATTTCTCTTGAGCAACATATTTCTTGAGAAGGGCCTAACTAATATTGTACCCTATTTTTTTTTTAATGATTGCTTTTCAAGAACTTCAAGTGCAATTATAAATAATTCAAAACATTCCAAAAAAATTGCATGCAAGGAGTAAGAAAATAGAGATATTGATAAGCGTATTTTACATGACATCTGAAGTTATAATTTGTGGGTTGTGATCTTTTTATCAGGGGACAGTGGCAGTGGAAATAATCTGGAATATTTATATTCCAAAACTATGGTTAAGCAAAGAGGCAATGAAAAATATGAAAAGACTTAGGATATTACATATAACTGACTGTCACGGATGGTTCCAACGGAACCTCGATACAGATTATAGCTCCGGGAGTGATTCGGAAGACAATGTTTCCTATGATTCCAATTGCCATGATGGTTTCATTGAGTACCTGCCCAACAACTTGCGTTGGTTTACCTGGTATCGCTATCCTTGCAAGTCATTGCCAGAAAATTTTAAACCGCAAAGACTTGTTCATCTTCAACTCCAGGATAGTTTGTTGCATGAATTATGGACTAAAAGAAAGGTACAATTTTATTTGAAATACTTTTTATTTTTGCAGAATAACAAAACTCCCCTAAAATGTAGACATAAATTTGTGTTGTTTTCATTATGGTCTTTGTCATCATTTGTCCTCCCCACGAGTCTCGGGTTTGAGATCCACACCGCCACATGGCATTATTGAAATAATATATTTTAGTCAACTGTCAAGTTTAAGGTGTTTTAGTTATTCTACCTTTTATTTATAGTACTAGCTAATGACAACACTTTTACATAGTTTATGAACATGTATTTTCCATAAAAAAATAAAAACAATTTCTAATTTCCCACAAAATATTTTGTTGACTCTCATAATCTAAAGCAGTAAATTAAACAATTTCCTTGTGTGTTTTGAATAGATACTTTTGCCGTCTCTACGAAGGCTAGATCTCCGGTTCTCTAGAAGCTTGATGGCAACACCAGATTTCACCGGGATGCCAAATTTGGAGTATTTGGATTTGGGTGGATGCAAGAGTCTTAAAGAGGTTCACCATTCTCTGGGATGTTCCAGAAAACTCATCTCTTTAGAGTTGTATGGTTGTATAAACCTTGAGAGGTTTCCATGTGTAAACGTGGAATCTCTTGAAAAACTATATATAATGAGTTGCTGTAGTTTAGAGAAATTTCCAGAATTCCTTGGAAGACTGAAGTTGAAGTTAGACATTGGTGAGAGAGGAGGCTCCAAGATAAGTAGTGATTTGCCTACAGAGCTAGATTTGAGTAATTTCAAAAACCTTGTAGCTTTTCCAAGCAGCATTGGCATGTTGAAAAGCTTGGTGAAGTTATATCTATCGTATTGCTCTAAGTTTGAAAGCCTTTCAGAAGGGATAGGGGATTTAGAAAACTTGGAGGAGCTTCATGCCGACTATACTCTAATCTCACAACCTCCATCTTCCATTATCCGGCTGAACAAACTTAAATCATTGAGTTTTGCAAAAAAAAAAACAGGAGGAGGTCTCGTAGATGGAGTGTTCTTTGTTTTCCCTCACGTGAATGAAGGGTTACACTCATTGGAAAAGCTAGATCTCCGTTACTGCAATTTAACAGATGGAGGACTTCCGGAAGACATTGGATGTTTATCCTCATTGAAAGAGTTGCATCTCCAGGGAAATAATTTTGAGCATTTGCCTCTAAGCATAGCCCAACTTGGTTCTCTTCGATCCTTGGACTTATCAGAGTGCAGGAGGCTTAAAAAGTTTCTAGGTGTTAATGTGGCTGAAGGGTTACGCTCGTTGGAAAATCTGAATCTCAGTTCCTGCAATTTAAAAGAGGGAGTACTTCCCGAAGACATTGGATGCTTATCCTCTTTGAAAGAGTTGAATCTCCAGGGAAATAATTTTGAGTATTTGCCTCGAAGCATAGCCCAACTTGGTTCCCTTCGATCCTTGGATTTATCAGAGTGCAAGAGGCTTAAAGAGTTTCTAGGTGTTAATGTGGCTGAAGGGTTACGCTCGTTGGAAAATCTGAATCTCAGTTACTGCAATATAATAGATGGAGGACTTCCGGAAGACATTGGATGCTTATCCTCTTTGAAAGAGTTGAATCTCCAGGGAAATAATTTTGAGCATTTGCCTCGAAGCATAGCACAAGTTGGTGCTCTTCAAACCTTGCACTTATCAGATTGTCATATGGCTCTGAAAATTATTCCTAATTTACCACAGAGGGTGATATTCACGCCATTGCCATATGGCTCTGAAAAGTACAATGATTTGATTCCTATTACGTTTGCCCGTGCCATGTTTCAGAATATCTCTGATTCACATTCCTTGTCATTTAGAATGTCTACCATTAAGCATGCTGGGAATAAAATCCCTAGTTGGTTCCACCATCGGGGGACAGATAAAAGTGTATCAGTCTATTTGCCTGAAAATTGGTACACATGTGATAACTTCTTGGGATTTGTTGTATGTTATTATGACAGATTAGTTGAAACCACAGCTCACTTGATTCCCTTATGCAATCGGATGCCGTGGATGATCCAGAAGTTATCCGTAAGCAATCATTCAAAGTGCTATAAATCTTATCTTGATGAACGATCTACTATTCATTTATTCTTTGTACCTCTTGCTGGCGTATGGGATACATCTAAGGCAAATGGAAAAACACCAAATGACTATGGATTTATTGGTGTAACTTTTTCTGGACAAATTAAGAATTACGGATTTCATTTGTTGTATAAAGATGAACTAGAGCATGAGGCGTTGTTACAAATGAGGGGAGATAATGATGAACCAACAGAGCACTCCATTTGGACAAGGAGGAACAGATATGATGATAGTGAACACCATAATGAAGCCAGTTGCTCGTCTTCTAAGAAACAAAGGTCTGGTACTACTGTTGATGTGCCAGTTAATCCAGAAACTTTGCAGCTCTTTCCGACAAGTCCTGGATTTTCGCTTAAATAAAGGTATATATAACTCATGATTTGGTAGTGAAATTGTGTTCTCTCTCTCTTTTTATGTAATTCTCTCCTGTTAGCTACAAATTAAAGTGTTCTTCCATAACCAATACATTTTTATTTTCATTCTTAGTCACCAAATTAGTGCCAGAAAGTGCAGGCAACACTAACAGTCCAAGACTGGGAATCTAAGGGGTTAAGAAGGAACTTTGCATGTTGAATATGAAATGGTAAGTAATAAGAGATGTAATTGCATCTTTATTGATCCTCAAGTAAAGCAACCTTCTGGAAATGTGTGTTTGAGAATCCCCGAGCTATTTTAATTCTCCTTAAAGTTTTGAATTAATATTCGTTCGATGGATATTTTGTCTCTGAATTAAGTTTCTAAGTTCCAAGACGAAATGAATATAGAAATGTGTATTAGAACGGATCAATTGGGGGAGTTAATTGAATGGATGGAGTAATTGATCCCTTGATCATGTCTTTCTGCAGATGCATACCCAACCGATATCGAGATTTCACTGTGTTATCACTTGAAGGTGTTATAAGTTCATAGAGAAAATGCAAAGTGTTATTCCCAAAGGTAGGATCCATGACTTACTACTCGTTCAGTTTGATGCTCTTGCTAATTGTTATATTCGGGGAAATTGCTGCTTTAAAAACAAAAACTTTATGTGGGATGCGTCAGTTTAAATACTTCTGCATCAAACATTTTCATGTCGAAAGGTATTGATCATGCTCTAACTTTAGCTATACACTTTTGTAAGTATTGCTCTGTCAGATTGTTATATTAGCAGAAATCTCTTTTTAAATGTAAATATCTCTTTTGTTATGGCAGGTAAGTTGGGAATAAGATGAAACAATCAAAGATGCAGTACAGCTTGAGACGGTGGGGGAAGCTGGAGTGAAGTTGAGGTTCGTCTCCCTCGTGATCTCTTTCAGTTCTCTTGTAATTCTTTTGTAGGAAAATGTGAAATATTGTGGATTAGGAATACAAGATATTTTAGGCATAATACATATATTGGACCTTAAACTTGGCTTCAAATTTTAACTTTGACCTCCAACTTTCAAAGTGCACAAATAGACACTTTAACTATCCAACCTTTTAATAAATAAACACGCGTGTCTTACCTGGCAAAACGCGTGATGTGCACGCATTTTGCGCGAGTGAGAAGTAATTTTTGAGGCACACAACAGTAAAAAAATGCTGAAATGACAATTCTACCCTTAATTAATTTTATTTTATTTTTGGTTCTAAAATGGACGTGTAAAGTGTTTAATTAGTTGGCAGCCATTGAGTGGCTCACTAATACACGTGGAAGCGTTTGCATTTCATGCGCTTTGCTCCAAAAATCACGTGTTTATTTATTAAAAGGTTGGATAGTTAAAGTGTCTGTTTGCGCACTATGAAAGTTGGAGGTCAAAGTTAAAATTTGAAGCCAAGTTTAAGGTCAAATATATGTATTATCCCTACTTTTTATCTTACTGTCTGCTCTCTAGCTTATTGACTTGTGATCTTTATGAAAGTAATAAGTAGCATCTGTATCTCGATTACTTATATAAGTGGTTTGTAACAAGTTTCACTCTTTTGGTTTTTGATTACTTCTGCAGTACAAACTGGGACCATGGTACTTTTAGAACAACGCTGGTAACTCCGCCTGTGAAGGACACATGTTTCCTCTGTTTGTAACAGAACAACTAGTAATTTGTTGTTTCAGAATATCTCATCATTGCGGCGTGACATTTCTGCTTCAGTCACTAAGAGTGTTTACCACAAGGGGGAATATTGCAAGTTGGTTCCACCATTCGGGAATTGGTAGAATTAATTGTATCAGTCAATTTGCCTGATAATCGGTACGTTAGTTAAAACTTCTTGGGATGACCCAGGAACTTGCCTAATCTTACCATTCAGAATGTGATACAAAGTTCTCCAACCATTCAGATTTTGAAACAGAATTATCCGACCATTCAGAATGTGATACAGAATTATCCGACCATTCAGAATTTGATATAGAATTTACTATTAATATTTTCTTTGGTACCTTTTGCTGTCTCCGCTCTATAAAACAAAGGTCACATGTGAATCTGCAGCCACAAGATTAGGCATAGTATGTAGAGCTGTTCTTAATCGACTCCTCATTTGATCCGATCTCCAAGGGAGAAAGTTAGAGGGAAAACGTCTGTATGGAGCTATGTTGCGCGGACTCTTCATTTTTGAAGGCGAACCCGTCTCGACGCGACACTGGTGCGGGTGCGGGTGCGGGGTGCGTGTCGGATTCGGTCAATCCGATCCGGACACTTTGACCAGAGCCGAGAAAAAATTGGGAAAGGAGATCAGTTTCCGGCGACCTCTTTAGAGAGGCGGCGGAGCTGTAAGAAGCTTAGAAAAAGTAAAAAGAAGAAGAAAGAGGAGTTTTTTCTTCATTGAAAAAGTAAAGAGAAGAAGAAAGAGGAGGAGTTTTTTCTTGAAAAAGTAAAAGTAAAGTCTTTTCAACTTTTCAACTTTTCAACTTTTCAACTTTTCTTGAAAAAGAGAACTAAAGGCTAAGGGTTTCTCTAGATAAGTGTGTC

>PGSC0003DMG400016882 LRR-RLPs
