## Supplementary material for "RLP/K enrichment sequencing; a novel method to identify receptor-like protein (*RLP*) and receptor-like kinase (*RLK*) genes": Note S2

>M6-PGSC0003DMG400009417 M6 C-LecRK

AACACAATTAGGAGTCCAAGCTGAATGAAAAGAGAAGAGAACACTGGAGCTTATTTTTTTATAGATTTTTTTTTTAAAGTTTCAGTGTTCTCTTCTCTTTTCGTTCAGCTTGGACAAGCTTAGTTTAATGCAAGTTTGGCTTTGTTTGTCAGAACTTGGTACTTTCATTTATCCTTCATACTTGAATCTAGCACACCTTCCCTCTCAACAAAGGAAAAGTGAAAACTGAGAACACACTGTCGATGTCAAACCTCCACTAATTTGAGTGATATCACACAATCCATAACACGGAGGCCAAGCAACGACTACTCCTATCAGGTTCATAAGACACAGGGGTAGAGGGCAATGACCTGGTGTCAACATCGTTCTTGGGCCTTCTTATTTCGTAAGCTTAGTTTGAGATGGCAGTGGAGGCCAATTGTGGTGAACCGTGAATCTTTCAATCTTCAAATCTTGGAGACCTTATCTGAAATGAATTCTTATTTTGAAGCTGATCCCTCAATTGTTAGCATAATCAAGTAAATAAATCCATTAATGAAGATAAAGATACTTTTAGGGTGAAAAGTTATCTTTAGTTGTGGTATTAGAAATAACTAGTTCAGAATACCCGAGCTTTATTGTCCATTATGAGGTATTTATTAGTTATAAACAAGTTCATGATACCTGAGTTTTACCTACCTAGATACAAACAACTATTCCTACCTAGCACTTTCCAGTCCCCGCAATCTTACTAGTATTATTTACAACTGAAGTCGATGTTCAAACTTAAGGAAACAACAGAAAGAAAAAACATTAAAACTACAAATATAGAAAGATGGTGAAAAGAATCAACAAATGTGTACAAGAGTTGGAGATCATGTGGCTAAATGGATGGGGTTAGTGCAGGAATGTCAAGTTTGAGCCACTTTTATAACATTAGGAACACATATAGACTGATAGAATAACATTAGGGGTATATATGTACTGATAGTTTAATGGAGGGTAAAATTAAACCTTTTCAATAGTACAGCTGTACAGGAAGTGAGGGACAAATATGACCCTTTTCCTAATTTCTTTTCTTCTTCTTAGATAAATGAAGAAGTTAGATCAAGTATTATTAAAACACTAGTGCAACTGAAACTTATTTGGACCTCTCAACATGGTTTTTCTTAGTGTATGTTTTTTATTTTTAATTGGTCAAGTTATTATGTCTCATCTCTTTGGCACTGTTGTTTATTGTTTCTCAAGCAGGAAGATGTGATCCTGGTTGGTTTAGCAGCCCAAATGGGAAAAAGTGTTTCAAATATAATGCGACCTCTTTGCAATGGAATGAATCAGAGACTTACTGCAATGGTTTTGGAGGAAACCTGGCTGCCATAACATCTTCAGAAGAACTGAACTTTGCCCAAAAGCTCTGTGGCAATGATAGTAACGGATGCTGGGTTGGCGGAAGGAGCATCAATACCACAGCTGGGTTGGGTTGGAAATGGTCTGATAATGCTTCTAGCTGGAATGAGAGTATTGCTCCTAAGACAACTTTGGACTCAAGTTGCAAAAATTTCTCATGCTATGGCTTTAAGTCGGTGGATTTGTGTACATCGATAACCAATGGTGCTATTATAGCTGAGAGATGCAATGTAACTCATGCTTCTATATGCACACTTGATGCTGGTAAGTGATTGCAGACCAATTTATTTCTAATTGTTTCTTTCACCCAGTGTAATTTTGAAAAGTAGGAGCAAAACTCTTTCCCTGATATAGTGTGGTTATAATATGTGATTAAAAGCCTCATAGTCCCCTTTTGTTTTTTCCATGAGAAGTGCATTTTAGCTAACCATTATACAGGCCTACATTGTTATCTGGATAAATCTATTACATTCCTTAATTGGTTTTTAAATAATCCTCTAAATTTTATATTAGGGAGCAAGTGTCGTCATATGCGTTGCCACAGGGAGTATCTCATTATTCTTGCAGTTGTGAGTGGGTTGATCCTTTGCACTACTTTGGCTATTGTGATATGGCTTCTTGTATTTCGGCGAAGTAAAAGACGCAAAAGATCATCGCGTGCATCATTAGCATTAGTTCCTCCGTCTTGGAAAATTTTCAATCGTGATGAAGTAAAGTCCATGACAAAGAATTTCAGTGAAGCAAATCGTCTCCTTGGGGATGCCAAAACAGGTGGTACCTATAGTGGGCTTTTACCTGATGGTTCAAGGGTGGCTGTGAAAAGATTGAAGAGGTGTGGTATTCAGAGGAAGAAAGAGTTCTACTCTGAGATTGGAAGAGTTGCGAGACTTCATCATTCAAATCTGGTGTCTATCAAAGGATGTTGCTATCATCATGGTGACCGCTATATTCTGTATGAGTTTATTATCAATGGACCCCTGGATAGGTGGCTACACCATATTCCTAGAGGAGGTAGGAGTTTGGACTGGAGCATGAGAATGAAGGTCGCCACCACCCTTGCTCAAGGGATTGCGTAAGTAGTGTTCCAACATTATCTTCATGTGCATTGTAGATTAGGAATAAATATTTTAACTTATCCGTGAGGTGGATTGATTTTAGTCAGAAATGTGAGTTTATTCTTTAAATTTTATCATACTTCCCTTCTTTGTCATGAGTATGAACTCATATATACATGTTTGGGTTGATGTCCAATGAAGATGTTATCGTGGAAGTCAAACAAGGGTGAAAATAGATGATAAGAATGCATTATCTATATTGTTAAGAAAAAAGGGTGTGTTATCTGTCTTGCAATATTCCTTTTCCTATTTTCTTTTTCCTTCCCTAGATTTTTCATTCAGAAGAAAGTCAAATTCTGCACATCTTGACCAAGATACTTCTTGTTAACATATAATGCAATTAATAAATGAAAACCTTTTTGGTTTACAACATCCGTTCCAGTACTAAGGAGAATTCACTTCCTAGGGATCTACCTATCTGCTTTTCCTACTCCAGCTAGATCTTACCATTGCCAAACGTGTTTTTGCAGCACTGGCCTTCTGGCTTCTCATGTTTCCTTTCAGGTTTGGGGTGGATCTTTCCTTTCTCAATTATCACATACTCGGTTAGGTCGCCCGTGTGTGTGTTTTTCAATTTAAGTGTTTGAACAGGAACTCTCTTCATGAATTTCACTAAAATATTGTAGGAAAGCTATGATTGATTTTCTATTCGGGGATTCATTATTAACTATTTATGACCCTATTTATTGGTGACAGAATTGTTGAACTTTTATCTAGGGAATATTTCCTTGACATGGAAATTATGGCAAAAATTTTTTTCAGGGGTTTGTGAGACTACTCACCACCTAACCCCCAACAACAAATATACATGGAAGAAATATTGAAGTGTACAAAAAGAAGCCAATATATTATTCCAATTATTCACGATGATATAGTGAAATAATTAGCATAAGATAGACCAAACCTTAGGCTTATCTACATCACCATATTTAATATATGCAATGTATTATGTATGGTCCCTGGTAGTTTTAACTTTTCATGGACTTCTTTGTTAAGAGGTATTATTTTCCATAATTTATTCAGTGTAAGTTGAAAAATATAGGCCACTTGAGTAAGCTGGGTTCATTTCATAATGTACATACTTTTAAAGATTTCGATAGTTTGCAAGTAGGTCTTTGTAATTTTTATATGCTACAAGGAATCCAGAGAGAAGCGAGTGAGAAAAGATGAAATGCTTAAGGAAATGGATACCAGCGACATCTTAGTAAGAAAATAAAAACTGCATATTACTTTCCTGTTGTCTTCATGAGCTTTTGTTCCAAGTGTCTTACCTTGTTCTAGGGCGGATGTCTATGGAAAGTGAAGTCATAGAAAATAGGATGTCTATGGAGGTCACTGTTATAAGCGAAGCAACCTAGATGCCTAATATATCGATGGATAGATACTTTAAGACCAGTCATGCCTATGGCATATTTGTTAAAAAAGGGAAAATAAACAGAGACAATTCAGGGTCTATTACTAGTTTATTACCTTTGTAATTTATGGAATTGAAGGATCTAGCATGTTAATCAACTGCATAAGGTGAAAATGAGTGGTCTGGTTGTGTTATGGTTGTATTTTTAACCATGAAAGGTGGTGATAGTTGTTAAAACTCTGTTTTGTGTCACTGTGTCAGCTAGTGCACAAGAGGAAGTGCGAGTGGACAAAAGCATGACATTGTTTGTTTGCTAATCTCTGTGATGTCAGTTTCTTTGCTTATATTGTTTAATGTTTACTTTCATCATATGTGACAGGTTTCTTCACGACAAAATGAAGCCACAAGTTGTGCATAGGGATATCCGTGCAAGCAATGTTCTTCTTGACGAAGACTTTGGCGCCCATCTAATGGGAGTTGGTCTGTCTAAGTTTGTACCATGGGAAATTATGCATGAGAGGAGAGTCATGGCAGGTGGTACTCATGGGTATCTTGCTCCGGAGTTTGTTTACAGAAATGAGCTTACTACAAAGAGTGACGTCTACAGCTTTGGAGTTCTCTTGCTTGAAATAATTAGTGGACGAAGACCAGCACAGGCAGTTGATTCTGTTGGATGGCAAAGTATATTTGAATGGGCCACTCCTCTGGTGCAGGCTCATCGATATGTTGAACTGTTGGATCCTCTCATATCATCTTCTTCCTCTTCTCAGATTCCTGAAGTTGGAGTCATCCAAAAAGTTGTTGATCTTGTTTATTCTTGTACACAACATGTACCATCAATGCGTCCGAGGATGTCACACGTCGTCCATCAGCTACAACAATTAGCTCAGCCCCCTATTGTAAAGTAAGTGGATAAGAATTATAGTTCAGCATTTCTTTTGACTGAAATCAGCCTGCAATCTTTTATCCTAGTTGCAAGGGCTAATTGTATTAGGCATTATTTTTGAACTTGACATGCAAGGCCTTGAGGCTTGAACCAGTCCATGATGTTTGAGATGTAATACAACAACTAATCTGTATCACTAAAGCCAGTATGCTTTTCATCTGCTGCTGCTTTGCATTGTTTACTTGTAACACCTGTATTTTTAAAATTTGTTTCTGTTTCCTGTGTTAAAATATCACTATGCATAACAACCAATTCAATAAATGTCCATGAAATTTT

>M6-PGSC0003DMG400000179 M6 G-LecRK

CAAATATTCTCTGAGCTTTTCAATGTTTCCTTATAAACCAAAATCTTTTTTTCTTTAGCTAAAGCAAAATTCATCACCATGATTCCACAGAAAGCAATTCTCATTTTACTTTTCATCACAATTTCTATGGCATTTGAGATCAGTTCAGAAACAGTAAAGTCAAGAATGCAAAATGTAAACATGACCCATGTAGCTGAACTCACTAAAGAGTCTGTTTCAGCTGAAAGAAAGTTGGGATCTTTAGAAGGGAGCATAAAGTTTGAATCTTTGAACAAATCTAGAGTTTTACTAAGTGGGAACTTCACTGTTTTGAGTGAAAACAAGACATTTGAAATGGGGTTTTTCAAGACTAATGATGAAAGTAAATGGTATTTAGGGATTTGGTTTGCTTCAATTCCTACCCCTACTTTTGTTTGGGTTGCTAATCGTGAGAAACCCATCAAGAATCCATTTTTAGCTACTTTGGAGTTTAGTGAGGATGGGAGATTAGTTTTGAAGGAAGATTCAAGAACCATTGTATGGGAAACAACCAATTTGGAGAAAGCTAGTGATGTGAAGCTTTTGGACCAAGGGAATTTGGTTCTTGTTTCTAATAAAGGTGTTTTGGTATGGGAAAGTTTTGATTTTCCTACAGATACATGGCTTCCTGGGATGAACTTGACTGCAACAAAATGGCTGACATCTTGGAGAAGTACTAATGATCCATCACCTGGAAGGTACTCTTTAAGGCTCCAGCCACATAGTTATGGTGAGATAGTTCTTTTTTATAATGGTACTTATCCATATTGGTCCACTGGGAAATGGAGTGAAAATGCATTTGTTGATGTACCAGAAATGACTGTTCCTTACATTTACAAGTTCAATTTTGAATCACCTTTTACCCCTATGGCATCATTTGGATACTCAGAGGTGACATTAGAGAATGGAATGCCTCCTCCTTTAACTAGGTTCATGGTTGATTTCACTGGACAGGTTAAACAGTTCACTTGGATCCAACAGGCGCAAAGTTGGAACATGTTTTGGTCACAGCCAGAGAATATGTGTAGTACTTATGGTCTTTGTGGGAATTTGGGGTTCTGTAATAGTAAGTCATTGAATCCATGTAAGTGTTTACCAGGGTTCACCCCTTTGGATGGTGATTCATGGGATGCAGGGGATTTTTCGGGTGGCTGTCGCCATGAGAGTAATGAAGTTTGTGGTGAGAATGATAGGTTTGAAGAGGTTGGAATGGCTAAATATGATGGAGCAAGGGTAGTTTCAGTTTCTGGAACTAGAAATGAATGTGAGAGAGAATGTTTAGGTAATTGTTCATGTATTGGTTTGTATCGCAATGAGAAGACTAATTTGTGCAAGAGTTTATATGGGTCATTGCTCAATCTTAGGAATCTTACTTCTGATGGCACTGTTGAGGATAAGCTATATGTTAGAGTTCAAGGAGGAGGGAATACTCGAAAGAATCAGATTCAAGTCAGATTACTTGTGATCGAAATGATCTGTGGATTTGTGGTAATTCTGTTAGTTGGAATTGGGGCTTTCTTGGTAGTTAGGAGGAGAAGAGTAAGGAGGAAGAAAAAAGATGAGGAAGATGTATTTCCTATAATGAATTTGAAGGTATTTTCATACAAAGAGCTTAATGCTGCAACAAAGGGATTCTCGGAGAAGCTTGGTCATGGTGGTTTTGGAACCGTGTTCCTAGGGGAGCTATCGGATTCTTCACTCGTAGCTGTGAAGAGACTTGAAAGACCTGGTGGTGGTGAAAAGGAATTTCGAGCTGAGGTATGCACAATAGGGAACATTCAGCATGTTAATTTAGTGAGATTGAGGGGATTTTGCACTGAAAATTCTCATAGGCTTCTGGTTTATGAGTACATGTCAAAGGGTCCTCTTAGCGCATATTTAAGGCAAGATGGTCAGAATCTAAGTTGGGATGTTAGATTCCGAATTGCTGTAGGGACAGCAAGAGGTATTGCTTATTTACACGAAGAGTGTCGCAATTGTATAATACATTGCGACATTAAACCAGAGAATATTCTGCTACATGAAGATTTTTCAGCAAAGGTTTCTGATTTTGGACTTGCAAAGCTCCTTGGTAGGGACTTTAGTAGAGTGTTGGCAACCATGAGGGGTACCTGGGGCTATGTTGCACCCGAGTGGATCTCTGGATTAGCGATCACAACGAAAGCTGATGTATATAGCTATGGCATGACATTATTGGAACTGATAGGTGGTCGACGCAATGTAGAGTCACCGCCTTCAGCTAAAGGTGAAGAGGGCGGAACAGAGGAGAAGTGGTTTTTCCCACCGTGGGCAGCACGTCAAATAGTCGAAGGAAACATTGCTGCAGTTATGGATGAAAGACTACATGGAACGTACAACCTCACGGAGGCAGAACGTGTGGGATTGATAGCTATTTGGTGCATTCAGGACGATGAATCAACGAGGCCTTCCATGGGAATGGTGGTGAAAATGTTGGAAGGAGTTGTCGAGGTGACAATGCCTCAACCTCCGAAGCTGCTCCAAGCATTAGTATCTGGTGAATCATTCCATGGGGTAGGAATTGATTCAAGTAACGGGACATCACGAGACGGAGGTTTCAGCTCGGGTTATAATCCTCAACGTTCAATATATTCAAAAGACTCTCAAGCTTCAAGTTAGGTCTGCTTCTCAACTAGTATGCAAGTTGATTTTTCCCAGCTTCACTGAATGTAGAGTTCTTTTTTATTGTCTTCAAAAATCCCACTGAGAATTAGTAGCAAAATGCTCATCTTGGTATTTTGTAAAATGTATCATGAATATAGAAAATCTTGTTTACTTTGCTTAAAAATTGTCACCTTTTTTTTTTTTTGTAAAGTCCAAAGAACCAAAGCAAAACAAAACATACATTTCACTTTGGATCAGAGGGTTGGGGCGGAGCTAGTACATTACACTATTGATATATGGTTAAAATTATTTTTTTATGTATATATAATAGACATTGAACCCTTTAACCTCTTCGTGTGTTTACTTCTTCATATTTTGAACTCCTTAATGAAAATTGTGACTTGCATAGGATTATAGGAGAGGGCTTCCTAACCTTAGAATGAGACAATAGTTGAAAATATCAAAAGTTATAATAGCAGAATTATTTAATGGATATATCTTTATTGGTAGATATTTCGTATAGTGAACTTAAAATGAGATATAGGTGTATAATATAAAGCATTAAAACATGTGTTTGGATTATACCAACCAAACATTATGCAAAATTTTATATTGAGATTATTTTTGTTAATATGGTAAACCAAATGACTAATAAGAATATACCCTCTACCTAATTTAATGGGAGAGTTAATTTTATATTTTATAAATGAGTGTAACGTTTGTACCTGCCCAATTATAACCACACTTTACTTTCTAAGTTTAATTTACATAACTGTCTTGACTACAACAATAACAACTAACCACTTGTCAATAGTTTCCTCTTTTGTAGCTAAATAATATAACAGAAATTGAAACAAGAAAGACCATATACTCTCACACAAACAAGCAAACAGAAAAAGGAGAAATGTTTTCCTAACTTGTCAACTTTAGTAACAAAATTTTCCCAAAATTAATTTTTTCAAAAGAGTAAAGATTGAGGGGGTTAAAGAAAATGGTCAAAGGAGAGCAGAGGTAGACAACTCTGCACTCTCACCAAAGAAGTTTGTTACCTTGGAAATGGAGACACCTGGCAGTTCAGGTGGGGAGGTGCCGATATATGGGGGCTCCTCGTCTATGGAGCCGACTGAGTTACACTCGGCTCCATCTACCATGCTGGCGGATTATCCGCCAGTACAAGGCTTTCAAGATTTGAAGAACTTGGTTAGCATGGAATCTGGCAAGTTATGGACTATTGCAGGTCCCATAGCCTTTAATATTCTGTGTAATTATGGGATTAATTCTTTCACAAGTATATTTGTTGGACATATTGGTGATGTAGAACTCTCTGCCGTTGCCATATCTTTATCGGTGATAGCAAATTTCTCATTCGGATTCTTGGTTAGGTTCTTTTTTTGTTCTTACTAATTCTCAAAGTTCTGTTCTTTTTTATATTATGTTATATGGGCATTGAGTATTCGTATAGCCGACCCCAACTTGCTTGAAATAGCGGGATAGTTGTTGTTATTGTTGTGATTGTTTGTAAAATGTAATACATCGAAGATATGTTGGATTTTGTCAGACTCTTTGATGAGACTTATGACTATAGATCACAAATAGTTTATTATGAAACAGAACTAGGATTTTGATTGTCAAATGATATAAAGTCTCAAAAAGGTTAGAATAAGCATTCTTTTAGGTTATTGGTTATGTTAGTTTTCTAATTCTTTATCTTATTTTTGATGGATTATCAGTTAGGCATGGGTAGTGCACTTGAGACACTTTGTGGACAAGCATTTGGTGCAGGCCAAATAGAATTGTTAGGAGTTTACTTGCAGAGGTCATGGATAATACTTGTTGCTTCTTGCTTCTGCATAATGCCACTCTACATCTTCTCCACGCCGATACTGAAGTTATTAGGCCAAAGAGATGACATAGCAGAGTTAGCTGGAAAATTTTCGATACAAATCATACCTCAAATGTTCTCCCTTGCTATCAACTTCCCAACCCAAAAGTTCTTGCAAGCACAAAGCAATGTCGCGATCCTGGCATGGGTTGGGTTCATGGCTTTAGCCATGCATATCGGGGTGCTCTTTCTTTTTATCAAAGTTTTCCGATGGGGCGTTACTGGTGCTGCTGCAGCCTATGATGTTTCTGCTTGGGCTATTGCTCTGGCTCAAGTGGTTTATATTGTTGGCTGGTGCAAAGATAGTTGGAAGGGGTTGTCTTGGTTGGCCTTAAAGGAACTTTGGCCCTTCGTAAAGCTTTCTGTTGCATCAGCAGTGATGATTTGCTTGGAGATCTGGTATTTTATGACTATAATTGTTCTAACTGGTCATCTTGAGGACCCCGTCATCGCTGTTGGATCTCTTTCTATCTGGTACAATCTTGCAAATATGCACTTAAAATTTATAATGTGTTCATAATTGTTCTGTTTTTCAAGCTTAGATTCTTTCTTCTTCATTGTTTCAGTATGAACCTTAATGGATGGGAAGGCATGTTGTTCATTGGAATTAATGCAGCAATAAGGTATACTACTATACTAGTTATAGCAGGACATTGTCAGTGCACAAAAGTTAAACTCTAACGAATAGTGTAACTGCTAACGGTTGTTTTCATTTGTTGATATTTCAGTGTTCGCGTTTCCAACGAGCTTGGATCAGGCCATCCTAGGGCTGCAAAGTATTCCGTCTTCGTCACAGTAGCTGAATCTCTCATGATTGGGATATTCTGCATGGTACTAATCATATTAACAAAGGACCATTTTGCATTGCTTTTCACAAGCAGTGAGAAAATGCAAAAAGCTGTTTCTAAGTTGGCATATCTTCTTGCTGTTACAATGCTGCTTAATAGTGTTCAGCCAGTTATATCAGGTACCAAAATATGTGTGTTTCTTACTTAAAACAGTTCTCCACACACACATCAGTTGATGTGCCAAACAAATAGTACTATTTGGTACAAATCTATGTTGCTCGGACTCGTCAAAAAATGTCAATGGGTGTGTGTCGGATTCTTGATAAGTAGAGTATTGTTGAAGAATCCGCGCAATCAAAAAGTTAAGAGTCTGCGCAACTTTAGGTACAAACTCAAAGAATAAATTAGATTGTTCCTAATATTGAAGGTGTTGCTGTTGGAGGAGGATGGCAAGCTCTAGTAGCATACATTAATTTGGCTTGCTACTATGTCATTGGCCTCCCACTTGGATTCCTCTTAGGCTATAAAACAAGTTTGGGAGTTCAGGTAACAACATAAATCTTTTTACTAGTATTAATTTAGCTCCGGTTTGATCATAGATTTTAGAGTAGATTAAAAAAAAAAAGGTATACTAACCTGTTCTTTTACTTGATCTCAACTAGACATCTATGCCCTCCAATTTTGAATCAAATACTTGAACTTATATGAAGTTGAACAAATAGACACATGCGTCCTATATGGCATAATGAAAATCAGTTTGGGTGTATCCCACATGAACAAATAGACACATGCGTCCTATATTGCATAATGAAAATCAGTTTGGGTGCATCTCACGTGAATTGGCACGTAGGACGCATTTATCTACTGGTACAACTTTAGACAAGTTTAAGTGTCTATTTGTGCACGCCCAAAGTTGAAGGGCAACTGAAGTCAAGTTAAAGAGCGTTTATGTATTATTTTGAACATTGTTTGTTCATGAAATTTGGTCAGTTTTCTAAAAATATTCAATTGAGCTAAGTTTTTGGATGAAATTTCACGTCTACTCACGAAACAAGTAAAATGCATGTCCAAACATAACTTCATATTCCAAAAACTATTGTTCAACACAACTTCAAAAACTCACTTTTTCAAGTTTCAACTAAATCTATGGTCAAAACTCAAACACTAGCTTGAGTTTGAGTTTTAGGAGGGAACTTTGAATTAACGATAAAGTTAATAAACTATTGATGTTTGCGTCCGGATAAATTGTCTACATTACACTAATTGTCACTTCATTTTTTTTCTCTTTTCTGTAGGGAATATGGATGGGCATGATTTTTGGAACTTTTTTGCAAACAATAATTCTCTGTGTTATCGTATACAAAACCAATTGGAACGAAGAGG

>M6-PGSC0003DMG400000915_ M6 G-LecRK

ATGGAGTTAATCTTATTCTTTCTCATTTTATCTTTGTCCGGTATTCCTGTCTTTGGTTTTTTATCAAAAGAATTTGTCTATCCCAACTTTACTGCTTCCAATTTCCAATTTGTCGATAGCACCGGAAGTTTCTTGGTTTCTCGCAATGGAACATTCAAAGCTTCTATCTTCAATCCCGGTCCTGAACAAGTTAATTTCTATTTGTGTGTTATCCATGCTGAATCCAATACCATTATCTGGTCTGCAAATGGTGATGCACCTATTTCAACTTCAGGAATTATGAGGTTGACCAAGAATGGTATTAGTATTACTGAAAAAGATGGCAGTTTTAAGTGGTCAACTCCACCATTAAAGTTAGCAGTATACGCGATGCAGCTAACTGAGACTGGAAATCTCCTTCTCTTTGATCAGTTTAATGGGACACTTTGGCAAAGCTTTGATCATCCGACAGATACAATTGTAATTGGACAGAAGTTGCGTGTTGGCATGATGCTTTCAAGTGCAATAGCAGGTGATGACTTGTCAAAGGGTCATTACAGGCTTTCTTTGTCTGCTTCCGATGCTATGTTACAGTGGCAAGGTCTGACATACTGGAAATTGTCCATGGAAACCAAAGCTTATACTAATTCAAATTATGCAGTGGAATACATGTCTGTCAATCAAACAGGTCTTTATCTTTTTGGTCAAGGTGGTTCTGTAGTTGTGATAATGGTGAACTTGTTACAATCCACTTTTAGAATTGCCAAATTGGATGGTTCAGGTCAGTTGATCGTTATTAGTTTTGTTGGTTCTGACATCAAACGAGATTTTGTTGGGCCAGTGGATGGGTGTAGAGTACCTTATGTTTGCGGAGGGCTTGGTGTATGCACCTCTGATGTATTATCAGATAATCCTAAATGTTCCTGTCCTGCTAACTTCAATCTTGGATCACATGATTCAAGAAGCTGTGTGCCTAGTGATAGTTCCTATTCTTTGCCAGTTTCTTGTAATTCCACCAACCACAGTAATCTGTCGAATTCTTCCTCCATGTCGTACATAGGACTTGGCTTTGGTATGGACTACTTCACTACTGATTTTACCAAGCCCTTCAGGGATGGGGTAAATTTGTCCGTTTGTCAAAATCTCTGCTCAGTAGACTGTTCATGCTTGGGGATATTTTATGCAAACTCGTCTGGTTCTTGTTACAAACTTGAAGATGAAATAGGGTCAATTATGTTAAGGACTAGTAGAAATGATGATCTGTTGGGATTTGTTAAGACCCTGGTAGGAGTTTCAACAACCTTAGGAAACAATGATAATTTTGATCAGGAAGCTGCAGACTTTCCTTTAGTTGCAACAGTGATCTTACCTTTCACAGGAATCTTCCTACTCATGGCGCTGGGTTTCATTATGTGGAGAAGATCTAGACCCCAAAGAACGGGAAAGATAGAATCCAAAAAGAGCCAACCCAATTCTCCTTTTTCAGAGGACCTGGATGCCTTCTCCATCCCAGGATTACCTGTACGGTTTGAGTACAAACAGCTTAAGGCTGCAACTGATAATTTCAAGACTCAGATCGGCACAGGAGGCTTTGGTGCTGTATATAAGGGTGTGCTCCCTGACAACACTCTGGTTGCAGTAAAGAAGATAATAAATGTAGGCATTCAAGGGCAGAGAGACTTTTGTACTGAGATTGCAATCATTGGCAATATTCACCACATAAACTTGGTCAAGTTAAAAGGTTTCTGTGCTCAAGAGAGACAACGCCTATTGGTTTATGAATTTATGAACCGTGGATCGCTTGATCGCACTCTTTTTGGTCATGTACCTGTATTGGAATGGCAAGAACGGGTTGAGATAGCTCTTGGTTCTGCCCGCGGACTTGCATACTTGCACAGTGGCTGTGAACAAAAGATTGTCCACTGTGATGTGAAGCCAGAGAACATTCTCTTGCACGATAATTTTCAGGCAAAAATATCTGATTTTGGCCTCTCCAAGCTGCTGAATCGTGAAGAATCCAGTCTATTCACAACTATGAGGGGGACTCGTGGCTACCTTGCTCCTGAATGGCTCACAAGTTCTGCAATCTCAGACAAAACGGATGTGTATAGCTTTGGAATGGTACTCCTTGAAATTGTAAGTGGAAGGAAAAACTGCTCTTTAAGAACACAAAGTCACAGCCTTGATGATAGTGCTACTGGAGATCACTCATCATCTTCATCTGCACAGGGACTTGTTTATTTCCCTTTATTTGCATTAGAGATGCATGAGCAAGGAAGGTACCTTGAACTTGCTGACCCAAAACTTGAAGGGCGGGTCAGTGGTGGAGATATTGAGAAGTTTGTGCGGGTAGCATTGTGCTGTGTCCATGAGGAGCCATGTCTCAGGCCAACTATGGTTAGTGTTGTTAGCATGTTAGAGGGTGAGATACCACCAACTGCGCCAAGAATGGAATCTCTGAACTTTTTGCGCTTCTACGGGCGACGTTTTTCAGAGGCATCAAACATGGAAGAAGCTGGAGGCCAGATTGATGCCATGTTATATCCGCAGGCAAATACTTCTCATACAACCTCTCGGAGCGTCTCTAATGCTTACTTCTCTTATATTTCGTCCCAACAAATCTCTGGTCCAAGATA

>M6-PGSC0003DMG400001059 M6 G-LecRK

AAAAATTTTATACTAGAACAATAGTTCTTTAATGGCATGTCTAAAAATCCTACTATTTTTCCTCTTCATGAACATTTCTTTTGCCCAAAAAAACACAATTTTTCAAGGCCAACACCTTAGAGTTGGAGAAAAATTGGAATCAGCAAACAAGGAATTCAGACTTGAGTTTTTTAGCCTTGATGCCAATAAAACACATTATATTGGCATATTCTACAATTTACCTTCTAACATGACCTTATTCCCCGATGATGATCGCCCCGTTTGGGTTGCAAATCGCGATATTCCGATACAATATGCATCAGGTAACACACTCACATTGGATGATGATGGAAAGTTGAAAATTTTCTATGGAGAGGATAGTTTTTTTATGTTAAGTTATTATAATGTAACAACAAGAAATACAAGTGCTACACTATTTGATAATGGAAATTTTGTGTTGGTGGAATTGAATACTAATGGTTCAGAGAATAAAACATTGTGGCAAAGTTTTGATTATCCTACTGATACACTATTACCTGGGATGAAACTTGGGAGAAATGTTAAAACAGGACAGATGTGGTCACTGGTTTCTTGGATAAGCAAAGATGTACCTGCCTCGGGGTCATTTACGCTTGGTATAGACATGAATGACATTGACCAATTGATAATATGGTGGATGGGAAGTGTGTTTTGGACGAGTGGACGCTGGGGAAATGAGACGTTTGGCAATGTATCGCGAGTATCACACTATGAGTATGTTAACTTGAGTTTTGTTTCGACAGAGGATGAGAAGTATGTGACTTATTCTGTCAGTGAGACTCGAACTTTGTCTAGATATGTAATGGATACATTTGGTTTTATTAAGGAAAGAGGAGCAGCAGGACCTTTTGGGGTGTGTTTTTACAAGCCTAGTGCTGGTTGTGTGACAGAAGAATCGATTGGGTGCTCGATCAGAAACGATTCGTGGTTCAAGAGGAGTCCAAATAGTGTCTCTGGTAATAGGTTTAGGTTTGAAGACAACCACATGAGCTTATTTGATTGCAAGAGACAGTGTGAGAAAAACTGTTCTTGTGCTGCTTTTGCCTCTATCACAGCTAATGGAACTGGTTGTGATATTTGGAGTAACGTATCGATTTTGAACTCACTTGAATCGGATGTTTTCATTCTTGGTGATGAAAGAGGTATGTTCATATTTATCTCAAATATCCTTTTTATGTTGTTGTCGTGTGACCAGGAGGTCACAAACTCACGGGATCCAGCTATGGAAACTGCCTCTTGCAGAAATGCAAGGTAAGGCTGCGTACAGTTGACTCTTGTGGTCCAGGGAAGTGCACTATTTCTATTGTCTAGCAAGTAACTTGAGTTAGTAAAGCGGACTCTAACGATGAATTGTGCAATGTGTATTCATAGATTTGGTGCCAATAAGAAATCGGAACAGCACAGATGTACCAAGCACAATCTCACTTCCTCCAGATTTAGCACCAGAACCATCTCCTTATTCAACGTCAGGATCATCTAAAGGAACTTGTATGTATTGTGTATAATCTTGCTCTTGAATTATTGTATATGATTGAAGGTGAAAACCTTTTTTTGATTGTTTCCTCGATATAGTCCATTTAGTAAAGAACTATACGTTTTAGGCACGTATTCTTTTGTAGTATCATCATTACACAATCTAGTCCTTGTTTCGAGTATATCCAGAAAAAGATCATTTACATCATCTGTATTGATGTTGTTACTGCTCCTTTCCTGAATGTTCTACACTGTTTTCCTTTTTACTTGTCATGTTTTCTGCACTGTCGTTTTTCTCCGTTTCCATAACTACTATGATTTGTTGCTCTTGAGCCAAGGGTCCTTCAAAAACAGACTCTCTACCTTCACATGGTAGGGGTAAGGTCTGTGTATACTCTACCCTCTCCAGACTCCACTTGTGGAATTTCACTCGTTATGTTGTTTTGTTGTTGATGCAGCAAGAAAATGGTGGGTATGGCTTATAGCTGCAAGCGGTTTGACACCATTTGTAGGATTTTTCGCCCTATGCTACTTTCTACACGGAAAAGGTGAGCGCTCAAAGAATTTATCTAGAAGCTTGCACCAGTAAAACTCAACTATGTTGCTCGGACTCTGCTAAAATGCTGCCACAGTACTCCTGTCGGATCCTCAAAAAAAAAAACGCACTACTTTTGAAGGATCTAACACACACTAAACAACATTTTTTAAAAGTCCGAGCAATATAGAAATGCTCTATACTATGGTTTCTGATCCCACAACTTCCTCTTTCACTATCAGGGAAAGCAAAAGCAACAGCTTTGCTATTGCTTAATCAAACAGCAAATATATCTAAAAAGAGAAAAATAGACAAGAAAATGAGCCATGAAGTGCAGTTATACAGCCTCGAGAGCCTAGCAATCGCCACAGACAATTTTTCACCAGGAAATAAGCTCGGGGAAGGTGGATTCGGACCAGTTTACAAGGTAAACATGAAACCTTTTATATGAATATGATTCGCGTAAGCATAAATACTTAAACGAGTCATTTTTTTTCAGGGAGAAATGCCTGATGGGCAAGAAGTTGCCATAAAAAGACTTTCGACTAGTTCAGGACAAGGCCTATTGGAGTTCAAGAATGAAATTCTATTGATTGCAAAACTTCAACATACTAACCTTGTTAGGCTTCTAGGATACTGTACTCTACGCGAAGAACGGATTATAGTTTATGAATACATGCACAACAAAAGCCTAGACTTCTTCCTCTTTGGTAAACATCATGTTTTTCATCTTCAACAATTAGAACAACTTTGTCAAGAAATTTCATCTTTTACAATGATATGAACAGATAGTAACAAAAAGGAGCTACTAAATTGGGACACACGATTCAGAATCATCGAAGGGATTGCTCAGGGTATACTGTATCTGCATAAATATTCAAGGCTGAAAGTGATTCATAGAGATTTAAAAGCAAGCAACATCTTACTTGACGCGGAAATGAATCCAAAGATATCAGACTTTGGCATGGCGAGGATATTTGGAACACAAGAATCTGAAGCAAACACGAAAAGAATTGTTGGAACACAGTAAGTAAATAAGCAATGTTTTCATAAACAAGTTTTTATATAATATTACTTGAAGTTCTGAATACATTTTTCGGATACTACTTCAGTGGCTATATGTCTCCAGAGTATGCCTTGAGAGGCATTGTTTCAACGAAAACGGATGTATTCAGCTTTGGAGTTTTACTGTTAGAAATTGTTAGTGGCAAGAAAAACAACAGCTGCTATGATTCCGAACATCCACTAAACCTCATAGGACTGGTAAGTTTCTAGATTATATTATGAACATATCTCGCGAAATGGAGAAACATATAACTATTAGTACTATGTTTTTAGGCATGGGAATTGTGGAGAGAAGAGAGAGCTTTGGAGCTAATAGATGCAACATTAATTGAATCATGCTCGCGTGATGAAGTAATGAGATGCATACATGTGGGACTCCTTTGTGTGCAAGATTATGCAAAAGATAGGCCTTCAATGTCAAATGTTGTTTCAATGCTTGTGAATGATACGAAACAACCACCACCACCACCAGAACGACCAGGATTTTTCATCGAGAGAGGGGATCAACGTGCAGAGATCTCCGAAGAAGTAGTCAGATACTCGATAAACGGGCTATCAATATCAGAGTTGAGAGCAAGATAAGATACAAGATGGTCAATTGGGAAGTAACTTATTTAACATTCTTTTGATGAAGTGTAAAAAGTTACCAACTTCAAGTACATACTTATAGGACTAATTTCGAGTCCTGTTTCTTTGATACTGTTGCATTATGATGCATTTTGTGAGTAGAATATTTGAGATGGATCCAGAATTTAAGTCAAAGATGGTAAACGAGATAAGGCAAGGCGAAGTTATGGCAATGGAAACTTCACCATTACCTTCCCCCTCCCGCCTGATAGCAACGTCACCAGTTACATTTAAAATGTTTTGAATACACTTGACACTCATCGATTATATAAGTTGAAAGAGAATATGAGAGGAGTTTCTGAACTATCAATATTTGTATATTAGACAAAAGAGGTAACTGAATGTACATATATTTTCTCATTCTGAGCTTTACATGTTCCACAAATGGGCTGTGAATTAGTTTACACTCAATTCACTACAAAAATAAAGAACAATTTTGGTTTTCACCTAAACTCACTACAATTCTGATGATTTAGCTTCATCAATATTGTTATCTTTATCGCTAACAAAAACTAAACGATCACCATCTCTGATCAACTCAATTTCGTCGATTTCAGCTCCATCTTTGCTTACAACTTTGGCCTGAGAGATACCATATCTACTAGAACCAAGTTGAAGTAGCTCTTTAAAACTCTGTGGGAGTAAAATGAGTTTCCCCCCGTTGTCTCCTTTCTCAGGGCAACACACTGTCACTCGGGGAGCATAAGTCTTGGTTGTCATGACACTTACATTACTGTCGTTTGCAGATAAAAGCACGTCATGCTCATTGGTCTGCTTTGCTGACATTATCCCGAATAATGAGTTGTGGAAGTTATTAGACCTACGTCTAGGACGCGAACGCCCTAATGATCCATCTAGTCCTAGAAATGAGACTCCATGGGATGCAGGGGTGATTGTTGGTTCACTTTTGAACCTCCCAAGAAATCTAACCCCTCGTTCCTCGGGGATAGGATCAACGGATCGAGTTCTCATCATTACCCTGGATTCGAAGAGTTCTTTGATGTCTTCATGTCCTTGTTGCTCAGCAAGGTCCCGGGGGGTCCAATTATGTTCGTCTGGTTGATCAACGTTAGCCCCTCGATCCAAGAGGTACTTTACTATTTCTATGTTATCTTCACAAACAGCAACATGGAGAGCTGATGAACCGTTGATCTTGGGAAGCGTGACATCCCCGCCATGACGGACAATCTCCTTGAGCAAATTCAAGTTGTTTTGTTCAACAGCAACGCAGGCGAAATGTCCCACATCACCAGCTGATAGTTTAGCACCGTTGTCAACGAGTAACTGGATAACTGACTCGTGCTTCCCCATGATAGCCTCCCACAATGGCACATTGCCTTCTGAATCTGTGTATACACAAGAAACAAAATCAAATTCTTATATCTGGCTTTGTGTCAGTGGACATGGTAGGAAAAGGAGACAACTGATAGTTAGAGTGTGAAACTCGCAAAAAAAAAAACGATAGTTGAGGTGTGTTTTTGACTATAAACTCTCTAATGTTTGGACCAAAACAAACTTTTGGCAACAACAGAATTTGAAAAAAGAAATGGAGCATACCTCTACTATTGGGATCAGCCCCAAAATCGAGCAGAAGAACCACACAGTTCTCAATGCCTTTAGACGCAGCGATA

>M6-PGSC0003DMG400002017 M6 G-LecRK

AATTCTGCTCACTAAGGCTTAGGTAGTTGTGAAAGAACACCTATGATGGTTTTGCTTCCATTGGAATCTGAACTTGAACCTCATGTTTTGGGCTTCATGTATTAGTTTTTTTCTTATCGAGTTGCCTATTGTCATGTAGATAAGAGGAGGATGCTAACGGGAGCCCATATTGCAATTATTGTCTCCGCGACATTCAGCCTGATTGTTATTGTTTGCACTATATATGTATTCAAGATCCTGAAAAAAAGGTAAAAAAAAAAAGTCAATTTTTACAATTCTTAGAGGCACAACTTTTCGCTATCTTTTGGAAAGACTTTAATTTTTACTAGATTTTATAAAGCATTGGGAGTTTAGTCCATTTTTGCATCGCTTCAAAACACTCTTCCGCAACAAATTTGCTTCATAACAGGTTTACTCAAGGTAACAGTTTAGAATTACCAGTTTTCCGCTTCAAGTGCATAGAAACTGCTACTAAGAACTTCTGCTAACGAAATAAACTTGGAGAGGGGAGATTTGGACCTGTTTACAAGGTATATATATATCACTCGTGTGCTCTAATCTTCGAACTTCACTAGAAAAAGTAGAGAAAATTTCACATAGTTTATACTCATTTGATTATTAGTTGAAGTCATGTGTAATTCATATATAGGGGATATTAGCTGATGGACAAGAGGTTGCTATTAAACAACTTTGGAGGAGCTTAGGACAAGACATTGAAGTGTTCAAGAATGAGGTTACACTAATTTCCGAACTTCAACACAGAAACTTAGTCAAACTGATGGGATTTTGCATTAACGGGGAGGAAAAGTTGATGATCTATGAATACATGTCCAACAGAAGCTTAGATGTTTTCATATTGGGTGAGCTAATTAAGAAGAGTACTTAAACAGTACTTTTCTTGATGAATGATGGTTAAATTACTCTTTGTGCTTGGAAGGAGGATCAAGATCTAATGATCAGGCGGCATTGAGTTGGAATATCGTTGAAGGGATAGCTAGAGGTCTTCTTAATCTGCATCGTGATTTCTGACTAAAAATCATTCACAGAGATCTTAAGACGAGCAATATTCTGTTAGACGAAGATGTGAATCCCAAAATCTCAGATTTTGGAATGGCTAGGATTTACGGAGGGAATCAAACTATAGCAAATACCAACAGGATTGCCGGTACCTAGTAGGTATTAATTGTTCATAGCAGTGAAATTGATCCTAACTAAATTGTATAATTGCATATCAATGCAGTGGTTACATGTCTCCTGCATATGCAATACATGGAATTTTTTCCGAGAAATTAGATGTCTTCAGTTTTGGTGTCATATTGTTAGAGATTATCAGCAGTAAAAGAAACAACAACTATTGTCACTCTGAAGACTCTCTCAACCTCCCGGACCAGGTAAAAACTGAAAAACAGAAAGAATTTCTCATTTTCTCATATATAGTCAAACCACTTTATAATAACGTTATCTATAGTGAAAATGTAGTATAGAGAACGTATAATATAACATAATATGGAAGTTCAAGAAAAACTTGGTTGAGTATGATTGTTATCGATAGGTCTGAATGGAGTTGAATATTCTTCTATCCTATACTACTACTATTTTCAAATGACAAAACTACTATAATTGGATCTTAGGCATGTCAACTTTGGAATGAAGGCAAGATACTCGAGTTTCCAAATCCAAATGTGTTTGTTTCACTTGAAACACCAACACAAGTTGTTAGATGCATCATTCTGATTGGTGGGATTGCTTTGTGTGCAAGAACATGCAAGTTACAGGCCAAATACAGCTACAATTGTTTCCATGCTCTGCACTGACACAGTTCTTCCTTCGTCTAAGCGATTGGCATATAGCAGCAAAGGCTCTTGTGATCCAAATATTTCACCTTGCTCTAGAAATTCCATTTCTATTTCAAATGTTCAAGCATGTCAA

>M6-PGSC0003DMG400003313 M6 G-LecRK

TTCAAAATTACATATTCCGAAACTTTTGTACGAATTTTGATTTCATACTATAACATGAATATAAAATTTTCAATTCTTCTTTTACCTTTGCTTGTTGTTTCATCTTATTGTTCAACAATCTTTAGTTTATCATCTGAAGGTTCACTTTCTGCTCCACAAGATTTAATTTCGTCTTGAAATGGAAAATTCACTGCTGGTTTCTACTCTGTTGGTGATAATGCTTATATCTTTGCTATATGGTTTACTAATTCATTGGCTGATGGAACCAACACAATTGTTTGGATGGCTAATCGTGATCAACCGATAAACGGACAAAAATCACATCTGTCGCTGTTTAAATCAGGCAATCTTGTACTGATTGACGCGAATCAGATCAATGTTTGGGAATCTGGTACACAATCATCAACTTCTTCTGTTGAGTTAAGATTGTTGGATAATGGAAATCTTGTTCTTGTAACTTATGAAGGTCAAAGATTATGGCAAAGCTTTGATTCACCAACAGATACACTTCTTCCTGAACAACCACTTACGAAAACCTCAAAGCTTGTATCGCGTAGAAGCTTGAATAATTTCTCCTCTGGATTCTATCAGTTACATTTTAATGAAGATAATGTCCTGCATCTTGTCTTTGATGGAATAGAGATGACAAGTGTGTTTTGGCCAAGCCCGTGGTTAATCGTTTGGGATGCAGGTTAGTAGAGACGGATTCAGAATTTAAATTTGATTCGTTAAATCCTTAGTTTGTTGTCACTCAACTCACATATATTATTTTTGCATGAAAAATACTAGTTTTAGTTGAACTCATTATATGGTTCATTTGAGTGTATTGGTTAGGAAAGACGGATTCAGGATTTAAATTTCGTTGGTTCAATATTTAGTACAACTCATTTTTGAAGTTTTCGGTTTTAACTGAACTCATTGCTGATATACTCCATCCATTTCTGCAGGTCGATCCACTTATAATGACAGCAAAACAGCAGTTCTTGATCGTTTTGGGAATTTTACATCGAGTGATCGATTTGGATTCCAATCTGCTGATTATGGTTTGGAATTGCAAAGAAGATTCAGCCTTGACTTAGATGGAAACCTTCGATTGTACAGCCTAGATAAGCTGAGCAACATTTGGAAAGTTTCTTGGCAGTTATTTCCAGCAGCCTGCAGGATTCATGGAGTTTGTGGATTGAACAGTTTATGTTCCTATGATCCTGTCTCTGGCAGAAAATGTTCTTGTGTACCAGGATACAGGATGAAAAATCCTAAAGATTGGTCCTATGGTTGTGAACCAGAATTCGATATCTCTTGCAACGATACAAGCTCGTTGGACTTCATTCCACTTCGCCATGTTGAGTTTTACGGGTATGATATTGCATATTTTCGCAATAAAACATTGCAAGAATGTAAAGATTTGTGCTTGAAACATTGTGATTGTAAAGGTTTTGAATACAAGTATGTTGAGGGTAATGGCACCTATGGTTGTTACCCGAAAACACTTCTGTTCAATGGCTATGTTCAAGCTAGTTGGCCTGATTTTGTTTACCTTAAAGTGCCTAAAGCACGACGTTCCTGGCAGGAGCAGTATGAAGGAAATCTTCAATGTGACGATAAAAAGGTGATGTTGGATAGAGCTTATAGAAGGAAAGAACATGGTGGATGGATGAAGCCATTCATTTGGTCAGTTGTTGTAGGTGGAGTCTTTGAAATTCTCTGTGTTCTCACTTACTTTATTAAAACGAGAAAAAGCTCGAATGAAACCACACAAGGCTACCTTCATCTTTCGACAAGATTCAAGAAATTCACCTATGCTGAGCTTAAAAAGGCCACTTCCAATTTCAGTGAAGAGATAGGTCGAGGAGGTGGTGGTATTGTGTACAAAGGGAAGTTGTCTGATGACAGAGTTGCAGCTATAAAGTCTCTCTATGGAGCTAACTATCAAGGAGAAGCAGAATTTCTAGCTGAAGTGAGCACTATAGGCAACGTTAATCACATGAATTTGATCGAAATTTGGGGATATTGTGCTGAGGGAAAGCACAGGCTTTTAGTTTATGAGTACATGGAGTATGGTTCTTTGTCAGATAATTTGAATGCCAACAATCTTGATTGGGAAAAATGGTTTGAAATCGCGTTGGGAACGGCTAAAGGACTTGCATACTTGCATGAAGAATGCTTAGAATGGGTTTTGCATTGTGATGTAAAGCCTCAGAACATACTTTTAGATTCCAATTACAAGCCAAAAGTTGCTGACTTTGGGCTTTCAAAGATATTTAACAGAGGTGGCTTGGATCATTCAAACTTTTCAACGATAAGGGGGACGAGAGGATACATGGCTCCAGAATGGGTATTCAAGATGCCTATTACATCCAAAGTCGATGTCTATAGTTATGGCATTGTTTTATTAGAGATGATCACAGGTAAGAGTCCAGAAGTTTGTGCTGATGGTGGCTCGGGGGATGATGATTCTATGGGACTAGGATTGTTGGTCACTTGGATTAGAGAAAAAATGCGCGAAAACAGTGAAAGAAAATCATGGATTCAACAAATTGTGGATCCTGCATTGAATGGTAAATTCGATTTGAAAAAAATGGAGATCTTATTAGAGTTAGCTTTGCAATGTTCGGAAGAAGATAGAGATGCAAGACCTACCATGTGTGAGGTGGTGGATAAAATGCTTCATCCAGAAAATTTCAAGTTAAAATTAGACATGTTAATGTAGTTTCTCAAGATGTAAAATCTTGCCAAAGGTGATATAGTACAAGCTTATATGTAATGTGTCTTGAGTAGCAAGTACTTTTGAATTCCAAAATGTAAATCCTTATAGAATGACATAGTTTCTTAACAAAATTGTGCATTATAGCAACTGTTATGGTCTTGTGCATCAGTGATAGATGGAAAGTGTGTATGAGTACAT

>M6-PGSC0003DMG400004928 M6 G-LecRK

GGATCATTCACTATGGAAATTGGAATGGTATCAACCAGCAGAAAGCAAATGAAAAATTTTCTGCTTATGTTGCACATATTGTTATGTGGTTTTGTATGCTCAATTTCTAGTGCAAAAGATACCATCACTACAACTGACTTTCTAATAGAGGGTAAGACCATTGTTTCATGTGATGAAAGCTTTGAATTGGGATTTTTTTCCCCAGGTAGTTTCGCGAATAATTGGTATATTGGAATATGGTACAAGCACGACGATGGGCCTGACAAGTCTGTTGTATGGGTTGCCAATAGAGAAACTCCACTCAACAATACATCAGGTGTTATGCTGAAAATCGTTGATCCAGGACGACTTGCTCTTGTTACTTCTGCTAATATGAGCATATGGTCTACTAATTCGTCAAGGATTTTGTCTGTGAAGAATCCAATTGCGCAGTTGTTGAGTTCAGGGAATCTAATCATAAGAGATGCAAATGAAACTAAAACAGAGAACTTCCTGTGGCAGAGTTTTGATTATCCGACTAATACGCTCTTACCAGGCATGCAACTTGGTAAGAACTTTGTGACTGGACAAGAATTTTACCTGTCCTCATGGAAGAACGAATATGATCCGGCTCCAGGGAAATATACGTATCATTGTGATCCTACTGGATATCCTCAGGTTGTCATGAGGAAAGGTAAAGTTAAGGTGTTCAGTGGCGGACCATGGAATGGACTCCGCTGGAGTGGTGTACTAGGATTGACAAAAAATACAATTTATACATTTAAGCTGGATTTTGATGAAAAGAAAGCAATTTACAGCTACGCGCTAGTGGATAATTCTGTTTTATCGAAATTAACCTTGAACAGTCATGGTATGCTTAAGCGTTGGACGTGGGATGAGAAGATAAAGGAATGGCATGTTTACCTGGCTTCACCAGCAGATGCCTGTGAAAATTATGGAGCATGTGGTGCCTATGGTAGTTGCAACATCATATTTTCTCCTGTTTGTCGTTGTTTAGACAAGTTTGTGCCTAAAGATCTGAAAAATTGGTCAGTGACAAAGTGGTCCGGCGGGTGTATTAGAAGGACACCCCTTAATTGCTAGAATGGAGATGGATTTCTCAAGTACTCTGTTATAAAGCTTCCAGATACACAGTATTCACGGTTTGATGTGAGAATGACACTCCGAGAGTGCGAGAAAGTTTGCCTAAAGAATTGTTCTTGTATGGCTTACTCAAATCTTGATATACGCAATGGTGGCAGTGGCTGTTTGCTTTGGTATGGCGACTTAATTGACATTCGAGAATTACCTGGAGGACAGGACATTTACATAAGGATAGCTAACTCAGAGTTAGGTAAAATTTATGCAACAAATTCATTACTATTTGATCAATATGAAGCATTAGCTGAATGAAGTAAACTTCATAATTCTTAGCTTTCGTTGTGTTTTGCTTATCTATTTGTTAAAAAGTTCACATAGCATATATTTATTTTAGTTTACCCTCCATTTTTTTTCAGGCTCGAAGAAGAAAATGAAAACACTTGTGCTGAGTCTATCATTGTTGAGCGGAGTGTCTGTCATTGGCCTGATCATAGGGCTGTATATTTTGATAAAGGCGAGAAAGAAGAAGAGGAAGATGATTCTGAAAGATGATCTAGAACTTCCACTGTTTTCCCTATCAACAGTAACTAAAGCGACCTCTAATTTCTCAGACAAGAACATGCTCGGAGAAGGTGGATTTGGATCTGTTTACAAGGTAGCATATCTTGTAGGTTTATTATATGTACTATAATTGACAGTAAACAGATAAGTAACACTTTGAATTATCCCTGTCAAAACATCCTTAGCTTCAATATGAGTCAGCTGTATGTTATGATATCGTAGGGAATCCTGGAGGAAGGACAAGAAGTAGCTATAAAGAGGCTCTCAAAATCTTCCTTGCAAGGAGTCAACGAGTTCAAGAATGAAGTTAATTGCATTGCCAAACTTCAGCATCGAAACCTTGTGATGCTTATAGGTTGCTGCGTTGAAGGAGAAGAGAAAATTCTGGTCTATGAATACATGCCCAATAGAAGTCTTGATTTCTTCATATTTGGTATGCTTACTCTTGCTTAACACATAAAACCTCAAGTACTGTTCATAGTATAGCATTGAGCTTGCAGCTTTCACCATGATTTTGAAAAAGGTTGTTAATTTCAAATGGAAGTAGATTCTGTTTAACATACTAAAAGCAGTTAGAGCATCAGTTACTATACTAATTGTTGGAAGATCTGTTAACAGATGAAGGGAAGAGCTCGATACTCGATTGGCCTAAGCGCTTCAACATCATCAATGGAATTGTTAGAGGACTTTTGTATCTCCATCAAGATTCTCGGCTAAGAATTATCCATAGAGACCTCAAAGCTAGTAACATTTTGCTAGATGCTGACATGAACCCCAAGATCTCAGATTTTGGCATAGCTAGAAGTGTGATAGGGAATGAGACGGGTGCTAATACACATCATGTGGTTGGGACACAGTAAGTGATTTCACACTGCGACCTTTTAGCATTTATCCCATGTACTACTCATATAATATCATCTGTCTAACACTATTTTCTGTCTAATATGTGATGCAGTGGCTACATGTCCCCAGAATATCTAGTACACGGTGTTTTCTCAATAAAATCAGATGTATTTAGCTTTGGCGTGTTAGTTCTAGAGATAATCAGTGGTCGAAGAAACAGAGGATTTGCCCGTGAAAGTCACAGTATCAACCTTCTCGGACATGTAAGTCTTGTATGTTAAAAAGTACTTACATATTCAGACCACTCGTAAGGTAATTACAAGTGATAGTGTAACTAAACTGATAGTGCAAAAAGTCTTACTAAGTGTCGGTGTATACAACTTAAATCTATCAAACATGCTTCACTTACTTATACATGTTCCTTATTACTATATACTGATTGTGTTATTAACAGGTATGGAAACTTTATAAAGAAGGTAGACCATTGGAGCTGATTGATGTGCATATTACAGATTCATGTTATTTTACTGAAGTGCTACGGTTAATTCATGTCGCTCTATTATGCGTACAACATAGTCCAGAAGACAGGCCAGACATGTCTGAAGTGGTTGTGATGTTGGCTAATGATACCATTTTGCCTGAAGCTAAGGAGCCTGGTTTTTTCACAGAAAGTAAATTCCCTGATTCTGAATATTCTACAAGCATGTATTCGAAAAATGAAATCACTATTACACAGTTAGATCCCCGGTAAATTAAATTTCATTTTCTTGTAGATCTTTAAGTGTCTAGAAGCTTGAATAACAGAGGAGAACATTTTATGAGTTGTATCATATTAGGAAAGAATTTACCTAACTCTTGAATTAAAAAAAGACAGTCCGGTGCACTAAGTTCCCAACATTCATGGATTCGGGTCAGAATTGCACACGGGTCTATTGTACGACAGTTTTACAGGAACTTGAGTGTTAGTTACAAATTGCCATTGTTGTGTATGTTACATGGTTTTGTTATACTTACAGATACGTATCCACTCTGGGTGTGTGTAAGAATCTAATATGAATACTACAGTTTTTTTTGAAATTTTAATTTACTCTCCA

>M6-PGSC0003DMG400004930 M6 G-LecRK

ATTAAAAGGCTTAGGCATCAAATACATTGATGGAATCTAAATACTTCATACTTTTCTTACTATCTTTCTACTCTTTTATACCGAAGATTAGTTCTACTACGACAAATGTAATCAATACTAATCGGTTCATAACAGATGGTGACACTATTGTTTCATCTGGTGGAACCTTTGAGCTGGGATTTTTCAGCCCCAATGGTTCCACAACGCGATATATTGGGATATGGTACAAGCAAATTCTTCCTTATGTCCAAACAATTGTATGGGTTGCAAACAGAGAGAAACCACTCACCAATACATCTTCAGTCGTTTTGAAGGTCAACAAGCCGGGAATACTTGCTCTTCTTAATGACAAGAATGAAACTATATGGTCCACGAACACCTCACGATCAGTCCAAAATCCAGTTGCAGTGTTGTTGGATTCTGGTAATCTTGTCCTAAAAGATGCAAATGATGACAATCCAGAAAATTTCCTATGGCAGAGTTTCAATTTTCCAACTGATACTTTCTTGCCTGATATGAAACTCGGAAAGAATTTTAAGACTGGTATTGAAGTTTACCTTTTAGCATGGAAGAACGATAACGATCCAACTCCAGGGGAATACAGTCTCCACATTGATCCTACTGGATATCCACAAGGCCTCATTAGACATGGGGCAAGGGTATCAGCTCGAGCAGGACCATGGAATGGTTTACGGTGGAGTGGAGCACCTGCACCACTACAAACACAAAGCAGCATATATACTTTTCAATTTGTTTTCAATGAAGAGGAGGTTTACTATAGTTTTTCTCTCATTAACAACTCACTGTTAACAAGGTTAGTCCTGACTAACAATGGTTTTATACAACGTTTGACGTGGGTGGATAGGACAAAGAGTTGGCATCTTTACCTCAATATTCCTCTGGATACTTGTGATACATATAGCTTATGTGGTGCATATGGGAACTGTGTCATAGATAGTTCTCCTGTTTGTGGATGTTTGGACAAGTTTGAACCTAAATATCCACAAAATTGGCAAACAGGAGACTGGTCACAAGGGTGTGTTAGGAAGACACCTATTGATTGCAACAAGGAACATGGATTTCTAAAATATTCTGGAATCAAGTTGCCAGAAACTAACAACTCTCAGTACAATAAGACCATGACACTCGAAGGATGTAGGCAAGTATGCTCGACGAACTGTTCTTGCACAGCTTATTCGAGCCTAGATATAAGCAATGGAGATAAAGGTTGCTTGTTTTGGTCTGGGGAGTTGATTGATATCAGAGAGCTCTCTGGAAGAGGACAAGACATTTACATACGAATGGATTCTTCAGACCTAGGTAAGAACTTATATTAACGAAATTTCTAATGATGTGTTTTTGAACTTTTAAATTTTTGAACATATGTAGACCATATTATATATGCAAAAGTAGAGGCCTATATCAATTCGAAGAAGCTAATTTAGAAAATGGTTGTAGTATCTCAGGCAAGTTCAAACAGAAAGAAGACAGGGACAGTACTCGCGGTGAGTTTCTCGTTGTTGGTGGCAGTGATTCTGCTAGGCCTGATTTTGTTCATGTACATACGTAAAAAGAAGAAACTAAAACTCAAAGAAGATTTTGAGCTGCCACAGTTTCAATTGTCGTTAATAACAAGAGCCACAGATAACTTTTCGGTCAACAACCAGATTGGAGAAGGTGGATATGGACCTGTTTATAAGGTATAAACTAAAAATTATTGGAATAGTGACCAGGCTAGAGATGACTTTCATTCTGTAATACTCGCTCGTCTTTGTAAAATATATTTCCTTCTGATATACTTTTTCCATTTCAAGGGAGTGCTCGAAGAGGGACAAGAAATTGCTGTGAAGAGACTTTCAAGGACTTCCATGCAAGGAATTGATGAATTCAAGAATGAAGTTACCTATATTGCCAAGCTTCAGCATAGAAATCTTGTGAGACTTCTGGGTTGCTGTATTCAAGGGGAAGAAAAGATGTTGATCTATGAATACATGCCTAATAAAAGCCTGGACTCATACATATTTGGTTTGGTTTTCTAAACTATTCTACAAGCTAAAATGACATGTTTTCAAAAGTTGCTAACAAGAAAGCTTTTCTTCTTTTTTTTTTGTAGATCAAACAAAGAAAAAGTTACTTGATTGGTCAAGGCGTTTCGACATCATTAATGGAATTGCTCGTGGCTTATTGTATCTCCATCAAGATTCTAGACTACGTATTATCCATAGAGACCTTAAAGCAAGCAATGTTTTGTTAGATACAGAAATGAATCCAAAGATATCAGACTTTGGGATGGCTCGAAGTGTTGCAGGAAATGATATGGGAGCAAAAACATGCCATGTAGTTGGGACACAGTAAGAATCGATCATACAGAAGTACTCTCATATGAAATCCCCCTTCTCATGGTGTAGTTCTTTTTTTGGAACAGCGGTTACATGTCCCCTGAATATGCAGTAGATGGAATCTTCTCAGTAAAATCAGATGTGTTTAGCTTTGGCGTATTGGTGTTAGAGATTGTGAGTTGCAAGAAAAATAGAGGATTTGTCCATCAAGATCACAACCTTAACCTTCTCGGTCACGTAAGCACTCTCTCTATACATTCTTGTGATCTATATTACATATAGAATTAATTTGTCATGATCTAAATTACTCAAACAGGCATGGAAGCTTTACAAGGAAGATAGGTCCTTGGAACTAATTGATGAGAAGCTAGCTGAATCTTGTCATATTTCTCAAGTTTTAAGGTCAATTCAAGTGGGACTATTATGTGTGCAACACTGTCCTGAAGATAGACCAAACATGTCTTCTGTGGTTCAAATGCTAGGTAATGAGAGTTTACTTGCAAAAGCTAAAGAACCAGGATTTTTCATGGAAAGAACGTTACATGATTCAGATAATTCAGGAATGCAGACAGGAAGTTCGAAAAATGAAATAACAATGACAATGTTAGATCCTCGGTAGAGTACTGTGATTGAGGCAACTACTGTTGATAAGCATGAGTTTTTAGAGGGCATGATAACAACATTGAAGCGCCTGAGAAGCGCTCAAGCATCACCTCAGGGATTAAGCGTGCCTTTGAGTATTTAGAAAAATGGTTCATAATTGCCATGCTAGCTATTATAAGTAGCTATTCAACTTGAATTTTCATTTGTATCTTATCTTAGAAAGAAATGTAAATTCTAGTAAGCAAAAATGTCCAAATTATATTGCCTATGACAATACATAAAATCGAATCAAACCGACAGATGCATGAGCTCTCCAATATCTGCATGGAATAGTCCAATATGTTGAAAT

>M6-PGSC0003DMG400005044 M6 G-LecRK

ATGTCACCAGAGTATGCAATGGGAGGACTATTTTCTGAGAAATCTGATGTCTACAGCTTTGGTGTATTGCTACTAGAGATTGTCAGTGGAAAGAGAAACAGCGGATATTATGACCACGAAAGGCATCTCAACCTTCTCAGTTATGTAAGTGCAAAATCAGCAGGATCTCTTGCCACAAGTTATTCACACAAACTTATAGTATAAATGATTTCAGGCATGGAAGCTACAGACCGAAAGCAAAGAGTTAGACTTAATGGATAAATCAATTTTGGACTCATATTCATCTGCAACAGTGCTGAGATGCATACATATTGGCCTGCTTTGTGTCCAGGATCATGCTGTTGATAGGCCATCAATGCCGTCTATCGTTCTTATGCTAAGTAGTGAGATGGATCTGCCTCAACCAAAGAAACCTACATTTATATTTCAAAGATGGTTGAATTCTAATACCCAATCACAATTCAGCAAAACTCAGTCTGTTAATGATCTTACTGTCTCTGTGGCAGAAGGCCGATAA

>M6-PGSC0003DMG400005689 M6 G-LecRK

AAAAAAACCTGGTCAACAAAAGAAATCAAAGCCAACAGCATGCTTGTCTCTGTTTTGGTCATACTATTTTGACTGTAAACATTCATGTGTGGATTAAGGAGCTTCTCTCATATGGCTACCAACAGAATTCTGATTGCTTTCATCCTCGTGTTATCCTGTCACTTTTTGCGTGCAGATTTTGCTAATTTCAACTCAATATTGAAGGGAGAAAGTTTAGCTGATTACCAACAGCTGGTCTCAGCAAATGGCTTCTTCAAAATGAAATTTTTCAGTCCAGGCAAGTCAAGGAACCGCTACTTGGGTATATTTTACACTCAGCCAAGTCTTGATATGTACTCTGATGTTAATGGAGGAGGAGATGAAAAAGCTTTATGGATAGCTAACAGAGACGATCCGATAACGGATACATCTGGAAGTTTTATGATTGCTCCTGATGGCAGATTAATAATTTCTCATAAGGAAGGTAATATTACTTTGTTCAGTGCTACACCAACAACAGCCACTAACTTAACTGCTATACTACTCGATAATGGGAATTTTGTGCTGAGGGAACTGAATACCAATAGCTTTGTAAATCGAACGCTATGGCAAAGCTTTGATTATCCAACTGATACACTTCTGCCTGGAATGAAGCTTGGGATTAACTTGCGAACGGGGCATAAATGGTCATTGACTTCATGGGTAAATGATCAAGCACCTGGTTCAGGGTCTTTTACCTTTGGTCTTGACCCTAATGGTACAAATCAGCTGATGATCTTGTGGATGGGCAAGGTCTATTGGAAGAGTGGCCCGTGGCCTACTGGCCATCTTGCTCTTAAATATGTCTCAAATGAGGATGAGAAATACTTTATGTATACTGTTGAATTGGAATATCAAAGATACTTTGTGAGTCCATTTGGAATAATTCAGGATACTTTTAAAGGAAATGCAGTGTTTGGTAATTGTAGTAATGAGACTCCCCATGCTGGCTGCGTAAAGCAGGTATTGCCCCAGTGCAGGGCAGCTAAAAAGTACTGGTTTGAGCTTAGACAAGTTTACATGTTTGGAGACAGCGTTAAATTTGATGAGAACTATACTTTGAGCCTCTCTGATTGCAAGGCCAAGTGTGAGAATGACTGCTCGTGTGTTGCCTATGCCTCAGTCGATAGTGACACCGGAATTGGCTGTCAAATTTGGGGGAATGACACGAGTTTTGTGACAGCACAAAACAGTCTAGCCAGAGATGTTTTCTTTCTTACAAGAAAAGGTAATTCTTGAGTACAAAGTTATTCAATAGAGTGAGAGGACATGTTAATAATGTTGCATCTCATATTATGATTACTTTATCCAGCTCAAGATAAATTGTTTTGTGAGGGCATCACACTAATATCACCAATCTGCATTACTCTCATTTGCACGTCATTCAAAAAATTACTAGTACTAGAACAAGCAATTTTCTCCGATCGCGATGCTGCAGGATTTGAGTTAATAAGAGTGTCATCTTCATTTCAAAAATACACTCGAGCAAATGAACAACCATACCATAGAGAATCTTGTTTTGGAGGCTATTAAATTATTATTTTTTGGTATTTCTAGAGTACCGACAATTATGATAGCATCATAGTGTATTCCCAAGGTAAATATGTGGAATTTTGGTCATAAACACTGACTTGCAATTTTGACAGATAGGAAGCGGAAATGGTGGATATGGTTGACAATAGCAGTGTCACTAATGGTATTTGCGTTTATTTTCTCTTTATTCTGTCTGATACGAAGAAAACTCAGAGCAAGAGGTGAGCAATACATCCGCTCTAGCATTAGTCTAAGGTGACATTTTTTTTCCTTTGAAATCTTCAATTTTGCATTTGTCACCTAGGCAAAGTCAGACAAATGGAGAAAATATTATACGAGCTTGAAGATCCTAAGACCATTTCTGGCCAGTACAATACTAACAAGACTGCTAGACTAAGGAAGAAGTTCAGGCATGATATACACATATTCAGCCTAGAAACTATGAATATGGCCACAAACAATTTCTCGTCTTCTAATAAGTTGGGACAGGGCGGTTATGGACCTGTTTATAAGGTAAACGTTTAGAAGTTTGTTGTCTTAAGCTGATTTTTAGATTTGCTAAGTAGCAAAAATTTCAGGGGATGTTACTCGACGGGCAAGAGATAGCCATCAAGAGACTTTCAAGAAGTTCAGGGCAAGGATTGGTTGAGTTTCAAAACGAAATTGTGCTGATTGCTAAACTCCAGCACACTAATCTTGTCAGACTTTTAGGATGTTGCATTGAAGGGGAAGAGAAGATATTAGTTTATGAATACATGATGAACAAAAGCTTAGACTTCTTCCTATTTGGTATGGATTTCCAATTTCATGCCTAAATTGCTTAATTAATCATATTTGATAACTAATACCCCTTCCCATTTTGATTGAACAGATCCTAGCAGAAAAGATTCACTAAAATGGAATACGCGTATGAACATTATTGAAGGAGTTGCTCAGGGACTACTTTATCTCCATAAGTATTCAAGGCTGAGGGTGATTCACAGAGATCTAAAAGCCAGCAATGTCTTGCTTGATGACAATATGAACCCCAAAATCTCAGATTTTGGGTTGGCCAGGATATTTGGAATGCAAGAATTTGAAGCAAACACGGAGCGAATTGTTGGTACCTAGTAAGTCCTTAATAGATATAATTATCTAACAATGGTGATTCACTGTGTCCTTTTTATTATGCGCTAAATTGCTACCTTTTCAGTGGTTATATGTCTCCAGAATATGCCATGAATGGCATTGTGTCCATGAAGACAGATGTATTCAGTTTTGGAGTTCTAGTACTCGAGATTTTGAGTGGCAAAAGAAACAACAGCTGCTATCATTTAGAACGCCCACTAAACATCATCGGATATGTAAGTTCCTCTCGTAAAATATCAATATAGCATGATCAAGAGGCGGATGGAGTGTTTATTCAATGAGTTCAACTAAACCCAATACTTTAGAAACGAAACATAAATATACAACATCAACATACCCATTGGAATCCCACAGGCCACAAGTTGGGTCTGGGAGGGCGGTGTGCACACAGACTTACCTACCTAATAAAGGCAGATAGGCAGTTTTCAATAGACCCTCGACTCAAGACACGAAACATGGATATACATGTGAAAATCACTTGAATTATAAAATTCTGAAGTATAATGAGTCTAGTTCTAAAAACTTTAAAGGTTCAACCTGTCAACTTTACATTGTTTTGGATCCGCCCTGGCATGATCACTTAGTTGAACAAGGAAATTTCAAACTTGATATGCTCAATTTGTCTCAGGCATGGGAATTGTGGAAAGCAGGCAGCGTCCTTGAAGAACTAACAGATCCTGTATTGACAAATGAATCAACACCAAAGAATGAAGTGATGAGATGCATTCATGTAGGATTACTCTGTGTTCAAGTCAATCCTATGGACAGGCCATCCATGTCAAATGTGGTTATGATGCTCACAAATGATAGCCTACAACTACCCGTCCCCAAGCAACCAGCATTTTTCATTGAGACAGCTATGACTGAGACTGAAACTCGCGAAGAAGTTGGACATTGTTCCACAAATGGGCTATCAATTTCAGATATGGTAGCGAGATGACTTTCATCATTGCACCCCTCAAAATTGTCAAAAATGGTAATATGCAATCTCCCTAATGATACAAAATAAAAACCACTGCTTGTCTCCTTCATCCAGATTTTACTTTATCTAGCATCGAAATTAAGTATGAGTTGTTGACAAAATAAAGTAAGCTTGTTAGAGCCTTTTGAAGATTATACTGTGTCTAAACAATTTTTACACATTTGTTTTGAATGCAAATGAGTTTTCAGAGTATCAATTGTTACTGAAATCAGAACTCATAAATGTAGTAATTGTCACTAGTTTTGCAAAGATGATTAGTATCTGTACACTAGAAAATCAAATGGATGTTTCCCACGTTCCTTTGTTGCTTG

>M6-PGSC0003DMG400007073 M6 G-LecRK

CCAATCTCAAGAAGCAAATGTATCATATAACAAACAAGCTAAAACAACAAACTAAACATGTCTCCTTCATGGCCTACTACACTTCTTGCCTCTCTCTTTCTTTTCTCCCAAATATTTTCTTGCATTGCTCAAGTTCCAGCTGAAAACACCTTTAAATTCGTCGATGAAGGCGAATTAGGAGATTATATTGTAGAATACAGAGCAAATTATCGCGTTATTAGTATTTTCAATAACCCATTTCAGTTCTGTTTCTACAACACAACTCCTAATGCATGGACATTAGCATTGCGAATGGGCACTGTCCGTTCTGAATCACTCATGCGTTGGGTGTGGGAAGCAAATAGGGGAAACCCCGTTAAAGAAAACGCGACATTCACATTTGGAACTAACAGAAATCTTGTATTGGCGGATGCAGACGGTTGAATCGCTTGGCAGACCAACACGGCCAATAAGGGGGTCACCGGGTTCAAGTTGTTACCTAACGGTAATGTTGTGCTACATGACTCCAAGGGTAAATTTGTCTGGCAGAGTTTCAACTATCCCACTGACACATTACTAGTGGGCCAAACTCTCCGATTGTCCGGCCCGAATAAGCTTGTGAGCTGGGCCTCGGTGAAAAAGAATGCGAACGGGCCTTATAGCTTGGTGGTCCAGCCGAGATTGTTCGCTGTCTACAAGGGAAACAAGGTAGACGTGGAATTAGCTTGGTTCGATTATGGGAATAACACGTTGGAATCGATAAAATTGAATAGTGGGAATCAAAGATTGAAATTGGATTATAGATTTGCAAAATCAACACAGAGAAGTTCACATGTGATGGCATTTACTAAATACAACACAACATTGACATATCTTAGACTTGAAATTGATGGAAATCTAAAGGCTTATACTTTCGTTAGAGATGAAGAAGCAGATGAATTCCGTTGGAAAGTAACTTACAGTAGGATTTAAAACGTTTCATATTTGAAACGTCTAAATCCATGTGCACATTATCCTTTACGAGCAAATTTCACGGGTTATATTATTATTGTTGTTGTTATATTGTTGTAAACGTAGCTCTATAAGATAATGATTAGTGTGTCCAGTACAATAAGTGGGTCCACAACTAGTAGTCCAATAAATGAGTAGTACTTCCTATTTAATGTCAATTATATTTTTCTTTTTCCCTTTTTGAAAAAATATTATGTTTTGGTTGAATTAAGA

>M6-PGSC0003DMG400007455 M6 G-LecRK

ATTTTTTAGAGGAATGAGGTGGAAAAAGGAAAAAAAAGAAATTGGGTCCTTTGTGATTTCTTGCATTGGCAGTAACCAACGTCCGAAGGCCAAATCTCATCGATTGTGACATTGTAACAAAATATACATAATTCATCAATCTTCATAAACCTATCATCATAATCATAATCAATACAACATTATTGTAAACAAAACAAATATTCCTTCTTGTATATACACACAATATTAAAAAATTATAAACTTTTTTACATTCTTGATTGTGTATGTGGGAAATTGGGCAGCAAATTTGCAATTTTTGATCCTGTATCTAATGCTTTGTGGATGTTTTCACAATTTGATTCACCTTTTTGGAGTGGGGTTTCCTTAGAAAGGGTAAATTCTACATAATCTTGAATTTCTATATCTTTTTCTTGATTTGGGTGTCTGGGTATTCGAATAAATCTGTTTGGTGTTGTGGATGTTTCTTGAAACTGCATTTGGTGATCTGGGGTACTTTTATTTTGGTCATAGTTGCTGGTGATAATCATTATTTGGCTGCTTTGTTTAGAAAGATTGAATCTTTTTTACTCTGATAAGGTTCTGCAGTGTGTGGAGGGATTGTTCTTATGAATTATAGCTAAAAAGGAGATATTGTTCTGCTTTAGACCTGTCACTAAATTCTGTGTCCTGTGGCAACATTTTGATTTCGCTTAAAATCTGATATGTTTAAGGTTGAGGTTGAGGTTCTTGTTGTGTTGAAAATCCTAGCTGTTTAACTTGGGGGTGGTGTGTGTGTATACTATGAGTGAATTGGTTGGTGAGTTTGGATAAATTTTGGAATAAGAGGAAGTAATTTACTGCATATTAGGGATATTAGTTTTATGTCAGCAAGTCTGCATTGTATAAGAAATGTTGTAAATGTAGCTAGCTTTGGTGGTGGTGAGATGTTAAAACTTTAATCATCCACCTCCAAGTTATAGAATTTGAAGTTCCATTATCTGTGACTTCTCTCCGGGATCGTTACTCTCTTTCGACAAATGCAAATATAGTCTGTCCTAGCCTCACTACCAATCTTACAGACTTATAATGGATTCTTCACCATCTCATGAAGAATCTGCTCCTCCAACTGATGTGGCACCTCCAAAGTTTAGTCCTCCGCCTACACCTTCTAGTCCATCTGATAGCTCACCACCACCTCCATCTTCTAGTCCACCTCCTAGTTCACCGCCACCTGTATCTTCTAGTCCACCTCCTAGTTCACCTCCACCTGAATCAACTAGTCCGCCACCTAGTTCACCACCACCTCAATCTTCTAGTCCACCACCAGCCTCACCTCCACCTGAACCAACTCCTTCTTCTCCTGTCAGTTCACCTCCACCAACTTCATCACCTCCCCCTGCATTGTCACCTCCGCCACACGTACAGAATCCAGAACCTCCTCCAGCACCACAATCACATGGAAATTCATCCCCACCACAATCTCCTGAACCTAAAAAAGGTTCATCATCATCTTCACCACCTTCACTACCTTCATCCAACTCACCTGGGGATGGAGGTGGAGGTGGAAATTCACCACCATCCCCAATATCTAAGTCATCAGATAATTCGCCTCCTTCACCAGCCTCAGGCCCTCCTAAAGATTCGCCACCTTCACCAGCTGCATTATCTCCAAGTAACTCACCATCTTCTGATTCACCCTCAAACTCATCAACACCAACATCTGTAATACAATGGTCACCACCACCTCCATCCTCTAATGGTGCACAATCTTCTCTTAATCCTCCTTCATCACAAGTTCCTAGTAACCATAGTTCCGGAAACCATTCAACAGGAAGTCCAAAGCCTGCTGGTGGCAACAGCAATGGTACTGCAGAGACAGTTTCCATTGGCGTTGTAATTGTTCTTTTTTTGGTTGGTATCATCGGAGCAGTTGCATGGTGCATACGGAAGCGAAAGAAGAAGAATTCTGTCCATTGTAGTGGTTATGTCAAGCCAATCTCTGCGTGCTCCTCTCCAAAATCAGGTAAATTTTCAGAAACTATAAAATCATTCCTTTTAGTTTGTTATTCAGTGTTCAAGATGCTTGCAAGAAAAATCAATGTAATATTTGTAGTTGAGACCTTTCAAGTGAAATCTGAGAAGATGTGAAAGTTGTGTACAGAGTCTGCTTTATTGAAGGTCAATGAATCAGCACCTGAGAGAAATGGAACGAGCTCTAATTTCCTCAATTCTCCAGCAGACCCTGGCGGATTCAACAGTTCAAAGGCATGGTTTACTTATCAAGAACTTGTTGAGGCCACTAATGATTTTTCAGCGAATAATGAATTAGGGAAGGGTGGATTTGGTTCTGTATACAAAGGCTATCTAGCAGATGGAAGATATGTCGCGGTAAAACAGCTAAATATTGGTGGGAGCCAGGGCGAACGGGAATTTAGAGCTGAAGTTGAAATCATTAGCCGGATACACCATCGTCATTTGGTTTCACTTGTAGGTTATTGCATTTCAGAGAACAAAAGGCTCCTCGTTTATGACTATGTATCCAATAATACCCTTTACTTCCATCTTCATGGTAAGCATAACATTTGTAGATTTAATTATTTCATTTGTTATGTTTGGTCACTAAAGCGAGCTGTTGATACTTGCAAGTTACAGCACAAGGGAGGCCTGTCATGAATTGGCCAACACGTGTTAAAATTGCTGTTGGTGCAGCTCGTGGAATAGCTTATCTGCATGAAGATTGTAATACTGCTCTCTTCCTCAATTCTTATGAGTTATCCATGTTTTATTCATGTGATATCTTTCATTGGTTGAACAAACGTCCCCCTTATAGTTAAAAGAGAATGTACAGTTAAAAACTTTACCACAACTTGGAATTAGTTTTAATGATCCCTAAAGCTGTGGCGGGATCATGGATGGAAATTTAAGCTCCACAACCACCTTCTTCCTACTCTTCCTAAAAGCATACCTTAACGAAAGGAAAATAAAAAAGAGAGAGAAAACAAAACATGTAGTAATAATTTTCAGCTCATAGAAGTAGAAGGCCTTCCAGTTGGCATTTCCTTTTTGTAATGTAATGATATTACTTATGCCATTTAGAATTGAGTTCTAAGGTCCTTTTTCATGCCATAAATTACCTCGTTATATAACTTTTTGGCATGAGTGTGGATTATCATATCTGCATTAAGATTGTAAGTAGTGTTCTGTTCCTTAATCCTTTTATGTTATTCATGTTTATTTTTATTTCAGTTCGTTTGTATGGCTGAAAAAAAATGTTTCTCCTCTTTCCTGCCCCAAAAGTTAAGAGAGAAAGCAGGTGGTAAAACTTTTTACCATATGACGAATCGGATTGTGATATTAGATTATGTGTTCCTATCATATCACTTACATTCTTTTTATTGATCCCTACTCCACTAGCTGAACAGCTGTAGTAAATTAAATGATGCAATCTGAGCTACTTCCAGAACCAACTTTCTCCTATTATCCCTATAAGAGGACTTGAAGTAAAAAGAGAAAATGGAACGTATAAATAGTTTAAGCTCAATGGAAGTAGCAAGATCTTGATATTGGATTTCCCTTTGGTAATGTTATAATATAACTTCCGCAATTTAGAATTGTGTCATCAATTTTTATGCATCTCGTAAATTATCTCGATTATATGACCTTTTGGCAGTATAAATAATGAATTCTAATCACTTCTCTAATTAGTTACTTAAAAGATGTTTTTGTAGCTGAATGTACCAAATGAGGCTCAAAATCGAACTAAGACTGAACAAAACATGATTGGTTCCATTGAGAGAACAGAAAAAATTGTATGACTAAGTTCTTTATACACTCTTGTATTGCACCATGTAGTTACATTCCCGAGAAAGAAATAATTTAATCCATGAGCAGGCATAGTCTCCTTGAACATGAGTATATCTTCCACACTGAAGTGAGCGCATCAGGCTCTGCATTTGTAGTGCATTATATGCTTTTCGCTTGCCAATATTCACTCTACGACAAGCATTTGAATAAATCTTCACTGATCGAGCCAGGTTGAAAAGTGAGAGAAACATGTTCAACTGCTCTTGTGAAGCATATAGTGAAAAGCAAGTGACAAACTGGTTTCTCCATATAACAAAGCTGAATGTGTTAATCATCACTGAGTTATTATGCTATACTGTTGAATAAGTAATGAAAAGTTAAGAAAAGTAGACCAGGCGAAGAGATTGTGATTTTCTATGCTCATTACCTTCATTCTTGTTACTTGAGCTACCTTTAATTATATAGTTAAAGTATGATACTCCTTTTTTATTTGGTTCTTAATTAATTCTTTTGGTGTGGATGACAGGTTATCCTCGTATTATTCACAGGGACATTAAGTCATCAAACATTCTTTTAGATGATAACTTTGAGGCCCATGTATGTCTAAGTCTTCGAACTCCTATATCCACATTCTTTGTCTCACGAGCCGCCTTTTACAATTAATTTACTTTCAACAGGTTGCTGATTTTGGACTAGCTAAATTAGCTCAGGATGCAGAAAGTCATATCACAACACGTGTCGTGGGAACATTTGGGTATGTTTCAGTGCCAACTTTTATATTAAGTTGACAGCATAAAATAAATATTTTATTCCATCTTTTTGAATTTCACATATTTATTCTAATTCCTTAGATCTGACTCAGATACATGGCTCCTGAATATGCATCAACTGGCAAGCTGACTGAAAAATCTGATGTATATTCCTTTGGCGTGGTGCTTCTGGAGCTAATTACTGGAAGGAAGTCAGTGGATACATCTCAGCCTTCAGGGCAAGAGAATCTAGTTGAGTGGGTATGTTTGTAGCCAATCTATTATTTGTCTTTCTCGAAATCCATATTTTCATCAGGTCTCCTTTATTACCTCTTTGTTAGAAATTTTGACATTAATATCCGACCTAAGGCAATGAAAGTGCATATCAGCATGCAGAGAGAACTTAATTTTTAGTCCTTTTAGCTAGGATAACCTACAAAATTTCATTTGAGTCAGGATTTGGTTGTTCCTCCGTCTTTTCCCATACTGCACAAGAACTTGGTAGCTGATATGTGAAGAAACCGCCATTTTGCTACTGTACAGTAGGCCTCTAGTTGGTCTGCTAGTGAAGTTCCTCTTCTTTATTTTTCCCTTTTTTACCATGAAATGTGCCTTATCCTGGGAAGAATGTGCTTGCAATGCTGTAAGCCTGAAACATTGTGTCTACATTTTTATTTTTATTTTACAACAATCAAACAGCTTGATCAAGTTATCCGCTTTAATTTCATTGATCCTTTAATGTTAGAAAACTGAAAAACCATTAACAGTAGCAGCCGAAAACTTGTATCTTTTATCATGCTGAATGTGTGAATTCCATTTACTATAGTTATTTTATTTTCGAACTCTTTTGTTTTCTTATCGTCAGTGCATAGTAGCAGTAGCCAGTTTACAAACTTTTAATGCGTTGTATTCTGGTACTAGTTTCCACTGCTAAATTTGAACCAGGCTTAAAAATAACTACTCCATTATATTGTACTGTAACTTGATTTTTTTGTTGTCCAGGCACGACCTTTGCTTAGTCGAGCACTTCAAAAGGAGGAATTTGATCTGTTAGCAGATCCGTGCCTTGAAAAGAACTACGTTGGCACGGAGATGTTCCGAATGATTGAGGCTGCTGCATCTTGCGTGCGTCACTCATCTGCAAAGAGACCAGCAATGGGACAGGTAACTCTTGTTCTGTTATATTGAAAACCTATGCAAAAGACACCTTCAAGACTCTTCTTGTGAAGGGCATTTACAACATTCTGAATTCTGATCATGGTTATGCTTCGTAGATCATGAGAGCTTTCGATGGTATGGCTATACAAGATCTGTCAAACGGGATGAAAGTTGGTGAAAGTGCAATACATAGCGCTGCACTACAATCTGCAGAAATAAGTTGGTTTAGGAAAATGGCATTTGGTAAGCAAGATTTTAGTTCGGATTTTTTCAGCGAAAGTAATCAGAATAGTAGAGAGTCAAATGAACATCCTTAGAGGATTTTATAGAACAAAAAGGCTTATTGAAGCTATGTATGCATCTGTTGATACATCAAAGGTACCTTGTTAGCTTAGAGATGCCTTGGTTTAGCTTTCATATGTATAAGTTCCCTTTTGGCTTTGAGTAATTTTC

>M6-PGSC0003DMG400008493 M6 G-LecRK

TAAAAAGTTGTATATAGCTAGTGTCCGGCATGCTTTTATTTCAACTATAATGGCTTTTGATTTTCAATCACAACAAGAAATCATGTACGTATAGATTATTATCTAAATTTTCATGTTTTTCCCATTTGTCTAAGTAGATTTTATAGTAGTTGATTATTTTTTTTTCGTATTTCCTTTTCATTAATTTGCAGAAATATTAGGTGCTACAAATTTAAAATTTTGAATATTTGAGCAATGGGTTGGAATTGCTTTGGAGTTTTGCAATCATACAAAAACAATAGAGAGACTCAAATCCAAGGTAAGTACACTCATAACTCTTATTTCTGTTTTCTCAATTACATGCTTAAAACAAATCGTTCTTATAAAATTCAGAATCCAAAAAATTTAAAAATTTGACGCATCTGCTCTACTTTTTGTAGTCCATCCAAATTTGAGAAGTACTTCACATAATTTTCATTTTCTTTTTGTGTTTTTCATTGTGTAGAGTTCACGACCAACAATGTGAGGGTTTTTTCCTATAACTCATTGAGATCAGCAACAAGTCACTTCCATCCATCTAACAAAATAGGAGGTGGTGGTTTTGGAGTTGTTTATAAGGTAATCATCTCTTGTTTATGGTGTCATGCCTTTTAATGGTCCTTGTGGATGCTAGTTATTAAGTACTGAGTTTAACATCTATACGTTCAAAATTTAGAAGAAGTCACCTAAAAGATAACTATAATAATTTAAATCATTTGAAGTGGGATTTGCTAGTGAGTAGAAAAATGAGAGCATGTAAGGATAGATGCAGCCTAATGATCAATGAAGTGGGATGGGAATAATGAGATTTTTAGGTTCAAATCTCAATGGAGGCCCAAAAAGGGTACCGCAGCCTAGATAGATTCCTTCCAATGTAACTAATCTTTGGTGGAATAGCGAGGTACACATAAGCTGATACAAACACCACCATCATAAAAATTAAATTAAATAAGGTAGATTACCTGCTACATTAATAATGTAAAAATATTTTACACTATCAATATATATAGGTTAAATCTTTTATGTATTTCCACTCTTTGTACTTATGCAACTTTGTTCTTTAAATGTAGGGAGTTTTGAGAGATGGAACTTGTGTTGCCATTAAATGTCTCTCTGCTGAGTCAAAACAAGGGACTAAAGAATTCTTAACAGAGATTAATATGATTTCAAATACCCAACATCCAAATCTTGTTCAGTTGATTGGTTGTTGTGTTGAGAGTGGCAATAGAATGTTGATATATGAATACTTGAAGAATAACAGCCTTGCCAGTGCTTTACTAGGTAATGTTCTCACAGATCAGTTTGAAAAAAAAAAACTTATTTTGAACCAAATGAACATGTCATAGTTGAAGTTAGGAATGTTAAATGAGCAGATTGGACTGAATCTGGACGGGTTAAAATTGTTATTTCAGGTTCTAATGGTAAACGCGTTGCCCTGAATTGGCCTCAGAGAGTTGCTATATGTTTAGGTACAGCATCTGGTTTAGCATTTCTTCATGAGGAAGCCTCACCACCGATTGTTCATCGAGATATTAAGGCTAGTAATATCCTTATAGATGAAAACCTTCATCCTAAAATTGGAGATTTTGGTCTGGCTAAGCTTTTTCCTGAAAATGTCACTCATCTAAGTACGCGAGTAGCAGGAACAATGTGAGTAGTTACATTGAATTTCAGTTTCATAACATTCGATATTCCTCTCCTGTCTCGTCTCAGTAAGGTATTCTGATGTGCAGAGGTTACCTTGCTCCGGAGTATGCCTTGTTTGGACAACTCACGAAAAAGGCTGATGTCTATAGCTTTGGAGTTCTTGTACTTGAAATCATAAGTGGAAGAAGCAGTAGTAAATCAGCATTTGGAGTGGATCTCTTGGTTCTGGTGGAATGGGTAGGATTATACTTTCAGCTTTATGATCTTCAAGATTCATCAGTAATCAGCATTTTGAAAGTTTACGCGTTGCAGGTATGGAAGCTAAGAGAAGAAGCACGGCTTCTGGAAATCATTGATCCAGAGCTAACAGAGTATCCAGAAACTGAATTGCTGCGGTTCATCAAAGTTGCTCTCTTTTGCACACAATCAGCACCTACCCAAAGACCTAATATGAAACAAGTGATTGAGATGTTGTCTAAGGAAGTCAACCTCAACGAGAAGTCGTTAACTGAACCAGGAGTTTATAGACCACATAGCTCGAAACGATCAAGTTATGGTAGTCTGCAGAAATCATCCTCTGCGGGGAAGAATGGGGTGCAATCTGTAAATCCTTCTGTAACATCAACAAAGTTTGAGAGCTTTCAGAGTGTCACCCAAATGCTTCCTAGATAATGTTAACTGGTGTTCTTATTCTTCATACTTGTATATTGTCGTTCTTCTAACGTGGGCTATCGAGGAGTAAATGTATTGTCTCTCATTAGACTTACCTTTTTCATACATTTTCTCATGTAATGTAAAGTTGGCAGTAGCTGATAAGGGAGAAAGAGAGACAAATCACACAGAATGAGCTACTATGTAACTACAAATTTCAGATATGAAGATGAAGTTTACAGCAGAGACTATGAACACAGTAAAGGATAGAAACTTTTCTCTATGCACATATTAGCTCCTAAAACTAGAAATGAAGGTAAAAACTTCCTGATAGAAGTTATTCTCAAACTCATAACATTTAAGTTAACGTATAAAGGATGACTGCTTTACAAGATCAACACAGTAGCATTTTCATTGTGCAGCAGAGAATAAACCTTCAAGATGCAGCAGTAGTAGCTTGAGGCTTTGCAGAGGCAGCAACAGAAGTTTCTATTCCTGTCTTTGCATCTTTTCCTGGTTCCGAAGATGGTTTTGCAGAAGGCTTCATATGCTCAGGGGGGTTGTGTTTGAGTTTCACAAGTATTTGACGCATTCTAGAGAGTTCAGTTGGCCTGCCAGGTTTGGGTCCTTGGATGACATTGATGGATGGCTGCTTCTCCTTATCCTTCTTTCCTTTCTTCTCTATGGTCGGGACCTCCTTTGGGATAAGCTCTCCAATTGCCCTCCAATAAGCCTTATTAGCTTCAGTATGGAACTTCTCTTGGTTGGCCAAGAATAACTGGAAAAAAAGAAAAAGATGACACATGAACTGCAACACATTGATTGACTAAGGAGGAACTTGCCGTACTGTGGAACTATGTTTAGAGATTTGACAAACCATAAAACCAAAGACATAGTTCTGTGGAGATTAGCAAATAAAACTCACCAAAGCACAAAAACTCAGCATATATTGTTCTAAAGGTTATCAGCACATTTTGATCTCTTGTGGAAAACTTCCACTTTCAAAAATTTTAATTTACACCTCTTACGCAGTATTAATGCTAAATCATATGAAATGTAAACAGACAAAATGTAGAAACTGCTTTTCTCTGTGTCCCTACCTTGGTTGAGTCGAAAGTACCAATACAATGATTTTTTCAAAACTAATAACTTAAAAAGGAGAGTAACAAGTACTATGATTAAAAATTGGATGGTACTTTTACTAGGTTAACTAGGTTAATCTCCCTGTCTCAACACTCATATCCTATTCCGAAGAGTTGAAAGCAATGATTAGGAAGAGAAGGTAACTAGATGCATGACATGTCAGACCTTCCAGCTAAAGCAACTGTTTGGAAAACAAACATCCAGCACAAGCAAATTTATGCCAAACTAGTTAACAAACCGTGACTTTAAGCAAAATGCCTCGGAAAGACTTTTAATTTATATAGCTATAGGTCTCACCTTCTCCTTCTCCCGGTTAGCAGCTATGTTGTTCTCACATCTAAGTTTCCATTTCCTATAGTACTCTGCTTTGTATTCCTCTGCTTCTCTGATTATTTCCCTAACAATTTCTTTCTCCCTTTTCTCCTTCTCCTCCAGCCGTATGGCGTTCAAT

>M6-PGSC0003DMG400008508 M6 G-LecRK

CATGGAAATGAAGAACAATCATTACTTTCTGCTACTCTTTTTATGCTTCAGTCTCAACAGTAATCTCTGTATCGGAAGGGACACCATTTCTGCGACTGAATCTCTTTCTTTTGGTGAAACTCTAGTCTCTTCAGGTGAGATATTTGAGCTTGGTTTTTTCAAACCAGGTAACTCTCTTAAATATTATCTAGGCATATGGTACAAGAACGTTATTTTATCGCAAACAGTAATTTGGGTAGCAAATAGGGATAAACCACTTGATTATGGTGCTGCTGGGATGATAATTAATCAAGGAAACCTGGTGCTTCTTGATAGATTTCAAGGCGAAGTTTGGTCGGCACTTGCTGGAAACATCAATCCAGATATTTCAGTTACTGCACTTCTTCGTGATGATGGGAATTTGATTTTGAGTGTTGTGTCAAATTCATCAACACCCTTACTACTTTGGCAAAGTTTTGATCACCCAACACACACTTTTATGCCTGGTGCTAAGCTTGGATATGACAAACGTACACAAAGAAAACAGGTTCTGGTTTCATGGAAGAATTCGAACGATCCAGCACCAGGATTGTATTCTATGGAAATGGACCCAAAGCATGCCCAGTTTGTTTTAAAATGGAATAGGACTACAGAGTATTGGGCAAGTGGTTCATGGGATGGCCAAAGATTCAGCTCAGTGCCTGAGATGAGGTTAAACTACATATATAATTACAGCTATATCGACAATGAGAATGAGTCATATTTTACATATTCCCTTTACAATAGTTCAATCTTATCAATATTCATTATGGATGTCTCGGGACAAATTAAGCAACTAACATGGTTGGATGATAGCAGTGTTTGGAATCTATTCTGGTCTCAACCAAGACCAGATTGTGAGGTTTATGCTAAATGTGGGGCATTTGGAGTCTGTGATGAAGCTAATGCAACCTGCAATTGTTTGAGTGGTTTCAAGCAAAGATCAGATACAGAATGGAATTCAGATGATTATTCTGGTGGCTGTGTAAGAGACCAAAAGGTGCAGTGTAATGCAATTACTGAAGACACAGATAGTTTATGGATAACTTCAATTGTGAGAGTACCTTCTTCTCAAAATACTAATATCACAGTTGGGGAAGCCTCACAGTGTAGATCTGCATGTTTCAACGATTGCTCTTGCACTGCTTATACTTATGATGGTTCCGGTACCTGTTCTATCTGGACAGGGGATCTGTTCAATCTGCAACAACTCAGCACAACTGAAACAGAAAGGACTATATTTGTCAAACGCAGCTCACCTGAAGGTTAGAATAGCATAGTATAACATGCTTTGATTCTTTATTTCATATTAGTACAATGTTATGAATAGAATTCCTTATTAACGTTAGCACTTAACAACAATGATTCTCCTTTCAGCTCAAACTAATGCTAAGAAATCAAAGAAACTAAAAGCAATCATTTCGTCAATATCAACTTTGATGTTTCTACTCATAGGCAGCATTAGCTACATTTACTATAGAAGAAGAATGGCAAAAAGAGCAGGTATTTGCTGAATGAATCTAACTTTGGAAGATATTTTCTCTTATACCATTTTGGACTTAATTCCAAAGGTCATCCTCAATCCAATTGAGGTAATGAAATCCCTCCCACTCCCCTTTTCTATTTCAGATAGTAGTAAAGGTACTCAAGGGGCACGTAAATCTCACTGGCACAAAGCTGAAGGAGAAGCAAAAGTATTAATGAATGAAAACAGTGATGAAGCGATTGATGTTCCATACTTTCACTTGGAGACTATTTTGGAAGCTACAGACAACTTCTCAAATGCGAATAAGCTCGGACAAGGAGGATTTGGTCCTGTTTACAAGGTTAAAACTTCATTCCTACATCTATGTGGACTGGCATTGACCCATAGTAATGAGATGAATTTTTTTTGCAGGGTATCTTTCCGGGTGAAAAAGAAATTGCGGTAAAAACGTTATCTAGTCAATCAGGACAAGGCATAGATGAATTTAAGAATGAAGTGACTCTAATTGCAAAGCTTCAACATCGAAATCTGGTGAGGCTATTGGGCTATTGCATCAATGCAACCGAACAAATATTACTTTATGAATATATGCCGAATAAAAGTTTAGATACATTCATATTTGGTGAGTTAGCCTTAATAAATTCTTTATTCTCAAAGTCATACATTTCTTATGATGATCTAATTTTATTGACGTACCAAACTTTGTTATGGTAACAATGCAAGATGGAACACTTTGTCAATTATTGGATTGGAAGAAACGTTATGACATCATATTGGGCATTGCACGGGGTCTATCTTATCTTCATCATGATTCACGACTAAGGATCATTCATAGAGATTTAAAAACCAGTAACATTTTATTAGATGAGGAGATGAACCCGAAAATTTCAGATTTTGGCTTGGCAAGGATTGTTGAAGGAAAAGTAACAGAAGCAAATACAAAGAAAATAGTGGGGACCTAGTAAGTATAATATTCCTTCCTGTCATTAATTTCTTACAAAAGGAGCTTGGGAGAAAATCTTATACATAATGTTTTGTTGACCTGGTGAAATCTAGTGGCTATATGTCTCCTGAATATGCACTAGATGGATTGTTTTCCATTAAATCAGATGTATTCAGTTTTGGAGTAGTCGTACTTGAGATAATCAGTGGAAGAAGGAATACAGGATTTTATCAGTCAGAAGAAGCCTTAAACTTGTTGGGCTATGTAAGTGAAGAAACTTCCCATGACTTAATTATTATATCTTCTTCTTAGCTTTGTCTCTCTGATATAAAAATTATCACAGGCATGGAAATTGTGGACAGAAAAAGCTGAGACACAATTAATAGAGAAGTCATTACTTGAATCATGCAACAGAAGTGAAGCAATAAAGTGCATAAATATTGCGCTCCTGTGTGTACAAGAAGATCCAAATCATCGTCCTAATATGTCAGATGTCATTTTGATGCTGGGAGGGGAAGGTACTAACCTTCCAACACCTAATAGACCAGCTTTTGTAATAAGGACGCATGCTTCTAGTACATCTTCTTCTTCCTCCGATAAGAAATACATCGTATCCAACAATCAGGTGACGATTACAGTTGAAGAAGGACGTTAGCAGGAATAGTGAAGGCATTTAATTCGTATGGAAAGGGCTAGGGTGGCACTAGCTAGGACTTATAGATGAGCCATATTATACTTTTTTAACCTCTACTTTGGTCCTTAAAAAGTCCAGGGATACTTGACCAAATGTCGTCTTCTCTATATATAATAGTTAGGACAAAAGTGGAAATGTTGGGAAAACTTTCAAGTGTTTGAACTCAAACATATACTATATTTTTGGTTTAGAATATAGTAGGCCTTAGCAATCAAGTACAAGAATAATTCGATTATTCCTGATTCCTTCAAGCTTCAAATATATAAGCAATGTAATTTTTTTTTTTT

>M6-PGSC0003DMG400008523 M6 G-LecRK

AAACAAGAAACTTGTTTTGACTAGTTTCATGGACTCCAAAGAAGAAAAATCTCCTTCCGTAAACACTTGCACTCCTTATTTTCTCTCATTCTTTCAGTCATCAATTTCTCTATGTACGTAGCATGGAAATGAAGAACAATCATTACTTTCTGCTACTCTTTTTATGCTTCAGTCTCAACAGTAATCTCTGTATCGGAAGGGACACCATTTCTGCGACTGAATCTCTTTCTTTTGGTGAAACTCTAGTCTCTTCAGGTGAGATATTTGAGCTTGGTTTTTTCAAACCAGGTAACTCTCTTAAATATTATCTAGGCATATGGTACAAGAACGTTATTTTATCGCAAACAGTAATTTGGGTAGCAAATAGGGATAAACCACTTGATTATGGTGCTGCTGGGATGATAATTAATCAAGGAAACCTGGTGCTTCTTGATAGATTTCAAGGCGAAGTTTGGTCGGCACTTGCTGGAAACATCAATCCAGATATTTCAGTTACTGCACTTCTTCGTGATGATGGGAATTTGATTTTGAGTGTTGTGTCAAATTCATCAACACCCTTACTACTTTGGCAAAGTTTTGATCACCCAACACACACTTTTATGCCTGGTGCTAAGCTTGGATATGACAAACGTACACAAAGAAAACAGGTTCTGGTTTCATGGAAGAATTCGAACGATCCAGCACCAGGATTGTATTCTATGGAAATGGACCCAAAGCATGCCCAGTTTGTTTTAAAATGGAATAGGACTACAGAGTATTGGGCAAGTGGTTCATGGGATGGCCAAAGATTCAGCTCAGTGCCTGAGATGAGGTTAAACTACATATATAATTACAGCTATATCGACAATGAGAATGAGTCATATTTTACATATTCCCTTTACAATAGTTCAATCTTATCAATATTCATTATGGATGTCTCGGGACAAATTAAGCAACTAACATGGTTGGATGATAGCAGTGTTTGGAATCTATTCTGGTCTCAACCAAGACCAGATTGTGAGGTTTATGCTAAATGTGGGGCATTTGGAGTCTGTGATGAAGCTAATGCAACCTGCAATTGTTTGAGTGGTTTCAAGCAAAGATCAGATACAGAATGGAATTCAGATGATTATTCTGGTGGCTGTGTAAGAGACCAAAAGGTGCAGTGTAATGCAATTACTGAAGACACAGATAGTTTATGGATAACTTCAATTGTGAGAGTACCTTCTTCTCAAAATACTAATATCACAGTTGGGGAAGCCTCACAGTGTAGATCTGCATGTTTCAACGATTGCTCTTGCACTGCTTATACTTATGATGGTTCCGGTACCTGTTCTATCTGGACAGGGGATCTGTTCAATCTGCAACAACTCAGCACAACTGAAACAGAAAGGACTATATTTGTCAAACGCAGCTCACCTGAAGGTTAGAATAGCATAGTATAACATGCTTTGATTCTTTATTTCATATTAGTACAATGTTATGAATAGAATTCCTTATTAACGTTAGCACTTAACAACAATGATTCTCCTTTCAGCTCAAACTAATGCTAAGAAATCAAAGAAACTAAAAGCAATCATTTCGTCAATATCAACTTTGATGTTTCTACTCATAGGCAGCATTAGCTACATTTACTATAGAAGAAGAATGGCAAAAAGAGCAGGTATTTGCTGAATGAATCTAACTTTGGAAGATATTTTCTCTTATACCATTTTGGACTTAATTCCAAAGGTCATCCTCAATCCAATTGAGGTAATGAAATCCCTCCCACTCCCCTTTTCTATTTCAGATAGTAGTAAAGGTACTCAAGGGGCACGTAAATCTCACTGGCACAAAGCTGAAGGAGAAGCAAAAGTATTAATGAATGAAAACAGTGATGAAGCGATTGATGTTCCATACTTTCACTTGGAGACTATTTTGGAAGCTACAGACAACTTCTCAAATGCGAATAAGCTCGGACAAGGAGGATTTGGTCCTGTTTACAAGGTTAAAACTTCATTCCTACATCTATGTGGACTGGCATTGACCCATAGTAATGAGATGAATTTTTTTTGCAGGGTATCTTTCCGGGTGAAAAAGAAATTGCGGTAAAAACGTTATCTAGTCAATCAGGACAAGGCATAGATGAATTTAAGAATGAAGTGACTCTAATTGCAAAGCTTCAACATCGAAATCTGGTGAGGCTATTGGGCTATTGCATCAATGCAACCGAACAAATATTACTTTATGAATATATGCCGAATAAAAGTTTAGATACATTCATATTTGGTGAGTTAGCCTTAATAAATTCTTTATTCTCAAAGTCATACATTTCTTATGATGATCTAATTTTATTGACGTACCAAACTTTGTTATGGTAACAATGCAAGATGGAACACTTTGTCAATTATTGGATTGGAAGAAACGTTATGACATCATATTGGGCATTGCACGGGGTCTATCTTATCTTCATCATGATTCACGACTAAGGATCATTCATAGAGATTTAAAAACCAGTAACATTTTATTAGATGAGGAGATGAACCCGAAAATTTCAGATTTTGGCTTGGCAAGGATTGTTGAAGGAAAAGTAACAGAAGCAAATACAAAGAAAATAGTGGGGACCTAGTAAGTATAATATTCCTTCCTGTCATTAATTTCTTACAAAAGGAGCTTGGGAGAAAATCTTATACATAATGTTTTGTTGACCTGGTGAAATCTAGTGGCTATATGTCTCCTGAATATGCACTAGATGGATTGTTTTCCATTAAATCAGATGTATTCAGTTTTGGAGTAGTCGTACTTGAGATAATCAGTGGAAGAAGGAATACAGGATTTTATCAGTCAGAAGAAGCCTTAAACTTGTTGGGCTATGTAAGTGAAGAAACTTCCCATGACTTAATTATTATATCTTCTTCTTAGCTTTGTCTCTCTGATATAAAAATTATCACAGGCATGGAAATTGTGGACAGAAAAAGCTGAGACACAATTAATAGAGAAGTCATTACTTGAATCATGCAACAGAAGTGAAGCAATAAAGTGCATAAATATTGCGCTCCTGTGTGTACAAGAAGATCCAAATCATCGTCCTAATATGTCAGATGTCATTTTGATGCTGGGAGGGGAAGGTACTAACCTTCCAACACCTAATAGACCAGCTTTTGTAATAAGGACGCATGCTTCTAGTACATCTTCTTCTTCCTCCGATAAGAAATACATCGTATCCAACAATCAGGTGACGATTACAGTTGAAGAAGGACGTTAG

>M6-PGSC0003DMG400008524 M6 G-LecRK

GTGAAGTTTCTCCACTTGCCATAATCTTGTCTCCAACCCCATTAAGAGAAAAAAAAAATAAAATTAAACAACAGAGCAAGAAAAGCTGGAGCTCAAGAAGGTGAATGATACTCAAAGAAGTGAAATACAGGAAGAAAAATTGTTTGGATTTTCTAAAATCCAATTTAATTTAAGGTAAATTAGTTTTAGGTACTTTTAAACTTTATTTTGGAGTAAATATAATGTGTATGTTATTCATTTTCTTTTTTATGGCTTTTATGGTTCTTGAGTGATTGATTTTTGGATTTTCATCTTGGTTGTTGCATGTTATTTGCTTTTTGCGTGTGTGTGTGTGGATTTCATGGTTCTTGAGTTTATTCTGATTCAATAATGAATTCTTGAAAGTTAGTATAGTTTTTCATATTGAATTTACATGTGGAAACTACTTTTGATTTTGACTTTTATGTTATAAGCATTCTAGGTGGAAAAGTTATTTAAAAATTGTTATATATAACTTAAATCCTTTCGACTTTTGTCATTTCAGGCATTTATGTTTGTTATTGGGTATATGTGTTATCAATTACTACAATTACCTAAAAAGAAAAATTGTGTCTTATTACTCTCAACATAATTTCGTTGTCTAGCAAGATACGTCTAGTTCTCCCTCTACTTACCTATCCATTAGGAAAGTGGAACTTTCTTGACCTTCGAAAAGGTATAAAATGGTTTTATAAATATGGTATGTTATATGGACTAATTTGTTAGGTGCTACTTGCTTGCTTCAAGCACAATACTTTGATTTCTTTTTAAAGTTTTAGACAAAGGCGGAGCCAAGATTTAAAATTTATGGGTTTTAGATTCTAATCTTTTATGTTACTGGGTTCTAAATTGATAAAATGTATAAATTCAAAAGATTTTAAAGACAATTACAAGATTTAAACTAAAACTACTTTATTCAGACTTAACCCTTAGCTGATATTCTAGCTCCGCCCCTAGACTTTTTTATTAGCATCCTCAAATACTAAAGTGGTGTTATTGCCCCTTCTTTTGTCACACTTTTTAGCTAGTTTTCTCAAAATGCATGAAAAGGAAGAGATGCTCTATTTCATCTTTTAAGTTTAATTCTTGAGGCTATTTCTTGAATTCATTTTTTTTTTTAAAAAATAAAATAAAATTGAGGGCCTAAGAAAGGGATTACAATGTGGGGAATCGAATACTCACACACGTAGACAACTAACTGAGTTATTGAGATTCTCCTTTATCTTGATTTCACATGATTTGCTTATATTTTGTTGTGATCCAAGTTGTCTATCTATTTTAGCTTTAGGTTATCAGTTCTGTTCCTATTTCAGGCTAATTGCAAATGATTTGAGGCTTTGAAGGTTTAAGAGGATGTTGGACTTAAACAAAATAGAACCTGAAGAAGGGAAGCTAGACTTGGATCTTCGTCGCGTTAACTATGATCTTGATGACAAATCTCATCTTAGATATGGTGATTGGGTCACTTTCAAGAGCCATTCATCTAAAAAAGTTAGAATCTCTCGACCTGAATCTCCGAGAGGTGAAAATTCAGATAGACAACCTCAGGATGCAGATTATCCCTCACTTAGCTATGAACTCGAAAGTGAGATTCTAGCCAGGTTTCCTAGGTGGGAGTATTGGAAACTTAGCCTCGTAAACAAGAGATGTTCAATGCTTCTAAAGAGCGGTGAGATATTTGAGATTCGTAAAGAGAATGGATTTAAAGAACCTTCGGTATATATGTTAGCAAGTGGGGAGACTAACTGGTGGACATTTGATCGAGAGTTTAAGTCCCGGAGGAAAATCCCTGACTTACCATCAGATGTGTGCTTCACATTTGGCGATAAGGAATCGCTTTGTGCAGGCACACATTTACTTGTTTCGGGTAGGGAAATCGATGGCCTAGTTATATGGAGGTTCGAATTAGCAACGAATTGTTGGTACAAAGGTCCCTCTATGGTTAATCCAAGATGTTTATTTGCATCAGCTACTTGTGGCACCTCTGTTTTTGTTGCTGGTGGTGTTGGTATTATGGCGAATAGTGAGGTATACGACACAGCTGAGAAATACAACCCCGATAGCAGATCGTGGGACCTACTACCGAGGATGAAGAGGAAGAGGAAGCTTTGTTCAGGATGTTACATGGACAAGAAATTCTACGTGATCGGGGGGAGGAATGAGAACGGAGAGCTTACCTGTGGAGAATTCTTCGATGAGGGTATGAACAAGTGGGAGCTGATTCCGGACATGTTGAAGGATGATCCAGTCCTAACATGCCACTCACCACCCCTCATTGCTGTCGTGAACAACGAGCTGTACTCACTTGAGGCATCTTCAAACCAGCTGAAGCTTTACCTGAAAAAAACCAACACTTGGAAAATGTTAGGACAAGTGCCAGTGAGAGCTGATTCAAACAGAGGATGGGGCATTGCTTTCAAGTCATTGGGAAATGAACTTCTTGTAATCGGAGCTGCTTCGTCTTCAGCCTCTTATTCGGGTAATTCCATGGCTATATATACTTGCTGTCCAGATCCTGATGTAATTGATTTGCAGTGGAAACCTCTAGACAGTGGTCGAAATCGGCTTAGTAGTTTCATCTTAAACTGTTCTGTTATGGTAGCTTGAATAATGTCATTTATCTAGCAAACAAATTATGTATTGTGATCAAAATATATAAGTGGCATATGGCCACTGGTTTCTGATATTTTCTTGCATATATTCACTACTGTTTGCTCTGCTATGTTTGATGTAATTTGCATTTATTGTTGCATTATTATGATTTATATTGAAGGTGTTATTCTA

>M6-PGSC0003DMG400010240 M6 G-LecRK

TCTTCTTTGCTTGTAATAACACTTTGTTCCTTATTCATAGTAACATTAGCTAATGTTCCTCCTTCCAAAACCTTCAAGTACATAAATGAAGGCACATTTGGCGAATACTCCGTTGAATATTTTGCAGATTATCGTGTGTTGGACATTGTCAACTTCCCTTTCACCCTTTGTTTCTTCAATTCTACTCCAAACACCTTCATACTTGGGCAACGAATGGGTCATCGTAGATCTGAGTCCATTATGCGATGGGTCTGGGAGGCTAATCGTGGTAAGCCTGTTCGCGAAAACGCCACTCTAACGTTTGGTACTAATGGGAATTTGGTACTAACTGATGTCAATGGCACGGTAGCTTGGCATACCAATACTGTCAACAAAGGTGTAACTGGGCTAAAACTTTTGCCAAATGGCAACTTGGTTTTACATGACAAAAATGATAGATTTGTTTGGCAAAGTTTTGATCACCCAACGGATACACTTTTGGTGGGCCAAAGTTTAAGGGCTAAAGGCCCATCTAAGATTGTTAGTAGAAAGTCCAGTAAAGAACCTGTTGATGGGCCTTATAGTTTTATCCTAGAAGGGAGACATTGGGCCATGTATTATACAAGTGTAAATTCACAAAAGCCTTTACCATATTTCAAATCAGATGATTTTGGTAATGGGAAAGGCTCTTTAGCCAATTTTGATTTCTACTGTGAGCCAGAATATGGTGAAAATTATGCTTTTGAGCTAGGATTTAGGATCAACATGACCAATTCTCCTAGCTCTGGCACATCTATATTTACTAGGCCCAAATACAATAGTACATACACTATGTTTAGAGTTGATATGGATGGTGATTTAAGAATGTACACTTATGAACCAAATGTTGATTGGGGAGCATGGGAAGTCACATATCGACTTCTCGATAGGGACAACGATGGAAAAAGTGAATGCAAGCTACCGAGGAGGTGTGGATCATTAGGCGTTTGTGAGGACAGCCAGTGTGTGGCGTGTCCGAGCCCTCAAGGCTTGATTGGGTGGAGTAAGGGATGTGCACCACCAAAGTTGCCAGTATGCAGGGGAGGATCAAATGTGGACTATTATAAGGTGGAAGGAGTTGAACATTTTACTAGTGATTATAATGAAGGAGTTGGACCTATAAAAATAAGTGAATGCAAAAATAGGTGCAGCAAAGATTGCAAATGTGTGGGCTTTTTCTACAAACAAGAGTCATCAAAATGTTTGGTTGTACCTGAGCTTGGAGCTTTGGTTAAAGTGGACAATTCCTCCCATGTTGGCTACATCAAGATCTCAAAGTAGATTAAGTTATTACTGATTTGTATTTATATCAAGCAAGCATTTTATATTTTTTTAAAATTTCAGTAATAATAATGTAACTTTTGTCAAAATATAATGATAAAAAATGCAC

>M6-PGSC0003DMG400011251 M6 G-LecRK

ATGTTAATTTCTCCAAAAAGGAAATTTATTTTGGGATTTTTTAGTCCAAATGTTTCTAATAATCAAAGGTTTGTTGGTATATGGTATGTTGATGGTCCAAAAAATTCATTTGTTTGGGTTGCAAACAGAGATAAACCAATTTTTGATAAAAATGGTGTTTTTACTATTGAGAAAAATGGGAATTTAGTGGTGAAAAATGGTCGTGGCGATTTAATGTGGACGAATAATGTTACTGCTATGAATAATAATTGTACAGCACATCTTTCTGATAATGGAAATCTTAGGATGTTTAATGGCAATAACAAGGAGTTGTGGCAGAGTTTTCAGCATCCTACAGATACATTTTTACCAGAAATGAGATTTTATTTGGATGAAGTTTTGAGGTCATGGACTAGTGATAGTGATCCTTCACCTGGACGGTACGTAATCTTGTATCGTTTGATCATCTTATCTGAAATCTTTTATTTACTTAATTTTCTTTTGTACATGCCCCCTCACGTTTGGTATGAATTTTTTACATAATGTGTGGTGGTGAGACTCTCTGATAGCATGTTGCGTTTGTGACATCTCATTTAAAAGCTTAATTGAGAATGTGCTTTTTTATTTTACGTAATTTTGTTTTGAACACAACCCCTCACGTGTTTGACTCATGGAAGTTGAAAGACAGAAGGGTTGTTGGTCAGTCACGGACTATTTGATTTCATTATGACAATTCAATGTTAATGACCTCATTCAATATATATATATTTTTTTGATCCTTCACTATTGTGGGCTGGATGGATAGCTAGTAGTTGTGAGTAGCAGTCCCCTTTTTATAGATTAAAGGTGATGATTTTTTGATATGGGGTGGTAGGGAGACTCGAACCTACAAACTTTGCTTGCCCTGATATGGTAATGCATTTGTGACATCTTATCGAAAAGATTAAGTTGAAAACATGTTTTTCTTTACTTAATTTTGTTTTTAACACGACCCCTCACGGTTTCTTGAATCTTTGATATTGGGTGATAGTGAGACTCGAAACCTAAGAACTCTACTTGCACTGATATGGTAATGCATTTGTGACATCTTATCTAAAAGATTAAGTTGATAACATGTGTTTCATAACTTACTTTTGTTTTTAACACAACCTCTCATGTGCTTCTTGAATCCTTGATATTGGATGGCAGTGAGATTCGAACCTAAGAACTATGCCTGTTCGGATACAATGTTGCATTTGTGACATCCCAAAGTCAAAGAGCATATTTTTATTTACTTAATTACGTTTCCAACAGGTACTCATTAGGAGTAAATACTCGCGGATCACCACAAATTGTGATTTTGGATGGAGAGAAGAGACGTTGGCGGAGTGGATATTGGGATGGACGCATATTCACAGGAGTTACAGATATGAAGCCCGAGTATCTACATGGTTTCAAGCTTTACAATGAAGGAGATATGCTATACTTCACTTACACTGTTGCAGATTCTTCTGATTTGGTGAGGTTCCATATTAGTCCAACCGGATACGAGCTGGAACAAAGGTGGGATAAAGATAACGAAAACTGGAGTATAATACAGTCTCATCCATCAGGTGATTGTGACTTGTATAACTTATGTGGGAATTTCGCGAAATGTGATGTCACATATTTGAAAAAATGCATTTGTTTAGTTGGATTTGTACCGAAAGATTTGGGGCAGTGGAATGCGAGGAACTGGTCAGAAGGGTGTGTTAGGAGGAAAGAACTGGAATGCAGAGGAAATAACAGTGCGTTGAAGAGTGTTAGTGGAAAGAAGGACGGATTCTTTGAGATTGAGAAAATCAAGTTGCCGGATTTTGCTGATACTGCATATTTACAAAATATCGATGAGTGTCGAAGCATGTGCCTTGAAAATTGTTCATGTACTGCTTATGCTTTTGTTAGCGGAATCAATTGCATGATGTGGAGTGGAGATTTAGTCGATATGCAGCAGTTTCAGGAAGGCGGAAACACTCTCTATGTTCGCCTTGCTGATTCTGAATTTGGTAAAAATGTGTACTAACTTTTCTGTTGGTATCATTTTAGATGTTCTTATTATAAACTTTCTTGAAACAGCTGGAAGCAACAGGACAGTCAAAATTGTGGTAATACCCGTTATGGTAGCCGGGGCTTTACTTGTTTGTATGGTAGTTTTTCTACTATGCAAGTACAAAGCCAAAACTCGAGGTAAGATTTCAGAAAGGCTGGTGTTTTACGACAGTCATTTGTTGTTATGCAATATACCTATATCTTCTTCAATGTCCCATTTTCAGTGTCCAATAGGATCAACCAAATGGAGACAAGCGATCCAACTAGGAGCGGAGAGTTTTCTATGAACACATCAGGAGCAGGAGACTTGAGTATTGAAGGGCATCAAGGGAGTGGTTCAGAACTTATATTCTTCAGTTTCAGCGGTGTAGCAGCAGCTACCGACAACTTTTCTAACGAAAACAAGCTGGGACAAGGTGGATTTGGCCCTGTCTACAAGGTTGGTTTTCATTCAATTGACAAGTGTGTCTTACTTTTCTAGAGAAGTATGACATTATTGTGTTAAACTAACGTCGATGTTTGTTTATTTAGGGAAAACTACTATGTGGCTTAGAAATTGCAGTGAAGAGACTTTCAAGAAAGTCAGGACAAGGTGTGGAGGAATTCAAGAATGAGATCAAGCTAATCGCCAAATTACAACATAGAAATCTTGTTAGACTTATGGGATGCTGCATTGAAGGAGAAGAAAAGATGCTTCTTTACGAGTACATGGCAAACAGAAGTTTAGATTCATTTCTATTTGGTTAGTACTCTAGATATGCTATAAATTATTTCATCGATAGAGAGATCGATCACTTACAATTATTTTGGTTGGTAAATTTTTTGACAGATCCTGTTAAGCAAGCTCAATTAGACTGGAGGAAACGCTTCAACATTATTGAAGGGATTGCACGAGGGCTATTGTATCTCCATAGAGATTCGCGACTCAGAATAATCCATAGGGATTTAAAGGCTAGTAACATTTTGCTAGATGAAGATATGAACCCGAAAATTTCAGACTTTGGTATGGCTAGGATATTTGGTGGAAATGAAAATGAAGCAAATACAAATAGAGTAGTTGGGACATAGTGAGTATCATTTGATTTTTATACCTTTAAGAGTGAGTACCTTTTAACTTGTTCAAAGAACTAACTCCATCAACGCGTTTCTTTTTTTGTTTTTACTTAAATCAGCGGATATATGGCTCCTGAATATGCAATGGAAGGCTTGTTCTCCGGAAAATCTGATGTTTATAGCTTTGGTGTACTATTACTGGAGATCATCTGTGGACGGAGGAACACAAGCTTTCGTTCAAATGAACACTCTGGCATTATTGGTTATGTATGTCCAACAACTACTTTTCAATATCTGTTGTTAAGTTAAATCATTTAACAACAATTCTACGTTTTTCTTTTTAGGCATGGCAGAAGTGGGATGAAGGTACACCAATGGACTTAGTAGACCGTTCTATTTGGGACGAGTGTCAATATGATGAAGCATTGAGATGTATACAGTTAGCACTAATTTGTGTACAAGACATGGCAGTTCACAGACCGAGCATATCATCCATCGTGTTAATGTTGGAGACGGACAATATACGATTGCCCTTGCCTAGGCAACCTACATATACCTCAATGAGAAAACATGAAAATGCAGAAACATGGAATGAGAAGCAGGATTTTTCTGCAAACAATGTCACAATTAGTGTTATATTTGGTAGATGA

>M6-PGSC0003DMG400013383 M6 G-LecRK

AGACTCCATAGCTGTAAAGCTTGCAAACGATGTCGCTTCATCCTTCTTCTTCATCCTCACTTCTCTTCTTCCTTCTCTTCTTCTTCTTGAACCCCTTTTTCTCTTCTTCACAAAAAATCCTCTCATCATTCAATTCCAAATCCCCTCCTTGGAATCCAACCCAAAACCAAATCCTACTTTCACCCAACTCCACTTTCGCTGCTGGGTTTCTTCAATCTTCCCAAAATTCCTTCAATTTCTCCATTTGGTACTACAAAATCCCAGTCAAAACCGTTGTTTGGTCAGCTAACCCAAATTCCCCACTTAATTCCTCAGCTACCCTTTTCATTTCTTCCTCCGGTGAACTCAAGCTAACCCCATCTACATCTTCCTCAGCTCCAAATTTATGGCCTTCAATTATTCGAAACACCTCCTCTGTTCTTTTCCTACAAGAAGATGGAAGCTTGGTTTATGCAAACTGGAACAGTTTCTTGAACCCAACTGATACTTTTCTTCCGACCCAGAACATCACCGGAACAAATTTAACTTCCGGTAATGGAAAATTCCAGTTTGATGGATCGAATTTGAATACCCTTTTCTTCGATGAAAATGATAGTTATTTCACTTTTCCCCAAAATGCCCTTCAGAGGTTGGAGGAAACAGGGGAAGTTACTCAAGTTAATGGAAGATTTTTTTCGTCGGATTTCGGGGAGAAAGGGAAACTGAGAAGGATGAAATTGGATGAAGATGGGAATATGAGAATCTATAGTTTTGATCTTAGTTCAAGTTTAGCTAAAAATTGGACAATTGTTTGGGAAGCAGTTAATCAATTATGTACAATACATGGTACTTGTGGTACAAACTCTATATGCTTGTATGATACTTCTACAACTCAAACTTCTTGTGTTTGTCCACCTGGTTTTAGGAAAGACGCTAGTAAATCTTGTGTGAGGAAAATTCCATTGATGACGAAGGATTCGAAGGCTAGTAAGTATTTACCTCTTGATTTTGTTAGTTTTACTGGTGTGGGTAACCAAACAGACCTTAAAGCTTTGAGCTTTTCGAGTTGTGAAAAGAATTGTTCAGATAAGAATGATTGTTTAGGTTTTTTGTTTAAGTATGATGGGACTGGATATTGTGTGCTTGTTTTGGAGAAATTGTTGTATGGATATTGGTCACCAGGTACTGAGTTTGTTATGTATTTGAGAGTTGATAGTAAAGAGAACGATATATCGAATTTCACAGGCCTGACTAGTTTGATGGAGACTTCATGTCCAGTGAGAATTAGCCTTCCATTTCCACCAGAAGAGTCTAAGACAACCACTAGAAATATTGTGATCATTTCGATGCTTTTTGCAGCTGAGTTGATTAGTGGAGTGTTCTTCTTTTGGGCATTTCTCAAGAAGTATATTAAGTATAGGGATATGGCTAGGACTTTTGGTCTTGAAGTCATGCCTGCGATTGGACCGAAAAGGTTTAGCTTTTCTGAGATCAAGAATGCAACTAATGATTTCACTGATAAAATTGGGAGGGGTGGATTTGGAGATGTTTACAAAGGGAAGTTGAGTGATGGACGTGTTGTTGCCGTGAAGTGCTTGAAGAATGTAAAAGGTGGTGATGCTGAGTTCTGGGCTGAGGTCACGATCATAGCACGTATGCACCATTTGAATTTGGTGAGGTTGTGGGGGTTTTGTGCTGAGAAGGGTAGAAGGATACTTGTGTATGAGTATGTACCTAATGGTTCATTGGGTGAATTCCTCTTCCAAAAGGCACCGATCCAGTCACCAGATGGGCAGAAACCCATACTCGATTGGAATATAAGGTACAGGATTGCGCTTGGTGTGGCACGAGCAATTGCCTATCTACATGAAGAGTGTTTGGAATGGGTATTACATTGTGATATCAAGCCTGAAAATATCCTCTTAGGCGACGATTTCTGCCCAAAGGTGTCTGATTTTGGGCTAGCGAAGCTAAAGAAGAAAGAGGAGATGATGACCATGTCAAGATTTCGAGGCACGCCAGGATATGTAGCACCAGAATGGACCAAGGCTGATCCAATTACTCCAAAAGCCGATGTGTACAGCTTTGGATTGGTGCTGTTGGAGATAGTTTCAGGTACAAGGAACTTTGAGCATCATAACTCAAAGGTGGAAAGTGATCAATGGTTTTTCCCTGCTTGGGCATTTGATAAGGTGTTTAAGGATATGAATGTTGACGATATTTTGGACCCTCAAATCAAACAATCGTATGATTCAAGGGCACACTTTGACTTAGTGAACAGGATGGTGAAGACAGCAATGTGGTGCATCCAAGATCAGCCAGATGCACGACCATCAATGGGGAAAGTGGCTAAGATGTTGGAAGGGACGGTGGAGATCATCGAGCCAAAGAAGCCTACCATTTTCTTCTTATCAGAGGAATAATATAAGGCTCAATCCTCTTCATAATTTCCTCAAAATGCTGTCTCAACAAGTTGCAGTAAGCATGTATTCCATGTTGTTTGAATTTGGATCATGTTGTGTTTTGTCTGTTTTTTTTTTCCAGTTTTGTAATCTCTTGTGTTGTAGTTGTAAAGTACATTTTGATCAATGAAGAAGCAAATGGTTGTTACATAGAAG

>M6-PGSC0003DMG400013882 M6 G-LecRK

TTCATTTTGGTCCAGAGTTTAAAGTTTGAGGAACCTAAACAGAGAAGATTACCTTGTTAGAAAGCATCAGCTATGTGTCAGAAGAAAGAGTTGTTGATAGTAATTGTTTCATTTCTTGTCCCATTTTGTAGTTCCATAGATACCATTTCCTTTAACCAATCTCTCAAAGATGGTGATTTGTTGATCTCTTCAGGGAAATCTTTTGCTTTGGGTTTCTTTGGGAATTCCCCTGGTAAACGTTACGTCGGAATTTGGTACAACAATGTTCCTGAACTAACTGTTGTTTGGGTTGCCAACAGAGACAATCCTGTTAATGGCACATCTGGGATTCTCACCATCGATTCAACTGGAAATCTTGTTGTACATCTTGATGCGGAAACAAAAACTACAGCCTGGAGAACAAATGTTTCTTCAGCTACAAAACGAGCAAACTCGTATACAGCTAAGCTGGAGGATTCAGGGAATTTTATATTGTTTCAAGACTCCAAAATGGATGTTACTGAGTGGCAGAGCTTTGATTATCCTACTAATACTCTGCTTCCTTCAATGAAATATGGAATAGACAAGAGGACGGGTTTGAACCGGTTTCTAACATCCTGGAAATCGTTGAACGATCCAGGCACTGGAGAGTATCGTTACACAATGGAGCTCAATGGGACACCTCAAGTTTTCTTGTACAAGAACTCTAGCAGGATTTGGCGGACAGGATCCTGGACGGGTCATGGTTGGAGTGGTGTACCAGAAATGAGCCCACGATTCATCTTCAGCCTTAGTTATGTGGACAATGATACCGAGGTTTCCTTGACATATGGCATACATGACAGTTCAATAATCTCAAGAATGGTCCTGAATGAATCAGGGATCGTGAACCGGTTAACTTGGCAAGAGAGTGAGCAAAAGTGGGTGCAATTTTGGTCTGCCCCTAAAGACCCTTGTGACAATTATGAACATTGTGGAGCATTTAGCAATTGCAATCTGTTCAATTTGGCTGAATTCGAATGCAGTTGTCTTCCAGGGTACGAGCCAAAGCTAAGCCGACAATGGTTCTTGAGAGATGGCACTCATGGATGTTTGAGGAAGAAAAATGAAGAGGTGTGCAACAGTGGTGAGGGATTCGTCACATTAAGTCATGTAAAGGTCCCCGACACAGGTGCAGCTCGGATGAACAAGAGCATGGGATTGAAGGAATGTGAAGAGTTGTGTCTGAAGAACTGCTCCTGCACAGCCTATGCAAGTGCTAATATTAGTGCAGGAGGAAGTGGATGCATCACTTGGTATGGTGAGTTGACAGACATAAAGCAATTTATAGATGGGGCCCAAGATTTATATATCAGAGTCTCTGCATCTGATTTAGGTAAGGAGTTAACTTAATATATTATGTGAGGATTTAGGTAATTCAATATTATGTGGAGAAGAATGTGCAAGTTGATCATTTTTGGTTAGTAGTAGTACACAGTATCTTACCCTGGTGTTTACTGGAAGTTCTCATTTTTTGTTTGTTCAGCGCAATTCTCGAAGAATTCAAGAGGCCATAATAGGAAAAGAATGATAGGGATTCTAGTAGGGTCTGCTGCAGCAATAATTCTTTCACTCTCTTTGGCATGTTGTTTGGTGATCAATATAAGGAGAAATGGTAAGGATATTATTCTACATCATACTTTAGTCGAATATTTTCAGTATCTGTACAGCAGCTACAATGGTGAAGCAACTATAGCAACCTAGACCTTCTACTACCTTTCTATATCGTGCAGCAAAAACAATAACATTACTGCAAATTTCATTTGATCCTAATCGGTGTTTGCTGATGTCACAGATAATGAGAGAAGTGAAAGTATAGCATCCTATGACGGCATGGATGAAAGTAAACATACAGAAATCTCTATCTTTGATTTAACCACAATAACTAATGCCACTGATAAGTTCTCTGATGCTAACAAACTTGGTGAAGGAGGATTTGGTAGCGTATACAAGGTAACAACTCCCAAACATGATTGAATTCCTAACAATCAACATTATCTTTATAGGAGGCTACAAATAGATTTAATAATACTTGTTAAACCAAGATTTTCTCCCTAAAAATAAGAAGAAGAAGCTATGTTTACCAAATAATAATCTGGAATCAATATCAGGGTCACCTGACAGATGGACAAGTAATAGCTGTCAAAAGACTTTCAGTAACTTCAGGGCAAGGGACCGAGGAGTTCAAGAATGAAGTCACACTAATTGCAAGACTTCAGCACAGAAATTTGGTGAGGCTTTTAGGATGTTGCGTTCAACGAGGAGAGAAGATGTTGGTTTATGAATATATGCCAAACAAATCCTTGGACAGTTTTATTTTTGGTAAGGAACATCTCCAGAACCGGAATTTCTTCCATTGACCTCCAGTCTCTACATGCTTTTACATGTTTACTGATGATTAATGATTACTGCTTTAACGTGTGTGTGTGTCCCTAAAAGATTTCTTATGCACAAATTTTTAAACTGTGATCCAGATAAAACAAAAGGGTCTTTGTTGGATTGGGGAAAACGGTTTGAAATCATACATGGGATTGCACGAGGAATGTTATACCTTCACCAAGACTCTAGACTAAGAATTATTCATAGGGATCTAAAAGCAAGCAATGTTTTACTCGATGCCTCCATGCAACCAAAAATATCAGACTTTGGAATGGCCAGAATATTTGGAGGTGACCAAATGGAAGCAAACACAAATCGAGTTGTTGGAACATAGTAAGTATGTCAAGTTGTGTATAAACATAAATGAGTTATGTTAGATATGTGCATGTGATCTTTAAAATGATAAATCTTGACATTGCTTATCTATTGCAGTGGTTATATGTCACCTGAATATGCAATGGTAGGGCACTTTTCAACAAAGTCTGATGTTTTCAGTTTCGGTGTTTTGTGTTTGGAAATTATCACTGGCAGAAAGAACAACAGTCATAAGGATCAGGAGCAATCTCGACATTTAGTTGGATATGTAAGTACAAACATATATATGCTGTTAAAATCGATAAATTGCATTTATTATCTTACAGTTGCCTTAGACATAAGCTGTGTCATGGATGGTAGTACTGCAAATAATAATTTGGAAACAATTCCCTTTCAGGTTGTTAAAAGTGCTGAATAGATCGAGATTTTGTATTTCAGAATTTCAATTTTTGGTGTTATAGTCGCTTCTCTTATAGAGAAGAAATTGTCTGAAATTTTGATTGTGATGCATTGTGGTGTTTCTATTTCAGGTTTGGGATCTTTGGAAGGATGAAAAGGCATTAGATGTTGTTGATCCATTGTTAGGTGGCTCATATGAAGCTTGTGAAGTTTTGAGATGCATCCACATTGGCCTTTTGTGTGTACAACCGTTTACAGATGACAGACCGACTATGTCAGAATTGGTCTTCATGCTGTGCAATGAAACAAATCTTCCTTTCCCTAAACAACCTAGCCTTATATTCAGATCAGAGAATCATAGTTCAGTTAAACACTCTTCATCAGCAAGTATAGCAACATCTTCTGTCAACGATATGTCTATTTCAACAGTTCATGGTCGCTAGAGCTTATGCTAACTATTCAAGAACTATTTTGAAAGAAATACAGAATTCCTTTTCTGTACTTTGCTTTCATATTATTAGAGAGATTTATTGGTGAGTTATACACAACATTTAACATGCTAGTCCATTCTTGATTGAAACTAGTGGGGTTGGGCTCTAATAGCACTGGAACCAAAACTGACTTCTGATTCCCCATGGCCCATGTTACAGACTTAAAGGTAGGTGGGTGGCAAAGGATTGGTAGGAAAGAAGCATAAACACAGAGGGTTTCTATTTCATCTGTGAATTAAATGGTGCTCTTTGTCAGCTGTATTTGATAATTGCCACACTTCATGTTTTCTTTAACAGTCTTCTCTGTGTTCATCAATGCTCTAAAAGCAACAAAGGCTAATCCTGCTAGCCACACACAACTTTTCAGTGTTTTTATTGATTTGCAGGATAGTAATTTGGTTGTGGAGTTTTGTCAATTAATGAGATATTTAACTATTATTAGTTGGTTATGTTTTTCCTCAAAAATTATTGATAAAAAAATACAAACATGTAAGTTATCTTATGAAAGATTTTAACTTTTGTTTCATAATTAGATCATGAGATAGACATGTATCAAATACAACAATAATATTATGATCATGAGTAAGAGCACTGTTGTCTCAATCTTTCTGTGATGAGTATCTGTAAAAGTAATATCAACTAGTCTATTGAAAATCTTTTAGCATTATTTGCATTGAAACATTGAAGACTATATCATACTTTGGGAAGTAATGAGCACTAGATACATTTGATAAACAGATAACAGGAATACATATACACATCAGACAACTAACGAACATAAGGTGCTGTAATTGTCACCTCGTTGATCGAAGATGATGCTGTTTCGGTAGTAATCAAGTCTGAGTTAGGGGTAATGAAGGGCTGCTTCGGGAAGGGAACTGTTGCTTCAATACTTAGCATGGACAGAATTGTTGACATGGTTGGTCTGTCTGCTGCGTTGAGTTGCAAACACAATAGTCCTACTAGAATGCAGCGTAATGCTTCCTCACTGGGACAAGAATCAACTATTGATGGATCTACCAGCTCAAAGGCCTTACCATCATTCCACAACTCCCAGGCCTGAAGCAGCAGTTCA

>M6-PGSC0003DMG400013885 M6 G-LecRK

GGGATGTGTACCAAAATACCATCTTCCTTCCTTCCCTCTTCCTATCAAACAAGAGAAGAAGTACACCAACAAATGTCTCAAACACCAACTCTGCAGCAAAACAAAATACTCTTTTCTTAAATTCCACAATGAGAAACACCAAAGATTACTACTTTCACCAATTCTTGTTCTTGATTTTGATTTCTCAATTTCTTCATCCAATTCTTGCTATACCCACTGATACAATCACCCCAACACAGCCTTTAACCAAAGATCAAACCTTAGTATCTTCCGATCAACACTTTGAATTAGGCTTTTTCTCTCCCGGCGGTGCAAATTCAGACAAATGGTATATCGGTATTTGGTACAAAGAGATTCAAGACAGGACTATTGTCTGGGTTGCAAACAGAGCTAAACCCCTTTCAGCTTCATCAACATCAGTCCTAAAAATCGCAGAAATTGGTACCCTTCTTCTCGTCGATGGCCAAACTGGAAATTCCGTCTGGTCTTCCGAGCAAACTCCGGCGACCAACGTAGTTGCCCAACTCCTTGACTCCGGGAACTTTGTCATTCGACCGGAAAACGATGACAGAGAACAGAGCTACCTGTGGCAGAGCTTTGATTATCCAACAAACACTTTATTACCTGGGATGAAACTTGGGTGGGATTCAAAATCCGGGATGAACAGAAACATAACATCATGGAAATCAGCAATCGATCCAGCTCCCGGAGATTACACCTTCAAGATCAACACCAGTGGGTTCCCGGAAGTTTACTTAACAAATAAACAACAAACTATCTACAGGAGCGGCGCTTGGAATGGAATTAGATTCAGTGGGGTACCGGAAATGAAGGCAAGTGAGATTATCTCATTCGAATTTCAGTTCAAATCAGATGAAATTACTTACATGTTCAAATTACACAACAAAACTTTATACTCAAGATTGTCTGTGAATCATACTGGTTATCTTGAAAGATTCACTTGGATCCCAACAAGCAACATGTGGAATCGATTTTGGTATGCTCCAAAAGATCAATGTGACGGATACACAGAGTGTGGGGTATCTGGAATTTGTGACACAAATATTTCACCAATCTGCAAATGTATGGTGGGATTCAAGCCTAAAAATCAAGTGGCCTGGGAATTGCGAGATGGGTCTGATGGTTGTGTTCGATTTCATAAATTGGATTGTAAAACTGATAAATTTAACGTATTGAGGAACATGAAGTTGCCAGATACGACGAATTCGTTTGTGGATACAAATATGAATTTGGATGAATGTGAAGCTATGTGTATGAAGAATTGTTCTTGCACGGCCTATACAAATTCGAATATTAGCGGAAGTGGGTCGGGTTGTGTGATTTGGAGCACCGAACTTATTGACATGCGGCAGTATACGGCGGCGGAAGGTGGCCAAGTTCTCTACGTTAGGGTGGCTTCTTCTGACGCAGGTATCATCCTATTTCTCTACTAGCTTCGCTTGCCATTTCTTTTTTCGCTTAGTACAATGATAACGATTAATAATCGATAAAAAATAGTTGTATTAAAAAGAATTTGTACAATTAATTTTAAGTAATTTGGGGTTGAAAATTGATGGTTAGATAATCCACTTCTCTGTCAAGATCAAATTTTGGGTCACCGGGATGATTTGAGTCGACTAGGTCTGCACGAGTGGAGTTAGTTGGAATTTTTTATTTACTTGTTAGGTTAGACGATATTTCAACACAGAGAAGTAGAGTAAAATAGAAAGGGTGATGAAGTTTTTGGAAAATATTTTTGTGGAAGATTGGAGAGTATTTCCATGATTGAAAATATGCAATTTAATATTCTCGTTTAAGTTTATGTGTCACCCAATGTTTTTAATTTAAAAATAAATAACATATTTTCATGTATATAGTAACAATTCAATTTTAAAATGCTTATTTTATTTCTAATGAGATAATTTATAGCTATACAAACGATGACTTGTATTTTACCACAAATTTCAAATGTCTTTTTTTAAAATTCTGTGTCAAATTAAATTAATATCACATAAATTGAGACGAAGAAATAGTAATAATATTTTGCATATTTTTGTCGGAAGTTGATTTATCTCATTCATTTGTAAACTTGCTATTCTCTGCATTATTCCACTATATACATACTGTTCTGTTTGAATCTATTTTCATAATTTCAAAGTCGAGTTCAAGACGAAAAATCTATTTGTCCAAAAATTGTTAGTACCAGTTTTGCATGTCATCCAATCCAAATGGACCAAATATTCCTTGAATAATATTTTAAATGTATTGTTTTGTCATTTTAATAAGAAGCATATTACACATTATCATAATTTTAATTTACTCTTTTCCTATTGAATTCCAAAGTTGGGAAGCATGCCGTTAGATTTGCCTCTAATAGATGAAATTACAAAAATATATTTATTTTTTTGAATTTTTCTTTTCTATTTTTTTTTTTTTGGGTATTGCTATTTTTTTTAAACCACCCCCCCCCCCCCCCTCCACAAACGTCATGCAGAAATTAAATTAGAGATTTTATTTTGAAAATAAATACTCTACCCCCTTTGAACAACACGTATAGTTGTCTTTTAATATCCTTGGTCCCCTTTTTAAATTCCCTTTCACGTTACCTACTTTCGATCTAATATTTTACATAATCAACTAGATTTCAAAATAAAAATAAAATTCATAATTGCTAGTTGAAAAAACAAGAAGTTAGTATAATAATTATTGAGTGCATTTAATCCTGAAAAGGACTTGTAAACAACCTACTTTTCAAAGTTATTCTGCTAATGAGTTTGCCAATATATTCTTTGTGGTATGCACGTTTTCAATATAATTTTTCTTTTTCTATGCTTTGTGTATTTTACTTTGAAAAATCGAAAAGTATAAAAAACCTAACCATATCTACCAATCCCCCATCAACAATTTGCCTTCAATAGGGACTTAATTTTATACAATTTGCTTTCAATAGGGACTTAATTTTATTTATTTACTATATTTGGACTTTCTTTTTCTCATTTGATTTGATAGGGATGATTAGTATTTGGAAAATATGTTACACGGCCAAGTATCTCAAGGCTCTAAAAGTAATCAGCCACCTAATTCCACCTTCTTTCCAATCACCTGTTCATAAATTTGAACAAACATTATCGATTTATTGTCACAATTCAGCTCGTAATAATAATAAATAATCGAAATGTGCATTTGCTTTTTCTTTTGTACTTTCCTCTTTAAATCTGCATGCTTAAGAGGAAACTTTTAATTATGGTGGGAAATAAAATTTCCAAATAAAGCTTATTTAGCTCTAAATTATTTTCCATTAATTGGTCCCACAATAAATGGGTCCCACAATAAATGGGTCCCACAAAGAGATACTCCTTTGCATTTCAAAACGGTGACGACGCAATTTCCTGGTCGGTGTAATGGGCCCTTACATGGCCGTCGATCTTGACTTTTATTGGGCCATCCCGGCCCAGTTGTTTTCCCTTTTTTCTCCTTTTTAGAGAAAAAAAGCATATGTTTCCTAATGTAATACAAGAAAATGATTGGACACCAATTCCTTTGACACGTTTGGGAAATGGTCATATACAGGTTTGAAAAGTAAGGTTTGGTGGGGTAAATTAATTCAAATATTACAAAGGGATTAAGAAGTATTTATTAAGCTAATTATTTTTTTTTACCAAGAATATGTCATGGATGATATTAATAGTTTTTTTTGGACACAACTTTGTGAAGTAGGTGAATTAGATCATGTTTAGTGACACCATTGGACTGTAGTCTTTTCAATTCAAACTTTGCTATTGTCACCGAAAATGATCTTTAACAACACTATAGGTTTTCATACATAAAGAGACATTCAAATTTGGCCTCAGCTTGCAAGTAAACACCTTAACTTTAGTATACATACCTAGACACCTCAAATTATTTCTTCTGTCTCAATTGAACTCTCCAACTTACAAAATGATAATCTAGACACCTTCAAAATTAATGTGTCAAGTCAGCATGGGTGTCCATGAGACATAGTGGGACGAGTTGGAGTGTCTAGTTGTTAGTTGAGGTGTCTAGATATGCTCTTTTGAAGTTGGAGTGCTTACTTGTCGGCTGAGGTCAAGTTTGAGTGTTTCTTTATGTATTATGCCAGTAAAAAATGATGAACTTCTTCTTGCAGAACTAAACTCTTGAATAAAATTAAGCATGATTAATTAACTCGAAAAACTTTTAATTTCAGTGCAAATTGGAGGTGTGGGATCTGGAAATAGTTCTGGCAAGACAAAAATAGTTGCAATAGCTGCTGGGGTTACAGTTGGTATAGCTCTTGTGCTATTTGGACTAACCCTTTGCTTCTTATCAAAGAGAAGAAAACATCAAGGTTCAATCCGAACTAAATCCGTAAACAGAGGTATTATATGAATCTATCATGATAAATAAGATTGTCTAATATAAAATGCTTCATTTTCTAATGAATGTTTCTTATGATCCTAAACCAGGCACTAGTGAACGAAGTCAAGAACTTCTGATGAATGCAACTATTATTCCAAGTAAAAGAGAATTTTCTGGTGAAACTTCCACGGAAGAGTTTGAATTGCCATTGTTTGATTACAGTACCCTAGCTACGGCAACAGAAAATTTTTCTGATGCGACTAAGTTGGGCCAAGGTGGATTTGGTTGTGTTTACAAGGTAAAATCTCAGTTATATGTAAACCATAGTGGTATAATAAAGAAGAAAAAATGATCGACAAACTCAATAATATAATCTTTTGATCAGGCAATGCTGGTTGGTCAAGAAGTAGCAGTCAAAAGGCTCTCGAAGAATTCCGGACAAGGAGTAGAGGAATTTAAAAATGAGCTAAGATTAATTGCCAGGCTTCAACACAGAAATCTAGTCCGGCTTCTTGGCTGTTGTGTTGATATGGAAGAGAAGATGCTGATCTATGAATACCTGGAGAATAAAAGTTTAGATTCAATTTTGTTCAGTAAGTCTCTTTCGTTGTATCGCCAAGAGAGTAGTGTTTCACATCATTTACTTACAGGCTTATATAATTTGTGACTCACATTTGAATATGTTCCTCAGATAAACAGAATAGCTCGCTGCTCAACTGGCAAAAGCGGTTCAATATTATTTGTGGTATTGCTCGGGGGCTTCTATACCTTCACCAAGATTCAAGATTTAGGATTATCCACAGAGACCTTAAAGCAAGCAATATTCTGCTTGATAAGGATTTGACCCCCAAAATATCAGATTTTGGCATGGCAAGAATATTTGGCGGAGATGAGACTGAAGGAAATACAAAAAGAGTAGTTGGAACCTAGTAAGCACTCAACATAATTTCTTCACATAATTCATAGAACCTTTAAAGCTGTTACTAAATCTGTTTTTTTGTTTTGATGTAGTGGTTACATGTCTCCGGAATATGCAATGGATGGCCTCTTCTCTGTTAAGTCCGATGTTTTCAGCTTTGGGGTTTTGGTGTTAGAAATAGTAACCGGGAAGAAGAACAGGGGATTCTATTATCAAAATAACCAACTCAACCTTCTTGGACATGTAAGCTTCATTATTGGTAATACATTCATCATTTGATTCATTTCTGATAAACAAAGTAACCGATTATGCAGGCGTGGAGACTATGGAAAGAAGGGAGCGGCTCAGAGCTACTAGATCCATCTTTTGGAGAATCGTTTTCTCCGTCTGAAGTAATGAGATGTATACAAGTTGGACTGTTGTGTGTCCAGGAACAAGCAGAAGACAGACCAAACATGGCCACTGTGGTTTTGATGCTAGGTAGCGAAAGCGCGTCACTACCTCAGCCTAAAAACCCAGGATTTTGCCTTGGAAGAAGACCCGTTGATTCTGATTCATATTCGACTAATTATGAAGAAACTTGCACTGTCAATCAAGTTACAGTTACCATGATAGATCCCCGGTAGTTGATGGTATATATCGCCCTCAAGAAGTGTATATTACTACAAGAGGTTCACAATTTTCTCTCGTATAATTTCTCAGTTTTTGCTTTTTGGGTAAGTTTTGTGTAAGATAAAGAGTATAATTAGGTCTTTTTTTCTCATTACATACAATTGTGAGTTCAAAAATCTGTCTTTTTTCATCATCAATGTAATAGTCCAATCATTCCTTTTCCATATCCTTGAAATCCATTGTAAATCACTGTGC

>M6-PGSC0003DMG400013886 M6 G-LecRK

GCCTCATCAGAACCTTCTTCTTTCCTTTTGTTTTCTATACCCTTTACCATTCTCCTTTCTAAATTCCCAATTTTCTCCATTCTTGATTTGTGGCATCATCATTTCCAAATTCTTGAATGGCCACTGGCTATAGATTTCCCCCCTTTTGGTTCTTGTTTCTTTTCATTCTTTCTTGTTTCTCCTCATTCTGTTTAGGTGGTGACAGAGTTACACTAGGAGAGATTATTAAAGATGGTGATAACATCACTTCTAAAGGAGGGGATTTTGTGTTGGGGTTCTTTAGTCCTGCAAATACTAGTAAAAGGTATCTTGGTATATGGTATGTTGACGTACCAGTAAAGACATATATTTGGGTTGCCAACAGGAACAATCCTGTTCATGATAAAAATGGTACCTTTTCAATTGATAAAAATGGGAATTTGGTGGTTAAAGATGGACATGGGGATTTACTATGGTCGTCTAATGTTTCTGTTAATACCACAAATTCTACGGCTTGTCTGAGAGACGAGGGAAATCTTGTCATCCTTAATAATGACAGGAATGCTACGAGGTTAAATTCTGAATTGTGGAAGAGCTTCTCGGATCCTACAGATACATTTTTGCCTGGAATGGAAGTTCTCATAGAAAGACGAGGCCAAGAGCAAAAAGTTTTCCGTTCTTGGACAAATGAGAGTGACCCTTCACCTGGGAGGTACTCAATGGGAGTTGATCCTCGCGGAACACCTCAGATTGTTATTTGGGATGGACCGAATAGACGATGGAGGAGTGGACATTTTGATGGAGCGGAATTCATTGGTGTTCCAGATGTGATTAGAACTACTTTTTTCTCTGGCTTCAGAATTCAAAACCAAGGAGACGATAAATTGTTACTCACTTACAGTGCCTCAAATACATCTTCTTTTGTGAGGTTTCAGCTTACTGTGACTGGAAATGAGCTGCAACAGAGGTGGAACGAAGACCAGGGTGAATGGAACACGTTGCAATCTAGGCCTGTAGGTGGTTGTGACTTGTATAACTTCTGTGGGAATTTTGCAGAATGTGATAAAGATGTATGCCAATGTTTAAAAGGTTTTGTACCACGTGTTCAGGAGCAATGGCATGCTGGGAACCGAACCGGAGGATGTGTTAGGAAGACAGAATTGGAATGCAGGAGAAATAGCAGTGTTTCAAGGAATGATAGTTCAAAAGATGATGGATTCTCGACTATTCGGAGAGTTAAATTACCTGACCATGCTGATGTTTCAGAAATAACCACAGACGAGTGCAAAATCAGGTGCCTGAATGATTGTTCCTGCAATGCTTATGCTTATGTCAGAGGAATCAATTGTATGATATGGCGTAATGATTTAGTTGATATAGAGCATTTTCAGGAAGGTGGGAACACTCTTTATGTTCGTCTTCATCCTTCTGATATCGGTAAGCACAATACTTTCTGCTGAATTCATAAATCTCTTACCAGTTTTCTTCTCATTCGTGCAATTATACAAATCTAAAACCAAAGTAGTTTGTGCTTTAGTACAAAGATCAATTTCTTGTTTGTCTTTACTCTTTATTTGTAGCAAGCTCAAACAATAAGCAATATATCTTTTTCTTCCTTGTGTTTGAAGAATGATTAGCCTTTCTAATTTCTTGATTTCTCCTTTGAATATCTTTTGAAACAGGTCAAAAAAAGAAGACGATCATAATTGTTGTAATATCAATTCTAGCAGCTCTGGCTTTGGTTGTTATGGTGGCCATTTGGCTAGTATGCAAGTACAGGGCCAGAAAACGAGGTAAATTTTGATGCAGTAACATTTCTGACTTGTTAGTTGCATTTATTATATGTTCCATATTGTACTTATACCTTTGACTTGCAATTTTCAGAATCCAAGAGGACCAGTGAAATTCCCAAAAACCATTTAGTTAGGAGTGGAGAGTTCTCCACAGAATATTCTGGACCAGGCGACATCAGTGCTGAAGGGCATCAAGGAAATGGTTCAGAATTAGCATTTTTCAGCTTCAGCATGGTGGCCACAGCTACTGATGACTTCTCTCTTGCAAACAAGCTTGGGCAAGGGGGATTTGGCCCCGTTTATAAGGTAAATTTTCAATCTTGATAGATTATTCTGTTCCTTTAGGAAGTATATGGCCATGTTTTGGACTAAAGTTGATGTTTTTCAAATCTTTAGGGAAAGCTACCTTGTGGTCAAGAAGTTGCTGTAAAAAGGCTTTCACAAAAGTCTGGACAAGGCGATGAGGAGTTCAAGAATGAGATCACTCTAATAGCAAAATTGCAACACAGAAATCTTGTTAGACTTTTGGGTTGCTGTGTTGAAGGAGAAGAAAAAATGCTGATTTACGAGTACATGCCCAACAAAAGCTTAGATACATTTCTATTTGGTAAGTTCAGGAAATTCTATCAAAATAATTTCTGCTGCTCCCACAGATGTACTATAGCAACACATCTCATTGAGATAGACACTAACAACTGTATTGGCTGTTAAACTTTGACAGATACTGCTAGGAAATCCCAGTTAGACTGGAGAAAACGCTTCAACATTATCGAAGGAATTGCCCGAGGATTACTGTATCTCCATAGAGATTCAAGGCTTAGAATAATCCATAGGGATCTAAAGGCTAGTAACATTCTGTTGGATGAAGAGATGACACCAAAAATTTCAGACTTTGGCATGGCCAGAATATTTGGAGGAAACCAAAATGAAGCAAATACAAATAGGGTCGTAGGAACATAGTAAGTATCATTTGATTTTAAATCTTTTAAGAGACTAATTTCAAAATTTCTTGTTCCTGGAGCTAAATCCATTTCATTTTGTACCTTGTTGCTAAACAACAGTGGATATATGGCTCCTGAGTATGCTATGGAGGGCCTGTTCTCTGGCAAGTCTGATGTCTACAGCTTCGGCATAATATTGCTTGAAATTATATGTGGACGTAGAAATACAAGCTTTCGGACAGATGAGCATTCAGGCATCATTGGATATGTAAGTCTGACTAATTTTCTCCAACTATTTCAATCTTTTTAGTCCAGAACAAGCTGTCTGAATAATGGGGCTAATTTCTTGTGTGATGAGCAATGACGGAGTGATCATCTTTCTCATATTTTTCTGGTGGAAAATATTTCAGGCATGGGAGAAGTGGGATGAAGGTAGACCAATGGACCTAGTGGATCGTTCGATTTGGGATGGTTGTCAACACAATGAAGCATTGAGATGTATACACTTGGCATTGCTTTGTGTTCAAGATTTGGCCGCTCACAGACCAAACATGTCTTCTGTGGTTTTGATGCTGGAAACTGACAATGTAAGGTTGCCCTTGCCTAGGCAACCTACCTATACTTCAATGAGAAGATCTGTGGATGAAGATATATGGCATGGGAATCAGGATTTAACATCTTCCAACAATGTCACAGTTAGTGTACTAATAGGTAGATGATTGGTTAATGCTTGTTCCTTCTTACATACCTGCTTCATTTGAAGAGAATGGGATCAAGTAGTGTAACTATGTAAAGTTATACTTGGAACTATGTTCATAATTTTTTGTTAGCTAGGTAAACTTTAGCACATCATGTGTATTGTAATTAGAAGGTGGTTCAGCTCTGTAAGCTGAAAGAGAGATATTTCACAACCTTGTTCAATCAGTATGCTTGAAGTTTTTTTCTTTACTGTTTGTAGATGGAGATGCTCCACCAATCGAGCAAACCCCACTTGTCATATCTAGTT

>M6-PGSC0003DMG400013887 M6 G-LecRK

AGTAATACTTTTAAAGGGGAACTGCACTTTTGTTGTAGTACACCTGTGTATATATAGACTAAAGTTACCAGTTTCAAAAAAAATAAAAATCAATTAGTGCTTTCACTTCCATGGAAAGAAAGATATGCTCACGAAGAACTGTGTATGGCTCATACATGTTTGCAGCTGCAAATGACAAGAAAGACATTCCTGTAGCAGCTATTGCAACCACAGTCTTAATAGGCTCAATAACAGTTATCTTATGTGGATATCTTTTCTGGAAATTGTTGGCTAAGCACAGAGGTAATTAATCAAATAAGATTTTTGAGAAGACAAACTAGAACTTCTCTGTCTGCAAATGGTTTGATTTCTGAATGTCCACTCTAAACCAGAAAGGAAGAGGAAAAATGAAGCACTCTTAAGAGAAGCATCTCCAAAATTTTACCAAGAAGGCATGAGTAAAGATGACATCAATCAAGTCAAAATTGAAGACATCACCTTGTACATATTTGATATGTTAGCGAATGCCACTGACAGTTTTCACCCAGCTAGCAAGCTGGGGCAAGGAGGCTTTGGTCCAGTCTACAAAGTAATGTTTCTTGATGCATCTCTGCAGTGTTCTGACAAATGAACACAATATAAAATAAATTAGATCAATTATCTGCAGGGACAATTGCCATATGGACAAGAAATTGTAGTAAAAAGGCTTTCACAGTCTTCCGGTCAGGGGCTACAGGAGTTTATGAATGAGGTCGTTGTGATTTCTAGACTACAACATCGTAATCTTGTTAGACTCCTTGGCTGTTGCACAGAGAGAGGGGAAAAGATGCTGGTTTATGATTTCATGCCAAATAGAAGCTTAGATGCATATGTTTTTGGTCAGTGCAGAATTTTCTTTCATTGGTTTTGCCTGTCCTTCAACTTAAGTACTTCCAGAATCAAAAGAAAGCTGTATTTCTTTAAGGAAACAAAATCCATCATACATTTCTCAGTGATACCTTTTCAGGTTCACACCAGGAAAAGGTCCTTGATTGGGGTAAACGAGCCATCATTATTGAAGGAACTGGTCGAGGCCTCCTTTACCTTCACAGAGACTCAAGACTATGAATTATTCATAGAGATCTGAAGGCAAGCAATATCCTGTTGGATGAATACCTAAACCCAAAAATTTCAAATTTTGGCATGGCGAGGATTTTTGGAGCAATCAAGATCAGGCCAACACTATAAGAGTTGTTGGCACATAGTATGTTTGCATTGCTTCCTTCATTTTTTCATGGATGAACTGCCATCATTAGTCATTGATTTTCTGTGTAATATGTTATGCAGTGGTTACATGGCCCCTGAATATTCAATGTACGGAAGATTCTCAGAAAAATCAGATGTCTACAGCTTCAGTGTGTTGGTATTGGAAATTGTCAGTGGAAGGAAGAACTCTAGTTTTTATGATGATGAACTGACTCTACTTGCATATGTAAGTTCATTCTTGCAACTATATATGAAGCTTTAATGTTCTGCTAGCTCATCAAACCAAAAAAATAGAGGAGAAAAATTATGGAACTTCTGAAAATAACATTTTTGCTGTTTGACAATGTCATTGTTCATTAGGCATGGAAGTTGTGGAATGAAAACAATATCATAAAATTGATAGACCCCAAAATATTTGATTCATGCTTTGAAAAACAGATGGTGAGATGTGTACATATTGGATTATTATGTGTTCAAGAATATGCAGAAGATAGGCCAAATGTCACCACAGTTCTGTTAATGCTCACTAGTGATATTGCTGAACTACCTACTCCTAAACAACCTGCATTTACCGGAGGACATGCTTCACCACAGCAAGTATCTTCTAAAAGCCAAGGGTCCGTGAATGCTGATACCATAACTGTATTGGAACCACGATAGGATGCAGAGTTTTCAACAGGCAGGCAAAGATCAGTACTTTCCCTAAAATGGTAACAGTAGCATAGTATTTCTTTCTTTTGAATGTGCATAGGGTAAAACACGCTCCTGCAATAACTCGCAAAACACACAAAAGAGGTAAAAGTGCACTAGCTATGTGATGAATTAATTACAGAGTGAGCTTGGAGAACATGAAACAGTTGATCAATTTTTAACTTTGCATTATGAAAATCCTTAGAAAGAAAGTAGACAAAAGGACAAAAAGAACCGCAAATCCCCAATTAACAAGTTCAAATAATGGCAATCAAAGACTTGAAAAATCATACCTGCTGTACTGATACACGAAACGGCGTAAGCGAAAAATAGCCACTCGTTTTACCACTGTTTTTTTTCTTAGAAAATTACAGAGAAGTGAATTGGTGAATTAATCAATGTTGAGGAAAAGTTAACTCTGCATTATTTTAGGTCATTCACATTTCACACAAGTTATTTGGGTGTTGGTTAAAGACTATAATTGTTTCATTTTTAAAATTTTTTTGCATAAACTAATGTGAAACTTACCTAAATCTCACTAGTTTAGGAGTTAATTACTTATATGTATTTTAGTTTGTAATATTACGTGTGTTACCCCCAATTTTGGTGCACTAAGAAACATCCAAATACATGTATTTCTGAATATATGGGCCAAAATTAGGTGTAAGTTGTTTTAGATACATTGTATCCAAGTAGATTTACATGTATTTCGTATACATAAAAAAATTGGTTCGCCTCCTTCACCTCTCTCCCATATCGCTCACCACTCTCCAATGTATTAGGGATACATAATAAATCTCGCTCACCTCTCTCCCTATTTTAGTCTATCCAGAA

>M6-PGSC0003DMG400015476 M6 G-LecRK

CTTGGTGGGTTCTCTAAGAGTCAGTTCACTTATGAGGAGCTAGCTAGAGCGACTTCTGGATTTTCTAAGGCCAATTTGTTAGGCCAAGGTGGTTTTGGGTATGTACATAAAGGTGTTTTGGTTGATGGAACGGTGGTAGCTGTCAAGAGTTTGAAGTTGGGCAGTGGACAAGGTGAACGAGAATTTCAGGCAGAGGTTGATATCATTAGCAGAGTTCATCATCGCCACCTTGTTTCTCTTGTTGGATATTGCATTGCTGATAGCCAGCGAATGTTGGTCTATGAGTATGTTCCCAACAAAACCTTGGAATTCCATCTTCATGGTACTTCCCCTGTCATTGAATACTCCCT

>M6-PGSC0003DMG400017034 M6 G-LecRK

TGGAAGAAAAGGATTCCAATTGTGGTAGGACAGGTTCATGGAGTTGTACTATTACTTGTTTTAGGAGGGATTGGTGATACTATGTAATCCAAAAGAGAAAATAGTTGTGCACAGATACTTGTCACAGCCATTGACATTGTATAGATGAGAATTTTTTTCAAGATAGCATTTTCTTCATGGAGATACTAATTGACTCTCTGAAAAA

>M6-PGSC0003DMG400017106 M6 G-LecRK

TCTCTTGTAACTACCCTTTTCAACCTCCTCAGCTTTATCTCGGTGACACAACTATTCAAATCACACGCATATCGCTCGCTGGGCAACTAACAGTCTTACAATTTATATCGAGAGATTGTTAAAAGTCCGGGATAAACACGGAGAATAACACACCATGGATAGCATTACCACTGTCGTTTTCTGTGAATGACACCACAAATAAGTTCATCGCAATTGGCTGTGACACATATGTTATAGTTCAGGGTTATTATTCCAAGTTTGAAAGTAGTAAAATCGAGGATTATTATTCCTACATAACAGGATGATGCATGTCCATGTGCAATAGTATTGATGATGATAAATGTTCTGGTGTTGGTTGTTGCATGACATCCATCCCTAATGGGGCCTGGAATGTTACTATAACACTAAGGAGCTATTATAATCATACTTATGTGAAAGATAACCCTAGTTACGTTGTAGCTATGGTTTTGTAGTTGAAGAAGCTAATTTTAGTTATTCTAAGAATTATCTAAGAAGTTTGCAGAATATGGAGGAGCTTCCTCTTGTGCTAGATTGGGTAATTGGTGAAGAAACATGTGAAATAGCTAAGAGAAACTCGACTACATATGTATGTAAGAGTA

>M6-PGSC0003DMG400017277 M6 G-LecRK

ATGATTGCAGAGCAGAGAGTGTATCTTCCTATCCTTCTCTTCATTTGTCTCCACTTTTTTTCATCCCGGTTTGCTTTCTCGCAACCTTTCTGGAACTATAATGTCAGTTTTTTGAATTCTACTACTGGACTTTCCAATTATTGGATCAATATTCCGTATGTTAGAGTTTACAATACCTCCGAGCCTTCTGAAATGACGCCTATTATTTGGAGCAGAAAGGATAGCCCTCAGTTTCTCTTTGCCTTCTATTGCCCCCTTCTAACTAACAGTACCACAGAATGCCTTCTTGGTATCCTTTTGTATCACAACAGATCCGATGGGGTTTCAAGTATTCCCCAATTAGTTTGGTCTGCTAACAGGAACCATCCAGTGATAGTTAATGCAACGTTGCAACTAGGTCAAAATGGCGACTTGGTCTTGACAGACTCTGATGGCACTCTTGTTTGGTCCACTAATACTACTGGAAAATCTGTTTAAGGCTTAAACATCACAGAAATGGGAAATCTTGTGCTCTTTGATAAGAGAAAGCAAGCAATTTGGCAGTCTTTTGATCATCCAACAGATTGTTTGCTTTCGGGGCAGAAGTTGATTTTGGGGCGGAAGCTGATAGCAAGCATTTCAGAATCCAATCAGTCCTTTTCTTTTACTATTCTTAATGGAAGGCTGGTGTCTTCCATAGATGCCAGCCCACCTCAGTATTATATCGCTTCAAGTGGGGATGATGCTCAAAATAGTCCTTTTTACAATTTTGACGGTTGAACATTTACTGCTCTACAAGACCCTTATACATCCCAATTCATAAAGCTTGGGCCTGATGGACATTTAAGGTTATACCAGTCGGGTGCATATGATTGGAAACAATTAGATGAGGTCATGGATTCATATCTAGGGAACTGTGGGTATCCAATGGTGTGTGGAAGATATAGCATTTGTACAAATGATGGGCAATGTAATTGTCTGGTAGAAGGAAACTTCTTTAGGCCAATAAATAGGAACCCGGATCTCGGATGTTCACAGCTAACATCCATTTCTTTCAACTCTTCACATTATCATAGTCTTATAGAGCTCAGCTACACCACATATTTTGCATTTGAGATCAACTTTTACGCAAGTTCAAACATGTGGCTTGAGGGAACAAAGATGGAAAATTGCAAAGCTGCCTGTTTGAGTAACTGTTCCTGCAAAGTTGCTGTTTGGTCTAAAACTCTTAGAAAAAACTGCTTATTACTGAATGAAGTCTTCTCTCTTAAGGACAACTGGTCTGGTAGTGACAACACAAAAGTTTTTCTTAAGGTGCAGAATTTCACAAAGGCTCAGTATCAGCCGACAATCGTTTCTCAAAGAAAGCAATCAAGACCTCTGAAAGTGATTGTAGCATCTGCTCTTGCTGCTTTAGTGGGAATAATTTTAAGTATTAGTGCATGGTTTGTTCTTTTCAAAAAGAGGGTTGTCTGTCAAGGCTAGGGATCTTCTGGATTTAGCACCAATCTTACCGGGAATCCTAACTCGATTCTCTTACAATGAGTTGAAAATAATTACACAAGATTTCAGCAGAAAGCTCAGGGAAGGAGGATTTGGCTCAGTATATGAAGGAACACTGAGTAATGGCAACAAAATAGCTGTGAAGCGTCTGAATGGTTTAGGTCAAGTAAAGGATTCATTCTTAACAGAGGTAAAGATAGTTGGTAGCATTCACCATGTCAACTTGGTAAAACTCATCGGCTTTTGTGCTGAAAAAGATCACCGGGTTGTGATCTATGAGTACATGGTAAATGGATCATTGGATAGGTGGATTTCTCATGAAAAGCAAGAAAATGGGCTTGGCTTACGAGGCAAAGGATAATATCGGATATTGCAAAAGGATTAGCTTATCTTCATGATGAATGCAGCTAGAAGATAATTCATTTGGACATCAAACCACACAACATCCTCTTAGATCAAAATTTCAATGCTAAGATATCTGATTTTGGGTTGTCGAAACTAATAGAGAAAGACAAAAGCAAAGTTGTGACTAGAATTAGAGGAACACCCTGTATTTAGCTCCTGAATGGTTGAGGTCGGTGATCACTGATAAAGTAGACGTATATGCCTTTGGAATTGTGCTCTTGGAAGTTTTCTGTGGGCGAAAGAATTTGGATTGGTCGCAGGCTGATGAAGATCATGTCCATTTGCTTAGTGTTTTTAAGAGAAAAGCAGAACCAGAGCAGCTCATGGATATGGTTTACACAAACAATGAGGATATGCAGCTCCACAGAGAAGAAGAGACAGAAATGATGAGCCTCACAACATGGTGTCTACAAGGCAATTTCAACAAGAGGCTGTCCATGTCAGTGGTGGTAAGTATTGGCAGGTTTAGTGTCTGTTGAAAC

>M6-PGSC0003DMG400017278 M6 G-LecRK

CAGAATTTGGTTTAAGGGCAGAAGCTCATAGCAAGCGATTCAACAACCAATCAGAGTCAAGGTTTGATTACTCTTACTGTTCTCAATGGAAGCTGGGCAACTTACACAGATACAGATCCACCTCAATAATACTACAAATCATAATATAAAGAAAGACCTTATTACAGTTTTGATGGTCAAACCTTTAATGCTTTACAAAATCCTAATAAATCAACAGCTCAATTCATGAAGATAGGGCCAGATGGACATTTAAAGGTATACCGATGGTTTATAGATGATTGGAACGAAGTATCTGACATTTTGTCGCCATATGTAGAAAACTGTGGGTAACCAATGGTATGTGGAAGTTAAAGCATTTGTACAAATAACGGGCAATGTACTTGTCCGCAACAGGAAAACTTCTTAAGACCATTTTCTGAGAGGAAACCAGATCTTGGATGTTCACAGCTAACTTCCATTTACTGCAACTCTTCGCAGTATCATAGTTTCGTAGAGCTCAAGAATACTACATATTTTTCATTTAACATTGATCAGGAACTAAATTCGAGCAGATTAAGGCTTGGGCAGACAAAGTTGGAAGATTGCAAAAGGGCCTGTCTGAATAACTGTTCTTGCAAAGCTGCTGTTTTAGAATATAATTGGAATGGGACTCGAAGAGGGAACTGTTTGTTACTGAATGAAGTTTTCTCTCTCAAAGACAGCGAAGAAGGAAAAGGCAAAACAGTATTTCTTAAGGTGCAAAATACCTCAAAGACACAGACTCAGTCTCTAATCATTCCTGGAGGAAAGAAATCAAGACCTTTCAAAGTGATAATAGGATCTACTCTTGCAGCTTTCTTTGGGATAATTTTAAGCATAGCTACTTGCTTTGTTATTTTCAGAAAGAGGACACATGAGTCCAGAAAGGCTGCGGATATTGTGGATCTAGCACCAATCTTACCGGGAATACTAACACGATTCTCTTACAATGAACTGAAAATAAATACAGATGATTTCAGTAGAAAGCTCGGGGAAGGAGGATTTGGCTGTGTTTATGAAGGAACACTGAGAAATGGCACTAAAATAGCTGTGAAGCATCTGGATGGTGTAGGTCAAGTAAAGGAATCATTCTTAACAGAAGTAAAGGCAGTCGGTGGCATTCACCATATCAATCTGGTAAAACTCATTGGATTTTGTGCCGAAAAAAGCCACAGGCTTCTAATCTATGAGTACATGGTGAATGGATCGCTGGATAGGTGGATTACTCATGAAAATAAAGAAAATGGGCTTACATGGAACACAAGGCAGAGGATAATATCAGATATTGCGAAAGGATTAGCTTATCTACATGAGGATTGCAGCCATAAGATAATTCATTTGGACATCAAACCACAAAACATCCTTCTAGATCAATATCTCAATGCTAAGATATCAGATTTTGGGCTGTCGAAGCTAATTGAGAAAGACAAAAGCAAAGTTCTGACTAGAATGAGAGGAACACGGGGTTATTTAGCTCCTGAATGGTTGAGCTCGGTAATCACTGAGAAAGTTGATGTTTATGCTTTTGGAATTGTCCTTTTGGAAATTCTCTGTGGGCGAAAGAATTTGGATTGGTCTCAACCTGATGAAGAAGATGTCCATTTGCTGAGTGTATTTAGGAGAAAAGCGGAACAAGATCAGCTCATTGATATGGTTGACAAAAATGAGGATATGCAGCTCCACAGGGAAGCAGTGACAGAAATGATGAGCCTTGCTGCATGGTGTCTACAGGGCGATTTTTCCAAGAGGCCTTCCATGTCATTGGTGGTTAAGGCATTGGAAGGTTTGGTGACTGTTGAAACCAACTTGAATTATGATTTCACACATGTACCTGAGGTTGGAGCAGGCAACCAAGAGAGGGAAGCCATTAGCAGTTCAAAATTTCCTTCAATTTTATCGGGACCAAGGTAAACAATATG

>M6-PGSC0003DMG400017282 M6 G-LecRK

GAAAAAACTACTCAAATAAAAGGATGAATTTTAAGATTGAGACTGATCAATCAAACAGAGCCTTCTTTCTGAAAAAAATAGTACAATGGCGCTTAATTATAGTGCCATTCTTTTCATGGCCAGTCTCCTCTTCATTTCTTCACAGCATCTAATTAGTTGTCAACCTCCTGAAACATTTATCACCAATTCCTCCATTTCCTGGATCAATAGCCGCGCCCGCTCCTATGTAGTGAATTCCACAGATGGAGCCACACTGACACCCATCCTATCGAGAGAAAGGAACGAGACTCTCTTTGTCTGTGGTTTCTTATGTGATTACAATGGTACTGCTTGCGTTTTTGGTGTCCTTTTATCCCCAAAGATCAGTAGCAACAATTCTTATCTATATTACGCTAGATTAGTTTGGTCAGCAAATCGGAATAATCCAGTTAGAGCTAATGCAACCTTGCAACTTAGGCAAGATGGAGGCTTATTCTTGATGGACTCAGATGGCACTTTGATTTGGAGTACAAGCACAAGCATAAGAGGAAAGGTTGTTACGGGATTCAAATTGACAGTACTGGGAAATATTGTTCTCTTTGACAAAAGTAACAACATCATTTGGCAGTCTTTTGATCATCCTACAGACACTTTGGTTCCTGGACAAATGACAAATGCTGGACAAAAGCTGAGATCCAGTGTATCAGCAACAAATTGGAGTGAAGGTTTGTTCACATTTGAAGTTCAACTATTTGGCTTTTTAGCATACATTGAGTCAAATCCACGTCAACTCTATGTCTCAAGACTTGGGGATAAAGGTTTCCCGTTTCCTGGACTATTCTTTGATTCGAAAATAATGATTTTCAACGGGACATTATTCCTAAATTTTGCATTTTCAAATACTAGATTCATGAGGTTTGGGGATGATGGGCATTTAAGGGTGTATCACTGGAGAGGATCTTGGTTAGAGGAGGCTGATCTATTGACAGACTACATTGGTGAGTGTGGGTATCCAACTGTGTGTGGGAATTACTCTGTTTGTGCAAATAGGCAGTGTACTTGTCCTCAAACCGCGGTTGATCAGACTAACTTTCTCCAGCAAAAAAATTACAGGCAGCCAAACCAAGGGTGTGTCCTTATCACACCTATCTCTTGTGAAAATTCCCAATATCATATTTTTATGGAGCTCAAGTACATAGCTTATATTCCCCTCTATTTGCAATATGGTACAAACAAAACAGATTTGGAGAATTGCAAAAGGTCTTGCTTAAGCAATTGTTCTTGCAAAGCAGCTCAATTTCAATTTAGTGGTCCTGGTAACACAATGGGTCACTGCGCATTGTTGAATGATGTATTTTCGCTCACATACGATTATGGAGCACCCAACAAGACCACTCTTTTCATCAAGGTGCAGAGCTCTTCTAACTTACAAGCCTCTCCATCACTTTTCTCTTCAGAGAAGAAATCAAAGCGTGTCCCAGCAATTGTAGGATCCACCATCGGAGCTTCATTTGGTTTGCTTCTCATGGTTTTAACTTGCTTTGCCTATATTTTCAGAAGGAGAAAAGGGCTCGAGGAAGACGAGGAGGAGTTCCTTGATCAAATACCAGGAATGCCTACTAGATTTTCCTATGAGGAACTCACTGTGATGACAGAAAATTTCAGTGAGAAACTTGGGGAAGGGGGATTTGGTTCTGTATTTGAAGGAACAATGAGTGATGGCACCAAAATAGCGGTGAAGCGTCTGCAAGGTTTTGGTAATGTGAAGAAATCATTCTTAGCTGAGGTAGCAACAATCGGTAGCATTCAGCATGTCAATTTGGTGCAACTTGTTGGATTTTGTGCTGAGAAATCACACAGACTTCTGGTTTATGAATATATGGCCAATGGCTCTTTAGATAGGTGGATTTTTCACGGAAAGCAGGAACGATCTCTCACATGGGATATCAGGAAAAAGATCATATCAGACGTTGCCAAGGGCCTAGCTTACCTACATGAAGACTGTAACAACAAAATTATTCATTTGGATATTAAACCTCAGAACATTCTTCTTGATCATAATCTCAATGCAAAAGTATCAGATTTTGGGTTATCAAAGCTTGTGGGGAAAGATGAAAGTAAGATAGTTACAACAATGAGAGGAACACCAGGCTATTTAGCACCTGAATGGTTACATGAGGTCATTACTGAGAAAGTGGATGTTTATAGCTTTGGGGTTGTGATCTTGGAAATCATCTGCGGCCGGAAGAATTTGGATCGTCATCAAGATGAGGATGATATGCATTTGCTTAGTTTGTTCATGAGAAAGGCAGTGGAGAGACAACTCTTGGAAATGGTGGACAAGAAGAGCGAAGACATGCAGCTACATAGAAAAGAGGCTGTGGAAATGATGAAGATTGCAGCCTGGTGTTTACAAAGTGATTATACTAAGAGGCCTTCCATGTCATTGGTAGTTAAGGTGTTTCAGGGTTTAGTGGCTGTTGAAACCGACTTGGATTACAGCTTCACATTTCCAACAATGACGAGAAGAGTTGCCGGAACTAATCAAGAAAGGGAATCTGTGGTTGGTATCAGTTTTCCACTTCCATCACAGTTGTCAGGACCTAGGTAAATTCTTAGTAGTATTTATCATCAAGACTGAATAGCTAATAAGTCACAGCCATATGTCCATACCTTTGAGTTATTACTTGTTATTTGCTGCGAAAAGACATTTTGTGTTTTTCATAATCCTTATTATATCAATACTATTTCAGAAACATAATTTAGAAAAGTAGAACACACTGTCTACTATAGAAATGTTAGTTTTCGTGCAGTCTAATTAACTTATTGAGTAT

>M6-PGSC0003DMG400017284 M6 G-LecRK

CCTCTGCTGTTCGTCTCGATTTGCTATATCGCAACCTTTACAGGACTACAACTTCACTCTTTCGAATTCTACTGCAGGGCTTGCCGGGCTTTCCTCTTTCTGGATCAACAGGCCCTCTCTTAGGCTTAATTCCACCGACGACTCCTTTGGTTTGGTGACACCCATACTTCAGCGAAGAAATGCTGGCACACGATTCCTCTGTGGCTTCTACTGCAAATACAATGAAAAAAAATGCCTTCTTGGTATCCTTTTGAACCACAACAAATACAACAAAAAAAACAGTATAATAAATAAACCCCAGTTAGTTTGGTCTGCTAACAGGAACCATCCAGTGAAATTCAATGCAACCTTGCAACTAGGCCATGATGGCAACTTGGTCTTGACAGACTCCGATGGCACTCTTGTTTGGTCCACTGATACAATTGGAAAATCTGTTTCTGGCTTAAACTTAACAGAAATGGGAAATCTTGTACTCTTTGATAAGAGAAAACGAACAATTTGGCAGTCTTTTGATCATCCTACAGAATCTTTGCTTCCAGGGCAGAATTTGGTTTAAGGGCAGAAGCTCATAGCAAGCGATTCAACAACCAATCAGAGTCAAGGTTTGATTACTCTTACTGTTCTCAATGGAAGCTGGGCAACTTACACAGATACAGATCCACCTCAATAATACTACAAATCATAATATAAAGAAAGACCTTATTACAGTTTTGATGGTCAAACCTTTAATGCTTTACAAAATCCTAATAAATCAACAGCTCAATTCATGAAGATAGGGCCAGATGGACATTTAAAGGTATACCAATGGTCTGTAGTTGATTGGAATGAAGTTTCTGACATTTTGTCGCCATATGTAGAAAACTGTGGGTAACCAATGGTATGTGGAAGTTAAAGCATTTGTACAAATAACGGGCAATGTACTTGTCCGCAACAGGAAAACTTCTTAAGACCATTTTCTGAGAGGAAACCAGATCTTGGATGTTCACAGCTGACTTCCGTTAACTGCAACTCTTTGCGGTATCATAGTTTCATAGAGCTCAAGAATACTACATATTTTTCATTTAACATTGATCAGGAAATAAATTCGAGCAAATTAAGGCTTGGGCAGAAAAAGATGGAAGATTGCAAAAGGGCCTGTCTGAATAACTGTTCTTGCAAAGCTGCTGTTTTAGAATATAATTGGAATGGGACTCGAAGAGGGAACTGTTTGTTACTGAATGAAGTTTTCTCTCTCAAAGACAGCGAAGAAGGAAAAGGCAAGACAGTATTTCTTAAGGTGCAGAATTCCTCAAAGCCACAGACTCGGTCTCTAATCATTCCTGGAGGAAAGAAATCAAGACCTTTCAAAGTGATAATAGGATCTACTCTTGCAGCTTTCTTTGGGATAATTTTAAGCATAGCTACTTGCTTTGTTATTTTCAGAAAGAGGACACATGAGTCCAGAAAGGCTGCGGATATTTTGGATCTAGCACCAATCTTACCGGGAATTCTAACACGATTCTCTTACAATGACCTGAAAATAACTACAGATGATTTCAGTAGAAAGCTCGGGGAAGGAGGATTTGGCTGTGTTTATGAAGGAACACTGAGAAATGGAACTAAAATAGCTGTGAAGAATCTGGATGGTGTAGGTCAAGTAAAGGAATCATTCTTAACAGAAGTAAAGGCAGTCGGTGGCATTCACCATATCAATCTGGTAAAACTCATTGGATTTTGTGCCGAAAAAAGCCACAGGCTTCTAATCTATGAGTACATGGTGAATGGATCGCTGGATAGGTGGATTACACATGAAAATCAAGAAAATGGGCTTACATGGAACACAAGGCAGAGGATAATATCAGACATCGCCAAAGGGTTAGCATATCTACATGAGGATTGCAGCCATAAGATAATTCATTTGGACATCAAACCACAAAACATCCTTCTAGATCAATATCTCAATGCTAAGATATCAGATTTTGGGTTGTCGAAGCTAATTGAGAAAGACAAAAGCAAAGTTGTGACTAGAATGAGAGGAACACGGGGTTATTTAGCCCTTGAATGGTTAAGCTCAGTAATTACTGAGAAAGTTGATGTGTATGCTTTTGGATTTGTGCTCTTGGAAATTCTCTGTGGGCGAAAGAATTTGGATTGGTCACAAACTGATGAAGAAAATGTCCATTTGCTAAGTGTATTTAGGAGAAAAGCGGAACAAGAGCAGCTCATGGATATGGTTGACAAAAATGAGGATATGCAGCTCCACAGGGAAGCAGTGACAGAAATGATGAGCCTTGCTGCATGGTGTCTACAGGGAGATTTTTCCAAGAGGCCTTCCATGTCATTGGTGGTTAAGGCATTGGAAGGTTTGGTGACTGTTGAAACCAACTTGAATTATGATTTCACACATGTACCTGAGGTTGGAGCAGGCAACCAAGAGAGGGAAGTCATTATCAGTTCAAAATTTCCTTCAATTTTATCTGGACCAAGGTAAACAATATGCTTAAGATTTTGTCTTTGAAGGAACCAACATATGGTTGCGTAATGTCAAATATCGCAACATTTTCAATTCTAGCATAACTTTTTTACCGTGTATGGGATTATAATTCATGAAATGTAATGTTCAGTTACTTCTATATTAAATTTTTGAGCTTCATGTTACTATCATGTAGAGGCTTTTCAATTCATGTGTGAAAATAAGTAATGATATTGTACGTGGTGGTGCAGCTACTGAATTCAAA

>M6-PGSC0003DMG400017286 M6 G-LecRK

TACAAGTACAGATATCACTTTGGATATTCTGTATGCTCATTTCTTTCTTCAACTATAGACAACAAAATTCCATATATTTTATTGATTTTGATTAGGAAACAATGTTAGCAGAAAAGAGACTGTATTTTCCTATCCTTCTCTTCATTTATCTCCTCTGTTGTTCGTCTAGATTTGCTATATCGCAACCTTTACAGGACTACAACTTCACTCTTTCGAATTCTACTGCAGGGCTTGCCGGGCTTTCCTCTTTCTGGATCAACAGGCCCTCTCTTATTATTAATTCCACCGCCGACTTCTTTGGTTTGGTGACACCCATACTTCAGCGAAGAAATGCTGGCACACGATTCCTCTGTGGCTTCTACTGCAAATACAATGAAAAAAAATGCCTTCTTGGTATCCTTTTGAACCACAACAAATACAACAAAAAAAACAGTATAATAAATAAACCCCAGTTAGTTTGGTCTGCTAACAGGAACCATCCAGTGAAATTCAATGCAACCTTGCAACTAGGCCATGATGGCAACTTGGTCTTGACGGATTTTGATGGCACTCTTGTTTGGTCCACTGATACAATTGGAAAATCTGTTTCTGGCTTAAACTTAACAGAAATGGGAAATCTTGTACTCTTTGATAAGAGAAAACGAACAATTTGGCAGTCTTTTGATCATCCTACAGAATCTTTGCTTCCAGGGCAGAATTTGGTTTAAGGGCAGAAGCTCATAGCAAGCGATTCAACAACCAATCAGAGTCAAGGTTTGATTACTCTTACTGTTCTCAATGGAAGCTGGGCAACTTACACAGATACAGATCCACCTCAATAATACTACAAATCATAATATAAAGAAAGACCTTATTACAGTTTTGATGGTCAAACCTTTAATGCTTTACAAAATCCTAATAAATCAACAGCTCAATTCATGAAGATAGGGCCAGATGGACATTTAAAGGTATACCGATGGTTTATAGATGATTGGAACGAAGTATCTGACATTTTGTCGCCATATGTAGAAAACTGTGGGTAACCAATGGTATGTGGAAGTTAAAGCATTTGTACAAATAACGGGCAATGTACTTGTCCGCAACAGGAAAACTTCTTAAGACCATTTTCTGAGAGGAAACCAGATCTTGGATGTTCACAGCTAACTTCCATTTACTGCAACTCTTCGCAGTATCATAGTTTCGTAGAGCTCAAGAATACTACATATTTTTCATTTAACATTGATCAGGAACTAAATTCGAGCAGATTAAGGCTTGGGCAGACAAAGTTGGAAGATTGCAAAAGGGCCTGTCTGAATAACTGTTCTTGCAAAGCTGCTGTTTTAGAATATAATTGGAATGGGACTCGAAGAGGGAACTGTTTGTTACTGAATGAAGTTTTCTCTCTCAAAGACAGCGAAGAAGGAAAAGGCAAAACAGTATTTCTTAAGGTGCAAAATACCTCAAAGACACAGACTCAGTCTCTAATCATTCCTGGAGGAAAGAAATCAAGACCTTTCAAAGTGATAATAGGATCTACTCTTGCAGCTTTCTTTGGGATAATTTTAAGCATAGCTACTTGCTTTGTTATTTTCAGAAAGAGGACACATGAGTCCAGAAAGGCTGCGGATATTGTGGATCTAGCACCAATCTTACCGGGAATACTAACACGATTCTCTTACAATGAACTGAAAATAAATACAGATGATTTCAGTAGAAAGCTCGGGGAAGGAGGATTTGGCTGTGTTTATGAAGGAACACTGAGAAATGGCACTAAAATAGCTGTGAAGCATCTGGATGGTGTAGGTCAAGTAAAGGAATCATTCTTAACAGAAGTAAAGGCAGTCGGTGGCATTCACCATATCAATCTGGTAAAACTCATTGGATTTTGTGCCGAAAAAAGCCACAGGCTTCTAATCTATGAGTACATGGTGAATGGATCGCTGGATAGGTGGATTACTCATGAAAATAAAGAAAATGGGCTTACATGGAACACAAGGCAGAGGATAATATCAGATATTGCGAAAGGATTAGCTTATCTACATGAGGATTGCAGCCATAAGATAATTCATTTGGACATCAAACCACAAAACATCCTTCTAGATCAATATCTCAATGCTAAGATATCAGATTTTGGGCTGTCGAAGCTAATTGAGAAAGACAAAAGCAAAGTTCTGACTAGAATGAGAGGAACACGGGGTTATTTAGCTCCTGAATGGTTGAGCTCGGTAATCACTGAGAAAGTTGATGTTTATGCTTTTGGAATTGTCCTTTTGGAAATTCTCTGTGGGCGAAAGAATTTGGATTGGTCTCAACCTGATGAAGAAGATGTCCATTTGCTGAGTGTATTTAGGAGAAAAGCGGAACAAGATCAGCTCATTGATATGGTTGACAAAAATGAGGATATGCAGCTCCACAGGGAAGCAGTGACAGAAATGATGAGCCTTGCTGCATGGTGTCTACAGGGCGATTTTTCCAAGAGGCCTTCCATGTCATTGGTGGTTAAGGCATTGGAAGGTTTGGTGACTGTTGAAACCAACTTGAATTATGATTTCACACATGTACCTGAGGTTGGAGCAGGCAACCAAGAGAGGGAAGCCATTAGCAGTTCAAAATTTCCTTCAATTTTATCGGGACCAAGGTAAACAATATGCTTAAGATTTTGTCATTGAAGGAACCAACACATGGTTGCGTAATATCAAATATTGCAACATTTCCGATTCTAGCATAACTTTTTATAATTCATGAAATGTAATGTTCAGTAACTTCTATATTGAATTTTTGAGCTTCATGTTACTATCATATAGAGGCTTCTCAATTCATGTGCAAAA

>M6-PGSC0003DMG400017315 M6 G-LecRK

ATGTTTGTAGAAACGAGACTGTATTTTCCTATCCTTCTCTTCATTTATCTCCTCTGCTGTTCGTCTCGATTTGCTATATCGCAACCTTTACAGGACTACAACTTCACTCTTTCGAATTCTACTGCAGGGCTTGCCGGGCTTTCCTCTTTCTGGATCAACAGGCCCTCTCTTAGGCTTAATTCCACCGACGACTCCTTTGGTTTGGTGACACCCATACTTCAGCGAAGAAATGCTGGCACACGATTCCTCTGTGGCTTCTACTGCAAATACAATGAAAAAAAATGCCTTCTTGGTATCCTTTTGAACCACAACAAATACAACAAAAAAAACAGTATAATAAATAAACCCCAGTTAGTTTGGTCTGCTAACAGGAACCATCCAGTGAAATTCAATGCAACCTTGCAACTAGGCCATGATGGCAACTTGGTCTTGACAGACTCCGATGGCACTCTTGTTTGGTCCACTGATACAATTGGAAAATCTGTTTCTGGCTTAAACTTAACAGAAATGGGAAATCTTGTACTCTTTGATAAGAGAAAACGAACAATTTGGCAGTCTTTTGATCATCCTACAGAATCTTTGCTTCCAGGGCAGAATTTGGTTTAAGGGCAGAAGCTCATAGCAAGCGATTCAACAACCAATCAGAGTCAAGGTTTGATTACTCTTACTGTTCTCAATGGAAGCTGGGCAACTTACACAGATACAGATCCACCTCAATAATACTACAAATCATAATATAAAGAAAGACCTTATTACAGTTTTGATGGTCAAACCTTTAATGCTTTACAAAATCCTAATAAATCAACAGCTCAATTCATGAAGATAGGGCCAGATGGACATTTAAAGGTATACCAATGGTCTGTAGTTGATTGGAATGAAGTTTCTGACATTTTGTCGCCATATGTAGAAAACTGTGGGTAACCAATGGTATGTGGAAGTTAAAGCATTTGTACAAATAACGGGCAATGTACTTGTCCGCAACAGGAAAACTTCTTAAGACCATTTTCTGAGAGGAAACCAGATCTTGGATGTTCACAGCTGACTTCCGTTAACTGCAACTCTTTGCGGTATCATAGTTTCATAGAGCTCAAGAATACTACATATTTTTCATTTAACATTGATCAGGAAATAAATTCGAGCAAATTAAGGCTTGGGCAGAAAAAGATGGAAGATTGCAAAAGGGCCTGTCTGAATAACTGTTCTTGCAAAGCTGCTGTTTTAGAATATAATTGGAATGGGACTCGAAGAGGGAACTGTTTGTTACTGAATGAAGTTTTCTCTCTCAAAGACAGCGAAGAAGGAAAAGGCAAGACAGTATTTCTTAAGGTGCAGAATTCCTCAAAGCCACAGACTCGGTCTCTAATCATTCCTGGAGGAAAGAAATCAAGACCTTTCAAAGTGATAATAGGATCTACTCTTGCAGCTTTCTTTGGGATAATTTTAAGCATAGCTACTTGCTTTGTTATTTTCAGAAAGAGGACACATGAGTCCAGAAAGGCTGCGGATATTTTGGATCTAGCACCAATCTTACCGGGAATTCTAACACGATTCTCTTACAATGACCTGAAAATAACTACAGATGATTTCAGTAGAAAGCTCGGGGAAGGAGGATTTGGCTGTGTTTATGAAGGAACACTGAGAAATGGAACTAAAATAGCTGTGAAGAATCTGGATGGTGTAGGTCAAGTAAAGGAATCATTCTTAACAGAAGTAAAGGCAGTCGGTGGCATTCACCATATCAATCTGGTAAAACTCATTGGATTTTGTGCCGAAAAAAGCCACAGGCTTCTAATCTATGAGTACATGGTGAATGGATCGCTGGATAGGTGGATTACACATGAAAATCAAGAAAATGGGCTTACATGGAACACAAGGCAGAGGATAATATCAGACATCGCCAAAGGGTTAGCATATCTACATGAGGATTGCAGCCATAAGATAATTCATTTGGACATCAAACCACAAAACATCCTTCTAGATCAATATCTCAATGCTAAGATATCAGATTTTGGGTTGTCGAAGCTAATTGAGAAAGACAAAAGCAAAGTTGTGACTAGAATGAGAGGAACACGGGGTTATTTAGCCCTTGAATGGTTAAGCTCAGTAATTACTGAGAAAGTTGATGTGTATGCTTTTGGATTTGTGCTCTTGGAAATTCTCTGTGGGCGAAAGAATTTGGATTGGTCACAAACTGATGAAGAAAATGTCCATTTGCTAAGTGTATTTAGGAGAAAAGCGGAACAAGAGCAGCTCATGGATATGGTTGACAAAAATGAGGATATGCAGCTCCACAGGGAAGCAGTGACAGAAATGATGAGCCTTGCTGCATGGTGTCTACAGGGAGATTTTTCCAAGAGGCCTTCCATGTCATTGGTGGTTAAGGCATTGGAAGGTTTGGTGACTGTTGAAACCAACTTGAATTATGATTTCACACATGTACCTGAGGTTGGAGCAGGCAACCAAGAGAGGGAAGTCATTATCAGTTCAAAATTTCCTTCAATTTTATCTGGACCAAGGTAAACAATATGCTTAAGATTTTGTCTTTGAAGGAACCAACATATGGTTGCGTAATGTCAAATATCGCAACATTTTCAATTCTAGCATAACTTTTTTACCGTGTATGGGATTA

>M6-PGSC0003DMG400017318 M6 G-LecRK

GAAAACTTCTTAAGACCATTTTCTGAGAGGAAACCAGATCTTGGATGTTCACAGCTGACTTCCGTTAACTGCAACTCTTTGCGGTATCATAGTTTCATAGAGCTCAAGAATACTACATATTTTTCATTTAACATTGATCAGGAAATAAATTCGAGCAAATTAAGGCTTGGGCAGAAAAAGATGGAAGATTGCAAAAGGGCCTGTCTGAATAACTGTTCTTGCAAAGCTGCTGTTTTAGAATATAATTGGAATGGGACTCGAAGAGGGAACTGTTTGTTACTGAATGAAGTTTTCTCTCTCAAAGACAGCGAAGAAGGAAAAGGCAAGACAGTATTTCTTAAGGTGCAGAATTCCTCAAAGCCACAGACTCGGTCTCTAATCATTCCTGGAGGAAAGAAATCAAGACCTTTCAAAGTGATAATAGGATCTACTCTTGCAGCTTTCTTTGGGATAATTTTAAGCATAGCTACTTGCTTTGTTATTTTCAGAAAGAGGACACATGAGTCCAGAAAGGCTGCGGATATTTTGGATCTAGCACCAATCTTACCGGGAATTCTAACACGATTCTCTTACAATGACCTGAAAATAACTACAGATGATTTCAGTAGAAAGCTCGGGGAAGGAGGATTTGGCTGTGTTTATGAAGGAACACTGAGAAATGGAACTAAAATAGCTGTGAAGAATCTGGATGGTGTAGGTCAAGTAAAGGAATCATTCTTAACAGAAGTAAAGGCAGTCGGTGGCATTCACCATATCAATCTGGTAAAACTCATTGGATTTTGTGCCGAAAAAAGCCACAGGCTTCTAATCTATGAGTACATGGTGAATGGATCGCTGGATAGGTGGATTACACATGAAAATCAAGAAAATGGGCTTACATGGAACACAAGGCAGAGGATAATATCAGACATCGCCAAAGGGTTAGCATATCTACATGAGGATTGCAGCCATAAGATAATTCATTTGGACATCAAACCACAAAACATCCTTCTAGATCAATATCTCAATGCTAAGATATCAGATTTTGGGTTGTCGAAGCTAATTGAGAAAGACAAAAGCAAAGTTGTGACTAGAATGAGAGGAACACGGGGTTATTTAGCCCTTGAATGGTTAAGCTCAGTAATTACTGAGAAAGTTGATGTGTATGCTTTTGGATTTGTGCTCTTGGAAATTCTCTGTGGGCGAAAGAATTTGGATTGGTCACAAACTGATGAAGAAAATGTCCATTTGCTAAGTGTATTTAGGAGAAAAGCGGAACAAGAGCAGCTCATGGATATGGTTGACAAAAATGAGGATATGCAGCTCCACAGGGAAGCAGTGACAGAAATGATGAGCCTTGCTGCATGGTGTCTACAGGGAGATTTTTCCAAGAGGCCTTCCATGTCATTGGTGGTTAAGGCATTGGAAGGTTTGGTGACTGTTGAAACCAACTTGAATTATGATTTCACACATGTACCTGAGGTTGGAGCAGGCAACCAAGAGAGGGAAGTCATTATCAGTTCAAAATTTCCTTCAATTTTATCTGGACCAAGGTAAACAATATGCTTAAGATTTTGTCTTTGAAGGAACCAACATATGGTTGCGTAATGTCAAATATCGCAACATTTTCAATTCTAGCATAACTTTTTTACCGTGTATGGGATTATAATTCATGAAATGTAATGTTCAGTTACTTCTATATTAAATTTTTGAGCTTCA

>M6-PGSC0003DMG400017327 M6 G-LecRK

AGAGACGCGTTCCCAAAAGAAAGGCATTATTAGAACACAGAAGGAGTAGATTGCAAACATTTTGCTAGCTTTAGTAAGAAATGATTGCAGAGCAGAGATTGTATTTTCCTCATGTCCTGTTCATCATTTTTCTCCACTTTTTGTCATTCAGGTTTGCTTCATCCCAGATTTCATGGTACCTAAATGGCAGTTATTTGAATTCTACTGCTGGACTTTCCACTTCATGGACCAATAGGCCGTTTTGGTTTGATGCAGAATCCACATATGGATTGTCTCTCTTGACACCTATCCTTCTGAGTGGAATAAATGGAAATAAATACCACTTCTGTGGCTTCTACTGCAATAGCCAAAGTGTTGAATGCCTTCTCGGGATCTTCTTTGCATTCCCAGACAGCTCCTCTACTGAACACAAATTAATGGATCCCCAGTTGGTTTGGTCTGCTAATCGAGACCGTCCAGTGAAAGCCAATGCAACCTTGCAACTAGGACAAGATGGCAACTTGGTCTTGGCAGACTCTGATGGCACTTTTGTTTGGTCCACTAATACTACTGGAAAATCTGTTTCTGGCTTAAACATGACAGAAACGGGGAATCTCGTGCTCTTTGACAAAGCCAATCGCACAGTTTGGCAGTCGTTTGATCATCCTACAGATTCTTTGCTCCCGGGGCAGAGCTTGGGTTCTGGGAGGAAGCTCATAGCAAGCGTCTCAGCAACCAATTGGAGTCAAAGTTTGCTTTCTTTAACTATTATCAATGGAAGCTGGGCCACTTACATCGAATCTGATCCACCTCAGTTTTACTATGCTTCAAGTAATCCGGATACTAGTACTTATTTCAGTTTTGATGGTCAAACACTTACTGTTCAGCAAGACCCTCCTAAATCACCAGCTCAATTCATGAAGCTTGGGCCTGATGGACATTTATGGGTATACCAATGGGACGAGAATGCATTAAATTGGAAAGTCATATCTGATGTTTTGTTTAATGTACATGCAGAGAACTGTGGGTACCCAACGGTGTGTGGAAGATATGGCATTTGTACAAGTAATGGGAAATGTAGTTGTCCGCCAGAGAGAAATTTCTTCAGGCCATTAGATGAGAGGAACATAGATGTTGGATGTTTACAGCTGACTTCCATTTACTGCAACTCTTCGCAGTATCATAGTTATGTAGAGCTCAAGAATACCACATATTTTGCATTTGAGTTCAGTGATGAACTAATTTCGGGCATATTACCGTTTGAGGGGAAAAAGTTGGAAGATTGCAAAAGGGCCTGTCTAAGAAACTGTTCCTGCAAAGCTGTTGTCTTCGGATATGATTCAGATGGGGCAAGAAGTGGGAGATGTTTATTACTGAATGAGGTCTTTTCTCTCATCGACACTGAGAACGGAATGGACAACAGAGTATTTCTTAAGGTGCAGAATTCCTCAAACACTCAGAATCAGTCTCCAACCATTTCTGGAGGGAAAAGATCAAGACCTCACATACTGATAATAGGATCCACTCTTGCAGCTTTCTTTGGGATAGTGTTAAGTATCACAGCTTTCTTTGTTATCTTTAAAAAGAGGACAAAAGATTCCAGAAAGGCTGGGGGTTTTCTGGATCTCGAACCAATCTTTCCGGGAATGCTAACTCGGTTCTCTTACAATGAGCTAAGAATAATAACTGAAGATTTCAGCAGAAAGCTCGGGGAAGGAGGATTTGGCTCAGTATATGAAGGAACACTGAGTAATGGCACCAAAATAGCTGTGAAGCGTCTGGATGGTCTAGGTCATGTGATGGAATCATTCTTAACAGAAGTAAAGATAGTCGGTGGCATTCATCACATCAATCTGGTAAAACTCATTGGATTTTGTGCTGAAAAGAGCCAAAGACTTCTGATCTATGAACACATGGTAAATGGATCACTAGATAGATGGATTTATCATAAACATGAAGAAAATGGGCTTAGATGGCATACAAGGCAAAGGATTATAACAGATATTGCCAAAGGATTAGCTTATCTTCATGACGAATGCAGCCAGAAGATAATTCATTTGGACATCAAACCACAAAATATCCTTCTAGATAAAAATTTCAATGCTAAGATATCTGATTTTGGGCTGTCGAAACTGATCGATAAAGTCGAAAGCAAAGTTGTAACTAGGATGAGAGGAACACCAGGTTATTTGTCTCCTGAATGGTTGAGGTCAGTTATCACTGAGAAAGTAGATGTGTATGCCTTTGGAATTGTGCTCTTGGAGCTTTTGTGTGGTCGAAAGAATTTGGATCGGTCCCAAGCTGATGAGGATGTCCATTTGCTAAGTGTTTTCGAAAGAAAATCGGAACAACAGCAGCTTATGGATATTGTTGACAAAAACAATGAGGATATGCAGATCCACAAAGAAGCAGTGACAGAAATGATGAGCATTGCTGCGTGGTGTCTACAGGGTGATTTCACCAAGAGGCCTTCCATGTCATTGGTGGTTAAGGCATTGGAAGGTTTGGTGACTGTTGAAACAAACTTGGATTACGATTTCACAAGCCTACCCGAGGTTGAGGATTGCAACCAACAGAGGGAAGACACTATCAGCTTAGTATTGCCTTCAATTTTATCCGGTCCAAGGTGAACAACTAGCTTAAGAGGTTGTAAGTTTTTTTTTCCTTTCCCAAGTCTGTCTTTCTTTTGTGAAAATGTATTTCCATGATTCATCACCAGATCCATCTTTCTCTTGTACATAAATTTGGCTAACAGGTAAACTATGTCATAATGTAAAAGGTAAACTTAAGTAGCTTTGATAGATAACCAATTAACGTCTTGTGGCAGTGAGTGATATTATTTTCTTTTTTAATTTGCATCTTTGTGGATCAATACATCTTCTTATGCAGCACATTTTATATTTCTAAGAGTAAGGGGTCTACTTATTCTGTGTTATAAACAGAAAAAACTGCAG

>M6-PGSC0003DMG400017330 M6 G-LecRK

AAACAACCCAAGATCTTAAATTTAAGTTCAATCATTTTTCATCTCAATGATGAGCTAAATGCCCTTTGCCTTCTTTGGAAAACCAAAAAATTGGTCTTTATGGCTTTATGCCAGATCTTACCTTTCTTTTTGTTCTTCATTTTTTGTTGTTCTTTTCATTTGATAACCAGCCAACCTTATGATTATCCATCTGCAAATCTTTCTACAACTTGGGTCAACAGCATTTCGGCAGACCATTCAGTGGATTTCACTGATGGCTCAACAGTAAGAGCCATACTACTCAGAGGAAGTTTTGGTCCAAAATATGCTTGTGGTTTCTACTGTAATGGAAATTGTGATACTTACCTCTTTGCCATCTTCATTTTGCAAACCAACAGTGGTTCCGGAATTACTCTTACCACGATTGGGTTCCCACAAGTTGTTTGGTCTGCTAACAGGAATAAACCTGTTAGGATCAATTCTACTTTGGAACTTACAGCACAAGGAGATTTAGTGCTTAGAGATGCTGATGGTACTTTAGCTTGGTCAACTAACACTACTGGGAAGTCTATTGCTGGCTTAAACTTGACCGATGAGGGGAATCTTGTTCTGTTTGATTCGAAGAATGCAACTGTTTGGCAATCGTTTGATCACCCAACTGATGCTTTAGTTCCAGGGCAGAGGTTAGTATCAGGGATGAAGCTGACAGCAAGTGTTTCAACGACAAATTGGACTGATGGAGGTTTGCTTACTCTCTCTGCTACCGATGATGGTTTGGTTGCTTTGGTAGAGTCAAATCCTCCAAAAACATACTTTAAAGCGACTATTGGTGGATTGAATGCAAGCAGAGGCTCAAATTATGTTGTGTATTTGAATGGTAGCCTAGCTTTACTCTCAAACTCTAGTAATTCCAGCAATTCACAGACACTGGTTCCACTTAATCTAGCGACCTCAGCTCAATATATGAGACTGGAGTCTGATGGACACTTGAAAGTGTATGAGTGGCAAAGTGGGTGGACTGAGGTGAATGATCTCCTCACAGGTTTTAACGGCGAGTGTTATTATCCAATGGCCTGTGGAAGATATGGCATTTGCTCAAGGGGGCAGTGTAGTTGTCCCAAATCGAGCTCTAATTCAACGACCTATTTCAGACAGATAGATGATAGGCAGGGTAATCTGGGTTGCTCTGAGGTCACAAGGCTGACATGTAATGCTTTAAGTAACCACAGATTTTTGGATCTTCAAGATGTGGACTATTTCACGTTTACGACAGATATCACAAACACTAGTATGAATACCTGTAAAGATGCATGTTTGAGGAACTGTTCCTGTAAAGCTGCTTTGTTTCGCAGTGGTTTAAACTCATCCACGGGAGACTGCTACCTACCATCCGAGATCTTTTCACTAGCGAATAATGAGAAGGACAAGACAAGGTATGATTCCCATGCATTTATAAAAGTACAGGTTGAACCTGAAGTAGCTGCTGCTGAAGAGAAAAAGCGCGTTAATGGTGCTATACTGGGCTCTGTCATAGTAGTTTCCATCTTAGGGATCATAATTGGAATTGCAGTTTTCATATTCTGGAAGAAGAAGAGAAAGGCCAATGAAGATGAGGAGGATTATCTAGATCACGTACCAGGAATGCCGACTAGATTTTCATACGATGATCTAAAGGCTGCAACGGAGAACTTCACCAAGAAGCTTGGTCAAGGAGGATTTGGGTCAGTTTTTGAAGGGTGCTTAGCAGATGGCACAAAGATTGCAGTGAAATGTCTTGATGGAATAGGACAAGTCAAAACGTCATTCTTAGCTGAGGTTGAAACTATTGGCAGCATACATCATGTAAACTTGGTGCAATTGATTGGCTTCTGCGCTGAGAAATCTCATAGGCTTTTAGTATACGAGTGCATGAGCAACGGATCACTAGAAAAATGGATCTACCATGGGAAACAGGAGCAAACTCTAGATTGGAACTGCAGGAGGAAGATTATTCAAGACATAGCCAAAGGATTAGCTTACCTTCACGAAGAATGCAGGCAAAAGATTCTGCATTTGGATATTAAGCCCCCAAACATACTCCTGGATGAGAAGTACAACGCTAAACTCGCTGACTTTGGGCTTTCAAAGCTAATTGATCGGAATCAGAGCCAAGTTATGACTCAGATGAGAGGCACTCCTGGTTATTTGGCTCCTGAATGGCTTAGTGGAGTAATAACAGAGAAGGTAGATGTCTACAGCTTTGGCATTGTGCTCTTGGAAATTTTGAGTGGAAGAAGACATTTTGAAGCATCGGAGTCTGAAGACCAACAGGTAATGGTGAACTTATTCAGGAAAAAGGCAGAGGAAGGGCAGTTGGTGGATCTTATTGATAAGCACAGTGAAGATATGCAGTTTTACAAAGAAGAAGTAGTAAAGGCGATGCAGATTGCTGCCTGGTGTTTACAAAGTGATTACACGAAGAGGCCATCAATGTCGATGGTGGTCAAGGCCATGGAGGGTGTTATAGATGTTGACAAGGATCTAGACTACAGCTTTAGGCCACAAACCGTTTCAGCAATACCAAATATCATTTTTGCAGATTCAACTCCTTTATTGCCTTCGATCCTATCAGGCCCAAGGTGAGGAGGAAGTTCCTTGGAATTTTTGTTAATTTCTTGTGTTTATGCTATTGCAGCTTGCTCGCTGCTGTGTAAATTTCTTTTCCAATGTTCACAAAGTTATGATTGTTTTCTTCAATTTATT

>M6-PGSC0003DMG400017331 M6 G-LecRK

TTTAAATCATCTTCATTCAGTGTTCTTAATGGCTTTATGCAAGATTTTTCCTTTTGCCTTCATCCTCCTTTGTTGTTCTTGTTATCTGATATCGGGCCAACGTTTCGATTACCCCACTGCAAATCTTTCTACAACTTGGATCAACAGTGTTTCTGCACCCCATTCAGTGGATTTCACTGATGGCTCAAGAGTAAGGGCTATACTACTCAGAGGAACTTTTGGTCCAAAATATGCTTGTGGTTTCTACTGTAATGGCAATTGTGATACTTACCTCTTTGCCATTTTCATTGTGCAAACTAATAGCGCGTCCCAAATTACTTCTCCCGCGATTGGATTCCCACAAGTTGTTTGGTCTGCTAACCGGAACAATCCGGTTAAGATCAATTCAACTTTGCAGCTAACAGCACAAGGAGACTTGGTGCTCAGAAATGCTGATGGTACTTTGGCTTGGTCAACTAACACTGCTGGCAAATCAGTTGCTGGTTTAAGCTTGACCGATGAAGGGAATCTTGTTCTGTTTGATTCAAAGAATGCAACTGTTTGGCAATCGTTTGATCACCCGACAGATGCTTTAGTTCCAGGGCAGAAGTTAGTATCCGGGATGAAACTAACAGCAAGTGTTTCAACAACGAATTGGACTGAAGGAGGCTTGTTTTCCCTCTCTGCTATCGATAATGGTTTGGTTGCTTTCATAGAATCAAATCCTCCCCAAACATACTTTGACGCGACTATTGGTGGATTGGATCCTAGTGGAGGATCCAATTATGTTAAGTATTTGGATGGAAGCTTAACTTTCTTCACAAATTCGAGTGGTACAACCGAGTTGGTTCTGGTTTCTATTACTCCAGCATCCTCTGCTCAGTATATGAAACTCGAGTCCAATGGACACTTGAAAGTGTACGAGTGGAAAAATCGATGGGGAGAGGTGGATGATCTCTTGACAGGCTTTCGTGGCGTGTGCAATTACCCCACAGTCTGTGGAAGATATGGCATTTGCACAATGGGGCAATGTAGTTGTCCCATAAGCTCTAATTCAACCACCTATTTCAGACCGATAAACGTTAGGCTGCCTAATCTTGGTTGCTCTGAGGCCAAAAAGCTAACTTGTAACAACTTAAAGAAACACAGACTTTTGGAAGTTGAAGAGGTGGACTATTTCACGTTTACTGCAGATATTAGTAACACAGATGTGAATACTTGTAAACGCGCATGTTTGAATAAATGTTCCTGTAAAGCTGCGTTCTTTCGTAGTGGTTTAAATTCTTCCAGGGCCATAGGGGAGTGCTACTTACCAACTGAGATCTTTTCACTAATGAATAATGAGAAGGACAAGACAAGGTATGACTCTGTCGCGTTTATAAAAGTACAGGTTTAAGCTAAACCAGCTGCTGCTAAAGGGAAAAAATCGCGTTTTATGTACAAAAATGAATCTATGGGAGGACATGGATGGAAACTTAGCAAATTATGCGCGGGATTATAAGTTCATGCTAGACATGAACTATATGAAGGTATTATTACATAAATAAATGGTTTGATAAGCCAAATTAGTGTTAAGAGTTCTCTGTATTCCTTCATTTCTCTGTGTTATCTGTAATTTCAATTCAATAGACTATAATAAAGACA

>M6-PGSC0003DMG400017946 M6 G-LecRK

AGCTATGCAAAATTTCCACGCGACTATGTTTTTATGTCAACCGTAGCAAATAGATTATGAAGCTTCATATTCTGAACATCTCCATGACTCCATCTGCAAGTGAAATCAAATTCCATCTGCTATCTCATTTTAATGGAGGCCATAAACATACTTTACTTCTTGTCATCCATATGGATTCTGTCGAGTGTGAGTGCTCAACATACCATACCTGCAAATCAACCTCTAACAGATGGAAGCACCATTATTTCATCAGATGGCAAGTTTAAGTTGGAATTTTTTTCCCCTGGGACATCCGGGAAACGCTACTTAGGAATATTGTTTAATAAAGTTACTGTACAGACAGTGGTATGGGTTGCTAATAGAGACAATCCACTCAATGACACATCGAGTGTGCTAAATTTTACCAGACAAGGAATTCTTACTCTTCTAAATGGTTCTGGCAGAGTCATTTGGTCCTCCAATTCCACTAGACATGGGCAAAATCCAGTAGCACAACTCCTTGATTCAGGCAATCTTGTTGTTCGAGATGCAGCCGAGGATAAGCCTCAGAACTACCTGTGGCAAAGTTTTGACAATCCAGGTGACACGGCTTTACCTGGAATGAAGGTTGGGATTAATCTGAAAACTGGTTTTCATCGTTCCCTTTCGTCATGGAAGAGTACAAATGATCCTTCCAGGGGTCAGTTTACCTGGACATTTGATACTGGTGGATTCCTACAAACATTCGTTATGAATGGTTCCATTGAACTCTACAGAGCTGGACCATGGAATGGCCGAGTCTTTCCTAATGCACCATCTCGTGACACATCGTGGAATGGTTATAACTATACATACTTATCTGATCCTAATGAAATATCATTCATGTATGAACTTACAGACAGCTCTATCATTGCAAGGGTAGTGATGCAACTAAATGGATTTTTACAGCTTTCAGAATGGAATAATCGGACCCAAAATTGGGACGATGTTGTCAGTGCACCAGCTGATAACTGTGACATTTATGGCCAGTGTCATTCATATGGTTTGTGCAACAGCGGCAATTCCCCAATCTGCAGATGTTTGGATAAATTTGAACCCAAAGATCCAACAGAATGGGCAAGGGGAAATTGGTCTGGTGGCTGTGTTAGAAAGACAGCAATATTGAATTGCCAAAAGGAAGTTAAATTCTTCAAGTATCAAGGCATCAAGTTGCCAGACACTCGTTTTTCCTGGTACAGTGGAGGAGTGACTCTTAATGCATGCGAAGAATTGTGCTTGAGAAACTGCTCGTGTATGGCTTATGCAAATCCAGACATAACAGGGACAAATGAAGACTGCTTGCTTTGGTTTGATGAGCTGATGGACATCAGAGAGTTTGGCGCTACTGGGCAAGATATCTATGTAAAGTTGGACTCTTCTGAGTTTGGTATAAATCTGTTTCTATACTTTTTCATGTTAGGCTGTTAAGTATGCCACATTAGATTTTGAGTATTCTAGACCAAATTAAATGCCATTTAGTGAGCAAGTTGAGGTTTCTCTACATTATTTGAAAACCAAAACAGTTTTTTTTTTTCGTCTTACGAATTTATTTCTGTCATCAGGTAACTCCAGTACAGGGAAAGTAAACAAACTGACAATCAGCTTGCCACTGGCAGCATTAGGTCTGCTCCTAGTACTATGCTTAATCATGTACATAAGGCAAAATAAGAAGAATCAGCAATATTTTAGAAAAGGTAAACCTCAGAGGTTTTGCTGCTTTTGAATTTGTGGTTTTGTTTTTTCTAGAGTTACATTTGGTAAATATCAGAATTTCTCTTTACATCCAATTAAGTCCTTATTTTAACAGGAAGAAGAACAATAATCTCTGAAATGTTCTGCACTAAAGAAAGGAAAGATAAAGAGCTAGATTTACCGTTGTTTGATTTTGAAACTGTATCTCATGCTACCAATAACTTTTCATTGAACAACAAGCTTGGAGAGGGTGGTTTTGGACCTGTTTACAAGGTAACTATTTTAACTTCAGCTCTTTCTATTTCATCCAAAATAACTGCCTAAAAAGGAATTTCCGAAACTTTAAAACGAAAACACATGTTTCGGTAAGAAAATTGATATGTCATTCTCCTATTTCATCTAAACAGCAATGTCAAAATCTAGTTTATTGTTTGTCAGTAAAATATTCTGGAGAAGCTCATTCAACTAATTTACTGGATATCTAGGGTATGCTGAAAGATGGACAAGAAATAGCAGTAAAGAGACTTTCAAGATACTCAGCACAGGGAGCTGACGAGTTCAAGAATGAGGTTATCTTCATTGCTAAACTCCAGCATAGGAATCTTGTGAAGCTTCTTGGTTGTTGTATCCAAGCAGAAGAAAAAATGTTGGTTTATGAGTACATGCCTAATAATAGCTTGGATTGGTTCCTTTTTGGTTGGTTCTCACTTCTCTCAACTCTTAACACATTTAGTTATCTGTAGTAATAGTTCAATCGTCAGCTGAATTTTGTTATCTGTTCTTTGATTTGAAATATCTGAAGTGTTCTACTATAAGCTCAGAATTTTTGCAGATACAGATAGGAGGTCACTGCTTGATTGGCCTAAGTGCTTCCATATTATTAATGGAATTGCTCGCGGACTTCTCTATCTTCATCAAGACTCAAGATTGCGGATAATCCACAGAGATCTTAAACCTAGCAATGTTTTGCTAGACATAGATATGAACCCAAAGATATCCGACTTTGGCATGGCAAGAAGTTTTGGAGGAAATGAGACTGCAGCCAAGACAAAAACAGTGGTTGGGACATAGTAAGAATCTTTAAACCGATGAAGTATTTGCAGTCTACTAAAAAGCTACTTACTTTGTTTATTTGACATTATCCTACAGTGGATACATGTCCCCGGAGTATGCAGAAAAAGGAGCATTTTCAGTGAAATCAGATGTATTTAGCTTTGGAGTTGTAGTACTGGAGATTTTAAGCGGGAAGAGAAATAGAGGGTTCTTGCATCCAGACCATAACCATAATCTTCTAGGACATGTAGGTGATCTTCTTGTCTATCTTATATGTTTAAGTAAAACTTATAGCATCTAACTGCATGATAGTAAATGTTCAGGTGTGGATCCTTTTCAAAGAAGGCAGGGTTATGGAAGTAATTGATACACAACTAAGGCAGTCATGCAATCAGTCTGAAGTTCAAAGATCAGTTCATGTAGGTCTGTTATGTGTGCAGCAGTGTCCAGAAGATAGGCCGAATTTGGCATCAGTTGTACTCATGTTGGGTAGTGATGTCGCACTACCCTCGCCTAAAGAGCCAGGCTTTTTCAATGGCAGGAACCAATTCATTGAAGCTGACAGTTCATCTAGCAAGCATGGAGAAACTTCAGTGAATGAATTAAGCATCACACAGTTGGATGCCAGATAGGAGATAAATTTTCAGGTCATGGATGTTTTCTGTACGCTATATTAACTTAATTGTTGCGGTCATATAGAACAAGAACTACTGGTTGTTCAACTAAATAGTTTTACTCAAAAATCTGTGGCATGGTATTAAGCATGACAACTTGACACTCAGTTTGCTGTATATAAAGATAGGAGCAGAGTTGTGAAAAGAGATAAAAATATTCTCTACATACAAAAGCACCTAGTACATTTGCTTATTTAAGAAGAACAGCTTATTCTGTAAAAAACTTTTAGCCTCCCTTGGGGAAGTAAGAACAATTTTTCCTTCGTTGGCTTGTATGAGTCCACGACTAGTCAGGAGACCAGGTCCCTTGACTGTAGTAATAACAGGAAGTAATGTAAATGCTCAAAGTAAAGAACATAGATGTTTACCTGGAAACTTCCTTGTTCAAGAGAGGAAAAATTACGACCTGTCACACAGGATTTCTCAAAACTCCAGTAACTTCAAGTAACTTGTGAGTACTGGCTCTAGTAATCACAAGGATTAATATCTTAATCCTACAAACCTTTAAACACCTTTATTACAAAACACAAAGCAATAACCCTATTGTCCACTTTATCAACTACCTCTAAGTAGTATAAACTTCAACAAATACCACTCAGCTAACTCAAGCCGCAAACTTAACTAACACTAATATGATATGGCATGATCTATATTCCCGTATAACATAGGAATATATTCTTAATAAAAGACCATAAGACACTAGACTCAAATAACAACTTCAGACTCTGCAAGACTAAACCGGGTCCCTCATTGTGTGTCAGAGTAGTTCTTTCTTTAAAACGGGTTTTGCTTTTGAATAATCCTAATTGCAAATAGTAGGTTGTTTCTTGTCAACATTTTCAATATATAAGAGACACAAAACAACTTAGCAGCATGTCATTGGTCGAGGACTGTTGTCCTTTCTGCGCCTCTACTATAGTGCCCACCAACCACTGTAGTTTCTACAACTTTACAACTACGCTTCATTGACTTGTCCTTTACGACAAAAACACAGAGGACCAGGTCCCTTTACTAGATCCTATTTCTTTATAGCTTGTTAAATCATCAAAATCCCAACATACAACATTTTCCCCCTTTTTGATGATGATAAACTATTGCCCATGTCTACATATCTTTTGCTTGCACATTCCCCCTATCAACATGCTTCCCTTGTCGGAAAGACCTGATCCCTTGATCTGAACTGGCATACATGTTATCAGTGTATCTTCCTCCTTTTGACATCATAGAAAAGTAAGAACAGTGAACCAAGCAAAATTACATGCATTCTGGGTAAATTCATGGCCACTAGGCAATACTAATATGAACAAACAGACAGAAAGAAAGGAGCGAGAAACAAACATTCTTGGTTTTCCAACTCCGCCCTCTCACTTTTCCATTGTTCAAACCAAAATTGAGTAACACACTTTGGTTGATAAGCGGTCAGCTCCGCCTTCTCCACCGTATAAAAAAATCCATAATTCCGACTATCTTATACACCTTATCAATGAACTCTTGAGGATCCTCATCAACCTTGAAAACATGGAACTTCGAGGGATTCAACCTAGTGAAGTCCCTTATCCTTGTAACTGCCGTACCCTCATTTGGATTCAAGGGAG

>M6-PGSC0003DMG400017949 M6 G-LecRK

TATATATGAGCTATTTCATCAATAATCCCTGCCGACTATTGAAGGAGAATCCTGCAATATTTTCCTTTAGTAAAACAAGGATGATGGAAAGCAGACATGTTCATTTGTTATTATTATTATTATGTTACATGTTTTGTGTTTTGATTCTTACTAGAGCTGCAGATAATATAGCTAGAGATGAACCTTTAAGAGATGGAAAAACGATTGTTTCGTCAGGTGGAAATTTTGAGTTGGGATTTTTCTCCCCAACTCCAACATCGAGTAAACGATACCTGGGAATATGGTTCAACAAAGTATCTGTACAGACGGTGGTATGGGTTGCTAATAGAGACACTCCACTGAATGATAAAAATGGTATGCTCAATTTTACAACACAAGGAAATCTTATTCTTGTAAATGGTTCTGGCAGTGTGGTTTGGTCTTCCAATACCACAAGAGGAGGCGTGCAAAATCCAATAGCACAGCTTCTAGATTCAGGCAACCTTGTTGTTAGAGAAGATGCAACTGAGAATTCGAATTATGTATGGCAGAGTTTTGACTATCCTGGTGACACAGCTTTGCCTGGAGTTAAGATTGGGATCGATCTTAAGACTGGTTTTCGTCGTTTCCTCTGGTCATGGAAGAGCAGAAATGACCCTTCTAATGGTCAATATAGTTTCATATTTGATCCTAATGGATTCCCACAGCCATTTCTCATGAACGGTTCCATCCAACGCTTCAGGGGTGGAGCATGGAATGGTCAAAGCTTTGCTAATTCACCATCTCTACTACCAAGTCCTTCTTATAAATACATATTTGCATCCGATCCAGATCAAGTATACTTTACTTATGAGCTCATTGACACTTCTGCTATTGCAAGGGTAGTGATGCAACTAAATGGGTTTCTAGAGCTTTCGACGTGGAATAATCTAACTCAGAATTGGGATAACTTTGGTAGCGCACCAGCAGACAACTGTGACATTTATGCCCAGTGTAACGCATATGGTTTGTGCAACAGTGGAAACTCTCCAATCTGCAGTTGTTTGGATAAATTTGAACCCAAAGATCCAACAGAATGGGCAAGGGGAAATTGGTCAGGTGGCTGTGTTAGAAAGACAACAATATTGAATTGCCAAAAGGAAGTTAAATTCTTAAAGTATTCAGGGATCAAGTTGCCAGACACTCGATTTTCCTGGTACGGTCGAGGAGTAACTCTTAATTCATGTGAGGAATTGTGCTTGAGAAACTGTTCATGTGTGGCCTATGCAAAACTAGACACAACAGGGACAAATGAAGATTGCTTGCTTTGGTTTGATGAGCTGATGGACATCAGAGAGTTTGGTTCTAGTGGGCAAGATATCTATATAAAGTTGGACTCCTCTGAGACAGATATAAGTACAGGTTGGTATGCTTCTTTACATTAGTCTTGAGTTTTCTAGACCAAATTGAACTGTCTATTAAAAATTTGATAAGACTAAGATGAATTTATATGAATCATCAGGAAACTCCAGTAAAGAGATATTAAAGAAATTGAGAATCTGCTTGCCATTGGCAGCATTAAGTCTTCTCTCAATGCTATGCTTGATCTTGTACATTAGGCATACGAAGAACAAGAAGAAGAAAGAAGAGGAGAATCAGAATCAACAACGTTTTAGTGAAGGTAAACCGGATGTTTTGCTACTTTGAAATTTGAAATAAGTGGCTTGAATTGATGGTAAATCTTAGAAATTCCTCTTAACAGGAAGCTCTGAAATGTTGTACATTAATAAGAGCAAAAATGATGATCTAGATTTACCATTGTTTGATTTTGCCACTATCTTGGAGGCTACCAATAACCTTTCTTTGAGCAACAAGCTTGGAGAGGGTGGTTTTGGACCTGTTTACAAGGTAACGCTGTTAACTACAACTCTTTCTTTTTCTCTAAACAAACTGCTTATCTTCAATATATTATATATGCTGGAACATTGACAAACTTTCGGAACCTTTAAAAGGAAACAAGTTCCCGATTAGAAAATATGTCTCCGATTCTCCAGAAATGTCATTAAAAGTTAATTCCACTTCAAACAAGCTATCTAAAAAAGTTTACTCAGAATTTTTCAATCGAAAAACTAGAGAAGCTAATTGCCTGAGGATATGTAGGGTGCGCTGAAAGATGGACAAGAAATAGCAGTAAAGAGACTTTCAAGATACTCTGCACAAGGAACTGATGAGTTCAAGAATGAGGTTATCTTCATTGCTAAACTCCAGCATAGGAATCTTGTTAAGCTTCTTGGTTGCTGTATCCAAGCAGAAGAAAAGATGTTGGTTTATGAGTACATGCCAAATAATAGCTTGGATTGGTTCCTTTTTGGTTGGTTCTCACTTCTCTCAACTCTTAATACATTTTGTTATCTGTAGTAATAGTTCAATCAGCTGCAGATGGAGTTGGTTATCACTTCTCTGAACTTTTGATTTTTGCACATACAGATAGGAGGTCACTGCTTGATTGGCCCAAGTGCTTCCATATTATAAATGGAATTGCTCGAGGACTTCTCTATCTTCATCAAGACTCAAGATTGCGAATAATCCACAGAGATCTTAAACCTAGCAATGTTTTGCTAGACATTGATATGAACCCGAAGATATCTGACTTTGGCATGGCAAGAAGTTTCGGAGGAAATGAGACTGGAGCCATGACAACAAGAGTGGTTGGAACATAGTAAAGATCTTTAAACCAAAGAGCTACTTACACTCTACTAAAAAGCTAACTTTCTTTATTTGTGATTTTTTTTAAAAAAAGAGATACTTACTTTCTTTGTTTGACATTATCCTACAGCGGCTACATGTCTCCGGAGTATGCAGAAGAAGGGAAATTCTCAGTGAAATCAGATGTTTTTAGCTTTGGAGTTGTAATTCTGGAAATTTTAAGCGGGAAGAGAAATAGAGGGTTCATTCATCCAGACCATCACCATAATCTTCTAGGACATGTAGGAACTCATCTTAGTACTTAAATAGTGTAAAAACCATGTTAGTTAAAATTATAGCATCTAACTGCATGATGTGAAATGTATGTTCAGGTGTGGATCCACTTCAAAGAAGGAAGGGTTATGGAAGTAATAAATACACATCTAAAGGAGTTATGCAACCTATGTGAAGTGCAAAGATCAGTCCATTTAGGTCTATTGTGTGTGCAGCAGTGTCCAGAAGATAGGCCTAGCATGTCATCAGTTGTGCTGATGTTGTCTAGCGATGTTCCATTGCCCTTACCTAAAGAACCTGGTTTTTTTACTAGCAGAAGCCGATTCGGTGAAGTTAATAGTTCATCTAGCAAGCTTGGCGAACATTCTGGAAATCAATTAAGCATCACACTATTGGATGCTAGATAGGATATATATATATATATATATATATATATATATATATATATTTATATATATATATTTTCGTTGTTTGCTTGTAGAGGCTAGTGTCACGGACTTAGAGTCTCACTAAAATTCCGTACGGTACTCGACAACTTACACAGGAATCTAATTATAGTGGCGGGACATGCCACTAGGCTTTATTTACTTAATTCTTTAGCTTCCAGTTGTACAACCAAACAACTTAAAGTGAATTTGTCAAATGTACAAGTGCTTTCTCAGTCTAATAATTCTTCATAATCTATATTGTCCTCACTCCTCAGATTGTGAAATGAGTTAAGACAATTTGTCATAATTGTACAAAGGGCATGGACAGCTCTTTTTCAAAGCAAGTACAGTTGATAGTATTTTAATTAACAACTTGACGGTGATCAACAGGCTGGTCAATC

>M6-PGSC0003DMG400018452 M6 G-LecRK

CTTCAACCATGTTAGCAGTTAGAACTTGTCCTTCTTTCTCCCCCATTGATATCATGACTCTCTATTTCTTACTTGTGCTGCTTATCTTTGGATCCGCTAGAGCTCAATCAAATCCACCTAATACAATAAGATTAGGATCTTCTCTTCATCCCACTGGACAATCTACAGCATGGTACTCAGCTTCAGGTCATTTCGCCTTCGGCTTTTATCCACAAGGAAGTGGTTACAAGGTGGGAATTTGGTTAGTGGGAGGCCCCAATAACACTATTGTGTGGACCGTGTCTCGTGATGATCCAGAAATATCTGCTGATTCTACCTTGGCTTTCATTGATGGAATTGTAGTTTTGAAAACAAATAACAGTGAGGATAAGACTGTTGCAGCTAGTTCATCAGAATCTGCTTCTTATGCCAACTTGCTTGATAACGGTAACTTTGTTCTTTACAATCAAGATCATGAAGCTATCTATGAGAGCTTCAGGTCTCCTACTGATACAATTTTAGGTGGCCAGAAATTAACAAGTGGGGGTAAATTGGTTTCTAGTTTCTCCTCCACAAATCATTCCTCAGGGCGATTTCGCCTCTTGATGCAGGATGATGGGAACCTTGTTGCATACCCCAGATACTTGTTTCGGCCAGTGGATGCCTATTGGTCTTCACAAATTTATTGTGATGGTTGCAGAAATCTCCTGCTTCTTAACAGCACTGGCATCTTATTTCTGATCAATGACACCGATTCATCAGTACTTCGACAATTATACACTCCACTTGTGCAGAATGACAGAGCTATTTATCGGGCCACTCTAGATTATGATGGGAATTTTCGACTCTATTCTCACGAATTCAACTCCAATGGGAATTCTAAAATGGTAATGGAGTGGGAGGCAATTGATGATATTTGTCTTGTGAAGGGTTTCTGTGGCCTTAATGGCTATTGTGTTGGAAGTAATCACAATGCACCGTCTTGTATGTGTCTACCAGGGTCATACTTGAGAGAGAATGATCCAAGCTTTGGTGATTGTCAAAGGAACTTCACCAGAGGAAAGTGCATTAATGGGAAAGAAGATGTAAGTGTCTACAAGATTACTACCACAACCAATTTGACCTGGGAGGATCCTCCTTACTTTGTTACATCACTTCTAGGAAAAGAAGATTGTGGTAAATCTTGTTTGGAGGACTGTGATTGTGATGCTGCTCTTTTTGATGAAACTGGCCAGTGTATGAAACACAAGCTGCCTCTGAGGTATGTCAAAGGTGCTCCTGAAGGATCTCACACAGCTTACTTCAAAGTCAGCAATCTAGCAGTGCAAAATCCAACAACGGATTCAGTGAAGCCACCATGGGTGGTGATCTTGGTTTTATCGATAAGTTTCGTTATGTATTCAGGTACTGCTCTTGCATTTTCAGGCTTTTATGTCTTCAAATTCCGGATAATAAAGTACAGGAAGCTACTTCAGACAGGAACCACAGGCTTAACGAAGGATTTCATACTCAAAACTTGCTCTTACAAGGAGCTAAAACAGGCAACGGATGGTTTCAAAGAGGAATTAGGAAAGGGTGCTTTTGGAGCAGTATATAAAGGGAGCTTTGACAAAGGTAAAAACCTTGTGGCAGTGAAGAGGCTAGAGAAGGTAGTGGAAGAAGGTGAAAGGGAGTTCAGAGCAGAAATGAAGGTCATTGGTAGAACTCGACACAGAAACTTGATTCGTTTGCTAGGATATTGTGCTGAAGGATCAAAAAGAGCTCTTGTTTATGAATATATGAGCAATGGTTGCCTAGCAAATCTGTTGTTCCGAGGAGCTACTCGGCCGGATTGGAATCTAAGAGTAAATATTGCACTAGATGTGGCCAGGGGAATCCTCTACCTCCATGAAGAATGTGAAGCTCCAATTATCCACTGTGATATAAAGCCACAAAACATCTTACTCGACGAGTTGTTGACAGCTAAAATCTCAGATTTTGGTCTGGCCAAGCTGCTAATGCCTGATCAAACAAGAACATTCACTGGTGTAAGAGGGACTAGAGGGTATTTAGCACCAGAGTGGCAAACAAATGCCCCCATCTCTGTCAAGGTTGATGTCTTCAGCTATGGTATTGTATTGCTAGAAATCATTTGCTGCAGAAGAAACATTGAAGTCTGTGTGACTAAAATAGAAGAAATCCAGCTTTCTATATGGGCATATAGTTGCCTTGTTGAAGGTGAAGTAGACAAGCTTGTTGGAAGTGAAGAAGTAGACAAGAAAACCTTGGAAGGAATGATCTCTGTAGCGATTTGGTGCATTCAGGACGAACCAGCTCTTCGTCCTTCCATGAAAAAAGTATTACACATGTTGCAGGGAATTATTGATATTCCAATTCCTCCATGTCCAAGTCCCAGTAGTTAATTTCAGTAGTTGTAAGTGGTTTGGTTTATTATATTGTTGTTTTCTTCAATAACATCTAGTGAGGATATATAGTTAATTCAACATACATGAGTTCAGATTATGTTTTATTTTAATGAGATCATTTGCAGGCCCTATGGGTGTAATCTTGAAAGATATTACTCAAAAATGAGAATGTTTTGATGTGTCAGAGTATCAAAATACTCAACTATTACACTAGGAGATTATCAAAATTATGCTTTCCATCATATGTTGGGAGTTGCGAAAATTTAGACATATATACATGTGGGAATATTTAAGGTGTTGAGAGCCACTAGATTTGGAGCTTCTAATTAAGGTGGATAACACTTCAATATAGTGCTGATATTATGCAGGGAAAGGGTATTCCTTCCTGGCACCTTGATAGTATTGTGGTTAAAGGCTTAAAGCAAGGAAGACTCTTTATCAAACCAATGGCATTATTTATTGCTGTAGAAAAGGAAATTAAATAGTGGATGCAATGGCTAACTGGAGTTTAAAGGGGCATACATCTATCAGTTCAATGATCTACTACAAGAAGCTAAAGCTGCATACAACAAATTGTACAAAATTCAGATGCTTAGTCAAAGGTGCAGAGTATGCAACAAGATTCTTGTAAGAACTTCATGCATCAAGACATGATAAACCACTTCATATTTTGTACGTTCTTATAAAATCAGATAGAAAAAGTAGTCGAAAGATGATTTTATTAATTCAGTAGTCA

>M6-PGSC0003DMG400018457 M6 G-LecRK

GGAGTAATACTACCATTTAAAATTGGAGATGTCCAAGTTAGTATACAAAAGAATACTTATCAAAGAGTGTATCCATATTCCCATCCTTGTGTCATGGCTAACTTAGCATTTACTTTCATGTTGTTGATGCTTCCATTTGTATTTGCAGCTATTCAAGTTGGAGCTCAGCAACAGCCCTCCAACATAACCTTAGGTTCTTTTCTTTATCCGAACCGAAATCCAAGCTATTGGCCATCTTCTTCAGGCCTTTTCTCATTTGGTTTCTACCAAAAAGGGGATGGATTTCAAGTTGGCATATGGCTACCTGATAAAACTGTTGTTTGGACTGCGAATAGAGATGATCCTCCACTTCATTCGGATGCATATATCGAGTTCACAAAGGAAGGCAAACTGTTACTATGGACAAAACAGAACAAGAAAGTCATTGCAGAATCTCCAAAGCCAGTAACTTCAGCATCCATGCTTGATTCTGGTAACTTTGTTATCTATAACAAAACAAATGTTGTTTGGGAAAGTTTTGACTATCCAACAGACACCATCCTAGCCGGTCAGCGTCTAACATTAGGCCAGAGCCTTGTCTCTAGTGTTACTGAAGTTGATCATTCAAGTGGAAGATTTTATCTTAGTATGCAGGCTGATGGGAACCTTGTTGCCTACCCAACCAATTATCTAAATGGACCTGAGGCTTCCTATTGGGGTTTTATGCTACAAACCAGAGGTGAATTTGGTGAGGTTAGCTCAGTTTTCATAGGTCTTACTCCAACAGGCCAACTCTTTCGGAATGCTTCTAATGGTTTTCGTTTCCAAGTCAATACCATAGCGAATACCTCTAATCAATCATCAACCAACAAAGCAGCAGCAATTTATCGCGCAACTCTTGATCCTGATGGGATTTTTAGGCTTTATACACATAACTTCGACAACTCAAGCTTGGAAATCAATTGGTCAGCGCTGGAAAATCAATGCAAAGCCAGGGGATTCTGTGGTTTAAACAGTTATTGTACTGCAAAAAATGGCAGCAGCATAGGCGATTGTTCTTGTTTTCCTGGATTTTCATATGTCAACCGTGCAATGAAATTCCAAGGCTGCTATCGGGGATTCGTTTATGAAGAGTCTTGTGGAAGCAGCATAGACTCAACATTGGCCTACAATGTCACGGCGATTCAACACTTGAAAGTTGGAGGATATCCATTTTCTCAGGTTTCAATGGTTGAGCAGGACTGTAGAAATTCTTGCTTGGATGATTGTACTTGTTGGGCTGCTCTATACATGAATGGATTGTGCAGCAAATTTAAGCTTCCATTAATTGATTCGACAATTGAACAAAATGATGAGTCCATCAAGAGTTTCGTCAAGCAGAGCTATAACATCTCCCAAATGGCAGGTCACTCTATTCCCTATCCTGGAAAAACAGGGAATAGAAATGAAAGTCGGAAAGAGATAATATTGATTCTATCCCTGGTTCTTGGTTCTGTAGCATTTCTGTTCACTGTTTTTGCAATTTTCAGTTATATTTTCTACAGAAGCAGAGCGGACGAATACCAAAAGCTATTGGAAAATCCATATTTGGGGCCAAATGAGGAGTTCACTCTTCGATCATTTTCTTACAGTGAGCTTGAGAAAGCCACTGAGCATTTCAAGGAAGAGGTAAGGCATGGTTCATTCGGGAACGTCTATAAAGGAATTCTATCAGAAGGTAACAGAACAGTTTCTGTAAAACGACTGGAGAAAATCGGGGATGAAGGAGAAAGGGAATTTAAAGCAGAAATGACTGCGATTGGGCAAGCTCGTCACAAAAACTTGGTTCATTTGCTTGGTTTTTGCTTGGAGGGATCAGAAAAAATTCTTGTGTATGAGTACATGAGCAACGGGTCACTCGCAGATATCATATTCAATTCAGAAACAAGACCTTCTTGGGAGCAAAGGATGAGACTTGCACTGGACATATCAAGGGGAATACTTTATCTGCATAAAAAATGTGAGACTTGCATAATTCACTGCAACATAAAGCCACACAATATCCTAGTAGATGATTCGTGGACGGCTAAAATCTCTGAATTTGGGCTTGCAAAGTTTCTGGTGCCTGATCAAGTAGGAAACCAACTTCAGCTTAAAGGAACAAGAGGATATTCGGCACCAGAGTTGCAGAATAGTGCATTGATCTCCGATAAGGTAGATGTTTATAGCTATGGAGTAATGCTCTTGGAGATCATATGCTGTAGAAGTAATATGGATGTCAACGTCTCAACCGAGGATGAAATCTTTCTTCCTAATTGGGTACACAAATGCTTTGTCGAGAATGATATCATTAAGCTAGTGGGAGATGAGGAAGTAGATGTGAAGTCCTTGGAGAGGATGGTGAAGGTTGGATTGTTATGCATTCACGATAATCCTGATCTGCGCCCTTCTATGAGGAATGTAGTTCTCATGTTAGAAGGTACTATGGATATACCATTTTTGGCATGGTATTAAGTCTTTTACACTTCGTTTGGCTGGTTGCTACATCTCATTCAAAATGTATCATATTATATTGTATTGTGTTGTGTACTTGTATTATATTGTTTTTATGAATACAAC

>M6-PGSC0003DMG400019062 M6 G-LecRK

ATGGTTCTGTGTGGTATTATTTATATAATTTATCTGAAAAGAGTGGCAAGAAGTAAAGGTAACAATGTAGAATTTTCATCTGCTTTTTCTCAACTACTCCAACTTTTGGATATCGTAATCACTTCAATTGTTTTGATTATAGAAGCCAGAGAAATTGTTTTGGGAATGCCTATGGAGTACTTGCCTCATAGAGATAGTTTTGATGAAGATTTAATCACGGAAGATGATAAGAAACGGATTGATGTCCCGTTTTTCAGCTTCAATAGCATACTAGTTGCTACCGAAAACTTCTCAAATGCAAGTAAGCTCGGGAGAGGTGGATTTGGACCTGTTTATAAGGTTTTTGCAACTTTCTTTTTTTTTCCTTCTTCTGTTCTAAGTTGTTTTACCATTCAGATTTTTTTCTTTCGTTTTCGTTATATAACAGTCTGGAGATATATTTATGCAGGGTACGTTTCTCGGAGGTACAGATATGGCAGTTAAAAGGTTGTCAAGTAATTCTGGACAAGGTGTTGAAGAATTTAAAACTGAAGTAATGTTGATTGCCAAACTGCAGCACAGAAATTTAGTTAGGCTTCTAGGCTATTGTGTTGAGAGAAATGAAAAGATTTTGTTATATGAATATATGCCCAACAAAAGCTTGGATACGTTTCTATTCGGTTAGTATACTTCCCTCCCTACCATGAAGCTCGTTGCAGTAATAGTTTTATGATTGATTCTCTCTGAATTTATCTTACAAATCTTGAAATTAGATCATGCATTCTGTCGATTACTGGACTGGAGGATTCGTTTTGAAATTATATTGGGTATTGCTCGAGGGTTACTTTATCTTCACCAAGACTCGAGGCTAAGGATCATTCATAGAGATCTAAAGACGAGTAACATATTGTTAGACCAAGAAATGAATGCCAAAATATCAGATTTTGGCTTGGCAAGGATCGTTGAAGGCAAAAATACAGAAGCCAATACTAACAAAGTCGTTGGAACATAGTAAGTCTCTCGGTTCTGTTTTCACCTTTGATTTGGTGGGTAAAAGATAGATATATATACATCTAAAACTTGGTATTAGTTCACTATATGTATTGGTTTACACAAATTAGATGCTAAAATAACCTATTCTGGAACACTAAAAACAGTGGCTATATGTCTCCGGAGTATGCATTGGAGGGACTCTTTTCAATCAAGTCTGATGTATTTGCCTTCGGAGTAGTCCTACTCGAGATCATAAGTGGGAGGAGAAACATGGAATTTTTCGAAGATGTAAACCTCATTGGTCATGTAAGTACCAAAATTTCCCCCCATTAATAAACATCTCATTCTGTTTAAAGCTTTCATTGACTCAAAATTACTTATCACACACAGGTATGGAGACTGTGGATGAAAGATAGTGCATTAGAAATGATGGATCAAACCATAGTTGAGTCGTGCGATGAAAGTGAAGTGCTAAAGTGCGTGAATGTGGCACTCCTATGTGTTCAAGAAGATCCAGCTGAGCGTCCTGTAATGTCAAATGTTGTTTTTATGCTTGGAGGTGAAAGCATAACACTACCAAGACCAAATCAACCAGCTTTTATAGCACGAAGAAATAGTGCTGGTGCAAGTACATCTTCTTCGTCTTCTAAGCTATACAACACTTCCAACAATGGGTTGACAATAACTCAACAAGTAGGCCGATGA

>M6-PGSC0003DMG400019237 M6 G-LecRK

GCCAAGTGTAATTCAAGAAGTGGACAACTTTGTCAAAATTCAAAATTGTCCAAAATCTTGCAAGCCCCACTTTGTCTTCCAAGTAATTCAGTCACAAAAATGAAGTCTTGGTCATATTGACATGTTATCACAGCACCAACACCTTTTAATATTTTTAAATTTTTTTTTGCCTTATTTTGTATAACCAAGAAATGAAAAAGGGGTAGTAGCTATGGAGAAATTTCTACCTTTTTTGCTTCTTTGCTTTTTAATTTTTCTTCGAAAAGTTAATGGACAAGGAGATATGATAACTACCACTAAGTTTGTCAAAGATGGTGAAACTATTGTTTCTTCTGATGGAACTTTTGAGTTGGGATTTTTCAGTCCTGGAAATACTTCTACAAATCGTTATGTTGGAATTTGGTACAAGAAAATTTCTGTTGTTACTCCTGTTTGGGTTGCTAATAGGTTGGTTCCACTTACTAATAAATCCGGGGTTTTGAAAGTTATTCAATCGGGCGGTGTTGCTCTTCAAAATGCTACAAATTCAACAATTTGGTCGACAAATTCATCGAGATCCGTGCAAAACCCCGTTGCGCAGTTGCTGGACACGGGGAATTTTGTGTTGAGGGATGCTAATGATACGAACCCAGAGAATTTCCTCTGGCAGAGTTTTGATTACCCTACTGATACGTTGATTGCGAATATGAAACTTGGAAGGGATTTGGTTACTGGTTTTGAAAGGTATCTTAGTTCATGGAAAAGCAGTGATGATCCTGCCCCGGGGGATTATACTTATCATTGTGATCCAACGGGTTATCCACAAGACGTTATGCGAAAAGGGGCTGATGTGATTTATAGAGCAGGGCCATGGAATGGTTTAAGGTGGAGTGGTGCACCTAATATGGTGAACAATTCAATCACTTCGTTTGGTTTGGTAATGAACAATAAGGAAATTTATTATAAATATGAACTTGTGAACAAATCTGTAGTTTCAGCTCTAGTAGTAAAGCCTAATGGGAACACTATGCGTTTGATTTGGATTGCGAAGACTCAGGGTTGGGTTAATTACCATTCTGCTGATGCAGATGACTGTGACACTTACAAATTATGTGGTGCATATGGGACTTGTAACATTTTGAGTGATCCTGTGTGTCATTGTTTAGATAAATTTGAGCCAAAGCATCAGGATGATTGGAATAGAGCTGATTGGACGAGTGGTTGTGTCCGGAAAACCCCATTGAATTGTACTGGAGATGGATTTATTATGTATTCTGGTGTTAAACTGCCTGATACACGAACTTCTTGGTTCAATGAGACCATGAGTTTAGATGAATGCAGGGCGGTTTGCTTGAGGAATTGTTCGTGTATGGGTTATACTAATCTGGATATACGCAATGGAGGAAGCGGGTGCTTGATATGGATTGAGGAGCTGGTTGACATCAGACAGCTATCTCAATCAGGACAGGATATCTACATTAGGATGTCTGCTTCAGAAATAGGTAATATGCAATTTACTGAGGTTTTCTTTATCACACAATGCTTGTTATCAAGAATGATGTATTGAATAGTTCAATTCTATAAGGCTGATTAACGAACGAAGAGTAATGCATTTACAGGGAATTTAGGTGTAAACATCCTCAAGCAATGGATCCTTTTGTTGAGAATAAGACTTATATGACCATCTTGTTCTACAATAAAATAATTTGCACAGGATTTAAAAAGTTTATCGGAAATTCTAATTCAAACAACTACCTGAAGAAACTTTTGGATGGAAAATGTTGCAGGTTCTGCTGGATCAAGTTCAAAGGGAGATAAATCTGTTATACTTGCAGTAGCTTTGCCACTATTGTTTGCGTTGATTCTTCTAGGTTTAGGTGTTGGCCTGATTCTTTATAAACGGAGAAGGAGAGAGGATCCAGTGGTTACGACAAGGGGTAAGTGAACATAACAAGTTTTTATCTGTTGAAAAGCCGGAAGTAATTAATGGTTATTAGTTCTGTATTCCATTAGTATCTCTTTTCAAGCTCGTTGTTTGAACAAAGTCTGCTTGGTTGTTCAAATTGCAGGGAGATTTAGTGGTCACAACAACAAGAATGATAACAGTAATCAAAGTCACCATGAAGATTTTGAGCTACCACTTCTTGATTTGCTTACATTAATCAATGCTACTGACAACTTTTCAATTGCGAACAAGATTGGAGAGGGTGGCTTTGGACTAGTTTACAAGGTAGAAACATTTGATCCTTGAGAGCACATTGAAACCTTAAATCGATTTAACTTTTTTGGTTGAAATGAAAGTAAGTTCTACTTTTACAGGGTGTGTTAGAAGGGGGGCAAGAAGTAGCAGTGAAGCGGCTATCAGAAACATCAAAGCAAGGATTTCACGAGTTCAAGAATGAAGTTAACTGCATTGCTAAACTTCAGCATCGAAATCTTGTAAAGCTTCTAGGATGCTGCGTTCAAGGAGAAGAGAAGATGTTGGTCTACGAATACTTGCAAAACAAGAGCCTTGATATATATATATTTGGTTAGTCTTTCACATCCAGTGTGCATCCATTTCTTTCAAGTGAACTTCTGATGTGATTTCACTGAGCTTTGTTTCGTGTTTAGGAGAAATTATAATAAGTTGAAAACATTTGGCAGATGAAGAAAGGAGCGCATTACTTGATTGGCCAAAGCGCTTCAACATTATCAATGGAATTGCTCGTGGACTGATGTACCTCCATCAAGATTCGAGGCTCAGAATCATCCACAGAGACTTGAAAGCTAGCAATGTCTTACTTGATACTGAAATGAACCCAAAGATATCAGATTTTGGGATGGCTAGAAGTTTCGGAGGAGATGAGACAGGAGCCAATACACGTCGTGTTGTTGGAACATAGTAAGATTTAAATCTTTGATTATACATATAGTTATGCACATTCACA

>M6-PGSC0003DMG400019238 M6 G-LecRK

TATATATATGTATAGCTAGGAACTAAAACTTTAGTAGAATAAAAGGCTACTACTTAGATTGAGGTGTTGAATTTTAAAGCTAAATCTATAAGCATAATGAGTTATTGCATCTGTCAGCTAAAAATCTAAGACGAAATAAAAGAGGAATTTAGTTGGAAAAACTCATTGAGAACTAAACATATGATTTGCAGTACTTCAAATGTTTGCAGAACACCTCTACTAAATGTCAACAAATTAAAAATGACGTACCGTACTTCAAAATCCAGCAGAATGTTGCAACTCAATATCAGTTATCAACTCCCCCAAGTGATCAACAGGTAGAGACAGTCAAATTTATTGATTACTTTGTTGGAATGGACCTTGCAGATAGTCTGGAGAAATCAGATGGTGAAAAAGGAAGAGCACTATGCTGGATTCTTCCATTATCCGTTGGTTTAGTTCTGGTAATCCTTAGCATATTGCTGATATGCCACAGGAGAAGGAAGAAGGCTTTAAAGCTTAAAACGAAAGGTAAGCTGGATAGGTCACTTTTTTGCCTGATCGTTGAGCAAAATAGAGGAAGTGAAATTCAATTAGACGGTCTTTGCTGCTTTAACACAATTCAATCAATTTCCTTTTAGCTGCAGAAAAATCGAGATACAGCGGAAGTATCAAGCAGGATTACAACAGTGGTAGTTGCACTGAAGAATTTGAGATACCACTGTTTGACTTATCTACCATAACTAAGGCTACCGATAACTTTTCAGTAAACAGAAAGATTGGAGAGGGGGGCTTTGGTCCTGTTTATAAGGTAAAAACACTTGTTTCATGGGCAATTTTGTACAAGAAAAATACCAGCTTATCACACAAAGTCTAGATGACTGACAGAAGCTTCCTGGTCCTCCAGAGCAAAGCCTTCATCTTTAGTAGGCAAAATGTAATGAATTAAGTATATGCTGGAGCGTTTTTTGACCTAAACTATACTTAAACTGATTCTCGTAATTCCTCCAGTCCTTTTTCGTGGGAAGCAGCTTACTTTGTACCTTATACTTTCAATTTAGATATTATTTATTGTTTTTGAGAAAAAAGATATAAAATATATCGGAAACAATCTCTCTACCTTCAAAAGGTAGTGGTAAGGTTGTGTACACACCAGTTACCACCCTCCCCAGACCCCACTTGTGGGATTACACCAAGTATGTTGTTGTACAATATGTAAAATATTTAACTTGCAGATTTTGATGTTTAGGGAATACTTGAAAAAGGACGAGAAATAGCTGTGAAGCGGCTGTCTAAAACATCCACGCAAGGAGAAGATGAGTTCAAGAACGAAGTGGTATATATTGCCAAACTTCAGCATAGAAATCTTGTGAAGATTCTAGGATGTTGCATTGAAGGAGAAGAAAAAATGTTGATATATGAGTACTTGTCAAACGGAAGTCTTGATTCATTTATATTTGGTAACAAATTCCCATACCAATATTGTATATACTGAATTATCTTCTCCCAGTTTCGAATCTCCCATCAAATAAGAAAAAAAAAAACATGTAAGGAGTGAACTAAGACCTCAGCTTTATTAGAGCCTAATAATAATGTACACAGTAGCCCAAATTGATCTTTACGTTCAATACGTGAATGGACTGATTTAGTTTGATGTTTTAGAACTACTTTCTATCTATAGCTTAGTGAACCAACAAAATTTGGCAGGTGGCAGACAAAGCAGGATATTAGACTGGTCTGAGCGATTTCACATCATCAATGGGATTGCTCGGGGACTTGTGTATCTGCATCAAGATTCTCAACTGCGGATAATTCATAGAGACCTGAAAGCTAATAACATCTTGCTAGACAATGATATGAATGCAAAGATATCAGACTTTGGCATAGCAAGATGCCATGAAGAGGATGGTAATGAAGCCATGACAAACCGAGTTATTGGAACATAGTAAGAATCTATTACTGGGTAGTAGGAAATGCTTGAGATAGTTATGTTTTCCAGCATTAGTAACTATATTTTGTGTACATTTTCTGCAGCGGGTACCTCTCACCAGAATATGCGCTATATGGGCAATACTCTGTAAAATCAGATGTGTATAGCTTTGGTATCTTAGTACTGGAAATTGTGAGTGGAAAAAGCAACAGACGATTCGCTCCTTCAAACCTTAACCGTAGTCTAATCGGGCATGTAAGATTTTTATGTATTTTTCTTATCTATTCATTGGTGTTATCAACTAACAAGTATCTTGTAAGGCTAGTGATGCTTATAAGCTTGTAAAACTGCAGGCATGGGAGCTCAACAAGGAAGGGAGGTCAATAGAACTACTTGATGAACGTGTAGGTGATTCTTGTTCTACACCTCAAGAAGTGGTACGATCAATTGGTGTTGGTCTATTATGTGTCCAAGAACGTCCAGATGATCGTCCAAGTATGTCTTCCGTAGTTCTAATGTTAAACAATGAAGGTACGCTGCCTCAGGCTAAACTACCAGCTTTTTACATGGAAGGAGATGTATCAGATACTGAATTACTTTCTACCCAATTTGCACATATTACAACGGATGACACCCCGTTAGAAATTAGATGA

>M6-PGSC0003DMG400019241 M6 G-LecRK

TTGATTTTTCACATTTTCCCAAATATTCCAAAAGGACTAAACAATTTCCATGAAACGCTTTATCTTTTTCTATTTCTGTTCCTCTAATATAGTGTCATTTCTATTCATATTTTCCACTGCTTTAGACACTATATCGAACGATAGCCCCATAAAAGATGGACATACCATAGTTTCAGCTGGTGGAAACTTTGAGCTTGGATTTTTCAGCCCTGGAAATTCCAAGAATCGGTATATCGGTATATGGTACAACAATTTACCTAAAGGCAGAGAAGTAGTGTGGGTTGCCAATAGAGTGAACCCACTGAATGACACTTCTGGCATATTAACAGTCAGTAGCAAAGGGATTGTACTACTCAATGGAAACCAAGATGTGATTTGGTCATCAAATTCATCGAAACGTTTGATGAATCCAGTGGCTCAGCTTTTGGACACTGGAAATCTTGTCTTAAAAGATGACAGTTTGGTAAATCAAGAAGACTATGATTGGCAGAGTTTTGATTATCCAGACAGTACTTTGTTACCTGGCATGAAATTAGGCCTTAACTTGGTCACGGGCAAGTATTGGACTATGTCGTCGTGGAAGAGCAGTGATGATCCTTCTCCAGGAGAATATTTGGATCGCCTCGACACCAGTGGTTATCCTCAATTCTTTGTGTGGGAAGGTCCTGCAATAAAGTTTAGCTCAGGAATATGGAATGGACATTTATTTGTTGGAGGACCAAATTTGAAACCAAATCCTTATTATACATTTGAGTTTGTGAACAATGACAAGGAGATCTACTATAAATATGAGCTTATCAATACTTCTCTTCCTACCAGGTTGGTCTTGAATCCAGCAGGCCTGCTTCAACGACTTTTGTGGATCGAACGGAACCAGAACTGGTTCCTTTACTCCACTGGACAGATGGATAATTGTGATCGTTATGCCTTGTGTGGTCAATTTGCGCGGTGCAACATCAATGACTCACCTCCATGTGATTGTTTGAGAGGATTTCAGCCTAAAAATCAACAAGGATGGGATGCAGCAGATTGGTCTAGTGGGTGTGTACGAAGAACTCCGTTGACATGTGGGACAAGTGACAGGTTTCTGAAGTATTCTAGTGTGAAATTGCCAGATACACGCCATTCCTGGTTTGACAAAAGCATTGGCCTCGAGGAATGCCAAAGGCTGTGTCTGAAAAACTGTTCGTGTACAGCATATTCCAATCTTGATGTTAAAAATGGCGGGAGTGGATGCTTACTTTGGTTTAATGAACTCGTTGATATCAGGGAATATGCTGAACTTGATCAAGATCTCTATGTAAGGATGGCTGCATCAGAATTAGGTAAGTCTCCTTATGCTAAGATGAATCATACCACAGGGAATCTTAGTAACTCAATTGATTGACTATCTGAACTTTCACCTTGTTAACGAGGGTTGGGTTTCCAACCTTGTTAACGAGGGTTGGGTTCCCAACCTTGTAATCCCCTTTCCCATTTCCCATTTAGCCTTCCCCTACCCCCACCCCCATTTCAAAAATAAGAATCATACCGGAAGGTTTTCACAAGTCATAATGAACTTTTCACAGGTTCAGGCTATATGGGAAACACAAGAATATCAGTCATTGCAATTATATTAACAGTTTCAGCAATCATCCTTGTTGGTTTTCTATTTTGGTTTGCTATGCAGAGAAAGAAAGGGGAAAGAGGTTAGAAAATTTGTCCGTGGCGTTGAAAGTAAAGTAATTTCATTCAGCTCAATATCTTTACAAAAAAAAGTTCAATTTTACAGGAGTAGGAGAAGGAGAAGGAAAGGAAGATATGGAATTACCTTTGTTTGATGTGATGACTGTTTCTGCTGCAACTAATAATTTCTCATCTGCTAATATCATTGGAGAGGGTGGTTTTGGATCTGTATATCGGGTAACTAGAATTATATTCTCTCTGTATCTTAGTCAAATATATGAATGATTCTCTTGTGGCTTAGACTTCGCGTGGATTATGTGACAGGGAAAACTATCAACAGGACCAGAAATAGCAGTAAAGAAGCTCTCCAAACATTCCGGACAAGGTTTCGAAGAGTTGAAGAATGAAGTTGTTCTGATTTCCAAACTTCAGCATAGAAATCTAGTTAGGCTTCTAGGCTGTTGTCTTGAAGGGGAAGAAAGGATGCTAATTTATGAATATATGCCAAACAATAGCTTGGACTTCTTCATCTTTGGTTCGATTCTCTTGACTCATTTCATCAAAAGATTCATCTTATCATATCAGCATTGTTCTGCTAATGCAAAATGAACTGATGTTTCTGCAGATGAATGCAGGAAGAAACAACTTCCCTGGGAAAATCGTTTTCGTATTGCAATGGGAATATCAAGGGGCATTCTTTATCTTCACCAGGACTCCAGATTAAGGATCATACACAGAGATCTAAAGACCAGCAACATTTTACTGGACAGTGAATTGAATCCCAAAATTTCTGACTTTGGACTGGCCAGGATTATTGGTGGTGACCAAAACGAAGCAAGAACAAAGCGAGTAATAGGGACATAGTAAGTTCAATTAGATAACTCATTGATGTTTTCATCTTTTCCTCTGTTAGAGTTCCACATTGGTTGGGGAATTGACTGGTTTATATGGACTTGGGCAATTCTCCCTCATAAACTAGATTTGAGTTGAGTTAGACTCAAGTGTCATATCCTTACATGGTATCAGAGGCAGGTAAATCCCGGTTGGGGCTCCTGGTCCATGTTGGTGGTTTGCTTATATGAAATTGGACAATCTTCCCCTTATGAGCTAGCTTTTGGGGTTGAGTTAGGCATGTGTCAAATCTTTACAACTTCCATAAAATCATTTAACCCATATTAAACATTATATGTCTTTGTGCAGTGGCTATATGTCTCCAGAATACGCAGTTGATGGGAAATTTTCAGTGAAATCAGATGTCTTCAGTCTTGGTGTTCTTCTACTAGAAATAGTCAGCGGAAGAAAGAACAGAACATTTCATCATCCAGATCACCATCATAGTCTTATAGGACATGTAAGCAAGATACAAATCCAGACATTTGCTTTTGTCATTTCATACTACATTTGTATTTCCTACATATCATTGAAAACTATACTTGATTTTTCTATAGGCTTGGTTATTATGGAATAAAGGGAAGGCTTTGGAAGTAATCGATGATTGTTTAAAAGAATCGTTTGTGGAATCCCAAGTCCTAAGATGTGTTCATGTGGCACTGTTATGTGTTCAACGACTAACAGATGAAAGACCAACAATGTCATCAGTGGTTTTCATGTTAAGCCATGAAGAAGTGGCATTACCTCAACCAAAGGAGCCTGGTTTCTTCATAGAGAGAAGTACAGCTGAAACAGATTATTCAAATGAGAAAAGATGTATCAGTGACAATGATTTGACATTAACAATTCTTCAACCAAGATAGCAATGGTGGAACCAGAAATTCAAGAACCTTCTATTGTATAAAGGTGATTCAACAACTAATGTATATACAATACAGAAATTCTGTTCATCCCTATATGTACAGCAAAATATTTTGACGACGAAGGGGCTTCAATTGAACCCCCTCTAGACCATGCAGCTCCGCCCCTGATTGTTATTGGACGAGCATTTACTAACATGCTTACACGGAACAATTGTGTATATCTTTCCGATATTTGATTAAGACTCAAGAGCCATCGGGACCATTGTGTCAAGTTGTGCAAATTGTCAGAACATGATGGAAATGGACATATAAGTTGTTACACAGTATAATGACTAATAGTAGATCTGAATGGCTAAATGATAGCCACTGACTTTGTTGAGCAATTATTTTTCCTCAGGACTACAAATCAACATCATGTCCTGGGGGAGTGTATCTTTGCAATCACATTCATCATATCTTTTTCCACATCTCTATCACCAAACTCAAGTTTTGTGCAAACTTTTTCCTGATTTTTCTACCCGGTGTTCAGTACCAGCTTTGGGG

>M6-PGSC0003DMG400019275 M6 G-LecRK

AAACTTGCTAGGACATAAAATTATTACAATTTCTAACAAGATACTAAAATATTATCTAATTCTTTCATGCTCAAAAAGTCAAGAACTATTGACCACCTATAATATTTCTTAATTCCACAACAGCTCCCGTTTAGTCTTGGTTATTATCCAGTCAGATCTTATAGCAATCTATGAAGACCTCATTATCTTTGAACTGCAGCAACATTTTCCTCATGTTTGATACTGCTCAGAGCAGATATGCCTTTTACTAAATTATTCGTAAAATATATACTCAAAAGTCCAAAGCAGCCATTAACTTTTTTTATAAGCAGCCATGACTTGAAGCAGCCTCAACTCTCATTACAAAGCTAAGACTTTTCATAGTCATAGACCTTTCACTCCTTTCAAATAATACTCTTTCATATGTTAGAAAGCTCAAGTGATGGCTATTTTAACACTCGTTTGTTCAGTTTCTTCTTGTAAGATTCACTGTCTGAAAACAAATTAAAGTGTCATTACTATGCAGCATTTGACATAGAAAAAAATTGGATCTCAAAGCATGAAGGCATTACTAAGGGAAAGTTTTTACTTATCTTTGCTCTTGTGTTTGTGTTTCATCAACCGAAGCTTTGGCGCTACAGATACAATAACTACACCTCATTTTCTGAAAGATGGTGATGAAAATGTTGCTTCAACTGGTGGAATATTCGAAATGGGGTTCTTCAGTCCAGGTAATTCGAAAAATCGCTATGTGGGTATGTGGTATAAGAACACATCTGTTAGGACAGTGGTGTGGGTTGCCAACAGAGAAGCTCCTTTGACTGGTGGATCAGGTATCTTGAAAATAATCAAGCCAGGAATTCTTGTCCTTGTCAATGGCACTAGCCATGTTGTGTGGTCAACTAATACCACTAGATCTGTGCAGAATCCTGTAGCGCAGTTGCTAGATTCTGGAAATCTTATTGTCAAAGAAGCTAGTGATGATAGCCCGGGGAATTTGATATGGCAGAGCTTTGATCACCCAACGGATACGCTATTAGCAGGCATGAAGCTTGGTAGGAACTTTGTAACTGGCAGAGAGGTGTACTTGTCATCATGGAAGAATGAGGAGGATCCAGCTCCTGGTGATTCTACATATCACTGCGATCCTTCTGGTTACCCGCAGAATATCCTGAAGAAGGGGTCGGATGTAGTATACCGCTCTGGACCATGGAATGGTCGCTCTTTCAGTGGGAACCGAAACTCAAGAGAAGGTCCTTCCTATACATATGGAGTATATTCAAGTAAGACAGAGGTGTATTTTGGATATAACCTCACAACTTCAATTCTCGTGAGGTTGACATTAAGTCACAATGGAGTCCTACAGGTTTGGACTTGGGGTGATGGAGACCAAGGTTGGATCCCTTTCCTCTTAATACCCGCTGATAATTGTGATATGTATAAGTTGTGTGGCGCATATGGCAGCTGCAACAGTCAAGATTCTCCAGAATGTGGATGCTTAGACAAATTTGTGCCCAACAATAGTGATGCTTGGAAGAGGAAAGACTGGTCAGGTGGCTGTGTTCGGAGAACTGAACTAAATTGCCTCCAAGAAGATGTATTTTTGAAGTATTCGCATATCAAATTGCCAGACACACGGAATTCCTGGTCCAATGTGACTATGACACTAGAAGAATGCAAGAACACCTGCTCTAAAAATTGTTCTTGTATGGCTTACTCCAATTCTGACATACTCAACGGAGGAAGTGGATGCTTATTGTGGTTCAAGGATCTGCTTGACATTCGGCAGGCGCCCAACGGAGGGCAAGATATCTACATTAGAATGGCTTCTTCTGAATCAGGTTGGCTTGTTGCAACTTAATTTATTTATTTTTTTGATAGATTGCCTGAAGCTAGTAAAGGAATTTAAGGTTGCAGGCACTTTAATAAAGCAGACTGTAGTTTTGTGCCTATGCTTACTATGATACTATCTACTGCCAACTCCTACCATATATAGCCCGTAACTACCAAATAATAGCAGTACAGAAAGCAATTATTTTTGAGTGAAGTTGGTATGAATTTACAAGCCAAATCTGCAAGCATGATGACTATAGAACAAGCATTTGATATGACTTGCATTACACAAGGAATTGAAGAACCTTGACAACTCAACCTCAACAAACTAAAAAGGATGTGCAGTACTCCAAAGGCTTGCAGAGACTTTTAAAACTCAAAATAAAAATCATCTCCGATGGATAGGATAATTGCAATTTATTATTTATGTTTGTTGAATTGATACTTGCAGATAGTCTGGAGCAATCAGATGGAAAGAAAAGGAAAGTACTTTTCTGGATTCTACCATTATCAGTTTGTTTGATTCTGGTATTCCTTACTCTGGTGGTCTACCATAGAAGAAGGAAGAAGGCTTTAGAGCTCAAAAACAAAGGTAAGCTGGATACCCCTTAATTTTTTCTTGACATCTAAAGTCTAAGCAAAATAGAGGAAAAGACATTTAAATTATACTGAAAAGAACTTTCACTTCATTCGCTTTCTCTTATAGGCAAAATTATGTCATTTGGTTTTGGCAGGAAGGTCAGGATACAGTGGCAATTACAAGATGAATTACAACCGTGGCAATTGCACTGAAGAATTTGAAATACCACTGTTCGACTTATCTACCATAGCAAAGGCTACCAATAACTTTTCAATTGACAGACAGATTGGAGAGGGGGGCTTTGGACCTGTTTACAAGGTATATACACTTAAGACCTGGGTCTTCAGAGTAAACCTTCTTCTTTAAATACAATATAGGAGTTGTATGATCAAAGGTATAAGCCTTTGCTTTGTGAAACCAAAAGGAGACCTATACAATAGCTAAGGGAAGAAAGGTTGGGATATGGGGGATATCCTTCTCAGAATGTATCCAAGAACCACTAGTCTAGCTTTTCAATAAGTGTTGTCCCAAGTTAGTCAATATAGAGCCTTTAGCATTTTCAAGAAAAAGCTGCAAGAGTAACCCATCACCCTTTGGGTGGAACTGCAGATTAAAGTGTTGAGGATATAGTTTGTTTAAGAAGAGAGTCATTTGTTCTTTGTAGTCATTTATTTGTTCATCTTTCCATGGTGACAAGTGGCAACTAATTCCCAAATCAAAAGGGCGAATATTTCTTATAGACTTTAGTTTTAGTTTTAAAGAAGATTATCATAGGGCTAAACTGAAAATATAATGAACTTATAATAGGGATTTGGCACATTTGACCTAAATTTGTTAACGGAGATTTTGATCAACTTAGGTATTGAAAATATTAACATGCAGAGATTGTGATGTTTAGGGCATACTTGAAGGACAGGAAATAGCCGTCAAGCGGTTGTCTAAAACTTCTACACAAGGAGAAAAAGAGTTCAAAAATGAAGTTTTATATATTGCTAAACTTCAGCAAAGGAATCTGGTGAAGATTCTTGGCTGTTGCATTGAAGGGGAAGAAAAAATGTTGATTTATGAGTACTTGCCAAATGGGAGTCTTGATTCATTTATATTTGGTACAACTTCTCAATCCAATATCATGCATTTGAATAATAATTGCATTTCAGTTAATTGATGCCTATAGGCATACACTGATTTTCTTCCAACTTCAAAATCCCCATAAAGCAGGGAAAAAAAATGTAAGGAGAGAACTAAGACCCCAGTTGTATACGAGCCTAATAATTACATACTTGGTAGTTCAAACTTCTCGTATTATGTTTCCATCAGCAAGGAACTGATTTTGTTTTAGGTTTTAATACTACCTTCCACTAAACCTATTTAGTGTGTTAATGACCCAACATTATTTGGCAGATGACATACAAAGCAAGGTATTAGACTGGCCTAAACGATTTCACATTATCAATGGCATAGCTCGGGGACTTATGTATCTGCATCAAGATTCTCAATTGAGGATAATCCACAGAGATCTGAAAGCTAACAATATCTTGTTAGACAAGGACATGAATCCAAAGATATCAGACTTTGGCATTGCAAAAATATGTGATGGGAATGACATTGGGGACAAGACAAACCAAGTTGTTGGAACACAGTAAGAACCTTATACTTGATAGCTAGAAATTTGAACCTAAGAAGAGTTTTGAGATATTAGGTACGTTTTCCAGCATTAGTAATTCTATTTTTTGTTTACATTTCTATGCAGTGGGTACCTCTCACCGGAATATGCATTGCACGGGAGATACTCAGTAAAATCAGATGTATTTAGCTTTGGTATCTTAATATTGGAAATTGTAAGTGGGAAAAGCAACAGAAGATTCTCTCATCCAGACCACAACCTTAATCTACTTGGACTTGTAAGATTTGATGAGTTTCATTTTACTTATCCTTTGGTCTTATGTTATCTTCTAGTAACAGATATCTGCAAATCTAGTCAGCGTTAATAAATCGTGTAATCTACAGGCATGGAAACTCTATAAAGAAGGCAGGTCAACAGAACTACTTGATGAATACCTAGGGAATTCTTGTTCAACACCTGAAGTGGAACGATCAATCTGTGTTGGTCTATTATGTGTCCAACAATGTCCAGAAGATCGTCCAAGTATGTCTTCTGTGGTTATGATGTTAAACAATAAAGGTGTATTGCCACAGGCTAAACGGCCAGGTTTTTACATTGAAACAGATGCACCGGATGGTGAACTCTCGTCTAGCCATACTGAGACCCCCATCACAATATTGGTAGCCCGATAAGAAAATATTGTTTCACTTGCTTTCAACAAGCTTGCAGTTGTAAAAAATCTTTTCTTTTATTCTTTGTAGAAATGTGGTAAAAAAATATTTGGTATATCAGCTATTTCGAAC

>M6-PGSC0003DMG400019276 M6 G-LecRK

AAAAAATGAAGTCTTGGTCATCTCTGAGACAATCTTACTGAAAAGAATACCACAAGAGATGGAATATGATAAATTCTTTTTAACAATTTTGAAGAAGAATGCAATGCAACAAAAGGTAAATATAACTATCATGGATATAACAATATTTCAATTTATTCTACTTCTCTTGTATTTTGTGTTTCTTCAAAATGCTAGTGGTCAAGGAGATACCATAACTACAACTCAATTCATTAAACATGGTGATACTATTGTTTCGTCTCGTGGAACGTTCGAGTTAGGATTCTTCAGTCCTGGAGATACTTCTACAAATCACTATGTTGGAATATGGTACAAGAAAACCTCTGCTACTACTCCTGTTTGGGTTGCCAATAGACTAGCTCCTGTCACTAATAAATATGGGGTGTTGAAAGTCATTCAGCCAGGTCTGATTGTACTTCTGAATGATACTAATGCAACTGTTTGGTCGACAAATTCATCCAAATCTGTGCAGAATCCTGTTGCTCAGTTGTTGGACACAGGGAATTTTGTGGTGAGGGATGCTAATGATCCGAATCCAGAGAATTTCCTCTGGCAAAGTTTTGATTATCCTAGTGATACGTTGTTGGAAGGTATGAAACTTGGGACGGACTTGGTGACTGGATTAGAAAGGTATCTTACTTCTTGGAGAAGTAGTGATGATCCTGCTCCGGGGGATTATACTTATCATTGTGATCCTACGGGTTATCCACAAAACCTTATGCGAAAAGGGGCTAATGTGACATTTAGAGCAGGGCCGTGGAATGGTATCAGGTGGAGTGGTGCACCAAATCTGGTGAACAACTCTATCATTTCTTTCGGTGTGGTTATGAATAGTAGGGAAATTTACTACAAGTATGAGATGGTGAACAAATCCGTTATTTCAACATTTGTTTTAGAGCCTTACGGGAAAGCTATGCGTATCATCTGGATTGGAAAGGTTCAGGGATGGGTAAATTACCACTCTGCAGTTGTGGATGACTGTGACACCTACAAATTATGTGGTGCATATGGGACTTGCAACATTTCGAGTGATCCTTTTTGTCAATGTTTAGATAAATTCAAGCCAAAACATCCTGATGATTGGGAAAGATCTGACTGGTCTAGTGGTTGTGTCCGGAAGATCCCACTGAATTGCACAGGAGATGGATTTATTAAGTATTCTGGTGTTAAACTGCCTGATACGCGAAATTCTTGGTTCAATGAGACAATGACACTAGATGAATGCAGGGTAGTTTGCTTGAGAAATTGTTCGTGTATGGGTTATACTAATCTGGATATACGCAATGGAGGAAGCGGGTGCTTGCTATGGATTTGGGAGCTGGTTGACATCAGACAGCTACCTGCATCAGGACAGGATATCTACATTAGGATGTCTGCTTCAGACATAGGTAATTAATAATTAATATGCGAGTTCATAAATCATAACCATGTCAATTGTACTTCAATGCCAGAAAGTCAACTCATATGAGGGACAAAGGTAATCCAAGTCCACATTATGGGAATAAGACTTCAAGGACCGTCCTGTTTTCCTCGACATAAATACAACAAATATAAAAAGAAAAAAAACTTCTAGCATTACTAGAAACAACTATTGATAATATTTCGGTGGGAAATGTTGCAGGTTCTGCTGGATCAAAAGGAAAGATAGCTGTCATACTTGCAATAGCTTTGCCACTATTGGTTGCATTGATTCTGCTCGGTTTAGGTGTAGGCCTGATTCTTTGTAAACGAAAAAGGAGAAAGGATCCAGTGGTTACAACAACAGGTAAGTGGAAGTACCAATTCTATCTATTGTAAAACTGGAAGCACCTAGTGGTCATTAGTTCTGTATACCACTAGTATTTCTTTTCGAGCTCCTTATTTGAGCAAGGTCTACTTGTATGTTCAAATTGCAGGGAGATTAGGTGGTCACAGCAACAAGAATGATAACAGTAATCAAATTCACCAGGAAAATTTTGAGCTTCCACTTTTTGACTTATTAACGTTAACCCATGCTACCAACAACTTTTCATTTGCAAACAAGATTGGAGAGGGTGGCTTTGGACAAGTTTACAAGGTAGAAACATTCCTCAGACCCCAAGTAATGATACTCGAGACTTAATTTCTTTGCTTTTCTTGAAAGGAATATTAACTCTTGGAGTCCGTACATACAGCAGTGTTTGCTAGTATGATGAGCGGAATGAGTGTTAAGTTGAGAGCCTTGTTTCAAGACCGATAATGTTTTATAAAATAAAGGTTTCAAATTCAAAGAAAAGGAATATCAGCTGGAAAGATTACCAACCATGGGAGAAGCTGTGGAATAAAATGTGCAAGTAGATGCTGCCTGAACAAGAACTTGATTGCAATAGTAAGGAAAAAATGACGATTCACTCCATCGAATTTCTAACAGGAGTGCTACTTTTACAGGGTGTATTAGAAGGTGGACAAGAAGTAGCAGTGAAGCGGCTTTCAGAAACATCAGAGCAAGGACTTCACGAGTTCAAGAATGAAGTTAAGTGCATCGCAAAACTTCAGCATCGGAATCTTGTAAAGCTTCTCGGATGCTGTATTCAAGGAGAAGAGAAGATGTTGGTGTATGAGTACTTGCCAAACAAAAGCCTGGAATTATATATATTTGGTTAGTCTTTCACATCTACTGAGCGGCCATTTCTTTCAATGCAATCAATTGTGATTTCACGAAGTTTGGTTTCGCTGTTATAGAAAACTTTAGTAATAAGTTGAGAACATTTGACAGACGAAGAAAGGAGTGCATTACTTGATTGGCCTAAGCGCTTCAATATTATCAATGGAATTGCTCGAGGACTGATGTATCTTCATCAAGATTCTAGGCTCAGAATCATCCACAGAGACTTGAAAGCTAGCAATGTCTTACTTGATATTGAAATGAATCCAAAAATATCAGATTTTGGGATGGCTAGAAGTTTCAGAGGAGATGAGACAGGAGACAATACACGCCGTGTTGTTGGAACATAGTAAGGTTTAATATTTGATTATATATGTAGTCATGAACACATGGTGCATTATATAATCCTGTAGTTTCTTTATTGCAGTGGTTACATGTCCCCGGAATATGCAGTCGAGGGGATATTCTCAGTGAAATCAGATGTCTTTAGCTTTGGGGTATTAGTACTAGAGATTGTGAGTGGAAAGAAGAACCGACGATTTGTTCATCCAGACCACCACCTCAACCTTCTTGGACATGTAAGTACAGTAACTCCTAACGGTAAAAACTCAGAAAAAGTGTTTGTTAATGCATTTAAGTTTTATGTGCCGACAGTAAATTACATGTAAATCTCTTTTATAAGAGCTTGTTAACTAACCTACTATCCCAAGTTAAACCACAATGCTAGAGTAAAAAGTCTTTGTACTATTACTGGTATAAGTTAAATCCTTTGTTGATGGAGGGATCGTTTCATGCAGGCATGGATTCTTCATAATGAAGAACGGGCATTGGAACTAGTAGATCCATATCTCGTAAACTCATACTATATATCAGAAATACTAAGATCAGTCCATGTGGGATTGTTATGTGTTCAACAAAATCCAGAGGACAGGCCAAGCATGTCAACTGTGATTATGATGCTGAGCAATGAAGGCATTTTGCCATTGCCTAAACATCCCGGTTTCTTCACGGAAAGGAAAGTTGAAGATGTTGATCAATTGTCTTGGAGTACACAAACACCAAGTTCTATAAACGAAGTTACCATCACGCAGTTGAATGCTCGATAGAAGCATGTCCATGTCCATAGTCAGTGGCAAAGCCAAGATTTTCACTAAGAGTTTCAAAATATGAAAGAAGTAAACACACAAAGTCAAGATGATTCAACAACTATTATATATACATAAACAATTAATTGTAGACCTTTACATATAGAGTGTAATTTTTTGGCGAAAGGGTTCGGATGAACCCCTTGCATCAACTTGGCTCCATCCTTGTCGATTGTGAGTGTCATTCATTTCTCAGAATTGTATTAAAATCCTCCCTTTTTTTCTTTAAGAAATTAAAAAGCAAAATCTGCAGGAATTGTATTGAAGTATGTCTAAATTTTAAGGCCTTGAATTCAATTTCAG

>M6-PGSC0003DMG400019278 M6 G-LecRK

TATGCCTTCACACGACTCCTCTGTCTGTTCCTATACCATGTTAATTCGTTAATTGTGTATTCACAGGTATCCTGAACAATGGTATGCAGTGGACTGGTCTAATGGTTGTGTAAGGAAAACTTCTTTGTCTTGTAACCAAGATGGTTTTCTTAAATTTAGGAATATCAAGATGCCGAATTCTAGACACTCCTGGTATAATGTAAGCATTAATCTTGAAGAATGCAAGAATATGTGCTTGGCTGATTGCAATTGTACAGCCTACTCAAATCTTGATATAAGAAATGGCGGAAGTGGATGCTTACTATGGTTCGGTGAACTCATTGATATTAGAGAGTACAACAAAAATGAGCAACACCTGTTTGTGAGAGTTGCTGCTTCAGAATTAGGTAATGACCCTACTCTATACCATTTTCGTGTTCTTTTGCACAATATAAGCACAACTCTTTAGAATTTAAGGAACTTTACTATCTGATGTCACTGATAGTAACATATGATAGTCGCGGAAGCTATTCATACTGATATTGCCAATTCATATGTCAATTGAAAACTGCTTATGGACTTTTTATTAAACAGATTCAGTCAGGAATCGGAGGGGAAAGAGGTCAGCCCTGATTGCGGTCATTGCAGCACTAGTAGCAACTTCTATCCTCATCTTTGTAGCTTGGTTTACCTTCCAAAGAAAGAACAAAAAAACAGATAAACATACTGGAGGTATTAGAAAATTGTTAACTGTGAATGCTTTTTTACCTCCATCCAGCTAACATTGTTCGACTTTATCAAATGATTGAGGAAAAGTTCAATTATATCATGTCTGTGATTGGAAAATATATGTTGACAGGTTCAGAAGTTGGAAACAAGGACCTCGAGTTGCCATTGTTTGATTTAGTTACTGTTACTACTGCCACTGAAAATTTCTCTTCTGCGAATGTGATTGGTGAGGGTGGCTTTGGACAGGTTTACAAGGTAAATCAACTTTGATCTTCAACTAAAAGCTAGAAGGGACATATTTGACCCTTGAGAGAAAATCGCAGGGTATTCTACCAGATGGACTAGAGATAGCAGTAAAGAAGCTATCAACGTATTCTGGACAAGGCGTTCAAGAGTTAAAAAATGAAATTGCTCTCATTTCCAAGCTGCAACATAGGAACCTTGTCAAGCTTTTGGGTTGCTGCCTTGAAGGAGAAGAAAATATGCTAATCTATGAGTTTATGCCCAACTCTAGCTTGGACTGTTTCATTTTTGGTCTGCTTTTAGTGATTCATTAACTAAATATTACTACTTGATAAGTCCTCAATATCTCTGTATCACTCAAGAGTTGAATTTACATTGTGCAGATCCAAGCAGAAAAGCTTCACTTGCATGGAAGAATCGCTTTGAAATTTCTGTGGGAATATCTCGAGGTCTTCTTTACCTTCACCAGGACTCAAGATTTAGAATTATTCATAGAGACCTCAAGACCAGCAACATTTTACTAGATGGCGGCATGAATGCCAAAATTGCTGACTTTGGCCTTGCCAAAATTTTTGGCGGAGAACAAGTGGAAGGAAATACTAAGAGAGTGATAGGGACATAGTAAGTTCTACAAATCTTTATGATAGACATTCATACAGTACATGCACTTAAATTTGATGTTGATGCAGTGGATATATGTCACCTGAGTATGCTGTTGATGGGAAATATTCAGTAAAATCCGATGTATTTAGTATCGGCGTTATCATTCTAGAGCTAGTCAGTGGCAGAAGGAACAGGAAATTTCGTCATTTGGAACATCATCACAATCTTTTGGGACATGTAAGCACATGCAACACTAAACTTCGTTCCCTTAAGTCACTTTTTCTTTAGTGAATGCATTTGTATAAATGTGTTCTGTTTGTAATTTAGGCATGGATACTTTGGAATGAAGACAAAGCGTTGGAACTGATGGACGAATGTTTGAAAGAATCATTTTCAGAATCTCAAGTGTTGAGATGCATCCAGGTTGGTTTGTTGTGCGCCCAAAAACACCCTGAGGATAGGCCTACAATGGCATCAGTAGTTTTCTGGTTGGGAAATGAAGGCCTGGTTCTTCCTCAACCAAAGCAGCCTGGATTTTTTATAGAAAGGAATTCAATGGATTCAACAGAATCAGCTCAATTGATCGTTAGCTCGTTGTAATCGGGCAAGGGCATTGTGAATTCAAGTTCAATCAACACATATAATATCTAATTAATGAGAAGAAAATCTGAAAGTGTGTATACACATTTCAGCACACGCTTTGTAGCACAATTAATCCGGATTCACACTGGAAACTACAACTTTTTGAGAGAAACTTAAGTCTCCTCATCAAAAGTGACTCCATTCAAGAAGTTGAAGATTCTTCTTGTAGCATGGCAAACTATTCTTTATATTTTTTCTCTCAGTTGAATATTAGATGGTGACTTCACAACTTTATAAAATAAATGCATTAATATTAATAGAAAATAAAAAAGACATCAGTTCCTTTTTTCTTAGTTCTGCTATTACTAGTCCAAGATGACACATCGTCATTACTACTAACTCATTTTAGGAAGACTTTAGGCAATTTGCTCCAATACATTTCATGTGTTTACACCAAGAAAATAACCGCATGCCATATGTTTGTATGAATAGTTCTATAGCGTGCAATTGCAAACACCAGGTTAATGGTGGAAATGGGAAGTCAAAGGCATACAACTTTTGACAAGTTCAAACTTTGACATCTAATTCTTGGCAGCAGTTGTTGGCCTTTCATTCAAAAAAGTATTTTTTTTCATTTTATTTATATAAAACATTTTTAGGCATGGTTGTTAATTCTAAACTGAGAAAACAAGAAGTTATCAGTATCTTTTCTGTGTCAAGGCTTTTTTGAACTTGTCAATCAAAAACATGAGTCAGAATCTATCAAAAGGGCTACTGTTTTTATTCTTTTGCTCACATTTTTTCTTCATCTTACTAATTTCCGCTGCATTAGACACGATCACAACAGATAAATCCATTAGAGATGGTGACACAATTGTTTCAGCTGGTGGGATTTATGAGCTTGGATTTTTTAGTCCTGGAAATCCGAAGAATCGTTACGTTGGCATATGGTACAAGAAGATATCACCTAGAACTGTTGTGTGGGTTGCAAACAGAGATATTCCACTTAATGACACATCAGGAGTTTTAACACTCAAACCCAATGGGATTCTTGTACTTGTTGATAATTCCAATATCTCAATTTGGTCATCAAACTCATCAAGATCCTTAAAGGATCCAAAAGCACGGATCCTGGATTCCGGGAACCTTGTTGTCAATGATGTAAATGAAAGAGACCTGGAAATTAATTTCGCGTGGCAGAGTTTTGATTATCCAGGAAATACTTTTATACCTGGCATGAAACTTGGACGTAATTTGGTCACGGGCATGGATTGGTACATGTCATCATGGAAGAGCATTGATGATCCTTCTCCTGGTGAATATATAAACCGTCTTGATTCTCATGGATACCCGCAATTGTTTGTGTGGAAAAATTCAACTATAGTGTCTAGCTCAGGGATGTGGAAAGGTAATGCATTTAGTGTTAGTGCAAATAGTAAACCAAATACACATTACACTTCCGAGTTTATTATTGATCAGCAGCAAATTTACTACCAATTCAAGCTTAAGAATGAGTCACTGCCCAGCAGGATGGTGCTTAACCCGGAAGGTCTGATAGAACACATAACATGGATTGAGAGCAGTCAGAGTTGGTTTCTCTACTCAACAGTACAATTTGATAACTGTGGTCGTTTTGCTTTATGCGGTCCTTATGCAAGTTGCAACATCAATAACTCCCCTCCATGTGACTGCTTGCAAGGTTTCAATCCTAGGTATTGTTAAAATGAGTGATCATCAAACCTTAAGCTATTAGAGATAACACACTTGATTACTTAGTTATGATTTTACACTCCCCCTCATGTGTGTGCTTGATTCTATTTCTTGGGCCAAGCACATAGATATTATATTGATTACTTAATAATGACTTCACAGGGTTCCTCAACAGTCTGCAGCAGATTGGTCTACTGGTTGTGTTAGGAGAACTTCTTTGGATTGTAACCAAGATGGTTTTCTTAAATTTACGGGCATCAAGATGCCTGATTCTAGAAACTCCTGGTTTAATGAGAGCATTAACCTTGAAGAATGTGAGAAATTATGCTTAGCTGATTGCAACTGTACAGCCTACTCAAATCTTGATGTTAGAAATGGGGGAAGTGGATGCTTACTATGGTTTGGAGAACTCATTGATATTCGAGAATTGAGCCAAAATGAGCAAAACCTGTTTGTGAGAGTTGCTGCTTCAGAAATAGGTAATAATCCTACTCTGCACCATTTCCTCATTTTGTTGTTCAATATAAATGCACAGCTCTTTAAAACTGAAGAAGCTCTGGTCAGCAAGTCACTCATTCAGTTCATGGCCAACTCATATGTAAATTGAAGACTGCTTATGAACTAATTATTGAACATATTCAGACAGGAAGCGGAGTAGAAAGATGTCAGTCCTGATTGGTGTCATTTCAGCAGTGGTAGCAACATTTATCCTCAGTTTTTTAGCTTGGTTTTCCTTCCAAAGAAGGAAAAGAAGAATAGGTAAATATAGTTGAGTCACTCTTGTCTACCAACTTTATAAGAAAATTGTCAAGTCTAAATGCTGTTCTGGCTAGGTGAATTTTCTATCTAGCTGACATTGTTTGCCTTTACATAATTCTCATTGATATCATGTTTGTGGTGGGCGACTGTATATTGACAGGTCCATCTGTTGAAAATGAGGACATGGAGCTTCCATTGTTTGATTTAGTTACTGTTACTAGTGCCACTGGGAACTTCTCTGCTAAGAATGTGATTGGGAAGGGTGGATTTGGACCGGTTTACAAGGTAAATCAACTTTGATCCTCAACGTCTAACTTGAAATGGCATCTTTGACCCTTGACATAAATCTCAGGGTATCCTACCAAATGGACAAGAGATAGCAGTAAAGCGGCTATCAAAGCATTCTGGACAAGGCTTTCAAGAGTTAAAAAATGAATTCGTTCTCATTTCCAAGCTGCAACACAGGAACCTTGTTAAGCTTTTGGGTTGCTGCCTTGAAAGAGAAGAAAGGATGCTAATCTATGAGTTTATGCCCAATGCTAGCTTGGACTATTTCATTTTTGGTCTGCTTTTCATGATCCATTAATAAATAGTACTTGATATAAGTCCTCAATATCTCTGCATCACTCAATGATTAGTTCTACATTGTGCAGATCCAAGCAGAAAAACTTCACTTTCATGGAAGAACCGCTTTGAAATTGCTATAGGAATATCTCGAGGTCTTCTTTACCTTCACCAGGACTCAAGATTAAGAATTATTCACAGAGATCTCAAGACCAGCAACATTTTATTAGATACTAACATGAATGCCAAAATTTCTGACTTCGGCCTTGCCAAAATATTTGGTGGAGATCAAGTGGAAGGAGAAACTAAGAGTATAGTTGGGACATTGTAAGTTTCTACACATCTTTAAATCGTATAATCTTTATTATATTAGATCAGTACATGAGCGTAAACTTATTGAATTGATGTAGTGGATATATGTCCCCGGAGTATATTGTTGATGGGAAGTATTCAGTAAAATCAGATGTATACAGCATCGGCGTAATCATTCTTGAAATAGTTAGTGGCAGAAGGAACAGGAAATTTCATCATTTGGAACATCTTCACAATCTTTTGGGACATGTAAGCATATACATTACACAACTTCATTCCCTTAATTCGCTCTTAGCTTGCTATTGCGAATACATTTGTATAAATGTGTTCTGTTTGTAATATAGGCATGGTTACTTTGGACTGAAGGCAACGCGTTGGAACTGATGGATGAATGTTTGAAAGAATCATTTTCAGAATCTCAAGTGTTGAGATGCATCCAGGTCGGTTTGTTATGCGTCCAAAAACTCCCAGAGGATAGGCCTATAATGGCATCAGTGGTTTTTTGGTTGGGAAATGAAGGTCTGGTTCTTCCTCAACCAAAGCATCCAGGTTTTTTCACAGAGAGGAATCCAATGGAATCTACTGATGAAGAATGTCTAAGTAACAACGCGACGTTAACAGTTCTTGAGCCAAGATAGACAAGTAATCACTATTGGATGTTTTGAAATGTATTTACTTTGTCTTGTTTGTCAAACAGTACTAGTGTATATATGATAGAATATACTAAAAGAGATAGTTTAATGCCAGTGTTCATATCAAGGCAGGGCAAATAATTGATATACAATTCTTGTCTTTGATAATTGAGCAATCAATGGACTAATTATTGGCAAAATTTGTAAGCATAAGTAGTTAGAGGAATCTATATCATGCATTACTTTTTTGGGGGACAATGAATATATTCATCAAACATAGTTTGCTATACATGATACATAAGCCCTATTCTATATGAAGTATTTTTTTTTGTTTCCCACTCGGTGTTTGGAGCCCACATTGGAGCTCCGGTTAAATTCGAGTTGCCCATAGCAG

>M6-PGSC0003DMG400021804 M6 G-LecRK

TATGCTGAACCGGTGACCATGATGTCTAGACAAATAAAAGAGCTCTAATGTTGTCCTGTCTAACAAGCTGATGGAGAAAGTCGAACTGATTACCTATATTTGGGTTGAATAGTTGCAGCTAGTCCGGTGAAATCAAATGGACACAAAGGAAAATTACTCTCCTGGATTATCCCGTTATCGGCTGGAGTGATTTTGATAATCCTAAGCCTAGTGATCTGGATTAGAAGAAGGAAGATATGTTCAAAGAAAAAGAAAGGTAACCTCGACAAGTCATTATGTTCGGAATATAATTAGTAGATGAACTCAAATTCAAAGGAAAATCTTTCTTGTCTAGTATTTGCCAAACTGAATTTGGGACATTGGTCTTTTGGCAGGATGCTGGGGGCGCAATGGCAATTACAAGATGGATTACCTCAACGGAAACCTCAGTGAAGAATATGAGTTACCACTGTTCGACTTATCTACAATAGCTAAATCTACGAACAACTTTTCAGGGATGAGCAAAATTGGAGAAGGTGGATATGGACCTGTTTACAAGGTAATCACTTATAGCATTTGCAGTTGTTTACATGAAATAAGCAATTCACAGCCCACATAATTGAGCATAGAGTGATTCGGGAAGACTGACAGTAAGGAATGATTTAATGACTTTATAGTTTTGGAAGAGTTTTATAAACATTAGCTAAACTTGATTGAATGTATAATAGTTGCTTCATAATTTGAGCTTCTACCTTTTGGTGGAAGTATAATTTAACTTTCATGTTCGGTGTTGAAAAAACTTAACGTGTAGATATATGATGTTTAGGGCGTGCTTGAACACGGACAGGAAATAGCTGTGAAGCGGCTGTCTAGGACTTCCACACAAGGACAAGATGAGTTCATGAATGAAGTGATGTATATTGTGAAACTTCAGCATAGAAATCTGGTGAAGATTCTCGGATGCTGCATCGAAGGAGAAGAAAAAATGTTGATCTATGAGTACATGCCAAATGGAAGTCTTGATTCATTTATATTTGGTAAGGATTTCACAAAAAAAAGAAAAAAATTGTGCTTAATAGTTACATTTCAGTTGATAGATGCCTTGTAGGAGTACCCAAATAATCCTCTTGGCCCAAAGTTATGTTTTTTATTGTTACTTTTCCGATTTAGTTCGATGTTTAAATCCTACTCTCAGTTTTACCTGATTTGTGTGAATTAATGCTCCAACTATGAAACTTTTAACAGATGACACACGAAGCACGGTATTAGACTGGTCTAAGCGCTTTCACATAATCAATGGGATAGCTCGGGGACTTATGTATCTGCATCAAGATTCTCAGTTGAGGATCATTCACAGGGACCTGAAAGCTAACAACATTTTACTAGACAATGACATGAATCCAAAGATATCAGACTTCGGTATAGCAAGAAGTTGTGAAGACGACAAGTTTGGAGCCAAGACACACCGAGTAGTTGGAACATAGTAAGAATCTTAAGATGATATTTTACTGAATCATTCAAAACTGGAATAGAAGAAGAGCCTTGAGATTATCGTAGTATGTTTACCCAGTATTAGTAACGCTAATTTCTTTTTACGTTTTATGCAGTGGCTACCTCTCACCGGAATATGCTGTACATGGTGTCTACTCGGTGAAGTCAGATGTATTTAGCTTTGGTGTCTTAGTTCTTGAAATTGTGAGTGGGAAAGGGAACAGAAAATTCTCTCATCCGGACCATAACCTTAACCTCCTCGGACATGTAAGGGTTGTGTTTTTCATTGTCTATCAGTAAATTATCTCGTAAACCTGGTCATGCTAACACGATACTAATATCTACAGGCATGGACACTCTATAAAGAAGGTAGATCAATGGAATTACTCGGGGATTTCCTCATTGATGTTCATTCTACACCTGAAGTTATACGATCAATCCATGTTGGTCTATTGTGTGTTCAACATCGTCCAGAAGATCGTCCAAGTATGTCTTCAGTGGTTATGATGTTAAACAATGAAGGTGTACTGCCACCTGCTAAACAGCCAGCTTTCTTCGTAGAAACAAATACACCAGATAATGAATTCTCTTCTAGTCAATATGCACATAGTACAGTGAATCAGATCACCATCACAACATTAGAAGCCAGATAGTAAATTCTTCATTGCAAATAGCAATCTATTATACTCTTCATTCTTTTCTTCTTTGTAGCA

>M6-PGSC0003DMG400021993 M6 G-LecRK

AAAAGTATGAAAATTTTATGATGATGAATTGATGGAGGACAAAAAAGTTTATTCCTTAGCCCTTTTACTTGTGATTTTATTGATCACACTCATCGTTATCGCACGTATAACGTTGAAATATTCGGAGACATTTTTTCTTATTTTCGGAGCTGATATCGCGTTGATTATTGGTGTTTTTGCTGGTGTTTTGATCCGAAGGAAGTTCAACAACAGGAGGTTATTGCTAGAAAATCAGTTAGACTCTGATGGACGCGAGCTGAGGATCGAGTATAGTTTCCTAAGGAAAGTAGCTGGTGTTCCAACAAGATTTAAGCTGAAAGAGCTCGAAGAAGCAACCAATAACTTTGGATCGTTGGTAGGACGTGGATCATCAGCTTGTGTTTTCAAAGGCGTGCTTAGTGATGGTGCATCAGTAGCTGTTAAAAGGATCGATGGAGAGGAACTTGGCGATAAGGAATTCAAATCAGAAGTTGCAGCTATTGCTAGTGTTCAACATGTTAATCTTGTACGTTTACTTGGATACTCTAGTGTTCCTCCATCAGGACCTCGATTTTTGGTTTACGAGTATATAGTTAATGGATCACTTGATAATTGGATTTTTCGAAAAAGGGGAATTCGAGGTTGTTTGTCTTTGGATTTGAGGTGTAGAGTTGGTCTAGATGTGGCTAAGGCACTTTCTTATTTGCATCATGATTGTAGATCGTGTATTTTACATCTCGATGTCAAGCCTGAGAATATACTTCTAGACGAGAATCATCGTGCTATTCTAGCTGATTTCGGGTTGTCTAAGTTGATGGGGAAAGATGAGAGTAGAGTTGTTACAACGATTAGAGGTACTCGGGGGTATTTAGCCCCCGAGTGGCTCCTGGAGAACGGGATTTCTGAGAAAAGTGATGTTTATAGCTACGGAATGGTGCTATTGGAGATGATTGGAGGGAGGAGAAATATTACTTTAGCTGAAAATGGAAAGAGTACCAATTCAAAGAGTAAGTTCAGCTATTTTCCTAAGATTGTGAGTGAGAAATTGGAGCAGGGAAAGATCATGGAAGTCGTGGACGAGAGGCTCGTCCACGAGGCAGCTACAGGAGGTGTAGCTGTAGAAATGCAAGTGAAAAGATTGGCGTGTGTGGCGTTGTCTTGCATCCAAGAACGGCCTAGTCTTAGGCCAACAATGGCACGAGTCGTGGAAATGTTGGAAGGTCGTGTGCCAGTAGAAGCGCCTACACAAACGACAATGTTGATCCTCGATCTATTAGACGAGGGTGTTGATCATCATCCCGGGATACCTAGGGTTGCTGCAGCCATGGCAAGATCTAGTGATACTAATTCGCTTCGCTCTTATACAATGTCTATCCTCTCAGGGAGGTAGTACCTACATAATACATAACGGATTTCGTTTGCTTTCTATCAAATTTTTCAATATCAATAGGTTGATTGTTTCTCTGGTGTTGTATCCTGAATCCGAAAGTGAGTACTTGAATTCTCCCCATAACAAAAAAGTGTAAGAATTTGATTTTGTTTACACCAAAATATA

>M6-PGSC0003DMG400022391 M6 G-LecRK

AAAGATAACATTCAAAGCATAACGAAAAGAGAAAGAAGTGTGAGCAAATAGTATTGCAGGCTGCAGCTATAGGTATTCTCCATTTTAGCTTATACAGAATTAATCCACATGCAAATTGGCAGAATAAAAACTTCATAAGCAATGAAATTAAGCACAAGAGAGATACTTCTATTTCCAGCTGTTCTTGTTCTGTTTCTTTCTATATCAACTGCTTCAGGTATCATTAGCACCAATAAATTCCTGCGAGATTCAGAAACTTTAGTCTCCAATGACAAAAGATTCATAATCGGGTTCTTTAGTCCTGAAAATTCGACGAATCGTTATGTTGGTGTTATGTTTAATGTCCAACCACCAACTGTTGTATGGGTTGCCAACAGAGAGAAACCTTTACAGGATTCTAGTGGAAGAGTGACAATATCTGATGATGGAAATCTTGTAATCTTGAACTCACAGAATAGGAGTATATGGTCATCAAATATTTCACCAGCTGTTAGAAATTCTACAGCGCAGCTCTTGGATACTGGTAACTTAGTTTTGAACGATAGCTCCAACGGGAGGGTTCTCTGGGAAAGTTTTCGGGATCCTTCAGATTGCTTCTTGCAGACAATGAAAATTGGCGTTGATGTAAGTACTAACACGACAAATCTGCTGAAATCATGGATAAGTCCTTCAGATCCATCTGTTGGGAGTTTCTCAGTTGGTATTCAACCTGAAACAGTTCCTCAGATTTCCATATGGAAGAATGGGAAACCTCATTGGCGTAGCGGTCCATGGAATAAACAGGTTTTCATTGGGGTACCAGACATGACTTCATTCTATCTCAATGGATTTGATCTAGTTAATGACAACAAGGGCACCGTGTACCTTACCTATTTATATGCAAATCAGATTGAGCTGACGTTTTTCACCTTGAACTCAACAGGGTTTTTGCAGCAGAAATATATGGATCCTAGTAAGAATGATTGGGAAGTAACATGGGAATTTCCTGCAACTGAGTGTGATTTTTATGGAAAATGTGGACCTTTTGGAAGCTGTGATCCTACAAGCTCACCAATCTGTTCTTGTTTAGAGGGATTTAAGCCGACAAGTGAAGAGGAATGGGGGAAAGGAAACTGGACTCGTGGATGCAACAGAAAGTCCACGTTAGCGAGCGAAAGAAACAGTTCTAACCTTGAGCAAGGGAAGCAAGATTGGTTTCTGAAGCTGCAGTCAATGAAAGTGCCGGATTATGCTATTTGGGTACCTTTTGTAGATGAAGATTGTGTTAACAATTGCTTGAGGAATACTTCCTGCATAGCTTATTCATACTACACAGGCATAGGATGCATGCATTGGGAAGGAAGCTTACTTGATGTTCAGAAATTCTCTATGGGTGGGGCAGATTTATTCCTTCGCCTTTCATATTCTGAGCGTGGTAAGACCTGCATTTCAGAACTATAATATAAGCTATAAAACTCTGCTTGCCAATGACATATGTATAATGGACACTTATTTCGTAATGCAGATCAAAAGAGAGAATATAAAGTAGTCATTGCCATCATAGTCCCAGTAGGCTCAATAATTCTTGCCATTTTCGGATACATTTCCTGCAGATATGTAGCTAAGCGCAGAGGTAACAAGCTTATCTGAAACATTAGAAAATTCATAGTAGCTTTAATCATACGAATCACTTTATCTCTAATGTCTGTCCTCTTGAATCAGGATGGAAGAGAATGAGTAAGATCTTATTAAGTGAATCATCGCCAAACTATTACAAGGAAGACAAGATAACAGAGGACATCAATCAAGCTAAATTGGAAGAACTGCTTGTATACAACGTTGATATCTTAGCAAACGCAACTGAGAATTTTCATCTGTCCAGCAAGCTTGGACAGGGAGGTTTTGGTCCAGTTTACAAAGTAAGGATTTGTTTGATTGAGTGTCTTGTAATTGTTTTTAAGAAAAGATATCTAAGTATATGATATGATTGCAGGGGAAATTGCCAGATGGACAAGAGATTGCTGTGAAAAGGCTTTCACAGTCTTCTGGTCAGGGGCTACAGGAGTTCATGAATGAGGTTGTGGTGATTTCAAAACTTCAACATCGTAATCTTGTTAGACTTTTTGGTTGCTGCATAGAAAGAGGGGAAAAGATGCTGGTTTATGAATACATGCCAAAAAGAAGCTTGGATGCCTATCTCTTTGGTTCGTTATGATTAGTTTTCTTTTATTTTCCATAGAGTCTAACTAAAACAAACTTTTCTTTGATTTAAACTTGTCAAATTCGCAATGTATTTTCAGGGTCACAACAACAAGAGGAAGAGTTCCTTGATTGGAGTAAACGTGTGATCATCATTGAGGGAATTGGTCGAGGCCTTCTTTACCTTCACAGGGATTCAAGACTACGGATTATTCACAGGGACTTAAAGGCCAGCAACATATTGTTGGATGAATACCTAAACCCCAAAATTTCAGATTTTGGGATGGCAAGGATTTTTGCAGGCAACCAAGATCAGGCCAACACAAGCAGAGTTGTTGGAACCTAGTAAGTTCTTGTATCTTTCCCTTTTATCTTTTCATGAATGAAAAGAACACAACAAGTAAACATTCAAAATTATAACATGATGCAGTGGTTACATGGCACCTGAATATGCAATGGAAGGAAGATTCTCAGAAAAATCAGATGTTTATAGCTTTGGAGTATTGTTATTGGAAATTACAAGTGGAAGGAGGAACACTAGCTTTCACCAAGATGATGGTGCATTAAGCCTGCTAGCATTGGTAAGTTTTTATGAAAATTAAGCTCCAATTCATATGAAATGCCAACTTTTCATAGTTCAAGTTTTCAATGTTTTGTCAATGTGCTTGCTCTTCAGGCGTGGAAATGTTGGATTGGAAACAAGATTGTGGAATTGGTTGATCCCAAGATAACTGACCTGCACCTTGAAAAGGAAATGGTGAGATGTGTACAAGTTGGATTATTATGTGTACAAGAATATGCAGAAGACAGACCAAATGTCTCCACAATTTTGTCTATGCTCACCA

>M6-PGSC0003DMG400022426 M6 G-LecRK

CACAAATTTGAAGAGAAGTCATTTTCCTTAAATTTAAAAAAAATGAGTTGATTTGAAAAACATTTAACCCAATCAAATATGAGAAAATTGGAAAACATTTTCCTTCGTACCAAACACACCCATTGTAAGCAAACATATTTGATTTCTAAAAAACTGCTACCTTCAATTGTTCTTTTGATCATCAATTGGTTTCGTTTTTTGCTAATGTTGTGATTTTCTACTTGTTTTGCAAGATCATAGTGTACGCCCATTGTTGAATTGGAATATCCGCTTCGAAATCATATTGGGCATTGCTCGTGGACTTCTTTATCTACACCACGACTCCAGATTACGTATCATTCATAGAGATCTAAAAACGGGCAATATTTTATTAGACGAAGAGATGAATGCCAAAATATCAGATTTAGGCTTGGCCAGGAATGTTGAAGGCAAAAATATTGATGCTAAGACGCAAAAAGTTGCCGGAACTTAGTAAGTATATATATGCGCTTTCAGTTTAACTTTCTGCAAAATATATAATCGCAAAAAAGCTTATGAAAATACAAGTACATGGTGCTAATCGGAAAATTGTGCAATCACCTGTTTATAGAGAGCAGCCTTGTTTCATGAACTAGCCCCTCTCAAATCCCCACTTGATCAGGAATTCTTCATTGACAGTGTCACCGGAGCTAGATGGATCTTAATTACTGGTGTGATATTATTGAATTTTTAGTCAAAATCATTCTTAAAATATAGAGAAAATACATTAAAAAAAACCCTGAATTTGTCCACTAAATTTACTTTAGTACTTTTAGCTTTATAAATAATTATTTACTTCTAAATCTCATTTCAAGTGAATTAAAATCCACCCAAAAAGCTGACGTCGCAAATAAAAAAAAGAAAGGCTAGTGAAAACGTAAAATATCAAAATAAATACTCTTTCTGTCCCTAATTACTTGTCCACTTTCTCTTTTATAGTTATTTCTAATTATTTGTCCATTTTGACAAATCAAGAAATGACAAATTTTTTTACCTATTATACCCTCAATTAATTACTTTGAAAAAGGTAGAATTTTTTGAAAATCTTAAATTTTAATTCATTTACTTCATAATTAATAAAGGTAACATGATAAACTCACTATGTCAATAATTATTTTCTTAATAGGTGTGTCAATTCAAAAGTGAACAAGTAATTAGGGACAGAGGAAATAAATAAATAAGCCTTTTTTTTCTTTCCAACCAATAAGGTTTCATCTTCCTCACACCCCACCACCTTTCTCTGCCACAACCCCACCCTTACGCCTCGAACCCCCCACCTGCACCTCTCTTTCCCCTATTTCGAACCCTGCCGCCACCCACACTTGCCTCACCCCTTTCTTCTTCTTCTTCGAGCACCCGTCCCTGCCACGTTTTTTCTTCTTTTTCAAAACACTCCCACACATTCCTTCATTTCAAATTATTCTTGCTTATTCTTAGAAATGATGAATAGGAAAAAAAATGAAAAAACCAACAACTTATTATTGCTTCAAGATCTTACTTTTGGAAGGGTTGTTGATTGTGATGAACTTGAAGTTTTTTTAATAAAAATAATATAATTTCTTCATCTCAAATTAGTTTTTTTAATTTGTACTCCTATTATTTATTTATTTTGATAATGTGCTTTAAAAATTGAATGTTTTCTAGTCTTGGATCCATAGAGTTTGATTTTATTTTTATAATCTTAGATTAATTCTCAATTTTTTTTCTATTTTTTACCTTTCTAAAATTGTATAAATAAAAAGAAACAAAAAAGAATTAAAAAAACTCAAAATTTAAATAAATTTCAAGGTGGCGCTGATGTGACAATGATGTGGCATTGATTTGGCAATGAAATGGAGCATGTGTAATACACTTGCCATTGTGAGACTGATATTGTAATATTAGATTTGAAAAAAAATCACTTCAAAAATGTTAAGGGGTATTTAAACATCCATGTAGTTTAAATGCTAAAGTAAGTTGTGCTGCCAAGTTTAAGGGGGTTTTTTTTATGTGTTTTCTCTAAAATATACCCTAAACTTAAACTATATCCTTAAAATATTATTCTTATCAAGAAAACATTTTTCAAGTATTTAAAAGTGAATAATTTTCATCCTTCTACTTGAATTTATGAGAAATCTAACAAATCCCACAAAAATCAAGTTATTCCTTCGTAATTATTATTCTTAGATACGCCAAAAATACTATCGTGAACTTTTATGGTAATTGAGTTTAGCATTATGCAAAATCTTTAAATTTGACAGTTTTTTTCATTTATTTTTCATGTAATTTCATATTTAAAATATTATTAGTTGGTATAAGTTTCTTCTCAATTGAATATACTAGAAGTGACGTTTGTCGGAAACTTCCCCTTCTAATAATGTAGAATGAAACTCAAATGCGAGACATGATCAATATTTTGATTACTTCAATATCAGATTGACCCAAATATTTATTGATCTAAATACAATTTGACTTTGAAAAAGGTAAAGGTAATTAAGCTTCAGCTTATTTCTGTTATTAAATGAGAAAAAGAGCTGATCATGTATCCTCACACCTTCCAAACATTTTTTGTTTCAGTAAGCAATGGTGTCTAAGATTCTTAATATTTATTTTCGAAACTTCTTATAGCAGGTAATGAAATTATGTAAGAAAACTTTTGTTGATAGTCACATGAATTGCTCATGTTTTGTAGGATCCCTTAGTTTTTTTACTATATCTAACATAAGGATGAAAATTGTTAACTTTAAAATATTTAGAGGATGTTTTTTTGCCAAAAAAGATATTTTAAGGATGTAGTTTAAGTTTGGGGTATAGCATAAGGATGTTTTTGACCAAAACTTTGATATTGTTTCCCGGTAGTATTTTTCCCCAATTGAGTTGTGCTATCATGGGCCGAGCCGCTTTTACCCGAAAATTATGCGGTGAATAGAGGTTAAATGTTAATTATTATGAATATATATTTAATTTTGTATATTATTCACCTAACTAAAAAGAAAATTTGCAAATCCTCGCCACAGGCGTTTACTGATCATGTATTCATTCATCTACTCAAGGCCGTTTTTATCTCATATACATGATGGTATACGTTATCTTGACTCATACATGGTGGGATACACTATCTAAGATAATTAATCCTGGGATTACTTGTTCGTTTAATTGGATTAATTCTTGTCTTGTGATATACAGCGGCTATCTTGCACCAGAATATGCATTGGAAGGACTTTTTTCAATCAAGTCTGATGTTTTTGCTTTTGGAGTGGTTGTACTGGAAATCATTAGTGGAAACAAAAACTTGGATGTTCTTGAAGACACAAACCTCTTGGGTCATGTAAGTACCAAAATACTTTCTCTATTTAGAACATGTAATCTTAAGACATAAACGAAATGTGAAGATTGACATATAAAGTAGGCCTTCTGTCCCAATTTATATGAATATTGGGCATCGTTGATAACTCGAAATTAAATATAATTACAGGCTTGGAAACTATGGATGGAAAATAAGGCATTGGACATAATGGAACAAAGTCTAGTGGATACATTCAATGAAATTGAAGTAGTAAAGTGCATAAACATAGAACTTTTATGTGTGCAAGAAGATCCAGGTGATAGGCCAATTATGTCAAATGTGATAATTATGCTTGGAAGTGATAGCATGCCTCTTCCAAGACCCAATCAACCAGCTTATGTCACAAGGAAAAATTACACAATAGGAGCAAGCACATCATCCTCCTCTTCTTATAATTATAATAATTACTATAATTATGATTCATTCACTGACTCCAAAAATGAGTTGACCATCACTGATATGCAAGGCCGATAAATATTATATAATAGTAGTAAGTTTTGCACGATATGGACATGTTAAATACGTATTCTAAGCTTAGTGTGAATGACAATATTTATGTAGTAGGCGGATCTACGTAGAGGAGAGGGAATGCAAGGGCACCCGATCGAACACTCGAGAAGAACTTCATGTATAAATATCTCGAGTATATATATATGTATGTACTCCATTAAAATCTATGTGTGTTAGCATATATTGAATGTAAATATAGTAATTTGTATATTATGATAAATATAGGGCTAATAGGTTAGGGTTAGCACTACTATTGTCGACTATTCTCTTGTATGGAGCACACAATTGAAGCAGATATGTTGATGCTCAGCTTTGTGCTTATAGTAATTCTTGGATAGTACTAATACTTTGTACTGGTAATAAAATCCTTGATAATCCTCTAAATGTTGCATTAGTTTTC

>M6-PGSC0003DMG400022787 M6 G-LecRK

CACACTTCGACTAATTCATAGGATACGTATCACCTCTCACCAGCAACAAAATCTTGATTTGTGGAAAAAAAAAAAACCAAAGTGATGATCACTAGTCTACAATATTTGCTTGTTCTTCATCTTTCTTACTTAGGACTTTGCTTATCAATATGTAACATTAATAGCACTCAATATCTAAAAAGATCCAGCAACTTTATGGTGTCCTGATAACATGTTCAAGTTGGGATTTTTCAGTCCTGGAAATACTACGAATCGTTACGTTGGTATTTGGTACAACTTCTCAGAAACAACTGTTGTTTGGGTAGCAAATAGAGTTAAACCATTGAAAGATTCATTTGGTGTTTTCAAAATTTCTGAGGATGGAAATCTTGTTGTGATGAATGGAATGAAAGATGTTATTTGGACATCGAACATTTCATCTACTTCACACGTTGTTACGAGTTCTATTGCTCAGTTACGAGGAGGGAACCTTGTTTTACTTGACAACTCGAAAAATAGGAACATTATATGGCAAAGTTACCAACATCCTAGTGATACATTTTTGCCTAAAATGAAAATACCTATTAACTCTGTTAAAGCAGGGGAGAAAACAGGGCTAAGATCTTGGAAAAGTCCTTTGGATCCAAGTTTT

>M6-PGSC0003DMG400024258 M6 G-LecRK

ATGGTGTCATCGACTCAATTCAGGAGGTTTACGTATACTGAACTTAAGAAAGCAACAAAAAACTTCAAGGTTGAGCTAGGAAGAGGAGGTTCAGGAGCTGTTTATAAAGGAGTTTTAGCAGATGGAAGAGCGGTAGCAGTCAAGAAACTTGCAAATGAGTTTCAAGAAGAATTTTGGGCAGAGATGACTACCATAGGAAGAATCAACCATATGCATTTGGTGAGGATGTGGGGATTTTGTTCTGAAGGTAGACGTCAGCTTTTGGTCTATGAATACGTCGAGAACTCATCACTAGACAGGCATCTTTCCAGGGCTGATATTCTTGGATGGAAACAAAGGTTTGCAGTGGCACTAGGGACAGCCAAAGGTCTGGCATATCTTCATCATGAATGTCTTGAATGGGTGATTCATTGTGATGTTAAGCCTGAAAATATACTTCTCGATAGTGAATTCGAGCCCAAGATTGCAGATTTTGGACTTGCAAAGCTGTCCCAAAGAGGTGGCCCTGGTTCAGATTTCACAAAGATTCGCGGTACTAAAGGCTACATGGCTCCTGAATGGGCTTTGAACCAACCTATAACAGCAAAAGTTGATGTTTATGCCTTTGGGATTGTAATACTAGAAATGATCAAAGGAAGTAGGCTGTCAAGTTGGGTGGTGGATGATAGTGAATGTGATCATGAACAAGATTCACAGCTCGGAAAATTTGTCAGGATGGTGAAGAGGAAGATTCAAAGTGGAGAAGATTCTTGGGTGGAGAAACTTGTCGATCCACGACTAGAAGGCAAATTTAGCAAGAACCAGGCAGTGACACTGATTGAAATAGGCCTTTCTTGTGTTGAGCAGGACAGAAATAAAAGGCCTACAATGGCCTCAGTAGTCCAAACATTGCTGGATTGTGAAGATGAAACAACACATACAACATAG

>M6-PGSC0003DMG400024259 M6 G-LecRK

ATGAAAAGTTTGAATCTTTACCATCTTTTTCCCTTTTTAGTTTTCTTATTTCTTGTTTTGTCTTTCCCTTTGATAACATCATCAAGAATTTTGCTTAAAGGTGATTCTTTTTCAGTTAAAGATGATTCACATTTCATTACTTCTCCACAAAAAACATTCACTTGTGGTTTTTATCCTATAGAAAGTACTAGTAATGCTTATTATTTTGGAATTTGGTATACAAATTCAAGAGATAAAACTGTTGTTTGGAATGCTAAACAAGATAGGCCTGTTAATCTCAAAGGGTCAAAGTTGACATTAAGAAAGAATGGAGCTTTAGTTTTAACAGATGTTGATGATACAATTGTTTGGCAAACAAACACAACATCATTTGATGTTGAAAAGGCTGAGGTTCTTGAAACAGGAAACTTAGTATTGAAGAATCCTAAAGGTGATGTTTTGTGGCAGAGTTTTGATTTGCCTACTGATACTTTGTTACCTTATCAAAGATTTACCAAGAATCATAGATTAGTTCCTCCTTTGAGGAAAGGGAGTTTATCACCTGGATACTTTAGTTTGTACTTTGATAGTGATAATGTATTGAGAATGATTTATGATGGTCCTTTAGTTTCGAGTATATACTGGCCTAATCGCGATAGGGATGTGTATGGGAATGGTAGAACTAGTCAAAATAGTAGTAGATTTGCTTATTTTGATGGAATGGGTAGATTTTTCTCAAGTGATATGTTGCAATTCAATGTTTCTGATATGGGGTTTGGGATATTGAGAAGATTGACTATAAATACTGATGGGAATTTGAGGATGTATAGTTTGGATAACTCGACAGGGTTATGGAAGATGTCGTGGCAAGCTATTGCACAACCTTGTATTGTGCATGGAATATGTGGAAGATTTTCGATATGTACTTATGTATCGAAGCCTAAATGTACATGTCCTCCTGGATATGAGATGGTTGATGGAAGTGATTGGAGTAGAGGTTGTAGGCCAATGTTTAGATCAAGAAGTTTGAAGTCTAGACATGTCAAGTTTGTGGAAGTTCCTAATGTTGATTATTGGGGTTTTGATCTCAATGCTACTACTCCTTTGTCATTTGAATCTTGTAGGGAATTGTGCTTGGGTGATCCACGTTGTCGTGCATTTGTCTATAGGCATACTGGGGAGGGATCATGCTATACTAAAGGCAAACTTTTCAATGGATACAGGTCTCCTGGTTTCCCTGGAAGTGTATTCTTGAAACTCCCTATGAATCTAAGTGCATCAGAATCAGGTCATTTGATGCTTGAAGGTACTGATGCAAAATGTGGATTGAGTCCCGAAAATGTTCTTTTTGGTTCTCCTTCTATGTATGTATTGGATTTTAAGAAGGTGAAGTGGATTTATCTTTACCTGTTTTGTTCCACCCTTGGTGGAATAGAGATTCTGTTCTTCGTGTTAGGCTGGTGGGCGTTGTTTAGTAAACATGGGATTCCTGCTTCCATTGAGGATGGATATAAAATGGTGTCATCGACTCAATTCAGGAGGTTTACGTATACTGAACTTAAGAAAGCAACAAAAAACTTCAAGGTTGAGCTAGGAAGAGGAGGTTCAGGAGCTGTTTATAAAGGAGTTTTAGCAGATGGAAGAGCGGTAGCAGTCAAGAAACTTGCAAATGAGTTTCAAGAAGAATTTTGGGCAGAGATGACTACCATAGGAAGAATCAACCATATGCATTTGGTGAGGATGTGGGGATTTTGTTCTGAAGGTAGACGTCAGCTTTTGGTCTATGAATACGTCGAGAACTCATCACTAGACAGGCATCTTTCCAGGGCTGATATTCTTGGATGGAAACAAAGGTTTGCAGTGGCACTAGGGACAGCCAAAGGTCTGGCATATCTTCATCATGAATGTCTTGAATGGGTGATTCATTGTGATGTTAAGCCTGAAAATATACTTCTCGATAGTGAATTCGAGCCCAAGATTGCAGATTTTGGACTTGCAAAGCTGTCCCAAAGAGGTGGCCCTGGTTCAGATTTCACAAAGATTCGCGGTACTAAAGGCTACATGGCTCCTGAATGGGCTTTGAACCAACCTATAACAGCAAAAGTTGATGTTTATGCCTTTGGGATTGTAATACTAGAAATGATCAAAGGAAGTAGGCTGTCAAGTTGGGTGGTGGATGATAGTGAATGTGATCATGAACAAGATTCACAGCTCGGAAAATTTGTCAGGATGGTGAAGAGGAAGATTCAAAGTGGAGAAGATTCTTGGGTGGAGAAACTTGTCGATCCACGACTAGAAGGCAAATTTAGCAAGAACCAGGCAGTGACACTGATTGAAATAGGCCTTTCTTGTGTTGAGCAGGACAGAAATAAAAGGCCTACAATGGCCTCAGTAGTCCAAACATTGCTGGATTGTGAAGATGAAACAACACATACAACATAG

>M6-PGSC0003DMG400024260 M6 G-LecRK

ATGAAAAGTTTGAATCTTTACCATCTTTTTCCCTTTTTAGTTTTCTTATTTCTTGTTTTGTCTTTCCCTTTGATAACATCATCAAGAATTTTGCTTAAAGGTGATTCTTTTTCAGTTAAAGATGATTCACATTTCATTACTTCTCCACAAAAAACATTCACTTGTGGTTTTTATCCTATAGAAAGTACTAGTAATGCTTATTATTTTGGAATTTGGTATACAAATTCAAGAGATAAAACTGTTGTTTGGAATGCTAAACAAGATAGGCCTGTTAATCTCAAAGGGTCAAAGTTGACATTAAGAAAGAATGGAGCTTTAGTTTTAACAGATGTTGATGATACAATTGTTTGGCAAACAAACACAACATCATTTGATGTTGAAAAGGCTGAGGTTCTTGAAACAGGAAACTTAGTATTGAAGAATCCTAAAGGTGATGTTTTGTGGCAGAGTTTTGATTTGCCTACTGATACTTTGTTACCTTATCAAAGATTTACCAAGAATCATAGATTAGTTCCTCCTTTGAGGAAAGGGAGTTTATCACCTGGATACTTTAGTTTGTACTTTGATAGTGATAATGTATTGAGAATGATTTATGATGGTCCTTTAGTTTCGAGTATATACTGGCCTAATCGCGATAGGGATGTGTATGGGAATGGTAGAACTAGTCAAAATAGTAGTAGATTTGCTTATTTTGATGGAATGGGTAGATTTTTCTCAAGTGATATGTTGCAATTCAATGTTTCTGATATGGGGTTTGGGATATTGAGAAGATTGACTATAAATACTGATGGGAATTTGAGGATGTATAGTTTGGATAACTCGACAGGGTTATGGAAGATGTCGTGGCAAGCTATTGCACAACCTTGTATTGTGCATGGAATATGTGGAAGATTTTCGATATGTACTTATGTATCGAAGCCTAAATGTACATGTCCTCCTGGATATGAGATGGTTGATGGAAGTGATTGGAGTAGAGGTTGTAGGCCAATGTTTAGATCAAGAAGTTTGAAGTCTAGACATGTCAAGTTTGTGGAAGTTCCTAATGTTGATTATTGGGGTTTTGATCTCAATGCTACTACTCCTTTGTCATTTGAATCTTGTAGGGAATTGTGCTTGGGTGATCCACGTTGTCGTGCATTTGTCTATAGGCATACTGGGGAGGGATCATGCTATACTAAAGGCAAACTTTTCAATGGATACAGGTCTCCTGGTTTCCCTGGAAGTGTATTCTTGAAACTCCCTATGAATCTAAGTGCATCAGAATCAGGTCATTTGATGCTTGAAGGTACTGATGCAAAATGTGGATTGAGTCCCGAAAATGTTCTTTTTGGTTCTCCTTCTATGTATGTATTGGATTTTAAGAAGGTGAAGTGGATTTATCTTTACCTGTTTTGTTCCACCCTTGGTGGAATAGAGATTCTGTTCTTCGTGTTAGGCTGGTGGGCGTTGTTTAGTAAACATGGGATTCCTGCTTCCATTGAGGATGGATATAAAATGGTGTCATCGACTCAATTCAGGAGGTTTACGTATACTGAACTTAAGAAAGCAACAAAAAACTTCAAGGTTGAGCTAGGAAGAGGAGGTTCAGGAGCTGTTTATAAAGGAGTTTTAGCAGATGGAAGAGCGGTAGCAGTCAAGAAACTTGCAAATGAGTTTCAAGAAGAATTTTGGGCAGAGATGACTACCATAGGAAGAATCAACCATATGCATTTGGTGAGGATGTGGGGATTTTGTTCTGAAGGTAGACGTCAGCTTTTGGTCTATGAATACGTCGAGAACTCATCACTAGACAGGCATCTTTCCAGGGCTGATATTCTTGGATGGAAACAAAGGTTTGCAGTGGCACTAGGGACAGCCAAAGGTCTGGCATATCTTCATCATGAATGTCTTGAATGGGTGATTCATTGTGATGTTAAGCCTGAAAATATACTTCTCGATAGTGAATTCGAGCCCAAGATTGCAGATTTTGGACTTGCAAAGCTGTCCCAAAGAGGTGGCCCTGGTTCAGATTTCACAAAGATTCGCGGTACTAAAGGCTACATGGCTCCTGAATGGGCTTTGAACCAACCTATAACAGCAAAAGTTGATGTTTATGCCTTTGGGATTGTAATACTAGAAATGATCAAAGGAAGTAGGCTGTCAAGTTGGGTGGTGGATGATAGTGAATGTGATCATGAACAAGATTCACAGCTCGGAAAATTTGTCAGGATGGTGAAGAGGAAGATTCAAAGTGGAGAAGATTCTTGGGTGGAGAAACTTGTCGATCCACGACTAGAAGGCAAATTTAGCAAGAACCAGGCAGTGACACTGATTGA

>M6-PGSC0003DMG400024426 M6 G-LecRK

CATTGTTTGGGAGAACGGCAGGGTTTAAACGACGTGTTTTGACATAAAGTGTGATCATCAAGACTTTATAAAAATGTCTTCATCATCTAAGGATGATCAATCATCCAATACACCATCTTCGGATTCTTCATCAAATTCACCATCATCATCTTCTTCGAATTCTCCACCGTCTGGCTCATCCTCAAATTCTTCGCCATCCCAAGACAATTCATCTTCATCCAATGATTCAAAATCCCAATCTCCTCCTCAAAGTAATTCATCATCTTCTCCGCCATCAGGGGATAATAATAACTCGTCTCCTTCCGGTGGTAACAACAACAATAATGACAATAACAACGACAACAACAACAACAATAATGGTAACAGCAACAACAACAATAATGGTAACAACAACAACAATGATGGTAACAACAACAACAACAATGGTAACAATGATAACGACAATGATAACGACAACAATAACAACATCGGTAACAATAAGAATGACAATAACAACAATAATGGTGGGAACAATAACAACAACAATAATGGTGGTAACAAAAATAATAACAACAACAACAATGGTGGTAACAACAATAATGGTAACAAATCTTCACCACCAGGAAAATCGTCAAATTCAAATGGAGGTTCATCACCTCCTCCACCCCCACCACCACCACCTCCTCCACCACCTCCTCCTTCTCCTTTTTCTCCCAACTCTGGGTTTCTTTCTCCTCCAAAGCATAAGCCACCACCAAGCAAGTCCCTAGATAATAATAATAACAACAAGCCTAATGATAATGGGACAGCCATCATTGTAGGAGTTGCTGCTGCAGGAGGTCTATTACTTCTTGTCATGTTAGTTTTCCTTATATCATGTTGTAAACGCAAAAGGAGAAGGCCGCGCGACCAAATGGGCTACTACAGGGACAATTCTCATGGAGGCAATAGTATGTTCTTTCTTCTCTTAACTTATACTGTATGTTTTATGTTGAGAGCATACTGTTTTCAAATAGAATTAGCTTCAAATGCAGTAGCTATGAAGAGAACTGGTAAATGTGCAGAGTAGTGAATATGTTTAGGTTTAAATAGTGTTAGTATTGAGAATGTATATATGGTGCAGGTACTGACTACTATAACAGTGGACCATATGGGAATTGGCACAACAACAACAATATTCAATCAACGGATCATATGGTGAAAATGCCTAATCCACAAGCACCGAGTACTAACGTAAGTTCAGAACTGAGTTGGCCAATAGCACCTCCACCACCACCACCAATGATGAGTAGCAGTGAATTAAGCTCTGCTGCTTTCTCTGGTCCGCAACAACCTCCATTGCCACCCCCACACCCATCAATGGCTCTTGGTTTTAACCAAAGTAGTTTTACATACGATGACTTATCTACCGCAACTGGAGGATTTGTTAAGTCTAATCTATTGGGACAAGGTGGTTTTGGCTTTGTACATAAAGGGGTCTTGCCTAATGGTAAGGATATAGCTGTCAAAAGTCTAAAAGCTAATAGTGGACAAGGTGAACGAGAATTTCAAGCTGAAGTTGAAATTATTAGTCGTGTACATCATCGACATCTTGTCTCTTTAGTCGGTTATTGCATTGCTGGATCTCAAAGGTTATTGGTATATGAGTTTGTTCCTAATGGCACTCTTGAGTATCACCTTCATGGTATTAACTTAATTTCCTTTTTGTTATCTAAATACTCATTCTACATATGTCGTATTTAACTCATTGATCCCTTATTTAATTAGGAACTGGTCGCCCTGTTATGGACTTCCCTACTAGGCTTAAAATTGCATTAGGATCTGCCAAAGGCTTTGCGTACCTTCATGAAGATTGTAAGTTCATATATACTATTCTCATGTCTATATATTCTCATCACAAATTAAAGTTCTATTTCATATATAAATTACAGGCCACCCTCGTATTATTCACAGAGACATTAAAGCTGCTAATATTTTACTGGACCATAACTGTGAAGCTAAGGTATTTGGTAAAAATAGCTGAAACATTAAGCAAATTGATTCCTCATTCAGGTTTCTAACAAAGTTTATTATTACTCCCTTTGTAAGGTTGCTGATTTTGGATTAGCTAAACTTTCCAACGACACTAATACACATGTTTCTACACGTATCATGGGCACTTTTGGGTGAGTTTAGTTTGATAATTTTTCAAATATATCCAATGGATTTGCATTAATCGAAAGAGTATATTTATTTGTGTAGGTACTTAGCACCAGAATACGCTTCAAGTGGCAAGCTGACTGAAAAGTCTGACGTCTATTCTTATGGTGTTATGCTGTTGGAACTTATAACTGGGCGTCGTCCTATAGACATCAGCAGTGATGATGATACTTTAGTGGAATGGGTATACCATGATGATTACTTTACATCTTCGAATGATAAAAATTAATTCAGTAGGACATCGATCAGATTATTTAAACATATATGTATATATGTGCAGGCTAGGCCAATTCTGATTCGTGCAACAGAGGGTGGAAACTACGATGAATTAATAGATCCACGTTTGGAAGGAAATTTTGATGCCCAAGAGATGCTATGTATGGTAGCTTGTGCTGCTGCATCCATCAGACATTCAGCAAGAAGACGTCCCAAAATGAGTCAGGTTAATTTCTTCTTAATATTTGTCAAAAGCTTTTCCTAATTCACAATTTTAAAAAGTATAGTGGTCCAGCGACTGACATATATATATATATATATATATATATATATATATATATATATATATATGTTGTAGATTGTACGTGCTTTAGAAGGTGATGTGTCTCTGGATGATTTAAATGAGGGAATGAAGAAGTCCCCGAGTACTGCAATGTTTGGTTCTAGTGAAAGTTCTGAATACGATGGAGGTTCATATGACCTTAAGAAATTCAAGAAATCTGGAATGTCGAGTCAAGAATTCACAAGCAGTGAACATGGTACCGGTGAATTTGTCCATTCCGGTGGTGAATCTCAAGAACATGGGCAGAAAAGGCACACCCCTTAAGGAAAATGGATCATACCTTCAACTTCTGATGGATCTATAAGTAATAATTACACACATTTGTTGGGCTTACAAGGATGCTTTAGGCATTAAAAAACTAGTTTTATGGGGGAAATATAATAATGTATTTAAACAACAATTGTGAAGCTTTATTAGAGATTTGAATGCGGGTTGTAAAGAGTATCACATCCTCCTAACCATTTTGTGGTTGGCTTTGGTTGTCAGTTTGAATCATTTAGTTTTGTTCGACTTTGGAAGTTGGGATATATATACTAATTCTGAATTGTTTTCAATATATTTCGTCAATGTGTATGCTAATTACTAATCTTTTCTTTGTTT

>M6-PGSC0003DMG400024998 M6 G-LecRK

TTTTTAACCATACTGATATGAGAAGAACTGCAATTTATAGTACTTTTCGTATAGTTATTTCTGTTGTTGTTTCTTTTACTTTGGTTATTCTTGGTATTTGTACTGTGGATACTTTTTTGTTCAATATTTTCATCTTAACATTGTATTATTGCTTTTCTTTCAAACCTTTTTTATAAACGTTTTTCTTGAGCTGAGGGCCTATGGGAAACAAACTCTGTTTATGTTGTTGTAGTAGTAGTACTTTTGAACTATTGTAATTTTAATTTGAAATTATTGAGTTAATCTAATGTAATCTGATATCAGGTAATCTTGTCATCAGGGAAGAAAATGAAAACAGGCCTGAGTTTTTCGCCTGGAAGAGTTTCGATTATCCAGGGGATACTCTGTTGCCTGGAATGAAGCTCGGGCGGAATTTGGTAACAGGCCTGGATTGGTACATGTCGTCGTGGATGAGTCCTGATGATCCTGCTATAGGTGAATTTGTTGACAGGATGGATGTTCAGGGGTATCCACAGTTATTTGTATGGAAAGGTTCATCAATTGCATTTAGTTCAGGTCCCTGGAATGGTTTAGCATTTAGTGGGAGTCCTAGTTTGCAACCGAATACGTATTTTACGTTCGGTTTTGTACTGAATCAGGAGGAAGTTTATTACAGATATGATCTTAAGAATGGATCACTGCTCACCAGGGTGGTACTAACCCCTGGAGGATTGATAAATCACTATACCTGGATCGATCGTACGCAGAGTTGGTTCCTTTATTTGACAGCACAGTTTGATAACTGTGATCGTTTCGCGTTGTGTGGTCCTTATGCAAGATGTGTTATTAACAACTCGCCCCCGTGTGACTGTCTGAGGGGTTTCGAGCCAAAGTATCCACAGGAATGGGATGCAGCTGACTGGTCTAGTGGATGTGTTAGGAGGACTCCGTTGGCGTGTCAGCAAGATGGATTTAGGAAGTTTACAGGGATTAAAGTGCCTGATACGCGGAAATCACGGTTTAACGAGAGCATTGGACTTGAGGAATGCAGAATGTTGTGCTTGGCTGATTGTAACTGTACAGCCTACTCAAATATGGATGTTCGAGATGGTGGAAGTGGATGCTTACTCTGGTTCGGAGACCTCATCGATATCAGAGAACTTAGTCCAAATCAGCAGGACCTGTTTGTGAGAGTGGCTGCTTCAGAAGTAGGTAATAACTACTTACTGTTTTTCGTTTCTCTATCTTGATCTTGTTGAATGTGGCGAAAAACATGTATGTAAAATTTCAGCAAGTATGTATCTAGATAGTCAAAAGTCTCGATAGTTAGCTCGTGGGGTCTTCAGTCAATAGAAAAATCATCTATGAATGTGTGTTGGTGCATTGTTCTCTCTCACTCAAAGACGGAACGTTGCCTGTTTCTTATGGTCTAAAATTTTTGTTGGGAACTCATGAATGTGAATCATATTGTTATGACAATGTTGGAATTTTCTGATCGATTATGTACTAACTTATCGACAGACAAGAAACGAAAGAATAAGAAGTCAAGGCTGATTGCAATCTTATCAGCAGTAGCCGCGACTTGTATCCTCAGCCTTCTAGCTTGGTGTGCATTATTCCACAGAAGAAAAAAGAGTAAAGGTAAACAACTGTTGTTCATTGTTATCAAAAGATCGTGTTGCTAACGTTCTATTTGTATCAGGCTAATGTATACATTTATACCAATTGATTGCTCATTTTACAGGTAGGGAAGTTGGAGCAGACGATATGGAGTTACCATTGTTCGATTTAGTAACTGTTGCTAACGCCACTAAGAACTTCTCTTCTGCTAATATTATTGGAGAGGGTGGCTTTGGACCTGTTTATAAGGTACAATAACTATCCCATCGTCGAATTTTAGCTAAAAAAAGGCATCTTAAACACTTTGTTTTGAAAAAAAATTCAGGGGAAACTACGAAATGGACCTGAGATAGCTGTGAAGAGGCTTTCGGAATACTCAGGACAAGGTCTTCAAGAGTTAAAAAATGAACTGATACTGATATCCAAACTACAACACAGGAATCTTGTCAAGCTTTTAGGTTGTTGCCTTGAAGGAGCAGAGAGGATGCTGATATATGAGTATATGCCAAACAATAGCTTGGACTATTTCATTTTTGGTCCGGTTCTCTCGACTAGTTACCTTTATACTATGTTGCTCGTATCCTCCAAAAATGATGTTTCACCAGTGTTGGATCCTCAAAAGATGCACTACTTCTGGAAGATCCAACAAGGGCGCGGAGGCAATTTTGGAGGACCTGAGCAACATAGCCTAACTATTTATTTCCAATGTGCTACTAATGAATGAATGAATGAGGATTCACTGACATTTTACATATTGCAGATCCAAGCAGAAAAGAATCACTTACTTGGAGAAATCGCTATGAAATCGCGATGGGAATATCTCGAGGCCTTCTCTATCTTCATCAGGACTCAAGATTAAGAATAATACACAGGGACCTGAAGGCCAGCAACATTTTGTTGGATACTGACTTGAATCCAAGGATTTCTGACTTTGGTCTAGCTAAGATATTTGGTGCAGACCAAATGGAAGGAAAAACAAAGCGAGTAATCGGAACATAGTAAGTCTTATAAACAAACTCTATGATAGATGTTTTTGTTTGTTGTTGATCAACTTAAATTTGATGCATTGTGCAGCGGATACATGTCTCCGGAGTATGCTGTCGATGGGAAATATTCAGTGAAATCTGATGTCTTCAGCCTTGGCGTGCTTTTACTCGAAATTGTCAATGGCAGAAAGAACAGGAAGTTTCATCATTTGAGTCATCATCATAATCTTTTGGGACATGTAAGTACATATAGCACTAAATTTTTACCCTTTAGCCTCTTTTTTTACTAAACACTCGACCCCGTTTGAGCTTACTCACATCACCGGTCCCAAGCTCCCATAAAGGAAGAGGGTTGCAGCAGGGTAGCAGCCAGTATAAAACTAGTCAGTTCATGTCGAATCCTCACAATACAGTCGGAACTGAGGCGTTGTAGTAGTTGTTATAGCCTGTTTTCCATTAAAAAAAACTTGACCTGCCTGAAGTTACTCATATTACCGGTCCCAAACCCTCATAAAGGAAGAGGATCACAGTAGGCTAGAAGCCTCTATTAAACTAGTCAATTCATGACGAATCCTCATAATACAGTTGGGGCCGAGGTGTTGTAGTAGTAGTTGTTATAGCCTTACATTTCCATTCCCAAGCCCCTACAGAAGTAAGAGGGTCACAATAGGCTAGCAGCCAAGCATAAAACTAGTCAATTCATGACGAATCCTCATAATGCAGACGGGACTGAGGCGTTGTACTAGTCCACATAGCCTTTCTTTTACTAATGCAAATGTATAATTCATGATGACCGAGGCATTGTAGTAGTCACCGTTGTAGCCTCTTTTTCCTAAATGCATTTGTATAACTGTGTTATCGTAAAAAAAATTTCAGGCATGGTTACTATTGAATGAAGGCAATGCATTGGAACTAATGGATGAGTGTTTAAAAGACTCATATGTTGAATCACAAGTGTTGAGATGCATTCAAGTAAGTTTATTGTGTGTACAAAAACTCCCAGAAGACAGGCCTATAATGGCATCAGTAGTGTTCTGGTTAAGCAATGATGGTGTGGAATTGCCTCAACCAAAGCAGCCTGGTTTCTTTATAGAGAGGGATTCAACTAATCAGTCAAATGAATCAACTGATGAAAGATGTGTTACTGACAATGAACTTTCACTAACAATTCTTGAGGCAAGATAGACAAAATATGAACTTGCAGTCGTGTGACTAGGAGGTCAAGGGTTCAAGTCATGACAATAGCCTCTTGCGG

>M6-PGSC0003DMG400025558 M6 G-LecRK

TGATGTTATTTTAGTTTTGATGATACAAACAAAGGAATCTTATAATAGTACATGATCCCTTGTGCCAATTCGTGGAAAATGCCAGCTGGAAAAAGTATAGAAAGTTGGTGAGGTTGGTACACGACTTCCAAGCGTGATAGAATCCTCGGCCAATACCATGGCTTCCTCATTTTTTTCATTTAGATAAAAGGCTTTGATCAATAGACATAATCTTAGTTTTTGCATTCCTGAAATCCTCAAGAACACAAGCAACAACCACTCCGTCCAAGAAGTATTATCCTCAAGAGCAACAGGACTTGATAGAGCTGCATGCTACCTAGTTGTCTTAGGATTATATCTTTTGTGCTTAATTGTAAATATATCCCTAATCTATAAAGGAATTAGATCGTTGTTCAGTTCTTCTAAAAGTCTAAGTCAGTTGATGTTAACTGATTTAGTGGGCAGTGTTGTATCTGGTTAGTCAAATTCTAAATCATCTAAAGTTAGGTGGTTTAGTGGACAGTGTTGTATCTGCTTAGTTGAAGTCTAAGTTATCTAGAGTTAGGTAGCTTAGTGGGAAGTGTTGTATCTTCTTAGTTGCTCTCTTTCTAGTAGAGTTACTACATAGGGGGGGGGAAGAGATTTAGAGGTAAATTCCTAGGTTGTAGCTGTTGTACTCTAAATTGTTGCTCCATTTGAGTGAAATAGAGTTGGATTGAAAGCTTGTGAGGACATGTCATGATTTTCGCCACCCTTGAGCAAGGGGGTTTCCACGTAAACTACTTGTGTTGATTTCTTTATTGTTTAAGCCTTCATAACTATTCATACTGAGTCAATGTACCTGGTCCGTTGGCTGATAGTGGACGCATGAATCTTAACA

>M6-PGSC0003DMG400026154 M6 G-LecRK

GTCTCACTGAAATGAAATAGAGGGCAAGAGCTACCTATTTGAATTTATTATGTTCTTCATTCAAATCATCAACAATTCCATAAAACAATGCAATTAAGTAATTTTCTCTTGTTGTTGTTGTTTGCAGCATTGATGATTTTACATCGATGTGATGCAAGGAGGAGCACCATTGATGGAAGCAGTAAATTAGTTGACAATGGCGAAACACTTGTATCTGCTGGTGAAAATTTTGAAATGGGATTCTTTTCTGATGATGACGGGTTAAACAAATATGTAGGCATATGGTATTACAAATTAAGTCCTAGAACTGTTGTATGGGTTGCTAATTGGAATAATTCGATTCAGGGTAAAAGAATAATGAATGAGGATAATTCGGTTGTCGTTGAAGATGGTAATCTCAAAGTTTTAAGTAATGGGAGTACTTATTTCTCCACACTATTAGGTGGTGGTTCTAACAGGAAAGTAGAGTTGTTGGATACAGGGAACTTAGTTTTAGTTGATGAATCAGGAGGTGAAATGTGGCAGAGTTTTCGCAATCCAACCGATACATTCCTACCAGGTATGAAAATGGATAGTATTTTGAATTTGATTGATTCAAAATATGTGAACTACATATTCCAATTAGATCAATCAAGTGACAAGGAATATGTTATAATTGCAAAGCAGGGCGGATTTCGCTGGAAAGGGTCTGCGAAATCAGGAAATTTAAGATTAGGTGAGATGCCTGGTTATGTTGCCTACATGTTATCCAACTTCACTAATAAAAACATTGTCAATAATTCATTGGAATCGATAGGAACATTTAACAAGAGTAGGCTCTTAATGAACTCCTCTGGGGAAATACAATTCTACGGTTGGGATAAGGAGATCAGTGGATGGTCTTTGAAGTGGTCAGCGCCAAAGGATACGTGTGATTTGTTTAAGTTTTGTGGGAAGTTTAGCATTTGTAACAGCAAACATGATCCAGTATGCAAATGTTTGCCCGGGTATAAGCTCAATCCTCCAGATAATTCAAAAGCAGGAGATTTTTCAGGTGGTTGTTCAAGGATGTCAGTTAGCTCTTGCAATGAAGACAATGTAGAAGTACTCGACACGTTCTTGGATTTGAGGTCAATGAAATTCAAGAGTCCAGACAAGATAATCAGTAATATTAGTACCCGTGAGGACTGCAGGAGATTTTGTCTAGGCAACTGTACGTGCCAGGCCTATACTTACAATGATTCTGTATGCTGGATTTGGGACACTAGTCTCATGTCTCTCCAGGAGAACTATGCTGGTGGCTTCAACATCTCTGTTCGTGTCAGCATTTCCGATATTGGTAATTTCTTTTTCGATTTAACTTATGTACACTGACAGTTTAAGCTAATGTGTGACCACAAATAGTACAGTAATTTGATTTTTGTGCATTGCAGAAGCAACAAGGAGAAATTGCAAGCCTTGTGGCATAAACCTAATACCCTATCCATTAAGCAGTGGACCAAATTGTGGTGACCCTCTGTATTATAGTTTCTCATGTGATGATTTAGCTGGTCAGGTAAGCTTCTACACATCGAATGGCAAGTATACTGTCGTTAACTTTGATAAAGATAACAAAACATTTGTTATAGAAGCAGCTCATAAAGAATCTGTTGGTACTTGTGATGACAAAGGACCGGTGACAGGAATTTCTTGGTTCAACCAGTCATCACCTTTTAAGGTGATAAACTGGTGTTACAATCCTGAAGAGAATCTTACTTCAGGACCTTTTAGTCGAGGGAAAGATTTAATATTGATTAGTTGGAAACCACCATTGGAGCCATTTTGCAAGACTTCAGAAGATTGCAATGATTGGCCAAATTCTAGTTGCAATATGACAAAACAAGGAGAAAGAAGATGTATATGCCAAACTGATTACAAATGGAATGGCTTGACTTTAAATTGTTCCTTGAGTTTAGGTATATGGATCAATGTTACTTACTACAAATGATTTTCTTGCCAATAGTTCAATGATACCTTTTTCTATGTTCTCCATTTGACCTTTTATATGACCAACACATATTTCATTTCTTAAATTTGCTTCAGAACTTGGAACACAAGGATCATTCATTACGAAGCTGGCATCCTCACGGAATCAGAGAACCCTTGTTATTTCCATCTCTGTAGTTCTTGGAGTTATAACTTTGTGCAGCATTAGTTATATAATTTATCAGAACACTAGAGTGACAAGAAGTAGA

>M6-PGSC0003DMG400026155 M6 G-LecRK

ATCTGATTGAAAGGAAAAAACCAAGTCTTACCATGACTATCAACATGATCTTAATTGATGATGTGCATCTTCTGTTAACAAATTTGGTGCAACCATGCAACTGCAACATTATTCGATTCGTCTACTTTTTGTGATCCTAACATGTTTATTTATCAGTATTCATCAGTCTCTCGGAGTTGAAATGATTTCAGTAAATCACTCTGTTTCTGGAAACCAAACTATCGTCTCTTCGGGAGAAAACTTTGAATTAGGATTCTTTTTTCCTGGTAGTTCGAAAAATTACTATCTTGGTATATGGTATAAGAATGTAATGCCACAAACAATAGTATGGGTAGCAAACAGGGAGAAGCCACTTTCTGCTACTGATATGAATTCTGTTGAGTTGAAAGTTTTGGATGGTAATTTAGTGCTTATTACTGAATCCAAAAACTTGTTTTGGTCTACAAATATCAGCAACACAATTAGTTCAAATACACTAATGGCAATTCTTTCTGATGATGGTAATTTGATTTTTAGTGATGGATCAAATTCAACTACACCACTTTGGCAAAGCTTCGATAACCCGACTAATACATGGTTGCCGAATGCTAAGGTGAAGTATGACAAGCGTACAAATACAACAAAGATTCTTACTTCTTGGAAGAACTCAGAGGATCCATCTCCGGGGATTTTTTCCGTGGAAATGGATCAGAGTAATAAGCAGTTCTTGATAAAATGGAACAGGACTGAGATGTACTCAGCTACTGGATCTTGGAATGGGCATATTTTCAACATGATGCCTGAGATGAGTTTGAATTCTGATAGATACAGATTTTCATATGTTGATAATGAAAATGAGAGCTATTTTACTTATTCCCTCCGTAACTCCTCCAAAATCAGATTAACACTAGATGTCTCTGGGCAAATCAGGCATCTGATTTGGGCCGAAAATTTAAAGGAATGGCAAATTTTTACATCTCAACCAAGACAACCCTGTGAGGTGTATGCTAGTTGTGGTGCCTTTAGCATTTGCAATAAGGAGTCCGCGACCTTCTGTAATTGCTTGACTGGTTTCACACCTATATCTGATGCAGAATGGGACTTGAATGATCATTCTGGTGGCTGTGTTAGGAAGGAAAGCTTGCAGTGTGGTGATGGTAAAATGAAGGGCGGTTTTTCGGAGAATCCCCATGTGACATTGCCCGGATATTCTCTAACTGTGTCGGCTGCTAGTTCTGAGGAATGCCAGTCGACTTGCTTGAGTAACTGTTCTTCATGTAATGCATATGCTTATGATAATAATGTGTGTTCAATTTGGAATGAAGTTGTGAATCTGAAGCAACTCTCACCAGGAGATGGTAGTGGAAGTGTGATTTATACCAGACTCGCTGTCTCTGACGAAGCATACGGTGAAGGTAAGTTTAAAGTTAAGAGTTGTATTTGATAGTGTTTGCCATTTCTATGTATGCAATGCTCATCTCCAAATAGCTACCGATAGAGCTTTCTATGGATGTTTGTTTTTTCTTGTCTGATGTTTCAGTTCTTTACTTCATGTCTTTCAACAGTACGAGTCTATACATTCTCTAAATCTTAGTGCTCATAAGAATTAACTAGGCAATTATAAGTTTACGCAAACACCTTCTATGTTTTAAACAAACATCAGATAGCCACTGTCCAAATTTAAGTTTACGCAACATCATCTACGTTATAAACACACATAAGATAGTCACTGACCAAATGTATGGTTTTCATGTTTTGATTAGTTACAACTAGCCAAAATGAGTTGAAGATCTTTCGCCAGTAAACATTACCAGTTCGTTGAATTCCCACATCAGCTCTTTAAGGATGGGATGGTCTTTTCATATGGTTTTGGACAGTTCTTACCTCATAAACTAGCTTTTATAGTTGAGTTAGCCTCAAGGTCCATTTCTTATCACGTTATTAGAATCAGACCCATCCCAAAATAGTCTATTTCTTGTCACAATTTCCTTATGTTTCTTCTTTCTGTCATCAGATGCAAATTCAGGGAAATTGTCCCTGAAGGTTAAAGTCATCATATCTGGCGTGGTAGCCGTTGCAGTTCTACTTCTATGTAGCTTTAGCTACATTTATCATAGGAGAATAATGTCAAAGAAAACAGGTACTTGTGTTGCTATTACAATCTCTGGAGTAGTTGTTTTACCACTACCAGAAAAATTTCTCCTTCCATTTTTTTTTTCCGAATCAGATCAAGAGAGATATCTCAAAGGGCTGCTCTTTTTTTTTTTTTTTTTATCTTAGCAAGCCAACAAAGCACAGGAGGCAATCCAGTTCCACATTGGTTAAACAGAGAAAGAGAAGCACAAATTCTGATTAACGAAAATGATAAGCAAAACATTGCCGTTCCATTCTTTAGTTTGGAAAACATACTAGCAGCCACAGATCATTTCTCAGATGTAAATAAACTTGGACAAGGAGGTTTTGGTCCTGTTTATAAGGTTAGCATTAATAATTCACTTTAACATGAGTTCAACTTTATGTAGTCATTTCATAGCATCCTAGTTTCCACATCGAAAATCTTTCAGGGTATATTTTCAGATGGACAAGAAATTGCTGTGAAAAGGCTATCAACCCAATCTAGACAGGGCATTGAAGAATTCAGGAACGAAGTCATACTGATCTCGAAACTTCAACATAGAAATCTAGTCAGACTTTTAGGCTACTGTATCACAGGGTATGAACAGATTCTACTTTACGAGTATATGTCAAACAAAAGTCTAGACACCTTCATATTCGGTTAGTATATATATCTGAACTTATCTTATTTCTTAATCTCTCTATTTACTTGATCAATGTTGCTAAATTTTCTCCTTGGTTACCAAAACAAGATCCTACACTTAGCAAGTCATTGAATTGGAAAAATCGATTTGAGATCATATTAGGAATTTCTCGAGGGCTTCTCTATCTTCATGAAGACTCAAGGCTCAGAATCATTCATAGAGATCTGAAAACAAGTAACATTCTTTTGGATCAACAAATGAACCCAAAAATCTCAGACTTTGGCTTGGCAAGGATAGTTGAAGAACAAACAACAGAAGCAAGTACCAAAAAAGTGGTTGGAACTTAGTAAGTAATACAAACATGACAATGTTATGAACTTTGCTTAATTTCATCGCACATCTGACTCATGATTATTCCATCATTATTTGATTTAGCGGATATATGTCTCCTGAATATGCATTGGAAGGAGTATTTTCGATTAAGTCAGACGTATTCAGCTTAGGAGTAGTTATACTGGAGATAGTAACCGGAAGGAGAAATACAGGATTTTATCAATCTAAAGAAGCATCTAACCTTTTGGTCCATGTGAGTGCCTTGAGTACACGTTATTTTTTTCCAATGTGTACGTTGATCTTCATATTCATTATTCTAAATCATGATATCACAGGCATGGAATTACTGGAAAGAAAAAAGAGCACTACATTTGCTGGATCATTCATTGCTCGAATCATGCAACCCGAAAGAAGCAATGACGTGCATAAACGTTGGACTCTTATGTGTGCAAGAAGATCCAGGTGATCGTCCTTCCATGTCAATTGTTGTTATGATGCTTCGTTGTGATAGTACGTCTCTTCCAAAGCCTAATCAACCTGCTTTTGTGTCTAGGTGAAATGTTTCTAATTCAACTTCTGTTTCATCTGGTACACCACCTAGCTTCTCCAAAATTGAGTTGACAATCACAGTTGAAGAAGGCCGATAACGTGATGAATTCATTAGTTTTTCAAATGAATTTAAACGCGAATTTAAATTATTTTTTCACCTTAGATTAAAAAACAAAATTACAGGAATACTTGATCCTTTTCTTTTTCTCAATTTTTTTGTTGTTTAGGCAGTAGAAGACTTGGGACCAGTACTGCTGATATTTATAGTGAAAACTGAAAGGTATATATGCTTAATATTGTGTACAACATAGAATATATTGTGTTGATAAAAAAAAGTTCATTTTCTGTTTCATTTTGGTCTCTTCTACCTTCTTAACTTGTCTTAAAACAGTAAGAATGGAAGAAAAAAAA

>M6-PGSC0003DMG400026156 M6 G-LecRK

TTAAAACAGGAGCAACTTACAAGGAGGTGTACTAGGAAAAATAATGGAGTTGGATTGTGTTGTAGCTATTTTAATTGTATTTTGTTGTTCTTTCGTTATGCTTCTGGCAAGTGAAGAAAGTATCAAGATGGGCAAATCAATCACTGGTAATCAGAAAATGATTTCAGCTGGTGGAAGTTTTTCCTTGGGCTTCTTCACTCCACGTAATTCTACATTTACCTATCTTGGCATATGGTACAACACGATCCCCGAACAAACAGTCATTTGGATTGCTAATAGAGAATCTCGAATTCCTCAGAATTCTACTGTTGTTTTCACTATTGGTGAAGATGGGAATTTAGTGATTTTTGAAGTAAAAGATCAACTTATTTGGTCATCAAATGTTTCATCCATAACTTCAAATTCTACTGTAGGAGCACTACTAGATAATGGAAATCTAGTGCTCTTACATGGTGAGTCTGATATATTGTGGCAGAGCTTTGATCACCCTACCGATACGTTTATGCCTGATATGAAACTTGGTTATAATAGAAAAACAGGACAACGGACTCTGATCAATTCGTGGACAAGTAATGAAGATCCTCGGCCTGGAAATTTCTCGTTTGGGATAGATCCTGAAGGGCAGACGCGTTTTTATATATTGAAGCAGAACAGTATTTATTTTAGGTTTGATGATAGTAATGAATACACTGGAGGGAATATCGGAGGAATTGCTTGGTATATTTCTGTTGTTTCGGGAAATGATGGAGTTTTTGTTACGTATGGTTATACCAAAGGGTTGGTAACATTAAGGATTGTTTTGCATCCTAATGGTTACATTCAGTTATTGGTATGGCGTCAAAATGCTAATAAGTGGATAGTTAAACTTCAGGCACCAGAATTTCTTTGTGAACGTTATGCGGAGTGTGGTCCATTTGGCAGCTGTGATGTTTGATCAAGTGGATTATGCAGATGCTTAACTGGATTTGAACCAAGATTTTCGACAGATTGGGAGAACGGGAAATGGAATGATGGTTGTGTTAGAAAAGTAGCATTGAGCTGTGATGACGGAGACATATTCTTGAAACATGACGAGGATATGAAGTTTCCGGATCATGCTATTTCCTTAGGACATATGAGCATTAAGGACTGTGAAACTCGCTGCATTAGGAACTGTTCTTGTTCAGCTTATGCTTATTCAAATAGCACCTGTTTAATTTGGTTTGGAGATTTACTGGACCTTTCCCACAACTTTCCTGGTGGCAGAGTTCTCAATGTTCGAGTTCGTGGCTCCAAGCCAGGTATTTTCACTTCTATCCTTACTAATTCTACTAACAAATGATTCTCTACACGTCTTTGGTCAAGAAAACTAAATTTATTTACTTTTTGTCTTCTGGATGCAGTTACTCATGGTGGATCTAGGAATTTGGCTCATAGACACAAAGTTTTCGTTGCGAGTATTGTTTCTGCAATATCTATCATCTTGGTTCTTATAAGTATTTTTTTCTTTATCTTCAAAAGAAAGCACTTGAGAAGACAAGGTATGTTTAGTGTACGTCTTAATTGTAAATACAAATACTGAATTCGAATTTCGTATAGATTACACTGATTTGCTTTACTGATTGCTGTTTGAAATTAGGCTGGAAGAATGGAACTAAAGCTTTCATTAGGTCCGTGTCTGGTTTATATTCGGTTGAAAAAAGCGATATGAAGTTGCTGCAGTACAACTTTCAGCAAATAAGAGAGGCTACTAACAACTTTCATGGAGAACACAAACTTGGAGAAGGTGGATTTGGTCCTGTGTTTAAGGTGCTAAAACTAAATTCATTGTTGATATTTTACTGCTGCTACTTTGAGAGAGACGTCATACGTCTCATTTTCGTTTCATGACTATGGTGGATGTGGATTTACATATTCTGTTTCAGGGGTTTTTAGCTGAATTCGGAGACGTAGCCATCAAAAGGATAAGCAAGAGGTCATCACAAGGACTGGAGGAGTTTATGAATGAACTGAAGCTAATCGCGAAACTGCAGCATAAGAATCTTGTCAGCCTCTTAGGATGCTGTGTTGAAGGGGAGGAGAAAATACTAATCTATGAGTATATGTCCAATTGCAGCTTGGACAAATTTCTGTTTGGTTCGTCTTCTACTCCGTTTTAAGACTTCACATAACTTAAATTTTTACTCCTCTCAATTCCCTTTGGTTAATGAGATGTTCAAATGTCTTTTTAGACCCTTCGTTGAAGGTTACACTCGATTGGGTTACACGTTTGGGGATAATAGAAGGAATTGCTCAAGGAATGCTTTACTTGCATAAGTACTCGAGATTAAAAGTCATCCACAGGGACTTAAAAGCAAGCAATATTTTGCTAGATCAAGAGATGATCCCTAAAATATCAGACTTTGGAATGGCAAGGATTTTCGGGACTGATCAAACACAGGCCAATACAAATCGAGTCGTTGGTACATAGTAAGTAATAACCCCTTATGATTTGTGGTTTCCAAGCTTTACTAGCTTTTATGTCATTTGTGCATATCTAATGGATACTTATTTATACAAATATGGCATAGCGGCTATATGTCTCCAGAATACGTAGTATATGGCCAATTTTCAGAGAAATCAGATGTTTTCAGCTTCGGAGTATTGCTCTTAGAAATCCTGAGTGGTGAACGTAACTCAGACTTTTTAATGACTGAAATTTCCGCGTCCCTCTTAGGATGGGTAAGAACACTCAAGACTTTAATTAATTCTGTTTTCTTTCACTTTTTTTCGTGTTTTATATATATGGTGCAACTCATTTGAACTTTTTTTAGGCATGGAATAAGTGGAAAGAAGGAAAAATCTTGGAGTTGATTGATCCGTCCATAAGAGAAACTTGCGACAATAACAAAGCAACAAGATGTATTATAGTTGCCCTTCTTTGTGTTCAAGAAATCCCAATTGATCGACCAACAATGTCTGATGTATCTTTCATGTTAAGCAACGAAACAACGCCCATCCCTGAACCAAAAGAACCTGCTTTTCGTAGTAGCTGGCAATCTCAGCAGTTAAGTGATGTATCCATAAATGAGATGACGATTACCTTGCCAAAACCGCGATGAAATAATTTAAGTTCTCCTATTTCTTGTAGTAAATTTTTCTTGTTAGCATAGAGCTGTCATAAATTTTTTTTTGTTGAACTTACAAAAGCGTAGCTCTGGAAGAAATATAGACTAAATTTCTCTGTTTAATTGATGTTTTCATGTTTTTTTTTTCTGTTTCTACAGTGTATCAAGTTTGTGAAAATGTTGTTTTCAACATTCACTTGGATT

>M6-PGSC0003DMG400026183 M6 G-LecRK

AAAACAATGGAGTTGGATAGTTTCTTACCTCTTTTACTGTTAATTTTCTCATGTTCTTGCTTACCCGTTCTTCCTTGCCAAAATTGTGTCAAAGTTGGCAAATCAATCACTGGGAATCAGAAATTGGTTTCAGCTGATAGAAGTTTTGCCTTAGGATTTTTCACTCCATCAAATTCAACGTACACCTATCTTGGCATATGGTACAACATAATCCCTCAACAAACAGTCGTTTGGGTTGCCAACAGAGAATCTCCCATCCCACAGAATGCTTCACCAGTTTTCACTATCGGCGGTGATGGAAATTTAGTGATTTTCGATGAAAAAGGCAAACTCATTTGGTCATCAAATGTCTCATCAACAGCACCGGAATCATTATTAACTTCAAATTTTACTATCGGAATATTGTTAGATAACGGGAACCTTGTTCTTCGACACTACGGTGAGTCCAATTATTTGTGGCAGAGCTTCCAGTATCCTACAGATACGTTATTACGCCATATGAGATTTGGTTTCAATAAAATTAGTGGCTTACGAAATGTGATCACTTCATGGACGAGTAATGAAGATGCTCGCCCCGGAAATTTCTCTTTTGGAATAGACCCTACTGAAGGAGTTATGCGTTTTGTCATTTGGAATCGAAACGGTATTTATTATAGATTTGACGATAGTTGCAGAGTGTATACAGTGTGGTGTTTCGCTGTTGTTACTGAAAATGATAATGTATATCTTAATTTAGATTACTACAGAATCTTTCTGAGATTAAGAATCGTTCTGAATCCGAGCGGGCACCTTCAGGTGATGGTGTGGAATCGGAATGGAAGTAACAAATGGAATGTTGAATTTGAAGCACCACAATCTAAGTGTGACCTTTACGCGCAGTGTGGTCCGTTTGGTATCTGTGGGATCAGATCAAGTGGATTATGCAGTTGTTTAGACGGATTTGAACCTAAATTTACGACAGATTGGGTAAATGAGAAATGGAATGGAGGTTGTGTGAGAAAAATAGCGTTGGGATGTGATGGTAGAGATCAGTTTTCGAAGCACGAGAATATGAAGTTGCCAGATCATGCTGTCTCTTTAGGAAATATGAGCACTGAAGAGTGTGAAACTCAGTGCATTAAGAACTGCTCCTGTTCAGCTTATGCTTGTACGAAAAACGCCTGTTTTATATGGCATGGAGATTTACTGGATCTTGGGCATGACACTACTAACGGCAGAGCTCTCTATGTTCGAGTTCACGTGTTAGAGCTAGGTATGTTTACTTGTATCATTGTTGGTTCTGTTAACAACAAGTGAAAGACTAGTATAATTTTCGTGTTGTGGATGCAGTTACTAATAATCTATCTGGCTCAGGGAATTCAGCTCGAAGATACAAAATTCTCACTGTGAAAATTGTTTCTGCAATTTTTGCAATCTTTCTTCTTGTTAGTATCTCTGCTTTTATCTTCAAAAGAAAGCGCTTGAAAAGACAAGGTGAATCTAGCATTTCTAGACTGTAATGATTTCAGATTTCATATTGTTTACGGTGCTTTGCCTTACTGATGGTTGTTTGAAACTAGATAAGATGGACGGAGATTCTGCTTTAGTTAGCTCCATGCCTAGTGGATCTTTAGTTGGGAAAGATGATATGAAGTTGGTACAGTACAGCCTGCAGAATATAAGAGACGCAACAAACAACTTCCATGAAAGTAACAAACTTGGAGAAGGTGGTTTTGGCCCCGTGTTTAAGGTGCAAAATTCAATATTTTGATGTTCGTATTTAATTTCATGAATATAGAGAATTGTGGCTTTAAACATTCTGTTTCAGGGATCTTTGGCTGAATTTGGAGACGTAGCCATCAAAAGGCTAAGCAGGAGGTCATCACAAGGACTGGAGGAGTTCATGAATGAGTTGAAGCTAATCGCGAATTTGCAGCACAAAAATCTAGTCAGCCTCTTAGGATATTGCGTTGAAGGGGAGGAGAAGATATTGATCTATGAGTATATGCCCAATTGCAGCCTGGACAAATTTATTTCTGGTTATACTCATTAGTATTATTTTCACGAGAATCATAACTTTGTTTTCCTCATTTCCGTTGGCTAATTGGATATTCATTTTTTCTTTAAGACCCTTCGTTGAAGGTTACAATCAATTGGGATACACGTTTTGGGATAATTGAAGGAATTGCTCAAGGAATACTTTACCTGCATAAGTACTCGAGATTAAAAGTCATCCACAGGGACTTGAAGGTAAGCAACATTCTGCTAGATCAAGAGATGATCCCAAAAATATCAGACTTTGGAATGGCAAGGATTTTCGGGACTGATCAAACACAGGCCAGTACAAAACGAGTGGTTGGTACATAGTAAGTATAGTAGCCGCATCTGTAACCAGTTTTTATGTTAATACCTCTTGTGTATAGCTACTTAAAACTGATTTACGCCAATATGGCATAGTGGCTATATGTCTCCAGAATACGTAGTATATGGCCAATTCTCGGAGAAATCAGATGTATTCAGCTTTGGAGTATTGCTGTTGGAAATCTTGACTGGTGAACGAAACTCAGACTTCTTCATGACTGAAATTTGTATGTCTTTATTAGGATGGGTAAGTACACTCGCGCATTTGAAAACTCAGTCCAAGTTTCTTTTAGTTTCTTTTAACATTTCACGTCATTTGGCATGTTAGGCATGGAAAATGTGGAAAGAAGGAAGAATATTGGAGCTAATAGATCCATCAATAAGGGCGACCTGCGACAGTAACAAAGCAACACGGTCTGTTCTGGTTGCTCTTCTTTGCGTTCAAGAAATTCCTACTGATCGACCAACAATGTCTGATATAGTTGTCATGTTAAGCAACGAAACAACGCCCATCCCTGAGCCAAAAGAACCGGCTTTTCGCAGCAGCTGGCGACATCAAAAGTTAAACAATTTTTCCATAAATGAGTTGACATTTACTTTGCCAGTACCTAGATGAAAGGATTTAGCTTTCCATATTTCTTATGGAATTAATCTTATTTTGAAAAGAACCATCTTTGAAAAAAATGTTTAGCACTGTAAATAAAAAATCAAAAACCAATTCAATTACTACCTCTGTCCCAGTTTATATC

>M6-PGSC0003DMG400026184 M6 G-LecRK

ATGGCGAAGAAAACAGGTTTGTGTTGCTATTACAGCCGTTGCAATTCTACTTCTTTTTTTTGCTCTATGAATCAGATAAAGAGAGAAATCTAAAAAGACTGCACATGTTTTCTGATCTTAGCAAGCCAACAAAGCACAGGAGGCAATCCAGTGCCACATTGGTTCAACAGAGAAAAAGAAGCACAAAACCTGATTAACGAAAATGATAAGCAAAACATCGCCGTTCCATTCTTTTGTTTGGAAAACATATTAGCAGCCACAGATGATTTCTCAGATGTAAATAAGCTTGGACAAGGAGGTTTTGGTCCTGTATATAAGGTTAGCACTATTTTTTCACTTCAATAGTTCCACTTTATGTAGTCATTTCATAGCATCCTAGTTTACACATCGAAAATCTTTCAGGGTATATTTTCAGATGGACAAGAAATTGCTGTGAAAAGGCTATCAACTCAATCTAGACAGGGCATTGACGAATTCAGGAACGAAGTCATACTGATCTCGAAACTACAACATAGAAATCTAGTCAAACTTTTAGGCTACTGTATCACAGGGTATGAACAGATTCTACTTTACGAGTATATGTCAAACAAAAGTCTAGACACCTTCATATTCGGTTAGTATACATATCTGAACTTAGCTTATTTCTTAATCTCTCTATTTACTTGATCAATGTTGCTAAATTTTCTCCATGGTTACCAAAACAAGATCCAACACTTAGCAAGTCACTGAAATGGAAAAAGCGATTTGAGATCATATTAGGAATTTCTCGAGAGCTTCTCTATCTTCATGAAGACTCAAGGCTCAGAATCATTCATAGAGATCTGAAAACAAGTAACATTCTTTTGGATCAAGAAATGAACCCAAAAATCTCAGACTTTGGCTTGGCAAGGATAGTTGAAGAGCATACGACAGAA

>M6-PGSC0003DMG400026185 M6 G-LecRK

ATGATCTCAGTAAATCACTCTGTTTCTGGAAATCAAAGTATCGTCTCTTCAGGCGAAAACTTTGAATTAGGATTCTTTAAACCTGGTAGTTCGAAAAATTACTATCTTGGTATATGGTATAAGAATGTAATTACACAAACAATAGTATGGGTAGCAAACAGGGAGAAGCCACTTTCTGCTACTGATATGAATTCAGTTGAGTTGAAAGTTTTGGATGGTAATTTAGTGCTTATTACTGAATCCAAAAACTTGTTTTGGTCAACAAATATCAGCAACACAATTAGTTCAAATACACTAATGGCAATTCTTTCTGATGATGGTAATTTGATTTTGAGTGATGGATCAAATTCAACTACACCACATTGGCAAAGCTTTGATAACCCGACTGATACATGGTTACCGAATGCTAAGGTGAAGTATGACAAGCGTACAAATACAACAAAGATACTTACTTCTTGGAAGAACTCAGAGGATCCTTCTCCGGGGATTTTTTCCGTGGAGATGGATCAGAGTAATAAGCAGTTCTTGATAAAGTGGAACAGGACTGAGCTGTACTCAGCTTCTGGATCTTGGAATGGGCGTATTCTCAGCTTGATGCATGAGATGAGTTTGAATTCTGATAGATACAGCTTTACATATGTTGATAATGAAAATGAGAGCTATTTTATGTATTCCCTTCGTAACTCTTCCAAAATTAGATTAACACTAGATGTCTCTGGGCAAATCAGGCATCTGATTTGGGCGGAAAATTTAATGGAATGGCAAATTTTTACATCTCAACCAAGACAACCGTGCGAGGTGTATGCTAGTTGTGGTGCCTTTAGCATTTGCAATAAGGAGTCCACGACCTTCTGTAATTGCTTGACTGGTTTCACACCTAAATCCAATACAGAATGGGACTTGAATGATCATTCCGGTGGCTGTGTAAGGAAGGAAAGCTTGCAGTGTGGTGATGGTAAAAAGGGCAGGTTTTTGATGAATCCCAAAGTGACATTGCCCGAATATTCCATATCTGTTCCGGCTGCTAGTGCTGAGGAATGACATTCTACTTGCTTGAGTAACTGTTCCTCATGTAATGCATATGCTTATGATAATAATGTGTGTTCAATTTGGAATGAAGTAGTGAATCTGAAGCAACTCTCACCAGGAGATGGTAGTGGAAGTGTGATTTATACCAGACTTGCTGTCTCTGACGAAGCATACGGAGAAGGTAAGTTTAAAGTTAGGAGTTGTATTTGATAGTGTTTGTCATTTCTATGTATGCAATGCTCATCTCCAAATAGTTACTGACAGCTTTCTATGGATGCTTGTTTTTTCTTGTCTAATGTTTCTTCAGTTCTTTACCTTATGGACTTCAACATTACGAGTTCTCTAAATCATAGTGTTCACAAGAATTAAATAGGCAATTATAACTTCACATAAACAGACCTTACGTTTTAAACAAACATCAGATAGTCACTGACTAAATATAAGTGCACGCAACAGCCTCTATGTTATAAACACACATAAGATAGTCACTGACCGAATGTATGGTTTTCATGTTTTCATTAGTTACAACTAGCCAATATGAGTTGAAGACCTTTGCCAGTAAACTATTACCAGTTGGTGAGTTCCCACATTGGCTCTTTAAGGATGGGATGGTCTCTTCACATGGTTTCGGACAGTCCTTAACTCATAAACTAGATTTTATAGTCGAGTTGCCTCAAGGGTCCATTTCTTATCACATTATTAGAATCAGACCCATCCCGATTCTTTGTTTACCCCATATTCTGCCCCATCATATTGTTCACGCTCCAGTTGGCAGGCCTAGGCATGAGCGATGTTGAGTTCCCACATAGTCTCTTTAGGGATGAGATGTTCTTTTTATATGGTCTCGGACAATCATCATATCATGAAGCTAGCTTTTAGAGACAAAAGTTTATTTCTTGTCACAGTTTTCTTATGCTTCTTCTTTTCTGTCATCAGATGCAAAATCAGGGAAATTTTCCCTGAAGCTTAAAGTTATCATATCTGGCGTGGTAGCAGTTGCAATTCTACTTCTATGTAGCTTTAGCTACATTTATTGTAGTAGAATGATGGCGAAGAAAACAGGTTTGTGTTGCTATTACAGCCGTTGCAATTCTACTTCTTTTTTTTGCTCTATGAATCAGATAAAGAGAGAAATCTAAAAAGACTGCACATGTTTTCTGATCTTAGCAAGCCAACAAAGCACAGGAGGCAATCCAGTGCCACATTGGTTCAACAGAGAAAAAGAAGCACAAAACCTGATTAACGAAAATGATAAGCAAAACATCGCCGTTCCATTCTTTTGTTTGGAAAACATATTAGCAGCCACAGATGATTTCTCAGATGTAAATAAGCTTGGACAAGGAGGTTTTGGTCCTGTATATAAGGTTAGCACTATTTTTTCACTTCAATAGTTCCACTTTATGTAGTCATTTCATAGCATCCTAGTTTACACATCGAAAATCTTTCAGGGTATATTTTCAGATGGACAAGAAATTGCTGTGAAAAGGCTATCAACTCAATCTAGACAGGGCATTGACGAATTCAGGAACGAAGTCATACTGATCTCGAAACTACAACATAGAAATCTAGTCAAACTTTTAGGCTACTGTATCACAGGGTATGAACAGATTCTACTTTACGAGTATATGTCAAACAAAAGTCTAGACACCTTCATATTCGGTTAGTATACATATCTGAACTTAGCTTATTTCTTAATCTCTCTATTTACTTGATCAATGTTGCTAAATTTTCTCCATGGTTACCAAAACAAGATCCAACACTTAGCAAGTCACTGAAATGGAAAAAGCGATTTGAGATCATATTAGGAATTTCTCGAGAGCTTCTCTATCTTCATGAAGACTCAAGGCTCAGAATCATTCATAGAGATCTGAAAACAAGTAACATTCTTTTGGATCAAGAAATGAACCCAAAAATCTCAGACTTTGGCTTGGCAAGGATAGTTGAAGAGCATACGACAGAAGCAAATACCAATAAAGTAGTTGGAACTTAGTAAGTAATAAAAACATGACAATGTTATGAACTTTGCTTAATTGCATCGCACAGCTGACTCGTGATAATTCCATCATTATTCGGTTGATTTAGCGGATATATGTCACCTGAATATGCATTGGAAGGAGTATTCTCGATTAAGTCAGACGTATTCAGCTTAGGAGTAGTTATACTGGAGATAGTAACTGGAAGGAGAAACACAGGATTTTATCAATCTAAAGAAGCATCTAATCTTTTGGTCCATGTGAGTGCCTAGAGTACATGTTAATTTTTTCCAATATATATGTACGTTGAACTTCATATTAATTATTCTAAATCATGATATCACACACAGGCATGGAATTACTGGAAAGAAAAAAGAGCACTACATTTGCTGGATCATTCATTGCTCGAATCATGCAACCCGAAAGAAGCAATGACGTGCATAAACGTTGGGCTCTTATGTGTGCAAGAAGATCCAGGTGATCGTCCTTCCATGTCAAATGTTGTAATGATGCTTCGTAGTGATAGTACGTCTCTTCCAAAGCCTAATCAACCTGCTTTTGTGTCAAGGAGAAATGTTTCTAATTCAACTTCTGTTTCATCTGGTACACCACCTAGCTTCACCAAAATTGAGTTGACAATCACAGTGGAAGAAGGCCGATAATGTGATGAATTCATTAGTTTTTACGCCAATTTTCTCCGCATGAAATTATTTTTTCACCTTAAGAGAAGATAAAATACATCAAATCAACTTGATAGCAAAACTTTTTTATCAGTTTAACTACAATGGTG

>M6-PGSC0003DMG400026810 M6 G-LecRK

ATTGGAAGAGACATTATTCAAAGTGTCTGAAACCTTTTCAGACCACTCCCTTTTACTCCCAGAAAAATTTTGCTTTGTTGATTTTCCGGCATTCCCTTCTCTTGCTTTCCAGGGAAAGATAAAGAGTTGAAAATTTACAATGTGAGTCATATCTCTTGTTAATTTAAGCTCACCACTCAAAACTTCCCCAAGATACTAACTTTCTGTTTCACTCTAGCTTTTTCACTGAGATATGTTGATTTTAAAGCTTTTTCACTTTTGGGGTTCTCATTTTCTTCAAATCTTGAAATACCCATTTAAGGATTTCAACTTTTGGAGTTATGTTAGACCGAGTACTTTTGCTCTACATTGTCCTTAAAATTGGGATGTAATTGTAAAGATTCATTCTTTCTTGCATGTTCACTTAGATTTGTTCTCTTATCTTTCCTTAAAAATAATTTTTTCTCTTACCTAAATTGCTCTTTTTTGCTTAGAAAAATTGAGATATATATGTGTGTATGTGTGGACTCACGGTTTCTTTGCAACGCGTATTTTGTCAAAGGCATGTTTGTTTCCAAGTCTATGCGTCTATATTGACTTTTAAATAGTTAATTTTTGGGGTATTAAACTACTCCCTGATTAGGTTTTCAACTTTGGATGCTTGCAAAACATATTTTAGTGTCACTTTAAAGTATAGAAAAGTGGTTGAATATGCATAAGGACCTAAATAGGGTCCCATTTGTAGTTTTGAACCAGGATTTCAAACAAAAGCTGTTTTCTTTAGCCATTTTCTTTAGCCACTTTTAATTAATTTTCTTTGTGTACTGTTTATTTTATGGTCCATCTGTAGGTCATATTTTTTTTAGTTCAATCATAAGGTGTGGATCTTTTGTTTTCTCACATGGCAAGTTTCTTATAGGGAAGCATTTGCTGTTTCAAGAAAGTGTCTGCAGGCAATATTCTTCCAATTGATGTGGTTTGGATGTCTTTTCTGAGGTTCAATGCTTCCCTTCTCTTTCAATCAATTTCCCACCTTTTGTTTGATCTTGGTGTAACCTTGAATCAATTGGTAGTTTAAGAGTTTTACCTTATAGTTGATTGTGATATTTCAATTCCTTTACACAAAAGAACAGTTTCTTGTTCATTGTAAATCAGTGGAATGACTGAACATATATTTTGCTGATATGCAGAAGCTGGAACCGGTTTTCTGTCTTTTCATCTGCTTCCGCAAGTAGTGACAGATATGGATAGAGGAGACAGAATAATAGCCTGTTATTCTAATTTGCTGCTTTCTGTCTCTATCGGGTTCTTGCTTTTTTCAGTTGTCAGTTCACAAATTCCTTTGGGTTCCAAACTAACAGTAGAAGAGAACAATTATTGGTTCTCATCCAATAAGGACTATGCAATTGGATTCTTGAACTTCTCCGACCAGTATAGCTTTGGTGTCCGTTTCAATGCAATTTATATACCTAGCAGTGAGCAAGCAGCCATTTGGACTGCAGGGTGGAATGTTAAAGTTAGTAGCAAAGCATATTTTGAGCTTTCCCTAACTGGGGAAATGATATTGTTTGATCCAGCAAAGGGAAAAATTGTTTGGCAAAGCAAAACTGGCAACACATCAGTTGAATCAGCTGTTCTTCTTGATGATGGCAACTTTGTCCTGTTAAATAGGAATAAGAGTGCTGTTTGGCAGAGTTTTGAAAGTCCATCTGACACAATGTTGTCTGGCCAGAGTTTATCTGTTGGCCAGTCTCTTCGAGCTTCTAGCAGGGGTTCTTTAACAAGTTATTACAGTCTTCATATGAATGTTTCTGGTGAGATGCAGCTTAGGTGGGAGACTAGTATCATTTACTGGACTGTTGGGGGTCCTAAATCTGCTGTACGAGCCATTCTTGGCTCTGACGGAATCCTCCAACTACTTGATCAGCAATCGCAAGCTGTTTGGTCAGTCTATGGTGAGGACCACAATGATTCTGATGTGAAGTTTCGGTTCCTCAGACTAGATTCTGATGGTAATCTTCGGATATACTCATGGGAAAATAATGCAACGTCATGGAGAAAAGTTTGGCAAGCTATAATTAATCAGTGTGATGTCTTCGCAACCTGTGCTACCAATGGTATTTGCACCTTGAATGCATCTGACTCTTATGTTTGTTGGTGTCCATTTAGATCTACAAGAGACTCAAACTCTGAATGTCTAATTCCATATAAACCAAGTTGTGAATCTGGTTCATCCATGATCCTTCATGAGCATATGTATTTGTACGGAATATACCCACCAAACAACACAGTTGTGCAAACCAATTTGCAGCAGTGTAGAAGTCTGTGTCAGAAAGATCCATCCTGCCATGCTGCATCCTTTATCAACAATGGCACTCCACAATGCCACACGATGAACTCCCGGTATGTTGGTGGTCAATCAGATCCTTCTTTGGGTTCCGTTACTTTTGTCAAAACTTGTTCAGACCCAATTGCTGTTTTACCTCCGCCTGTACCTGCTCCGAGAAAGGTATCACAGAAAATCTGTGTTGCTTGTCTGTTAGAAGTTGCTGCTGCGTCAATCGTTGTGTTTGTCATGCTTCAATTCAGCATTGGAGTTTACCTTTTCAGGAGAAGGAAACATATGATGCAGAAATCTGCTTTACCCCATATAGTGCCTAATGCTACTGGCTGCATCGTGTTTTCTTACTCTGAAATCAAAGATCTGACTGATAATTTTAAGCAACAAATTGGGCAGAATGTGTTCAAAGGTCTACTTCCATATAATAGGCTTGTTGCAGTTAAAGATCTCAATGCCTCTATTGATGAAAGGAGGTTCCGGGCTGCAGTTCTAAAAATAGGAAGCATATACCACAAAAATCTTTTGAGGCTGGATGGTTATTGCTGTGAGTCCGGCCGTAGGTTATTGGTTTATGAGCTTGCCAAATATGGTTCTGTTGACAAATGTTTAGAAGAACCTAGAATGTGCAAGAGATTGACGTGGAGAAAAAGAATGGATATATGTTTGTCAGTGGCAAGGGCCATATCTTACTTGCATACAGGATGTAGGGAGTTTATTAGTCATGGAAATCTTAAATGTGAAAATGTTGTCTTAGATGATGAATTAGAGGCCAAAGTTAGTGAATTTGGGCTGAGGACTATTCAAGCTGAGGCTTCCAGTAGCGGGGGGCTAGCAGAGACAGATGTAAGAGATTTTGGCAAAGTGATGGTGGTATTAATCACTGGATGCCAAAATGCGGATGAAGCTTGTTATTGGTCCTATGAAAAATGGGTGAAAGCTGATAAGGAAATGGCCATGGATCAGCGAATTGTGGTTGGCGCCGATTCAGAAGAGTTAGAACGAGCATTAAGAATTGCATTTTGGTGCTTACAAGGTGATGAACGCATGAGGCCATCAATGGGAGAGGTAATCAAGGTACTGGAGGGCACTCTGACTGTCGATCCTCCACCACCTCCTTTTGCTCATAATCAACAATGGCCACCTGATGACGAATCACCATCTGAATCATATTAAAGTTCTAATTCGTATATTTCATTCCCATTAAGTCATCTGTTTCTGTACAGAACTTACCAGGCACCTTCGACTTGGCATGATGTAGAATGCCTAGTTTTTCCAGTATGGCATGCTTAATCTGAAATTCAAGACGAGCATCCCACTCTGCCTCTTCTGAAACTGTAACCTTGGTATGTAACCCTCCAAGTACAGTTTTGTGTTAGAAAATCCTGATTCAGAAATCCAGGTTTCTTTTTAAAGTTTTTGATTCTGTTCATGTTAATTATCGCTTCTTATCACATGTGTTTACATTTGATATTTTCTTGGAGGTGACTAAACTTCCTTAACTCTGCCCAAAATGTCTCCTTGGCTTGTGCTATCAGTTGGCACTGGCTTTTTTACTAGGCAGTGGTCCAAGAGATCGCGGATGGGGTTTGAACCTGGACATCTCTATTCTTAGGATTACTCTGACCATTAAGTCGATCCTAACTTAAAATGGAAATGGAAGTTTGTGTCTCTTTTTCTTAACAGAGGTGGTGTCCATGCTATTTGCACCTCGACCATTTCACCAGGTACTTGGTATCTCCACCAACCTAGGTATCAGGTAACTCTATCTCTAAAGGTTAGAAAAATGGAAAGAAATCACCTAGCATTTTGAATTTCGTTGATAGGTCACACTTGTGTTTTATATATGTTCTGTTTTGATTTTGGCTTGCTGCACCTTTGTAGCCTGGTAAATAGGGTCATTTTCGGTGAACATCCATGATCAGTATTTCAGCAATGAACACAATTTCATTATTTTTATTAGTGAAAGAGTTG

>M6-PGSC0003DMG400027233 M6 G-LecRK

AGTAAGCCTTGCTACCAAGATCAACTGAGATTCAGCTTTTTGTTGCTCTGATTCGTTTTCTGTTCTCCTTTTTCAAGTAGGTAAAGTAGGTTGACTCGCTACTTTACCTACTGTGTATAAGGAGACGGAGCAGAGTAAGAAATGCAACTTAGTATGGTATGCAAAAAGATTCTTGATTTCAAGAACTGATTTTTTTTTCAAGTTTTCATTGTTGTTAAGAGCATGGGATCATCTTTCTTGTTATTCATTGTTGCCCTTTTGTTATCTTGTTTTGTTGTACACTCTGGTCCTCTTTCTCTTCAACCATTAACTCCAAATTTCACTGCTTCAAATTTCAAGTTCATTGATACTTCTGGATCATTCTTGTCATCACCAAATGGAACTTTTAAAGCAGCAATTACAAACACAAAATCCCAAGAAAGATCCTATTACTTTGTCATCGTTCACTCGGAATCTCATGTAGTTGTTTGGTCAGCAAATCGCGATATGCCTGTATCTGATTCAGGTGAGTTGCACCTTTCTGTTGATGGCCTTGCTCTTTTTGATGATTCAGGTGACACGGTTTGGTCAGCGAAGCGTAGTAGTACTAGTAGTTCTGTTACTTCAATGCAGCTTCTTGAATCAGGAAACTTTGTGTTGGTTGATGCTTTTAATAATATTGTGTGGGAAAGTTTTGATTCTCCTACAGATACAATCGTTGTAGGCCAACGATTGCCTGTTGGGAAATCGTTGGTTAGTTCAGTGAACGAAGATGAAATTGCTAAAGGCGATTATAAACTTGTGGTTGTAGAGAATGATGCAATGTTACAATGGAATGGAATGACTTACTGGAAGTTAAGTATGGAACCTAAGGCTTTTACTGATGCATATACTCTGGTGGAGTATATGATGATCAGTAGCAACGGGTTGTTCCTTGTAGGGGCGAATGGGACGGAGAGAGTGATTCAGGTTATTTTGGATGAGGTGAAAGATCCTGACTTTAGGATTGCGAAATTGGAAGAGAATGGACAGTTCAGTGTCAAACGATTTAGTAACGGAAATTGGATATCTGAATTCGATAGTCCTATTGATAGTTGTCATGTTGCTTTCACTTGCAAGAAGCTTGGAGTTTGTGATGAGGGGAGATGTTCTTGCCCACCGGGATTTCGTGTTAGTTCTGAGGTGAATGGTAGTTGTGCTCCAGTTGATAGAAATTTGGTAATGCCAGTTTCTTGTAATGCATCTTTGAATATGAATGTAACTGAATTGGGAAATCGTGTTTCGTACTTGAGGCTAGAGAATGGTATGGACTATTTTGCTAATGATTTTATCGAACCTGTGAAGCGTGGTGTTAATGTATCTGCTTGTCAAGATCTTTGTTCTAAGAAGTGTTCTTGCTTGAGTGTTTTTCATGATCAGTCGTCGGGTTCATGTTATATGATTGAAAATTTTCTTGGTTCGATTTTGAGGGGTTCGGATTCTGGTAACGGAAGAGGTAGATTGGGATACGTTAAGGTCATTTCTGAACCGTCATCATTTGATCCAAATGACAACTCAAGTGACAATAGGTCAAGATTACCCGTCGTTGCTTTGGTTTTACTTCCTTCTTCTGGGCTTTTACTGATCATTGTGATGATGGCTGGGATTATGTGGTTGATGAGAAGAAAAAGATTGATGCAGATTTCAGGGAAAGAGTTTGGACGGACGGATTCTTCTTCGTTTGCTGAGCTTGATAACATTTCCATACTGGGGTTGCCTGTGAAATTTGACCATGAGGAGATTCGTGTAGCCACGGAGTATTTTAGAAATCAAATTGGTACTGGTGGATTTGGAACAGTATATAAAGGTACTCTTTCTGATGGAGCTGTTGTTGCTGTCAAGAAGATGAATGCTCTTGGGGCTCACGGAAATAGGGAGTTTTGCACTGAGATCGCGATTATTGGGAGAGTTCACCATGTTAATTTGGTTAGTTTGAAAGGATTTTGTGCACATAGGGGAGAAAGGTTTCTGGTGTATGAATATATGAATCGAGGCTCATTGGATCGAACCCTTTTTGGACTTGGACCTGCCTTAGATTGGCATACCAGATATGAAATCGCGCTTGGAACAGCACGTGGGCTTGCTTACCTGCATGGTGGTTGTGAGCAGAAGATCATACATTGTGATGTGAAGCCAGAGAACATACTCCTCCATGACAACCTGCAAGTAAAAATATCAGATTTCGGGCTTTCCAAGCTTCTTAATTCTGAGCAGTCTAGTTGGTTTACCACGATGAGAGGAACTCGAGGCTACTTAGCTCCTGAATGGCTGACTAGCTCAGCCATAACTGAGAAATCAGATGTCTACAGCTACGGAATGGTGCTACTAGAAATAGTACGAGGGAAGAAGAACTCCTCGTTCCAACCACCAAATGACACTACTTCTCAAAGCGAAAGTAGTGAAAGGAATAGGCTTCCTCCATCGTCTCTTGCCTCAGCAAATCAGCCAATCTACTTCCCATTATTTGCGTTGGAAATGCATGAACAGAAAATGTACCTGGAGCTTGTGGATCCTCGTGTTCTGGGGAGTGTCAAAATTGAGGAAGTCGAGAAGCTAGTACGTGTGGCTTTGTGTTGTTTGCATGAAGAGCCAACTCTGAGACCAACAATGGCCAATGTTGTTGGCATGTTAGAAGGTGTGTTGCCTTTGGCTATGCCACAGGTTCAATCACTCAATTTTCTACGATTTTATGGTCGTAGATTCACTGAAGCATCAACGATCGATGGGGATCAAGGGGTTAATGTGTTTGAGTTACATCAACAAAACAGGAACCTTAGCAGTACTACTAGTAGCTCTTACAATTCCTTCTCTTATATGTCTTCACAGCAAGTCTCAGGTCCAAGATAGCACTAGTAAATCTTATGTTGTTCGAACCCTCCAAAAATGTTGTCACATTCATGTCTGGTCCTCCAAAAGTGCATTGCTTTGGGAAGATCCAACATAATTTTGACATTATTTTTGAAGAGTCGGAACAACACGGAGTAAATCTGCTTTGTTAGTTCAATTTGTTATGCTACTGTTTGAAAACTTTAGTTGTGATGTGATGTTTGCACCTCTTCTGTGAGGCAGTAGAAAGAGTATAAATATAAGTTGTAATTTATTCAGTTTC

>M6-PGSC0003DMG400028461 M6 G-LecRK

AAATACAACATCAGCTTCAAATCATCTTTACCCTATCTAGACAAGTAGTTAAGCAGATGGGAATAGATTCATTTCAATGATCGTCAAGTCATTGATATAAATTAGAGGTTTCACAGATGAACTATTTGTATGTATAAGAATTCTTTTCCCTTCTAATATGTTTTTTGGTCAATGAACATTTTAGTTTTTTTGTCTAAACAGAATCATGGCTTTACCCTCTGTCAAACATTCGTTGACGCTTTTGATCATTGTGCTCCTTGAGTACCTTACTTTGGCTCAATTTCCTGATTATCACACGGTGAAATTACCCACAACATGGATAAACAATAAAGTCTCGTCTATCAATGTTACAAAGCCCTTGACATGGCCTTTTTACTCATTCAATCAATCATCTGGACCTTTCATGCGAATTCTGATGCTCAAAGAGATAGAAAAGGACATGGGGTATGCTTGTGGTTTTGCTTCAAATGGAGAGGACGATTCCTTTGTGTTTTCCATTGGAATTATACGTCTTCGTCAGGACTTTCTAGGAGCTCAGCGCTGGACTGACCTTGAACTGGTCTGGTTTGCAAACAGAAACCATCCAGTTCGTGAAAATGGAACGCTGCAATTGCTGCAAGATGGAGATTTAGTGTTAAAAGATGTGGATGGAACTCTTGTATGGTCTACAGGCACAGCGAATAAGTTTGTTTCGGGCATTAAGATGATGGAGACTGGGAACCTTGTGCTTCATGACATGTATAATCAGACTGTATGGCAGTCTTTTGATCATCCAACAGATGCATTGCTTCCAGGCCAGAAATTAAGGTCTGGTCAAAGACTTGCGGCTAGAGTTTATTCGTCCAATTGGAGTGAAGGGAATTACTATATCTCTGTAACCAATCAAGGCTTGTTTGCATTTTACATATCAGACAAGCCACAGATGTATTTTAAGTTTTTGGTCAGTGGAGAAAGGGATAGCTTTGATGAAAGTTATGTCAAAGCAGTGAATGGTACTCTTGCCTTGTACATATCTTCAACCGAGCCGAATGAGCCAAATGCTGTCTTCTCAAGACCTTCAAGGATGAAATATTTGAGATATGATTATGATGGACATCTTCGAGCCTACACCGAGGGAAGTGATCAGACAAATGATCTTTTGGCAGATTTCATTGGGCTTTGCGTTACCCCACAGCTTGTGGCAGTTTGGAGCTTTGCTCAAATGGAATTTGTTCTTGTCCAGGTCCATTAATAAAAAGTAATGATCGACAAAACAACGGATGTGTGGAAGTTAGTCCGGTGGAATGTGATGGACAACGCTCCCACAGAATGGACCGTGCTTCGGATGTCTATTACTTAAACTATGTCGACACTGATGCAGCAGCTCTGAGAGGAACAGATGAGGAAAGTTGCCAAGAGTTGTGCTTGAAGAACTGCTCTTGTAAAGTAGTTCTATTCCGGTATTTCACTAACTTTTCAAGTGGTGACTGCTATTTACCCTCTCCTGTTCTGTCACTTATAATTGATGGAAAGGGAAGAAGTGTCTATGAATCATCTGCATTCATCAAATTACCAAATGATGCAGAAAAAGGTAAGTCTACAGCTACAAGAAGAACTGGCATAATAGCTGGATTAACTGGTGGAACAGTTCTGCTGGTAGCATTAACTGTTGGCATCTTGATTGCTTTCAACAGAAAACACACATTACAAGAAAATAATGATGATTACTCCGGGGAGATATCTGGTCTTGTAAGATTTTCTTATGAACAGCTGAAGATGGCAACTTGGAATTTCCAGAAAAAGCTTGGTCAGGGAGGTTTTGGCTCAGTTTATGAAGGAGTTATTCGAGATGGTCAGAAAGTAGCAGTGAAGGTTCTTGATGGTTTTGGCCAAGGAAAGAGGGAATTCTTGGCAGAGGTCCAGACCATAGGAAGCATCCATCATGTTAATCTTGTTAGACTAATTGGAGTTTGTGCTGAGAAAGAGCACACAATATTAGTTTATGATTTCATGAGTAATGGATCATTGGATAAATGGATTTTTGGCACAACCTCTACTCAATTTACCCTTGATTGGAAAATAAGAAGAAAAATCATCCATGATATAGCAAAAGGATTGGCTTATCTTCATGAAGAATGTATGCAAAGAATAGTCCACTTAGATGTTAAGCCACAGAACATTCTGCTGGACGAGAATCTTTGTGCAAAAGTATCTGATTTTGGCTTGGCCAAGTTGGTGGATAAAAATCAGAGCCACATAGTGACCAGAATTAGGGGAACCCCTGGATACTTGGCTCCTGAATGGTGTAGTGCATTCATTACAGAAAAAGCAGATGTCTACAGCTTTGGAATTGTTGCAATGGAGATCCTCTGTGGACGCAAGAACGTGGACTATTCACACTCACTCGAACATCCACATTTGCTTAGCTTATTTATGGAGAAAGCCGAAAACAATCAGCTTATCGATATGATGGGAAACTACAGCGATGATCCACAATGCAATACATCAGAAGTCATCCGTATGATGAAGCTTGCTGTCTGGTGCTTACAGAATGATTTTACTCTGAGGCCTTCCATGTCAATGGTGGTTAAAGTAATCGAGGGGACAATGGACATTGAATCTCACTTAGACTATAGTGTTCCAAATTCACAAACAATTGCAGCAATCAAAAGAGTGGCAGACACCACGACTAACTTGTTTCCTTCAATTCTTTCAGGACCTAGGTAAGCATTTCTAAGCATGATGAAGAGTCAATAGAGAAAAAGAAAATCCCCCCTCCTTTATCCTATTTCTTGTCTGATCCCATGATCATCATTGTTGTATAGCTATTTATTGGTTTCATTTGTGTTGTTTTTCTATGACTTAGTAAGATACTACAATTAATTAAGAACCATGGCGTGATTTCCA

>M6-PGSC0003DMG400028617 M6 G-LecRK

GTCACAGTAAAAGTTCTACAAACAATATCATCGAAACTTGGGAACACACATTGGAATGTCAGCCGAGCGTCTTGTATGCAGGAGAGGGATTTTAACATGACGGATCCTAATTATAAAGAAGTTTTCAGCAATGTAAAATGTAATTGTTCATTCAATAATGGCTCAGTTTGCCATGTTGTAACAATGTAAGCTCTGCTTTGATCTTTAAAACTTCACATTTTACTCTTAATCAGAAAGTATTACAGTTAAACAGATGCCATCTTCTGCATCATGTAATGAGAATTTCTTACGCTTTTTTTGTTGATACAGCCAATTGAAGCGTCTTAATATGACAGGAATTCTGCCTCCAGAATTTGCAAATCTTACTCATCTGCAAGAATTGTGAGTATCAGTTATAGTTTTTTCTCCCCTTTCCGATTACAGAAACTTTTATTCAATTGTTTTTATAAAATCGTAACAGTAGTAGCAAGTTAACTGTTTGTTGCTTTTATTTATACCCTTCGAGGTGGCCTAGTGGTCTGGGCTTGGGAATTCCATGTCGGAGGTCTCAAGTTCAAAACCCTTTGTCAAGCGAAAACTGACAAGGGGTTTGCCTTCTGCTTGCCTAGTGCGGGTTACCTCTCCTGTGTGCATAAGAGTGGGGATTTTACCTGTGGCTTAATCAATTTTTGCATTTCCAGAGATCTGACTCGTAATTACTTAAGTGGATCGATTCCCTCAAGCTATGGCCAGCTTCGCGTCACTATTTTGTAAGTTCTTTTCTTGTTTAATCTTTTCATTGAAGAATTTTTGACATTTTGTTCAAACTCAGGTCACTCTTGGGTAACCGTATCAGTGGACCGATTCCGAAGGAGTTAGGTGACATTTTCACACTGGAAAAACTGTAAGTTGAAGATATTGCATACTTCCTTTAATTTTCTACATATGTCACATTCATGAATTCCATATTTTCAGAGTTTTGGAAAATAATCTACTTGAAGGACCTCTTCCTCCGAACCTTGGTAGCTTGAACCGCCTCAGAGAAATGTAAGACATAGTAACCATATATTCAACGTTTAATATCTACTTATTGTTATATCTATTCCAAATTATTTGGCACTTTTCGATTTCCACTTGTCAAACTAAATTGCATTTCTTTCACCGTTTTTATTTGAAAAGAACTGTCAAGTACATGAGGTTAATGAAGTATATCTTACCCATGATAGAACGATTTTACTCACTAGTCTATTGGTACCTAAAATAGTGGTGTCAACGAACCACTCAACAAGAGCAAAATACGCTCGAATTTATTTGTGCAAAAGAAGAAGAAGTTCAGAACATTCGTAAGGAATAATCTGAGGAAATAGATTTATTTATAGCCATAGAAGTACTGTTTCTGAAAGTTTGCAACCTTCCAGAATAACTTAAGAGAAGTACTATTTCGGAAAGTTTCCTTAAGATTGTTGTTTATCTGTATTAATAGGATACTAAGAACAATACTTTGCAGAACTTCTTAAGAAACATCCCTTCATTTCTCATTTATTATTATTATTATACCCAACACTTTTAATATGAAGAAAGAGTTAATCTAGTTTAATTTCGCTCCAAAAGTTGGTCAAATTGTATCAAACATAAGTAAAAAGTGTTGATCAAATTGGGATGAACAGTGTGATTTTTCAGTTATCGTGGTTAATTGATGGATCATTTTGCAGGTTTCTTTCTGCCAATAATCTCAATGGAACTATACCTGAGAACTTCAGCAATCTTAAGAATATGACAGATTTGTAAGCAAGAAAATGTTTTAAGGTACATTTGACTTTTGTGCTTCAATCTCTTAATTCTTTCGTCTGTTTGCAGTAGGATCGATGGGACTAGCATATCTGGAACCATACCAGATTTTATTGGCAATTGGACCGAAATGGATAGATTGTAAGTATGCATCTTGTAGTTGTAAATGTTGATGGAATATTCTTCTCCCGGAAGGATTCTAAACTTTAAAAAAAGCAAAAAATTGGTATCATACATTGTATTTAATAGCAAATATCATGAAACTTTTTGCAGAGACATTCAAGGAACGTCAATGGAGGGACCTATTCCTCCTACAATATCCCAGTTGAAAAACATGAAAGAGTTGTAAGTACACACGTCTCTCGTACTACCTGCTATAGCGGAAGTTCTCATGTGCATTGTTGTTTTCAGGACAATATCTGATTTGAGAGGAGAACAAATGCAATTCCCAAATCTTAAGGGCTTGACGAATATGGAAAGGCTGTGAGTAGTTGGAAATTTAGCTATTACATCTCTTCATTTTAATTTTGTATTCTCATGGTTCTTTAGTTTGACTCAGGACTTTGAGAAATTGCTCAATTTTTGGTCCGATTCCGAGATATGTAGGTGACATGCCTTTGAAACTTTTGTAAGTTTGTCATCTCCTCTTTCGAAAAAAAATTAAATTTGTCTGTAATGCAAACCTGGAATTACCCTATATTGTATAAAAGGAAAATCTTCTACATACCAATGCATTCGTCGTATTTGCATGATGGAGCAAAGATTTTCACTAATTGTCTGGATACCACAGTGATTTGCTTTGATGTCTGGTTTGATGCTAATCCTACCTGCTTCTGATAAGAGTTATTTACTTGCAGGGACCTGAGTAACAATATGTTGAATGATACAATCCCAGATACATTTGAGAAGTTGGATTTTGATAACATGTAGGTACCTTGCAAGTGAAACTTTTACAAGTTGAAGCATAAATCTTGTACATGTTTGTTTTTTTTCTTTTCTCTCAAAATTTTCATTCGGTTATAACAAGCAAAAGTTTTATCTGCAATGTAGGTTGCTTGGTAACAACACATTAAGCGGAGCAATTCCCAGCAGGATGTTGAGTATGAGTGGTAGAAAGAACATGTTGGTATTTGCTCAATTTGTTCCGTTTTCCTCATCTTAGTAGTAAATTTTGGTTATTATTATAGTTAGCCACACTTCAAAGTTAACAGTTATTTTCCCATAGCTTTTTGTTTACTGCAAGGACACATTTACAAGAATTTGATGGCGTCGACAACTATAATTTTATGCAGTGATTTATCATACAACAATTTCACCCATACATCTACTCCAGAATGCCAGCACTCTACCGTGTGAGCATTCATATATTATTGCTTTTTTTATTAGTGAGAGTTTCGGTCCTTTAACTGCATAACTTACTAACATTCTTTCTTTAATAGAAACATAGCCGCCAGCTATTCAGGTATAATGAACACGTTCAAAAACACGTAAGTCTTCCTAAATGTTGTCTTATGCATCAGGTGCTAGTTTGTTTGATGCCTTCTATCTATTGTCGGGCTTAATCTATTCGAAATTGTAGGGATGCCTGGTGTTCCATGAAACCCCTCATTTGTCCCACAGAAGCGAAATGTAAGAGTTACATCCTATTAACCTTTTCTTTTTTATACCTGAAAATTCATAAATGAAGTAGCAGTTTGTCAAAGGACCTCAGGGGTTATGTCTGGATCGTTTACCACTGAATCGACTCTTGGAGTGGCAGACTGTGGAGTTTGAAGTACAAAAGCATAGCCTTGGAGTTGAGAGATAATTTCACAACCCTTCCTCATGTTATGGTCTTTATGTCAATTGCAGATACTTCTCTATTCATAAACTGTGGAGGAGGTAGAACTACATTTGAGGGAAATGATTATGAGGAGGACAAGATCAGTTCGGGTCCATCATATTTTTCTTCGTCCTCCGATAAATGGGCATTCAGCAGTTCAGGAGTGTACATGGGTCGACAAAATAGCTATATTGCAACAAATACATTTTCACTGGATGTGTCCGGTCCTGACTTTTACAACACAGCACGCCTTGCCCCGAATTCTCTCAAATATTATGGGCTTTGTCTACGAGAAGGTATCTACAGAGTGCGCCTTCACTTTGCTGAAATAATGTTTTCTAATGACTCAACATATAGCAGCCTTGGAAGGCGGATATTTGATGTAGCAATACAAGTAAGTTTGAGGCCTAACTTCTTTTGTAAATAATAAATATGTAATCTAAAATGTCGATTTATTCAGTTCTGATATGATTTGGCCACTTTCTGGTACTTGATGTAGATATACCGAATTCTGATATGATTGGCAACTTTCTGATTTAATATTGAATGTTGGAGATAATTTTGATTTCTTACACAGGGGAGAGTAGTTTTGAAGGATTTCAACATCATGGAGAAAGCTAATGGTGTTGGAAAAGGTCATACCGAGGACTTTGATGATATTACTGTTAGTAGCACTCTGGAGATTCACTTGTATTGGACCGGGAAGGGGACTAACGCCATTCCAGAAAGAGGTGTATATGGACCTCTGATTTCAGCTATCACAGTGACACCAAGTCCGTTTATCTCTCTCTGCCTATAGACTTTACACTTTTCTTCCCTCCCACAATCTTATCCCATTTTCTTCTTGCAGACTTTAACACAGGCTACGGGTTACCTGTCGGAGCTGTTATTGGCATTGTACTTGCTTCAATTGTGGTGGTTCTGATAGTGTTATTTGCTCTGTGGAAGAAGGGCATTTTTGGAGGGAAAAATAATGAAGAAGAATTAGGTAAACATACCTCAAAGAATTAATTAATCACTCTCTATATTTGATATGGTGATACTTTTGTTGTGATAATCAGAATTATTTACGAGATTTCTAGCTGCGACATACAAATAATAATCTGCCTGTAATTGACCAGCATTGAGATGATTGTTCTGTTGGTGGATCGACATCACGACATATCATAGATAGTTAGCACACAACTAAATGCTCATGACTGCTTAAACTATATATCTTCGTCATACAGATGTTCATTGTTGTTCATCATACAGATGTTTTAGAAGATCTATATATACTGACGTTTGCAATATTCTTCTCGTTTACAGAACTCAGAGCCCTTGATTTACAAACAGGTCATTTTAGACTAAGACAAATTAAAGCTGCTACAAATAACTTTGATCCTGCCAACAAAATTGGTGAAGGAGGGTTTGGACCAGTTTACAAGGTATCAAATCTCTATCTATCTCTACCCATACTGATTTTGTGTAAGGACAGCTTGTGGTTCTGTGGTGATATTTCCTACTATGCGTAACGCTTATAGGAAAATAGCTGATATAATCCATATCCTACGTTCTTCCTTCCGTGTGTGATATGATCGTTGTGAATTAATCTTGCTATCTGTAGGGTGTACTCGCGGATGGTGCAGTGATAGCTGTTAAGCAGCTGTCTTCCAAGTCAAAGCAAGGAAATCGTGAATTTGTCAATGAGATAGGCATGATTTCAGGTCTACAACACCCAAACCTTGTCAAGCTTTACGGTTGTTGTATAGAAGGGAACCAGTTATTAGTGATATACGAATACATGGAAAATAACTGTCTTGCTCGAGCTCTCTTTGGTAATGCTTTGCAATACCATTTTGAAAATTATTAATCATGTAATGATAATTTTATGCTTATTAGTCCTCGTATTCTCCTTTCTTAGGCCGTGACGACCAGAGGTTGAACCTAGACTGGGCAACCAGAAAAAGGATATGCTCAGGAATAGCAAAAGGGTTAGCTTACCTTCATGAAGAATCAAGGTTGAAAATCGTACACCGAGATATCAAGTGTACCAATGTGCTTCTTGATAAGGATCTTAACGCTAAGATTTCCGATTTTGGATTAGCTAAATTAGATGAAGAAGAGAACACTCATATTAGCACCCGAATTGCTGGAACAGTGTGAGCCTTCTTATCATAACTATCTCTGTACTATCATTTGAATTTGTTCGTCATGGTCAGTTTGAAGTATCTTACATTTCTTTTTCCGTTTTTGAAGAGGTTATATGGCTCCCGAGTATGCAACGAGAGGCTACTTGACAGATAAAGCTGATGTTTATAGCTTCGGAGTCGTTGCATTGGAGATTGTCAGTGGGAAAAGCAACACAAACTACAGGCCAAAAGAGGAATTTGTTTACCTTCTTGACTGGGTAAACTATCTTTCTACTAATATCTGCAACATTATATCTTTAGTTTAAGACCACAAGATGCAAATGTCTTTGTTACTTTCTTAAACTCTGTACGGAGGGATCTTTGCATATTTAGTTTTTCTTTTCTGTAGCTAATTTCATGTTTGCTATGTATGTGAATGTTTGTTTTAGGCTTATGTCCTACAAGAACAGGGAAATCTGTTAGAACTAGTGGATTCACATCTCGGTTCCAATTACTCCAAAAAAGAGGCAATGCAGATGATAAATCTTTCTCTTTTGTGCACAAATCTCTCTCCTACTCTCCGACCATCCATGTCTTCTGTGGTGAGTATGCTCGAAGGGAAACTCCCCGTGCAAGCACCAATAATCAAGCTGACTACATCAACCAACGAAATGAGATTTAAATCGTTTGAAAAAATTTCACATGACAGCCAAACAACATACTCTCAAGAGTCTCAAGGACAAGCACCTTGGAATGATTCGTCAGGGTCAGGGTCTCTGCCTGCTAAAGATGAAAAGTTTAGTTCCTCGAGTACACTTCTTCCAGATCTTTATGATATAAACATAGATTGATTTTGATTTTGATTTTCACACAGGATCAAATATTTATTTTCTTGTGTATACTACTTTGTAA

>M6-PGSC0003DMG400029520 M6 G-LecRK

CCCCACGCCACCCCCTCCTTTGAATTTTCTGAATATTCCACATAAACACGGTTCCTTAATGTACCAAAAAACCTTAAAATTTGTAATTTTTCTATTGCATATACTTTGTACAAAGTCCTCTTGAGTTTTGTAATGGACTAATTATTTTTGTGTACCAGCTATGACCAAACCCAAAAAAATTCTTGAGAACTTTTTGAAACCCTTTTTCTCAAGCTCCAACAAAGGTAAGCAAAAAATCTTGTGAGGGTACCCTTAATCTTGATTTTTTTTTTTGTTGTTCTAATGGTGATTTTGGGTTTTTCTCTTGTTGTGAATTGACTAATCAAAAAAAGGGAAAAATGAGGAGGACTTAGAGAAAATTGCAGCACAAGAACAGAAGCAATTCCCTTTTGAAGTTCTTGTTTCTGCTACCAAAGATTTTCACCCAGATAACAAGCTTGGTGAAGGTGGATTTGGGCCTGTATTCAAGGTAAAAAAAAAACTGAAAATCAACAAAATTGAGTCTTTTTGTGATTTGGGTGTTAACAAAAACTCAAGCTTTCCTGAAAAGATTGAGTCGTTTTGTGATTTGGGTGTTAACTGAAACTCAAACTTTCTTGAAAAAATTGTGTCTTTTTGTGATTCTTGAAGAAAATGAATCTTTTTGTGATTTGAGTATGAACAAAACCTCAAGCTTTCTTGCAAAAATTGAGTCTTTTTGTGATTCTTGAAAAAAAAGAAGAAGTCTTTTTGTGGTTTGGGTATGAACTGAAACTAAAGCTTTCTTGAAAAAATTGAGTCTTTATGTGATTTGGGTATTGAACAGAAACTCAACATTTCTTGAAAAAATTGAGTCTTTTCTTATTTATAGGGGAAATTGAGTGATGGGAGGCAAATAGCTGTGAAGAAGCTTTCACATAGTTCAAGGCAGGGGATGAAGGAATTCAAGAATGAAGCTAAGTTGTTGGCTCGTGTACAACACAGAAATGTGGTGAATTTGTTAGGGTACTGTGTACATGGGGTAGAGAAGTTGCTTGTCTATGAGTATGTTGCTAATGAGAGCCTTGACAAAATCCTCTTCAGTAAGTTTTATTAGTTTTTATCTTCTTAGATCTACTAGCTTTTGTATGTTCTTTTAAAATGAAATGAAATGACATTCATTGAGAGGCGTAGCCAGGATTTTTAGTTTATGTGTTCTAGGTCATAATTGTGTTTGTTTACTGAATTCTGGATAGATTATTCGTACTTATTAGATGAATTTTTTAATGCAATAACAAGGTTTGAGCCAAAGTTACCGGGATCTGCAGAACTAGTAGACACCTGTGGCTCTGCCCTTGTCTTCACTATCCAAAGATTAAAACAAATTGAACTGCAGATGGATGCTCTTGTCAGTTTTTTGAGGAGTGAGATACTGGAAATAGCCTTGGGATATGCATTTTCATTTTGCTATTGGGTCCAGAAATTTTTCTTTCAGTTACGTTGTCAGTAATCTAGTTCAATGATCTCTGTGTCAGTCCGTGAAACCACTAGGGATCGTCATATTGTTCTGTATTCGTGTGTTTTAGTTGCTAAACAAACTCTAAGCTCAAATGTTCCTTGAACTCCAGTGATGGACATAACGTTTATTGTTTTCTCGACATCCAGCAACAAACATGTGGCTGTTCTTTCTGTTGTCTTTCTTCTTTTTTGATAACTTTGATGTTTGGACCAGCATGTGTGCACCTCAACTACTTCATGGAATACTTGCATCCTCCTATCAATACATGTATCATGTAAATCTGTTCACCAACGGTTAGACAGATGAAAAGAAATTACTTAGTTTGTTTTTTTGCCTTTGCTAGGATTCGTACCCCACCCATTGTGTTCATTCCAGTTTCATCAACCTTAGGCGGCCGCAGTACGTTATTGAAACTTCTTCTGTCGGAAACAACCTCCCATTCCCCACTAAGGTACATCATACACTCCCTAGACACCACGTGTGGGATTACACTGGGTATGTTGTAGTAGTATTTGTAGATATTGAAACTCCTTGTACTTCTCAAGAATTCTGTAACCCTGAAACATCATTCGACGACATTTGAGGACCTTAACTTCCTCGTGTAAAGTTAAACTTATCAGTAGAAGCAATTTGTGATTCCGAGCCCGAATATAATATAATTCTTCAAGCAAGCAATTTTTCTCATAGTGGTCTTAGAAAAACCAGACTCTTTACTTTATGAATCTTTTTAACCTTTCCGAGTTTTGAGCAAGAAGAAACATGTTTAGTAGTTTTAGAAACTAGTACTGACTCTTAAATTTTTGTATTAGAATCTGTTAACAGAGATGCACTCATCTGGAAGCAACGGCATGACATAATTGTTGGCATTGCTAGAGGGTTGCTATATCTTCATGAAGGTTCACATACTTGCATTATCCACCGTGACATAAAAGCTAGCAACATCCTACTTGATGACAAATGGGTGCCCAAGATTGCTGATTTTGGAATGGCTAGACTCTACCCTGAAGATAAAACACATGTTAACACCCGTGAAGCTGGTACAAAGTATGATTCTATAGAACCTTATAAACTGTGTTTAGTGATGACTATGTAGAAATTTCGTTATATTGCACAAGTACACTCACTCGATATTGCTAACAACTCTCTTTGATAATTTAACTTGGAAGCCAGTGGCTATATGGCGCCAGAGTACGTTATGCATGGACAACTATCAGTGAAGGCTGATGTATTCAGTTTCGGAGTTGTAATTCTGGAGCTCATAAGTGGCCAGAAGAATTCTACGTTTAACAGAGATCCTGATTCTCGAAGCCTTCTTGAATGGGTAAGAACTTGATTTCTCTGCACTTTTCAGTCTTGATATTCCTTGAAATTTCTGGATTACAAGAAGAAGAATAACCTGTCAATGCCTTCATTCCTGTTGAGTATAATAGGAGAGTCCTCGGAATCAATTCGAGCTTCCTATAAACATGTAACAGAAAAAATAATATAGCCATATCGTAGAACTTTTTAACTACTTGAAAAGTAAGATCGGACTACTTTTATCTATGTTGAGGGTGCAAAACATAATCAAATCGGTAGATGGAGGGAGAAGGACTTTACTTATAAGGTGTAGGTATACTCTTAATGGTGTGAGACCTTTGGGAGAAAACTGTGTGGCTGTGACTGTAAGTGGATAGTATCACACCATGTAAAGGTATTTTTTAGGCCAATTTAGCCTAAAAGTGGGTAGTAGTGCCAATTATTTGACGGATAGAGTATGGAGATGAAGGCATTGGGTGAGTTTGCTGCTAGTCGTTGGAAATGCACAAAGGAAAAAAACTATCAGTCTTGGTGA

>M6-PGSC0003DMG400030209 M6 G-LecRK

TTTTGATATTTTCTTTCAATTATTAAACCAAAAAAAACACCAATTCTTCTGGCCAAAAAAGTATTTCAAAAATGAAAATTCAACTTAAAAGAAGGCCAAAGTAGAAATGTAAAGTCAAAGTGGACTCTTCTTCTTCCCACCTTCCCCTGTTTTCTCCACCTCCACAACAATGGCACACAACTCATAAATCCACACCAAATACCTCATTCTTCTTCACCCATTTCCTCCAAAAATGAGACTACTATTCATACTTTTCACCATTCTTATCATCACCATAGAAGCTGCAGATATTCTTCCTAATACTACCCTTTCAGCTTCCAACCCAAACTCCAGATGGTCATCTCCAAACAACACCTTCTCTCTTTCCTTCCTACAGCTCGACCCCACAAACCAGAGTTCCTATTTTGCTGCTATTTCTTACAACAATATACCCATATGGAAAGCCGGCGCCGATACCGTTAACGGTGGAGCAGTTGATATCTCTGGTGAACTACGGTTCCTCTCTAACGGTGACCTTCAGCTCATAAGTGGATCATCTGGTTCTGTTGTTTGGAGTTCAGGAACTGTAAATCGTGGTGTTTCTTCAGCTTCTCTTGATGATAATGGAAATTTTCGTCTCAAAAACGGTACGGTATCGAACATTTGGTCTACTTTTGATAACCCAACTGATACTATTGTTCCTGGACAGAATTTCACTAATAATCATGTTTTGAGATCTGGGTTGTATTCTTTTAGGCTTATGAATAATGGAAACTTGTCACTTTTGTGGAACGCTTCTATAGTGTATTATAATTCTGGGTTGAATTCTTCTGCAAATGTTAACTTGTCTTCTCCTAGTTTGGGAATGCAGCCTATTGGGATTCTTTCACTTTCTGATCCTAGTCTTTCGAACCCTTTGAATGTTGTTTATAGTAGTGATTATGCTGATGAAGGGAACATTCTAAGGTTTTTCAAGCTGGATGATGATGGAAATCTTAGGATTTATAGTTCTACACAGGGTAGTGGGACTCAAAATGTGAGATGGGCTGCTTTAACTGATCAGTGTCAGGTTTTTGGTTATTGTGGGAATTTCGGGGTTTGTAGTTATAATGAAACTGATCCAACTTGTGGATGTCCTTCGCAGAATTTTGAGCTTACTGATCCGAATGATAGTCGAAGAGGGTGTAGGAGGATAGTTGAACTTAGTAATTGTCCTTCCAATGCAACTATGTTGCAATTGGATAATGCTAAGTTCTTTACTTATCCTCCGGAATTATCTGAGCAGATTTTTTCTGCTGGTATTTCAGCATGTAGGTTTAACTGTCTTGTTAATGGTGCTTGTGTTGCTTCGACTTCTTTAGCAGATGGCACGGGGATGTGTTATATTAAACAGCCAAATTTTGTTAGTGGCTATCAGGCGCCGACCCTTCCTAGCACTTCATTTCTTAAGATATGTGGGCCTGCAATGCCTAATCCTACTGCAGTTTCAGAAACTGTTCAGGAGAAGAATGGAGGGAGGGTTCCTGGCTGGGTTGTCGCAGTTGTGGTTGTGGCTAGTTTCTTGGGTTTGATCCTGTTGGAGGGTGGTTTGTGGTGTTGGTGCTTTAGGAATAGTTCTAAATTTGCATCACTGTCATCTCAATATGCACTTCTTGAATATGCCTCTGGTGCCCCTGTGCAGTTCACATTCAATCAGCTTCAGCGGGCAACCAAGGGGTTTAAGGAGAAGCTTGGTGCAGGAGGATTTGGGGTTGTTTATCGGGCAGTTCTGGCTAATAGGTCTGTTGCTGCAGTGAAGCAGCTTGAAGGAATTGAGCAAGGAGAGAAGCAGTTCAGAATGGAGGTTGCTACTATTAGTAGCACACACCATTTGAATTTGGTGAGATTAATTGGGTTTTGTTCTGAAGGGCGCCACAGGCTACTGGTTTATGAATTCATGAAAAATGGTTCTCTTGATAAGTTCCTCTTTTCAGAAGATCATTCGTCGGGAAGGCTTTTGAACTGGGAGCAACGGTATAATATTGCTTTGGGGACAGCAAGAGGTATCACATACCTTCACGAGGAATGCCGAGATTGTATAGTTCACTGTGACATAAAGCCTGAGAACATACTGTTGGATGAGAACTACATTGCCAAAGTATCAGATTTTGGCCTCGCCAAGCTCATTAACCCAAAGGACCACAGACATAGAACCTTGACAAGTCATGTGAGGGGTACTAGAGGATATTTGGCTCCTGAGTGGCTGGCCAATCTTCCTATAACTTCCAAATCTGATGTGTACAGTTATGGTATGGTATTACTGGAAATAGTGAGTGGAAAAAGGAACTTCGAAGTCTCTGAAGAGACTAACCAGAAGAAGTGCTCTTTGTGGGCATATGAAGAATTTGAAAGGGGTAATATGGAGGCCATTATGGACAAAAAGCTTTCCAATCAGGAGATGGATATGGAACAAGTTATACGAGCAATTCAGGTGAGCTTTTGGTGCATCCAAGAGCAACCATCTCAAAGGCCAACGATGGGCAAGGTGGTTCAGATGTTAGAAGGCTTTTTTGAGATTGATCGGCCACCAGCACCTAAAGCTACTGAAGGGTCCTTTGCTGGAACCAGCTTAACCGCCAGTAGCACAAGCGGTCTCTCCACCTTTGCAGTCTCGGCCCCAGCTCCCTCTTCATCATCATCATTCCAAACTGCAGGATTCCTGTCTTCATCTTCTGCAATGAATGTCGAGAGGCAAATCATCATCACTTCTACACTCCGAAATTAAGTGAGTCTAAACCATACACTTTAGATTCCCTACAACAACTGATGGGAGTTTGGAGTTTTAGTATCTCAAAGATCATAGTTTGACTTGTGTCTTACAATTCATATTGTATAATAGATAGGGTTTGCATATATATATATATATATATATATATATTGATATGTTTCATATTAGAAAATACCATTCACTTATTTTCCATATCCTATTATTCACCAGAATTCGACGACTCTCTCATGTCCTGTCCTGTCTGTGGTGCTTGTCCAACTTCTTTTTATACATGAAGCTAGGGGTATGCATCAGTTGGTTCGGTTTATCGTTTTTCGGGGTCAGGGATGATTTGATTAGGGGGTGGTTGATGTGAACCAGAAACATGACTATAATATCAT

>M6-PGSC0003DMG400030225 M6 G-LecRK

ATGTCTGGTAAAGCTGGTCATAATGCACCTGTATTTGAGAAGTACCCGCAGAAAGCAAAGAACTTTATGTGTTCATGTCTTGCAAAGGGTGACAGAAATACCCAGAAAACTCCTGGAGGTCTCATCTATAGGCAAAGATGGAACAATATGTAGTTTGTTACTAGTGCTGCATTCCTTGCCACTACTTATTCTGACTATTTGGCCTCTTCTGGCAATACCTCAAGTGTTCTTCTGGTTCTGTTTCTCCGAATGAGCTTCTCTCATTCGCTAAGTCTCAGGTAAAGATTTTAACTTTGATAGAGTGCTACAAGTCTGTATTATGACGATTCATCATAAATTGAAGCATGGAACTAGCAATGTGTTCTTGGTGAACTAATCATCATGTTTGATGTTGTTCTAGGTGGACTACATTCTTGGTGACAACCCGAGAGCAACAAGTTACATGGTGGGATATGGAAACAATAACCCAAGACAAGTACACCACAGAGCTTCCTCCATTGTCTCGTTTAAGGTTAATCCTAGTTTTGTTAGCGCCAGGGAGGCTATGCTACCTGGTATAGTAGAAAGGTGAGCGATCCTAATCTCCTTACTGGTGCTCTTACCTGATGCCTATGACAACTTCGCTGATCAAAGAGACAACTATGGGCAAACTGAGCCTGCCACTTACAACAATGCTCCGTTGATCGGTGTGTTGGCAAGACTTCATGCTGGTCACAACGGTTACAATCAGCTCCTCCCAGGTAAATTTTAGTGAATTAACATCTTTATATGAGCATTCCGAGGATTCATCATAAAATGACTAGTGTTAGCCCACCTACTACAACTTGCTTGGGATTGAGACGTAGTAGTTGTTGTAGTACTAAAATTTTCGTATAACTCATTTAGTCTCAAGTTCTAACCTGTAATAGAGAAAACCCACTCAATGACACATCCAGTGCGCTAAATTCTACCAGACAAGGAATTCTTACTCCTCTAAATGGTTCTGGCAGAGTAATTTGGTCCTCCGATTCCACTAGACATGGGCAAAATCCAGTAGCACAACTCCTTGATTCAGGCAATCTTGTAGTTCGAGATGCGGCCGAGGATAAGCCTCAGAACTACCTGTGGCAAAGTTTTGACAATCCAGGTGACACAGCTTTACCTGGAATGAAGGTTGGGATTAATCTGAAAACTAGTTTTCATCGTTCCCTTTCGTCATGGAAGAGCACAAATAATCCTTCCAGAGGTGAGTTTAACTGGACATTCGATACTGGTGGATTCCTGCAAACATTCATTATGAATGGTTCCATTGAACTCTACAGAGCTGGACCATGGAATGGCCGAGTCTTTCCTAATGCACCATCTCGTGACACATCTTGGAATGGTTATAACTATACATACTTATCTGATCCTAATGAAATATTATTCATGTATGAACTTACAGATAGCTCGATCATTGCAAGGGTAGTGATGCAACTAAATAGATAA

>M6-PGSC0003DMG400030553 M6 G-LecRK

AAAAAATGTCTCCTCTCTTTGTTTTTCTCTTTGTACTCTCTTTTTTCTCTCTAAAAAATTTAGCAAAAGCTAACAATTTCACAAGCCAAAATAGTTCAATAATTGGTAACACAATTGGTGGAAACAAATGGATTTCAATTGCCTCAACAATAGTTTCTTCAAGTGGTAATTTTGAATTGGGATTCTTCACACCAGGTAACAATTCCAATTATTACATAGGTATATGGTACAAAAAAATAAGTAAACAAACCATAATTTGGGTTGCAAATAGAGAGACACCAATTTCCATATATGAAATGGATTTTGCACAATTCAAGATTGATAATGGTAATTTAGTACTCATAAATGGTACTAGACACATAATTTGGTCCACAAATATCAATTTTACAACAAATTTCAACAATTCACAAGTTGTAGCAACACTATTTGATGATGGCAATCTAGTATTAAGTAATCAGGATATTACCAATTCGACGAATTATCTATGGACAAGTTTTGATCATCCCTCTCATACGTTCATGCCTGGTTCAAAATTCGGTTACAACAAACATACAAGATTTAAACAGGTACTTACTTCATGGAAAAACGCAAACGATCCATCACCAGGACCCTTTACACATGAAGTTTACATGGAAAATAAGTATATTGGACAAGGTGTTAATATGTGGAACCATAGTGTTGTTTATTGGAATAGTGGACCTTGGACAGGTAACAATTTTACAGGGGTACCATATCAACCAAATCCTATGTTTAATTATACATATGTTAATAATGATGATGAAGTTTATTATATGTATAATTTCTTTAACCCTTCATTAATATCAAATTTTATTATGGATGTTAATGGACAAACAAAACAAGTATTGTGGATGGATAGTACTAATGATTGGAATGTATTTTACACTGATCCAAAACAAGTTTGTGATGTTTATTCTTATTGTGGAGGTTTTGGTATTTGTAATGAGGTAAACTCTACTTCAACATGTGATTGTTTGAATGGTTTTAAGCCTAAGTTTGAGAAAGATTGGAAATTGAGTAGTTTTTCAAGTGGATGTATGAGGAAAACAAGTTTGAATTGTGGTGATTTTGGTGAAAAGGATAGGTTTTGGATGTATAAGAATATGAGATTACCTACAAATAATGAGAGTTTGGGAGTTGGAAATGAAGTAGAGTGTGAAAGTGGTTGTTTGGAGGATTGTGATTGTGTTGGTTATGCTTATGGTAGTGGAAATATTGGGTGTTTGATATGGAAAAGGGAGATGTTGAATTTGCAACAACTTTCTCAAGATAATGTTAATGGAAGCACCATTTATGTTAGGCTTGCAGCTTCTGAGTTTTCAAGTAACCAAGGTAATGACTTGCAATTTCTCCTATTTTGTTGAATTTTAGATGCATGATGTACTTTATTTTGTTTTTTTGGTTCTCGTTTGGTGTATGATATCTGTTTTGGGATCTGACTAAGATGTATTCGCACCACTTTTAGGGATGGGGTGGGGGAAGGGGAAGCTTTTCCTACCGAAAACGACTTCAATCTCAAAACTTGAACTAGAGACTAGAGACATCTAGTTGAGGATAGAGGAGTACTTACCAGTTACCACTTCACCACAACCTTTGATAATACATATATGATATATACTTCAAATTGTTACCTCTAACCAGTTACCATTCCATCACAACCTTTATACATATATACTTCAAATTGTTACCTTTGTAGGTAGAGTGGATTTACCTAGTTTATAAAGTTTCTTTTGAGTCCAAATTTGAATTTCATTAATTACTTCTTTTTTTTTCTCTATTATCAAGAAAATGAAAACCAAAAACACATATTTTCTTGGTTTAACTATCCAGTAGCTGAAGTTTACTGCCTTGATTAATTTAGGTTTACATCAGGTAAGCCTAATTAAGGTGTTAAATCTTTTTCTATTAATTCATTATTGAATTCGAGACGTCTGGTTAAAGATGGAGGCCATCACTCACAATTAGTGACACATTGAAACATACTAACATTTTTTTTTCTCTTTTGTTATTTGTCTATTTATGAACTTGTCATTAATCTTTTCATTTTACTTGGTAATAATGTGTTCTTTGTTCTCTTCATAAACAGATCAAAAGCAAACTTCAACAAAGCTGAAAATTGCAATTCCTATTGGTGTAATTGCTGCACTTCTTATTCTATCATGCTTTACCTACATATACTTTAGAAAAAGAAGAAACTCAAAAGTCAAAGGTAACAACAAATGTCTCTTTTTTTTAATGACATTTTTCTCCTTTTTATTAATAAAAATATACAAAATTTAGTCTTATCAAGACTCTAAACTCATTCATTTTTTTTTATTTTTAAATATAGAGAGTACTACAAAATTTCACAGGCAGAATACAGAAGGGGAAAGACTAGAATTGATAGATATTCAAGATGATGATATTGAGGTTCCATTTTTTAGCTTTGAAAGTATATTAGTGGCAACAGATGACTTCTCTGAACAAAATAAGCTTGGTCAAGGAGGATTTGGTCCTGTTTACAAGGTATTTATGCAATATCATTTTGCGATTTATTTTTATAAGTGGTGCACGGTTCAAAACTCGATTTGTACATGTTTGGCAGAACCCAATAATTTTGATTCATACGTTATATTTGTGTTTAAAAAATCGATTAATAAGTATAAATATTTTATCCACATCCCAATAAGCTCTCTTTTCAAGAATCTGAAATCTATACACTCAAAATTCTAGTTTGCTGCATGTACTACTTTTCTTATTAGTTGTCATATCCTTTTTATTGCCACTTTCCTTTTTTTCAAACTACTTTGAAATGCTTTTCTCGAGTTTGATTGTCTATCAGAAACAATTTCTCTATTTCCATGAGGTAGGAGTAAGGTCTATGTACCATCTACCCTCTCTATGCTCCACTTATGAAATTACACTACTGAATATGTTAGTAGTATTATATTGTTTTTGATGTGGTGGACGAACCTGTGACGTGTGCCTAACCAACATAATCACTTTTTAAAATAAATAAATAAATTGTATCACAGTATTTATTTTTAAAAATAAATAATTTTTTTGTAGGGAATATTTTCTGGAGGAAGAGAAATTGCATTAAAAAGGTTATCAAGTCATTCTGGACAAGGTATAAATGAATTCAAGAATGAAGTGATGTTGATTGCAAGGCTTCAACATAGAAATCTTGTTAAACTTTTGGGCTATTGCATTCAATCAAGTGAAAAGATTTTACTCTATGAGTATATGGCAAACAAAAGTCTAGACACCTTCATATTTGGTTAGTATTTTTATTTTTTAACTAATATCTTTATACTGTCGATGTCTACAAGTTAAATTTATTTGATTGTTTTTCTTGCATAAGATCGAAAACGAAGCGTGGTGTTGGATTGGAGAAAAAGATTTGAGATTATAGAGGGAATTGCTAAAGGACTTCTTTATCTACACCATGATTCAAGATTAAGGATTATTCATAGAGATCTCAAAACAAGTAACATATTATTAGATGAGGAATTGAATCCAAAAATATCAGATTTTGGATTAGCAAGAGTTGTTGAAGGGAAAATTACACAAGCAAATACAAATAAAGTGGTTGGAACCTAGTAAGTACTCATAACTCCCTATTGTGATGTGACCCTTCTCCAGAATCTGCTAACGTAGGATGTTTTGTGCATCAGACTGTCCTTTTTGAAATAATTTGCCAATTGTGTTGATTTTCTTGTTCATTAATATGTTTTGTTGATACTATTAATGCAGTGGTTATATGGCTCCGGAGTATGCAATAGACGGATTATTCTCAATTAAATCAGATGTATTTAGTTTTGGAATAGTTATACTCGAGATCATCAGTGGAAGAAGAAATACTGGATTTTTTAATCAAGAAGAAGCGTCGAATCTCTTAGGCCTTGTGAGTGAATCATTACTTAAATTCTTGCTTCTTTTTTTTCAACACTGTCACAGAGTTTCAAACTTCATAGGTTAAACTCCAGTACATTGTCACGGAATTTCACCTATGAAATTTGAATCGGTCTCTATGACATTTATTTCTCAATTACATGACTGAAAAGTGACTAGTCCGCGCTATGTATTACAAAACTTTATGCTAACATAAATATGTGTACTATATTTTAGGCATGGAGATTGTGGACAGAAGATAAGTCAATGGACTTAGTGGATCAATCACTACATGAATCATGCAACAAAGAGGAAGCAATTAAATTGATAAATATTGGGCTTTTATGTGTACAAGAAGATCCAAAAGATAGGCCCAATACATCAAATATAATAATGATGCTTGGTAGTGAAAATTCCAATATTATTTCACTTCCAAGACCAAATCAACCAGCTTTTATGACAAGAAAATGTGGTAATAATAACACATCTTCTTCTTCTAATGCTAAGTCTGATGGTGTCTCCAACAATCAGCTTACAGTCACCATTGAAATGGGTCGATAACGCACTACTAGAAATCAGTCAATTTTTCATCATGAGCATGTTCATTAAAAAATCCATCGAAAATATTTTTGATAGAACGTCTGTTGAAAATTGACTGATTTAAGAGTGAGATGCCAGCAATTTTGACATAGCAATATTGTAGTTTTATCTTTACTTATTCAATGTATATATTCAACTTAAGTATGTACTTTAAAAACATCTTTAGTAATTAAATTATGTATTTTT

>M6-PGSC0003DMG400031860 M6 G-LecRK

CTTTGATAGTACAAACAATTTCTGGTTAGACCAAGAAATATAGTGAGCCATTGTGTATATGAAATCATTTGTTACTTCCTATTGTCTAAACCAAGGCCTTCTGATTTCATACAAGATGAGAATATCTCTTAATCTTCTTGCATTCCTCCTCCTAGTCCTTCATCATCAGTCAAATGGCCAAAATAATCCAAATATTAGCATAGGTTCTACTTTCACTGCTGGAGAAAATGTTTCTTTGTTATCCTTGTCTGGGGACTTTGCCTTCGGATTTTATCCTGTTTCAGATACTCTTTTCCTAGTTGGAATTTGGTTCAACAAAATCCAGAAAACAACACTTGTATGGTCAGCAAATCGCGACAATCCAGCTGAAATCAGATCAACTATTAGTCTAACAAATGGTGGTCAGCTTGTGCTCAACTATGCTAATGGCACAGTTCAACAGATTTACAATGAGTCTGCAAGGTTGGGAACCATGCAAGATGATGGAAATCTTGTGTTGAGAGACTCAAGTTCAAATAAAGTTTGGGAAAGCTTTAAATCTCCCACTGATACACTTTTGCCAGGACAAACTTTATTATCAGCAGGAAAGCTCTATTCTAATTCTATTTCGGATGGGAATTCCAAGTACTCTAAAGGGAAATTCATGCTAGAAATGCAAGCTGATGGAAAATTGGTACTATCTGCTTATCATTTTGGAGACACAGGATATTGGCTGAGTAGTCAAGATGATAAAGGAAACCGTGAAGTGAAATTAGTTTTCGACCAGAGGAATGCATCTTTGTACCTTGTGAAAGATAACAATAACACAATTTTTTCTTTTCCATCGAATGTGCCAACCCCAGTTGAAGACTACTACCACCGTGCAACTATAGACAGTTTTGGCAACTTCCAGCAGTATGCTTATCATAAGACTAATGGTAGCAATTGGATTAGAGTTTTGAAGATCCCTTCAGAGCCATGTATGGTTAATGCTGTTTGTGGGGCTTATGGATTCTGCAGCTCAAATGATAATGAAACTGTAAATTGTGACTGTCTTCCGGGGTATGTCTTCTTGGATCCAAGTAATCCTACCAAAGGTTGTCATCCGGAGACAATGATCAACTTCTGTGCAGGTCATCTATCTAGCGGAAATTTCACAATAGAGGCTATTCAGGATAGTGACATGCCTTACAATGAGGTTGGGGATTATGAAATTTATCGTGATGTTGACGAGGAGGAATGCAAGAAAGTTATCATGGAAGATTGTTATGCCATGGCTGCTTCTCTAATCAATAGGGAATGTCACAAGAGGCGTACTCCTATACTCAACGCCCGTAGAATTTCTATGACCACAGGCAGCGTTTCCTTCATCAAGGTGCCTATTAAGTCAGCTAAAGATGACATATCAATAAAAAAGAAATCCAACACCAGAGCCCATCTCACTGCTGGCCTTATAACTACCAGTTCACTTACTTTCCTATTTGGAGCCTTGGCGTTTTACTATCATCCTGCTCCTCGGAGACTTGTAAAACGAGAGTGGAATCCAAATTCATCAAGAATAGGTATCAATTTCCGCGAGTTCACATACAAAGAACTGCATGAAATAACTAATGGCTTCAGCAAGCAACTTGGAAAAGGAGCTTCTGCAAAAGTTTACTATGGTAATCTCAAGTTAAAGGATGTTCAAGTAGAGATTGCGGTGAAGCTGATGCAAGACGTTGCAGAGCCAAGTGAGAAGGTATTTATGACAGAGCTAAAAATCATAGGCAGAACTCATCACAAGAATTTGGTAAAGCTATTAGGATTTTGCATTGAGGATAATCACTTCATTTTGGTTTATGAGTTGATGAAGAACGGAGCTTTATCAGATTTTTTGTTCAAGGAAGAAATACTGCCTACTTGGAGTCACAGAACTGAAATGGCACTAGGAATTGCAAGAGGACTGCTTTACTTACATGAAGAATGTGATTCTCCAATCATCCACTGTGACATAAAGCCTCAAAATGTACTTCTTGACAGTAAATATAATGCCAAGATTTCAGATTTTGGGCTATCGAAACTCTTGAAGAAGGATCAAACAAGAACAGATACTTGCGCAAGAGGGACAGTGGGATATTTAGCACCAGAATGGCTAAAGAATGCTCCAATAACACCCAAAGTTGACATTTTTAGTTTTGGTGTAATGTTACTGGAGATAACATGTGGTAGAAGGCATATTGAGCTTAGCAGGGTAGAGGAAGAGAGCGAAGACGATGAAGGTGATGATTTATTGCTTGTAAATTGGGTTTCAGGCTGTGTGAGAAGTGGTAGAATTGACAAACTTGCAAGGAGTGATCCTGAAGTATTGAATGATATTAAAAGACTTGAACGATTCGTGAAGGTGGGATTGTGGTGTGTCCATCCTGATCCAACTGTTAGACCTTCAATGAAAATAGTGATGCAAATGCTTGAAGGCATAACTGAGGTTGGAGTTCCTCCAATGTTCTATAACTAAATGATTGTTTAAATCATCAACAGGGAGGAGCATCGGAACAAGACGAAAAACTGGAAGGAAAAAGGCCTCCTTGGCTTTGCAAACTGGAAGTGAATAATGAAAATTGTCATAATGTTAATATTTAATACTTGTGCTATTGTAAAATTATTCTAAGTCTCAGAGTTTGATTGTTAAGTTTTTTCTTTTTTTGTATGAAGTCCTGAATTTGAATTGTTACTTTTGCTAGTTTATAATCTGCACTTCAGTTTCTCCCTGGTCA

>M6-PGSC0003DMG400031861 M6 G-LecRK

GAAAAGAATTCCACACATTCCAGTGACCTCATAATCATAGCCTTCTGCATTATAATATGCAAGTCTTTGAATCATGGCATTTTGACTAATAGATTTTTACTCTCCATTTGTATCAAATAAGTTTATTTGTTGTGAAAACAATTATAATATCTCAAAAAGAATAAAGTTTTTTCCGTAAGATTCCAACAATTCATGGTGTAACAATGGTTTCTTGGAAAATCCTGTTATGGTTTGTCTCAATTTATCTCACATTTATTAGCTCTTTCTGTCAAGAAACTACAAATATTGTCCCTCTTGGTTCCAATATCATAGCTGGAACTGAATCTTCACACTGGCTTTCATCCTCTGGAGATTTTGCTTTTGGTTTCTACCTTCTTAACAGTACTGGATTATTCCTTGTTGGCATATGGTTTGATAAAATCCCATATAAAACTCTCGTTTGGTCAGCCAATCGCGATGATCCAGTTCATGCTGGATCAAAGGTTAATCTAACGTTGAGTGGACGCCTCGTGTTGACAGATACTAATGGGAGAGAGTTTGTTTTGTACAATGGCACAGGCACTAGTCATGCTACTATGCAAGATGATGGAAATTTTGTCCTGAAAAATTCATTCTCAGGTGTCTTGTGGGAAAGCTTTGATTTCCCAACAGATACAATTTTGCCTGGTCAATATCTTGATATGGGGCAGGCGCTGTTTTCCAGTGCCAATGGGACAGTAGATTACTCGACGGGGAAGTATAGACTACTGGTGCAGACTGATGGGAATGTGGTCTTGTCTGCATATAGAACTGCTGATTTTGGCTACTGGAATTCTGTAACTATAAACAATAATAATGTAAGACTGGTTTTCAAAAACACCAGTCATACTTTGTTCTTACAAATAGAAGCTCAATCTTATCCAACTTGACAGCGAACTTGCCTGATTAGTTCAAGATTATTACCACAGAGCTATGATCACTGATAAAGGAGATTTCCAACAGCTCTTCCACAGAAAAGTAAATGGGAGAGGTTGGAATGTTGCGCTGCAAGTTATCGAGAAGCCTTGTATTGTGAATAACATTTGTGGTGTATATGGCTTTTGCCAATCACCAGACAATAAGGAAGTCAACTGTAGTTGCTTGCCAGGCTATTCTCCAAGGGATCAGTACACTCCATCTAAAGGATGTTATCCGAATGAGATGAAAGATTTTTGTGATCTGAATTCTTCCCTTTCGGATGTCTACGTTGAAAGGATCTCTAATGCTGATTTCCCTAATAGGATGTATGCTGAACTAGAAAGGGTATATGATACTACAGAGGAGCTTTGCAGGCAGGAAGTGTTGAATGATTGTCTTTGTGAGGCAGCTGTGTTTAATGATTCAACATGTTTCAAGAAGAGGATGCCTATACAGAATGCCAGAAGTATTAATCCTGATACAAATAACATGGTTGCATTTCTTAAAATTAGCAACTCTTCGATAACTCAAAACAAAAAGCACTCTCTATCAAAGGGTGTTTTGGTAGCTGGTGTGACAGTATGTTCAGTACTCGCTCCGCTTTTTGCAGCAATTGCTATATATTATCACCCTTTTGTGAAGAAACATAGAGATGCCAAAACCCCTCCAAAAAGAAAGGCAATTGAGATGAATTTAAGGGCATTTTCGTTCCAGCAGCTGCACGAAGCCACCAATGGTTTCAAGAATAAATTAGGCCAAGGGGCTTCTGGTGCTGTTTATAGCGGAATTCTCAAGTTAGAGGATGAAGAAGTTGAAGTTGCTGTGAAGAAATTAGGAAATGGAATTGAACAAGGTGATGATATAGAATTTTTGGCAGAAGTAAGGGTGGTAGGCCTAACACATCACAAAAATCTGGTTCATTTGTTAGGCTTCTGCAATGAAAAAAGCAACAGGCTTGTTGTTTATGAATTAATGAAAAATGGGGCTGTGTCAAATTTTATCTTTCGTGATGGACAAATACCAATCTGGAAGCTAAGAAGTGACATTGTACTTGACATTGCCAGAGGCCTGTTATACTTGCATGAAGAGTGTGAGAACCAGATCATTCATTGCGACATTAAGCCACATAATATCCTTCTTGACTAGAACTACTCTGCTAAAATTGCTGATTTTGGTTTTGCAAAGCTTCTTATGAAAGACCAGACGCTAACAAACACTAATTTTAGAGGAACCATGGGTTATATGGCACCAGAATGGCTAAAAAATGTGCCCGTGACAACCAAAGTTGACATCTACAGCTTTGGTGTTTTGCTGCTTGAGATTATTTTCTGCCAAAGGCACATGGACTTGAACCCAATCGGCCAAGAAAATGAAGAGCCCGAGTTGATCTTAGTCGATTGGGTTCTTCATTGTATGAGAAACAAGAAGCTGAGAGCTGTTGCTAGTCTTGATGAAGAAATTATGTTACATTTTCACAATTTTGAGAGGATGACTATGGTTGGCCTATGGTGCTTGTGTCCTGAGCCAAATCTTCGACCATCTGCAGCAAAGCTTGTTCAGATGTTGGAAGGAACTATAGAACTAGGTGTTCCTCCTATGGTTGATACAACTATATGAGTCATTTTCTGTATTATCTTGTAATTTTTCCAACAATTATCATTCTATTTTATTAGTCATTAGACCTGGTTAAATTTCAGTGTTTTTTCTCATAGTTCTCTGGTGCAAAAGTTAGTGTTT

>M6-PGSC0003DMG400034859 M6 G-LecRK

ATGGTACAAAGATGTAGGTCCCGGTACCGTTGTATGGGTTGGCAATAGAAGAAATCCAGTTAATTGTGGTGTATTGCTAGGACTATACACAGAAGGAGACATTTTTCTCCACGATATACTGGGAACAACGGTATGGATTCATAAGCCCAACAGTACAGTTCCCTTACCAGTACTGAAGCTCCTTGATTCAGGTAATCTTGTGTTTGGAGATTCTAGCAACTAACGGCTGGAGAGTACCTATGGCAAAGCTTTGACCACCCAGTTGACACATTGTTACCAAAAATGAAGCTTGGTTGGGACAAGAAAACTGGCATCGATCGATCCATGAGGTCCTGGAGGACAGAAGGTGATCCAGCTCCAGATGATTATTTGTTCAGGTTAGACTCAGGTGACTCAGGTCAGTTACCCCAACTAGTTCTTGAGAAAAACCAGCGAATACATTCAAGATGGGGTCCTTGGGATGGACAGATCTAGAAATTCAGTGGCGGTTATGCCCTCATGGATAACCATGCATTCAAGCCAATATTTCATTCCGATACAAATGCATTTTTTTTCACATTCGAAGCCAAGAACGATTAGAGTTTGAAACTTTCACTGAATCCAGATGGAAAATACAATTCTTGAGGTGGAATAATGCTACCAATTACTAGGAGGAAGTGAATACACTGAATAAGGACACATGTGACCAGTATTGCACTTGTGGGCCTTATGGAGTTTGTAGTGATGGAGATCTCCCCTGCGGGTGTCCTGATGGTTTCACAGCTGCATCACCAGATGAATGGGGCATGATGAATTTTACGGAGGTATGTAGGAGAAACACCTCACTTAACTACACCAATAAAGACGTCTTTGTGGAGAATACTGGATTGAAGTTGCTTGATAATGCTACTTACTAGGGAATGTTGTACCACCAGGAGTGTGAACAGAAGTGCTTAAATGAAAGATCGTGTATGACTTATACTATCATCAATATCAATGGAAATGGAAGCAAATGTGTCGTTTGGTTGGGAAATTTACTTGATATGAGACATTCACAGATAGTTGGGAACAACATTTTTATACGGATGGCACCTGAAAAACTAGGTAAAAATGATTGCTTGCTTTAACTGATTTAATTCCTAGATTATATTATTAATATGGTGCAAAACAATGTGTGCAAGAATGACATTTGAAAGGTGAAGTAGTTAATTGTTGTGTAGGTTTGATTATTTACACAGAAACATCAATTAATAGAATTTCCATATTAATTCTTGATTAAGTGGTTCATCGCAGATCGATGTTTGAAGCATTAGATTGAATGTTGTCACATAACTGCTATATAATCGAGCATAAGATGAAACAACCCTGCATGCATCTAGTTATTCTCATTGCAAAATCTTTAACATAATGAGTAAATACCTCTAGAAATTCCTATAAAGTTTATGTCAAATTTGAAGCCTAATTTTATGAGTAAGAAACAACTAACACTTTTAACTTTCTTATGGACAGATGCACATGTTTCCGACATCAGATGGAAAAATGAATGGACAGCTTTTGCTTCTTTAATCTCAGAGTTGGCACTACTTATTATTTTTCTTATAGCTGTTGTGGTTCTATGTCAAAAAATGAGGACAGAGGTGCTAGGTAAAAAGAATCACCTTGCGCTGCTATTATATTATTTTCATAGTTCTTTATTACAAACTAAGCTTAGTTTCACTAGAACACTATTATTTAGCAAACACAAGACCTAAGTAAGAAAGCTAGCCTAATACTCATATTAAGGTATGATTCACATCCATTATTTTCTGATTAATTTGCTTTCTCATAGCAGATGAACAAGAAATCAATCAGCCGACAAATTCTCAAGGAGCTTTGCTGCAAGGCACTAATGTCATTGCATATGATTCCAGTGATCTAGCTGCAGCCACTAACAATTTCTCGCTGGCTAAAAAGATAGGGCATGGTGGCTTTGACAATGTTTGTAAGGTTATTATTTCTTTCGACAAGCATGAAAATGAATATTTACAGTAACTCAGACAACGTAATTAATACCTAAATGATACATTTGTAGGGTGTACTAGAGAATGGAGTAGAAATAGCAGTGAAGAAGCAGGACGTGACTTCATGTCAAGGATTTACTGAGTTCGAGAATGAAGTCAAGTTAATAGCAAAGCTACAACATCGTAACCTAACCAAATTGTTGGGATACTGCATTAATGGAACAGAAAAATTTCTAGTCTATGAGTTCATGGCGAATAATAGCCTAGACAAGGTCATATTTGGTATGACATCTTAAATAATATTTTTCTTTTTTTACTATTGTCTGTGTTGTTTTGATATTGGAAAATCCACAAAAGACAGAAATTTAGGAGTATCCGAGTGAAATTGAGTAACTGTTTTCTGTACATGTCTAGATCCTGCAAGAAGGGGTACAGTAACATGGCCAATGCGCTTTAACATTATCAAAGGAATCGCAAAAGGATTAGTTTACATGCATCATGACTCAAGGTTGACAATAATCCACCAGGACTTAAAAGCTAGCAATGTCTTGCTAGACAGAGAGATGACACCCAAGATATCTGACTTAGGGTTGTCAAGGGTATTCGAAGATGATGTTGAAGAAAAGACTCAGCATTTAATTGGAACACGGTTAGTATATTATCATTAAGACTACATGTTTATGTATATTTTAGAGTTTTTACAGATGGTTTTATGTTGTGGCTGTAGTGGTTATATGCCTCCTGAGTACATGAAAAATGGACATTATTCAACAAAATCATGACGTTTTTAGTTTTGGAATCCTGGCATTGGAGATTGTGAGTGGCCAAAAGAATTATGATTATCGGCACCCAGTCTATGATATTGGTCTAGTAGACTATGTACGTAAGTACATCATCGATCCCCTGGTTATCGTCATTACTTCACTTAAATTGGTTACATAGAAGTTTCACCTTGTGCTAAAACAATTTACATGATTGATGTTCGATCTATTGTTGGTCAAGCTTGGAGAATTTGGAGTAAGGGGAATGCCATAGAACTACTGGATCCAATGATTGAAAAACCAGGTGATGAGAATGAAGTTTTAGGATGCATCCTTGTTGGGGTCTTGTGTTGTCAACGCCGTGCCCAAGACAGACCTTCAATGATTCAGGTGGTTTCCTTACTGGAGGAGAATGAAATGTCAAGGTTAAATTTTGTGCCTCAGGAACCTTAA

>M6-PGSC0003DMG400035841 M6 G-LecRK

GGAATTCTTACTCCTCTAAATGGTTCTGGCAGAGTAATTTGGTCCTCCGATTCCACTAGACATGGGCAAAATCCAGTAGCACAACTCCTTGATTCAGGCAATCTTGTAGTTCGAGATGCGGCCGAGGATAAGCCTCAGAACTACCTGTGGCAAAGTTTTGACAATCCAGGTGACACAGCTTTACCTGGAATGAAGGTTGGGATTAATCTGAAAACTAGTTTTCATCGTTCCCTTTCGTCATGGAAGAGCACAAATAATCCTTCCAGAGGTGAGTTTAACTGGACATTCGATACTGGTGGATTCCTGCAAACATTCATTATGAATGGTTCCATTGAACTCTACAGAGCTGGACCATGGAATGGCCGAGTCTTTCCTAATGCACCATCTCGTGACACATCTTGGAATGGTTATAACTATACATACT

>M6-PGSC0003DMG400042814 M6 G-LecRK

ATGGAGATCAAACACAATTATTTTCTCCTCTTTTCTTTGATGTACCTATGTTTCTCTCTCAAAACCCATCTCTCCATTGAAGCTGCTGACACCATTTCTGCAAATCAATCCCTTTCTGGAGACCAAACAATTATCTCTTCAGATGGAAAATTTAAGCTTGGTTTCTTCAAACCAGGTAATTCTACCAACTAATACATAGGCATGTGGTATGATAAAGTGGTTGAACAAACTGCTGTTTGGGTTGCAAACAGGGAAAAACCAGTTCTTGATAAGAATTCTGCAGAGTTAAAGATCTTAGATGGTAATTTAGTACTGGTGGATGAATCTCAAACTCCAATTTGGTCCACAAATATCAGTTCCAGCAACTCGAGTTCTGTTGTAGCAGTCCTCCAAGATGATGGTAATTTGATCTTGACAGATGGGTCTAATTCAACACCACCACTTTGGGAAAGTTTTAATAACCCCACCAATACTTGGTTGCCTGGTTCTAAACTTTCTTACAACAAAGTCACCAGAACAAAGCAACTTCTTACTTCATGGAAGAGTGCAGATGATCCTGCACCTGGGCTGTATTCACTTGAACTCGACCCAAATGAGAAGCAGTATATTATAAGGTTCAAAAGAAGTGAGCAATATTGGAATACTGGACCATGGAATAATCGAATTTTCAGGGATGTGCCTGAGATGAGGACTAACTATATTTACAATTTTAGCTATGAGGACAATCAGAATGAAAGCTATTTTACATATTCCCTTTATAATGATTCTATTATCTCAAGATTCATTATGGATGGCTCTGGACAGATTAAGCAACTCACATGGTTGGATAATACAAATCAGTGGAATCTGTTCTGGTCTCAACCAAGACAACAATGTGAAGTTTATGCATTTTGCGGGCCATTTGCTACCTGTCAGGAAACCAATCCCTTCTGTAATTGTTTGGATGGTTTCAAGCAAAGTTCAGAGACTGATTGGAATCAAAATGATTTTTCATGTGAGAGGCAAACGAAGTCGCAATGTGGCAATGGTAATGGGGAGAAAGATGACTTTTGGATGCATCCCCAGATGAAAGTACCTGAAAATGCTCAAAATATTTCTGCTGGAAGTGCTGAAGAATGCAGATCTACCTGTTTGAATAATTGCAATTGCACTGCTTATACTTATGAAAGTTCATGCTCAATATGGAATGGTGAACTCTTGAATATGCAGCAACTCCCACAAAATGATGGCAGGGGCGAGTTGATCTATGTCAGGGTAGCTGCATCTGACATCCCAAAATCAAAGAGTAAAAAGGGAATTCCAATTGGTGTTTCTGTGGGGTCTGCTGCTGCCGTACTGATCCTTCTGGGAATTCTTTTTGTTGTATTCCGGAGAAGGCGTAGGCATATTGGAAGTGGGAAAATAGTGGAGGGTTCATTGGTTGCATTTGATTACAAAGACTTGCAACATGCGACTAAAAATTTCTCAGAGAAGTTGGGTGGTGGAGGCTTTGGTTCTGTTTTCAAAGGGAAACTATCTGATTCATCTGTTATTGCTGTTAAAAGGCTCGATAGCATCAGCCAGGGAGAGAAGCAATTTCGATCAGAAGTCAGCACAATCGGGACAATCCAACATGTTAATCTTGTACGCCTTCGTGGATTTTGCTCTGAAGGTAACAAGAAACTACTGGTTTATGATTACATGGAAAATGGATCCTTGGATTCACATATTTTTACCGAAAAGCAGTCAGATGTTATGGATTGGAAAACGAGGTACCAGGTGGCACTAGGGACAGCTAGAGGCTTGACTTATCTCCATGAGAAGTGCAGGGACTGCATCATACATTGTGACATAAAGCCTGAAAACATCCTTCTTGATGCACAATTATGCCCCAAAGTAGCAGATTTTGGCCTGGCAAAGCTTGTTGGGCGCGATTTTAGCAGAGTCCTTACTACTATGAGAGGAACAAGAGGTTATCTTGCACCAGAATGGATATCAGGGGTGGCCATTACAGCCAAAGCCGATGTTTATAGCTACGGCATGATGCTTTTGGAAATTGTATCAGGGAGGCGAAACTCAGAATACTCTCACGATGGGAAAGTGAAATTCTTTCCTAGTTGGGCTGCACGTGTAGTAGTTGATGAAGGCGATATCCTTAGCCTATTGGACAATAGGCTGGACAGAGCTGCTGATGCTGAGGAGGTGTCAAAAATATGTAAAGTTGCATATTGGTGCATCCAAGATGATGAATTCCAAAGACCCTCAATGGGTCAGGTTGTTCAGATTCTTGAAGGAGTTTTGGATGTGAATTTGCCTCCTCTCCCACGCTCTCTTCAAGTTTATGCAGACAATGAGGAGCACATCGTTTTCTTCACCGAGTCATCCTCGAGCCAGACTAGTTCACAAGCTCAGAGCAAGACTTCAAGTACAACATCTCAATCTAAAAGCATCACAGAGTCTACGAATTCCAGGTCTTGA

>M6-PGSC0003DMG400042886 M6 G-LecRK

ATGGACTTAGTTAACTCATTAACTCCTTTGAAATTAACTCAGTTTTTCATTATTTCCATCTTTTTCTCAATTTTTTTTCAAATTGAAGTTGAATCTGAGACACCCAATCAATTTTTCAAAGGATTTAAAGCAAACCCAGATCCCCAAATTTCGAATTTTCAGCCTCTTCTTACTGATTCTACTGGAAATTATTCACTGAGTTTTCTTCGAGTTGAAGAAAATCAACTCAGTCTTGCTATTATCCATGTTAAATCTTTCGAGTCGATATGGGTTGCTAATTTGACCCGTTTAGCTAAATGGGCTGACCCGACTGAGCTCTACTTCAATGGCAGCCTTGTTCTGTCTGATTCACGTTCAGGGGTATTTTGGTCAACTCACACTGATGGAGACCGTGTTTGGCTTTCAAATACATCGAATTTGCAAGTTCAGAAGGTTGATAATGGAGTGACGTCACCTTCTGTTTTATGGCAAAGTTTTGATTTTCCCTCAAATACCCTTGTGGAAAATCAAAATTTCACTAGTGCCATGACTTTAGTTTCTTCAAATGGGCTTTATTCCATGAGATTGGGCTTTGATTTTTTCGGGTTGTATTTTAAAGGTGAATCGGGTTCGGGTCCGGGTCGGATCTATTGGAGGCATAAGGCATTAGAAGCTAAAGCTGATGTGATGGAGGATCAAGGACATATTTATGTTGTATTAAAGTCAAATGGATTTTTGGGAATGTACCAAAATGAGTCAGTCCCAGTTGATGTTGAATCATTTAACAGCTTTCAACAACCCTTATCCGGTGTCCGTCGGGTCCGGGTCGAACCGGATGGTAACTTAAAAGGGTATTTTTGGGACGGGTCGAGTTGGGTATTGGACTATCAAGCGATAAAGGAGACATGTGAATTGCCTAGCCCCTGTGGGGCATACGGGCTCTGTCATCCAGGCAAGAGTTGTTCTTGCCTGGATAACAACACGGATTATAGTATCGGCGGGTGTGTCTCGCTGAACACTCAGGAGCCCCGCGATTTTTGCGGGGCCTCCTATGATCTTGATCACAAGAAGTACAAGGGGTTATCAAGAAATGGTGTTGATTTGCCTAACACTGAGTTAATGGACTATCAACATATGGTTTCTTTTCAAGAATGTCAAAGTGCATGTGAAGGAAATTGTACATGTTGGGGTGTGGTGTACACTAACACATCTGGATTTTGCTACATACTAAATTACCCCATACAAAGTTTAGTAGGGGTGAGGGATGAGTCCAAGATGGGGTTTTTCAAAATGAGGGAGGGTGTAGGAAAAGATAAGATGGGGGTGGGGTTTGGTGTAGGGGTAGGGTTATTATGTGGGGCCATCTTGGTATTAGGTGGGGTCATAGGTTTAGGGTATTATAGATATATAAAAAGAAAAAGAGAAAGAGGAGTTAGTGGTTATGTTGATGATGATGGAGTAGTGGTTGGACCATATAAAGATTTGGGAAATGCAAGTTTTAGGTCAATTGAACTAATTGAAAGATGA

>M6-PGSC0003DMG400044943 M6 G-LecRK

ATGGCTTCCATGCTTTCTTTGTTTACTATCTGTCATCTTTTTATCTCCATAGCCCTGTATGCATGTTCCGCTGGTGCTACTGACTCCTTGTTTGCATATCAATTCATCACCAACTTGAACCCTTTAGTTTCATCCAACCATAATTTCGTAATTGGATTTTTTGGTTATTATTCATCGACTCTACGTGTATGCAAGACGTACCTGGGATTGGGGTACAGGAGTGTAGATCCCAGGGCTATAGTATGGGTAGGTAATGAATTGAATCCACTTGGATGTTTTACAGCGCTAGCACTAACCAAAGAAGGATTTTATGTCAAAGATATAGTACTGGGAACAACGGTATGGATTCATAAGCCCAACAGAACAGTTCCGTCACCAGTTCTGAAGCTCCTTGATTCAGGTAATCTTGTGTTTGGAGATTCTAGCAATCTGACGGCTGGAGAGTATCTGTGGCAAAGCTTTGACCACCCTCTTCACACATTGTTACCAGGAATGAAGCTTGGCTGGGACAAGAAAACTGGCATCGACCGATCCATGAGGTCCAGGAGGACAGAAGATGATCCAGCTCCAGGTGATTATTTGTTAAGGTTAGACTTAGGTGACTCAGGTCAGTTACCCCAACTAGTTCTTGAGAAAAACCAGCAAATACAATCAAGATGGGGTCCTTGGGATGGAGAAAAATTCAGTGGCGGTAATGCCCTCATGGATAGCCATGCATTCAGACCAATATTTCATTCCGATACAGATGCGATTTTTTTCACATTCGAAGCCAAAGAGGATTCGAGTTTGATACTTTCACTGAATCCAGATGGGAAACTACAATTCTTGAGATGGGATAATAATGCTACCAATTTCTGGGATGAAGTGAAGACACTCAATATGGCCATATGTGACCAGTATAGCACATGTGGGCCTTATGGAGTTTGTACTGATGGAGAATTCCCCTGTGAATGTCCTGATGGTTTCACAGCTGTATCACCAGAAGAATGGGACAAGATGAATTTCATGGAAGGATGTAGGAGAAACACCTCACTTAACTATACTGATAAAGACGTGTTTGTCAAGAACACTGAACTAAAGTTGCCTGATAAAGCTACTTACTGGGGAATGTTGTACCCTCAGGACTGTGAACAGAAGTGCTTAAACAAAAGATCGTGTATGGCTTATACTAACATCAATATCAACGGAAATGGAAGCAAATGCGTTGTTTGGTTGGGAGATTTACTTGATATGAGACGTTCACGGATAGCTGGGAACGACATTTTTATAAGGATGGCAAATGGAAAACCCGGTAAAAATGATTGCTTACTTGTGATATGATTACTTGCTTATGACACGACAACTAATTCAATGCCTAAATTATTTTTAGTATAGTGCAAAGTTATGTGTGCATGAATGACATTTGAATGGTGAAGTAGTTAATTGTTGGTGCATAAACATCAATAAATAGAATTTCCATATTAGTGCTTGATTAAGTGATTCCTGCAGCTTTGTCTTTTTTTAAAAAAAAATAAAAATTGTGTCTATAACTTGATTCTTACATGAACCACTCATCCCTCATCTCTATCAGAGGGAAACAGTTCTATAACTTTTCAAAATGCATATGTATCTCCGTTATATAGAAACATTAGATGTGTTTCAAGCATTTTACTGAATGTTGTCACATAACTGCTACACAATCTAGCAAAAGTTGACACTAACCTGCATGCATCTAGTTGTTCTCATAGCAAAATCTTTAACTTTGTATCAACAGAATATTATGAGTAAGAAAACAACTAACACCTTTTACTTTCTTATGAACAGATGCACCTGTTTCCCATGGGAAAAGAAAATTGAATGAGGGATCGATAGCAGTGATTTCTGTATTAGTTCTGGAAGTACTTATCTCGATTTCATATATTGTGTATACACATCGAGCAAAGAGTACAGAGGTGCTAGGTAAAATGAATCACCTCTCGCTGCTTATATATTATTTTCATAGTTATTTTTTTCAGCCCAACCTTAGTTTCACTCAAACACCATTCATGAGCAAAACCTCCTTTCAACTCTTGAGACAACAAGAATGAGCAGAGACATTGTTCAAAAGCATCTTGTTCTGCCAATACCCTTTAGTTTGTAATGAAAAACAAAATTTACTTAGAGCACACATGACTCTGTAACCTAGCTTGTGGTTCGCTTCCATTATTGTCTGATATATTTGTTTTTGAAAATGAATATCTTCATCTGAGCAGATGTCATTGCGCCAGAAAATTCTATAGGAGCTTTGCTGCAGGGAACAGATGCTATTGCATATGATTCCAGTGATCTAGCTGCAGCCACTGACAATTTCTCGCTGTCTAATAAGATAGGGCATGGTGGCTTTGGCAATGTTTATAAGGTTAGTATTTCTTTTCAACAAGCATGAGCATGCATATTTACAGTAACTCAGACAACACAATTAATACCCAAATGATACATAAATCTGTAGGGTGTACTAGAGAATGGAGTAGAAATAGCAGTGAAGAAGCAAGACGTGGCTTTGCGTCAAGGAGTTGAGGAGTTCGAGAATGAAGTCAAATTAATTGCAAACCTACAGCATCGTAATCTAACCAAGTTGTTGGGATACTGCATTCATGGAATAGAAAAATTCGTAGTCTATGAGTTCATGGCGAATAAAAGCCTAGACAAGGTCATATTTGGTAGGTGATCATCTTAAATAACAACTTTGTTTTACTATTATTTGTGTTATTTTGATATTGGACAAGCCATAAGAGAGAAATTTTATTAGGCCTATCCGAGTGAAATAGGGTAACTGTTTTCTCTACATGTCCAGATGCTGCAAGAAGGGCTACAGTAACATGGCCAACGCGCTTAAACATTATCAAAGGAATCGCAAGAGGACTAGTTTACATGCATCATGACTCGAGGTTGACAGTAATCCACCGGGACCTAAAAGCTAGCAATGTCTTGCTAGACAGCGAGATGACACCCAAGATATCTGACTTTGGGTTGGCAAGGGAATTTGAAGACGATGTTGAAGTGAAGACTCACCGTGTAGCTGGAACATAGTAAGTAGACTAGCACTCATTATCATTAACACTACATGTTTATGCATATTTTACAGATAGTCTTATGTTGTGGCTGTAGTGGCTATATGTCTCCTGAATACATGCAAGCTGGACATTATTCAACAAAATCAGATGTGTTCAGTTTTGGAATTTTGACATTGGAGATTGTGAGTGGCCAAAGGAATTCGATTTATCGGCACCCAACCTATGATGTTGGTCTAGTAGGCTATGTAAGTATATCATCCCCTGATTATGGTCACTACTTGTACACCTAAGTTTCACCTTGTGCTAAAACAATTTACATGCATGATTGATCTTTTGTTGGTCAGGCTTGGAAAATTTGGAATGAAGGGAATGCCATAGAACTACTGGATCCACTGATAGATAAACCAGATGATTTGGATGAAGTTTTAGGATGCATCCATGTTGGGATCTTGTGTTGTCAACGGCGATCCCAAGACAGACCTTCAATGGTTCAGGTGGTTTCCTTACTGGAGGAGAATGAAATGTTAAAGTTTAATTGTGTGCCCCGGGAACCTTACTTCTATAGGGAGACGAGTCAGAGTAGCAGGAGTCGCGAAAGTATGAATGGGCTAAGTATTACTGAATTCACCGGAAGATCATGA

>M6-PGSC0003DMG400045997 M6 G-LecRK

ATGACTTTTTTAGCTATGCTTCTTGTTGGTCTCTTATTGTTTCCCCTTTCAGCCATTGCTCAATCTTACCAAAATGTTAGTTTGGGTTCATCACTCACTACAAGTGATGTTACTAGCTTTTGGTCATCCCCATCTGATGAATTCGCTTTGGGCTTCCAAAAAATTGGAAATGAATCAAGGTACTTACTAGCCATATGGTTCAACAAAATCACAGACAAAACCATAGTTTGGTCAGCCAATAGGAATAATCTTGCCTTAGATGGATCCAAAGTTCAGCTCTCCGCGGATGGAAGACTAGTCCTCACTGATCCAAATGGTCAAGAGATGTGGGCTCGTGGCATGGCGAATGCTCAATTGGCCTATGGAGCCATGCTTGATAACGGAAACTTTGTGCTAGCAACTAGTAGTTCAGACACTCTATGGCAGAGTTTTAATGAGCCCACCGATACGATTTTGCCTGGTCAAGTACTTAACCAAGATCACAGTCTTGTTTCAAGTTTCTCTGACACAAATGTATCGAGTGGGAGGTTTGAGTTCATACTTCAAACTGATGGAAATCTAGTGCTTTACACAGTCAATTATCCAGCTGAGACAACAAATGCTGCATATTGGTCATCTATGTCAGTTGGCAGTGGTTATCAGGTGATCTTCAATCAATCAGGCTTTATTTTTCTTCAAGCAAAGAATGGAACTTTGATAAATTCGATATCTTCTAATGTGGAAAATTCAAGAAGCCAATCAATGTATCATCGAGCGATTCTTGAATATGATGGAGTTTTTAGGCACTATGTCCATCCGAAATCTTCTGGTAGGGAACCGATGGCATGGTCTAGTTTGTATAACATTCCTGATAACATCTGTTTGTCTATCAGGCAAGGTACAGGAGGGGGTGCCTGTGGTTTCAACAGCCTATGCTCAATAGGAACTGATCAGAGGCCGAGATGTGATTGTCCACTCGGATATATCTTGGATGATCCAAATGACAAATTAGGCAGCTGCAGACAAAATTTCTCTGAGCAAAACTGTAACCATGAATCGCGAGAAGTTGAGTCTTTCACATTCCATGAAATGTTGGATACTAACTGGCCTGACTCTGATTATGAGTCTCACCGCGATGTTTCAGAGGATTGGTGCAGACAGAATTGCTTGAATGATTGTTTTTGTGATGTTGCCATTTACAGTGACGACAACATCTGTTGGAAGAAAAGGTATCCTCTTTCAAATGGGAGAGTAGGTCCAACGATCGGAGGGAAGGCTTTGATAAAAATAAGGAAAGACAACTCCACAGTGGGAACACCAAATGTTGAAATAAGAAAAAAGAAAAATCAGTCTACTCTGATCATTTCTGGTTCTGTGTTGTTGGCCAGTTCAGTTTTCATGAATTTGTTCTTAATTCTTTTGGTCCTATTGTACATTTTTAAGTTTATAGGGAAAAAACGAAAGAGGACTGCACTGTATTCAGCTGTTCCGGGAGTGAACTTAAGAAGTTTTAGCTACAAAGAGCTAGAACAAGCTACAAATGGATTCAAGGAAGAATTAGGAACTGGAGCTTTCTCAACAGTTTACAAAGCGGTTCTTGATGATGAAAATGGGAAAGTCGTTGCTGTCAAAAAACTCCGAAACATGGTAACAGAAGGAGAAGGAGAAGAAGTATTTGAAGCTGAAGTGAATTCAATCAGCAGGACTAATCACAAGAACTTGGTCCAACTCCTTGGATTCTGCAACGAAGGCCAACACCGCCTTTTGGTATATGAACACATGAAAACCGGCTCAATAGCACACTTGCTATTCAAAGATTCTAGGCTTAGTTGGTCCAAAAGGGTACAAGTTGCCATCGACACTGCCAAAGGACTTTGTTATTTGCATGAAGAGTGCAGTACCCAAATTATACATTGTGACATCAAGCCCCAAAATGTGCTTTTGGATGAAAATTTGACAGCAAAAATAGCAGATTTTGGTATGGCCAAGCTACTGAAAAAACATCAGACTCAAACAACCACTAGGATACGTGGAACCAAAGGGTACGTAGCCCCAGACTGGTTCAGAAGCATGCCTGTAACTGTGAAGGTTGATGTGTACAGCTTCGGGGTCTTGCTCTTGGAGTTGATTTGCTGCAGAAAAAACTATGAGCAGGATGTGGCAAACGAGAACGAAATGATACTACTGGAGTGGGCTTATGATTGCTATAGAAGAAATAAGTTGCATTTACTTGTAGGTGATGATGAAGAGGCATTAGAAGATATCAAGAGATTCGAGAAGTTCTTGTTGGTTGCAATTTGGTGCATTCAAGAAAATCTAGCATCAAGGCCTAACATGAAGAAAGTCATGTTAATGCTTGAAGGTTCTGTTGAAGTATCAATTCCTCCTGACCCTTTTTCGTTTGTCGATTCTATTTAG

>M6-PGSC0003DMG400046324 M6 G-LecRK

ATGGTTGTTAATTATAGTGCCACTCTTTTCATGCTCAGTCTCCTCATTCTTTCTTCACAGCATCTGATTAGTTGTCAACCTTCTGAAACCTTTATCACCAATTCCTCCATTTCCTGGATCAATAACCGCGCCCGCTCCTATGTAGTGAATTCCACAGATGGAGCTACACTGACACCGATCCTCTTGAGAGAAGGGAACGAGACTCAGTTTGTTTGTGGTTTCTTATGTGATTACAATGGCACTGCTTGCGTTTTTGGCGTCCTTATATTACCAAATATCTATGATCCTTATCTATATAACCTGAGATTAGTTTGGTCAGCAAATCGGAATAATCCGGTTAGAGCTAATGCAACCTTGCAACTTAGGCAAGATGGAGGCTTGTTCTTGATAGACTCAGATGGAACTATGATTTGGAGTACAAGCACAAGGGGAAAGGTTGTTGCGGGATTCAACATGACAGTACTGGGAAATCTTGTCCTCTTTGACAAATATAACCACACCATTTGGCAGTCTTTTGATCATCCTACAGACTGTTGGCTTCCTGGTCAAATGACAAATGAGGGACAAAAGCTGAGATCCAGTGTATCAGCAACAAATTGGAGTGAAGGTTTGTTCACATTTGAAGTTCTACGATTTGGCTTTTTAGCATACATTGAGTCAAATCCACGTCAACTCTATGTCTCAAATTCTGATTGGGAAAGAGCTTTCATGTTTCCTCGACTAAGATTCATTGAAGGGATGTTAATTGTAAATTTTGAATTTTCAAATGCTAGATTCATGAGGTTTGGGGATGATGGGCATTTAAGGTTGTATCACTGGGAAGGATCTTGGTCAGAGGAGGCTGATCTACTGCCAACCTACATTGGTGAGTGTGGGTATCCAACTGTGTGTGGCAATTACTCTGTTTGTGGAAATGGGCAGTGTAGTTGTCCTCAAACTGCGGTTGATCAGACTAACTTTCTCCAGCAAATCAATTACAGGCACCCAAACCAGGGGTGTGTCCTTATCACACCTATCACTTGTGAACATTCCCAATATCATATTCTTATAGAGCTCGAGGACATAGCTTATATTCCCCTCTACTTGCAATATGGTACCAACAAAACAGATTTGGAAAATTGCAAAAGGTCTTGCTTAAGTAACTGTTCTTGCAAAGCAGCTCAATTTCAATTTAGTGGTCCAGGTAACACAATGGGTGACTGCGCATTGTTGAATGATGTGTTTTCGCTCACAAACAATGATGGAGCATCCAATAAGACCACTCTTTTCATCAAGGTGCAGAACTCTTCTAACTTACATGCCTCTCCACCACATATCTCTACAGAGAAGAAATCAAAGCGTGTCCCAACAATAGTAGGATCCACCATCGGAGCTTCATTTGGTTTGCTTTTCATTGTTTTAACTGGCTTTGCCTATATTTTCAGAGGGAGAAAAGGACTCGATGAAGACGAGGAGGAGTTCCTTGATCAAATACCAGGAATGCCTACTAGATTTTCCTACGAGGAACTCACTGTGATGACAGAGAATTTCAGTGAGAAACTTGGGGAAGGGGGATTTGGATCTGTATTTGAAGGAACAGTGAGTGATGGCACCAAAATAGCGGTGAAGCGTCTGCAAGGTTTTGGTAATGTGAAGAAATCATTCTTAGCTGAGGTAGCAACAATCGGTAGCATTCATCATGTTAATTTGGTAAAACTTGTTGGATTTTGTGCTGAGAAATCACACAGACTTCTGGTTTATGAATACATGGCCAATGGATCTTTAGATAGGTGGATTTTTCACGGAAAACAGGAACAATCTCTCACATGGGATGTCAGGGAAAAAATCATTGCAGACGTCGCCAAGGGCCTAGCTTACCTACATGAAGATTGTAACAACAAAATTTTTCATTTGGACATTAAACCTCAGAACATTCTTCTTGATCAGAATCTCAATGCAAAAGTATCAGATTTTGGGTTATCAAAGCTTGTGGGTAAAGATGAAAGTAAGATAGTTACAACAATGAGAGGAACACCAGGCTATTTAGCACCTGAATGGTTGCATGAGGTCATTACTGAGAAAGTGGATGTTTATAGCTTTGGGGTTGTGATCTTGGAAATCATCTGTGGCCGGAAGAATTTGGATCGTCATCAAGATGAGGATGATATGCATTTGCTTAGTTTGTTCATGAGAAAGTCAGTGGAGAGACAACTCTTGGAAATGGTGGACAAGAACAGTGAAGACATGCAGCTACATAAAAAAGAGGCTGTGGAAATGATGAAGATTGCAGCCTGGTGTTTACAAAGTGATTATACTAAGAGGCCTTCCATGTCGTTGGTAGTTAAGGTGTTTCAGGGTTTAGTCGCTGCTGAAACCGACTTGGATTACAGCTTCACATTTCCAACAATGACGAGAAGAGTTGCCGGAGCTATTCAAGAAAGGGAAGCTGTGGTTGGTATCAGTTTACCACTTCCATCACAGTTGTCAGGACCTAGGTAA

>M6-PGSC0003DMG401004946 M6 G-LecRK

ATGGGATTTTTTTGCCCGGCTAGTTTCACGAATAATTGGTATATTGGAATATGGTATAAGCACGATGTACCTGATAAGTCTGTAGTATGGGTTGCCAATAGAGAAAATCCATTCAACAACACATCAGGTGTTATGTTGAAAATCATCGATACAGGACAACTTGCTCTTCTCACTGCTGCTAATAAAATCGTGTGGTCTACTAATTCATCGAGGCCTTTGGCTGTGAAGAATACAGTTGCACAGTTGTTGAACTCGGGGAATCTTGTCATAAGAGATGCAAATGATACCAAGCCTGAGAAGTTCCTGTGGCAGAGTTTTGATTATCCTACTGATACACTCTTACCAGGCATGAAGATGGGTAAGAACTTTGTAACTGGACAAGAATTTTACCTGTCCTCATGGAAGAACGAATATGATCCAGCTCCAGGGGAATATACGTATCATTGTGATCCGACTGGATATCCTCAGGATGTCATGAGGAAAGGTAAAGTTAAGGTGTTCAGTACTGGACCATGGAATGGACTCCGCTGGAGTGGTGTACCAGGACTGACAAAAAATACTATTTATACATTTAAACTGGATCTTGATGAAAAGAGAGCATTTTACAGCTTCGCGCTTTTAGGTTCTGTTATGACCAAATTAACCATGAACAGTAATGGTGTGCTTCAGCGTTCAATGTGGGCTGAGAATAGACAGGAATGGCATGTTTACATTTCTTCACCGGAAGATACTTGTGACAATTATGGAACATGTGGTTCATATGGTAGTTGCAACAACATACTCACTCCTGTGTGTAATTGCTTAGACAAGTTTGTGCCTAAAGATCCAAGAAATTGGGCAATGACAAATTGGTCCAGCGGGTGTATTAGAAGGAAACCGCTTAATTGCCAGAATGGAGATGGATTTCTCAAGTACTCTGGTATAAAGCTACCAGATACACAGTATTCATGGTTTGATAGGAGTATGACACTCCACGAGTGCAAGCAAGCTTGCTTGAGGAATTGCTCGTGTATGGCTTACTCAAATCTTGATATACACAATGGTGGGAGTGGCTGTTTGCTTTGGTGTGGCGACTTAATTGACATTCTAGAATTACCTGGAGGACAGGACATTTACATAAGGGTTGCTACCTCAGAGTTAGGTAACATTTGTGAATTTGATCAAAGTTGGAGTATTAGCTGAATGATTTAAACTTCCCAATTCTCAGATTTAGTTGTATTCTACTTAGATATCTGTTAAAAGTTCACACAGTATATGTTCACTCTTCCATGTCTATTTAGTTTATTTCACCCCTCCATGTTTTGTTCAGGCTCGAAGAAAACAAAACTACTCGTGTTGAGTCTGTTACTGTTGATTGGAGTGACTGTGACTGGCCTGACCATAGGGCTGTATATTTGGAAAAAGAAGAAGAAGAGGAAGATGAATTTGAAAGATGACCTAGACCTCCCACTGTTTACCTTATCAACTCTAACTAAAGCATCCTCTAATTTCTCAATCGAGAACAAGATTGGAGAAGGTGGATTTGGATCTGTTTACAAGGTAACATCTTAAGTTTATGAATCATTCTTTCAGTAAATAATATCTACTATTATTGACAGTAAACAGATGACACTTACTGAATAACTAAGCCAGATATATGTTGATATTATAGGGAATCCTGGAAGGTGGACACGAAGTAGCTATAAAGAGGCTATCCAAGTCTTCCTCGCAAGGAGTTAACGAGTTCAAGAATGAAGTTATTTGCATTGCCAAACTTCAGCATCGAAATCTTGTGAAGCTTATAGGTTGCTGCATTGCAGGAGGAGAAAAAATGTTGGTCTATGAATACATGTGCAACAGAAGTCTAGATCTCTTCATATTTGGTCTGGTTACTCTCTCTTAACTCGAATAACCTCTATTAGTACAAATAGTAAAGTTGATAGTTTAATCACATTATCAGTGTTCTAACAGAGGACCTGTTAACAGATGAAAAGAGGAGCTTGTTACTCAACTGGCCTAAGCGCTTCGACATCATCAATGGAATTGGTAGAGGACTTCTGTATCTCCATCAAGATTCTCGGCTAAGAATTATCCACAGAGACCTCAAAGCTAGTAACATTTTGCTAGATGCTGACATGAACCCCAAGATCTCAGATTTTGGCATAGCTAGAAGCGTTGTAGGGAATGAGACTGGTGCTAATACACACCATGTGGTTGGGACACAGTAAGAAATTTAATACTACTCACTTCTTAGCATTTATATTATGTATAATATCCTTTATTTTACATTACTTTCTGTTTGCAGTGGTTACATGTCCCCAGAGTATGTAGTCCATGGTGTTTTTTCAGTAAAATCAGATGTATTTAGCTTTGGCGTATTAGTGCTAGAGATAATCAGTGGTCGAAGAAACAGAGGATTTGCCCGTGAAAGTCACAGTATCAACCTACTCGGACATGTAAGTCTTGTTTGTAATTAAAGACGAGTTTAAATTTTGTGCACTATATGGTAGCTACAGATATTTCTCTCTAATAATTAATTGATAACACCTCGTACAAATGATAGTTAAACTGCATTGAACTTAAATCCATCAGGCACATCAACCGCTGCATGTTTCTTGATTATGTACTGATTGTGTTATTGATAGGTATGGAAACTTTATAAAGAAGGTAGACCATTAGAGCTAATTGATGGCCATCTAATGGATTCGCGTTATATTTCTGAATTGCTACGCTTAATCCATGTCGCTCTGTTATGTGTACAACAATGTCCAGAAGACAGGCCAGATATGCCTACAGTGATCTTGATGTTGACTAATGATGCTATTTTGCCTCAAGCTAAGGAACCTGGTTTCTTCACAGAAAGCAAAGTCGCGTCTGAATGTTCTACAAGCATGTGTTCAACAAACGAAATTACCGTCACACAGTTAGAACCCCGGTAAATTACATTACATGTTCATTCAGATCGTTCAATGTCAACAAGCTTGAATTACAGTGTTTTGTGCTGTAAAGCCATAATTTCCTATGGTATCTGTTATTAGTAAATATGGGTGTGTTGTAGTTTTTATTAATTCTAATGTATTTTAAAGTATTCGTACAAAGTACCTATTCAACAGTACATCAAAATAATCGAAAGATCCATATAAAATGAATCAACACAATGAGAGTTCACAATAACAACAGATGCCATTGTTGATGGCTTTTAGAAGCTCCAGTCCCATAGTACTCCATTAAGTACAGAGTATCAGTTTTGTTGTTCCCATTCTCCGAGTTTAGGATATTCATTAACTGTGGGGTTCCAGTTATACGATCAAACATCAGGCAGTCGCTACGCTAACAATTTGAATGATAGCAGAATGGTTGATTAAAGTCATGAAATTGCCAAGACTACAGTGAATGGAAGAATTGCAATTCTAATTTCTTATTTTCTTTCTAGGAAATGTACAA

>M6-PGSC0003DMG401005047 M6 G-LecRK

TTCACGTAAATTCAATAGCTTTTATTTAGATCATCGTTTTCTTTCGGGTGAAGCTATAATAAGTATGTAGGCAGGTTTGATAGAACTTAGTAGCTTTGATCTATACTCTGTATTTTTGCTTAAAAATACTCTAATCATGTATAAATTATTTATCAAGAACTCAGCAAGCTAATTTTTTCTAGAGTTTATAATTTATAAACTTAAAATTTTAGCTCAGACTCTTATTATAAATATTCGTAAACTTCAAATCTTAAATCCGGCTCATTATAAGTTAATGCTTACCTAAAAAAAAGAAGGTGAGAAAAGGGGTTTAAAATTAAAACATGGTCATAAAGTGAACAAAGCGTGTGTTACATTTTTTTATTTTGTATTTTGTATTGCTAACGCGAATTGTTGCGTGACATAATGAAAAATACATTAATTTTTTGCTGCTTTTATTGTTATATTTTCATGAGGATATCGTATCTTTTTTTAATTTTAAAATTTTTGACAATAAATGTAAAAACTAAAATTAGTGGACAATTGGTCCTAAAAAGTTAGGTGTAAGTTCACACAAAAGATTCTTTTTGAAATGGATGAGAAGAAATCATTTTTTTGGACCCACTTTATATTCCAAAAATAGTCACTTGGCCGATTATTTTGACTTTGATTAATTTCAAATTTACATATAGTATGAGTTCTACTTTTAATTTTATTTTGACTTTGATTAATTTCAAATTTTACATATAGTATGAGTTCTACTTTTAATTTTTCATGAGGTTTACTTTCATATTGTAAATATAAAAAATAAATACTTTTTGACAAGGACAAAGTCTTTGGGTCATTTGAATGGTGTACAAGTTACATTTAAATTATAATATTAAAAAATAAATAATATTATTTAATATATATGTATTTGACTTTACATTGGATAAATTAGTATTAACTATTATATTTTCTATTTTAACCTTAGATTTAATCTAAAGGACCTTGGAAGAAGAGTCTCAAGAAAGGCCATTTTGGTCATTTTTTGTATTTTATCTAGAGATTATTATCCGTATTAAAGGTTGTATAAAAATAATATCACAATTACGGTGTAATGGCTAGCCCCAAAATCAAACTTTCAATCACACGGAATAATATAATCTCACATTTCATCTCGGGATTGTTATCTATTATTCCAACTATCAAACGACTATAGTAATTACTAAGTTATGTAATGTGCACTAATGTTTTTGGATTCATGACAAAAGAAACTTGATCTTCCATTCATTTGTATTTTTCGAGCAATTAACACCTAGTTCAAGTTACTCAATCGTGATAAAAAATAATTTTGTGACTTTACTGTTAAAAAAAAATACAACCTTTGATGGAAGACTCATTTTTATGTGTTAATTTTCAGCAAAAAGTGGAAATGTTGGATCTGAAGATGGTTCTGGCAAAACAAAAAGAATTGCTATGGCTACTGGAATTACAGCTGGTGTTGTTCTTGTTCTAATTGGAGTTGTAAGCATTTGCTTTTTATCAAAGAGAAAAAAATTATTGGAAGATCCAATAAGAAAGAAAACAGAACAAAGAGGTATGTTATAATATAAGCCTATAATTTCGATTTCAGTATGGTATTCGATATTTACTTTAAGGCATAATACATAAAGAATGCACATCTATAGACACCTCGACTCGTCTTCACTGTGTCTCATGGACACCTAATACGTTATTGTGACACATAAATTTTGGAAATGTCTAAATGATCATTTTGTAAGTTGGAATGTACGTTCACTAGACATGATGGAGACGAGTTGAGGTGTCTAACTATACATTACTCAAAGTTAGAGCCTTCAGTTGTTAGTTGAGGTCAAGTTAAAGTGTCTATCTGTATATTATACAGTAACCATCTCATAATTCATACTTCTTACACTATTTTTGAACCAGGCTCTATTGAAAGAAGTCAAGATCTTCTAGTGAACACAGCCATTATTCCAAGTAAGAGAGAAATTTCTGGCGAAACTGTCGCGGACGAGTTTGAATTGCCATTATTTGATCTCAGTACCTTAGCTGTGGCTACTGAAGATTTTTCTGATGCAAACAAGCTAGGCCAGGGCGGTTTTGGTTGCGTTTACAAGGTAAAGATCATCTAGACGTTAAAAAAGCAAAAGCTTATGTTGCTCGGATTGGTCAAAAATATCATCATATCCATGTCAGATCCTCCAAAAACACATTACTTTGGAAGATGTAACACGTGCAGTGACATTTTTCGAGGATCATAGCAATACATCAATTAAGTAGTACTTTTTTGCATTTACAAGTTAAAGCTGATCATATATCAGTAAAAAGAACATATGCTGCTCAGACAGGACAAAAATGTCGTTGCATGCACTCGTGTCAGATCCTCCAAAAATGCATCACTTTGTAGGATTGATCTAACACGGGTGCAGTGACATTTTTCGAGGATCTGAGCAATAAAAACTAATCCTTTTTTTTGCATTTACAATGTAAAGCTCAACTAGATGTGAAAAAAACTTTTGTTGTTCGGATTCTACAAAAATGTCATTGCACCCTTGTCTGATCTGACACTGGTGTTACAACATTTTCAAGGATATGAGCAACATATGAAAAACTAGCCCTTTTTGCCTTTACAAGGTAGACGTGAAAACAACCTATGTTGTTCGTATCCTCCAAAAATATCATTGCACCCTTGTCGGATCTCCAAGAAAATGCTTCATTTTTGGAGGATCTGACACGAGTGCAGTGACATTTTTCAAGATTCGAACAACATTGGGAAAGACGGACTCTGTTTTATTGGAAAATGTGAAGACATTCAACAAAAGAACATCTTTTTTTTTTTATTATCAGGGAATAATAGATGAAGGTCAAGAAATAGCAGTGAAAAGGCTGTCAAAGAATTCAGGCCAAGGAGTAGAGGAATTCAAGAATGAGCTAAGATTGATTGCAAGACTTCAACACAGAAACCTCGTTCGCCTTCTTGGATGTTGCGTTGAGATGGAAGAGAAAATGTTGATATACGAATACATGGAAAATAAAAGCTTGGATTCAATGTTATTCAGTAAGTCGTTTTCCGTTCAATATATTCTTATGATCAGTAACTATGTTTATGTGTGCTATTGGACTTCAAAATTAACAAATATACGCGTCTCAGATAAGCAGAAAAGCTCATTGCTAGATTGGCAAAGACGATTCAGCATTATCTGTGGGATTGCTCGGGGGCTTCTGTATCTTCATCAAGATTCAAGATTTCGGATTATCCACCGAGACCTCAAAGCAAGCAACATTCTCCTTGACAAGGAAATGATCCCGAAAATATCAGATTTTGGCATGGCACGGATTTTTGGAGGCGATGAGACCGAAGGAAATACAAAGAGAGTTGTTGGAACCTAGTAAGTAGAATCACATTATTCCTTATAAATCATTCATAAAAAACACTTTGAAAAGTGTTGGTAAATCGACTTTTTTTCTCTCTTAATGTAGCGGTTACATGTCTCCTGAATACGCGATGGATGGTCTATTCTCTGTTAAATCAGACGTTTTCAGCTTCGGAGTTTTAGTGTTGGAAATAGTAACAGGGAAGAAGAACAGAGGATTTTATTTTCAGAACAATGAAAGAAACCTTCTTGGCCATGTAAGTACACCAAAATTATCACAACTTAGCCATCATTTCTAACAAAGTCAGAACAGAGAATTTTGAGTTTATGCGTTCTGAATTCTTAAAAAAATATAGCTAATTAGTAGGTTCTTGATAAATTATTTATGCATATTAAGTAGATTTCATAACATAAATACAAGGTTTTTACTAAAGTTGCTACTGAGTTCTGTCGAACCTGTAACTTTCTCTCTAGTTCCGCCCCTAACTATGTAATCACTCTGCAGGCGTGGAAACTATGGAGAGAAGGAGGAGCCTCAGAACTACTCGATTCATCAGTTGGTGAATCATTTTCTCCGTGTGAAGTAATCAGATGCATACAGGTCGGGTTATTGTGTGTTCAGGAACAAGCAGAAGATAGACCGAACATGGCAACCGTTGTATTGATGTTAGGAAGTGAAACCGCAACAATGCCTCAGCCTAAACACCCTGGTTTTTGCCTCGGAAGAAGACCTGTTGATGAACACTCGGAAACTATTTACGAAGAAACTTTCACTGTGAATCAAGTAACGATTACCATGCTTGATCCCCGGTAGAGTTAAACGTTCAGACAGATCATAGGCTAGCCCGCTTCAACGTGACAGAATAGAAATCAAGGCAAATTCCGCTGATGGGAGTGTGAACATAGAGATTCCACGACGAATGAATGCAATACATGGAGGATGGGTCCCTGCTGGCAAAAATGTATAGATAGTACATTAGTGTTTTTTCTCCACCCGACCATATCAGGAGGATCAATAGTGTTTTAACTTACCCTCGTATAGTACATTAGTACTACGAGTGGTTAACAATTTTCTCTTTGTACACTTACCTTGATTCTTTTTACAAAAATAATGTTGTTTTTTGGGTAGTTAATTATAGAA

>M6-PGSC0003DMG401009323 M6 G-LecRK

GCTGCCAAAAATGAAATCTTTAGCCAATTCCATCTTCATCTTAATATTGACCTTATTCTTCAGCCATACATGTATCATTAACCTTCCATCTGTGGCTGCAATAACCTCTATGAAGCCTGGTGATGAACTCAATCACACTCAGGTTCTTGATTCTGAAGATGGAATATTCAAGTTAGGATTTTTTTCTATCTCACAAACAAATAACTTTTATCTTGGTATATGGTATGCAGGTGATCCAAAAGATAAGAAACTATGGATAGCCAATCCAAATACGCCTTTATTGAACAATTCAGGATTGCTCACTATAGATACTACAGGAACATTGAAAATCACTAGTGGAGGGAAAACAGTTGTGAATATTACCCCTCCTTTATTGACAGGAAGTTTAATAGCTAGGCTTCAAGATTCTGGAAATCTTGTGCTTCAGGAAGAAACTCAGAACAGGACTTTATGGCAAAGCTTCGATCATCCTACCGATACTCTTTTTCCAGGTATGAAGTTTGGTTATAACCTTACAACAAAGCAGAATTGGACTTTAACTTCTTGGTTGAGTAGTTTTATTCCGGCTTCTGGGGCTTTTACTTTAAGTTTGGAGTCCATTCAAGATGCATTTCAGCTGGTTATACGTCGAAGAGGAGAGGTTTATTGGACAAGTGGCGCTTGGAACAATGAAATTTTTCCATTCTTAACTGCATTGTATGATTCTTCTAACAGATATCAGTATAATCTCAACTTGGTGTCTGAAAAAGATGGTGTGTTCTTCCAATTTGATGCTCCAGATGGAAGTTTCCCAAGTCTTGTGTTGATTTTTAATGGGGCTATTCTTGGTGGTGGTGAAGACAGTCGTGTATACGCCCTTTATAATGAGTTCTGTTATGGTTACGAAAGTCACGATGGTTGTGTCTCTAATCAGTTACCTGAGTGCCGAAAGGATGGGGATAAATTTGAGCAAAAGAGTGGAGACTTTATCGATAGATCTAATTCTAACAGTTATGATAATGCAAGCACTAGTCTTGGTGATTGTATGAAGAGGTGCTGGGAGCACTGCAGTTGTGTTGGCTTCACCACCACCAGCAATGGAACAGGCTGCATAATTTGGAATGGTGATGGGGAGTTTCAAGTTGATGAGAGTGGTAATACAGTGAAAAAATATGTTCTGGTTTCATCAAAATCATCTAATGGTAAATCCTTATAGCAGGAATGCACCTTGCTTATGAACTAATATATTTTGAATAGATCAAGAGTCACCTGTATAACACTCAGCAACCTATGTGTGCAGGAAAACAAAAGAATTGGATATGGATAGTAATTGTTGTAGCTATTGTTGTCCCTATGCTGATATCTGGTTTCATCTGCTACATTATAGTGAGAAGGCGCAAACTACAAGGTAAATCTCTGAAAAAGTGAAAATTGATTAACTAGTAAGCTTTTTTATTAATTATGTTAAGCATGTTTAAATATCTACAAACTTTTCTTGCACAGCTGAGAAAAGAAGGGAGGAAGAGTATATACGTGAGTTGACAGCATCCGATAGCTTCAACGACACCAATATGAAAGAAAAGGATGGAAGGGAAGTCCAAGATTTGAAGATATTTAGCTTTGGATTTGTATTGGCAGCCACAAACAATTTCTCAAGTGAAAACAAGCTTGGAGAGGGTGGTTTTGGACCTGTTTATAAGGTATAATTACATTATATGCCTTCGAAAAACTTCAGTGCCATCATTGCATTAACTGAATTACCTTTAAAATTTTATGACAGGGAAAATTTCCAGATGGGAGAGAGGTGGCAGTCAAGAGACTTTCACGAACATCAGGTCAAGGGCTTGTAGAGTTCAAGAATGAGCTTATACTCATTGCAAAAGTTCAGCACACAAATCTAGTTCGAGTTCTAGGATGCTGTATTCACGAAGATGAGAAAATTTTGATATATGAGTATATGCCTAACAAGAGCTTGGACTTCTTCCTATTCGGTTCGTTACCATTTCAAAACAAAGAATTAATAGCTCAAATAAGTTCTGAAAATTTTGACCTGAAATGCTTTCTGTCTTTGACAGATCCTGAAAGAAAGAAGCTACTGGATTGGCAGAAACGTTATGAAATTATCGAAGGATTTGCTCAAGGATTGCTCTATCTTCACAAATACTCAAGAATGAGAGTGATACATAGAGATCTTAAAGCCAGTAATGTGCTATTGGATGAGAACATGAATCCTAAGATAGCTGATTTTGGTATGGCAAGAATTTTCAAACAGAACGAGACGGAGGCAGTGACTGCCAGGGTTGTTGGAACATAGTGAGTAGAACTTATTTCATGATATAAGTTTTATTCATTTAGACATTTTCATATTGACTAAACGGGTAATGTTTCCTGTCCTCTATACAGTGGTTACATGGCTCCCGAGTTCGCAATGGAAGGAGCATTCTCAATCAAGTCTGATGTCTTCAGTTTTGGAATCTTAATGCTGGAAATTGTGAGTGGTAGGAGAACTACTAGCTTACAACAATTCGATCGTCCACTCAATTTTATAGGTTATGTAAGTTTTCCTTGACACCCCTTTTTCCTTTTAAGTACTATAATACATAAAGTTTGAGAATACTGATCATTTTGGTTTAGGCATGGGAACTATGGAAAGAAAAGTGTGCACTGGAATTGAAGGATCCAGCTCTTGGAGATTTATGTGACACTAAACTGTTGTTACGAGTTATTCATGTTGGACTATTATGTGTACAAGAAGGAGCAACTGATAGGCCAACAATGTCTGATGTTATCTCATTGCTTGGTAATGAAAGTATGCCACTGCCTACGCCAAAACAACCAGCATTTTTCACAGGAAGAAATGAAGCAGAATCAAATTCATCGGGTATCAAAGCAGAACAATGTTCGGTTAATGATTGTTCCATCACTGTCATTGAAGCAAGGTAGCTTAGAGACATTATCTTGCTGTATTCTGTTGTGCAACACTCAGTAGCTATACATGAAAAGCATGTTTCCTTATGGTATAGATTAGTCAATTTGTACTGTTAACACTTGTATAGCTGTCTTTAGCTGACTTTGTAACAATATATTAAATCTCTCTCCTACAAAGACTAGAATTATCATTAACTTTTCTGTTTTTTTTGCATTTTCTCACTTAAATGTAGTAACATTTATTAGTATAACAACCTTAAGATTGTTAGATTCTCTTTATTTATTAGCTCCTTTATATTTTTAAAATTAGGAGACTCATCAATGATGATATTTTTTTGTGGTCCACATAAATGGATAACTTATCTGAGGATCATAATTCATAAGGATGACTGTAAACTAATCTAGTCCAAACAACTTACTCGACTAGTTAAGGACATTCTTAATCTTGATTCAACAATGCATCAATGAAGTTCACAGTGTATTTCAACAAGTAGATATGATATCTTTTTGAATTATATTAGCAAGCAGCAAAATGAAATAGAGAAATATGAGGAAGAAAAAAGAAATTAGAAAAGTCAAAGAAAGAACAGAAACAAAATGCAAATTGATTTCACATTTCAACAAGAAAATTAAAGATTGTCTCTTTCAACAAATTTTCTTACAAGTAAGGTAATATTTCTACTTAGAAAAATCTTCAAAATTTCTGGTTGGAAAATGTTTAATATAATTGTCTATTCTCTAATGCATTATTGGCTTTGAACAGTAAACAACCTAAAAAGGAATAATCTATTATTATCTGCAGCAACATTTCTTTCTTTCTGGCTGCCAAAATGAAATCTTTAGCAAATTACATTTTCTTCCTCATTTTGACTTTATTCTGTAGCCAAACATGTATCATTATCTTTCCATCTGTAGCTGCAATAACCTCAATGAAGCCTGGTGATGTACTTAATCATTCACAGGTACTTGATTCAGAAGGTGGAAGATTCAAGTTGGGATTTTTCTCTATCCCACAAACAAATAAAACTTATCTTGGAATATGGTATGCAGGTGATCCAGTAGAGAAAAAATTATGGATAGCCAATCCAAACACACCTATATTGAACAGCTCAGGATTGCTCACATTAGATTCTTCAGGATTATTGATAATCACCAGTGGAGGTAAGACAGTTGTGAATATTACCACTTCTTTATTGACAGGAAGTTTAATAGCTAGGCTTCAAGATTCTGGAAATTTGGTGGTTCAGGATGAGACTCGGAACCGGACTTTATGGCAAAGCTTTGATCATCCTACTAGTTGTCTTTTGCCTGGTATGAAGCTTGGTTATAACCTTACCACACGGCAGAATTGGACTCTAACTTCTTGGTTGGTTATTAGTACAATTCCGGCCTCAGGGGCTTTTACTTTAAGTTTGGAGGCAATTCAAGATGCATTTCAGTTGGTTGTTAGACGAAGGGGTGAGGTTTACTGGACTAGTGGGGCTTGGAACTATCAAGGTTTTCCATTCTTACCTTCATTCCGTGATTCTGCTACTACTTATCAGTACAATCTCAACTTGGTATCCGGTACTGATGGGATGTTCTTCCAATTTGAGGCTACAAAGGGAAGTTTCCCAAGTCTTGAGTTGTTTTCAGATGGAGCTATTGCTGCTGGAGACGGAAGTATATACACCCGTTTTAATAAGTTCTGTTATGGTTATGGAGGTGATGATGGTTGTGTCTCTAGTCAGTTGCCTGAGTGTCGAAAGTACAGTGATAAGTTTGAGCAAAAGATAGGAGACTTTATAGATTTATCTGGTACTACTACTAGTTATTATGACAATGCAAGCATTAGTCTTGGTGATTGTATGCAGAAGTGCTGGGAGCACTGCAGTTGTGTTGGATTCACCAGTCTTAACAGCAATGGGACAGGCTGCCGAATTTGGAATGGAAAGAGGGATTTTCGAGTCGATGAGAGTGGTAATGCAGTTCAAAGATATGTTCTGATTTCACCAAAATCATCTAAAGGTTAGTATTTATGTCATAAATGCACATTGTTTATGAACTAAATTTTTGGTCGAACGAACACATTTTGTTGCAAACATCCAAAAACAGAATACAGTATATCCTAAGTAAAAAGCTACAACTGACCACAGCTAAACAAATTTAAAAACAGCCCCTACTCTTCCTCCCTTCTATACAGTTTTTCTGATTGTAAAGTTCTT

>M6-PGSC0003DMG401013876 M6 G-LecRK

GAAAAACTTGAGACTTTAAGAAAGGGGATGAGAGGAGTACACATTTTCGTCACAAGGCGCATAATTTCCATAGCAGCTGACGACAACCCTTTTAACCCTTTGTCACGTAATTCCTACTCAAGCAAATCTTTCATTGGGCTGTCTAATTCTTGTGCCTCTTCGTAGTTTAACCTTAGGCCCAACATCTTAGCCAGCCTCAGATAATTGGTCCATAAGCATTATATGGGCTATCCATGTTCACAACTTTATTAGTACTTATTTTTTAGTGGAATGATTGCACTACTGCAGTTTGATGTCATTTTCATCAATTGGATATTTTCCTTACGTAGAATTTTCAAATATTTTGACTAAGGTGTTTGATTAGTTGCGTAACTTTGGTTTGGTAGAATTCTCATAGACTAATTTTATTTTTTCTGTTTCTCTGAGATGATAGCATTGATATACTTATGGATACTTGAATGTTTGTTTGTGTTAATTCGTCTATTCCCTCGACTTGTCAAAACTTTTTAAAAACAGGTATTTGGGAATGAAGATATTGTTGGTTGATTTAGTTCTCTTGCTTTCTTGTTTTCATTTAGTTAGGTGCAATAATACGCGTTCAGACAACTTATTGCCTGGACAAATCCTTTCCTTTTCTGCAGCAGATACTTTAGTGTCCAAACAAGGGAAGTTTGAACTCGGTTTCTTCTGTCCAGGTAATTGTGAAAAGCTTTTTATTGGTATATGGTATACAAACATAAAGCCCAAGACCGTCGTTTGGATAGCAAATAGGGACAACCCTATCCGCCCTCCGTTCAATAATTCTCATCTAGAGCTGTCTGATGGAAATCTAGGAATCTTTAATGCACGTAAACAGAGAGTTTGGGACTCTAATTTATCCACAGCAGATGTCAATAGGGCAGTACTTCTTGATTCTGGAAATTTCATTTTGACAAATGGCATTGAAATACAGTGGCAGAGCTTTGATTATCCTACCGATACATGGCTGCCTGGTGCGAGGATTGGATTTGATAAGTCCAAGAACACACTCCAGAAACTTACTTCTTGGATGGATACGAATGATACCGCTTCAGGACACTTCTCTTTGCAATTAGACCCTAATCAAAATGGTGACTTAGTCATACTACGTAACCTCGTGGAAATGTGGATCTGGGTGCCTTTGAATCTCGTGGTATCCAACTTCAGTTATAATCCAACTGAACATAGTGCATTTATTACATACAATGTCTTTGGTGAATCTGTCATTTCTCGAATTGTCCTAGATCATAGCGGGTTCATGAGGGAACTCTTCTGGTCTAATGAACGTCAGAATTGGATGGTTGTGTGGTCTGTACCTGTTGATATGTGTGATGCAACGAGTTTATGTGGTGCTTTTGGAATTTGTGATATGAATACCAGTCCAATCTGTGGATGTTTACAAGGTTATGAACCCAAGTTTCGATTAGGTTGGGAAACTAATGATTATTCTGGTGGGTGTGTGAGAATAACTCCGCTGCAGTGTGGCAGCAATAACTATGGTTTTGTTCGTATGCAAAATGTTGAACTTCCTGCATCATCTGAATCAGTGCAAGTGGGGAATGATCAGATCTGTGAATATATATGTTCATCCAATTGCTCATGCAATGCTTATACTTACAGCAATAGTGGAGTGTGCTTATTATGGAATGATGATCTTATCAATCTGAGAAGATTATCTCATAATTCACGTCAAGCAGTGTTAAGCATAAAAATGTTTGAGCGTTCAAACAAAGGTTAGGTGCTTCTAAACTGCATCTTGTTCTAATTGTGCTTACTTGAAACTGACATTATGTTCGTTTTCTCCTCTCCCCATCTATCTTATTGCTCAGGCAATAAAAAGTCATCGTTTGTAGTGCTAGTAGCATCCATCACTTCAGCAATTTTTGTCTGTGGTACATGCTGTTACTTTCTATGGAGACGAAAGCTCAAGAAAAAAGGTACTTCTTTATACTTCCAATAGCTATGCCAGCGCTTTATGAATGTTAATGTGGCCTACATTGGTTGAGGCCATGAGGGTATAGGATCTTGTCTCCTTATATTGTCTTGGCTATCCTCACCTCATGAACTAGCTTTCAATCGAGATTGACATTTTCTTAACAATGAACATATGTGGGATGCCAATAAACGAACCTAGTCCTTCTTGATTCATTATTAAACTGTGCAGTCTTCGTGTGATATTCTAGTTCTTACTAGTAGTAGCAGGTTATCTGACTTAACTGTGTTCTCGCTGTCACACTCACTACAATCATGGATTAGATTATAAGTTAAAATTACCAATGTACAGCTTCATCTATAGACTTCATGTTTCTATTGCCTTTTCAGGGATTTTGAGAAAGATGAAGTTCAGGGAAATGTTACTTTCTGACTCAGCAACTAACATGTCAAAACCTGGAACTTCCATTGGAAAGGGACAAGAAAAGAACGGTGACATTGAACTGAAGTTCTTTGAACTACATGACTTGAAAGCCGCGACAGACAATTTTTCCCCTGACAATAAGCTTGGAGAAGGAGGTTTTGGTCCTGTTTTTAAGGTATTTTTACTACAACCTGAATACCTGCAAGTCTGCAACACAAATATGCGCTCACACCTAAAGAAAATGGAAAGAAATCCCTCCGGAAATAGCCATAGTACATATCCTGATTTTAGGTAATATCTTTGCTAGCTGTACTTATGTTTTGGAGGTGTGACAAATGTTTTTTTGCAATTTCTAGGGTCAATTGCCAGATGGGCAGAAAATAGCTGTGAAAAGGCTGTCAACTCAATCAAGGCAAGGCATTAGTGAATTCAAAACTGAGGCCCTTCTCATTGCAAAACTCCAACATAGAAACCTTGTTAGGTTCCTTGGCTGCTGTGTAGAAGAAGAGGAGAAAATGTTGGTATATGAATATATGCCAAATAAAAGTCTAGACTATTTCATATTCGGTATGATAATTTACTACTATCTGTTTTTGTGTTAACCCTTTCCTCCTGTCAAATGTGTGCTTACTGGATAAATTTGATATGTACAACTTAGACAAGAGCAGGAGGTCGTTGCTTGATTGGAAAAAGAGGCATGAGATCATTATTGGTATAGCTAGAGGGATACTCTATCTGCATCAGGACTCAAGATTGAGAGTAATCCACCGTGATCTGAAAGCCAGCAACATTCTTTTAGATGAAGACATGAACCCCAAGATATCAGATTTTGGTACAGCAAGAATATTTAGCGGAAACCAAGATGAAGCAAATACTCTTCGAATAGTTGGCACATAGTAAGCAAGCAAATACTGAAAGGTCTTTTCCCATACTACTTACCAATATTTTGCCTACTCATTATAGCTCCTTTTCTATGCAGTGGCTATATGTCACCTGAATATGCTCTTGCTGGTCTCTTCTCAGTAAAATCTGATGTCTTTAGCTTTGGTGTCATATTATTGGAGATCATTAGTGGAAAAAAGAACAGAATCTCTTATAATAGTGATTCCCCTCCAAATTTGATACGACAGGTAAGTTGTAAAATGCAAATGTTCGAATCTTTCTTTGATTAAGAATCAAATTTATCCATTTCAATGAACTAATGATTCAGGCCTGGGAGCTTTGGAATGATGGCAAGGCCTTCGAGCTGATAGATCCAACCATAGTTGATTCTTGCCCCAGTGAGGAAGCACTACGCTGTATTCAAGTAGGACTTTTGTGTTTGCAAGTGAATGCAGGGGACAGACCAACCATGTCATCAGTTCTATTCATGCTAAGTAATGAAGCAACGATTCCCTCCCCAAAGCAGCCATTGATTACCCCTAACTCTGACTCGGGTACAACAGAAACAACACCATCTTCGATCAATGAGGTGACTATTACTGTACTTGATGGTCGTTAGTCAGCTGATGAGTGATGAATCATCAGTCATCAGCCAATCCGTTCGAGTGTGGGAGGGTTATTGACCCGTCCATTTACTAGCTCAACGAATTGGGTTGATATACTGTCCAAATTGATTCAGGAGAGACTGTCAAAATAATTTTCAGAAGACATTTTTTTATTTGATATGTTATATATATAGTTACCATAAAGAAACCAATAATTTTAGGTACTAAAAATTCATAAAAGAACAAATAAAATAATTAAACCTTACTAAGAATTGAACAAATTGAGTTACAACTCACATTGCAGTCCTGTTTCAGTCAAATTTAATCCACTCAAATTCTAATCAACCCGTCTAATTGACACGTTAGATC

>M6-PGSC0003DMG401019239 M6 G-LecRK

CAAATAGCCGTACCACTAACTGATTAAATTAAAAATAGTTACATTGGGATAAAACGCTTCCTAACAAAGGCGACTTCATAAGACTGAAACCCGAGACTTTTAATTAAGAATGAAGGAGTCATATCACCACAACTGTGTCAGTGATTTGATGTTGTTGTAAGCAAATAAGATAGAGCATGGCCCCAACATCCATGTGTGGCTGTAATTAAATTCAACTATTCATTATAAACAAATAATCTCAAAACCAAAGAATACGATGAGTATTAATCAAACCAGCCAAGCTACGGTGTGTTCAAATAGGACTCAGGTCAAGTGTCATCAATATGCTGCATGAACTGGTCATAAGAGTCGGAAGGCAGACTTGCTAAGCACTTCGACATAGAGGAAAGACAACACAACGTCTATTGGATCGCGAAGCATGAAGGAATTACTAAGGGAGAGTTTTCACTTATTCTTGCTCATGTATTTGTATTTTGTTCAACGCAGCATTGGCGCTACAGATACCATAAGTACAACCCACTTTCTGAAAGATGGTGATGCCAATATTACTTCACCTGATGGAACCTTTGAAATGGGGTTTTTCAGTTCAGGTAGTTCCAAAAATTGCTACGTGGGTATGTGGTACAAAAGCTTATCTGTTAGGACAGTGGTGTGGGTTGCCAACAGAGAAGCTCCCTTGACCAGTGGATCAGGTATCCTTAAGGTTATTGAGCCAGGACATCTCGTCCTTCTCAATGACACTAACAATGTTGTATGGTCCACCAATACCTCTACATCTGTGCAGAATCCCATTGCACAGTTGCTGGATTCTGGGAATCTTGTTGTGAAACAAGCTGGTGATGATAATTTCCTTTGGCAGAGTTTTGATCACCCAACTGATACACTATTACCGGGCATGAAGCTTGGTTGGAACTTTTTAACTGACAGAGAGGTGTACTTGTCATCGTGGAAGACCCAAGAGGATCCAGCTCCAGGTGATTATACATATCACTGTGATCCTTCTGGATATCCACAGAACATCCTGAAGAAGGGGTCAAATGTTGTCTATCGATCAGGACCGTGGAATGGTCTTCATTTTAGTGGAACAAGAAACCCAAGAGGCAGTCATTTATACAAATATGAGATATTCTCAAGTAAGACACAGGTTTATTTTGGATATAAGCTCATATCTTCAGTTATCACAAGGTTGATACTAAATCAGAATGGTGCCCTAAACCTTTGGACTTGGGGTGATCATGTATCTGATTGGATCCCTCAACTCTCAATACCCGCAGATAACTGTGATGCTTACAAACTGTGTGGCGCATACGGGACTTGCAACAGTCAAGATTTTACTGTATGTAGATGCTTAGACAGATTTGTGCCCAACAATAGTGAAGCTTGGAAGAAGAGAAACTGGTCAGGTGGCTGTGTTCGGAAAACAGAACTAAATTGCCTTCAAGGACATAGATTTTTGAAGTATTCGCATATCAAATTGCCAGACACACGGAATTCCTGGTCCGATGTGACTATGACACTTGAAGAATGCAAGAATACCTGCTCTAAAAATTGCTCTTGTATGGCTTACGCAAATCCTGACATACGCAATGGAGGAAGTGGATGCTTGTTGTGGTTCGAGGATCTGCTTGACATTAAGGTGGTTTCCGATGAAGGTCAAGATATCTACATAAGAATGGCTGCTTCTGAACCAGGTAGGTTTTGTTGTCACTTAAAGATTTTTGATGGATTACCTGAAAATAGTAAAAGAATGTATGGTTGCAGTCACTTCTTTGGAAACAATTGATAGGTATTTAGTTTTCTGTAGAACAAGAAGTTGAGATGCATGTGGCTTGAATCAAAGAAAAAATCTTCTCAAAATTAAAGAGATTTATAGTAACATTCAAAAATTAAAAATGACTTGCATTATATTGAATTCTAGAACCCTTACAACTCAATATCAACAAATTCAAAATGATGTGCAGTACTCCAAAGCTTACAGAATCCTTTTTAATTCAATATAAAAATTATCTCCCCCGTGGATTGGACAGAAAGTGAAATTGATTATTTTTGCGGAATGGATACTTGCAGATAGTCTGGAGAAATCAGATGGAAAGAAAAGAAAAGTACTTTTCTGGATTCTGCCATTATCAGTTGGTCTGATTCTGGTAATTCTTAGCATGTTGATCTTCCATAGAAGAAGGAAGAAGGCTGTACAGCTCAAAAGGAAAGGTAAAATAGGATAAATCTCATTTTGTCTTGACATCTTAGCAAAATCAAGGAACATAATTCAAAGAAAAGTGAAGAATTTTAATATTTGGTCTGGATCACAGCTTAAACAAAATTCACTTCGTTTGCGGGCAGGTAGGTCGGGACGCAATGGCAACTACAAGATGGATCATAGTGGTAATTGTGAGGAAGAATTTGAGATACCAATGTTTGACTTATCTACCATAATGAAGGCTACCAATAACTTTTCAATAGACAGAAAGATTGGAGAGGGGGGCTTTGGACCGGTTTACAAGGTATAACTGGTGCACAGGCAAAAACTTCCTGATCCTATAGAGTAAAGTTTCATCTTTAGATTCAGTACAGAAGTTGTATGATGAAATGCATAATGAAATAAAGCTTTGGCTTGTGAAACCAAAGGAGACATATACAATAGCTAAGGAAGAACAAGGATATGGTGGAACACAACTGATTGTCTCTTTCTAATTGGAACACAATTGACTCTGTGTACTTCCTACATCAAGTAAGGTAATGAAAATATTAACATGCAGAAATTGTGATGTTTAGGGCATACTTGAAGAACGGGAAATAGCTGTCAAACGGCTGTCCAGAACTTCCAAACAAGGAGAAGGTGAGTTCAAGAATGAAGTTTTATATGTTGCAAGACTTCAACACAGGAATCTGGTGAAGATTCTTGGTTGCTGCAGTGACGGGGAAGAAAAGATGTTGATCTATGAGTACTTGCCGAACGGGAGTCTTGATTCATTTATATTTGGTACTATTCTCCATCCAATATTGTGCAAATTAAATAAATAATTGTGTTTCAGTGATGATGCCTATAGTAGTACTTCATTAATCTTTTTTATTTTGAAGTAAAAGTAACTAGTAGTACTTTGGTAATTTTGTTCCAATTATGAAAGCACCATAAAGCAGGAAAAATATTGATAGACCAAATCTGTTGTGGGAGTATTTAATACTACTTTCTCCTATAACCAATTAGTCAGTTAATGACCCAACAAAAAAAATTTGGCAGATGACACACAAAGCAAGGCATTAGACTGGCCTAAGCGATTTCACATTATCAATGGTATAGCTCGGGGACTTATGTATCTGCATCAAGATTCTCAATTGAGAATAATTTACAGAGACCTGAAAGCTAACAACATTCTGCTAGACAAGGACATGAATCCAAAGATATCAGACTTCGGCTTGGCAAAAATATGTGAAGAGGATGACATTGGAGCCATGACAAACCGAGTAGTTGGAACATAGTAAGAACCTTCCACTGGATAGTATTATTTTGATAAGATTTACTGGATAATAATTTGAACCTAAGAAGAATCTTCAGTTAGTTAAGGCATGTTTTCCAGCATTACTAATTATAGTGTTGTGCACATTTTAATGCAGTGGGTACCTCTCACCGGAATATGCATTGCACGGAAAATACTCAGTAAAATCAGATGTATTTAGCTTTGGTATTTTAGTATTGGAAATTGTCAGTGGGAAAAGCAACAGAAAATTCTCTCATCCAGATCATAACCTTAATCTACTCGGGCATGTAAGATTTTACTTGTTTTATTTCTTATTTTTTTGATAACCGAGGTGTTGGAGCCAACATACTTGAGCCGAGGGTCTATAAAAAATAGTCTCTCTACCAACGAGGTAGTGGTAAGGCCTACATACACACTACCCTCCCCAAACTCCACTAATGGGATTATACTAGGTATATTGTTGTTGTCGTGGTGCCGAGGCCAGCTTGCGCACACCTCAACTTATTTCACGGGATACCTGCAAACTCCCACCACCAATAGGTACCGGATAACTCTGTCTAACAAGGTTAGGACTGATGGGAAGAAATTGCCTAGTGTTTTTGTCTCTGCTGGAATTTGAAACAGATCATCGACCACTAGGTCACACCCTTGGGTGCAGTTGTTTAGTTTTATTTATTCCTTGGTGTATGTTATATACTAGTAACAAATATCTTGTAAAACGACCCCTGCTAATAAGCTTGCAATATCTACAGGCATGGAAACTTTATAAACAAGGTAGGTCAATAGAACTACTCGATGAATGCCTAGGTGATTCTTGTCAACATCTGAAGTGGTACGATCAATTAGTGTTGGTTTATTATGTGTCCAACAATGTCCTGAAGATCGTCCAAGTATGTCTTCCGCAGTTCTGATGTTAAACAATGAAGGTGTGTTGCCGTTGGCTAAACAACCAGGTTTTTACATAGAAGGAAATGCACCGGGTGGTGATTTCACTCCTAGCCAACCTGCACAGATTACAGTGAGTGACACCCCCATCACAATACTAGCTGCTCGATAGAAGATATTGGTTCACTTGTTTGCAACAAGCTTGCAGTTGCAATGATCTTTTCTTAATTCTCTATAGTAATGTGGTGAAAAGAATTTGTGCATGATTGTGCATTTTTAGGTCTATCAACTATGTCAAACATATGTAGGAAGCTTGCTGACTGGCTGTTACTTTGATTTGCGCAGTATATCTGAT

>M6-PGSC0003DMG401019242 M6 G-LecRK

ATGTTACTAACTTCAGCAGCATTAGACACAATCACTACAGATAAATCCATTAGAGATGGTGACACAATTGTTTCAGCTGGAGGAATTTATGAGCTTGGATTTTTCAGCCCTGGAAATTCTAAGAATCGTTACGTTGGCATATGGTACAAGAAGATATCACCTAGAACTGTTGTGTGGGTTGCAAACAGAAACATTCCACTTAATGACACTTCAGGAGTGTTAACACTTAATCCCAATGGAATTCTTGTACTTGTTGATAAATCCAATGTCTCAATTTGGTCATCAAACTCATCAAGATCCTTAAAGAATCCAAAAGCACAGCTCCTGGATTCCGCGAACCTTGTTGTCAGTGATGGAAATGATAGAGGCCTGGAAAATAATTTCGCGTGGCGGAGTTTTGATTATCCAGGAAATACTTTGTTACCTGGTATGAGGATAGGAAAAGATTTGGTCACGGGAATGGATTGGCATCTAACGTCATGGAAGAGCACGGATGATCCAACCCCTGGTGAATATGTAGATCGTGTTGATTCACATGGATACCCTCAATTGTTCTTGATGAGAAATTCATCTGTAGTGTTTAGCTCAGGGCCATGGACTGGTGCTTCATTTTCTAGTAGTCCTAGTAATAAACCAACATTGTATTATACGTTCGAGTTTGTTATCACTCAGAAGGAAATTTACTTCAAATACGAGCTTAAGAATGACTCTATGCCCACCAGGGTGGTGCTCAATCCGGATGGGGTGATACAACACCTAATATGGATTGAGCATACTCAGAGCTGGTTTCTCTACTTGACAGCACAACTTGATAATTGTGATCGTTTTGCTTTATGCGGACCTTATTCAAGTTGCAACATCAATAACTCCCCTCCATGTGACTGTTTGAAAGGTTTTGAGCCTAGGTTTGTATTTGTTAAACTATGTGATCATCTCGCGTTAAGCTATCAGAGAGAGTGCACTTTTGAATTTTGATTACTAAGTTATGTTTTCACACACCCCTCATGTGTGTGCTTGATTCTTTTTCTTGGACCAAACACATATAATTATTTTTTGCCTTCACAGGTATCCTCAAGATTCTGCAGCAGACTGGTCTAGTGGTTGCGTAAGGAGAACTTCTTTAAATTGTAACCATGATGGTTTTCTTAAATTTACGGGTATCAAGATGCCGGATTCTAGAAACTCCTGGTATAATGAGAGAATGAACCTTGAAGAATGCGAGAAAATGTGCTTAGCTAATTGCAATTGTACAGCCTACTCAGATCTTGATGTTAGAAATGGCGGAAGTGGATGCTTACTATGGTTTGGAGAACTCATAGATATACGAGAGTTCAGCCAAAATGAGCAAAATCTGTATGTGAGAGTAGCTGCTTCAGAATTAGGTAATCATCCTACTCTGCATAAATGCACAACTCTTTAGAATTGAAGGAACTCTACACCTAATATACTTACATGATGGACATTAAAGTTGATTAGGTGCTTTAGTAGTTGGTAATATGGCGATTTCATTTTCTGTGGTACTGGAACGGAAGTAGCTTGCTGCAAAGAGCATTTGAATAATTTCATATCTTACTCTACGACTATATGGATACTAGGACATAACCGGTTAGTAGGATTGCCAAGGGATGAGACTTCATTCCCTTTAAAGGCTTAAAAATATGCTAGACCTCATGACAGATCAAGGCTGTTGACAGCAGGTGGTAATCATATGCATTCTGTCCTTAGTAGTGCCGCTTGTAATCAGTGACACTGTACTTTAATGTGTCATCTTTCAAAGAAGATTTCCGATTTAACAATGCAGCTTGAAATCAATTAAACATTGACTACTTTTTCAGTAATCTTGTTGATTGAATGCCCTGGTCAACAAGTTACAAAGTCTGCAGGGCTTCATAACAAAATAAGAAGTAATACAATCAATGCAATTAACTACTGATATGCCTAACTCTTATGTAAATTGAATACTGATTATGAACTAATATAGAACATGTTCAGACAGGACGCGGAGGAGAAAGAGGTCAGTCATGATTGGTGTCATTTCAGCAGTGGTAGCAACACTTATCCTCAGCATTTTAGCTTGGTTTTACTTCCAAAGAAGGAAAAGAAGAAGAAGAATAGGTAATATAGCTGAGGTACTCTTGTCTATCTGGCTGACATTGTTCGCCTTTACCAAATTCTCATTGATATCATGTTTGTGGCAGGCGAATGTATATTGACAGGTTCAAAAGTTGAAAATGAGGATATGGAGCTTCCATTGTTTGATTTAGTTACTGTTACTAGTGCCACTGGGAACTTCTCTTCTGCTAATGTGATTGGGGAAGGCGGCTTTGGACCAGTTTACAAGGTAAATCAACTTTGATCTTCAACTTCTAGTTAGAAGTGTCATCTTTAATCCTTGATAGAAAATTGCAGGGTATCCTACCAAATGGACAAGCAATAGCAGTAAAGAGGCTATCGGAGTATTCTGGACAAGGCATTCAAGAGTTAAAAAATGAAATCGTTCTCATTTCCAAGTTGCAACATAGGAATCTTGTCAAGCTATTGGGTTGCTGCCTTGAAGGAGAAGAACGGATGCTAATCTATGAGTTCATGCCCAACGCTAGCTTGGACTATTTCATTTTTGGTCTGCTTTTTCGTGACTTATTGGCTATACTAGTACTTGACAAGTCTTCAATATCTCTGATCACTCAATGATTAGTTCTACATTGTGCAGATCCAAGCAGAAAAGCTTCACTTGGATGGAAGAATCGTTTTGAAATTGCTATGGGAATATCTCGTGGTCTTCTTTACCTTCACCAGGATTCAAGATTAAGAATTATTCACCGAGATCTCAAGACCAGCAACATTTTATTAGATAGTGACATGAATGCCAAAATTTCTGACTTCGGCCTTGCCAAAATTTTTGGTGGAGACCAAGAGGAAGGAAAAACTAAGAGAGTAATAGGGACATAGTAAGTTTCTACACATCTATAAATCATAGAACCTTTATTACATTTAATCAGTACATCATGAGCATAAATATATTGAGTTGATGCAGTGGATATATGTCCCCGGAGTATGCTGTTGATGGGAAATATTCAGTGAAATCAGATGTATTCAGTATCGGCGTAATCATTCTTGAAATAGTTAGTGGCAGAAAGAACAGAAAATTTCGTCATATGGAGCATCATCACAATCTCTTGGGACATGTAAGCACATATATATGGAACAAAACTTCGTTCCCTTTATTCACTCTTAGCTTTCTATTGTGATTAATTACAACTGTATAAATGTGTTGTTCTGTTTGTAATTTAGGCATGGTTACTTTGGAGTGAAGGCAACGCGTTGGAACTGATAGACGAATGTATCAAAGAATCATTTTCAGAATCTCAAGTGTTGAGATGCATCCAGGTTGGTTTGTTATGCGTCCAAAAACTCCCCGAGGATAGGCCTACAATGGCATCAGTAGTTTTCTGGTTAGGCAATGAAGGTCTAGCTCTTCCTCAACCAAAGCAACCTGGTTTTTTCATCGAGAGGAATTCTATGGAATCAACAGAGTCATCAGCTGATGAAGTATATGTAAGTAGCAGCGTGTCGATAACAGTTCTAGAGCCAAGATAA

>M6-PGSC0003DMG401029444 M6 G-LecRK

TGGAATAAAATGAAAAAGCACTTAGGTTTAATGTCAATTTGTGCAACAATACATTTTTCATTTCACTAGGTGCATTATTAGTTGACAATCATGTTGATGATGAGGTGCTAGCTTATTGTCAAGGTCCCCACTCAATTCTCACACTAACGTGGCTTTTGGATTTTCACTCATTTTACTTTTTCGAAATAGTTTAAATATATCAATCATTATATTTCTGGTCTAACGTTGAGTTAAGCTTAGATGTCATATCTTTACATCAATACCCATTTTATTTTATTTTCTTCTTCTTTTTTCAATGAGCATTTAATAATTTAAGGAAAAGAAATGTGTTATTTCAACCAAGTTAAAACTTGTCAATTCACAATGTTATTTCAGGAACATACTACACAATGATAGTGATGGGAGAAAAGACTTATATTGTCAGCGCGTATAGAAGTTAAACTCTACATTTTTCATTATTCGTTTTTAAGTGTCAATATTTCCATTGGCCTAAAAGTCTCTGATTGAGAATGGTATGTGGTTTGTGATATTTCTACAGAGTCAACAAATTAAAAATCCTATTTGGCTGCTTGTGCTATTAGACTTTGACAACATATTGTGCTTTTGTTCATATAGTGTGGTAATAAGAAAAGTCTTAGAAAACCAAAAAGATGAGCAACAGCAGCAGCAGAAGAAGAAATTTGCAATATTTTGTTCATATAATTCTTGTTATTCTTCGTTTTATCGATACAGGATTATGCAGTGAAGTGGATAACATTACCAGCATTCAATCTCTGAGAGATCCTGGAATTTTATCGTCTCCAGGAGGCTTATTCAAGTTAGGATTTTTCAGTCCTCAAAACAGTACTAATAGGTATGTTGGTATTTGGTACAATTTTTCTGTAACAACTGTCATTTGGGTGGCTAATAGGGACAAACCTTTAAGAGATTCTTCTGGTGTTGTAAAGATATCTCGTGATGGAAATGTCGTTATAACGAATGGAGAGGAGGAGATTCTTTGGTCATCAAATGTTTCTACTTCTCAGGTAAACTCAATTGGCCTTCTCCAAGATTCTGGGAACTTTGTACTCGTAGATTATCGAAACAATATGAGCACAATATGGCAGAGTTTTGAACATCCTTCTGATTCAACTATTCCTAGAATGAGAATAAGTGAAAATACAAGGACAGGGGAGGTGGTAGAAGCAAGATCTTGGAGAAGTCCTTCGGATCCTAATATCGGAGACTTTTCTTTAAGAATGAACTCTGGAGTCATTCCTCAGGTGTATATATGGAAAGGAAACCGTCCCTATTGGCGAACTGGTCAATGGAATGGACAGATCTTTATAGGAGTGCAGAATATGTATGCTGTGGTATCTGATGGATTTAATGTAGTGAATGATCGTGAAGGTACTGTATATTTTACTGGACCTATTCGTGATAATTTTTCAAGGATATTAGTCTTGGATTGGAGAGGGAACTTGGTTCAATCATATTGGAATGTGACTGAGACAAACTGGAAGATAATATGGTCAGCTCCCAGTAACGATTGTGAAGTTTATGGAACGTGTGGTCCATTTGGGAGCTGCAATCATTTGGAGTCGCCGGTCTGTTCTTGTCTGAAAGGTTTCGAGCCAAAGCATAGGGAAGAATGGGAAAAGGGGAATTGGACTAGTGGTTGTGTTAGGAGGAGTGCTTTGCAATGTGAAGTTAAGAATAACACCGCGAATTCAAGTAAAGAAGACGGATTTCTAAAGATGGAGTTGATGAAATTGCCTGATTTTGCTGAAAGGTCATCTACTACAGAAGACTTATGTAGAAGCCAATGCTTGGGGAATTGTTCCTGCATGGGGTATGCATTTGATTCAGGTATCGGCTGTATGTCGTGGAGTGAAATGATTGACATTCAGCAGTTCCAAAGCTCGGGGAAAGATCTCTTTATTCATGTGGCACATTCAGAGCTTGGTAAGTATTCTGGTGCTGAATAAAAGACTCATTGTCAGTCATTTGAGATAGACATAATGCATAAACGTGCCCTTTTACTTGGCCTCAATTCACATCTATGCTCTCATGCACACTCAAAGTTAGATGGCATGGATGTCAGCAAAGGCCAAGTCAAAAGACACGTTTATGTATTATGCCTTTGAGATATGTTCGACTTAATTGAATTCCATGTATTCTGTAAAGAACTGATTTTTGTTCACCTTTAGTGTTTTCTGCAGATCATGGTAAAGACATAAAGAAAATCGTAATTCCAGTAATCGTTGGCTCTCTTACACTCTGTGTTTGTCTGTTCCTTTGCTATATAATGGTGATCAGACGTAGAGGTAAACGTGCTATTAAGTTTTTGTATTATACATTTAGAAGCACTTTGTTCGCTATAGTTAACTTGTATTCCTGCTGATATTAGGAGTGAAAATAGAGGAGGTAGCTTTACTTGGTAACAAAAGTTCAGTTAATATGGAAGAATTACCAGTCTTCAGCCTCGATACGATTGCAAATGCAACATCCCAATTCAATGAGGATAATAAGCTTGGTCAGGGCGGTTTTGGTCCAGTTTACAAGGTACGAGTTTCATCTCTCTCGATCACTTCCCATATTTCAAGTTACTGATATTGGTGATTCCTTGTAGGGGAAATTGGAAGATGGGAAAGAAATAGCTGTCAAGAGGCTTTCAAAAGCCTCGAAACAAGGGCTAGAGGAGTTCATGAATGAAGTGTTGGTGATCTCGAAAGTCCAACATAGAAACCTTGTTAGACTCTGTGGATGTTGTGTTGATAAAGAGGAGAAGATGTTGATTTATGAATATATGCCAAAGAAAAGCTTAGATGTGTTTCTCTTTGGTTAGTCCTTGTTACTTCTCATCTGCATTACTCCACGTCTATGTTGCTCGGACTCTTCAATAATGTTGATGGATGCTTGCTAGATCCTCCAAAAGTACTGCATTTTTGGAGGATCTGACATGATTACATCAGCATTCTGAGCAACATTACTCCACATTTCTTCTGATCCTTTTGTTTAAATTCTGCATTCATATTTGACAAACATCTAGATGAAGGACATCGAGACATTTTGGATTGGACAAAACTTTCCATAATAATCGAAGGAGTTGGTCGAGGACTCCTTTATCTACACAGAGATTCAAGATTGAAGATAATTCATAGAGATCTAAAGCCAAGTAACATCTTGCTCGATAATAACTTCAATCCAAAGATTTCAGATTTTGGCATGGCTAGGATTTTCGGATCTGATCAAGATCAAGCAGACACAATGAGAGTAGTTGGTACTTAGTAAGTGATCATTTATCTATATACAATACCATAACTTTGCATAATTTTCCACCATATTTACCAACTAAACTACTCGAATCGTACTGACATTTTTACAGTGGATACATGGCCCCCGAATATGCAATGGAAGGAAGATTCTCAGAGAAATCCGATGTTTTCAGCTTTGGAGTTCTAGTCTTAGAGATCATAAGTGGTCGAAAAAGTTCAAGTTCTTGGACTGAAACATCCTCTTTGAGCCTTTTGGGATATGTAAGACTCCTCTTGTAGTTGCTATAGTGAAATGTTTGTTATAAAGAATGAATAATATAACATAACATGAAAAATCAGTTCCAAAAAAGAACTTGGCTGTTATATAATGAAATGATATTATAAAGGACGACTGTTATAAAGAGATGCGACTGACTATAGGTGCCTTATATTTAGTTTGATGATCAACTTTTCATTGTTTTCAGGCATGGAAGTTATGGAAAGAACAAGATTTATCAACTTTTATCGATCCGTTTATATTGAATACGAGCTCGGAAATGGAGATCAGAAAATGCATACAGATTGGTTTACTATGTGTTCAAGAATTTGCTGAAGACAGTCCAAGTATTTCATCTGTTCTTGTTATGCTTACTAGTGAAACAAGTAGTCTTCCAGCACCATCACAACCTGCTTTTACTGAAAGAAGACACTTCAGAATGTGCAATGAAAATAGAGAAACTAAGTTTACTTTGAACAAAATGAGTATCACAAACCTTACTGGTAGATAACATGAGCATATAAAAGTACTTTTTTAAGATTTGCACAATTGTTATAGTATGATCCATTTTGTCTCAATTTGTATGATACCTTTTGTTTCAAAATGTTTGACAATACTGGATCCGTGTAAAGCACGGAGTTTTTGTCACTAGTTTCAAA

>M6-PGSC0003DMG402003983 M6 G-LecRK

GAGCTAAAACAGGCAACGGATGGTTTCAAAGAGGAATTAGGAAAGGGTGCTTTTGGAGCAGTATATAAAGGGAGCTTTGACAAAGGTAAAAACCTTGTGGCAGTGAAGAGGCTAGAGAAGGTAGTGGAAGAAGGTGAAAGGGAGTTCAGAGCAGAAATGAAGGTCATTGGTAGAACTCGACACAGAAACTTGATTCGTTTGCTAGGATATTGTGCTGAAGGATCAAAAAGAGCTCTTGTTTATGAATATATGAGCAATGGTTGCCTAGCAAATCTGTTGTTCCGAGGAGCTACTCGGCCGGATTGGAATCTAAGAGTAAATATTGCACTAGATGTGGCCAGGGGAATCCTCTACCTCCATGAAGAATGTGAAGCTCCAATTATCCACTGTGATATAAAGCCACAAAACATCTTACTCGACGAGTTGTTGACAGCTAAAATCTCAGATTTTGGTCTGGCCAAGCTGCTAATGCCTGATCAAACAAGAACATTCACTGGTGTAAGAGGGACTAGAGGGTATTTAGCACCAGAGTGGCA

>M6-PGSC0003DMG402004946 M6 G-LecRK

ATTTTAAGGCCTTCAGAACTAGCTAGATTTACACTGAAAAGGTCAGGTTACCTCCAGATATCTAAACCTGCATAGGATAGGTAGCATGTGATAGAGCAGTTCTTAATGGAAAGTTTTGTGCTTATATTGCAGATATTGTTATGTTGTTTTGTATGCTCAATTACTAGTGCAAAAGATATTATCACTACAACTGACTTTCTAACAGATGGAAAGACCATTACTTCATCTGATGGAAGTTTTGAGATGGGATTTTTTTGCCCGGCTAGTTTCACGAATAATTGGTATATTGGAATATGGTATAAGCACGATGTACCTGATAAGTCTGTAGTATGGGTTGCCAATAGAGAAAATCCATTCAACAACACATCAGGTGTTATGTTGAAAATCATCGATACAGGACAACTTGCTCTTCTCACTGCTGCTAATAAAATCGTGTGGTCTACTAATTCATCGAGGCCTTTGGCTGTGAAGAATACAGTTGCACAGTTGTTGAACTCGGGGAATCTTGTCATAAGAGATGCAAATGATACCAAGCCTGAGAAGTTCCTGTGGCAGAGTTTTGATTATCCTACTGATACACTCTTACCAGGCATGAAGATGGGTAAGAACTTTGTAACTGGACAAGAATTTTACCTGTCCTCATGGAAGAACGAATATGATCCAGCTCCAGGGGAATATACGTATCATTGTGATCCGACTGGATATCCTCAGGATGTCATGAGGAAAGGTAAAGTTAAGGTGTTCAGTACTGGACCATGGAATGGACTCCGCTGGAGTGGTGTACCAGGACTGACAAAAAATACTATTTATACATTTAAACTGGATCTTGATGAAAAGAGAGCATTTTACAGCTTCGCGCTTTTAGGTTCTGTTATGACCAAATTAACCATGAACAGTAATGGTGTGCTTCAGCGTTCAATGTGGGCTGAGAATAGACAGGAATGGCATGTTTACATTTCTTCACCGGAAGATACTTGTGACAATTATGGAACATGTGGTTCATATGGTAGTTGCAACAACATACTCACTCCTGTGTGTAATTGCTTAGACAAGTTTGTGCCTAAAGATCCAAGAAATTGGGCAATGACAAATTGGTCCAGCGGGTGTATTAGAAGGAAACCGCTTAATTGCCAGAATGGAGATGGATTTCTCAAGTACTCTGGTATAAAGCTACCAGATACACAGTATTCATGGTTTGATAGGAGTATGACACTCCACGAGTGCAAGCAAGCTTGCTTGAGGAATTGCTCGTGTATGGCTTACTCAAATCTTGATATACACAATGGTGGGAGTGGCTGTTTGCTTTGGTGTGGCGACTTAATTGACATTCTAGAATTACCTGGAGGACAGGACATTTACATAAGGGTTGCTACCTCAGAGTTAGGTAACATTTGTGAATTTGATCAAAGTTGGAGTATTAGCTGAATGATTTAAACTTCCCAATTCTCAGATTTAGTTGTATTCTACTTAGATATCTGTTAAAAGTTCACACAGTATATGTTCACTCTTCCATGTCTATTTAGTTTATTTCACCCCTCCATGTTTTGTTCAGGCTCGAAGAAAACAAAACTACTCGTGTTGAGTCTGTTACTGTTGATTGGAGTGACTGTGACTGGCCTGACCATAGGGCTGTATATTTGGAAAAAGAAGAAGAAGAGGAAGATGAATTTGAAAGATGACCTAGACCTCCCACTGTTTACCTTATCAACTCTAACTAAAGCATCCTCTAATTTCTCAATCGAGAACAAGATTGGAGAAGGTGGATTTGGATCTGTTTACAAGGTAACATCTTAAGTTTATGAATCATTCTTTCAGTAAATAATATCTACTATTATTGACAGTAAACAGATGACACTTACTGAATAACTAAGCCAGATATATGTTGATATTATAGGGAATCCTGGAAGGTGGACACGAAGTAGCTATAAAGAGGCTATCCAAGTCTTCCTCGCAAGGAGTTAACGAGTTCAAGAATGAAGTTATTTGCATTGCCAAACTTCAGCATCGAAATCTTGTGAAGCTTATAGGTTGCTGCATTGCAGGAGGAGAAAAAATGTTGGTCTATGAATACATGTGCAACAGAAGTCTAGATCTCTTCATATTTGGTCTGGTTACTCTCTCTTAACTCGAATAACCTCTATTAGTACAAATAGTAAAGTTGATAGTTTAATCACATTATCAGTGTTCTAACAGAGGACCTGTTAACAGATGAAAAGAGGAGCTTGTTACTCAACTGGCCTAAGCGCTTCGACATCATCAATGGAATTGGTAGAGGACTTCTGTATCTCCATCAAGATTCTCGGCTAAGAATTATCCACAGAGACCTCAAAGCTAGTAACATTTTGCTAGATGCTGACATGAACCCCAAGATCTCAGATTTTGGCATAGCTAGAAGCGTTGTAGGGAATGAGACTGGTGCTAATACACACCATGTGGTTGGGACACAGTAAGAAATTTAATACTACTCACTTCTTAGCATTTATATTATGTATAATATCCTTTATTTTACATTACTTTCTGTTTGCAGTGGTTACATGTCCCCAGAGTATGTAGTCCATGGTGTTTTTTCAGTAAAATCAGATGTATTTAGCTTTGGCGTATTAGTGCTAGAGATAATCAGTGGTCGAAGAAACAGAGGATTTGCCCGTGAAAGTCACAGTATCAACCTACTCGGACATGTAAGTCTTGTTTGTAATTAAAGACGAGTTTAAATTTTGTGCACTATATGGTAGCTACAGATATTTCTCTCTAATAATTAATTGATAACACCTCGTACAAATGATAGTTAAACTGCATTGAACTTAAATCCATCAGGCACATCAACCGCTGCATGTTTCTTGATTATGTACTGATTGTGTTATTGATAGGTATGGAAACTTTATAAAGAAGGTAGACCATTAGAGCTAATTGATGGCCATCTAATGGATTCGCGTTATATTTCTGAATTGCTACGCTTAATCCATGTCGCTCTGTTATGTGTACAACAATGTCCAGAAGACAGGCCAGATATGCCTACAGTGATCTTGATGTTGACTAATGATGCTATTTTGCCTCAAGCTAAGGAACCTGGTTTCTTCACAGAAAGCAAAGTCGCGTCTGAATGTTCTACAAGCATGTGTTCAACAAACGAAATTACCGTCACACAGTTAGAACCCCGGTAAATTACATTACATGTTCATTCAGATCGTTCAATGTCAACAAGCTTGAATTACAGTGTTTTGTGCTGTAAAGCCATAATTTCCTATGGTATCTGTTATTAGTAAATATGGGTGTGTTGTAGTTTTTATTAATTCTAATGTATTTTAAAGTATTCGTACAAAGTACCTATTCAACAGTACATCAAAATAATCGAAAGAT

>M6-PGSC0003DMG402005047 M6 G-LecRK

CAATGGCTATCTCATGAAAAGAACCATAGCAATTGTGACAATCTGCATTGGTGCAATACTCATCGCGCTCTCGTTTGCTTGTTGTTTGGTAATAAGGAAAAGGAGAAGAGGTAAAGGTACCCCTTAATTTCTTCGTGTGTTTACTTCTTTATTGAATTCCCTTAGTGAAATCTGGCTCCGCGAATTACAATAGTACTAAGTTACTTCATGTACAGATAAGGAGGACCAATTCACCAGTTTGAACACTTTGACAAGGAATTTAGCATCCTATGAGAACTCTTCAAGAGGTAATGAGATGGATGGAAGTGAACACGTTGATGTGTTAATCTTTGATCTAAGCACTATCATTTCTTCCACTGATGATTTCTCTGATGCTAACAAACTCGGAGAAGGAGGCTTTGGTAGCGTTTACAAGGTACCTCCCGATATTTTACAACTTCTTACTCATTTGTTGAGCCAATTTTACTTGACAACAAGCTTACATTTTGATTTCAGGGACAGCTAAACAACGGCCAAGAGATAGCGGTCAAAAGACTTTCGAAAAATTCAGGGCAAGGAGTGGAGGAATTCAAGAATGAAGTAACGTTAATCGCGAGAGTACAACACAGGAATCTTGTGAGGCTTTTAGGATGTTGCATAGAAAGAGGAGAGAAAATGTTGATCTATGAATACTTGCCAAACAAAGGCTTAGACTGTTTCATCTTTGGTACATATATACATGTCCAGATTGTGTAAAAAATATAAATAGGCATGTTACTGATCCTTTCGAATCGTGTATTGCAGATAAAACTAAAGGGTTGCAGTTAGATTGGAGAAAACGATTTGAAATCATCGTTGGAATAGCACGAGGATTGTCATACCTACACCACGAGTCACGAGTACGAATCATACACAGGGATCTAAAAGCTAGCAATGTTTTACTAGATGCCTCAATGCACCCGAAAATATCAGATTTCGGAACTGCTAGGATATTTGGTGGTGACCAAGTTGAAGCTAACACGAATCGAGTCGTTGGAACATAGTAAGCATCACTTTTATGTATGAACATATGTGTATTTTTCTCAAAGGCACTTAAACTTGTATAAAGTTGAACAAGTAGACACACTCGTCCTACATGACGATTCGCATAAGATACACTCGAACTGATTTGTATTATGCCATGTAAGACGCATATTATGTCATGTAGAACGTTTGTGTCTACTTCAACTTTATATAAGTTTAAGTGTCTACTTGTGCACACCTCATAGATGTCAGCTGAGACTAAGTTAAATGGCACGTTTATTTATTATGCCAAATTAAAATTGTGGGTCTTGACATAGCTATTGGTTCATTTCAGTGGTTATATGTCACCAGAATATGCAATGGAAGGGCATTTTTCAGTGAAATCTGATGTTTTCAGTTTTGGTGTTCTGTTGTTGGAGATCATTACTGGCAGGAAGAACACTACACATTATCAGGATCATTCATTAAATTTAGTTGGAAATGTAAGATATCTCTTCATCAAGAACTCACCTTTAGTTAAGAAGATGTATATATCTTGTTTTGCTTTAGCCCTTATACCATTGCTCTAGAGAAGGATATGTGGAAATGTTCATCTCTTCTTTTTTTTCTTATACGATGGAGTCGGGGCTAGCTTGTGCATACTTTGACTATTACACTGTACTTGCTACCCTCCACCAGCATAGGGATGGAAACAATGGCGGAGCTACATGCATTGGAAGGGTATTCAAATTGAATCCCCTTCTTTAGGAAGCTATGATAGAAACTATTGGGTGTGTCTAAACAAAGTCCCACATGAAAAGTAAAAAAAAAACTACTTTAAGATGTCGGATAATCTTAATGTTATGAAGTCTTCTGAAAAAAACTATGCGGGTTTGGTCCAAGGGGACAATATCATGCCATGTTCAGAGTATCTTATTGGATTATTTTAGCCTAACAACTGATATCAGAGCCAGTGGTTTGGCAGGACAATTATTGAGATGGTAGAGTGATGGAGTGGGGTCCGACTTGGTGCCTTTGACCATCTATGGGCCGATTTACTATCTTTACCTGTAGCCTTAGAGATGCTTACACAAAAAGAAGGTAGTGGCCGTTTGTGACCATAAATACTCTATTCATATGGTGGCACGCAGTGAAATCTCCAGATAAGCCTATAAAGTGATAGGTCATGTGGAACTCAGTTTAAGGAGAAGACTGTTGAGTGTACGTAAACAAAGTTTCACATGAAAAAATAGAAAAAAGATAACTTATAAGGTATTGGATACTCTTAATATTGTGAGGCCTTTTGAGAAAACCGTTGGGATTTTGTCCGCAAACTGACAATGTCACATCATGTTAAAAGTATCTTTGAGTTATTTTAGCCCAGTAGAAACACATTCTTTCATCGAACATATATGAAATTTTGGACCCCCTTAACCCCAGACACGACATTTTTGTATTTACTTGAATTCCCTAATCGAAAATCCTGGCTCCGCCACTAACCAGAAATGTTGACTTTGCTCTCCCTTGTTTCAGGTTTGGGATTCATGGAATGATGACAGAGCTTTAGATGTTGTTGATCCCTCATTAGGGGATTGGTATGAAAGTAGTGAAGTTTTGAGATGCATCCAAATTGGACTATTATGTGTACAATCATATGCAAATGAAAGACCAATGATGTCTCAAGTTGTATTCATGCTTTGCAATGAAACAAATTTGTCTAATCCTGGACAACCTGGTTTTGTGTTCAGATCAAGAAATTCTTCTTCACTTCCTTATTCATCATCAGCTAGTATTGGAAATTCTGTAAATGACATTTCTATTACTGCACAACATGCTCGCTAAGTGTGTGTCGGATCCTTCAAAAGTAGTACCATATATTTTTGTGGATCTGACATTGGTGCATCAATATTTTTGAAGAGCACTGAGCCATATGGTCATTAGGTAATTTATACTTGATCTATTTATTGATTTAGTTTAAGATTTTTTACATTTATCAATACTGTATTTTAACGTGTTATAACAGGTAGCGTATCTT

>M6-PGSC0003DMG402007101 M6 G-LecRK

CAAGAAGCAAATGCACCATATCAAACAAGCTAAAACAATAAACTAAACATGTCTCCTAATTCATGGCCTACTACACTTCTTGCCTCTCTATTTCTTTTCTCCCAAATATTTTCTTGCATTGCCCAAGTTCCACTTGAAAACACCTTCAAATTCGTCAATGAAGGTGAATTAGGACCTTATGTTGTAGAATATCAAGCAGATTATCGCGTTCTTAGTATTTTCAATAATCCATTCCAGCTCTGTTTCTATAACACAACTCCTAACGCATGGACTCTAGCTTTGCGAATGGGCACTGTCCGTTCTGAATCACTCATGCGGTGGGTGTGGGAAGCAAATAGAGGAAACCCCGTTAAAGAAAACGCGACCCTCACGTTTGGAACTAACGGAAATCTCGTATTGGCTGACGCTGACGGTCGAATTGCTTGGCAGACCAACACGGCCAATAAGGGGGTGACAGGGTTCAAGTTGTTACCTAGTGGTAACATGGTGCTACATGACTCTAAGGGTAAATTCGTTTGGCAGAGTTTCAACTATCCCACTGACACATTACTAGTGGGCCAAACTCTCCGATTGTCTGGCCCGAATAAGCTCGTGAGCCGGGCCTCAGTGAAAAAGAATGTGAACGGGCCTTATAGCTTGGTGGTGCAGCCAAAATTGTTTGCTATCTACAATAGGACTAAGTTAAGCGTGGAACTAGCTTGGTTCGATTTTGAAAACAGTACGTTGGAATCAGTAAAATTGAATAGTGGGAATCAAAGATTGAAATTGGATTATAGATTGGCAAAATCAACAAAGAGAAGTTCACATGTGATGGCATTTACTAAATACAACACAACATTAACATATCTTAGACTTGAAATTGATGGAAATTTAAAGGCCTACACTTTTGTTAAAGATGAAGAAGCTGATGAATTCCGTTGGAAGGTAACTTACCAAATGCTATAAGAAAATGTTTTTTTTTTTAAATAATAATATTAAAAACTAACTTTATAAGATAAGGATTTCATTTCCAGTAAAATAAGTGGAGAAAACGAGTGGGTCCACAGCTATTATTTTCCTTTTTATTTTGGTTGAAGGGTGTATTTTATTATTATGATTTCTATTTTTCATTTCTGTTAAAGGATGTATTTAATTATGCTTTCTACTATTTAAGATGCTATGACAAATGTGAA

>M6-PGSC0003DMG402009323 M6 G-LecRK

ATTAAGCTTGGTTATAACCTCACCACAATGCAGAATTGGACTTTAACTTCTTGGTTGAGTTTTAGTGTTCCTGCCTCAGGGGCTTTTACTTTAAGTTTGGAGGTTTTTGAAAATGCATTTCAGTTGGTTATTCGTTGAAGGGGTAAGGTTTATTGGACTAGTGGGGCTTGGAACAATCAAGGTTTTCCATTCTTACCTTTATTCCATGATTCTGCTACTATTTATCAGTACAATCTCAATTTGGTATCCGATAAAGATGGGATGTTCCTCCAATTTGATGCAACAGAGGGAAGCTTTCCAAGTCTTGCATTGAATTCAGATGGGGATATTCTTGGTGGATACCGGTTTACATACACCCTTTTTAAAGACTTCTGTTATGGTTATGAAAGTGAGGACGGTTGTGTCTCTACCCAGTTGCCTGAGTGTCGAAAGGATGGGGATAAATTTGAGCAAAAGAGTGGAGACTTTATACATTCATCTAGTATTACTACTGAGTTTTATGACAATGCAAGCATTAGTATTGTTGATTGTATGCAGAAGTGCTGGGAGCACTGCAGTTGTTTTGGCTTCACCACCGCCAGTGGCAATGGGACAGGCTGCCGTATTTGGAATGGAAATGGTGATTTTCGTGTCTATGAGAGTGGTAATGCATTGCTAAGATATGTTCTAGTTTCACCAAAATCATCTAAAGGTAAGTCTTTATATCAGGAATACTCATTTTTATGTACTCAATTGTCAACTGTATAACTCTCAGCATGCTATGGGTGCAGGAAAAACTTGGATATGGATAGTAATTAGTGTAGCTTTTTTTATCTCGTTGCTATTATCTGGTTTCATCTGCTACATTATAATGAGAAGACGCAAACTACAAGGTAAATCACCAAACTGAGGATTTAGAAGCTAGTAGGTTGTTTAGTCATTTATTTTAAGCATGAATGTATATCTACATACTCTTCTTACATAGATGAGAAAAGAAATGAGGAAGCATATATACGCGAGTTGACAGCAGTAGACAGCTTCAACAACGCAAATATTAAAGAAGAAGATGGAAGGGAAGTACAAGATTTGAAGATATTTAGCTTTGGATTCATATTGGAAGCCACAAACAACTTCTCTAGTGAAAAGATGCTCGGAGAGGGTGGTTTTGGGCCTGTTTATAAGGTATATTTGCACTATGTTATATGTAGCTCAGAGATTTGCTTACTGCTCTCACTACATAACGAATCAACTGTAAAAATGGATGTCAGGGAAAATTTCCAGATGGTCGAGAGGTGGCAGTCAAGAGACTTTCAAGAACCTCGGGACAAGGGCTTGTGGAGTTCAAGAATGAACTCATACTAATTGCCAAAGTTCAGCATACAAATCTAGTTCGAGTTCTAGGATGTTGTATTCACGGAGATGAGATAATGTTGATATATGAGTACATGCCTAACAAGAGCTTGAACTTCTTCCTCTTTGGTCTGTTACCATTCCACAAAAAAAAGACAGTAATTTTTCCGGTATTTAAAAAAAAAATTGATGTAAAATGCTTTCTCTTTAACAGATCCTGCAAGAAAGGAGCTACTGGATTGGCAGAAGCGATTTGCAATTATCGAAGGAATTGCTCAAGGATTGCTCTATCTTCACAAATACTCAAGAATGAGAGTGATACATAGAGATCTAAAAGCCAGTAATGTGCTTTTGGATGAAAACATGAATCCTAAGATAGCTGATTTTGGTCTGGCAAGAATTTTCAAACAGAACGAGATGGAGGTAGTGACTGGCAGGGTTGTTGGAACATAGTGAGTTGAGCCTAGTCATTTCTTCGATATAAATCTATGGAACATGTGTACTTATTCATACTTACTGAGTAAAAAATGTTTTCTATCCTCTGCACAGTGGTTACATGGCTCCCGAGTTCGCCATGGAAGGTACTTTCTCAATCAAGTCTGATGTCTTCAGCTTTGGAGTCTTAATACTGGAAATTGTGAGTGGTAGGAGAAATACTAGCTTACAACAAATTGATCGTCCACTTAACTTAATAGGTTATGTAAGTTTTCTTTAAACCTTTCTTTCTTTAATGTACTAAAGTTCACAAAGTTCAACAATACTGATCATTTTGGTTCAGGCATGGGAACTATGGAAAGAAGGGTGTGGACTAGAACTGAAGGATCCATCTCTTGAAGATCTTTGTGACACTGAACAATTTTTACGAGTTATTCATGTCGGACTATTATGTGTACAAGAAGGTGCAGCAGATAGGCCAACAATGTCTGATGTTATCTCAATGCTTGGTAATGGAAGTATGTCACTGCCAATTGCAAAACAACCTGCATTTTTCACGGGAAGAGATGAAGCTGAATCATATTCGTCCACAATCAATACAGAAGAATGTTCGGTTAATGATTGTTCCATCACTGCCGTTGAAGCAAGAT

>M6-PGSC0003DMG402013876 M6 G-LecRK

TTAGTCCTCTATCAGACTTCCCCCAAAGTTTCATTGCCCATCACCTAGGAACTTTCTTTATTCTTCTCTTTCGTTTTTATGTAGTAGTGAGAAATGAAGATAATATTGGTTCTCTTGCTTTCTTACTTTCATTTAGTTAGCTGCAATAGTACACGTTCAGATACCTTGTTGCCTGGCCAGTCCTTTTCCTTTTCTGCAGAAGATACTTTAGTATCCAAACAAGGGAAGTTTGAACTCGGTTTCTTCTGTCCAGGTAATTGTGAAAAGCTTTTTATTGGTATATGGTATACAAACATAAAGCCCAAAACTGTGGTTTGGATAGCAAATAGGGACAATCCTATTCGTCCTCCGTTCAATAATTCTCATCTAGAGCTGTCTGATGGAAATCTACAACTCTTGGATGCACGTAAACAAAGAGTTTGGAGTTCTAGTTTATCCACACCTTCAGCATATGCCAATAAAGCAGTACTTCTTGACTCTGGAAATCTTGTTTTGACAAATGGCATTGACAGACAGTGGCAGAGCTTTAATTATCCTACAGATACGTGGCTGCCTGGTGTGATGATTGGATTTAATAAGTCCAAGAACACACTCCAGAAACTTACTTCTTGGAGGGATCTGAATGATCCAGCTTCAGGAAGCTACTCTTTGCAATTAGGCCAGAATCAAAATGGTGAGTTAGTTGTTCAACATAATTTCCAGGAAGAATGGAGGTCTGGGCCTTGGAATGAAGGAGGGTTTGCTATTCTAGGACAACAAAATTATTATAAGGATTTATTCGACTTCAGTTATGATCCAGCTGAATATAGTAAATATATTACTTACAATGTCTTTGATGAATCTGACATTTCTCGAATTGTCTTAGAATTTAATGGGTCGATGAAGCAATGGTTCTGGTCTAAAGATCATCAGAGTTGGCAGTTAGTGTGGTCTCGATCTACTGATAATTGTGACCAAATAAATTTCTGTGGTGCTTTTGGAATTTGTGACATGAATGCCAGTCCACTCTGTGAATGTTTAAAAGGTTATGACCCGAAGTTTAAATCAGATTGGGATATTAGAAACTATTCAGGTGGGTGTGTGAGAAAAATTCCAATGCAGTGTGGCAGCAATAACAAAAAATTTGTCCGTATGCCAAATGTTAAACTTCCTGTGTCATCTCAATCAATGAAAGGAGTAAATGATCAGATCTGTGAATATATATGTTCATCCAATTGCTCATGCAATGCTTATGCTTACAGCAATAGTGGAGAGTGCTTATTATGGTATAGCGATTTGATTGATCTGTCAATATTACCTAATAATTCAAGTCAAGTAGAGTTCAACATTAAGCTTTTTGAGCGCTCGAACAAAGGTTAGGTGCTTCCAAGCTGCATCTTATTTCTAATGTGCTCCTTTTCAACCAAAAATATGTTATTTTCCTCCTTTCTTGATCTTCCTTAATGTTTAGGAAAGAAGAAACTACTACTGGTAGCGCTAGTAGTGTCCATCACTTCTGCAAGTTTTGTCTGTGGTACATGCTGTTACTTTCTATGGAGAATAAAACTCAAGGAAAAAGGTACATTCTAATTCTTGCTAGTAGCAGTTGAGTAACTTAAGTTTCACTTCTATTTGGCATGACTGAAAGAGGTCAATCAAAATGTAACCTTTGATTTAATTAGTCATGGAAAGGAAATTATATTCTCTGAATCATGCTTAGATAAAAGTTAAAATTGCCCATGTCCAGCTTCTTCTAAACTCATCAATAAACTTCCTGTTTATATTGATTTACATCAGGGATTTTGGGAAGGAAAAAGTTCAGGGAAATGTTACTTTCTAAATCAGCAACTAACTTATCAAAACCTGGAACTTCCAGTAAAATGGGACAAGAAAAGAAACGTTCCATCGAGCTGAAGTTCTTTGAATTACGTGAGTTGAAAGCTGTGACAGACAATTTTTCACCTGACAATAAGCTTGGAGAAGGAGGTTTTGGGCCTGTTTTTAAGGTATTGTTACTACATAAGAAAATATATTGACAACCTGAATAACTGCAACACTAAATTCTACTCAGGCCCAAAGAAAATGGGAAAAAGTCCCTCTTGAGGTGAACAGGTAGCGTAATGGTTTTAACTAAAATCTTTTCCATACTTGTGTTTTAATGGTGTGATTAATTGCATTGCAATTTCTAGGGTCAATTGCCAGATGGGCAGAAAATAGCTGTGAAAAGGCTATCAACTCAATCACAGCAAGGAATTAGTGAATTCAAAACTGAGGTCCTTCTCATTGCTAAACTTCAACACAGAAATCTTGTTAGGTTTCTTGGCTGCTGTGAAGAAGACGAGGAGAAAATGTTGATCTATGAATATATGCCAAACAAAAGTCTAGACTATTTCATATTTGGTATGATAATTTACTACTATCTGTTTTTGTGTTAACCCTTTCCTCCTAAGTAGTGCTTACTGGATAAATTTGATATGAGCAACTTAGACGAGAGCAGGAAGTCATTGCTTGATTGGAGAAGGCGGTATGAAATCATTATTGGAATCGCTAGAGGGATACTCTATCTTCATCAGGACTCAAGATTAAGAATAATACACCGTGATCTGAAAGCCAGCAACATTCTTTTAGATGAAGCTATGAACCCCAAGATATCAGATTTTGGTACTGCTCGAATATTTAGTGGAAACCAGAATGAAGCTAAGACACTTCGAGTTGTTGGAACATAGTAAGCAAGAAAATACCAAAAGGGATTTCTCTGTCGCATCCTTACCATTTTTCAATACTTGGCTCATTATTGCCCTTTTCCTTTGCAGTGCCTATATGTCACCTGAATATGCTCGAGCTGGTCTCTTCTCAGTAAAATCTGATGTCTTTAGCTTTGGTGTTATATTATTGGAGATCATAAGTGGCAAAAGGAACATGGTCTCTTATAATAGTGATTCCCCTCCAAATTTGATACGACGGGTGAGTTTTAAGATACAACATTTGAAACTATCTTTTGATTTAATAATCATACTTATTTTCTAAATGAACTGCTGCTTCAGGCCTGGGAGTTGTGGAATGATGGTAAGGCCTTTGAGCTGGTAGATCCATCAATAGTTGATTCTTGTCCCAGTGAGGAAGCATTACGCTGCATTCTAGTAGGACTATTGTGTTTGCAACTCAACGCAGCAGACAGACCAACCATGTCAACAATTCTGTCCATGCTAAGTATTGAAGCAACAGTTCCCTTCCCGAAGCAGCCCTTCATTACCCCTAACTCAGACTTGATTACTACCGAAACAGCATCATCTTCGATCAACGAGGTGACAATTACAGCACCTTATGTTCGTTAGTTGTCTGATGTGTATATGTATTCCTGTTATCTGTTTATCAAATGTATCTAGTGCTCATTACTTCCCAAAGTATGATATAGTCTTCAATGTTTCAATGCAAATAATGCTAAAAGATTTTCAATAGACTAGTTGATATTACTTTTACAGATACTCATCACAGAAAGATTGAGACAACAGTGCTCTTACTCATGATCATAATATTATTGTTGTATTTGATACATGTCTATCTCATGATCTAATTATGAAACAAAAGTTAAAATCTTTCATAAGATAACTTACATGTTTGTATTTTTTTATCAATAATTTTTGAGGAAAAACATAACCAACTAATAATAGTTAAATATCTCATTAATTGACAAAACTCCACAACCAAATTACTATCCTGCAAATCAATAAAAACACTGAAAAGTTGTGTGTGGCTAGCAGGATTAGCCTTTGTTGCTTTTAGAGCATTGATGAACACAGAGAAGACTGTTAAAGAAAACATGAAGTGTGGCAATTATCAAATACAGCTGACAAAGAGCACCATTTAATTCACAGATGAAATAGAAACCCTCTGTGTTTATGCTTCTTTCCTACCAATCCTTTGCCACCCACCTACCTTTAAGTCTGTAACATGGGCCATGGGGAATCAGAAGTCAGTTTTGGTTCCAGTGCTATTAGAGCCCAACCCCACTAGTTTCAATCAAGAATGGACTAGCATGTTAAATGTTGTGTATAACTCACCAATAAATCTCTCTAATAATATGAAAGCAAAGTACAGAAAAGGAATTCTGTATTTCTTTCAAAATAGTTCTTGAATAGTTAGCATAAGCTCTAGCGACCATGAACTGTTGAAATAGACATATCGTTGACAGAAGATGTTGCTATACTTGCTGATGAAGAGTGTTTAACTGAACTATGATTCTCTGATCTGAATATAAGGCTAGGTTGTTTAGGGAAAGGAAGATTTGTTTCATTGCACAGCATGAAGACCAATTCTGACATAGTCGGTCTGTCATCTGTAAACGGTTGTACACACAAAAGGCCAATGTGGATGCATCTCAAAACTTCACAAGCTTCATATGAGCCACCTAACAATGGATCAACAACATCTAATGCCTTTTCATCCTTCCAAAGATCCCAAACCTGAAATAGAAACACCACAATGCATCACAATCAAAATTTCAGACAATTTCTTCTCTATAAGAGAAGCGACTATAACACCAAAAATTGAAATTCTGAAATACAAAATCTCGATCTATTCAGCACTTTTAACAACCTGAAAGGGAATTGTTTCCAAATTATTATTTGCAGTACTACCATCC

>M6-PGSC0003DMG402014514 M6 G-LecRK

TTTCTCCTATTGTCCTGACTCGTGCTTGGTACTATCATACATGTCCTTACTTTCCCTAATATAAGCCTCAGGTACACCTTTAGCCTCCAAACATCACCAAAGAACCCTCTAATGACGTTGTTGTAAGATTTTTCTACGTCAATGAACATCATATGTAAGTCCTCTTTCTCTCCCTAAACTGCTCGATTAATCGCCTAACGAGATGTATGGCTTCTGTAGTTGAAAGCCCTACGATAAGTCCAAATTGGTTCTAGCTGATAGATACACCCCTCCTCAACCTCATCTCCACCACTCTCTCCCAAACTTTCATAGTGTGGGTCAACTAACAACTTGATATCCCTATAACTGGTACAATCCTGAATATCACCCTTGTTCTTGTACAACAAGATCAAAGATGATTTAATTACTAATTAAATTTATCATTATTTTGTTTTATGATTTAGTAGGGATTATTAATCCTTTTTACTGATTAAGTGGTGACTTTATCTCTAAATTCAACTATTGAAAATTAGTGATCCAAATTGCTATTTTTTCCTCATACAATTTAATAGTAACTTCTTTAAAAATTTACTAGTAATGAAAATAGTGACTTATGTTTCTAAATTCATTGTGACAGACAATTATCATTTCTTAAATTTTATACTTCTTATGTTCATTTTATATGATTGTGTTTGAATGTGTATGGACTACGGAGTTTTAAGAAGAAAAAGGGACTTTTTGAGACATAATCGAGTATATGTCACATACCAAAAGTATCTATAGGATCTTATGCTTTTAAACATGCCATGCCATTCTGGAAATGTGACAGTTTGGACATTAATCTCTTAAATGTGAAACTTTCTCCGTCTGACCAGATATAATGCTTAAATCTTGTTATAAGAGGCTCCTTGTTCAGAGCTTGTTTTCATTGATTGCAATTTGCATTTAGAATGGGGTCCAAATTGTTGATATTATGAAATTTAGCAGTTTAATCTATCAACATAAGTTCAAAAAAGCTGTTTACTAACCAGTATGAGTTCAACTTGGTATATTCATTTGCTAATTGTTTCGGTCGATTCATCAGATGACTTGTACCATGTTCTGCGTCAAAGTTTTAGACAGCAGAATGTAGAAAATGGTTCTCCTGTTTCCCCAAAGCAAGTGACACCATTCCGCCCACTTCACTACTTTGACAACAATGCTCATTCCAATTCCCTTAATTCAGAAGCTGGCAGTCATTTTTCACCAAAAGACCACATAGTGGATTTAGCTGTTTCGGGTGGCAATTTCACATATGAAGAGCTGTGGTTAGCAACTAGTGGATTCTCTACATCCAACCTTCTAGGTGAAGGTGGATTTGGGTATGTACACAAAGGAGTCCTCCCTACTGGAAGAGAAATTGCAGTTAAACAGTTGAAGGTTGGTAGCCATCAAGGGGAGCGTGAATTTCAAGCAGAAGTAGAGACTATTAGCCGAGTGCATCATAAACATCTTGTGTCTTTGGTTGGATACTGCATGAATGGAACAAAGAGATTGCTTGTTTACGAATTTGTTTCCAACAGGACCTTAGAGTATCATTTACATGGTATGTTCTTTCTCTTTTTCCTTAATTCTTTGGTCGAATCCTGTATCAATAAGATTTGTGACCAGCTTAAGTCATTTTTTGAATTATGCTAATATTTGTACAGCAGAGAGTTGATAATTAAGGTTAAAATAAATCTACCTTAACGGGGGCTTCCTTAATGCAGCTTAGTTTCACTGTATTGATTTTGAAAAGCATGTAAAATACACCTCACGAGTTATGAGTTCCATGTTAAATGGACAATCTTTAAGGGGGTACTTAACCAAGCCTTTTAAGTAAGATTCCTCGTGGTATATGCCTATACCACAAAAAGGTGGCCTAGAATTTCATCAAACCATGGTTTTTAACATGAGAAGGTAAAAGAAGAGCTTGCAAATACATGCTACTTCTGTAACCCAAGCCCTTTAATCTTGCATGGTAAAGCACTGCTATTTCTATGCTTTTGGAAAGACCAACATTTAATATCTTCTGACTTCTGCTGTCCCAGGCACTTCATTTGCAATTTTCCACATACTTTTCTATACCCTTATGGTGTATTGGTTTAACACACTTCATGTGAGGTACATTTGATGGAGGATCTTTCATAAGTACCACATTTTAGATGTGTGATTTGTTTCATCATCTGTTTTAAGGTTTGTATAAAGAATGTTCCTATCCACCCTGATGCATGTTTTCTTTTGAACTTTTGTTTTACAGAAATGCTCTTGTATCAGTTTATACTTCTCTGATCACTTCTAAATATCCGATTGAATAGAGTTTCCATAAGTTCCTCTGATATAAAGGTCTTTCTACCAAGATGCTATTTAGAACAGATTGTATGATATGCAGGGGAGGCACAATCCACGATGAAATGGGCATCTAGAATGAAAATTGCTATAGGGTCAGCCAAAGGATTGGCTTATCTGCATGAAGACTGTGAGTGCGCTTCATCTCTGTCTTGGCTTCCTATGTTTGTTCTTAATTTGCATGTTCGTCCATTAGTTGACTAGAGGTGTCCAAAGGGGTAAATTGAATATTAAGTGGCCTGGATGCTTTTTAACAACGTTGAACATGTTTCCAAAGGGAACTCTTGTTAGTGTGTTTGTAGATTGTGACAAGTTGGGTAACATAATGGTTTTCATCACATATCTATCTTCACTTTTTCTCCTGAAAAGGATGTAAATCTAATTTTCTCCCTCCCAATAGATACCATTGATCCTCACTTCTATAATACCATTTCTGAATCTCCTCGAGATTTGAGAGCTGAATACCTCGGTTCTAGGGTCTGCTAATGGATAGCAGAAGGCCATAATAATTCCTAAATTGTGAAAGATATAACATGAATATGCATCAGGCAAACTGTACTTTCCCCATATAATTTTATATGCCATGTACTTTCCTGTCAAATATGAAATCAAATAGTTTTAGTATATTTTCTTCCACTACTACCAATTTAATATAGATGTCAACGTATCTTTGCGCTCTGTAGGACAATTTAATTGTGGTTGCTTGAATTAGCTTGAATAGAATAAATATCACTTCTTTTGATGTAGCAACAACAGGTAATCGTTCTCTGAAGATACTAACTTGTGGCTGCAGGTAACCCCACTATCATTCATCGTGATATCAAGGCAGCTAATATTCTTCTTGATTCTAACTTTGAAGCTAAGGTAACTTTGTGTTTCGCTTTTTGGTATTGCCACTATATTTTCTGTTCTATTTTTTGAGAACTGTTTTTTTATTTTTTTTTGAGAACTATTTTCTGTTCTATCTTTACTATCAAAGAAAGGTCGGGCTGTTTTCTATTATCGACAACTGGCTTCTGAGTACACCTAGAATCCGCTGTTGATGTAACTAGTTTTTTGTTTTTAACAATAAAATCCATATGAAGAGATTAATTTGAACCCAAATGAGCATCTACTCAGGTGCAAATGTTAACTGCATTATTGATTCTGATTTTTGTTTCTTTAATTTTCATGTGACAACTAAGAACATATTTCCTCGAAAGTTATTTAACATGAATGGGAACTTTGATTTGTATCCAGGTTGCAGACTTTGGACTTGCCAAGTTCTTATCTGATTCAAACCATCATGTCAGTCATATCTCCACTAGAGTGGTGGGAACTTTTGGGTGAGTTTTATTGAACCTGTCTCTGCTACCATGAATTAGATATCAGAAGGAAAAAAATTGACTAGATCAAGCAAAATTACTATATACAGGTCACCATACTGAGTGGACCAGTCTCTTCTACTTTTATCAAGTAAAAAGTGGTGCTTCCCTCGTTCTAGCGTTAATAATATTCATTTTTTTGGGCTAACCTCATACGTAGCATAGGCTATGCCCTTCCCATGTTAACTTTGTTTCATCCAACTTGCAAACTGATATCTATATCAGATACCTATAGAAAATCGAGCTAGAATGGGACTCTCAAATCTGGGGATTTTAAAGACATTGATGACTTGCATGACCTCGATTTTAGGTCAACCTGCTCATAATGCAGTGATTCCTCTCAGAGTTATAACCGAAGTTTTCCTGCTAGTCGGAGGATGAGAAGAGCGCATGGGATGTTACTGCTGTTTTCCCGTGTCTGATAGTGTTGGGTTTCTTAGTCCTTGATTTTCCCTTACCAACACTGATTGGGGAAAGAGGGGGTTTGGGGGGTTAAGGAGAGGGGCTTGGATTTTGGAAACTGTAGAACTTAAGCTACTTCATGATTAGTAATGATTTCTGGATTACACAGCTTCAGGTAGATGGACAACGTAAAAACTAACAGGGTAAAACCTACATTTCCTGATCACAATTTCTTCATGTATTTGATAAAACTGACGTCTTGCAGGTATCTGGCCCCAGAGTATGCACAGAGTGGAAAGACCTCAGATAAGTCCGACATCTTTTCTTTTGGAGTCATGCTCCTGGAGTTAATTACAGGGCGTCCACCTATAATCTCAACTGAATCCTCCGCATGCAGCAGTTTGGTCATATGGGTATATTAACTGTTCCACTTTCTTGACCGCTATTCAGCATTCTTAAGTGCCTAAAACTAATGTTCAACTAGATGCATGAGTAAAACGTTTGTGTGCAACCTTTATAACTGGTTAAATAGCATAATGCTTAAATTTACTTTCTGTCCAACTACACTAATCAAATCCTCAATATGGAATAACATTCATCTTACTATAATCTTATCCTTAATTAGCTATGTATGCACTGTATTTAATCTGAGGCAGAACATGCAAGCACAGCTTATGCAATTCTCTGAGGTGGCCTTTCTATTTATAAGAGATCTTGATCCTCCGGAAGGTTTATCATTTGTGGTTTTAGAAGTACTGGGTGGTTTTATCCTTTATTGCTTTTCTTTTTTGCGAGTGATAAATGGTTGAGAAATTTTATCACGGCAGTAATCAATATCACGGAAACAACCTTTCTACCTCCACGAGGTAGTGGTAAAGTCTGCGTATACTCTACCCTCCTCAGACCCCACTTGTGGGACCTCACTGGGTATGTTGTTGTTGTTGTTTGTAATAATCAATCACTTAAAAAAAGAAACATGGAAAAGCATTGGCTTAAAATCAGAATTTGGCAGACCACCGTCCTTGTACAAAAGAAATGTGCAAATCAAAAGGATCTGGTGTTCTTGAAAAGTTCATCTCTGTCAAGCCAAAGGCATAAAAATTCACGTCCATCTTACAAATTTTTAAAAGAAATTTGTCATCCTTGTACCAAATGATTTGGCTTTTGGACACCCGTTTTGATGATGGAACATATCTCAAGTTTTCTGTTGCATCTTTTACAACTATTAGTTTGACCAAAGTTTCAGTCTTAAAGCATGGAGTTCACCATGAATTTGATATCATAATCTATCAAATTTTCCCAAAATAAGGCTCCCTTTTTAGGAAATCTTGGAGATACGGGTTCCTGGAATCATTTTGTTGATCTCAAAGATTATTACTGTTGGTTCTACTTAAGTCCCACAATTGATCAGAATTAATTAACATAAGTTAATTCGAAATTGAGATCAGAAAGACGGATTTCAGAGTTAGCAGCTTCTTGCATGCAGCCTACTCACAGACAGAATTAGTTATGTCACTTTAGTTAATCAGAATTAACTTTACACCTCTTCCGGGGCATTTTTTTTCATAACCTCGTCATCAAATACAATATTTTGTATTAGGAATCATTACCGTAGTTTCTGGTTTTCATATTCTTGGTTTTTGATTAAGTACTTTTTGAGAGTCGTTTATTAACTGGATGTACACAAATTTCTTGATCAACATATTGGTTTCTTGTAGCAAAATAACAAAATATTTTGGGAACTAGAAGCATCTCTAGGCCGATGCTTAAATGCATTTTCAAAGATATGGATATCATTGTTGTCAACCCTCTTCTTATAGTATACCACTTGTACATAGCATAAGAAGACTCTTTCTGATACACAGGCAAGGCCTTTGCTTAGAAGTGCATTGGATGATGGAAAATTGGACGCCCTAGTAGATCCATGTTTGGGACAAAATTATAACAGTGAAGAGATGGCTAACATGATTGCTTGTGCTGCTGCTTGTGTGCGTCATTCATCACGGAGAAGACCGCGGATGAGTCAAGTAATAACATCTTCTCTTAGTTCTTTTTCTGTTAGTGCTGGCGGATTTCTTTAAAGGGGTTTATAAACCAAAACCTTGCTTCTTATTTCTTCCTTCTTATTGTTCAGGTAGTTCGTGCTTTAGAAGGCGATGCCTTTATCCTGGACCTTGATGAGGAAAATAGACCAGGACAGAGTACCATTTGTGACTTTGATTCTGATGGAAGCAACTATTTTGAGAAGTTCAAGAAAATCCAGCGGAGCACTCTGAAATCAAAAGAAGAATGGTAATACCACCGTGCACCGGGGGCTGTCATGGAAGTACACCAACTGAAGATGGAGTGAGATGAACGAAGCTCTTCTCCCCTTTTTATCTTTCTTTTCACTGATGTAGATGTAAATCGAGGGTGAAATATATGATCACAACCATCAAAGTTGTCTCACTTAGTGAGAGGATTGATAGTTTGTAGTTACTTGAAAGTGCAAAAAGAGAAGAAAAAGGAATTGGCATGAGATATTCAGTATATCACCAAGTAAAAATCTGTGTATAAAGATGCATGTTCAGATGTACATATAGATTTCTTCCCCTCTCTCTTTTCTTATTCCTTTTTGC

>M6-PGSC0003DMG402019242 M6 G-LecRK

AAGTCTGCAGGGCTTCATAACAAAATAAGAAGTAATACAATCAATGCAATTAACTACTGATATGCCTAACTCTTATGTAAATTGAATACTGATTATGAACTAATATAGAACATGTTCAGACAGGACGCGGAGGAGAAAGAGGTCAGTCATGATTGGTGTCATTTCAGCAGTGGTAGCAACACTTATCCTCAGCATTTTAGCTTGGTTTTACTTCCAAAGAAGGAAAAGAAGAAGAAGAATAGGTAATATAGCTGAGGTACTCTTGTCTATCTGGCTGACATTGTTCGCCTTTACCAAATTCTCATTGATATCATGTTTGTGGCAGGCGAATGTATATTGACAGGTTCAAAAGTTGAAAATGAGGATATGGAGCTTCCATTGTTTGATTTAGTTACTGTTACTAGTGCCACTGGGAACTTCTCTTCTGCTAATGTGATTGGGGAAGGCGGCTTTGGACCAGTTTACAAGGTAAATCAACTTTGATCTTCAACTTCTAGTTAGAAGTGTCATCTTTAATCCTTGATAGAAAATTGCAGGGTATCCTACCAAATGGACAAGCAATAGCAGTAAAGAGGCTATCGGAGTATTCTGGACAAGGCATTCAAGAGTTAAAAAATGAAATCGTTCTCATTTCCAAGTTGCAACATAGGAATCTTGTCAAGCTATTGGGTTGCTGCCTTGAAGGAGAAGAACGGATGCTAATCTATGAGTTCATGCCCAACGCTAGCTTGGACTATTTCATTTTTGGTCTGCTTTTTCGTGACTTATTGGCTATACTAGTACTTGACAAGTCTTCAATATCTCTGATCACTCAATGATTAGTTCTACATTGTGCAGATCCAAGCAGAAAAGCTTCACTTGGATGGAAGAATCGTTTTGAAATTGCTATGGGAATATCTCGTGGTCTTCTTTACCTTCACCAGGATTCAAGATTAAGAATTATTCACCGAGATCTCAAGACCAGCAACATTTTATTAGATAGTGACATGAATGCCAAAATTTCTGACTTCGGCCTTGCCAAAATTTTTGGTGGAGACCAAGAGGAAGGAAAAACTAAGAGAGTAATAGGGACATAGTAAGTTTCTACACATCTATAAATCATAGAACCTTTATTACATTTAATCAGTACATCATGAGCATAAATATATTGAGTTGATGCAGTGGATATATGTCCCCGGAGTATGCTGTTGATGGGAAATATTCAGTGAAATCAGATGTATTCAGTATCGGCGTAATCATTCTTGAAATAGTTAGTGGCAGAAAGAACAGAAAATTTCGTCATATGGAGCATCATCACAATCTCTTGGGACATGTAAGCACATATATATGGAACAAAACTTCGTTCCCTTTATTCACTCTTAGCTTTCTATTGTGATTAATTACAACTGTATAAATGTGTTGTTCTGTTTGTAATATAGGCATGGTTACTTTGGAGTGAAGGCAACGCGTTGGAACTGATAGACGAATGTATCAAAGAATCATTTTCAGAATCTCAAGTGTTGAGATGCATCCAGGTTGGTTTGTTATGCATCCAAAAACTCCCCGAGGATAGGCCTACCATGGCATCAGTAGTTTTCTGGTTAGGCAATGAAGGCCTGGTTCTTCCTCAACCAAAGAAGCCTGTTTTTTTATCGAGAGGAATACAACAGAATCAGCAGAGTTATCTGATGAAGGATATCTAAGTAACAACGTATCGATAACAGTCCTAGAGCCAAGATAGACATAAAATCCAACAATTAGAACTTTTGAAATGTATCCACTTTGTCCTGTTTTTGTCAACAGTATTAGTGTATTTATGGACTATATCATGAAGTGCATTGAGGAAGATAGTCTAGTATTTCTGTTTTTCATTGACATGTGTATTCTTCGATACTATAATAAACGTAACGAATTGGCCTATTCTGAGTAACTTGTGAAAATTCTGTCTCCACTTCTGGCCATCAAAAT

>M6-PGSC0003DMG402026153 M6 G-LecRK

CAGAGAAATTGTTTTGGGAATGCCTATGGAGTACTTGCCTCATAGAGATAGTTTTGATGAAGATTTAATCACGGAAGATGATAAGAAACGGATTGATGTCCCGTTTTTCAGCTTCAATAGCATACTAGTTGCTACCGAAAACTTCTCAAATGCAAGTAAGCTCGGGAGAGGTGGATTTGGACCTGTTTATAAGGTTTTTGCAACTTTCTTTTTTTTTCCTTCTTCTGTTCTAAGTTGTTTTACCATTCAGATTTTTTTCTTTCGTTTTCGTTATATAACAGTCTGGAGATATATTTATGCAGGGTACGTTTCTCGGAGGTACAGATATGGCAGTTAAAAGGTTGTCAAGTAATTCTGGACAAGGTGTTGAAGAATTTAAAACTGAAGTAATGTTGATTGCCAAACTGCAGCACAGAAATTTAGTTAGGCTTCTAGGCTATTGTGTTGAGAGAAATGAAAAGATTTTGTTATATGAATATATGCCCAACAAAAGCTTGGATACGTTTCTATTCGGTTAGTATACTTCCCTCCCTACCATGAAGCTCGTTGCAGTAATAGTTTTATGATTGATTCTCTCTGAATTTATCTTACAAATCTTGAAATTAGATCATGCATTCTGTCGATTACTGGACTGGAGGATTCGTTTTGAAATTATATTGGGTATTGCTCGAGGGTTACTTTATCTTCACCAAGACTCGAGGCTAAGGATCATTCATAGAGATCTAAAGACGAGTAACATATTGTTAGACCAAGAAATGAATGCCAAAATATCAGATTTTGGCTTGGCAAGGATCGTTGAAGGCAAAAATACAGAAGCCAATACTAACAAAGTCGTTGGAACATAGTAAGTCTCTCGGTTCTGTTTTCACCTTTGATTTGGTGGGTAAAAGATAGATATATATACATCTAAAACTTGGTATTAGTTCACTATATGTATTGGTTTACACAAATTAGATGCTAAAATAACCTATTCTGGAACACTAAAAACAGTGGCTATATGTCTCCGGAGTATGCATTGGAGGGACTCTTTTCAATCAAGTCTGATGTATTTGCCTTCGGAGTAGTCCTACTCGAGATCATAAGTGGGAGGAGAAACATGGAATTTTTCGAAGATGTAAACCTCATTGGTCATGTAAGTACCAAAATTTCCCCCCATTAATAAACATCTCATTCTGTTTAAAGCTTTCATTGACTCAAAATTACTTATCACACACAGGTATGGAGACTGTGGATGAAAGATAGTGCATTAGAAATGATGGATCAAACCATAGTTGAGTCGTGCGATGAAAGTGAAGTGCTAAAGTGCGTGAATGTGGCACTCCTATGTGTTCAAGAAGATCCAGCTGAGCGTCCTGTAATGTCAAATGTTGTTTTTATGCTTGGAGGTGAAAGCATAACACTACCAAGACCAAATCAACCAGCTTTTATAGCACGAAGAAATAGTGCTGGTGCAAGTACATCTTCTTCGTCTTCTAAGCTATACAACACTTCCAACAATGGGTTGACAATAACTCAACAAGTAGGCCGATGAAATCAATATACATATAAGTTAAATGCATAACGCACACAAGCCAATAATGTGCCGGTATATCTTTTCTTCAATTTCAGTTTGAATTTCTGTTAAATTATTTTGGCACGCAAAATAACCGATTGTGTTGTAGTCTCGGA

>M6-PGSC0003DMG402029906 M6 G-LecRK

AGTTTTTGATATGTGAAAATGACATACTTCACAAAATATCTTCTTGTTCTTGTTCTTCTCATCTTTCCCCTGCATATTTTAGCACAAACTATACCATTAGGCAGCTCTCTCATAGCAAATGATGAAACAACACCATGGCTTTCTCAATCTGGAGATTTTGCATTTGGATTCAAGAAAATTCAAAGTCAAAATCAGTTCTTGCTTTGCATATACTATGCCAAGATTAAAGATGCCACCATAGTTTGGTATGCAAATGGTGGTAACCTCGTGCCACGAGGCTCTATCGCGGAGTTAAATCCTCAAAAGGGACTAATTCTCCGCGATCCTAAAGGGAAAATGATCATTTGGAGCACTGGTCGGATTGGTTCTAATGTTGCTTATGCTGTCATGAATGATACTGGAAACTTTGTTCTTGTTGGGGTTGATTCTTCAGTTTTATGGGAAAGTTTTCAATATCCGACTGATACCCTTATGCCTACTCAAATACTAGAGATTAACAACAAGCTTGTTGCTCGAAAATCGGAGTCTTTCTTTGTTCCTGGTAGATTTTATCTTCGTATGTTGAGTGATGGGAATTTGGTTCTTGTTACTCAATCCAAGCCAACGAATTTCGATTATGATGCTGAGTATTACAACAGTCATACTTCTGATTCAGGAGATGAAGCCAATTCTGGTTATGGATTGGTATTCGATGAGTTTGGTTCAGTTTACATTTTAAAAAGGAACAATCAAAGACTTGTGTTGACACCTCCCAATGTTCCTTCAATTTCAGAAAACTATCACAGGCTAAGTCTTGATTTTGATGGTGTTTTAACTCATTATTATCATCCTAAGAGTACGGGGGATCAGAAGTGGAGTACTCTTTGGTCTTTGCCTGATAACATATGTCTAGCAATTCTTGAAGATGTTGGAAGTGGAGTTTGTGGATTCAACAATGTATGCAATCTAGGTGAAAATCAAAGACCATATTGTGAGTGTCCAAAAGGGTATTCGTTGATCGATCCAAATAATAAGTATGGAAGCTGCAAACCAAAATTTGTTCATAGTTGTGATGAATTTGGACAGGGAAATCCTGAAGAATTATATGATTTTGATGTGGTGACAGATGTTGATTGGCCGTTATCAGATTTCGAGAGGATTTATCCTTCTGCTGAAGAAGAATGCAAGAAGGCTTGTTTAGAAGATTGTTTCTGTGCTGTTACTATTTATAGAAGCAATAGTTGTTGGAAGAAGAAACTTCCATTGTCAAATGGGAGAGTAGATACCTCTTTGAATGTAAAAGTTTTCCTTAAAATAAGGAAAGTTTGAACATAAGAGCTTCTTAAATTAGATATAATGTATGTCTGCTATAGGTTCATGGTTGAATTCGTCGATCCAACTTGCAATTTTACACAATTCAATAGTTTGGATTTGTAATAATTAATGGACCATATTGTATAACAAACTTTGATTAGCAAATAATGAAATATTTCATACATTTCAAGAAACATATTGCACTGAGATACAGATACAAGTTATTGATGTACTCTTTAGGTCCAAGCAAATGAATAATTGAAAAAAAAAAACATGGTTGATCTAAATAGAATCGACAAACGAAAAAGGGTCAGGAGGAATTGATACTTCAACAGAACCTTCAAGCATTAACATGACTTTCTTCATGTTAGGCCTTGATGCTAGATTTTCTTGAATGCACCAAATTGCAACCAACAAGAACTTCTCGAATCTCTTGATATCTTCTAATGCCTCTTCATCATCACCTACAAGTAAATGCAACTTATTTCTTCTATAGCAATCATAAGCCCACTCCAGTAGTATCATTTCGTTCTCGTTTGCCACATCCTGCTCATAGTTTTTTCTGCAGCAAATCAACTCCAAGAGCAAGACCCCGAAGCTGTACACATCAACCTTCACAGTTACAGGCATGCTTCTGAACCAGTCTGGGGCTACGTACCCTTTGGTTCCACGTATCCTAGTGGTTGTTTGAGTCTGATGTTTTTTCAGTAGCTTGGCCATACCAAAATCTGCTATTTTTGCTGTCAAATTTTCATCCAAAAGCACATTTTGGGGCTTGATGTCACAATGTATAATTTGGGTACTGCACTCTTCATGCAAATAACAAAGTCCTTTGGCAGTGTCGATGGCAACTTGTACCCTTTTGGACCAACTAAGCCTAGAATCTTTGAATAGCAAGTGTGCTATTGAGCCGGTTTTCATGTGTTCATATACCAAAAGGCGGTGTTGGCCTTCGTTGCAGAATCCAAGGAGTTGGACCAAGTTCTTGTGATTAGTCCTGCTGATTGAATTCACTTCAGCTTCAAATACTTCTTCTCCTTCTCCTTCTGTTACCATGTTTCGGAGTTTTTTGACAGCAACGACTTTCCCATTTTCATCATCAAGAACCGCTTTGTAAACTGTTGAGAAAGCTCCAGTTCCTAATTCTTCCTTGAATCCATTTGTAGCTTGTTCTAGCTCTTTGTAGCTAAAACTTCTTAAGTTCACTCCCGGAACAGCTGAATACAGTGCAGTCCTCTTTCGTTTTTTCCCTATAAACTTAAAAATGTACAATAGGACCAAAAGAATTAAGAACAAATTCATGAAAACTGAACTGGCCAACAACACAGAACCAGAAATGATCAGAGTAGACTGATTTTTCTTTTTTCTTATTTCAACATTTGGTGTTCCCACTGTGGAGTTGTCTTTCCTTATTTTTATCAAAGCCTTCCCTCCGATCGTTGGACCTACTCTCCCATTTGAAAGAGGATACCTTTTCTTCCAACAGATGTTGTCGTCACTGTAAATGGCAACATCACAAAAACAATCATTCAAGCAATTCTGTCTGCACCAATCCTCTGAAACATCGCGGTGAGACTCATAATCAGAGTCAGGCCAGTTAGTATCCAACATTTCATGGAATGTGAAAGACTCAACTTCTCGCGATTCATGGTTACAGTTTTGCTCAGAGAAATTTTGTCTGCAGCTGCCTAATTTGTCATTTGGATCATCCAAGATATATCCGAGTGGACAATCACATCTCGGCCTCTGATCAGTTCCTATTGAGCATAGGCTGTTGAAACCACAGGCACCCCCTCCTGTACCTTGCCTGATAGACAAACAGATGTTATCAGGAATGTTATACAAACTAGACCATGCCATCGGTTCCCTACCAGAAGATTTCGGATGGACATAGTGCCTAAAAACTCCATCATATTCAAGAATCGCTCGATGATACATTGATTGGCTTCTTGAATTTTCCACATTAGAAGATATCGAATTTATCAAAGTTCCATTCTTTGCTTGAAGAAAAATAAAGCCTGATTGATTGAAGATCACCTGATAACCACTGCCAACTGACATAGATGACCAATATGCAGCATTTGTTGTCTCAGCTGGATAATTGACTGTGTAAAGCACTAGATTTCCATCAGTTTGAAGTATGAACTCAAACCTCCCACTCGATACATTTGTGTCAGAGAAACTTGAAACAAGACTGTGATCTTGGTTAAGTACTTGACCAGGCAAAATCGTATCGGTGGGCTCATTAAAACTCTGCCATAGAGTGTCTGAACTACTAGTTGCTAGCACAAAGTTTCCGTTATCAAGCATGGCTCCATAGGCCAATTGAGCATTCGCCATGCCACGAGCCCACATCTCTTGACCATTTGGATCAGTGAGGACTAGTCTTCCATCCGCGGAGAGCTGAACTTTGGATCCATCTAAGGCAAGATTATTCCTATTGGCTGACCAAACTATGGTTTTGTCTGTGATTTTGTTGAACCATATGGCTAGTAAGTACCTTGATTCATTTCCAATTTTTTGGAAGCCCAAAGCGAATTCATCAGATGGGGATGACCAAAAGCTAGTAACATCACTTGTAGTGAGTGATGAACCCAAACTAACATTTTGGTAAGATTGAGCAATGGCTGAAAGGGGAAACAATAAGAGACCAACAAGAAGCATAGCTAAAAAAGTCATGTAAGAAAGAGGCAAAGTTATAAATGGATATTATGTACACC

>M6-PGSC0003DMG403026153 M6 G-LecRK

ATACCATAAAAAATGTTACTGACTCAAGTAGTTGCTGGAAAAGTTTATGTTATACAAATGAGTAATTTTGTCTTGTTGTTGTTTGCAGTGTTGGTGATTATATGTGGATGTGATGCAAGGAGCACGATCGATGAAAGGAGTGAGTTAGTAGACAACGGGGAAACACTGGTATCTGATGGAGAAAACTTCGAAATGGGATTCTTTTCTGAAGGGTTAAACAGGTATGTAGGTATATGGTATTACAAGTTAAGTCCTCGAACTGTTGTATGGGTTGCTAATTGGAACAATTCGATTCGAGCTTCAACAAAAAATAAGGAGAGTGTTTCCATTGTCGTTCTTGATGGTAACCTCAAAATTTTGAATTCAAGTAATGGCGAAACTTATTTCTCCTCAGGATTAGATGGTAACTCTAGCATTAGAGCAGAGTTGCTGGATACAGGGAATTTGGTTTTAGTTGATGAATTAGGAGCTAAAATGTGGCAGAGTTTTCGGAATCCAACGAATACTTTCCTTCCTGGCATGAATATGGAGAGTCTGGAATTAACTTCATGGGAAGGATCTGGAAATTGGAAGTACAAATTCAAAGTAGATCAAGGAGACAACAAAAAATATGTGATAGTTGAGAACGGAGTGGGTACTCACTGGCAAGGGTCTGTAGGAGGGTTTGCCAAATCAACTGCTTATGAATTCTTCAGATTCAATGATATGCCTTTTTTTGTAACCTTCTTTTTATCCAATAATTCAGTTGAAGCGAGAAGAACATTATTCAATAATTCAGTGGAATTGACAAGAACATTATTCAATAATAATCGTAATTTTACTCCGATTGATTTTTCAAGGTTAGTCAATCATAGTAACAAAGATTCTATTTTCAAAGGTACTAGACTCCTGATGATCTCCTCTGGAGAAATACAATTCTACTTTTGGGATTACCAGAGTGGATGGTCCTTGGTCTGGTCTGCCCCAAGAGATAAATGCGACCAGTTTAGAAAATGTGGGAAATTTGCGATTTGTAACAGCAATGAAAGTTCAGAATGCAGATGTTTGCCTCGTTTTCAGAATGATCCTCCAGAAGATAATGGAGACGGGGAATATTCAGCTGGATGTTCGAGACTATCAGATAACTCATGCAAAGAAGACACATTTGTGCATCTGAGCTCAATGAAATTTGGAAATTCAAATTGGACAGATAAAAAGATTAATACCAGTGAGAAATGCATGAGTCATTGTTTAAGCAACTGCTCATGCAAAGCATATACATATTATAACAATTCTGGATGCTGGTTTTGGGGGTCGAATCTCAATAATCTCCAGGAGAACTACCCTGGTGGCTTCAACCTCTCTGTCCGCGTGCCCCCTA

>M6-PGSC0003DMG400000578 M6 L-LecRK

CAATCATTAAAGCATTAAAATTTACATTCCATCTGAATTACTATATTTTGTTTACTGAAACTGAAATAGGACAAAAAATTGCATTATGTACGTGAGGACTTTAAGTTCTCTTGTGTTTGGAAGAAAGTGAAATGAAGTTATCTATAGACCACATCCTTATGTACAACCTTGACAATTGATAATTTGCTACTCAAAAAATTCAAATCCCAAAAAACTTTATCACTAGCCAAGAATTAGCTCATCATGGATTACTTCTTCTGGTTTTTCTTCTCGTTTGCCATACTAGTTAATAGTAATTTAGGTTCGTCCGCGGACGGACCTTCAATTAACGTGACGAAGCATATCTCTTTTCAAGATTTCAGTTCTAGTAATCCCAGATTGAAGCAAGATCTTACACTTCTCGGTAGTGTTATCGTCTCGGATGAAAAAAAATCCGTTCAAATTCCTGATCCTGAGCGAGAAGGTGATGATCTAAAGCATCTAGCAGGACGAGCTATTTATTCTTCACCTATCCGTTTCTTTGATCCTCGGACTCAAACACCGGCTTCTTTTGAAACGACTTTCTCGTTTCAATTTCAAGTGAAGTCCAATTCAAGCAATGAGACGTCAGATCAGGGGAAATCTGTTGGTGGTAGTGGCCTTACTTTTATAATCGTTCCGGATGAATTGACTGTTGGCCGTGCTGGTCCGTGGCTTGGGATGCTGAACGATTTGTGCGATGAGGATTATAAAACAGTAGCTATTGAGTTTGATACTCGGCAGAACCCTGAATTTGGTGACCCGAATGATAATCACTTGGGCATTAATTTGGGTAGCATAGTATCAACTGCAGCGATTAATGCTTCGGATGCCGGAGTTCAATTGAATGATGGATCAATTCACAGAGTATGGATATCTTATGATGGGCGAAAGCGATTTGTTGAAATTCGTCTTGCTCCTGATGGCAGAGGATATCCTTCTAAACCAGTTTACTCTGGTTTACTTGATCTTTCACCTTACTTGAATGAGTATATGTTTGTTGGATTTTCAGCTGCTACTGGTAACCATACACAAATCCACAACATTCTGTCATGGAATTTCACCTCAATTAGTCAAGCTTCGCTTCGAATTCCTTCAACAGAGACGTGCCAGAACAAAATCATGCTTCAAAATAGCACACAATCTGAAATTGATCATCGGAAAACTCCTAATAGTTTCTTTATTTTCCTTGCTGTTGTCATTCTTCTGGTTATCGTTCTTATTAACCTCTATTTCAGTAGCTACAAGAGAGATGACAATTCCGACGAAGCATTCACTCTGCCCCAGAAAAAGCAGAGACCGCGGCCACCAAACAAGGCACGTCGCTTCACAATAGCTGAGATCTCCATCGCAACAAGGAATTTCAGCGAGTTACAAATATTAGGCAGTGATGAGAAGAGCATTACGTATAAGGCCACGATCCTGAACGGGTGCAACGTTGTTGTAAAACGATTTTTAACTCAATTTTTCAACACACACGGGTTTGAAAAGCATCAATTTCACAAGGAAATCAAGGCTATCACCAGAATTCGTCACCCGAATTTGGTCCCAATTAGAGGATGGTGCTATGACAACCAAGAAACAATAGTTGTGTACGATTTCATCCCCAATGGAAGCCTCGATAAATGGCTGTTCGGCGTTGGCGTCTTGCCTTGGACGAGACGTTTTAAGGTCGTTAAAGATTTAGCAGATTCTCTTGTCTACCTTCATTCAAAGCAACTCGCTCACAAAAACGTTAAAAGTAGCAGCGTGTTTCTTGACGTGAGATTCAGAGCAGTAGTGGGTGATTTCGGGTTTGTGCTTACTTCAGCCGGGTCAACCCGGTTTGAGGCCATGGTGAGTCAGACAGCTGATGTGTTCGAATTCGGAGTAGTCGTGCTAGAGATCATTGCGGGTCGGAGCAGAAAGTCCAACCCGGGAGAGCGAGACTTGTTGGATCTTGCATGGGCAATGCATGAGGTACAACAGAAGGAAACTCTAGTGGATCGAAGAATGGGCGCAGTCGTGAACCTGGAGCAGGCGATTCGGGCTTTGGATATCGGGTTGCTCTGCACGTTGAACGAGAACAAAGGAAGGCCTACTATGGAAGAAGTGGTGGAGTTCCTGAATATGGAGAAACCGATACCGGAGTTGCCATCGGGTCGACCCGTTTGTTTGTTCCCGTACAGCAGCACCACAGGCTTATGCAGTGGGTACGCTTGTACAGCATTCAAATGAAGTCATGTTTTTTGCTTGACCAATTGGTTCTTCGGCGTTCCACGTAAGCAGCAATTATTCTTATCTGTTATACTACAAAATAATGACAGGGAAGCACATGTATATGCTATGTGACGTTATTTTCATGACATAGACAACTATTGTCTGATCTGTGGTCGGAGTTAAATATCACAAGAACTCCTAAAACGATTTGAAATTATTGTAAATTATGTACTGCAATTCAAATGTTCTATATATAAAATT

>M6-PGSC0003DMG400000789 M6 L-LecRK

ATGCGTTTTTGTTTGCAAAATCTCATAAGGGTCACTCTGTTCTTCATCTTCACCATAAATTCACCTTCCCTTTCAGCTCCTTTCTTGCCTATAAACAATGTAACTCTTTATGGTGATGCTTCATTCACCGCAAAATCAATCACCCTCACCCAAGAACGCAACTGTTCATCATCATCTTCAACACCTCCCATTTCTGGCATTGGAAGAGCTTTCTACACATACCCAGTTCGTTTTCTTGATTCTTTAACCAATAACACTGCTTCTTTCTTGTGTACTTTCTCTTTTACTATACTTCCAACCCCTTCTTGCCCTTTTGGTGATGGCATGGCGTTTTTGGTCACTTCTGATGTTGATTCTTTGAGCATCTCTGATGGGTATATGGGTCTTCCGAATCCCGACTCGGAAGATTCGTTCTTGGCTGTGGAATTCAATGCCAATGATAACCGTATTGGTGTTGATACTAAAGAAATTAGGTCTTTGGCTTCTACTAATGTTTATTCAGCTGGGATTGATTTGAAAAGTGGGAAAGAAATGGCGGGTCGGATTGAGTATAAAGATTCAGAGAAAATGATCAGAGTTTGGATTGGGTATGAACTGCAAATTAGGCCTCCTAATCCTGTTCTTTCTACCAGAATTGATGTTTCCAATCAGTTGAAGGAGTTTATGAGGATTGGTTTCACTGCTAAAGGGTCTGCAGTTTATAGCTTTAGTCGTTGGCGATTTAGAACGTTTGGATTGATTTCGTCTCCAATATCATCGTCTTGGGATCAATCTGATGAAGGAAACTGTTTGATGTGTTTCCCTGAGGAGGAAATTGGTGGGCATATTTCTGATTCTCATCATAGTAGTAGTAGTAGTGGTAGTAAAAGTTTACTGAAATTGACTTATGGAGGGTTAGCTGCAATTGTCACACTTGTTGGTTGTGCTTTGTCTGTTGTGTTTGTTCTTGCGTTGAGAAGGAAAAAACGCGATAGAGTAGGGGAGACCAATGAACGGCAAATGTGTAGATTACAAGGAAATAGAGTGCCTCAAAGATTGTCATTATCTGAAATAAAATCAGCTACAGAATGTTTTAATCATGAAAGGATAATCGGAGAAGGAGCATCTGCTGTTGTGTATGAAGGGGAAATTCCTTCTAGGGGATCTGTGGCTGTTAAGAGATTTGTCCAAGGGAGTAGATTGGGTCCTTCACATATTCCTTTTAATACTGAATTTGCTTCTATGGTTGGCTGTTTAAGACACAAGAATTTGATTCAGCTTCAAGGGTGGTGTTGTGAGAGAAATGAATTGGTGTTAGTGTATGAATTCATGCCTAATGGTAGCCTTGACAAAATCCTCCACGAGCGATCGCATTTGACCAAGTTTCTGACTTGGGAGAGAAGATTGAACATAGTTATTGGTGTGTCATCTGCTCTTATGTATCTTCATGAAGAGTGTGAGAATCATATAATCCACAGAGATGTTAAGAGTTGCAATATAATGCTTGATGCTGAGTTCAATGCCAAGCTTGGAGATTTCGGTTTAGCAGAGGTGTTTGATAATTCTAAGACAAGGGATGCTACTGTACCAGCTGGAACAATGGGATATTTTGCACCTGAGTATGTCTATACTGGTATTCCAACTGTTAAAACAGATGTGTATAGCTTTGGTGTTGTGGTACTGGAAGTGGCATCAGGTAGAAAGCCTATCGATGAAGGTGGAGGTTTGATTACTGATTGGGTGTGGGACTTGTGGGAGAAAGGGAGGATAACTGAGGCTGCTGATCCTAAACTAATGGGGCGGTTTCAGAAGAACGAGATGGATAGAATGCTCATTGTGGGACTTTCTTGTGTGCATCCCGATCATGAGAAGAGACCGCGAATAAGAGATGTTTTCCGTATGTTTAAAGATGAAGCTCCACTTCTCATTTTACCTCCAATGAAGCCCACTGTGAGAATTCAATCTGTTTTACCAGAGAGCTGTGAAGAAATCATGAATTGGGCTGCAAGAATGGATGATACACCATGGTCCACTCCTAGAACTCATTTTAGCAAGAACTAG

>M6-PGSC0003DMG400001773 M6 L-LecRK

CCAACGGCTAGTTTGATGAACCCCACGGCGGATATAACTACATCTGCCACATCGCCCCCTCCGGAGCAACAGATTGCTCCGAAGGAGGCTAATGGTACTGCAGTACAGAAAAACAGCAGCAAATGTCATAATAGTTTTTGTAAGCAGGGTGCTGGTGCTGTTGTTGGGGTTGTTACTGCTGGAGCATTCTTTCTAGCTTTTGCTACATTGGTACTTATTTGGTTGTATTCGAAAAAATTCAAGAATGTGAAGACTTCAGAGTCTATGTTGCCTTCTGATATTATCAAAATGCCAAAGGAATTTAGCTATAAGGAGCTTAAAATCGCGACGAAAGGGTTTGATTCTACTAGGATAATAGGGCATGGTGCATTTGGTACTGTCTATAAAGGGATTTTGTCTGAGACTGGTGATATTGTTGCGGTAAAGAGGTGTAGTCATAATGGACAAGGGAAAGCTGAGTTCTTTTCTGAATTGTCGATAATCGGAACACTTAGACATAGAAATCTTGTTAGGCTACAAGGATGGTGTCATGAGAAAGGTGAAATCTTGTTAGTTTATGATTTGATGCCAAATGGTAGTCTTGACAAGGCGTTATTCGAGTCCAGAATGATCCTCTCGTGGCAACACAGGCAAAAAATCTTGTTAGGTGTTGCTTCTGCATTAGCATATTTACATCAAGAATGTGAAAACCAGGTGATACATAGGGATATCAAGAGTAGTAACATTATGTTGGATGAAGGGTTCAATGCAAAGTTAGGAGATTTTGGACTAGCACGACAAATCGAACACGACAAGTCCCCCGATGCAACGGTAGCAGCCGGTACAATGGGGTATTTGGCACCAGAATACTTGTTGACAGGTAGAGCAACTGAGAAAACTGATGTGTTTAGTTATGGGGCAGTGGTTCTTGAAGTGGCAAGTGGGAGGAGGCCAATTGAGAAAGAAACAGCAACTAGTGGTGTTGGATTAAACAGCAATTTGGTTGAATGGGTGTGGGGATTACACAAAGATGGGAGTTTACTATCAGCAGTTGATTCAAAATTGAATCTTGAATTTGATGAAAAAGAAATGACAAGAGTGTTGTTAGTTGGTTTGGCTTGTTCACATCCTGACCCTATAGCTAGGCCAACAATGAGGGGGGTGGTACAAATGTTAGTAGGTGAGGCTGAAGTACCAATTGTACCAAGGGCTAAACCATCAATGAGTTTTAGCACATCTCATTTGTTAATGACATTGCAAGATAGTGTTTCTGACTTGAATGGTATGATCACACTTTCAACTTCTTCATCAGAAAACAGTTTCAATGGTGTTGGTGGACATGATGGCATGGACCTAGTTTAG

>M6-PGSC0003DMG400004396 M6 L-LecRK

ACATTTCTCTAACTATGGCTGTTTTTGCATATTTCACATACCTTATTTTGATAATCCCCTTTGTCACTTCATTGTCCTTCAATTTTGATAGTTTCAATCCCAATGATATGAATGTAACATATGAAGCAGATGCTTCTCCAGCAAACGGTGTAATTCAGCTCACCAAAAACCAGCTTGATAGTAGTGAAAGCTTTAGCGTAGGCCGAGCTACATATTCTAAAGAACTCTATCTTGGGACAAGGCCTCTGGGAAAGTCACAGATTTCTCTACGCACTTCTCCTTTGAGATCAATTCACAGGGCAGAAATTATTATGCTGATGGTCTTGCCTTCTTCCTCGCTCCTGCAGGTTCAGTAATTCCTGAAAAATACTCAGCTGCAGGAGAGGGCCTTGGCCTTGCAATTGATAATGGACTAAATACATCGAGAAATCAGCCTTTTGTTGCTGTCGAGTTTGACACCTTTTAGAACGCTTATGATCCACCAGGTGATCATGTAGGCGTTGATATCAACTCCATGTTATCTCTTGTTAATGTGGCCTGGCTTAGTAGAATTCCCTATGGTACGAAAACTGATGCCTGGATTACTTATAATTCAACCTCAAGAAATCCTAGTGTTGTCTTCACCGGTTTCAGACAACAAGGGAATACCACAGTCACTGTCCTGCAGAACCTATCTTACAATCTTGATCTGAGGGAATATTTGCCAGAATGGGTAACTTTTGGTTTCTCAGGTGCAACAGGAACCTTCTTTGCAATACAAAACATATATTCTTGGAATTTTACTTCTTCTTTAAAATATAGTGACAGCATAACAGATCCAGATGTACCCCAACCAAGCCCTGTGCCAGAAGATACTTCAAGCAAAAAGAAGTCAGGACTAGTGATTGGATTGATTTCTGGTGGTTGTGTTTCGGTAGCAGTATCTGTTTTTATATTGTTTGCATTTTGGAGAAAGAGGAAGGTCAGAGAAGATGAAGATGATGATATTATTGATGGTTCCATGACTAATGAATTTGAAAGAAACACAGGACCAAAGAGTTTTTGTACGGTGAGTTGGCTAGATGTACAAATAACTTTTCGCCGGAAGAGATGCTTGGGCATGGTGGGTTCGGAGGTGTTTATAAAGGATATCTCAGGGAATATGACTCCTATATTGTTGTTAAGAGGATTTCAAGGGAGTCAAAGCAAGGAATAAAAGAGTATGCATCAGAAGTCAGGATTATCAGCCGGTTAAAGACATAAACATTTGGTGCAACTCATTGGTTGGTGCCATCAAAAAAGAGAACTTCTACTTGTCTATGAGTTTATGCCTAATGGAAGCTTAGATCATCATCTTTTCAAGGGAAAAAGCCATTTGACATGGCCAATAAGATTCAAGATTGCTCAAGGCTTGGCCTCAGCGTTGTTTTATCTACACGAAGAATGGGAACAATGTGTGGTGCATAGGGACATAAAGTCGAGCAATATTATGTTGGATTCCAATTTCAATGCTAAACTTGGGGAAAGCTTTGGCAGGCACGATGGGTTACATGGCTCCTGAATGTCTCACCACTGGCAAAGCTAGCAAGGAAACTGACTTCTATAGCTTTGGTGTTGTCGCGTTAGAAATAGTTTGTGGAAGGAAACCTATTGATCCTAAATCTGAGGAGCATCAAGTAAATATTATCGAATGGGCTTGGAGACTTTACGGGATGGGAAATCTTAGTGAAGCAGTTGATCCTAGACTCTCATCAGAGTTGAATGAACAGGAGGTGGAGCACTTGCTAATTGTTGGCTTATGGTGTGCTCGATCATCCAGATAACAATTGCAGACCTTCTATTAGGCAAGCAATTCAGGTGCTTAATTTTGAAGCTCCATTGCCCACACTCCCTCCAAACATGCCTGTGCCGATATATTGCAGTTATGAATCAATTACTTCACTTGCATCACCATACGACTCCAATGGTCCTCAGATTTCTGAAATAAACTCTTCAGTGACTAGAGATCATTCGACACAGCAGCATCTTCGCCTTCCGCATCACTTCTGTACACACGTTGATATTGTCTTGTAAAGGAACTGGTTTTTTATTTCCCTGCAGTGTCCAACTTTTGCTCTTCGTATTCTTTATTATTTATACTGTTA

>M6-PGSC0003DMG400004397 M6 L-LecRK

ATGGCTGTTTTTGCATATTTCACATACCTTATTTTGATAATCCCCTTTGTCACTTCATTGTCCTTCAATTTTGATAGTTTCAATCCCAATGATATGAATGTAACATATGAAGCAGATGCTTCTCCAGCAAACGGTGTAATTCAGCTCACCAAAAACCAGCTTGATAGTAGTGAAAGCTTTAGCGTAGGCCGAGCTACATATTCTAAAGAACTCTATCTTGGGACAAGGCCTCTGGGAAAGTCACAGATTTCTCTACGCACTTCTCCTTTGAGATCAATTCACAGGGCAGAAATTATTATGCTGATGGTCTTGCCTTCTTCCTCGCTCCTGCAGGTTCAGTAATTCCTGAAAAATACTCAGCTGCAGGAGAGGGCCTTGGCCTTGCAATTGATAATGGACTAAATACATCGAGAAATCAGCCTTTTGTTGCTGTCGAGTTTGACACCTTTTAGAACGCTTATGATCCACCAGGTGATCATGTAGGCGTTGATATCAACTCCATGTTATCTCTTGTTAATGTGGCCTGGCTTAGTAGAATTCCCTATGGTACGAAAACTGATGCCTGGATTACTTATAATTCAACCTCAAGAAATCCTAGTGTTGTCTTCACCGGTTTCAGACAACAAGGGAATACCACAGTCACTGTCCTGCAGAACCTATCTTACAATCTTGATCTGAGGGAATATTTGCCAGAATGGGTAACTTTTGGTTTCTCAGGTGCAACAGGAACCTTCTTTGCAATACAAAACATATATTCTTGGAATTTTACTTCTTCTTTAAAATATAGTGACAGCATAACAGATCCAGATGTACCCCAACCAAGCCCTGTGCCAGAAGATACTTCAAGCAAAAAGAAGTCAGGACTAGTGATTGGATTGATTTCTGGTGGTTGTGTTTCGGTAGCAGTATCTGTTTTTATATTGTTTGCATTTTGGAGAAAGAGGAAGGTCAGAGAAGATGAAGATGATGATATTATTGATGGTTCCATGACTAATGAATTTGAAAGAAACACAGGACCAAAGAGTTTTTGTACGGTGAGTTGGCTAGATGTACAAATAACTTTTCGCCGGAAGAGATGCTTGGGCATGGTGGGTTCGGAGGTGTTTATAAAGGATATCTCAGGGAATATGACTCCTATATTGTTGTTAAGAGGATTTCAAGGGAGTCAAAGCAAGGAATAAAAGAGTATGCATCAGAAGTCAGGATTATCAGCCGGTTAAAGACATAAACATTTGGTGCAACTCATTGGTTGGTGCCATCAAAAAAGAGAACTTCTACTTGTCTATGAGTTTATGCCTAATGGAAGCTTAGATCATCATCTTTTCAAGGGAAAAAGCCATTTGACATGGCCAATAAGATTCAAGATTGCTCAAGGCTTGGCCTCAGCGTTGTTTTATCTACACGAAGAATGGGAACAATGTGTGGTGCATAGGGACATAAAGTCGAGCAATATTATGTTGGATTCCAATTTCAATGCTAAACTTGGGGAAAGCTTTGGCAGGCACGATGGGTTACATGGCTCCTGAATGTCTCACCACTGGCAAAGCTAGCAAGGAAACTGACTTCTATAGCTTTGGTGTTGTCGCGTTAGAAATAGTTTGTGGAAGGAAACCTATTGATCCTAAATCTGAGGAGCATCAAGTAAATATTATCGAATGGGCTTGGAGACTTTACGGGATGGGAAATCTTAGTGAAGCAGTTGATCCTAGACTCTCATCAGAGTTGAATGAACAGGAGGTGGAGCACTTGCTAATTGTTGGCTTATGGTGTGCTCGATCATCCAGATAACAATTGCAGACCTTCTATTAGGCAAGCAATTCAGGTGCTTAATTTTGAAGCTCCATTGCCCACACTCCCTCCAAACATGCCTGTGCCGATATATTGCAGTTATGAATCAATTACTTCACTTGCATCACCATACGACTCCAATGGTCCTCAGATTTCTGAAATAAACTCTTCAGTGACTAGAGATCATTCGACACAGCAGCATCTTCGCCTTCCGCATCACTTCTGTACACACGTTGA

>M6-PGSC0003DMG400004401 M6 L-LecRK

TCTCTCAAATACTTGCTTCTTATCATAATTATCCCTCTAGTCACTTCACTCTCCTTCAATTTTGATACTTTTACACCCAATGATCACAATGTAACCTATGAGAGAGATGCTTCTCCAGAAAATGGTGCAATTCTACTCACCAAAAACCTCCTCAATCGCGATATTAATGCTAGTATAGGTCGAGCTATATATTCAAAACCACTGCATCTATGGGACAAGGCCTCTAGAAATCTCACTGATTTCACTACACATTTCTCCTTTAGCATTAATTCACAGGGAATAACAAAATATGGAGATGGTTTTGCCTTCTTCCTTGCCCCTGCAGGTTCAACAATTCCTTATAATACAACAAGAGGTGGAAGTCTTGGCCTTACAAGTAATAATGAGCGACTAAATTCTTCGAGCAATCATTTTGTTGCTGTGGAGTTTGACACCTATCAGAACCCCTATGACCCAAAGGGTGATCATGTAGGTGTCGATATCAACTCTATGCAATCTGTTGTTAATGTGACTTGGTTTAGTAGCATTCCAAATGGTACGAAAACTGATGCCTGGATCAGTTATAATTCTAACTCAAAAAATCTTACTGCTGTCTTCACTGGTTTTCAACTCCAAGGGAACAATATCACAGTCACTGTCCTGCAGAACCTATCTTACAATCTTGATCTGAGGGAATATTTGCCAGAATGGGTCACTTTTGGCTTCACTGGTGGAGCGGGAAGGAGATTTGCATTTCAAAGAATCTATTCTTGGAATTTTACTTCTTCTTTAGATAATACGACAGATACAGGGGTAACCTTAACAAACACCATGCCAAAGGATGCACCAACAAAAACTAAGTTAGGACTTGTGGTTGGATTGATTTCTGGTGGTTGTGTTTTGGTTGCAGTATGTGCTTTGGTGTTATTTGCATTTTGGAGAAAGAGTAAGGTGAGAGAAGATGATTATGTCTTTGATGGTTCCATGACTAATGAATTTCAAAAAAGTGCAGGACCAAAGAAGTTCTTGTACAGTGAGTTGGCTAGATGTACAAATAACTTTTCACAGGAAGAGATGCTTGGGCGGGGTGGATTCGGAGGTGTTTATAAAGGATATCTCAGTGAATCCAACCCTTATATTGCTGTTAAGAGGGTTTCAAGGGAATCACAGCAAGGAATAAAAGAGTATGTATCAGAAGTCAGGATCGTCAGCCGGTTAAGACATAAACATTTGGTGCAACTCATTGGTTGGTGCCATCAAAAAAGAGAACTTCTACTTGTCTATGAGTTTATGCCTAATGGAAGCTTAGATCATCATCTTTTCAAGGGAAAAAGCCATTTGACATGGCCAATAAGATTCAAGATTGCTCAAGGTTTGGCCTCGGCGTTGCTATATCTACACGAAGAATGGGAACAATGTGTGGTGCATAGGGATATAAAGTCTAGCAATATTATGTTGGATTCTAATTTCAATGCCAAGCTTGGGGATTTTGGGTTAGCTAGGCTAGTTGACCATGATAAGGGATCCCAAACAACAGTTTTGGCAGGCACAATGGGTTACATGGCTCCTGAATGTGTCACCACTGGCAAAGCTAACAAGGAAACAGACGTCTATAGCTTTGGTATAGTCGTGTTAGAAATAGGTTGTGGAAGGAAACCTATTGATCCTAAAGCAGAGGAGCATCGAGTAAACATTGTCGATTGGGTTTGGAGACTTTATGGGATGAGAAGTATTCTTGAAGCAGTTGATCCGAAACTATCATCAGAGTTCAATGAAGAGGAGATGGAGCATTTGCTAGTTGTTGGCTTATGGTGTGCTCATCCAGATAATAATTGCAGGCCTTCTATTAGGCAAGCAGTTCAGGTGCTTAATTTTGAAGCTCCATTACCCACTCTCCCTCGGAACATGCCTGTGCCCACATATTGCAGCTTCTCACGCCATTTACCAACTATTTCAGTTTCATCAACACATGAAACCAGCGGTACAGAGATTGCCCTTACCCAAATCCAGCATTCAGTAAACGGAAACACCACTGATGGTTCAATAAAAAGAACATCTTCTTCAGCATCTTCATCTTCAACATCAGTTTTGTATCCAAGTTAATATTGCCCTGTTATAACCCTGCATTTTGTCTTCCATTAGCAGTGTCCATCTTTTCCTTTCTTACTTTTTGCTGTTTCTCTTCTAATGAAGAGGATTCTTTGGATTACATTTCCTGTAATTTTAACTGTTCTATTAAAC

>M6-PGSC0003DMG400008822 M6 L-LecRK

ATGTTCTTCAATCTTTTCATACTAGTAGCACTCTTTTTAGTTGATTTTTCACCAGCTTCTTGTGAAGTTGATGCATTCATTTACAATGGATTCCAATCAGCTAATCTTAGCTTGGATGGCATAGCACAGTTCACATCTGATGATCTTTTGTTGTTAACAAATTCAGGAACACAAAATCAGGGCCATGCTTTCTATCCAAATCCAATTCATTTCAAGAATTCATCAAATGGTACTGTATTTTCTTTCTCCACAACTTTCGTGTTCGCGATAAGGTCTGATTATGGAAATTTGAGTGGTCATGGACTGGCTTTCGTTATCGCGCCACATAAAGGACTTCAGGGGTCTTTGGCAAATCACTATCTTGGTCTTTTTAACTCTAATAATAATGGGAATAGATCAAACCATGTCGTTGGAGTTGAGCTCGATACAATCATTAGCGAGGATTTTGGTGATATCAATGACAATCATGTTGGAATTGATATAAATGGATTGAGGTCTGTAGCATATCATACAGCAGGTTATTTTGATGATACTGGTTTTTTTCATAACTTGACTTTAATTAGTGGCCAGGAAATGCAAGTTTGGATTGATTATGATGGGAGGACTAAGCAAATGAATGTAACAGTAGCTCAATTACATATGGAAAAACCAGTTAGACCACTTTTGGCTTTGAAATATGATCTTTCGTCTATTCTTGAGCAGAGTATGTATGTTGGTTTTTCATCGTCAACAGGTTCAGTCCCGACACATCATTATATCTTGGGATGGAGCTTCAAGACAAATGGGAAAGCTCAAGAACTCTCGCAACTTCCTAAGCTTCCTCCTCTTGGGAGTAAAGGGACGTCAAGATTTGTAACGATTGGTTTGCCTATAATCTCATTGGTTTCACTTGTTGCAGCTGTGTTAGCGGTGGTTTTTTATGTAAGAAGGAAGAAGTATGAAGAAATTCATGAAGATTGGGAACGCGAGTATAGACTACAAAGGTTTAAGTATAAAGAATTGTACATTGCTACTAAGGGATTCAGAGAAAAGGAGGTATTGGGAGCTGGAGGATTTGGTAAAGTTTATAAAGGAGTGATGCCTATCACTAAACTTGAGATAGCTGTAAAGAAGATATCTCATGAATCAAGACAAGGTATGAAGGAATTTGTTTCGGAGATTGTAAGTATTGGTCGCATGCAACACAGGAATGTAGTACAACTTTTGGGTTATTGCAGGCGAAAAGGGGAGTTGATTTTGATATATGAATACATGTCTAATGGAAGTCTAGACAAGTATTTATACGATCAACCAAGATACACCCTCGATTGGAACCAAAGATTCAAAGTCATTAGAGGTGTAGCCTCAGGACTATTTTTCCTACACGAAGAATGTGATCACGTAGTGGTTCATAGAGATGTTAAGGCCAGTAATGTCTTGTTAGATGGTGAACTAAATGGCAGGTTAGGTGATTTTGGACTCGCAAGGCTGTATGGTCATGGGACCGATCCTCAGTCTACTCGTGTTGTTGGTACTCTTGGTTACCTTGCACCAGAGCATACTAGAACTGGCAGGGCAACACCGAGTAGTGATGTATTTTCATTTGGGGCATTTCTGCTTGAAGTTGCATGTGGTAGGAGGCCGATACAGCCAAGACAAGACGGTGATGATCTGATTTTGGTCGATTGGGTGTTCTCGTGCTGGAATAGGGGTAATATACTTGATGCTGCTGATCCAAACATAGGCATTGATTTTGTTCCAGGGCAAGTGGAGTTGGTCTTAAAGTTAGGCTTGTTCTGTTCTCATTCAGATCCTTCGTGTAGGCCAACCATGCGACAAATCTTGTTGTTCTTGGATGGTGTTGTGGCCTTACCAGAGTTATCAGAACTCGGTGTTTCATCAGCTAACCTAACATTTGAACATCGCGGAGGTTTTGATGATTTTGTCAAGTCATATCCAGCTTCTTTGGGCTATGCATATTCCTGCTCTCCATCTGTAACAGACTCCTTTCTCTCTGGTGGCCGTTGA

>M6-PGSC0003DMG400008897 M6 L-LecRK

ACTAGATTTGACTATATTGTGTGATTATTTTTCATATATACATATTGAAGTTTGACTGGCCTCAAGAAAATGACATTACAGAGTTTACATTGTAGACTTGAGCATTAGGAGATAGATAGTCATATTTGTGATTGTTATCAGCACTAATCCTAGGAGAGTTCTGAGCAGACATGGAGAAATTGTTTTTAGCCATGATTTTTCAGTTTCTTGGTTTGCTTTTTCATGCTCCATTAGGCTATGCAGCTGCAGAAGATCTCGATTTCACCTACAACGGGTTCCACTCAACAAACTTGAGTCTTGATGGAATAGCAGAGTTCACATCAGATGGCTTGTTAAGGCTAACTAATGCTTCTCAATTGCAGGAGGGTCATGCTTTTTATCCTAGACCTATCAATTTCATGAATTTAGCATCCAATGGTTCAGATTTCTCTTTCTCAACCACATTTGTCTTTGCCATAGTGCCTCAGTACTCGATTTTAGCTGGTCATGGTATGTCTTTTGTGATTGCACCAGCTAGAAGGCTTCCAGAAGCACTTCCAAGCTCTTATCTTGGCTTGTTTAATGATTCTAGTAATGGAAATGCCACCAGCCATGTTGTTGCGGTGGAGTTTGACACGCTACAGAACCGTGAGTTTGATGATATTGACAGCAACCATGTTGGCATTGACATCAATGACCTAAAATCTGTAATAGCCAAGCCGGCCAGTTATTATAACAGTAATGAGCAATCCTTCAATAACTTAACTCTAGCTAGTGGCCAGCCAATGCAAGTTTGGGTGGAATATGATTCTGTGGATAAGCATATGAATGTCACAATGGCTCCACTGCACGTTGCAAAACCAGACCGTCCTCTCCTGTCCTTAGTCTACGATCTTTCATCTGCCACGGATGAAAACGTGTCTGTTGGCTTCTCAGCATCCACCGGTGCAGTTGTTTCAACACATTATATCCTTGGATGGAGCTTCAAATTGAATGGAATGGCTCGAGGGCTTGATCTCTCTCAACTTCCTAAGCTTCCTCGAGTAGTACCTAAGAAAGAATCCAAGCTTTTATTGCTTGGCTTGCCCGTGATTTCTTCAGTTTCATTTGTGATTGTAGTCCTGGTAATAATTTATTGCATAGGAAAGAAGATGAAGTTTGCTGAATTGCAAGAAGAGTGGGAGCTTGATTATGGACCACACAGGTTCAAGTACAAGGATTTGTACACTGCCACAAAGGGCTTTAGCAGGAAGGATTTGTTGGGATCCGGCGGATTTGGTAGAGTCTACAGAGGGGTATTGCCCATATCCAACATTCAGATAGCAGTCAAGAGGGTCTCTCATGAATCAAGACAAGGGATGAGGGAATTTGTTTCAGAAATTGTTAGTGTTGGCCGGCTACGTCACAGGAATTTAGTACCACTTTTGGGTTATTGCAGGCGTAAAGGAGAGTTACTTCTGGTTTACGAATGCATGCCTAACGGAAGCCTAGACAAGTATTTGTTTGACAAACCAAGCTGCACTCTTGATTGGAACCAAAGATTTCGAGTCATCAAAGGTGTAGCATCAGGACTAGTCTATCTTCATGAAGAGTGGGAACAAGTAGTGATTCATAGAGATATAAAAGCGAGTAACGTGTTGTTAGATGTTGAATTAAATGGAAGATTAGGAGATTTTGGGCTAGCAAGATTGTATGATCATGGGACAGATCCTCAGACTACGCATATCGTTGGAACTCTCGGATACCTTGCCCCAGAGCATACAAGAACAGGCAAAGCCACGACTAGAACTGATGTATATTCATTTGGAGCATTTTTACTTGAGGTGGCTTGTGGGAGAAGGCCAATAGATCAAAGGGTATCGGACGAGGATCTCGTTTTGGTTGACTATGTATTCTCTTGTTGGACTAGAGGTGATATTCTTGAAGCCATTGATCCAAACTTGGGCAATAACTATGTAAAAGAAGAGATTGAGTTGGTACTAAAACTTGGACTATTGTGCTCTCAGACAGAACCAGCAGCTAGGCCAAATATGAGGCAAGTGTTGATGTACTTGGAAAATGCCCTGCCTTTGCCAGAGTTATCACTGATGCAAACTTCAAATTTTAGCTTAACTTTTGCAGGCTTTGATCATTTTGCAATGTCATGGTCTTCTTCTGTTACTGACTCCCTTCTCTCCGGTGGTCGATGACTCGGGCGAAATTTTAATATATTCCTAGTAAAAATAAAACTTGTTATATAAGCTATATTTCTTCCACCTAAGGTTGTATACTTGGTTCATTTAGGCATCTTGGAG

>M6-PGSC0003DMG400010466 M6 L-LecRK

CGTGTTTCCTTTGGATTCTCTGCTTCATTCACACCCCTCTTCTTCACCCCCTGTTTCCAGTTTTCTTCAAAAAAAATCACTATTTTCTAAGCTCAAAAAATAGCTAAAAATGAAATCTTTGTGCAATTTCATCTTCATTGTGTAATGGGGTTTTGCTAAAATCTAGCAATGAAGTCAATTCTTGAAGAACTTTCCAGCAACATATTCCTGTAAAAGTCATCCTGTTGTAGTGTTGTTCTCTCCTGTCTACTTTCTTGCGTTTTCACTGTTTCTAAGCTAAAAAAAAAGCTAAAAAACAAATCTTTGTGCAATTTGATTTTCATTGTGTAATGGGGTTTTGCTAAATCAATTCTTGAAGAAATTTCCTTCTGTAAAAGCCATCATGTTGTAGTGTTGTTCTCTCCTTAATCCTCACTCTACTGTGTCAGTTTTCTACTTTTTTGCATTTTCACTGTTTCTAATCCCAAAAAAAAGCCTACCCTTTTTGTATCTTCACCAAATTCTTGTGATTTGGTGTGAAAAATCTTACCTTTTGTACACTTTGATCTTCATTCTCTGATGGGTTTTTTCTAGAATCTAGCAATGACATCAAATCTTGAAAAAAAATTCAGGGGCTTTCTTGTATGGCTCATCTTATTGTTCTGTTTCTGTAGTATTAGTTCAACATTTGGTAAAACTGATTTCGATTTTGAAAATTTAACACTAAGTAGTTTAAAACTTCTTGGTGATGCACACTTGAGTAACAACACTGTAAGGTTGACAGGTGATCTTGCTGTACCAAATTCCGGTGCCGGAAAAGTTCTGTATTCAAAACCAGTAAGATTCCGGCAACCGGGGTTTGATTTTCCGGCGAGTTTCTCGACATTCTTTACATTTTCAGTTACTAATTTGAACCCATCATCGATTGGTGGTGGTCTTGCTTTTGTGCTTACGCCGGGTGATGAGTTAGTAGGTGATGCTGGTGGGTATATGGGAATCTTGGATGCTAAAGGGACACAAATTGGTACAATTGCTGTGGAATTTGACACCCTTATGGATGTTGAATTCAAAGATATTAATGGTAATCATGTTGGTTTGGATCTAAATTCAATGATTTCAACTGAAATTGGTAATTTGGATTCCATTGATATTGATCTCAAGAGTGGTGATTTGGTTAATTGTTGGATTGAATATTCTGGTTCTAATGGAGAGATGGATATATTTGTGTCATATACTAATCTCAAGCCAAAAGAATCATTTTTATCAGTTAACATCAATCTTGCTGAGTATGTAAATGATTTCATGTTTGTGGGGTTTTCTGGTTCAACTCAAGGGAGTACTGAGATTCATAATATTGTGTGGTGGAGTTTCAGTTCATCATTTGATGCAACTCCAAACTCGGGGGAGGTGGTGGCTGAGCCACCACCGCCAACGACTAGTTTGATGAACCCAACGGCAGATTCTGTAACGTTGCCAACACCTTCTATGGCTCCATCAGAGTCTAATAGTAGCACAGGTGTATTGCAAGAAAAGAGTAGTAGGAAATGTCATAGCAATTTTTGTAGACAAGGTCCTGGAGCTGTTGTTGGTGTGGTGACTGCTGGTGCATTTTTTCTTGCATTTGCTACATTGGTACTTATTTGGTTATACTCCAAAAGATTCAAGAGAGTGAAAAATTCTGAAATTGTGGCATCTGATATCATCAAAATGCCTAAGGAGTTCAGCTATAAAGAGCTTAAATTGGCCACAAAAGGTTTCATTCCAACGAGAATTATAGGCCGTGGTGCATTTGGGACTGTTTACAAGGGCATTTTATCGGATACCGGTGGCATTGTTGCAGTCAAGAGATGTAGTCATAATGGACAAGGGAAAGCAGAGTTCTTATCTGAATTATCTATAATTGGAACCCTTAGGCATAGAAATCTTGTTAGGCTTCAAGGATGGTGCCATGAGAAAGGTGAAATTTTGTTGGTTTACGATCTTATGCCTAATGGGAGTCTTGATAAGGTACTATTTGAATCAAGAATGGTTCTTTCATGGTCACATAGGCGAAAAATCTTGATAGGTGTTGCTTCTGCCTTGGCATATTTGCATCAAGAATGCGAAAACCAGGTGATTCATAGGGACATTAAGAGTAGTAACATCATGTTGGATGAAGGGTTCAATGCAAGATTAGGAGATTTTGGATTAGCAAGGCAAGTTGAGCATGACAAGTCCCCCGATGCAACGGTAGCAGCCGGGACAATGGGCTACTTGGCTCCAGAGTACTTGTTAACCGGTAGGGCAACCGAAAAAACTGATGTTTTCAGCTATGGAGCAGTGGTTCTTGAAGTGGCTTGTGGAAGGAGGCCAATTGAGAGGGAAACAACTAAAGTTGAGAAAGTTAGAGTGAATAGCAACTTGGTTGAATGGGTATGGGGGGTGCATAGAGAAGGGAATTTACTAAATGCAGCTGATTCAAGACTTTGTGGTGAGTTTGATGAACAAGAAATGAGAAGGGTTCTAATGATTGGACTAGCTTGTTCACATCCCGACCCTACGGCTAGACCAACAATGAGAGGTGTAGTTCAAATGCTGGAAGGTGAATCTGAAATCCCAATTGTCCCAAGAACTAGACCTTCTATGAGTTTTAGCACATCACATCTTCTAATGAGTTTGCAAGATAGTGTTTCTGACTTGAATGGTTTGATCACACTTTCCCCTTCATCATCCGAAAGTAGCTTCACCGGTGGTCGCAATGGCAATGGCATGGAGTTGGTCTAAATGCAAGACTAAAAAAGACTTGACTTTTCGAGTTAATGCACCTATAAGAGCCCAACAATCCCACATATGGGGCCACCATCAAGGCGGAACAGACACAATGACTACTCTGATTGGGCGTGGACAAGGAAAAAGTGGCGTCTTTTTTGTTTTGTTCATAAATTCATTAATTTTATTCATAATATATATGTACCAAATAAGCTTTAAGATTTAGTAATGTAATTTTCTCCTTTTCCTTTCATATACTATAGTCCATTGAGTGCAAGATTAGAGATATTTAGTGGGGGAAAAATAAA

>M6-PGSC0003DMG400011071 M6 L-LecRK

AAAAAAAAAAAAAATCTCACTATCCATCACCAAATCTTGAAAAAAAATCTTGTGTATTGTCTTTTTTTTAAGGCAAATCGACTTCGATTCTTGAATAATTCACATCACTTATGCTTCTATATATCTTGCTTTAAAATGTTCAATTTTCTCACATTCCTCTGCTTTGTCATTTCCTTCTCAATTCTTGCTTCATCTCAACTTGATGAATTCATTTTCACAAAATTCAATCAACCAAACAACAACATAACCTTAAGTGGAGTAGCAAAAATAAGCCAAAATGGGTTTATTCAATTAACCAATGACACAAGTAGACTAATAGGACATGCTTTTTACTCTTCACCTTTCCACTTCAAGACTACTAATGCCTCTACCTTCTCTTTCTCAACGTGTTTCGCGTTGGCCATAGTCCCCGAGTATCCAAAACTCGGGGGCCATGGCCTAGCGTTCACAATTTCTCCGTCAAAAGACTTTAGTACAGCTCTTCCGAGCCAGTACTTAGGTCTGCTCAATGCTAGTGATATTGGTAACTTCTCTAACCATATTTTTGCTGTTGAATTCGATACGGTACGTGATTTTGAGTTTGGTGATATTGATGATAACCATGTTGGTGTTAATATTAACAGCTTAGAATCAAATAAATCAGCTGCAGCTGGCTATTTTATTAACGATCAAAATGTGAAACAAGATTTGAATCTTAAAAGTGGGAAAGTCATTCTTGTTTGGGTTGAATATGATTCTGTTACAAAATTAATTAATGTAACTCTTTCACCAAATTCTTTAAAACCAAAAATTCCTCTTTTATCTTATAATTTTGATGTCTCTACAATTTTTAAAGAAAATATGTATGTTGGCTTCTCTGCTTCTACTGGTTTGCTTGCTAGTTCACATTTCATTTTGGGTTGGAGTTTTAAGTTAAATGGAGAAGCAAAATCACTTGACTTGGATTCTCTACCATCACTTCCTGGACCTAAGAAGAAGCACAGAAGACTAATTATTGTCATCTCGATTATAGTGTTTGTGTTCGCGTTAATTTCAATCTTGATTGCTTTTTATACAATACGGAGATTCAAGAATGCTGACGTGATTGAGCAGTGGGAGCTTGAGATTGGTCCTCATAGATATTCGTATCATGAACTCAAGAAAGGTACTAGAGGTTTTAAGGAGAGTGAACTACTCGGGCGAGGTGGATTTGGGAAAGTTTACAAAGGTGTTTTGAGAAATTCAAAAGCAGAAATTGCTGTGAAACGAATTTCTCACGAATCTAAACAAGGGTTGAGGGAATTTGTGTCTGAGATTGCGAGCATTGGAAGACTCCGTCATAGGAATTTGGTTCAATTGTTAGGATGGTGTAGACGTCGTGGTGATTTGTTGCTTGTGTATGAGTTTATGCCTAATGGAAGTTTGGATAACTTCTTGTTTGATGAGCCAAAAATGGTGTTGACATGGGAACAAAGGTTCAGGATTATTAAAGGGGTTGCGTCTGGTTTACTTTACTTACATGAAGGCTATGAACAAGTCGTGCTACATCGAGATATTAAGGCAAGTAATGTGTTACTAGATGGTGAATTAAATGGTAAGTTAGGGGATTTTGGACTTGCTAGATTGTATGAGCATGGGTCTAATCCGTCCACGACTAAAGTTGTAGGTACATTAGGGTACCTTGCACCAGAATTACCAAGAATAGGACGAGCCACTACAAAGACTGATGTGTTTGCCTTTGGTGCTTTGTTACTCGAGGTGGTTTGTGGACGTAGGCCAATTGAACACAAAGCAGGACCCGAGGAGTTTGTTCTCGTGGACTTGGTGTGGGATAAATGGAGAGAAGGTAAAATACTTGACATTGTGGACTATAGATTGAAAGGCGAATACAATGAGAGTGAGGTTTTGATGGTGTTAAAATTGGGACTAATGTGCTCAAATAACAAGCCATTGGCGCGACCAAGTATTAGGCAAGTTGTAAGATACTTTGAAGGTGAAATTGGGATGCCTGAGGCTCCAACAACACAAGGTTCTGATGATCTTGAAGGATTTGGATTCGATGAAAGTGAACGTTGGAATTCCTTAGCATCGACTAGTATTATGATCAAGTCAAGTTCATTTACATGTTTGTCTAATGGAGATGAAAATAGTACTAATTATGTTTCCTCTTCTACTTCACCACTTCCACTTCTATATAGTAGTGAATGTACCTAGATACTAATGTGAACTAGCATTGCAAACTTTTCTATATTTTCCTGAAGAACACTAATGTACATTGTTTTTTCCTTTTCTTTGTGTTACCTCTTTGTAGTAGTTTTTTTCCCACAGAAAACTTAGGAAGCTTCCCGGTATTTCAGTGAGAACCTGTTTTCCACCATGTGTTTTCCCAGTTACCTGGTTTGATGGGAAATGGGTGGAAAAATCATGTTTTATAATACAAATTTTTCATTGATTCGCGATGGAA

>M6-PGSC0003DMG400018184 M6 L-LecRK

TAGGAATTAAAATAATTAGTCAAAGGATGAGTCATCTGCATCATTTTCAGAGAAACAGCACTCATCTCTAATTCCCTTCAAAGTACCATGTAATTAACACATAATTTCCCAGTTTCCTTCACAAAATTCTCTTCTTCTTCTTCATGCAAACCCCAACAATATGCCACCATTATCACTGTTCTTCATTCTAATCAACCTTACAATCTTCCCATCCATTTCCGCCGTCGAATTTCTCTTCAACTCCTTTACTGCTAACACCACTCCTGCCCTCAACCTCATCGAGGATGCCCGCCTTGAGCCACCAGTTATCCGCCTCACAAATGACTCCAACCAATTTTCCCTTGGCCGAGCTTTTTACCCATCTCAAATTCCTATAAAATCCACTTCAAATTCCACTTCAATTTCTTCCTCTTTCTCTACTCAGTTCATCTTCTCTGTACTTCCTGATGATTCCTCTAGTCCCGGATTCGGAATTGCTTTTGTTCTCTCTGCTTCAACATCCCCACCTAATGCCCTCTCCAGTCAATACTTTGGACTTTTCTCTAACGCCACCGTACACACGGTGGCCCCACTTCTCGCCGTCGAGTTTGATACGGGTCGTAACCCGGAATTCAATGAACCAGATAGAAATCATATCGGGATTGATCTTAACAGCATTGAATCTATTGTTACCCAAACGGCTGGGTATTATAGTTCTTCTGGTAATGATAATGATTCTTTTGTGCCTTTAAATTTGCGGTCTGGGGAAAATATACATGCTTGGATCGAGTTTAATGGACCTGAATTCGAGATTAATGTTACTATAGCTCCGGCTGGTATGTCACGGCCTGTTAGGACTTTGTTGAGTTATAGAAACCCCATAATTGCGAATTACACGTCTGCCCAGATGTATATGGGGTTCTCTGCTTCCAAGACTCAATGGGTTGAGGCGCAAAGACTTCTAGCTTGGAGTTTCAGTGATTCTGGAGTTGCAAGGGATATTAATACTACGAATTTGCCGGTTTTTCAGCTGGAAAATTCCGGGTCTTCGCTATCTCCCGGTGCTATTGCTGGGATTGTTATTAGTTCTGTGTTTGCTGTGTTGGGTTGTTTATTTATATTTTATTGGTTTTGGTGGCGTAAGAAGGAAGAAGATGATGTTATTGAAGATTGGGAACTTGAGTACTGGCCTCATAGATTTTCATATGAAGAACTTAGCCAAGCTACAAAAGGGTTTTCTAAAGATGAGCTACTTGGGGCTGGTGGTTTCGGTAAAGTATACAAGGGAACATTTGCTAATAACACAGAAGTGGCAGTGAAATGTGTGAACCATGACTCAAGACAAGGAATTAGGGAATTCATGGCTGAGATATCTACTATTGGAAGGCTTCAACATAAGAATTTAGTACAAATGAGAGGGTGGTGTAGGAAGGGGAATGAACTTATGATTGTGTATGATTACATGCCTAATGGGAGTTTGAATAAATGGATATTCGATAAGCCGGAGAAGGTTATGAATTGGGTAGACCGGAGGAGGGTCCTAGCTGATGTTGCAGAGGGGTTGAACTATTTGCATCATGGTTGGGAACAAGTTGTTGTACATAGGGATATTAAATCTAGAAATGTTTTGTTAGATAGTGAAATGAGAGGGAGATTGGGTGATTTTGGCCTAGCTAAGTTGTATACTCATGGTGGTGTCCCTAATACTACTAGGGTAGTAGGTACATTAGGGTACTTGGCACCTGAAGTGGTGACAAGGGCTAACCCAACTGCTGCTAGTGATGTTTATAGTTTTGGGGTGGTGGTTTTGGAGGTGGCATGTGGGCGAAGGCCGATTGACTTTGGGATTGTGTTGGAGGAAGAGGAAGTGTTGATTGATTGGGTTAGACAAAAGTACAGGGAAGGGCGATTATGTGAGGCAGCAGATAAAAGGATTAAGGGGCAATGTTCAGAGGAAGAAATGGAAGCTATGTTGAAACTAGGACTTACATGTTGTCACCCTGATCCTCTCCGTCGACCTACTATGGGAGAGGTTGTTGCTGTATTGCTTGGTGAGAATGTGGATGCAACGCCAAATGAGTTGATCGTTGAATTAGCACCTAGTGAAAGGAACACGAGAGATAGAAGCGACTGGTCAACTGAGGAATCAGAACCACTTTCGGCTGTGTAGATATTATACTTTTGTTTCCTTTATTGTTGAATCTTTTTCATCCCTTGAAATTATTGTTTAAGGTTTGTTCTTTGTATTGTTCTTACATGCTTTTGAACTCTAAACTTAGCATTAGTTGCAAATAGTTATGGATATGCTGCTATGCCTTTGCGCTACCTTAAAGGGTCAACATCCATGGCAATTAGCAAACGTCTATACGCATTAAGATTCACCTCGGTGGAATAACG

>M6-PGSC0003DMG400018677 M6 L-LecRK

ATTTTACACAACTCCAACATGCTTCAAAATAGCTTCTTTTATTGACCCTTCCTCTTATTAAAATCCCTTACATCTCTATACTTCTTTTCACTACACAATATGGAATTTTTGACACCAAAAATCTTCATATTTTTCTCTTGTCTTCTACTCATATCACAAGCCAAAATCATAAAATTTGACCAACAATATGGTGACCCTTTTGATCATACATATGTTCCCTTACTTGAAATCAAACATCCTGCACAAATAAGCAATCAAGCACTTCAAATCACCCCAGACTCAGCTTCCTCTGCTTACCAAATGTTCAATAATTCAGGCAGAATCCTATTAAAGCAGCCTTTCAAACTATGGGTAGACAATGATATTTCAAGGACAGCGTCTTTCAACACATCATTCCTAGTAAACATTTTTAGACCGGACAACAATACACCAGCTGAAGGTCTAGCTTTCTTGATATGTCCAAATTTTGATTTGCCACTAAATAGCCAGGGCCAGTATTTGGGCCTAACGAATGCTACAACAGATGGTGAATTATCCAACAGAATAATCGCGATTGAGCTAGATACATTCAAACAAGAATTCGATATTGATGATAACCATATTGGGATTGATATAAACAGTGTTAAATCCGTTAAGTCGGAACCTTTAACACCACATGGAATTGAACTAGCTCCAATAGGTGAAAGAGTTTACAATATATGGATACACTATGACGGATTCAAGAAAGTACTTGATGTGTACATTATTGAACAAATGAGGAAAAATGGGACAACACCAACAAGGCCAAAAGTACCAATACTAACACACAATGTTGATTTGAGAGAATTTGTAAATCAAGAATCATATTTTGGATTTTCAGCTTCAACAGGACATTTCAATCAGTTGAATTGTGTGTTGAGATGGAATTTAACAGTTGAGTATTTTCAAGAAAAGAATGATCAAGAAAAGGTACTAATAATAAGTCTAAGTGTTGGTGTATCAGTATTGGTTGTTTTATTGATTTTGTCAGGGTATTTTGGGTATTTTTTTTATAAGAAAAAAAGGGATGATAGGTCACAATCTAATATATTAGGGGCATTAAAGAGTTTACCTGGAATGCCTAGGGATTTTGAGTTTAAAGAATTGAAAAAAGCTACTAATAATTTTGATGAAAAGAATAAACTTGGTGAAGGTGGATTTGGAGTTGTGTACAAAGGGTATTTAGTTGGTGAAAATTTGGAAATTGCTGTGAAGTGGTTTTCTAGGGAAAGTATCAAGGGTCAAGATGATTTTTTGGCTGAGTTGACAATTATCAACCGTTTAAGGCATAAACATCTTGTCAAACTACTTGGTAAGTATTTTGAACTCTTTACTCTTATGGTTTTTTTTTTGAGTATTTGTTTTTTGTTCAAGCTCGGATAAAGGAGGAATGTTATGACTACCAGCAAAAGAATAATCAAAGTTTTTTGAGTGGATATAAAGATTTTGATATATATGCTTATTGAGTTGAAGTGTAATTGGTTGATAGATTTGTCCTGTCTTTGGAATTTTATCCATGTTTTATGTATAGGTAGCAAAATCTACTAAGATTTTATGGAGAAAAAAAAAAGAGACAGGTTTTACACAGTCATTTCTCCGTTTTGAGGACACTTTTTATTTATTGAGTCGTGTTTCTTATCCTTCCTCGACATAGTAACCGCTTTCATATTTTTTATAACTTTCGAGTACAAGTGCATGTTCATCCATCTTCATAGTCATTTCTTTGTTCCTTTAGTCGTTTTGATTGCACTTTTTTTATTGAGTCGTATTTCTTCTCCTTCTTCAACATAGTGATTTGTTTTATGTTTTTTTTTTTCAAATTTCATAACTTTCGATCACAAGTGCATGTTCATCCATCTTCGTTTTGAACTAAAGCTTGAATTAAATTTTATTAATTAAGGTACCAAGAATATTTTCTTGTGTTCTAAGGAGTTCCATTTTGACTCATGTTAGGCCCATCTCCATCTTGAAAACGGTCCCAATAATTATATCCAACTACTATAAGATAAGTGAAAAAGTTAAAAGCTTTGATGCAACTTGCAAGTGTCAAAAATGGTGTGCTCATGTACCTTTTACAATAGTTTGAAAGTGATAAACATATACTTTTTCGTCTCAAAATATTTATTTATATATTTTTTATACGCTTTTTGAGAAAACATTAATTAGTAAAGATATTTAATTATTTCCCCCTTATTTATATTTTAAAACATAATCTCTATTTATATGTCATATTTGAGGTATCTGTAAATTTCAAAAATAATTACTAACTACTAAATATAAAATGAGGTAGGGGTGATTAATTTTAACTTGAATTTTTAAAATATAATCCAAAGTTTCTTGGAGGACCTACTAACTAATAATTTTAATGCCCTCTGGATGAAAAATCTAAAGTTAATAAATATTAAATTCTTAGCTGGAAGACAGAAGAGCACTGATTCCATTTCCTTTGTGTCTTTTTGGTTTTCTATACCACCAATTTCGAACATAGGGTCCCACACTCTTTTGTTCTTTGTTAGCTTCGGTATAAATATCGAGTAATTCTGATCATTTAAATTAATTATATGAAAAAAATTGTCTAGTATTTTTATTTCGATTAAATTTGAATCATTGTTTGAACAAACTTTTCTTTTTTAGGGAAGCTTCCTGTCTTTGAAAATCACTTTCATTCAAATGTCACCTATTATGATAAGTTTGGTACTTTCTAATATTAATATTAATATAATATAATTGGACAATTGCTTCAAGATGCACTCAACTCCAACTAAGACTAAGATTGTAACCCTACAACATGTCATCATAATTGAGAGAGGTCCAGATATTCCAAAATTAGGTCCTACAAAAAAGTTGATTGGAACACATACAACCCAACTTTTTTTGCCTTGTAAAGATTTGATATCCTCAAAATAGTATTGATTAGATATATTTATAGATTATTATTATTGTAGTAACAATTTAGCTTATATATGCATTAACAATTTTAACTCCACCCATGTGAGCTCGCTCAAATTGTCTGTCCTTAATCCGAATAAAGAAGGAGGGTTGTTGAGGATCACTCAGAAACACCATTCTCTATCCCATAAGAGTAGGAGTAAGGCTGCATCAATTCATAGATCATTATTATTGTAGTCATAATTTGGCCTATATATATGTACTAATAACAATTTCAACTCTGTCATCAGTGTGATCTCGCTCACATTGTCGGTCTCAAGATTGAATATAGAAGGAGAGTTGTCGAGGGTCAACCGAAAACGGTCTCTCTGCCTTATAAAGATATGAATAAGGATATTTACATCTTACCTTTGTCAGACGTACCCCACTTGTAGGATTTCACTGAATTTGTTGTTATTATTGTTGCACTAACAATTTCAAGTGATCTAAAGTAATTTTTGAACCTCCTTTACCTTCTAATGAGAACGTAACCTAGATGTTACATGTAGTTATCATTTAGGTGACCTAATGGTATAAAAGTTCTTTTACGCTGTCAGTGTATATATATATGCTAATCACATGCATATTATTGTGACAGGATGGAGCCATAAGCATGGTAAGCTACTACTTGTATATGAATACATGCCAAATGGTAGCCTGGACAAACATCTCTTCTCATCAGAGCAAGAAAAAGAGCCACTCAGCTGGAGTGTGAGGTACAACATTGTATCGGGAGTCGCGTCAGCTCTGCACTATCTTCACAACGAATACGAGCAGAAGGTAGTCCATCGCGATCTCAAGGCAAACAACATCATGCTCGACTCAAACTTCAATGCACGTTTAGGTGATTTTGGATTAGCACGAGCAATTGACAATGAAAAGACCTCATATGCTGATGAAGCTGAAGGTGTACTTGGCACGATGGGATACATCGCGCCAGAATGCTTCCACACAGGAAAAGCCACTCAACATTCTGATGTCTATGCATTCGGAGCTGTGTTGTTGGAGCTAATATGTGGCCAAAGACCTGGAACTAAAGTTAATGGCTTTCAACTCTTTCTTGATTGGGTTTGGTTCTTGCATCGCGATGGGAGAATCCTTGAAGCCGTTGACAAGAGGCTCGAGGATCAATACGTGGTTGAAGAAGCAGAGAGATTGTTACTACTTGCTCTGGCTTGCTCCCATCCAATTGCCAATGAAAGGCCTAAAACACAAACAATAGTTCAAGTTATATCAGGATCAGTACCAGCGCCACAAGTACCACCATTCAAACCATCATTTGTATGGCCTTCTATGGTTCCAATTGACATAGATTCATCGAGCATTGTCGATACAATATCCATCACAACTCCTCAGTACAGTTCAGAGAACAACAGTATTGAATATCAAAGCAAGTAGAGATAGCAATAGCTTGATTTAGGCCACTAAGAGATTTTTTTTTGCTTATTGTGTAATGACAAGAATAGCAGCTCATGTATATATTTCTCAGCCTTTTATGTTAAATTCCATCCTATACATTGTTCCTTCTACGGTTTGACTGTTTGGTTTGTTGTATTAAAAATGATGTGTATTGCATAATTTATAAAAAGAAGTTGTTCG

>M6-PGSC0003DMG400020326 M6 L-LecRK

CAACAATAGAACAAATTAGCATATTTTCTCTCCACCAAGAAAATATTACCAATGTTGTTAAATTTTGTCATACTAGTGTCTCTCTTCTTTGCACCAACTTCTTGTGAAGTTGATGGATTCATTTACAATGGATTCAAATCAAGTAATCTTAGCTTTGATGGCATTGCTAATATTACATCCAATGGTCTTTTGTCGTTAACAGATTCCAAAACACAAGATCAAGGCCATGCTTTCTATCCAAATCCAATTCATTTCAAGAATTCACCTAATGGTAATGCATTTTCATTCTCTACAACATTTGTGTTCGCGATAAGGTCTGATTATAGACCATTGGATGGTCATGGACTAGTTTTTGTTATCGCGCCACAGAGAGGAATTCAGAAGGCTTTGGCAAACCACTATCTTGGCCTTTTTAACTCAACTAATAATGGGGATAAATCAAACCATGTTGTTGGAGTTGAGTTTGATACAATAATTAGCGCGGATTTTGGTGATATGAATAACAATCATGTTGGAATTGATATAAATGGATTGAGGTCTGTGGCAAATCACACAGCAGGTTATTTTGATGATACTGGTTTGTTTAACAACTTGATTCTAGTTAGTGGCAAGGCAATGCAAGTTTGGGTGGATTATGATGGCAGAACTAAGCTAATCAATGTGACAGTAGCTCCATTACATATGGAAAAACCAGTTAATCCACTTTTGGCTTTGAAATATGATCTTTCGCCGATTCTTAATCAGATTATGTATGTTGGTTTTTCATCATCAACTGGTTCAGTCCCAACACATCATTATATCTTGGGATGGAGCTTTAAGGCAAATGGGAAAGCTCAAGAACTTTCTCAACTTCCTAGTCTTCCTCGTGTTGGGCGCAAAGGAACTTCAAGATTTGTAACGATTGGTTTACCGGTGATCTCTTTGGTGTCAATTGTTGTAGCAACCTTAGTGGTGGTTTACTATGTAAGAAGGAAGAAGTATGAAGAAGTTCTTGAAGATTGGGAACGTGAGTATAGACCGCAAAGGTTCAGTTATAAAGATTTGTACACTGCTACTAAAGGATTCAGAGAAAAGGAGCTGTTGGGAGCTGGAGGATTTGGTAAAGTTTATAAAGGAGTGATGCCTGTAACCAAACTTGAGATAGCTGTAAAGAAGATATCTCATGAATCAAGACAAGGGATGAAGGAATTTGTTTCGGAGATTGTAAGTATAGGCCGTATACAACATAGGAATGTAGTACCACTTTTGGGCTATTGTAGGCGAAAAGGGGAATTACTTTTGGTTTATGAATTCATGTCTAATGGAAGTCTAGACAAGTATTTATACGACCAACCAAGACTCACCCTCGATTGGAACCAAAGATTCAGAGTTATTCGAGGTGTAGCATCAGGACTATTTTTCCTACATGAAGAATGTGACCATGTAGTGGTTCATAGGGATATTAAGGCTAGTAATGTCTTGTTGGATGGTGAACTAAATGGCAGGCTAGGAGACTTTGGACTCGCGAGGCTATATGGTCATGGGACCGATCCTCAATCAACTCGTGTCGTTGGTACTCTTGGTTATCTTGCACCAGAGCATACTAGAACTGGCAGGGCAACACCTAGTAGTGATGTATTTTCCTTTGGGGCTTTTTTACTTGAAGTTGCATGTGGTAGGAGGCCGATAGAGCCAAGACAAGCCGGTGATGATCTGATTTTAGTGGATTGGGTGTTTTCGTGCTGGAATAGAGGTAATATTCTTGAGGCAGTTGATCCAAACTTAGGCAATGATTTTGTTCCAGGGCAAGTGGAGTTGGTCTTAAGTCTAGGCTTGTTCTGCTCTCATTCAGAGCCCTCATTTAGGCCAACAATGCGACAAAACGTGTTGTTCTTGGATGGTGTTGTGGCCTTGCCAGAGTTATCAGCCCTCGGTGTTTCATCTGCTGGTCTAACATTTGACCATCGCGGAGGTTTTGATGATTTTATCAAGTCGTATCCAGCTTCTTCTGTTAATGCACATTCCCCCGCTTCATCCGTAGCAGAATCACTCCTTTCTGATCCCCGATGAGTTGGACATATCTTCTTTGGTGTATTATTGATATACTGATCCTGGTAAATATAACATGATTTTAGTGATTATGTACTTCTGATAAATTATCGCAATTCATACACCTGAAATATATTGTGACATATCTTAAGATATCTGCTATGGTATCCTGATCTTATTGTTTACTTCCAATGCTAAATAAATACTAATCTGAGATTCAGCAATGGC

>M6-PGSC0003DMG400020327 M6 L-LecRK

AAAAAATGGATAAAAAAGATCATTTTTTTGTCATATCAAGAAATGTATAAACAGTAGAACAAATTAGCTTTATTTGGTCACAAATCACAATATTATCTCTCAATCAAGAAAATATTGGCAATGTTGTTCAATCTTGTCATACTAGTGTCTCTTTTCTTAGTTGGTATTGCACCAGCTTCTTGTGAAGTTGATGCATTCATTTACAAAGGATTCCAATCAAAGAATCTTAGCTTGGATGGCATAGCAGAGTTTACGCCCGATGGTCTTTTGCTGTTAACAGATTCTGAAACTCAAGATCAGGGCCATGCTTTCTATCCGAATCCAATTCATTTCAAGAATTCAACTAATGGTACTGTAGTATCATTCTCCACAACATTTGTGTTTGCTATAAGGTCTGATTATGGAATTTTGAGTGGTCATGGACTGGTTTTTGTTATCGTGCCACAGAGAAGAATTCAAGGAGCTTTGGCATACCACTATCTTGGCCTTTTTAACTCAACTAATAATGGGGATAGATCAAACCATGTTGTTGGAGTTGAGCTCGATACAATCTACAGCGAAGAATTTGGTGATATCAATGCCAATCATGTTGGAATTGATATAAATGGATTAAGGTCTGTAGCAGTTGAAACAGCAGGTTATTTTGATAATACTGGCTTGTTTCATAATTTGACTTTGATAAGTGGCCAGCCAATGCAAGTTTGGGTGGATTATGATGGCAGCACTACGCAAATTAATGTAACAGTAGCTCCATTACATGTGGAAAAACCAGTTAATCCACTTTTGTCTTTGAAAACTGATCTTTCGCCGATTCTTGATCAGACCATGTATGTTGGTTTTTCATCATCAATTGGTTCAGTCCCAACACATCATTATATCTTGGGATGGAGCTTCAAGATAAATGGGAAAGCTCAAGAACTCTCTCAACTTCCTAATCTTCCTCGTCTTGGACGCAAAGGGACATCAAGATTTGTAAAGATTGGTTTGCCATTAATCTCTTTGGTTTTACTTGTTGTAGCAACCTCAATGGTGGTTTATTATGTAAGAAGGAAGAAGTATGAAGAACTTCTTGAAGATTGGGAACGCGAGTACAGACCACAGAGGTTCATGTACAAAGATTTGTACACTGCCACTAAAGGATTCAGAGAAAAGGAGCTGTTGGGAGCTGGAGGATTCGGAAAAGTTTACAAAGGAGTGATGCCTATCACCAATCTTGAGATAGCTGTAAAGAAGATATCTCATGAATCAAGACAAGGGATGAAGGAATTTGTTTCGGAGATTGTGATCATAGGTCGTCTACAACACCGGAATGTAGTACCACTTTTGGGTTATTGCAGGCGGAAAGGAGAGTTACTTTTGGTTTATGAATACATGTCTGGTGGAAGTCTAGACAAGTATTTATACAACAAACCAAGATTCACCCTTGATTGGAGCCAAAGATTCAGAGTTATTAGAGGTGTAGCATCAGGACTATTTTTCCTACACGAAGAATGTGACCATGTAGTGGTTCATAGGGACATTAAGGCCAGTAATGTGTTGTTGGATGGTGAACTAAATGGTAGGCTAGGAGACTTTGGACTTGCGAGGCTATATAGTCATGGTACCGATCCTCAGTCTACTCGTGTCGTTGGTACTCTTGGTTACCTTGCCCCAGAGCATACTAGAACTGGCAGGGCAACGCCTAGTAGTGATGTATTTTCCTTTGGTGCTTTTCTGCTTGAAGTTGCCTGTGGTAGGAGGCCAATAGAGCCAAGACAAGACGATGATGATCTGATTTTAGTCGATTGGGTGTTCTCATGCTGGAATAGGGGTAATATTCTTGAGGCTGTTGATCCAAACATAGGGATTGATTTTGTTCCAGGGAAAGTGGAGTTGGTCTTAAAGCTAGGCTTGATCTGCTCTCATTCAGAGCCCTCGTATAGGCCAACCATGCGACAAATCTTGTTGTTCTTAGATGGTGTTGTGGCCTTACCAGAGTTATCATCACTCGGTATTTCATCAGCTGGCCTAACGTTTGACCATCATGGAGGTTTTGATGATTTTGTCACGTCCTATTCATCTTCGATTAATGCACATTCCCGCTCTCCATCTGTAACAGACTCCTTTCTCTCTGGTGGCCGATGATTCTTACATTAAATATGTTAGTAACTTTAACTTGTAAGTTGTAATCATGATTAGCACCTTATCTTAGTTTTAATTGAATCTCTATACTACTGCACCTCAAATAAGATGGAAATGCA

>M6-PGSC0003DMG400020328 M6 L-LecRK

AAAAAAAATTAGCCATGTTTTTCAAGATTGTGTCCTTGATTGTGTCTTCTCTTTTTGTTGTTACAAATTCTGAAGACTTTGGATTCATTTACAATGGGTTCAATAGAGCAAATTTGAGTCTTGATGGCATTGCACAATTGACATCAAATGGCTTATTAGAGTTAACAAATACTTCTAGGCTACAAAAGGGACATGCTTTTTATCCTACACCAATTAATTTCAAGAACTTAACAAATGGTTCAAATTTCTCTTTTTCTACCACTTTTGTCTTTGTTATTGCGCCTTCTGTTTTGCCTGGTCATGGTATGGCTTTTGTAATTGCACCTGTTGGTGGTCTAGTAGATGCACTTCCAAGCCCATTTTTAGGCCTTTTCAATGATAACACTACTGGGAGAGTCAGTAACCATGTTTTCGCGGTCGAGTTTGATACTTTACAGAACAGAGAATTCAATGATATTGATGGTAACCATGTTGGAATTGATATCAATGGATTGAAATCCGTTGAATTGAAAACAGCAGGTTATTATCATGGCGTTAAATTCGATAACATGACTCTTGCTAGTGGTCAGCCAATGCAGGGTTGGGTGGACTATGATGGTGTGGCTAAGCAAATTAATGTAACTCTGGCTCCAATGAATGTAGCAAAACCAAATGCACCACTTTTGTCATTGTTGTATGATCTTTCACCAATCTTGAATGAAACCATGTATATCGGCTTCTCTGGATCGACCTGCTCGGTTGTTTCAACACAATATGTTCTAGGATGGAGCTTTGAAATGAATGGAATAGCTCAAGGTCTTGATCTTGCTAGACTTCCTAAGCTTCCTCGAGTTGGACCGAAGAAACAATCTAAACTTTTGTTGATTGCTTTGCCAATGATATCGGCAGTTGTTGTAGTAATAGCCTTTTCTGTGTTGATTTACTATGTCGGAAGGAAGAGGAAGTTTGCGGAATTGCTTGAAGATTGGGAGCTTGAATATGGACCTCATCGGTTCAAGTACAAAGATCTTTATATCGCGACAAAGGGATTTGCTAACAAAGAGTTGTTAGGTTGTGGCGGTTTTGACAGAGTTTACAAAGGAGTATTACCAACTTCTTCAATTGAGGTAGCAGTCAAGAAGGTGTCTCATGACTCAAAACAAGGATTGAGGGAATTTGTAGCAGAAATTGTTAGTATTGGTCGTCTACGCCACAGGAATTTAGTGCCACTTTTGGGTTATTGCAGGCGTAAAGGAGAGTTGCTTTTGGTTTACGAATACATGCCTAATGGAAGTCTTGATAAGTTCCTATATGACAAACCAATATGTGCTCTCAGTTGGAACCAAAGATTTCGAGTCATCAAAGGCGTAGCATCAGCACTAGTCTATCTACATGAAGAATGGGAACAAGTTGTAATACACAGAGATGTGAAGGCTAGTAATGTGTTGTTAGACAGTGAATTGAACGCGAAGTTGGGAGATTTTGGACTAGCAAGATTGTATGATCACGGGAGCGATCCTCTCACTACACATGTAGTTGGAACTGTAGGTTACCTTGCCCCTGAGCAAACAAGAACAGGCAAAGCCACAACCATTAGTGATGTATATGCTTTTGGTGCATTTTTGCTTGAGGTGGCTTGTGGGAGAAGGCCAATAGATCCACGAGTATTGGATGAGGACATTGTGTTGGTTGATTACGTGTTTTCTTGTTGGAGTAGAGGTGATATTCTTCAATCCATTGATCAAAATTTGGGGAATGAATATGTTAAAGAGGAAGTTGAATTGGTACTAAAACTCGGACTTGTTTGCTCTCAGACAGAACCAACAACGAGGCCAAGTATGAGGCAAGTTTTAATTTACTTGGAAGATATCGTGCCTCCACCATTGCCAGAGTTGTCATTGTTACAAAGTTCTGCCCATTATTTTTCATTTCCAGGTTCTGATCATCACTTGTCTATGTCTTCATCATTTTCTCCAGAGAAAGTACTTTCATATCCCTCCACCGATTCGAACTCTCTTCCATTCACATGTCCACGATTAGAACTCTTTCTGTATTGCCTTGATGAATTGTGATTCCAATCCCAACTGCAACAACAACTGTACTTCAATTCCAAACAAGTCCGTGTTAGCTATGTGAATTCTTACTGTCTATATCGTTTCAATTAAGCATGTAATATTTGTTACGATACAAATAATGGACTAATGTACACTTAGAATGATCCGATTTCTTTTGATCTATAATGTTGTCGACATGTATTGCTA

>M6-PGSC0003DMG400023671 M6 L-LecRK

AAAAAACATAGTTTTTTGTTATATGTTGATATGTATATTCTAGGCAAGATTAGTTATTTAACTAACTTTGTTCATCTTTTGATCATATTCATATTTTTGGTTCTTAGAGTGGATTGTATGAGTTTCAGCTACCAAAATTTCACCAAGACAAACGAAAACGATTTCATTGCTACTGAGCATTCCTACATAGCCGCTGATGCTATCCAAGTAACTCGCGATGTTCGTGGTGCTTCCATCAGTAATTTGTCTGGAAGGATATGGTATTCACAGCCTATGAATCTATGGAACAGAAGGAAAAACAGAACAGCCTCGTTTAATTCAACGTTTGTGATCAATATCAAACCTGAGAGTGATCCATGGGGTGAAGGATTGGCTTTCATATTGACTAAAGAAAGTGGTACTGACTCAATCCCGGATAACAGTTATGGACAATGGCTTGGGATTGTGAATGAAAGTACTAATGGATCTTCTTCAAACAACATATTTGCAGTTGAATTTGATACAAGGAAAAGTTACCCTGATGATCTTGATGATAATCATGTTGGCATTGACATAAACAGCATCAATTCTGTCAAGCAAGTCTCGTTGATTGATCGTGGTGTTAACCTTTCACTAGCTGTTGATGTTATAGCAAGTGTCCGGTACGATGGAGAATCCAAGATGTTGAAGGTTTATACATTCATGAGCAATCAAACCGGAGACAATGAGAGAAATCCCATCATTTCCATGCCTCTTGATCTTTCTAATTATCTCCCCGAGGATGTTTTCGTTGGATTTTCAGCCTCAACTGGGATATATAATCAGCTGAACTGCATAAAGGCATGGAATTTTACAAGCACAGATATTGGGAACAGTAATGAAGTTAGCCTGTTATGGTTGTGGATCTTGATACCAGTAATGTCAGTTCTTGTTATGTTCTTAGGTGGGGTTTTCGTATATTTTAGCTGGTGGAGAAAAAAGAAGAGGATGCAAGTGTTAGACCAATCTGAGAATATAGAGATTCAGATTCAGAATTCAGCTACTGCACCTCAGAGATTCCAGCTGAAGGACTTGAAACGAGCAACGGGGAACTTTGATCCGAAGAACATTCTTGGAAGAGGAGGATGTGGAGTAGTCCTCAAAGGGATGTTAGATCAAAAAGAGGTGGCTGTGAAGAGATTCTTTAATGATTCAAGTCAAGGGGCAAAAGATCTCATAGCAGAAGTCACAACTATAGGCAATCTGCACCATAAAAACCTTGTCAAATTGATAGGATGGTGCTATGAGAGCAATGAACTCCTTGTGGTGTATGAGTTCATGCCAAATGGGAGCTTAGACAAGTTGATATCTTGCGAAGAAAACGGAAAAGGTTTGAGTCTGAATTGGGAAATAAGGCATGGTATCATCTGTGGGATTGCTCAAGCACTTGATTATCTGCACAATGGATGTCAAAAAAGAGTACTTCACAGAGACATAAAAGCCAGCAACATCATGCTTGACTCAGAACTCAATGCTCAGCTAGGAGATTTCGGATTAGCTAGGACAGTTCAAGTGAATGGAAAGACTCACCATTCAACAAAAGAGATTGCTGGTACTATAGGCTATATGGCACCAGAGAGTTTCCATATAGGCCGAGCCACAGTTGAAACAGACGTCTTTGCATTTGGAGTGCTAGTTCTTGAAGTTGCTTGTGGTCAAAAACCTGGAAATCGATATGAGGAGAACAATTATACTAACAGAGTCATTGAATATGTATGGGACTTGTACAAGATTGGAAGGATCATTGATGCCATAGATGTCAGATTGGATAGAGACTTCAATGAAGAACAAGCCGAATGTGTGTTAATATTAGGATTGGCTTGTTGCCATCCAAATCCTTACGAAAGGCCAAGCATGAAAACTGCTTTACAGATTCTTACAGGGGAATCAGTTCTACCAAACATCCCAACTGAGAAACCTGCATTTGTATGGCCAGCCAAAGCTCCATCGTCCAACGAAGCTGCTAGTGGCTCTCTCCTGGATGACCAACTTACACAGATTACAGTACTCAGTGGCAGATGACTGCAGAGTATGAGGAACCTCAGCAAGGTTAGGTAACTCTTTAGGAGGCAAAAGATACCTGTAAATTCACATATGTAAAGGCATGAATGTCTCTTTATTTTTTTTCTTTCTATGTATGTGGTGGGGAGTGGGAGATATCTTCTTCCGAGTGTTCGTTTTTCTGCATTATAAATAACTTTCTATTTAGTCAATGTCATAATCAACTTTACAAGTAAATACTGAAAACAAAAGCAGCAAGCATTGCAACAAAGGCAGCTAGCTACATGAGTTACTCCTACAAAACATACATGTTCTACCAAAATCACTAGAAGATCCATAATCATCTTCATGTTCTACCACAACAAGAACAGTTTCTCTGGATTGTTCATGTCTGCTACGTCCATCGTT

>M6-PGSC0003DMG400026646 M6 L-LecRK

CACAAATACTTAACAATTTGTACTTCCAACAACTCCATGGACAATCCCTATAGCCTAGTTATTCTTGCATTGATCTTTTGCACTATTCTTGTAAGAGTTCAATCTTTTGATTTTGCTTACAATGGATTCAATAACTCAAATATCATTAGAGATGGAGTTGCAATTATCAATCCAAGTGGTGCACTTAAGCTCACTAATAGATCATATAATGTTATAGGTCATGCTTTCCACCCCAATCCTGTACCAATTTTCAACTCCAGTACTAAAAATGTTACTTCATTTAGTACATACTTCGTCTTCGCCATTTTCCCTCTTTAGAAAACCTCTGGTGGCTTTGGTTTTGCTTTCACCTTATCACCATCCCCAGGTTTTCCAGGTGCTGAGGGAGATCACTATCTTGGAGTCGTCAACAGGATGAACGATGGCAATGATACTAACCACATCTTTATGGTTGAATTTGACACTGTGAATGGTCACAATGAAGGTGTGGACACAGATGGAAATCACATTGGGGTCAACAGAAATGGCATGAGGTCTATAGCGTCAGAATCAGCTGACTACTATGTTAATGATTCTCTTGTAGCAGAGCAGGTAAATCTACAAAAGGGAGAATCAATTCAGGCTTGGATCGATTATGATGGAATGAGCAAAGTGGTGAATGTAACTATATCTCCCATAACACTTCCAAAGCCAAGTAGACCTCTGATCAGTGGGGTCGTAGATTTATCTACTGTGTTGGACGAAACTATGTATGCTGGTTTCTCAGCAGCCACGGGAAGAAATAAAGCGAGCTCTCATTACATCTTGGGTTGGAGTTTTCGGTTGAATGGAGCTGCTGATCCTTTAGATCTTGATAACCTACCTGTTGCCCCACCTGAAGTGGTAAGACCCTCTAAGAATTCCCATCTCAAAAGGGCCTTGATAGGAACATTTTGCTCCGTTGTTATCCTGCTCCTTGGTGCATTTTATATCTATAGAAGGATGAGGCTTCATGAAGTTCTAGAGGACTGGGAGCTGGATTGTCCTCACAGGTTCCGATTTAGAGATCTTTATAAGGCAACTAAGGGATTCAAGGAGAGTGAACTAATTGGAGTTGGGGGATTTGGTGCTGTTTACAAGGGTGTTTTTCCTACTAATGGAGCCGAGGTTGCAGTGAAGAAGATATCGAGCAATTCTCTCCAAGGAATGAGAGAATTTGCAGCGGAGATCGAAAGCTTAGGCAGGTTAAGGCACAAACACTTGATCAACCTTCAAGGCTGGTGCAAGAACAAGAATGATCTTCTCCTAGTGTATGACTTTGTCCCAAATGGAAGTCTTGATTCATTACTTTACAGGCAGAAAAGGGACATTGTTCTAACATGGGAACAGAGATTTAACATCATCAAAGGGATTGCTGCAGGGCTTCTTTACTTGCATGAAGAATGGGAACAAGTGGTAATACATCGCGATGTGAAGAGCAGTAACGTCCTAATTGATGGGGAAATGAATGGTCGGTTAGGGGATTTTGGGCTAGCAAGATTATATGACCATGGCAAGAATTCACACACAACAAATGTTGTTGGCACAATAGGGTACATTGCACCAGAATTGACACGGACAGGGAAGGCCTCAACGAGCACGGATGTGTATGCATATGGTGTACTACTTCTTGAAGTAGCTAGTGGAAGACCTCCTATTGTCTATGAACCAGGGAAAGGAGCTTTAGTACTAGCTGATTGGGTGATAGAATGTTTACAACTAGGTAATATTCTTGATGCAGTTGATCAAAGGTTGAATTCAGTCTATGTAGATAAAGAGGTTCAAATGATTTTCAGTCTTGGGCTTGTTTGTTCACACCCAAGACCAGAAGCTAGGCCAACAATGAGACAAATAATGAAGTATCTCAATGGAGATGAATCACTTCCAGTTTTTGAGCAGCAGTTGAGCTGTATTGGGTCTGATAGGGTTGATGAAATCACAGCTAAGTTCTTGGAAGTGTTTTCTACTGATACAATCAATATATCACGTCGTTCGCTATCCGTCGGACAGATATCATCTAGTTCACTCCATTCTGGTAGATGAAAACTCTGTATAAAGGACATACACTCTTCTTTTGTTTTGTTTAACCACTTTGTTGGGATTGATTAAGTAGCTTCTATTTACCCAAATCCTGTTTTTCTTCCTCATAATTCTGCAATTTAGATAGGTCATTTATTTTTAGGCTCATTCAGCATTCACAAGTTAACATAATCTCACTTCTATGTCTGCAATATCACTTATCACATCCTATATTTTTGTTTATGTACAAAC

>M6-PGSC0003DMG400026647 M6 L-LecRK

TAATTGTCTCTTTAAATTGAATATAAATATTGCACAAAAACTTAACAATTTGTACTTCCAACGACTCAATGGACAATTCCTATAGCCTAGTTATTCTTGCATTGATCTTTTTCCCTATTCTTGTGAGAGCTCAATCTTTTGATTTTGCTTACAATGGATTCATTAACTCAAATATCATTAGAGATGGAGTTGCAATAATCAATCCAAGTGGTGCACTTAAGCTCACTAATAGATCACAAAATGTTATAGGTCATGCTTTCCACCCCAATCCTGTACCAATTTTCAACTCCACTACTAAAAATGTTACTTCATTTAGTACATACTTCGTCTTCGCCATTTGTCTCTCTTGAGAAAACCTCTGGTGGCTTTGGATTTGCTTTCACGTTATCTCCATCCCCGGGTTTTCCGGGTGCTCAGGGAGCTCATTTACTTGGAGTCGTCAACAGCAAGAACGATGGCAATGATAATAACCACATCTTTATGGTTGAATTTGACACTGTGAATGGTCACGATGAAGGTGTGGACACAGATGGAAATCAAATTGGGATTAACAAAAATGGCATGAAGCATATCGAGTCAGAATCAGCTGATTACTATATTTATAGTACTTCTGAAACAAAGCCAGTAAATCTTCAAAATGGAGAATCAATTCAGGCTTGGATCGATTATGATGGAATGAGCAAAGTGGTAAATGTTACTATATCTCCCATAAAAGTTCTTAAGCCAATTAGACCTCTGATCAGTTTGGTTGTAGATTTATCTCCTGTGTTGAACGAGACTACGTATGCTGGTTTCTCAGCAGCCACGGGAGATAAAGCGAGCTCTCATTACATCTTGGGTTGGAGTTTTCGGTTGAATGGAGCTGCTGATCCTTTAGATCTTGATAACCTACCTGTTGCCCCACTTGAAGTGGTAAGACCCTCCGAGAATTCCCATCTCAAAAGTGCCTTGATAGGAACATTCTGCTCCTTCTTTATCCTGCTCCTTGGGGCATTTTATATCTATAGAAGGATGAGGCTGCATGAAGTTCTAGAGGACTGGGAGCTGGATTGTCCTCACAGGTTCCGATTTAGAGATCTTTATAAGGCAACTAAGGGATTCAAGGAGAGTGAACTAATTGGAGTTGGAGGATTTGGTGCTGTTTACAAGGGTGTTTTGCCTACTAATGGAGCCGAGGTTGCAGTGAAGAAGATATCGAGCAATTCTCTCCAAGGAATGAGAGAATTTGCAGCGGAGATCGAAAGCTTAGGCAGGTTAAGGCACAAACACTTGATCATCCTACAAGGCTGGTGCAAGAACAAGAATGATCTTCTCCTAGTGTATGACTTTGTCCCAAATGGAAGTCTTGATTCATTACTTTACAGACAGAAAAGGGACATTGTTCTAACATGGGAACAGAGATTTAACATCATCAAAGGGATTGCTGCAGGGCTTCTTTACTTGCATGAAGAATGGGAACAAGTGGTAATACATCGCGATGTGAAGAGCAGTAACGTCCTAATTGATGGTGAAATGAATGGTCGGTTAGGGGATTTTGGGCTAGCAAGATTATACGACCATGGCAAGAATTCACACACGACAAATGTCGTTGGCACAATAGGGTACATTGCACCAGAATTAACACGGACAGGGAAGGCCTCAACGAGCACGGATGTGTATGCATATGGTGTACTACTTCTTGAAGTAGCTAGTGGAAGACCTCCTATTGTCTTTGAACCAGGGAAAGGAGCTTTAGTACTAGCTGATTGGGTGATAGAATGTTTACAACTAGGTAATATTCTTGATGCAGTTGATCAAAGGTTGAATTCAGTCTATGTAGATAAAGAGGTTCAAATGATTTTGAGACTTGGGCTTGTTTGTTCACACCCAAGACCAGAAGCTAGGCCAACAATGAGACAAATAATGAAGTATCTCAATGGAGATGAATCACTTCCAGTTTTTGAGCAGCAGTTGAGCTGTATTGGATCTGGTAGGGTTGATGAAATCACAGCTAAGTTCTTGGAAGTGTTTTCTACTGATACAATCAATATATCACGTCGTTCGCTATCCGTCGGACAGACATCATCTAGTTCACTCCATTCTGGTAGATGAATCTGTATAAAAGGACATACACTCTTCTTTTGTTTTGGTTAACCACTTTGTTGGGATTAATTAAGTAGCATAGAATAATATATAACTAGGAAAAAGGTAAGTCTCAAGTCAACATTTTCACTTAGCTGGAGTGCAAAAACCATGTAATTTTGAGAAAGAAAAGTTATAAAATTAGTCTAGAAAGTAAATTTGTAATTATAAACAAAAAAAAATAAAAAAATCTATGTAGAATATTCGTTATAAAATCTATTTGTAATAAAAATGGTGACATGATACAGA

>M6-PGSC0003DMG400027140 M6 L-LecRK

CCCCACTCTTTCTTCATTCCTTTGTAATTACACAGACAAATCACGTGGCAAATCAAAGTTGAACTCTACAAAAAGAATGTCTAGATTCAGCTATAAATCATCACCCCACGCCGTTCTTCAGCTCTACAAACAAATTTCAATCTAACTATGCCATGGCCAAACAAATTACACCAACCCTTTTCTTGATTCTCTTTATTTTGCTGTCAAGAACAGTTTTTTCTGATGAATTTGTGTATAATGGCTTTAATGGGGTTGAAGCTAGTAATCTTACTGTAAATGGGGTTGCTCATATTTTAAAGAATGGTGCTTTAAAGCTTACAAATGAAACATCTAGAGTTGTTGGTCATGCTTTTTACAAAAATCCAATTAAATTCAAGAATTCCCAATCTGGTAAAGTTTCTTCCTTTTCTACTGCTTTTGCTTTTGGGATTGTTCCTGAATATGCTAAGTTAGGTGGGCATGGCTTTGCTTTTACAATTTCAAGGACTAAAGAGATGAAAGGGGCTCTTCCTAGTCAGTATTTAGGTCTTTTAAATTCAAGTAATATTGGGAATTTTTCAAATCATGTATTTGCTGTTGAATTTGATACTGTTCAAGATTTTGAATTTGGTGATATTAGTGATAATCATGTTGGTATTGATGTGAATGATTTGAAGTCTAATGCTTCAGTAAATGCTAGTTACTTCTCTGAAGGAAGTTTTAGTAAACAAAAGCTTTTTCTTCAGAGTGGAAAGACAATTCAGGCTTGGATTGATTATGATTCAAGTAGAAATGTGTTGAATGTTACACTTTCACTTTCTTCAACAAAACCAACTTTTTCAATTTTGTCTTTCCCTTTGGACCTTTCTCTAGTTTTTGAGGAATTCATGTATGTTGGATTTTCTGCTTCAACTGGTTTGCTTGCTAGCTCACATTATATATTTGGTTGGAGTTTCAAGATGAATGGAAAAGCTCAATCTTTGGATCTTGATTCATTGCCTTCTCTCCCTAAGCCTAAGAAGGATCAAACAGTCTTGATTGTAGCTACCACGATATCGGCTGTTGTGTTTCTTGTATTCGGACTCATTTTAGCCTTGTATGTCATTTGGAAAATCAAGAACATAGATGTGATTGAGCCTTGGGAACTTGAGATAGGTCCACATAGGTTTTCTTACAAAGAGCTAAAGAAGGCTACTAGGGGTTTTAGAGATAAAGAGTTGCTCGGGTTTGGTGGATTTGGTCGAGTTTATAAAGGAACACTGCCTAAGACGAATACCGTTGTAGCTGTGAAGCGAATACATCATGAAGCAAAACAGGGTCTTCAAGAATTTGTGTCTGAAATCGCGACTATTGGTCGCCTTAGACATAGAAATTTGGTTCAGCTCTTAGGATGGTGTCGGCGAAGAGGTGATTTGTTACTTGTTTATGACTTTATGCCTAATGGAAGCTTAGACAAATACATTTATGATGAACCAAGAGTTACTTTAACTTGGGATCAGAGGTTCAAGGTTATTAAAGGTGTGGCTTCTGGTCTGTTGTATTTACATGAAGAATGGGAACAAACTGTGATTCATAGAGATATTAAAGCAGGGAATGTGTTGTTAGATTCCGATATGAATGGACGACTTGGAGATTTCGGACTTGCTAAGTTATACGAGCATGGAGCAAATCCTAGCACAACAAGAGTAGTCGGTACGTTGGGCTACTTAGCTCCAGAATTAACCAAGACCGGAAAGCCCACACCAAGTTCAGATGTTTTTGCCTTTGGTGCATTGCTGCTAGAAGTTGCTTGTGGAAGGAGGCCAATTGAGGCAAAGGCATTGCCTGAGGAGTTAATCCTAGTGGATTGGGTATGGGATAAATGGAAGGAAGGTGCCATTCTTGAAGTGGTTGACTCGAGATTAAATGGCGAGTATGATGAGATGGAGGCCATAATGGTGCTTAAATTAGGACTCATGTGTTCAAATAATACACCATCTAAGAGGCCTAGCACGAGGCTAGTGGTGAGATACTTGGAAGGGGAGGTGACACTCCCAGAGACGCTGGCAGCTCCCGATGAATATGATGGAAAGAAGGGCGGTACTAGTGGTATGGAGTTAGAGGTCTTCATAGATTCCTATCCGTCTTCGTCTTATTTTGAGAAAGTAAGCACATGGTCATCACCTTATGACGGTGAAGGCGATATTGATATTGAAGCTAATTCAGTAACGCCATTAACGGATTCTGGCAGAGAGGACAACAGGTAGCCATGCAAAATGGAGGGACCATTCTACTTGCCCTCCCAAACTCCCTCTATATTTTAGTACTTCTTTTTACGGTTTGATTTTTGACTTAAATGTAAATTATTGCTTTTGAAATGAAAATGATATTAGTATCAACCATTCCTATGGTTTTGAGAATATTGGTGTTAGTAGGATGTGAATATATTTTTGCAATCAAATCAAGTTTGAATCTTGATGAAGCTT

>M6-PGSC0003DMG400029735 M6 L-LecRK

GAATGATATTTCCCCATTTTCTCTGCTTCCCATAATTCTTTATCCATGTGAATTAGCTCTGCTTCTTTCTTAGAATGAACCAATATTATCCAAAAATTCTTCTTTCTTACCTGTTGTTAACCCTATTTTCTTGTAATCAATCAGTTGCAGAAACTGATTTTGTCTTCAATGGCTTTAAACAATCAGATGTGTCAATTTTTGGGAATGTTACAATTGAATCTAGAATTCTTACTCTTTCAAATGACTCAACTTTTTCAATTGGTAGAGCTCTGTACCCTTCAAAGATTGTCACAAAAGAAGCCAATTCATCTAAAGTTCTTCCCTTTTCAACATCTTTCATTTTTGCAATGGCTCCTTACAGAGACAGGCTTCCAGGACATGGTATAGTTTTCTTGTTTGTGCCACACACAGGTATTGATGGTACTACTTCTGCACAGAATTTAGGTTTCTTGAATTTTACAAATAATGGGAATCCTGATAATCATGTTTTTGGGGTTGAGTTTGATGTTTTCAAGAATCAAGAATTCAATGATATAAATGAAAACCATGTTGGAATTGATGTTAATTCTCTTGCATCAGAGTTTGCTCATGAAGCAGGATATTGGCCTGATGAGAAAAATAAGTATAATAGTGATGGTAGCCTAAATGAGGAGTCTTTAGAGACTTTTAAGTTGAATAATGGAAGAAATTATCAAGTTTGGATTGATTATGCTGATTTCAACATTAGTGTAACTATGGCACCAGTTGGTATGAAAAGGCCTAAGCAGCCTTTGTTGGATTTTCACCTAAATCTTTCCCAGGTTTTTGAGGAAGAGATGTATGTTGGGTTTACAGCAGCCACAGGAGAACTTGCTCAAAGTCACAAGATTTTAGCTTGGAGTTTTAGTAACTCGAATTTTTCTATAGGTGATGGTTTGATCACACAAGGGCTGCCTTCATTTGAGCTTCCTGAGGATCCAGTTTATCGATCGAAGGGATTCATTGCAGGTATGACAGTGTTAGTCTTGTTTCTTGTTGTGGTCATTGCTGTAGCTTCATGGCTTTTGATTAAGAGAAGAAGGAGAATGAAAAAGGAAAGAGAGGATATGGAAGATTGGGAATTGGAATATTGGCCACATAGGATTACTTATCAAGAAATTGATACTGCAACAAAGGGTTTTGCTGATGAAAATGTGATTGGAATTGGAGGTAATGGGAAGGTTTATAAAGGGGTTTTGGCTGGGGGTTCAGAGGTTGCAGTAAAGCGAATTTCTCACCAAAACAGTGAAGGGGCAAGGCAATTCTTGGCTGAGATTTCGAGTCTTGGTAGGCTAAAGCAAAGAAATTTGGTGTCATTAAGAGGGTGGTGCAAGAAAGATAGAGGCAGCATGATTGTGGTTTATGATTATATGGAAAATGGGAGCTTGGATAAAAGGCTGTTTGAATCTAATGAGAGAAACATGCTGAGTTTTGAAGACAGAATTAGGATTTTGAAAGACGTGGCATCAGGGGTTCTATACTTGCATGAGGGATGGGAGTCAAAAGTGTTGCATAGGGACATTAAGGCTAGCAATGTTTTACTTGACAGGGACATGAATGCAAGGCTAGGTGATTTTGGTCTAGCTAGAATGCATGATCATAGTCAAGTGGCTAGCACGACTCGAGTTGTTGGCACGGTGGGATACTTGGCGCCAGAATTTGCCAAGACTGGCCGTGCCTCCACACAAACCGATGTTTTTGGATATGGAGTGCTAATTATGGAGGTGATGTGTGGAAGGAGGCCTATAGAGGAGGAAGGCAAGCCACCTTTAGTGGATTGGCTGTGGGAATTAATGAGACGAGGTGAATTGATTAACGCCTTTGATAGTCGATTAAGGACTAATCAAGATTTCAATGAAGAGGAAGCTTTAAGGGTGTTGCAATTAGGCATGATATGCGCGAGCCTAGATGCCAAAGCTAGGCCAAGCATGAGACAAGTGGTGAAATTCTTTGACAGGAATAGTGATATTGATGAATATGAAACTGAGGATATGGATGCTTACCTTCTAGAGAGTTTGAGATCTAATACCATGTTGTCCAATTTCTCCTTAAGTTTAAGCCATGGTTCACATCCAACATTTGAAGAAATTAGAGAAGGTTTTTCTTCTTCCATGTCAATTTCTTGGACAAATTCTCTGGTGGATGGTAGATGAGTTTAACATTTGGTTGGTTTATGTCCATCACTCTGATTTAACTTATGTTACCTTCCAGGGCCTTGCTTTGATTGAATTTGCATTTGGCAATGCTGCTTCATGTACAAATTTATATGATGGAGTAGGAAAAAGGATTTGGTTATGACTTATGAGATATACATGTATTCATTTCTTTATTTGTACCTTGATACTATTTGTAGTAGAAGAGTTTGTTCTCTTTTTTTGTTTTGTTTTTACTATAGAATTTGTTCTTTATCTTTTTCATTTTACACAAGATAAG

>M6-PGSC0003DMG400030144 M6 L-LecRK

GTATATATAAAATTTATTTTTTAGCAAAGCACTAAACAGAGTGTTTGGTGTTTATTGTATATATTCAGTTGAATGTCTAACAATGTGATTAACATATTGTCCAACTATGGTTGTTTTTGTGTCTTTATCTTCTCTCAAATACTTGCTTCTTATCATAATTATCCCTCTAGTCACTTCACTCTCCTTCAATTTTGATACTTTTACACCCAATGATCACAATGTAACCTATGAGAGAGATGCTTCTCCAGAAAATGGTGCAATTCTACTCACCAAAAACCTCCTCAATCGCGATATTAATGCTAGTATAGGTCGAGCTATATATTCAAAACCACTGCATCTATGGGACAAGGCCTCTAGAAATCTCACTGATTTCACTACACATTTCTCCTTTAGCATTAATTCACAGGGAATAACAAAATATGGAGATGGTTTTGCCTTCTTCCTTGCCCCTGCAGGTTCAACAATTCCTTATAATACAACAAGAGGTGGAAGTCTTGGCCTTACAAGTAATAATGAGCGACTAAATTCTTCGAGCAATCATTTTGTTGCTGTGGAGTTTGACACCTATCAGAACCCCTATGACCCAAAGGGTGATCATGTAGGTGTCGATATCAACTCTATGCAATCTGTTGTTAATGTGACTTGGTTTAGTAGCATTCCAAATGGTACGAAAACTGATGCCTGGATCAGTTATAATTCTAACTCAAAAAATCTTACTGCTGTCTTCACTGGTTTTCAACTCCAAGGGAACAATATCACAGTCACTGTCCTGCAGAACCTATCTTACAATCTTGATCTGAGGGAATATTTGCCAGAATGGGTCACTTTTGGCTTCACTGGTGGAGCGGGAAGGAGATTTGCATTTCAAAGAATCTATTCTTGGAATTTTACTTCTTCTTTAGATAATACGACAGATACAGGGGTAACCTTAACAAACACCATGCCAAAGGATGCACCAACAAAAACTAAGTTAGGACTTGTGGTTGGATTGATTTCTGGTGGTTGTGTTTTGGTTGCAGTATGTGCTTTGGTGTTATTTGCATTTTGGAGAAAGAGTAAGGTGAGAGAAGATGATTATGTCTTTGATGGTTCCATGACTAATGAATTTCAAAAAAGTGCAGGACCAAAGAAGTTCTTGTACAGTGAGTTGGCTAGATGTACAAATAACTTTTCACAGGAAGAGATGCTTGGGCGGGGTGGATTCGGAGGTGTTTATAAAGGATATCTCAGTGAATCCAACCCTTATATTGCTGTTAAGAGGGTTTCAAGGGAATCACAGCAAGGAATAAAAGAGTATGTATCAGAAGTCAGGATCGTCAGCCGGTTAAGACATAAACATTTGGTGCAACTCATTGGTTGGTGCCATCAAAAAAGAGAACTTCTACTTGTCTATGAGTTTATGCCTAATGGAAGCTTAGATCATCATCTTTTCAAGGGAAAAAGCCATTTGACATGGCCAATAAGATTCAAGATTGCTCAAGGTTTGGCCTCGGCGTTGCTATATCTACACGAAGAATGGGAACAATGTGTGGTGCATAGGGATATAAAGTCTAGCAATATTATGTTGGATTCTAATTTCAATGCCAAGCTTGGGGATTTTGGGTTAGCTAGGCTAGTTGACCATGATAAGGGATCCCAAACAACAGTTTTGGCAGGCACAATGGGTTACATGGCTCCTGAATGTGTCACCACTGGCAAAGCTAACAAGGAAACAGACGTCTATAGCTTTGGTATAGTCGTGTTAGAAATAGGTTGTGGAAGGAAACCTATTGATCCTAAAGCAGAGGAGCATCGAGTAAACATTGTCGATTGGGTTTGGAGACTTTATGGGATGAGAAGTATTCTTGAAGCAGTTGATCCGAAACTATCATCAGAGTTCAATGAAGAGGAGATGGAGCATTTGCTAGTTGTTGGCTTATGGTGTGCTCATCCAGATAATAATTGCAGGCCTTCTATTAGGCAAGCAGTTCAGGTGCTTAATTTTGAAGCTCCATTACCCACTCTCCCTCGGAACATGCCTGTGCCCACATATTGCAGCTTCTCACGCCATTTACCAACTATTTCAGTTTCATCAACACATGAAACCAGCGGTACAGAGATTGCCCTTACCCAAATCCAGCATTCAGTAAACGGAAACACCACTGATGGTTCAATAAAAAGAACATCTTCTTCAGCATCTTCATCTTCAACATCAGTTTTGTATCCAAGTTAATATTGCCCTGTTATAACCCTGCATTTTGTCTTCCATTAGCAGTGTCCATCTTTTCCTTTCTTACTTTTTGCTGTTTCTCTTCTAATGAAGAGGATTCTTTGGAT

>M6-PGSC0003DMG400047213 M6 L-LecRK

ATCTTGCTATTTTTAGTAATTTTTGTTATCCCTTCTTTATATTCATTGAACTTCAATCTCACCAATATCAATCCTTCTTATGTTAATCTCTCTATCAATGTCACTGGAGACGCCTATGTAACGAAAGAGGGCATACAAGTTACGCCTAATGAACGTAACATTACATTAAATGGGAAAACAGGACGCGCCACGTACATTGAGACACTGCAGTTATGGGACAAAGCTACAGGGGAAATGACAGATTTTACTACACATTTCTCTTTTGTTATTGATTCAAATGGTAACGATTGTTTTGCTGATGGACTAGCCTTTTTCTTAGCTCCTGTTGGTTCCAGTATTCCAGTTGGTTCGTCTGGCAGTGGCCTTGGTCTTGTTGATGCTGAGACAGAAAACATGTCATCACATGAGTCATTTGTTGCTGTAGAGTTCGATACATATGTGAATAACTGGGACCCCTCAGACATACATGTAGGCATCAATATAAATTCTATGGAATCTGTTACTACAACAATATGGAAAAATGACATTAAACTTGGGAAGAAAAATGATGCTTGGATTAGTTACAATGCTAGTTCCAAGGTTCTTGAAGTTGTTTTCACAGGATTTTGGAAAAAATATAATCGAGACAAACTAAGCTACCCGGTTGATCTGAGGGATTATTTGCCTGAGAATGTTAGTTTTGGCTTCTCAGCATCAACTGGACAATTGTTTCAGAAAAACAATGTCAAGTCTTGGGATTTCAATTCAAGTTTGAGTTTTGATGCTATCAAAGTTCCACAACAGCCTAGTGCAAGTCCTCCTATACAAGCTCAAGATCCAAGAAATCATCCTATTCAAGGTCATAATCCAAATGATCCTCCAATTACTCATGTTCAAGAACCAAAGGGTCCTCCTCAAGAAGCACTCAGCACCTCAGGAAAAGGAAGCAAGGGACTAGTAGTCGGATCAAGCATAGGTTTACCTATTCTGATTCTTGGTTTGATAACTGCCAGTTGCTTTTTGTGGAGGAAAAAGAGGAAAGGAGATAAGAAATTAGACCATGTCTTCATTGATCTAGCTATGGATGATGAATTTGAAAAGGTTACCGGTCCTAAGAAGTTCTCGTATAGTGAATTAGCTCGTGCAACAAACAACTTTGCTGAGGTACAGAAGCTTGGAGAAGGGGGATTTGGTGATGTTTATAAAGGTTTCTTGAAGGAATGTAACTCATATGTTGCTGTTAAGAGAGTGTCAAAAGGGTCTAAACAAGGGATAAAGGAGTATGCATCAGAAGTGAAGATCATCAGTCGATTAAGGCATAGAAATTTGGTTCAACTTATCGGTTGGTGTCACGAGAAAGAGAAGCTGCATCTTGTTTATGAATTGATGCCCAATGAAAGCCTAGATAAACATCTTTTCAAAGAAAAGTCATTGTTGGTCTGGGAAATAAGGTGGAAATTTGCTCAAGCAATTGCCTCAGCTTTGCTGTATCTACACGAAGAGTGGGAACAATGCGTAGTCCATAGGGACATAAAAGCAAGTAATGTCATGTTGGATTCAAACTTCACTGCTAAATTAGGTGATTTTGGGTTGGCTAGGCTTATTGACCATGACAAAGGATCTCAAACAACAATGTTAGCAGGGACGGTGGGGTATATGGCACCGGAATGTATCATGAATGGAAAAGCTAGCAAGGAGTCAGATGTATACAGCTTTGGAATAGTAGCATTGGAAATAGCAAGTGGAAGAAGATCAATAAATATTAAAGCACCAGAAGATGAAGTGAGATTGGTTGAATGGGTGTGGAGCCTCTATGGAACGGGAGAGCTAGTCAAAGCAACTGATCCAAGATTGAATAAAATGTTTAATGAGAAAGAAATGGAACGTTTAATGGTCATCGGTCTATGGTGTGCTCATCCAGACAACAAACTCAGGCCATCAATAAGGCAAGCAATCCATGTTCTAAATTCTGAAGCCCAATTACCAAATCTTCCATCAAGAATGCCAGTTGCAACATTTTCACCCCCTCCATTGAATATGTTTTCGTCTCCATTTTCAAATACCTATGAAGTTAATAAGCAGGAGCAGACTATGACTTACACTATTTCCTCAAGTGTCTATACTTCGTCAGCTGCTTCATCGACAAAATCACTTTTGTAA

>M6-PGSC0003DMG401009415 M6 L-LecRK

AAAAAAAATGGCTTTTACTATTGTTTTTCAACTTTACAACTAATATGTTTGAGATTATAAGATTAAAAAATATTTTGATATACATATGTTTAAATTAAGACCATAAAAACCTAAAATCTTCTTTATTATCATAAACTCTGCAAGTCAAAAACAAGACAAACATATTGAAACGGAAGGAGTTTATGTAATATTATTATGTTGTTCAGACTCTCCAAAAATGTTGTTGCACATGTATTGAATTCTTAAAAAATAATACTACTTTTGAAGGATCCGATCGGTATCCATCCACATATTTAGAGAGAGCGAAGCAACACTAATTCAATAGACTCATTAACCGCGTGAGCACATGCCCTTATCTTTTTATGCTTTTCTTTTTCTAGTAGCTAGTAAGTGATCAATGTCTTATATGCTTATTCAATTAATTTATGTAATTTTAACAAGGGACTAGTCTTTCTATTCTTACATTTAGTATGCTAGGAATTTTGACATGGGACTAGTTTTTCTAGTCTGACATTTAGCGTGCTAGGAATGTACTAGGTTGGATCATCTTCTGACAAGCAAATTTCCCCTTTGGAAGAATCAAAGTCTTCTTGAACATTTAATAACTTGGTCCTTCTTTTTGTCTTCAATCTCCTTTTCCTTTCTTAGAATTCTCAAGTCATCCCAAGGGTTTGTCAAAGTTTCTTTTACCCTGACTAACTAGATTTGCGTAGCATAGGGCCCAACAAAGTGAAAAACGCTATATATCAAGACCATTTTTTAATACTTGAAGTCGAGACCTCTAGTTAAGGAATCACACTCCCCGTTAGGCGATAGTAAATGTTTTAATAGTTGATTCATACCATTTGGTAAGTAAGAGTTTAAGCCAAAAATATCAAGGATGTAAGAAAAACTAATGTAATGCCTTAAAGTTTTGACCAAGAACCAACTTGCTACTATGGTCCCTCCAATTTAGAAAAACAAGGCACTATCTTGAAGCCCTTCATTTTGATTTGCTTAAAAGAAGATGATCATCCTTTGCTGACCATGGAATAAGCAGCCATTTGTCAAAGTTCAAAACAATTTTTTATGTCAAAATTTTGGGAAGCTTCCTTCTTGTCCTAACAGTCCTAACATCCCTTTCATTCATTGTCAACGATTAAGACAAGTTTTGGATGTTAGTTGAATGATGGTAAATTAAATCCAATGAAAAGGACATTAATTCTGTATTTTAGTATACATTCATCTTCAAACACCTAATCTTTGTGTCACCTACTCTTGTCATCTGACCCTACAAAGAGATAAGGTTTGTAAAATACAAAAAAAAGTTGACCATTTGGGTATTCTGCAACTTCTGTGTCTAGCTGTTTTTACTGATTAAAAAACAAACATATGCATCTTACAAAATTACATGTTATGGTAACAGGATGGTGTCACAAGAATGGAAAGTTACTGCTTGTATATGAATACATGCCTAATGGCAGTCTAGATATGCACCTCTTTGCGGGCCCAGATAAAGAGCCGCTGAGCTGGCTCGTCCGCTACAAGATTGTGCAAGGTGTTGCCTCAGCATTGCACTATCTGCACAATGAGTATGAGCAGAGGGTGGTCCATCGCGACTTGAAGGCAAGTAACATCATGCTTGACTCAAAGTTCAATGCGCGCCTTGGGGACTTTGGCCTTGCACGGGCACTTGACAACGAAAGGACCTCGTATGCTGAGGCAGAAGGAGTGCTTGGTACAATGGGATACATTGCACCAGAGTGTTTCCACACAGGGAAAGCCACACAGCAATCTGATGTGTACGCGTTTGGGGCAGTGTTGTTGGAAGTTGTATGTGGCCAGAGACCTGGAACCAAGATTAATGGCTTTCAATTCTTTGTTGATTGGGTTTGGTACTTGCACCGCGATGGCCGTATCTTGGAAGCTGTTGATCCAAGGCTTGGAGGTGACTATGTTGCTGATGAAGCAAAGAATCTTCTACTACTTGGTTTGGCTTGTGCACATCCTATAGCTACTGACAGGCCTAAAACACAGGCAATAGTTCAGATTATATCAGGGTCAGCACCAGCACCAGAAGTTCCACCATTCAAGCCAGCATTTGTGTGGCCATCAATGATGCCAATTGACATAGACTCAAGTGTTATGGACACAACATCCATTACTACTTCTCAATTCAATTCAGGATGGAGTCTTGACTATCAAAGCAGGGAGACCCCAACATATGCAGACCACTCTTTGGTGTAGTTACTTTGATGGTAAACAACTATTACCAAAAGGAAAATTGATTTTTTTGGCCTTCTATTTTTTAAAAATATTTCCTGTAATTATGTTTTCTTGGGAGGAGGTTAAAATTTCCTTCAGTTTTTTACTTTTTTGTTTGATGTTTTCTTTTTCCTTTTCAATTTGTTATTGGGGCATGATGAAAAGATCAGGCAGCCTATGTACATTCATATAACTCAGTACTGTTACATTATGTCATATTTACTGTTTTCCTGTCTTTGTCAACCAAAATTATGTAAAAGAAACTCATGTTGAGCCAAATTGGAGGAGACAGAAAATGAATAGTGGCGAAGTATCTTTTTTGCCTCAATTGTACTACTACATACAACTTAAAGAATTCCAAGATCTCAAATTGTAATGAAAGATAAGACCTGATCAATTCTTATAAAAGCAATAGATAACAGAAGAACACAATACAAAGATAGATCATAGAAGCTTTGTTGATATGCATAAATGCTCTGCCATTGTAGTATTAGATCGTTGACAATATATAGTAGATGAAATCCCACTAAGATTTATATTCTTTCTTACTTTCAAGATCATTTCTGTAAATTAATATAATGGAGGACCTGAGATGCTTGCCAAAGTTTTACAAGTTAGTGCGCATAAGATCAAGAGTTTCCCCCGCCTGCAACTTTGTTGATAAACGGAGCTTTCTATAATCAGTGAAAGTGAAAGGAACATAGATACGGGGGTGTTCAGCATCCACCAGCTCAGGTGGTGGTTCCACTGCTGTGTCCCTTGGTCCCAAGAGGAATGATGCAATAGACACCCGTAGTTTTGCTTCCTTGCATATCACTCTATGCTTGACTGTGTGTAATCTTCCATTGCTCCATACCTTCACATAGCAAAAGCAAGTATCATTGTCTAGTTCTTTTTTCTCAATCTTAGAAGGGGTAACGGAGTAGGAATTATCTAACTTCCTGCTAAACAAACCCCGCCAGAGGCTAGCATAAGATTGTTGGAACAATTCAAATCTTGAAAATGAACTAGGTAGAGTAGCAGCATAGTTACCGTGCCTATGTCTCCAAGATTCACAAGGAGGGTGTTTGGCCAAGGATCAACTGCTACAAATTCACCAGACTTCTTCATGACCTCAAGACCGCCAACGCTTTCATCATCCTGTAGAATAGTAAGGAATCCTGAGTCTGTGTGTATCTGAACCCCAGAGGAACCTACAGCTTCTGGGGTGAAGTGGTACTTGTTAATCCTAAACTGACAAGGCCAATCCTCAAAGGAAACATTTGGCAGTCCCAACCCTTCACTCATCTTTCGCAGAATGTCCGTCATTAGTCCATTAATTGCTGTAGCATACTTCTTGATTGTCTCTCTGTTCACAGTAGGAATCAAACTGCTTTTAAGCATTCTAGTAAAGCAGTCTTTT

>M6-PGSC0003DMG402009415 M6 L-LecRK

TGCAACTTCTGTGTCTAGCTGTTTTTACTGATTAAAAAACAAACATATGCATCTTACAAAATTACATGTTATGGTAACAGGATGGTGTCACAAGAATGGAAAGTTACTGCTTGTATATGAATACATGCCTAATGGCAGTCTAGATATGCACCTCTTTGCGGGCCCAGATAAAGAGCCGCTGAGCTGGCTCGTCCGCTACAAGATTGTGCAAGGTGTTGCCTCAGCATTGCACTATCTGCACAATGAGTATGAGCAGAGGGTGGTCCATCGCGACTTGAAGGCAAGTAACATCATGCTTGACTCAAAGTTCAATGCGCGCCTTGGGGACTTTGGCCTTGCACGGGCACTTGACAACGAAAGGACCTCGTATGCTGAGGCAGAAGGAGTGCTTGGTACAATGGGATACATTGCACCAGAGTGTTTCCACACAGGGAAAGCCACACAGCAATCTGATGTGTACGCGTTTGGGGCAGTGTTGTTGGAAGTTGTATGTGGCCAGAGACCTGGAACCAAGATTAATGGCTTTCAATTCTTTGTTGATTGGGTTTGGTACTTGCACCGCGATGGCCGTATCTTGGAAGCTGTTGATCCAAGGCTTGGAGGTGACTATGTTGCTGATGAAGCAAAGAATCTTCTACTACTTGGTTTGGCTTGTGCACATCCTATAGCTACTGACAGGCCTAAAACACAGGCAATAGTTCAGATTATATCAGGGTCAGCACCAGCACCAGAAGTTCCACCATTCAAGCCAGCATTTGTGTGGCCATCAATGATGCCAATTGACATAGACTCAAGTGTTATGGACACAACATCCATTACTACTTCTCAATTCAATTCAGGATGGAGTCTTGACTATCAAAGCAGGGAGACCCCAACATATGCAGACCACTCTTTGGTGTAGTTACTTTGATGGTAAACAACTATTACCAAAAGGAAAATTGATTTTTTTGGCCTTCTATTT

>M6-PGSC0003DMG400000295 M6 LRR-RLK

TTTGCCAAATGTTCATACAAACTTAGAAAAAATCACCAAAAAAAATGTCCAAAATTATTTTTTCACTTTTCATTTTTCTGTTTTTCATCATTTCTCATGGAAAATCCCAAGAATCTCCAAATCAAGAAAAGGCCATTTTACTCCAACTAAAACAACACTGGTTTACTTCACCAAATGTCACAAAATGGATTTCATCATCAAATCATTGTTCTTGGGATGGAATAATTTGTACCCAAAATTTAGTTTCTGGTATTCAAATCCCTTTTGGAAATATCTCAAAACCAATTCCAAATTTCATTTGTGATCTTAAAAATCTCATCTTTCTTGATTTTAACCATAATTTCATCCCTGGAAACTTCCCTGATATTTACAGTTGTTCAAATCTTGAATTCTTAGATCTTTCTTATAACTACATGGTTGGTAATCTCCCTGATGAGATTAACCGTCTTTCGAGTAATCTAAAATACCTAAACTTAACTGCCAACAACTTCAATGGTGATATCCCTAAAGGAATTGGTGGATTGAGTCATCTCAAAGTACTTGATCTGATGGTGAATTTGTTTTATGGATCTTTTCCTGAAGAGATTGGTGAATTGTTGAATCTTGAAGTACTTATGTTGAGCTTGAATGGTTTTGCTCCACATGCTATACCATCACTATTCACTAAGTTGAAAAATTTGAAAATTTTTTGGATGACAGAAGCACATTTGATTGGAAATATCCCGGAAAATATTGGTAATATGACAAGTCTTGAATTTTTGGATTTGTCGAATAATGGATTGAGTGGTAGTATCCCTGATGGTTTGTTTCATCTCAAGAATTTGAGTATTGTGTATTTATTTACTAACAAATTGTCAGGGGAAATACCACAGTCAGTTTCTTCAATGAATTTAGATGTTGTTGATTTGTGTAACAACAGTTTGACAGGAAAAATACCAGAAGATTTCGGAAAATTGACTAAAATGACTGGATTGTCTTTGTTTTTCAATCAATTATCAGGTGAAATACCATTGAGTATAGGCAAGTTATCATCATTGTTATTTGTTAAGTTGTTTGGGAACAAGTTATCAGGTGAAATTCCACCTGATTTTGGTCGATTTTCGAAACTTTTTGACTTCCAAGTTTCAGAAAATCAACTTGTTGGAAAGTTACCTGAGGGTATTTGCAATAACAAGGCTTTAGCTAGGATGGTTGCTTTTGGCAACAATCTAACAGGTGAACTGCCAAGTTCACTTGGAAGTTGTGACAGTCTGAGGTATCTTCGAGTCGAAAATAATCGACTTTCTGGTGAGGTTCCTGATGGATTATGGACAGGTAAGAGTTTGTCAATGCTTTTAATGAAGAATAACTTGTTAACTGGTCAGCTTCCACATAGAGTGGCATCAAATTTGTCACAAGTTGATATCAGCAATAACAAGTTTTCGGGGGAATTACCAGCTGGTATGGGTACTTGGCACAATCTAAGTGAGTTCAAGGCAAGTAATAATCTCTTAAGTGGTCAAATCCCTCAAGAATTGACTCTTCTTCCAAGGATAACTAAACTTTTTCTTGATGGTAATCTACTATCTGGAAATTTTCCATCGAATATATCGTCGTGGAAGACTCTTGTCACATTAAATAGCAGAAAAAATCAGCTTTCAGGTCCTATACCTTCTGCACTTGGTCTTTTACCAAATCTCATTGATTTGGACTTGTCAAGTAACCAATTTTCAGGTGTTATTCCAACTGAATTAGGTAACTTGAGGTTAACTTCACTTAACCTGTCATCCAATCGCCTGTCGGGTGAAATCCCATCTCAGTTGGAAAATGCAGCTTTTGGTAAGAGCTTCTCGGATAATCCTGGTCTTTGTGCAAGTAATCCATCAGTAGAAGTCGCGTCTTGCAAGAGGGAAACTAAGTCAGATATGTTTCCTGTTGGACTTGTTGCTGCTCTTGCTAGTGTAGCAGCAGTTACTTTCCTTGTGGCAGTTCTGTATGGTCTGTTTGTGTTGAGAAGTCATAGAAAAAGTAAACAAGAATCAGTTTCGACATGGAAACAAACATCATTTCACAAATTAGACTTCACAGAATCAGACATTGTATCCAATCTGACCGAAAACAACATAATTGGAAGTGGAGGATCTGGACAGGTTTACCTCGTGCCTTTAAGCCGATCAGGTGATTATGTTGCTGTCAAGAGGATTTGGAGAAACCAAAGGTTGGATCACAAGCATGAGAAACAGTTTCTTGCTGAAGTTCAGATATTAGGCACGATTCGACACTCCAATATAGTGAAACTCCTCTGCTGCTTCTTCAGTGAAGAGTCGAAACTTCTTGTCTACGAATATATGGAGAACAGGAGCTTGGATATATGGCTTCACTCAAAGAATAGGTTGAACAATGCCTCAAGATCAACTCCACATCTGGTCTTGGAATGGCCTAGGAGGCTGCAAATTGCTATAGGAGCTGCTCGTGGTTTTTGCTACATGCACCATGACTGCTCACCTCCTATTATTCATCGCGATGTGAAGTCAAGCAACATCCTATTGGATTCTCAATTCAATGCAAAAATTGCAGATTTTGGCCTTGCCAGGATGTTACTCAAGCCTGGAGATAACACAGTGACAGCAGTAGCTGGCTCTTTCGGATACATTGCACCAGGTATAAGACCATCAACTTATCTATAACATCATTATAAAGTTTAGAAGTTGGTTTAATCTCTTTCATGGAAAAAAGTTTCCTATGCTACACAATTCAATAACATATCAAGGGGTCGTTTTGTTCACGTACTAGTTATGCATGAGTTGTAACACGGGTTGTTTATGTAGTAAATAGTAATGTAGATATTAGTACGCGAGTAGTTATTCTTAGTACACTAATTCTCGCGTAACTCATATAAATGCTAATATATTCATGATGGTATTTCAGAGTATGCGCGAAAAACTAGAGTGACAGAGAAAATTGATGTGTATAGTTTTGGAGTCATACTATTGGAACTGGTGACTGGAAAAGAAGCCAATCTTGGAGATGAAGATTCTTGTCTTGCAGACTGGGCATGGCGCCACCTTCAAAAAGGGAAACCGATGGCTGATGCATTAGATGAGGACATCAAAGAAACGCGATACTTGGAAGAAATCTTCGTTGTGTTTAAGCTTGGAATCTTTTGCACCAGCACATTTCCTTCATCAAGGCCAACAATGAAGGAAGTTCTGCAAATCTTGATCCAATGTAGCAACAGTTCACCTACATCTGGAGAAAAGAAAAATGAAACAGAACAGGATGTTTTGCCACTCCTCAAGAACTCAAGGAGTGAGAGGATTGAAGAAAATGATGATGTTGGTTTAACATCAATTATTTGATGGCATTACAATATGATTAGAGTGAAGGAGAGCCTTGACGTAACTGGTAAAGTTGTTGTGGAAATAGTCTCTCGCAGAAATGCAGGGTAAGGTTATGTACAATAGATTCTTGTGGTCCGGTCCTTCCCAGGACCTTGCGCATAGTGAAAGTTTGTGCACCGGGCTGCCTTTTTTTAACAATATGATAAGAGTAAGAAATAGCAAAGAATAAGACCAATAGATAATGTATCAGGTAGTGAAAACTAGCATTGTACATACACTGTATTGGATCCATAGTGTTGCTTTGCATTGTTTTGCTACATAAATTTTGCTTTTCCAATGTTTTCTTTACTCATAAAATAACTGTCAAAAAGGTACATTAAAACCATAAACTATTGAATGGAGGATAGATTTCCTAAAAATCCAGAAAAATAGAAACAAGCTTGAACTACCTCAGTACTTTAACTTTTGGCAACCATACAACAACATATCCGATGTAATCCCATGAGTCCCAAACTTTTGGCAACAATACAACAACATATCTGATTTAATCCCACAAGTGGCTCTGGGGAAGGTAGTGTGTACGCAAATCCTACCCCTATCTCGTAGAGATAGAGAGGTTGTTTCCGATAGTAACTTTTTGGCAACAATAAATCTTGTAAAGATCACACAAAAATGTGGCTACTAGAAGAACTTTATTTGTATCCTAGCTTTATTTCTGTCTAGTGAGGAAAAAATTTAAGTTTACACTGGGGATAAAGATGATCCTTAGAGGCACCAGGAGTGAAAAGGTCAGCATATTGATGTTTAGCTGAAGAGAGATGTGCTCATTTGGTACATTGAACTATGGCTCACTTCAAAGCTCACTCTTGCATGACGCGGTGCACAAGGAAGCAAGAAATATTTCAATCATCTTGGAGAAGAAAGTAAGAAGAGGGTTCTCTTTAGCAAGAAGAGGGGTGATTGTAAACGTACAAACTTTGACTAGCTAGGCACTTTTGGTTGTGTCACTCTGACCATGATCATGATTCCTCGAGATCTGTGTATTGGGAATGCGAGGCAGCCAAAAGTGCAGCTCCAACGCCTGAACCATCATTAGAAAGCTCAATGACTATGCTGTCCGCAGCTTCTCCCAGTAACTCTTTCATAGTTTCCTCCAAGCAATTTCTGAACTTGGTGTAACGCTCAAACAATGCACCGTCCACAGCTATGACAGACCTCTGGTTCTCTCCGTCCTTCAAAGTGTCTTTTCCCAATTTCTTGAGGATGCCCACGATTCCTGCTGCAGAAAGACGAGCTCCATGAGAGGTGATAATGTCGCATAGCTCCACAACTATTTTCCTCATTTTCAAGGTAGAATTAGGTACCTGTGAAAAAAATGTATATATAGCAAATAGGATGAGAAAGACGTTCTCCACTTAATTAAGCCCAGCCAACATTGTGATGTTCAGACAGGAGCTAGGAATTTGAGTTTATGGGTTCCGGATAAATTATTTACACATATTAAGTGGGTTTTTTAGTACAAATACAAAGCTACTGGTTTTTTGTCCTATCATATATAGGACAAAAAAGATGAGAACTTCAAAGACTAGTAATGTTTCACTTTAAGTCAAATGTTTGATCTTAATACCTCTAAGATATCCTTCAGTTTATTGCCAACCACCTTGAGATCAGCCGACTCGTCGTGATGCATAGCAGCCATATCCGGAGTCCTGCATGTGTGGCAGTAAATAACCTTTTTCACAATATTTGACAATTCAAAGCTTACATATGCTTAATAAACGTACACGAATCTAGTCCAACGAGAAGCATAACTTACAGTGTAAGAGATTAGTGCAATTACATAAACCTTAAGTTGCACTAATTGCATGGAGAAGAATAAGGTTAAAATTGATGAATGCAAATAGGGCCTGCCAAAAAATAAATTACTGCCGTAATAAATGTGTTGGTCAAAGCAATACCCTACCTCAATATGAAAGGAATTTTCAGTTTGGATGGGACATAATCACCGAATAATGAAGCTTCTTTAGCCATTCTACACAATACTCTACGCAAAATTTCTCCAAGATACATCCCGGAGATCATCTTTTCATAAATCTGTGCAGTTTCCGAAAGTTCAACAACAGATGTAAGATAGAAAATGTAGAAAGGAAGGGGATTGTCTTAGTTACCTGCTCCCCGGGGTTTAAACTCTCAACATCAAGATTTTGGTCATATTCTGTTACTGGAAGATGTGATGAGCGGAAATTACCCCATTCCATGTTGATAACCTTTAAAGGCCAGTCAAAATTACGTTTAAGTAAACAAACGTGTATGAATCTACCAAGGAAAATATTTTTAGGAAATGAATTAGGCACATATCAAGTCATGAAGGTGTTATGAAATGTCATAAGCATCTACCATCTCTCCGGATTTCGGCAACAGACCATGCCATTTGGGAATCACATGAGCCCGTTCAACATATGCTGCATTGGTTCCAGTACCCAAAATTACTGCAGCAATGACATCAGGATTATTGTACCGACCTCCTGCTAACGTTCCAACAGTATCATTGACCTAAACAACCAAGTCAACTAAGAAATTCAGGCTCAGGAATTGCTTAAGATTAAAAATCAGAAATATCACACATACAAAAACCTAATACATGAAAATCATCAGCTTACTTATATTTTATCGAATTTGATTGGGAAAAAAACTTACTAAAGCAGCCACACGCACATCAAGGCCGGCCCTTTCCATTGCTTTTGCTAATTCTCCAACCACATCTTGGCCAACCTATGTTAGATTCCATGAGGAGTAGTTCAGAATTGAAATACATTAAAACATTACAAAAGGAACTCTTTATAACGTACTTTAAATCAAATTGGTGAAAATGCAATGAAACAATGATCAACTTTGCTTGTATATAGTTGTATGCATTAAATCCTCATCAATGCCAACCTATTTTAGGCCATGAGGAGTAGTTCATCATTGAAATACAACATAACGTTGGAAAAGGAAATCCTACAACGTATTTTAAATCAACTCAGTGAAAAAGCAATGAAATATACGTTGTCGGACTCTCCAATAATATTGCCGCACCTTTGTCAGATCCTTCAAAATGTACTGCTTTTGAAGGATCCTACAAGCACCTGGCGAGTGGCGAAATTTTTGAAGAGTCTGAGCAACATAGATGAAATAATCATCAACTTTGCATGCATAAAGTTGTATGCATTAAATCATCATCAATGCAAAACCATACATTGAAATGAATGGACTAAATCTAAGGTTGATTGTAGGCATGATCAATCGGAAGTGAAAATCAGAAACTCTGTATGACTAACCACGTCTTCTATGGAGAAGCCCTTTGTCCATTTAATAAGAGTTCCTGATGCAATTGACAATTGTTTCACAGGGAAAGAGAAGGTAAAGCCTAACTCCCTTTGTCTACCAGGTGGAAGATGAAAATCATCACCTTCTGTAGCTACAAATTCTGCAAGTGCTGTGGCAATAAAATCAAATAACACCTGCAATTAACAGTTTGAAGTTTAATAATGTGAGTACAACCTTGATTGGATTAAAAAAGTTGCAGGATGCATGGCTGATACAGCATAAGACTTACTTCAGATGACGATCCAGCCATCACATTCTGTGGAATTGAAACTTCTTTAACTTCATGTTTAACTATACGCTTTTCTTTCCCACCCAACTGCACACGCATCACACGAAAGTTTGTGCCGCCTAGATCCAATGCATAAAACAGACCTTCTTCGTCCCTAAGATGAAGGAATAAAAGTAATACCAGTCATTAAACTGAAGATACAATCATACATGGAAAACATACCCCACCCCTAATGCACTACAGAGGTCTATTGGCAGCAGCTTTTGAAAATTTGTCGATACATGTGAATTTTTATACCCAATTTTAGTAAGTTTATCACAAGAACATGGAATTGTTACCAAGTCCCTACCTCCGCCTACTACTATAGGTATGCCATTTTTAAACTTTAGAGAATGTCAAACTGATTCATATGTCATATATTCTTTTCTTATGGATCTAACTTCATTATGTTGATCACAGAGATGTCTGTGTAGACAGCACAGTAACAAATAATGAGCAGACAGAAGAAGATGGAACTGAAAAATTTACACGCATGATAATTGTTAAAGAATGGAACTGGCAATGTAAAATTCAAAAAAGGTGGTCAAGATACAAAACAGTAGAAGAGAATATAGAATAAACATCAAACTCAGGATCCATAGATAATGAGTGACCTGATCACTTGTCAGTTAGAAACTTCTTAGTGGACTCCAAGTTCAACAAATACTTCAAATAATAGAGTTTCATTTCTGCTCTTTGTGATTTATAACGATGCTCAAGGTAATCACCTTAAGAACATCAAAAGAATCATACTTTAGGACTAAATAATAATTTGAAAAGTACTTAATCAATTATCCAAACATTACGAAAGTGAAATCAATCTCCATGATCACAGCTTTATTCAAAAAAGAGAAATTCTTGACACCCAAATTAGCAAAAATCAGTTTGTGCACATATACACACACAAATACTCATCCAATGTGGAGGCTATCAACTTAGCTAATAAGCATCTTTAGTTTGCCAATGAATCGTTAGTAAAACACACAATCTGATATAACTCTCCACCCCATAAGCTAAAATCAAGAGACAACACTACCAAGATCACAGCTTTATTGAAAAAAGAGACAAATTCTTGACACCCAAATGAGCAAAAATCAGTTTATACACACAAAAATACTCTCCCAGTCGGATGGTTATCAACTTAGGTAATAACCATCTTGAGTATGTCGATGAATCAATTGTAAAAACACAAAAAAGAACTGACATAACTCCAACCCAGAAGCTAAAATAAAGAGACAACACCTCCAACCGAGTTGGAGCTAGAATTTGATGTTTATTGGTTCTGAACTTGTCGAGAAAAAACTATTAGGTTCATCCGAACCTGTATCTAATATGTAGCTCCACCCCTGCCTCCAAAATCACAACTTTACTGAAAAAGAGACAAATTCTTGAACACCCAAATAAGCAAAAGACAGTTTATGCACATTCTTTTTTTGAGAAGGAGTTTATGCACATTCTTAAACACCCAAATTAGCAAAAAACAGTTTATCCACATTCATAAACACCCAAATAAGCATAAAAACAGTTTATCCACATTCTCAAACACCCATATAAGCCCAAAAAAAACAGTTTATGCACATTCTTGAACACCCAAATAAGCCAAAAACAGTTTATCCACACTCTTAAACACCCAAATAAGTAAAAAACAGCTTTACTTAAACACCCAAATAAGGCACTCAAGGGTGTGGCCTAGTAGTCAACAACGTATTGAGCACCATGAAGTCTCAGGCTCAATTCCCAGCTCCTATCTATTCTAGCCTTGGTGGACT

>M6-PGSC0003DMG400000358 M6 LRR-RLK

TTTGCCAAATGTTCATACAAACTTAGAAAAAATCACCAAAAAAAATGTCCAAAATTATTTTTTCACTTTTCATTTTTCTGTTTTTCATCATTTCTCATGGAAAATCCCAAGAATCTCCAAATCAAGAAAAGGCCATTTTACTCCAACTAAAACAACACTGGTTTACTTCACCAAATGTCACAAAATGGATTTCATCATCAAATCATTGTTCTTGGGATGGAATAATTTGTACCCAAAATTTAGTTTCTGGTATTCAAATCCCTTTTGGAAATATCTCAAAACCAATTCCAAATTTCATTTGTGATCTTAAAAATCTCATCTTTCTTGATTTTAACCATAATTTCATCCCTGGAAACTTCCCTGATATTTACAGTTGTTCAAATCTTGAATTCTTAGATCTTTCTTATAACTACATGGTTGGTAATCTCCCTGATGAGATTAACCGTCTTTCGAGTAATCTAAAATACCTAAACTTAACTGCCAACAACTTCAATGGTGATATCCCTAAAGGAATTGGTGGATTGAGTCATCTCAAAGTACTTGATCTGATGGTGAATTTGTTTTATGGATCTTTTCCTGAAGAGATTGGTGAATTGTTGAATCTTGAAGTACTTATGTTGAGCTTGAATGGTTTTGCTCCACATGCTATACCATCACTATTCACTAAGTTGAAAAATTTGAAAATTTTTTGGATGACAGAAGCACATTTGATTGGAAATATCCCGGAAAATATTGGTAATATGACAAGTCTTGAATTTTTGGATTTGTCGAATAATGGATTGAGTGGTAGTATCCCTGATGGTTTGTTTCATCTCAAGAATTTGAGTATTGTGTATTTATTTACTAACAAATTGTCAGGGGAAATACCACAGTCAGTTTCTTCAATGAATTTAGATGTTGTTGATTTGTGTAACAACAGTTTGACAGGAAAAATACCAGAAGATTTCGGAAAATTGACTAAAATGACTGGATTGTCTTTGTTTTTCAATCAATTATCAGGTGAAATACCATTGAGTATAGGCAAGTTATCATCATTGTTATTTGTTAAGTTGTTTGGGAACAAGTTATCAGGTGAAATTCCACCTGATTTTGGTCGATTTTCGAAACTTTTTGACTTCCAAGTTTCAGAAAATCAACTTGTTGGAAAGTTACCTGAGGGTATTTGCAATAACAAGGCTTTAGCTAGGATGGTTGCTTTTGGCAACAATCTAACAGGTGAACTGCCAAGTTCACTTGGAAGTTGTGACAGTCTGAGGTATCTTCGAGTCGAAAATAATCGACTTTCTGGTGAGGTTCCTGATGGATTATGGACAGGTAAGAGTTTGTCAATGCTTTTAATGAAGAATAACTTGTTAACTGGTCAGCTTCCACATAGAGTGGCATCAAATTTGTCACAAGTTGATATCAGCAATAACAAGTTTTCGGGGGAATTACCAGCTGGTATGGGTACTTGGCACAATCTAAGTGAGTTCAAGGCAAGTAATAATCTCTTAAGTGGTCAAATCCCTCAAGAATTGACTCTTCTTCCAAGGATAACTAAACTTTTTCTTGATGGTAATCTACTATCTGGAAATTTTCCATCGAATATATCGTCGTGGAAGACTCTTGTCACATTAAATAGCAGAAAAAATCAGCTTTCAGGTCCTATACCTTCTGCACTTGGTCTTTTACCAAATCTCATTGATTTGGACTTGTCAAGTAACCAATTTTCAGGTGTTATTCCAACTGAATTAGGTAACTTGAGGTTAACTTCACTTAACCTGTCATCCAATCGCCTGTCGGGTGAAATCCCATCTCAGTTGGAAAATGCAGCTTTTGGTAAGAGCTTCTCGGATAATCCTGGTCTTTGTGCAAGTAATCCATCAGTAGAAGTCGCGTCTTGCAAGAGGGAAACTAAGTCAGATATGTTTCCTGTTGGACTTGTTGCTGCTCTTGCTAGTGTAGCAGCAGTTACTTTCCTTGTGGCAGTTCTGTATGGTCTGTTTGTGTTGAGAAGTCATAGAAAAAGTAAACAAGAATCAGTTTCGACATGGAAACAAACATCATTTCACAAATTAGACTTCACAGAATCAGACATTGTATCCAATCTGACCGAAAACAACATAATTGGAAGTGGAGGATCTGGACAGGTTTACCTCGTGCCTTTAAGCCGATCAGGTGATTATGTTGCTGTCAAGAGGATTTGGAGAAACCAAAGGTTGGATCACAAGCATGAGAAACAGTTTCTTGCTGAAGTTCAGATATTAGGCACGATTCGACACTCCAATATAGTGAAACTCCTCTGCTGCTTCTTCAGTGAAGAGTCGAAACTTCTTGTCTACGAATATATGGAGAACAGGAGCTTGGATATATGGCTTCACTCAAAGAATAGGTTGAACAATGCCTCAAGATCAACTCCACATCTGGTCTTGGAATGGCCTAGGAGGCTGCAAATTGCTATAGGAGCTGCTCGTGGTTTTTGCTACATGCACCATGACTGCTCACCTCCTATTATTCATCGCGATGTGAAGTCAAGCAACATCCTATTGGATTCTCAATTCAATGCAAAAATTGCAGATTTTGGCCTTGCCAGGATGTTACTCAAGCCTGGAGATAACACAGTGACAGCAGTAGCTGGCTCTTTCGGATACATTGCACCAGGTATAAGACCATCAACTTATCTATAACATCATTATAAAGTTTAGAAGTTGGTTTAATCTCTTTCATGGAAAAAAGTTTCCTATGCTACACAATTCAATAACATATCAAGGGGTCGTTTTGTTCACGTACTAGTTATGCATGAGTTGTAACACGGGTTGTTTATGTAGTAAATAGTAATGTAGATATTAGTACGCGAGTAGTTATTCTTAGTACACTAATTCTCGCGTAACTCATATAAATGCTAATATATTCATGATGGTATTTCAGAGTATGCGCGAAAAACTAGAGTGACAGAGAAAATTGATGTGTATAGTTTTGGAGTCATACTATTGGAACTGGTGACTGGAAAAGAAGCCAATCTTGGAGATGAAGATTCTTGTCTTGCAGACTGGGCATGGCGCCACCTTCAAAAAGGGAAACCGATGGCTGATGCATTAGATGAGGACATCAAAGAAACGCGATACTTGGAAGAAATCTTCGTTGTGTTTAAGCTTGGAATCTTTTGCACCAGCACATTTCCTTCATCAAGGCCAACAATGAAGGAAGTTCTGCAAATCTTGATCCAATGTAGCAACAGTTCACCTACATCTGGAGAAAAGAAAAATGAAACAGAACAGGATGTTTTGCCACTCCTCAAGAACTCAAGGAGTGAGAGGATTGAAGAAAATGATGATGTTGGTTTAACATCAATTATTTGATGGCATTACAATATGATTAGAGTGAAGGAGAGCCTTGACGTAACTGGTAAAGTTGTTGTGGAAATAGTCTCTCGCAGAAATGCAGGGTAAGGTTATGTACAATAGATTCTTGTGGTCCGGTCCTTCCCAGGACCTTGCGCATAGTGAAAGTTTGTGCACCGGGCTGCCTTTTTTTAACAATATGATAAGAGTAAGAAATAGCAAAGAATAAGACCAATAGATAATGTATCAGGTAGTGAAAACTAGCATTGTACATACACTGTATTGGATCCATAGTGTTGCTTTGCATTGTTTTGCTACATAAATTTTGCTTTTCCAATGTTTTCTTTACTCATAAAATAACTGTCAAAAAGGTACATTAAAACCATAAACTATTGAATGGAGGATAGATTTCCTAAAAATCCAGAAAAATAGAAACAAGCTTGAACTACCTCAGTACTTTAACTTTTGGCAACCATACAACAACATATCCGATGTAATCCCATGAGTCCCAAACTTTTGGCAACAATACAACAACATATCTGATTTAATCCCACAAGTGGCTCTGGGGAAGGTAGTGTGTACGCAAATCCTACCCCTATCTCGTAGAGATAGAGAGGTTGTTTCCGATAGTAACTTTTTGGCAACAATAAATCTTGTAAAGATCACACAAAAATGTGGCTACTAGAAGAACTTTATTTGTATCCTAGCTTTATTTCTGTCTAGTGAGGAAAAAATTTAAGTTTACACTGGGGATAAAGATGATCCTTAGAGGCACCAGGAGTGAAAAGGTCAGCATATTGATGTTTAGCTGAAGAGAGATGTGCTCATTTGGTACATTGAACTATGGCTCACTTCAAAGCTCACTCTTGCATGACGCGGTGCACAAGGAAGCAAGAAATATTTCAATCATCTTGGAGAAGAAAGTAAGAAGAGGGTTCTCTTTAGCAAGAAGAGGGGTGATTGTAAACGTACAAACTTTGACTAGCTAGGCACTTTTGGTTGTGTCACTCTGACCATGATCATGATTCCTCGAGATCTGTGTATTGGGAATGCGAGGCAGCCAAAAGTGCAGCTCCAACGCCTGAACCATCATTAGAAAGCTCAATGACTATGCTGTCCGCAGCTTCTCCCAGTAACTCTTTCATAGTTTCCTCCAAGCAATTTCTGAACTTGGTGTAACGCTCAAACAATGCACCGTCCACAGCTATGACAGACCTCTGGTTCTCTCCGTCCTTCAAAGTGTCTTTTCCCAATTTCTTGAGGATGCCCACGATTCCTGCTGCAGAAAGACGAGCTCCATGAGAGGTGATAATGTCGCATAGCTCCACAACTATTTTCCTCATTTTCAAGGTAGAATTAGGTACCTGTGAAAAAAATGTATATATA

>M6-PGSC0003DMG400000728 M6 LRR-RLK

ATCAGTGTATTTCGTGTCTCTGAATTAGGTGGGACAAGATTCCCATAAAAGGCTCTCAAATTTTCAACCAAAAATTCAGTGCAGCACAAACACTATGCACGTTAAAGGTGCTTTGCAACTTTATGGATTGTGCCCATGTCTGGTTGACTAAAAATCATCTAAAGCCTATTTATAAATGATAATATTAGATGTCTGCTTCGCTCTTAGCTTCTCAGGATTTATAAAAGGAGGCAATGCATGGTTGATGGTTTTCTGATTTGCTATCAGTTGGAGATCTGTAATGTTTATTGCCTCGTGAGACTATCATTATACAGAGTTCATCACATAAGGAATAGCCTCCACAATATCTTGGACCTGGAGCTTGACTCTATCAACTTGAGGGCTTCACTTGCTATCTTTATGAGCTACACCTGCATGTAATAAAAGTAAGAGTTTAGGAAACCTAAAACCTGTCCTTTTAGGGCTTAGCAATGTAACGTCAAGAGCCATGCACTTCTAAAATTGCTGTAAGCTCAGCTTCTATAGTTCCATGCATGGCTGCGATATCTCTAATCTAGGCAGTTTCAATTGTTAGATTCGGTATGAGGGATGTGGCCTTAGAAGATCTTGATGACATTATAGTTCTACTGTGCAGAGATATTGTGAATGCAGAAATAGAAAATTTCTCAAACTAAGAACAGTTATGTTTTTTTTTAAAATTTTTTCTTTTCTTATTATTGAACCTTCTCCATACATCAATCCTTTTTTTTTTGATAACCGTGGTGCCCGGGCTAGCTTGTGCACACCTTGACACAATCCCATCAATCTTAATCAACTAAAGGTAGTCTGCATATGACACTCCTCATCTTTTGTACCTTTTATCTCCATGGAACCTAAGCCAGTCAGTGAACTATCTAATTTCTCTAAAACTTTTCATCTGCATGTTTTGGGTTCAACCAATATTCTTTCACCGGGCAACCCCTAGGGATTTGGGTTAGTGGCAAGAGCACAACTCATTATGTGTGGGTTAAGCTCACGTTATCAGTTTGAATCCTACCATAGACTAAAGCTTGGTATTTAAGTGGAGAAAGGTAGAAGGCTGGGCCCATTATCCACCGACTTTTGGACCGTGCACCGAAAGAGCTGGGAGGATTTCTCAGTTACAAACAAAAGATCCTTTCATGGGGCATCTAATAGACTTTCTTGAACCGAATATCCATTCCAGAAGGAGTTTCCAAAGGAGATGCTTCTCTAAATGTCCGCATAGTTTTTCTGCTGCAGTTCACCATGAAATTCTCAATGATTTAATTGCATGAATTAGTTTCTTGGATTCAACTCTTCCCTCAACAACCCACCCCCCACCCGCCCAAAAAAAGAACTTTATTTCTGTCAGAAGTTTGCTAGTGCATCATGTAGTGACATCTTCTTTTCCTATGCCTACTATTCTCACAGAGATGTTCAAGCAAGCAGCATACTTCTTGATGATAAGTTCGAAGTACGGCTAGGGAGTCTAAATGAAGCCTGTGCACAAGAAGGAGAGAGCCATCAAAATCGAATTTCCAGGCTGCTTAGATTTCCACAGTAAGTACTTGTTGGTAAATTCTGTGTTCTAACCATCCTTTTCTTTTCGAAGGCTAACCACTATTACTACTAGGTTGTTATGTAATGTTCAATAATTTTGTTTCAATTTTTGTCTCCCATGATGGATCTTAAAATCAGGCAGTTTTATAGTTAATATGTCTGTATCTCCAATTTGATGTTATAGAAACTACTTTGGCCTGGTTTCTAAATTGGATTTGAGTTCCACAGTTTTGCTGCCACTCAAGTCTTTCTGGTGTATATTTATTAGCTCTGATATAAATGAGAATCCTAACGTTCATGCAGCAACCTTACTCTTAACCTTGTCATCTCGTTATCTGTGCTTATACAAAAGGGCTTTAGCTTTGTTTCTAAATTAGAAAAACTTGTAATATAGGCTCTGAATAGTTTAGTAGTCCTGATGATAGTTATCATGCAGTTGCATATAATCTTATTTCTTCTCAGTATTCAAGTTACAAGGAACACTGTAAAGAACAGTGGGTCTAGTCTGAAAGACCATCACCCACTTGTTAATTACTTGTTTTATCCCTCAGTAAATTATTCCTTTTCTATGTTGTATGAATCTTTTTGCCTTGATTTACCTATGTCTAGCGGGTGTCACATGGGTGTTGTTGTTTAAGTAGAAATTAGCAAAAAACTGTATGTACCCTTACTGGATACTTAAAACTCATGTATGGGATATGCATGCGTAGCCGAATCCATGAAACATAGTCCTTTGATGTACATGTTTTCACTTAAGGAGAATTTGTTGGTTATGCTATTTGGTAGGGAATGCATAGTTCTCATTCTTTTAGCTTTCCATCCTGTGCTGAAGTAAGTTTAACTATTTCTCACTCTCATTGTCTAGCTTTAACTATTACCTATTCTCGTTATGCCTACACATTGAATGCAGGAAATTATTAAATTTTGTATTGTGGTACTAAATACTTCTGGAACGACCTAGATATAATGGCATGTGAAAAATTGGTTTGAAGTCCCATGTTAGATGCCAAAGATTTTAGGCCTTCTATTCTATTATTTTAATCTTCATGGCTTCTGTTAAAACTTTCCAGGTTCTCTTATTTTAGAAATTGAATTTACGACCACTGATCACAGTAATCCAGTAAGCTTTTGTTTTATAGTGAAAACTTTTCCATATGTATTTCCTGTCTTACAAATGGTTATCACTGTTTTTCCAGCTTTTCGTCGAAGCTTGTTGCATGATCTCAAAAGTTTAGTTATATTCTTTCAACCAGTAACTGTTTAATAGTTTATACTGCTTATTGACATAGTTCTCCCTTTCTTATTTGCAGGACGTCGGAACAAGGTGCATCAGGTATGTGCAATTTTATTTAGTTCTACTTTCTTTAAATAGAGTTTACTTGGATATTCTTAATATTTATTAAATTGGTGTAAGTAACTATTTAAAATTATTTGCGTCACATTTCCAGTATTGTTTGTTGGTGGGTAAAATTTCCTTAATGTCTAGGCAACAAATCAAAACTATAGATTCTCATTAGAAGTAAGAATAAGTGAGACTATTAGGTGGAAATTTACGATATAAAGCTGATTCCAAAGAGTGTAAAGGGCTGGACAACTGTATATTGAACTGTACTGCCTAAAGAACTGGAGATTTTGATGGCGGTGTTTTGCTTTACTTTTTGTGGGATCTTTGTAGCTTGAACTATCACTAGGGAATACTTCAAAGTCTACTCCTCTCGTCCTAATATTCTTTCTTACCTGAATGGACAGGTACACCTAGTGCCACTTGTGCGTATGATGTCTACTGTTTTGGAAAGGTTTTGCTCGAGCTAGTAACTGGCAAGCTAGGTATCAGTGCATCCAATGACGCCAGCATGAAGGAGTGGTTGGATGGTACACTTAAGTACATTAGTATATATGATAAAGAACTTGTCACAAACATTGTGGATCCTTCCCTAATCATAGACGAAGATTTATTAGAGGAAGTATGGGCGATGGCTATTGTTGCCAGATCCTGCCTCAATCCCAAACCGTCAAGGAGACCCTTAATGAGATACATCCTTAAAGCTTTGGAAAACCCTTTGAAGGTAGTTAGGGAAGAACACACTAGCTCAGCAAGACTTAGAGCAACTTCATCGAGAAGTTCATGGAACGCTGCACTGTTTGGCAGCTGGCGTAGCTCCTCAGATGTAGCAGCCGTACCTGCTGCAGCTTCAGCACATAAGTTGGAAGGAACAAGTAGTTTGAAGCAATCCGGAACAACAGGCTCTCAAGGAAGTGGTCCTAATGGGGACAACAACGGTCATTCTTCATCAACTAGACGACAGTCCAAAGAGATTTTCCCTGAGCCATTGGAGGAGCACGATGTAGAGAGAGGACCACTGGCGGAGTAGAAGGGAGGGTGATTGTCATTAATTCTTTGTTAATAATGCAGCAGGATCTGCATTAAGGTTGCATTTTTGTTTGGGGACACCTTGTACATTCTCCGTTTCCGGAATTCTTGAGTTTGCAAGCCATTGTCTTGCCCTGATCTTGAAAATGGAGATGCATAGTTTACCTGGAGACAGCAAACATCAAATTGTTTATTGGGGTTTTGGGATTTTAAGTTCTTTTTCTTTTTCTGGGGATAGATTTGGAAAACAGGATTTAGTAGTGGTGGTGGTTCTTTTTGTTAATTTTGTTTTTGGGGGGTTTATGTTTGCTTCCAAATTCCAATTCTTTTCTTATAATTCTTCTCTTTTACTAGTCTCTCCATGGAATGTAAATTATACACAATCTTTCATTTTCCAATTGGAGGAGAGGGAAAAAAAGAAAAAGAGTATCCC

>M6-PGSC0003DMG400000996 M6 LRR-RLK

GGCTGTTGTCTCTTTTTTGCCTTAAAACCTGAACTTCTCCTGAATTTTTCTACTCTACATATTACTGCAGGCATGAAGTCTTTTAATGTTTTCATTTGTGTTATGCTTTCTTGCTTATTATGTGTTCTTTCTGCAGATGCATCTTCCAGGCTTGCTGGAAATGAAACGGATCACTTAGCCCTCCTCAGTATTAAAGCACAAATAACATATGATCCTTCTGGTGTTACAAATTCTTGGAATGATTCCTTCCATCATTGTAGTTGGCAAGGCGTTACATGTAGTGCTCGACATCAAAGGGTGACCGTGCTGGATTTAAGCTCTAAGCAGGTTGTAGGAACATTAGTTCCTCAGATAGGAAATATGAGTTTTCTTAGAGAACTAATTCTTCAAAATAACACCTTCAATAGTGAGATACCACGCGAAATCGGAAGGCTATTCAGGTTGAAGAATTTAGTGCTAAAGGACAACTCATTTACTGGTGAAATTCCAGTGGAACTATCAAATTGCTCTAGACTCATTTATCTTGATTTGGATGGTAATTCACTGACCGGGAAAATCCCTGTGGAGCTTGGTTTGTCTTTGAGAAATCTTCAGGTACTTTTTCTTAGATCAAACAATTTAACAGGTGAACTACCATACTCTTTGGGTAATCTTTCATCCCTTATTGCCTTAGCAGCAACAGAGAATAGATTAGAAGGAAGTATACCGTATTCGTTAGGCCAGTTGACTAACTTGAGTTATATCTCTCTTGGTGGTAATATGCTTTCTGGTTCAATTCCCATTTCAGTTTTTAACTTATCCTCTCTGTATCATCTTGCTGCACCAGTTAACCAACTTAAAGGAAGTCTTCCCACTGATATTGGATCCACACTTCCGAGTCTTCGCATAGTTTTTTTGTTCTCAAACTTATTAAGTGGAGTGCTTCCAAGTTCCATTTCTAACTTAACCAATCTTGAAATTCTCTCATTATCAAGAAACCAGTTGTCCGGGAAAATCCCTTCTCTGGAAAAACTCCGTAATCTTCAAGGACTAGCTATGCATTTCAATAATCTTGGAACCGGAAGAGAAGATGACATGGATTTCTTTTCTTCATTGGTTAACATTACCAGTTTTAAAGAACTGAGTTTGAGTGTGAATAATATAGCAGGCCAGCTGCCTAAGAATATTGGCAACTTAACCAATTTCCGGTCTATTGGTTTTGCTAGGAACAAACTATTCGGACGAATACCTGATGGATTTGTTGATCTCAGCAACATGGAAGTGGTAAGTTTGGAGTATAACCAATTGACTGAAGAAATTCCAGCTAGTCTTGGTAAACTTCAAAAACTCAAGTATTTCTATGTTAACGGAAACAAACTCTCGGGGGAAATCCCTTCCTCTATTGGAAACATAACATCTTTATATGGACTTAATCTAGCTCAAAACAACTTAGAAGGGACTATTCCTTCAGTTCTTGGAAACTGTCAGTTGTTGCAAATGTTGTATCTTTCTCGAAACAGACTCAGTGGGACGATACCAAAAGAGGTTCTTAGTATCTCATCCTTGTCTATCCAACTAGACCTGTCTGGAAACCAGTTAAGTGGATCACTTCCCCTTGAAGTAGGAAGTTTAGTCAATCTTGGGTTTCTTGATATCTCCGAGAACAAGTTATCTGGAAAACTTCCGAGCACTTTAAGCAGTTGCATCAAGTTGGAAAACCTCTACGTTCAGGAAAACATGTTTGAAGGTGTGATCCCATCATCTTTAAGTTCTTTGAGAGGGATGGAATATTTGGACTTGTCGCATAATAATTTCTCTGGTCTTATTCCAAAGTACTTTGAAACTTTCATATCATTGAAAAGCTTAAACCTGTCATTTAACAATTTTGAGGGTGAAGTACCTCGAGAAGGAGTTTTCAGTAATGCAAGTGCAGCAATAGTCAACGGAAACAGAAATCTTTGTGGAGGTTCTCAAATGCTAAAGCTACCACAATGTAAGTTTCCAATGCCGAAAAAAAGAAGATTGTCATCAACCTTGAAGATAACTATATCTGTTGCTAGTGCATTATTTGGAGTAGCACTTGTTCTGATGCTCCTAATTTTGTGTTTCTTAAAGAGGAAACGTAGCCCTTCACTCGATTTATCAGATGATTCGTTCTTGAAAATCTCCTACAGGGAACTTCTGAAAGCAACTAATGGATTCTCCTCAGATTATTTAATTGGTAAGGGTGGTTTTGGTTCTGTATACAAGGGAATTCTTGGTCCAGATGAAAAAACTGTCGCGATCAAAGTACTCAATTTGCAGCATAAAGGAGCTTTGAAAAGCTTCATTGCAGAATGTGAAGTCATAAAGAACCTCAGGCATAGAAATCTTGTCAAGTTAGTAACTGCATGCTCAGGTACTAACTTTCAAGGCAATGATTTTAAGGCCCTCATTTACAAGTTCATGGTTAATGGTAGCCTAGACGATTGGCTACATTCATTTTCTAATGATGGAAGTCTGCACGTTCGATATCTAGATCTTTATCAGAGAGTAAACATTGCATCAGACATTGCTTTCGCGCTAGATTATCTTCATCATGGAACCCAAACACCAGTAGTTCATTGTGATTTGAAGCCAAGCAATATCCTTCTAGACAAGGACATGACTGCCCGTGTTGGCGATTTTGGTTTATCAAGATTCCTCCAAGAAACATCTCAAAGGGAAACTAGCACTATTGGTAGTAAAGGATCAGTTGGCTATGCAGCTCCAGGTAAACATCTCTTTCCCTATTTTAGCAACTCTCACATTCCAACTTTCAGCCTGACATGTTTAAGATCATACTATGTTTAAGATCGTAATATTCAAAAGTCTTACATACTTTCTTAAACTTCGTGGCAAGTCCAAACCAGACAAATATAAAATGGAGTGATTTTGATATGGACTAACGATTTTTTTTCTGCAGAATATGGAATGGTAAGTGAAGTATCAACTTATGGTGACATATACAGCTATGGAATCATCATACTGGAGATGTTAACAGGGAAAAAACCGACAGACGACGCCTTTTCAAATGGAATGAACCTTCACAACTATGCTAAAATGGCTTATTCTACTGGTAGAGTCATGGAAATTGTTGATCAAATGGTTTACCACAACTTGCAAGAAATGAAAACCAAAGATTACATTGAAGAGTGTTCAATTTGTATGTGTAAAATTGGGATTGCTTGCACTATAGATTCACCAAAAGAAAGAATGAGAATTAGTGATGTTATTAAAGAGTTGCAATTAGTCAAAGAAACTTTATCTTAATGTGATAGAACACTATGATATTAGAAGTAAATCATAAAACATTATCTTTTTTGTTCTATATATTGTGAGTTGGTAAAATTTC

>M6-PGSC0003DMG400001185 M6 LRR-RLK

TCAATGGTAGTTTCTCTTGTCTCCTTGTTTTACAACTTTTTCCTCTTTCTCATACTTTCGTCCTCCAACTTAACTGCTCAACCATCTAGCACAGACACAGACTTTTCATGCTCGTTTAACTCTTCCTTTTCTTGTGATACTTTTATTTCATATCGTGCCCGTCCACCAAATTATTTAGATGTCGGAAGTATATCTGATCTTCTTGAAGTCAGTCGTTTAAGTATAGCTACGGCCACAGGTCTAGCCTCAGAGGATACCGAGCTATTCCCTGACCAGCTTTTACTTGCACCTGTTAAATGCTACTGCAATAGTTCTCATTATTTCTCCAATGTCACCTATCAGATCATGAAAGGCGACAGCTTTTATTCAGTTTCAATAGGAGCTTTTGAGAACCTTACTAATTATCACGTGGTGCAAGATATGAATCCAACACTAGATCCAACCAACTTGACAATTGGTGCAGAAGCTGTTTTTTCTTTGTTTTGTAAATGCCCCACACATTCTGATCTCGAAAAGGGACTTCAGTACCTGGTCACTTATGTTTGGCAGCCTTGGGATGATGTGTTGCCTGTAAGCTATATGTTTGGAGCATCTGCAGCTGACATTCTGGCTGCCAACAATTACAGAAATTTCACTGCTGCAATATATTCTCCTGTCTTGATACCAATAAATCTTCCTATTATTCTGCAATCTTACCCCTCTTCTGCAAGCAGTAGAAAATCCAAACACGGATGGATAGTAATTACTGTTTTAAGTATTATGGGGTTGCTTGCAGTTTTTTCATTTTTCTTGATGGTATACATGCGTCATCTAGACGAGAAGAGGAGGGCTAACTTAGCCCATAACTCTTCTACTCTTGAAACTTCTGATCTCTTTCATACCAAGAAAGCCTCAGAAGGTGAAATTACGGACCATAAGAACATTCAAGATAAGCTTCTTCCTGGAGTTTCTGGCTACATTGGAAAGCCAATAATATATGACCTGAAGATCATCATGGAGGCAACTGTCGACCTCAGTGAGAGATACAGGTTAGGAGGATCAGTGTACAAGGTTACAATGAACGACCAAGTTGTAGCAGTCAAGAAAACAAAACAAGCCTCGGAGGAAATGACAATACTTCAAAAACTACATCATGCAAATCTAGTGAAACTAATGGGCTTTTCATCAGATTATCTTGGTAATTCCTTTTTAGTTTATGAATATGCTGAAAATGGCTCACTGGATCAATGGCTGTTTCCCGGATTATCATCATCTTCTTCTGCCTCTGTCTTAGTGGCATCACTTGGTTGGAGTCGAAGATTACACATTGCCTTGGATGTTGCAAATGCTCTGCAATACCTGCATGAACATACTCAACCTAGTATAGTTCACGGGGACATCCTAACGTGTAGCATACTTCTTGATTCAAGGTTTAAAGCCAAAATTGCAGGTTTCTCTACAGCCAGACATGCCACCAACTCAATGATGCTGAAAGTTGACGTGTTTGCTTTTGGGATTGTGCTGTTGGAGTTGCTTTCAGGGAAGAAAGCAATGGAATCAAAAGATTATAATGATGAGACTCTGATCATGTGCAGAGAAATTAGTGGAATCTTGGAAGTTGAAGAGAACAGAGAGGAGAAGTTTAGAAAATGGATGGATCCAAAGTTGAGCTTTTACCCTATTGATGATGCTCTAAACTTGGCTGCCTTGGCGACAGCATGCACATCGGAGCAATCAGCAGAGAGGCCTAAGATGACAGATATTGTCTTCAACCTCTGTTTTCTTGCTCAATCATCTTTTGAAATGTATGGACCGTCTTGGACTTCAGGTGAAGCTGAGGAGATTGTCCAGATTGTTAGTCCTGTAATAGCACGTTGAGTTAAAGATGGAGCGTAAATATAGTTCAAGAACTTTTGTCAGTTCGTCTCAAATTTGTAACATATTCTGGATTGTAAAGCCACAGGAACGATAAATAAGCTGATCTCCAGAATATATTGCAAATTTGAGTTTTTTTGTCCAGTTTTTCTTTGTATTTCTCTTGCTGCTAATTTAGTTGCTATTGTATATTCTTGTATTGGATGTTTTTCTCAACTTTGATATAAGTGGAGTGCAATTTGACCTGTATTTCTTTCGTCCTCTCAAAAGAAAAACAGAAAAAATTAAGTTTCTGAAAAGACACGGCAAACTCGTTCTTATTTATATTTATTTAAGATCCAGACGAGTACATGTTTA

>M6-PGSC0003DMG400001301 M6 LRR-RLK

TAATTTCTTAGTAGCTAGCTTCTAAAGGCACCACTCATATTTGTAAAATAACAGACTGGGTTTTCATTTGTCAAGTCAAATGCCTCACCAGTCTTCATCTTCCTTCTCCTTGCTATTTCCTACCTATTTCTCTTGCGCATATTATCAAACACCTTGTGCCAATTACCCAAAACTGCTGTTGGATCCCTAAACCCCCCAGCTAACGATGGCTCGTTTCAGATTTCTCCTTTTTCTTCTGTTGATTCTGATTTCTCTTGTTCTTCCGTGCCTTGATGCTGCTGTAGTTGAAGATGATCTCAAGTGTTTAGAAGGTTTCAAGAAGTCTTTGGAAGATCCAGCTGGGAATCTGAATTCATGGAACTTTAAAAATACTACTATCGGAGCTATCTGTAAGTTTACGGGTGTTGATTGCTGGAATGACAACGAAAATCGCATCATCAGCCTCATTCTTTCGACTATGAACCTCGGAGGGAAGGTTACAGAGCCTGTACAGTATTGTGCCAGCTTGACAACCCTTGATCTCTCTGGTAATCGCTTTTCTGGTCCAATTCCTTCTCAAATTTGTACTTGGCTACCTTTTTTGGTAACCCTAGACTTGTCCGGTAACGATTACTCCGGCCCTATCCCTGCTGATCTTGCTAAATGCACCTTTTTGAATAAATTATCGCTCAGTAATAATAAGCTTACTGGAAATATTCCCCCTGAATTTTCTAGTTTAGGTAGGCTTAAAAGTTTTTCTGTGGCAAATAATCAACTCTCCGGTAGAATCCCGGCAGCTTTCGATTCTTCTAACTTTAATTTTGAAGGAAATAGTCTCTGTGGTGGACCTTTGGGGAAATGTGGAGGACTAAGTAAGAAAAGCTTAGCTATTATTATTGCAGCAGGAGTCTTTGGTGCTGCTGCTTCTATGTTGTTGGGTTTTGGTGCGTGGTATTGGTTTTTTACCAAGTCTGGAAAGAGGAAGAGAGGGTATGGAGTTGGGAGAGACGACTCTGATAGTTGGGCTGATAAGCTGAGGGCTCACAAGCTCACTCAGGTTATGTTGTTCCAGAAACCGCTCGTCAAGGTTAAATTAGCTGATTTGTTGATTGCCACAAATGGTTTCAGGACGGACAATGTTATCAACTCAACTAGAATGGGTACTACTTATAATGCTGTGTTACGTGATGGTTCTGCACTTGCCATTAAGCGGCTTAATACTTGCAAACTGAGCGAGAAGCACTTTCGTGATGAGATGTATAGGCTAGGACAACTTAGGCATCCTAATTTGGTGCCACTGTTGGGGTTTTGTGTTGTGGAAGAGGAGAAGCTTTTGGTTTATAAACACCTATCAAATGGTACTTTGCACTCATTCTTGAATGGGAATGCAAGTGAACTGGATTGGCCTACTCGGTTTAGAATTGGTTTCGGTGCTGCTAGGGGCCTTGCTTGGTTACATCATGGTGTCCACCCACCTATCTTGCACCAAAACATATGTTCTAATGTTATTTTCCTCGATGAAGACTTTGATGCTAGAGTAATGGATTTTGGGCTGGCAAGGTTGATGACATCAGATGCAAAAGAGTCTAGTTATGTGAATGGGGAGTTGGGTGAATTTGGCTATGTAGCTCCAGAGTACTCTAGCACAATGGTGCCTTCACTGAAAGGGGATGCTTATAGTTTTGGGGTAGTGCTTTTGGAGTTGGCTACTGGGCAAAAACCTCTTGAAGTTATTGCTGGTGAAGAGGGATTCAAGGGTAATTTGGTAGACTGGGTGAATCAGCTCTCTGCTTCAGGTCGCATTAAAGATGCAATTGACCAAAACATGCGTGGGAAGGGGCATGATGAAGAGATTGTGCAATTCCTAAAAGTTGCTTGTAATTCTGTGGTTTCTCGGCCCAATGATAGGTGGTCTATGTATCAGGTTTATGAAGCACTTCAGAGCATGGCTGAAAAACAGGGTTTCTCCGAACAGTATGACGAATTCCCGTTACTATTTGGCAAAGAAGGTGCTACTAGAAGTCCTGTTTGAAGCCGGAATTGCGAAATTATTACTAAATGTTCCACTCCAAATGAACATGATGCATCCTGTTGTAAGTTTTCAGTTCCTATTGGTGTAAGTGGCATGGCACTGTTTTGATTGGCTCAGAAGTACACCCTTTAAATATTTCTGTATTATATTTGCAAAATGCACAAATCATTCTTAAGTATTTGCTTGCTTGATATTTGTAGTATAGGAAGCAACTGATAATTATTGTTCATGTTTGTGATTACCCATTGCCTATAGCATTAGATGTTATGTATTCAAAGGATTCTCAACTGAACGTCTCATTTACTGGAAACTGGTCTATATTACTATTTACTGCCTGTCTAGCTCTTGTTTGGAAGCCAATGGGGACG

>M6-PGSC0003DMG400002012 M6 LRR-RLK

TGTAAGTAGTTAGTCAAAATTGTGTCTAAATGAAAGGTTTTTGGTGGTCTACTGTTTCTTCATTTATATAAGGAAAGTTGGTTTGACTAAAATAACATCCCCTATTGACTTGCTTATGGTATAGGGAAATGTAGGGTCTCTTGCTTTTGATTTTGATGTATTGGTTATTGGTGAGTGAATTTAATGGTGTCAATTTGCATTTTGGGACATTATTGGTGTTGCTGTTGGACTTCAATGAATTTAGTGACTGGTGTTGTTGCTGCTTGTTCTTTTATCTAGGTTTATATACTCTCTTATTTCACTTACATAATGATTAGTGATGATTGCAGTTTGTGTTTAAAAATTTTGCTAATTCATTTTCTGGGATTCTTGAAATGATTGTGCAATCTTGTCTTTCTACTTGGTGGGGTTTGTCTACTTTCTTGGCCCATTCTGAATTATTAGCTTTGTCAAAATGCTGTCTTTTTTCCTTTTTAGTTCAGTATTATATGTAACTCTCTGCTAGTGTCACACGGGTTTTTCATTGGGAGGAAAATAAGAGGAGGAAATGCAATCTACCATTTGTTTAGTGCTGCTATTTTTAGTTGAGTTAGCTAAAGGAAGCTTAGATTTGGATGCTCTTTTGGAGTTTAAGAAGGGAGTTTTGAAAGACCCTTCTGGGAAAGTCCTTTCTTCATGGGATTCTAAGTCATTGGGACCAAATGGTTGCCCCCAAAACTGGTTCGGTATCGGTTGCAGTGATGGCCATATCACTTCAATTGAACTAAATGATGTAGGTCTGGTTGGAGTTTTGGATTTTGCAGCTATTAGTGGCCTAAAAATGCTGCAGAATTTGTCTGTTGCAAACAACCAGTTAAGTGGGAAAATTACTGAGGAAGTTGGGTTGATTATGTCATTAGAATTTTTGGATCTTTCCAAGAACATGTTTAGGGGTTTTATACCCTCTAAGCTGACTAGTTTAAAGAACTTAGTATCTCTTAATCTTTCTTTAAATAGTCTCGATGGAATGGTTCCCACTGGTTTTGCTAGTCTTGAGAAATTGAAGTATCTGGATTTGCACTCTAATGCCTTCTCAATTGATATTATGCTTCTTCTGGCCTCATTGGGGGATGTATAGTATGTAGACCTCAGTAGCAACAAGTTTGTTGGATCACTTGACCTGCAAGTAGGAAATTCGAGCTTTGTCTCGTCAATTCAGTATCTGAACATTAGTCATAATAACCTGGATGGAGAGCTATTTCCTCACGATGGAATGCCATATTTTGACAGTTTAGAGGCTTTTGATGCAAGTGACAATCAGCTTACTGGTACTATTCCGTCCTTCAATTTTGTAGTCTCCCTTCGAATTCTTAGGCTCGGGAACAATCAGTTGTCTGGATCACTCCCCGAAGCTCTTCTAGAGGATAGCTCAATGATTTTATCAGAATTGGACCTAAGCCAGAATCAGCTTGCAGGTGTGTTGATCCTTTCTTCTTTAGTTAATTTGTTTGGTTGAAGTCAATCTAAGTAATTTATGGTTAATTACTACTCCCTTTGTCCCAATTTATGTGACACTTTTTGTTTTTCGAGAGTCAAACAATTTAAGTTTGACCGGAAATTTGCGTGTGGAATCTTCAACTTTTTTGAAATGAAATTTATATATTTGTAAACTACGTAAAAAGTACTATAAGTCACAATAATTGGCAATTCAAAATAATTAAAAGATATATTATAAATTTACGGTCAAAGATAGAATTGTTTGAATCTCGAAATCCGAAAAGTGCCACATAAATTGGGACGGAGGGAGTACTATCTATGATCTTAGGTTGTTGTCAGTTAGATAACAAATGAATAATATTTAGATTGCTATCTGCTGGAAAGCATGCTTGTTTTGGTTGCAATCAAATGTAATGTATAGAACACAATTCCTGAGCACTTCGGAATTGTGTTCTAGAGGAATGCTGTATAATTCTCTGCTTATGTGTAATTATCATACACATTGTTCCCAATTCTTCCCTAGGTTAGAGATATTTCATACTTAGATTCAACTCTAAGGCAATTATTGCTGTAAACTTGCATTTTAATTAGTGGAATGTTTCAGTGTCTTTTTGATTATTTTTTTTTGCTGCATTATTAATTTCATCTCTCCTTGAGCCTAGGGTCTATTGGAAATAGCCTCCTTACCCTCACAAGGTAGGGGTAAGGTTTGCATTCATGTTACCCTCCCTAGACTCCACCTATGGGATTATACTGGATTTGTTGTGGTTGTTGTTTGCTGCGTTATAAACAAGCTGCATAAAGATATTCTTCATGCGGTTGGAGTTCTGATGTATTTTAAGTCTTTTGTTGTTCTTTTAATTATTATATTTAGTTGTTCTTACCACAGGCCTAAACCAAGCTACTTCCTTTCCAAACTATTTCATGAAGCCATTTGCTATTTTGGCAGGTCCCATTGGAGGTATAAGTGCTGTGAATCTCAAGCTTCTAAATTTATCATATAACCAGCTCTCAGGCCCCTTGCCTTTCAAGGTTGGGCGCTGTGCCATCATAGACCTGAGCAACAATCGGCTAACTGGAAATGTGTCCCGGATCCAAGGTTGGGGAAATTATGTTGAAGTTATTGTGTTAAGCTCAAATGCATTGACAGGAACCTTTCCCAACCAAACTTCCCAGTTCTTGAGGCTTACTTCGTTAAAGATTTCAAACAACTCACTTGAAGGTTTGTTACCAACTATGTTAGGTACATACCTCGAACTTAAAACAATTGATCTTAGCATTAACCAACTTAGTGGTACTCTCCTTCCCAGCCTTTTCAACTCTACCAAATTGACTGATATTAATGTTTCTTTCAACAAATTTACCGGTAGTGTACCTATCATGGCATTTAACTCAGAGAATCTAAGCTTGATTTCTCTAGATGTTTCACATAATGCATTAGCTGGTCCGTTACCTCCGGGGCTGGACAAGTTCCCGGACATGGTAAATCTGGATCTCTCTGATAACAAATTTGAAGGTGGCCTTCCTAATGATCTCTCGGACAAATTGGAATTTTTTAATGTGGCTAATAACAATTTTTCTGGACCTGTGCCACAAAATTTATGGAGGTTTCCTGACTCCTCATTCCATCCGGGAAATCCCTTGCTTGTACTTCCAAAGCACGCCAAAGCTCCTTCAGAGGGAGATTCAACTTTAAGCTTGAGAAGTCATGGTTCTCGCATGAAATCTACCATCAGGGCTGCCCTCATTGCTGGGCTGATCTGTGGTGTATCTGTTATATCTTTGTTGACATTAATAATCTACCGCAAGGCCCATCAGCGAGATGGCGGGAAGGACGATATGAAAGGAACCAAGGAAAAGAAAGGTATGGAACTATTCTTCTTCTAATAAGACAGGTGATCAGATCAACTGATACTAATGCCTTTTCATTTCCACAGGTTTGTCTTTATCTGATATAGAATGTGGACATGACACTAGAGAACACAGCGTGCCAGTATCAACAGTTCAAAATGAGCCATTATCTTCACCTATATCTGTCATGTCATCCGCCAATCTCTCTCCTTCTAAGGTTCAAGATCAATCCAAAAGCCCAAACTCTCTCAGGGTTTCTTCTCCTGATAAACTGGCTGGAGATTTACATCTACTAGATAACGCCTTGAAGGTCACTGCTGAAGAATTATCGTGTGCTCCTGCCGAAGCTGTGGGCAGAAGTTGTCATGGAACATTGTACAAAGCTACGCTTGGTTCAGACCAAGTTCTTGCTGTCAAATGGCTTAAGGAAGGGATAGTGAAAGGCAAGAAGGAATTTGCTAGAGAAGCTAAGAAACTGGGGAGTATTAGGCATCCTAATTTAGTTTCTCTACTGGGTTACTACTGGGGTCCCAAGGAGCATGAGCGGCTTCTCATATCAAATTACACCGATGCACCATGTTTAGCTCTTTATCTTCTTCGTAAAGGTCAGTCTCCCCTTTTCTATCTATAGAGTATATCACCTGCATTTATTTCTGTAGCTTTTCTGATTTTTGTATGGTGCTGAAGTCGTGATCCAGTTTTACCAAATAAATTAGTAGTACTCCTTCAATGAAATCAAACATAATGGACCGTGAATGCGCAATACCTTGAATGTGTATATTGTTATTGTAGTTGTCAAGATTGCTCATACCTGATATGCTGAAGTTTAATCACCATCTTGCCACTTGATTTCCATTCCGGTGCCTCAAGTGTACTTTTCTTCCAAAAGATATTCTAGCTTGCTGGTTGAATATGTATCTGAGACTATTAATACACATTTCACATAGCAACTTTCATGAAATTTATAAATGGAGCTATCTTCCAAAATCTGAACTTTTCAATTCATATATATTAGCAATGTTAGAATTACACATAGCAGAAAAGATACACACCTGTTTTTTGCTTTCATGGTAATACTAAAAGGTAAAACAGTTCCACCTTCGCTAATTTTGCCTTGATGGTGATAGTGGGAGAGTCAAAAGTCGAAAAGAGGATTCCTCTCTTGTTTCTCTCATTCAAAACTGTGCATGAGAATTTTATCTCACGTGGCTCATTTTGTATTGATTACTTTGGTTGGGCTCGATTGATGTCATTTTGATAAGGGCATCCCGGTGCACTAATTAGAGACTGCTACTTTGAATTCCGTTTATTGCATATAGCACTGTACTTTGTTTTCTAGTTTTGAGTGATGATATGGATGCCGGTTTCATTCTTTGATGTATGCAGATGCAGAACCCTGTAAGTTACATCCTTTATCTCTCGATGATCGACTCAAGATCTCTGTAGATGTTGCTCGTTGTCTAAACTACCTTCACCATGAAAGTGCCATACCTCATGGCAACCTGAAATCCACAAATGTACTTATAGACACGTCTACCGTGAATGCCCTACTAACTGACTATAGTCTTCATCGGTTGATGACATCTGCTGGAACAGCTGAACAGGTTCTAAATGCCAGTGTACTCGGTTATCGTCCACCTGAATTCGCGAGTACAAGTAAACCTTGCCCGTCATTGAAAAGTGACGTCTATGCTTTTGGTGTTATCTTGTTGGAACTTTTGACTGGAAGAAGTTCTGCAGAAATTGTTCCTGGGAACTCAGAAGTGCTGGATTTAACCGAATGGGCGAGATTATTGGCTTTTCAAGACCGTTCAATCGAGTGCTTTGATCCATTCTTATTAGGAAAGCAAAGCAACGACGAAGACATGCATACGATTCTTGATAGCATGCTTCAGGTTGCACTAAGGTGCATTTTGCCAGCTGATGAAAGGCCTGATATGAAATTGGTTTTCGAGCAACTTTGCTCGATAGCGCAATAACACAAGTTGAGGGGAGAGGGGTAAATTTGAATACTATGTAGTTGTATATTTGTATCTTAAAAGTTGATATTTCATTCATGTTCCATTTTTAATTCAATTGTTTTAGAAACAAATTCATTTGTGAATTAGCACATTAAGTGATGAATGAAGTTGTATTCTTCACATTTATTGACAAGAAATAAAATTTTCTTGTAT

>M6-PGSC0003DMG400002689 M6 LRR-RLK

TACACAAATTCAACTCTTTTGTTCCTTAAATTCAACTTGTGTTTTTTTGTTCTCTTCCTAACTTCCCGCGCTTTATGCCTTAAACCCAAAAAAATGAACCGTTTTTTCTCCTTCATTCTCCTCCTCACACATCTCATCTCCGCCGTAACCGCCACCGGCGACGGCAACGACACCACCGCACTCCTCCTCTTTCAGTCACAAACTGACATCCACGGCACTCTCCTTCACAATTGGACACTTCCGTTGAACTCCACCACCGCTTGCACAGCTCAATGGCTTGGTATAAAATGCATAAACAACCGTGTCTCCGCCGTAATCCTCCCGTCGTTCAATCTCCGGGGACCCATTACCGCTCTCTCTTCTTTACCTTTACTCAGCCTTCTCGACCTCCGTAACAATCGCCTTAACGGAACCCTCACACCCATTACCCAATGTACTAATCTTAAGCTCATTTACCTCTCCGGCAACGATTTTTCCGGCGAAATTCCGCCGGAAATTTCTTCTCTCCGTCGTCTCCTCCGCCTTGACGTTTCAAATAACAACCTTGAAGGTTCAATTCCGACCCAGATTGCGAATTTGACCCGATTGCTCACACTTCGTTTACAGAACAATGAACTTTCTGGAACAATCCCAAAATCCCTAAATTCTCTTCAAAATCTTAAAGAATTGAACTTTTCGAATAATGAGCTTTATGGGTCTGTTCCAAATGGTTTATATAATCATTTTGGTGAAAATAGCTTTGATGGAAATGAGGGTTTATGTGGAATTGGGAATTTACCTCAATGTTCATATACAGGGGAAATAGTTCCTTCAAACCCTAGCTCTTTGCCTACAACCTCCACTGCAACAATCGAAGAACCCCACGAAAAATCGCGAAAAGGGTTGAGTAGAGGAGGGGTGGTAGCAATAGTGATGGTGAATGTGGTGGCATTGTTAGTTGTAGTGTCATTTATGGTAGCATATTACTGTGGGAAGTATTCAAGAACTCAGAGTTTTTCAATGTCAGGAAGTGAGTGTGGGAAGAGGAGGAGTAGTTATTCGAGTGAAAAACGAGTTTACGCTAATAACAATGGTGGTGGTGGTGGTGATAGTGATGGTACAACAGCTACTGATAGAAGTAAACTCGTGTTTTTCGATAGGAGGAAGCCATTTGAGCTTGAGGAATTGTTAAGAGCATCAGCTGAGATGCTTGGTAAAGGGTGTTTAGGTACTGTTTATAAGGCAGTTCTTGATGATGGGATTACAGTTGCTGTAAAAAGATTGAAAGATGCAAATCCATGTCCAAGAAAGGAGTTTGAACAGTATATGGATGTTATTGGAAAGCTTAAGCATCCAAATATGGTGAAACTTAGAGCTTACTATTATGCCAAGGAAGAAAAATTGCTTGTTTATGACTACTTGCCTAATGGGAGTTTGCATTCACTTCTTCATGGTAAACATTAGTAAATGTTCATGTCAAAGATTAAAGATTGTTTTGTAATTCAAGATTGTCTCTTTTGGGAGTTGTTAAGGTCTATATTGCTCAGACTCTTCAAAATGTCGATGGGGTGTGTCGGATCCTCCAAAAATAGTGCATTTTTGGAGAATTCTGACGTTGTAATTCAAGATTGTCTTTTTTGGGGGTTGGTAAGGGTTATATTCCTTGGACTTTTCAAAAATATCGATGGATGTGTGTTAGATTCTCCAAAAATCTCTAGAAATAGCGCATTTTTGGAGAATCTTACACGGGTATGACATGTTTTGTAATTCAAGATTGTCTTTTTTGGGGTTGGTAAGGGTTATATTGCTCGGACTTCTCATAAATGTTGGCAGGTACATGTCGGATCCTCTCGAAATCTCCAGAAATAGTGTATCTATGGAGGATCTGATGCGGGTATGGCAACATTTTTGGAGAGTCCGAGCAAGTTGTAGGTATGATACTAATATTGATGGGATTTTTTTTTCTTGATTGTAATAGGAAACAGAGGACCAGGGAGAATTCCATTGGATTGGACAACAAGGATTAGCTTGGTGCTAGGGGTAGCTAGAGGGCTTGCTCACATTCATGAAGAGTATGCAGATTCAAGAATCCCACATGGGAATGTGAAATCATCTAATGTGTTGCTTGACAAAAATGGTGTGGCTTGTATATCTGATTTTGGGTTGTCATTGCTTTTGAATCCTGTTCATGCCATAGCTAGATTAGGAGGATACCAAGCGCCCGAACAATCTGAAATCAAGAGGCTATCTCAAAAATCTGATGTTTATAGCTTTGGAGTACTGTTATTAGAAGTTCTCACTGGTAGAGCTCCTTCAGAGTACCCTTCACCGACTAGGCCTCGCGATGAGGAGGAAGAATTGCCAGTGGATTTGCCTAAATGGGTTCGATCAGTAGTGAGGGACGAGTGGACTGCTGAAGTGTTTGATCAAGAACTGTTGAGGTACAAAAACATTGAGGAGGAATTGGTGTCAATGCTACATGTAGCCATGGCATGTGTAGTGCCACTACCTGAAAAAAGGCCTACTATGGTTGAAGTTGTTAAGTTGATTGAGGAGATTAGAGTTGAACAATCTCCATTAGGTGAAGATTATGACGAATCGCGCAATTCTCTTTCCCCCTCCCTTGCCACCACTGAAGATGGACTCCCCGGTTACTAATTTTCGATCATCGTGGCATTGATGTTTCATAGTCCTAGTAGTATATGTTCAAATTTGCAGCATCTTACTGCAAATTGGGAAGTTCAAAAGCTATGGACTTATGTTTAGTAGTTATTTGATTGCTTTGTGTTATTCCAAATTACTGCACTTTTGGAAAATCTGATGATCCAACAAGCATATAAGAAGTGCAGCAATCAGTCCATTATGTATTTTTTTTAATGTAATGTTAATTCTTTGTATTTAATTCATTTTCAAGATTTGTGTTCTAAGGCAAAAC

>M6-PGSC0003DMG400002870 M6 LRR-RLK

TTTTGCATTATTTCGTTGTTGTTACCGGAGGTGGAGCTAGCAAGGGCCGAGGCCTCTTCCCCGAAAAGTTACACCGTTTATACATGGTTTAAATTATTTTTATGTATACATAATAGATGTTGAATTTCTTTGACTTCTTTGTAGTGAAAATTCTGGCTACGGCACTCATTGTTTTTGCTTTCATTTCTGTATTCTCTTTTTCAAATTGCTTGGAAATACTTTTTTTAAAAAAAGTTAAAGGGTTTATTGAAAACAACCTCTTTGAGGTAGGGTAAGGTCTGCGTATACTCTACCCTCTTCAAATCACGCTTGTGAAATCACATTGAATATGTTGTTGCGGATTTTGAAATTAAAAGCTCAAGAATTTTTTTTCTTTTTTTCTTTGGCAGTGCAAGCTTTAATGGAGATGAAGAAGAATTTAGATGATCCACTTAATGTTTTAAACTGGGATGGAGATGCTGTTGATCCTTGTAGCTGGAATATGGTCACTTGTTCAAATGATAAATTTGTCACTAGCCTGTAAGTAGAACAATTTTCACTTATTCTAAAAATAGTATTCCTAACAAGATTTCTTAAAATAGTTGTATTATTGATTATGCAGGGAGTGTCCCAGCCAAAATTTGTCTGGCAAAATATCACCATATATCCACAACTTGACACACCTTGAACTTATGTGAGTACTTACTTAACCAAAGTAAATAAAAAACGTAATACATAAACGTGTCTTTTAACTTGGTGTCATCTATCATCTACGCCTCCAATATGGATGTGCAGAAGTAGGCATACACGTCCTACGTGAAAATTCTCATCCTACGTGTATTATGCTACGTAGGATGTGAGTGTGCCTACTTGTGCACCACCAAAGTTGGAGGATAGATGTCAGGTGACACCAAGTTAAATGACATAATTATGTATTAAGCCTGAATAAGATATCAAGACATTTGAGTTTTGATTCTTATTCTTAGATGTTAATATGAGTGTCCTGTTTTTTACAGACTTCTACAGAGCAATAATATATCAGGATCACTTCCTATGGAGCTTGGAATGCTTAAAAAACTTAAGACAATTGATCTTTCTGATAACAAGTTAACCGGCGAAATCCCGTCATCTTTAGCTCAGCTGAAAAGCCTTCAATACTTGTAAGCAAAGAGAATTCTGAGATCTTATACAAACAAGTTTTTTTGGGGTTAAAAACTAATGATGTTTGTTCTTGAATTTTGATGATTATAGGAGATTAAACAACAATAGTCTAACTGGAGCTATTCCTTTGGATTTGGCCAATATGACCCACCTCTCACTTATGTAATTTTCTAATAACATTTTCATGTTATGTTATGTTATATGTTCTTTATAACAACATTTCGATATAGCAGCTAAAAAATATCCAGACAAATGAAGTACGCAGTTATAGAGAGGTTTGACTGTATTTTTCACTTATGGAGTTGTACTGTTTTGCATGTTTAGGGACTTGTCTTTCAACAATCTCAGTGGTCCTGTACCAAGGCTTCTAGCCAAAACATTCAAGTAAGGTTCTTGAAGAATTCTTTGTGTATCACTTTTTCCTTAATCAGTAAATGAAACCAATTTTTTGTATTTATTTGTCTTTTTTTTAACCTTATTGACACAGTATATTGGGAAATCCAATGATATGTGCAACTGGAAAAGAATCAGAGTGCAATGGAACAACACCAATGCCCCTCTCCTTCTCTTCAAACAATCCACAGAGTAAGAACTAACCTTATTTTAACAGCCAAATGATAGTTTTTTTTTTTGTTTTTTTTTATGACCGTGGTGTCCGAGCTAGTTTTCTTACACCTTAACTAATTCCACGAGATACCTACCACCTCCCACCAACAACAGGTACCAGGTAACTCTGTCCATCAATGCTCGGATGGAGGGGAGAATCACCTAAGACCTCATGACTCTCAACCACTTCATTGGCCAGTAGGCCACACCCTTGGGTGCACAACCAAATGATAAATAAGCGCCTCTCTACCTTCACAAAGGTAGGGGTAAGGCTACATACACACCACCCTCTCAAGACCCCATATATGGGATTACACTGGGTATGTTGTTGTTGTTGTTGACGGTCGATATAATGATTATGATGTTTTTACTAGTCATACAGATTTATGAAGTAAATATGCTTCACTGATTTTCTCTACTGATAGATTCAATATTTATAGTTCATTCCCAGCTAAAAATATCCTGTGTTATCTCAGATGTTCAGCCTTCTGGAAAACCGAAGACTCACAAGGTTGCGTTAGCCTTTGGAACAAGTCTAGGATGCATCTTCCTACTAATTGTTGGGTTTGGATTCTTTCTATGGTGGAGACAAAAGCACAATAAGCAAATTTTCTTTGACAGTAATGGTGAGTATTGTTTGATCAGAACAATTTGAAGTTACGGTTTAGCTCATCGATCCTCTCTTATAAACGTTATTGACATAAATTGCAGAACAACATATTGAAGAAGTGTGCTTAGGCAACTTAAGGAGGTTCCAATTCAAGGAACTTCAGTCTGCCACAAACAATTTCAGCAGCAAGAACATACTAGGGAAAGGTGGTTTTGGTAATGTTTATAAAGGACGTCTTAGTGACGGGATTATTGTAGCTGTGAAAAGGCTCAAAGATGGAAATGCAGTTGGAGGCAATCAACAATTTCAGACTGAAGTCGCGTTGATCAGTCTGGCAGTGCATCGGAATCTCCTCAGGTTGTATGGATTTTGTATGACACCAACTGAAAGGTTGCTGGTGTACCCTTACATGTCTAATGGAAGTGTAGCTTCGCGTCTCAAAGGTAAACATATCTAAAAACTGTCTCCTTTGGCCTATAATTACATTGTCCTTATTTGCTCAGCTCATAACGTATCATTTCTTGCAGCCAAACCAACCTTGGATTGGGGTACAAGAAAAGGAATAGCTTTGGGAGCTGCAAGGGGCTTATTATATCTACACGAACAATGTGATCCAAAGATCATTCATAGGGATGTGAAAGCTGCAAATATATTGCTTGATGATTACTGTGAAGCTGTTGTTGGAGATTTCGGATTAGCAAAGCTTTTGGATCATCACGATTCACATGTCACAACCGCGGTTAGAGGAACTGTAGGACATATAGCTCCAGAATATCTCTCCACAGGACAGTCCTCTGATAAGACGGACGTCTTTGGTTTTGGTATTCTCTTGCTAGAATTAATCACTGGACAAAGAGCTCTGGAATTTGGTAAAGCAGCAAACCAGAAAGGCGCTATGCTTGATTGGGTAAGTACGACAAATCCCTCGAATTGTAGTTACATTTCTCTTTTTCAGCATGCTTTGACATGACTTCTTGAGCCAAGGGTCTATCAGAAACAACATAACCTCTCTACATCCCAAGGTAGAGTTAAGGTCTGCGTTCACTCTACCCTCACCATACCTCATTAGTGTGATTACACTAAATATGTTGTTGTGCTGTATACTTGTAAGGTGTGTAGGCGCACGTGCATTCACACTATCTAAAATATGGACTCTATAGGTATTAAGAGATGCAACGAACAAACCATCTTTAAATCCTGGATCCGTCCTTGTTATTTTTTCACTTGCAGGTAAGAAAGATTCAACAGGAGAAAAAGCTAGACATGTTGGTGGACAAAGACATGAAAAACGAGTACGATAAGATAGAGTTAGAGGAAATGGTTCAAGTAGCATTACTATGCACTCAGTATCATCCAAGTCATAGACCAAAAATGTCTGAAGTTGTAAGAATGCTTGAAGGAGATGGACTAGCAGAGAAATGGGAAGCATCTCAAAGAGCAGAACCAACAAGATGCAGCAGAGCCAATGAATTTTCCTCTTCAGAAAGATACTCTGATCTTACTGATGATTCTTCATTACTTGTACAAGCAATGGAACTATCTGGTCCAAGATGATAATTGAATTAAGACACAATGTCACTTTTGCTGGTGGGAGGTGACAGGTCTCTCGTGGAATTAGTCGAGATGCGTGAAAACTGGCTCAGACATCACATACATAAAAAATAAACATGTGTCATATTGAGCTTTGTTTATAGGTTTATACAGAAGCTGCAAATAGAATATCATACATTCTGATTGTAAAGGACAATATAATAACTTTGTAAGGCTGAGAGTTGTTTTTCTGTTTTTCCTTTTTTGGTTCTTGTTTGTTTGAGTGGAAATGTTTATGGGATGCTATAATTTGTTTATGTATATATATAATTAGAAATTGAA

>M6-PGSC0003DMG400002902 M6 LRR-RLK

AGAAAAGAAACAGAGAAGCATTTGCAGATTGTTTCTTTAAGGGGAAGGTGGACATGTCTAGTAACAAACATGGCCGCCGTTCCCCTAATCCTTCATCTCCTTCCTCAAGCATCGTCTCCGAAAAAACACCATCTTTTCCTTATTCATATCTCCATTTTCTTCCTCATTTCTCTAGCTAATCCACAATTTTCTTTCTCCATTTCCGAGGACGATGCCTTGATCAAATTCAAAGAATCTTTAAAAAACACAACATCTTTAGATTCATCTTGGCTTAAAGGTACTAGTCCATGTGACAAAAACAAGAAATGGGATCGTATTCAATGCGAAGGGAACGTGGTTGAAGGCCTCCTTTTAGGTGAAGCTGGCTTATCAGGAGAACTAGATATTGATCCATTGGTTGCACTTCCAGGTCTCCGAGTTCTTGAAATTGCGAATAATTCATTTTCAGGCACAATCCCTGAATTCTTCTTGCTTGGTGCACTTAAATCTATATATATAGATGGAAACCAATTCTCAGGTGAGATTCCTAAAGATTTCTTTGCCAAAATGGGGTCTCTTAAGAAGATTTGGTTTGCACGAAACAAATTCTCCGGTTCAATTCCTGAATCTTTAACCAACTTGAAGTACTTGATGGAACTTCGTCTTGAAAGTAATGCATTTTCAGGACACGTTCCGTCATTATCACAAGTGAGTTTAACATCAATAGATCTTTCGAATAACAAGTTGCAAGGCGAAATCCCTCAGGGGATGTCGAAATTTGGTGCTGATCCTTTTAAAGGAAATGCAGACTTATGTGGACAACAATTAGGGAAAGAATGTAATAAGGGAATCAATAAGACTCAAGGTGCACCAATGTCAAAATTGAAATGGATCATTCTTGGTATAGTAGTTAGTCTTTTATTGGTGACCATTCTTTTTCGCGTCAAGCGCAAGGAAGATCATTTCGACAAGCTTGGGAAAGAGAATCTTGATGAGGGGTTGCACGTTGGTAGCTCGATTAGGAAGAGCACGAGCATTCATAGCAAAGGAGGTGATTCTGTTCGTGGATCAACACGAAGGGGAGGATCACAAGGTGGTAAGGCTATGGGAGATCTTGTGTTGGTGAACAACGAAAATGGTACATTCGGATTGCCTGATTTGATGAAGGCAGCTGCTGAGGTACTCGGAAACGGAGTGTTAGGATCAGCTTACAAGGCAAAAATGGGAAATGGAATGTCTGTTGTGGTTAAGAGATTGAGAGAGATGAATAAAATGAATAGAGATGTGTTTGATGCAGAAATGAGAAAACTTGGAAAGTTAAGGCACAAAAATTTACTTCAATTATTGGCATATCATTACAGAAAGGAAGAGAAATTGATGGTATCTGAATATGTTCCCAAAGGCAGCCTATTATATCTGTTACATGGTATGACAATTTCTTTCATCTACAGTCAAACCTCTCTATAATAATGTCGTTTGTCTAGATAAGTTTTTGGCTGCTATAGCGAAATGTTGTTATCTAGAACATAAGTTTCATATCTCAACTGTTACTCCCTTTGTATTGAAACATACCTCGATGATGAGTTGGCCTACACACGCCTGTTGTTGATTGAGAACAAATATTTGACCAACTCAACATCGAGGTGTGTTTTGATACAAATGGAGTGAATAGTTAAGGTAGGTTAAGAGAAAAATGGATAGTTAAAGTATGAAACTCGTGAAAAAAGGATAGTTTAGGTGTGTTTTTGACCATTATGGTAAAAATCAGTTCTGAAGAAAACTTGGTCACTATTATAGTGAAATGTTATTATAAAGGATGATTGTTATAGAGAGGTTTGACTGTATATCAAATCATTAAGGTCTTCAACTTCTTTTCAACATTTGCACGAGTAGATTTTATCCGATAATGTGATCCGATCTATACTTACCGGTCCCTTCCAAGACATCCGGTCACAATTGGGACGATACAAAGAAGATTAGCATGATCCC

>M6-PGSC0003DMG400003002 M6 LRR-RLK

ATGTTTGACACAATTGTATGTGATGAAACAGTGACCAGAGATGCATTGCTCAAGTTTCTTGCAGAGGTTTCTAACAATAATAGTACAATTCTTGGTACTAATTCGGGCTGGAACTCGACATCTGATCCTTGCAAGGACAAGTGGCTTGGTGTTACATGTAACGCGAAGACTCTCTCTGTGGAGAAAATCATTCTTGATGGTTATGATTTTTCATCTGGAACGTTTGATGCTACCACGATTTGTAATGTGGAGCCTATCGCGGACACCCTTTTCCTCCTGAGTCTAAAAAATGACCTACTTCAGGGGAAAAACTTTGATAGTATAGATAAATGTAAGCATCTAAGACATCTTGTTCTTGGTGGAAACCGGTTCTCAGGTAGCATTCCACAGTCATTTTCAAGATTGAATAACTTGAAAAGCCTTGACATATCCAACAATAGATTATCAGGTGTTCTCCCTGATTTATCTAGGATTTCAGGCTTGATTGAATTCGAGGCTCAGAATAACCAGCTATCAGGAGAGATACCTGAATTTGATTTCTCAAACTTTGTTGCATTTAATGTCTCTTTCAACAATTTCAGTGGATCGATTCCGCCTGAGAGTGATAGTTTCCCTGTTAGTAGATACTTGGAAAATCCTCTGTTGTGTGGCTCCCCGTTGGCTGGAAAATGCTCATCACTGTCCGAAGATTCAGAATCAGCTGATCATGACAAGCCAAAGAGCAGCTTGTCAAGAGATGAAATTCTCATGTTTTCAGGTTATTTCCTCATAGGCCTGGCTCTGGTTCTTCTTATCCTACTATGCGTCTATAGAAAACGGAAGCAGAAGCAGCAGAAGAAGAAAGAATCAAGAATCGAGCTGATGAAAAAGGTTGCTACAGTTGATCATGACAGTACTACTACATCAATGGATCAATCAAAGAAAGGCGCGAGCAATTCAGATATCTCATCAGCTGTCATTTCTTCCGATGACAGCAACACCGCGGTTTCACAGTCACTCGTGGTTCTTACGAGTCCTGAGGCGAATAACGGATTGAGATTTGAGGATTTGCTTAAGGCTCCAGCAGAGTTGCTTGGAAGAGGGAAACATGGTAGTATGTACAAAGTGATGTGTGACAATCCCAAAATGACACTGGCTGTTAAGAGGATAAAAGATTGGTCAATAACTGGTAGTGAATTCAAAAAGAGAATGCAGAAACTAGACAAAATCAGGCATCCAAATGTGTTGCCTGCTGTTGCCTTTTATTCTTCTAGACAAGAGAAGCTTTTGGTCTATGAGTACCAAGACAATGGCAGTCTTCTCCTACTTCTACAAGGTAATTGACTCTACTAGGCTATTCTTCTGTGTATTGAATTGTGCGACCATTTCATCTAGGCATATAAACAGACACTCGAACTTTGCCTTAGCTGGCAAGTAAGCACTCAACTTTGAGAGAGTGCACATCTAGACCCCTCAAATTGGTCTCAACTG

>M6-PGSC0003DMG400003175 M6 LRR-RLK

ATGGGTGGTAGATGGAATACTTATGGATTCAAGTTATCATATCTTGCACTTCTGATTATTCTTTATGACGTTCACGGATGTAGCTCACTCAATTCTGAAGGTAATTCTATTCTATGTTTGGACTGTCCTACTGGTTTTGGTGTTTATGTCTTAGTTATATCTAGTTTGTTATGTCCATTTGAATATTGGATTGGTCTACTTTGCAAAACAAGGATTGGCATTGTTCGGACTACGATCTAAAGTGGATTGTGATCCATATGGAGTTCTTGCAAACTGGAATCTTGATCATTGTGACCCATGTATGTGGTTTGGAGTACAGTGTCTGGCAGGCAAAGTGCAAATGTTGTAAGTCCCTTGTTACATACGATCATATCTGTTAATGAGTATTCAATTCCATAAAACTTACTTTATAATTGTAATGCTTTGTTTCTTTTTTAGTTAAGACCAAATCCGTTAGATTTATTTTAATGACAACAAATGTGCAAAAGATGAACTCATATGTTTTGACTTTTGAGCTATCTCTCATGTTGTTAATTATTTTCTTTTTCTTCTCTGTTTAGGGATCTTCATGGACATTCTTTAGAAGGCACATTGGCACCAGAGCTTGGGAATCTCACTCACTTAAGATCAATGTAAGTTTTTACCTACGACAAAATGTTTTTTGCTTTTTTAGTTCATAAGTGAAAATTTGTGGGGAATTAAAGAGAGATTGTTGCTTGTCATGGTTGAAAACTGATTAAAACTTAGTATTGGTTGAAAATAGCCTTGCGCATTGAAAATAAGCACGGGTTAAAAATGACCCTAATGAGTTAAATGGATTGAAATAGCCTGGCAGCTTAAAAATACCTCCTTCTTTTGATTTCTTTTTTAACTTGCAATTTTTTAAAAAAAAAATTGATCCTCCTCTTCTTCTTTTGGTTTTTTGGTTAAAAAAACTCCCTAAATAATCTTGTCTTGTTTGAAAGCTCTTTTGAAAGTGCTTGCCTCTCGACAATGATAGCATTTGAAAGGAGTGTTTCTAAGGATCGTTCTTCCGTTACAATATTCCATAAATCATAGGTTTCAAGATATGAATTCATTTTCACTGACCAAATCTGATAGTTTTTGGTAGTGAAAATTTGTGGGACAATGAAAGAGAGATTGTTGCTTGTCATGGTTGAAAATTGGTTTAAAATAGCCTTGAGCATTGAAAATAAGCACAGGTTGAAAATAACCCCTAATGAGTTCTAAAAAATAGATTGAAAATAGTCGTGGTGGGTTAAAAATAGCCTTGACAAGTTAAAAATGGGTTGACAATAGTTCTTTAGAAGAACTCACATATCCCATAAGATCAATGGGGCTTTGGATACTACTGTTGCTGTTTTTTTAAAGATGGTGCAGCTAAATAATGTTGAAAATGTCTCTCAATTTTTATTAAGCATTAGAGGGGTTTAATAAACAACTTGAAAACATAATTTGCAGAAAATCTGCATAACAAAAAGTAGTAGATTAGTAAATTGGGTGGAATAGACTAAAGATAATTAGTTCCATGTCTAATAATCATAGTTTTCATCATCTTGCATGTTGTATCTTCCTGTGTGTTGTGTATGACTCCTTTTTTAATCTTAACAATTTAAACGTTTTTTTCTCTATGATTGGCTCACAGTGTGCTGTCAGAAAACAATTTTTTTGGTGCCATTCCAAAAGAATTTGGACGACTTCAAAGGTTGGAAGTGCTTGACTTGACATATAATAATTTGAGTGGGAGAATTCCAGCAGAAATAGGAGACCTGCATGCGCTAAGAAGCTTGTGAGTTCATAATTTAATTTTTGTGCATGTGAAGATGCGTGCTGCCTATATTAGTTTATATATGTGAGTATTCATGAGGAAGAACCATTAAAAATATTCTTTTGAACTGTGTCTTATACCTTTTCACAGGTTGATTCGTAACAACAACTTGGAAGGAAAAATTCCATTTGAAATTGGGAAGCTTCGTTTGCTCTCCGAATTGCAATTTGACGAAGAACTCATCCCTGCTGCTGCTGGTGGAACATGCTGTTTAAATAGAAAATTTGTACACTGGTAAGGTTCTTTTGGTTTTATTGGACCTTTTATATCCGTGTTCTGGTTCTTGTTACATTTATGTAACCAAATATCAACATGCAAAGATGTCTGAAATGTATGAAATTTGTATTTAAGCCATTTCAGGCTGCCATTATTTTCAGAATATGGTCATAATTCCTGTCATAAGGTTTTCTTGGTGGGAGTGAGGGGGTGGGTGAACACAAGTTGATAAAGTTGAGTATTAATGACAACAAGGGATATTGTCGTTTATGTCAAATCAAATAAAGCCTGGGAAATAGGTGATTTCATAGATGTGGCGATGCAATGACTTGTGTTGCTAATGCTTGAAATTTATTGTCTAGTTTGACATCTAGTAGTAGTACTTGACCATACATCAACTCAAACAAAAAAGCTTTGTAAACTACAAGGAGGTGATTGTCCAGTCATTGCATCCATCCAAGGCTAGGGTTGAGGATGAGGGATGCAATATATTGGAGATCTTGGGATTGATCCTGGAATCAAGAGAGCATCAAGTTATCATATTGGTTCTAATTTATTTTAATTTTTTTATCAAGAAACTCTTCTCAGATATCTTTAACTCTTACATAGAATACTGACTATCAAAACTAATAGCATGGTTACTTTCAATAACACTTCTCAATCAGATATTTTCATTTGCCTGCTTCTCTCTGGTAATTTGATGTCTGTTTCTGGCAGCATCTGTCATGGCAGTCTGATACCATTAAAGACCGTAGACTCCTTTATCAAACCACTTAAAGGGACACTTCTGCGTTATTTCACTATGTAAGTAACTAAATGGTTTTCAATCACCATGTGTTTCTACTTACATTTTATCACTCTAGAAGCGTCTTTGTGTCAATAAAGGTTCTCCAAACAACTTGGAAGTGGCTCATTGGATAATCAGACAGGGTATTCCTCTGACAACCTACCAAGTGAGTGAAAAAGTTTTACCCCACATCATTAGCAGGAATTATGTTGAAAGTGCACAAAAGATTGTTTTCACTTTTCTTTTTTAGCAGGTTCATCTAGGCCTCACATCATCCATACTATTCAGAACCATGCGAATATCGCACGCCGCAAGTTAGTAGAACAATCCAGCAACCTTGCTGCTGTTCCTGCCAATGCTGGGAAACCTTTGGGTCCTATTGTTTCAGTGCCAAGCACCAGAAGTAGTGGGTCATTTCATGTTGTGCCAAATACCCAAGAAACACCTCCTACACCCTTACCTGCCTCGCAACGACAGCCTGAGTCTCAATCCAAACCAAATCCAATGGGACGGTCAAGTAGTGTTGTTAAACAGCCAACTAATAGACAAGCTCCACCTGCCCAAAAATTTGGTGATACATGGAAGTATATAGTTGGAGGTTTTGGGTTTTTTCTGCTCATTGCTGTTGTAGCCGTGTTCTTAGTTTACAGAAGAAGAGCTGCAAAAACAATAGGCCCTTGGAAGACTGGGTTGAGTGGACAGCTGCAAAAAGCATTTGTCACAGGTATTTCAATTAAAATTTGTGACAAGAATTACTGTTAATATTCTGTCCACATTCTGATATTCAATGACAAAATCATTGTTGGTTGATAATAGTCTTAAATCACAGCCATGAAAGAAGGTATAATCCTGATGTTGATTGCAAAAAATTGTTTCTCATTGCAGGGGTCCCTAAGCTTAACAGGTCTGAGCTTGAAACAGCATGTGAGGATTTTAGCAATATAATTAGTAGTCATGATGCATATATAATCTACAAGGGAACTTTGTCTAGTGGAGTAGAGATTGTTGTTGCCTCGACTACTCTAACTTCTCTCGAAGATTGGTCTGATAGTTCAGAAATGGCCTTCAGGAAGAAGGTTTGTTGTTTCCATTTACATCAAAAATAAAATATTATCTCTCCATAGATCTCACAGTCTTACACTTCACATAATGACATTGTTGTTATCTCTTCATCCTTTTCGGTTGTAAAAGACATGACATTTTGTGGTTGTGCTACTGAACAGATTGATACACTTTCAAGAGTGAACCACAAAAATTTTGTTAATCTTATTGGATACTGTGAGGAGGATGAACCTTTCACTAGGATGATGGTCTTTGAGTATGCTCCAAATGGAACTCTTTCTGAGCATCTACACAGTAAGATTCTTTGTTGTCCACAGTTCAGCTTTAATTACTCACAATGATGGCTTGATTTAGTTCAGAATAACACTTCTTATTTTATTTTGAAAAAAAAAAAAGATGTTCAGAATGACTTAAAATATAAGTTTAATTGTGTCAGTTGAAGAAACCGATCCTCTTGACTGGTCTGCAAGGATGAGAGCCATAATGGGGACAGCATATTGTCTTAAGTACATGCATGATCTGAATCCTCCAGTGCCCCATTCCAGCTTAGATTCAAATGCCATATTTTTGACTGACGATTATGCTGCTAAGGTCTGCTCCGTCTTCTTCAATATGTATATATAATTTATCTACCATATTGAACTAAGAAACATTTTTCATTTGTATTTCATATTGAAATAGAATTTTTTATATTTTCACAGATTACAGAGATTGGTTTCTGGTCAGAACTAATAGCCAAGTCTAAGGACTCTAGGGATGATTTAGAAAATTCTGAGCTGCCAGCACTTGCTGATCCTGAAACAAATATCTATAGTTTTGGGATTCTATTACTAGAAATAATTTCTGGAAAGTCACCTTATTCAGAAGAACAAGGATCTCTTCTGAACTGGGTAAATGTCCTTCATATGTTTCCCTTATTTTCTTATACTTGATCGTTACAATGCGTTCCCTAGTCTGCATGCATTTTCTTGTGTGAATGTATTTTCTTTCAGACTTATACCAATGCAAAAGCATATTGATAACTATTATTTTATTTAGGCGGAGCAATACCTTAAAGATAAGATGAATGTGAGCTCCTTGGTTGATCCAACACTCAAGTCCTTCAAAAATAATGAGCTTATGGCTATCCGTGTGGTAATTAAAGAATGCCTTCAAGAAAATATAAGGAAGAGACCAACGATAACCGAAGTCATTGCAAAATTAAGGGAAGCGATTGATATATCGCCTGATGCTGCAGTTCCAAGGCTTTCTCCCCTTTGGTGGGCTGAACTCGAGCTCTTATCTTCTGAGGCAGCTTAG

>M6-PGSC0003DMG400003210 M6 LRR-RLK

TCGCAGTATGGATCTAAAAACGTGCTGTATTATACTATTAAAAAAAACTGAGAGAGACTGGCTCCCTCATTAAGAATTATTGAACAACCAGCCAAACCAAAGCATTAAATATTCTCCTATTTTGGTTGTTTCCTAACTCATTCATTACCAAATACCACACTAATTAAGCATATGATTAGCTGAGGACACATATTATGAAACAACATTCTCAGGCGCCTCAGCCCAAAAATTAGTTCTTGAACTAGCAGCTCAGTGTTTTAAGAGGTCAGAGATGTTCTTGTTTCTGGTTAATTGCCTTCTCTTCTTTGCTAATTTATTTGCAGTCTTTTCGATCTCATGGCCTCAAGATTCTGTTCATCTACTTGGGTTTCGGAGCAATCTTCCAGAGCTCTCTCAACAATTGCTCCCTTGGAATCAGTCTGTCTCACACTGCCAATGGAAAGGTGTTTCCTGCTATTCAGACATAACTTCTCATGTCAAATCATTGATTCTAACAGACTTATTGCTACCTGGAACCTTGGACAAGGCCTTACCTAATCTCTGCCTTCTTCCTCGCTTGGTCTCTCTTGATCTCAGTGGCAACCATTTCACTGGTGGCATCCCTGATATGCTCGCAAACTGCAGCCAACTTGACACGATTCTACTCAACGACAACAGATTTTCAGGATCCATCCCACCAGAGATTTTCAAATCAAGCAAACTCATAAAAATTGATTTGGGTTTAAATCAGCTCACTGGCACTATTCCCTCTGAGGTAATCCTTTCTACAAACCTTCAACATCTCGGGCTGTGGAACAATTTCCTCAGTGGAAATATTCCAAAAGAGCTATTCAATTTGCCAAACTTGACACATCTCCACCTCAACACAAATGAACTGACTGGACCTCTACCTGACTTCCCATCCTCGTGCTCACTTTCCGAGTTTTTTATTTATGAGAACCGTTTGTCAGGTTCTTTGCCAATTACCCTAGGCAATTGTCACAACCTTACTTCATTCTCTGCATCATCTGCTCATCTTGGAGGAGTTATTTCCCCAGAGGTTTTAAGGGACCTTTCAAATCTTGAATTTCTCTATTTAGATGATAACAACTTTGAAGGGGAAATTCCTGAGACCTTATGGAATGGGAGCTTACAAGAGTTAGCTCTTTCCTTGAATAAATTCAATGGTAGTATATCAGAAAAGATTGGTGGTTCTCATCAGATAAATTACATTGATTTATCAGTTAACAAATTAACTGGTCAGCTTCCAAGATCAGTTGGACGTCTAAAGAATCTGAACAAGCTTTTTCTGTATGATAACATGTTTAGTGGTTCATTACCAGCAGAACTTGGGAACTGCACTTCTCTTGTTGCGATAAGTCTCGTCACTAACTTCATTGGTGGGGAAATTCCTCTAGAGCTCTGCAACCTTCACTCCCTAATAAAATTTCAAGTGTACGGAAATCAAATTCAGGGCCAAATTCCAGAATGCATTGGCAGAATAAGTGGACTGGAGGAGCTAGATCTTAGTGAGAACCGATTGATCGGCAAGATACCACCTGGAATCACAAATATGACGAAACTTGTGTTACTCTCTTTGGCTCATAATAATCTTACAGGGGAGGTACCACCTAATCTTGGAAAAAACAATTTTCCTGGCTTATTTAAGGTTGATTTAGGTTATAATATCTTCAGCGGACCTATTCCTTCTGAACTATGTAATGGTAACAGTCTTGGAGTCCTGGTTCTTGAAAACAACAGTTTTAATGGCAGCTTCCCAACATATCTAGCAAAATGTGAATCCCTCTATAGGGTAAAACTCCCCAACAACAATCTGCAGGGGAGTATTCCTGACTACATTGAAAAGAATGAGAAGATTTCTTACTTGAATGTTCGAGGAAATATGCTTGCAGGAAGGATTCCAGCAGCATTTGGTTATTGGACCAATCTTTCAACAATTGATCTTTCAGAGAATATGTTTTCTGGTTCCATACCTGCCGAAATGGGGAAGCTGCAAAATCTTGTAAGATTGAACATTTCTTCAAACAGACTGACAGGAAAAATTCCTCTTCAGTTGAGTTACTCTGCAAAACTGGCAGAGTTGGATCTCAGCAACAACAATCTTTCGGGTAGAATTCCGAAAGAAATTGCATCATCTTTAGTACTGACAAATCTTCTACTCCAGGACAACAAACTTTCTGGTGCTCTTCCAGACACTTTCTCCTCCTCACAAAAGCTTGTAAAGCTGCAGCTCGGGGACAACTTGCTTGAAGGCTCAATTCCATGCAGCCTCAGTAAACTTAGGGAACCTAATGTTGCACTGAATCTGAGCATGAATAAGTTTAGTGGTCAAATTCCTAGGTGTCTGAGCAATCTAGACAATTTAGAAATCCTTGATATATCAAGCAACAACTTGTCTGGCGCAATACCTTCAGAAATGGACAAGATGAGGTCCCTTTCGTTTCTCAACATATCATTTAATAACTTTTCAGGGGAGGTACCCATTTTGTGGGGAAAACTATTAAGTTCACATCCAGGAACTTCCCAGGGAAATCCAGGACTCTGCTTATCAGACACTGAAAGTAGCAGCTGCAAGCATGTGAAAAAATCACAAAGAAACTGGAGGACCTTGGCTGGTGTAATTAGTGGTGTGTTTTCCGTGGCAGTCATAGCTGCGGCAATATACTTGCTAGTGACTCGGATTCAGCACCCTTCCCTGCTGAATAAGCATCGGCTCGTCAAGTATCAGGCAGAATTTGAAGATCTGCCAGATCGTATAAATTTTGAGGACATTGTACGTGCAACAGAAGGCTGGAGTGAAAAATATGTAATTGGAAGAGGCAAGCATGGAACTGTTTATAAGATGGAATCTGCGAAATCCAAAAAGCTTTGGGCTGTCAAGAAGGTTGACTTAGCTCAGAGAGCATTCAGCAATGAAATGAGAAGCCTCAATTCTGTCAGACACCGCAATTTGGTAAGATTGGGAGGACACTGCACCAGACATGGATATGGCTTCATTCTAACAGAGTTTATCCCTGGAGGAACTCTTCATGATGTCCTTCACCAGAGAAAACCACCTGTAGTCCTGGACTGGGAGTCCCGCCATCGTATTGCTCTTGGTGTTGCTCAAGGTCTGTCTTACCTTCATCATGATTCTGTCCCTCAAATCATTCACAGAGATCTCAAATCAGATAATGTAATGCTGGATACTGAGATGGAGCCAAAGATTGGAGACTTCGGGATTGCTAAAACAGTATCAGACTCTGATGAAAACTCGACAAACTCCAAAATTGTTGGGACACTAGGATACATTGCTCCAGGTTAGCCACACGTTACTATGCTAGTCTTCAGTAATAATTTAGTTGGCAGATTACTTTTGATGATTTACTAAATTTTATTACTTTCTCCCCCGGTCCCCCCTCCTTCTCCTTGTCTGTCCATCAATTCAGAGAATGCGTACTCAGTTCAATTGACAGAGAAAAGTGATGTCTATAGCTATGGAGTTCTTCTACTGGAGCTCTTCTGCAGAAAAATGCCAGTAGACCCCAGCTTTGAAGAAGGATTAGACATTGTTTCCTGGGTGAGGAAAAACTTGCACAGAAGTGACAACAATTTCCTTCACTTGCTTGATGAAGAAATTAGTTTTTGGTACATAGAAGAACAATGGAAGGCCCTGAAAATGGTGTATCTAGCGCTTCAATGTGCTGAACTCGAGGCAAGCACCAGGCCTGCAATGAGGGATGTAGTAAGGTCTCTTGTCGAGTTAAATCATAGATGTAAAAGAGAAGAAATGAAACTTTGAATAGTGAAAGCCATAATCAACTTTTCCAAATTCCAAAATCTTTTTTTTCTTTGCTTACATAAAAATTGAACCATTTCTCGATTTGTGTGTGTCATCCTTTCATTCAAGGGCTAAATTCTAACATAATCGATCCTGGGTGTTTCTGTTTAACTTTGTGACATCCACTGCAATAATATAGACTAAGAAAATGCATGGCGTTTCAAATCAATATATGGTAAAAGCAGTTTTCTCGTTCTAGCCATAGCTCTTTTCATAATGGGCCACTGTCCTTATCAACTTTTGCTTGTTACTTTCTCCAGCTGCTCGAAAATCTGAACAGTTTTGAG

>M6-PGSC0003DMG400003211 M6 LRR-RLK

TAAAATAATCTCCTATATTCATTGTTCCCTCGGACAGTCGTTGCCTACTACCCCCACACGTATCTTTAAAAAACAGGCTGACCTCAACTTAAAACCTAAATAATAGTCTCAGGTTCCAGCCCAAAAATTAGTTTTTGAACTAGCTCCTTATTGGATGCAGCTTCTTTAAAGAAGTGTTGAAGAGTTCAGAAATGTTCTTGTTTGATCTTGTTCACTGTTTTATTTTCTTTGCCAGCATTGTAGCGGTTTTTTCGGCATCATTGCCTCAAGATTCTGTTCATCTACTTGGGTTTCGAAGCAATCTTCCAGAGCCCTCTCAACAATTGCTCCCTTGGAATCAATCTGTCTCACACTGTAAATGGAAAGGTGTTACCTGCTATTCAGACAAAACTTCACAAGTCAAATCATTGGATTTCAGAGACTTCCTTCTATCTGGAATCTTGGACAAGGCCTTTCCAAATCTTTGCCATCTTCCTCGTTTGATCTCCCTTGATCTCAGTGGCAACCATCTCACTGGTGGCATCCCTGCTATGCTTGCTAACTGCAGCCAACTTGACACGATTTTCCTCAACGACAACAGGTTTTCAGGATCCATCCCACCAGAGATTTTCAAATCCAAAAGGCTTGTATACCTTGAGTTGGGTTACAATCAGCTCAATGGCACCATTCCTTCTGAGGTGGGCCTTTCGACAAGCCTTCAATATCTTGGGCTGTGGAACAATTTCCTTAGTGGAAATATTCCAAAAGAGCTATTTGGTTTGCCAAACTTGACAGATCTCTACATCCACACAAATAATCTAACGGGACCTCTCCCTGACTTCCCGTCCTCATGCTCACTTTCCCAGCTTCTTATTCATCAGAATCGCTTCTCAGGTTCTTTGCCAATTTCCCTAGGTAATTGTCACAACCTTTCTGCATTCTATGCAACATCTGCTCATCTTGGAGGAGTTATTTTGCCAGAGGTTTTTAGGGGCCTTTCAAATCTCGAATTACTCTATTTAGATGACAACAACTTTGAAGGGGAAATTCCTGAGACCTTATGGAATGGGAAATTACAAGAGTTGGTTCTTTCCATAAATAATTTTAATGGAAGTATATCAGAAAAGATTGGTGTTTGCCATCAGATGACTTACATTGATTTGTCAGCTAACAAATTAAGTGGTCAGATTCCCAAGTCAGTTGGACGTCTGAAGAATTTGTACAAGCTTCTTCTGTATGATAACATGTTTAGTGGTTCATTGCCAGCAGAAGTTGGGAACTGCACTTCTCTTGCTGAGATAAGTCTCGTCAGTAACTTCATCAGTGGGGAAATTCCTTCAGAAATTTGCAGCCTTCAAAACCTAGAAACATTTAATGCATTCTAAAATAAAATTCAGGGCCAAATTCCAGAATGCATTGGAAGAATAAGCGGGCTGCAGGAGCTAGCTCTTTATGAGAACCGATTGACCGGCAAGTTACCGCTTGGAATTACAAATATGACGAAACTTTCATTTCTGTCGTTGGCTCATAATAACCTTACGGGGGAGGTACCACCTGATCTTGGAAAAGATAATATTCCTGGCATAGTTAAGGTTGATTTAGGTTATAATAACTTCAGTGGACAGATTCCTTCTAAACTGTGTAATGGCAACAGGCTTGCAGTCCTGGCTCTTGAAAACAACAGGTTTAATGGCAGCTTCCCAACCTATCTAGCAAACTGCAAATCTCTCTATAGGGTAAAACTTTCCAACAACAATCTGCAGGGGAGTATTCCAGACGACATTGAAAAGAATGAGAATATTTCTTACTTGGATGTTAGACAGAATATGCTTGTAGGAAGGATTCCAGCTGCATTTGGTTACTGGACCAATCTTTCAATGATTGATCTTTCAGAAAATATGTTTAATGGCTCCATACCCACAGAACTTGGAAAGCTTCAGAATCTTGTAAAACTGAGCATTTCTTCAAACAGACTGACCGGACAAATTCCCTTTCAGTTGAGTCACTTTGAAAAACTGGAGGAGTTGGATCTCAGCAACAACAATCTTTCAGGAAGAATTCCTAAAGAAATTGCATCATCTTCAGTATTGACAAATCTTCTACTCCAGGACAACAAACTTTCTGGTGCTCTTCCAGACACTTTCTCCTCTTCACAAATGCTTGTAAAGCTGCAGCTGGGAAACAACTTGCTCGAAGGCCCAATTCCATGCAGTCTGAGTAAGCTTATGCAACCTGTTTTTTCACTAAATCTGAGCATGAATAAGTTTACTGGTGAAATTCCTAGGTGCCTAGGCAATCTTGACAAATTGGAGGTCCTTGATATATCAAGCAACAACCTGTCTGGTGCAATACCATCAGAAATGGAAAAGATGAGTTCCCTTTCGTTTCTCAACATATCATTTAATAGCTTGTCAGGGAAGGTACCCAATACATGGGAAAAACTATTAAGTTCACGTCCAGGATCTGCCTTGGGAAATCCAGGACTTTGCTTAATAGACACGGATGGTAGCAACTGCAAGCATGTGAAAAAATCACAAGTAAAATGGAAGACTTTGGCTGGTGTAATTAGTGGATGTGTGCTTTCCATGGCAATCATAGTTGCTGCAATGTACTTACTAGTGACTCGGATTTGGCACCCATCCCTGCTGAATAAGCATCGGCTCGTCAAGTGTCAATCAGGAATTGAAGATCTACCTGATGGTATAACTTTTGAGGACATTGTACATGCAACAGAAGGCTGGAGTGAAAACTATGTTATCGGAAGAGGCACGCATGGAACTGTTTATAAGTTGGAATCTGCCAAATCCAATAAACTTTGGGCTGTCAAGAAGGTAGACTTAGCTCAGAGAGCATTCAATGATGAAATGAGAAGCCTAAATTCTGTCAGACACCGCAATTTGGTAAGATTGGGAGGATATTGCATGATACATGGATATGGCTTCATTCTAACAGAATTTATACCTGGAGGGACTCTTCATGATGTCCTTCACCAGAGAAAACCACCTGTAGTCTTGGACTGGAAGTCCCGCCATCGTATTGCTCTTGGTATTGCTCAAGGTCTATCATACCTTCACCACGATTCGGTGCCTCAAATCATTCACAGAGATCTCAAATCAGATAATGTAATGCTGGATTCTGAGATGGTGCCAAAGATTGGCGATTTCGGGATTGCTAAAATGGTATCAGATGAAGAAAATTCGACGAACTCCAATATTGTTGGGACTCTAGGATACATTGCTCCAGGTTAGCCACACGTTACTATGCTAATTTTCAGTATAATTCAGTTGGCAGATTACTTTGATAAACTAATAACTTTTACTACTTTCTTCCCTGGCCCCCTTTCTCTTTTTGTCACTCCATCTATTCAGAGAACGCATACTCAGTTCAACTGACAGAGAAAAGTGATGTCTATAGCTACGGAATTCTTCTTTTAGAGCTTTTCTGCAGAAAAATGCCAGTAGATCCCAGCTTTGAAGAAGGATTAGACATTGTTTTCTGGGTGAGGAAAAACTTGCAGAGAAGCAACAATTTCCTTAGTTTTCTTGATGAAGAAATTAGGCTCTGGAATGTAGAAGAACAATGGAAGGCCCTGAAAATAGTGGATTTGGCGCTTCAATGTGCTCAACTCGAGGCAAGCACCAGGCCTGCAATGAGGGATGTAGTAAGGTCTCTTGTCGAGTTATGAAGAGAGGAATGCAAGATGATCTGAAGGCAGGTGCAGTTCAATCCAGAAGATCAAGAGCTCCTGCATTTCTCTTCTTTATTTTTGGTGTAAAGAACAGTTAGTTGATGATGTTGAATCATTACTAGACTCTATGGCCTCTTTGTAAGTTTTGTTTGAGGCTGTATCTGTAGATATAATATACCAGGTCTTTATTAGGGTGCTGATGAATATAACATGTTACCAAATTCAAAAAGAGAATAGATGTAAAGATAAGTAATGAAACTTTTAGTAGTGAAACCTGTGATCAACTTTCCAAATTCCAACATCTCATTTTTTTCTTTTTTTGCTTACATAAAAAAATTGGACAAAGAAAATACATGATGTTTCAGTTTAACAAAATCGTGATCTTGGGTTTTTCAGTTTAACTTTGTGACATCCAATGTAGCAACATAGACTAAGAAAATGCATGATGTTTCAAATCAATTTTTGGTAAAAGCAATTTTCACCTTCTAGCCATAGCTCTTTTCATCATGGATCACTGTCCGTAATCAGTTTTTGCTTATTACTTTCTCCAGCTTCTCCAAAATCTGAACAGTTTTCAGGTATCACAATTAAAATGAGAGTGTGAAGGAACAAGGCTTTGCTACAAGCTAGTCCTTATCTTATGGACTATAACATTCCGGTTTATAAAAGAAATCAAAAGAACAACCCCCACCCAAAAAGATAAGCTTTTGTTGAATTTGGTTGTACTAATGATGGTTCCTGCGAAAGAGTGCAGAAGACCTCCGCCTTACCGGTCTTTCTTTGTGTTCCAAACTTGCAGTAGCAGAATTATTTGCAGAAAATATAGATATGATGAATAGAAATGGCTTGTGGACAATCATTGTATTTCAAACAAATACAGATATTCAAATAAGGTAGATATAACTCAGATATGAGTTACCAACAGGACTTCTAACAAGACTTAACAGACGAATGGCAAATACATAGGAATATCAGATCAGGAGAGGAAAATAATAGTCCTTGAAGAATTGATCAGAATTCAGTAGAAAGGAAATATGAAAGAAGTGCAAGAATAAACTGGAAGAAATACTCTGCAGGTGACAAATAGATGAAAGACATACCTCAGTTACGAAAACTGCAACACCCTAGCACAATACGATGCGCACCTAAATAATCATCCAAGTAGGAGAAGCCCAACTTAAATGTCCCAGATTCCTTTTTCATATTTTTTATGTAGTTGACTTGGATGTGCATTACAAGACAACTGTCACAAGATTTTGAGAATTTTTCATGTTAAAATATCCACCAGAATTTGCATTGGATATATTCACATGGCACCACATATGCTGACAAATCTCAGACTTCTAATAATAGAGATTCCATTCTGAGACAGATCCACTAAATTCCCAAGTTTCTGCTCTTAGCAAGTGCAGCACTATCACATATGATGATAAGTCTAGGACCTCTAGTAATAAAGCTTCCCTTCTGAGCCAAATCCACTAAATTCCCTAGCTTCTGCTCTTAGCAAGTGCACTTTTGTGCAAGACGATTAGAGACAAACTCCAGGAATTTGGTGCTTGCTCTAATACCAAAGTTACAAGCAGCTCAAAAGAGAAAATATAAAACACTAAACCACATTAAAGAAAAGCTAGGAAGCTAAAACCAAAATTGAAGAAGAAAAAGTTCCACACTGTTTCTCTTAACAGAACAGGTGAAATTTCTCATAAAGAGCTGATCATATCCTACATCCAATCACTTTACTGAAGTTAGAGTACAGATAAATGTTTGAAAACCCTAGTGTCCAAAGATAGGGACAATAGAAATATGAAGCTGTAAAATGCTCCAAGAAAAAATAATTAAATTTCTGAGAGGATTTCCAAAATAGCCTCGAGGTAAAATATTTATACAAGGTTAAACTTGTCTGCTTCCTCAGTCCAATAAAGCAGAGTTCAGGAGTTTTACTTTGAAGAGACCGAACAAATTTAGAGAACTACTCCATTACTGGCTATGTAACCTATGGCTAACTAGATTAAATTAGAAGCTCAATGTGCATAGACGTGGTGTTCTTGGGTTATACTTAAGACCAGCTAACTTGAAGAAGAATTCATTAGTATACATGGTTACGGACTGAGGCACAGTGCAACAGGATAATATTTTCAACTTAAGTAAGAAATTACCATTATAAAGATAGGATGTCCTTGATCACATCATACAGAATTAGATACCTAATGGAGCTTGCATGTCTGTTTAATTAACACTAAACAATGTTAACTATTGCAAAATGCTCAAGTTCACAATTTTATTTATATCCTCTCCACTTGAATATTTTCATGCAACATATTTTTGTCATTTGGATATATATATGTTGAATGAAATTAAGGAACCTAAAAGTCAGTTCACCAGACAATCTATAATCAGTTATGTACCACATCTTTTTACCTAATACATTTTTTGTGAATCAACACCACTATTTATACCCTATTCTACATATCAAAGGGACAAGATGAAGTTTGCAAGGATCAATGCCTAGTTTATACTCAGCATACAAATAAAGCTCATATAAGATACAAAACTAACTAAACCTCTTTTCATCTCCCAAATATAGTTTGAGCAATAAAATAGAGAGATGGAAAAGTATTATTTGATCGCGAAATTATTAACCACAATAGATTGTGGCAGCAACACCAGACCTCAGTCACAGTCATACTAAGTCATCTACTTTGTTGTTTTTCACATACTTTTCCGCTTCAATTTTAATCAAAAGCCGAATGCTGATAGCCTGATATGCATTCAACTACAGATCCTTTTTTTATTTTTGATGACAAGGGAAACCGCAGCCGCTACCCTTTGGGTGCGCACAGGGTAAAACTCCCGCTCATATGCAATAGCTCGCAAACCACATAGGAGAGGTAACATGTGGGTGCGCACAGGGTAAAACTTCCGCTCCTATGCAATAGCTCACAAACCACATAGGAGAGGCAACCCACACTAAGCAAGTCTGGTGTGACGAGCTCGACCCAAAAGGCAAACCCTTGCTTTCTCTGGCAAGGGGTTTCGAACTTGGGACCTCCAACGTGAAAGTCCCAAGCCCAAATCACTGGGCCACCCCGAAGGGTCAGATGTATGGATCCTTTTCACCTAGATATTCAACATCAAGGCATCTCAGAATTCTCAAAAGCAATTTACTCCTGGCAACTTCTAATTTTGAAATCCCTAAGTTCATAAAGGGGCTGTGCCCAAGCATTAGAAAACTAGCATTGCCTTCGAAGGTCAAATTATCGATTTAATTAATGGTACAGAGGGCTTTTAGAAGCGGAGAAATTGGGAACAGCATTGTACCTGTTAATTGTTGATGGAGTCATCGGAATCAAAAGTAACTCAAATTAACGAAGACAAATAAATAAAAATCTCATTTCTCAACTCAAACTATATTGGATGGGCATAAATTTGTGGAACCCATTAAGCTTTTTTCTCCAACTTGC

>M6-PGSC0003DMG400003280 M6 LRR-RLK

TTTCAGGTGATCGAGGGATATCACACGGTGAGCTAAACTGGCCTACTCGTCTAAGGATCATCCAGGGAGTTGCTGGTGGAATGTGTTATCTTCATTCCGAGTTTGCATCGTATGCAGTGCCTCATGGAAACCTCAAGTCTAGCAACATTCTACTCAATGAAAAATACGAGCCACTTCTATCAGATTACGCCTTCTATCCACTGATCAACAACACGCAAATTGTTCAATCCCTTTTCGCGTATAAAGCACCAGAAGCCATACAACATCAACATCTCACTCCAAAATGTGACATATATTGTCTTGGAATCATAATACTTGAAATCCTTACAGGGAAATTCCCTTCTCAGTATCTCAACAATCAAAAAGGTGGAACGGACGTTGTTCAATGGGTGCAATCAGCTATAGCTGACCAAAGAGAATCCGAATTGATTGATCCAGAGATAGCGTCTGCAACAGATTCCGTTGAACAAATGGTGAAACTCCTTCATGTAGGAGCTGCTTGTACTATAAGCGATCCGGATAAGAGGATTGATATGAAGGAAGCATTAAGGAGGATAGAAGAGATAA

>M6-PGSC0003DMG400003429 M6 LRR-RLK

AAGCAACACGTCCACACTTCCAAACCACAAAAAAAAAAAAAAATGTCTTCTAGTCATTTTCTTTCATTCCTTTTCCTTTCTCTTCTAATTTCCGGGATCTTTTCGGACCTTAACGCGGATAGAGCCGCTCTACTCCACCTGAGTGCGGCCTTCAAAGGCCGCACTCTTGGGTGGAACACAACCAATTCAACCCCATGTTCATGGCAAGGTGTCAAATGTGATACTACAATCAACCGTGTAATTGACCTCCGTCTCCCCGGATATGGTCTATCCGGTGAGATGCCCTTAAATTCCATCGGAAACTTAACTGAACTCCGTACTCTCAGCCTTCGGAGTAACTCTTTATCCGGTCTTCTTCCTTCTGACATAGGATCATGTACAGAGCTACGCATACTTAATTTGGAAAACAACGACTTCTCCGGTTCAATTCCAACGACTTTTTTCAATCTGAACAATCTTATACGTGTGAGTTTTTCAGGGAACAGATTCTCCGGTGAGATATCGGACGCATTTAACAATCTGACTAGGATGAGAACTCTGTATTTGGAGAATAACAACTTCTCTGGCTCACTTCCTGATTTGAAAAACCTCTCACAGCTTAATGAATTCAATGTGTCTTTCAATAGATTAACCGGTTCGATCCCTTCAAGTTTAGGTCAATTCGTAGCTTCTTCATTTCTCGGAAATTCACTCTGTGGCTCTGCTTTCTCTCTTAGTCCTTGTCCTGAAAACAATAATATCACCAATCAATCGGATAAACTTTCCGGTGGAGCAATAGCTGGAATAGTAATTGGCTCGATTATCGGATTCTTCATACTACTATTAGTTTTGTTTATGTTAGTGAGATCAATTTATAGGAGTAAAAAAAGTTTCCGTCAAGTGAACGTCTCTCCTACGCCAAATCAAGTACTTTCGAGTCCTCGTGATTCAATGGTGACAGAAAATCATGATATCGAAAATGTGTATTCGGAGAGGAAAGATAGCAAGAGGAAAGTTTGTGATGATAGGACAAAAGGGATGGTGTTTTTTGGGGAAAGTTTTGAGCTTTTTAGTTTGGAGGATTTATTGATGGCGTCCGCGGAGGTTTTGGGGAAAGGGTTAACTGGAACTACTTATAAAGCATATTTAGATAGAGATGTTGAAGTTGTTGTGAAGAGGTTGAGAAATGTTTGTGTTTCAGAAGAGGAGTTTAGGGGAAAAGTGGAGGAATTGGGAGGAATTAGTCATGGGAATTTGGTGCCTCTTAGAGCATATTATTATGGAAGAGATGAAAAGCTTCTTGTTTATGATTCTATGCCTACAAGTTTGTATGCTATTTTGCATGGTAAGTATCTTATTTCTAATAAACTTTAGCTTTTGTTGTTTTAATAATAATATTTGCTACTTGCTTCTTTTTGGTAGTTCTAATTGGTTTGTTTTCTCTGGTTTTGGTTTGGGAGTCGATCAACCTAATTAAAATTCTTAGCTTAGATTAGATTAGCTTCTAGTTAATTGTTTTTTAGCTTTAGCTATGAAAACTGAAATTATGTGGAAGAGATGGTATGAAAACAGGGGAAAGGAAGAAGATCATTTGGTAGTTCTCTTTTACATTAGTTAATTGTCATCAGATCCACTGTCGCTACCCACTATAGGCTTAGTGGAATTTACAAAGAGTGTTAAAATGTAAAAGTCACTAGTAATTGCCTCTAAAAATATATTTAGCAGCAATTAAGCTATTGTCGTAAATTAATAGTGGCTAAAGATAATTTTTGATGTTGTGTATACTGATGTTCACATCACAACTAATGAAGGACCCATGTAATATTATTAGAGAGAGAGGGACGTACAACAAGAACTTTCTGTGTAGAGGGAAAGCCTCCTTAATTAGTATTTAGTTGAATAGGCCTTCTATCAACTCACATTCTATTAAATGTTATGAAAACGAACTATAGAATTTTACTAGTATTAATTATTTCGTATTCTAACCAATTACTATTACTATGAGTTAGTTTTAACCTTTCTGATTTTGTTAGGCGAAGGAAAAGAAGCATTGACATGGGTAATAAGGAGCAGAATTGCATTAGGAGTTGCCAATGGTATCGAATATCTCCATTCACTAGGCGCTAAAGTCACTCATGGCAACATCAAGTCATCTAACATCCTCCTCACACACTACTACGATGCATGTCTATCTGAATTTGGCATAACACAGTTGATTTCCTCAACATTGAACTCCAAAATGACCGGATACTACGCGCCTGAGGTGACAGATATTCGAAATGTTTCCCAGAAAGCTGATGTCTATAGCTTTGGCACGATTCTGTTGGAACTCCTAACGGGTAAAAATCCTAGTAGTGTCATCAATGACGAGGGAATTGACCTGCCTAAATGGGTGAAATGTATAGTTGAAGAGAGGGGGACAACACAAGTGTTTGATCCTGAGTTGATTAGTTTTCAGAATTGTGATGAAGAGCAAATGGTTTCATTGCTGCATCTTGCTATTTCTTGTACATCCCACCACCCCGAAAGGCGGCCGCCTATGGCTGACACTACCAGGAGGATCAAAGAAATTGTTATGTGAGTTTTGTTAATGTTTGATCTTTCCCCCTCATTCTTGTCCAAATGAAGGTGAAAATCATGGTTTTATTCCTTTCTTGGTTCTACAAGAGGGGTGAATTCATACCTTAAAAATTGGGCACATGAATATGTGATCTCTCTATAAAACTAGATATTTTATGTATATATATTTTGAATTAATATAATATTATTTATTGCCTC

>M6-PGSC0003DMG400003551 M6 LRR-RLK

GTTAAAAAGTGCTGGCCCTTGTTTTGAAAAGTGTTCATTTGCACATTCTCCACTTCCTCCCTCACTCTTCTTCTCTGTGTCACTCTGTGCCTCTGTGCAAAGTGAGGGGGCGTCTGCCATGGCCATTAACACTCATTTTCTCCCCTTTATCATGCACCCATTTTAACCTTCACTATAAACCAAGCCAACTCCTTTTTCTTATGTTCTTGATTTCTTCTTCTTTCATGGCTGCTTATCTACTTCTGTTTCTTGTTTTTCTCTCTACATTATGCTCTGCTCAGAGAAACTCACAAACCCTTTTGGAAGTTCAAGCGCTTACTTCTTTTAAACTTAGTATTCATGATCCACTCGGTGCACTCACTGACTGGGATTCTTCTTCCCCTTTTGCTCCCTGTGATTGGCGAGGTGTTTTCTGCGTCAATGGCCGGGTCGGTGAACTTCGTCTCCCTCATTTGCAACTCAGTGGGCCACTCACTACCCAAATTGGTAACCTGCGCATGCTGCGCAAGTTAAGCCTCCGGTCCAACTTCTTTAATGGCACTGTTCCAGCTTCGTTATCCAAATGTACCCTTTTGCATTCTGTTTTTTTACAGGGTAACGCATTTTCCGGTAATCTTCCACCGGAAATATTCAACCTTACTGATTTACAAATCTTTAAAGTTGCCGGAAATCAGTTGTCTGGTGAAATCCCCGGAGAGCTTCCTCAGAGTCTACGATACTTTGATCTTTCGTCAAACTTGTTTTCAGGAGACATTCCGAGGAAATTTTCTGACCGGTCTCAGTTACTTCTGATTAATCTTTCTTATAATCGCTTTTCCGGCGAAATTCCGGCGAGTCTCGGCCGGCTTCAACAGCTTCAGTACCTGTGGCTTGCGTACAACAACTTGGTAGGAACTTTGCCCTCTGCAATTGCTAACTGTTCATCGTTGGTTCACTTGAGTGCTGAAGGAAATGCTATAGGGGGTCTTATTCCGGCAGCAATTGCTGCTTTACCGAAGCTTCAGGTGATATCTTTATCACATAATAATCTCTCTGGTTCCTTACCAGCTTCATTGTTCTGCAATGTTTCCATTTATCCTCCCTCACTTAGGATTGTTCAGTTGGGTTTCAATGCATTCACGAATATTGTCAAGCAAGAGTCTTCTAAATGTTTTAGTTCTTTGCAAATTTTGGATCTTCAACACAACCAAATACATGGTGAATTTCCTTTGATTTTGACGAACATTTCTGCATTGACGTCGCTGGATGTGTCATGGAATTTGTTCTCTGGAAAAATCCCAAGTGCCATTGGGAACTTGTGGAGATTGGAGGAACTGAGAATGACAAATAACTCATTTGAAGGTGCTCTTCCGTTTGAGATTACGAACTGCAGTGACTTGAAAGTTCTTGATCTTGAAGGGAATCGAATGACGGGTGACATTCCAATGTTTTTAGGTGATCTTAGAAGCTTGAAGACATTGTCACTGGGAAGAAATCAGTTTTCGGGTTCCATTCCTTCTAGTTTTAGAAATTTGAGTAATCTCGAAAATCTCAACTTGGGAGGAAATGGTCTCAATGGAAGCTTGCCCGAGGAGGTAATGGGTTTGAGCAATTTAAGCACACTGAATCTCAGTGGAAACAAGTTTTCTGGCAGTATGCCAGTTGATATTGGGAATCTTCAGCAGCTCTCGGTTCTGAATCTGAGTAAAAATGGCTTTTCAGGCACTATTCCTTCTAGCATTGGGACTTTGTATAAGCTAACGGTTGTTGATTTGAGCGGACAGAATTTCTCAGGTGAAATACCATTTGATCTTGCTGGTTTGCCTAATCTACAGGTTATAGCTTTGCAAGAAAACAAGTTGTCTGGTAATGTTCCTGAAGGTTTCAGTAGCTTATTAGGCATGCAATATTTGAATCTTTCTTCCAATTCCTTTTCTGGCCATATACCATCTACATTTGGGTTCTTGACCTCATTAGTTGTACTTTCTTTGTCTAACAACCACATAAATGGTTCGATTCCTCCCGACTTGGGCAACTGCTCTGCTCTGGAGAATTTGAATCTTCACTCAAATTCATTGAGTGGCCAAATACCTGCTGATCTCGGGCGTCTTTCCCACTTGAGTGTGCTAGATTTGGGTAGAAACAACTTAACAGGTGAAGTCCCAGTGGATATTTCCAATTGTTCATCCTTGACATCACTCGTACTCGACTTGAACCATCTTTCAGGGAACATACCAGAGTCACTATCCAGGTTATCAAACCTAACTGTCCTTGACCTCTCTACTAACAACTTCACTGGAGAAATCCCAGCTAATCTTACCATGCTCTCTAGCTTGGTGAGTTTCAATGTGTCAAACAATAATTTGGGCGGCCAGATTCCGGAGATGTTGGGCTCCCGTTTTAATAACTCATTGGATTATGCTGGCAACCAAGGTTTGTGTGGGGAGCCATTGGAAAGAAGATGTGAGACATCTGGTGATGGTGGGAATAAATTGATTATGTTCATTGCTGTGGCTGCCAGTGGAGCTCTTCTCCTTTTATCATGCTGCTGCTTATATACTTACAACTTCTTGAGGTGGCGCAGGAAGCTCAAAGAGAAAGCAGCTGGAGAAAAGAAGCATAGTCCTGCAAGGGCTAGTTCAAGGACCAGTGGTGGTCGTGGCAGTGGTGAGAATGGTGGACCCAAACTTGTTATGTTCAATAACAAAATCACACTCGCTGAAACAATTGAAGCAACTAGAGAATTTGACGAGGAACATGTGCTAAGCAGAACACATTATGGGGTGGTCTACAAGGCCTGTTACAATGATGGAATGGTGCTCTCAATTCGTAGACTTTCAAATGGAACACTAGGTGAGAACATGTTCAGAAAAGAGGCAGAATCACTTGGCAGGGTGAAACACAGGAACCTAACAGTTCTGCGTGGGTACTATGCTGGCCCGCCTAACCTCAGACTACTTGTCCATGATTACATGCCTAACGGGAACCTTGCAACACTGCTCCAAGAAGCATCCCACCAAGATGGTCATGTCCTCAACTGGCCAATGCGCCACCTCATTGCACTTGGCATTGCTCGTGGCCTGGCTTTCCTTCACTCATCCTCAATGGTCCACGGAGATGTTAAGCCGCAGAATGTCCTATTTGATGCAGACTTTGAAGCCCATCTTTCAGAGTTTGGCCTAGGCAAGCTTGTAGTAGCCACACCAACCGAGCCCTCCACATCAACTTCAGTTGGTACCCTAGGCTATATATCTCCAGAAGTAGCATTAACCGGAGAAACCACAAGAGAATCAGATGCTTATAGCTTTGGCATTGTTTTGCTGGAACTGCTAACAGGCAAAAGACCCTTAATGTTCACACAAGATGAGGACATAGTGAAATGGGTGAAGAGGCAATTGCAAAGAGGGCAAATTTCCGAACTATTAGAGCCAGGATTGCTCGAACTGGACCCTGAATCATCAGAATGGGAAGAGTTCTTGCTAGGAATTAAAGTAGGCTTGCTATGCACAGCACCTGACCCACTTGATCGTCCCACCATGGCCGACATTGTGTTCATGCTCGAAGGCTGTCGTGTGGGACCTGATATCGCGTCTTCAGCTGATCCCACATGCCAACCTTCACCCGCCTGAATCCCCCGAGGTTCATTCTAATTCTTTTCCCTTTTTTATTATTAGCTCTGTATAAGTTGTGAGAGTAAATTTGCAATGCAATGCAACGCAGCAGTAGCTAGCTAGCTAGCAACTTAGTTCGTTCAATTTCATGATTTTTTTTTTCTTGTAGATTTTATCAAGGTAAAAGAGAAACAATATTTCTCTTTTATTGATTCTTGTTTTATGCTATTATTCAAACACCTCACTCCCCATGGTGCCGGGATTTTTCTCCTTTGGCGTGGGGACCAAAATGTAGTGCTATCAATAAGTCAAAGAAAGCTAA

>M6-PGSC0003DMG400003714 M6 LRR-RLK

GCTCAACACATGGCGTTAGTCTTTGTCAGTGCTGCTTCTTCTTCTCCTTCTGAAACTGAAACTTTTTTTTTTTTCATCTCATACGCTCCTCTCTTCCCCACTTCCATAACAAACTCCATTGACACTCACCATTCTTGATTCTTCCCCCATTTGATCTTCTTCTTCTTCTTCTTCTCTCTACCCTTACCCACCACCACTCTTTCTCATGTCAACGGCTACTTTTCTTCTCTCTCCGCCATTGGTGGTGGTGGCACTTACATTGCTATTTATCAGTGTTTTCTCCATGGCTGAAAACGTTACAGAGACGGAAATGGCAGCTTTAATGTCGTTCAAAAGGAATTTGGAAGACCCACTTGGGGTATTAGACGGTTGGGATTTTTCTACCACATTAGCTCCGTGTGATTGGCGTGGGGTTCTGTGTGATGCCGGCAGAGTTCGTGAACTCCGCCTACCTGGTTTGCAGCTGAGTGGACGGCTAACTGACCAGGTTGGTAATTTACGCCAGCTACGGAGGTTAAGCTTACATTCTAACAACTTTAATGGCTCTGTCCCACGCTCTTTAGCTCAATGTGCTTTGCTACGCGCTGTTTACTTCCATTACAACTCTTTTTCCGGTGAACTCCCGCCGGCTATCTCGAACCTCACTAATCTTCAAGTTTTAAACCTTGCTCACAACTTCCTCTCCGGTCATGTCTCTGGTAATGTTCCAGCAAGTCTTCGGTTTCTCGATCTTTCTTCGAATCTCCTCTCCGGTGGCATTCCGGGTAACTTTTCAGCTGGTTCTCAGCTCGAACTGTTAAACTTGTCGTTTAACCGGTTCTCCGGTGAAATTCCGGCGAGTATCGGAGCTTTACAGAAGTTAGAGTATCTCTGGCTTGATTCTAACCAACTTTACGGCACTATACCTTCAGCAATCTCTAACATTTCCACATTAATACACTTGAGCACTAGTGATAATCATCTTCAAGGACTAATCCCAGCAACAATCGGTTCACTTTCCAGTTTACAGGTAATCTCATTGTCGCAAAATCAACTATCTGGTGTGGTCCCTGCATCGTTTTTCTGTAATGGGTCAGTAAATGCTCATGCTATTAGGATCATTGAGCTGGGTTTCAATGCAATCACGGGTTTAACTAAACCTGGGAATGCAACATGTTTGAGTGCTTTAGAGGTCTTGACCCTTCACGGGAATCGCATTAATGGTGTTTTTCCTGAATGGTTGACGAATTTTTCGTCGTTGAAGGTTCTTGATATCTCCGGGAATGCAATTTCTGGAACTCTGCCAAGTGATTTAGGGAATTTAAGGTTTCTTGAGGAGCTTAGACTGGGTAACAATACATTAACTGGTGATATTCCTTCTAGTATTGTGAACTTTGGTTCATTGGGGGTACTTGATTTAGGTGGAAATCGGTTTTCTGGATTGATTCCACAGTTCTTGGGTAAATTGACAGGGTTGAGAATGCTGTCTCTTATTGGGAATCGCTTTTCTGGTTCGATTCCGTATAGTTTGGGTAGTCTTTATCAGCTTGAGTTTTTTGATTTGAGCATGAATGTTTTGAATGGGAGTTTGCCTCAAAACCTTATGTTGCTTAGCAATTTGACTAGTTTGAATTTGTCCAGCAACTTGTTTTATGGTGAAATTCCAAGAGGGATTGGAAGTTTGCATGGGTTAGAGGTTTTAAATGTTAGCAATTGTGGGTTCTCAGGGAACATTCCGACTAGTATCGGAAGTTTGTTGAGGTTGACAACTCTTGATGTGAGCAAGCAAAACCTATCTGGTGAATTACCCTTTGACATTTTCGGGTTGCCAAGTCTACAAGTTGTGGCTTTACAAGAGAACATGTTAACTGGGGATGGTCTTGAAGGTTTTAGCAGTTTATCTTGTCTGGAATATTTGAATCTGTCATCTAATGCCTTTTATGGGCAAGTTCCGAAGACATATGGTTTCCTTACATCCTTGAAAGTTCTTTCTATGTCTAAAAGTGGCATTAATGGCTCAATTCCAGCAGAGTTGGGTAACTGTTCCGGCCTTCAAGTGCTTGAGCTTCGAGGAAATAAGTTAACAGGTCAAATTCCGAAGGATTTCTCACATTTATCTCATCTAAGGATACTTGATTTGGGCCACAATGGGTTAACCGGGGAAATCCCTGAAAACATCTCTAATTGTTTGTCTTTGGCTACTCTCTTGTTGGATTCTAATAATATCTCAGGTCACATACCAGAGTCGTTGTCGAAATTATCAAACTTGGAAATGCTGGATCTCTCCTCAAATAATTTGAATGGATCAATTCCTTCAAGTCTGTCCCTGATTTCCAGTCTGAAGTATTTGAATATTTCCCATAATCATCTCGAGGGTGAGATTCCAGAGGCGTTGGGTTTTCGGTTCAAGGACCCTTCTTTGTTTGCTGCGAATAATGCTTTGTGTGGAAAGCCATTGAAGGAGTGCAATGACGGTAAAAGGAAAAGAAAGAAGTTGATCTTATTCATTGTTTTGGCTGCTGTAGGGGCCTTTTTCGTTGCAGCGTGTTGTTGTGGGTACATCTACAGCCTTATACTTTGGCACAAAAAGTTAAGAGGTTCATCTGAAGGGAAGAAGCGAAGCCCAGGCCGTGCGAGTTCAGGAGGAGAAGGGGTTCGTGGAAGCGGAGAGAATGGAGGTCCAAAGCTTGTTATGTTTAACAACAAGATCACTTATGCAGATACTTTAGAAGCGACACGACAATTTGACGAGGAAAATGTGTTAAGCCGTGGAAAGTATGGCCTAGTATTCAAAGCCACTTTTGCAGATGGAATGGTCTTGGCTATCCGCCGCCTTCCTGATACATCAATCGAAGTGAACACATTCCGCAAGGAAGCTGAATCCCTCGGCAAGGTGAAACATCGCAATCTGACTGTTGTTCGTGGATACTATGCAGGACCTCCACCTGATGTTCGATTTGTTGTGTACGATTACATGCCTAATGGAAATCTTGCCACACTTTTACAAGAGGCATCTCATCAGGATGGCCATGTGCTCAATTGGCCAATGCGCCACCTGATTGCTCTTGGCATTGCTCGTGGCTTAGCTTACCTTCATTCAGTTTCATTGATTCATGGGGATGTAAAGCCACAAAATGTATTGTTTGATGCAGATTTTGAAGCTCATCTTTCGGATTTTGGTTTAGATAAACTAAGCCTAGTAGCAACCCCAGCAGAGACTTCAACATCTTCCATCCCAGTTGGAACTCTAGGCTACATAGCACCTGAAGTGGCATTAACTGGACATCCCACAAAAGAAGCTGATGTATACAGTTTTGGGATTGTATTGTTAGAGATACTAACAGGAAGAAAGCCAGTAATGTTCAATGGGGACGAGGACATTGTCAAGTGGGTGAAGAGACAACTTCAAAGAGGCCAAATTTCAGAATTGCTCGAGCCGGGGTTGCTTGAACTGGATCCTGAATCATCAGAATGGGAAGAGTTCTTGTTGGGAATCAAAGTTGGTTTGCTTTGCACCATGCCTGATCCTCTTGAAAGACCATCGATGACCGATATTGTATTCATGCTCGAAGGCTGCAGAGTTGGTCCTGACATTCCTTCTTCAGCTGATCCTACAACTCTTCCTTCACCTATGTGATATGAAAATTCAATTCTTTGCAATGAAATCAATTTGCAAGAAAATGTCCTTCATTTTTCTCAGTTCAATGTTGTTCATGCATCTCTTCAATGTATTGTTACTCCATTCATTATCATTTTGCTTGTACATTGTTCATTCAAGAGTTGGTTGGTTTTGTGCA

>M6-PGSC0003DMG400003790 M6 LRR-RLK

TTGAAAACCAAATATTACACATGACAACATTTCATACTACCTTAGTTTACATTTTGTATGTGCTACTTCTTTTTTCACTTCCATTAAGTATCACATCTTCAGCAAGAACAGAAGCTGAATCTCTTGTCAAATGGAAGAGAAATTTACCTTCTACGTCTTTCTTGGACTCATGGTCCATTTCGAATCTCGAAAACTTGTGCAATTGGACATTTATTGTGTGTAATGATGGTGGAACAATTTCTGAGATCAATCTTTCTGATGCAGCCCTCTCAGGCTCACTTGATCACCTTGATTTCACTTCATTTCCGAGTCTCGTGAATTTCAATCTTAATGGAAATAACTTCAGTGGATCGATACCTTCGAGTATTGGCAATGCCTCATTGCTCACTTTCTTGGACCTGAGTAACAATATCTTGTCAGGTGTCATACCTGAAGAGATTGGAAAATTGAATCAACTTGAGTATCTAAGTTTTTATAACAACAATATCGAAGGTGTGATTCCGTATCAGATTAGCAATCTGCAGAAGGTAATGCACTTGGATCTTGGATCAAATTTCTTAGAAACTCCTGATTGGTTGAAGATGAAAAATATGCCTGTGTTGACGTATCTAAGCTTTGGTTATAATGAACTGAGGTTAGAATTCCCTGAATTCGTACTTCGTTGTCATAATCTAACATATCTTGATTTATCGATAAATCATTTCAATGGTTCGATTCCTGAAACAGTATTTACCAACTTGATCAACCTTGAAAGACTTAATCTTTCATCTAATTCATTTCAAGGTTCATTGTCACCAAATTTTACCAAGTTGTCTAAGTTAAAAGAACTTCAGCTAGGTGTTAACATGTTTTCTGGTTTAATTCCTGATGAAATTGGCTTGATTACTAGTCTTGAAGTTCTTGTACTTTTCAACAATTCATTTGAAGGAAAGATTCCATCTTCTATAGGTAGACTCATAAATCTTCAACGGTTGGATCTTCGAAAGAACGATTTGAATTCAACTATTCCTTCTGAGCTTGGTTTTTGTACTAAACTCACTTTCTTAGCTCTAGCAGAAAATGACCTGCAAGGGTCATTGCCTCCATCATTCTCCTCTCTTACTAAGTTATCTGATGTGGGGTTGTCTGATAATAGTCTTTCTGGTGAAATCTCATCGAATTTCATCACCAATTGGACTGAGTTGACATCTTTGCAGCTTCAAAACAATATGTTTACAGGGAAAATCCCACCCGAAACCAGTCAATTGACAAACCTTGTATATCTCTACCTCTTTCATAACAATTTCACTGGTTCCATTCCCTATCAGATTGGGAACTTGCAAAACTTGTTGGAATTAGACTTTTCTGATAACCAGCTTTCAGGCATAATACCTCCAACCATTGGAAATCTGACCAATCTGAAGACGTTGCAACTTTTTCGCAACAATCTCAGTGGAACCATTCCTCCTGAGATTGGAAAATTGATATCCCTTGTAACCATCGACATCAACACCAACCGACTAAGTGGTGAGCTGCCGGACAGCATTTCTGACCTCAGTGAGCTGAAGATTCTTTCTGTATATACTAATGATTTCTCAGGCATTGTCCCTAAGGACTTTGGAAAGAACAGTCCTCAATTGTCAAGTGCCAGCTTTGCAAATAACAGCTTCACTGGTGAACTACCTGCTGGATTGTGTAGCCCGAATCTTGAGGAGTTGACGATAAATGGGAACAAATTCAGTGGGAAGTTACCAGATTGCTTGAAGAACTGCACACTGCTAAGACGAGTACGGCTTGAAGGCAACAATTTATCTGGTAATCTAGCTGATGCATTTGGAGTGCACCCGAATCTTGTTTACCTTTCTCTTAGTGACAACCAACTTTCAGGTGAACTTTCACCTGACTGGGGGAAATGTGAAAATCTCACAAGTCTAAGAATGGATGGAAACAAATTTTCTGGTGTAATCCCAAGTGAGCTAGGGAACCTGAGAGCGTTGCGCGTGTTAGCTTTGGAAGGAAATGAATTGACTGGTGAAATTCCTTCTGAACTGGGGAGGTTAGACCTGCTGTACAATCTCAGCTTGAGCAAAAACAATCTTACTGGAGGCATCCCTCAGTCTGTTGGCAATTTAACTAAGCTCCAGTATCTTGATTTGTCAACAAATAAGTTAAGTGGTAACATACCAGTAGATCTTGGGAAGTGTGAGAGCCTGTTGAGCTTGAATCTTGGCAACAACTCATTATCAGGTGGGATTCCATCTGACCTTGGAAATTTGATGCAGTTGAGTATTCTTGTGGACCTAAGTAGCAATTCGCTAACGGGAACCATTCCACAAAACTTGGCCAAGCTCACCTCATTAATGCATCTCAACCTCTCACATAACAACCTCTCAGGTAGAATTCCTCCAGCATTAGCTCAGATGATCAGTCTTCAGGAGATGGATTTTTCCTACAATGAGTTTTCTGGTCCAATCCCAACCGACGGAGTATTTCGAGGAGCATCCGCTAGTTCTTTTCTTGGAAACTCTGGTTTGTGTGGAAATGTAGAAGGATTATCCTCATGTAATTTGGATACCCGAAATGACAAGTCCAGAAATAATAATCAAAAGATCCTCATTGGGGTACTTGTACCGGTGGTCAGCCTCATACTTTTGGCAATCCTGTTTGTTGCATGTCTCGTGTCTCGAAGGAAGGCTAAGCAATATGATGAGGAGATCAAAGCTAGCCAGATACATGAAAATACAGAATCTCTGATTTGGGAAAGAGAAGGGAAGTTTACATTTGGTGACATAGTGAAAGCTACTGAAGATTTCAGTGAAAGGAACTGCATTGGAAGAGGAGGCTTTGGAAGTGTCTATAAAGCTGTTTTGCCCTCTGGGCAGATTGTTGCAGTCAAAAGACTCAACATGTCAGACTCAAGTGACATTCCATTGACTAATCGTCGAAGCTTTGAGAATGAGATTAGAACTTTGACAGAGGTGAGACACAGGAATATAATTAAGCTCTTTGGTTACTGTTCCAAGAATGGGTGCATGTACTTGGTTTATGAGTATATAGAAAGAGGTAGCCTCGGGAAAGTTCTATATGATAATGAGATGGGAATGGAACTTGGATGGGGTACAAGAGTGAAAATTGTGCAAGGAATAGCTCATGCACTCGCTTACTTGCACCATGACTGCTCTCCACCCATTGTGCACCGTGATGTATCATTGAACAACATTTTGCTTGAATCAGAATTCGAGCCACGACTCTCTGACTTTGGCACAGCAAAGTTGCTAGCTTCAGACTCATCAAATTGGACCACAGTTGCTGGTTCTTATGGCTATATGGCACCAGGTATGTTACATACTCATTACCTAATGCATCCATATTTCTGTTGATTAAGTAGTATATTCATTGGAAGGATACTTCAATAATAGTGTCTTTCTTCTTGTGCAGAGCTTGCACTTACCATGCGTGTTACAGAAAAGTGTGATGTCTATAGTTTTGGAGTCATAGCAATGGAGACTATGATGGGAAGGCATCCAGGGGAGCTTTTAACTTCATTATCAGCATCAACAACATTGTCTCCAGAAATACTTTTGAAGGATGTTCTTGACCAAAGACTTCCACCTCCCACCAGCCACTTGGCAGAGGCAGTGGTTTTTGTGATCACAATTGCCTTGGCATGTACACGTACCACTCCTGAATCGCGACCAACCATGCGTTGTGTAGCACAAGAATTGTCTGTTCAGACTTTGCCTTACCTTCCACAGCCATTGGGGACAATAGAAATGAGCAAACTAACAAGTTTTCAGAAATAATGAGAAGATAGAACCTCATGTACAGGTGTGAAATGTACAGACTGTAGAGTTTTAGTGTTGTAATGCAACTTCCTCAAGATCTAAGGTCTATATAGTATCAAGGATGAAACCTTGGATCTATCCAAATTACTATGTTTCTTTTGCACTTTTCCCTAGAAAAAAAGAAATAATATCATCCTGTTAATCATGAATGTTACAAAGTTGGTATATTCATTTGGACTAGTATGAATGTTACAAAGTTGCACAATTCCCTTATCCTAGTCTTTACGTTATTGCTAGGCAAAACACTAGCAGAAAGATACAATGTGTTCACTTGCAATAAAAATAGCAACCTAACAAAAATTTGGCAAAGAATACATCACACAGATACCAGGCATGGTACTTAGCATTCCACTCCACACTGCTTGAGTATAGGTCCCTTCTGCTTTGGTAGGAATGGGTCTATTCGCACCCAAACAAGTGAGAAGATTGAAGCAAGAAGAATTGACCAGAGGACCACAATGGTGGGAGTCCTGTTCTGCCTGCCCATCAATCCCTTTAGGAAAGGATAGAGATGCACAATAACCCAAAAGGCGAAGAATAACTTTCCGAAGAGAGGACCCCATGAACCGTAACCATTGTTGATGGCATCCGAAACACCAGCTACAACACCGACCATATTCAAGATGATCAGAGTGGTTGGTGGGATAAGAAGTGTGGTCCATTTGAAGAGGTAAAGCTCCCCAAATTCAGCATCATCAGCTGCTTTTGATGTTACAGTAAAGTTTGTGTCTACACCAGCAAGAACTTTAAGAAGGCCTTGGAAAACTGCAAACAAATGTGCTGAAACACCACCGATGACCCAAAATTGCTCGTTACGCCACCAGGCCTCAATGCTTACTCCACTCCATCGCAGCTCGAGTACACTTGTGGCTATGATGGAGATAAAGAGAGCCATGAACCAAATGCTTGCAAAGTTGTTAAGCTACAAAACAAACATGGATTTTGATTTATAAAATCTAAATCAAATCTTTATATTACCGAACCAAATGACTTCAGTTCACTTACTGTGGGGACGATAAATTTTCCAGTGAGAAGGCATACAGCTGGAAGGGTGCAGTAAGCTAGCAGAGCAATGGAAGTGAATGGGTAAACAATGGTGTTAATGTATGCTAGCCTTTCCAACCATTTGAGTTTTCCACCCCAGGCATACCAAAGGGGACAATGACGACTCATGAAGATCTCGACAGAACCAAGAGCCCATCTCAGGACTTGGTGCAATCTATCTGATAGATTGATTGGGGCTGATCCCTTAAAAGCTGCTCTCTTCGGACAGCAATAAACTGATCTCCATCCTCTGCAGTGCATCTTAAATCCAGTCAAAATGTCCTCTGTGACAGAACCATAAATCCATCCAA

>M6-PGSC0003DMG400003918 M6 LRR-RLK

TCTATCTGGGAATCTTTTAGTAGGAGAAATACCACAAAACCTGTACAACTTGTCAAACCTTGTATATCTTGACTTACATCATAACCAACTGAATGGAAGTATTCCATCAACCATTGGAAATTTGTCTAACTTGCACTTTTTGGATTTATCACAAAATCTGCTATCTGGGTCAATCCCTGTTGCTCTTGGGGATTTGCAAAATTTAACTCATTTTAATGTATCTTACAACCTTTTATCTGGTGCTATTCCCTCAATTGAGAGCATCAAGAAGTTTGGACCCTCTGCCTTTTTTCATAATAGTGGTCTTTGTGGTGATCCTTTAGAAGTTTCATGCTCTGCCAGTGGCACTACTCCTGCAAAAGGGAAACCAAACTTGAGTGTCTCAGCTATTGTTGCTATTGTCGCTGCTTCAATAATCCTTACAGGCGTTTGTCTGATAACCATCATCAATATGAAGGCTCGACGGAGAAGGAGGAGAGAAGATGAAACATTCGTTGTAGAGAGCACGCCACTGGCTTCAACAGATTCAAATGTCATTATAGGAAAGTTGGTTCTTTTCAGCAAGACTTTGCCTTCAAAATATGAGGACTGGGAGGCTGGCACAAAAGCCTTGCTAGACAAGGAGTCTTTGATTGGTGGGGGGACAATTGGATCAGTTTACAGAACTTCCTTTGAAGGTGGTGTATCAATTGCAGTGAAAAAGCTCGAGACATTGGGGAGAATCAGAAATCAAGATGAATTTGAGCATGAAGTTGGCCGGTTAGGGACCCTTCAGCACCCCAATCTTGTTGCTTTACAAGGGTACTACTGGTCCTCTAGCATGCAGTTAATATTGTCGGAGTTTGTTCCGAATGGAAACCTATACGACAATTTACATGGACTTAGTTATCCTGGAACCAGCACTGGTGCTGGTAACCCAGAATTGAATTGGCCAAGAAGGTTTCAGATAGCAGTTGGAACAGCTCGAGCACTCACATATCTTCACCATGATTGCAAACCTCCAGTCCTTCATCTTAATGTCAAGTCCACTAATATACTCCTGGATACAAACTACGAAGCCAAATTGACTGATTATGGCCTGAGTAGGTTCCTTCCCCTGCTGGATAACTATGGACTAACCAAGTTTCACAATGCAGTTGGTTATGTTGCACCAGAGTTGGCGCAAAGTCTGAGACTAAGTGATAAATGTGACGTTTACAGTTTTGGGGTGATTCTGCTGGAGTTGGTCACTGGGAGGAAACCAGTAGAGAGCCCAGTAGCAAATGAAGTGGTGGTCTTGTGTGAATATGTTAGAGGATTGATAGAAAGGGGTGCTGCTTCAGACTGTTTCGATAGAAGTTTGAGGGGCTTTGCAGAAAATGAGCTGATTCAGGTAATGAAGTTAGGCTTGATTTGTACTTCTGAAATATCCTCAAGGAGACCTAGCATGGCTGAGGTAGTGCAGGTTCTTGAGTCCATCAGGAATGGCTTGGGATCATAGTAAAGAATCAGTAGAGCCATAAGACTATTTGAAACAGCAGCTGTTCATAGAATGAGAAGAAAGCAGTCTAGCTGTTGTGCTAGACTCGCTTCTCATGAGATTCGATGAGGGAATCAGAAAGGGTCTCAAATAAGAGAGACGCGCAGACCAATTAATAGTATCAGTCATATTCTTGGATTTGGAATCAGAGTGATTAGGAGGTGACAGCTTTTGGTACAGAGAGAAGATTCTTTAGTATTTGATTTGTGATTCTTTTAATCAATTGGTCATTGTAAGTTTACAATAAAGCAATTCTTTTCCTGTAAGATGCAAAGGTATTTTGATGTCTCATATGCAGCTTGGCAACTGCTTTTTATAACCATTCAATATTGCTCTTCATTTACTGTTAAGAATTACCTTTGACATGCATATATCAACAACCTTTGAAGGGATGAAAGAAATCGATTCATTTCTCGAATTACATATCTAGTGCTAACACAAGAATGAATGTGAATAAATAATGAGTCCATACATCTACCTGTAACTATACAACAATGAGATAATAAGAAAATATCAAGATTTAAGACCTTACTTATTGACACAAAGAATTTGGAAATGGATTTCATTTGGAGCCATTTCTACATGCTATATGTTTTTCCTTGCATCAACTTTGACTGCTAATTTAGCTATCTTCAAAATCCATAAACTGGGAGCTGGAATGCATTTGTGAAAATGACAGTTGGAGAGGACCAAGAACACACTTCTGGAATCATCTGCATACAATCTGGCCTTTTTTTGTTTGTTTTCTCTGCATGTAGTTCTCAGTGCTGCTGCATTCTGATCTCATTATCCGAACATTGATAACTTATGGATCCATTAAATCACATAACACACGACCCTTAAAATCTCTGCAGAAGTTTTGGACCTTTGGTTACGGTTTACTGTAAGGGATCATATCATTAACTCAGAGACTTACGTAACTCCAAAACTTTCTGGATACGCTGCATAACTGTGGAAATGTCCTTCTCACTTAATGTCGAAAAACATAACCTGAACCACCCAGGTTCAACACAGTGGCAGGAAGAACCAGGAGTTGCATTGATCTTAGCTACATTTAGTAGATTATCCCAAAGCTCAATCTCACCCTTCTCATTATACGACCGAATTAACCTGCTCATGTCAGCCCAACAATAGAAACCCCCACTACTTTTGGTGCACTCAATTCCCAGCTGCTTCAATCCACTGACAAATAGAGAAGACATCCTTCTTAGCCTCTCTCTGTTCTTTTTGATAAATTGTTGTACAAACTTTGCATCTGATAGCATCTGGATGATTAAATGCTGTGTTGGGGCTGAAATGGATGAGAATCTTGTCAACTTTTTCGCAGCAGCTAGAACATTTTCATTGCAAGAATAGATGACCCCCACCCTGAAACCAGGAAGAGAAAGGTCCTTAGAGAGACCATACACAATGTGTACCCTGCTCCTGTCAAAATCTTCAGAATCAATTATCTCTGCTATGCTGACAAATTCTTCACGTCCATAAGTTGACCCTGCCAAGATTTCATTGGATATTACATGGATGTTCTTCTCAGTTGCAAAGTCTAAGAGGTTATAAAGTGTCTCCCTAGAGAAAATATTTCCCACAGGATTTGAGGGGTTAGAAATGATAATCCCTCGTACTTTAAGACCACGTTTCTTAGCTTGGTTAAAAGCTCGGTCAAGAGCATCAATGCTCAAGTTAAAGTTGTCTGCACTGCGGCAAGGAACAGGTATAATCTCCACTCCAGTTCTCCATTTGACATCCCTATCAAGACTGCAAACGCATTGAGCTCATTTTATACACTATATACGTAAAGATGACCAAAAGATTGAAATTTATTTGGGAAGGAGAATAAAACTCACTCAGGATAATATGGTGAGGGAACAAGAAAAGCATTTCCAGGATCAGCTAGGCAGAAGCTCAGTATCTCAAGCGCAGGAGCGGCACCACCAGTTAGAACCATCTGCGAGGGGCTGAAGGACACTGATCTCTCCAGAGCTTGAGACATAAATTCTCCCACAGCCTGCATACAGGTGAGGAAACATGAAAAGCAATTCAAAGAGATACCATAAATAATGTAGAATCACACAAAACACCTAATTGTTTTTATGAATCGATCTTCATAGTGGCTCAAGGATGATTTCATATATTGCTGTTATACAATTCTCAAACCCACAATTTAAAAAGGTAAATCCATCAGATCATCTCCCACCCTAGGATACCCAATGACAAAGGTAGCCACTCATATTTCAAAGAGGCCAAAATTGCCTTCCATCTTCAACAGATACTGCAAATCTAGCAATAGATCCTTCAACAGGTTACACAATGATGAGTATTGAGGTGAGTATCTGTCAAGGAAATGATAGACCACCTCTCAGAACTACTGAGTCCATTATTCGCTATTAACTTGCTTATAACGAAAACCATGCAAAGTAAAGCAGGTTAAACTATTCTCTTATGTGCTCAAACTAGATTTTTTATCATATCACACTGCAAGGGATCCTATTGTTCCGGTCAGGACAAACATCACCACAGAAAATCATCAAATGCTGAAACAGCATAAATGACTGAGCTAGATCAACAATAATTATGAATTTCCAACTCAAGTTTTACCAATGTAACGCTCAAGTAGGAGAAGATATTTACCACAAAAAAACACCCCCATCCCTCTGGATACTGTTTCAATGCTTTGGCTGGTCAAACACATTTTCAGGTTTCAAAATTTTCCTATCATTTCTCTTCTTTTGTATTTTAACCTCCTTTATTAGACAGTCATGGGCTCCAAAAGCTCAAGAAAAAAATCAAAACATTCAGAACCGTGTTTATTTTTCAAAGTGTTGCTTAGTGTCGATAGCGTAGAAGTTAGAGAATTGATGAACTTTTCAGAAGTTAATTGGCACTGTAAAATGAACGTGAAGCTTATTCAATGCTTTTCCTTATGTTTGTTGCACTTTATAGACTTAGAAAGCTAAAGACGGGACATACTAAAACCTTGTATACAATAGGTAATTTTGGGATTGTCAAATGCCATAACCAATCCCGCATCCAGGGGTTCACATTTCAAATTAAACCAGCTATAAAGCACACTGGACTAGCGGATGGGTGACTGATATGGAGTAACTGTTACACAAGCATATGAACATAACATTTTTGAAATTATATGGGCATGACATGAGCCCATCCTAGGTTACAACTATAAACTAATACCACCCAAATAGGTATCTCGTTGATTTAATCCTATAGTCTAAACTCAAAGTAGCAACTTTAAATAATACGTACAAACTTTTTCAAGTAAATTTTTTGTTCAATTTCCTTCTCTAGCCTGTAAAAACTATAGCCATATGAACAACTACTAGTACTTTATTAGTTCAAGAAGAACATATTATATCCAATACATGTCAACATTATATCAACAAACAAGTCACTAGTAGTGTGCAAGGAAAAATCAAAAAGCTTACCACTTTCAACTCCAACAACCCATCAAATGGCTGATAAGTAGCTATTCCGGTTATACTCGAATCTTGAGTCATCCATCTTGACACATTTTCTGCTAGCCATTCTTGAACTAAATCCAATGACAACTATAACCATACAAATAGAGACAAAATTCAAAATCATTGAAACAACAAACAAAAGTAGATTCCCCAACAAGAAATGTACTGCATTTGGTACATTTCTCTAGAAAATGATTCAATATATCCAAACATTCAAAATTATTAGTACCTTGTTTTCTGCTAATCCTAGCTGAATAACACCATCTGGATTACTTGACTCATCATAAGGGTCTTCAGAAGCTCTTTTCAATCCAACATAGAAAGGGCTACTATCAATTTGTTTTGAAATAATCGAATTAGCTCTAGCAGATACCTGAGATTGCCCAGTAGTTCCCTTCGGTGACAGTAAAACACGTGAGTGAACCCTTTGTAACCCAGACCCAGAAGACACTTCCGGTAAATGGGTCGACGACGTTGACCTCGCCGGAGCCGTAGATTCGCCGTTATTGTCACCACCGGAGGAACGATTTCGTTTTAGGTAGAGTTGCCAAAAATAGAAAAGAGCACAGGGTATAACTGAACCTAAGAAAAGACCACCCCGACCTTGAACTACACCTTGTAAGGGTACTATCACCCTCATCGCCGTCGTGGCTCCTCCTCCGTCTCTGCCGCCGGCGCCGCCAGTTGAAATGGTTGTTGTTGTTGTTCTAGTTGGTGAACGGTCTCGTGACCTTGTCATAACTGTGACATATTGGGGGCAGAATTAAGACGTCACCTTATCGGAGTATGACGTCAGTCAGCTATCTGGCCGGCGAGACGGTTGACGGAGAGTACGAAGGGGGGTGAGTGATGGAGAAGAGATTTGGAGAAAATGGTGGCAAATAGTTGGTCGAATTATTATTTATCTTTTTGTCCGCCTAAAGTTTTTTGTTTT

>M6-PGSC0003DMG400003958 M6 LRR-RLK

TATTTTCTAACTAACTCTCTTTTCTCCTTTGACAATTCACACCAATAAACTCCCCATTTTTCCTTCTCTCTACTGTGAAGCACCATTGAAAATCCAATGCAAAAGCTCACTAGTACTTCATAAAAAACTCAACTTTTCACTCAAATCTGAAAGTTCACAAAGTTCAGAGAAATGGGGGTTAATTATTTCCTCTTAACTTTCATTTTTGCTCTGTTTGTTCTTGAATTTTCTTGCATAGAACCTGTTTGTGGAAATGCTGAGGTGAGAGCTTTAATGGAGATTAAAACAGCTTTGGACCCAGAAAGCAAGAAACTTTCTTCATGGACAAGTGATGGTGATCCATGTAGTGGTTCTTTTCTTGGTGTGATTTGTAATGAACATCATAAAGTAACAAATATTACTCTGACAGATAAAGGACTTACTGGAAAGTTGTCACCAGCAATTGCCGAGCTGAAGTGTCTGTCTGGTCTGTTCTTGCATTATAATTCTTTAACAGGGGAAATACCAAAGGAACTTGGGAATTTGACTGAGTTGAATGATCTTTATCTCAATGAGAACAATCTTTCTGGGATTATACCACCAGAACTTGGAAACATGGCAAGTTTACAAGGTTTGTAGCCTTTGTAAACATCTTTCTGTTTAATTTTGTGGGAGATTGTTTCTCTTCTTTTCTTAGCTCCCACCTTTTTGCTGCTTTGGTGACTCAATCTTAAATTATCTGTTGTGATTTCTAAGAAATAGTTGTATTAAATGATTTGTCATTTTAGAAGTTTTAAGATAAAATAATTATCCTTTTCTCATTCTACTTTTAATAGTAATTGTTCTTGAAGATCCTGTTTTGGTGACTCGAACCGACAACCGTTGGAGGTAGAGAGTGCTTACAACTTGAGCAGCCCCTGCTATCAGGGAAATTGCTTGCTGTTTCATTCTACTGTTTTTTGTGTGTATTAGTGAATAATGAGTCTACGTGAATTTAGGTGACAGAGTAGACAAGTTGCAAAATTTCTTAGAGATCTATGCAGATTTTTTGGTTCTGTAAAGGTAGGACTTTGTGGGGATTGATGGTTTACTCGGTTTTTTGAATTCCTTCATCCATAAATTAACAGTGCAAGTTTGAGAGAAAATAAAGTGTTACTAGTATATATATGAATTGTTGAATTTCCTAACATAAGGAAAGCTTTAAGTACAACAGTAGAAGTGGCTCAAAATTGCTTTAAGTAGCAGATTTGAATTCCAAGTATCCCGTGTTAGAATTTCTACTTCCGCCACTGTTGCACCCACACCCCCTTTTCTGGTCTTTATTCTTTCTTGTTAATTATATTTTTGGTCTGAACAGCATACTGAAATTTCTTTTCCAAATAAGGTGATGAAAGAAATGCAAGAACTTCTTTTTTTTTCCCTTTCTACAATGATATGTGCTCTGAGATGATTTGTGCATTTTGTGGTTCCACAGTGCTGGACTTGAGTTGTAACCAGTTGACAGGGAGCATACCAACAGAGATTTGGTTCTTGAAGAAGTTGAGTGTTCTCGAATTGCAACGTAACAGGCTGACAGGTAAAATTCCAGAGAACGTAGGAGTCCTCGGGACGTTGAAAAAGCTATATTTGGGATTCAATCAGCTATCTGGTCCGATTCCATTAAACTTAGCTGCTGCTCCCCAGTTGGAGGTTCTAGAAGTACAAAATAACACCCTCTCTGGAGTTGTTCCTCCAGGTTAGTTTCCCTTTTTGAGTTCTTAACTTTTTATAAATCTTTTTTTCCTAGCTTGAAGTTATATCTTTTGATTTCTGATTTTAAACCTCTTTTTTCTCCTTTTTATCTGGACTGATTTGTAGCTCTAAGGAGATTAAATGAAAAATTCAATCATGAAAGCAATCCTGGTTTATGTGGTACTGGATTTACTTCATTGAGAGTTTGCACTGCCTGGGATCATGTCAATGTGAATCAAGTTGGTTCTGATGTACCTGATTCCAATAATAACGGCATCCCTATAGATGTCCCAGGAGCTGCTCATATTAGTCCGTTGCCTTGTAATCAAACTCATTGTTCAAGATCATCAAAATTTCCTCAAGCAATTATCGTTGCTTCTGTGATTACAGTCACTGTAACTTTGGTAGTAGCTGTTGTTTTCGGTATATTTAGACGAAGAAGGCTAAAACAAAGAGTAGGGAACACAAGTGATGCATCTGATGATAGGCTTAGCACTGATCAGGCAAAGGAAATATACAAGAGAAGCCCTTCTCCACTACTTACTGTTGAGTACTCAAATCGTTGGGACCCTATGACCCCTGAAAAGAGTTGCAATAGCATGTTCAATGAGTTCTTAAATGGATTCAAGTTCAATTTGGAAGAGGTTGAGTCTGCAACTCAGCATTTTACAGAGACAAATCTATTGGGACGGAGTAATTTCTCTGCCGTCTATAAAGGAATTTTAAAAGATGGATCTACGGTAGCTGTAAAAAGCATCAGCGTGACAAGCTGTAAGTCGGAGGAGACGGAGTTCATGGAAGGATTGAGCTTATTAACCTCCCTGAAGCATGAAAACCTTGTCAAGCTAAGAGGTTTCTGTTGTTCAAAGGGAAGAGGCGAGTGCTTTTTAATCTATGATTTGGCCTCTAAAGGGAACCTTTCTCAGTATCTTGATGTTGAAGTAAACAGCAACCATGTTCTTGACTGGTCCAAAAGAGTTTCGATCATCAAGGGGATTGCAAAAGGTTGGTCTTCTTTGCTTTTGTTCATCATTACATCTTCAACATTCTCTACAAATTTCTCAGAATATTTTAAAGTTCGGAAATATTATGGCTGTATAATATTTTCCCCATAATGTGGAGTCTTCTAGGAGTTCCATTGCATAGGCAGTCTATTACCACTAAAGTGCTTCTCACAGGATTCAGGAACAGGATGTGACTGTAGATTCATTTGCATGGTAATTAGTGGCAGAGTCTGGACTTTCAATAAGGGGTTCAAAGTATGAAAAAGTAAACACACGAAGAAGCCGAAGGGGTTCAACATCTACTATATATACATAAAAATAATTAAAACCATGTATAAACAACGTAATTTTCTGCCGACCACCTCGGCCTACCTAGCTCCACCCCTGTGGTTAATATATTATCAAATTACTCCTCCATCCCAAAATAAGTGTCGCTTTAGTTCATTTCACGCCCATTAAGAAAGGTAATAAATGCAAGGCATAGTTTACTAAGTCACCCCTAGTTATTAGATGAAGAGGAGTAGTTCTTTAATATAAAGGTATAGTTGGAACCAAGAAGTACTATTTTTTGTCTTGACTTCTAAATAGGCAATTATTTTGGGATCAACTTTTATGCTAAAGCGACACTTATTTGGGACGGAGGGAGTAATATGAATACAGAGAGAGACTGGAATTATTG

>M6-PGSC0003DMG400004146 M6 LRR-RLK

CAAACCATGAAACAAATTCATCTGTTTTGCTTGTTATTTTTGAGTGGTTTTTTCTTTGTTTCATCAAATAAGGATGAAGTTTCTATATTGTTAGCAATAAAGTCTAGTCTTGTTGATCCAATGGATCATCTCAAAGATTGGAACTTTTCAAACAATGAAAGTTCTATTCATTGTAAATGGAATGGTGTGTTTTGCAACTCAAAAAGCTATGTAGAGAAGCTTGATTTGTCAAACATGAATCTAAGTGGCAGAGTTTCAGATCAAATTCAAGGACTTCAGAGTCTATCTTTGCTTAATCTTTGTTGCAATGACTTCTCCACTTCATTGCCAAAGTCATTAGCTAATCTGACATCTTTGAAGAGCATTGATGTGAGCCAGAACAACTTTGTTGGCGAATTCCCTGATGGTATTGGAATGTCGAATCCTGGTTTGAAATATGTCAATGTTTCAAGTAATAACTTCGAGGGGTTTCTCCCTGAGGATCTTGGCAATGCTATGTTGCTCGAAGTCATGGATTTTCGAGGGAGTTTCTTTGAAGGTTCAATCCCTGGATGTTACAAGAATCTGAAGAATTTGAAGTTTCTTGGCTTATCAGGGAATAATCTTTCCGGAGAGATCCCTCGTGAGCTTGGTGAATTGAAGGCTATGGAGACTATGATTCTTGGATATAATCAGTTTGGAGGTTCAATACCAGCTGAGTTTGGTAATATGAGTAGCCTTAAGTATCTTGACTTGGCTGTTGGCACGTTAAGTGGACAAATTCCAGCTGAGTTAGGTAAGTTGAAGAACCTGACTACAGTGTACTTGTATCAAAACAGTTTTGAAGGTAAGATTCCAGCTGAAATTGGAAATATAACATCTTTAGTTTACTTGGATCTTTCTGATAACAAGATCACTGGGGAGATTCCGAATGAGATAGCAGATTTGAAGAATTTGCAGCTGCTAAATCTCATGTGCAATAACTTAACTGGTCCTATTCCTACCAAACTTGGTGAGTTAGAAAATCTTGAGATACTTGAACTATGGAAAAATTCATTAAATGGCTCTTTGCCAATGAATCTTGGGAAAAAATCTCCCTTGCAATGGTTAGACGTTTCTTCGAATATCTTGACAGGTGAGATCCCTCCTGGCTTGTGTGATTCTGGCAATCTAACAAAGCTCATTCTGTTTAATAATTCCTTCTCTGGATCAATCCCATTAGGCCTCTCTAACTGTTCTTCACTTGTTCGAGTTAGAATCCAAAATAATCTGCTTTCAGGAATGATTCCGGTTGGGTTTGGAACACTTCCTATGCTCCAAAGATTGGAATTGGCTAAAAACAATCTCACTGGTGAAATTCCTGTGGATTTCACTCTCTCCACCACTCTTTCTTTTATCGATGTCTCTTCAAACCATCTTGAATCATCTTTGCCTTCTAGCATTTTGTCCATTCCTAGCCTGCAAACCTTCATAGTCTCCAATAACAATCTTAAAGGCAACATTCCTGACCAATTCCAAGATTGTCCTTCCCTTTCTTTGCTTGATCTTTCAAGCAACCATTTCTCGGGGAAAATTCCTCAGAGTATTGCTTCATGTGAAAAGCTGGTGAAGCTTAATCTAAGAAACAACCAATTCAGTGGTGAAATTCCAACTCACATTGCCACATTGCCAACTTTATCTATTCTTGATCTATCAAACAACTCGTTAGTTGGAAAAATACCAATGGACTTTGGTAGCTCTCCAGCTCTAGAAATGCTTAATCTTTCCTACAACAAGCTTGATGGTCCAGTTCCAAGAAATGGGATACTCATGACTATTAATCCCAATGATCTGATAGGAAATGCTGGTCTTTGTGGTGGCATACTTCCACCGTGTTCTCAAAGTCTCACCATAACTTCAAATGCGAGAAAGAATCGTGTCAATCACATAATCGTGGGATTCATTGTTGGAATCTCAGTTATCTTAGCTGTTGGGATCATGGTTCTAGCAGGGAGATGGATGTACAATAGATGGTACTTGTGCAACAGCTTTTTCAAGGAATTTCGGTTCAACAAGAACAACTCAGAATGGCCTTGGAGGCTTGTAGCATTCCAGAGGCTTAACTTCACAAGTACTGATATTCTGGCTTGCCTAAAGGAGTCAAATGTGATAGGCATTGGTGGCAATGGAATTGTCTACAAGGCTGAGGTTCTGAGGCCACATTCTGTTGTAGCAGTTAAAAAGTTGTGGAGATCAGATGGAGACATTGAAGCAGGAGATGATCTCGTTGCAGAGGTTGATCTTTTAGGGAAGCTTCGTCACAGGAACATAGTTAGGTTACTAGGCTATCTTCACAATGAGACTGATATCATGATGTTATCTGAATACATGCCTAATGGAAATCTTGGAGCTGCTTTACATGGAAAAGAAGCTGGAAAAATGCTCGTAGATTGGTTGTCGAGGTACAACGTTGCCCTTGGTATTGCTCATGGACTCGCTTATCTTCACCATGATTGTCATCCACCAGTCATACATCGCGATGTCAAGTCAAACAACATTCTCCTAGATTCAGATTTTGAGGCAAGAATTGCAGATTTTGGCTTGGCAAGGATGATGCTTCACAAGAATGAGACTGTTTCTATGATAGCAGGATCATATGGCTATATAGCACCAGGTTAGTTACATTTCCTGATAATGCCACTAACAAGAAAATTCATTTCATTTTATATAGTCTCATTTTCCTTTCTCTGTTGTTTTTGGCAGAATATGGATACACATTGAAGGTTGATGAGAAGAGTGACATATACAGCTATGGGGTGGTGCTTTTGGAGCTTGTAACAGGAAAAATGCCATTAGACTCTTTGTTCGGAGAGTCGATTGACATTGTAGAATGGGTTAGAAGGAAAGTAAAGAACAAGGCATCAGAAGAAGCATTAGATGCTGATGTAGCTGGCCAATGTAAACATGTCCATGAAGAGATGCTTCTTGTCCTAAAAATTGCACTTCTTTGCACTGCCAAGCTTCCCAAAGAAAGACCATCAATGAGAGACATTATAACCATGCTTGGAGAGGCAAAACCAAGAAGGAAAAGTATTTGCCAGAATTGGGGTTATAGTACTAGTGCTAACAAAGACAAGTTAATCTTTGCTCATTCACCAGTTATAGGCCTTCTATAAGTACCAAAACCTTTTTTTTTCCACTTCAATATCTAGTATATCTTCATTTTCATTAGTTCATATGTAAA

>M6-PGSC0003DMG400004667 M6 LRR-RLK

ATCTAAAGCTGTCACTTAGCTTACTGCTGAAAAACTCATAAACCCAAAATTTTCAACAAACAATACCAAAAAAAAAAAAAAAATATCTTCAAAATTTTGAAGAATCATAGCAAAAAAAAGTAATCTTTGCCAAAAAAATTCAAGAACAATAGCAAAAAAAAAAAAAAAAAAAAAAAACTCCATTGTTGATTGTTCTTAGATCTCTAAAACCTCCATTAATGGCACCCCACAAGCTTTTTTTTCTTGTTAACTTCTTCTTCACCTTCATTTCATTCACATTTTCCGATGACAGCACGGTCATGTCAAATTTTCTGAAGGGTCTGTCTAAACCACCATCTGGTTGGGATGCTTCAAAGTCTTACTGTGAATGGAGTAAAGTTACTTGTGATAAATCGAGTAATACTGTTGTTTCCATTAATCTTGAATCTCAAGGTCTTTCTGGGGTTTTGCCTCCTGAGCTTAATCAACTGTCAAATCTCAAAACTCTTTCAGTTCAGAGGAATTCGCTTTCTGGGATTCTTCCATCTTTCGCAAACATGTCGAATTTAGCTGAGATTTACTTGAATGACAATGGATTCACATCTGTTCCTCAAGATTTCCTTTTGGGTCTTACTAGTTTGCAGACTTTTAGTATTAGTGAGAATGGGAAGTTGAGTCCTTGGCAAATTCCTAGTTATTTGATTGATAGCAACAATTTGGCTAATGTTTATGCTAGTAATGCTAGTTTGTTTGGTGTTATACCTGATTTTCTTGGTAAATTCCCTAATCTTCAGGATTTGAGATTGTCTTATAATAATTTTACTGGTTCTTTGCCTGCTAGTTTTGGGGATTCTGAGATTAAGAATTTATGGCTGAATAATCAGGGTAAAGGGTTATCAGGAACTATTGATGTGATTTCAAGTATGGTTCAGTTGTCACAAGTGTGGTTACATGCTAATTCTTTTACTGGTCCTATACCCGATTTGTCGAAATGCGAAAATATATTTGATTTGCAGTTAAGGGATAATGATTTTACTGGTGTTATACCTGATTCGGTGATGAAACTTCCTAAGTTGTTGAATATATCTTTGCAAAACAATAAGTTGCAGGGTCCAATGCCACAGTTTGGACAGGGGATTAAATTGGAGATTGAGGGTACTAATAGTTTTTGTTTGGATACTCCTGGACCTTGTGATTCGCAGGTTATGACGCTTTTAGATGTGGCGGGTGGTTTTGGATATCCGGGGTTTTTGGCTGATTCTTGGAAAGGAAATGATGCTTGTAATGGATGGTCACATGTAAGTTGTGATGGATCGAAAAAGAATGTGGATGTTGTAACATTTGGTAAGCTACGTTTATCGGGTTTTATATCTCCTGCTTTTGCTAATTTGACTAGTTTGAGGAATTTGTTATTGAATGATAACAATCTTACTGGATCTATACCGGAAAGCTTGGCAGCATTGCCTCTGCTTCAAACACTTGATGTTTCTAATAACAATTTATCTGGTCCTGTACCTGCATTTCGATCAAATGTGAAGCTTATTAATGGTGGTAACGTGTTTCTTGGAAAGAATATTAGTGATGGAGGGGGTAGTGGTGGATCACCGGGGTCTGGGTCGAGTAGTGATGGTGGAAATTCATCTGGTGGTTCGAAGGGATCCTCAGTTCAACCTGGAGTGGTTGCTGGTGTTGTTATATCTGTCGTTATATTTATCCTTGTTGTGTTGTATGTGTCTTACAAATGTTACATAAGGAGACGGCATAAGAGATTTGGAAGGGTTCAGAATCCGGAAAGAAGCAATGATATGGTTAAGCCTGGTGTAGTGATAGGTAATGGATATGCAGGAGTCCCTAGTGAATTACAGAGCCAGAGCAGTGGGGATCATAGTGAGATGCCGATATTTGAAGGTGGAAATGTTGCCATTTCCTTTCAGGTGCTTCGACAAGTGACAAACAACTTCAGCGAAGAAAATATCTTGGGTAGAGGAGGATTTGGAGTCGTTTATAAGGGTGAATTACATGATGGGACTAAGATTGCTGTGAAGAGGATGGAATCTGGCGCGATGGGTACTAAAGGAATGAATGAGTTCCAAGCTGAAATCGCGGTCCTTACCAAAGTTAGGCATAGGCATCTGGTCGCTCTTCTTGGTTCCTGTGTTAACGGCAATGAAAGACTTTTGGTGTATGAGTACATGCCACAAGGAACTTTAAGTCAGCATTTGTTTGAATGGCAAGAACACGGGTGCCCAATTTTGACCTGGAAGCAAAGGGTCACAATAGCATTAGATGTTGCAAGGGGAGTCGAGTATCTCCACAGCTTGGCACAGACAAGTTTTATCCATAGGGATTTGAAGCCCTCTAACATACTTCTTGGGGATGACATGAGAGCCAAGGTTGCAGATTTTGGATTGGTTAAGAATGCCCCCGATGGGAAATATTCTGTTGAGACACGGTTGGCTGGAACTTTTGGCTATCTGGCGCCTGAATATGCTGGTAAGTCTGTCTTTCCTCTGCTGATCTTTAATCCTTTTAGTTGTAAAACTGGTAGAAAACTAAAACAATATCCTATCATTAGCTTTCATGTTCTTTTTAATCTTCACTATATTAGAGATAAATGGAGAGGTGTTAGCTTCGTTGAAGGTCATTTCCATGTTCAAATGCTAGTTTGTTAATCTCGATCTGATTCAAGGATTGATACCGCAAACGTTGTCCAGTAAATTAACTAATGTTAGCAAAAAAGAGTACGATTTGAAAGACAAACATAAGGTAGAAATCTAGAAGCCCACTCATCCGGTTTTCAAATTGAAATTCGGATGTTGGTACAGCATGTGTGACTGATCCCATTGTATCTAAATAGAAATTTGGCTTTCAGAATATCACTCTTCTTCAAGACATTAAGTTCAACATATCTAGATTTAGTGTTGGCTGTCAATAGTTTTCTGCTCGTAACATGTTTAATTTCCTTTAGAGATCTTGTTCTGGCCTGTTTCTTGCATGTCTTAGCAAGGTTGTCTGTGATCATTATTAAATTTGGATTAAACGTGACTGCTCCTCTTTATTTACAGCTACTGGACGAGTCACGACCAAAGTAGATGTGTATGCATTTGGAGTTGTTCTAATGGAGATCATCACTGGTAGAAAAGCATTAGATGAGACCTTGCCAGATGAGAGGTCTCATTTGGTCACCTGGCTTCGCAGGGTCCTTGTCAACAAAGACAACCTCCGAAAGGCTATTGACCCAACTCTTGATCCTGATGAGGAAACATACGAGAGCATTTGTAAGGTCGCTGAGTTAGCAGGCCATTGCACTGCACGAGAACCATTCCAGCGCCCTGACATGGGACATGCAGTCAATGTCCTTGCCCCCCTCATAGAACTGTGGAAGCCTACAAGAAATGAAGACGAAGACAGTGGCATCGACCTCCAAATGAGCCTTCCTCAAATCCTTCAAAGATGGCAAGCTGATGAAGGAACTTCAAGAATGTTCGATGACATCTCCTTCAGTCAAACACATTCAAGTATACCCTCGAAACCTTCTGGATTTGCAGATACTTTCAATTCAACAGATTGCAGATGACATTAAGGCAACAATCAAGAAAGGCACACAATTCATTTGGTTAATTGGTTTCGTCTTTAACAACGCTTTTTACTTCGTCTTTTCATAGCCCTTTTGGTGATAGATCATTAGACCAAAGAAGCCAAGATGGATAGCAGAGTTTATTTATGTGGTTTTGTTAGAAGAGAAAAATTGTATTAAATTCTTTTAATCTTACATAGTTGATGTAAATAGATTTGAGTTCTTGTGTTAAACATGCAAAATATACTCTTGAAGTTTTTTACTACAAGATTTCATTATTTGTGCCACCAAGTGTGTA

>M6-PGSC0003DMG400004963 M6 LRR-RLK

CTTTTATTTCCTTCCTCTCTAAGAAAAAAACAATTTAATACAAATGACTATTCAAAAAATTATCACCCTTGTGTTATTTTTTCTTACTTTGACGTCCACTTTTGTTCAAGGTGATCAATCTCAGTTCTTTGTACTCATGAAGAAATTTGTCACAGGAAGTTCTTTGTCTAATTGGGATATTGGAAAACCAATTTGTCAATACAAAGGTGTCGGTTGTGATGAAAGGGGAGACGTTATCAAAATTAACATATCGGCCTGGTATTTATCTGGCCAATTTCCAAGTGATGTGTGTAGTTATTTGCCAAGATTGAAAAGTCTTCATATTGGTCATAATAATTTCCAAGGTGGTTTTCCTAAGTACCTAACAAATTGTTCTTTTTTGGAAGAATTGAACATGACTAAAACATCACTTACAGGACAAATTCCTGATTTGTCACCATTACAATCTTTGAAACTACTTGATCTTTCATGTAATTAATTGACAGGGGATTTTCCTTTGTCAATTATTAATCTCACCAACTTAGTTATCCTGAATTTCAATGAAAATCGCCATTTCAATCCGTGGCGATTACCAGAGGACATTTCCAGGCTTATAAATCTCAAGTGGATGATTTTAACAGCATGTAATATGCATGGGACAATTCCAGTGTCAATAAGTAACATGACTTCTCTTGTTGATCTTGAATTGAGCGCGAATCGCCTTGTTGGTAAGGTGCCTAGAGAACTCGGGAAGCTGAAGAATTTGAAACTCCTTGAGCTTTTTTACAACTTACTTGATGGTGAAATCCCCGCGGAGCTAGGGAATTTGACAGAACTTGTGGACTTAGACATGTCCGCGAATAATTTCACAGGGAGAATTCCAGAGTCTATAAGCCGGCTTCCTAAGCTGGAAGTGTTGCAGCTTTACCATAACGCGCTATCTGGAGAGTTTCCAGCAGCACTTGCTAATTCAACAACTTTAACTATCCTGTCTCTTTATGACAATCTTTTTACAGGAGAAGTTCCACAACACTTTGGTCTTTCATCAGCTTTGTTGGCATTGGACTTGTCAGAAAATCGATTTTCTGGAAAGCTACCGCCTTTTCTGTGTAGTGGAGGTAAATTGAGTTATATTCTTTTACTTCAAAACATGTTCTCAGGTGAACTGCCTGATGGATATGTGAAATGTCAGTCTGTTCTTCGCTTTCGAGTGAATTACAATCAGTTGGAGGGAAGAATACCACAAGAGCTTTTTACTCTTCCACATGTTTCAATCATTGATTTGAGTTATAATCATTTTAGTGGTCCAATTCCAACAACAATTGGAAGTGCTAGGAATTTATCAGAATTGTTTATGCAAAGTAACAAGCTTTCAGGTTTGCTTCCTCATGAAATATCTACATCTTCTAATCTTGTGAAGCTTGATCTTAGCAATAACCTTTTGTATGGTCCAATTCCTTCTGAAATTGGTGGTTTAAAAAGCCTCAATTTGTTACTCTTGCAAGGTAACAAGTTCAATTCTTCAATCCCCGAGTCACTTTCTTCGCTTAAATATCTCAACTATCTTGATCTGTCTAACAATCTTTTGATAGGAAAGATCCCTGAGAGCCTAGGTGAATTGTTACCAAACTCCATGAACTTGTCAAACAACTTACTTTCTGGTGCTATACCTCTTTTGTTTATAAAAGGAGGTGTTTTGGAAAGTTTTTTAGGCAATCCGGGACTTTGTGTCCCTACCTCTTTGAACTCAACAAACACAAGTTTTCAAACATGTTCACATAGCTATAACCATAAGAAGAGGAACAACATTGTTTGGGTTATTGGGACATCGGTGGGGATAGTCATTGTTGGATTAGTCTTGTTTATCAAACGATGGTTTGGTAACAAAAAGGCGGTGATGGAACAGGATGATCATTCATTGTCATCATCATTTTTCTCCTATGATGTTAAGAGTTTCCATCGATTGAGTTTTGATCAACGTGAGATCTTTGAAGCTATGGTTGAGAAGAATATTGTTGGATATGGAGGGTCAGGGGCTGTCTATAAGATCGAGTTGAGTAACGGAGGAGTAGTTGCTGCAAAGAAGCTGTGGAGTCACAAACATAAGCATTCTGTTTCTGAGGATAAATTGGTTTTGGATAAGGAACTTAAGACAGAAGTTGAGACTCTTGGTAACATAAGGCACAAAAACATTGTGAAATTGTACTGTTATTTCTCAAGTTTGGATTGCAGCTTGTTGGTTTATGAATACATGCCTAATGGAAACCTTTGGCATGCTCTTCATGGTGGGAAGTTCGTGTTGGATTGGCCTATTCGTCATCAAATTGCACTTGGAATAGCTCAAGGATTGGCCTATCTTCACCATGATCTTATGCCACCGATTATTCATAGAGATATCAAATCCACCAACATCCTCCTGGATATTGATTATCAACCAAAAGTCGCGGATTTTGGTATAGCCAAAGTGTTGCAAGCCAGAGGAGGCAAAGATTCCAGCACCACGGTCATCGCAGGAACTTATGGTTACTTGGCACCAGGTATAATTTTCCCAATTCATATTTGTTACTTCAATCTTTGAGCTTAAGAGTTATAATACATATCACAATCTTGATTGAGTAGATTAAATTAAATCCTTTATGTGCAAATAACATCCCATAGTTGAAGTGGCCTAATGTTACTCTAAATTGGCCATTTAATATAGTCATTGCCTAACGTTAAGAGTATATTTGACCCTTTTCCGTTCTTAATTATGGCTTTTTATGTGTAAAAACCATCACATATGCTCAAGTTTCTTGATTGAACTTTCATTTAACACATGTTTAAGTGTTCAAATTAGGCTAATGTTACTCAACTTTGGTGTAACAGAGTATGCATATTCTTCAAAGGCAACTACAAAGTGTGATGTGTATAGTTTTGGAGTTGTTCTAATGGAATTGATTACTGGAAAGAAGCCAGTGGAGCCAGAATTTGGAGACAACAAGAACATTGTTTATTGGGTATCAACAAAAGTGGAGACTAAAGAAGGTGCATTTGAAGTATTGGATAAAAAAGTCTCAGATTCTTTCAAAGAGGACATGATTAAGGTTCTACGTATAGCTATACGTTGTACGTATAGTACACCAACACTTCGTCCTACCATGAATGAAGTTGTTCAGTTGCTAATCGAAGCGGATCCTTGCAAGTTCAATTGTTGCAACATGTCAAATAAGAAGAAGAATGACACACAAGATCAACTGATCAATAAGCCACCAAAGAACATATATGAGTTGTAAAATATTTAGTGTATCAAATTAGTATACAAGGTATATAGAATATGTGTACTCATGAGATATTACACCCTCAAAAGAATCAGTCTTTATGTTTTCTAATAAATGCTACTACTTTTTATTAGTATATAGATTCTCCAATTAGTGTTACTTTGCTTTTTTATATTTTTTTGGCCTGTTGTAAGTATTTGGGAATTTTATGAA

>M6-PGSC0003DMG400004966 M6 LRR-RLK

GCGCTCTTCATAGTCGTCTCCTTCACACGTTTTTCTGAAGGCGGAAGAAAAAAACTTCTCTTCGTCAACTTCTTCGTCAGCTATGGCGGCGCTATTCAATTGCCGGCCGGAACTTATCCACTTCTTCTCTATTTTCTCCTTCTTCCTTCTCTTCCCCGTCGCTTTTTCCGACGAGCTTCAAACACTTTTATCCATCAAATCCTCTTTATCAAACCCCATCACCAATGTATTCCAAAATTGGGAACCCAATACTCCTCTCTGTAAGTTCACTGGCATCACATGTAATTCCGATGGGTCAGTCAAAGAAATCGAGCTTTCGAGTCAAAAAATTTCTGGGTTTGTTCCTTTTGATAAAATATGTTCACTTAATTCATTGGAAAAGCTCTCTCTGGGTTACAATTCATTTTCTGGGGAATTTACTGATGATTTGAACAAGTGTGTTAGCTTGAATTACTTGGATGTTGGTAATAATGAATTTACTGGGTATTTCCCTGATGTATCTTCACTCAGTGAGTTAACCCATTTTTATGCAAATAACAGTGGGTTTACAGGAAAATTTCCATGGAATTCTTTTGCGAGTATGAGTAATTTGATTGTGCTTAGCCTTGGTGATAATCAGTTTGATCGGACACCGTTTCCTGAAGTGATTTTGAAGCTTAATAAATTGAATTGGCTCTACTTATCGAATTGTGAGCTTGAAGGGGAAATTCCTGAAGAAATTGGAAATCTTACAGAGCTGATTGATTTAGAGCTGTCGATGAATCACCTTACTGGTGAAATTCCTTCGGGAATTACGAAGCTGAAGAAATTATGGCAGCTTGAGTTGTATGAAAACCAGCTTACAGGGAAGTTGCCGGTTGGATTTGGAAATTTGACAAGTCTTGAATATCTTGATGCTTCGACTAATAATCTGTACGGTGATTTGTCGGAAATTCGAAAGTTGAATCAATTGGTAAGTTTGCAGTTATTGCAGAACCAATTTTCAGGGGAAGTACCAGCAGAATTGGGAGAGTTCAAAAAGCTGGTGAACATCTCTTTGTACACTAACAAACTCACTGGTCAGCTTCCGCAGAAGTTGGGTTCTTGGGCGAATTTTGATTTCATTGATGTATCAGAGAATAGTTTCACTGGTCCAATTCCACCGGACATGTGTAAGATGGGGACGATGAGAGGATTATTGATACTTCAGAACAATTTCACTGGCGGAATCCCTGAAAGTTATGCTAATTGTACAACAATGACGCGGATTCGAGTGAGCAAAAACTCACTCTCCGGTGTGATTCCTGCAGGAATTTGGGGGCTACCAAAGCTCGAAATCCTTGATGTTGCAATGAATGAGTTTGAAGGTACTATTACTTCTGTTATTGGAAATGCAAAATCTTTAGGTGAAATTGATGCGGCTAATAACAGATTCTCTGGTGAATTGCCTTTTGATATTTCGAATGCTTCTTTCTTAGTGAAAATTGATTTTAGTAACAATCAGTTTTCCGGTGAAATTCCGGGAACAATTGGGGAGCTTAAGAAGATTGTTAATCTTAATTTACAGAACAACAAGTTTTCTGGTTCAATACCAGATTCTTTAGGCTCTTGTGTTTCACTAAGCGACATTAACATGGCTAACAATTTGCTTAGTGGCTCTATTCCTGTTTCTCTAGGATCATTACCGACTTTAACTTCATTAAACTTGTCGGAGAATCAGCTTTCAGGGAAAATTCCGACGAGTTTATCAAATCTGAAGTTAAATCTTCTTGATTTTTCCAACAATCAGTTGACTGGAGCAATACCAAACTCTCTATCAATTGATGCTTATAAGGGTAGCTTTGCTGGGAATAATGGTCTTTGTAGCCAAAACATTAAGAACTTCCGTAGATGTTATGGTGAATCTGGAAAGCCTAGAGAATGGTACACTCTGTTGATCTGTTTATTGGTGGCTGTGATTGTGGTGCTCGTTTCATTTGCGGGCTATTTGTTCTTGAAGAAGAAGAGTCATAAGGAACATGAGAGGTCTTTGAAGCAAAATTCTTGGAATACAAAGTCATTCCATATATTGACCTTCACAGAGGATGATATTCTTGATGGAATCAAACACGACAATTTGATTGGAAAAGGTGGTTCTGGTAGTGTATACAGAGTGCAGCTTTCAGATGGAACAGATTTCGCGGTGAAACATATTTGGACTTCTGATTCAGGAAACAGGAAAATATCAGGAACAACATCTCCAATGTTGGGAAAACCAGGGAAGAAATCGAAAGAATTTGAGGCTGAGGTTGAAACACTGAGCTCAATTCGACATGTGAATGTGGTGAAATTGTACTGTAGCATAACGAGTGATGACTCGAGCTTGTTGGTGTATGAATATATGCCAAATGGAAGTTTGTGGGATCGATTGCATACTTGCAAAAAGATGTCACTTGATTGGGAGACGAGGTACGAGATAGCTCTTGGTGCAGCCAAAGGACTGGAGTATTTACACCATGGTTGTGATAAGCCAGTGATTCATAGAGATGTTAAATCAAGTAACATCTTGTTGGATGAGTTTTGCAAGCCAAGAATCGCCGACTTTGGTCTTGCTAGGATTGCTCAAGCTGATTCAACCAAAGATACAACTCATGTCATTGCTGGAACACATGGATATATTGCTCCTGGTAAGAAGCTCATTATATTGTAGTTGATTTTATACGTAGAATTTGAGTATATCTACTTAACACTAACCATGTATTCATTTTTCACCATCGGTAAAATTGACTTACGAAATAGGCAACTCCCCATATCAAGCAATATGGTATCTTGAAAGATATAGTGAAAACTTGCTATAAACAGTTGGATCACTATGTTGCTCGGATCCTTCAAAAATGTTGCTGCACCCGTGTTGGTTCCCTCAAAAATAGCGCATTCTTGGAGGATTTGACATGAGTGTGACAGCATTTTTGAAGGATCCGAGCAATAAGTGAATTGCACTAATAAGAGTGTAAAACAATATTAGTGTTAATTAACTTAGATCCTATCTTGATTATCTATTCATTTGGTCATAGTGAGTGTCTCATTTCTGGCATCCTTTTTGAACTTATTTGAATTGAATTGGTGTTGCAGAATATGGATACACACACAAAGTGAACGAGAAAAGTGATGTATATAGTTTCGGAGTTGTATTGATGGAGCTTATATCTGGGAAACGGCCAATAGAATCCGAATACGGAGAGAATGGCAACATAGTTACATGGGTGAGCAGCAAATTGAAGAGCAAGGAAAGTGTATTAAGCATAGTGGATTCAAGAATTCCTGAAGCTTTCAAGGAAGATGCAATCAAAGTCTTGAGGATTGCTATTGTATGCACATCTAGGCTTCCCACATTAAGGCCAACAATGAGAAATGTAGTCAAAATGCTGGAAAATGCAGAACCTTGCAGATTAGTTGGGATAATTGTAAGCAAAGATGATGGTAGTATCAAGACAGAACAATTGAAGGATCACACCAAGATGTAATTGGATCCCCCGTCTTTGCTCGTTTCACTCTTCAACCTGATCTCTGTCCTCCATGGAACATCGAATCTTGTAGCAGTATGTAAAGTTCTGAAGATGGCCTCTTGGCATTGTAACATATAGTAGAAGCTAGATCCATACTAGTATGTGGAATTAGATTGAATTTAGAGATGCATGGCATAAGTGCTATAAGATTGGTTTAGTTATTCAAATCCCTTTTCAACCCTGCA

>M6-PGSC0003DMG400004972 M6 LRR-RLK

AGCTTGAAATCTTTTGTTTAAGCCGTTAACTATACCCTTACTAAATGATTCTCTATAAAATGTAATGTTTCCAGGACTTAAGAGCATTATACATTCAAAATTTTTGGAATATTTGAAGCTTGATTAAATTTATATATCTTTTTAATCTTCTAAGGTGCCCATAGTATCTCAATTAGCATATGATTTGCTGCAACATTTTTGGCTGCAGTTTGTCTGGACATTTACAGAGTCTTAAAATTGAAAGATTATGGCTAATGTTTAATCTTTCTGTCGTCTTGATTGATCGTGCTGGACTTCTGTTGACATGTATGTCTGTTGTACTTGAACTTAATTTAATACATTTGTTGTTCTCCTAGAAACTTCACACGATTACCGAGTGTAATGAAATCTTAAAGGGTTCAACATGTATATTCAAAAGTCATCACTGACGTTATCGGCAATATTCTGTATAGTTTCTGGAAAAATCAGTAAAAAGCCTCCTATGCTTGCTTGAACTTGTAATTGTTCTACATAGATCCCTTTTAAGTTTTATCTCACAGTTGGTTTCTTAGTAATCGTCTTTTATTTTTGTTTAGGCATGCCTCTGGTGCCAAGCCTACTGGTCTTTGTCGTTTATCTCAACTGAGAGTTGCTGATTTCTCCTTCAACTTCTTCTTTGGTAGCATACCGAAGTGTCTGGAGTACCTTTCAAAGTATAGTTCTTAATTCCGATCCACAAGCATTTTTTATATTCTTTATTTCAATTTCTGAGTGCCTATTATTTTGTTCTTTCTATAGAACTAGCTTTCAAGGAAATTGCCTTCAAGCAAAAGATCCCAGGCAGCGTTCTACTGCTTTATGTGGTATGTGTAATCTGCTAGACAACTCTTACACTTTAGCAGAATCATTAACCAAGCCTTTGAATCTTTTATGCATACATTAGTGTACGAAAATGCAAGCTGGTCATGCTTACTTTTGATTGTATATCCTAAAGAGGCTTCTGCGACTAAGTTCGCCCATTTCACTAACAGTCGCAGTTGAACAGATCAACTTTAAAATAATTGAATTGATTATTGTAACTTTTGCATTATAGGAGGCACTCCACCTGCCAAAAGCAATCCAGCGCAGAGCAATAATAAGCACGGGACTGTTGAAGGAAGAGCCAAACATAAGTCTGCTTCAAAACCTACCTGGCTTTTTGTGTTAGAAGTGGTAACTGGAGTTATTGCAGGTTCTCTCTTCCTTGTGGCTATTTTGGCTGCCATTCAGAAGTGTAAGAACGAATCTTCTATTATCATCCCTTGGAAGAAATCAGCAAGTATGAAAGACCATTTGAAAGACCATATGGCAGTTTATGTAGGTGAGTACCATGCAATTTCAGTTGGGGGATTACAGTTAAGAATTGCCTTTAAACATTTGTATTACTTGATTCTGTATTGACTTAGGGTGTGTTCGGTATGAAGGAAAAATGGTTTCCTTGGAAAACATTTTCCAGAAAATATGTGGATTCCTTACTTATTTTCTTCTGTTCGGTATGTAAGTAAAAAACACCACCCTAATAGGATTTGTATATAACCTAGACAAATACTATGGGAGGTGGGGGTAGGGGTAGGGGTCCCAAATGGTGGGGATGGGGCTTACGGGGTGGGGGTGAAGATGAGGTGTGTTGGAGGGTGGGGAGGACACAATTGATGTGAAATACCACTTGTGGAACTTGTTTTCCCTACCTCCATACCGAACACACCCTTAATTCTTTCTCATATAGGATCTGATTCTTCAATTTAGTACTACAAATAGGACACCACGTAGGACGCAAATTGCCATGTAGGATGCCACTTAGGACGTGTGTATTTATTTGTTCAACTTTATACAAGTTTAAGTGTCTATTTGTGTATACCCAAAGTTGGCGGACATAAATGTGAAATGAGGCCAAGTTAAAGGGCACATTAATGTATTATGCCTTTTTCTTTTTTTATGATATTTTCACAATAGCTTCCTTACTCTTGTATTTTTCAAAACCTGTTTTGGAAAACAACTTCTCTACCTCCACAAAGGTAGGTTTAAGGTCTGCATACACCTCACCCTCCCCAGACCCCACTTGTGGGATCACACAGGGTATAGGATCACAAGTTCAGAAGCATTAATTATAATTCACATTTCTATTAGTAACAAATGTTTCTTTGTCAGATACTGAGATGCTAAAGGACGTTGAAAGATATAGCAGACAAGAACTTGAGGTAGCTTGTGAAGATTTCAGCAACATTATTGGATCCTCACCAGACAGCTTGGTCTATAAAGGAACTGTGAAAGGCGGACCTGAATTAGCTGTAATTTCCATCTGTGTCAAAGAAGAGCAGTGGACCACCTATCTAGAGCTTTATTTTCAGAAAGAGGTGAAGCCTGTTCTCTCCTTAAAATACATGGCTACTGCTGAGCTCTCGTGCCCAATAAAAATTTCATCTTTGCTGCAATGCTCTAACTTGTATGGCTTACAAACTCAGGTAGCAGAATTGGCCAGAATAAATCACGAGAACACTGCAAAACTTCTAGGTTACTGCAGAGAGAGCAGTCCCTTCACAAGGATGTTGGTGTTCGAGTATGCATCTAACGGGACTTTATATGAACACCTCCATTGTGAGTGGAAAAAACTATTTTTTATACATTTGCTACTGCATGTTTTAGAGGCTGAAATCCAAACTCTTTGAACTTATCACAGATGGAGAAGGATGCCAACTTTCTTGGACACGGCGAATGAAAATAGTTATTGGCATCGCTAAAGGACTGAAATATCTCCATTCAGAACTTGACCCGCCATTCACTATATCAGAACTGAATTCAAATGCTGTATATCTCACAGAAGATTTTTCTCCCAAGGTAAGCGAAAGATTTCTGTATCACTGCTGCAAAACCAATTCACTGTTAAACTTTCTGGTTCATCTTACTGTCAATTGTGCTCATCCATAAGACAATTTTAGTTTACTCTCCACGGTTTCTGCTCATTTTGTCTTTGAACTCGCCGAGTTTCCTTACAAACAATTTGGGCTTACATTTTTCCATTTTGGTTTCCTTCAAGCTCAACCCTTTGTGCTTTTTAACCCGAGCAATAGAAAGCTGATTTTCCATACCCTTTTCTCTCAAAGTTTAATTTATCTGGTGCAGCTTGTTGATTTTGAAAGTTGGAAAACAACATTTGCCAGATCAGAAAAGAATTCCGGTGCTATCAGTAGTGAAGGAGCAATGTGTGTCCTTCCAAATTCTCTACAGAGCCGTCATCTTGACATGCAGGGTAACATCTATGCATTTGGAGTACTACTGCTCGAAATAATCAGTGGAAGACCTGCAAATTGCAAAGATAAAGGGTGCTTAGTAGATTGGGTAAGCATTTCTTTTCATGAAGTTTTTGTATAGGCATAGTACATAAATATACACTCAAACTTGGCCTCAGATGGCAAATAAGTAGTCCAACTTTGAGAGTGCACGTCTAGACGAGTTGAGTTGTTTAGATGTGCACTCTCAAAGTTGGGAGTGCTTACTTGCTAGCTTAGGCCAAGTTTGAGTGTCTGCTTGTGTATTATGCCTTTTGTATAATGATAACTCTCTCTGTGGAACCTGGACTCGTGCTTTATGGAATCTCAAGTTGACTCTGATATGGCAGGCCAAAGAATTCCTTGAATTGCCAGAGGTATTGCCATATGTTGTAGATCCCGAGTTGAAACATTTCAGATACGAAGACCTCAAAGTGATTTCTGAAGTGGTCAATCTCTGTATCCACCCAAGTTCGAGTAGCAGGACTTCGATGAGAGATTTGTGTGCTATGCTGGAGAGCGACATTGACACATCCATAACTGCAGAGCTCAAGACATCGTCTTTAGCATGGGCTGAGCTTGCACTCTCATCCTAA

>M6-PGSC0003DMG400005103 M6 LRR-RLK

CTCATGGCCACCATCTTGAGAAAAAATTCTCCAATTTCTCTCGCGTGGATTTCATGTTCCCACAGAAACCAAACATCAAACATAGCCATACCAAACTTTCTTCTCTATTCTTCTTACTAGCGAAAAAAAAAAATGTCAGCCAGTGCCAACTTTTTCCGGCCATGCCCAATTTTATTGATCTGTTTTTTCATATTCTTGATTTCTCCTTCTCATCAACAAGATGAGTTACAACTCCTTATGAATTTCAAATCCACTCTGAAAACAACACAAATTTCACATCTTTTTGACACGTGGACACCTCAGAATAACATTTGCAACTTCACTGGAATTTTCTGTGATTCTGACAGCAAATTGGTTAAAGAAATCAATCTTTCTGAACAGAATCTATCTGGGGTTGTCTCTTTTGATTCACTATGCTCTCTGAAATCACTAGAAAAGATTTCTCTTGGTACTAACTACTTGTATGGTAGAGTCAGCGACCATTTGAAAAACTGTACAGAGTTGCAGTATTTGGATTTGGGTAACAATTCTTTTTCAGGGGAAGTTCCAAATTTGTCTTCTTTAAGCCAATTGGAGTTTTTGAATCTCAATAGAAGTGGATTCTCTGGTTCATTTCCATGGAGTTCACTTGGAAATCTTACTAGTTTAACTTTCTTGAGTTTGGGTGACAATTCATTTAATAAAAGTCCATTCCCTCTTGAAATCTTGAATCTTGATAAGTTATATTGGGTTTACCTTACCAATAGTAGTATTGAAGGCCAAATTCCAGAAGGTATTGGAAATCTCACTCTGCTTGAAAATATTGAGCTTTCATATAATGACTTGTCAGGCAAAATCCCAGATGGAATAACCAAACTTTCTAAGCTTAAACAGCTTGAGATCTATTCTAATGGACTCACAGGAAAGTTCCCTGTGGAGTTTGGGAATCTCAGTAGCCTTGTCAATTTTGATGCTTCAAGTAACAATCTTGAAGGTGATCTTTCAGAGCTCAAGTCTTTATCTCTTCTTGAATCTTTGCAGCTGTTTGAAAACCATTTTTCTGGTGAGATTCCTGTTGAATTTGGAGAGTTCAAGTTTACAGAACTGTCATTGTACAGAAACATGTTTTCTGGTTCTCTTCCTCAAAATATTGGATCATGGGCAGAACTTCAGTACATTGATGTTTCTGAGAATATGTTCACTGGTTCAATACCACCTGATATGTGCAAGAAAGGGAGTATGACTGATCTTTTGCTTCTCCAGAACAACTTCACTGGTGGGATACCTTCGAACTATGCTAATTGTTTGTCGTTGCAGCGTTTAAGGGTTAGCAACAATTCACTTTCAGGAGTAGTCCCTAGTGGAATTTGGAGTTTGCCAGATTTGGAAATCATTGATCTCACATTGAATCTATTTGAAGGCCCAGTGACATCAAATATTGGTGAAGCAAAGTCTTTAGCACAGTTGTTCCTAGCCTACAACCGGTTCAATGGTCAATTACCACAGACAATATCAGAAGTTTCTTCATTAGTAGCCATTAATCTGAGCGCGAATCAGTTCTCTGGTGATATTCCAGCAGCAATAGGTGAACTTAAGAAGCTTAATACTCTTCATTTAGAGTATAATTTGTTTTCTGGAAGTCTTCCAGATTCTATTGGATCATGTGTTTCTCTCTGTGAAATTAATCTTGCTGGTAATTCACTTTCTGGTGCAATTCCCGAAAGTCTTGGCTCTTTGCCTAGCTTGAACTCTCTCAATCTATCTGATAACAGCCTCTCAGGTCAAATTCCAGCGACCCTTTCGAGTTTAAGATTAAGCCTTCTTGATTTGTCAAACAATAGGTTGAGTGGTAGCATACCGGATTCTTTATCAATAAAAGCTTTTAGTAATAGCTTTTTGGGAAATCCAGATCTTTGTAGTGAGAATTTTGGTAGTCTCAGGCCATGTTCATCAGATCCTCACACATCTAGAGACCACAGGACAGTCATGTTGTGCTTGATAGCTGGGGTAGTTGTTCTTGTTCTGTCACTTACGTGTTTCGTGTATGTGAAGTTTAAGCACAATAATCAGAACACTCCCGTAAAGAGACTTGATTCTTGGGATATCAAACAGTTTCATGTATTGAGCTTTAGTGAAGATCAAGTCTTGAAGGCATTGAAGCAAGAAAATTTGATTGGTAGAGGTGGTTCAGGGAATGTGTACAGATTAGTCTTGAATTGTGGAAAACAGTTGGCTGTGAAACACATTGTTAAATCCGATTCTGGTGACCAAAAAAGTTACCGGGACAGCTCAGCCATACTGGTGAAGGAGAATAGAAGATCAAAGGAGTATGATGCTGAGGTGACCACGTTAAGTTCCATTAGGCATGTCAATGTTGTCAAATTGTACTGTAGCATCACAAGTGAGGACTCAAATATGCTGGTTTATGAATACTTAACTAATGGAAGCTTATGGGACCGATTGCACACGTCTCAGAAAGTCAAGATGGATTGGTTGGTGAGATATGATATTGCATTAGGTGCTGCTCAAGGGCTTGAGTATCTGCATCACGGATATGACAGCCCCGTGATGCACCGGGATGTCAAGTCTAGCAACATCTTGTTGGATGAACAGATGAAGCCAAAAATTGCTGATTTTGGATTAGCCAAAGTTTTGCATGTTAATGGTAATAAGGACTCTTCTCAGGTTGTAGCTGGGACACATGGCTACATTGCTCCTGGTACATATTTTTCTCTTCCCTTTTCCATTTCATTGTCACTTAAAAATAACTTTGGCAAATTCTAAATTTTTTCTTTCTTCATTGTTTGACACAACAGAGTATGCTTACACAACCAAGGTGACCGAGAAGAGTGATGTCTATAGCTTCGGGGTGGTGCTAATGGAATTGGTGACTGGAAAGAAGCCAGTGGATGCAGAGTACGGGGAGAACAGTGACATAGTCCAATGGGTTTGTAGCAAGATAAGAAACAAAACTAGCATGATCGATTTGGTGGATTCAAGCATTTTCGAGGGGTTCAAAGAAGATGCTGTTGAGGTTCTTAAAATTGCAGTTCATTGCACATCAAGGACGCCAGCATTAAGACCTTCCATGAGGATGGTAGTTCATATGCTAGAAGAAGCCGAGCCTTGCAAGCTGACAAATGTAGTCGTAAATTCACCGAATGAGGATGGCAGGAAACAAGGACTTGCTAACTAATGGCAAGTCATGATGTTGAACTATCTCGTTGAGAGCAGAGGAATTTTGCAGAATATTGAACTATGTTGTAAGTTCACAAAGAATGCTTAGTGTACATATGACACTTTTGAAGAAATAGTGTATTTCACAGCTGCATTAATAAATCCAACTAGCAAGTAATGTATTTCACATCCAAATTACTTATAGAAACTGAGCAGTTGATCTACCCTACAACCAATTTTATTTTTCT

>M6-PGSC0003DMG400005254 M6 LRR-RLK

AAAGATTTAAAAAGTAATTGGCACCTTGCTTTTTAGTGGCCTACAATGGAAAGAATGGCTTTTATTGCCAAAAGGACTTTAGGCAAAGTATGCAATAAATCCTCTTCCCACTCACTTTGAAAATCTGTCTTCAGTAGTTCAATTAGTATCTGTCATAATTCTTGCCTACTGGATTGATCCATTTTCAATATGAAGAGGTAGTAAATCAAATGAAGTTTCCCTTTCATTGAACAACACAGCTCTCAGATCCCTTTCATGCATATCATATAGGTAAGTTAACTTGTGTGCCATATATTTTAGAACTAAAAGAGTAAATCTTGGCTTTCACAATAACCAAATGAATTTTTTCTATTCTGTCATTCTGAGATAGAGTTCTTCCTTTTTCGTTACTTCTTTAGCTGATATTATTTCTGGATTGTATAGTTTTCTTGCATTGTGTTAAAACTTTTTCATTGTATTTGTTAATGGTTTCTTATAGCAATCTGATTTTATCTCTCTTTCACAGTGGCTAATGAGCTGCTTCCTCTAAGCGGAACCAACCGGCTGATTTTGTCTAATGCTCTTCAAAAGTAGTCAAGGAGAGAGAGGGAAGAAGTGAAGGACTGCTGTAACTCGGATCTTGCTTTTGCATTTTTGTCAGCCAGATCAGTGAAAACATAATGAAGCTTCAGCCTTTACTGGCAGCTATTGCTTTTCTTATTCTGCTGCTATCAGTATTTCTTCATGTTATAGCTGATCTTGATTCCGATAGACAGGCGCTACTCGACTTTGCAGAATCTGTCCCTCATATCCGAAAACTCAACTGGAATTTGGCTCTCCCTATTTGCAAATCTTGGGCTGGTATAGCTTGTAACAAAGATGGAACGAGAGTCATTGCAATCCATCTCCCTGCTGTAGGACTTTTTGGTCCTATCCCAGCTAATAGTATAGGCAAGCTAGATGCTCTTAAAGTCCTTAGTTTAAGAGCCAATTATCTTAATGGAAGTGTTCCTTCTGATCTCCTTTCGATACCGTCTCTCCAATTTGTTTACCTTCAGCATAATAGCTTCTCTGGTGATATTCCTGTTTCTTTATCTCCTCGTCTTGGTGTACTAGATCTCTCCTTTAACTCTTTTACTGGAGAAATCCCCCCCACAATCAAGAACTTGACGCGATTATCTGTGTTGAACCTTCAGTTCAATTCATTGACAGGAGAAATCCCCAGCCTTGACACCCTCAGGTTAACTAACCTGAATTTGAGTTACAATATGCTAAATGGCTCCGTTCCAGACCCCCTCCAGAAGTTTCCACTTACATCATTCGCAGGAAACTCTCATTTATGTGGTACACCTCTAAATAGCTGTTCCTCGACCCCGTCACCTTCTCCAGCAGCTGATGGCTCTGCAATCCCTGAAAAACAGAAGGCTGTCCACTCTAAGAAACTAAGTACCGGAATCATCATTGCCATTGTGGTAGTGGTTTCTTTAGTAATGTTTCTCCTAGTTTTAGTAATCTCCTTCTGCTGCTTGAAAAAGAAAGTGAGTCATAGCACTAGCATAATTAAAGAGAAGGTGGCTAATGGGGGAAGAAGTGAAAAACCCGAGGACTTCGGAAGTGGTGTGCCAGATGCTGAGAAAAACAAGTTGGTGTTTTTCGAAGGTTGCTCTTACAGCTTCAATCTGGAGGATCTTCTTAGAGCTTCTGCTGAGGTTCTTGGAAAAGGGAGTTATGGTACAGCTTACAAAGCTGTCTTGGATGAGGCAACGATTGTGGTGGTGAAACGGCTGAGAGAAGTAGGGGTTGCCAAAAAGGAGTTTGAACAGCATATGGAGATAGTTGGAAGGGCTGGGAGGCACCCTAATATCGTTCCACTCCGCGCATATTATTATTCCAAGGATGAGAAACTTCTTGTTAATGAATACATGCCTGCTGGTAGCTTGTCAGCAGCCTTGCACGGTATAATTGCTAACCTTTATCTAACCTGTGTTGCTCAGACTCTTCAAAATGTTGCTGCACCTGTGTCGGATCCTCCAAAATTTACTACTTTTGGAGGACCGACATGTCCACATTTTTGAAGAGTCCGAGCAACACAACCTCTAAACTTTTTTCTTGCACATTAATTGCTTTGATCATCTGTAATGATTCATATTTTTACGCAATCTTGATAATGAATGACAGGTAATCGAGGTATTGGAAGAACTCCACTAGATTGGGATTCAAGATTGAAGATATCTCAGGGAGCAGCAAAAGGAATTGCCCACATCCACACTGAAGGTGGTGTGAAATTTACTCATGGAAACATCAAGTCCTCCAATGTACTCCTGACAAGAGACCTAGATGGATGCATTTCTGATTTTGGCTTAACACCTCTGATGAACTACATATCATACAAATATAGATGTGCTGGTTACCGCGCACCAGAAGTTATTGAAACACGGAAAGGGACACAAAAATCCGATGTTTATAGCTTTGGAGTGCTGCTCCTTGAGATGCTAACAGGCAAATCCCCTCTCCCATTATCAGGGCAGGATGAGGTAGTTGATTTACCTAGGTGGGTCCGGTCCGTTGTAAGAGAGGAATGGACTGCTGAAGTTTTCGACGTAGAGCTGTTGAAGTACCAGAATATTGAAGAAGAGATGGTGCAGATGCTTCAGATTGGACTAGCTTGTGTAGCAAAGGTGCCTGATATGAGACCTGCTATGGGAGAAGTGGTTAGAATGATTGAAGAAATTTGACAACCAGAAGGTGAAACGAGACCTTCCTCGGAGGACAGCAGGTCCAAGGACTCTAATGCACAAACACCTGAATGAAAAATGCAGTTTTTAGTCATTCAGCTTTCAATCTTGATCATTTTAAGAATGCAATTTATCTCCATTGCCAACTTTTGCATGTTCTTGGCCATACAAGTAGTGTTATTTTTATTTTATTTTTTTTATCTTAATGTTATATCCTATTCTGGCCTCTATAGAAACTTATCAGCTCAACATAAATGAGAACCTTTGTTTTCTCTATTTTTAAATCTTTG

>M6-PGSC0003DMG400005485 M6 LRR-RLK

CTAACAACGATAAATCTTAGGAATAATAGCTTCCATGGTGAAGTCCCTATGGAAATTGGCAACCTCTTGCAACTGCAACATCTTAATCTCACTTGGAATTCATTTACTGGAACCATTCCTGCTAATCTGAGTTGCTGTAAAGAACTTAGATCACTTGCTCTAGAATATAATAGTCTGGTTGGTAAAGTATTACCTGATCAGTTCAGTTCACTTTCAAAGTTGAATTATTTAGGCCTTGGAAGCAACAATCTAACAGGAGGTATCCCATCTTGGATAGGAAATTTTTCGACTCTTCGAGGTCTTTCCCTTGCAATCAACAATCTACAAGGACCAATACCTCGAGATATTGGACGTTTGTCAAACTTGCAAATATTCCAAGTTTATGGAAATCAGTTGAATGGTACAATTCCTCAATCTCTTTTCAATATATCCTCTGTTTACTATTTCTCTGTTACTCAAAACTTGTTGTATGGAGAGCTTCCATCAGATATAGGCCTTACTCTTCCGAATCTTGTGGTGTTTGCTGGTGCTGTGAACGATTTCACTGGACCAATTCCTGTTTCATTATCAAATGCAACAAAACTTGGTGTACTTGAATTGTCTCAAAATAAGCTCACCGGAAATGTCCCTACGAGTTTAGGACAATTGCAAGGGTTGTATAGGATGAACTTTGAGATCAATAGTCTTGGAAGAAACACAAGTGGGGATTTGAGGTTTCTTGATTTCCTAGTTAACTGCACAAGTTTGCAAGTTCTTAGCTTTGAGGACAACATTTTGGGAGGTGAATTGCCTAAAACAATCGGTAACCTTTCCACGAGGCTAGAAATATTTGCTCTTGGTTATAATATGATAGTTGGTTCTCTTCCTACAGGACTAGAAAATCTTGTTAATTTGACCCTTCTATCACTGGATAATAACTACTTGAGAGGCAGTGTTCCTGAGTCTTTAGGTAAGCTTCGACGCCTGCAAGGACTGCTGTTGAATGGAAACAAGTTATCAGGAAGGATTCCATCTTCTATTGGTAACTTAACATCTTTGTCTACTTTACACATTGAGGACAATGAATTAGAAGGAAACATACCTCTGGAGCTCGGGCAGTGCATCCGCTTATCAAGGCTAAACCTTACAGGAAATAATCTTGTTGGTTCCATACCAAAGGAGCTTGCAGGTCTTTCTTCTCTTTCGATTTCCTTAGCCTTAGCAAACAATTCGTTGACCGGTTCCTTGCCAGCTGAATTTGGAAAGTTGATAAATCTCAAGGAAATGGATATTTCACATAACAAATTATCAGGTGAAATTCCAAGTACTCTAAGTAGCTGTGTCAGCTTAGAGCGCTTCATAGCGAATAACAACCTGTTTCGAGGAGAAATTCCTGAATCCTTGCAAGGTTTAAGAGGTTTAGAAGAAATTGATTTGTCACACAATAACATCTCAGGAGGAATACCTGAGTTTCTTGGGAAACTTCCATATCTTAGGAAGCTTGATCTTTCATTCAACGAACTTGAAGGCGAAGTACCAACTGAAGGGATCTTTGCAAATGAAACCGCCGTCTCAATCTCAGGGAATGAAGAACTATGTGGAGGTCCTCCAAACTACAATTTTCCTACATGCCCTAAACAAAAAGATGCATCATCAAAGAAACACATTAGTTCAAGGATAAAAGTAGCAATTATTATTTCGGTTGCATTTCTATTTTTACTTTTGTGTTCTTTTGCAGCTTGCTATATAGTAATTAGGAAGTCAAGAAAGAGAGATCTTACTGGACGGTCCTCAAGAGAAAGGCAGTCTGATCATTTTGATGATGAAGAACCTACTTTATTTAATGATCCAATTTTGACAGCAAAGATAACTTACCAAGATATATTCAAGTCAACCAATGGATTTTCTGAGGATAATCTAGTCGGTACAGGAAGTTTTGGTTCTGTTTACAGAGGAAAGTTTCAGGTTTTTGATAAAGTTATGGCAGTGAAAGTATTGAACCTTCAACAAAGAGGTGCTTTGAAAAGCTTTTCGGATGAATGTAGAGCTTTGAAAAGCATAAGGCATCGAAATCTACTTAAGATAATAGCTGTTTGTTCAAGCATCGATTACCAAGGTAATGATTTCAAATGCAAAGTCTTTGAGTTCATGGAAAATGGAAGCTTAGATGACTGGCTGCACTCGAAAGGTGATGAGCAATACCTCAATATTATCCAAAGACTAAACATCGCAATAGATGCTGCTTCAGCACTTGATTATCTCCACAACAATTGCCAAGTATCAATTGTTCACTGTGATTTAAAACCGAGCAACATACTCCTCGATGAAGAGATGACTGCCCATGTTGGTGACTTTGGATTGGCAAAATTTCTCTTCAAATCATCATGGAACAAATATACATCTATTGCACTAAAGGGTTCTATTGGTTACCACCAGGTATGTCATTAATCCATGTTGATTTCTCTTGCATATCTATTTCCCTTATCAAGGGTCCTAACATGTAAGATGTTTGTGTACCTCAATCTCATATACTCCCTCCCTCCGTTTCAATTTGTTTGTGTGGTTTTGACTTGACACAAAGTTCAAGAAAAATTAAGAATACTTTCGAATCTTGTCGTCTTAAACTAAAGATACATGGAATGTAAGAACATGCCCTTTAATCTGTTGAAATTTAAAGAGTTGTCAAAAAGCAAAGAGACACTCTTTTATAAATGGACTATAAAGGAAAGTAAGACAAATGAATTGAAACGAAGGGAGCTTATCTTTCTGATGCTTAAGGTGTCGAATGAACAAATCAATAACTAAGCATTCATATGAGGTTGTCATTTTCATGTGAGTGAACATATGTTCAGTGTCCCAACATAATTCATCATTATCTTGCAGAATATGGATCAGGTGTAAATGTGTCCACTCTAGGAGATGTTTATAGCTTTGGTATCATGTTGCTAGAGTTGTGTACCGGTAGAAGGCCAACCGATGAGATCTTCAAAGATGGCCTGAATATTCACCAATATGTCAAATCCCATCTGCCTAGACGTGTCACGGAGATTGCTGATGCATCATTGCTTTTAGCATATGAAGAACACAATATATATGAGGACAATGCAAGTGATTTGGAAGAAAAGGCAATACTTCAAGATGATGAGTATATATCCAAGCTAAACACCAGCACTATTATACAAGAATGCTTGATCTCAATCATGAAAATAGGACTTTTATGTAGTAGCTCATAGCCAAGAGATCGAATGCCAATAAGCATAGCTTTGAAAGAGATTCACACAATCAAGAACTTATTTCTAGAGTCTAAGAGGATAAATAATAGTATAGATAGATCAGATGGTAATAAGCTAATCAAATTATCATCTTCAGAATTTTATTGATGAAGGCTAATATACATAGAATTACAACTTTGAAATAGATAATATTTACAATTAGTACTATAGATAGCCTCACAAAATGTGAATTGTGACCAAGAAATTAAGGGATATGGTCGAGTTTTTTTTTTTGAGTAATGGATGGAGTTCCTTTTTCCATCCTATCCCATTCACC

>M6-PGSC0003DMG400005896 M6 LRR-RLK

GAGAAATCAAAGGACAGAAGGAGACAGAAAAAAAGGTGTCATGAATTTGTACTTGTCTTTAGGGTTTGATCCACACTGCACCACAACTCTTTGCTAAACTCATCTCTCTTTTTTACAAAATTCACTCACATTATTAAAAAAAGGTACCATTTTTACAACAATTTATAGTTATTATGTACTTGCATCCTTATATGTTACTGTTATGTACTTTGTTCATTTGGGGTTCACAATTCTTGAGAAAAAAGTATGTAATTTTATGATAATCTTGAATTTTCATCATGTGTGTATATAAGTATTGTGATTTCAGAGGTGTTGTTATATACTTATGTACATTAGGTGTATTTCTTGTAAATGTTGTTTCTGGTGTAACAAATTTGAATGATTTCAAGATTTTGAGTGATTTTAGAAATGGATTAGAGAATCCTGAACTATTGAAATGGCCTTCCAAGGGGAATGATCCTTGTGGTCCTCCTGCATGGCCTCATGTGTTCTGTTCTAGTGACAGAGTTACCCAAATTCAGGTTCAGAGTTTGGGTTTAAAGGGTCCTTTACCTCAGAACTTGAATCAACTTGATAAGCTTCAAAATCTTGGCCTTCAGGGGAATGGTTTTAGTGGGAAATTACCTACTTTCAATGGATTATCTGATTTGAAATATGCATACTTGGACAACAATAAATTCGATACGATTCCTGGTGATTTCTTTAATGGTCTTAGCAGTGTTCAAGTTTTGGCTTTGGATTATAATCCGTTTAATGAGAGTGGATGGTCTATTCCCATTGAGTTGCAAGATTCAGCTCAGTTGACGAATTTTTCTTGTGTGCAATGTAATATAGTTGGACCTGTGCCTGATTTTTTTGGGAAATTGCCTTCTCTAAGTGCGTTGAAGCTGTCGTATAATCGTTTGACCGGTGAGATACCGGACAGTTTTCGTGATTCAATGCTTCGAATTTTGTGGTTGAATAATCAAGATAGTCCTGGAATGACGGGTCCAATCGACATAATTGGTACTATGGATCAGCTTATGTTGTTATGGCTTCAAGGAAACAGTTTTAGTGGACCAATTCCTGATACAATTGGTGATTTAAATGACTTGAAGGAGCTGAATTTTAACGGAAATCAACTAGTTGGTCTGATTCCGCAAGGTTTGGCAAATCTAGACTTGAGTGTGTTAGACTTGAACAATAATAAGCTCATGGGTCCTATCCCAAAGTTTAGAGCTGCGAATGCTACATATTCCTCAAATTCTTTCTGTCAGCCAACTCCTGGTGTTCCTTGTACTCCTCAAGTTAATGCACTTTTAGATCTTCTTTGTGGTTGGAATTATCCAGCCAACCTTGCTCCTGAATGGAGTGGCAATGATCCATGTACAGGTCCTTGGTTGGGAATCAGTTGCAATCCTAAAGGCCAGATTACAATTGTTAATCTACAGAACAAGAACCTTACGGGTACGCTTAGCCCTTCACTTGCAAACTTGGATTCACTGCTTGAAGTTCACTTAAAAAGAAATAGTTTGCACGGTCGAGTTCCTGCAAACTTAACAGAGCTCAGATCCTTGAGATTGTTGGACTTAAGTGGGAACAATTTTGATCCACCATTACCGAAATTCCGGGATCGTGTGAAAGTCATTACAGACGGAAATGCTCATCTTGTTGCTAATGTAACTGCGGCTCCTCCTCCTCTGAGTATTAGCCCATTTCCTCCATTGAGCCACAGTCCTAAATCAAGCAAGGAGTTACCATCGAAAAGCCCGTTTCCCGGTGATAATCAGCCAACATTATCAGATACACCGCCCTCCCCAGAGAAAAGTTCAGGTTCAGACTCAAAGTCTCCCAGGGGAACCAAAGGGCAGACCACATCAGAGAACCATGATAAGACCATGATCATTGTAGTTGTAAGTGCAGCTTCTTTCGTCTTCACTCTTCTTGCCGTTGTTTTGTATTTCAAGAGCAAGAGAAAACGAGAAAAGGATTCAGGTACTATTGTTATCCACCCCAAAGAGCCATTCGATCAAGATAACATTGTTAAGATCACTATCTTAGAAGATCCCATGATGTACAGCTTACAGAGTGGTACTACAACAACTACTACAACTAGTGGTGGAACAAAAGGAAGTCGAGTGATTGAGATTGGAAACCTTGTCATTTCAACCCAGGACCTTCGGAGGGTAACCGATAATTTTGCACCAGAGAATGAGCTTGGTCGTGGTGGCTTTGGAGTTGTTTACAAAGGCGTAATTGAAGATGGAATTCAAATTGCAGTGAAGAGAATGGAATCTGCCATTATCAACAGCAAGGCATTGGATGAATTTCAAGCAGAAATAGCTGTGCTTTCTAAGGTTCGACATCGCCATTTAGTTTCACTTTTGGGTTATTCCATTGAAGGAAATGAGAGGCTTTTGGTATATGAATATATGTCAAAGGGCGCTTTGAGTAGACATCTTTTCCGCTGGAAGATCCTGAACTTAGAGCCTTTATCATGGACAAAGAGACTCAATATTGCTCTAGATGTCGCTCGAGGGATGGAGTACCTTCATAATCTGGCACACCAGAGCTTCATACATCGCGATCTCAAATCATCAAACATTCTTCTAGATGATGATTTTAGAGCAAAAGTTTCTGATTTTGGATTAGTGAAACTCGCTCCTGACAAAGAAAGGTCCGTTGCAACAAGGCTTGCAGGAACATTTGGATACCTTGCTCCCGAGTATGCTGGTATGTGTGCATTTTTTTCCATTGGTATCTCTTTCTTAGACACTCGTTTTGACTGTTCTGGTAAATATTATCATTTAAGCATCAATTGTATTTTTTACGCTCAGGTAAGCTGTTACATGAACACATCTCTACCTGAACGAAACCAATACTCTTTTAGTCCCTCTATTTTAAGAAATGCATCAGTTCTATAATGAGAATCCAGACTGCTGAATTGCGCTTCTGACCTTCCAAGCCCTTCCTTCTCCAGCCAAAATAGAACAGAGTAACTAATGGAGCATTTACGTTTTTCCTTTAAGTTGTCGGTGTGCCTACTAAATTGATATACCTCAGACATTGATGTATCAATCTCGCATCATTCAGTGATATTCTCATTCTGTTCACTATCTTTGTTAAACAATTCAAGTAGAAGGTTTCACATCCTCTCTCCATATGATCATTCAGAGCATAGTTATCATCTTGATTCCATAAAAAGAAACAGAAACTCTATCCCTTTAAAGATACTTTTCCCTTGAGGCATGGTACTGAAATATGTTTGTGTAAAGCTGTTCAACAAGCATGTCATACCTGTTAATTTTGGTGTAGAAACTATTGCTTACTTTATCTTTATGTTACCCGTATATTTTAGTTCTCAGTCTTGTTGAACGAATTACCTAGTCACCTTTCATATATCCTAACTCACATTTTACCTCTCCAGTGACCGGAAAAGTCACAACAAAAATAGATGTCTTCAGCTTTGGGGTGGTGTTGATGGAACTTGTGACTGGACTAACAGCCCTTGATGAGCATCGGTCCGAGGAAACTCGATATCTAGTTGAGTGGTTCTGGCAGATAAAATCGAATAAAGAAAACCTCCTTGCTTCCGTTGATCCAGCTCTTGACGTGAAGGAAGATATCCACAAGAGCATTTGCACTATGGCTGAATTAGCTGGACATTGCACTGCAAGAGACCCCAATCACAGGCCTGACATGTCACACGTTGTGAATGTGCTTGGCCAACTAGTTGAGTCTTGGAAACCCGTAGAAGAGTTAGATGAATATTCAGGAATTGACTACAGTCCACCTCTGCCAGAAATGTTGAAGGACTGGCAGGATGAAGACACTGGAGACTTCACCAGTACTAGTCAAGATAGCAGAGGAAGTATTCCTGCTAAGCCAAATGGATTTGCGGATTCTTTTACATCCAACGATGCTCGATAGGTGAAGTTGGTTTTGGCTACCAAAGTCCATTGCTTCCTTACTATTAGTTTTGCTAGCATTTTCGACATGAATTTTGCTTTTGGCTCGTGAAAATAATCTATGGAGGCATTGCGTCCAGTCTATATATGGTGAATATACAGATACCGAGTTGTCATGAATGATTGATACAGAGCAAAAAACAGAAAAAACTTTTCAATTTAATGTTGTTTACACAGTTATTTGTTTTGAATGTTGCTTGTAAAAGTTATATTTATAATAGGTTGGTCCAATTTGCTGTATAAGTTATTTACATGACACTCTTGAGAATCTTACGCATGAATTTGAGAGTGTTAATAATGCAAAAAAAG

>M6-PGSC0003DMG400006065 M6 LRR-RLK

GAAGAACATTCTCTTATTGATTCTTCTCTTTCAAATTCAATTTTCTATGTTAACTGCTTCATCAAATGAGACTGACCAACAAGCTTTACTAGCTTTCCAAAATCTTATTACAAGTCCGAGTCATTTTTTGGCCAGTAATTGGACCAAAAATATTTCTTTTTGCTCTTGGTTTGGTGTCACTTGCAGTTCCAAAAGGCAAAGGATTGTGGCCTTGGATCTTCCTAATTTGCAACTTCAAGGACAAATTTTCCCATCTTTGGCCAATTTGTCCTTTCTCAGAGAGCTGAATCTTGGGAACAACAGCTTTCACGGTGGCATCCCTTATGGACTTGGCCACTTGCCTCACTTGCAAGTGATTGATATTCAAAACAATCAGCTTAACGGAAGTATACCAACAAGTCTATTTCAACACCGGAGAGTTCAGAAGATTTCATTGGCTTTCAATAAACTCGGTGGTGAAATGTGGAAAGGTCCATGGTATGTACCCAAACTCAAAGTCTTTGATCTCAGGAACAATAGCCTCACAGGTATGATCCCTCCTTCTGTTGGAAATGCAACAAAGTTGCTGAACTTCAGTGGGAATAGAGTCTGCGGCAACATTCCAAAGGAAATTGGTAATCTCAGCAAACTTGCATATTTGTCCTTGGCCGATAATCAATTAACAGGTTCCATTCCCGCAGTACTGTTTAATATCTCGTCGCTACTTGAAGTGCATCTGGGAATCAATAGCCTTTCTGGTCCTCTCTTGCTAGATGAAGGGAATGCTGTGTCAAATCTGAAGTTTTTAAGTCTATCTAACAACCAAATTTCTGGTTGCATTCCTTCCAACATATGCCAACTCACAGAGCTCGAAGTTTTGTCCATATCTTACAACAACATAACTGGAGAAATACCCAGAAATATTGATTGTTTATCCAAGCTCAAAAAGTTCTTAATTGGTGATAATCCAACAAAAGGGACTATTCCCACATCATTGGGAAATATTTCCACTCTGCAATATCTTTATTGTGGAAACAATCGCATAGTGGGGCAAATTCCTCCGGAATTAGGGAAGCTATCAAATTTGCAGCAATTATGCTTTAGGAATAATTATAATCTAATTGGTCAAATTCCAGAGGCTATTTTCAACATATCTTCTTTGGAAGTTATTGATTTCAGTTTCAATAACCTCTCGGGGAGAATTCCAACCACTACAGGTCTTCATCTTCCAAACGTTAAGGAACTTATCTTGGCAGTCAATCAGCTGAAAGGGGAAATTCCATTGTTCACAACAAATGCTTCCAAGCTTGAGAGATTGGGGCTAAATAATAACTTTCTCACAGGAACTATTCCTAATAACTTGGGAAATCTTCATGAGCTGCAAATACTGCTCCTACATACTAATCAACTTGCCAATGAACCAAGACAGCGTGAGTTGCTATTCTTCAATTCTTTGGCGGACTGTAGGATGTTGCGATATCTACAAGTGGGTAACAATCAGTTGAATGGTTTTCTGCCCAATTCTATTGGGAATCTTTCATATACTATTCAAAACTTTTATATAGGAGATGCACACATCAATGGCCTAATCCCCACAAGTATAGGCAACATGAGCGGTCTTACATTCCTAAGCCTTGGAGGAAACAACTTGACAGGGAGTATTCCTCCTGAGATTGGTAAGCTTAAACAACTCCAAGGGTTGCATCTAACTAACAATAAATTGCAGGGACATATTTCTGAGGCAGTATGCAATTTATCTAATTTGGTTCAATTATATCTGGAACATAATGAGCTCACTGGAGTGATCCAAGAATGTATAGGAGATCTAAGCATGCTACAACTGCTTTATTTAAATTCTAATAGATTTTCATCAAAATTTCCGTTGAGCCTTTGGAAAATGAGTGGTCTTCTCTTTCTAAACGTGTCACAAAATTCTTTAGAGGGAGAAGTTCCATCAGATATTGGAGAACTTAAAGTCATTATAGCGCTAGATCTTTCCAGTAATCACTTTTCAGGAATGTTACCAAGCAGATTGGGAGAACTCCTAAACCTGCAGTCTCTTGTCCTATCAAACAATTCATTTTTTGGACAAATTCCATTATCCCTTGCCAACTTGATAAGCTTAGAATTCTTGAATTTGTCTTTAAATGCATTGTCAGGTACTATTCCGAAGTCTTTGGAAAAACTCTTATACCTTAAAAGCATTAATGTTTCATTTAATGATTTAGAAGGTGTAATACCGAGTGGTGGTGTGTTTGCAAATTCCACTCTGCAATCATTTCTTGGGAACAAAGGTCTATGTGGAGTCCACATATTGGAGATTCCTGCTTGTGCTATCACTTATCCTGGAAAACAATCAAAGCTTAAGGAGGTTGTGCTAAAAATTGCTACTCCAGTGGTTATTTCATCCTTGATGATATTCTTGTTGGTTTCAATTTGGATAATGAAACGACAGAAGAAAGGAAAGTCCAAAGATGTTGAAAAGGTACCGGAGATCAGGACTCATCAATTAGTTTCTTATCACGAGATTCAACTAGCAACAAATAATTTTGATGAATCAAATTTAATTGGTGTGGGAGGTTCTGGCTCTGTGTACAAAGGCACATTATCTGGTGGAACTTTGGTGGCCATTAAGGTTCTGGATTTGCGAAGTGAGGAAGTATGCAAAAGGTTTGACACCGAATGCGAAGTGATGAGAAATGTTAGACATAGAAATCTTGTTCCGGTGATTACTACTTGTTCTAGTGACTACATAAGAGCCTTCGTTCTGCAATATATGTCAAATGGGAGTCTTGAGAATTGGTTGTACAGAGAAGAGTTCAACTTAAACCTTCTTCAAAGAGTCGCCATAATGCTTGATGTGGCTGTGGCAATTGAATATCTACACCATGGGCATAACACTCCAATAGTTCATTGCGACGTAAAGCCAGCCAACGTTCTTTTGGATGAAGATATGGTGGCTCATGTTGGTGATTTTGGTATCTCTAAAATTTTAGCGATAAGCAAGTCCATGGCTTATACAGAGACATTGGGCACTCTTGGATACATTGCACCAGGTATAAAAAAAATCTTTTCTCTTTGATTTTCTCTTATCATAATTAAGCCTCTCCAAGTTCTAGAAGTAAAAAGAGCAAGTCTTTATTTATGCAGAATTATTGTTGTATTTCAATTGAGTAACTTTTCTTCAATCCTTTCCTAAGAATATGGCTTGGAGGGAATAGTGTCTGCTAGTGGTGATGTTTATAGCTACGGCATCATGTTGATGGAGATTTTGACCAAAAGAAGGCCAACAGATGAAGAGATATGCAACCAAAATCTTGACTTGAGGAAATGGATCACACAATCATTTTCAGGGACTATGATGGATGTTGTGGATGCCAATCTTTTTTCTGAGGAAAAACAAGTCACCTTGTAAAAGTGAAATATGCATAGCCTCCATGATAGAATTGGCTTTAGACTGCACAAAGGAAATGCCAGTATCAAGAATAACCATGAAAGAAGTAGTCAAGAGGCTTAACAAAGTCAAGAACACATTTCTGGAAACAT

>M6-PGSC0003DMG400006280 M6 LRR-RLK

CCAAAGCCAAAAATTACTATCATATTTTATCAGTATCAATGGAATCTTTCAAAATACTTGTTGCATCTTCTTCTTCTTCTTCTTACCTTTGATGTTTTCTAAAGTATTTGAACTTCAATTACTAGTTATCTAAATGGCTACTTCTAACACAAGTCTCTTCTTTTTCGCGTATTTCCTACTCGTGTTCCTCATTACTCCATCTCAATCGCATAACCTGTCTCTGAGACGACAGGCTAAAACTCTAGTTTCATTGAAACAAGCATTTGTAGTATCCTCTGTTCCTAGTACTCTGTCTACTTGGAACATGTCAAATTATATGTCTATATGTTGTTGGACCGGTATCACGTGTGATGATACCAAATCAGTAACTACCATTGATATATCCAATCTAAACATTTCTGGCTCTTTATCACCTGATATTCATGAGCTCACCAGACTTCGCATCCTGAATATTTCTAACAACTTGTTAGGTGGGAACTTGAGCTGGGAGTATCGAAAGTTTAACGTACTTCAAGTGTTGGATGCTTATAACAACAATTTCACTGGTCCACTCCCCTTGGGAGTTACTCAACTTCTGCAGCTGAAGTATTTGAATTTCGGGGGTAACTACTTTTCGGGGAAAATTCCTCTGAGTTATGGTAGTTTTAATCAGCTTGAGTTCCTGTCTCTTGCTGGGAATGACTTGCACGGTCCTATACCAAGGGAGCTGGGCAACGTTACGAGCCTCAGGTGGTTACAATTGGGTTATTATAATCAATTTGATGAGGGGATTCCGCCAGAGTTGGGGAAACTTGTAAATTTGGTTCATCTAGATCTTTCTAGCTGTAACTTGACGGGTTCGATTCCAGCAGAATTGGGCAATCTTAATATGTTGGATACTCTTTTCTTGCAAAAAAATCAACTTACTGGTGTATTTCCTCCTCAGCTAGGGAATTTGACAAGGTTAAAATCTCTTGATATATCCGTCAATGAACTCACAGGAGAGATACCAGTTGACTTGTCAGGACTCAAGGAGCTCACATTGTTGAACCTCTTCATCAACAATTTGCACGGTGAGATTCCTGGATGTATCGCGGAGCTGCCAAAGTTGGAAATGTTGAATCTTTGGAGGAATAATTTCACTGGATCGATTCCTTCTAAGCTTGGGATGAACGGTAAACTAGTTGAAATTGATCTTTCTAGTAATAGACTCACTGGCTTGATACCAAAATCTCTATGCTTTGGGAGGAATTTGAAAATCTTGATTCTGCTTGATAATTTTCTGTTTGGACCTTTACCTGATGATTTTGGGCAGTGTCGAACCTTGTCCAGAGTCAGAATGGGACAGAATTACTTGAGTGGATCAATACCAACAGGGTTTCTTTATTTGCCTGAGTTGTCACTGGTGGAACTGCAGAACAACTACATCAGTGGACAACTCTCGAACGAGAAAATCTCAGCGTCTTCTAAACTTGAAGGGCTGAACCTGTCGAACAATCGCTTGTCTGGTGCACTTCCCAGTGCTATTGGAAACTATTCAGGGCTGAAGAATCTTGTGTTAACTGGAAATGGTTTCTCAGGAGATATCCCTTCTGATATTGGCAGACTAAAAAGCATCTTAAAGCTGGACCTGAGTAGGAACAACTTCTCTGGCACAATCCCCCCTCAGATTAGTAACTGTCTTTCCTTAACTTACTTGGATTTGAGCCAAAATCAACTTTCTGGTCCTATCCCAGTTCAAATTGCTCAAATTCACATCTTAAATTACATCAATATTTCCTGGAATCACTTCAACGATAGCCTTCCCGCGGAGATTGGCTCGATGAAGAGCTTAACTTCAGCAGATTTTTCCCACAATAACTTATCTGGATCAATACCTGAAACAGGACAGTATTTATATTTCAACTCAACTTCCTTCATTGGCAACCCTTATCTCTCTGGATCCGACTCGACTCCTAGCAACATTACATCCAACTCACCGTCAAAACTTGGAGACGGAAGTGACAACAGAACTAAGGTTCCAACAATATACAAGTTCATATTTGCATTTGGGCTCTTATTCTGCTCCCTCATTTTTGTTGTCTTAGCAATAATCAAGACAAGAAAGGGGAGTAAGAATTCAAATTTGTGGAAGCTGACAGCATTTCAGAAGCTTGAATTCGGAAGTGAAGACGTCTTGCAGTGCTTGAAAGACAACAACGTCATAGGGAGAGGTGGAGCAGGGATAGTGTATAAAGGAACTATGCCAAATGGTGATCATGTCGCGGTGAAGAAGTTGGGAATAAGCAAAGGCTCACATGATAACGGCCTATCTGCTGAACTTAAAACATTAGGAAAGATCAGGCATAGGTACATTGTGAGACTGCTCGCGTTTTGTTCAAACAAGGAAATCAACTTGCTAGTTTATGAGTACATGCTAAATGGAAGCTTAGGTGAAGTGCTTCATGGGAAGAACGGCGGGCAACTCCAATGGGACACTAGGCTAAAAATTGCCATAGAAGCTGCCAAGGGCCTTTCTTATTTGCACCATGATTGCTCCCCTATGATAATCCACCGCGATGTCAAGTCCAACAATATATTGTTGAACTCTGAACTTGAAGCTCATGTTGCAGATTTTGGCTTAGCCAAGTACTTTCATAATAATGGTACTTCTGAGTGCATGTCTGCAATTGCAGGGTCTTATGGCTACATTGCTCCAGGTAACTAGTTTGTTCATACATTAGTTGATTTAATTTCTTCTCGACTGGCCTATTATGCACTCGGACCAATCTACAACCTACTAATAATGGTCTTTACATGTTATCTTTTTGTATAGAATATGCATACACGCTGAAAATTGATGAGAAAAGCGATGTGTATAGCTTTGGAGTGGTGTTGTTGGAGCTTATAACAGGACGAAAGCCAGTTGGAAATTTTGGAGAAGAAGGAATGGACATTGTCCAATGGGCGAAAACGGAGACAAATTGGAGCAAAGAAGAGGTGGTGAAAATCTTGGATGAGAGGCTAAAAAATGTTGCAATTGTAGAAGCTATGCAAGTATTTTTTGTAGCAATGCTTTGTGTTGAAGAGTACAGCATTGAGAGGCCTACAATGAGGGAAGTAGTCCAAATGCTTTCTCAAGCTAAACAACCAAATACTTTCCAAATCCAATGATCTAATTGTGGCTCTACTAATTCTTATAGTATATCTTTTTTTGTTTCTTGTGTGCTTGGGAAGACCCCTTTTGTTAGCTTTGCAAACAAATTACAATTATGTTAGATCTTAGGGCATGATATCTAACTATCCTCTCAACTTGGAATAGTGTATTGGATGTGTAGAACTAGTATTAGTATCCGCGTGATGTGTGGTGAATATCAAAAGAAATTGATTACATAAAATGAATCTCTCTGTTTC

>M6-PGSC0003DMG400006364 M6 LRR-RLK

ATGTATCTTGCAAATTATGGGATTAGAGGTAACCTTCACCATCTAAATTTCACTTCTTTTCCCCATCTCCAATTGATTAATTTTTCTAATAACTCACTCAATGGTACACTTCCAAAAAACATGTTCAATCTTTCAAGATTAAGTTTCCTATATTTGAGTTACAATCATTTTGATGGTGTGATTCCACATGAAATAGGGCTATTGAAGAATCTCAAGCACTTAGACTTGTCGGTAAATCTTTTTATTGGCTTTATCCCTTCATCTATTGGTAACTTGATTAAACTTGAGTTGTTATTTTTATCAGTTAATGAACTCTATGGCCCTGTCCCTTCAAGTTTTGGTAACTTAAAATCCCTTGTGCATTTGTGTTTGCTTAGAAATAAACTCAATGGATCACTTCCAAAAGAGTTTGAAAATCTTACACATATGCAAATTTTTCAAGTAGCAGAGAACAATTTTTCTGGTCATTTGCCACAAAATGTGTGCCTTGGTGGATCACTTGCTAAATTCATAGCATATGACAATAAGTTTATTGGTCATGTCCCAAAAACCTTGAAAAATTGCTCTACGCTATCAAGAGTTAGGCTTGATGGTAACCAATTGTCTAGCAATATATCAGAAGCTTTTGGTGTCTATCCAAGTCTAGTTTACATGGATTTGAGTCACAATAAACTATATGGCGAGTTATCGTCACAATGGGGCTTTTCTCATAATCTAACAAGTTTGAAGATCTCCAAGAACAACTTGTTTGGTGCCATACCAGTTGAAATAGGAAATTTGATCAAACTAGAAATGCTTGATCTCTCTTCAAATCACTTGACCGGCGAAATTCCTAAAAGTTTAGAGAGGCTAACTCTCTTGCTGGTACTTGATTTACATGAGAATATAATTTCAGGAAAAATACCTATAGAAGTGGGGAAGTTGTCTAAACTTACAAGACTTAACTTAGGTGCGAATAATATGAGTGGAACGATTCCAGGAGAAATAGGAGATTGTAGGCAGCTTTGGTACTTGAATTTAAGCAAGAACATGCTGAACACAACTATTCCTTCTAATTTAGGTAACTTGCACTCTCTAGCGTATCTTGATCTCAGTTATAACATGCTCAGTGGTGAGATACCGTGGCACATAGGGAGTTTGAGAAGTTTGGAAAGAATGAATCTTTCGCGTAATAACTTGTCTGGTTCAATCCCGCCTAGTTTCAATGAACGTGTTAGTTTGAGGTCAATTGATATATCTTGCAATCAATTGGTTGGTCCCCTTCCCAAGATCGTGGCATTTCAAAATGCATCGCGTGAAGAATTAAGAGATAACAAGGATTTATGCAGCAATGATCATATTGGTATGAGACCTTGCTCTTCATTGAGGAAAAAAGAGAGAACAAGTAGAAAGCTTATACTGACCTTAACTCTGTCTCTAATTTTGGTTGCCTTGCTTCTACTTGTGATTACTTGTATACTTTTTCGAATGAAAAGGAAAAGAAATAGAAGTATTCAGCCTAGAGAACCATCCAGCAATTCTTTTTCAGTTAGGGACTTTGATGGGAAAATAGCACATGAAAACATAATTGCAGCAACAGAAAACTTTGATAGCAAATATTGCATTGGGAAGGGAGGACATGGAAGTGTCTATAAGGTTGAGTTACCATGTGGTCAAGTTGTTGCAGTGAAAAAAGTTCATGCTTTAGAAGATGAAGAATCTGATGACAACTTGATCAAAAGCTTTTCAAAAGAGATACAAGCTCTAGTAAACATCCGCCATCGGAATATTGTGAAGCTTTATGGGTTCTGTTCACACGCGCGCCATTCATTCTTGGTTTATGAGTTGTTGGAAGGAGGAAACTTGTCTCAAAACCTTAGCAATGAGGGTAGAGCAAGAGATTTAAATTGGCTCAAGAGGGTTGATATTGTCAAGGGGATAGCAAACGCGTTGTGTTACTTGCATCACGCGTGTTCACCTCCTATAGTTCATAGAGATATATCAAGTAACAATGTTCTCTTAGATGATGAAGGCAATCCCCATGTTTCTGATTTTGGTACTGCTAAACTCTTAAGGCCTAACTCCACCAACTGGACATCGTTTGCTGGAACTCTTGGCTATGTAGCTCCAGGTATTTAGTCTTCTTTATTTTCTTAAACTTCGTGTTAAGTCAAAATCAGACAAATAAATTGAAACAAATGGAGTATATTTTTAACTAGGAATTCATTGCTTCAAAATTGTTCACTCTTTCACACAATTTTCTAATTTGTTTGTAGAGCTTGCTTACACAATGAAGGTAAATGAGAAGAGTGATGTTTACAGCTTCGGGGTGATATCACTAGAAGTTATTATAGGTCATCATCCAGGTGATATTACTCAAGCTATATCATCATTTTCGCCAGCATCAGGAGCGAATGGAACGCTGTTAAAGGAATTGATTGACAAACGGACACTAGCTCCTGGAAAACAAGAAGCAGAAGAATTGATCAAGATCACAAAACTAGCATTCGCGTGTCTTCATCAGAGTCCTCTAGCTCGGCCAAGTATGAAGCAAGTTTGTGCATCTTTATCAAAAGAAAATTGGCCTTCAAAAGGCTTGTTCTCCACTGTTACATTAGGCCAGTTGATTGAACCAACACTACTAAGATGTTGA

>M6-PGSC0003DMG400006502 M6 LRR-RLK

CCATGGCCTATTATCTTGACAAAATCTTCCTTCTCTTATTTCTCTTCACACTTAGTACTCCCTCAAAATCAGATAATATGACAGATATTGAAGCACTCTTTGCTATCAAGAATGAAATAATTGATCCCTTTGAATCATTAAGTTCATGGAATGAGTCTGTTCCTTTATGTCAATGGCGTGGCGTTGTATGTGGAACGCGAAATCAACGAGTCATTGAACTAAACCTTCTTGACCATAAGCTAGCCGGTGTGTTATCACCCTTTGTTGGAAATTTGAGCTTCCTCATACGACTAGACATCGGAAACAATACCATTTCTGGCAATATCCCTCCTGAATTAGGCCGTTTGACAAGCTTACGCCATCTATATTTGCAAAACAATTCATTTTATGGTGAAATTCCTGTCAATCTATCGTTTTGTAGTAACCTCGTTGAGTTATGGGCATACAAGAACAATTTTGTAGGTGTTCTTCCAAGGGAGCTTGGTTTACTTACCAAACTCAAATATTTAGATTTCGGTTACAACAAGCTCATTGGTGAAATCCCAAAATCTTATGGTAACTTTTCAAGTCTATTGGAAATGTACCTTTTGGATAATGATCTCGAGGGTAAAATCCCCGATGAGTTTGGGAAACTGAAGAGTTTGGAAATATTTGACGCGGACTTTAATAGGTTATCTGGTAGAATTCCGAGCTCATTGTACAATCTTACCTCATTGAAAGTCATTGATGTGTCATTTAACCAACTCGAAGGTACTCTGCCTCGGGATTTAGGGATCAACCTTCCTAACCTGGAGTGTCTCAGCATTAGTGAGAATCAATTCACTGGATCGCTTCCTCATTCACTTTTCAACCTCACAGGACTCGCGTACCTTTATGTTGGAAAAAATGACTTAGGAGGACGTGTTCCTAGGTTCGATAAGTTGCATAATTTGTATATTTTAAGCATGGAAAATAACCAATTTGATAATGACTTGACCCTCATGTCCTCCTTGACAAATGCAACTAACCTCAACCGTCTGTATATGCAAAACAACGCGTTTGGAGGGACATTTCATGAGTTTTTCAGCAACCTTTCAAGTAATCTGGCGTTTGTAGACTTGTCAAGCAATCATTTTTCAGGACATATCCCGATAGACATTGGGAAATTCGTCAACTTGGAAGAGCTTAGGCTGAGCGGAAACCAACTCACAGGTATCATCCCTTCGACTATTGGGAAGCTTTATAAGCTACGATATTTACACCTGTCCCAAAACAGAATTTCGGGTAGTATACCCTTTTCCATTGCAAATCTGACCCTTCTAACAGAGTTAAATCTTGATCATAACAATCTTAGAGGTAGAATACCTTCAAGTGTTGCAAAATGTCAAAGTTTGTTGTACCTTAACCTGAACCAAAACAATTTAATCGGTTTGATACCTAAAGAAATTTGTCTAATCTCTTCCTTAGTTGTTCTAAACCTTTCCAGTAACAACTTGACTGGCGCCCTTCCCAAAGAAATCGAAAGCCTGAAAAATCTCTTATCATTAGATGTCTCTTTGAATAATCTCAATGGTGAAATTCCTAGTACACTTGGGAGTTGTGTGACAATTGTTAACTTAACAATGAAAAGGAATTTATTCCGCGGGATTATTCCTTCCACTTTCAGTTCACTAAAGAGTCTTCAAGTTTTAGATCTTTCGCGGAATAAGCTATCAGGTATGATTCCAAAGTACTTGGAAGGATTTGCCTTGCATTTCTTAAACTTATCGTTTAATGATTTTGAAGGTGGCCTTCCACAACGAGGCATCTTTGAGAACGCGAGTGTAGACTCTTTCGTTGGAAATCTTAGAATCTGTGGTGGTGTACCTGGCTTGAAACTTCCCAATTGCAACTTCAGTCATTCAAAGAAAATGAACTTTAAGCTTGTTATTTTAGTAATATTAGGAATTTTGGGGCTAGTTGTTATGGTTTGTGCTTTTTTCTTCTATCGATTTGGAAGGTCCAAAAGAACAATTCCTTCATTAGATAACAATCTAAACCAACTCATAGCTATGTCCTATCAAAGTATTCTCAAAGTTACCAACGGGTTTTCTACGAGCAACTTGATTGGTGTTGGGAGTCATGGATATGTTTACAAGGGAATTTTAGAAACGGATGGTAAACACGTTGCTATAAAAGTGTTGAATCTTTTACAATATGGAGCTATCAAAAGTTTCATAGCAGAATGTGAGGCCTTGAGAAATGTTAGACATCGGAATTTAGTTAAGCTACTTACTGCATGTTCTGGTGTTGATTATCATGGCAACGAATTCAAGGCACTGATTTACGAGTTTATGGCCAACGGAAGTTTGGAGGACTGGTTGCATCCAGATAATTCAAGACCGAATGTTCAGCCAAGAAGGTTAGGATTTCTACAAAGGCTGAATATTGCTATTGATGTTGCTAGTGCAATTCATTATCTTCATAATGATTGTCAAATATCAATAGTTCATTGTGATCTCAAACCAAGCAATATTCTCCTTGACAATGAATTGGTTGCCCATGTTGGGGATTTTGGATTGTCAAGATTCCTACATCTTACTGACGAAACCACTTGTAGAATTCAGACAAGCTCTTCAACTTTTAAAGGATCTATTGGCTACATAGCTCCAGGTAAATTTCACTCTATATAAATGCCTCTAAAGACTTTAGTGACTCTGGATGCAATGACATTAACTCAATTGCCTTTATATTTTCATTGCAGAGTATGGTATGGGAAGCGAGGCATCAACGCAAGGTGATGTGTATAGTTTTGGTATCGTCTTACTTGAAATGTTAACTGGGAAAAGACCAACAGATGACACGTTTGGCAGCGATTTAAGCCTTCATGATTTTGTTAGGAATGCCATGCCTGATGGAGCACTTGAGATAGTAGATCCAGTTCTTAATTTTGAAGAGGAAGAGATAAGCACAGAGAGGTCTCAAATTCCAAGGTTCATGAGAAGGCAAAAAATGGTAGAGGGTTTAATTTCCTTATTTGGAGTTGGTATTGATTGTTCAATGTATGATTCAAGTAAAAGAAAAAACATGAAGGAGGTTGTTAGAGAGTTGTGTTCTATAAGAGATTCTATTGTTGGTTGTACAATATGAAACACTTTTCTTGTTTTGTTT

>M6-PGSC0003DMG400006563 M6 LRR-RLK

AGTAAACATACAGAGCTAAGGAGATCAAGTGTCATTACATTTCTGTATATTTAGGTCTCTGAAATGCATAGCATAGTTATTTAAAATTTATGTTTACTTGTGAGTAAGATTTTGGAACTTGCGTTGTATTTTTAAAGGTGTGAAATTATGAACCAATGCAGGTATTCTTTAAAGCCAATAATTGAAAAGTTGGATAAACATATGTTTTTTGCTCATGGAGCTGTTAATTGTATAACGCCTGGCCATTTCTTTCGTGATTAAGGTGGTGTCTTGTGAGGACCTTGATTGGTCCAACATATACCTGCATTTACCTCTAAGCATTAGGCATATAAGGAAGAAGATGCCTAACTCTTTTTTTGTCAGCTAGGATTCAAACTCTAGATTTCATGATTCTATATTCATCGAACACTAGACCACACCCTTGGGTTTGTATCCCTTTTCGACTTTCAGCTTGTGCATTTCTTCATCTGTTGTATTCCCTTTCTCTCTTTTGATAATGTTTGACCCTTTATTTGCATTGTGGTTTCTCTTTCCTATTCAGTCTATCAGGAAAATGGTGTGAGGGTTCAATAAGTTCACTAAACATCTGAAGACTAAGATGCTAATATTGATAGTGATCAGTTCAATTTCCACCTTTAACTTTTTTAAATTGCATCGATCCTAGCAGAACTCTTATTCTATTTTTACTTTTCTTTACATGTTAGATGATGATGGAGAGAAAGGATATGAAGGTATAACAGATATAATGAATTCATGATACATGTTGTGAAAAGACAACTTACCAAAAAGTTATATATGAGAAAGGAATAAATCGAAACATATTTGGACTCCCACATTATGTTTGAATTTTTGAAAGACAGATTATGGAATATTTTATCATGATAGAACATGTTGGTAGAAATAAACATGAAGTTAGGTCCTCATCTTGCACACTTATTGAAGATCAAATGCTGATTCTGATATTGCAGAAGTGTCAGAAATTGCTCTTTGCAAGGATCCATTCCTGATTTCAGCAGAATTGCAAACCTAAGCTATCTGTAAGTGCGAGTTTGTTATTGTTTTGTTCTTCTTTTGTTAAGCTTTATCTGTATATTTTTACTTCAAGATTGTTCAACTAGTAGATCTCTGAAAATGTCTAGCTTCTGCTTGCTATACAGTGATCTCAGCTGGAATCAACTTTCTGGATCTATACCACAGAATAAGCTTTCCAACAATATGACAACTATGTAAGCCTTTTGATTTGGCCTCGATAGTTTCTTCATATAGTTATATAAATTGACAGATGTTGGCTTTTGTTTCCACAGCATTTGAACTGGATTGCCTGGGTTCCTTTTACTTATTAGCATATTCTAATGTATATTTCAAATGCATGATGCAGTATTCTATCACATAACCGTCTTAATGGATCTATACCCAAGAGCTTTTCTCTCCTTCCTTCCCTGCAGAAACTGTGAGAATCTGGACTTAGTCTTGCAATATCTACTATTTCACTGAAGGAAATGTACCATTAACGAATTGGTGGGACAAATTTGTCACAGGTCTCTTGAGAACAACTTTCTGAATGGTTCCATTTCCACTGACATTTGGCAAAACAAGATCTTTAATACAACTTCCAGACTTGTGATGTAAGATGTGAAGCAGAAAAATAACAGATCTTATTTCAATAGAAGTATTTACTTCCAGAGTCAAACCTTTGAATGAGGATTTCTTTCTAACAACTCTTTCATTGATTTTGCAAACCTAGTGATCTGCGGAATAATTCAGTTTCAAACATTTCGGGTGCTTTTGAACCTCATGTAAATGTCACACTAAGGTATGATTTCTTCTTTCAAATTCAAAAATAGAACAATTTGAATGTCGGTTTATGTCCTCATGGGCCACATAAATAGGACAAACAAATGGGTAAAGAAACTAGAATAACAGAAGCAATATGCCATACCAAAAAGGGAAAGTACGTCGATTTTCCTCGTGTGTGTGACAACAATTACTTAAATTACTTGAAGAAGCACAAATGCATGGAATCACTTCTGTTATGTTTTATCCACACACACCGATACACATGCTTTTAGCAAATTCATATATGGTGCTTTATAAAATGTAAACAAAATGGAAAGAACAAGGAGTTGAGAGAACCTTTGTGGTTATCAGTTTTTTCATTGTTTGAAAAATTTAAGTATGCTTAAAAAGGAGGTATAGATAGAAAAGGGAATTCTGAAATTCTTAGTCACATATTTTCATCTAGTTGCAGTGGAAAAGGATGAAATGATGAAGGGCCCTATAATATTGGTTATGGTCCTGATCAGCGTAAATCTTGAAAAAGTACAAAGTTTTACCCAATAATATAAATGAGAAGTGTGTGTGTGTGGGGGTGGGGGGTAAAAAGATATAAATAGAGGGAAACTATGAGCATAAAAGGAGTTTTTGTCTTTTGTTGAGTCTCTCATTTATTCGTTGAAAGGAGGCCTGAACAAATGGAGAGGAAGGAAGATTCTGTAAAATTTGAGTCCACTACATGCCTTATGATGATTTTTTAATGTGACACAATGTAATTCTAATCTAAGTTGCTCGGACTCTTCACTTTTGGTGCCGCACCCGTGTCGACAGGATATGGGTGTGGGTGTGGGATCCGTGTCCGATTTTGGGTATTTGGACCAAAATTGATGGAGAAAGTCTGGACAGTTTTAATGATTTTTATAATCAATACAAAAGCTAAGGTGAAATTGAAGAAAATGGAATACCTTTTATATAGAAATTTCTATGTCACTCATTTTCCTTTGATCTCCTTCCAAGATTCTCTTCAAGTTCTCCACATAATATCTCATAATATAGGTTTTATAACTCTTTTAGATATTTTAATTATATTTAGGCGAATCCCTGCACCCGTATCCATTCATGGATCCATATCCCTGAATCTTAAAAGTTATATCATATCATGAAGGATCGGACCTTTAGATCCGCACACGTATCGGACACCCACACCCTAGTCCAAGCAACTTAGATTCTAATAGTTTGTGAAACTGAGGGAGATTTTCTGCAGTATATCACATCAAGACAAGTGATATAAACTGGTATCATTAAATTCTTGAAGAAGGAAAACATCACATTTAGGAAAGTGGAGAATGGAAATTGTGCCAAACAAAAAGAGTTATTGGTCTAAAATTACTTGTACATCTTTGTTACTGTCTCTGGAGTTCATGTTTTCAGTTATGCACTCATTCTTGTTGTCCAATTATTCATAAATATCTGATTTTGACAGTGAATTGAGGTCATCTTTCAAAAATTACTTTCTGATCTCTTTCTATCTCATGAGGTCAATTCATCTTGAATTCTTAATTGAAAATCTGCTCAATGTTTCTCAAAAGAAACATGTCAATAAAAATGTTTTAGATTTAGTACTATGTTAAAATTCATTTATTATTTTCATAGTCTTCCCCTAACTTTAGCAGACGATTTCCAGGTTCCAAAGCAATCCCATCTGCAGCAGTACAAGTATCAGAAACATTGGTAAATACTGTGGCCCTGACTTTGGAGCAGATGACGACGAACCTAGTAATTCAACAAATCCAAGTGGTGTTTGTCCTATTCATGCTTGTCCAACTGACAACTATTATGAATATGTTCCTGCATCACCTAAACCATGCTTTTGTGCCTCACCTCTTAGAATTGGGTACAGGTTGAAAAGTCCTAGTATTTCTTATTTTGATCCGTACAAGCAACTATTCGAATCTTATGTGACTAGTTCCCTTCAGTTGGACCTTTACCAATTGTGGATCGATTCATATTTTTGGGAAAAAGGACCTCGTCTGAGGATGTATTTGAAGCTTTTCCCGGTGGTTGGGGACAATACATTTAATGAAAGTGAGATTCTTCGTATCAGCGAGATTTTTGCCTCTTGGGAATTTCGAGGAAGTCACCTATTTGGACCATATGAGCTCCTGAACTTCACACTCCTCGGACCTTATTCACACTGTAAGTACTCACAGCTTAAATTGATTCTAATTCAACTTGTAACAACTTAGGCTGTGTTACTACAATTGCATGACAATTGAATCATTGACATGCAGCATGTGGATAAGATTGAAATATTTGACTATCATTGCCACTTCTAGCCAACAAGTATAAGGATCTTAATTCATATAATCAAATAGTTTAGATAAGCAACAAAAGACTATGATCCATGATCTTCTAAGGACTTCATATATATTGGGTTGAAAAAGACAATGAAGTCTTTGTTTATATCCAAAGATTTTTTAAGACTTCAGATATTTTACGTTGAGGAACCTTACTTGATTAAAAGGTTGAAACTTCAAGGGGTTCAAACTTAGATAGTTTGGTCCAGTCTCTCTTCAAGAGCTTCAATTCTGGAGGTCTCAGCCCTTACAGTTTTTGTTGGGATGAAAGTGGATAAAAAAAAATGGACCTTAGGCCTCCCAAAAAGCTAGCTCATGAGGAGGGGATTGTCCAAAATCATATAAGGAGAATAACCCTTGTCACTCCAACACCTTGCTGCACTCCCAAGAATTGACATTGGAGCGTGGACAATATAATAAGGGGGGGTTCAACACCGGAAACAATAAATTGGGATCAGTGTGGATCTGAGACCATGTGGGAAATAAAGAAGAAAAATATTACTCCCTTTATCCAGTTTATATGACTTACTTTTCTTTTTAGTCAGTCTCAAAAAGAATAACACATTTCTATATCAAGTAATAATTTAGTTTTAAAATGTCAATTTTACCTTAATGAAATGATTTATAGCCACACAAATATCTATCACTTTTTTTTAGACCACAAGTTTTAAAAGTATTCCTTTCTTTCTTAAACTCCGTGCCAAGTCAAACTAACTCATATAAAATGGGATGGAGGGAGTATTAAAATTATGTATTCTTTTATTACATTGAGCCCTATTTATAGATTCTACATTACAATTCATTTCCAAGTAGGACAATATACACAATCCTATTCGAACACTCCCCCCTTAAGCTGGTGCATACAAGTCAGATGTACCGAGTTTGTTAAGATGTAATTAATATGAGGATCAGTGAGGGACTTGGTGAAAATATTTGCAAACTCATCACTCGACTTCACAAATTTTGTAACAATATCTCTTGGGATCATCTTTTCTCTGACAAAGTACCAGACAATCTCAATGTACTTAGTCTTCTCATGAAATACCACCGATTTGATGTGATATCAAGGGCTGCTTGGTTATCACAGACAAGATAATGGCTCAATTGTTTGATTTCCTTTGTACAAGCTTTGATTTCTTATATTATCTGCAGTGAATCCTGAAATGAAAGGTAAAAAACAAAGCAAAGGCGTTGTGATAGCTTGCATTGTAGCAGCAGGTGCTTTTGCTGCCTTTGTTTCCTCAATTGCCACTATATTAATCACAAGACGACGTGCTAAGTATCAAAAGATCTTGTCAAGAAAACGTTTGTGTAAGTCCTCTTATTAGATTAGCTATAATTTTACTTTATCAAGTCTGCTATTTTTAATTCTTTCTGCAAAATTGATGACTATGGGTTTTTGTTTGCAGCTTCAAGCCTCTCTATAAAAGTAGATGGCGTTAAAAGTTTCACCTTCAGAGAAATGACTTCGGCCACTAACAACTTTGATACCTCAACTCAAGTTGGTGAAGGAGGATATGGTTCCGTCTTTAGGGGAATTTTAGCTGACAAAACAGTTGTTGCTATCAAGCGAGCAAAAGTAGGATCCCTGCAAGGGCAAAAGGAATTCTTGACCGAAATAGAACTATTATCAAGGTTACATCACCGTAATCTGGTTGTGTTGCTTGGATACTGTGACGAAGAAGGAGAGCAGGTATGACAAAAGCTGAAAACCTTTTCAATTCCAACTGTTTTAGTTACTAATAATTCACACCATTCATCATTTTTAATCATAGTTAGTCATATTTATTTAGTGTAGCATCATAAACCTTCTTGTATTCTTTCTCACTTCAGTTGCTAATGATCAGACTGAATGTGTAGCAGACTAAATGCTTTCTTGAGCTACTGAAGTCTTCTTGTTTAGCACAAAGAGTGATATAGTCGAGCAATTATGAGTCTTCTGTATACATTTGCAGATGCTGATTTATGAGTTCATGCCCAATGGAACTTTGCGAGACTGGCTTTCTGGTAAGAACTGTTTCCTGGTTGTTTCTAGCTTTCATATGATTAATCCTCCTCAATTTCTTTCTTTTAGTTTAGTAGTGAAGATCCAATGCGGAGTTGCAATGTTTATAGATGTTCCTTGCACATGTTGTGTCATTAGTAGTACTATGGTGAATTGATAATACAGGCTTCAGTATTTTATTGTGTTTGTGGAGAAACCATTGTCAAGGTGGTGATCGATCTAAAAGTTGCATAATGGAAAAGATATCGCAGGAAATCTAATAGTCAGAGAAACATAGATTCCATCTTCTTTTTCCTTATCAGAACAAAAAAATTTGTTAGTTTCTTTATGCGGCTGTGATAAACTTCAAGAAACTTTTGTATTGCTATCTAAGTAGGTGATTTCCATCATTTGACAGCCAAATGTAAAAAAATTTTGAAGTTTGGAGCAAGGTTGGGGATCGCACTGGGGGCATCTAAGGGCATCCTTTACCTCCATACTGAAGCTGATCCACCCATATTCCAAAGGGATATTAAAGCCAGCAACATTCTTTTAGACTCTAAACTCACTGCCAAAGTTGCTGATTTTGGATTGTCTCGGCTTGCACCTGTCCAGGATGATGAAGGTCTTCTGCCTAATCATGTGTCAACAATGGTGAAGGGGACACCTGTAAGTCTGTCTATGCTCTAGATTTAAGGAAATAATTGTGCAATGATCTTACTGATTTTCTTATGTATATGCACAGGGTTACCTTGATCCCGAATATTTCTTGACCCGTAAAATGACAGATAAAAGTGATGTCTACAGCATTGGGGTTGTATTTCTGGAGATCTTAACTGGCATGCATCCTATCTCACATGGAAAAAACATTGTCCGTGAGGTATTTCTGTCGGCTGAACACCTATTGGAATACAAAAACAGTGAATATTTGTCTAGCATTTTAGCAATAAAAGCATAGGTGTATGTATATAACCATTAACTGTCCATGACAGGTAAAAATGGCTCATAAATCGGGAGTAATGTTCTCCATTATGGATAAGAACATGGGATCTTACCCCTCCGAATGTGCGGAGAAGATGGTGGAGTTGGCCCTGAAGTGTTGCCAGGACAAACCAGAAGACAGACCCTCAATGCTGGAGGTGGTTAGGACGCTAGAAACCATACTCCAAATGATGCCATATACTGACGCCGATCCTTTGGATGATAAAGCCAGTTTCAGTGAAACTACATCATCAGCTTCATTTTCTAATACTAGAAGTGGAGACCTATTTATATCATCATCCAATGTATCAGGTGGTGATCTTATTAGTGGTGTTACTCTTAATATTACACCTCGTTAACATTAGCTTGTCTTCCACTGCATAAACATTGGCTTGTACACTATGACTAGTAGGAAGTCTATACGACTTGTCTTTTATGATGTATACGAGTGTACAATAATACCATCTTACTGCTTCTAGATTTATGTATATTCCTCAAGAATCTGTTGTGACAGACAACCCTT

>M6-PGSC0003DMG400006923 M6 LRR-RLK

ATTTTGAAGTCACTGTTAAATGGGTTTCTTCAAGATTTTTGTGATAGTGTTGGTTATTTGCGCTTTCTTTATAGCAAATACTTATCAGTTACAGTCCTATGAAATTCAAGCTCTTCTTCAGTTGAGGAAGCACCTGGAATATCCAGTTCAGTTGGATGTTTGGGAGAATTACCATGGAGATTTTTGCAGCTTGACTTCAACACTGCATATGAGCATTACCTGTCAAGATAACTCTGTTACTGAACTCAAAATCAAAGGAGATAAGTTAGTTAAGTTGAATGAATTTCATGGATTTGCAATTCCAAACCAAACATTATCTGAGGGTTTCTCCATTGATTCTTTTGTTACCACATTGACAAGGCTAAGTAGCTTAAAGGTTCTTACTTTAGTTTCTCTTGGTATTTGGGGACCCCTTCCTGATAAAATTCATAGGTTAGCTTCACTTGAACTTTTGGATATGAGTTCAAATTTTCTATTTGGTTCTGTTCCTTTTCAGATGTCAAGATTGATTAAGCTTCATACTTTGACATTAGATGCCAATTTTTTCAATGAAACTTTCCCTGATTGGTTTGATTCCTTACACAATCTTACTATATTAAGCTTCAAGAACAACAGGTTGAAGGGTCAGTTTCCTTCTTCAATGTCTAAAATCACAACCCTCACTGACATTGTTCTGTCACACAATGTGCTTTCTGGTAAATTGCCCGATTTAACGGCCTTGTCTAAGTTGCTTTTGTTGGATTTAAGGGAAAATCATTTGGATTCTGAATTGCCACCCTTGCCAAAAGGACTTACCACAGTTCTTCTTAGCAGTAACGTTTTTACGGGTGAAATCCCAGTAGAATTTGGCACACTAAATCAACTTCAACACCTTGATCTGTCGAACAATTCTCTCAGTGGAATGCTGCCTGCTGGCTTATTCTCTTTGTCTAGCATTAGTTACTTGAATTTAGCGTCTAATGTTCTTAGTGGTTCACTTCCTAGCCATCTCAATTGTGGGGGTGAACTTGGTTTTGTTGATATTTCTAATAATAGATTAGTTGGTGGGCTTCCTTCTTGCTTGAACACTATTTCAGATAAGCGAATTGTTAAGGTTGGTGGAAATTGCTTGTCTCTTGACACTCAATATCAGCATTCAGAGGGCTATTGTAAACAAGCTAATTCGGATAAGAAACAGATCACAGGAAAACAAATAGCATTGTTGGCAGGTGTTATTGGGGGAATTGTAATTGTTGTGGTGTTTCTGTTAGTTGTCCTTTTCATCTTCCGTAGAAAACGACATGCGCGTAACATGGTGGATAAGCACATTCCTCCGAAAGTTGTGCAAAATACACAGCCCGGGGTTCCCTCTGAACTCCTAGCAAACGCCAGTGAGTTGATGTTTTTCTGATTTCATTTTTAATCTGTTTAATTTATATGAAGGAAATGTTCTCCAGAAAAGCTTGATCTCAATCTTCGAATTCTCAGGAGTCATTTCTCAAGTGGCAAACACAGGATCACAAGGTGCCCCATCATATCGAGTGTTCTCCATGGAAGAATTGCTAGAAGCAACAGAAAATTTTGATCAGTCAGCATTACTTGGTGAAGGGTCAGTTGGAAAAGTATGGCTTCAAAACAATACTTATG

>M6-PGSC0003DMG400006942 M6 LRR-RLK

GGTGCTTACTCAGGGAGTCCACTCCATTAACAGGATTAACAATAAATTAAACTTTACAAATTCTCTAAATCCATAATTCCAAGGTTCCACTTCATCATTCAAAGTCAAATGTCATCTTCTATTCTTTTGCTTTTTTCATACACTTATATTTCTCATGTAAACAATTCATCAAACACCTTCTATGCATCCCTACTCCCTAAAGCTCAAAGATGAAACTTTTCAGATTTTTCAATCTTCCCCTTATAGTATTCATCTTATTGATTCTTGTATTTCAACCCCTTCTTCATTGTACTGCTGTTGCTGGTGCTGTAGCTGAAGATGATATAAAGTGTTTAAAAGGAGTCAAGAACTCCTTAACAGATCCAAAAGGGAACTTGAATTCATGGAATTTTGCTAATTCTACAGTGGGGTTTATATGTAAATTTGTTGGTGCTTCTTGCTGGAATGACCGTGAAAATCGTCTCATAAATCTTGAACTTCGGGATATGAATCTTGGAGGTAATGTTCCAGATTCGTTAAAGTATTGTCGAAGCTTGCAATATCTTGATCTTTCTGGTAATCGGATATCTGGTTCAATTCCTTCTGATATTTGTACTTGGCTACCTTACTTGGTAACCCTTGATTTGTCGAACAATGAATTTACCGGTTCTATTCCGTCTGATCTTGTTAGTTGTTCCTATTTGAATAAATTGATGCTTAATGATAATAGGCTTTCTGGGAATATACCACCTCAGTTTTCTAGTTTGGGGAGGCTTAAAATATTTTCTGTGGCAAACAATGATCTTTCTGGTAGGATTCCGGAAGCTTTTGATTCGGCTGAGTCGTTTGATTTTGGAGGAAATGATGGGCTTTGTGGTGGTCCGTTGGGGAAATGTGGGAGGCTTAGTAAGAAAAGTTTAGCTATTATTATTGCTGCCGGTGTGTTTGGTGCTGCTGCTTCTCTGTTGTTGGGTTTTGGGGCTTGGTATTGGTATTTTACGAAGGCGGGGACAAGGAGGAAGATGGGGTATGGATTAGGGAGAGTTGACTCGGAAAGGTGGGCGGATAAGTTGAAGGCTCATAGGCTTACTCAGGTTACATTGTTTAAGAAACCGCTTGTGAAGGTTAAGTTGGCGGATCTGTTGGCTGCCACGAACAATTTTAGTACGTCGAGTGTGATAAACTCGACTAGGACGGGGACTACATTTAGAGCTGTTCTACGTGATGGTTCTACGCTTGCTATTAAACGGCTTAAAGCTTGCAAGCTGAGTGAGAAGTTGTTTCGAATGGAAATGAATGGACTAGGACAAGTTAGGCATCCTAATTTGGTACCACTTTTGGGCTTTTGTGTTGTTGAGGAGGAAAAACTTTTGGTTTATAAGCACCTGTCAAATGGTACTTTGTATTCATTATTGAAAGGGAGCGCAAGTGTGTTGGATTGGCCTACTAGGTTTAGGATTGGCTTGGGTGCTGCAAGGGGCCTTGCTTGGCTTCATCATGGTTGTCAACCACCAATATTGCACCAAAACATATGTTCTAACGTTATTTTCCTTGATGAGGATTTTGATGCTAGAATAATGGATTTTGGGTTGGCAAGGCTGGTGACACCTCCAGATGCAAAAGAGACTAGTTTTGTGAATGGGGAGTTGGGTGAATTTGGCTATGTTGCTCCGGAGTACTCAAGCACAATTGTGGCTTCACTGAAAGGGGATGCTTACAGCTTTGGGGTTGTGCTTTTGGAGTTGGCTACTGGGCAAAGACCTCTAGAAATCACTGCTGCTGATGAAGGTTTTAAGGGAAATTTGGTGGACTGGGTAAATCAGCTCTCTGTTTCTGGTCGGATAAAAGATGCAATTGATAAGCATATTTGTAGAAAGGGCCATGATGAAGAGATTGTAAAATTCCTCAAAATTGCTTGCAATTGTTTAATTTCTCGGCCCAAGGAGAGGTGGTCTATGTATCAGGTATATGAAGCACTGAAGAGCATGGCTGAGAAACATGGTTTCTCTGAACATTATGATGAATTTCCCTTACTATTTAACAAACAAGAGACCAGCAGTCCCATTTGATACATGAATGGGAAAGCATTACGAGATGCTTCACATCTGGTGAACACAAATGCATTATGGTACGTTCTTCACTTCTCTTAGGTGCAAGTGGAATCAGGCACGATACATATGGTTCTGTGAAGTACTACCACAGATATTGCTGTATTGTACTGTGAAGTGTCAAATCATTCTGAGGTGTTCTGCTCGAAATTTATGTTACAGATAGCTAATATTATTGGTATCTTAGGTTTATGTTTCTCTTTGCTCATACTTATGCATTAAACAAGGCAGTAAGCAAAAGCTGTCACAGTTTATTTGAAAGCTGTAATACCACTAATTTAGCTGTATATAACTGACCTGGGCATTGGCCTTTTTAGGGTCGAGATATCTTCAGTATGTCCCTTACAGTCATTTATGGTTTGTAAGATCAAATCACACTAGATTTCCCTGATAGATTGGTTCGGTAGAATATCTGCCTCCACACAGATATTAATTCATCTAACCGGAATGCTATCAAGCAGAGTTATTAGTATTTACTCCTTCTGATTAATTTTGGATGATGTTGATCAGTAATTGTAGGTCCTTTTGCAAAATTTGTGTCTGAATGGGAATAATTTTCACTAAAATGGAAGTTGAATGGGCTTTCTAAAAGTTATGTGCCCTCTTGAATTTGATTTGGCAGCTGAAATTTTCATTCACTTGATTCTAATCTGCTTCTTTACAATCTCAGAAACATAATGCACAATTAAAGTTTCCAAGTAGGAAAAAAAATAATTCTTGTTGACAGTTGAAACTCTTTGCAATAGTAGTTATCAAACCTCTTTACTTTTTCCCTCAAGCTCAAGCTCGGATGCGTGTTTCCATGTGCTCTCCTGCTTCCCTGAAAAGTCATTTTGTAATATTTCAAAACAATAACTTGTAGTCTAATAGTAAATGAAGGGAAAAGGTCTGTTAGCAAGTTATC

>M6-PGSC0003DMG400007234 M6 LRR-RLK

CTCATCTTTACTTGCCTTATTTTCCTCAACCAATTTTTTCTTACAATCTCATAACCATGAACCATTTCTATCTCACTACAATTTTCTTCTTTCTCATACAATTTTCACTTGTTTTCTCTACTTCAAATTTTCCTCTTCAACTCATTTCACTTTTATCGTTAAAATCTTCATTTCAAGATCCAAACAATACATTTCAAGATTGGAATCCTACATCAACTTTCTCAAATTTTGGTTCACAACCCTTTTGGTGTTCATGGTCAGGCATCAAATGTGACACAAAAACAAGTCACATTACATCACTTAATCTCTCAAAGAGGAATCTTTCTGGTAAAATACCTCAAAATATTGCATCCTTGATACACTTACATCACTTGAATTTAAGTGGAAATTCATTTGATGGACCTTTACCATCCATCATTTTTGAATTTCCATTTCTTAGAACACTTGATATTAGTCATAATTATTTCAATTCAGCATTTCCTCTTGACGGGCTAACGAATCTCAAGTCCCTAGTACACTTGAATGCCTATAGCAACAGCTTTACAGGCGCGTTGCCTATAGAGCTTGTTGGGCTACAAAATCTTGAGTATTTGAATCTTGGTGGCAGTTACTTCACTGGTGAAATTCCGGCAAGTTATGGGAATTTTAACAAGTTGAAATTCCTACATTTAGCCGGAAATTCACTGAGTGGGAAAATTCCCGAGGAATTGAGATTGTTGAACCAACTCGAACATCTTGAAATTGGATACAACAATTATACTGGAAATGTTCCTCCTGGATTTTCCTCGTTATCGAATTTAGCCTATCTTGACATCTCACAAGCTAATCTTTCAGGTGAAATTCCAATTCAGCTTGGAAATTTAACAAAGATTGAATCCTTATTTCTTTTCAAGAACCATTTTATCGGTACAATTCCCTCGAGTTTTTCCAAACTCACATTGCTGAAATCATTGGATTTATCGGATAATCGTCTCTCAGGCAACATTCCTTTTGGATTTTCAGAGTTGAAAGAGCTAATCATGTTGTATTTGATGAACAATAACTTATCAGGTGAAATCCCTGAAGGGATTGGTGAGTTACCAAATCTCGAATTGTTAGCTCTTTGGAACAATTCACTCACTGGAATTTTACCACGAATGTTAGGATCAAATGCTAAGTTACAAAAACTTGATGTGTCCTCAAACAATTTGACTGGTCCAATTCCCCAAAATCTCTGCCTCAGCAACAACTTAGTTAAACTCATTCTGTTTTCGAACGAGTTTACTGGTGAGATTCCTTCATCCTTAACTAACTGCAACGCTTTATCCAGGTTGAGAATCCAAGACAACAAACTCAATGGCACGATTCCATCAGGATTCGGTTTTTTGCCTGATTTCACTTACATGGACATCAGCAAGAACAAATTTTCGGGGCAAATCCCAAAAGATTTTGGAAATGCCCCGAAAATGATGTACTTGAACATTTCTGAGAATTTTTTCGGTAGTAACTTGCCTGATAATATCTGGAACGCCCCGAATTTACAGATATTCTCAGCAAGTAAGAGTGGCCTTGTTGGAAATCTCCCTGATTTCAAAGGATGTCAAAGTCTTTACAAAATCGAACTCGAAGGAAACAATCTCAATGGTAGCATTCCATGGGACATTGAACATTGTGAAAAGCTTATTTCTCTAAATCTTCGACGAAATTCATTTACAGGAATCATTCCTTGGGAAATATCTGGTATTCCTTCGATAACAGACGTTGATTTGTCACATAATTTTCTCACAGGAACAATCCCTTCCAATTTCGAAAAGTCTAGCACGTTGGAACATTTCAACGTGTCGTATAATCAGCTCACTGGTCCATTGCCTTCTTCAGGGTCTATGTTCTCAACGTTCCATCCTTCGTCCTTCGTAGGAAACGAAGGTCTTTGTGGTACTATCATACATAATCCGTGCAGGACTGATGAACTCAAAGATGGAAAAACAGATTTTTCGAACCATTCAAAGAAAACAGATGGTGCATTGATAGTATGGATTGCGTCTGCAGCATTTGGAGTAGTAATTATCATTCTTATTGTGGCAATTCGGTGTTTTCATTCAAATTACAATCGAAGATTTCCATGTGACAAGGAAATCGGACCGTGGAAGTTAACAGCATTCAAAAGGTTGAATTTCACAGCAGATGATGTACTCGAAAGCATAGCAATCACAGATAAGATCATCGGTATGGGGTCTGCAGGTACTTAACGGATCTATATTGTTAGTTATTATGAATTCAATTGAACCCGCAATTTTTCATTAAGTATATGTGGGGGGGAAATCATACAAAATTAGTGTACATATGTTTTTTAAAAAATGCTAACATCACCTAGACAAGTTGGCAATGTGGAATTTATTAATAAAAATCTC

>M6-PGSC0003DMG400007517 M6 LRR-RLK

TTACTCCTGGTGGTGTTAATTATGAAGGTTAGTTCTTGTTATTTTTAATAAAGATTGAAACTTTGAGACATTATTGGAATGAGATTGATCATATTAGAAAATATCAAGAATTTAAAAGTATGATTTACTCCTGTTGGTGTTAATTATGAAGGTTGTTTCTTGTGGTTTTTCTCATTAAGATTGAAACTTTATGACTTTTTTGTTGAAAAAATCATATGAGGTGTTTGAAAATTTTGAAAGGAATGAAATTTTGATCAAATTTTAGAAATATCAAGCATTTTGAAATATGCTTACCCTTGTTGGAGTTGATTATGAAGGTTAGTTCTTGTGGTTTTTCTCATGAAGATTGAAACTTTATGACTTTTTATTTGAAAAAATCATATGGGGTGTTTGAAATTTTTGAAAGGAATGAGATTTTGATCAAATTTTAGAAATATTAAGAATTTTGAAATATGTTTACCCCTGTTGGAGTTGATTATGAAGGCTAGTTCTCATAAAGATTGAAACTTTGATACTTTTTTGTGAAAAATTCATAAGGGGTTTCTGATTTTTTTTGAAAGGAATGAGATTTGAACAAATTTAAAATATTTCAAGAATTTCATCTATTTTAACTTTCTTGTTATGTTGATTTTGGACAGTGCAAGCTTTAATGGAGATCAAGAACTCATTACATGATCCACTTGGTGTTTTGAATTGGGATGAAAGTTCTGTTGATCCATGTAGCTGGAATATGATCACTTGTTCAAATGATAAATTTGTCACTGGCTTGTAAGTATACCCGATTGTCTGGTATCTGCATTGGAGCCCGACTAACAAAACCTTTTCTATTCCTAATACTTGAACCGGAGACCTCTAGTTAAGGATGAAAGTGCTTACAAGTTTTCTTGAATTGTGCAGGGCTTGTTCAAGCCAAAGTTTGTCTGGGAAAATATCACCATACATCCACAACTTGACTCACTTGGATCTACTGTGAGTACTTTATATGAAGAAAAAATAATGAAATTTCAAATGATTGATTATTTGAATCAAGAATATAGAAAATAGTAGTACTAATATTACTTTTTAACTTTTTTTTTTTTTTTAAACAGGCTTCTGCAGAGCAATAATATAACAGGACCAATTCCAACTGAACTTGGAATGCTTCAGAAACTTAATACAATTGATCTTTCTGATAATCAACTCACTGGGAAAATCCCTTCATCTTTAGCTCAATTAAAAAGCCTTCAATACTTGTAAGCAAAGAAAATTTTAAAGATTTTGATATAAAAATGAGTTCTTGAATTATAAAAAGCTAATGATTATGGTTTTTTTTTCTTTTTCAATTGTGTTATAGGAGACTAAACAACAATAGTCTTAGTGGAGCTGTTCCTTTGGAATTGTCTAATATGACTCAACTTACTCTTCTGTAAGTTCCTTAGCTTCATTTAGTTTTGTTTATTTTCTCATACTTGAAGGGGAGCCTCGGAGCAATAGTAAACTTGTCTCCTTGTTATCTATATAGGTCACGCGTTCGAGTTAGGGGTAGGCTGCCTATAGTATAGCCTTTCCCCGGACCTTGCGTGAACGTGGAATGCTTTGTGTACCTTTTTCCTAATTGCTTTCTGTTTTCATGAGTATATTTTGATGGGATTTTTGTTGTTTTTTTCTAGAGACTTGTCTTTTAATAATTTGAGTGGTCCTGTGCCAAGGCTTCTAGCTAAAACATTCAAGTAAGTTTCTTGAACAATTCTTTGAATTTCATTTTTTTTCCCTTTGCTAAAGATTAGAGAAAATAAATCATTTTTTCTCTCATTGATGCAGCCTTTTGGGAAATCCTATTATATGTGCAACTGGAAAAGAACCAGAATGCAATGGAACAACAACATCCATGCCTTTGCCCTTCACTTTGAACAACTCACAAAGTAAGAGTCTTGAATCATCCTCCTAAGTATTTTTTCGAGCTAGCTTGTGTGCCTCTCGACTATTCCACAAATCTTGGTTGTTGTCAAGTCTCCTGCTTTCAATGTCGTTATGATTAACTACAACTCTATTATAAGGATTCATCATGAAGTCTTGGATAGTTTCTGTGTTGTACTATTCACAACTGAAAAGTCTCTTGTTATATCTCAGATGCTCAGCCTTTTAGAAGATCTAAAACTCACAAAATTGCCTTAGCCTTTGGGACAAGTCTTGGATGCATCTGCCTACTAATTATTGGGTTTGGAATATTTCTGTGGTGGAGACAAAAGCACAACAAGCAAATTTTCTTTGACATCAATGGTGAGTATGATATGCAGCTAAAACCTATATTGCTCGAATCTCGAGATGCTAGTAATGTCTTGCTTTAACCAAGAGGATCATGCTTGTTGATAATTTCAGAACAACATCATGAACAAGTGTGTTTGGGACACTTGACGAGGTTCCAATTTAGGGAACTTCAAGTTGCCACAAACAACTTCAGCAACAAGAAGATACTGGGAAAAGGCGGTTTTGGAAACGTGTACAAAGGTTGTCTTAGTGATGGCACGATTGTTGCTGTAAAAAGGCTCAAAGATGGAAATGCAATCGGAGGAGACAAACAATTCCAAACAGAAGTTGAAATGATCAGCTTGGCAGTGCATCGCAATCTCCTACGTTTGTATGGGTTCTCCATGACACCAACTGAAAGGCTTCTGGTTTACCCTTACATGTCTAATGGAAGCGTTGCTTCTCGTCTCAAAGGTAAAAAGAAACAACCTTAGAGAACTAAATATTCTGCAAGTATTCTTAAATATGGGGGATGGTTGAGATCCCGTCATCCTTAACCATAGTAGGCTTACCTGTGTGAACCGGAATAGTCGAACTAGTGGACTTCAGATACCTGATGGTAGATTTCTCTATTTCTGTTGTTTCTTGCAGCCAAGCCACCATTGGACTGGACTACAAGAAAAAGAATTGCATTAGGAGCTGCAAGAGGTTTACTATATCTACATGAGCAGTGTGATCCAAAGATTATCCACAGGGATGTGAAGGCAGCAAACATATTGCTCGACGATGACTGTGAGGCTGTCGTGGGAGATTTTGGATTGGCAAAGCTATTGGATCATCGCGATTCACATGTCACAACTGCAGTTAGGGGAACTGTAGGACATATAGCTCCTGAGTATCTCTCTACAGGACAGTCCTCTGATAAAACAGATGTTTTCGGGTTTGGAATTCTCCTGCTAGAGTTGATATCTGGCCAGAAAGCTCTAGAATTTGGTAAAGCAGCAAATCAGAAAGGCATTATGCTTGATTGGGTAAGTATTACACATTCCAAGAAGTGACAAAACCCTCAGTTTC

>M6-PGSC0003DMG400007634 M6 LRR-RLK

TATATATTCATTTTTATCAATTTGGCATTTTCTTCACACCTCCAATGTTGACCAAAATCAACTCATAATTCTAACCCCACCTTTTTTTTTTTTTCATGGAGTTTTCCCATTTTCCGACCGGCATCTTCGCCGGAAAACAGTCCCTTTATCTCATTCTTCTACTAAACTTCATACCCATTTTCGTAACTTCACAATCTCCGGCCACTACTGAACGTGACACTCTACTGAAGATTAAGCGTCAATGGGGAAACCCAGTAGCACTTGACTCATGGAACTCTACTTCTTCTCCTTGTAGCTGGCCGGAAATTGAGTGTGATGATGGGAAGGTTACCGGAATAATAATTCAAGAAAAAGATATTACTGTTGAAATTCCAAGTTCTATTTGTGAACTTAAAAATCTTACTTTTCTTAATCTCCGGTTGAATTATCTTCCCGGAAAATTCCCAACTTTTCTTTACAAATGTTCCAATTTGCAGCATTTGGATCTTTCTCAGAATTACTTTGTGGGTTCCATTCCGGAGGATATTCACCGGCTTGGGAAACTCAAGTATCTTAATCTGGGTGGTAATAATTTCACCGGAGATATCCCGTCGTCAGTCGGGAATTTGACGGAGCTGGAGACTTTGTGCATGCATTCGAACTTGTTTGATGGGAGTTTTCCGGCGGAAATCGGAAACTTAACCAATCTTGAAAGTTTGGGGTTGGAGTTTAATGGGTTTTCTCCAATGGCACTACCGCCGGAGTTTGGGAAGCTGAAGAAGATAAAGTATATTTGGATGAGAGAGACGAAATTAATCGGTGAAATCCCTGAAAGTTTTGGGGAATTTCAAAATCTTGAGCTTATTGATTTTGCTTACAACAATCTGGAAGGAAAAATCCCTTCTGGGTTGTTTCTGTTGAAGAATTTGACGATGATGTACTTGTTCAGTAATCGGCTTTCGGGTCGAATCCCTGAAACTTTTGATTCATCAAAGTTGATGGAACTTGATGTTTCCAACAACAATAATTTAACTGGAACAATCCCAGAAAGTTTTGGAGGGTTTAAACATTTGGAAATTATGAATCTGTTTGGTAACCAATTGTATGGAGCAATTCCAGAGAGCATAGCCAAGATTCCATCTTTGAAGGTATTCAAAGTGTTTAGGAATAAACTGAATGGGAGCTTACCATCTGAAATGGGACTTCATTCGAAACTCGAATCTTTTGAAGTTTCACTGAATTCCTTCACTGGAAACTTGCCAGAGCATTTGTGTGCTGGAGGAACTCTTTTTGGTGCTGTAGCTTATGTTAACAATTTGTCTGGTGAAATACCAAAATCACTCGGAACTTGTTCTAGCTTACGTTCGATTCAGCTCTACAAGAATCAGTTTTCGGGTGAAATTCCATCTGGGGTTTGGACTTTGGTTGACATGACAAGTTTGTTGTTGAATGATAACTCATTTTCCGGGGAGCTACCGAGCAAAATTGCATACAATTTTACGCGGTTAGAGATCAGTAACAACAAGTTTTCTGGTGAAATTCCTGTGGGGATATCTTCTTGGAGGAGTTTAGTGGTGCTTTTGGCAAGTAACAATTCATTTTCAGGTCGGATTCCAGTGGAATTGACTAGTCTTTCTCAGATAACTCAGCTAGAGCTTGATGGTAATTCGTTATCAGGAGAACTTCCAGCCGATATAATATCGTGGAAGTCGTTGACAATATTGGATCTTGCCAGGAACAAACTCTCTGGCAAGATTCCAGCAGCTATAGGTTTAATCCCTGATCTTGTTGCTCTAGATTTGTCTCAAAATCAATTTTCAGGTCCAATTCCACCTCAACTAGGTGTAAAAAGGATCACTAGCCTTAACGTGTCGTCCAATCAACTTATCGGGAATATCCCTGATGCATTTGCTAACCTAGCCTTTGAAAACAGTTTCTTGAACAATCCCAGTCTTTGTACCACCAACTCACTTCCATATCTTCCAAGTTGTAATAATGCGAAAGTCGCCAATTCCAAAAGATTGTCCCATAGAGTCCTAGCCCTGATTCTAGTCCTTGCATTCGCTGTTTTCCTATTTAGTGTTATATCGACCTTGTTCATGGTTAGAGACTACAGGAGGAAGAAGCATAAGCGCGATGTTGCAAGCTGGAAGTTGACTTCATTCCAGAGGTTGGATTTCACAGAGGCGAACATTTTGTCAAGTTTGACGGAAAATAACATGATTGGAAGTGGTGGATCGGGTAAGGTGTACAGAATTTCCGTTGGCAGACCAAATGAGTATGTTGCTGTGAAGAGGATTTGGAGTGATAGGAAAGTAGACTACATTCTGGAGAGGGAATTCTTGGCTGAGGTTCAGATATTGGGATCAATTAGACATTCCAATATCGTGAAGTTATTGTGTTGTATATCAAGCGAAGATTCAAAGTTGCTCGTTTACGAGTACATGGTAAATCACAGTCTTGATGGATGGCTTCATGGGAAGAAGAGAGTATCCTTGAGTAATAAAGTCATGGATTGGCCAAAAAGATTGGAGGTTGCCATTGGAGCTGCTCAAGGACTTTGTTATATGCACCATGACTGCACTCCACCAATCATTCATCGCGATGTAAAAACAAGCAATATCTTGTTGGATTCTGATTTCACAGCCAAAATAGCTGATTTTGGGTTGGCCAAGATTTTAGAGAAGAAAGGAGAGCTTAACACCATGTCTGCTGTTGCGGGTTCCTTTGGTTACATAGCTCCAGGTAAAAGCAAAATCCAAGCTTGAACTTCGTATTACATCAATTTCGTTAACATTATATCAAAAGTATGACTAACTAATAATTTACTGTTTGGTTACAGAGTATGCTTATACAACTAAAGTGAACGAGAAGATTGATATCTATAGCTTCGGGGTGGTGCTATTGGAGCTAGTGACTGGAAGACAACCAAATTTTGGGGACGAGCATACAAGCCTGGCAGAATGGGCATGGAAGCAGCACGGAGAAGGGAACACTGCCATCGATAATATGTTAGATACAGATATCAAAGAAACATGTTACTTGGAAGAAATGAAGACTGTGTTCAGACTCGGGCTTATATGCACGAGCAACTTACCTGCAAGCAGGCCTTCCATGAAAGAGATTCTGCAGATACTTCATCGATGCAAGTCCTTTAGGTATTCTGGAGGAAAATCACCAGACACAGAGTATGATGTTGCTCCTCTCCTTAGCGGAAACAACAGTGAAAAATACATTGCTAGTTACAAACGTATTAATTCCAATAAGGTAATAGATGATAGTAGTGATGATGGTCTAATAATTTCCAGTGTGTAATATTGGAGATGCCGAAGTAAACTGAGTTGTTGTAAAATGATCATAGAATAGTGCCACTTAAGTAAAGTACAAGTTTGAATTTCTAGTGTACTCTTGAATTTGTTCACTGCTTGTAGATTGAGTGAAATGCATCATTTAACTTTTGCTAGTTATGTAGATTGTAA

>M6-PGSC0003DMG400007709 M6 LRR-RLK

TTTTCTCCATCTGTCTCTAACTATTCCTCAAAGTTTTATCTGAGTAACGACTTCTCAAATGCAGATCATTGTTATCCAGAGGTATGCTTGGGTCACCTGAAAAGGTACACCCTTAAGGAACTGCGGACTGCCACTGACCATTTCAGCTCAAAACGTATTTTGGGAAAGGGAGGATTTGGGGTTGTGTACAAGGCCAGCTTAAACAATGGTACTGTTGTGGCCGTCAAACGATTGAACGACTACAATGCTGTTGGTGGTGAAAATCAATTTCAGACTGAAGTAGAATTGATTAGTTTGGCTGTCCATCGGAATCTTCTCCATCTTCTGGGGTTTTGTTCAACTGAAAGTGAACGGCTTCTTGTTTACCCCTATATGCCAAATGGAAGTGTAGCATCACGATTAAAAGGTTAGCTCTTGCCTCCATGTGTT

>M6-PGSC0003DMG400007862 M6 LRR-RLK

AGTTGAGGCCACTGTTATGATATGTGAATACTAATTGTGGTCATCTCCTGTGGCAGAGATCTGAGCAACAACCATATTGGCGGAAGCATTCCGTCTAATCTGCCAGTTACCTTGCGGAATATGTAAGTTCTACCATTTGACTTATGAAGGATCCCCAGAAACACGGGAAAAAGAAAGAAAAGAAAGAGGACATATTGCTAAACCTAGCTTTACTTATTGACTAAAACAGTTGCTATTCTTCACCTTGGACAGTTTTCTTTCAGCTAACGCATTCACTGGAAGTATCCCTACTTCTTTGTCTTCCCTGAGTCAACTTTCAGCTATGTAAGTATCATTTATAGTGTAGATTAAAAGTTGCTTTCTTCATGAAGAATCAAACTCAGGTATTTCGTTGTGTGTCAGGTCTCTGAATGGCAACCATTTGACTGGAGAAATTCCTGATTCCTTTCAAGGCCTTACCGCTTTGGTCAATCTGTATGAAAGCAAACAAAAGCATTTACATACTTCCATTTAGTTTTTTTCATATGAATAACTGGTTTGGGTGTTTATCTTTCAACAGTGATTTGTCAAGTAACAATTTAAGTGGATCATTGCCACCATCTGTAGGAAATTTGTCTTCATTGACCACTCTGTGAGTAATTCAACATATTATACAGTTTGTTTGTAACTTTATAGTTTGATCAATATGATAAAACTAACTTCTTGATTTCCAGATATTTGCAGAATAATCAGCTTTCTGGAACTCTTGATGTTTTGCAGGATCTTCCCCTTGCAAATTTGTACTCTTCCAAACTAGATTTTACTCAAATATAATTCAGTTCACTAGAATGATAGAATCTTACTAACATCTCTCCACCCTCCCTGTGTCCTTTCTAGGAATGTGGAAAACAACTTGTTTTCTGGGCCCATACCTCAAAAGTTGTTCTCTATTCCCAATTTCAAGTAAGTACATTCATTTTATGCTTTCATATACCCGTTTCATTTTTGTTTATGTTGAGTCAACATCTTGTATCTTAAAATGTAATCTTGGTCACATTTAATATCACTGTATCCCTAGTCGTTTGTTGGAGATATCATTTGACATGTAACATATCCCACAAAAAAGATATATAATCATAGAAGACAATAGGCATGTATGTCGAAACCGCGGAAACAAATATCTTTTCAAGTTCAATGTTACTTAATACTGCTCCTTATTTTCCAATAATGCCTCACTTTTGTACTATCTACCAGTGGTTATAACCAAACATATAACTTTTCTAGCTCCCAATACCCCCCTTTAAAGCTTGAGAACTTGAATCGTCTGTGCTATGCTTCATACACACAAGTAAACCTTTCAGAAACTTCTTGTTTACAAGTTCCTGAAATCTTACTTAGTAAAAAATGGATGTACTTGTATGCTATGTTGCTTGGACTCGTATTGGATCCTCCCAAAGTAATCCGACACTGGTGCAACAACATATTTGGAAAGTCCGAGCAACATAGCTTGTATGCTGCATGACCAGAAATATAGACTTAACGTCTGACATGGATATGAATCCTAAACTTGCCCTTAGCATAAATGAAAGCTCAATGAGTAAAAGTAAGAATAAAACAAATAAGAAATTGACCTGAAAAAAGAAAAATAGAATCCTGACAAGCTTGTCATTTAAATACATGAATTCTATCATATGGCCAGTCCCTCAAAACTTGGAAGTAAACAACTCATATAGACACAATAAATAGACTGCAGATATAGGTATAGATTTACAAACCGTAGTACAATATGCATATGAAAACATACAAACAGTTAAGTTATGATATTCATGACCTGTAAAATAGACAAAATTGCAAGTTATTTAGTTTAGCAAGCTGGCTCCTGGTCTAAACTTGATCATAATTTAAGTTCACATTCCATAGTGGCTATACTCTAGTTAGGTCACCCACCTTAATCATATATCGAGCCCTTAACTTTCAGGAGGCAAATAACATGGAAGTCAGGCAGTCTTATTTATTAAGCTGACTTAAATGGTACAAGTTGGTAGATATATTTGATATATGCAAAGATCAATGTATCAGTTGTTGGATCATTTTGTTACCATCTGATACATGCCTTGTCGATTATGATCAGTGATGGTTTGTACTGTCTTAGTTACCGGTTTTGTTTTTCGTGCAGAAAAGATGGAAATCTTTTCAACAGTAGTACTGCTCCTTTACCTCCACCCACATCATCAACAGCACCTCCCATTTCTGGGTCTCCAACTTCAGCACAGATGCCATCTTCTCAAAGCAAACCTAAAACACAGGCCGATGGACCATCAGCATCTGCAGAATCAAGTTCAACAGGAAAAAAAAAGTCTTTATCTGTTAAAAGGGTGGTCTGGATATCAATTGCAGCAGTTTTGTCATTTATCATATTGGTTTTAGCAATCCTTCTCTGTCTGCCAAAGTGCCTCAGGAAATGGCACGAGACTCAAAGAACCCCCAAACACCATGAAATAGCCCCTTATATGGGTGCTAGAGAGAATCTTGGAGATAGTGGTTCATTAGTCCAACCTGGCCATGATATAGAAAAAGGTGCTACAAAATACTCAATATCTATGTTATTTTATGATATCTTTTATCACTGTTTCTTAAAGTTCAACTTTTTTCCCTTGGGCTTCGTTTGGTAGGATGTATTAGAGAAATTAATACATGTATTAGTGATTGTATTACTTAATCCTTTGTTTGCTAGGATTTTCAGCATATGTATAACTAATATTATTCAATATTATAGGGTGTATAACTTATACCTAACAAACCATGGTATTAGTAATACAATGGTTATTAATACATGGATAAGTATGGTTAAAGACAGAACTGTCCTTAATATATCAAACTAAACCGTGGATAAGAAACTATATCTAGCATAACTAACCCCAACATAATTAATCCCAACATTACCAATCTCGGCACTACTAATACACCCTATTTCAATACTATTATACATTCTACCAAACGACCCCTTAGTTACTTGGAAAAAAAAGATACTGGGTCTGTTGTGTTGGTGCACTTGAGCCAAGGGTCTTTCGGAAATAGCCTCTCTACCACCACGAGGTAATGGTAAGGTCTGCATACATTCTACCCTCCCCAGAACCTACTTAGTGGGATTTCACTGGGTATGTTGTTGTATTGGATGTCTGTTGTGCAGCATTTAGAACTGAAAAGTTAATGAATCTCAAAGTTTAACCTTCTACTCTTTAGTCTTTAGTGATAAGTGGCTTTCTTATGTGTTAAACAGCTCCCCTAGTTGTGACGCCAAAAGAAAAGCAACCAAGAAGACCGGCTCCTATTCCAAAGCCACAAGATGAGCAGCAGGTAAATGTGCAAACCATGAGTGCTGTGCCAAAGAAGGATGATAGTGAGATAAACCTTGGCAGATTGGATATTGATTTGACGCCACCACCTCCGCCTCCTCCTCCTCCGCCTCCGCCTACTCTTCCACAAGAGAGAGTTATTGTCGAGCCGATTGCGCCTGCTGAAGATACAGCAACGAGGCTTCCAACCAGGCCACTTCCACTAACTTCTGTGAAATCTTACACAATTGCATCACTTCAGCAATACACTAATAGCTTCTCTCAAGATAATCTGTTGGGATCAGGAATGCTGGGTAGTGTATACAGGGTGGAGCTTCCTAATGGAAAGGTATAACTTTCAATCGGCATGAATATATTATGATGTACCAATTTATTAACCGTGGTTTCCTACTTCCACTAGATTGCGAGAGTCTTGACTTACCACTTTTTCTCTCCTTGCTTTTATTTCTCCATCATGTTTGGTATTCTGGAATGCTAAATTATCAGCAAGTAGCATCAAAATTGCCTCTTTATCTCTAGTATAAATTGTTACATTGCTTAGATTTATCAAGAATGGTTGTGATACAAGCACAATTATATGCAAAAAGGCTGTTTTTAATGGAAAATTCACCTTCATATATCCTTAATTAGAGCACTCCTTAGCAAATCTATTAAATGAGTTCTACCTATTGTATCTGATATAATGTTTCTTGAATTACTTTAGCTAGTTGAGACCTGCTTAAGGACAAAAGTTGTAATTGCAATATTTTTCCTCTAGGTTTCTTCTAGTTCTTTTCTTTTCCTCCTTGTTCCAAATTAATCTTCAACTAGTATTTTTTTTTTCCAGTAACCAACATCATCGACGAAATCTTGTCCTAGAAGTGTATTACAAATTTGTTAAAATGAAAAACAGCCTTCTAGCTATATGATATTACTATTTTTCTGCCTTGACTTTTGTCTCACTTTATAAGAGGAACCGCCAGATACTGCTGCATCTTTTTTCTGGTCATGATTTTAACTATTAAAGTAAGTATATGCATGCTGTTTCATTTATATAAATTCCTTGGGGATGTATATGCATTTCTCTGGCATGTAAATTTTACCGCTCTTCCTTCTCTTGCACTATGCACCACATCATTCAAGTGGTTGCAACATTTCACTTCTGACACATGCTCCTTTTGGATGCTTCTGCAGTTGCTTGCTGTCAAAAAACTGGACAAAAGAGTTTGTGATCAACAAAAGGATGATGAGTTTCTTGACCTAGTAAACAATATTGACAGAATTCGTCATGCTAATGTTGTTGAGCTGATGGGCTATTGTGCAGAGCATGGTCAAAGGCTTCTGGTCTATGAGTATTGCTGTAACGGGACACTACAAGATGCACTGCATTTCGACGATGAATTCAAGAGACAACTTTCATGGAATACTCGCATGAGGATGGCTCTTGGAGCTGCAAGAGCTTTAGAGTATGTCAAATAACATTACATACATTTAATTATGCACATCATGTTTGTACACGCTCGCGTTCATGCGTGTGTGTATTAGTGTTTTGGGACCTATAATCAGTGTTTCTGCACATACCATATGATAAATCAGTGCTATTAAACACTAAACAATAAAATGGTTCTGATATAGTGGCAAATTCAATAACAAGCTCATTTTAGTCCTAACTTTCTTGTGTTGTGTATATTACATGATAGAAAGCCACACAAACGACTTGGAATGGTCTTAACTATATGTAAAATGTTAGATGTAGCTGTCTTTTGCTTCCTCTCCTTTCTTTGCTTCTGAATGTTCAAGAAAAAGATGAAGAAAATTAAATTAATTGGTTTGCCACGCTATGCTGTAAAAGGGAAGCCCTATGCACAAAACATCCCACATTAGCAGTCCGGGGAAGGGCCGTACCCCAAGGGTTGTGATGTAGACAGTCTACCCTAATGCAAGCATTAGTGGCTGCTTGCAAGGCTCGAAACCGTGACCTATAGGTCACACAGAGACAACTTCACCGTTGCTAAGCAGTCTGATTCTCATTTCAAGAATTGCATGAAAAAGTGAGGTAGACTGCTAGAGCATGTTTGTAATTAATAATTATTCTCGTTTGATGCAATGCAGGTATCTCCATGAAGTCTGTGAGCCACCTATTATTCATAGAAACTTCAAGTCTGCCAATCTTCTGCTTGATGAGGAGTTGGCTGTACATGTTTCAGATTGTGGTTTGGCTCCACTTATGTCATCTGGTGCTGTGAGTCAGGTAACTTGTGACCCTAATGTTATGGAGTAGAAAGAATTGAAATGCTTCCTTGGATTATTAGTTTCCTCTATGAGCTGCAAGTGGAAAGAAAAATGGATTCCTAGTGTTTGATAGAAGTGGAATGGGAGCTGAATAAATAATTTAAAAAAGGAACAGTAGTGATGGTCAAAGTGTTTTCAACAAGATGTTGGAATAAGGAGCAGTAATAATGAATTCCTGTAAATGTGTAAAGTATATTCTTTGGGAAATGGTTGGTTCTCACCCTGTAGGATAGGGTGAAGTGTTGTCAAAGGTGCGCTTAAAGCACGCTTGAACCCTGAAGTGAGGCTCAAAATGTGTTGAGCATTTCACATCGCTTAATGTGAGATTAGGGGTCTATAGGAAACAACCTTTCTACATCTGAGGTAGGGGTAAGGTTTGTAGACACTCTACCCTTCCCAGACTCCACTGGGTATGTTGTTGTAAGTAAAGACAAATGGTGAAGGTGCTCTTCTCATGATGTGTTTTATATGAACAGAAAATTTCTCTAGGATGCATGTGGATCATCTATCTTGCTGTTATTTGTTGTTCTTTTGATAAAGTAGTCTTGGGATTGATTACTGTTACGTTCAATTTAAGGTATGTTAAAAACTATATCAAGTTGACATTACACTGATTTTTCAGTTGTCGGGACAACTTCTCACAACATATGGCTACGGTGCTCCCGAATTTGAGTCAGGGATTTACACTTCCCAAAGTGATGTTTACAGTTTTGGTGTGGTGATGCTGGAACTAATAACAGGCAGAATGTCATATGACCGGTAAAAAAGCCTTCTGCATCTTTTTGAGCTAATATAGTTGTGCCTCATTTCTCAGTTAACACACCTGCATATCATATTCAGGACACGAAGTCGAGGAGAGCAGTTCTTGGTCAGGTGGGCAATCCCTCAGCTACATGATATTGATCTCTTGACAAGGATGGTTGATCCTGCTCTCGATGGAAAATACCCCATTAAATCACTATCACACTTTGCTGACATAATTTCCCGCTGTGTTCAGGTGAGTTATTCTGTTGTTCCTTCATGTGAAACACATAAATGCTTACTAATATGCATTGTCTAATCGCTGAGTGTACCGAATAGTCTTAGATTGTTCGTCATCTCAATCGGTTGGAAGGATATTTCTAGGATCTAGTTGTGGAATAGGTTTGCCTTCAAATGTCACTTAACTTTTAAACTCTTATACAGCATGAGCGAGAATTCAGGCCACCAATGTCTGAAGTTGTTCAGGACCTCATACAAATGATAAGAAGAGAGTCTCACAAAAGATCTGATGGTGAATGATTTGAGTTAGACGCGCATACTGGGAATTCAGCCGATGAGAAACTAAGATTATTTCTTCTTCGAAGGAGCTGAGATACTCGGAAAAAGAAAGGCGGTTTGCAGGCCAAGCACCGTGCTAAAGACAACTTCAAACTGAAGCAAAATGCACCTCAAAGAAGCGCGATTCATGAAAATACCAACACATGCAGGCGACTTAATGACCTGTAAAAACAGCTTGCTCATCTGATTCTTGTCAAGCTGCATCATTTTAATGTTCATTTAGGCTTACATGTGAGCACCAATGCATATAATTCATTGATCTTGACAAAATGGTATATTTTCTTGAAGTTGTATATCTTTACTTGCTGCAACATGTACAAACTTCATTTGGTTGTACAAGGATACCTTCACTCTTTCTCATGTACTTGTTGCACATGCCAAAGTTATTTACTACTAGTCATTACCTAAGGAGTTGGAGACCTACAACCACTCAACCATGTAACAATATGTAAA

>M6-PGSC0003DMG400007948 M6 LRR-RLK

TTAATAATTAATTCTAAAATAACTTATTATTTTCGACCGTGAAACATTATGGCCATGGCTGACACAGAAACAGAGAGAAAAATAGCCAGACTAAATATTGGAAGTCCCCACTTCCCTTTTTTCTTTCTCAAAAATCTTCACGTGAATCTGACTCTGCCTTTTGCCATATCTTCTTAGACTGCTGCTTTTCTCTTCACTCTTCCACCTCTTTTGTCACGTATTGCACCTCTACACATATTCCACAACAATTCAATCTGGAAATTTTTTCTTAAACAAGATGAAAATACAACAATTCTTACTACTAACCATTCTTCTGTTTTTATCTGTTTCTTTTCATTCCACTGTTTCGTTGATTCCTTCAGATGCATCAGCATTGTTAGCATTCAAATATAAAGCTGATTTGGACAACAAACTTGCATTTTCTGCTAATACAAGTTTCAGATTCTGCAAATGGAAAGGGATACAATGCTCGGAGAAGAAAGTTATTCGTATTGTTATCGAAAGTTTTAGCTTACGAGGTACGTTTCCAGCTAATACATTGTCTATGCTTGATCAACTGAGAGTATTGAGTTTACATAACAACTCACTTACCGGTCCGATTCCGGACCTTTCTCCACTTATTAATCTCAAAGTTCTTTTCCTGGACCATAATTTGTTCACCGGTTCAATACCATCTTCCATTTTCACTCTTCATAGGCTTAAAACCCTTGATCTCTCTTACAATAACCTCACCGGTTCAATATCCGTTGCAATCAACGGGTTGAACCGGTTGTACTATCTCCGGCTTGACTCGAACCGGTTAAACGGATCAATCCCACCGCTGAACCAATCTACCCTCCATATCTTCAACATTTCTCACAATGCCCTTTCTGGTCCTATTCCGGTAACGAAAACGTTATCCCGGTTTAAAACGGCGTCGTTTTCAGATAATAAAGGACTTTGTGGTGAGATCGTACATAAAGAATGCCGTCCAATACAACCGTTTTTCAGTCCATCCACCGCTGCCGCCACTAAAATTACGCCGCCGCCGTCTAAAACTCCGGCAGAACTCGGCCAAAACGAAGAGTTACGACAGGGTAGCCCGTTAAACCGTAAAGAAAACAAATTACATAAAAGATCCTTACTCATTATTGGAGTTTCAACAGCTTGTTTAGTCCTTTTATGTTCAGTTATACTATTAGCATTAGCGTCGAAGAAGCACAGGAACTCGAAGAAATTAGGAGAAACTAAGAAATCCGCATTCGATCCAAGTGTTAGCGGCAATGCAGAAGCAGTAATAAGAATTGAAGAAGACAACAATGAACTGGAAGAGAAGGTGAAGAGAGTACAACAAGGAATGCAGCAAGTGATGGGTATAAGTGGGAGCTTAGTTTTCTGTGCAGGTGAGGTGCAGGTGTATACCCTGGAGCAGCTGATGAGAGCTTCTGCTGAGCTTTTAGGTAGAGGAACAATGGGGACTACTTACAAAGCAGTGCTTGATAACCGTCTGATTGTATGTGTTAAGAGATTGGACGGTGGGAGATTGGCAGGTACAAGTCAAGAGGAGTTTGAGCAGCATATGGAATCAGTGGGCGGGCTTAGGCACCCGAATTTGGTTCCTTTTCGGGCTTATTTTCAGGCCCGACAAGAAAGGCTTTTGGTTTATGATTATCAACCCAATGGCAGCCTGTCTTCTCTTATTCATGGTACGTTCTTCTCTGTATATCTTGTACTTATTATTTATTAAGATGCGCATAAACTATACGAAAAATATTATTAATGCAATTATTTTGACCTTTGATAAGTAAAATATAATATAATGAACGGAGTATTTTGAAGGTGTTGTCCAATCAAATAGGGTCCATACACCATACCCTATTGTTACAGTTTCTCAACCCCAATGTTCTCACAAACACAGAGAATCTTGCAGCTGACTCTGATGACATTTTTGGACACATATTTCTGTCATCGTAAAATACTTTTATGTTTGTTTAGTGTCCTAATCATAGATTATAGTATTACTTTATTGAAGCTAGTTCTGTCTGCATTAAGAATGTTATTAATTTTGTTTCACGTGTGGTTGCGTAATGTGAAGTTAATAAGAGTACGGTGGTTAGCATTAATAATTCATAAAGTAATATAAATAATAGCTTTATTTCATATTTAATGCCACGACGGTACTGAATTTTTTGGTCAATCTGACGTGGCATATGGACTGGTCAACCTGCCACGTGTTCAGGTTCCAAGTCATCAAGAGCAAAGCCACTTCACTGGACTTCATGTTTGAAAATAGCAGAGGATGTAACTCAAGGACTTTCCTACATCCACCAAGCATGGAGGCTTGTCCATGGCAATCTCAAGTCATCTAACGTCCTCCTCGGCTCCGATTTCGAGGCGTGCATAACTGACTACTGTCTCTCCATCCTAGCTGTCCCCTCAGACGATGACAATCCTGATTCTGTAGCCTATCAGGCTCCAGAAATTCGGAAACTGAATCACAACAACCACAACCACCACCGCCAAGCTAGTGCCAAGGCTGATGTCTATTCGTTTGGCGTTCTTTTGCTTGAACTTCTTACGGGGAGGCATCCTTCGGAACATCCATATCTAATGCCAGATGATATGATACATTGGGTGAAGTCTACTAGAGAGGATCATGATGGTAGTATTGGAGAAGATAGTAAGTTGGAGATGCTTCTTGAAGTTGCTATGGCTTGTAGAGTGAGCTCACCAGAGCAAAGACCAACAATGTGGCAAGTGTTAAAGATGATACAAGAGATTAAAGAAGCTGTTGTTATGGAAGATAGTCATGAAATGGACCTCCTCACTGGCCCTCCTAAATCCTAATTCTTAGTTATTATAAAATTGAATTTGTATTATATTCACTACATGAGACTTTTCAATATTAGTAATTGCAAAAGAAAGGAACAAAAGGGGAAAGACTT

>M6-PGSC0003DMG400008146 M6 LRR-RLK

ATGGATTCTTGATTGTTCTATCTGGTTGTTTTGGAGGGAAAAGGAGGAAGAAATTGAAGAAAAGGTGTTTTCTTGAAAAAAAAAAAAACTCAAGAATGGCAATTTTGAGGTTTCTGTTAACTGTGTTTTTTCTAGTTGGTTTGTTATCAAGATTCCAACTTGTGAATTCCCAGTTGGTATTTCATGATCAAGATGTAGTGGAAGCAATTGGGAAAGAGCTATAAGTACCTGGATGGGGTGTGATGAACACAACTGATTTCTGCTCTTGGCATGGAGTTATTTGCAGCTCAAACAGTTCAATGGTGGAAAGACTTGATCTTTCTGGTTTTAGATTACAAGGTAATGTAACTCTAATATCTGAGCTCAAAGGTTTAAAGTGGCTGGACCTTTCTAATAATAACTTCCAAGGTTCTATTCCATTTGCATTTGGGAATTTATCTGAGCTTCAATATCTTGATTTGTCTTTTAACAAATTTGGAAACTCAATACCTAGTGAATTAGGTAAGCTCAGAAGCCTAAAGGCATTGAACCTTTCGAAAAACTGGCTTACAGGAGCTATACCTGATGAGCTTGAGGGGTTAGAGAATTTGCAAGATTTTCAAATATTTACTAACAACTTGAATGGTTCTATCCCAATGTGGATTGGTAATTTGACCAATCTTAAGGTTTTTGCTGCTTATGAGAATGAATTTAGTGGTGGTGTTCCTGTTAACTTAGGATTGTACTCTGATCTTTCTTTGTTGAACCTTCACTCAAACCAGCTTGAAGGTACTATTCCTGAGAGCATTTGTGCTAATGGGAATCTTGAATTTCTTGTTCTAACTCAGAATAAGTTGACCGGAAAGATTCCTGATTCGATTGGGAATTGTAAAGGACTTTCTAGTATTAGAATTGGGAATAACAAGTTGATTGGAGGTATCCCTAAATCAATTGGGAATATCAGAAGTCTTACTTATTTTGAAGCAGATAGTAATAGTTTGTCTGGTGAAATTGTCACCGAGTTCGCAAAATGCTCAAATCTCACTCTACTTAATTTAGCTTCAAATAAGTTTAATGGAACTATTCCTCCTGAGTTTGGTCAGCTTAACAGTTTGCAAGAATTGATTGTTTCTGGTAATAACCTTTATGGAGAGATTCCAATATCAGTTCTTAAGTGCAAGAATCTCAATAAGCTTGACTTGAGCAACAACAAGTTCAATGGTACCATACCGGCAGATATCTGCAATACGTCCAGATTGCAATATTTGCTGTTGGGGGAGAACTCGATTCGAGGGGAGATACCTCATGAGATAGGGAATTGTGTAAAGTTGCTTGAGTTGGAAATGGGGAGTAATGAACTTACTGGAAGTATCCCTCCTGAGATTGGTCATATGAAGAACTTGCAGATTTCGTTGAATTTGAGTCATAATCGTCTCCACGGAAAATTGCCTCAGGATTTAGGAAAACTTGACAAGTTGGTTTCTTTAGATGTATCTGATAATCAGCTCTCGGGGAATATTCCGCCGGCATTGAAGGGCATGCAGAGTTTGATAGAGGTTGATTTTTCGAGCAACCAATTCACTGGTCCAATACCTACTTTTGCACCATTTCAGAAAAGTCCTAATTCAAGTTTTCTTGGAAACAAAGATCTCTGTGGTGATCCTTTGAGTGTTGGTTGTGGGGATTTAAATGGTTATGACCACAGCGTTTACGGTCACAAGGTCTCTTACCGAATCGTTTTGGCTGTTATCGGTTCTGGTTTGGCAGTTTTTGTATGTGTCACAGTGGTTGTTTTGCTGTTCATGATGAGGGAGAAACAAGAGAAAGCTGCCAAGGAAGCAGCTGGAAAAACCATTGATGAAATTTCTAGCAAACCAGTGATACTAGCGGGAAACGTCTTTGTTGACAATCTAAAGCAAGCAATTGATTTTGATGCAGTTGTAAAGGCAGTCATGAAAGACTCGAATAAGATTTGTGTTGGTACTTTCAGCACTGTCTATAAAGCAGACATGCCTTCCGGGATGATTTTGTCGGTGAAAAAGCTAAAGTCAATGGACAAAACTATAATTCATCACCAGAGTAAGATGATCAGAGAGCTTGAAAAACTTAGTAAGCTCGTTCATGATAATCTTGCTCGTCCTATCGGATTTGGAATCTATGAAGATGTCGTTCTTCTGTTGCATGAATATTATGCCAAAGGAACGTTGACTCAATATCTTCATAACTCTAGCCAGAAACCTGAATATAAGCCCGACTGGCCAACAAGACTTGCCATTGCCACTGGAGTTGCAGAAGGATTGGCATTCCTTCATCATGTTGCTATCATCCATCTTGACATTTCTTCCGGGAATGTGCTTCTTGATTCTAATTTCAGGCCTTTGGTTAGTGAAGTTGAAATCTCTAGGCTTCTCGATCCATCTAGAGGGACTGCAAGTATTAGTGCTGTTGCTGGTTCATTCGGATATATTCCTCCAGGTACAT

>M6-PGSC0003DMG400008296 M6 LRR-RLK

TTATGTACTCCTAGGTTTCGAGTAGGTGACGAAGTTCTTGTTGTCTTTTCTTGTGTGGGGGTACATACAACTTGTTCTTTAACACGGAGCATCCTAGATTAGAGGCATGCTTGGTATTTTGTAACAAATTCATAGAATGCAAGTCATTTTCCTTAGTGTGAATGGATGACTAGTTGACTGCTACTGAAAACGGTGATTCACTGTACGATTCACTAAGAAATGTGTATAAAGGCTTCATTGTCAGCATAAGTATTTTCAGTGAGTGAAGCTTTTTTCTTTTGAAACAATGATGTCAAAGACAGTTGTATATGCATTAGCCATATTTTCTATTACCTTTCTCATACCTCTGTCATCTGGGCAAAATCCAAGCTTTGAAGTTGAAGTTGCTGCTTTGAAAGCTTTCAAGAGCTCAATCTCTGACGACCCTTTCAGTGCACTTGTGGATTGGACTGATGCAAATCACCATTGCAACTGGTCTGGAATCATATGTGATCCTTCTTCAAATCATGTCATCAACATTTCGCTTATTGAGACACAGCTCAAAGGCGAAATTTCTCCGTTTCTAGGAAACCTCTCCAAACTCCAGGTTCTTGATCTAACTTTGAATTCATTTACTGGAAATATCCCACCCCAGCTGGGGCATTGCACAGACCTGGTTGAGCTTGTTTTTAACCAAAACTCTCTTTTTGGTGAAATTCCTGCCGAGCTGGAAAACCTAAAAAAACTGCAGTTAATAGATTTTGGAAATAACTTTCTAAATGGGAGTATACCTGATAGTATATGCAACTGCACTGAATTGTTGTTAGTAGGCCTCAACAACAACAATCTCACAGGTAAATTACCATCTGAAATTGGCAATTTGGCTAATCTTCAGTTGTTTGTAGCTTATACAAACAATTTGGTTGGTTCTATTCCTACCTCCATTGGAATGCTGACAGCATTGCAAACCCTTGATCTGAGCGAAAACCAGTTATCTGGACCTATACCACCAGCAATTGGCAACTTGTCAAGTTTAGGAATTCTTCAGTTACATCTCAACTCTCTTTCTGGGAAAATCCCATCTGAACTTGGCCTCTGTATCAATCTTTTTACCTTGAACATGTATACTAATCAATTTACTGGAAGCATACCTCCTGAGCTTGGAAATTTAGAAAACTTGCAAGTGCTCAGATTGCATAACAATAAGTTGAACTCCAGTATACCTGCCTCAATATTCCACCTGAAGTCATTAACTCATTTAGGACTCTCACAGAATGAACTAACTGGTAATATTCCTCCCGAGTTGGGATCTTTAACGTCACTAGAAGTGCTTACCCTTCACTCAAATAGGTTATCGGGGGAGATCCCATCAACTATAACAAACCTGGCAAACTTGACATATTTGTCTTTGGGCTTTAATTTATTGACTGGATCACTTCCATCGGAATTTGGGTTACTCTATAATCTGAAGAATCTAACTGCAAATGATAACCTCCTGGAAGGATCTATTCCATCGAGCATAACAAATTGTTCTCATCTTCTTGTTTTAACCCTTGCTTATAATAGAATAACGGGGAAAATACCAAATGGGTTGGGGCAATTGTCCAATCTTACATTTTTGTCTTTGGGATCAAACAAAATGATGGGGGAGATTCCTGAAGATCTCTTCAACTGTTCAATGCTTGAAATTCTAGATCTGAGTGACAACAATTTCAGCGGGAAACTCAAATCAATGATTGGCAGACTCTCTAAACTTCGAGTTTTAAGAGCTCGTACTAATTCATTTCTTGGGCCAATCCCACCAGAGATTGGTAAACTGAGTCAACTCATGGATATAGTGCTTCACGAAAACAGTTTCTCAGGTGCGATACCACCAGAAATTTCAATGCTTTCAAACCTTCAGGGTCTTTTGCTATCTGAAAACAAGCTGGAAGGTGAACTTCCTGTGCAACTTTTTGAGCTCAAACAGCTCAATGAACTCCGGCTGAAGAACAATAATTTCTTTGGTCCACTACCGCAACATATATCCAAACTAGAATCACTGTCGCTCATGGATCTGAGTGGAAATAAGCTTAATGGCACAATCCCAGAGAGCATGGCGAGCCTCCGCAGATTGATGACCGTAGATCTTTCACACAACCTTCTTACCGGAACTATTCCAAGAGCAGTACTTGCCAGCATGAGAAGAATGCAACTTTACTTGAACGTTTCCAGCAATTTGTTGCACGGAGAGATCCCAGATGAGATAGGCGTGTTAGAAATGGTTCAAGAGATTGACATGTCAAATAATAAACTGTCAGGCAGCATTCCCAAATCCCTAGAACGCTGCAAAAACTTATTTTCACTGGACCTATCAGGAAACATGCTCTCTGGTCCTGCTCCAGGTGAAATTCTCACCAAATTAAGAGAGCTTGTGTTCTTGAACCTCTCAAGGAACAGATTAGAAGGCAGTCTTCCTGAAATAGCAGGATTATCACATCTTCGCTCTCTTGATGTTTCACAAAACAAGTTCAATGGAATCATTCCCGAGAGATTTGCCAATATGACTGCATTGAAATATCTCAACCTTTCTTTCAACCAACTTGAAGGTCACATTCCAAAGGGGGGTGTATTCAACAATATAAGGTCGGAGGATTTACTGGGAAATCCATCTCTAAGTGGAACAAAGTTTCTCAGTCCATGCAACATAAAAAGAAACGGAACAAGTTCTCATGGTTTTTCCAAGAAAACCTGGATCATACTTGCAGCACTTGGATCTGTTTTCAGTCTCATTCTTCTTGTCCTGGGAATATTTTTATTTCATCGGTACATGAAGAAGAAGAAAAATGTGAATGAAGCGGAGGATACAAATCCAAAGTATACCGCAGCACTGAGCCTCCAGAGATTTTATCAAAAGGATTTGGAACATGCTACCAATAATTTCAGTCCGGAAAACATTATTGGAGCCAGCAGTTTAAGTACTGTGTACAAAGGTACACTGGAAGATGGGAAGATTGTAGCAGTTAAGAAGCTGAATCACCAGTTCTCAGCAGAATCTGGTAAATGCTTTGATAGGGAAGTCAAGACTCTGAGCCAACTCAGACACAGGAACCTAGTTAAGGTGCTAGGTTACGCTTGGGAAAGCAAGAAGCTAAGGGCTTTAGTTTTAGAATACATGGAGAATGGGAACTTGGACAACATGATTTATGGTCAAGTGGAGGATAACTGGACGTTGTCCAATAGGATTGACATTTTAGTTTCAGTTGCAAGTGGACTGTCATACCTGCATTCAGGCTATGATTTTCCAATAGTGCACTGCGACATGAAGCCTTCAAATATTCTTCTGGATAAAAATATGGAGGCACATGTGAGTGACTTTGGGACAGCTAGGATGTTGGGTATTCATCTCCAGGATGGGAGCAGCATATCATCAGCATCCGCATTTGAAGGAACGATTGGTTACATGGCGCCAGGTAATAATTCTTACAACTTCTTGACCCACACTTTACTAACTAGATTATGTGTTACAATATCATGAATACAAGATGTCTACAATGTAAACGAGCTTATATGGGTACCACAAATAATATTTTGTCCATGATGAATGTTTCAAATACATTCCATTTTAGCATCATTCAGTAGTAACATAACTTCTGTCAATCACAATACGCAGCACAGCAGATTTTTTTTTCCTTTCAATCACAAAATACTGATTTAACTTTTATGTGGAAGAATAGACAAGCAAGTAAATGATGCAATGTTTAGAACATATAATCTGTAAAGTATTTTTTTAAATGATAATGTCAGTTTTAATCTATCCAAAAATCTAGCGCAAGCTTTGAAAATTTGAATTTCTTGGACATTTTACATGCTCCTAATTTAATTTGAGACCTCACTTACATGAAATGTTATCTATGAAACAGAGTTTGCATATATGAGGAAAGTGACCACAAAAGTAGATGTATTCAGCTTTGGTGTCATTGTGATGGAGATCATTACAAAAAGAAGGCCAACAAGTCTTACCGGAGCAGATGAATTACCAATGACTTTGCATCAAATTGTTCAGAATGCCCTTGCGAATGGCATAAACAAGCTAGTTCAGATTGTGGATCCTAATCTAGCTTCGTATGTCTCCAAGAAACAGGATGTAGTAGAGGGACTTCTTAACTTGGCTTTGTCCTGCACCTGTCCTGATCCTGAAGATAGACCTGACATGGAACAAATTCTGTCTTCTCTTTCAAAGCTAAGTAAGATGGACTGCATGCCTTCGCATTTGGTAAAAGATTAAATTTGATGTAGAAGAAGCTGGAATATCATCACTCTTAATGTGAACCACCACCAAGTTTCCCATGGTGGCTAATGGAACTAGTTCATAACTTAAAATGTATTTACTATTTAAATAAATTTTCCATATTTCAAGCCACAAATGAACTCTCGAGTGTTCCTGCTTTGCTAGAGAGATATAATGGAACTGTCCCATCAGATTATGTTGAAGTTGACATGCATCATGAAATTGAACAA

>M6-PGSC0003DMG400008403 M6 LRR-RLK

AGTGACCATATGCATGAAAAATCAAATTAGTTATTTGCTGATCTGAAGCAAATTTTTGAACAATGGCAGCCACTCGGCTGTTACTCTGCTTTATCGTCTTCTCGTCTGGCATTGATTTCATATACACTGTTACAGATCCCAGAGATGGTAAGAACTTTGTATAAAATTGTCGGATCCAATATTTAAACTTTATGAGTTTTACTATTTGTTGCAGTTTTTGTGAACTTTTACTTATAAATTTATGTTCTGCGTTGAAAGTACTGAGTTCAGATAATAATGTTTTGTTATGAATAAGTGTGAACTTGGTTGATTGCAGTAACGATACTTCGATCACTGAAAGATCAGTGGGAAAATACGCCGCCTAGTTGGCAGAGATCGGATGATCCTTGTGGAACATCTTGGGAAGGTGTAACCTGCAACAACTCAAGGGTAACTGCATTGTGAGTATGCTGTTTTCTTCCAATATATCTAGGGATCGTTTGATTTACGTTTTAGTTATACGATGACTAATAACACATGAATTGTTATGCATGATTGCGTAATTTAAGTGCATTTCCACCTTCAGATGCTCTTATCCACGTATTACTTGTAGATATATTCTTGTTCCACATTCTATCTGTGAAAAATAATACATAGATTCTCGTCTAAGTTATACGAGGTTTAGTTTCCAATATTAAACTCCATATGACTAGTGCAGAGTTTAATACCGGAAACAAAACATCATGTTAGTTATATGGAAACTATATTAGCTTACACGTTTTTTACATTGAAAACCAAATGTTGTATAAGTAAAATGAAAATTACTTCTTTTTTTTTTAGGAAAAACTACATAATTAGGACATAGGTCGCTATCGTAAATACTATTATTATCTATACTTTCAAAATACTATGTATCTTACCACTAATTAAGAATTTCTTACCATATACCAAGGAAGATACACTTAGATACATGTATTTTTGCTACCATATACACTAAAATAGGGAGAGAGGAGAGCGAAATGGGGAGGAATGCGAGCGAGATTCGCTATGTATCTCAAATACATAGGAGAGTGGCGAGTGAGATGGGAGAGAGGTGAGAGGGACAACCGAGATTTGTTATGTATCCCAGATACATGTGAATCCACTTGGATACAATATATCTAGAACAAATTACACTTAATTTTTACCCTATGTATCTTGAAATACATGTATCTGGATGTATCTGGACGCACCAAAATATGGTAAGATACGTAATATTGCAAACTAGAGTGTATCAAAGTAATTAACTACTAAACTAGTGGGATTTACGTAAGTTTCTCACTTTTTTATACTTCAAACCAAGCCACAACTTAGTTTTTGAAATTGTTGTTTCACCAACGAAGTAGTTGTTAACAATTGATTGTGTTTATTGTTTCTTTCAGGGGATTATCAACAATGGGACTGAGAGGAAAATTAAGTGGTGACATTGGAGGGCTCACTGAATTAATTTCCTTGTGAGTTCGGTTTTTAAATCCAATAAACTTCTGTATATGACACCAACAATTATTAGCAATAAGCTTGTATTTGTCACCATAATTTGCTCACTTCAGTTGCTAAACTTGCACTGCAGAGACCTGTCGTTCAACCGAGGACTTACAGGTTCCCTTTCTCCGCGGATAGGAGATCTGCAGAAACTAAACATATTGTAAGACTACGAAAACTTGATCATTTTGATTGTATAGTGTATACCTGCTATGTACACAAAATTGATATTTATAATATTTTCTTCATAGAATGATCATTTGATCAGCTTTCTTCTTCGTAATTTGTATCATGATTCAATGACATACATAACTTACTTGTACGCGAATGAATTTATTTACTTTTCAGAATCCTCGCGGGCTGTAGCTTCAGTGGCAGTATACCGAGGGAGTTGGGAAGACTTGCTGAGTTATCCTTCTTGTAAGAATTTCTCAGCAACATCAACTTGTGATCGTTTTTGGTGTATCATCAACTCATTTAGTTCCATGTAAATTTCTTTGACAGGGCACTTAACTCAAATAATTTCACCGGTGAGATTCCACGAACTCTTGGCAATCTGTCAAAACTGTACTGGCTTGATCTAGCAGATAACCTGTTAACAGGACCAATTCCTGTTTCAACCTCCTCAAGTCCGGGGTTAGACCTTCTAAAAAAAGCAAAACACTTGTGAGTAAACTTTATATGTCCATATTACATGCTGTTCTTAAGTTGGTATATATACCTCTTAAGGTCTGACTTCTACGTTATGTCATTATCGCAGCCATTTTAACAAAAACCAACTTTCAGGCTCCATTCCGGACATTCTTTTTAGTGCTGACATGGTGCTAATACATGTGTAAGTCAAGAAACTAAAGGGTGATTCGCGCTATTCTTCCTGCCCTCCTTCTCACTTTGTGAACTTCTGATATGGCAGACTGTTTGATGGAAATCAACTTAGCGGGAGCATTCCATTTACACTAGGACTTGTTCAGACACTTGAGGTTCTGTGAGTAGAATTGACCGTCTTATTTATCCTATCTTTCGTGCTACTTCACCTTCTGTTTTGGATTATATGCTTGTACTCTTTTATCTTTTGCTGGTATTCATGTAACCTAACATATACAGTCGGCTCGATAGAAATGCCTTAAATGGTAGCGTCCCATCAAACCTCAACAATCTCACAAGCGTTGTAGAGTTGTAAGTATACTTTTTCCCATCATTGCTCACTCAATTGTCTGCTTGAAATCAATAACTTGTAGATTGAAGTAGTCCTTTTGTGCTCTTATTTCTGTCTTCAGGAACTTAGCTCACAACGAACTGTCCGGTCCACTACCAGATTTAACTGGAATGAATTCCCTTAATTATCTGTAAGGAAAATATCCAGAATCCTGATTTTCCCTAGTTGTTTCTTAATGTTGAAGTGATACATCATATTTTGTTTCTCTTCCAGGGACTTGAGCAATAATTCCTTCCATAAATCGGAGGCGCCAATATGGTTCTCGACTCTAGAGTCTCTCACCACTCTGTGAGTATTAATCAGTTATGAATGCAACTATGAATTTCCCAAAAAGTTTGATGAATCTAAGTAGGAAGAACTGATGTAATGCTCAGAGTCATCGAGTATGGCTCACTCCATGGATCAGTGCCTCAGAAACTTTTCGCTTTACCTCAATTACAACAAGTGTGAGTCTCGCGATCCTAACTGGATATTCCATGTCTATTACCTGCAATGAAGAGGTCGTAATGATCATAATAACTTATGCAGGAAACTGAGGAACAACGCATTGAATGACACATTGAACATGGGTGGCATCAGTGGCAGGCAACTCACACTTGTTGATTTACAGAACAACGAGATTTCCTCAATAACACTTGGCTCTGGATACAAAAATACCTTGATGTAAGATGGTTCATTTCTCCTTTTCGTCGATTAGATGATTGTTTTTTTTCTTTCTCCTTGATTGCTTTGTCGTTGTCCTCAGACTGATCGGAAACCCTGTGTGTGACACGGCCCTTGGAAACACGAACTACTGTCAGCTTCAGCAGCAATCTGCAAAGCCTTACTCAACCAGCCTAGCTAACTGTGGGAGAAAATCATGTCCTGCTGATCAAAAAGTTAGCCCACAAAGTTGTGATTGTGCTTATCCGTACCAGGGAACATTCTACTTTAGAGGACCCTCTTTTAGAGAGTTGTCCAATGATAATACGTTCCATTCACTCGAAATGAGCCTGTGGGTGAAGTTGGATCTCACGCCTGGTTCGGTTTCTCTGCAAAACCCCTTCTTCAACATTGATGATTATCTTCAGGTGCAGTTAGAATTATTTCCACCAACAGGAAAGTATTTTAATAGATCAGAGGTTGAGAGGATTGGATTCTCCTTGAGTAACCAAACTTATAAGCCGCCTCATGAGTTTGGACCGTACTATTTTATAGCATCTCCTTACACTTTCCAAGGTATAATTACTGAGTTTCTTTACAACTGTAACTTCTGGACCAATACTGACTTAAAACTAAAACTGTTTCAGCCGAACATGGAGAAACTTCTATTAGCTCAAAACAAGTTATAGGAATAGCTACCGGGTGTACCATCCTGGTCCTCTTGCTCGTTGCTCTTGCAATATATGCTATTCAACAAAAGAAACTTGCTGAAAGAGCTATTGGATTGAGTAGACCATTTGGTAATGTCATCTTTTACTGAAATAAACCGATGAGTTATAACTGCATATACATTCATGCCTCTCTATAACAGTTATCCTCTATGATTACATTTCACTATAACGACCAAGTTTTCTTTGGAACTGATTTTTCATGTTGTATTGAATTATATGTTTTCTATAACAACATCTCGCTATAGCAGCCAAAGAAATATCAGAACAAATGACGTCTTTATAGAGAGGTTTGACTATATGTCCATAATGCAACTAGGATTAAACTAAAATGTCTGATTACTTTGACCTTCTTTGTCCAGCATCGTGGGCTCCAAGTGGAAACGACAGTGAAGGTGCACCACAACTAAAGGGAGCCAGATGGTTTTCTTATGATGAACTTAAGAAATGCACGGGAAACTTCTCGGAGAGAAATGAGATTGGTTCCGGTGGCTATGGGAAGGTTTGCTACCAAAATCCACAGTAGATATAGAATAAAGGGATAAACTCTTAGTTACTTTGCTTATTTGCGTCTGTTCTGTTTCCTAGGTATACAGAGGAACACTCGCAAATGGGCAAGTAATTGCTATCAAAAGGGCTCAACATGGATCCATGCAAGGTGGACAAGAGTTCAAGACAGAAATTGAATTGCTTTCACGGGTTCATCACAAGAACCTAGTTGGTCTGGTTGGATTCTGTTTTGAGCAAGGGGAACAAGTACTGGTATACGAGTATATGCCAAATGGTTCACTCAGGGAGAATTTGTCAGGTAAGTTACATCCTCCTATACATTAAAGACTGCTTCCTATTTATAGTAATTTACACAAAAAAGTTCCTAATATCTCCATTTCGCTGTAGAAGTTAATTTCCACTATGCTCTACAGGGAAAACGAGCATTTATCTAGATTGGAAGAGGAGACTTCGTGTTGCGCTTGGTTCAGCCAGAGGATTAGCTTATCTGCATGAGCTTGCCAATCCTCCTATAATTCACAGAGATGTGAAATCAACTAACATTTTGTTGGATCAGAACTTAACAGCAAAGGTCGGGGATTTTGGCTTGTCCAAACTCGTATCAGATAGCTCAAAGGGTCATGTTTCCACTCAAGTTAAAGGCACATTGGTCAGTACCACATCCTGCACATTTTCGCGTTCTATTTCAAGAAAAAGATTGAATTTTGTTGAACTCTTGAAGCATGAAATTGAATTTAAGACTATTGATTACATTGCAGGGTTATCTAGATCCTGAATACTATATGACTCAACAACTAACCGAGAAGAGCGATGTGTATAGCTTTGGTGTTGTAATGCTAGAACTCATAACTGCAAAACAACCAATAGAGAAGGGGAAATACGTTGTTCGAGAGATGAGAACAGCAATAAATAAGAACGACGAGGAGCATTATGGCTTGAGCAATATGATCGATCCAGTCATTAGAAACATGCCAAATCTAATTGGTTTTACAAGGTTTGTGGACGTGGCGATGCAATGTGTTGAAGAGGCTGCTGCTGATCGTCCAACGATGAGTGAAGTGGTAAAAATGCTTGAAAGTATTCTTCAAAATGATGGTCTGGAAACGAATTCTACGTCTGCATCATCATCGATAACGGATTTTGGGACAGCAATAGCTGCTTCTAGACATCCTTATAACAAGGAGGCTCTACAGAGAAAGGAGATCAATGATACTCATGCGTTTGATTACAGTGGTGGATACACTCTTCCAACAAATGTGGAACCAAAATAGTGACAACCAAAAAAAAAAGGACTTGTCTCTACTGTCTCTCATTCTTTAATTTATTCTCTAGAAAAAAAATTGGAAAATTGGTGAAAGGGATAAGAGAATTTAGTTGTTGTATTGTCCCTATATTGGCTACTTTTCGCGAAACAATCACGTATAGTATCAGGATCGAAGTTCAGATTTTCTATTGCCAAGACTTACTGTCTAGTGGTACTAGTGAGAGTCTCCTATACACTAAGTGTGAGTTCGATTTTTACCACCCCCGATCCTCAAATGTAATAGAAAAAAGTTAAAAGGAAAAAATTTAGATTTTGTACCATCAAGGCTGCAGTCTAGTGGTATACCTAAGAGGTCTTCCATACACTAAGTGTGAGTTCGATTTTTTACTATCTCCTTTCCCATTCCCTGATTCTACAATGTAATAGAAAAAAATCAAAATTAAAAAGTTCAGATTTTC

>M6-PGSC0003DMG400008581 M6 LRR-RLK

ATGAATTTATTAACCATTTTCATACTCTTCTTGTTCTACTATTACAGTGTTTTTGCTAACGGAGTGGTTGCGAAAGTTGATTTGAATGATGAGGTGTCTATTCTACTTTCTATTAAAGAAAGTCTCGTTGATCCATTGGATCATCTTCATGATTGGACAGTGCCCAATAATGCAGCTGCAGGGAACAATAGAAGTATTATTGTTCCCTGCAGCTGGACGGGAGTTGAATGCAACTCCCATGGGGCAGTCGAGAAGCTTGATCTGTCCCACATGAATCTCACTGGCACAGTGTCTAATGATATCCAAAAGTTGAAGAGTTTGACTTCTCTAAATCTATGCTGCAATGAGTTTTCATCGCCATTGCCGAAAAGTTTATCAAATCTAACTGCATTGAGAAGCATTGATGTGAGCCAGAACTATTTTGTCTATGACTTCCCTGTAGGTCTTGGAATGTCAGAAGCACTTATGTATTTGAATGCTTCGAGTAATAACTTCTCCGGTTATCTTCCTGAGGATATTGGAAATGCTACATTGCTCGAGACATTGGATTTTCGAGGGAATTTCTTCGAAGGTTCAATTCCTAAATCTTATAGGAATTTGGGGAAATTGAAGTTTTTAGGTCTTTCAGGGAACAATTTGACTGGACACATTCCAGGAGAGTTGGGGCAGCTTTCGTCGCTTGAGACTGTAGTACTGGGATACAATGTTTTCGAAGGTGGAATTCCAGCTGAATTTGGAAATCTGACAAATCTCAAGTATCTTGATCTTGCTATTGGAAATCTTGGAGGTTCAATTCCTTCTGAGCTAGGAAAGCTGAAGCTGCTCGATACGATTTTCTTGTACAAGAACAAGCTTGAAGGCAAGATTCCGCCAGAGATTGGCAATATGACTTCATTGCAGTTGCTTGATCTGTCTGATAACATGCTTACGGGTGAAATTCCAGCTAAGATTGCTGAACTTAAGAATTTGCAGCTTTTGAATATAATGTCAAACAAATTATCAGGTTCAGTTCCCCCTGGGATTGGAGGTTTGACTCAGCTGGAAGTAGTTGAGCTGTGGAATAACAGCTTTTCAGGTCCTTTGCCAAGTGATCTTGGAAGAAATTCGCCATTGCAATGGGTGGATATTTCATCCAACTCATTTACCGGTCCAATTCCAGCAGGATTATGCGCCAAAGGTGACCTCACTAAGCTTATCATGTTTAACAATGCTTTCTCAGGTCCAATTCCAACTGGTTTATCAACCTGCACGCCCCTTGTTCGTGTAAGAATGCAGAACAATCTTCTTTCGGGGACGATCCCAGCAGGTTTTGGTAAACTTGGGAAACTACAGAGGCTGGAACTGGCAAACAATAGTCTGACAGGACAAATCCCAAGTGATCTTGCTTCTTCTACCTCTCTTTCTTTTATTGATTTCTCTAGAAACCACATTCAGTCTTCTATTCCTTCATTCATTCTTGCAATTCCAACTCTTCAAAATTTCATTGCCTCCGACAATAAGATGACAGGCGAAATCCCGGATCAATTCCAGGATTGTCCTTCTCTTACTGTTCTGGATCTATCCACAAACCATTTCACTGGAGATCTTCCAGCTAGCATTGCTTCCTGTGAGAAGTTAGTAACTTTGAACCTTCGAAATAATCAATTAAACGGTCCAATTCCAAGAGCGATTTCCATGATGCCAACATTAGCTATTCTAGATCTATCCAATAATTCTTTAACTGGTGGGATACCTGAGAACTTTGGCAATTCACCAGCACTAGAAATGCTGAATGTTTCTCATAACAAACTGGAGGGTCCTGTACCGGAAAATGGCATGTTGAGAACAATCAATCCGGATGACCTGATTGGAAATGCCGGTCTATGTGGTGGAGTGCTTCCTCCTTGTTCTCATAATGCAGCATACACATCAAAGCAGAAGAGCTTGCACGTGAAGCACATCATTACAGGATGGCTAATTGGTGTGGCAGCTGTTCTGCTTTTTGTGACAGCAGGTCTTGTAGCTAGATCTCTATACAAGAGATGGCATGAAAATGGGAGCTGCTTTGAACCAAGCTTTGAGATGAGCCGTGGTGAATGGCCATGGAGGTTGATGGCATTCCAGAGGCTAGGATTCACTAGTAATGACATCTTGGCTTGCCTTAAAGAGTCAAATGTCATTGGAATGGGAGCAACAGGGGTTGTATACAAAGCTGAAATGCAACGCGAGAATATGGTTGTTGCAGTAAAGAAGCTGTGGAAATCTGGAACTGATATTGAGATGGGGCACAGTGATGATCTCGTAGGTGAGGTGAAAGTTCTTGGAAAATTGAGGCATCGAAATATTGTCAGGCTTCTGGGGTTCCTTCACAACAAACGTGATGCAATGATAATATACGAGTATATGCAAAATGGTAGCCTTGGAGAGGTTTTACATGGTAAACAAGCAGCAGGGAGACTGCTTGTAGATTGGGTCACTCGATATAACATAGCCCTTGGAGTGGCACAAGGCCTTGCTTATCTCCATCATTACTGCCATCCACCAGTTATTCACCGGGATGTCAAGTCAAATAACATACTTCTTGATGCAAATCTAGAGGCAAGAATTGCAGATTTTGGTTTGGCAAGAACGATGCTCAAGAAGAATGAAACAGTTTCCATGGTTGCTGGATCTTATGGATACATAGCCCCCGGTAAGTCCTTCAAATTTTTCATCAATGTTTACTCTAGAGTGATTTTTGTTACTAATATTTAACAATCATTTATCTGGATTTTGACTTTGCAGAGTACGGATACACGTTGAAAGTGGATGAGAAGAGTGATATTTACAGCTATGGAGTTGTTCTCATGGAGCTCCTAACGGGTAAACGACCTCTAGATCCTGAATTTGGGGAATCTGTTGACATTGTAGAGTGGTTTCGGATGAAGATCCGAGACAACAAGTCTTTAGAAGAAGCACTTGACCCAAATGTAGGAGCAACACAACATGTTCAAGAAGAAATGTTGTTAGTTTTAAGGATAGCAATTCTATGTATAGCCAAACTCCCCAAGGACAGACCATCCATGAGGGATGTTTTGACGATGCTCGAAGAGGCAAAGCCTCGGAGAAAAAGCAATAGCAACAGTGGAGGTAGCCATGATGCTACTACTAACAAAGATAAGCCAATATTCAGTACATCACCTGTAAATGGTCTTTTGTAA

>M6-PGSC0003DMG400009068 M6 LRR-RLK

CGTGATTAAGAGTGGTTAATTTCCTTTCCCTTTTTCGTAATTGCAATCTGTGTCTTCCTTTAGTGAAGAGCTTCAAAAAATGGCGGAAATGGTGGATAATGCTCTTCTGGTGCTCCCTTGCCTTACTGGCACCACGTGCACCCACTTCTCCTCTTCTTCTAAATCCAAGCAAATTGTTAAAGAATTGATTTTGTGTATTACTGTTAACTTCTTCATAGCATAGTAGTAGTAGTTCCACTTTTGTCTCCACGTGCTTTTCCCACTCTTTTCGCTTCTTTGTTAACCAAATACATACATATACCCCATTTCTTTATTTGCTTTCTTCTTCTTCAAATTAACTGCCTTTCAATCATTTATTGCCTCTCACCCACTTCCCCCCCTTCTTCTCATTTAATCATTGTTTCTCTCTAGCTACTTACTTAATTCTGCATTTACTTCCCATCATTTTCCTCTTCATCTACTCCTTCTTTTCAGTTTACTCTGCCAAGTGTTTGATTTTTTGCCTCACTGGTACTTGCTTGTATACTATACAGTTTCTGAATTTATGGGGTTTTCCGTATTCTCCATTTTGATGTGATTTGGCTATCTGGGTATTGCTATTCAGGTTCATTTCAATGTTCTTTTTGTTATTGCCGACAGTTTTCAAATTATGCATACTTTGCTTTAAGGTCAAGAAATGTGTTAAAAATTCAATCTTTGTATGTGTGTGTGCTTGGAAATAGGTGAAGAACGTGAAATGCCTGTTTGTTCCTTTTTCTTTTTCTTTTTTTTTCATTTTTTGGCAGCCCCAAAAGCTGACGGAAGAAATTTGTAGCTGGTTCTTGTTTTTACTTTAGCTGTTCTGTTCATTGAAACCTAACTTTTACTTAACTTGCGTGGTTGACCCAAAAACAAAATGTGTTCTGAGTGTCTACGTAACTTTAGTTGCCCTATAAATTGTATATTTATGGTTAAGAAACGGTTCTCTAGTTTAAATGGTAATTATTTATAGTGAATAATGAACATCATGCCGTTTCAATGTCTGATTCCATTTTTATTTTTTTTTGAAATTCTTGTAGAGATGAAGGGTGCTGTTACTAGAAGTTGATGCCTTTGTAGTTTAGCTCAGAGGAGGAGAAGGATGGCTACAAAGTTCCTGTTTTTTTCAGTTTTATTCTGCACAGCTCTGTTCTTGCTAGCAAGTAGTGAACCATATGAGGACAAAGAAGCGCTTCTTGATTTCCTTAATAATGTTAATCATACCCGATACCTGAATTGGGATGAGAGAACTTCTGCTTGCAGTTCTTGGACTGGAGTCACTTGTAATCATGACAAATCAAGAATTATAGCAATTAGGTTGCCTGGAGTTGGATTCCGTGGTTCGATTCCTGGAAACACTCTTAGCCGCTTGTCTGATCTTCAGATCTTGAGTTTAAGGTCTAATAGTTTTAGTGGCAGTTTACCTTCTGATTTTGCAAACCTTGGAAATCTTACTTCTATTTATCTTCAATCCAATAACTTTCAAGGTCCATTGCCTGCTGATTTTTCTGCTTGGAAAAGCCTTTCGGTTTTGAATTTGTCAAATAATGATTTTAGTGGGAGTATACCTTCTTCAATATCGAACTTGACTCATTTGACTGCTCTGGTTCTTGCTAATAACTCGCTTTCTGGCAATATTCCTGATCTCAATCTCCCTAGTTTGCAAATATTAGATCTATCAAACAATAACTTCACTGGAAGTATCCCCAATTCTCTTCAAAGATTTCCTGGTTCAGCTTTTGCTGGTAACCGGCTTTCTCCTGCAAATTTTTCACCTTCATTTCCTCCAGTTCCTCCTCCAAGTGTCCTACCGAAAAAGAAATCGTTTAAACTCCGTGAACCTGCAATACTTGGAATTGTAATTGGAGGGTGTGTTTTAGGTTTCTTGGTAGTAGCTGCAGTGCTGATTATGTGCTTTTCGAAGAAAGAGGGTAAAAGTGGGGCCACAGAGAAATCAATCAAGAAAGAAGATTTAGTCAGGAAGGGAGTTTCCAGCAGTCAGCATGGAGTAGGGAATCTCGCATTCTTTGAGGGTTGTAATCTTGCATTTGACCTTGAAGATCTGTTGAGAGCTTCAGCGGAAGTGCTTGGAAAGGGAACATTTGGCACTACATATAAGGCAGCTTTGGAGGATTCTACCACAGTTGTGGTGAAGAGATTGAAGGAGAGTGTGGGAAGAAAAGATTTTGAGCAACAGATGGAGGTTGTTGGCAATATCAGGCATGAGAATGTGGCTCCACTGAGGGCATACTACTATTCCAAGGATGAGAAGCTCATGGTGTATGATTTCTACAGCCAAGGGAGTGCATCTTTGATGCTGCATGGTATGTTAAGCACCCAATTTCTACTAATCTTGCTGTTTTATTCTTGACCGATCCATAGAGAACTGGTTATCAGGACCAAAAAGATAATGCAGATCATTTTTAGCTATGATGTGTCTTGTGCAGTAGTATTTGAAATTGCTTGTAGGCTAATTCCAACATAACCAATTACCTTTGAATTAATAGAGTATCTGCTGTTGTTGTCCTCTTGATAGTTTTTTTTTATCATCAATAGCTTATAGGACAGCATAAGAGTCTCGGAGTCATGTCTAAGTGCCATTTTTGTTTTAATTTGTCAAAATTCAATAACATCATGTTGCTCGCGAACTTTGCTTTGGCGCTATAGTCAACTTTCATCATTGTTAGGTGGATAGTCTCTTAACTGCACCTTCAAGTATTCCTATCATTATATCTGATGCACTCTGTAATTGCAGCAAAGAGGTCTGCTGATCGAATTCCTCTGGACTGGGAAACCCGACTACGCATTGCCATTGGTGCAGCTAGAGGCATTGCTCAAATACATGGACAGTCTGGGGGGAGGCTTGTCCACGGGAACATAAAATCCTCAAACGTATTCCTCAACTCCCAAGGTTTCGGTTGTATTTCTGATCTTGGCTTGGCAACTATAATGGGTCCAATTGCTACACCAATTGTGCGAGCTGCTGGTTATCAACCTCCAGAAGTTACAGATTCAAGAAAAGTATCTCAAGCCACTGATGTCTACAGCTTTGGTGTCCTAATACTTGAACTTCTCACTGGAAAGTCACCTACACATGCCACAGGTACAAGCGACATTGTCCACTTAGTCAGGTGGGTACATTCTGTCGTTCGTGAGGAGTGGACTGCTGAAGTATTCGACGTGGAGCTTTTGAGGTATCCAAATATAGAAGAAGAAATGGTAGAGATGTTACAAATAGGTTTGACTTGTGTATCCAGAATGCCAGAGCAGAGACCAAAAATGACTGAAGTAGTAAAGATGGTAGAGGGTGTTAGGAGAGTGAATACGGGGACTCGAACATCCACTGAAGCTTCTACACCGAACCTGACACCACCGATGACTGAAATAGGGTCGTCTTCCATTACACACTGATATTTCTTTGTTATTTAGTGATTAGTTACTGTTCTATATATGCTTAATTGTTTCATTGTTATAAGTTTTTTCCTCAGACTTCTGTCAAACATTATTTCAGCTACAAAATTGAATAGAAATATTTACTTGATC

>M6-PGSC0003DMG400009180 M6 LRR-RLK

ATTAATCAAAATAAAAATAGGAAAGTTCTGACAAAGCCTTTCTTCTGTTCCCGAGCTGAAGCTTGGGGTTTTGTGTTTTGTGGTGCCCGGTACTGGTGGTGAAAGAAAACTCCGACGACTGTGACTGAGGTGGAAGAGGAGAAAACGACGCCGTGTAGATGAGGTTGCTGCTGCTGTTGCCGATACTCATCGGCATTTTGCTTTATTGTGCTAATTATGCCATTGCCGACACAGATCCAAATGACGGTCAGCTATTGCTTTTAATCATTCTCATTTATTGTTTCAATCTGTTTGTCTGTATGCTTACTGATCATTAATGATTTAATGGAGGTTGATTGATTTGGAAAGTCAATGAGTATCAATCTACCTGACAATGAACTCGGCAACCGTCACAAATTTGTTGATATTTTCCTTACCTGGATAGTATATTGTCTTATTGTATATGTAAAAATGCGACAAGTTCCTTTCTAATCAAGGCATGAATCATATAGAGGGAGCTCTCTTGGATTCTTATATATGTTATGTATGTATTGGCCATGTCGGTGGACTTTTAGTACTGTTGAATCTTTGCATTATTACTTCCATTCATTCTGAGAACTGACATATTGAAACTTCGTCAGACATATTTGACTTAGTTACCATTTTCAGCCAAGTTGCTCCAGAAGATTTTGGCTTTTCTTCTTTCTGCTGCTTGGAAGCGGGACAGGGTAGTGGGGGTGGGACGCCAAAACAAGTGTGATCAAGGGCATTATAACCTAAAAGACTCTTCTCCCAGTGATAATACACAAATAGGTTCATGGCTTCATCAACGCACTTGTTGCGTTTCACCATGTTTATTGATGTCTTGTAGATTTACCACTCGTTAACAGTTTGTCCACCGTTTGATTACTCTTCAAGCTATTATTAATGAAAAGAAACTTTTTGTTTGAACAGTATAAGAATATGTTGACCTGTAAGTATGCCTAGCTGATTTTTTTGGTGGCGCTTTGATTTTCAGCTTCTGCTCTGAGAGTTCTGTATAGCAGCCTAAATTCACCTGGACAGCTAACAAAATGGAATGCCAATGGTGGTGATCCTTGTGGAGAATCCTGGACAGGTATTACTTGCTCGGGCAATAGAGTTACTGAAATGTGAGTTCCTCTCTTCCTTACTGTTGTAAGTAATTATTGTTGATGTTTTCCTTCTTTATCTTCTTCATCATGTTTCAATACTTGGAATTGTTGACTTGGTGATTTCTGATGCTTTCTTCCCTTTTCATTTTCCTCCAGCAAGATATCACGTCTTGGACTCTCTGGTTCACTGGGCTACCAATTGGCAAGTCTTACATCGGTGACTAATTTGTAAGATAAATGGCCTTCAATAATGTCCTCATAAATGTTTCTTCGATCTATATCTATATCAAATTCTTACACACTTCTAATTTTGCAGTGACATAAGTAATAACAATTTGGGAAACCAGTTACCTTATCAGCTTCCTCCAAATGTGCAAAGATTGTATGTCTCGTAGTTAGTATATTTTCATCATTCCAGATGCTATTTTTCCACCTATCTTTCTTAAACCTTGCCTTGCAATGCAGAAACCTTGCTGCTAATGGTTTTAATGGTGCCCTTCCATATTCCGTTTCGCAGATGACTTCCCTCCGATACTTGTATGAAAAATCTTTGAATTTGACTGCAATTTGCTTTCCAGCAATAAGCTTTTGGTTTAAGGTTGCGCTTTTCAATGACTTTAATATTTTTATCTGGCTGTAGAAATGTCAGTCATAATCAAATTCAAGGACAAGTGACTGTCGCGTTCGACTCTCTGTCTTCTCTCGATACATTGTAAGCCCTATCTTCTCTCATGCCATTTCTTGAGTGTTTTAATATTACAAAGTTGTGGTCATAACTCATATATGAGGTGTTGTAGATAAAACATGTCATTCTTACTTCATTAGTATAAACTTTTCTTCTCGCATCCGAGTTGTTGACTTGTTGCTTTGGGAAAGAAAGTCAATATCATGAAAGACAGTATCTGATCCTTGTTGTAATCTGTAAACTTATGTGATTGTCCCAGTTAAGTCAACTAAGAAGCTGTATACATTTCGAGATATTCTTAAGTCAAGTCTTAGAAACCTCTAAATGACTAGTTTATCATTCTGTCTCATAGTTTTTGATTGATTGATCTGTACGCTTCAGGGATTTCTCATTCAATGCTATGACGGGTGATCTTCCTCAAAGCTTCAAGACACTGACTAGTATGAATAAAATGTGAGCTAATAAATTTTCTGCTTCGTCCTTGTAGACAATTGGTTGAGTCCTTCAGTGGTGAAGTTTACATGTTGAAGTATGCAGGTATTTGCAGAACAACCAGTTTACAGGAACTATTGATATCTTAGCCAATCTTCCTCTTGATGATTTGTGAGTTTTCAGTCTATGCTTCTTTTTAGATGTTATCCCTGATGTTTTATCTTATTTATATAGACTCGATTGAAAGCATGGTTGAAAGTCATGTGTATTCCTTTATGCTTTAGGAATGTTGAGAACAACCGTTTTACTGGCTGGGTTCCTGATCACTTGAAAGGGATCACGAAGTAAGTATATCATGTTAGAGTAATAAATTTTGAAACTTATATCAGAAGTATTTTGAATTTATTCAGGGTTTGTGACATTTCTGTTACAGATCAAATGGTAACTCATGGAACTCAGGATCTGCACCCCCTCCTCCACCTGGCACACCCCCTGCAAGCAGGCCGCACCACAAATCTGGAGGCAATAATAGCCCTACTGATGGTGCAAGCAGTGGTGATGGTGGTGGTAAATCGGGGATAGGGGGTGGAGCCATTGCAGGCATAGTGATATCTGTTCTAGTTGTTGGGGCAATTGTAGCATTCTTTGTTATCAAGAAGAGATTGAAGCGGTCATCGACGGACATTGAGAAACATGATAATCAGCCATTTGCTCCTCTTGCTCCCCCCCAGGAAGTACATGGTACTTCTTTTCATAATTTCTTGGTTTTATCAGTTAAGATGTCTTTTAGACCCTTTCATGGCCTCTGTGATTGTTATTTAACTGTGGAACATATCAATCAACCACACTATCATCAACCTCATTCCTGTATCCTGCTTAAAAAGGTGTAAAATAAACCTGCTGCATAAAATGGAAATTTTATGAAAGTGAATTTGGAGAATCAATCTTATCCTTTGTGCTTTATCACTGCCTGTGGTGTATGCCTAATAGTGGATCAACACAGCCACTGTCTTCGCTATAAAAGAAAAACACCAAAGTAAAAACAAACTTTATATCGAAATGTAAAACTGCTAAGGCTGATAGGTCCTATCTTCCATGGAAGCAAAACTTCTTGCTTACTGAAAAGTTCCAAGAACTTGATTGTAGAAATCTGAACTGATGTTTATACTCTGTATTGCCTGTAGATGCTGATAGTGGGATAAAGAGAACTTCAGATTTTTTGATAATGAAGAAAACTTCAGCTTTATGATCTCTTTCAGTGGATCTGGTCCTATGCTTGTTATAAGTTAGCTCCACTTGATGGTTTTCCTTCCTATGAAACTACTATTCATAAATCTTATTGACCTTTATAGAACAACTATTGCTTCAAAATGCAACTTTCAGCTAAGTAAAAGTTTCTCCCATTCTAAACCATTTGTGCTCACCACCAAATTCACCTCTTTGCAAATTCTAATTCTTTGCTTTTCTCAACTCCATCACTCGTGGTGCATAGTTATGCCTTTTAAGCCTCATCTCACTTCCGTATCTCATTAATCCGAACCTGAGTATGCATTTGCCTGTTTGTAACACTTACAGTCTTGTGCTTCTTCTCTGTCTGGATTTTCTGTTCATTCATTGTCCTTTTTTTTTACTGCAGAATTGAAAACTAATGAAGCTTCCTCTGCGCCAATAGTGAAAGTATTTGAAGCTCCTGCAGTAGTAAATCTAAGACCTCCACCTATTGAACGCCACAAGTCATTTGATGAAGTTGACATAGCAGCAAAACCAATTGTCCCACCCAAGAAAGTTAATACAGCTAAAATAGATGCAAGACAATATTCAATCGCAGACCTACAGATGGCGACTGATAGCTTCAATGTTGATAACCTTATTGGCGAGGGATCATTTGGGCGTGTTTATAGGGCTCAGTTCGAAGATGGAAAGGTATGATTTCTCTCTTATGACTGACTGACTATCTATGTGAAGAAAATGGCTCTCTCTAATTGGATAAAAAAATCATGAATTTTTTATGTGCTGAGAATGAATGTTTAAGCAAGATAGTTGAAATGACAGATAAACGTTCAATATGAAAAAGTACCTATCTAAAGAAAATGAAAGAAGTTTGATCATTAAAAATTAGTTGAAAAATTATGTTTGTTCAAAACTCAGTTCGGAGCTACTATCATCTTGTCCTGTGTAACTGAACCATCCTCATAAGTTCCTTTCTAATTTTTGAAATGCCATAGGTTCTTGCAGTCAAGAAAATCAACTCTTCTGCACTCCAGAATCCTGAAGATTTTCTCGATATTGTTTCTGAGATATCACGGTTGCACCATCCAAATGTTACTGAGCTGGTTGGATATTGTTCAGAGCACGGGCAGCATCTGCTGGTTTATGAATTCCACAAAAATGGTTCTCTGCATGATTTCCTGCATCTGTCAGATGAGGAAAGCAAGCCTCTAACATGGAATAGCCGAGTTAAGATTGCACTAGGCACAGCAAGAGCACTAGAGTGAGTTTCCATGATCTACAAAGATATTTGATGCCATTTTATAATGTTCACGGAGCTCATAACAAGGATTTATAATTTCCTTGTACCATTGGTTGAAATACTAGTCTCTGTCATATATTTTCTGGTATGTGTAGTCAAACAATAATGAAGTTTCTTTTCCTGGGGGGGGAGGGGGGGG

>M6-PGSC0003DMG400009181 M6 LRR-RLK

GAGGAGAGGAGAATAGTATGGTAAAGTCTCTCTCTCTCTCCCACGTATATTACACGCTTTCAAATATCCATAGAATAAACAAAGTCCACCAGCCATCTATCTATCTAGCCGTCCCAAAAGTAAACGCTTTATTTCAAAAACTCATACCTCACCTTTTTGCTACATAAAAATTGCACCTTGTCAACATCAACTACACTCTCTCTCTTCTCTTCATGGGGTTAGACTAAAGTCTGCACGATCAACAGGACAAAGAATTGCCAAAAAAATACCCACATGGTTGATTGGAGTATCTTGGCATCTTCAACTGTCTTTCTTCTTCATCTTTAGTTGTCTAAAAATAATAATGATCATGAATAATAAAGACAGCCGCCACCAATCTGGTCTGACATATCATCAATTTCGAGCAGCCATGCTTCTTACTTCCTTTCATTCTTCTTCTTCTTTACCCCACCATAAATCTTTCAATCCAATAATATTGATAGTTGTTCTACTTCTTTCTTCAATTGCTACCATCTGTCATGCTTCTTGCAATCAGCTTGATCGTGACTCTTTGTTATCATTCTCTGTTGGTATCTCTTCACCTTCTCCTTTGAATTGGTCCTCTTCCGTGGATTGCTGCACCTTGTGGGAAGGTGTTGGTTGTGATGATAATGGCCGAGTAACCACTCTCTGGCTTCCCTCAAGAAGCCTTTTTGGAAACATAACCCCTGCTATTGCAAATTTGAGCAAACTCTCTCAACTCAGCCTGTCAAATAATCGATTTATTGGTCCCCTCCCAGATGGATTTTTCAAGTCATTTAGTAGCTTGCAGATCATTGACTTGAGTTATAACCGTTTATCAGGACGATTGCCGCTATCAGATAGATTAACATCACCCATCAAGACAGTAAATCTCTCTAGCAACCACTTCAATGGGACAATTCTATCATCATTTCTCGAGCCAGCAATTAATTTAGAGAGTTTCGATATTAGCAACAATAGCTTCTCTGGTCCAATACCCTCCTTTATTTGCAGCTACTCAGCCGCTGTCAGAGTTCTTGATTTCACTTCCAATGACTTCAGGGGCCAGATGCCCCAAGGGTTTGGGAGTTGCTCCAGTTTGGTTACTTTGAGAGCAGGATTCAACCATCTTTCGGGATTCATTCCTGATGATATTTATAGTGTGTCAACACTGCAAGAAATCTCTTTACCTGGAAATAAATTTTCTGGACCCATCCCAGAAAGCATTGTCAACCTTGTCAACCTCAGAATCCTCGCACTCTATGGCAATGAATTGACAGGTTTGATCCCTCAAGATATTGGAAGGCTCTCTAGATTGGAACAGTTGCTCCTCCATATAAACAATCTAAATGGCACTGTGCCGCCATCGCTGATGACTTGCACCCGTCTCACTGTGCTCAATTTAAGGGTCAACTTCTTGGAAGGTGAACTCTCAGGTCTTGATTTCTCCAACCTTAGTCGACTTGGCATAATCGACCTTGGAAATAACTTCTTCACTGGAAGCATTCCACAAAGCCTTTTTTCATGCAGGTCATTAACTGCAATCCGTCTGGCTACTAACTATCTGACAGGAGATATCTTGCCTGGCATAACGTCTTTACAAGCTCTGTCCTTCCTCTCGGTTTCCAATAAAAGCCTAACCAATTTTGCAGGGGCAATAGAAGTCCTTAAGGGTTGTAAGAATCTTACCACACTAATCCTCACCAAAAACTTTTATAATGAAACGTTGCCTGATAATGAGAACTTGATTGGGTCTGAAGATTTTCAAAATCTCCAAATTCTGGGTTTAGGTGGTTGTAATTTCACCGGACAGATACCCACATGGCTGGTTAAACTTGGGAGGGTAGAGGTTCTAGACCTTTCTATGAATCAAATCACAGGCAAAATTCCAGGTTGGTTGGGGACTTTACAGAATCTATTTTACCTGGATTTGTCTCAGAATTTTCTATATGGAGGCTTCCCAGGTGAACTTACTCAACTGCAAAGACTCGCATCACAAGAGGCTGCTGATCAGGTCGACAGAAGTGCTTTAGAATTGCCTGTGTTTGTTCAGCCAAACAATGCCTCAAATCAGCAATATAATCTACTGTCAAACCTGCCACCAGCTATTTACCTTGGAAACAACAACCTTGATGGCAATATTCCAACTAAGATTGGCCAGCTGAAGTACATTCATGTTCTTGATCTGAGCAAGAACAACTTTACAGGAAACATTCCAGAAACAATATCCAACCTCACTAATCTGGAGAAACTAGACCTCTCTGCCAACAACCTCTCAGGTGAAATTCCTTCTTCACTCAAGGGTCTTCATTTTTTGTCTTCTTTCAGTGTTGCTCACAACAATCTTGAGGGACCTATTCCAACAGGAGGTCAGTTTGACACATTTCCTATCACTAGTTTCTTAGGTAATCCAGGACTGTGTGGTCAAATCCTGCAGCACCCTTGCCCTGATCGATCAGGAACTACACAACCTTCTGCAGTTAGAAAAACCGCAAAAAGGAAAATCTTAATTGGACTCATCCTAGGGATCTCCTTTGGCATTGCCTTCACAGTCATCATCATAGCATTCTGGATATTCTCCAAGAGGAGGATCCTTCCAAGAGGTGACGCCGAGAAAAATGACATGGAAATTGTGTCCTACAACTCCACTTCTGGATTGTCTGCTGAGATTGGAAAGGATAACAGCATGCTTGTTATGTTTCCAACCAATAAGGATCAGATTAATGATCTCACAATTTTTGACATTCTGAGAGCCACGAATAACTTCAACCAAGCAAACATAGTAGGTTGTGGAGGCTTTGGCCTGGTCTATAAAGCAACTTTAGCAGATGGAACTACACTGGCTGTTAAGAAACTTTCAGGAGATATGGGTTTAATTGAAAGAGAATTTAAAGCAGAGGTGGAAGTTTTATCCACTGCCCAACATGAGAATTTGGTCTCTCTTCAAGGTTACTGTGTGCATGATGGCTGTAGGTTGCTGATCTATTCCTATATGCAAAACGGAAGTCTGGACTACTGGTTGCACGAGAAAACTGATGGAGCATTCCTACTTGATTGGCCAACTCGACTGAAGATTGCACAGGGGGCAAGTTGTGGGCTGGCTTACATGCATCAGATATGTGAACCACATATTGTTCATCGTGACATAAAGTCCAGTAACATCCTCCTTGATGAAAAGTTTAAAGCACATGTGGCAGATTTCGGATTGTCCAGAATGATTCTTCCTTACCAAACTCACGTCACCACTGAACTGGTTGGTACCCTCGGCTACATCCCACCAGAGTATAGTCAATCATGGATAGCCACTTTGAGAGGAGATGTCTATAGTTTTGGAGTTGTGATGTTAGAACTCCTGGCTGGCAGAAGACCTGTGGACATGAGCAAACCAAAGATGTCAAGGGAATTGGTTGTATGGGTGCACCTAATGAGAAATGAGGGTAAACAAGAAGAAATATTTGATCCTATACTGAGAGACAAGGGTTTCGAAGAAGATATGCTTCAAGTGCTTGATGTTGCATGTATGTGTGTCAGCCAGAATCCTTTCAAAAGGCCAACTATAGCAGAAGTGGTTGAGTGGCTCAACAGAGTAGTATCCAACCAGGGAGCACCTAAATAAGGTGGTGGTACACATAAATTTGATGTAGATGTGAGTGTCTGTATAGATGAACTTTTGCTTTTTACACGTAGGAAGTTAACATACAGGAATAAATGTATAAAGTGAACAATTTCCTCAATAAATTACATTTACGATTTTGCGTCATGGCTAACATTGGATGACAGGTAATG

>M6-PGSC0003DMG400009196 M6 LRR-RLK

TTCTTGAAAGATGACATTGAGAAATTTGACTTACCTGATTTGTTGAAAGCCTCTGCTGAAGTATTGGGAAGTGGAGTGTTTGGTTCAACTTATAAAGCTGCACTTAGTACTGGTCCTGTCATGGTTGTTAAGAGGTTTAGACATATGAATAAAGTTGGTAAAGAAGATTTCCATGAACATATGAGAAGGCTTGGAAGATTGAGTCATAAGAACTTGCTTCCTGTTATTGCTTTCTATTATAGGAAAGAGGAGAAGCTTCTTGTCTTTGAATATGTCAACAATGTCAGCCTTGCTGTTTATCTTCATGGTA

>M6-PGSC0003DMG400009238 M6 LRR-RLK

ATGAAACACAAAACATTTGTTCCTTTGTTCCCTCCCCTTATTTCACCTGAGCTCAGAGAGTATATGCAAATCTTGTACATAGATTTTCTTGAAACCCCTTATCATTTTCTTGAAGTTTTCAGCTTTCTGACTGTCAAAGCCAAAAACAAAAAAAAAATGAACTTAAGGTATTCTCCCCATATAGTTATTACTATATTCTTGTCTTTTTCTCTTCAAATTGTGTTTTCTCAATTGCCCCCAAATCAAATTAATGACATGAGAAGTGTCTATGATCTGTTTCAGAATGAAACTGGTGCTTCTTTTGTATGGAATGGAACTGATAAAGCTTCAACCCCATGTTCTTGGAAAGGGGTTTCTTGCAATTCTGATAATTCTTCCCTTACCAAAGTTACCTTTTCATTGTTCACTATTTCTAGCTCTGAATTCTTGCCTGTTATTTGTCAAATTGATACTTTGGAGTCTATTGATGTTTCCCAAAACTTTTTGAGCTCTATCCCAAATGAGTTCATCACTGTTTGTGGGGGAATTAGTGGGTTGAAATTGTTGAACTTTAGTGGGAATAAATTGGAGGGTTTTTTGCCTACTTTCACTGGTTTTGGAAAGTTGGAGTCTTTGGACTTTTCTGACAATAAGATGAATGGGAAAGTTGACTTGCAGTTGGATGGATTAGACTCACTCAAGAGTTTGGACCTTAGCTTTAACATGTTTAATGGTTCAGTTCCTACCAGTCTTGGAAAATTTAATCTTCTGGAGGAGCTTCATCTTTCTGCAAATGCTTTTGGAGGTGAATTCCCCACTCAAATTGTGAACTTTGGCAACTTAACTTTGATTGACCTTTCTCTAAACAACCTATCTGGTGTAATTCCTGATAGATTAGGTGAACTTTCAAAATTGCAAGTCTTGATTTTGTCAACCAATAGATTGAGTGGCACAATCCCACAATCCCTTAGGAATATCACAACGCTGACGCGTTTTGCTGCAAATCAGAATAATTTTGTTGGAAATATACCCTTTGGTATAACCACATAACTGAGGAATTTGGACCTCAGTTTTAACAGATTAAATGGTACAATTCCTCAGGACCTCTTATTCCCAATGAAATTGCAGTTTGTTGATCTCACTTCCAATAAATTAGAGGGACCTGTTCCTTCAAACATGTCGATAAATTTGATCAGGTTGAGATTGGGGCAAAATGCTTTGAATGGATCATTTCCATCTGCTTCATTTGAAAGCCTTCAAAGTTTGACCTACTTGGAACTGGACAACAACCAGTTAACTGGACCAATTCCTTCTGAATTGGGAAAGTGCCAAAAGATGGCCCTTTTGAACTTAGCCCAGAATAAGCTGAGTGGTGTTATTCCTGTTGAGTTGGGTGATATTTCTAATCTTCAGGTACTGAGTCTTCAGTCCAATAACCTAGTTGGAGAAATTCCAAGTAACATTTCACAGTTGAACAGATTGCAGAGGATCAATTTCAGCTAGAATTAGTTGACTGGTTCCATTCCAAGTTCATTATCAAGTTTGAGAAATCTCACAAACTTGAATTTGCAGGGGAATAATCTAAGTGGTCGAATTCCGGTTGACATTAGTAACTTAAATATGCTATTAGAACTCCAACTTGGAGGGAACCAACTCGGTGGGCCTATTCCTGATATGCCTTTGAGTTTACAGATTGCTTTGAATCTGAGTGGCAATCTTTTTCAAGGACCTATACCAAGTAGTTTTTCAAGATTGACTTCATTGGAAGTTTTGGATCTCTCTTACAACAGGTTCTCGGGTCAGATTCCAGACTACCTGACGGGAATGAGAGGATTGACTAGGTTGGTGCTTTCAAACAATCAACTGTCTGGAGTTATTCCAAAGTTTGAAAGATTTGTCAGTGTTGAGACAGATGGAAATGGGGATCTGATATATCCATACACAGTTAATCCACCACAAGCTGCTGCAAAAAAGAGAAAATCAATAGTTGTTGCAGTTGTTGTCCCAATTGCATGTGTAGCAACAATTGCTGTATTAGTGGTGATTGCTATTTCAATCTCTAGAAGATATTACAGGATCAATGATGAACATTTTCACTCAGGATTAGAAATTTCTCAATCCCCTGTCGTTCAGGGAAAGGTTTTAACTGCAAACAGCATTCACAAGTCAAATATAGACTTCACAAAGGCCATGGTAGCAGTGTCTGAGCCCTCAAATGTTGTGTTCAAGACAAGATTTTCAACCTACTATAAAGCTGTTATGCCGTCGGGTACAACCTATTTTGTCAAGAAGCTTAATTGGAGTGACAAAATATTTCAGTTGGGAAGCCATGAACTATTTGGGGAAGAGCTAAAAAATATTGGAAAGCTGAATAATTCAAATGTTATGATTCCTCTAGGCTATCTTTTGGCTGCTGATAGTGCTTATCTCTTCTACGAATTTGCCCCCATTGGCTCGTTATATGATGTTCTTCGTGGTAGCTTGGGCTACTCGTTAGATTGGGCAAGTCGCTACAGCATAGCTATTGGAGTAGCTCAGGGTTTAGCTTTTCTTCATGGATGTGAAAAGGGTCCAATCCTCCTCCTCGACCTTTCCAGCAAGAGCATTCTCTTGAAGTCCCAGAATGAGGCTCAGATAGGAGACATTGAACTCTACAAGGTGATGGATCCTTCGAAGAGCACAGGCAGCTTTTCTGCTGTTGCTGGTTCTGTCGGTTATATTCCTCCAGGTAATGAACTATTACTTGGTTAATGGTAAAAGTTTAAGATCCTTCAACATTGATTTATACTCCTTAAGGGTAATGTGACGTTTTTTGTCCTAACTGTCGTACCATGCTTTAAACTCATCTTTTTGTGTTTAATCTTCATATAGTTATTGTTGTTTTCAAACTTAAGACTTCATTATATCCTCCCACCATGAAACGGTGTTGAACTATCTTTCGATATCTAAAGTGAAACGGTGTGTCAGTATAGCTTAGAGAGGGTGGGGTGCATTAAAATGTAACATGAGAAGCCTCAAATTTGAGCATATATTTAGTGATGCATTTCCTTTTTGTGTTACAGAATATGCATACACAATGAGAGTCACAATGGCTGGAAATGTATATAGCTTTGGGGTTGTCCTGCTGGAATTGTTGACAGGAAGGCCTGCAGTTAGTCAGGGAACCGAGTTAGCCAAGTCAGTGCTGAGCAACTCTGAGAAGCACAGCAAGTGGGATCACATTCTGGATTCTAGCATTTCTAAAACATCGCTCAACGTTAGGAGCCAGATGTTAGCAGTCCTCAAATTGGCTCTAGCTTGTGTCAGTGTATCACCAGAAGGCCGGCCAAAGATGAAGAGTGTGCTTCGAGTGCTTCTCCATGCAAGGTAATCGGACGTCAAAGGAAAGTTAAGGAGAAAACTGCCAAGGTGGTTTTCTTCTTTTTCTTTTGGGGAGGAGCTGCAAGTGGTGAAGGTGTTGTGAGCTATGCAGAATATGCCAGTTTTGCCAGTTTATACATATTATATTAATGTAAACATATAGATATATGATGATAGTAGTTACATAACTTTGAATATGATGTTAGTTCTCAGTGTAACATAGTCCATTTTTCTTGTTCATGTGTAATATATATAATATATAAAAGGTCTTTGGGAGTGAATCCAATTT

>M6-PGSC0003DMG400009380 M6 LRR-RLK

AAAGCTTACACCATTTTAACAAGCTCAGCAAACACCCATTTCACCATTATTACAGAATAATCTCTGTATCTTCTTCCTGTGCATGAAAAACCATGCAAATGAAACTATTACTCTTCTTTCTGAGTACATTCCCTTTGATTTTTGCTTTAAATCAAGATGGGTTGTATCTACAAAAACTGAAACACTCTTTGTCAACCTCAGATCAAGGGGTATTTTCTACTTGGTATGAAAATGATCCTACCCCATGTAACTGGACAGGTGTCACCTGTAATGACGCCGGAGATTCTCCCTCCGTCGTCGCTGTTAACCTCTCCGGTGCTTCTCTTGCCGGACCCTTTCCGGTTTTCCTCTGCCATCTTACTTCGCTTTCGTCACTCTCGCTTTCGAATAATCTTATTAATTCTACTCTTCCGGTTTCTATTTCTGAATGCCGTAGCCTCACGTACCTTGACCTTTCTCAGAATCTCATCGGTGGAACTATCCCCGACACTATTTCCGATCTTCCTTACCTCAGGTAACTCAGTTTGAACCTCAATTTTGCATTTCCCCTGTTTTTTTTTTCTTTCAATTTTTCGAAAAATATCAACTAATTGTGACTTACATTTAGCCATATATGCGTATTTGGGTCTGTTTTAAGGTGAATATAAGATCTATAGTTAGTGTATATAACATAAACATCTCGCATTAGCAAGGTCCGGGGAAGGGGCTGTAAGCAATGATTCAGCTGAACCCACATAACCTGCCCGGATTTGCTTGCTGGTGTTATAGTAACTCTAAATTACTATTCATTGTTTGAATAAAAATGCAGATGGAACGAAATGACTATTATGATCTTTTATTGCAGATTTTGTTATTCTTCTTGTTTCATTAGTATATATTATGATCTTTTATTGCAGGTATCTTGATCTTAGCGGATGCTATTTTTCGGGGAATATTCCGGCAAGTTTCGGAAGATTCAGGCAACTGGAGACTCTTATTCTGACTGAGAACATACTTACTGGTGAAGTTCCTGCTGCATTAGGTAATGTAACGAGTCTCAAGACACTTGAACTTGCTTACAACCCTTTTGCATCGAGTCAGTTTCCTCCTGAACTCGGTAACTTGACTAATCTTGAGACATTATGGCTAAGCATGTGTAGTCTTGTTGGTTCAATTCCAAAAAGTATTGAGAAATTGAGTCGATTGACTAATTTTGATGTGTCCAATAATGGACTGGTTGGGTCAATACCAAGTGCAATTTTCCAGCTTAATAGTATTGTTCAAATTGAGCTCTACAATAATTCCCTCACCGGAGAATTGCCTTCGGGATGGTCTAACTTGACCAGGTTGAGAAGATTCGATGTGTCGACTAACAAGTTAAATGGGACTATTCCTGATGAGTTGTGCGAGTTGCCACTTGAGTCACTCAATTTATTTGAGAATCAATTTGAGGGGTTTCTTCCAGAAAGTATAGCTAACTCGCCGAATCTGTATGAGCTTAAGTTATTCTCAAACAGATTTTCTGGTTCATTGCCTAGTGAACTGGGGAAGAACTCGGCTTTACAGTATCTTGATGTTTCATACAATACATTTTCTGGTAAAATTCCTGAAAGTTTGTGTGAGATGGGAGCTTTAGAGGATATTATAGTGATATATAATTCGTTCTCTGGGAGTATTCCGGCTAGTATTGGCAACTGCCGGAGTTTACTTAGGGTCAGGTTTCGGGCTAATAAGCTATGCGGGGAAGTCCCAACTGAGTTTTGGAGTTTGCCTCAGGTTTATCTCTTAGACCTTTTTGGCAATGCATTTTCAGGAAATATATCACACATGATTTCTGGTGCCAAAAATTTGTCTAACCTACAGATATCAAGAAACAAATTCTCAGGGGTAATACCTAGTGAAGTAGGAAAGTTGAAGAACTTGGTTGAGTTTTCCGCGAGTCATAATGAGCTAACGGGAGAACTTCCAGACACATTAGTGCAGCTAGGGCAGTTAGGAACCCTTGATCTTAGTTTCAATGAGCTATCAGGGAAAATCCCCTTCGGAATTCACACAATGAAGCAACTCAGTGAGCTTGACTTGGCAAACAATGGATTTTCTGGGGAAATTCCGGAGGAAATTGGGACTTTGCCCGTGCTTAATTATCTTGATCTTTCTGGGAATTACTTCTCAGGGGAAATCCCACTCAGTCTGCAAAGCTTGAAGCTTAATAAGCTAAATTTGTCTAATAATCAGCTGTCAGGGATGATTCCTGCAGTTTTTGATAAGGGTGTTTATAGAGACAGCTTTCGAGGTAATCCAGGTTTGTGTCAAGGTGTTGCTGGTCTTTGTCCTACGAAAGGTAGAGGACAGCATGAAGGATACTTATGGACTTTGAGAGCTATCTACACAGTTGCTGGCTTCGTTTTTCTTGTCGGGATTGCTATGTTCATTTGGAAGTATCAGAAATTCAAGAAGATTAAGAAAGGAAACACTATGACAAAGTGGACATCATTCCATAAGCTTGGATTTAGTGAATTTGAAATACCCGTTGGCCTAGATGAAGCTAATGTAATTGGCAATGGAGCTTCAGGAAGAGTGTACAAAGCTGTCCTAAGCAATGGTGAGGCAGTAGCTGTCAAGAAGCTATGGGAGAGAACAGTTAAAGATGAAACCCCTTATGGTGCTCTTGAGTCTGATAAAGACGAGTTTGAAATTGAAGTTGAAACTCTGGGTAAAATTAGGCACAAGAATATTGTGAGATTGTGGTGCTGTTGCGATACTGGGGATAGCAAGCTCTTGGTATATGAGTACATGCCAAATGGAAGTTTGGGCGATTTGCTGCACAGTTGCAAGGCCAAATTGTTGGATTGGCCGTTGAGGTTCAAGATAGCTTTAGATGCTGCTGAGGGGCTCTCTTATTTGCACCATGGTTGTGTTCCTCCAATTGTTCACCGTGATGTTAAGTCAAACAACATATTGCTGGATGATGAGTTTCGAGCCAAAATTTCAGATTTTGGTGTGGCAAAAATTGTTAAAGCAGGCAGCAAAGGTGACGTTGAATCCATGTCTGTAATTGCTGGTTCCTGTGGTTACATTGCACCAGGTATTGATTTGTTCAAAACTATATCACTGGCTTTAGTCATTTTTACCGATATATTGGCTCATCTAGCTTATATTTTCGGAAAGATTGCAATTTCATTTTAAAGTAGCCAAAACTAACATTAGTGAAATTCATTGATTGCAGAGTATGCATATACTCTTCATGTGAATGAAAAAAGTGACATATATAGCTTTGGAGTGGTCATTTTGGAGCTGGTGACAGGCAAAAGACCAGTCAGTCCAGAATTCGGGGAGAAAGATCTAGCTACTTGGGTACACACGACGTTGAACGAGAAAGGAGTTGATCAGTTGCTCGACCCAAATCTAAACTCCAGCTTCAAAGAACATATATGCAAGGTTCTTGATGTTGGTCTACGTTGTCTTAACCAAACTCCAGCTAATCGCCCCTCAATGCACAGAGTGGTGAAAATGATCCAAGAATCAGTTCCTTATAATGTGCCAGAAATGGAAAACAAGAATGGTAAACTTTCCCCTCACTTTCCAAAGTCTGTCTAGTTAATATATTAGATCAAGTAATAGAGCAGGAGTGTTTCTTGTCAAGAATTGGCAAGGAAGAAGCTGCATAATGAAAAATTAGTAGTCTTTTTCAAGAATTAAGCATATTGTACTCTTTGAATTCTACAACCCTTTATGAATCTATATTACGAGGATTCATCATGAATTAACAAATTTTACGCTAGCTGTCAGCCTACCGCAACACAGATGGGGTACTGAAAGTTGTAAGTATTCTTAACTAATGCCTTTGATTAAGTTTGTTAAACTTAGGTCCTACTTTGCTGTCCATCATTCACTGAGCTGGGGCCTTTAGAAGTAAA

>M6-PGSC0003DMG400009561 M6 LRR-RLK

AAAACTACAAAAACAAAAACAAAAATCTTTCATTTGTAGGAGCCCTTTGAGAGGGCGCGCACCTGCAAGTGCGCGCACCAGAAAAAAAAAAAAGTTTAAAAAAATAAAGAAGACGGTCTCTCTTCTCTATGCAATCATGGCATCACAAAAAAACTACAAAAACAAAAACAAAAACAAAAACAAACATGTCCTCCTCCTTGCTATCATCATGTGTTCATTAGCATTTGTGACAGAGGGAAAATTATCAGAGCCTGAGGTTCTTCTCAAGTTTCGTCAATCATTAAAGTTCGATGGAGATCCCTTCTCGACGTGGGATATTAATGTACCACCTTGCATCAAAGAGAACAATAAGCCTAAATGGAACAATCTATTTTGCGAAAGTGGAAAGGTTTATGGTTTGAATCTTGAAAATCTTGGGTTAAGTGGTACCATTGATTTGGACATACTTAAAGACTTGCCAAATTTGAGAACCATAAGTGTTTTCAAAAATAAATTTGAAGGTTCCTTACCAATATTAAATAAATTGCCTGCATTAAAGAGTGCTTATTTCTCAAATAACAAGTTTTCTGGTCCAATTGATCAAAAGATATTTGAAGGGATGAATTCATTAAAGAAACTTCACCTAGCAAATAATGAATTTACAGGACCACTCCCTCCTATTTTTGGTGATATGCCAAATCTAAGGGAGTTGAATATTCAAAACAATAAGTTTGAGGGTCCAATTCCTCCTTCCTATTCACATCTATATTTTCCTGCTTATGATGGTAATTCCTACCTCGATCTTTTCCTTTTTTATATTTCATTAATTATTGTGTCCAATGAATTAAACACTTTTTTTGTTTATCATTATGCCATGTTTGTCTTCCAATCCTAGATCATTAAAAAGATCATATCATTGGTTACATAATAAACAAAAGATCATTAGATTCGGGTCATCCATGTAAATAATAAGGGAAGAGACACAACTATTCAGATTATTTCTGAAAAGTAGTAGTACTCCATCTCTTTAGAATTATAAAAGAATTGTCTTAATAGAAACCAACAACACTTTTTCTTTTTTTTCTTAAAAAACAATCCTCTCTTCTTCTTTCAAGTTCTTGTGAAAGGATTTCATAGAACAAACTTCTGTAACAGGTACGCGATTTTAGACTATGGCTTAATATGTAGTTCGTCTTTATATGTGATTACATCTAGTGAAACCAAATTCATTATTGATTTATTAACTAACTAAACATAAGTATTTTTCTTCTTATAAAATAGGCAATGATGGTCTTTGTGGTCCTCCTTTAGCAAAATCATGCAATAAGGAAGATGAACAAAAGAAGCAAGAATCATCATCATCGTCATCGTCATCAGGTTGGAAGATTGCCCTTATTGTGGTTATAGTAGTTGTAGTGATAGGCATTATTGTGTTTGTACTAATCAATCGTCGCAAGAAGAATCACCAACAAGAAGTAGTACTTGGAGGATCATCATTAAGTTCTTCCTCACCAACAAGTCAAGACCAAAAACTAATCCCTCAATCTCATGATCATCTAAACAAAATGGAACAAGGCCAATCATCAGCTGCATCTACTCCTGATCGCGCGTCTAATGATGGCGGTAAAAGAGCTGAAGTTGCTGGACAAAAACTTCTATTCTTGAAAGATGACATTGAGAAATTTGACTTACCTGATTTGTTGAAAGCCTCTGCTGAAGTATTGGGAAGTGGAGTGTTTGGTTCAACTTATAAAGCTGCACTTAGTACTGGTCCTGTCATGGTTGTTAAGAGGTTTAGACATATGAATAAAGTTGGTAAAGAAGATTTCCATGAACATATGAGAAGGCTTGGAAGATTGAGTCATAAGAACTTGCTTCCTGTTATTGCTTTCTATTATAGGAAAGAGGAGAAGCTTCTTGTCTTTGAATATGTCAACAATGTCAGCCTTGCTGTTTATCTTCATGGTACCTATCTCTACTCTCTTTTGTTTCAATTTGTTTGTCTCTTTTTAACTTGACATGCACTTTAAGAAAAAAAAAGAAAACCTTTGAGACTAAAGATACTTAAAATGTAGTAGAATATCCTTTAATTTAATTTTGTGGTATGAAAAATTTCATCAGTAAAGTTGGAATTAAAAAGTTCCCTAAAAGAAGAAGAAGAAAGAAACATTCTTTTTGAAACCAACTAAAAAGATAAAATAGAGTATTAATTATCCTTTCTGTCTCAATTTAAATGTCTTACTTTCACTCTTGTTCTTTTCTAAAAAGTGTGTCGGTTTCTATATTTAGTAAGTTTTCAAATTGTGACATTCTACCTGCTTAGTTTAAGACCACAAAATTAAAAAAAATTCTTTATTTTTTTTAAAACTATTTACTGACTTAGATTAAGACACTTAAATTGAAAGGAAGGATGTACTCCTGCATCTATCTTTACTTGTCTAGTATATGGGATGTCCAATAATACTTGTCCAGTTTATAAAACTAATGAATAATTTTTCATTTTATGTCTATTTTACTTTTGCTATTAAATACTATTCATTTTTTAATGTTTAAATGACTTATATTTAAATAAAAGTGATTTAATAAAACTACCCCTATTATTTACTGCTTCATAATCCATGTGCCAAGTCAGTAGTGTACGACGGACAGAGGGAGAGGGAGTAATATTTTTTTCATAGTTTTGACCACCTTTTCTTGATTTTTCAGGTAATAGCAAATCACGTGGTAACCAAAGTCTTGATTGGCCAACTCGTTTGAAAATTGTGAAAGGAGTAAGCAAAGGAATTTTATACCTTTACAATGAGCTACCAAGCTTAACATCACCTCATGGTCACCTCAAGTCTTCAAATGTTCTTCTAACTGAAAACTTTGAGGCAGTTCTCACAGACTATGCCTTACTACCAGTAGTAAATGCTGAACATGCACATGAACACATGATCTCATACAAAGCACCAGAGTTAAAACAATCAGGCAAAATCAATAGAAAAACTGATGTTTGGACACTTGGTATGTTAATCCTTGAGATCTTGACTGGAAAATTCCCTTCAAATCTTTTAGGCAAAGGAACACAAGATAGCGATGATTTAGCGACATGGGTTAATACTACTCTTGGAGATGAATCATCTGAGAAAGAAGTGTTTGATAAAGAAATGAAAGGAACAAAAGATTGTGAGAGTGAAATGATGAAGTTATTGAAGATTGGACTAAGTTGTTGTGAGGTTGATGTGGAGAAGAGATGGGATATAAAAGAGGCAGTTGAAAGGATTGATGAAGTAAAAGAGAAAGGTGATTTCAGCTCTAATGTTGCAACTAATGAAGTTGACAATATGCATACTTCAAGATAAAAAAATATTTTTGATTGATAAGCTCATGATCATAATATGGAAATTTTGATTATACTGGATGCATATTGCATGACTATATATACTTTGGGTATATGATTTTGTTTTGTCATGGTGGAGGATAAATTGAACCATGATGTGGGGTTACTACAAAAGAAAAAAAGAAAAAAAAAAGAAAAAGAAAAATAAGTGATAGAAAAATGAAGTTACTTAATTTTTGTTTGCATATTGTAAATTTGTCCTTTATGAATCTATAGTCTAGGTGTGTTTCTTTTATTTTTTTTATTATTCATTCATCAAAGCTTATTTGGGACTTGCTTTGCAAAATTGGC

>M6-PGSC0003DMG400009843 M6 LRR-RLK

TTAGAGACTATTTTGACCCTACGGCCAAAATTAATGGACCATATTTGTCATTTTCTTTCTTCTTCACCCTACCCCTCGTTTTCCCACTTCACTCTTCTCCATAGTTCCTCACTGTCTCCGAAAATCCAGTCCATTTTTCAGACCTCCTTTAACGCACAACATGACAACAACAAACACTAAACTTGGCCTCTAGTCACTAAACCAAATCACTCGACATTTTTAGTTGCATAACTTGCAATGTCCCCATTGATGCTTCTGCTCTGTTTTCTTCTTCAACTAATCTACAGTTTCTACTTTTGTTCTTCTTCTTCTTCTTCTGCTGCTGCTGCTACCTCTACCCCTTCTGCTCCATTCAACTCTTTGCTTCCCTCCGACGCTGTTTCTTTATTGTCTTTCAAGTCAAAAGCTGACCTTGACAACAAGCTTCATTACACACTTAACGAGCGTTTTGATTACTGTCAATGGCGAGGAGTGAAATGTGTACAGGGTCGTGTTGTGCGTTTTGTTCTTCAAGGGTTTAGTCTCCGGGGTACTTTTCCGGCGAATAGTTTAACTCATCTTGACCAACTCCGAATCCTTAATCTTAGGAATAACTCACTTTCCGGTCCTATTCCTGATCTTTCTGGTCTTCCCAACCTCAAAACTCTGTTTCTTGATCATAATTTTTTCTCTGGAACTTTTCCCCTTTCTGTTCTTTCTATTCGCCGCCTTGTTATTCTTGATCTTTCTCGGAACAATCTCACTGGTTCACTCCCTGTTAGGCTTACTGTTCTTGACCGGTTGAACTATCTCCGGCTTGATTCCAACTGGTTTAACGGTTCGATTCCGCCGTTGAACCAGACCCAACTCCAAATATTCAATGTGTCTAATAATAACCTTACTGGTCCAGTACCTGTTACTCCAACTCTGAAGAAATTTAACATACGCTCGTTCTTACGGAACCCTAGTCTTTGTGGTGAGGTTGTCGACAAGCCGTGCCGTTCGGCTCCATTCTTTGATTCGCCTTCCTCTGCTGCCTCTCCGCCCACACCGTTATACCAAAACGCACAGTCGCAAGGTATTCTTATTACTCCTCCACCTCAGCACAAACACAAAAAAGTTGGTGTTGTTTTGGGTTTTGTAGTTGGAACATTGATCTTGATTGCAGCTGTTCTCTGTCTTTTTGCTTTTGTGAAAAAGAGAAGAGAATAAACTGAAACTGAATCAAAAGCAACAAAATGTACCATTGAGACCATCACTAATAGTGCGGCGAACGCAACGGTTTCTGAGCCAGATGATAGTAGCCAAGAGATTAAATTAGAAAAGGAAATGAAAGTGCTTCAAGCCCCCAAACAACAGATGAAGAGTGGAAATCTGATATTTTGTTCAGGGGAAACAGAGTTGTATTCTTTAGAACAACTAATGAGAGCATCAGCAGAGTTGCTTGGAAGAGGCACAATTGGAACAACGTATAAAGCTCTGATGGCTAGTCAGTTAATCGTTTCAGTAAAGCGATTGGATGCTGGTAAAACTTCGATTACGAGTGCAGAAGCGTTTGAGCAACACATGGAGTCAGTTGGGATGTTGAGGCACCCAAATTTGGTTGCAGTCAGAGCATACTTTCAGGCTAAACAAGAAAGTCTTGTTATCTATGATTATCAGCCTAATGGCAGTCTCTTCAATCTAATTCATGGTAAAGTTTCTAACTTTTTAATATATCATTACTAGTTCATCTGAAATGGATCAATGTTTATTTTTCATTTTTGGTAATTCTGCTAATTGGTCAATGTGAAGTCAATTTGGTGGACCTGTGATGGAGAAAAAAGTGGAAGAGTGAGTTGTCTGTGGCTACTGGCTAATCACTATCTAGCTTTTTTTTTTTCTTTACAGAAATTAGTTGTAGTGTTTTGGATCTTTCAGTGGTTGCTATGGTTGTAGTAGCAGGGGACTTTTTTAGCTTTTAGTGTGGATGCCTAACATGGCTCCCATTGAGTTCACTACCGACAATTGCAAAGTTTGTGTTTACAGGGCAGCCAAATAGTAATGTGTTTTAGTTTCAGGCTGTCACTGACTGACCGGTATGGGTTGATAGCATTATATTCCTCAAATATTTTTATCATGCATTGAGGCAAATGGTCCTTGGCTCTTGCGTCTGTCTTTTTGGTATTTATGAGTATGTTTGCTTCTTCTTTAAGTTGAATAGGATGTTGACAGTAACTTTTTGGTGTTTTCCTTTTACTTGTCCAAATGTTCTCTGTTTCTTAAATATTAAATTTACTTTAAGTGAACGTTAACGTTGTTAGATAGAAGGATTAAGGGAAAGATCTAGAACATCCTTGGGAACTCTTAGTGTAGTGACAGAAATGGTATGAATTTGCTTAATTTACATTTTTTTGAATCTTAGGTATTGCATTTTTTGTATCCCCTTGTGATACGCTATCAAGAAATTGGGATTTCATTTTCATAGATTACTCTGCTCAAGAAGATTGATTCCTCTATATAATTGTGTGTGTTTTTTAAGTGGTTGGTCTTTTCACCTAAAACCTGATAATCTGTTGTTCGTTTACTTATATATTGTGTTCAATTTTCTATTGTTCGTGGCAGGTTCAAGATCAACAAGGGCAATGCCTCTTCACTGGACATCGTGTCTAAAAATAGCAGAAGATGTGGCTCAGGGCCTTGCTTACATCCATCAAGCGTCAAATCTCACTCATGGCAACTTGAAATCCTCCAACGTCCTTCTAGGGTCTGATTTTGAGGCCTGTCTAACAGACTACTCCATCATCGCGCTTGCAGATATTTCTTCGGAGGATGATCCTGATTCAGCATGCTATAAAGCACCAGAAGTGCGCAAATCAGCACGAAGAGCCACACCAGGGTCTGACGTATATGCCTATGGCATTCTGTTATTAGAGCTTTTAACGGGTAAACCTCCATCTCAGCATCCCCATCTCTCTCCCCCTGATGTGCCAGATTGGGTTAGAGCTATGAGGGAGGATGATAATGAGGAGGATAGATGGCTTGCAATGCTAGTTGACCTTGCTAGTATATGTAGCTTGACATCACCAGAACAGAGACCAACAATGAGGCAAATCTTGAAAATGATACAAGATATAAAGGATAGTGCAATGGTGGAGAATAACAAAAGAGATGCACATAATGGATACTCATAGCTTCTTCTGAATTGCAGTTACACCATTAACCAGCGAGGGTACATGGTCAATTGTAAGTAAGTGTCATCTCCCCCTTGGCAAACTGCCATTGATCGACAGCATATCTGTAATTGAGGTTTGCTCAGATAGACCAAATGACTCATCAACTATAATGGTGCTTGTACCTTTACTATTCAGCAGTAAACAACCTAAAACTGACAGGTTAAGGTCTATCTGCTAGAACATGGTCAACTTAGTCACTGAAGATTACGACACATTTTGTTCGAAATCCCTCTCTTTTTTGGTCACACAAACATATGGTGCCTATATTTTTAGGGCATTTTGTAGTAGGTGGTTTTGGTTGATTCTTTTTCTAAAGAACTTCAAAATGTATTCGCTTTACCTTAGTGGCAGAAGAATATTGTATAATCTTGAATGGTAAAGCTAACCCAAATTCTGAACTTTCCCTTCTCTGCATGGTTGGTTAGTATCAACTTAGCAGGAGAAGAGTTGTTTAATG

>M6-PGSC0003DMG400009941 M6 LRR-RLK

CTTCAAATACCCTTAAACCTATCCTAAAATTTGTACAAACCAGACTGAACTAACAGTGTAACAATGTCACTCCCCAAAAAAATATCCCTTTTCCTCCAAATTTTCATCTTTTTGGTTTTCACCATTAATGCAAACTCTGATCTTGAAACCCTTTTGAAGCTCAAAGAATCCATGGTTGCTCCTGGAACTTCTGCACTTCTTGATTGGAACAACAACACAAATTACCCTTTTTCCCATTGTTCTTTTTCTGGTGTTACATGTAACAATAACTCTCATGTTATATCTATAAACATCACTAATGTTCCTCTATTTGGTACTATTCCACCTGAAATTGGTCTTTTACAAAATCTTGAAAATCTTATTATTTTTGGTGATAATATTACTGGTACACTCCCTTTAGAAATGTCACAACTTTCTTCTATTAAACATGTTAATCTTTCTTACAACAATTTTTCTGGTCCTTTTCCTAGAGAAATCTTGTTGGGGTTAATAAAGCTTGAATCTTTTGACATTTATAACAACAATTTCACTGGTGAACTTCCTAATGAGTTTGTAAAGTTGAAAAAGTTGGAAACTTTACATCTTGGTGGAAACTATTTTCATGGTGAAATACCAGAAGTTTATTCTCATATTGTAAGTTTAAAGTGGTTGGGTTTAGAGGGAAATTCACTTACTGGGAAAATACCAAAGAGTTTGGCTTTGTTACCAAATCTTGAAGAACTTAGATTGGGTTATTATAATAGTTATGAAGGGGGTATTCCATCTGAGTTTGGTAATATTAGTACACTTAAACTTCTTGATCTTGGAAATTGTAATCTTGATGGTGAAGTTCCTCCAAGTCTTGGAAATTTGAAGAAGTTGCATACTTTGTTTCTACAAGTGAACAGACTTACAGGTATTCATGAACTCACTATTTGTTTAGTTATACAATCAAATCTCTCTCTAATAACGTCGTTTGTCTGAATATATATTAGTTGACATGGTTAAATATTGTTATAGGAAACATATTTGTTACAAATCATTTTCCAAGATGTAAGACTGAACATCAAATTGTTTTAAAATACTCAATAGTAATAGTTGTTTCTTGAGTGACGTGCAAAGTCTAGAGTTGATAAGTAAAAGTGAATGGAGTGAGAATATCATAAAAAGTCTCTTTGAGAAAAACTGTGTGTTTATAGTGAAATGATGTTAGAGTATGAGTGTTAATAGAGAGGTTCTTGTGTAACATATATGTATATGTGCAAAATATTATGCATATAAAGTTTTGAAGTCATAATTCTGATAAATTTTGAATCTTGAATTCGTAAAATTTAAATCTTGGATTCGTTTATGATTTACAGGTCGCATACCATCTGAACTATCTGGTTTAGAGAGTTTGATGTCGTTTGATTTGTCTTTAAACCAACTGACGGGAGAAATACCAGAGAGTTTTGTGAAGTTGCAGAATTTGACATTGATTAACTTGTTTAGAAACAACTTGCATGGTCCAATTCCCCCTTTTATTGGTGACCTTCCAAATCTTGAAGTGTTGCAGATTTGGGGAAACAATTTTACTCTCGAATTACCGGAAAATCTTGGGCGCAACGGGAGGTTTTTGTTTCTTGATATTTCTGTTAATCATTTTACTGGAAGGATACCACCTGATTTGTGTAAAGGGGGGAAGTTAAAGACACTTATTCTAATGGAGAATTACTTCTTTGGTCCTATTCCTGAACAACTTGGTGAGTGCAAATCGCTTACTCGAATTCGCGTTAGGAAGAATTACTTAAATGGTACTATTCCAGTTGGTTTTTTCAAGTTACCTGCATTGGATATGCTTGAACTTGACAACAACTATTTCACTGGTGAGCTGCCAACGGAGATAAACGCGAATAATCTCACTAAACTTGTACTTTCCAACAACTGGATCACGGGGAACATTCCTCCGTCATTAGGGAACTTGAAGAATCTAGTAACTCTATCACTTGATATGAACAGATTGTCCGGTGAAATTCCTCAAGAAATTGCGAGTTTGAATAAACTCGTGACCATCAACTTGAGCGGCAACAATTTAACAGGTGAAATCCCGAGTTCAATTGCGCTTTGTTCAGAGCTAACATTGGTTGACTTGAGCAGAAATCAACTGGTTGGTGAAGTGCCAAAAGAAATCACCAAGTTAAATAGCTTGAATGCGCTGAACTTGTCAAGAAACCAACTGAGTGGCGCCATTCCTGGAGAAGTAGGAGTGATGAATGGCTTGACAGTTTTGGATCTTTCTTACAATGATCTTTCTGGAAGGAGACCGACCAACGGACAACTAAAGTTCTTCAATGACACTTATTTTGTAGGAAATCCAAAACTCTGTTCGCCTCATGCTACTATTTGCCCGTCAGCTTCCAATTCACCACAAAATGCGCTCAAAATCCATGCTGGGAAGTTCACAACTACCCAATTGGTGATTACAATAATCATCTTAGTCACTGTTGCATTGCTGTTGGCAGTTACCGTGTTGTTCATCAAGAAGGAAAAGTTCAAGAATTCGAAACTTTGGAAGTTAACAGCATTCCAGAAACTTGATTTCAGAGCTGAGGATGTTTTGGAGTGTTTAAAAGAGGAGAACATAATTGGGAAAGGTGGAGCTGGCGTTGTGTACCGAGGGTCTATGTCAAATGGCATCGACGTTGCAATTAAGAAACTTGTAGGCCGAGGAACCGGACACCATGATCACGGATTCTCAGCTGAAATCCAAACACTAGGAAGGATCAGGCACAGAAACATCGTACGATTACTAGGATATGTCTCAAACAAAGACACAAACTTGTTGTTGTACGAATACATGTCGAATGGGAGCTTAGGTGAAATGTTACATGGTGCCAAAGGAGCACATTTGAAATGGGAGACGAGGTACCGTATTGCTGTGGAAGCTGCAAAGGGATTGTGTTATTTGCACCATGATTGTTCGCCTTCGATTATTCATAGAGATGTCAAGTCCAATAATATTCTGCTGGATTCTGATTACGAGGCTCATGTTGCTGATTTTGGTCTAGCCAAATTCTTGCAGGATGCTGGTGCATCAGAGTGCATGTCCTCTATTGCTGGCTCATATGGTTACATTGCTCCAGGTAAATTAATACAGTTACAGTATTTTTCATATTTATACAATTATGAATTTTCCGTATTAGTATTCTTTTATTGCTGGCTCATATGGTAATCATATTTATACAATTAGGTGGCATATAATGTAGTTACTGGTAAAATTTAATGATAATCTTAAAATGTGATACTAATTAACTTTCTATAATAGATTAAATTATGATGATACTAATTAACCTTAAGCTATGTTGCTCAAACTCTTAAAAAATGTCAGTGGGGTACATGTCGGATCCTCAAAAAGTAATGCATTTTTGGAGGATTCGACACGGGTGCAACATCATTTTTGAAGAATCAGAGAAAAATAAACCCTTAAGACCTTTTATATACCTGTGTTATTGGTCAATATTTTCCTATTTCATGAAAGATGCACTTCACCAACTTTTAGTACCAATTACCAAGACCTATGAACTTTTAGAATTTACTGAACTGGGGTTTAAAGTAGTAGGTATTTAAAAGCTATGCTCCAGGACAAAATCTATTTAAAACATTTGTCCAAAACATTAAATTGTGCAATTAATGTTTTTCTGAGGTGCATTTTTGTTTGGATATGAAACAGAGTATGCATACACATTGAAAGTTGACCAAAAGAGTGATGTATACAGTTTTGGAGTAGTACTGTTGGAACTTATCACAGGTCACAAGCCAGTTGGTGAATTCGGAGACGGTGTAGATATAGTCAGATGGGTAAATAAAACAATGTCCGAATTATCTCAGCCATCTGATGCAGCCTCAGTTTTAGCAGTCGTTGACTCGAGGCTACATAGTTACCCTCTTGCAAGTGTTATCAATTTGTTCAAGATTGCTATAATGTGTGTTGAAGAAGAGAGTTGTGCTAGGCCTACTATGAGGGAAGTTGTTCACATGCTTACAAATCTTCCTCAGTCTACTACTACTACTCCTACTCTCCTTGCCCTTTAAAATTGTGCGGATATTAAGTGTCTGGTTGAAAACTCGTGGAGTTTGAGGCCGGGAACACGAGCCTCATGAGTCTATTTGGGTATGGGGAACAAAAAAATTGTGCAAGTAGAGATATGTATTTGAAATGTATTGCTTGTTTTTATATTTTTGAAGTAGTTGTATAAAAAGGGGCATGAAAAAAAGAAGGAAAAAGACTCCTTTAATTACTTATTATGGGTATCGAGTATAATTATGTGGCCTTAGGAAATAAATATCAATATGGCAATTAGTCAATATTTGCTTTTAATAACATGGCCTATGTAATGTTTTCAAGAGTTTGCGAAGTTAAAATTTG

>M6-PGSC0003DMG400010055 M6 LRR-RLK

TGAAGAAGTGTGCTTAGGCAACTTAAGGAGGTTCCAATTCAAGGAACTTCAGTCTGCCACAAACAATTTCAGCAGCAAGAACATACTAGGGAAAGGTGGTTTTGGTAATGTTTATAAAGGACGTCTTAGTGACGGGATTATTGTAGCTGTGAAAAGGCTCAAAGATGGAAATGCAGTTGGAGGCAATCAACAATTTCAGACTGAAGTCGCGTTGATCAGTCTGGCAGTGCATCGGAATCTCCTCAGGTTGTATGGATTTTGTATGACACCAACTGAAAGGTTGCTGGTGTACCCTTACATGTCTAATGGAAGTGTAGCTTCGCGTCTCAAAGGTAAACATATCTAAAAACTGTCTCCTTTGGCCTATAATTACATTGTCCTTATTTGCTCAGCTCATAACGTATCATTTCTTGCAGCCAAACCAACCTTGGATTGGGGTACAAGAAAAGGAATAGCTTTGGGAGCTGCAAGGGGCTTATTATATCTACACGAACAATGTGATCCAAAGATCATTCATAGGGATGTGAAAGCTGCAAATATATTGCTTGATGATTACTGTGAAGCTGTTGTTGGAGATTTCGGATTAGCAAAGCTTTTGGATCATCACGATTCACATGTCACAACCGCGGTTAGAGGAACTGTAGGACATATAGCTCCAGAATATCTCTCCACAGGACAGTCCTCTGATAAGACGGACGTCTTTGGTTTTGGTATTCTCTTGCTAGAATTAATCACTGGACAAAGAGCTCTGGAATTTGGTAAAGCAGCAAACCAGAAAGGCGCTATGCTTGATTGGGTAAGTA

>M6-PGSC0003DMG400010230 M6 LRR-RLK

AAAAAAGAGAAAAAATCTTGTAGAGAGAGAAACAAGAGAGAGGAAAAAACTTTTTTTTTCCCACCATTTTCCCATATTTTCCCTCTTGTTCCTCCTCAACCATTTTTCTGAGAAAAAAAAAATAGTTAAAATGAAGCCTCTGTTTTGAAATCACCCTGCAAACTGCAGTCAAGAAAAAAAAGAAAAGATATTACACTATTTTTTACAAAAAAAAGAAAAAAAAGATTGATTTTTTCTTGTGTGATTTCAATTTTCTTCTTCAATCTCTGTTATTTCTTTAATTTTTTTCAAAAAAGAAAAAAATAAATTCTGTCCCACCGCCTGCATTTTCCTATTGCAGTTATTAATTTTTGGCTTTTTTTTCAAGAAATGTGACGAATTGGCATTTCCATTTCTGCAATAAAAAGCAGAAAAAGATTTCATTCTTGTCTGGTAAGTTTTACTCTATACATTTTACCATCTTTTGGGTTTTTGGATTTTTATTTTTCTTTTGATTGCTTGCAAATTCAGTTATTCCCTCATCATCATCTTTTTCTTTTTCTTTTTCTTTTTCTTTTTCTCTCTCTCTCTTTTTCTTTTCATGGTTTGGCTTGTGAAGTGTTCTTAGGTTTTTCACCAACCCCTTTTCAAGAAACATATTTTTACTAATTTTTTTTCTTTTGTGTTTATTCATTTTCCAGATAAAAATTGAATCTTGGAGATTAATCAATCGAGTTTGTGGAAGTGTTTGGTGAGGGAGATGTTGTTATAGTCTAAATTTTAGAAACTGTGAAAAATGAGAGACCCCAATTTTGTCCATTGTGTCAATAGGGTTAATAGTTTTTCATTTAACTTGTAATATAGTTTTACATTGTAAAAATTGAATGATAAGCAAGCAGTGGTGTATTTGCTCTCCTTTTTTTCAACTTGTTTATATTTACGAGTGAATTCAATTGTTTGGAGAGTAATTTTGTACAAGAAAGCTGGCGATTTTGTATAGAGTCCTGGCTGTTGACAGATATTGTTACTGCCTCAACAACTGATTCATATGGTTGTGCAAGCTATGGGTGGTAGATGGAACACTTATGGAATCAAATTGTCGTATCTCGCTCTTTTGATTATCATTTTGGATATTCGCGGTTGTTGTTCTCTTAATTCGGAAGGTAATTCTTATGCTTGAGTACAGTTCTATTGGTTTTGTAGTTCCGTCTTAGGCTAGTTTTAAGTGTTTGGTATGTTTTCTTGTTTGCTAATTATTGTTGGTGTGAGAAACAGGATTGGCACTATTGAGATTCCGCGTGAAAGTGGATTCTGATCCGTATGGCATTCTTGAAAACTGGAATTCAGATCATTATGACCCGTGCTTGTGGTCTGGTGTCCAGTGTATGGACGGGAGAGTGCAAATGTTGTAAGTCTCTTGTTAAATATACCCAAAAATTTCCTAGTGTGAATCCTTTTCTTATAAATTTATATGTGTTGATTTGCTATCGACAAGCCGGTTGAAATGATAATGACAGACAAAGTGCAAAAACTGAGTTGTGAGTTTTTCTGTTTGTGGTTCTTCGGATATTGTTAAATCATTTGTCTGTTCTCTATTTAGGGACCTTCATGGATATTCGTTGAAAGGAAAACTCGCAGCAGAACTTGGAAATCTTACTCACTTAAAATCACTGTAAGCTTCAGCTATACTTATATACTTCTCATTTGTTTCTTTAGCTCATATTCCTGTCCTCACCATCTGAAAAAAGATTGTTCCAATCATTGATGTTTTTTTCTACTTACAAAATGCTTTCATGATTCTGGTCTTAGTCTAGAAACAATATTTAGTGTGCTATGTCATGGAACTAATACTTCCCTATCGTGGTAGACACTATCTTATCACTAAACAATTTGAAATTGCACTATACTTACATGTCTGGTCAGTACAGGTTTCCATTTTTCGTTCCCATTTGTTACTTAGGAGATGTCTCAAAATATAAAGAGCGTCATATACATTAGGATATTTAGCATATTTGGTCAGGCAAATTGCACCAAAGAAAAACTTTACCTTTTTTTTTTCCAAAGAAAAATGTATAAAGTGAATTGGGTAGATTTATGGTAATCAATGTTTCACCTTCTGTTCTGCATCTATGTACTATGTATTAATAAATTTTCTTCCTCTTTGAATGCTTTTCTTCCACAGTGTGCTTTCTGAAAACCGTCTTTTTGGTGCTATTCCTAAAGAATTTGGACGACTCAGAATGCTGGAAGTGCTTGACATTAAGAACAACAATTTGAGTGGAAGAATTCCAGCAGTAATAGGAGATTTGCAATCACTTAGAAGCTTGTAAGTTGATAATTTGTTGTGTTTCCGTCTTATAATATATTTGTATGTATTTATGCTACAACATTGCTAACAAAATGTTCTTGAATTGTGCTTGATACTTCTTTTCCAGGTTGATTAGGGACAATAACTTCAAAGGGAAGATCCCTTTGGAGATCAGAAAGCTTCATCTGCTCTCGGAATTGCAATTTGATGATTACCTCACCTCTGGTGTTGCTGTCGGAACTGGTTGCATAAATAGAAAATTTGGGCGTTGGTAAGACTTCCGTAGTTGTTTGGTCCTTTTATCTTCATGTTTCAGCTGTGCTGCATTTTAAAGTCGCAGACTTGTTTCTATGTTCAATTATATTGAAAATGCATAAAATCCATCATTTGAGGAACATATACAAATATACTACTTTCATACGAACTACCTCCTGCTATAAGTTTGGTATTAACATTAAATTAAGTGCAAACAATTATCCCATGGAGTTGCACTGAAGATCTTTGAGACATTTTCTATGCATAGTGTGTCCCCACTAAACTCTCATACTTCCCTTACAATAGTGAAGCAACTATGATGGTGATTGTCGAAAGCATTGCCATGGGGCACTTGATACTTTCCTGTCTCCAACCCTAATATTCAGAGGGGATTAGACTTTAGTGTTTCTAAAGTTGTGATTCATCTTCTTATTCATCAAGACCTGTTTATGTGGAAATTTGACTGATCTATTGCATAATATTTGTTTATTTTGTCGACTTTTCTGTACATTAACCAATAAAATTGCTGGAAATGTGGTTATTCTTTCCCATGAGTAGAAGAACACTTGGATATTATCGTATTCCTGTTTTGTAGCATGAAACTTCAGCCAGCTATATGAATAAATTTATTCTATTCATCCCGTTGACTATGTTTGAATTCTGGATCGAGATACTCATCTTGGACAAGTACCTTCAGCTTTACCCTTATAACTGATGACAGATTACTGACTACCTCTTAATATGCTAATGTTTTATTTCTCAATCACGCACTTTTAAGTTTCATACTCCGCTCTGAAAATTTGGTGACTTTTTCTTGCAGCATCTGGCATGGCAGTTTGAGGCCATATAAGACTATAGGCTCCTTTCTCAGACCAATTAAAGGGATGCTAGCATGGTATTTTAGTTTTTTCACACTGTAAGTAACTAAATACGTTTTTTTCACCTTATGCTCTCTAAATAAGTTTAATGGTTCAGAAGCAACTTTTTTGTTTCCTAAAAGGTTCCCACGTTTTTTGGATGCTCAGGCAGACTTCTGCTCAGACAACCTACCAAGTGAGTGAAAATATTTTGTGACATATCTTTTATTCGATTTCAAAGAATTGTACACAAAAAAAAGAAAGTATTTTATTCTCTTTTGTTTTTGCAGGTTCAACTAGGCCTCACAATATCCACGCTGTAGAGAATCAAGCAAACATTGAACGCCGTAAGTTAGCTGAACAATCCAGTAACTTTGCTGCTGCTCCTGCTAACGGAGGGAAACCATTGGGACCTGTTGATCCTATTCCAAGCAGCAGAAGTAGTGGTTCATTTCGTGCTGTGCCAAGTACTGAAGGAGCTCCTCCACCTCCTCTGACTACACCATCACGGACCACACATGATCATCATCCGCCTCCAAATCCTGGGGGACACTCTAATGGTGCTGTTAAACAGCCAATTGCTAGTCAAGCTCCACCTGGTGGAAAGTCTGGAAGTACATGGAAGTATATAGGAATTGGCATTGGGGCTTTTCTGGTTGCTGTTTTTGTGTGCCTGATAATCATCTGCAAGAGCAAAGCTGCACAAACAATAGGCCCTTGGAAGACTGGATTGAGTGGACAGCTGCAGAAAGCATTTATTACAGGTACTTTTGATTTAATATTGTGTCAAAGCTGATGAATCCATATGTTCTAGTTCAACTGTTGGATTCATTTTTCTCTGGTGGATATAATCCGAATCTTCAACTCCTACAATAGTTTTTGTTTCATACTCCAAGATGATGTTAAAAGCATGTATTAATACTCCTGTGCTATATTGTTTCTCATGTCTTATGCATGAAATCAATTTTGCAGGGGTCCCAAAACTAAACAGGCCGGAGCTTGAAAATGCATGCGAGGATTTTAGCAACATAGTTTGCCATCAAGATACATTTACAGTCTACAAGGGAACTATATCCAGTGGAGTAGAGATTGCTGTTGTGTCAACTGCTGTAAATTCTTTGAAAGATTGGTCCAAGCATTCTGAATTGGCCTTCAGGAAGAAGGTTAGTAGTTGTCATCTCCTTCACAAATAAAAAAACTGCTTCATAACTTATTACCATCTCACTCTTTCTGTTGAGATATCCTAGTCGTCTCGGTTTTTTTACTTATAAGATTTCTACTTATTGTGCACTTGGGTAGATCGATTCACTCTCAAGGATAAACCACAAGAATTTTGTTAATCTAATTGGATATTGCGAGGAAGATGAACCTTTCACTAGGATGATGGTCTTTGAGTATGCAACAAATGGAACACTTTTCGAGCATCTGCATGGTAAACTGGTTAATTGTTCAAGTTTCAGTCTAATCATTTTTGTGATTTCGTTTAGATTAATGCAGTCTACATGTTTTGTTGTTTCAGATGAAGAACTTGAACCTCTTGACTGGCCTGCAAGGGTACGAGCTATAATGGGAACAGCATATTGTCTCGAGTACATGCACAATTTGAATCCTCCTCTGTCGCATTCTGATGTGAATTCACACTCCATATTTTTTACGGACGATTATGCTGCTAAGGTATGCTTCATTTTGTATACCACATTGAATTTAGAAACGTTTTACTGTCTTGAATTAAACGAATCGTTTTTTATTTACACAGATTACAGAGCTTGCTTTCTGGTCAGATATAATGGTGAAGTCAAAGTCCTCTAGTGGTGGTGATGTAGAGAACTCTGAGCTACCACCACTTTCTGATCCCGAAACAAATGTCTACAATTTTGGGATTTTGTTGCTGGAAGTAATTTCTGGGAAGTCACCTTACTCGGAACGAGATTCTCTTCTGAGCTGGGTAAAGTCTCAATCTATGTAACTCTAATTTTCTTCGGTGTTTAATATAGACTAGTGAAACGTTCTTCTCTGCTTTTGTTTAGGCGGAGCAATGCGTTAACGATAGACAGTTGGTTGATCCAAAATTGAAGTCGTTCAAGAATAATGAGCTTATGGTTATGTGTGAGGTAATGAGAGAATGTATTGGAGAAGATTCAAGGAAGAGACCAACAATAAAAGAAGTGATTAAAAAGCTAAGGGAAGCGATTGATATCTCCCCTGATGCTGCAGTTCCAAGGCTATCTCCCCTGTGGTGGGCTGAACTTGAGATCCTATCCTCTGAGGCAGCCTAATATACTGACTAATGTGTCTTCTACTTTTGTAAGTAGATAGTTGCTTTGCTATGCATAAATGTTCTGAAATATACTTCACTTAATGTCAACTGTGTACCACATTTTTCTGTTCTAAATATATTAGTTCATTCTTTT

>M6-PGSC0003DMG400010348 M6 LRR-RLK

CTCTGTCTTCATCTCTTCGTCATTAAATGCAACGTCGCCACAACTAACCAGAAGTGATATGACATTTAGGTTGACATTTTTGGCAAACATCTCATAAAGGCAATATTAGCACTGAGCCACTGATGAAGCAAGAAATGTGATAACTTCTCGTTGTTGGTTAGGATGTTGCATAATTTGTCACCTCTATATTGATTCTACAGTTACTTCGCAGGGTGAATACTACTGGATATATGGATCAGAAGACAATTGTGTTTTACGGTTTAGTGTCGAGGCGTGTGTCATGATGTCAGATTTTGTCAGTGGCTTACACTTACATACATACGATTTTTTACCCTGCAGAGATACTTGTTTTAGAGCATAAGCTACTATACAATGAAGTAAATGATGTGCTAAACCTATTAATTATTACCAATCTCAAATAGCATTGATGAAAAAATTCTCTCAAGTTTCATAAACAAACATTACAATGTTTTTGCACTGTATTTTACTGACAATAAGAAGATACTCCTATGTGAAACCGTCATCAGCAATGAAGGCTCTGTTGTTAAGAGAATTTGTGCTTTCCCGATCTAAAGAGAAACTTGGACCAATATGTTGATTTTGCCATATCAAGCCATAATTTATACTGTAAACACCCAAATAAAGCCACAGGAACACCAACACAAGCAGCAATGAGCTTTGGCTCCCAACCAATATGATTACTATAGACCAAAAAGAATGTTGCTGCAAAAGCCAAGAGCATGCTAACTATTGAGACAAACAGCATTGTGAGTCCAAACAATAACTTGGCTGGCAACGACACAAGAAAATCACCTTCAGCATAGTGGGATGTCACAATGGACAAGAACATAATGATGGAAACTATTGAGGTGAATAGTGCCACTGCATCTGAATTGACAAAGATCCTAAATCCATACAATTTTAGCAATATCAGAATGCAATTATCACTGTTATTACCGCCTGGCACAGTGAATCCAGCAGCAAAGACCATGGTGGTAATCAAAGTAGCGATAATCATACAAGAATTTGCAGTGTCTTTCATCCACCTTTACCCTTCTTTCAGTAATAACTTATGCTCTTCTGTGAATATTTCTCTAGGGGTTTTCCCATCTTTGTTTTGTTAGTGCAAAGTTTTAATGATAAACAACTTATTCTTGCATCAAAAAGGAAAGAATCAAGAATGTGTACATCCTTGACTCTGTAAAAAAACAATGGAGGACATATAAGCCTAGCTTCAATGACTGATGATCCCTGAATTTGGCACAAGAATTTGGCCATGCAAGCATGCGTCTAATAGAGAAATTAGCTAAAAATGAGCTTGTAATTGGACACCACCATCACTTTTGTCTTTTTAGTTATTCCAAAACTATCCTTGAAAATAAAAATAATTCAAAGTAAAAATATAAATATGAACAATTAACTCTCTTGAACTTATCACTTAGAAAAGGTAAATGAAAGACAAAAATGGAAGAAATTTTTAACAACTCTCTTGAACATTGAATAATTGAAAAAATGTCAAAAGTTCAATTAGTATGGACTAGATGGAGTAAAATATATAGTTTATATTTTATTAGTCCATATATCATAAAACAATTTTTTTTTTGTTTCTAGTCAATAGTAATATGAATTTTACACATCATATTAGGCCTCTTATTGTAATTTGAAGTAATTCCATTGACCCATCCCTGTGTAACACACTTGTGGAGATGTGTAACACATTATGAAGGGTAGAGAGGACTATTGATAGACACAAAAATCACGCCAAAACTAATGGCAGTATTCAAAATGCAAGGCCACAAGTAGTGATCAGAAAGACCAAGATTCTTGCACATTTCATTGAAGGCAAAAAACATTTTCCCTGTAGGTCACTAATGCCATTTTCTAGTCTTCACTAAAAGAAGTTGTTGAACTCCAAATTGTCACTCTAAACCAAAAAATACTCCAGTCATCAATTCTTGTTTTAGGAAAAAGAGAAACCTGTCAGTTATAGTTATGGGAAGAAGTTGCGATCTTCTCTTTGCTCTTTCGGTTTTCATTCTGCTCCATACTTCACTATCTATTAGTACTGATGAAGCTGCTCTTCTTGCCTTCAAATCTCACATTTCTTCTTCTGATCCTAACAACATCTTAGCAACCAACTGGTCTTCTTCCACCCCTGTTTGCACCTGGATTGGAATCACTTGCACCTCCCCACACCATCGAGTCGCAGCTTTAGACATTTCTAGTATGCAACTTCATGGTACCATTTCTCCACACCTTGGAAACCTCTCATTTCTTGTTTCCCTCAACATTAGTAACAACTCTTTCCGTGGAGATTTGCCCAAAGAGTTGGCTCATTTGCAGAGGTTGAAATTGATTGATGTCTCAAGCAATAACTTCACTGGTGCCATTCCATCATTTTTAAGTTTGCTAGTAGACCTTTGCATTTTGCGCCTGTCGAGCAACCAATTTTCAGGAAAAATCCCATCTTCACTTTCCAATCTAACAAAGCTGGAGGTGTTGAAAATAAAGGGAAATCTTCTCGAAGGAGAGATCCCTCGAGAACTTGGTGATCTTCATTACATGACTGCTTTAAACCTGGGAAGTAACCACCTTACTGGCTCTATACCACCATCAATTTATAACATTACAACCATGCGAATCATTGCTCTTACCAACAATAATCTTAGTGGTGAGCTTCCAACAACTATATGTGACCATCTTCCAAACTTGGAAGGGCTTTACCTCTCAGAAAACATCCTAGGTGGCGTTATTCCACCAAACTTGGAAAAATGCAAAAACCTCAAAATTTTGTCATTGTCTGTAAACGAGTTCACTGGAACCATACCAAGAGAGTTAGGCAACATAACAGATCTTACAGATATAGCTCTTGGAATACAGCACTTGGGAGGTAGTATTGCATCTCCTAGCAAGAAATTTTGAATTTTCTTCTTTTTCACTCTTCCACAAATTTGGTACTCATATATATATATATATATATATATTTCCCCACTTGACAGGAAAGATACCAGTGGAGCTAGGTAATCTTAAGAAACTACAGATTCTGTCATTAAACGACTGTAAGCTTACTGGTTCCATCCCCGCAAGCATTTTCAACATGTCAGCACTGCAGCTCCTAGGAATATCAGAAAACAGGCTTTCAGGTACTCTACCTTCAGATTTAGGTCGTGGAATGCCCAACCTAGAAGGATTCTCTTGTTTTCAGAATAGTTTGAGTGGTTTTCTCCCTGCTTCTATCTCAAACTCTTCTAGACTCAGAGTACTTGAACTCTCATATAACAGTTTTACAGGCCCAATTCCTGAATCAGTTAGTGACTTAGAATACATTGAGATTTTGAACTTGGGGGGGAATAATTTTTTCAGCAATTCAGCATTGAGCTTCTTGACATCTTTGACAAACTGTAGGAAACTGAAAGAAATCACTTTTGCCGAGAATCCCTTGGACGGTTTTTTGCCTGCCTCCATTGGGAATTTCTCTGATTCTCTGCAAATTTTCCAAGGATGGTATTGTAAACTGAAAGGCTTCATTCCTGGAGAAATTGGCAATCTTACCGGTGTGATAAAGATGGATTTGTCACAAAATGAGTTGACTGGATATATTCCAAAAACCATCCAAGGCTTGTCAAACCTTCAAGAACTTTACTTAGGTGGCAACATGATAAAAGGAACCATACCAGATGTTATGTGCAATTTAAATAAACTTGGAGCATTAGATTTGTCAAAAAATCAAGCTTCTGGTTCCATACCACCATGCTTAGGGAACGTTACCAATTTGAGGTATCTTTATCTAGCTAACAACAGGCTGAACTGGACCTTACCTTCAAGCTTGGGGAGCCTTCAAGATCTCCTAGAATTCAATGTTTCGTCCAATTTATTAACTGGGGAAGTTCCTATAGAGATTGGAAATTTAAAGGTTGCAACACTAGTTGATCTGTCAAAAAATGATTTTAATGGTAAGATCCCTAGCACACTAGGGGGTCTGGATAGAATGCTTAGTCTTTCTCTAGCACATAATAAATTATATGGGCCTATTCCAGATTCATTTGGCAAAATGCTGGCCTTGGAATTCTTGGATTTGAGCAATAACAAACTTAGTGGTGAAATTCCAAAGTCATTAGAAGCTCTCGTGTATCTCAAATACTTGAACTTCTCATTTAATGAACTCAGTGGAGAGATTCCCACGGGTGGTCCTTTTGCAAATGCCACAAGTCAATCTTTCTTGAAAAATTATGCCCTTTGTGGCGACTCTAAGTTTCATGTTTCACCATGCATCATCAAATCTCCCAAGAGGTCAAAGAAAAAAAAGGCAATATTGGTTTTGGGAGTGGGTATGCTTTTTCTTGCATTAGCCCTCACATATGTATTTTTGAGATTGAGAAAGAAAAAAAAGAATGCAGGTCAATCTGATATGTCTCTGATAAAATGTCATGAAAGAATTTCCTATTATGAACTTTAACAAGCAACCGAAAGATTCAGTGAAAGCAACTTGCTTGGTAACGGGAACTTCTGCAAGGTCTACAAAGGGAGACTAAAGGATGGTACTCTTTTGGCAGCAAAGGTATTCAATGTGCAATTGGAGGGTGCATTCAAAAGTTTTGATACAGAATGTGAGATGTTGCGCAACCTCCGCCACCGAAATCTGACCAAAGTCATCACCAGTTGCTCCAACCTTGATTTCAAAGCCTTAGTGTTGGAATACATGTCCAATGGGACACTTGATAAATGGCTATACTCTCACAACTTGTTCTTGGATTTATTTCAGAGATTAGATATAATGATAGATGTTGCATCTGCAATGGTATATCTCCACAGTGGTTGTTCAAATCCGGTGGTGCATTGTGACTTAAAGCCAAGCAATGTCTTGCTAGATCAAGACATGGTTGGCCATGTCAGTGATTTTGGAATTGCAAAATTGTTAGGTGCAGGGGAGACTTTTGTCCAAACAAGGACAATAGCAACCATTGGATATATTGCTCCAGGTATTAAAACTTTTACATCTTCCTCATGTTTTTAGAATA

>M6-PGSC0003DMG400010350 M6 LRR-RLK

TTTCTTTCATTTTTATCTTCCGCCAAATTGGTACTAATCATGTATTTTCCTCACTTAACAGGAGAGATTCCAGCAAGCATTTCTAACATGTCAGAACTGCAGAACCTAGGATTTGCAAGAAACAGGCTTTCAGGTAATTGTTGTGTAACAAAAAATCTGAAGTGTTTATGTAAATATATGACACGTGAGCTAACTCAACCCCAAAAACTAGCTCATGAGGGGAGGATTGCCCAATTCCATATAATCTGACCACCTGTCTATTCCCCAAACAATGTGGGACTTTTAGCCACTCTAACTCCCCTTTCACACCCAGGTCCAACTGGAGCGTGGACCGGCAGCCCAAAGGGGATGGGATGACCCAGATACCATGTAAAGATATGACATCCTCTCCTCATGAACTAACTTTTGGAGGAGAGTTAGACTCATGTGTCATATCTTACATGGTATCAAAGCCACGTCCATCCTCGTTTGGACTCCCGCTCATCACTCCAGTTAGGCCTAGGCAAGAAGGGGTTGTTAGAGTGGTTAAAAGTCCCACATTTGTTGGGAATGAACAGGTGGTCTGCTTATATGAACTTGGGCAATTCTCGCTCATGAGCTAGCTTTTGGAATTGAGTTAAGCCCACATTTCATATTTTTACATTCTCAAGTACTAACATGTCAACTCATATATTTCCCCCTCTTGACAGGAGAGATACCAATGGAACTAGGTTATCTTCAGAAACTACTGTTTCTGGCATTAGACACAAATGAGTTGACTGGTTCTATCCCTGCAAGCATTTTCAACATGTCAGCACTGCAGATCTTAGGAATTGCAGAAAACAGGCTTTCAGGTACTCTACCTTCAGATTTAGGTCGTGGAATGCCCAACCTAGACGGATTCTATTGTTATCAGAATAGTTTGAGTGGTTTTCTCCCTGCTTCTATCTCAAATTCTTCAAGACTCAGAGTACTTGAACTCTCATATAACAGTTTCACAGGCCCAATTCCTGAATCAGTTAGTGACTTAGAATACATTGAGGTTTTGAACTTAGGGGCGAATAATTTTGTCAGCAATTCAGCATTGAGCTTCTTGACATCTTTGGCAAACTGTAGGAAACTGAAAGAAATCACTTTTGCCGAGAATCCCTTGGACGGTTTTTTGCCTGCCTCCATTGAAAATTTCTCTGATTCTCTGCAAATTTTCCAAGGATGGTATTGTAAACTGAAAGGCTTCATTCCCGGAGAAATTGGCAATCTTACCGGTGTGATAAAGATGGATTTGTCACAAAATGAGTTGACTGGATATATTCCAAAATCCATCCAAGGCTTGTCGAACCTTCAAGAACTTAACTTAGGTGGCAACATGATAAAAGGAATCATACCAGATGTTATCTGCAATTTATATAATCTTGGAGCATTAGATTTGTCAGGAAATCAAGTTTCTGGTTCCATACCGCCATGCTTAGGGAACACTACCAGTTTGAGGTATCTTTATCTAGCTAACAACAGGCTGAATTTGAGCTTACCTTCAAGCTTGTGGACTCTTCAAGATCTCATAAAATTCAATGTTTCGTCCAATTTATTAAGTGGGAAAGTTCCTATTGAGATTGGAAATTTAAAGGTTGCAACACTAGTTGATCTGTCAAAAAATGATTTTAGTGGTAAGATCCCTAGCACGCGGGGGTCTGGATAGATTGATGAAACTTTCTCTAGCACATAATAAATTAGATGGGCCTATTCCAGATTCATTTGGCAAAATGCTGGCCTTAGAATTCTTGGATTTGAGCAATAACAATCTAACCGGTGAAATTCCAAAGTCATTAGAAGCTCTTGTGTATCTCAAATACTTGAACTTCTCATTTAATGAACTTAGTGGAGAGATTCCCACGGGTGGTCCTTTTGCAAATGCCACAAGTCGATCTTTCTTGAAAAATTATGCCCTTTGTGGTGACTCTAAGTTTCATGTTTCACCATGCGTCATCAAATCTCCGAAGAAGTCAAAGAGAAAAAAGATAATCTTGGTTTTGTACATCCTTTTGGGAGTGGGTATGCTATTTCTTGCATTAGCCCTCGCATATGTATTTTTAAGATGGAGAAAGATAAAGAAGAATGCAGATCAAGCAGATGTCTTTCCCGTAAAAGGGAAACGTGAAAGAATTTCTTACTATGAACTTGAACAAGCAACAGAAGGATTCGATGAAAGCAACTTGCTTGGTAGTGGGAGTTTCAGCAAGGTTTTCAAAGGGATATTTAAGGATGGTACTCTTTTGGCAGCAAAGGTATTCAATGTGCAATTGGA

>M6-PGSC0003DMG400010351 M6 LRR-RLK

ATCAATTCTTGTTTTAGGAAAAAGAGAAACCTGTCAGTTATAGTTATGGGAAGAAGTTGCGATCTTCTCTTTGCTCTTTCGGTTTTCATTCTGCTCCATACTTCACTATCTATTAGTACTGATGAAGCTGCTCTTCTTGCCTTCAAATCTCACATTTCTTCTTCTGATCCTAACAACATCTTAGCAACCAACTGGTCTTCTTCCACCCCTGTTTGCACCTGGATTGGAATCACTTGCACCTCCCCACACCATCGAGTCGCAGCTTTAGACATTTCTAGTATGCAACTTCATGGTACCATTTCTCCACACCTTGGAAACCTCTCATTTCTTGTTTCCCTCAACATTAGTAACAACTCTTTCCGTGGAGATTTGCCCAAAGAGTTGGCTCATTTGCAGAGGTTGAAATTGATTGATGTCTCAAGCAATAACTTCACTGGTGCCATTCCATCATTTTTAAGTTTGCTAGTAGACCTTTGCATTTTGCGCCTGTCGAGCAACCAATTTTCAGGAAAAATCCCATCTTCACTTTCCAATCTAACAAAGCTGGAGGTGTTGAAAATAAAGGGAAATCTTCTCGAAGGAGAGATCCCTCGAGAACTTGGTGATCTTCATTACATGACTGCTTTAAACCTGGGAAGTAACCACCTTACTGGCTCTATACCACCATCAATTTATAACATTACAACCATGCGAATCATTGCTCTTACCAACAATAATCTTAGTGGTGAGCTTCCAACAACTATATGTGACCATCTTCCAAACTTGGAAGGGCTTTACCTCTCAGAAAACATCCTAGGTGGCGTTATTCCACCAAACTTGGAAAAATGCAAAAACCTCAAAATTTTGTCATTGTCTGTAAACGAGTTCACTGGAACCATACCAAGAGAGTTAGGCAACATAACAGATCTTACAGATATAGCTCTTGGAATACAGCACTTGGGAGGTAGTATTGCATCTCCTAGCAAGAAATTTTGAATTTTCTTCTTTTTCACTCTTCCACAAATTTGGTACTCATATATATATATATATATATATATTTCCCCACTTGACAGGAAAGATACCAGTGGAGCTAGGTAATCTTAAGAAACTACAGATTCTGTCATTAAACGACTGTAAGCTTACTGGTTCCATCCCCGCAAGCATTTTCAACATGTCAGCACTGCAGCTCCTAGGAATATCAGAAAACAGGCTTTCAGGTACTCTACCTTCAGATTTAGGTCGTGGAATGCCCAACCTAGAAGGATTCTCTTGTTTTCAGAATAGTTTGAGTGGTTTTCTCCCTGCTTCTATCTCAAACTCTTCTAGACTCAGAGTACTTGAACTCTCATATAACAGTTTTACAGGCCCAATTCCTGAATCAGTTAGTGACTTAGAATACATTGAGATTTTGAACTTGGGGGGGAATAATTTTTTCAGCAATTCAGCATTGAGCTTCTTGACATCTTTGACAAACTGTAGGAAACTGAAAGAAATCACTTTTGCCGAGAATCCCTTGGACGGTTTTTTGCCTGCCTCCATTGGGAATTTCTCTGATTCTCTGCAAATTTTCCAAGGATGGTATTGTAAACTGAAAGGCTTCATTCCTGGAGAAATTGGCAATCTTACCGGTGTGATAAAGATGGATTTGTCACAAAATGAGTTGACTGGATATATTCCAAAAACCATCCAAGGCTTGTCAAACCTTCAAGAACTTTACTTAGGTGGCAACATGATAAAAGGAACCATACCAGATGTTATGTGCAATTTAAATAAACTTGGAGCATTAGATTTGTCAAAAAATCAAGCTTCTGGTTCCATACCACCATGCTTAGGGAACGTTACCAATTTGAGGTATCTTTATCTAGCTAACAACAGGCTGAACTGGACCTTACCTTCAAGCTTGGGGAGCCTTCAAGATCTCCTAGAATTCAATGTTTCGTCCAATTTATTAACTGGGGAAGTTCCTATAGAGATTGGAAATTTAAAGGTTGCAACACTAGTTGATCTGTCAAAAAATGATTTTAATGGTAAGATCCCTAGCACACTAGGGGGTCTGGATAGAATGCTTAGTCTTTCTCTAGCACATAATAAATTATATGGGCCTATTCCAGATTCATTTGGCAAAATGCTGGCCTTGGAATTCTTGGATTTGAGCAATAACAAACTTAGTGGTGAAATTCCAAAGTCATTAGAAGCTCTCGTGTATCTCAAATACTTGAACTTCTCATTTAATGAACTCAGTGGAGAGATTCCCACGGGTGGTCCTTTTGCAAATGCCACAAGTCAATCTTTCTTGAAAAATTATGCCCTTTGTGGCGACTCTAAGTTTCATGTTTCACCATGCATCATCAAATCTCCCAAGAGGTCAAAGAAAAAAAAGGCAATATTGGTTTTGGGAGTGGGTATGCTTTTTCTTGCATTAGCCCTCACATATGTATTTTTGAGATTGAGAAAGAAAAAAAAGAATGCAGGTCAATCTGATATGTCTCTGATAAAATGTCATGAAAGAATTTCCTATTATGAACTTTAACAAGCAACCGAAAGATTCAGTGAAAGCAACTTGCTTGGTAACGGGAACTTCTGCAAGGTCTACAAAGGGAGACTAAAGGATGGTACTCTTTTGGCAGCAAAGGTATTCAATGTGCAATTGGAGGGTGCATTCAAAAGTTTTGATACAGAATGTGAGATGTTGCGCAACCTCCGCCACCGAAATCTGACCAAAGTCATCACCAGTTGCTCCAACCTTGATTTCAAAGCCTTAGTGTTGGAATACATGTCCAATGGGACACTTGATAAATGGCTATACTCTCACAACTTGTTCTTGGATTTATTTCAGAGATTAGATATAATGATAGATGTTGCATCTGCAATGGTATATCTCCACAGTGGTTGTTCAAATCCGGTGGTGCATTGTGACTTAAAGCCAAGCAATGTCTTGCTAGATCAAGACATGGTTGGCCATGTCAGTGATTTTGGAATTGCAAAATTGTTAGGTGCAGGGGAGACTTTTGTCCAAACAAGGACAATAGCAACCATTGGATATATTGCTCCAGGTATTAAAACTTTTACATCTTCCTCATGTTTTTAGAATA

>M6-PGSC0003DMG400010400 M6 LRR-RLK

GGCATTTAAATCGAGCCTTCGATTGGAACCTCACCACATTTTGTCCAGAAATTGGAGCTCAGACACACACTTCTGCAGCTGGTATGGAGTTTCATGCTTGCACCAGAGAGTTGTAGCTTTGCAACTTCCTAATTTGGCCATTCAAGGCAGAGTTTCTTCAGAAATTGCCAACTTGACTCAACTTGCTCTTCTTGACCTTAGTAGCAATGATTTGAATGGCAATTTACCAAGTGAGTTGGGCTTCTTGCAACAACTGAGACTTTTTAATGTGACTGGGAATTCTCTTGATGGAACTATTCCTCTGAATATAAGCCGTTGTCGTGTACTGCAAGAACTTCATTTATCAGATAACATGATCAAAGGAAGCATACCTCAAGAACTTGGTCGTTTAAGTCAACTTAGAATTTTGCGTCTCAACGACAATAATCTGACAGGAAAGATTCCACCTTTATTAGGAAACTTATCAAAGCTGGAATATTTTTATCTGCATGAAAATGATTTGGAAGGTGAAATACCTGATGAAATTGGTGATCTTACCAATTTGAGGCTCCTTTCCTTTCGTGGCAATGACTTCACTGGATCGATACCTGCTTCAATTTTCAATATTTCAAGATTACAAATTGTGGATCTCTCGATAAATGCCTTATCAGGGGAACTTCCAAGTGATTTGGGTAATTATCTTCCTATCCTTGAACATCTTTTTCTGGACAGCAACAGGATCACGGGGAAAATCCCGGTTTCTTTATCAAATGCATCAAAGATTACTCATCTCTTCTTTACTGAAAATGATGTGCAAGGAAATATTCCAAATGAGTTTGGTAAGTTGCATGAACTAGTCTGGTTTGAATTTGAATATAATCGAATTTCTGGTACCATTCCATCCAGCTTATTCAATATTTCGTCTCTTGAAATCCTCAAAGCCAGGCATAATTACTTAAATGGGCATCTTCCATATGATCTTGGTACTTGGCTACCTAACTTGCAGGAGATTTTCCTTTCACACAACCAATTCAGTGGAGATCTACCCGCTGCAATATGCAATGCTTCAAAGCTTGAAAATCTTGAAGTTGCAAACAATAGTTTTACCGGACCGATTCCCATGATGTTAGGTAACCTGGTGGACCTTATAACTCTAAATCTGCAAAATAATCTGTTGGAGAACAATCCTGGAGCCACACAACTTGACTTCTTGAATTCATTGGTAACCAGCAGGAATCTGGTGTACCTGATTCTTGCATCTAATCCTTTGAATGGAATTTTTACGGAATCAATAGGAAATTTATCAAGCAATATCAGGGTCCTTGATGCATTCGACTGTGGAATTAGAGGCAGTATGCCAACAAACATTGGAAACTTAAGTGGCTTGATTTTCCTGGGATTAGCAAGTAATAACATTGTCGGTAATGTGCCCCCCTCGTTTGTTGGCCTGCAAAACCTGGAAAGGCTGTATCTTACTAGAAATAAATTAGAAGGGACATTTCCAGCTGAACTGTGCAGCATAGGAAGATTAGGTCTACTGCATTTGGGAGAAAACAGACTGTCAGGAGAAGTACCCTCGTGTATCGGAAATCTGACTGAATTAAGAGTAATGTCAATTGCTGCCAACAATTTCAGCTCAAAGTTACCTTCAAGTTTCTGGAGATTAGTAAAGCTTGATGATCTGAACTTATCGAGAAACCTGCTGCATGGTTTCTTCCCTTCTGATGTGGGAACTCTGAACGCAATGAGGATCATGGACCTCTCTTTCAATAGGTATTTTGGTGAAATACCCAATTCCATAGGATCTCTTCAATATCTAGTTTCACTAGATATGTCAAGAAATGCATTACAAGGTCCCATACCGGCCACATTTAGCAATCTAATAGTTCTTGAGGCTCTGGATTTGTCTACTAATGCTCTATCAGGTGTGATTCCCAAATCTCTTGAGCTATTGCGAGATCTCAAGTATCTCAATGTTTCTTTCAATAACTTGCAAGGAGAAGTTCCAAAGAAAGGGGTATTTGCAAATGTTAATTACCGGTTCTTGATGGGAAATCCAAGACTTTGTGGAGCACCTGACTTATATATTCCTATATGTCCAGCTCAAGGTAGAAAGACAACAAGGAAAAAGTCCATTATTCTTAGAACAGCAGTACCTGTTGCTGTAACATTTCTGATATTGGTCGCGTTGTTTTTCACATGGATGATTTGGTCAAGAAAAAAACATGCGATTGGAAACAATGAATCGGATTACCCTCCTAGAATTGCACATCAGAAAATCACTTATTATGAGCTTCTCCAAGCTACAGAAAATTTCAGTCAGTCCAAGCTAATAGGAAGTGGAAGTTCCGGTACAGTCTACCAAGGTACATTTTCTGGTGGTGCTGTCTTTGCAATCAAAGTTTTTGATATGCAATGGCAACGGGGTTTAAGAAATTTTGATTCAGAATGTGAAATTTTGAGCAATGTAAGACACAGAAACTTGGTCAAGATAGTGTCGACTTGTTCAAATTTGGATTTCACAGCAATTGTGCTGGAGTACATGCCTAATGGAAGCTTAGACAAAAGATTATACTCTGATGAAAATTGCTTGAGCCTGGTTGAAAGACTCAATATAATGATCGATGTAGCACTAGCCATGGAGTATCTTCATCATGATTATACAGTGCCCATTGTGCACTGTGATCTGAAGCCAGGTAACGTGCTTCTTGATCAGGATCTGACTGCTCATGTTGCAGATTTTGGAATTGCAAAAATGTTGGCACAAGAAGGAAACATGGCTCAAACCAAGACTCTCGGAACTATTGGTTACATTGCTCCAGGTATGATAATGTCTTGAAATGAAGGTGTCCTATGCTGGTTACTTTTTTCCTTTATGTCTACCATAGCTTTATACTGTAACATGGATAACTACACATTTTAACTGGGAATTTGTTGAAACAGAATATGGTTTAGACGGGCAAATTTCCACTAGCAGTGATGTTTATAGCTTTGGGATTTTGTTGCTTGAGACATTCACAAGAAAGAAGCCTACAGATGACTTGTTTAGCGGAGACTTAAGCTTGCATAAATGGACATCTCTCTCCTTTCCTGATGCTGTACTGGATATTCTGGATGCTGACCTTGTTAGCGACATTGGCTTTACAATTGATGAAAACCAAAGCCAAATCAAACAATTGCTGGTTTTGATAATAAATGTTGCATTCCTGTGTTTGAAAGAGTTGCCAGAGGAAAGGATAAACATGAGAGAGGTTGTGGTGCAGCTAAACAAGATAAGAGCTGAATTAACCAAGCTTTTAACTCAATCCCATTCAAAATTTGAGAAAGAGGATTGATGCAGCAACAACCAAATTCAGTTTTCCACCTTCTATGGCCTGTCAAAACATCTGTTATTTCTTAACATTTTCTTTC

>M6-PGSC0003DMG400011160 M6 LRR-RLK

AAAAAATCTCTCTTCCCTCTCTCTGAGTCCTCTCACTTTCACTCATTTTCTTCATATCATCTCCACACCTGATTCTCTTACACCAAAAACACAGTACAAAAAAAATAGTAAAAAAAAATTTGAAAAAAAAAAACACAAACAAACTCAGATAAAGCAAAAGGAGAAACAGAGTTGGTACAAATACCAACAAAATAGACAGACTGATAAAGAACCTGTTTCTTTTCCTTTACACTCTCTCACTTTTTTTTTTTTAGTTTTTGACTGAAAAAAGACACAGTTTTTACTCATTTATATGAACCAAAATGTTAAATAAATTCATGGGGTTTGCTTAATATATGTGTTGAATCATTTGATTTTACTTTCTTGGTTTTTTTTTTTTATCATCAATGGCTTCCATTTTCTTGATTTTTGTTCTCTCTTTGCTTCTGTTTTACCCTTTTCTTTGTTTTTCATATGAAGCTAGGAATCATGAAGGTAAACAAAAAAAAAAAAGAACCCCATTTGCTTATTTTTCAGCTTTTGATTTTCAAAATTTGATCTTTTGTTTTGATGTATAGTGGAAGCTTTGATTGCTATAAGGAGAAGTTTGGATGACCCAAAAGGAGCTTTAAGTAATTGGGATGAAGATTCTGTTGATCCTTGTAGTTGGGCTATGATCAGTTGTTCTTCTGAAAATCTTGTTATTGCACTGTATGTTTAATAGTGTGTTTTTTTTTGGAATTTTATGGTACTATATGATAATATTGGTTGTTAAAGATTGAAACTTTTACTGTTTTATCAACTAAGAAAGTATTTTTTTTTTGTTGTAGAGGAGCTCCAAGTCAAGATTTATCTGGGGTTTTGTCTGGGATGATATCAAATCTTACAAACCTTAAACAAGTGTAAGAGACAATTTTTTAATGTTTTTGTTGTATGTAATTTCAAATTTTGATGACCACATTTGGTTAATTTTCTTGTATGTTGAAAATTGTGTAGTTTATTGCAAAACAACAACATTTCTGGGCATATTCCAAGAGAGCTAGGGAAACTTCCTAATCTTGAAACATTGGATCTGTCTAATAACCATTTCTCTGGTCATGTTCCTGATTCTTTGGGTCTGTTAAACAGTCTTCAGTATCTGTAAGTGTATTAAACTTTCAAAATTTTCATTTTTGTGTGATTATCAAGATTATGTTTTTAGAGTTGAATGGTCTAAAACACACCTGAAGTATCACGTTTTTGCGAGTTTTCTACGAATTATCACCTAAAACATATCTCACCGCCATCTCATAGTCGATGTGTGTTTTGATAAATAGTTGGTGATGGTTTAAGAAGGAAAAGTGAACACATGATAGTTTAGGTAGAAAATGTGCGAAATCGTGATACTTCAAAGTGTGGTTTTTACCATTATCTCTTGTTTTTCTGAGTACCCTTTTTCCTTTAATTGTTGTTTTCTCATGTAGACAATGTTTAATTGCTATCTTATTCTTTGACTATATGTTTTTGTTTTTTTTTGATGTCTTGTTCTGGGGCTTGTCTTGTAATGGAGTAGAAGGTTGAACAACAACAGCTTGTCTGGTGCTATTCCTCTTTCTTTAGCCAGGGTACCTCAGCTTGCTTTCTTGTGAGTATTGCTTCTTTAAATCTATGACTTTTGTTTGATTTTGTTTTATTCTTTTGTCTGATTGTCCTGAGAATTGGAATTATCTAGTCACTTTCTTGAATCAGCCTTATGTTAAAGCTTTTTTGGTTTAAAGGAAATTGAATAGTTTGTTCTGTTTAGGGGTATTATTATCTTTTAACTTATTATTCATTTTTATTTTATTTAGTATGACGATGATATGCAATGAGCTTAAATGAAAGTGGGGATTCATATTGTTGACCCCAACTTGTTTGGGACTGAGGCGTTGTTGTCGTTGTCGGATAGTGTTGTCTTTTTTTTGTATAAAGAAATGAATAGTGTGTTCTGATTAGGGGTATTATTATATTTTAACTTATTTTTAACTTTTATTATATTTAGTATGACGATGATATGCAATGACCTTAAATGAATGTGGGGATTCATTATGCGGACCCCAACTTGTTTGTAACTGAGGCGTTGTTGTTTTTGTTGTCATCGAATAGTGTGTTCTGTTTTTGTGTAAAGGAAATGAAGAGTGTGTTCTGATCAGGGGTATTATTGTATTTTAACTTATTTTTTACTTTTATTTTATTTAGTATAATGATGATATGCAGTGAACTTAAATGAAAGTGGGGATTCATATTGAACTTGTTTGGGATTGAGACGTTGTTGTTGTTGTCGAATAGTGTGTCTGCTTTGTGTAAAGGAAATGAATAGTGTGTTCTGTTCAGGGTTATTATTATATTCTATACTTATATTTCACTTTTATTTAATCTAGAATGACGATGATATGCAACGAGCTTAAATGAAAGTGGCTATTCATATTGCCGACCCCAACTTTTTTTCTTGGTTGAGACTGAGGCGTTGTTCGCGTCGTTGCTGTTGAATAATGTGTTCTGTTT

>M6-PGSC0003DMG400011205 M6 LRR-RLK

ATGTCTATGATTGTTTCTGATCCATTTCATGTACTTGAAAGTTGGAATTCTTCCAACGTTCATGTATGTAATTGGACAGGAATTGTATGTGACAAGAAAATGCATAGAGTTGTTGAGCTTGATCTTAGTCATCATTCGTTACGTGGAACGATTTCTCCTTCTCTCTCTGGGCTCTCTTTCTTGCAGATTCTTGATTTGTCTGGTAACTTGTTAGAAGGACAAATTCCAGCTGAATTAGGCAATCTTGTTAAGCTAAATCAGCTTAGTTTGTCTTCTAATCTCCTCGAAGGGGAAATCCCAAACGAGTTAGGATTCCTTCATGAATTGAAGTATCTTGATTTGGGAACAAACAATCTTTCTGGTGCAATTCCTCTGTCTTTGTTCTGTAATTGCTCTGCTTCATTACAATATATGGACCTTTCTAACAATTCGCTTCGCGGTGAGATCCCTATGAATGATCATTGTGAGTTGAGTGGTTTGAAGTTTCTTCTTTTATGGTCTAATGAACTTGTTGGTGAAGTTCCTATTGCTCTTTCGAAATCTACAAAGCTAGAATGGCTTGATCTTGAATCGAATTCGTTGAGTGGTGAACTTCCTTCTGATATTGTAAGTAAAATGCCAAAGTTACAGTTTTTGTATTTGTCTTACAACAACTTTGATAGTCATAGTGGTAATACTGACCTGACGCCTTTTTTCGCGTCTTTAGTTAATTCTACTAACTTGCAAGAACTTGAGTTAGCTGGGAATAATCTTGGTGGGGAGTTACCTCCTATTATCGGTAACATCTCCAAGAACCTTGCACAGATTCATCTTGACGATAATCTTATCTATGGTCCAATCCCAACACAAATTTCGAGTCTTGTAAATATTACTCTATTGAATTTGTCTAGTAATCATCTGAATGGTACAATTCCTCCTGAGTTGTGTCGAATGGGGAAACTTGAGAGGCTTTATCTGTCGAATAATTCACTCTCTGGTGTCATTCCTTCTGCTTTTGGAAATGTTTCTCATTTAGGCCTTCTTGATCTGTCGAAAAACAACCTCTCTGGATTAATCCCAAATACCTTTGCTAATCTTCCGCAACTGAGGAGGCTTTTGTTGCACGATAATCACCTCTCTGGAACGATTCCCCCGAGCTTAGGTGATTGTGTTAACTTGGAAATACTTGATCTTTCTCACAACAGAATCACGGGGACGATTCCAAGTGCAGTTGCTGGATTGAGTAGTTTGAAGCTCTATCTGAACTTGTCTAGTAATTACCTACATGGTCCAATTCCTTTAGAGCTCAGCAAAATGGATATGGTGTTGGCAATTGACTTGTCTTTGAATAATCTCTCGTCTAATGTCCCTTCACAGCTTGGCAGCTGCATTGCTCTCGAATACTTGAATCTATCGCGTAATTCCTTAGAAGGACCTCTTCCATCTTCGATAGGAAGATTGCCTTACCTTAAGGAAATCGACGTTTCATTCAATGTGTTGAGTGGGGAAATACCACAGACATTTCAAGCATCGTCGACTTTGCAGAACCTCAACTTCTCCTATAACCGTTTCTCGGGGAACGTGACAGATACTGGTGCATTTTCATCACTAACCATCAGCTCATTCATGGGAAATTTCAAGCTTTGTGGCTCAGTACAAGGGATGAGAAGTTGCCATAAAAAGAGGGGTAGACATTTCATTATCACAATCCTCCTTTCATTGCTTATAACTCCTATATTTTGCATAGTCGGGTACCCTCTCCTCCTAAGATCAAGATTCAGGAACCAATTCGCGAGTTTTAATAAAAGCAAGGTAATTGAAGATGTGGAGCAAGGTGGTGGGAGGAAAGAGCAAAAGTACCCGAGGATATCTCGTATGCAGCTGATTGAAGCAACGGGAGGTTTTAGCAGCTCGAGTTTAATAGGAGCAGGAAGATATGGGCGTGTTTATAAAGGAGTCCTGAAGAACAACATGAGCATAGCTGTTAAAGTAATTGACACAAAAGGAGGTGGAGAGATATCAGGTAGCTTCAAAAGGGAATGTCAAATCTTGAAAAGGACTAGACATAGGAACTTGATAAGGATCATAACGACATGTAGCAGACCGGACTTCAAAGCTCTCGTGTTCCCTCTGATGTCAAACGGAAGCCTAGAGAACCATCTGTACCCAAGCCACAGGCTGAGACATCAACTTGACTTAGCTCAATTGGTTAACATTTGCAGTGATGTAGCAGAAGGAGTGTCCTATCTCCACCACTATTCGCCTGTTAAAGTCGTGCATTGTGATCTAAAACCAAGCAATATCCTTCTTGATTACAACATGACAGCTTTGGTAACAGATTTTGGGATTTCAACATTAGTAAAAGCTGTTGAGGAAAATACTAATCCTATCGACGAATCAGTTAATTACAACTCAACAGATGGATTACTTTGTGGATCAATTGGTTACATTGCTCCTGGTTAGTAACAAATAATAGTTCTATTCTATCTAATTCATTTCAATTTATATGACTAAAGTTTCTGTTTCTTTGATTGTAGAATATGGAATGGGAAGGAGGGCTTCAACAAAAGGAGATGTATTTAGCTATGGAGTTTTGTTACTTGAGATTGTGACAGGTAAACGTCCCACAGACGTTCTATTTCAACAAGGCTCGAGCTTACACGAATGGGTGAAGAGTCAGTATTCACATAAACTCGAGGCAATGATTGATGAAGCATTACAAAGATGTACAACAGTTCAACGCGATAGCAATATATGGCACGATATCGTGTTAGAGTTGATTGAACTTGGCTTAATGTGCACTCAGTACAATCCATCAACTAGGCCAACAATGTTGGATGTTGCACTTGAAATTGGAAGGTTAAAACAGTATGTTTCAACTTCTTCAAACACTGTTACTGAAGATGTCACAACAAAAAGTTGAACAGTTTTTTTTTTTATATATCTAACAATGTACCTTTTAACTTTATTCATTGTACTTCTATGTTCTTTTTTTTTTTTACCATTTTAATTGTTCCTTCTTGGGGTGCGGACCTTCTCTGGATTCTGCGTGAATGTGAGATGGTTCGTGCACCAAGACTACCCTTC

>M6-PGSC0003DMG400011509 M6 LRR-RLK

ATGAAAACATGGAATGCATCCACACACTTTTGCCATTGGTCTGGAGTAACTTGTGGCTGTAAACATGTTAGAGTCATTCAACTAAACGTTGCGAACCAAAAACTAGATGGCCCCTTGTCATCCTTCATTGGGAACATGAGTTTCCTCAGGTCTCTATACCTGTCAAACAATAGCTTCCGTGGTGAAATCCCTTCTGAGATTGGTCGTTTGAGAAGGCTACACCGATTGTATCTTGGTAACAACTCATTTCATGGAGAGATCCCCTCAAATTTATCAAGGTGTTTGAATCTTGTATCTCTTGTCCTTGAAGGTAACAAGTTAGTAGGAAGCCTTCCGCAGGAGCTAGGTGCCTTGTCCAAACTCGAGTACTTGTTACTCACTCGAAACAACTTGACAGGTGAAATCCCTTCTTCTTTTGGAAACTTAACATCATTGATAGGATTCTATGCACCATTGAATAATCTTCAGGGGAAAATCCCTGACTCTTTTGGGCAGCTGAAAAACTTAGAAATTATCGGAGTAGCTGCAAACCATTTGTCAGGTACTATCCCTTCTGAAATCTTCAACATATCTTCCGTCACAACATTTGATGTTGGAATGAACCAAATCCAAGGTACACTTCCATCAAGTTTGGGAATCACCCTCCCGAATCTTGAATTATTTATCATCGGTGGGAACAATATTTCTGGATCGATCCCATCTACATTGTCCAATTCATCAAAGCTAGTGTATTTTCTTGCTGAAAGAAATCAACTTACTGGAAGTGTACCAAGTCTTGAGAACCTGAATGGGCTTCAACAGTTGACAATTCCTGGGAATTATCTTGGAACTGGAGAACCAGATGATCTAAGCTTCATAGCCTCTCTGACTAACGTGTCCCGTTTCAGAATTTTGGAGATTCAGTTCAATAGTTTTGGAGGGGTTCTGCCTGCATCCTTTAGAAATCTTTCAACTGAGCTTCAGGTTGTGCAGTTGTCATACAACAGAATACGTGGTAACATACCGTCTGAGATTGAAAATTTCGTCAACGTGGAAGAATTTCAGGTGAGGGAAAACTTACTGGTACTATTCCAATCAATTTTGGTAAAGTTAAGAAACTGCAAATTTTGGATTTGTCTCAGAATAGATTTTCAGGGAAAATTCCGTCATCTCTAGGAAATCTTAGTGTAGTATCCATCCTTCTTCTGCACGATAAAAATCTCACCGGTGAGATCCCAGCAAGTTTAGGAAATTGCAATTACGTGATTGAAATATATGTTGCTAAAAATAACCTGTTAGGACAGATACCTAAGGACCTTTTTGCTCTCTTCCTTAGTTGCTGTGGACATTTCTGAGAACCATTTAGATGGTTTCATTCCTCTTGAAGTTGGAAATATGATAAATCTTGAATATCTTAATGTTTCCGTAAATAACTTGACCGGTAAAATCCCAAGTACTATTGGTAGCTGTGTGACACTAGAAGCCTTAGACATGAAAGGGAACTTCTTTCAAGGGATTATACCTCCTTCTTTCAGCTCTTTACGAGGTCTCCGTGTTTTAGACCTTTCTAGAAATAATTTGTCTGGCCAAGTACCAAAATACTTGGAAGATTTCAAGTTTCAACTCTTGAATTTAGCGTTCAATGATTTCGAAGGAGCGTTACCAAATGAAGGCGTCATTAAGAATGCAAGTGCTATATCAGAAATTGGAAATCCAAAACTTTGTGGGGGTGTGCCTGACATACATCTTCCGGGATGTGACATCAAAAGATCGAAAAAATTTGGTTCGAGGTTTATCCTCAAGATTGTCATCTCTGTTGTCTTTGGTATACTAGGCCTTGGTATGCTTATCACTTTACTATTTTGTTTCTTGTTAAAGAAACCGAAAAGGCCGCCTGTTTCAAGTTCATTGGGTGAGTCTCTCATAAATGTGTCTTACCGAAGTTTGCTTCAAGCTACTACTGGATTCTCTGAAGACAACTTGATTGGTGCTGGGAGCTAAGGATCTGTTTATAAAGGAACTCTCGATGGAGGAATGGTTGTCGCAGTTAAAGTGCTAAACCTCTCGCGACATGGAGCTTCCAAAAGCTTTATGGCTGAGTGTGAAGTCTTGAGAAACATTAGACATCGGAATTTGGTCAAGGTTCTCACAGCATGTTCAGGAGTTGACTATCGAGGGAACGATGTCAAGGCTTTGGTTTACGAGTTTATGGTTAATGGAAGCCTCGAAGACTGGTTGCATCCTAGTCCAAGTGAAGATACAAGTCAGGCAGCTGAAACAAAAAAGTTGAATATTCTTCAAAGGCTCAACATTGTTATAGATGTTGCTTCTGCAATCAATTATCTTCATCTCCACTGCGAGACTCCGATAGTTCACTGTGATCTCAAGCCAAGCAACATTCTTCTAGACAACCAATTGGTCGGGCACATTGGTGATTTTGGTTTGGCAAAGTTCCTTCAACCAACTGCACAAAATTCTTCTATTTCAGAAGGAAGCTCTGGTCTTGTTAGAGGAACAATTGGTTACACTGCTCCAGGTAACAATTAATACATAAACAAACACTCGAACTTGGTATATGCCCTTTTTTTCTCTCTCATTTCTTCTATTTTTTCTTGAAAATGATATATTCATAGCAAAACCACACTGCTTAACCATAACTTGGTTCGACAGAGTTAATTTATCAATATCATAAACTATCTCTTAATTGTCACACACATACTTCCATTACTTTTATAAAATCAAGTCATACTAAGTTATGTCCCTATTAGGAAAGTTATTGTTCCTCGTCTTTTTTTTTGTTTTGTGAATTTTAGAATATGGGATGGGAAGCGAGCTATCGACTTGTGGTGATGTGTACAGCTTCGGGATCCTTTTACTAGAAATGTTCACCGGAAAAAGGCCTACTGATGGAATGTTTAGGGATGGGTTAGACCTTCCAAGCTTCGCTAAACATGCATTGCTTAATGGAGCTATGGAAGTCATAGATCCATCCCTTATTTATGGAAGTGAAGAAGATGAGAAAGGTAAAAACACAAATATCTACCAGAATAAGGTGTATCTGGTTTCTGTTCTTCGAGTCGGGGTTGCTTGTTCTGCATATTCTGGAGCTGAAAGAATGAACATCACTGAAACTGTTTCCCAATTGTACTCTATCAAGGAAGCTCTTCTGAAATCAGAAAAAAATTCAGGAGCACGTTGA

>M6-PGSC0003DMG400011580 M6 LRR-RLK

AATAAAGGTTGAATCTTTCTCATTTTTATCCAACCCCTTTTCAAAATCTTGCATGTATATAGGAAGATCCCCACCATTATTGGAAAAATGTTATAGGTTCTTGGTTTTAGATATAGTATAATCTTTGATTTTATCTTTTCCTCAATTCATGAGTTTTATGGTCTTTGCTCTGTCATCATCTAAAGGAATTACTTGAGAGTTTTTTGCTTTTTCTGGGTTTTTGGTGGAATTCAGCTCTTTTTGACAAGGTGATGGACTCAATAGCCTAGAGGTATGCTAAAGCTTAGATCTTTAATGAGTTTCTTATTTATTCCTGAATTATTCCTTTTATATAGTTGAGTTTCTGGGCATTTCTGAGTTGATTGGATTCTTTGAAGTTAGTTGATTAATTGGTATGCATAGAACTTTTTCTGGTTTGTCTAGTTTATCTAGTTAGTATTGTCTCTTCTTGATGATTGACCTTAAGGGTGTGGAGGGTAGGACTTGGATTAGGGTAGTAGGTGGTAATGCGTTGTCTTGTTGTCCGTTCATGTTAGTAGTCATACTATTGCTCATTCTGTTACTATATTTTGGTTCCTTATACCTCGATTGTGTACTGTTTTGTTGTAGTTTCTGTTCTGCTATTCTCTATTGTTTTGCTATTATCTGTTGATTCTTGTACTTCCGTTACTTCATTTTCTGGACTGTTTTTTCTTGAGCCGAGGGGCTACCGGAAACAACCTCCCTACCTCTGAGATAGGGGTAAGGTTTGCGTACACTATAGTCTATACCCTCCCGAGTCCCCACTTTGTGTGACTACACTGGTTATGTTGTTGTTGTTGTTGTATTTTATAAGGATTGACACATAGTTGGATTAACAGGTCAAACTGAGACTGTCTTATTGGAAGTTTGTTATGATGTCTGGAGATTTTGAATCAAGAAGTGGTTTGGTTCTTCTAATTTGGATAAGTGCTTTGGTGGCTGCTGTACTATTGCTGGTTTCTCCTGCCGAAGGATTGAATCAGGAAGGAATGTACCTTCTTGAATTGAAGAAGAATTTCCAGGATTCGTTTAACCATTTGGGGAATTGGAATCCAAATGATGAAACTCCTTGTGGCTGGGTAGGTGTGAATTGCACTTCTGATTATAATCCAGTTGTGCAATCTCTTTATCTGAGCTATATGAATCTTTCAGGCACGTTGAGTTCTAGCATTGGTGGTCTTGAATATTTAGCTTATCTTGACCTGTCTTACAATCAATTCACCGGGAACATTCCCAAAGAGATTGGAAATTGCTCAAAATTGCAGTCTCTCCAGCTCCATTTTAATACTTTTTACGGTCCGATTCCTGCTGAACTGTATAACCTGTCTAATTTGAAGGATGTGAACATGAGCAGCAACATGATCTCCGGTCCTATTGCTGAGGAGTTTGGGAAACTCTCGTCTCTAGTTACTTTCGTTGCATACACCAATAATCTTACCGGTCCAGTGCCTCGATCTATTGGAAGTTTGAAAAATTTGACAATATTTCGAGTGGGGCAGAATGCACTTTCTGGAAGTTTACCTGCAGAGATAGGTGGTTGTGAGAGCTTGGAATCTCTTGGTCTAACACAGAATTGCCTGGAAGGAAACATACCAAAAGAACTTGGCATGCTTAGTAAGTTGAAAGAGCTTGTGCTTTGGGGAAATCAGTTCTCTGGTTATATTCCAAAGGAACTTGGAAATTTAACACAAATTCAATTGCTTGCTTTATATCAGAATAACCTGATTGGAGATATTCCTGCTGAAATAGGTAAACTCAAAACTTTGACGAAGTTATACCTATACCGGAATGGATTAAATGGTAGCATTCCAAGGGAGATTGGGAATCTTTCTATGGCAACTGAAATTGATTTCTCGGAAAACTTTCTGAAAGGAGAAATTCCAGTTGAGTTTGGCCAAATAAAAAGTCTGAAATTGTTGTTTCTTTTCCAGAATCAGCTCGAAGGCGTTATACCAGATGAACTTACTACTCTAAAGAACTTGATTAGTCTTGATTTGTCAATTAATCACCTAACAGGTCCTATTCCATTTGGTTTTCAGTATCAAAAGGAATTGATTCAGTTACAGCTTTTTGAGAATTCTTTAACTGGTACTATTCCTCAACGTCTTGGCATTTATAGTCGACTTTGGGTGCTTGATTTAAATAACAATCAACTCACAGGAAGAATTCCTCCATTCGTTTGTCAGAATTCCAATTTGATTTTGCTGAATCTTGCATCAAATAAATTGCATGGATATATTCCATCTGGTGTTCTTAAATGTGATTCATTGGTGCAACTTCGTCTGAACGATAACAGGCTAACAGGGACATTTCCTTCTGAGTTGTGCAAATTGATAAACCTGTCTGCTGTTGAACTGGGGCAGAACAAGTTCACTGGTCCAATACCTCCTGATATTGGATATTGCCAGAAGTTGCAGAGGCTTGATTTCTCAGGCAATAGTTTCAACCAGTTGCCAAGGGAGATAGGAAATCTTACCAGGCTTGTGACATTCAATGTCTCAGCAAATTCGCTCACTGGACCGATACCTCCAGAAATTCGGAACTGCAAGGCACTACAAAGACTGGATCTCAGCAAGAACAGATTTACTGATGTTATACCTGATGACATTGGTAGCTTGTCGCAGCTAGAACGTCTTTTGCTTTCAGAGAATAAGTTATCTGGAAAGATACCAGCAGCATTGGGGAGCCTCTCTCATTTGACTGAGCTGCAGATGGGCAGTAATCTATTGTCAGGTGAAATACCATCAGAGTTGGGTAATCTTAGTGGCTTACAAATTGCAATGGATCTTAGCAACAACAAGCTCTCTGGCAGTATACCACCCAACCTTGGTAATCTTATCCTATTAGAGTACCTATATCTCAACAACAATCATTTGAGTGGTGAAATACCAAGCACATTTGGAAACCTGACAAGTCTCTTGGGCATTGACTTCTCATACAATGACCTGACTGGACCACTGCCGGATATACCACTCTTTCGGAACATGGATATCAGCAGCTTCATTGGAAACAAAGGTCTTTGCGGTGGGCCTCTAGGGGAATGTAATGCATCTCCAGCTTATGATGCCAATAATCCTCCTAGAGTCGAGAGTGCAGATTCTCCTCGAGCAAAGATTATTACTGCAGTTGCCGGTGTAATTGGTGGAGTTTCTCTTGTTTTGATTGTGGTAGTTTTGTACTATATGAAGCAGCATCCAGTTGAGATGGTTGCAACCCAGGATAAAGACATGGCATCTTCAGATCCAGACATTTATTTCCGTCCCAAAGAGGGGTTCACTTTCCAGGACTTGGTTGAGGCTACAAACAATTTTCAAGATTGTTATGTTCTTGGAAGAGGAGCCGTCGGAACAGTTTACAAAGCAGTCATGCAGTCTGGACAAACGATTGCAGTCAAGAAGCTTGCTTCTAACAGAGAGGGTAATAACATCGACAATAGCTTCCGTGCAGAGATCTTAACTCTAGGAAAGATTAGGCATCGTAACATCGTAAAGCTATATGGCTTTTGCTATCATCAGGGTTCTAATTTACTTCTCTACGAGTATATGGCCAGAGGTAGCTTGGGTGAACTGCTTCACAGTACATCTTGTAGATTGGATTGGCCAACTCGCTTTATGGTGGCTGTTGGAGCTGCTCAAGGACTTTCTTACTTACATCATGATTGCAAGCCCCGGATCATCCACCGTGATATAAAGTCCAATAATATTCTGATTGATGAGAAGTTTGAAGCTCATGTTGGTGATTTTGGTTTGGCAAAAGTTGTTGATATGCCTCAATCTAAGTCGATGTCTGCAGTTGCTGGCTCATATGGCTACATAGCACCTGGTATGTCTTTCTGTTTTCTCCTGGACAACTTCTTATGTTCCCTTGTTATCTATAGAAATATGTCGAGAAATGTACTGCCATTTGATCCTTATTCCCACATATTTATCTGTATCTTCACTCTGAAAGTGTTACCTTTTTCTGTTGATATGAATAGGATTTTCTTGTTGAGATTAGATAGTTAGAATTGTATGCAATGTCTATTTCTTCGATTATCTGAAGACAACAATGAAATGCTTTTAACTTTTATCTGTGTATACTTCTCATTTTATATGTGTTCTATGCCAGGAGAAACCATACAAACTGCTCTATGTATCTTAGGCAAACTTTGGGGACTCTTCCTGATGCATATGGAGCATTCTTGTTCCCACATATTTATGTTCAGTTTAGATATTCGACCATGTGTTGTTCACTTTGAAAGTCTAATTCTTCTCTTTGTGATATGAAATGCATATCTTGTTGAAGTTTGATAGTTAAAATTGTTTACGAACAAGTTAATTTCTTTGATGAAATGACAGAATACAAGAAAAAATGCTTTTAACTTTTCTCTGTCTATGTAGCTTTTAATATCCCTCTCGTAATTTTCATTCTGTGTTATAAAACCATTTTTTCCTTTTCAATTGATGTTTTCTACCTTCTCTGTTGCAGAATATGCTTACACCATGAAAGTTACGGAAAAATGTGATATCTATAGTTATGGGGTAGTCCTTTTGGAGTTGCTAACTGGAAAAGCACCCGTACAGCCCCTAGAACAAGGTGGTGATCTTGTCTCATGGGTGAAGCATTATGTTAGGAACCATTCATTGACACCAGGGGTACTCGATAGCCGATTGGATTTGGAAGATGTAATTACTGTTAGTCATATGCTTACAGTTCTTAAAATTGCTCTAATGTGCACTAGCATGTCCCCTTATGATCGTCCTTCAATGCGCGAGGTTGTTCTGATGCTGATTGAGTCTGAAGAGCAAGAAGGGAACTTTCTCTCATCTCCAGTTTATGATCCCCCTCTGAAAGATAATTCCTTCTAGATGCATGTAAGAAGCACATCTCAATTAACTTTGATAATTTTTTAGTACTAATTCTCCTTGTATTTTTTTTCTTAATGGCTAAATTATAGATATTCCCATGTATATTTCTCATAGTGAAAATTTCATCAATCACTTTCTAGAACTAGTTTGGAGGTGATTGATGTTATTTCAATAGTTCTCCATCTGATGGTTATGGAAAATAATATGCTTTATGCTTTTCTCACTATATATTGAGGGCTACCTAGAT

>M6-PGSC0003DMG400011833 M6 LRR-RLK

TTTTAATACATAGATGGTGATAGTTCAGGTAAGAAAAGGGAACAACTAATAGTTGGAGTGTGAAACTCGCGAAAAAGCAATAGTT

>M6-PGSC0003DMG400011932 M6 LRR-RLK

ATGTCACTACTAGCATTCAAGAATATGATCATTGATGACCCTTTCAAGATCATGGACTCCTGGAATGAAACTATACATTTTTGTGACTGGCCTGGGGTTTCATGTGGTCGACGTCATCCCCGAGTTACAGTTTTAAATCTTACTTCTCTGAAATTGAGGGGTTCTCTGTCACCAAGCATTGGTAATCTGAGCTTTCTTTATGTCCTTAAGCTCCAAAATAATAGTTTTTCAGGTGAAATTCCATCAGAGATTGGCTACTTACATAAGTTAAAGTTTTTACGCCTTGATAACAACTCGTTCACTGGTCATATTCCATCAAACATTTCAGGATGCTTCAATCTTGTTTCTGTTGTTCTTTCGTATAATATGATGGTAGGCAAAATTCCAGCAGAATTAGGCACATTGTTGAAACTCAAACAACTTTCTCTTGTTTCTAACTATTTTACAGGAGGAATTCCGCCTTCATTTGGTAATCTTTCCTTACTTGATACCTTTTCCGCAAGCAAGAATAATTTGTTAGGGAAAATACCTGATGAATTATGTCAATTGCTCAACTTAAAGTACTTTGTAGTGAATGAGAACAACTTGAGTGGTACCCTGCCTCCTTGTCTTTTCAATCTTTCCTCTATCGTGGCCATAGATGTTGGAACAAACCATTTAGAAGGAAAATTGCCTCCATTGCTTGGTATTACTCTTCCTAATTTAGAGTTCCTTAGCATTTATAGAAACAATGTAACTGGAAATATTCCGGTAACATTGTCAAATGCTACGAATCTTCAGTCTCTTATTGCTAGTAGAAATGGACTCACAGGAAAAGTACCACCCCTTGGAAATTTGCTGAAAATGCGGAGATTTTTAGTTGCTTTCAATTATCTTGGGAAGGGGGAGGATGATGACCTTAACTTTCTCACCACATTGGTGAATGCCACCAGCTTGGAGCTTTTGGAGCTCAATACAAATAATTTTGGTGGAGTCTTACCTGCATCTGTTAGTAATCTTTCGACTGAGCTTATAGAGCTTTCGTTGTCTTATAATCAGATTTCTGGAGAAATCCCAAGATGGATATCCAATCTCAAGAAGCTTCAGGCTTTCTTTGTAGCTTACAACAGATTTCACGGTGAAATTCCGTCTGAAATTGGTGATCTGATATATTTGCAAGAGCTGGCTTTACTTGGAAATCAGTTCTCTGGCAAAATTCCAATCTCATTGGGAAATTTAGCTTCTTTAACCAAACTCAGTTTAAGAGAAAACAATCTCCAGGGGAGAATTCCTTCCAGTTTAGGCAAATGCCACAATTTGGAACTGTTGGATCTTGGTTCAAATAACCTTAGTGGGTTCATACCATCAGAAATTCTTGAACTCTCATCCTTGTCGGAGGGCGTGGACCTATCTCAGAACCAATTAACTGGTTTCCTTCCGATGGAAATTGGGAAGTTGAGAAATCTAGGCTACTTAAACCTGTCTGATAACAAGTTGCAGGGCCAGATTCCTACCAGTATAGGTACTTGTGTGAAACTTGAAGCACTAGACTTAAACAACAACAACTTTCAGGGTAGTATTCCTTCAACCATGAATAGTTTGCAAGGCCTGGAACTTTTAGTCCTTTCTCACAACAATCTGTCAGGTGGAATCCCGGGATTCTTGAAGGACTTCAAATTTCTGCAGATTTTAAATCTGTCTAGCAATACTTTGGAAGGTGCTGTCCCAACAGGAGGCATCTTTAGTAATGCTACTGTGGTTTCCATCATAGGAAATAGAAATCTCTGTGGGGGTGTACCAGAGTTGGACCTTCCGGCTTGCGTTGTAGAGGTAAAGAAAGAAAGGAAATCAGGTTTTCCCTTGAAAATAGTAATTCCTGTAGTTAGTGGGCTTATAGGATTGACATTCATAGTTTGTTTTCTTGGCATACGACAGTTTAGTAGATCGAGAAAACCAACACCTACATATCTACCTGAAAACTCAACCTTAAGAGTATCTTATCAGTGTCTACTTAGGGAAACTGATAGATTTTCTGCATCAAATTTGCTTGGAATGGGTGCTTTTGGGTCTGTATATAAAGGAATTTCTGAACACGATGGTACGTTTTTTGCTGTTAAGGTACTGGACCTTTCACACCATGCAGCTTCCAGGAGTTTTTTAGCTGAATGTGAGGTTCTAAAGAACATTAGACATCGTAATCTTGTAAAGGTTCTAAGTGCATGCTCAGGTATTGATTATGAAGGTAATGAGTTCAAGGCCATAGTTTATGAATACATGGATAAGGGAAGTCTACATGACTGGTTACATTTCACTCCTCAGGAGAAATCTGAGTCACAGGAAGAGCACAAGAAGCTCGGGTTTATTCAAAGATTAAACATTGCCATTGATGTTGCCTGTGCCCTGGATTATCTCCATAATGATTGCCAACCCCCAATAATTCACCGTGATCTGAAGCCAAGCAATATTCTTCTCGATGAGAACATGACTGCACATGTTGGTGATTTTGGTTTGGCAAGATTTGTTCCACCAGCAATTCCAAACTCATCAGCAAATTCTAAAAGCTCTACAGGTGTAGGGGGAACTATTGGTTATACACCTCCAGGTAATAATAAATCAAGATTTCTGCTGTCATATATGTCTTTTGATAAGAACATCTCTAATTCTGTAGTATTTCTTGTTGACAGAATTAGGTATGGGAAGCGACGCCTCAATCTATGGAGATGTGTACAGCTTTGGCATCCTACTCTTGGAAATGTTTACCGGGAGGAAACCAACTGATGAAATGTTCAAAGATAATTTAAACCTTCATAAATATGCCAATGCTGCTTTGCCAGACAGAGTGATGCATATTACAGATCCCATTCTTCTCCAGGAAAGAGATGAACTTGGAATGAAATACAATGTTGATGACAATACAAGCAGTGCTGGCGACATATTTCTGTCATTCTTAGTTAAGGTGATTCAAATCGGAGTTTCCTGCTCCGTTGAATCTCCAAAAGAAAGAAAGCAGATCAGTGATGTCGTTGGAGAACTCAATTCATTAAGGAAGTTGTTCTTGGAGCAAGCATACCGGAAAGAAAAGGTTATAAGCTTCAGCCTGCATTGA

>M6-PGSC0003DMG400011989 M6 LRR-RLK

CACAACTTCATCACAAAACCAAAAACAATCAATCATTTAGCCTTGATGATGGTTCCCACAATATTTTATAACCTTCCCTTCTTATATCTTTTCATAGTTATAACTTTTGCTTCCACTGAGGAAGCAACTGCTCTTCTTAAATGGAAAGCAACATTCCTAAACCAGAACAATTCCTTACAGGCTTCATGGACACTAAGTAGTCCTGCTGATACCAAGAATTCTTCCAGAATTGGAGCGACGAGTTCTGATGCATGCAAGGGCTGGTATGGAGTTGTATGAGTTAATGGTCGGGTTAACAGGTTGAATATTACAAATGCTAGTGTCATTGGCACACTGTATGATTTTCCATTTTCATCACTCCCATTTCTTGAATATGTTGATCTTAGTATGAACCAGCTCTCTGGCACCATCCCACCTGAAATTGGTAAGCTCGCTAATCTTGTCTACCTTGACTTGTCTATCAATCATATTTCGGGCACAATCCCACAACAAATCAGCTCACTAATAAAGCTTGAGACCCTCCACATATTTGACAACCAATTAAATGGTTCCATTCCGAGGGAAATAGGTCGCCTAAGGTCTCTTACTGATCTAGCTTTGAGTTCTAACTTTCTTAGTGGTTCAATTCCTGCTTCATTGGGGAATTTGAACAATTTGTCTTTTTTGTATCTTTATAATAATAAGCTTTCTGGCTCCATTCCTTCAGAAATAGGAAAACTTGTCAATCTTGTTGAAGCTTATATTAGTAGAAATCAATTAACAGGTCATATTCCTCCAGAAATAGGTAATTTGATCAATGCAAAAGTGTTGTATGCTTTTTCCAATAACTTGTCTGGTCCCATTCCTGATGAAATTGGGAAAATGAAGTCACTTGAGAACTTAAGCTTCCAGACAAACAATCTTTCTGGTCCAATTCCCAAAACAATAGGTGACTTAACTGAGCTCAAACTCTTGCACCTTTATTCCAACCAACTTTCTGGTCCCATTCCTAGTGATTTTGGGAATCTGAAAAAATTGAATGATCTGCAATTATCCAATAATCAACTTACTGGTCCTATTCCTGGTTCATTTGGCAATTTGAGAAACTTGCAATTTCTGTATCTCCGTGCCAACAAACTTTCTGGTTCTATTCCAAAAGAACTTGCATATTTGGATAACTTGGTTATAATGGAAATGGATGAAAATCAGTTTTCAGGCAATTTGCCTGAGCATCTTTGTCAAGGTGGAAAACTTGAGAACTTCACGGTGAATAGTAACAAGCTGACAGGGCCAATACCAAGAAGCTTTAGCAAGTGCTCGAGTTTCAAAAGAGTTCGCTTGGATAACAACAGTTTTACTGGGAATTTATCTGAAGCTTTTGGCGTCCATCCACACCTTTACTTCATTGATTTGAGTGATAATGATTTTCATGGTGAACTCAGCAGAAACTGGGGGAAATGCAAGAAATTGACTGATTTTCAGATAGCCAGAAATAACATTAGTGGTAGCATACCACCAGAGATTGGAAACATCAAAGGTCTTTTGGGACTTGATCTTTCAGCAAATCATTTGATTGGGCGGATACCAAAGGAATTTGGAAAATTAACCTCTCTGGTTAAGCTTTCTGTGCAAAATAACTATATTTCTGGCAATATTCCTATAGAACTTGGGGCACTGACAAAGTTAGATTCCCTAGATCTGTCAAACAATAGATTGAATGGGTCGATCCCAACGTTTATAGGAGATTACCAGCACCTATTTCACTTGAACCTGAGTTGTAACAAGTTTGGGCAGAAAATTCCAAAGGAGATAGGGAGGATAACTCATCTTAGTGTACTTGATTTGAGCTATAATCTTCTTGATGGAGAAATACCTGCTCAGTTAGCTAGTTTGCTGGACTTGGCAAACTTGAATCTTTCCCACAACAGCCTTGCTGGGCGTATTCCTGAAGAATTTGAAAGTTTGACCAGTTTGCAGGATGTTATCTTGTCATACAATGAGTTGGAAGGTCCAATCCCTAATAATAAAGCTTTCATGAATGCCTCATTAGAAGGTAATAAAGATCTTTGTGGCAATGTGATTGGATTCCAACCTTGCGAAAGGCCATCTTCTATTGTGAAGAAGCACTCAATGGCGAAGGGACATAAACTCATCCTCATCACTGTACTTCCTATTATGGGAGCACTAGTACTACTCTGTGCTTTCATTGGTGTTCTCTTTATGTGTAATAAAAGAAGGAGAGTTAGAGACGTTGAAAGAAGGGATTCCAGTGATGATGTTGATGGTTTGATTTACATATCCACATTAAATGGAAATGCATTGTACTGGGATATCTTAAAAGCCACAAAGGAGTTTGATGCAATGTTTTGCATCGGAAAAGGAGGATCTGGAAGCGTTTACAAGGTAAAACTTCCAACAATAGAAAATGTAGCTGTGAAGAGACTTCATTCTTCATTTGAGGTTACCCATCGCAAAAGCTTCATGAATGAGATAAGGTCATTGACAAGTATCAAGCATCGCAACATTGTGAAACTCTATGCCTTCTGTACAAATGCACAACACTCATTCTTGGTTTACGAGTACATGGAGAAGGGGAGCTTGTCTAGTATTTTGAGCAATGAAGTAGAGTCCAAGAAATTGGATTGGCTTACAAGGGTGAATATCATCAAAGGTGTTGCTTATGCTTTATCTTACATGCACCATGATTGCTCACCCCCGATTGTTCATCGAGACATATCAAGCAGTAATGTTTTGCTTGATTCCGAGTATGAAGCTCGTGTTTCAGATTTTGGCATAGCTAAGATTCTAAAGCCAGACTCATCCAATTGTACTGCACTTGCAGGCACATATGGCTATGTAGCACCAGGTAAGATCTAACTTCTTATCCTTATATTAGTTCATGAAATATATCTGCAAAATCTATATTGTTTCTATCTCATTTGAAACTTTATTGTTTCAGAGCTTGCCTATACAATGAAGGTGACGGAAATGTGTGACGTCTATAGCTTTGGAGTATTAGCATTGGAGGTAATCAAAGGAAAGCATCTTGGGGAATACATTACTGTGCTAGCAAATTCATCGACAAGAGATCATGTGCAGCATAGCGATTTGCTAGATGAACGCCTTCCATATCCTGAAGATAGAGTAAAAGAGGTTTTGGTTTTTATCATCAAGCTAGCAAGCTCATGTTTGCTTGAAACTCCAAAATCAAGGCCAACAATGCACTTCATCTCCAGTATGTTGTCAATGGATCCACCTGTTTCTCAAATAACTTCTATGAAGGCGAAAAGAGTCCCCGAGATGTAAGGTGTCAGTGTATGAAAAAAAAAGAAGCAAGTTTAGACACCAAGTGGTTGTGAAGTAAGAATAAGAAGCTATTATTACTGGTACTAGTATTTGTGTATTTTATTGTTAGTTATGCTTATCTAGTTTAGTGTTGTACTAGAGTGGATACAATATCAAGTAGAAGGTAATATTTTAATTTTCTTGTGCCAATAAGTGCTAATAGAAAT

>M6-PGSC0003DMG400012218 M6 LRR-RLK

TAGCTATTGTGTATTTATGAGAAAATATTCGAAGTTTGATCAATCATAGCGACGCTGGTAAAGGCCCACACTTCCTTTACTCCGCCGCTCCTTCTATTATCTCCTTCTCCATTTTCTGCTTGATTTTTTGTTCCCTTATAATTTTAACATATAGTTTTTATAACCCCAAAATTCTTCTTCTTCAGCTCATAAATTCTCTGTTTCTAGACACCAAATCTCTCTTTGAACCCAAGTTTTATTTTTTTTAACTAAGTTGGTGACACTTAATAGTAGATTCCACTCAAAAGGGTTGTTACAGATTTAGATTCTTGAATTCACTCTTAAGCTTTGTTAAGTTTTCTTTAGTGTTTATCTTTCTCCCCTCACTTGTAAAAAAGACAACTTTATCTTTTTCCAGCTTTTTTTTTTGTTTGTTTTCATCTACTGTCTTGGGTCTTTTCTTGTTTTCTCTTTCAATTTTGTTTGGATTATTGCTTTTTGAATCTTTTGCTTTCTGGGTTTTTCTTCTTTTTGTTGATTGCTTAAGTTCTTGGAGACTGTTTCTTTAGATGGGTTCTTTCAGTTCTGGATTTTTGTAGAAACTTAGTGTGTTCTTGTGGTGTGGAATTATTTGGGGGAAAAGATGAGACTTTGTGTTGTATTTCAAGTTTTAGTTACTGTTGGACTAATTCTTGGGATTTGTGATGCTTTTCCATCAAATGAAGGTGTGTATTATTAACTTAAGTTCAGATCTTGATGTAGTTACTGCTGCAAAGATTCAATTTTTGTCTTTTAATTGAGCATTTAGTGTTGTAATGTAATGTTGTTTAGGGAAGATCTTGTATTTTCTTGTTTTGAGATACTTGCTATATTTTGCTGTCATATTTTCAAATTGTCTATCAGAAATAGTCTCTCTACCCTGCGAAGGTAGGTGTAAGGTCTGCGTACTATGTTGTTGTTGTTGTTGTAGATAGTTGCTGTATTTTATGAGATTGTAATTCAAAGAGTAGGTAGTTTATTCTGAGTGAATCTGATTGGGGACTCTTTGTTGATACAGTTTATGCTCTCAATACCTTCAAGGAAGCTATATATGAAGACCCTCTTCTGGTTTTGTCAAATTGGAATGGTTTAGATTCAGATCCTTGTGGTTGGGCTGGAGTCTTTTGTTCTATGGCTCAAGATCATGTCATGAAAATGTAAGTTAATTGTCATCAATTTTATGTGCAATACATGGATAGGTTGTTTGGTTAATGATTTTTCTCAACTAAAATGCAATCTAGTAAATCTTTTATGGACTCAAGTTTAGTGCAAATGAGTCTCTAATACGCATTTCCACTGGATTGTGTGACAGAAACATTTCTGGCGCTTCGTTGAAGGGGTTTCTTTCACCCAACTTGCATTTACTCTCCTCTTTGCAAGAACTGTAAGTTATTTCCGAATAGATTTGTAAGGATTACATAGTTCTGGTTGTTCTTTTATGTGGTTTTGTTAAAGCTTATGTGATTCCCATGCAATATGCAGAATTTTGCATGGAAATGTGCTGATTGGTACAATTCCTAAGGAGATTGGTCTATTGAAAAACCTGAAGGCCTTGGATTTGGGTGCTAACCAGCTAACAGGACCAATTCCTTCCGAGCTTGGCAATTTGACAAACATTATGAAAATGTTAGTCTAACTTGAATTTTGTAGGTTCATTTGTACTTGCCTTTACCTATCCTTGTTGAATCTCCCTTACTTTTCTAGCTTCCATGTGTCTCTGCAGAAACCTACAATCTAATGGATTAACAGGGAAATTGCCTGCTGAGCTTGGTAATTTGAAATACCTTGAGGAACTTCGGTTGGACAGGAACAAACTCCAAGGACTAGTGCCTGCTAATACTGGATCAGATTTTACATCCGGTGTACGTGGAATGTGAGTTGATTAGCTACGTCTTGTACTTTATCGTTTTTCTTTTGAATTTTCTGGTATGTGGATAAGGATTAATGCACGATTCTCTGTTTACTTGCAAGATATTTTTGGTTGTCCTTAGAGCGTTTTCCGTCTTGGCCCTCTAGGATTCTGTCAGATATTTGAAGGATTGAGTGACCAAAGAATCTAATTTTCTGATCTTTATGACAGTCATTGTCTGGAATGATGATGACAGAATCCAGGAAAGAGATGCTTATGGGATAATGTCTGTTCTTTCTGTCAGCATTGTTTGATATGTACTGCCTGCAGATTGACACTGATGGCTGTTTTACTTGAGTCTAGTGTAGTACATTCATTGTCTGTTGTTTGATGGGAGGATGGTGAATTGGGGGACCCCAATGGAGTGTGTTTTTGCTATAGCCTTTAATATTTGACCAAATGAGAGCAATGTTACAGAGTTCAGAATGTATGTTTCAAAAGTTTAACGATGATGTTCTTGGAATTGATCTGTATTTCTGTGAAGAAAAATAAGTAAAAACTGTTTGATGTCCTGTATATTTGACCCAACTGATTTTGCTATAGCCTTACGTCCATCACATGTATCTTTATCTGATTGTTTTTTTTCTTTTCTGTTGTACCCATTTAGGTATGCCTCTGGTGCCAGCGCTACAGGTTTTTGTCGCACATCCCAGCTCAAAGTGGCTGATTTCTCATTCAACTTTCTTATTGGAAGCATACCAAAGTGTTTGGGTTACCTTCCAAAGTCAGTTCTTAATTCTACGCCATAATTCTTGTTTTACCTCATCTTCTCTATTCCTGAATGCCTATTAGTTTTGTTGCTGTAAATCTATATCCAAAGTAGCACTCTGATGCTCTACCAGTGTTGTTTCTCTTACCTTTTCTGTTTCTTAATGCCTGTTATTTTGTGATTCTATAGATCTAGTTTTCAAGGAAATTGCCTTCAGCACAAAGACCCGAAGCAACGGTCTGCTGCTTTATGTGGTAAGTAAACAATCATAGTTACAATTTTCAACACTATGCATAGGAGAATTGATGTCTCAACCTTTTTTCTTTAATAGTTGCAGTCAAATAGATTTATTTAAGAAACTGAACTGGTTGTTTTGACATTGCATTATAGGTAGCACTCCGCCTCCCACTAGCCATGCATCGACACTCAGCAATAAGCACCAGCCTGTTGAAGAACCGAGACATCAGACTTCTTCCAAGCCCACTTGGCTTTTGGCTTTAGAAGTGGTAACAGGAGTTATGGTGGGTTCTCTCTTCATTGTAGGTCTTGTGACTGGCCTTCAGAGGCTTAAGATCAAATCCTCAATTATACCTTGGAAGAAATCAGGAAGCGAGAAAGATCATATGACAATTTATGTAGGTTAGTACTATCATGCACTTGAACTTGAGCACTTAGTACTTTCAAAAGCATTGAGCCTAATCTTCGTTTCTGTTTACCGCTGATGATCTTTGTAGATACTGGTATGCTGAAAGATGTTGTAAGATACAGCAGACAAGAACTTGAAGTGGCTTGTGAAGATTTCAGCAACATCATTGGATCCTCACCAGATAGCATAGTTTACAAAGGAACCATGAAAGGTGGACCTGAAATCGCTGTAATATCCCTTTGCGTCAAAGAAGAGCACTGGACAGCCTATCTAGAACTTTATTTTCAGAAAGAGGCAAGACTCAATTCTTTCTTCATCATCACTATTTATCTCTTCCAAAAGTTTTCATCTTTAGTTCCATGCTCTAACTTCTGTAAATTTACATTTAGGTGGCAGAATTGGCCAGAATAAATCACGAGAACGCTGGGAAACTTCTAGGCTACTGCAGAGAGAGTAGTCCCTTTACGAGGATGTTGGTGTTTGAATATGCATCTAATGGGACATTATATGAACACCTTCATTGTGAGTAAAGTGAATTAATTTTCTCATGCACTTCATTTTATGTTTTCCAAAGCAAACTCCTAACTCTTTGAATGTATCATAGATGGAGAAGGATGCCAACTATCTTGGACACGCCGGATGAAAATAGTTATTGGCATCGCTAAGGGACTGACATATTTCCATTCAGAACTTGACCCTCCATTTACTATATCAGAACTGAATTCAAATTCTGTATATCTCACCGAAGATTTTTCTCCCAAGGTAAACGAAAGACCATT

>M6-PGSC0003DMG400012307 M6 LRR-RLK

GTTTTTTAGAAGTTTTGAACTTTAGATTCTTGATGCTCTTTTTGTTCTTCTTATGTGACTACAAAAGCTAAGTTTTTGAGGGGTTGAGGGTATTCATAATCTTTGAAATGGTTAAAGTTTTGAGCTTTAGGGAGTGGTGGGGGATTGTTTCTTTGGGTTTCTTGGTGGGGTTGTTTCTGTTGTCACAAGACACTGTTGCTTTTCTTGAAAATGATAAGCAAGCATTGTTGGATTTTGTTAATCAGTTGCCTCATTTGCACACTTTAAACTGGGATGCAAATTCTTCCGTTTGCAAGAACTGGACTGGAGTGGGGTGTAGTGAAGATGGTTCTAGAGTGATAGCACTGAGGTTACCTGGGGTTGGGTTCAATGGTCCAATTCCTAATAATACATTGAGTCGTTTGACTGCATTGCAGATCTTGAGCCTTAGATCTAATGGTATCAATGGCACTTTCCCTATGGACTTTGATAACCTGAAAAACTTGTCTTATCTTTACCTTCATTACAACAATTTCTCAGGTCCTCTGCCTTTTGATTGCTCTGTTTGGCAAAATCTCACCAATTTGAACTTGTCAAACAACAGATTCAATGGCACCGTTCCTAGCTCGATATCGGGTTTAAGTCATCTTACTCCCCTTAATCTTGCTAACAACTCACTTTCTGGTACCGTTCTTGATTTTCATTTTCCCAACTTGCAGCTACTGAATTTGCCAAACAATAACCTTATAGGTATTGTGCCTAAATCACTTCAAAAATTCCCAAAGAATGTGTTCATTAGTAATGATATATCTTTACTTGACTATCCTGTTTCGAATTCTTCTATTGTTTCCCTGCCTCAGCAACCAAATCCAAAACTCAAGAATGATGGTAAATTGAGTGAGCGG

>M6-PGSC0003DMG400012443 M6 LRR-RLK

CTTTGTCTAATCTTAATATAACTGTTCCTTTGATTTGAAATCTCAATCTTTATCAGCTTAAATAACAATTGTATAATCTTTTTTTTTCTTTATTAGATTCTGTCCCTCTTTCTTGAAATTAACTAAAAGAGTTTGCTTGAGTTGAAAGTTGGGTTCTTGCTTCTCTTACAAGTGATGGGGTGTTGAAAAATCTTATCTTTTTCATTTTTTCTTCCTAGTTTTAGGCTTTGAGTTTTGTAATTTAGTGTATTTATTTATGATTTGGAGTTAAGGGTTTGTTAATTCATACAAGTAGTTATAGGTTTGCTTTAATTACAAGTAGTTGGTAGGCTTTTTTCTTTGAGATTTTTGAGGTGTTTTCTTGGGATTTGTGTTATTTGAAGTTTCTGCAACTGGGGTTCTCTTCTTCCTTTACTATAATAATAAGGGTGGTGGTGCCCTGCTGCTACCGTTAAGTAGTCTTGATTTTTTCAGTCTTGGGATCTGATTTGATTTTTGGTACTAGGGGATCTGAGGAAGGTTCTTTCTTGGTGCTAATTTGCATCAAAGGTGGAAACTTTTCTTCATTTGGAACTGTGAATTTGGAATACATGGTGGTGGAATCTTGATTTTGAGCTGTTTATATAAAGATTTGTGTGTTTTCATCAATGGTGATTTGGGAGTTTCTGCCAATGAGTTTTCTGTGTTGGGTGTTTTTGGCTTATCTTTTTTGTACATCTTTGAGTCTTGAAACCCCAGTTCAAACCTGTCATCCATATGATTTGTTGGCATTGAAAGAAATTGCTGGCAATCTAACAGATGGGGTTATTCTATCAGCTTGGTCTAATGAACCTAATTGCTGTAAATGGGATGGGGTTGTCTGTGGTAATGTTTCTGCTCAAAGTAGAGTGATCAGGCTAAATTTGTCAAGAAAAGGTTTGAGAGGTGTGGTTTCACAGTCCTTGGAGAAATTGGATCAGTTGAAATTGCTCGATCTTTCGCACAATCACTTGGAAGGTGGATTGCCTTTGGACTTGTCCAAATGGAAGCAGTTGGAAGTTCTTGATTTGAGTCATAATGTGTTGCTTGGACCAGTGTTGAGGGTGTTTGATGGATTGGAATCAATCCATTCTCTCAATATATCAAGCAATTTGTTCACTGGAAATTTCAGTGAGTTTGGTGAATTCCCTAACCTTGTTGCATTTAACACAAGCAACAATTCGTTTACTGGTAGTTTCAAGTTTGAAATTTGCAGTTTTTCCAAGAAGCTTAAGGTTCTGGATATATCACTTAATCATCTCACTGGTGATCTTGGAGGACTGGATAATTGCAGTTCATTGCTCCAGCAGCTACATGTGGATTCTAATGATCTTGGGGGTCACCTTCCGGACTCGTTGTATTCAATGACATCTTTGGAGCAACTTTCACTGTCTGCCAATAATTTCTCAGGCCAGCTAAGCCCACAGCTTAGTAAGCTTTCCAAACTGAAATCCTTAGTTTTATCAGGAAATCGCTTTCATGGTTTGCTTCCTAATGTGTTTGGTAATTTGACATTGTTAGAACAGCTAGCTGCACATTCTAATAGATTTTCAGGACCATTGCCCTCTACGATTTCGTATCTTTCTGTGCTTAGGGTGCTTGATCTTAGGAATAATTCTTTGTCTGGTCCTGTTGATCTTGATTTTACTAAATTGACAAGTTTGTGCACTCTTGACCTTGCAACTAACCATTTCAAAGGTAACCTTCCTGTATCGCTCTCTAGTCGGGAGTTGAAAATCATGAGTCTTGCCAAAAATGAATTCACCGGGCCAATTCCTGAGAACTATGCAAACCTTTCATCCCTTGTGTTCCTCTCATTGTCCAATAATTCCCTTTCGAATTTATCTGGAGCTTTATCTGTTCTGCAGCACTGCAGAAATCTTTCAACTCTTATTCTCACCAGGAACTTCCGTGGTGAAGAGATTCCGAAAAATGTGAGTGGATTTGAGAACCTGATGATATTTGCATTGGGGAACTGTGGTTTGGAGGGGCGAATTCCGATATGGTTATACAATTGCAGTAAATTGCAAGTGCTTGACCTGTCATGGAATCATTTGGATGGCGAAATTCCTCCTTGGATTGGTGAAATGGAAAAATTGTTCTACTTGGATTTCTCGAATAATTCACTCACAGGTGAAATCCCAAAAAATTTAACTGATCTAAAGAGCCTCATTTCCCCACACAACTACGCATCTAGTCTGAATTCTCCCACTGGTATACCGTTGTTTGTCAAGAGAAATCAGAGCGGTAGTGGTTTGCAGTACAATCAGGCTTCAAGCTTCCCTCCGTCTATCTTATTGAGTAATAACAGACTAAACGGGACAATCTGGCCCGAAATTGGTCGTCTGAAACAACTTCATGTCTTGGATCTCAGTAAGAACAACATTACGGGGACTATTCCTAGCTCGATTTCAAATATGGGGAACCTGGAAGTTTTGGATCTTTCATGTAATGATCTCAATGGATCAATTCCTGCTTCATTCAATAAGCTCACATTCCTTTCAAAGTTCAATGTAGCTAATAATCACTTGCAGGGAGCGATTCCAACAGGGGGGCAGTTCTTGAGCTTTCCCAACTCGAGCTTTGAGGGTAATCCTGGACTTTGTGGAAAAATCATTTCTCCTTGTGCTGCCAGCAATTTGGACCTCCGACCGGCTAGTCCTCTTGCTTCTAGTAGTAGTAGGCTTGGCCGAGGCGGAATTATTGGAATTACAATCAGCATAGGGGTAGGAATTGCACTTCTTCTTGCAATAGTGCTGCTTAGAGTGTCTAGAAGAGATGCTGGTCACCAGATTGGGGACTTCGAGGAAGATTTCAGCAGACCACCTCGATCGTCTGATACTTTTGTTCCTTCTAAGTTGGTACTTTTTCAGAATTCTGATTGCAAGGAACTGACTGTTGCAGACTTGCTTAAATCTACAAACAACTTCAACCAGTCGAACATTGTTGGATGTGGAGGATTTGGTCTTGTTTACAAGGCGGAACTTCCTAATGGCATAAAGACTGCGATCAAGAGGCTTTCGGGGGATTGTGGTCAGATGGAGCGTGAATTCCAAGCTGAAGTGGAAGCCCTCTCGAGAGCTCAGCACAAAAACCTTGTATCCCTTCAAGGTTACTGTCAGCATGGGAGTGATAGATTGCTGATATATTCTTACATGGAAAATGGAAGCTTGGACTATTGGCTACATGAAAGAGTCGACGGGAGCTCATTAACATGGGACATTAGGTTAAAGATTGCACAAGGAGCTGCTCGCGGATTAGCCTATTTGCATAAGGAACCAAATATAGTTCATCGCGACATTAAAACCAGCAACATTCTTTTGAATGAGAGATTTGAAGCTCATCTAGCTGATTTTGGACTATCAAGGCTGTTGCATCCCTATGATACTCATGTCACGACAGATCTCGTTGGAACCTTAGGATACATTCCTCCTGAATATAGTCAAACATTAACTGCTACTTTTCGGGGGGATGTTTACAGCTTTGGTGTTGTTCTACTTGAGCTATTGACAGGCAAGCGTCCCGTGGAAGTATGCAGGGGGAAAAACTGCAGGGACTTGGTATCATGGGTTTTTCAACTGAAATCTGAGAACAGAGTGGAGGAAATATTCGATACATCGATATGGGATACAAGTTACGAGAGGCAGCTTTTGGAGGTGTTAACTATAGCTTGTCAATGCATAGTGCAAGATCCACGCCAGAGGCCCTCGATCGATCAAGTTGTCTTATGGCTCGAGGCAATCGGAAGTGTAAAAGAAAGATGAGATTCAACTAGTTTTTCCTTGTAGACTTGGCTTCTTTTGCTTACAAAGTGAAAAATATCAGTGTAAAGCAACAGAAAATTTCAATTTTATAGAGTTAGTTATAAAACTTATGAACAGCTTTGTTTGGTTCACTTTGTTTCATACACAAAAAATAGTGATAGTTGTGTTTGTATACATATGGTTAACTTGGGGACAAGTCCTTTACCAATATACAACTAATTACTCCC

>M6-PGSC0003DMG400012514 M6 LRR-RLK

GAAATCGGGATTCTTAATAATTTGGTGATTTTAGCACTGGATACTAACCATATTTCTGGTAATATCCCAATCTCCACATTCAATATCTCCTCATTACAGGTTTTGGTACTGAATCACAACAATCTAATTGGATCCTTTCCAAGGGAGGTTGGCAATTTGACAGAATTGCAAATTCTGGATCTTGCCGATAATAATCTTACAGGTATTCAAAAGGGGTATGCAAATCATTACTAGCAAGATTTATAATTTCTTAAATCTTAGTAATTACTAACTATACATCTACATGCATACATTGACAGGTGAAATACCGAAAAAATTTAGCAATCAGATTGAGTTGTTTGATCTTAGTTTCAACAGGTTTACTGGTTCACTACCAATGGAGATTTTCAACATATCGGGTATGAGAGAGATTCAAGTTATTTTCAATAATGTGACTGGAACTCTCCCATTAAACATAGGTTCTACGTTACCCAACATTGAAGTTATTCATCTGAGAAACTTAAATCTTTATGGGACTATACCTCATTCTCTCTCCAATTGTTCCAAACTCACTAGTCTAGACCTTTCTCTCAATAGACTAACAGGTTTGATTCCCAACTCTCTTGGAAGTTTGACTCATCTACAGTCCCTAAGCTTGGCAATTAACAATTTAATGGGCGATAGTAGGTTAAGCTTCTTGACTTCCTTGTCAAATTGTAGAGATTTAAAAATTATTTTCCTCTCTGCGAACCCTCTATATGGTGTGCTTCCAGAATCCATAGGGAACCTTTCCAGTACTTCTCTTCTAAGGTTTAGAGCTATTGATTGTAAAATTAAGGGGGAAATTCCAAAAGGATTTGGGAACTTAAGCAGCTTATTAGAACTCGATCTTTCTAGAAACGGTTTCACTGGACCAATTCCCACAACAATCAGCAACTTGAGATTCCTTCAGAGCTTGAAGTTGAGTGTCAACAAACTTTCTGGATATATTGGAGATGATCTATGCAAATTACAGAACTTGGGTTATTTAAACTTGACTTTGAATCAACTTTCAGGATCTCTTCCGAATTGTTTGGGAAACTTAACTTCCCTTAGGGAGATGATTCTAGGTTCCAACAAATTGCGTTCAATCATTCCAAAAAGCTTAGGAAACCTCATAAACCTCTTGAAGCTAGACTTATCCTCAAACAACTTGGGTGGCTCTATACCTCTGGAAATTGGAAATCTAAAGGCTGCGATATATATGGATCTATCAATAAATACACTCTCAAATGGAATTCCTAAAGAAATTGATAGCTTGCAAAATCTTATATACCTTTCTTTGAGAGACAACAAATTGCAAGGATCAATTCCTGGCTCAATGAGCAGCATTTCAGCTTTGGAATTTCTAGACCTTTCTCATAACAATGTTTCAGGATTGATTCCCAAGTCTTTGGAGAAGCTTCTATATCTCAAGTATTTCAACGTTTCATTCAACAAGTTGGTTGGTGAAATCCCCTCGAATGGTCCTTTCAAGAACCTCTCCGGTGAGTTTTTCATGTCTAATGAAGCATTGTGTGGATCTCCAAGATTTTGTGTTCCCCCATGCCACAGTAGCACATCAAAGCATAAATCCAAGAGGAAAAAAGTGATTGTTCTTGTTCTACCGGGTACACTTGTATTAGTCTCCATCGCCTTTGCCCTTTTATGGTTAAGGTATACAAGAGGTAAAAGAACTGATCCTCAGCAAGCAGTTGATTCATCCTTTGTCATCTCAACAAAAGGAAGAATTTCATACTATGAACTACTTCGAGCAACTGATTCACTTAGTGAGATCAATTTAATTGGTTCTGGGAGTTTCGGCTCTGTTTACAAAGGCATCCTCAGAGATGGGACTTTCATTGCAGCTAAAGTGTTCAACCCACAATTGCAAGTGGCATTTAGGAGTTTTGATACAGAATGTCAAGTTTTGCGCAACCTTCGTCATAGGAATCTCACCAAAGTCATCACCAGTTGTTCCAACCTTGAATTTAAGGCTTTAATACTCGAGTACATGCCCAATGGGAGCCTCAATAAGTGGTTGTATTCACACAACTACTTCTTAGACATCAAACATAGGCTAAGTATAATGATAGATGTAGCATGTGCATTGGAATATCTACACCATGGATGCTCATTACCCGTGATTCATTGTGATTTGAAGCCTAGTAACGTCTTGTTGGATGAGGAAATGGTTGCCCACCTAAGTGACTTTGGAATTTCAAAGTTGCTTTCTGAAGATGAGAGTTCTTTATACACTAAAACATTAGCAACGTTTGGTTATATTGCACCAGGTACGTGATCTCTTAATCAATTGCTTTTGATTATTTCTCTTTCCTTATTTTCCACTAGTAAATGTTGAATTTCAAAATTGATTGATTGCAGAGTATGGACTGGAGGGATTGGTGTCAATAAAATGTGATGTTTATAGTTAAGGGATCATGTTGATGGAAACATTTACTAGGAAAAATCCTAATGATGAAATGTTCGATGGAGATTTTAGTTTGAAGCAATGGGTGAGAAATTCACTCCCACAAGCAGCAATGGAAATTGTTGATGCCAACTTAATATCGCCACATGATAATCAATTAATGAAAAAGCTAGATTGTGTGGTATCCATCATGAAAGTGGCAATAGATTGTTGTGTTGAATCTCCAAAAGGAAGGATTGACATGAAAGATGTTGTGGGGAGGTTAAAGACGATCAAGATTCAACTTCTTGCATGTTGA

>M6-PGSC0003DMG400012588 M6 LRR-RLK

TGAGAGCTTCAGCGGAAGTGCTTGGAAAGGGAACATTTGGCACTACATATAAGGCAGCTTTGGAGGATTCTACCACAGTTGTGGTGAAGAGATTGAAGG

>M6-PGSC0003DMG400012594 M6 LRR-RLK

ATGCTGTAAGCAGTAACGAAATACTATTTCCATGGACAGAATCCCTCTGCCGCAGATGTTCCCATTGACAATTTAAACACATTTTGAACCTCCTTTTCTTTTCTCTCAAAAGGTGTAGCTTGTGGATGCACTTATTAATTTTTAATGTGCTACATAGCTATGTGATGACTAATGTATAGCCCCTTAATTTTTACAGATTGTGTGGTCCCTAAGTTTTAGAACTGATTTGCTTACGACCTCTTTTTCCTAATTGCAGGGAACTTTATAGTAATAATATAAGTGGAAGAATTCCTTATGAACTGGGGAATTTGACGAATTTGGTGAGCTTGGATCTTTACCTGAACAAGTTAGTTGGTCCAATCCCCGACACTTTGGGCAAGCTTCAGAAACTACGTTTCCTGTATGGGTTACACATCCTTTTTCCTCTTCTATATGTTTGTTTTTCTTTTTCTTTTGCTTGGCTGAATAGTTTCTTAAGCCATGAGCTTTTATTTTCTGAACATTTTGCTTAGTCAAGAAAGGCCACCTTCTATGTTAGTGGCTCTGTACTCTGTAGTACTATTTCTCTTGTTTAGTTATTTGGAAATAGGGATAGACTTTGCTCCTTATTGGCCAAGTGGCAGACATGAAGATAAAAACATTTCAATCACCTGGAGTATCATTACCAATGAGACAGCATCATAATTTTCTATGTGGCTTAAATTATTCATAAAGAGAGAGAGAGGAAGAGGAGGCGGAAGAAGAAAACCTGAAGCAGCTCATAGAAGTACCCAAAGTTGGTGTCTTAGGGTGTCCAAAGTGCTGGACATGCATGTCTGAAAATTCAAAAAATATACAGGAAGTTAGACACACACATTCTCAATGTGTGTCCAACACCAGTACTTGACAATGAAATTTGTCCATCATCCTTGGCAGTCATCGTTCCTATTTCCCTTATATAGGCTTGCAAAAGGCTAGGTGGATTTGAGTGGTTGATAAAATTGTGGTTGCTGCTCTCTATAAAGGTTCATTGGCGTGCTGCTGAACAGGATAAAAAATTTACATGCTATTCTTGTTATTTCCCGAAGCTCTGAAAACTAGATACTTCTGATGATCGTATATATGAGCATAAATCTCAACCTGCTAACAGTGTGAAGAAGCTTTCTTAGATATTGGGCATGAATTTTGTAGCATCTTCTAATACTACTATTCTTCTTCATTAGTCAAGTATTTACTACAATGTACACATGGACTGATTAATTAGCCGAGTTCTGTCTGTTTAAGATTGATCAAACTTTTTTTTTCTTATTGTTATTGTTTTCTTTTTGCTAAATTGAAGGAGGCTTAACAATAACAGTTTGACTGGGCAAATTCCGATTCTTTTAACCACAGTCACTTCACTTCAAGTGCTGTAAGTATAAAGCTGGCACTTTTTTATCATGTGGTTTTTTTCGCCATCAGTTTTTTCTAATTCTTAGGTACTAATGTTCATTCCTCGCAATGCAGTGATTTGTCAAACAATAATTTAACAGGACCTATTCCAGTCAATGGTTCCTTCTCGCTTTTTACTCCTATCAGGTCTGTGAAAAGCACTAGTTTATATTTTAAGTTCTGTCACTGGAATCATGATAATGAAATTCTTACATCTGAATGTCTATATTTCAGTTTTGCTAATAATCCATTGGATACTCCTCCACCTTCTCCACCTCCACCTATTTCTCCAACACCCACTTCTCCAGGTTTGTTAAAATAATTTATGTTTTATGGGTTAAGACACTGCCCCCCCCCCCCCCCCCCCCCCCCCAAAGGAAGAAGGCAAAAGGGCTGACTGTTTTAGGGGAAGAGATTTGCACATTATAATGAGTTTTAGTAGTGTATGTGAGATAAATTTTACAGGTGATTTTGGGGTTATCTACTGTTTGATAGGTTTGAATAAATAGATTAAACTTACAAGTGATATTGAAGAGTGGATAGAAGTGGCCTACATCAGGAAATGCAACTGTATGCATTGGATATTGTGACTATAAAGGACCCAAACTTTCAGAATTTATTGTCATACATTCTCTCAATTCTTTCCCCATGTGTCTTATTTCTCACATGGTATCGGAGCTTTGGTGAAAAATTAGTTTCAGTGCATTTACCAGCAACACGCGGGAGAAAAATCTCCTTGCTGTGCAATTCTCATTGACCATGAATGCTGTCAAAATGTATGTATCCTTGCCGTCTCTGACTACTGTACATAAATTAACACCACCAAAAGGCCGAATAACAAACAAGTACCTGTTGTTATGAATAAACTGAAGGATCTTTACCTATTTCTGCATCTTTTCTATTTTGTTAGCACTTTAGCTTTACAACATAGTCCTCGGTGCGAGTGTAAATGGTGAAACTACCATATCCCTTTTCTCAATTTTTTTTAACTACTTGTGTTAAGTTTTTATCATCCATCAAAAAAAGGGAATTAAAACTTAAATTTTGAAATTCCTAGGTATGGGAAACAGTGCAACCGGGGCGATTGCTGGCGGTGTTGCTGCAGGGGCTGCTCTTCTGTTTGCAGCTCCTGCGATTTTGCTTGCCTGGTGGCGCCGAAGGAAACCGGAAGACCACTTCTTTGATGTTCCAGGTGGGAAGCCTCAATCAAACACCTGAATTAGTTTTTTGTTTTCATTTAATTTTAATTTAATGAGTTCTTGCAATTTTACATTCTTTTGCAGCTGAGGAGGATCCAGAAGTTCATCTTGGACAACTCAAGAGGTTCTCCTTGCGCGAATTACAAGTTGCAACTGACAGTTTTAGCAACAAAAATATACTTGGAAGAGGAGGATTTGGCAAGGTTTACAAGGGCAGGTTGGCTGATGGCTCTTTAGTTGCTGTTAAAAGATTAAAAGAGGAGCGTACTCAAGGCGGGGAGTTACAGTTTCAGACAGAAGTTGAAATGATTAGTATGGCCGTGCATCGAAATCTACTTCGTTTATGGGGCTTTTGCATGACTGCAACTGAACGGTTGCTTGTTTATCCATACATGGCGAATGGAAGTGTTGCTTCACGATTAAGAGGTATCTCTTAGTCTTGAATTTTCCATCCTTCTTTTTCTTTGCTCCATTGAGTCGAAATACATCAGATCAAGCACTTGAAAAAGCTGAAAAATGATAACAACGTTTTTTTTGAACTTTTAAGTTATGGTCTGGTAATATAAGTTATCTGGAAAAGTTACTGCAGATGTGTACATCATTGTGTAAACTTGGCATTATTGGATATTGTCGTGGTTGTATTAGGAGTTAGTTTGTTATAGTTAGAATATTCAATCCCGTAGTTAGAAGCTGTTGTTCTTTGTGATTTTACCCGTGGTCTATCAAGCAATTTCAAGAGCGTTGCTATCCTTGATGTCCTACCAAATATTAAGATTTCCGTTCAAGTTTTATGTTCTGAATTCATAGTGTCAGATATGGAAAGTAATCTTCATTTATGCATGAACCGCATATATCAATTAATTAAATTGCGTTCAAGCTAAAATGGGAACTTGAACCAGTTTACTTGACATCTTGAACTTGTCCTTACAGTAATTTATTTTTCTTGTAGAGCGGCCAGAGTCAGATCCACCACTTGGCTGGCCAATAAGGAAGTGTATAGCACTTGGATCTGCAAGAGGCCTTGCTTATTTGCATGATCATTGCGACCCTAAGATCATCCACCGTGATGTCAAAGCTGCAAATATATTGTTGGATGAGGAGTATGAAGCAGTTGTTGGGGATTTTGGGTTAGCCAAACTCATGGACTACAAGGATACTCATGTTACTACTGCTGTACGTGGTACGATTGGGCATATCGCCCCGGAGTATTTATCCACGGGTAAATCTTCAGAGAAAACTGATGTGTTTGGCTACGGAGTTATGCTTCTAGAGCTCATAACTGGGCAAAGGGCATTTGATCTTGCTCGACTTGCAAATGATGATGATGTCATGTTGCTAGATTGGGTAAGTATTTACTAGCCTACATGCTCTAGTTATTCTACTAGCTTATGTCTTTTTCTCTTTTCTTTATTCGTTGAAACTTTATACTACTTACAATTGCTAGTTTTTGTCGTGCAACTGACTTCAAACATGGAAGTGAGCACGGGTGAGAAGATCCCACTTTCCTGCATGCATCTCCCTATAAGCAGAACTCTATCATTGCTGTACCCAGCATGATGAAAAGCTGATTTGTAGACCCACTACAAATTAGGACTGGCGAATAAGACACTGTCATGTCCCTTCTCTTCCCCTGCCTTTTCCTTTAATATCCCCAATCCCGTTCCTGTTCTGTTATCGCTTCCCTACCCCCGTATGTAACCCACATCCCATATTCTCGTTTTATATCCTCATCCATCAATGAGAAATCTCAGCTTCATCTCATCTCCCTCACTCTTTTGCACAGATTACACACACTCATAGCTCTCTTTGTTCTTTACTCAAGTCTCAGGGCTGCAGTTCTGTTTAAAGTAAAAAGTTTTATATTGACCCCCCCCAAAAAAAAAATAATAATAATAAAAAAATTGTTAGCTCTAGTAGGTTTTGCATTGCAGTTTGGCATGTATGATTTCTGTAATCTTTGATACTACATCATACGAAACATCCAAATTTCCAAAATAATTTGATTCCTGGGACTTGTATTTGTGCACATTCTTCATGTAATCTTTTTGTTTCTCTTTCTAGAACTTAACCATCTTGCGAAAACATTTGTTGCATTAGCTGTATGGTCAGGGTGGATCACTTTCAATATCGTTTAGTAGCAGATTGTTCTGGAATTGCTTTTTAGTGTTAAAATCTTTTTCCCTTTGGTTAGCAAAAGCACTCATATAGCATATTGCTATATCAGAAGATTTAGATACTTTTTCCCTGCCATATTGACTGTCGCCTTCCGTGCAGATAAAACTACATCTCTTTTTTTTTCCTTTGTATGGAATTGCTAGGCTATTTCTCTGTATTTGCATCACAGAGTAGGTGAGATAGAGAATACACTTTACGAGAGGGGACTTCTTTGGGAGGATCCTCTAGAAAGGGAACCCACCCTTTAGGACTTGGGAGTGCAGAAGATGGTCAATTCTTTCACTATGGTTTAAGTATAGAAAATTGAAAGACAAGTGGCTATCCGAACTTGGAAAGTAAATAAAGTGGCTATCTGAACTTGGAAATTCAGTATCTCCCTATAGGGCTGCTTATCAAAAATCTCAACAACTTCTATGGTGAATGCTTTGTTATTTTAGTGATGAAATTAATTATTTGCTGACATGATGAGGAAAGTCCGCATGCACTGTGATTTCTTTTTAAAAAATGGCACTCAATTGAAAAAAATGGTTATGTACCTTATTTGACCTCAAAAATTGTGTTAAAATTTCAGGTAAAGGGACTTCTTAAGGACGAGAAGTATGAAACATTAGTAGATGCAGATCTTCAGGGTAATTACAATGAAGAAGAGGTGAAACAGCTTATTCAGGTAGCTCTACTCTGCACACAGAGCTCTCCTATGGAACGTCCAAAGATGTCGGAAGTGGTCAGAATGCTTGAAGGTGATGGCCTTGCTGAGAGGTGGGAAGAATGGCAGAAGGAAGAGATGTTCCGTCAAGACTTCAACCATGCTCACCATCCTCACACTGATTGGATTATTGCAGACTCCACTTATAATCTACGACCCGATGAGTTATCAGGGCCAAGATGA

>M6-PGSC0003DMG400012817 M6 LRR-RLK

GGCTTAATCACTTTTCCAGAAAAAAATCCACTAATCCATTCATCTCCTTCAACAATGCATCTTCAAATCTTGCTTATACTTCTATTACCTACATTGATTCTCTCAATAAACCAAGAATCCCTTTATTTACACACCATAAAGCTTGGATTTGATGACCCAAATGGTGTTTTTTCAAACTGGAATCTCCATGATAACTCCTCACCCTGTAACTGGTTTGGAGTAAAATGCGACTCTTTAACTCGTTCTGTTACATCTATTGACCTCTCCAACACCAATATCGCCGGCCCATTTCCGGCTTCTCTTCTTTGCCGGCTCAAGTATATCAAGTACATTTCATTCTACAATAACTCCATTAACTCGACACTTCCGGTGGAGGAGTTATCCGCTTGTAAATCTCTTGTCCATCTAGATTTAGCTCAAAATTTGTTAGTGGGTAGTCTTCCATCGAGTTTGGCGGAGCTTCCTGAGCTGAAATACCTTGATTTGACCGGAAATAACTTTACCGGCGAAATTCCGGCGAGTTTTGGGGCATTCCGGCGACTTGAAGTTCTGGGTCTGGTTGAAAATTTGTTAACTGGGACTATCCCGCTGGAGATTGGAAACATTTCGAGTTTGAAGCAGCTGAATTTGTCGTACAACCCGTTTTCTCCGGGTCGGATCCCGCCGGAGATTGGGAACCTCACGAATCTAGAGGTGCTATGGTTAACTGACTGTGGGTTAATCGGTGAGGTTCCGGGTACGTTAAGGGGATTAAATAAGCTTGTTAACTTGGACCTTGCGTTAAACAACTTGTACGGTCCTATTCCGAGCTGGCTCACTGAGTTAACTAGTGTTGAGCAAATTGAGCTGTACAATAACTCGTTCTCTGGCGAGTTTCCGGTGAATGGGTGGTCGAATATGACCTCGTTGAGGCGGGTCGACCTGTCGATGAATCGGGTTACCGGGTCGATCCCGAGCGGGTTGTGTGAGTTGCCACTTGAGTCGCTCAATCTTTATGAGAATCAATTGTATGGTGAGTTACCTATAACCATTGCAAATTCACGCAATTTATACGAATTAAAGCTCTTTGGTAATAGTTTAAATGGAACTTTACCTAAAGATCTTGGTAAATTTTCTCCATTGGTATGGATTGATGTTTCAAACAATGAGTTTTCGGGTGAAATTCCGGTGAATTTGTGCGGAAATGGGGTTTTAGAGGAGGTTTTGATGATAGATAACTCGTTTTCCGGTGGAATTCCGGTGAGTTTAAGCCAATGCCGGAGCTTATTACGTGTGAGATTAGCTCATAATAAGTTCTCAGGTGATGTCCCTGTGGAATTTTGGGGGCTGCCACGCCTTTCGCTGCTTGAGTTAACGGACAATTCATTTTCTGGTGGAATCGTGAAAACTATAGCTGGTGCATCGAATTTATCCGCTTTGATTTTGTCAAAGAACGAATTTTCGGGTACTATTCCTGAAGAGATTGGCTTTTTGGAAAGTTTGGTTGATTTTGTGGGAAATGATAACAAGTTTTCAGGGTCATTACCGGTTAGTATAGTGAATCTTGAGCAATTGGGAAGAATGGATTTTCACAACAATGAATTAAGTGGTAAGTTTCCTAGTGGGGTTCATTCTTTGAAGAAATTGAATGAATTGAACTTGGCAAACAATGATCTTTCTGGAGAAATTCCCCGGGAAATCGGGAGCTTGTCTGTTTTGAACTATCTTGACCTATCAGGAAACAAGTTTTCGGGGGAAATTCCAGTTGCGTTGCAGAATTTGAAGCTCAATCAGCTGAATTTATCGAATAATGGCCTTTCGGGTGGTATTCCTCCTTCATATGCAAAGGGAATGTACAAGAATAGCTTTCTGGGGAATCCCGGTTTATGTGGAGATATTGGAGGCTTATGTGATGGAAAAGATGAAGGTAAAACTGCTGGTTATGTATGGTTACTGAGATTGCTTTTCGTACTTGCTGTTTTGGTGTTTGTAGTTGGGGTAGTTTCGTTCTATTGGAAGTATAGGAATTACAAGAAAGCAAAAAGGTTGGATAGATCGAAATGGACCTTGACGTCGTTTCATAAGTTAGGTTTCGATGAGTATGAAGTACTGGAAGCTCTAGATGAAGACAACTTGATTGGTAGTGGTTCTTCCGGGAAGGTTTACAAGGTCGTTTTGAGTAATGGTGAGGCTGCTGCTGTGAAAAAACTTTCAAGAAGTTTGAAAAAAACAGATGAGAGTTGTGACATTGAGAAAGGTAACTATCAGGATGATATATTTGAAGCGGAGGTTGAGACATTGGGCAAGATTCGACACAAGAACATCGTTAGGCTATGGTGTTGTTGTACAACAAGGGGTTGCAAACTTTTGGTTTACGAGTACATGCCTAATGGAAGCTTGGGCGATTTGCTACACAGCAGCAAAAGTGGGTTGTTGGATTGGCCTAAGAGATTTAAGATAGCTATGGATGCTGCAGAGGGACTCTCATATTTGCATCATGATTGTGCTCCTCCGATTGTTCACAGAGACTTGAAGTCGAACAATATCTTGTTGGACGGGGAGTTTGGAGCTCGAGTTGCTGATTTTGGTGTGGCAAAGGCGATGGATGTTGATGACAAGGGAACCACGTCTATGTCAGTCATTGCAGGGTCTTGCGGTTATATTGCTCCAGGTTAGAGGATTAATCCTTCACTTTGGGAGTTCTACTAACATGATTTTGTTGAATTTGGTTTATGCCTTCCATTGCAAATTGTAGTTAAGATGTATGCCTAAATATAACATATTTCATATATTGTTATATTATAATTAAGCGTCGAATAAGGTCTGCTGTACACTCTACCCTCCTCAGGCCCCACTTGTGGGATTACATGGGGTATATTGTCATTGATGATTGTTAAACGTCCAATATCTATAATCAGGCAGACAGCTATATATGAGTAATTTTGGTCCATTCTTGAGCTATGTTGCTCAAATCCTTCAAAAAATGTGGCATGTCTTTGTCGGATTCTCCCAAAGTTATTCAATTTTGGAGGATCCAATACAGGCGCAACAACATTTTTGGAGGATCCGAGCAACATAGTACTTGAGTGCCCTTGTACATTAACACTAAGGTTTGCTCCAAGAACAGGGCATTCTTTGTTTACATTTGCTTTCTGGTTCTGGTAGTGCTCCTCCGGTTGAACTAAATTCTGTTCGTCTTGCAGAATATGCTTACACACTTCAGGTGAACGAGAAGAGTGATATATACAGTTTTGGCGTGGTAATCCTCGAGCTAGTGACAGGGAAACTCCCTGTAGGTCCTGAATACGGGGAAAAGGATTTGGTGAAGTGGGTTTGTGCTACTCTAGACCAGAAGGGTAGAGATCATGTTATTGACCCGAAACTCGACTCTTGTTTCAAGGAGGACATAAGCAAAGTCCTAAAAATTGGCCTCCTCTGCACTAGCCCCCTCCCAATTAACCGACCCTCGATGAGAAAAGTCGTAAAAATGCTGCAGGAAGTTGGTGGCGGAGACCAGCTCAAGACAGCATTAACAGATGGCAAGTTGACCCCTTATTACCACGAAGACGCATCAGATCAAGGAAATGTAGCTTAAAGATAGTCAACATATCTTTGCTTAGATGTAGCGACATGGACAGTGAAGATTCATAGGCATAGTCGTTGTATCGTGAAAGTGAAGATGTGTGAATTTTTTTATTTGCTAATTGCTTAATACTTTGTCTATCAATACGTGATTTTTGGTTTGATTTTGGAGAGAAGGGGAAAACAGAATTGAGGTGAAATGTAAAAGGA

>M6-PGSC0003DMG400012929 M6 LRR-RLK

GCTGCCTGCAGTAACACCTTGAAAACCATATCTCATAGAAGCAAAAAACAAGAGTCTGGTATTGTTGCTGGTTAAATGATGAATGTTGTTGTAGTTGTGATCCTTAGTTTTTTCACATTTTCAATCTCTGTTATATTTTCATTGAATTCAGATGGTGTTTCTCTGTTAGCTTTGAAATCTGCAATTTCTAATGACCCATCTGGGTTTCTTTCTTCTTGGTCGGAGTTAGACTCATCAGCATGCCATTGGAATGGAATCACATGCGATGAAAATCAGAAGGTGACTTCTATTTCTCTGTCGTCGAAGAATTTGAGTGGTTACATTCCTTCTGAAATCGGAGCACTTTCTTCTCTATCAATTCTTTCTCTTTCCTACAACAACTTTTCAAAACCAATCCCTATTCGTCTCTTCAATGCTACTTCCCTTCACTCCCTTGACCTCTCTAACAATGCTTTCACAGGATTCTTACCTCAACAAATCACATCCCTTATAAACCTTAAACACCTTGATCTTTCTTCTAATTTTCTCAATGGCTCACTTCCCCAAGAGCTAACTCATCTTACACTTCTCACTGGAACTTTAAATCTTTCATACAACAGATTTACCGGCCAACTTCCGGCGAGTTATGGTAAGTTTCCTGTAACATTGAGTTTAGACCTTCGGCATAACAATCTGACCGGAAAGATACCTTCAATCGGGTCGTTGTTGAATCAAGGGCCTACTGCATTTTCCGGTAACCCTTTTCTTTGTGGGTTTCCATTGGAGACTCCGTGTGCAGAACCAGAAGCTCAAAAACCACCACCGAACCCGAACCCTAATGAGGAGGAGGTGGGGTTTGTAGGAAAAGGGAAATCGGGAAATGGGTCAGTGGTGGTTTCTTTGATCTCCGGCGTATCGGTTGTGATTGGGGTGATGTTTGTTTCAGTGTTGGTGTTGAGGAGAAAATGGAAGTTGGATGAGGGAAAAATGGGAAAAGAAAAGTTGGAGAAGCCAGAGGAGGGGCAAAAGGGTAAGTATGTAGTGTTAGACGAGGGGTTTGGGATGGAACTAGAGGATTTGTTAAGGGCTTCAGCTTATGTAGTAGGTAAGAGTAGGAGTGGGATAGTCTACAAGGTGGTGGCTGGCGCTACGGCTGTTGCTGTTCGACGATTAAGCGAGGGGGATGTCACGTGGAAGTTCAAGGAGTTTGAGACGGAGGTGGAAGCCATTGGGAGAGTACAACATCCAAATGTGGTGAGGCTTAGGGCTTATTACTATGCAAGTGATGAGAAATTGCTGGTAACTGATTTCATCCGCAATGGCACCTTGCACAATGCTTTACATGGTAATCTCACTTCATTCTTACTTGGAATTTTCTTTACTGTTAAAAGAAAATTCACTCTAAACATACTACTTACGGAGTATTATTCTAAATCACTATAAGGATAAACTTTTAGTAGTGCTGGAGTAAGTTATCCTGCCTATGCGAGTGAGAAAGGGTGTACTTTTTTGGAGAAAATTAGATCAAATCCTCATAAGCAGTATTATCTCTGTGTCCATGGTACTTCAGGGGTTTTATTAATATATCAGACAACCTTGAAGGTGGCATTGTATTTGATATTTATGCAAAAAAAGAAGTTTAATGTCTACTCCATTGAAGTAAGTGTTTACAACCTAAGATTCCTTAAAGCTGCTCATGATTATTCGTTGAAGTATTCTCTAGGCACATTTAAGAATTTCTGTCATGGTTTCAAGCATTCTCTGGTAAATATTAATGATCCCAAATTATATGCTTTTAAGTCGTTAGCTGCCTACTGTCCTTATCTTCTTTTGATCAAGAATAGGCAACCAAATGTAATTATGATGAACAAGAACTAGTTTGGGGTAGTGGGGTGGAGAGGGGAATAATACGTGCATGGTTCATTTGTTTTTATCTGAGCAATTGGCATTTGTGTTTTCTGCTTCCTGGAAAACATCTATGCAAGTGTTAAGCTATTGAGACCTTTTGGGCTCCACAATTCAACATCTGTTGGTTTATTGTAATTCACTAGTTTCTGGTCCCCATTATGGCATTTTCCCCTTGTTTTCTGCTTCCTAGGGAGTACATTTTATGTGGCCTAAGATACATTGAGAGAGGCTTTAAAGTTGAAGTAAACATTCGGAAGTTACACTAAGGAAAAAAAACATTAGATCTATCAATTGGATGTTAAGCTATTCGTTCGATTCACGTTCATTAGGAATTGAACTAAGCATGGTTACACATAAAATGTGAATCCAAAGTGTGAAATTTTTGCAGAACAAAATATCGACTAATCATAAAATGTTGTAGTTCCTCCTTGTAACATCTATCATTGTCAAGTTCACTTTAGGATGTTGGACTTTATGTAAAAGAGGCATTCAGCTGACACCGAGAAGAAAACTATTACACCACAGGTTATTTTTGTCTTCAAATTCTTTTTTAACTAAGTGGAGAAGTCCACGACGTTAAGCTTTCTGATAGGTAAATATAAAACAAAAAAGATAATAAGCAAGACACGAAAACGAAATTTCATCTGATTCCAAGACTTGGTAAGTCATTCCCTGTTTCCTTTAGTCTGTTTGTCTTGTTATTCAGAGGTGTCGTTTAATGTACCTATAATTTGTATTGTGGTTATAGAATCCCTGAATAGGCAGTTACTGGACTTGTGTTAGCCTTGTATTTTGCTCTCTTTATTTTCAGAACCTTGCACTTCACAAAAACATTAATGTTCTTTTGAACTCCAGTTTGTTTTTCTTAACTTCAATCTTTGTGTACAAGTCTTCTCCACATAATCCTTCCCTTTTGCTCGTACTTTGGTATTGTCAATTGTGAACCAATCCTATATAAACACAAGTGGAAATGAAGATTGCACCTACTTTGGGGTGGCTTCTTGAGAATGTGGTTCCTCTAGTTCATATTGGTAATTTTTTGTCTCGACAAAATACCAACTGACCTATTTCTTGTTTTCCTTGACTCAGACTTCCATTGTTTCAAGCACATATTTTTGTACCAATTTCAGGATTTGAAGTGAAAAATGTTCTATTTGTTATTCTGAAAAGCTGTCATCGATTTGGCATGAATCACTTTTTAATTTATCTTCCACAAGTCCAAAAATATTTTTTGGATTTCAAAACTTGCATTTTCCGGCAAAAAATTTCTCATGGAAAAAAACACATACACAACTAATTTCTCTAGAGAACTGGGTGATTTTCTTTTTATGTTGGAGCATTATCCTTGTTAATCAATCTTTTTATGTTTATTTTTTACAATTGCAGGAGGACTTGGTAATAACTTGCCACCTCTGTCATGGGCAGCTAGATTGAAGATTGCTCAAGGCACAGCTCGGGGTCTAATGCACATTCATGAGTGTAATCCCAGGAAGTATGTTCATGGGAACATAAATTCAACAAAAATCCTTCTTGATGATGACTTGCAAGCCTACATATCTGGTTTTGGATTGACTCGCCTTGTTTCAGGTTGCTCCAAGCCCATAAACAGTAGTACCAAAAAGCTAAGTACAAGTCAAATCATAATAAGTCCTCAAAACTCAACTCCGTCTTGTACTATGTACATGGCACCTGAGGGTCGAGTGCCTGGCTCCAAGTTCACCCAGAAAAGCGACATCTACTCATTTGGTATGGTACTTTTGGAGATTCTGACTGGTCAGTTGACAGATGGAGGATCGGAGGTGGAGGATGATGGGAAGGGGCTTGAGAGTTTTGTTAGAAAGGTTTTTAGACAAGAACGACCCTTGTCTGAGATCATAGACCCTGCTCTCTTGAATGAGGTTCATGCTAAAAAGCAAGTTGTTGCAGCATTTCATATCGCTCTCAACTGCACAGAACTGGATCCTGAGCTTCGGCCAAGGATGAGAACAGTTTCTGATAACCTTGATCGCATCAAACTGCAGTAAAAGAAACTTGATCAATGTAAAGGCTTTCAATGTTTTGCTGACCGCCTATTGTAGAGCTGTTTAGTAATTGTGAATCAGAATTTTCAGTTTGTGTTGTGCCAAATCCTAGATTTGGCGTTTCTATTGTAACTCTACTAATTCAATGTTCTATTTGTTGAATCATCAATTGAAAATTGCTTGTGCTAGCGAATCTTACAAGGCTATTTGTGTTTGATGTACGGACATGATCTGAATTCCAGATAACATGCAAAGGAGCATGGTCAGACCCTGCCTATTCAAATGATGTAGTTCAATAGTTGGAAAGTTCTGTATGAATTCTTGATTGCCCATTACCCTATTCAATCTTTTGAAACTACACTCCACCTCTATTCTCTCATTCTACCAAGTGAAGTTGCTAC

>M6-PGSC0003DMG400012955 M6 LRR-RLK

CACCCCATTCCAACAGAGAAATACAGAGAGAAAGAGCAAAATACAAAAGAGACACCAACTCATTTGCCACTCTACCCCAAATATAAAAACCCTAATTTCCCTCTTTTTTTCTCCTTTCCCAACTCTTTCCCCACACCCCAACCCCTCTCTTACCACCCCACCTACCCTAATCTCTCTCTCTGTCTCTGAACTCTTTCTTGGATTTCCATGGTGCTCTCTCCATAGTCATCCTTAAGCTTTTCTTCCTTTTACCCTTTTTCAGACTTTCCTTCTTTTTCTCAATGCCTCATTTTTAGCACTTAACTCACATTCTCAACCCAATGACTCTCTCTCAATCTCTCTCTTTACTCCTCCTTGTACTTGTGTTTCGGCTCAATGGGTCTTCTGCTGCAGAAGATGAAGTTCGATCACTTTTGGAGTTCAAAAAGGGTATTAAAAACGACCCATTAGGTAAAATCTTCAGTTCATGGAGTCAAACGGGTCTTTCAAATCCATCAGCATGTCCAAAGTCGTTTCATGGTGTCGTTTGTGATGAAAACTCTGATTCTGTTTTCTCCATTTCTCTTGACGGGCTTGGGCTTGTAGGTGACTTGAAGTTCTCAACTCTTAGTGGGTTGAAGCAGCTCAAAATTCTTAGTCTTTCTGGTAACTCTTTTACGGGTAGGGTTGTTCCTGCTTTGGGTTCAATGTTAACTCTTCAGCATTTGGATCTCTCTGGAAATCAGTTTTATGGAACAATTCCAGCACGAATTAATGAGCTTTGGGGTTTGAATTACCTCAATTTGTCCAATAATAATTTTACATTTGGTTACCCAATTGGAATTAGTAATCTTCAGCAGTTGAGAGTACTGGATTTGCATAACAATGGGATTTGGGGTGATATCGGGGAGTTGTTTTTGGAATTGAAGCGTATTGAGCATCTTGATTTGAGTAACAACTCGTTTTTCGGGTCTCTTCCTACGAGCCCGGAAAATGTATCACTCTCTTCGACAATTCAGGTCATGAATTTGAGTCACAACAACTTGGGTGGTGGTTTTTTTCCTGGAAAATTGTTGGAGGCATTTGAGAATTTGATGGTTTTGGATTTGAGGAATAATGCTATAATGGGGCAGCTTCCTTCTAACGGGTTTATGCATAATTTGAGGGTTTTAAGGCTTGGGAATAATAAGCTTTATGGATTGATTCCCGATGAGCTCTTGCAGGGAACAGGTCCATTGGAAGAACTGGATCTTAGTGGCAATGGATTTTCAGGTTGGGAAATGGATATCAGTTTTTTGGAATATTTACATTTCTTAGATCAGTATATATTGTTGTTGTTGAATGATCATAATCAAATACACCTTTTATTTGTATAGTGTTTTTGTGGATGCCTTTAACTTCAAGCCTCTTAATTTTTGGAGGTGGACTAAAATTGGAAACTTTCTTCTGTTTTAGGCATTGAAACTTTTACAGTTATGCAAAGTTGCGCGGACTCTTCACTTTCGATGCCGCACAGTGTCGGATTTTCCAAATATACACTACTTTGTAGAATCCGACACGCACCCGTTGACATTTTTGAAGAGTCCGAGCAACATAACAGTTGTGAATGTAAAATCTCGTTCTTTTTGTTTTCTTGTCAGGTTATAATTAGTTTTCTTGTGTGAGTAGTGGTTGACAATCTATATAACAGAGAATAGAAATCTTCAAGGTCTTTCAGCTTATGCCAAATACAGAAAAGTTTGCTTTTTTGAGTAAGTAGGTTTTTTAAAAAATGTTTTAACAAAACCAGATTTTGAACAAACTTTTACCGCAAGCTTTTTTTGTTTGCCTCAACAGAACGTGAGCAGCGTGGACTGTATAGACTAGTTTGAATTGCAAAGCATGTTCAATAGAACTTGCCCATAATAAATAGTCAACAAAGTGTTTCCTAATTGTAGTAAATTTAATAAAAGCCCCTAATGGAATAGTTGCATGAATGATTGCATCATTAGTAGCCATAAATAATCCTTTTTGCAATGGTATAACTGTATGCTCAAAGTTCATGGCAAGCTAGCTGGAGCAATATTTAGCCCAGTGTCATCAAAGGCGCTTAAGCCCTGAAGCAAGGCTCAAATCGTGTTGAGCGCTTCGCTTCGCTTAATGTGCGCTTTAGTGTCGTCATCAAGCTTCTAAGGCATACTTTTCCTTAAGTTTATATATTTGTAATTCATGCTTAGGACTAAAAATAAATATATTTATTTTTTCTCCCTTTTCGCCTTTTTCCATTAAAGCCCATACTTGTGCTTAAAGCCCAATCAAACCTTAGAGCTTTTTTACTCTTTTTGCTTTCGATAAGACTGATTTAGCCTTTATCTTGAAAAATTGAAAATGTTGAATTTTTTCTTCTGCTTCCAATAGCCAATGTTATAGCACAATATTTGTACTTACAGTAAAATGAGTTTTATTTCATTAATGAGCTATTTGCTCTTAATATGCTCATCTTTCTATTTTGTTATCATTGCATTACACCGTCCAGGGCTTCCCCCTTCCCTCTAGTATAGGCCGCTCATGATGACGTTGGGCCTTGGAATTGGAAAATAATTGATGACAACATTTCTTTTTTGATAAGGAATTGATGACAACATTCCTGTCTTCATTATTATCAAATCAGTACTCAGATTAGAGAAAAGAATGGAGTCGGTGAGACTTGATTATTGAAGAAAAGCAGAGCAAATTTACAAGGGTGATAATTTGGGTCTCTGAGCTTAAAACTCCTCGGTGACCCTCTGATAGGGAAGTTTGTTCTGTTTCATGAGCTACAGAGAGGGTTCAAAATTTCATATTTCTTTAGTTACACTTGGATATCTTTCATCATAGGACCAGAGCGTGGTTCCATTTAGAGCCTAAATGTGGAGATCCTCTGTTCAGAAGTTGAATGGTGCCGTTGACTGATGACTGATATTTCATTCTTCTCATCTTGCATTTATTCTTTGAGGAAATGACTCTCAGATCAGCTAGATAGTTTAGTATACTTGTCTCACTAATATACTTTTTTGCCTTCTAATTTTTTTAGGCTATATAGTATTTCTACCATATTCTTTAGCTAGAAATAGTTGCACTTGGACCTGAAAGTTAAATTGGATGTTTAATAACAATGCACATTTTTTTTTAATAATATGAGCTCTTTTTCTGTTGGACTTGTAAACTTATTCTAAAGAGATATTTAAAGTTTAATATTTAGTTTCTACGCATGCAGGTTCAATTCCGATTGTGAATTCAACAAAGTTGAGAGTTCTAAATATTTCATCGAACCACCTATTAGGTTCACTGCCATCCTCCATAGAAAATTGTGCAGTTGTGGATTTGAGCAGAAATATGCTTGTGGATGGTATATCAGTTATTGAAAGCTGGGAAGTCAATTTGGAAATTATTGATTTGAGTTCAAATAGGTTAACTGGAAATATTCCCAATATAACCTCTCAGTTTCAACTATTGACTTCACTTAATTTTGGAAACAATTCTCTTGAGGGAACCTTACCTTCAGCACTGGGCACCTTGCCAAGACTGGTTAAACTTGATCTCAGCACCAATAAACTTGGTGGACCGATTCCATCTACATTTTTTACTTCGACGACATTGATGAACCTAAATATATCTGGAAATCAATTGTCAGGATCAATTCCTCTTGAAGGCTCTCATGCTAGCGAATTAATTGTTCAATCACCATACCCTGCACTGGAGTCTCTTGATCTTTCAGAAAACAATTTCACAGGTAATTTGTCTTCTGCCATTGGTAATCTGCGGAGGCTTCAAGTGCTAAATCTTGCTAAGAATCAGTTATTAGGCATGCTGCCTACTGAATTGGGTGACCTTAGAAGTTTGGAATTCCTTGATATATCTAACAACAATTTTAGTGGTATGATCCCCGAAAATCTTTCATCAAATTTGAGGGTGTTTAATGTGTCTAACAATGAATTAAGTGGTGCTATCCCTGATAACTTGAGAAACTTTAATGAAAGTTCATTTCGTCCTGGAAACTCAAATTTGGCAATCCCAAGAAACTGGTTACATGATAATCATGGTGATCCTGATCAAAATAGTCAGCACCATCATAACTCGAAATCCAGTATCAGGGTGGCTATTATTCTTGCCTCAGTTGGAGCTGCTTTAATGATTGGTGTTGTTCTCCTGGCTTACCATCGACAACAGTTCCAAGATTTCCACTTACCAAGTGGATTTAACAGCCAATCTGCTGGAAGGGATGTTAAACTCGGGAGGTTCAGCCGGCCTGGAATTTTCAAGTTCCATGGCTCCTCAGAACCACCACCAATTTCCTTGAGTTTCTCTAATGATCACTTGCTCACAGCGAACTAGAGATCTCTGTCTGGGCAAATTGAGTCTGGTACAGAAATTGATGAGCAAGTTTTTCCAGAAGGAGTAACTGCAGTTTCAGCATCTACGCATCTTGGCACTGTGGGTAATAATCCTGCAACATCTGGCCGGAGATCCTCTCCAGGCTCTCCAATAGCTTCCTCCCCTCGTTTCGTCGACACAGTTGAACAACCAGAGACATTGGATGTGTATTCACCGGATCGATTGGCTAGGGAATTGTTCTTCCTGGATGGTTCACTGTCATTTACTGCTGAGGAGTTATCTCGTGCCACTGCTGAAGTTCTGGGTAGAAGTAGCCATGACACATTGTATAAAGCTACTCTGAATAGTGGGCATGTTCTTACTGTCAAGTGGTTGCGGGTTGGGTTGGTAAAAAACAAGAAAGAGTTTGCAAAGGAAGTTAAAAAGATTGGATCTATTAGGCACCCAAATGCTGTTCCTTTACGAGCATATTACTGGGGACCTCGGGAACAAGAAAGGCTTATTTTAGCAGACTACATACCAGGAGATAGTTTGGAGTTGCATCTCTATGGTAAGATTTTCTTACGAATAACTTCAACAATGATATTCCATGTCTTCCATTGTTGTTTGGTTACTGTTATTAGAAAGTTCCCTTTTGCTGGTCTGGTGGGTGATTATCTTGTGTTAGGACCAATATTGATATCATTCGCATAAATAGTTTCTATAGCAACATAGCTTATTTCTAGTGGTGGTTTTCTAATTGCTGAGATCTCTTGTGTTTTGTGGGATTGTTTGATAAAAATTATGGAGGATAAAGTTGACTTGGCCCTATTATTTATTTTTGGTTGGCCTCTCTTGCAAATGTCCCTCTTTTACATGTCTTACAATATGCAGATAGTGATGAGATCTTATTTGGATCTGTCGTCTCTTATGAATCCAGTGGTAGAGTTGTTGATCAAGCGGAAAAGGAAAAAAATAGGACTGCAATTGTAGAATCTGAATGATTTTTTACAAAATTATGTTTGTTAGAATGTATGTGCAGATCTTTATGTGTCATCTGGTAACCTTCAAAGGAGCTCCAAAAAGGTACAATAGATCTTTGGTTCACTAATTGCCGTACCCAACAGCCACACATCAATAGCCCATTAAAATTAGGGAGTAGACCTATTCTAAGATCTCGTTAACATGACCTAGAATGTAGTGATAGTTGCTCAACAATCCGCTTTAAATAGCATTTCCATTGTTTCTGAATCATTCATGATGGTAAAACTTGATAAATATGGAGAGTTGGCTTTGTCATAGTATCCCCTGTGCAGTGAGGACTGCTTACTTGTAGACATGTTGGGGGTTGTTGGAAACTTTTTACTTTTTGTAATTTTTTGCAGTTTTAAGTTTTTAATGAATGTTTACTTTTTACAGAGACAACCCCTCGGAGGTACTCCCCGTTATCCTTCAACCAAAGGTTAAAAGTTTCCATTGAAGTTGCTCGGGGTTTGACATATCTGCATGAGAGAGGGCTGCCTCATGGAGACTTGAAACCTACAAATATAATCTTGGTTGGTGCTGATTACAGTGTCCGCCTCACTGATTATGGTCTCCACCGCCTGATGACTCCAGCAGGGATTGCAGAGCAGATACTGAACCTAGGGGCTCTAGGATACCGTGCTCCTGAACTTGCTACTGCGACCAAACCTATCCCATCATTTAAGGCTGATGTATATGCTCTTGGTGTGATTCTAATGGAATTGCTGACAAGAAGAAGTGCTGGAGACTTAATCTCAGCACACTCTGCTGCAGTTGATTTAACAGATTGGGTTAGGTTGTGTGATCAAGAAGGTAGGGGAATGGACTGTATCGATAGGGACATCGCAGGAGGAGAAGAACACTGCAAAGCGATGGATGATCTGCTAGCTGTGTCGTTAAGGTGTATTCTTTCGATAAACGAAAGGCCAAACATCAGACAAGTTGTTGAAAATCTGGGTTCCATATCTGTGTGAAATTTTTGTACATGTTGTCTTGAAATTACAATTCTTTTTTCTTTTCCTTCACATTTTTTATTATGTCCTATTGGTTGCCTTCTAATGTTGGTCAATCCTTACATATTCCCCATTTTCTTTCCCCTCCCCCCTTCAATTTTTTTGTAAAGGGATGATCAGTCCATTGATTGATTGGTTGACATTTTTCTCTAATTAGAAAAGATTGCAATTATTTATGTGTAATTTATTTTTTGC

>M6-PGSC0003DMG400012992 M6 LRR-RLK

CTTACTGGTTCAATTCCGATTACTATAGGTCACTTAACTGAGCTCAAAATTCTATACCTTTTTTCTAACCAACTTTCTGGTTTTATTCCTAGAGAGTTGGGAAATTTGAAAAGCCTCGCTTATTTGGATTTATCCACTAATCAACTTACTGGTCCTATTCCTAGTGAGTTGGGGAAGCTGATAAATCTCATAGAGATAGATTTATCTAAAAGTCAACTTACTGGTTCAATTCCGATTACTATAGGTCACTTAACTGAGCTCAAAATTCTATACCTATTTTCTAACCAACTTTCTAGTTTTATTCCTAGAGAGATGGGAAATTTGAAAAGCCTCACTGATTTGGATTTATCCACTAATCATTTTATTGGTCCTATTCCTAGTGAGTTGGGAAATCTGATAAATCTCACATAGATAGATTTATCTAAAAGTCAACTTATTGGTTCAATTCCGATTACTATAGGTGACTTAACTAAGCTCAAACTTCTATACTTTTTTTCTAACCAACTATCTGGTCCCATACCTAGTGAGTTGGGTAATTTGAAGAACCTCAATGGTCTGTCCTTGTACGAAAATCAACTTACTGGTCCAATTCCTGCTTCCATTGGCAATTTGAGAAGCTTGCGATTTCTGTATCTCAGTGCCAACAAACTTACTGGTCCAATTCCCGCTTCCTTTGGCAATTTAAGAAACTTGCAATTTCTGTATCTGAGTGCCAACAAACTTTACGGTTCTATTCCAAAAGAACTTGCATATTTGGATAACTTGGAAGAGCTGATAATAAGTGAAAATCAGTTTTCTGGTCAAATGCCTGAGCATCTTTGCCAAGGTGGGAAACTTGAGAACTTTACGGTGAATAGTAATAAGTTGTCAGGGCCTATACCAAGAAGCATGTCCAAGTGCTCGAGTTTCAAAAGGGTTCGCTTTGATAACAACAGTTTTACTGGAAATTTATCTGAAGCTTTTGGTATCTATCCGGAGCTACAGTTTATTAATTTGAGCAAAAATGATTTTCATGGTGAACTCAGTAGCAACTGGGGGAAATGCAAGAAGTTGACTAATTTTCGAGTAGCCAGAAATAGAATTAGTGGTCACATACCACCAGAGATTGGAAATCTCAAAGGTCTTCAAGGACTAGATCTTTCAGCAAATCATTTGGTTGGGAAGATCCCTAGGGAACTTGGAAAATTGACCTCTCTAGTTAATCTTTTTTTGCAAAACAACCAGATTTCTGGCAATATTCCTATGGAACTTGGATCAATGACAAAGTTAGAATCCCTTGATCTATCAAACAATATGTTGAATGGGTCGATCCCAACATTTATAGGAGATTACATACACTTGTTTCAGTTGAACCTGAGCAACAACAAATTTGGCCAAAATATTCCAATGGAGATAGGGAAGATAACTCAGCTTAATGTACTTGATTTGAGCAATAATCTTCTGGTTGGAGATATATCCCCTCAGTTAGCCAATTTAAAGGTCTTGGTAAACTTAAACCTTTCCCACAATGGCCTATCTGGGCGCATTCCTAAAGAAGTTGAAAGTTTGACTGGTTTGCAGTATGTCGTATTGTCATACAATGAGTTGGAAGGTCCAATCCCCAATAATAAAGCTTTTATGAATGCTTCATTAGAAGGTAACAAAGGTCTTTGCGGCAATGTAACAGGACTCCAGCCCTGTGATAGGCCATCTTCTGTGGTGAAGAAGCACTCAATGGCAAAGGGATGTAAAGTCATCCTCATCACTGTACTTTCTGTTATGGGAGCACTACTACTGCTCTGTGTTTTCATTAGTGTTCTTTTTATGTGTAATAAAAGAAGGAGAGTTAGAGACGTTGAAAGAAGGGATGGTGATGGTTGGCTTTCAATATCTATGTTAGATGGAAAGGCATTGTACAGGGACATCTTAAATGCCACAGAAGAGTTTGATGTAAAATTTTGCATCGGGCATGGAGGACACGGAAGCGTTTACAAGGTAAACCTTCAATCATTAGGGAATATAGCTGTGAAGAGACTTCATTCTTCATATCAGAATACACATCCCAAAAGCTTCATGAATGAAGTAAGGGCATTGACTGGGATTAAGCACTGGAACATTGTGAACCTCTACGGCTATTGTTCAAAAGCACAACATTCACTCTTGGTTTACGAGTATGTGGAGAGGGGGAGTTTGTCTAGTATTTTGAGTGAGGAAGTTGAGTCCAAGAAATTGGATTGGTTTAAAAGGGTGAATATCATCAAGGGTGTTGCTTTTGCTTTAGCTTACATGCACCAGGATTGTTCACCACCCATTGTTCATCGAGACATATCAAGTAGTAATGTTTTGCTTGATTCAGAGTATGAAGCTCGTGTTGCAGACTTTGGCATATCAAAGCTTCTCAAGCCAAACTCATCCAATTGCACAGCGCTTGCAGGGACATATGGTTATGTTGCACCTGGTAAGTTTAAAATATTTCATGAACTAAAGCCAAACATTCTATTGTTTATATCTCATTTACATGTTTATATTGTTTTGGAAATTGTATTGTTCAGAGTTTGCATATACTATGAAGGTTACTCAAATGTGTGATGTTTATAGCTTTGGAGTATTATCTTTGGAGATAATCAAAGGAAAGCATCTTGGGGAATACATTACCGTGCTAGCAAATTCATCTACTATAGATCATGAGCATCTTAGTGATTTGTTGGATGAACGCCTTCCATATCCTGAAGATAGAGTAAAAGAGGTTTTTATTTTTATCATCAAGCTAGCATGCTCTTGTTTGCTTGAAACTCCAAAATCAAGGCCAACAATGCACTTCATCTCTCATAAGTTATCATCTATGGATGCACGTTCACATACTCATGTAAAGCGAGTTC

>M6-PGSC0003DMG400013198 M6 LRR-RLK

GATGGGATAATTATTACTGCAGCAGATTTTCAAGCACTTCAAGCTTTTAAACAAGAATTGGTTGATCCAAAAGGGTTCTTGAAAAGCTGGAATGATAGTGGATTTGGAGCTTGTTCTGGTGGATGGCTTGGTATTAAATGTGCTCAAGGACAAGTTATTGTTATTCAGCTTCCATGGAGAGGTTTAGGTGGAAGAATTACTGAAAGAATTGGACAATTTCAGTCTCTTAGAAAACTCAGCTTACATGATAATGTCATTGGTGGTTCAATTCCTTCTACTTTGGGACTCATACCTAATCTTAGAGGACTTCAACTTTTTAACAATAGGTTATCTGGTTCTATTCCTGCTTCTCTTGGTTTATGCCCTCTTCTTCAAACACTTGATCTCAGCAATAACTCATTCTCTGGTGCAATTCCAGACAGTCTTGTTAATTCAACTAAGCTTTATAGGCTTAATCTTAGCCATAATTCTTTGTCTGGTTCAATTCCCACAAGTCTCACACAATCTCCCGCACTCATCTTTCTTGATCTCAAGTATAACAATCTTTCAGGCTCAATACCAGATACTTGGGATGGAAATGGAAATGGAAAAAGACTCTTCCAACTTCAGTCTCTTACACTTGATCACAACTTCTTTTCTGGAAGTATTCCTGCTTCCTTAGGCAAGTTGAATGAACTTGTTGAGCTTTCACTTAGTCATAATCAATTGACTGGAGTTATCCCAAGTCATATTGAGGGACTCTCTAGGCTTACAACACTTGATTTGTCTTATAATGCCATTAATGGAAGCTTGTCTGATAGCTTCTTGAATCTATCCTCACTAGTGGTCTTGAACTTGGAAAGCAACCAACTTGATAACCAAATCCCAGCAGCCATAATAAAATTGCAGAAACTCTCAGTTCTCAACTTGAGGAGTAACCACTTTAGTGGTGATATTCCTGTCACTATAGGAAACATCTCTACTCTTAGACAACTAGATTTAGCTCATAATAATATAAGTGGAGAAATCCCAGCTTCTTTAGATACTTTACCAAATCTTAGTGCCTTCAATGTCTCATATAATAATCTGTCTGGTCCTGTTCCAACTCATCTCGCACGGGAATTCAACTCGAGTGCCTTTGTGGGTAATCTTCAGCTATGTGGTTACAGTGCATCAACCCCATGTCCAATCTCTCCAGTTAGCCCATCCCCCGAAACACCAAAAAAGCAACGTCGTAAGTTAAGTACTAAGGACATAATTCTCATAGCAGGAGGGGCACTTCTCATAATCTTGGCTCTGCTATGTTGCATTCTCCTATGCTGCTTGATCAGGAAAAGATCTGCAGCTGAAGCAGGGAAGGATGGTCAAGGCACGAGTAGGGTGGCTGCAGCAGCCAGAGGTGAAAAGGGTGTTCCACCAACTGCTGGAGAAGTTGAAGCAGCAGGAGGAGGAGATACAGGAGGAAAGCTTGTCCATTTTGATGGACCTATTGTGTTCACAGCAGATGATCTTCTTTGTGCAACTGCCGAAATAATGGGAAAGAGCACTTATGGTACGGTGTACAAGGCCACGTTAGAGGATGGCGATCAAGTTGCTGTGAAAAGATTGAGAGAGAAGATCACAAGAGGTCAAAGGGAATTTGAGAGTGAAGTCAATATTCTTGGCAAGATCAGGCATCCAAATCTTTTGGCCCTTAGAGCATATTACATGGGACCTAAAGGAGAGAAACTTCTTGTTTTTGACTACATGCCTAAGGGAAGTTTGGCAACTTTTCTTCATGGTAAGCACAAAATCAGTTTTCCATTTCACTTTGTATTTGTGAGAACTTCAGATTATAAATCTTTCTGTTTTTTGCAGCTCGCAGTCCTGATACACCAATTGATTGGGCAACAAGGATGAGAATAGCAAAGGGTACAGCAAGGGGATTGCTCTTCCTTCATACAAATGCCAACATCATTCATGGGAATCTTACATCGAGCAATGTTTTACTTGATGACAACACAAATGCAAAAATTGCAGATTATGGTCTTTCGCGCCTAATGACTGCTGCTGCAAATGCAAATGTTATTGCAACTGCTGGAGCACTTGGGTATCGGGCACCTGAACTTTCAAAACTCAAGAAAGCAAACACAAAGACAGATGTATACAGCCTTGGTGTTATTATACTGGAACTTCTAACTGGGAAGTCACCAGGAGAAGCAATGAATGGTGTGGATTTGCCTCAATGGGTTGCATCAATTGTGAAAGAGGAGTGGACTAATGAGGTATTTGATTTAGAACTAATGAGGGATGCATCCGTTATCGGTGATGAATTATTGAATACCTTGAAATTGGCTTTGCATTGTGTTGATCCGTCACCATCAGCTCGACCAGAAGTTCAGCAACTTCTCCAGCAACTAGAAGAAATTAGACCTGAAACAGCTACCAGTTCTGGGGACGACGG

>M6-PGSC0003DMG400013489 M6 LRR-RLK

AAAGAAAGAAACTTGGCATTTGGGTGTTTTGAGGTTTTTCCAGTTCTTGAAAATGCTAGTTGTGTAACGACTGAAACTTTGTCATTTGGGTGTTTGAGGTTTTTTCAGTTTCTTGAAAGAGCTAGTTGTATAAAGACTGGAACTTTATCATTTGGGTGTTTGAGCTTTTTCCAGTTCTTGAAAAGCTAGTTGTGTAAAGACTGAAACTTTATCATCTGAGTGTTTGAGGTTTTCAATCTTGAAATGAGCTAGTTGTGTAAAGACTGAAACTTTGTCATTTGGGTGTTTGAGATTTTCCAATTCTTGAAAGCTAGTTGTGTAAAGATTGAAACTTTGTCATTTGGGTGTTTGAGGTTTGCAATCTTGAAATGAACTAGTTGTGTAAATACAGAGACTTTGTCATTTGGGTGTTTGAGCTTTTTCCAGTTTCTTGAAAGAGCTAGTTATGAAAACACTGAAACTTTATCATCTGTGTGTTTGAGATCTTCAATCTTGAAATGAGCAAGTTTATCAGGTGTTTGTGGAGGTGTTGAAAATGGATAAAAAGGGCATTTTTTTTAATGATAAATTATGGGGTTTTGTTAGAAAAATTACTTTGTTGAGTAGGTGAGATTGTAGAAATCTGTACTGGAAAGAATGTTGTATCTGTTTTGGAATGTGGGGCTGCTACTGTTTTGTACATTGATGGAGAGTTCTTCTGCTACACTTTCTCCTGCTGGTGTAAACTATGAAGGTTAGCTTCTTTCTTGAAATTTCTCACTTGTACTGAAAATTTCCTTTTATCAAATTAAGCTAAGAGTAGAAGAACTTGTTAATTCCTCTGGATCAGAGTTAGAGCCACATGGGTGCAAGGGGTTCATCTGAATCCCCTTCGACGGAAAATTAAACTGTATATCTAAGGTTACGAATATTTTTATGTATATATAATAGACATTGAATCCCCTTGTATTCGTCCTGTGTTTACTTCTTCATAATTTGAACCCCTTAGGCCTTTGCAATGTATATGAGTATGTGACTATGTGTTCTATCTTTTTTCTCTACGATTTCCTCTCGTTATATTGAATCTTTGATGCAATGACAAACTGAAAAGTGAAAACAGTGGAAGTAATCTAATAGAGTGACTGTTTTTCCATATAATAATGTGCAGTTGTAGCTTTGACGGAAATTAAGAAGGCCTTACATGACCCCTATAATGTTCTGGACAATTGGGATGTGAATTCTGTAGACCCCTGCAGTTGGAGAATGGTCACTTGCTCCTTGGATGGCTATGTGTCTGCTCTGTAAGTTGTTTGGATGAAAGAGTAATTTAGTAGAAACTTAGTTGGCTCATGAAAAGGCCTGATTTAATCCAGTAAATTTATCGCTGAATATTTTCTCTTGCATCAGAGCACTACCTAGTCAAAGTCTAGCTGGCACCTTATCGCCTGGAATTGGCAATCTTACTAACTTGCAATATGTGTAAGTTCTTGACTATATACATCTCTTAGTTGCTGCTACTTGGTTCCGTAATACCTAAGGTCACTTCTTATATGATTTTCTTAGTGTTATTAAGCCATTTTCTCGCAAACTGTTCTTAACTTGTTCATGTAGATTGTTGCAAAACAATGCCATTTCTGGTCATATTCCTGATGTGATTGGGAAGTTACAAAAGCTTCAGACACTTGATCTTTCCAACAATAAATTTGAAGGCGAAATACCCACTTCCTTCGAAGACCTGAAGAATTTGAATTATTTGTGAGTTCATGAATTTTCACTTATTATTATTCACTTTCAAGACTCCAACCATATATTTCACTCTGTCAAAGATTGAATACATCATGCTACCTATTGACTCAAATCTTCTGTTTCGTCCTCATCCTTGTGATTGACTTAGCTGATCATTGCTTGCAGGGATGTCTCTTATAATAATCTAAGTGGTACCCTTCCTAAAATATCAGCGCGAGCTTTCAAGTAAGTCGGCCTTTTCTATTGCCGTAATATACTCAGTTATTTCATTTTCAAGTAGTTATCTTAATATTATTGAAGTTAAACTATTGCTGTACGGCTGTACATGTAAATCTCGATGCTTAGTTGATAAAAATCCATTATTAAAAATACATTGACAAATGCTTTTGGGTAAACTGGTACTTGAAAAGTATTTCATTAGTATTCCACACTATAGTGAGATTGACTTCCTCTTCATGGCTTTCAGAGTGCTCGGGAATCCTTTAATTTGTGGTCAAGGTTCTGAGAATAATTGCTCAGCAGTTTATCCAGAACCACTGTCCTTCCCACCTGATAGTCTGGCAGGTACAGATAGCATTTTCATGCTCAAGTCCCTTCCCTTGCACCTGAAAATGGAAAACTGTTAAAAACATATTATACATACTGAGGAAGTTTTGCTATTTGTGTTATTGAAAATGTGTTGCATCTATTATTTTTGATATCACAGATCAATCTGGTGCTGCACGCAAGAGCCGTCGTGTGGTTGTTGCGTTTGTTGCAAGTTTTGGTGCTGCTTTCTTGGTCATAATAGCCCTCGCTTTGCTTCTTTGGTGGCGCTACAGAAATAATCAGCAGATCTTCTTTGATGTCAATGGTTAGTTGATTTACTTTCCCCCCCTCTTCTCTATGATAATTTGAGCTTTTATTAGTTCTGCATTTCTGATTCTTTTCCTGCCACGGATATATAAGTTGAGCCAAGTGAGCTTTTGCATTATTGTCTGAGTGATTGGTGTTAGGTATCATGTGAATATTTTTTAAGCTAGGATGTTACAGAGTCACCTGCTTGTGGTTTGAATCTTTATTTTAATCTCTAAAAAGATTTTTGTGATTGTTGATTTTAAATTCATTTGTTTAGTTTTAGAGAGGACATGAGTGGTCAAGGAATAGGAATTGGGGATAAATTCTGTGTTTACCATATTCAATAAACCTGTTGAATATGAAGAACTTTCGAACCCCGTCAGTTAGAAAATTATGCCCTTTGGCATACTCAGTAGTTCTGGTCAATTTTCCTAGTAAGGACTCAGAAAAGCTGTTTGAATTTTATTGTTATCGGGACAAGTCCTGATCCACTACAGGCTACCTTAAACATTTAGTTCCAAACATTGACAGAATAATTTGTTAATCACATATCTGAATAATTACACTTGCATTTTCTTGTGCAAGTTGAAGTAAATATTTCTGTAAAAAGGAAGATGGTGTTATGTTCACTCTCTCGGGCCACCATGCCTGGTGAATTGCCTTGCCACATTCATTGCGTTGAAAGTTACTCTTGATAATCTTGTACTTTGGTTGTCTGATTCATATATGTGGATCAGTTTTGGGACGTTTTTGCCAACCTCTATTATTTTCTCCCTGTTTCACTTTTCAAAAGATTTTCTCCATCTGTCTCTAACTATTCCTCAAAGTTTTATCTGAGTAACGACTTCTCAAATGCAGATCATTGTTATCCAGAGGTATGCTTGGGTCACCTGAAAAGGTACACCCTTAAGGAACTGCGGACTGCCACTGACCATTTCAGCTCAAAACGTATTTTGGGAAAGGGAGGATTTGGGGTTGTGTACAAGGCCAGCTTAAACAATGGTACTGTTGTGGCCGTCAAACGATTGAACGACTACAATGCTGTTGGTGGTGAAAATCAATTTCAGACTGAAGTAGAATTGATTAGTTTGGCTGTCCATCGGAATCTTCTCCATCTTCTGGGGTTTTGTTCAACTGAAAGTGAACGGCTTCTTGTTTACCCCTATATGCCAAATGGAAGTGTAGCATCACGATTAAAAGGTTAGCTCTTGCCTCCATGTGTTAATACATAATACTCACTATTGATTTCAGATAATGAGTCCAACATTTCCATTTATAAAAGTTAAATATTCTCACATGGTGTTTGTTTGTTACTACTCCCTTCATTCATTTTGTTTGACAGATAATTGATAGGATACAGAGTTTAAAATTTTAATTTCTTTCATTTATTACTTTGTAGTAGAATAAATTTATTAAATAGTGGTTATTATACTTGTTAAAGAGAACATCATATAAATTTTAGAATGTGGATAGTATGAATTTGAGTTCTTAAAGCTGTAAAAGGTGTACGGACAAATATAATACTAGAATAGTGCCAAATTTTGGACCATGTCTAATTTAAAAGAAGTCATGTTAGTGAACAGTCCCTACTTCCCAATTTATTGTCATTAAACATTAGTGAGTGCTCTGAAGACCAGCAACCGTTGTCATTGGTGAATTGAGGCCATAAGAGTCTGGTTAAATACAAGCTTAGATGTTATGACCCACCCTTACGTCAGTCCATCCTTGAGCCCGCGGTGGGCTGCCCACCTTCACATACTCTGTACTGCTGTAGTTATATGATAAGACATTATCAGTTTGCTGCTGGCTGAAACCATTGTTGTGTGTTTCTGCATGTTCGAAAGCCAAGTTCATTTTACTTTTCTGGAACTTTATCAGTTACTTTTTAATAGTTTCCAAGTAACTGAATTTTGTAACAGTTGAAGCTGATACATATATTGGCTTGATTGAAAATGCATTGTGGTTTCTAAAATTTGCAGATAGCATAAATGGTAAACCAGTTTTAGACTGGTCAAGGCGGAAAAAGATAGCACAAGGTACAGCAAGAGGGCTAGTTTACTTGCATGAGCAGTGTGATCCCAAAATTATCCATCGTGATGTCAAAGCCGCAAACATTCTGTTGGATGAAGAATTTGAGGCAGTAGTTGGTGATTTTGGGTTGGCAAAGCTCTTGGATCACCACGATTCCCATGTGACAACTGCTGTTAGGGGCACCATTGGCCACATTGCTCCTGAATATTTGTCTACTGGCCAATCATCTGAAAAAACTGACGTTTTTGGGTTTGGAATCTTGCTACTTGAGCTGATTACTGGCCAGAAAGCTATGGACTTTGGTCGAGGGGCCAACCAGAACGGTGTTGTTATGCTTGATTGGGTGCGTCTCTCCTCTTTCTCTTTCTCTGGATGTTGTACCAAGTACTTTTTGAAATATGTACTGTGAGTTGTGAGAGGGAATAAGCTATACAGAGGCGGGTTATGCGTCTCTCCTCTCTCTCTTTCTCATAGCTGTTCTCTTTTCAGTTATTAGGAGTTGTAGGTTCAAACCTCAAATATAGCTAGCTAGTGGTGTGTACATGTAGTTTACAAGGCTCTTTTCCATTAGCATACTTCAACGAACTCTCTCGTTTAATTACTATTAATGATATGCAGGTGAGAAAGCTGCATCAAGAGGAGAAACTGAACCTTATGGTAGACAACAATTTGAAAAATAACTTTGACCAGATTGAGCTAGAAGAGATGGTTAAAGTTGCCCTTCTATGCACCCATTTCAACCCTTCTCATCGCCCAAAAATGTCCGAAGCATTAAGGATGTTAGAAGGCGATGGATTGGCTGAAAAATGGGAGGCATCACAAAAGCTCGAGACTCCAAGAAACCGAGCTTTTGAACATACACCACAAAGATATTCCAACTACGTCGAAGACTCTTCACTAGTTGAACCAATGGAACTTTCTGGACCTAGATGACATCCATTTTTCGTCCTCGATTCTTATCATCCTTGAGTTGCTAAAGTTGAAGTGAGAAGTATTTCATTGCTGTTGAATGTATGTATTCATTTTGTAAAAGAAAAGGGAATAAGTCATGTGTAAGCTGAGTTCATATATGATATATTCCCTTGTTCTTTCAATTTTTGAAATGATATGCTATGTAATTATGATTTCATTTGTTCTTTTTATAAGAAAGATTTTGAGATTTATGAGTAATAAAACGCCATAGGCTTGCCATTCTCATTAAGT

>M6-PGSC0003DMG400013500 M6 LRR-RLK

ATGGTTGGTTCTTTACCTCTTGAAATTGGAAATTTAAAGGCTGCGACACTGATAGATCTGTCAATGAATCAATTCTCAAATGGAATTCCTATAGAAATTGGAGGATTGCAGAATCTGGTGCACCTTTCTTTGAGGCACAACAAGTTACAAGGATCTATACCTGACTCATTGAGCAACATGGTAGGTTTGGAATTCCTAGACCTTTCTCATAACAATATATCGGGAACCATTCCTATGTCTTTGGAGAAACTTCAAAACATGAAGTATTTCAATGTTTCTGTCAACATGTTGCATGGTGAAATTCCCTCGGGGGGTCCTTTCAAGAACCTCTCGAGTCAGTTTTTCATCAACAATGAAGCATTGTGTGGTTCTTCAAGGTTTAGTGTCCCGCCATGCCCCACATCATCAAAGCATAGATCAAATAGGAAAAAAGTGCTTGTTCTATTTCTTATGCTGGGAGTTGCACTTGTATTTGTTCCTATTACCCTTTTGTTTGTATGGATAAAGCATAGAAGAGATAAGAAATACCCTCAACAAGATGATTCATTGTCTATTTCAACGAGAGGAAGAATTTCGTACTATGAATTGCTCCAAGCAACTGATTCTCTCAGCGAGAGTAATCTGATTGGTTCCGGAAGTTTTGGCTCTGTCTACAAAGGCATTCTCAGAAGTGGGACTGCTATTGCAGTTAAAGTGTTTAATCTGCAACTGGATATGGCATTCAAGAGTTTCGATACAGAATGTGATGTTTTGCGCAACCTTCGCCATAGGAATCTCATAAAAGTCATCACTAGTTGTTCCAACATTGATTTTAAGGCTTTAGTGCTCGAGTATATGCCTAATGGGAGTCTTGAGAAGTATTTGTATTCTCACAACTACTTCTTGGACATCAAGCAAAGACTAAGCATAATAATAGATGTGGCATGTGCATTGGAATATCTCCACCAT

>M6-PGSC0003DMG400013525 M6 LRR-RLK

GTAAACTTCCATCCATCAGCAAGTTGTAGCTCATAATAAACCTCTAAATTATGTCATAAATTTCAATTCTCTTAGCTTAATTCCTTCAAATTCTTCATTAATGGGGTTTTCATCGAAATGGGTTTTTGTATTTACGTTATTTTCATTCATAAATCTTTCAATTTCTGATCCAAGAGCAACAGAAGCAGCTCTAATTTGTACAAACAGAACAGCTGCACAATCCAATCGACAAGTCTATGTAGCAACTTTTCTAGAAGCCATGGATTCCATTACGCCATTAGTGGGGCGAGACAAATTTGGAGCAGTAATTAATGGAACAGGGAATAATACAGTTTATGCGTTTGGTGAATGTATGAAGGACTTATCACAAACAGATTGTAATCTCTGTTTTGCTCAGTGCAAAACTCAAATCCTACGATGCTTACCTTTTCAAAGATTAGTTACTGGTGGGAGATTATATTATGATGGGTGTTTCCTAAGGTACGATTATTATAGATTTTTTAATGAAAGTTTGAGTTCCGTTGATAGGACGATCTGTGGGAACACGAGTTTTTCTGGGAATCAGAGTTTGTTTAGTGAAAATGTTGGGAAATTGGCGAGGGATTTGGAGTTTGGGGGTTTGAGGAATGATGGGTTTTTGACTGCAGTTGTGAGTACTGGGAATTTGAGTGTTTATGGGTTAGCTCAATGTTGGGAATTTGTGAATGGAAGTGCTTGTCAAAGGTGTCTTTCTGATGCAGTTTTGAAGATTAGTTCTTGTTCGCCTAAGGAAGAAGGGAGGGTGTTGAATACTGGTTGTTATGTTAGGTATTCTACTCAGAGGTTCTTTAATAATTCTGCAAGTGATACTCCTCCTGCTGGAAATGGAGGTATGTTCTTTCTTTCTTGAATGTATGAAATGAATAAATTGTTTCGTTTAGTTGTGCGATCATTTCGTCTAAAAGTGTAAGCTATTAGAGATAGTACTTTTTGCTTGACATGTCATATCTTTACATGGTGTCATTGGGTCTGACTGTGAAGGGGTGTTAGAATGGTTAAAGTTTCACATTGGTTGGGGACTGACTTGAGCAATTCTCTTGTGATGAGCTAGCTTTTGGAGTTGAGTTAGACTCAAATGTCATATATCTTTACGGATGTCATGTTTAGTTGTGTAATCATCTTATTTAACATATGCTTGCGATGTTTCTTGGTTTCATGAGTTATTTCAATCTTATCCAATATTCAATGATTTTAATGTTTAATTTTGTTTTTGTATGATTTGCAATGATACTTGCTGCAAGCCTAACACTATTGGATGTAGAGCAAGAAAAAGTTCAACTTGTTCTGCCCAATTATTTTTTTTACTAACAAAACAGATAACTACTTTGTCATTTGGAATGATTTGAAAAACGGCGGTTGGAAATCGAGGTGATTTCGACATTTCTTGAAAGTTGCTCTTGTTATCTGTTTAAAATGTGGTAGTTTGTATTTCTTGGATTTTAGTTCCATTCCAGGATTTAATAACTATCTAGACATTACTTGTGATGATGTTTCTTTTTGTCATCTGATAAGTTGAAATGGTTGACTTATATATATACATGAAATGAATGAAGAGTGAATCAGCATTGAGGCAAATGCAGACAATGAATCTTTTAGAAGCTGTGACAAAATAAACTGTTTATACAAAACTATGTCATTCATGTTGTCGTATTTTGATTAATTATCAGGAGGATCGAATCGTTTGGCTGTTATTCTTGCAACAACTCTGAGTATTTCTGCTTTCTTGCTATTTGTATCGGCTGTTTCATTCTTTGTGCGGAGGAAGGTTCTGAAACAAAAGAGAGGTACTAATTTTTAAGTCAAATCATTTTCATTCTTTCCGAAAAAAGAAATCTCATTTAAGTCTGATTGATCAAATGGTAATTTGACTTGAATTTGCTTCGATGATGTAGAGAAGAAGCAGCTAGGAGCTCTCATAAGAACGGTTAATAAATCAAAATTGAATATTTCATATGAAACTCTTGAGAAGGCGGCAAATTACTTTAACAACTCGAATAAATTGGGACAAGGAGGTTCTGGTTCTGTTTACAAGGTAATTCTCTTTGAACATTATATTCATTTTTCTTATACTAATGAGATTTGCCTTACGATAGTCGTGTTTACACATGAAATAGGGGATCTTGCCAGACGGGCAGGTTGTTGCTATAAAGAGGCTCTTCTTCAATACGAGGCAGTGGGTTGATCATTTCTTTAATGAGGTCAATTTGATTAGCGGCATCCATCACAAAAACTTGGTAAAGCTATTGGGTTGCAGCATTACAGGGCCGGAAAGTCTTCTAGTGTATGAGTACGTGCCCAATCATAGTCTTGCGGATTACGTCTTTGGTAAGTATCCAATCCACCTTAATTTATGAAAATTTTTAAATTGGTGTAGTCTTATCTCGTTATGTAAATGCCTTTGTTTTGCACTGAATCAGACACAAAGAACCTTCAACCACTTACATGGAGTCAAAGATACAAAATTGTATTGGGTACTGCTGAGGGATTGGCCTATCTTCATGAAGAATCGAAATTAAGGATCATCCACCGAGATATAAAGTTGAGCAATGTTCTTCTAGATGAAGATTTCACGGCAAAAATAGCTGATTTTGGTCTAGCCAGGTTATTTCCTGAAGACAAGAGCCACATTAGCACAGCTATTGCTGGCACTCTGTAAGTTCAATAACTTTTTCCGGTTACTTGTTACTTTTTCTTTAATCAATACACTAAGAAAGTGCTGAATTAATAAGTTTTGGCATTGCAGCGGTTACATGGCTCCAGAGTACGTTGTACGTGGCATACTGACCGAAAAAGCTGATGTCTATAGTTTTGGAGTTCTTGTTGTCGAAGTCGTGTGTGGAAAGAAGAACAATTATGTTTTCAAGAACACTAATTCTCTTCTGCAGCAGGTACTTTCTTGTGATATTTTTTCTTTCCTTAACGGACATTAATAGTATTTACTCAACTCATGTCATTGTGGTTTCTAGTTGTGGAACTTGTATGGGATGGGGAAATCTTATGAAGCAGTTGACCCTTTATTAGAAGGTAACTTCAATAAAGAGGAGGCATCTAAGCTGCTGCAAGTAGGTCTTCTTTGTGTACAAGCATCAGCAGAACTACGACCCTCCATGTCAAGTGTCGTGAAAATGCTCACCGAGAACCATGAAATTCCTCTGCCAACACAGCCACCATTCCTCAATTCTGGTAGTTCAGAATTTAGTCCATTTAATCTGCATGGCAAAAGATTTTTCGGTCAACCAAGCTCCTCTACACAATCTTCTGGGAATAAATTGACGGAGAGTTGGATTGAGCCTAGATAAACTAAGTTGATCATCTGGAATTGGTATTTGATTTTGTTCTCAGCAAAGGCCAACTTCTTGAAGGTGAATTGTTCAGCCAACTCAAGTTTTTTCTGTGATTAACTAGTACTCACTACTCACTCTCCATTTTTTTTTACTGATATTTGTTTAAAGTCTTATTTAGTCAACTTCCTTTTGTGAATACCCCAGTGAGCTATTTCATTTCTAATGTAAAAAAATTGTATAGTATCTGCAGCAGTTGCCATTAACTATATCTACTGTTTTCTATTTTCGATTTTCTATTTAGTAATTTTCATCAAGGATTATGCTTCACAATTGTCATCATCTTGTCTTAGAAGTGGCTATTCAGAATCCCCTGCCTAAAGTAACAGGCATTGGTGAAACTACACATTACAATTACATCTACA

>M6-PGSC0003DMG400013682 M6 LRR-RLK

CTAAGCTTCTCTTTCCCTCACCTCCCTCCCATCTCTCTCCTCTTTGTGCTTCCCCATGCATGCCCACACAATCCCATTTCCCTAGAGATGTTGAAATTCTTTTTGCATGCCCTCTAAATCATAGTATTTTCTCATTTATAATTGCCCTTTGTAAAGGGCGCGCACCCACAAGTGTGCGCACCCCTACAACTAGTCTATGCAATGTCGGTGGCTTATCGTTATAGCAACCACAATCGCCAGTATCGTTATCACCACCACCATCTCTTAATCCTTGTCGTGTTATTATTACTAGTCGTTGTCCCGATCAAATCGGATAGTTCTGAGGCAGAAATCCTCTTTCGATTTAGCAAGTCCTTGCAGAAAAATGGTGCAACCGCTAATTGGAATACGAAAGTATTGCCTTGCGATAAAAATACAGATAAACCTAATTGGGATAATGTTATTTGCGAAAACGGATTCGTTTTCGGGTTGCAATTGGAGAGTAAAGGCTTAAGTGGGGCAATAGATGTGGATGCTCTTAAAGACTTGCCTAATTTCAGGACCATAAGTGTTATGAACAATAATTTTGAAGGACCTTTACCGAATTTAAGTAAACTTGCGGGGTTAAAAACGGCTTATTTCACAAATAACAAATTTTCTGGCCAAATTGACAATACTCTTTTTGAAGGGATGCATTGGTTAAAAAAACTTCATCTTGCTAATAATCAATTAAGTGGTAAAATCCCATCTGTTTTGGGTCAATTGCCTAAGCTTACAGAATTGAGACTTGAAAACAACAAATTTGAAGGCCAAATACCTGATTTTGTTCAAGAGAGACTTATGGATATGAATTTTGCTAACAACTCTCTTGAGGGTCCAATTCCTCATGGCCTTACTAGTCTCAAACCTTCAGCATTTGAAGGTAACTTTTTTTTCCATGTGTAGGATAAATTCCATTTTCTGAAATTATTGTAACACTTATCTAGCTACGACGTACACAAATGTCTACATTTTAAAATGTAGTCATAAAGATATGTCATGTATGCCTAACTCCACCTTAGAAGCTAGCAAGGATTGATCAAGTTCATATCATCAGTAGATCATCAATCCATTTCCCAATCAATAGGAGACTCTTTAACACTTCTTTCACACGCATCTCCCAAAACAGAGATGAACCTGACTCTGACACCATATAAAGATATGACGTTTGCGTCTAACTAAATCTCAAAGTTAACTCACGAGGAAAAAATTGTCAAAGTCCCAATCAGCACATTCCCCATATGGCGCTTTAACCCACATTCTAACAATAGTTCTTCGTCATATTGACAATAGATCAAGAATTGTAGTTCATAACAATTTTTATATAGTTTTTAAATATCTAAATTTTTTAATTTTAGAATATTGAGTTAATCTAAACATGCTCTCATTTTTGTTCATTTGGTAAAATAATACAGGTAATGATCTTTGTGATGGTCCATTATCGAAATGCACAAGTGAACCAAAAGTAGCTCTCTGGACAATTATATTGGTGGTTATCGCAGTTGCAGCAGCAGTAGCAGCAATTGTTGTTGTTATCATCATTCTTCGTCGGGGCAAGCAGACACCCGAAACTGAAACAAGACCAACACCTACACCCTCTGGAGCTGCAGGGGGCGCTACTAACCAAACAGGAGCGCCCTCAGCAGCAGATCTAAACAAAATGGAACAGGGCAGCAATCAAGCAATGGCTGCTCCTGCTCCTGACCAGTCTCCAGAGGGGACCGGTGGACTAAATAATAATAAGAGGCCTGAAGTACAAGCAATACAAAAACTTTTATTCTTAAAAGATGATATTGAAAAGTTTGATTTGCCAGATCTACTAAAGGCTTCAGCTGAAATATTGGGTAGCGGTGTATTTGGTTCGACTTATAAAGCTGCACTTAGCATGGGACGTGTTATGGTTGTCAAAAGATTTAGGCAAATGAATAATGTAGGCAAGGAAGATTTTCATGAGCATATGAGAAGGCTAGGTAGGTTGAGTCATAAGAACTTACTTCCTGTTGTAGCTTTTTACTATAGGAAAGAGGAAAAGCTTTTGGTCTCAGAGTATGTCAACAATGTCAGCCTTGCAGTTCATCTTCATGGTACGTATTTCATACTTTTAAGTTTCCGCGCTAAAATTAGATCATAAAATCATGATTTGCGCTACCAGTTTATTATATTAACTCAATCTATTTTAACCTTTGTTTTATTCAATTCATTTCAAAACAGAGTTGAGATATGCAACTCAAATCTTTTGAAGAGGGCTCATTAATTTGTTGTTCTCTCTACCTACTCTACTAGCTAGAGTGATGATCTGACGTGTGATCATCTCATCTAAGATATTAAGTTTTTACTATGATGAACTCTGTGATCATCTCATTATACATATTGTTTATTTTTTTCAGGTAATAAATCGCGGGGGCAACCAAGTCTTGATTGGCCAACTCGATTAAAAATCGTAAAGGGAGTAGCCAAAGGACTATTGTACCTCTACAACGAGCTACCAAGCTTGACAGCACCCCACGGTCACCTCAAGTCCTCGAACGTGCTTCTGAATGAATCCTACGAGCCCCTCCTCACGGACTATGCATTACTACCCGTAGTAAACCTAGAACACGCTCAAGAACACATGATTGCTTACAAATCTCCTGAATTCAAACATAGTGGTCGTATCACACGTAAAAACGATGTGTGGACGCTTGGAATATTAATCCTCGAGATCATAACGGGTAAATTCCCGTCTAATTTCTTGCAACAAGGCAAGGGGAGTGACACGGATTTGGCTACGTGGGTAAGGTCCGTGGTAAATGAGGACATGACGGAGGTAGATGTATTCGAGAAGGAAATGAGAGGTACGACGAATTCAGAGGGGGAAATGATGAAGTTATTGAAGATAGCTTTGGGTTGTTGTGATCTTGATATGGAAAAGAGGTTTGATATGAAGGAAGCAATGGAAAGGATTGAGGAAGTAAAAGAGAGAGATGGGGATGATGATTTTTACTCTACAAGAGGATTATCTGATGATTATACTCAAGTTTCTATGAACATTTGAAGAGTGCTGGTTATACATACTCATAGTTCGTGTGTAACACTTTTTTATTTGTTCAAAAATCATCATGGTAAATATGGAATGGGACTTTTATTTTGTTTGATGCATGGTTGGGGGTCGTATATATATTGGTCAATGAAACCACAAAATAAATAAATATGAAAACAAAAAAAAAAGAGATAGAAAAAATGAAGTTAAGTGTTTTTTTGTTGTGATTCAATTGAAGCAAGGGAACATGTAAATTGTCCTTGATGAATCTAGTTTAGCTAGTTGCCTTCACTTTTTTTCTTTTCTTTTCTTTTTGAAACTTTATCTAGTTTATTATATTTGGGGCTTGTAAAAATTAAGTAAATCCAAAAGCATGCAAGAACAAAGAATTATAAATCTTTCAATTGC

>M6-PGSC0003DMG400013953 M6 LRR-RLK

GGTTATGGCCTTGGCTCTTCCTGATTTGCAACTTCAAGGCACAATTTCCCCATCTTTGGCAAATTTGTCCTTTCTCAGTGTTCTCAATCTCCAGAACAACAGCTTCCACAGTGGCATCCCTTATGGACTTGGCCACATGACTCGCTTGCGAGTGATTGACGTTAAAAACAATCAGCTCAACGGAAGTATACCAACAAGTCTATTTCAAAACCGGAGAGTTCAAGTAATTTCAATGGCTTTCAATGAACTCGGTGGTGAAATGTGGAGAGGCCCATGGTATGTACCAGAACTCAGAGTCTTAAATCTCGGGAACAATAGCCTCACGGGTATAATCCCTCCTTATGTTGGAAATGCCACAAAATTGATGAACATCAATTTGTCTGGGAATAGAATCAATGATAACATTCCAATGGAGATCAGTAATCTTAGCCAACTTGCATTTTTGTCCTTGTTCAATAATCAATTAACAGGTTCCATTCCTCCAACACTGTTTAATATCTCGTCACTACTTGGCATATATCTGTCATTCAATGGCCTTTCTGGTCCTCTCTTGCTAGATGAAGGGAATATTGTGTCAAATCTGAAGCATTTAAGTATGTCTAACAACCAAATTTCTGGTTGCATTCCTTCCAACATATGCCAACTCACAGAGCTCAAACTTTTGTCCATATCTTTCAACAATATAACTGGAGACATACCTAGAAATATTGGTTGTTTATCCAAACTCGAAAGGTTTTATATTGGTGAAAATCCAATAGAAGGGACTATTCCCACTTCATTGGGAAATATTTCTACTCTGCAATATCTTGATTGTGTAGACAATTGGATGAAGGGGCAAATTCCTCAAGAATTAGGGAAGCTATCAAATTTGAGGGAATTAAGCTTTGCCAATAATTCTAATCTTATTGGTCAAATTCCAGAGGCTATTTTCAACATATCTTCTTTGGAAGTGATCAATTTTAGTTTGAATAACCTCTCGGGGAGAATTCCAACCACTACAGGTCTTCATCTTCCGAACCTTAAAGAACTTTACTTGGGAGACAATCAGCTTGAAGGGGAAATTCCATTGTTCATAACAAATGCTTCCAAGCTTCAGACACTGGCACTAAATGATAACTCTCTCAAAGGAACTATTCCTAACAATTTGGGGAATCTTCGTGAGCTGCAAATACTGCTCCTACACACTAATCAACTTACCAATGAACCAGGAAAGCATGCGTTGAAATTCTTCAATTCTTTGGCGAACTGTAGGATGTTGAGATATCTAGAAGTGGGTAACAATCCGTTGAGTGGAGTTCTGCCAAATTCTATTGGGAATCTCTCATCTACTATTGAAAAAAATTATTTAGCAGATGCACACATCAATGGCCTCATCCCCACAAGTATAGGCAACATGACCGGTCTTACATCCTTAAGCCTTGGCGGAAACAACTTGACAGGGAATATTCCTTCTGAGATTGGTAAGCTTAAACAACTCCAAGGACTCGATCTATCTAACAATAAATTGCAAGGACATATTCCAGAAGTGGTATGCAATTTGTCTGATTTGGTTCAATTATATCTGTATGGTAATGAGCTCTCTGGATTAATTCCAGAATGCTTAGGAAATCTTACCATGCTTCAACAACTTTATTTGGCTTCTAATAAATTTTCATCAAAATTTCCATTGACCCTTTGGAAAATGAGTGGTCTTCTCTATCTAAACGTGTCACAGAATTCAATAGAGGGAGAACTTCCATCAGATATTGGAGGACTGAAAGCCATTGTAGAACTATATCTTTCTGGTAACCACTTTTCAGGCATGATATCAACCAGATTCGGGGAACTCCAAAATCTGCAGTATCTCGACCTATCGAAAAATTCATTTTTTGGCCGAATTCCATTATCCTTTGCCAACTTGATAAACTTGGAATTCTTGAATTTGTCTTTAAATGCATTGTCAGGTACTATTCCTAAGTCTTTGGAAAAACTCTCATACCTTAAAAGCATTAATGTTTCATTTAATGATTTAGAAGGTGAAATACCAAGTGGTGGTGTGTTTGCAAATTCCACTCTGCAATCATTTGTT

>M6-PGSC0003DMG400013956 M6 LRR-RLK

ATAAGAGCCTTTGTTCTGCAATTTATGCCCAATGGAAGTCTTGAGAATTGGCTGTACAAAGAAGATCGACACTTGAACCTTCATCAAAGAGTCACTGTAATGCTTGATGCAGCTATGGCAGTTGAATATCTACACCATGGTCATAACACTCCAATAGTTCATTGCGACCTGAAGCCAGCCAACGTTCTTTTGGATGAAGATATGATGGCTCATGTTGGTGATTTTGGAATATCTAAAATTTTAGCCATAAGCAAGTCCATGGCTTATACAGAGACATTGGGCACTCTTGGATATATTGCACCAGGTAAAAGAAAATCTACTCTCATTGATTTTCTCTTATCATAATTAAGCCTCTCCAAGTTGTAGAAGTAAAAAGAGAATTATTGTTGTATTTCAATTGAATTAACTTTTTTTTAAATCCTTTTTTAAGAATATGGCTCGGATGGAATAGTGTCTGCTAGTGGCGATGTTTATAGTTACGGCATCATGTTGATGGAGGTTTTGACGAAAAGAAGGCCAACAGATGAAGAGATATGCAATGAAAATCTTGACTTGAGGAAATGGATCACACAATCATTTTCAGGGAGTATGATGGACGTTGTGGATGCCAATCTTTTCTCCGAGGAAGAACAGATCACTTCAGAAAGTGAAATCTGCATTGCGTCCATGATAGAATTGGGTTTAGACTGCACAAAGGAAATGCCAGAATCAAGAATAAGCATGAAAGAAGTAGTGAAGAGGCTTAACAAAATCAACAACACATTTCTGGAAACATAGTAGTGATCAGTATCTATTCCGGTGGTGTTCTTTTCTGAGCTGCAAAT

>M6-PGSC0003DMG400014281 M6 LRR-RLK

AAGCCAAAGGAGCATCTGAGTATCAACATTGCAACCTTTGAGCAGTCTTTGTTACGGATTAACCCTACAGCTATTCTCTCAGCCACTGAGAACTTCAGCAAGAGTTACATCATTGGTGATGGTGGTTTTGGTACTGTATACAAAGCCAAACTTCCTGAGGGCCAGAACATTGCTGTCAAGAGGCTGAACGGAGGCCACATGCACGGTGATCGCGAGTTCTTTGCTGAAATGGAAACAATTGGTAAAGTAAAACATGAAAATCTGGTGCCATTGCTTGGTTATTGTGTCTTTGCAGATGAGAGGTTCTTGATATATGAATACATGGAGAATGGAAGCCTTGATTTCTGGTTGAGAAACCAAGCAGATGCAGTAGAAGCACTTGACTGGCCAACTCGTTTCAAGATTTGTCTAGGGTCAGCTGTCGGACTTTCCTTTCTACACCATGGTTTTGTTCCACACATCATCCACAGAGACATCAAGTCAAGCAACATACTCCTAGACAAGAACTTTGAACCACGAGTCTCTGACTTTGGCCTGGCTCGAATAATTAGTGCTTGTGAGAGCCATGTTAGCACAATCCTTGCTGGTACATTCGGGTACATACCTCCCGAGTATGGACAAACCATGACGGCCACAACAAAGGGTGACATCTACAGCTTTGGAGTGGTGATGTTGGAACTGGTAACAGGAAGGGCACCAACAGGACAGGCTGATGTTGAAGGAGGCAATCTTGTTGGCTGGGT

>M6-PGSC0003DMG400014576 M6 LRR-RLK

TTGGTTTGTGTCTTGTTTTTTTTTGGTTTTTAGTTGCTTTTTTTGGCTGCAACTGTTAATAAGGACTGGTCTCTAATGCATATAAGAATAGAAACCCAAATTTGGATAGTTTGAAAGCATATTACTTTGCTAAAGATTGAGTAGTTGTTGGTGAGTTTGGTGTATGATTTCATCACTTGTTTTGGCTTATTCATGGTAAATAGTATTCAACCTTTTGTTCTTTAAGAAATTCATAGTACTAAAATATTTGTGAATTACTTGTTTCTTAAAGAAAAATGTGATCTTTAGCTTTTTGTTGTAGAATAGATTATTGATTGACCAATAAGGATTATAGTGGAGTTGTAAGTACTTCGTCATCATTAATCTGAGGTTTCAAGTTCGAGCCTTGAGTATGGACTTGCCCTTGTTAGGTAGCGGTTTGTCCTACGTGGGACTTCCTAGCGCGAACCCGTATCCGGCTCCAATGCGGTTGCCGTACACCGGATGGGAAAAAAGGAGAAGATTATTGATTGATTTGGTGGTAAAGTTTGGCAAGTAGTATAGCATAATCACATGGGGTTTGAGTCTGGCTTTTTGTCTTTAATTGCTTTTGGTCTGCTACTGTTAACCAGGACTAGCCTTCTGATGCAATAGGAATAAAGATGGGATGCCTCAATACTAGACAGTATTTGCTTTAGTTAATGTTTCTATTTACATAATTGAAACTAGCCATGTGGGAATATTGAAAATCTATGTTGCTCGGACTCTTTAAAAATATCGAGGGTGCGGCAACATGAGTAAGTCTTACATAGTCTTTTCAAGCTAAAACTATAGAATAGTATGTTACTTTATTAACAATCTTTGTCACATAAGCTATTCTTGATTTTCAAATGTAGGGAACTTAAATTATTTAACAATCTTTATCATCTTATTCTTGAAAAAGTTTTTAAAGGAATCTAATTTACTCATGATGGTCCTAAAGTGACAGAAGATTAGAGCAAAGGACTTGTTGTACCAGATGAATAGAATTATGTGTTCCCTCTGATTTTTGGGCCAAAGCAAAAGAAAAAAATTCACACTTTTTTGGCAAGAGATGCCCTACTGTCCCATTGTGCTCTACAGCTTTTTTGCTTATCTTTTTGACCTCACTTAGTATTCATATCTTGCAACTGTTTTGGTAGGTGGATAGTGAAATATAACCTCTTGTTTTTTTTCTGGTTCAAATGGTAAAGAATCAACTTCTTGTTGAATTGCACCTGTCATTCATGTAACAGGAGACAGCTGCTTCATATGTTAAATTTTCTCATTTTAGAATTATGAACAATTATGTGTTACTATTCCTACCATCAAAGTCTGCTATGTCAACTTCAAACTCCTTGTCCTCTCTGCCCAGGTTTTTGTTCTTTTGTCTAGTTAAAGTTGCATCTTTATCTTTCTATAACAAATACTCCCTCCCCTTCAATTTGTTTGTCTGGTTTTGACTTGGCACGGAGTTAAGAAAAGTAAAGAACACTTTTGAATCTTTTGGTCTTAGTTCTTAAACATGTCATGTGGAAAGTTGAAATTAATAGTTGCCAAAAATGGGAAAGGGGCAAATAAGTGAAAATGTGTAAGTGTTCCATTTTATAGTAGTTTGAGTAAATGTTTTGATGGACATGAATCTTCTTGCTACAGGTAACAGAGGGCCTGGAAGAACTCCTTTAGACTGGACAACAAGGTTAAAGATTGCAGCCGGAGCAGCTCGAGGGCTAGCCTTCATCCACAACTCATGCAAGTCCTTGAAACTCACTCATGGCAACATCAAATCAACTAACATCCTCATTGACAAGAGCGGCAATGCACGTATCTCCGACTTCGGTCTAGCCATCTTCGCCACACCATCTTCCGTCCCAAAAACCAACGGCTACCGAGCCCCTGAAGTCGCATTGGATGGTCGTAAAATAACTCAAAAGTCCGACATCTACTCATTCGGGGTCCTATTACTCGAGCTGCTAACCGGTAAATGCCCCTCAGTCGTGGACAACGGTGGTGGCCTCACAACCGGCTACGGAGGGGTTGTGGACTTACCGAGATGGGTGCAATCCGTGGTGAGGGAAGAATGGACCGCGGAAGTGTTCGATTTGGAGTTGATGAGGTATAAGGACATTGAGGAGGAAATGGTGGGGCTATTGCAGATAGCTATGGCATGTACTTCAACATCACCGGATCAAAGGCCTAAAATAAATTATGTTGTGAAAATGATTGAAGAGTTGAGAGGTGTTGAAGTTTCACCTTCTCATGACACTGCTACTGATTCTGTTTCTGACTCTCCTGCTGTTTCTGAAGATAATACATGTGGTGCAGCTAGTCAGTAGTGAGAGTTTTTTTTTTTCTACTAGTTTGTTATTCTGAGTTATGTAAATTAACCTCTTAGTTTTAATCCTTTTATCTATAAGATATAATTTAGGAGAAGTTTTATA

>M6-PGSC0003DMG400014917 M6 LRR-RLK

GTTCATTACTACTATTTTTATTTGTATCTCAATTTTGTCAAAAATAAAAAAATCCCCAACAAATGATGTGGAGGATATTCGTTTTTCTTGTTCTTTTACTCAATTCTCATACCGGATTTTGTCTAACTTCCGACGGACTTTCACTTCTATCTCTAAAATCCGCCATGGATGATGGCGGTGGTGGTACAGTTTTCTCCGATTGGAATGAAAACGATGATACTCCTTGTACTTGGACTGGCATTTCATGTGCTAACATTTCTGGTTCTTCAGAGCCACGTGTCGTTGGGATTACTCTTTCCGGTAAGAATCTCCGGGGATACTTACCGTCTGAGCTGGGAACGTTACTCTATCTCCGGCGACTGAATCTTCATGGAAATAATATTTACGGTTCTATCCCTGACCCTTTATTCAATGCGACGTCGCTTCATAGTATTTATTTGTATGATAATAATATCTCTGGTATACTTCCTCCTTCTGTTTGTAATCTTCCTCGTTTGCAGAATTTGGATATTTCTGATAATTCACTTTCTGGAACTTTCTCTAAGGATCTACGTAATTGTAGGCAGTTACAGAGGCTTATTCTAGCTCGGAATAAATTCTCCGGTGAGATTCCGGTGGGGGTGTTCCCGGAGTTGGCGAATTTGGAACAACTTGATCTTTCGTCGAATTTGTTCAATGGTTCGATTCCTGAAGATATTGGTGAATTGAAGTCACTATCTGGAACTTTGAATCTTTCTTTTAATCATTTTAGTGGAAGAATTCCCAAATCTGTAGGTGATTTACCTTTAACTGTTAGCTTTGATCTAAGGAATAATAATTTATCTGGTGAAATTCCTCAGACTGGTTCTTTTGCTAATCAAGGTCCGACGGCTTTTTTGAATAATCCCTTGTTATGTGGATTTCCTTTACAAAAAAACTGTAAAAATAGTTCAAACAATTCAACACAAGTTCAGGGTTCGTCTGATAACGAGGGAACGAATTCGCGCAAGGGGTTAAAACCTGGATTCATTATATTGATATGTTTGGCTGATGCATTTGGAGTGGCATTTATAGGACTAGTGATTATTTACTTGTATTGGAAAAAGAAAGACTCTGGTGGTTGCAGCTGTACTGGAAAAGGGAAATTTGGTGGAAATGAGAAGAGAATGCTTTGTAATTTCCCATGTATTAATGGATTTCCGAATAATGATTCGGAAGTTGAGTCTGAGAAGGGAGGTGGCGGTGGGGCTAGTGGTGGTGGTGTTAGTGGTGGGGAGGGTGATCTTGTGGCTATTGATAAAGGTTTTAGCTTTGAGCTTGATGAGCTGTTGAGGGCTTCTGCTTATGTGTTGGGAAAGAGTGGGTTGGGGATAGTGTACAAGGTTGTGCTTGGGAACGGTATCCCAGTGGCAGTGAGGAGGTTAGGGGAAGGTGGGGAACAGAGATATAAGGAGTTTGTTGCTGAGATTCAGGCAATCGGAAGGGTTAAGCATCCTAATGTTGTGAAGTTAAGAGCTTATTATTGGGCTCCTGATGAGAAACTTCTCATTAGTGATTTCATTTCCAACGGCAACTTGGCTTCTGCTCTTCACGGTGAGCTTTTCATTATAGTAAAACATCTTCAGTTTTGTTTGTCTAGTAGATATAACCTGTTGAAAGAATCATTAACACATTCATATTTTAGATATCGGGGTATACTATATTCTTATCATTTAGTAGCATGATACACTGATGGATGTTTCGTACTTTCCAAGATCGATATGTGTAAAGTGTAATTCTTTTGCTGGATTAGTATTTTACTTATTGATCACTCTTTTGAGGAGTGGAAATACAGTACAAGACTTTTGGTATTAGGTTAGTTAGGAGTTTGGGATGTTCTAGTATTTATTCACTAGTATAACTGTTTAATATATTCCCGATGACAAAATAATGACTTTTAGCTGTTGTGTGTTCCTTGATGATTTTGAATAGCAATTATTTATATGCCATCTAGTGATAATTTTATATTGAATGTGTCCCATGTGCTTGCAATCCTTGAGTGAAGGATGAGTTTTGTTTGCGACTTGATAATTTGCGTAGGATTTTTGTTTAATTGTATCAGCTGATAGGATAGTCTTTATTATGTATCATAGATGAAAAGTTATCTGCATAAAGGGTAAACCAACAACCTAACATTTTCCATCCATTTCAATGGATGTTAAAGCACATAGTAGTATATTAACTTATAATGATCCAGCTTAAGTCAATTTGCAGACTTTAACGATTTACTGCCGATGCAATAATCATATGTTGACATCATATCGAACCTGGTAGAAAGGGCATTTGGGCATGTTTTGAAGAATTTAGTCTCTCTTTTTTCATGTGACATATGGTTGACTGCCCCGATAAACCTTGCGGAAAATGATTTATCATCAATTAGAAGCAAACCAATAAGTACAAGGGGCCAAGAACTGAATTATGTCATGTACTAGGTCAATTCCTTAAGTTGTGGAGTTACGTATTCTGTCTCTCAAAAGAGTCTAGATATTACATTGTTTTATTATGTCTTTTAGAGATGTATTACTGATTTGCATATGCTAATATTCAAGAATTTTCGAATCGCTCCGCCTGGTGCAGTTTGACTTTACAGATGTTTACTTTTTCGAAGCAAACTGGAATCTGCAGTATCTGGTCTGGTAACTACCATCTAGCATGAGCGGGACTTATACTCCCTTGAGATGGATTCTAGACTTCGAATTTTTAAAACTTAGTTTTCTGCAATTGAATTTTGCACTGTTGCTGTTGGTATAGCTGCTTGAATGTTACATTTTTAAAAAGAATTAGATTTTTTCCCCTGATACAAATCGTTGTTTACATTTATGAATATCTACTTCACTGCCTCATATACATGTCACTATGAATCTATAACACATTCGTTACCCAAGAGTAACTGCTTGCTTTCTGAGACAAATATATTTGATGACCTTTGGTTCTATGGTACCTGATGTTCCCAGTTTCTTAATTTACCTGTTTTTAAAATTATGTAGCTGGAATTTGACAGTAGAGCAACTTACCTTTTGCATTCTGAAGAAGTTGAGGAGTGTTTTCTTTCAGCATCTCTTTTTCCAAATTGTTTTGCCATGCTAGATCTGTATGTATTGACGTAGTGACAGTAGTTTCATGCTCTAGCTGATTGAGATTAGCCAAATTAGTGTGAACACCGATTTGCCTATTGAGCTGAAAAGTTATTAACTTGTATCTTGTTCGTCATACTGTGGTAAATGTTGTGATGACAATCTCGTTTGCCTTTATACAGGGAGAAATGGGCAACCATCACCTAGTTTAACATGGTCAACGAGACTGAAAATTGCTAAGGGAACAGCGCGAGGTTTAGCCTATCTCCATGAGTGTAGCCCAAGGAAATTTGTCCATGGAGACGTCAAACCTTCCAATATTCTCCTTGATACTGAACTGCAGCCATATATATCTGATTTTGGCCTCAATCGATTAATCAATATCACGGGAAACAACCCCTCCTCTTCTGGTGGATTTATGGGTGGAGCTCTTCCTTACTTGAAGCCTGCTCAACCAGAACGTCCCAATAACTATCGAGCACCAGAAGCTCGAATCACTGGTAACAGACCCACCCAAAAATGGGATGTGTACTCATTCGGGGTGGTCTTGCTTGAACTGCTGACTGGAAAGTCTCCTGACCTCTCAGTTCCTACAACATCCACTTCAACGGAGGTTCCAGATCTTGTAAGATGGGTAAGAAAGGGATTCGAGGAGCAAAATCCACTGTCGGATATGGTAGAACCAATGTTGCTTCAAGAAGTCCATGCTAAGAAGGAAGTACTAGCTGTATTTCATGTAGCTCTTGCATGCACCGAGGCAGATCCAGATATTCGACCAAGGATGAAAACTATCTCAGAAAACATTGAGAAAGTTGGAGCTTAAACTTTTCGGTACACTCATGTCAAGGAGTTCCTGAGTTTCATGGACTTCTTTAATCCTTTTTTTTGCAGTAATTGCTCCTATTATTTTTTTTCTTTTAGAGGTTAATCAAATCTTTACTTAACTACAGTACAAAATACTAGTGTGCTGCAACAAAATTCCTGTCATTTTTGAGAAGTTAATTAGTTGCTTATCTCTTTAATCTATTTCTGTAATTCTGATGTTAAAACAACTTTTCATGGCCAA

>M6-PGSC0003DMG400015081 M6 LRR-RLK

CACAACAAGTTGCTAGGATCTATTCCTGACTCAGTGAGCAACATGGTAGGTTTGGAATTCCTAGACATTTCTCATAATAATATATCAGGAACCATTCCTATGTCTATGGAGAAACTTCAAAACTTGAAGTATTTCAATGTTTCTGACAACAAATTGTATGGTGAAATACCCTCGGGGGGTCCTTTCAAGAACCTCTCGAGTCAGTTTTTCATCAACAATGAAGCATTGTGTGGTTTGTCAAGGTTTAGTGTCCCGCCATGCCCCACTTCATCAAGGCATAGATCAAATAGGAAACAAATGCTACTTCTTTTTCTTTTGCTTGGAATTGCACTTGTATTTGTTCCTATCTCCTTTGTGTTCCTATGGATAAGGTATAGAAGAGGTAAAAGATCTCCTCAACAAGCTGATTCATTGTCTATTGCAACAACAGAAAGAATTTCATACTATGAACTACTCCAAGCAACTGATTCGCTTAGCGACGGTAATCTAATTGGTTCTGGAAGTTTTGGCTCTGTTTACAAAGGTGTTCTCAGGAGTGGGACTGCCATTGCAGTTAAAGTGTTCAATCTGCAACTGGAAGCGGCATTCAAGAGTTTTGATATAGAATGTGAAGTTTTGCGTAGCCTTCGCCATAGGAATCTTGTGAAAGTCATCACTAGTTGTTCCAACCTTGATTTTAAGGCTTTAGTGCTCGAGTATATGCCTAATGGAAGTCTTGACAAGTATTTGTATTCACACAACTACTTCCTAGACATTAGGCAGAGACTAAGCATTATGATAGATGTAGCATGCGCATTGGAATATCTTCATCATGGATGCTCCTCTCCTGTGATTCATTGTGACCTGAAGCCTAGTAACATCTTGCTGGATGAGGATATGGTTGCCCACCTAAGTGACTTTGGTATTTCAAAACTGCTTGGTGAAGATGAGAGTGATTTATACACAAAAACCTTAGCAACATTTGGTTATATTGCACCAGGTACGTCTCTTTGTTTTTGCTAATTTAAACATTGATTTTTTGTCTTTTTTATTTCCAATCAATAAAGTATATTGCATCTTTAAATCAATTGTTTGACTGTAGAGTATGGACTGGATGGACTGGTGTCAATAAAATGTGATGTCTATAGTTACGGGATTATGTTGCTGGAAACATTTACTAGGAGAAAGCCTAGTGAGTTTAAGGGAGATCTTAGCTTGAAGCAATGGGTGAGTTATTCACTCCCAGAGGCAGTAATGGACGTTATGGATGTCAACTTGGTTACACCAATGGATAATCGCTTACAGAAGGAGTTAGATATCGTGGCATCAATCATGAAAGTAGCATTAGATTGTTGTGCTGAATCTCCGACAAGAAGGACAAACATGAAAGATGTTGTAGGGATGCTACAGAAGATCAAGATTCAACTTCTTGCATGCTGAGCATGTCCTCTGTATCTCTTGTTCTGAATCGATCACTTTTTTTTGTTTTGTTTTAGCACTTGATTTGTGCATGATTCTGTTTTCAAAAATG

>M6-PGSC0003DMG400015157 M6 LRR-RLK

TTCAGAACCACCAAGCAACTAAAAGAGACATCCTACTCCTAAAGCACCACCAGTGTGGTGCCTCATCCAACTCCCAATTCCACCATAATACTGCACCAATCCGATGGCCTTCACAGCCTCACATATACACCTTTCTTTTTTGTCTTTTTTCACTGTCATCCTTCTTGTTCAAGCAAAACTCAACCTTTACTCACCTGATCACAGTGCTCTTTTGCTTGTCCAAAAAGGCTTAGGCATCCGTGCTCACCGCAATGCTCTTGAGAACCCATGCAACTCTGTTGGAATAACATGTGAAAAACGACTCACAAACAATTCATACGTGCTAAGAGTCACAAGGGTTGTCTTCAAATCCTACGGGTTGAAGGGTACTCTGTCTCCTGCCATTGGGAGGCTTTCTGAGCTCAAAGAACTGTCCCTCCAAAACAACAAACTCTTTGACAGAATACCAACTGAAATCGTTGACTCGAGGAAATTGGAAATCTTGAACCTTCAAAACAATCAATTTTCTGGCAAAGTCCCACCTGAATTATCATCTCTAGTCCGCCTTCGTACCCTTGACCTTGGCTCTAATGAGTTATCTGGGAACCTGAACTTCTTGAAATACTTTCCTAACCTTGAAAAACTCTCCCTTGCTGATAACATGTTTACTGGCAGAATACCTCAATCCTTGAAATCTTTCAGAAATCTCCGGCTCCTCAACATTTCAGGCAATAGATTCCTTGAAGGTACGGTGCCTGTCGTGAGTCAAGTAGAGCACTTATCAGCAGAATTAAATCGAAAAGATTATGTTCCTAAGCGTTACATTCTTGCTGAGAACGAAAGAAGGCCAAGGCACAGCCCTGCAATGGCACCAGAGTCCAAATCAGGATATGCCCAAGCTCCAGGAACTAGTACAGTTGTAGTACCAGAAAAAAAAGACAAACACAACAAGATAAGGAAGGTAGGCGCCTGGATTCTTGGATTCTTTGCTGGAGTTTTCGCTGGGGGGTTATCGGCGTTGGTCTTCTCCGTCCTCTTCAAGGTGCTAATGTTTTTAATCAGAGGGATCAGAAATGATTCAGGCTTAACAATTTTCAGTCCATTGATTAAGAAAGCCGAGCACTTGGCCTTTCTGGAGAACGAAGATGGATTGGAATCGCTAGAACTCATTGGGCAAGGTGGATGTGGAAAAGTTTATAAAGCTGCGTTACCTGGAAGTGACGGAAAGATTATAGCCATAAAGAAGATCATACAACCACCAAGGGATGCTGCAGAACTCACTGAGGAAGATAGCAAGGCAATGAATAAGAAAATGCGCCAGATTAAATCAGAAATCAAAATTGTAGGTCAAATCAGACACCAGAATTTGCTTCCCCTACTGGCACATATGCCAAGACCAGACTGCCATTACTTGGTCTATGAGTACATGAAAAATGGGAGCTTACAGGATACCCTCCAGCAAGTCAGAGAGGGGACACGGGAACTAGATTGGTCGGCACGTCACCGAATTGCAATGGGAATAGCTGCTGGACTCGAGTATCTCCATATGAACCATACTCAGCGTATAATTCACAGAGATCTAAAGCCAGGCAATGTCCTCCTTGATGATGACATGGAAGCTCGAATTGCAGATTTTGGCCTTGCAAAGGCTGTACCAGATGCTCAAACACATATTACAACTTCAAATGTGGCAGGAACTATAGGATACATAGCTCAAGAATATCATCAGACACTAAAGTTCACTAATAAGTGTGATATATACAGATTCGGGGTGCTGCTAGGCGTGCTAGTTATGGGAAAGCTTCCATCTGATGAGTTCTTCCAGAACACTTCTGAGATGAGTTTAGTGAAATGGATGAGAAATGTCATGACTTCTGAGGATCCAAACAGAGCAATTGATCCAAAGCTGATGGGTAATGGAAATGAGGACCAAATGCTTTTGGTTCTCAAGATTGCCTGCTTTTGTACCCTGGAGAATCCTAAGGAGAGGCCTAATAGTAAGGATGTTAGGTGCATGTTGATGCAGATCAAGCATTAAGTATGTAGTGGATAACTTACAGTAAATAAAAGTTTGTAATGAAACTAAAAAAATATTTCTTGTGTATGACGGTCCTTGTGTATTCTTGTTACATATCCAATATGAGAGACTGTCCTTGTGTGTTTTGTTTCCTCTCCAAAATGAGAAGTTTATATTTAGCTGTATGAAAGTTCTTTGCAGAAGTTTCCTCTGCTATATTATGATACCTTAAAGCATGGTACGAAATGTGAATCGAATTCTTTCCACTAGA

>M6-PGSC0003DMG400015162 M6 LRR-RLK

TTATGACACCTTGGTCCCCACATTCCACACTCACTCTTTCCTTCCCCTTTAAACAGTCATGCATTTACATAAAGAAAGTGACTGTTTTTTCTCTCTCTAAAATTTCATCTTCTTTCATGGCTTCTCGCCGGAAAATCAAGAATTCCCGCCGGGAACCACATTCTTGATTCGCATTCTTCTTTAAAAGGGTCGGAACAAGAAAAAAAAGGGTGTTTATAAATGGCGGTGTTTTTAAGGTTTGTTTTTTGTGTTTTATTCTTTGCTTTACTCGGATTGAGTCGTTACCGGGTTTTTTCAGAGCCTACACAGGATAAGCAAGCTCTCCTTGCTTTTTTCTCTCAGATCCGACATGCGAATCGGGTCCAATGGAATTCTTCTGCTTCTGTTTGTACTTGGTTCGGAGTTGAATGTGACCCGAATAACTCTTTCGTTTATTCACTGAGACTTCCTGCTGTTGGTCTCGTCGGAGAAATTCCTTCTAACAGTTTGGGCCGGCTGAGTCAACTTCGAGTTCTTAGTCTTCACGCCAATCGTCTATCTGGTTCAATTCCTTCGGATTTTTCTAACCTTAAGCTTCTTCGTAGTCTCTATCTTCAAAAAAATGAGTTATCCGGCGAGTTCCCGGAGAGTATTCCCGGGTTGACCCGGTTAAACAGGCTGGATCTTTCGTCGAATAACTTCACTGGTACTATCCCTTTCTCTATCAATAATCTGACCCATTTGACTGGACTTTTATTGCAGAACAATAGCTTTACTGGTACTCTTCCTAGCATTAATCCGCCTGGTTTAGTGGATTTCAGTGTTTCGAATAATCAGCTAAATGGGTCAATTCCTACTGCACTTTCCAAATTCCCAGCTTCCTCTTTTGCTGGTAACATTGATTTATGTGGCGGCCCATTACCCCCATGTACCCCATTTTTTCCTTCTCCTTCACCTTCCCCTGAAACAGAACCAAAAACCCCTCCGTCAATTAAGAAATCCAAGAAGCTTTCAACAGCCGCCATTGTTGGAATAGTTATAGGTTCAATAATTGGGGTTCTCTTACTCTTGTTACTCCTTTTCTTCTGTTTAAAACGAAGAAAAACCGACACATCAAAGGTACAAAAACCTCCAGTAGCATCAAGAGCTATCGGGGCAGTAACAGGGGCAGCGGCTGAGGCAGGGACATCGTCGTCTAAAGACGATTTAACCGGTGGTTCAGGTGAAGGGGAAAGGAATAAACTTGTGTTCTTTGATGGGGGTGGGTATAGTTTTGATCTTGAAGATTTGTTAAGAGCTTCTGCTGAGGTTCTTGGAAAAGGGAGTGTTGGAACAAGCTATAAAGCTGTTCTTGAAGAAGGTACTACAGTTGTTGTTAAAAGGCTTAAAGATGTCGTTGTGCCCAGAAAAGAATTCGAGCAACAATTGGAAGTAATAGGGAAGATGAAACATGAAAATGTGCTACCTTTGAGAGCATTTTACTATTCAAAAGATGAGAAATTGTTGGTATCTGATTATATGCCTGCTGGAAGTTTATCAGCTCTTCTCCATGGTACGCTGCTCTTATTATTTTTAGTTAAAACTTGTTCGTTTTAGTGTATAATTTGCTCTATTATATGTTACTGCAATTCTGTGTCTAAAATTTGCTCTCAACACTTAAGTTCCATTTTTCCACCAATGTTTTAAAACTTAAATGGCAATAAAGATAGTAGCATTCAATTCTTGCTAGATTTTAGTGACTTTGTTCTGATTCTTGAAGAGTCTTGAAGGGGGATAGGTGGGTTATAGGTATGAAAAAGATTGAATTTTTATTTTGTTTCTTTTTTTGGAAGAATAGATGGGGCTTCAATCTAATACACGTGCAGATCATGTACACATTGTAGCCATAGTTTTACAGCCCTTCTGAACTGGCATTGCATTATTGTTTCACTCAATTATTGAAGTTTGGTACCACTTAATTTACTTCTTTAGTACATTTATTGTCAATTCAACCGCAAATTGCGCATTTTGTTTCTATAATTCCTTAAGCTACAGAACATTCTTTTTTGAGAGAAATGGAACAATGACATTTTTATTATGATTTATTCGAATCGAGAGCATATCGGATACAATCTCTCTATCTTAACGGACACACTTGTTAGATTACACTTTTACTCTTATTTTTTTTATGAAACGAACGTTACAATGCAACAGACCACCATGATGGTCAACATACCTTCCTTTATAAATGATGAAATGAGTTTAAAAAGTGGAAAAAGAAGGCTTCTTTCTTGTTCAAAGATTTTCCTTTTATTTTGCTACATTGCTTGCGTCTATTTATCAACAAATAACTTGTATTATTATAGTTATATATATTTTAAAATATTTATTATACATAGGGGAAATGGGGGAGGGAATAACAAAGTGTTGTGGAGTTTGGTGATATCAAATATATTAATTAAATATTAATCTTTTGTTTTCTAACCCCACAATTTTCAAATCATGATTTAATTAGTAGTAATTGCTGTAAGTCTAAACAATACTTATATACTTAAAAAGCTTATATTACTTTGGCTTTAAGCTATTTGGTGGTTATAACTGTCTGATTATTATTCATCTAATCTTGATTCTTTTGCTTTACTTTATTCATCATTGTTTCTTGCTGTCCCCCATTTTTCTTCTTTTGTTGAAATAACGTGCTGCTTTACTCTATTGTCCATTGATTCTTGCTGTCACCCATTTTCTTCTTTTTTTGGTGAAGTAAAGTGCAAAAGAAAAGCTAATAATACAATGTGTCCTGTTGAGTCTTGAGTAGTTCAACTGCTTTGAATTTGAATCCAACTTATTAATTTGCATTGACATTAGTTTGTTAACAACATAACCAGTGTAATCCTACGAGTGAGGTTTGACTTTTGAGTAGGGTAGGATGTACACAGACCTTATCCTTACCTTTGATTGTCGTTGGTTTGGTTGTCTAATTAAAATGTTAACTTTTGGTTAATTCTTGTAGGTAGTAGAGGATCAGGGCGTACGCCACTTGACTGGGATAGTCGAATGAGAATAGTGTTAGGTGCAGCAAGAGGCATTGCGTACCTCCATATTTCAGGAAAAGTTGTCCATGGAAACATCAAGGCTTCGAATGTTCTCCTTAAACAAGATAACCAAGATGCATGTGTATCTGACTACGGATTGAACCCTCTATTCTCCACCTCTGCACCAGTAAACCACCGTGTTGCAGGATATCGTGCACCTGAGGTTCTTGAAACTCGAAAAGTAACATATAAATCCGATGTGTATAGTTTTGGAGTGTTAATGTTGGAGCTCTTAACGGGGAAAGCTCCAAACCAAGCTTCACTAGGTGAAGAAGGCATTGATTTGCCTAGATGGGTCCAAAGTGTTGTAAGAGAAGAATGGACCGCGGAGGTGTTTGATGTTGAGCTAATGCGATACCATAATGTGGAAGAAGAGATGGTACAACTACTTCAGATAGGTATGGCATGTGTTGCTACCATGCCAGACCAACGACCTGCCATGACCGAGGTGGTTAAGATGATCGAAGAAATGAATCGTGGTGATACAGATGATGGTCTTAGACAATCCTCCGACGATCCTTCTAAAGGTTCTGAAGGCCAAACACCTCAGGAGTCTAGAGGTTCACCTCATGGTGTAACACCATAAAATGAAATTAATCATAAAAATGGATGGCTTGATTTGTCAAAAATTCTCAGAAAAGCCTAAAGTAAATTGTATTAGATTCTTAGGGGGGGAGCCTTTTATGTGTTTTTATTTTATTTCATTTTTTTTTTCTTCTTGAAATTATTATTTTGAGAAAATTTGTGGAATGATTTTTACAGATGGTTAGTTGAGAATTTGAATATTTCAGTGAAGTGATTGGGAAATTGAAATTTG

>M6-PGSC0003DMG400015276 M6 LRR-RLK

CTATAACTATTTTTTCAACTTTTTGATGTGTTTGACATTGTTTGAATTATACCCCCAATGTTCAATTTTACTTCACCTACTACATACAACTGGTAAATTCTCAATCTAATATTACTTATCTATTATTCTTCTAAGTCAAGGTTGCCACGACCAGAAATGCGTAGGACGATCCTATTTTTCGCTTTGTTTCTGGTGGTGGAATCAGCTGAATTTTTCCCAAATGAAAGAGATGCTTTGTTGCAGCTGAGAGATAGTGTAAATTCAAGTGTTCTTGATTTACATTCACATTGGACAGGGCCTCCTTGTTACAACAACCAGAGTAAATGGGCTGGAATTTCGTGTTCTAATGGGCATGTAAATCAACTTGTTTTAGAAGGAATTCAGTTAACTGGCTCACTTCCACCAACATTCTTGTTGCACATAACTTATTTGACAAAAATCAGCTTTAGGAACAATTTATTGTATGGTCCTGTTCCCAATCTCACAGGACTAATCTACCTGGAGTTTGTTTTTCTGTCAGATAATCGATTTTCGGGTTTGATTCCATTTGGGTACACTGAGTTAAAGAAACTAACAGAATTAGAGCTCCAGGGGAATGATCTTGAAGGTTCAATTCCACCATTTTATCAACAGACTTTGATTGGTTTCAATGTGTCGTCGAATAAACTGGAAGGACCTATCCCTGAAACTCCTGTCCTGAAGAGATTTTCGAAGAGTTGTTATGAGAATAACGCGAATTTGTGTGGTGAAGTTGTAGGAATACCTTGTTCAATATCACCTGCAAGTCCACCTAAAGCACCAAGTGAAAAGAAAAAGAATGGCCTTAAAGTATGGAGTATTGCTTTGATTGCAGCAGCAGTAGCAGTACTTGTTCTAATTTCAATCATGTTCATTTTGTTCCGTTACTTTAGAAGATCTCAAAGAATGGAAACACAAGGAGAAGACAAACAAGGTAACAGTTTTATTGTATCATCTATGTTATCCTAGCTAGTGCCAGTTATTCTGGGGGTAAAGTTATTACTACTTAGATTGAATTAGATGATGAGTTTATCTGGTGAGACACATTTTTCGAAGGAGAAATATGTATTTTTTCGATATGGTTTATGTTAATCGCGAAAAGGTCATTCAGAACTCGTTGGCTCCAGAGGCATTAAAGATGTTTGTGATTGGGAAAAGGTCATTCAGAACTTGTTGGGGCTGCTCCAGAGGTGCTAAGATGATTAGTTCATGATAGGAAGACGACCAATCCATGTCCATTATGAAGTCGAATTCAACCATTTCTAGCTCAATAAGATGGGATTTAGTCTCTCCCCCCAGAATTGTGGCTACACAACAACAATAACATACTAGTATAGACCCATAAGTGAGGTCAGGGAAGGGTGAGGTGCCCCTACTTTTGTAGAGTAGAGAGACTGTTTCCGATAGAATTGTGGCTATACAACATAGAACTAGATTAGCTAATTCAATTAAACTTAAAGATCCATGTGATGAAGGAGCAGTTTATATTGTTATTATTTCCTCTTTTTAACATTTCCAGTTTCTATTTTGACAGGTAATATTTCAATAGAGAATGTGGTAAAGAGAAGTTATTGGTCCAAGAGTACAGAAGATCCAGAAAAAACATTGGACTTAATATTCTTTGAAAAGGATATGTCAGTCTTTGACATGGATGACTTGTTAAGAGCATCAGCTGAAGTTTTAGGAAAAGGAAAGCTAAGCACAACATATAAAGCAATCCTGGAATCTGGTTCTGTTGTTGCTGTTAAGAGACTCAAAGAAATGAATTCTTTGAGCAAAAAGGAATTCACACAACCGGTGCAGTTGCTTGGGAAATTGAAACATGAAAACCTTGTGGAGATGTTGTCCTTCTACTATTCCAAGGAGGAGAAGCTCATTGTTTATGAATATGTTCCACAAGGAGACTTGTTTGGGCTCTTACATGGTTGGTTCATCAACTTTCTTTTACTTTTGGGAAAATGTCTAAAGTTAACCCCTCCTGAATGAAATACCTTAATTTTATCTTTTGTTATACTTTGAGGTTATTCATATCCTTATCCTTAGTAAAACGTCTCCTGTTATACTTCGAGGTCATTCTGATTCTAAACTATGAACTCATGATCAAATGACTTTTGAGGTGGGCTTTCCCTTCTCTATAGTGCTACTGTGAAAATTGAAATTGCTTAGAGGTAGGTGTAAGGTCTGCGGACACTCAACCCCTTGTGGGATTATACTGAACATGTTGTTGTCATCTTATAGTAGTAATAGAGGAATATCTTTTACCATTTTCCCATCTTTTTTCCCCTTAAGATTCAACCTTGTAAAAACTTATGATTCTTGATTTCAGAAAACAGAGGAATTGGGAGACAACCACTAAACTGGACAGCAAGAATATCCATCATAAAAGATGTAGCAAAGGGTCTAAATTTCTTGCACCAGTCCTTACAATCACATAAGGTGCCACATGGGAACATCAGATCAAAAAATGTTCTAATCCAGCAAGACCAAAGCAAGAACTACCATTCTAAGCTGACAGATTATGGATTCTTTCCTTTGCTACCATCTAAAGAATCCTTAAGAAAAATAGCAGTAGGAAAATCTCCAGAGTTCTCTCAAGGGAAAAAGTTAACAAGAAAAGCTGATATCTACTGCTTTGGTATTCTTCTGTTGGAAGTTATCACTGGAAAAGTACCAGGTGAATTGTTTTCACCAGAAAATGGGGTTACTAGTTATGTTGATGATGATTTGTCAGAGTGGGTAAGGACAGTGGTGAATAATGATTGGTCAACAGACATTTTGGATATGGAAATACTTGCACAAAAAGAAGGGTATGATGAAATGTTGAAGCTTACTGAATTGGCTTTGCAGTGTACAGATGAGGCACCAGAAAAAAGACCTAAGATGAGTGAAGTGCTGAGAAGAATTGAAGAGATTGAGCTTGAACCAAAGAATTTAGTCACTGATGTGGAAGATGATTCAGTCTCTTACTTGAAGACTTGAACTTTCCCTCCTAAACTGTAATCAAGCCGTAGCAACAACAACAATATATCCAGTATAATTTCATCAGGTATAGCCTGAATAATATATTCAGTATAATCTTATCTACTGGAGTCGGATTCTGAAGAGAATTTCAAGTTGTGTCTGAACAATACATCCAATATAATCCCATCAAATGGAGATTACCGTAACCCAATCCTAACATGGGAGTAAAGAAAGGGTAAAATATGCACATACTTTACTCTGATTCCAACCGATGGGATCTTTGAGAGAATAGAGTATGCGCATATCTTACCCAACAAGGATAGAGAGGCTGTTTCCTAAATTGTAATCAAGAAAATAGATTAGAAATTTAGGAACAGGCTTTTAG

>M6-PGSC0003DMG400015391 M6 LRR-RLK

CAAGAATTTGGTGTCTTCTAAGTGCATGACAGCTATAATCTTTTTTATGCAGGGATATATCATACAACAATTTCACACAAATATCTACTTCAGTATGCGATTCCTCTACCTTGTGAGCATTGTTCCTCTTCATTTGGAACCTACATTCATCTACTATTTCTTTTTTTTTATAAGTGAAAATTTTGGTCCTTTAACTACATTACTTACAAACCTTCCCTCTTTAACAGAAACTTAGCTGCCAGCTATTCAAATACAATAAACACGCTCAACAACTCGTAAGTCCTAAATGTTGTCTTATGCATCAACTGCTAGTTTGTTTGATGCTTTCTGTCTATTGTCTATAATTATGTGGATTAATCTTTTTGAAATTGTAGGGATGCCTGGTGTTCTATGAAACCCCTCATTTGTCCCACAGAAACGAAATGTAAGAGTTACAGCCTATAAACCTTTTTTTTATACTTGGAAATTCTTAACTTGACCCTTGAGAGGGCCTGAAGTAGCAGTTTGTCAAAGGACCTCAGGGGTGGCAGCCTGTGCACTTTTAAGTACAAAAGCATAACCTTGGAGTTGAGAGATAATTTCACAACTCTTCCTCATGTTATGCTCTTTATGTCAATTGCAGATACTTCGCTATTCATAAACTGTGGAGGACCTAGAACTTCATTTGAGGGAAATGATTATGAGGGGGACACGACCGACAGAGGTCCATCATATTTTTCTTCACCCTCCGACAGATGGGCTTTCAGCAGTTCAGGAGTGTACGTGGGTCTTCAAGCAGCCAGTTATGTTGCAACAAATACATTTTCACTGAATGTGTCCGGTCCTGACTTTTACAATACAGCACGCCTTGCCCCTAATTCTCTCAAATATTATGGGCTTTGCCTACAAGGAGGTAGCTACAGAGTGCGACTTCACTTTGCTGAAATAATGTTTTCTAATGACTCAACATATAGCAGCCTTGGAAGGCGGATATTTGATGTAGCAATACAAGTAAGTTTGAGGCCTAACTTCTTTTGTAAATAAGGAGTATGTAATCTAAAATGTTGATTTATTTAGTTCTGATATGATTGGCAACTTTCTGATTTAATATAGAAGGTACCAATTTGTTGGAGATAATGTGGTATATTTTGATTTCTTACACAGGGGAGAGTAGTTTTGAGGGATTTCAACATTATGGAGGAAGCTAACGGTGTTGGAAAAGTCATTACCAAGGACTTTGCTGATGTTACTGTTAGTAGCACTCTGGAGATTCACTTGTATTGGACCGGGAAGGGGACTAACGCTATTCCAGACAGAGGTGTATATGGACCACTGATTTCAGCTATCACAGTGACACCAAGTCCGTATATCTCTGCTTATAGACTTTACACTTATCTTCCCACAATATTATCTTAATAAAATCCCATTTTCTTCTTGCAGACTTTGATGTTGATACAGGCAGCGGGTTACCTGTCGGAGCTGTTATTGGCATTGTACTCGCTTCAATTGTGGTGGTTCTGCTAGTTTTATTTGCTCTGTGGAAGAAAGGCATTTTTGGAGGGAAAAATAATCAAGAAGAATTAGGTAATACATACTTAAAAGAATTTTTTCACTCTTTCCATTTGATAAAATGGGGATTCTTCTGTTGTGATAAATAGAATTTTTTACGAGATTTCTAGCTGCGACAGACAAATAATTATCTCCCCGTAATTGACCAGCATTGAGATGATTGTTCTGTTGGTGGATCGACATCATGACATATCATAGATAGTTAAACTGTCTTCGTCATACAGATGTTCATTCTTTTAAAAATTGTTGTTCATCATACAGACGTTTTAGAAGATCTATATATACTGACGTTTGCAATATTCTTCTCATTTACAGAACTCAGAGCCCTTGATTTACAAACAGGTCATTTTAGACTTAGACAAATTAAAGCTGCTACAAATAACTTCGATCCTGCCAACAAAATTGGTGAAGGAGGGTTTGGACCAGTTTACAAGGTATCAATTCTCTATCCATACTGATTTTGCATAAGGACAGCGGGTCTTTGGTGATATTTCCTACTGGTTATCCTTGATGTCACGTTTATAGGAAAATAGAGTAGTAATAGTTGAAGTAACCCGTATCCTTCTTCTGTCTGTAATATGATCGTTGTGAATTAATCTTTCTATCTGTAGGGAGTACTCGCCGATGGTGCCATCATAGCTGTTAAGCAGCTGTCTTCCAAGTCGAAGCAAGGAAATCGTGAATTTGTCAATGAGATAGGCATGATTTCAGCTCTACAACACCCAAACCTTGTCAAGCTTTACGGTTGTTGCATAGAAGGGAACCAGTTATTAGTGATATACGAATACATGGAAAATAACTGTCTTGCTCGAGCTCTCTTTGGTAATGCTTTGCAACAACATTTTAAGAATCATTAATTAGGCAGTGATAGTTTATGCTTTACATTCCTCTAATTCTCGTTTCTCAGGCCGTGACGACCAGAGGTTGAACATAGACTGGGCAACCAGAAAAAGGATATGCTCAGGAATAGCAAAAGGGTTAGCTTACCTTCATGAAGAATCAAGGTTGAAAATCGTACACCGAGATATAAAGTGTACCAATGTGCTGCTTGATAAGGATCTGAATGCTAAGATTTCAGATTTTGGATTGGCTAAATTAGATGAAGAAGAGAACACTCATATTAGCACCCGAATTGCTGGAACAGTGTAAGCCTTCTTATCATAACTGTCTCTGTACTATCATGTGATGAGAGAGTTACAGTTCTGTGCGTTTAAGTTTGTTCATCATCAACATTACTATCGTTACTTTGAAGTATCTTACGTTTTGTTTTCTTTTTTTGAAGAGGTTATATGGCTCCGGAGTATGCAATGAGAGGCTACTTGACAGATAAAGCTGATGTTTATAGCTTTGGAGTTGTTTTATTAGAGATTGTCAGTGGGAAAAGCAACACAAACTACAGGCCAAAAGAGGAATTTGTTTACCTTCTTGACTGGGTAAACTATATCTTTTTACTGTACTGGCATTCCCAAATACTAATATCTGCAACATCATATCTTTAGTTTAAGACCACGAGACAAACAAATTGAAACGAAGGGAGTATTTACTTTGATACCTCCAAAGAAAATATTAGAATCAGAGTTGTTTCATTAGTTTTCTTTTCTCCAATTTCATCCTGATACGTGGATGTCAATGTTTGTTTTCTCTTTGTCATGCAGGCTTACGTCCTACAAGAACAGGGAAATCTGTTAGAACTAGTGGATCCACGTCTCGGTACCAATTACTCCAAAAAAGAGGCAATGCGGATGATAAATATTTCTCTTTTGTGCACAAATCCCTCTCCTACTCTCCGACCATCCATGTCTTCAGTAGTGAGTATGCTCGAAGGAAAACTCCCCGTGCAAGCACCAATAATCAAGCGTACTACATCAGACGATGAAATGAGATTTAAATCGTTTGAGAAACTTTCACATGACAGTCAAACAACACAAGTGTCAACATACTCTCAGGACAGTCAAGGACAAAGTATGAATGCACCTTGGAGTGATTCCTCAGTCTCAGTGTCTGTCCCTGGTAAAGATGGAAATGTTACTTCCACAAGTAGGCTTCTTCCAGATCTTTATAATGTAAACTTGGATTGATTCATTGACACCTACATTTTGATTTCCATCATGTAATTATCTAACTCTATCTTCACACAGGTCCCAGATAATTTCTTTTTGTGTATACTCCAATTTGTTCAATTGTGGTCCATTATGTTTTTGTAAAATGTATGTTACAATTCTTATTAATTAGTTCAATTGAAAT

>M6-PGSC0003DMG400015458 M6 LRR-RLK

AAAAAATGAAGACATAAACTTCACAACATATCAACCATAGTCCATACACCTAATATCCAAAAAAAATATGTTAAAGATTACTCTTTTCTTCTTCTTCCTCTTCTTTTACTCAGTTTTATCTAATCAAATCTCCAGTTTAGAACACAAAGCTTTGCTTGATCTTAGATCATCTTTACACATCAGAGCAAAAAACTGGCCAAGAAAATCCAACCCTTGTACCAATTGGACTGGAATTCATTGCAACAATGACAAAGTTGTGGCTATTAAGTTGACAGGACTAAGAAGAATTCACAAAGGGAACATGTCTCCACAGTTTGCTGTTGAATCTTTAGTCAATTTTACTCACTTGACATCATTCAACTCATCAGGTTTTGTTCTTTCAGGGCAAATACCTGATTGGTTTGGCCAAAAGTTAACTCAACTTAAACAACTTGATCTTAGTTCAAGTTCAATTCTTGGTTCACTTCCATCCTCATTAGGGAGTTTGAGTAATCTTGATTTTCTGTCTTTGTTTAACAATTCAATTACAGGAACTGTTCCTTTAGAACTTGGAAAGTTGAGTTCTTTAAGGGTTCTTGATCTTTCTTGGAATTCATTTACAGGTGAAATTCCAACTTCTGTTTCAGATATTAAGAATCTGTCTGTTCTTGATTTGTCTTCAAATTTCTTTTCTGGACAAATCCCAGTTGAGTTTGGTTCATTTCTTGGACTTAAGTTGCTTAAATTGTCAAATAACAGTTTAAGTTCTTATATTCCTTCTGAAATTTCAAATCTTTCTGCATTAGTAGAACTTGATCTTGGCTATAACTACTTGTCAGGGTCATTGCCTGAGGGTTTTTTCAGAATGAGAAATTTGAGTAAATTGTTAGTTGGAAATAATGAGCTAGAGGGTACATTGTCAAGTACTATGTTTACTTCAAATGATTCTGTATTGGTGTATATAAATCTTAGTTGGAATCGATTCGAAGGCAAAGTTCCTGATGTTTTGTGGTCATTGTCTGATCATTTGAGGATTCTTGATATATCAGGGAACAATTTTACTGGTGTGGTTTCAAATAGTACCTATTTTAGTGCTACTGGTGCTTCATATAACCTTTCTAATAATCTTTTCTATGGAAATGTGACGTTTGATGTTCGGAGTTTTCAGTCGATTGATCTGTCTAAGAACTATTTTCAAGGACTAGCACCTGATAAATCTGGTGTTCTAGTTATGGATAATTGCTTTGCGGGGTTATTGAATCAGAGGAGTTTAGATGATTGTAGTAAGTTTTATGCTGATAGGGGTCTAGTTTTTGTTGATGGTGGAATCCAAACTCCGGTGCAAAGAGCCTCGAAGGGCAGGCATAAACGTTTGTTTATCATGGTTGGAGTTTTTGGTGGAGTTGGTGTTCTTATGCTTATGGCATTAGTCATTTTATTGTTTTGGAAGGTATGCTATAGAGGGAATTCAAACGAAAGAGGCATATCGAATGTTAGTCCTGTTGTGGAAGGAGGAAATAATCAGGCACCACCAGTAAAGTTCTCTCGTGATACATCTGGTTCGATAGAATCATTTACTTATGATCAGATACTAAAAGCCACAAGCAACTTCAATGAAACAGATCTTATCAAGAATGGTCATACAGGAGACATTTTCCGAGGAATATTAGAAGGCGGTGTCACCGTAGTTATCAAGAAAGTAAATGTACACCATATCGAAAAAGAAACATACAAGTTGGAGTTGGATTTGTTAAATTGGACAACACACCATAGGCTTGTCCCTTTTTTGGGACATTGCTTAGATCATGAGAATGAGAAATTCTTGGTCTATAAATATATGCCAAATGGAGATTTGTTCAATTTGTTGGCCCGGGATATGGATTTAGAAGACGAAAATGGACAGTCTTTGGATTGGATAACAAGATTAAAAATTGCAACAGGAGCTGCTGAAGGCTTAGTTTATTTGCACCATGAGTGTAATCCTCCCCTTGTTCATAGGTATTTTTTTCTTTCACTTTTTTGCATTCTTGTTTGATAATGTTGTACACCAAGCTAATTATATTCTGCATGTTCTTGTTCTTGAAGAGATGTTCAAGCTAGCAGTATCCTTCTCGACGATAAATTTGAAGTAAGGCTGGGAAGTTTGAGCGACGTATGTGCACAGGAAGGTGATAGCCAGAATTTGATCTCAAAGTTTTTCCGCACGACAAAGTAAGTATTACTATGATCATAGTTTGATCCTAATACCAGATTTTTCAGAAATGTAATAGACATTTAGTTAGATTTATGTGTTTACATTTCTGTATATTGCTTTTTGTTCAATGTTCTAAGATATATTATGTTTCTTGTTGACAATAAACAGGAATCCTGAGGGAGAGCCTGCTGGTATGTACTATTCTTTTCGTAAAAAAACGCATAGTTGAATAGAGTTATTTTTCTGAATTTCTCATATGCTGCCTTTATACATGAACAGGATCTTCATTTACAAGTTGCAGTACTTATGATGTATACTGTTTCGGGAAGGTCTTACTCGAGCTTGTAACTGGTAAGGTTGGAGTAAGTCACTTAGATGATGCCTCAACCAAGGAGTGGTTGGACAAGCTTCTTCCTTTCATCACTATACGCGAGAAAGAACTAGTGTCCAAGATAGTCGATCCATCTCTAATTATAGACGATGACTTACTTGAAGAGATGTGGGCTGTAGCAATAGTAGCCAAAGCATGTCTTCATCCGAGGCCTTCTAGACGTCCTGAAATGAGACGTGTCCTGAAAGCTCTTGAGAACCCGTTCAAGGTTGTTAGGGAAGGAAGTTTCAACTCTATAAGGCTACGAAATACTTCATCGAGGCGATCATGGAGTGCTGCTTTTCTTGGTAGTTGGAATCATAGCTCACAGGATAGTGTTAATGCTAGTGGCCAAACAAGCAAAGAAGCTGGCATGAACCTGAAACAGCATTCTATTAGCAGCGGGAACGAACATTCATCATCGTGTAAGAGATCATCGAGTGAGATTTTCCCCGAACCTCAGGACACACTAGATATAGAGAGACAAGACATGAGCTAGCCGTCTAGCACGAATCTATGCCAAATGCATTTCTGCAGGCAGGGGTTTTGTAAGTATAAGGAACTTTGATTTTTTATTAGATGAGCGGCGGTACTATGATCATTCTGAAGACATGGGGCCTACACACTACACACATGTTATGCGGTTGGTAAACTCTTTCCTTCTGCAACAGCCTATCAAATCGGATCAGTATTTTTGAATATGTTGTGTTACATCATGTACATGTATTCTGTTTTCCTCTTTTCTTTTAGTGTGTCACAAATGCTCATGTGCTTTGTAAATTTTCAAAAAGTTTACTCTTGTTCCTGTGATGATAGAACAAGTATATAACTGTGTCAAGACTCAAGAGTGTACACTATCTAATCCAGATTCACATTTTCATGATCTCATATTTTGCAATTGCATGAAAAAGTATGATATGAATGAATAATAGAACCTGCACTTTCTTGAGCCGAGGGTCTATCGGAAACAACCTCTATACCTAAGGTAAGGTCTGCGTACACCTGCTCGTCTCAACACCCTTGCACCAGGTATGTTGTTGTAATTGGATAAATAGGGCCTCAAAACTGCTATAGTGTGTAAATCCCTTGGTGGCCTAAATGTAACTAAATACATGTAGATGCACCTGACAGTTACAATGAGC

>M6-PGSC0003DMG400015666 M6 LRR-RLK

CAAATTCCCAACTAGTATTTCTTTATCTTCTCCAGCTCAGACACAGACACACAGTCCATTTTTCTCTCTCAAAATCAACCAACAAGAAGATGAACAACATGTTTACCACTCCTTTGAACAAGAATTGAATCAAGAAATCAAATTACCCAGAAAAAATCCTGTTTGGGGGTTTTGGGTTTACCCAAAATCTTCAAAAAATGAGCTCAAAAATCTTGTTTTTATTCACCATTTCTGTATCAATTTTCACTCTTTTCTTGCCGACAAACTCCGACATTGTTTCCGATCGAGCTACCCTTTTATCTATCCGATCTGCTCTTCGTGGACGTTCACTTTTATGGAATATCACAAGTCCAACTTGTTCATGGCCTGGTGTTATTTGTTCACCTGACAAATCTTCTGTTTTAGAGCTTCATCTACCTGGAATGGGACTTTTAGGCCAAATCCCACCTGGGCTTTTCTCAAATTTGACTAAATTGAACTTTCTTAGTCTCCGGTATAATGCACTTTCCGGTGTAATTCCGCCGGACTTGTTTACTTCACTTCATGATTTGCGTAACTTGTACCTTCAAAATAATCTATTTTCCGGTCCGATTCCTGATTCAATCTTCTCTCTTACCAATCTTGTCCGGCTTAATCTTGCTCATAACAATTTCTCTGGTTCAATACCGGAATCTTTCAACAATTTGACTAGTTTGGGTACACTTTACTTGCAGGGAAATGGGTTTTCGGGTCAGATACCGGATCTGAATTTACCTGGTATGGTTCAGTTCAATGTATCGAATAATCAGCTAAACGGGTCAATTCCTAGTAAACTTGCGGGTCAACCCAAGGATGCTTTTTTGGGTACTTCACTTTGTGGTAAGCCACTTGATTCATGTGATGGTAGTAGTAGTAGTAGTATTGGAGAAGGTAAAAAGAAGAAACTTTCTGGTGGAGCAATTGCTGGTATAGTAATTGGTTGTGTTGTTGGTTTGCTTTTACTTCTTTGTTTGCTGTTTTTCTGTTGTAGAAAAAGGGGTAAAAAAGAAACAAGATCTGCTGATGTTGGTGCTGTTAGTAAGCAGGTTGAAGTTGAAATGCCTGAAGAAAGAGGGGTTGAGAGTAATGGTGGAAAAGATGGATTCTTGGGTAGTGCAATTGCAGCTATTGGTGTTGGAGGGGGTAATAAGGATAAAGGTAAAGCTGAGGCTGTTGTAAATGATGGAAAAAGTTTGGTGTTTTTTGGGAAAATGGCCAAAAATTTCAATTTGGATGATTTGTTGAAAGCTTCTGCTGAAGTTTTGGGAAAAGGGACATTTGGTACTGCTTATAAAGCTGCTTTGGAATCAGGGATTACATTAGTTGTGAAGAGATTGAGAGATGTTACTGTACCTGAGAAGGAGTTTAGAGAGAAAATTGAAGATGTTGGGAAAATGAATCATGAGAATTTGGTTCCTTTGAGAGCTTATTATTATAGCAGAGATGAGAAGCTTCTTGTTTATGATTACATCTCAATGGGAAGCCTTTCTGCACTTTTACATGGTGAGAAAAATAACTTTTTTTTTTCTGTCATTCTTTTTCATGTCATTATATTGCTGTTATGCTGTTCCTTGAATCGAGCGTCTATTGAATCATGTCATTATAGTGCTGTACTGTTGTTTACCATTTACATGATCTGCAAATTTTAGTTGTCTGATTGACACTGATACTACTGCTTTCTATGTTGCCTAGACTCTTAAAATGTGGTGCTTGTCGGATCCTGGATCCTCCAAAAGTAGTGTATATTTGGAGGATCCAAACACGAGTCCGAATAACATAGACTGTTTTTGCTGATTTTATGAGGCTAGTAAGTTTGGATGATCATTTGTTTTCTTGGATCTTGTTATCTGTTACTGAATTGGTGCTGTTGTGTAAAAGTTGCTGCATTTGCATTATTGTGCCCAATTGTGTTCAGCCACTAAAATTTAAGTTTTTTTTTTTACATTATTGTGCTGTTGTTGCATTGTTGTTCATGTTTAATCAGATTTTATCGTAATCTAATGTGATGAACATTGTGTTTAGTGAATATGATATCTGACATGTTTTATATATAACTCAACCTGGAATGGCTTTACTATGTCCAATTGTTGCTGTTCAAACTCGTGTTCATCATTTAGATTGAATGTATACTTTATCCACAAGGTGATCTGCACAATTTTTAGATTCCTAAGTCCGAGAGTCTATCAGAAACTACCTCTCTACCTACAGAAGGTAGAGGTAAGGTTTGCGTGCATCCCACCCTCCACAGACCTCACTTGTGAGATCTCATTCGGTATGTTGTTGGAAGAGTTTAGGAAGAAGTCGTAAAATTGAATATATGTGTAGGACAGGTTCACTGCGTTCAACAGGACGAAACAAGATCATTTAACATTTAATCTGTTTGTATGGATTTAAGCCAAATGTACCTTTGTTAACTTATCGACTATATGTTGGGTAAAAACTATGAATTATTTGGGTGCTTTATTCGGCATATTCCACTGCTACTATGTCATGTCACAATAGTTTTGTCATTCAACTTGTCTTTTCTATAGGCAACAAAGGAGCAGGTAGGACACCGTTGAATTGGGAAACAAGGGCTGGCATTGCCCTTGGAGCTGCACACGGGATTGCGTACCTTCATGCACAAGGGCCCTCAGTATCACATGGCAACATTAAGTCATCCAATATCCTCCTCACCAAATCATATGAGGCCCGTGTTTCAGATTTTGGCCTTGCTCAACTCGTCGGTCCATCATCAACTCCCAACCGTGTTGCAGGTTACAGGGCACCAGAGGTAACTGACCCTCGTAAAGTCTCCCAGAAAGCAGATGTCTACAGCTTTGGCGTCTTGTTGTTGGAACTCCTAACAGGGAAGGCTCCCACACATTCAGTAATGAATGAGGAAGGTGTCGACCTACCAAGATGGGTGCAATCAGTGGTACGAGAGGAATGGACAGCCGAAGTCTTTGACCTCGAGCTCCTTAGGTACCAAAACGTCGAGGAGGACATGGTCCAGCTATTGCAAGTTGCAGTTGATTGCACTGCTCAGTACCCTGACAGACGCCCTTCAATGGCCGAGGTCACTACTCGCGTTGAAGAACTCTGTCGTATGGATTCTGGAGGTGGTGACATAATTGACAATGATGCAGAAGTGCAAACTGCATGAAATTGAACAAGAAAAAATAGTTCCATCACATTATTTCAATTTTTTTCTTTTTCGTTTTCGATTGAGCATTTCGATAACTTTGTTCATTCTTGGAGTTCTTCAGTGAAGATAAATTGTTCATTTTATTTTATTTTTTTTGAATTTATTTTCCAGTAGCAACAGAGTTTTATTAAAGATCTGGTTGGTAGTTGAAATGGAGCTGTTTTTTTTGTTTTATAGATATTGATGTATTCTTTTTTGAAGGTGTGATGAACTGTATAAATAATTGAGATGACCTGTAGTTACA

>M6-PGSC0003DMG400015806 M6 LRR-RLK

CATGAGAATTTGGTTCCTTTGAGAGCTTATTATTATAGCAGAGATGAGAAGCTTCTTGTTTATGATTACATCTCAATGGGAAGCCTTTCTGCACTTTTACATGGT

>M6-PGSC0003DMG400015814 M6 LRR-RLK

ACCCCATGTAAATGGTCTTTCATAAAATGCAACTTTCAAGGTTTTATTACAGAAATCAATATTCAGTCTATTCATCTTGAACTTCCTTTGCCTACAAATCTTTCTTCATATAAGTACCTGAAGAAGCTTGTTATTTCTGATGCTAACATCACTGGTGCAATACCCTTTAACATTGGTGACTGTTCGTCACTTGTTACTATTGATTTGAGCTCAAATGGTCTTGTGGGTACCATTCCTTTAAGCATTGGAACCCTTGTAAACCTTCAAGATTTGATTTTGAACTCAAATCAGCTTACTGGGAGAATCCCAGTTGAGATTGGGAACTGTAGAAACTTGAAAAATCTTGTTCTTTTTGATAACAGGCTTAGTGGTGGTTTACCAAGTGAAATAGGGTTGTTGTCAAATCTTGAAGTGTTAAGAGCTGGTGGGAATAAAGATGTTACTGGTAAAATCCCAAATGAGTTTGGGGATTGTGGGAATTTGACTGTTTTAGGCTTAGCTGATACCAGAATATCTGGTTCTTTGCCTGTTTCTTTGGGTAAGCTTAAGAATCTTGAAACTTTGTCTATTTATACTACAATGTTGTCTGGTGAGATACCTTCTGATTTAGGTAACTGTACTGAGCTTGTGAACTTGTATTTGTATGAAAATAGTCTTTCTGGTTCAATCCCATCTGAGCTAGGGAACCTTAGAAAGTTAGAAAAGTTGCTTCTTTGGCAGAATAATCTTGTTGGTGTTATTCCAGAAGAGATTGGGAATTGTACTAAGCTGACTATGATTGATTTGTCTTTGAACTATTTGTCTGGGAGTATACCATTATCTTTTGGTGGTCTTGTTGTGTTACAAGAACTTATGCTTAGTAACAACAATGTTTCTGGTTCAATCCCTTCTGTTCTTTCACAATGTACAAGTCTTGTGCAGTTGCAGTTTGACACAAATCAGATATCAGGTTTGATTCCTTCTGAGTTGGGGAATTTGACAAGTCTTGTTGTGTTTTTTGCTTGGGATAATCAACTTGAAGGAAGTGTGCCTTCAACTTTGGGTAGTTGTAGTAACCTTCAGGCATTGGACTTGTCACATAACTCGCTTACTGGTAGCATTCCTCCAGGCTTGTTTCAGTTGAAAAACCTGACTAAGCTACTGTTGATTTCTAACGATATTTCGGGTACTATACCTCGAGAAATTGGATATTGTAGCTCATTAGTGAGGTTAAGACTTGGAAACAACCGGATTGCTGGTGGAATTCCTAAAGAAATTGGAGGTCTTAAGAGCTTAAACTTTCTTGATTTGTCCGGAAATCGCCTTTCTGGACCAGTTCCTGATGAAATAAGTAGCTGTACTGAGCTACAAATGGTAGACCTCAGCAGCAATACACTGGAAGGTCCCCTGCCTAACACTCTGTCTTCACTATCAGGGATTCAAGTCTTGGACGTTTCGAATAACCGGTTTGGAGGGCCTATACCGGCTAGTTTTGGGCGTCTTGTGTCATTGAACAAGCTGATTCTCAGTAAGAACTCGTTCTCAGGGTCTATACCTCCATCTATTGGCCTGTGTTCGAGTCTTCAATTGCTAGATCTTAGCAGCAATGAGCTCTCGGGAGGCATACCGATGCAGTTAGGAAAGATTGAGTCCCTTGAAATCACTCTCAACCTTAGTTTCAATGAACTAACAGGTCCAATTCCTGCTGAAATTTCTTCACTAAGCAAGCTGTCGATACTTGACCTTTCGCACAACAAGCTTGAAGGGAACTTGAATCCACTAGCGAGGCTCGATAATCTTGTCTCACTGAATGTCTCATACAACAACTTCACTGGTTATCTTCCGGACAATAAGCTCTTTCGACAGTTGCCATCATCAGACCTAGATGGAAATGAAGGTCTATGTTCATTCGGAAGGCCTTCGTGTTTTCTCAGCAATATTGATGGAGTAGGAGTAGCTAAAAATGGAAATGATGAGGGAAGGTCAAAGAAGCTCAAATTAGCCATTGCATTGCTAGTCATCATGACAATAGCAATGGTGATCATGGGAACTATTGCTATCATACGCGCACGAAGGGCGATGAGAAGGGATGATGATTCAGAGATGGGAGATTCTTGGGCTTGGCAGTTTACTCCATTCCAGAAGCTCAATTTTTCAGTGGACGAAATACTCAGATGCCTTGTGGATACTAATGTCATTGGAAAAGGGTGCTCGGGGATGGTCTATCGTGCAGTTATGAACAATGGGGATGTCATAGCAGTGAAGAAGCTTTGGCCAATAACCATGACTACAACTAATGGAGGCAATGATGAGAAATGTGGGGTACGTGATTCGTTCTCTGCAGAAGTTAAGACGCTTGGTTCAATTCGACACAAGAACATTGTCCGATTCTTAGGGTGTTGTTGGAACAGAAGCACAAGATTGCTCATGTATGATTACATGCCTAATGGGAGCTTAGGAAGTCTTCTTCATGAGAGGGGTGGTAACCCTTTGGAATGGGAATTGAGATACCAAATATTGCTCGGGGCAGCGCAAGGGCTCGCCTACTTGCACCATGACTGTGCTCCTCCAATTGTTCACAGAGACATCAAGGCCAACAACATTCTCATTGGTCTTGAGTTTGAGCCTTACATTGCAGACTTTGGCCTCGCAAAACTAGTCGATGATGGTGATTTTGGTCGTTCCTCCAATACAGTTGCTGGTTCTTATGGCTACATTGCTCCAGGT

>M6-PGSC0003DMG400016015 M6 LRR-RLK

TTTTTTTCCAATATATATCCTTTTTCCATTTCTCTTCATAGTAGTATGTTTCAAGTGCGTAGTAACATCATGGAGAAACACATTTTCTTATTGATACTTGCTATCTTAGTTCAATTTTTCTTTGTTTCTTCTATATCAGCTACTATTTCCTCAAATGAGACTGATCAAGAAGCTCTACTAGCTTTTCGAAATCTTGTTACGAGTGATTCTAGTCATTTTTTATCCAATAATTGGACCAAAAATACTTCATTTTGCTCTTGGTTTGGTGTCACTTGTAGTCCAAAAAGGCAAAGGGTTGTAGCCTTGACTCTTCCTAATTTGCAACTTCAAGGCACAATTTCGCCTTCTTTGGCCAATCTATCCTTTCTCATAGAGCTAAATCTCTCGAACAACAACTTCCATGGTAACATCCCTTATGGCATTGGCCACTTGCCTCGTTTACGAGTGATTGATATTCAGAACAATCAGCTCCATGGTAGTATTCCAACATGTCTATTTCAACACCTGAGTGTTCAAATCATTTCATTGGCTTTCAATAAACTCGGTGGTGAAATGTGGAACTGTACATGGTATGTACCCGAACTCAGAGTCTTAAATCTCAGGAACAATACCTTCACAGGTATAATCCCTCCTTCTATTGGAAATGCCACAAAGTTGATGAACATCAGTTTGAATTGGAATAGAATCAACGGCAACATTCCAATGGAGATCGGTAATCTAAGCCATCTTGTAGAGTTGTCGTTGTCTCGTAATCAATTAACAGGTTCCATTCCTTCAACATTGTTTAATATCTCCTCCCTTCTCGTCTTGTCTCTGGCATACAATAGCCTTTCAGGTCCTCTGTTTCTTGATGATCGACGTAATGTTCTTTCATCAAACCTCGAGCATATAGGTGTATCATACAATCAAATCACTGGTCACATTCCTTCCAACATCTGCCAATTCAAAGCTCTCAAAGTCTTGTCCATATCTTACAACAACTTAACTGGAGAAATACCGAGAAATATTGGTTGTTTAGCCAAGCTCGAAGAGCTTTATATCGGTTATAATGCAATAGATGGAACAATTCCTACTTCATTAGGCAATATTTCCACTCTTCAAAAACTTCATTGTGGAAACAATCACATGGAGGGAGAACTTCCTCTGGAATTAGGAAAGCTATCAAACTTGAGACAAATCAATTTCGAAGAAAATTATAATCTTATAGGTGAAATTCCAAATGCTATTTTCAACATATCTTCTTTGGAATTCATTGCTTTCACTTTCAACAACCTCTCAGGTAGAATTCCAAATCTTCTTCATCTTCCAAACCTTATACAACTTCTCTTAGCAATCAATCAGCTCGAAGGTGAAATTCCTCGGTACATTACAAATGCTACCAATCTTGAGCTATTGGAACTATCAGATAACCTTCTCACAGGCAGTATTCCTTATGATTTAGGAAATCTTCGCGAGCTGCAAGAACTTTTCCTTCATCATAATCAACTTACTGAGTTGTGATTCTTTGATTCTTTGGTTAAATGTAGGATGTTGAGATATGTACAAGTGGGATCGAATCCGTTGAATGTTGTTCTGCCAAGTAGTATTGGCAATCTTTCATCTACTGTTGAATACTTTCATATTGGAGATGCACAAATCAATGGATTCATTCCCACTAGTACAGGCAACATGAGCGGTCTTACAACGCTAGTTTTTCAAGATAACAATTTTACAGGAAACATTCCTCGTGAGATCGGTAAGCTTAAACAACTCCAAGGTTTATTTCTAGTTAACAATGGACTACAGGGGGATATAACAGAGGTAGTATGTGATTTATCGAATTTGGTTCGATTAGCTCTGTCTGAAAATGAGCTCTCGGGGGTGATTCCGGAATGTCTAGGAAGTCTTTCCATGCTACAACACCTTTTTTTAGGTTCTAACAAGTTTGAATCAAAGCTTCCTTTAAGCTTTTGGAAGATGAGTAGTCTTCTCTATGTAAACATGTCGCGTAATTCTATAGAGGGAGAAGTTCCATCAGATATCGGAGAACTTAAAGCTATTGTAGCAATTGATATCTCTGGTAACCACTTTTCGGGGATGATACCAAGCAATTTGGGGGAACTTCAAAACTTGAAGTTACTTTCCTTATCGAACAATTCGTTTTCAGGTCCAATTCCATTATCCTTTTCAAACTTGAAAAGCTTGGAATTCTTGGATTTGTCTTTGAATGACTTGTCAGGTACTATTCCTAAGTCTTTCGAAAAGCTTTTGTACCTTACAAGCATCAACGTCTCGTTTAATGTTTTAGAAGGTGAAATACCTAGTGGTGGTGTGTTTGCAAACTCCACCCTGCAATCATTTCTCGGGAACAAAGGTCTATGTGGAAGGCAAATATTGGAGGTTCCTGCTTGTGCTGTCACTACTCCTGAACAACAACAATCAAAATCTAAGAAGCTTGTGCTAAAAATTGTCACTCCGGTGGTTATTTCATTTTTTCTGATATTCTTGTTGGTTGTCTCAATTTGGATAATGAAACGAAAGAAGAAAGTTAAGTCCAAAGATTTTGAAAAGGTTCCGGAGATGAGGACTTATCAATTGATTTCTTATCATGAGATTCAACGAGCAACAAATAATTTTGATGAATCCAATTTGATTGGCGTGGGAGGTTCTGGCTCTGTGTACAAAGCCACATTACCTAGTGGAATTGTGGTTGCAATAAAGGTACTGGATTTGGAAAATGATGAAGTATGCAAAAGGTTTGATACTGAATGTGAAGTGGTGAGAAATGTTAGACACAGAAATCTTGTTTCAGTGATCACTACGTGTTCTACTGATCACATAAGAGCCTTCTTTCTGCAATATATGCCCAACGGAAGTCTTGACAATTGGTTGTACAAAGAAGATCGCCACTTAAACCTTCGTCAAAGAGTCACCATAATGCTTGATGTAGCTATGGCAATTGAATATCTACATCATGGTAATGACACCCCTATAGTTCATTGTGACCTCAAGCCAGCCAACGTTCTTTTGGATGAAGATATGGTGGCGCGTGTTGGTGATTTTGGCATCTCAAAGATTTTAGCTGTAAGCAAGTCTATGGCACATACAAAGACATTAGGCACTCTTGGATATATTGCACCAGGTATACTACTATACTCTATACTAGTCTCTTTCCATTTCACTTATGTATCGAAGACCTTTCATTATTGTTATATCAGATGAGTAATTGAGTAATACTTTTTTTCATTTTCGTTCTTTTAAGAATATGGCTCGGAGGGAATAGTGTCCACTCGTGGTGATGTTTACAGTTATGGCATCATGTTGATGGAGGTTTTGGCAAAAAGAAGGCCAACAGGTGAAGAGATATTCGATGAAAATCTTGGCTTGAGAGAGTGGATAACGCGAGCATTTCCAAGAACTGTGATGGAAGTTGTGGACGCGGATATTTTACATGATGTAGAAAAAATCACTTCCAAAAGTGAACTCTGCATACTTTCCATGATAGAACTTGCTTTAGATTGCACAAAGGCAACACCATAATCAAGGATAACCATGAAAGATGTAGTCAAGAGGCTTAATAAAATTAAGAACACATTTTTGGAAACATAGAAGTTAGTTAGAATCTCTTTTAATGTTGTTTTTTCTTATAAGTTGTAAGTTGATACTCAATCTGTTTGTCATGCTTTTCATTCTAGTCCGTTATAAAAAGAATGTCTCTTTTCTTTTTTGGCAACTGTTAAATTCTAACTTTCTACATGTCTATG

>M6-PGSC0003DMG400016164 M6 LRR-RLK

ATGTGCTACTCTAGAAGTATATATTTATCGATTTCCTTCATCAAGCTGATATTAATCTGCTCATGATATTGGCACATTGCTAACTGCTCTTATTTGGGGGACTGGTGATTGCTTGTATCTTTGCCCGTGCATGGTTTGGGATCTTAGCCTTTTCCGATTGTACTACTTTCCTACCCACACAAGCATGTGTAGAAATCTACTCGCAATTTTGGTTCTCAATATTCCAACTAAAGAGTAGTATTCTTTTAAGTTAATAATTAGGAAACAACATTCTTGAGTGTTTGGTCATGCTTTTGCCAGAGTGTTGGATATTTCATCTTCATTTCTTGTCCGTTATCACATTATTCTCGTAGTGCCTCTGATAAGCATTCTAAGGTCATAACTGGTAACATTTGCTAAGAAGATCTCCTAAAAATAAGTTTTCATCCTTGTCAATAGGAAGTTCTACTTTAGCGAGATTTGTGAACTCATCAATTTTCTTGTCCTTAGTTGTGTTATGTATTGAACTGACTGGGCGGTCAACATCCTGGCAGTTCTTTGCAAGGAAATGGAATAACTGGTAAAATTCCTGAAGCATTAGGAAATTTAACCAGCTTGACCATGTTGGATCTGGAAAAAAACAGTTTGTCTGGAGAAATACCAGCTTCCCTAGGCAATCTAAAAAAGCTTCAATTTCTGTAAGTAATATTATTTCATTGATTGTTGTTTTTTTTAGAACTGAGTTTTACATTTGGTGTTTGACTTGGGAGTCTATAATGATGTTTCTTTGAACAGGTTTTTGAGTCAAAACAATCTGTCTGGGACTATCCCTGAGTCACTTTCAGGCCTTCCCAACTTGATTAACCTGTAAGTGATTCATTTAGCCTGATCTTTGATGTGCTTACAACGTGTGCCAAGTGATACTGACCTGACCCGGTGATATCTCTTTGCCAGTCAGCTTGCATTAAATGATCTTACTGGAAAAGTTCCTGACAGTTTGTTCCAAGTCCCGAAATACAAGTATGCAGATCTCAATTATTGCATTTTTTAATTTGATGATAGATTGAAGCTGCTTAATAATTTGAACATTTTAAAAAGTTATAATACATAAATAGACACTCAAACTTAGCCTCATCTGGCAAGTAAGCACTCCAACTATGAGAGAGCACATCTAGACACCTCAACTTGGTCCCAACTGACAACTAAACACTCTAACTCGTCCCCACTGTGTCTAGTGAATACCTGACGCTGATGTGGCATATAAATTTTGGTGGTGTCTTGATGATCTTTTTGTAAGTTGGAGTATTCAACTGACACAGTGAAGACGAGTTGAGGTATCTAGATGTGCATACTCAAAGTTGGAGTGTTTACTTTCCAGCTGAAGTCAAGTTTGAGAGTGTGTTTATGTATTATACCCTTTTTATAATAGCCCTGCCTCTTGCATGAGTCAGAACTGTTGATATTAGTTTAACTCAGTTTGCCTACTAGTTTTTCAGAGAACCATTTGAATTGTGGCTTAAACTTCAGTCGTCGTTGTGTATCTGACAGTGAAGGTAATTAATTAAAAGAGCAAATACCACATTTTTAAACTACAAGTATTTACTTTAAAAGTTAGATGTTGCTGGAGATGTTTGGCAGTATTCTAAATTCTAAAGTTGATATCCTGGCAGGTTCTCCCAGTAAATCAAAGACTGGTCTCGTAATTGGTATCGTTGGTGGATTTCTTGGAATTATCCTTCTTGGGGGATTAATGTTATTTTTCTGGCGGGGGAGGAACAAAGGCTACAAACGTGAAATCTATGTTGATGTTCCAGGTCCTTATCAGTGTTTTACACAGTATTAACTTCAATATATGTTGTTAGCTCATCAAAAAAAAAAATACTCCCTCCATTTCAATTTGTTTGTCTTACTTTCTTTTTTAGCTCGTTTAAAAAAGAATGTCCCTTCCCTTTTTGGCAACTCTTCAATTTCAACTTTTCACGTGACATGTTTAGGACCACAAGATTCCTTCTTTATTTTTTTAAACTCCATGTCAAGTCAAAACGAAACAAACAAATTAAAACGGTGGGAATATGTGTTAGCTCTCCTGAAAAAAAAGAAGAAGGCTGAAGTAGTTACCTATTGCTACATTTCTTAGTTCCATCTTACATTGGAGACTTGTTCTTTTATCAGCAGTACCTTAGTTCACTTGCACTGTAAGATGGTATAGGCTCATTGAGTATGTTGTGACATTTTTCTTGTGTCTGTTCGATGTGCCCAGGTGAGGTTGACAGACGAATTCCATTTGGTCAATTGAGGAGATTCGCATGGAGGGAATTGCAACTTGCTACTGATAACTTTAGTGAAAAGAATGTTCTTGGACAAGGAGGTTTTGGCAAGGTTTATAAAGGAGTGCTCAGTGACAGCACTAAAGTTGCTGTGAAACGCTTAACTGACTATGAGAGTCCTGGGGGAGATGCTGCATTTCAGCGTGAAGTTGAGATGATTAGTGTTGCTGTTCATAGAAATCTTTTGCGGCTTATAGGATTTTGCACCACACCGACTGAACGCTTGCTAGTGTACCCGTATATGCAGAACCTAAGTGTTGCCTATCGTCTACGAGGTACTTATTTTTCCACAATAGTGCTTAGATCTTGGTTTGCAGTTTTCCCAAGGCTTTCCTTGTTTATCTCAATATCAAATTTTCCTGGGATCACTTATTTCTTTTGCTCCAAACTCAGCTGAATTTCTATAGAAGTCTGGAATTTTATGGTCAAATGCTTATGTCAATAGTCTAACTGTCTAATAGCCTATTTGGCCAAGCTTCTAAAATCAGCTTATTTTGATTTTAGGCTATTAGACTCCTATCAAAGGTCTAATCACCTACTTTTTCCTAGTGATCTTTAGTAGAGTAGACAAGTTGAACAGATAATAAAGGAGGGTATTTGCGTAGGTACTCGTATTCCTAATATTCTTAGAAGTGTGTGAGTTCTTGAAAATTTGGTTCTCTTCAGTCTTCAAGAGCCATAGTTGGATTGTACCTCGTCACGAGGGAACTTCCATCACGTATGAATTGACCGATATCCATAATGAGAGCTTAAATGAGTTCATGAATTCATTCTTGCGCATTCATCATTCCCCCCGAGGTGACTTTTTTGAAAACCTTGCATGAATTGAAGATGACACATGCAGTAGTCGGTCAAACTTTAAAATTCAGCATTTATCTCCCAGAGATGATGAGATTTTGGTGGCTTCTTATTTGTAGTATTTTGTTAAATGCAGAACTTAAACCTGGGGAGTCTGTTTTAATTTGGCCAACCAGAAAGCGTGTGGCACTGGGCACTGCACGTGGACTTGAATACCTGCACGAACACTGTAATCCAAAAATTATTCACCGTGATGTTAAAGCAGCCAATGTTTTACTGGATGAAGATTTCGAGGCTGTAGTTGGTGATTTTGGCCTGGCAAAGTTAGTTGATGTTAGGAAAACCAACGTGACAACGCAAGTTCGGGGTACTATGGGCCATATAGCTCCTGAATATTTGTCCACTGGGAAATCATCAGAAAAAACTGATGTTTTTGGATATGGAATCACGCTTTTGGAGATTGTAACTGGTCAACGCGCTATAGACTTCTCACGCCTAGAAGAAGAAGATGATGTCTTGTTACTTGACCATGTGAGTGTACATAACTCAACGACTGATATTAACTTGTGATTCTATTGGTCTGCATATTACTTTTAACGTGTCCTCTAAAATTATTCAGTTTCTTCTAAGTTTATTTTTTTATCTTATACTTTCACTCTTCTCGAAGCTGCATATTGGTGGATGGCTCATTGTCCATCACAAGACATTCTCATGAATTATATACGTAGTTCTTCTTTTTTATGTTGCTTTCTTGGTATATGTGGTAATTTTATATTCAACAGGAAGGAGGGACAAAAAAGAAGTTTCATTTTCCATATAGCACTCTTACGCAAATTCTTTTGCATGTCAAACATAGGTCTTAATACTCAGCCTCTTTTGTCGGATACCCTTTTTAGGTGTAAAGTAAATTAGTGAAAAAAGGTGAAATAACATTGAGAGGACGCAAAATTTCTTAATTAGGGTAGAAGGATAGTAGATAGTGCATTTTCTCGCTATCCGTCCATTCTAGTAGTTGTAGTAGTGCTCCTTTAGTTTCTTGTCCTATGGTTTTTGTTACCATCTGTTGTTTCTTGTATTTCGCTTGTCGTATTATTTTGATGTAGTTATTGTTCAGCATGCTTTGCTATGCTTTCACATCCATTACTTTTTCTGGACTGCTTTTGTTTGTTTTTCCTTGAGCCGAGGGTCTATCGGAGACAACCTCTCTATTCTGAGGTAGGGGTAAGGTCTCTATGTAGACTCTATCCTCCCAGATTCCACTTTGTGGGAGTACATTGGGTATGTTGTTGGTTACTGAATGGCTGCATGTGTCATCTTGAAGGTCAAAAAACTGCAAAGGGAGAAAAGGTTGGATGCTATTGTCGACCGCAACCTACATAAGAACTATAACATGGATGAAGTAGAGATGATGATTCAAGTTGCGTTGCTGTGTACTCAAGGATCACCAGAGGATCGTCCAGCAATGTCAGAGGTAGTACGAATGCTTGAAGGAGAAGGACTTGCTGAGCGGTGGGAAGAGTGGCAGCATGTCGAAGTTACACGTAGGCAAGAATACGAGAGACTACAACGACGATTTGATTGGGGCGAAGATTCAATCTATAATCAAGATGCAGTTGAACTGTCTGGTGGAAGATAACAAAATCTGTATCTCCAAACAAGTACATAATTTCCTGTAATAGTCTCTCCTTGATACATATAACTTGTCGCTAAGTGGCAATCATTCCACGTTTTCATATTGCTTGTGTCATATCTTGATTGTATTGTTTTTCGTATGCTTCTCTTATTTAATCGATTTTTTCCCTGTATGATTGGCCTGTGCAGCGACATAATATATATCGACTTTCTTTTTGTTTCAATTGTAACATCAATCTTTGTTATAGTGATTTGAGTTTTGC

>M6-PGSC0003DMG400016685 M6 LRR-RLK

GAACATTTCTTCTTTCTTTCCACCATTTCTGCTCTGCTCACTCTTTCCCCTCCACCATTGAAGAAACATGCGTTTTCTTCTCCTCTTCTTCCTTTCCCTTATTCACCATTTCCATCTCCTCCACTTCACCACCGCAAAACCACCTTACGTGCCGGAATACCGGGCATTACTCTCCCTGAAAACTGCCATTACCGATGACCCACAATCTGCTCTTCTTTCATGGAATATCTCAACAAGTCATTGTACATGGAGAGGTGTCACGTGCGACCGGTATCGTCACGTGACTTCTCTCGACATCTCTGGTTTTAATCTCACCGGTACTCTCACGCCGGAAGTTGGTCATCTCCCTTTTTTGTTCAATCTTTCTGTAGCTGTTAACCAGTTCTCTGGACCCATTCCTATAGAGCTCTCGTTTATACCAAATCTGCGTTACCTTAACCTCTCTAACAACATTTTCAATTTGAGTTTCCCTTCCCAGCTTACCCATCTCCGGTACTTGAAAGTTCTCGATATTTATAATAACAATATGACCGGTGACCTTCCGGTTGGGGTTTACAATTTGACAAATCTTCGACATCTTCATTTGGGTGGCAATTTTTTTAGTGGCAGTATTCCACCGGAGTATGGTAGATTCCCATTCCTAGAATACCTTGCAGTTTCTGGAAATGCGCTCGTCGGTATGATACCTCCGGAGATCGGGAATATTACCACACTTCGTGAGCTTTACATTGGATACTACAACACGTTTTCCGGTGGGTTACCGGCGGAAATAGGGAACTTGTCGGAGCTCATTCGGTTAGATGCTGCAAACTGTGGACTTTCCGGTGAGATTCCGCCGGAGATAGGGAAGCTTCAGAAATTAGATACACTGTTCTTGCAAGTGAATGGTCTTTCTGGGTATGTTACACCGGAATTGGGGAATTTAAAAAGCTTGAAATCTTTAGATCTATCAAACAATATGCTCTCCGGTGAAATACCGTTCACATTCACAGAGCTGAAGAATCTAACTCTGCTAAATCTTTTCCGTAACAAGCTTTACGGGTCGATACCGGAGTTCATAGAAGATTTGCCGAAACTGGAAGTATTGCAGCTTTGGGAAAACAACTTTACCGGAAGTATTCCACAGGGTTTAGGCAGAAACAGTAAGTTAACAAACGTTGACATCAGTACCAACAAATTAACCGGCAATTTGCCCCCAAACATGTGTTCCGGCAACAAGCTACAGACGTTGATCACTCTCGGAAACTTCTTGTTTGGCCCAATTCCAGAATCTTTAGGTGAGTGTCAATCACTTAATAGGATTAGAATGGGAGAAAATTTTCTAAATGGGTCTATTCCAAAAGGGCTATTAAGTTTGCCGAAGCTTTCACAAGTAGAACTTCAAGATAATCTTCTCACTGGTACATTTCCAGTGACTGGTTCTGTTTCAGCTAGTCTTGGACAGATTTGTCTTTCGAATAATCATTTTACGGGGCCTCTACCATCGAGCATTGGTAATTTTATTGGTGTTCAAAAGTTGCTTCTTGATGGGAACAAGTTTTCTGGTCAAATTCCAGCTGAATTAGGGAAATTGCAGCAGCTGTCGAAAATGGATTTTAGTGGTAACACTTTTTCAGGCCTGATTCCACCGGAGATAAGCCGGTGCAAGGCTTTAACTTATGTTGATCTTAGTAGGAATAAGTTATCTGGTGAAGTTCCTACTGAGATCACTGGTATGAGGATACTGAATTACTTGAATGTATCTCGGAATCAGTTAGTTGGGAGTATTCCTGCACCTATTGCTGCAATGCAGAGTTTAACTTCGGTTGATTTTTCGTATAACAACTTATCTGGATTAGTTCCGGGTACTGGTCAGTTCAGTTACTTCAATTACACATCATTTATTGGTAATCCAGATCTTTGCGGACCCTATTTGGGTCCTTGCAAAGAAGGTATTGTTGATGGGGTTAGTCGACCTCATGAGAGAGGTGCATTTTCGCCTTCTATGAAGCTTTTACTTGTTATCGGATTGCTTGTTTGCTCGATTGTGTTTGCTATTGCTGCAATTATTAAAGCTAGATCGTTAAAGAAGGCGAGTCAGGCTCGTGCCTGGAAGCTTACTGCTTTCCAACGCCTGGATTTTACTTGTGATGATGTATTGGAATGTTTGAAAGAGGATAACATTATTGGTAAAGGAGGTGCTGGAATAGTATACAAGGGGGTAATGCCAAATGGTGAACTTGTTGCTGTTAAAAGATTGCCGGTTATGAGCCGTGGTTCTTCCCATGATCATGGGTTTAATGCCGAGATACAGACACTTGGGAGTATTCGACATAGACATATTGTTAGATTATTGGGATTTTGCTCAAATCATGAAACAAATCTTTTGGTTTATGAGTACATGCCTAATGGAAGCCTTGGTGAAATGCTTCATGGAAAGAAAGGGGGTCACTTGCATTGGGATACCAGGTATAAGATAGCACTGGAAGCTGCAAAGGGACTTTGTTATCTTCATCACGATTGCTCGCCTTTGATCCTCCATCGTGATGTAAAATCAAACAACATTCTTCTGGATTCCAGCTTCGAAGCTCACGTTGCTGATTTTGGGCTTGCCAAGTTTTTGCAAGACTCGGGAACATCAGAATGCATGTCTGCTATTGCTGGTTCTTATGGCTACATTGCACCAGGTATGCTCTCAAATTTTTACTTCTTTGTAGTCTTGGTTTTGATGTGTGTTGTTCCTTTCAAAAGAAGTTTATTATTGTTGTAAGTTGTCAACTAAAATTGTGATTTATCAGTCTCTTCAAGTGTTTATATGGTACTTCTTGGTGCTAGTAATGATCAGTGTCCTTGTTATAACACGGAAGAACAACATGACACGAATATGTCCTTGATTGGTAATCTACAAGAACAACAACAACAACAATGATAACTACGTGTTGAAAATGAATGCCTCTGTTCATAGATGAGCCTTGTAACTCTTTCTTTAATATGATAAACAGCCGTCAACTGGTGCTTTGATGAATTCAAATTTGGTTTCTTGCTGTTTGGCGATAAGTTTTTTCATTTTGACTTACAAAGTTTGCTTATTGCTTGAAATACAATCTCTATTTTTTTTGTATGCGCCTAATTTCGTTTTCATATAAACTGTGCAGAATATGCATACACACTCAAGGTAGATGAGAAGAGTGATGTATACAGCTTTGGTGTGGTTCTGTTAGAATTGGTGAGTGGGAAAAAGCCGGTTGGAGAATTTGGTGATGGCGTTGACATAGTCCAATGGGTAAGGAGAATGACAGATGGGAAAAAGGAAGGAGTTCTCAAAATCCTTGATCCAAGACTCTCAACAGTTCCCCTTCATGAGGTGATGCATGTGTTCTATGTCGCAATGCTGTGTGTCGAAGAGCAAGCTGTTGAACGCCCCAAAATGCGCGAGGTTGTGCAAATGCTAACTGAGCTTCCCAAGCCATCTGGTCCAAAAACAGAAGATTCAACAATCACTGAGTCGCCCCCATCATCAGGTCCTGCATTAGAGTCTCCCACTTCGACTCCTGGAGACACGAAAGACCAGTACCATCATCAGCCATCACCTCAATCTCCTCCACCTGACCTACTCAGCATATGACCTACAATGTTCCCTTCTAATAGAGGGTGTAATTCTCTTCCTTTAGTTTTATTATCAGTTTTATTGGGGGTTGGTTTATTGTTTAGGCAATAAGCTTTTACTAGGAAGTGAGTTTCATTGTACAGTAACATTGGCTAAGGTATCTGAGTAACATTATCCTTCAAATTTTCCCATTTGCTGTTTAGCTTTTTGATATTGAAAGAATC

>M6-PGSC0003DMG400016709 M6 LRR-RLK

CGACACCTGTAGATACACACTGCCAGTGGAGTCTTCTTATATAAATTTCTCTCTCTCTCCATTAAAAAGAAAGCTTGGAATCGGAGAGTCTCTTAACGATGGTGACACCTACCACTAGTGACGACGTTACCTTCTTGTCATTTCCCTGTCTTTCTTTCGCCATGCTTCTTCTTCTTCCTCTTACCTCTGTCTCTGCACAACCCCTTTAATTCTTGACTAACAAGAACAAAACAGAAAATGAATAGAAATAGCTGTAAGCAAGACACCCTTTACCCCCTTTTGCCAGTAGTGGTGATTGTGGCAGTGGGGTTTTACTCATCTTTTGTTGCGGAAGGAAACACTGATCCGAATGACTGTGAGTTTCTTCACTGGTTGAAAATCTTGGATCCGTTTTTAGAGCTTCATTAGTGATTTTGTGCTTAGTGGGTTTAGGTGTTTTTCTGTTTTCTTGAATGTGTTGAAGGAGGAGGCTTTTTTTGTTTTTTGTTTTTGTTACTAGGTGAATCTTGTGATATAAGAACGCCTTTCTTGTGATTATTTGAATGCCTGATTTGGTTAATGGGGTAATGTATTTGTTTGCCCTTCTTCTGGGGTTCTTTTTCTTCCCTTTTTGTTGGGTTTCCCAGTCAATGCAAGTCAAAGATTGATACGTTAGTACAAAAAATATGGATTCTTGATCTTTGTACTAAAAATTATATGACTGGGATTTTCTGTGATTATAATGCGAGGACGCATGCAATTGAATTCGAATGCTAAAACTGTTTGATTTGATAAGATAATATGTTTTCTTGCCCTTTCTTGGTGTCTTTCTTGTATTTAATTAGTTTCAGAGCCAACACTAGCTTAAGATTGAGGCTGTACTTTATTCTCAAAAAAAAAAAAAGATTGAGGCTGTACTTGTTGAATCTTGATAAGATATATATTGAATCATGGTTTTCTTGTTGGAGAGATGTTCTGGCTTTGGCTTTGTCGTTGCCTGACTCTTGTAATCTGCAGTGCAAGCTCTTCAGGTCTTGTATGCTTCACTGAATAGTCCTTCACAGCTCTCTAATTGGAAGTCAAATGGTGGCGATCCCTGTGGTGAGTCTTGGAAAGGAATCACTTGTCAGGCCTCTTCTGTTGTTTCAATGTAAGTTTGCATAGCATTAACTTTGTGCGCATCATTATACATTTTAGTTATGATTTGATTGATATAATGGTTGGTTCTTGCTGAAACAGTCAAATTCCTGATTTGGGACTCAATGGAACGTTGGGATACATGCTAAATGGTCTTACCTCATTAAAGAACTTGTAAGAGCTTGTCCTTTTAGTGGTAAAGGGTAATTTAGAGGACCTACTATCACAATAAAATTGAACTTCTGCTATCTGTTGTCTTCAGGGATATGAGCGGCAACAACATTCATGATTCTATTCCATATCAGCTTCCACCCAATCTCACTAGCCTGTAATGTTGACTCCTTGTTATGTGTTTGGTTTGTTTGAAAACTTCCCGGTTTATGATTGTCTCACTGGAACTATAATCATTGTCAATGCAGGAATCTTGCAAACAACAATATAAGTGGCAACATTCCGTATTCCTTTGCTGGCATGGTTACACTTAGCTATCTGTGAGTGGCCAGATTCACTAGGCACTTTCCAAAACAAAAATAGCATTTGCTTGGTCTTCTCTTTATTCTATTGAATCATTTCTGGCAACTATTTAAAATAGCTTACCCTACGATTCATTTTTCAGGAATCTTAGCCATAATTCACTCTCTCAAACAATTGGAGAATTGTTCACCAACCACTCCGATCTGGGAGCCTTGTAAGTGTTCTACTTAATCTTTACTTGATTGATCCTTTCTGGTAATTTTTTTTTTCCTTTTCTTTTTTTCTAATAGAGTTCATTCATTGCCACACTTGAAGGGCTTCTTTTCCTACACCAGTCCAATTAGTTACTAGCTTGAATTCATAACACCATGGTGTCTCCGACCCACAGTAGCTGGCCCTTCATTTGTTTTGACGTGCCTGCAATGTACAGACATGACTGCTTCCAAATATGTTCTCAAAACAAATATAGGATCTAATCATACACAAATGCCTTTTCAGAATCATTATCTGCTTAATTTTAATGAACAATGAAGTCTGGAGGCTTGTAAAGTCCTACATTTTATTTTCCTTCGTAGCTTTTTTATGTGGAAATTCGTTTTGATTGTATTAGGATAGGTGCCCTGCTCCTCTTTGAAGCTTTGTAAACCTACATTAGAGCTCAGTACCCCTGCTACAAAGAGCTTTCTGGCTATATTGATTCCTCATGTCTTTTGAACATCTGTTTCACTAGTTAATAGTTATAACTTTTCACTGATTTGCTGCTACTTCAGGGATCTGTCCTCGAATAATTTTTCTGGAGATCTCCCCTCTTCATTCAGTTCGTTGACCAATCTTTCCACACTGTAAGCATAATAACTTTCTACTAATAGGATTTGCTGTTCCGCTACTTTACTTACCCATGAAATTGGCATCCCCTGTAGTCATCTGGAAAACAATCAGCTAAGTGGCCCTCTCAATATCCTGGTTGGTCTACCTTTGACTAATTTGTAAGTTTCAATACCTCTTCTTGAATTAATGGGCTTAGCTAGTTTGATTCAATGCCAGTAGAGTGTTTTATGGGCAAAACAAAACCTTTTTGCAGAAATGTTGCAAAGAATCATTTCAATGGATGGATCCCGAATGAGCTCCAATCTATCCCTAACTTCATGTAAGTAATGTCACATCCTATATATTCTTACATGGTTTAGAATGTTACGGACTTACAGTTTCAAGGGGTGTGACATAACATCTAATATGACAGATACGATGGGAATTCCTTTGCAAATGGTCCAGCTCCTCCCCCACCACCATACACTGCACCTCCTCCTGGAAGATCTCGCAACAATCGTAGCCATTCTCCTCCTAGTTCACGTGCAAGACCAGGTTCTGATGGCCAGCCATCTCCAGAAAATGGTGATAAAAAAAAGGGATTGACAACTGGAGCTATCATTGGCATAGCTCTTGGATCTTCTCTTGCAATTCTTTTTGCTCTGGTAGCGCTTGTATTTTGCTTCCGGAGGGGCAAGGGCAAGGAAATTGCTACAAGACCCTCTACAGGAAGTCTTCCAGTTGGCACCGATAAAGGTGTGTTCCCTTAAACTTTTTTCCTTCCTGACTCAATATTGTCATTGGTGCTAATACCCTCAGATGACTAACAGTTGTGGTGTGATGTGGCATCTTAATTTGGAAATTGGTACGAACTTGGTTGATTTCTCATGTACTCTGTTTAATCTTAGTGAATATGGAGGTACAAGAGCAAAGGGCGAAGCCTGTGGCAGCTGTTGCTGACTTGAAGCCCCCACCTACCGAAAAGGTGACTGTTGAAATGATACAAGGGAAGAATGGATCTGTCAGGAGAGGGAAGTCTCCAATAACTGCATCTGCTTATACTGTTGCTGCATTGCAAACAGCAACAAACAGCTTCAGTCAAGAAAATCTCATTGGTGAAGGATCTCTTGGGCGTGTTTATAGGGCAGAGTTTCCTAATGGGAAGGTAATTTCAGATGTTTAACGTTGTAACTTCTTACCTTTCCTACTCATGATGTGCATTCAAAACTATTTGAAGCTGAACAATATTGAATAAATTTAATCTGTCTCTCATGGACATCTTTCATTTTTGATGCTTCAGAGAATCACATACTTGTGAGTTGTATAACGTTCATCAGATACATAAGGACATCAAAAGGGGTATATCAAGATGTTGGCCACTCCACCTATCACCCTATTGCTAAGACGCTATAGCATAGTATCAAGCCGTTACACGTGTGGGACAGTGTTGCATTACCTAATGTCCAACCTCGGGTAAATGAGGCCATAAAGATATGGTAACTCAACCCCAATAGCTAGCTCATGAGGGGAGGATTGCCCAAGTCCATATAAGGAGACCACCAGTCCATTCCCTAACCAATGTGGGACTTTTACCCACTCTAACACCCCACTTCACGCCCAGGCCCGCTGGAGCATGGACCGGAGCCCAAATGGAGATGAACCTTGATACCATGTAAAGATATGGCACCTGGGCTTAACTCAACCCCAAAAGCTTCGCTCATGAGGGGAGGATTGTCCAAGTCCATATAAGGAGACCCCAGTCCATTCCCCAACCAATGTGAGGCTTTTACCCACTCTAACAAAGGCATCCAAAGTAGTAAATAAGGTTCTTGTCTTACTATATCCATTGCCTTAACGTTATGGTTGAACCCTTAGATCCTTTCACAATATTTGTCAAACCGGGCAAACACTAAAGGCTACATACTTGATCTATGTGTTACTAAACCTGAGTAGAAGTTAGTAGACTTCTTGATACTCTGCTCGCATGGGAGTTAAAGTATTTCCTCTTGCTAGTACATAATCTGTTTTGAGTGATTCTCTTAACTGAGATTAAGTCCTTGATACTCTTTGATTCAACTTGCTCTTTTAGATCTTTCACAAAGTTGATTGCCCACAGTAGGCTCACATTCTGATGAACACATGTCAATTGTTCACTCCTCTGGTTGGTCTGGTCACATCTTTGCTTTTATCTGTCTTGTGATGTTTGATGGGTGTAATTATATGGTCACTTTGATGGTCATCTCACTTCTCTTCTCAGCCTGAATTAACTATGATACTATAATCTTGTCCTTTGGTTACATGATGCAGTCCTAAGCAAAAATATTCTTAGCAGTTTCAAAGTTTGACATTATATATCTTCTACTTTGTGGTTTATTATTGGATGTGCTACTGGTTATTTCTTTTTGAATGAACCAATTGTGTTGACTATTTCTCTGCTCCAAACTTGATGCCAACTGTGTTCCAATGAATGATTTAAGAATTTCATGTGTCAACAACATTTTTGATGAGGAAACTTTTAGTTTTTCATTCATGTAGTTTCTACGGAAGGTGCCAACTGAAAATCTCAATTGATCTTTTGAACTGTCCTTACCAGATAATGGTCATCAAAAAGATCGACAGTGCAGCACTGTCGCTGCAGGAGGAAGATAACTTTCTTGAAGCTGTCTCAAATATGTCAAGGCTGAGACACCCAAACATTGTTCAATTGATAGGATATTGTGCTGAGCATGGACAGCGTCTTCTGGTTTATGACCATGTTGCAAATGGTAGCTTGCATGATATGCTGCACTTTGCTGATGAGAGGAGCGAATTGCTTACTTGGAACGCCCGGGTTAGAGTAGCGCTTGGCACTGCTCGAGCTTTAGAGTATGTTATAGCTGTATAGTATTTTACCTGTCTTGCCCGTTTCTTGTTGGGTTAGCTTAAAACTTGACCGTCTTCTTTTCAGGTACTTGCATGAAGTCTGTCTACCTTCTGTTGTACATAGAAACTTCAAATCCGCAAATATTTTACTAGATGAAGAGCTCAATCCCCATTTATCAGATTGTGGGTTAGCAGCTTTGACTCCAAACACAGAACGCCAGGTGAGTGAGTAAAGAAGATTTAATCTACGTCGTCGTTTTAACAGAGGGTGTTGATAATATTGGGTGGTACGTGATGCATATCATGTTTATGAGATGATGTCAATCAACTTACCTAGTACTGTAGTAGGAGCAATATTAATGAGCAACTATTTCTAGGTAGATCTTTACAGACTCGGTCATGAAATTGGAGATTCTGATTGTGGCAGCTTAGTTTTGTTTTCTAATTCTTCTGGAGCAATCTTGTTAGATGCTTAAACACTTGGCTTACCATAGTGCACAAACTTTAAGGAAGACCCCTTCCCCCTGCCTGAAATTCGATTTAGTTCTGTGAATTATTTTTGTGAAAAGAGAACCTTCTTGTACTGAAGTTCGGGTGAAGGAAGCTAATGTGGGAAGGCTAATTGACGCTGTTAAGGGCATTGCCTAAAATTGTAATTACTTCTGAATGTTGACAGGTGTCATCCACACAAATGATTGGTTCATTCGGTTATAGCGCTCCTGAATTTGCTTTATCTGGAATATACACGGTAAAAAGTGATGTATATAGTTTTGGTGTGGTGATGTTGGAATTGTTGACAGGAAGGAAGCCACTTGATAGGTAAACTAGATCTTTTTACATTGCTAGTCAACTTTTACACTATTGAATTCTTTTAATGACTTGATTCTTGTTAACAATCATATAGTTCAAGGGTGAGATCGGAACAGTCCCTTGTGAGATGGGCTACGCCTCAACTCCATGATATAGATGCCTTGTCAAAAATGGTTGATCCTTCACTAAATGGCATGTATCCTGCAAAGTCTTTGTCCCGTTTTGCTGATATTATTGCACTCTGTGTTCAGGTGAGCTTTACTTGCATGTTCTTTCATTTGGAGCATAACAGTTGTTAACATGTGAAGCTTAAAAGATGAAACCACTTCTAAAATATCTTTGATGGAAGCGAAAGTCAATGATAATAGGGAATTTACACACTTAAGGACCAAACTCGACATTCTAACCCGTCAAATAAATGATACTCTACCAGTTTTTGCATTCTATATAAGTCCAAATTCAGGGGCGTGGATAGATTGGGACCACCTGCACTCTGAGTTTGCTCATGGCTCAGTGCTGAAGTGAATTGATGTGCTTCAGCAAATAGATGACCTTTCTTACACATACTCGGTGAGTCTTGGAATGGCGAGAGCACCCATCTTGGTTTAAGTTAGTTTGGGAACTTCCAAGGCGTAAGTCCTTGCACACCATGGTCCATAGTGGCTTTGGGGGCTTAGTCTTTAAATGAGAAGTTCTTAATTAAGCTTTTCTTGATCTAAGGTGACTTCAAGACTCACTCAACTCACGAAGAAAAGTCAGATGGACTCAACAAAATTATAATGAATACTTGCTCATTGACATATCTGAAGTGAATTACTTTAATTGCAATAATTTGAATGCGAGTGTTTTCTCCTTGGTAATAACATTTAACATGTATTTCCTGTACAGCCTGAACCAGAGTTTCGACCCCCAATGTCTGAAGTTGTGCAAGCACTGGTTCGGTTAATGCAAAGGGCGAGTGTGGTCAAAAGGCGATCCAGTGATGAATCAGCTTTCACATACAAGACTCCGGATCATGAGACGGTTGACATGCCATAGATAAACATACACGACGAAGCAGCCTTTGCAATTTTGTAAGTAACTTCTGTGAAGGTCAAGAGCAACTATGGCAGATAAAACTCGGAGGTGTCAATACCAGGGACATTAAAAATGTAATTATTTTTCACATCTTCCTCGTTAAAAAAAAAAGAAGCTTTTTTGCATCCCAATTTGAGGTACATAATTCACTGACAATGTCCTCCAGAGTATATTTACAAACCATCTTGGGATACTGGATATAATTTATTGTCAATAACAACATTGCTACACATTATCCTTCTGTGTTGTTTATATTACAGTTTTTCCAATAACTTTTGGCTAGCCTAAGAGCTAACTGTAGTGTGCATCGGTTGATTTGGTT

>M6-PGSC0003DMG400016872 M6 LRR-RLK

ATGTCTGGAGTATATGACTCAAGAACTGGTCTAATTCTGATTTGGATAGTTATCTTGTTGGCTTCCGTGACGATGGTTTGTCCTGCAGAAGGATTGAATGCTGAAGGAATGTACCTACTTGAGTTGAAGAAGAACTTAAAGGATGAGTTTAACAATCTTGAAAATTGGAATCCTAGTGATGAGACGGCGTGTAGATGGAAAGGCGTAAATTGCACTTCTGATTATAATCCGGTTGTGCAATCTCTTGATTTGAGTTTTATGAATCTGTCAGGCACTTTGAGTTCTAGCATTGGTGGTCTCGTAAGTTTAACCGTTCTTGACTTGTCTTTCAATGGATTCACAGGGAATATTCCGAAAGAAATAGGGAACTGCTCAAAAATGCAGAGTCTTCAGCTCCATGATAATGAGTTTTATGGGCAGATTCCTGATGAATTGTATAACCTTTCTCATTTGAAAGACTTGAATTTGTTCAATAACATGATATCTGGTCCTATATCAGAGGAGTTTGGGAGACTCTCTTCTTTAATTTCTTTTGTTGCATACACCAATAATCTCACTGGTTCACTCCCTCGATCACTCGGGAAGCTAAAAAAATTGGAAACATTTCGTGTGGGGCAGAATCCGCTTTCTGGAACCTTACCTGCTGAGATTGGTGATTGCAAGAGTTTACAAGTTCTTGGTCTAGCACAAAACAATATAGGAGGAAATATACCCAAAGAGGTCGGTATGCTTAGAAGATTGAAACAACTTGTACTTTGGGACAATCAACTCTCGGGCTATATCCCGAAGGAACTTGGAAACTGTACAAAACTTGAACTGCTTGCTTTATACCAGAACAACCTGGTTGGAGAAATTCCTGCTGCAATAGGTAAGCTCAAATCTCTCAAGAGGTTATACCTATACCGGAATGGATTAAACGGTACAATTCCAAGGGTCATTGGGAATCTTTCGTCAGCGATAGAGATTGACTTCTCAGAGAATTATCTAATAGGAGACATACCAATTGAGTTCAGTCAAATAAAGGGACTGATGTTGTTGTATCTGTTTCATAACCAGCTCAATGGTGTGATACCTCGGGAACTTTCTAGTTTAAGAAAGTTGGAACGTCTTGATCTGTCGATTAACTACTTATATGGCTCTATTCCATTTTCATTTCAGTATCTCACTGAATTGGTTCAGTTACAGCTCTTTCAGAATTCTTTAAGTGGAACTATTCCTCAAGGTCTCGGTAACTATAGCCGACTTTGGGTGGTTGATTTTTCTGACAACTACCTCACAGGAGGAATTCCTCCTAATATTTGTAGGAATTCAAATTTGATTTGGCTAAATCTTGGTTCAAACAACTTGCATGGAGTCATTCCATCTGGTGTTATTAAGTGTGACTCATTGGTACAACTTCGTCTTGACGGTAACTGGCTACAAGGGAGTTTTCCTTCTGACTTGTGCAAACTGAGTAATCTATCTGCTCTTGAATTAGGACAGAACACGTTCGGTGGTCTAATTCCTCCTGAGATTGGAAACTGTCAAAAGTTGCAAAGACTTGATCTTTCAGGCAATTATTTCACACATGAATTGCCTCAAGAGATAGGGAATCTTGAAACACTTGTGACGTTTAATGTCTCGTCTAATTTGCTGACAGGTCAAGTACCACAAGAAATTCTGAAATGCAAGGCGTTGCAACGACTTGATCTCAGTAGGAACAGCTTTAGTGGTGCTATACCTGATGAAATTGGAAAACTAGCACAGCTTGAACGTCTTTTGGTTTCGGATAATAAGTTCTCTGGAAAGATACCAGTAGCATTGGGGAGGCTTTCTCGTTTGAACGAGCTGCAGATGGGTGGAAATTCATTTTCAGGTGAAATACCATCAGAATTGGGTGATCTTACTGGTCTACAGATCGCGATGGATCTTAGTGACAACAATCTCTCTGGTTCAATACCACCTAAGCTTGGAAATCTTATTCTACTAGAGTGTCTCAATCTCAACAATAATCATCTGAGTGGTGAAATACCAATCACATTCGGAAACTTGACGAGTCTAATGAGCTGTAACTTTTCATACAATAACCTCACTGGACCATTGCCGGATATACCACTCTTTCAGAATATGGATGTCAGCAGCTTTATTGGAAACAACGGGCTTTGTGGTGGTCGTTTAGGTGGATGTAACAAATCTCCTCCGTTCAATTCTGATCCTCCAACCAAAAATGCAGGTGCTCCAAGAGGGAAGATTGTTATTGTTGTTGTGGCTGTTGGTAGTGGTGTTTTTCTTGTTTTGATTATGGTAATTTTGTATGTGATGAAGCGGAAACCAGTTGATCAGATGGTTGCATCAGTAAAAGATAAGAATGCGTCATTTCCAGCTTCGGACATATATTTCCCTCCCGAAGAGGAGTTCACTTTCCAAGACTTAGTTGAGGCTACAAACAGTTTTCATGACAGTTATGTTGTTGGAAGAGGAGCTGTTGGGACAGTATACAAAGCAGTCATGCAATCTGGACGAAAAATCGCTGTTAAGAAGTTGGCTTCTAACAGAGAGGGTAATAACATAGAAAAGAGTTTCCGAGCAGAAATTTCAACTCTCGGAAAGATTAGGCATCGTAACATTGTAAAACTGTATGGCTTTTGCTATCATCAAGGTTCTAATCTGCTTCTTTACGAGTACATGGACAAAGGTAGCTTGGGTGAATTGCTTCACGGTGCATCTTGTAGCTTGGATTGGCCTCAACGCTTCATGATAGCTCTTGGGGCAGCTGAAGGACTTTCTTACTTACATCATGATTGCAAACCACAGATCATCCACCGCGATATAAAGTCCAATAATATTTTGCTCGATGAGAAGTTGGAAGCTCATGTTGGTGATTTTGGTCTTGCTAAAGTTATCGACATGCCTCAAACTAAGTCGATGTCTGCAATTGCTGGATCATATGGCTACATAGCCCCTGGTATGTACATTTCATTATTTTTTCATTAGAACTTGTGTAGTATTGCCTGTCTACTGTCTGCGAATAATGTCAAGAAACATATGGCCGGCTAGATTGCTCTAACAAGACTAATATATTTTAGATAAGCAATGACTATAACTCTTCATCGAAAAATCAGTGAACTTACACGAAAAATCTCAAGTATAGTTCCTCCCATGAGCTGACTAGTGTATGTTAGTTTTCCTCACACTTTGACTAGCAAATAGTACTAGTGTTTCCTTGGTGAACGTTTTTGCTTCTTTTATTTATACTCAACAACAACAACATACCCAGTGAAACCCCACAAAGTGCCATCTGGAGAGAGTAGAGTGAACACAGACCTTACCCTTACCTCGTAGAGGTAGAGAGACTGTTTTCAAAAGACCCTCGACTCAAGTGAAGCGAAGAGAACAATACAACTAGTAGAAAAGAGTGCAGAGCATACATGCAAGGAATAGTAACATCAACGACATAATACGATCACCTAAACATAGTAAACAACAGATGATATTTGAAGTAAGGGTAATACTATTTGATGAACTATTTGAAGAGAGTAAGCAGAAGCGTCTACTTTGTTTAACGGTTCTGCTTTTCCCATTGCAGAATATGCTTACACCATGAAGGTAACAGAAAAATGTGATATCTATAGTTACGGAGTAGTCCTTTTGGAGTTGCTAACAGGAAGAACACCGGTGCAGCCACTCGATCAAGGAGGTGATCTTGTCACTTGTGTAAGGCATTATATCCGGGACAATTCGTTGACACCAGGGGTACTCGATATCCGATTGGATTTGACAGATAAAACTACTGTTAGTCACATGCTTACAGTCTTGAAAATTGGTCTAGTATGCACTTGTTTGTCCCCGGCTGATCGCCCTTCAATGCGCGAAGTTGTGTCAATGTTGATGGAGTCTGATGAGCAAGAAGGTAACTTCATCTTGTCTCAAAGTTGA

>M6-PGSC0003DMG400017291 M6 LRR-RLK

TAAATGATCATCTCTAAGCTTCAAAAAATTATTAATATTATGGACTTGCTAAACTTCTAGTCTAATTTTGTTGACTATTTTGCTTCACAGAGATGTTTCAAATAACAATTTTACATGGGAAACCTCACCTTTTGAATGTCCCCGTGGAAATCAGTAAGTTGTTTGATAAAATTGTTTTTCTTCATTTGACTTATTTGTTTGTGATAATTTTCTAACATCAAGACGATTTATTCAGAAACTTGGTGGAGAGCTATTCTGCATTGGGACAAGAATCGTAAGACTAATATGCCTATAATAAATTTATCTGTTGCACTGATCTTACAGAGGAAATTATTGTTCCTTGAGTTCACCTATAACTAAGAGTGAACTCTTACCTACCATAGAAAACTTTTGTAGCACTCTTAACAACTTTTTCTTTTGTTTTCTGTGTTGCAGAAACAAAATTCATCCTTGCCTGAAGCAGAATTTCCCTTGTTCTGAGCCAATAGATCGGCGTAAGTAAACTTTTATGTGTCCATTTCTGTAATTGATAGCACAACAAAATCTGTCTCCTGTCACATAAATAGGTGTACTTGTAGTGTGAGGTCAGCTAGTGGCAGAAAATATAAACAACAACAACATAGCCAGAGTAATCCCACAAGTGGGGTATGGGGAGGGCGGTGTGTACGTAGTCTCACCCCTGTTAGAGTGGGTTAGAGTCCTATATTGGTTGGGGAATGGATTGGTAGTCTGCTTATATTGACTTGGACAATCCTCCCCTCATGAGCTAGCTTTTGGGGTTGAGTTAGGCCCAGGTGTCATATCTGACTCCTAGTAACAACAAATAATACAATTTAAGAGTTTGGGACTGACTGCTGTCTTCTGATCTGTCCATTGTGATTTTCTGCTGAATTCTTGTTAAGAGTGCCCTTGTCTACTGGTTTGCAGAACAATATTCTATGCACATTAATTGTGGAGGCAAAGAGGTAACCATCAAGAATGGTACAAAATACGCAAACTATGAAGCTGATTTAGAACCCAGAGGTGCTTCTATGTATTATTCAAGACATAACTGGGCGTTTAGCAGCACAGGAAATTTTATGGACAACGACGTTGAGTCTGATGTCTATATCAATACCAATGTCTCTGCCCTGCAAAATGTCAAGGTCCCTGAACTGGAACTATACACTACTGCACGTATATCTCCCCTCTCCCTCACATATTATGGACAATGTCTAATGAATGGAAACTACACTGTCAAGCTCCATTTTGCGGAGATAATATTCACAAATGATACTTCATTCAACAGCCTTGGAGAGCGTATATTTGATGTCTACTTGCAGGTAAATTCATGCTCTGGGAGAAAGTGAATTTCATGTTTGCTGCCACCAGTGCATTTATTCAATCTTCTTGAGACAGGAAAATTTGGTGTTGAAGGATTTTAACATTGCAAAAGAGGCTGGTGGGCCTGGTAAGCCAATTGTGAAGACGTTCACTGTGAATGTGACAAGCCACACGCTGAAGATACATTTTTACTGGGCAGGAAGAGGGACTACAGGAATTCCATTTAGAGGGGTTTATGGTCCTCTAATTTCAGCTATATCAGTTGTTAATAGTATGTTCCTTTACTCTCTTCAAGTTTATATCTGTCTTGCTTACAGTTGTATGCTCAATTCTTCTTTAATGAATGCCATAGGGAACATAGGACATTATTCTGACGTACTAGTCCTTAAAAAAACCTTTCCGAGACCAAATGACTCTACATACCAAGGCTTAATAAGTTTTGTTAACCATCTTGATGTTATAAATTCTATTCCTACTGATTCATTATGCATCTAAAACTGCTGTCAACATATTCTTTCAGGAAGTTAATGTGTATCAGAACCTAAGAAACTTCATATACCCTTAAATTTCATGATGCATTTCAAATACAGTGGGGAGACCACTTATAGTGTCCCCAGTTCAGTTCCTTAATCTTGTCTGGTAGTGGAAGCATCAAGACTTATACTAAGACTTCAAAAGAAAATGTTTCAAATTGCATATGCAAGTCAATTAAAGCTTGCTAAAATCATCACAAGTTTTGAGATGTCATATTATTTTTTCCTTTCCGTGCAGATTTTCCCCCTCCTCTGCCTGCTCGTCTGCCTGCTCATCTGCCTGCAGATCAAAAGAGTAAAATTAATGTTGGCATAATTGCCGGCATAATCGCTGGATCAGTATTTCTTGTCCTTCTGATTATTGGTATCTTGTACAAGGGAGGCTATCTAGGCAAAAATGTATCTACTGATAGAGGTAATAACTACAAGTATTTTACTTTTGAAAGTTTAGTATCCATTTATTTGACCTATGAGTCAGCAATAATGTTCTCAAGTTAAATGCTAAACTACATAATTACTTTCTAGCCAGTATTTCAGTTCATACCAAATTATATCAGCTCAAATACTTGAAGTTATGCTTTACGTTCTATTGACTTGCTTGTTCATCTCATCAATCATTTATTCATTTTAGGCTTTCAAAAATCATTAATGGTCTGAAGCACTTGACCAAAGTTTATATGAACCTTGAAGAAACATAGTGTTATAGCCATTAGCTTCCACTGAGATACTTTTCCTCTTGAACAGAACTTAAAGGACTGGATCTACAAGCTGGACTATTTACTCTCAGACAGATCAAAGCCGCGACAAAGAATTTTGACCCAGCAAACAAGATAGGGGAAGGTGGTTTTGGTTCAGTTTATAAGGTAATCATCTTTTCCTTGTCTATTTTAAATCAAACACATAACTATAAATGTCATTATATTAGCATGAGAAGGATCATAATGGCATAGTGCAGATGTCTCTTAAGAGAGAAAATGAAAGACTAAAATCTTAAATTTCATAAACCTTCAAAGGAAGTGTGAATTTTCACCTGTAATGAGGAGACTGCACAGGAAATGCTCTATTACCATATGTGTTTATATATATCTGAGAAGTGGAGTAATTTACTGATCTTTTTCCATTCAACTAAGATGGAGATTTCTTATCTTCGCGACATAGGATTCAATCTATTATAATCTTACTTGCTGATCATAAGCTGCAGAAGTCTGAATAGAGTTTTGAATCATGACTTTTGTACACAGGGTCTATTATCTGATGGAACAGTAATTGCGGTGAAGCAACTTTCTGCAAAGTCAAAACAAGGAACTCGAGAATTTTTGAATGAAATAGGGATGATCTCTGCTGTACAACATCCAAATCTTGTAAAGCTTTATGGATGCTGCATTCAAGGAAACCAATTATTGCTAGTCTATGAGTACATGGAAAATAACTGTGTATCTCGTGTTCTTTTTGGTAAATACTTCCTAACTCCGATCAATCAAAATATGTCTGCTTCTTATGCACTTGACATTGTTTATCTAACGGAGCTGGTATATACAACTTTTTCTTTTTGTTTGTCTTTCTCTTTCAGGCAAAGGTCCAATTGGCAAAATGAAATTAGACTGGTCTACTCGGAAAAAAATTTGCCTAGGAATTGCAAGAGGTTTAGCATATCTCCATGAGGAGTCGAGCCTGAAGATAGTCCACAGAGACATCAAGACCAGTAATATTTTGCTTGACAAGGACTTCAATCCAAAAATATCTGATTTTGGCTTGGCAAAACTACATGAGGATGATACTACCCACATTAGTACCAGGATTGCAGGAACTGTGTAAGTATTATGAAACTAGTTCCTCTTGTTCCGATCCCATTAGCAACAACATTTGGTTGATATGTTTCATAGAAAATGGCTCAACATTTAATCCACTCATTTATCTTGAAACACTCGTGGTACTGTTCTCATCCGAAAGCAACTTTGCAGGTATAATCTGCCAAAAGTAAAGCAAGCCAAGAACTTATTACCTGCCTAAAGAAACTTAACTTTTATAGATATTAAACAGCCTTTGTAGAACAGTAACATAGAATTTTATTCGTCAATGGTAAAACCACCTCTCTTTCCTTACCATAGTTGTAACTCATTGGTTTGCTAATTATTAGCACCTCAACGTCTCACTTCGGTCATAATTTTACATTTTGCAATTCAACATAGCTATAGGTGAACTTTCAAGAAAATATTTTCTTGCTACTTCAAATTTTAAGAAATGTGAAGATTGTTTGTATCAGAATTCTCTGTAATTAGCAAGTAACTGTATTAATGATGAGGAAACCAAGATCTAATAAGTTGGAACATCTTTGGTATTTGGTGGTACAGGGGTTACATGGCTCCAGAATATGCAATGCGAGGCTACTTAACTAGCAAAGCAGATATTTACAGCTATGGAGTTGTTGCATTGGAGATTATTAGTGGGAAAAGCAATACAAATTATAGGCCAACGGAAGATTGTGTTTATCTTCTTGACTGGGTAATTTTAATATCTCTCTACAACAATAAACCCAATGTAGTCCCAAAAGCGGGGAATTTCACTACTTCTCTCTATTCGACCAAATATCTGGGCTTCCATCATCCTTAAATCACTTCCCCTTCTAGTTTCATCTCAACCAGAGAACGTTTTATCAACATTCCGAGTAATGCTAATAGCATCTTATTAATCTTCCCCTGCATCAACTTGTACATTTAGAGCCAAAGTACTTTAAAATTGGCTTTGGCAAATGAACATGTCATACTTTTTTAGTACATTAACTGTTCTTCATTTTCTGTTTTGATTAGGCTTATGTTTTGCAAGAGCGCGGGAGTATATTGGAATTGGTTGATCCAGACTTGGGTTCTGACTACTCGTCACAAGAAGCAATTGTGTTACTGAATGTTGCCCTTTTATGCACTAATGCAGCTCCAACACTTAGACCAATAATGTCTCAAGTGGTAAGCATGCTCCAAGGCCAAACTCTTGTTCAGGATGTCCTTTCTGATCCTGGGGTGTCAACAAGCGGTTCGGGATTCAGATCAACGAGAAGTCACTTCTGGCAAACTCAAAGCCTAACTGATGGTACGCTTACTGATTCCACTCTCTCAACTGGAAATGCGGAAGAAATTGGCATTCTTCAAAGAGTAGAGCCTAATGTATCTAACAGTTAAACAAAATGTCTGTTAACACAAGCAAGTGTATATATGAACATCAAAATGTTTAATGAAGATTAAGTTTAGTCTTTACAAAACTGCTAACTGGTCCTCCTTTTATAGTACTTCATTTGTTTATTTTCTAAGCCAACCTCATTGTTTCATAGTTCTATCAATTCCTTCTTTATTTTTC

>M6-PGSC0003DMG400017645 M6 LRR-RLK

TGTCACAAAAATTGTACAACTAGACTAGCTATAGACAAACACATCTGTACTCCATTATTTAATCTCATCATTTAAAATCACTTTTCTCTATGTATCATTACTCACTCTATTATAGATTCTTGAATTATTTCATTCCACATTTCTCTTTAATCACTTCAAAAAAACAAAAAAAGTGCTCACTCTTTTGTGAGCTCAAAAGATTTTTTTGGGATTTTGAGTCTTTTAGCTATGAAGGGTTTCTTGTTTCTTGGTTTGTTTTTGTTTTTTCTAGTTCCATTTTCATCATCTCAGTTAGTTCCTGCAGAAAGTAGAATCCTTTTTCAAATTCAACAGTTTCTTGAAAATCCACCTGTTCTTCAAGAATGGAATAAATGGACAAATTTTTGTTTTCTCCCACAATCTCCTTCTCTTGTTATCACTTGTTCTGGTAATCATATAACTGAGTTAACAATTGTTGGTAACAAAAAATCCCCTTTTGAGTCTTTAAAATCCTCTTCTCCACAAGCCCTTTCTGGGAAATTTTCAATTGATTCATTTTTCACTGTTGTTACTAAGCTTTCTAGTTTAAAAAAGTTGTCTTTAGTTTCTCTTGGTTTGTGGGGTTCATTGCCTGCTAAGATTAGTAGGTTACATTCACTTGAGGTACTTAATATAACTTCAAATTTCATTCTTGGTGAGATTCCATCATCTATAGTGAATTTCAAGAACCTGAAATCTCTTGTTTTGGCTAGAAATTTGTTTAATGGAAGTGTACCAGATCTGAAAGGGTTAAAAATACTTGAAGAGTTGGATTTGAGTGGTAATAATTTAGGACCAAAGTTTCCATCTTTAGGTGATAACAACAACAATCTTGTTTCACTGAATTTGAGTAACAATCTCTTTAGATCAGAAATACCTAATGGACTCAACAAGTTTACACATTTGCAAAATCTTGATTTGTCTTCTAACAAACTAGTTGGTCCAATGCCTTCTTTCCTGTTTTCACTTCCAGCAATTCAGTCTATTAGTATTGCTAAGAATCAGCTTAGTGGTGCCCTTCCAGGAAGTGTGTCTTGTAGCAACAATCTGAAATTTGTCGATTTTTCGAGTAATTTATTGATGGGAAAGTTGCCTGCTTGTCTTGGATCAAGTTCCAGAAACAGGACAGTGATTAATGTTTGGAATTGTTTGTCAAGTACTAGTACTAAGTCTCAACATCCACATACATTTTGTGAGAAACAAGCTATAGCTGTTAAGCCCCCTCCAAGAACTAGTGATGAGAAGGAACAATCCACAGTTAAGCTTGGTGTTGTCTTAGGACTCATTGCTGGAATTGTTGTAGTTGTGGGTGCAGTTGGTTTCCTGATCTTTTTCATTGTCAAGAAAGTTGTGAGAAATAGAGATCAAGGATATAAAAATGACAGCTTTGCTTTTGAGAAGAATTCAACCCTTTCCAAAACTGCTGATGGAGGTAAATTTCCTTAATGATTAGGGTCTATTGGAAACAGTCTCTTTACCGGTCTCTTTACCTGTCAAGGTAGGGGTAAGGTTTGTGCACACTCCCCCACTAGACCCTACACTGTGTATATTGAATCTTGAATAGTGATAGGGGTCTATTGGAAACAGCCTCTCTACCTCCCGAGGTAGGGTAAGGTCTGTGTACACTCTATCCTCTCCAGACCCCACTTGTGGAATTACGCGGGGTTTGTTGTTGAATCTTGAATAGTGATAGTGCATCATTATGACAATTTTATTGATAACTTGATTTAATCAGGGAATGCAAGGAGGACAATGAGGATGGTATCATTAGGACTTCCACCATATCATGTTTTCACTTTAGAGGAAATGGAAGAAGCAACAAACAGTTTTGATCCAACTAATCTGGTTGGAGAAGGTTCCCAGGGTCAGGTAAATATGTTGATTCATTATAAATAGCAAAAAAAAAAGATATTTATTACTATATCCTGTCCATATTATTCGTCATTTGGGACGCCGAACAATTCCATTGTAGTCAAATACATAATCTACATGAGTATACGAGTCTGTAGATACTCATATTGCTATACTTCTAGTCTAGACCCAAAAAGTCAACCCGAACAACACTTGACATTTTAGAAAAACTAAAGATAGTAATATGGATATGCTTTTCATAAATCTTGGTGAATACAAACTTGTGTTTTGAACATCGGGATTGATGTTATTATTAATGGCAGCTATATAGAGGTTGGCTTAGAGATGGATCAGTGGTCCTAGTGAAATGTCTGAAATTGAAGCAGAAGCATTCACCTCAAATTCTGCAGCAGCACATGGAGATGATATCAAAGCTAAGGCATCGACATTTAGTAAGTGTACTTGGACACTGTGTTGTCACTTACCAAGATCATCCAAATACAGCTAGCACTGTATTTATTGTGCTTGAAAACGTCGTCAATGGATCACTAAAGGATCATCTTAGTGGTAAGCTATTACGAGTTGATTTCTCTCATGGCCACTCAACTAATAGTTATTACCTCAGAAGTCACTTTCGAAAAGGTGACTTCTGAGATAATGACTATTAGTTGAGTTACTATCTAATAGTCATTACTTTCTTCAGTCGGAACGATGAAGGACTGAATTTTATATTCTGGGATTCGTTCTTTGGCAGATTGGAGGAAAAGGGATGTATTGAAATGGCCACAAAGAATGGGAATAACGATGGGCATTGCAAAAGGAATTCAGTATTTGCACACTGGAGGAGTCACTGGAAATGATATAAAGCTTGAGAATGTTTTGTTGGATGAAACTCTTACTGCTAGAATAAGCAGCTACAATATATCGTTACCACCTAAGGTAATGCTAGTAAATCACAACAAACAACACGTTAATCTTGGAGAAAATCTCATTTTTAACCATCCAAAACATTCGCAGGTTGGTTCGGAAAGTCTTCTTGCTGGACCAGACAATTTTACCAGGTATTATATTACTCTGCAGCTACATTCACCTTAATTTCTCTGTTTCCTGGCTATCATTTCTGAATTGTCTGCTAATATAAACTTCTTATTCAGCGCGAAAGAAGCAGAAAAGGAAGACATTTATCAGTTGGGAATTATTCTACTAGAGGTTATTATTGGTAGACCAATCAACTCTCGAAGTGAAGCAGAAGACCTTAAGCTTCAGGTATCAATAACAATATCCTAATAACACTCTCAAACGATAGAAATATATCAGATGGTCACTCAACTAATAGTAATCATTTCAGAAAGTCACATTTCTTCTTGATTCGGACATTCTTTTACAAAGAAAGTGACTTTCTGAAATAATAACGATTAGTTGAGTGACCATCTGAGAAATCAACTCAAAAAATAAGAGATACGCTATAAAGAATTAGAAGAATCACGAATTACGTGATATGAAAGAACACACATTTCTTCACATAATGCATAAGAGCCTAGTCTATTTACACAATCTATCAAACAACCTTGAACCAAATTGCCTTTTATGACATTCAGGTGGAGACTGCCTTAGCGGAATCACCATCAAAGCTACGAGATTTAACAGATCCTTGTATACGAGGTACATTTGCATATGACTCATTGAAAACGACAGTCCAGATAGCAATAAACTGCCTTGAAAAAGAACCAAGCAGGCGGCCATCAGTCGAAGATGTTCTCTGGCATATGCAATACTCAATTCAGGTTCAAGAAGGAACAACCAACAGTGGAAACCTCAGTGGGAACCAAAGTGGGAAACTCAGTGGAAAAATAAGTGGAAATCTTAATAATAAGTTTTACTGATAATAGAGATCTGTCATTATATTGTTATATCTTGTTGTTTAGTCATTTGTATGCATCCATCTTGTATAGCAAAGTTCAGTAGCTTACTGAATAAATGTTTTTGCTTTCTACATCACAAAAATGTACCCAAGTTTACAATACAAAGCAATTTTCATGTCGCGAAAAACGGTGTTTACATCAACTCAATGAAGACATAAAGTGTATCTTCATGTGCTTTGTTATCGCTGAGTGTAATCACCCCCAACAACTTCATCATCAATTATTTTCAAAATAACTAGCTGAAGATATAATTATAAGTAAACAACACTGCAGTCTCTATGGTCATGATATGATTCAATATCGTATCAGAGCAGGCAAAACCTCCCGGTTCATGTCTCACCACCAGAAATTTCCATGTGCTTGGCCTATTAGGCCTTTACACAGAGTTATGTTGAGGATATAATTAAGTAAAAACTAAAAGTATTGTCTCTCCAGCAACTTACTTAAACTTTTAGATGAAATGCTCACAACGTATCTTTGTTGACTTAAAAGGGTACTTTATACCATTTTCAGCAGCCGCGCCTGCCACGGTGGCTGCAGCAACAGCAGAATCTGCTGTTGCATTCATCGTTGGTGCTCCAAGAGCACCGACATCAATTACCAATTCCAGCCACAACAGCTGAATTGGCTGATCAGGCTGCTTCTTCTATCTCTGCATTCTCCAGCACCCAAAGAACAATAACAGAGTAGTTAAGTATTCCCAAGTATCTTTCTCTCCAGCCATTACTTTGTCGACACTACTAAAAAATCACTATTTTCTCCAAGAAATTTTTCACTGATATTTTGATTGAATCGGTGAGAAAAGAGTAATGTTCTCGTACAAAAAAACTAGAAAGTTTACCACTATTTCATTGAGAATCTGTTTTCCACCATGCATATTCCCAGTGAACTGATTTGGTGGAAAAACATGTTCTATAACACAAATTTCTCACTAATTCACCGTGGAAAAAATCTATTACTACTACTCATCATTAGTACTTCATCCGTTTCCTCTTATGTTGAACTTGATTGATTAGGCGCGGCTAAGAGCAACATGGGAGGTTTTAAAGTTCAATTTTTTCCAAATATAGAAAAAGACACTTTTATAACGTAC

>M6-PGSC0003DMG400017690 M6 LRR-RLK

CACATGTTAGTTGGCATGTCTCATCTTAAGCGGCATGGTCTGCAAATTCTCAGAATAACTTGCTAATGATCTCTTTTTTTAATTTCAGGGAACTTTATAGTAATAACATAAGCGGAAGAATTCCAAATGAATTGGGGAACTTGACAGAGTTGGTTAGTTTGGATCTTTACCTGAACAACTTAAATGGTCCTATCCCTCCCTCATTGGGCAAGCTTCAGAAACTACGCTTCCTGTACGAGTTATACATTCTTTTTCCTTCCGTTTTTTTTCTTCTCTTTTTGTTTCTTTTTCGGTTTGGTTATTTTTTTTCAATGTTTCAAATCTTTAGCATATATATCTGAAGTTTAGTTGAATCAGAAATGATCACCTTCTATTTTTATTTTTATCAAGAAAGCAGCGTCTATGTTAAGGATCCGTTTGGCTATAGATTTCCCAAGTAAAATTTGGGGGAAAATTTGGCATCTAGTGTTTGTCCATACAATTTGCCATTATTTGGCAAATATTTTTGGCAAATAACCCAAATTCCCAAATACTAGAAAAAAACTAGTATTTGGGCCAAATCTCATTATTTGGGATCTTTTAAAAATTTAAATGCTACACCAAACTTTTATCTTTTACAAAAACACCCTCTATAGTAGTTGTTTGCGTTGTATTACATAATTTTTTACGTAAACACCAAAGTAATGAAATATTCAGTGAATATGAAAGTGATCATATGGTTATTGATGAAAATGATGAACAATCGGCTTAGGATAACAAAGTCACATCTTTGTCTACTTGACGATGTATGGAATGATGCTTGTTGCACTCACTCTGAACTACCATATTGCTCCAATGTCATGGACACTGCTTGTTATTTGTTGCAATGAAGATCCAATTGACTTGTAATATAAACTTATGGTTAGTTTTGATAGTTTTTAAAACTTTTATAAATCATATTTCCTAAAAAAGTGAAATATGTTTCCCAAATACTATGGCCAAACACATGGTGAAATTTCACCCAAATTTTAACACAAATAATATTTGCCAAAAATATTTGGAAATTTATGGTTAAACGCTAGCTAAATCATGCAAATGGGGTTGTAGTTTCTAATTCTCTAAGCAAATCTGTATGATGTTCTGTAACTTTTCTGTTAATTATGTTTACATTGATGGGTTCTGTCCAAAAAAAATTGATGTGTGGAAGCATTATGAAAATCACTACTCCTGTCAGCGAATAAACCTTTGTAGTTTGAGAGGTCAACAGGGAATATATCTGTATGATTCATTCTGTCTTTACTCTGTAGTGGAAAGTTAGTTGATATGCTTGGCAAACCTTGTTCTTGTTTCCCAAATACTGTATGGTTGCTGAGTGAAGACCAGGTGACTGGACTGGACTCTTTTTTTGTACGACAAACAGAGATTAATAAGTAATAATATAAAATGTTAAGAATAATATGAACCAACTTGAATAAAAATATATTCTAAAAGTATTCAATTATTATCTGATTAACAAAGTTGTGCTTGCTTGGTCTCTCTCGATATAGACTCACAGCCGTGCAGCTCAACTGAAAGTGACAATTCTGAGATACTTCTCATCTTATCAGGAAAATGGCCTGTGGATTTAGTTCATTGCTAAAATACTTTATGTATGAACGGTGATTCTCAACCTGCAAACACTATCAGCAAGTTTTTCCTAGCATATTGGAATTAATCTTTCTGGAGTATCTTGTCCTATATATATAAATGTATTTGTCTTTGTCTACCTTTAGCACATGTACGCTACATTTAGCACATGTACTGAGTAATGAGATCTAGACCAGTATTCGCTTTGACGAACTTCTTTTCCTGTCTTGCTTACTCTTGATCAACTTCCTTTTATCCTTTTTTTTTCTTTGCTGAATTGAAGGAGGCTCAATAATAACAGTTTGAATGAAGGTATTCCCGTGTCTCTAACCACCATTGTTGCACTTCAAGTACTGTAAGTATAAAGTTGTTCCTTTATTTATCAATGCTGTATGTTGTCTGTGTTGTCTAGTTCTTCTGGGTTACTTATTGTCATTGCCTTTCAATGCAGTGATCTCTCAAACAACCATTTGACAGGACCAGTTCCAGTCAACGGTTCCTTTTCACTTTTTACTCCTATAAGGTGTGTTGGAAGCATAAATTTGTAGTTCCTTCTGTTCCTGAAGTTACAATAGTGAATATCTAAAGATTGAGTACCTATATTTCAGTTTTGCTAATAATCAGTTGGAAGTTCCTCCAGTTTCTCCACCTCCTCCTCTTCCTCCTACGCCCTCATCGTCATCTTCAGGTTTGTTGACATAATTTGTGTTTATGCTGTTATGAAATTGTGCTACCTCTCCTTTTTCCCTATTCCAATTTAGTTTTCATGAAAATCCCTTAAGTTATTCACAATAAGGGGAGATGGCAATGGTAAGAAAGGAAAGGCCTGGTTGCCTGGGGAAAGCTGAAGAGATGAGCCCCAGACTGGATTCCCTTATGAAAGCACTTTCTTTATGAAAAAAACTACCATTACTGTAGGCACTGTAAATTACTCTTTTTTAGATCAATTATACAGGTATTGGGGTCTTGTGGTTATCCAGCATTCATGACTTTCCAAAATAAAAAATTCAAGTAAGCTTGGAGATACTATCAACATTTTTTTCTTTGTTGGAAATTAGATTTACTGTAAACTCTGAAGTTCGAGTGATACGTGTGTCAAAAGTACCATATATTTTTTAGAATTGCCTGTATTTTAAAATTTAGGTCATACAAATAAAAAGTTAACTCAATCATCTTTTCAGTGGGCAACAGCGCAACTGGAGCTATCGCTGGAGGAGTTGCTGCAGGCGCTGCTCTTCTATTTGCAGCTCCTGCAATTTTTATTGCTTGGTGGCGTCGGAGGAAACCGCAAGACCACTTCTTTGATGTTCCTGGTTTGGCAAAACACATTGAATCTGAGTTAATTGTACTTTCAACTTGATGTGTTCTCACTCATTATAACTGCTTTTGCAGCTGAGGAGGATCCAGAAGTTCATCTGGGACAACTCAAAAGGTTTTCCTTGCGTGAACTACAAGTTGCCTCGGATAATTTTAGCAACAGAAATATACTCGGTAGAGGTGGATTTGGTAAGGTTTATAAGGGCCGGTTAGCTGATGGCTCTTTAGTTGCAGTGAAAAGACTAAAAGAGGAACGTACTCAAGGTGGAGAGTTACAGTTTCAGACAGAAGTAGAAATGATCAGCATGGCTGTACACCGAAACCTACTTCGTTTACGGGGATTTTGCATGACACCCACTGAGCGGGTGCTTGTTTATCCGTACATGGAGAATGGAAGTGTTGCATCACGTTTAAGAGGTATCTCTTTTTTAATTTTCATGTGCTTGCCTCTACTTTTTGCTCACCTTTAGGCACGCTCACAACAACGACAACAACAACATACCCACAAAGTGGCGTCTGGGGAGGGTCGAGCATATGCTGACCTTACCTCAGATGTCAGGTAAAGAGGTTGTTTCCAATAGATTCTTGGCTCAAGACAAAAATAGTCTAGAAAAAGACATAATTGAAGTACAAGAGAAAATAGATAGCAACAAAAACAACAGATAGTATCAAAACAGTACTGCAACAAAGTAATACGGTAATAGAACTACATAAAACATCAAATATCAACAAAAATCGAAGAACAAGAAACTACAAGCATAGTACTACAACTACTAGTAAGGGCAACAATATAATGCATTCCTACCTACTAGTATGGATGTACTTTTACCTACTAATCTTCTACCCTAATTTGTGTCCTCCACAGCTTTCTATCTAAGGGCATGTCCTCGGTAAGCTGCAATTATGCCATGCTGTATCTAATCATCGCTCCCCAATACTTCTTCGGTCTACCTCTACCTTTCCTAAATCCATCCATAGCCAACCTCTCTCACCTCTGCCCTTGGCATCCGTGCATCTTTTCATCACATGTCCAAACAATCTCAACCTCACTTTCCGCATCTTGTCTTCCACCAATGTCACTTCCACCTTGTTTTGGATATCCTCATCCTTAATCCTATCTCTCCTAGTAAACCCACACATCCATCGCAACATCCTTATCTCTGCCACTTTCATCTTCTAAATACTGGCCAACACTCTGACGTTACAACATAGTTGGTCTAACCACAGAAAAAAAGCCAATAGCTAATGTTAATGATAAGCCTTTAGTCTTTTAACTTTTCTTTCATAAATTTGTAACAAATTGAAAAAGTTACTGCAGAAGTGTAAGCCAGTGGTGGAATCTTCACATCAATTTCATTGTGCTATTAGTTATTATAGTGATTGAGTTTTTTATGAGTTCTCATGATGGGTATGTATATTTAACAAGATGTCAATGTTGTTTATGATTTTGCTTGTGTTCTATGGACCAATTTCACAAGTGGTGCTATCCTTGATATCTTACTGAATATAGAGACTGCAGTTCATGATAAAATTTATTTAATGAGCACATCTGAAAGTCAAAAAGAATTAAATGTCTTGAAAGAGAATTCATATTTTTATTATTGTATGAAACTGGATAAGTAATGCCTCATGACTTGGCGTTTCATGATTTAGAGTGATATTGACCCAAAAAGATCAGTTTTCTTTTTATAACTTATTCAAGCTTAAATAGGAACCTGTTTCCAGTTTGCTGAATAGCTTGAATTTATGCTCATGGGATAATTTACATTTTTCTAGAGAGGCCTGAATCAGAGCCCCCACTTGACTGGCCAAAAAGGAAGCGTATTGCACTTGGATCTGCAAGAGGCCTTGCTTACTTGCATGATCATTGTGATCCTAAAATTATTCATCGTGACGTCAAAGCCGCAAATATCTTGTTGGATGAGGAGTTTGAAGCAGTTGTTGGGGATTTTGGGTTAGCTAAACTCATGGACTACAAGGATACTCATGTTACCACTGCTGTACGTGGTACAATTGGGCATATTGCCCCTGAATATTTATCTACTGGTAAATCTTCTGAGAAAACTGATGTGTTTGGCTATGGGGTTATGCTTCTAGAGCTCATAACTGGGCAAAGGGCTTTTGATCTTGCTCGACTTGCGAATGATGATGATGTCATGCTGCTAGATTGGGTAAGCATTTATACTGTCCTGTATGCTTAAGTTATTTTGCTTCTGCTATCAATCAGCAATTTGGATTTGCTTAAAGATCAAGTCACTTCTGCTATCATTGAACAGTCTTGGAATGTTCTGAAAAGATCAAGTCAACTCTCTCTTCTCTTCTATAAGTCAAAGTTTGATGTCTTTTTCTACTGTAGTGTTATTTGTTGAACACTGATGTGCTCATCCTGAAGAAATGGTGTGGTACGAGACCTGCATCGTTGCTTATTATTTATTGTGGTTTTGTGGTGTGAAAGTAACATTCGTTGCAGTTCCCTGTGAGTAATTTTTTTAAAAAAAGGTTTAGATGTTTCAGGAGGATTTAACCATTCATTACTAAAATTGCACGAGCATGTAGGTTATAATTTGAGTGGATTACTTTCATTATCTTATTCTTGCAGGTGCTTCAACAACTAGGTTGCCAAATTCGTCCCCTCAATTTGTGACTAATCTAGAAAATGTGCATCTCCACTTCTCATTTCTGATGCTATCTGAAGATTAGAAAAATATACTTTTTTCATTTTCATTCAAATAAGCTAAGGTGTATCTGCAACTGCCATATTAATGTCACTTCAGTCATGTTGTATTGGGGGTAGATATAGTATACTTTAGGACAGGTACTCATTCAGGAGCATCCTTTAGAAAGGCTAGTCTTACGTGGGATCGCCATCCTGGAGAAGGAAAAGGAGATGGTCAAACTGTTAGGAAGTGACTATTAAAAATAAAAGCTTTTCTGAACCTCTTGTGTCTCTGAAACTGTAATAGCAACATGAACACTAATAGACAGCAATGCCACTAATGTCAATCAGAAACACTTGGCTGTGTTTGAGCCTCATGCACTGATGTGCTGTTTTTTTAGTCTCCCTTCAGATCTTAGTAGCTTTTCTAGTGACTGTGCTCTTAGAGATGGCACATATGGTTGCATGCATGGCTATTATTTGACTTGTAACATTCCACTGGATATTCAGGTGAAGGGACTCCTGAAGGACAAGAAATATGAAACATTAGTTGATGCAGATCTTCAAGGTAATTACAATGAAGAAGAGGTGGAACAGCTTATTCAGGTAGCTCTACTTTGCACGCAGAGTACGCCTACCGAACGTCCAAAGATGTCAGAAGTTGTAAGAATGCTTGAAGGTGATGGCCTTGCTGAGAGGTGGGAGGAATGGCAAAAGGAGGAGATGTTCCGGCAAGATTTCAACCATGTCCACCACCACCATACTGATTGGATAATAGCTGACTCCACTTCAAATATCCGACCGGATGAGTTGTCAGGGCCAAGATGATCTTTCATTTATTCTGCCGGTAGCACATATAGGACCGACTCTTTTGAGGGAAAATCCGCTTTGTATCCTTGTATCTGTAATCGGTATATGCCATCTTCTTTTCTTTTTTTTTTTTTTTTTGGGTTAATTGGTATATGTCATCTTTGTGCATACCATTTTCCTTCTTTTTTTGTTTCAGTCAGTTATATTGTATTTGATTATGTCCAAATGTGTATAGGTGATTGGGTCATTACAACAGGTCTGAACATGAAAATCAACTTATAGCAGGAATGGCTGTAGAAAGCATCTTTGCACCTTCTCTTTTTCTAATGTATCAGATAGAGTGGCGTTTGTTAAAGAATAAACACTTTGTAAACTTAAAGTAATCATAGTTTCTTTCATGTTCAGTGACAGTTTGTCACCTCCTCTTGATGGCTGAATTTTGTAGTGTTATAG

>M6-PGSC0003DMG400017713 M6 LRR-RLK

CAAAAACAATCAATCATTTAGCCTTGATGATGGTTCCCACAATATTTTATAACCTTCCCTTCTTATATCTTTTCATAGTTATAACTTTTGCTTCCACTGAGGAAGCAACTGCTCTTCTTAAATGGAAAGCAACATTCCTAAACCAGAACAATTCCTTACAGGCTTCATGGACACTAAGTAGTCCTGCTGATACCAAGAATTCTTCCAGAATTGGAGCGACGAGTTCTGATGCATGCAAGGGCTGGTATGGAGTTGTATGAGTTAATGGTCGGGTTAACAGGTTGAATATTACAAATGCTAGTGTCATTGGCACACTGTATGATTTTCCATTTTCATCACTCCCATTTCTTGAATATGTTGATCTTAGTATGAACCAGCTCTCTGGCACCATCCCACCTGAAATTGGTAAGCTCGCTAATCTTGTCTACCTTGACTTGTCTATCAATCATATTTCGGGCACAATCCCACAACAAATCAGCTCACTAATAAAGCTTGAGACCCTCCACATATTTGACAACCAATTAAATGGTTCCATTCCGAGGGAAATAGGTCGCCTAAGGTCTCTTACTGATCTAGCTTTGAGTTCTAACTTTCTTAGTGGTTCAATTCCTGCTTCATTGGGGAATTTGAACAATTTGTCTTTTTTGTATCTTTATAATAATAAGCTTTCTGGCTCCATTCCTTCAGAAATAGGAAAACTTGTCAATCTTGTTGAAGCTTATATTAGTAGAAATCAATTAACAGGTCATATTCCTCCAGAAATAGGTAATTTGATCAATGCAAAAGTGTTGTATGCTTTTTCCAATAACTTGTCTGGTCCCATTCCTGATGAAATTGGGAAAATGAAGTCACTTGAGAACTTAAGCTTCCAGACAAACAATCTTTCTGGTCCAATTCCCAAAACAATAGGTGACTTAACTGAGCTCAAACTCTTGCACCTTTATTCCAACCAACTTTCTGGTCCCATTCCTAGTGATTTTGGGAATCTGAAAAAATTGAATGATCTGCAATTATCCAATAATCAACTTACTGGTCCTATTCCTGGTTCATTTGGCAATTTGAGAAACTTGCAATTTCTGTATCTCCGTGCCAACAAACTTTCTGGTTCTATTCCAAAAGAACTTGCATATTTGGATAACTTGGTTATAATGGAAATGGATGAAAATCAGTTTTCAGGCAATTTGCCTGAGCATCTTTGTCAAGGTGGAAAACTTGAGAACTTCACGGTGAATAGTAACAAGCTGACAGGGCCAATACCAAGAAGCTTTAGCAAGTGCTCGAGTTTCAAAAGAGTTCGCTTGGATAACAACAGTTTTACTGGGAATTTATCTGAAGCTTTTGGCGTCCATCCACACCTTTACTTCATTGATTTGAGTGATAATGATTTTCATGGTGAACTCAGCAGAAACTGGGGGAAATGCAAGAAATTGACTGATTTTCAGATAGCCAGAAATAACATTAGTGGTAGCATACCACCAGAGATTGGAAACATCAAAGGTCTTTTGGGACTTGATCTTTCAGCAAATCATTTGATTGGGCGGATACCAAAGGAATTTGGAAAATTAACCTCTCTGGTTAAGCTTTCTGTGCAAAATAACTATATTTCTGGCAATATTCCTATAGAACTTGGGGCACTGACAAAGTTAGATTCCCTAGATCTGTCAAACAATAGATTGAATGGGTCGATCCCAACGTTTATAGGAGATTACCAGCACCTATTTCACTTGAACCTGAGTTGTAACAAGTTTGGGCAGAAAATTCCAAAGGAGATAGGGAGGATAACTCATCTTAGTGTACTTGATTTGAGCTATAATCTTCTTGATGGAGAAATACCTGCTCAGTTAGCTAGTTTGCTGGACTTGGCAAACTTGAATCTTTCCCACAACAGCCTTGCTGGGCGTATTCCTGAAGAATTTGAAAGTTTGACCAGTTTGCAGGATGTTATCTTGTCATACAATGAGTTGGAAGGTCCAATCCCTAATAATAAAGCTTTCATGAATGCCTCATTAGAAGGTAATAAAGATCTTTGTGGCAATGTGATTGGATTCCAACCTTGCGAAAGGCCATCTTCTATTGTGAAGAAGCACTCAATGGCGAAGGGACATAAACTCATCCTCATCACTGTACTTCCTATTATGGGAGCACTAGTACTACTCTGTGCTTTCATTGGTGTTCTCTTTATGTGTAATAAAAGAAGGAGAGTTAGAGACGTTGAAAGAAGGGATTCCAGTGATGATGTTGATGGTTTGATTTACATATCCACATTAAATGGAAATGCATTGTACTGGGATATCTTAAAAGCCACAAAGGAGTTTGATGCAATGTTTTGCATCGGAAAAGGAGGATCTGGAAGCGTTTACAAGGTAAAACTTCCAACAATAGAAAATGTAGCTGTGAAGAGACTTCATTCTTCATTTGAGGTTACCCATCGCAAAAGCTTCATGAATGAGATAAGGTCATTGACAAGTATCAAGCATCGCAACATTGTGAAACTCTATGCCTTCTGTACAAATGCACAACACTCATTCTTGGTTTACGAGTACATGGAGAAGGGGAGCTTGTCTAGTATTTTGAGCAATGAAGTAGAGTCCAAGAAATTGGATTGGCTTACAAGGGTGAATATCATCAAAGGTGTTGCTTATGCTTTATCTTACATGCACCATGATTGCTCACCCCCGATTGTTCATCGAGACATATCAAGCAGTAATGTTTTGCTTGATTCCGAGTATGAAGCTCGTGTTTCAGATTTTGGCATAGCTAAGATTCTAAAGCCAGACTCATCCAATTGTACTGCACTTGCAGGCACATATGGCTATGTAGCACCAGGTAAGATCTAACTTCTTATCCTTATATTAGTTCATGAAATATATCTGCAAAATCTATATTGTTTCTATCTCATTTGAAACTTTATTGTTTCAGAGCTTGCCTATACAATGAAGGTGACGGAAATGTGTGACGTCTATAGCTTTGGAGTATTAGCATTGGAGGTAATCAAAGGAAAGCATCTTGGGGAATACATTACTGTGCTAGCAAATTCATCGACAAGAGATCATGTGCAGCATAGCGATTTGCTAGATGAACGCCTTCCATATCCTGAAGATAGAGTAAAAGAGGTTTTGGTTTTTATCATCAAGCTAGCAAGCTCATGTTTGCTTGAAACTCCAAAATCAAGGCCAACAATGCACTTCATCTCCAGTATGTTGTCAATGGATCCACCTGTTTCTCAAATAACTTCTATGAAGGCGAAAAGAGTCCCCGAGATGTAAGGTGTCAGTGTATGAAAAAAAAAGAAGCAAGTTTAGACACCAAGTGGTTGTGAAGTAAGAA

>M6-PGSC0003DMG400017864 M6 LRR-RLK

ATGAATATGAAGAAGTTGGAAAGAAGAAATACTATTGTTATTGTTATATTAGTTGTTGCATTTGATCACTTTACTGCTTGTTTGGCTAACATTGCCACAGATGAAGCTGCTCTTCTTGCCTTCAAATCTCACATTTCTTCTTCTTCTTCTGATCCCAACAACGTAGTTGTCTTGCAAACCAACTGGTCTTCTTCAATTCCATCATCAGTTTGCACCTGGATAGGGATCACTTGCAACGGGCGCCACAGAGTCACTGCTTTAGACATATCTAGCATGCAACTTCATGGTACCATTCCTCCGCACCTTGGAAACCTCTCATTTCTCGTATCCCTTGACATTAGTAACAACACTTTCCATGGAGATCTCCCCGAAGAGTTGACTCATTTGCGGAGGTTGAAATTGATTGATGTTACAAGAAACAATTTTAGTGGTACCATTCCATCATTCTTAAGTTTGTTACCTAACCTTCGATTCGTTTACCTTTCAAACAACCAATATTCAGGGGAAATTCCATCTTCCCTTTCCAATCTAACAAATCTTCAAGAGTTAAGAATACAGAGGAATTTTCTTCAAGGAAAAATCCCTCCTGAAATTGGCAATCTTCATTACTTGACTTTTTTAGACCTGCAAGGTAATAGACTTACTGGCTCTATACCATCGTCAATCTTCAACATGACTTCCTTGACAACACTTGCTATAATACACAATCGTCTTGTCGGTAAACTTCCTCTAGATATTTGTGACAATCTACCAAATCTTGAAGTTCTTTTACTATCAACAAACAACCTGGATGGCCTAATCCCACCAAATTTACAGAAATGCTCAAACCTACAACTGTTGACATTGTCTGGCAATGAGTTCACTGGACCTATACCAAGAGAACTTGGGAACTTAACTATGCTCACAGTCTTGCATCTTGGAGAAAACCATTTAGAAGGTACGTCGTATGAATTACTCCCTCTGTTTCAATTTGTTTATCTAGTTTTGACTTGGCACTGGTTTAAGGAAGTAAAAAAGGACTTCTGAATCTTATTGTCTGTAGAATGTACCAAAATACCCTATAATCTTAGAGTCTTAAACAGGTCACGTGAAAAGTTGAAATTAAGAAAAAAAGACAAACTGACTAAAAAGGGAAGTAAGACAAACAAATTGAAACGGAGGAAGTGTATCTTTATGCTTTTGTGTTTATTTTAAGATTTCTCAAAAGCTAAGCAATATCAATATGTACTAATGTTTATGCATTTTACTCTAATTGACAGGAGAGTTATCGGTCGAGATAGGTAACCTACAAAACCTCCAAGTACTTGGGTTAAGAAATAATAAGTTAAATGGTACTATTCCTGCAGAAATTTTCAACATTTCAGCATTGCAGATATTGACAATGTATGGAAATCAGTTATCAGGTAGTTTACCTTCAGATTTAGGCGTTGGAATGCCTAGTCTTGAAGAACTTTACCTTGGATCAAATGAACTCAGTGGACGAATTGCTCCTACCATCTCGAATTCTTCAAAGCTAACACTTCTAGACCTTGCAGACAACAAGTTTACAGGTCCAATTCCTGATTCACTTGGTTCATTAGAGTTTCTTGAAGTCCTATTCTTGGGAGGTAACAATTTCATTGATGAGCCATCTTCTTCTGAGTTAAGATTCGTCTCGTCCCTGACTAATTGTAGATATTTAAGAGAAGTAGTCATTGAAGATAATCCTTTGAATGGTTTTCTTCCATCTTCTATTGGAAACTTTTCTGATTCTTTTCGGATGTTTGTTGCACGTAGAACCAAACTGAAGGGTACAATCCCGGAAGAAATTGGAAACCTGAGTAGCTTAGGTGTGCTAGCTCTCTCTCATAATGACTTCACTGGTTCGATACCTGAGAAGTTGAGAACCATGAAGAATCTCCAAGAATTTTACTTGGAAAATAACAGCCTAAGTGGAACAATACCTGATGATATCTGCAGTTTACGAAATCTTGGAGCTTTAAAATTGACTGGAAATCAAATATCTGGTTCAATTCCAGCATGTTTAGGAAATGTTTCTACTTTAAGATATCTTCATTTAGCTTTCAACAGATTGACTTCAACCTTACCTGAAACCTTGTGGAGCCTTCAAGATCTCTTGGAACTCAATGCATCAGCAAATCTGTTTAGCGGGCGTATACCTCCTGAGGTTGGTACTCTAAAGGCTGTGTCACTAATTGAAAAGTTGATTAGTCTTTCCATGGCACACAATAAATTGGAAGGGCCTATTCCGAGTTCATTCGGGAAAATGGTGAGCTTGGAATTCTTGGATTTTTCTTACAACAATCTTACTAGTGAAATTCCAAAGTCACTTGAAACACTTTCACATCTCAACTACTTTAACATCTCTTTTAACAAATTAAGAGGAGTAATTCCTAGTAGTGGTCCATTTGCAAACTTCACCAGTCAATCTTTCATTTCCAATTCTGCACTCTGTGGTGCTCCTCGCTTTAATGTGTCGCCATGCCTCATCAAATCCACCAAGAAATCAAGGAGACACAGAGTTCTAACAAGTCTGTATATCATATTAGGTATTGGATCGATGATTCTCACATTGGTTCTAGGATATGTTTTGTTAAGGTGGCAAAAGAGAAGGAAGAATCCAGGTCAAACAGATGCATCTTTGGTTAAAAGACATGAAAGAATCTCTTATTATGAACTTCAGCAAGCAACGGAAGGATTCAGCGAGAGCAATTTGCTTGGTACCGGAAGTTTCAGTATGGTATACAAAGGGATACTTAAAGATGGGAATCTTTTGGCAGCTAAGGTATTCAATGTGCAATTGGAGGGAGCATTCAAGAGCTTTGAAACAGAATGTGAGATACTACGCAACCTTCGCCATAGAAATCTAACAAGAGTGATCACTAGCTGTTCCAATCCAGACTTCAAAGGCTTAGTATTGGAATACATGCCCAATGGAACACTTGATAACTGGCTACACTCTCATGATTTGTTCTTAGACATGTTGAAGAGGTTGGACATTATGATAGACGTGGCATCGGCATTAGATTATCTCCACAACGGGTATCCAACACCCGTGGTGCATTGTGACTTGAAGCCAAGCAACGTCTTGCTTGATCAAGACATGGTTGGTCATGTGAGTGATTTTGGCATTTCAAAATTGCTAGGTGATGGGGAGACTTTTGTACAAACAAGGACTATTGCAACCATTGGATACATTGCTCCAGGTAATTGAAATTTCAAACTTGACATCCTTTAGATTTTCACATTTCAAACTTTATCTAACAAGATTTCTCCCATGATTGCAGAGTATGGACAGGACGGAATAGTATCCAAATGTTGTGTTGTTTATAGTTTCGGTATCATGATGATGGAAACATTTACAGGTATGAGGCCAAGTGATAAAATGTTTACGGGAGATTTGAGCCTAAGATGTTGGATTAATGATTCTTTTCCTAATGGAGTTGTGGATGGCAATTTGTTAAGGCCAGAGGAAGAACATATTAAAGAGAAAATGCAATGTGTATTATCTATCATGGAATTGGCTTTGAGTTGCACTTTAGTGTCACCTGATGCAAGAGTTAACATTGAAGATGCTCTTTCAGCCCTCCAGAAAATAAGACATCAGTTTGTCAATAGTTGTGCAAGGACTTAA

>M6-PGSC0003DMG400018024 M6 LRR-RLK

GTTTCACTATGGAGGAACAATCTCAAGGGATCCTTACCACGCGAGATTGGCAACTTAACCAAGATGCAAGTTTTAAATCTAGAGGAGAATAGGTTTACTGGTACGTATACATTCAAATTAAGTGATAGCAATCTGAACAGTTTAACCAGTGATTCATTTTAGTTAAACTCATTCTTTCTTCAGGTGAAATACCCAAAGAGATAAGAAATCTCGTTGAGTTGGAGGAACTTGGTCTTGCGTTGAATAGTTTTAGTGGTTCACTTCCAATGGAGATCTTCAACATATCACGACTGACAATAATTTCTCTTTCAGCCAATAATCTATCAGCAACTCTACCACCAAACATAGGTTCTACCTTACCCAACATTGAAAGACTTTATCTGCATAGCTTAACCAATCTTGTTGGGACTATTCCTCATTCTATCTCCAATTGTTCCAAACTTACTAGACTAGAGCTTTCAGATAACAAACTCACAGGATTGATTCCCAATTCTCTTGGATATTTGACTCATCTACAATTCCTAAACTTGGCCACAAACAATTTAACCAGTGATTCATTTTTAAGCTTCCTCACTTCCTTATCCAATTGTAGAAATTTAATAGATCTTTCTCTATCTTTGAACCCTCTAAATGCCATGCTTCCGGTCTCCGTAGGGAACTTCTCCATATCTCTTATAAAGTTTGGTGCCAGTGCTTGCAACATCAAAGGCAAGATTCCAAATGAAGTTGGAAACTTAAGCAACTTATTAGACCTTGATCTTTCTAATAATAACTTGATCGGATCAATTCCTACATCAATTCACAACTTGAGAAACCTTCAGCGCTTGTACTTGAACAACAACAAACTTACAGGATTTATTGGAGATCATATCTGTAAATTGCAGCATTTGGGTGATACTGATTTGGGTCAAAATCAACTTTCAGGATCTCTTCCTAATTGTTTAGGGAATGTTACTTCCCTTAGATTGATATATATGGATTCCAATAAATTGAATTCCAATATACCAACAAGCTTAGGAAATCTTAAAGATCTAATGGAGCTTGACTTATCGTCAAACAACATGGTTGGTTCTTTACCTCCGGAAATTGGAAATCTAAAGACTGCGACATATATGGATCTATCAATGAATCAATTCGCAAATGGTATTCCTAGAGAAATTGGAGGATTGCAAAATTTGGAGTACCTTTCTTTGAGGCACAACAAGTTGCTAGGATCTATTCCTGACTCAGTGAGCAACATGGTAGGTTTGGAATTCCTAGACATTTCTCATAATAATATATCAGGAACCATTCCTATGTCTATGGAGAAACTTCAAAACTTGAAGTATTTCAATGTTTCTGACAACAAATTGTATGGTGAAATACCCTCGGGGGGTCCTTTCAAGAACCTCTCGAGTCAGTTTTTCATCAACAATGAAGCATTGTGTGGTTTGTCAAGGTTTAGTGTCCCGCCATGCCCCACTTCATCAAGGCATAGATCAAATAGGAAACAAATGCTACTTCTTTTTCTTTTGCTTGGAATTGCACTTGTATTTGTTCCTATCTCCTTTGTGTTCCTATGGATAAGGTATAGAAGAGGTAAAAGATCTCCTCAACAAGCTGATTCATTGTCTATTGCAACAACAGAAAGAATTTCATACTATGAACTACTCCAAGCAACTGATTCGCTTAGCGACGGTAATCTAATTGGTTCTGGAAGTTTTGGCTCTGTTTACAAAGGTGTTCTCAGGAGTGGGACTGCCATTGCAGTTAAAGTGTTCAATCTGCAACTGGAAGCGGCATTCAAGAGTTTTGATATAGAATGTGAAGTTTTGCGTAGCCTTCGCCATAGGAATCTTGTGAAAGTCATCACTAGTTGTTCCAACCTTGATTTTAAGGCTTTAGTGCTCGAGTATATGCCTAATGGAAGTCTTGACAAGTATTTGTATTCACACAACTACTTCCTAGACATTAGGCAGAGACTAAGCATTATGATAGATGT

>M6-PGSC0003DMG400018171 M6 LRR-RLK

TAAACAAATCCGAAAAAAACACCCATCTAAAATTCCTCCCCACTCTCTCTTACATAATGGCCTAGCTGTTCTTGGATCCGGAGATTCTGGCAACTTTTTCAAATTTTTATATGTAACCTACATAAAATCCATTAGCCCACAAGACATGATTTGGTTTTCCTCTCTGTCTCTCCTCCCTAGGCTTAAACCTCTACACCAAAACCAAGAACAAGGCTTCCTTTCTCTGCTTAATCTTGAAGTTTGTAGCAGCAGTCACATATTTGTTCTAATCTAATGCCTGTTTATTCATGGACCCTCTTTTTCTTTTTCTCCTTCTTTCTCTTTAATACCTCCTCTGCTCTCAACCCTCAAGGTCAGGCTCTTCTTTCATGGAAGACAAGCTTAAATGGATCATTAGATGTGTTGAGTAATTGGGATCCTACTGATGAAACTCCATGTGGTTGGTTTGGTCTTACTTGCAACTTCAATAAGGAAGTTGTTGAATTGGAATTGAAGTATGTGGATTTGCTTGGGATTGTTCCTTCTAATTTTAGCTCATTGGTGTCATTGAACAAGCTTGTTTTGTCAGGGACTAATCTTACTGGTGTAATTCCAAAAGAGATTGGGATGCTTCAGGGACTTAAGTTCTTGGACTTGAGTGACAATGCATTGACAGGTGAAATTCCAAGTGAAATTTTTCACTTGCCAAAGTTGGAGCAACTTCATATCAATTCAAACCGGCTGGTCGGATCGATCCCAGAGGACATTGGTAACCTCACAAGCTTGGTGTGGCTTATATTCTATGATAATCAGCTGAGTGGTGGAATTCCAAGTAGTATTGGGAATTTGAAAAAACTGGAGATTATTAGAGGAGGTGGAAACAAGAATCTTGAAGGTCCATTGCCTCAAGAAATTGGCAACTGTAATAATTTGGTTATGTTAGGATTGGCAGAAACCAGCATTTCTGGTTTTCTACCTTCTTCTCTTGGCCAGCTGAAAAGACTCGAAACGCTTGCTGTTTACACCTCCTTGCTCTCTGGACAAATCCCACCAGAGCTCGGGGATTGCTCCAAGCTCCAAAACATTTATCTTTATGAAAATTCACTCACTGGTTCAATCCCAGCTCGGCTTGGTAACCTCAAGAACCTTCAAAACCTCCTTTTGTGGCAGAATAATTTGGTTGGAACAATCCCTCCCGAGCTTGGGAACTGTCAACAATTACAAGTCATTGACATTTCAATGAATTCATTAACCGGAAGCATTCCGGAATCATTTGGGAGATTGAATTCAATGCAAGAGTTACAGCTGAGTGTTAATCAGATTTCAGGTCGGATTCCAGCACAAATTGGCAACTGCACTGGTCTTACTCACATTGAGCTCGACAACAATGAGATAACCGGTTCCATACCATCGGAATTCGGAAACTTGTCGAACTTAACTTTGCTGTTCTTGTGGCAAAATCGTCTTGAAGGGGAAATTCCTTCTTCAATTTCCTTGTGCCATAATCTCGAGGCTGTTGACTTGTCACAGAATGCATTGACTGGTTCGATTCCTAAAGAGATATTCGATCTTCAGAAGCTGAACAAGCTACTCCTACTGTCTAATAATTTCTCTGGTCCCATTCCACCAGAGATAGGGAAATGTTCATCGCTGATTCGACTTCGGGCAAATGATAACAAGCTGACCGGATCACTGCCACCGGAGATTGGGAAATTGAAGAACTTGAATTTCCTTGATGTCGGATCCAATCACCTCACCGGGATCATTCCACCGGAATTATCTGGATGCCGGAATCTGACATTTCTAGACTTGCATTCCAACTCAATCAGTGGCAATTTACCGGAGAATTTGAATCAGCTTGGAATCCTCCAGTTCATTGATGTTTCCGATAATTTAATTGAAGGCACTTTGAGTCCGAGTTTCGGCTCACTGACTTCACTCACCAAACTTGTTCTGGGTAAGAACAGGTTTTCTGGTCCTATTCCAACTCAACTCGGTTCCTGCATGAAGCTTCAGTTGATTGACTTGAGTGGTAACCAACTCTCCGGCAAGATTCCGGCGAGTGTTGGTAAAATCCCTGGTCTAGAAATTGCACTGAACCTCAGCTGGAATCAACTTTCCGGTGAAATTCCGGCAGAGTTCGCCGCATTGGACAAGCTTGGAGTGCTTGATCTTTCTCATAATCAGCTCTCGGGGGACCTCCATTTTCTCGCGGACCTACAAAATCTCGTGGTTCTGAATGTCTCGCACAACAATTTATCGGGGCACGTGCCTGACACCTCATTCTTCTCTAAGCTCCCACTCAGTGTACTTGCTGGCAATCCAGATCTATGCTTTCCCGGCAACCAGTGCTCCGCCGATAAGGGCGGCGGTGTCCGTCGTACCAAAGCGGCGAGGGTGGCGATGGTGGTGTTGCTTTGCGCAGCTTGCGCTCTCCTTGTGGCAGCTCTCTACATCATCCTCAGCGGCAAAATACGGAACCGCAAGGCCCACGATTACGATCTAGATGGCGACAATGACGTTGAACTTGGCCCACCATGGGAGGTCACAGTATACCAAAAACTCGACTTATCTATAACTGACGTCGCAAAATGTTTAACCGTTGGTAACGTCCTCGGCCGTGGTCGGTCCGGAGTTGTGTATAAAGTGAATATTCCGTCAGGATTGACGATCGCTGTGAAAAGATTCCGAGCATCGGACAAGCACTCGATGTCGGCATTCTCGTCGGAGATTGCCACGCTGGCACGAATTCGACATCGGAACATTGTTAGATTACTTGGATGGGCAGCAAATAGAAAGACTAAGCTAATGTTCTATGATTATTTACCAAATGGTACATTAGGTTCATTTTTACATGAAGGTTTTGGAGGGTTAATTGAGTGGGAAACCAGATTCAAGATTGCATTAGGGGTGGCTGAGGGCTTAGCTTACTTGCACCACGACTGCGTTCCGCCCATCCTCCACCGCGATGTCAAAGCTCAGAACATTTTGTTAGGTGATCGTTACGAGCCGTGTTTAGCTGATTTTGGACTTGCTAGACTCATGGAGGAGGAAAATTCTTCAGTTACAGCAAATCCTCAGTTTGCTGGATCTTATGGCTACTTTGCCCCAGGTAGATATTTCTCTTCTTCCTAGTTCAAGTTCATTTCTTTAAAAAATCGACTAGTTATGATTCATTTATTACTTTAGTTATTAGAGGAATTCTCCTCTAATGCAATTATGTGAGGTTTCATTTATCACTTAGTTCAAGTCCATGGTGGAGCTAGAGGGTGGAAGGGAGTTCCTTTGAACCCTCTTCGTCACAAAATCACACACATATATAAGAACTATTTTTTCAATATATGTATGGAAATTGGAAGGAAGTAATCTCAAATACGCTTGATATCAATACAGACTTAGGTAAAGATAGTACAAATGGAAGAAAAAGATCCATATATAGCTTAATGCTAAGGATAGGTTAAAGACTGGTATGAAACTTTTATCATGATTATTAATAGTATTATATATTTATATTTTTAACATATTTTTTTTACTTTATGATGATGATATGATGTGAGCTTAAATGAAATTAGGATTCATATAGCGACCCAAGTTTGCCTTTGGATTTTTTCCGCTTTTTTCTTTATAGAAAAAAAAGAACTGAATGCGAGGTAGCAGAAGAGAATAGATATAAAGCACTGCAAACACAGAACAAGAAGGAGAATTCAGATGAAGTTGCTTCCATACGGATGGAATTGGGTCATTATTTTGACAATATATGCATGTGAAGCCATGTGCTAGCTTTAACTATGTGCAAAACCAAGTTTGTCATGACATACTTGGCAGCTACAGTGCTATTTCCCCATCTTTTTGCCTGTGTATGGCCCCCCACTAAAGATACTTAGGATGGGGGATACTCTAAAAAAACCATAACCTAAGTTAGTGGTCACACTTTCACCAGCCAAGCATGGACACATTGCTGCTGTCTCCATATCCCTTTGGCCATTGCTTGTTGTCCGATTGCGATATGTGAAAAATAATCGCCCAACAAATGCCTTTGTTATGTTTCAAAAAACATCAAGATCAAATAGGTGATAAATATCTGGCATGAAAAGGACTAAACTCTGTTCTATTGTTTTGTCTTTTGCAGAGTACGCTTGTATGCTAAAGATTACAGAGAAGAGCGATGTTTTTAGCTTTGGAGTTGTGTTGCTAGAAATCATTACTGGGAAAAAACCAGCCGACCCATCCTTCCCTGATGGCCAACATGTTATACAATGGGTTCGCGACCACCTAAAGAGCAAGAAAGATCCAGTTGACGTAATTGATCCAAGGTTACAAGGACATCCTGATACACAAATTCAGGAAATGCTTCAAGCCCTTGGAATAGCCTTGTTGTGTACTAGTAACCGGGCAGAAGATCGTCCAACAATGAAAGATGTGGTGGCATTACTCAAGGAGATTATCCATGAGCATGCAACAGGAAATGAAGCTAAAAAGACATTGAACAACTCCTCAAAAATGTCCGATACTTCATCATATTCCTACTCGTCTTCATCTGTCACTCCAGCTCAGTTGCTACGCCTCCAAGGGTCGTTTAATTGCTCTCTGACTCACTCATCCTCCTCCGTTAGCTATAGCACCAGCAACCAATAATAATGTAGCGATAGTCATGATAGATCTGTTATTCTTTTGAACTAGATAGTATTAGCAACCGTAAAGAGCTAGGTAGAGGGAACCTTATTGTCAATTTAATTGTTAAGCAAATGAAAATGAAATCCATTGTACAGCAAAGTCAAAATTTTCTTTATCTCCC

>M6-PGSC0003DMG400018253 M6 LRR-RLK

ATCTGTTCTTGATCTTTCGCGTAATAACTTAACTGGAGAACTCAATGATCTTGATTTTGGGACTTGTCAAAATCTCACTGTCCTAAATTTGTCTTTCAACAACCTTACCTCAGTTGAGTTTCCACCCTCTTTGGCAAATTGCCAGAGCCTCAATACACTGAACATTGCCCATAATTCGATCCGAATGGAGATTCCTAGCGAGTTATTGGTGAAGTTGAAGAGCTTGAAGCGATTGGTGCTAGCCCACAATCAGTTTTTTGATAAAATTCCATCAGAATTGGGGCAGAGTTGCAGCACACTTGAGGAGCTTGATCTTTCTGGAAACCGGCTGACTGGTGAACTTCCTTCAACTTTTAAATTGTGCTCCTCACTTTTCAGTCTCAATCTTGGTAACAATGAGCTATCGGGTGATTTCTTGAATACTGTTATCAGTTCATTGACAAATCTTAGGTATCTTTACTTGCCATTCAATAACATAACTGGTCATGTACCAAGGTCTTTGGTGAACTGTACAAAACTTCAGGTACTCGACCTTAGTTCCAATGCGTTTATTGGGAATGTACCTTTTGAGTTGTGTTTAGCTGCCTCAGGCTTTCCACTGGAGATGATGTTGTTAGCCAGTAATTATCTTACTGGAACTGTGCCTAAACAGCTTGGACATTGTAGGAACCTCAGGAAGATTGATCTCAGCTTCAATTACCTGACTGGTTTGGTCCCATTGGAGATTTGGACCCTTCCAAATCTTTCGGAATTGGTAATGTGGGCTAATAATCTCACAGGTGAAATTCCTGAGGGCATTTGCATCAATGGAGGAAACCTTCAAACACTCATTCTCAATAACAATTTCATCTCAGGGGCACTTCCACAGTCCATTTCCAATTGTACCAACTTGGTTTGGGTTTCGTTGTCTAGCAATCGACTGTCTGGAGAGATACCACAAGGAATTGGGAATCTTGCAAATCTTGCTATCCTTCAGTTAGGTAATAACTCACTTACTGGATCGATCCCTCGGGAATTAGGTTCGTGCAGGAACTTGATATGGCTAGATTTGAATAGCAATGCTCTAACAGGTTCAATCCCTCTTGAGCTTGCTGATCAAGCTGGCCATGTTAATCCGGGAATGGCTTCTGGAAAACAGTTTGCTTTTGTGAGAAATGAGGGTGGAACTGAATGCAGGGGTGCTGGTGGACTGGTTGAGTTCGAGGGTATCCGCGAAGAGAGGCTTGCAATTTTACCTATGGTTCACTTCTGTCCATCAACTAGGATCTACTCTGGTAGGACGATGTACACCTTCACGAGCAATGGAAGCATGATTTACCTTGATCTTTCCTACAATTCTTTTTCAGGAACCATTCCTGATAATTTAGGTTCATTGAGCTTCCTTCAAGTTCTGAATTTGGGACACAACAATTTTACCGGAACCATTCCATTCAACTTTGGAGGTCTAAAGATAGTTGGAGTTCTTGATCTATCACATAACAGCCTACAGGGATTCATTCCACCCTCGTTAGGAGGCCTTTCATTTCTTAGTGATCTTGATGTTTCAAACAACAACTTATCGGGAACGATTCCTTCTGGTGGACAGCTGACTACATTTCCAGCTTCAAGATACGAAAACAATTCAGGCCTTTGTGGCGTCCCCCTTCCTCCTTGTGGCTCGGGCAATGGACACCATTCATCGAGTATTTACCATCATGGGAATAAGAAGCCTACAACTATAGGAATGGTGGTTGGTATTATGGTCTCTTTTGTATGTATTA

>M6-PGSC0003DMG400018263 M6 LRR-RLK

GAGATATTATCCTCTTGCTGTTTAGAGAACAGTTTAATCCTCAAAACAAACCAAACACTTGTGGTTTCTCTTTCTTCCCATTTACTCTTCTTTTCTCTTTCTTTTTAGTACACATATTGGCCTCTAAGAAGAAACTAAGGTTGCATGGTGGTGGATTTAAACAATGGTCCAAACAAGAACAATCCAACTGATCATCTTTTGTCTGCTTATTATTTCTATGCAAGTTTTGGTGCAGCTGGGAGCTTCAACAAATTCTGGTGACTGTAAGTCCTACTTATCACATCTCTTTTGTTGTTTTTTTAACTCTATAATTATGAATTTAACTTATATACACCATATCGCGAATAAAAAATAAAGTTATACTAGTACCTAGCACAGTTTAACATGTTATATCGTAAGTTACTTATTGTAAATAGTTCTTTTTTTAGATGATCTGAGTTATGAAAGCAAATTGTTTTCGTTTTTTTTTACTAGGGAATATATTAGTGTCCACTATTTTAACTCCAATTTATTTTGATCCCTTTTTCAAGTTCATTACTTAAAAGGTCATTTCAACTATTGAACATAGTAATGCTTTTCATTGTTTTGTATCAGAAAAGTCAAATATGATTTTCTTCATCAATTTCATTCCGTATGTTCCGAATTATTTTAAAAATAATTAGTGTTTGGTATTTTTGTTCAAAAATAAAATGATTGGTGCAGATGTTGTGTTAGAGTCACTGAAGGAAGAGTGGGAAAACATACCTCCAAGTTGGGATGGGTCTGATCCTTGTAACGATCCATGGGAAGGCATCGACTGCAACAATTCACGTGTTATTTCTATGTAAGCTTTTTTTTTTGTGATTAAATTAAAATTGTTAAAAGAGAATCATATTAATCGGAAATAATAATAATATTACCACTCTTTAAATTATTCCTAAATTTATGAATAAAATGTTGAAATTAAGAAATAAAAACACAAGTGCACTCTGTCAAAAATTCCATTTTTCCTTAGCTCGACAACGTCACGTGTTTAATTCAATGTTCACCCACCTCATTTATGCCAGCTAAATAATATCTTCGATTTTTTAAAAAAAAAAAAAATCTTCAATTTTTTTAAACACTAGAATAAATAAGGAATTATTTCCAAAAAAACAGCAAAATATTTTTTTTCTTAAAAATTCTTAATTAGTAAGAAAAGTACAAAAATATGTAACTGTTACTGTATTTACTCTTCCATTTTGGTCAGTTCATGTGTATATATAATAGAAAATAAAACTAAAATGCTTATTATCTAATGAAACTAGGTTAAAATTAAATCATATTAATTTTGTTAAATGACATTGCTAATTGTAATTTGAGGTACTAATTAATGATTGTGAATTTCAGAAAATTATCAAGCATTAATCTGAAAGGGGAGCTATCTGGAGATATCGAAGGCTTGTCTGAATTGCAGATATTGTATGTCTTCTACTCATCATAATTGAAAGTGTTAAAATTTGGAGAAAAAAAAATTGTCTTAATTATGTTCGCTCTGTTTCAATTTATTTGTCTGATTTGGATTTGACACATAATTAATGAAAGTAAATAAGATTTTTGAAATCTTGTGTGGTAATATGACCATTTTTGTTTTTGTGCTCGCCAAAGTGTTCCTAAAGGATTCATTACAAAAAATAATAGTTTAATGATTTTATTTGGTGCGATAAAATGAAAGTTGAGTGATCATCTGTGAAATTATTCCTGTTCTTGAAGCACCATTTTGCTTGTGATCATGCATTGCATGCACACAAATATTTTTGACTGTTGAAAGTTTTAAATTTATGTTCTGATTATCTGTTTCCTTAATAGGGATCTATCATACAACAAAGGCATGACAGGTTCACTCCCTCAATCAATTGGAAATTTGAAGAGTTTATCAATTCTGTAAGGGTATTATTATATGATAATGATTAAATGTTGTTAAATTCACCAGCTGTTTTCATTAGAGTAATTGCTCCGAGCTGCCTCGCAGGATCCTTGTTGGTTGTGGGTTTTCTGGACTGATACCAAACACAATTGGATCCCTTAGCCAGCTGGAATTTCTGTAAGGATATCAATTTCTTCTGTTTGCAACTTTTTATGCTTAACGAATTCAAGATATGGAGTATTTGAAGAACTTTTAGTCCTTTATGTTTTGTATAAAATTGGAAATTTGGCATGAGATGAGTTGGATGGAGTAACTTGGTCAGTGAGGATTTATATAGCTGACCTCAACTTGTTTGGGGACTGACGCTTAGTAATTTTTGTTGTTGTAGTAATTCTTGTATCCCTTTACTTTGTCAAACTGTGAAGTCTTAGAGCTTCAGGTCACTAGATATGTTATACTAGATTGAAATATTTTTCCTGTTTCTTCTACTGGGCTACATTTCTAAAATACTATCTGAATCATGCAGATCTTTGAATTTAAACAACTTTATCGGACCGATACCAGCTTCTGTTGGTAATTTATCAAAGCTACGTTGGCTGGACTTAGCTGATAATAAGCTCAGTGGACCTCTTCCAGTTTCCCATGGAAGCACACCTGGTCTAGACATGCTTGTTCACACAAAGCATTTGTGAGTAATACTGATTACTGTCAGTTCCTTAAGTAACACCTTCGTCTCTCATTTTCATCTGATATACCACTTACCTCGCAGTCACCTGGGGAGAAATCAGTTCTCAGGTGAGATTCCAGATCAGCTATTCAGCTCAAATATGACTCTCAAACATTTGTAAGTTCCACTAAACTGGAAGTGTATATATTTTTGAGAATTATTTGATGGATCAACTTCATTATTGTCCCACTCCAACGCTATGGTGCTAGCTCTGACACTTGTTGCAAACTACACCCGTTTGGCTAATGGTTTCTTCTGTTTGGGTCTTGTAATTGCACAGGCTGCTTGAGCAAAACCAGCTTACTGGAAAAATACCCCCTTCATTGGCACTTGTGCAGACTCTGGAGGTGGTGTAAGTATTTCCAAAATCCATCATGCTCTTAAAAAATTCTAATAAGTGTCCATCTTTCTTGCTTTCGATTACCTGGTGAAGTTTTATTAACTAGATACCCATGTCCACTGTAGTCGTCTAGACAGGAATTCATTAGACGGATCGATTCCATCAACCCTCAAAAATCTCACACTTATGAGTGAACTGTGAGTAGACAAGCATTATGCTAATATAAAAAATCCTTTTTATGAGTGTATCAGACCATCTATAAATTTTCCGTATCACTTGCAGCTTCTTATCAAACAATAAATTCACTGGCCCCTTGCCCAATCTTGCTGGCATGAATGCCCTCTACTACTTGTAAGTTCCTTGAAACTCTTTTCTGCTAAGAAAATGCAGCAATCTTGATATTGTGATAATTACAGATTATGTCTTTCAGTTACTTTAATTTGATTCTGTGGCCTATGGTCATGTTTTGCAGAGATATGAGCAATAATACATTCAGCTCAGCTGATTTTCCGCAGTGGTTTTCAACTTTGCAGTCATTGACGACTCTGTATCTATCTCTCTCTCGCTTTCTCTTTATCTCTTAACATGCATGTGTGTATCTGTCCCTGAGTTCACGTGCCTCAGCAGAACTCACATGCGTCTGAGTGTATATGTTATCCTAATGAAGTGGTTCATAGAATTTGTTATGCACCTCATTTGTATTTGTTCTGCTCATAGGGTAATGGAGAACACTCAACTTCAAGGAGAAATTCCGCAAACCCTCTTCAGTCTTTTTCAGTTGCAGACTGTGTGAGTGTTCTGTTACTTTTAGTACATGGTTCATGTCATGCAACTATTCTCATTTTCGCTGTGTGCTAGTCTTGTACAAAGTTTGAAAGGTAGTACAAAGTGACCTAGTATATAATTCTATGTTTTGCATTTCGATTTTACGACAATGAGGTGGTTATTTATGACAGCAACTTGAGGGGAAACAAGATCAATGGAACCCTGAATATTGCATCCAACTATAGCAGTCAATTAAAACTGATTGATTTGCAGAACAATTCCATCGATTCATATACAGTAAGACCTGGATTTCCTTTTCAAATAATGTAAGTCCTCAAGCCACTTTTTCTCCACTTTAGCAATGAGAAAGAAATAATGGATCATTCATGTTAGGACATTTGATTCGTCTGAATACAATTTAGTCATTTAACAATCTCAGTTTTCTACTTGGTCACTTGGCAGACTTATGCATAATCCAGTTTGTTATGAAGAAGGATCAGAAGACTATTGTGGCATTTCCCAAACGAACTTTGGATATGCAACTCCACAAGATAATTGCCTTAAAACACAATGCAGTTCTGATCAAATTCTTAGCCCTACCTGTAAATGTGCTTATCCGTACACAGGCGACTTATTCTTCAGAGCTCCTTCCTTTTCTGACTTGACAAACACAAGCATTTATGAGTCACTTCAGAAGTCTATGCTATCTTCTTTTAGCCCAAATCAGGTGCCAGTAGATTCAGTTTCCTTGAGTAATCCAAAAAAGAATTCAGAGTACTACCTTGTGTTACACCTGCAAGTTTTTCCATTTGGCCAAGATCATTTCAACCGTACAGGGATTGCCACAATTGGATTTGCACTTAGCAACCAGACTTACAAACCCCCACCAAATTTTGGACCATTCTTTTTCAGTGGTGAAAGCTACAAATACTTCGAAGGTGATTGGTCATTTCATTTCATTGATATGTAAATGTATTTCTTATGCACAAGTTTCATCTGTTAGCCGTACTATTAGTCGTGGTATTACTCTTGTAGTTTCTTGTCCTTTAAGTTAGGTTATCGTCTATAGTTTTTTTACACTTCGATATATTGTTTTCAGTTGGATCAAGAGGAACTCACAAGTCAATCACTAAAGGAATTATAATTGGAGCAGCTGCTGGTGGTGCTGTCCTTTTAATTTTATCAGTAATAATAGGAGTTTTAGTTTGCCAAAAGAAAAGAGATCAAGAAGCTGTTAGAAAGAGTAACCCTTTCGGTGAGTTCTGTTACTGTTCTTTAGTGTTGCTTCTAACTTTTTATTTCGCTGATGGTTATCGTCTTTTTTATCTGTCCAGCTACATGGGATTCTATAAAAGACAGTGGTGGTGTTCCACAATTAAAAGGTGTCAAATGTTTCACATTTGAAGAGCTGAAAAAATACACCAATAATTTTTCAGAAAGCAATTACATTGGATCTGGTGGTTATGGGAAGGTTGGTTTCTATTGACTTTTGATAATACAATGTTATATTATAATGACTGGTTAAAAAGCTGTTGCTTTAATTCTATCCATACTATCTGCACCATCTTTCGGGCAAGCTAAAGTTAGTCGATCACTTAAAACTTCTCGCGAACTTACAGGTATACAAAGGCACACTTCCAGATGGACAACTTATTGCAATTAAAAGAGCAGAACAAGCATCTAAGCAAGGCGCCCTCGAGTTTAAAACTGAGATTGAGATTCTTTCAAGGTTTCATCACAAAAATGTTGTCAGCCTTGTAGGATTTTGTTTCAGGCAAGGTGAACAGATGCTGGTCTATGAGTTTATACCAAATGGATCCTTGAAAGAGAGTCTTTCAGGTATTTTGAACTCTCATCGACTGTTGTTTTGAAAATCAGCTGTAGGTTGGAATGTCTGGGCATTGATAAGTTAGTGTTAACTTCTGCAGGAAAGTCCGGGATTAAGTTAGATTGGAAGAGGAGACTTCGGATAGCCCTTGGAGCAGCCAGAGGTTTGCAATATCTTCATGATCATGTTGACCCTCCTATCATCCACAGAGATATCAAATCGAATAACATCTTGCTGGATGAGCGCTTGACTGCAAAAGTTGCTGATTTTGGTCTTTCCAAGACGATGTATGAGCCAGACAAGGGTTACGTCAGCACTCAGGTTAAAGGAACAATGGTAAGTCCAGACTTTACAAATGTTGTCATATGGTAGCATTTCTCAGTTTCTATATTTCTTAAGTCACTGGAAACTCCTGTCCTTTAATAGTTATAGACTGGTGATCGTCTTGGTTCAATTCTTAAAAGTTAACACTGAAGTTGCGTCCTGATCAATTACACATTGTGCTGAACTTATGTATGTAAGTTCATACAATGAAAGGAGTTCATTGGTTAAACTGCTTAAAAAGACTAAAACAACAGTTGATTTGTAATTTTCGCAGAGTAGAAAACTTGAAACCCTTGAACACTTGCTATCAACTTTGTTTGAAGAATCTGTAACTGAACATTTAATTAATACGAGCTGCTTTGTACATTTATCGGCTTAGGGCTACATGGATCCAGAGTACTACACAACACAACAGTTGACTGAGAAGAGTGATGTCTATAGCTTCGGAGTTCTGCTGCTAGAGCTTATAACAGCACGGAGTCCTATTGTGAGAGGAAAATACATTGTGAAGGAATTGAGGCAAGCGATAGACAAGTCTAAAGACATGTACAACATTGATGAGTTTGTTGATCCTGCTATCCCCTCAAACATGACACCTATAAGCTTCAGGAAGTTTGTGGATCTAGCTTTCACATGCCTCGAAGATGCAGGAGCTAATAGGCCAACAATGGCTGAGGTAGTGAAAGAGATTGAGAATATCATGGAAATAGATGGATTGAACAGTTGTGCTGAATCCAGATCAACTTCTGTGAGCCACGAAGGAACCAGCGCAACCTTTGATCATCCTTACAGTGAGGAAAGCCTAAATCGTTACAATGGGGTCACCAATTCGCGATAAACACCAACTCTTCTCCTGAAACCAGGTACATTGTCATAGAAATGGAAAGTGTATTCAAGCAACTGTTGTGCTTTTGTGTTAGCTGTGTACATTCTGTGAACCTCTGTGTGCGGCGTAAATTGCCTTATACTAGAAATGATTCAAAAGTCCATCTAAGGATTTTTATAAACAGCCTTGTATGTTTCAACTGTTTGCTTCTGATAATCCAGGAGCTAGACTCCGTGGATTCAAAATGAGTGATTTTATTGTACGTATACATTTTAATTTTGATTTTTTTTTTACCTATGTATACATAATACGAATCAAAAAGTAATGCTTCAATTGAACCTACACAGA

>M6-PGSC0003DMG400018264 M6 LRR-RLK

ATGGTAGTTTCTCTGGAAATGACACTGCAAATTCTAGTCTATTTTCTGGTTGTTCTGATTCATGTTCTGTCAATAGCAGCATATACTAATCCTGTTGATGGTAAGTTGCTTTTATTCTTCTGTATCAAATGCTTCTTAACCAGTGTACACTAATATTTGATTTTCTGTGAGTACTAGTGTTCAAATCTTAGGTTTCCTTTTTTGTAATAATATGAAGAAGTTGTCTCTTAAAAAAAGTACTCCAATAAATAATCCAGAGATTATGGAAAATATTCAATAATCTTGAGGGAACATTATATTAAACATTTCCTGCTCTCTTCTAGATATGGTCAATTAAGTGTTCTAGTATAGAAGAACTGATTCAATATTTTATACAGCTGCTGCTTTAAAGTCTCTCACGAATTACTGGTATGGTCCTTCCGATTGGAATGGAGAAGATCCCTGTGGGAGTAGTTGGGAAGGAGTTGGATGCAGAGATTCACGAGTGATTTCCATGTAAGAATTCGACGTGTATCCCTGCTTAAAATTACAGAAATCATATGGTTGATGCCATCTTTCTCAATTTCAGAACATTGTCAAGTATACGTTTGAAAGGTCAGCTATCTGGAGACGTCCAAGGATTATCTGAACTGGAAACTTTGTAGGTTTTCCTTTAACATTTCATGTTTCTCCTGCACACCCTTTTTTCTTATGACTGAGCAATACTTTGGGATAATTTTTCAATTCTAATTATTGACTGTGTCTTATAGGGATCTATCATATAACAAAGATTTGACAGGATCTCTTCCTCAGTCTATTGGAAAATTGACAAAGTTATCAAATTTGTAAGTGAAGGTGAATAATCATCGCTCAAGTATTCACTTACCAGCTAATCTCATGATAATATGCTGGTGTTTTCTTAATTTTATGACAGGATCCTTGTTGGTTGTGGATTTTCCGGACCAATACCAGAATCCATAGGAGATCTGAGGCGGCTGGCTTTTCTGTAAGCATTCTGTGGAACTCTTTGTACTTTGAACGCGCGACCAGTACGTGACATATTTTACCAGAACTCCAAAGGTCATATATGAGCTTCTTAAAATGGTTAACGTAATGTGTGGCTTTTGGTGCTAGCTGTACTCAAATTTTGGTCCATCTCCAGGATAGGCAATGCAGTTCAGTTCATGAGGAGTTTAATCCATGCTGCTATACTTGTGTTTTATACGTTATCAAAATCTGAATAGTTGTGTTTGAACCATATCTACCTGTATATGATTTCTTTCAATCCTTCTTCTGGTATATGCTCTAAAACAGCCCCATTCTGTATTATGCAGATCTTTGAACTCTAACCGATTTACTGGAAGCATACCTGCTAGCATTGGCTATCTGACAGAGCTTCATTGGCTCGATTTATCTGACAATCAGATAGAAGGAAACATTCCAGTCTCTAATGGGAGCAACCCTGGTCTTGATAAGCTTGTTAACACTAAACGCTTGTACGCATCTCCTTTTTCTTGATGTTTTTTTGTCTCTCAACTGCATTTTCTAAGATTATTTACATTTCTTCAGTCACTTTGGGAAGAATCAGCTATCCGGTGAAATTCCAGCTAGTCTTTTCCATTCTAATATGACTCTGACACATCTGTAAGTTCCATCGAATTGGAGAATGGAGTGAATGAATAATTTTGTCCGTAATGGAGACCTCACTTGTAGTTGAGCTTGCCTTTTGATATTCTTGTTAGTGCATGTTTTTACTTCTGAGAGTAGTCATGTGAAAGAAGCTTGATAATTATATCACAAACAACTTTCCTTTTGATCATGCTTTACAGTTTCCTCATGCTTGTATATTCTAACTTGTAGGCTAGTTGAGAATAACAAATTTACCGGGGTAATCCCGGATACATTAGGATTTGTCCAGACGATGAAAATGCTGTAAGTATTGTTATCTTGTGCTTTCCCTTTATAATCTAGTAAAATTTCTTGCTCTCTATGCCTTATAAATTGTTTCTGGTATTGTAATAGTCGTCTTGACAGGAATT

>M6-PGSC0003DMG400018367 M6 LRR-RLK

CTCTGTTAAAACAATGCTTTGCTTGCATTTTTAGTTTTTGTTCTTTTGTTTTTCAGCTCAATTATATGAAATCATGATGTTTTTTCTCAGCTTCTTCCTCATTATTTCTTCATAATTCACTTCATATTTGGTGGGTTTTTCCTTCTTGTTACTGTACTTCAATTTTTACTTCTCCTTCTTCAACTGTCCAGAGTTAGTTTTTTTTTTTTTTTTTTTTGTTTCTAGATTTGTTCTATCCTTTTCTCAATTTTTGAATACCCATTTTGTTATCAGTTGGGTGTCATTGATTTGCTTCTTTTTGGTTGTGTTATGATGAATTTTGGGTTTGATTGAAGTACTGATTTTTATTTTTGGGCTCTTTCAGATTGGAGGTTTATGTATTGGTTATGCAGAGTGTGTTTCCAAAGAAAGAGTAGATTGAAGATAAGGGTTACAGTTTGGATCAATGGGTTGGAAGAGATCTGTGTTAGGGTGCTTGAATTTGGAGATCTTGGTGGGGATTTTGTTAATTTTTGCTGTTCAGCTTTCAGATGAAACTGACCCCGGAGATGGTAAGGAACAACAAATATTAGTATTATCTTTCGAGTTGTTTTGCGTTGAAGGAGAGGAATGCTAATTTTCATGTTTTGGGCAATGCTGGTTATGAGGGACTGAATTATGAGCTGAGCTAGTTGTTAACTTGAACTTTATTTTGGGCAAACGTTTTCAAGTCGGGGGGGAGCTGGAGTTATCGAAATTGGTCGGTTAAATGAACAGTCTCTATCCATTTCCGGAATTAAGTTAGGGCATGACTTTAGATATGAGGTTGCGTAGGAAATAGATTAGGTAGAAGATTAGTAGGTAATGGAGTGTTGTCTTGCTTATCCTTTGATGCTGGTAGTCCTAGTATTTCCCTTGGTAGTTTCTTGTCCTTGGATTTGTGTTACTATTATTGCACTGTTTTGTTGCGGTTACCGTTGCTTGTTTTTTTCTTCTCAGAATGCTTTGTCATGCTTTCTACAGTATTTCGCCACTATTATTTCACTTCTGTTATTTCTTTTTTCGATATGCCTTTTCTTGAGCTGAGGGTCTATTAGAAACAACCTCTCTACCTCTTAAGCTAGGGGTAAGGTCTGCGTACTCTACCCTCCCCAGACCCCACTTGTGGGATTACATTGAGTATGTTATTGTTGTTTGAAGTAATTTACAAACAACTCCATCATACACAAACAAAAAACGTTAAAAATAACCTAATCGTTTCACTTACATTCTCAAAAAACTTGCATGCCATAGTTACCAATTCTACTAACAGTGGACAAGCTGTCATTATTTCAGAAAAAAGAAAAGAAAAAGAGACAGTCCCAGGAAATTGAAGGGGGGAGAAGAGGAAATGAAGTACTGACTCCTAGGATGCTAGTCAAGTGTAGGGGGCTATGGCCTCGTCAAGTGATATGTGAATTCAACACATATTATTGTCCTAAATATCTACTAAATTGCCATCTTGATCGTATAAACTTTGCTTGGTTTGAAGTTAGCATGCTTTAAATGTTTGGAATCAAATTTTCTAGTTCATTCATGATCAATATCAACAATATTCTTATATGGTTTGAAATTTCTAACTCTGGCATTTTCCTTTCTGCTGGTCTAGTTGCTGCAATTAATGCCTTACATGCTTCCCTGGGATCACCTCCCCTTCCTGGCTGGGGTGTCAGTGCAGACCCATGTGATGGACAATGGCAAGGGGTCGTATGCCAGGAGACAAATATAATATCCATGTAACTACCTCTTATTCGTTGCAGCATTTCATGGATATTTGGAAGTCGTAATGTTAATCCTTCTAATCATCTTCTTTGCAGTCAATTAAATGCTGCTAACTTGGCAGGTGAACTAGGGGATAACTTAGCATCATTTTCTTCTCTCAAAACGATGTACGTGTACCCCACAGATATATCTCCTTTGACATATATTTGGTTGTGTTTTCTGTGCATGTGCTGCATATATTATGTGGATGCTTCCTTTTCTAGAGGTTTCTTTTATTGTGTGACATGTGAACAATAATTGTGGAAACTTCTTGTGGCAGAGATCTGAGTAACAATCACATCGGTGGAACCTTCCCATCCAGTCTGCCAGTTACATTGCAGAATATGTATGTATTATCATTATCCTAGAAGGGGAAGCAGAGAATGAAGCTTGAACTATCACAGCTTACCTTTTGAGCAAACCATTTGATAGTCTTTACCTTTTGGCAGTTTTCTTTCAGATAATGATTTAGCCGGAAGTATCCCCAGTTCTTTATCTTCCCTAAGTCAACTGTCAGCAATGTGAGTTTCAGCTTATATTTTAGATTGGAGATTTCCCGTTACCGTAATCATGGTTCCAGTCAGCTATACTTCTCTTTTGTTTCAGGTCTCTCAATGGAAACCAACTGACTGGAGAACTTCCTGATTCCTTTCAAGGCCTTACAGCATTGGTTAATCTGTATGACACCAAAACAAAAGCATTTCAATATTTCCATAGAGGTTTTTAATTGTCCATAACTGGTGGATTATTTGTATTCTTGCAGTGATTTGTCAAGTAACAGTTTCAGCGGAGCTTTGCCGTCATCAGTAGGAAACTTGTCCTCCCTGACCACTCTGTGAGTGATTCTACATATTGTTGAATTGGTTTGTAATTTATGACCAATAGAAAAGCAGCTTATTTCTTTATCTCCAGACATGTGCAGAATAACCAGCTTTCTGGAACTCTTGATGTTTTACAAGATCTTCCCCTCGCAGATTTGTACTCTTCGTCACCATTTTTTACTCAAATATTAATTCCAGGTCCATAGAATGATATAGTATCGTACTAATAGCTCCCATCTTTTGTGTCCCTCCAGGAATGTGGAAAATAACCTGTTTTCTGGGCCCATACCTCAGAAGTTGTTATCTATTCCCAATTTCAAGTAAGTACATTTCACTTTCTGCTTCTAATATTGTTGTTTATCTTCCAGCATGTAATCTACAATATATTCTTGGCCGCCTATTACATTCTTGTGCAGTAGTGCCTCTTTGTTCGACACAATCTGCTAGACAGATATTCAGAAACTATGGAGACAGATTCCTTGTTGAAGTTCGATTATTACATGCCAAACTGCTCTGTACTTTTTGTTAATACTTAATACCCTCTTTCTTAAATCTGAGTGACAACCTTGATCAGCTTGTTATTCATGGACAATTAAGGCAGATAAGATCTGAAGGGCTACAATCGTTAACATAAACAACAATTGTACTCCACACTCCTAAAGCTGCAGCACATGTATCATCTGTACTCAGCTTCATATTCCCTACCCACTTCAGACCATACTAAACTTTTAGTTAACATACTGGTGAATATCAAACTTCACTAAAGGGGAGAGTTTCTGTGCAAAATGACAAGACTTGAATAGGCTTCATCATCTCATTAAGTATAGATCAGCGGAAGAGTTCAGAGTGGCTTCTTTATTACACCCAAGTTTGTTGGTTGGCAAAAAAATGCAATTGCTTTTGGCAAACTTCTCAATTTTCTGAACCTATATGTTTAAACATTTTATGTGCGTAGAAATTTGTGTGTGAGTGTGTGTGTGTGTGTCTGTCTTTATGTAAGTTTGTGAATGCAAGACCTCGTTAAATCCTATAAGCTAGAAGAAACCTTGATCTGAACTACATTACTCTTTGTCATAAGCAGGAAAAATATTATGAAATGTGTGTCGTGCTAGTCTGTATGTTAGCTGCACCATCTTTAACAATAGTATTTTAGAAAATTGTTGAGTCCCACATCGTGTAGCATGAGGAAGTTGACTCCTTTATATGGTCTTGGGCAATCTTCATCTCTTGAGCTAACTTTTGGGGTTGAGTTAGGCCCAAAGTCTTTTTTATACTATCGGAGATCTGATATGTGGAGGAATATATTTATGTTGCAAAACCAGATTATGCTTACAGGTTTATGGCACTGACTTAGATTAGAATTGCTAGTGGGCACACTGCATATCAACAGTTTGAGTCCTAAATGTTAGAATAATTTCAAATTACATTAGGTAAAGTAGCCAAATTTTGATATCATCAGCACAATTACCTAAAAAAAGAATAATCCTAACACTATACTGCTCAATTTGCCTACACAATAAATTGGAAAAAATGTGACGTATACAAGCTAAAGGATTATAGTAGCACAAAAACCATATGACACCGATATATGATGTATATGTCACGCTGGACCATGGACTGCCAAGCTGGATCATGGACTGCATTTTGGTCATATTTAATAATTTAAGTTCACTAGCGGGCTAACATTTATTTAAGTTTTGAACACAGGCGTGATTTTCATATCATATTGTTCTATAAGATTTGTTTGCCGGGAAGTTTTATTTATTGAACAATATTGGTGAGGGTACATATTGATCCATGCTAGGAAGATGTATCAGTTGTTTCATTATTTTTTTCAAGTGTCTTATATAGGTTTTGACTCTTGATTGACTTTAATTATGAGAATTGATGGCTAATGCTTTCATGTACTTACTAATTTTGTTCTTGACGTGCAGAAACACTGGAAATCCTTTCAACAGTGTCTCTCCTTTGTCTCCACCCAACAGTAGTACCGCCCCTTTGTCTCCACCTACTTCATTTACGAGACCCCCTCCAGCACCACCAGGACCACCCTTTTTTAAGCCTCCAACTTCAACACAGACACCGCCGACTTCAGACAGAAAACCTGGACAAAAAGCTGATGGACCATCAGCAACTGCAGAATCTAGTTCAAAAGGAAGCAAAAAGTCTGTTAAGAGGGTAGTCTGGATATCCATTGCATCAGTTTTGTCATTTATAATATTGGTAATAGCAATCCTGCTCTTTCTGCCAAGATGCTTTAGGGAAAGGCAAGAGACTAATTGGTCCAGACGGCACGAAATAGCTCCTGATGTGGGCTCCAGGGAGAATCGTAGGGATAATGGTTTATTGGTCCTACCTGGACATGATGTAGAGAAAGGTACTGCAAATTATTAAACCTCTCAACTTTTATATCGTGTGTATTGTTGCTAGTATTTGTTAGTCTGGCTTTTTTCCTTAGTTCTTACAGGGAAAACTAGTACTCTTTAATTTAGGCTCTATTTAAATCTGCAGTCTTGACATCTAAGTATGCAGCTAAATATATAGTGGTTATCCATGTGCCCAGGGGCGGACCTACCCTTACCCAAGGCCTTCGTCAGAAAATTACATGGTATATAGAGGTCAAATTTTGATTTAATAAAATATATACAAGTATTGACACCTCTTGACACAAGTTAAAGGTTCAGTCTAGTGGTTAAGACATTGCTTAATGACTACAACCATTGACACCCCTTAATGACAATGGCCGCTACAGTATTGTCAAATTCTCTATTACAATCATAAATGCCCTACAAAAGATTCGTGAATTTGATATCTGTTTTTGCAACATTCATAACTAAAAAGCTATGACTCTTAACCTCTTCTACTCTTTGATGATAAGTGAACTTCATATGTTTTGAACAGCTCCTCCTCTGGTGAGGCCAAAGGAAGAGCAACAACCAAGGAGACTGGCCCGCACTCCCATGCCACAGCAGGAGCAGGAGTTAAATGTGCAAAACCTGAGGGCAGTACCAAAGAAGGATACTAGTGAGATAAACCTAAGCAGAATTGATATTGATTCGATGTTGCCACCACCTCCACCCCCACCACCACCTCCGCCACCGCCACCACCCTTTTCTCAAGAGAGAGTTATTGTGAAGCCAATTCCACCTGCCGACAATACTTCAATGAAGTTTCCCCACAGGCCACTTCCACTAACTTCTGTGAAATCTTACACAATCGCATCTCTTCAGCATTACACCAACAGCTTTTCTCAAGATAATCTGATAGGATCAGGAATGCTGGGAACTGTGTACAGGGCAGAGCTTCCAAAAGGGAAGGTATAAGTTATGAATGATATATAATGTACTGTAATTTGATACTCGAAATTACCAACTGTGTTTTCCTACCCTGCTGTATTCTCTTGAACCTAGATTGTCAGCTTTTCTCTTCTTATAATTTCCTTTTTTGGGGGTCTTTACGGTGTTCTGTGAGCACTAAAGATGCCTCTTATCTGTTGTTGCATTACGTATCTTGCTCTTGCATTTCTATGCTTAATTGCTTAAGAATGATTCACGGTCTGATCGAATCGTATGTAAATATGGTTTTGTTGGTGGAAATCCAAAGTCATATATATTAGGGTAAACTGTATTTCATAAATACAGATAAAAGCAAAATGACTTCCAATTGCTGCATAAGATATTTCTTGTAACTTAAATTTTCTTTAGGACGTTTTTTGGTTCTTATCCTATATAATCTTTAAGTGGCATTTTCTTCCAAATAGCCAACACCATTGATCAACTGTTGTAATAGAACAATCAACATTCAAGTTGCATGAGCTTATCATTCTGATTTGTGCCTTAGTTTTGTCATTTGCTGTATAGCAGGATCTGCCAAATACTGATGCATCATTTCTAATGAGAAACTATTTTATAATCAGAATATACAATGTCCCTGTTTTATTTTCAGTGTACATTTATTTATCTGGCATGCAATCTTAACTCTCATCCCATTGCTAACATTATTTGAATCATTTGGTGGTCACATTGTGTTACTTCTGACACTTGACCCTTTGAATGCTTGTGCAGTTACTTGCTGTCAAGAAGCTGGACAGAAGAGTTTCCAATCAACAGAAGGATGATGAGTTTCTTGACCTAGTAAACAATATTGACAGAATTCGTCATGCTAACGTTGTCGAGTTGATGGGCTATTGTGCAGAGCATGGTCAAAGGCTTCTGGTTTATGAGTATTGCAGTAGTGGGACACTACAAGATGCGCTGCACTCTTATGATGAATTCAAGCAACAACTTTCATGGGATACACGGATCCGGATGGCTCTTGGAGCTGCAAGAGGCTTGGAGTAAGTAAAATAACCAGTCACATATACATAGAAACCACATGCAAATATACGAGTATATACATATGAGCTATGGAAAATCTGATGTTGAAGAGTTATTTGGCAGTGATCTATAAGAAATTTCTGTAAGCAGTACTAATAATAATTTAACACTGGAAGGTAGCAAAGTACTCAAATCACAAACTCGTCCTGAACTTTCATGCTATGTACATACTACATGACATAAAATGATAAAATACTGTTGGGAGATGGTCTTTATAAGTATATGTTAACATTATATTTGGTGCTGTATTTCGGTTTATACTACTTTCCATTCCTCCTCCACATGAGTTCATACAAGGGATGCTGAACACGATTGTTGAAAGAAAATATCCTCATAGTCATCCCAGCCGTGAATTATTATTAGTGGGATTGCTAGGTGATCTAATTCTTAGATTTGCATGACAAGCGAGTTAAAGCAAATTTTAAATCTCATGTATTAATTATTTGTTCTTTTATGCAATGAAGGTATCTCCATGAGGTCTGCGAGCCACCTATTATCCACAGAAACTTCAAATCTGTCAATCTTCTCCTTGATGAAGAGTTGGCTGTGCATATCTCTGACTGTGGTTTAGCTCCACTGGTATCATCAGGTGCCGTGAGCCAGGTAACTTACCATCCAAATATTCTGGATTAGAAGTGTGGAACAAGCTGACATTTACCTCTTGGGGTGGGATGGTCATTTTCTGCTTGCATTTCCAGTTGAAAGAAAAGCAAGTTCCAGGTTTATGATGGAAGTGAAAATGTGAACTAAGTAGTAGTTGTTAAGAAAGGAGTGTAATGATGATCAAAATGCTTCCAAGTAACACAACGGAGTAGAGCAGTAATTAACTACAAATGCAGTTTCTCTTAAATGGATAACTTATTGTTTAGAAATAGTTGATTTTGTAGTAGTGCCAGAAGTTATACGGTCAATTTCAGAAGGGTTAAGTCTTTTTCCCTGTAAATCTGGAAATCAGGAAACAAATCTAGCTAGTCTGAGACTACCTGGTTGCCCAGCTAGTCTGAAGTATTTGTCATGTTTTAGGCTTTTATAACATTTCAATGCTTCACGAGATCTAATAGACACTAGTAGAATTTGTAAGCGGGAGAGTAGATCCTTTCCAACATATGCTGGACTTGCCTTCTCATCTTCACCCCTTGACAAAGAGATGAAATTTAGTAATCCTTGAAACTGTTCCCAACATTCTTCAGAAGTAGTTTTGCCGCTGCACATTTAGTATGATGGTGACTAGCTCCTGTATTTTTTTCCCTGTTGGAGAATTTTGACTTCTAAAGATGTGGGAAGAACAGTTTCACTATGTTAGTTTGACCTAGTCATACATAAAGACACTCTACAGGTGCTTTTGTTAGATGGGTTTTCAGAAACTGAAAATTCTATTTTATTTTTATGAAGTCCACATTGTCTCCTCTCATTAATAAGAAACAGTAAAACACAAGATTGAACTAAACAAGACATCACGGATATGACTGTTTTTTTCACTTGTACTTCATATGATTATAGATCTGCTACTCTAGGAGCATGTGGATCATCAATCTTTGCTGAAACCTTACCTTTTTTGTTTTTCTTATGCACTGTGCTGAAGAATTTGAAATTGAATAAGTGAAGTTGTACTTGATTTTCAGTTGTCAGGACAACTTCTTACAACCTATGGCTATGGTGCCCCAGAATTTGAGTCAGGGATTTACACTTCTCAGAGTGATGTTTACAGTTTTGGCGTGGTGATGCTGGAACTCTTAACGGGCAGAATGTCATATGACCGGTAAAGCCATTTGCATCTCATTGAATAAATTTTGTTGTCATGTTTTCTCCCAGCTTTATCATTACCTATATATCATATTCAGAACTCGAAGTCGAGGAGAGCAATTCTTGGTCAGATGGGCAATCCCTCAGCTACATGATATTGATGCCTTGACAAGAATGGTTGATCCTTCTCTCAAAGGAAAGTACCCACTTAAGTCATTATCACATTTTGCTGACATAATTTCCCGCTGCGTTTTGGTGAGTTTTTTAATCCTTTCTTTTATTACAAGCGAGTAATTTGTCCAAATCTCCATTTTCTTCAATCCTTCATCAAACTTTGGGTACAAATGAGGAAAAAGGTAACATGGAAAATGATTTATTTCTTGTCACAGAAATTAAAATAAGAAGTAAGTAAAAGGAAGGAGCTGTACTGTGTCAAAATAGGAACTCTACACATGGTGTCTTTTGTTTTTCATAGCCACTAGATTATGAATTATTTACCAGTTTAAGGACATCTGTGAATCTGTAGTATGAGATGTACCAGGTTTATAGGTATAGTTAAAAGTTACAGGGAGCGGAAACATCCCATGTCGATGTTCCGATATCGTTTTCTGCTCAGGTGTCTGAATGCCTTTTCATGACTCGTTGAAGAGTGTGTGTATGTTGTAGTTGAAGCAAACAACACTAGGAAAGTCTGGATCAATGAGCAATTTACCCATAATACCCTATTCACCAAATTAAATCTTGTGGCAGAATACGCGTGGTTAACTAGCACACAATCACACACCAAACAAATAAATAATAAAAATAACACAGGGAATTTACTAGGTTTGTCAAAACCCATTTCCTCGGGGAAGAAGCGGAGAGAATTTTCCACTATAAAAGATAAGAAAACAGTACCAAGAGATCCCCTATTACAACCCTTTTAAATACCCAAGTGTTTGCCCAATCCAACTGCCCCAGCCTAACAATATAAGCAAATAATAGTGGTTCAAAGCTTTTCGGGGTTGGGTCACAATTCGGATCTGCCAGAATCCAGACACAAAGCTGTAGAAGCAACAGATATATTATATTTGCTTTCCTTCTTCTGCTTATTGTTCTCCTGTCTTTTTGTTTATTCGGTGGGTGTTGTATTTTTCACTAGGTAATATGCAATTTTTTGTACAACCAAATGCTTGTGTATGTGGGGGAGAAGAAGAATGAGGCACACTCTATTGATTGGATATTTGATTGCAACTGATAGATCCATTGTATTTATATAATGGTCAAATTTTCAATGTTTCAATGAACAGAAATCGATACTGATAAGTTAAAAGGAACCAAGTAGTTGGTGTAACTCCAATGTGAAGCCTGTTCACAGTGTCCTCCCCATTATCATCTTTCAGCTACCCTTCTAGCAATGACAGAGAAGAGATTAAATGCATCATATGATTTCTTAGGGACTATTTCCATTTGGCTCCACCTTCCATATCCTATTCCCGCTATCTTATAGATTGATCGTACTCACAATTGTGTGCAGGACAACTAGTACAGAGTACTATATTACTGTTAATCAAGGAAGAGTGAAGTAGTTTTCGTGTGGCTGTTTTCTGAAGCCCATCTCATATCTCTTCCTAGCATGACTAAGATTTAGCTGTTGTATAGGATTTGCCTTGCGCATTTTGTTGTAACTATCTTTAATATTTCCTTGCAGCCTGAGCCTGAATACAGGCCACAAATGTCAGAAGTGGTTCAAGACCTCATACAAATGATAAGAAGAGAATCTCCCAGTAGATCCGATGAAGAATGATTTGAACCAGACATGCATGCTATAGCAGATTCAGGCATGGAGTAACCAAGACTTCCAAGATTTTTTTACTTTTGCAAAGAACTTAACTCGAACTGAGATACTTGGAACCAAAGATTTTGAGGCTGTAGATGCAGTGCCCTGCCAGCAAACCTTCATACCTCCAAGACAAAGGGCTGGCTAGAAACTGCGCGATTTCAACCAACACAGCTCGGGTTATCCTGATTCTTAGTACAATTTTTTGCAATCATTCTAATGTTCATTTATAGTCCAAACAGAAACCACAATGCATAAAATGCACTAACCCCATTGTTTTGAAAGCTTATTTATGGTTGTTAATTGGTATATCTGAAGTTGTAAATCTTTATTTGTTCTGCTGTACATTTTCAATTGTTTGTACAAGAATATTCTTCACACTTTCAACATGGACTTTCTGTAGCACATTTTTCAGCCTTCAGTATCATCATGGATATATTGAATAATCACTTGAAG

>M6-PGSC0003DMG400018752 M6 LRR-RLK

TTAAGAAGTTGGTCAATCTGGTTCAAATCACGAATTTCTTGAGAAAAACAATCCTTGACAAGATGGAGAAAAGCTATGTTCTTTTCCTTATCTTATTAGTATTTCAAGAATCAATTTTTTCCCTATATGCAACTTCCACAAATGAAACAGACATAGAAACTTTGTTATCTTTTAAACAACTCATTACAAATAACTCTTCTTTCCTTTCCAAGAATTGGACATCAAATACTTCATTTTGCTCATGGTTTGGTATCACTTGTAATCCAAAAAACCAAAGAGTCATAGCTTTAAATCTCCCAAACATGAATCTTCAAGGCAAAATTTCACCATCAATAGGTAACTTATCCTATCTTACTATACTTAACTTTAGTAACAACATTTTTCAAGGCACTCTTCCTTATGAACTTGGAAATTTGCCTCTTTTGGAAGTCATTGATGTTCATAACAATCAACTAGAAGGAACTCTTCATCCTTTTGTTGGTAACATTACAAAGTTAGAGAGATTGAGGTTAGATGGAAATAGATTTAGTGGTAAAATTCCTACTGAGATTGGGAATCTAAGCCAACTTGTTGAGTTAGATTTGTCACATAATCAGCTTTCAGGTTCAATCCCTGGATTGATCTTCAGCATGTCATCGTTACGTGCTGTGTATTTGGTTAACAATAGTCTTTCTGGTTCATTCTTGGTTGATGAAATAAATGGTGTGATGAATCTTGAAGTTATAGACCTTAGTTATAATCGAATCATCGGTGAGATTCCTTCAAGACTTTGTCAATTCAGTAAGCTAAGAACTCTTGTTCTGTCTTACAACAATTTAACAGGTCAAATTCCGAGAAATCTTGGTTGTTTATCAAGGCTTGAGAGTTTTTATATAACTCAGAATGCAATAAGTGGTACAATTCCACTTTCATTGAGTAATATTTCCACTTTTCAATTTCTTGGTTGTGTGAACAACCATATAAGTGGTACAATTCCTCAAGAACTTGGAAATTTACCAAACTTGAAGATGTTGGGATTTGATTTCAATAATCTTACTGGTGTAATACCAGAATCTATCTTCAATATATCTTCTTTGGAGTACATTGCTTTCAGTGATAACGATCTCTCAGGGCGAATTCCAACCACTCTTGGACTTAAACTTCCAAACCTTAAAGGAATTTTCTTGCCAGAAAATCAACTTGAAGGACAAATCCCAATGTATATCACAAATGCTTCTAACCTTATTGAGCTAGAGCTGTCCTATAACTTGTTCACAGGCATAGTTCCAAGTGATTTGGGAAATCTACACCAGCTCGAGTTCTTGAATCTTGGTGGTAATCAGCTTACAAATGAACCAGGACAACAAGAGCTTGGATTCTTGAACTCTTTGGTGGATTGTAGGAGGCTGCAGTTTCTGATATTGGCTAACAACCCCTTGAATGGCATCTTGCCTGATTCGGTTAGTAATCTTTCGTCTACTATTGAAATGTTTAACATAGAGAATGGTCAGATAAATGGACAAATTCCAAGAGGAGTAGGCAATATAAGCAGCATGTTATCGCTAGTCTTGAACGGTAACCAATTGACAGGAACCATTCCACCAGAGATTGGTGAGCTAAAGCAACTTCAGAGGCTTTATCTAAGCAAAAACAAGCTACAAGGTTCAATTCCAGAGGAAATATGTGACTTGGTAAACTTGGGAGATACATTTCTACATGAAAACGAGCTCTCTGGGGCAATTCCAAGTTGCATTGGAAAACTCACTAGGCTGCAAAGGCTTTCATTTGGCTTCAATAAGTTCACCTCAAGCCTACCTTCAAGTCTATGGGAAATGGATAGCCTGATTTTTCTAAACGTGACGCAAAACTCAATACAAGGGGAACTTCCTCTTGACATTGGAAAATTGAAATCTATTGAAGGTATAGATTTTTCAAGCAACCAACTTTCCAGTGTGATTCCAAGTACATTTGGAGACCTTATAGGCCTTAGTTATCTTTCACTCTCAAACAACTCGTTACGGAGTGCAATTCCTTCATCATTTGGCAGTTTGTTAAGTTTGGAGTTCTTGGATTTATCTTCAAATGCGTTATCAGGAAGCATCCCAAAGTCCTTAGAAAAACTTCAGTTCCTTAAAGAAATCAATTTTTCCTACAACCATTTAGAAGGTGAAATACCTACTAGTGGTGTGTTTGCAAACTCTTCTTCTCAATCTTTTGTTGGAAATAGAGGTTTATGTGGAAAACCTATATCAGAAGTTTCTCAATGTGCTACGGATAGTGCAACGAAACGATCAAAGTCTAGAAAACATGTGCTAGTAGTTGTCATTCCTGTGATTGCATCAATTCTCCTCATATTTGTTGTTCTGTTCGTATGGATTAAGCGAAGATCAAGAAGAACAAAACTTCAAGGCCATGATCAAGAGTTAACAGAAATCACAACACACCAGTTGATCACATACCGTGAACTTCAACAAGCAACGGATAGTTTCAGTGGCTCTAACATGATTGGTTCAGGAGGTTCTGGTTCTGTTTACAAGGGCATATTGGCTAACGGAACAATGGTCGCGATAAAGGTTCTAAATATGCAAAATGAGGAAGGATGCAAGAGGTTTGATACCGAATGTGAAGTGATGAGAAGCATTAAACATAGAAATCTTGTCAAGGTGATCACTACATGTTCCAATCAATACGTTAGAGCCATCGTTCTCGAGTATATGCCAAATGGAAGCCTTGAAAGTTGGTTGTATGATAAGGAACATCAAGTCTTAGACATGTTTCAAAGAGTTAGCATAATGCTTGATGTTGCAATGGCATTGGAATATCTTCACTATGGTTATGATACACCTATAGTCCATTGTGACTTGAAGCCTCAAAATGTTTTATTAGATGCTGACTTGGTGGCACATGTTGGTGATTTTGGCATTTCAAAGATTTTAGGTGAAAACAACTCCATGGCACAAACTAACACATTGGGTACTATTGGCTACATTGCACCAGGTAAACAGTTCTTCTCTGACTACTCTTCCTAATATAAGGCATATTTTCGGTCATCTTCTCTCTGAAAAATAGAATATTGGTCAATTAAGTTGATTTAGGTGCTACTTACTATGCCTGAATCTAGTTTCACTAACGCTAGGCTAAGTCTTGAACCTTTGAGCTGACCGTTCACTTTCTTTGTAGCAAGAGGGCTTCTCTTTGCCCCTTTATTGGTTGTATCAACTAAACTGCCCTTTCATATAGATCGTCGCAGCATGAAACTATTAATAGATTCCTAGCTTCCTATTAATCTTGAAAAGTTCTAGCTTAGTTCTTTTTTTTTGTTAGAATTATCTTTTTAATTAGACCGGACCCACTTTTATACTTATCCTACAAGTGCTATTGGGGACCCACTCAAAAAAAAAAAAACTTAAACTAAGAAAAATCCACCCTTTAATAGATTAGGGCAGAATCAACTGGATTTAACTTATATCTACTGATAGTATAGAAACTATGTTTAATTCGTTATAACATATTAATTTCCTTTCCTTTTTGTGTTTTTAGAATATGGCTCAGAGGGAATAGTGTCCACGAGTGGTGATGTTTATAGTTATGGTATAATGTTAATAGAGATATTGACAAGAAGAAGACCAACAAATGAGTTTCTCAATGAAAACATGAACTTGAGGCAATGGGTCAGTGAATCATTTCCAAGTTCTTTGAAGACAGTTGTTGATGAAAATATTTTATTTGGAGAAAATGAGATTTTCATATTTAGTATGATGGAGTTGGCTTTGGAATGCACAAAAGAAAGACAAGAAGAGAGGGTAAATATGAAAGATATAGTGAATAGACTTGGTAAAACTAAAGAGGACCTCTTCTTGGAAACAAAGAAGAAAAAATCTTATGAGTAAAATTTTATTTATTTTTAGTTTATGGTTTAGTTATTCTTT

>M6-PGSC0003DMG400018811 M6 LRR-RLK

TAAGGGGTCGTTTGGTAGAGTGTATTAGCAAAAATAATGCATGCATTAATTATATCTATTAGTAATCCCTTGTTTGGTACATTTTTTCATGCTATGCATTAGTTATACACTCTATTGTGTATTGAAGAGTGTATTACTAATACCATGGATTTCTAGGTATTAGTAATGCAAAGGATTTAGTACATGCATTAATATGGTTAAAGACTCAATTACCCCTCAAAAGTACTTTTACATCTTTTCCACCATAATTGTGAAGGATACTTTTTGTAAATAAATATTTTTATACAATGCATACTATTTTTAATACACCAAACCAAACAATGCATAAGAAATAATATATGCATAACTAATACAAGTATAACTAATACATGCATTACTAAAGCTTGCATTACTAATACACTCTATTGGGTATTATTTTTATACACTCTACCAAACGACCCCTAACTCTATAAA

>M6-PGSC0003DMG400018849 M6 LRR-RLK

ACTTAACCCACTAAAATCATCATGAAACACACTCTATTTTTCCAGCACTCCTAAAAAGCATTTTCTCTCAAATTTACCTCTCTTTACATACCCCATTTCTCAACCATTTCCACCATAAACAACATCACCATGAGACACTTGTTTCCTTTCTTTCTTACTACAGTTTTCTTCTTCCTAACAAGAATTTCACTTGTTTTCTCCATTACCAATCTACCTCTACAACTTATATCTCTTTTGACTTTAAAATCTTCTTTGCATGATCACCACAATACCTTCAATGACTGGGATCCCACCCTCGCCTTCGCTAGACCGGGTTCTCACATTTGGTGTTCCTGGTCTGGTATCAAATGTGACAAGAAAACTAACCAAATCACCTCTCTTGATCTATCGAAGCGCAATCTTTCTGGTACAATTCCAGAAGATATCAGAAACTTGGTACACCTTCATCACTTGAATTTGAGTGGAAATGCTTTGGAGGGTCCTCTTCAAACAGTCATTTTTGAATTGCCTTTTCTCAAGACACTTGATATTAGTCACAACTTGTTTAATTCAACTTTTCCCTCTGGTGTGTCAAGGCTCAAGTCTCTAACATACCTAAATGCTTATAGTAATGACTTCATAGGCCCTTTGCCTGAAGAAGTTGCTCAAATCCCAAATCTTGAGTACCTTAACTTTGGGGGAAACTATTTCAACGGTGTGATTCCAAAAAGTTATGGTGGCTTAGCAAAACTGAAGTTTTTGCACTTGGCTGGAAATTTATTGAATGGTCCTGTTTTGCCCGAGTTGGGTTTCTTGAAACAGCTTGAACATGTAGAAATTGGTTACCAGAATTTCACTGGAGTTGTCCCTGCTGAATTTTCTTCTTTGTCAAAGTTGAACTATCTTGATATCTCATTAGCCAATCTTTCTGGTAATCTCCCTGTTGGACTAGGCAACTTGACAAACCTTGAAACTCTGCTTCTTTTCAAGAACCATTTGTATGGTACAATTCCATTGTCTTTTGCCAGACTGACATCCTTGAAATCATTGGATTTGTCTGACAACCATCTCTCCGGAACAATCCCCGAAGGGTTTTCAGGATTCAAGGAGCTTACAGTGTTGAATTTGATGAACAATAACTTAACAGGTGAAATCCCACAAGGCATTGGTGAGCTTCCTAATCTTGAATTATTGGCTCTATGGAACAACTCACTCACTGGGATTTTACCACAGAAGTTGGGTTCAAATGCAAAGCTACAAAAACTTGATGTCTCTTCAAATTATCTCTCAGGTCCCATTCCACCAAATCTTTGTCTCAGCAATAGCTTAGTGAAACTTATCCTGTTTTCAAATCAGTTTATTGGTGAGCTCCCTTCTTCCTTAGCAAATTGCACTGCATTATTCAGGTTTAGAATTCAAAACAACAGACTCAACGGGTCAATACCATTAGGCTTCGGTATTTTGCCCAATTTAGCTTACTGGGACTTGAGCAAAAACAATTTCACTGGTCCAATTCCAGAAGATTTAGGGAATGCCGCGACATTGGCGTATTTGAACATATCGGAGAATTCATTCAACAGTGAATTACCGGAGAGTATCTGGACTTCACCATCTCTACAGATCTTTTCAGCTAGTTACAGTGGTCTTGTCGGGAAAATCCCCAATTTCAAAGGGTGTAAAGCTTTCTACAGAATCGAACTGGAAGGAAATAATCTCACAGGAAGCATTCCATGGGATATAGAACATTGTGAGAAGCTCATTTGCATGAATTTTCGACGAAATTCGCTAACGGGAATCATTCCTTGGGAAATTTCTGCAATTCCTTCCTTAACAGAGGTTGATTTGTCTCACAATTTTCTTACAGGAACAATCCCTTCGAATTTCGCCAATTCCACTACCATAGAAAACTTCAACGTATCCTACAATCAGCTCACTGGTCCAGTTCCTTCTTCAGGCTCTATTTTCTCCAGCCTACATTCCTCTTCCTTCATCGGAAACGAAGGCCTGTGTGGTACTGTCATTCAAAAGCCTTGCGGGACTGACGGACTCGCAGCCGGGGCAGCGGAAATCAAGCCACAGCCGAAAAAAACAGCTGGCGCTATTGTCTGGATCATGGCAGCAGCATTCGGTATTGGTTTATTCGTCCTCATCGCCGGTAGCCGCTGTTTCCACGCGAAATACAGCCAGCGATTTTCAGTAGAGAGAGAAGTTGGGCCATGGAAATTAACCGCATTTCAAAGGTTAAACTTCTCGGCGGATGATGTACTTGAGAGTTTGACAATGACAGATAAGATCCTGGGAATGGGTTCAACAGGGACAGTGTACAAGGCGGAAATGCCAGGCGGAGAGACTATAGCAGTAAAGAAATTATGGGGAAAACAGAAGGAAACGATCAGGAAAAGGCAAGGCGTTTTAGCTGAGGTGGATGTGTTAGGCAATGTGAGGCACAGGAACATCGTGAGGTTGTTCGGCTGTTGCAGCAACAACGAGTGCACCATGTTGCTGTATGAGTACATGCCTAATGGTAGCCTGGATGATCTGTTGCATGGTAAGAATAAGGATGATAATTTGGTGGCTGATTGGCTCACTAGGTACAAAATTGCTCTTGGGGTCGCTCATGGAATTTGTTATCTTCATCATGATTGTGATCCCGTCATTGTTCATCGTGATCTTAAGCCTAGTAATATTCTTCTGGACGGTGATCTGGAGGCTAGAGTAGCTGATTTTGGTGTTGCCAAGTTGATTCAGTGTGATGAATCTATGTCCGTGATTGCTGGATCCTATGGCTACATCGCACCTGGTAAGTTAATACTCCCTCCGTTTTAATTTTACATGAAATTTAAGAAAGTAATTCGTAATATTCAACTTTTTATCGTGTTCTTTAATTTAGTTAGTTATTAATGGCAGGAATATTTGTCTGTTTGCTTGATGGAATTACATGATTGTTTTGGAGTTCAAGTTTTTTTCTTGTGCGGCTTATTGCTATCCAAAATTATTTTGTTCAGATCAGTTTAGTTCGTTCTGTAGGGTGACGAAAAAGAGTTCACTTGCGTGCTAAAACGTGGTGTCATGTCAATAGCATGATGCATTTTTTTAGTGAAAGCATGATGTATATTGTTTTTTTTTTCGTCCACTATCGTCATTAAATCTTCTTTGCCCACACAATTGTCAAATTCCTCAACTGATTCATACTAGTGGAATCTTCTTTTAATCATAAACAGATCTTGAAGGCACTAATTCATTTTTTGAATTTGAGCGCCACAATATGATATAAATATCTAAATGTTGAAACTTTTATTAGGAAGTTAAAATAATATTCTTATGAAGATTCTAATATTAGTGTAGTGAAACAGTTAATATAATTTTCTTAATAAAAGGTATCAAGTATTTCTTTTCTCATACTTCTTTATGACAATTGGTGCAAATATTTATTCTCTTTGTTTTTGTCCATGATTTTGTGGTGCTTAAAGAGGGCATGGCGCACTAGCTGCGCGTGAATAGCTCCATTAGCCTTTAAGCTTTTATTTTACACTTTTTGTTATTCTTTGGACCCACCAAGATTTTGTCTTAGTTACTTATCTGTGATTAACCCTTTGATTTAATTAGTATATAATATAGTACTAGTACTACATTCTAATGCTTAAAGTCAACCATTTTGGGCACTAGTAGACAGAGTGTTATGATAACAAATTAAAATGATCTTATCTTTTTTCTTTTTTTTAATATAGTTATTGTACTAAATATGTTTGTTGATACAGAATATGCATACACATTGCAAGTTGATGAGAAGAGTGACATCTACAGCTACGGGGTTGTGTTACTTGAAATTTTATCAGGGAAAAGATCGGTGGAACCAGAATTTGGCGATGGAAACAGCATTGTGGATTGGGTTAAGTCGAAGATCAAGACTAAAAATGGCATAAATGATGTGTTGGATAAAAATGCTGGCGCTTCATGTCTTTCAGTTAGGGAGGAAATGATGTTGTTGCTGAGGGTGGCATTGCTGTGTACTAGTCGTAACCCGGCTGACAGGCCATCGATGAGGGATGTTATATCTATGTTGCAAGAGGCTAAGCCCAACAGGAAGTTGCCTGGAACTGGTGATAATGCAATTGCTGCTATTCCTTTGGCACAAAATGCTAATGTGGAGTGTTGATATTATAGGAGGAGATTTGGGGTTGTTTTGATTTTACAACTTACTAATGAATTGTGGGGGAAATTGGTATAGGAGAAGATTAAATAGTTTTATTTGCTTCTTAAATTTTTGCATAGTAGGAAATTCTCCAGTGTTGTCATTCCAATGTTGTGATGAAATGGGTATCTTTTCCAACAAATTAATCTTCAATAGCGTACTTGT

>M6-PGSC0003DMG400019051 M6 LRR-RLK

ATGAGCGGTCTTACAGCCCTAAGCCTTGGAGGAAACAACTTGGCAGGGAATATTCCTTCTGAGATTGGTAAGCTTAAACAACTCCAAGGGCTGGCTCTATATAACAATAAATTGCAGGGACGTATTCCAGAGGTGGTATGCCATTTATCGAATTTGGTTGGCTTATTACTGGATGGTAATGAGCTCTGTGGATTAATTCCAGAATGCTTTGGAAATCTTAGCATGCTACAAGCGCTTTATTTGGATTCTAATAAATTTTCATCAAAATTTCCGTTGAGCCTTTGGAAGATGAGTGGTCTTCTCTATCTAAGTGTGTCACGAAATTCAATAGAGGGAGAACTTCCATCAGATATTGGAGGACTGAAAGCCATTGTAGAACTATATCTTTACAATAACCACTTTTCAGGTGTGATACCAACCAGATTGGGGGAACTCCAAAACCTGCAGCATCTTGACCTATCCAACAATTCATTTTTTGGCCAAATTCCATTATCCTTTGCCAACTTGATAAGCTTGGAATTCTTGAATTTGTCTTTAAATGCCTTGTCCGGTACTATTCCTAAATCTTTGGAAAAACTCTCATACCTTAAAAGCATTAATGTTTCATTTAATGATTTAGAAGGTGAAATACCAAGTGGTGGCGTGTTTGCAACTTCCACTGTGCAATCATTTCTTGGGAACAAAGGTCTTTGCGGAATGCACATATTGGAGATTCCTACTTGTGCTATCACTAATCATGGAAAACAATCAAAGCTTAAGGAGGTTGTGCTAAAAATTGTTACTCCAGTGGTTATTGCATCCTTTCTGATATTCTTGTTCGTTTCAATTTGGATAATGAAACGACAGAAGAAAGGGAAGTCCAAAGATATTGAAAAGGTATTGGAGATCGGGACTCATCAATTAGTTTCTTATCACGAGATTCAACAAGCAACAAATAATTTTGATGAATCCAATTTAATTGGTGAGGGAAGCTCTGGCTCTGTGTACAAAGGCACATTGTTTGGTGGAACTGCAGTGGCCATTAAGGTTCTGGATTTGGAAAACGAGCAAGTATGCAAGAGGTTTGATACCGAATGTGAAGTGATGAGAAATGTTAGACACAGAAATCTTGTTTCAGTGATTACTACATGTTCTAGTGACTATATAAGAGCCTTTGTTCTGCAATTTATGCCCAATGGAAGTCTTGAGAATTGGCTGTACAAAGAAGATCGACACTTGAACCTTCATCAAAGAGTCACTGTAATGCTTGATGCAGCTATGGCAGTTGAATATCTACACCATGGTCATAACACTCCAATAGTTCATTGCGACCTGAAGCCAGCCAACGTTCTTTTGGATGAAGATATGATGGCTCATGTTGGTGATTTTGGAATATCTAAAATTTTAGCCATAAGCAAGTCCATGGCTTATACAGAGACATTGGGCACTCTTGGATATATTGCACCAGGTAAAAGAAAATCTACTCTCATTGATTTTCTCTTATCATAATTAA

>M6-PGSC0003DMG400019415 M6 LRR-RLK

ATGGCCTATTATCTTGACAAAATCTTCCTTCTCTTATTTCTCTTCACACTTAGTACTCCCTCAAAATCAGATAATATGACAGATATTGAAGCACTCTTTGCTATCAAGAATGAAATAATTGATCCCTTTGAATCATTAAGTTCATGGAATGAGTCTGTTCCTTTATGTCAATGGCGTGGCGTTGTATGTGGAACGCGAAATCAACGAGTCATTGAACTAAACCTTCTTGACCATAAGCTAGCCGGTGTGTTATCACCCTTTGTTGGAAATTTGAGCTTCCTCATACGACTAGACATCGGAAACAATACCATTTCTGGCAATATCCCTCCTGAATTAGGCCGTTTGACAAGCTTACGCCATCTATATTTGCAAAACAATTCATTTTATGGTGAAATTCCTGTCAATCTATCGTTTTGTAGTAACCTCGTTGAGTTATGGGCATACAAGAACAATTTTGTAGGTGTTCTTCCAAGGGAGCTTGGTTTACTTACCAAACTCAAATATTTAGATTTCGGTTACAACAAGCTCATTGGTGAAATCCCAAAATCTTATGGTAACTTTTCAAGTCTATTGGAAATGTACCTTTTGGATAATGATCTCGAGGGTAAAATCCCCGATGAGTTTGGGAAACTGAAGAGTTTGGAAATATTTGACGCGGACTTTAATAGGTTATCTGGTAGAATTCCGAGCTCATTGTACAATCTTACCTCATTGAAAGTCATTGATGTGTCATTTAACCAACTCGAAGGTACTCTGCCTCGGGATTTAGGGATCAACCTTCCTAACCTGGAGTGTCTCAGCATTAGTGAGAATCAATTCACTGGATCGCTTCCTCATTCACTTTTCAACCTCACAGGACTCGCGTACCTTTATGTTGGAAAAAATGACTTAGGAGGACGTGTTCCTAGGTTCGATAAGTTGCATAATTTGTATATTTTAAGCATGGAAAATAACCAATTTGATAATGACTTGACCCTCATGTCCTCCTTGACAAATGCAACTAACCTCAACCGTCTGTATATGCAAAACAACGCGTTTGGAGGGACATTTCATGAGTTTTTCAGCAACCTTTCAAGTAATCTGGCGTTTGTAGACTTGTCAAGCAATCATTTTTCAGGACATATCCCGATAGACATTGGGAAATTCGTCAACTTGGAAGAGCTTAGGCTGAGCGGAAACCAACTCACAGGTATCATCCCTTCGACTATTGGGAAGCTTTATAAGCTACGATATTTACACCTGTCCCAAAACAGAATTTCGGGTAGTATACCCTTTTCCATTGCAAATCTGACCCTTCTAACAGAGTTAAATCTTGATCATAACAATCTTAGAGGTAGAATACCTTCAAGTGTTGCAAAATGTCAAAGTTTGTTGTACCTTAACCTGAACCAAAACAATTTAATCGGTTTGATACCTAAAGAAATTTGTCTAATCTCTTCCTTAGTTGTTCTAAACCTTTCCAGTAACAACTTGACTGGCGCCCTTCCCAAAGAAATCGAAAGCCTGAAAAATCTCTTATCATTAGATGTCTCTTTGAATAATCTCAATGGTGAAATTCCTAGTACACTTGGGAGTTGTGTGACAATTGTTAACTTAACAATGAAAAGGAATTTATTCCGCGGGATTATTCCTTCCACTTTCAGTTCACTAAAGAGTCTTCAAGTTTTAGATCTTTCGCGGAATAAGCTATCAGGTATGATTCCAAAGTACTTGGAAGGATTTGCCTTGCATTTCTTAAACTTATCGTTTAATGATTTTGAAGGTGGCCTTCCACAACGAGGCATCTTTGAGAACGCGAGTGTAGACTCTTTCGTTGGAAATCTTAGAATCTGTGGTGGTGTACCTGGCTTGAAACTTCCCAATTGCAACTTCAGTCATTCAAAGAAAATGAACTTTAAGCTTGTTATTTTAGTAATATTAGGAATTTTGGGGCTAGTTGTTATGGTTTGTGCTTTTTTCTTCTATCGATTTGGAAGGTCCAAAAGAACAATTCCTTCATTAGATAACAATCTAAACCAACTCATAGCTATGTCCTATCAAAGTATTCTCAAAGTTACCAACGGGTTTTCTACGAGCAACTTGATTGGTGTTGGGAGTCATGGATATGTTTACAAGGGAATTTTAGAAACGGATGGTAAACACGTTGCTATAAAAGTGTTGAATCTTTTACAATATGGAGCTATCAAAAGTTTCATAGCAGAATGTGAGGCCTTGAGAAATGTTAGACATCGGAATTTAGTTAAGCTACTTACTGCATGTTCTGGTGTTGATTATCATGGCAACGAATTCAAGGCACTGATTTACGAGTTTATGGCCAACGGAAGTTTGGAGGACTGGTTGCATCCAGATAATTCAAGACCGAATGTTCAGCCAAGAAGGTTAGGATTTCTACAAAGGCTGAATATTGCTATTGATGTTGCTAGTGCAATTCATTATCTTCATAATGATTGTCAAATATCAATAGTTCATTGTGATCTCAAACCAAGCAATATTCTCCTTGACAATGAATTGGTTGCCCATGTTGGGGATTTTGGATTGTCAAGATTCCTACATCTTACTGACGAAACCACTTGTAGAATTCAGACAAGCTCTTCAACTTTTAAAGGATCTATTGGCTACATAGCTCCAGGTAAATTTCACTCTATATAAATGCCTCTAAAGACTTTAGTGACTCTGGATGCAATGACATTAACTCAATTGCCTTTATATTTTCATTGCAGAGTATGGTATGGGAAGCGAGGCATCAACGCAAGGTGATGTGTATAGTTTTGGTATCGTCTTACTTGAAATGTTAACTGGGAAAAGACCAACAGATGACACGTTTGGCAGCGATTTAAGCCTTCATGATTTTGTTAGGAATGCCATGCCTGATGGAGCACTTGAGATAGTAGATCCAGTTCTTAATTTTGAAGAGGAAGAGATAAGCACAGAGAGGTCTCAAATTCCAAGGTTCATGAGAAGGCAAAAAATGGTAGAGGGTTTAATTTCCTTATTTGGAGTTGGTATTGATTGTTCAATGTATGATTCAAGTAAAAGAAAAAACATGAAGGAGGTTGTTAGAGAGTTGTGTTCTATAAGAGATTCTATTGTTGGTTGTACAATATGA

>M6-PGSC0003DMG400019698 M6 LRR-RLK

TCTTCACATTCTACTACTCCTTTCCCTTTCTCTATTTGTCTGTTTAAAACTTCTACTAATCCTATCTTTCTTTTCAGAAAAATATGGAACCCCTTCAAGAACTCAAGCTATAGATTCAAGAAAATCTCCATTTAAGCTATAAAGTTTCAATCTTTGAAGATGAAAGCTCACAAAACTGTGTTTTACCAACACCCTTTAAGCTTAAACAAGCTTTTCTTTGTTCTTCTTCTTATCTTCTTTCTTCCTCCAGCTTCACCAGCTTCTGTTAATGGTCTTTTTAAAGACTCCCAACAGCTTATTTCCTTTAAAGCTGCACTTCCACCAACCCCAACTCTGCTTCAGGACTGGTTGTCATCTACTGACCCTTGTAGTTTCACTGGTGTTTCATGCAAGAATTCTAGAGTTTCTTCTATAGATCTCAGTAACACTTTTTTAAGTGTGGATTTCAGTTTGGTCACTTCTTATTTGCTTCCTCTTTCTAATTTGGAGTCTTTAGTGTTAAAGAATGCTAATCTTAGTGGTTCTTTAACTTCTGCTGCTAAATCCCAATGTGGGGTTTCTTTAGACTCCATAGATCTAGCTGAAAACACAATTTCAGGACCTATTTCTGATATCTCTAGCTTTGGTGTTTGTTCAAACCTTAAGTCTCTTAATCTTTCTAAGAATTTCTTGGACCCTCCTGGTAAAGAAATACTTAAAGGTGCAACCTTTAGCCTCCAAGTTCTTGATCTTTCTTACAATAATATCTCAGGTTTCAACTTGTTTCCATGGGTTTCATCCATGGGGTTTGGTGAACTTGAGTTCTTTTCTCTCAAGGGTAACAAACTAGCTGGAAGTATTCCTGAATTAGACTTCAAGAATTTGTCACATTTGGATCTTTCTGCAAATAATTTCTCAACTGTTTTTCCTTCATTCAAAGATTGCTCCAATTTGCAGCACTTGGATTTGTCATCCAACAAGTTTTATGGTGATATTGGTTCTTCACTTTCTTCATGTGGAAAGCTTAGTTTTCTCAACCTTACCAATAACCAGTTTGTAGGTTTGGTCCCTAAACTACAAAGTGAAAGCCTACAGTATTTGTACTTAAGAGGGAATGATTTTCAGGGTGTGTACCCAAACCAGCTTGCTGATTTGTGCAAAACTGTGGTGGAATTGGACTTGTCATACAATAATTTCTCAGGTATGGTTCCTGAGAGCCTTGGTGAATGTTCAAGTTTGGAACTTGTTGATATTTCCAACAATAATTTCTCTGGTAAGTTGCCTGTTGATACTCTCTTGAAGTTGAGTAATATGAAGACTATGGTCTTATCATTCAACAAATTTGTTGGTGTTTTGCCTGATTCTTTCTCTAATTTGCTGAAATTGGAGACTTTGGATGTGAGTTCTAATAATCTCACAGGGGTTATTCCATCTGGGATTTGCAAAGATCCTATGAATAACTTAAAAGTGTTGTACCTTCAGAATAACTTGTTTGAAGGTCCTATACCTTCCAGTCTAAGCAACTGTTCACAGCTGGTGTCACTTGATCTTAGCTTTAATTACTTGACTGGGAGAATACCATCTAGTTTGGGGTCATTGTCAAAGTTGAAGGATCTCATCCTTTGGTTGAATCAGCTTTCAGGGGAAATCCCACAGGAGTTGATGTACTTGCAGGCTTTAGAGAATTTGATTCTTGATTTTAATGACTTAACTGGACCAATACCTGCAAGTCTTAGCAACTGTACCAAGTTGAATTGGATTTCATTGTCAAATAACCAATTGAGTGGTGAGATACCGGCTTCTCTTGGTCGTTTGTCAAATCTAGCCATTCTTAAGCTTGGAAACAACTCAATCTCAGGGAATATACCTGCTGAGTTGGGTAATTGCCAGAGCTTGATATGGTTGGATCTCAATACTAATTTCCTGAATGGCTCCATTCCGCCGCCTTTGTTCAAGCAATCTGGCAATATTGCAGTGGCATTACTGACCGGGAAGCGATACGTGTATATCAAGAATGATGGGAGTAAGGAGTGCCATGGAGCAGGGAATTTGCTGGAGTTTGGAGGGATTAGACAAGAACAGCTGGATAGAATCTCAACAAGGCATCCTTGCAATTTCACAAGAGTTTATAGAGGTATCACTCAGCCAACATTTAACCACAATGGCTCTATGATATTTCTTGATTTATCTTATAATAAGTTGGAAGGTAGTATCCCAAAGGAATTAGGGACCATGTACTATCTGTCTATATTGAATTTGGGGCATAATGATCTGTCTGGTATGATTCCTCAAGATCTTGGAGGCTTGAAGAATGTTGCAATTCTTGATCTGTCATATAATAGGTTAAATGGCCCGATCCCGAATTCCCTCACCAGTCTTACATTGCTTGGAGAGATTGACCTGTCAAACAATAATCTCAGTGGAATGATTCCTGAATCTGCACCATTTGATACATTCCCTGATTATAGGTTTGCAAATAATTCTCTCTGTGGGTATCCTCTCCCCCTACCTTGTAGCTCGGGGCCGAAATCGGATGCAAATCAGCATCAGAAATCTCACCGCAGACAAGCATCGTTGGCAGGGAGTGTGGCGATGGGTTTGTTATTTTCCCTCTTTTGTATCTTTGGTTTGATTATTGTTGCCATAGAGACGAAGAAGAGGAGGAAGAAGAAGGAGGCTGCTCTTGAAGCTTATATGGATGGTCATTCACATTCTGCAACTGCAAACAGTGCCTGGAAGTTTACGAGTGCTCGTGAGGCGTTAAGCATTAACCTCGCAGCATTTGAGAAGCCTCTCAGGAAGCTCACATTTGCTGATCTTCTCGAAGCCACCAATGGTTTCCACAACGACAGTCTTGTAGGCTCTGGTGGTTTTGGTGATGTCTACAAAGCTCAGTTGAAGGATGGCAGTGTTGTGGCCATTAAGAAGTTGATACACGTCAGCGGACAGGGTGATCGAGAATTCACTGCTGAAATGGAAACCATAGGGAAGATCAAGCACCGGAACCTTGTCCCTCTTTTGGGCTACTGCAAAGTAGGGGAAGAAAGACTACTGGTTTACGAATACATGAAGTATGGAAGTCTTGAAGATGTCCTGCACGATCGGAAGAAAATCGGGATCAAGCTGAATTGGCCTGCAAGAAGGAAAATCGCCATTGGAGCTGCGAGAGGGTTGGCTTTCCTACACCATAACTGCATTCCACACATCATTCACCGGGACATGAAATCAAGTAATGTCTTGCTTGATGAAAATTTGGAAGCCAGAGTATCTGATTTTGGAATGGCAAGGTTAATGAGTGCTATGGACACTCATTTGAGTGTCAGCACTCTTGCAGGCACTCCCGGATAAGTTCCTCCTGAATATTACCAAAGCTTTAGATGTTCGACAAAAGGAGACGTTTATAGTTATGGTGTCGTATTACTTGAGCTTCTAACCGGCAAACAGCCAACAGATTCAGCTGATTTTGGTGACAACAATCTTGTCGGATGGGTAAAGTTGCACGCTAAAGGAAAAATAACTGATGTCTTTGATCGAGAGCTATTGAAAGAGGATCCAAGCATTGAGATTGAACTTCTACAACACTTAAAGGTAGCTTGTGCTTGCTTAGATGATCGACATTGGAAACGTCCCACGATGATACAAGTTATGGCTATGTTTAAGGAGATTCAAGCAGGGTCAGGCATGGATTCGACATCCACAATCGGAGCTGATGATGTCAATTTTAGTGCAGTTGAAGGAGGGATAGAAATGGGGATAAATGAAAGTATAAAAGAAGGCAATGAGCTAAGCAAACACCTTTGAATGCACTAAATCAAGAGTTTGTTGAAAGCTCACAAATTTTCCAAAATCATATGCAAAGAATTGCCAGTGTAATTTTTAGCCAATTATTGTATGTACCACTAGTTCCCATCCATAAAATCTTGTGTTATTATACATAAAAGGAAGGATGTTTTTCTTTTTTAAAACATGTAACAGTGTTGTATATAAATAGTGTATGAGTTTGTTTTTGC

>M6-PGSC0003DMG400019801 M6 LRR-RLK

ATGGTTTCCATGACTATTATTATAGAATTGCTAGTTTTGTTAACGTACCTCTTGGATACTAATGCATCGAACATTTCAACTGATGAGGCTGCTCTTTTAGCTTTCAAAGCTCATATAACTTCAGATCCAATGGAATGCTGTCGAAAAACTGGACAAAAGGGACTCACGTTTGCAATTGGATAGGCATATCTTGCAGTAGAAGGCATCAAAGAGTGACATCACTAGACCTCAACGGCCTTAGACTAAGAGGTTCAATCGCCACAGAAATCGGGAACCTCTCCTTTCTTAGTTTCCTGGACATTGGCAACAACAGCATCCACGGCGAAATCCCTGATGAACTCGGCTATTTGAGGAGGCTGAAATACTTGTCCTTGTCAATGAATAATCTCACAGGTCAAATTCCAGCTAGTCTAGGATTACTGAGAAGGCTCGAATCACTTGATCTTTCTGAAAATGATCTCCTTGGATCTGTTCCTTCGTCTATCTTCAATGTCTCTTCTTTGAAGGTCATCAATTTGATTTATAATGCACTGTCTGGGACTTTGCCGAATGACATTTTCCATAATCTTCCGCGGCTGGAGGGCTTACAAATAGCAAGAAATCGACTCTCTAGCCAACTACCAAGCAACTTAGACAAATGCAAAGAGCTTACATACTTGTCCTTGTCATATAACCAGTTCACAGGTAAAAATACTGAACCAATTTCTTTTTCTGCTACTTTTGGTTCATATTGACAATAAAGATTGCAGCAATTTTTAGTGAAAACAATGATAGGCTGTTGAGAGGTATAACTTATTGTAACGTCTATTACAGGAAGCTTTCCTAGAGACATGTGGAACTTGACAAAACTTCAACATCTATTTCTTGGATGGAATAACATAACAGGTACTTATAATGTTCTAGTTTAACTGAATCTGCCTACAACTGCATTTATTGCTGGACTTCTCACCTCATAGCACTAGAGATCTTTGACATTTCCAAACTTGCTCTTCAAGTCATTTATATGAAATTAGTATTTGAATTTCAAAGGGTAGTAGCAGTAAATGTTAGTATAGCTCTTTGTTCTTCATTTTCAAACAACAACAGCAACTACTCCATCGGTTCAATGCTAGTTTGTGATGTGAATCCTCATTATCCATGTTGATCCATTTAGTCCCATCTTCAAGAGGACGTTAAAACAGATTGCGAGGATTGCTGAACTATTTAAAAGGCATACAACTAACAGTTTTCTGTTATTTATAGTAACTTAGTGACTGTAAAGCTGTTATACAGTATTGGACATTTCAGTGTAGTAACCACACAATTCGACATATATTTTACAGGACATATACCAAGTGAAATTGATAACCTATCAGCAATTCGAAGATTAAGTCTACGACGGAACAATTTGGTGGGAAGTCTTCCACCAACTATTGGAAATTTGTCAAATCTAGAGATGATAGACCTAGGTGACAACAGACTATATGGTGGCATTCCTCAGGAGTTTGAACATCTTGTAAATTTGAAAGAAGTGTATCTTGGTACAAACAGATTATCGGGTGAAGTTCCAAGGCGTATGTACAACATTTCTGGACTAAAAAAGATTTCATTTGTAGCAAATACGCTTTCAGGAACACTTCCCCCAAACATAGGCCATACTCTTCCAAATCTTGAAGGCCTTTACCTTGGGTAAAACCAGTTCACCGGGCTGATTCCTACCTCCATTGTGAATTCCACCAAGCTTATCCAGCTTGATTTTGGTCGCAATCTGTTTAGTGGACCTGTACCTATGAATCTTGGAAAGCTACAACAGCTTCAATTCATCAACTTGCAAGTAAACCAATTGAGGAATGATCAATCTACAGGCGAGTTAAGTTTCCTGACTTCATTATCTAATTGCAAGTATTTGAAAACTGTTCAAATAGGAGGTAATCAATTTAATGGAAGTCTTCCAAAATCTCTAGGTTCAGGAAACTGGTCCTTTTCCCTTGAATATTTCATTGCCCTTCAAAGTGGTATCACAGGAAAAATACCAACTAACATCAGTAACTTTAGGAACCTGGAGTGGTTAAGCTTGGGAGAGAATAAGCTTATAGGTTCTATTCCTCCAGATTTGGGGAACTTGAGGAATTTGAAAAGATTTAGCCTGGAGAAAAACAATTTAGATGGAATTATACCAACTAGCTTGTGTAACATGGAAAACTTGTACCAAATTATCTTGGGGAAAAATCAACTTTCTGGCGAATTACCAAGTTGTTTTGGAAATCTTTCTTCTTTAAGAGAACTCTACTTAGATTCCAATGCACTAATTTCCCATATTCCATCAACTTTTTGGAGGAACAGAGATATTTCAATTCTCACTTTGTCCTCCAATTTGCTGAATGGGTCTCTTGCAGTAGAAATGGGGAATTTAAGGAGTTTGCGGATATTACATTTACATGGTAATCAGTTCTCTGGTCAGATTCCTAGCACAATTGGCCAACTACAAAATTTGGTTAGCCTTTCATTGTCAAAGAATATGTTAGATGGTTCCATACCTGAACTATTTGAAGATTTGGTTTCATTGGAACACTTGGATCTATCTAGCAACAATCTATCAGGCATGATCCCCAAGTCCTTGAGGAATCTTGAACATCTCATGTATTTCAATGTCTCGTTCAATGGGCTCATGGGTGAAATTCCAGATGGAGGGCCATTCGTAAATTTTACAGCTGAATCATTCATGGGTAACCCTGCATTATGTGGATCATCACGCTTCCATGTGATGCAATGCAGAGTCACAAGTCTTGAAAGGAAAAGAAAGAGTAGAGTCTTAACTTCTGTTCTTGCATCAGTTTCCTCAGGAGTTGTAGTCACAACCATTTTCATCATTTGGTTTCTGAAATGCCGAAAAAGGAGTACGGAACTTCCTCTAGTTGATACATTTGGTCAGGTACATAAGAGGATTTCGTACTATGATATTTCTCAAGGGACAAACAACTTTGATGAAGCAAACTTGATTGGAAGGGGGAGCCTTGGTTTGGTGTACAAAGGGACACTAGCAGATGGAATGGTTGTGGCAATAAAGGTATTCAATACAGAACTGCAACATGCATTTAGAAGTTTCGAGGTGGAGTGTCAAGTTTTACGTAGCATTCGACACAGAAACCTTGTTAAGGTGATAAGTAGTTGTGCCAATTTTGATTATAAAGTGTTGGTGCTAGAGTACATGCCCAATGAAAACCTCGAATGTTGGCTTCACTCTACTGACAAATTTTTGGATTTAACTCAAAGACTAAAGGTGATGATTGATGTGGCTTCCGCTGTGGAGTATTTACATGGAGGTCATTTGTTTGTAGTCGTCCATTGTGATTTGAAACCAAGTAACATACTTTTGGATGGAGATATGGTGGCAAAAGTGAGTGACTTTGGGATATCCAAACTTCTAACAGCTGAGACGCTAATAGCACATACCAAGACTTTGGGTACTATAGGCTACATGGCACCAGGTAAATGTTATTATCAAGTTCATTTGAAGATTTTAAATGACGTTAATTATTTTCACTAATGATTGCGTGATTTTTTGCAGAGTATGGATCAGAAGGCAAAGTATCAACCAAGGGAGATGTTTATAGCTTTGGTATTTTGTTAATGGAGACTTTTACAAGAAAGAGCCCCGTGGATGATCTATTCGTTGGAGACTTCACCTTGAAAAGATGGACATGCCAATCATTACCAAACCGATTGGTGGATGTAGTGGACATTAATCTATTTTCACTGGACGAAGAAAATTTTACTTCTAAAGAGAGATGCTTCAAATCAATCATGGAATTAGCTCTTGAATGCACAAACGATTTGCCTGAAGAGAGAATTCGCATAGAAGATATTACTCTGCGACTCAAGAAGATCCTAACTCAATTTCTGTAAAATGTTGTAACAAACTAA

>M6-PGSC0003DMG400020594 M6 LRR-RLK

CCCAACCAACCCCACCCCACATCTCCATTCTTCCTTATTTATGCACCCACCACCCTAACATCTCCACCCCACTCACTCACAATCAATGGCTTTCCACCACCATCACTACCTCCTCCTCCACCTTCTCCTTTTCTTCTCTTCCCTTTCTTTCACCTTCTCAGATGATGCTGATGTTATGTCCAAGCTTCTTGCAGCTCTTTCTCCAACTCCTTCTGGTTGGTCAACTTCTAAGCCTTTCTGTTCTTGGACTAATGTCATTTGTGACAAGTCTTCAGCTACTGTAACTTCAATTAATCTTGATTCTCAATCTGTTTCTGGTTCTTTACCTTCTGAAATTACTCAACTTTCCAATCTTAAAACCCTTTCTCTTCAAAAAAACAAACTTTCTGGCCCCTTACCTTCTTTTGCCAACATGTCAAAATTAGCTGAACTTTTCTTGGACAATAACCAATTCACTTCTGTTCCTCAAGATTTCCTTTTGGGGGTTCCCAATTTAGTAACTTTAAGCATTAGTGAAAATGGGAGACTCTCCCCTTGGCAAATTCCTATGTATTTGACTGAAAGTACAAATTTGGGATCTTTATATGCTAGCAATGCTAGTATTGTTGGTGTCATTCCTGATTTCTTTGATGTTTTTCCTAATTTACAGAACTTAAGGTTGTCTTATAATAATCTTACTGGTTCTTTGCCTGCTTCTTTTGGTGGTTCTGAAATTATGAATCTTTGGTTGAATAATCAAGTTAAAGGGTTATCAGGTAGTATTGATGTGATTGGAAGTATGACACAATTGTCTCAAGTTTGGTTACATGCTAATTCTTTTACTGGTAGTATTCCTGATTTGTCAAAATGTGAGAATATATTTGATTTGCAGTTGAGGGACAATCAGTTTACTGGTATTGTTCCTGAGTCTGTGATGTCTCTTCCCAAATTGTTGAATATTACTTTACAGAACAATAGGTTGCAGGGTCCAATACCTCAGTTTAAAGATGGGGTTGAGTTTAAACTTGGAACTACTAATAGTTTCTGTAAGGACACTCCGGGACCGTGTGATCAACAGGTCACTGCACTTCTTGATGTGGCTGGTGGTTTTGGATATCCATTGTCTTTGGCTGAGTCTTGGAAAGGGAATGATGCTTGCAATGGTTGGAGTTTTATAAGTTGTGATACAACAGGGAAGAATGTAGCTGTTGTTACCTTAGGTAAGAGGGGATTTTCGGGTACTATTTCACCTGCTTTTGCGAATTTAACTTCGTTGAGGAGCTTGTTTTTGAATGACAATAATCTTACTGGTACAATCCCAGAGAGTTTGACAACTTTGCCTAATTTACAAGTTCTTGATGTGTCTAATAATAATTTGTCTGGACCTATACCATTGTTTCCACCTAGGGTGAAGTTTACTCATACTGGTAACTTGTTTCTTGGGAAAAATATAACTACTGGTGGTGATGGTGGCGGAAGTGGTTCGGATGGTTCTGGATCGAATTCCAGTGGTCGAGGTGGAAGTTCATCTGGTGGTTCGAAAGGGCCTTCAGTTGGAATGATTGTTGGTGTTGTTATAGCTGTTGTTCTTTTTGTTTTAGTAGTGTTATTTGTGTCTTACAAATGTTACATGAAGAGACGCCATAAGAGATTTGGAAGGGTAGAGACTCCGGAGAAAAGCAATGAAATGGTTAAGCCTAGTCTACCTAGTGTCGTCGGTGGTTCAAATGGATATACTGGTGGCACCAGTGAGTTACAAAGTCAGAGCAGTGGTGATCATAGTGAGATACCAGTTTTTGAAAATGGAAACGTTGCCATTTCCATCCAGGTCCTTCGACAAGTGACGAACAACTTCAGTGAAGAAAATATCCTGGGGAGAGGAGGGTTCGGAGTTGTTTATAAGGGCGAACTACACGATGGGACTAAAATCGCGGTGAAGAGGATGGAATCTGGGGCAATGGGTACCAAAGGAATGAACGAGTTCCAAGCAGAAATTGCGGTCCTTACCAAAGTCAGACATAGGCATTTGGTTGCTCTACTTGGTTCATGTGTTAATGGCAATGAGAGACTTTTGGTGTATGAGTATATGCCGCAAGGAACTTTGAGTCAGCATTTGTTTGAATGGCAAGAATTAGGCTACAAACCGCTTACTTGGAAGCAAAGGGTAACGATAGCATTGGATGTGGCTAGAGGGGTTGAATACCTACATAGCTTGGCACAACAAAGTTTCATTCATAGAGATTTAAAACCATCAAACATCCTTTTGGGAGACGACATGAGAGCCAAGGTTGCAGATTTTGGTTTGGTTAGAAATGCCCCTGATGGAAAATATTCTGTGGAGACACGTTTGGCTGGAACTTTTGGCTATCTTGCACCTGAATACGCAGGTAAGCATTTCTACTCCTTGTATCATTACGTCTTTTTTTAACTGAGAACTGATTTGTATGCCCCTAGCTATAAACTTTAGATTTTTGCAAATTTTGAAATTATTCGATCATCTGCATTGCTAGGCATTGTTATCTTCATTACATTAGAGAGACATTTGGGTTATAATGTTATTACTGAGGCAATCTCCACGCATAAATGCTGCTTTGTCATTTTTATCGAACCCATGAACTATATACTACTATTATAAAAGATAAGTTTCCATTTCGTGTGGATTAGCATCTGAATTATCCTATTACATCTCCTCTTTGTTACAGCTACTGGACGAGTCACAACCAAAGTAGACGTTTACGCCTTTGGCGTTGTTTTGATGGAGATCATTACTGGTAGAAAAGCTTTAGATGAAACAATGCCAGATGAGAGGTCTCATTTGGTTACATGGTTCCGCAGAGTCCTTATCAACAAAGAAAACCTCCGAAAGGCTATTGACTCGACACTTGATCCTGATGATGAAACATATGAGAGCATTTCCAAGGTTGCTGAGTTAGCTGGCCATTGCACTGCTCGGGAACCTTTTCAGCGACCTGATATGGGACATGCTGTTAATGTCCTTGGTCCCCTTGTAGAGCAGTGGAAGCCTACAACAAGTGAAGATGATGACGGCTATGGCATTATTGACCTCGACATGAGCCTTCCTCAAGCCCTTCAAAGGTGGCAAGCTGATGAAGGAACTTCAAGAATGTTCGATGACTTCTCGATCAGTCACTCGCAGTCGAGCATACCCTCAAAACCTTCCGGATTTGCAGATACTTTCAGTTCAACAGATTGCAGATGATATCCAAAAGAATTGACAACAGTTCAAGCCACGCCACGCCACACGCGGTTTTAATCCTTTGTGGATCAACATAGAGTATGCCAATATTCATTTGATTAATTGGTTTCTTCTTGTCAATGTCATTTACTTTTTATGTGATTACATATTTAAGTTAATATAGTCTTGATGTAAGGTGTGATAACAGTTAAAAAGTCAGGTGAATGTCACATTTTTTCTTTTGCCTCTTTGGAGACAACAGAAAATTGTAACAAATTCTTTCATATTACATTGTTAATGTAAATAAAATTTTGAACTTCCCC

>M6-PGSC0003DMG400020697 M6 LRR-RLK

ATAATGGAGAAAAGCTTCACATATTTTCACTTAATAGCACTCATGTTGCTTTACTTGTGTGTGGCTAGTTTAGCCATGACTCAAACGAACATTACAACTGATCAATCCACTCTTCTTTCCTTAAAATTCCAAATCATTTCGGATCCCTTTCACTTATTGGATGAAAGTTGGTCTCCAGCTATTTCTGTTTGTCATTGGGTTGGAGTCACTTGTGGTTCTCGTCACAAAAGAGTGACATCATTGAATATTTCCAACATGAATCTTATAGGCAGAATTCCTTGTGATTTTGGGAATCTCACATTTCTTGTTTCTCTTGACTTGAGTAGCAACAATTTCTACGGAAAATTGCCTCAAGAAATGGCACATCTTCGTAGGTTAAGGTTTGTTAAACTTAGTTTTAACAACTTTAGCGGAGAGGTTCCTTCCTGGTTCGGGGTCTTACACCAACTTCAAGTTCTGACTCTAAGGAATAACAGTTTCAATGGTCCCATCACCCTCTCTTTATTTTCTAATATCTCCACACTTCAAACTTTGGATTTGACATACAATTCCTTAGAGGGCCAAATACCAAAAGAAATAGGAAATCTTGAAAACCTGAGGGTTTTAAGATTATCTGGCAACAAGTTTATAGGTTTTATCCCTCCGTCTCTCTCGAACGCCTCAATGTTGATGACATTAGAACTATCTCTCAATTTCCTTGGAGGAAACATTCCAGAAGAGATTGGAAAACTTCAAAACCTAAAACTTTTGTCAATAGAATCCAATCAACTTACAGGCTCTATACCATTCTCGATCTTCAACAATATTTCTGGAATTGAGATCATTGCATTTACAAATAATACCTTATCAGGAAATCTTCCTTATGATATGTGTAATTGTCTTCCAATGCTCGAAATGCTCTATCTAAGTGACAACAAGCTATATGGTCATATGCCTGTAAGCTTGCCAAATTGTTCGAACCTTCAAATATTGTCTTTATCTAATAACAAGTTTGACGGACCCATACATAGTGAAATCGGGATTCTTAATAATTTGGTGATTTTAGCACTGGATACTAACCATATTTCTGGTAATATCCCAATCTCCACATTCAATATCTCCTCATTACAGGTTTTGGTACTGAATCACAACAATCTAATTGGATCCTTTCCAAGGGAGGTTGGCAATTTGACAGAATTGCAAATTCTGGATCTTGCCGATAATAATCTTACAGGTATTCAAAAGGGGTATGCAAATCATTACTAGCAAGATTTATAATTTCTTAAATCTTAGTAATTACTAACTATACATCTACATGCATACATTGACAGGTGAAATACCGAAAAAATTTAGCAATCAGATTGAGTTGTTTGATCTTAGTTTCAACAGGTTTACTGGTTCACTACCAATGGAGATTTTCAACATATCGGGTATGAGAGAGATTCAAGTTATTTTCAATAATGTGACTGGAACTCTCCCATTAAACATAGGTTCTACGTTACCCAACATTGAAGTTATTCATCTGAGAAACTTAAATCTTTATGGGACTATACCTCATTCTCTCTCCAATTGTTCCAAACTCACTAGTCTAGACCTTTCTCTCAATAGACTAACAGGTTTGATTCCCAACTCTCTTGGAAGTTTGACTCATCTACAGTCCCTAAGCTTGGCAATTAACAATTTAATGGGCGATAGTAGGTTAAGCTTCTTGACTTCCTTGTCAAATTGTAGAGATTTAAAAATTATTTTCCTCTCTGCGAACCCTCTATATGGTGTGCTTCCAGAATCCATAGGGAACCTTTCCAGTACTTCTCTTCTAAGGTTTAGAGCTATTGATTGTAAAATTAAGGGGGAAATTCCAAAAGGATTTGGGAACTTAAGCAGCTTATTAGAACTCGATCTTTCTAGAAACGGTTTCACTGGACCAATTCCCACAACAATCAGCAACTTGAGATTCCTTCAGAGCTTGAAGTTGAGTGTCAACAAACTTTCTGGATATATTGGAGATGATCTATGCAAATTACAGAACTTGGGTTATTTAAACTTGACTTTGAATCAACTTTCAGGATCTCTTCCGAATTGTTTGGGAAACTTAACTTCCCTTAGGGAGATGATTCTAGGTTCCAACAAATTGCGTTCAATCATTCCAAAAAGCTTAGGAAACCTCATAAACCTCTTGAAGCTAGACTTATCCTCAAACAACTTGGGTGGCTCTATACCTCTGGAAATTGGAAATCTAAAGGCTGCGATATATATGGATCTATCAATAAATACACTCTCAAATGGAATTCCTAAAGAAATTGATAGCTTGCAAAATCTTATATACCTTTCTTTGAGAGACAACAAATTGCAAGGATCAATTCCTGGCTCAATGAGCAGCATTTCAGCTTTGGAATTTCTAGACCTTTCTCATAACAATGTTTCAGGATTGATTCCCAAGTCTTTGGAGAAGCTTCTATATCTCAAGTATTTCAACGTTTCATTCAACAAGTTGGTTGGTGAAATCCCCTCGAATGGTCCTTTCAAGAACCTCTCCGGTGAGTTTTTCATGTCTAATGAAGCATTGTGTGGATCTCCAAGATTTTGTGTTCCCCCATGCCACAGTAGCACATCAAAGCATAAATCCAAGAGGAAAAAAGTGATTGTTCTTGTTCTACCGGGTACACTTGTATTAGTCTCCATCGCCTTTGCCCTTTTATGGTTAAGGTATACAAGAGGTAAAAGAACTGATCCTCAGCAAGCAGTTGATTCATCCTTTGTCATCTCAACAAAAGGAAGAATTTCATACTATGAACTACTTCGAGCAACTGATTCACTTAGTGAGATCAATTTAATTGGTTCTGGGAGTTTCGGCTCTGTTTACAAAGGCATCCTCAGAGATGGGACTTTCATTGCAGCTAAAGTGTTCAACCCACAATTGCAAGTGGCATTTAGGAGTTTTGATACAGAATGTCAAGTTTTGCGCAACCTTCGTCATAGGAATCTCACCAAAGTCATCACCAGTTGTTCCAACCTTGAATTTAAGGCTTTAATACTCGAGTACATGCCCAATGGGAGCCTCAATAAGTGGTTGTATTCACACAACTACTTCTTAGACATCAAACATAGGCTAAGTATAATGATAGATGTAGCATGTGCATTGGAATATCTACACCATGGATGCTCATTACCCGTGATTCATTGTGATTTGAAGCCTAGTAACGTCTTGTTGGATGAGGAAATGGTTGCCCACCTAAGTGACTTTGGAATTTCAAAGTTGCTTTCTGAAGATGAGAGTTCTTTATACACTAAAACATTAGCAACGTTTGGTTATATTGCACCAGGTACGTGATCTCTTAATCAATTGCTTTTGATTATTTCTCTTTCCTTATTTTCCACTAGTAAATGTTGAATTTCAAAATTGATTGATTGCAGAGTATGGACTGGAGGGATTGGTGTCAATAAAATGTGATGTTTATAGTTAAGGGATCATGTTGATGGAAACATTTACTAGGAAAAATCCTAATGATGAAATGTTCGATGGAGATTTTAGTTTGAAGCAATGGGTGAGAAATTCACTCCCACAAGCAGCAATGGAAATTGTTGATGCCAACTTAATATCGCCACATGATAATCAATTAATGAAAAAGCTAGATTGTGTGGTATCCATCATGAAAGTGGCAATAGATTGTTGTGTTGAATCTCCAAAAGGAAGGATTGACATGAAAGATGTTGTGGGGAGGTTAAAGACGATCAAGATTCAACTTCTTGCATGTTGAGGAAGATCTTATTAGAAACTTTTCTTCCAAATCATCTCATTGTAACAAACATTTGCAATAATAGTTAGCTTCAAATAAGTTATAGCATTATGTCTTTCATAACATAGAGAATTGATGGGCATGTATACTTCTTAGGTCTTGAATTTGTAAAGTGGTTTTGGCCTAATACCTCTACTTTTTTCCATCTTTTAACTTTAACAATGTAAAACATAGAACTCTCAATATTGAATAACAGAATTTCG

>M6-PGSC0003DMG400020744 M6 LRR-RLK

ATGTCACTACTAGCATTCAAGAATATGATCATTGATGACCCTTTCAAGATCATGGACTCCTGGAATGAAACTATACATTTTTGTGACTGGCCTGGGGTTTCATGTGGTCGACGTCATCCCCGAGTTACAGTTTTAAATCTTACTTCTCTGAAATTGAGGGGTTCTCTGTCACCAAGCATTGGTAATCTGAGCTTTCTTTATGTCCTTAAGCTCCAAAATAATAGTTTTTCAGGTGAAATTCCATCAGAGATTGGCTACTTACATAAGTTAAAGTTTTTACGCCTTGATAACAACTCGTTCACTGGTCATATTCCATCAAACATTTCAGGATGCTTCAATCTTGTTTCTGTTGTTCTTTCGTATAATATGATGGTAGGCAAAATTCCAGCAGAATTAGGCACATTGTTGAAACTCAAACAACTTTCTCTTGTTTCTAACTATTTTACAGGAGGAATTCCGCCTTCATTTGGTAATCTTTCCTTACTTGATACCTTTTCCGCAAGCAAGAATAATTTGTTAGGGAAAATACCTGATGAATTATGTCAATTGCTCAACTTAAAGTACTTTGTAGTGAATGAGAACAACTTGAGTGGTACCCTGCCTCCTTGTCTTTTCAATCTTTCCTCTATCGTGGCCATAGATGTTGGAACAAACCATTTAGAAGGAAAATTGCCTCCATTGCTTGGTATTACTCTTCCTAATTTAGAGTTCCTTAGCATTTATAGAAACAATGTAACTGGAAATATTCCGGTAACATTGTCAAATGCTACGAATCTTCAGTCTCTTATTGCTAGTAGAAATGGACTCACAGGAAAAGTACCACCCCTTGGAAATTTGCTGAAAATGCGGAGATTTTTAGTTGCTTTCAATTATCTTGGGAAGGGGGAGGATGATGACCTTAACTTTCTCACCACATTGGTGAATGCCACCAGCTTGGAGCTTTTGGAGCTCAATACAAATAATTTTGGTGGAGTCTTACCTGCATCTGTTAGTAATCTTTCGACTGAGCTTATAGAGCTTTCGTTGTCTTATAATCAGATTTCTGGAGAAATCCCAAGATGGATATCCAATCTCAAGAAGCTTCAGGCTTTCTTTGTAGCTTACAACAGATTTCACGGTGAAATTCCGTCTGAAATTGGTGATCTGATATATTTGCAAGAGCTGGCTTTACTTGGAAATCAGTTCTCTGGCAAAATTCCAATCTCATTGGGAAATTTAGCTTCTTTAACCAAACTCAGTTTAAGAGAAAACAATCTCCAGGGGAGAATTCCTTCCAGTTTAGGCAAATGCCACAATTTGGAACTGTTGGATCTTGGTTCAAATAACCTTAGTGGGTTCATACCATCAGAAATTCTTGAACTCTCATCCTTGTCGGAGGGCGTGGACCTATCTCAGAACCAATTAACTGGTTTCCTTCCGATGGAAATTGGGAAGTTGAGAAATCTAGGCTACTTAAACCTGTCTGATAACAAGTTGCAGGGCCAGATTCCTACCAGTATAGGTACTTGTGTGAAACTTGAAGCACTAGACTTAAACAACAACAACTTTCAGGGTAGTATTCCTTCAACCATGAATAGTTTG

>M6-PGSC0003DMG400020832 M6 LRR-RLK

AGATACCAATGGAGCTAGGTAATCATAAGAAACTTCAGGTATTGGTATTATCACAGAATGAGTTTACTGGCTCTATCCCTGACAGCATTTTCAACATGTCAACACTGCAGTTCCTAACATTGTCAGAAAACAAGCTTTCAGGTACTCTACCTTCAAATGTAGGTCGTAGAATGCCCAACCTAGAAGTATTTATTTGTGAAGGAAATAATCTGAGTGGTTTTATCTCTGATTCAATCTCTAATTCTTCAAGACTAGCAATATTGGAACTCTCATTCAACAGTTTCACAGGTCCAATTCCTAAATCATTTGGTAACTTAGAATACCTTGAGGTTCTGAACTTAGGGTTGAATAATTTTATCAGTGATTCATCATTGAGCTTCCTTACATCATTAACAAACTGTAGGAAACTAAGATCACTCTGGTTTAATGAGAATGCATTGGATGGGGCTTTATCAGCGTCTGTTGGTAATTTCTCAAAGTCGATGCAAAATTTTCAAGGAAATGGCTGTAAGCTAAAGGGTGTCATTCCTCGAGAAATTGGTAATCTTACTGGAGTGATACATATGAGTTTGTATAGCAATAAGTTGACTGGACATATTCCAAATACTGTCCAAGACATGTTGAACCTCCAAGAATTTTATCTACATAGTAACGAGATTAAAGGTACCATACCAAATGTTATATGCAGTTTAAAGAATCTTGGTGCATTAGACTTGTCAGGAAATCATTTTTCTGGTTTGGTACCCTCATGCTTAGGGAATGTTACAAGTTTGAGGTATCTTAATCTAGTTTACAACAGGCTGAATTCAAGATTACCTGCAAACTTGGGAGCCTTCAAGATCTCATTGAATTTAATGTTTCATCCAATTCATTAAGTGGTCATATTCCTCCGGAGTTGGGAAATTTAAAGGCAGTGGCACTAATTGATTTGTCAAAAAATGATTTTTCTGGTAAAATTCCTAGCACTTTAGGAGGTCTAGCTGAATTGATTAATCTTTTTCTAGCACATAATAGATTAGAGGGGCCTATTCCAGATTCATTTGGCAAATTGTTGGCATTGGAATTCTTGGATTTGTCCTATAACAATCTTAGTGGTGATATTCCAGAGTCATTAGAAGCTCTTGTGTATCTCAAATACATGAATTTCTCATTCAATAAACTCATTGGAGAAATTCCCATTGGCGGCCCTTTTGCAAATGTCACTAGTCAATCTTTTTTGTCCAATGATGCACTCTGTGGTGACTATCGGCTTAACGTAAAACCATGTCGAACCAAATCTACAAAGAAAAAGAAGTCAAGAAGAAAAAGAGCGCTGATAGGTTTATATACGCTGTTAGGGATTGTGTCACTCCTTGTGTTGGTAGTTGGATATGTGGTGTTAAGATTGAGAAAGACAAAGAAGAATGCAAGTCAAGCAGATGTGTCTCTGGTAAAAGAACATGAAAGAATTTCCTATTATGAACTTGAACATGCAACAGAAGGATTCGACGAAAGCAACTTGCTTGGTACAGGGAGTTTTAGCATGGTCTACAAAGGGATACTTAAGGATGGTACTCTTTTGGCAACAAAGATATTCAATGTGCAATTGGAGGGTGCATTTAAAAGTTTTGACACGGAATGTGAGATACTCCGGAACCTTCGCCACAGAAATCTGACCAAAGTTATTACAAGCTGTGTCAACCTTGATTACAAGGCCCTGGTGTTGGAATACATGCCCAATGGGACACTTGATAAATGGTTATACTCCCACAACTTGTTCTTGAACTTATTACAGAGACTGGATATAATGATAGATGTTGCATCTGCAATTGACTATCTCCACAATGGCTATACAATGTCTGTGGTTCATTGTGACTTGAAGCCAAGTAATGTCTTGCTAGATCAGAAAATGGTTGGTCATGTCAGTGATTTTGGCATTGCGAAATTGCTAGGTGAAGGGGAGGCTTCTGTTCAAACAAGGACTATTTCAACCATTGGATATATTGCTCCAGGTATATTAGAACTTCTTAAAGTTTTTCTTGTTTCTCATTTCAGTTTTGGCATATTCCATTTTAACTAGGAAGCTTTTATGATAAATTTTTAAAAATAATCGTAGAGTATGGACAAGATGGAATAGTATCCACGAGTTGTGATGTTTATAGTTTTGGCATCCTGATGATGGAGACGTTCACAAGAACAAGACCAAGTGATGAAACATTTACCGGTGAATTGAGCATACATCGTTGGGTTAGTGATTCCTTTCCTGATGAAATTCATAAGGTGGTGGATTACTATTTGGTTCAGCCAGGGGATGAACAAATTGATGCAAAGATGCGATGTTTGTTATATATCATGGAATTAGCTTTGTACTGCACTTTAGTGAGACCTGATGCAAGAATTAGCATGAAAGATGCTCTTTCAACACTCAAAAAGATCAGGCTACAACTTGTCAATAATCGGCACTAG

>M6-PGSC0003DMG400020848 M6 LRR-RLK

ATGGGAAGAAGTTGCAATCTTCTCTTTGCTCTTGCAGTTTTCATTCTACTCCATCACCATGCTTTACTTTCTACTGTTCCTAATATAAGCACCGATGAAGCTGCTCTTCTTGCACTGAAATCACACATTTCTTCTCATCCTAACAACATCTTATTAAGCAACTGGTCTTCTTCGACCCCGGTTTGCAGCTGGATTGGAATCACTTGCAGCTCCCGCCACCATCGAGTCACTGCTTTAGACATTTCTAGCATGCAACTTCGTGGTACCATTCCTCCACACCTTGGAAACCTCTCATTTCTCGTCTCCCTTGACATCAGTAACAACACTTTCCATGGAGATTTGCCAGTAGAGTTGGCTCATTTGCAGAGGTTGAAATTGATTAATGTCAGAAGCAATAACTTCACCGGAACCATTCCATCATTTTTAAGTTTGTTACCAAACCTACGCTTTGTGTACCTCTCGAGCAACCAATTTTCGGGGAAAATTCCTTCTTCTATTTCCAATCTGACAAAGCTGGAATTGTTGACAATACACACAAATTTTCTTGAAGGAGAGATCCCTAAAGAAATCGGTGATCTTCGTTACTTGATTGTCCTAAACCTGCAAGATAATCAGCTTACTGGCTCTATACCACCCTCAATCTTTAACATTACTACAATGCAAGTAATTGCTCTTACTGACAACAATCTGACCGGAAATCTTCCAAAAACTTTATGTGACCGTCTTCCAAACTTGGAAGGGCTTCACCTCTCAAGCAACTCCCTAGGTGGAGTTATTCCACCAAACCTGGAGAAATGCCGAAAACTTCAAATATTGTCTTTGTCTTACAATGAGTTTATTGGAACTGTACCAAGAAAGCTAGCCAACTTAACAGCTCTTACAGAATTATATCTTGGATTGCAACAGTTGGAAGGTAGTATGCTATTTCCTCACAAGATATCTGAAATTTTCTTCCTTTCCTCTCTTCACCAAACTTGTACTCATATATTTCCTGAAATTGACAGGAGAGATACCAGCGGAGCTAGGTAATCTTAACAAACTACAGCTGCTGATGTTAGACCAGAATGAGTTTACAGGTTCTATCCCTGAAAGCATTTTCAATATATCAGCAATGCAGAGCCTAGATTTTTCAATAAACAAGCTTTCAGGTCCTCTACCTTCGGATTTAGGTCGTGGAATGCCCAATCTAGAAGAACTTTATTGTGGAGGAAATAGTCTGAGTGGTTTTATATCTGATTCAATCTCAAATTCTTCAAGACTCAGAATGGTTGAACTCTCAGACAACAGTTTCACCGGTGTAATTCCTGAATCACTAGGTAACTTAGAGTACCTTGAGGTTCTGAACTTGGAGCTGAATAATTTTATCAGCGATTCATCATTGAGCTTCCTTACATCATTAACAAACTGTAGGAAACTAAGAGCACTCCGGTTTAATGAGAATGCATTGGATGGGGCCTTATCGGCGTCTGTTGGTAATTTCTCAAACTTGCTGCAAAATTTTCAAGGAAATGGCTGTAAGCTAAAGGGCGTCATTCCTCGAGAAATTGGTAATCTTACAGGAGTGATATATATGAGTTTGTATAAGAATGAGTTGACTGGACATATTCCAAATACTATCCAAGACATGTTTAACCTTCAAGAATTTTATCTACAAAGCAACGAGATTGAAGGAACCATACCAAATGTTTTATGCAGTTTAAAGGATCTTGGTGCATTAGACTTGTCAGGAAATCATATTTCTGGTTTGGTGCCCTCATGCTTAGGGAATGTTACTAGTTTGAGGAAACTTAATCTAGCTTACAACAGGCTGAATTCTAGATTATCTGCAAACTTGGGGAACCTTCAAGACCTCATAGAATTCAATGTTTCATCCAATTCATTAAGTGGTCATATTCCTCTGGAGTTGGGAAATTTAAAGGCAGTGACACTCATTGATTTGTCAAAAAATGATTTTACTGGTAAAATTCCTAGCACTTTAGGGGGTCTAGCTGAATTGACTGATCTTTCTCTAGCAAATAATAGATTAGATGGGGCTATTCCAGATTCATTTGGCAAAATGTTGTCACTGGAATACTTGGATTTATCGTATAACAATCTTAGTGGTGAAATTCCAAAGTCATTAGAAGCTCTTGTGTATCTCAAATACATGAATTTCTCATTCAATAAACTCAGTGGAGAAATTCCCACTGGTGGTCCTTTTAAAAATGTCACCAGTCAATCTTTCCTATCCAATGATGCACTCTGTGGTGTCTCCTGGTTTAACGTAAAACCATGTCAAACCAAGTCAACAGAGAAGCCAAGAGGAAAAAGAGTGCTTATCGGTTTATATACATTGTTAGGGATTGGATCACTCCTTGTGTTGGCCGTTGGATATGTGGTGTTAAGATTGAGAAAGACAAAGAAGAATGAAAGTCAAGCAGATGTGTCTTTAGTAAAAGAGCATGAAAGGATTTCCTATTATGAACTTGAACATGCAACAGAAGGATTCGATGAAAGCAACTTGCTTGGTACCGGAAGTTTTAGCATGGTCTACAAAGGGATACTTAAGGATGGTACTCTTTTGGCAGTAAAGGTATTCAATGTGCAATTGGAGGGTGCATTCAAAAGTTTTGACACGGAGTGTGGGATACTCCGGAACCTTCGCCACAGAAATCTGACCAAAGTCATAACAAGCTGCTCCAACCTTGATTTCAAGGCCCTAGTGTTGGAATACATACCCAATGGGACACTTGATAAATGGTTATACTCTCACAACTTGTTCTTGAATTTATTGCAGAGATTAGACATAATGTTAGATGTTGCATCTGCAATGGACTATCTCCACAATGGCTATTCAACGCCTGTTGTGCATTGTGACTTGAAGCCAAGCAATGTCTTGCTAGATCAAGAAATGGTTGGTCATGTCAGTGATTTTGGCATTGCAAAATTGTTAGATGCAGGGGAGGCTTTCATTCAAACAAGAACAATTGCAACCATCGGATATATTGCTCCAGGTATATTAGTTAAGCATTTCTTTCCCTTAGATATGGATTGATGCTTTCAGTTTTGGCATATTCCATTTTAACTAGGAGGCTTTTATGATCATTTTTTACTATGATTACAGAGTATGGACAAGATGGAATAGTATCCACAAGTTGTGATGTTTATAGTTTTGGAATTCTGATGATGGAGACATTCACAAGAAGAAGACCAAGTGATGAAATATTTACCGGAGAATTGAGCATACAACGTTGGATTAGTGATTCCTTCCCAAGTGGAATTCATAAGGTGGTGGATTACTATTTGGTACAGCCAGGGGATGAACAAATTGATGCAAAGATGCAATGTTTGTTATCAATTATAGAATTAGCTTTGAGTTGCACTTTAGTGACACCTAATGCAAGAACTAGCATGAAAGATGCACTTTCAACACTCCAAAAGATTAGGCTCCAGCTTGTCAGTAGTCAACACTAG

>M6-PGSC0003DMG400021008 M6 LRR-RLK

TATTTGTAGTTGTTGTAATCAAGGTGTACTGATATTGATTTCTTGTTGTAAGTGAACCCCATTTTACATTGTGGTGTTAGAATGAAGTTACTCAAGTTCTTGATGGTTTTTGTTCTTTATTTTGGCTCTGTCATTGGGCAGCTTCCGTCTCCGGATATATTGGCTTTGCTCGAGTTTAAGAAAGGTGTCGAATATGATCCAACGGGGTTTCTTCTTGAATCTTGGAATGAGGAGTCTATTGATTTCAATGGATGTCCTTCATCTTGGAATGGGATAATGTGTAATGGTGGTAATGTTGCAGGAGTTGTTCTTGACAACTTGGGTTTAATTGCTCATGTTGATTTGAGTGTATTTGCTAATCTTACAAAGCTTGTTAAGCTTTCAATGGCTAACAATTCAATTGCTGGGAAAATGCCTAATAACATAGATAGATTTAAGAGCCTTCAGTATCTAGATCTCTCAAGTAATCTATTCAACTCTTCTTTACCCCCAGAGGTTGGACAGTTAGGGAGGTTAAAGAATCTGTCATTAGCTGGTAATAACTTCTCTGGTATGATACCGGAGACAATTTCAGGGCTTGTGTCTATGCAATCTTTGGACTTGAGTCACAATTCTTTTTCTGGGGCTCTATCTTCATCTCTGATGAAATTGACTGGTTTGGTATACCTGAATCTATCTGTTAATGTATTCACAAAGGAAATTCCTAAAGGATTTGAGCTGATGGAAAATCTTGAAGTGCTTGATTTGCACGGAAATATGTTTGATGGCAATTTGGATGCAGAAATGTTGCTGCTTACCACTGCAACTCATGTTGACTTGAGTGGAAATTTGCTAGTGAACACTGCTTCTCAGCAGCAGAAATTTTTGCCAGGTTTATCGGAGTCGGTGAAGTATTTAAATCTTAGCCATAATCAGCTAAAGGGTTCACTTGTAAGTGGAAATGAAGCTCAGACTTTTGGCAACTTGAAGGTCTTGGATTTGAGCTACAATCAGCTCTCAGGGGAATTGCCTAGTTTCAATTTTGTTTATGATCTACAGGTGCTTAAACTTGCCAACAACTTATTTTCCGGATTTATTCCGAATGATCTTTTGAAAGGAGATGCATTGCTTCTCACTGAGTTGGATTTGAGTGGAAACAACCTCACTGGTAAATACATGAAACCCTTTGGTAAATAACTCTTTTCACTGTTTTTCTCAATCTCTTCATTTTGCAGGTCCAATAAGCATGATAACATCCACAACATTGCGAACTCTTAACTTATCATCCAATGCTCTTTCTGGTGAACTTCCCTTTGTGACTGGAACTTCTGCGGTGCTTGATCTCTCGAAAAATCAGTTCAAAGGTAATCTAACTAGAATGCTGAAATGGGGGAATATTGAATTTCTTGACCTTAGCCAAAACCGTTTGACTGGAAACATCCCTGAGGTTACGTCTCAGTTTCTGCGTTTATATCACCTCAACCTTTCGAGCAATACCTTGACAGGATCTATCCCGAAAGTTATTTCACAGTTTCCTAAGATCACTGTCCTTGATCTCAGTTTCAACCAGCTGGATGGGCCTCTTCTTACTTCTCTACTAACTATGCCCACTATTGAAGAACTTCACCTACAAAACAATGCATTTGTTGGAAGCATTGATTTCTCTCCTCCGGCTAGTACTCCTAAGCTTCGTGTTCTAGATCTTTCTCATAATCAGCTTGCTGGTTACTTTCCTGATGGATTTGGGTCACTGATTGCACTTCAAGTACTCGATATATCAGGAAATAATTTCTCTGGATCTTTGCCAACTTCCATGGGTAATGTTAGCTCTCTTACTTCTTTAGATGTTTCAGAGAATCATTTCTCAGGTGAACTGCCAAAAAACCTGCCTAACAGCCTCCAAAGCTTTAATGCATCACTCAATGACTTCTCTGGTGTCGTGCCTGAGAATTTGAGAAAATTTTCCTTATCATCTTTCTACCCGGGAAACTCTCAACTTCAATTTCCGAATCCTCCTTCAGGATCTGGTCAAGCTTCAACAGAGAAGCATAAGAGCAGGCAACTTAAAACTATTATTAAAGTGGTGATCATTGTTGCTTGTGTGGTTGTTCTTGTTGTTTTAATCATGCTAGCCATTTTTGTATTTTACTTGCGGGCATCGAAGAACTCTCAGGCACATGTTACTGACAAAGACATCCGCCATCAAACCCAATCAAATCCTTCTGGTTTTAGCAGTAGAGAAGGTGCAGTAGTTTCAGCACAAGATGTTACAGCTGCTTCACGGAAAGCATCATCATCAGAAGTAATTAGTGCTGATGAGAAAACGACAGGTTTCTCCCCTTCGAAAACAAGTCATTTCACTTGGTCACCAGAGTCAGGGGAATCATATACTGCAGAAAGCCTTGCCGATAATTTGGCTGGAGAGCTGTTCTTCCTTGATGATACAATCTCTTTTACAGCTGAGGAACTATCTAGGGCCCCAGCTGAAGTCTTAGGCAGAAGCAGCCATGGGACATCTTACAGGGCAACACTAGATAACGGGTTGCTCTTGACTGTGAAATGGTTGAGAGAAGGAGTGGCCAAACAGAGAAAGGATTTCACAAAGGAGGCTAAAAAGTTTGCCAACATTAGACATCCAAATGTGGTGGGATTAAGAGGTTACTACTGGGGTCCCACACAACATGAGAAGCTTATTCTTTCAGATTACATATCTCCCGGAAGTCTATCAAGTTTTCTATATGGTAAACTTTATCTTCTCGCATCAACCTTGCAGTTCGTCTTATGCATATCGTTTGCTATAACACTTTGAACTCATAAAACACTTATAATCTTTCCTTCTTTTGATATTTTTTAGTATTTTATGGCAAACAATTTGTCAGTTTGATATCTTGACTGTTCTTGTGGGGAATGAATTTTCTCAGTCTGTGATACACGAATTAAGGATAAGAAGCCATTTCAATTTTTAGTTAGCATAGTGAGTCCAGCTAATTCATTCATTCATTATACAGCTTGTAATCAGTTCATTCACAATCATATATGAAACTACACTCTTTCTCTTCTGAATCTATTCTCTCAATATTTCTCCTTCTTTAGCTGTTCTTCTTGAGGGAACTCCTGAAACAGTCACTTGATCCTCAAATTCCTTGTGAATTCTTGCAATCAAGTAACAGTTTGTATTATCATGAAATATAGTATCACTTTCACTGTGGAATTCTTCAACTTAAACTGCTGATGTCTAGATTCTGATTTATTGTTGGAAAACTTTAAACTTGTCCAAGGCCCTATACACTACGTATGTCTTTGTTACCTGGTCCGACTGGACGTGTAAGCAGGCCTTTACTTGGTTAATATACCTGCACTGGCAGTAAGGTTCAGCAATAGTTGTGGTACTTTGCTCATAAAAAGGTCACAATGAACTGCAAATTCTTCAGTGCTCAAATACCTTATTTTCGATTGGTGATTCTACCTGCTCAACTTCTCTACTATTTTGTATGCTTTTGTGTAGTTTGTATAGTGTAAGATTTCTGTATATACATTTAACGTGCGACACCTTTGCAGATCGACCTGGAAGAAAAGGTCCACCACTAACCTGGCCCCAAAGACTCAAAATATCAGTTGATATTGCGCGTGGCCTGAATTATCTCCATTTTGACCGCGAGGTTCCTCATGGAAACCTTAAAGCAACCAACATCTTGTTGGATGGTCCTGATCTTAATGCACGAGTTGGTGATTACTGCCTTCACCGTCTTATGACTCAAGCGGGGACAATTGAACAAATTCTTGATGCTGGAGTGTTAGGTTATCGTGCACCAGAGTTGGCTGCATCGAAGAAACCAGTGCCTTCCTTCAAATCAGATGTATACGCTTTCGGAGTCATTTTGTTGGAGGTCCTAACTGGGAAATGTGCAGGTGATGTTGTTTCTGGTGAAGATGGTGGTGTAGACTTGACTGATTGGGTAAGGTTGAAAGTGGCAGAAGGTGGAGGATTCGATTGTTTTGATAATGCATTGTCATCTGAGATGGGAAATCAAACAATAGAGAAGCAAATGAAGGAGGTTCTTGCAATAGCTCTAAGATGCATACGTTCGGTATCTGAGAGGCCAGGTATCAAGACTGTATATGAGGACCTTTCATCTATTTAGTTTTCCTCATTTTGGCTTTTGCTTGGGAATCAAATAGCTCATTGGTTTCCCTTGAGTTTGTAGAATTAGCAAGTTTAGTGTGTGAATGGTCTAGTTTTATTAACCATTGTTTGTTGTAAATTGGGGCATATTTATAATGTATACAACTTTCACCCCCCACCC

>M6-PGSC0003DMG400021017 M6 LRR-RLK

ATGGAGAAGCACATTTTCTTATTGATTCTTCTATTTTTTATTCAATTTTCTGTATCATTTGCTACCTCAAATGAGACAGACCAACAAGCTCTACTAGCTTTCCAAAATCTTATTACAACTCCCAATCATTTTTTGGCCAATAATTGGACCAAAAATACTTCATTTTGCTTTTGGTTCGGCGTCACTTGCAGTCCAAAAAGGCAAAGGGTTGTAGCCTTGACTCTTCCTAATTTGCAACTTCAAGGCACAATTTCCCCATCTTTGGCCAATTTGTCCTTCCTCAGTGTTCTCAATCTCCAGAACAACAGCTTCCACGGTGGCATCCCTTATGGCATTGGCCACATGCCTCACTTGCGAGTGATTGATGTTAAAAACAATCAGCTCAACGGAAGTATACCAACAAGTCTATTTCAACACCGTAGAATTCAAGTAATTTCATTGGCTTTGAATGAATTCAGTGGTGAGATGTGGAGAGGGCCATGGTATGTACCCCAACTCAGAGTCTTAAATCTCAGGAATAATAGCCTCACCGGTATAATCCCTCCTTCTGTTGGAAATGCCACGAAGTTGCTGAACTTCAGTTTGTATGGGAATAAAGTTGGTGGCAACATTCCAAAGGAAATTGGTAATCTGAGCCAGCTTGCATTTTTGTTCTTGTTCAATAATCAATTAACAGGTTTCATTCCTGCAACATTGTTTAATATTTCATCACTTGTTGCCCTATCTCTTGGAACCAATAGCCTTTCCGGTCCTCTCTTGCTTGATAAAGGAAATAGTGTCTCAAATCTCAAATTTTTAAGTATATATGACAACCAAATTTCTGGTTGCATTCCTACCAACATATGCCAACTCGCAGAGCTCAAATTTTTGTCCATATCTTTCAACAACATAACCGGAGACATACCCAAAAATATTGATTGTTTATCTAAGCTCAAGGGGTTCTATATTGGCGATAATCCAATAAAAGGGACTATTCCCACTTCATTGGGAAATATTTCCACTCTGCAATATCTTTATTGTGGAAACAATCGCATAGTGGGGCAAATTCCTCCGCAATTAGGGAAGCTATCAAATTTGAGGCAATTAGGCTTTGACCATAATTTTAATCTTATTGGTCAAATTCCAGAGGCTATTTTCAACATATCTTCTTTCGAAAGGATTGATTTCAGTTTCAATAACCTCTCGGGGAGAATTCCAACCACTGCAGGTCTTCATCTTCCGAACCTTAAAGGACTTACCTTGGGATTCAATCAGCTCGAAGGAGAAATTCCATTGTTCATAACAAATGCTTCCAAGCTTGAGATACTGGGGCTAAATAATAACTTTCTCACAGGAACTATTCCTACTAATTTGGGAAATCTTCATGAGCTGCGATATCTGTTCCTACATGATAATCAACTTACCAATGAACCAAGAGAGCATGAGTTGCGATTCTTCAATTCTTTGGCAGACTGTAGAATGTTGCGATATCTACAAGTGGGTAACAATCCGTTGAATGGCACTCTGCCCAATTCAATTGGGAATTTTTCATCTACTATCGAAAACTTTAATATAGTAGATGCACACATCAATGGCCTCATCCCCACAAGTATAGGCAACATGAGTGGTCTTACAGTCCTAAGCCTTGGAGGAAACAACTTGATGGGAAATATTCCTCCTGAGATTGGTAAGCTTAAACAACTCCAAGGTCTGTATCTACGTTACAATAAATTGCAGGGACATATTTCTGAGGCAGTATGCAATTTATCTAATTTGGTTGAATTATATCTGGAACATAATGAGCTCACTGGAGTGATCCCAGAATGTATAGGAGATCTAAGCATGCTACAACTGCTTTATTTAAATTCTAATAGATTTTCATCAAAATTTCCATTGAGCCTTTGGAAAATGAGTGGTCTTCTCTTACTAGATGTGTCACTAAATTCTATAGAGGGAGAAGTTCCATCAGATATTGTAAGATTGAAAGCCATTGTAGAACTATATCTTTACGGTAACCACTTTTCAGGCATGATACCGACCAGATTGGGGGAACTCCAAAACCTGTTGTCTCTTGACCTATCAAACAATTCATTTTTTGGACAAATTCCATTATCCATTGTCAACTTGATAAGCTTGGAATTCTTGGATTTGTCTTCAAATGCATTGTCAGGTACTATTCCTAAGTCATTGGAAAAACTCTCTTACCTTGAAAGCATTAATGTTTCATTTAATGATTTAGAAGGTGTAATACCGAGTGGTGGTGTGTTTGCAAATTCCACTCTGCAATCATTTCTTGGTAACAAAGGTCTTTGCGGAATGCACATATTTGGGATTCCTGCTTGTGCTATCAATAATCCTGGAAAACAATCAAAGCTTAAGGAGGTTGTGATAAAAATTGTTACTCCAATGGTTATTTCATCTTTTATGATATTCTTGTTCATTTCAATTTGGATAATGAAACGACAGAAGAAAGGGAAGTCCAAAGATGTGGAGAAGGTACTGGAGATCAAAACTCATCAATTAGTTTCTTATCACGAGATTCAACAAGCAACAAATAATTTTGATGATGAATCAAATTTAATTGGTGAGGGAAGCTCTGGCTCTGTGTACAAAGGCATTTTATCTAGTGGAACTGTAGTGGCCATAAAGGTTCTGGATTTGGAAAATGAGCAAGTATGCAAGAGGTTTGATACCGAATGCAAAGTGATGAGAAATGTTAGACACAGAAATCTTGTTCCAGTGATCACTACGTGTTCTAGTGACTATATAAGAGGCTTTGTTATGCCAATTATGCCCAATGGAAGTCTTGAGAATTGGCTGTACAAAGAAGATCGCCACTTGAACCTTCATCAAAGAATAACTGTTATGCTTGATGCAGCTATGGCAGTTGAATATCTACATCATGGTCATGTCGCTCCAATAGTTCATTGCGACCTAAAGCCAGCCAACATTCTTTTGGATGAAGATATGGTGGCTCATGTTGGTGATTTTGGAATCTCCAAAATTTTAGCTATAAGCAAGTCCATGGCCTATACCGAGACATTGGGTACTCTTGGATACATTGCACCAGGTATAAAAAATCTACCCTCTTTGATTTTCTCTTATCATAATTAAGCCTCTCCAAATTCTACAAGTAAGAAAAAGCAAGGATTTATTTATGCAGAATTATTGTTGTATTTCAATTGAGTAACTTTTCTTCAATCCTTTTCTAAGAATATGGCTCGGAGGGAATAGTGTCCGCTAGTGGTGATGTTTATAGTTATGGCATTATGTTGATGGAGGTTTTGACCAAAAGACGACCAACAGATGAAGATATTTGCAATGAAAATCTTGACTTGAGGAAATGGATAACACAATCATTTTCAGGGAGTATGATGGACATTGTGGATGCCAATCTTTTTTCTGAGGAAGAACAAATTACTTGTAAAAGTGAAATGTGCATAGCCTCCATGATAGAATTGGCTTTAGACTGCACAAAGAAAATGCCAGAATCAAGAGTAACCATGAAAGAAGTAGTCAAGAGGCTTAACAAAATCAAGAACACATTTTTGGAA

>M6-PGSC0003DMG400021018 M6 LRR-RLK

ATGAGTGGTCTTCTCTATCTAAACGTGTCACAGAATTCAATAGAGGGAGAACTTCCATCAGATATTGGAGGACTGAAAGCCATTGTAGAACTATATCTTTCTGGTAACCACTTTTCAGGCATGATATCAACCAGATTCGGGGAACTCCAAAATCTGCAGTATCTCGACCTATCGAAAAATTCATTTTTTGGCCGAATTCCATTATCCTTTGCCAACTTGATAAACTTGGAATTCTTGAATTTGTCTTTAAATGCATTGTCAGGTACTATTCCTAAGTCTTTGGAAAAACTCTCATACCTTAAAAGCATTAATGTTTCATTTAATGATTTAGAAGGTGAAATACCGAGTGGTGGTGTGTTTGCAAATTCCACTCTGCAATCATTTGTTGGGAACAAAGGTCTATGTGGAGTGCACATATTAGAGATTCCTGCTTGTGCTATCACTAATCCTGGAAAACAATCAAAGCTTAAGGAGGTTCTGCTAAAAATTGTTACTCCAGTGGTTATTGCATCCTTTATGATATTCTTGTTCGTTTCAATTTGGATAATGAAACGACAGAAGAAAGGGAAGTCCAAAGATGTGGACAAGATACTGGAGATGGGGACTCATCAATTAGTTTCTTATCACGAGATTCAACGAGCAACAAATAATTTTGATGAATCCAATTTAATTGGTGAGGGAAGCTCTAGCTCTGTGTACAAAGGCACATTGTTTGGCGGAACTGCAGTGGCCATTAAGGTTCTGGATTTGAAAAATGAGCAAGTATGCAAGAGGTTTGATACCGAATGTGAAGTGATGAGAAATGTTAGACACAGAAATCTTGTTCCAGTGATTACTACATGTTCTAGTGACTATATAAGAGCCTTTGTTCTGCAATTTATGCCCAATGGAAGTCTAGAAAATTGGCTGTACAAAGAAGATCACCACTTGAACCTTCATCAAAGGGTCACTGTAATGCTTGATGCAGCTATGGCAGTTGAATATCTACATCATGGTCATGTCACTCCAATAGTTCATTGCGACCTAAAGCCAGCCAACATTCTTTTGGATGAAGATATGGTGGCTCATGTTGGTGATTTTGGAATCTCTAAAATTTTAGCCATAAGCAAGTCCATGGCTTATACCGAGACATTGGGCACTCTTGGATACATTGCACCAGGTATACTAGTCTCTTTCTATATATTTTCACAAAAATCAGAATTTAGCCCTTCCAAAATCATAAAAGCAAGTGTCTGTTAGTACATAATTATGTTATATTTAAATGAGTAATTCTTTTTCATTTTCATTTTCATTCTTTTTTTAAGAATATGGCTCGGAGGGAATAGTGTCTTCTAGTGGTGATGTTTATAGCTATGGCATCATGGTGATGGAGGTCTTGACGAAACGAAGGCCAACAGATGAAGAGATATGCAATGAAAATCTTGACTTGAGGAAATGGATCACACAATCATTTTCAGGGAGTATGATGGACATCGTGGATGCCAATCTTTTTCGTGAAGAAGAACAAATCACTAGTAAAAGTGAAATGTGCATAGCGTCCATGATAGAATTGGCTTTAGACTGCATAAAGGAAATTCCAAAATCAAGAATAAGCATGAAAGAAGTAGTGAAGAGGCTTAACAAAATAAACAACACAT

>M6-PGSC0003DMG400021184 M6 LRR-RLK

TTTTTTTTAGAGAGAGAAGATGAGTTGTAAGAGAGAGAAAATATATGTCTGCAAGAAGACAAATGTTGAGTGTCACTGCCATTTCTGGGAGCTGTGAGGTTTCTCTTTCATTCATATACAAAACAAGACTAGGGTTTTACCATTGTGGTTTGGTTTTTTACCTGAGCAGATAGAAATTTTGAACTTCATTTATCAAGATTTCTAAAGGGCTAAATACTTTGCTGGCCCATTTCATTTTCTTGTACTTTAATGCAAATTTTTTTTGAGGCCACCAGTTGAGTGCATTAAAGAACTTGAGGTTCCAAGATAACATGAAGCTGTAATCATTACAAGAAGTTGAATTTCTTCAGTAAATCTTTGAAAAAGACTCTTCTTTTTTTGTTTTTCTTCTAGTTGAGCTGATGTTCTTGGTAAGGGAATACTGAACTTGTTTTACTATCTTGAATATTTGAAAGTTCAGTCTGTACTTTGTTTTTGTCACTCATTGTCTCCTTTCAGCTTGTATACCTAGAAAAGTTTCTATTTTTAGTTGTATCTGTCTTGTATTGCTGTTGAAGGAGGACTCTGGTGATTTTGGGTTGAAGTTGTGTACTTTTTTCCCCTGTTGTTTTACTGCTTTGAACTGTGATGGTTGTTGTGAAAGATGGTTTTCATATGTTGAGGTTTTGCTTAAAGTGATTCTCATACTTAAAGCTGCTCCTTTTGAATTTCTTGGAAGGTTGAAGATTTAGTTGTATTTTTGGGAGTGAGTTTGTTGTCACTTGGGCTAGTAAAGCTTTTTCTGGTGTTGCAATGAACCTATTTAGGTTTTTGATGCTTGTGCTTTGTTTTGGTTCTGCAATGGGGCAGCTTCCTTCCCAGGACATTTTGGCCCTGCTTGAATTTAGGAAGGGTATCAACCACGATCCAACTGGTTATGTACTGCAGTCATGGAATGAGGAGTCCATTGATTTCAATGGTTGCCCTTCATCTTGGAATGGTATAATGTGCAATGGTGGTAATGTTGCAGCAGTTGTCCTTGATAACATGGGTTTATCTGCTGATGCAGATTTGAGTGTGTTTGCTAACCTCACAATGCTTGTGAAACTCTCTATGGCTAACAATTCAATTGCTGGACAAATGCCAAAGAAAATTGGTGATTTTAAGAGTCTTGAATATCTGGATATTTCAAACAATCTATTCAACTCCTCTTTACCACCAGAGATTGGGAAAATAGGAAGCTTGAAGAATCTGTCACTTGCTGGTAACAACTTCTCTGGCCCAATCCCTGATACTATTTCAGAGCTCATGTCAATTCAATCTTTGGATTTGAGTCACAATTCTCTTTCTGGGCTACTTCCTTCTTCTTTAACCAAGTTAAATAATTTAATATACCTCAATCTTTCTCGTAATGGATTTACAAAGAAAATCCCTAAAGGTTTTGAGCTAATGGCTAACCTTGAAGTTCTTGACTTGCATGGAAATATGCTTGATGGTACTTTGGATCCAGAGTTCTTACTGCTTACTACTGCAACTTATGTTGACCTAAGTGGAAATTTACTCGCGAGTGCCGCTTCTCAACATGAGAAATTTTTACCAGGCATGTCCTCGTCAGTCAAGTACTTGAGTCTTAGCCATAATCAGCTTACGGGGTCACTAGTAAGTGGCGGTGAAGCTCAGGCATTTGGAAACTTGAAGGTTCTGGATTTGAGCTATAATCAGTTGTCCGGGGAATTACCTGGCTTCAATTTTGTTTATGATCTTCAGGTCCTGAAACTCAGCAACAACAGATTTTCTGGTTTCGTTCCAAATGATCTTTTGAAAGGAGATACTTTAGTTCTCGCAGAATTGGATTTAAGTGGAAACAACCTCACAGGTATATTCCAAACTGCTCTCTGAAATTCATTTTGCAAATTAAATAGTATTTCGTTGTGTTGAACTTGAGTTACAACAAAGTAGTTAATTTCTGATCATTTTGTTAGTGTGCTTTACTCAGTCCCTTCATTTTACAGGGTCGATAAGCATGATTACATCAACAACTCTGCGCGTTCTTAACTTATCCTCCAATGCTCTTTCTGGTGAACTTCCACTGGTGACAGGTAGTACTGCAGTTCTTGATCTCTCAAAAAACCAATTAGAGGGGAATCTAACTAGAATTCAGAAATGGGGAAATGTTGAATTTCTTGACCTCAGCCAGAACCAATTGACTGGAAACATCCCTGAGGTTACAGCTCAGTTTCTGCGATTAAATCGTCTAAACCTTTCTCACAATGCTCTTACTGGATCTCTCCCAAAAGTTATCACACAGTTTCCCAAGATCACAGTCCTTGATCTTAGTTTCAACCAGCTGAATGGACCTCTTCTCACTTCTCTGCTAACATTACCCACTATTGAAGAACTTCACCTTCAAAACAATGCACTTGTTGGAAATATTGATTTCGCTGCCCCCTCTGCTACACCACCCAATCTCCGCGTTCTTGATCTTTCTCATAATCAGCTTGCTGGTTCTTTTCCTGAAGGGTTTGGTTTTTTGACTGCGCTTCAAGTACTTGATATAGCTGGAAATAATTTCACTGGGTCTCTGCCTACATTGATTGGTCAAGTTGGTTCGCTCACTTCTCTAGACATTTCACAGAATCATTTTACTGGTCCTCTGCCGATGAACCTGACTGATGGCCTCCAAAGCTTCAATGCATCACTCAATGACTTATCTGGTGTTGTCCCTGATAATTTGAGAAAGTTTCCTTTATCATCTTTCTACCCTGGAAACTCTGAACTTCAATTTCCAAATCCTCCCTCAGGATCTGGCCAAGCTTCACCAGAAAACCAGAAAAGTCGATCACTTAAAACTATAATCAAGGTGGTAATAATAGTTTCTTGTGTAATTGCTCTTATTATTTTGGTCTTGCTAGCCATTTCCTTTTATTATATACGAGCTTCAAGGAAGCGTCATCCTCGTGTTGCTGAAAAAGTTGTTCATCGCCAAGCCACATCAAATCCTTCTGGCTTTAGCAGTAGGGAGGGTGCCGGTGGTGCAGTTGTTTCAGCTGAAGATCTTATGACTTCACGGAAAGGATCATCAGAAATAATTAGTCCAGATGAGAAAATGGCTGCTATAACTGGTTTCTCCCCTTCAAAAGGTAGCCATTTCTCTTGGTCACCAGAGTCTGGAGATTCGTATACTGCAGAAACTTTTGCAAGATTGGATGTGAAGTCTCCGGATCGTCTGGCTGGAGAGCTGTACTTCCTTGATGATACAATCTCTTTTACACCTGAGGAACTTTCCAGGGCTCCGGCTGAAGTCTTAGGCAGAAGCAGCCATGGTACATCATACAGGGCAACACTGGAGAATGGGTTGCTCCTGACTGTGAAATGGTTGAGAGAAGGAGTAGCGAAGCAGAGGAAGGATTTCGCAAAGGAGGCTAAAAAGTTTGCCAATATTAGGCATCCTAATGTAGTAGGATTAAGAGGTTACTACTGGGGTCCCACACAGCATGAGAAGCTCATTCTTTCGGATTATATATCTCCTGGAAGTCTTGCAAGTTTCCTCTATGGTAAGCTTCATGTATTCTAAACTCTCATATTATGTTTTGCTAATACTTTGTAAAGGCTTTGTGTCCCTGTTCTTTAATCTAGTCATTTTCCTCTGACTGTGTGATATATCTTACAGTTCATATGGTTTTTCCATGCACATTATATATGAGCACTTTAAGCTGTGTGATCCTGTAGTGGGAATTTGTGTGAACGCAGTATTTAGTCCTTGGTTTGATGTGGTAATACTTTAGTACATCTAGAGCCCAATATATTATGCATGTCTTTTGTTTATGTAACTTTCACACCAAGTAGTAATACACCATCACCATGTGCTCTTCGTATCAGGTCACTACATAGGCTGTGAGGGATTTGCAGTATGGTACTTATTATTTCCAGTAGCTAAAATTCTGTATGTTTGAGTTCTTAATGGATCTGAAAATTGTGATGTTTAGCTTTTGTTTCATGATTAACTTTGTTAGAAATCAGCTAGAATATGATACATTTTCCTTCTTATATGAATCCTTCAAATGGACGTGCTTTCTATTCATTTGACTAACCAAATGTTTTCATTCAGTTGATAATTATATATGTGGTCATTCTCAATATACTTCTCTAATGTGCTGCACGTTTGTAGATCGGCCTGGTAGAAAAGGTCCTCCACTAACCTGGCCCCAAAGACTCAAAATATCAGTTGATGTTGCACGTGGCTTGAACTACCTCCATTTTGATCGTGAGGTTCCACACGGAAACCTTAAAGCAACCAACATCTTGTTGGATGGGCCTGACCTTAATGCAAGGGTTGCAGATTACTGCCTTCACCGCCTCATGACTCAAGCCGGCACAATAGAACAGATTCTTGATGCAGGAGTGTTAGGTTATCGTGCCCCTGAGTTGGCTGCGTCGAAGAAACCACTGCCTTCCTTCAAATCAGATGTATATGCTTTTGGAGTTATTTTGTTGGAGCTGCTAAGTGGGAAGTGTGCAGGTGATGTTGTCTCTGGCGAAGATGGTGGAGTAGACTTGACTGACTGGGTAAGGTTGAAGGTGGCAGAAGGTCGTAGTGCTGATTGTTTTGATAGCGTGTTGTCACCTGAGTTGGGAAATCCGGCAATGGAGAAGCAAATGAAGGAGGTTCTTGGTATAGCCGTGCGATGCATTCGTTCTATATCTGAGAGACCTGGTATCAAAACTATTTACGAGGACCTTTCTTCTATATAGTTTGTACTCATTTTGGTTTTGGTAAACCATATGTTTTGCTCTACTTTGGGAATCAAATAGCTCATTGACTTCCCTTTTGAGTAGTAGAATTAGCACAGTTAGTGTGTGAATGGGGCTAGTTTTATTGACCATTGTTTGTTGTAAATTGGTGCATATTTAATGTGTAAACAAACTTTTCTCCCCTTTTGGCCTATGAGCTATCAATGTTTGTCAGTTTTTTGGAGTAAATACTTGTGACTTTATTTTCCAAGTTTTGCAGCACAGTTTCACAGAACATTCTCAGATGGTCAATCAAACACAATGCTGACATCAACATTGCAGTGCAAACTGACCGAAAAAGGTGTTCAAGAGAAAGTGCCAGGTTATGAACAAATGATACTGGTTTTTCTTGTCCACTTTTCATAAAAAGTTAAATTGCAACATCATTATAAGACACCTAAGGTGGAAAAAGGAAATGTTACAAAAACCTACGGATGCCACTAGTTACTATGCAAGTGAGGAGAGGGGTGGGATAAGGGTATGAATTTAAACAACAGGAACTCTAAATTACAGGAAAATATATCATAAGATGCAACATATTTTATTGTTGCAACTTTAACAATGTATGTTCGTCAAACCTGAATCCAGCCATCTCTGCCTCCTTGCAGATATCCTTGCACTTAGCCACCCTTCCAGAAGCCATATAAACTTTGATCAAGCAGCAGTATATGTCAAAAGTTGGACTAAAACCTGCTTCCTTAAGTTTAGAGAAATAGCGTACAGCCCTTGAGAGATCGTTCAAAGCAACAAAAAGCTTAATAACTACACGATAAGCTGGAAGATCAAAAAGGCAATTTGAAGCCTCCATTTCCCATAGCATGTCTATTGCCTCTCTATATTTGTTCGCTGAATAACGACAGTGAATCAGAGTAGTATACATAACTACACTTGGATTACCACCAGATTTCCTGAACCAGTCATATAGTCCCTCCACGATCTCAAATCTTCCCAGCCTAATGCAAATCTTCATGATAGCTGTACAGTCCTGTGGTGTCAAATTCAGATCATCTCTCACAGCAAGCTCCTCTAATAATACCAAGACAAGTGTGCTTTTATCAGGATCCTTACCAAGCTCTAAGATTAGCTTAGCATACACCCTAGTATCAAGCACTCTGTTAGATTCCTTAGCCATTGAAAGTAGCTTCAAAGCAAGCTTCAGGCTTCCTCCTTTAATATAACCCCTTAACATTGCTTCATACACACCCCGACTAGCCAAGCCAGTGAACTTGTCCAAATCAAATGGCAATTTCAACTCATTTGACCCCGCCAAAACTTCAATGGTAGAACCAAGAATATGATCATCTGGGAAAAGATGGGGTTGTTTCTTCACCCAACAGAATGTTTCAAGAGCTCTCTCCGGAAGCCCCAAGTAACCCAACTCTCGGATTGTCAACGACAACGATCCTTTCCTAAGAAATGGACTCCATTTACTGAGAACCTTTGATACATTTTCCTCAGTTGAAAGGCCTTGTATATCTTTCGCCAAACCAATCAAGAAAGTAGGATTCTTGTACACTTGCGTTGATATGGATTGCCTTGAACGATTTGTGAATCCCTTTTGTGTAGGAAGTCCTAAAGGTCGAATCTTGTAAGGAAGGGGCAGAGGTAGAGGCCTCTTTCTATCCGATTTTCCAGGCTTCTGAGGGATCCTCCCTTTGAAAAGAGATGAAATTGCTTCCATCTCATCAGACTCCAATATAATTCCCTCATCATCATCACACATCCTTCCTTTTTCTACTTCCAGCTCCAATTCTTCATCCATTCCATCAGAACAAGCCTCAGTTGCAACTCCCTCATCCTCCTCGTACAAGACCCTTTTAAAACTATAGTCCACATCTGGTGGTAGCTTGGGTCGTCGAGGATTTCTAAGGTTCTTTGGAAGTTTTCTATGATTAGCTCTGTTATTAGATACAGCCATGAAAACAACTCTTCCCTTTCTTCTTCCAAAACTTAAAGGTGTTCCAGAAGCAGCTGTACTGGCACTACTAGACTGCGAGATAGAATATGTGGGTGAAGACCAAGTAGAGGACTCATAAGTGGCCGAGAACATGCAATAACTCCTGCTGAGCATTTGGTCGAGCAGCTGGTCGGCACAAATTTGTAAACAAAGGGATATGAGAGACCTTATAGTGTGACGGAGTATAAAACCAACTGTTAAATGAAAGTAGAGCAGTTTTTTTGACATATAAGTTACTAT

>M6-PGSC0003DMG400021353 M6 LRR-RLK

AGCATTCCTCATATGAAAAATCCACTCTTTATGGTCATTTCTTGCTACTAAATCTTCACTTATAAAAAAAAAGATTACATTTTTTGGATTACTCTGTGAAAACTACTGTGTTGTCAGCAAGAACAACACCATAGTAGTAGCAACTAGCAAGCATTGTGCTTTTTCTCACTCCAATTATCACTCTCTTTCTCCTTTTTTTGTTTTGGGGTTTAGTGATCTGAACCCCATGTTTTCCACCTCTTATTGTCTCAAGATTTGACATGGGGGTCTTCAAGAATCAACCTCATTAGGTGGCAAAGATTGTAGCTTTCTTGATTTTTGTTGTAGTGGGGGTTTGTTTTTTTTTTTGTTGAATGTTAATGCAAATGCTCAAATGTTGTTGCTTTGTGGATCATCTATGGAGTTTTATTTTCACTACATATGGTTTGTTGTAGCTCTGTTTCTTGCTTTTGCTTCACTTGTTTTCACTGTCACTGACCCTAATGACTTGTCTGTTATTAATGAGTTGAGAAAAGGGTTGGAAAATCCTGAGGTCTTGAAATGGCCTGAAAATGGTGGAGACCCTTGTGGTTCTCCAGTTTGGCCACATATTGTGTGTAGTGGTAGTAGAATTCAGCAGATTCAAGTTATGGGGTTAGGGTTAAAAGGCCCTTTACCTCAAAACTTGAATAAGTTGTCTAGGTTGACACATTTGGGGTTGCAAAAGAATCAATTTAGTGGAAAGTTGCCATCTTTTAGTGGTTTGTCTGAATTGAGCTTTGCTTACTTGGATTTCAATCAGTTTGACACAATCCCACTAGATTTCTTTGATGGACTTGTGAATCTACAAGTTTTGGCTTTGGATGAAAATCCTCTAAATGCCACTAGTGGTTGGTCATTGCCAAATGGGCTTCGAGATTCAGCTCAGTTGATCAATTTAACAATGATAAACTGCAATTTAGCTGGTCCTTTGCCTGAATTCCTTGGAACTATGTCTTCTTTAGAGGTTTTGTTGTTGTCTACAAATAGGCTTTCAGGGCCTATTCCAGGTACTTTCAAGGATGCAGTGCTCAAGATGCTTTGGTTGAACGATCAGTCGGGTGATGGTATGAGTGGTCCAATAGATGTTGTTGCAACAATGGTGTCACTTACACATCTTTGGCTTCATGGGAACCAATTTTCAGGTAAAATCCCAGTAGAGATTGGTAATTTAACAAATCTGAAGGATCTCAATGTGAATACAAATAACCTTGTTGGATTAATCCCTGAAAGTTTAGCAAATATGCCATTAGACAATCTTGATTTGAATAATAATCATTTTATGGGACCAGTTCCTAAGTTCAAGGCTACTATTGTTAGTTTTATGTCCAACTCTTTTTGTCAAACCAAACAAGGAGCAGTATGTGCCCCTGAGGTTATGGCACTTTTAGAGTTTCTTGATGGTGTGAATTATCCTTCTAGGCTTGTTGAATCATGGTCTGGAAACAACCCTTGCAATGGACGTTGGTGGGGAATTAGCTGTGATGATAACCAAAAAGTTAGTGTCATAAACTTGCCTAAGTCTAATCTTTCCGGGACCTTGAGTCCTTCAATCGCGAACCTTGAATCCGTTACTCGCATTTATCTTGAATCAAATAATCTTTCTGGTTTTGTTCCATCTAGTTGGACTAGTTTGAAATCTCTGTCTATTCTTGATTTGAGTAATAACAATATTTCTCCACCTCTACCTAAATTTACCACCTCTTTGAAACTTGTTCTAAATGGAAATCCAAACCTGACTTCTAGTCCTCCTGGAGTAAACCCTTCACCAAACAACAGCACAACTCCCGCAGCTTCACCCGCGTCGTCTGTACCATCTTCACGACCCAACAGTTCAAGCTCTGTGATCTTTAAACCCGGTGAACAGTCACCCGAAAAAAAGGACTCAAAGTCAAAGATAGCCATAGTTGTGGTTCCTATTGCTGGTTTTCTACTTTTGGTTTTTCTTGCTATTCCACTGTACATTTATGTCTGTAAGAAGAGTAAAGATAATCATCAAGCTCCAACTGCTCTTGTGGTTCATCCTAGAGATCCATCTGATTCGGATAATGTGGTCAAGATTGCGATCGCCAATCAGACTAATGGAAGTCTTTCCACAGTAAATGCAAGTGGTTCTGCTAGCATACACAGTGGTGAATCCCATATGATCGAAGCTGGGAATTTGCTTATATCGGTTCAAGTACTTAGGAATGTGACTAAGAATTTTTCTCCGGAAAATGAACTTGGGCGTGGTGGTTTTGGTGTGGTTTATAAGGGAGAATTAGATGATGGGACACAAATAGCTGTCAAAAGAATGGAGGCTGGTATCGTTAGCAACAAAGCTTTGGATGAATTTCGATCAGAAATTGATGTTCTCTCAAAAGTCCGGCATCGGCATTTAGTGTCTCTACTGGGATACTCTGTTGAAGGTAGTGAAAGAATTCTGGTTTATGAGTACATGCCACAAGGTGCACTTAGTAGACATCTTTTCCGCTGGAAAAAGTTCAAGTTGGAGCCTCTCTCTTGGAAGAAAAGGCTTAATATTGCCTTAGATGTTGCTAGAGGTGTGGAGTATCTACACACACTAGCACATCAGAGCTTCATTCATAGAGATCTCAAATCCTCAAATATCTTGCTGGGTGATGATTTCCGAGCAAAAGTATCCGACTTTGGACTCGTGAAACTTGCTCCTGATAAAGAGAAATCTGTGGTCACCAGGCTGGCCGGAACTTTTGGATATCTGGCACCTGAATATGCTGGTAAGCAATTTTTTGCTTACTTTGTCAAAGAGAACAGATGATTTTTTAAAATGCATGTTTATATGACAATTTTTACTTTTGCAGACACCTAGAGTTTGCTCAACTCAATTCTGATGTATAGCATTTTATTTTTTTGCATACCAATGTACTGTTTTCCGAACTAACATGAGATGTTTATTTGGTCCAGTTACGGGTAAAATCACCACGAAAGCAGATGTCTTTAGCTTTGGTGTTGTGCTAATGGAGTTGTTAACTGGGATGATGGCACTTGATGAGGATAGACCTGAGGAAAGCCAGTACTTGGTTTCGTGGTTCTGGAATGCCAAATCCTCTAATGAGAAGCTTATGACAGTCATTGATCCAGCTCTAGACGTGAAAGATGAGATAACTAAGAGCATTTCCACCTTAGCTGAACTTGCTGGTCATTGCACTGCAAGAGAACCTGGTCAACGGCCAGATATGGGCCACGCTGTAAACGTGCTCTCCCCACTTGTTGAGAAATGGAAGCCTCTTGAGGATGATCCAGAGGACTATTGTGGTATCGACTACAGTCTTCCCCTCAATCAAATGGTCAAGGGCTGGCAAGAATCGGAAGGAAAAGACTTAAGTTATGTGGATCTCGAGGACAGTAAGGGTAGTATCCCTGCAAGACCAACTGGATTTGCAGATTCATTTACATCAGCTGATGGTAGATAATGGAGGTACTTCTATGTAGTAGATGTAGATATCAATTTTCTTTGTATTGTGTGAGATTTTGATCGTATTTTCCACGTGCCTTCGTTCATTTCTCCCCCTTCAATGTGAATGTATTAGAAATTCAAACTATGTGTTGCCTCAGTTCCTTCTGTAGATACAAAATAGTGATGAAGGAGAACTATACATCGGTTAAACTCCTCGCTCAGTCACTATTTTCATCTTCTACTATGGTGAGATTTAAGATCATTTCATACAACTTTGCTTATTTTCGTCGTTTAGTTTGCTGCTTCAGAAGTTCTGTGTTGAGAAACATACTGACGTTGAAATAGCCCGGTAATAGAATATGTCGTTTGATTCGA

>M6-PGSC0003DMG400021394 M6 LRR-RLK

ACTTCTAATTGAGTTGTTGAATTTTTGTTGTAGATAAGGACAAGGTAAAGATTCTAAGTTGGGAAGAACGTTTCCGTATTATAGTTGGAACAGCACAAGGCATTGATTTCCTTCATGGAGGTTCAGAGATCAGAATCATCCATAGGGACATCAAGAGCTCCAATGTACTTCTTGATGAAAATCTTGAAGCAAAAATTGCTGATTTTGGACTTGCTCGGTGTTTTGGAGCTGACAAAACTCATCTCAGTACTGGAATTGCTGGAACATTGTAAGTCAACAATTCACAAATACGTATCAATACAGTTCACAAACACTTGTCCGTATCTTAAACTTAATCTGTGATTTTCGGTGTACAGAGGATATATGGCTCCTGAATACCTCGTAAAAGGACAGCTAACAGAGAAGGCTGATGTCTATAGCTATGGGGTGCTTGTTCTTGAAATCGTTAGTGGCAGAAAAAGTATTGCCTTTGCAGAGGACTCTGGCTCTCTACTACAAACAGTAAGAATGCTATAGATTTCTCTTTATCATGGTCATTATATGAATATCATTGAACTTTAGTAGTAAAGTAAAAAAAAACTATTTTTATCATATAACTCAGTATATTGTCATTTTGGTTTCAGGTGTGGAAGCTGTACACAACAAATCAGGTAACTGAAGCATTAGATCCTCTATTGAAAGGCGATTTTCCACCAGAAGAGGCATCAAAGGTACTAAAAGTGGGGTTGTTGTGCACTCAAGCTTCTGTCGCTTTAAGACCATCAATGAGTGAAGTTGTCCAAATGCTAACGTGTGAGGGCTATCAAATTCCAGAACCATGCCAACCACCATTCTTA

>M6-PGSC0003DMG400021609 M6 LRR-RLK

CACAAAACACACAAGCCCACTACACTACCAATCTTAGTCATTATTTCTCTATCTACTATCACTCCTGTGAATGCAGTTACTTCATTTTCCTTTCCACCTTGTACACATGAATATCTAACAATAATCCCAAACTCAACATTCATTCAATGAAACTTGTACTCCATTACCTAAAATTCACACCTTTTTCCCCTTTTCATTTCCATTCCCTTCTCTAACTACTCTGAAAAAAACATTTGTTTGCTTCTCTTGTTGTAGGAACTAAAAAAAAACACCATCATAAAAATGAGATCTTTTAGTCCTTTTTTAGTTCTAATACTTCATTTTGGCTTATTGTATGGATCTTTATTAGCTGATGATAATACTATGCAGCTTAATGATGATGTTTTAGGCCTCATTGTGTTCAAATCAGCTCTTCTTGATCCTTACTCAAAACTCTTATCATGGAGTGAAGATGATAACTCTCCTTGTGCATGGGAGTTCATTAAATGCAATCCAATGAATGGTAGAGTCACTGAGCTCAACTTAAATGGCTTGAGTTTATCAGGAAAGATCGGTAGGGGGCTCGAGAAGTTGCAATCTTTAAAGGTACTATCTTTGTCAAACAACAACTTTACTGGTGCAATTAGCCCTGAGGTAGCCTTGTTGACAAATCTTGAAAATCTCAACTTTAGTCATAATGGCTTATCAGGGAATATCCCTGGATCTTTTTCGAATATGTCATCTTTACAGTTTCTTGATCTGTCTGAGAATGCGTTATCTGGACCTGTCTCTGATACCATGTTTGATAACTGTGGTGATTCGTTACGTTATCTTTCTCTATCCGGGAATTATCTTGAAGGTTCATTTCCTAAAACAGTTTCAAAATGCACTAGTTTGAATCATCTCAATCTTTCAAGAAACCATTTTTCTGGTGACCCTGGATTTTCCGGGGGTATCTGGGGATTGACAAGGCTTAGAACATTAGATCTTTCACATAATGAACTCTCTGGATTAGTACCAATTGGTGTTTCAGTTTTACATCAGTTGAAAGAGTGTTTGTTACAAGGGAATCATTTCAGTGGAGATTTACCAGCTGATATTGGATATTGTCCACATTTGAATAGATTAGATTTGAGTAATAATCAATTCACAGGACAAATTCCAATGTCACTTCAAAAGGTGAATGTTCTTTCTTTTCTCAGTTTATCAAACAACTTGATAAATGGAGACTTCCCTCAATGGATAAGTAACATGAGCAGCTTGGAATACTTGGATTTTTCAGGCAATAGTTTAGAAGGAGCATTGCCTGATTCAATAGGGGATTTGAAAATGTTGAAGTACTTGAGTTTATCTGGTAACAAATTGAGTGGAAACATTCCAAAGTCTATGGTGTATTGTACTAGTTTATCGACTATTCGGATAAAAGAAAATGCATTGACTGGTTGCATTCCTGAAGGATTGTTTGGTATAGGATTGGAAGAAGCAGATTTTTCAAGAAATGAGTTAACTGGTTCCATCCCTCCTGGTTCTGGTAAATTCTTTGAATCACTTCAATTTCTTGATCTGTCAGGAAACAATCTTACTGGAAACATTCCAGCTGAGGTGGGACTTTTCTCCAAGTTGAGATATTTGAATCTTTCTTGGAACAATTTTCAATCAAGATTGCCTCCTGAAGTTGGATACTTTCAGAATCTAACGGTGTTGGATCTTCGACACAGTGCTTTAGTTGGATCAATTCCTGGTGATATATGTGATTCTGGTAGCTTGGGAATTCTTCAACTTGATGGAAATTCATTCACTGGACCTATTCCTGATGAGATTGGAAATTGTTCATCCCTCTACTTGTTGTAAGTATATTTATTCTGATTTATCTCATTATAGTTTGGTCCAGCCTCTCTATAACATTCATTTCACCATGAGAACCATGTTTTCTGTGGAACCGATCTCTCATGTTATGTTATATTACATGTTATATTATATGTTATCTATTACAACATGTTGCTATAGCAGCAAAAATAATATTGGAACAAACAATGCTGTTATAGAGAGGTAGTAAGGGATATGTTTATTTGAAAATGAGGTTCTGTATCACTTTTGCTTTGTCTATATTCCCTGTAGTTCTTGAATTGAATGTCAAATTTGTTAAATGCAGGAGTTTGTCACATAATAATTTAAGTGGGTCAATACCAAGGTCCCTTTCAATGTTGAAGAAGCTCAAGATTTTGAAGCTAGAGTATAACCAATTGAGTGGTGAAATACCACAAGATCTTGGCAAGTTAGAAAATCTTTTGGCTGTTAATATATCCTACAACAGGCTCGTTGGACGCCTTCCAATGAGTAGTATATTCCAGAATTTAGACCAGAGTTCATTGGAAGGGAATTTGGGTATTTGTTCACCCTTGTTGAAAGGTCCTTGTAAGATGAATGTGCCAAAGCCTTTGGTTCTTGATCCTTATGCCTATGGAAACCAGAGGGGAGGTCAAAACCGGGACGATGGAACTTCAAGAAGCAATAACAAAAGTTTCAAGAACCATAGATTCCTCAGTGTTTCATCCATTGTTGCAATCTCTGCTGCAGCTGTGATCGCGGTTGGAGTCATGATAATAACCTTACTCAATGCTTCTGTTCGAAGGAGGATCACATTCGTTGACAATGCCTTGGAAAGCATGTGCTCAAGTTCTTCTAAGTCAGGGAACCTAGCCACTGGAAAGCTTGTGTTACTGGACACCAAATCGTCCCCGGATTGGACTAGTAGCAGTCTTGAATCAATCCTAAACAAGGCATCTCAAATCGGTGAAGGTGTCTTCGGAACTGTTTACAAGGTTCCATTGGGAGGGGAAGGAAGAATAGTAGCTATCAAGAAGCTTGTAACATCAAAAATCCTCCAATATCCTGAGGATTTTGATAGAGAAGTTAGAGTTTTAGCAAAAGCAAGGCATCCAAATCTAATATCCTTAAAAGGGTATTACTGGACTCCTCAACTTCAGCTTCTAGTATCAGATTATGCACCAGAAGGAAGTTTACAAACCATACTGCACGAAAGGCCTTCGTCTTCAACTTCCCTTCCCCTGTCTTGGTCCACCCGATTCAACATTGTGCTTGGAACAGCCAAGGGACTTGCACATTTGCACCACGCGTTTAGGCCAGCAATCATACACTACAACATAAAGCCTAGCAACATCCTCCTTGATGAAAACTTCAACCCGAGAATATCAGATTTCGGGCTGGCAAGGCTAGTGACAAAACTCGATAAGCACATGATAAGCAACAGGTTCCAGAGTGCACTAGGCTATGTAGCACCTGAATTGGCATGCCAGAGCTTAAGGGTGAACGAAAAGTGCGACGTGTATGGTTTTGGAATGTTGATTCTTGAAATTGTCACAGGTAGAAGGCCAGTTGAGTATGGAGAGGACAATGTGTTTATACTCAATGATCATGTGAGAGTTTTGCTTGAGCAAGGCAATGTGTTGGAATGTGTTGATCCAACTATGAATACTTATCCTGAAGAGGAAGTTTTGCCTGTTTTGAAGTTGGCATTGGTGTGCACTTCTCAAATACCATCAAGTAGGCCTTCAATGGCTGAAGTGGTTCAAATCTTGCAGGTCATCAAAACACCTGTTCCTCAAAGAATGGCAGCATACTAAAACATGTCACAAACTTGATGAATTTCCCTTTCTCTTTGGCTTTCTATCATCCTATTGTTGTTATGTTAACCTCCTAATTTGATGTTTTTGGACTAGATAGTGATAGAGAGAGTTTTGTACTTCTCTGTCATTTGATGAAATATATGAGAGGTTGGTTCAATGACTTCAATCATGTTTGTTG

>M6-PGSC0003DMG400021789 M6 LRR-RLK

CTTTAATGTTTTGTAGACATATTTTCATGGAAGATGGTGTCTTATGTCACAGTGATAAATTGCAACAAGGATACTTGACATAAAGTGCTATATCCGAAAGATGGTACTGTTCTTTGCAATTGTGATATTGGCATGTCTCCACTCATTTGTCTTACCTGATGGGCAAGGTACCTGGTTCTTCCAGTTTTACATCATAAAAACAAACTTTGTTTTTATTTATCTACTCAAATGGATAAATAATAATTTAGATGTGACCACTATATCTTGTTTACATTTTGCTTCTGCCGGTGAAATGATGTCTTTGTTCTTCTAATAAAGTAGGGTGACATTTGTTTTTTTATACTCTGGCCCATTTTAGGAGATGCACTGTACACATTGAAGAACTCACTAAATGCTTCAAATGGTCAACTTGCTGATTGGAACCAGAATCAAGTTAGTCCTTGTACTTGGTCCAAGATCACTTGTGATGATAAAGGCAATGTCATTATGGTGTAAGTACTATTGTTAGAGCCATTAGAATCACCGTGTGCATATACTATGATGATACTTGTTTTTATGCACTAGAAAGGTTAATCTTTGGTTGGTCAACTTCCTTTACTGGATGTGATTTTGATAATGTCTGAATCTGTATTTGCAGGTCATTATCAAGTTTGGGATTCTCGGGCACTTTAACTCCCAGAATAGGTGTTCTAAAGAATGTGAATACACTGTACGTTTAAAGTACTCGGGCTGAGTTATTATTGATAGAATGATGTCTTGAATGCTGCCCACTTATAATTTTCTAGATGCTATTTATTGCTACTTTAAAATGTTTATTTAAATTTTGTTTCTGAATATACAAGTTTGATGTCATGGCGTTGCACTAGAACAGAAATCCCTTCAAATCATCAGATGTTAACTAAAAGTTGAGAACACAATGATTAATATTTTCAAGGGAGCTTCACTTAGCGGAAGGGGGGAATAAGAAAAAGAAAGTACCTACATATTGTGGTGTTATAAATGAACAAGGAGAAATATTTGGCCATTTCACAAAGGCAAACAGCTTTAAGAGAGATGGAAGCTGTATGAACTGATCTAGCAAAAAGGGAGTCTCTGTTCGGATGGCTGTCTTTTTTCTTGTGGTAACTGTGTGGTGTCTCTACACCTGTAACTGCTGGTTCCGTTGGACTAAGAGCAACCTAAGCCATTTCACGAAGACTAGAAATGTGAACTATACAAGATTAGTCCTCAGTATCAGTGACTTTATCTTTTTTCCTGAGTGGTCATTCCGTCACATTGGTATTTGCACTTTGTCTACTTCTTAGGGTCCTATTTTTGTATTTGACTTATATATATTTCAAATCAATAATTTGGATGCATAACCATACGGAAGGGTACATAGTTCTTTGATGTAGCTCTTGATGAACTTTGATTTGTTAGTATGTTGTTGAGCTCAAAATTGTTAGGTCTACGACTAAAATTTTTGGGATGATTTCTTGTCTTCTTAGATCCACGACTAAAATTTTTGGGATGATTTCTTGTTTTCTGCTACTTACCTAGCTCACTATGTCGTATACACTCCCTGCAGGTCCTTGCAAGGAAATCGTATAACTGGTGAAATTCCCAAAGAGATTGGAGATTTAACAAGCTTGACAATGTTGGATTTAGAAAATAACCGTCTTTCCGGAGAAATACCAGCTTCGTTGGGCAATCTTAAAAAGCTTCAGTCTCTGTAAGTCATATTATTTCATTTAATCATTTGAGGATAGAACTTAGTTTACTTGGTGACTGGTATAGGTGTGAACTCTATTATTCCATTGAGCAGGTTTTTGAGTCGAAACAATCTCACGGGGACTATCCCTCAGTCACTTTCGGGTCTTCCAAACTTGATGAACCTGTAAGTTATTTATCTTGCCTTTTCCTTATTCCAAGTATAAATTACTTCTCTGAATAGGAGAATAATACATTTTCTTTTGGGATGTCAATTCAGACTTCAGTGATCCTTAACCTGTTACCTGTATCATTCGATATTTGACTTGATTCTCTGATTCCGCTTTGCAGTCAGCTTGGTTCCAACGGTCTTACTGGCCAAATTCCTGAACAGTTGTTCCAAGTTCCAAAATACAAGTACGATTACAGTTCTTACTTTAGACAGTAGCTCAAGGCTTCTTAGTTTTTGCCTTCTTTACTAATGCATAATATTTAATGTTAGTATAACTTCGTTTTATTCTAGTTTTACAGGGAATCATTTGAATTGTGGCTTAAATCTCACTCATCATTGTGAATCTGCTAGTGGAGGTAAGTTCTACCAGCAAATATGTGAATTTTTGAACTGTATATACTTCATGCTTCTCTGTATTCTGGAAGAATGAAAGTAATGTAGCTAAAAATTTTGAATTTTACTACCATTATCTTGATTTTATTTATTATTATCTTTTAGCTATAAAGGAGGCTTTATCTTTTAGTTTACTTTATTAGAGAAAAAACTTTAGTATTTTCAAGAGGTCTAGCTTGTTAGGGGTTTCATATACTATATCTAGTATAAGTGTGTACTTTGTTTGGAACGACACTTGATGAATAGACTCTATTAATTTCCTTAATATGTCATCTATTTTTTCTTGTTTCTTTCCGTTCCCATTCTTACTTAGTTCCTTCTCCCTTTTCTTGGTGCTATTTCCTCCTTGTACGCAAGACATGGAACAAGATTAAAGTTATTAGTGTTATTTAGAGTTATACTTGCCATTGGCTGTGGGATATGGTTGTTGTGTGGTATACTGGTATCTAATTTAATGGATGGAAGAATCAAGCACCTCATGGTGCTGCCTGCACTAAAAAGGATGAGTTTCTTTCCATTAGTCCTAATCTTGGAGAGGTGTAACTTATAACACTGTCTTACTTGTAGGTGCCAAGCATTCTTAACGGAGGGATAATGATGTAGCTGAAAAATATAGCATTATTTACTACCTTTACTGTTGATCTATTGTCGTCATCTTTTAGTTAAAGTTTGAAGTTTGATATATATTTTTTTAGAAATACTATCATATTATAGTAGGAGAGAATTTTAATAATTTTAGGTGTTGTAATGCTTAAAAAAAAGAATTTAGTAATTCTAAGAGTTGTAATGATTCTAAAAAAAGAATTTTTTTAACTCTAAGAGTTGTAATGATTAAAAAAAAGAATTTTAACAATTCTAAGAGTTATAGTTATCTAGGTTCCCTAATCTATAATTAGAGTCCATAATTCCTTTGCTTGGAACAACATTTGATGAGTAAATATTGTTGATCTCTTATTATATCTCTCGTCCCCGTTTCTTCTCGTCCTCTCCCTTAATCATCATGCTCATTCTCTCTCCTTTTGGTGCTACATCTCAGAAAGTTTCTGAAGTTGATAAAGCTGTTGATTACCTATCCGATGTATTACTATTCATTTAGCAAGGAAATCGTCATTTAACATATGCACCTTGTTGTACTTTCTGGACAAACCAAATTCTAAAGTTGAATTTCTTGCAGGTTCTCCTAGTAAACCAAAGACTGGTCTCATAGTTGGTATTGCTGTTGGATTTCTTGGACTTCTCGTTCTTGGAGGATCTGTGTTATTGTTCTGTAGGAGACGGCACAAAGGCTATAGTCGTGAAGTCTTTGTTGACGTTGTAGGTTTGTGTCACATTTTTTTGACAATAATTTCATCAGTTTAAATCACCTGTTATTCTTCGTAAGGGAAGAAGATATTCTGGCCATGCATTGTTGTATCTTGTAATTCCATCTTAAGTTAGTGACCTCATTCTTTATACACCAGTACTTAGTTTTCATTTACCTTCAAGATGGTATGATTCCTTTGAGTATATTATGATATTATATTTTATGTTTTATTTGGCGCACTCAGGTGAGGTTGATCGACGAATTCCATTTGGTCAATTGAGGAGATTTTCATGGAGGGAATTGCAGCTTGCTACTGATAACTTCAGTGAAAAGAATGTTCTTGGACAGGGGGGTTTCGGCAAGGTCTATAAAGGAGTGCTCAATGACGGCATTAAAGTTGCCGTGAAACGGTTAACTGATTATGAGAGTCCTGGGGGAGATGCTGCATTTCAGCGAGAAGTTGAGATGATAAGTGTGGCTGTTCATAGGAATCTCTTACGGCTTATAGGATTTTGCACCACACCGACTGAACGCCTGCTAGTATACCCATATATGCAGAATCTAAGTGTTGCTTATCGTCTACGAGGTATTGTTTCTTCATAGCATTGCTTTTATTCATTGTAGTTTCTTTTACATCCTGAGCATGCTATTTTCCCTTACTGTGCCTGTTTCATCTAATTTATTTTATCCCAATCAATAGATCCTTCATTTCATTAGTCTTACATCTCCTGCGGCGCATATTTTGATTTTGTCAGCTTGATTGTAAACAATCATGAGTCATTTCCCCGTTCGATCTGTGAAAAAGTGTGCATTTGTCATTGTACGGTCTAAATTGACTATTAAAGTACTGTAGAACTGCTGATCTGGTAAAAGAACTTGTACATGGCATCAAGTTCGAGGGGCATGCTTATAGAGCTAATGTGTGTAGCTTAATTGGCAGAAGCATAAACCCTTTAATTTGTGTTTTCATGCTTCAGAATCTTATTCTTTTTCCTTCTCACAAAAGAATCTTATTCTTTCAACTTTGAATGAAGTGCTAGAAGTTTAATGCCTTAGGGTTGCATCACCATGAGTAAAATTTGCATGCTTCCTTTTTTCTCCCAGAATATGGTAGGGAAAGACAAGTTCTGAGCAGATTTAAGGGTTCTGTAGAATTATTTTCTGATATTCATTAAAGGGGTTGAGTGTTCGGAGATTGCAATTTTCTTTGGTCAAATGGCAATACTCAAATCATACCTCATTAATCATTAATAGATGTATTCAGTTTTTTTTTTAAATCATTAAAAGATGTATGTGCGTTAAGTTGGTATCACAGATCGTGATAAGAAAGAGCTATGTGACTTCATAAATTGATTCTTGTGATCCTGGCTTGAACTAGAAGGTGTTACATGAAGTAGTTGGTGCAGCTTAAGATTTCAGCAGTTTTCTGCTGGAGTTGATCTGTCTGTTATTAGTTTTCAGTGGTGTCTAATTAAGTTTTTAATCACTTACAGAACTTAAACCTGGGGAGCCTGTTTTGAATTGGCCAACTAGGAAGCGTGTGGCAGTGGGTACTGCACGTGGACTTGAATACCTACATGAACACTGTAACCCAAAGATTATTCACCGTGATGTTAAAGCAGCTAATGTGTTACTAGATGAAGATTTTGAAGCTGTAGTTGGTGATTTTGGCCTGGCAAAGCTAGTTGACGTTAAGAAAACCAATGTGACAACTCAAGTTCGTGGTACAATGGGCCATATAGCTCCTGAATACTTATCCACTGGCAAATCATCAGAAAAAACTGATGTTTTTGGCTATGGAATCATGCTTTTGGAAATTGTAACCGGCCAACGTGCAATAGACTTCTCACGCCTAGAAGATGAAGATGATGTCTTGTTGCTTGACCATGTGAGTGTACTTAACTCAATGATCTTCATATATTTTCAAATGGGCATCAAATTATTGCATCAAACTTTTCATATATCTTATCTCTACTGACCAATCTTTGTTTCTCTCCATATTCTATTGGGACGACTCATTTGTCCTTTGCATGAGAGCCTCTTGGATGGATATACATTCGTTTATCTTTCTCCTATTGGTGAAACTGCTGCAATTATTCTATGATAATTGAAATGGTGTAAACAATTATTATTTTTCACGTAGCACCAGGGTACAAATGCTTTGAATATGAGCTAGGTCTTAATATTGAGCCTTTTAGGCACAAGAAATATGATGACTAATATATATTTGATGCTAATTTTTCTCTTTCAAAAGTGTGGATATGTGGACCCTGAGTCTCTCTTATAACTTTACATGATTATTCTTTAAGCTTCATAGGACCTTTATGTAGTATTAGTTGCTAAAAAGAGATCATCTTGAAGGTCAGAAAACTGCAAAGGGATAAAAGACTGGATGCAATTGTAGACCGTAACCTGAATAAGAACTACGAGCTGGATGAAGTGGAGATGATGATTCAGGTGGCATTGCTGTGCACTCAGGCATCACCAGAGGACCGTCCAGCAATGTCAGAGGTGGTGAGGATGCTGGAAGGAGAAGGGCTTGCTGAGAGATGGGAAGAGTGGCAACACGTTGAAGTCAACCGTAGACAAGAATACGAGAGACTTCAGAGACGATTTGATTGGGGAGAAGATACAATGTTTAATCATGATGCTGTTGAGTTGTCTGGTGGAAGATGACACATGAGTAATAACTTTGAACCACAAAAATGTTTCTTGATATATAGATGCCTGGTTGCTAAATGTTTTTTTTTTTTTACTTTTTATGGAAAACCCCTTGTTCTAACTAGGCAGATCAACAGTGGACTCTGTATTTATGTAAGTTTTAGTCATTATTTTCATGTTCCATCAACTCCTGGGAGTGGGAATTTGATTCCTCAAATTGTTACATATTTTTCCTCACTAAACTATTTTT

>M6-PGSC0003DMG400022000 M6 LRR-RLK

ATGGTTTCCAGGACTATAATTATAGCATTCCTAGTTTTGTTAATGTACCTCTCAGTTACAAATGCATCGAACATTTCAACTGATGAGGCTTCTCTTTTAGCTTTCAAAGCACAGATAACTTCAGACCCCAATGGAATATTGTCAAAAAACTGGACAAAAGGAACTCACATTTGCAATTGGATAGGCATATCTTGCAGTAAAAAGCATCAAAGAGTGACATCATTAGTCCTCATAAGCTTTGGATTTAGAGGTTCAATAGCAAAAGAAATTGGGAATCTCTCCTTCCTTAACAAGTTGGATGTTGGAAACAACAGTTTCCACGGCCAAATCCCCGATGAAATTGGGAGTCTGAGGCGTTTAAACTACTTGTCTTTGCAAATGAATAATCTCACTGGTCAAATCCCAGAAAGCCTTGGATTTCTCACAAAGCTTCAACTTCTTGATCTTTCTGAAAATGATCTATTTGGAAATGTTCCATTTTCCATTTCCAATGTGTCTTCCTTAAAGATCATTGATTTTGGATTTAATCGGTTTAGTGGGAATCTCCCAAGGGGATTTTGTACTAGGCTTCCAAATCTGCAAGGTTTATTTCTCTCAAAAAATCAATTAGCAGGCCAAATCCCAAGTGGGCTGAATCAATGCACACAACTTATTTACCTTTCCTTGTCATATAATCAGCTCACAGGTAATTAGTAATGTTAATTCAATTCTTGGACTATATATACTTGATTCAGAATCATTTAATCCTTCTACAATGTTATGATGCTATCTGATTGTGTTCATATATGTATTAACAGGAAGTTTGCCAAGAGACATGTGGAATTTGACAAAGCTTCGAGAATTATATCTTGGATGGAATAACATAACAGGTATCTTGGTTCTAAGTCTAATAATTTCTAGTTAATTAGAAATTGTAGTAGCAGCAAAAAATGTTCATATAGCTCCTTGTTATTCATTTTCAAACAACAATGACTACTACTACATCGGCTCAATGCTAGTTTGTGTCAACTTTATGAATCCTCATTATCCGCGTTAATCCATTTAGTCCCCATCTCAGTCCAAGGTTCAATAATTTAAGTTTTCTAACATTAAAATTTCTCTATATTTTCTGTTAACATACAAATATCTAACAAGATTAAAAGCATCCTAGCGAACATAGAGTAAGTTGGAGGTACCAACATGATTAATCTTTAATATCAACTAGTTGGTATCACTTTTATGGATCTTACTATTCCATTTTGCTCTACTTTTCGCTGAGTCAAAATGGATTCTAACATATAGTTGGTCTTTCTTTCAACTTCCTCCCATGTGATTTTAGATCTACCTTGTTCCTTTTTTAAGCAAATCCAATCACCATATATAGTTTCATACCTGCAGACCAGTGCATCTAGAGATCTATGTAAGACTGAACTATCTCAGAGCTACTTAACCAATTGCTAACCATGGTAGTTTCTAATGTCTTATCTCAAGTGACTCACACATCCATCTTAATATAGATAATTGACTGACGATTTTAAGTTTCCTTTCTTCAGTTACCATATAAGAAGAGAGGAAAAGAGAATAGCAGAAGTAAATGAATAGTTCTCATGTTGCCAATATTTAAAATTTAAATGCCCTAATTAATAGAAAACCTTTCCTAGGCTTGTTATTGTTCGTCTTCAAGAGGACGTTGAAACAGAATACAAGGATTGCCGAACTATCCGGGATTCTAATTCTCTTACACAATATTGTACGTTTTAGTGTTGTAACCACACAATTTAATATATATTTTACAGGACATATCCCAAGTGAAATTGATAACCTATCAGCAATTCGAAGATTAAGTCTACCACGGAACAATTTGGTGGGAATTTTTCCACCATCTATGGGAAATTTGTCAAATCTAGAGTTGATAGACCTAGGTCTGAACAGTCTACATGGTGGCATTCCACAAGAATTTAAACATCTTGTAAATTTGAAAGGATTGTTTCTTGGTCAAAACAGATTATCGGGTGAAGTTCCAGGGCCTATGTACAACATTTCTGGACTAGAAATGATTTCATTTGTAGGAAACGGGCTCTCAGGAACACTTCCCTCAAACATAGGCCATACCCTTCCAAATCTTGTCGGCCTCTATTTTGGGAACAACCAATTTACCGGGCTGATTCCTACTTCCATTGTGAATTCCACCAAGCTTATCCAGCTTGATTTTGGTCGCAATCTGTTTAGTGGACCTGTACCTATGAATCTTGTAAAGCTACAACAGCTTCAATTCATCAGCTTGCATTTTAATCAATTGACGAATGATCCATCTACGGTTGAGTTAAGTTTTCTTACTTCATTATCTAATTGCAAGTATTTGATGACAGTTCAAATAGGAGGTAACCAATTTAATGGAAGTCTTCCAAAGTCCTTAGGATCAGGCAACTGGTCCTTTTCCCTTGAATATTTCTTTGCCTTTTATAGTGGTATCACAGGAAAAATACCAACTAACATCAGTAACTTTAGGAACCTGGAGTGGTTAAGCTTGGGAGAGAATAAGCTTATAGGTTCTATTCCTCCAGATTTGGGGAACTTGAAGAATTTGCAAAAGTTTTACCTGCAGATAAACAATTTAGATGGAATTATACCAACTAGCTTGTGTAACATGGAAAGCTTGTACCAAGTTATCTTGGGCAAAAATCAACTTTCTGGGGAAATACCAAGTTGCTTTGGAAATATTTCTTCTTTAAGAGAACTCTACTTAGATTCTAATGCACTAGTTTCCCATATTCCATCAACATTTTGGAGAAACAAAGATCTTTCAGTTCTCTATTTGTCCTCCAATTTGCTGAATGGATCTCTTGCAGTAGAAATAGGAAACACAAGGAGTTTGTGGATGTTAGATTTATCTGGCAATCAGTTCTCAGGTCAGATTCCTAGCACTATTGGCCAACTACAAAGCTTGGTTAGCCTTTCATTGTCAAAGAATATGTTAGATGGTCCTATACCTGAACTATTTGAAGATTTGGTTTCATTGGAATATTTGGATCTATCTAGCAACAATCTAACAGGCATGATCCCCAAGTCCTTGAGACATCTTGAACATCTCATGTATTTCAATGTCTCGTTCAATGGGCTCATGGGTGAAATTCCAGATGGAGGGCCATTCGTAAATTTTACAGCTGAATCATTCATGGGTAACCCTGCATTATGTGGATCATCACGCTTCCATGTGATGCAATGCAGAGTCACAAGTCTTGAAAGTAAAAGAAAGAGTAGAGTCTTAACTTCTGTTCTTGCATCAGTTTCCTCAGGAGTTGTAGTCACAACCATTTTCATCATTTGGTTTCTGAAATGCCGAAAAAGGAGTACGGAACTTCCTCTAGTTGATACATTTGGTCAGGTACATAAGAGGATTTCGTACTATGATATTTCTCAAGGGACAAACAACTTTGATGAAGCAAACTTGATTGGAAGGGGGAGCCTTGGTTTGGTGTACAAAGGGACACTAGCAGATGGAATGGTTGTGGCAATAAAGGTATTCAATACAGAACTGCAACATGCATTTAGGAGTTTTGAAGTGGAGTGTCAAGTTTTACGTAGCATTCGACACAGAAACCTTGTTAAGGTGATAAGTAGTTGTGCCAATTTTGATTATAAAGTGTTGGTGCTAGAGTACATGCCCAATGAAAACCTCGAATGTTGGCTTCACTCTACTGACAAATTTTTGGATTTAACTCAAAGACTAAAGGTGATGATTGATGTGGCTTCTGCTGTGGAGTATTTACATGGAGGTCATTTGTTTGTAGTCGTCCATTGTGATTTGAAACCAAGTAACATACTTTTGGATGGAGATATGGTGGCAAAAGTGAGTGACTTTGGGATATCCAAACTTCTAACAGCTGAGACGCTAATAGCACATACCAAGACTTTGGGTACTATCGGCTACATGGCACCAGGTAAATGTTATTATCAAGTTCATTTGAAGATTTTAAATGACGTTAATTATTTTCACTAATGATTGCGTGATTTTTTTGCAGAGTATGGATCAGAAGGCAAAGTATCAACCAAGGGAGATGTTTATAGCTTTGGTATTTTGTTAATGGAGACTTTTACAAGTAAGAGCCCCGTGGATGATCTATTCGTTGGAAACTTCACCTTGAAAAGATGGATATGCCAATCATTACCAGACCGATTGGTGGATGTAGTGGACATTAATCTATTTTCACTGGGTGAAGAAAATTTTACTTCTAAAGAGAGATGCTTCAAATCAATCATGGAATTAGCTCTTGAATGCACAAATGATCTGCTGGAAGAGAGAATCTGCATGGAAGATATTACTCTGCGACTCAAGAAGATCCTAACTCAATTTCTGCAGAATGCTGTGACAAACTAA

>M6-PGSC0003DMG400022139 M6 LRR-RLK

GTTTTCTGGAAGATTTGGAGATTGTAGGATGGAGGAGTGTTCTTAACTAGTTGAAAGAATTGATTTTTATGCAGTTGGTTGGTGGTGGTGTTTGTTTTCAGCAAGATGAGCTATTAAAGATTCTAAAGAATTGATTTTTATGCAGTTAGTGGTGTTTTTTACAGCAATATGAGTCATAAAAGATGCCAAGAATATTTTTCACTAAATGGGGTGTTGAAAAATGTTGTCTTTATGATAATACTTCTGAGTTGTAGTTTTTTTGTGGTGGAGTCAAATGCAAGAAAACTAGCTGAAAATGATCAAGTAGGGAAGTTATTAGCCTTCAAGAAGTCCTCTGTTGAATCTGATCCAAATGGATTCTTGAATGAATGGACTTCATCTTCTTCAAGTCCTTGCACTTGGAATGGGATTTCATGTTCAAATGGTCAAGTTGTTGAGCTTAATCTGTCTAGTGTTGGGCTCAGTGGTCTACTTCACCTAACTGATCTCATTGCCTTGCCTTCTTTGCTCAGAGTTAATTTCTATGGCAACCATTTCTATGGCAATCTTTCCTCAATAGCTAGCTCCTGTAGCTTTGAGTTTCTTGATTTATCAGCTAATAATTTCTCTGAGGTTCTTGTTTTAGAGCCTTTGTTGAAGTCTTGTGATAAAATTAAGTACCTAAATGTCTCTGGGAATTCAATTAAAGGGGTTGTTCTAAAGTTTGGACCTTCACTATTGCAGCTTGACTTATCAAGTAACACCATTTCAGATTTTGGCATTTTAAGTTATGCACTGTCCAATTGCCAGAATTTGAATCTCCTTAATTTCTCCTCAAATAAGCTTGCTGGCAAGCTAAAAAGCTCCATTTCTTCTTGCAAGAGTCTATCTGTTCTTGATCTTTCGCGTAATAACTTAACTGGAGAACTCAATGATCTTGATTTTGGGACTTGTCAAAATCTCACTGTCCTAAATTTGTCTTTCAACAACCTTACCTCAGTTGAGTTTCCACCCTCTTTGGCAAATTGCCAGAGCCTCAATACACTGAACATTGCCCATAATTCGATCCGAATGGAGATTCCTAGCGAGTTATTGGTGAAGTTGAAGAGCTTGAAGCGATTGGTGCTAGCCCACAATCAGTTTTTTGATAAAATTCCATCAGAATTGGGGCAGAGTTGCAGCACACTTGAGGAGCTTGATCTTTCTGGAAACCGGCTGACTGGTGAACTTCCTTCAACTTTTAAATTGTGCTCCTCACTTTTCAGTCTCAATCTTGGTAACAATGAGCTATCGGGTGATTTCTTGAATACTGTTATCAGTTCATTGACAAATCTTAGGTATCTTTACTTGCCATTCAATAACATAACTGGTCATGTACCAAGGTCTTTGGTGAACTGTACAAAACTTCAGGTACTCGACCTTAGTTCCAATGCGTTTATTGGGAATGTACCTTTTGAGTTGTGTTTAGCTGCCTCAGGCTTTCCACTGGAGATGATGTTGTTAGCCAGTAATTATCTTACTGGAACTGTGCCTAAACAGCTTGGACATTGTAGGAACCTCAGGAAGATTGATCTCAGCTTCAATTACCTGACTGGTTTGGTCCCATTGGAGATTTGGACCCTTCCAAATCTTTCGGAATTGGTAATGTGGGCTAATAATCTCACAGGTGAAATTCCTGAGGGCATTTGCATCAATGGAGGAAACCTTCAAACACTCATTCTCAATAACAATTTCATCTCAGGGGCACTTCCACAGTCCATTTCCAATTGTACCAACTTGGTTTGGGTTTCGTTGTCTAGCAATCGACTGTCTGGAGAGATACCACAAGGAATTGGGAATCTTGCAAATCTTGCTATCCTTCAGTTAGGTAATAACTCACTTACTGGATCGATCCCTCGGGAATTAGGTTCGTGCAGGAACTTGATATGGCTAGATTTGAATAGCAATGCTCTAACAGGTTCAATCCCTCTTGAGCTTGCTGATCAAGCTGGCCATGTTAATCCGGGAATGGCTTCTGGAAAACAGTTTGCTTTTGTGAGAAATGAGGGTGGAACTGAATGCAGGGGTGCTGGTGGACTGGTTGAGTTCGAGGGTATCCGCGAAGAGAGGCTTGCAATTTTACCTATGGTTCACTTCTGTCCATCAACTAGGATCTACTCTGGTAGGACGATGTACACCTTCACGAGCAATGGAAGCATGATTTACCTTGATCTTTCCTACAATTCTTTTTCAGGAACCATTCCTGATAATTTAGGTTCATTGAGCTTCCTTCAAGTTCTGAATTTGGGACACAACAATTTTACCGGAACCATTCCATTCAACTTTGGAGGTCTAAAGATAGTTGGAGTTCTTGATCTATCACATAACAGCCTACAGGGATTCATTCCACCCTCGTTAGGAGGCCTTTCATTTCTTAGTGATCTTGATGTTTCAAACAACAACTTATCGGGAACGATTCCTTCTGGTGGACAGCTGACTACATTTCCAGCTTCAAGATACGAAAACAATTCAGGCCTTTGTGGCGTCCCCCTTCCTCCTTGTGGCTCGGGCAATGGACACCATTCATCGAGTATTTACCATCATGGGAATAAGAAGCCTACAACTATAGGAATGGTGGTTGGTATTATGGTCTCTTTTGTATGTATTATACTACTTGTTATTGCCCTTTATAAGATTAAAATGACTCAGAATGAGGAGGAGAAGAGAGATAAGTACATTGATAGCCTTCCAACTTCTGGCAGTAGCAGCTGGAAACTATCAACCGTTCCCGAGCCTCTAAGCATTAATGTGGCAACATTTGAAAAACCATTGAGGAAGCTGACCTTTGGTCATCTTCTAGAAGCGACCAACGGGTTCAGCTCTGAAAGCATGATTGGCTCTGGAGGTTTTGGTGAAGTCTACAAGGCTAAGCTGAGAGATGGCAGTACTGTTGCAATCAAGAAGCTCGTGCACGTAACAGGTCAAGGAGACAGAGAATTCATGGCTGAAATGGAGACTATTGGAAAAATCAAACATAGAAACCTTGTCCCTTTGTTGGGATATTGCAAGATTGGGGAGGAAAGGCTTCTTGTATACGAATACATGAAATGGGGAAGTCTTGAATCTGTTCTTCATGATGGGGGGAAAGGAGGGATGTTTCTTGATTGGCCAGCTAGAAAGAAAATCGCGATAGGATCAGCGAGAGGATTAGCATTTCTTCACCACAGCTGCATACCACACATAATTCATCGCGATATGAAGTCCAGTAATGTACTTTTAGACGAAAACTTTGAGGCACGAGTTTCTGATTTTGGAATGGCAAGGCTGGTAAATGCTCTGGATACTCATTTAAGTGTGAGTACACTTGCTGGTACTCCAGGTTATGTTCCTCCAGAGTACTACCAAAGTTTCAGGTGCACCGCAAAAGGGGACGTGTACAGTTATGGTGTCATACTGTTGGAGCTTCTTTCAGGGAAACGGCCAATAGATCCCCGTGTATTCGGTGATGATAACAATCTTGTTGGATGGGCAAAGCAACTTCATAACGAGAAACAAAGTCACGAGATTCTTGATCCTGAGCTAATAACAAATCTATCTGGTGATGCTGAACTCTATCATTACTTGAAAGTTGCATTTGAATGCCTTGATGAGAAATCTTACAAAAGGCCAACAATGATACAAGTGATGACCAAGTTTAAAGAACTACAAACTGATTCAGAAAGCGATATTCTTGATGGCATTTCAGTGAAAGGTTCTATACTCGAGGAATCGCAAGAAAGGGAACCGTAAATCAATGATATTTCATCTTTTTGTGGTTAGAATTCTGTTCTTTTTTTCTCTTTTTTTGGATTGCCTCATGGATAGTAGTAAAAACTTTTATCTTTAATGGTTCTTCACCATTGAGATTGATAAGAGTTTATACTTCAGATTTTGCCTACTGTGTAAATATTAGAACACGACGATGAATGGGAGATGTGAA

>M6-PGSC0003DMG400022189 M6 LRR-RLK

ATGGACAATCTTCTCAAGTTCATTGTAAGTGTTTTACTTTCTCTTCTTTTTATCAGATTCTGTAATGGAGGCGAGTTTATCGAATCCAAATCATTTATCAGTTTTCTTGTTGCTATTGATCCAAACAATGTACTAAACGTTGATCGAAATGCTTTGTTTTCTTACCCTTGTTACTACAAATTACATGGTGTTAAGTGTGATTTACGTTCCGGTGAAGTCATAGAAATAAGACTTGAGAACATGAATCTTAGTGGTGTAATTGATGCAGAGTCACTCTGTAAACTCTCAAACTTGAGAGTTCTTAGCTTAGCTAGGAATAAGATTGAGGGAACTGTTCCTGAATCAATATCATCGTGTAAAAGTTTAACTGTTCTTGATCTAAGCAGCAACTTTCTTAGTGGAAACTCATTAGTACTAATGCAGTCATTGACAAGGATCAAGAATCTTAAAAAACTTAACATTTCTAAAAACAATTTCACTTACTCACAACGTATGATGAGTCTACACGCGTTGAGACGTGTAACTATTAAAGAATCTTCATCTTCAACTACAGTAAGTGATGATAAGAAGACAAAGAATCATCATGTGAGTCTGATGGTTTGGGTTCTCATATCTCTTGGAACTGTGATTGTGTTTGTAGTTTTAGTATTCTTGTATATGAGATGTGTTGGGACAACTAAGGAGAAGGAGGTTATAAAAGAAGTAGTTGATTACCCGTCTCCTCAAAAAACTCCCTCAGATGAAGTTGTGGATGGAGTGTGTAACACTGAAGAGAAAAGCTCAGAACTTTACTTCTTCATGGAAGATGTTGAGAAGTTTACGATGGACGATCTTCTTGAAGCAACTGCTAACTTAACAAAACAAGGCATTTGCAGCAGCCTTTATAAGGTTCATATCAGTAGAAGTGGTGTGTTTGCTGTCAAGAGACTGAAGAAACTACAAGTGGGGTTCAAGGAGTTCTCGCGAACGATGAAAAAGATAGGGAACTTGAAGCATCAAAATGTACTTCCACTTGTTGCTTATTACTCTAGCAATGATGAAAAAATGCTCATCTACAAATATCAGAACAGTGGAAGTCTACTAACACTTTTCGAAAGTAAGTAGTCAAGAACTTCAAATTTCTCATCTTGTAATTCAGAATTTTATATCATATTGTTGTTGTTTATTGTTGCAGATTATGCAGAGGGGAAATGGAACTTCCCATGGAACTTAAGGCTGTCGATTGCAGTTGGCATTGCAAGAGGTTTCGCGTACTTATACAGAAGTTCAAAGAAAGGCAATGTCATTCCTCATGGCAACATTAAGCCATCAAACATTCTATTAAACGAAAACGGGGAGCCATTGATAAGTGAGTATGGATATTACAAATTTCTAGACCCTGACAAAAGTTGCCTCTATAAAGACAATGGTTATACAGCCCCTGAGAAAACGTTGACAGAAGAAACTGATGTCTATAGCTTCGGGGTGATTCTGCTGGAACTGCTAACCGGTAAAGTTGTGGAGAAGACAGGACTAGACCTTGTCAAATGGGTGAAATCTATAGTGAGGGAAGAATGGACTGGAGAAGTGTTTGATAGTGAAGTTTCCAACTATGAAATGTATGCTTTCCCTCTACTAAACGTTGCACTCAAGTGTGTGGATCGTTTGCCTGAGGAACGACCTACTATGACAGAGGTTTTGGAGATAATCGAAGAAATTGTGAATGATCAAGAAGACATTTCTCCTTCTTCTATGACTTCTTTTGAATCCACCCCTGGTTCAATGAAGCATGTGTATGCTCTTTAG

>M6-PGSC0003DMG400022264 M6 LRR-RLK

GTTCAATAATAATAATAGAAAAATTCAAAGTGCTCTTGGATCCAGATCTTCTTCTGTTTTTCCTCTATAAACAAAACATGGGGTTCCCCCATTTACAATCCCAAAGTCCTCAACTTTCTCTCTTCTTCCCTCTAAGCAGCAGCAAACAATGCCTGCTCCTTCAACATATCCCTTTCTTCTTCTTCTTTTACTTTTCCTCTCTTCTTGTTATTCACTTGATGTACAAACTCAAACACTTCTTGCATGGAAAAAAACATTAAACATAACTTCCAATGATGCTTTAACATCATGGGATTCTTTAGATAAAAGTCCTTGTAATTGGTTTGGCATAAGTTGCAATTCCAATGGTCATGTTGTTAGTATAAATCTTAAATCAGTTGATTTACAAGGTTCATTGCCTTCAAATTTTCAGCCTCTTAAATTCTTGAGCAGTCTTGTTCTTTCATCTGTTAATCTTAGTGGTCCAATCCCAAAAGAGTTTGGAGATTATCTTGAACTTAAGTTCATTGATATTAGTGATAATTCAATCACTGGTGTCATTCCTCAAGAACTTTGTAAGCTTATAAAGTTACAAACTTTATCACTTAGTTCAAATTTTCTTGAAGGTGATATCCCTTCAGACATTGGGAACTTGTCTGATCTTAAGAAGTTCTTGATTTATGACAATCAACTTAGTGGTGAAATTCCAAAGGGTATTGGAAAGTTGAAGAATCTTGACGAATTTAGAGCTGGAGGGAATCAAAATCTTAAAGGGGAACTCCCTTATGAGATTGGAAACTGTTTGAACTTGGTTTTTTTAGGCCTTGCTGAAACAAGCATTTCAGGGAATTTGCCAGTGACAATTGGAAACCTGAAAAAGATTCAGACAATTGCAATATATACAGCTCTTTTGTCTGGACCAATTCCAGAAGAGATTGGGAATTGTAGTGAACTACAGAACTTGTATTTGTATCAGAATTCGATCTCCGGTTCGATTCCAAGGAGTATAGGGGAACTCAGGAAGCTTCAAAGTTTGTTGCTTTGGCAGAATAGTATAGTTGGTGTGATTCCAAATGAGCTTGGGAATTGTAAGGCTATTACAGTTATTGACTTGTCTGAGAATCTCTTAACAGGTAGTATTCCTACAAGCTTTGGTGAACTTTCAAGTCTTGAAGAGTTGCAGTTGAGTGTCAATAAGTTATCAGGTACAATACCTACTGAAATATCAAATTGTACTAAATTGTCTCATTTGGAAGTTGACAATAATGGCATATCAGGGGAAATTCCTAATGAAATAGGTAAGTTAAAGAGCTTGACTCTATTCTTTGCTTGGCAAAATAATTTAACAGGTGAAATTCCTGTTACATTATCTAACTGTGAGAATCTTCAGGCTATAGACCTTTCTTATAACAATCTTTTTGGTTCCATACCAAAAGAAATCTTTAGTTTGAAAAATCTAACAAAGTTGTTGCTTCTTTCCAATGATTTGTCTGGTTTTATACCAACTGATGTTGGAAACTGCACAAATTTGTACAGATTTAGAGTGAATAGGAATAGGCTAGGGGGTACTGTTCCATCAGAAATAGGGAAGTTGAGTAATTTGAATTTTCTTGACATGAGTGGAAATCATTTTATGGGAGAAATTCCATCATCAATATCAGGATGTCAAAATCTTGAGTTTCTTGATCTCCATTCGAATGCGTTTACTGGTTCTTTGCCAGAGAAACTGCCAGGGAGTCTACAGTATGTGGATATTTCAGACAACAGGCTTTCGGGTTCGTTAAGTTCAAGTGTTGGTTCTTTAACAGAATTAACAAAATTGAATCTTGGGAAGAATCAACTTTCTGGTAGAATTCCAGCTGAGATAGTTTCTTGTAGTAAGCTTCAGCTGTTAGATCTCGGGTACAATGGCTTATCAGGTGATATACCTAAAGAGTTGGGTCGAATACCATCCCTTGAAATCTCTCTTAACCTTAGTTGTAACCAGTTTACAGGAGTGATTCCAAGTGAGTTTTCAGGTCTTAGTAAATTAGGAAACCTTGATCTTTCCCACAACAAACTCACCGGGAATTTGGACGTTCTCACGAACCTTCAAAACCTTGTTTCACTCAATATATCATTCAATGACTTCTATGGAAAACTTCCCAATGTTCCATTCTTTCATAAGCTTCCTTTGAGTGATCTTACTGGAAACCAAGCTCTATACATTTCTGGTGGGGATGTGATCCAAACGGGGCCTGCTGGACATGCTAAAACCACCATGAAGCTTGCTATGTCAATCCTAGTCAGTATAAGTGCTGTGCTGGTGCTGCTAGCCATCTACACATTGATCAGGATGCGAATGGCGGCTAAGTATGGACCGGAAGTTGATACTTGGGAAATGACTCTTTACCAAAAGCTGGACTTTTCGATAGATGACATTGTTCATAATTTAACATCAGCTAATGTCATTGGCACTGGGAGCTCCGGGGTTGTCTACAGAATAATGACCGAAAATGGGGCGACTTTAGCAGTCAAGAAAATGTGGTCATCAGAAGGATCAGGAGCATTTGGTTCAGAAATTCAAACACTTGGTTCAATCCGGCACAAGAACATCGTACGCCTTCTCGGTTGGGCTTCAAACCAGAACATGAAGCTTCTGTTCTATGACTACCTTCCCAATGGGAGTTTGAGCTCATTACTCCATGGTGTTGGCAAAGGAGCAGCAGAATGGGAGAACAGATTTGATGTGGTCCTTGGAGTAGCTCATGCTCTTTCATACCTACATCATGACTGTGTACCCCCTATAATGCATGGAGATGTCAAAGCAATGAATGTATTGTTAGGCTCTAGAATGGAGCCTTACCTGGCCGATTTCGGGTTGGCAAGGATTGTCAACACTGATGTTGATGCAGACTTGTTGAAGGAGAGCCAAAGGCCTCATCTTGCTGGTTCCTATGGATATATGGCTCCAGGTAATTATTTTTACTGCATTTTTACTCACTATCTTAGATTGATGACGGAGAACGTAAAAGTAGTCAATTTATGATGAATCCTCACAATACAAAAGATATTGTTGGTGTTGTACTCATTAGTCCTTATATCAGCTTGATTAGTTGCATCAACTTTTCTTGAATTGACAAAGACTCTTGCTTATTATGTGTCCACAGAACATGCTTCGATGCAACGGATCACAGAAAAGAGTGATGTATACAGCTTTGGTGTGGTCCTCTTGGAGGTACTAACAGGAAGGCATCCACTGGACCCAACGTTGCCCGGGGGTGCACACTTAGTACAATGGATACGCGATCACTTACAAAGCAAGCGCGATCCAAATGACATTCTTGATCCAAAGCTTAGAGGCAGAGCTGACCCTGAGATGCATGAAATGCTGCAAACTTTAGCTGTTTCATTCCTGTGTGTGAGCACGAAAGCTGATGATCGTCCGATGATGAGGGATGTTGTAGCCATGCTCAAGGAAATTCGGAATGTTGATCCTGTTGTGTCCGAATCCGACTTGTTGAAGAAAAATGCAAGTGTAACACCTCTTCCAAAATCACCTGGCACCAAGAATGTGGATTCTCAAATGTCTTGTAGCTGCTCTTTTGTATTTTCTGATAACTCTATCTCCAAATAAGGCATCAAAATAGTAAACAAATTTGTGAATATAGCATGGTTTATTAGCAGTATGAGTTAGAGCAGATTATAGATTTTAAAAGGAAATCATGTATTCTATTAAAGAAGTTTTCAATTTAGCATAAGCATTATAGAGTGAAA

>M6-PGSC0003DMG400022425 M6 LRR-RLK

CTTTTGGTTGTTGGAGATTTGTCTGTTGTTTATGTTAAAGTGAAGATCTTAGTGTGTTTTGCAGAGTGTTTTCTTTTTGCTGGAACTGTGAAGCAAAGATTTGAAGGAAAAAGGACAGAGGAAGAGGAAGCAGGAAATTTGTGTTCTTGGAGTGTCCCTTGGTGATGATTGGTCACAAGATGTGTGGACTATATATGAAGATTTTGTTTGTATTTGCAGTTTTTGCTGTTCAAAATTGCCAGGGGTTTACTGATCCTCGTGATGGTATTAATAATCTCTCTGTGGTTTGAATTTTGTGTATTATACGGTATAGAATGTCTCAATAGTTTGAATTTGTTATCTTTTCTATTAGCTTTAATGAAATTACAGTTTGATGTTTTGTGTAAAAGAAAGGTGAAGTTTATGACTTTGTGTACTTTTCTTGCTCTCCAGTATTTGCCATAAATAGTTTATATACTAGTCTGGGCTACCCCTTGCTTCCTGGATGGGTGCCATATGGATTAGACCCTTGTGGTGATCTATGGCATGGTGTACTCTGTGTCAATTCCAATGTAACTGGAATGTAAGATTTCCTTCTTTTTTTTTTTTTGCTTGTGTTTCTCACATGGTTTGTATACTTTATAATAGATTTCTTTTCTTTTTTTGGCAGAGTACTAAATGATTCAAATTTGGGAGGTGAATTAAGTGAGGGCATAGGAAGCTTTGCTTCAATCATACAAATGTTGGTACCATCTTTATCTTCATCATTGCATTGATTTAACTTATGTTGCTGAAATTTGCCTTGTCATTTGGTTCTGTGCACTTTGTCATTCCCTGCATCATTTGGCACCACTATTACATTGTTTAATTCTATGGATTTAGTTAACTGCTACGTATTAAATGTCTACGCTTTGTGTAATGATCAAACCAGAGATCTCAGCAACAACCATATTGGAGGCAGTATACCATCCAATTTGCCTGTCACCCTCAAAACCTTGTAAGTTAATAATCTTTGTCCTTACTGAGCTTCTTCACCACTTTATAAGTTATTTGCTCCACATAAATCCACATTTTTAATCCTATTATGGTATTTGTGTATATACAGGTCTATCTCTGGTAACCAGCTTACTGGGAGCATTCCAGAAAACATATCCTCGTTAGGGCAATTAACAGACTTGTGAGTTATATTTGACTTATCTTGATATACTTTTCTATGACCATGCTTTTGTTGCCATTACATTTAATTCCTTCCTTGTAGGTCGTTGAACAACAATCATCTCAATGGAGTCATTCCAGATGCCTTTATGCAACTTAAGACTTTGATTAACGTGTACTCATACTTCTTTTAGCTTTTTACCTTTTCGCAGTGATGAAACTACTATTCTTGGTGTTGTTTGTATTTATGGCTTCATGCTCTTATTTTGCAGGGACTTATCTGGGAACAGTTTAAGTGGTCAGCTGCCTCCTTCAGTGGGGACTTTGTCGTCACTAACTACCTTGTATGTATTTCTCCTTGTTGGTATGATAATTAGGGAAATATTAGGAAATGAAACTACTTTTCTGGCTATGAGGAATGTGAGTGCTTTCCTCTCTCTAGACACTGAATAGAAGTTCAAATAAGTCCTTAATTCCTTCAGAAATTTAACATAAGTTCTGCTCATACATAATTTAAACTGATAAATTTTGTAAAGATAAATATAGTTTTTGACATTTCATTGCATATGTAACTATTGTATGGTTGTGCTGTATGTTGGTTTGTCAAGATGTGTTTTCTGTGAACAAGCTAGAGGAATGATATTTGAGGATAATACTTCACAAAGTTGATTAATTTTGTCTAGAATCTCAATTTGCAACCATTCAACTTAATATACTTGGATGGTAAAAGGATGTTAACATAATTTCTCAACTGAACTTGTGAATATACTTGTATTTATGGTGAATTGTGATACTCTAGTTTTGATTTTTGAATATATACTTTTGGCTCTATGCAGACACTTACAGAACAATCAGCTTTCTGGGATCCTTGACGTTTTGCAAGATCTTGCTCTAACGGATTTGTAATCTCCTGGACTAATAAATTTTCCATCATTATAGATAATGTCATCAACTATGTCATTTTCTGCTCAATGTGTGCTTAAATTTGGTCACTTTCTTGTTCTGCAGAAACATAGAGAATAATTTGTTCTCTGGGCCTATACCACCAAAATTGTTGAGCATCCCGACCTTCAGGTAAAAACCTAGGAATTTTTTTGCCCTCTTTAGTTTGAGTTTTCTTTTAAGCTGATGAAATCCATTTTATTCTATCTTCAACTTCTGTAATATGAAATGGAGTATCTTATTTCTCTTTGTTTTAATTTGGCTTCATCTCTCTGAGTACTTCCATCAATTTAAAATTGGATAAGCATTGGTCACTGTGGTAATGGGACAATGTGACTTAAAAAATTTAAAACAAAAGGGAGTTGTCTCTTTGGCTAGACTCTAAAGATTGCACTATTTAGATCTAATTTCATTGCCAATTGTGCTTGCAATGAATTTTTATGGGAAGCCAGTATAACTGTTGTTTCTAATAAACTTGACACCATTCCATGAGCTGACCATGAAAATGTTGGTTTTACAAGTTTTAGTTTCTATAACTTTGTCATGCATCGATCTGATAAATAAATCAGTACACTGGCTGATAAATGGGTAAAGTAGCAAACTTGAAGATATGGCATGTAACGATATTACTAAATAGTATGGTTATATGGCATGTAATGATATTACTAAATAGTATGGTCTTTTTGATGAAGACTGCATAGTACACCTTATGAACCAGTTCTTTACTCCATCAATCAACCATCCTCAATCCCAAATTAATTGTGGATGACTATATGAATCCTCTTTATCCGTTCTACCTTATGCATCATCCTACATGTCACTCTTAACTTTATCTAAAATTCTATTTTGTGCAAGATTGATAATAGATATTTAGTTGATGTTTGTTGTGAAAGCTCTTTTGGAGTTCTTTTAAAGACCTTATTGAAACCATGTTTGCTTAAGTTCTTATAAGAAAATTTTCATTTCATCAATGAAGAAGGGGGTTTGAGGGATTTTCATTAAAAAACAATTCTTCTTCCACTTATAAAAGTGCTCACAAATCTTCAAAAGATCAACTGTGTAAACACATTTGTTTTGCACATCTACAAAAATATTTCTCCAGAACCATTTGATTGGTTAAAATGAAATTACAATGGATAACCAATCAACATGATGCATTTTTGTGATAGACTAGCCTTAGTCATGGCCTCAATTGTTTAGTTGATACTCTTTGGGGGAAGCGAACTATGGTAATCTTTTCTTCTTCATGAAAGGTTCAGTTGTATACTTGTAAATTTTGTTGCCTTACATGGTGAAATCTATGTTTTTTATCAGACGTGCAGGGAATCCTTTTAATACTACTATCATCCCTTCACCACCCATGATAACGCCTTCTCCAGCACCATTTGCGCTTTCTCCAGATTTAGCCCCTTCACTGCCATATATTGTACCGCCAGGACAGGAATTACAGCCCTCTGGAAGGCAAAAGAGTAGTAATTCAACCAAACATGTCAAATTGATTGCTATAGCAGGATTAGTATCCCTTGTCATACTTGGATTGGGTGTTTGCCTATTGATGTCACGATTCTGCAAAAGAAGAAGAGAGACCCAAAAAGAAACCAAGAGACATGAAACATATGATCATAGTTTGCCCAAGGCCAAGGGTAACCCTAAGCATGATCAATCCCTGCAAAAACCATATTATGATGCAGAAAAAGGTAAATAGTTCATATGTGTACTTTCTTCTTATAGATTTATGTTTATCGTCAGTTACCTGTTGTTTTTAATGGCATTATGATGACAAAATGGATATGTCCAAAAGAAGCAGGTTTGAGGCCAGTGGCTGGAAATGGGAAAGCTCAGGGTAGGAACATGTCAATGAATACTTCAGAGGCTGTGCAGCAGAAAGACGTGAAGAGCACTATATCATATACAGACGATGAGCTGGGCTCGGAGATGGAGTCAATGATATCAGATGTTCTTCCACCACCGCCACCACCTTTTCTTCAAACTCTTTCTGCTGAGAGGATGGTTGTAAATCCAATTGTTCCCACAATTACCTCAGGCAAGCATGCTATGAAGAATGTAAACTCTGCAGAGTTCTTCACCATTGCCTCTCTTCAGCAGTATACAAATAGCTTTTCTCAAGATAATCTTATTGGAGGAGGCATGCTTGGAACAGTCTACAGAGCTGAGTCTCCCAAAAAAGTAAGCATCTTTATTTTCACGAGCTTCAAATCAGTAAATACTGTGTGCTTGCATGCATTGTGACTGCATCAGTACCACATCTTGAACTTCATTTCACTTTTTTAAAGGTGGAAATTACTCGTTATTTTTTTATTAAAAGCATGGTTTTCTGTGACAGACATTATATCTATTTTACATTGAACTTGTAGATGTTTTGTGGGTCTGTAATGTTTGACTCGAGTTTTGCATAATAGAAGTAGTCCAACCTTGTAGATGGCAACAGTTTCCATTAGTAGACTGTGTAGATATACCTGACATGCCTCCTGATTCATTAAATTAGTTTCCTGTTTTCACAATAACCACTTGTACTTGTCATGTTTGTATATAACCTATTTAGTTGCCGTACAAGGGTAGAAGGTGCTGAGACTTTGCTATTTCTTGATGCCTTTTGTTTTGTTAGCTATTGGCGGTCAAGAAACTAGACACTGCTATGACCAAGCGTCAAAGTGATCAAGAATTTGTTGAAATGGTTTCCAATATCTCTAAACTTCAGCATGAAAACATTGTCAAACTTGTGGGTTACTGTTCAGAGCATGGACAACGGTTGCTTGTCTACGAGTATTGTAGAAATGGTACCCTCCACGAAGCATTACACTTAGATGATCAGATCCACAGAAAACTTTCCTGGAGTACACGTGTACGCATTGCACTTCAAGCAGCAAGAGCATTAGAGTAAGTACGAAAAAAAGCTAAATGATATTAGTTTAGAAGTGCTTGTAACTTGTTTTAATGAATTGTGATGGTTAAATAAATTATAAATTGGCTGTTAAAGTCAGCTCTGCATGTTTCCCGGGCTTAATGTTTGCTACACCACAATTAATTTTGTAGTTTTATCCTTAGCTTGTAACCGTTTAGAGATTGAGTTGCTTCCAGTTTCACTGAAAGGTACTTGCATGAAGTTTGTCAACCACCTATTGTCCACCTGAACTTCAAGTCCACTAATGTTCTCCTTGATGATGAACTTGCTGTACATATGTCAGATTGTGGTTTGGCACCTCTGATGTCATCTGATTCCATAAGACAGGTAATCCCTTCAATCCTCAAACTAGAAACAAGAATAACAGGAGATTTGACTTGCAAACTGTAGTTATTCTGAATTCGTAATGGGAAAATTTATCCTTTTGGACTGTGATCTTCATTGAACTTGTAAACACTTCCTTCTCAACTTCTAGAAACTGCATCATTATTCATTAGTATGAAAACTAGGTGCCAATTGGCTGCTAAAAATTGGTGCATTCTGTTTAGAAATGGTTGTATGTTTCTATTTCGAGGGGATTAAAATGAAAGGACAAGACCTGGTATTTCTGCTAATTTTGGTTTGCATCAAAAAGCTGTCCATCTCATATTGCATTTATCGATGAATAAAGTGTTGATAACTTCAGTATAACTTTCAGTTGCAAGGATCCGGCTATGGTGCTCCTGAACTTGAATTGGGAAACTATACTTGTCAGAGTGATGTTTATAGCTTTGGAGTTGTTATGCTAGAACTTCTCACAGGAAGACAATCTTAAGACAGGTTAATTTCTTTTCTGACAGATATATTTTAACACCTTGCTAGTTGTTATTGTTGCAACTACATATTTTTCTAAAATTATACGACTCCTGACTTTGAGCCCATTAAAATATGATTTCCAGGTCCCGGCCTAGAGGGGAGCAGTTGCTGGTTCGATGGGCTATTCCACGGCTGCATGATATAGATGCATTATCAAGAATGGTTGATCCTTCCCTAAATGGTTGTTATCCATCCAAGTCTTTGTCTCGGTTTGCTGACATAATTTCCTTGTGTATTCAGGTTAGTCTCTTCCTTTACAGGTTCCTCATAGCTCTTTAATGAAGTATAAAATCCCACTTACAGTTTTGAGTTACCACGTGTGAGAAACTTTACTGGTTATTCTTCATAAAATGAGACTTTAATCCAGTTAAATTCCAGGGATTTGGTAAATGTCTGTTTTGGATAAACCTACTTTACAGGCTGCATATAACTATGTCCAACAAGAAAACAAACTCACATTTGAGCTATTTAGGCTTAAGTAAATTTTAGAATCAACTCAGTCTAGGGAGAGAAGAGGTTCTTTGTTAGTAAATTCGCCATTCTGCTATAGCTAAAAAACACACACATAATATTTGAACTGATGTTTTTCGAGCTGTCGCCTTTCATACTTTTCTCATGTCTTCTTACTTTTTTGTTGGGAAGTCGGAACCTGAATTCAGGCCACCAATGTCCGAAATCGTGCAGAATCTTCTGCAGATGATCTAAAGATATCATCAAGACTGAGTCTTTTAGCATAGATGGCAAGGATTCAAGGAGAAATCTTTACACAACATTTTCATTAAAGCTCAAATATTGTGTGCTAACATGAGTAATGTAATGTAAATAGCATTATTATGTTTTGCAACATTTTCATGTGTGATTCTTTTTGACATATCTATGAAAAAGTAGAAAAAGACTCTCATTCTTTTAGTGTAATCGAGGAGTTTGTAATATATTATCAGCCAATTCTTTATGCAAGTTTTGTGATCATTTGAAG

>M6-PGSC0003DMG400022565 M6 LRR-RLK

CCAAAACTAGCTGCATCCCTGCCAATAACACTCATCTTTCACTGTCTCACTCTCACAATGATACCCTTAATCTCCATGGCTACTTCTTCATCTTCTGTAAATTTCTTCTTCTTCTTCCTCTTTCTCTTTACCCATTCATTCTTTTGCTTTTCTCAACAAAACCCAGAAATTCAAGCACTTCAATCCTTCAAACTTAGTATTCATGACCCACTTGGAGCACTCACAAGTTGGGATTCTACATCTCCTTCTGCTCCATGCGATTGGCGAGGAATTTTTTGTTCAAATGATTCCTATGTGAGTGAAATTCGTCTTCCACATTTACAGCTTAGTGGGTCTCTCACTACCCAGATTTCTGATTTGCGAATGTTGCGTAAGATTAGTCTTCGTTCAAATTTCTTTAATGGAACTATACCTGCTTCAATCTCCAAATGCAAACTTCTGGATACTGTTTTCTTACAGTATAACTCTTTTTCCGGTGAAATCCCGCCGGAGATTATGAATTTGACTGAGCTTGAAACGTTTAATGTGGCTGGAAATCAGATGTATGGTGAAATACTCAGTGAGTTGCCGGTGAATCTGAGATATTTTGATGTCTCGGAGAACTTGTTTTCCGGTGGTATACCGGAGAAAATATCGGACCTCTCTCAGGTTATTCTGCTTAATCTGTCGTACAATCGATTCTCCGGTGATATTCCGGCGAGTTTAGGTAGGCTTCAACAACTTCAGTATCTTATGCTTGACTACAATGAGTTAGAAGGAACAGTTCCTTCTGCAATTTCGAACTGTTCGTCGTTGGTGCATTTGAGTGCCGAAGGGAATGGTATTACTGGTGTTATACCGGCGGGGATTGCTGCTCTTCCGAAGATTCAGGTGATAAATTTATCGCATAATAAGCTTTCTGGATACTTAGCAACTTCATTTTTCTGCAATGGTTCTGTGTATCCTCCTTCCCTTCAGATTGTTCAGTTGGGTTTCAATGCCTTCGCTGAGATTCTTCACCCACAATCGTCTAGATGTTTTAGTTCATTGCGAGTTTTGGATATTCAACACAATCAGATACATGGAGATTTCCCTTTTTTCTTGACAGACAATTCTGCATTGTCGTCCTTGGACTTGTCAGGGAATTTGTTTTCGGGTACAATCCCGAGTTCCATTGGGAATTTGTTGAGGTTGGAGCAACTGAGAATTGGAAATAATTCATTCGAAGGTAATATACCGGTTGGGATAACGAAATGTAGTAGTTTGAATGTTCTTGATCTTGAAGGGAACAGATTCATAGGGGAAATTCCAGCTTTTATAGGTGAACTTAAAAGCTTGAAGATTTTGTTCATGGGAAGAAATCAGTTTTCGGGTTCAATCCCTTCTAGTTTTAGCAATATCACTAGTCTTGAAAGTTTGAATTTGGAAGGAAACCATCTTACGGGAAGCTTGCCTGAGGAGTTGATGTTTTTGAGCAATTTAAGCACATTGAATCTTAGTGGGAACAAGTTTTCAGGTAGTATTCCTGTTGGTATTGAGAACCTTCAGCAGTTATCGGTTCTAAATCTGAGTAAGAATGGCTTCTCAGGGACTATCCCGTCTAGCATTGGAACTTTGTATAAGCTGGTAGCTCTTGATTTGAGTGGACAGAATTTATCAGGCGAATTGCCATCTGTTCTTGGTGGTTTGCCTAGTTTACAAGTGATTGCTTTGCAAGAAAACAACTTGTCTGGAAATGTTCCTGAAGGTTTTAGTAGCTTAATGGGTTTACAGTTCTTGAATCTATCTTCCAATTCATTTTCAGGTCATATACCATCTACGTTTGGGTTCTTGACATCGTTAGTTGTCCTCTCGTTGTCTAAGAATCACATATCTGGTTCAATACCTCCTGACTTGGGTAATAACACTGCTCTGAAGATTTTAAATCTGAGGTCAAATTCATTGAGTGGCCAAATTCCTTCTGATCTTGCACGTCTCTCCCGTTTGAGTGTGTTGGATTTGGGTAGAAATACCCTGACAGGTGAAATTCCAGAAGTTATTTCCAATTGTTCATCCCTCACATCAGTTGTGATGGACACGAACCATCTTTCGGGAAACATTCCAGCATCACTCTCCAGCTTATCAAGCTTAATTACCCTTGACCTCTCCGGAAACAACTTGAGTGGAAAAATCCCAGAAAATCTGACAGTGCTCCCAAAGTTGGTGAACTTCAATGTATCGAACAATAAGTTGGAAGGCCAGATACCAGTGAAGCTCGGTTCTCATTTTAACGATCCATCGGATTTCAGTGGCAACCAGGGTTTGTGTGGGGAGCCATTGAATAGAAAATGTGAGCGGACTGGTAATGATAAGAATAGACTGATTATGTTTATTGCAGTGTCTGCTAGTGGAGGTCTTCTCCTTGCATCATGCTGCTGTTTTTACATATACGCCTTGCTGAGATGGCGTAGGAAGCTCAAAGAAAAGGCAGCTGGAGAGAAGAAGCATAGCCCTGCAAGGGTTAGTTCGAGGACCAGTGGTTCTCGTGGCAGTGGCAACAATGCTGGACCAAAACTTGTTATGTTCAATAACAGAATTACAGTTGCTGAAACAATCGAAGCAACTAGGGAATTTGATGAGGAAAATGTTCTTAGCAGAACACGCCACGGAGTGCTGTTCAAGGCCTGTTACAGTGATGGAATGTTGCTCTCAATCTGTAGACTACCAGATAGATCACTAGATGAAAACACATTCAGAAAAGAGGCTGAATCGCTTGGCAGAGTGAAGCACAGAAACCTAACTGTTCTCCGTGGGTACTATGCTGGACCTCCAGACCTTAGACTACTTGCATATGATTACATGCCAAACGGGAACCTTGCAACACTTCTTCAAGAAGCATCCCATCAAGATGGTGGTCATGTTCTCAATTGGCCAATGCGACACCTCGTTGCACTTGGCATTGCCCGTGGGCTTGCTTTCCTCCACGCAGCCTCTATCATTCATGGAGATGTTAAACCACAAAATGTACTGTTCGATGCAGACTTTGAAGCCCATCTTTCAGACTTTGGCTTAGACAAGCTTACTGTAGCCACACCAATCGAACCTTCCTCTTCAACATCAGTTGGGACTATAGGCTATGTAGCGCCAGAAGCAACATTAACCGGAGAAGCCACAAGACAATCAGATATTTATAGCTTTGGCATTGTATTACTGGAACTGCTAACAGGGAAAAAGTCGTTAATGTTCTCGCAGGATGAGGACATTGTGAAATGGGTAAAGAGGCAATTACAGAGAGGGCAAATTTCAGAATTGTTAGAGCCAGGATTGCTTGAACTGGACCCTGAATCATCAGAATGGGAAGAGTTTTTATTAGGAGTAAAAGTAGGCTTGCTTTGCACAGCACCGGACCCCCTTGATCGTCCCACCATGACCGACACTGTGTTCATGCTCGAAGGTTGTCGTGTTGATCAAATTTCACTCTCCTGAAGAACCACGACCGGCAATGGTGACACGTAAAACCCAAGGTCTCATCCTAATTCTTTGCCTTTCCTATTGACTCTTTGATATTTTGAGGAGTTGCATTGCAATGCAACAATAGCTAGCACTAGCACTTAAGTTTTTTGTTTAAGGAAAAGAACAAAAAGAAAATATTCTCAATTTTTTTTTAGCTATAGCTATTTTATTGATTTTGGTTTTCTATTACAAAAGTGCCTCTTGAGTTGAAAACAATGAAATGAATTCTTTTTAAGTAATTCAAG

>M6-PGSC0003DMG400022632 M6 LRR-RLK

CAAACCTCAGTCAAATGACAACAAGCACAGCCTCTTCATTCTTGTTTTTGCTCTTCCCCCTCTGTTCCTCTTCTTCTTCCTAACCCCCTTCTCACTTCACTTCTCCACACTCAACCCACCCACCATGGCAGTAAAATCAACTAACCCCATCTTCTTCTTCACTCTTTTCTTCCTCATTTTCACATTAACCCCATCTTTTTCCATTGAAAATGATGTTAAATGTCTTGAAGGTATCAAATCTGCACTTTCTGACCCTTTAAACAAGCTCTCATCTTGGTCATTCTCAAACACTTCTGTAGCTTCAATTTGTAAACTTGTTGGTGTTTCTTGCTGGAATGAGAAAGAAAACCGCCTTCTTTCTCTTCAACTTCCTTCAATGTCTTTATCCGGTTCACTCCCACCTGCCCTTCAGTTCTGTACTTCTCTTCAATCTCTTGATCTCTCTGGTAACTCTTTTTCCGGTCCAATCCCGGTTCAAATCTGTTCTTGGTTACCTTATCTCGTCAATCTCGATCTCTCCAGCAACTCTTTCTCCGGTTCCATACCCCCTGAGTTTATTAACTGCAAATTCTTGAACACCCTTATTCTAAATGACAACAAACTTACTGGGTCGATCCCATTTGAGATCGGCCGGCTCGACCGGTTGAAACGGTTCAGTGTGTCCAACAATGGCCTTTCCGGTTCAATTCCTGATGATTTAGACCGGTTTTCGAAAGATGATTTCGATGGGAATGATGGGCTTTGTGGGAATCCAATTGGATCTAAATGTAGTAATTTGAGTAACAAAAATCTTGTTATTATTATAGCTGCTGGTGTTTTTGGTGCTGCTGCATCACTGATTCTTGGATTTGGGATTTGGAGATGGTTTTTAGTTCAGCCTAGTAAGAAAGATAGAGAATTCGGTGACGGTAAAGGTGGTGGTGGTATCAGTGATTATTGGGTCGATAAATTGAGAGCTTACAAGCTTGTTCAGGTGACATTGTTTCAAAAACCTATTAATAAGATTAAGCTGAATGATTTACTAGTAGCTACAAATTCATTTGCTAGTGAAAATATTGTTATTTCAACTAGAACTGGTATTTCTTATAGAGCTATGTTGATTGATGGATCAGCATTGGCTATTAAAAGGTTGAGTAGTTGTAAGTTAAGTGAAAAACAGTTTAGGTCTGAGATGAATCGATTAGGACAGCTTAGACACCCGAATTTAGTCCCGTTATTGGGATTTTGTATTGTTGATACTGAAAGGCTTTTGGTTTATAAGCATATGCAGAATGGAAGTTTGTATTCGCTTTTACATGGTAATCTTAGTACTGGTATTAGAAATGATAGTTCTGAGCTTGGTTGGTTAGCAAGGGTGCGGGTTGCTGCTGGTGCAGCTAGAGGATTAGCTTGGCTTCATCATGGGTGTCAACCGCCATATGTGCATCAATACCTTAGCTCGAATGTGATTCTTGTAGACGATGATTTTGATGCTAGGATCACGGATTTTGGGCTTGCAAGGCTTATTGGTTCTGCTGACTCGAATGATAGTTCGTTTGTGAATGGGGATTTGGGAGAGTTTGGGTATGTAGCTCCTGAGTATTCGAGCACGTTGGTTGCTTCAATGAAAGGGGATGTGTATAGTTTTGGAGTAGTGCTGCTTGAGTTGGTGACAGGGCGGAAGCCTCTTGGTGCGGGTAATGCAGAGGAGGGATTCAAAGGTAGTTTAGTCGATTGGGTTAATCAGCTTTCGAGCTCAGGTCATAGTAAGGACGCGATCGACAAAGCTTTTGCTGGAAGTGGTCAAGATGATGAAATTTTGCGAGTTCTTCAAATTGCATGTTCGTGTGTGGTTTCAAGGCCGAAAGATAGACCTTCGATGTACACAGTGTATCAGTCCTTGAAGAGCATGGTGAAAGAACATTGTTTCTCTGAACACTTTGATGAGTTTCCAATCAACTTGACCAAAGAAAACCATGATCATAAAGATTAGGTTATAGCATTGAAATGATTAGACAAATGTTTATGAATTTTGAATATTCTAGTTATGGCATTTGTATTTCTTGATAACTCTGTATAGTTTGAGAAGTAAATGCATGATCTTTGTACTTTCTTGACACATTTT

>M6-PGSC0003DMG400022900 M6 LRR-RLK

AAAAATTATACTAATTCTACCTAACAAAAAAACAGAAATCAACCAAACAAGTAAATTTTTCTTTTGTTATCTCTTAGTTAACAAATGAAAAATCCAGAAATTTACTTCATAACGGGGACGAATATAGTAATTTTGAAGAAATAACATGGACATTTTTCGTGTTATGATCAAAACATTCCTTCTTTGCATCGTCTTCTTGATCAATTACCCCGTGAAAATAATTTCAGTTTCCGATACTGATGCTTTGTTGAAGCTGAAACAATCATTTACAAATGCAGCGTCATTGGAATCCTGGAAACCTGGAACGGATCCGTGTGACAAAAATATACGTTGGTTGGGTGTTTTTTGTCAGAAAAAAATGGTTACTGGACTTCTTCTTGCAGAAACGAATTTATCTGGAATTATAGACGTTGAGGCATTATCACAAATGCCTGCTCTGCGTACACTTAGTTTTCAAAGCAATTTATTTTCTGGACCAATTCCGGAATTCAATCGAATGGGTGCTTTAAAAGGTCTTTATTTGTCGGAAAATCAATTTTCTGGTGAAATTCCATCAAATTACTTCGAAAAAATGTTGTCGTTGAAGAAGTTATGGCTCTCTGACAACAAATTTTCGGGTGAAATTCCATCTTCATTAATGGAGCTGCAATATCTCATTGAATTGCACCTGGAGAACAATCAGTTCACAGGTCCAATACCGCCAATCTCGAAGTAGTCAACATTAGAATCAATCGATCTTTCTAATAATAACTTGAAAGGGGAAATCCCTGTTTCTTTATCAAGATTCAATGAGAGTTCGTTTAAGGGGAATTCCGAGCTTTGTGGTGAAAAATTAGGCAAAGCTTGTAATCCAACCATAAATAGCAGCAACGACAACAACACTAATAATAACGCAAATAATGGCACTCAACAAAGTCCGGATAGTAATTCGAATAAATGCATCTCAATGTGGATTATGATATCATCAGTTGTTCTTTTATTGATCATGATCATAGTAATATACCTCATATGTCGCTACCATCAAAATCGTCTTGCTTCAATTGAAAGTTTTGATGAGCCATCACTTGGGAGGCGAATTTCGAGTGATAGCAAGAAGTCATTTGAACTAAGTAGACGTGGCAGTTCGATTAGAAAAGGATCGATAATGGGAAAACGTGTTGGAGATTTGACAATGGTGAATGATGATAAAGGTGAGTTTGGTTTAGCAGATTTGATGAAAGCTGCTGCAGAGGTGTTAGGGAATGGGCCATTGGGTTCTTCTTATAAGGCTATGATGTCAAATGGATTGACAGTTGTGGTAAAGAGGATCAAAGAAATGAACAAGATCGGAAAAGATGGTTTCGATGCAGAGGTTATACGTTTAGGAAGATTGAGACATAAAAATATTCTAACACTTTTGGCTTACCATCATCGCAAGGAAGAGAAGTTGTTTGTGTATGAGTACATTCCTAAAGGAAGCTTGCTATACATATTGCACGGTAAAGTACCTAGCTCTAATTTTATTATGTCATAAATAGAAATGCATGGTAAGACTGCGTACAATAAACTCATTTTTGTTAATTTTGTGCATGTGCTTAGAATAGGAGGCCAAAAACTTCTACAAGTCGTATGTTTACATTGTATTAGTAGTCTAATCGTCAAGTGTTTCATATTAGGTGAGGAACAGAAAGTAGTCTCCTTGTATAATCTTGGCAAACTATACGTCTATATCCATGAACTAGCTTTGAAGTTGTGTGAGGTCTAAAGGTCCGTTTGCTTCACAAGTTATCAAAGCCAGACTCATCTGTACTCTTGACATGTAGGCCCCCATGTTATATTGGGATTATTCTCACCTTATGGTCTTTTTCAAGGCCACTTTTTGAATTATGTTCCCGTTAGCCCAGTGGGTTAAGGGGAACATAAGTTAAATAGAGACCTCAAGAAAGGTCATTTCTATAGTTGTCCTGTTATTTTCTCCATGTTAATTAGTTGAGGGACGGATTTGGGCTACGCTTAACTCATGAACTAGTTTTCGAGGTTTAATTAGGCATAATATTCATTTTCTTCGCGCTAATTAGTCTGTGCATGATTACTTATAAAGATATGAATGATCACAGGTGAAAGAGGGCTGCCCCATGCTGAGCTCACCTGGCCAGTGCGTCTAAATATTATACAAGGAGTAGCTCAAGGACTAAATTATCTTCACACTGAACTTGCTTCTTCTGATTTGCCCCATGGCGATTTGAAATCGAGCAATATACTCATAAGCACAAACCACGAGCCATTACTCACTGGTTACGGATTCTGCACCTTAATGAACAACGCGCATGCAGTTCAAGCACTAATCGCGTTCAAGTCACCAGAAGCAGTACAAAATAACCAAGTTACACCCAAATGCGACGTTTATTGCCTTGGAATTGTGATTCTTGAGATAATAACAGGGAAGTACCCGTCTATATATCTTAATACCGGTAAAGGAGGGATAGACATAGCACAATGGGCAAAATCCGCGATTGCTGAGGGAAGAGAAACTGAATTATGTGATCCACATATAATAGCAAGCGCGAAAGATTCAATGAGTTCAATTAAGGAACTAATTCACATTAGTGCAGCATGTGTAGAGAGTAATCCGCAACAGAGGATAAATATTAGAGAAGTCATTAGGAGGATTGAAGAAATACAACAACAAAACGGAGTAGGCCAAACAGGACAAACAACAACAACAACGCCTCAGTCTCAAACGGGACAAGCAATTGAGATAATGCCGTCTCCAAGAGATGGAGATGCTGAAATACAAGTGTCAGAATCCTAACAAGGAACAACAAATCATTAGGAATTTTAGCAGACTCCGGTAATTCAAGGCGTGGTTAATATCACAGCTTCTCCTATGGTCATCAGACTAATGATAGTGCTTTTTTTTTCTTCATTTTTTAAGGTTCGTTAACTAGGGATAGTACATATATTACTGACTAAGTTTTAGTTTCATTTTGTGTATTTTCCCCCTTCTTTTAGATGCCCCATTATCTCCCCTAGTAGAACAAGAAATGAGAGGAGAAGCTGTATCCTGTGTAGAGATGAAAGAGCTTAATGCTTGCTTTAGAATTTTATATGCAATACAATTTCCTGTTTCTACCATGTTGAGTAGTACTTAACTCTTGTAGACCAAAAATTCCTCAATTACAAAATACAATTATTGACCATGTCACAAGTACGCTAGCTTGTCCGTGCTTTGACTAATTCCACTGGATATAGTCATTTCTAACCAACAACATGTATCAGGTAACTCTTTCTTACTTATTGTTTTTATCTCTGCTAATTTTTGAACCTTAGACCTCATAATTCTCGATCCACTTTGTTGATCACTTAGATCACTATTGGACGTTGTATGTTAATATTTTTTCACACAATAATAAACAGAGAAGTATGGAAGGCCCTATCAGATAACTATAAC

>M6-PGSC0003DMG400023004 M6 LRR-RLK

TTATCTTGAGCCAAGGGTTTATTGGAAGCAAACTCTCTACCTCCAAGGTAAGGGTAAGGTTTGCGTATACTATACCCTTCCCAGAGCCTACTTTGTGGGATTACATTGGGTATGTTGTCGTTGTTGTAGTTCATCCCCGTTGCTCTAAAGGTGGCTTTAAGTTTGTTTTTGGGTGTCTATTTATAACCTCATTTTGATAATGGAGTCTGTCTCACTTTAGTTTATTGTCTTGTTCCTTATTGTACAGTTTTGCAACTATGCTATAACCATTTTACTGGTAGCATACCCACTCAACTAGGGACTCTAACAAAGCTCAATGTCCTGGCTCTGCAATCCAATCAGTTAACTGGTGCGATCCCCGCTAGCTTGGGTGACTTGGGAGTGTTGATGAGGGTGGACTTGAGTTATAATAGTCTTTTCGGTTCAATTCCTTCAAAACTTGCGGATGCTCCCCTACTTGAAGTCTTGGATATCCGAAACAACAAGATTTCTGGCAATGTGCCTCTTGGTAATGAATTTTCTCGCCCTACATTGTACTCTCTTCATTGGAATTTTGAGAGTTAACCAATTTTTTTTTTTTTTTTTATGACATTTCCGTATATGCCTTTTAGATAATTTAGTTGTTAATTATTGTGTTTCATAGTACTTATTACGTAGTTTTCAAATATATAAGTCTTTTTCAAAAACCTTAAAGCTTCTATGTCCGATTCATGGTCAAAGTTTAGACGTTTGAATCTTGAAATTCCAACTGTGCCACATAAGTTTGAGACAAAGGGAGTTATGTATATGTAGCCCCATAAGAGTCGTACGTATAGTTTCTTGGTAAATTATATCTTATCTGGTCTTTTGTTTTCTGTCTTATGAAGCTTTGAAGAGATTAGTTGGAGGTTTCCAGTATGAGAACAATCCAGGACTATGCGGTGTGGGTTTCCCATCCTTGCGAATTTGTACATCTTTGGATCGCTCGAATCCTAATAGACCAGAGCCCTATGGAGTTGGCTCAAGTGGTTTATCTACTAGAGATATTCCAGAAACAGCTAATCTCAATCTAAATTGCAGCGGAAATGCCTGCACAACGTCGTCTAAAACCTCACAGGCATCTGTTGTTGTAGGGGTTATTGTGGTTACTATTATTGCATCTGCTATAGGAATTCTTTCCTTCACACACTATAGAAGACGAAAGCAGAAACTTGGAGGTGGACTGGAAATGTGTGATAGCCGTCTCAGTATCGACCAGGCCAAGGAAGTTAATAGGAAGAATGGTTCTCCCCTAGTCAGCCTTGAGTACTCGATTGGCTGGGATCCGTTGGCTGAGGGCCGGCGTTATGGTGGAGTTTCCCAAGAGATTCTGCAGAGATACAGATTCAATTTGGAAGAGGTGGAGTCAGCTACACAGTACTTTGCTGATAAGAATCTATTGGCTAAGAGCAACTTTTCAACTACATATAGAGGGACTCTAAGAGATGGATCACTTGTTGCTGTTACGAGAATTGCCAAAATTAGCTGCAAGTCAGAAGAAGCTGAGTTTTTAAGAGGACTAAACATTTTGACGTCTCTGAGACATGATAATTTAGTTAGGCTGAGAGGTTTCTGTTGCTCTAAAAGCCGGGGAGAATGTTTTCTTGTTTATGATTTTGTGCCAAGAGGAAACTTGTTGCACTACCTAGATTTGAAGGAGGACGAAACTCGTGCTCTTGAATGGTCCACGAGGGTATCTATCATTAGTGGCACTGCAAAAGGTCAGTTTTTGAATCAACTCTATCTTGTAAGATATAAAGTACAGTAATTTGACATTATATTACTTTTTCTGAGCTTACTGTCAACAATTCAATCATGCCTTTTTATCTATCTTGAGCAGGCGTTGAGTATTTGCACGGGTGCAAGGTGAATAAGCCAGCTCTAGTGCATCAGAATATCTGTGCTGAAAACTTGCTCCTTGACCAACGATTCAAACCATTGCTTTCAAATTCAGGCCTGCATAAGCTCCTCACAAATGATACCGTCTTTTCAACACTCAAGGCAAGTGCTGCCATGGGATATTTAGCTCCAGAGTACTCCACTACAGGCCGATTTACTGAGAGGAGTGATATTTATGCCTTTGGAGTGCTCATTTTCCAAATTTTATCTGGCAAGCGAAAATTCACCAGTTCAATGCGTGCTGCTGCTGAATTGTCCAAATTCCACGACTTGATGGACGTGAATCTCCGTGGAAGGTTCTCTGAGTCTGAAGCAACAAAACTAGCGAAAATCGCTTTGTTATGTACACATGAATGTCCAGAGGAGAGACCAACCATGGAAACAATCGTACGAGAAATTGGTAATTTAGGTAGTCCTTCCTAATGCTAATGATCTTCTTCCATCAACTATTGCCTGTGGCAAATTTTCAATCCTCCTAACTTATTAGCTGGAGATAGATTCGCACTAACCTCTCACGAGTATTGGAGCTTCAAATATTTAGTCAGTCTGAGAACTTCTTCGACTGAGGACTTGACCATTTTGTGAATGTTTGCATCTCGTATGCCATTTAACCTTAGGATTTCTCATGTTGGAAAATTTTCCTCTGTTTATCTCATGGATTCTAATTAATCTGTTTGTAAAAAGATGAATGTTCTTCCATTTACCTAGGATCATTGCTTCGTTTTCATTTTGTGCCT

>M6-PGSC0003DMG400023057 M6 LRR-RLK

ATGTCGATGTTTAACAAATTGTTGTTGGTTTGGTTTTTTGTTGTAACACTGGTTCTTGGATATGTGTTTTCTGAAACAGAGCTTGAGGTTCTGTTAGAGATAAAGAAGTCATTTCTTGATGACCCTGAAAATGTTTTGTCAAATTGGTCTGATAAAAATCAAAATTTCTGTCAATGGAGTGGTGTTTCATGTGAAGAAGATACATTGAAAGTTGTGAGATTAAATCTTTCTGATTGTTCAATTAGTGGTTCAATTTCACCTTCTATTGGCTTTTTGCATGACCTACTCCATCTTGATCTTTCTTCTAATCTATTATCAGGTCCCATTCCACCTACTCTGTCTAATCTTTCTTCTTTACAATCTCTTCTTCTTTATTCTAACCAACTCACTGGCCCCATTCCCAATGAAATTGGCTTGTTGAAGAATCTCCAAGTTCTTAGAATTGGTGATAATGTTGGCTTAACAGGGCTGATTCCAAGTACTCTTGGTGATCTTGAAAATTTGGTTACTCTTGGTTTGGCTTCATGTAGTTTGAGTGGTATGATTCCTCCTGAATTGGGAAAACTCGGGCGGGTTGAGAATATGAATCTGCAAGAAAATAATCTTGAAAATGGAATTCCTGCTGAGATTGGTAACTGCTCTAGTCTTGTTGCGTTCAGTGTTGCAGTTAACAATCTCAATGGATCAATCCCGGAGGAATTATCCATGCTGAAAAACCTTCAAGTTATGAATTTAGCGAATAATAGTCTTTCAGGACAGATTCCTACTCAGCTTGGTGAAATGAATGAGTTGCAGTATCTCAATTTGCTTGGTAATCAGTTGGAAGGTTCGATACCAAAGTCGTTGGCGAAGTTGAGTAATGTTCGGAATTTAGACTTGTCTGGTAATAGACTTACTGGTGAAATTCCTGGAGAATTTGGGAGCATGAATCAGCTACAGTTCTTGGTTCTGACTAGCAATAATCTTTCGGGTAGCATACCGAAAACAATATGTAGTTCTAATCGAAACAGTAGTTTGGAACACATGATGCTATCTGAAAACCAACTTTCTGGTGAGATTCCTGTGGAGTTGAGAGAATGCATTTCACTAAAGCTGCTGGATTTGTCTAATAACACGCTTAATGGTTCGATACCAGTTGAGTTGTATGAGTTGGTTGAGTTAACTGATCTCCTACTTAACAACAACACATTGGTTGGTTCAGTTTCTCCGTTGATTGCAAATCTTACCAATCTTCAAACATTGGCATTGTCGCACAATAGTCTTCATGGCAATATACCGAAAGAGATTGGGATGGTCGAAAATCTTGAGATTCTGTTTCTATATGAGAATCAGTTCTCTGGAGAAATACCAATGGAGATAGGCAATTGCTCCAGATTGCAAATGATCGATTTTTATGGGAATGCATTCTCAGGACGCATTCCCATTACAATTGGAGGGCTGAAGGAGCTGAATTTCATTGATTTTAGACAGAATGATCTCTCTGGTGAGATTCCTGCCAGTTTGGGAAACTGTCATCAGCTGAAGATCCTTGATTTGGCGGATAATCGTCTATCTGGTAGCGTTCCTGCTACGTTTGGTTATCTTCGAGCATTGGAGCAGCTTATGCTTTATAACAACTCACTTGAAGGTAATCTTCCTGATGAACTGATCAATCTTTCAAACTTGACAAGAATAAATTTTTCTCATAACAAACTGAATGGTAGTATAGTTTCCCTGTGTAGTTCGATGTCTTTTCTTTCGTTTGATGTTACAAATAATGCATTTGATCATGAGGTTCCACCTCATTTAGGATATTCACCATTTCTTGAAAGGTTAAGGCTAGGAAACAATCGTTTCACCGGAGAAATCCCTTGGACATTGGGATTAATCCGTGAGTTATCACTGCTCGATCTCTCTGGAAATGAATTGACTGGTTTAATACCTCCACAGCTTTCACTATGTAGAAAGCTCACACATCTCGATCTCAATAACAACCGGCTTAATGGATCAATTCCATCTTGGCTCGGAAACTTGCCACTCTTAGGTGAGCTTAAGCTGTCCTCAAATAAGTTCTCGGGACCTCTTCCTCGAGAATTGTTTAACTGCTCAAAGCTTTTAGTGCTATCTCTAGAAGAGAATTCAATCAATGGAACTCTGCCCCTTGAAATAGGTGAACTCAAGTCCCTAAATATCCTAAATTTCGACAAAAACCAACTCTCTGGTCCCATTCCATCTACCATTGGCAACTTGAGCAAGCTATATATACTCCGTCTCTCGGGAAACAACTTAACCGGTGAAATACCAAGTGAGCTTGGAGAACTCAAAAATCTTCAAAGCATACTGGATTTGAGTTTCAACAACATCAGTGGTCAAATTCCTCCTTCTGTTGGGACACTCACGAAGCTTGAAGCACTTGATCTATCTCATAACCATCTCACCGGAGAAGTTCCTCCTCAAGTCGGTGAAATGAGTAGCTTGGGAAAGCTTAACCTCTCCTACAACAACCTTCAAGGCAAATTGGACAAACAGTATGCACATTGGCCAGCTGATGCATTCACGGGGAATCCACGTCTTTGTGGAAGTCCTCTACAAAATTGTGAAGTCTCTAAGTCCAACAATCGGGGTTCAGGACTAAGTAACTCGACAGTTGTGATCATTTCAGTGATATCGACTACTGTGGCGATCATACTTATGTTGCTTGGAGCTGCACTCTTTTTCAAGCAAAGGCGAGAAGCCTTTAGAAGAGGAAGCGAAGTAAATAGTGCTTACTCATCCAGCTCTTCCCAAGGACAAAAGAGACCACTTTTCGCAAGTGTGGCTGCTAAGCGTGATATCAGGTGGGATGACATCATGGAAGCAACAAATAATCTAAGCAACGACTTCATAATCGGATCTGGTGGTTCAGGAACTGTCTATAAAGCTGAGCTGTTTAACGGAGAGATCGTGGCAATCAAGAGAATACCAAGCAAGGATGATCTCTTGCTAGACAAAAGCTTTGCAAGAGAAATCAAGACACTTTGGAGGATAAGGCACAGGCACTTGGTGAGGCTACTTGGATACTGTAACAACAGCGGAGAAGGTTCAAATGTATTGATCTATGAGTACATGGAGAACGGAAGTGTGTGGGATTGGCTTCACAAGCAGCCAGCTAATAATAACAAGAGGAAAACATGCCTCGATTGGGAGGCAAGGCTGAAGATAGCGGTGGGATTAGCACAAGGAGTTGAGTACCTTCATCATGATTGTGTACCCAAGATCATTCATAGGGACATCAAATCAAGCAACATTCTACTAGACTCAAACATGGAGGCACATTTGGGAGATTTCGGGCTGGCTAAAGCCGTTCATGATAACTATAATTCCTACAACACTGAATCAAACTTGTGGCTTGCTGGTTCATTTGGTTATATTGCTCCAGGTATATATCATTCCTGAATTCTCGAGTATTAGCATTCGAAAATATTTTTTTCTTTTTTATACTAAGTTGCTTATTTGTTCGCAGAGTATGCTTATTCTTCGAAGGCAACTGAGAAGAGTGATGTTTATAGCATGGGGATTGTACTGATGGAGCTAGTGAGTGGAAGGATGCCAACTGATGGGAGTTTCGGAGAAAACATGGACATGGTGAGGTGGGTTGAATCATGTATCGAAATGTCTGGGACAATTAGGGAAGAGCTCATTGATCCTGTTTTGAAACCATTGTTACCTAATGAAGAAAGTGCTGCACTTCAGGTGCTGGAAATAGCACTTGAATGCACCAAAACTGCACCTGCTGAAAGGCCATCTTCGCGTAAAGTTTGTGATCTGTTGCTACATGCATTCAACGACAAGGTGGTTCATTCTGATAAGATGAGTCCAGACAATTACGTCTAG

>M6-PGSC0003DMG400023192 M6 LRR-RLK

ATGACAACTTTGTCATATTTTTTCATTATATTCTTCTTGTTAAGTTGTTTGAGTCCAAGAAGAGTGAACTCTCTGTCTTCTGATGGAAAAGCTTTGCTCTCTCTTCTCAAAGCAACTTATGACCCTTATGCTAAATCATCATCATCTTTTGTACTTTCTTCTTGGAATGCTTCAACTTCAACTCCATGTTCTTGGCAAGGCATTTCTTGTTCTCCTCAACAAAGAGTTATTTCTGTTTCTATTCCAAATACATTCCTCAATCTTTCTTCTTTTCCCTTTGAGCTTTTTTCACTTTCATCTCTTCAGCTTCTAAATCTTTCTTCTACTAATATATCTGGCTCAATACCTTCATCTTTTGGTTTGTTTACACATCTTAGGCTTCTTGATCTTTCTTCTAACTCTCTTTCAGGACCTGTTCCTTCAGAACTTGGTGGACTTACCTCACTTCAGTTCCTTTTCTTGAATTCAAACAGATTGAGTGGAAGAATCCCATATCAACTTGCTAATCTTTCTTCACTAGAAATCCTCTGTCTTCAAGATAATCTCCTTAACGGGTCAATTCCGAAAGATTTAGGCTCATTGGTATCACTCCAACAGTTCAGGATTGGAGGGAATCTAGAGTTGAGTGGTGAAATACCTGCAGAGCTAGGAATGCTCACCAATCTTACGACATTTGGTGTCGCGGCTACTGGACTTTCAGGGGTTATTCCACATACATTTGGTAATTTGATCAGTCTACAAACTCTGGCAGTGTATGATACTGAAGTGTTTGGTTCAATACCACCTGAACTTGGGATGTGTTCAGAGCTCAGGAACTTGTATTTGCACATGAATAAGCTCACCGGTCCAATCCCTCGTCAGTTGGGTAAGTTACAAAAGATTACTAGCTTGCTTTTATGGGGAAATTCGCTTACTGGCCCTGTCCCGGCTGAGTTATCGAATTGTTCATCCCTTGTGGTTCTTGATGTCTCTGCAAATGACTTGTCTGGCGAAATTCCGGGTGATTTAGGGAAGCTTGAAGTTCTTGAACAGCTTCACTTATCTGATAATGCACTTAGTGGGGCTATTCCAATGCAGTTGAGTAATTGTAGCAGCTTAACAGCTCTTCAGTTGGATAAGAACCTTTTCTCCGGCACGATTCCTGAGCAAGTTGGTGAGTTGACGTACTTGCAGATTTTCTTGTTGTGGGGGAATTCGGTTTCAGGAACTATCCCGGCTGCTTTCGGAAACTGTACTGAACTGTATTCGCTTGACCTCTCAAGGAACAATCTCACTGGTTCAATACCTGAAGAGATATTCAGTTTGAAGAAGCTGAGTAGGTTGTTGCTTCTTGGGAATTCGTTAACGGGAAGATTGTCTCCAAGTGTGGCAAAATGCCAGTCTCTGGTCAGGTTGAGGATTGGCGAAAACCAGTTTTCTGGACCAATACCTGATGAGATAGGACAGCTGCAGAATCTTGTGTTTCTTGATTTATACATGAACCATTTCTCTGGTGAATTGCCTTCTGAAATTGCCAACATTACGGTTCTTGAGCTGTTGGATGTGCATAACAATTACCTAACCGGGGAAATACCATCATCGCTGGGAGAGCTTGTGAATTTAGAGCAGCTTGATCTCAGCAAGAACAGCTTCACTGGTGAGATTCCGTGGAGTTTTGGTAATCTCAGTTACTTGAACAAACTTATTCTTAGCAATAATCTGCTTACTGGTCCAATTCCAAAGTCATTTAGCAACTTGCAGAAGTTAACTCTACTTGATTTGAGCTCCAATAGTTTTTCTGGTGCAATTTCACCTGAGATTGGTTACATGACAAGCTTAACAATCAGTTTAGATTTGAGCTCAAACCGATTCACAGGTGAACTCCCTGAGACGTTATCCGGATTGACACTGTTGCAATCTCTTGATATTTCTCATAATATGTTGAGTGGAAGAATCACAACTCTAAGCCTCTTAACCAGTCTCGCTACTCTAAATGTATCGTATAATAATTTCTCAGGGCCTATTCCTGTAACACCTTCCTTTAGAACTCTAACATCAAACTCTTTTCTTGAGAACTCACTCCTTTGTGAATCAATTGATGGCTTCACTTGTTCTGCACATATAACTAGAAGGAACGGATTGAAGTCTGCAAAAACCATAGCTTTGGCTGCAGTGATTGTGACTTCTGCATCCATAACAGTCGTGGCAACGTGGTATCTTGTGACAAGGAAGCATAGATATGAATTTGAGAAATCACCTGGCATGTCGGTCTCTGTAATAGGGACAGAAGATTTTTCCTATCCGTGGACTTTCATCCCTTTCCAGAAGCTCAATTGCACCGTTGACAACATCTTGGATACCTTGAAAGATGAAAACATTATAGGGAAAGGCTGTTCGGGGGTTGTTTATAGAGCAGAAATGCCAAATGGCGAGTTGATTGCAGTGAAGAAGCTCTGGAAAACGAAGAAAGATGAGGAACCAATAGATTCTTTCGCTGCTGAAATTCAGATTCTCGGTCACATTAGGCATCGAAACATAGTGAAGCTCCTTGGATATTGTTCAAACAAGAGTGTTAAGCTTCTTCTCTACAACTACATCTCAAATGGTAATCTTCAACAGCTCTTGCAAAGTAACAGGAACTTGGATTGGGAAATCAGGTACAAGATTGCGGTTGGATCAGCTCAAGGCCTTGCTTATCTTCACAATGACTGTGTGCCAGCAATTCTTCATAGAGATGTCAAGTGCAACAACATACTCCTCGACTCTAAATTCGAGGCTTATTTAGCAGATTTTGGACTTGCAAAGCTGATGAATTCTCCAAATTATCATCAAGCTATGTCCAGAGTAGCAGGCTCTTATGGCTATATAGCTCCAGGCAAGTATCTTTCAACTTTATTTTCTCATCTTAACCTAAATGTTGTGCAGCTATATGATTCTTTGAATCCTTGTGTTGAATCGTGTGATCATCGCATCTCTAAAAGCTTAAGTTATTAGAGAAAGGATACGCTTTTGTATACACTTTTCTCTACTAAATCATATCTTCAACACATGTGGCTCTGATTCTTGCCAAGCACGTGGATAATGGGTGACGCTTAGACTTCAACCTAGGATACCATGTTGAAATGTGTGACTCTCTCATTTAGTTTAAGTTATTAGAGATGGTACACATTTATTTACTTCATTATATCGTCAACAGTTCCCAGATATAGGTCAAAACGAGAGTCACAACGTTATGAGCTATAGTCAAATTTGGAAGGATAAACTAAACAAGTTATGCTTCATAGATGTTAACACTATATTTTACCACTATGGTTGCAGAATATGGATACACAGTAAATATAACAGAAAAGAGCGATGTCTACAGTTATGGAGTGGTGCTGCTGGAAATATTGAGTGGACGTAGTGCGATTGAGCCTCAGATTGGTGATGGACAACACATTGTCGAGTGGGTGAAGAAGAAGATGGGAAGCTTTGAACCAGCTGTTACAATTCTTGATTTGAAGCTACAGAGTTTACCGGATCAAATGGTGCAAGAAATGCTTCAAACACTTGGAATAGCTATGTTCTGTGTGAATTCATCACCCACTGAAAGGCCAACTATGAAGGAAGTAGTGACACTTCTAATGGAAGTGAAGAGCCCAACTGAAGAATTTGGAAAAACTTCTCAGCCTTTAATCAATCAGTCAACTCAAAGTTGA

>M6-PGSC0003DMG400023282 M6 LRR-RLK

TCATCTCTCACCAATAATAAATATCATGTAACTCTATTTACCAAAGTTAAGGTAGATGAAAAGTAATAACCTAGTGTTTTTTTGTCTCTACTGGGATTTTAGCTTACCTCATAGTTATCACTTACTTCATTGACCAATAGGCCACAACGTTGGTGCAATAAGTTTAGAACTATTCAGTTCAAGACTTTAATCTTTTCTGGTGTTATTTTCTATCCCACTATAGTCAGATCCTTCCTGTGCTTCAGATATGGAGAATTTGATTGTAGGAAAGATGAAAAAACAAAGGCCGCTCTTTAGTATCTTTCCTTGTAACCAAGACTATAAAGTCAAGTGAAACATTATATACTTGAGTCACTTCAAAGAAAGATGTAAATTCATTGGCAATATAGTAAATCCACAACTAAATACAAATTGACATGTTAAACACATCAGCTAATCACCCAACATCCTGTATGTGTTTGTGTATATCTGTTTAACATCTTTCATATGTGCTTTACTGAATTCCTAAGTTATCTTATTCTTTAAGTGAAATTTTTTAGTTTCTTATAATAGATAAATTATACAACAAACATTTGATTAGGTATACATTGCTTTTAAGTAAAAGGGGTTTGTTATATGTATATTCTTGTTTCTATACTTTATTGTTTGGTTACTGATTGCTACTTTTCTTCTTTCAGTTTCTCATTAAGAATGGGCATCCATCATATCTGTTATGGAGTTTGTCATCCCCTTGTGAAATGACTGTTTAGCCCTTCCCATAATCAATCATGTCTAGAATCCAAACTCTCCATTCCCTCCAAATCGTTGTTATTTTCTTGAAGTTTCTTGTTTTCTCTAATTCATTGTCTGTTGAGACAGAAGCACTTCTTGAATTTAAAAAGCATCTTGTTGACCCTTTGAATGTCTTGGAATCTTGGAAATATTCAGATTCCCCTTGTAAATTCTATGGCATTCAATGTGACAAACACACTGGCTTGGTCACTGAAATTTCACTTGACAATAAGTCCCTTTATGGATTTATTTCCCCTTCAATATGTGTCCTCCAAAGCATTACTTCTCTAGTGCTTCCTTCAAATCACCTGTCTGGAAATCTTCCAAGTGAATTGGCGGATTGCGCCAATCTCAAAGTATTGAATGTTACTGACAATAACATGAACGGTGCGATACCTGATTTATCTAGCTTGGCTGAGTTGGAGGTGTTAGACTTGTCAGATAACTGTTTTTCTGGAAAGTTTCCAGCTTGGTTTGGAAAGTTGACTAGTTTGGTTGCATTAGGCCTTGGTGGAAATAAGTATGATGAAGGTAAACTTCCTGATTTGTTTGGGAAGTTGAAGAAAGTTTATTGGCTTTTCTTGGCAGGGTCAAATTTGACAGGACAAATCCCGGAATCTATTTTCGAAATGGAGGCACTAGGTACATTAGATATTTCGAAAAACCAGATATCCGGGAATTTTCCAAAGTCGATAAACAAGCTGCGAAATCTCTTCAAGATTGAACTTTATCAAAACAATTTGACCGGGGAATTGCCAGTAGAACTTGTAGACCTTATACATCTGCAGGAAATTGATGTATCTAGAAACCAGCTGCATGGGACTTTGCCTAAGGGGATTGACAACCTGAAGAATATAACGGTTTTTCAAATATTCAAGAACAATTTCTCTGGGGAAATCCCTCCAGGATTTGGAGATTTGCAGCATCTTAATGGCTTTGCTGTGTACAACAACAGTTTTACTGGAGAAATTCCAGCTAATCTTGGCCGGTTTTCGCCACTGAATAGCATAGACATATCTGAGAACAAGTTTAGTGGTGCATTTCCAAAATATTTATGTCAAAACAACAACTTGCAAAATTTGCTTGCTGTTGAAAACAGTTTCACAGGGGAGTTTCCAGGTAATTATGCTTCTTGCAAGACTTTAATGAGATTGAGGGTGAGTCAGAATCAACTCTCTGGGAGAATTGCTGAGGGTTTATGGGGGCTTCCAGAAGTCACAATGATAGATTTCAGTGATAACAACTTCACTGGAACTGTATCTCCTGGAATCGGCGCTGCGACTAAGTTGAATCAGTTGGTATTATCAAACAACAGATTTTCAGGTGAGCTTCCAAAAGAACTTGGAAAGCTCACACAGTTGGAAAGGCTATATTTGGATAACAATGATTTCTCTGGCATAATACCATCTGAACTTGGCACATTGAAGCAAATTTCATCTTTGTATTTAGAGAAAAACTCACTTTCCGGGTCGATACCATCGGAGTTGGGTGAGTTCCCAAGGTTGGCAAACTTGAATCTTGCTTCAAATCTTCTCACAGGCAACATACCGAATTCGTTGTCAATGATGGCCTCGTTGAACTCTCTGAATCTTTCGAGTAACAAACTCAATGGCTCAATTCCAACAAGCTTGGATGACCTGAAGTTATCTTCATTAGACCTCTCTAACAATCTGCTAACTGGAAGAGTTCCTACAGATCTTTTGACGGTGGGAGGAGAAAAGGCATTTGTAGGAAACAAGGGGCTTTGTGTTGATCAAAGCATCAGGAATGTCCGAAGGAACTCAGGCATTGGCGCTTGTAGTGGAAAGGCTGCTCAGGAAGTGTTCATGAAGAGCAAACTAGTTGTGCTTTGCATTGTATTACTTTCTTTGGCTGTTCTAATGGGTGTTTTCATGCTTTTGAGTTACTGGAAGTACAAGTGCAACGCTGAAGCTGATTCAGAAAAGTGTCTGGGACATGCAAATGGAATGAATCCAAAATGGAAGCTTGAAAGCTTCCAACATGTGGAACTTGATGTAGATGAAATATGTGATGTTGGTGAGGACAAGCTGGTAGGAAGTGGAGGCACAGGAAAAGTTTATCGGTTAGATTTGAAGAAAGGTTGTGGCACAGTGGCAGTAAAGCAGCTTTGGAAAGGAAATGAGGTGAAAGTTTTGACAAGGGAAATAGACATTCTAGGAAAGATCAGGCATCGAAATATAGTTAAACTGTATGCCAGTCTAATGAGAGAAGGTTCGAATATGTTGGTTTTCGAGTACTTGCCAAATGGTAATTTATTCGAGGCACTACACCGGGAAGTCAAGGCTGGGAAGACAGAATTAGATTGGTACCAGAGGTATAAAATTGCAGTTGGAACTGCAAAGGGAATTGCTTATTTGCACCATGATTGTGTCCCTCCTATTATTCATAGAGATATCAAGTCAACAAACATTCTACTTGATGAGGAGTATGAAGCGAAAGTTTCTGATTTCGGGGTTGCAAAAGTTTCAGAAATTTCCTCTAGAGTCTCCGAGTTCAGCTGCTTTGCAGGCACTCACGGCTATTTAGCTCCTGGTATGTTTCAAAGAGGAAAAACCAAATACTATGTTCCCACTTCTTTCTATTTTCACTCTTGCTAGTTTCCTAACATCACATTTGAATTTGTTTTATTGACAGAGCTAGCATATACTTCCAGAGTGACAGAAAAGAGCGATGTATACAGCTTTGGAGTTGTTCTATTAGAATTAGTAACTGGAAGGAAACCAATAGAGGAAGCCTATGGAGAAGGCAAAGACTTGGTTTATTGGGCATCAACTCATCTAAATGACAAAGGAAGCGTTCTGAATATTCTTGATCAAAAGGTGGTTTCCGAGCTTGTCCAAGATGACATGATCAAAGTGTTGAGGATTTCTGCCCTATGTACGACTAAGCTTCCTCATTTACGCCCCAGCATGAAAGAAGTAGTAAAGATGCTTGTTGATGCTGAACCTTGAACATTTAGTTTTCAAGCAAATGTGAGAAAAATGAGAAAGATCCTTTGTAGTTTTCTCCTTTTCCCTCTTACCATATAGTTTAAATTCCAATCTTCAGTATAGTTTTTGTTTTATGATTATGCATATGCCATGTAAAATTGTATCCAACTTTGTCTGTACAAGTTCATTTTGTAAAATGGAATGCATTTTTCTGGTTGAGCTTATAA

>M6-PGSC0003DMG400023283 M6 LRR-RLK

ATGATTGTCTTGAACTCTAGTGAATTGTTTGGTCCATTGTCACCAGCAGTTGGTAATCTTAGTTTCCTCAGGGTACTTTGGCTCAGCAGAAACAGTTTCACTGGCAAAATCCCTGGAGAGATAGGAAAGCTATCGAGATTACAGAGGTTGAACTTAGCAAACAATTCATTTTCTGGTGAAATACCAAGAAACATATCGCGGTGCTCCAATCTTAATTATATTCATTTAGGTAATAACAATCTGAGTGGAAGCATTCCAACAGAGTTGAGATCATTGAATAAACTTGAGAATTTACTACTTCAGTTGAATAAATTGACTGGAGAAGTTCCAGCTTATCTAGGTAATCATTCTTCACTCTTGGCAATTGCATTAGGAGATAATCAATTACATGGAAAGATACCTGATATTTTTGGTGGACTCAAACATTTGGTGTTTCTTGATTTTGCTTTGAATTATTTGTCTGGAGAGATTCCTATGTCGATTTTCAACTTGTCATCATTGCACACACTTCAGATAACTTTTAATAAACTTGAAGGGGGTCTTCCATCAGATATGGGGTTCAATCTTCCAAGTTTAAAGTTCTTTCTCCTTGTGGGAAATCAACTGACAGGAAAGGTTCCTACTTCAGTACTGAATTCTACATCACTTATGGAATTTGGTATCGACGGGAATAATTTCACTGGACAAGTACCAGCCTTTGGAAACCAAAAGGATTTGTACTGGCTTGGACTTGCTGAAAATCATTTTGGAAATGGAATGTTGGATGATTTGAAGTTTATGTACTCCCTACAAAATTGTACTAGTTTAGAGCAGCTAATACTGGAAAACAACAATCTTGGAGGAGTACTGCCAAGCTATATAGGCAACATGTCCAATCTTTTGCGGCTTTCTGTAGGGGGGAATCTCATCCAAGGAAACATTCCCACAGAGATTTTACAGCTTGAAAACCTGCAAGTACTTGGTCTCGATAAAAATAATTTTACTGGTATGATTCCTGAGTCTATTGGACAACTAAGCCAACTTAGTAGGTTATACATTCACCACAACAGATTTTCAGGTGAGATTCCACATTCTCTTGGAAATTTGACGAGGTTGATTGAACTCGATTTGGGATCAAACAACTTGCAAGGTACTGTGCCTTCAAGTCTTGGAAGCTGCAAGTTCTTATCACTCTTGTATCTGAATGGAAATCAACTTAGTGGTCTTATACCAAAAGAACTGTTTGGGCTCTCCCTTATAGAGTTTGACCTCTCAAACAACCACTTGACGGGGTATTTTCCTGTAGGGATTGGCTCTGGAAACTTGACTGGACTGATAAACTTGATCTACATGAACCATTCATACAACAATTTATCAGGTGAAATACCAAGCAGCTTTGGCACTTTGACTAGTTTGAGGGAGCTATACCTTGGTAACAATGCCCTTCAGGGAGTTATTCCTGCATCCTTTAGTTCTTTGAGAAGTCTTGAATATATTGATCTTTCTCATAACCACTTTGTAGGGAGGATTCCAAAGTTTCTGGATGAACTTGTCTCCTTGAAGTTCTTAAATTTGTCTTATAATGATTTGGAGGGTGAGGTACCTTTGAAAGGAGCTTTCAGAAACATGAGTGCTGTTTCACTTGTTGGGAACAGTGAGCTCTGTGGGGGTATTCCAGAGTTCAAAATGCCAAAGTGCAGCAACAAAGTAGCGTCAAGAAGACGAAGGTTATCTCATAGATTGATAATAGTAATGCTGGTTATTGGAGGATTATTGGCAGCAACAACAGTGGCCATTCTTATTTTCTTGTGTGCTAGAAGAAAAAAGAGGTCTACTTCATCAGAGAATTCATCGTTGGACGTGATCCCAAGAGTGACTTACAATAGTCTTTACAAAGAAACTAATGGCTTCTCTACGTCAAACATGATTGGTTCGGGTGCATTTAGCTTCGTGTACAGAGGCATTCTAGAGGAAAATGGAAAATTTGTTGCCATAAAGGTGCTCAAACTCCAAGTTAGGGGAGCTTCTAAGAGCTTTCTTACTGAATGTGAAGCACTAAGGCATATCAAGCATAGAAACCTGGTAAAACTCTTAACATCATGTTCAAGCATTGATTACCAAGGTAACGACTTCAAGGCTCTCATTTATGAATACATGGCCAATGGTAATTTGGCAAATTGGTTGCACAATAGAAGTACAGATGGTGAAGAAAATCATGAGCCTAAAACTTTGCACATGCTACAGAGACTCAATGTAATAATTGATGTGGCTTCTGCATTAGATTATCTTCATCATCAATCTGGAACGCCATTGACACATTGCGATATCAAACCAAATAATGTGCTTTTGGATGAAGATTTTGTTGCTCATTTAGGTGACTTTGGACTGGCAAGGTTTCTGCCTGATGCTGCAAATCTGCTCTCCTTAAGCCAATCAGCCAGCTCTCTCAATATAAGAGGAACTATTGGATATGTTCCACCTGGTAATTAACTCTTCTCCCTCTCTAAACATCATTTTATATTTATGAAGGTCCTCTTACTTATCAAGTTTGTCAATTTATGTATGTGCAGAATATGCTACAGGTAATACGTTTTCGACCTATGGAGATGTCTATAGTTATGGAATTCTACTTTTGGAAACATTTACTGGAAGAAGTCCAACTGATGAGATTTTCAAGGATGGTCTAAACCTTCATGATTTTGTTAAGAGAGCCATACCTGAGCAAGCGAAAGATGTTTCTGATCCTAAACTTGTCTATGATGAAACGGGCAGACTGATCAGCAACAATAAAACAATCGAGTGCCTCACATTGATTTTTCGAGTTGGGATTGCTTGTTCAGTGGACTCTGCAAAAGATCGCATGGACATTGCCAATGTTGTTAATGAACTGAATGTAATCAAAGATGCCTTCTTGAGGAACTAGCAATATTGGCTATGAAGTTTTTGAGTTTCATAATAAAGATACTACTTGTATAAGTTATATTCTCACTATCCACTTTTCCCTTCCCCTTTTCCTGCTTCTTCTTATGTAATGGAAA

>M6-PGSC0003DMG400023329 M6 LRR-RLK

CAAAATTGTTCTATTCCTACTTTCTTGAATGCTTTGTGTGTCCCACTTTACACATTACACTAAACCCCCCAATTGCCATTTGAATCTCACCTATTCATCATCATGTCATCAATTATCTACTATTACTCTTCCTTGTATATCTTAATCATTCATTCTACTCTTCAAAATCCATGGCTATTAGTCCAGGTCACTTTTCTTACTAACTTGTATATTTACATATTACTATTTGCATATATACATGTTGTGAACCTCATATAGTCGACCTCAACTTGTTAGGGACTGAGGCGTAGTAGTTCTTATACATGTTGTGAAGCTTATATAGTCGACCTCAACTTGTCAGTGACTGAGGCGTAGTAGTTGTTATACATGTTGTAAAGCTTATATAGTCGACCTCAACCTGTCAGGGACTAGTATTTGTTATACATGTTGTGAAGCTTATATAGTCGACCTCAACTTGTTAGGGACTGAAGCATGGTAGTTGTTATACATGTTGTGAAGCTTATATAGTCGACCTCAACTTGTTAGGGACTGAAGCATGGTAGTTGTTATACATGTTGTGAAGCTTATATAGTCGACCTCAACTTGTTAGGGACTGAGGCGTAGTAGTTATTATACATGCTTTTATATAATTTTCTTGATATTTTTTTGGGTATATTGAACTAAGTTAATTTTGCCTTAGCTTTTAAATGAACTTATGTTAAGGGGATTCAACAGTCTAATATATATATATGTGCCTATCATTTAACATACGTATGTCATGTATTTTTTGTACCTTTGTCGTCCCTCAACCCCTCTCCAATCTTTTCTATCTATCATTAGAAGTAGGACGAGTAATATTCCGGTGAAGGGAATAAAAATTAGTTTAAAGCCTTTATGTTTTGATGATCATTTGAGAAATTAACTCCTAGGAAGACTAGTCACTCAGAAATAGTCATTTTGGTTCTTTTGTTCTTGCTGAGTATGAGCATGAAATATATTGGTAGCACAAAAAGAAGTGGATACTTCTTGAACTAATTGGTAATATTTTCATTGTGTTGCAGCTACAAGGCCTATTTCTTTGACATCATATTCTCCGACCTAGACTTGTTTGAGATTGAGGCGTAATTGTTGTTGTTATATAGAGTATCGGAAACAAGGAGAAGGATCAACTGGAAGCTCTCCTCTGGTTGATAGAGAAAATTCAATGTTTACAGCCTTTTCGTTCCGGAATTTTCACTTGAACTTGCTGATATTCCTACTTTTATGCAGCAATGGTCTCTCCTCAAATGATTATGATGTTGAAGGTATGGAGTTATAGGTTTTGGTTACTGTACCTGTGAAAATTTCATATGCTTGCAAAATAATGTGCTAATAAAATCGCGATAACTGTTATTCTTTATTGAAATCATATATCATATATACATCATTTCAGGTCATGCTTTAATTGAGTTGCTGAGGGCTCTGAATGATTCACACAATAGAATCAAAGATTGGAACATTGATTTTGTGACTCCATGTTTCAGTTGGTCTCATGTTACATGTAAAAATGGAAATGTCATATCCCTGTAAGACTAATTCTTTCCTTTCTGTTATAAGTAAATCAAGTAGATTCATATAGTCGACCTCAACTTGTTTAGGACTGAGACATAACTATTGTTGTAATCAAACAGGAACTTAGCTTCTAATGGATTCTCAGGAACACTTTCACCGTCGATAACCAAATTGAAATTTTTGGTTAGCTTGTAAGTATACTGCACTTTGTATTAAAATTAGAGTTGCTGGTAAAAATCACACCTATAGTATCCCCTTTTTTCGAGTTCCCTATCACCAACTATTTATCAAAAACACACCTCAACTATTAGTTGTTCCCTTTTTCCTACCTGAGATATCACCATCGTTCAGGCAGGAAAAATGAAATGTTGCTATTACTCTATGTCTCGCGAACCTAGGAAAATTTTGTGAAGTGTTGCTAAAATGTTAATGGAACTGTGCAGGGATTTACAGAACAATGATCTATCTGGTTCCTTACCTGATTACCTTAGCAGCATGTCAAATCTACAAAACTTGAATCTTGCAAACAATAATTTCAATGGCCCTATTCCACCAGATTGGGGTCAGCTCTACAAACTTAAATATTTGTAAGTACTTGTTTTTCTTTGTTTCTTTGTTCTCTCCGTTTTCCTAAAATATCAAGGAATGTCACTCGGACTCTTCAAAAATGATGACAGGTTTGTGTCAGGTCCTCCAAAAGCAGTGCATTTTTGGAGGATCCAACACGGGTGTGACAACATTTTTTGAGAGTCCTAGCAGACCTTAACATAGCTCTTATTAAATAGTTCTTGTAACAAAGTCTTTCTATGATATGTCCTTTTGTCCTGCATCAGGGTCATGAGAGGAAATCACATAACTGGAACTGTTCCAAATTCACTTATAAACATCACAGGCTTAAAAGAACTGTAAGAACTATGTTTACCTTACGGGCGTAATTTACTGTTCGTTTGAAATTTTCGGTTGGATTGAGGCCTATGTTGCTCAGACTCTTCAAAACTACTATAGTATGTGTGTCGGACCCTTCAAAAGGTACCATTTTGGAAAGTCCGAGCAACATAGAATGAGTTGTTATTTGATGTTTTCATTTATTTCTGGTTGTAGTTCTTACTTGCTTTAATTCTTCACAGGGACCTGTCCTCCAATGATCTGACTGGAGGTATCCCACTGCAGTTGTTTTCAATACCAACATACAGGTAATGTCATCATTTATTCTTCTTTCTTGCCAACTTGTTTAGCCTTTCCATTTTGATGTTGAGCTCTCAAAAAGGGCTACTCATGCACAAAACAGACATGGGGGTGTGACATAGGCAGTCTACCCTGGTGCTAGCATCAGTAGCTTATTCCACAGCTCAACCTCATGGCCTATAGGTCATATGGAGACAACTTTACGTTGCTCCAAGGCTCCCTTCAACTGATAAAAAAAAATGAATTTGTTACTCATGACTGCATTAATCAACTTGAAGCTATTATTATTGACTTAGTTGCTTAACTATGTTGTTTTAACAATCATTCTATACGTCGTAATCAGTTTTTCAGGAACTCATCTTGCTTGTGGCAATAGATTTCAGAAACCCTGTGTTTCTAGTTCCTCTGCTCCAGGTTAATAGTCTTGTCACCAATGCATCTTTTAGTTCTTAAATCTTAGCCAACCAAATGATTATTGCTCAAGTGATATTTTTTTCAACAGCTTCTAGCAGAAGACCAAAACTTGAGGTTGCTATTACTGGTGCAAGTTGTGGTGCATTCCTTCTTCTCTTAGTTGGGGCTATCATCACATATCGTTGTAATTATAAACGTAAACTCAAACAAGATCAATTTTTTGATGTGGAAGGTATATGCTTTACTCCTATTTCTTCAACAGAAAGAGTACATGGTTTTTCTGTACTGAACAAATTTCACTTCAAATGCCATCTACTCATATGGCATTTTAGGTGACATTGAGCGCAACATTTCATTAGGACAACTAAGAAGATTCTCATGGCGCGAACTTCAGATTGCAACAGACAATTTCAGTGAAAGCAACATTATAGGACAAGGAGGTTTTGGTAAAGTTTACAAGGGCTATCTTTCAGACAACACAAAAGTGGCAGTGAAAAGACTCACTGATTACCACAATCCTGGTGGAGAGGCTGCATTTCTAAGGGAAGTGCAGCTGATTAGTGTTGCAGTTCATCGAAATCTCCTACGTTTGATCGGGTTCTGTACAACCTCTTCTGAGAGAATTCTTGTTTATCCATTTATGCAGAATCTTAGTGTTGCATATCGGTTAAGAGGTATGCCCCCTATCACTCAACCACTTTTTTTGACATTCTTGATACTATTCAGTCTGTTAAGTCTGTGTTTGTGTTTTTTTCCTCTCTTTTCGACAGATTTAAAACCTGGAGAGAAAGCTCTGGATTGGCCAACGAGGAGACGGATAGCATTTGGAGCAGCACATGGTCTAGAGTACCTACATGAGCATTGTGATCCTAAGATCATTCATAGAGATCTAAAAGCAGCAAACATCCTCCTAGATGATAACTTTGAACCTGTTCTTGGAGATTTTGGACTAGCAAAGCTAGTTGATACAAAACTGACTCATATTACAACTCAAGTTCGCGGAACAATGGGCCACATTGCTCCAGAATACTTGTCTACCGGAAAATCATCTGAGAAAACAGATGTATTTGGATATGGTATTACTCTTTTAGAACTTGTAACAGGGCAACGTGCAATTGATTTTTCGCGTCTTGAAGAAGAGGAAGATGTCTTGCTACTGGATCATGTAAGTCCCTACTTCCACACTTTCCAATTTTTGAAGAGTCATAACGGATACCTTTTTACTTCTACAGATCAAGAAACTACTAAGAGAGAAAAGGCTAGGTGACATTGTGGATGGAAACATGAAAACTTATGAACCCAAAGAAGTTGAGACAATCCTTCAAGTTGCATTGCTTTGCACCCAAAGTTCACCTGAGGAACGTCCGAAAATGGCACAAGTAATCACCATGCTAAAGGGTGTAGGTTTAGCTGAAAAATGGGCAGAGTGGGAGCAACTTGAAGAAGTAAGAATTCAACAATTTTCGCTCATGTCGCGACAGTTTATGTGGGCTGAAGACTCAACACATGATCAAGAAGCTATACAATTGTCTCAAGCAAGATAAACAAAAAGAAGATGCAGAGAAATGTATGCATAAGTACTTCTGCAATAAAATTTAAGAGTAGTTTTAGCTTCTGCTGCAGCTATAGCTAGTTATGTAACATGTAAGAGTAATTTTATCATACATCTAGCTAGTTATGTATCATGTAAGAGTAATTTTATCATATACGTATATAAGGCACTAAGGTGTCTTGTTTAATTTTACAATGTATATATATATAGAGAGATAGAGAGACCATCTATTACTGAATTTACTCGATTCCTAGCTCGTTGTACTAAAATTCCTGCTCGATATTTATATTTAAGTTCC

>M6-PGSC0003DMG400023550 M6 LRR-RLK

ATGTTGAGATATCTAGAAGTGGGTAACAATCCGTTGAGTGGCGTTCTACCAAATTCTGTTGGGAATCTCTCATCTACTATTGAAAAGATTTATATAGGAGATGCACACATCAATGGCCTCATCCCCACAAGTATAGGCAACATGAGCGGTCTTACAGCCCTAAGCCTTGGAGGAAACAACTTGGCAGGGAATATTCCTTCTGAGATTGGTAAGCTTAAACAACTCCAAGGGCTGGCTCTATATAACAATAAATTGCAGGGACGTATTCCAGAGGTGGTATGCCATTTATCGAATTTGGTTGGCTTATTACTGGATGGTAATGAGCTCTGTGGATTAATTCCAGAATGCTTTGGAAATCTTAGCATGCTACAAGCGCTTTATTTGGATTCTAATAAATTTTCATCAAAATTTCCGTTGAGCCTTTGGAAGATGAGTGGTCTTCTCTATCTAAGTGTGTCACGAAATTCAATAGAGGGAGAACTTCCATCAGATATTGGAGGACTGAAAGCCATTGTAGAACTATATCTTTACAATAACCACTTTTCAGGTGTGATACCAACCAGATTGGGGGAACTCCAAAACCTGCAGCATCTTGACCTATCCAACAATTCATTTTTTGGCCAAATTCCATTATCCTTTGCCAACTTGATAAGCTTGGAATTCTTGAATTTGTCTTTAAATGCCTTGTCCGGTACTATTCCTAAATCTTTGGAAAAACTCTCATACCTTAAAAGCATTAATGTTTCATTTAATGATTTAGAAGGTGAAATACCAAGTGGTGGCGTGTTTGCAACTTCCACTGTGCAATCATTTCTTGGGAACAAAGGTCTTTGCGGAATGCACATATTGGAGATTCCTACTTGTGCTATCACTAATCATGGAAAACAATCAAAGCTTAAGGAGGTTGTGCTAAAAATTGTTACTCCAGTGGTTATTGCATCCTTTCTGATATTCTTGTTCGTTTCAATTTGGATAATGAAACGACAGAAGAAAGGGAAGTCCAAAGATATTGAAAAGGTATTGGAGATCGGGACTCATCAATTAGTTTCTTATCACGAGATTCAACAAGCAACAAATAATTTTGATGAATCCAATTTAATTGGTGAGGGAAGCTCTGGCTCTGTGTACAAAGGCACATTGTTTGGTGGAACTGCAGTGGCCATTAAGGTTCTGGATTTGGAAAACGAGCAAGTATGCAAGAGGTTTGATACCGAATGTGAAGTGATGAGAAATGTTAGACACAGAAATCTTGT

>M6-PGSC0003DMG400023554 M6 LRR-RLK

GGTTCCATTCCTGTAGGACTATTTAATATCTCGTCGCTACTTGCCATAAATCTGCTATACAATAAACTTTCCGGTCCTCTCTTGCTAGATGAAGGAATTTTTTTGTCAAATCTGAAGTTTCTAAGGATATCTCGCAACCAGATTTTAGGATGCATTCCTTCCAACATATGCCAACTCAGAGAGCTCAAAGTTTTGTCCATATCTTTCAACAACATAATTGGAGACAAACCCAAAAATATTGATTGTTTATCCAATCTCCAGGAGTTTTATAATCGTCATAATTCAATAAAAGGGACTATTCCCACTTCATTGGGAAATATTTCAACTCTGCAATATCTTTATTCTGGAAACAATCTCATGGTGGGGCCAATTCCTCCAGAATTAGGGAAGCTATCAAATTTGAGGCAATTAATTTTTTCCAATAATTCTAATCTTATTGGTCAAATTCCAGAGGCTATTTTCAACATATCTTCTTTGGAATTAATCAATTTCAGTTTCAATACCCTGTCGGGTAGAATTCCAACCACTACAGGTCTTCATCTTCCGAACCTTAAAGTACTTGACTTGGGAGTCAATCAGCTTGAAGGGGAAATTCCATTGTTCATAACAAATGCTTCCAAGCTTCAGATACTGTCGCTAGAACATAACTCTCTCTCGGGAGCTATTCCTACTAATTTGGGAAATCTTCATGAGCTGCAATCACTCTTACTACATACTAATCAACTTACCAATGAACCAAGAGAGCATGAGTTGAAATTCTTCAATTCGTTGGCGAACTGTAGGATGTTGAGATATCTAGAAGTGGGTAACAATCCGTTGAGTGGCGTTCTACCAAATTCTGTTGGGAATCTCTCATCTACTATTGAAAAGATTTATATAGGAGATGCACACATCAATGGCCTCATCCCCACAAGTATAGGCAACATGAGCGGTCTTACAGCCCTAAGCCTTGGAGGAAACAACTTGGCAGGGAATATTCCTTCTGAGATTGGTAAGCTTAAACAACTCCAAGGGCTGGCTCTATATAACAATAAATTGCAGGGACGTATTCCAGAGGTGGTATGCCATTTATCGAATTTGGTTGGCTTATTACTGGATGGTAATGAGCTCTGTGGATTAATTCCAGAATGCTTTGGAAATCTTAGCATGCTACAAGCGCTTTATTTGGATTCTAATAAATTTTCATCAAAATTTCCGTTGAGCCTTTGGAAGATGAGTGGTCTTCTCTATCTAAGTGTGTCACGAAATTCAATAGAGGGAGAACTTCCATCAGATATTGGAGGACTGAAAGCCATTGTAGAACTATATCTTTACAATAACCACTTTTCAGGTGTGATACCAACCAGATTGGGGGAACTCCAAAACCTGCAGCATCTTGACCTATCCAACAATTCATTTTTTGGCCAAATTCCATTATCCTTTGCCAACTTGATAAGCTTGGAATTCTTGAATTTGTCTTTAAATGCCTTGTCCGGTACTATTCCTAAATCTTTGGAAAAACTCTCATACCTTAAAAGCATTAATGTTTCATTTAATGATTTAGAAGGTGAAATACCAAGTGGTGGCGTGTTTGCAACTTCCACTGTGCAATCATTTCTTGGGAACAAAGGTCTTTGCGGAATGCACATATTGGAGATTCCTACTTGTGCTATCACTAATCATGGAAAACAATCAAAGCTTAAGGAGGTTGTGCTAAAAATTGTTACTCCAGTGGTTATTGCATCCTTTCTGATATTCTTGTTCGTTTCAATTTGGATAATGAAACGACAGAAGAAAGGGAAGTCCAAAGATATTGAAAAGGTATTGGAGATCGGGACTCATCAATTAGTTTCTTATCACGAGATTCAACAAGCAACAAATAATTTTGATGAATCCAATTTAATTGGTGAGGGAAGCTCTGGCTCTGTGTACAAAGGCACATTGTTTGGTGGAACTGCAGTGGCCATTAAGGTTCTGGATTTGGAAAACGAGCAAGTATGCAAGAGGTTTGATACCGAATGTGAAGTGATGAGAAATGTTAGACACAGAAAT

>M6-PGSC0003DMG400023587 M6 LRR-RLK

ATGGAGAAGCACATTTTCTTATTGATTCTTCTACTTGTAATTCAATATTACTCTGTTGCTTCCTCAAATGAGACTGACCAACAAGCTCTATTACTTTTCCAAAATCTTATTACAAGTCCTAATCATTTTTTGGCAAATAATTGGACCAAAAATACTTCTTTTTGCACTTGGTTTGGTGTCACTTGCAGTCCAAAAAGGCTAAGGGTTGTAGCCTTGACTCTTCCTAATTTGCAACTTCAAGGCACAATTTCTCCATCTTTGGCCAATTTGTCCTTCCTCAGTGTTCTCAATCTCGAGAACAACAGCTTCCACGGTGGCATCCCTTATGGCATTGGCCACATGCCTCGCTTGCGAGTGATTGATGTTAAAAACAATCAGCTCAATGGAAGTATACCAACAAGTCTATTTCAAAACCGAAGAGTTCAAGTAATTTCATTGGGTTTCAATGAACTCAGTGGTGAAATGTGGAGAGGGCCATGGTATGTACCTGAACTCAGAGTCTTAAATCTCAGGAACAATAGCCTCACGGGTATAATCCCTCCTTCGGTTGGAAATGCCACAAATTTGTTGAACTTCAGTTTGTTTGGGAATAGATTCAGCGGCAACATTCCAAAGGAAATTGGTAATCTGAGCCAACTTGTACAAATGTCCTTGACTGATAATCAGTTAACAGGTTCCATTCCCGTAGAACTGTTTAATATCTCGTCACTACTGTCCATACATCTGCGATACAATAGCCTTTCCGGCCCTCTCTTGCTTGATCAAGGGAATATTGTATCAAATCTGGAGCATTTAAGTATTTCTTTCAACCAAATTTCTGGTTGCATTCTTTCCAACATATGCCAACTCACAGAGCTCAAAGTATTGTCCATAGCTTACAACAACATAACTGGAGAAATACCCAGAAATATTGATTGTTTATCCAAACTCGAGATGTTTCTTATTGGTGTTAATCCAATAAAAGGGACTATCCCCACTTCATTGGGAAATATTTCCACTCTGCAATATCTTTATTGTGGAGAAAATCACATAGTAGGGAAAATTCCTCCGGAATTAGGGAAGCTATCAAAGTTGAGGCAATTAGTCATTTTGGATAATTCTAATCTTATTGGTCAAATTCCAGAGGCTATTTTCAACATATCTTCTTTGGAAGTGATTGATTTCAGTTTCAATAACCTCTTGGGGAGAATTCCAACAACTACAGGTCTTCATCTTCCAAACGTTAAGGAACTTAGCTTGGCAGTCAATCAGCTGGAAGGGGAAATTCCATTGTTCATAACAAATGCTTCCAAGCTTGAATTACTGGAGCTAAATGAAAACTTTCTGACAGGAACTATTCCTACTAATTTGGGAAATCTTCGTGAGCTGCGAGCACTGCTCCTACATACTAATCAACTAACCAATGAACCAAGAGAGCGTGAGTTTTTATTCTTTAATTCATTGGCTGACTGTAAGATGTTGAAATATCTACTAGTGGGTTCCAATCTGTTGAATGGTGTTCTGCCAAATTCTATTGGGAATCTTTCATCTACTATTCAAAACTTTTATATAGGACATGCACACATCAACGGCCTCATCCCCACAGGTATAAGCAACATGAGCGGTCTAATGACCCTAGACTTTCAAGGAAACAACTTGGCAGGGAGTATTCCTCCTGAGATTGGTAAGCTTAAACAACTCCAGGGGCTGTATCTAACTAACAATAAATTGCAGGGACATGTTCCAGAGGCGGTATGCCATTTATCTAATTTGGTTCTATTATTTCTGAATGGTAATGAGCTCTCTGGATTAATTCCAGAATGCTTAGGAAATCTTAGCATGCTACAAATGCTTAATTTGGGTTCTAATAAATTTTCATCAAAATTTCCGTTGCTCCTTTGGAAAATGAGTGGTCTTCTCTATCTAGACATGTCACAAAATTCAATAGAGGGAGAAGTTCCAACAGATATTGGAGGACTGAAAGCCATTGTAGCTCTAAATCTTTCCGGTAACCACTTTTCAGGCATGATACCAACCAGATTGGGAGAACTCCTAAACCTGCAGTATCTTGACCTATCAAACAATTCATTTTTTGGCCAAATTCCATTATCCTTTGCCAACTTGATAAGCCTGGAAATCTTGAATTTGTCTTTAAATTCCTTGTCAGGTACTATTCCTAAGTCTTTGGAAAAACTTTCATACCTTAAAAACATTAATGTTTCATTTAATGATTTAGAAGGTGAAATACCGAGTGGTGGCGTGTTTGCAAATTCCACTCTGCAATCATTTCTCGAGAACAAAGGTCTATGTGGAATGCACATATTGGAGATTCCTGCTTGCGCTATCAATAATCCTGGAAAACAATCAAAGCTTAAGAAGGTTCTGCTAAAAATTGTTACTCCAGTGGTTATATCATCCTTTCTGATATTCTTGTTGGTTTCAATTTTGATATTGATACGACAGAAGAAAGGAAAGTCGAAAGATGTTGAAAAGGTACTGGAGATCGGGACTCATCAATTAGTTTCTTATCACGAGATTCAAAGAGCAACAAATAATTTTGATGAATCCAATTTAATTGGTGAGGGAAGCTCTGGCTCTGTGTACAAAGGCACATTGTTTGGGGGAACTGCAGTGGCCATTAAGGTTCTGGATTTGGAAATTGAGCAAGTATGCAAAAGGTTTGATACCGAATGCAAAGTGATGAGAAATGTTAGACACAGAAATCTTGTTTCAGTGATTACTACTTGTTCTAGTGACTATATAAGAGCCTTTGTTCTGCAATTTATGCCCAACGGAAGTCTTGAGAATTGGTTGTACAGAGAAGATCTCCACTTAAACCTTCATCAAAGAGTCACTGTAATGCTTGATGCAGCTATGGCAGTTGAATATCTACATCATGGTCATGTCACTCCAATAGTTCATTGCGACCTAAAGCCAGCCAACATTCTTTTGGATGAAGATATGGTGGCTCATGTTGGTGATTTTGGAATCTCTAAAATTTTAGCCATAAGCAAGTCCATGGCTTATACCGAGACGTTGGGCACTCTTGGATATATTTCACCAGGTATAAAAAAATCTATTCTCTTTGATTTTCTCTTATCATTATTAAGCCTCTTCAAATTCTAGAAGTAAAAAAAGCAGGTCTTTAATTATGCATAACTATTGTTGTATTTCAATTGAGTAACTTTTCTTCAATCCTTTTCTAAGAATATGGCTCGGAGGGAATAGTGTCTGCTAGTGGTGATGTTTATAGCTACGGCATCATGTTGATGGAGGTTTTGACCAAAAGAAGGCCAAGAGATGAAGATATATGCAATGAAAATCTTGACTTGAGGAAATGGATAACACAATCATTTTCAGGGAGTATGATGGACATTGTGGATGCCAATCTTTTTTCTGAGGAAGAACAAATCACTAGCAAAAGTGAAATGTGCATAGCCTCCATGATAGAATTGGCTTTAGACTGCACAAAGGAAATGCCAGAATCCAGAATAACCATGAAAGAAGTAGTGAAGAGGCTTAACAAAATCAACAACAAATTTCTGGAAACATAG

>M6-PGSC0003DMG400023612 M6 LRR-RLK

ATGGTGGGGCCAATTCCTCCAGAATTAGGGAAGCTATCAAATTTGAGGCAATTAATTTTTTCCAATAATTCTAATCTTATTGGTCAAATTCCAGAGGCTATTTTCAACATATCTTCTTTGGAATTAATCAATTTCAGTTTCAATACCCTGTCGGGTAGAATTCCAACCACTACAGGTCTTCATCTTCCGAACCTTAAAGTACTTGACTTGGGAGTCAATCAGCTTGAAGGGGAAATTCCATTGTTCATAACAAATGCTTCCAAGCTTCAGATACTGTCGCTAGAACATAACTCTCTCTCGGGAGCTATTCCTACTAATTTGGGAAATCTTCATGAGCTGCAATCACTCTTACTACATACTAATCAACTTACCAATGAACCAAGAGAGCATGAGTTGAAATTCTTCAATTCGTTGGCGAACTGTAGGATGTTGAGATATCTAGAAGTGGGTAACAATCCGTTGAGTGGCGTTCTACCAAATTCTGTTGGGAATCTCTCATCTACTATTGAAAAGATTTATATAGGAGATGCACACATCAATGGCCTCATCCCCACAAGTATAGGCAACATGAGCGGTCTTACAGCCCTAAGCCTTGGAGGAAACAACTTGGCAGGGAATATTCCTTCTGAGATTGGTAAGCTTAAACAACTCCAAGGGCTGGCTCTATATAACAATAAATTGCAGGGACGTATTCCAGAGGTGGTATGCCATTTATCGAATTTGGTTGGCTTATTACTGGATGGTAATGAGCTCTGTGGATTAATTCCAGAATGCTTTGGAAATCTTAGCATGCTACAAGCGCTTTATTTGGATTCTAATAAATTTTCATCAAAATTTCCGTTGAGCCTTTGGAAGATGAGTGGTCTTCTCTATCTAAGTGTGTCACGAAATTCAATAGAGGGAGAACTTCCATCAGATATTGGAGGACTGAAAGCCATTGTAGAACTATATCTTTACAATAACCACTTTTCAGGTGTGATACCAACCAGATTGGGGGAACTCCAAAACCTGCAGCATCTTGACCTATCCAACAATTCATTTTTTGGCCAAATTCCATTATCCTTTGCCAACTTGATAAGCTTGGAATTCTTGAATTTGTCTTTAAATGCCTTGTCCGGTACTATTCCTAAATCTTTGGAAAAACTCTCATACCTTAAAAGCATTAATGTTTCATTTAATGATTTAGAAGGTGAAATACCAAGTGGTGGCGTGTTTGCAACTTCCACTGTGCAATCATTTCTTGGGAACAAAGGTCTTTGCGGAATGCACATATTGGAGATTCCTACTTGTGCTATCACTAATCATGGAAAACAATCAAAGCTTAAGGAGGTTGTGCTAAAAATTGTTACTCCAGTGGTTATTGCATCCTTTCTGATATTCTTGTTCGTTTCAATTTGGATAATGAAACGACAGAAGAAAGGGAAGTCCAAAGATATTGAAAAGGTATTGGAGATCGGGACTCATCAATTAGTTTCTTATCACGAGATTCAACAAGCAACAAATAATTTTGATGAATCCAATTTAATTGGTGAGGGAAGCTCTGGCTCTGTGTACAAAGGCACATTGTTTGGTGGAACTGCAGTGGCCATTAAGGTTCTGGATTTGGAAAACGAGCAAGTATGCAAGAGGTTTGATACCGAATGTGAAGTGATGAGAAATGTTAGACACAGAAATCTTGTTTCAGTGATTACTACATGTTCTAGTGACTATATAAGAGCCTTTGTTCTGCAATTTATGCCCAATGGAAGTCTTGAGAATTGGCTGTACAAAGAAGATCGACACTTGAACCTTCATCAAAGAGTCACTGTAATGCTTGATGCAGCTATGGCAGTTGAATATCTACACCATGGTCATAACACTCCAATAGTTCATTGCGACCTGAAGCCAGCCAACGTTCTTTTGGATGAAGATATGATGGCTCATGTTGGTGATTTTGGAATATCTAAAATTTTAGCCATAAGCAAGTCCATGGCTTATACAGAGACATTGGGCACTCTTGGATATATTGCACCAGGTAAAAGAAAATCTACTCTCATTGATTTTCTCTTATCATAATTAAGCCTCTCCAAGTTGTAGAAGTAAAAAGAGAATTATTGTTGTATTTCAATTGAATTAACTTTTTTTTAAATCCTTTTTTAAGAATATGGCTCGGATGGAATAGTGTCTGCTAGTGGCGATGTTTATAGTTACGGCATCATGTTGATGGAGGTTTTGACGAAAAGAAGGCCAACAGATGAAGAGATATGCAATGAAAATCTTGACTTGAGGAAATGGATCACACAATCATTTTCAGGGAGTATGATGGACGTTGTGGATGCCAATCTTTTCTCCGAGGAAGAACAGATCACTTCAGAAAGTGAAATCTGCATTGCGTCCATGATAGAATTGGGTTTAGACTGCACAAAGGAAATGCCAGAATCAAGAATAAGCATGAAAGAAGTAGTGAAGAGGCTTAACAAAATCAACAACACATTTCTGGAAACATAGTAGT

>M6-PGSC0003DMG400023614 M6 LRR-RLK

ATACCCAGAAATATTGATTGTTTATCCAAACTCGAGATGTTTCTTATTGGTGTTAATCCAATAAAAGGGACTATCCCCACTTCATTGGGAAATATTTCCACTCTGCAATATCTTTATTGTGGAGAAAATCACATAGTAGGGAAAATTCCTCCGGAATTAGGGAAGCTATCAAAGTTGAGGCAATTAGTCATTTTGGATAATTCTAATCTTATTGGTCAAATTCCAGAGGCTATTTTCAACATATCTTCTTTGGAAGTGATTGATTTCAGTTTCAATAACCTCTTGGGGAGAATTCCAACAACTACAGGTCTTCATCTTCCAAACGTTAAGGAACTTAGCTTGGCAGTCAATCAGCTGGAAGGGGAAATTCCATTGTTCATAACAAATGCTTCCAAGCTTGAATTACTGGAGCTAAATGAAAACTTTCTGACAGGAACTATTCCTACTAATTTGGGAAATCTTCGTGAGCTGCGAGCACTGCTCCTACATACTAATCAACTAACCAATGAACCAAGAGAGCGTGAGTTTTTATTCTTTAATTCATTGGCTGACTGTAAGATGTTGAAATATCTACTAGTGGGTTCCAATCTGTTGAATGGTGTTCTGCCAAATTCTATTGGGAATCTTTCATCTACTATTCAAAACTTTTATATAGGACATGCACACATCAACGGCCTCATCCCCACAGGTATAAGCAACATGAGCGGTCTAATGACCCTAGACTTTCAAGGAAACAACTTGGCAGGGAGTATTCCTCCTGAGATTGGTAAGCTTAAACAACTCCAGGGGCTGTATCTAACTAACAATAAATTGCAGGGACATGTTCCAGAGGCGGTATGCCATTTATCTAATTTGGTTCTATTATTTCTGAATGGTAATGAGCTCTCTGGATTAATTCCAGAATGCTTAGGAAATCTTAGCATGCTACAAATGCTTAATTTGGGTTCTAATAAATTTTCATCAAAATTTCCGTTGCTCCTTTGGAAAATGAGTGGTCTTCTCTATCTAGACATGTCACAAAATTCAATAGAGGGAGAAGTTCCAACAGATATTGGAGGACTGAAAGCCATTGTAGCTCTAAATCTTTCCGGTAACCACTTTTCAGGCATGATACCAACCAGATTGGGAGAACTCCTAAACCTGCAGTATCTTGACCTATCAAACAATTCATTTTTTGGCCAAATTCCATTATCCTTTGCCAACTTGATAAGCCTGGAAATCTTGAATTTGTCTTTAAATTCCTTGTCAGGTACTATTCCTAAGTCTTTGGAAAAACTTTCATACCTTAAAAACATTAATGTTTCATTTAATGATTTAGAAGGTGAAATACCGAGTGGTGGCGTGTTTGCAAATTCCACTCTGCAATCATTTCTCGAGAACAAAGGTCTATGTGGAATGCACATATTGGAGATTCCTGCTTGCGCTATCAATAATCCTGGAAAACAATCAAAGCTTAAGAAGGTTCTGCTAAAAATTGTTACTCCAGTGGTTATATCATCCTTTCTGATATTCTTGTTGGTTTCAATTTTGATATTGATACGACAGAAGAAAGGAAAGTCGAAAGATGTTGAAAAGGTACTGGAGATCGGGACTCATCAATTAGTTTCTTATCACGAGATTCAAAGAGCAACAAATAATTTTGATGAATCCAATTTAATTGGTGAGGGAAGCTCTGGCTCTGTGTACAAAGGCACATTGTTTGGGGGAACTGCAGTGGCCATTAAGGTTCTGGATTTGGAAATTGAGCAAGTATGCAAAAGGTTTGATACCGAATGCAAAGTGATGAGAAATGTTAGACACAGAAATCTTGTTTCAGTGATTACTACTTGTTCTAGTGACTATATAAGAGCCTTTGTTCTGCAATTTATGCCCAACGGAAGTCTTGAGAATTGGTTGTACAGAGAAGATCTCCACTTAAACCTTCATCAAAGAGTCACTGTAATGCTTGATGCAGCTATGGCAGTTGAATATCTACATCATGGTCATGTCACTCCAATAGTTCATTGCGACCTAAAGCCAGCCAACATTCTTTTGGATGAAGATATGGTGGCTCATGTTGGTGATTTTGGAATCTCTAAAATTTTAGCCATAAGCAAGTCCATGGCTTATACCGAGACGTTGGGCACTCTTGGATATATTTCACCAGGTATAAAAAAATCTATTCTCTTTGATTTTCTCTTATCATTATTAAGCCTCTTCAAATTCTAGAAGTAAAAAAAGCAGGTCTTTAATTATGCATAACTATTGTTGTATTTCAATTGAGTAACTTTTCTTCAATCCTTTTCTAAGAATATGGCTCGGAGGGAATAGTGTCTGCTAGTGGTGATGTTTATAGCTACGGCATCATGTTGATGGAGGTTTTGACCAAAAGAAGGCCAAGAGATGAAGATATATGCAATGAAAATCTTGACTTGAGGAAATGGATAACACAATCATTTTCAGGGAGTATGATGGACATTGTGGATGCCAATCTTTTTTCTGAGGAAGAACAAATCACTAGCAAAAGTGAAATGTGCATAGCCTCCATGATAGAATTGGCTTTAGACTGCACAAAGGAAATGCCAGAATCCAGAATAACCATGAAAGAAGTAGTGAAGAGGCTTAACAAAATCAACAACAAATTTCTGGAAACATAG

>M6-PGSC0003DMG400023638 M6 LRR-RLK

ATGAAGAAGCACATTTTTATATTGACTCTTCTCTTTCTAGTTCAATTTTCTCTATCAATTGCCTCCTCAAATGAGACGGACCAAGAAGCTCTACTAGCTTTCCAAAATCTTATTACAAGTCCGAGTCATTTTTTGGCCAATAATTGGACCAAAAATACTTCTTTTTGCTCTTGGTTTGGTGTCACTTGCAGTCCAAAAAGGCAAAGGGTACAAACCTTGACTCTTCCTGATTTGCAACTTCAAGGCACAATTTCCCCGTCTTTAGCTAATTTGTCCTTTCTCAGCGTGCTCACTCTCCGGAACAACAACTTCCATGGCGGCATCCCTTACGAACTTGGCCACTTGCCTCGCTTGCGAGTGATTGATTTTCAAAACAATCAACTCCAAGAAAGTATTCCAACAAGTCTATTTCAACACCAGAGAGTTCAAATCATTTCATTGGCTTTCAATAAACTAGGTGGTGAAATGTGGAAAGGGCCATGGTATGTACCAGAACTCACAGTTTTAAATCTCAGGAACAATAGCCTCACAGGTATAATACCTCCTTCTATTGGAAATGCCACAAAGTTGCTGAACTTCAGTTTGTATGGGAATAGAGTCGGCGGGAACATTCCAAAGGAAATCGGTAATCTCAGCCAACTTGAATTTTTCTCCTTGTTCGATAATCAATTAACAGGTTCTATTCCTGCAGCACTATTTAATATCTCCTCGCTTCTTGTCACATCTCTAGCATTCAATAGCCTTTCCGGCCCTTTCTTGCTTGATGAAGGGAACGTAGTGTCAAATCTAGAGTCATTAACTATATCTAACAACCAAATTTCTGGTTACATTCCTTCCAACATTTGCCAACTCACAAAACTCAAAGCGATGTCCATATCTTTCAACAAGATAACTGGAGGAATACCCAAAAATATTGGTTGTTTATCCAATCTCGAGGAGCTTTATATTGGTGACAATCCAATAAAAGGGAATATTCCCGCATCATTGGGAAATATTTCCACTCTGCAAAATCTTTATTATGGAAGCAATCACTTGGAGGGGCCAATTCCTCCAGAATTGGGGAAGCTATCAAATTTGAGGCAAATTAGCTTTGTCCAAAATAATAATCTTATAGGTCATATTCCAAAGGCTATATTTAATATATCTTCTTTGGAAATAATCGATTTCAGTTACAATAACCTCTCGGGGAGAATTCCAACCACTACAGGTCTTCATCTTCCAAACCTTAAAGAACTTTTCTTGGGAGTCAATAAGATCAAAGGGGAAATTCCATTGTTCATATCAAATGCTTCCATGCTTGAGATACTGGGGCTAAACAGGAACTTCCTCACAGGCAATATTCCTACCAATTTAGGAAATCTTCGTGACCTGCGACGCTTATACTTAGAGGGTAATCAACTTACCAATGAACCAAATAATCATGAGTTGCAATTCTTTAATTCTTTAGTGGACTGTAGGAAGCTGCAATATCTAACAGTGGGTAACAATCCATTAAATGATATTTTGCCCGATACTATTGGAAATCTTTCATCTACTATTGAATCAATTCACATGGGAAATGCACAAATCAGTGGCCTCACTCCCACAGGTATAGGAAACTTAAGCGGTCTAATGTCACTAGCTTTTGTGGAAAACAACTTGATGGGAACCATTCCGTCTGAAGTCAGTAAGCTTGAACATCTCCAAGGTCTGTATCTATATAGCAACAAATTGCAAGGGAATATTCCAGAGGTAGTTTGTCATTTACCTTATTTGGTTACATTATCTTTGCATGTTAATGAGCTCTCTGGGGTGATCCCAAAATGTTTAGAAAATCTTACCACGCTACGAGTTCTTTCATTGAGTTCTAACAAATTTTCATCAAAGCTTCCTTTGAGTCTTTGGAAGATGAGTGGACTTCTCTATCTTTTCATGTCACGAAATTCTATAGAGGGAGAAGTTCCACAGGACATTGGAGGAATGAAGGCCATTGTAAAACTAGATCTTTCTGGTAACCACTTTTCCGGCATGATACCAAGCCAATTGGGGGACCTCCAAAACATGAAAGTTCTTGACCTGTCAAACAATTCATTTTCAGGCTCAATTCCCTTATCCATTGCCAATTTGATAAGTCTAGAATACTTGAAATTGTCTTTCAAAGCATTGTCAGGTACTATTCCTAAGTCTTTGGAAAAGCTTTCATACCTTAAAAGCATCAATGTCTCATTTAATGATTTAGATGGTGAGATACCTAGTGATGGTGTGTTTTCAAATTCCACTTTGCAATCATTTCTTGGGAACAAAGGTCTATGTGGAGTAAACATATTGGAGATTCCTGCTTGTGCTATCACTAATCCTAGAAAACAATCAAAGACTAAGGAGCTTGTGCTGAAAATTGTTATTCCAGTGGTTACTTCATCCTTTCTGATATTATTGGTTGTGTCAGCTTGGATAATAATGAAACAAAAGATGAAAGGAAAGTCCAAAGAAGTGGAAAAGGTACCAGAGATCAGGACTTATCAATTGGTTTCTTATCATGAAATTCAACGAGCAACAAAAAATTTTGATGGATCCAATTTAATTGGTACGGGAGGTTCTGGCTCTGTGTACAAAGGTACATTATCTAGTGGAACTGTGGTGGCAATAAAGGTTCTCGATTTGCAAAATGAGGAAGTATGCAAGAGGTTTGATACTGAATGTGAAGTGATGAGAAATGTTAGGCACAGAAACCTCATTTCAGTGATTACTACTTGTTCTAGTGAATACATAAGAGCCTTTGTTCTACAGTATATGCCCAATGGAAGTCTTGATAATTGGTTGTACAAAGAAGATTGCCACTTGAACCTTCTTCAAAGAGTCAACATAATGCTTGATGTGGCTGTGGCAATTGAATATCTACACCACGGTCATGATACTCCAATAGTTCATTGCGACCTAAAGCCAGCCAACATTCTTTTGGATGAAGAAATGGTGGCACATGTTGGTGATTTTGGCATCTCTAAAATATTAGCTGTAAGCAAGTCCATGGTACATACTGAGACATTAGGCACTTTTGGTTACATTGCACCAGGTACGTACACTAGTTTCGTTCTTTTTCACTTATCATTAACCCTTTCTCTTATCAGAACTAAGCTCTTCCGAATTCTGGAACTAAACAGAAACCAAGTCTTTCATTGTACAAAACAATTATTATCTTCCGATTGTAATTTATTTTTCATTTTCATTCTTCTAAAGAATATGGCTTGGAAGGAGTAGTGTCCATTAGTGGTGATGTTTACAGCTATGGCATCATGATGATGGAGGTAATGACAAAAAGAAGACCAACTGAAGACGAGATATTCAACGAAAATCTTGGCTTGAGGCAGTGGATAAGACGAGCATTTCCCAGGACAATTATGGAAGTCGTAGATGCCAATCTTTTTCAAGAGGAAGAATCTGTCAATTTAAAAAGTGAAATATGCATAGCCTCCATGATGGAATTGGCTTTGGATTGCACAAAGGAAATGTCAGAGACAAGGATAACTATGAGAGACGTAGTCAAGAGGCTTAACAAAATCAAAAACACATTTTTAGGAACATAG

>M6-PGSC0003DMG400023639 M6 LRR-RLK

GGAGAAGCACATTTTTTTATTGACTCTCCTCTTTCTAGTTCAATTTTCTATATCTTTTGCTTCCTCAAATGAGACAGACAAAGAAGCTCTATTAGCTTTCCAAAATCTTATTACAAGTCCTAGTCATTTTTTGGCCAATAATTGGACCAAAAATACTTCTTTTTATTCTTGGTTTGGTGTCACTTGCAGTTCAAAAACGCAAAGGGTTGTGGCCTTGGCTCTTCCTAATTTGCAACTTCAAGGCACAATATCCCCGTCTTTGGCCAATTTGTCCTTTCTCAGAGAGCTCAATCTCGAGAATAACTTATTCCATGGTGGCATCCTATACAGACTTGGCCACTTGCCTCGCTTGCGAGTGATTAATGTTCGAAACAATCAGCTAGAAGGAAGTATTCCAACAAGTCTATTTCAACACCAGAGAGTTCAAATCATTTCATTGGCTTACAATAAACTCAGTGGTGAAATGTGGAAAGGGCCATGGTATGTACCCGAACTCAGAGTCCTAAGTCTCAGGAATAATAGCCTCACAGGTATAATCCCTTCTTCTGTTGGAAATGCCACAAAGTTGCTGAACTTCAGTTTGTCTGGGAATAGAATCAATGGTATCATTCCAACTGAGATTGGTAATCTTAGCCAACTTATAGAGTTGCATTTGTTCAATAATCAATTAGCAGGTTCCATTCCCGCAACACTGTTTAATATTTCGTCGCTAATTAGAGCATCTCTGGCAAGCAATAGCCTTTCTGGTCCTCTCTTACTTGATGAAGGGAATATTGTGTCAAATCTGAAGTATCTAAGTATATCTAAGAACCAAATTTCTGGTTGCATTCCTTCTAACATATGCCAACTCACAGAGCTCAAAATTTTGTCCATATCTTATAACAATATGATTGGAAACATACCCAGAAATATTGGTTGTTTATCCAAAATCGAGGAGTTTTATATTGGAAATAATCCAATAACAGGGACTATTCCCACTTCATTGGGTAATATTTCCACTCTGCGGAATCTTTATTGTGGAAACAGTCGCATAGTGGGGCAAATTCCAAAGGATATTTTTAACCTATCTTCTTTGGAAATGATTGATTGCAGTTACAGTAGCCTCTCAGGTAGAATTCCAACCACATCAGGTCTTCATGTTCAGAACCTTAAAGAACTTTTCTTGGGACACAATCGGCTCGAAGGGGAAATTCCATTGTTCATAACAAATGCCTCCAAGCTTGAGAGATTGGGGCTAGAAAATAACTTTCTCACTGGCACTATTCCTACTAATTTGGGGAATCTCCGTGAGCTGCAAGAACTGTTCCTACATGATAATCAACTTACCAATGAACCAAGAGAGCATGAGTTGCAATTCTTCAATTCTTCGGCGGACTGTAGGATGTTGCGATATCTACAAGTGGGTTCCAATACGTTGAATGGCATTCTGCCCAATTCTATTGGGAATCTTTCATATAGCAGATGCACACATCAATGGCCCCATCCCCAGGGGTGTACGCAACATGAGTGGTCTAATAGCCCTAAGCCTTGGAGAAAACAACTTGGCGGGAAGTATTCCTTCTGATGTTGTTAAGCTTGAACAACTCCAAGGGTTGTATCTACATAACAATAAATTGCAGGGACATATTCCAGAGGCGGTATGCAATTTATCTAATTTGGTTCAATTAAGTCTGGCTGGTAATGAGCTCTCTGGATTAATTCCAGAATGTTTAGGAAATCTTAGCAAGCTACAAGCCATTATATTGAGTTCTAACCAATTTTCATCCAAGATTCCTTTGAGCATTTGGAAGATGAGTGGTCTTCTCTATCTAATCATGTCACAAAATTCTATAGAGGACTGAAGGCCATTGTAGGACCAGATCTTTCTGGTAACCACTTTTCAGGCATGATACCAAGCCAATTGGGGGACCTCCAAAACATGAATACTCTTGACCTGTCGAACAATTCATTTTCAGGCTCAATTCCATTATCCTTTGCCAACTTGATAAGTTTAGAATTCATGGATTTGTCTTTAAATGCGTTATCAGGTACTATTCCCAAGTCTTTGGAAAAGCTATTATACCTTAAAAGCATCAATGTCTCATTTAATGTTTTAGAAGGTGTAATACCGAGTGGTGGTGTGTTCGCGAATTCCACTCTGCAATCATTTCTTGGGAACAAAGGTCTATTTGGAATGCACATATTGGAGATTCCTGCTTGTGCTATCACTACAACTGGGCAACGATTAAAGTCTAAGAAGCTTGTGCTTAAAATTGTTATTCCAGTGGTTGCTGCTTCTTTTCTGATATTTTTGTTCTTTATAGTTTGGATAATGAGACGACAGAAGAAAGCAAACTCCAAAGACGTGGAAAAGGTACCAGATATCAGGACCTATCAATTGGTTACTTATCATGAGATTCAACAAGCAACAAATAATTTTGATAGATCCAATTTAATTGGTGCGGGAGGTTCTGGCTCTGTGTACAAAGGTACATTATCTAGTGGAACTGTGGTGGCAATAAAGGTACTCGATTTGCAAAATGAGGAAGTATGCAAGAGGTTTGATACTGAATGAAGTGATGAGAAATGTTAGGCACAGAAACCTTATTTCAGTGATTACTACTTGTTCTAGTGAATACATAAGAGCCTTTGTTCTGCAGTATATGCCCAATGGAAGTCTTGAGAGTTGGTTGTACAGAGAAGATCGCCACTTGAACCTTCTTCAAAGAGTCACCATAATGCTTGATGTGGCTCTGGCAATTGAATATCTACACCATGGTCATGAAACTTTGATAGTTCATTGCGACCTAAAGCCAGCCAATGTTCTTTTGGACGAAGAAATGGTGGCACATGTCGGTGATTTTGGGATCTCTAAAATACTAGCTGTAAGCAAGTCCATAGCACATACTGAGACATTAGGCACTCTTGGTTACATTGCACCAGGTACGTACACTAGTCTCATTCCTTTTCACTTATCATTAACCTTTTCTCTAATCAAAATTAAGCTCTTTCGATTTCTAGAAATATTTAAAAATCAAGTCTTTGATTGTAAAAAACAATTATTATCTTTCGACTATAATTTATATAATTTTCATTCTTCTAAAGAATATGGCTTGGAAGGAAAAGTGTCCAGTAGTGGTGAAGTTTACAGCTATGGCATCATGATGATAGAGGTATTGACAAAAAGAAGACCAACTGATGACGAGATATTCAATGAAAATCTTGGCTTGAGACAGTGGATAAGACAATCATTTCCCAAGACAATTATGGAAGTTGTGGATGTCAATTTTATTCATGAGGAAGAACATTTCAATTCAAAAAGTGAAATATGCATAGGCTCCATGATGGAATTGGCTTTGGATTGCACAAAGGAAATGCCAGAGTCAAGGATAACAATGAGAGACGTAGTCAAGAGGCTTGACAAAATTAAGAACACATTTTTGGGAACATAGAAATTAGCATCTTTTAAAGTGGTGTTCTTTTTTAGTTGCAAGTTAATATGTTGATCAGCTTTTTCTGTTTAAATGGTTTATAAGGAATGAAGTTCGAGGATTAAAATGGAGCAAGGTTGGGATGGAGTACCAAAATGAAATTTTGGTGTAAGTTAGAGGGGCTATCTATGTATTTGACCTTGTGTGAATGTACG

>M6-PGSC0003DMG400023643 M6 LRR-RLK

ATGGAGAAGCACATTTTCTTATTGATACTTCTCTTTTTAGTTCAATTTCGTATATCAATTGCTTCCTCAAATGATACTGACCAACAAGCCCTACTAGCTTTCCAAAATCTTATTACAAGTCCCAATCAATTTTTGGCTAATAATTGGACCAAGAATGCATCTTTTTGCTCTTGGTTTGGTGTCACGTGCAGTACAAGAAGGCTAAGGGTTGTGGCCTTGGCTCTTCCTAATTTGCAACTTCAAGGCACAATTTCCCCATCTTTGGCCAATTTGTCCTTTCTCAGAGAGCTCAATCTCGAGAACAACTTATTCAATGGGGGCATCCCTTACGGACTTGGCCAGTTGCCTCACTTGAGAGTCATTGATTTTCAAAACAATCAGCTCCAAGGAAGTATTCCAACAAGTCTATTTCAACACCAGAGAGTTCGAATCATTTCATTGGCTTTCAATGAACTCAGTGGTGAAATGTGGAAAGGGCCATGGCATGTACCAGAACTCAAAGTCTTAGATCTCGCGAACAATAGCCTCACGGGTATAATCCCTCCTTATGTTGGTAATGCCACAAAGTTGATGAACTTCGATTTGTCTGGGAGTAGAATCAATGGTACCATTCCAATGGAAATTGGTAATTTGAGCCAACTTGCATCTTTGTCCTTGGTTGATAATCAGTTAACAGGTTTTATTCCTGGATCACTGTTTAACATTTCATCACTACTTGGCGTAACTCTGGGATTGAATCACCTTTCTGGTCCTCTCTTGCTTGATGAAGGGAATATTGAGTCAAATCTGAAGATACTAAGTATATCTTTCAATCAAATTTCTGGTCGCATTCCTTCTAACATATGCCAACTCACACAGCTCAAAATTTTGTCCATATCTTTCAATAGCATAACTGGAGACATACCCAAATATATTGGTTGTCTAGCCAAGCTCGAGATGTTCTTAATTAGTGAGAATGCAATAAGTGGGACTATTCCTGCTTCATTGGGCAATATTTCCACTCTGCAACATCTTGATTCGGGGGACAATCGGTTGGAGGGACAAGTTCCTCAAGAATTAGGGAAGCTATCAAATTTGAGGGTGTTAAACTTCCACAAGAATTTTAATCTTATTGGTCAAATTCCCAAGGCTATTTTCAATATATCTTCTTTGGAAGCGATTGATTTCAGTTTCAATAACCTCTCAGGTAGAATTCCAGCCACTACCGATCTTCATCTTCCGAACCTTGAAGTGTTTATGTTGGCATTCAATCAGCTCGAAGGAGAAATTCCTCAATACATAGCAAATGCTTCCAAGCTTGAGATACTGGCGCTAAATTATAACTTTCTGACAGTTGCTATTCCTACTAATTTAGGAAATCTTCGCGAGCTGCAATCACTGTTCCTAGATCATAATCAACTTACCAATGAACCAAGAGAGCCTAAGTTGTTATTCTTGAATTCTTTGGTGAACTGTAGGATGCTGAAATATGTAGATGTGGGTTTCAATCCGTTGAATGGCGTTCTTCCCAATTCTATTGGGAATCTTTCATCTACTATTGAAGTATTTGATATAAGAAATGCACACATCTATGGCCTCATTCCAACAAGTATAGGCAACATGAGTGGTCTTATATCCCTAGTCTTTAATGAAAACAACTTGACAGGATATATTCCTCCAGAGATTGGTAAGCTTAAACAACTACAAGGTCTGTACCTAAATAACAATAAATTTCAGGGACATATTCCAGAGGCGGTATGACATTTATCTCATTTGGTCGAATTAGCTATGGATAGTAATGAACTCTCTGGATCAATTCCAGAATGTTTAGGAAATCTTGGGATGCTACTTTATCTTTCGTTGGGATCAAATAAATTTTCACCAAAATTTCCCTCGAGTCTTTGGAAGATGAGTGGACTTCTCTACCTAAACATGTCACATAATTCTATAGAGGGAGAAGTTCCACCAACTATTGGAGAACTGAAAGCCATTGAAGAACTATATCTTTCCGGTAATCACTTTTCAGGCCAGGTACCAAGCAGATTGGGAGACCTCCAAAGATTGAAGACTCTTGACCTATCAAACAATTCATTTTCAGGCTCAATTCCATTATCCATTACCAACTTGATGAGCTTAGAAGACTTGGATTTGTCTTTAAATGCATTGTCAGGTATTATTCCTAAGTCTTTGGAAAAACTCTCATACCTTACAAGCATCAATGTTTCATTTAATGATTTAGAAGGTGAAATACCCAGTGGTGGTGTGTTTGTAAATTCCACTCTGCTATCATTTCTCGGGAACAAAGGTCTATGTGGAATGCACATATTGGAGCTTCCTGCTTGTCCTATCACTAATCCTGGACAACAATCAGTGTCTAAGAAACTTGTGCTAAAAATTGTTACTCCAATGGTTATTTCATTGTTTCTGATATTCTTGTTGGTTTCAATTTGGATAATGAAACGAAAGAAGAAAGGGAAGTCCAAAGATGTTGAAAAGGTTCTGGAGATCAAAACTTATCCATTGATTTCTTATCATGAGATTCAACGAGCAACAAATAATTTTGATGGATCCAATTTAATTGGTGAGGGAAGCTCTGGCTCTGTGTACAAAGGCACATTCTCAAGTGGAACTACAGTGGCCATAAAGGTTCTGGATTTGGAAAATGAGCAAGTATGCAAGAGGTTTGATACAGAATGCGAAGTGATGAGAAATGTCAGACATAGAAATCTTGTTCCAGTGATTACTACTTGTTCTAGTGACTATATAAGAGCCTTTGTTCTGAAATATATGTCAAATGGGAGTCTTGAGAATTGGTTGTACAGAGAAGCTCGCCACTTGAACCTTCTTCAAAGAGTCACTGTAATGCTTGATGCGGCTATGGCAATTGAATATCTACATCATGGCAATGACACTGCGATAGTTCATTGCGACATAAAGCCAGCAAACGTTCTTTTGGATGAAGATATGGTGGCGCATGTTGGTGATTTTGGAATCTCTAAGATTTTAGCCGTAAGCAAGTCCATGACACAAACCGAGACATTGGGCACTCTTGGATACATTGCACCAGGTATACTAAAATTATAACCTTTCTATTTAATTTTTCTCTTATCAAAATAAAGCCCTTGAAAATTCTAGGACTAAATAAAAAGCAAGTCTTTGTTAGTATGAGCATTATTGCTATATCCAAATGAGTTAGTTCTTTTTCGTTTTCGTTCTTTTAAGAGTACGGCTCAGAAGGAATAGTGTCGGCTAGTGGTGATGTTTACAGTTACGGCATCATGTTGATGGAGGTTTTGACGAAAAGAAGGCCAACAGATGAAAAGATATGCAATGAAAATCTTGACCTGAGGAGATGGATAACACAATCATTTTCAGGGAGTATGATGGACGTTGTGGATGCCAATCTTTTTTCTGAGGAAGAACAAATCACTAGTAAAAGTGAAATGTGCATAGCCTCCATGATAGAATTGGCTTTAGACTGCACAAAGGAAACGCCAGAATGCAGAATAAGCATGAAAGAAGTAGTGAAGAGGCTTAACAAAATCAACAACACATTTCTCGAAACATAG

>M6-PGSC0003DMG400024485 M6 LRR-RLK

ATGTTGCTAGTACTTGCAGGTTCAGAGACTCTGGAAGTGAGACGAGCATTAGTTAAGTTTATGAACAAGATATCACTTGGGAACATTTCAAAAGAAGTAAACTTTGGATGGGATTTATCATCTGATCCATGTACCAACAAATGGGAAGGGATAACTTGTGATTCAAGAAACCAACATGTAAAAAATATTGTTCTTGATCAGAAGAATCTCACTGGAATTCTTGATGCTTCTTTTGTCTGTGAGGCTACTTCCCTTGCTGTACTGAGCTTGAACGAAAACGAAATTGTAGGAACATTACCTCAAGAAATATCAAACTGCAGACGTCTCACACATTTGTATCTGCGTGGCAACAAATTATCAAGCAATCTACCAAGCTCTCTTTCGCGGTTGAGCAATTTGAAAAGGTTTGTAATTTCAGGCAATGCCTTCTCGGGGCAGATACCGGATATGTCAAGAATCTCAGGGCTGATAACTTTTTTGGCTGAAAGAAATCAGCTAACAGGGCAGATACCAGAATTTGATTTTTCCAATCTTGTAGAATTCAATGTCTCATACAATAATTTAACTGGTCCAATTCCTGATGTCAAGGGCCAGTTTAGTTCCAGCAGTTTTTCGGGTAATCCCGGATTATGTGGAGTGCCATTGCCAAGCATATGTCCACCCTCGCCACCTCCACCTCCCCATCCAGTTGCTAAGGAGAAAAAAGTTTCATACTTTATTTATCTTGGCTATGCTATTCTTGGGCTGATTATCATACTTCTACTTGTATGGAAGCTTTTTAAGTTCATAAGGAAGAAAAAGAGCAATAGTACTCCAATGGATCACAACAAGACTATTAGCTCTTCTGGTGTTACAAAGACTCCTAAAAATAGGTCAGAATATTCAATAACATCTAGTCCCGAAAACTCTATGCTCTCAGCATCATTTGAAATCCTTTCAAGTCCCCTGGCTAACAAATTGAGATTCGAGGATTTGCTTCGAGCTCCAGCTGAATTGATTGGGAAAGGAAAACATGGAAGTGTTTACAAGGTCAATGTTGATGGAGTGACATTGGTAGTAAAGAGGATCAGTGGTTGGAATATTTCGAAAGACGATTTCAAGAAGAGGATGCAGAGAATACATAGAATGAAGCATCATCATGTATTGCCCCTTGTTGCCTTCTATAGTTCCAAACAAGAGAAGCTAACAGTTTACAAGTACCAACAGAATGGCAGTCTATTCAAGCATCTCCATTGTAAGTTTTATTAGCGCAAACTTAAATATTCTCACTTGAATATCAAACCATAGAATGATACATGAGTAATAGGCCTTCCTGTAAGAACTATCAAGAATTGATATCAGTAAAGTAAAATGTTAGATAGAACTATAGTTCACTTACCATATTCAGGGTCTAGCTAATTCAAAATCTTAAAGTATTTTATCAGTCTAAGCTGCTCTGACTCTCCAAAATATTGCCACACCTGTGTCAGAACCTCCAGTAAAGCATTACTTTTGGAGGATCCGACAAGCACCTGTCTGAAATTTTTGAAAAGTTTGAGTAACATAGTTGTCGGTTGATGAAGTAAGAAAGAAACTATAGTAATTAGTATTATTAGTTTGGGTAGCGAGATA

>M6-PGSC0003DMG400024661 M6 LRR-RLK

TTTGTATTAAAAAGTTTGGTTTGGTCCATTTTGCAGGCCCTCTCTTTCTGAAATTCAACTCCATTTTCCCATGTTTATACCCTGCACAGAGATATTGTTTCTGAACTACCAAAACCCCTTTTCTTAATTATAGTAAAATACCATTTCTTTCTTGTTTTGCACAAATTTCTATAACAACAACAACTCATCTGTCACACAACAAAAAAATAAATAAAAATAGAACAAATAGTACTGGAAGAAATCAAGATTCAAGATTGAGTTATGTCAGAAGAGGAATCTGATATTCTTCTTCCTGTTGTATTATTCCACTTCTTGCTTCTAATCACTACTGGTAAGTCTGAGTTTTTTTTTTTTTCGAATTTTCAAGATTGTACTCAGAATTCAGATTGTTGTTTTATGAGGTGTTTTAGCTGATTTGTAATACTAAAGTTTTGATTTTTATTTATAGTTTGTGGAGATACCCTGGAGAGTGACAAGCAAGTGTTACTGAGTTACAAGGATTTTCTTGAACATCAAAATCCGGTTAATAAAGGATACAGGCATACAGAATGGAATGCTTCTGATTCGTCACCGTGTAATTGGCGTGGGGTTTTTTGTGATGGCGGAGTTGATCGTGTTACTCGAATTGATCTCTCTGGTGATAGTTTGGCTGGAAATATGTTTTACAACTTCTCAGCTATGACAGAGTTGACATATATTGATTTGTCTATGAATACAATTGGAGGGTCTATTCCTGCAGACTTAGGCCAATGTAAAAACCTGAGATTCTTGAATTTATCGCATAATATTATTGATGGGGAGCTTAATTTGACTGGTTTGAACAATTTGCAAGTTCTTGATTTGACCATGAACAGGATTCATGGTGAGATCAGTTTAACTTTCCCTGGAATTTGTGATAGTTTGGTGGTTGCAAACATTTCGAATAACAATTTTACTGGTGAGATTGGAAGAACTTTTGATCAGTGCTGGAATCTTAGATATCTTGATTTGAGCTACAATAACTTGACTGGAGGATTGTCACTTGGTTTTGATAAGCTTAAGGAGTTTTCGGTATCTAAAAACAAGTGTAATGGCTCTCTGACTTCGTCGTTTTTCACTCCAAACTGCACTTTGCAGGTGTTGGATTTATCAGAAAATGGATTCGTTGGAGGAGTGCCTAAAGAGATATCGAATTGTAAAAACTTGGAGGACTTGAATTTGTCTGGAAATAACTTTTCAGGGCCAATTCCTGAGGAAATTGGATCAGTTATGAGTCTTCAAGCACTTTACTTGGGAAGCAACAATTTTTCAAGGGATATTCCAGAGAGTCTATTAAGTTTAAGTAACTTAGTGTTTCTTGATCTTAGTAGAAACAACTTCAGAGGAGAAATACAAGAAATTTTCAGGCAATTTACACAGGTAAAGTTTCTTCTGTTGCATGGTAATTCTTATACTGGAGGTATAGTTACCTCTGGAATTCCCAACTTAGTGAACCTATCTCGATTGGACTTGAGCGATAACCAATTCTCCGGTCCATTGCCAGTTGAACTTTCCAAGATGAAAGGCTTGAAGTTTCTGATTCTTGCATATAACCACTTTAATGGAAGTATACCATCAGTATACGGGGATATTCCGACACTTCAGGCCCTTGATCTTTCATCTAATAAGTTAACTGGCGAAATACCACCAAGTTTAGGGAAACTAAGCTCACTTTTGTGGTTAATGCTTGCTAACAATTCATTGACCGGTGGAATCCCACCCGAGTTGGGAAACTGCAGTAGCTTATTGTGGTTGAATCTTGCAAACAATCAGCTTTCAGGTCCAATTCCACCTCAGTTAGCAAGAATTGGCTCAAATCCTATGCCTACTTTCTTGTCGAATAGGGCTAAGGATAAGGTGACTGCTGGCTCGGGGGAATGCTTTGCTATGAAGAGATGGATACCAGCTGATTATCCTCCATTTAGCTTTGTATATCCTCTCCTTACCAGGAAGAATTGTAGAAGCCTATGGGATAAATTGCTCAAAGGGTATGGTTTATTTCCAGTGTGTGAACCTGGTAGTAATGTTCGTTCAAATCAGATATCAGGCTATCTTCAACTTAGTATGAACAAATTTTCTGGTGGGATCCCTCCTGAAATTGGCAACATGCAGAATTTCAGTATGCTTCATTTGGGTGTAAATGAATTCGCTGGCACGCTCCCTTCAGAGATTGGAAAAATGCAGCTAGTAGTACTTAATATTTCACAAAACAGAATATCTGGCGAGATTCCAAGCCAGATTGGCAACATTAAGTGCTTACTGAATCTTGATCTGTCCAGCAACAATTTCTCTGGTCTGTTCCCAGCTAGTTTTAGTAAATTGACTGATCTAAGCAAGTTCAATATCTCTTACAATGCGTACATCTATGGTACTATACCAGAAAGTGGGCAATTAGCCACATTCGAGAAGTCATCATACCTTGGCGTTCCATTGCTGCACCTTCCACCTTTCATTGATAACACTAGGAATAATACTATAAACAAAGGTGGAAGTTTCAAAAGGCCTACAAAGGTTGGTACAGTTTTGGTAATCATGGCTTTGCTACTGGCTTTGCTAGTTTGTGGACTTATGACACTTGTTATCTGCCTCGTTCTAAAATCTCCGATAGATAAACCAGGATACCTACTGGAGGATTCAAAGGGCCGACACGATCTTGCATGGAGTTCAGGTGCATCCTCACCATGGTTGTCTAATGATGTTAAGGTTATCCGCTTGGACAAAACAAGCTTCACACATTCTGACATATTGAAGGCCACAGGCAGATTCTCGAATGACAGAATTATAGGGAAAGGAGGATTTGGGACAGTTTATCGAGGAGTCTTGCCTGATGGAAGGCAAGTGGCGGTGAAAAAGCTACAAAGAGAGGGAATTGAAGGGGAAAGAGAGTTTAGAGCTGAAATGGAAGTACTAAGTGGAAATGACTTTGGTTGGCATCCGAATCTTGTAACACTTTACGGTTGGTGCCTTAATGGATCAGAGAAATTGTTAGTCTATGAATACATGGGAGGTGGAAGCTTAGATGAGATCATCACAGATAGAACCAAATTCACATGGAAGAAACGACTCAACGTGGCAATTGATGTTGCACGTGCTTTAGTCTTCTTACACCACGAGTGTTACCCTTGTATCGTCCACAGAGATGTCAAGGCAAGCAACGTGCTGCTAGACAAAGACGGAAGGGCAAGAGTCACAGATTTTGGCCTAGCTAGGGTTATGGATGCTGGAGATAGTCATGTTAGCACAATGGTTGCTGGTACAGTCGGGTACGTTGCACCAGAATATGGACAGACATGGCAGGCCACAACAAAAGGCGATGTCTACAGTTATGGAGTGCTGGCAATGGAGTTAGCCACAGGGAGACGCGCTGTAGATGGAGGCGAGGAATGTCTAGTTGAATGGGCGAGACGTGTGATGGGAGACGGAAGGCAAGGATTCACCAGAGCCATAATACCAGTTTCTCTTTTGGTATCAGGCCTAGCAGAGGGAGCAGAGGAATTGTGTGAGTTGCTTAGAATAGGAATAAGGTGCATTGCTGATAATCCTCATGCTAGGCCTAACATGAAGGAAGTATTGGATTTGTTAATCGCGATTTCGCGCAGCCAAAGATCAGGATCCAGTCGTAGCACTTCTCCTTCATTTTGATCAATTTTTGTTTCAAATTACACAGCAAAGTGTCAATAGTTTTTTTTTCTTCACATTTGTAACATACACACATTACATTCTTCTTGTAAACAGTTCTTTTTAGATTCTTTGTTCAGGCACTCTTGTTTGTTAGGATCATAAGAG

>M6-PGSC0003DMG400024795 M6 LRR-RLK

CCTATGGCCAACCCTGTCCTACAACACTGTGTCAAGTGTCAATCTTGACAATGAATCTAATTGAAGACAGATATATTTGGAGGAAAATCAAGATTTCTGAAAATTCCATTTGTTTCCTTTCTTTTACATAAAATTCTCTGTTGTGCATTTCCAACACCAGTAAACCATGATAAAGAAATCATTGCAGATAACCTCTTTCTTATGACTACTACATTTGGTGCAAATGAAATAGAAGTACAGTAGATCAGAGAGTACATACACACCCTCATTGTCATCATAATCATGTCAGGCAAACTAACAGTCATCTTATTCTTCTTGTTTCTCAACATATCTTTTTTATTCCCAACATCCATTTCTGGTTTAAATCAAGAAGGGGTTTCTTTAATGTCATGGCTTTCCACTTTTAACTCTTCTGCTTCTGTTGCACCTTTCTCCTCATGGAATCCAAGTCATGAAAATCCATGCAAATGGGATTATATCCAATGCACCAGTAATGGATTTGTTTCAGATATCAAGATCAGGTCGATCAATCTTCCCGCGATATTCCCAAGTCAAGTTCTTTCTTTTCACTTTCTGGAAGTGCTAGTTTTATCAAATTGTAACCTCACTGGTGAAATTCCTGCTTTTATTGGAAACTTGTCATCGTCTCTGAGGATTTTGGATCTCAGTTTCAATGCTCTAACAGGAAGTATTCCACCTGAAATAGGAAGATTATCACAGCTGAAGCAACTTTTGTTGAGTTCTAATTTCTTTCAAGGTCAGATACCAAAGGAGATAGGAAGATGTTCTGAATTGCAGCAGCTTGAGCTCTTTGACAATCAGTTTTCTGGAAAGATACCTGAAGAGATCGGTCAGTTAACGAGTCTTGAAATCTTTCGTGCTGGTGGGAACGTGGGAATTCAAGGAGAAATCCCAATGCAGATATCAAGCTGCAAAGAGTTGGTCATTTTGGGTCTTGCAGATACCGGTGTTACGGGTCAGATTCCACATAGTATAGGTGAACTCAAGAAGCTGCAGACTCTTGCTGTTTACACAGCTAATTTGACTGGTGAAATCCCTCCAGAAATTGGGAATTGCACTTCTTTAGAAGAATTGTTTGTCTATGAAAATCAAATTACAGGAGAAATCCCAAGTGAACTCGGTCTCTTGAAGAATCTTAAGAAGGTTTTGCTTTGGAAGAACAATCTGACGGGGGAAATTCCAGGAAATCTTGGAAACTGTTCAAGTTTGAAAGTGATTGATTTCTCTGTGAACTATCTATATGGTGAAATCCCTCCGTCTTTCGAGAATTTAGGCACATTGGAGGAGCTTCTATTATCAGAAAACAGCATTTCTGGAGAAATCCCATACTATATTGGCAACTTTTCCAGCTTGAAGCAGCTTGAATTGGACAACAACAACATTTCAGGTGTGATTCCACCTACCATTGGGAAACTAAAGGAGCTAAATCTGTTTTTCGCTTGGCAGAATCAACTTCAAGGGAGCATACCGACTGAGCTAGCAGATTGTCAGAAACTTCAGTCTTTGGATCTTTCTCACAATTTCCTTACAGGGTCCATCCCAAACACTTTATTCAATCTCAGGAACTTGACTAAGCTGCTGCTGATATCAAATGTTCTGTCCGGTGGAATTCCACCTGATATTGGAAATTGCACCAGCTTGTCCAGATTACGCCTTGGATCAAACAGGTTAGATGGTCCAATTCCTCCAGAAATAGGGCGTCTACCGAGTTTGAGTTATCTTGAACTGTCGGAAAATCAATTCACTGGATCAATCCCTCCAGCTATTGGTAACTGTCCGCAGCTAGAAATGGTTGATCTTCATGGAAACAATCTTCAAGGAACAGTTCCTTCCTCTTTTGTGTCCCTCACTGGTCTTAATATATTAGACCTATCCATGAACAGAATATCAGGTAGCATTCCAGAAGATATAGGGAAACTCATATTACTGAACAAACTCATACTGAATGGAAACAACATAAACGGAACGGTTCCTAAATCACTTGAACTCTGCCAGGATTTGCAGTTGTTGGATTTAAGCAGCAACAGACTCGCGGGCTTAATCCCTGAAGAGATTGGTAACCTCCAAGGACTAGATATTCTGTTCAATGTAAGTCGTAATTTTCTGACCGGACAGATTCCTGAAAGCTTCTCTAACCTCTCCAAGCTAGCCAACATGGATATTTCTCATAACATGTTGACTGGAAGTCTTAGAGTACTCAGTAACCTTGATAACCTTGTTTCTCTGAATGTTTCATACAATAACTTTTCTGGTGTTCTTCCTAACACAAAATTCTTTCAAAGTCTCCCTCCTAGCGCTTTTATTGGTAATCAAGAGCTCTGCACTGACAGAGCTGCATGCCACTTAAGTGGTGATCACCATGGATTAAAATCCATAAAGAAGATCACCATTGCTATTGTACTCAGCATTTTCATGACCATGCTAATTGTGACAGCTTCTATTGCCATCTTTATCCGAACACAGGGAGAGATATGTCAGAAGGACGACGGAGAAAATGGTTTGCAGTGGGAATTCACCCCATTCCGAAAGCTTAGCTTCTCTGTAACTGATGTAGTGCCAAGACTTAGTGAATCTAACATTGTTGGAAAGGGCTGCTCAAGCTTTGTTTATCGCGTTGAGACTCCATCAGGACAGGTGATCGCGGTGAAGAAACTATTGGCTAAGAAAATCGGTGAGGTTCCTCAGAGGGACTTTTTTTCTGCAGAAGTTAGAACACTGGGATCAATCAGGCATAAAAACATAGTGAGGCTTCTTGGATGTTGTAATAATGGCAAAACAAGATTGTTGTTGTTTGATTACATTAGTAACGGGAGTTTATCGGGACTTCTCCATGAGAAGAGGGTATTTCTGGATTGGGATGCTAGGTTTAATATCATATTAGGAGCAGCTCAGGGCTTAGCTTATCTTCACCATGACTGTAGTCCTCCTATAGTCCATCGCGATATCAAAACCAATAACATTTTGGTAGGTCCACAGTTTGAAGCTTTTCTTGCAGACTTTGGACTTGCAAAGCTATTAAACACATCATCAGATACTTCAAGAGCTTCCACCATAATAGCTGGTTCCTATGGGTATATAGCTCCGGGTGAGTACATTTCCTCAAATTTCCGCCCTATATATTCAAATAATTATTAAGTGCTCAACATTTTGATAACATGATTATGCTAGTACTATTGAATATTAGCTTTAAAGAAACACTTGAAAGTTCACTATATTAGTCTTGTACCCCGTGAAATTCGCTGTGCAAGACATATAGTAGGTATATATACTATGTTTGGACAGCATTCTAAATGTTTTGGTCCTACATTGCAGAATATGGATATAGTCTAAGAATAACTGAGAAGAGTGATGTCTATAGCTATGGCATTGTGCTTCTCGAGGTCCTAACGGGGATGGAACCAACTGATTCCCGGATACCAGAATGCACCCACATTGTCACTTGGGTCAATCAGGAGCTAAGAGTAAAACATAAGGAATTCACAACAATTCTTGATCAGCAATTACTTTTAAGGTCTGGTACACAAACTCAGGAGATGCTTCAAGTTCTTGGGGTGGCACTACTTTGTGTTAACCCATGAGCGAATGAAAGACCAACAATGAAGGATGTAGCAGCAATGCTAATGGAAATCAGGCACGAAAATGAAGATCTTGAGAAGCCTAATCGAGGAGTGGTACCAAATCCTAAAGAAGCAATTAGTTGTCCTAGTTTCTCTGGTTCATCTCAACCTCTCATCAGATCCCCTCCTAAGTAACTCTAGTTAACTTAGCCTCTCTCTAGTTCTAACTAGTTGATTCTTTCAAGTAACTAATAAGGATGTATAGATTATCAATATTCATTTCACAAAATAAGAACATTACTCTTTCTAGTCCCGACTATAAGTCGTTTTGACA

>M6-PGSC0003DMG400024890 M6 LRR-RLK

ACCATTTAATTTAGGCTAAAAACTAACTTTGATCTTCTATGTAATGTGTTGTTCTTCAGTGAGAGATAAAAGGAAAAAAAGCAAAGAGAGGAAAAGGGGAGGCTTAGCTTCTTCCTTGATCTTGAAGACTCACTTTCTTTCTTTTGAAGATGTTTGATTCTTTAAATCTATCTATACCTGAAGTTTCTCAGATTCCCTTAACTTTTTTCTCCTTTTACATTGCCTCAAAGTCTCATAATAACTCACTTTGTTTCTTCTTCAAGTCCAAACAAATTCACTTTCTTTGAAAAACTTCCAAGAAAATATGTTGCCACTTATTCTTGTCACACTCTTTACTCTTGTATGCACTTCCATTTCCTCCTCTTCCTCTTCTATTAGTACTGATTTTCATGCTTTGTTGAGTCTAAAAAAAGGATTTGACTTCTCAAATTCTGTTTTGAGTTCATGGGATAGTTCAAATCCTAGCTCAGTTTGTTCTTGGATGGGAATCAAATGCTTACAAGACCGAGTTGTATCGATAAACTTATCCAATATGGAGCTTTATGGTTCTGTTTCCCCTGCTATTTCAAGGTTAGATAAGCTTGTTGAGCTTTCAATTGATGGAAACAACTTCACTGGTGAAATCAAGATTGAGAACATGAGGAGTCTTCAGTCACTCAACATTTCAAACAACATGTTTAGTGGGAGTCTTGATTGGAACTACACAAGTTTGGCTAATTTGGACGTTCTTGATGCTTACAACAATAACTTCTCTTCTTTTCTTCCTGTTGGAGTTGTTAGCTTGGAGAAACTCAAGTACTTGGACTTAGGTGGCAATTATTTCTATGGACGAATACCGGAGAGTTATGGGGATTTGATTGGGTTGGAATACTTACAACTTGCTGGGAATGACCTGCATGGAAGAATCCCTAGAGCTTTGGGGAATCTTACTAACTTGAAAGAGATTTATTTGGGATATTTCAATGTTTTTGTAGGTGGGATTCCTAAGTAATTTGGAAAGTTGGTGAATCTTGTTCACATGGATATTTCTAACTGTGAATTAGATGGTCCGATTCCTCCTGAATTAGGGAATTTGAAATTGTTAAACACACTTTTTCTACATATAAATCTTCTTTCAGGTCCAATTCCAAAGGAACTAGGCAACCTTACTGGCTTAGTCAATCTTGATCTTTCAGCCAATGCACTTACTGGTGAAATCCCATTTGAGCTCATTAACCTTCAGCAGCTAAGTCTTTTCAACCTGTTCATGAACAAATTACATGGTTCTATACCGGATTTCATCGCTGATTATCCTGACCTGAAGATTCTTGGTCTCTGGATGAACAACTTCACCGGTATAATCCCGGAGAAGTTAGGCCAAAATGAGAAGCTACAAGAGCTGGATTTGTCTTCAAATAAGCTCACTGGTACAATTCCTAAACACTTGTGTGCTTCTAAGCAACTCAGAATCTTGATTTTACTCAAGAATTTCCTCTTTGGTTCAATCCCAGAGGATTTAGGCACATGTTTAAGTCTTGTTAGAGTGAGATTGGGACAGAACTACTTGAATGGTAGTATTCCAAATGGATTCATTTACATGCCTGACCTAAACTTAGTTGAGCTTCAGAATAACTACCTGTCTGGTACCTTATCAGAAAATGGTAACACTTCTTCAAAGCCAGCCAAGTTAGGTCAATTGAATTTATCAAACAATCAACTTTCAGGTTCTTTACCATTTTCACTTTCTAATTTCTCTTCCCTTCAAATCCTTTCACTTGGAGGGAACCAATTCTCTGGTCCTATTCCTACTTCTATAGGTCAACTCTCACAAGCTTTAAAAATTGACCTCAGTCAAAACTTTCTTTCTGGTGAAATCCCCCCTGAAATAGGCAACTGTGTCCACTTGACTTATCTAGACTTGAGCCAGAACAACTTCTCTGGCTCAATTCCACCCCGAGTATCCGAAATCCGAATCTTGAACTACTTGAACTTGTCAAGAAACCACTTGAATGAGACCATACCAAAGTCAATTGGCACCATGAGGAGTCTCACTACTGCTGATTTCTCATTCAATGATCTATCAGGCAAGCTCCCAGAATCAGGCCAATTTGCTTACTTCAATGCTACTTCTTTCGCGGGCAATCCTCAACTCTGTGGTTCTTTATTGAACAACCCTTGCAATTTTACACTAATCACTGATCCACCAGGAAAGTCTCATGGAGATTTCAAGCTGATCTTCGCCCTCGGATTGTTAATCTGTTCACTAGTCTTTGCAGCTGCAGCTATCATAAAGGCCAAGTCTTTCAAGAAAACAGGCGCGGACTCTTGGAAAATGACAGCATTTCAGAAGGTTGAATTCAGTGTTGCTAATGTTCTTGAATGTGTAAAAGATGGCAATGTGATAGGCAGAGGAGGAGCCGGGATTGTGTACCATGGGAAAATGCCAAATGGAGTTGAAATTGCAGTCAAAAAACTGCTTGGCTTTGGGAACAATAGTCATGATCATGGTTTCAGGGCGGAGATTCGGACACTAGGCAACATTCGACATAGAAACATTGTAAGACTAGTTGCTTTTTGTTCCAACAAGGAAACGAATCTCCTTGTGTATGAGTACATGAGGAATGGTAGTTTAGGAGAGGCGTTACACGGGAAAAAAGGCGGATTCCTTAGCTGGAATTTGAGGTATAAAATTGCTATAGAAGCTGCTAAAGGATTGTGCTATCTTCACCACGACTGTTCGCCATTGATCGTTCACAGAGACGTCAAATCCAACAACATTTTGCTGAATTCAAACTTTGAAGCTCATGTTGCTGATTTTGGACTAGCAAAATTTTTGGTAGATGGTGGCGCCTCAGAATGCATGTCTGCTGTTGCTGGCTCCTATGGTTACATTGCCCCTGGTACATTACTTTTTATTTTTTTTTTTTTTTTTTTTATTATTTATTATTATTTATTTTTTTATTTTTTTTTTTTATTTTAATTTATTATCTTTTAATTTTAAATTTTATTTTGTTTTATTTTTTTAAATTTTTTTTTTTTTTTTGGGTTTATTTTTTTTTTAAAATCATATATGTACGTTGGTTAGTTAATTTAGTTAGGTATGTAGACATTACCTGACAAAGTGATTTTTGCCTGGTCGTACATACAACAACAATTAGTCTTATCAATTGGTCATAAAATCAAACTTTTAAGATAAATATTTTATCGCATTTTGTTTTTTTTAGTTTGGAGTATGTTGAATTAGAGAGAAATTTTGGGCTAAGGCCCAAGATCACCCGTGATTGAAGGCATAGAGTTTAGGGCTTTGTTGGAGTTTTCTGGGTAATCGTAAATCGTATCATGGGCCCGTTTTACGAGAAAGCACTTAATTTCTGTGCCATAGAGAGAGTTCTTAGTGGGGGCCCAATAGAAGATCACTCTGCTAAAGTGGGTCAAAAAACTGACTTGACTGACACACAAAGATAATTGGGCCCCCTCGATTGTCGTGCTTGCCAACTCAACATGCACTTGTTCAAAAGAAATTAGGCATCAACTAAATGGAATAGAGTATATAATTTATAGTCACAATATTACATATTTTACTGTATAACTCTTTAAAGGAGTCAACTTTTTTAAAATTAAAGGAGGGAGAGAAAGCAATAGTGTGAATAGCAATGAAGTCAAGATATTCATTAAGGAGATTCGTCACATATATTTAAGGGGTGTTAACTGACTCCAAATATTGTTTAATGCATATATATTAAATTTGCATATACATTGTCATAAATATTAACCTATTTTTTTAAAAAAATGTTGCAGAATATGCATATACATTGAGAGTAGATGAAAAGAGCGATGTATATAGTTTTGGTGTGGTTCTATTGGAGCTGATCACCGGGCGTCGTCCGGTGGGTGAATTCGGTGACGGAGTGGACATAGTACAATGGAGCAAGAAAGTGACAAATTGTAAAAGGGAACAAGTGACGCACATTGTTGATCCAAGGTTAACCACTGTGCCTGAAGATGAAGCCATGCATTTGTTCTTTATTTCTATGCTATGTATTCAAGAAAATAGCGTAGAACGTCCAACAATGAGGGAAGTCATTCAAATGTTGTCCGAATTTCCACGACAATCATCAGAATATAATCAACCTTCGTCTTCAAAAGTTGTTTTACGGAAGCTAAAAAGCCTTGAGAATGACCAAACTACTTGTCCGAAGATTCGGAAAGAACTTTTGGTCTAACCAATTTTCATTTTTATATAATATGGTCCTCTTGGCAAGTTGTTTTAATTTTAACTTTGTATTATTCTTTATGCTTATTTAAGGTTAGGGGTTTTGTGACAAAGAGAAGTAGGAATATAATTAGTACTATGTGTTATTAGTGGTTAATTAGTTGCTTTTGATTAGTTGTTAGGGTTGGGGGTGCTTAAGTTCTTGGCAAAGGCAGACCAGGCCCAATTAAGTTGGTTCTGAATGCTTTTTGTATCATATGGTTATATACAGGTATATATTCTTCCTTTTACGGGTGAATTGATTCTCGTTCCAATAATGTTTAATTGTTTATTGTTGGTCATGTTGTTGCGGGTGAATTGGTTCTC

>M6-PGSC0003DMG400024954 M6 LRR-RLK

GGAAGGTTGGTTTTCGATGAGCTGTGTCTTCTTCTTCTCTCAATATTTTATAAACTTCCCTTCACATGGTCCCCTTTTTATCATAAATCTACACTTATTCTCCTAACTTTTTTTCTATTCAATTCCTAAAAAAAAACAGAGATGTGGGAACATTTAGAGACGCCATGAAATTCCAAACCTTTGATCTTGTTCTTCTTCTTTTTCTACTACTTGAATATCAAGCTTCTGGGCTTAACAATGATGGACTTCTGTTACTGACATTCAAGTACAATATTCTGAGTGACCCTTTAAGTGTTCTGCAGAACTGGAATTCTTGGGATGATACACCATGTTCATGGAAAGGGGTAGGTTGTGGAAATCCAAATGCTTCAGACCCAAACCTTCGAGTGACAAATTTGTCTCTTCCTAATTCTCTGCTTCTTGGTTCTATTCCTTCTAGTCTTGGTATGATTCAATATCTTACAAATCTTGATCTTTCTAATAATTCCATTAATGGCTCTATTCCTCTTACTCTCTTTAGTGCCCCTGAGCTTCAACGCCTTGATTTTTCTAACAACAGAATATCCGGCCAGTTGCCTGAGCTTGTTGGACATTTAAATAATCTTCAGTTTCTTAACCTCTCCGGTAATGCTTTGACGGGAAGATTACCGGCAAATCTCACTAGGCTTAGTAATCTAACTGTTGTTTCTTTGAAAGATAATTATTTTTTTGGTGCTGTTCCTTTTGGGTTTGATTCAGTTCAAGTTTTAGATCTGTCTTCAAATTTGATTAATGGGTCATTGCCTCCTAATTTTGGGGGCAGTAATCTCCGTTACTTTAATGCTTCTTTTAATAGACTTTCTGAAGATATCCCACCGGAATTTGGCAGCAAAATCCCTCCAAATGCAACTATAGATCTCTCTTACAACAATTTCAGCGGTGCAATTCCAGAGTCAATTGTTTTTATCAACCAAAATAGAAAAGCTTTTTCTGGTAATCCAGAATTATGTGGTGCGCCGCTGAAAAATTTATGCCCAATTCCGTCAACAGTAACAACTCTGCCAAATTCCCCTGAACCAACTTCTTCCCCAGCAATTGCAGCCATTCCGAAACCCACAGACTCAAACCCAGCCGCCGAATCTCCAAAAGAGGGCAAAAATGGGTTGAGAACTGGCACAATTATAGGCATAATAGCAGGGGATATTGCAGCAGTAGGAGTTTTAGCACTGATTTTCATGTACGTATACCGAGCCAAAAAGAAGAAAAGGAACATAGAAAATAGCATTAAAAAAGAAGCTGAAACTGCAAAAGATTTTGACTGGGCATCGTCAGCATCAGAAGAAGAATACAACTGGTTAAGGTCATGGACTTGCTTGAAAAAACAAAGGCACGGAGATGATGACGAATTATCAGAAGCTTCCCATTCAGAAAGTGAAAAATCCCAATTGGGTCATCCTCAGCAGAATCACGTGCATACAGAGCAGAAAACAGGTGAATTAGTAACGGTTGATGGAGAAAGGGAACTCGAATTAGAAACACTACTAAAAGCATCTGCATACATTTTGGGAGCTAGTGGGTCGAGTATAATGTACAAGGCAGTGCTAGAAGACGGTACTACACTGGCGGTGCGACGAATTGGTGAAAGTGGTGTGGAACGGTTCAAAGATTTCGAGAATCAAGTAAAGGTAATTGCTAAATTGGTGCACACAAATTTGGTTAAAATCCGTGGCTTCTATTGGGGCGCTGAAGAGAAATTGGTCATCTACGATTTCGTTCCTAATGGTAGCCTTGCCAATGCACGTAACAGTAAGTCCACTTTTTCCCATATACATTATGTTTTAGATCAAATAACGTTACATCAACTTTTTTATTGAGTAGAATCGCGTTTTTTGTTATTGTTTTACTATTTTGCTCCTGTTTTACAGTGGAGAATAAATTGAAACAAAATTGACTAGTCATTATGTGTGTACATTGTACCTAATCATCTCTAATTCTTAATTTGTTTCAATAAGTATCTTTTGCATAATTGCGAACATTTTGGAATATAAAAGTTCGGAGTAGGTTTGTACTTTAAGACATTTGGTTATGCTTAATATTCAAAAGAAAGACGTAACAAGTCACTCAAATTAAGTTCACACAAGTATAAATTTGTGCGCAAATTAGAAAAATATTATGAATTTCTGTTAAAATGCGCGGGAAGTAAAGGAGATAGTGGACCTTGGGGTGGAGTTGTTGAAACTTTAGTGTTTGTTTTTAATGTGGAAAATAGCAGTAGTTCCCTCACTGCAAGTCAACCGTGTCTGTTTCTAAATGGGCGCCTCACAAATTCATAAATACCGCCAAACGCCCCCTCTTCTGCTGTCACTTACATGGAAAATGACTACCTTTTTTTTAAAAAAATATTGTACTCTGTAGGCCATAAGTATTCTCTTTTGTTATGTCAAATTCATGCATTTCTTTTGCCCACAACACCAATAATGTTAAAAGAATCTTCACCTTATTTACCATTTATAGTATGTTGTTTATTTTATAGATATTATTATTGTGAGAATCTTAGCGCTCAGTTGATTGACTAATTATAACGTGCGCATTTTGATGGTTTCAGGAAAGGCTGGTTCTTCGCCATGTCACGTACCCTGGGAAATCCGGCTGAAGGTTGCTAAAGGTGTGGCTCGTGGGCTCACTTACATTCATGAAAAGAAGCATGTCCACGGCAACTTGAAGCCCAGTAACATTTTACTAGGGGAAGATATGGAGCCCAAGATTGGTGATTTTGGAATTGAAAGGCTTGTAACAGGAGACAGTAGTCATAAGACCTATGGATCTGCTCGTAATTTTGGAAGCAAAAGGTCCACAGCATCCCGTGAGAGCTTCCAGGACTTTACATCTGGGCCTACTCCAACCCCAAGTCCAAGTGCATTGGGCATATCTCCTTACCATGCACCCGAATCGCTTCGTAGCCTCAAGCCCAACCCGAAATGGGATGTATTCTCATTCGGGGTAGTCTTGCTGGAGTTGCTAACTGGAAAGGTGATTGTCTCGGATGAAATGGGACCCGCATCGGTCATCGGTGCTACCACATCAGCAGAGGAAGAGAAGAGTAAAGTGTTACGGTTGGCTGATGTGGCGATACGTGCTGATGTGGAAGGGAAAGAAGATGCTTTGTTGGCACTAATGAAAGTAGGATATAGTTGTATATCACCAATACCTCAGAAGAGACCAGGCATGAGAGAGGTTGTTCAAGCATTGGAAAAATTTCCAACTAATTCAACCAGTTCATCATACTATTATGGACCTTAATTAGCACTGATTAAAAGATGGACACAATTGTAGAGACAAGAGACTTTGTATGTAATGAGACTGACATCTATGTTATTTTATTTTATTTTCAATTTTTTTTGGTTTGTCTCTTAATATAGTATTAGCCATTGGTTTTCTTTTTTAACAGTGATTAGTGTAATTGTCCGCGTTTTGTTGTGTTGATTTTTCCTAAACCTTATGGTGACTTTGGATTACCAATATATATGACAACTTATTCTGTG

>M6-PGSC0003DMG400025076 M6 LRR-RLK

TAATCTCTTTGCTATATTTAATGTATAGTTTCATTTGTTTGCTCTTTAGTTTCTCAAGAACTGTCACTCTCACTCAAAATCCTCTCCGCCACCGTCAACCGCCGTCAACCACCGTCAACCGCCGTCATGAAAAAATGCAAAATGACAAATGTGTTTGGTTCATGCTAGTGACCATTTATCTTCACCACCTTCATTAGTGAGCTGCTAGCTAGCGCCGCCGTGTACGGAGGAGCAAAGAGGAGCTGGTTTTCTCCAGTAAGAGAACTTTAATAGTAGCAGTTTAGTAAATGTTTCATTAAAGATAAACAAGAACTTTTTAGTTCACATGCATGGTAAGTAAATTTTTTGAGTTAGTGTGAGTTTGAAGCAAACCTCCATTAACACACCACATGTATGCTGTCTTGTTAGAGTGAACTTTCAAACACTATTATCATTTTTTTCAACAACCCCATTTCTTGACTTTGATTTTCACTCAATAATGGAGAAGAAAAGCTACCATTTTTACACTTTCCCTTCATGTTATTTCTTGTTTTTGTTGATTTCTTGGTTAAGTTTGAGTCCAAAGATTGTATTTTTAGTGAGCTCTGTTTCTTCAGATGGACAAGCGTTGCTTTCTCTGCTTAAAGCAGCTGACCCTTACACTAAATCATCATCTTCTGTGCTTTCTTCTTGGAACCCTTCAAGTTTAACTCCATGTTCTTGGCAAGGGATTACTTGTTCTCCTCAAGAAAGGGTTATTTCACTTTCTATCCCAAATACTTTCCTAAATCTCTCTTACTTGCCTTCTGAACTCTCTTCCCTTTCATCTTTGCAGCTTCTTAATCTTTCTTCTACTAATATTTCTGGAACAATCCCTCCATCTTTTGGTTCTTTTTCTCATCTTAGGCTTTTGGACCTTTCTTCTAATTCACTTTCAGGTTCTATTCCATCAGAACTTGGTGGGCTTAGTTCACTTCAGTTCTTGTTCTTGAATTCAAACAGATTGACTGGTAAGATCCCACCAGAGTTAGCTAATCTTTCTTCATTAGAAATCTTTTGTCTTCAAGATAATCTCCTTAATGGGTCAATTCCATCACAGTTAGGGTCCTTAGTGTCACTCCAGCAGTTTAGGATTGGTGGAAATCCATATTTGAGTGGTGAAATTCCTGCACAATTAGGCCTGCTCACCAATCTTACCATGTTTGGTGTTGCGGCGACTGGTCTTTCGGGTGTTATTCCGCCAACATTTGGGAATTTGATCAATCTTCAAACACTGGCAATTTATGATACTGAAGTATTTGGTTCGATACCGCCTGAACTTGGGATGATTTCGGAGCTTAGGTACTTGTATCTGCATATGAATAAGCTCACTGATTCGATCCCTCCTCAGTTAGGTAAGTTGCAAAAGCTTACTAGCTTGCTTTTATGGGGAAATTCATTAACTGGACCTATACCAGCTGAGGTTTCGAATTGTTCGTCCCTTGTGATTCTTGATGTTTCTGCGAATGAATTGTCTGGTGAAATTCCTGGTGATTTGGGGAAGCTATTGGTTCTTGAACAGCTTCATTTGTCTGATAATGCACTTACCGGTTCTATACCGTGGCAGTTAAGTAATTGCACTAGCCTGACAGCTCTTCAGCTAGATAAGAACCAATTATCAGGGCAAATTCCCTGGCAAGTTGGTAAGTTGAAGTACTTGCAGAGTTTTTTCTTGTGGGGGAATTCGGTTTCAGGAACCATTCCAGCTGCTTTTGGAAACTGTACTGAGCTATATGCACTTGATCTTTCGAGGAACAAGCTCACTGGATCGATCCCTGAGGAGATATTTGATTTGAAGCAGCTGAGTAAATTGTTGCTCCTTGGGAACTCGTTAACTGGACGATTGCCTCGAAGCGTTGCAAGATGCCAGTCTCTGGTGAGGTTGAGGCTTGGTGAGAATCAGCTTTCAGGACAGATTCCTAAGGAGATTGGTCAATTGCAGAATCTTGTGTTTCTTGATTTGTACATGAACCATTTCTCTGGAGGCTTGCCTTCTGAAATTGCCAACATTACAGTTCTTGAGCTGTTGGATGTGCACAACAATTACCTAACTGGGGAAATACCACACCAGATGGGGGAGCTTGTGAATTTAGAGCAACTTGATCTTAGCCGGAACAGCTTTACCGGTGAGATCCCTTCGAGTTTTGGCAATCTCAGTTACTTGAACAAATTTATTCTCAGTAACAATCTGCTTACTGGTTCAATTCCAAAGTCTTTTAAGAACTTGCAGAAGTTAACTCTACTTGATTTGAGCTCGAATAGTCTTTCTGGCGAGATTCCATCTGAGCTTGGCTATGTCACAAGCTTAACAATCGGTTTGGATTTGAGCTTGAATTGTTTCACGGGTGAACTTCCTGAAACATTGTCCGGTTTGTCACAGTTGCAATCTCTTGATATTTCTCATAATTTGTTGAGTGGAAGGATCACAATTCTAAGCTCCCTGACCAGCCTTACTTCCCTGAATGTTTCAGACAACAATTTCTCGGGGCCTATTCCTGTCACACCATTCTTTAGAACTCTCACTTCAGATTCTTTTCTGGAGAATTCACTCTGTCAATCAGTTGATGGTTATAGTTGTTCTTCACATATAATGGGAAGGAATGGGTTGAAGTCACCGAAAACCATAGCTCTGGTTGCGGTGATTCTGACATCTGTAGCCATAGCAGTCGTGGCTACATGGATTCTTGTGACGCGAAACCACAGATATGTGTTTCAGAAGTCCCAAGGCGTGTCTGCCTCTTCGGTAGGAGCAGAAGATTTTTCCTACCCTTGGACTTTCATCCCCTTTCAAAAGTTCAATTTCACCATAGACAACATCTTGGATTGCTTGAAAGATGAAAACATCATTGGGAAAGGATGTTCCGGTGTTGTATACAAAGCAGAAATGCCTAACGGTGAGGTGATTGCAGTGAAGAAGCTTTGGAAAACAAAGAAAGACGAGGAGCCAGTAGATTCTTTTGCAGCAGAAATTCAAATTCTTGGACATATTAGGCATCGAAACATTCTGAAGTTATTGGGATATTGTTCAAACAAAAGCGTTAAGCTTCTTCTCTACAACTACATTTCAAATGGCAATCTCCATCAGCTCTTGCAAAGTAACAGAAATTTGGACTGGGAAATTAGGTACAAAATAGCCATTGGATCAGCTCAAGGCCTTGCTTATCTTCACCATGACTGTGTTCCAGCCATTCTTCATAGAGATGTCAAGTGCAATAATATACTGATCGACTCCAAGTTTGACGCTTATATTGCAGACTTTGGACTAGCAAAGCTAATGAATTCGCCAAACTATCACCACGCTATGTCCAGTGTAGCAGGATCTTATGGATATATTGCTCCAGGTATGCATGCACTTAGAATATTTTCAATTACTAATATTTCAACATAAAAGCCAAATCGATCATAACATCTTTCAAATATTTACTTTATGAGTCATAGTTCGTTGAGTGTGACACAAATATTTGTCAACTATGGTTGCAGAATATGGATATACAGCGAATATAACAGAAAAGAGCGATGTCTATAGTTATGGAGTGGTGCTACTGGAAATACTTAGTGGGCGTAGCGCTGTTGATTCTCAGATAGGTGATGGGCTTCACATCGTTGAATGGGTAAAGAAGAAAATGGGAAGCTTTGAACCAGCTGTTACAGTTCTCGACACTAAGCTACAAGGTTTACCCGATCAAGTAGTCCAAGAAATGCTGCAAACGCTTGGAATAGCGATGTTCTGTGTCAACTCATCGCCGGTTGAAAGGCCAACAATGAAGGAAGTTGTGGCACTTCTAATGGAAGTAAAGAGTTCACCTGATCAAGAATTTGGGAAAACTTCTCAGCCTCTAATAAAACAGTCCTCAACTCAAAGTTGATTAATTTTGTTATGTTTTTTTTTTTTTGCTTTTGACTAAGAGGAGTAGGAATTAGTTTTCTTTAGTTCCCAAGATATGATCAATTCATTGTTCCAAGCATTGGATAATGTCTTGGGGTCCATTGTGTACATATTTCAGAAACATTTATGTAGTTTATTTGCTTCTTCATTTCTTCCTATCCATCTCTGTGCTGAGATTTTAGCTTTCAAAATGCTTGCAGTTGCACCATATAATATTATAGTTTACCATAGTTTCAAAAAAAGAGAAATATAGTTTGACCTATTCAGAAAAATTCTATTGTTGGTGTACGAATGTGGTTTCAAACCTTCCGGTGATGCTAAAAGTAGTTTCAGAAAAGAGAAATATAGAAAGAAGAAATCAAAGAAGGGGAATATTGAAGCAACGGTAAAATTATCTCATGGGTTCAAACCGTGGAAGCAGGCATTAATACATTAGGTTAAATTGTCTACCTCATATCCTTTGAAGTGTGGCCTTTCTCTGAGCTTTGCTAACACATGATGTTTTGTTCATTGAGTTGTTTGTAGAAAGAAGAAAACAAACAACTCTTAAAAACAAATATAATACTCATGTAATATCATTAGAGACTATTGGTCTAGTGGTAAGAGTGTAGCGCATAATGCGTGGGTAGTTGTGTC

>M6-PGSC0003DMG400025330 M6 LRR-RLK

TTTTGAATTCTCCACTGGTGTCTGGACAACGGTTCTTCTCTCTTTTCTTTTTTTCTTTCTTTCTATTATACACAACTTTCTAGACTACTTTCAAAGTAAAGTCTTCTCTTTTTCAAAATACACACCCCCGCCCCCCCTCTTCCTCTTTTTCTCACTTCGTGCTTGAAATTTCATTATTATGGATTTTCCTTCTGTAAGAACTGTAATCATACCAATTGTTTCTATCAAAACCCCACCATTTTAGCTTTACTCTTTTTTTTTTCAAGATCTGTATTGATTTTTGAACTTGCTACTACTCCACATTGCACTTGATATTTCTAGTCCAAACTGAAAAAGTAAGCTCAAATTTGCATTTTTCATCGAGAAATCGAAATGGGTGTCTGGGAGTTGATCATTTTGGGACTATTTACAATGACCATTTTCAGTAGAAGCTGTTTTTCACTAACCCCAGATGGTAAAGTTTACTTCTTTGTCTAAGGTTTTCAGCATTGTATGTTGTTCTTGAGCTGAAAAGATTCTACTTTTTTAGGTTCTGCTTTGTTGGAAATGAGAACCCTTTTGAATGATACAAAAAATGTTTTGAGCAACTGGAATGATTCGGATGAGTCGCCTTGCAGGTGGACTGGCATTTCTTGCTATCCTAATGACCAAAGAGTTCAATCTATGTATGTTTAAATCACTTTTTTATTTTCACTATCAGTGCTTTATTCACTTTGTTTGCTTGGGTTCATTTTCTGCATTTTGTTTTTTAATTTTCAGCAACTTACCTTATATGGAACTTGGAGGAATTATATCTCCTAGCGTTGGTAAACTCACCAGATTGCAGAGATTGTAAGTCCCATAGTCATTGAGGCAATGCAACAAAATTGTATTTGGGCTAAAGTTTCTGTCTTGCTACTTGCCTTGATAAACTATGTGTTCTTTCCTAAAACCAGGGCACTTCACGAAAACAGTCTGCACGGTGTGATTCCTAATGAGATTGGCAATTGCCCTGAACTCAGAGCTATGTAAGTAGGAGTTTCATATATAACTGAGTGATAACATGTTTTCTTGGAGATTCCACATGTTTCTTGATGAATTTGTCACAAGCTAGGCATTATTTATGTGTACAGATACCTCAGGGCGAATTTTCTTCAAGGAGGCATACCATCAGATATTGGAAATCTTTCAATACTGACCATATTGTGAGATATATAAATTTTTTAATTCACCATCTTTTTATATAGTTAATGAGGTTGTTGTTTTAATGATTGGTACCTTTTAATCCTTTTTAGAGATTTATCAAGCAATTCACTTAGGGGTGCTATACCTTCTTCTCTTGGTCGTCTCAGACATCTTCGTCTTCTGTAAGTTCAATTTCTGTTCATTGTAAATTTTTAATTTGATAACTTAATGCATTACACTGATGTGTTAAACATGAAAATCAATTCAGGAACTTGTCAGCCAATTTCTTCTCGGGTGAGATCCCAGATGTTGGAATACTGAGCACTTTTGGTAACGATTCGTAAGTTCTCCTTCCATTGGAATATAATCCCACATTTGATTTTGCCCTCTGTATTCATCATTTTCTATATACTGTAAATAAATTTAGATGGCTCAAAACTCAGAAAATACAATTAATCGTGAATAATGCTTCACATTATATGTATAGAAGTGCAGCAAAATATTCACATTCCATTTGGCACTACAATCACACTTATAGGAAATACAATACAAGATAAATCAGGTCTTTGCACACCAGTTTTCTGGTGATTAACTGCTGTTTGCTGCATAGGCTAGGTTTGTGGCAGTGTTACAATGACACCCAGCAAATACAAACTAAGCAGAATACCCATCAAAAGATCGATGTATTAACATCACAAGTGCAGTTGAGATTTTTTACAACTGAACAAACCGAATCTATCAATTGATAGTCCTTTAATCATGATCCTAGAAGTTAAATACCATGTGATAATTGAGATGGGAACTACATTAGCAAATTGATGGTGCTTCAATTTTAAAGATGGTGATTAATTTGTAAGTATACCTGACTTAAATTTATGGTACTCCAATCATGCATATGTGTTGCATATGTTGAGATAGACTTTTTGGTGTTAAATTGCCTAGTGGGTGATTGGCTGCTACATAGTTGAAACTAGGAATCTAGTCCTGTATCATGATCAGTTTTACTTGTGTACCTTGCAAAAATTATTCTTCCCCATCATCTCAAAAGGGAATTTGAAGCTTGATGCTCTCAAAGTCTCAAATAGTTAAGCTTATAAAGTATTTTCAAATGTCACAAGGGCAAGAATGTATGTGATGGGTCAATTCACGTAGGTTTCTTTCTGGAGTAAATTCTTAGTCATCCATTGGTAAGGGTGTTGCATCACCAACACTAGCTAGAAACAACTTGTTTAACTGGAGTCTCGTGAACTGACCTGCCGTTTACCTTTCTTCTTCATAGTCTTGGGAGATAATATTTTCAGTGTTAAATTTTTCTATTAATGGCTATCTTATAGCAGACTATTCTGATGAGGATAATGCTTATGCTTAGGTTTATTGGCAATTTAGATCTCTGTGGTCAACAAGTCCGCAAGCCTTGCAGAACATCTATGGGATTCCCTGTAGTATTGCCTCATGCTGAAAGCGATGAAGCTGCAGGTTGTTATTTGCTACTGCTTCATATCTATTTCCTTTTAGAAGTTTACTGGCTTTTGGAACTTCCCTTTCCTCCACCAACTGAATCTATTTTTTTTCTCTTTTACTTCTATGTAGAAAGCTTGATTTGTTGCTAAATTAATGTGATTTCTGCAGTCCCTACGAAGCGATCCTCTCACTATATTAGAGCAGCAGTTATTGGTGCAATAGCAACATTGGGTTTTGTACTCATTGTGTTCTTTATTTTCCTCTGGGTTTGGTTGTTATCTAAGAAGGAAAGGACAGCAAAAAAATACACAGAAGTCAAGAAACAAGTCCACAAAGAGCCAAGTAAGTGGGTGAATAGTACAATACCATTTCTTTTCCTTCTCCCAATTTTTGTCTTAATTTGTCGAGGCTTCAGGTGCCAAGCTTATTACTTTCCATGGAGATCTTCCTTATCCTTCATGCGAGCTAATAGAGAAGATTGAATCCCTTGATGATGAAGATGTTGTTGGGGCAGGAGGGTTTGGTACTGTATATCGGATGGTTATGAATGATTGTGGAACATTTGCTGTTAAAAGAATAGACCGGAGTCGTGAAGGCTCTGATCAAGTCTTTGAGAGAGAGTTAGAAATCTTGGGTAGCATCAAACACATTAATCTGGTCAATCTACGAGGTTACTGCAGGCTTCCTACTGCAAGGCTTCTCATCTATGATTATTTGGTCATGGGAAGCTTGGACAATTTCCTACATGGTATGTTTCCAACTTTTCAAAATTTGTCCCCAGATCCTTCCTCCTCTCCACATCTCCTCTTAAGCTTATACAAGACATCAAATATGGAAAGTGTGTATATTTTTGCCATACATTTTTTATTTGGTTTATCCCGAATACAACCATTATTTTCAGTCATCTGTTTTTAAACTGGAGAGGGGGAAAAGCGCTATTGCTTTCATTTAAACAATCTGTTTTTCTTGGCTATTCATTGCATGCATCTTTATTGTATTTTTGCAGAACGTGTGGATGATCGGTTGTTAAACTGGAATGCACGTCTGAAAGTAGCTCTTGGTTCTGCAAGAGGATTGGCCTACCTACATCATGACTGTTGTCCTAAAATAGTTCATCGAGATATCAAATCAAGCAACATACTCCTTGATGAGAATCTAGAGCCTCGTGTTTCAGACTTTGGCCTTGCCAAGCTTTTGGTGGATGAGGAAGCTCATGTCACGACTGTGGTTGCGGGTACTTTTGGTTATTTAGCACCAGGTATGCATATTCACTTAAATCAGCAAGGCATAAAGATAGATTCACATATATGTTGCCTTTTAATTGCAAGAATGGTTTATTTTCTTATTTTTTCGCTTGGATAAACCTTAACCTATTCTTCTTTTTATAATCATCTGTATGTAGGTTTGATTTGCTTAACCTCTATAAATTGAAATTCCAAACTCACCTCATCTTCTAGTGAAATGTCTGACCAACGATTTCTTGTTTGTAAATTGTAATATAACCAGAGTACTTGCAAAGTGGAAGAGCCACAGAGAAATCTGATGTGTACAGCTTTGGGGTTCTCCTTTTAGAGCTTGTGACTGGAAAAAGGCCAACAGATCCATCCTTCGTAAACCGAGGCTTAAATGTGGTTGGCTGGGTATGCATGCTTCCCTTGAAACTCACCAAATTTTAGAATTTCATGAAGACTCGTTAATTTTTTAAACTTCATAGGAGACACTAGTTCGGAAATTATGGGTGATTGTGATATGGAACTTCCCTTTTCTGTATTTAGCTGTACAATTTCCACATATTGATAGTGTCACAATTTGAAAAATGAACAGATGAATACCTTGCTGAAGGAGAATAGACTGGAAGACATCCTAGATGCGAGTTGCAGAGATGCCGATGCAGAGACGGTAGAAGCAATTATAGAAATAGCTGCAAGATGCACTGATGCAAATCCAGATGATAGACCTACAATGCAACAGGTGCTGCAATATTTGGAGCAAGAGGTCATGTCACCATGCCCGAGTGATTTCTATGACGAGTCTCATTCAGATTATTCTTGATGTATGGAATAATTTCCTGGGACAAATATTTAGAAACAATTTGCAGCCTGATACTGAAATGTGTAATCTGCCATTGCAGATACTCTGTATGCTCCTTGCTTTTGGAGCCCTGCTGCTTTTCTTCCTAATGTCTTTGTGCATCTCTTGTGCCTTTTTATGTAGATACCTGCAGTTAGTTTTAAAGCTACAATTATGAGTCCTAACTAGCCATGGCATTATGTAGTATTTATAATCTATTCATCACTATTGAATTTTCCTTCACCTATAATAGAGTTTAGGTTCCTCCTAAGAAAGTTCTTTAAGAAAATTTGATGAAATATTTGAGGACACTCAACTACTGGTGGATGTCATATCTGCAAAGCATATTGTATGCCACTCAATTATCA

>M6-PGSC0003DMG400025635 M6 LRR-RLK

ACCAAGTTGAGGCTTCTTGGATTGTCAGTAAATAGCTTATCCGGGGAGTTCCCTCCTCCATTGTACAATTTGTCATCCCTGGAACTAATATCACTTTCCTTTAACAACTTTTCTGGTAATCTCAGATCCGGTTTAGGCAACTACTTCCCTAATCTCCAAATACTTGGCGAAATGTCAATTCATTGGTTCCATACCGTCCTCTTTGGCCAATGCTTCAAAATTGCTAGAGCTTGATTTTCCTGTAAACAATTTCACTTGAAATATACCCCAAGGGTTTTGGTAACTTGCAGATTTTCTTGTGACTCAACGTTTGGAGTAACCAACTTGGATACGGTAAGCATGATGACCTTGACTTTGTAATTTCTCTTACCAACTGCCAGCAGTCTATAAATGCTCCATTTTGGAGACACCAGTTTGTACGTGCATTACCTCATTCAACAGTCAACCTTTCTAGTCAGTTACAACGCCTACTCTACTACGGAAACAGAATTACTGGAAGCATACCCAGAGATATCTCCAACCTGGTGAATTTGATCTACTTGAGATGAGTAACAACAATCTTACAGGTAGCACTCCAGGTTCTATTGGAAGACTGACAAACTTGGGAGGTCTCAACTTGGGTAACAGTCTCTTGACAGGGGTCATCCCTTCTTCAATAGGAAACCTCATTGAACTCGTCTATCTTTATTTGGGACTAAACAGGTTATAGGGTAACATACCTTCAACATTGGGAAACTGTAGCCAACTGCTAAAATTGGGTATTTCTGAAAACAGCCTAACAGGAACTATACCACAACAACTTTTTTCTCTCTCATCCCTCACAAATATTTATGCATCTTATAACTCTTTGACTGGTCCATTACCGGTCTATATTAGTAACTGAAGTTATCTTACTTATCTTGATTATTCTCACAACAATTTTTCGGGGTTGATTCCACGAACTCTTGGAAAATGCTTGGCCTTGGGGGATCTATTTGAAGGGAAACTCCCTCCAAGGGACCATTCCCAATTTAGAAGATCTGCCGGATCTCCAATCCTTGGATCTTTCCCAAAATAATCTATCAGGGCCAATTCCTCACTTCATTGCAAGTTTTACTTCCTTGCTCTACTTGAACTTATCTTTTAACAATCTAGACGGTGAGGTTCATGTCACCGGAATATTTTCAAATTTGAGTGCAGATGTATTAAACTGGAACTCCGAGCTTTTTGGTGGGATTCAAGAGCTGCATTTACGACCTTGTGTTTATCAAAAGACACGGAAGAAGCATGTACTTCCCTCAAGCATGATACTTCACAGAATGAATATCATAATATGTGGCCTCTGCACTTCATTATCTTCACCATGAGTGCCAAACACCCATGATCCACTGTGATATAAACCACAGAACATTCTTCTTGATGAAGATCTCACAGCTTATTTGGGTGATTTTGGTTTGGTGAGACTCATTCTTGATGAAGATCTCACAGCTCAGTTTAGCTCACTTGGAGTTATGGGAACCATTGGCTATGCAGCTCCAGGTAGTGTTCCTAGGTTAACAAAAACTGTGATATGAAATTTCTTCAAACAATAGTTGTTTATTTATAAGCTAAATTTGCTTGTTGCAGAATACGGCATGGTAAATAAGGTCTCCGCTTTAGGAGCTATGTACAGTTTTGGGATTCTCATATTGGAGATATTCACTGGAAGAAGACCCACAGACACTTTGTTCCAAGCAAGCTCTAGTCCCCACCACTTTGTCGAAACAGCGTTGCCAGAGAAAGTTATGGAGATTCTAGACAAAACAACTTTTCATGGTGAGATGAGTAAAGCAACCAACGTCAAAGAATACTGGGGCAGTATCAAGAAAGAAGAGATGGAATGCTTGGTTAGCATACTTGAGATTGGTGTCGCATGTTCAGCTGAATCCCCAAGAGACAGATTGACCGTGACACAAGTTTACAGTAAGTTGACACTAATGAGAGAACAATTTCTACAAAGGGGAGGTTGAGGCCACAGGCTACAAGTGAAAGCATTGGATTTCATCTAAAATGGTTTAAATTCTGTCTACTTGGTAAGAATGGTAACATTCTGTCCAGCATGCAGAAGAACTAAGGTAGTGTCAGATTTCTTGTTACTTAATGGATTTAGTCTAGAAACATCAGATATTAGTTCTTAATGTCTACATCAGATAATACGTTTCATTCATGAATGTAAAGAATCAGATGTTTTAGGAGATCATAAGGCAAGATCAAATGGTCATGGTGAGAATGTTCCTTTAAGGTTGTGATTCTAGACACAGATGAGGTATTCTTGTCCCAAAGTTGGAGCATCATGATCAATACTTGGAGGGTTAGTGAGAACACTGTTGATCTTTGCATTGCTTCCTTAGCTTTGATTGTCGTTTGTCCTTGTCCTTTATTTGAATACCAATGCCATGCCCACATGACATTGTTTAGCCTTCATCTTGTCAAGTCCACACCAAGTTGCCTTGCCCTCAAGTCCTTGGTCCTCACTTGCCACTTTGGAATCAAGCCTTGGTGCCATTGTGCACACCTTAGCTTTGTCAACCGTCCGTCGCCAACTTAG

>M6-PGSC0003DMG400025771 M6 LRR-RLK

CAACACTTTCTTTTTTTTGCCTTAACAATTCTCCATCTCTTGCTGCCTTTTTGGTCACCAAAACAACTTCTCTAATAGTTGGTTTACCTCTTCCTCTTTCTTCTTATGTATATAGAAAGTAATCATGAATAATCACTTCAAGAAATGATGACTATCTCAGAACATTACTATTCTTTCTGGTCATTTGGGGTCTTTTTCTTTCTTCTTTTATGTCCTTCATTTGGTCTCAACATAGATGGAACTCTCTTGTTGTCTTTCAAATACTCTGTTCTTGATGACCCTTTATCAGTTCTTGATAATTGGGACTAAAATGATGCAACCCCATGTTTATGGACTGGTGTAACTTGTGCACCAGACATGTTTAGAGTCATTAGTTTACTACTCCCAAATTCTAACCTTGTTGGTTCTGTTCCACAAGAATTAGGCTATATTCAACATCTTCATACTATTGATCTTTCAAATAATTTCTTGAATGGAACACTTCCTGTTTCACTTCTCAATGCTTCTGAACTTCAAGTTCTTTCACTGTCAAACAATGCTATTTCTGGTGAACTTACTGAATCTATTGGAGTGTTAAAGAGCTTAAAGGTTCTAAATCTTTCTGTCAATGCATTTGTAGGAAATATCCCACAAAAGCTAAGTTCTTTGCAGAATCTAAAAGTTGTTTCTTTGAGTAAAAACCTATTTTCTGGTGCAATACCAACTGGATTTCAGTTTGTTGAAGTGTTGGATCTTTCTTCTAACTTGTTGAATGGAACATTACCTGATGATTTTGGTGGTGATAGTTTGAAATACTTGAACTTTTCATCAAACAAGTTATCTGGTTTAGTGTCACCACAATTTGCCAAGAAAATACCAACAAATGCTACAATTGATCTTTCTTTCAACAGCTTTACAGGTGAAATCCCAGAGTCATTAGCTTTATCAAACCAGAAAACAGAATTCTTTGCAGGAAATACAGATCTTTGTGGCAAACCACTCAAGAAACTCTGTACAATCCCTTCAACACTTTCATCTCCACCTAATATAACCACTAATCCTCCAGCAATTGCAGCCATTCCAAAAGAAATTAACTCAACCCCACTTCAAGATTCTGATGGAACAACTCAAACTGCAGCTCAGAATCAACAACAGCATGGGCTAAAGCCAGGTACCATAATAGGAATCATAGTTGGTGACTTAGCAGGTGTTGGTGTTCTTGCTATTGTCTTTCTTTATGTCTACAAATTGAAGAAGAAGAATAAAGCAAGTGAGGCAACCATAGAAAGCTCAATTGACAAAGATCAAAAATACAACAAAACACCAGAACCATCAATACTTGTTGTAAAAGAAAAAGACACCATTTTTCCTAGTTGGCCATGCTTGACCATAAATCAAGAAAGTTCAGATACTGATGACAGCAAAAACCAAGAAAGTGAAGACCAAATTGAGTATGAAACAGAACAAAAGAATGAGAAAAGCAAGAAAACAGAAAGGTCATTTGTAATGGTGGATGGTGAAACAGAATTAGAGCTAGAGACTTTGTTGAAAGCTTCAGCTTATATTTTGGGGTCAAGTGGAGCTAGCATAGTTTACAAAGCTGTGTTAGAAGATGGAACTGCATTTGCTGTTAGGAGAATTGGCGAGAGTGGTGTTGATAAATTGAAGGATTTTGAGCAACAAATTAAAGGAATTAACAAGCTAAGGCATCCAAATCTTGTTCGTGTTAGAGGATTCTATTGGGGAGATGATGAGAAATTAGTCATCTATGATTATGTTACTAATGGAAGCTTAGCCAATATTGGTTACAGTAAGTTACTTTCCTTTCATCTTCATTATCTTATCATAATTATAGTTTTGATAAATCTTTTCTGTCCTTATCATAATTACCTCCAAATATTCGGATTAGTCAAGTGTACTTGACATTTAATGTCACTGATATATTTGCAGCAACATTCCCTGATTTGTCACTTAACAAAAACATTAAAGCCTATAGTTTAGCGGTAGTAATAAATACTGTACTAAGGGGTCGTTTGATAGATGATTAGAATTATGATGATATTAGTAATATAGGAATTGATATTATTTCATGCTCATATTAATTATGCAGTGATTAGTTATTCCACCATGTATTTGTCATGAGGGTTTTTAAATTGCAAACTAGAGCTACCAAACTATTACTTTTGCAGAACTTAATTAATACCTGCATAACTTACATCTAGACCAACTACCAAACAACTCCTTTTTTAAGTACACTGTCTGTCATTAGAAAAACAAGTAGTTTAATTTTAGTGTACATGTACTACACAGGCATGGATTTTCGTTACTCAATTTTGCAGATACTATACTTCATATGTGGTTGTGTTTTGAGGCTCCCAATGGTCCTCCATGTGACATTGTCATAATCCGTGAAAGTCCCCAAACTATACTACAATACATATAATACGTCAAAATACATGTGCACCCCTTCTCCTCAATATTGTACTACCACAAACGTTTCGGAGACTTGATTTTTTCAAACTCCACGTTTAAGATTATATTGAGTGTATTATTATTGTTT

>M6-PGSC0003DMG400026004 M6 LRR-RLK

TTCTTCACAAAAGCCAACACTCCTTTCTTGAAATTTGTCACTAAAACCAACTTTCCCTACAAACTTTTCCATCAATTTTTCTTGCAAAAACCAAAAATGGGTTTTCTCTCTGTTCTCTCTTCCATTTTCCTTATCATCCTTATATTAACATTCCCACAACAAAGGGTTTCTTCTGAAGCCATTAATGATAGTACTAAAACTTCTTTATCTTTGCTTATGACTATCAAAGCTTCTTTGGACCCAAAAAGCCTCATTCTTTCATCATGGTCACCTACTTCTACTGACCCATGCAATGATTCTTTTGAAGGTATTGCTTGTAATGAGCTTGGTCAAGTTGTTAACATTTCTTTGCAAGGCAAAGATCTTACTGGAAAAATCCCACCTGAGATTGGTCAGCTTCAAAGCCTGTCTGGATTGTACTTGCACTTTAATAAGCTACATGGGGTTGTACCTAAAGAAATTGCCAACTTAACTCAGCTCTCAGATTTGTATCTTAATGTGAATAATCTCTCTGGTGTTATTCCTCCTGAGGTTGGCAACATGTCTAGTCTTCAAGGTAGGTTCTTTTTTGCTTTCCTGTTTTGGCTAGTTATGGTAAAAAGCCTCTGTCTTTATGCTTAGTTTTGACTTAATTTGAATGAATGTTTGAACTTTGCCAAATAGATCTATACCTCTATTAATGAAATGGGTAGCTCAATAAAGTGTCCAATAAGTGATAATGCTGTTAGTTATTGAGTAATTTCACAATGCCTATGGCTTTTCTTGAAAAAAGATTGGATTTTTAGTACTCATTAGTCATTGAAAGTCTATATTAAAAAGGTTCTTGAAGTATCTTTTATGAGGTGATTTTTGTTCCTATGTAGCTTACAGCTAGGTGCTTAGAACATGAAACACTTGAAAAAGCCTCCATCTTTGGTTTTATGTTAAGCTTTTCTCTGTCATTTGTCTCTTTCCCCATTGATTTGAGCTTCAATATGGGCTAATGCTTGAATAAAGTTTCAACTTTCCTGGCAAACTTGCAAATTTCTTAGTCTCTTGTTTGGGTGGCTTTTTCTGCATTTTTGGTGGAATAAATCTCAAATAAAAGTAACCGAGCTCCATTTGTTTTCTGTTTTTACACAGTACTGCAACTATGTTATAACCAGTTGACTGGAAGTATACCTAATCAACTTGGAGCTTTGAAGAAACTGAGTGTTCTTGCTTTGCAAGTCAATCAGTTAACTGGAGCAATTCCTGCAAGCTTGGGGGATTTGGAGATGTTAACCAGGCTTGACTTGAGCTTCAATAATCTTTTCGGGTCGATTCCAGTGAAACTCGCTGATGCTCCAAAGCTAGAAGTTCTTGATATCAGAAATAATACCCTTTCTGGCAATGTCCCTCAAGGTAGTTCTATTTGATTTTCTTGATAGTACCTTTAATCCTATTTCATTGCTTAATATGCTTAAGAAATGAAGAAATCTTAGGTTCTTTGATTCCCCCTAATATCAATCCAAATGTTTACTAATATTAGACATTATAGTAGAACTAACTCTCTTTTTGGTCTTTTTGGTTTGGGCAATGGATGAAATGAAGCCTTGAAGAGACTGAATGAAGGATTTCAGTATGCAAACAACCCAGACTTGTGTGGAATTGAATTTTCTTCCTTGAAACTCTGCACTGTATCTAGTCTAAACCAAAACAGACCACAGCCATTTGAACCTGGTTCGAATCGTCTTCCCACAAAAGACATTCCAGAGTCTGCAAATGTACAAACAAAACAAATGAGTCAGTCAAGAAAATCACAAACTGCTGTGGTTGTTGGTGTTATTGCACTTTTTGTTGCAGTAGCAGTAACTGGACTTTTCACATTTTCGTTGTACCGTCGAAGGAAACAGAAAATTGGAGGTACTCTTGATACCTCCGATAGGCGACTCAGTACTGATGAGGTGAAGGAGATCTCTAGAAGAAGCGCGTCGCCCCTCATCAGCCTGGAATATTCCAATGGCTGGGATCCTCTAGGCAAAGGCCGAGGTGGAAGTGCTTTCTCTCAGGAAGTATTTGAGAGCTTTATGTTCAATTTGGATGAAGTTGAGTCAGCTACACAGTACTTTTCAGAAGCAAACTTGTTAGGCAAGAGTAATTTCACAGCTGTTTATAAAGGGACATTGAGGGATGGGTCTGTTGTTGCTATTAAGTGCATTTCCAAGACTAGCTGCAAGTCAGATGAAACTGAATTTCTCAAGGGACTAAAACTCTTAACCTCATTGAAACATGAAAATCTTCTGAGGTTGAGAGGCTTTTGTTGTTCAAAAGGTAGGGGGGAATGCTTTCTGATCTACGATTTCGTTCCAAATGGAAATTTGTTGCAGTACCTTGACGTGAAGGATGGAAAAGGAAAGGTCCTTGATTGGTCTACCAGACTTTCTATAATCAAAGGCATCGCCAGAGGTGAGCTACTGTTTGGTTATTGTCCCATGTTTGATGTTTTATTTATAGCTCTTTGACGACCAGATTTGAATAACAACATTTGTTTTCTATTGTTTTACAGGTATTCATTACTTGCATGGAAACAAGGGAGGCAAACCTGCTCTGGTCCATCGAAATATATCAGCTGAAAAAGTACTTGTCGATCAACACTACAACCCATTGCTCTCGGATTCTGGTCTGCACAAACTACTAGCAGATGACATTGTCTTCTCAACACTCAAGGAAAGTGCTGCAATGGGCTACCTAGCTCCTGAATATACAACCACTGGACGGTTCACCGAAAAGAGTGATCTTTATGCATTTGGCATGATCATATTTCAGATTCTCTCCGGAAGACGTAAAATCACCCCTTCTAACCATCAAGGGGCAGAGTTGTGCAGATTTGAAGACTTCATTGATCCGAATCTTTGGGGCAACTTTGTTGAAGCGGAGGCAGTTCAGATGGCAAAGGTAGCTCTACTTTGCACGCACGAGTCACCAAATCAGAGGCCAGACATTGAAATCGTCATGCAAGAACTGAATGATTTAACCTCCAGCAAGTCGTGAAAGCTATTGCAAACAACATCCAAGTTCTCCAATGACCATTAGGAGCCTGATACTAACTTCTTACTGATGCCAGGAGTAGGCACCCTTTTCCCATCCTTTTGATCCTGCATGACTTGTGCCTGAGGACAACACATGAATGTCAGGTGCCTGTTCCACATAGTTTAGTACTAATGCACCTCGACGTAGGTAGCCTCAAGTGATCAGTTCTCTTGTATTATGTAACTTAACCATAATTCTAAGCAAGTTTTCTCCACTTGCAAACTAGTGTACCAGAAGTGCAATTTCTATTTAGTATCATATATGACCTCTTGC

>M6-PGSC0003DMG400026072 M6 LRR-RLK

CTAAGACCTCATTTTCTATCATCACTAAATGGACTAAAAACTAACAAACTCTGCATTCAGAAACTTCTTTTTCTTTTATCTATTTCTGGTGACTGACACTGCAAATTCCCATTTTCCCTTTAGAGGGGTCTCATTCAAACTTTTGAATTTTTTTGGGTATTCAAGAAAACAAAAAGTTTGAAGCTTTCTGCCTCTTTATGTGTGAATAAATAAGAGAAACAGAAAATCATCTTACTATTTCAGGTATAACTCATGTTTTCTTTAAGACTTGTTCATTTGCTAGTCTTTTAATTACATAAAAATCCAAACTTTATGTGTTATATGAGTTATCAGATGATTCTTAACAACTAAGCTAGCATTTAGCCATAGATTTTAGATGATATTTTGAAAATTTATATTCCTATCTGTTTGGCCATGAAATTTGACATTTTTGGAATTTTGGGACTTTCGCTCTTAAAACTTCAAAATCTATAGCCAAACAGGACCTAAAGGATGCATGTTACTTATGAGTTCTTAAGAAAGTTTACTTTTTTGTTTCTATGTATATCTCTGTTTCTGTTAAATTATGAAAGCTGTTTGTTTTCTGTCAATTAGTTGATTTCCATGTTCCTTGTTTGCCTATTGTAGTTTGCTACAACTTTCTTGGAAAAAAGTTGTTTTGTTATTGTTACAAATATGGAAATCCTAGATTTCAATCTTCTTGCTTATATTCCTATAATTCTTGCAATTATGTTTCGAGGGGTCTATCGAAAAAAGTCTCTACCTACCCCAGATTCCACACTTGTGTTTTGGGGTAAATGAGTATCTTCTTTCATCTGGACAGTCTGTAAAATAAGGTGAAACTATTATATTTTGAGTCACTTTAAAGGGAGATAATTTAATTTTTATATAGTGTAATTCACAACTAAATGAGAAATTGACATCTTAACATAAGTTTATTTCCTTCCCAACCCCACCCAACAATTCCATCTTTTATAATCTAGACTTCCTGTATCTGTAAGTTTTCTTTTTCTTGAACTACAATTTTTATTTCATGTTGGAAAGTTATATGAAGTATATGACAAACATTTTGTTAGGTATACATACAACACTTTAAAAAAAGACAGCTTTTTTTGAATTGGATATGAGTATTATCTTGTTTTTTTTAATATACCACTTTATCTTAGTGTTTGATATCTGAGTTTCCACTGTTTTTCTTTCAGCTGCTTCTTGGATGGACACCCATCATGGCAATAATGGAGTTCTTGATTGGATTTGTGTCCCCTTTATGAAATGACCCTTTTGCCCTTCTCAAATTTCAAGAATGGCCAGAATCCAAAAACTTCAAAACTCTCTCTCAATTCTTGCTGTTTTCTTGTTTTTGAATTTTTTTGTCCAGACATGTAAGTCCTTAACCTCAGAAAGTGAGGCTCTTTTGCATTTCAAAGAGCAGCTTAATGACCCTTTGAATTACTTGGATTCTTGGAAGGACTCAGAATCTCCTTGCAAGTTCTATGGAATTACATGTGATAAAAATACTGGTCTTGTTATTGAAATTTCCCTCGATAATAAGTCGCTTTCCGGGGTGATTTCACCTTCAATTTTTTCACTCAAAAGTCTCACTTCTCTTGTGCTACCTTCAAATGCACTATCAGGGAAACTTCCTAGTGAAGTTACCAACTGCACCAGTCTCAAAGTCTTGAATGTCACTGGAAATAACATGAACGAAACGATACCTGATTTGTCTAAGTTGACTAACTTAGAAGTGTTGGACTTGTCGATTAACTATTTTTCCGGGGAATTCCCATCTTGGGTTGGAAATATGACTGGTTTGGTTGCGCTAGGCCTTGGAGATAATGACTTTGTTGAAGGTAAAATTCCTGAGACTCTTGGGAATTTAAAGAAAGTGTATTGGCTTTACCTGGCAGGTTCAAATTTGACAGGAGAAATCCCAGAATCCATTTTTGAAATGGAGGCACTAGGTACATTGGATATTTCAAGAAATCAAATTTCTGGGAATTTCTCTAAGTCAGTAAACAAGCTGAAAAATTTATGGAAAATTGAGCTGTTTCAGAATAAGTTGACAGGGGAACTTCCTGTAGAACTGGCAGAGCTCTCTCTTTTGCAGGAATTTGATATCTCTAGCAACCATATGTATGGGAAGCTGCCTCCAGAAATCGGGAACCTGAAGAAGTTAACAGTTTTTCATGTATTCATGAATAACTTCTCTGGTGAAATTCCTCCTGGTTTTGGAGATATGCAGCATCTTAATGCCTTTTCTGTGTACAGGAACAATTTTTCTGGGGTATTTCCAGCAAATCTTGGTCTGTTTTCCCCCCTGAATAGCATAGACATATCTGAAAACAAGTTTACTGGTGGATTCCCAAAGTACTTGTGCCAAAATGGGAACCTGCAGTTTTTGCTAGCTATTGAGAACAGTTTCTCAGGGGAGTTTCCGAGTACTTATTCGTCGTGCAAGCCTTTACAGAGATTGAGGGTCAGTAAGAATCAACTTTCTGGGCAAATTCCTAGTGGTGTATGGAGACTTCCAAATGTATTGATGATGGATTTCAGTGATAACGAATTCAGTGGAACTATGTCCCCGGAAATTGGGGCTGCTACTAGCCTGAATCAGTTGGTATTATCAAATAACAGATTTTCAGGTGAGCTTCCAAAAGAACTTGGGAAACTCACACAATTGGAAAGGCTTTACTTGGATAACAATAACTTCTCGGGCGCAATACCTTCTGAACTTGGTAAACTAAAGCAAATTTCATCTTTGCATTTGGAGAAAAACTCATTCTCGGGCACTATCCCATCAGAATTGGGTGAATTTCCTAGGCTTGCAGACCTGAATCTTGCTTCAAATCTTCTTACAGGTAACATTCCCAATTCCTTATCAACAATGACCTCTTTGAACTCTCTGAATCTTTCACACAACAGACTCACTGGTACAATACCAACTAGCTTGGACAATCTGAAGCTGTCTTCATTGGATCTCTCAAACAACCAGCTATCAGGAGAGGTATCATTAGATCTTTTAACACTGGGAGGAGACAAGGCATTAGCTGGTAACAAAGGACTTTGTATCGACCAAAGCATCAGATTCTCGATCAACTCAGGCTTGGGCAGTTGTGGTGGAAAGGCTGCTAAACATAAGTTAAACAAACTAGTTGTGTCTTGCATTGTGTTGCTTTCCTTAGCGGTTCTAATGGGTGGCTTATTGCTCGTCAGTTACCTAAACTACAAGCATAGTCATGAAGTAGATCATGAAGAAAAACTGGAGGAAGCCAAGGGAACAAATGCAAAATGGAAACTTGAGAGTTTCCACCCTGTGGAATTTGATGCAGATGAAGTTTGTGATTTCGATGAGGACAATTTGATCGGAAGTGGAGGTACAGGAAAAGTTTATCGGTTAGACTTGAAGAAAGGTTGTGGAACAGTAGCAGTGAAGCAGCTGTGGAAAGGGATTGGTGTGAAAGTTTTGACAAGGGAAATGGAAATCTTGGGAAAAATCAGGCACAGAAATATAGTTAAGTTGTATGCCAGTCTAATGAAAGAAGGTTCAAATATCTTGGTTTTCGAGTACATGCCAAATGGTAATCTTTTCGAGGCACTACACCGAGAGATCAAGGCTGGCAAGCCAGAACTAGATTGGTATCAGAGATATAAAATAGCCCTTGGAGCTGCAAAAGGAATTGCTTATCTGCATCATGATTGTTACCCTCCTATTATTCACAGAGATATCAAATCAACCAACATTCTACTTGATGAGTATTATGAAGCAAAAGTTTCAGATTTCGGGGTTGCAAAAGTTTCAGAAATTTCTTCAAGGGGATCCGAGTTCAGTTGTTTTGCAGGCACTCATGGTTACATGGCTCCTGGTAAGTTTCAAGAAAACCTTAAAATGAGTGACCATTTTGTCTCTCCTTGTTACTTTCCATTCTTGAATTTGCATTTACTTTCCTATATGTTTGCTTGACAGAAATGGCATACACTCTCAGAGTGACCGAAAAGAACGACATTTACAGCTTTGGTGTCGTGCTATTGGAGCTAGTAACTGGAAGAAAACCAATAGAAGAAGCCTACGGAGAAGGAAAAGACCTGGTCTACTGGACCTCAACTCATCTAAATGACAAGGAAAGTATAAACAAAGTTCTTGATCAAAAGGTGGTATCTGAACTTGTTCAGGATGAAATGATCAAAGTGTTGAGAATTGCCACACTTTGTACTACTAAGCTTCCTAATTTGCGTCCCAGCATGAAAGAGGTTGTCAATATGCTCGTCGATGCTGAACCTTTGACGTTTAGGTCTTCAAGCAAATCTGAGAAAAAAGGGAACAATTTTTCTGAGGTCTAGTTTCTCTTCTTTTTTTATTGGGTTTTCCCTGTTGATAGTTGCTCTGTAGTATAATTCTCAAGTTAGTTTTAGGTATCTGATATACATAAATTTATACCATGTGATGTGATATTCCACTTTCTTTATGTAAAAGTTCGAATTGTGTAAGATAAAATGTACTTTTTTCTGCCGTGATTGGAAACTTCAGAAGTTGGATTAATCAGAAATGTGGGTAGTTAATGTCCATGATTAAGAGTCCCACATCGGCAAAAATACATTAGATATGTTGATTATATAAATTAGCAGGCCTTACATTCTAATAATACTTCTTAAAGCTGCATAGGTGTAGACTCAAAGCTAACAATATCACTAGCGAGCTGGGCCTTACACGAAAACTTGTTTTGTTGTAAACTTTGAATCTGATTCAATGAATATCCACATAGATAGTTGGAAGCAAAAATGATATACATAAACATTTCATACATTTGTATAAGAGGGATGGAAACCAACAATATTACTCTTTTTATCTTTTGTGATCCTCCTCAGTATTACATGGTACATTCTGCAGATTTTTCCTGCTCTCTATGACATTACGGTTCCAAATCTTCACTCCTTTGCTTCAAGAAGTTGAGTTAATGCACCATCAATTTGAGCAAATACCTCCTCTTTGGAAACGCTCCCGTCCACCTGAAATTAAAATAGCTCAGAGAAGATGGAGAAAGAAGATAGAAGTTTTAACACTGAGAAATGATAAGAACACGTTCATGCACGCGTATGGAAAGAAAGAACATACAGTATCCTTGTTCAGTGAAGAAAAAGAGAAGGGTAAGCGAAAGGGGAGTCCTTGACTAAGTATATTTTCCCCCAGATTGTTTGGTATGAACGTAAAACTCAAAGGATTTGCTCTTCCTTTATCTAATTGAGGAGAGATGAATAATTTTAAGATAAAAGAGGATGTATGGCCCTTTTGCTTTGGCTTCGCTTTGTTTTACAGCTGTTGAGAGGAAGGATCAGCCTGGATGAACTGGAAGATAATTTAAATTAATCATTTTGTAACCTACATGTCAAACAACATAATTTTATAACCATATCATTGTTCTTTTGCCAATTAGAATACCAAACTTCTTTTTCTTCAAACCATTTCCACCCAGTCCCTTCCACATTGTCTTTTTT

>M6-PGSC0003DMG400026749 M6 LRR-RLK

AGTTTTATCTCGTAATTTACTTCAAGGGAACATTCCAGAAGGGATTGGCAATCTACACAAGATGAAATTGTTGTCCATACAACATAATAAACTTGCGGGTTTTATACCGTTCACAATTTTCAATATTTCTAGAATCGAATTCATTGCATTTACAGACAATAGCTTATCAGGAAATCTTCCCAATGGTTTATGTAATGGTCTCCCAATACTCAAAGGGCTTTATCTATCCACAAACAAACTTCGTGGTCATATGCCTACAAGCCTGTCAAATTGTTCACAACTTCAAATATTGAATTTATCAGGCAATGAGTTTGGCGGACGAATACATAGTGAAATTGGAAGATTGAGTAATTTGCTGATATTGTATCTTGGATCTAACCATTTCACAGGTACGATCCTAGCCACAGAATTATTAGTTGTTTTATGGATTCTTTTATGAACACTTGTAATATTGAACTTGAACTGCAGGAATAATTCCACAAGAGTTAGGAAATCTTGCTAATTTGGCGAATTTAGGCATGGAGGATAACCAGATTAGCGGCTCTATCCCGATCAACATATTCAATATCTCCTCACTGCAACGTTTTGGACTATGGCGGAACAATCTCAAGGGATCCTTACCACAAGAGATTGGCAACTTAACCAAGATACAGATTCTAGATCTTAGAGAAAATACATTTATTGGTACGTATTCGAAGTGATAGGAAAATTATAAACTTATACTAGCTTGAACTAACTCATTCTTAATTTCTTTAGGTGAAATACCCAAAGAGATAAGCAATATGATGGAGTTGGAGATACTTAGTCTCGGGTTAAATAGTTTCAGTGGTTCACTTCAAATAGAGATGTTCAACAGTACTTCACGCTTGAGGATAATTTCTCTTACAAACAATAATCTATCAGGAACTCTACCATCAAACATAGATTCTGTCTTACCCAACATTGAAGAGCTTTATTTGGGTGTCTTAACCAATCTTGTTGGGACTATACCTCATTCTATTTCCAATTGTTCAAAACTTACTAATCTAGAGCTTTCAAATAATAAACTCACTGGCTTGATTCCCAATTCTCTTGGATATTTGGCTAATCTACAGTTCCTAAACTTGGCAGCAAACAATTTAACAAGTGACTCATCGTTTAGCTTCCTCACTTCCTTAACCAATTATAGAAATTTAACAATTCTTTTTCTATCTTTGAACCCTCTAAACGGCATGCTTCCAGTATCTGCAGGGAACCTTTACACTTCTCTTACAATGTTTTATGCCAGTTCTTGCAATATTAAAGGGCGAATTCCAAATGAAGTTGGGAACTTAAGCAACTTATTAGGGCTTGATCTTTCTGGAAACAACCTGGTTGGATCAATTCCCACATCAATTGGCAACTTGAGAAACCTTCAGCGCTTCAACTTGAGTTACAACAAACTTACAGGATTTATTGGAGATAACATTTGTAAATTGCAGCATTTGGGTGCTATTTACTTGGGACAAAATCAACTTTCAGGATCTCTTCCTAATTGTTTAGGGAATGTTACTTCCCTTAGATTGATACATCTTGGTTCCAATAAATTGATTTCCAATATACCACCAAGTTTAGGGAATCTTAAGGATTTAATGGAGTTTGACTTATCCTCAAACAACATGGTTGGTTCTTTACCACCAGAAATTGGAAATCTAAAAGCTGTGACACATATGGATCTGTCGATGAATCAATTATCAAATGGAATTCCTAGGGAAATTGGAGGATTGCAAAATCTGGTGCATCTTTCTTTGAGACACAACAAGTTGCAAGGATCTATACCTGACTCAGTGAGCAACATGGTAGGTTTGGAATTCCTAGACATTTCTCATAATAATATATCAGGAACCATTCCCATGTCTATGGAGAAACTTCCAAACCTGAAGTATTTCAATGTTTCTGTCAACAAGCTGTACGGTGAAATACCCTCGGGGGTCCTTTCAAGAACCTCTCGAGTCAATTTTTCATCGACAATGAAGCCTTGTGTGGTTCTTCAAGGTTTAGTGTCAGACCATGCGCCACATCATCAAAGCATAGATCAAATAGGAAAAAAATGCTTGTTCTATTTCTTGTGCTGGGAATTGCACTTGTATTTGTTCCTATTACCTTTGTGTTTGTATGGATAAGGTATAGAAGAGGTACAAGTGATCCTCAACAAGCTCACTCATTGTCTACCGTAACAAGAGAAATAATTTCATACTATGAACTGCTCCAAGCAACTGAGTCGCTTAGCGAGAGTAATCTGATTGGTTCTGGAGGTTTTGGCTCTGTTTACAAAGGTGTTCTCAGAAGTGGGATTGCCATTGCAGTTAAAGTGTTCAATCTGCAATTGGAAGCGGCATTCAAGAGTTTTGATACAGAATGTGAAGTTTTGCGCAGCCTTCGCCATAGGAATCTCATGAAAGTCATCACTAGTTGCTCCAACCTGGATTTTAAGGCTTTAGTGCTTGAGTATATGCCTAATGGAAGTCTTGACAAGTATTTGTATTCGCACAACTACTTCCTAGATATCAGCCAGAGACTAAGCATTATGATAGATGTGGCATGTGCGTTGGAATATCTCCATCATGGATTCTCCTCACCTGTGATTCACTGTGATCTAAAGCCTAGTAACATCTTGTTAGATGAGGATATGGTTGCCCACCTAAGTGACTTTGGTATTTCAAAACTGCTTGGTGACGATGAGAGTGATTTATACACAAAAACCTTAGCAACATTTGGTTATATCGCACCAGGTACGTCTCTTTGTTTTTGCTAATTTGAACATTGATTTTTCGTCTTTTTTTTTTCCAATCAAGAAAGTATACTGCATCTTTAAATTAATTGTTTGACTGTAGAGTATGGATTGGATGGACTGGTGTCAGTAAAATGCGATGTCTATAGTTACGGGATTATGTTGCTGGAAACATTTACTAGGAGAAAGCCTAGTGATTTTGAGGGAGATCTTAACTTGAAGCAATGGATGAGTTATTCACTCCCCGAGGCAGTAATGGATATTGTAGATGCCAATTTGGTAACACCAACGGATAATCGCTTACAGAAGGAGCTAGATGTTGTGGCTGCAATCATGAAAGTTGCATTAGATTGCTGTGCTGAATCTCCGGCAAGAAGGACAAACATGAAAGATGTTGTAGGGATGCTACAAAAGATCAAGATTCAACTTCTTGCATGCTGAGCATGTTTTCAGAATCTCTTGTTCTGAATCGATCACTTTTTCTGTTTTGTTTTAGCACTTGATTTGTGCATGATTCTGTTTTCAGAAATGCTAGTTGGTTGCAATAAATGTCACTTTTTCCTCTT

>M6-PGSC0003DMG400026760 M6 LRR-RLK

TGAATTTTCTTGTTTCCATTCATCAATGGTGACACCCAACCAAATCAAGAATCCATTTTTGCAGACCTCTTTAGTCTCAAAACTCCATTTTTAGCTCTGCACTTTGGACTTTCTTGTTTCCCATTCATCCATGGTGACACCCTACCAAATCAAGAATCCATTTTTGTAGAGACCCCTTTAGTCTCAAGACTCCATTTTTAGCTCTGCATTAGATCCCTTTAGTGTTAAGACTCCATTTTTAGCTCTGTAAGTTGAGTATTTTAGTGTCAAGACTCTATTTTTAGCTCTGTATTAAGCATTTCAGTGCTCAAAAATTGTCAAGTTCAATGAAAATCAAGAACCCATTTTTCCTCTAGAGTTGAACAACTTAAATGCTCAATCATTCAACAATGGAAATGTCAATTTCAATGGAAATCAAGAACCCTGTTTTGTTCATAAGGTACTGTTCTAGTCACAAATTCTTGTTTTTCATTTTCTCAATATTTTCCATAGTTTTACCACTTGTAAGATCAGATGATGCTGAAGCACTTTTAGCATTGAAATCCACAATAGACCCCTCAAATTTTCTTGATTGGAAAAAAGGAACTGATTTTTGCCAATGGCAAGGTGTTAAAGAATGCATGAAAGGTAGAGTTACTAAGCTTGTTGTTGAAAATCTCAATTTGAGTGGTTTATTGGACCAAAATGCTCTCAATCAACTAGATCAATTACGAGTTTTAAGCTTCAAAGAAAACTCCATTTCCGGCCAAATCCCAAACCTTGCCGGACTTTCCAATCTGAAATCCCTTTTCCTCAACAACAACAACTTTTCCGGTGAGTTTCCGGCTACCATTACCCTCCTCCACCGTATCAAAGTTGTGGTTCTCGCCGGAAACAAACTCTCCGGTGACATTCCTCCGGCGTTAACAAATTTATCCCGGTTGTATGTTCTTTATCTAGAAGACAATCACTTCACCGGAGAAATCCCACCGTTGAATCAGCTCAGTTTAAGGTTCTTTAATGTGTCAAACAATCAACTTTCAGGTGAAATCCCGAATACTCCGGCGTTATCTCGGTTCAATGAATCATCTTTCGCCGGAAATGTCAATTTGTGTGGAGTACAATTGCATAATAACTGTCCTTCAACTAGTCCAGCGTATCCCATCATTCCAGAAAATGTATCTCATCATAAAAAAAGCAAAAAGCTTATCATCATCATTGTCCCTGTGGTCGCCGGAATCTTGATCCTCTGTGTTTTGATGGCGATTTTTATAGCATGTGTTCGAAAACGAAGAAGAAAAACTCAAGATGGAAAGAGTAAAGGAGTTGAAGCCGGTGGAGGCGGGGGAGGGGGAGGGGAAGAAGAGGGGACTAGTCGCGGTGGCAATGGCAGTGGTGGTGGTGGTGGCAGGGATGGTAATAATGGTGGAAAAGAAGGGGCATTTTCATGGGATCAAGGAAGTGGAGGACTAGGGACATTGGTGTTTTGTGGACCAGGTGATCAACAAATGAATTACACCTTAGAGGATTTGTTGAAAGCATCAGCCGAGACATTAGGGAGGGGTATAATAGGAAGTACATATAAAGCCGTGATGGAATCTGGCTACATTGTGACAGTGAAGCGGTTGAGGGATTCGAGGTTTCCGAGGTTAGAGGAATTCCGGAGGCATGTGGAGATTATTGGTAGGCTAAGACATCCTAACTTGGTCCCACTCCGAGCATATTTTCAAGCTAAGGAAGAACGTTTGCTTGTTTATGATTATTTCCCTAATGGCAGCCTCTTCTCCCTCGTTCATGGTATGTATTCCGTCGTTCATTTATTTATTTATATTAAATTTTGCTTTTATCCATTTTTGTGTACTGAACTCTAGTGAGTAGTGAAAGGAGGAAAAAACAGGTATAGTTGAGCTCATGAATAGTGAAAGGAAGTTAAAAAGGGATAAATTTGGCAGGATGAACTTGTACTTGAGTGGTATAAGGTGGAATAAGATAAGGGGAGGTAACTTATTTACGTCTAGTATATGCATCGATCGAATAGATGCATTACATGTGTTTTTTTACTAAATTAGTTATTGAATAGTTTATCCTCTCTTGAGGTTACACTAGTTTAGTTTTGTGCAAATACCAAGTGGGGACAAATTCTTTTCATGAGATCACTAGTTTATAACTCAATCTCTCGAAGATTCTGCTCCTTTATTGACATGAGTCATTTTCTTATTTTGAATTCAAATTTGTTACCTCTTTATGCTTGGCTGTTTTGTGGGGGAGCATTGGAGGACTTTAGTGTCACTTTATTTTCCTATTTTTTTGTGTTTATATTTGTGTACTTTGCACATTTTCCTTGAATTGTTTATACTTTTTAATATGTTTCTTTATCTTCTTCAGTATTCTGTCCCTTCTTAGTTTGTTTTTTTGTTATTTTTGAATAATTTCCATATAATTGGTTTGGTAGTGTATCACTTTCTGGTCTCTCCTGTGGGTTCTGTTTGGTGATGGTGGGGAAGAGAGGGCCAACAAAGGAAGCTTGTAGGAGCCAACAAAGAATTGAGATAGCTGGCCAAGTAGGACAAATATGTAAATTTGTAAAGGAAGTTCCAAACTTGCTTTGGTATACTGATGTGAAGATTATAGAAAAAAATTGTTTTGTTTAAATAATTAAGTTAAAAAAAAATCAATAATCAAATAAGTAAACTCTTCTCATCTTGTTCTCTACGACTACGGATGTTATACAAAAAGAGGATAAAGAACAAAAGGGGAGTATAGTTATATATTGGCCTTTTGGAATCTTAAGAAGGATGATATTTCATTCAAATGGATTCTGAAAAAGATGTCTATTGTGTTTCGGTTATTGCACTATTTTGTTGTTGTTACAGTTAATTTTATGAATGCTTTGTGCTTTCCTTATTATTTGCCATATTTCTTCAATGCCATTTTTCCTTTCATAACTACTTTGTTTAGCTGCACAAGAGCCTAGGGTCCTTCGGAAACAACCTTTCTATCTCAACGAGGTAAGGTCGCGTATACTCTACCCTATCTAGATCCCACTTGTGGGATTTCACTTTGTTGTTGTTGGATTGTGAAAAAGATGGCCCTCTGGGAGGAGGAAACTCAAATTAGGAACCAAAACAATGATGTCACCGCTACACTCGCATATTTTGGCCTTGTCATTTATTGCATAGAGCCAAAACAGTTGTCCACCATATATGGAGATAAGGCATGATACTTTGAAATTAGAAAGAC

>M6-PGSC0003DMG400026792 M6 LRR-RLK

TTATCACATAACGGTCTAACTGGTCCTGTACCTGAAGCCCTAGCAAAACTGACTTATCTTCAAAGTGTTGACTTATCCTTTAACAAATTAACGGGAGTGTTGCCAAAGCAGCTTGGAGATCTTGGCCATCTCTCCTTATTCAACATTTCACACAACCAGCTACAGGGCGAACTACCTTCTAATGGTTTTTTTAATACTATTTCTCCTTATTCCGTGTCAGCAAACCCATCTCTATGTGGGGCTGCTGTCAATAGGTCTTGTTCTACAGTCATGCCCAAGCCAATTGTTCTGAATCCCAACTCTACTGATTCTTCTCCCAACACTGTCCCTCAAAGTATTCGCCATGAGAAGAAAATCCTAAGCATCTCTGCCCTCATTGCTATTGGTGCAGCTGCTGTTATTTTTGTTGGTGTGATTGCCATTACTGTGCTCAATATTCGTGTCCGGTCAGCGGCACCTCGCTCTGCTGCTGCAGCCCTTACATTTTCTGGTGGAGATGATTTTAGCCATTCTCCATCCACGGATGCGAATTCTGGTAAGCTTGTCATGTTTTCTGGTGATCCCGACTTCAGCACAGGGGCACATGCTCTGCTTAACAAAGATTGTGAGCTTGGGCGAGGTGGTTTTGGAGCAGTGTATCGCACTGTCCTTGGGGATGGGCATCCAGTGGCCATTAAGAAGCTTACCGTTTCCAGTCTTGTTAAGTCTCAGGAAGATTTTGAGAGGGAGGTTAGGAAATTGGGAAAAGTACGTCACCATAATCTGGTCACCCTTGAAGGTTATTATTGGACACCATCTTTACAGTTACTTATATATGAATTTGTGGCTGGTGGAAATTTGTACAAGCATCTCCATGAAGGATCTGGTGGGTGTTTCCTCTCATGGAATGAGCGGTTTAACATCATTCTCGGGACAGCAAAAAGCTTGGCTCACTTGCACCAAATGAATGTAATCCACTACAACCTAAAGTCCAGCAATGTCCTAATTGATAGCTCAGGTGAGCCTAAAGTTGCAGATTATGGCTTAGCAAGACTATTACCCATGCTTGATCGATATGTCTTGAGTAGCAAAATTCAGAGTGCACTCGGTTACATGGCACCTGAATTTGCATGCAAAACTGTGAAGATAACCGAGAAGTGTGATGTGTATGGATTTGGAGTTCTTGTTCTGGAGGTAGTCACTGGAAAGAGGCCAGTTGAGTATATGGAAGACGACGTGGTAGTACTGTGTGACATGGTACGTGGAGCATTAGAAGAAGGCAGGGTAGAAGACTGTATAGATGCAAGGTTGCAAGGAAAATTTCCAGCAGATGAGGTGATTCCTGTGATGAAGTTAGGATTAATCTGCACATCACAAGTCCCATCAAACCGGCCTGATATGGGAGAAGTGGTTAACATTTTGGAGCTGATTCGATGTCCTTCAGAA

>M6-PGSC0003DMG400027071 M6 LRR-RLK

CAACCATTTTTGATTCATATAACCACCAATACACACTTATAAAAAAATCCATTAACCACCACAGCCTGGTTACCTCATTCACTTCTTTCCTCCCTCCTCCCTCCGTCCGGCCTACCGGAAAAACAAGAATCTCAATGGCTTCAAATTTCCACTTTTTTCTCTTATACCTTGTGACCCTTTTCCTTTTTGCTCAAGCAAGACTGAATCTTTATCCACCAGATCATGCTGCACTTTTGCTTGTACAAAAGGACTTAGGCATCATTTCTCTTAACAACCCCTGTACGCTCGCAGGAATATCCTGCGAGCGTAGACCGGGTAACAGAACACAAGTTCTGAGAGTTACCCGTATTGTATTCAGATCCAATGGATTGAAGGGAACTTTGTCTTCTGCTATTGGCAAACTCTCTGAGCTCAAAGAGCTTTCTCTTTCCGACAATCAACTATCTGAACAAATCCCGATTCAGATTCTTGATTGTCGGAAATTAGAGATTCTTGAACTTCAAAGAAACAGATTTTCTGGGAAGATTCCATATGAATTGTCATCTTTAACCCGTTTAAGGCTAGTTGACTTTTCATCGAATGAGTTTTCTGGGAATCTTGATTTCTTGAAGTACTTTCCTAATTTGGAAAAACTGTCTCTGGCTGATAATATGTTCACTGGAAAAATACCCTTTTCATTGAAATCTTTCAGGAATCTTCGTTTTCTCAACATTTCAGGGAATAGTTTCCTTGAAGGTCCAGTACCTGTCATGAGTCAAATTGAGCATTTATCAGCAGATTTGAATCGAAACAATGCCGTTCCCAAACGTTACATTCTTGCTGAGAATTCAACAAGGTCAAATCAGATATCTGCAATGGTGCCTGCTCCAGCTCCAGCACCGGTCAATCGTGTTGTACCTGTGGTGCATAAACGTAACAAGAAGAAAAAGAAGTTAAGATCTTGGTTTCTTGGTTTCCTAGCTGGAACTTTTGCTGGGGGTATATCTGCTGTGATCTTTTCGTTGCTATTCAAGCTTGTAATGTTCTTCGTAAGAAGGGGAAAGAACGATAATTCAAGTTTAACGATATTTAGTCCGTTGATCAAGAAAGCGGAGGACTTGGCTTTCTTAGAGATGGAAGATGGAGTAGCATCACTTGAAATGATTGGAAAAGGTGGATGTGGAGAAGTTTATAGAGCCGAGTTACCAGGGAGTAATGGGAAGATTATAGCTATAAAGAAGATTATACAATCCCCAATGGATGCTGCAGAGATCACGGAGGAAGATACTAAGGCGTTGAACAAGAAAATGCGTCAAGTTAAATCAGAAATTCAAATTGTAGGTCAAATCAGACACCGGAATCTGCTTCCATTACTGGCGCATATGCCAAGGCCAGACTGTCATTACTTGGTGTATGAATATATGAAAAATGGGAGTTTACAGGATATCCTGCAGCAAGTCACAGAAGGCACAAGAGAATTAGATTGGTTGGGACGACACAGAATTGCAGCGGGGATAGCTGCTGGTCTTGAGTATCTCCATATAAACCATACTCAACGCATAATTCACAGAGATCTAAAGCCAGCAAATATCCTACTTGATGATGACATGGAAGCTCGAATAGCTGATTTTGGGCTTGCAAAGGCAGTTCCAGATGCTCATACACATATTACGACTTCAAATGTTGCAGGAACTGTGGGATATATTGCACCAGAATATCATCAGACACTGAAGTTTACAGACAAGTGTGATATATACAGCTTCGGTGTTGTGCTAGCTGTGTTGGTTATCGGAAAGGCTCCATCTGATGAATTTTTCCAACATACTTCTGAGATGAGTTTAGTTAAGTGGCTGAGAAATGTAATGACTTCTGATGATCCTAAAATAGCAATTGATCCTAAGCTGATAGGAAATGGATATGAGGAGCAAATGCTTTTGGTTCTCAAGATAGCTTGCTTTTGTACTCTCGACAATCCAAAGGAGAGGCCTAACAGTAAGGATGTTAGGTGCATGTTAACTCAGATCAAACATTAGCTAAGTCATACGGGGTATACGTTGGTTATGTAACTAAGAATGTGTTCTGTATTTCAGTTTTTCAAGTTGATTTTTTGTTGTAATTGTGTTATACATGTCAAATCTTATATGTTTACTGCAGTGTTAATAATTATGTTCTTTCCACTTTGCTACTTAATTTACTAAAACTCTA

>M6-PGSC0003DMG400027115 M6 LRR-RLK

TTCTTGTGGTGTGTGTAAAAAAAAAAAAAAACTCTCCCTATCCCATTTCTATACTTATCAAAGCCAAATAAAAAACAAGAATGTCAACTGAAAAATCCCATTTTTTCTAACAAATAATGAAAAATCAAGACCAAGTTCAAGATTTTAGCTGTTTTTACCAGTACCCATTTGGGGTTTTCATGTTTTTAATCCAAGAAAGAGATGTGGGGTGTGTTAAAAAGAAGGAAAAATGGATACCATTTATCCAAATTTTGTTGTTTTTGCAGCTGATGATTTGTGGAGCTCAGTTAGCTTTAGGTCAAGTTTGGGATGGTGTAATTATCATTGAGGCAAATTATCAAGCACTTGAAGCGCTCAAACATGAGTTAGTTGATCCTAGAGGGTATTTAAGTAGCTGGAATGATAGTGGTTATGGAGCTTGTTCTGGTACTTGGGTTGGCATAAAATGTGCTCAAGGACAAGTTATTGTTATTCAGCTTCCGTTAAAAGGATTTGGGGGAAGAATCACTGAAAGTATTGGTCAATTTCAAGCACTCAGAAAACTTAGTTTTCATGATAATGAAATTGGTGGTTCAGTACCTTCATCTTTGGGTTTTTTACCCAATCTAAGAGGTGTTCAATTGTACAATAACAGGTTTTCTGGTTCTATTCCTCCTTCACTTGGTTTATGTCCAGTTCTTCAAACACTTGAACTTAGTAACAATTCTTTATCTGGTGCAATCCCTGATAGTCTTGTGAATTCATCTAAGCTTTATAGACTTAATCTTAGTTACAATTTATTGTCTGGTTCCATTCCTGTTAGTATAACTCAATCACGCTCTCTAGTCTTTATTGACTTGACATATAACAATTTGTCAGGGTCTATTCCAGATTCTTGGGGTGGAAATGGAGATAGACAGTTTAAATTAGAGTCGTTGACGCTTGATCATAACTCTTTGTCTGGTGGAATTCCTGTTTCTTTTGGGAAGTTGAGTGAGCTTCTTGAGATTTCGTTGAGTCATAATCGTATCGTTGGTGTCATCCCAAATGATATTGGAAGGCTTAGTATGGTTAGGAATCTTGATTTTTCTTATAATGAGATTAATGGTAGTTTGCCAGAGAGTATAACCAATCTGTCCTCCCTCATGGTTCTCAACTTGGAAAGTAACAATCTTGATGGTGAAATCCCGTTGGATGTAAAGAAGTTGCAAAAGTTGTTGTTTCTTAACTTGAGGAATAACCGGTTTAGAGGTGATATTCCAGCTGCTATCGGGGACATTTCTGGTCTAGTCGAGATAGATTTGTCTCGTAACAATCTAACCGGAGAAATCCCAGAGTCTGTTAGTGAGTTACCTAATCTTAGTTCCTTTAATGTTTCATATAACTCTCTGTCTGGTCCTGTTCCTACTTATCTTGCTAAGAAATTCAATTCAAGTGTTTTTGTGGGTAATGTTCAGCTATGTGGCTATAATACTTTGAATCCATGTCCTGTTACGCCTTCTCCAGTTACACGAGGTAGTAAACGAAGTATTAAAGACATTATTCTCATAGTAGTAGGGTGTCTTATAATAGTCTTGTTTCTACTCTGTTGCATTCTCCTCTGCTGTTTGATCAAGAAAAGAGGTGAAGCCAAAAAAGCAAAAGACGTAAAGGATGTTCCTCCAACTGCCGGAGAAGTTGAGGCGGCTGGGGATAACGGAGGAAAACTTGTCCATTTTGGTGGAGGTATGATGTTCAGTGCAGATGATCTTCTATGTGCTACTGCAGATATATTGGGAAAGAGTACTTACGGTACAGTGTATAGGGCTACATTAGAAGATGGAAGTCAAGTTGCAGTGAAAAGAATGAGAGAAAAGATCACGAAAGTTCAGAGGGAATTCGAGAAAGAAGTCAACTTTCTTGGAAAGATAAGGCATCCGAATCTTCTGGCTATTAGGTCTTATTACTTAGGTACGAAAGGCGAGAAGCTTCTTATTTTCGACTACATGCCTAAAGGAAGTCTCTCAAATTTTCTTCATGGTAAGCAGAAAACAGATTTCCATTTGACATTCTTTTGCTTCTGTTTTTTTGAGAATCAGATACTTTCTCTGTCTCATTTTAACGTAATGAAAATTGGATTATGTATCATATCGGAAACAACTTCTCTAACTCACACAACGTAGGTAAGGTCTACTTACACTCTACACTCCCCAGACCCCACTTGTGAGATCGCATGGGGAGGAAATTGTTATGGAGGGGGTAATATAGCTTTATTGTTTTTGTTGCAGCTCGTGGACCGGATACACCAATTGATTGGCCAACAAGGATGAGAATAGCAAAAGGGACTACGAGGGGATTGCTGTACCTTCACACTAATGCCAACATCATTCATGGGAATCTTACATCAAGCAATGTTCTACTTGATGAAAACACAGATGCAATAATTGCAGATTATGGCCTTTCACGCCTTGTAACTGCTGCTGCAAATTCAAATGTTATAGCCACAGCAGGGGCACTTGGTTATCGCGCACCTGAGCTTAATAAGCTCAAGAAAGCCAATGTGAAGACGGATGTCTACAGCCTCGGAGTCATTATACTTGAACTCCTAACTGGCAAGTCTCCAGGTGAGGCAGTTAATGGTGTAGATTTGCCTAGAAAGGTGGCGTCAATTGTGAAAGAGGAGGGAACAAAAGAGGTATTCGATTTGGAACTAATGAGGGATGCATCCATAATTGGTGATGAATTGTTAGTTACACTAAAATTGGCTTTGCACTGTGTTGATCCCTCGCCATCAGCTCGACCAGAACTTCAGCAAGTTCTCCGGCAACTGGAAGACATCAGGCGTGAAACGCCAACTGCAGGTCCCTCGACAAGTTACTAGATACTCAGTTTTTGATTTTATTAACATGATTCTTTCATTCTTGAATTTCATTGCTAGAGTACTAGATCTGTTTCAAGTGTCTTTAAATAAACGTGTGCAACATTGTTAAAAGTGTTTGAGCTTAATGGAGTTCTAAATACCATCCCCTTTTGGCCTAAAGAACATAACTTGAGATTGAG

>M6-PGSC0003DMG400027467 M6 LRR-RLK

ATGTGGTAATGGTAGATATGTTTATCTCCTCAGATTTTAAAGCTGGAGAAGATTGTTACAAGAATGAATTTTATTGAACTATCAAAAGCGACTTCGAATTTCAGCCAAGACAATGAAATCGGGAATGGAATGCTGGGGAAAGTGTACAAAGCACTAGTCCCAAATGGATGGACTGTTGCCATAAAGAGGCTCCAAAGTTCTGAGGATTTGGAGGAAGAGTTTGTTTCCGAGATCACAACTCTAGGTATTTTGAGGCATCCGAATTTAGTGCCCCTAATTGGATTTTGTTATGAGAGGGACGAGCGGATTCTGGTTTACAAATACATGCCTAATGGAAACTTACATGAGTGGCTACATTCAACCGACGACAGGGCAAGACTCTTGGACTTCCCTCTAAGGGTTAAAATTGTTGTTGGTGTCGCGAAAGCCCTGGCTTGGCTTCATGATGGTGGAAATTTCCATGTCGTGCATAGTAACATTAGCACACAATGTATCTTGCTCGATGAAAATTTCGATCCCAAGTTATCCAATTTTGGGAGGCAACATTTGTAGAACCAAATGACATAGGTTCAAACTTGAGTCTATCGCCAATGGTCGAGTTTTTAGACTTTACAACATACAAAAAAGATGTCTATAAATTTGGGGTTGTGCTTCTCGAGATTTTAACGGGGAAGGAATCTTACGAATTGAGCTGCTCGAGTCTAAATTTATCGAGCAGCTCTTTTGCCAGTCCTCTTGATGTGGATAAACTACTGTTAGGCCAAGGATTTGATGATATGGTAATGCAACTACTTGAACTTGCAAGTAACTGCATGAAATTCATCCCGGATCAAAGGCCAACGATGCAACAAGTGTATCAGACTGTTGCTGCGATTGCTCGAGTCCATGACCAAACAGGGGACTCTGAAATACAATTGCACTTGGACTAG

>M6-PGSC0003DMG400027468 M6 LRR-RLK

TTTCAACAATGTGTAACTGCAACATTCTTTACTTAGTACTACCATTTTTCCTACTATTTTGCAGCTTTTCTCTAAGATGCAACGCGGTGCAGAGTGATATCGATTGCTTGAAATCGATAAAAGATTCGTTGGTAGACCCTTTAAACATGTTGAATTCTACATGGAAGTTCGATAATCAGACAGAAGGTTTCATATGCAGATTTGCAGGAATACAATGCTGGCATCCTGATGAGAGCAAAGTTTTATCCATTAGTCTTCCAGACATGGGACTAAAAGGTGAGTTTCCGCGAGGCATACAAAATTGCACTTCTTTAACTTCTCTAGATCTTTCAAGCAACGAGCTATATGGAACCATCCCTTCGGATATATCAAGATTAATTGGATTTATTGTAACGGTTGATCTCTCAAGCAACCTATTCTCAGGCGCGATCCCAAGTGACATTTCAAATTGCTCTCTACTCAATTCCCTCAAATTAGAAAACAATAAACTAGAAGGACTGATCCCACCTGAAATTCGCTTGTTAAAGCGACTCAAGACGTTTAATGTTGCAAATAATATGTTGACAGGGCCAGTGCCAAATTTTGTCACTTTTCCAGTTGAAAGCTATGCAAATAATCCAGGACTTTGTGGTAGTCCTCTGGAATTATGTGTTACAGAACGCGTAGTAAGCCGTCATATTCTGTTTGCTTGTGGTTTTGTGACGGGTTGGTCGCTTTCTACATTACTTGGTGTATATCTTTTTTTCTTTGGTGTAAAGAAGATGTTTCTATTGATCAACAAGAGAACAAAAGTAATGGTGATTGATGGAAATAAAGTAAACAACAACCAAAAGGTAATTTTG

>M6-PGSC0003DMG400027482 M6 LRR-RLK

ATTTGGTGCCTATAATAGGCTTTTGTTCTGAGAGGAAAGAGCGAATTCTAATTTACAAATACATGCCTAATGGAAACTTACATGAGTGGCTGCATTCAACCGACGACAAGGCAAGACTCTTGGACTTTCCTCTAAGGGTTAAAATTGTTTTAGGTGTCGCGAAAGCCCTGACTTGGCTTCATGATGGTGGAAATTTCCATGTTGTGCATAGTAACATAAGCACACAATGTATCTTGCTCGATGAAAATTTCGATCCTAAGTTATCCAATTTTTGGGAGGCAACACTTGCAAAAACAAATGACAGAGATTCAAACTTGAGTCTATTTCCAATAGTCGAGTGTTTAGACTTTACAACATACAAAAAAGATGTTTATCGATTTGGGGTTGTGCTTCTCGAGCTTTTAACGAGGAAAGAATCTTACATATTGAGCTGCTCGAGTCAAAATTTGTCGAGCAGCTCTTTTGCCAGTCCTCTTGATGTGGATAAACTACTGCTAGGCCAAGGATATGATAATATGGTAATGCAACTACTTGAACTTGCAAGTGATTGCATGAAATTTATTCCGGATCAAAGGCCAACGATGCAAGAAGTGTATCAGACTGTTTCTGAGATTGCTAGAGTCCATGACCAAACAGGGGACTCTGAAATACAATTGCACTTGGACTAGAAGCAGGGGCGGAGCCACTAAAGGGACGTGTAGCTTTGGCTCAAACCATGTATTTG

>M6-PGSC0003DMG400027499 M6 LRR-RLK

GTGACAGTATTCTTTAAAAGTGCATGCAGCTTCTTCAACCCCAAGCATTTCTTGAAGTTTACAGCTTTAATTCCTGAAAAAAACCAGAAAAAAGTTGCATTTTCCATGGGAATGGCAAAAACAGTGCAGGGCTCATGTTGGCTTCTTGTATTCATGGTGTTTCTTCTGTTTAATTTATCAGATGAGTTCTTAGAACCTTCACAATCTGAAGCTATAGTGAAAATCAAGCAGCTTTTGAACTTCCCTCAAGATTTGAGTAGTTGGAGTGACAATACAGACTTTTGCAATAGTGAGCCAAACACTGCCTTAACACTCATGTGTTATGAAGATAACATAACACAGCTTCATATTTCTGGCTATAACTGGTTTCCTAATTTGCCTCAGGGGTTTTCTACAGATACCCTTTTCTCTAATCTTGCTCTTTTGCCTAATTTAAAGGTACTTTCTTTGGTTTCTTTAGGTTTGAGAGGAACTTTACCTAAAAAGATTGGATTTTTATCTTCTTTGGAGATTATTAATATCAGTTCAAACTTCTTTTATGGTGAGATTCCAGGAGAGATTTCATATTTGAAGAGTCTTCAGACACTTATATTGGATGATAATAAGTTTACAGGTCAAGTTCCTGAAGGGGTTGGTTTACTTCATTCTTTGTCTGTTTTGAGTTTCAAGAACAATTCATTTTCAGGTTCATTGCCTAATTCTTTGTCAAATTTACAAACCCTAAGAATCCTTTCTATTTCTGGTAATAATTTTTCTGGGGTTGTTCCAAACCTTCATAATCTGTCAAATCTTCAAGTTCTTGATTTGGAAAGTAACAATCTTGGACCTAATTTCCCTAATATTCCAACCAAGTTGGTATCTTTGGTACTTAGGAAGAACAAGTTCAGTTTAGGGGTACCAAAAGAACTGAGTTCATGTTATCAGTTGAAGAAACTGGACATTTCTTCCAATGAGCTTGTTGGACCCTTTTCACCAACAGTTTTATCATTGCCATCACTCAGCTATTTGGATATTTCTGGAAATAAACTAACTGGGAAGCTTCTAAAGAATGTGACATGTAGTCAAGACCTTTCATTTGTGAATTTATCATCAAATTACTTGACAGGGGAGTTGCCTGATTGCTTGAAACCTAGTTCTAGTTCCAAGATTGTATTGTTTTCTGGGAATTGTCTGTCGAATAAAGAGCAATGGCAGCATCCTTATTCATTTTGCCATAATGAAGCTTTGGCTGTTAGTATTGAGCCTCATAAGGGAAAGGTTAAAGGGGGAAATGGTAAAGCTGTTCTTGCATCAAGTATGATAGGAGGTTTTGTTGGAGTAGTAGCCATTGTCGGTTTGGCTTTTGTGGTTGTTAGAAGAGAGTATGCCAAGCAGAAAGCTTGCCAAGCCCCTCAAACAAGATTGATACTTGAAAAGGTCTCTCCTGCTCACACTCTTAAGCTGCTTAATGATGCAAGTAAGCTCTCTCTTTCTCTCACACAAACATATTATGTTAATGATTTCAACTTATGAAACTTTTTAGGTCCTGATAATTGTAAATCTAATGTTCTGTAAGGTCTGTGTACACTCTACCTTCCCCTAATCCCGCCGGTGGGATTACACTGAGTATGTTGTTGTTGTTGTTATCATTGTACTTGAAAAGTCTATGTAAACATTTGAATTTAAGCATTATTCCACAAAAGGAGTTTTCTTTATATATAGAATGAAAATATCAATTTGCTTCAGGAAATTGTCCAATACTGTGTTATTGGAATAAACTCTGCTCTTGTCTAAATTTCAAGATGTATCTTGATCTTTGATTCTTGATAGGGTATCTTTCTGAAACGAGGAAGTTGGGATTGCTTGGGGCCCCTCCTTATCGAACGTTTGTCTTGGATGAGCTTCGAGAGGCCACTAATAACTTTGATATATCAAATCTAATCGGTGCAAGCTCCAGTGGGCAGGTATGGATTTCCTTACTAAATACTGTAAGACTGTTTGAACAATGTTAGTCCACATGGACTGTTTTGTAGTTCATTCCTCAATTCTCTTGCATCTATGGTGAATGCTGATTGTGCTTTTATATATTTTGAACAATAGATTTACAAAGGGCGTCTCACAGATGGCACTGTGGTTGCTATAAAAAGCATAAAAATGAGGAAGAGGCATAGTGTCCAGTCCTACACTCACCAACTCGGGCGTATTTCAAAGATCAGATACTGCCATTTAGTTAGTACTATTGGACACTGCTTTGAGTGCTACCAAGATGACTCAAGTGTTAGCCGAATATGTTTAGTCTTTGAATTCGTGCCAAATGTGACGCTGCGGGGGGTCATATCAGGTATATAGTCTAGATAACTTCATCTGCTTCGCGAGTTATCATTTGGCACAACAACTGCAACAGTTTATAATTGAGCATTTAATTTACTTATTTGACTATTTCTACCATTTTACATAAATTATACTAAGTCTAAGTAATTGACTTCTTGGTATGTAGACGTTATAGAAAACAAGCATCTTTGCAACAATCTGTTCCATGTTGTTTTGATATATATTACTAGATTGTCCTCAATCACGACAAGTACCTACTACATTGTACTTCTTGTCAAAAGTGCTGATGAATTTTAAGGAGACATGAAATGCTTTTAATTACTTTCTCCTAGATTATGCAACGCGTTGTTTCTTCTTTGCCCGATGCTTAAATGTACAGGTGTTTATTGTCTTTAAGACAGTTGTTCTGAATTCTTTGCAAATGCTCGGTTTTGTAGAAGCAAATTCAGCTCAAAAGTTTACATGGACGCAACGGATGTCAGCTGCAATAGGGATAGCAAAGGGAATTCAGTTCCTTCATACTGGAATAGTGCCAGGGATATTCTCCAATCAATTGAAGATTACAGATGTATTACTGGATCAGAATTTCCACGTAAAGATTAGCAAATACAATCTGTCTTTGTTAATTGAAAATAAGAAAATGGTATGCAAACCTCTCTTCTCTACTATATTCTCTATTAATCACTCATTTCATTCTTGAACATTCACATTTCGTTATTGCAACTGTTCTTAGGATGCTGGACCGTCTTCTAGTGGATCAAAGGGAAATGATGGGCAAAGGTAAATGATCTCATCATTCTCAGCTGCTACTGCTACTGCTACATTTTCGGTCTTCATTACTCCACTTTTATGGTTCGATTTCAGAAGCACCTATTGGAAACTAATCTTTTTATACTTCTAAATGCAAATTGGTCTAAAAAGGTTTTAGAGAATCTTAAGTGTATCTCGACCAATTCTAGACAACACCTCTCCAGCCACTGCTATTGGGGGTAGGCACCGTTATAGCATCATGCCATTCATTGGTTATGATTCTTGAAGACAATATATAGAGCCATAGACTTGGACCGTCTTAGGAGATTCCGAGATAGTACTCTACCGTTTAGTAGACTATTAATGAATTCCTTCTTTTGTTAATATCTTCCTGAATGCAGGTTAAAATACGAGGAGAAAGATGATGTGTATGACTTTGGAGTAATCTTACTCGAAATCATATCGGGAAGGACAATCGATACAAAAAATGATATAGATGTTTCTAAAGACATTGTAAGAGTCACTTATGACATTTATGAGAAAATTCTTTCAAGTCATCAAGTTGAAGAACTATTTTGTTAAAACTTTTTCTTGTTGTTCTTATTGCAGCTAATAGTTAGCTTAACAGCTGACGAAATAGGTCGGAGGAACATCATTGATCCAGCTGTTCGGAAGGAATGCTCAGATAGTTCTCTTAGAACATTAATGGAACTATGCATAAAATGTCTTTCAGATGAACCATCGCAACGTCCTTCTGTAGAAGACTTGATATGGAACTTGCAGTTTGCAGCTCAGGTCCAAGACCCTTGGAATAGAGACACTTATGGCAACCAAGAGTCCCCTGGTCATGTGTAAGAACTTGTGCAAATATCCAATGATCAATGGACCTTTTCTTGTACAAAATGTAACATTATACCACTAGGATAGTAGTTTCAATTATTTAGTAGTCTCTATTGCAACACTTTTGGCTATATGAGGTTTCAGTACTCTTTTGAGGTTAACTCTACATTTTAGCCAAATTTTTTAAGCAAATGGCTATTTCTTGCACTTTTTTTTTTAACGAGCCTAAAATTTAAAAGGTGGGCCTTGCACCGACCATCCAATGGACCATGGACTAAACGGCACCCAAAAATG

>M6-PGSC0003DMG400027536 M6 LRR-RLK

CTCTGCGCACACACAACTCAGTAGAGAGAGAAAGAGAAGCAGTACAAAAATGTGAAAAGTTGTAGCAGATAAAAAAAAAAAGGGGTTCATCAGAAGAAGGAGAAGAACTAAACTGAACTGAGAAATACAAAAAAGGTTTGTTCAGTGAAGAGTAAAAGGGGTAGCAAAATCTTGGAAATTAAGTGTAAAAATTTGATCTTTTTTTGTTGGGTTATGAGATGAGATCGATTCTTGGGTTTTGAGCTTTGTGGGTGTGTTTATCTTGGTTCATGGTAATGAGGGGTAGGGTTTTCTTGAATTTTTGGGTAAATGCTGGGTTTTTTGAGTTTGCTTTACTTGGATCTACGAGATTGTTTAGTCAGCTGGCTGCTTGATTTGTAAAAAAAAAAACAGTATTTTGATGGTGAAGGTGATGGAGAAGGATGCTGTGGTGGTGTCACTGGTGGTATGGCTAATCTTGGTTGTACATCATCTTAAGCTCATTTATGCTAATATGGAAGGTTTGTGGGTCCATCATCCTCATTTAGTTAATTTTTAGGTCTATATTTGTTGTTCTGTTTTGTGAGATGAGTTTCAAATGTCGAAACTTTTCGTACTAGAAATTAAATATTTGCTGAAAGTGAAATGAGTATGTGCAAAATTGGATTTTGAATTTGACTAATGTGTGAGTTGTTGAATTTTCCCAAAAGGGTAAGCAAGATGTGCAGTGAGCTCCCTGCATTGTTACGGTGGCAAGCAGTTTAGTTTTAATGTCAGTTAAATGTGAAAGTTTTTAGTTATCATTGCCATTCAACCTTAGTGCATTATGTTTCCTATACAAGCTTGAATAATTTGATGTGATTTAATTTGGTTGTTGCCAAAATCTTGATAAACTGTTTTTTTCAGTGAATAGATTAAATTTTTATGTGCATTTTGGTGGATTAAATATCGTCTATGTTTTCCTAAATGGATACACGAACAGGTGATGCATTGCATAGTCTACGCGTCAATTTACAAGATCCTAACAATGTGCTACAGAGCTGGGATCCTACCCTTGTTAATCCTTGCACATGGTTTCATGTTACCTGCAACAATGACAACAGTGTTATAAGAGTGTAAGAATCTGTTGTCTGGTTTACCTCCGTTTTAATGGATTTGCAATTCCACTCTCTTGTGGTGGTGAGCAAATATAATTGCAGTTTGTGCTTCCATAACACTTGTTATATGATCAACCTTTCCAGTGATTTAGGAAATGCTGCTTTATCTGGTCTGTTAGTCCCACAGCTTGGCCTTTTGAAGAATTTGCAGTACTTGTAAGTCTCACTTCATGAACTATGTTTGGAATTATTTTATGAATTTAGACTTGGAAAACTGGCATTAAGTGTATTGGTTTTTCCAGATGTTGAAGATTTTTCATATTACCTCTAGAATTTGCTGCATGGAAATACCATAGATGGCTTGCATATTAACTTTTGCATAAAACAAAGCTGTTGTAAGTGGCAAAAGCGAGCAGAGAACACTTTCTCCTCTCCATCTCTTGTTTCACTTGGTGTTGTAGTCTATTGCATAGTTGTTAATGTGTTCCCTGAACAACACAAATTTGAATGTATACTTCATAAAAAAAAGATTTAAATTTGTGTTTCACAATACATACTAGTAAATTAAGGAACTCGGATACTTGGATGAGACTAGCAGTGAATTAACTTATGTGGCAAAGTAGTGTCTGTAATTGTTTGTGAAAACATTGTAGTTGAACTTAAATTTCTGTCATACACTCCCCTATCAAAAGATAAGCATAGATTCTAAATATCATTACATTAGCCAGTACCCTAAGCACTCTTTTTTTTCTTTTTGGTTTCCCACCCGGTGTCCAGTACCCACATTGGAGCCCGACTAAATGTGGATTTGTGTTGGGAAGTCCCACATTGGGGGGTAAAGCGCTCCCTAACAAAGGCGACTCCGTATCCTAGAGGGCTCGAACCCGAGACTTCTTGGTTAAGGATGAAGGAGTACTTACCACTCCACCACAACCCTTGTTGGTACCCTAAGCACTCTTAATTTGGAATAACAAACTTCAACTCATATCAAATCCTTTATTCCCTCTTTCCTTGTCCCAGGTAGCGTCTAATCATGATTTAGGATTATGGAATAGGATAAAGCAAACAAGAGAAACAAGTCCATTGTAAAAGTAAAGCAAAATCTAGACTTAGAAAGAGGAAATATGGCTTGAGGGTAGGTTGATATCTTGGAGGAGAAGATCTGAAGTTTTGTGGTCATATAACAGATGCTATTCTGGTTTTCACCAAGAGTTTGAATCATGATTCAATTAAGCATAGATTTTGTACATGTGTTAAGCTTAGTTTGGAGTTTGATAATATACCGTTCACTTAAATTGTCAACGTGCTATAGTCATATTCTGTCTCAAACTCGCTTTCATAGTTCTTATTGTAACCCATGCTCTTTTGAAGATGCAACTGTATGGAAGTTCACAGAGACAGTAAGTGCATGTGCACCTGGTGAAGCTGGGCATTACATGATGTATACCCATGCATCAGACAATGTTTTAATGATGGGCATAATTATTACCTAGTGACTGCTTCATTCTGCTCACACATTCATATATTCCTATGGATGCCTAGATTTTCTGGAGGCTTGTGTAGTATGTGTGCTAAGTATTTTCCAAGTGTATGCAAGTGACAATCACATCAAGAATGAAACAATATAAAAAGAACATCAAGAATGCTGATGGAATTTAACGATCACTGGGCATACACACTGTTTAGATGAAGTTCGAAATTAACATAATGACAAACGTTAAATTTTTATTGATGAGCTATAAAGATTGTATGAAATGTTTGATGAGCTGATCCTTTGCAGCACATGGGATGACAGATTTTTATCTAGCAAAATGCTTGTAACCAAATATAAGAAAATTCCTCCATGACATAAACATTGGTAACTAGTTATAATTGTTAGCTTGGAACTCCCAGACCTTTCTCATTTGTTGACATTTAACTGTTGCTTGATGCCAGTTTTGATGAATGTGTGAACTTTGCTTCATTATCTTTGTTGCCCCATTATCCTAGGGATGTCATTGGTATGATATATTGAATTGATTCATTCTCTGGTTTTCTTGTCACAGGGAGCTTTACAGTAATAACATAAGTGGTCTAATACCTAGTGATCTTGGGAATTTGACTAATTTGGTCAGCTTGGATCTCTACTTGAACAACTTCGTTGGTCCCATCCCAGATTCCTTGGGCAAACTGTCGAAATTGAGATTCCTGTATGTATTTTTATTCTATATTAGATTTATGTGGATATTGTGATGCCAAATGTGTATGTCATCAACCAAGATGGACCTTCTTTTGCCGGCATGTTGAGAAAGGAAAGGAAGCCTGAATTCTTTTCTAGGTTTCACATAAGATTACCTCTATTAGTATTAGGTTTCACACTAAGTTTTGATGGTAGAAAGGTATTTGAGGACTTTATGGGTCTGATATCATTTATAAAACAGATCTTACATATAAGTCTTTTTATTTTCTCCCTTTGTTCTAATCCTTATATAAAGGAAAAGAATAGAAGCCTATGCTTTGTCTGATATTTGGCCCTGATATTGTAAAATTAATCAGAGTTTAGTAATTGGTGTTTTATTTTTCTCCCATTGTTGGGATCCTTTTCTAGAAGAGAAAAGAAGTTACGAATCCCATGCTTCATCTTGCATTTTTCACTTCTGAAGTTTCTCTTGCTTAATGCTCCACAAACTTCTTTCTTTTGGCTTGACTGTCTAAGAAGTGTACCAAATAGGTTCTATTATGCAGAATCAGCATGAAGCCTCATTAACTGCATCTTGGGACTTACTATAGAACTATTGCAGTACTGCCAATGTTGACTGACATGATTTAGTCCAAAAAGTGTTTCAAGTTTCTGTTTGTCAGGGGGGAAATGTTTTGAGGCTGTTTATTGTTGTTCTGAACTGACAACTTCATTTCCAGCTGAAGTGCCGGTTCTGGCTTGAAGTTCGAATATGTTTTACATCCACATATAAATAATTTATTAAGATGATTCTCCTTGTCAATAAATAAATAAATTAAGATGATTCAATTAGTAACCTTTTCTTCTGTTTTATTCTCCAATTATTACTTTTACCAAAAGTGAAGCTCATTCCTGTACACCTTTGTGATTTTTTTGCAAGGATAGTCGGCTCAACAATAATAGCTTGACTGGTAACATCCCTATGTCACTGACTAATATCTCATCACTACAAGTGTTGTAAGTACACTAGTTATTTTGTGACTTAACTTAGATAATTCTTTCTGTCTTCCATATCTTTTCATTCATTTCCTTTTCCTTCTAGTGCATATACAGATATACTCATACTCAATTCTTGTCTCATCTGTGTATAGGGATCTGTCAAACAACCGTCTCTCAGGTGTTGTTCCAGATAATGGTTCCTTTTCTCTATTCACGCCTATCAGGTATATTTCATTTAAGGTGGTGCACCATCAAACTGGCTGGTGGTTTTAGTATGCTGCTATTTGCTAATATATTTTGCTTGAATTTACAGTTTTGCGAATAATTTAGATCTTTGCGGGCCTGTAACTGGACGTCCTTGTCCGGGATCTCCTCCATTCTCCCCTCCACCTCCATTTGTTCCACCACCACCAATTTCTGCTCCAGGTGGTTCTCTAAATTGGTGTGGATAAATTGTTTTCCTTTTTTTTTTTGCTTGTTTGAGTTTGGTGATTTTTGTTGGTCCATCCATCTTGCCTCTCTTAGTTTTTCCTAACCTAAGTGAGAATTTGTGCTATAGAAGTACTGACAAGAAAGGGGCAGAAGAGGAAAGCCCATCTTAGCTAGTCAGGCATTTTGGTTAAGTAGTTCTTATGGGAAACTGGTATGCACGAGACTACATTTAGTCTTTAAGCATTCTGGTCTTTATACAAATTTAATTCAGCATTGTGGAGATCTTTTCTTTGGTCCCCTTGTAAATTATTATCTGTGGTATTTGAAGTTGAGTATGTCTGAAATTAATCATATTTATGCCAGAACTTGTGAAGATTTTCTTTGCTTTGTAAAAACTGTCCTGGAAATTAGATCTGAATGATAGATACAAATTTGTGTACTTCTTTCTTTTTGAAGTGATTTAAATAATGAAGGGCCAAAATTGTGTTAAATATAGTCTGGATCATATGAAGGAAATTAATCAAATTTATGGTAACTTCAGGAGGAAATGGTGCAACTGGAGCAATTGCTGGAGGTGTAGCTGCTGGTGCTGCTCTACTATTTGCTGCTCCTGCCATTGCATTTGCCTGGTGGCGCCGTAGAAAGCCACAAGAATATTTCTTTGACGTACCAGGTTAGCAGTATTCACAAATCACAATACCCAACCATAAGTCCATAACTCCTACTTACTCCCTCGCGTGTTTGTGGTTTCTCTTGCATGTTTATTTTTTGGCTCCATAATTACCGTTTTTGCTTAAACTTATTGCAGCTGAAGAAGATCCTGAAGTTCACTTAGGTCAACTGAAAAGGTTCTCCCTCCGAGAGCTACAAGTTGCAACCGACAGTTTTAGCAATAAAAATATTCTGGGTCGAGGTGGATTTGGTAAGGTATACAAAGGACGCTTAGCAGATGGATCATTGGTGGCTGTTAAGCGGCTTAAGGAGGAGCGTACTCCTGGAGGGGAGTTGCAATTTCAAACAGAAGTTGAGATGATTAGCATGGCAGTGCATAGGAATCTTCTACGATTGCGTGGTTTCTGTATGACACCAACTGAAAGACTGCTTGTCTACCCCTACATGGCGAATGGAAGTGTTGCATCATGCCTTAGAGGTGACGCCTTCTGAAATCTATCACTCCGTAACATATTTGGATGCTCTCACCTTTAATTTAGAGGGTATTATTACATAATGCAAGAACTTCTTTCGCTGGTTAACGTTCTATCTGGCATAACATACTCTTTATAATAAAACATATTCTTGTATGTTATTCCTGTAACTTGGTAAAAGGTAGAAATATAATTTATTTTGTATTCTCTTGACAAATTAATTTATTTTATCAGAACGACCGCCTTCTGAACCGCCACTTGATTGGCCAACGCGAAAGCGCATTGCTTTGGGGTCTGCCAGGGGATTATCGTATTTGCATGATCATTGTGACCCTAAGATTATCCATCGTGATGTGAAGGCTGCAAATATATTGCTAGATGAAGAATTTGAGGCTGTTGTTGGAGACTTTGGTTTGGCTAAACTTATGGACTACAAGGATACACATGTTACAACTGCTGTGCGTGGTACAATTGGGCATATAGCTCCAGAATACCTGTCCACAGGGAAGTCTTCAGAAAAGACTGATGTTTTTGGGTATGGGATCATGCTTCTGGAGCTAATCACCGGCCAACGTGCTTTTGATCTTGCTCGGCTTGCAAATGATGATGATGTCATGTTGCTTGACTGGGTATGTTGTCATCCTGCTTTACATGTAACATTACACGAGTACCATAATGTGCTCATTTTTTAATCTGTGCATCACAACACTAGCTGACTAATAAGTGTTTCTGCCTTTAGCAGGAATCTTTATGTCTATGTCTGAACTTGTTGTGGTTCTTGTTTCAGGTGAAAGGACTCCTCAAAGAGAAGAAATTGGAAATGCTGGTTGACCCTGATCTTCAGAACAAGTATGTGGAGGCTGAGGTGGAGCAACTGATCCAAGTAGCATTGCTTTGTACACAAAGCAACCCAATGGATCGGCCTAAGATGTCGGAAGTGGTGAGAATGCTTGAAGGTGATGGTTTGGCTGAAAGATGGGATGAGTGGCAGAAGGTAGAAGTTCTCCGGCAGGAAGTGGAACTTGCACCACATCCTGGTTCTGATTGGCTTGTTGACTCAACAGAGAATTTACACGCAGTTGAATTATCGGGTCCAAGGTGACCGACCTCGTACTACCCCAAGTAAAGAAAATCAAATCATTAGGAACTCTTTATTTTTGTTGACCTTGTGAAAGTTTTTCTTTGTGAAATTTAATGAAAATCTCCAGTTGTAAGCCCATGAAGTGTCTGTTACATTGTGCATACAAATTTGTACTGACAGGAAATTCCAAATCCATTCCCTGACAAGTTGGCACATAGGCGTAGTAGTTAAGTGCTTTAAAATTGTCCAACACAGTAAACTACTGTCAAGCAGGCTCTTGTTTTCAGTGTCATTGAAAATGCTATCAATTTGACAAGGTTATTTACAAGATTTTGGATTTTACAATCAAAGAATAGTGCTTAACATTCTCTTACGGTTTTTATCATAATCTCGTGCTACAAGATAATGGCAAATATTTCATCAATAGGCCACTTGCAGAGTTTGTTTTGCCGCTCTATGCCCAAATTTTGGGCTCTTTCTGACAATTCAGATTAGTACCAGATGCTCTGATACAAGTACAAGATCTGCTCAGTGATACTTTCGATCATTTCTGTCATTACTTTTTACTCCAAAGAAAATTTCTTACATGCAAATTTTTTGAACTGCATACTCTTCAACATACAAGGATCCATCGAATGTACTACAGACCAAAGGGCAATAGAATAAACAGAAGAATCTCAAAACTCACAATCAACAAACAAACCAGAGCCTGCAGAAGGTAGAGAAATACACTTCACTTATCATGGCTGGACCTGATATTATCTTTGAATGACTGTGAACGAGAACCAGGATTAGAGCCATTTGTTGCATTAAACAGGTTTGGATATGAATAATATGTTTCATCGTCCATGGAAGGCAATCTCTGTGGTTGGTGAAGACCCTGACCAGGATGATGAATATGGGAGATGGGAGACTCAACACCCACTGTTGAATCATCGTATGAAGGAATATTCGACACCTGATGTTCCAATGGACTGCCATGAAGTGGGCTGGGGATACCATCAGAGTTTGACGATTGGCGTTGGAGGTCTAAGTGAGGCTTTTCTAGGATGTGAGATTCTTGTTTCTCAGCATAAAAGGGGTTCTCACTGGGGCTTACAGGCACCTCTTCGCTCCTTTTCCTGGTAAGCTCGCTGATTCTAACCTGAGCAAGGCTAGCTGCTGAGCGAGCAGCAGCAGCTGCACGTTCGGCTGAATCAGCAGCTGCTTGGGCAGCAGCCAATACATCTTGCAAATCTACATCAGTCTCAACTTTGGTCTTTGTAGTAGAGGGTGGTTGAGTACTTTGGGTTGTGGTTCTTTCTGTTGAGGAGAATGGAGGTGATTTTGGTGGGGGAACCATGAATGGCAAAAACTGTTTCTCTTCTTCCGGTGCAGATAAACTATCAGCTGATCCAAAATCTTTTGAAACTACCTTCTCTTGCAAAAGTTCATCTCGCTCCATATCTTGTTTGTGAGTTTTGAGCTCACCGTCTTCAGAAGGCTTAGCTACCTCCTCATCGGAGTCAGATAGTGCAGAGGCTGGGAAAGGAAGCATTTCAGGGGCAGAAACTCCACCGGTGCTAGGTCGTAGTGGCTGCTTAGGGACTTCTGGAAAATCTAAGTCAAAATCAGCCTCAGATTCCGTTTGTTTATCAAAGACTTGATCTGATGCAGAGTGCTGCACCTCATCCAACCTTTCTTTGGGAAGTGGCACCTTAGCCCCACTGACAAATTGTGTGGGCCCATTCTGCAAAGCAAAGGGAAGAAATCATGATGATTGAAAAGAGGACAAAATCAAAGCCAGTACAGACTGAAATATACCAGCAAGTCCTCATGTGACTTCAACAGCTCAGTTTCAGAAGCACTTGGATCCCAATCTAGCTGATGCTCCTCCGCAATTTCCTTTAGCAGCTTCATTTTCACATCAGGAGCAGGAGCTCGAATAGACAGCAACTCTATCAACTGTCAGAGACAAGAAAAAAGAGAGAAAAATCAGAAACAAGAAACAGTTCCTCTTTCTCGTTTAGCTATGGCATAAAGGAGATCTAAGGTAACCTGGCGATTGACTCCGCATTCTGGCATGAGCTCTGTAGCAGCAGCAATGAATTCCTTCCCATATTTACCAGCAAACATCATCTGAACCTGCAGCAATTCCGGAAGATCTGCACATCTTGGTGCAGCAAAACAAACACTTGAAATAGCTTCTTTCAGATCTAGAGGGCATTCCCTGGACATTATAGAGTTTGGGGGTTCAGCTCTTTCCAATTATTCAAGTTAAGAAAATTATCAACAAAATTCAACTAGAAGTCTAGAATATTAAGACTTGGAAAGAAAACCTCCACATTGCAGGACAGTCAAGTTCAAGAGTATCAGTTTGCTTTAAAACAAATATCGAAGATGAAGCGCTTTGGGTCCATTTTGGAACTAGAGATCCACATTTATGGTCTCATGTAGATGATCAAATTATTTTAAGGTTTCACACTCCCATAACACAAGCAAGAATGTATTGTTCATTCTATGCCCTAGATACTTCTCGGTCATATAAGCAGAAAGTTAAACGTAGTTAATCCTTGTTTGGCTAAAATTGACTTTATGCAAAAGGATCAAACATGCTGCTGCTCGGATACATGCACATACTTCCATTTAATCGACCTACAACTTAGACATTTCAAAGTTGGTAACTTTTTTTCAGAATACATGAAATTTAGCAAATTAAAGTACTGTGATTCATTAATTGAACTAGAGTCGCTCCATGAAAAAGTATCCATATCTAGTGCAGATAATTCTCGCAAAACAACAGCTAGGGTGGAGTATAGTAGATCTGAAAGTTTGGTATTGAAGGAGATATGGAACTAAATGTTAACAGCAGCATAGACAGCAAAGGCTAGAGGCAAGCAAGGAAGACAGAGAGGAACATTTGGGACCCGTTTGGCCATAGATTTCCCAAGTAATATTTGGGGGGAAATTTGGCAAATAGTGTTTGTCCATACAATTTGCCATTATTTGGCAAATATCCCAAATTCCCAAATACTAGTTTTTTATAGTATTTGAGCCAAATCTCATTATTTGGGATCTTTTAAAAATTGAAATTTTATACCAAACTTTTATCCTTTACAAAAACTCCCTCTATAGTAGTTGTTTGCGTTGTATTACATAATTTTTTACGTGAACACCAAAGTAGTGATGAAATATTCAGTGAATATGAAAGTGATGAACAATCGGTCCAGGGTACCAAGTCATGTGCTTTGTCCACTTCATGATGTATAGAATGATGCTTGTTGCACTCGCTCCACACACCACATTGCTCCAGTTTCATGGACACTACTTGTTATTCGTTGCAACGAAGATCCAACTAACTTGTAATACAAACTTATGGTTAGTTTTGATAGTTTTTAAAACTTATGGTTA

>M6-PGSC0003DMG400027555 M6 LRR-RLK

CAGCTGAAGAAGATCCTGAAGTTCACTTAGGTCAACTGAAAAGGTTCTCCCTCCGAGAGCTACAAGTTGCAACCGACAGTTTTAGCAATAAAAATATTCTGGGTCGAGGTGGATTTGGTAAGGTATACAAAGGACGCTTAGCAGATGGATCATTGGTGGCTGTTAAGCGGCTTAAGGAGGAGCGTACTCCTGGAGGGGAGTTGCAATTTCAAACAGAAGTTGAGATGATTAGCATGGCAGTGCATAGGAATCTTCTACGATTGCGTGGTTTCTGTATGACACCAACTGAAAGACTGCTTGTCTACCCCTACATGGCGAATGGAAGTGTTGCATCATGCCTTAGAGGTGACGCCTTCTGAAATCTATCACTCCGTAACATATTTGGATGCTCTCACCTTTAATTTAGAGGGTATTATTACATAATGCAAGAACTTCTTTCGCTGGTTAACGTTCTATCTGGCATAACATACTCTTTATAATAAAACATATTCTTGTATGTTATTCCTGTAACTTGGTAAAAGGTAGAAATATAATTTATTTTGTATTCTCTTGACAAATTAATTTATTTTATCAGAACGACCGCCTTCTGAACCGCCACTTGATTGGCCAACGCGAAAGCGCATTGCTTTGGGGTCTGCCAGGGGATTATCGTATTTGCATGATCATTGTGACCCTAAGATTATCCATCGTGATGTGAAGGCTGCAAATATATTGCTAGATGAAGAATTTGAGGCTGTTGTTGGAGACTTTGGTTTGGCTAAACTTATGGACTACAAGGATACACATGTTACAACTGCTGTGCGTGGTACAATTGGGCATATAGCTCCAGAATACCTGTCCACAGGGAAGTCTTCAGAAAAGACTGATGTTTTTGGGTATGGGATCATGCTTCTGGAGCTAATCACCGGCCAACGTGCTTTTGATCTTGCTCGGCTTGCAAATGATGATGATGTCATGTTGCTTGACTGGGTATGTTGTCATCCTGCTTTACATGTAACATTACACGAGTACCATAATGTGCTCATTTTTTAATCTGTGCATCACAACACTAGCTGACTAATAAGTGTTTCTGCCTTTAGCAGGAATCTTTATGTCTATGTCTGAACTTGTTGTGGTTCTTGTTTCAGGTGAAAGGACTCCTCAAAGAGAAGAAATTGGAAATGCTGGTTGACCCTGATCTTCAGAACAAGTATGTGGAGGCTGAGGTGGAGCAACTGATCCAAGTAGCATTGCTTTGTACACAAAGCAACCCAATGGATCGGCCTAAGATGTCGGAAGTGGTGAGAATGCTTGAAGGTGATGGTTTGGCTGAAAGATGGGATGAGTGGCAGAAGGTAGAAGTTCTCCGGCAGGAAGTGGAACTTGCACCACATCCTGGTTCTGATTGGCTTGTTGACTCAACAGAGAATTTACACGCAGTTGAATTATCGGGTCCAAGGTGACCGACCTCGTACTACCCCAAGTAAAGAAAATCAAATCATTAGGAACTCTTTATTTTTGTTGACCTTGTGAAAGTTTTTCTTTGTGAAATTTAATGAAAATCTCCAGTTGTAAGCCCATGAAGTGTCTGTTACATTGTGCATACAAATTTGTACTGACAGGAAATTCCAAATCCATTCCCTGACAAGTTGGCACATAGGCGTAGTAGTTAAGTGCTTTAAAATTGTCCAACACAGTAAACTACTGTCAAGCAGGCTCTTGTTTTCAGTGTCATTGAAAATGCTATCAATTTGACAAGGTTATTTACAAGATTTTGGATTTTACAATCAAAGAATAGTGCTTAACATTCTCTTACGGTTTTTATCATAATCTCGTGCTACAAGATAATGGCAAATATTTCATCAATAGGCCACTTGCAGAGTTTGTTTTGCCGCTCTATGCCCAAATTTTGGGCTCTTTCTGACAATTCAGATTAGTACCAGATGCTCTGATACAAGTACAAGATCTGCTCAGTGATACTTTCGATCATTTCTGTCATTACTTTTTACTCCAAAGAAAATTTCTTACATGCAAATTTTTTGAACTGCATACTCTTCAACATACAAGGATCCATCGAATGTACTACAGACCAAAGGGCAATAGAATAAACAGAAGAATCTCAAAACTCACAATCAACAAACAAACCAGAGCCTGCAGAAGGTAGAGAAATACACTTCACTTATCATGGCTGGACCTGATATTATCTTTGAATGACTGTGAACGAGAACCAGGATTAGAGCCATTTGTTGCATTAAACAGGTTTGGATATGAATAATATGTTTCATCGTCCATGGAAGGCAATCTCTGTGGTTGGTGAAGACCCTGACCAGGATGATGAATATGGGAGATGGGAGACTCAACACCCACTGTTGAATCATCGTATGAAGGAATATTCGACACCTGATGTTCCAATGGACTGCCATGAAGTGGGCTGGGGATACCATCAGAGTTTGACGATTGGCGTTGGAGGTCTAAGTGAGGCTTTTCTAGGATGTGAGATTCTTGTTTCTCAGCATAAAAGGGGTTCTCACTGGGGCTTACAGGCACCTCTTCGCTCCTTTTCCTGGTAAGCTCGCTGATTCTAACCTGAGCAAGGCTAGCTGCTGAGCGAGCAGCAGCAGCTGCACGTTCGGCTGAATCAGCAGCTGCTTGGGCAGCAGCCAATACATCTTGCAAATCTACATCAGTCTCAACTTTGGTCTTTGTAGTAGAGGGTGGTTGAGTACTTTGGGTTGTGGTTCTTTCTGTTGAGGAGAATGGAGGTGATTTTGGTGGGGGAACCATGAATGGCAAAAACTGTTTCTCTTCTTCCGGTGCAGATAAACTATCAGCTGATCCAAAATCTTTTGAAACTACCTTCTCTTGCAAAAGTTCATCTCGCTCCATATCTTGTTTGTGAGTTTTGAGCTCACCGTCTTCAGAAGGCTTAGCTACCTCCTCATCGGAGTCAGATAGTGCAGAGGCTGGGAAAGGAAGCATTTCAGGGGCAGAAACTCCACCGGTGCTAGGTCGTAGTGGCTGCTTAGGGACTTCTGGAAAATCTAAGTCAAAATCAGCCTCAGATTCCGTTTGTTTATCAAAGACTTGATCTGATGCAGAGTGCTGCACCTCATCCAACCTTTCTTTGGGAAGTGGCACCTTAGCCCCACTGACAAATTGTGTGGGCCCATTCTGCAAAGCAAAGGGAAGAAATCATGATGATTGAAAAGAGGACAAAATCAAAGCCAGTACAGACTGAAATATACCAGCAAGTCCTCATGTGACTTCAACAGCTCAGTTTCAGAAGCACTTGGATCCCAATCTAGCTGATGCTCCTCCGCAATTTCCTTTAGCAGCTTCATTTTCACATCAGGAGCAGGAGCTCGAATAGACAGCAACTCTATCAACTGTCAGAGACAAGAAAAAAGAGAGAAAAATCAGAAACAAGAAACAGTTCCTCTTTCTCGTTTAGCTATGGCATAAAGGAGATCTAAGGTAACCTGGCGATTGACTCCGCATTCTGGCATGAGCTCTGTAGCAGCAGCAATGAATTCCTTCCCATATTTACCAGCAAACATCATCTGAACCTGCAGCAATTCCGGAAGATCTGCACATCTTGGTGCAGCAAAACAAACACTTGAAATAGCTTCTTTCAGATCTAGAGGGCATTCCCTGGACATTATAGAGTTTGGGGGTTCAGCTCTTTCCAATTATTCAAGTTAAGAAAATTATCAACAAAATTCAACTAGAAGTCTAGAATATTAAGACTTGGAAAGAAAACCTCCACATTGCAGGACAGTCAAGTTCAAGAGTATCAGTTTGCTTTAAAACAAATATCGAAGATGAAGCGCTTTGGGTCCATTTTGGAACTAGAGATCCACATTTATGGTCTCATGTAGATGATCAAATTATTTTAAGGTTTCACACTCCCATAACACAAGCAAGAATGTATTGTTCATTCTATGCCCTAGATACTTCTCGGTCATATAAGCAGAAAGTTAAACGTAGTTAATCCTTGTTTGGCTAAAATTGACTTTATGCAAAAGGATCAAACATGCTGCTGCTCGGATACATGCACATACTTCCATTTAATCGACCTACAACTTAGACATTTCAAAGTTGGTAACTTTTTTTCAGAATACATGAAATTTAGCAAATTAAAGTACTGTGATTCATTAATTGAACTAGAGTCGCTCCATGAAAAAGTATCCATATCTAGTGCAGATAATTCTCGCAAAACAACAGCTAGGGTGGAGTATAGTAGATCTGAAAGTTTGGTATTGAAGGAGATATGGAACTAAATGTTAACAGCAGCATAGACAGCAAAGGCTAGAGGCAAGCAAGGAAGACAGAGAGGAACATTTGGGACCCGTTTGGCCATAGATTTCCCAAGTAATATTTGGGGGGAAATTTGGCAAATAGTGTTTGTCCATACAATTTGCCATTATTTGGCAAATATCCCAAATTCCCAAATACTAGTTTTTTATAGTATTTGAGCCAAATCTCATTATTTGGGATCTTTTAAAAATTGAAATTTTATACCAAACTTTTATCCTTTACAAAAACTCCCTCTATAGTAGTTGTTTGCGTTGTATTACATAATTTTTTACGTGAACACCAAAGTAGTGATGAAATATTCAGTGAATATGAAAGTGATGAACAATCGGTCCAGGGTACCAAGTCATGTGCTTTGTCCACTTCATGATGTATAGAATGATGCTTGTTGCACTCGCTCCACACACCACATTGCTCCAGTTTCATGGACACTACTTGTTATTCGTTGCAACGAAGATCCAACTAACTTGTAATACAAACTTATGGTTAGTTTTGATAGTTTTTAAAACTTATGGTTA

>M6-PGSC0003DMG400027581 M6 LRR-RLK

ATGAAACTACAAGATTTACGTGCAACTATTGTCTTTTTTCTATCATTGCTAGCAATTCTTCCTCACATTATTGCTAATCTTGATTCAGACAAACATGCTCTACTTCAATTTGCTGCATCTGTCCCACATCTTAGAAAACTCAACTGGAACTCTGCTTTGTCAATCTGCAATTCTTGGATTGGGATCACTTGCAACAAAGATGGAACGACGCGAGTCGTTGCAATTCATCTGCCTGGTGTTGGACTTACTGGCCATATCCCAGCAAACAGCATTGGCAAACTAGATGCTCTCCAAGTCCTTAGCCTAAGAGCAAACAATCTAAATGGAAATCTTCCTTCTGATATTCTTTCAATCCCTTCTCTTCTCTCTATCTACCTTCAACATAATAACTTCTCTGGTGATATTCCTGTTTCCTTTTCATCAACACTTGGTGTAGTAGATCTCTCGTTTAACTCTTTTACCGGAGAAATTCCCCCCACGATCAAGAATTTGCCTCGATTATCCATGTTGAACCTAAAGTTCAATTCATTATCTGGATCAATTCCCAATCTTGATGTGTCAAGGTTGAGTTTTTTGAATTTGAGTTACAACATGTTAAATGGCTCAGTTCCATATTCCCTCAGGAAGTTCCCACTTTCTTCTTTTGTGGGGAACTCTAATTTATGTGGAACACCTTTGAGTAGCTGTTCTTCGAGATCTCCATCGCGTAAAGGTGACAACTTTAAGAAACTTAGTAATGAAATTATCATTGCAATTGCAGTAGGTGGTCCTTCTGTTATAATTCTCCTTGTTTTATTCATCTACTTCTGTTATTTTAACAAGAAGGTTGATAACAACACAAGCATGGTTGAAGAGAAGAATGAGAAGTTAGAAGACTTTGGAAATGGGGTTCAAGATTCTGAAAAGAACGAGTTAACGTTCTTTAAAGGGTGCTCTTATAACTTTGATCTTGAGGATCTGTTGAGTGCTTCTGCTGATTTTCTTGGTAAAGGGAGTTATGGAACTGCTTACAGAGTTTCGTTGGATGAGGTATCGATGATAGTTGTGAAAAGGCTGAAGGAAGTAAGGGTTGTTAAAAAGGAGTTTGAACAGCATATGGATATTGTTGGAAAGATTACAAGGCACCCAAATATAGTGCCATTTCTTGCTTGTTATTACTCAAAGGATGAGAAACTTCTTGTTTGTGAATACGTCCCTTATGGTAGTTTGTCATCCGCCTTATATGGTATAAGCTGCTTAACTCTTCTTAGATAAAACT

>M6-PGSC0003DMG400027586 M6 LRR-RLK

GCTAAACAAGGTATATAATTAAAAGAACCCTTGTTGACATATATAGTTATGGGCAGAAGTTGCATTGTCCTCTTTGGTCTTGCAGTTTTCATTCTGCTTCATAACCAAACTTCACTTGCTACTGTTCCTAATATTAGCACTGATGAAGCTGCTCTCTTTGCACTGAAATCACACATTTCTTCTCATCCTAACAACATCTTAGAAAGCAACTGGTCTTCGTCCAGCCCGGTTTGCAGCTGGATTGGAATCACTTGCAGCTCCCGCCACCATCGAGTCACTGCTTTAGACATTTCTAGCATGCAACTTCATGGTACCATTCCTCCACACATAGGAAACCTCTCATTTCTTGTTTCCCTCAGGATCAGTTACAACGCTTTCCATGGAAATTTGCCAGCAGAGTTGGCTCATTTGAAGAGGTTGAAATTGATTAATGTTAGAAGCAATAACTTCACTGGAGCCATTCCATCATTTTTGAGTTTGTTACCAAACCTACGCACTGTGTCCCTCTGGAGCAACCAATTTTCGGGGAAAATTCCATCTTCCCTTTTCAATCTAACAAAGCTGCAAGTGTTGAGTTTACGGAGTAATTTTCTCGTAGGAGAGATCCCTCGAGAAATTGGTGATCTTCGTTACTTGACTGTCCTAGACTTGCAATTCAATCAGCTTACTGGCCCTATACCACCATCAATATTTAACATTACGACAATGCGAGTGATTGGTCTTACCAACAACAATCTTACTGGAAAGCTTCCAACAACTATATGTGACCATCTTCCAAACTTGGAAGGGCTTCAACTCTCAAGCAACTCCTTAGATGGAGTAATTCCACCAAACCTTGAGAAATGCAGAAAGCTTCAAAGATTGTCATTGTCGTCCAATGAGTTTATTGGAACTGTACCAAGAGAGCTAGGAAACTTAACGGCTCTTATAACATTATTTCTTGGACAACAAGTACACTTGGAAGGTATATAGTATGATATTTCCTTACAACATTATATCTAAAAATTTCGATCTCTCTTCCGCCTAACTGGTAATATTTCAAACAATTGACAGGAGAGATACCTGTGGAGCTAGGTAATCTTAAGAAACTACAGATGCTGGGATTAGCTGAGAATGAGTTTACTGGTTCGGTCCCTACAAGCATTTTCAACATATCAGCACTGCAGCTCCTATCACTTTCACTAAACAAGCTTTCAGGTACTCTACCTTCAGATTTAGGTCCGAACCTAGAAGAATTTTACTGTGGAGGAAATAATCTGAGTGGTTTTATCTCTGATTCAATCTCAAATTCTTCGAGACTCAGAATGCTTGATCTCTCATTCAACAGTTTCACAGGTCCAATTCCTAAATCACTCGGTAAATTAGAATACCTTGAGGCTCTAAACTTGGGAAACAATAGTTTTATCAGCGATTCAACATTAAGCTTCCTTGCATCATTGAATAACTGTAGGAATCTAAGAAATCTCTGGTTCTATAATAATCCTTTGGATGCCATTTTTCCGGCATCTGTTGGTAATTTCTCCGACTCCTTGCACGGTTTTGAAGCAGATGGTTGTAAACTTAAGGGCATGATTCCCCAAGAAATTGGTAAACTTACTGGTTTGACAAAGATGAGTCTATCTAACAATGAGTTCACCGGATTTATTCCAAATACCATCCAAGGCATGTTGAGCCTTCAAGAACTTTACCTAGATAAAAACAAGATAGAAGGACCCATACCAGATTATATGTGCAGTTTACAGAATCTTGGTGCATTGGACTTGTCACAAAATAAATTTTCTGGTTCCGTGCCACCATGCTTAGGAAAATTTACCAGTTTGAGGATACTTCATCTAGATTACAATATGCTGGATTCTAGATTACCAGAAAGCTTGGGGAGCCTTAAAGATCTTTTAGAATTTAGTGTTTCATCCAATTTATTAAGTGGGCAAATTCCATTTGAAATTGGAAATTTACAGGCGGCAACAATCATTGATTTGTCAAAAAATGATTTTTCTGGTAAGATTCCTAGCACTCTAGGGGGTCTAGATAAATTGATTTATCTTTCTCTAGAACATAATAGACTAGATGGGCCTATTCCAGATTCATTTGGCAAAATGTTGGCATTGGAATTCTTAGATTTATGCTATAACAATCTTGTTGGTGAAATTCCAAAGTCATTAGAAGCTCTTGTGTCTCTCAAATACCTCAACTTTTCATTCAATAAACTAATTGGAGAGATTCCCACTGGTGGTCCTTTTTCAAACGCCACAGATCAATCCTTCTTGTCTAATGATGCACTATGTGGTGATGCCAAGTTTAATGTGACAACATGTGTCATCCAAACACCCAAGAGGAAAAAGGCTAACTTAGTTTTGTACATCAGTTTGGGAGTGGGTCTGTTACTTCTTGTATTAGCCCTTGCATATGTATTTGTAAAATTGCGAAAGACAAAAAAGAATACAGGTCAAACAAATGTGTCTCCGTTAAAAGAGCATGAAAGAATTTCCTATTATGAAATTGAACAGGCAACAGAAGGATTCAACGAAACCAACTTGCTTGGTAATGGGAGTTTCAGCATGGTCTACAAAGGGATACTTAAAGATGGTACCCTTTTTGCAGCAAAGGTATTCAATGTGCAATTGGAGCGTGCATTCAGAAGTTTTGACATAGAATGTGAGATACTCCGAAACCTTCGCCACAGAAATTTGACCAAAGTCATCACCAGCTGCTCCAACCTTGATTTCAAAGCCCTAGTGTTGGAATACATGCCCAATGGGACACTTGATAAATGGTTATATTCTCATAACTTATTCTTGAACTTGTTGCAGAGATTAGATATAATGATAGATGTTGCATCTGCAATGGACTATCTCCACAATGGTCATTCAACAGCTGTGGTGCATTGTGACTTGAAGCCAAGCAATGTCTTGCTAGATCAAGAAATGGTTGGTCATGTCAGTGATTTTGGCATTTCAAAATTGTTAGGTGTAGGGGAGGCTTTTGTTCAAACAAGGACAATTGCAACCATTGGATATATTGCTCCAGGTATAGTACAACTTTTCAAAGTTTTCTCATATCGTTTAAATACTCAAAACAATTCTTCCTCCCCTAAATATAAATGGATTTTCTTTGTTTTTTGCATATTCCATTTTAACTAGGAAACTTTTATAATTGTGATTTCAGAGTATGGACAAAATGGAATTGTATCCACAAGCTGTGATGTTTATAGTTTTGGCATCATGATGATGGAGATGTTCACAAGAACAAGACCAAGTGATGAAATATTTATCGGAGGCTTGACCATACAACGTTGGGTTAGTGATTCCTTTCCAGGTGAAATTCATAAGGTGGTGGATTCTAATCTGGTACAGCCAGGGGATGAGCAAATTGATGCAAAGATGCAGTGCTTGTTGTCTATCATGGAATTAGCTTTGAGCTGCACTTTCGTGACACCCGATGCAAGAATTAGTATCGAAAATGCTCTTTCAACACTGAAAAAGATTAGGCTCCGGCTTGTCAGTAGTTGGCACTAGCTGGTGGAATTGGACCGGCTACTCTTGTATGCTATTTGATTAACTATGCCAGTAACATTAAACTTTCAATTTTGC

>M6-PGSC0003DMG400027683 M6 LRR-RLK

GTCGTCGCCTGTCCCCGTTTTAACACTCTTTTTCTTTTTTGCAATTTCTTTGCAAGTTTTCAGTTTTAGTTCTCACTGTTAGCCTTCTTTCTTTGACCAAAGAAAAACTGTATCATCAAAAAAATACTATCTTTTTGCCCCTTTTCTTCTTTGTAATTGACCATCAAGATCCAGCATTTGATATCCTCAACTGGGTTGTAAGTATCTTGCATCTGTAATTTCTTGGTCTTCTTTTGATATTTTTAATGCTCAATTTGTTTATTGGTAGAATTTCTTATTAAAAATTATTTCTTTATTATGTTGTTGCTTGCCTTTGACCAAGGACTTGGTAATTTGTTGAATTTTGAAAGTTGTGTAGAAAGGGTTGGACTTAATTTATGGAAATTGAAGTTTTTTTTTGGTATTTCTTCCTAGTTCTTATAAAAGATTTTTTTTTTTTTATTGTTTTGGTAGCTGCTTTGAATAAGGACTTGGTAAATTTGTTGAATTTTGAAAGTTGTATAGAAAAGCTCTGGACTTGTATTTCTGGAAATTGAAGTTGAATTTGGCCTTTTGTTGTTTGATCTTAAGAACCCTTTTTGGGTTTTGTGGATTATGTGAGATGGGGTTTGTTTAATATCTGGTAGGGGTCTTTTTTCTTTTTTGAACTTTCTTCTTGAAGGTGTGAACTTTTGAGCTTTTGGATGTGGATGTGGAAGATCAAGTAAATTCTGTATTTCGGAAAGAATCTCTCTACCATCTCAAGGTAGTAGGGGTAAGGTTTGCGCAGTACCCTCCCTAGACCCGACCTGTGGGATTACACTAGGTTTGTTGTTGTCATTGTTGTTGTTGACCTTAGTAATTCTGACTAACCATTTCTTTAATACATTGGTTGCCTTCTTGAATACTTTGAAAAAGGACAGTGGTTTATTATTTGTTGAAATTTGGAAGTCGTGGTCAAAAAGCTCTCAACTTTGAAACTTTCTGGAATTGGAATTCACTCTGTTGTTGTTTGATTTTCAGAACCTTTCCTGGGTTCTACTAGCCTTGCTTGTTGGGATTCGTTGTATAGCAAGTAAGGAGGGTTTCTTCAACTTCCATCTTGGAGGTGTGAAGTTTGTGCTTGTGGCTTTGTAAGGTCAGATAAAATCATCTTTCAATAAAAAAATTTCGTGTGTTTTGTTTATATCAAGTAGGGGTGTTCTTTTTTCTTCTTTTCTTTCAACTTCCGTGTGAAAGGCGAAAACTTTTATGGTTGTCGCTGTAGATGATCAGGTGAAATCATCTTTTGAATGAAGTTTTCCTTTTGTTGATGATAATTTTATGTAAGTGCATCTCTTAGATTGCTGTAATTGTGCTTTTTCTCTTCCCAGGTCTCTTTTGCCTCCTTGTTTGCTGTTTTTCCCATTAAAATCTGCAAGGAACAGTTTTCTGCTTGTATCTGACTTGTTCGATAATCTTTATTTTGGTTGTCCTGTAAGTGTGGGACTTGAGTACCTTTACTTCTGGAGAATCACTAGTTTGCTTAAGAAGAAAAGTTGTCTAGTGTTGATATGAAATGAGTCCTAATTGAGCGGATGAAACGTAAATTCGTATAGTTTTTTCCGACTAGTTTGGAAGTGAGGCAGCGTTGATTGATTGATAAAACTGATTCATCGTCTTATTTTTTTCCTTGCAGCCTTACATTTTGCTTTTTAACTCGGGAGAAATTGGATTTGTTCTTGCAAAGGATTTTTCGGGGTTGCAAACTAGTCTATACAGATCTTAAATGTTTCAGTCAACTGCTTAATTAGGACCTTTGATTGTTGCGGTTATGACTGGGCCTCTCACCAAGATGGCTTTGGATAACAGAGCTCTTACTACCTTTGCTGCTATTTTGATCTACTTGGCCTTGAGCTGTGCTGTTTGCTCTGCTGTTCAAAGCGACATTGACTGTCTGAAATCTATAAAAGATTCATTTGAAGATCCATTGAATTTCTTGAACACAACATGGAGGTTCGATAATCAGACCGAAGGTTTCATCTGCAAGTTTGCTGGGATTCAGTGCTGGCATCCTGATGAAACCAGGGTTCTGAGTATCAGTCTTCCTGACATGGGTCTCAAGGGCAAGTTTCCACATGGTCTTGAGAATTGCACCTCTATTACATCATTAGATCTTTCGAGCAACAATCTCCATGGAAGTATTCCAAATAATATCTCCAAAATTATAGGTTTTGTGGTAACGCTGGATCTCTCATCCAACCATCTATCAGGGGATATCCCAGTTAATCTTGCAAACTGCTCTTTTTTGAATAGCCTTAGACTTGACAACAACCAGTTTACGGGTCAAATTCCTCCAGAAATTGCCCTGCTTGGTCGGCTCAAGAATTTTAATGTAGCAAACAATCAATTGTTCGGACCAGTCCCACGTTTCATAAATTCAACTTTTCCTGTAGAGAGCTATGCCAATAATGCCGGACTGTGTGGACCACCATTACCTCTTTGTGAGGGTTATGCTAAGAAACCCCGTACTGGAATCATTGTTGGTGCAGCTGTTGGTGGGGTGACTTTAGGTGCTGTATTATTGACCATTGGTATGTTCTTCTATATGAGAAAGATATCCAGGAAGAGGAAAAAGGAAGATGATCCTGAAGGGAATAAATGGGCGAAGAGTATTAAGGGTTCCAAGGCAATCCAGGTGAGAATTAAAAAACTGTGTCATTACTGAAACTTCTGTTCGCGCTTTGTTGATAGGTCATAAATCAATTCTAGTGGGTTGTGGTTTGTGTTTACGGGTTCTACTGTGGCTTTGCGATCATATTTAGAACCATAAATTCGTGTTTTCATATAAGCATTTAACTAATGAGTTTTTCACCATAAGCTCTTGTGTGAGTTTTCTCTTGCTCAAGCATTCTTAATTGACTTAAGGTGCCTTATTTTCAGCTTTCAATGTTTGAGAAATCTACTTCAAAAATGAGACTAAGTGATCTCATGAAGGCCACCAACAACTTCAGCAAGAACAATATTGTTGGGTCAGGAAGAACCGGGACATTCTACAAAGCAGTACTTGATGATGGCACTTCACTTATGGTTAAGAGGTTGCAGAATACTCAACACTCAGAGAAAGAGTTTACATCCGAGATGGCTACGTTGGGAAATGTAAAGCACCGTAACCTGGTGCCTCTTTTAGGTTTCTGCATGGCTAAAAAAGAAAGGCTGATGGTCTACAAAGACATGCCAAATGGCACCTTGCATGATAGGTTGCATTCGGTGAGTGAAGGGGAAAAAACACTTGAGTGGCCTATAAGAATGAAAATAGCCATTGGAGCAGCCAAAGGATTTGCATGGCTTCACCACAACTGCAATCCTCGTATCATTCACAGAAATATTAGTTCTAAATGCATCTTGCTGGATGTGGAATATGAACCTAAAATATCGGATTTTGGACTTGCCAGGCTCATGAATCCAGTTGATACTCATTTGAGTACCTTTGTAAATGGGGAATTCGGGGACTTTGGTTATGTTGCTCCTGAGTATGCACGAACTCTCATGGCTACACCAAAAGGTGATGTGTACAGTTTCGGTGTTGTACTTCTCGAGTTAGTTACAGGTGAGACACCAACTTCTGTCGCCAAAGCTCCCGAGACCTTTAAGGGGAATTTGGTGGAATGGATCACACAACTCTCTGGTGAATCTAAACTTCAAGACGCGATTGACCATTCATTGTCTAGTAAAGGTTATGACAGTGAGATCTTCCAGGTCCTTAAAGTTGCTTGTCGATGTGTGTTGTCTGCTGCTCCTAAGGAGAGGCCAACAATGTTTGAACTGTACCAGCTTCTGAGAGCCATCGGAGAGCGATATCATTTCACAACTGATGATGACATTATGATGCCTGAGAGTGATAGCGGATTCCAAATGGACGAGCTCATTGTTGCTCAATAAGTAAACGAGGCCCTTCGATAATGGTTGAGTAAGATACAATAATTACTAGATCATTTTCTATCCTGTACAAATACCACGGTTCACTATCTATATAGATCATTCTATTTCTACTGTAAGTCCCTTATTCATAACTACCCAACTATGTAATTTGTGCTTCTTTTTCTGTATTTTTAATGGTGAAGAGATTTCTACAGATTGGTGTTAGTATTTTTCCTCTTTTCAGATTTACTGCTTGAACAGCGTAGTCTCATCTTCCAGTAATTATAAAATGCAAAGTTGGAAAAAATCCACCACTCAAAGCTGAC

>M6-PGSC0003DMG400027703 M6 LRR-RLK

TGAATCCTGATATTCAAGCTAAACATAAAAACAAAGGTGCTATCATAGCTATCGTTGTATCAGGTGTTTTTGCTGCATTTGTTTCAGCAATTTTAACTGTGTTAATCAAAAAACGACATGCCAAATACCAAAGCATCCTATCAAGGAAACGGCTATGTAAGTTCTCTCATTTTGGTTATAAGCTACCTTTACTTCATAAGAATATGTTTCTTTCAATTCTTTCCGCAAAATTGACGAGCATAAAATGTTTAATTTCATAATGCCAGCCGCAAAGCTTTCTATAAAATTGGATGGCCTTAGAAGCTTCACTTTCAGAGAAATGACCTTGGCCACTAACAACTTTGATCATTCAAATCAAGTTGGTGAAGGAGGATATGGGGCTGTCTTTAAGGGAATTTTAGCTGACAAAACAGTTGTTGCTATCAAACGAGCAAAAGAAGGATCAGTACAAGGGCAAAAAGAATTTTTGACTGAGATATCTTTGTTATCAAGGGTACATCACCGGAATCTAGTTTCATTGCTAGGATACTGTGATGAAGAAGGAGAACAGGTATGATAAACCTCAAAACTTCAAAATTCCAAATCTTTTTGAATCTACTCATTAAGATGGAATTCATCATGTTGATCCTTTCCTCGGCCATATGAGTAGTTCAGGCTCACATTGTATGCTATAGCTTCTTATTTTGCTCCGTAGCAATCTTTTGTCATTCGTTCTGACATCTCTTTGGTATATCATCTATGTGAAGCAATTAGTGTGTTCCTGTTTATGTATTTGCAGATGCTGGTTTATGAATTCATGTGCAATGGAACTTTGCGAAACTGGCTTTCTGGTAAAGAAACTGTTCCCTGGTTTATTCTGATCCATCATCTTCCTCTGCTTTATTTTTGGTTAGGGAAGCTATAGAGCCAGCAGAGGAATTTGACATTTATCATTCTTCTAATAATTGTGTTGTGTAAGCTAGCTTGCTTGCGCTTTGACTAATCTATTAAATACGTGCTACCTCCCACTAGCACAAATATCAGATAACTCCGTCCTTCAAGGTTTAGGAAGATGGATCACCTCTGTTGGCGTTCCAACTTTAATTACCAAGTTTATTACTCCAAATCTACCACCACTAGACCACCTTTCAATGTTCCTAAACAAATAGTAGCAGTATCATTGAACTGGTACAGTAGACGGGTAATATATGCTCTTAGACATTGTTTTTTAAGTCTATAGATGGAGATCAGATATCAAGAAGCAAGCTTTGATCTAATAACTGTATGTCCACATGGTTATCACCTAATTTAGATTATCTTGTTGTAGTTACTGCTCCATTACTATCTGCTGTTTTTGTTACTATGTGTTGTTTCTGGTACTTCTGTTACTTCTTTTTCTGGACTGCTTTGGACTATTTTTCCCTTGAGCCAAGGATCTATCAGAAACAGCCTCTCTACCTCTAAGGTAGGGGCAAGGTGTACACTCTACCCTCCCCAGACACCACTTTGTGGGAATACACTAGGTATGTTTTTGTAGTTGTAACACTTGATATCATTTTTTGACAGCTAATTGTAAAGGAACCTTGTCGTTTGCTACAAGATTGGAGATTGTACTAGGTGCAGCAAAGGGCATTCTTTACCTCCATACAGAAGCTCATCCACCTATATTCCATAGGGACATCAAAGCCAGCAATATTCTTCTTGACTCAAAAATGATAGCCAAAGTTGCTGATTTTGGATTATCTAGGCTTGCACCTGTTCAGGATGATGAAGGTGTATTACCTAATCATGTTTCAACTATTGTGAAGGGAACACCTGTAAGTCCTCCACTGCATCATTATGTTCTATTTGACCTCACTTGTATCATTAAAATTTAATGTATTGAGACTCTGCTTGTATATGCAGGGTTATCTTGATCCCGAATACTTCTTGACCCGTAAATTGACAGATAAAAGTGATGTTTATAGCCTTGGAGTTGTGTTTCTAGAAATCTTAACCGGAATGCGTCCCATCTCACATGGAAAAAACATTGTCCGCGAGGTATTTCTCTCATTGTCAACTTCCCGATTTCCATGTCTTATCGCGTGCATATGTTTGATAGATGATTTATCCGCACACACAAAGTGAAGTACTCCTGACTGATGTAACTGTAGGGATAGAGGCAGATTCACGAATTAAACTTGTAATGCTCTTAGTTGTTACCATTGAACACACTACATGTCGCACTTTTGAAATGATGCTTATAGAATTTACTTTTTGTTGATACATTTTAGTGATTTTACATATGTATGTTATAGAACCCAACAATTTTATGGTGCATCCTCCTCTGGGCTGAGAAGGGCTAAGGGGAAATGTGAACATAATTTTTACGCAATCTTTCCTTGCATTTCTGCTATGAAGCTTCACCCTTCGAATCTAGCTCCACCATGTTTAATAGTCCCTTGTACCATCAATGGCAGGTAAATTTGGCTCATGGCTCTGAGAAGATGTTCTCAGTTATGGACAGTACAATGGGATCTTATCCTTCTGAATGTGTGGAAAAGTTTGTAGCTTTGGCCCTCAAGTGTTGCGAAGACAAACCAGAAGACAGACCTTCAATGTTGGATGTGGTCAGGGAGCTGGAAACTATTCAGTCTATTATCAATATGATGCCAGATATTGACGCTGATTCAGTAGATTCTAAAGCAAAATTTAATGAACCAAAGACATCGTCTTCATTTTCTGATTGCACTAGCAAGGACGCGTTTTTATCGTCCTCCAATGTATCGGGAGCTTACAGTATCAGTGGTGTCAGTCTTACTATGCCTCGCTAACAGAATAGACAATAGTTGCTCATCTTTTGTACTAGAGAGAAAATAGCATGTAGAATAGAGAGTTAGCTTCTGTTTTATATATGTATATTTCAAGGATTTGTTAACAAAAACCTCTTTATCATAGCCTTCTGAGTTCTGACAGTATGTGTATGTAATCAGAATCCAAAACTTTTCTACAAAATGGACTGAGATAGGTACTGCTATTTGTTTGCAAGGAGTTGCACCAACCAATTCCTCAAAACAACTGACAACAGCGGAAGGGGGTTCATCGCCAGAAAATCACACTGTATATATAAAAGGTCAATCTTTTTATCTATATAAATGATGAGAACCCTCTTGTAATCCAACATCATTACCAGAAATAGAAATATTATCTAAGCTATGGCAACAATGAGAGTCATACAACAAAACAGGCAAAACTCGTTAGTTTTGAAGTAATGTTATCTTTAGATCACATAACATGTCATTGTAAACAACACAAAGGAACAGAAAACAAATCCTCGAATCAATATGAGCAAACTCCTATTTTGTTCTATGGAACCAAAGTTAGACATGGTTGACACACTCCAAGCTCCTACAAGAAGTCAGAAAGACTATTTCTTGGACTAAAAAGACAGCAAAGTTCATTTTTATTTATAGATCATAGTAAGAGAACAGCGGGTAAATGGCTCTACCGGGAGGGAGGGAGGGAGCCGGTTTTGGATTGGTGTGTATTTGAAACAAATATTGACAATCACTTTGAAGCAGCATCAGCAGTATGCATACTGTCGTCATTCTGGTCCAGCAACCTCTCTTTCTCTCCAACTAGTGTGTCTTCAACACGAACAACCTTTCCTTCATCTGGTTTATCTGCTTTCTGTGATCCTGTGAACTTTGCTAACATGACGAATGTCATTCCCCCAAGAATGTTATTTGCGCATCTCAAAGCAATAACCGAGGTCTTAAATACCAAAGGAGGAAGACCTTTAGCTAGCACAAACTCAACACCGTTAAGTGTTTGATATCTTAAATTACTACTGAGACCCATATGTATTGCCCATGTTGCTGCATTCAAAAGTGTAGGAGGTGGCTTGTTTGGTGTTTCGAAATTCGGATCCATCTTCTTCCTCAACTTGATTAAACCATTAGATAATGCAGTCCCAACAAGCCCTGCTGCAAATCCAACTGCAGCAAACTGAGTTCCTTTGTACACAAATGTACCAAGCCTACTCAATAAACCATAAGAACCAGACTGAAACATGTGGCTCGGTGGACAACTGGCGAACATCAAAGGGAGAGTTTGGATCGACGCAGATGCCGTTGGTGCCAACAGATACATCAACACAAAATTCATAATCGAACCAACAACAAGCGTCGAAAACACAAAATCTAATTCATTAAGCCCAAAATTAGGCCTTGATGCCATATCTCCAAGAACATTAGCACTTACACCAACCAATTCCTCCATAAGAACCTTAAAAGGAAACTGTGGATCTGCAGCCACCCTTGATCTCCATCCATTAACAAAAGCACCAATTGGTCCGATACTATCCCAAGAAGACTTTGAATCATTTGAATCTCCACCTTTACTCCAATCACCACTACCCCCACCCCCACCACCCCCACTACCCCTTCCCATCCCAATACTTCCTCCATCACCTCCACCAACACCAACACAAGAAACTTGATACTTCAGGTGATCCAATTGTTTAAACACAGTATCCCCAGTTTTCAAAGGGTTAAAATAGACCCTATTATCACAAAATCTGGCTGAAACACCAGAAAGATCACGACTTTGATTATGATTCTGATAGTAACCCGTAAGAGGAGAACAACGAAGCTGAGACATCGCCACCATTTTCTACCAACCAACCTAATAAAATGGGAAAATAGAAAGTAATGGCCATATTATTCATCACATCAACAAACAAAGAATCAGATTCACTAAAAGATCAAGAAAGACTCTTTCTTTGACACAAATAATATATATACACACAACAAAAAAGATGCAACAGATTTAATAATAAATCAAGAAAAGGAAAAGGGGTTACCTTTTATGAAGAGATGTGTTCAGGGGCTGCACATCAAGTGATCTGAGCAGAGCAAAAAAAGATGGCAGCAGCGGCTAAAGTGACTAAAATGGGGATCTGCATGAAAGTGGGAATCTTTATAACCTAAATAAAAATACGGGCGGCAGGGCCGAAGTTAAAGAAAAAC

>M6-PGSC0003DMG400027710 M6 LRR-RLK

ATGAAGTACCTGGAAGATTGGGAGAAGGGTGATCCATGTACATCTAACTGGACAGGAATTCATTGTTTTAACAAAGTTGGAGCCAATGGCTACTTGCATATCAAGGAACTGTATGTTATTATATTCTTCTCTCTTATTAGTTGTTATGTTTTCTTCTTCTTTATACCTGGAATTGCTGCACTTGAGTAGAGGGTCTTTCGAAAACAGTCTCTCTACCTCCACAAGGTAGTGGTAAGACCTGCACACACTCTACCCTCTCCAAGCCCCGGGATTTCACCAAGTATGTTCTTGTTGTTGTAATTAGGTAGATTCAAAGCCAATTTACTCTAGCGAAACTTGGGTTGGTGTTTGAAAAACCCGAACCAACCTTGCCTATGTACACCTCTACTTTGGGCCATAAAAATACCCTTGATGTTAGAAAATCTTTGTATCTTTAATCGAGTTCTAGTACGGTGGATTTCACGTGTAAGAATATATCAGGAAAAAGGTCTAAATTTACCCCTTTGCTTTTGGTTGTGGTTAAGAAATTACCTTTCATTATGCTTCAAGTTGGAGTTCCTCTAGCATTAACAGCAAAAGAAAGAGACTTGGTTCTTTCGATATCTGCATAATACTCCTGCAATCCAGTTAGCAGCATAACTAGTATCTCCAAGGGGGGACCTGGTTCTTTTGTAGTTGTTTGTTGATCATCAAAATCAATTTAAGCTCATCTCAACAGTTTGCTAACGACAAGGGCATGAATAACTAAGTTAACTCATAAATTAGAGCACTTAGTTGTAAACTCTAGCAAAAGATGTGTCTTTACTACTTTGAGTTATGACTATCTTGTATTCTTGTGTGAGGGTGATGCATGACGCATTAAGCATATTATATTTCCCCTTTGTGCAGCCGTATGATGGCTATGAATCTTTCTGGAAGTTTAACACCTGAGCTTGGACAACTATCTCATCTTCAGTTTCTGTGAGTTGTTACACATATTACTACATTGTACAAATATTTTGATTTTAAGATATAGTAGAGTTTTACTGATTTTTCTATCTGTTTTACTTATTTTTATCTTTATAATTCACTTTCTGAAAGGAATTTCATGTGGAATGATTTGACTGGGAGTATACCGGAAGAAATTGGAAACATCAAGTCTTTGAAACTTTTGTAAGTTGAACGAAATTTGATCTGTTTTTTTTGGTACCATCTTCTTGATTTTGCAGTCACCCTACATGGTTCCCCGAGTGCTTTGAACCCTGAACTATCTTATCTAGGAGATCTAGATAATCATTTGGCTATGATGGTGATATACTTTATGTTGGATAGAACAGGTGGATTTATGATTTTAAGTTTGATGGTTTCAATCTTTACGTTCTTATCACGGAGTCCATTACATTTTCAAAACTAGGGGTTCAAAGTTCAATATTTTTAAATTTATAGTGATTTGTCATATATGTATCTGTGCTTCCCGTCGAAAATGTTGAGTTTAGTTGAACCTATTAATTAGATCCTTCATCCGCCACTGGATAGAATTGAGGAACATTTTTTTTCTTGAATTTCTTATGAAAATTGTGCTTAATGTAAGATTGTTTCAAGAAGAAGTAAAACCTAATGCTTGGAAAGATCATGACTATGTCGATTGGTTTAATCCCTTCAGAATTATGTTCATGCAGGCTCTTGACTGGAAACAAACTTTCAGGATCTTTATCTGATCAGCTTGGGTATCTTCCGAACCTGAGAATATTCCAGATAGATGAGAACCAGATTTCCGGAAGAATTCCTAAATCCTTCTCCAATTTAAACAGTGTCCAACATATGTAAGTGCAACTCATTTGATTGTTTTAAACAGTGGAACAAGTCTAAGGTTCTACCTTTGTAATCTTATGTGATATATCATGTGTCTACAGCCACTTCAACAACAACACACTGATTGGCCAGATCCCTCCCGAGCTTTCCAATTTGTCCACTCTGCTTCACATGTAAGTTTCTTAGTTTCTCTTCTTCAATAGACATGCATACTCTCAGTAGAAGTTAGAGCAAAGCAAGTTTTTGTAACTTCTAAAATGACTTTTATCTCTACTTATGTCTGTAAACTAACGGAGTAATGTTTTTGTGATGATACTTTCTTTTTGTCAACAGGCTTATGGACAGCAATAATCTTTCAGGATACCTTCCTCCTGAGTTTTCAATATTACCACGTCTTCGAATAATGTATGTCCCTACTCACACACACATATATGATATGCATATATTGAACCTCATATGCAACTTTCATTAGCTGTATTTTGCTCAAGAGGTATAAACTAACTGTTACAAAAAAACAATTGTTATTCAGAGGCAACTTTAAACTGAACTTTCTGAAGTAACTAGTTATAAATTTGAAGGAATGCTGATTCCTTACTACCTATGAGTGAGACTGTCTATGTTTGTCTTAGTTGAAGTGGCGTTATTGGCCAAATGATCGAACAACTGAAGCTGAGAACATTTCGTTACAGACCTATCCTCAGAACCCTCTAATTATCATCTTCAGATCTTTCTGTTGATTATAAAATTCATCGATAATCTTTGCTTGTAATATCCAATGATAAAAAATTGATCTTATATAAGACAAATATAGTAATCCTTGTCATACAATAACTGCTATGCCATTTGCAGTTCCACTGAGAGGTGCATGGGTTGAAATTTATGATGAAGTTCTGAATTTTCTTATGCAGACAACTTGATAACAATAATTTCAGCCGTTCCGAGATTCCAGCTTCGTATGGGAACATGTCATCATTAGTGAAACTGTAAGTGATCCTTGAAAATGTTCCTCTCTTTTGTTCTTCTCCTATTCTCTATGACTCAGCATTATAAGTTATACATTTGAAGGACAACAATGGTAAATAATATGTACAAACATAAGCATATATTCATTGTGTACAATGTATATGAAAGAGGGAATGTTCAGGACATCTTATTATGACGATTAGGGTAGATAGAATAAGATGCAATAGTTGAATACTACTGTCACCTAATCAAATGAAGATTCAATGCTTCTGATAATATTGCAGAAGTCTCAGAAATTGCACGTTGCGAGGATCCATTCCTGACCTTAGCAGAGTACCGAGTCTACACTATCTGTAAGTTCAAGTTTTGAAATTGTTTCTTCTGTTCCCGTTGATTTGTGACCTTTATACTTCATGCAACATGATTGTAAGATCTTCTAAAATGTTAACACTTCTTGCTTACTGTTTGTGCAGTGATCTCAGCTGGAATCAACTGTCTGGATCTCTACCAAATATGCTTTCCAACAATATGACTGCTATGTATGCTTTCTACAGATTTCTTCATATATCTAGCATCTTAGTTCAGCCATATTTTGTGTTAGTTCATTATCTCATGTTGTGAAGAGTTCTACATCAGCCCTTTAAGAAACATTACTAAAGAGCTAGCCTTTGGGGTTGAGTTAGGCCCTACATCCATAACACAAGTTAAACATAATTTTTATCATGATTGCACATTTCTCTGTGTTTTTCCTGTCATTTTCCCTAATCTAATGTGTTGTTGTACATTCTCATGCAGTATTCTGTCACACAATCGTCTTGATGGATCTATTCCGAAGAGTTTTTCATCTCTTCCTTTGTTGCAGAAACTGTAAGGATCTTCACTGTGTCTTGTAATACATTCTCTATAGTACTGGTTCTTTAACATTTCAAAGCTCAATTTCGTGTATAACAAATTGATTTGACGGACTTGTCACAGGACAATGGATAACAACCTTTTGAATGGTTCCTTTTCTACCGACATTTGGCAAAACAAGTTTACTTCAACGTCGTCACTGTTGATGTAAGACCTGAAGCTAAATTGTAACATCAAATATCTGATATGAAATGTACTATCATTTACACAGAAGTCTGGGAGTCTTTTTAACCTTTCTATCTTCTTTGCAAAACTAGTGATCTTCAGAATAATTCACTTTCGGACATTTCCGGTACTCTTGAACCACCTCTGAATGTCACTCTAAGGTAAGGTTTTTCTTTCCATATTCAACGACAAAACCTAAGAGAAGCCATTGAATTACTAATTTTATGATCTGGCTATCACTTCATGTCCACATTTTCAATAATAAAAGACATTAT

>M6-PGSC0003DMG400027967 M6 LRR-RLK

CAAAATTTTGTACCTTTATTCTAACCAACTTTCTGGTCCCATTCCTAGTGAATTGGGGAATTTGAAAAACCTCACTGATATGGAACTATCCCATAATAAGCTTAGTGGTTCAATTCCAATTACTTTAGGTGACTTAACTGAGCTCAAAATTTTGTACCTTCATTCTAACCAACTTTCTAGTCTCATTCCTAGTCAGTTGGGGAATTTGAAAAACCTCACTGATCTGGAACTATCCCGTAATAATCTTACTGGTTCAATTCCAATTACTTTAGGTGACTTAACTGAGCTCAAAGTTTTGTACCTTTATTCTAACCAACTTTCTGGTCTCATTCCTAGTGAGTTGGGGAATCTGAAAAACCTCACAGAGCTGGATATATACCATAATCAACTTAGTGGTTCAATTCCAATAACTCTAGGTGACTTAACTGAGCTCAAAATTTTGTATCTTTATTCTAACCAACTTTCTGGTCCCATTCCTAGAGAGTTGGGGAATTTGAAAAACCTCAATGATTTGGATCTACAAGAAAATCAACTTACTGGTCCAATTCCTGCTTCCTTTGGCAATTTGAGAAAGTTACAATTTTTGTATCTCCGTGCCAACAAACTTTCTGGTTCTATTCCTAAAGAACTTGCATATATGGATAACTTGGTAGAGCTGATACTAAGTGAAAATCAGTTTTCAGGCCATTTGCCTGAGCATCTTTGCCAAGGTGGGAAACTTGAGAACTTCACGGTGGCCAGTAACAAGTTAACAGGACCAATACCAAGAAACTTGAGCAATTGCTCCAGTTTTAAATGGGTCCGCTTTAACAACAACAGTTTCACTGGGAATTTATCTGAAGCTTTTGGTATCTATCCGAAGTTGCTGTTCATTAATTTGAGCGATAATGATTTTCATGGTGAACTCAGCAGCAAGTGGGGGAAATGCAAGAATTTGACTGATTTTCGGATTGCCAGAAATAACATTAGTGGTAGCATACCACCAGAGATTGGAAATGTCAAAGGCCTTTTAGGACTTGATCTTTCATCGAATCATTTGATTGGGCAGATTCCAAAGGAATTTGGAAAATTAACCTCTCTAGTTAATCTTTTTCTGCAAAACAACCGAATTTCTGGCAATATACCTGAGGATTTTGGGTCACTAACAAAGTTAGAGTCCCTCGATCTATCAGACAACAGATTGAATGGGTCAATCCCGATGTGTATAGTAGATTTCGTACACTTGTTTCAGTTGAACCTGAGCAACAACAAGTTTGGGCAGAAAATTCCAAAGGAGATAGGGAGGATAACTCAGCTTAATGTACTTGATTTGAGCTATAATCTTCTGGTTGGAGATATACCCCCTCAGTTAGCCAATTTGAAGGTCTTGGTAAACTTAAATCTTTCCCACAATGGCCTATCTGGGCACATTCCTACAGAACTTGATACTTTGACTGGTTTGCAGGATGTCATATTGTCATACAATGAGTTGGAAGGTCCAATCCCTAATAATAAGGCTTTTATCAATGCCTCATTGGAGGGTAATAAAGGTCTTTGCGGCAATGTAGCAGGACTCCATCCCTGCGAAATGCCGTCTTCTGTGGTGAAGAGGCACTCAATGGCGAAGGGACGTAAACTCATCCTCATCACTGTACTTCCTGTTATGGGAGCATTAGTACTGCTCTGTGTTTTCATTGGTGTTCTCATTATGTGTAATAAAAGAAGGAGAGTTAGAGATGTTGAAAGAAGGGATGGTGTTGGTTGGCTTTCGATATCCATGTTAGATGGGAAGG

>M6-PGSC0003DMG400027971 M6 LRR-RLK

ACAGGGCCAATACCAAGCAGCTTAAGCAAGTGCTCGAGTTTCAAAAGGCTGCGCTTCAATAATAACAGTTTTACTGGGAATTTATCTGAAGCTTTTGGCATCCATCCACACCTTCATTTCATTGATTTGAGCGACAATGATTTTTATGGTGAACTCAGCAGCAACTGGGGGAAATGCGAAAATTTGATTGATCTGCGTGTAGCCAGAAATAACATTAGTGGTAGCATACCACCAGAGATTGGAAATGTCAAAGGGCTTTTAGGACTTGATCTTTCATCAAATTATTTGATTGGGCAAATTCCAAAGGAATTTGGAAAATTAACCTCTCTGGTTAATCTTCTTTTGCGACACAACCAAATTTCTGGCAATATTCCTATGGAACTTGGGTTATTGACAAAGTTAGATTACCTTGATCTATCAGATAATAGATTGAATGGTTCGATCCCACCGTTTATAGGAGATTACCAGCACCTGTTTCACTTGAACCTGAGCAACAACAAATTTGGCCAAAAAATTCCAATAGAGATTGGGAAGATAACTCAACTTAATGTACTGGATTTGAGCCGTAATCATCTTGTTGGAGAAATCCCCCCTCAGTTAGCCAATTTGAAGGTGTTGGTAAACTTAAATCTTTCACACAATGGCCTATCTGGGCGCATTCCTCAAGAATTTGAAAGTTCAACTGGTTTGCAGGATGTCGTATTGTCATACAATCTGTTGGAAGGTCCAATCCCTAATAATAAAGCTTTTATCAATGCCTCATTGGAGGGTAATAAAGGTCTTTGCGGCGATGTAGTAGGAATCCATCCCTGCGGAATGCCGTCTTCTGTGGTGAAGAAGCACTCAATGGCGAAGGGACATAAACTCATCCTCATCACTGTACTTCCTGTTATGGGAGCACTTGTTCTACTCTGTGTTTTCATTGGTGTTCTGTTTATGTGTGATAGAAGAAGTAGAGTTGGAGATGTTGAAAGACGGGATAGTGATGGTTGGCTTTCGATATCCATGTTAGATGGGAAGACATTGTACAGGGACATCTTAAACGCCACAGAAGAGTTTGATGCAAAATTTTGCATCGGGCAAGGAGGACACGGAAGCGTTTACAAGGTAAACCTTCCATCATTAGGGAATATAACTGTGAAGAGACTTCATTCTTCATTTGAGAATACAGCTCCCAAAAGTTTCTTAAATGAAGTAAGGGCATTGACTGGGATTAAGCACAGGAACATTGTGAACCTCTATGGCTATTGTTCGAATGCACAACACTCGCTCTTGGTTTATGAGTATGTGGAGAGGGGGAGTTTGTCTAGCATTTTGAGCAACAAGGTTGAGTCCAAGAAATTGGATTGGTTTAAAAGAGTGAATATCATCAAGGGTGTTGCTTTTGCTTTATCTTACATGCACCAGGACTGTTCACCCCCGATTGTCCATCGAGACATAAGTAGCAGTAATGTTTTGCTTGATTCTGAGTATGAAGCTTGTGTTTCAGACTTTGGCATAGCGAAGCTACTCAAGCCAGATTCATCGAATTGCACTGCACTTGCAGGCACATATGGCTATGTTGCACCTGGTAAGTTTTAAATATTTCATGAAAGCCAAACACTATTGTTTTTATCTTATTTACAAGTTTAAATTGATAATATTGTTCTGGAAACTGTATTGTTCAGAGCTTGCATATACTATGAAGGTTACACAAATGTGTGATGTCTTTAGCTTTGGAGTACTATCATTGGAGATAATAAAAGGAAAACATCTTGGGGAATACATTACCGTGCTAGCAGATTCATCGACTATAGATCCTGAGCAGCTTAGCGATTTTCTGGATGAACGCCTTCCATATCCTGAAGATAGAGTAAAAGAGGCTTTGGTTTTTATCATCAAGCTAGCATGCTCTTGTTTGCTTGAAACTCCAAAATCAAGGCCAACAATGCACTTCATCTCTCATAAGTTATCATCAATGGATGCACATCCACCTATTCATCAGAGAAACCCCCATGTAAGGCGAGCAATATAA

>M6-PGSC0003DMG400028123 M6 LRR-RLK

CAGTTACAAGAGAGCATAATGTCTAATGCAATGTCAAGAAAGAATGTGGTTGATCCAGCAATTAGAAATTCGTGCTCGGATGAATCATTGAAGACGATGATAGAAATATGTTGCAGGTGTTTGGAACAAGATACAGAAGATATGCCCTCAGTTGAAGATGTAATATGGAATTTGCAGTTTGCTGCTCAAGTTGAGGATTCATGGAGAAAAGACTCTTCAAGTAGTGATGCCTCTCCAATTTCACATCTCTACAGCTTCAAACCCAACCTGTCGCGCAATAGCAATAGCTCGTTTGTGTGACAAGCTATTTCTAACGTGGATAACTAATCTCGATACTATTATTCCATCCTGTGACAAGAATAACTAAACTAACAATTTATGTAATAATTTTCTTTGCATAGCTTATACAAAAAAGTAAGTTTTCATACAATCTTTTTCCTAACAAGCAGATGCCATAGCAATTTCCTTGTGTATATACCAACCCAACCTTAATAGTACCTAGATATTACAAGAGCAACATATAGTGCAGAAACCTATATTAGACGTGGACGATGAAAAACAAAAACAAAAGAATGCAAAGAGGGACTAGTTAATGGAAAAGGGAAGCATTGCAAAATGATTTTTCGGACAGTAGAAACAGACTATCTATTCGTTTGCAGGAGGAGTTTCCATTTTCTGAACTTGAGGGGCTGTCACTGCCACTAAGCTAAGATCAGATGAGTTCGAATCTCCAGATCTTTCACGGATCTCCCGGTACTCCTGGCATATGGCACATAAGTGGCAGAAAAAGTGTGTGGCGAAGTCTCCACATGGAGCTTCCTGCATATATATCAGGGCACACAAAATTGTTAAATGCAGAAATACTGATGTAAGCAGTTTGCATCTTTCTATTCAGGCATGGAACCATAGTATTTCGTAGAACAGAAAAGGGGGCAGGAGCTAATGGGAGGAGAGGAAGATGAAGAATTAAGAATTCACGGTAACTGGTTATAAATGCTGCAAACTCATCCAATAACAAACTCAAAACTGAAATTTCTAACTCAGACATCATCTCAATGTAAACCAAACAGAATAGGAAACTGAAGTCAGTTCTCTTTCTAGATGGAGTCCTAATTTTCCCGATCTGAAGTTGATAGTTTATCGAACTGAAGTCCACCACTACTAATTCAAATGAAGTATAATCATCTCTCCCGTGGAGTCGATTACACACTGTAACTTTTTAATCTTTTGTAACTTCTACAGATAACATCCAATTCAAAAGAATGTATGTGGAATTTTAAGACATTTCTGATTAATTTTTCTTTCTAATGAAAAGGTTAACTTCAAACTATCATCATAGAAGTCTTAATATAGTCATTAATATTAATATATATCACACCATAATGGGGAGTTTATGCATAGACGTAAAGAAACATCCTCCCATGCATAGCCCATGTTATAGGAGATACAACTATGAATATGGAAGGTTAGATTGAAGGTAAATTTTCAAGCAATCTAACTTCACTCATTTTTTCAAATGTCTTTCCTGTTGAATCAATAGAGCTACTGATAATATGCTAAACCACATTCTCATCCATGTGCACTATAAACAGCAAAACAACAGAGATGGCTACCTGAAGATTGTACTTTGATCTCAATGTCTTTCGATAGCCACAGGCATAACATGCAACAAAGCACCCAGGCAAACCCAGAAGAATGCCTTCAGTCAAGAAGCAGCAAATTGTGTTGAAAAAAGCCCACAAAATAAAGTGAGTTGTACATGATCCCATCAAGGTTCCTGACCCCAAGAAGTCCGCATTTTTTCCAAACAGAAAGCAAGGGCAAAGAAGTCCAATAAAACCTGTGCTCAAAATTTTGCAATTAGTGCTAACTCGAGGTTGTCAACCAACATCTAGTGTCATTCCTCTGGGATGACAGAAGAGCCATTGCTTAGGAACAGGTGTTGGAATTTTAAACATAAACACACATAATTAACCCTCAAAAGAGAAGTTCCACAATCACACTTTCATCTTATTATATGTTCAGAAAGGAATGATTGATCTAGTCCCATCGGCACTGCTGAAAACCAGTTTCAGGAAATTCCGTCAGTTGTATTATTTATAACTTCATGAATATGACAAACAGTGAGACCACAAGGGGGAGCAAATAGCAAATAATCTGGAAACAGAATGGAAGCAGTGTTGACTGTTGAGAAAAGAAATGCATGAATTTTATCCAGAACTCTATGGAATTACACTACACTGGGTATGTTGTTGTTCATAAGAATATATACATAAATCCTGTGGGATTACACTACACTGGGTATTATTTTGTTGTTGTTCATAAGAATATATACATATATCCTTATGGGTATACTTTTGTCCCAGCAAAAGAGCACCAAGAAGGTTCTGTATATGTACAGTATACAAAAACTCAGATACCCCTTTATAAATGCAGGGCACTGAGGAACTCTAGCAAGGCACCTAAATCATGCACACTTAAGCATGTTACACCATAAATCAAAAAAGTACAGGCACTTCTAAGTTCCTAAGTCTGGATAGGACTAGTTTTGCAAATTTGGTAAGAAACTGTATTTTCAAAGGCATACCAACTTCAAGCTAAATTTTTATCTAAGAATCATATAAGCAGAGAAAAATACTTCATTCCATTATAGAGCCCGAGGAAAACTCAATTAAATGTATGGCTTATAATAGCTTCCTTAGTGGTTCACTCAGAGATTTATCATGTTTTCCATCTAATAAGATAAGAGGATAATTAATCATGAATCTTATTTTACTACCCAAAATTGCACCTTTCATTCACTTTTTTCCCTTCTCATAAAGCAAGATCACAGGGACGAGAGTAGCGTACAGCTCTGTGGATCATCAAAACAAGCACAGATTCCAGAACTCCACTGAGCATGGTCATCTTTTTTGGTCAGTTGAATTTCTTCTGTAGTAGAAACTTGTGGTACTTCTGCATCGGACTGATCCAGAGGAATATAAGGTGGAGGTGAATAATTCTTGTGCCCTACCATTATTAGTCAGGTGCAAACAGCAACCTGCAGCACAAATTCAAGAGAAAGCAAAAAATATCAATTAAGAACAAGCGAGTTACACATAAACATCAAATCAGTTCTCTGATATCATGAAGGACGGTTACTTTTTGACTAAGTTGCTCGGACTCTCCAAAAATGTTGCCGCACCAGTGTTGGATCCTTCAAAAATACATTATTTTTGGAAGATCCGACACACACTTGTCAACTTTTTTGTAGAGTCCGAGCAACATAGCTT

>M6-PGSC0003DMG400028147 M6 LRR-RLK

TCTCACTCCAATTATAACTGTATATTACTGTTTCTCCATTATTCTTGGCTTTGTAGTGATCTGAATCAACCCATGTTTCCCACCTCTCCTTGTCTAAATTCTAACATGGGTTTGTACAAAAAACAGCCATTAGTTGGCACAAAGAAATCAACTTTCTTGATTATGTGATTCTTGTCAATACATGCTGTCTAGTGGTTCTTGGTGAAATCATCATCAATGGAGGCTGTTCAGCTAATGGGGTTTGCTCTAACACTCTTTATTTCACTTTTTTCAGTTGTTTACAGTGTTACAGACCCTAATGACTTAGCCATTGTCAATGAGTTCAAGAAAGGGTTAGAAAATTCTGAGCTTTTAGAATGGCCAGTTAATGGTGATGACCCTTGTGGTCCTCCAGCTTGGCCTCATATAATTTGTACTGGAAACAAAATTCAACAGATTCAAGTTATGGGGTTGGGGTTAAAAGGCCCTTTGCCTCCGAACTTTAATAAGTTGTCTAAATTGACTAATTTGGGGTTGCAAAGGAACAAATTTAGTGGAAAGTTACCATCTTTTAGTGGTTTGTCTGAGTTGCGCTATGCTTTCTTGGATTTCAACATGTTTGATAGTATTCCATTAGATTTCTTTAATGGGCTTGTGAGTTTAGAGGTGTTGGCATTGGATGATAATCCTTTGAATGCAACTACTGGTTGGAGTTTGCCTAATGAGTTGCAAAGTTCAGCTCAATTGACTAACTTGACTTTGATGAACTGCAATTTGGTTGGTTCTTTGCCTGAGTTTCTTGGAAATATGTCTTCTTTAGATGTATTGTTGTTGTCCAAGAATCGACTTTCGGGGACTATACCAGGTACTTTTGAGGATTCTGAGCTCAAAATGCTTTGGTTGAATGATCAGACTGGTGATGGCATGAGTGGTTCAATTGATGTGGTTTCAACAATGAGATCACTTACTAGTCTTTGGCTTCATGGGAACCATTTTTCAGGTAAAATTCCACAGGAGATTGGTAATTTGACATATCTGCAGGATCTTAATGTTAATAGCAATGATCTTGTTGGTTTAATTCCTGAATCTTTAGCAAATATGCCATTAGGCCATCTTGATTTGAATAATAATCATTTTATGGGGCCAATACCAAATTTCAAGGCTATAAATATTAGTTATCAATCCAATTCTTTTTGTCAAGCCAAAATTTGTGCACCTGAGGTTCTTGCTCTTTTAGAATTCCTTGATGAGTTGAATTATCCATCTAAACTTGTTGAATCATGGTCAGGAGATAATCCTTGTGACGGGCCGTGGTGGGGATTAAGTTGTGACATTAACCAAAAAGTTATTGTGATAAACTTGCCTAAGTCCAATCTTTCTGGTACCTTGAGTCCTTCAATTGCAAAATTAGATTCTCTTACTCATATTTATCTTGGATCTAACAATATTTCTGGTTCTATTCCATCTAGTTGGACTAGTTTGAAACATTTGGTTCTGCTTGATTTGAGTAATAACCACATTTCCCTTCCTCTGCCTGAATTTACTCCCCCCTTGAAACTCGTCCTGAGTGGAAATTCATTACTGAACTCTAGTCCTCTTATAGCAAGCCCTTTGCAAAAGAATAGTACATCCACTGCTGTTTCACCCTCTTTGCCGACCAATAAGTCAAGCTCTTCCAAGTCTAACTTAGTCATATTTGTGGTTCCTATTGCAAGTTTCGCACTTTTAGTTTCTTTTGCTATGCTACTATATGTTTACGTTCGAAAGAGGAGTATGGATAGGCACAAAGGTCCAACTTCTCTAGTGGTTCATCCTAGAGATCCTTCTGATTTGGATCGGATGGTCAAGATTGCAATTTCCGATGAAACTAAAGGAAGTCTTTCGATACTGACTGGAAGGGGTTCTTCTAGTATACACAGTGGTAAATACCCTGTGATGGAAGCTGACAATTTGGTCATATCAGTTCAAGTACTTAGGGATGTAACCAAGAACTTTGCTCCAGAAAATGAACTCGGGCGTGGTGGTTTTGGAGTGGTTTATAAAGGAGAATTAGACGATGGAACAAAAATAGCTGTCAAAAGAATGGAGTCTGGAGTAATTAGCAGCAAAGCGTTGGATGAATTTCAATCTGAAATTTCTGTTCTTTCCAAAGTCAGACATAGGAATTTAGTGTCTCTATTGGGATATTCTGTTGAAGGCAATGAACGGATTCTGGTTTACGAACACATGCCACAAGGTGCTCTTAGCACACATCTTTTCAACTGGAAAAACTTGAACTTAGAGCCTCTTTCATGGAAGAGGAGGCTGAATATTGCATTAGATGTTGCTAGAGGAATGGAGTATCTACATACGTTAGCTCATCAGTGCTTCATACACAGAGATCTCAAGCCCTCAAATATCTTGCTGACTGATGATTTCCGAGCAAAAGTATCAGACTTTGGACTCGTGAAACCCGCCCCTAATGGAGAGAAGGGCTCTGTGGTCACTAAGCTAGCTGGAACTTTCGGATATTTAGCACCTGAATATGCGGGTAAACATTTCATTTCCTGTTGTTTTATTCCTAACATGACATTTTATTTTGTAAAACAGGAAACAGATGTTCTATTTGCCATTTGATGTTCTCTTTGGTATGATTACAGTTACATAAAACTCAAATAGTTTAGTTACTGTTGTCTGTTTTGGGGTGGATAAGTTAAGGACTATTTTGCATAGTCTAATTAAATCTTATGTCACCTAATAAATCTGTTCACTGATAACTAGTTAATGACTATTCTTGTAGACACCTAGGATTTTCTTTAGATGATTCTGTCTTTTCCGTCTGCTGGTTTCCCAACTCATTTTGCCTGCTGTTTTAATACAGTTACTGGTAAAATCACCACTAAAGCCGATGTCTTTAGTTTCGGTGTTGTCCTTATGGAGTTGTTAACTGGATGGATGGCACTTGATGACGACAGACCAAATGAGAGCCAATACTTAGTTGCATGGTTCTGGAACATCAAATCCTCTAAAGAGAAACTTATTGCAGCAATCGATCCAGCTCTGGATGTGAAACAAGAGAGTACATTTGAGAGCATCTACACCGTTGCAGAACTTGCTGGTCACTGCACAGCAAGAGAGCCTGGCCAACGCCCCGACATGTCCCATGCTGTGAACGTGCTCTCCCCGCTCGTTGAGAAATGGAAGCCTCTTGAGGAAGATTCAGACGATGATTGTGGTATCGATTACAGTCTTCCACTCAATCAAATGGTTAAGGGTTGGCAAGAATCAGAAGGAAAAGACTTGAGTTGTGTCGACCTTGAAGACACTAAGGGTAGTATCCCTTCAAGACCTACTGGATTCGCCGAGTCCTTTACATCAGTTGATGGTAGATGAAAAAAAAGGTACTGCACTACTTCTAGTGTAGTTGATCTGATATTCTTATTCAGTAAATGTGGATAGCATTTTTGTTTGTTTTTGTGTGATATTTTGATCCTGTTTTTTATCTATGCTTTCTTTCACTTTTATTTGTAGAAAATTTTACTCCAATGTATAGTCTTTTGAAGTTATGTGGGATGTAGATATGGTATAGTGATGATTGATGAATGAAAACTGTTTCCA

>M6-PGSC0003DMG400028429 M6 LRR-RLK

TGAATTCTAAAAAGGACTTTTTTCTGAAAGCTCTGCACCAAAATCTTTCTTTTGGGTAATTTAGCTTTTTGGATCCAAATGGTAACTTTGCTACTCATTTTCTCATTTTTTCTTCATTGGGTGGCCACCACTCCAGCTTCCATGGGTTTTCTTTCACCTAAAGGTGTCAACTATGAAGGTAAAGATTAAAACTTTACTATAATCCAATCATTTCTTGTTTTTTTTTTCCAGTGAAAAATCCAAACTTTATTGCTTAGCACTTGTTTATTTTTGTTTTCATAGTTTAGAGTGGTAAAGATGAAAACTTTACTACAATCCAAAAATTTGTTGTTTTTTTTTCCAGTGAAAAATCCAAACTTTGTTGCTTAGCACTGTTTACAAGCATTTATTTGTTTATGTTTTCATAGTTTAGAGTGGTAACTTTACTTAAAGATTTGTTGAATTTATTGTGGGGTAAGCAGTGGCTGCATTGATGTCTATGAAGAATATAATGAGAGATGAATACCATGTGTTAGATGGATGGGATATAAATTCTGTTGATCCTTGTACTTGGTACATGGTTGGTTGTTCTTCAGAAGGCTTTGTTATTTCTCTGTGAGTTTTCTCCCTTTCTGTTACATCTTGTCCATTTCTTTTCCCTTTCTGTGAAAATAATTTATTATTATTCTTTGGTTATATATTCTTGGCAGTGAAATGGCAAGTATGGGATTATCTGGCACACTTTCGCCTAGTATTGGGAATTTAACTCATCTTCGGACAATGTAAGACTAATTTGACATGAATTTCCCACTTTATTCTTATTTTGATGTGATGCTGCTCACATATATCTTGGAGATACTTGAAGTTAAAATCACATAGTACTTTTAGTTTAATAATTACTAATACTATTAAGAGTGACATTAAGTGATTTAATTGCAAGATATGTTGAAAAATATCTTTTAATATGATCTGTTTAAGTCATTTTTGCATTGGTTTTTGTTGTAGGTTGTTGCAGAATAATCATTTATCTGGTCCAATTCCTACTGAAATTGGACAGCTCACTGAACTGCTGACCCTTGATTTGTCTGGTAATCAGTTTGATGGAGGAATTCCTAGGGCGTTGGGTCGCCTGATTTATTTAAATTACTTGTGAGTGTAGGCTAAACTCTTGATTTTGTTCTATCTTTATTACTCGTATGTTACAGTTCCCCTTGTTGGCTGTGTATGTAATTCTTTGGTGTGTCAAATTCTCAGGAGGCTTAGCAGGAACAGATTGTCTGGACAAATTCCTAAACCTGTGGCATACCTTTCAGGTCTTTCATTCTTGTATGTCTTCCATCCATGACATTTATATGTCCAATACAAGGGCTGTATGAGAGTTAAATAAACTGTGGATGGTGTTGTGTAGGGACTTATCATTCAATAACCTTAGTGGTCCAACTCCAAAAATTTTGGCCAAAGACTACAGGTAAGATCCAAGGAAATATTATTATATCTGTCTCAATGTTTGGTCCTCAATAGTCGGTGAACTGGAAGAATTCTTCGGGTCTCTAGTTTGATGAATCAAGGAATTCAGGGTTTACACTTTTACTATGAGGAGGATGGTATATGCCACAGTACTGTGCCAGAATCTTATTCATTAATGGAGTTTAAGATAAACACCTTGATCTTAAATTCAACTGCAAAATAGAAGCTTGATTCAATATTGTTGCATCTGCGACATGCCATATGGCGCAGTGCCACACATAATGTTTCTCTCGTAATAAAGTTATGCTTTCTATAGTTTTAGTAACTGTTATTTTGCATTCCAATTTGTTGTTGTGGTATGTCCCTTCACTGGAAACCAGCAGTCAGTTCTTGCAGACCTGAAAGTAATATCTCAAATCTCATTATCAAAAAAAAGAAGAAAAGAAGAAGTAATATCTCAAATCTCCTTATCAAAAAAAGAAGAAGTAATATCTCAAATCTTATCATTGAAGTATTGTCTTCTTACCTGGTCTAATGGAACTTCCATGATGTTTGTTTGATATAATTCCTATATTGCTAAATATGCTAATGTCAAGCAGAATATTTGCTTGAGATAATTTCTGATGTTATAGAAGTTTTCTATTCTACTTGCAATGCTTGTTGTTTACCTTCCCCCTGTCATCATACTAGCAACCCTTCGGGTTCTTTTGTATTCTATTTCAGCTTATAATTTGGTTGTTGCCTTAGTATTGCAGGAAACAGATTTCTTTGCTCTGCATTGACCACACAAATTTGTGGGGGTGTGCCAAAACCAGTAAATGGTATTGTTGAAATTTTACCAGAACTTGTCATTATATTGAGGCTGATGTTTTTAATCAAACTCTATTTTTACCATGCAGAAACAACTTCAGACAAGAGGATTAGCAATCATCATCGATGGGTTGTTTCTGTTAGCGTTGGGGTCAGTTGCACATTCATAGTTACAGTCTTGCTTCTGGTTTGTTGGGTGCATTGGTACAGATCTCGTGTCCTCACAGGATATGGTAATTACAAGATTAGAAACACATCTCCGATTTCTTGCACATTAAGACTAGATACATTATGAGTCTCTGATCATTTGATCTGCAGTGCAACACGATTGTGAGTTCGCCACTGGCCACCTGAAGAGATTTTCATTTCGTGAACTGCAAATTGCCACTGGAAATTTCAGCTCTAAGAACATACTAGGACAAGGTGGATTTGGAGTGGTTTACAAAGGATACCTTCCAAATAGGACAGTAGTGGCTGTCAAAAGATTAAGAGATCCAAGTTTCACTGGAGAAGTGCAGTTTCAAACAGAAGTTGAGATGATAGGCTTGGCTGTGCATCGGAACCTTCTACGTCTGTATGGATTCTGTACGACTCCCGAGGAAAGATTACTTATTTACCCTTATATGCCAAATGGGAGCGTTGCCGATTGCTTGAGAGGTTTGCTTTGAAGTATATTTGCTTGGTGCTTTTGCTGAAGCATTCTTTTTTCTTTTCACAATGGATTTCGTAAACTGATCAAGTTTAACATATGCATTTCAATTATTCCTAGTGCTCTCAATGTTAATCTGGCAAAAACTCTTGAAAGGTCATGGTTTATGGAAATCGAGGACGTTGTGCTTTATTTTTACTGTACATCTGAAAACACCAGGGCCCTATGATGAAAACAACTTTTGATTGATTATCTTAGATAGTTAAATTGGATTTACATTGTCAATTTTTCTTGTGAAAGTGGAATGAAATCTATTATCAACATATCAATCAACAATGCATCAAACCCAATCTAGTTGGGGTTGACTATATGAATTTTTTGTATCCATTGCAATTAAGACTCCTGATGTAGTGCATTACTTAAGTAGGCATGCCTCTAGCAATCTATTGCGACCTTACGTGCTATTAAGCTTAATTTTTATGTGACAAAGTTTCTATCCATCTTCTGAAACTCTAATGCCATTTATGGTATCGACTTACATTATCCATGAATAAAATAAATTGTTATTTGGCAGTAAGGTAACTGTCGTGTTGTCATTAATTTTTAGCATGACCAATTCAAGTCAATTTTTAGGTATATGAACACAAAAATCAAGTTGGAATTATTGGAATGTAAACACATTAGCTTGATGTAGTGACTGTGTCATCTTTTGCTCTGTTCTCACATAATCTTTTGTTATCTACCAGACAATGGCCGGGATAAACCATTTTTGGATTGGAGAAAGCGCATGCATATAGCTCTTGGAGCTGCCAGAGGTCTTCTGTATTTGCATGAACAGTGTAATCCTAAAATAATTCACAGAGATGTTAAGGCGGCTAATATTCTGCTTGATGAAAGTTTTGAAGCTGTTGTTGGTGATTTTGGTCTTGCTAAGCTTTTAGACTGTAGAGATTCTCATGTTACAACTGCAGTTCGTGGTACTATAGGTCATATAGCACCAGAGTATTTATCAACTGGTCAATCATCTGAGAAGACAGATGTCTTTGGATATGGGATACTACTTTTGGAACTCATTACAGGACACAAGGCATTAGATGCAGGAATTGGTCAGGGTCAGAAAGGAACTATTCTTGACCGGGTGTGTAGCTTCTAACCCTCCTACTGTTTGAAGCTTTATATAGTTGTGCATATAATTTCTAGACTTTATTTGATTAACCTGTTAATAGTTAATCTTCTTGCAGGATTTTGTTTGAGTCATAATCAATATCAATATTTGACTTGAAATTTTTCACTAAAATTGACGCAATAACTAGTGTCTAACGCAGAAGTTGCTCCACATCTTATGAGAGAAGTACAGTTCCAAAATTCTGAATCATTGCTCGAAAACTTGTTCTTTTTACTGACTGCATAATCTGGTTCATGATCTGTAGGAATTATGGACAACATATTCAAGGTTTTTGAATCACTTACTGATCAGTGCAATAAAATTATTGCTGTGTATAACCTTACTGTTGCATACTTTAAGCCCTTAAATATAAATTATTTGGTTGATTTTTTATTACATATAATGTTACACTTGCATACTTTAAGCCCTTGATTAACTTATTTGGTTGATATTTATTACATTATCATGTAAGTCTGAACAACCATGTATTATTCTAGAATAGTTGTGTGAAAAAAGGTACCCGTTCTATGCTTCTTCAATTATGATTTTTCTTCCTTCTCAGGCTGTTTTATGTGGAGTAAAGATTTTCAGAACATGATAGAGTTCAGAGTTTGTTATGTCTATTATTCGGGCCTATTCATTCTGACAACCTGTCCAATTTACAGGTCCGGAATTTATTTGAGGAAAAGAAAGTGGAAATGCTTGCGGATAGGGATCTAAGAGGATGTTTTAATGCAGAGGAGCTAGAGAAAACAGTTGAAGTGGCTCTACAGTGTACTCAATCTAACCCCAATAATCGGCCTAAGATGTCGGAAGTTTTGAGAATCCTGGAAGGCGTTACAGAACAAATGGGGCATGTGGACGAATCACAAGGTGGAAGCAACACATGTCAAACAAGGGCTTTCAGCTTCTCTAGGAATTTTAGTGATTTTCATGAAGAATCCTCATTTACTTTTGAACCAATTGAGCTTTCAGGACCCAGATAACAAAAGCAGCCTGCACTGGCATCGTCATAAAAAGCATGCAATCTTGATATATCTAGTGTAGCCATACATACGAGTATATTGTTGCTTTAAGTCAATGTCCTAGTACTCTGTAAGTTACTGTAATTTGTAAATTTTGTTGCAAAAACATTCATTATACTAGTTCTTGGTGGTTCACAATCTGTGTTAGAACTAGCTAGACTAACTTCATTTTGGAACTTCATTCTTTGGAAATGACAAACTTC

>M6-PGSC0003DMG400028435 M6 LRR-RLK

TTAGCATGAGGCTAAGCACCAAAGTTTGTTAATTGAGAATTAACACCTTGAGCAACATCATAAAAATTTGCAAATTTGCTTGCCGTTTGTCAAAATAATTTGCAGGCACAATTCCAGCAAAGATTGGTAATTTAGGAAAGTTGCAGCAATTACAATTGGAGAACAATCGCATCGTGGGTTCAATACCGCGCACCATATCGAACTTATCATCACTATGGATGCTTAATTTGAACACAAATAATTTATCAGGTACCTTGGTCTTGTTTGTCTTGGAATTAATTGTTGTTTTTTATCAACTCTTATATATACTGATGGGCACAATCAAATAAAAGTAATTTAGATTTATTCTTTCGTGTGGCAGGTATTATACCCAGAGAGATTGGAAATCTTCACAAACTGGAGATACTTTATTTGCAAATTAATAAGTTAAGTGGTTCCATACCGGAAGGGCTCTTCAATATCTCTACTCTAACAGAGGTGTCAGTTTTCTTCAATAACCTTTCGGGTAGTCTTCCATCTGCTTCAGGGTACTGGCAAACAAATCTACAGCGTTTGAATCTTGGTCAAAACTACATTGGTGGAGTTATACCTACTTCAATCTCCAATTCATCAAATCTAGAGATTGTAGATCTTAGTATAAACAAATTCAGTGGCCAAATTCCTAACTCGTTGGGGGATTTGAGGAATCTTGAATATCTGGACTTGTTCGTTAATAATTTATCGTCTCCGCACCTAAGTATCTTAACTTCATTGGTGAACTGCATATATTTGAAAGAAGTAATATTAGGGGATAATCCTCTGAATGGTGTTCTTCCAGATTCCATTGGCAATCTCTCCACTTCTATCATGGAACTTGAGTTACATCATTCTGAAATTAGGGGCCAGATACCATTAGGAATTGGGAATTTAAGTAACTTAAACACTTTGACCCTATACGGCAATGACTTGACTGGATCATTACCAACAACATTATGTGATTTACACATTCTTCTAAGGCTAGGACTTGGTGGTAATAGGTTGAGTGGATCCTTGCCAGAGTGCCTTTGTAAAATGTCATCGTTGGGCTTGCTTTCTTTGTACTATAATCGGATTTCTGGTCCGATACCATCTTGCATTGGTAATGCAACCTCTTTGAGAAATGCTTATCTAAATTCAAATAGGCTCACTAACATACCCATAAGTCTGTGGAGCCTCAAATATCTTTTGGTGCTTAACTTGTCAAATAATTCTTTGATTGGTTCTATACCTCCAGAATTAGGGAACTTGAAAGCCATAACGTCCTTAAATCTGTCCTGGAATCACTTTTCAGGAAGTATTCCCTCCACAATTGGAGACTTGCAGAGCCTACTTTATCTTTCCTTGTCTTATAATGAGTTACAAGGATCTATTCCAGAGTCATTTGGGAAAATGATAAGTTTGGAATCGGTTAATCTATCCAATAACATTCTTTCAGGTACCATTCCAAAATCATTAGAGTTGCTTCGGTATCTGAAGAATTTCAATGTATCATTCAATAGATTAGAAGGTGAAATCCCAAGTAAAGGACCTTTTCTCATTTTCATCTCTCAATCTTTTATGGGAAATGGAGGGTTATGCGGCGGTTTAGTTTTCCAACCTTGTATGACTAGGGCTTTTCATCATTCAAGGAGAATCAAACTGCTTCTGATTTTACTTGTCTCACTGGGAGCTGCTGTAATGGTACTTGGCTCAATCGCTGTGTTTATGTTAAGGAGACCTCGGAATAGAAATATTCCAACTCAAGCTGAATCCTTTGCTGCAACCACACTAGCCAGGATTTCATACATTGAAATTGAAAGGGCAACTCAAGGGTTCGACCAATGCAACTTGCTAGGCTCTGGAGGTTTTGGCTCTGTTTACAAGGGCATGTTTGCAAATGGGATGACTTTGGCAATCAAAGTGTTTAATTTACAGGTTGAAGGTGCGTTAAAGAGTTTTGATGCTGAATGTGAAGTTTTACGCAACCTTCGTCATAGAAATCTTACCAAAGTTATCAGCAGTTGTACTAACTTGGATTTTAAAGCATTACTTCTTGAGTATATGCCCAATGGAAGCCTAGAGCTGTGGTTACATTCTGATGATCGCTTCTTGAATATGATCCAAAGATTAGACATAATGATCGATGTTGCATTTGCTTTGGAATATCTCCATCATGGTTATGAAACAGTTGTTGTACATAGTGACCTGAAGCCTAGCAACGTCTTGCTAGATGAAAAGTTGGTTGGACATGTGAGTGACTTTGGCCTGACCAAGTTATTAGGAGAAGGGGAATCAATTGCTCATACTAAAACACTTGCCACAATGGGCTACATTGCACCAGGTAACTTTCTTTAAGTCCTTTTATCATCGCCTATGATGTGTTATTTTACATCAAATAATTAGTTGACTGGTTCTAATATAATGCATCACATTAGAACTGGGTGAACGCCTAGGTGGATGGCTTATATTAATGGGAAGAAATCACCTAGTGTTTTTTGTCTCTTTGTTGGAATTTGAACCTGAGATCTCATAGTTCTTAATCCACTTCATTGGCCATTAATCCACTATTAGCCTTGGGTATAAATATAGATAGTATTTTGATTTCTTTATTTGGCTCGTCAAGAATCTAATTACTCTGGTAACATTGGATCAGAGTATGGATCAGTAGGATTAGTTTCTACAAGATGTGATGCGTACAGCTATGGCATTATGCTCATGGAAACTTTTACAAGAAAAAGGCCATATGATGAAATGTTTCAAGAAAATTTGAGCATGAGGAGTTGGGTCTGCAATTCACTAACTGCAATGCCAGAGGATATTATTGATGCCACATTAATGGAACCAGAAAAGACTCATTTTCAGGAAAAGTTGCATTGTGTGTCTTCTATTTTGGAGTTGGCATTGCATTGCACAACTGAATCTCCCAATGAAAGGTTGAACATGATAGAAGTCCTTGCAAATATCAAGAAGATCAAGTTGGAATTTCTTAGCAAATGATGTACGTCGCACTGGTAACTATCCCTTTTTTAAAATATTCTAATGTCGTCTAGATATATTATTTCCTTATTCAGGAATAGAAAAAATTATTGTCTTGTCTTTATCGTACATAAATATTGAATTGGATATTCTGGTTTTCCATACAAATATATCCACTTGTATAAGAAAAAACAAGAACTAATTGAAACATAACTTTAATCATGATTTTATAAGTAGAAGAATCCAACATTTTTGCCCATGGAAATTTGAATTGTGATGAATGTTGAAAACTGAAGTAAACAACTCAAGTCTCCGTTACTGTTAGTTTGTTTTTGCAGTGTCAACTCAAGTTTGATTCCAATTGAACATTTGTGCAGGTTGTGCTGCCGTTTTCCTGGCTTCTAGCAAATTTGGGTGACACCAATTGAGATGAAAAATCAACATTTGTTCCCAGATTTGAAATTTTCTTGTTTCTGTTGTGTTTTTGTCTTGTGCATATTTCGACATTTGGTAAAATAAGTTTTAATATATAAGATGTGTACTACAGTTGGAATGGAAGAATGCCCTGTATAATATTGGATTGT

>M6-PGSC0003DMG400028436 M6 LRR-RLK

TATATACCTTGATATCACCTTCTTATTTATCCTTTCCTGCCTGAAAACTTAATTTGTTCCCCATGGAGAGATCATACTGTTTCTTTCTTTCACTACTACTATTGAAACTCTACTTGTTCGAGGCCACAGACATAAAAACAGATCAATCAGCTCTACTTGCCTTAAAAGCTAGCTTTACTCTAAATTCTTCTCATCCCTTGACTCAAAATTGGTCTTCTCAATCTTCCGTTTGTGATTGGATTGGAGTCACTTGCGGCTCTCTTCACCATAGGGTGACTTTACTAAATATATCCAACATGGACATTCTTGGAACCATTCCCCCAAACTTGGGAAACCTCACCTTCCTCGTTTCTCTTGATTTGAGCAACAATAATTTACATGGAGACCTCCCTCAAGATTTTTCTTGTTTACGTAGACTGAAGGTGATGGATTTTGCGTTCAATAATTTCAGCGGAGAGATACCAGTGTGGTTTGGTTTCATTTCAAAGTTAGAAACCCTGAGTGTGAGAAACAATCATCTTCAAGGCAATATCCCTAAGGAGATTGGGAATCTTGAGTATCTAAAGGAGTTGATTTTATTTGAAAACCAATTTACTGGCTCTATCCCGCTCTCCATTTTCAATATGTCGTCGTTGGAAATTTTAGGTCTAACATATAATCAATTAACTGGGAGTCTTCCTGTAGATATATGTCTTCGTCTTGCAAAAATAAAGAGCATTTCAATCATTTCCAACCACTTGAGTGGTCAAATTCCAACAGGTTTGTCTAACTGCACAGAATTGCATGATTTTTCATTGTCATACAATAACTTCAGTGGAACCATTCCACCAGAAATTGTAAACTTGGTGAGGCTTGAGCTCTTAAATCTCGGGGGAAACAACTTGCACGGTACTCCTTTCGTCCACTTTTACTTGTTCAATTTAGAAAATAATTGTATTTATATTGCTGCTTAAGGATGGCCATCCCATTTGGCAAAATACATCTAACTTGTCTTTATTTTTCATTTTGATACTTCATCTTAACCTTGTATGTTTCATTTTAACCCTCCCAATTCTATCATTTTATTTTTACTTTATAAAATAAGTTTATCCATATTCAAAAGCCTTCTAGCATGAGGCTAATTTCGAAGTTTGTTTATTGAGCAACAGGATAAAGGTTTACAAGTTTGTTTGCCGCTTGTCAAAATAATTTGCAGGCACAATTCCAGCAAAGATTGGTAATTTACGAAAGTTGCAGCAAATACAAGTGGAGAACAATCAAATTGTGGGTTCAATACCGCGCACCATATCCAACCTATCATCACTATGGGTGATTAATTTGAACACAAACTATTTATCAGGTAGCGTACGTGGTCTTGTTTGGTTATCCGGAGGCAATAAAATAAACTAATCGAGATATATATACTCTTTCATGTGTCAGGTGTTATACCCAGAGTGATTGGGAATCTTCACAAATTGGAGACACTTTATTTGCAATTTAATCAATTAAGTGGTAATATACCAGAAGAGCTCTTCAATATCTCTACACTTACAAGGGTTTCTCTTCTCTTTAACAACGTTTCGGGTAGTCTTCCATATGCTTCAAACTACTGGCTAACAAATCTAGAGATTTTGAATCTTGGTGTAAACAACATTGGTGGAGTTATACCCACCTCAATCTCTAATTCCTCATATCTAATGAGATTATATCTTAATGGTAACAAATTCAGTGGTCAGATTCCTAACTCATTGGGGGGTTTGAGGCATCTTGAACATCTGGACTTGTCCAACAATAACTTATTCTCTCTGCACCTAAGTATCTTAACTTCATTGGCAAACTGCAGATCATTGAAAGAAGTTTACTTATACCATAATCTTCTCGACGGTGTTATTCCAGATTCCATTGGCAATCTCTCCACTACCATTATGGAATTTTATTTAGATAGTTGTGAAATAAGGAGTCAGATACCATTAGGAATTGGGAATTTAAGTAACTTAAACACTTTGTCCCTATCCCGCAATGATTTGACTGGATCAGTGCCAACAACATTATGTAATTTACATATTCTTCAAAGGGTAGCAATTGATGATAATAGGTTAAGTGGACCGTTGCCCCAGTGCCTTTGCAAATTATCATCCTTGGGCTTGGTTGATTTTTCTAATAATAGGATTTCGGGTCCGGTACCGTCTTGCATTGGTAATGCAACCTCTTTGAGAAATGTTTATCTAAATTCAAATAGGATCACTATCATACCCATGAGTCTATGGAGCCTCAAAGATCTTTTGGATCTTAACTTGTCAAATAATTCCTTGGTTGGTTCTATACCTCCAGAGTTAGGAAAGTTGAAAGCCATAACATCCATAGATCTTTCCAGGAATCACCTTTCAGGAAGTATTCCCTCCACAATTGGAGACTTGCAGAATCTATTTTATCTTTCGTTGGCTTATAATGAGTTACAAGGATCTATTCCGGAGTCACTCTAGAATACGATAAGTTTGGAATCAGCAAATCTATCCAATAACTTTCTTTCAGGTATGATTCCAAAATCATTAGAGTCACTTCGACATCTGAAGGATTTCAATGTATCATTCAATAGATTAGAAGGTGAAATCCCAAGTAAAGGACCTTTTCTCAATTTCACCTCTCAATCTTTTATGGGAAATGAAGGGTTATGCGGCGGTTTGCTTTTCCAACCTTGTAGGACTAGGTCTTTTCATCATTCAAGGAAAAGCAAACTGCTTTTGATTATACTTGTCTTCCTGGGAGCTGCAGTAATGGTACTTGGCTCAATCGTTGTGTTTATGTTAAGGAGACGTCGGACTAGAAATGTTCCAACTCAAGCTGAGTCCTTTGCTGCAACCACACTAGCCAGGATTTCATACATTGAAATTGAAAGTGCAACTCAAGGATTCGACCAATGCAACTTGATTGGCTCTGGAGGTTTTAGCGCTGTTTACAAGGGTATGTTTGCAAATGGGATGACTTTGGCAATCAAAGTGTTTAATTTACACGTTGAAGGTGCGTTAAAGAGTTTTGATGCTGAATGTGAAGTTTTACGCAACCTTCGTCATAGAAACCTTACCAAAGTTATCAGCAGTTGTACTAACTTGGATTTTAAAGCGTTACTTCTCGAGTATATGCCCAATGGAAGCCTAGAGATGTGGTTACATTCTGATGATCACTTCTTGAATATGATCCAAAGATTAGACATAATGATAGATGTTGCATTTGCTTTGGAATATCTCCATCATGGTTATGAAACAGTTGTTGTACATAGTGACCTGAAGCCTAGCAACGTCTTGCTAGATGATAAGTTTGTTGGACATTTGAGTGACTTTGGCCTGACCAAGTTATTAGGAGAAGGGGAATCAATTGCTCATACTAAAACACTTGCCACAATGGGCTACATTGCACCAGGTAACTTTCTTTTAGTCCTTAAATATCGCATATGATGTGTTATTTACATAATAATTAGTTTACTATTACATCACATTAAAACTAGGTGAAGGCCTGAGTGGATAACTTAATTAAAGAAAATGTAATATATCACAGATTAAGTATGAACAAAAATGGTATAGAAAACATTGAAATATGACTGTCAAACTCTATATGCATGCATCTACGAATTTTTTTTTATCAAAAGTTCTCTCAGAGGATATTAAATAGATAAGCCAAAGGGGCATAGAGGGGGACTATGCCTGACCTCAGCAATACAAATTTACGATATTAGGAGTGAGGAAACGAAAAAGTAGGGGGGGGGGGGGGGGTTAAGGGTTGCCCCACATCAATTAAGCATACAAAAAGATAAATTCTAGGATTTTAGAGCAAGCAAAGAAGACTCAGCCAGGATCAAAAAGAAAGTTGTTCTCTTCACTTAGATTCTAATGTTAGGCTTGCCAACTACATCATTCTTCAACGTGGATATAATCTTAAACGGAAGATTTATAGGCCTAGTCTCAGTGAAGAAGGTAGTCATTTTGCACTATTTAGCCAAGTTAGCAAGCATGTGAGCAGGAGTGTTACCTTCTCTTAAGGTATGTGTGAAAGTGAAACGGCTAAGTGAACTCATAGATTGGATCTGTCTAATATCATCTTTTATCTGCTAGAAGTCCATATCGTTTCAATTGACTGCCTTACACTTGTTTCATTCCAATTATACTTTCAGTTTTTAAAGTCCTTATCTGCTCAAGGTAAATTCAACTATTTTCAATATCTAGCAAATATATATATATATATATATATATACAGACACAAACACACACATATATATGTATAATATTTTCATTTCTTTATCGGGCTCGTCAAGAAACTAATTACTCTGGTAATATTGGATCTCAGAGTATGGATCAGTTGGATTAGTTTCTAGAAGATGTGATGTGTACAGCTATGGCATTATGCTCATGGAAACTTTCACAAGGAAAAGGCCATATGATGAAATGTTTCAAGGAAATTTGAACATGAGTAGTTGGGTCTGTAATTCACTACCTGTAGCACCAGATGATATTATCGATGACACCTTATTGCAACCTGAAGAGATTGATTTCAACAAAAAGTTGTGTTGTGTGACCTCTATTTTGGAGTTGGCATTGAATTGCACAGCTGAATCTCCTAATGAGAGGCTCAACATGAAAGATGTCCTGGCAAATATCAAGAAGATCAAGCATGATTTTCTATGCAGATGATGTAACAAGGTTTATTCAATGGTAGGTATCCAACTTTTTAAAACATTCTAATGTCATATGGAGAAATTATTTCCCATTTCAGGAATAGAAAGATTACTGTCTTGTCTATACCATGTAGAAATGGGTTTTTTCCATACAAATAATAATAATAAAAATAAAAAATAAAATAATAATAATAATATAATAATACTTAAATCAAGGAGAACTTCATTAGTTTTGAAATTGGTCACATAATCAAGGAGAACTTCATTAGTGAGTAAAGTAAATCAAGAATTCATGAGGACCAACATTTGGAAATTATGATGAATGTTGAAAAGGGAAGCAAACAACTCATGAATCCGTTACAATTAGTGTATTTTTGTGTGTGCTATGAGTTTTTATCTGCAATGACTGCCAACTCAAGTTTGATGCCATTTAAATTTTTGTACAGGTAGTGCTGCCTTTTTTGCCTGTCGTTGGCAAATTTGGGTTACACCAATTGAGTTGTCAACAAAGATTTGGTCTGAGATTCGGTCTTCTCTTATTTCTTGTGCCTTCCTCTTGTGCATGTTTCAAGTTTGAGTATGTTTATGTAATAAATTTAATATATAAGATGTGTATTAAAGTATCATACTGGTGATCATGAATTATTATTGAGATAGCAGCTTCAATAGTTTATGAAGA

>M6-PGSC0003DMG400028437 M6 LRR-RLK

AATTTTCAAATGGTATTCCAAGAGAAATTGGAGACATGCAAAATCTGATACATCTTTCTTTGGCACAAAACAAGCTACAAGGATCTATTCCTGACTCAATTGGTAGCATACCGAGTTTGCAATTTCTAGACCTCTCGAATAACAATCTATCTGGATCAATTCCGATGTCCTTAGAGAAGCTTCAGTATCTCAACTATTTCAATGTTTCCTTTAACAGCTTGCAGGGTGAAATTCCCTTTAGTGGTCCTTTCAAGAACCTTTCCAGCCAGTCATTCATGTTCAATAAAGCATTGTGTGGTGCTCCAAGATTTCGTGTCCCTCCGTGCCCCACTTCTTCAAACCATAGATCAAAGAAGAAGAAGTTACTTCTGATAGTCTTTCCTCTACTGGGAGCTGCTGTGATAATTGTTTTTGTAACATTGGCATTTATGTGGATGAGGTACAGAAGAGAAGGAAATGTTCCAGTTGAAGCTGATTTGTTAGCTACAATGGAAAGAATTTCATACTATGAAATCATACAAGCCACTAATGATTTTAGTGAGAGTAATTTCATTGGTTCGGGGAGTTTTGGTTCTGTTTACAAAGGAATTCTCAGAGATGGGACTACCATTGCAGTTAAAGTGTTTAACCTACAAGCTGAAGGTGCATTCAAGAGTTTTGAAATAGAATGTGAAGTTCTTCGCAACCTTCGCCATAGGAATCTCACCAAAGTCATTAGCAGTTGCTCCAACCTTGATTTCAAGGCTTTAGTGCTCGAGTACATGCCTAATGGGAGCCTTGAAAAGTGGTTGTATTCGCACAACTACTTCTTAGACATCCTGCAAAGACTAAGTATAATGATCGATGTGGCACGTGCATTGGAATATCTCCATCATGGTTGCTCAACACCTGTGATTCATTGTGATCTGAAGCCTAGCAACGTTTTGCTAGATGAAAACATGGTAGCTCATGTAAGTGACTTTGGCATTTCTAAACTACTCTGTGAAGATGACAGTGATTTACACACTAAAACCTTAGCAACATTTGGCTACATTGCACCAGGTAAATCTCTTATAATTCTTGTTTCAGTATCCTATCTTCATTTTCCTTTTCTCGGTTTAAATGATAATCCGTAAGTGTGATTTACTGTAGAGTATGGAAGGGAAGGATTGGTGTCGCTAAAATGTGATGTTTATAGTTATGGGATCATGCTAATGGAAACATTTACGAGGAGAAGGCCTAACTATGAAATATTTGATGGAGATCTTAGTTTAAAGAAATGGGTAAGTGATTCACTCCTAGAGGCAACAATCAAAGTTGTAGATGCAAATTTGCTAAAGCCAGAAGATAAGAAATTCATGGACAAGGTAGATTGTGTAGCATCCATCATGAAAGTGGCACTATATTGCTCTGCTGAATCTCCTGAAGAAAGAATCAACATGAAAGATGTCGTAGGAATGCTACAAAAGATCAAGATTCAACTTCTTTCATGCTCTACAAGTACATAAAAACATATCAGTCATGCCACTTAAGGGTATAGCAAGTGTTTGGTAAACTTCGTATCTGCATACATAGACTGGTTTCCTTTTTAGCTGCATAAGGTTATTTTCTGCAGCACTATGTAAAATCGG

>M6-PGSC0003DMG400028438 M6 LRR-RLK

ATGAAGAGAAAATTAAATTTCCTTTTCCCTCTTGTAGTTTCAGTAATATGGCATCATTTTCTGATATTATCAGCCACTGCAGAAGTCAATCTCAAAACTGATGAAGCTGCTCTACTTGCTTTGAAATCCTATGTTACTTCTGATACTTACAATATCTTGACAAGCAACTGGACCTCAACTACTTCCGTTTGCAATTGGATTGGAGTCACCTGTGGCTCTCGCCATCAAAGGATCATAACCTTGGATATTTCCAATATGGGATTAATAGGTACCATCCCTCCTCACTTGGGAAATCTGTCCTTTTTGGTTTCTCTCGATATCAGTAGCAACGGTTTCCAAGGAATTCTGCCACGAGAGTTGGCTAATTTGCATAGATTGGAGTTAATAAATGTCACTTCTAATAAGTTCACTGGAGATATTCCCTCGTGGTTCTCGCTTTTGCCAGAACTCCAGCACTTGCATCTAGCTTTTAACAGTTTCACAGGTATTATTCCACCAGATAGTTGTAACGCTTCCAAACTTGAGAGTTTAGTCTTGGGATTTAATCAGTTACAAGGGGAAATTCCAAATGAAATTGGTAATCTTCAGAACTTGACTTGGTTGAGCTTGGGATCAAATCAGCTAACAGGCTCCGTACCACTTAGCCTGTACAACATTTCATCGTTACAACGTTTGGTTTTGACAAAAAATAGTTTGTCTGGAAATCTTCCTGTCGATATATGCTTGAACCTTCCAGAACTTATGGTGCTTGCACTTTCTGACAATGAATTTGATGGTCAAATTCCTTTGGGGATAGATAAATGTTCAAAGCTTCAGATTTTGTCATTGTCATTCAATAAGTTCAGTGGATTGATACCTAAACAGATTGGGAACTTGAATGTGCTGTCTATTTTATATCTTGGTCGTAATGACTTGAAAGGTGAGTTATACATATTTGCTTTTAGGTTCCTTCTTTGTTCATACATGAAACATATTTTTTGAGTCATGTATGCACTCCTTAATTGATATACTGCAGAAAGTAGACTTTCTATTTATGATTTAATTTGCAGGTGAAATTCCAGAAGAGATTGGAAATCTTCGTAATCTAGAGATATTGGATGCACAGAACTGTAGCCTAAGTGGTCCTTTACCATCTTCTATAAGTAACTTAACTTCACTTCAATCACTCAATCTTTATGGCAATAATCTCTTTGGCACCTTGCCCCGGTATATATGTTTGAACATGCCTGATCTCAGTGCATTCGATCTTGGGAATAATTTATTCAGTGGAAACATTCCAAAAGAATTTGGAAACTGCAGTTCACTTTCTGACCTCTTCCTAAGAGAAAACAACTTAACAGGTATCATCTTGTCATTTTCTTTTTCCATTATTTGGTGTATGTTCCTATGTTTTTCATATCCTAATAAGTCACTCAAATGTCATCAGGTGAGTTACCAAGGGAGATTGGTAATCTTTTCAATTTGGGAAGATTAGATCTACATTACAATTTCTTAACAGGTCCTATTCCATCTACAATATTCAACATGTCAAACATAAGGGGCATTTCATTTTTAGGGAACTTCTTTACTGGAAGTCTACCATCAGATATAGGACTTGGTCTTCCTAATCTTGAAGAACTTTATCTTGGTTACAATAACCTTACTGGAGCCATCCCTAATTCTCTCTCAAATGCTTCCAATATTTTCCGGCTTGGAATTGGATATAATGACTTTTCTGGTCATTTTCCGAGGTCGTTTGGTAATCTAAGACGTTTAGAGTACCTCAACGTGAACGGCAATCATTTCACAAGAGATCCTTCATCTCCAGGATTGACTTTCTTTGATTCCTTGACAAATTGCAGACATTTGAGAAAACTGTGGATTGGTTATAATCCACTCAATGGAAATCTTCCGGCTTCTATTGGAAATCTCTCAAGTTCTCTCGACTACATTTATGCTGTTAATAGTGAAATCAGAGGCTACATTCCCAGTGAAATAGGAAATTTAAGTGGTTTGTCTTTCTTGTTACTTCAAGGCAATTATTTGAGTGGGTTCATTCCAAGAACAATAGGGAATTTAAAGAATCTTCAAGCTTTGAATTTGTATGACAACAAAATGATAAGTGGACCTATCCCTGAGGAACTCTGCAATTTAAAGAAATTAGGATTCTTGAGCTTGGGAAATAATGAGCTTTGCTGTTCTATACCATCATGTTTAGGGAACATTACATCTCTTAGATACATTTACCTTGGTTCCAACAAATTGACTTTTAGCATACCTCCCAGCCTGTGGAACCTCAATGATCTCTTGCATCTTGATGTATCCTCAAATTCCTTGAAAAGCTCTCTACCTCCAGAAATTGGAAATCTGAAAGTCGCAACATTGCTGAATATATCGAAGAATCAAATCTCAGGAAGTATACCGAGCACAATTGGAGGCATGCAAAATATGGCTGAACTTTCTTTTGCAGAAAATAGACTAGAAGGGTCAATTCCAGAGTCAATGGGGAACATGGTCGCCTTGGAATCGCTAGACTTGTCTCATAACAAGCTCTCTGGTGGCATTCCCAAATCATTGGTAGCTCTTTCACATCTCAACTATCTTAATGTATCTAACAACAGATTGAGTGGTGAAATTCCAATTGGAGGTCCTTTCGTGAACTTCTCTTATGATTCATTCCTTTCAAATGAGGCATTATGTGGTGCTGCTAGGTTGCAGATACCAGCATGTAGGTCTAATTCTCCTATTAGGAAAAGAAAAAAGAAGGTGCTTCTTATTGTGTTGATCTTGTTGGTGGCTTCTTCAGTAATAATGCTCTCAGTCATGCTCACCATCTTTTTGGTGATTAGAAGTCGAAAGAGGAAGACAAACATTGCTACTCATCAGGCTGATGCCTCTCCAGCCACGGTACATGGAAGAGTTTCATACCATGACCTTCAACAAGCAACCAACAGATTCGGTACAAGCAACTTGCTCGGTTCTGGAAGTTATGGCTCTGTTTACAAAGCAACATTTGGGAGTACTATAGTGGCCGTTAAGGTATTCAACTTGCAAACAGAAGGCGCATTCAAGAGTTTTGATACAGAATGTGAAGTCTTACGCAACCTTCGTCATAGAAATCTGACCAAAGTCATCAATAGTTGCTCCAGCATTGATTTCAAAGCCTTTGTATTAGAGTACATGCCTAAGGGAAGCCTGGATGATTGGCTTCATTCTGAAACTTGTTCCTTAGATATAATGAAGAGAGTAGACATAATGATTGATGTGGGATCAGCATTAGACTATCTTCATCAAGGCTATTTTGTGCCTGTAGTTCACTGTGATCTAAAGCCTAGCAATGTCTTGCTCGACGAAGATATGGTTGCACATGTGAGTGACTTTGGCCTGGCTAAGCTATTAGGGGTAGGAGAAAGTATTGTACAAACCAAAACACTTGCAACCATTGGATACATTGCACCAGGTAATAATCATGTCAACAAGCTTCATTTCTCCTTCAAATTTCAACCATTTTAAATATACCATATTCTCTTGATAAATTGTTTTATTTCAATGTAGAGTTTGGATTGGAAGGACTTGTATCTACAAGGTGCGACATTTACAGCTACGGTGTCATGTTGATGGAAACCTTTACGAGAAAGAAACCAACGGATTGAATGTTTGCTGAAAATTTAAGCCTGAGGGAGTGGATAAGACAATCATGGCCACGCAGAATGGACGAGATAATAGACCCCGAGCTTATGATACCAGTGGAGAAGAACAAAACAGGAAAGGTGCAGTGCTTATCATCCATCATGGAATTAGCTCTGAGGTGCACTGCAGACTTGCCAGAGGAAAGAATGAATATCAAACATGTTCTTGCTCAACTGAAGAACATCAGAACTATGTTGGAAAATGTAATAAATCACTAA

>M6-PGSC0003DMG400028473 M6 LRR-RLK

ATGAATATTAGCACTGATCAATCTTCTCTTCTTGCCTTGAAATCCCACATCACTTCAGATCCTTTTCATGTTCTTTCGACGAACTGGTCTTCTTCTACCTCTGTTTGTGATTGGATTGGAATCACTTGTAGCTCTCGCCATCAACGAGTGATTGCACTGAATATTTCAAACATGGGTTTTTCGGGTACTATCCCGCCACAACTTGGCAACCTCTCTTTTCTTGTTTCACTTGATCTAAGCAAAAACGATTTACAGGGTGAACTTCCATCAGATTTTTCTCATTTAAGGAGATTGAGAGTCATTAATCTTAGTTACACCAACTTCACCGGGGAAATTCCAATAGGAATAGCCACTCTTCCGAGTTTGATACGGTTTTCCATGGGATACAATAAATTACTCAATGGCTCCAATGTGCTCTCTAAATTCAACATCTCAACCCTTGAATATTTGGATCTCAGAAATTCTGGTTTGACTGGTGATTTCCCTTCTGATTTATGTCATCGTCTTCCTAGATTGCAAAAGCTTGCACTTGGCGACAACATGTTAAGCGGAGAGGTACCAAGAAACATATCTGAATGCTCAGAGCTTCAACTCCTAATGTTGTTTCAAAATAACTTTGTTGGAACAATTCCAAGAGAAATAGGTAACTTGCAGCTGCTGCAACATTTAGATCTTGCAGATAACACGTTAAAAGGTATGGCTTTACCCTTTTTTTCTCAGGACATTAGATTTAACTTATTTAAAAGAAAATGTATTTTAGAGAGTTGAAGTGTTTGGACAATCTTTAAAAAAATGTAAGTATTTTTTTAATAGTAGCATAAGAAGATTTGTTTGTTTTCACTAATTGCTCTGTCACTTCTCCTCTCGATTTGTATACTTGTTGCCGGTTTGACCAAGCTTCTGGCATACCCATTTGGAAGTTAAACTCTTGTTAAATATATGTTTTTTATGTGATATTTCTGTTTAACAATATGACAGGCACAATTCCTGATGAGATCGGCCATCTTTATAACTTAAAGAACATATTCATGGATAAAAATTATTTAACAGGCTCAATCCCTTTAACCTTATTCAACATTTCATCACTTGAAATGCTATATATGAATGATAACAAGCTTGAAGGACCTTTACCGAGACAGGTTGGAAATTTGACTATGCTTGCCTGGTTTGATCTTTCCAATAATTACCTGGCAGGTACGGAAAAAAAATCTAATTCAAGTCATGTTAGAATTTTTAGTTTTTTGGGATATGTCTACTGATCATAATCATACTGAATTTTTCAGGTGTAATTCCACATGAAGTTGGTAACCTTCAAGGGTTGACAGATCTTCCATTGTCTTACAATGAATTTAGTGGGTCGATCCCTATTGGTATCTTTAATATCTCATCTCTTGTAACTATTGGACTCACAGAAAACCACATTTCAGGTAATCTTCCGTCCACTATAGGCAATGGGTTACCTAATGTTGAAAAATTATTTCTAGGTGGAAACAACATTAATGGTGTTTTACCTGGATCCATCTCAAATTTGTCTAAGCTCACCTTTCTCGAACTCTTTAGTAATGAATTTACTGGTTCAATTCCTGACTCTCTGGGAAATTTAAGACAACTTGAACTTCTCAACTTGTACGGTAACTCCTTTACAAGTGAATTGAGCTTCATTACTCCTTTGGCCAATTGTAAACACTTGAGAATATTGATATTGTCTTTCAATCCCCTAAATGCAATGCTCCCAAGATCCATTGGCAATATCTCTTCTCTTCAATCATTTATTGCTGTAGGCTGTAACCTCAAAGGCAATGTTCCAAGTGAGATCAGAAATATGAGAAATTTATCCACTTTGTCTCTGGGAGGTAATAACCTTACTGGAATTGCCCCGACGACATTAAGTTCTTTGAAAAACCTGCAACGGCTTTTTCTTGATGCAAACAGAATAACTGGTCCTTTCCCAATTGTTTTATGTGAGCTGCCCAACTTGGGGATACTAAGCCTTTCACAAAATAAAATTTTGGGACATATTCCTAGCTGCGTGGGGAATATGACATCCCTAAGGGAGGTTTATCTTGATTCCAATAACTTTACTGCTAGCATACCTTCAAGTCTATGGAACCTCAAAGACATTTTGAAGCTGAACTTGTCTTCTAATTCCTTCAATGGATCTCTACCACTAGAAGTTGGAAACCTCAAGGCTGCACTACTTCTGGATCTTTCTAGGAACCAAATATCAGGCAACATTCCAACTACATTGGGAGGTCTACAGAAATTGATTCAACTATCTTTGGCTCATAACGGAATTGAAGGATCTATTCCTGAGACATTTGGGGAACTCCTAAGTTTGGAAGCATTAGATCTTTCATATAACAAAATGTCTGGTGTGATTCCAAAGTCATTAGAGGCACTTAACCAGCTAAACTCCTTTAATGTCTCATTCAATAGGTTACACGGGGAAATTCCGAGTGGAGGACCTTTTCTTAATCTACCTTACCAGTCTTTCATGTCGAATAAAGGATTGTGTGGTAACCCTCAAAAGCATGTCCCAGCTTGTCCTTCTTATTCAAAGAATCAGTCTAAGTCGAAGAAAAGAAATCTGATATGGATTAAAGTTACCTCTTCAGTTATCTCTGTAACAGGGCTTGCTTCAGCAATAGTTTTCATGTTGATGAGACGTGGGGGTAAAACGATCAATGCTGAAGTTGAGTGGTCTCCTGAAGTAGCACCACAAAGATTTTCTTACTATGAACTTCAAAGAGCAACTCAGAGTTTTAATGAAAATAACTTGCTTGGTAGTGGAAGTTTTGGTTCGGTTTATAAAGGGATGTTGGCTGATGGGATGATTATAGCTGTTAAAGTTTTCAATGTGCAGATGGAAGGAACATTTCAGACCTTTGACAGAGAGTGTGAAATCTTGCGGAATCTTCGTCACAGAAATCTCACAAAGATCATTAGCAGCTGCTGTAACTTGGATT

>M6-PGSC0003DMG400028474 M6 LRR-RLK

AACACATTTTCTCCCATTCACTCAGAAAAAATGAGGATTTAATTAATGGAGAAAAATCCCTCTTTTCACCTTCTAACAATTTTGTTCGTTCTTCTTCATGCTCTAGCAACTTGCTTAGCCATGAATTTTAGAACTGATCAAACTTCTCTTCTCGCGTTGAAATCCCAAATCACTTCAGACCCTTATCAAATTCTTTCAACAAACTGGTCTTCTTCTGCCTCTATTTGTAGCTGGATTGGTGTCACTTGTGGCTCTCGTCATCAACGAGTTACTGTACTCAATATTTCAGACATGGGCTTTTCGGGTACCATACCGTCACAAATTGGCAACCTCTCTTTTCTTGTTTCACTTGATCTAAGCTACAACTATTTCCATGGTGAACTTCCACTTGAGTTTTCTCGTTTGCGAAAATTGAGAGCTATTAATCTTAGTTTCAACAACTTCACCGGAGAAATCCCTAAATTCCTCGGTGATTTTCAAGACCTTCAAATATTGTCTCTTGAAAACAATAGTTTCAGTGGGTTCATTCCTTCTTCTATCTCAAATATGAAGAATCTTGGATTCCTGAATCTGAGGTACAACAATTTAGAAGGAAATATTCCAGCTGGAATAGCCGCTCTTCGGAGTTTGAAATGGTTGAGCTTTGGATTTAATAAACTCAATGGTTCCAATGTGCTCTCTATGTTCAACATCTCTACACTAGAATATTTGGATCTCAGAAATGCTGGTTTAACTGGTGATTTTCCATCTGATTTGTGTGGTCGTCTTCCAAGATTGCAAAAGCTTGGACTTAATTTCAATATGTTAAGTGGAGAGATCCCAAGACGTATATCAGAATGCTCGCAACTTCAAGTCTTACTGTTGATGGAAAATAAATTGATTGGAACAATTCCAGGAGAACTTGGGAAATTACAACTGCTGCAACAATTAGCTCTCGGAAATAACAAATTAGAAGGTATGTTGCTTCCAATCTTAACAATATCTACATATATAGTAAGAAATCGAAGATATTGTTTTGTGCTGAGCACATGCATTCCCTTTAATTTTCAGTTTCGACAAGAGGTTAAGTGTTCAATTTTCCTCTCCTAAACAAGCATATGTATGGTTTTGACCTGGTGTGAGAATTCGTTACATCATAAAGTAGACCTTTGCTTATACCTGTCTGATAAATTATTTAATGAGGTATTGCAATTTGTTCAACTTAATTCCTTAGAACTTCATTTGATACTTTTGCATAACAATTTGACAGGCACAATTCCCAATGAGATTGGTCATCTTCATAACTTGAAGCAATTGGGATTGGAACAAAATGCATTAACAGGCTCAATCCCATTAAGCATATTCAATATTTCATCACTTCAAGTTTTATCAATGTGGGACAACAAGCTTGAAGGAACTCTACCAAGAGAGGTTGGAAATTTGACTACGGTTAACGTACTTGATCTTGGAATGAATAGCCTGACGGGTATGGAATAATATTTGACCACAGGATTTCAGAATGCTAGTTCTTTCAATACTTCTTCTAATGGTAATCTTACTTCATTCTGCAGGCGTACTTCCAGATGAAATCGGTAACCTTCAAGAGTTGCTGATGCTGAAGTTGGATTTCAATAACTTTAGTGGGTCGATCCCTATTGGTATATTTAATATCTCAACTCTTGTATCTATTACACTCACACAAAACCGCATTTCAGGTAATCTTCCTAACACTATAGGCAGCGGGTCACCCAATCTTGAACGAATTTTTCTAGGTGCAAAAAACATTGATGGTCTTTTACCTAGTTCCATCTCAAATTTGTCTAAGCTTACTGTTCTCGAACTTAGTGCGAATGAACTCACAGGTTCAATTCCTGACTTTCTAGGAAACTTAAGACTTATTGAAATCCTCAACTTGCAAGGTAACTCCTTCACTAGTGACTCTTCAATCTTGAGTTTTATTACTCCTTTGGCCAATTGTAAATACTTAAGAGAACTGATAGTGTCTATCAATCCCCTAAATGCAATACTTCCAAAATCCATAGGGAATCTTTCGTCTCTTCAAACGTTTGAGGCAATAGGTTGTAATCTCAAAGGCCATATTCCAAATGAAATTGGAAATTTGAGAAATTTATCTTATTTGAAGCTGGATAAAAATGACTTCACTGGAATTGTCCCGACGACAATAAGTTCTTTGGAAAAACTTCAACAGTTTTCACTTGGTGCAAACAGAATTAGTGGTCCTTTCCCAATTGTTTTATGTGAGCTACCCAACTTGGGCATGCTAAACCTTTCACAAAACCAAATGTGGGGAAGTATTCCTAGTTGCTTAGGGGATGTGACTTCTCTAAGGGAGATTTATCTTGATTCCAATAAATTCACTGCTAGCATACCTTCAAGTCTATGGAACCTCAAAGACATTTTGAAGCTGAACTTGTCTTCTAATTTCTTCAATGGCTCTCTACCATTAGAAGTTGGAAACCTCAAGGCTGCAATACTTCTGGATCTTTCAAGGAACCAAATCTCAGGCAACATTCCTAGTACATTGGGAGGTTTGCAGAAATTGATTCAGCTATCTTTGGCTCATAACAGAATTGAAGGATCTATTCCTGAGACATTTGGGGAACTCGTAAGTTTGGAAGAATTGGATCTTTCAAATAACAATATTTCTGGTGTGATTCCAAAGTCATTAGAGACACTCAAGCAACTACATTCCTTTAATGTCTCATTCAACAGGTTACATGGGGAAATTCCGAGTGGTGGACCATTTCTTAATCTCCCTTACCAGTCTTTCTTGTCGAATGAAGGATTGTGTGGTAATCCTCAAAAGCATGTCCCAGCTTGTCGTTCAAATTCAAAGAATCATTCTAATTCAAAGAAAAGAAGAATGATATGGATTGTAATTGTCTCTTCAGTTATATCTATAATAGGGTTTGCTTCAGCAATAATTTTCGTGTTGATGCGACATCGGGGTAAAGTAATCAAAGGTGGAGATGAGTGGTCACCTGAGGTAGCACCACAGAGATTCTCTTACTATGAACTTCAAAGAGCAACTCGGGGGTTTGATGGAAACAACTTGCTAGGTAGTGGAAGTTTTGGTTCTGTTTTTAAAGGGACGTTGGCAGATGGGATGATACTAGCTGTCAAAGTTTTTAATGTGCAGATGGAAGGTACATTTCAAACCTTTGATAGAGAGTGTGAAATCTTGAGGAATCTTCGTCACAGAAATCTCACAAAGATCATTAGCAGCTGTTGTAACTTGGATTTTAAAGCATTGGTACTTGAGTACATGCCAAATGGGAGCTTAGACAAGTTGCTATATTCTAGAGATTATTCTCTAAGTATAATGCAACGATTGAATATCATGGTCGATGTTGCATCTGCTCTGGAATATCTCCATCATGGTTACTCTGTACCTGTTATTCATTGTGATCTGAAGCCTAGCAACGTCTTACTTGACAAAGACATGGTGGGACACCTAACTGACTTTGGCATAGCAAAACTTCTAACAAAAGAGGAATCTATTGCTCACACTACAACCTTTGCCACAATTGGTTACATTGCTCCAGGTGAATCATTTATTTAACTTGTAACAAAGTTTTTGAGTAACAGTTGATCAGATTTATGCCTTATAAGTTCAGTTGCTGATGTATTTCAATTTTTCTTTTCTAGAATATGGATTGGAAGGACTTATATCCAAGAGGTCTGATGTTTATAGTTATGGTATCATGTTGCTGGAAACTTTTACAAAGAAGAAACCGAATGATGAAATGTTCACGGGCGATTTGGATTTGAAAAGCTGGGTACATAGTTCACTTCCTGATAAGCTGGATGAAATCATAGATGCCGACTTACTAACACTAGATGAACAAAAGCTAAATGAAAAGTTACAGTATGTGTCATCTATCATGGAGTTAGCTATGAATTGCACAGCTAAATCTCCGGTTGAAAGGATGAACATGACTGATGTTGTAGCAGCATTGGAAAAGATCAAGCAGCAGCTTTCTTCATGTTACTGATTGAATGGTTTAATGACATGACATCAGGTACTTTTTCCATATCACTTTTGGGAGGCAAAAAGGAAAGTTGTAAAACAGTTCACTAGCATTTTAATCAGTAATTGTTCTTCAAATGCTTTTGCAAGAGACTGTTTGGCTATCGTAATTGTCCTTCGTTGAGATGAGTATTCTGTACAATGTATTGCTAATTAAGGATCAATAAAATACAGTGCTTCTTAAATTCAAGTGAAGTGATCCTCATCTGGCCTTAGTAAATATATCCAAAAGAGTCACTTACCCTTGCTTTAATCTTGTATCCAGTGGAAA

>M6-PGSC0003DMG400028478 M6 LRR-RLK

CAATTCTATTGGACTTACTTACCTCCACTTGAAATCCCCTTAATACATCTACTCTTATCAACTCTATATGGCATTTACATCAACTACTTCCAACTGAAAGAAACAATCACAATTCCCACAAACAAAAAATGAAAATCTTGTTCACACTCTTCATCTTCCTTTCTCTTTTCACATTCTCAAAATCAGAATCCCCTTGCAAAAACAACACAGATTTAGATTTATCTCTTGTTTTCAAGGCTTTCAGTTCTGTTTATGGTTTCAACATCTCTTGGTTCAACTCAAACTGTTCTTCTCCAATAACTGAAATTAACTTATCTTCAAGAAACCTCACTGGCATAGTTTCATGGAAGTACTTAACAAATTTGTCTCATCTTCATACTATAGATCTATCAAATAATTCCTTAAAGGGTTCTGTTCATCCTCTGTTTTGGTCCATTTCATCTTTGGTTCAAGTCAATTTGTCAAAGAATAAGCTAGGTGGAGCTGTTGCTGTAGCAAGATCATCATCATCTATTCAAAGACTTGATCTTTCTTTCAACAGGTTTACCAACTTAGGCTCTGTTTTTTATGGTTTTCCTAATCTTACTTCTCTTGACCTTTCACATAATGATATCAAGTTTTTACCTTTTTGGTTTACCAACCTAACTAAGCTTGAAAATTTGAGCATTTCTAGCTGTAACATTTATGGTAATCCAGAACCCCTTTCACATATCAAGTCACTGAAGCATTTAGATGTTTCTGTCAACCATATGAATGGTAAATTTCCTAATGATTTTCCTCCTCTGTCTGGCCTCAATTTCTTGAATATCTCATTCAATAACTTTACTGGGGAAATACCCAAAGACCAGTTTGCTAAATTTGGGAACTCTTCTTTCATTCATGCTGGGCACTTGCAAATCAAGAATCTCTTACCAAATCCCAAGAATTCCTCACAGTTACACATTAAAAATCATAACTTTACTACCCCAACACACAAAAAAATGTTGCCTTTGAGACACAAACCTATCAAACCAAACACACACAAGAAAAAGCCCAAATCAAGAAAGAAAGTTTTAACCATAGCTATTTCCTCTGCTTCAGCATTTCTAATCTTGGCCATGGGATTAGTCATATTGTGTCTATACAAGAGGAGAAAACAATTGGCTAGGAAGAACAAATGGTTAATTTCAAAGCCAATACAAATACCCTTTAGAATGGACAAATCAGGGCCATTTTCATTTGAAACAGAGTCAGGGAATTCATGGGTTGCTGACATAAAAGAACCAAGTTCAGCAGCTGTAGTTATGTTTGAGAAGCCATTGATGAACCTAACATTCAAAGACCTCATAGCAGCCACATCTCACTTTGGAAAAGAGTCATTGTTAGCTGAAGGAAGGTGTGGGCCCGTTTACAGAGCCGTACTTCCAGGTGACCTCCACGTGGCAATCAAGGTCTTGGAACATGCTAGGGAACTAGCCCATGAAGATGCTAATACCTTGTTTGAACAACTCTCCAGGCTTAAGCATCCTAATCTGTTGCCTATTTCTGGTTACTGCATCGCAGGTACAAATCTTCTTTCACTTATTACTGTTTCCTCTGTCCACCGTCGATCGATGTGACACAATTCAAATTTTAAGGGGAGAAAAATATCTAGAGGTGAGCATATTGAATGCATTCACGTGGGAGGATGGGATGAGAAATAGGACAGACAGTAGTAAATGAGACAGAAATCTTTGCAACCATAGAAACGTAGGCAGTAGTTATGATTACATACAACAAGATGAAGCAGTAAATAGGAATGCGTGACTTATTTGGATGTTAGTACTATGACTACATCATTCAATTTATATGTGCTTTTAATGCTCTTGGTAAATTGTGTCCATTTTTTACCCTACAAAGTCAAACTTTCAAATCTTATGCCAGATTGGGATTTGAAAAATACTGAAACTGCAGTATGGTTTGGTTCGCAGAATTACGATAACCAATGATATTCTATGAATTGAGATATTTCACTACATTTATCTATCGTGAAATTAAAATGGTGGAATAAAAGAAATTTGTGATTAATTAATATCTCAAACTAAATATGAAAAAAAATAATTTTTAATTTTATTTCGAGATTATAATTTGTATCACATATATCACCACGACATTTATGATGATGCTCTTGGTAAATTGTGTCCATTTTGCACGATAAGGTCAAATCTTTAAAAATCTTACCCGATTGGGAGTTCTACTGTTCAGCCTTGATTTATGATTTTTGGAAGAATGTTGCACAAATATTATCCATAAAGTATTATTTTATGAAACCGTTATTGAATTAAAAAGATAAGAGGATATTTTAAAAGCATTAATGGTTCAGAAGAATGTCATTTTCAATTGAAAATGCTTAAAGTTGGAAAATTATTTCATGACATGCATTGATTGATCAATTGTAGTATCAAAGTCCTTGTCAAATAATTTTGGAGGACTTTTCTAGCTAGGAACCAAATTGCTCCATTTGATAAGAACAGAATCCAATTATATTTTCTTTCAATTAAGTCATATTTAAAGCTAATTTAATACTTCCCTTTGATGCAGGGAAAGAGAAGCTAGTCTTGTACGAATTCATGTCCAATGGTGACCTCCACCGATGGCTACACGAGCTTCCAACTGCTACCACCAAAGTGGAAGACTGGACAACCGACACGTGGGAGTTACAAAACGGTTCCCAAATTACGTCACCCGGGAAAATGGAATGGCACACACGCCACCGTATTGCGGTTGGCGTAGCACGTGGACTAGCCTACCTCCACCACGCGCAGTCAAAACCCGTGGTCCACGGTCATTTGGTGCTATCTAATGTCTTACTAGCCGATGACTTTGAGCCTCGGATAGCCGATTTTGGATTGAGCCAAAACCAAGCCGAGGGATCTAGTGAGATGGATGTGTACGATTTTGGAGTCGTATTAGTAGAGATATTAACAGGGAAAATTGGCTCGGATGAGACGATTAAATGGGTGAGGAGGCTAGTGAAGGATGGACATGGAGCTGACGCGCTTGATTCGAGACTAAGACTCGGTGGTGACTCGGTGAGTGAGATGGTGGAGTGCCTCCGAGTTGGGTACTTGTGTACGGCGGAGACCCCAAATAAACGGCCTAGAATGCAACAAGTTTTGGGTTTGCTAAAAGACATACAGCCCCACTACCAACTCGGGAAAACGCAAAATTTATAATCTTTAGGTAAACCACTCTCACTCCAGCGCGTGATTTACACGACGAGGAAGATTGATTGGCAGCATTGCATTATTGTACTCCCATGTGTTTGGTTGACCATTGAAAAGTCAAAAGTACAAGGCATCTATTCCACTTCATCAATGGCATTTAGTCACGTGCAGATCACGTGATGAAGTGTAGTGAGAGGAACATGTTTGCTTGCTTGTCTCATTTTATTCTCTCTTCTTTTTTTTAAACCTTTTTTTTTTTTTTTGTAGTTTTGTATAGTATTAAATTAGAAAAAAGAAAAAAAATTGTGGGTAAATCTTTTTTTTTAAATATATAAA

>M6-PGSC0003DMG400028688 M6 LRR-RLK

CTGTAAAGAACACTTGGTTTCTTGTTTTTCTTCATTGTCTTGTCAAAAAGTTTCAATCTTTGCAAAAGAAATGAACACCCTTCTTGTGTTAAAGCTTGTATCTTTACTTGTTCTATAAAGAACACTTGGGTTCTTGTTTTGTCAAAAAGTTTCAATCTTTACAAAAGAAATGAACACCCTTCTTGGTTTAAAGCTTGTAGCTTTACTTGTTCTATAAAGAACACTCGGGTTCTTGTGTTTTTTTCATTGTCTTGTCAAAAAGTTTCAATCTTTACAAAAGAAATGAACACCCTTCTTGGTTTAAAGCTTGTATCTATACTTGTTCTTGGTATTTCTGCATTTTTTTTGGGTGTTGAGTCTCAAGATGATGATGTTTCAGTGATGTTAGCTTTGAAAAAGAACTTAAACCCACCTCAAGAAGTTGGATGGTCAGACCCGGATCCATGCAAATGGAACCATGTGGGTTGTTCGGATAAACGGGTCACCCGGATCCAAATTGGGCGCCAGAACATTCAAGGTACCCTCCCTCCAGAAATCTCTAAACTTACTGAACTTGAACGTTTAGAGCTTCAAGGGAACAATATTTCTGGCCCTTTGCCAAGTTTGAGTGGTTTGAGTTCATTGAAAGTGTTACTGCTTGGTGAAAATCAGTTTAGTTCAATTCCTGCTGATTTCTTTACTGAAATGTCTTCATTACTCTCTGTTGATATTGATAAGAACCCTTTTGTTGGTTGGGAGATCCCTGAAAGTCTTAGAAATGCTTCTTCACTTAAAAACTTTTCAGCAAATTCAGCAAATGTAATTGGGAGAATTCCTAACTTTTTTAGTCCTGATGAGTTTCCTGGATTGGTAAATTTACATTTGGCTGGCAATAACTTGGAAGGTGAATTGCCCTCCAGTTTTTCTGGCATGCTGCTTGAGTCTTTATGGTTAAATGGACAGAAGCTTAATGGAGGAATTGATGTTCTAACAAACATGACATTCCTGAAGGAGGTTTGGTTGCATTCCAATAACTTTTCGGGGCCGTTGCCGGACTTTTCAGGATTAAAGGCTTTGGAGACATTGAGTCTTAGAGATAATGCCTTTACAGGTCCCGTGCCAAGTTCTTTGATGAATCTTGAGTCTTTGAAGTTCGTTAATTTGACGAATAATTTCTTTCAAGGACCAATGCCGGTGTTCAAAGGTTCTGTTGTTGTTGATTCGGCAAAAGGTACAAATAGCTTTTGCTCGTCACAACCAGGTGATTGTGATCCTAGAGTAAATACATTGCTTTCAATTGCAAAAGCAATGGATTATCCAACAAATTTTGCGAAGAACTGGAAGGGAAATGATCCCTGTGCTGATTGGTTTGGTCTTACTTGCAGTAATGGAAACATTACTGTCATAAATTTCCAGAAAATGGGGCTTTCAGGAACAATTTCTCCTGAATTTGCTTCACTGAAGTCACTACAAAAAATAGTTCTTGCAGATAATAATCTGACTGGTACAATTCCAGAAGAGCTAACCACATTGACAAGCCTTACAGAGTTAGATGTGTCAAACAATCAAATTTATGGAAAAGTACCAGCTTTTAGGAAAAATTTGATTTTGAAATTTAGTGGTAACCCTGATATTGGAAAGGATAAGAGTGATGCACCATCCCAAGGCAGTTCTCCAGGTGGTTCAACAGGCTCTGATGATGGAAATGCCCAAGCAGCTCGTAAGAAGTCCAATCGCCGGGTTGGAATTGTGGTGTTTTCTGTAATTGGAAGTGTATTTATGCTTTGTTTGATTGGAGTGGCAGCTTTCTGTCTGTACAAAAGCAAACAAAAGCGTTTTAGCCGAGTGCAAAGTCCAAATACCATGGTGTTGCATCCACGTCATTCTGGTTCAGACAATGATAGTGTGAAAATCACAGTTGCAGGATCTAGTGTCAGTGTTGGTGCAGTTACTGAAACTCATACTGTTTCTGCTAGTGAAGCTGGTGATGTTCAAATGGTTGAAGCAGGAAATATGGTGATTTCCATCCAAGTACTGAAGAGTGTGACCAACAATTTTAGTGAAGACAACATACTAGGCCGGGGAGGCTTTGGCACTGTCTACAAAGGTGAATTGCATGATGGGACTAAAATTGCAGTCAAAAGAATGGAAAATGGAATCATCACTGGGAAGGGACTAGCTGAATTCAAATCCGAGATTGCTGTATTGACTAAGGTCAGACATAGGCATCTTGTTGGACTACTTGGGTACTGTCTTGATGGAAATGAGAAACTTCTAGTCTATGAATACATGCCTCAAGGAACATTAAGCAGTCATCTCTTCAACTGGGCAGAAGAAGGGCTTAAACCCATGGAATGGACAAAACGATTGACCATTGCATTAGATGTTGCTAGGGGTGTAGAGTACCTCCATAGTTTAGCACACCAGAGTTTCATTCACAGGGATCTGAAGCCTTCAAATATTCTTCTAGGGGATGATATGAGGGCTAAAGTTGCAGATTTTGGCCTTGTCAGACTTGCCCCAGAAGGGAAAGGTTCTATAGAGACAAGGATTGCTGGGACATTTGGATATCTGGCACCGGAATATGCAGGTATGCTCTTTGCTTACTTATATGTGTATGCTAAATTGCTGATAAAATACTTGTATGCAATTTGTGCCTAGATAAACTTGGTCTTAAGTCCAATACCTCTAAAGTATGAATTATGTGAACCTTAGGGGTTTGATCTCAGGATGCTTCCAGAGAATTAGCATTCATTTGACTGATAAGAGTCGGATGATAATAACTGTACTCCAAGAAACTTAGCCTGTTTTACATTAATCAACTTTTGTCCTATAATAGGTTTATAACAGTATATTTTAATATTTTGTCTTTTTTCTAAGTCAATTATACTTGTGCTCATGGTTAAGAACCTGCCATTCATACATTGTCATATATCAAATATCCTTATTGATTCTGTCATCCCCTTTAATGCTATGATTTCTTGGTTTCTAAAAGTGGATACCCATCCTACTGGAAACACTTTTTCCTGATGCATTTGCTTAATGCCACCAGTGTCTACTTTGGCCTAATCGCACACTTTAAACATGTTATTAATCTTATGACAGAAGATCCTCCTTTTTACTTTTCCGCTTGGTGGAAATTGGAATTCAAGGATAGCCAAATTCGTTATTTGCTCACTTTCCTGATCGTGATATGTGCATAGGAGTTTTTGGAATGTAGATACATCATTATCTAAACTTGTATTTTATCTTGTTTCGTCTGCCAGTAACAGGGAGAGTGACAACCAAGGTGGACGTGTTCAGTTTCGGTGTGATTTTAATGGAGCTCATCACAGGAAGAAAGGCTCTTGATGAAAGCCAGCCTGAGGAGAGAATGCATCTAGTAACATGGTTCCGAAGAATGCATCTGAATAAGGATACATTCAGAAAGGCAATTGACCCCGCAATTGACCTTAGTGAGGAAACACTCGCCAGTGTCAGCACTGTAGCTGAGCTAGCTGGACACTGTTCTGCAAGGGAGCCATATCAAAGGCCTGACATGGGCCATGCTGTCAATGTTCTTTCTTCTCTTGTGGAGCTCTGGAAACCATCCGACGAGTGCTCAGAGGACATATACGGCATTGATCTCGATATGTCATTACCCCAAGCGCTTAAAAAGTGGCAGGCTTATGAAGGTACAAGTCACATGGATTCATCCTCATCTTCTTACCTTCCAAGCTTGGATAACACTCAAACTAGCATACCTACTCGCCCCTACGGATTTGCCGAGTCATTTACATCTTCTGATGGTAGGTGAGAAGTTGACACTCTTTGATCTAGGTTTGGTTGTTCATTCTCTCCGTTTTCTTCCTCATCGCGAGTCACACATCATTTTTTTTTTGCTTTTAACTCTGATGCTTTTACCTTAACTCATTGTTAGTTTTGGCACTGTGGTGAATAGGGATTCTTTTTAAGATTTTTGTTTTCTTTTGACTCTTATTGTATGATGAATCTCTCTCTCTTTTCCATGTGAATACATTGTGGTTTGGTTGGAAATGAAAGGGATGCCATCAATTACCAGTAGATCATAGTACTTTTGTATTCTTGAATTGTTTTTTTGTTAGAAAAATTTGTAGGTGCAAGCTGTATTGAAATGTAAAGTGGCATTGAAGTGTGCAAATCATAATGTTAGCTTTTTGCTTATGATGCTTTTTCAAGCAAAATGTAAAGTTTTTTGTTATGATTACAGATCAACTGCTG

>M6-PGSC0003DMG400028801 M6 LRR-RLK

AACACATCCTAGAACTACTATACATTTTTCATTACTCCTACAAAGAACACCATTCTTTCTTCTTACTTCTGCTACTACATTTCTCTCTGCTATGAATTTCTAGCAGTTCTTGGTTCTTGCGCCATATTCAGCTAAAACAAGTAGGCACTGCCCATTTCAGTTGGCTAGTTCAATTTCAGTTTCACTGTCAAATTTGACAATGGTGTTGTTAATCATCTTCATAATTCATGTTTTCATGTTTGTAACAGCTTCGTCGGATCTTAGTTCCGACCGTGCTGCACTTTTAGCTTTTCGCTCCGCCGTGGGTGGCCGTACATTTCTCTGGAACACCACCAGTACCTCCCCATGCAACTGGGCTGGTGTACAATGTGAAAACAACCGTGTTACAGTTCTCCGGCTACCGGCGTCAGCTCTCTCCGGTACCCTTCCGGTGAACACAATTTCCAACCTCACTCGTCTTCGTACACTTAGTCTTCGTCTCAATCGCCTTTCGGGTCCTCTCCCTTCTGATCTCTCAAAATGTGTTGAGCTCCGTAACATTTATCTCCAGGGTAACTTTTTTACCGGAGCTATTTCTTCTTCTTTCTCTGGGCTTCACTCTTTAGTTCGGTTAAATCTTGCTGAAAACAATTTTTCTGGAGAAATTCCATCTGGGTTTAACAGTTTGACCCGATTGAGAACATTTCTACTTGAGAAAAACCAGTTTTCTGGGTTAATGCCTGAGCTAAAGTTTTTCCCAAATCTCGAACAGTTTAATGTGTCTTTCAATAGATTAAATGGGTCTATTCCGAAAAGTTTGGAGGTAATGCCAGTGAGTTCTTTTACTGGGAATTCTCTATGTGGAAAGCCTATTAATGTTTGCCCAGGAAGTAAGACTCAGCCTGCTATAGCTACTGATGGAATCGAAATTGGGAATTCAAATAACAAGAAGAAGAAGCTATCTGGTGGAGCTATTTCCGGTATTGTCATTGGGTCTGTTGCTGGATTTTTCATTCTTTTGTCGATTTTGTTTGTGTTAGGTAGGATGAAAACCGGTGATAAAACAAGGTCTCTTGATGTTGAGACAATTAAGTCACCGGAAACAGAAGTTCCCGGGGAGAAACAAATTGAGAAACCCGAAAATGGAGGTGTAAATAATGGGAATTCAGTTGCTGTAGCAGCACCTGCTGCTGCAGTATTGAATAGTGGTGAAGAGAATTGGGGAGAAAATGGGGTTAGGAAAAAGCTTGTGTTTTTTGGAGATTATTATAAAGCATTTGAATTGGAGGATTTGTTGAGAGCATCTGCTGAGGTTCTTGGAAATGGGACATTTGGAACAGCTTATAAGGCAGTTTTGGAGATTGGTACAATTGTTGCTGTTAAGAGATTGAAAGATGTGGCAATATCTGAAAGGGAGTGCAAGGAGAAAATTGAAACAGTTGGAGCAATGAATCATGAGAATTTGGTGCCTCTCAGAGCTTATTATTTCAGCAGGGAGGAAAAACTTCTTGTCTTTGACTATATGCCAATGGGAAGCTTGTCTGCTCTTCTTCATGGTAAGTTATAGTTACACTCTTTTACTGTCGATTCATCTTGTTCGGACTCTCTAAAAATGTTATCGCATCTGTGCCAAATCCTCGAAAAATGCATTACTTTGGAGGATTCAACACGCACTCAGCGGTATGGATCCTTTAACAATACACTACCTTTTTGGAGGATCTCACATGCACTTAGCAGCATGGGTCCTCAAAATATGCACTACCTTTGGAGGATCCAACACACACTCAGTGGCATGGATTCTCAAAAAAAAAAAAATCACTATTTTTGGAGGGTCGACATGTATTGGTCGATATTCTTAATGAGTTCAAGATACACTATATTTTGTTCCCTTATTCAGTAGATACACTATATTTTGTTCCAGATTTTGACATTTTGTTAGAGTAATTAGTTTGACTTGTTGCTAACAGTGTAACATGCAATGTCTGTAGTATTTGGCCCCTCAATCCAATGAAAATGTGATCCTAGAATTAAGCATTAATTTGTTATCATTACCTTAGTCTTCATTATGTAAAGCATGCTTCTAATATAAGAAAGTATGTTAAATCTAAAGTTGTTTGTAATTAAGAGCAATAAGTAGTACTCCATTTCATTTAGTGCTTTCATTATAGTAAAAGGGTCCTTTGTTCTTTATAGATACCCTGTCCATATACAGGGGAAAGGTGAACAAGCACACATATAATCTTTCACTTGGCCCTACAGTAGTAGGGTCTATCATTTTATGTTCTTTTTCTATCTAGAGCATCTGTATAGGACTAGAATTTTGAATCTCCGAAACTAATTATTCAACTGGTGATTAGTTTCCTGGCATGTGAATCTTGCATGAACATGTACATTTTTCGCCTTTGGTATACTTGAAACTTGTTAAGTGACTAAGAAGCTGTAATGAGCTGTATGCTGCAACTCTACTTTCTATTCACTACAATTTTTTTAATTTACTAGGAAGCAAGGGGGCTGGCAGAACACCATTGAATTGGGAAATCAGGTCCAGTATTGCCCTCGGAATTGCACGTGGCATCAAATACCTTCACTCTCAAGGTCCTGATGTCTCCCATGGGAATATCAAATCATCCAATGTTCTTCTCACCAAATCATACGAAGCACGTGTCTCAGATTTTGGCCTGGCTAACTTGGTTGGATCGCCATCCTCCCCTACTCGAGTTGTTGGATATCGTGCTCCAGAGGTAACTGATCCTCGCAAGGTTTCCCAAAAGGCTGATGTCTATAGCTTTGGAGTATTACTCTTGGAGCTTCTTACTGGAAAAGCACCTAGTCATGCCCTCTTAAATGAAGAAGGTGTTGATTTGCCAAGGTGGGTTCAATCTGTCGTTCGTGAAGAATGGCCTTCTGAAGTTTTTGATATTGAGCTCCTCAGGTACCAGACTGCTGAAGAAGAGATGGTCCAACTCTTGCAGCTAGCAATAAATTGTACAGTTCAATATCCTGATAAGCGACCTTCAATGGCTGAGATCTCTAAACAGATTGAAGAGCTTCGACACCCAACCTCACATGATAAACAAGAACTCTCTGATCTTGCCAATGCAACAGAGTGAAGTGGTTATTATTAGTCCCCCTCGAGATGTGCAGTAATTAAGCTTAAACTCAGATTCATCACCAATCTTTTTTTAAGCATATTAAACTTGTTGTTTGTTGGGGGAGGGGTGTCTCATCTTCTTTCTGTTTGAATTAACTGATCATGGCCCCTTTTTCTCTATGTTTCTGTAGTAGTCTAGTGGTTACCCCTTTCATTTTCCTTTAGATGAGGGGCCAAAGCCAATATCTGTACCATTTATTCTTGTTGGATGTATAATGCTTTCTGTAATTATTACTGCTTCTGCTTTTACTTCTGCTTTGTCAGTTGCAACTGTCGATGGCTATTCTGTGTGTATATGAGGCATGCATGCATGTGGTTCTTGATATACTGAAGCCAGCAGGGAGATTGCATTGAGAGTTTTCATAGATGCTGAGTTGGCCAAATATTG

>M6-PGSC0003DMG400028845 M6 LRR-RLK

TGTGTGGCTTTGGATTCTTGTAGCTGAATAGTGGTTTTTCTTTAGAATGGCAAAAGCATCTGACCCTTTTCTTCTTTTAGTGTCTTTTTCTCACTCAAAAAAGAGTCTTTCTGTGTAATTTTGTGTTGTTGGTGTGGGAGAGATGAGGTTGTTGTTGCTCCTGGCAGTAGTACTGTTGGTGGCCATTTTGGCGAATTTTTCAAGCTTTGTTAATGGTGATACAGACCCATCTGATGGTTAGCCCCTTTTGATTTATTCATCATTCTCTTTATTTATTTATTTGTTTTTTAGTATGTGAAGTAGTAAGTACAAGTTCCAGTTTTCTTGTTCTCAAGGAAAAATTTAGCTGAGAGTGGGCTTGAGTCTTGACAACTCATGAAATGAACAAACCCAACTGAAAAAACTTGTTTCTTTGAAGCCCCATGATATTGAAAAGAAAATAGCATTTGAAAACACTAGGTCTAGGCTTGGTAGATAGAGTTATGTGGTACATGTTGTTGGTGGGAGGTGGCAGGTATCCTTGGAATTAGTTAGTTGCACACAAGCTGGCTCGGACACTGTGGTTATCATAAAAAAGATCTTTGAAAATGATGGATATAATAAGTCTTGCTTGGGCTGCATCAATGGTAGTCTTCACTTTCGAGTAAGGGAAATACTCATTACTTATGGCTGAACAACCCCTTAGTCTTGTCGACTAGATTGAGTCATAGTTATTGTAATTGTTGTATTTACTGTTGCCGATGCATTTAATAGTTACAACAACATATCCAATGAAATCCTGTAGCTGGGGTTTGGGAGGGTAGAGTGTATGCAGACCTTACCATACCTCGTGGAGGTAGAGGGGAGACTCTTTCGGGAAGATGCACTTAATAGCTAATAGAAAACGTCAACCTTCGCACAATAATTTTCTTTCATTAAAAATTGAACAAGTCTGGGTAAAAGGTAAATCCTTACAGAAACTTGTGGTTTGTGAATGTGATTGGTTTTAGTTACTATTGCCAGCCTTATATAAGCAATTACTGGATAATGTTTTCTACTTTATGTGGTAAGGGGTGACAAAAACCAATGAGTTTGAAGTTGATCAAAAACTTAATAAAGCTATATAATTTACTTTTGGACTCCTGATGATCCAAAGAGTTTCACAGTCATGGTTTGGTTAAGTTGCTCGCTACTTTGTACCATGCTGGTCAAGTTGCCGTAGCTGCACATTCTTTTAACAGATTGTCCTATGTAAAATCTGTCTCTTGAGCCGTTCTCTGTCCATATATAACTATTTCTTTATCATTAAAATTTCTTACACAATCACATATGTCTTGATTTTAAAATAATCTAGTTGACTTCTGTTTGGTTTTCAGCATCTGTTCTGAGAGTCCTGTATACCACCCTAAACTCACCAGGGCAGCTGACAAAATGGAATGCCAATGGCGGTGATCCTTGTGGAGAGTCCTGGACAGGCATTACCTGCTCGGGCAATAAAGTTACTGAAATGTGGGGTTCGGTCTTCCTTTTTGTCTTCATCTCCATGTTGTTTCAACTTTTTGGAATTATTGACTAGGTAAACTGATGATTAGTTTCTTTTCATTTTCCTTCAGCAAGATGTCAGGTCTTGGGCTCTCAGGTTCATTGGGGTCTCAGCTAGATAGTCTAAAATCTGTGACCAATTTGTAAGATAAATGGCCTTCATTTGTTCCTTATTAATTGGATTTATGTTGAATTCCTGAATGTGGTCATCTTTGTGAATGTTGCAGTGACATAAGCAATAACAATATGGGGAGCCAGTTACCTTACCAGCTTCCTCCAAATGTGCAAAGATTGTATGTTTGTTTATTTGCATATCTTCCTCCTTGCAGGTTCTATTTTCTTCCTCTTATCTTTCTTAGAACTTGCCTTGCAGAAACCTCGCTTCTAGTGGTTTTAGTGGTGGCCTCCCATATTCCATTTCACAGATGACCTCCCTCCAGTACTTGTAAGTGAATGCTTAAATTTAATATTTGCTTAAATACTTGGTCCTCTTCATTCTCTTTGGGGCTCCTTATGGTTCTTTGCAATGTTTATGCCTGGTTGTAGAAATGTCAGTCATAATCAAATTCAAGGATCAGTGGTTGTCTCGTTTGACTCGCTCTCTGCACTCAACACATTGTAAGGCCGATTCATCTCCTTCCATTACACCAGAGTTTTCATATTGCCTAATTATGATCTGAATGAGAATAGTTTGCTGTTCTCAAATGTTATGATGAAAGTACACTTATTCTATTATTTAAGTTGTCTCGATTTGAGAAAGAGGTTAAACACACTTGTTCCATTTCCCACATTACTGTGCAGAAGTGAGTCATGTCTTTGGTCTGCTGCCAAAGTATTGAATTATGCCAATTGAGCTTAGTTGCTACTTAGATTGTACTTATTCTTAAGCAAGTATTAAAAGCCTCAAAGGTGCGCTTAAGCCCTAAAACGAGGTTTAAAACGTGTTGAGCGCTTTGCCTTGCTTAATGTGTGCTTTAGTGTCATCATCAAGGTCCTAAGGCATACTTTCCCTTGACTAAACAAGTGATATTTCACTTTATCGTAAAAAAATCCAATTTCTTTGTTCATATATTTGTCATGCATGCTTATAATTATTAGTCTTGGACTAAACATATATATACTTGTATTTTTTTCTCCCTTTGCACCTTTTTTTCATTAAAGCTTACACTTTATTTGCGCTTAAAGCCCCAATGGACCTTAGAACTTTTTTGTGCTTTTTGCTTTTGATAACGCTGGTACCTTATCATATTTTCTTGTTTAAAACATAATAGTTTGAAATTTATCATTCTCGGTCACAATTTTGATTGATCTTTTTAATGCTTCAGGGATTTCTCTTTCAATTCTATGACGGGTAATCTTCCTCAAAGCTTCCAGTCACTAACTAGTATGAACAAAATGTAAGAAGAGTTTTCATACTTCATCTATCTATTTGCAAACTTCCATCAGATATAGAAGTTTTGTTGTTATAATATGCAGGTATTTGCAGAACAACCAGTTTAGTGGAACTATTGATGTCCTGGCCAATCTTCCCCTTGGAAATCTGTGAGTTGTATGCCACATGTGTAGATATTATGTCTTATGTCTACATCTCACTTCAGTTTCAGTTAAATCCATTGGTTGAAAACCATGTTGATTCTTCTATTCTTTAGGAATGTTGAAAACAACCAATTTACTGGCTGGATTCCAGATCAGCTGAAAGGGATAGTGCAGTGAGTGTCTTAGATTGCAATAATTTTCATTCTTCTTTGCAAATTCTTTGGGGAGTTCTTTCTAATTGTTTTCTGCATTTATGAATCTTTCATAACAGATCAAATGGTAATTCATGGATCTCCGGACCTGCACCCCCACCTCCACCTGGCACACCTCCTGCAAACAATCCCAATGGAGATCACAAATCCAGAGACAACAGCAGCAATTCGGGCAGTAGGAAATCAGGTATAGGGGGTGGAGGTGTTGCAGGAATAGTGATATCTATTCTAGTTGTTGGAGCAATAGTAGCATTCTTTATTATCAAGAAAAGATCAAGAAGGTCGTCCACAGACATAGAAAAACTTGATAAGCAGCCACTTGCTCCTCTTACTTCACAGAAAGTAGAAGGTATTTGGCTTTCATAATTCTTGATATTGTTAGAAATGTCTTGAATTCTCTCTCAACTCTTTGGATATTTCTTAACGAATAATTATAATCAATTGCGGTGTTACTATTTACTCAGTTGCTTGGGAATATGGAAAATGATGTCTAACCTCTATCCTTACCTTGCTAATGAGTGGTCATAAACCATGCAAGTAGTATCAAAATTTTGCAAGGGGTTGCCATTGATACAGAAGGAAGCAGCAAAGCTGGAAAATATTTATCTTCGAAGTGGAAAAACTGTATGGATTTGAAAGATCCTAAGTTCTGAACAAACAAAAGTTTCATACTTTATTGGTAGTCCCGTAGGTATAATCTTTAGAAATTCAAACTGATCTTTTGGAGGAGTTCCTAAATTACGTGGTGCTTTTAAAGCTTATGATTATTTTTTGACAAATTAACATCAGCTACCCTTGATGGTTTAACTTCCTGGGAGACAAAAACTCTTGAGAATATAGTTGACCCGAAGAACTCTTGCTTCAGACAAATATGAAGTTGCTTTCTCCTTTTTAATCCATCAAAAGCCCACAAAACAAACCGTCTCTTACAATACGAGTTCCTTCTCATTTGCAACTTCATCAGTAATTTTCCAGTTCATGCTTATAACTTTCATGCCTCATTTCATTATCGACTTCAATGATGTCATGGAATCCAAATTGGAACTTGTATTTGCTTATGTCTAACTTTTTCATTGTTGTGTTGCTGTTTTTTTAAAAAGGTAACATCACTGTTATGTTTTGTTTGTTCCCGGAATTCCTACTCATTGTCTGTTTTTGGTTGCAGAGTTGAAACCTTATCAAACCTTCTCCACAGTAAGCACGGGCACATTTGAAACTCCCACTGCTGTAACTCTAAGACCTCCACCTATTGATCGCCTTAAGTCCTTTGATGAGGTTGACATTTCACCGAAACCCATTGTTCCTCCCAAGAAAATTAATACAGCTCAAATCAGGGCTACACAGTTTTCAATTGCAGACCTACAGATTGCTACTGATAGCTTCAATGTTGAAAACCTTATTGACGAGGGATCAATTGGGCGTGTTTATTGCGCTCAGTTTGATGATGGCAAAGTAATACTTCTCTCTATGACTACCCATTAAAAAAACTTTTTTTCTTTCTTATGACTATATAAGATGATTTAGTGATGCTAACACATGTTTAGGCAAAAATGTATTATCAGAGAAACTACATACTTATTGTGTTCCTTTCTGTTATTTGAAATGTTGTAGGTTCTTGCTGTCAAGAAAATCAATTCTTCTGCCCTCCAAAATCATGAAGATTTTCTCTATATAGTTTCTGAGATATCACAGTTGCACCATCCAAATATAATTGAGCTTGTTGGTTATTGTTCAGAGCATGGACAGTACTTGCTAGTTCATGAATTCCATAAAAATGGTTCTCTGCATGATTTCCTGCACCTGTCAGATGAAGACAGCAAGCCACTGGCATGGAATAGCCGAGTCAAGATTGCACTTGGTGCAGCACGAGCACTAGAGTGAGTTTTTGCAATCTCAGAATTATTTTATCATATTTTATATTCAATTGGTTACAGTTATGCGGGACTATTTCCTTATTGATTCATTCCAGAAAGGATGACACGTTTCTGTATTTGGAAACAATTCACTTTATATTTCCCATTTTTACCATTAGTGAGAAGTTTTATAGCAACACAAATGTTATGGTATGTTTAAGGCCACAAGTTTCAAAAGTTTTATAGTCAGACAAATGCTATGCCATGTTTAAGATCGATCACAAGTTTCAAATTTTTTTATCCCTTTCTTAGATTCAATGCCGATTGGAACTATGTCACATAAATTGGAAGGGGAAGTAAAATTCATGGAGCATGCAGCAATGCTATGTTTTTCTCTTTATTTATTGATGGCCAATTACTAGTAGTTTACATGAATCTTCTGCTTATCTATACCGGAGATTCATAGAGTAGATCTTCTGGGTAGAAGAAAAGGGGATTCTGTGAAAGTCAATACCAAGCACGTAGTCGTGACTCGTGATGAACTTTACCAGAGGATCCATAGTAATATTGCATTTAGAGGTCATACTTGTTGGACTAAAAAGACCACTATAGAGTTTAGTCTTTTTTAAACATTGATGACCAGTAAGTTAACATAAAGAAGAGTTCATTTATTCCTTTTAGTTAGAGCCTAGTATCCTAAATATGCCGGTCTTGCCAATATAAAGCTAATCAAAACTTCAATGCTCCCATCATTTCGTTTTATGTGGTGGTGTTTGAGTGGGGACAGAGTTCAAAAAGAAAGAAAGACTTCTGGAACTTGTGGTCTTAACCTTGCCATGTCATTTGTGTGGCTATAAAAGCATGTCATTAGGGCTAACAATGAAAATGTGAAGTTAAAATTGTTTCCAAATATAGGAAGGCGTCAATCTTCTTGAAAAGGCTTAAAAGGAAATCCGCCATATAAAATGATACAAAGGGTGTAACAAATTTCCAATTTATGTGGTTTTCATATCTGTTCATTTCCTGTTTTCCATACACAAATTTCTTCACTAGCTAATATAAGCATCCTTTGCAGGTATCTGCATGAAGTTCATTCACCATCATTGGTTCATAAAAACATTAAATCGGCAAATATTTTACTTGATCTGGAACTCAATCCTCATCTATCAGACTCTGGATTAGCAAGCCTTATTAATGATGCAGATCAGGTGGCTTTAAGCTCCTTACTTTATGCATTGTTGCAAAGTGTTTCGGTGGATTCGATATAGAAGCATGCATGCACATAAGTACACATGTATTATATAGGCATGTTTTGACATACTATTGGCTGTGTATGTGTCCTTTTCAAGGAGATAAGAAATATCAAAAAACCACATGGTTGAAACGTCATTTGTTTCAGGAGAAGCTTTGTTTTGTGCACAAACCTTTGAAACATGAAATCATTTACAACAGTTTCTGGGGTGCATAAATTGGATTCCAAGAGAAGTTGTCAAGTAAAAAACAGAAAATTTTATCCAAATCTAGAGCATGGAATTATCCAGTATATTCTATAGAAGTAACACTACCATATAAGTCATTTTAATGAACTAGTCTCTCTATGGTCCCTGTTGCTTTATCTCTTGCTCTTCTCTACTTACTCAATTGGTAAAATTGTAGCCGTTTCTTCTTCTTCTTAAAAGATTCACATAACTATCATCTTCTCAAGATGTTTGAAAGTGATCATCTTTGCAGGCACTAAACCATAATACAGGGTCTGGATATGGTGCACCTGAGGTAGCTATGTATGGGCAATGCACCATAAAAAGTGATGTCTACAGCTTTGGAGTTGTTATGTTGGAACTTCTTACCGGACGGACACCTTTTGACAGGTAAAGTTACTTAGGCATCATGATATTGTGTCAAATTACTAAGAAACAAACAGGAAGAAATTCAGAAAGTCTTAACTTTCTTGATCAATATGTATGCTCTTGATGCAGTGAGAGACCAAGATCTGAACAATCTTTAGTTAGATGGGCAACACCTCAACTCCATGATATTGATGCCTTAGCTAAGATGGTTGATCCAGCACTCAAAGGGCTCTATCCTGTTAAATCCATTTCCCGCTTTGCTGATGTAATTGCTCTTTGTGTCCAGGTAATATTTTGTATAGACAGTGTGCCAAATTCATAAATCTGCTGTGTTAATCAAGAACCTAGTTCTGTTAGTTCAGTCAACACTATCTCATTGTCCTGTATGCTATTGTTGACAGCCTGAGCCTGAGTTCCGGCCACCTATGTCAGAAGTGGTTGAAGCCTTAGTTCGGCTAGTTCAACGAGCAAATATGAGCAAGAGGACATATAGTATTGATCGGGGTTCAAGGGGTGAAACAGATGCTCAAGACAACCAACCTTAGTTCCGATGTAATGTAGCCTTTTCTACCAGAATGTCTCTCTTTATAGTTTGTTGTTTATGGGATCTATCAGCTTATCTGTTCAGCAAGATCTTGTTGGTTTCATTGCGTTGTACACACTGCTAGATACTAAAATCTGTTCATAGTGCATATTCAATTTTCATTTTTCCCTGTATCTGCT

>M6-PGSC0003DMG400029146 M6 LRR-RLK

GGAAAATGAAGGATCATTCATATTAGAAAAAAGTTGCTCAAGCTTTATTTAGTTCCTCTTTTTCCATGCAACTTATGATATAGAACTTTCATATCTAGCTAGAGAGAAAAAATGGCCAGCATAGGATCAAAATCGATCATGTGTTTGATCGTGTTCGTGCTATTTTACATGCAGCAAGTCAAAAGTATCTCTGAGCTGGAGCTACTTTTATCGATGAAAACATCGATGAAGGATCCATTAGGATCTCTTCATGATTGGATTCCGCGGTCTCAATCTTTTTGCCATTGGAATGGTGTTGTTTGTGATGACTTGTTGCATGTTGCAAAAATTGAGCTATCAGGGAAGAACTTATCTGGTAAATTATCCGAGACGATTTTCAACTTTCCACACGTTGAATCGATTGATCTCTCGAATAATCAACTCTATGGAGAAATTCCAAGCAATATCTCAACTTGTTTGGTGTTGAGATTTCTCAATCTTAGTAACAATAACTTCACAGGTCCACTTCCTCAAGGCTCGAGAATCCCACTTGTCGAGACATTGGATCTCTCAAACAACATGATTTCGGGGAAAATCCCCGAAAATATTGGTTTATTCTCAAGGCTCAAAGTTCTTGATTTTGGTGGAAATGTGTTGGTTGGAAGCATTCCAAAGTCTATTGCAAATATTTCTAGTTTGGAGTTTTTGACTCTTGCTTCAAACCAACTAATCGGAGAAATCCCTCGAGAACTAGGTCTTTTGAAGAACTTGAAGCTTATTTATTTGGGATACAACAATTTCTCTGGTGGAATACCGGAGGAGATTGGTGAATTGTCTTCTTTGTATCATCTCGATCTTGTGTACAACAATCTCACAGGTGAAATCCCTTCATCTTTAGGTAATCTCACCAATCTTGAATACCTTTTTCTTTACATAAACAAGCTCACTGGCCCAATTCCAAGATCTCTATTTAATCTCAAGAAAATCATCTCGCTTGATTTAAGTGATAATTTCTTGTCTGGTGAAATTCCTGAGCTTATTTCCCAATTACAAAATTTGGAAGTTCTACAATTGTTTGCGAACAATTTCACTGGTAGAATTCCAAATACTTTATCTACCTTACCTCGACTTCAAGTTCTTCAACTATGGTCTAATAAATTATCTGGTGAGATTCCCAAGGACCTTGGGAAACACAACAATCTCACAATTCTAGACCTTTCCACCAATAACCTCACGGGTAAAATACCCGAAACCATATGTTACTCCAACCATCTTTTCAAACTCATACTCTTTTCAAATTCACTTCATGGCGAAATCCCTGTAAGCTTGAGACATTGCAAAAGCTTACAAAGGGTCCGCCTTCAAAACAACCACCTCACCGGTGAATTGTCCTCAGAATTCACAAAGCTCCCACTTGTTTACTTCCTTGATATCTCAGGCAACAATCTATTTGGCTCCATTAGCGAAAGACGATGGGATATGCCATCACTTCAAATGCTAAATTTGGCTAGGAACAAGTTTTTTGGTACCTTGCCTGATTCTTTTGGTAGCAAGAAACTCGAAAACTTGGACTTATCGGAAAATGATTTTAACAGTACTATTCCTAAGAATTTCGGAGAGTTATCAGAGTTAATGGAGCTCAAACTAAGAAGCAACAAGCTTTCAGGCGAAATCCCCAATGAATTGTCTTCATGCAAGAAGATTGTGAGTCTTGATCTGAGCCATAATCGATTTTCAGGTCAAATTCCAACTAGTCTCTCCGAAATGCAAGTTCTTAGCCTTCTTGATTTGTCAATGAATGAATTATCAGGTGAAATCCCGCCTAATTTGGGGAAAGTTGAATCACTTGTCTTGGTTAACATCTCTCACAACCATTTCAGTGGATATTTACCATCCACCGGAGCTTTCCTAGCCATAAATTCAAGTGCGGTGGTTGGAAACCAACTATGTGCCCGTGGGGACGACGACATTACAAGCGGCTTAACGCCTTGTAAATCGCTCAAGAAAAGCTCAATTTGGTGGTTCTTCTTGACATTTCTTCTTGGAATACTAGTTTTGCTAGTATTCTCGGCTTTAGTTGTTGTATTTATTCAAAGAAGAAGAGAATTGAAGCTAAAGAAAGTGGAGAGTACTACTCAAGATGGTAATAATTGGGAAATTCAATTCTTCGATTCAAAAGCCTCAAAATCCATCACCCTAGATGACATCTTAGGGATTGGAGTTTCGTACAAAGGGTTTTATTCTGAAATAAGTAACATGCAAGTATTCGTGAAGAAGTTGAATGTGAATATTCCAACAAGTTTTTGGACAAACATCCAAGAACTAGGAAATATTCGACATCCAAATGTTGTGAAGATATTGGCTGCATGCAAATCGGAGAAAGGAGGTATTTTGGTTTATGAGTATGTAAAAGGGAAAGATTTGAGTGAAGTGATTCGAGTCATGAGTTGGGAACGTAGACAAAAAATTGCTATTGGCATATCTAGAGCATTGAAATATTTGCATTGTTCTTGTTCACAAAGCATTTTCATCGGTGACCTTTCTACCCGGAAAGTTATCATTGACGGAAAAGATGAACCTCGCCTGAGATTGAGTCTTCCCACTACGTCCTACGTTGGTCCAGGTAAATTTACCATTTTACCCTTGTCATCTACGGTTACTTCTCTTGTTTTGACATTTCTTAAGAGTTTAAGTAATATACAACCATAAGCTAAAACATCATATATTCTATCAGTTAACATGCATGTCTTATTTGCAAGATTATAATTAAGCTTACCTACATACATCTTTCAATTGACTAGATCGTGTAAATTAATTTATATTGATGATGCATATAACTTAATTAACTCGTTAGTTTAATAATTAACAACCTTTGTTATAACTTAAACAAACCACATATAGTTTCTAAAGAGTGTAACGTAGCTATGATGACTTTAATGAAAATTGATTTTTTCAGCTTTAGCATTGATGTGATAGGAATGTCATGTTATGTTGAAAATTATTTGATCTAACTAAAAGAAATTTTGTATTTTTCTAATTATTAGAATAGAAATAATTCTTAGAATTTACGTACTTGGTCTTTATTTGGTTTCTATTTTAACTTTAGTTCTCATAGTCTCTAGGAGTAATGATAAGGTCAAGGAATTAAGCAAACAAATGTCATGGAGGCAATCATGAATGTTTTTTTCTCTTAAAATAAATGTAGAGAAATATAAATCTTATCATTCCTTTCTTTACGTAGAAAACAATAAATCTAGTCTAGTGTAACAAAAGTCATGACCTTTTATGCCATGCAAGTACAATACTGTTACTTAATTACGAGTTACAACTTTTCTCATAAATTGTAGAATACAATGGAATATCGGAGAAGAGCGATATACATGGATTTGGGCTTGTTTTGATTGAGTTATTGACTGGAAAAAATTGCGATGATGCTGAATTTGGGAAGCGTGAGAGTATAGTTGATTGGGCGAGATATTGTTACTCAGAATGTCATTTGGAGACATGGATTGAACCACTATTGAAGAGTGATGCAGTGAATAATCAGAATAAAATGGTGGAAATGATGAATGTGGCTCTTCAATGTACTGCTAGTGAGCCAGCAGCAAGGCCTTGTGCTAGTGATGTTGCAAAGACATTGGACTCTTTTGTTAGATCAAATTCTTGTGGTTTAGGGTTGAAAAGGTGTGGTAGTGTTTAGATTTTGGGATTTCTAACTTCAAGTCCAAGATTTTTGTTTTTATGTTAATGTTACTTTTTTTTTTTGGTTTACTTTGTATTGGGCATAAATTAATGAGTGTTGCTCAATGTTTTGGATGTTATTTGAGTGTAAAAATGTAGATATGAAACTAAATGCTAATATATAT

>M6-PGSC0003DMG400029442 M6 LRR-RLK

TTCCAACACTCACTAAAAGCTTACTCTCTAAGCTCTCTTCTTTCATTTTTTTTTCTATCTATTTTCTCTATGCATCATGGACAACATGAACACCAACTCTGCTATAACATTGCTCATTGTCTTGTTTTTTTTCTTGATTCTTGCATCCGTACACACGGTATCTTCCTCCATCAAAACTGATGCTGAATCACTCTTGCTATTCAAGAATATGATACAAAAAGACCCTTCTGGTGTTTTATCAGGATGGCAACTTAAAAACAATCCATGTTCATGGAATGGTGTAACTTGCAATAGTCTTGGTAGAGTAACAAATCTTGATCTTCAACAATCTGAACTTGTTGGAGAGGTTTCTTTTTCTCCTTTCAATTCTCTAGATATGTTAACTGTCCTTAATCTTTCATCGAATTCGTTTTATGTAAATGCATCAACCTCTTTAACTCAGTTACCTTATAGTTTGAAACAACTTGAACTTTCATTCACTGGCTTAGCAGGCTATGTACCTGAGAATTTCTTTGCAAAATGTCCAAATCTTGAATATGTTAGTCTTTCTTTCAATAACATAACAGGTTCTTTGCCTCAGAATTTCTTGTTGCATACTGATAAGTTACAGTATCTTGCAATGGACTATAATAATCTTACAGGATCAATTTCTGATATCATGATTGAGAGTTGTAATTCATTGTTACGACTCGATTTGTCAGGGAATCAGATAATGGATTCAATTCCTTCTGCTTTATCTAATTGTACAACTCTTCAAGAATTGGTTTTGTCTGAGAATTTCTTCTCTGGTTCAATTCCAACTTCTTTTGGTGAGCTTATAAGTTTGCAAAGATTGGATCTTTCTAAGAATCATTTATCTGGTTGGATCCCATCTGAATTAGGAAACTCATGTAGTTCACTTGTGGAACTCAAGTTTTCTAATAACAATATCACTGGTTCAATCCCGAATTCATTCTCATCATGTTCTTCTCTTCAAAATCTTGATTTGTCGAATAATAATCTAACCGGTCCATTCCCTGATTCCATCCTACAGAATTTAGCTTCTTTAGAGTCTTTGCAAATGAGTAGTAACAAGATTTCTGGTTCATTTCCTGCTTCTCTTTCATATTGCAAGAAACTCCGCGTTGTAGACTTTAGTTCCAACATGATCAATGGGATGATTCCACCAGATTTATGCTCTGGAGCTTCATCATTGGAAGAGCTGAGAGCACCTGATAACTCACTTTATGGTCCAATTCCATCTCAATTGTCTCAATGTTCACAGCTCAAGACTATTGATTTTAGCTTGAACTATTTGAACGGTTCGATTCCATCAGAGCTAGGAAAACTTGAGAAACTTGAGCAGCTTATTGCTTGGTACAATAGTTTGGAAGGGAGCATACCAGCAGAGTTGGGAAAATGCAGTAACTTGAAGAATCTTATACTTAATAACAACTACTTGAGTGGGAAAATTCCAGTTGAATTGTTTAATTGTGGTAATCTTGAGTGGATTGCTCTCACAAGTAATGGACTTAGTGGTGAGATTCCTAAAGAATTTGGCCATCTTTCAAGGCTTGCTGTTTTGCAACTAGCAAATAATAGCTTGAGTGGTCAGATTCCAAGTGAATTAGTCAATTGTAGTAGCTTGGTTTGGTTGGATTTGAGTAGCAATAGGTTAACCGGGGAGATCCCACCGCGACTTGGTAGGCAGCAAGGGGCTAAAGCATTGAGTGGGATTCTATCAGGTAACACATTGGTGTTTGTCCGAAATGTAGGTAATTCGTGCAGAGGCGTTGGAGGATTGCTTGAGTTCTATGGGATACATCCTGAAAGGCTTTTACAGGTTCCATCATTAAAGAGTTGTGATTTCACTAGGCTGTATTCTGGTCCAGTTCTTAGTGCTTTTACTCGGTATCAGACTATCGAGTACCTTGATCTTTCGTACAATGAGCTTCGTGGGAAAATCCCTGATGAATTTGGTGATATGATAGCTTTGCAAGTTCTTGTTATATCACATAATCATTTGTCAGGAGAGATTCCTTCATCACTTGGAGGGCTAAAGAATTTGGGAGTGTTTGATGCATCACATAACAGATTGCAGGGTCAAATTCCTGATTCGTTTTCACTTCTTTCTTTCTTGGTGCAGATTGACTTGTCTAATAATGAATTGACAGGACAAATTCCACAAAGGGGTCAGCTTAGTACACTTCCTGCAAGCCAATATGCAAACAATCCAGGACTCTGTGGGGTTCCATTGTCTGAATGCCAGTATAACTCACCTGCCACGAATACTGGAGACGGAGGAGGAGGAAAAAGAAGTTCAACTGCATCATTGGCTAATAGCATTGTTCTTGGAGTTCTCATTTCCATTGCCTCAGTATGCATTTTGATTGTTTGGGCTATTGCTATGCGTGCAAGGCGTAGAGAGGCTGAAGGAGTGAAAATGCTTAGTAGTTTGAGTACTAACTATGCAGCTTCATCATGGAAGATTGACAAAGAGAAGGAGCCATTAAGCATCAATGTAGCAACTTTCCAAAGACAACTTAGGAAGCTGAAGTTCTCACAACTTATTGAGGCCACAAATGGATTCTCAGCTGCAAGCCTCATCGGTTCTGGAGGATTCGGAGAAGTCTTCAAGGCAACATTGAAGGATGGATCAAGTGTTGCAATCAAGAAACTCATCAGGTTAAGTTGCCAAGGCGACCGAGAGTTCATGGCTGAAATGGAGACATTTGGAAAGATCAAGCACAAGAACCTTGTACCTCTCTTAGGATACTGCAAGGTTGGTGAGGAAAGACTTCTAGTTTACGAGTTCATGGAATATGGCAGCCTAGAAGAAATGCTTCACGGGAAGACAAGAACGCCTGATAGAAGGATCCTAACATGGGAGGAAAGGAAAAAAATAGCCAGAGGAGCAGCCAAAGGACTCTGCTTCCTTCACCACAATTGCATCCCACACATCATACACCGCGACATGAAGTCAAGCAACGTGCTATTAGACAATGAAATGGACGCCAGGGTGTCTGATTTTGGCATGGCAAGGCTGATAAGCGCACTGGACACACACCTCAGTGTGTCCACACTAGCAGGAACACCAGGATATGTACCACCTGAGTACTATCAAAGTTTTAGGTGCACAGCAAAGGGAGACGTGTATTCGTTTGGAGTCGTGCTTCTTGAACTCTTGACAGGTAAAAGGCCAACAGATAAGGAAGATTTTGGTGACACAAACTTGGTTGGATGGGTGAAAATGAAAGTAAGAGAAGGGAAATCAATGGAAGTAATTGATCAAGAATTGTTATCAGTGACTAAAGGAAATGATGAAGCAGAAGTTTTAGAAGTTAAAGAGATGGTAAGGTACTTAGAGATAACAATGCAATGTGTAGAGGATTTTGCATCAAAAAGGCCTAATATGTTACAAGTTGTGGCTATGTTGAGAGAACTAATGCCTGGAAGTAGTTCAAGCAACAGTGGCTAAACAGCAAACCAGAAACAAAAAAAAGGAAAAAAAGAAGAAAAAAACCAAAAGAATGTAAAATTCTTTCTTGTTTTGTTTGTGCAAGTATTCATCTGCATTTGTATTTAGTCAACAACCTCTCTTATTTTTTTGCAATATTATTCCCATGTTTTAAGAATCTTGTGTATGGATATTGATAACAT

>M6-PGSC0003DMG400029644 M6 LRR-RLK

AAAAAACAGAGCAAAACCCAAGAAACAAAAAAGAAGGGATTTTTACTTAGTTTTTTGATATGCAATTGCAAACCCAAGAAGAAATTTTTGGAATTTGAGGCAAAGATGTCTTCTTCTTCTTCAAATCCCAACAAAAAATACCATTTTTTTCTTTTCTTGGTTACCCTTTTCTTCACTTCTTCTTCTTCTTCTGCTGCTGTTACTGATGTTGTTGCACCATTTTCTAATTGGGACATTCTTGATGATACCCCATGTAAATGGTCTTTCATAAAATGCAACTTTCAAGGTTTTATTACAGAAATCAATATTCAGTCTATTCATCTTGAACTTCCTTTGCCTACAAATCTTTCTTCATATAAGTACCTGAAGAAGCTTGTTATTTCTGATGCTAACATCACTGGTGCAATACCCTTTAACATTGGTGACTGTTCGTCACTTGTTACTATTGATTTGAGCTCAAATGGTCTTGTGGGTACCATTCCTTTAAGCATTGGAACCCTTGTAAACCTTCAAGATTTGATTTTGAACTCAAATCAGCTTACTGGGAGAATCCCAGTTGAGATTGGGAACTGTAGAAACTTGAAAAATCTTGTTCTTTTTGATAACAGGCTTAGTGGTGGTTTACCAAGTGAAATAGGGTTGTTGTCAAATCTTGAAGTGTTAAGAGCTGGTGGGAATAAAGATGTTACTGGTAAAATCCCAAATGAGTTTGGGGATTGTGGGAATTTGACTGTTTTAGGCTTAGCTGATACCAGAATATCTGGTTCTTTGCCTGTTTCTTTGGGTAAGCTTAAGAATCTTGAAACTTTGTCTATTTATACTACAATGTTGTCTGGTGAGATACCTTCTGATTTAGGTAACTGTACTGAGCTTGTGAACTTGTATTTGTATGAAAATAGTCTTTCTGGTTCAATCCCATCTGAGCTAGGGAACCTTAGAAAGTTAGAAAAGTTGCTTCTTTGGCAGAATAATCTTGTTGGTGTTATTCCAGAAGAGATTGGGAATTGTACTAAGCTGACTATGATTGATTTGTCTTTGAACTATTTGTCTGGGAGTATACCATTATCTTTTGGTGGTCTTGTTGTGTTACAAGAACTTATGCTTAGTAACAACAATGTTTCTGGTTCAATCCCTTCTGTTCTTTCACAATGTACAAGTCTTGTGCAGTTGCAGTTTGACACAAATCAGATATCAGGTTTGATTCCTTCTGAGTTGGGGAATTTGACAAGTCTTGTTGTGTTTTTTGCTTGGGATAATCAACTTGAAGGAAGTGTGCCTTCAACTTTGGGTAGTTGTAGTAACCTTCAGGCATTGGACTTGTCACATAACTCGCTTACTGGTAGCATTCCTCCAGGCTTGTTTCAGTTGAAAAACCTGACTAAGCTACTGTTGATTTCTAACGATATTTCGGGTACTATACCTCGAGAAATTGGATATTGTAGCTCATTAGTGAGGTTAAGACTTGGAAACAACCGGATTGCTGGTGGAATTCCTAAAGAAATTGGAGGTCTTAAGAGCTTAAACTTTCTTGATTTGTCCGGAAATCGCCTTTCTGGACCAGTTCCTGATGAAATAAGTAGCTGTACTGAGCTACAAATGGTAGACCTCAGCAGCAATACACTGGAAGGTCCCCTGCCTAACACTCTGTCTTCACTATCAGGGATTCAAGTCTTGGACGTTTCGAATAACCGGTTTGGAGGGCCTATACCGGCTAGTTTTGGGCGTCTTGTGTCATTGAACAAGCTGATTCTCAGTAAGAACTCGTTCTCAGGGTCTATACCTCCATCTATTGGCCTGTGTTCGAGTCTTCAATTGCTAGATCTTAGCAGCAATGAGCTCTCGGGAGGCATACCGATGCAGTTAGGAAAGATTGAGTCCCTTGAAATCACTCTCAACCTTAGTTTCAATGAACTAACAGGTCCAATTCCTGCTGAAATTTCTTCACTAAGCAAGCTGTCGATACTTGACCTTTCGCACAACAAGCTTGAAGGGAACTTGAATCCACTAGCGAGGCTCGATAATCTTGTCTCACTGAATGTCTCATACAACAACTTCACTGGTTATCTTCCGGACAATAAGCTCTTTCGACAGTTGCCATCATCAGACCTAGATGGAAATGAAGGTCTATGTTCATTCGGAAGGCCTTCGTGTTTTCTCAGCAATATTGATGGAGTAGGAGTAGCTAAAAATGGAAATGATGAGGGAAGGTCAAAGAAGCTCAAATTAGCCATTGCATTGCTAGTCATCATGACAATAGCAATGGTGATCATGGGAACTATTGCTATCATACGCGCACGAAGGGCGATGAGAAGGGATGATGATTCAGAGATGGGAGATTCTTGGGCTTGGCAGTTTACTCCATTCCAGAAGCTCAATTTTTCAGTGGACGAAATACTCAGATGCCTTGTGGATACTAATGTCATTGGAAAAGGGTGCTCGGGGATGGTCTATCGTGCAGTTATGAACAATGGGGATGTCATAGCAGTGAAGAAGCTTTGGCCAATAACCATGACTACAACTAATGGAGGCAATGATGAGAAATGTGGGGTACGTGATTCGTTCTCTGCAGAAGTTAAGACGCTTGGTTCAATTCGACACAAGAACATTGTCCGATTCTTAGGGTGTTGTTGGAACAGAAGCACAAGATTGCTCATGTATGATTACATGCCTAATGGGAGCTTAGGAAGTCTTCTTCATGAGAGGGGTGGTAACCCTTTGGAATGGGAATTGAGATACCAAATATTGCTCGGGGCAGCGCAAGGGCTCGCCTACTTGCACCATGACTGTGCTCCTCCAATTGTTCACAGAGACATCAAGGCCAACAACATTCTCATTGGTCTTGAGTTTGAGCCTTACATTGCAGACTTTGGCCTCGCAAAACTAGTCGATGATGGTGATTTTGGTCGTTCCTCCAATACAGTTGCTGGTTCTTATGGCTACATTGCTCCAGGTAAAAAAAAATTTTATTTCATGTCATCATTTTCCTTACACCTAGCATATTATTCTGAGTATACATTTTGCTTAGTTATAGTATAAGCATTGATTGCGTATTGCAACCACAACAACATACCCAGTGTAGTCCCACAGAAAAAGAAGTAACGGATGTGAAAGCATAACAAAGCATGCTGAACACTAACTTCATCAAAATAATATGATAATCGAAGTACAAGAAACAAACAGTAGTAACAAAAATCGAAGGACAATGATTGCACATTGCATTTATCTACAAAGAATATTAGTAGTTCTTGATGATCATTGATCCAAGGGTACCAATATTGTTCATTTTTCATTACAGAATATGGCTATATGATGAAGATAACTGCAAAGAGCGATGTCTACAGCTACGGAGTGGTTGTATTAGAAGTCTTGACAGGTAAGCAGCCAATCGATCCAACAATTCCCGAAGGAGTGCATCTAGTTGATTGGGTAAGACGGAAAAGAGGAGGAATTGAAGTCCTCGATCCAAGTCTACACTCGAGACCAGAATCAGAAATCGAGGAAATGTTACAAGCATTAGGAGTAGCCCTGTTATGTGTCAACTCCACTCCTGATGAAAGGCCTACAATGAAAGATGTAGCAGCAATGCTAAAGGAAATCAAACATGAAAGGGAAGAATACGCCAAAGTCGATGTTTTACTCAAAGGCTCTCCAACAACAACAACTACTACTGATCATAATCAAGAAAACAACAACTCATCAAAGGGTGTTTTAGCAACATCTTCATCAGGACAGAAAGCAAGAAGTATGTATACAATGAGCAATAACACAAGCTTCTCTGCTTCATCATTGCTTTATTCATCATCATCATCATCAAATGCCAAAAGT

>M6-PGSC0003DMG400030113 M6 LRR-RLK

GAAGTTGACTTCATTCCAGAGGTTGGATTTCACAGAGGCGAACATTTTGTCAAGTTTGACGGAAAATAACATGATTGGAAGTGGTGGATCGGGTAAGGTGTACAGAATTTCCGTTGGCAGACCAAATGAGTATGTTGCTGTGAAGAGGATTTGGAGTGATAGGAAAGTAGACTACATTCTGGAGAGGGAATTCTTGGCTGAGGTTCAGATATTGGGATCAATTAGACATTCCAATATCGTGAAGTTATTGTGTTGTATATCAAGCGAAGATTCAAAGTTGCTCGTTTACGAGTACATGGTAAATCACAGTCTTGATGGATGGCTTCATGGGAAGAAGAGAGTATCCTTGAGTAATAAAGTCATGGATTGGCCAAAAAGATTGGAGGTTGCCATTGGAGCTGCTCAAGGACTTTGTTATATGCACCATGACTGCACTCCACCAATCATTCATCGCGATGTAAAAACAAGCAATATCTTGTTGGATTCTGATTTCACAGCCAAAATAGCTGATTTTGGGTTGGCCAAGATTTTAGAGAAGAAAGGAGAGCTTAACACCATGTCTGCTGTTGCGGGTTCCTTTGGTTACATAGCTCCAGGTAAAAGCAAAATCCAAGCTTGAACTTCGTATTACATCAATTTCGTTAACATTATATCAAAAGTATGACTAACTAATAATTTACTGTTTGGTTACAGAGTATGCTTATACAACTAAAGTGAACGAGAAGATTGATATCTATAGCTTCGGGGTGGTGCTATTGGAGCTAGTGACTGGAAGACAACCAAATTTTGGGGACGAGCATACAAGCCTGGCAGAATGGGCATGGAAGCAGCACGGAGAAGGGAACACTGCCATCGATAATATGTTAGATACAGATATCAAAGAAACATGTTACTTGGAAGAAATGAAGACTGTGTTCAGACTCGGGCTTATATGCACGAGCAACTTACCTGCAAGCAGGCCTTCCATGAAAGAGATTCTGCAGATACTTCA

>M6-PGSC0003DMG400030117 M6 LRR-RLK

GGAATTCTTGGCTGAGGTTCAGATATTGGGATCAATTAGACATTCCAATATCGTGAAGTTATTGTGTTGTATATCAAGCGAAGATTCAAAGTTGCTCGTTTACGAGTACATGGTAAATCACAGTCTTGATGGATGGCTTCATGGGAAGAAGAGAGTATCCTTGAGTAATAAAGTCATGGATTGGCCAAAAAGATTGGAGGTTGCCATTGGAGCTGCTCAAGGACTTTGTTATATGCACCATGACTGCACTCCACCAATCATTCATCGCGATGTAAAAACAAGCAATATCTTGTTGGATTCTGATTTCACAGCCAAAATAGCTGATTTTGGGTTGGCCAAGATTTTAGAGAAGAAAGGAGAGCTTAACACCATGTCTGCTGTTGCGGGTTCCTTTGGTTACATAGCTCCAGGTAAAAGCAAAATCCAAGCTTGAACTTCGTATTACATCAATTTCGTTAACATTATATCAAAAGTATGACTAACTAATAATTTACTGTTTGGTTACAGAGTATGCTTATACAACTAAAGTGAACGAGAAGATTGATATCTATAGCTTCGGGGTGGTGCTATTGGAGCTAGTGACTGGAAGACAACCAAATTTTGGGGACGAGCATACAAGCCTGGCAGAATGGGCATGGAAGCAGCACGGAGAAGGGAACACTGCCATCGATAATATGTTAGATACAGATATCAAAGAAACATGTTACTTGGAAGAAATGAAGACTGTGTTCAGACTCGGGCTTATATGCACGAGCAACTTACCTGCAAGCAGGCCTTCCATGAAAGAGATT

>M6-PGSC0003DMG400030119 M6 LRR-RLK

GGAGGAAGAAGCATAAGCGCGATGTTGCAAGCTGGAAGTTGACTTCATTCCAGAGGTTGGATTTCACAGAGGCGAACATTTTGTCAAGTTTGACGGAAAATAACATGATTGGAAGTGGTGGATCGGGTAAGGTGTACAGAATTTCCGTTGGCAGACCAAATGAGTATGTTGCTGTGAAGAGGATTTGGAGTGATAGGAAAGTAGACTACATTCTGGAGAGGGAATTCTTGGCTGAGGTTCAGATATTGGGATCAATTAGACATTCCAATATCGTGAAGTTATTGTGTTGTATATCAAGCGAAGATTCAAAGTTGCTCGTTTACGAGTACATGGTAAATCACAGTCTTGATGGATGGCTTCATGGGAAGAAGAGAGTATCCTTGAGTAATAAAGTCATGGATTGGCCAAAAAGATTGGAGGTTGCCATTGGAGCTGCTCAAGGACTTTGTTATATGCACCATGACTGCACTCCACCAATCATTCATCGCGATGTAAAAACAAGCAATATCTTGTTGGATTCTGATTTCACAGCCAAAATAGCTGATTTTGGGTTGGCCAAGATTTTAGAGAAGAAAGGAGAGCTTAACACCATGTCTGCTGTTGCGGGTTCCTTTGGTTACATAGCTCCAGGTAAAAGCAAAATCCAAGCTTGAACTTCGTATTACATCAATTTCGTTAACATTATATCAAAAGTATGACTAACTAATAATTTACTGTTTGGTTACAGAGTATGCTTATACAACTAAAGTGAACGAGAAGATTGATATCTATAGCTTCGGGGTGGTGCTATTGGAGCTAGTGACTGGAAGACAACCAAATTTTGGGGACGAGCATACAAGCCTGGCAGAATGGGCATGGAAGCAGCACGGAGAAGGGAACACTGCCATCGATAATATGTTAGATACAGATATCAAAGAAACATGTTACTTGGAAGAAATGAAGACTGTGTTCAGACTCGGGCTTATATGCACGAGCAACTTACCTGCAAGCAGGCCTTCCATGAAAGAGATTCTGCAGATACT

>M6-PGSC0003DMG400030169 M6 LRR-RLK

AAGCCCCAAGATAAATAGTATTAAGCAGGGTACATGATCACACGAACTTCCATTAATACAAGGGAAACTAGGTAGGTAGAGCATCCTACCAGGATGTTTTTTGTGGTCTCTTTAAAGAGGTAAAGACGGGGAAAAAGTATTTGCAGGTTTAAATGGTTAGTTTCAGTTTTCGGGTACAAAATTTTCCTGTTAGTTCAAACTTGTTTTGTAGATTGACATGACCAAGCAAAACAATTTTATGCATGATTAGTAAAAGTTTTACAAAAACAACAGATCTTCTCAAAGAAATTTTGGATGTGGTAACACAGCTGGAGCATAAAAGAAAATAAAGACTTGTTTTTTCTGCTTAGATGCTTCATGAGATTGAAGTTTGTTAATTAGGACTAAAAGATATGCACAAAATTTTGTAAGTTGTTTATCGTGAGTTTAGACTTTAGCACTCTTCAAAGGGTATCAGTTTCTACTCTAGGACCATACTAAGTTCAACTAGGATATGATACAATATTAGGATTAGTCGATAGTACATAGCATAACTAGCCCTTGGTTAAGGGTCGTAAAGAGAGTTATTTTTTAACTATTTATTTGTGTTTCCTTTAATTGAAATTTTTATTTTTGGTTTATGGGATTTCTTATCCTATAGTGCATCTTAGTTAGGTTAAAATCATCTTATTATTTTACCTCATGTAATTTGACAGATGAAGAGGTTATATGGTTTGCCTTTATAATGGATAAATTTATTATTTTGAGGTTGGCATTATGCAGCCACCAGAAGACTTTTCCCTTTATCTCCGTCTTGTTGTTCCTTGCAACTACAACGAGTTAAAGATTTTGAGACATTAAAAATCTGGCTGGTTAATCAATTTGCAGCATATTGGTTGCTGCTGTCTCCTGGCAATTATACCCTCTCAAGGTCCACATTTTGTTTGAGAATTATCTTTTGGTCCCTGAAGACATCATTCTACTTTTGATTAGCAGGTGTTATGAAAAGCCTATGCACGAGTACATCACATAATTTTAAAGTTATAGACGTCTTAAATCTAACAGGTCTCCAATAGGAAAAAAAGAAAAGGGAAGTATATGAAGTGACTTTGTTCGCATGGAGTATGGAATATAGAATACGCTGCCTTAAGGGGTTTTAGGTGCAAAAGGTGTCCACACACTTCACACTTCAGAATTGCGAGCTTGTTGCTGTCACAGTAGTATGCACTATAGCTAATAGCAGATTAAGTGAACTTTTGCTTAGCATGCGTAATTTATCAGTTGTATATCATCGCTTAATGTTGGTAGGACTGGAAATTAGTCGTTTTTGCAGTTGGAAGGAGGATTCAAATTGGAAATAGGTGTTAGGATTTGTATTTTATAGTGTTCCTTTTATTGGGTGCAGTTACGCCAGAATATTGACTCTTCAGATAAAAGTCATGTTTAAGTTGTTTGACACTTGATACATTTTGTTAGATGATGTTTGACACTTGATACATTTTGTTAGATGAGGTCTGATATCTAAAAATATCTGTACATCACATTTTATTGTTTTGATATCTGAAAGGCTAAAACTTCATTCTGCTACTCTTTTGGCAAGCAGTGCTTTACTTTTGGTTGCTAGAGGTTGAAAGTTGTGAACCATAAACTTGTAGATTCAGTTCATATCTATAAACCATTTTATCTTTTTTTATCATTGTACTCCATTATAATGCCATTGTTCCTTGCTGTTCTTGCAGATATTTACCTCCGGAAATAGCCTGCCTCACTAAATTGGAGTGCCTTGATCTTTCATTCAATAAGATGAAGAATTTACCTGTTGAGATAACTCATCTTAACTCCTTGCTGTCACTGAAGGTTGCTAATAATAAACTGATTGAAGTACCTCCGGGGTTGTCTTCTCTGCAAAGATTAGAACGTTTGGACTTTTCTAACAATAGATTAACATCCTTGGAGAATCTTGATCTCCTTTCAATGTATAATCTGCAGAGTCTCAATCTTCAGGTATGTGCTACTGAAATTACAGTTTCTTTCCAGTCTTTTTGCTTTTAGGCTTTTTCTCTAGTTTCTTTTCCTTCTCTGATAAATGTTTATAACATGAGAGGCATCTACTCTTAATGTGGATGAAATTCTTGCATCCCACTGGAAAAAAACCCGAAATGTATTTGTCTCAATTGTCTAGTCAAGAAACTTGATATATTATAATTACCTTATCCTGACCCATATATCTTCATTGAAATTTGCTGAAGCACAATAAGCTTCTTAGGTGTTGTTCGATTCCTTCATGGGTATGCTGCAATTTGGAGGGGAATTTCATAGATTTGTCCAAGGATGACACCACCAGTTCATCTTCAGAAATGGACGTGCTTGAAAGTTATGAACAAGAAACTTCTGAGAATACCCAAAATGGTGAGTTCCTCGGAATCTCTTGTCTCCCCTCCTTATATCTATTGCTATTTCTGCACATTCTGATTCCTTTGGATCTTGTTTGTTGCCCTTCTAAGAAGTGTTGTTCGGAAGGGTAGTACCTTTGAGAAGTCTGATGTGTGAACTTTTAATTGTGTAGGAGTTTCTACCAAATTGTCAGGTCATTTATGTGGCTCATCACCCAGTCATCGATGTTTTCGGCCTCGAAAATCCAAGAAATGGAAGCGGCAGTATTATATGCAACAGAGGGCTCGTCAAGAGCGATTAAAAAACAGCAGGAAATGTGTTGCCTGCAAACATTCTAAGCTAATTGATGATTCTCTGGTGGAGGCCTCATCAAGCATAGTGGATGATGATACACATGTTAAAGAACTGATTACTGAAGAAGCTGAGTGCAAAGGCTCTTTAGCTAGTGGTATTGATGAACATATCAGATTAAAGGAGGACAATTATATAGGAAGATCTTCATATGTTGCTTCTGACTCCATTGAAACATGCATAGATATTCAGAATTGTAAAACATGTGATGCTTCAGTAGGTTCTGTTTCTGATGCTGCTGATGTGGCTGAGGAAAGTTTGTCTTCAGAGGTGACTAATAGTCCTCCAAAATCGAAAAGGCACCTGGATGGGGTCATTGATAATCCAAAACCATGCAAAACCCGCAGACCAACAGATCATTCAGAGGTATCCTGCAAATACAGCATGATGTCTTTCTGTGGTATTGATGATTATTTACCAGATGGCTTTTATGATGCTGGGCGAGATCGGCCATTTATGACGCTGAGGAGTTATGAGCAGAATTTGCATCTTGATTCACGGGAAGTCATTATAGTTGATAGGTAACTTCCATGTTTGCATGCCATTAATAGACTAATAGTTATGGACCTCAACTACTAACTTCAGGGAATGCTTTGCCTCTCTACTCATTTTGATTACTTATTTAATCTGTTTAGGCAAAGAGATGAAATGCTGGATGCCATTGCTCTACGTGCTCAAGCTCTAATCTTTCATTTTAATCAGATAGATGGTCTGTTTAAAGATAGAGAACATGTTGCTGTAGACAACTTGCAAATTGCATCACTGCTTGCACTTTTAGTGTCGGATCATTTTGGTGGAAGTGACAAAAGTAATATTGTACAGAAGGCAAGGAAAGATGTATCTGGGTCGAACTATAGCAAGCCTTTTGTCTGCACATGCCCTACTGGAAATGATGACACTACCAGCATGGTCACAAAAGAAAGTCCTAGCATCTTAGAAGATATACTATTCCTTAATCTTTGTGAGAAAGCACTTCACTCAATAAAATCAAGGCAAAATTCTATTGTTGTTCCTATTGGGAGTTTGCAGTTTGGTGTATGTAGGCATAGGGCCTTGCTTATGAAGGTATTTTGTCATCAGACATGTTTCTAATTAGATATGGTGGTACTGAATTTAGTTTATTATTATGAACTGATAAATTTCGTTTGTCTTCTGCAGTACCTGTGTGACCAAATAGAACCTCGAATATCTTGTGAGCTTGTTAGGGGTTATTAGGATTTTTCACCGCATGCGTGGAATGTGATCGTGGTAAAAAGGGGTGAATCATGGGTTCGAATGATAGTTGATGCGTGTCATCCCCTTGATATCAGAGAAGAGACAGATCCAGAATACTTTTGCAGGTCAATGATTTTCCAATTCTGTGTTACTTTCTTCATGACTAATTTCTGGTACTACAAGCATTTATATCAATACACGTGTATGACGATATTGCATTTTCAATTTAGCCAGTGTGATTATTGGTTATTTTGGGGTGAAATTGCCAGCTTCTTGAGTAGTTCCTTAGTTACTAAAAATCTCTAGTTTACCTGAGTTATTTCGGAATGAAAATGGCCAGATAACTGAGCTTGCTGTTTCTTTCAGGTATATTCCACTGAACGGGGATAAATGTTCCTGTTGTGCCAGATGCTAGTCCTGGTCAAGTTAGTTCTTTTCCTTCTCTAACCGGTGATGATAAAATTCATAAAGCTCCCTCCTCAACCCTCGTCCCGTGTAAATTGGGAATTTGGGATCCCTTGAGACTTTGGCAAAGGTGTGGTTCCTGATTGCTTTTAACTTCTCCAGTTATTATTATGCAAATATACTTAAGTTCTTATTCCAAACTTCAATGACTTGAATTAAATATGTTTTATTCTTAGAGTGACTGAACTACATGTATATGTGTTAACCCACAACAATTAGAAAATGATTTTTCTGGCTTTATTGTACTTTCCATTATTGACTGGAATTTACCAGTCTATCTGGTCTTGTGCTTTTGATTTGTTTTGGAAATCTGAAATAGATAAGAAGCAAAGCAATCCTTTTTGACTTATTCTGCTATTCAGCAGGAATCCATTATATTGCAGGCTTATGCTTTTCATTTTGCAAATTTTTCATTTATCTTCTGATAAAACTTGATTAGTGACAAGTATTATACCACTAGATGTGGGAATTTTTCTTGGCTAGCATCTTTATTGACGAGGCATAGTTGATTTTTTAAAAAATTTATTCTGCTCAATTAACCTGAATTGAGCAGAATGAATATAAAAGAGTCATATGACTGATACCGACTAGTTCAGGATTGAGGTTTAGATGATTGAATGAACATTTTTGCCACTGCAGGTGCGAACTATAGAGATGTCCAAGTCCACGGCGGATGAGATTAAGAATTTTGAGTTTAACTGTATTGGAGAAGTTAGGGTGTTGGGTGTCTTAAACAGTTCGTGCATTGTCAAATATTATGGTCATCAAATATCTTCAAGATGGGTTCCTTCATCAGATGGCAGTTCAGAAAGTCGTACTTTGCAGTCTGCCATTTTGATGGAGCATATTAAAGGGGGTTCACTAAAGGTATATGTTTTCCTAGAACTCTTCTCGTACCTGACTTTCCTCTCCTTTATTTCCCTTCCCTTTTGAGGTGGGAAGGGTATAAATAACACCTGATTGCCTTTTGTATTCAGCTCATTTTACTTTCTTATCCTTAACTGGCAGAAACATGTAGATAAGCTATCTAATGCTGGTGAGAAGCGTCTTCCCGTTGAGCTGTCGGTATTTATTGCCCGAGATGTAGCAAGTGCATTGACAGAGCTGCATTCTAGGCATATCATACATCGTGATATAAAAAGTGAAAACATCCTGATAGATCTGGATAAGAAGAGAGCTGATGGTACCCCTACAGTAAAGCTATGAGACTTCGATATGGCAATTCCCCTTCGTTCATACTTACATACTTGCTGCATTGCCCATGCTGGAATTCCTCCACCAGATGTTTGTGTTGGTACACCTCGTTGGATGGCTCCTGAGGTCTTTCAGGCGATGAATAAGCGCAATATCTATGGGTTGGTAAGTCACCAATCCTTCTGACCAAAAAAAAAGTCACCAATCCTTGAAGTTAGAATCCTAACCATTGAATTCTCTACTAACAAGTACCAATTAAATGCCCCACCGGGTGCGTTGATCTGCTAAAACAATGTGGTTCTTTCTGATTTGCCAATTTGTCACCCTGGGGTAAAATCAATTCAACACAATCAAGAACTTACCACCTTCAAAAGACACACATTTTCTACTAAAAACTGATATATCACTCATTCCTCTATTAAAATTCAGCTTTATCTTGTCATTATTAACTTCAAATGTTTTAAGCTAGAAAAGAAAGCATGTTTTGTCTGCTCTATTGCTGCACTGCTACAGATTAAAGAACAATAATATGTACTCATTGACAAATTCAAATTTTCACCTTTCTTTCTTGGAAATTTGTTTGTTTAAATTTCCTCATTTTTGATGCTTGAATCTATTTCGGAGATGAGGATTTTTGTTTTTGAGAGACTATTTTTATATGACGGCTAATCTCAGAGGATGTATCCTGTGGCAAAAGGGGTTGGGGAATATTATCTCTTACTTCTTTGAAGTTGAAAAGGTCTGATGAGTTGCTTCAAGGTAAAAAGGTTTTTCTTGCTTTTACTTTGCAGGTTCCCCTTTTCAATAGGTAGATGAAAGCTAATGGACCTAAAATAAGTCCACTATGGCCAAAAAATATCAGGAGGGGCGGGGAGCGGTTGGATTATTTCACCCTTAACCCAAATTCAATAGATGGGTAAATTGGCTAGAATGGGATATTGAACAACAAAAAAGTATTTATAGGTGCTAATTTCCTGTATTATCTTTTTCCTTTTGAAGATGAACAAATGAGTTTAATTGCTTTGAGTTGGTGACTTGAAAGATAAAGTGAGTTTTTGTCATGTCTAAACTGCAATTCTAGCTTAATGTATTCAAAGGTTTTCCAAACTAGGTTTTTACTTGTCTTTCTTTGGATGTAGGGTGCTGATATATGGTCGTTTGGGTGCGTCCTTCTGGAATTGTTAACACTGCAACTTCCTTACTCAGAGTCATCCGAGTTAGATATCCACCATTCTCTTCAGGTATTTCTTGATCTCTAACCTTCGATCTATTAATGCTCTAAATACCATGATGAGTACCTAGTGTGCTTGAAATATCATTGTGTGATGAGGACTTCAGAAGTTCCAAAGTCCCCTGTTGGACAATCTTTATTATGGCTGCCTTTGGCTGCTCAAAGGCTAATGGTTCTAGCATCTTGGTTGTTGGATATTTCGGAGCTCTCATGGTTTTCTTCCTTATATTGCTTTCGTTCATAAGAACTGCCTGCAGAAGTACACTTGTCCCATGCACATTTTTAAAAAAATAAGTTTTTGTTTGAATTGCATTAAATGGTAGAAAACAATATAATTCATGTTTAACACATGCAGGAAGGTAAACGGCCACAATTAACCGAAAAATTAGAGGCAATGGCTGCATCCAAAGCAGAGCTAGAGGATCTAGCGAAGTCTTGTTCTAGCTCGGACCTTGACAAGAAACAATCTGAATCTCGTATCCTGAAACTTCTTGTTTCTATTTATCGTTGGTGTACTGAGAAGGATCCAAACGACCGTCCCACAGCAGAGAACCTCTACAACCTCTTACTTACCTGTGCTAATTCCCTCCCTTCACAACAAAGTCAAGAAGATTCTTGATCTTCCTCAAATTTTGGTAGCCTTAGGATGTTGTAGATTAGCATTAGATTATTCAGCAGGAGTTACAGACCCAATACTTGGGTCTGTTGACCCCCTCATTGAGCAATGTAACATGTTCTTAGTCTTATTTGAAATGGAAAACCATACGGACGTATTCGCGTCCATAGGACATTTATGATGGTCTTGTTGTGTGTGTGTGTTGTGTATCAGTTTCTTTAGTCCTTTTCCAGCTTGTAGTTGTATACATTTGTACCCATCTAATGTATTCATCTTCACTAATTCCAATTGATGTTAATTATGAATGTATTCTGAGTTTCTAAGACTTGGATTGTGC

>M6-PGSC0003DMG400030298 M6 LRR-RLK

TGTAACATCCGACAATTTGAAGTGACTTTGAAGGAGCTTGAAACTTGGAAAATAGCCATTTTTTTGGAAAAATTTAGAATCTGGAAATTTGGGTTAAATATGGAAAAAAAAATGAGTTTTTGGGCAACTTTGAGTAGTCGTAACTCCTAGTTTAGGATGAGTTAGGAGTAGTTTCAGTTATGGTTGCGAAGCTCGTGGAATGACCTTTTCAACGCCACCGAGTTTGCTCGATTCTGAGTTCGTATGAGGGAGTTATGCCCTTTAGAAGTTGGGCAGTTGGAAGGGAAATCCGTCCAGAAATTTAAGGGCATTTTGGTCTTTTCACAAATTATTTTTTTGAAGATTACATTGTTGTATTAGGCTGATCCTTGGATCATTTTTGTCCCATTTTGAAAGAGAAAGAGTTAGGGTTCTTGAGAGTGAAGAAGAGAAGAAGGAGAAGAAGAAGAAGAGAAGAAAGAGGAGAAAGGAGATCAAAAAGTTTCCGTCGTGGATTATCGTCGGGGGTGATCCCTATCAAGGTATGTGAGTTTTCGTGTGGGTTGATCCTTTCCCTCACACACACCAAGCTTATTCTATGATTTGAGAAAAGATTGGAATTGTTGTTGGTGGTGTGGTTGAAGTCATTGTTCGTTTTGAGACATGATTGGGTTATGTATTTACGTTGAAACCTATTGTATGTTGAGGGAATTATGATTCTAAGAGTTATGACCCTTTTGAAGTTGATTCACCATCAATTCTCAGCAATTTTTTTTGGCTGTCGATCCCCATCTACGAGCGTGACCTACGGACCGTAGATCGATTGATGGTCCGTACTGGTCAACCGTCAATCGGGACAGGTTTTTGGGGCATCGATCTACGGGCACGACCTACGGTCCGTAACCTGACCTACGGACCGTAAGTCCGGACCGTTGGTCAACACTTAGTCATTTTTTTAAATGAATTCGGGGGAAGTTTCTGACGCGATCTACGGGCCTGCAGGACGGACTGTGAGTCCATCTATGGTCCGTCGATGGCGTCCGTAAGTGCCATCTGTGTCACCAGAAATCTGAAGTCTCAGCCAAGTTTCCCCATTTCTTTTCTCACTTTCTCAAGCAAGTTCCAAAGTATTATACCTATGTTTTATAGTTGTTTCCGACACTCCAAAGTACGTCTAAGCGTTCAAGAATCCCTCTATAATATGTGATCGAACACCTTGAATTCATAATTCCGATTCAAGGAAAGATAGTTCCAAGTCAAGTGAAGTTAGAAGTCAAGAAAAAGAGTATAGAGTTTCAAGCTAAGTAAGAGTCTCAAGTTTCAAGTCGAGTAAGAGTCTCAATTTCAAGTTAAGTCATGAGTTTAAAGTTGAGTTCTCTTTTTAAAGGCTATTAGGGAACTAAGTATTCCCAAAGGAGGTTATAAATGCTATTACATTTGAGTAAGAGGGGAACCATGTTTTCTTCTTTTTCCAAGAGAGCCTTTGGGCTAAGTTTTGAGTAATTATCTCAACTAAAGAAAAATGTGTTTAAAGATAGTTATGAGCCAAAGTATTTTTTTTTGGGAGTAGTATTGAGCACCGAATTGGGGACACGAGTTCTGATTTACTCAACTGTCCATAAGAACCATGCGCCAACATGGGATTTCTCGGATCATACTTTTTAGATGATCACGAAAGTTAAGCTAGTGGATCCACTAAGTAAAGTTAGCTTCCTATATCACAGCAAGGTATAGGATGGTCCTTGGGCAACGCGAGGTAAAGCGTTGTATCATCACTAGGGCTCATAGTGGTGGTTGTCGGTTAAAGAACCTCCCACATAAGTTATATCACTTTTATATAAGTACAGTTGAGTTGTTATTGTTTTATCATTAACAAGCTAAGTTGTTTATACTGTTTTACAAGCTATATATATTGCATGCGTCTTATTGCTTTATATATTGAGTTCAGTTAGTCATGAGTCGTACAGAGCCAAGGTAAGTGTTCTCTTGTATTTCCTTTCAAGCTTAAGTTGTGTTTTGGCTTTCCCGCTCGCATACTCGTACATTCCATGTACTGATGCCAGTTGTCCTACATCGTTTGATGATGCAGATTCAGGTACCCCGGATCAGCATCCTGTACGTCGTTGATCCAGCTGAGCACTCCAGAGTCAGTGGTGAGCCTCCTTGCATTCCGGAGGACTCAATTATTTTGTGTTCTTAATTTTTGTCTTATTAGGATGTTGCGGGGTCTGTCCCAACATCCATCTCAATATTTTAGAGGCTTTATAGACAAATAGTCAGTCAGTTAGTTTTGAGTCTCTTATCTATGTATATATGTAAATATCCTATTTTGAGATTCGAGTTGCCTTATGGCCATATTTTATAGGTTAAGTTGTTTTGTAATATTGCATTGCATTGAGCTATTCTGTTGAGTTAGGTTTCCGCTGAGTTAAGAAAGCCAGGCCAAGGGTTCGCTTGGGGCCAGAAATGGTCTCCGGGTGCCGGTCCCGCCCAGGGTGTAGGCTCGGGGCGTGACA

>M6-PGSC0003DMG400030443 M6 LRR-RLK

ATTTGATCCATCCCTAAATTTATATTCTTTCTTTCTCTCTTTATAAACTTTGTGTGAAGTTAATTAAATTTGGGCGAACGAGAGAGTACTAATTCAAATCGGCGAGTTTTGAAACCAGAACGCACACTACAACATGTTATCCCTGGGATTCTCACTTCCTCATCCTTCCAATAATAAACTAATTATTACCAACAAAAACCAATTTAATTTTCGTACCAATTTAGCCAAGGTTTGTTCTAATAATGTTAATGCAGCTGGGGATCAATTAGTGACACTGCCTGAGGGCCTCAAACATGTTGCAGTGATCATGGATGGCCATAGAAGGTGGGCAAAAAACAAAGGCTTGACTGTAAAGCAAGGTCACCGCGCTGGCGGAGAGAAAATTCAAGTCCTCACCCGCCTTTGTAGCCAGTGGGGAGTCAAAGTTCTCACCATTTTTGCATTCTCTACAGAAAATTGGGTTCGCCTCAAAGTAAGTCATTGTATTCCTTCCATCCTAATTTACATGACACAGTTTGAATTTCGAAAGTAAAATAGTATAATTTTGATTGTGATTCCGGGGAATCTCTAACTTTTGAAATAGAATTTACATATTTGAAAACTAGGTTAAAACTACTATAAGTCACCATAATTCATAATTCAAGATGTTTACATGAAAAAATCCGGTCAAATAAAGATTTATTTGAATTTCTAAATTTGAAATTAGGTGCACCATAAATTGGGGAATGAGGGAATAACTTTTATGACGATCTATCAATCATGTTAAAATAAATCATCAAGAAAATAAATTAAATAAAGAACAACAATAACATATTCAGTGTAATTTTATAGGTGAGGATCTGAAAATGATATAATATACGCTGAACTTACCCCCACCTCGAAAAGATATTAAAAGTTGTTTTTAAAGATCATCGACCCAAGTAACTTGTACAAAATCAACGACTATATATATATATATATATATATATATATATATATATATATCAGAGAAAGTGACTTTCTTTTCTAACAATGAAATTAATTATACAAGAAATGAATCATATGTCTGATTATATTTAACTATATATAATTATAATTACATATGATTTATTAATTTAATTTCTAATACACCAATGTTTTGTTTTGTTTGTTAACAACGCAACTCTAAACTTGAGTGCCGCAGGAGGAAGTTGACTTTCTTATGAAGTTGTTTTTAGAGCTGATAGGCTCACAAGAAAACTTGGATGAGTGGACAAGGTATTCTATATATTATCTATATGTGATAAATAAAAATATCTTGTGTGATCAGCTATCCGACTATCTATATAAGAAGATCGTTAATTCCACCATTTTAATGCAAAATGGCTCCCGAAACTAACTCCTCATGGTTTAGTAAATGAAAGGTGGTGTTTGATATTCATTTTGAAATTCAACTTATTCGAATTAAATTATATTATGTGATAGGGATGGTAGACGAGTATCATTTATTGGGGATAAATCAATCCTTTCCAAGTCTTTACAGAAAGCTCTTGCGGTAATGGAGGAGAGAACAAAATGTAATTCTGGACTACATGTTATTATAGCAATCAATTACAGTGGACGACATGATATATTACAAGCAACTAAAAGCATTGCCAGTAAAGTGAAAAATGGACATTTAACTGTAAAAGACATTGATCAGAGCCTATTTGAACAAGAACTGGAAACTTATTGCGCTGAATTTCCTGAACCAGATTTGTTAATTAGAACAAGTGGAGAGAAAAGATTTAGCAATTTCATGTTATGGCAATTAGCTTATACTGAATTCTATTTCGCGAATAAGTTATTTCCTGACATGGAAGAAGCTGATTTTATGGAGGCTTTAATGTCGTTTAAGTCGAGACAGAGACGTTATGGTGGACAAAAAATCTAGTAATATTTTCAACAACTTCCATTACTTAGAATTAAACCTAGCTCGACATATGAAGAATTATTGATTAGATCCCTTGAATTTCCTCTTGCTATTAGTACTATTAAATAATTTTATGGTGGTCGATTGTATCTATACCAGCACTATATAAGCATGAAATATCGTCATGTGGTTTGTGCTGAGTTTTTTTTTCTTTTCCCACCGGCCCAGTTAAATTTGAATCACGCACTTTGAAGGACTTGTAGCACTCCCAACAAGGTATTCATATATTCATAATTAAAAAAAAAAAAAAAAACAAGGAGTGGGGCAACGCCGGAAAATGAGACTATAAAAAGTCTTCAAAGTCATATATGCTTCCAAGATGTCAATCACATCACATCATTCATCAATTTGTTGGTTCAAAGTGATGAGTGGATATAAAATTTTCATTTTTTTACATCACATTTGATTTTTGCTAGACTATAATATAGTAAATTAAATTATAAAATTGAGGCGAAAAATATAATATATTAATTAACTATCAATAATAATATCATATATTAATTGAATTAGTATTAAATCGTAACTTGTCTTTTTAGCAAATCCTTAAATTTTTAGATGAGGGAAGTTCACTTTCTTTAGTCAAATGTCAACTCATTAGTCATTACTCATCTCATTATTCAAAGATCCTTGGAAAATAAAAGATTTGACAAACAAATAACAAACTCTTTGGTCTTCTCCTTTCTCTTAAAGGGAGAGTCGTTTACCTATACAAGTCTATATTTGTCGAACACTTAAAGTCATTGGCCATACTTTGACTCAAGGGATGGTATCCTGTTTGATTGGTAAATCTAGAACCTTTTTTTTTTTTTATTATTATTATTATTATTATTATTATTATTATAGTTGTAGTATTCAAATCAACTTATAAGCATCTTGATTAATTTCACAAGATATGTTAGCTTCCATCAACACAAGTACTGAGTAAGATTTGCTAAGAAAGTGTCCTTTCCCTTCACATGATCTTCTTTTTTTATTACAAGTTATATGCACTAATTGAGTCAAAATAACTACACAACCATGTCACTTGTAAAATAATTCATGTAAATATCACAATATATATATTGATAACCCGATAAAAATAATGTCTAACCTGATATAATAGGTTAAATTACATTGAAAATGGTTGTAAAAAATTTGTACACTACCATTGTGTACAACTTTAAGCCCTTTTTCTCAAACACTTTTGTACCCCACATAAGGAATAATGAACTCATTATAATTAGCAGAATCAAATGTGACTCAGTATTATTCCCTTTCTCCACATATCACCTAACCTTCTCAATCCCCAAATGGAGCTTCAAACTACTTTTGTTCATTCAAGCTCTCTTTTTTTTCTGTCATTCCATGCTATTCTCTTCATGTTCCTATACTTTTCTTTTCCAGAGTGTGCTTCAGCAGGCATACTTGGGAATCAAACAGACAAACTTGCATTACTTGATCTTAAGTCTCAAATAACTGAAGATCCACAAGGACTCATGGATTCTTGGAATGCCACTCTTAATGTCTGCCAATGGCCTGGTGTTACATGTGGCCATAAACATCAAAGGGTCATTTCTTTAGATCTCAAAGAACATAGATTGGCTGGCACTATATCACCTAGCATTGGCAATCTTTCATTTCTTCGAATTCTTGACATCTCAGATAACTCCTTTCACGGTGTAATCCCTCCAGAGCTTGGCCAATTGATCAGACTTCAAACCATGAACTTAAGCTTCAACTTCTTGAGAGGTGAAATCCCGTTTACCCTGTCTCGTTGTGTTAACGTTGTTAACCTTATTCTAGACCATAATATCCTTGAAGGACACATTCCTACTGAACTTGGTTCATTGACAACGCTGGAAATGTTGTACCTCAAGAACAACAATCTCACAGGAAATGTTCCGAATTCTGTTGGAAATCTTACGTCTCTAAGAGAGCTTTACATTTCCTACAATGACTTGGAAGGTGAATTACCTGAAACAATGGCCAATATGAGAAGCTTGATTGAGTTAGGTGTTTCAGTGAACTCTCTGTCCGGAGAATTTCCCCCTGCACTCTACAATCTTTCATCTCTTACACTTTTAAGCTTGTCATTTAACAAGTTCAGAGGCAGACTTAGACCAGATATTGGCCTTGCCTTCCCAAATCTTCAGAGACTGTATTTGGCCAACAATTACTTCACTGGTTCAATACCAGCTTCATTGTCCAATTGTTCAGATCTTCTGCGGCTTGATATTCCTCTAAACAATTTCACAGGAAATATACCGTTGAGTTTTGGAAACCTTAAGAATCTCTTATGGCTAAATGTTCTCAATAACCAGCTTGGAATTGGTGCACCTGATGATCTAAATTTCATAAATTCTTTAACCAATTGCAAAATGCTTGAGTTCCTGGACATTGCAAACAACAAATTTGGAGGTATGTTGCCCTATTCTATAACTAATCTCTCAACAACTTTGACAAAGCTACTGATTGGATACAACAGGATAAGTGGAACCATACCTAGAGAGATCTCCAACCTTGTCAACTTGGATATGCTTAGCATCCAAGGAACTCTTATCAATGGTAGCATTCCTGATTCTATTGGAATGTTATCAAACTTGAAAAATCTGCACATGGAATCGAATCAGTTGACAGGAAATATCCCCTCTTCCTTAGGAAACATTAGAGGATTGCTGTACATTTATTTGCAAGATAATAGCTTGGAGGGAACCATACCTTTAAGTTTAGGAAATTGTACATCTCTCCAAACCTTAGACATCGCTCAGAATAAACTCAGTGGTAGTATACCCAAACAAGTCATTGCACTCTCCTCCCTTTCGGTACTCCTTAATATGTCTTACAACTCTTTGAGTGGTTCCTTGCCTGTTGAAATAGGTAACCTGACCAATCTTGCTGCACTTGACATATCGAACAACAAACTGTCTGGTGAAATTCCACACTCACTGGAGAGTTGTTCATCCCTTGAAATACTCTACCTGCAAGGAAACATCTTTGAAGGAACTATTCCTCCTCTGGATGATTTGAAAAACATTCAGTATCTTGATCTTTCACGTAATAACTTGTCAGGCAACATTCCTAGGAGTATCACCAAGCATGTTTCATTACAGAACCTCAATTTGTCGTTTAACCACCTTGACGGAGAGGTGCCATTTCAAGGTGTTTTTTCTGATGCAAGTAGAATTCAAGTGATGGGAAATATGAATCTTTGTGGGGGAATTGAAGAGCTGCATTTGCATCCATGTCTCAAGCATGCAAATAAAAGACCCCAGAAGCACATTGCCCTCATTTTGGTGCTGGCACTGGGTACTAGTGCTGCATGTTTAACTTTGTTGCTGCTAGTTTCGTATTGCTGCGTGAAAAAGGGGAAGCATAGGCCTAGCACAGCATCTTCATTCAGGAAAGTATACACTCAAGTTTCCTATGAAGATCTCTTAAATGCAACTGGTGGGTTCTCTTCAAATAACTTGATAGGATCAGGAAGTTTTGGCTCCGTTTACAGAGGAAATCTTTCTCCAGAAGGAACAATTGCAGTAAAGGTACTAAAACTTGAAAAAAAGGGTGCTTCTAAAAGCTTTTTGGCTGAATGTGAAGCCTTGAGGAACATAAGGCACCGGAACCTGGTGAAGATCTCAACTGTTTGCTCAAGTGTTGATTTTGATGGCAATGATTTCAAAGCTCTCATTTATCCATTTATGGAAAATGGGAGCCTAGACGAGTGGTTGCACCCAAAGGAGGGACAGATGCTGCAGAAGAGATTGAGTATCTTACACATATTAAACATCACAATAGATGTGGCTTCAGCATTGCATTATCTTCACAGTCAATGCCATACATCTATAGTTCACTGTGATCTTAAGCCAAGCAATGTTCTACTGGACAATGATCTGACTGCTCTTGTTAGCGACTTTGGTTTGGCAAAGTTCCTGTCTGACTCGGGTCAAAATGCCGATGTAAATCAGTTCAGCACAACTGGGATTAAAGGCACAGTAGGTTATGCTGCTCCAGGTAAATATTATAGTGCGGATCTTAGTCTAAATATTGTCTGAACTCTGAAATCCATTTTTTTCTCCATAAGACCATAACATATTAACAATTTAAGCAATTACATACTTCCTTTAGAGACTCCGGATGCTTCTTGTCCTAAGCCCCAGGCATCCGTTACCTTTGCAGATCACTACCTCATGATTTTGCGCTATTGATTCTTATAACCAGAGGCAGAGGAAGCTTTAATTTAATGGATTCAACCAAATCTAGTATTTTCGACACTGAATATGGATATATAAGTTAAAAAAACTACTTAAATTTCAATAAATATCATATTTTGAACCCATAATTTCAGAAGTACAATGATTTCAGTTCTAAGAATCTTAGATGTTGAATCCATCAAATTTAAATCCTGGATGCCTCTTCTCTCAGATATAATTCCTGACTAGTCATTGTTTTTCTAAGGACCTTAATAAGAGGTTTCAGCTTTCTGGGGCATTTTACATAACATTCTCTATTGGAAAATCGATACAAGTGATCTGTGTTGCTTAGGCATTTTTGACAAATGTTTAGGATCCCTAATATTACTATGAAACACCTTCTCTGATACATCAAATTTTTCAGAATATGGCATGGGTGGTCAGGTATCATGTCAAGGAGATGTCTATAGCTTTGGGATTCTTCTATTAGAGATTTTCACTGGACGAAGACCAACCAGTGAACTATTTGAGGACAATGAAACTCTTCACAGTTTTGTTAAGCAGGCATTGCCCGGTCAAGTGATGGATGTTGTGGATCAATCTGCTTTGTACGAGACAGAACCAGGGGATTTTATGGATATACTCAGCTGCAGAAGTGACTTCTCGGATGAATTTGTTGAATGCTTAGTTTCTATTCTTACTGCTGGAGTTTCTTGTTCAGAGGAAACTCCACAAGCCAGAATAAGCATGGGTCAAGTTATTTTGGATCTCATTTCCTTCAGAAACAAATTGAGCAGAATTTTAGTTCATTCAGAGAAAGTGATCTCAAGAAAAGGTGAATCATAA

>M6-PGSC0003DMG400030474 M6 LRR-RLK

AAAAAACAGATTCATTACAAAAAAGCTGAAATTCCTTTTACCATTTCTCCACCTCTCACTCTTCACTTTGTACTTGGCACTAAACACTCCTTCTTCTTCTCCATTAACATAGATCCAAATCTCTTACTACTTCAGCTCAAGCTAAACGGGTCGGGTCGGGGTTTTATTCTGCGCAAAATGTGGTTACTGCAACTCGATGAATTCGATTTGATGTTCATGTTTACTGTTAATTTCACCTCAGAACCTCAGTATTACTCCTAGAACTTCAAGTGCTTTCTCTATTTTATAGGTAATTACACTGCGATTTCATATTGTTTGTTGAAATTTATAATTTTGTATTTGTTTAGCTTGATTTCTTTTGTGTTTTCTGCTAATTTTGGTGGCAGATCTGTTTGCTTATTTGTTTCCTATTTGTTTGGCATTGTTAGATTGAGCAGTCAATTTTTATCTATTTTGAAATATGATTTTTGAAGGTTTTCTGATAGTATTTTTTTTTGTAGCTTTAAATTTGATTTTTAGAAATTAATTAGGAAATCCGTGACGAATTCATGCAGAAAATTAGACTCATTGGAGGGAGGAGTATTAAGTAGCGGCAGAGCTAGGATTTTCAGTAAAGTGATTGAAAATATAAAGAAAGAAATATAATGATAAACTATGTATGATTTTTTTCTATTTCTTTTTTTTTGAACTTGTATATATGGTATGATTTTCCAGCTTGCACCCCTGGCTCCGGCGGTGGACTATTAAGTAGCAAGACCATGTTAGGTTCTGTATGTTGTTTAGTCTTCAGTGTTTGCTTCTTTGATAATGTGGAAGTATAATGAATTGAGAAATTGAAAATGTTGTTAGTCATTATGAACTATCATTGGATTATATTTGTGTAAAGGAAGCTTAGGATTCCCTATTACTGGGACCTCATATGAGTTGGGGTTGTTCAATAGTCTTAATTAACCCAACTCTCTTTGACATTTACTTGAGAGAAGAATTATGAATGTGGTAATAGGTAACCCCCAACCCCCAATAATAATAATAATAAAAATCCCATGATCTGGTAGATTTTTTGATGTAGTTCTTCTTCATCAACTGTTTCAAGCTTCAACCTTGAATAAGAATTATCATGAGGATTGAAAAACTTGTGATTAAATGGGTTAGTCTAGTCCCGGGTTGACTTGAAGGAGTGGTTATTTTCCTTCCCGTTGCTTATTAGATACTATCATGTCAGAATACTTAAGTGCTAGCTATTTGTATGTATTTTTTTTTCCTCAAAACATTTCTTGTTTATTAATAATATAATTACAAAAGAGGGTGACATAAGGCCTAACACCTCTTCTCACACGTAACACCAAAAAAAAAAGAGTTTCCATCCCGGAAGGCTTCTTCTTCAATTAGAGTTTTTACCAAAACCGAGGGTGGCTCATAATGCCAACCAGAATGGTTTTCTTTTTTTGAACCTAAAGGAGGCAAGGTTAAGTTTATCCATCCTCAGAAGGTTTATTTGTATGTATTAGCTCTTGCTCTTCTACTTCAAGACTCTTTTTTTCATGAAATTGAGAATGGAATATTTGTGAACCAAAGTTTTTTCTGTTAAATGTTTCTTCTTCTACTATATTCTTTTCTAATCTCGCTAATTCTTCTGTTTTGAGCTGTTTAATATTGTACATGGTTTGGTATCTCTTAGCATCTTTTTATTTTATTTTGGTGTAATCATTCATCCATCTGTAACATTTTCACCCACTTCGTTGTTAAGGTTTTCGTTATTTTTGGCTTAGAAAGTGATATGGCAAGGTTTCTGGTTTTTGTTTCCAGCATCTCTTTTGCATTTTGTTTATTTCTTGGCATCAGTATCTATGAGATTGAATAGCTTTTGAAGTAATTATAAGTTGTTAGTATGGTGGATAAGATTTTTAACCAAGTCAGCAGAGCTTTCATCATACTATGTAATTTGACTGACTGGTTTTTTTCTTTTTTTATTCTTCTTTTTATGGTTCCATGAGAACATGATTAGTTTTATTAATCTTCTTCTGCAGATGGTGAAAAATTAGTTAAATGTAGAACAATTACTATTTGGAATTCATAAGCAATTGTATAAGTCTAGTTCATTGCTAAGTGCTTTTGGAACGTATGTGCAAAGAGAAACATTAGTTGGACACATAATAATTGCTAATTGGAATCCACAAAAAGTCATCCAGTTTTAGTTCATGAAAATTTAGTGCTGATTCCAAACATATTTCTTGAGGATATTCTTATAACTTTGACTGCATAGATGCTTGCTATGTCATAATGTCTTTTTTAATCCTTCCTCTCTCACCTTTGGTGTTTGTTTGGTTCTCAAAATGTTTATCTACTGAATTATTCCAGGACGATCACAATACTGGCAGCATAGTGGCTTTCAATCTGTTGAAGGTTTCCTCATGAGAATCTACTACACCAATCAACTGCTACATTCTCTTTTTGTTGTAGCAACTCTCCTTTCGCTGTGCCGTGGTGATCTAAACTCAGACAAACAAGCCCTTCTTGACTTTGCTGCAGCAGTCCCACATGGCAGGAAGATCAACTGGGATCCTGCCACTCCTATCTGCTCATCTTGGGTAGGTGTTACTTGTAGTAATGATGGAGCCCATGTGGTGGAAGTACGACTTCCAGGAGTTGGACTCATTGGTTCACTTCCGAAAAATACTTTTGGGAAGATGGATAGCTTAAGGATTATCAGCCTTCGATCCAATCGTCTCAGTGGAAACCTTCCTTCCGCTATCACTTCACTTCCTTCTCTCCAATCCCTGTTCTTACAGCAAAATAACTTATCTGGACCTATTCCTTCTTCTCTTTCAAATAAACTGAATGTTTTGGACCTATCACATAATTTCTTCACTGGAAAGATACCTGAAACAATTCAGAATTTGTCTCAGCTAACAGGTTTAAGCCTCCAGAATAACTTACTTTCTGGACCCATACCCAATATTACTCTTCCAAGGCTAAAACATATTAATATAAGCCACAACAATCTTAATGGTTCTATTCCATTATCCCTCCAGAAGTTTCCAAATTCCTCCTTTGTGGGGAACTCACTCTTGTGTGGACTACCCTTAAATCCTTGTTCACCTGTTCTACCACCATCCCCTCCTTCCCCTTCCCCTTCCCCACCTGCACCCTCCCCAAACTCTCAAGAAAAAAGTTCAAAAAAGAAACTTAAACTAGGAGTTATTCTTGCTATTGCTGCTGGAGGGGTTGTGCTGGTCTTGCTGCTAGTTGTGATCCTGGTTTTGTGCTGCCTGAAAAGAAAAAGGGGTGATGGCAGAGGTGTGCGGAAAGGGAAATCTCCGGGTGGAGGAAGGAGTGAAAAACCTAGGGAAGAGTTCGGTAGTGGGGTGCAAGAACCTGACAAAAATAAGCTTGTTTTCTTTGAAGGCAGCTCTTACAATTTCGATCTCGAGGACCTATTGAGGGCCTCAGCTGAAGTGCTAGGAAAGGGGAGTTTTGGAACTGCTTATAAGGCTATTTTGGAGGAGTCATCAACAGTGGTGGTTAAAAGACTAAAAGAAGTCATTGTGGGGAAGAAGGACTTTGAACAGCAGATGGAAATTATTGGTAGAGTTGGACAGCATCCAAATGTGGTTCCTTTACGTGCTTATTATTACTCCAAGGATGAGAAGTTACTGGTGTATGGTTATTTCTCAATGGGCAACTTGTCAATGCTCTTGCATGGTAAGGCATCATATGCTCTCATCGTCATTTCCTTTTTCTATTTCTAGCGTTTGGTGGAGTAACTTGATTCGCCCTTTGTTATTGAATGCCTAATAGTAAGCTTCTAACACAATACAAAATGCACTCAAGTAATGTCCTAAACGAATCTAAACAATGTGAGTTTGAACTTTTTAATTTGTGAACTGGTGTGAAAAGAAGCTTTCTTTCTTGTAGCTTATTGCAAAGTTCGAGAAGAAAAAGTCTTCTTTAGAATATATGCATCAGTTTGATCTTTATAGGTCTATTATATGAGTATAAACAGCTCTTATGCTATTCTGTATCTATTGTTTCTTTAGAACTTGCTGGCTTTCTTTTACTTATCACTGGTAGTTTATCACTCGCACTATCT

>M6-PGSC0003DMG400030572 M6 LRR-RLK

CTATTTTCCCAACGTCTCTTTATTTTCTTAGCTGAAATCAACTTTACCTTTTCTCACTCTCAAAAGTCTCACCTTTCACACTCTTTTTTCTCTCTTAAGCTCCATTCTTGGTTATCTGGTGTACAAAAGGTAAGGTCTTTGAGACACCCTTTTGCTCGTTTTTCATTTTTCAGTTGTCACTGTTCTTGATTTCATCATATTTGTAGATAAAAATGCAATCTTTTTTGTTTTTTTGCATTTTCTTGATTCTGTGTAGTTGTGTTTTTGTTACTTCTTTGAATGATGAAGGAATAGCTCTTTGGTCATTCAAAAAAGGTATTGGACAAGACCCTGAAGGGTCACTTAAAAATTGGAACTTTTCTGATGAAACCCCTTGTTCTTGGAATGGTGTTACATGCAAAGACCTAAAAGTTGTATCTGTTAGTATACCTAAAAAGAAACTCACTGGGTTTCTTTCTTCTAGTCTTGGATCTCTTACTGAGCTTAGACATGTTAATTTGAGAAGTAATTTGTTTTCTGGTAGTTTACCTGTTGAGCTTTTTGAGGTTCAAGGTTTGCAAAGTTTGGTTCTTTATGGGAATTCCTTTTCTGGGGTTATACCTATTGAGGTTGGGAAGCTAAATTATCTTCAAACTTTGGATTTATCACATAATTTCTTGAATGGGTCTGTGCCTATAACATTACTTCAGTGCAAGAGATTGAAGTTTCTTCATTTAAGTCATAATAACTTTACTGGTGTTGTACCTGAGGGGTTTGGGGGTAATTTGTCTGCATTGGAAGAACTTGATCTTGGATTCAACAAATTTGATGGTAAAATTCCAAGTGATTTGGGAAATTTGTCTAATTTGAAAGGAACTGTTGATTTGTCTCATAATATGTTTAGTGGTTCGATTCCGGCTAGTCTTGGTAATCTTCCTGAGAAGGTGTATATTGATTTGACTTATAACAATTTGAGTGGTCCAATTCCACAAAATGGTGCTCTGATTAACAGAGGACCTACTGCTTTTATTGGAAATCCTGGGCTTTGTGGACCTCCGTTGAAAAACCAGTGTTCTGCGCAAAGTGACGCAAGTTCACCGTCCTCGGAACCTTTCTTGCCTAATAACGTACCTCCCTTAGATGGTGCTGGAGGGGATGGGAATGGTAGAGGTCTGAGTAGAGGTGCTGTAATCGCGATAATTGTGGGGGATGTGGTTGGAATTTGTGTTATTGGGTTGTTATTCTCGTATTGTTATTCGAGAATCTGTGGTTGTGGGAGGAAAAAGGATGAATCCAGATTTGGTTTTCAGAAGGGGGGAGGGAAAGGAAGGAAAGAGTGTTTATGTTTTAGGAAGGATGAGTCGGAGACGTTATCGGAAAATGTTGAACAATATGATCTAGTGGCACTTGATAATCAAGTGGCATTTGATCTCGATGAACTTCTCAAGGCATCTGCTTTTGTTCTTGGTAAAAGTGGTATCGGCATTGTGTACAAAGTTGTGCTTGAAGACGGACTTAACTTGGCTGTGAGGAGATTAGGCGAGGGTGGTTCGCAGAGATTTAAGGAATTCCAGACGGAGGTCGAAGCAATTGGAAAACTGAGACATCAAAATATCGTGACTCTTCGTGCCTATTATTGGTCTGTCGACGAGAAGCTACTGATATATGACTTCATTCCAAATGGAAACCTTACCACTGCAATTCACGGTACGCTTCAAACTCAGGCTGTTGCCTTCTCGTTTATCCTTTTTTCTTTGATTTCCACAACTGTTTTAAACTTTCGTGATTATTAAGTTGTTAACTTGTCTTTTTGCTTGTGTAATCAATTCTTAGGAAAACCTGGTATGGTGTCATTTGCACCGCTTTCGTGGTCCATTCGACTGAAAATAATGAAGGGTACTGCAAAAGGTTTGGTTTATTTACATGAGTACAGTCCCAAGAAATATGTTCATGGTGATCTAAAGCCATCTAATATATTACTTGGACACGACATGGAGCCCAAAATCTCCGACTTTGGACTTGGACGTCTTGCTAATATAGCCGGAGCATCTCCTACGTTGCAATCCAACCACATGACCTCTGACAAACCACAACAAAGTAAGCAAGGCAGTGCACCATCAGAGTCCGGGACAGTTACATCAACTACAACATCTGGTTCTTGCTACCAAGCTCCCGAGGCATTGAAAGTGGTGAAACCATCACAAAAATGGGACATTTACTCGTATGGAGTGATCTTGCTTGAAATGATCACGGGAAGAACCCCGATTATCCAAGTAGGTTCTACAGAAATGGATCTAGTAAACTGGATTCACTGGTGCATTGAAGAGAAGAAGCCACTTTCGGATGTATTGGACTCATATTTAGCACAAGATGCTGATAAAGAAGAGGAAATGATCGCGGTTTTGAAGATTGCAATGGCGTGTGTTCACAGCAGCCCGGAGAGGAGACCTTCAATGAGACACATATCTGATGCATTAGAAAGACTGCAAGCTTCCTCTGAATGAACGATCGAAACAGAACCTGAAGGCTTCACATTGCAGTAAGGATCACAGCATCTAACCGATAACTATGGAGCACGATCACTGCTTGGCGCGCGATCATATTGACTAACAGTAGTAATCTCCACTTGTCTGTTTGAATATTGGCCTTCTTGCTCTCTAATTGCAAGGTTTTTGTTGTGTCTATTTGTTGTTTCCTTCTATTGATGAACATAAGTTATTTGTTCTCTTTATTACTACTACTATTCATCTTTGTGTGTCACAAATGAGTTCATGTAAATAAAAATGATTGTCAAAATAATAAATAAGATAGGTGTAAGCAATAACTATATGTCTGTTGTGAGTCTGCAAG

>M6-PGSC0003DMG400030625 M6 LRR-RLK

ATTTACTCAAAACAAAAGAATATACACCAAACGGATTACCCACCCCCAAACCAAATCCTCATTTTTGCTTTTCTCACTCTCACTGAGTGAAACCGCAAACCAAACAGTTGGTGGACATTAGATTAAGGAAGGAAAAATGCGTCTTCTTTTTCTTCTTCTTCTTGTTATGCATTTTACTGACTTTTCTGCCGGTCGGTTACCGGAATACCAGGCATTGCTTGCCCTGAAAACTGCCATTACTGATGACCCACAGTTAACTCTTGCCTCATGGAACATCTCCACCAGTCACTGTACGTGGAATGGTGTCACGTGCGACACGCATCGTCACGTGACCTCTCTTGATATCTCTGGGTTTAATCTTACCGGTACTCTTCCGCCGGAAGTTGGGAATCTTCGTTACTTACGAAATCTGTCTGTCGCTGTTAACCAGTTTACTGGACCCATTCCTGTTGAAATCTCCTTTATTCCAAATCTTGGTTACCTTAATCTTTCTAATAACATATTCGGGATGGAATTCCCTTCGCAGTTAACCCGTCTGCGTAACCTCCAAGTCCTTGACCTTTACAACAACAATATGACCGGTAAACTTCCCCTTGAGGTGTATCAGATGACTAACCTTCGACATCTACACCTCGGCGGGAACTTTTTCGGTGGCCGCATTCCTCCGGAGTATGGAAGGTTCCCGTCTCTAGAGTACCTCGCAGTTTCAGGCAATGCACTGGTAGGAGAGATACCACCGGAGATTGGAAACATCACTACACTTCAGCAGTTGTATGTAGGATACTACAATACCTTCACCGGTGGGATTCCGCCGGCAATAGGGAACTTATCGCAGCTCCTCCGGTTAGATGCTGCTAACTGTGGACTTTCGGGGGAGATTCCACCGGAGATTGGGAAGCTTCAGAACCTTGACACCCTCTTCCTGCAAGTGAATTCTCTGTCTGGGTCATTAACTCCGGAGATAGGTTATCTGAAGAGCTTGAAATCTTTGGATCTGTCGAATAACATGTTCTCTGGCGAGATACCGCCAACATTTGCGGAGCTTAAGAATATCACTCTTGTTAATCTTTTTCGGAATAAGCTTTATGGGTCAATACCAGAGTTCATAGAGGACTTGCCGGAGCTAGAGGTGTTGCAACTTTGGGAAAATAACTTTACTGGAAGGATTCCACAGGGGTTAGGCACAAAGAGCAAGCTCAAAAAAGTTGATCTCAGTTCCAATAAATTGACTGGAAATTTACCCCCAAACATGTGCTCCGGTAACAATCTGCAGACAATTATCACTCTTGGGAACTTCTTGTTTGGCCCAATTCCTGAATCTTTGGGTAGGTGTGAATCACTTAATCGGATTAGAATGGGAGAGAATTATCTGAATGGGTCAATTCCAAAGGGGCTCTTAAGCTTGCCACATCTGTCACAAGTTGAACTTCAGAATAATATTCTCACTGGTACATTTCCTGATATTTCTTCCAAATCTAATAGTCTTGGGCAGATTATCCTTTCAAATAATCGCCTAACTGGACCTTTGCCGCCAAGCATTGGAAATTTTGCTGTAGCCCAAAAATTGCTTCTTGATGGGAACAAATTTTCGGGACGAATTCCAGCAGAAATAGGAAAGCTTCAACAGCTATCCAAAATTGATTTCAGTCACAACAACTTTTCTGGACCCATGGCTCCGGAGATTAGCCAGTGCAAGTTGCTGACTTATGTTGATCTCAGCAGGAACCAACTTTCGGGTGAGATTCCTTCTGAGATCACAGGTATGAGGATACTCAACTACTTGAACTTATCGAGAAACCACTTAGTTGGGAGTATTCCTTCCCCTATTTCTAGTATGCAGAGTTTAACTTCTGTTGATTTCTCATATAACAACTTTTCTGGTTTAGTTCCTGGAACCGGGCAATTTAGTTATTTCAATTACACCTCATTTCTGGGCAATCCAGATCTTTGCGGACCCTATTTGGGCCCTTGCAAAGAGGGTGTTGTTGATGGGGTTAGTCAACCTCACCAACGAGGAGCCTTATCGCCTTCGATGAAGCTTTTACTTGTTATTGGTTTGCTTGTCTGTTCTATTGTGTTTGCTGTTGCTGCAATTATAAAGGCCCGATCTTTAAAGAAGGCAAGTGAAGCTCGTGCCTGGAAGCTCACTGCTTTTCAGCGCCTAGATTTTACTTGTGATGATATTTTGGACAGCTTGAAGGAGGATAACATTATTGGAAAAGGAGGTGCTGGTATAGTCTACAAGGGGGTAATGCCGAGCGGGGAACATGTAGCGGTTAAGAGGTTGCCAGCTATGAGCAGGGGTTCCTCTCATGATCATGGGTTCAATGCAGAGATACAGACTCTTGGGAGGATCCGACACAGGCACATTGTTAGATTATTAGGATTTTGCTCGAATCATGAGACAAATCTTTTGGTTTATGAGTACATGCCTAATGGAAGTCTTGGGGAAATGCTTCATGGAAAGAAAGGCGGTCATCTACTTTGGGATACCAGGTATAAGATAGCCGTGGAGTCTGCAAAGGGTCTTTGCTATCTCCATCACGATTGCTCTCCTTTGATCCTCCACCGTGATGTGAAATCAAACAACATTCTGCTAGACTCCAGCTTTGAAGCTCATGTTGCTGATTTTGGACTTGCTAAATTCTTGCAAGATTCAGGGACATCAGAATGCATGTCTGCTATTGCTGGTTCTTATGGGTACATTGCTCCAGGTATTTTGTAAAAGCCTTCGTATCTTGAAATTCTTAGTTTCTAACATGTGTTCATATTTATTTTTCTTGTCACATCATGCTATGCGAGTTTAGTGCCGAAGTTACAAACTTACTGATGAACTAAAGTTTATCAGCTGAGACTAATTATCAGTCTGTTAAAAATGAAGCTTAGGTGTTCTAAACATCATAAATGTGCTATCCTATAGTCTTGTATTTCCACTACTAATAATGTTACGATGAAGGACAGCATTAAGACATTGATAGTGGTGGTCCTTGACTGGTAGTAAACTAGAGGATTGGGCCTTCTTACTTTTGAAAAGGATCTCTAACTACTTCGGTGATGGTTAGGTTGGAATTTATAAATGCTTATGTAACTTGGTAAGTAAGGCAACTGTGTTGCATGCTCCTGAAGCAAAGCAACATTAAATAGTGGTACTGGTACTTGATGAACTATTAAAACTGTTGGAACGGGTAAGGGGATGTAAAACTCTGGTTTCTTGTTTAGTTATCTAACAAACTTACTGATTGCTAGCAATTATTTGTTCTGCTCTATTGTCTACTTCTCATTAGCATCTTATTTTCAATTTTGCAGAATATGCTTACACGCTTAAGGTTGATGAGAAAAGTGATGTATATAGCTTAGGTGTGGTGCTATTAGAACTGGTAAGTGGCAAAAAGCCAGTTGGAGAATTTGGTGATGGTGTTGACATAGTCCAATGGGTTAGGAAAATGACTGAAGGGAAAAAGGATGGAGTTCTCAAGATCCTTGACCCAAGACTCTCAACGGTTCCCCTTAATGAGGTGATGCATGTCTTCTATGTCGCATTGTTGTGTGTTGAAGAGCAGGCTGTGGAACGCCCCACCATGCGAGAGGTAGTGCAAATACTAACTGAACTTCCCAAGTCACCAGGTGCAAAATCAGATGACTCAGCCGTCACTGATCAGTCACCCCCACCATCAGCCTCTGCATTAGAGTCCCCAACCTCAATTCCCGAGGAAACAAAAGACCACCATCAACCAACACCTCAAGCAACTCCACCTGACCTACTCAGTATCTGATTTGCAATGTTCTTGAAATAGGAGTGTTTTATTTAGTTTGATTCTCTAGTTCTATTATGATGAATTGTGCTAGCTTTGTTCTTTTTTTAAAAAAAATTGGGTCTATTTAGGCTCAGGGGTTTATTTTAACTCCAAGATGGGTTTAATGCTCAGAAGTTTTCCTCTTGTACAGTAAGATTGGTAAGGTTTTCTCAAGTGTATTAATGGAAAAATTGCCCTTCATTTGCTA

>M6-PGSC0003DMG400030626 M6 LRR-RLK

TTATGCAATATTATTTTTGAGTCCAAAAACCTTATCATTACTCTCTGATTCAAGTTACTGTAACTCTTGTCTTGTTCTTCGCCATTAGAGTCCAATACTACTAGCTTCTTCTTTTTTCTAATGTGTGAACACTCTTCACGCTTTCAGTACTATTTCTTCTCCTTAAACTCTGCCTCCGTTACACTTGTAATCCATGCAAAAATTACCCCCACCTCTTTCTAACTCTTATCTTCCCAACATTCAACCTCCATTTACAGATGTTTATGATTAATCGATTCTTGCCTTTTTAAGGACTTGGAGGTAGGGGAGATTCATTCAGTCATCAATGGATTACATGAAGCTTCAATTTCTGATACTCGTTTGTTTTTTTCTCTTCATTGTTCCGGCGAGTTCATCGCCTCGGGATATCGCTATTTTACTCCGGGTTAAGTCCGATCAACTCGAAGACCCGAATGGGTTGCTTGATGATTGGAACGGGTCTGCTCCAAATGCGCCTTGCAGCTGGAACGGGATTAAGTGTGATCGGAAAACCGGTCAGGTTGTGTCCATTGATTTTGCAAGTTTTGGAATCGCTGGTCGTTTTCCTGCTGATTTCTGCCGGATTTCGACTTTGCAGAAACTCAATCTGGGTGATAACAGTTTTGGTGAGTCCATTTCCTCTGACTCATGGTCACTCTGTTCGCATCTACACTTTTTGAATATTTCTTTAAATTTCTTTGTTGGCCGGCTGCCGGAGTTTATTACCAAGTTCGATAACTTGACCATCCTTGATGTTAATTCAAACAATTTCTCCGGTGAAGTTCCGGCGAGTTTAGGCCGTTTACCCAAATTACAAGTGCTGAATATAGCTAACAATCTCCTCAATGGTTCAATTCCTGAGTTCTTGACGAATCTTACCGAGTTGACTCGATTGGAAATTGCTGCAAATCCGTTTAAGCCAGGTCCATTGCCTTCCTCAATCGGCCGACTCGGTAAGCTTCGAGTTTTCTATGCTCGGTATGCGAACCTTATTGGAAATTTTCCAGATTCTATCAAAGACCTGAAATCTATTCAGAATTTTGATGTGGCAAACAACAATCTCTCCGGAAAAATTCCAGAAAGCTTCGGAGAACTCAAAACCATACAACAAATAGAGCTCTTTGGAAACCATTTTTCAGGCGAATTACCGGACATGTTTTCCGGTCTTGGTTCTCTTTCCAGGTTTGACGCTTCTGAGAACAATCTCACCGGGAAAATACCTGAAACCCTTGCCCATTTGCCATTGGAATCTCTAAATCTCAATGATAACCAATTAGAAGGCGAAATTTCAGAAAATTTAGCTCTTAACCCAAATCTTAGTCAGTTCAAGCTTTTTAACAACAGATTTTCAGGTACTTTACCTCAAAATTTTGGTTTAAGTTCAGATTTAGATGAGTTTGATGTCTCCGGCAACAATCTAAAAGGTTCTTTACCTCCCAACCTATGTTCTAGAAAGAAACTTAGGATTTTGAACCTGTTTGATAATAAGTTCAATGGGCCAATCCCAGAATCCTATGGGCAGTGTTATTCACTATCATATGTGCGTATCTATAACAATCAATTCTCTGGTGAATTACCAACTGGTTTCTGGGGATTTGATGGATACACATTTCTTGAACTGCGAAACAACAACTTTCAAGGTTCAATTCCAGCTTCAATCTCCAATGCTCGAGGCCTGACACAAATTCTTATCTCCGGCAACAAGTTCTCCGGGGAATTGCCAACAGAAATATGCAATTTGGAAGAGGTTGTGATCATGGACATTAGCAAGAATCAATTATCAGGGGAGTTGCCTTCGTGTATCACAAGATTGAAAACATTACAGAAGCTTGATCTTTCACAAAATAGGATCAAGGGTCAAATTCCCAAATCAGTTAGTTCTTGGAATGACCTGACTGAATTGAATTTAGCTAACAATCAATTGACAGGTGAAATTCCAGGTGAGCTTGGGACGTTACCGGTCTTAACCTACTTAGACCTTGCCACAAACTTACTTTCTGGCGAAATTCCATCTGAGCTGAGCAAGCTCAAGCTCAACAAATTCAATGTATCAAATAACAGGCTGGAAGGGAAAGTTCCACTTGGGTTTGATAACGATTTTTTCGTCTCAGGTTTACTGGGCAATCCGGATCTTTGAAGTCCGGATCTTAAGCCTCTGCCCCAATGCCGAAGACCCAAAAGTGTAAGCTTGTACTTGGTGTGAATTTTATCAGCTTTTGCCTTCATACTTGTCGGGTCACTTGTTTGTGTCTTACTCAAGGCCAGTAAGCTGCTACCAATCCGTAGTAAGCGTAAAAGTGTGTGGAGAATTACTGCATTCCAACGCGTCGGTTTCACAGAGAGAGAAGTGTTAGATGCACTGATAGAGGAAAATCTCATTGGAGCTGGTGGGTCGGGTCGTGTGTACCGGGTCAAATTGAAAAACGGACAGATGGTAGCAGTGAAGAAACTTTGGGCGGCTAAACGGGAAAGAGAATCAGAGGAGGTGTTCAGGTCTGAGGTGGAGACATTAGGGAGAGTCCGACATGGAAACATAGTCAAATTACTATACACTGGCATTGGTGATGACTTTAGGATATTGGTGTACGAATACATGGAAAATGGAAGCTTAGGAGATGTATTACATGGGGAAAAAGGTGGCTTGTTATTGGATTGGCAGAGGAGATTTGCCATAGCAGTTGGAGCAGCTCACGGATTGGCCTATTTGCACCATGATTCTGTGCCAGCAATAGTGCACAGAGATGTTAAGTCTAATAACATTTTGTTGGACGAGGATTTCAGGCCCAAAGTGGCTGATTTTGGGCTAGCTAAGGCAATGCAACGGGATGCTGAGGAGAGTGAACAAGCCATGTCCCACATTGCTGGTTCCTAAGGCTACATTGCACCTGGTAAGCTTAAATTCCTTATTGTTTTTCACCATTTATTTTAGGTCCAATGACTGTTAGAAATTTTCTAGTCTTGTTAGAAATTTATTTGCCAATAAAAAATGTTGAAAAAATCTAGTGCTAAGGAGACTATTTTAAAAAAAAAAATATTGATCCAAGATAGGCATAAATGAAGTGGTATGGACAGCAGGAATTCATATGCTAACCTCTACTTGTTTGGGATCGGGGCCTAATTATTGTTGTCGTTCAACTTTTTTGTTTACTTGTTGTGAAAAGATATTTAATATCTTGCGTTTGGCATACCAAGGACATCCAAAAATAGTTTTGTGGAAGTTTTGGTTAACTTCACATACTATCTTAAAATTTTGATTTGATTTGATTTGAAATTCTTTAATATGAAGCCAGAGAAGATTTTAGGAGATGATTCAAACATATAAGTAGATTGGAAACCTTAATAATATCCATTTAATTGTAGCATCTAGTTTCATTCCTCTAACTGGACGTTGTTTCTGAAGTCTGCATATTCTTGTTTGTCTTGCTAATGCTTGATTATATGTTTAACAAGGGGATTTGTCAAAGTAAAATCAAAGAATGATTGGGCACCCCATTTTTGAAGTCTGCATGACTATTTAGTTTTAACTACCTGTAATATTTAATCCTGAGTTTCGTTTGGGATTTGTCAAAGTAAAATCAAAGCATAGTAGGGCGCTCCTTTTTCTGAAGTCGGCATGACCATTTATTCTACACCGGTAGTAATATTTAATTCTGAGATCACATCAAATTATTGAATGATTTTGTCACCCAATGTTTGTTACACAGTCAAGATTTGTTTGAAAGAATAAAAAATAGGAGGGTCATGTGTGTCCTGTCACCTAGCATTTGTTAACATATTCAAGATTTGATCTAAAAGAATAGGATTGAAAGAGACATGTGCCTAAGTATTGTTACTTATTTATCTAATTTGTCGATATCTTAAATTCTATTTACTCCCTTAAGTCCTCTTTGATTGGACATACAGTTTCATGTGCCCTTCCTTTGCAAAACTGAGAGAATACAGAAACCAATATCCTTGATGGAAATTTTTTGTGGCCTTTTGATACTAGAAAACATTCCGGTTGCAATGACAAACAGTTTGTTAAGTTGATCTTGGCTACATAACGATTAGGAAGATGCTAAATATCTCATGGGATGGCACATGTGTTATCCAAGGTTTTCAGCTGCACGATTAAACTTAACCTCGATTCCTTTGTTTCAATTGCAGAATATGCATACACTCTGAAGATCACTGAGAAGAGTGATGTTTATAGCTTTGGTGTGGTACTGTTGGAACTAATAACTGGTAAAAGGCCTAATGACTCCTCTTTTGGAGAGAACAAGGACATTGTCAAGTGGGTGTTAGAGGTTGCAACATCGTCTAAGAAAGATGAAGGAACTGGCCATATTGTTACGTGCGCAAGTGGTATTCTTGATTTGAACCAGCTAGTTGACCAGAGAATGAATCCATCTGCAAGCAATTATGCAGAGATTAAAAATGTTTTCGATGTGGCTTTGCTTTGCACTTCAGCATTGCCTATTAATAGGCCCTCCATGAGAAGAGTTGTTGAGTTGCTGAAGGGTATCCCCTCCGCTCGCACTAAAACAACCCATTAGCTTTTCACAGTTTCAACATGTCCTTGCTCAAGTAACTTGATACTATTGTAAACTGTAAAAGAATGTGTGAAATTAGGTGTGTTTAGGTCTGCTTCATAGGTCGATTTAGTTTGCTTGCTGTGGTTTAACAAGCACCTTGTTAGATACTCCAGTACTACACAGAGTAGCTTAACTAAAAGTTTGCTGACCCTGTAAAACTCATGTTGTAACGACTAAACCTTAGAGTTTTTGTTGCAATATATGAACTTTACTATCATATACAAGTATTCTATTCTGTATCCATG

>M6-PGSC0003DMG400031028 M6 LRR-RLK

ATGGAGGTGAGCGTGAAGATGAAATTCCCCTCACAAGCACTACTGTTGGCTCTATTGCTTGTTTTACCGATCGTTTTAGCTCTCACCGAAGAAGGTAACTATTTCTTTGAACATCTAGAAGCTGTTTTGGTGTTTTCTAGCTCCTCTGTTTTGCTAGCAGAAGGTAGTGCTTTGTGTGCTATCTAGGAAATACGAAAGTTATTGTTCGAATTTAGTTTTTATTTTCAGTTTCTGGAGCAGCATTCCATGCTATTTAACTCTGATAACGAAAAATCCGTGTTATTTAAGATATATTTAGCCTCCAGTTTATGGCATAACAACCGTAATACTAGTTGATGAAAACTGAAAATGGTTATGACTGCTTGCGCAGGCGAAGCATTAATGTCGATCAAGGCATCGTTTAGCAACGTGGCAAACGTGTTGCTAGATTGGGATGATGTCCATGACGAGGATTTTTGCTCATGGCGAGGCGTGCTGTGTGGAAATTTCTCCATTTCCGTCGTTGCCCTGTAGGTTTTCCATCCTTGTCTCCTAACTTTTACTGATACACCAGAACAAAGCAGTCTGATGACCTGCTAGCTATTGTATAGCTCTTTGTTAATTTAGCTAAAACTTATACGTCTTTTATTTCAATTTGCCTTTTCTGCTTGCGAAGGAATCTGTCCAATCTTAACTTGGGCGGGGAAATTTCACCAGCCATTGGAGATTTGAAGAATCTAGCGGCATTGTATGGTCCATGTTCTCTTTTACTGTTCTTGGTCCATTGACTGTTATTTTTCCATGCTGATATTATATTCTAATGTTAATGACAGAGACCTTCAGGGAAATAAATTAACTGGTCAAGTCCCAGATGAAATTGGCAACTGCATTTCACTGATCTATCTGTAAACTGTTTTTAACCTATTCTTTTAATATCTTTTTCTTGCGTGTCAGTAAATTTAAAGTGTTCACACTAGTTGATTTGCCATTTTCTGCAGTGATTTGTCTGATAACTTGCTCTATGGAGATATACCTTTCTCAATTTCTAAGCTCAAGCAGCTAGAGTTATTGTGAGTTATTTAATCACGTTAAATTGATATCTTCTCATATCTGATATGGTTATGATGAATTAATTCATATGTGGTACAGGAACCTGAAAAACAATCAGTTGTCTGGCCCAATCCCATCCACATTAACTCAAATCCCTAATCTAAAGACGCTGTGAGTTCCATGACTTTAGTTTATCTCACTCAAAATTTAGTCCAATATACATGATTAACTAATGGTTATTTGATGGTGAAGTGATCTGGCTCGAAACCAGCTCATTGGTGAGATACCAAGGTTGATCTATTGGAATGAAGTTCTGCAATACCTGTGAGTGCATGTTCCTGGTGTTTTGGAGGTTTTCCTTTTTGTTTGTGAAATTTACAATGTTATAATTACCTTCTGTTTTGCAGTGGCTTAAGAGGCAACATGTTGACAGGAACATTGTCCCCTGATATGTGCCAGTTGACTGGTTTGTGGTATTTGTATGTGCTCTGCTACAAGCTGTCTATACGGAATGCTCTGTTGTCTGTTTGGTGTTATGTTTATTCTAACATAAGAAGTTTCATATTATTTCAGTGATGTGCGGGGCAACAACCTCAGTGGAATAATTCCAGATAATATTGGGAATTGTACAAGCTTTGAGATACTGTCAGTGTCCTCTCCTCGCTCTTGTAGCATTATGCTACGGTTTTTACTCCTACTGCTGCCCAATTATAACAAAGTATTTTTTTCGTTATTTCAGGGATATCTCATACAATCAGATAACTGGAGAAATTCCCTACAATATTGGATTTTTACAAGTGGCTACCTTGTTAGTAAATTCAACTTTTAAATGATTTTTAGCTGTTGTCATTTTTACCTTGTCTGTTCTGTTATGGGGTTCATTTCTGTGAATTGTAAATGGTGATTGTGGTCCTTTCAACAGGTCTTTGCAAGGAAATAGGCTAACTGGGAAGATCCCAGAAGTGATTGGTCTAATGCAAGCTCTTGCTGTTCTGTGAGTATTCATACTACTACATATTTTAATTTTTTTTATCAACTCCATTTCTTACTATTTACTACTTGCAAGTTAGAAGGTTTCATGATGTCTGTTTCCTCTGTAGGGACTTGAGTGAAAATGAGTTGGTGGGACCAATTCCTCCAATCTTTGGCAATTTATCCTACACTGGGAAGCTGTAAGTTTTCACCCCTATTTTAATGCATATACTTTTCTACGTGTGGCTCGTTTGTCTGATTCACTTGTACATTCAACAGGTACCTGCACGGCAACAAACTTACGGGGCCAATACCACCGGAGCTAGGAAATATGTCTAAACTTAGTTACTTGTAAGAGGATTAATTCACAATTTCAGATACATTATGCAGTCACCTTTTTTTTTGCTTCATGCATTGAGCTATCTTATTTACAGGCAATTAAATGACAATCAGCTAATGGGGCGAATTCCCTCCGAACTTGGCAAACTGGACCAGTTATTTGAATTGTGAGCCTTTTATTCTTGTAGTATATAAATCTGGTTCCATTTCTCTTAGAATATTGTGTTCTCAGCTTTCCTTTACCAGGAATCTTGCAAATAACAAGTTGGAGGGACCAATTCCTGAAAATATCAGCTCCTGCTCGGCATTGAATCAACTGTAAGATTGTTGCGTATCCTTTAGGAACTTCTTTTTTTCTCTTTCTCTTGCTATTTTCATTCATTCTGAGTCTAAACATATTTGTTTCTTTCTTTCAGTAATGTTCATGGCAACAACTTAACCGGGTCCATTCCTTCAGGGTTTAAGAATCTTGAGAGCCTGACGTATCTGTATGTTATATGCTTCTGGTTTGTCTAGTGATCATGAAAATGATTTTGAATTTCTTTCATATGGTACAAACTCATTGGTGACATTTTGTTGATATATACCTGGCAGAAATCTTTCAGCAAATAAATTTAAAGGTCACATACCCTATCAACTAGGGCGAATCATCAACCTTGATACATTGTGAGTTCTTGACTGTCAAGGTCAATAAAATTTATTAGTTCAAGTGATTTCTTGTACTAAAATTTGTACTCTGTAGGGATCTATCTGGCAATGAATTTTCTGGGTCTATCCCTGGTTCTATTGGAGATTTGGAGCATCTCCTCACACTGTAAGATATTAAACATTTTGATGTACAAGGAGCTTGTATCATATTAAGACTTGAACATAATCACAATATCATTGCACATTGTAGGAATCTGAGCAGCAATCATCTTGATGGACAAATTCCTGTAGAATTTGGAAATCTGAAAAGTATACAGACCATGTGAGTTTCTGCTGAGAGCTTAGGTTACACTGCCTTCAAAATGTGTACTTCTTAGTATACTGATTTTCTTGTTCTGTGATTGGTTTCTATTCTTTCAGTGATATGTCGTGCAACAAGATTTCTGGTGGCATCCCAAAAGAGCTGGGACAGCTGCAGACCATGATAACTCTGTAAGTACATTTAGTTTTTGTCAAGTAAATTGATGTAGTGTTTAATCAGTTGTCCTTAAAATCAGTCAATTCTAATGAATTTTTGATTGGCTTTGTTTCTTTATGAAGTACTTTGACAGCTAACGATCTTAGTGGAGCAATCCCTGACCAATTGACCAATTGTTTCAGCCTCACTAGTTTGTGAGTTCATCTATCTTTGTCTTAACATCATAGACATTCTAATTCTTCCTACAGTTACTGATCTTATGCCATTTTCCTTTAGGAATATATCCTACAACAATTTTAGTGGTGTTGTTCCTCTTTCACGGAATTTCTCGCGGTTTGCACCTGACAGGTATTTTCCATATAGAATGAAGCTTCATCATTATATATGAACATTAGTACAAGTTAATGACCACCTTACTATGGTCATCAGCTTTTTGGGGAATCCATTTCTCTGTGGCAACTGGAAAGGCTCAATATGTGACCCCTATGCACCAAGGTCTAACGGTATGACTTCATTTTTGTGCATTGGTTAGTGAATCTTTTGGTATGCTAGAGTCATGTGCATTAAAATGACTTGTTACTTTTGCAGCCTTGTTCTCTAGAACAGCTGTTGTTTGCACAGCATTGGGTGTCATAGCACTCTTATCCATGGTTATAGTGGCAGTGTACAAGTCAAACCAACCACACCAGTTTCTGAAGGGGCCTAAGACCAATCAAGGTAAAAATTAGTACATGTCCATTCTGTTCTTTTGTTTTTCAGTACTTTCAGGTATTTATGTTAGCTTTAATCTTGTTTCCCTCTAATTCCCTAGGTTCCCCCAAACTTGTGGTTCTTCACATGGATATGGCCATCCATACATATGATGACATTATGAGGATTACTGAGAACTTCAATGAGAAATTCATCATAGGATATGGTGCTTCCAGCACTGTATATAAATGTGTTTTGAAAGATTCCCGACCGATTGCCGTTAAGCGACTTTACACTACACATCCGCACAGCTTGCGAGAGTTTGAGACTGAACTGGAGACAATTGGAAGCATCAGGCATAGAAACCTTGTTAGCTTGCATGGTTACTCCCTTTCCCCTCATGGGAATCTCCTTTGTTACGAATACATGGAGAATGGTTCACTCTGGGATCTACTTCATGGTTAGTAACCCTCTTTTCCTTGTAATCTTTTATAAAGCATCTTCGTGTAAGATATTTATTACTGTTTTCCCTGTCGGAGAATGGTGCTATAAGCCAGGATTGATTTGAAAATGGCTAATTATCAACTGAATACATGTCATAGGGCCTTCCAAAAAGGTGAAGCTTGACTGGGAAACACGTCTGAGGATTGCTGTTGGTGCTGCTCAGGGTCTTGCTTATCTTCACCACGATTGCAACCCAAGAATCATCCACAGAGATGTGAAATCTTCAAACATTCTAGTTGATGAAAATTTCGAAGCTCATCTTTCTGATTTTGGGGTTGCAAAATGCATCCCTACTGCAAAAACTCATGCATCAACTTTGGTGTTGGGTACCATAGGTTACATTGACCCTGAGTATGCCAGGACTTCCAGGTTAACTGAAAAGTCAGACGTCTACAGCTTTGGCATTGTTCTCCTAGAGCTTTTGACAGGAAAGAAACCGGTTGATAATGACTTGAACCTGCATCAGCTGGTATTACTCTCCTCTTATAGTCTACGTACGTTGTTACTTGTAAAAAAGATTTAACTCAGACTGGATATAGAAAAGAACAACTTATCTCCAATTTTCCTACCTTCCTCTTAGCATTTGCGATATTATCATTCCATCAAATGTCTTTTGTCTTTTAACTCCTGTATTATATTTGATGTCTTTATAGAGTACTTTGATTTGTGTTACAGATAATGTCAAAGGCGGATGATAACACCGTGATGGATGCTGTTGATCCTGAGGTATCTGTTACATGTATGGACTTAATGCATGTGAGGAAAACTTTTCAGCTTGCGTTGCTGTGCACAAAAAGATTTCCATGTGAGAGGCCAACGATGCATGAGGTAGCTAGGGTACTTGTTTCCTTGCTTCCTCCCCCGCCAACCAAACCTTGTTTAGACCCACCTCCCAAATCCATTAATTATACAAAGTTTGTGATTGGGAAAGGACTACCGCAAGTTCAGCAGGGTGATGATTCCTCCGAAGCACAGTGGCTTGTTAGATTTCAAGAAGCTATATCCAAAAACTCCCTTTGACTTGGGTTCATAGACACTAAATATCAAATCAGAAGAGAATATGGAACACATATATATGTTGGCCAAAGGTTTTTTGATGTTAGAGATTTCCCCTATAAGATGGTTAGGTACACGGTAATAGCTTAGTGTATTTCACATTTGTAGATGGAATGTTTTATGGTGTGGATGATATCCACAGTAGGTTAGATCTACAACCAGTGCAGTTTAACTAAATCTTTGTTGA

>M6-PGSC0003DMG400031103 M6 LRR-RLK

CAATTTTATCATCACCATATAGTTGTATTCATTCACATATCACACTCACATGTTCTAGTACTCCTTCCATTCTGTAAAGTTCTAGTTGACTCTCTCCCTTTGTGGTCCTTTTCCATTTTCTTGCCTTATCAATCAATCAACTTTCTTAGCCAATTTTTTTCTCTGTTTTCACATCCTTTTTTATCCTCTTATTTAAGTTTTAGCTAAACCCCATTTCTTGATTTTTAGTCTTTTGCATGTCCTGAGTGGCATTATTTTTCAAGATTTCATATTATAGCTAAACCCCATTTCTTGATTTGTAGTATCTTGTCTTTTGCATGTCTTGAGTAGCATACTATCCAAGATTTCATATTATAGCTAAACCCCATTTCTTGATTTTGTAAAACCTTTGGTGATTTGCATGTCTTGAGTGGCATGTTTTTGTGGTCATTAAGGTTAAAACAATTCAAGATTTCATTTATTCCTTTGCTTCTTGAAATTGGGTTGAGTTGTATACTGAGAGTTATGGGCCCTGGAATCTCAAAACTTGCTTAATGGTATCTGTTTCTTGGTCTGCCATTACCTCTGTTACAACTTTTGGGCATTGCCATACTCATAAGTTTCCTAATTTTCATCCATAAAAAAAATATTGACATGACTTTAGTGTAAACTAAACAGCATGTCATTCATGCTCTTTTATCCTTGGAAATGGCAAGTTATAGTAACTCTGCAATCCAACAGCATTTGCTTTTTGTGCTGTTGATTCTTCTTTTCTTTACTGGTTTTGCTGAATCTTTAAATGAAGAGGGACTTATTCTTTTGGAGTTCAAAGAATCCCTTAATGATCCTGATAACAATCTTGAAAGTTGGAATTCTTCAAACTTGAATCCTTGCAAATGGGATGGTGTTAAGTGTAGCAAAAATGATCAGGTGATTTCTCTCAATATCGATAATCGTAACTTGTCTGGTAGTTTTTCTTCAAGAATTTGTGAACTTCCTTACTTGACAGTATTGAATGTGTCATCAAATTTCATTTCTGGTCAAATTCCTGATGATTTCGCGTCGTGTCATAGTTTGGAGAAATTAAACCTTTGTACCAATAGGTTCCATGGAGAGTTTCCTGTCCAATTATGCAATATTACCTCACTTAGACAGCTTTACTTATGTGAGAATTACATTTCTGGTGAAATCCCTCAGGATATCGGAAACTTGTCACTTCTTGAGGAGCTTGTTGTTTATAGTAACAATCTTACTGGAAGAATACCTGTTTCAATAGGTAAATTGAAAAAACTTAGGATCATCAGGGCCGGGCGAAACTATCTGTCTGGCCCTATTCCTGCTGAAGTTAGTGAATGTGATAGTTTACAAGTGCTTGGTGTAGCGGAAAATAGGCTAGAAGGGTCTTTTCCAGTAGAGCTACAAAGGCTTAAGAACCTCATCAACTTAATTCTTTGGGCTAACTCATTCTCTGGTGCAATTCCTCCTGAGGTTGGGAACTTCAGTAAGTTGGAATTACTCGCGTTGCATGAGAATTCATTCAGTGGACAAATTCCTAAAGAGATTGGGAAGTTAACTAATTTGAGAAGGTTGTACATTTACACTAATCAATTGAATGGAACAATTCCATGGCAAATGGGGAATTGTTTGAGTGCAGTTGAGATTGATTTATCCGAAAATCAGTTGAGAGGAAGCATTCCGAAAAGTTTGGGGCAACTTTCTAATCTGCGGTTGCTTCACCTTTTCGAGAACAGATTACATGGGAAGATTCCAAAGGAGCTTGGAGAGTTGAAGCTGCTTAAGAATTTTGACTTGTCTATAAACAATTTGACAGGTAGAATTCCAGCAGTGTTTCAGCATCTTGCATTCTTGGAGAATTTACAACTTTTTGACAATCACCTTGAAGGTCCTATTCCGCGATTTATTGGCCTCAAAAGCAACCTCACTGTAGTGGACCTGTCCAAGAATAATCTTGAAGGAAGAATACCTTCAAATCTTTGTCAGTTTCAGAAGTTAACATTCCTGAGCCTCGGGTCTAACAAGTTGTCTGGCAATATCCCTTATGGCCTAAAGACTTGCAAGTCCCTTGAGCAGCTAATGCTAGGAGATAACCTGCTCACTGGGAGTTTTTCTTTTGATCTTAGCAAGCTGGAGAACCTTTCAGCTCTTGAGCTTTTTCACAATAGATTTTCTGGATTGCTACCACCAGAGGTGGGCAACCTTAGAAGATTAGAGAGGTTGCTCTTGTCAAATAACAACTTTTTTGGGAAAATTCCTCCTGATATTGGAAAACTTGTGAAGCTTGTTGCTTTTAATGTTTCCTCCAACCGGCTCTCGGGTGATATTCCTCATGAATTAGGGAATTGTTTAAGTCTCCAAAGGCTTGATCTTAGCAAGAACTCGTTCACTGGAAATCTCCCGGATGAACTTGGTAGGCTAGTCAAGTTGGAACTGCTTAAGCTATCTGATAATAAGTTTAATGGACAGATACCTGGTGGCTTAGGTGGACTAGCAAGACTAACAGATTTGGAAATGGGAGGCAATTTCTTCTCTGGTAGTATTCCTATTGAGCTTGGCTACCTTGGAACTCTTCAGATTTCTCTGAATCTGAGTCACAATGCTCTCAATGGATCGATTCCTAGTGCCCTCGGGAACTTGCAGATGCTTGAAACATTGTACTTAAATGACAACCAGCTTATTGGTGAGATTCCAACTTCAATAGGGCAGTTGATGAGTCTGATAGTATGCAATCTTTCTAACAATAACCTTGTGGGATCAGTACCAAACACGCCTGCCTTTAAGAGGATGGACTCGAGCAACTTTGCTGGAAATGTTGGGTTATGCACGTCCGATTCCATCCACTGTGATCCTCCTCCGGCCCCTTGGATTGCTCCAAAGTCAAACTGGTTGAAACACGGTTCTTCTAGGCAAAAAATAATTACTGCTGTTTCTGCGACTGTTGGGATGATCTCTTTGGTATTGATTTTAGTTATATGTAGGATCATTAGAGGCAACAAAGCAGCTTTTGTGTCCGTGGAAAATCAAGTAAAGCCGGATGACTTGAATGATCATTACTTTCCACGAAAAGGGTTCACGTATCAGGACCTTGTCGATGCTACTGGAAATTTCTCAGACAGCGCGATCATAGGAAGGGGAGCTTGTGGCACTGTCTACAGAGCTCATATGGCTGATGGTGAGTTTGTTGCAGTTAAAAAGCTGAAGCCACAAGGAGAAACCGCGAGTGTTGACAGCAGCTTCCAAGCTGAATTATCTACACTTGGTAAGATAAATCACAGAAATATTGTCAAGCTATATGGTTTCTGCTACCATCAAGATTGCAATCTTCTCTTGTATGAGTACATGGGAAACGGAAGCCTCGGGGAAGTACTTCATGGGAATAAGACAACTAATTTACTAAATTGGAATAGCAGGTACAAAATTGCCTTAGGAGCTGCTGAGGGACTGTGCTATCTGCACCATGACTGTAAGCCACATATCATTCACAGGGATATAAAATCGAACAACATTTTGTTGGACGAATTGCTTGAAGCACATGTAGGAGACTTTGGCTTGGCAAAGCTAATTGACTTCCCTTACTCAAAATCCATGTCCGCGGTTGCTGGTTCATACGGCTACATTGCCCCTGGTAAGACCGGTGTTTAGGCAATTAAAACGATACTCTTTATTAGGATTACTTACATAATACAGAAACAAACTAAGTTCTAGTTTTTTCTTTTTTCAGAATATGCATACACAATGAAAGTGACAGAGAAATGCGACATCTATAGTTATGGCGTTGTTTTGTTGGAATTGATTACTGGTAGGTCTCCAGTTCAACCCCTCGAACAGGGAGGCGATTTGGTGACTTGGGTAAGGAGATCAATTCATGAAGGAGTGGCACTAACTGAGCTTTTTGACAAAAGACTGGATTTTAGTGTTGCAAGAACCAGAGAGGAGATGTCTCTAGTTCTCAAGATTGCTATGTTCTGCACCAATACATCCCCTGCCAATAGGCCTACAATGCGAGAAGTCATCGCCATGCTGATTGAAGCCCGGGAATTTGTGTGCACTTCACCACCATCACCATCCTCCGAAACTCCTCTGAGTGAAGCTGATGCTAATAAAGGTGCATAAAAACAACTTCCTCATCTTTCAATCTTTTCACTACAGTCAAACATCTCTATAACAGTCATCCTCTATAACAACATTTCAATATAACAAACGGGTTTTCTATGGAACCAATTTTCATGTTATGTTACATTACATGTTCTCTATAACAACATTTCGCTATA

>M6-PGSC0003DMG400031228 M6 LRR-RLK

CAAACCTGCAGGAGTCTTAGGAACTGCAACTTGCAAGGAACAGTTCCTGATCTAAGCACAATACCACACCTTCTTTACTTGTAAGCATACTTCATTGAGAAATTCCTGTGTCCTGTTAAACCACATTTTGAAGCTTATGTTCTTGTTGTTCTAACCTCTTGGTAATGTTGATATCTGCTTAATAATTCTTATATGCCATCTGCTTAATAATTCTTATATGCCAGATGTAGTTGAACTCATCTGCCACTCAATTTGTGCTAAATATAGAAAGTTAAATTTACAGCCAGTGGCAATTCATCTGATATAAATTTCTCATGTCACAGAGACCTTAGCAGAAACCAACTCACAGGAAATATACCCTCCAATAAGCTGTCTGATAATATCACAACCATGTAAGTTTCTGGTTAAGTGTGATACTGCTGAAATCTTTCCTGGTTTGATTCAGTCATCTTTAATCACACTTCCTTCCTCTTCCCCCCCACCCCAACTCTCTATCGATATCTGTGGATAGGTGGTTGAATGGACATTAGTGTCTTAACACGCACTCTTTTCTTTTTGATGTTTGAGTAACTCGGTCTTGGTATCAGACCCTCAATAGTGAACTAACTACTTTGATGATGATTGCAGCATTCTGTCAGGAAATATGCTTAATGGTTCTATCCCTTCGAACTTTTCTGTCCTTCCAAATCTGCAGAGGCTGTAAGGAGCTTCATTTGCTCAATCCTTGTTTCTTGATCCTCTGTCTGTTTCAAATTTCATGTCGATACTCTAATAAATGCAGTATTGACACTACCTTTCCTTAAATTATTTTTCTTGTAGGTCACTCAACAATAATAGGTTAAGTGGTTTTGTGCCAACCACCATTTGGGAGAATAAGACATTTTCCCCAGGCGCAAAACTTAGATTGTAAGTGTCTCATTTGTACCTTTTCCTCTTATCAGTAATTTGATTTTCTTGTTTCTGTTGTCTGCTGCCCCACAATCTATTGGCCAATTAAAGTTTTTAACTGTAAGATACAGACAAAAACTTTTTGTAGAGAGTCGAGCCAAATTTGTGTGCCTATTCCATTTTGTTAGCTAGTCAGTACACATTTTGGATATGCATAGAAGGTATCCTGGAAAAAATAGTACAGTCAAGGAATAGTGTGTCCATTGTCAACAAGGGTTTCCTATCTCTATACAGTTAATATATAACAAGGGATAATATGTCTGATGTTTGTTTTGTTCGAGGTAATGTAGCATGTATTCTTTCCTCTTACTAAACCATGCCATGTCTGACTAGAAAAGGCCCCTTAGATGTTAACATATCAGATTGCCAGGCTGATCCAGTTGTTCTACTTTGTGCAGGAACTTTCAACGTAATTTTCTCTCAGATATTTCGGGTAATCTTGATCCACCTCCAAATGTCGATATTATGTAAGACCTTATATCCGGTAGTCTTGTACATACATATTTGCGGACTTACTTTCAGTTCTTGGCTTTTATATTTAGATGTGGCGAATCAAGCTCCCCTAATTCAATTGCTATCTAACTTTCATTATCCTGTTTTTGATAAGCTTCAAATTACAAAGTAATTCTGTGGAGTCTTAGTTTCTATTTCAGATTTCTGTTATTCAACAGAACTACTATCATGAGGAAGGCAGGAAAACTTGCTATAGAATATCCCCATGACAACATTTAGGATAGTTGCATAACACTGCCTTCATTTGCTGTTTTCCTGCTAAACTCTTTCCATCAGTGTAGAGATTTACTTCGAGTATCCATTAATGAAGCTAGAGTAGTGAACCAACACAGTTTCAACTTCTTTTATTTTTTTCATCGTTTCAAAACTTCATTTTTTGTCAGAAGATGGGTAGTTTTCTTGCTAGCAGTGCACTTTATATGGAGCTCAAAAACAATGAGGTCAAATTCATATATGTGAATTTCTTTCCATATGATATGGCAATATGACAAAGCTAGGGTTATGGGGGAGAATTTGGTGGCAATAGGGGCCTGGGGAAGCAGTGGGGATGCGACCAATATGTGGGATAGGACCTCTAGTTGCATAAGGGAAGCGACTAGAGAGGTCTTGGGGGTTTCAAAAGGTAACTCTAGTGGACACGGAGGAGATTTCTGGTGGATTGGCAAAGTCCACGTAAGGTAGAGGTAAAGAAGGTGGCTTATGTGAAGTTGCTGGAGACCAAAAGGATGAGGAGGATAAGAGGGCGACGAGAGAAGTATATAAAGTGGAGATGAAGGAGGCAAAGTTGGCAGTTATGGAGGCAAAAATGGCAGCTTTCGAGCGCTTGTATGAGGAACTAGGGGACAAAGGTGGAGATAAGAAGTTGTACAGGCTACCCAAGGTGAGAGAGAGGAAGACTCGAAACCTGAACCAAGTGAAGTGCATTAAAGATGAGAAAGGCAGAGTATTGGTGGATGAGAGGCGAAGTTGGCAGTTATGGAGGCAAAACGGCAGCGTTCGAACGCTTGTATGATGATCCCTGCATTATGATCTTTTTATTTTTATTACAATTCTGGCATAACTTGCCTCTAAACCAGACAACTAACTAGTTACATACTGAAGTGGTAACAAACTTTCCAATTTGATGGAGAGATCATGCCTCAGCAATATCTTTCAAGCTGTATATATTTAATTCTAAAAGCAAATTGTTCAACTTCTTTAAATAAATTCCGGCTTCACCTACTAACTTTATTTTTTTGAAGAGCAGCTTCACCTACTAACTTTATACATGTTTGTGTGCAGCTGTGTGCGCCATATTTGCTGATTTCATGAATAACATAAATAATGAGTTTGAGTTTTGTTTTTAACCCATTTACTGGTACTTTAACGTGCAAATATTTTCAATGGCATAAAATCCCTACATACATTTTGCTTTATCACTCCTTTATATTTCTGATTAAGTATCCATTAGGTGCGACACTTTAACTGAATAACTTTCAGGCTTTATGGAAATCCTGTTTGTGGAAATGCAAATGAGCGCCAGATTACCCAATTTTGTAAATCAAGAGATGGAGGTGAAGAGTATGGAGGCTTGAACAACTCTATTCCAAGTTGTGCAGCCCAGCTGCCTTGCGACATGGATTTTGAACATGTCCCAGCATTAATGGATGGCTGCTTTTGTGCAGCACCTTTTGGAGTTGGTTTGCGTTTAAGAAGCCCTAGCATTTCAGACTTTCCTCCACATTATAGTGACTTTGAACAGTGGATAACAAAAAGTGTTAATTTGAATGATTATCAACTGCACATTGATTCGGTTGCATGGCAAAGCGGTCCTAGGCTTAGATTCTTTTTGAAATTCTTCCCTCCACGTGCAAATGACTCTGGAACCAATGATTTTGGCAAATTTAATGACAGCGAGATAGTACGGATCGCAAACAAGTTTGCCTTTTTTAATCTTACTGGAAGTGATATCTTTGGCCCATATGACCTGCTCAATTTCACTGCCATGGCGTATAATTCTGGTATGTCTCCAAATCCATTATGTTCTTGCATAAGTAGGATATAATAATACATTAACATGTGGACATTTTCAGTTGTGGTTCTGTAATGATTTCCTGAGGAATTTAAAGTAAAACCAATGTGAACAATAATAGAAAAAAACAACGTGAATCTCGTTTGTTTTTTACAACTTAGGTCATGCACGGAGAGTGGCAAAAACTCAATTTTTTTTAGTTTAAATCTCTCTAGGGAGGAACAAGAACCCAAGGTTGAGATTAGCATGCAAGCCGAGAGAAACACCTTTATTGTAAAACTTGTTTCTTCAGTTCTAATTTTGTGTTTAATGCCAAATATTGGATACTTTTCAACTAGACTGGAAATCTGGAACTCTTGTGCACCCGCATGACTAATTTTTCTTTTGTTCTTGCTGATTCTTGAATATAATTATTATGATTATTATATTTTTTTTTATCTTTACTGGTCATGCTAAGATAAAAGTTCTCTTTCCTATTGTGCTCCATTTCGCTTCTGAAAATGTCTAGTTATATCGTTTACCCTGTGATATAGAACATGTTTTTCTTCTTGTCATAGAAGTATGCGAGTATGAAAATTCAATGAAATGGTCTAGATATCATGCATGCTTTGAAGCTTAGCATCCTCATGGACATGGTAACCTGACAGAGATGCTAGTAATCTCTTAGATACTCTGTAAGGGTCAAGAGACTAAATTGACTACCTTAGTACAAAATCTTGATTAAGTAAAATCTTAGGTGAACCATATGGTACTTTTAAAATATGTGAATATCCCCATTTAGAAACTCAAGATGAATTGGCTACTTTTGTTTTATTTTTGCTGTAAAGAAGAATTTGTTAATGACGCAGAATCTGTGTGAAGAAGCCTTATAGAATCTTTGTAAGAAGAACAAGGATTTACTATTTTTTTCTTTCTATTTGAGATGTCTCACAACACAGTTCTTGTCCTGAATATAAATATACCATTGTTACCTTTCTTCAAAAAAATATATGCTATCATATAAGAGAGCCATCAACTTTTTTGAAACTGATAAGAGAGCCAATAACTTGCTCTGTTCTTGTCTTTCTTTGTGCTTTCTCAAGATCAAATAACAAATGTTTTTCTTTCTTGCAGTCCTTTTTCCTCCGTTGGGAGGAGAGGGAAGCAGAAAAAACAGAGGCACTGTGGTTGGAATTGTCTTAGGGTCCATATTTGCTGCAGCTGTGCTACTTATGGCCATAATATTTGTGTTGCTTAAAATGCGGCGGAGGTTACACGTTTCAAAAGATCAACCATGTGAGTTAATCTTCAGTTATTTTATGTGACTTTTCCTCAGCTATGGGGAGTTTCTTCTCCATTTCTCTCATTCACCCCACTAGGACCTGCTGGTTTAGTAGACAATAGGTTTAACTTTTCTTTGGTAATCTGTTGAATCTCCAAAATAGAGGACATGAAACCACATTCTTTGAGTGCTTCATGCATATAATGCAAACTTTGCACCTGATAATTTCCAATTTCCATCTTCTGATAAACAAAAAGATGCCAAAACCATAAAATCTGTCTTCCTTCCTTTAATTTTCTTGTTTAAGTTCTCTATTTAATCAAGAAATAACAAGAATTGCCAGTGTTAACCATCTTTTTCTATGTGGCAAATTATCTTTCTTAGTATGTATATTTGTAATTTAGCTATGATACTGCTTGGTGTTTTACTTTTAAGTAGAGAGTATGGTGCTTTAATTCCTCACATGGACAGATAACTTTACGGTTTTATGGGAAGTTTGTGCATATTTATGAGTAAACTCTCTGGCAATGCCCCATCACCCTTAAAACAAAAAGTGGAAAAGAAATAAATAGGAAGTGTGTGCTGGGCAAGGGGCATGTATATCTAAGTAATACTCTTGTTGTAGTGCTTCTGCTATTTGTACATTAATTAGACGATTATTTCTGTTCTTTTCTTAGCCAGATCAATTGATATTCTTAAAGTGAACAAAGTTAGCTGAAGCCAACTATTGTATCAAAGGTTCATTTGCATTAAAGTCATGCTACCATCACCTAAGCTTTTTCCTTTCATTCTTTTTATGTCAACTTTTCATGCTAAAGTTTTCAATGAATATCTGATTTTTAATAAATTGAACCATGTGGCGATGATGTATTTCTTGCTTAAACAAACTTAGCCAGGACCACGTGGAACTCATAGTGATATTCCTTTCATTTCTTTTGTAGTACCAAAATTTCCAATGCGAATAGAAGGTGTTAAGGCATTGGGTTTTAAGGAATTAGAGGCAGCAACTAACAGTTTTAGCAGTGCTACTGAAATTGGCCAAGGAGGCTATGGGAAGGTCTACAAAGGCACTTTAGCAGAGGGAACAATAGTGGCAATTAAACGTGCACAGCAAGGCTCATTGCAGGGTGAAAAAGAGTTCTATACTGAGATAGAATTACTTTCACGTGTGCATCATCGAAATCTTGTTTCCTTGGTGGGATACTGTAATGAAGGAAGTGAACAAGTAAGATTCTCCAAACTGGAAAAATAATACACACACACAGACTAGTTCATGTTGCTATTCCTGCATGCAGAGGGGAGAGGGGAGTATGTTCGGTGGAATAATTTCTATGATGTATCAGTTCAGCCACTTGTTGAATTCTGATTGCCCAGTTCTGGTTGAATACCTAGGTCAAATTGAATTTTTTTGCATATCCTTGATCATTTAGGATGCCTCATTGCCGCTGTGCAGAAGCAATTAATCATAATGTCAATTACCTTGTACTACTTAGAGAAATCCATTTGATATCACTCTAGATGCAAATGAAATGACCAGAATCACACAAATGATAGTTGAATTGCAGTGCAAAGTTCAGACTTACTGATTCATCTGTTAAAGTAAAATCAAGCAAAAGCCTTCTGAGATATTATTTTCATCATACTTCTTTTTTTGTCACCAATTATTATCACACGTTAAGTGATTCAATTTGTGAGCATATCCAGTCTACAGTACTTGGATCACTGAATATTAAGTTCAACCAAATTCCCATCTATCAGTAGGCAGTGGCTCATCAGTTACATCCCTCTTATACTTGCGTAGTCTCTAGTCTGGTCACATTAGTGAAATGCATGTCTTCTGAAAAAGTAATCAAATCAAAAAGCAGATGGAAGGATAAGAACAGATATTATGTTTTGTGGGTTTTTATTCTACTTTTGTAATAGTTGGTTACATGTGCATGCAGATGTTGGTGTATGAGTTCATGCCAAATGGTTCCTTGCATGATCTTCTAAAAGGTATGTGATGTATCCTTACTTTGCCTATGTTGTCATTTGACTTTCTTTTGTTAGAATCAAGCACATCTATTCTCCTCTTATCACTTTTTTTTTCCTGGTGTAGTAGTTCTTTTCTGAAAAAGAGGGTGAAAGATATGTTCCTCTAGTAAAGAATACTTGTATAGATGAAAGTATTTCTAAATTATCTAATCACTCAGAGATTTTTTTTTCTGAAGTTTTTACTTCATTTTGACTCCTGAATCCTGACAGTCACTGTTAGTTTTCCTACTTTCTTCACTTGTGTGGTCATCAACAAGCTTATCACATGGAACTTTATATTCTTAATTTTAGTCATGTGTCAGTTCCAATATGAATATCAAACAGGTATCTCCAAAATAAGATTATTAAAGATATTATACTTATTTTTACAGCTAGATATGGGGAACGTCTGAGTCTTGGAACAAGGTTATACATTGCTTTAGGTGCTGCCAGGGGAATCCTCTACCTCCATACTGAAGCTAATCCTCCAATAATCCATCGTGATGTCAAAGCAAATAACATACTGTTGGACTCTAAGTTCACCGCAAAAGTCTCTGATTTTGGAATCTCAAAGCTTGCACCATTACCTGATGCTGAAACGAGTGGAAATGTATCAACTGTTGTGAAAGGAACTCCTGTAAGGCTTTACTCATCTTATATAAACCTGCAATAGAAAATCTAGAAAATCACAATGAGATAAGACCTGCAACTTGAGTAATCTAGGTAGCATGGCTACAGACCAGGAGAACTTCATTCTCT

>M6-PGSC0003DMG400031752 M6 LRR-RLK

TTCAACAATGCATCTTCAAATCTTGCTTATACTTCTATTACCTACATTGATTCTCTCAATAAACCAAGAATCCCTTTATTTACACACCATAAAGCTTGGATTTGATGACCCAAATGGTGTTTTTTCAAACTGGAATCTCCATGATAACTCCTCACCCTGTAACTGGTTTGGAGTAAAATGCGACTCTTTAACTCGTTCTGTTACATCTATTGACCTCTCCAACACCAATATCGCCGGCCCATTTCCGGCTTCTCTTCTTTGCCGGCTCAAGTATATCAAGTACATTTCATTCTACAATAACTCCATTAACTCGACACTTCCGGTGGAGGAGTTATCCGCTTGTAAATCTCTTGTCCATCTAGATTTAGCTCAAAATTTGTTAGTGGGTAGTCTTCCATCGAGTTTGGCGGAGCTTCCTGAGCTGAAATACCTTGATTTGACCGGAAATAACTTTACCGGCGAAATTCCGGCGAGTTTTGGGGCATTCCGGCGACTTGAAGTTCTGGGTCTGGTTGAAAATTTGTTAACTGGGACTATCCCGCTGGAGATTGGAAACATTTCGAGTTTGAAGCAGCTGAATTTGTCGTACAACCCGTTTTCTCCGGGTCGGATCCCGCCGGAGATTGGGAACCTCACGAATCTAGAGGTGCTATGGTTAACTGACTGTGGGTTAATCGGTGAGGTTCCGGGTACGTTAAGGGGATTAAATAAGCTTGTTAACTTGGACCTTGCGTTAAACAACTTGTACGGTCCTATTCCGAGCTGGCTCACTGAGTTAACTAGTGTTGAGCAAATTGAGCTGTACAATAACTCGTTCTCTGGCGAGTTTCCGGTGAATGGGTGGTCGAATATGACCTCGTTGAGGCGGGTCGACCTGTCGATGAATCGGGTTACCGGGTCGATCCCGAGCGGGTTGTGTGAGTTGCCACTTGAGTCGCTCAATCTTTATGAGAATCAATTGTATGGTGAGTTACCTATAACCATTGCAAATTCACGCAATTTATACGAATTAAAGCTCTTTGGTAATAGTTTAAATGGAACTTTACCTAAAGATCTTGGTAAATTTTCTCCATTGGTATGGATTGATGTTTCAAACAATGAGTTTTCGGGTGAAATTCCGGTGAATTTGTGCGGAAATGGGGTTTTAGAGGAGGTTTTGATGATAGATAACTCGTTTTCCGGTGGAATTCCGGTGAGTTTAAGCCAATGCCGGAGCTTATTACGTGTGAGATTAGCTCATAATAAGTTCTCAGGTGATGTCCCTGTGGAATTTTGGGGGCTGCCACGCCTTTCGCTGCTTGAGTTAACGGACAATTCATTTTCTGGTGGAATCGTGAAAACTATAGCTGGTGCATCGAATTTATCCGCTTTGATTTTGTCAAAGAACGAATTTTCGGGTACTATTCCTGAAGAGATTGGCTTTTTGGAAAGTTTGGTTGATTTTGTGGGAAATGATAACAAGTTTTCAGGGTCATTACCGGTTAGTATAGTGAATCTTGAGCAATTGGGAAGAATGGATTTTCACAACAATGAATTAAGTGGTAAGTTTCCTAGTGGGGTTCATTCTTTGAAGAAATTGAATGAATTGAACTTGGCAAACAATGATCTTTCTGGAGAAATTCCCCGGGAAATCGGGAGCTTGTCTGTTTTGAACTATCTTGACCTATCAGGAAACAAGTTTTCGGGGGAAATTCCAGTTGCGTTGCAGAATTTGAAGCTCAATCAGCTGAATTTATCGAATAATGGCCTTTCGGGTGGTATTCCTCCTTCATATGCAAAGGGAATGTACAAGAATAGCTTTCTGGGGAATCCCGGTTTATGTGGAGATATTGGAGGCTTATGTGATGGAAAAGATGAAGGTAAAACTGCTGGTTATGTATGGTTACTGAGATTGCTTTTCGTACTTGCTGTTTTGGTGTTTGTAGTTGGGGTAGTTTCGTTCTATTGGAAGTATAGGAATTACAAGAAAGCAAAAAGGTTGGATAGATCGAAATGGACCTTGACGTCGTTTCATAAGTTAGGTTTCGATGAGTATGAAGTACTGGAAGCTCTAGATGAAGACAACTTGATTGGTAGTGGTTCTTCCGGGAAGGTTTACAAGGTCGTTTTGAGTAATGGTGAGGCTGCTGCTGTGAAAAAACTTTCAAGAAGTTTGAAAAAAACAGATGAGAGTTGTGACATTGAGAAAGGTAACTATCAGGATGATATATTTGAAGCGGAGGTTGAGACATTGGGCAAGATTCGACACAAGAACATCGTTAGGCTATGGTGTTGTTGTACAACAAGGGGTTGCAAACTTTTGGTTTACGAGTACATGCCTAATGGAAGCTTGGGCGATTTGCTACACAGCAGCAAAAGTGGGTTGTTGGATTGGCCTAAGAGATTTAAGATAGCTATGGATGCTGCAGAGGGACTCTCATATTTGCATCATGATTGTGCTCCTCCGATTGTTCACAGAGACTTGAAGTCGAACAATATCTTGTTGGACGGGGAGTTTGGAGCTCGAGTTGCTGATTTTGGTGTGGCAAAGGCGATGGATGTTGATGACAAGGGAACCACGTCTATGTCAGTCATTGCAGGGTCTTGCGGTTATATTGCTCCAGGTTAGAGGATTAATCCTTCACTTTGGGAGTTCTACTAACATGATTTTGTTGAATTTG

>M6-PGSC0003DMG400033049 M6 LRR-RLK

GAGGAAAACAAGATTGTAATTAAGTGACACTAATTATCATGTCTGAAGTGGAATCTCATGTTCTTCTTCCCATTGTATTACTCAATTTCTTCATTCTCATAACAGGTATGTCATATTTGATAGCTTTAATTTGATCAAGTTTAGTATTAGAGTCTCATTTTGTACATAATTAAAAAGAATCAATTTTTTAAACAACATTTATTGCAGTAAGACTAATGTTTTTTTTAACTTAGCAAAATCATAGGACCATTCTTGATTGTGTTACTCCAACACCAAGCATTAAATGTTGTTCTGTTTTGTTTTTGGGAGAATTGACTTGTATTTATTGTAGCATGATTAGTTTATGCAAAATATTATATAGGTAGGCTTGTTTGTGGAGACTCAATTGAGACAGACAAGCAGATGCTGCTGAACTTGAAGTCATTTCTTAAGGAACAAAATCCTGTTGACAAAGGATTCAAATATAACCATTGGAATCCTACAGATTTGTCACCATGTAGATGGCCTGGAATATCATGCAGTACCTCCATCAATCGTGTCACTGGAATCGATCTCTCGGAAAGTAATCTGGCTGGGAAATTGTTTAATAAATTCTCAGCCATGACAGAATTGAACTCTCTTGACCTGTCCAAGAACACATTTTCAGAGTCCATTCCATCAGACTTAGGCTGGTGTAGAAATCTGAAGTTCTTGAACTTGTCACACAATATCATTGTTGGTGAGCTCAACTTGACAGGCATGAACAAGCTGGAAGTTCTTGATTTGACGATGAACAGGATCCACGGGCTAATGATCCCTGAGATTTGAGACAACTTAGCTGTTGCAAATATTTCTAACAACAATTTCACTAGTGGGAGTGGATTTGAATTTTCTCACTGCAAGAAACTGAAGTATCTTGATCTGAGCTACAATTATTTGACAGGGAATCTGTCGTTCGGGCTTGATATGCTGAATATGTTTTCAGCATCTCATAACAACTTAAGTGGCTCTCTGCCTTCCTGGATTTTCACTCAAAACTGCTCTTTGCAAGGTTTGGACTTATCAGAAAATATGTTATTTGGTGAATTGCCTACGTCTATCGCGAACTGTAAACGATTAGTAGAGTTGAATTTGTGGGGAAATAGCTTTTCAGGGTCAATCCCTAGAGAGATTGGATCAGTACAGAGTCTTAAAGAACTATGCTTGGGAAGTAACAACTTTTCAAGTGATATTCCAGACACTCTATCAGGTCTAAACAAATTGGTATTTCTGGACCTAAGTAGAAACAACTTTGGAGGAGAAATACAAGAAATTTTCGGGCAATTGACACAGGTAAAATTTCTCGTGCTGCATGGAAACTCATATATTGGAGGCATAGTGTCATCAGGAATTCCAAACTTGGTGAACCTTTCGCGGTTGGACTTGAGTGATAACCAATTCTCTGGTCCATTACCAGTTGAAATTTCTCAGATGAACGGTTTGGAGTTTTTGATTCTTGCCCACAATCAGTTTAGTGGCAATATACCTTTAGAATATGGAGATCTTACTGCACTTCAGGCTGTTGATCTTTCCTCTAATAGGATCAATGGTTCAATACCACCAAGTTTCGGGAAGCTAAGGTCACTATTGTGGTTGATGCTTGCAAGCAATTCGTTGAGAGGTGAAATCCCATCAGAGTTGGGGAACTGCAGCAGCCTGTTATGGTTGAATCTCGCGAATAATCAACTTACTGGTCCAATTCCTCCTCAATTAGCAAGTATTGGTGCAGATCCACTGCCTACTTTTTTGTTGAATAGGGGAAAGGAGAAGCTCACTGCTAGCCCAGGGGATTGTTTTGCCATGAGGAGGTGGATACCAGCTGACTACCCTCCATTTAGCTTTATATATCCTCTATTGACAGGGAAGAGTTGTAGAATCCTGGGGGATAAGTTGTTTATGGGAGATGGTTTAATGCCAGTCTGTGAACCTGGTAGTAATGTCCGAAAGAATCAGGTACCGGGCTATATACAACTTAGTGATAACAAATTGTCTGGTGAGATCCCTCCTGAGATTGGCAACATGAAGAAAATGAGTATGATGCATTTGGGTGCAAATGAATTTTCTGGCAGGCTCCCTTCAGAGATTGAACAAGTGCACCTAGTAGTCCTTAATGTTTCACAGAATAAATTTTCTGGTGAAATCCCAAAGCAGATTGGCCATCTTAAGTGCTTGCTAAACCTTGACCTTTCATTTAACAATTTTTCTGGTCCATTCCCAGTTAGCTTTAGTAACTTGCATGATTTGAGCAAGTTCAACATCTCTTACAACCCATACATCTATGGAACTGTACCAGAAATCGGGCAACTGCCTACATTTGATAAGTCATCATTTCTTGGTAATCCGTTGTTGCGTCTTCCATCCTACATGCATAACTCTAACAACAACACAGAACGAAACAAGAATGACAATAACAAAAAGCACAAAAAGGTGGGTGCACTTTTGGTAATTGTGGTTCTGGTACTAGCTTTCCTAGTCTGTGGAGTCATGTCACTGCTTGTCTGCCTCCTTATCAAAGTGCCAAGGGGTTCCCCAGGAATCTTACTGGAGGATACAGAGGGCAGACATGATTCTCCATCAAGTACTAATGCATCCTCCTCACGGTTGTGTGCATCCTCATCACGGGGTTCTGATGATGTTAAGGTTATCCGTTTGGACAGAACAAGCTTCACACATTCTGATATACTGAAGGCCACATGGAACTTCTCGAATGATAGAATTATCGGGAGAGGAGGATTTGGGATAGTCTATCGTGGAGTCTTGCCTGATGGAAGGGAAGTAGCAGTGAAGAAGCTACAGAGGGAGGGAATCGAAGGAGAAAGAGAGTTCAGAGCTGAAATGGAGGGGCTCAGTGGGAATGGCTCCGGTTGGCCTCATCCCAACCTTGTAACTCTTTATGGGTGGTGCCTTGATGGATCAGAGAAACTTCTAGTCTATGAGTACATGGAAGGTGGTACCTTAGAGGACTTCATTACAGATAGAACAAGGTTTACATGGAAGAGCAGAATTCAAGCAGCAATCGATGTAGCACGTGCTTTAGTCTACTTGCACCATGATTGCTATCCTTGCATTGTCCACAGAGATGTCAAAGCTAGCAACGTGCTTCTTGACAAGGACGGAAGAGCAAAAGTCACCGACTTTGGCCTTTCTAGGGTCATGATTTCTGAACATACTCATGTTAGCACAATGGTGGCAGGGACTATTGGTTATGTTGCACCAGAATATGGGCAGATAATGCAGGCCACTACAAAAGGAGATGTCTACAGCTATGGGGTGCTAGCAATGGAGCTAGCAACTGGGAGACATGCTATAGATGGGGGCGAAGAATGTCTAGTTGAATGGGCGACAAGGGTCATGGGAGACGGAAGGAAAGGGTTCACCAGAGCCATTATACCAGTTGCTCTATTGGTACCTGGCCTAGTAGAGGGAGCAGAGGAAATGTATGAGATGCTTAGGATTGGGATAAGGTGCACAGCTGAGACGCCTCATGACCGGCCTAACATGAAGCAGGTATTAGATATGTTGATTAGTATTCCTAGCAGCCAAAAGGGATCCAGCCGTAGCTTTGGATCAAGTCGTAGCACTTCTCCATTATTATGATAGACTTTCTTGGAACCCAAGAGATAAAAACGTCAATAGTTTCTTTACAAGATGTGTAACATACCTAGCAGATTCTTCTTGTAAAACAGTTTTTCTTTTTTCACATTTGTTATTCTAGGAAAATGTAAGTCAATTTTGTTTGTTTGGATCACAAAATGTTCTCAAGGATATCAATTTTTGCTATGAAATTGAAGCACTCAAATATTTGACCCACCTACAAAGTTACACTACAGACTACAGCAGGTGGCAAGATTGTTCCCCTTATCAGGTACATTATTTGTCTGGTGCAATTCAAAGTCTCTATTTCAAACAAGATTCCTCTACCAGGGACTGTGCAATAGCAACTACAAATAGGAGTGTCCAAATTTCAACTTCTGGACCACATCTGATGGAAAATTTTAGTATACTTCCACTAAATCAACACACCACCAATTGTCAACGCCACAATACTCATTTTCATATCAGGCTAGCTTCATCATCCGTGGGAATAATGTGGATATGCACTCAAGCATCATACTGATTTCAAGTTAAGGCTCAAAATTTTAAGTCACTCTCTGATGTATAAACACTGCTAATAGAAGAGTACAAATTCTTGTCTACCTGCTTAACAAAAAGAAACTACAGATAACGTTTAGTGAAACTCTTAATGGTGTGAGACCTTTTAAGAAAAACATCGCGAGGTTGGATGAAATCTACCAGTATCCTACCATGTTAAGAATGTCTTTGGGCTAAACGTAACATTAATCTAGAGGTCACCAATG

>M6-PGSC0003DMG400033667 M6 LRR-RLK

CAATTTGGCATTTTCTTCACACCTCCAATGTTGACCAAAATCAACTCATAATTCTAACCCCACCTTTTTTTTTTTTTCATGGAGTTTTCCCATTTTCCGACCGGCATCTTCGCCGGAAAACAGTCCCTTTATCTCATTCTTCTACTAAACTTCATACCCATTTTCGTAACTTCACAATCTCCGGCCACTACTGAACGTGACACTCTACTGAAGATTAAGCGTCAATGGGGAAACCCAGTAGCACTTGACTCATGGAACTCTACTTCTTCTCCTTGTAGCTGGCCGGAAATTGAGTGTGATGATGGGAAGGTTACCGGAATAATAATTCAAGAAAAAGATATTACTGTTGAAATTCCAAGTTCTATTTGTGAACTTAAAAATCTTACTTTTCTTAATCTCCGGTTGAATTATCTTCCCGGAAAATTCCCAACTTTTCTTTACAAATGTTCCAATTTGCAGCATTTGGATCTTTCTCAGAATTACTTTGTGGGTTCCATTCCGGAGGATATTCACCGGCTTGGGAAACTCAAGTATCTTAATCTGGGTGGTAATAATTTCACCGGAGATATCCCGTCGTCAGTCGGGAATTTGACGGAGCTGGAGACTTTGTGCATGCATTCGAACTTGTTTGATGGGAGTTTTCCGGCGGAAATCGGAAACTTAACCAATCTTGAAAGTTTGGGGTTGGAGTTTAATGGGTTTTCTCCAATGGCACTACCGCCGGAGTTTGGGAAGCTGAAGAAGATAAAGTATATTTGGATGAGAGAGACGAAATTAATCGGTGAAATCCCTGAAAGTTTTGGGGAATTTCAAAATCTTGAGCTTATTGATTTTGCTTACAACAATCTGGAAGGAAAAATCCCTTCTGGGTTGTTTCTGTTGAAGAATTTGACGATGATGTACTTGTTCAGTAATCGGCTTTCGGGTCGAATCCCTGAAACTTTTGATTCATCAAAGTTGATGGAACTTGATGTTTCCAACAACAATAATTTAACTGGAACAATCCCAGAAAGTTTTGGAGGGTTTAAACATTTGGAAATTATGAATCTGTTTGGTAACCAATTGTATGGAGCAATTCCAGAGAGCATAGCCAAGATTCCATCTTTGAAGGTATTCAAAGTGTTTAGGAATAAACTGAATGGGAGCTTACCATCTGAAATGGGACTTCATTCGAAACTCGAATCTTTTGAAGTTTCACTGAATTCCTTCACTGGAAACTTGCCAGAGCATTTGTGTGCTGGAGGAACTCTTTTTGGTGCTGTAGCTTATGTTAACAATTTGTCTGGTGAAATACCAAAATCACTCGGAACTTGTTCTAGCTTACGTTCGATTCAGCTCTACAAGAATCAGTTTTCGGGTGAAATTCCATCTGGGGTTTGGACTTTGGTTGACATGACAAGTTTGTTGTTGAATGATAACTCATTTTCCGGGGAGCTACCGAGCAAAATTGCATACAATTTTACGCGGTTAGAGATCAGTAACAACAAGTTTTCTGGTGAAATTCCTGTGGGGATATCTTCTTGGAGGAGTTTAGTGGTGCTTTTGGCAAGTAACAATTCATTTTCAGGTCGGATTCCAGTGGAATTGACTAGTCTTTCTCAGATAACTCAGCTAGAGCTTGATGGTAATTCGTTATCAGGAGAACTTCCAGCCGATATAATATCGTGGAAGTCGTTGACAATATTGGATCTTGCCAGGAACAAACTCTCTGGCAAGATTCCAGCAGCTATAGGTTTAATCCCTGATCTTGTTGCTCTAGATTTGTCTCAAAATCAATTTTCAGGTCCAATTCCACCTCAACTAGGTGTAAAAAGGATCACTAGCCTTAACGTGTCGTCCAATCAACTTATCGGGAATATCCCTGATGCATTTGCTAACCTAGCCTTTGAAAACAGTTTCTTGAACAATCCCAGTCTTTGTACCACCAACTCACTTCCATATCTTCCAAGTTGTAATAATGCGAAAGTCGCCAATTCCAAAAGATTGTCCCATAGAGTCCTAGCCCTGATTCTAGTCCTTGCATTCGCTGTTTTCCTATTTAGTGTTATATCGACCTTGTTCATGGTTAGAGACTACAGGAGGAAGAAGCATAAGCGCGATGTTGCAAGCTGGAAGTTGACTTCATTCCAGAGGTTGGATTTCACAGAGGCGAACATTTTGTCAAGTTTGACGGAAAATAACATGATTGGAAGTGGTGGATCGGGTAAGGTGTACAGAATTTCCGTTGGCAGACCAAATGAGTATGTTGCTGTGAAGAGGATTTGGAGTGATAGGAAAGTAGACTACATTCTGGAGAGGGAATTCTTGGCTGAGGTTCAGATATTGGGATCAATTAGACATTCCAATATCGTGAAGTTATTGTGTTGTATATCAAGCGAAGATTCAAAGTTGCTCGTTTACGAGTACATGGTAAATCACAGTCTTGATGGATGGCTTCATGGGAAGAAGAGAGTATCCTTGAGTAATAAAGTCATGGATTGGCCAAAAAGATTGGAGGTTGCCATTGGAGCTGCTCAAGGACTTTGTTATATGCACCATGACTGCACTCCACCAATCATTCATCGCGATGTAAAAACAAGCAATATCTTGTTGGATTCTGATTTCACAGCCAAAATAGCTGATTTTGGGTTGGCCAAGATTTTAGAGAAGAAAGGAGAGCTTAACACCATGTCTGCTGTTGCGGGTTCCTTTGGTTACATAGCTCCAGGTAAAAGCAAAATCCAAGCTTGAACTTCGTATTACATCAATTTCGTTAACATTATATCAAAAGTATGACTAACTAATAATTTACTGTTTGGTTACAGAGTATGCTTATACAACTAAAGTGAACGAGAAGATTGATATCTATAGCTTCGGGGTGGTGCTATTGGAGCTAGTGACTGGAAGACAACCAAATTTTGGGGACGAGCATACAAGCCTGGCAGAATGGGCATGGAAGCAGCACGGAGAAGGGAACACTGCCATCGATAATATGTTAGATACAGATATCAAAGAAACATGTTACTTGGAAGAAATGAAGACTGTGTTCAGACTCGGGCTTATATGCACGAGCAACTTACCTGCAAGCAGGCCTTCCATGAAAGAGATTCTGCAGATACTTCATCGATGCAAGTCCTTTAGGTATTCTGGAGGAAAATCACCAGACACAGAGTATGATGTTGCTCCTCTCCTTAGCGGAAACAACAGTGAAAAATACATTGCTAGTTACAAACGTATTAATTCCAATAAGGTAATAGATGATAGTAGTGATGATGGTCTAATAATTTCCAGTGTGTAATATTGGAGATGCCGAAGTAAACTGAGTTGTTGTAAAATGATCATAGAATAGTGCCACTTAAGTAAAGTACAAGTTTGAATTTCTAGTGTACTCTTGAATTTGTTCACTGCTTGTAGATTGAGTGAAATGCATCATTTAACTTTTGCTAGTTATGTAGATTGTAAGGAAGTGGGAAATTGGAAAATGAAAATGCCAATTTATATAATCAAAGAGAAATGACAGTAGAGTGTGAAATTGGAAAATAAAAATGCCAATTTCCAGCTCAACACGTAGTTCATTAAGAAGATTTGGTTTCGACTGAAAATGACTCAATAGGAGGGTGTGGTGTGGAGGCGAT

>M6-PGSC0003DMG400033914 M6 LRR-RLK

TTTAATCTTTCTCATAATACGTTTTATGGAAATTTGCCATCCTTAAACAGAAGCTTCAGTTTTCTAGATCTATCAGGAAACTATTTTGAAGGCAAACTCCCAGATTATGCACAACGTAACGTGTCAATTAGTAGCAATTGCCTCCAAAATGCGACAAGTCAAAGGAATAGGAGCGAATGTGCTTCTTTTTATAGCGGGCGTGGCTTGTTGTTTGATAATTTTGGAGAGCCAAATGCCACAGAGCCTCTTCCAACTTCCAAATCCAACAGAAAGAGTCACAGAAATAAAATCATATTGGCGGCTGTTCTGGGTAGCGTTGGATTACTTGCACTTGTCTTTATCTGTATATTTCTGCTGATTTTTTGCACCCGCAAACGGGGTGCCACGAATCAAAGAGCCACTGAGGTTGGGCCTGGTCCTGCCAGTTCAAGTCCACCACCTCCTGCACCTGGAGCATCATTGAATTTTTCAAGTTTAGGAGATGCATTTACCTACCAGCAGATTCTGCAAGCCACAGGTGAGTTCAATGATGCAAATCTTATGAAGCATGGTCATTCAGGTGATCTCTTCCGTGGCACCCTAGAAGGCGGGACACTTATAGTTGTCAAAAGAATAGATGTGCAGTCTGCAAGGAATGAAGCATACTTGTCAGAATTGGACTTCTTCAGCAAGGTTTCTCACTCCAGATTAGTACCTTTTATGGGGCACTGTTTGGAGAATGAGAATGAGAAGTTCGTGGTGTACAAGTATATGCCAAATGGAGATCTTTCCAGTTCTTTGTTTAGAAAAAACAATTCAGATGATGATAGTTTGCAGTCATTGGATTGGATAACACGACTGAAAATTGCTATTGGAGCTGCTGAGGGTTTGTCTTATCTACATCATGAATGTAACCCGCCTCTAGTGCACAGGTATCAAGGACTTGTCATATTACATTTATCTTACTTCCA

>M6-PGSC0003DMG400038191 M6 LRR-RLK

TTGATTTTTCTTACAACAATTTTTCGGGTATGATTCCACTAACTCTTGGAAAATGCTTGACCTTGTGGGAGATCTATATGAAGGGAAACTCCCTCCAAGGAACCATTCCCAATTTAGAAGATTTGCCGGATCTCCAATCCTTGGATCTTTCCCTAAACAACCTATCAGGGCCAATCCCTCACTTCATTGCAAATCTTACTTCCTTACACTACTTGAACTTATCTTTTAACAATCTAGAGGGTGAGGTTCCTGTCACCGGAATATTTTCTAATTTGAGTGCAGTTGTATTAAGCGGAAACTCCAAGCTTTGTGGTGGGATTCAAGAGCTTCATTTACAACCTTGTATTTATCAAAAGACACGGAAGAAGCATGTGCTTGCCCTCAAGTTTATTTTGATAATTGTTTTTGCTGCTTCATTTTCAATCCTGGCCTTGTTAGTAGTTTTCCTTTGTTGGAGAAGAAATTTGAACAATCAACCAGCACCAGAAGATAGTTCTAAATCTGCGCATTTCTACCCCAATATTTCGTATGAAGAGCTCCGCACTGCAACAGGTGGATTTTCTTCTGAAAATCTTATTGGCTCTGGCAGTTTTGGAACTGTTTATAAGGGAACTTTTCCCTCCAATGGAACGGTCGTTGCTGTGAAGGTACTCAAACTCCAGCTTGAAGGTGCTTCAAAAAGCTTCTTAGCAGAATGCCAAGCATTGAGAAACACAAGGCACCGGAATCTTGTCAAGGTCATCAGTGTATGCTCAAGTTCTGATTTCAAGGGAAATGACTTTAAAGCTTTAGTCTTTCAGTTCATGCCAAAAGGGAACTTGGACGAATGGCTACATCCAGAA

>M6-PGSC0003DMG400039680 M6 LRR-RLK

GAATTTGCAGAATTTGCTTCATGGCTTGCCACTCGGGGTTTGAACTACTACTAAAGACTATACCACGGACATGTAGGAAGAGGAGGATGATAATAATAATAGTATTCATAATGTTTGTTCTAAAGGGTTGTTAATCACTTAGAGATATAGGCACAAAATTGTGCTTGGCACTGCTAGAGCACTTGCATTTCTACACCATGGTTGTTCACCTCCCATTATACATAGAGATGTCAAAGCTAGCAGTGTTTATTTTGGCACAAGACCCGAGAATCATGAACTTGGATCCTAGACTATCCAATTTTGGATTGGCCAAAATCTTTGGCACAAGACCCGAGAATCAGATCAATCGC

>M6-PGSC0003DMG401000515 M6 LRR-RLK

TTTTATCATATAACTCAGTATATTGTCATTTTGGTTTCAGGTGTGGAAGCTGTACACAACAAATCAGGTAACTGAAGCATTAGATCCTCTATTGAAAGGCGATTTTCCACCAGAAGAGGCATCAAAGGTACTAAAAGTGGGGTTGTTGTGCACTCAAGCTTCTGTCGCTTTAAGACCATCAATGAGTGAAGTTGTCCAAATGCTAACGTGTGAGGGCTATCAAATTCCAGAACCATGCCAACCACCATTCTTAAATTCCAACTTATTAGTTGGCGGTTCAGTCAAATCCAGTATAAGAAGTTTAGTCTCAAATGCACTCTATAAACTTGATGAATCATCATCTTATGCCACCACCACTGGGTCCTCAAGTATGCAAAGCTCATCAACTGGACCACACAAAAGTGACGAATTCCTTTTAAAAGAGTCTGAAAATAGGAAATAATTGCAAAGACAAATATAGAAATGGAAATTTTGAGAGATTTTTGATCAGTACATTTTTTGTACAGTAATAGAATTGCTGTAGTTGTGCATTTTTATGATTTTATTTTTGCATTTTTGGTGAGTGATGATACAAGAAAGTGTAGGGGTGGCAATTTCAGCCCATATAAGTGAAATGAAACTATTTTGCCCATATTAAATTAAATGGATCACTCATATT

>M6-PGSC0003DMG401002925 M6 LRR-RLK

TGGACTTGTAGACTTGAATCAAAACCTGTCGCAAAATTTTTATCATAAGCTTGGTGAGGGGTCAAACATGTCTATTGAAGGTTATGGGATGAGCAATGCTGGTGGTTCTGGTGCAATGGCGGTGGATAACAGCAGTGTTGGATCAAATGATTCACACACTCATATTCTAAACCAGCAGGGTAATCACATCCACAATAACGATTCTGTTGCTGCTAGCGTAGTCAGGGGGAGAGTTACTAGGTTGAGCGATGATGCCCTTGCTCAAGCCTTGGTTGATCCTCAATTCCCCACTATTGGGCTTGCAATTTATCATGAGTGGACAATTGACTTGAGGAGGCTTACCATTGGACAAGCATTTGCTCAAGGATCTTTTGGGAAACTCTATAAGGGAACTTATAATGGTGAAGATGTTGCCATCAAGCTTCTAGAGAGGCCAGAGCATGATCTTGAGAAGGCTCACTTTATGGAGCTGCAATTTCATCATGAGGTCAAAATGTTGGCAAATTTGAAGCATCCAAATGTAGTACGATTTATTGGTGCATGCCGTAAAGCCATGGTGTGGTGTATTGTGACTGAATATGCAAGAGGGGGTTCAGTGCGTCAGTTCCTCACGAGGCGGCAAAACCGAGCTGTGCCATTGAAGTTAGCAGTTGTGCAGGCCTTGGATGTGGCTAGGGGGATGGAGTATGTGCATGGCCGGAATCTGATTCACCGAGATCTGAAATCCGACAATCTGCTGATTGCTGCTGACAAGTCAATCAAGATTGCAGACTTTGGTATTGCTCGAATTGAGGTGCTGACTGAAGGAATGACACCAGAAACTGGAACTTACCGCTGGATGGCTCCGTAAGCATCTAGACTCTTTAGCAGCCTTGTCAAATCTGGTGTCTTGATCCAGTTTGATTTTATTGCCAAACCCTATAATAGAGCTCTTAAGCTTTTTATCCTATTCTTGTGGGCTATTTTTTCTCTGTCTGCATTACTTTCTGTATTTTTTTTCAAAATTGCTTTGATATGTGTTTCTTTAGCCAAGCATCTTCCGGAAACAACCTCTCTTCTTCCACGAGGGCAGGCTAAGGTCTGCTTTCTCTCTACCCACCCCAGACCCCACTGAGTGGGATTTCACAGGGTATGTTGTTGTTGTAGCATGCTGGATTCCCTTTATGGAATAAATTTTATTAGAACTGAAGTGAAGAGCATGGTGGTTGATGGGTTCAAGTCCTGTATGTCCTTTTGATGCCACCTCACTGATTGTGATTGTATTTTCTATTGTATACCTAGTGTTTCTTCCAGTTAACTTCTTAATTTAGTTCTGTACCTGTTAATGAAGGATAAAGAGTTGATTGACTTTACTTTCTCTTTCAATTCCTTTCCCTCTTTTTCCTTTTCCTTCTTGTCCATGACTTGTAACTTGATTCAAGTTAGACACACTCAAGGGTGTCACTTAGTTGTCAATGAAGTAGGAGCAAACCTTTGGAGATTAGGGTTCAATCATCCCAGAGACAGAAAACACCAAGTGTTCTCTTCCCATATGCCTAAACCTTAAACCTTTGTGGGTATAGTTACCCGAAATCGAGATGCATGCAAGTTGGATTGGATACCATTTTCAGAAAGAGTAAAATTGCATATATCTGAAAATGGGTCTAAGACATTAACTACCACACTGGGTTCACTTTCAGGATGTGTCTGAGAGATATGAACCACAATTCTATTCTTATTGAATATGTAGATGAAAGTTCTATGGGAAGTATATTATCTCTTTCTCTCCTTTGTGAGTTTCGACCTTCCAGCATCATCTTTAACATCACCTCTTTCTTCAAAGCAGTGAGGCTATTTTGGAAATAATGTATTGGTAAGAAAGAGAATGGGGAGGTGCGATTCACCAGAGATCTCTCTCTTTATAGGATGACATACATCCACAAGAGTCGCATTGTTTGTTGCTGCTGGAGAAACTTAAAAAGAGCAAGGGAATGACATGTCTACCAGAAACATTTCTTGTTTAGAGTGGAAGTTGGTTGCCGAATGAAAGTTTTTAAGTTGTTAGTAGTTGAACAGCATTCCTGTTTGCTTGGCTCGTGCTAACTTTCTCAAAGCACTGAAATGAATCAGTTAAAGAAGGATTGTTTAGGACATGTTATTTGGACATACACTGGTGTCTCTGGAATCTGCAGGGCTCTTGTTAACTTTCTCAAAGCACTGAAATGAATCAGTTAAAGGAGGACTGTTTAGGACAATTTAATTGGACATACACTAGTGTCTCTGGAATCTGCAGGTAGATTTTAAGTTGTGCAGGTGGAAGTCATTGCTCTGTTGGAGGATGGCAAAACAAACATCGATTTTTTTTTATGACAAAGGAAACCCGCAGCTGCTATCCATTGGGTGCGTACACGGTAAAACCCCCGTTCCTATGCAATAGCTCACAAAATTCGTAAGAGAGGTAACCTGCACAAGGCAAGCTCGGTGCGACGAGCCTGACCCAGAAGACAAACCCCTTGCTTTTGCTGACAAGGGGTTTTGAACTTGAGACCTCCAAAATGGAAGTCCCAAGCCCAAACCACTAGGCCACCCCAAAGGGTAACTGACATATCTATTCTGATGTTTCTGGAAGGAATTCTAGGAGGGCCTTACAGAAACTAGACTCTACAACGATAATATGATTCTCCGCTGAGTTACTACATAGACTTTTTTCATCATTAGACAATCTCTACGGAAAGGTAAAAGTGGAATATTTGGCTCAATCTTCAATGAATCTAAATCATTTATTGGAACCTTCACTATTCTTTAGGAAGAGGCCTCTTTAAAAGCATAAAAAGCGACAAGGTGATGTTTCTATAATCTACAATGATTATGTGGATTCTGCATTGAGTTGCTACATGGACTTGTTCCGCCCATAAGACTATTTCAATCTCTATGGAAAGGTAGAAGTGGAATACATGGCTTAATCTTCTCATATTAAAAGCAAAAAACGATCCATGGAGTAAAAAGCTTGGAGTGAAGCATACAAGTTATGGAAAAGTATAATGAAGTACCATCCACAAAGAAATTTAGTAATCTTCTCATGAACTTGAAACATTTGTTGAAACCTCCAATTATTGAGAGTACTGTTATTAGAAGCGAGAAACACAAAAAGCGACAAAGTCCATGGAGTTTGAAGCAAATATCGAGAAGTGAAGCACAATGGCTATGAGAAGTAAGTAATGCCGAACATAAAAGGATTTAGTACAATAAAGATGCAAATAAAATAAATGGTTTTTGCATCTAGTTTACTTAATATTGATATTGATCTTGCTCTTCAAAGTCGAGATTCAAAAAAGTAGATCCTTTCTTGTTAGGATCATTTCGCACGTCATTGTTTAAAGCAGTGCATGGGTTCAAGAAGTAATCCCATTCTTCAAATTGCTTTAAGTGATGGCTTTGAATAATAATGTGGGAGAGATGTGACACATGATGAAATCAATGCCCAGTAAGAAGAATGATAAGAATGCCTTATTTGGTATTAAGTCTTACAACAGGTGTTGCTAGACCTGATTTGACAAATGCCTTGTATAAGTTGAAGTAGATCGTTTAGAAACAAAAGGAGCATGAAAACCTCATAATGATCAATCAATTAGTATAACTTGTTAAATTACGGAGATCTTACGTCAGGCCGATTATGACTCAGTATTTCTGACATCTAACTTTTAGTGCCTCACACACTATGTTGTATCGGTCCTTTGCATTACATACATGTTGAAATGGGTGTGTCATGGTTGCGCGTACTCTTCAAGATACACAAATAATTTACAAAATGATACTTACATCTGATAAAATTCATACTCACAAACGAGTCAAGAAGATTTAGGATTTAAAGTTAATCGGTTCCCAATGATATCAAGTTTAAAAAGTAAATTATTATATGTGCCCAAGGGGAATCGAGCCTTCAATGAGGAGGGCAATAAAGCTCAAGCAAGTTCCCATTTATGCCACTGGACCAAGAGCTTACTTTGTTAGTAGATCCCAACAAATACTTTATATATTTAATTGACTGGATTCCACAAAACCTACACATGATACTTTAGGTCCGCCCCTGTTCAATAGCAACAACAACATACTCAGTGCAATCCCACATGTGGGGTTTGGGGAGAGTAGAGTGTATGCAAATGTTACCCTTAACTTGTTGAGATAAAGAGGTTTGTTCTGAAAGACCCTCAACTTTAAATAAACTTGCCAAACAGTTTGCAAAAGGGAATACAAATATTCGGAGCAATCACAATGACACATTTATGTAAAATGGCAAGTGTTACATAACAAACAAGTGAAAGAACAATAACAAAAGTGAAATAATGCAATAATCGACACACAAAAAACTTCAAGAGTAGTGCTAATACTACTGATAAGAATACTAGTGAAAGCCTATCTCGCAAAGAAGCAAGAGAATGCTCAATTACCAACTAACCATTTACCCTAAAACACGACCTCTATACTCTCCTATCAAATGCCATGTCCTCGGTAAGATCTAGATGCACTCCGTCTAATCACTTCCTCCCAATACTTCTTCGGCTCGATTTTACCTCTCTTGATGTCCTCCCTTGCCAACAACTCACACCTCCTCACTAAGGCATCTATACTCCTCCTCACGTGTCCAAATCATCTCAGTTTCACTTCCCTCATCTTGTCCAACATGGAGGCCACTCCTACCTTGTGTCGGATATCTCCATTCTTAATCTTGCCACTCTTAGTATGCCCACACACCCACATCAATATCCTCATTTGAACGATTTTCTTCTTCTGAACTCGGGATTTCTTAATAGGCCAACACTCCGCCTCATAAAATGTGGTCGGTCTAACCACTACTCTGTAGAACTTACCCATCCATACTCCATATAATATGGATTTCATGTTATTTTAGTTGTGGAAGAATAAGTATATTTGAAACATTTTGGATTTCTCCACTAAAATTCATCACCTCTAAAGAATGATAGAAAAAACGAAAGCATCAAATCTATTTAGGTGCCACGGGTGTCCACATGGGCTTGATTTGCCAAGTTTGTTAATATGTACTATGAAACGCCAAATTAATTTTAGAAGTATGCATTAGGTATGTAAAAATATATTTTACTATTTGTTTTATGTATTACGGTGCATGCATAGGCTGACTGTTGTATTGATTCATGCATTAGCGCCTGATTGTTAAACTGTTGAAAATTTATTGACATCTATTGGAAGATCTCCTTGGTTGGCACAATTTGACCTGGAGGCTGCAAAAATTATGACAACTAAGAAAATTACTTCATTTTGCTCAGTTTATTGTTCTACTGATACATGGGGTTTTCCTGTATTTAGGGGTCGTTTGGTGTGAGGGATAAAAATAAATAGTCCTGGGATAAAAAAATAGTCTTGGGATTAAATTTTGGTGTCTTGTTTGGTTGCCATGTTTGGGATAACTTATCCCACCATTTATACCATAGTGATGGGATAAGTTATCCCATATACATGGCAGAATAAGTAATCTCAGGATAATTAATCATGGGATAACTTGTTCCCAACCAAATGACCCCTTAGAGTCTTATTTAAGAGTTGGTAATATTTAGTGACTCTTTTGAGAAAAAAATTTCAAATCACGATTTCCCTTATGATTCTGCTCTTAAAAGTTCAGTTATTTGTGTCTCAGGCATATATGGCAGTTAGAATTTAAACCTTGAATTAGATCATCAATTCAGGTTGGTTTAGAGAATATGGTATAGAGTTAAATAAATCCCTTTCAATCTGGTGGAGATTGATTGCTATATCCAGTTGAGGGTAGATTACTGATGATAGAACTGGCATTTTTTTATTTCTTTTGTCCTGGACAGAAGTATCGCCATCAGTTTCCGTTTGGGTTTGTACCTTACACAATGTTAGAGAGGAAAGCATATTCCTTGTAAGCTAAGTGAGTTGGAGATTAAGCTTTTGTTTTCTTGAGATAATGTGACAACTGGAGAAGTTTTGATTTCTAATAGGACATTTCTTTCAGACTTGATAAAAAATAAGTGTTCATGTCTGCCTGAAATATCTTCAAAATGGTTGGGATCAGAAAGGATTTAGGAGGTTGACCCGGTGAGTGGTGGTAATTTTATAGTCTTCTCTGTGTGATAATAGGCTAGTATGATGAATGGCCTTCCTTGGGGAAATAGACAAAATAATATGTTTGCTTTTGTTAATTTGATGTGATAATGAAAGATCTTGCCAAAAGAGGTGTGGATCGGATCTCAATATCTGGTAAAACTCGAATACGTGCAGCCAGTCTGACTCTTTGTATAATTCTAGTATACTCAAATTTTTATTCTGATTAACTGAGAAATGAAGTTACTTTTCAAGTTTACTCTTGAGAAATTAGTTATTCACATTCTGCAGATCCTATCCCGTTTAACTTTGTGAATTATTGGCAAGTGTGTCTGGTTTCTTCTCTTTTCAGAAAAAAGAAAAATGGATGTTCAAAGTTGTTAGCAATAGCATCTGTGTTTCGTTGTGGATCGCTAAGCGTTTGACTTTGTTTTGTAGGGAGATGATCCAGCACCGGCCATACACCCAGAAAGTTGATGTTTATAGTTTTGGCATTGTTCTGTGGGAGCTCATAACAGGGTTGCTTCCTTACCAGAGCATGACTGCTGTGCAGGCTGCTTTTGCTGTTGTCAACAAAGGCGTCCGTCCAACAATCCCCATTGATTGCTTGCCTGTCTTATCTGATATCATGACCTGCTGCTGGGATTCTGACCCTGATAATCGGCCAACTTTCTCTCAGGTGGTCAAGATGCTTGAGGCGGCTGAGACAGAAATCATGACTAATATCAGAAGGGCACGTTTCAGGTGCTCCCCTCATCCTACGGCTACAGGGTAAGAGGAAACTGGAGGCAACTGATTTTGAGTCTGCCATGTGTATCTATGAGGCACAATTTCTTTGCAAATAGAAAAAGGGCTATAGTTATGAGAAAAAACTAAACATTTCTCTGCTGTATTTTGTTTCTTTTCTTTAATCCGTGTATTTAAGTGGTGACTAGTTGGTAATACAGGATATCTGATGTGCATGTCAGATTATGGCTGTATTTACTGAATTTGTTCAAAGTTATATCAGGTTCTGAATTA

>M6-PGSC0003DMG401006401 M6 LRR-RLK

AACCAACTTTCTGGCCCCATTCCTGCAGAAATAGGAAAACTTACCAGCCTTATAGAAGTGGATCTTGATACAAACCAGTTAACAGGTCATATTCCTCCGGATATAGGCAACTTGATCAACGCAAAGCTGTTCAGTGCTTTCTCCAATGAACTGTCTGGTCCCATTCCTGCAGAAATAGGGAAGATGAAGTCACTTGAGGAGTTAATGTTATACAGAAACAATCTTTCTGGTTCGATTCCAAAAGCTTTAGGTGAATTAACCAAGCTCAAAATTTTGTACCTTTATGTTAACCAACTTTCTGGCCCCATTCCTGCAGAAATAGGGAAACTTGTCAATCTTGTTGAAGTTGATCTTGATACAAACCAGTTAACAGGTCATATTCCTCCGGAAATAGGCAACTTGATTAATGCAAAATTGTTCTATGCTTTTTCCAATGAATTGTCTGGTCCCATTCCTGCAGAAATAGGGAAGATGAGGTCACTTGTGGATTCACTGTTCCAGAGAAACAATCTTTCTGGTCCCATTCCTAGTGAGTTGGGGAATCTGAATAACCTCAATGATCTGGTTTTATCCGAAAATCAACTTACTGGTTCAGTTCCTACTACATTTGGCAATTTGAGAAACTTGCAGACACTGCATCTCCATACCAACAAACTATCAGGTTCCATTCCAAAAGAACTTGCATATTTGGATAACTTGCGAGTGCTGACAATGACCAAAAATCAGTTATCAGGCCATTTACCGGACCAGCTTTGTCAAAATGGGAAACTTGAGTTCTTCACTGTGGCTAGCAACAAGCTGACAGGGCCAATACCAAGAAGCTTGAGCAACTGCTCGAGTTTCAAATGGGTTCGCTTTAATAACAACAGTTTCATCGGAAACTTATCTGAAGCTTTTGGAATCTATCCAGAGCTTCTGTTCATTGACTTGAACGACAATGATTTTCATGGTGATCTCGGCAGCAACTGGGGAAAATGCGAAAAATTGATTAATCTGCAGGTAGCCAGAAATAACATTAGTGGTAGCATACCACCAGAAATTGGAAACATCAAAGGGCTTCTTGGACTTGATCTTTCATCGAATCATTTGATTGGGAAGATTCCAAACGAATTTGGAAAATTAACCTCTCTGGTTAAGCTTTCTGTGAAAAACAACAATATTTCTGGCAATATACCTGAGGAGCTTGGGTCACTGACAAAGTTAGAGTCGCTTGATTTATCAGACAACAGATTGAATGGATCAATCCCAACGTGGATAGGAGATTTTGTGCTGTTGTTTCAGTTGAATCTGAGCAACAACAAGTTTGGACAAAAGATTCCAAAGGAGATAGGGAGGATAACTCATCTTACTGTACTTGATTTGAGCCATAATCTTCTTGATGGAGAAATACCTGCTGAGCTAGCCAGTTTGCTGGACTTGTCAAACTTGAATCTTTCCCACAATGGTCTATCTGGGCGCATTCCTAAAGAACTTGATAGTTTGACTGGTTTGCTGGATGTTGTATTGTCATACAATGATTTGGAAGGTCCAATCCCTAATAATAAAGCATTTATGAATGCTTCGTTAGAAGGTAATAAAGATCTTTGCGGCAATGTAACAGGATTCCAGCCCTGCCAAAAGCCATCTTCTATGGTGAAAAAGCACTCAATAGCAAAGGGAAGTAAACTCATCCTCATCACTGTACTTCCTGTTATGGGAGCACTAGTACTGCTCTGTGTTTTCATTGGTGTTCTCTTTATGTGTAATAAAAGAATGAGAGTTAAAGACGTTGAAAGAAGGGATGGTGATGGTTGGCTTTCGATATCCATGTTAGATGGAAAGGCATTGTACCGGGATATCTTAAACGCCACAGAGGAGTTTGATGCAAAATTTTGCATTGGGCAAGGAGGACATGGAAGTGTTTACAAGGTAAACCTTCCATTATTTGGTGATATAGCTGTGAAGAGACTTCATTCTTCATTTCAGAATACACATCCCAAAAGCTTCATAAATGAAGTAAGGGCATTGACTGGGATTAAGCACCGGAACATTGTGAAACTCTACGGCTATTGTTCAAAAGCACAACACTCGCTCTTGGTTTACGAGTATGTAGAGAGGGGGAGTTTGTCTAGTATTCTGAGCAATGAAGTTGAGTCTAAGAAATTGGATTGGCTTAAAAGGATGAATATCATTAAAGGTGTTGCTTTTGCTTTATCTTACATGCACCAGGATTGCTCACCACCAATTGTTCATCGAGACATATCAAGCAGTAATGTTTTGGTTGATTCTGAGTATGAAGCTCGTGTTTCAGATTTTGGCATAGCGAAGCTTCTCAAACCAAACTCATCCAATTGGACTGCACTTGCAGGAACATATGGCTATGTTGCACCTGGTAAGGTCTAAATTTAATTTCGATTTTATTCAGACTTCGTGAATTATAGCCAAACACTCTATACTATTTTTATCTCACTTACACATATAAATTGTTATACCATTTTTGGAAATTGTATTGTTTCAGAGCTTGCCTATACAATGAAGGTTACGGAAATGTGTGATGTCTACAGCTTTGGAGTATTAGCATTGGAGATAATCAAAGGAAAGCATCTAGGGGAATATATTACTGTGCTAGCAAATTCATCGACTATAGATCATGTGCAGCTTAGTGATTTGCTAGATGAACGCCTTCCATATCCTGAAGATGAAGCAAAGGATGTTTTGGTTTTAATCATCAAGTTAGCAAGCTCTTGTTTGGTTGAAATTCCAAAATCAAGGCCAACAATGCACTTCATCTGTCATAGGTTATCATCAATGGATGCACGTCCACCTACTCATGTAAAGTGACCTTTG

>M6-PGSC0003DMG401009161 M6 LRR-RLK

TGTACTTTCTTCTTGGAATGCTTCAACTTCAACTCCATGTTCTTGG

>M6-PGSC0003DMG401013731 M6 LRR-RLK

GCTTTGCTGTCTGGTGATGCTAATCCTGCCTGCTTCTGATAAGAGTTATTTACTTGCAGGGACCTGAGCAACAACATGTTGAATGGTCCAATCCCAGATACATTTAGGCAGTTGGACTTTGATAACATGTAGGTACCTTGAAGTGAAACCTTTACAAGTTGAAGCATAAATCTTGTAATTTTTTTTTTTTTTCCTCAAAGTTTTCTTTCAGTAACAAGAAGCAAAAGTTTTATCTGTGATGTAGGTTTCTTGGTAACAACGCATTAAGCGGAGCAATTCCCAGCTGGGTGTTGAGTAGTGATAAGAACATGTTGGTATTTGCTCAATTTGTTCCGTTTTCCTCATCTAAGTAGTAAACTTTGGTTATTATCATGCCGGCACACTTCAAAGTTAACAGTTTTTTCCCCAAAACATTTTGATTGCTGAAAGGACGCATTTGCAAGAATTTGGTGTCTTCTAAGTGTATGGCAACTATAATTTTTTTTGCAGTGATATATCATACAACAATTTCACCCATACATCTACTCAAGGATGCCAGCCCTCTACCATGTGAGCATTGTACTTCTTTGTTTGGGAACCTACATCCGTGTATTATTTCTATTTTTATTAATTAGAATTTCGGTCATTTAACTGCATAGCTTACAAACCTTCTTTCTTCAAACAGAAACTTAGCCGCCAGCTATTCAGATACAATGAACACGTTCAACAACACGTAAGTCCTAAATGTTGTCTTATGCATCAGGTGCTAGTTTGTTTGATGCTTTCTGTCTACTGTCTATAATTATGTGGCTTAATCTATTCGAAATTGCAGGAATTCCTGGTGTTCTATGAAACCCCTCATTTGTCCCACAGAAGCGAAATGTAAGAGTTACAGCCTATAAAAGGACCTCAGGGGTTATGTCTGGATCATTTACCACTGAACCAACTCTTGGAGTGGCTGCCTGTGGACTTTTTAAGTACTAAAGCATAACTTTGGAGTTGAGAGATAATTTCACAACTCTTACTCATGTTACGTTCTTTATGTCAATTGCAGATCATTCGCTATTCATAAACTGTGGAGGAGGTAGAATTACATTTGAGGGAAATGATTATGAGGCGGACTTGACCTATAGGGGTCCATCATATTTTTTTTCATCCTCTGATAGATGGGCATTCAGTAGTTCGGGAGTGTACGTGGGTCTTGCAAATGTTAGTTATATTGCAACAAATAGATTTTTACTGAACATATCTGGTGCCGACTTTTACAACACAGCACGCCTTGCCCCTAATTCTCTCAAATATTATGGGCTTTGCCTACTAGGAGGTAGCTACAGAGTGCGGCTTCACTTTGCTGAAATAATGTTTTCTAATGACTCAACATATAGCAGCCTTGGAAGGCGGATATTTGATGTAGCAATACAAGTAAGTTTGAGGCCTAACTTCTTTTGTAAATAATAAAAATGTAATGTTGATATACTGAGTTCTGATATGGTTGGCAACTTTCTGATTTAAGATTGAAGGTACCAATTTGTTGGAGATAATGTGGTATATTTTGATTTCTTACACAGGGGAGAGTAGTTTTGAAGGATTTCAATATCATGGAGGAAGCTAACGGTGTTGGAGAAGTCATTATCAAGGACTTTGATGATATTACTGTTAGTAGCACTCTGGAGATTCACTTGTATTGGACCGGAAAGGGGACTAACGCTATTCCAGACAGAGGTGTATATGGACCTCTGATTTCAGCTATCTCAGTGACACCAAGTCCGTATATCTCTGCCTATAGACTTTACACTTATCTTCCCACAATCTTATCTTAAGAAAATCCCATTTTCTTCTTGCAGACTTCAAAAAAGGTCGCAGGTTACCTGTCAGAGCTGTTATTGGCATTGTACTCGCTTCAATTGTGGTGGTTCTGATAGTTTTATTTTCTCTATGGAAGAAAGGCATTTTTGGAGGGAAAATTAATCAAGAAGAATTAGGTAACACATACCTCAAAGAATTTATTCACTCTCTACATTAGATATGGCGATATTTCTGTTGTGATAATCAGAATTATGTACGAGATTTCTAGCTGCGACATACAAATAATTATCATAGATGATTGTTCTGTTGGCAGATAGACATCATGACATATCATAGATAGTCAGCACACAACAAAATGTTCATGACATATCATACAGATGTTTATTCTTTTAAAAATTGTTGTTCATCATACAGATGTTTTAGAAGATCTATATATACTGACGTTGATATTATTCTCGTTTACAGAACTCAGAGCCCTTGATTTACAAACAGGTCATTTTAGACTTAGACAAATTAAAGCTGCTACAAATAACTTTGATTCTGCCAACAAAATTGGTGAAGGAGGGTTTGGACCAGTTTACAAGGTATTAATTCTCTACCTATATCTATCCGTACTGAGTTTGCGTAAGGACAGCTTGCGGTTCTGTGGTGATGTTTCCTACTGGTTATCCTTCTGTCTGTAATATGATCGTTGTAAATTAATCCTGCTGTCTGTAGGGTGTCCTCTCCGATGGGGCAGTCATAGCTGTTAAGCAGCTCTCTTCCAAGTCAAAGCAAGGAAATCGGGAATTTGTTAATGAGATAGGCATGATTTCAGCTCTACAACACCCAAACCTTGTCAAGCTCCACGGTTGTTGCATAGAAGGGAGCCAGTTACTGGTGATATACGAATACATGGAAAATAACTGTCTTGCACGAGCTCTCTTTGGTAATGCTTTGCAACAACATTTTGAAAATCATTAATCATGCAGTGATAATTTTATGCTTAATACTCCTCTAATTCTCCTTTCTTAGGCCGTGACGACCAGAGGTTGAACCTTGACTGGGCAACCAGAAAAAGGATATGCTCAGGAATAGCAAAAGGGTTAGCTTACCTTCATGAAGAATCAAGGTTGAAAATCGTCCACAGAGATATAAAGTGTACCAATGTGCTTCTTGATAAGGATCTGAACGCTAAGATTTCAGATTTTGGATTGGCTAAATTAGATGAAGAAGAGAGCACTCATATTAGCACCCGAATTGCTGGAACAGTGTGAGCCTTCTTATCATCACTGACTCTGTACTATCATGTGATGAGAGAGTAGCAGTTCTACCCATTTGAATTTGTTCGTCATACATTAACATGGTCAGTTTGAAGTATCTTACATTTCTTTTTCCTCTTTTGAAGAGGTTATATGGCTCCGGAGTATGCAACGAGAGGCTACTTGACAGATAAAGCTGATGTTTATAGCTTCGGAGTAGTTGCTTTGGAGATTGTCAGTGGGAAAAGCAACACAAAATACAGGCCAAAAGAGGAATTTGTTTACCTTCTTGACTGGGTAAACTATCTTTTTACTCAACTGTCAATATTCCCAAATACTAATATCTTTAAAAACTGCTGCAACATCATGTATTATAAGTCTTGCATGTACATGTTTAATTTAATTGCCCACTCTGAATTATAAGCCAGAAAAAGCTTCTCTTGTTAGTGTTTTGTGCAAATATATACCCCATCAGTTTCAATTTGTTTGTCTTACTTTAGTTTATTTTTGTTTGTTTCAAAAGGTCTCTTTGCTTTTTTGCAAACTCTTTATTCTAACTTTCCACGTGACATATTTAAAACCACAAGATTTAAGGGTATTTTGGTACATTCTACATATCTTTAGTTTAAATTAAAACCAGACAAACAAAGTGAAACGAAGGGAGTATTTGCTATGATACCTCCAAAGAAAATATCAGAATCAGAGTTGTTTCATTAGTTCTGCTCACATTCTTATGGATGTCTTTTCTCCAATTTCATCTTGCTTTGTATGTCAATGTTTGTTTTAGGCTTACGTCCTACAAGAACAGGGAAATCTGTTAGAACTAGTGGATCCACGT

>M6-PGSC0003DMG401017953 M6 LRR-RLK

TCTCCGGGAAATTTTCATTTATCATAAGAGTTGTTGAGCCAGTATTGTGGTGATGGGAAAAGAAATGGGAGTTGAAAAGCTTCTGGTTTTTTTTTTGGTACTAATCTCAATCAGATATTCAGAGCAATTACAATCATCTCAAGTGCAAACACTTTTGAGAATTCAACATCTTTTGAACTATCCAACAGCTTTGAGTAGTTGGAATAATGAGACAGATTTCTGTAAGATTGAGCCAAGTTCAACTGTTACTGTTGTGTGCTACGACGAAAACATAACTCAGTTGCATATAATAGGGAGAAAAGGGACTTCCCCATTACCTAGAAAGTTCTCTATTCGTTCATTTCTCAACGCGCTTGCCAAGCTTCCTAGCTTGAAAGTCCTTAGATTAGTTTCTCTTGGTTTATGGGGGCCGTTGCCTGACAAACTTTCAAGACTTTCAGCTCTCGAGATACTTGACTTGAGTTCAAATTACTTTCACAGTGGCATACCTGAAGCAGTTTCATCCTTAACGGACCTCCAAACTTTAGTGTTAGATGGAAATAGGTTGACTGGCGGAATTCCAGATGGCCTTGGTTCACTTTCAGTTTTGGCAGTTTTGAGTTTGAAGATCAATTCATTAGATGGACCTTTGCCAGATACACTGCGTGATTTGAATAATCTTCGTGTACTTTCACTTTCGAAGAATAATTTTAGTGGTGATGTCCCTGATCTTAGCAGTTTACAAAATCTTCAAGTTTTAGAGTTAGAGGACAACTCTTTTGGTCCAAAGTTTCCTCAAACTGGAAGTAAAATTGAAAGCATTGCTCTAAGGAATAACAAATTCACTGCAAACATACCAGAGAAAGTTCAGTCCTATTATCAGCTTGAACATTTTGATATTTCTTCCAACAACTTTATGGGGCCATTTCCTCCGTCGTTGCTATCTTTGTCATCCATCACATATCTGAATGTTGCAGGAAACAAGTTAACAGGAATGCTCTTTGAAGATAATCCATGTAACACAGCATTAGATTTTGTTGATCTGTCTGCTAATCTGTTGACAGGATCATTACCTAATTGCCTTCTGTCTAGTTCTAGGAACCGGATTGTGCATTTTTCCAATAACTGCTTAGCAACCGGAGATGGTACTCAACATCCATTTTCATTTTGCCGGAATGAGGCTTTGGCTGTTGGTGTCTTACCTCAGCATCAAAGACAAAAACATGCTTCTAAAGTGATTCTTGCCTTGACCATATGTGGTAGCATCACCGGTGGAGTCATACTCGTATGCCTGACCATCTTGATTGTCAAATCTTTTCTTGCAAAGAAGGCTTCTGCCCAAAATACCCCAACCAGATTTATTTCAGAAGATGCACCATCTTCATATACCTCCAAGTTATTCACAGATCCAAGTAAGGTTCCTCTCAATAACCAAATTACTGTTTAAAATAGAAGCTTAGGGCAAATGTATCTGCTCTCTGCGCCCTAGATGCCTGAAGGCGAGGGCTTTCCTTCAATTCCCAAGTGCCAAGTATACTAACTTGATACCTTTGTATTTCTTGTATAAGGATACATTACTCAAGCAATGAAGTTAGGAGCACTTAGTCTTCCATCTTATAGAATATTCTCTCTAGAAGAACTTGAGGCGGCAACGAACAACTTTGATACAACAACTTTTATGGGTGAAATTCCTAATGGCCAGGTATGGTATATGAACTCTATTATTCTTTCCTGCTATAGCATTGAGGTCTTTGGGTGTTATACTTATGGCCATATTAGTAATGTAGCTCACGTGGCAGATGTGGACAAGTCATTCCCAAGAAAGGGAGTTTCAAGTATTTGAGGTAATAATCCATGGAAATGGAGAGATTGATGATGATGTTAAACATCGTATTGGAATGAGGTGTATGAAATGGAAGCTTGCATCTAGTGTCTTGTGTGATAATAATGTGCCACCATGTTTTAAAGGCACGTTTTATAAAGTGGTGGTCAGACCAATCATGTTGTATAGGGCAGGTGTTGGCTGTTCAAGAACTCGCATGTCAAGAAGATGAGAGTAGCGGAAATGAAAATGGTGAGATGAATGTGCGGACACGCTAGGAGAGATATGATAAGGAATGAATCTATTCGGTACAACGTGGGAGTGGCCAAGATGAGGGAGGCAATATTGAGATGGCTCGGGCAAGTGAAGAGAAGATGCACAGATGTTCCATTGAGGAGGTGCGAGAGGTTGGCTATAGTGGTATTAAGAGAGGTAGAGGTATGCCAAAGAAATTTTGGGGAGAGGTGATTAGACATGACATGACTTGCGTGTAGCTCATCGAGGACATGACCCTAGATAGGAAGATATGGAGGTCGAGGATTAGGGTAGAAGGTTAGTAGGTCAAGAGTGTTGTCTGGTTTCCCTTTTTAGTATTGTTATTACTACCCTTCTATGATCCTATTCTTCGATATTAGTTTTACTGGTTTTTTCCTTTGCTTCAGTTATCGTATTATTTTGTTGTTGTTGCTACTAATCTTTTCTTCATTGTTTTCAGCATGAATTCTTCGACTATGTTTGCTTTACACACTTGTTTTCTATATGTTTTACTTGAGCCGAGTATCTTTCAAAACAACCTCTTTACCTTCACAAGGAAGGGGTAAGTAAGGCTGCGTACACCCCACCCCACCCCACTCTCCCCAGATCCCACTTGGGATTATACATAGTATGTATCAGTATGGGTGTTATATTAGAGAAGTGGTGTGGACTTGTAGTAGTAATGCCATCTGTTTGTTCTCTATCATTTTGGCACAAAAGGATGAGTTTGACTCTTGAAGAGTAAACCTATTAATATTACCAGCTGTTCTCGAGGACTAAGGCTCACAAACGTTTTCTGTTTTGGATTCAGATGTACAGAGGGCAGATGAGAGATGGTTCCTACATTACGATTAGATGCCAGATAATGAAAAGAAGCAACACTAGGCAAAATTTTATGCATCACATAGAGTTGATATCCAAGCTCAGACATCAGCATTTGGTCAGTGCTCTTGGACACTGCTTTCAGTTTTACTCAGATGATTCAAGTGTCAGCAGAATTTTTCTCGTCTTTGAGTACGTACCTAATGGGACCCTAAGGAGATGGATTTCTGGTAATTACTTGACAAGCAGATATCTACCCCTTCCCTATTGTTGGATATAAATGTGCTTTAGTAATTTGATGCTCTCTGTTGCTTAATTGCAGATAAACATGCTAGACGTAAACTAACCTGGACACAACGTATAGCAGCTGCTACAGGGGTAGCAAAGGGACTACAGTTCTTGCATAATGGAATTATCCCTGGCCTCTTTTCGAATAATTTGAAGATAACCGATATTTTGCTGGATCAGAACCTTGTCGCCAAAATTAGCAGCTACAATCTGCCTGTTCTGTCGGAGAATATCGGCAAGGTGTGAC

>M6-PGSC0003DMG401021514 M6 LRR-RLK

AACAATCTTTCTGGTCCCATTCCTAGTGAGTTGGGGAATCTGAATAACCTCAATGATCTGGTTTTATCCGAAAATCAACTTACTGGTTCAGTTCCTACTACATTTGGCAATTTGAGAAACTTGCAGACACTGCATCTCCATACCAACAAACTATCAGGTTCCATTCCAAAAGAACTTGCATATTTGGATAACTTGCGAGTGCTGACAATGACCAAAAATCAGTTATCAGGCCATTTACCGGACCAGCTTTGTCAAAATGGGAAACTTGAGTTCTTCACTGTGGCTAGCAACAAGCTGACAGGGCCAATACCAAGAAGCTTGAGCAACTGCTCGAGTTTCAAATGGGTTCGCTTTAATAACAACAGTTTCATCGGAAACTTATCTGAAGCTTTTGGAATCTATCCAGAGCTTCTGTTCATTGACTTGAACGACAATGATTTTCATGGTGATCTCGGCAGCAACTGGGGAAAATGCGAAAAATTGATTAATCTGCAGGTAGCCAGAAATAACATTAGTGGTAGCATACCACCAGAAATTGGAAACATCAAAGGGCTTCTTGGACTTGATCTTTCATCGAATCATTTGATTGGGAAGATTCCAAACGAATTTGGAAAATTAACCTCTCTGGTTAAGCTTTCTGTGAAAAACAACAATATTTCTGGCAATATACCTGAGGAGCTTGGGTCACTGACAAAGTTAGAGTCGCTTGATTTATCAGACAACAGATTGAATGGATCAATCCCAACGTGGATAGGAGATTTTGTGCTGTTGTTTCAGTTGAATCTGAGCAACAACAAGTTTGGACAAAAGATTCCAAAGGAGATAGGGAGGATAACTCATCTTACTGTACTTGATTTGAGCCATAATCTTCTTGATGGAGAAATACCTGCTGAGCTAGCCAGTTTGCTGGACTTGTCAAACTTGAATCTTTCCCACAATGGTCTATCTGGGCGCATTCCTAAAGAACTTGATAGTTTGACTGGTTTGCTGGATGTTGTATTGTCATACAATGATTTGGAAGGTCCAATCCCTAATAATAAAGCATTTATGAATGCTTCGTTAGAAGGTAATAAAGATCTTTGCGGCAATGTAACAGGATTCCAGCCCTGCCAAAAGCCATCTTCTATGGTGAAAAAGCACTCAATAGCAAAGGGAAGTAAACTCATCCTCATCACTGTACTTCCTGTTATGGGAGCACTAGTACTGCTCTGTGTTTTCATTGGTGTTCTCTTTATGTGTAATAAAAGAATGAGAGTTAAAGACGTTGAAAGAAGGGATGGTGATGGTTGGCTTTCGATATCCATGTTAGATGGAAAGGCATTGTACCGGGATATCTTAAACGCCACAGAGGAGTTTGATGCAAAATTTTGCATTGGGCAAGGAGGACATGGAAGTGTTTACAAGGTAAACCTTCCATTATTTGGTGATATAGCTGTGAAGAGACTTCATTCTTCATTTCAGAATACACATCCCAAAAGCTTCATAAATGAAGTAAGGGCATTGACTGGGATTAAGCACCGGAACATTGTGAAACTCTACGGCTATTGTTCAAAAGCACAACACTCGCTCTTGGTTTACGAGTATGTAGAGAGGGGGAGTTTGTCTAGTATTCTGAGCAATGAAGTTGAGTCTAAGAAATTGGATTGGCTTAAAAGGATGAATATCATTAAAGGTGTTGCTTTTGCTTTATCTTACATGCACCAGGATTGCTCACCACCAATTGTTCATCGAGACATATCAAGCAGTAATGTTTTGGTTGATTCTGAGTATGAAGCTCGTGTTTCAGATTTTGGCATAGCGAAGCTTCTCAAACCAAACTCATCCAATTGGACTGCACTTGCAGGAACATATGGCTATGTTGCACCTGGTAAGGTCTAAATTTAATTTCGATTTTATTCAGACTTCGTGAATTATAGCCAAACACTCTATACTATTTTTATCTCACTTACACATATAAATTGTTA

>M6-PGSC0003DMG401023552 M6 LRR-RLK

ATGTTGAAATATCTACAAGTGGGTTACAATCCATTGAATGGCGTTCTGCCCAATTCTATTGGGAATCTTTCATCTACTATTCAAAACTTTGAAATAGGAAACACACACATCAATGGCCTCATCCCCACAAGTATAGGCAACATGACCGGTCTTACATCCTTAAGCCTTGGCGGAAACAACTTGACAGGGAATATTCCTTCTGAGATTGGTAAGCTTAAACAACTCCAAGGACTCGATCTATCTAACAATAAATTGCAAGGACATATTCCAGAAGTGGTATGCAATTTGTCTGATTTGGTTCAATTATATCTGTATGGTAATGAGCTCTCTGGATTAATTCCAGAATGCTTAGGAAATCTTACCATGCTTCAACAGCTTTATTTGGCTTCTAATAAATTTTCATCAAAATTTCCATTGACCCTTTGGAAAATGAGTGGTCTTCTCTATCTAAACGTGTCACAGAATTCAATAGAGGGAGAACTTCCATCAGATATTGGAGGACTGAAAGCCATTGTAGAACTATATCTTTCTGGTAACCACTTTTCAGGCATGATATCAACCAGATTCGGGGAACTCCAAAATCTGCAGTATCTCGACCTATCGAAAAATTCATTTTTTGGCCGAATTCCATTATCCTTTGCCAACTTGATAAACTTGGAATTCTTGAATTTGTCTTTAAATGCATTGTCAGGTACTATTCCTAAGTCTTTGGAAAAACTCTCATACCTTAAAAGCATTAATGTTTCATTTAATGATTTAGAAGGTGAAATACCGAGTGGTGGTGTGTTTGCAAATTCCACTCTGCAATCATTTGTTGGGAACAAAGGTCTATGTGGAGTGCACATATTAGAGATTCCTGCTTGTGCTATCACTAATCCTGGAAAACAATCAAAGCTTAAGGAGGTTCTGCTAAAAATTGTTACTCCAGTGGTTATTGCATCCTTTATGATATTCTTGTTCGTTTCAATTTGGATAATGAAACGACAGAAGAAAGGGAAGTCCAAAGATGTGGACAAGATACTGGAGATGGGGACTCATCAATTAGTTTCTTATCACGAGATTCAACGAGCAACAAATAATTTTGATGAATCCAATTTAATTGGTGAGGGAAGCTCTAGCTCTGTGTACAAAGGCACATTGTTTGGCGGAACTGCAGTGGCCATTAAGGTTCTGGATTTGAAAAATGAGCAAGTATGCAAGAGGTTTGATACCGAATGTGAAGTGATGAGAAATGTTAGACACAGAAATCTTGTTCCAGTGATTACTACATGTTCTAGTGACTATATAAGAGCCTTTGTTCTGCAATTTATGCCCAATGGAAGTCTAGAAAATTGGCTGTACAAAGAAGATCACCACTTGAACCTTCATCAAAGGGTCACTGTAATGCTTGATGCAGCTATGGCAGTTGAATATCTACATCATGGTCATGTCACTCCAATAGTTCATTGCGACCTAAAGCCAGCCAACATTCTTTTGGATGAAGATATGGTGGCTCATGTTGGTGATTTTGGAATCTCTAAAATTTTAGCCATAAGCAAGTCCATGGCTTATACCGAGACATTGGGCACTCTTGGATACATTGCACCAGGTATACTAGTCTCTTTCTATATATTTTCACAAAAATCAGAATTTAGCCCTTCCAAAATCATAAAAGCAAGTGTCTGTTAGTACATAATTATGTTATATTTAAATGAGTAATTCTTTTTCATTTTCATTTTCATTCTTTTTTTAAGAATATGGCTCGGAGGGAATAGTGTCTTCTAGTGGTGATGTTTATAGCTATGGCATCATGGTGATGGAGGTCTTGACGAAACGAAGGCCAACAGATGAAGAGATATGCAATGAAAATCTTGACTTGAGGAAATGGATCACACAATCATTTTCAGGGAGTATGATGGACATCGTGGATGCCAATCTTTTTCGTGA

>M6-PGSC0003DMG401027271 M6 LRR-RLK

GTTTTCTTCATTTCTTGATATGGCTTCTTCTTAGATAGTACTCACTACTACTGTCTACTACATTTTATTTCCTAAATTTAAGGGCTTCCTAGCTTTTTGGTTATTTTCCATTTTCAAGATTCCTCAAATGGAATACTCCCTCACATTTCTTGAGCATTTCAAGACTCAATCTTTTTCACTCTAAACACCCTTTTGTCTTCTACACCCCACCCCCACAATCTCCAATTCTGCAGAAAGTTTCATGTGATGGATTCTTGATGGGAAAAGGAGGAAGAAGTTGAAGAAAAGGTGTTTTCTTTGTAAACAACCTCATAACCATTTCAAAGATTCGATCTTTTTCACTGTAAACCCCTTTTTTCACCCCCTTAATCTCATCAATGGATTCTTGATTGTTCTATCTGGTTGTTTTGGAGGGAAAAGGAGGAAGAAATTGAAGAAAAGGTGTTTTCTTGAAAAAAAAAAAAACTCAAGAATGGCAATTTTGAGGTTTCTGTTAACTGTGTTTTTTCTAGTTGGTTTGTTATCAAGATTCCAACTTGTGAATTCCCAGTTGGTATTTCATGATCAAGATGTAGTGGAAGCAATTGGGAAAGAGCTATAAGTACCTGGATGGGGTGTGATGAACACAACTGATTTCTGCTCTTGGCATGGAGTTATTTGCAGCTCAAACAGTTCAATGGTGGAAAGACTTGATCTTTCTGGTTTTAGATTACAAGGTAATGTAACTCTAATATCTGAGCTCAAAGGTTTAAAGTGGCTGGACCTTTCTAATAATAACTTCCAAGGTTCTATTCCATTTGCATTTGGGAATTTATCTGAGCTTCAATATCTTGATTTGTCTTTTAACAAATTTGGAAACTCAATACCTAGTGAATTAGGTAAGCTCAGAAGCCTAAAGGCATTGAACCTTTCGAAAAACTGGCTTACAGGAGCTATACCTGATGAGCTTGAGGGGTTAGAGAATTTGCAAGATTTTCAAATATTTACTAACAACTTGAATGGTTCTATCCCAATGTGGATTGGTAATTTGACCAATCTTAAGGTTTTTGCTGCTTATGAGAATGAATTTAGTGGTGGTGTTCCTGTTAACTTAGGATTGTACTCTGATCTTTCTTTGTTGAACCTTCACTCAAACCAGCTTGAAGGTACTATTCCTGAGAGCATTTGTGCTAATGGGAATCTTGAATTTCTTGTTCTAACTCAGAATAAGTTGACCGGAAAGATTCCTGATTCGATTGGGAATTGTAAAGGACTTTCTAGTATTAGAATTGGGAATAACAAGTTGATTGGAGGTATCCCTAAATCAATTGGGAATATCAGAAGTCTTACTTATTTTGAAGCAGATAGTAATAGTTTGTCTGGTGAAATTGTCACCGAGTTCGCAAAATGCTCAAATCTCACTCTACTTAATTTAGCTTCAAATAAGTTTAATGGAACTATTCCTCCTGAGTTTGGTCAGCTTAACAGTTTGCAAGAATTGATTGTTTCTGGTAATAACCTTTATGGAGAGATTCCAATATCAGTTCTTAAGTGCAAGAATCTCAATAAGCTTGACTTGAGCAACAACAAGTTCAATGGTACCATACCGGCAGATATCTGCAATACGTCCAGATTGCAATATTTGCTGTTGGGGGAGAACTCGATTCGAGGGGAGATACCTCATGAGATAGGGAATTGTGTAAAGTTGCTTGAGTTGGAAATGGGGAGTAATGAACTTACTGGAAGTATCCCTCCTGAGATTGGTCATATGAAGAACTTGCAGATTTCGTTGAATTTGAGTCATAATCGTCTCCACGGAAAATTGCCTCAGGATTTAGGAAAACTTGACAAGTTGGTTTCTTTAGATGTATCTGATAATCAGCTCTCGGGGAATATTCCGCCGGCATTGAAGGGCATGCAGAGTTTGATAGAGGTTGATTTTTCGAGCAACCAATTCACTGGTCCAATACCTACTTTTGCACCATTTCAGAAAAGTCCTAATTCAAGTTTTCTTGGAAACAAAGATCTCTGTGGTGATCCTTTGAGTGTTGGTTGTGGGGATTTAAATGGTTATGACCACAGCGTTTACGGTCACAAGGTCTCTTACCGAATCGTTTTGGCTGTTATCGGTTCTGGTTTGGCAGTTTTTGTATGTGTCACAGTGGTTGTTTTGCTGTTCATGATGAGGGAGAAACAAGAGAAAGCTGCCAAGGAAGCAGCTGGAAAAACCATTGATGAAATTTCTAGCAAACCAGTGATACTAGCGGGAAACGTCTTTGTTGACAATCTAAAGCAAGCAATTGATTTTGATGCAGTTGTAAAGGCAGTCATGAAAGACTCGAATAAGATTTGTGTTGGTACTTTCAGCACTGTCTATAAAGCAGACATGCCTTCCGGGATGATTTTGTCGGTGAAAAAGCTAAAGTCAATGGACAAAACTATAATTCATCACCAGAGTAAGATGATCAGAGAGCTTGAAAAACTTAGTAAGCTCGTTCATGATAATCTTGCTCGTCCTATCGGATTTGGAATCTATGAAGATGTCGTTCTTCTGTTGCATGAATATTATGCCAAAGGAACGTTGACTCAATATCTTCATAACTCTAGCCAGAAACCTGAATATAAGCCCGACTGGCCAACAAGACTTGCCATTGCCACTGGAGTTGCAGAAGGATTGGCATTCCTTCATCATGTTGCTATCATCCATCTTGACATTTCTTCCGGGAATGTGCTTCTTGATTCTAATTTCAGGCCTTTGGTTAGTGAAGTTGAAATCTCTAGGCTTCTCGATCCATCTAGAGGGACTGCAAGTATTAGTGCTGTTGCTGGTTCATTCGGATATATTCCTCCAGGTACATATCATTTGAACACACATACTCCCTCTGTTTCAATTTGTTTGTTTGATTTTAACTTGACACAAAGTTTAAGAAAGTAAAGAAAACTTTTGAATCTTGGTGTCTTAAACTAAAGATATGTAGAATGTACCAAAATGCCCTTTAATCTTGTGGTCTTAAACATGTCACGTGAAAAGTTTGAATTTAAAAGTTGCCAAAAGAGGAAAAAGGCATTCTTTTTTAAGTATGGCAAACAAATTGAAACTATTATCTTTCTCTTTTCGCGTTTGTAGAGATTATAAATCTGTAATTGACTTGTTTTTTTTTTTCTAAACAGAGTATGCATATACAATGCAAGTAACAGCTCCTGGAAATGTTTACAGCTATGGGGTTGTTTTGCTCGAGATCCTAACCACCCGACTACCTGTTGATGAAGCTTTTGGTGAAGGAATAGATTTGGTTAAGTGGGTACATGGTGCACCTACAAGGGGGGAAACACCGGAGCAGATACTCGATGCAAAGCTCAGCACCATTTCATTTTCTTGGAGAAAGGAAATGTTAGCAGCACTAAAGGTTGCGTTGCTGTGCACTGACATGACACCAGCAAAACGACCAAAGATGAAGAAGGTGGTAGAAATGCTGCAAGAAATCACGGAAAGCTAAACAAAGTTCGTATCATCAAATCTAAGTATGTGTTGATCTCTGTCTCTTCATATGTGAATGCTCCTGTAAAAATACAAGTGTACGACATTCTAAGTCGAGAGGACAAAATGGTCGAACTCTTAAGGGTATGATGTGTCTTACTTCAATCATATGATCACCAGTTCTTTTTCTCCAAAACATGAAGATGGAGATGATGTAGATTAATTTCTCACAAGGAGAATATGTTTACAAATAGCATTATCTTTCATTTTCCAGTTCCCCTTTCTGGTACATTGCTTCCAAATCAAGTATAGATTAAATCATAGTGAAAGATTAAGGGAATGCTACTATTCAAC

>M6-PGSC0003DMG401029358 M6 LRR-RLK

CATTTCTTGGTTTTGTATTGATATTACTAGTTAGCAGGATTTGTATCTGTTGTGCTTCAGAGAGTGATGTTTATTGCTTGAAATCAATAAAAGAATCATTACATGACCCTTTCAATTCTTTAAACTCTTGGAATTTCAACCAAAACTCCATCTGTCTTTTTGAGGGAATTGACTGCTGGCAATCTGATGAGAATAAGGTAATGAGCATCACACTTTCAAACTTTGGATTAATAGGTGAGTTTCCTCGTGGCATTCGAAATTGTACAAGCTTGACAGGTCTAAATCTTTCTGGTAATCATTTGTATGGAACCATCCCTTCTGATATAGCAGTGATTCTTCCATTTGTTACAATACTTGATCTCTCAAATAACACCTTTTCGGGCAATATAACACCTGGTATAGGTAATTGTCAATTCATAAATGTTCTAAAGTTGGACTATAATGATTTAGAAGGTGAAATTCCAAGTAGAATAGGCTATTTGCCTCGCCTTAAGACGTTCAGCATAGCTAACAATTACTTGACTGGGCCAGTGCCATTGTTTTTTAGCAAGAATATCACGGCTGAGAGTTTTGCAAACAATCCAGAGCTTTGTGGGAAGCCCTTGAAAGGATGTGAGGATTCTTGGATATGGAAACATGTAGATCGTGCTTCGTTCATCAAAGCGTTTGTGATTGGTTGGGTACTTTTGTTTACAATGGTTTTAGTCCTTTGCTTATTTAAATTTCCTACCAAGGCTATCAACAAGATAGTCTCATTAAACATATGGAAGCTAAGGAAAAAGAAACATTTAACTTCAGGAAGTGAAGAGGAATTAAGCAGCCAACATAAGGTAAAGATCTTGCTAAATTTGCTGCACTGCTTATAGTCTGCAAAGTTACCTTATCTTCTGTAGTTCCAATTTTGTGATTAGTCCTTGCTAGTCCATTATGTTATAAATTCTTTATTCGATCGATGATCAAACATGATAGCAATAAGGTTCTCAAGTTTCTTTCCTGTTTTGACTTCTGTGTATATTCAAATGATTTAACTTGCTGTTTTATTTACAGATATTAAAGCTGGAGAAGTTTGTTATGAGAATGAGCTTCACGGAGCTTGAAAACGCGACTTCTGGTTTTAGTGAAGACTATCTAGTAGGAAATGGAATGTTGGGCAAAGTGTACAAGGCAATTCTTCCAAATGGCTGGATACTTGCCATCAAGAAGCTCAATGACTGGGAGAATTTGGAGGACGAGTTTGTCTCGGAGATTACAACTCTTGGTGGTCTGAGGCATAGGAATCTATTGCCTCTAATAGGTTTCTGCGCCGAAAAGCAAGAAAGGCTTCTTGTTTACAAATACATGACTAATGGAAGTCTTGATGAATGGCTGCACTCGAATGAAGTTAAGGCCAATATTTTGGATTTCCCTCGGAGAGCTAAAATTGCACTTGGGATAGCGAAAGGCCTGGCTTGGCTTCATCATGGTTACGAATTGCACGTTACACATGGAAGCATAAGCACGCGATGTATCTTGCTAGATCAAAATCTTGAACCAAAAATATCCAACTTTTGGGAAGCAAAATTTTGGAGTAAGAATGATTCTGCATTAAGTTGGAGTCTTTTCCCAG

>M6-PGSC0003DMG401030114 M6 LRR-RLK

TGGATTGGCCAAAAAGATTGGAGGTTGCCATTGGAGCTGCTCAAGGACTTTGTTATATGCACCATGACTGCACTCCACCAATCATTCATCGCGATGTAAAAACAAGCAATATCTTGTTGGATTCTGATTTCACAGCCAAAATAGCTGATTTTGGGTTGGCCAAGATTTTAGAGAAGAAAGGAGAGCTTAACACCATGTCTGCTGTTGCGGGTTCCTTTGGTTACATAGCTCCAGGTAAAAGCAAAATCCAAGCTTGAACTTCGTATTACATCAATTTCGTTAACATTATATCAAAAGTATGACTAACTAATAATTTACTGTTTGGTTACAGAGTATGCTTATACAACTAAAGTGAACGAGAAGATTGATATCTATAGCTTCGGGGTGGTGCTATTGGAGCTAGTGACTGGAAGACAACCAAATTTTGGGGACGAGCATACAAGCCTGGCAGAATGGGCATGGAAGCAGCACGGAGAAGGGAACACTGCCATCGATAATATGTTAGATACAGATATCAAAGAAACATGTTACTTGGAAGAAATGAAGACTGTGTTCAGACTCGGGCTTATATGCACGAGCAACTTACCTGCAAGCAGGCCTTCCATGAAAGAGATTCTGCAGATACTTCA

>M6-PGSC0003DMG402002925 M6 LRR-RLK

GTTCACTTCTCTTTAGTTTTTACTTCGATTTTTCTCCGGAGCTCCGGTGGTCGGAAAGTTGATCGGCGTCGGAAAACAAGTAGAATTATAAATTTATAATCACAATCGAGGTAAATTTGCTTTTTTCTCCGTAATTTTGCCCGTTGATCATTCATTTTTTCTGTGTTTTAATAAAAGTTTAGTTTTTTTCTGTGATTTTGGAGTTAAAGTTTGAAGCTTTTTTTCTTAGGGAAGTAGTTTGAGTTGGAGATGAAAAATGGTGAACTTTTGGTTTACTTGGAATTGAACCCCCACCCCACCCCACAATTGGAAGAAAATTGTAGGTGTAAAAGTTGAATTTGGAGTTTTTTGTGATTTTGGAGTTGGTTTGAATGTGTATCTTTCTGGTTTTTATGTTTAAAGTTTGGAGTTTTTTTGCTTAGGGAAGTAGTTTGAGCTGGAGATGAAAACATGTTTTGTTTAATTTAGAATTGAAACAGAAAAATTCGTCTTGTTTGTTCCATTTTTGATAAGTTTAGATTATATTTTTTTCCTGTGATGTTTGAAGTTTTTTCTCTTAGGGAAGTTTCAGCTGGAGATTAGAATGATGACATTTTTGTTCTATCATAATTGGAAGAAAGAGGTAGATGTAGGGGTGAAAGTTGAATTTGGAGTTGTTCATCTCTTTCATTTTATTAGTTTTTTTTTTATTTTTTGTTTTTTTTTCTGTGGTTTTGGGAAGTTTATCAAATTGGTATATGCTTAAAGTTTGGATTCTTTTGGCTTATGGTGATGTTTTGTTTTGCTTGGAAATTAAAAACCCATATTTGGAAGAAAAATGTAGAGGGTGAAAGTTGAATTTGGAGTTTTTTTTCCTTTTTTGTCTTATTAGTTCTATTTTTTGACAAGATTTTTTTTTTCTGTGATGTTGGTGTTGGTTGAAGGTTTATCTTGATGATTTATGTTTAGTGTTTGGAGCTTTTTTCTCCCAAATTTTAAACATAAACAAGGAAGATAAACTTTCAACCGACTCCAAAATTACAGAAAAAAAGAGAAGCTAATCAAATATAGAAATAATAAGAAAACTGAGAAAACAAAACTCCAATTTCAACTTTCACCACTACATTTTTTCTTCCAATTATAGGGGGTTCAATTTCAAGAAGAACAAAAGGTCACCGTTCTTATCTTTGTGCTAGAGATAAGAATGGTGACCTTTTGTTTACTTGAAATTGACCCCCTATAATTGGAAGAAAAATGTGGGGGTGACATCCATGAGGTCTCAGGTTGAATAAACACTAGGTGATTTCTTTCCATTTGTCCTAGCCTTGTTAGATAGAGTTGCTTGGTACTTGTTGCTAGTGGAAAGTGACATGTATCTTGTGGAATTAGTCGAGGTGCATGCAAGCTGGTCCAGACACCAAGGCTATATAAAAAAGAAGAGTTGAAAGTAGCTGACCCTGTCAGTCTACTTAGAAAATTGATTTTTGATGAACCTAACCTTTTACATTTTTCATGTTTTGCAGTGTCTCTGGAGGTGTGGTTTGATTCTAGAAATCCTCGTGCAAAAATAATAAGTCGGATTGAAACTTTTCTGCTGTAGCTGCTTTGTCAGAAAAAATGATGGAGGCTCCAAAATCTGTTGGACTTACAGGCTTGAATCAAAACCTGTCGCAGAGCTTTTACCATAAGCTTGGCGAGGGGTCAAACATGTCTACAGAGAGTTATGGGATGAGCAATGCTGGAGGTTCTGTTGCAATGTCGGTGGATAACAGCAGTAGTGGATCAAATGATTCTCACACTCATATTTTAAACCACCAGGACAACCACATCCACAATAACTATTCTGTTGCTGCTAGCGTATTTAGGGGGAGAGTTTCTAGGTTGAGCGATGATGCCCTCGCTCAAGCCTTGGTTGATCCTCAGTTCCCCACCATTGGGCTTGAGATTTATGATGAGTGGACAATTGACTTGAGGAGGCTTACCATTGGACAAGCTTTTGCTCAAGGATCTTTCGGGAAACTGTATAAGGGAACTTATAATGGTGAAGATGTTGCCATCAAGCTTCTGGAGAGGCCAGAAAATGATCTTGATAGGGCTCATCTTATGGAGCAACAATTTCAGCAGGAGGTCATGATGTTGGCAAATTTGAAGCATCAAAATATAGTACGATTTATTGGTGCATGCCGTAAACCCATGGTGTGGTGTATTGTGACTGAATATGCAAGAGGGGGATCAGTGCGTCAGTTCCTCACAAGGCGGCAAAACCGAGCTGTGCCATTGAAGTTAGCAGTTAAGCAGGCTCTGCATGTGGCAAGGGGGATGGAGTATGTGCATGGCCTGAATCTGATTCACCGGGATCTGAAATCCGACAATCTACTGATTGCTGCTGACAAGTCAATCAAAATTGCAGATTTTGGTGTTGCTCGAATTGAGGTGCTGACTGAAGGAATGACACCAGAAACTGGCACTTACCGCTGGATGGCTCCGTAAGCATCTAGACTCTTTAGCAGTCTTGTTAAATCTGGTGTCTTGGTCCAGTTTGGGCTTATTGCCAAACCCTATAATAGAGCTCTCTCTGTCTTCATTACTTTCTGAATTTTATTTCTTTTTGAAAACTCCTTTGCTATGTGTTACTTGAGCCGAGCATCTTTCAAAAACAACCTATCTACCTCCACAAGGGAAGTGTAAGGTCTGCTTTCTCTCTACCCAACCCAGACACCACTGAGTGGGATTTCACTGGGTATGTTGTTGTTGTAGCATGCTGGATTCCCTTTATGGAATAAACATTTTTAGAACTAAAGTGAAAAGCATGGAGGTTGATGGGTTCAAGCCCTGTATGTCCTTTTGATGGCACCTCACTGATAGTGATTATATTTTCTATTGTATATGTGCTGTTACTAAGAGGGAGAAATTATTTGTTCCAGTTAGCTTTTTAACATAGTTATGTGCCTGTTAATGAAGGTTTACGAGTTGATCGATTTTATTTTCTCTTTCAATTTCTTTCCCTCCTTTTCATCTTCCTTCTTGTCTGTGATTTGTAACTTGATTCAAGTTAGTCACATCCAAGGGCGTGGCTTAGTTGTCAATGAAGTAGGTGCAAACCATGGAGATTAGGGTTCAATCACACCAAAAGTGCTTTCTTCCCATATGCCTAAGCCTTAAACCTTGGTGGGTATCTATTTATGTCTGTTTATTGATCCAAATAAATGGTAAAGTGCCTTAGTTGATAACTCCCAGCACCAGCCACGTCATATTCCATGAACTTATCAGCCCAAAGTGATTATTAACAGGAAAAGAACTAATCTTCAAAAAAATCAGTTCTTTTTCTGTTAATAATCACTTTGGGCTGATAAGGTTCATGGAATATGACATGGCTGGTGCTGGGAGTTATCAACTAAGGCACTTTACCATTTATTTGGATGCACTCCCCGATGAAGATGATAAGTTCTACATTGTGGTGGAAATTTTGGTCTTTTAGCAGGCATCTTAGAGAAGCATGGGAGACATTAGTTCAAATCCTAGAAGAGGCACAAAAACACTAGGTGATTTCACCTGATACTTATGTTAGTGGGGTATTGCAAGTACCTCATAAAATCATCGTCGGCATGCACTAGTGATCTTCGAGGATTTCTCGATTATCAAAAAGAAAAATCTAAATAAGTTCTATTCTTAGACATAATTAAGTTTCAACACTTTTATGCCTCCCAACTTTTTTATGCAGTTGTTATAATTGTTATATATAAGCGACTAATTAATGGATTCAAAAACAAAAACTGGTTACACCATAACTCAAGTATGGCTCTTTTAGTTTCTTTGCTAGCTCTGAGTTGGAAAGAAGTTCGTTCTTCTTTGGCTGTTTCCGAAAGGAATTCCAGGAGAGGCCTTACAAAAACTAGACTCTACAACGATAATGTGATTCTCCACTTACTACATAGACTTTTTCCAACATAAGACCATCTCAATCTCTACGGAAAGGTAGAAGTGGAATATTTGGCTCAATCTTCCATGAATCTAAATCATTTGTTGGAACCTTCGGTCATTCTTTACGGAGAGACCTCTTTTTAAAAGCATAAAAAGAGACTAGGTGATGTTTCTAGAATCTACAACGATTATGTGGATTCTCCATTGAGTTGCTACATAGACTTGTTCCATCCATAAGATTATCTCAATCTCTATGGAAACGTAGAAGTGGAATACGTGGCTCAATCTTCTCATGACCCTAAATCATTAGTTGTAACCTCTGAGCATTCTTTCTGGAGAGAAGTCTTATTAAAAGCAAAAGGTGATTCATGGAGTTTGAAGTGAAAAGCTAGGAGTGAAGCATACAAGTTACGGGAAAGTAAAAAGAAGTACCAATCACATACAAATTTAGTCATCTTCTCATGAACCTGAATCATTTGTTAACCTCCAATTACTGAGAGTAGTGTTTTTAGAATCGAGAAATGCAAAAGATGACAAGGTCCGTGATTTTGAAGCAAAAAGAGAGAAGTGAAGCGCACTAGCTATAAGAAGTAAGTAATGCTAGACATAGAAGGATTTACTACAATAAAGATGCAATTAGAATAACTGATCTTTTCATCCAATTTACGGTAATATTGATATTAGTCTAGCTCCTCTAAGTCGAGATTCAAAAAAGTAGACCCTTTCTCATTAGGATCGTTTTACATGTCATTGTTTAAAGTCCACACTTCTCTGAAATGCATGGCTTCAAGCGGTAATCCCTTACTCAAATTGCTTAAAGTAATGGCTTTGAATAACACTGCAGGAGAGATGTGATACATGAAGAAATCAATATCCGGTTAAAAGAATGATAAGAATGCCTTATTTGGTATTTAGTCTTACAACACGTGTTGCTAGACCTGATTTAACGAATGCCTTATATAAGTTGAAGTAGATCTTTTAGAAACAAAAGGAGCATTAAAACCCCTTAGTGATCAATCAACATGAACAACTTGTTAAGTCAGGGAGAGCCTACGTTAGGCCGGTTATGACTTAGTATTCCCGACGTCTTACTTTTTGTGCCTCACATACTATGTTACACCAGTCCTTTGCATTACACACCCGTTGAAATGGGTGGGTCGTGGCTGCGTGTACTCGTTAAAATACACAAAAAAAATTAACAAAAAAAGAAAAATGCACGTACTTATAATGATACTTGCGTACAACTGATAAAATTCATACTCACAAATGAGTCCATACTTAGAAGCGGATCTAGGATATGGAGTTTATGGGCTCCCAACGATCTCAAATTAAAAATAAAAATTATTATAAGTGGTCAAAGGGAATCAGACCCTCAACGAGGAGGGCAACAAAGCATAAGCAAGTTCCATTATGCCGCTGGGGAAAAAGCTTACTTTGTTGGTAGATCCCAACAAATAGTTTTTATATATTTAATAGATTTTTTCAATATAAATACAAAAGTTAGTGGATTCCCCAAAGCCTACACCCGATACTCTAGGTCTGCCCATGTCCAACAACAACAACAATATACCCCGTGCAATCCCACAAGTGGGATTTGGGGAGGGTAGAGTGTATGCAAATCTTACTTTTACCTTGTAGAGATAGAGAGGTTTCTGAAAGACCCTCGTTTTTAATAAACATGCCAAAATAGTTCAAAAGTGGGATAAAAATATGGGAGCAATCAGAACGACACGTAATGCAAAATGGGAAGTGTTACACGACAAACAAGAGAAAGAACGGTAACGAAAACAAAATAATGCATAATCAATGCAGAAGAAACTACACGAGTAGTGCTAATACTATTGGTAAGAACACCATTGAAAGCCTATCTCGCAGGGAAGTGAGAGAATGCTTAATTACCAACTAACCTTCTACCCTAATATGCGACCTCCATACTCTCCTATCTAACACCATGTCCTCAGTAAGATCCAGATGCACTCTATCTAATCACTTAATCCTAATACTTCTTCAGCCTGCCTTTACCTCTCTTGGCGTCCTCCACTGCCAATTGCTCACACCTCCTCACTAGAGCGTCTACACTCCTCCTTCTCTCATAATCACTCAGTCTCACTTCCCTCATTTTGTCCAACATGGAGGCCACTCCTATCTTGTATCAGATAGCTCAATTCTTAATCTTGTCTCTCTTAGTATGCCCACACATCTACCTCAACATCCTCATCTCAACGACTTTCTTCTTCTGAACATGAGATTTCTTAATAGGACAACACTCCGCCCCGTAAAACATGGTCGGTCTAACGGCTACTTTGTAGAACTTGCCCGTCCATGCTCCATATAATATGGATCACATGTTACCTTTAGTTGCTGAAGAAGTATATTTGAACCATTTCAAAATTTTGGATTCCTCTACTAGAACTCATTACCTCTAAGAATGATAGGAAAAACGAAGCATGGGTGTCCACATGGGCTTGATTTGCCAAGTTTGTTCGTATGTATTATGAAACGCCAAATTAATTTTAGAAGTAATGCATAATTAGTATGTAAGAATATATTTTACAATTTGTTTTATGTATTACGGTGCATGTATAGGCTGACAATTGTATCAATTCATGCATTTGCGCCTGATTGTTTAACTGTTGAAAATTTTTGTTTTATTTTGATATCTATCAGAAGATCTCCTTGGTTGGCACAATCTGACATAGAAGGTGCAAAATAATGACAACGAAGATAATTACTTCAATTTGCTTAGTTTATTGTTCCACTGATACATGGGGTTTTCCTGTATTTAGAGTCTTATTTAAGAATGGGTATATTTAGTGAATCTTTCAAGAAATTTTTCTTATCATGATTTTCCTGGAGTCTTTAGGATTCTGCTCTTGGAAGTTCAGTTATTTGTTTCTCAGGCATATATGGCAGTTAGAATTTAAACCTTGAATTAGATCATCATTTCAGGATGAGAGAGAGAATGGTATAGATTAGTTGCTTTATCCAGTTTATGAGTGTAGATTACTGATGATAGAACTGGCATTTCTTGATTTCTTTTGTCCTGGACAGAAGTATCACCATTAGTTTCTGTTTAGTTTTGATCCTTACATAATGTTGGAGATGAAAGTATATTCTTTATAAGCTAAGCGAGTTGGAGATTAAGCTTTTGATTTCTTGAGATAATGTGACAAACTGAGAAGTTTCCATTTCTATTAGGACATTTCTTTCTGACTTGAATCAAAAGTAAGTTTTCATGTCTGCCTGAAATATTGTTGAAATGCTTGCGATCAGAATAGGAGGTTGACCTTGTGAGTGGTGATAATTTTGGTGACCTAGTGCAGTCGTGAAACACCACATCCCTGTGATGGTGGATGATGTAGCAGTGCAATATAGTTCTTCTCTGTGTGATAATAGGCTAGTATGCATGTTAGACTTTCCTTGGCGAAATAGACAGAAGAATCTGCTTGCATTTGTTCATTTGATGTGATAATGAATGATCTTGCCAAAAAAAAAATGATATTGAATGATCTTTGCCAAGACAGTTGGGGATCGGATCTCAATATCTGGTAAAACTCGTATACGTGCAGCCAGTCTCAGACATTTGATTCTGATTAACTGAGAAATGAAGTTACTTTTCAAGTTCACTCTTGAGAAATTAGTTATTCACATTCTGCAGATACTATCCTGTTTAACATTGTGATTTTTTTGGCTAGTGTGTCTGGTTTCTTCTCTTTTCAGAAAAAGAAAAACGATGTTCAAAGTTGTTAGCATCTGTGTTTCGTTGTGGACCGCTAAGTATTTGACTTTGTTTTGTAGGGAGATGATCCAGCACCGGTCATACACCCAGAAAGTTGATGTTTATAGTTTTGGCATTGTTCTGTGGGAGCTCATAACTGGGTTGCTTCCTTTCCAGAACATGACTGCTGTGCAGGCTGCTTTTGCTGTTGTCAACAAAGGCGTCCGTCCAACAATCCCCATTGATTGTTTGCCTGTCTTATCTGATATCATGACCTTCTGCTGGGATTCTGACCCCGACAATAGGCCAACTTTCTCTCAGGTGGTCCAGATGCTTGAGGCAGCTGAGACAGAAATCATGACTAATATCAGAAAGGCACGTTTCAGGTGCTGCATTCATCCTACGGCTACAGGTTGATTCGGGAAAGAGGAAACTGGAGGCAAAAAAATGTTGAATGGCAAAACTGATTTTGAGTCTGCCATGTGTATCTATCAGGCACAATTTCTTTTCAGATAGAAAAAGGGTTATAGTTATGAGAAAAAACTAAACATTTCTCTGTTGTATTTTGTTTCTTTTCTTTAAACTGTGGATTTAAGACAACGACTAGTTGTTAATACAGGTTATCTGATGTGCATCTATGCATTTCAGATTTTGTCTCTGTGTTTGCTGAATTTGTTCAAAGTTATATCAGG

>M6-PGSC0003DMG402004902 M6 LRR-RLK

TCCTATTAACTATTCCGATCCGCCATTAATCTAATTCTTCAGTCAGCTTTACTGACCATATATTTATAGATCATCCATTGGTATCATTGATCATGTACAAGAAGATTTGCAGATCGTCCATTTCTTTCTCCTATTAATTTGTCAAGAGGATTACTCTTTTTTTTCTAATAAAGCCTTGATTTTTTTTTTGAGTTCAGAGACATAGACGAATTTAGAATTTAAATTTGATGGATCAACCTTTTAAGTTATGACTATTGAATCCATCTGATCACATAATGAGTTCGAAATTTAATGTTACAGTACTATTTGATAAAATTCCGTATTAAAATATTGGGTTCAATTGAACGCGAAGAAGGTGGGGGCAGGATGAGCTTGGTGGTAAAGGTGGCCCTACTCTTCTTCATTTTGCATGGCTGCAGCTTCATCAATGCAGACACAGATCCATCCGAAGGTTAGCTTAATTCTACGTGCTACGTAGCTGAATCATAAAGAAATTAATATATTCACGGCACTCATGATTAATTTGGAAGTTACTAACAAAGTTTCTCTGTAAATAAGAAAGAATTTCGTAGTTAACTTATGATAAGGCGTGGAACACTTGGTGTACTACTTTTCTTTTCCAGTATTCGATATCCGCTTTGTTGTCTCCAATTAATTCGGACTCATTACAGGGGAAGCACTTCCTACATGTAATTTTTCATTCTCAAAGCTCAAACCCGAGACTTTTTATAGCTAAGCTAGGTTGCTAATTTTCACTAAAAAGCACGACGATAAGCCAAAGAGTTCACAACATGCAAAAAAAAAGAAAAAATTGTTATAAATTTAGTAACAAAACAGTGGAGTTTGATTTATTTATTTTTTGGAGCTGTTCTTAGTCTCTGTGTGTGTATTTATATGTTCTCAATAATTTCGTTGGTGTCGTGCAACTTTGATTATCAGGTTCTGCCCTGAAAGTGATGTATAGCTCCCTAAATTCACCGGGGCAGCTTACAAAATGGAGCTCATCAGAAGGTGGTGATCCTTGTGGAGAATCCTGGAAAGGAATTACTTGCTCGGGCAATAGAGTAACTGAAATGTAATTATGTTCTCTTCACCATCCTCCTATAGTTTTTCGATATACACTTTATTTTTGGTAACTTCTCGTTAGTTAATTGATTTGGCAATTGCTGATTTTCAGTTTTCGTTTCATTTTCCTTCAGCCAGATATCCGGTTTAGGACTATCCGGATCAATGGGATACCAACTAATGAGTTTAACATCAGTAACCAACTTGTAAGATGATCAATGGCTTCAATTACTATTTCCTTAATTCGTTCTGTTATTTGTATTTACATTGTAAGATGTTGTAATTTGCAGTGACATAAGCAACAATAATCTCGGAAACCAGATACCTTACCAGCTTCCTCCAAATGTAAAGAGACTGTATGTGTTTTTATGCCTAATTTCTTCTTTCTTTGCTCAATTCCAGGCTCTAGTGTTTTTTACAAAGTCACTTACAAAATCTTTCTGCCATTTGCCTCGTATTGCAGTAACCTTGCTGGTAATGGTTTTACCGGTGTCCTACCTTATTCAGTTTCACAAATAACTTCCCTTCAATACTTGTAAGTGAATTTTTAAAAATTAAATTTACATGTATAGTATTATTACACTCGTTGTTCTGATTTTCTCTCCGTTGTATGTGCCTGACTGTAGAAATGTCAGTCATAATCAAATTCAAGGAGAACTAAATGACATGTTTGGATCACTTTCTTCTCTCAACACATTGTAAGGCCTTGTCTCGATTTCTTCCTTTTCTTATACAGATTGATATTACCCTGTGATGATTGAGATTTTCAATGTATCTTGTGCTATTCTCGATAGATAAATACAGTATCTCTTCTGTTTATCATGTGAACTATTGTCACGGAGTGAAAGTATCTTTTCTTCTTCTTGGTTGATTATATTATGTTATATCCCGGGTTTCTTACTTTGATTGTTTACATTTATGCTTCAGAGATATCTCTTTCAATTCAATGACCGGTAAACTTCCTCAAAGCTTTAAGTCACTGACTAGCATGAAGAAAATGTAAGACACGTATCTTCTGATTCATTATTGTACTGTATTGGTCGAGTGTCATTGAACTTTCTAGTATTGTTGTGTAGTGCAGCTACCTGCAGAACAACCAGTTCACTGGAAACATTGATGTCCTCGCCAATCTTCCTCTTGACGATCTGTGAGTTATGCAACATCTTTGTGTGTGTGAATGTTATCGATCTCTGAATGTTCCATTTCAAGACTTTCTCAGTTACTATATTGTTGATTCCTTTGTGCTTTAGGAATATTGAAAACAATCAGTTTACTGGTGGGATTCCAGAGAAGTTTAAAGGGATAATGCAGTGAGTGTTTCAGTCTGCAAATTATTTTTCATCCTTGAATATTTGTAGAAAATAACTTTCTAACTGAATTCACGATTTGTGATACAATTGTAGAAAATCTAACAGTAACACAGGGAGCTCAGGGCCTGCAGCCCCTGCGAACAAGTCCACTCAAAGTCACAAATCAGGAGGCAATGGCAATTCTTCCAGCGGTGGAGGTAAAAGTGATAGTGGTGAGAAATCTGGTATCGGTGGTGGAGGCGTTGCAGGCATAGTTATATCAGTTTCAGTGATTGGAGCAGTTGCAGTAATCTGTATTATCAAGAAAAGACGCAGGAAGTCGTCAACAGACATAGAAAAACTTGACGTTCAGCCATTTGTTCCTGAGTCACATGAAGTACAAGGTATATAATTTTATGACTTCTTTTTTGGGTCAGTTGAGAAACATAGCTGGTCAGTATACTTACCTCAATATCTCAACATATATACACTCTTATGGTGGATAGGATCCTTTCTTGACATGTTATCTAATTTGAATATAGAACTGAATTTATTGTGGTACACAAAGGGGAGCCCTGGTGTAACTAGTAAAGTTACTTCCATGTGACTAGGTTCGAACTGTGGAAACAACCTCTTGCAGAAATGCAGGGTAAGGTTGCATACAATAGACCATTGTGGTTCGACCCTTCCTCGTATCCCGCATATAGCAGGAGCTGCTCCAGGCTGTCCTATTGCAAATGTAGTTTTCTTTATGTAAACACACTTTTAACTCTGGATGCAACTTGCAAGTTCCAGCAAAGGTACTGAATCACCTCAGGTGAATTTTGGAAAATCAGTTTTCCTTTTGAACTCGTCGTCCATCATTTCATGTTCAAGTGAGCATGTCTATAATGATGAATTATCTTTCCCTTTTGCAGCTTCATTTGTAACCTGGTATTTCATACATCTTAAGCCTTACCTCAATTGTTTTAAGTTCTGTAGCGTCATTCCATCAGAGTTTCAGCTAACATTTTCCCTTTTCTGGTTACTAGTATTATTGATTTTTAATCATGTGTTAGTTTTGGAACCTATATTTTTCTATTCATCACCTATTTTGGCTGCAGATACGAAACTCAGTCAAGCTCCGTCCACAGTAAACATGGAAGCTACTGAAACTCCCACTGTAGTATCTCTAAGACCTCCACCTATTAATACTCATAAATCCATTGGTGGAGATGACATTTCAGAAAAACTCATTGTTCCACAGAAGAAAACTCGTACAGCTGAAAGAAAAGTTGTGCAATTTTCAATTGAAGACGTACAGATAGCTACTGACAGCTTCAGTGTAGAAAACCTTATTGGTGACGGATCTATTGGAAGTGTATATCGAGCTCACTTTGACAATGGCAAAGTAATATCTATTAGTTCTGTTTGTCGTTTTAACTGTTTCTTGTTGCAGCTAACATTACCTATGCTGATGTAAGTGATTCTGATCGTTTAAGTTCCTCTTCCATAGGTTCTTGCTGTCAAGAAACTTTATTCATCTGAGCTCCGGAATCCTGAAGATTTTCTAAAGATGATTCCTGAAATATCCCAGCTGGATCATCCAAATGTAACAGACCTAATTGGTTATTGCTCAGAAGATGAGCAGCACCTGCTAGTTTATGAATTCCACAGAAATGGTTCTCTGCATGATTTCCTGCATCGAATGGATGAAGACAACTCGCGACTAACATGGGATAGCAGAGTTCAGATTGCACTTGGCACAGCTAGAGCACTAGAGTGAGTTTCATAATTTACAAAATTACTTTCATTATGCATCACATAAGATTAATGCAGCTTGGAGAAAACTTTTGTGTGATTGGTTTCAGATATTTTTTTTATGGTAAACTACCTATCTATCATAAATCTTCTAGTTTTCCTTAAAACATTGATGACTAGTAAGCATGTAAAAGTGTTTCAATAATATTTCGAATAGAAAAGCAATTATATTATTTTAAGTTGTAGAACACCGCGTATGTTGGCTTTCCACTAGATCTAATTAGAATAAGTTTTCTATTTTGGTCGCGACCTCTCCAATTCTTATGAGTAGGTTTATTCTAGTAGCTAAATCTAAGCTTCCTTTAATTTGCAGGTATCTACATGAAGCTTGTTCTCCATCTTTAGTTCATAAAAATATAAAATCTGAAAATATTTTACTTGATGCTGATCTCAATCCTCATCTATCAGACAGTGGGTTAGCAAACCTTATGGCTGATGCAGATCAGGTAACACTTTATAAATTTGTTTTTTGCCTTACTCTAGATTATAATTTCCTAAAAATCCACATCAGTATCTTCTAAATGGCTTCAATTTGGACCCTTTATTTTTTCTCCCTACAGGGATTGAACCATAATATAGAGTCTGGATATGGCGCACCGGAGGCTGCTATATCTGGGAAGTGTACTACCAAGAGTGATGTATACAGCTTTGGAGTTGTTATGTTGGAACTTTGTACTGGTCGAAAACCTTTCGATAGGTAAATGTTAATTTAATGATTTTGAACCCAAAATCTGTTTAAGACAAATAATAATATACGTGTGTTCTTCTCTTCTCCACTCAGCTCAAGACCAAGATCTGAACAGTCTTTAGTTAGATGGGCAACCCCACAGCTTCATGATATTGATGCCCTGGAAAAGATGGTTGATCCAGCTCTATCTGGACTATATTCTGTCAAATCTCTTTCCCGGTTTGCTGATGTTATTGCTCTCTGTGTTCAGGTAATACTCTCGTTGTTTCAATTCATGTGACTAGACACAAAGTTTAAGATAGAAATGCAAACAATTGAAACGTGTGGTCTTAAACATGACATAACATCTCATTAGAGTAATCTGAGAAGTAGTTCATTCATTTTGTACTGAAAGCTGCTACATGCCATGTGTTCAATTTTCAATAGAGTACTACCCTAAATCGCGCTCACTATCACAATGGTTTGTGTATGTTGTGTCAGCTTGAGCCTGAGTTCCGGCCACCTATGTCTGAAGTGGTTCAAGCATTAGTCCGGCTAGTGCAACGAGCAAATATGAGCAAGAGGTTACACAATAATGATCAGGACTGAAGAGATATGATCAAGACCTGCTAATTGTTACACCAAATGGTTCTCTTAAACTGTTTGCTGCAATGGAGATCATCTGATGCTGATCAGCCCAGCAATATCTTCTTCGGTGCAGTTGTATTTGATTAAATACCATATGCTAAAATA

>M6-PGSC0003DMG402005482 M6 LRR-RLK

AAATGCCAAGTGAATAAATAGAGTACAAAGGATAAAAAGAAAAAAAGAAAGGATTTTTCACACCAAAAAGCATAGAATAAACCAGCAAGCTACCTCATTCTTGAATACATTGTCTGTAACAAGTATGGTTTTTTCTCTAGACTTCAAGTTTCCAATGCTATTATGTTCTTTTGTTTTCTATCTAATTACCCTCCTTTCATTCATGTTCAATATCTCACATGCCTTGAAGTTACAGAATGGTAATGAAACTGATATCCAAGCCTTACTTGCCATCAAAGAAAAGGTAACTCAAGATCCTCATGGTATTTTTACTTCATGGAATAATTCAGTGCATTTCTGCAGCTGGGAAGGTGTCACTTGTGGCCATCTTCATCAAAGAGTCACTAAACTGCACTTAACATCACTTGTCTTAGTAGGTACTTTGTCACCTTTCATAGGAAATCTTACTTTTCTTACTAGTCTCAAACTTGAGCTCAACAATTTCCATGGAAAAATCCCACCACAGGTTGGTGGTTTGTTAAGGCTTAAGCATCTTAGTTTGACTAATAACTCATTTTCTGGAGAGATACCGATAAATCTTTCTCGTTGTTTGAATCTTGTTATACTTGGTGTTGGTTGGAATCAGTTGTCTGGAAAGATTCCATTTGAACTTGGTTCATTGCAAAAGCTTGAGAGATTGCAAGTTCATAACAACAACCTCAATGGACCAATCCCGGAAACCTTAGGTAATCTTTCCGCAATCAAATCACTTTCCTTATCGTTTAACAATTTGGAAGGAACTATTCCAAGTTCTCTTAGTCAATTGAAGACATTGAATGTTCTTGGTTTGGGTATAAATAAGTTATCTGGTATTGTACCTGCTGAAATTTTTAATCTTTCATCTCTTGAGATATTCACAGTTTGTTATAATCAGTTGTATGGAACTTTGCCATCAGACTTTGGTTTAAGTCTTCTTAAGTTGAAAGTTCTTAAGATTGGTCATAACTGGTTTACTGGACCCCTTCCAAAGTCATTATCTAATGCTTCAAATCTAGTTGAACTTGATGCATATGGTAGCAATTTCACTGGCAAAGTGTCAATTGACTTTGGGGGGTTATCAGATCTTTGGTGGCTGATTCTTGCATCAAATTCCATTGGAACAGGAGAAGTTGATGACTTGAGTTTTTTCAATTCTTTGAGTAGATGCAGAAACCTGAAAGTTCTGGATTTGAGTGATTGCAAATTTGGAGGTGAGCTACCTGATTCTATTGCAAATTTATCGACAACTCTTCTGTCACTAAGATTGGGAGGTAATCAACTATTTGGTTGTATTCATTCAGGAATAGGGAATCTTGTTAACTTGACTGAATTGCAACTGCAAAAGAATGATTTTTCATGTAGCATTCCTGAAGTCGTTGGCAATCTTCGTAGGCTTCAGTTAGTGGATTTGTCCGAAAACAAGTTTTCAGGAAGTATTCCATCTTCTATGTCTAACATGACTCGATTATATTCACTTCATTTGGAAAAGAATGAGTTAACTGGTAATATTCCCCTTAGTTTTGGCGATTTCCGGTACTTGCAAGATTTAGACCTTTCTCAGAATCATCTTAGTGGTACCATACCAGATGGATTTATGAGTTTGTCTTCCTTAACAAATTCTCTTAATCTTGCTGACAATCAATTGTCCGGTCCGCTATCAGTTGAAATTGGAGCTCTCAATAATCTTGGAAGATTGGATATTTCTAATAACATGTTGTCTGGGAAAATTCCTAGTAGTATAGGGAGGTGTGTAGGTTTGGAAAGCCTTGTTTTAGCTGGCAACTTATTTGAAGGTATAATTCCTTCATCCATTATTTCTTTGAAAGGTCTTGAAGAATTGGATCTTTCGCGCAACAATTTATCTGGGCAGATTCCAACATCCCTTCAACTCATTTCCTTAAAAAAAGTGAACTTGTCTTTCAACCAGTTTGAAGGTCAGTTGCCAACTGAAGGTGTTTTCAGGAACGCAACTGCAATCTCCATATCAGGGAATAGGAAACTTTGTGGGGGTATACCGGAACTGGAGTTGCCAATATGTCCGAACGCTGACCCTGATGGTTGTGATAAGTCCAGAAGCATTAAGCTAATGATCCCTCTTCTTTCTGGACTTGTTGCACTGGTTTTCATCATGTCTTTAGTCATAATCATTCGGTTAAGAAAGGCAACGGGAGAGCCTTCTTTAACATCTTCGCCGGTTACTTATGAGAGCCTATATAGAGCGACAAATGGATTCTCTTCTGCCAATTTGATTGGAAATGGTAGCTTTAGTTCTGTTTACAAAGGGGTACTTGATCCAGGTCAATGTATGGTTGCGGTGAAGGTGATAAACATTGACCAACAAGGGGATACTTCTACGAGCTTCATGGCTGAGTGTGAGGCCTTGAGAAACATAAGGCATCAAAATCTTGTTAAGATCTAATGCCTGTTCAACTAGTGATTTTGAGGGTAATCCATTTATAGCTTTAGTTTACGAATACATGCCTAATGGGAGCTTAGAAAGTTGGTTGCATCCTATTCCGGGAGCTGATGCCTCAACTAATGAGGTGAGGATACTAGGTCTAGTTGAAAGACTAAGAATTTCTATTGTTGTGGCTTGTGCACTGGAATATCTTCATAACCATTGCCATAATCCAATTGTGTTGTGTGATCTGAAGCCGGACAACATTCTTCTCGACAATGACATGACTGCTCATGTAGCTGATTTTGGATTAACAATGTTTTTCTCAGAAACCATGAGCAAATATAGTTCAATTGGATATGCTGCACCAGGTAACGTAGTTGATAGTCCCTTGAATTTTAGACTCAAAGGCATCCTTGTTCAGAGCACTAACAGTCCTCCATGTTTACAGTATTGCTGCAGTTTTAGTCCGCCCTCAAACTTGTCTCTTTTTTTATTGGGATTGAGAGGTTCAGTGTGTTAGATTTCTCGTGCAACAGTACAAAATTTTCTTTCCTTGCAACGTTATTTGTCTACAATAAACTGATATAGTTCGAAAAACAAAGGTAGATTCTTGGTTTAGCTGCAATGATGCTTGGACTAGTTGTACAGAACGAATAGCACATATGCTTCGTCTCACCTTCTCTCTCATGGTGTTATTGTTTCTACTAAACATAAAGGATGATGACTGAACATATACACTAAATTATGGACAAAACCAATGTATTAAATAGACGGACATTTGAGGGAGGACCAAAAGTGCATGCACCAATATTGCAAACATAGGGACTACAAGTACTCCGTGTGAAAAAACAAGGATGACTTTTAAGTCTTAGTCCAAAGATAAGGGACTATTTTGGGTCTTTTCTCGTGTCTTCTTGAGAAAGTTTTCATATTGATTTTACATGTTTTAAGTTGATTACATGTTTTCTTTCCTCTGTGGGCAGAGTACAGCATAGGAGGTAAGGCTTCCGAATTTGGTGACGTTTACAGCTATGGGATCCTTCTACTAGAGATGTTTACTGGAAAGAGACCTACTGACAGCAGGTTCGAAAATGGACGGAGTCTTCACAGTTTTGCTAAGACGGCTTTGCTTGATGAAATAGTAGACCCCATGCTTCTACCTAGCAACAGCAGAGAAAGACAAGAGGCTGAGGAAGAAGGAGTACTTATCAATCAAGATGATACTAGTATAAAACAAGCGCAGGAATGCTTGATCTCGATAATACAGATTGGAGTTGCATGTTCAGCTGAATCCCCGAGAGAAAGGATGGATATCGGGGATGTTGTCAAGGAACTTCAACTAATTAGAGACATTCTCCTTGCTTCTCATGCAATTCACAGCTCAACCAGTGGGTAAATTCGCTCTACTAGCTTTGTTTTCAATCGTGATATTGAGTTCCTCTGCTAGATATTGAGAAACTTGTTTCTAAACACGTGTTGCACGTGTGTCCTGTGTAAATTCTTGTAGATCCATTAGAACGATGGAAGTTCTACGTAGACATGAATGTGTTTCCCATGTAAATTTTCTTCTTGTTGTTAATCGTTGTCTTTCTTTCCTTGGTTTCAGGAGCTTGAGGTTTGAAGGGTCTTCCAGTCGTTCTGTTACTAGTAATTGGCAAAATTTCACATCATTTCGTTTGCCTTGATGTTTTTTGCTCTTATATAACAAATTACTTTAGCAGATTGAAGATGTTAATCATCAATTCAATTATTGGTTCAGTTTGCCTAATGCTGGTTAGTTCAGTTGAGCTTAAGCAAATGGAATTGGAATTGTCTATGTATATGATGTTATGACTTGGCCAAACACATACACGACCCCTTTAGCTCGTCTCGAAATCTC

>M6-PGSC0003DMG402008032 M6 LRR-RLK

AAAAAAGTTCTCTTTCAATGCATTGCCTTCATTAAAATCCCATTACTCATTTCTGGGTAGTCTCATTTTTTTTTTTTCTCTGTATTTGTTTGCCATGAAAAATGCCGTCTTTATGGAAAGCATTTTCACTATCTTCAAGTTCTCAACACTCTTCTTCTTCTTCTTCTACTACTGCTAGTACTACTTCTTCTCCGGCCGATTCACCGAATACCCGGCGGATTTACGGCGGCGGAAGGAAGCTTACTAGGCAGAGGAAGTTGAGACATGTTAGTGATGAGGATCTTGGTTTGCGGAGACCTAATATTCAGGCACTGATTATTGATGAACGTTCAAAGTCTTTGCCGGGTTCTCCTGATTCTTACGCTGATTTCGGGTCGGGTTCTAGGTCGTCTCACCACCACCTCCGTCACTGTTCTAACTCCTCTGCTGTGCCACTGCCGTTGCCGCTCCCTGAGCTCAATTCTCTGCCTAAACAGAACTCGGTTGACTCTAATTTGCCGGGTAGAGTTGACAGGGAGCTTTTAAGTCCTCCTCTTGCCAGGTTAGTCTTGAAAATTGAACCCATTGTTCAGAATTTATACTTAATTTCAAATTTTAGTCTTACTTTTCGTTTTAGTCTGTTTCGAAATGAATGAGTCTTTCCATTTTTGGCAACCCTTTGATTGTAACTTTTCACCTTACATATTTAAGAGATTAAATGATATTTTGATATATTTTATGTATTTAGTTTAAGACCACGAGATTCAAAAGTGGTTTTTACTTTGTTAAACTTCTTGTCAAGTCATAACCAGTCAAACAAATTGACATGTTGGGTATGTTTATCAAACTTTTGAGTTTTTTCTTACATGTATGTTTTGTGCTTTAAGTAAAAAAAATGCTACCTTTCTGAGTGAAGTATTGTTTTGCCGATTCATACTGGTTTGGGATTGAGGGGTAGTTATTTTCGGTTTGATGATGAAAGCTTTTCACCATAGGTAACTCTGTATGTAAAGGGATTTAGTAGTAGTAGGTAGTTGGTTCTAATTTGAAGTAGTTTTTGGGGTGGTTTGATTGCAGTAACACTTTATAGGTCATTACCATAAAAGAACCATTTTTTCCTTTTTTCGGTTGTGGATTAACTGCAGTAACCTTTATAAGGTTCTTCACCATAAAGGTAGGTTTGGAATTGTGAAAGAGCATGTGGTTTGTAGTTGTTTCAAATGATGATTTGTAGTGTGCTACTTTGTTTTTGGGGGAAAATGTTTGATTAGCTATGATGATTTTGTAAGAACAAAAACCGAACGCGGAAATGTTAACCTGTTGCGAGTTGCAGGGAAACTTTTGGTCGAACTCCAGCTGAAGTGAAAAGTTCTCAGCATCCAAGATCATCTACTCCAACTTATCTTCGTAGGGGGTTCCCTCAGGATCTGAATGCTGAGGGCGTTGAATTCAGGTTGAATGTTCCTGTTAGAAGTGCCCCGAGTAGTGGTTTTACAAGCCCCGTCCTAAGCCCAAAAAGGTTTAGTACACAGGACCTTTTTCATCATCCCTTGCATCAGGCCTCATCATCACCTTCAGAGGCTTACTCTTTCCAACTTTCACCCACAAGGGTTATAAATAGCGCGGATCATTCTCCACTGAGTAGCCCAATTTTACCAAGTTCAGCTAACAGAATCAGGAACAGCAGAAGTGGTGCTGTGCATTCACATCATAAATCTCTTCCGGAAAGTTCTCTAGGTTGGAATGAAGCTAACAACAACAATGTGCATCCACTACCCCTTCCACCAGGAGTTCCACGACAACCAGAATCATGTATCATGCACAGCAATATGGATAAACCATGTGTATCACCAGCAAAGGGTCAATGGCTAAAGGGGATGCTTTTAGGTCGTGGTACATTTGGAAGTGTATATGAAGCTACTAACTGGTAAGGGATAGAGCTTGCTTAGTAAGTAATTCGGTAGGAATGTAACATTTGCTTCTTTTGTTTTTCTTCCTTTTACAGTGAAACCGGTGCTTTATGTGCAATGAAAGAAGTTGATCTTACTCCTGATGATCCTAAATCTGCTGAATGTATAAAGCAGTTAGAACAGGTGACGTCCTCATGTCTTCTTAGGCTACATAATCAAAATTTGTATATAAACTGCAATTGTATGAGATACATGGTGAGAGAAAACATATTTTAAAGGCCAAATCTTTGTTAAGTGAAGTGAACTTTAAAACAAGTACTTACTAAGCATGTTATATAATATGTCCTGGTTGTCTAACTTGCATTAGCAAATTAGTCAACTTAAGGTTATGTCCAACTTTGAGCTAACTATCATGTGCCTGAAGATGCAAAAATCAATATCGTATCCATGGAGCTAATTTTATTGATCATATTCTTTGACTTCATAAGCCTCCTTCATGTTTTGCATCATTGAGAAAATATTTTCGGAATGTTGGTAACTGACTTGGGAAAAAAGTTTGAGGGTTTCATAATGTGGAGTTTTTTATTCTACTTGCTTATACATAGTTTGTGATGGATTGCAAAGACTATGACATGTTTTTCACTTGTTTATGCTATTCTAAATCGATTCAGTTTCTTCCTTACTTCTGTTGTAGGAAATAAGAGTTCTCCAGCAGTTAAAGCACCAAAATATTGTTCAATATTATGGCAGTGAATTAGTAAGAACATTTATCTTTCTCTTGCCATGTTCAGGTATTTCTTTAGAATATTGAAAGATAAATGTTGTTGTTGTTGTTCAGATGGATGACCGCTTTTGCATATATTTGGAATATGTGCACCCCGGTTCAATTAATAAGTATGTTCGGGAACATTGCGGAGCTATGACAGAGTCAATAGTTCGAAACTTTACTAGGCATATTGTATCAGGGTTGGCTTACTTGCATAGCACTAAGACAATACACAGGTAGATACTGGGTTACATCGATGTGGTATATGTCACTTGATTTAAAGATTCATTAACGGAACACTAAAGCGTCTGTGTTGCAATTCCTTGTTATAAGCAAGGAAGAAACTATTTTCTTTTACCATTTTTTTTTAAAAAAAACTTTTCGTTGACAGTGATGCTCAGTCTAATGGTTGGTTTCAATGATTTTTTCTGAGTAGGGACATTAAGGGAGCAAATTTGCTTGTAGATGCATTTGGTGTTGTCAAGCTTGCAGATTTTGGGTTAGCAAAACATGTAAGCTTTCAGAACTCTAATGCCAAACTGAAAATGCTACTTTCTATGTTCGTTCTAATTTAGTATTTTTCTTGTAAACCCTGATCTGTGGTGAAAATGAATGAGCTTGATCTTTCATCGATTATTTTTCATGCATGTCTTACCACTGTAATTTATAACGTTTCTCTTTTTCCCAGCTTTCAAGCTGTGCAACTGATCTTTCTCTGAAAGGGAGCCCTCACTGGATGGCTCCAGAGGTGGGCATTGTTAAGCTATTTCGCGGCTACTGCCTCTGCCTGGCTTCAGTTTTCTTTCCCCTTTATTAATTGGCTTGTACTTATGTCTGGACTTAACTTTATATGCAGGTAATGCAGGCTGTGTTGCGCAAAGATGCCAATCCAGAGCTTGCCTTAGCTGTTGATATATGGAGCCTGGGTTGTACTGTTATTGAGATGTTTACTGGACAGCCCCCTTGGGGAGAACTTAGTTGGGTAAGTGTTTAAAGCATCTCATTTTGAGAACAATGTAATCGCATTACAAGCTTAAATATATTTGTGAGCTTTTAGGTTGTCAATATAGCATCATAGCTTGTTAATATATGAGCAAGGCTGACTGATTGGCTTAATCTCAATAAAGCCAGAGCTATGTTCTTACTTTTCAGGCTTTCGTGAATTGGTTTCTGTAAGTGTCTTAATGTTCATGCTTTGGCTTTATTAGGTGCAAGCAATGTTCGGTGTCTTGAACAAATCACCACCTATACCAGAAAAATTATCACCAGAAGGAAAAGATTTCCTTCAGTGCTGCTTTCGGAGGAAACCTGCAGATAGGCCATCAGCTGTGATGCTACTTGAACATGCTTTTTTACGTAGTACTAGTTTCCTTGAGCATAGTATCAATGTTGCTGGTTGCTCAGAGGATTCTCCGGGGAAGAAATTCCATGTGAGTGGTACATGCTGCTAGCCTGCTACTGATGTGACATTAAGGGTTAACGTCACTGTATTATTTGTGAGATTCTCTAAAAAAAAAAATCCAAATGATTTTAAACTCTGAATTCATTACATTTCAGGATACCCTGAGTCCTAAGAACCCAATCAACCATAAAAAGGAACAAAAACCACTGTTACCAGGAACATCTGGCAGGCATGCAAAATCACCATGTAGCAGGTAAATAACTGTCTTGTGTTTGTCATGTTATATACCATACACTTTATTCTGTGAGATTATTTTTTTCGCGTATTAGTATTTACAATTGCAGCTAGATACTATATCTTGTGTTGTGCCATTGACTGTCACGTCCATCTTATGCAAGTTAAACTGTGTCTCTTCCAGTGAAACTTGCCGGCAAACTCAACCTGAAACCTGTGAATATGGAGCAGCTTCTCACCATTCTCCACATTCTGCACTTGAAGTCTTTCCCTGTATTTCTTCAATGGAGCTGAACTCGAGTTCACGTGCAGCCAGCCAATCAAGCGTCCCTAGTATCTTTCGCCTGGGACCTGAAAACAGAAGTCCTTATAGAATCATAGGAAAGGAAATTTCAAACCTCTGTATAAGATCATAAATGAATGGAGAGGGGGAAAACTCATGTACAGATCAGGAGAATTAGATACAGAGAACTTCAGTTGGATGATTAGTCATAATGTTGCCATATGTGGGAGAGATCTCAATTCCCGTATGCATTCACATCTTCTGTGTAGTATTGATGTTTGTATTTGAAGGCAAACATATCTTTGTAACAGATTCATGAAGAACTGAAGATGATTCTGGAGTTAATAGTGTATTTTAGATTGAAAGCAAGATTTAGTGTTCACAATTAAATGATTTATGCTAAATTCAATATGAGTATCTTTTTCCCTCATTTTTACAACCAA

>M6-PGSC0003DMG402014186 M6 LRR-RLK

GAAACTTGTTAAGTGACTAAGAAGCTGTAATGAGCTGTATGCTGCAACTCTACTTTCTATTCACTACAATTTTTTTAATTTACTAGGAAGCAAGGGGGCTGGCAGAACACCATTGAATTGGGAAATCAGGTCCAGTATTGCCCTCGGAATTGCACGTGGCATCAAATACCTTCACTCTCAAGGTCCTGATGTCTCCCATGGGAATATCAAATCATCCAATGTTCTTCTCACCAAATCATACGAAGCACGTGTCTCAGATTTTGGCCTGGCTAACTTGGTTGGATCGCCATCCTCCCCTACTCGAGTTGTTGGATATCGTGCTCCAGAGGTAACTGATCCTCGCAAGGTTTCCCAAAAGGCTGATGTCTATAGCTTTGGAGTATTACTCTTGGAGCTTCTTACTGGAAAAGCACCTAGTCATGCCCTCTTAAATGAAGAAGGTGTTGATTTGCCAAGGTGGGTTCAATCTGTCGTTCGTGAAGAATGGCCTTCTGAAGTTTTTGATATTGAGCTCCTCAGGTACCAGACTGCTGAAGAAGAGATGGTCCAACTCTTGCAGCTAGCAATAAATTGTACAGTTCAATATCCTGATAAGCGACCTTCAATGGCTGAGATCTCTAAACAGATTGAAGAGCTTCGACACCCAACCTCACATGATAAACAAGAACTCTCTGATCTTGCCAATGCAACAGAGTGAAGTGGTTATTATTAGTCCCCCTCGAGATGTGCAGTAATTAAGCTTAAACTCAGATTCATCACCAATCTTTTTTTAAGCATATTAAACTTGTTGTTTGTTGGGGGAGGGGTGTCTCATCTTCTTTCTGTTTGAATTAACTGATCATGGCCCCTTTTTCTCTATGTTTCTGTAGTAGTCTAGTGGTTACCCCTTTCATTTTCCTTTAGATGAGGGGCCAAAGCCAATATCTGTACCATTTATTCTTGTTGGATGTATAATGCTTTCTGTAATTATTACTGCTTCTGCTTTTACTTCTGCTTT

>M6-PGSC0003DMG402015976 M6 LRR-RLK

ATTGCAACAACAGAAAGAATTTCATACTATGAACTACTCCAAGCAACTGATTCGCTTAGCGACGGTAATCTAATTGGTTCTGGAAGTTTTGGCTCTGTTTACAAAGGTGTTCTCAGGAGTGGGACTGCCATTGCAGTTAAAGTGTTCAATCTGCAACTGGAAGCGGCATTCAAGAGTTTTGATATAGAATGTGAAGTTTTGCGTAGCCTTCGCCATAGGAATCTTGTGAAAGTCATCACTAGTTGTTCCAACCTTGATTTTAAGGCTTTAGTGCTCGAGTATATGCCTAATGGAAGTCTTGACAAGTATTTGTATTCACACAACTACTTCCTAGACATTAGGCAGAGACTAAGCATTATGATAGATGTAGCATGCGCATTGGAATATCTTCATCATGGATGCTCCTCTCCTGTGATTCATTGTGACCTGAAGCCTAGTAACATCTTGCTGGATGAGGATATGGTTGCCCACCTAAGTGACTTTGGTATTTCAAAACTGCTTGGTGAAGATGAGAGTGATTTATACACAAAAACCTTAGCAACATTTGGTTATATTGCACCAGGTACGTCTCTTTGTTTTTGCTAATTTAAACATTGATTTTTTGTCTTTTTTATTTCCAATCAATAAAGTATATTGCATCTTTAAATCAATTGTTTGACTGTAGAGTATGGACTGGATGGACTGGTGTCAATAAAATGTGATGTCTATAGTTACGGGATTATGTTGCTGGAAACATTTACTAGGAGAAAGCCTAGTGAGTTTAAGGGAGATCTTAGCTTGAAGCAATGGGTGAGTTATTCACTCCCAGAGGCAGTAATGGACGTTATGGATGTCAACTTGGTTACACCAATGGATAATCGCTTACAGAAGGAGTTAGATATCGTGGCATCAATCATGAAAGTAGCATTAGATTGTTGTGCTGAATCTCCGACAAGAAGGACAAACATGAAAGATGTTGTAGGGATGCTACAGAAGATCAAGATTCAACTTCTTGCATGCTGAGCATGTCCTCTGTATCTCTTGTTCTGAATCGA

>M6-PGSC0003DMG402020263 M6 LRR-RLK

CTTTAGTGGCGTAGGGGTTAAGACGACCGGTGCCACAGTGCTTTTCTGAGACTCTGTGAAAAGAAAACTGTCACCATTAAAGCCAGAAACTTGAATCGGTCCCCTGTTTTTTCTCCTTGACTTGCTCTGCATGCAACACTAAATTCTCTCTCTTTTGTCAACTACCCTCAATTTTGCTCTCTGTTTTTTTATTTTCAATATTGTTCTTCTCCCAAGAAAAAAAGAACATTCCTTTAATTTGCAATTACAGGTCAATTTATTCGCATTTTTAATGGGGTTTTATCAAGATTGTAGCAGCATTAACTGTTTGGTAGCTCAATCTTGACTTGTCTGCACATTTAGACTTCAATGCTTTTCATTTTCACTGAACTAGAAAGGGTCCACCCATGTAATTTTCTTATTCATGATTAGCCAAGAACGTAAAATGGGTTGCTTCAAGATTGTTGGAAACTTGGCTCCATTTTGAAGTCACTGTTAAATGGGTTTCTTCAAGATTTTTGTGATAGTGTTGGTTATTTGCGCTTTCTTTATAGCAAATACTTATCAGTTACAGTCCTATGAAATTCAAGCTCTTCTTCAGTTGAGGAAGCACCTGGAATATCCAGTTCAGTTGGATGTTTGGGAGAATTACCATGGAGATTTTTGCAGCTTGACTTCAACACTGCATATGAGCATTACCTGTCAAGATAACTCTGTTACTGAACTCAAAATCAAAGGAGATAAGTTAGTTAAGTTGAATGAATTTCATGGATTTGCAATTCCAAACCAAACATTATCTGAGGGTTTCTCCATTGATTCTTTTGTTACCACATTGACAAGGCTAAGTAGCTTAAAGGTTCTTACTTTAGTTTCTCTTGGTATTTGGGGACCCCTTCCTGATAAAATTCATAGGTTAGCTTCACTTGAACTTTTGGATATGAGTTCAAATTTTCTATTTGGTTCTGTTCCTTTTCAGATGTCAAGATTGATTAAGCTTCATACTTTGACATTAGATGCCAATTTTTTCAATGAAACTTTCCCTGATTGGTTTGATTCCTTACACAATCTTACTATATTAAGCTTCAAGAACAACAGGTTGAAGGGTCAGTTTCCTTCTTCAATGTCTAAAATCACAACCCTCACTGACATTGTTCTGTCACACAATGTGCTTTCTGGTAAATTGCCCGATTTAACGGCCTTGTCTAAGTTGCTTTTGTTGGATTTAAGGGAAAATCATTTGGATTCTGAATTGCCACCCTTGCCAAAAGGACTTACCACAGTTCTTCTTAGCAGTAACGTTTTTACGGGTGAAATCCCAGTAGAATTTGGCACACTAAATCAACTTCAACACCTTGATCTGTCGAACAATTCTCTCAGTGGAATGCTGCCTGCTGGCTTATTCTCTTTGTCTAGCATTAGTTACTTGAATTTAGCGTCTAATGTTCTTAGTGGTTCACTTCCTAGCCATCTCAATTGTGGGGGTGAACTTGGTTTTGTTGATATTTCTAATAATAGATTAGTTGGTGGGCTTCCTTCTTGCTTGAACACTATTTCAGATAAGCGAATTGTTAAGGTTGGTGGAAATTGCTTGTCTCTTGACACTCAATATCAGCATTCAGAGGGCTATTGTAAACAAGCTAATTCGGATAAGAAACAGATCACAGGAAAACAAATAGCATTGTTGGCAGGTGTTATTGGGGGAATTGTAATTGTTGTGGTGTTTCTGTTAGTTGTCCTTTTCATCTTCCGTAGAAAACGACATGCGCGTAACATGGTGGATAAGCACATTCCTCCGAAAGTTGTGCAAAATACACAGCCCGGGGTTCCCTCTGAACTCCTAGCAAACGCCAGTGAGTTGATGTTTTTCTGATTTCATTTTTAATCTGTTTAATTTATATGAAGGAAATGTTCTCCAGAAAAGCTTGATCTCAATCTTCGAATTCTCAGGAGTCATTTCTCAAGTGGCAAACACAGGATCACAAGGTGCCCCATCATATCGAGTGTTCTCCATGGAAGAATTGCTAGAAGCAACAGAAAATTTTGATCAGTCAGCATTACTTGGTGAAGGGTCAGTTGGAAAAGTATGGCTTCAAAACAATACTTATGGTTCTCATTGATAATATATATCTTGATGTTTTCCATTCAATGGTCACGCTGAATTTTGGCTCAAACTGATATACTTGCTGTAAAACTTGAATAAATGAAGCAGGACGTAGATTTTGTTGCGTAGACCATGTCCAACAATGATAATGCAATGAATGCACTAGTTCACCAAAGTACAAAATGGATTTGGGTGTTGTTTTTTTTTAAAAAAAAGTGCTTGCCTATCATATTCTCATTCTACACGTGTATCTGTGACTTCTTTTGATGATAAAAACTGGTGTCACCAATAATGCTTTTGTGCATTTGATTTCATATCTAATGAACACATGACAGTTCAACTAAATATATCGAATGCCCTTAAAGGCAAAATTAGATAGTCTCACACGTTTGAGTGCCTATGCAGTGAGTTGATATCTCTGCTATTTTGATTGACATAAACAGGTGTTAGCCTTATTATGTCTTGCTTGAATAAGACAGTTACTTTAGTTATCAGCAATTCCCTCAATATATATCTCCATTTCAGATTTACAAGGGAAGATTAGAGAATGGAGCTTACATAGCTGTAAGGTCACTAAATGTGTACA

>M6-PGSC0003DMG402026747 M6 LRR-RLK

GACTTATCATCAAACAACACGGTTGGTTCTTTACCTCCGGAAATTGGAAATCTAAAAGCTGTGATACAGATGGATTTGTCGATGAATCAATTCTCAAATGGAATTCCAAGAGAAATTGGAGGATTGCAAAATCTGGTGCATCTTTCTTTGAGACAAAACAAGTTGCAAGGATCTATACCTGACTCAATGAGCAACATGGTAGGTTTGGAATTCCTAGACCTTTCTCATACTAATATATCATGAATCATTCCTATGTCTTTGGAGAAACGTCAAAACTTGAAGTATTTCAATGTTTCTGATAACAAATTGCATGGCGAAATACCCTCGGGAGGTCCTTTCAAGAACCTCTCGAGTCTGTTTTTCATCAACAATGAAGCATTGTGTGGTTCTTCAAGATTTAGCGTCCCTCCATGCCCCACTTCATCAAACCATAGGTCAAATAGGAAAAACAATCTTGTTCTATCTCTTATGCTGGGAATTGCACTTGTATTTGTTCCTATTACCCTTGTGTTTGTATGGATAAGGTATAGAAGAGGTAAAAGATCTCCTCAGCAAACTGATTCATTGTCTATGGCAACAACGGAAAGAATTTCATACTATGAACTGCTCCAAGCAACTGATTCGCTTAGCGAGAGTAATCTGATTGGTTCTGGAAGTTTTGGCTCTGTTTACGAAGGCGTTCTCAGGAGTGGGACTGCCATTGCAGTTAAAGTGTTCAATCTGCAACTGGAAGCAGCATTCAAGAGTTTTGATACAGAATGTGAAGTTTTACTCAGCCTTCGCCACAGGAATCTCGTGAAAGTCATCACTAGTTGCTCCAACCTTGATTTTAAGGCTTTAGTGCTCAAGTATATGCCTAATGGAAGTCTTGACAAGTATTTGTATTCGCACAACTACTTCCTAGACATCAGGCAGAGACTAAGCATTATGTCTAGATGTGGCATGTGCGTTGGAATATTTCCATCATGGATGCTCCTCTCCTGTGATTCATTGTGATATAAAGCCTAGTAACGTCTTGCTAGATGAGGATATGGTTGCCCACTTAAGCGACTTTGGTATTTCAAAACTGCTTGGTGAAGATGAGAGTGATATATACAGTAAAACCTTAGCAACATTTGGTTATATTGCAC

>M6-PGSC0003DMG402029358 M6 LRR-RLK

TACAAGAAAGTTGAGTTCCTTTGCCTTACTAGTTTGACTGACCCCATTGTTGCTAAAATAAATTCAAGATTTTGAAAGAAGTCTGTCAAGTATCAAAAAATGGAGAAAAAAAATTTCAAAACATTTCTTGGTTTTGTATTGATATTACTAGTTAGCAGGATTTGTATCTGTTGTGCTTCAGAGAGTGATGTTTATTGCTTGAAATCAATAAAAGAATCATTACATGACCCTTTCAATTCTTTAAACTCTTGGAATTTCAACCAAAACTCCATCTGTCTTTTTGAGGGAATTGACTGCTGGCAATCTGATGAGAATAAGGTAATGAGCATCACACTTTCAAACTTTGGATTAATAGGTGAGTTTCCTCGTGGCATTCGAAATTGTACAAGCTTGACAGGTCTAAATCTTTCTGGTAATCATTTGTATGGAACCATCCCTTCTGATATAGCAGTGATTCTTCCATTTGTTACAATACTTGATCTCTCAAATAACACCTTTTCGGGCAATATAACACCTGGTATAGGTAATTGTCAATTCATAAATGTTCTAAAGTTGGACTATAATGATTTAGAAGGTGAAATTCCAAGTAGAATAGGCTATTTGCCTCGCCTTAAGACGTTCAGCATAGCTAACAATTACTTGACTGGGCCAGTGCCATTGTTTTTTAGCAAGAATATCACGGCTGAGAGTTTTGCAAACAATCCAGAGCTTTGTGGGAAGCCCTTGAAAGGATGTGAGGATTCTTGGATATGGAAACATGTAGATCGTGCTTCGTTCATCAAAGCGTTTGTGATTGGTTGGGTACTTTTGTTTACAATGGTTTTAGTCCTTTGCTTATTTAAATTTCCTACCAAGGCTATCAACAAGATAGTCTCATTAAACATATGGAAGCTAAGGAAAAAGAAACATTTAACTTCAGGAAGTGAAGAGGAATTAAGCAGCCAACATAAGGTAAAGATCTTGCTAAATTTGCTGCACTGCTTATAGTCTGCAAAGTTACCTTATCTTCTGTAGTTCCAATTTTGTGATTAGTCCTTGCTAGTCCATTATGTTATAAATTCTTTATTCGATCGATGATCAAACATGATAGCAATAAGGTTCTCAAGTTTCTTTCCTGTTTTGACTTCTGTGTATATTCAAATGATTTAACTTGCTGTTTTATTTACAGATATTAAAGCTGGAGAAGTTTGTTATGAGAATGAGCTTCACGGAGCTTGAAAACGCGACTTCTGGTTTTAGTGAAGACTATCTAGTAGGAAATGGAATGTTGGGCAAAGTGTACAAGGCAATTCTTCCAAATGGCTGGATACTTGCCATCAAGAAGCTCAATGACTGGGAGAATTTGGAGGACGAGTTTGTCTCGGAGATTACAACTCTTGGTGGTCTGAGGCATAGGAATCTATTGCCTCTAATAGGTTTCTGCGCCGAAAAGCAAGAAAGGCTTCTTGTTTACAAATACATGACTAATGGAAGTCTTGATGAATGGCTGCACTCGAATGAAGTTAAGGCCAATATTTTGGATTTCCCTCGGAGAGCTAAAATTGCACTTGGGATAGCGAAAGGCCTGGCTTGGCTTCATCATGGTTACGAATTGCACGTTACACATGGAAGCATAAGCACGCGATGTATCTTGCTAGATCAAAATCTTGAACCAAAAATATCCAACTTTTGGGAAGCAAAATTTTGGAGTAAGAATGATTCTGCATTAAGTTGGAGTCTTTTCCCAG

>M6-PGSC0003DMG402030114 M6 LRR-RLK

TTTCCCCATGTGTTTAATACCTAAAACTATGTCGTTTTGACCCCTATTTTAAGCCCTATATAGGTATTTTAAGTCCTAAAACTACCCGTTCTAAACCGTTCTCCTAAAATCACAAATTCCCTCCAAGTTCTTTCCCCAAAGTTCTCTCTAAAGCTAAAGAAGAAGAAGCTAGGGTTTTACTTCAAGATCTCCAATTCCCCATAGCTTTTGGGCCTTGATCACTTAGGTATGTTAGTATTCACCAATGGAGTCCTTTTCTCCATTGGGTGCCCAAAGATCCTAATTTTTACAAGTTCAATTCTCAATTCAAAGTTAGAGTTTTATTCAATTTTCATGGTATCTTTTCAATTATGATTCAATTAGCTATTTTCTATCTTCTATTGCATGACTTTAAGTAATTACATGTTTTTGAGTTGAATTCATATGAACCATGCATGATCCATGATTTTTTACTATGAACACAAGATGATGTTTATGAACTATGAATTATGAATTTATGAAGATATGCATACTTTTATGAAATTGATGTAAATGACTTGAGGATGCTTTAAGATGAAAGTATCAATGAGATTGCGAAAGGTTTTCTTACATACTACTATGATCATGATTGTGAAAGGTTTTTTCACATAAGGATTCATAAGGTTGAAAGGTTTTCTCAACTAAGTCGAATAAAGATTTAAGGAGCTATTCTAAATGACTTTTAGCTTGGATTGGTAGCTTAATTGTGCCTTTATATGGATAATGATATGAATATGATTAATGACTATTTCGTGGGATTTTACTTTGCACCAAGAGGATATTGAGATGGAGGCTCTTTTATTAGTAGAGGTCAGGTCTCTAATAGTAATCTCCTTGTCCCTAACTACGTGCCCCCGTGGGATGTCTTAGCTAGTGGATCCACGTAAGCTATAAGTTTATGGTCCTTACCTTGGCAAGTAGGACATCTCTTTTTGGTATGGGATACACATCGGATTCCATATTTTATCTTACATGGTCTATGTCGGTCAATGGTAAACATCCCACAATAATGAACTAGGTTACTTTCAAAGTATCTCACAAAGTGTTTTAACATGTTAAGTTTGCATTGACCATAATTATGACTTGTGTTTTCTAACCTCTTATCTTATGTTTTAGTTTGGTCATTGCATGTCACGTAAAGTCCTTTTAGCATGATTTCATATCGTATATGCATTGCTATCATACTTAGTACATTACATGTACTAATGCATATTTGTGCCTACATTGTTTCACTAATGTAGGGTCCGATGCTCCTACCTCGCATACTTGTGGCTAGTTGATCTCTCCTTGTGGATTTGAATATTGGTGAGTCCTCATGTTCCGAGGACCAAGACACTTTATAGCTTTTCATTATTTTAGTTTTCAAATATTGTATGTGGTGTATGTGTTACGTCCTAACCAATTCTTTGATGTATTAGATGGCTTGTTGAGACTATGTTAGACTTCCGCTTTGTCAATCTTCTTTATGATATGAGAATATTTCTTTTGAAATATTTATTTCTATTATCTTGTATGATGTATAAGTGTCTTGTGTAGGGCCTTCCGGGGTTCGATACGCCATGTTACGCCTAAGGTATACTTTGGATCGTGACAAACTTGGTATTCAGAGCACAAGATTTAGAAAGGTCCTATGAAGTCTCACAAGCCACGTCAAGTAGAGTCTTGTTCAAGAGTGTGAAGTGCACCACATTTATGAATGAGAGGCTACAAGATGTTTAGGAAACTCCACTTCTTTCATTACTCTAAAGTCGTGCGCTAGAGTTTAACTCTATAAGGTCTCTCTCCTAAACCTTATTCGATGCTTTACAGGATATGGCTACTCGAAGAGCTTATGCTAGAAGGAATGTGAGGGAGAATG

>M6-PGSC0003DMG400000386 M6 LRR-RLP

ATGGTGTTTGAGAATGATAGAATCAAGAATGCTTATATTGCTTTACAAGCATGGAAACAAGCTATCATTTCAGATCCCTTTAATATCACTCTCAATTGGGTTGGTCCTAATGTATGCAGCTACACCGGAATCTTCTGTGCTCCGGCGTTGGATAACCCCAAAATCCGTACAGTTGCCGGCATTGACCTTAACCATGGTGACATTGCAGGGTACCTCCCGGAAGAGCTTGGGCTTCTTACAGATCTTGGGATTTTCCATATCAATTCAAACCGTTTCTGTGGTACAATACCAAGAAAATTCAGCAAGTTGAAGATACTGTTCGAGCTGGATCTGAGCAATAACAGGTTTGCTGGGAAGTTTCCTTATGTTGTTTTGAGTTTACCCAAGTTGATATTTTTGGATATTAGGTTCAATGAATTTGAAGGTAATGTACCTTCACAGCTTTTTGATAAGCCATTAGACGCCATTTTTATTAATCATAATCGGTTTGCGTTTGAGTTGCCGGATAATTTTGGGAATTCGCCGGTTTCTGTGATAGTTCTGGCGAGTAACAGTTTCCATGGGTGTCTTCCGGCGAGTATTGGAAATATGAGTAATCTTAATGAGGCGATTTTAATGAACAATGGGTTGCGTTCTTGTTTGCCGGCGGAGATTGGGTTGTTGAAAAATCTGACTGTATTTGATGTGAGCTTTAATCAGTTGATGGGTCCGTTGCCGGAGAATTTTGGTGATTTGGTGAATTTGGAGCAATTGAATGTGGCACATAATATGCTTTCTGGAACAATTCCGAAAAGAATTTGTCAGCTTCCTAAGCTTGAGAATTTTACGTATTCTTATAATTTCTTCACTGGTGAACCGCCGGTGTGTTTGGGGTTGCCGGAGTTTCATGATGAACGGAATTGTTTGCCCAACAGACCGGTGCAACGCTCTCAGGGACAGTGTAAGGCCTTTTTGTCTAAAAAAATTCATTGTAGTGCTTTCAGGTGTCATAAATTTGTTCCTGTTTTGCCACCTCCTCCGCCACTTTCACCCCCCTTGCCTGCCCCTCCGCCACCTATTTATTCGCCTCCGCCACCAGTTTATACTCCTCCACCTCCAGTTTATTCACCACCGCCACCACCACCCTCTCCACCTCCAGTTTATTCACCACCGCCACCACCACCCTCTCCACCTCCTCCATCACCTCCTCCGCCTCCACCACCAGTCTATTCACCTCCACCCCCACCTCCATCTCCTCCTCCACCATCACCTCCACCTCCACCCCCACCATCTCCACCTCCACCATCACCCCCACCACCAGTTTATTCACCACCACCACCGCCACCATCTCCCCCTCCACCATCACCTCCACCTCCTCCCCCACCCATTTATTCGCCACCACCATCTCCACCTCCTCCATCACCATTGCCTTATTGTGTACGCTCACCTCCACCTCCACCACCAAATTCCCCACCTCCTAATTCTCCACCTCCACCGCTTGCTCACTCGCCTCCACCCCCTTCACCTTACTATTACAATTCACCACCACCTCCGCCCCCAAATTCGCCGCCTCCAGCACCAAATTCGCCTCCTCCTCCACCAGTATACATATACTCATCACCTCCGCCACCAAATTCACCTCCTCCTCCACCACCAACATACGTATATTCTTCACCTCCTCCACCTCCTTCCCCATTGCCTTGTATAGAACCCCCTCCACCACCTCCTCCTTGCATTGAACCACCCCCACCACCCTCCCCTTCACCATCACCTCCTCCACCACCAGTTTATCATTACAATTCTCCGCCCCCACCATCTCTAGCACCTCCACCACCAGTTTATTTTTACAATTCTCCACCTCCACCATCACCACCACCTACTCCAGTATATGAAGGGCCATTGCCACCAATAATAGGAGTTCAATATGCTTCACCACCTCCACCACCCTTTTATTAA

>M6-PGSC0003DMG400000819 M6 LRR-RLP

CTCTTTTTCTACCTTAGCTTTCTATAAACATTTTTAACAAGCACCTTCCTTCTTTACTTCTTTGTTCCCTCCATTTCCCTGTTTGATTGGATACTTTTACCATTTTCCACCACTGCCCCAAAAAAACAAAAAAAGTACACAATTTTTTTTTTTATATTTATGAAATGAAAGCATTTTAAGGTACCCCTCTTGGAGTTTCTTCAATGTTATAAGTTCTATTTGGCTTTAATTGACAAAGTTTATTACTTTCAAGATTTTGTCAACCTAACTCATTTTCAAGCTTTTTTTGGCTTCAAAGAGTGATGGGGGTCTTCAAGAACATGTTTGTATGAAAAAGTTGTATTCTTTGAACACTTTGTGAACTCATTGTGTGATAAAGTGTTATCTTTGAACACTTTGTGTACTCATTTTCAAGAAAATTTAGCTTCAAAAGAATGATGGGGTTCTTCAAGAATTCTAACTAGTTTGTTTATGTGACAGAGTTTTAAACTTTGAAGATTTTGTTATCTCTTTTTCAAGAAAATTTAGCTTCAAAAGAATGATGGGGTTCTTCAAGAATTCTAACTAGTTTGTTTATGTGACAGAGTTTTAAACTTTGAAGATTTTGTCATCTCTTTTTCAAGAAACTTTTTAAGCTACAAAAAGTGATGAAGGGTTTCAAGAATTCAAGATTGTGTGTGTTAGTTATCTTTTATCTCCTCCATTTTGGTGTCTGGTGCAATGGTGATACTGAAAACATGGGAGCTCAAATGGAGAAAAATGAGATAAATTCACTTTACTCAACCATTCAAGGCTTTGTTGGTAAATGGTGGAATGGTTCAGATCTTTATCCAGATCCTTGTGGTTGGACTCCTATACAGGTAACACAAATTTTTATATAAATTTTTCACTTTTTTAGTTACTTTTCTGCTTTATGGATTCTGTTTAGTAAAAGGTAAAAAAAAAAAAAAAACTAACTTTTTTGTGCATATGGTTATAATTGTCCTTTTTAATATTTTGTGGTATACATGACAAGATATGACTTCTCATTGTATATAGATTGTTTGAAAGCTTTAAGTGGAAATTTTATACTATGACAGATAATCAAAGAAATTGATAAGTTCTTTTTTCAGATTTAGTATATTTGGAATGATAGCTAATTAGCCATAGAAGTTTGATGAACCAAACTTCAATAATGAATATCAGTACATTCTTGGCTTGTTTTGTTTAGCAGTTTTGACAGACCAACAGAGACAGAATATGTTTTAAAACACAATGATAATACAGGACAGGACAGAGCTTATACGAAATATAATCAGTTCGTTTCAATTTGTTTGTCTGATTTTAACTTGACACGTAGTTAAAGAATGCACCATAGTTTTCTTTAATCCTGTGGCTTTAAACATATCATGTAGAAAGTAAGACGAACAAATTGAAACGGAAAGTAGTATAGTTGATGCATTATACTCATATTTCTGATTTGTTTTGTTTCTATTTCCTCTGTCCAGGGTGTATCTTGTGACCTGTTTGATGGTTTTTGGTATGTTACTGACTTAAACATTGGACCTATACATGATAATTCTCTAAGTTGTGCATCAAATGTGGAATTCAGCTCCAATTTATTCACATTAAAGCACCTAAAATCTCTCACATTTTTCAATTGTTTCGTGTCGCCTCGCCATCATCCAACTTCAATCCCAACAGAGAGGTGGGAGTTTCTTGCTAGTAGTTTGGAGTCACTCGAGTTTCGATCAAATCCCGGTCTCATTGGCCAAATCCCAAGTACATTTGGTAGACTCATAAAGCTCCAATCTTTAGTACTAGTAGAAAATGGACTCTCTGGTGAAATGCCACAAAGTCTTGGGAATTTAGTGAACTTGAAAAGACTAGTTCTTGCTGGAAACAAACTCAACGGCGAAATCCCGGATACTTTTGGAAGGTTCACTCAACTTTTGATATGTGATTTAAGCAGAAACAAACTATCTGGTGTCTTGCCTAAATCATTATTTGGAGGGTTGATCTCACTTTTGAAACTTGATTTGAGTTACAACAACTTACAAGGCAAAATCCCACAAGAGATCTCACAAATGAAGAATTTAACACTTTTAGACCTTAGTAACAACAAGTTATCAAGTGGATTACCTAAGTCAATTCAAGAAATGACATGTTTGGAGGAGTTAGTCTTGTCAAACAACCCAATGGGTGGATTCTTGGATGTTCTTGATTGGTATAACATGATAAAGTTAACAACTTTGGATTTATCAAACATGAATTTGACAGGTGAGATACCTAAGTCTATAGTAGAGTTAAAAAAGTTAAGATTTTTGGGACTTAATGACAATAAACTCATTGGTAAAATCCCAAAAAATCTTGAAAATTTGCCTAATGTAAGTGCTATTTACCTATATGGTAACAATCTGACAGGTGAACTTCAATTCTCTCAATGGTTTTATAGGAAAATGGGAAGGAGATTTGGGGCATGGGGGAATCAAAATTTATGTTACACTTTTGGTTTGGTGTCAAGTGATGTTCCATTTGGGGTAAAACAATGTCAACAAGAACTCAAGTTTGTTATTGATGTTGAGATGAACTCAAGTTTTGTGGATTTTCAATCTTCAATGGGATATTTGAATGATTGTGGGATTTGGAGATGTTTTGTAGTTGAACTATTTATGGTTATTCTACTAATGTTGAAATGAAAAGAAGGAATTTGACAATAGTGATTTACCTTTTTTTTTTGATATTTTGTTGTTGTTTTACTTGAAATGTACCACACATTTTGGTTTGTCTTTTAACATATGAATGAATGAATGTACATATTTTACC

>M6-PGSC0003DMG400000888 M6 LRR-RLP

TTATTGATTCAGTCTGTTGAAGACTGAGCAATATTTGTCTTGATTAATTCCATTGTGTACATATTCTGATGAAGTTTTCGTATACGGAGGGTGGAGGGGATTGATCTCTGAAGGAATTTCCTAATACAGATGGTTCAAAGTAAACTCAATTACAATAATTCTAAACAATTGGAGTAGACATTAAGAAGCTTCACAGAATTTTCCTACCTATTACTTCTATCAATATATTCAATGAAAATATAAGTTGACTAGATCCAACAACAATTATTTATTCAACTGGAGTAAAGCGAATGTTCATAGAAGCTTTCATGTTTCAAGTTCTTCTTTCCTCTATATAACTAATTTGTTCATCAATTGATCTCATCCATACACTTAAGTTTTTCTTTTCAGACAATTAATTGAATCTATGGCATCATCTTCTTCTTATGCGAGTAATTCACAGTACTGTCCTCGATGGAAGTACGATGTCTTTCTAAGTTTTAGAGGTGAAGATACCCGGAGAAATTTTACGAGTCACTTGTACCAAGGTTTGGACAATAGAGGAATATTCACCTTTCTAGATGATAAAAGGCTAGAGGAGGGCGATTCCATCTCAGAAGAACTCGTGAAAGCAATAGAAGAGTCTCAAGTTGCCATAATCGTTTTCTCAAAGAATTATGCTACGTCAGGGTGGTGCTTGAATGAACTGGTGAAGATTATGGAATGCAAGGAGAAAGAAAATGGACAAACAGTCATACCGGTCTTTTGTTATGTGGATCCATCACATGTTCGACACCAAAGTGAGAGCTTTGCAGAAGCATTTGTCAAACACGAATCGAGGTATAAGGATGATGTTGAGGGGATGCAGAAGGTGCAAGGATGGAGGAATGCCCTAACAGCTGCCGCAGATCTTAAAGGATATGATATCCGTGACGGGTGAGTTAAAACACATATTTGCTTTTTATGAAAAAGGAAATGGGATAATTTCCAGTCAAATTAGGATGCTTTTGTGAAACCTAGACATATTCTACTATAGAACAAGAATCTATACCTCCAATATGAGTTAATTTAGTTGGAGAGAATACATAATAATCCTTCTAAACTATTCAAGTTTTGCAACTTTCATAATTCAACTATAGAGTGTTCCTATTACCCTGAACTACAATGTTTCCTATTTGATTAATGGTATGAGATACTCACAAATACTTCGACTTTCTCACATTTTCCGCTATCAATCTTCTTATTTATATAGGATCGAAGCAGAGTATATTCAGCAGATCGTCGATCACATCTCTTCCAAATTATGTAAGAGGGCTTATTCTTTATCTTCGTTGCAAGGTGTTGTGGGAATAAATGCTCATTTTGGGAAACTAGAATCCCTACTTCAGATAGAAATCAATGATGTTCGGATTGTAGGGATCTGGGGAATAGGCGGAGTTGGTAAAACGACTATAGCAAAAGCCATCTTTGATACTCTATCTTATCAATTTAAAGCTTCTTGTTTTCTTGCGAATGTCAAAGAAAATGCAAATTATAATCAACTGCATTCTTTACAAAATACCCTTCTCTCTGAACTGTTAAGAAAAAAAGATGATTACGTCAATAATAAGTATGATGGAAAGTACATGATTCAGAGTAGACTTTGTTCTAAGAAGGTGCTAATTGTTCTTGATGACATAGATCATGGTGATCATTTGGAGTACTTAGCAGGTGATGTTGGTTGGTTTGGTAATGGCAGCAGAATTGTTGTAACAACTAGAAATAGACATTTGATAGAGAAGGATAATCCAATATATGAAGTGTCTACACTGCCTAATGAAGAAGCTGTTCAATTATTCAATCAACATGCTTTCAGAAAAGAAATTCCAGATGAACGTTTTATGAAGTTCTCATTGGAGGTAGTAAATCATGCTAAAGGCCTTCCTTTAGCCCTCAAGGTGTGGGGTTCTTTGTTGCATAAGAAAGACCTAACTCAGTGGAGAAGAACTATAGACCAAATAAAGAAAAACCCTAATTCAGAAATTGTTGAAAAACTCAAAATAAGTTATGATGGGTTGGAACGTGAAGAAAAAGAGATATTTCTAGATATCGCATGTTTCTTCCGTGGATTTGAAAGAAAAGAAGTCATGCAGATCCTTGAGAGCTGTGATTTTGGAGCTGAATGCGGATTGGATGTCCTAGTTGACAAATCTCTTGTGTTCATCTCCAAATATAATGTGATTGAAATGCATGACTTGATTCAAGATATGGGTAGATATGTGGTAAAAATGCAAAAGGATTCGGGAGAACAAAGCAGACTCTGGGATGTTGAAGATTTCGAAGATGTGATGGTCAACAATACAGTAAGTAGGCTAAACAATGCTATAATATTCAGTTTTTAGTCTCCATCTTCTATTTTTTATTTAAAAGATGAAAATTGTTGGTACCCTATCAATTACAACTTCGTTCTCTTACAACATGAAAAACTTTTCGATTGGACTTTAAAGAACTAGTTGAAAATTGTATTACTAGAAATTAGAAATATAGAATTAATTGCATAAAGTATTTCTCTTTAGCGACATTTTTCTTAACTAATATTGTACCTTAATTTTATTTAATGGTCTCTTCAAATGCAATAATAAATAATACAAAACATAACGAAATTTGGCATTGCAAACAGTTAGAAAGTAGTCATATTGGCATGCAAATTTTATACAGATATTTGAAGTTATACTTTGTAAACTAATGAACCTTTTAACAGGGGACCAAGGCAATGGAAGCAATCTGGACACATGGTGTTCAAAAATTATGCTTTAGCAAAAAAGCCATGAAAAACATGAAAAGGCTTAGATTATTATCCATCATTAGTTTTAGTACACATGATGACTCCATTGAGTACCTGCCCAACAACTTGCGTTGGTTTGTCTGTCATAGTTATCCTTGGGAGTCATTATTAGAAAATTTTGAACCTAAGAAGCTTGTTTGTCTTTATCTCCAGAGGAGTTCCCTGCGTCAATTATGGACCGGAATAAAGGTACAATTGTTGAATTCAATTTTCATTATTTTTCTTTTTGTCAAATAC

>M6-PGSC0003DMG400001016 M6 LRR-RLP

CATACATTGACAGGTGAAATACCGAAAAAATTTAGCAATCAGATTGAGTTGTTTGATCTTAGTTTCAACAGGTTTACTGGTTCACTACCAATGGAGATTTTCAACATATCGGGTATGAGAGAGATTCAAGTTATTTTCAATAATGTGACTGGAACTCTCCCATTAAACATAGGTTCTACGTTACCCAACATTGAAGTTATTCATCTGAGAAACTTAAATCTTTATGGGACTATACCTCATTCTCTCTCCAATTGTTCCAAACTCACTAGTCTAGACCTTTCTCTCAATAGACTAACAGGTTTGATTCCCAACTCTCTTGGAAGTTTGACTCATCTACAGTCCCTAAGCTTGGCAATTAACAATTTAATGGGCGATAGTAGGTTAAGCTTCTTGACTTCCTTGTCAAATTGTAGAGATTTAAAAATTATTTTCCTCTCTGCGAACCCTCTATATGGTGTGCTTCCAGAATCCATAGGGAACCTTTCCAGTACTTCTCTTCTAAGGTTTAGAGCTATTGATTGTAAAATTAAGGGGGAAATTCCAAAAGGATTTGGGAACTTAAGCAGCTTATTAGAACTCGATCTTTCTAGAAACGGTTTCACTGGACCAATTCCCACAACAATCAGCAACTTGAGATTCCTTCAGAGCTTGAAGTTGAGTGTCAACAAACTTTCTGGATATATTGGAGATGATCTATGCAAATTACAGAACTTGGGTTATTTAAACTTGACTTTGAATCAACTTTCAGGATCTCTTCCGAATTGTTTGGGAAACTTAACTTCCCTTAGGGAGATGATTCTAGGTTCCAACAAATTGCGTTCAATCATTCCAAAAAGCTTAGGAAACCTCATAAACCTCTTGAAGCTAGACTTATCCTCAAACAACTTGGGTGGCTCTATACCTCTGGAAATTGGAAATCTAAAGGCTGCGATATATATGGATCTATCAATAAATACACTCTCAAATGGAATTCCTAAAGAAATTGATAGCTTGCAAAATCTTATATACCTTTCTTTGAGAGACAACAAATTGCAAGGATCAATTCCTGGCTCAATGAGCAGCATTTCAGCTTTGGAATTTCTAGACCTTTCTCATAACAATGTTTCAGGATTGATTCCCAAGTCTTTGGAGAAGCTTCTATATCTCAAGTATTTCAACGTTTCATTCAACAAGTTGGTTGGTGAAATCCCCTCGAATGGTCCTTTCAAGAACCTCTCCGGTGAGTTTTTCATGTCTAATGAAGCATTGTGTGGATCTCCAAGATTTTGTGTTCCCCCATGCCACAGTAGCACATCAAAGCATAAATCCAAGAGGAAAAAAGTGATTGTTCTTGTTCTACCGGGTACACTTGTATTAGTCTCCATCGCCTTTGCCCTTTTATGGTTAAGGTATACAAGAGGTAAAAGAACTGATCCTCAGCAAGCAGTTGATTCATCCTTTGTCATCTCAACAAAAGGAAGAATTTCATACTATGAACTACTTCGAGCAACTGATTCACTTAGTGAGATCAATTTAATTGGTTCTGGGAGTTTCGGCTCTGTTTACAAAGGCATCCTCAGAGATGGGACTTTCATTGCAGCTAAAGTGTTCAACCCACAATTGCAAGTGGCATTTAGGAGTTTTGATACAGAATGTCAAGTTTTGCGCAACCTTCGTCATAGGAATCTCACCAAAGTCATCACCAGTTGTTCCAACCTTGAATTTAAGGCTTTAATACTCGAGTACATGCCCAATGGGAGCCTCAATAAGTGGTTGTATTCACACAACTACTTCTTAGACATCAAACATAGGCTAAGTATAATGATAGATGTAGCATGTGCATTGGAATATCTACACCATGGATGCTCATTACCCGTGATTCATTGTGATTTGAAGCCTAGTAACGTCTTGTTGGATGAGGAAATGGTTGCCCACCTAAGTGACTTTGGAATTTCAAAGTTGCTTTCTGAAGATGAGAGTTCTTTATACACTAAAACATTAGCAACGTTTGGTTATATTGCACCAGGTACGTGATCTCTTAATCAATTGCTTTTGATTATTTCTCTTTCCTTATTTTCCACTAGTAAATGTTGAATTTCAAAATTGATTGATTGCAGAGTATGGACTGGAGGGATTGGTGTCAATAAAATGTGATGTTTATAGTTAAGGGATCATGTTGATGGAAACATTTACTAGGAAAAATCCTAATGATGAAATGTTCGATGGAGATTTTAGTTTGAAGCAATGGGTGAGAAATTCACTCCCACAAGCAGCAATGGAAATTGTTGATGCCAACTTAATATCGCCACATGATAATCAATTAATGAAAAAGCTAGATTGTGTGGTATCCATCATGAAAGTGGCAATAGATTGTTGTGTTGAATCTCCAAAAGGAAGGATTGACATGAAAGATGTTGTGGGGAGGTTAAAGACGATCAAGATTCAACTTCTTGCATGTTGAGGAAGATCTTATTAGAAACTTTTCTTCCAAATCATCTCATTGTAACAAACATTTGCAATAATAGTTAGCTTCAAATAAGTTATAGCATTATGTCTTTCATAACATAGAGAATTGATGGGCATGTATACTTCTTAGGTCTTGAATTTGTAAAGTGGTTTTGGCCTAATACCTCTACTTTTTTCCATCTTT

>M6-PGSC0003DMG400001072 M6 LRR-RLP

ATCTCTTCTTCTCCACAAAACTTCATCTCTCTCATCTCACATTTTGCTTAAAGATTCCATTTTTAAACCAAAAATTAAAAAAAACTCCACAAACCCCATTTCTTAAATCCATTTTTCTTCATAGATCTTCACTTTTTTCATCTCACATTTGGTTAAAGATTCAATCTTTGAACAGAAAAAAGCAAAAAAAAACCAAGCATTCTTGATTTTCAGATGACTTATGTGAGATGAAGGAGAAAACTCAAACACCCATTTGCTCTCTGTTTTTTCTTGTGCTCTGTTTTTTGTGTTCAGTTTTTACTGAACACACAAATGCTGAGTTTACTGTCTCTGAAAATGGCCCTTTAACTGATTCTGAAGCTCAGTTCATCAAACACAGGCAGCTTTTGTATTACAGAGATGAGTTTGGTGATAGAGGTGAAAATGTGAAAATTGATCCATCTATGGTGTTTGAGAATGATAGAATCAAGAATGCTTATATTGCTTTACAAGCATGGAAACAAGCTATCATTTCAGATCCCTTTAATATCACTCTCAATTGGGTTGGTCCTAATGTATGCAGCTACACTGGAATCTTCTGTGCTCCGGCGTTGGATAACCCCAAAATCCGTACAGTTGCCGGCATTGACCTTAATCATGGTGACATTGCAGGGTACCTCCCGGAAGAGCTTGGGCTTCTTACAGATCTTGGGATTTTCCATATCAATTCAAACCGTTTCTGTGGTACAATACCAAGAAAATTCAGCAAGTTGAAGATACTGTTCGAGCTGGATCTGAGTAATAACAGGTTTGCTGGGAAGTTTCCTTATGTTGTTTTGAGTTTACCCAAGTTGATATTTTTGGATATTAGGTTCAATGAATTTGAAGGTAATGTACCTTCACAGCTTTTTGATAAGCCATTAGACGCCATTTTTATTAATCATAATCGGTTTGCGTTTGAGTTGCCGGATAATTTTGGGAATTCGCCGGTTTCTGTGATAGTTCTGGCGAGTAACAGTTTCCATGGGTGTCTTCCGGCGAGTATTGGGAATATGAGTAATCTTAATGAAGCGATTTTAATGAATAATGGGTTGCGTTCTTGTTTGCCGGCGGAGATTGGGTTGTTGAAAAATCTGACTGTATTTGATGTGAGCTTTAATCAGTTGATGGGTCCGTTGCCGGAGAATTTTGGTGATTTGGTGAGTTTGGAGCAATTGAATGTGGCACATAATATGCTTTCTGGAACAATTCCGAAAAGTATTTGTCAGCTTCCTAAGCTTGAGAATTTTACGTATTCTTATAATTTCTTCACTGGTGAACCGCCGGTGTGTTTGGGGTTGCCGGAGTTTCATGATGAACGGAATTGTTTGCCCAACAGACCGGTGCAACGCTCTCCGGGACAGTGTAAGGCCTTTTTGTCTAAAAAAATTCATTGTAGTGCTTTCAGGTGTCATAAATTTGTTCCTGTTTTGCCACCTCCTCCGCCACTTTCACCGCCGTTGCCTGCCCCTCCGCCACCTATTTATTCGCCTCCGCCACCAGTTTATACTCCTCCACCTCCAGTTTATTCACCACCGCCACCACCACCCTCTCCACCTCCAGTTTATTCACCACCGCCACCACCACCCTCTCCACCTCCTCCATCACCTCCTCCGCCTCCACCACCAGTCTATTCACCTCCACCCCCACCTCCATCTCCTCCTCCACCATCACCTCCACCTCCACCCCCACCATCTCCACCTCCACCATCACCCCCACCACCAGTTTATTCACCACCACCACCGCCACCATCTCCCCCTCCACCATCACCTCCACCTCCTCCCCCACCCATTTATTCGCCACCACCATCTCCACCTCCTCCATCACCATTGCCTTATTGTGTACGCTCACCTCCACCTCCACCACCAAATTCCCCACCACCTAATTCTCCACCTCCACCGCTTGCTCACTCGCCTCCACCCCCTTCACCTTACTATTACAATTCACCACCACCTCCGCCCCCAAATTCGCCGCCTCCAGCACCAAATTCGCCTCCTCCTCCACCAGTATACATATACTCATCACCTCCGCCACCAAATTCACCTCCTCCTCCACCACCAACATACGTATATTCTTCACCTCCTCCACCTCCTTCCCCATTGCCTTGTATAGAACCCCCTCCACCACCTCCTCCTTGCATTGAACCTCCCCCACCTCCCTCCCCTTCACCATCTCCTCCTCCACCACCTGTTTATCATTACAATTCTCCGCCCCCACCATCTCTAGCACCTCCACCACCAGTTTATTTTTACAATTCTCCACCTCCACCATCACCACCACCTACTCCAGTATACGAAGGGCCATTACCACCAGTAATAGGAGTTCAATATGCTTCACCACCTCCACCACCCTTTTATTAATTCTTTCAGAAATTTCACTAACTCTTATCCTCAGAGCACAAAATAGTTCCTACAATTATTATTTCCTCAACTCTTCAAGCCATCAACAAAATCGAGCTAATATGTGATCACGAACCTTAAATGGTACGACAGAAAGAAAGATAAGAGGAAGGTAAAGAGGAGAAAGTTTCAATCTCCAATGATTTTATTTTTGAATTGTTATCAAATAGATTGAAGTTGTGGAGAGAATAAATTGGCAAAGTAAAGATTGAAACTTTGCTGGAAAATGTATGTATAGAGAGATCATAATCTTGAATTAGAAACCAGTTTACAGTGCTTGCTGCAACTGGTTTTATTTAATGATCATACTTTGATTTCTTTGTAATTTTTTCTTTTTTTTGGTTCTTCCAAACTATAGTATAAGATGAGCTTTTTATTTTCTTTTAAAACTTGTATTGATTC

>M6-PGSC0003DMG400001252 M6 LRR-RLP

AATTTCAGAGAGAGAAACAAAACAAAAACTCCTCAGGATGGGCAGCTTCATGTACATTGCGATCTTACTTCTTGCCATAATCTCAACCGTTCATTCCTGCCCGCCATCAGATCGGGCAGCATTATTATCTTTCCGAGCTGCCCTAAATGAATCTTACTTAGGAATTTTCAACTCGTGGAAAGGGTACGACTGTTGCCATGAATGGTACGGGGTGAGTTGTGATCCCACAACTCACCGCGTAGCAGACATAAACCTCCGTGGTGAATCCGAAGATCCACTCTTCGAAAAAGCTCATCGTACCGGTTACATGACCGGTACGATCTCTCCATCTATATGCAAATTAGAAAGACTATCAAGTCTCACTATAGCTGATTGGAAAGGCATCACTGGACCTATACCATCATGTATTACGTCCTTACCCTTCCTTCGAATCATCGACTTAGTCGGAAACAAGCTCACCGGCTCAATCCCTTCAACTATAGGTCGATTAAGTCGACTCACAGTGTTAAACGTAGCTGAAAATCAACTTACTGGCTACATACCTCGATCATTAACCAACTTATCATCTTTAATGCATTTAGATCTCCGAAACAACTTCATCTATGGAAACATCCCAACAAATTTCGGAAAATTACGGATGTTGAGTCGAGCTTTACTGAGTCGAAACAGGTTAACAGGTCATATACCGTATTCGATTTCGTATATTTACAGGCTTTCAGATTTGGATCTGTCATTGAACAAACTTTCTGGTAGCATACCTCCGTCTCTAGGTAAAATGCACGTACTTGCAACGTTAAATCTTGATGGTAACAATATTACTGGTACTATACCACCTAGTTTGATTAATTCACGTATCAACATTTTGAATTTGAGCAAAAATACTATTGAAGGTTATATTCCTGATTCATTTGATGAAAGATCTTATTTTATGGTAATGGATTTATCTTACAACAAATTAAAAGGAAGGATTCCAAAATCGATATCGTCGGCGACGTTTATCGGTCACCTTGACGTTAGTCATAACCATCTTTGTGGCCAGATTCCGGAAGGTTCTCCCTTTGATCATCTTGAAGCTTCATCTTTTACTTACAATGATTGTCTATGTGGGAAGCCTCTTAAACCTTGTCACTAAAGTGTGACTATTATTAGGTAAATTTAATTTTACTTTTCTTTTTTGAGATTTTTATTAACACTATATATTGGTGAATATGTTTGTATTATTATGTGATACTGAAAAGTAAAGTGTTTTTTTTTAGCATATATTTGTAATTTTAAATGTATAATTGTGCCTTGAGATATATGAAGGCTTTCTTA

>M6-PGSC0003DMG400001829 M6 LRR-RLP

TATATTTTTTTTCCAGCAATCTCAGCTCACATATGCTGGGCAACATCTCTGTGCCTGCTATGACGCTTTTTATTTGCTTCAATTTAAGAAAGGCCTCACAGTTGTTGATCAAAATATACATGACTGTGACAATGAGACCCGAGCCAAGACTTTGTCCTGGAATGTTACAGGAGATTGTTGTCAATGGGATGGTGTAACTTGCAATGGATTTACGGGTCATGTCATAGGCCTTGATCTTTCTTGCAGCATTCTTAACCAGACCATCAATGCTAACAGCAGCCTCACTAAACTTGGTCATCTCCAAAGACTCAACCTTGCCTTCAGCCAGTTTGATGACGACTTTCCACTTGGATTAAGTGAACTCAGTAGTTTGACACATCTCAATCTTTCGAATACTGGATTTAATGAAGGGGAGATACAGATTCCACCAGGATTATCCAAGTTGTCTAAGTTGGTTTCACTTGACCTCTCCTCGAATGATTTCCAAGTTGGCCGAACAACCTCTAGAAGTTTGCTTCAGAACTTAACTAATTTGGAGGTACTGTTGCTTGACAGCGTGGATGCACCATTTGAGTTACCTAAGAATTTTCCTTCTTCCCTTAGGTACTTAATTCTTCAAGACACCAACATGTGTGGGAACATAAGTGATTCTCAGCTTTTTCATCTACCAAACTTGCAAGTGCTGAGAATGGGATGAAATCCTTTATTAACAGGCACTTTGCCTAATTTCAATTGGAGTTTCAGTACAAGTATTTTAGAGTTGGACTTTTTCTATACTGATATTTTTGGTAAGGTACCTGATTCAATCGGCAACCTCCGTTCACTTTGGTAATTAGATCTCTCTTTAAACCGTTTATCCGGCTTGATTCCAGTATCCATTGGCAACCTCAGTGGAATTAGAGTGTTGAGACTTTCAAGTAACAGCTTCACTGGTAATGTTCCCTCATCTATCTCAAAATTAAACAAACTCTTTGATTTAGATCTCTCTTTCAACGATTTTCAAGGCTTGATTCTAGAATCCATTGGTAACCTAACTGCAATCACAAAGTTGACACTTTCAGGTAACAACTTCACCGGAAATGTTCCCTCAACTTTCAAAAAGCTGAAAACACTTAACTACTTATCTCTCTCTTTCAATAATTTTGAAAGCTCAATTCCAGACAACTTCGCCTACTTTTCAGAGCTATCTTTTTTATACTTGCAAACTAACAATTTCACTTGCCCACTTCCCTATTTGATTGCAACCTTGACTCGCCTTCAAGAATTAAACTTGCAAAACAATTTACTAACTGGTTCACTTCCCTCTAATTTAAGCAGACTTCAAGAGTTAGAAACACTTGATTTTTCCTTCAATTATTTCACTGGCACAATACCCTCTTGGTTATTCCGTCTTCCATCCTTGGTTTCTTTATCTATTCAATACAATGAATTAACTGGGAAATTGCCAAATGAGCTCAAGAGCATTACGTTGGCAACGCTAGAATTGATCTTCGGAACAACAAGCTGCATGGAGAAATCCCTGATTGGATATTATCTATGAGCATGTATATCCTGGATCTCTCTCATAACTTGCTCACAGGCTTTGAAAAACAAGTATGACATTCAAAGAATTTGGTGTGGCTTAATTTGGAGAACAATTTTCTTCAAGGGACTTTGCATCAATCTATTTGTGACATGTTTAACCTTGAAGTCCTCATTTTGTCTCGGAACAATTTCAGTGGTTCAATCCCAGGATGTTTGGGTAACTCCATTCTTGTTTTAGACTTGCGAATGAACAATTTTTATGGAGAAATACCAAAATTCTTACCTATAGGGTTACAATATCTTGGTGTGTATGACAATCAGTTGGGGGGGACAAGTCCCACGATCCTTGGTTAACTGTACAAGCTTGGAAGCTCTTGATTTGGGGAACAACAATATAAATGACACGTTCCCGTATGGCTGGACAAACTTCCAAACCTACAAGTTCTGATTCTGTAATCAAATCTCTTTCACGGTTCAGTTGGTGATTTGGATTCCGAGTTTCCATTTCCTGAGTTACGGATCTTTGACCTTTCCGTTAATGGGTTCACTGGAACTTTTCCATCTAATATTTTCAAAAGTTTCAGAGGTATGATGGATGTGGATGAAGGAAAAACAGGAATTACTCTAGCTATTAATAGGACTCACAGAGATTACCTTTACCATGTTAGTTTGGTGATAAAAGTCAATGAATTTGATATGAGAATCACGTCAATTATGACGAGTGTTGATTTATCAAGCAACAGGTTCGAAGGAGATATTCCAAATTCCATTGGAAGTCTGAGCTCACTTGTGTTACTCAACCTGTCTCGCAACAACTTCCGTGGTCATATTCCTGCAGAAATTGCAAAGTTGCAGCAGCTCGAAGCATTATAGATCTCTCGTGGAACAGACTCATTGGAGAAGTTCCTGGTCAATTGTCGAGTTTGACATTTCTTGAGGTCTTAAACCTTTCATACAATCATCTGGCTTGGCGCATTCCTATTGGGAAACAGTTCAATACATTTCCAGATGATTCATATTGTGGAAATCTTGATTTATGTGGATTTCCACTATCAAAGAAATGTGGAAACAATAATAGGCCAGATGAGTCAACACTTGAACAAGATGATGAAGATTCATTTTTTGCGAGCGGGTTCACATGGGAAGCTGTGGTAATAGGTTATGGTATGATATTTGGATTGTTGATGGGAGGCCTCATGTTCCTATTGGAAAAACCAAAATGGTTTGTCAACTTTGCTGAAGATATTTCTCAACAAATTGCTGCTAAGAAGCGAAGAAATCAAAAGAAGAGACGTCAAGGACGTGGTCTAAGAATGAATTAG

>M6-PGSC0003DMG400001831 M6 LRR-RLP

ATGGAGTACCAAAAATTGCTAATAGCTTTCTGCTTCTATTCACATTTTATACTTTTTGTCCATCAATCTCAGCTTACATATGCTGGGCAACATCTCTGTGCCCGCTATGAAGCTTTTTATTTGCTTCAATTTCAGCAAGGCCTCACAGTTGATCTTATTGGTTATGACTGTGACTATGAGGCCCTAGCTAAGACTTTGTCTTTGAATGTTACAGGAGATTGTTGTCAATGGGATGGTGTTACTTGTCATGGATTTACTGGTCACGTTATAGGACTCGATCTTTCTTGCAGTTTTCTTAACGGAACCATTAATTCTAACAGCAGCCTCACAAAACTTAGTCATCTTCAAAGACTCAACCTTGCTTTCAACAAGTTGAGTAACTTTCCACTTGTGGATAGCATTAGTGAACTCGAAAGTTTGACATATCTCAATCTTTCGCATTCAGTACATATGATAGGGGAGAAGCAGATACCACAAGTATTATCCAACTTGTCTAAATTGGTTTCACTTGACCTCTCCTTGTATTACACTGAACTCAAAGTTGATTGAAAAACCTTCAGAAGTTTGCTTCAAGACTTGACCAATTTGGAGGTACTGTTTCTTGACAACTCGTATGCATCATTTGAGTTACCTAAGAATTTTTCTTCTTCTCTTAGGTACTTAAGTCTTGAAGGTATCGGCATGTTTGGGATCATAAGTGAGTTTCAGTTATTTCATTTATCAAACTTGCAAGTGCTGAGATTTGGACAGAATCCTTTACTAACAGGCTCTTTGCCAAATTTTAATCGGAGTTCCAGTGAAAGTATTTTAGAGTTGAACTTCTTTTATACTGGTATTTCTGGGAAGGTACCCGATTCAATTGGTAATCTCCATTCTCTTTGGTATTTGAATCTCCAGATTTGTCATTTATCTGGCTCGATTCCAGAATCCATTGGCAATCTCACTGCAATCAGAGAGTTGATACTTTCAGAAAACGTCTTCATTGTTAATGTTCCCTCAACTTTCTCAAAGTTAAACAAACTCGTTCGTTTAGATCTCTCTTCAAACTATTTTCGAGGCTCAATTCTAGAATACATTGACAATCTCACTGCAATCACACAGTTGACAATTCCAGATAACAGCTTCACTGGAAATGTTCCCTCAACTATCGGAAAGTTGAACAAACTTAACTACTTATCTCTCTCTTCCAATTATTTTGAAGGCTCGATTCCAGACATCTTTGCCAACTTTTCAGATCTATTTGCTTTATTTTTTGACACTAACAATTTCACTGGTTCATTTCCCTGTTCAATTGCAACCTTGATTTGCCTTGTGAATTTAGAGTTGCAAAACAATTTACTAACTGGTTCACTTCCCTCTAATATAAGCTGATTTCAACTTCTACAATCACTTGATTTGTCCTTCAATTATTTCACTGGCACAACACCCCCTTCGTTATTCCTTCTTCCAAGATTGGTTTCTTTACATGTTCAACTCAATCAATTAACCGGGAAATTGCCAAATGTGCTCAAGATGAGTTATCTAGGATACGTACTATACCGTAGCATTGATCTTTCATACAACAAGCTGCAAGGTGAAATCCCCAATGCGATGTTATCTATGAGCATGGGTAAGCTGGATCTCTCTCATAACTTCCTCACAGGCTTTGAAAAACAAGTATGGCACTCAGAGTACTTACTATACCTTAATTTGGAGAACAATTTTCTTCAAGGGCTTTTGCATCTATCCTTTTGTGACATGTATAGTCTTGTATTCCTCATTTTAGCTCGGAACAATTTCAGTGGTTCAATCCCGGTATGTTTGGGTAACTCCAGTATTTCCATTTCTTTTTTAGACTTGCGAATGAACAATTTGCATGGAGAGATACCAAGATTCTTATCTACAGGGTTTCAATATCTTGGTTTATATGGCAATCAATTGGGGGGACAATTCCCGTGATCCTTGGTTAACTGTACAAGCTTGGAAGCTCTTGATTTGGGGAACAACAAGATAAATGACACGTTTCCGATATGGCTGCAGAAACTTCCAAACCTACAAGTTCTGATTCTGAAATCTAATCTCTTTCATGGTCCAATTGGTTATTTGGAGTCCGAGTTTCCATTTCCTGAGTTACGAATCTTTGATCTTTCCTTCAATGGGTTCACTGGAACTTTGCCATCTAATTTTTTCAAAAGTTTTAGAGGTATGATGGATGTGGATGAAGAAAAAACAGGAATTACTCGCTCCTACAAAGGGTCGACCAGGCCTTGTACAAGCATAAATCGATGTCGAAGAGATTACCTTTACCACTTTAGTTTGGTGATAAAAGGTAATGAGTTTGATATGAGAATCACGTCAATTATGACGAGTGTTGATATATCAAGCAACAGGGTTGAAGGATATATTCCAAATTCCATTGGAAATCTGAGCTCACTTGTGTTACTCAACCTGTCTCACAACAGCTTCCGTGGTCATATTCCTGCAGAATTTGCAAAGTTGCAATAGCTCGAAGCATTAGATCTCTCACGGAACAGACTCATTGGAGATATTCCAGATCCATTGTCGAGTTTGACATTTCTTGAGGTCTTAAACCTTTCCTACAATCATCTGGCTGGACGCATTCCTCTTGGGAAACAGTTCAATACATTTCCAAATGATTCTTATTGTGGAAATCCTGATTTATGTGGATTTCCACTATCAAAGGAATGTGGAAACAATAAAGAGTCACCACTTGAACACGATGATGGTGATGATGATGATTCATTTTTTGCGAGTGGGTTCACATGGCAAGCTGTGGTAATAGGTTATGGTTGTGGTCTGATATTTGGATTGTTGATGGGAGGTCTCATGTTCCTACTGGAAAAGCCAAAATGGTTTGTGAACTTTGCT

>M6-PGSC0003DMG400001832 M6 LRR-RLP

TTGGCAATCTCACTGCAATCAGAGAGTTGATACTTTCAGAAAACGTCTTCATTGTTAATGTTCCCTCAACTTTCTCAAAGTTAAACAAACTCGTTCGTTTAGATCTCTCTTCAAACTATTTTCGAGGCTCAATTCTAGAATACATTGACAATCTCACTGCAATCACACAGTTGACAATTCCAGATAACAGCTTCACTGGAAATGTTCCCTCAACTATCGGAAAGTTGAACAAACTTAACTACTTATCTCTCTCTTCCAATTATTTTGAAGGCTCGATTCCAGACATCTTTGCCAACTTTTCAGATCTATTTGCTTTATTTTTTGACACTAACAATTTCACTGGTTCATTTCCCTGTTCAATTGCAACCTTGATTTGCCTTGTGAATTTAGAGTTGCAAAACAATTTACTAACTGGTTCACTTCCCTCTAATATAAGCTGATTTCAACTTCTACAATCACTTGATTTGTCCTTCAATTATTTCACTGGCACAACACCCCCTTCGTTATTCCTTCTTCCAAGATTGGTTTCTTTACATGTTCAACTCAATCAATTAACCGGGAAATTGCCAAATGTGCTCAAGATGAGTTATCTAGGATACGTACTATACCGTAGCATTGATCTTTCATACAACAAGCTGCAAGGTGAAATCCCCAATGCGATGTTATCTATGAGCATGGGTAAGCTGGATCTCTCTCATAACTTCCTCACAGGCTTTGAAAAACAAGTATGGCACTCAGAGTACTTACTATACCTTAATTTGGAGAACAATTTTCTTCAAGGGCTTTTGCATCTATCCTTTTGTGACATGTATAGTCTTGTATTCCTCATTTTAGCTCGGAACAATTTCAGTGGTTCAATCCCGGTATGTTTGGGTAACTCCAGTATTTCCATTTCTTTTTTAGACTTGCGAATGAACAATTTGCATGGAGAGATACCAAGATTCTTATCTACAGGGTTTCAATATCTTGGTTTATATGGCAATCAATTGGGGGGACAATTCCCGTGATCCTTGGTTAACTGTACAAGCTTGGAAGCTCTTGATTTGGGGAACAACAAGATAAATGACACGTTTCCGATATGGCTGCAGAAACTTCCAAACCTACAAGTTCTGATTCTGAAATCTAATCTCTTTCATGGTCCAATTGGTTATTTGGAGTCCGAGTTTCCATTTCCTGAGTTACGAATCTTTGATCTTTCCTTCAATGGGTTCACTGGAACTTTGCCATCTAATTTTTTCAAAAGTTTTAGAGGTATGATGGATGTGGATGAAGAAAAAACAGGAATTACTCGCTCCTACAAAGGGTCGACCAGGCCTTGTACAAGCATAAATCGATGTCGAAGAGATTACCTTTACCACTTTAGTTTGGTGATAAAAGGTAATGAGTTTGATATGAGAATCACGTCAATTATGACGAGTGTTGATATATCAAGCAACAGGGTTGAAGGATATATTCCAAATTCCATTGGAAATCTGAGCTCACTTGTGTTACTCAACCTGTCTCACAACAGCTTCCGTGGTCATATTCCTGCAGAATTTGCAAAGTTGCAATAGCTCGAAGCATTAGATCTCTCACGGAACAGACTCATTGGAGATATTCCAGATCCATTGTCGAGTTTGACATTTCTTGAGGTCTTAAACCTTTCCTACAATCATCTGGCTGGACGCATTCCTCTTGGGAAACAGTTCAATACATTTCCAAATGATTCTTATTGTGGAAATCCTGATTTATGTGGATTTCCACTATCAAAGGAATGTGGAAACAATAAAGAGTCACCACTTGAACACGATGATGGTGATGATGATGATTCATTTTTTGCGAGTGGGTTCACATGGCAAGCTGTGGTAATAGGTTATGGTTGTGGTCTGATATTTGGATTGTTGATGGGAGGTCTCATGTTCCTACTGGAAAAGCCAAAATGGTTTGTGAACTTTGCTAAAGATATTGCTCAACAAATTGCTGCTAAGAAGCAAACAAGGCAAAAGAAGAGACGCCAAAGACATGGTTTAAGAATGAACTAGCAATATACCAGGTATGCATTAAAACCTTTAGTTTTGAATCAGGAACAACACTTCAGAGGTTGTTGGATAACTATATGTCAACTGTTTTTATTTTTTCCTAAATATTCAAAACTTAATTTGTCTCTGCAGAGGTTGCTGGAGAGTTGGACAAGTTTATCAGCAAGTCAGTGCTGATGTTTGGCAAATAGCAATGGATGCATCTGTTTGTTTACTTTATGTCCATGGTTGCCAAATACGTATGTAAATGTTACAGTTTAGTTCT

>M6-PGSC0003DMG400001870 M6 LRR-RLP

ACTTTCCACTTGGATTAAGTGAACTCAGTAGTTTGACACATCTCAATCTTTCGAATACTGGATTTAATGAAGGGGAGATACAGATTCCACCAGGATTATCCAAGTTGTCTAAGTTGGTTTCACTTGACCTCTCCTCGAATGATTTCCAAGTTGGCCGAACAACCTCTAGAAGTTTGCTTCAGAACTTAACTAATTTGGAGGTACTGTTGCTTGACAGCGTGGATGCACCATTTGAGTTACCTAAGAATTTTCCTTCTTCCCTTAGGTACTTAATTCTTCAAGACACCAACATGTGTGGGAACATAAGTGATTCTCAGCTTTTTCATCTACCAAACTTGCAAGTGCTGAGAATGGGATGAAATCCTTTATTAACAGGCACTTTGCCTAATTTCAATTGGAGTTTCAGTACAAGTATTTTAGAGTTGGACTTTTTCTATACTGATATTTTTGGTAAGGTACCTGATTCAATCGGCAACCTCCGTTCACTTTGGTAATTAGATCTCTCTTTAAACCGTTTATCCGGCTTGATTCCAGTATCCATTGGCAACCTCAGTGGAATTAGAGTGTTGAGACTTTCAAGTAACAGCTTCACTGGTAATGTTCCCTCATCTATCTCAAAATTAAACAAACTCTTTGATTTAGATCTCTCTTTCAACGATTTTCAAGGCTTGATTCTAGAATCCATTGGTAACCTAACTGCAATCACAAAGTTGACACTTTCAGGTAACAACTTCACCGGAAATGTTCCCTCAACTTTCAAAAAGCTGAAAACACTTAACTACTTATCTCTCTCTTTCAATAATTTTGAAAGCTCAATTCCAGACAACTTCGCCTACTTTTCAGAGCTATCTTTTTTATACTTGCAAACTAACAATTTCACTTGCCCACTTCCCTATTTGATTGCAACCTTGACTCGCCTTCAAGAATTAAACTTGCAAAACAATTTACTAACTGGTTCACTTCCCTCTAATTTAAGCAGACTTCAAGAGTTAGAAACACTTGATTTTTCCTTCAATTATTTCACTGGCACAATACCCTCTTGGTTATTCCGTCTTCCATCCTTGGTTTCTTTATCTATTCAATACAATGAATTAACTGGGAAATTGCCAAATGAGCTCAAGAGCATTAC

>M6-PGSC0003DMG400002118 M6 LRR-RLP

CAATCCTTCAGAGATGTCTGAAGATGAGGAGCGTCCTTGTCCTTCAGATCTCACCGGTGGCGCCACTGCAAAAGCAAGGAACTGTTGCTTTAACGCCGCCGTTACCGGTGGCGGAGGGGGGATTTTTAACTTTTCGCCACACCCAGATCAAGTTCTTGAAAATGTGCTGGAAAATGTTCTTTGCTTCTTAACTGATCGCCGTGACCGTAACGCTGCATCTCTTGTAAGCAAATCTTGGTATCGGGCTGAGGCTTTAACCAGATCCGAAGTGTTTATTGGCAACTGTTATGCTGTTTCGCCGACCCGGGTTACGACCCGTTTCAAATGGGTGACCTCTGTGGCCATTAAAGGGAAACCTAGGTTTGCTGATTTCAGTTTGCTTCCTCCAGATTGGGGTGCTCACTTTACTCCTTGGGCTTCTGTTTTGGGTGATTCTTATCGTGGGCTTGAGAAATTGTACCTCAAACGTATGTCCATATCTGATGATGATCTAGGTTTGTTGGCGCGTTGTTTCCCCAGTTTCAAAGAGCTTGTTCTTGTATGCTGTGACGGTTTTGGGACTAGTGGACTTGCTATTGTAGCTCGTGATTGCAGGTTAGTATTGTTTTGTCCGGATATAGATCTGTTTATGTGATAACCCTTTTGGTAAAAAGTTTAGATTTTTAATTTCTAGAAGCTGAATTCTTCTTTCTGTAATGGGAAATTAACAGCTTAAACTAGTGTAGATATATTGTGTTATTTGAATTTGTGCTTCATGTGTGCTGGTGTTTGAATTAGTTATGTGATTGAGTTGTTCTTTAACTGTTTTATTCTGCATTTGAAACTTAATGATACCTTTTGAGGACACTAGCATTGAAGTTGTAATATGTGGTGACTTGTTGTATTGGATTTGTGAGTTTGCAAGCTGTGAAATATGGTTTGGTAGTGTGATTTTCTTGTGCATAATAGAAGGTTTTTACTGCAGGCAAATTAGAGTGCTTGATCTGATTGAGTCTGAGGTATCCGACGATGAAGTGGACTGGATTTCCTACTTCCCTGAGAACAAAACGTGTTTGGAGTCTTTGACCTTTGATTGTGTTGAATGCCCTATTGACTTTGAGGCATTGGAGAAGCTAGTAATCAGGTCACCTAGTTTGAAGAGTCTAAGGTTGAATCGGTTTGTTTCTATTACTCAGCTGTACCGTTTGATGATTCGAGCTCCACAGCTTACCAATCTGGGAACAGGCTCTTTTGGCGCCTCAACGGTCATTGATGAACCAGATCCTGATTATGCTTCAGCATTTGCTGCCTGCAAATCCATTGTCTGTCTCTCTGGTTTCAAGGAAATTGCTCCTGAATATCTGCCTGCAATCTATCCAGTTTGTGGCAATCTGACCTCTCTTAATTTAAGCTATGGTGCCAACATTAATACTGAACAATTCAAGTCTGTCATCAGCCGTTGCCATAAGCTCCAAGTATTGTGGGTATGTGGTTTGAATTTTCGCTATATGTTCTTTTAAAGAAAATTGTACTCTGTAATAGCGTTATATTGAGTTCAATTTTGATGTTTCAGGTATTTGATTCTGTATGTGACGAAGGACTTGAGGCAGTTGCTGCAACATGTAAGGATCTGCGGGAGATTCGAGTTTTCCCTTTTGAAGCTCGGGAAGATGCAGATGCCCCAGTTTCTGAAGTAGGTCTCCTTGCGATTTCTGAGGGTTGCAGTAAACTTAAGTCCATCTTATATTTCTGCCAACGAATGACAAATGCGGCTGTTATAGCTATGTCAAAGAACTGCCCAGACCTTGTGGTATTCCGACTATGCATTATGGGTCGGCACTTGCCTGACCATGTTACTAATGAACCAATGGATGAAGGCTTTGGGGCTATTGTCAAGAACTGTAAGAAGCTTACTAGACTCGCTGTATCTGGTTTACTGACTGATAAGGCTTTCAGTTACATTGGACAATATGGAAAACTAGTCAGAACCTTATCAGTTGCTTTTGCTGGAAACAGTGACTTGGCTCTGAAGTATGTGCTCGAGGGCTGCCCTAAGTTGCAGAAGTTGGAGATCAGGGATTGCCCATTTGGAGATTTGTCTTTGCGTTCTGGTGTACACCACTATTACAATATGAGGTTCCTTTGGCTGTCATCTTGTAGAGTAACTCTACAAGGTTGTCAGGAGATTGCTGGCCAATTGCCCCGTCTAGTAGTGGAAGTGATAAGTGGTGATGAGGAAGGGAGTGAGACTGGTGAGCATGTCAATACCTTGTACATGTACCGATCTCTTGATGGACCGAGGGCTGATGTACCATCATTTGTGCAAATACTGTGACCGACCTTTTTAGTAGATAGGAATTTTCTGCAGTAATTGATTGGGACCTTTGAGAAATGATTAACTATTTTCTTGGGAAGAGATGCTCTCCAGAAATCTGTTGTGACCCATATGAAGGTGCAAGTTTCCTTCTTGAGAAAACGAATGGTGCACCACTTGCAAAGGCAGTGAGTAATCTATAATGTATTCAATGCTGGTCTCCATGGATGTCATTGTATATGGTTCAGTACAGAGATTGTTTTTTTCCAGCAAAGCAATCTAGAAGACAAATGGTTCTTTGTATCAGTAATCTTTTCTATTCACTTTCTTTTCTTCATATTCATGGAGCCTTCAATTAGTAAGTGGAGTCATTGGTAAGGCCACAGCAAGGGTAGATGTTCCAAACTGCACGGTGGCGGAGCTCAATGAGATTTCTCAAAGGCTTTCGGCAGAACCAACACTTGTGGAATACACCTGCATGCTCGGTCAGGGTACTGGCTCGTCAAGTAGTATATTTTCATACAATTGAGGCCCTTCTTATTTTTTGTTTCATTGTTAGTTTAGAGTTGGAAGTTTTTTATTTTTAGTTGGTGATAGCCTAATGGTCAGAAGGATACCTACCGCTGTTGGATCTCTCCATCTCGCTTTCTAGAAATATTTGTTCTTTCCAGGATGTGTTGTTTCGTAGCACCTGACTGTTGAATTTATTATTGTTATTTTTAGAAGAAATGTGAGATGGGTAATAAATGCCAAATATGCATGGACCAGATGATGCTCTAGTGTGTGTGTGTTTGTGTTTGTCTTTGCCTTTTCTTTATGTGGAATGATCAGCTTGATAGTGAAGACCCATAAATGATGTGTGGAAAGCCTTGTACAAGTACAAAAATGCAGGACCCTCCTCTCCTTCACTCACTCTCTTTGTGGGATCCATTTATGAGTCTTATCCTCTATTATCTTTCCAGCTGATGAATGGCCCTTCTCTGTCTCATCTCTTGTATCTGAAAGAAGAGCCAAGAGCAGGGAGTAGACTGTAGTTGTGGAAAGTATAGATGGATAGTACAGCACACCCCTTATTTTTTAGCTGATGATGCCAACTTTATCATTATTCTTTTGGATAACAGTCTTGAGGAGCCTGCCTGTTGTGTTATCCTGTTTACTTTTCATATGCGGCTGCTGCTGCAGCATTCTTCATCTCTATCATTGACTACAACTCAC

>M6-PGSC0003DMG400002426 M6 LRR-RLP

ATGGCATCTTCTTCTTCTTTTGTGAGTGATTCACAAAATTGTCCTCTATGGAAGTACGATGTCTTTCTAACTTTTAGAGGTAAAGATACCCGTAAAATATTTGCGAGTCACTTGTACGAAGGTCTGAGAAACAGGGGAATATTCACCTTTCAAGATGATAAAAGGCTAGAGCAAGGCGATTCCATCTCACAAGAACTCTTGAAAGCTATCGAAGAGTCTCAAGTTGCACTCATCATTTTCTCAAGGAACTATGCTACATCGAGGTGGTGCTTGAATGAACCAGTGAAGATCATGGAATGCAAGGAGGAAGAAAATAGACAAACAGTCATACCGATCTTCTATGATGTGGATCCATCACATGTTCGACACCAAAGTGAGAGCTTTGCAGAAGCATTTGCCAAACATGAATCGAGGTATAAGGATGATATTGAGGGGATGCAGAAGGTATAAAGATGGAGAACTGCTCTAACAGCTGCCGTAAATCTAAATGGATATGATATTGGTGACGGGTGAGTTGAAAACACATAATATGTTTCTATTACCCTAAGCTATAATGTTTCCTATTTAATACGCTCTTGAACATTAAAAATATGAAAATACTTTCTAATACTTGACTTTCTCACATTGCCAACAATCAATCTCTTATTTATGTAGGATTGAATCAGAGAAAATTCAGTAGATTGTCGACCACATCTCTTCGAAATTTTGCAAGAGTGCTCGTTTTTTATCTTATTTGCAAGATGTTGTGGGAATAAATGCTCATTTAGAGGAACTAAAATCCCTACTTCAAATAGAAATCAATAATATCCGGACTGTAGGGATCTGGGGAATAGGCGGAGTTGGTAAGACGACGATAGCAAAAGCCATCTTTGATACTCTATCTCATCAATTTCAAGTAGCTTGTTTTCTTGCGGATGTTAAAGAAAATACAAGAAAGAATCAACTACATTCCTTACAAAATACCCTTCTCTCTAAATTGATAAGAAAAAAAGATAATTACGTCAACAATAAGTATGATGGGAAGTGCATGATTCCGAACAGACTTTCTTCAATGAAGGTGTTGATTGTGCTTGATGACATTGATGAAAGAGATCATTTGGAGTATTTAGCAGGTGGTGTTGGTTGGTTTGGTAATGGCAGCAGAGTTGTTGTGACAACTAGAAACAGACAATTGATAGAGAAGGATGCTGCAATATACGAAGTGCCTACACTACCTGACCATCATGCTATGCAATTGTTCAATAAGAATTCTTTTAGAAAAGAGGTTCCAAATGAGTGTTTTATGAAGTTATCGTTGGAGGTAGTGAATCAGGCTAAAGGCCTTTCTTTAGCCCTCAAGGTGTGGGGTTCTTTGCTGCATAAGAAAGGCCTAACTCAATGGACAAGAACTGTAGACCAAATAAAGAAAAAGTCTAATTCAGAAATTGTTGAAAAACTCAAAATAAGTTATGATGGGTTGGAGCCCGAAGAGCAAAAGATATTTCTAGATATCGCATGTTTCTTCTGTGGGCATGAAAGAAAACAAGTCATGAAAATTGTTGATAGCTGTGATTTTGGAGCTGAATACGGATTAGATGTTCTAATTGAAAAACCCCTTGTGTTCATCTCTAAATATGATAGAACTGAAATGCATGACTTGATTGAAGACATGGGCAAATACATAGTGAAAATGCAAACAGATTCCGGAAAACCTAGCAGAGTATGGAATGTTGAAGATTTCGAAGATGTGATGATGGACAATATGGTAAGTTAGCTAAACAATGCAATAGTATTCATATTTCAAACATATATATATATACACTCTGTTCCTCTAGCTTCATATTTCTTAAGAAATACTTAACTAATATTTGTACCTTAAATTTTGTTTAATGATGCTAACTCTTCAAGAACTTCTAAGTATAATAATAAATAATATATAAAACATGACAAAAATAGCATTTGGGGATTGTACATGACATGTGAAGTTATGATTTGTTGACTCATTACCTTTTTATAGGGGACCATGACAGTCGAGGCAATTTGGTTTACATATTCGGAAAAACTATGCTTTAACAAAGAGGCAATGCAAAATATGCGAAGGCTTAGGATATTATGCATGCAGACTTATTCATGGACTTCCCAAAGTGATTCAGAAGATGGCTCTATTGAGTACCTTTCCAACAACTTGTGTTGGTTTGTCTGACATCATTATCCTTGGAAGTTATTGCCAGAAAATTTAATTCCAGAAGGCTTGTTAATCTTGATCTCCGATGGAGTTCATTGCATTATTTATGGAATGAAACAAAGGTACCATTCTATTTCAGATACTTTCTATTTTCTATTGTCCTAGAATTGTATATGATTCTTGTTTTTAAATGAAATAAG

>M6-PGSC0003DMG400002478 M6 LRR-RLP

CTTCCACTTGATTTGAGACAATGCTTTGTGTATTCTGCAGTGTTCCCAAAGGACACCAAAATGGAAAAGGAAAATCTAATCTCTCTTTGGATGGCACATGGTTTTCTTTCATCGAAAGGAAACTTGGAGCTAGAGGATGTGGGTAATGAAGTATGGAATGAATTATATTTGAGGTCTTTCTTCCAAGAGATTGAAGTTAGATATGGTAAAACTTATTTCAAGATGCATGATCTCATCCATGATTTGGCTACATCTCTGTTTTCAGCAAACACATCAAACATCAATATCCGTGAAATAAATGCAGAAGTTTACTTCATTAGGGGGCACGGTAGTTTCATTGATGAGAAAGATTATATACCTAGGAAGCCCATTGGTTTTGCTAAAGTTGTGTCTTCTATCTCTCCTTCACTCTTGAAACAGTTTGTCTCGTTGAGGGTACTCAATCTAAGTTACTTAGGACTTCAGCAGTTACCTTCTTCCATTGGAGATCTAGTACATTTAAGATATTTGAACCTGTCTCGCAATGCAATGCGTAGTCTTCCAGAAAGGTTGTGCAATCTTCAAAATCTGCTGACTCTTGATCTACGGTTGTGCTCAAACCTTTGTTGTTTGCCAAAACAAACAAGTAAACTTGTTAGTCTCCGGAATCTAAAACTTGATCACTGTTCATTGAAATATATGCCACCAAGGATAGGATCTTTGACAGGCCTTAAGACTCTAAGTAACTTTGTTGTGGGCGAGAGGGAAGGTTGTCAACTTGGGGAACTAGGAAGCCTAAATCTCTATGGCTCATTTGAAATCACAAATCTTGAGAGAGTGAAGAATCATAAGGATGCAAAAGAAGCCAATTTATGTGCAAAAGGGAATCTGCATTCATTAAGCATGAGTTGGAACATGTCCGGACCACATAGATATGAATCAGAAGAAGATAAAGTGCTTGAAGCCCTCAAACCACACCCCAATCTGAAATATTTAGAAATCAATTGGTTCAGAGGAATCCGTCTCCCAGAGTGGATGAATCACTCAGTATTGAAAAATATTGTCTCTATTATAATTATCGAATGCACAAACTGCTCGTGCTTACCACCCTTTGGTGAGCTGCCTTGTCTAGAAAGTCTAAAGCTATGGCACGGGTCTGTGGATGTGGAGTATGTTGAAGAAGTGGATATCGATGTTGATTCTGGATTCCCCACAAGAATAAGGTTTCCATCCTTGAGGAAACTTGCTATATGGGACTTTCGTAATCTGAAAGGATTGCTGAAAAAGGAAGGAGAAGAGCAATTCCCTGTGCTTGAAGAGATGGAGATTTGGTGGTGCCCTATGTTTGTTATTCCGACCCTTTCTTCTGTCAAGAAATTGGAATTGGATCCAACACCTTTAAGGTCCATGTCTAATCTTAGGAATCTTACACCTTTCAGGTCCATATCTAATCTTAGGAATCTTACTTCCCTCAACATTAGCTATAGCAATAGCAATGAAGCAACTTCACTCCCAGAAGAGATGTTCAAAAACCTTGCAAATCTCAAATACTTGAATATCTCTAAGTTCGAGAATCTCAAAGAGCTGCCTACCAGCCTGGCTAGTCTCAATGCTTTGGAGAGTCTGAAGATTGAATATTGTGGTGCACTAGAGAGTCTCCCCGAGGAAGGGGTGAAAGGTTTAACTTCACTCACAAAGTTATCTGTTCATGATTGCAAGATGCTAAAATGTTTACCGGAGGGATTGCAGCACCTAACAGCCCTCACAACTTTAACAATTTGGGGATGTCGAGAAGTGTTCAAGCGGTGTGAGAAGGGAATAGGAGAAGACTGGCACAAAATTGCTCACATTCCGTATCTGGGTATTCATTACTATTAGTATTTATAATTGTTTCTTTGTTTGTGAGTCTTTTTGGTTCTTGCCATTGGATTGCATGTAATTTTTTCTAGGGTTGTTTGTTTGTAAGTCTCTCATTGGATGTAATTTTATTTTGGAAACAAATTAACAATCTATTTGTATTATACGTTTTGAGAATCTATTACTTATTTGTAATTGT

>M6-PGSC0003DMG400002618 M6 LRR-RLP

TAAAGTAAAATAATGGGGAAAGGAGCAATTTTGAAATTGAAAAAAAAAAAAGAAAAAACAAAAGTGATCTCTTATTCCGGAAAAGCAAGGTGGGAAATCTTTACAAGACTCGTCTCTTTTTCCTTATCAGAATCAGAAGAAGAAAGAAGAAGATTCGTCTCTCAGGTATGGTGTTCCTCTTACAATCAATCTATAATGCTTATGTTTGTTTATTTGAGAGGCCGATCCGCCCACCCTCATGATGGATGGGTCAAACCTGTTTTTTTTTTTTTCCATAAAAACAAACCAGATCTTTTCCGTTTACCCTTTTTGAGTTTTGACCATGAAAATCAATTTTTGAGTTTTGACCCATTGATGAAAATCAACATTAGGTTAGGGTAAGAAAAGGGAGTTTGATTGAATTATGCTGGAATATTCTGTCTTGTTAGCATGTGTATTTGTGAACAAGGTTCACGCACGCGTTTACTTTGACCTTTTCCGTATTACATTGGGTCTTGTCTTCTAATTACGTAATTAATCTCGTGCGTTTTTGTGACAGCACAAAATTTGACAGCTATATAGTCTTTGCTTGTGGGGCTGTCATTATAGTTCATCTATATTGGCAGCCCAATTTCTGAGGTGTGTGTGTCTTCTTTCTCTTCACCGGCAAAACAAATATATCATCATGGCTTACTTGTCTGGAATCCTCCAAAGACCTATAGTTGCAGCATCTGCAGTTGCTATAGCTTCTGTATCTACTGATCTTCATGAAAAATTCTGGCCATCTAAGTCATTACACAACAACAACAACAACAATTCCCAAGAAAAGGAAAAGATAACATCTCACATATCTCTTTCTAAGCTACCCCATTTCTCCTCCTACCTCACCACAATACCTGTTCCCCTGCCTATTCTAAATAACAGTACCACTTTTACTTCACTTTCAATCCTTCCTTTTCCTACTTCTTCTCTGTTGCTCAACTCTTATCAATCAGCAGCATTGCCCAACTCTGCTAATCCCTCCACATATACATATACACTTCCTTCTTCACCTTCCGAGGTGTTGTATACATGGCACTTACCTGAGCCAAATGCATCACGGAATTCTGATTCTTCACCACTAAAGTCTACCACTCTGGTAGTATTGTTGGGATGGTTAGGTGCAAAGCAAAAGCATCTAAAGAGATATGCAGAGTGGTATGCATCAAGAGGATATCATGTCATTACATTTACTTTCCCAATGTCTGAGATTCTAAGCTATCAAGTCGGGGGAAAGGCAGAGCAAGATATTGAAATGCTTGTGAATCATCTAGCTGATTGGTTGGAAGAAGAGCATGGAAAGAACTTGGTCTTCCACACTTTCAGCAATACTGGATGGTTAACGTGAGTTCTTGATGATACTCATTAGTCTAAGATGAGAACAGAAACATAACTATTTACATTTTCCTTAATACTAGCTTGATAATTATTGATTGACTTCTTTTATTCTTTTGGCCTTACAGTTATGGTGTCATTTTGGAAAAGTTTCAGAAACAAGATCATGCTTTAATGAGAAGGATTAAGGGTTGTATTGTTGATTCTGCCCCTGTCGCAGCTCCTGATCCACAGGTATTGCTGACTCAAAAATTGTGATGTTAAGATATCATTTTTATGTGCAAGTGTCAGAAGAGTGTTTCTTGGAAAGTAGTCTTATTTATGAAAGTTAATGCAGAGCACTTGTTTAGAGCTATACATTTCTATTGAAATCCAACCTTTTTCATGCTTTTATTGGACAACTATTACAAGTGTCGTATATAGTCAGGTTTTAGCCATTCAGAATTGCACAGTTGTTTGGATTCTAATCAGGACCTGCTGTTCTCGTATTTTTATCCTGTTGCTTTTTTTTATCTTAGGGCCTTAACCATGCAGCCTGCTTTTCGAAAACTATTCTTATGCTTGATAACTTGAAAGAATCTTGAGAGTTTAGCTCTCTCTGACAACTTTTTAACAGGAAGCATTCTATGCAACTTTTGTTTCACGGTTCAAGCTTTAAGCCTACTTTTTCTGGCTGAGAATAAGCCATCGAGGAATTTCCCCTTGGAGCTACTAAACTGCACACGTCCCTGCGACTACTGGATCTCTCTAGTAACAACTTTGGAGGAATGCTTCCATGAGGCATCGACAGGCTAGAAAGTCTCACATCCTTTTCAACAATAGCAACTTTACTGGAAATATACTGCCTCAAACAGGAAACTTGACGAACTTGGAAGATTTGTACCTAATCCATAACATGCTCTCCGGAAGAATTCCAGTGGAAATAGGAAAGCTTCAGAGGCTGTGTGAACTACCTCTATGGAAACCAGTTGTCAGTAACTACCCCAAGAGAATTGACAAATAGCTTAGTACTCATTGATTTCTTCGCTAATCAATTTTCAGGCTCTATTTCTGATGCCATTGGTAGGCTTAAGAATCTTGTTATCTTCAGCTAACTCAGAATGAGTTGTCTGGTCCAGTCCCATCAAGCTTAGGCTACTGCAGGAAGCATGAAAAACTGGCCTTTGCTGATAACAAACTTTCAGGATAATTGCCACCAACTTTAAGATTTCTATCAGAGCTGGACCTGACTACTCTTTACAACAATTGCTTTGAAGGTCCACTTCCTGAATATCTTTCCCTTCTGAAAAATCTCAGCAAAGTTAACTTTTCTCACAATAAGTTCAGTGGAAACATTTCTCCGCTAGCTGGTTCAAATTCTCTGACAGTTGACATGACAAACTACAGCTTCTCTGGTCCTATCCCATCTGAGCTAGTGGTGTCCAAAAATCTGACCCATCTTCACCTTTCTAACAATTTTTTCACTGGAAAAACCCCTTCTGAATTTGGACAACTTGAAGATCTCAGATGTCTTGATTTATTGTTCAAAAATTTGACTGGAGATCTAGCACCTTCCCTTGCTGGCCTGAAAACCTTGGCCGTTTTCTTCTTATATCTAATCAATTTTCGGGGAAAATTTCCACATGGTTTGTGGAGGAATGGAAGGTCTTGGTGAGTTCGATCTTTCATTCAACAACTTCACTGGAACAGTTCCAGCAGAACTTGGAAACTCTCCAAAATAACTCAAAACTTTTTCTCAGTTACAACAGATTATTAGGTGCCATACCACCAGAATCAGGAAATCTCACAAGTTTAAATGTCTTGAATCTTCAATGAAACAATCTTTTTGGCTCCATTCCTTCAACTCTTCAAAAATACCAGAAGCTCTATGAGCTGAGGCTCTCTGAGAATAATATAAGTGGCTCAATCCCTTACGAAATTGGCTCTTTATCCGAATTACAGGTGATATTAGACCTGAGCAAAAATCATTCTATTTTATGTGCAAGTGTCAGAATAGTGTATCTTCAAGCATAGTCTTATTTCCTAAGGTTAATGCAGTGCAATTGTTAAGAGCCATACATTTCTATTGAAATCCAACCTTGTCATTCTTTTATTGGATAAGTATTACAAGTGTCATATATAGTCCGGGCTTAGCCATTTAGGAATTGCTCAGTTGCTTGTACTCTTATCAGGACCTGCTGTTCTCTTATTTCGATCATGTTTCTTTCTTCTATTTAGGCCTTAGACATGCAGCCTGCTTTTCAAAATATATTCTTATACTTGATAATGTGGAAGTTTTTCATTCTTGATTTAGTCTTCAAGGGAAGGTGGATTTCCTGGGGGTCAGCTGTGGACATTAGGAGGAGGGAGGGGAAGCTAGTACTTGTTTAGTTGCATGTAATAAATGTTAATCTGTGCTTTGGTAAGGTTAAACTAATTTTGTCAAGTTCCCTTCTTTGTTTGTATCAAGGTCTGGGCTTCTGGATTCTCTGCTGCCTTTTTAAAGAAGAATAGTGTTGCAACAAAACGCATCATGACTTTAAACGACAGAGATGTGACAATAGAAGCCAAAACTTCTGTTGAAACTAAGCCTGCAGTAACTGAAGCAGCGTTGCTATTAGTACTGGAAAAGTTCTTTGAGGTGGTTTTGAACCTTCCCACCGTAAATAGGTAAGGAGATGTCTTCCAGAATCACCTTTTCTGCAAATGTTTATCAGCCTGCAGTATTTTTAAACAGAAGTACCAATATTAAATCTACTTTTTGTCTTTGTACTCTACTCAATGTGCGAAGTAGACCTTGTTGATATCATGTCCATTCACCAATGGATTGTAATTACATTTTCAAGTCATTAGTGTGTATAGCTAGTGCCAAAAATAGTTTCGTGTTTGTACTTACATTGATGGTTGGATGAAATTGCAGGAGGCTTTCTGACGTTCTAGATCTATTGACATCCCGGCAACCAAGTTGCCCTCAATTATACATATACAGCAGTGCCGACAGAGTCATTCCTGCAGTTTCGGTTGAGTCCTTTGTAGAAGAGCAACGAAGAATTGGTCGCGATGTTAGAGCTTGCAACTTCGTTTCTACACCTCACGTTGATCATTTCAGAAATGACCCAGAATTGTATACTTCACAGCTTACCCAATTTCTAGAGGACTACGTCCTAAGCTCTTGCAAACAGCCTTCTTGAATAATTCCTAATCACAACACATTGATTCTTTCTGCCCTTTTTTTCCCTCACTTTAACCAAATACTTGGTAAAGCTTCTGTTTTCCAACTTTAAAGCAGATATACATTGATGTAGATTGTTACGTATACAGAGAATTAAGTTTTTTTTTCTTTTCCATTCTTATTTAATTTAGTTGTATCTATGGTCTCCCAGCAACAGAATTGTTGCATTGTACCCAAAATATACTCTACTTTACTACATATATATA

>M6-PGSC0003DMG400002885 M6 LRR-RLP

ATGTGGCCTCTCTTTTTCATTTATTCACTTTTCTGTTGTGTATTTCTATGTATTTGTTCTTCCTTTGAGCATCATCTTTGCTCTCCCAGTGAAGCTTCTGCTTTGCTTCAGTTTAAGCAATCCTTTAATATATTGGACTTCTCCCAGTGTGATACGTCTTTCCCAAAAACAGTTTCATGGAATGAGAGTAACGATTGCTGCACTTGGGACGGGGTCACTTGTGACATGTTAACCGGTCATGTTATCGGGTTGGATCTTAGTTGTAGTCAGCTTAATGGAACTATTTATCCCAACAGCAGCCTCATCCAACTTCATCATCTTCAGACACTAAACCTTGCTAAGAACTATGTTTATCCTTCTACGATCCCAAAAGACTTTAGCCGGTTGAGGAATTTGAAGCATCTCAATCTTTCTGATGCTTACTTTCAAGGGGAAATCCCAACAGAAATCTCATACCTTTCCAATTTGGTTTCACTTGATCTTTCTGTTCCTTCTTATGTGAACGGATTACAATTTGATCAGAGAGCATTTGAAGCAGTGCTTCAGAACTTGACAAATCTAGAGGTACTTTCTCTCTTTGGTGTCAACGTTTCATCTCCGATACCTATGAATTTGTCTTCCTCCTTAAGGTATGTCGATCTTAAGGTAACCAATTTGGAAGGTGTTCTCACGGAGAGCTTATTCCTTCTGCCAAAATTGGAAACACTCTATTTGAGTAATAACTATCTTCTCAAAGGAGTTTTACCAAAGATTCATCCAAGCAGCACTCTGTTGGAGTTGGATATTTCATACACAGGAATCTCCGGTGAGCTGCCTGATTCAATTGGCACCTTGAGTTCCTTGACTCGCTTGTACCTCTATGGATGTCAATTCTCTGGTCGCATTCCTGATTCCATTGGCAATCTTACACAAATTAGGTATTTGAATTTTGGTAATAATCATTTCACCGGCAATATTCCTTCCACAATCTCTAAATTGAAGCACCTCGCAGTTTTAGCTCTTTCATCTAACTCCTTTGGAGGTGAAATTCCTAATTTTTTCTCTAACCTCCGAGAACTACGTTATTTAAGTCTTTCTAATTGCAGCTTCATCGGTCCGTTTCCTTCTCCACTTTTAAGCTTGACACAACTTGAAACCTTAGACTTGTCGAGAAATTCCCTATCAGGACTACTTCCTAGAAATGCAAGCATGCTTCAAAAGCTAGCCGAGCTGGATTTGTCTTACAACTCACTGAATGGTACCATACCATCTTGGGTGTTTAGCCTACCTTTGCTATCTTCAGTGTCGCTCCATCATAACCGGTTAAGTGGTTCTTTTCCTCAATCACCTGTGAATCTCACAAACCTTTCTACCCTTGATCTTTCATCAAATAACATCACCCTCGATGCGGGAATACAAATCACTTTACCTAGCCTAGAAATTATGCAGTTATCATCTTGTGAACTGAAGGATTTTCCACACTTCTTGAGAAATGTAAAGACACTTATGGTGTTGGATATTTCTAACAATAAGATTCGTGGTCAAATCCCTAATTGGTTTAGCGGCATGAGGTGGGACTCCTTGTTGCACCTAAACCTTTCACATAACTCATTAACAGGCAACCTACA

>M6-PGSC0003DMG400002886 M6 LRR-RLP

CAACTCCTTTTCAGGTGAAATTCCTGCTGTTTTCTCTAACCTCCAAGAGCTACGTACTTTACATCTTTCTTATAACAGCTTCATCGGTTCATTTCCCGCTTCAATTTTAAGTTTGACACATCTTCAATACTTAGGATTGTCGAGTAATTCCCTATCTGGCCCACTGCCTAGCAACCAAAGCATGCTTCAAAAGCTAACTTTTGTGGATTTGTCATACAACTCACTGAATGGTACCATACCATCTTGGGTGTTTAGCCTACCTTTGCTATCTTCAGTGTCGCTCCAACATAACCAATTCAGAGGACTAGCCGATGAAGTGATCAAAACAAACCCAACATTAAAACAACTGCATTTAAGCAATAATCAACTCAGTGGTCCTTTTCCTCAATCACTTGTGAATCTCACAAACCTTGAAACCCTTGGAATTTCATCAAATAACATCACCATTGATGAGGGAATGAATATCACCTTTCTTAGCCTATCATCTTTATTCTTATCATCTTGTGAACTGAAGGATTTTCCACACTTCTTGAGAAATGTAAAGACACTTGTCTACTTGGATATTTCTAACAATAAGATATGTGGTCAAATCCCTAACTGGTTTAGCGGCATGAGGTGGGACTCGTTGCAGTTCCTAAACCTTTCTCATAATTCATTAACAGGCCACCTACCACAATTTCATTACTATAACCTACAGTATCTTGATCTGAAATGTAACTCCCTTCAGGGTCCACTACCTTCATCCATTTGTAACATGAGTTCGCTTATCTTATTAGATTTATCACGCAACAACTTCAGTGACTCAGTTCCACATTGCTTGGGAAGCATGGCTAGTCTAACGGTGCTAGACTTAAGAAGGAACAATTTCACAGGGAGTCTTTCCCCATTATGTGCACACAGCACTTCGTTGAGTACCATTGTCATAAATGGTAATCAATTTGAAGGACCTGTACCTGTGTCGTTGCTCAAGTGTGGTGGTCTAGAAGTACTTGATGTGGGGAACAATGCTATAAATGACAGGTTTCCATCTTGGCTCGGAACTCTCGAATACCTGCAGGTCCTTATATTAAAGTCGAACAAGTTCCATGGACCTATAAGTACGTGTCTGACTAAGTTTTGCTTTCCCAAGTTGCGAATTTTTGATCTTTCTCGTAATGATTTCAGTGGCTCACTTCCTGCAAAAGTTTTTGGAAACTTCAAGGCAATGATCAAATTAGATGGCGAAGACACAGGAAATATAAAGTACATGGACTCTGTGTTGTATTCGCCATTTGTCACATCGTATGAGGATTCAGTGAGTTTGGTAATCAAAGGGCAGGATATTGAGCTATAAAGAATCAGCACAATTACGACAACCATAGATCTCTCAAGCAACCATTTTGAAGGTGTCATTCCGAAAAC

>M6-PGSC0003DMG400002887 M6 LRR-RLP

AATTCCTGATGTTTTCTCTAACCTCAAAGAGCTAGTTGAATTGAATCTTTCTTTTAACAACTTCACCGGTCCATGCCCCTCTTCAATTTTAAGCTTGACAAGTCTTCAATACTTAGATTTCTCTCACAACTCACTGAATGGCACAATACACTCTTGGGTGTTTAGCCTCCCTTCCTTACGTGATTTGAAGCTCCATCATAACCAATTCAGTACAGTGGCTGATGAGATCAAAACAAACCCAACATTAGTGACTTTGTATTTAAGTCATAATCAACTCAATGGTCCTTTTCCTCTATCACTTGTGAATCTCACAAACCTAGCCCTCCTTGACTTTTCATCAAGTAACATCACCGGCGATGTGGGAATAAATATCACCTTTCCTAGACTATCAGCTTTGTTCTTATCATCTTGTGAACTGAAGGATTTTCCACACTTCTTGAGAAATTTAAAGACACTTCAGTTCTTGGATCTTTCTAACAATAAGATTCGTGGTCAAATCCCTAACTGGTTTAGAGGCATGAGGTGGGACTCTTTGGAACAACTAAACCTTTCACGTAATTTTTTGGCAGGCCATCTAGAACAATTTCATCATTATAGACTAGAGTCTCTTGATCTTAAATTTAACTTCCTTCAGGGTTCACTACCTTCATCCATATGTAACCTAAGTAGCCTTAGAATTCTGGATTTATCACGCAACAACTTCAGTAACTCGATTCCAAATTGCTTGGGAAGCATGGCTAAGCTAACGGTGTTGGACTTGAGAAGGAACAATTTCAGTGGGAGTCTTCCACTGTTATGTACTCAAAGCACTTCATTGACGACCATTGTCTTGAATGGTAATCAATTTGAAGGATCTGTCCCTATGTCGTTGCTCAACTGTGTTGGTTTAAAAGTTCTTGATTTAGGGAATAACGCTATAAATGACACATTTCCAGCTTGGCTTGGAACTCTTGAAGATCTGCAAGTCCTTATATTAAAGTCGAACAAGTTCCATGGACCTATAAGTTCTAGGAAGAAGTTTTGCTTTCCTAGGTTGCGGATTTTTGATCTCTCTCATAATGCGTTCAATGGCTCACTGCCTGCAGATTTTTTTAGAAACTTCAAGGCAATGACAAAAAATGGCACAGACAAAAGTGACATTAGCTATATGGAAACACCAGTTTACAGATCATTCCTTGAGTCAGTGAGCTCAGCCGTCATTCCTATATATAAGCTAGAAATGGCCATCAATGAAGTATACAAGGATTCAGTGAGGTTGATGATCAAAGGAAACAATATTGAGCTAGAAAGAATCAGCACAATTGTTACAGCCATAGATCTCTCAAGCAACCATTTAGAAGGTGACATTCCGAAATCACTAAAGGATCTCAGCTCACTTTGGCTACTCAATTTATCCCATAACAATCTCAAAGGTGATATTCCAATGGAATTGGGGCAATTGAATACACTTGAAGCCTTAGATCTCTCTTCGAATCGGCTCACTGGAAAGATTCCACAGGAATTGACAACAATGAACTTTCTGGAGTTCTTAAACCTCTCTCAAAATCATCTCGTTGGACGCATTCCTCAAGGTTCACAATTCAGCACATTTGAAAATGACTCGTATGGTGGCAACCTTGATTTATGTGGTCCTCCTTTGTCAAAGCAATGTGGAACGAGTGATCGCTCGCATGTTCCTCAATCACTGGAGGAAGAAGCAGAAGAAAATGAGTCATATTTTTTTAGTGGATTTACCTGGGAATCAGTAGTAATAGGCTACTGTTTTGGACTAGTTGTTGGAACAATCATGTGGAGCCTCATGTTTAATTATCGTAAGCCAAAATGGTTTGTGGAATTCTTTGACGGCATCATGCCCCACAAAAGGAGAA

>M6-PGSC0003DMG400002906 M6 LRR-RLP

AGTGTCCAACCTTGTCAAGCCCACTACTAAAAATGGAAAAAACAACAAATAGACCAGGCCCTTTCGTAAACCTGATATTCTAAGATAGGATTTTCGCCGCTATCTCCGCCTCTTCGCCTTAGCTCTCTTCACTGCTTCTGTACATGAACAACAATGTTGCCTCCATGTTACTGACCTTTATGCTTTTTTCCAGTTGTGTTCAAAAGTTTCTTCCTCTTGCTTTCTTTTCTTCTTTTCTCCCTCTCTTTGTCTCTGATTCTCATCATTTTGCAGGTGATTTTGCGATGGAGTGCAAGGGTAGTTCAATTCCTCCATCTCCAGGTTAAAAGCTCCATCTTAGAATAACAATAGCATTTTCCCTATTTTGTTGCCTTTAACTTCTATGTAATTTTGTTGTTTGTGGAAAAATATATGTTGTAGCCTTCCCTTGTCCCCTATCCCCTTAAGGATGATACTCTTAAATATGGGCATAGTAATTTGTCATTATTTATGAGTCTCCTCGCAGGTGTCTTACTGTTAAAATTTGTGAATGTGCAGCTCCCTCGGTCAATATCAAAGTTTGCCACATGATCTGCTAGCTTGTTTCCTCCCCTTAGAATATGTTGGAAAATAACTTGTTTATCTTGTAGACATGATTGAATCCCTAATAATAAAAAAAAAAAAATGGTTGAATCCCTGTCACCATTTCTGCAATTATCCATGGACATGCCCATTTCCTTTCTAATACTTTGTACAAAAGCATTGAGTCAGTCTGTATGATTATTTGATTATGTTGTGACTTCTTGCAATGCTTGGACGCCTTCAGAATAGCTTTAGCTACCGCGACTGTATTAGTAGTATTTTCTATACTTGCTCCTTCCACGTATTCTACATCTCCTTTAGCATTCCTTAAAACGAAGGCATAGGAACTAAGACCCGGATTTCCTCTAGAAGTTCCATTTGTATTGTATTTTACCCACCCATTCGGTGGGAATTCCCATAACCCTTTTCACTTTCATTTTAGGTTTGTAGCTTGTCCATTCTTAGAGCATATCCGGCCAGTTAACAGGACATGTCATAAAGGGTTTCCTGAGTCTTGCCAGCATATACATGTTCCTGGTGATATTGTGAATGACTCTTGCTACTGATGTCTTTTTTCCTTCATGCTTCATCCTGTTTCTCTTTCTCCACAATTCCCATATTATAAAACTAGAGCTCTGTAAAAAGGTCTCATATTTGTCCCAATTTTTGTCCCCCACCACAACATTATAACTTCTCTTACTTAGAGACCCTCCATGTTAAGACCTGCAAAAGAACAAAAATAGGACCAAGTCCTGTTAGCTGAATCAGATTTCAGAAACACATGGGCAAGTGTCTCATGTTGAGGTCTTTCACAACACCAGCATTTGGATGGGCCTTCAAGACCCCATCTTCTCATTATGTCATCAACGGGGATTTTAAATTTCCATAATCTCCACATTATGAAAGCCATTTTAAATGGTACACCACCCTTACTCCACATCCCTTTATAAATTTTGCTTTCCTCCGCCTTATGTCTGATATAATGCCAAGCAGATTTAACTGAGAAACTTCCCCTTGTCTCTGGCATCCAACAAGGTTTATCCATTGTTGCTTCTTCTTTTGGAGGATAGGTTTATCCATTGTTGCTTCTTCTTTTGGAGGATGAATATTCTGCAGAATATGGTTCACCAATTCCTGTGGGAGTATCTCATGTAGTACCTCTGTATCCCACTCCCCTTGTCTAGTTAAATCCCTAACTGTTTTATATGTATCATCACATTGAAATTCTTCTCCAGTAATTGTGTACAAATCCCTTAGGCCCGTCCAGTTATCATGCCAGACTGATACTGATCCTTTCTCCATTTTCCATAAAATTTGGTGTTCAATGAGATCTCTAGCTTGGAGCATTTTTT

>M6-PGSC0003DMG400002908 M6 LRR-RLP

ATGAAATTAGGACAATTGAGTATGCTTGAAGCTTTAGATCTCTCTTGGAATCGGCTCACTGGAAAGATTCCACAGGAATTGACAAGAATAAACTTTCTCGCCTTCTTAAACCTCTCTCAAAATCATCTCGTCGGACCAATTCCTCAAGGTCGGCAATTCAACACATTTGAAAATGACTCATATGGTGGCAACCTTGATTTATGTGGTGTTCCTTTAACAAAGAAATGTGGAACGGGTGATTCATCGCATGTTCCTCAACAATTGGAGTCCGAAGAAGAAGGCGAGTCATATTTTTTTAGTGGATTTACGTGGGAATCAGTAGTCATAGGCTACAGTTTTGGACTAGTTGTTGGAACTGTCATGTGGAGTCTCATGTTTAAATATCGTAAGCCAAAATGGTTTGTGGAATTTTTCGATGGACTCATGCCTCACAAAAGAAGAAGGCCAAAGAAGAGAGCTCAGAGACGATGGACTTAA

>M6-PGSC0003DMG400002910 M6 LRR-RLP

AACCTACAGTATCTTGATCTGAAATGTAACTCCCTTCAGGGTCCACTACCTTCATCCATTTGTAACATGAGTTCGCTTATCTTATTAGATTTATCACGCAACAACTTCAGTGACTCAGTTCCACATTGCTTGGGAAGCATGGCTAGTCTAACGGTGCTAGACTTAAGAAGGAACAATTTCACAGGGAGTCTTTCCCCATTATGTGCACACAGCACTTCGTTGAGTACCATTGTCATAAATGGTAATCAATTTGAAGGACCTGTACCTGTGTCGTTGCTCAAGTGTGGTGGTCTAGAAGTACTTGATGTGGGGAACAATGCTATAAATGACAGGTTTCCATCTTGGCTCGGAACTCTCGAATACCTGCAGGTCCTTATATTAAAGTCGAACAAGTTCCATGGACCTATAAGTACGTGTCTGACTAAGTTTTGCTTTCCCAAGTTGCGAATTTTTGATCTTTCTCGTAATGATTTCAGTGGCTCACTTCCTGCAAAAGTTTTTGGAAACTTCAAGGCAATGATCAAATTAGATGGCGAAGACACAGGAAATATAAAGTACATGGACTCTGTGTTGTATTCGCCATTTGTCACATCGTATGAGGATTCAGTGAGTTTGGTAATCAAAGGGCAGGATATTGAGCTATAAAGAATCAGCACAATTACGACAACCATAGATCTCTCAAGCAACCATTTTGAAGGTGTCATTCCGAAAACACTAAAGGATCTCATATCACTTTGGCTACTCAATTTATCTCATAACAATCTCAGAGGAGATATTCCAATGAAATTAGGACAATTGAGTATGCTTGAAGCTTTAGATCTCTCTTGGAATCGGCTCACTGGAAAGATTCCACAGGAATTGACAAGAATAAACTTTCTCACCTTCTTAAACCTCTCTCAAAATCATCTCGTTGGACCAATTCCTCAAGGTCGGCAATTCAACACATTTGAAAATGACTCATATGGTGGCAACCTTGATTTATGTGGTGTTCCTTTAACAAAGAAATGTGGAACGTGTGATTCATCGCATGTTCCTCAACAATTGGAGTCCGAAGAAGAAGGCGAGTCATATTTTTTTAGTGGATTTACTTGGGAATCAGTAGTCATAGGCTACAGCTGTGGACTAGTTGTTGGAACTGTCATGTGGAGTCTCATGTTTAAATATCGT

>M6-PGSC0003DMG400002914 M6 LRR-RLP

AAAAAAGAACCATTCTTTGTTTCTTCTCTCTCTAACCTCAAATCTTTCTCTCTCTATTTTTGCCCTACTTTGTTGGGGTTCTTGAAAAAAACTGGATTTTTGAGGGTTCTTGAAAAAAATTGGATTTTTTCATCTGGAATGCAAATCTTGGTTAAAGATATAATCTGGGTTGTGTTTTTCTTGGTTCAAAATCTGATCTTTTTTGCTTTACAGTTTGGGATTTCTTCTGAAGAACAGACATGCCTACTCTTGTTAATTACAGTGGTGAGTTTCAAAAACTTCATTTCTTGGTATTTTAAGTTATGGGGTTGTTCATATTTCAATGAATTTTGAAGTTCAGTTATAATCTTTTGTTTTTTCATCATGAATTACTGAGATGGAGATGGATTTGTTATGTTTAGATTCATATTAGTTGTTTATCATAAATAAAGTTTATAACTTTGCATAGTTACATGATTGTTGAGATGAATAGGAGCAATAATTTATTGAGATGAAGATGAATTTTTTAAGGTTTAGATCCATGAATATAGTCTATAACTATGTATAGGTGCATGATTGTTGATAACTTGAGATAAATATGGTTTATTAATTTACTGAAATGAAGATGATCTTTTAGGTTTAGATACAATCATGTTTTTTTTTTTTAAAGGAAGTTGGTTATGAGATAATAACTAAAAGTTGAGTGACTATCTAAGAAATTAACTCATATAACTTTGTATAGGTGCATGATTGTTGAGGTGAATATGAGCATTAATTTACTGAAATGGATAAAAAGGGAAAAAAGTTATGATAAGACTAGATCCTTTGGCTTTGAAGATTAATTTCTGTGTTTGATAGATCTTTTTTTGTCTTATGTTCTGAATCTATATTAGAGATTGGTAATATATTGAAGAAACTGTGGATTACTCTTTTGGGTTCTGTATCTAGAAACGTGTATCTATCTGATCTGATCTGAAGATGTATGTACATAATAGGTAGATCTAAAGGTTCAATTCACATGTGATAGTTGCTTATCTGTTATACTTGAAGTTGAAGTGAAAATTCATAAGAAATGGGCATATGGATGAATACTTGTAAAAAAATTTCATTAGACAAAATTTAGTATGCTTGCTTGTTGATTTAAGGGAAAATATAGCTGACAATGTATTTGCTCCTTGGATTTCAGGTGATGATGAGTTCTATTCTGGTGGATCGTTTTGTTCTGCAGATTTGGGTCTCATGTTGTCTTTGGGTCATGCTGATGTTTACTGTCCTCCTAGGAAGAGGGCACGCATTAGTGGCCCGTTTGTTGTCGAAGACCGGTCAAAGGATCCATCTCTTGAAATTCTTCCTGATGAATGCCTATTTGAGATCCTCAGACGATTGCCTGGTGGCCGAGAAAGGGGTGCAGCAGCTTGTGTTTCTAAGCGCTGGCTTACAGTGTTGAGTAGCGTGAGGAACTCTGAAATCTGCAGGAGCAAGAGCTATAACAATCTAAATGACACAATAATGATCTCTAAAGATGAAGATCTTGAAGTTGAATGTGATGGATACCTTACTAGGTGTGTGGAAGGGAAGAAAGCTACTGATGTTAGACTTGCTGCAATTGCAGTTGGGACGTCTACCCGTGGTGGACTAGGAAAGCTTTCTATCCGAGGCAGTAACTCAGTTCGTGGTATTACTAATGTTGGTCTATCTGCTATTGCACATGGGTGCCCTTCTCTTAGGGTCCTATCATTGTGGAATGTTCCTTCTATTGGGGATGAAGGTCTCTTGGAGGTTGCAAGGGAATGCCGTTCATTAGAAAAGCTAGATCTGAGCCACTGTCGTTCAATCTCCAACAAGGGTCTTGTTGCAATAGCAGAGAATTGCCCGAGCTTGACTTCATTGACCATTGAATCCTGTCCAAAGATTGGAAACGAAGGCTTGCAAGCTATTGGAAGATGCTGCACCAAACTACAGTCCCTTACGATTAAGGACTGCCCACTTGTTGGAGATCAGGGAGTTGCTAGTCTTCTGTCATCAGGTGCTTCAATGTTATCGAAAGTGAAACTACATGGTCTAAACATCACAGATTTCTCGCTTGCTGTCATTGGTCACTATGGCAAGCTGATTACCAGTCTTAATCTTTGTTCACTTCGTAACGTGAGTCAGAAGGGGTTTTGGGTCATGGGTAATGCTCAAGGCCTCCAGTCTCTGGTTTCTTTGACAATCACCTTGTGCCAGGGAGCCACAGATGTCGGTCTTGAAGCAGTGGGAAAGGGATGCCCAAATCTCAAACATATGTGCATTCGCAAGTGTTGCTTTGTTTCTGACGGTGGACTTGTTGCTTTCGCTAAAGAGGCTGGATCTCTCGAGAGTTTAATCTTGGAGGAGTGCAACAGAATTACCCAAGTAGGTATCCTTAATGCAGTTTCAAACTGCAGGAAGTTAAAGTCTCTTTCCCTAGTGAAGTGCATGGGAATCAAGGATCTAGCTCTACAAACTTCCATGCTGTCTCCATGTGAATCTCTTCGGTCCTTGTCTATCCGAAGCTGTCCAGGATTCGGGAGCACTAGCTTAGCTATGGTCGGTAAGCTCTGCCCCAAGCTGCATCAATTAGATCTCAGTGGGCTTTGTGGAATAACCGATGCTGGTCTTCTCCCACTCTTGGAGAACTGTGAAGGACTTGTTAAGGTAAATCTTAGCGACTGCCTGAACTTGACAGACCAAGTGGTGCTTTCATTGGCTTCGCGACATGGAGAGACCCTTGAATTGTTGAATCTCGATGGATGCAGGAAGGTTACTGACGCAAGTTTAGTGGCAATTGCAGATTATTGCCCATTACTTATTGATCTTGATGTTTCTAAGTGTGCAATAACTGATTCTGGTGTAGCTGCTTTATCCCGTGGAGTGCAAGTGAATTTGCAGGTCCTTTCTTTATCAGGTTGCTCCATGGTATCAAACAAGAGTGTCCCTTCTCTTAAAAAGTTGGGAGAGAATCTACTCGGCTTGAATCTCCAACATTGCTCCATTAGTTGCAGCTCAGTCGAGCTTCTTGTGGAGGATTTGTGGAGGTGTGACATCCTCTCCTAAACCAAAAGGGCACAAGGAAATGCATCAAAATGGTCAAAAAGAATAAGCTGAAAGCTATAATGGTTCATTCAGCCTGGCAATTACAGAAGCAAACATGAGTGTTCTTTTTTTAGCCCGTATATTCGTGGCTATCTAATACTTCAGTGTCTCGGTTGTTTTCCAGGAGGCTATGGCATCACTTTTTTCCTCCATTTTTTGCAGGCTTCGTCAAATCGAAGCCTGTTTCATCCATTTTTGGAGACAACTCCATCATGCCATGGTTTCCTGAGAATACAGTCACCTAGTACTGAGATTAAAATTTTCACTCATGTTGCTCCTTGAAAGAATCAGGCCGTTTGAATCATTCTCAGCTACAACATCGACGGACAATATCTATCGCGCATTTGCCTCCTTGGTTAGCTACATTGCAGAAATCAGACAGTAGTACTGTAGTTTTTCATGTTTTAGTGTTTCATTGTAAGTATTCTGTTGTGTGTCTTTTAGACTTTAGTTGCTTTGAGCAAATTTCTGTTGTTGTGGTGCCTGTTATTCCAGGGGCTTGGCCATTACAACCTTATGTAGGTTGTTATTTTTTAACCAGTCCCTAGTTTTTTGATAGGATTATCAATGTCCTATTTTTTACCAGGTTCCACATATTCAGATTTGTACTCTTGCAACTATGTTTTATATATGATGGAAGTACTAGTTTGTTTTGAA

>M6-PGSC0003DMG400002937 M6 LRR-RLP

GATTGATTTGTAAGAAGAGAACATCCATCCTGTTTTTGGCTGAATACGAAAAAGAAAGATGAGACGTTTCTTTTTCTTTTATTCATTTCTCTGTTTTGTGTTGCTAATAAGTCGAAGTTTTTCTTCTTCCTTTGAGCATCATCTTTGCTCTCCCACTGAAGCTTCTGCTTTGCTTCAGTTTAAGCAACACTTTGAGATTATATCCGACCCTTATTCCCGATGTCCTATTCATTTCCCAAGAACAATGTCTTGGAATGAGAGTAGGGATTGCTGCATTTGGAATGGAGTCACTTGTGATATGTTAACCGGTCGTGTTATCGCGTTGGACCTTAGTTGCAGTGAGCTTTGGGGAAGTATTCATCCCAACAGCAGCCTCTTCCAACTTCATCATCTCCACACACTAAACCTTGCTTACAATGACTTCAATTATTCTTCAATCCCACATAACATTGGCCGATTGAGGAATTTGAGGCATCTCAACCTTTCTTGGTCTCGCTTTGATGGAAAAATCCCAATAGAATTCTCATATCTTTCCAATTTGGTTTCACTTGATCTTTCTGCGAATGGATTACAACTTGATGAGAGAACATTTGAAACAATGCTTCAGAACTTCACAAATCTGGAGCTACTTTCTCTCTTTCAAGTCAACATCTCATCTCCGATACCTGTGAATATGTCTTTCTCCTTAAGGTACGTGGATCTTGGAATCACTAATCTGCGAGGTGTTGTCCCAGAGAGTCTTTTCCTTCTGCCAAACTTGGAAACGCTCTATTTGAGTAATAACTATCTTCTCAATGGAGTTTTACCAAAGATCCACCCGAGCAACACTCTGTTAATGGAGTTGGATATTTCATACACAGGAATCTCTGGCTCCATTCCTGATTCAATTGGCACCTTGAGTTCCTTGAATATCTTGATCCTCCAAGGATGTCAATTGTCGGGTTCCATTCCTGATTCCATAGGCAACCTAACACAAATTACGGAGTTGATTTTATCTTTTAATCATTTCACTGGCCATATTCCTTCAACAATCTCTAAATTGAAGCACCTCACACGTTTGGATCTTTCTGACAACTCCTTTTCAGGTGAAATTCCCAACGTTTTCTCTAACCTCCAAGAGCTACGTTATTTAAATCTTTATGATAACAGCCTCATCGGTTCATTTCCTTCTACAATTTCAAACTTGACACATCTTGAAAGCTTAGCCTTGTCGAGTAATTCCCTGTCTGGCCCACTTCCTAATAATGCGAGCATTCTTCAAAAGCTAACCCACATGGAATTGTATAACAACTCACTGAATGGTACCATACCGCCTTGGGTGTTTAACCTCCCTTTAGCATCTACATTGTGGCTCCATCATAATCAATTTAGTGGACTACCCGATGAAATCAAAACAAACCCAACATTAGAATACCTGTATTTAAGCAATAATCAACTCAGTTTTTTACACCAATCACTTGCGAATCTCACAAACCTTTCTACCCTTGACCTTTCAACAAATAACATCACCGGTGATGCAGGAACAAATATCACCTTTCCTCGCCTACAAGTTTTGCAGTTATCATCTTGTGAACTGAAGGGATTTCCACACTTCTTGAGAAATGTAAAGACACTTAAGGTGTTGGATATTTCTAACAATAAGATTCGTGGTCAAATCCCTAACTGGTTTAGTGGCATGAGGTGGGACTCCTTGTTGCACCTAAACCTTTCACATAACTCATTAACAGGCCACCTACAGCAACTTCATTATAATAGTCTAGAGTCTCTTGATCTTAAATTTAACTTCCTTGAGGGAGCACTACCTTCATCCATTTGTAACATGAGCAGTCTTAGCCTTTTGGATTTGTCAAACAACTACTTCAGTGACTCGGTTCCACATTGTGTGGGCAGCATGGTTGGGCTATCTGTGCTAGACTTGAGAAGAAATAATTTCACTGGGAGTCTTCCACCGTTTTGTGTGCAGAGCAATTCATTGAGAACCATTGTCTTGAATGGTAATCGTTTTGAAGGAACTGTCCCGATGTCGTTGCTCAAGTGTGATGGTCTAGAAGTCCTTGATGTGGGGAACAACGCTATAAATGACACGTTTCCAGCTTGGCTCGGAACTCTTCAAGAGCTGCAGGTACTTATATTAAAGTCCAACAAGTTCCATGGACCTATAAGTACTAGGCTGAAGTTTGGTTTCCCTGTGTTACGGATTTTTGATCTCTCTCATAATGAGTTTGGTGGCTCACTGCCTGCAGAAGTTTTTGAAAACTTCAAAGGAATGATCAAAACAGATGACAGAGACAAAGGTGAGATTGAATATATGCAACCACAGTCTTACAGCGGATTTAGTAACGTAATGTATGAGGTTTCAGTGAGGTTGGTGATCAAAAACCAGGAAATTCAGCTAGAAAAAATCACCACAATTATGACAACCATAGATCTCTCAGGCAACCATTTTGAAGGTGTCATTCCGAAAACACTAAAGGATCTCAACTCACTTTGGCTACTCAATTTATCTCATAACAATCTTAAAGGAGATATTCCAATGGAACTGGTGAAATTGAATATGCTTGAAGCGTTAGATCTCTCTTGGAATCGGCTCACTGGAAAGATTCCACAGGAATTCACAAGAATGAACTTTCTGTCATTCTTAAACCTCTCTCAAAATCTCCTTGTTGGACGCATTCCTCAAGGTTCACAATTCAACACATTTGAAAATGACTCGTATAGTGGCAACCTTGATTTGTGCGGTCTTCCTTTATCAAAGAAATGTGGAACGAGTAATCCATCGCATGTTCCTCAACCATTGGAGGAAGAAGAAGAAGATGATGAGTCATATTTCTTAAGTGGATTTACGTGGGAATCTGTAGTCCTAGGCTACAGCTTTGGACTCGTTGTTGGAACTGTCATGTGGAGCCTCATGTTTAAATATCGTAAGCCAAAATGGTTTGTCAAATTTTTTGAAGGCATTTTCCCCAAGAAAATGAGAAGGTCACAGAAGAGAATGCTGAGAC

>M6-PGSC0003DMG400002954 M6 LRR-RLP

TTTTCTAATAGATTCTAAAAGAGTTGATGCTGATATTGAAGATAGATTCAGTGCTCTACCAATAAACCTTGTAGATGATATTCTAAGGCTCTTACCAGTTCACGACGCAGCAAGAACTAGTATTTTGTCAAAAAATTGGAGATATATTTGGGCCGCGCTTCCCAATTTAATGCTTGATAAGACCTTTTGCAGTAAATTAGCTGTAAAATCTCAATTTGTATTGAAAGAAACAATAGATATGATTCTCTTACAACATATTGGAGATATTGTGAAGTTTGATCTTGATGTTTCAGGAGTACAATTGTGTTCATATGCAGACATCGATAGATGGATGCTTTATGTCACGAGATATGGTGTCAAGGAATTAACCCTTTTCATGTCAAAGAATATTAACAGTACTTATAAAGTGCCTTCTTATATATTCAATTGTCAAACCCTGACACAATTGAAACTCTTCAATTGCATCTTCAAATTACCAAATTCTTTTCTTGGGTTTCAAACTCTTACCATCCTTCAGTTGAAAGAAGTAACCTTCGCGCCTGCAATAGAGTTTTATGCTATCAAAACGCCACTTCTTGTCGAATTAATCTTGACACATTGTGAGGGCACTCAATATCTTAACATTGAGTTCTCATCAGGATTGAAGTACTTGTTTGTTCGTGAGAGTCACTATAATCTTGATTTAAATAGCAATTACTTTAAGAACGTCAAAAAATTGACATACTTATACCTTGTGATTGATAATCCAATACGTGTTGATGAAAGATTAACTTTGGAAAAGCTTCTTTTTAGTTCGCCTACACTTGAGACACTTACTTTGGGTTTTTATGTACTTGAGGTAAGAAGCTAACATGTTGTTTTACTTAGATATCTACGTTATCCTTCTGTTTGTATAATTTTTCGTTAAAGCTTTTGAATGCAGGTACAGTCCCAAAGGGGCTTCATTATACACTCAACTGCTTGTGGCATCTATAGTTAGGTGTAAACTTCAACAAATTGGGTCAGACTTGTTACACTCTGGAGTTGATTAAAAGTTTCCCCAATTTGAGTAAACTTGAGATTTGGGTACGAAATAGCAAATATATATGTATGATCTTGTTCGTTTTTGCCCTTGGCATACTTAAGTATGAGAACTAAAATGACGTTGTATTCTTTTAGGATAATGCTTTCGGTGACCCTGATGAAGCAGTTATGAAATATCTAGACGCACCATCATGTTTGGACCAAACACTCAACAAGCTTGAAGATGTTTCCATCCATATTTTTAGGCGATCCAAAGTTGTACTATCTTTCATAAAGTTATTGTTTGCTCATGCTCCATCTTTGTCAAGGATGAGCATTAAGCCGAAAAATGCTTCTAGTTCCAAAGAAGAATTGAATGTTGCTACAGAGTTGATGTGTTTCCCCAGAGTATCTCCTAAGGCAAAGCTATTCTATCATCCAACAGTTGGATTAA

>M6-PGSC0003DMG400002960 M6 LRR-RLP

ATACACAAGAGTAGTACATTATTTGATATCTTGAGTTACATCTAAAAATTATTTTTGTTGTAATTCCTCTCCATATTGACTTGATGGCAGCTTATACTGCTGTAATTTCTCTTCTTCAAACTCTTGATCAATCAAATATTTCAGAACTCTTTCATGATCACAGTACTGAAATGCTCGATTCTCTTCGGGCTACAGCTGAATATTTTCAAGATGTTCTTGACAACAATAGCAATGAGAAAATCAAATCTTTTGAGGAAGATATTAGAGTTGTTGTTAGTGAGGCAGAGGATGTTGTTGAAATGAAGATTTCTGAAATCATCAAAGAAGAAAGCTGGACATTTGGAATTTTACAACACCATGATATGCTACCAGTTGTTGAAAAAATGGATACGACAAAGAAACAAGTGATGGAGCTTCTTTCTCATGATGCTGATCAAATTCTTGAATTAACCGCGGATTCCTTGATTGACGCTTCTTCTATGAGTGATCCAATGCTGTCAGATCACTTGAAAGATCATATTGTGCAGGGACTTGATGGTGACTTGGAGATAATAGATAAGAGATTGAGAGGACCAACGCCTTATCTAGACATTGTCACCATATCAGGTATGGGTGGCATTGGCAAAACAACACTCGCTAAAAAAGCTTATGATCATCTTCCAATTAGGTATCACTTTGACATTTTTGTTTGGGTTACAATATCTCAAGAATTTCGATATAGAAATGTATTGTTAGAAGCTTTACATTGCATTTCAAAGCAAAGAGTTATTGTGAACACAAAAGATTATGATAAGATGAATGACAGTGAGTTAGCTGATCTGGTGCAAAAGAGTCTAAAGGGTCCTAGATACCTTGTTGTTGTTGATGATATTTGGAGTACGGATGCTTGGGATAGCATATCACAATTATTTCCTAATCGCAACAATGGGAGTCGAGTCTTATTGACTACTAGGGAAACCGATGTAGCGATGTATGCAAATACTAGTAGCCCTCATAAGATGAACCTCTTGGATTTAGATAACAGTTGGAAGTTACTTCGTGATAAGGTGTTTGGACTAGAACACGATCACCCTCCCGAGTTGGAAGAAATTGGAAAAACAATAGCAGGAAAATGCCAAGGACTTCCCTTAATAATTTCAGTGATTGCAGGGCATCTCTCTAAAACGGTCCCCAAGACATTAAAAAGTTGGAAGTATGTTGCCAAAACCTTAAGTAAAATCATTGCTAGTCATCCAAATAAATGCTTAGGAGTGCTCGGTTTGAGTTACCATCACTTGCCTAATCAACTCAAACCATGCTTTCTTTCTATGGGTGATTTCCCAGAGGATTTTCAAGTTGATACTTGGAGATTGATCCAATTATGGATCGCAGAAGGTTTCATAAGGACGTCTTCTGGAAGTCATGAAAGCTTGGAAGAAGTGGCTAAACATTATTTGGAGGATCTTATCAGCAGGAACTTGATAATGGCAGTAAAAAAAAGATTCAATGGTGAGGTAAAAGTATGCGGAATACATGATCTGCTGCGTGAGTTCTGTTTGATAGAAGCTGAAATGACAAAGTTTATGCATGTTGTGAGAACTCCATTGTCACCAGCACAAAAGCCTAATTTTCGTCGCTTCAGTATTCAAGAGTAATATTCTGTTGAGGATTCTTATAAGCTCTTACCCCCTGTAGCCAGATCTATCTACTGCTTTTCTTACTTGGATCTACCTTATGAACCCCGTATTCTTCCCCCTGTATTGTCCATCTACCATCATGATCCTATAATACATGGATTTTTCTGTTGTTTCAACCATCTCAGGGTATTGGCCATCTTTCCTTTCTCGTTCAGCTCATTTCCACTTGTGATTACAAAGTTATTTCATTTGAGATATCTCCAAGTTCGATGTCTTGGAAATATTCCTGAATCAATCTCAAAGCTTCTGAATTTGCAAACTCTAATTTGTGATGGTAATTCTTTAGAGATAACTTTAACTAGGAAGATATGGATGTTGAAGAACTTGAGGTATATACGTGTAGGGGGAGTTACTTATTTACCGAGTCCTAGAATAGATAACAATCTTGTGATAGGGATGCCAAATCTAGAGGAATTTTTTGGTCTTAGTTACTCTAGTTGTACAAATGAAGTCTTTTCTAGGATCCCCAATTTAAAGAGATTGATCATTCCTGCACATTTTAGCATACTAAACAATATTCTGTATCGTCCCTTCGATATGTCCAGCTTGACAAAACTTGAGACATTCAAGTTTTACTGGAAAGATTATTTGGAAGACCCCATCAAGAAATTTGTTTTTCCAGAATCACTTAGAAGAGTGTCTTTAACTTGGTGTTGTAAGTTTATTTGGGAAGAGATATCATCAACTTTTATCATGTTGCCACATCTTGAAGAGCTCAAACTTAAATATTGTCGAGCCGAAGATGATGTATGGAGATTGAGTGACAAAGACATATTCAGAAGCTTGAAGTTGTTGTTACTGAGCGAGCTAAATCTTAAGGGTTGGGAAGCAAGCAGCGATAACTTCCCTAATCTAAAACGCCTTGTTCTGAAGAAATGCAAGGACCTGCAAGAAATTCCAACAGATTTTGGGGAAATTTGTACTTTGGAATCGATTGAGTTACATGATTGCAGCACTACTTCTGAGGAATCTGCCAGAAATATCGAACAAGAACAAGAGGTCATGGGAAATAATATCCTTAAGGTCTACATC

>M6-PGSC0003DMG400002961 M6 LRR-RLP

ATGACAAAGTTTATGCATGTTGTGAGAACTCCATTTTCACCAGCACAAAAGCCTAATTTTCGTCGCTTCAGTATTCAGAAGGCATTTTCAGATGTGTATGCTTATAAAGTACTACCCCCTGTAGCCAGATCTATGTACTGCTTTTCATTCTTGGATCTACCTATTGAACCCCATATTAAGCTTCTCCCTAGATTGCCCATCTACCGTCATGATCCTATAATACATGGATTTTTCTCTTGTTTCAACCATCTCAGGGTATTGGTCATCTTCCCTACCTGGTTCGGCCCATTTCCACTTGTGATTACAAAATTATTTCATTTGAGATATCTCCAAGTTCGATATTTTGGAAATATTCCTGAATCAATCTCAGAGCTTCAAAATTTGCAAACTCTAATTTATGATTGTCATTCTTCACATATGACTTTACCCGGGAAGATATGGACGATGAAGAACTTGAGGTATATACATTTAGGGAAAGCTAGTAATTATATACCAAATCCTAGAATAGATAACAATCTTGTGACGGGGATGCCAAATCTAGGGGAATTTTTCAATATTTGTTACTCTAGTTGTACAAATGAAGTTTTTTCTAGAATTCCCAATCTAGAGAGATTGAGCATTCGTGCATCTTCTAAAGTCAGAGAAAATTTTCTCTATCGTCTCTTGGATATGTCCAGCTTGAGAAAACTTGAGGTATTCAAGCTTTCCTGGGACAATTATTCGGAAAAGCCCATCAAGAGATTTGTTTTGCCAGCATCACTTAGGAAGTTGACTTTAGATTGCTGTAGTCATTTTATTTGGGAAGAGATATCGTCAACTTTTATCATGTTGCCAAATCTTGAAGAGCTCAAACTTAAAGATTGTGAATCCGAAGATGATGTATGGAGATTGAGTGACAAAGACATATTCAAAAGCTTGAAGTTGTTGTTACTGAGCTACCCAAATCTTAAGTGTTGGGAAGCTAGCAGTGATAACTTCCCAAATCTAAAACGTCTTGTTCTGAAGAATTGCCAGGACCTGCAAGAAATTCCAACAGATTTTGGGGAAATTTGTACTTTGGAATCGATTGAGTTACATGATTGTAGCACTACTTCTGAGGATTCTGCCAGAAATATTGAACAAGAACAAGAGGACATGGGAAATAATATCCTTAAGGTCTACATCCATAACAGCCACAGTAAGTCATAA

>M6-PGSC0003DMG400002980 M6 LRR-RLP

AGTTGGTGGTGCATTTCTCTCTTCAGCTTTGAATGTTCTGTTTGATAGGCTTGCTCCTCAGGGTGATCTGCTCAACATGTTTCGGAAGCATACGGATCATGTTAAGCTCTTAAAGAAGCTGGAGGACATTTTGCTTGGTCTTCAAATTGTGCTAAGTGATGCAGAGAATAAGCAAGCATCAAATCAACATGTGAGCCAATGGCTTAATAAGCTCCAGAGTGCTGTGGATGGCGCTGAAAACATGATAGAAGAAGTCAATTATGAAGCTTTGAGGCATAAGGTGGAAGGCCAGCATCAAAATCTTGCAGAAACAAGCAACAAACAAGTAAGTGACCTCAACCTGTGCTTGACTGATGATTTTTTCCTTAACATAAAGGAAAAGTTGGAAGAAAACATTGAAACATTGGAGGTGTTGGAAAAGCAAATTGGTCGCCTTGGCTTAAAGGATCATTTTGGTTCAACTAAACAAGAAACTAGAACACCTTCAACGTCTTTGGTTGATGATGCTGGTATCTTTGGAAGGCAGAATGAAATAGAGAATTTGATTGGCCGTTTGTTGTCTATGGATACACAGGGAAAAAATCTGGCTGTAGTTCCTATTGTTGGAATGGGCGGCCTGGGTAAGACAACACTTGCTAAAGCGGTTTACAATGATGAGAGAGTGCAGAAACATTTTGGTTTGAAAGCTTGGTTTTGTGTTTCTGAGGCATATGATGCTTTCAGAATAACAAAAGGTTTACTTCATGAAATTGGCTCAACTGACTTGAAGGTTGATGACAATCTTAATCAGCTACAAGTCAAATTGAAGGAAAAGCTGCATGGAAAAAAGTTTCTTGTTGTCCTTGATGACGTGTGGAATGATAACTATCCTGAGTGGGATGACTTGAGAAATCTTTTTCTACAAGGAGATATAGGAAGTAAGATCATTGTGACGACACGGAAAGAGAGTGTTGCCTTGATGATGGGTAGTGAGGCAATCAACATGGGAGTTCTGTCTAGTGAAGACTCTTGGGCTCTATTCCAACGACATTCACTAGAAAATAGAGATCCTGAGGAACATCCAGAATTTGAAGAGGTTGGAAAACAAATTGCAGACAAGTGCAAAGGGTTGCCTTTAGCTCTAAAAGCACTTGCTGGTGTTTTACGCGGCAAATCAGAGGTGGATGAGTGGAGAGACATTTTAAGAAGTGAAATATGGGAGCTGTCAATTTGTTCGAATGGTATATTACCAGCGCTGATGTTGAGCTACAATGATCTTCCTGCACATTTGAAGCAATGTTTTGCTTATTGTGCAGTATATCCCAAAGATTATCTATTTTGCAAAGACCAAGTTATTCACCTGTGGATTGCTAATGGTCTGGTACAGCAGTTTCATTCAGGTAACCAATACTTTCTCGAGTTGAGATCAAGATCGCTGTTCGAAATGGTCTCGAAGTCTTCTGAACGAGACATAGAGAAATTCTTAATGCATGACCTTGTCAATGATTTGGCCCAAATTGCATCTTCAAATCTTTGTATAAGGTTGGAAGAGAACAAAGGATCGCACATGTTGGAACAATGTCGGCACATGTCCTATTCAATAGGACAAGATGGTGACTTTGAGAAATTGCAATCACTCTTTAAATCAGAGCAGCTGAGGACATTACTTCCAATCAATATTCAGCTTCAGTACCAAACTAAGCTAAGCAAGAGGGTGCTGCATAACATATTGCCAAGACTAACATCCTTAAGGGCATTATCGTTGTTAGGCTATAAGATTAAGGAGTTGCCAGATGACTCGTTTGTCAAATTAAAGCTCCTCAGATTTTTGGACCTTTCTAAGACAGAGATTAAAAAGTTGCCAGATTCAATTTGTGGATTGTATAACTTAAAGGCACTTCTCCTGTCATCTTGTGCTGATCTTGAGGAGCTACCACTGCAGATGGAGAAGTTGATCAACTTACATTACCTCGACATAAGCAACACTTTTCACTTGAAGGTTCCCCTACATCTGAGCAAGTTGAAAAGCCTGCAAGTGCTAATGGGAGCCAAGTTTGTTCTCGGTGGTTTGAGAATGGAAGATTTGGGTGAAGCACAGAACTTGTATGGATCTGTATCAGTTGTAGAGTTGCAAAATGTAGTTGATAGAAGGGAAGCTGTGAAGGCAAAGATGAAGGAGAAGAATCATGTTAAGAAGTTATATTTGGAGTGGAGTGAAAGTAGTAGTGCCGACAATTCACAAACAGAAAGAGACATACTTGATGAGCTATGCCCACATAAAAACATAAAAGAAGTCGAAATCATCGGATATAGAGGGACAACATTTCCGAATTGGTTAGCTGATCCTTTGTTTCTTAAGCTAGAACAGTTGTCTATTGCTAACTGCAAGAACTGTTTTTTCTTGCCAGCACTAGGACAGCTTCCTTGTTTGAAAATCCTTTCCATTAGAGGGATGCATGGAATAACAGGGGTGACGGAAGAATTCTATGGCAGTTTGTCCTCTAAAAAGCCTTTTAATTGTCTTGAGAACCTTATATTTGAAGATATGCCGGAGTGGAAGCAATGGCACGTACTAGGAAGTGGAGAGTTCCCTATACTTGAGAAGCTTTTTATTAAAAATTGCCCTGAACTCAGTTTGGAGACACCCATCCAACTTTCAAATTTAAAAAGGTTTGAAGTTGGTGGTTCTCCAAAGGTTGGAGTTGTTTTTGATGATACTGAGCTGTTTAGATCCCAACTTGAGGGAATGAAGCAGATTGTTGAATTAGATATACATGATTGTAACTCTCTTACCTTCTTTCCTTTTAGCATACTGCCCACTACCTTGAAGAGAATAGACATATCTGGTTGCCAGAAATTGAAATTGGATCCGCCAGTTGGTGAGATGAGTATGTTTCTGGAGTGTTTGAGTCTGGCAAAATGTGATTGTATAGATGATATATCACCTGAGTCGCTCCCAAGAGCACGCAGATTGTATGTATCTGATTGCCACAACCTTACTAGGTTTTTGATTCCTACTGCCACTGAAAGTCTCTGTATTTGGTATTGTGAGAATGTTGAAATACTCTCGGTGGCATGTGGGGGGACCCAGATGACGAAGTTGAATATTCAGGGCTGTAAGAAGCTGAAGTGGCTGCCAGAACGTATGCAGGAACTCCTTCCATCTCTTAAGGAACTGCATCTGATTAATTGTCCAGAAATAGAGTCCTTTCCTGAAGGAGGATTGCCCTTCAATTTACAACTCCTTATGATCAGCGAGTGCAAGAAACTGGTGAACAGCCGAAAGGAGTGGCGTTTACAGAGACTCCCCTGTCTTAGAGAGTTAACGATCGAACATGATGGCAGTGATGAAGAGATTGTTGGTGGTGAGAATTGGGAGTTGCCTTCCTCTATTCAAACACTCTATATAAAGAATCTGAAAACATTAAGCAGCCAACATCTCAAACACCTCACCTCTCTTCAATATCTAGACATTGCGGGTAATTTACCTCAAATTCAGTCAATGCTGGAACAAGGCCAGTTTTCGCATCTCACTTCGCTTCAATGTCTAGTAATCATGAATTTCCCTAATCTCCAATCACTTTCCGAATCAGCACTGCCCTCCTCCCTCTCTCAGCTGACCATCAAGGATTGCCCTAATCTCCAATCACTTCCTGAATCAGCACTGCCCTCCTCCCTCTCTCAGCTGGTGATCTCCTATTGCCCTAATCTCCAATCACTTCCTGAATCAGCACTGCCCTCCTCCCTCTCT

>M6-PGSC0003DMG400003195 M6 LRR-RLP

GAGCTTTAAATACTTCTTAATTACAAGTTCTAGAAGGAGTCAGAGAAATTTATGAGCCATCCGGGATTGAATCCTTATCTCTCATGCACTTTATCGACCACTAGACCAAATCACTGAGGTGACGGGTGTCTAGTCATTTATATATAAAGTCGCAACCATCTGAATTTTTCCTGTTTGAATGTCAAAGATACTCATAGCACGATCAAACTGTCAGTCAAACAAAAACAATAGAGTTGACATTTAACATATATGCACACAAAATTGCCTAACTATGATGCAAGCTGCATAATGATTTGACAATCAATGCGGCTGGATTTTAATTACTTGTTCTAGAAGGAGTCAAGGAGATGTTCATGAGCAACCCAGGATTTGAATCCTGATCTGGCATGCACTTTATCGACCACTAGATCAAATCACTGGGGTGACGGATGTCTAGTCATTTATATATAGAGTCGCAATCATCTGAATTTATCAAGTTTGAATGTCAAAGATACTCCTAGCGCGATCAAACCGTCAGTCAAACAGAAAAAATAGAGTTGACATTTAACATATACAAACACAAAGTTGCCTAACTATGAATGCAAGCTGCGTAATGATTTGACGATCAATGATGCTGGATTTTGTTACTTGTTCTAGAAGGAGTCAAGGAAGATATTTCATACCTCAAGTATATAAAATCTTGTGAATTATTCCTTCATTCCTCAAGTACTTATTATTTTTGTGCAGATCATGGGCTGTCTATTTCTTCTTCTTCTTCTTCTTGTGTTGCAATATCCACTTGTTATCTTTTCAAGTCATCTATGTCGACGAGACGAGAGTTTGGTTTTGCAAAACCTGAAGCTTAAACTGTCCATAGATGACTCTGCTTCTGCTGATTGTGAATCCAATGGTTACTTTCCCTATCCAAAGACACTTTCTTGGAATAGAACCACTGATTGTTGTACTTGGGATGGTGTAACATGTGATAAAGTTACAGGTCGAATAACTCGTCTTGATCTTAGTTGTAGCCAACTTCATGGAACAATTGACTCCAACAGCACCCTCTTCCAACTTTCTCATCTTCAAAACCTTAATTTAGCTTACAATGATTTCTCACCTTCCCAAATTTCAAGCAAATTTGGATGGTTTCCGAGTTTGACACATCTTAATCTATCTCGTTCTGGTTTTTCAGGGACAGTACCTTTACAAGTCTCATATCTATCTAAGTTGATATCACTTGATCTTTCTGCAGGTATTGAGGAGTTGAGAGTTGAACCTCACACTTTCAAAATGCTCCTTCAAAACTTAACTCAACTAAGGGAGCTTTACCTCACCTCAACATATATATCTTCATCACTTCCATCAAACTTGTCCTCTTCCTTGCAAGAGCTGAATCTTGTCGGAACGGAATTGTCTGGGAAAATCCCTGATGACATTTTCCATTTACCAAGATTGCAAATGCTTAACTTAGGGAGCAACTTGTATCTCACAGGGCATTTGCCAACAACTCAATGGAACTGCAGCAACTCTTTAAGGGAATTAGATCTTTCTTCCTCAGGCTTTTCGGGGAATATACCAGATTCTATTGGTCATCTCAACTCTTTAAGGAAATTAGATATATCAGGCTGCTACTTCTCTGGTAACATTCCTCCAACTATCGGTGATCTCACCGAGCTCACGAGTTTGCAACTCTTCTCCAACAACTTCAATGGTCAACTACCATCAACAATCTTAAAACTGGTGCAACTTGTTGAGTTTGATATCTCATCTAACAACCTTACAGGCAACATTCCAAATATTTTCGGTAACTTCACCAAGCTCAAATCTTTATCCTTTTCATATAACCTCTTTACGGGCCTTTTTCCATCCTCTGTTACAAACTTAACAGAGCTAGAGAGTTTAGACCTCTCAAATTGTTCTATAACTGGTCCTATTCCTTCTAGCGTCAGAACCGGATTCCCTAACCTCATCTTGCTCTTCCTACCATATAACTCCCTTAGCGGAGAAATACCATCTTGGATATTAAATCTTCCTTCCTTAAAATACTTAGTTTTGAGAGGCAATCAACTTACTGGTCAACTTAAGGAAGTTAGATACAACTTGTTAGAAGTTGTTGATGTCGGTGACAATAAGCTGAATGGACCTATTCCAACATCATTCTCTAAACTAGTGAACTTGACAACACTTGATCTTTCTACAAACAACCTTTCTGGTGGTCTAGATATTGGATTGTTTTCAAAATGCAAACAACTTCGACGTCTTGGCCTTTCTTTTAACAATCTGTCTGTATTTTCCAGTCAAAAAGATGCATCCTTGCCTAGTTCTCTTGGAAGTTTGTATGCATCTTCTTGCAACATAAGGGAGTTAAATTTCCTACGAGCTGCAAAATACATTGGTAACTTAGATCTTTCCAACAACAACATGCATGGAAAAATTCCTGATTGGGCATGGTCTAATTGGCAAGTCTCAGTGTCTTATTTGAATCTTTCTAGCAACTTTCTAACGGCTATTGACCTGCTTCATAACTTTGAGAGTCTTTACAATCTTGATATTGGATCTAACTTGATTCAAGGAGAATTACCTGCTCCACCTCCCTTATTGTTCCTCTTCATAGCCTCGAACAATAACTTCACCGGGAAGTTACCTAATTCACCATTATGCAAAATGAGTTCCCTTTTGATTCTTGATTTGTCCAATAACAGCCTGAGTGGAGTAATTCCAAAATGTTTAGTGAACATGAGCACAAGTCTCTCAGTGTTGGACCTGCACAACAATCATTTTAGTGGGACAATTCCAAATAACTTTGGTAAGGGAAGTAGTCTGAGAAGTCTCAATTTGCGGAAAAATAAACTAGAAGGAATGTTACCAAGAAAGCTTACCAATTGCAGAGGGCTCGAAGTAGTTGATCTCGGAGAAAACTTGTTGAATGATACATTTCCCAAATGGTTAGGATCACTTCCTAGGCTGCAAGTTCTCAGTTTGAGATCCAATAAATTACATGGCCCGATTACCATTTCAAGAAACCAAGTCTTATTCTCAAAGCTGAAAATCCTTGATCTCTCTTATAATGACTTCACTGGAAACCTGTCAGAAAGGTTTTTCAACAACTTGAAATCCATGATAATAGAAGATCAAACGGGAATACCTTTAACGTATATTGGAGAAGTTCGATACCATGATTCCGTGACATTGTCTATAAAAGGACAGCAAATTGAACTTGTGAGGATTCTTTCTATCTTCACAACCATTGATTTCTCAAATAACAGATTTAAAGGTAATGTACCAAACTCCATTGGAAATCTTGGTTCACTTCGCGGGTTGAATTTGTCACACAACAGCCTTATTGGACTTATACCTCAGTCTTTTGGAAATTTATCTGTTCTTGAGTCGTTAGACCTTTCATGGAACCAACTTTCTGGGAACATTCCTCAAGAAGTTGCTTCACTGAAATCTCTTGCTGTCATGAATCTCTCACAGAACCACCTGATGGGACGCATACCTAGAGGTCCACAGCTTGATACATTTGAAAATGACTCATATTCAGGAAATGCTGGATTATGCGGATTTCCCTTGTCAAGAAACTGCGGAGATAATGAGATGCCTCCTTCTACTGTGACTGATATTGAAGACGAAGAAGATTCAGGTTTTATGGATTGGAGAGCTGCGATAATAGGATGTGGTTGTGGATCAGTGTTTGGGATGTTTATAGTATATATCATTTTCTTAACAGGGAGACCAAAATGGTTTGTGAGAATTGTCAACCAACAGATCAACAGGATCCTTTTGAAAATTGAGGCAAAGAGAAGAATTGAGGCAAAAAAGAGAAGACGGCAGAAAAGAAGAAGGATGAAGTTTTAGTATATACTTCATGCATTATTGTCTTGCAACATACACAGTTTCAGCTTACACCTTGTTTTGGC

>M6-PGSC0003DMG400003380 M6 LRR-RLP

TCCATAATAGTTCTAATTCTTGTGTCACAAGGATGAATTTCAACAATGAATTGTCTGATCTGAGAGTCCCGAAATACTCGGATGTTCCAAGGGATCGAATAGATTTCTTTATATGGGAGTTAAAATTCCTTAATTGTTTTCTCCATTTGCAGAAATTCACTTTTGCAAGGGAATGTGGTATGCTAGATATCTCACAGAAAATGCTAGAAATTTGCAAGAGGTTGTGTCCATCACGTCGACATAATGCATTTGCATACTGGAAGGAGGTAATTTGCAAGAGGCTGTGCGCTATTATCATCCGGCCAGATGATGGATTTGCATACTGGAAGAAAGTAATTTGGAAGACTAAGCAAGAATTCAGAGCTAAATACTCCTTTCCAAAAACACTACTTGCAAACAACAAGGTAGATGATGATGATATTACTAATCCCAAATTTGTGATGGAATTCATCGATGCTGTTGTGGGGAATCTCAATGTTCTAGTCAAGATCAATGATCCATTTTCATTGCTTTTTGTTCCAGGACCCAAGGAACAAATAGAACAAGTGTTAAAGGAGTTGAAGTTATTGAGATTTTTTGTCTGCTTTGTTTCAAACAAATGTATAGAGCCTCAATACCAACATGCTACTTTTTATACTCACGCTTTAATTGAGGCTAGCCACATCGCAATGGTTGTGTGGTTGAATTTGCCAATCTATGGAAACAGAAATCAAGACTTGGCTTCAAATGAAGTCAGTTGTTTGCTTTCTGATTTCATGGAAATGAAGATTAAGTCCATTCAGCCAGGCATCAGCCGCAACAATATTTATATTGATGTCTTGAGGGCGTTGAAGTCAACCATACCACATGCTCAAGATAAGCATGCAGCTGAGAGTGGTATTGTGGAGACTCCAACACACAATCTGATGGTTGGTTTGAGTGATCAAATGGCAAACCTTCAGGAGATGCTCTGCCTTCTAAGAGACAATCTCATTCATCTGCCAATACTAGATCTGGAATTTCATCTACAAGATATGGATTCTGTTATTGTTGATGCAGGACTTCTTATTTATTCATTATATGATATCAAGGGGCAGAAGGAAGACACATCATTGGAGGTTATCAACCAGGCACTTGGTTTTGATCTTCCGAGAAACATTGAGTCTATTAAGGCGATGGTCTACCTTGTCATGCAAAAGGCATTTCATTGTAACTTGCCAAGGGTTCATGGACTAGGCTATGTCGATTTTCTGTTGAAAAACCTGAAGGATTTCCAAGGCCGTTATTCAGATTCACTCATTTTCCTCAAGAATCACCTTCAAGTTATTCAAACGGAATTTGAGAGCTTGCAACCTTTCTTGAAGGTTGTCGTAGAAGAGCCACACAATAAGCTCAAGACACTGAATGAAGATTGTGCTACACAGATAATTAGGAAAGCATATGAGGTGGAATATGTAGTTGATGCTTGTATTAACAAAGAAGCTCCTCATTGGTGCCTCGAGCGTTGGCTCCTGGATATCATAGAGGAGATTACTTGTATCAAAGCACAGATTCAGGGAAAGATCACAGTTGAGGATTCAATGAAGAGTGTCATTGCTCGTACATCATCACAATTGGCAAGGACTCCAAGCATGAATGAAGAGATTGTTGGGTTTGAGGATGTCATAGAAAATTTAAGAAAAAAACTACTGAATGGAACCAAAGGGCAAGTTATCATTTCAATTCACGGCATGCCAGGTTTAGGTAAGACGACTTTAGCCAACAGACTCTATTCTGACAGGTCAGTTGTTTCTCATTTTGATATTTGTGCACAATGTTGTGTGTCTCAAGTATATTCTTATAAGGACTTGTTATTGGCCTTGCTACGTGATGCTATTGGTGAGGGTTCTGTGCGTAGAGAACTTCATGCCAATAAATTAGCTGATATGCTTCGCAAAACTTTATTGCCCCGAAGGTACCTTATCCTTGTTGATGACGTGTGGGAAAATAGTGTTTGGGATGATTTAAGAGGTTGTTTCCCAGATGTCAATAACAGCAGCAGAATCATTCTGACGACAAGACATCATGAAGTTGCCAAATATGCTAGTGTTCATAGTGATCCCCTTCATCTTCGTATGTTTGACGAAGTTGAAAGTTGGAAGTTGCTTGAAAAGAAAGTGTTTGGTGAAGAAAGTTGCTCCCCTCACCTAAAAAGTGTTGGGCAGAGAATAGCAAAAATGTGTGGACAACTCCCTCTTTCAATTGTTTTGGTGGCTGGTATTCTGTCAGAGATGGAAAAGGAAGTAGAATGTTGGGAACAAGTGGCCAACAATTTGGGTACCCACATTCACAATGACTCAAGAGCCATTGTAGACCAAAGTTATCATGTTTTACCCTGTCATCTTAAGTCTTGCTTCCTTTATTTTGGAGCATTTTTAGAGGATAGAGTGATTGACATTTCAAGGTTAATAAGGTTATGGATATCAGAATCATTTATAAAAAGTTGTGAAGGCAGGAGGTTGGAGGATATAGCAGAAGGTTACTTGGAGAATCTTATTGGAAGAAATCTAGTAATGGTTACTCAGAGGGCCGATTCAGATGGTAAGGTCACAGCATGTCGCCTTCATGATGTATTACTTGATTTCTGCAAGAAAAGAGCAGCTGAGGAGAATTTTCTACTATGGATAAATAGGTAATATGATAAGTAATGGTAATTTCAATCAATCAAGTATTTGAAGTTATATCTGAAAATTAATGATATGCTTTTGCTAATTGATATATTCACAGGGATCAGATTACCAAACCTTCTTCCTGTGTTTACTCTCACAAGCAGCATGCTCACTTGGCCTTCACTGAAATGCATAATCTTGTAGAATGGAGTGCGTCTTGCTCATTTGTTGGCTCGGTAGTACTTTCCAAGAAATATGAGCCATACTTGGCTGGTCGTCCCTTAACCTCTCGTGCTTTATCAATTTCATGCATTTTATTAAATTTCAAGTTTCTAAAAGTATTGGATTTAGAGCACCGGGTTGTTATTGATTCTATTCCAACTGAGCTCTTTTACTTGAGGTATTTATCTGCACACATTGAACAGAATTCAATTCCTTCAAGCATATCCAATCTTTGGAACCTTGAAACTCTTATATTAAAGAATACATCAGTTGGGAGAGATAACACGCTATTACTACCTAGTACAATTTGGGATATGGTTAAATTGAGACATCTGCATATTCCTAACTTCAGCCCAGAAAATGAAGAAGCATTACTGAAGAACTCTGCAAAACTTTATGATTTGGAAACCCTTTCCTCTCCATATTTCTCTCGTGTTGAGGATGCAGATTTGATGCTGAGAAAAACACCTAATCTTCGAAAACTGGTATGTAAAGTCCAGTGCTTAGAATACCCCCATCAGCACCATCTTTTTAATTTTCCAATACGGCTTGAAATACTAAAGCTTTATGGATCAGAAGCCTTTAAAAACATCCCCTTTTGCATCTCTGCACCAAATCTCAAATGCTTGGAACTCTATGGCTTTTACCTGGATTCTCAGTACTTATCAGAAACTGCTGATCATCTGAAGAACCTTGAGGTACTCAAACTGTCCAGTGTTGAATTTGGTGATCATAGGGAATGGAAAGTGAGCAATGGCAAGTTCCCTCAACTCAAAATCTTGAAACTAGAATATTTGTCCTTGATGGAATGGATTGTAGCCGATGATGCCTTTCCTAACCTTGAACAATTGGTTTTGCGTGGATGTCGAGATCTTATGGAGATCCCTTCTTGTTTCATGGACATCCTTTCTCTCAAGTACATCGAGTTAGGAAACTGCAATGAGTCGGTTGTCAAGTCAGCCATGAATATACAAGAAACACAAGTCGAAGATAATCAAAATACTAATTTCAAGCTCGTTCTCATCTAGGTACACTACTGAAAAAAGCTTTATTCTGCATGATTTTGATGACTCCGAAATAATGGCCTAAATTGTATGAACTGTTTTCTCTGTTATCTTTACCTGGTGGCCTCGTTTTACATTCGGGTTTTTCTCTTGGTTTTCTTGCAGAAAAAGACATACCCCCAAGGCATTTAAAAGATTAGTTTTTCTTCCAGGTACTACTTATTATTTTTTCTTTCTTTCCTTTTTTTGAGTACACCAAATAGATAGATTCATCTTTTTTGTCTTTTCGATATGAAAGGGATATAATCAATTTCAACTGATATGAAAGAGAAGAAACTTACTGTGACCAGAGATGTGGATGCTGATGAAGTTCAATTAGTTGTGGAGAAACTGAGAAAGTGTGGCATGCTCTAACAAAATGCTGAGCTCTGGTGAACTCAACCATGATGGATTTTCAAATAATGACAGGGTTGTAGTCCCAACTTGTCAACACAAATGTGCTATACTCATTTTGCTTACTGTAATACCATTTCATGACACACACACAAACATTAACTGTAGTAAAGTTTTGATGGATCAGTAAATCTGAGTTCAACCCATTGTAATCCGTTCAAATTCAACTCAAAGGTATCCAGAGAGTGTAGCTGGAGCAATTTGGAATATCACATGTAATTTCTTTATGAGTTATTTCGTTTAATAAAGATACTGTAAAACGTCCAACGGCTGTTGCATTCATTGTAAACTAAATACATCTCACAATGTAACTATTTGAACAATTTTTTCATTTAACTCCTTGAGGTTTGATGTAAGTCATTAGATTTTAC

>M6-PGSC0003DMG400003543 M6 LRR-RLP

CTTTCTTTGTTTGTTCCCTTCTTCGGACACTCAGAATATATGCTCACGCATCCATTTAAAAACCAATCTTGAAGGCTCTTCTTCTTCATCAATTCTCATTACGATTAAACAAAATAATATAACCACACATAAAAAAATCCAGAAAATGGGTTGTTTTTCAATTTCTACTGTTCTGTTCTTTATAGTTCTCTGTGCTCTGTTCTTGCATGCGCATTCAACTACTCACTGGCAAGACATTGAAGTCTTGAAGCAGCTGAAAAATAGCGTCGACCCCAATTCAGTGTCACCTGGCTCATGCCTGAATTCATGGGATTTCTCTGTAGACCCTTGTGATAATCTCGGGGGTGAAAAATTCACTTGTGGTCTCCGTTGCGACCTTGTCGTTTCCTCTGTTAGCCGAGTCACTGAACTCGCTCTAGACCAGTGGAATTACTCCGGTTCAATCACTTCCGTCTCTTGGAATTTGCCTTATCTCCAAAACTTAGACATCACTAATAATGCCTTCTCCGGTTTAGTACCTGAGTCACTCTCGAATCTAACTCGGGTTCAACGAGTCGGTTTATCAGGTAACTCGCTATCCGGGTCAATTCCGAGTTCACTGGGTTCACTTTCAAACCTTGAAGAGCTTTACCTTGACAATAATTTCCTTGAGGGAGAAATTCCCCAAAGTTTCAACGGTCTCAAGAATTTGAAACGGCTCGAATTTCAAGGCAATCAGATTACTGGCGAGTTTCCCGAGTTGGATCAGTTAATTAATTTGAATTTCATCGATGCTAGTGATAATGCCATCTCCGGCGAACTCCCGGCGAGTTATCCGGCTTCTCTTATCGAGCTTACGATGAGAAACAACAGCATAGAAGGTAACATTCCGGCGAGCCTCATCGGACTGAATTTCGTACAAGTTATCGATCTGAGTCACAACAAACTCAGTGGGTCAGTGCCAGCGAGTCTCTTCACTCACCCATCCCTCGAACAGCTCACCCTATCATACAATCAATTCGGGTCAGTTCAGGAACCCGGGATCTCGTTTCAGAATAGTCAGCTTATTGCAGCAGACTTAAGCAACAATGAGATCCGTGGGTTATTACCCGGGTTCCTGGGTTTAATGCCCAGATTATCATCACTTTCACTAGAGAACAACAAGTTATCGGGTATGATACCGACCCAATATGCATTGAAAATGGTGTTTCCGGGTGAAGGGGTGTCTCAATTTGAGAGGCTACTGTTAGGAGGAAACTATCTATTCGGACCCATACCGGGTCCATTGTTGGATCTGAAAGCGGGTTCAGTCACAGTGAGATTAGGAGACAACTGTTTGTACCGGTGCCCGTTGAGATTATTTTTATGTGAAGGTGGGGAACAGAAATCATTATCGGAGTGTCAGGCTTTTGGTCCCATCATTCCTTGAGTAATGTGATAATGTCTTTACTTTGAGGTCCTCATTTTCATTTTGTACTAGTAATTATTGTGTTGTTTCAATTTTATCTGAAGGGACTAGTAAGTATTGTATAAATATATATTTTTTCCTGCAACCTTACCCCATGTATGGGGTTTTAGATATGATCTAGATGGATCTATGGTAGAGTTGTTATTGTTGTGAAGAGTGAGACAGAAATATAAGATTTCTGACTTAATGTGTTGAAGTAGTAATTGTGGGACAGTTGAGGAAATTTTATTAGCGGAT

>M6-PGSC0003DMG400003544 M6 LRR-RLP

CAAATATGGGAATTTTGGGTAGGAAAAAATCTTTCTTGTTCATCTTATTTTACATATTTTTTTCTGTTTTTGTTCCAAGGCCATTAGAATGTTGTCATTTGGTTGATAAAACATCTTTGTTAGACTTTAAGCAAAAAGTCACTTCTGATCCTTCAAACTTATTGCAAACATGGACACTTTCAACTGATTGTTGCAAATCTTGGGAAGGTATTGCTTGTAATTCCAATGGAAGGGTTGTTAACGTCTCTCGCCCCGGCCTTTCTTCAGGGGACGATTTCATTATCGATACCTCAATATCTGGAACTCTTTCTCCTTCTCTTGCTAATCTTTCTTTTCTTGAATTGCTTGATCTTAGTAATCTTAAGGACTTAACAGGGCCAATTCCACCTGAATTTGGCAAGTTATCGCGCTTGAGTTACCTCTTTCTTGATACTAATAAGCTCTCAGGATCTATACCTGTCACATTTAAATATCTATACCAACTTAAGAAGCTTTATCTCAGTGATAATACTCTATCTGGTACTATCCCTTGGAGTATTTTCGAAAGCTTCCTTTCACTCTCAGAACTTGGTTTATCAGCAAATCAACTTTCAGGTCCAATACCATCTTCAATTGGGAATTTAGTATCATTGACCAAGCTTGATATGAGTCGAAATAAGTTCTTTGGAAGCATTCCAGAGAGTATAGGAAGATTAAAAAATCTTGCATATGTTGATTTGTCTGAGAATCAGTTATCTGGAAAGATTCCAAACTCTATTGGAAACCTTTCCCAAGTATCTCTAATGTATCTGAACCAAAACCAATTGAGAGGAAAAATTCCTTCATCAATATCTGGTCTTAGCTCATTGGTGTTCTGCCGTTTATCCGAAAACCAGCTGAGTGGCAGTATTCCACCTTCTATTGGCAATCTTCCAAAGATCGAAAGGCTGATATTTGAGAACAACAAGTTAAGTGGAAAACTGCCTGCAACACTTGGACATCTTGTAACTCTCACTGATATGTACTTTTCCAATAATCTATTTACAGGGAAAATCCCATCAAGTTTTGGTAATCTAAGGAATCTACAGATACTAGATTTATCGAAAAATAAGCTCAGTGGTGAAATACCTCATCAGCTTGTGAAGTTAGTGAGTTTACAGACATTGGATCTGTCATTTAATCCTTTGGGGCTTTCAAGAATACCAAACTGGTTCAAGAAACTAAGACTGTTTCGTCTCCTTTTGGCGAAGACTGGGATCAGAGGGAACCTCCCAAGTTGGTTGGCTTCATCGTCATTATCAACACTTGATTTATCAAACAATGGTTTGACAGGAAAATTACCTACATGGATTGTTAACATGACCAACCTTTCACTCTTGAATTTATCAAACAACGCCTTGTATTCATCAATCCCTGATGAGTTCAAGAACCTGACACTTTTGATGGATCTTGATCTTCACTCAAACAGATTTTCAGGCAACTTGAGAGCAATTTTCTCAAAGAATTTTCAAGACCCCCTTGGTCACTACAATTCTATTGATCTTGCATACAACATGTTCACTGGACCATTAGACGAAAACACCGGAAATGAACCTGTCATGGATTCTATTTTGTCACTAAGTTTGTCCCACAATCCATTGACAGGACACATACCAAAATCACTGGGAAACTTGACAAGCTTGCAGGTAATTAAACTAGCCGAAAATGGGCTAACAGGTGGAATTCCAACAGAACTTGGGAATGCCAAGGAACTGAAAACAATTTTACTTTCAAATAATAATTTGGAAGGTGCAATACCAGCACAAGTTCTGAATTTGAAAGACCTTCAAGAATTTGATGTATCGGGAAATCGACTAAGCGGTAGTATCCCTCCTCATAAAGCTAATATCCCAAAGTCTGCATTTTTGGGAAATTATGGCTTGTGTGGAGCTCCACTTCCTCCTTGTAAGCACACATAGTGGAAACCTTCAAACACAATATTTCCCTTTTCTTTATTTATTTTTCATTTTCGACATGATTGTTCGTCATAATGTATTGTATTTCAGTTGTAATCACTCAAATGTTATTAGTAGATACTGAACAAAGTTGATTTTTCAAAAGCAACCTTGCCGAATTCTACATCAATATCCTCAATGGTTTAAAC

>M6-PGSC0003DMG400003710 M6 LRR-RLP

ATGAAAGTGCAGTTTTTTCTTCATCTTTTTTCATTTTTCATCTTCTTTGTATTATTAGCACAATGCAAAACTCATCCTATTGATATACAAGGACTTCAAAATCTCAAAAATGGAATAAACCCAAAATCCATTAAACCCGGTTCATGCTTAAACTCATGGAACTTCTCTATAGACCCATGTGATCATATTTTCACCGACCGGTTCACCTGCGGTTTCCGATGTGATCTCATCGTCTCCGGTTTTTATCGAGTCACTGAAATCAGCCTAGACCAAGCCGGTTACTCCGGTTCATTGCCTAATTCATTCAACCTCCCTCATCTCGAGATTTTAGATATGTCATATAATTCCCTCTCCGGTTCAATCCCGAATTCATTTTCAAATCTAACTCGGCTCCGGCGGCTCAGTCTTTCGAAAAACTCGTTCACCGGTCTAGTTACTTCTTCTATCGGTTCTCTATCCCGTTTACAGGAACTATTTCTCGATAACAACAAATTCACCGGTTCAATCCCGGAAAGTTTTAATGGTTTAATCAATCTGAATCGATTTGAGCTTCAACAAAATAATATTTCCGGCGAGTTACCCGATTTGAATCAGCTCATAAATCTGATCTACTTAGATTTAAGCAATAACAGAATCGCCGGAAATTTCTTCACGACAAATTTACCTGATTCGATGATTGAACTCTCAATACGCAACAATTATTTACACGGCGAGTTTCCGTTGAACTTCGAAGAGTTGAAGTTCCTACAGGTGTTAGATCTGAGCCATAATTTGTTTTCCGGGATGATTCCGGCGGCGCTATTTCATCACTCATCTCTTCAGCAGCTTACACTTTCTTACAATAACTTTACGTTTTTGCAAGTGCCGGAAGATTTAGTGCTTCGAAGTAAGCTTATAGCTATTGATCTGAGTTACAACAATTTGCATGGCTTATTGCCGGCGTTTATGGCGTCGATGCCGGAGCTGTCGGCTCTGAATTTGGAGCATAACAAGTTTACCGGCGTGATACCGATACAGTTTGCAGTAAAAGTTGTGGCTCCGAGGAATCATACGGCGTCGTTTGAGAGGTTATTGCTGGGTGGAAATTATTTGTTTGGGCCGATTCCTGGGCCTTTAATGGGCCTAAAGCCTGGTTCTGGAAATGTTAGTTTGGTGGATAACTGTTTGTATATGTGTCCAGATTCGGAATACATTTGTCATGGTGGGTATCAGAAGTCTTTATTGGATTGTAAAAAGTTTGGGCCTAAGATTCCTTGA

>M6-PGSC0003DMG400003802 M6 LRR-RLP

ATGTTTGAATTCGCCCGGAAATTGCAGAAGCGAGGCGGAGAGTTGATTAGCGAGTTTTGCAGTGAAATTGGAGTCGAAAAATGGGGTTTTAAGGTACGGTTGCACAATAGGACACCTTCAATTGATATATTGTCGGAATGTCTCATTCATAAAATACTCTGTTGTCTTAGTTTTAAAGAAGCAACGCGGATGAGCATTCTTTCCAAAACATGGCTGCAAGCCTGGTCGACTCTTCCAAACTTGGATTTCACAGTTGATTTTTCTAAAGGCAACGATATGAAAATAGTGGATGATATAATGGAGAGATATAGGGACGGAAAAATCCCTATAGAAAAGTTTGAATTATCAGATGATCGTTTGTTCTTTGAAAATCCTCATGGAGTTTTAGTTGTCCCTATGATTGATAAGTGGCTTGGCATTGTACTAACAAATGGTGTAAAAGACCTTACCATATCATCATACCCCTTGTTTATTTTCGCAATCTTGGCGGTAAAATCTGTGAGAAAATTGGTTCTGGAATCTTGTATTCTTTTGCCTAGTGTTGTTGTGAATTGCAATTCTTTGAGGAAACTTTCTCTATCTTATGTAAGATTAGATGAAAACATGCTTCAGGCTCTACTTAATAGCTGTCCGTTGATTGAATCTTTCATATTTGAGTATTGCAAGGGCTTGGAAACAATGAATCATCAAAAGATTAAGTCGGTTTCCCTGAAGTTCTTGAAGATTCGGCATTGTGGGGGAATATGGGAGATTGATGCTCCAAATTTGGTATCACTTAAGTACACTTGGGAAGACATTCCTCAAATTAAAATTGTGAGAGAGTCAAGACAAATGAAGTACTCAAGAATCATTCTTGAATGCTTAGACAGTGACAATGTAAATGCTGCATGGTTTTGTAAGTTGAGGAAGTTTCTGTCAAATTCGATCTCTTGGTCTGAAGTTCGCATTGAATCTTATAGATGCAATGACATCAACATTCAAAATTTGCAACTGGACCATAAAGATTCTAATCCCCACGTGGACGTTTTAAATGTCAACATTATATGGAAGAATGGGGAGAGCTCAACATTTGTGGATGCTTTGGTATGGAGTTGTCAGCCTAGGAGACTCAACGTGCATTCAACAAGAGAAAAGATCACATTTTTCATTGATCATTTGATGTATATGAAGAATTCGAGTCAGTCTACTTCTCATGGAAGCACGCCTCTTCTTAGTCAATTAAAAGGAGTAAAAGCCTACAAATTTGATAGGAAGAAGGAGAGTTGGCATCCTGTGGAACACAAAAGTAGTGAGCTGGTAACAACAAACTCTGAAATGTGGGATTGGTGTTATTTTTTATTAGATTGGTGA

>M6-PGSC0003DMG400003814 M6 LRR-RLP

ACCCAATTGATTTCTTAGCTTTACAAGCGATTCGAAAAGGCTTGGATGATTTGCCCGGTTCCAACTATTTTGCTTCCTGGGATTTCACTTCTGAGCCGTGCAACTTCGCCGGAGTTTACTGCGATGGAGATAAAGTGATCGCCTTGAATCTCGGTGATCCCAGAGCTGGATCACCGGGTTTGACCGGAAACTTGGATCCTTCTATTGGGAAACTCTCTGCTCTTGCTGAGTTCACGGTGGTTCCGGGTCGGATAACCGGTGCATTGCCGGAAAGTTTTTCTCAGTTGAAGAACCTGAGATTTCTAGGAGTTAGCCGGAACTTTCTCTCCGGTGATATTCCGGCGTCATTAGGCCTGCTCCGGGAGCTACAAACTCTTGATCTTAGTTTCAATCAGCTGACTGGGAGTATCCCTTGGGCTATTGGAGCTCTTCCGGCATTGTCCAACATTATACTCTGTCACAATCATTTGTCCGGTTCGATCCCACCATTCGTATCCCAAAGATTGACCCGTTTAGACCTTAAACACAACTCACTTTCAGGTACCCTTTTGCCTAATTCACTTCCTTCATCGCTTCAATACTTGTCATTGTCATGGAACCAGTTAACCGGTCCAGTAGATTATCAGTTGAGTCGGCTGAACCAGTTGAATTACCTCGATCTTAGCTTGAACCGGTTCACCGGCTGCATTCCGGGAATCTTATTTACTTTCCCACTCACAAATCTTCAGCTACAAAGGAATCTATTTTCGGGTCCGGTTTTACCCATTTCAGAGGTGACAATTCCGACGGTGGACATTAGCTTTAACAGGTTTTTCGGGGAGATATCTCCATTGTTGTGTAATGTGGAGAATTTGTATCTGAACAATAACCGGTTCACGGGTAAGGTGCCGGCGGTGTTGGTGGACCGGTTATTATCTGCCGGAATACAAGTTTTATATTTACAGCATAATTTTCTGACGGGGATTGAGATAAATCCGAGGGTGGAAATTCCGGTGAGCAGTTCATTGTGTTTGCAGTATAATTGCATGATTCCGCCGGTTGAGACGGCTTGTCCGTTGAAAGCCGGGAAGCAGAAGAGTAGGCCTACAAATCAGTGTGTGGAGTGGAAGGGGCAAAAGAGTGAAAAAACAA

>M6-PGSC0003DMG400003822 M6 LRR-RLP

TTGAAAACCAAATATTACACATGACAACATTTCATACTACCTTAGTTTACATTTTGTATGTGCTACTTCTTTTTTCACTTCCATTAAGTATCACATCTTCAGCAAGAACAGAAGCTGAATCTCTTGTCAAATGGAAGAGAAATTTACCTTCTACGTCTTTCTTGGACTCATGGTCCATTTCGAATCTCGAAAACTTGTGCAATTGGACATTTATTGTGTGTAATGATGGTGGAACAATTTCTGAGATCAATCTTTCTGATGCAGCCCTCTCAGGCTCACTTGATCACCTTGATTTCACTTCATTTCCGAGTCTCGTGAATTTCAATCTTAATGGAAATAACTTCAGTGGATCGATACCTTCGAGTATTGGCAATGCCTCATTGCTCACTTTCTTGGACCTGAGTAACAATATCTTGTCAGGTGTCATACCTGAAGAGATTGGAAAATTGAATCAACTTGAGTATCTAAGTTTTTATAACAACAATATCGAAGGTGTGATTCCGTATCAGATTAGCAATCTGCAGAAGGTAATGCACTTGGATCTTGGATCAAATTTCTTAGAAACTCCTGATTGGTTGAAGATGAAAAATATGCCTGTGTTGACGTATCTAAGCTTTGGTTATAATGAACTGAGGTTAGAATTCCCTGAATTCGTACTTCGTTGTCATAATCTAACATATCTTGATTTATCGATAAATCATTTCAATGGTTCGATTCCTGAAACAGTATTTACCAACTTGATCAACCTTGAAAGACTTAATCTTTCATCTAATTCATTTCAAGGTTCATTGTCACCAAATTTTACCAAGTTGTCTAAGTTAAAAGAACTTCAGCTAGGTGTTAACATGTTTTCTGGTTTAATTCCTGATGAAATTGGCTTGATTACTAGTCTTGAAGTTCTTGTACTTTTCAACAATTCATTTGAAGGAAAGATTCCATCTTCTATAGGTAGACTCATAAATCTTCAACGGTTGGATCTTCGAAAGAACGATTTGAATTCAACTATTCCTTCTGAGCTTGGTTTTTGTACTAAACTCACTTTCTTAGCTCTAGCAGAAAATGACCTGCAAGGGTCATTGCCTCCATCATTCTCCTCTCTTACTAAGTTATCTGATGTGGGGTTGTCTGATAATAGTCTTTCTGGTGAAATCTCATCGAATTTCATCACCAATTGGACTGAGTTGACATCTTTGCAGCTTCAAAACAATATGTTTACAGGGAAAATCCCACCCGAAACCAGTCAATTGACAAACCTTGTATATCTCTACCTCTTTCATAACAATTTCACTGGTTCCATTCCCTATCAGATTGGGAACTTGCAAAACTTGTTGGAATTAGACTTTTCTGATAACCAGCTTTCAGGCATAATACCTCCAACCATTGGAAATCTGACCAATCTGAAGACGTTGCAACTTTTTCGCAACAATCTCAGTGGAACCATTCCTCCTGAGATTGGAAAATTGATATCCCTTGTAACCATCGACATCAACACCAACCGACTAAGTGGTGAGCTGCCGGACAGCATTTCTGACCTCAGTGAGCTGAAGATTCTTTCTGTATATACTAATGATTTCTCAGGCATTGTCCCTAAGGACTTTGGAAAGAACAGTCCTCAATTGTCAAGTGCCAGCTTTGCAAATAACAGCTTCACTGGTGAACTACCTGCTGGATTGTGTAGCCCGAATCTTGAGGAGTTGACGATAAATGGGAACAAATTCAGTGGGAAGTTACCAGATTGCTTGAAGAACTGCACACTGCTAAGACGAGTACGGCTTGAAGGCAACAATTTATCTGGTAATCTAGCTGATGCATTTGGAGTGCACCCGAATCTTGTTTACCTTTCTCTTAGTGACAACCAACTTTCAGGTGAACTTTCACCTGACTGGGGGAAATGTGAAAATCTCACAAGTCTAAGAATGGATGGAAACAAATTTTCTGGTGTAATCCCAAGTGAGCTAGGGAACCTGAGAGCGTTGCGCGTGTTAGCTTTGGAAGGAAATGAATTGACTGGTGAAATTCCTTCTGAACTGGGGAGGTTAGACCTGCTGTACAATCTCAGCTTGAGCAAAAACAATCTTACTGGAGGCATCCCTCAGTCTGTTGGCAATTTAACTAAGCTCCAGTATCTTGATTTGTCAACAAATAAGTTAAGTGGTAACATACCAGTAGATCTTGGGAAGTGTGAGAGCCTGTTGAGCTTGAATCTTGGCAACAACTCATTATCAGGTGGGATTCCATCTGACCTTGGAAATTTGATGCAGTTGAGTATTCTTGTGGACCTAAGTAGCAATTCGCTAACGGGAACCATTCCACAAAACTTGGCCAAGCTCACCTCATTAATGCATCTCAACCTCTCACATAACAACCTCTCAGGTAGAATTCCTCCAGCATTAGCTCAGATGATCAGTCTTCAGGAGATGGATTTTTCCTACAATGAGTTTTCTGGTCCAATCCCAACCGACGGAGTATTTCGAGGAGCATCCGCTAGTTCTTTTCTTGGAAACTCTGGTTTGTGTGGAAATGTAGAAGGATTATCCTCATGTAATTTGGATACCCGAAATGACAAGTCCAGAAATAATAATCAAAAGATCCTCATTGGGGTACTTGTACCGGTGGTCAGCCTCATACTTTTGGCAATCCTGTTTGTTGCATGTCTCGTGTCTCGAAGGAAGGCTAAGCAATATGATGAGGAGATCAAAGCTAGCCAGATACATGAAAATACAGAATCTCTGATTTGGGAAAGAGAAGGGAAGTTTACATTTGGTGACATAGTGAAAGCTACTGAAGATTTCAGTGAAAGGAACTGCATTGGAAGAGGAGGCTTTGGAAGTGTCTATAAAGCTGTTTTGCCCTCTGGGCAGATTGTTGCAGTCAAAAGACTCAACATGTCAGACTCAAGTGACATTCCATTGACTAATCGTCGAAGCTTTGAGAATGAGATTAGAACTTTGACAGAGGTGAGACACAGGAATATAATTAAGCTCTTTGGTTACTGTTCCAAGAATGGGTGCATGTACTTGGTTTATGAGTATATAGAAAGAGGTAGCCTCGGGAAAGTTCTATATGATAATGAGATGGGAATGGAACTTGGATGGGGTACAAGAGTGAAAATTGTGCAAGGAATAGCTCATGCACTCGCTTACTTGCACCATGACTGCTCTCCACCCATTGTGCACCGTGATGTATCATTGAACAACATTTTGCTTGAATCAGAATTCGAGCCACGACTCTCTGACTTTGGCACAGCAAAGTTGCTAGCTTCAGACTCATCAAATTGGACCACAGTTGCTGGTTCTTATGGCTATATGGCACCAGGTATGTTACATACTCATTACCTAATGCATCCATATTTCTGTTGATTAAGTAGTATATTCATTGGAAGGATACTTCAATAATAGTGTCTTTCTTCTTGTGCAGAGCTTGCACTTACCATGCGTGTTACAGAAAAGTGTGATGTCTATAGTTTTGGAGTCATAGCAATGGAGACTATGATGGGAAGGCATCCAGGGGAGCTTTTAACTTCATTATCAGCATCAACAACATTGTCTCCAGAAATACTTTTGAAGGATGTTCTTGACCAAAGACTTCCACCTCCCACCAGCCACTTGGCAGAGGCAGTGGTTTTTGTGATCACAATTGCCTTGGCATGTACACGTACCACTCCTGAATCGCGACCAACCATGCGTTGTGTAGCACAAGAATTGTCTGTTCAGACTTTGCCTTACCTTCCACAGCCATTGGGGACAATAGAAATGAGCAAACTAACAAGTTTTCAGAAATAATGAGAAGATAGAACCTCATGTACAGGTGTGAAATGTACAGACTGTAGAGTTTTAGTGTTGTAATGCAACTTCCTCAAGATCTAAGGTCTATATAGTATCAAGGATGAAACCTTGGATCTATCCAAATTACTATGTTTCTTTTGCACTTTTCCCTAGAAAAAAAGAAATAATATCATCCTGTTAATCATGAATGTTACAAAGTTGGTATATTCATTTGGACTAGTATGAATGTTACAAAGTTGCACAATTCCCTTATCCTAGTCTTTACGTTATTGCTAGGCAAAACACTAGCAGAAAGATACAATGTGTTCACTTGCAATAAAAATAGCAACCTAACAAAAATTTGGCAAAGAATACATCACACAGATACCAGGCATGGTACTTAGCATTCCACTCCACACTGCTTGAGTATAGGTCCCTTCTGCTTTGGTAGGAATGGGTCTATTCGCACCCAAACAAGTGAGAAGATTGAAGCAAGAAGAATTGACCAGAGGACCACAATGGTGGGAGTCCTGTTCTGCCTGCCCATCAATCCCTTTAGGAAAGGATAGAGATGCACAATAACCCAAAAGGCGAAGAATAACTTTCCGAAGAGAGGACCCCATGAACCGTAACCATTGTTGATGGCATCCGAAACACCAGCTACAACACCGACCATATTCAAGATGATCAGAGTGGTTGGTGGGATAAGAAGTGTGGTCCATTTGAAGAGGTAAAGCTCCCCAAATTCAGCATCATCAGCTGCTTTTGATGTTACAGTAAAGTTTGTGTCTACACCAGCAAGAACTTTAAGAAGGCCTTGGAAAACTGCAAACAAATGTGCTGAAACACCACCGATGACCCAAAATTGCTCGTTACGCCACCAGGCCTCAATGCTTACTCCACTCCATCGCAGCTCGAGTACACTTGTGGCTATGATGGAGATAAAGAGAGCCATGAACCAAATGCTTGCAAAGTTGTTAAGCTACAAAACAAACATGGATTTTGATTTATAAAATCTAAATCAAATCTTTATATTACCGAACCAAATGACTTCAGTTCACTTACTGTGGGGACGATAAATTTTCCAGTGAGAAGGCATACAGCTGGAAGGGTGCAGTAAGCTAGCAGAGCAATGGAAGTGAATGGGTAAACAATGGTGTTAATGTATGCTAGCCTTTCCAACCATTTGAGTTTTCCACCCCAGGCATACCAAAGGGGACAATGACGACTCATGAAGATCTCGACAGAACCAAGAGCCCATCTCAGGACTTGGTGCAATCTATCTGATAGATTGATTGGGGCTGATCCCTTAAAAGCTGCTCTCTTCGGACAGCAATAAACTGATCTCCATCCTCTGCAGTGCATCTTAAATCCAGTCAAAATGTCCTCTGTGACAGAACCATAAATCCATCCAATCTGTCAAAAAAAGAAAAATGAAAACCAATTTATTAGAAGTTAGCTAAGCGAACACCCATAACTCATAATTTATTCATGAAATATAATGCTAACCTCTTTGCCCCAGTCAGTTTTCTCTTCATACCCACAGCTAATAACATGAATAGCTTCCTTTATGAGTGTTGTCGGGTTTGTCCCTTCAGGAAGCCCACCATCTTCCTTAAGTGTGGAAGCTATGAAAACGGGGGACTGCCCGAATCGTTTCTCAAAGTTCTTTTGGGACATGAGAGATGACTTCTCAAGCTCATCATATCCTTCAAGGCCTTCTTCAATTTCCTCAAGATCAAAGACAGGTCCAGATGGCTTCCTAGTGTAGTTCTTTCCATTCATTGTTTTCTTTTTCTTGTTGTATAGTCCTCCAAGTCCAAGTAAGGCCTTGATGCTCTTCTTCTTAGACTTGGACTTGGACTTTCTTGAACCACCACAACAGCAACTGGGCCAGCAATCACATGTCATCTTTTGACGCTTCTCAGAAACAGGTGGATCATAGCCGTATAACGCCTGCCTGTTAAAGACAGTACCTGTTCCAACATAAACTGGTCCTTGAATTCCATCCAGGCCTTTCATGTTGATCTGCCACCATGAAGGGAAAAAAGCATATTAATACCGGGAATAAATCCTTTGTCAACGCATTGCTGTATATATAGATCAACTTACATCAAAAAAGACAACATTACGATTGGCATATCGATCGTGGCGATCAATACCATCAAATCTCTGGGGAAATTGGACATAGCATAGCTTCTTTCCAATTTGTGGGTCCATTAAAAAGCACATGGCTTCTCTAACTGCCTTGCTGTTGTTAAGGTAGTGATCACAATCCAAGTTCAACATAAATGGTGCATTGGTCAGGACTGCAGCGACTCGAACCTATTTAGAAAAACAGCACAAATTACATAACATTAAGAAAAGGACTTAGCAGAATGTCGAGGAAGCAAGCCAGAAACTAATATACTAATTACTAACCAAAGCATTCATGGCACCAGCTTTCTTGTGGTGTTGATAGCCCGGTCGTTTCTCACGAGAAACATAGACAAGGCGGGGAAGCTCCTTACCTTCAATGTCAAGTGCACCTTCACTACCAAGATAAACTTGAATCATACCAGGATGGTCTCTGGTGTTGTTACCGGGCCATGGAGTTCCATCTTGCATTACCCATCCCTCTTCTGGTTTTTTCTGAGCTTTTGCTACTAATGCATTAATTCTTACTTTGAATTCTTCATATTCTCTCTGCCACAAGAGAGTGAGTTTTCAATTTGCAGAAAAATCTTATTTCAAGCCAAAGAAACTTAAGCCCCCGATAAATGATACTCCTTCTATCCCAACTTCTATGACATTCTTATATTTTGGGCACGTCCTAAAATGTTTGACATGATTCCTTCTACAACCTTCACAATAAACTATTTGAATTTTAAACCTTGATAGGATTCAGTCAACATTTTAGGCACTGCATCTCCTCTGATATAGTCGTATAAAATGAGACGAAATGAAGTAATAGATACATACCTTCATGGCCCTGCGATCCTTGACAAAGGTAGGCTGAACTTTATCTTTCAAGTAATCAATCTTTTCATTGAAGTAGAAATCGGGTGCTCTAGGCTCTACACTATATTTCTTACAAAATGGCACCCATCTCCTGGCAAATTCAGCTGTTTCTGCTAATGAATCGAACAACAGCATGGAAGCACCATCATCTGAAACATAACAGCTAACCTTCTCGACAGGATAATCAACAGCCAAGATTGACAGAACAGTGTTTGCAGTGATAATAGGAGGCTCCTTAAGAGGATCAACAGTACTCACAAACACATCCACTGGGGCTAATCGATTTGGCTCGCCTTCACGTTCAAACCTCAATGTCAATCGATCTAAATAAGTTTCACGGTTTATGGGTGACCATTTGGGGAACTGATCAAGTATCCATGATAATGCAAACCATATTTCACATATAACAGAGATAATCCATAATGGATAGGCATCATAGGCTGGTGTTAAGATCCGGAAATGGAAAAAGAATCCAAGAATTACAAGCCTGAGGATAATGACTATCCGATATGGACTGATTAGACTTGATGGAAGTGGAACTTTTCGCCAGAGTGGTTGCCGAGCTTCAGCCAAACTGCTTGGAAATCAGTTAAGTGGTTAGAATATTGAAATACTGCAAGTTTTATATGCTTAAACTTGATCTGTTTCAATTCATACACACTCATTCCATGTTGAAACATTATAGTATTTGATTAATTTTGTTAATAATATTGCATGTAACTATCTAACTTTCACAATTATAAAGTGTTCAAATTATTTAATTATACCAGTCAGTCACTGAGTGATATATAACTACAAGTGACAGAGACGATATATAACCGAAAATCCAATATTTAAAAGTTGTAAGGAGAATGAGGGGAAGGTGGAAACAAGACCAGATATAATTTGATCGCTATGAACAAATCCAAAGACCAGGTTGATTGATTTGTTAAAAATTTAAAGTATTTCTATAGGGGTCGTTTCCTGTTCCTTCTATGCTGTTTTCTCGATAATTCCTCACATAAAACCTTCTCGTATTAGCAACGATATAACTTCAACAAGATCAAACCAGAGGCGGGAATTAGCTTGAATAATCATAATTTTCAATATCTTCTTCCACTACTAAGAATATATGTCAAATAACATGTTAAATTCCCCTTCTAAGTCGAAGGTCTATCGAAAAAGCAAATCTAGTAGGGGTAAGGTGTGGTATATCATACCCTCCCAAACCTCACTTGTGGGATTACACTGGGTGGTGTATATAACATGTTAAATTTCCCTTCTAAACTGAGTTGTTAAATTACCCTTCTACGCCGAGGGTCTATCAGAAATAGTAAATCTACCCCACAAAGGTAGGCGTAAGATGTGGTAAATCACTATATCTTACCCTCCCAAACCTCACTCGTGGGATTACACTGAGGGTAAACAAAGAGAGTGAAAAGAAAGAGTACATACAGATAATCATCATCTTCATCAGGTTGGTCATTTCCACCATTATTAACAAGACCTCTCTTTTCTTGACGATTCTTCCATTTCTCAACTCTTTCCTTCCATTCTGCATCACCATAACTTTCTTTCTGCTCATCCTCCAATTCCTTCCCTGCAACTGTAAGTGATTTTCCATTGAAATAAAACAAAATCATGACCTTGAGTGACACATATATATAAAGTGCAACAATTAATCATTTCGAAATTAAATAATATAGCTGTACCGCTTCCAACAACTGATTGATTAAAAGCAGCACCACTATTGGTGTTGTGATTATACTCTCCATTCTCCTGCCAAACATACAACAACTTGCTTTCAGTTCAAATTAG

>M6-PGSC0003DMG400003834 M6 LRR-RLP

ATGAGCATTCTCTCCAAAACATGGCTACAAGAATGGTCGAATCTTCCCAACTTGGAATTCACTATTAATTGTTGGGAAGGCAATATAAATACAGCGAACACAACCATGGAGAGATATAGGAAGGGGAAAATCCCTATACAAAAGTTTGAATTATTAGAGACCTTTGCTAATTCTCATGAAGATTTCCCTCTCATTGATAAGTGGCTCGACATTGCACTTCAGAATGGTGTAAAACATCTTTTTCTTAACTTTAAATCATACCCCATGCCTATTTTAAGAATCTTGGCAGCAAAATCTTTAAGAGAATTGGATGTGCAGGGTAGTATGCCTGATTCGTTATCTAGTGGTGTTGTGAATTGCAAATCATTGAGAAAGCTTTCTCTATCTCATCTAAGATTAGACGAAAACATGGTCCAGACTATACTTACTAGTTGTCCCTTTATTGTCTCTTTAATCCTTGAGTATTGTACTGGGAAGCTTCAAAAGATCAAGTCGGATTCCTTGAAGGTCTTGAAGATACATCATCGCGGGATAGAGGAGATCGATGCTCCAAATTTGGTATCACTTGACTATATGGGGAGTCAAATTCCCGAACTTAATAAAATTGCAAGAGACTCAAAAATCAGGCTTGGATGCATCAACAATTTAAATGCAGCATGGTTTTGTAAGTTGAGAAAGTTTCTATCAAATTCGTCCTCGTGGTCTCATGTTTCCCTTCAATTTATGAATTGCTGTGAGATCAAGATGAAAGATTTGCAAATGGACCACATAGGTTCTACCCCTCGGGTGGACATTTTAAATGTCGATATTCAATGGATGAATCAGAATTTTGTGGATGCTTTGCTATGGAGTTGTCATCCAAGGAGACTCAACCTACTCTCAAATATTACTCCAACGATTACACATTTCATTGATCGTTTAGTGTTTTTGACGAATTCGAGTCATTCTACTTCTCATAAAAGCTTTCCTTGGCATAGTCAATTAAAAGAAATAAAAGCTTTTGATGGAAAAAACCAGTCGCTACAACTCATAAATGGGGAGCTTGCAAAAAGGACCCTTATGGAGGGGGAGAAAGTTTATTTTTTATTAGATTGGTGATGTAGTTAATTTGTTTCTATTTCAAATTGAATGGACTTATTCTTTTCTGCATTGGTGTCAGAGGTATTAGATATTAATAAAACAGAACTAAGAGCATAATTTTAAACTTGGATCTTATAGGCTTGCAAAAGCAAAAACAAAATTTTTAAGAAAGTAATGAGTATGGCAAATTTCTCAAAAGGAAAGAGACTTAATTTCTACATAATTACTAGTGTTTCAACGTAACTACAAATGCCTTCGATGTCATACTTTCTCGGTAGCCCGTTGATTTCTAATTTTCTCAAACTCTGTCTCGTGACATCATAAGAGAAACATAAAGTATCCGACATTAAGTCCAGGATGACTACAATCTCTTCATCACAGGACGTGCATGATGATATAACTAAGGGTTTGTCCTTCCACCACGTCGAAGGGAAGTCAATGATTTGAATTTTCCATTCTTCTTCTTTCTGAGTTTGTTCTAAAATCCACAAGTTGAACTATCCATGTTGCCAATATTCATAATCCATGATCGCTAGTTTACCTTTCACCTCTAACAACTCGTAATACCACCACGAGTTAGATGCATTATATAAACCAATATTTTTTGAATTTTCGGTTTTAACATCAAATGCATCTATAGTAAGTCTGTCCTGGAAAACAAATTTATAGATAACTCTATTGATGCAAACCCCGGGCTTGCACAGAATAGGAGGGGTTATGCTTTGAGTCTCTCTCCATGATTCGTCTACTCCTAAAGTGAAAATCCAATGTTTTATGTACCCTTATATAATATGCTTTGTTGTGAAAAGAACTTTGTACTTCTTTTCTTCTGGGTCAAAACCGAATGAACAGAACCACATGGAAGAACCATCATTTAGGTCAGGAAGAAATCTTACTTCTCTTGTACTAGGATTGAAAATTGCAGCAAAAAATAATGGTTCCCAAATATAAAATAAACCA

>M6-PGSC0003DMG400003835 M6 LRR-RLP

ATGGCTGACATGTTGCCTGAATGTTTCATTCAAAAAATACTCTCTTCTCTTAGTTTTGATGAAGCAGCGCGGATGAGCATTCTTTCCAAAACATGGCTGCAAGCCTGGTCGACTCTTCCCGACTTGGAATTTAATGTTAAGTTTTGCAAAGGCGATATAAAGATATTGGACACTATTATGGAGAGATATGGAAACGGAAAAATCCCTATAGAAAAGTTTGATTTATCAGAGCTCTTAGGCGATTCTCATGAAGTTTTCCCTCTGATTGATAAGTGGCTTAACATTGCACTTCAAAATGGCGTAAACAATCTTGTCCTTAACTATAAATCATACCCCGTGCCTTTTTTAACAATCTTGGCAGCAAAATCTTTAAGAAAATTGGTTCTGAAGAGTTGTACTCTTTTGCCTATTTCTTTATCTGGTGGTGTTGTGAATCACAATTCTTTGAGTAAGCTTTCTCTATCTTATGTAAAATTATACGAAAACATGCTTCAGACTCTACTTAATAGTTGTCCTTTGATTGTCTCTTTCATCTTTGAGTATTGTTCGGACTTGGAAACAATTGAGCTAGTGAATCTTCAAAAGATCAAGTCGGTTTCCCTGAAGGTCTTGAAGATTCGGTTTAGTGGGAGCATATGGGAGATTGATGCTCCAAATTTGGTGTCATTTGAGTACACAGGGAATCAAATTCCTGAACTTCAAATTGTGAGAGAGTCAAAACAACTGAAGCACTCAAAAATCATTCTTCACCGCTTAGACAATATAAATGCTGACTGGTTTTGTAAGTTGAGGAAGTTTCTATCAAATTCGACCTCTTGGTCTCAAGTTTCCCTTTATTTTTATGAATGCACGGAGATTAAAATGAAAGATCTGCAACAACACCACAGAGTTGCTACGCCCCAGGTGGACGTTTTAGATGTCAACATTCTGTGGCAGAATCTCGAATACCCTTCATTTGTGGATGCTTTGGTATGGAGTTGTCAGCCTAGGAGACTCAACTTGCAATTAACAAGAGAAATGGTTACAGTTTTCATTGATCGTTTGATGAATATGAAGAATTCGATTCAGTCTACTTCTCATGGAAGCAACCCTTGGTATAGTCAATTAAAAGAAGTAAAAGTCCACAAATTTGATAGGAAGAAGGAGATTTGGTATCCTGTGGAACACAAACTTGGGGAGCGGGCAACAATGAACCTTGATAGATATTATATTTTATTAGATTGGTAA

>M6-PGSC0003DMG400004056 M6 LRR-RLP

ATGTACAGAGGTGTAGCAGATGCTTCTATATCTCCAATGGAGAAACATGAGAAAGAGGCATTGTACTCTGCTATTCAAGGTTTTGTTGGAAATGAGTGGAATGGTTCATTTCTTTATCCTGATCCTTGTGGCTGGACATCAATTGAGGTACTAATCTTTTGCTCTCGATGTTCTTTTGCTAATTTTTAATGCTAAATTAGATTACATTAATTTGATATTTTAAACTAAACATTTATTTATTCAAAAATTATATAAAAAATACTATAAATTGTAATTTTTTGTATTTTAATATGATGAAATTGGTCAAAGTTACTCTAAAGAAAACAGGAAAACATGACAACTAAAAAGGAACGAAGAGTATATGATTTCATAATTTGCTAGTTGATTCTAGTCTTATGTTTTTTTACCATTATTGACTTTTCTTGCTGTAGTTCAAAATCAAGAAAACAGTAATGTCACAATGCCATGCATGGATGTCCTTAAATTTTTTATTTTATTTTAATATAAAGAAATGGATCTTGGGTCTTACTCAACCACATAAACTACATCATAGGAATTAGGATGAGTATTGTTCAAGACATTTGAGATCAATTACCCATCCCATATTCGACTCTTGTGGGCAATATAATATGGAAGCACAACATAGGTGAACAAAGAATTCGGATGAGCGGACTTTGGTAACATGATAAAAAAAAATGGATCTTGGCCTAACTCAAGACAAAAAATATCTCATAGGGTGAGGATTATTCAAGACCATATAGGAGATCAATTACCCATCTCATATCCGATGTGGGACTCTTAATAGATCAAACTGTCCTCTCAATGGTTGTTAATTAAAGTGAAAAAGACTTTCTCAAATTCTTACTTTTTCTGTTCTGGATTTGCAAACAGTGTTATATGAGTGTGTGTAAGAGAGACATCACTAGTTAGGAGCTTTCCAATTTCTAACACACCTTTTCTGGTAGTACTAGTAAATATATATGTTAGTATTCTATTGGTAATTTGCAATATTTATCACTTTCTTATAGTGATTTAGGCTTTATTTTGATGTAATTCATTCCTAAAACCAAGTAGTTATATGCATTGAAATTATCGTTCTTGTATGTTAACTTGACTTAACAGTGTAAAAATTCTTTACACCGTAACATAAACTCTTGTCTTAACACTTTAATCAACTCTGTGAACTCTTATTTTTACTATAATTCACAGGGAGTATCATGTGATTTCTCCAATGGCTTCTGGCACGTTACGGATTTAACTATTGGAGACCTATATGACAATTCCCTTCGTTGCTCCCAAGCTGCAGAGTTCACAGAACACTTGTTTAACCTCAAGCATCTAAAAAGACTTTCCTTCTCAAATTGCTTCCTCTCAGATCAGCACAAATCAATTCCCATTTCAAATTGGGATAGTCTCGCAAACAGCCTTGAATCGTTAGAGTTCCGATCAAATCCTGGTCTTATTGGGACAATTCCCACAAGTTTTGGCTACCTTAAGAATCTTCAATCTTTAGTACTGCTGGAAAATGGACTGAAAGGAGAAATACCAGAAGCTTTAGGCAATTTAACTAAACTAAAGCGCCTCGTTTTGGCAGGCAACAATTTTATTGGTCCTATTCCAACTAGTCTAGGAAGATTGACAAATCTTTTGATACTTGACACAATCAGGAATTTTCTATCTGGTGCATTACCTGTGACTATTGGTGATTTATCTTCGCTCATAAAGCTTGACTTGAGCAACAATCGATTAGATGGAAAATTGCCTCGAGAAATTGGAGGACTAAAGAGTCTAACACTACTTGATCTCAGCCACAACAATTTCTCTGATGGTTTGCCAGAGACATTTCAAGAAATGGACTCCTTAGAAGAACTTGTGTTATCTGACAATCCGATAGGCGATTATGTTACAAGAATTCAATGGAGGAACATGAAAAACTTGGAAATGTTGGATCTTTCAAACATGGGATTGAAAGGTAATATACCAAATTCCATGACAGAAATGAAGAAACTGAGATTCCTCAGCCTTAGTAATAACATTCTTTCAGGAATTATTCCGTCAAAATTCGAGAAAATATCAACTATTAATGCTCTATACTTGGATGGAAATGATTTCACAGGGAAACTTGAATTTTCACAAAGGTTTTATACAAAATTAAGGAGTCGTTTTCGAGCTTCAGGTAATGTGAAACTCTGCTTGTCCCTCGAGTTAACGTCAACAAGTCATGTTCCAGAAGGTGTACAACAATGTGAACAAGAAAGCACAGTAGATAGTAAAGATTCAGACTCCAATTTAAAGAACAAATATTTGAGTCAGAATTCCCACAATATTATAGCTTCTTTGGGGCATTCAGGACATGTTGTTAGTCATTTTACATTTTGTTATGTCGCAAGAATAATACTTATGTCTCTTCCATGGATCGTATTTTTATAA

>M6-PGSC0003DMG400004190 M6 LRR-RLP

AATCCACCGACGCGCAGTGCAAGCTAAATAGCGCGTTGTATGAAAAAAACTAATTTTGAGGAAAATTCCCAAAGTGAGGATTCATTTCACACACAGATCTTCTTCCTTCACTTCCATCATCTATCTCAATCTCGACTATAGTAATGTACGTATAGCTCTGCGTTTGGATTCCGGTTGATTTAATAATTCTACTCCCCTGAAGGTGATTAACCCTATTCAAATGGCTCGCCATGGGCATCTACCTTCAGTATTTCTCCTGATCTTCTCGTTATCCTTGATCCAAAGCGCTCTCGTTCACTCTAAAACCCTCAAACGTGACGGTACACATTTCGATCCATTTACTTTTTAAACCTAATTTATTAGCAATTTAGGTGAGAAAATCAGTTGAATTTGATTGATAATAGTTTGGTTTTGTTATTTTGTAGAGTATGTATGCAGCTTCAAGTGAAATTAATTTTGTAGTTACAGAGAAAATGTAACAATGTTGAACTTTGTAGATCTTATGTATTCTAGGTGCAAGGTTATAGTTTGAGTGGAGTTTAATTGAATTTTAAAAGTTCGAGTGAAAATAGTTAGATTTTGTAGATTATATGCAGTTTGTGGTGAAATTTTTGTAGTTTCAGCGAAAACAATATTGAACGTTGTACATCTTATGCAATTTCTAGGTGTAAGCTTATGTGTGAGTGGAAGTAATTGAATTTTATAGTTCGAGTTGTAGTAATTGGATATTATCCAGTTTTTAGATGAAAGCTTATAGTTTGAGTGGAAGTAGCTGGATTTAATAGGTTATGTGCAGTTTGTGGTGGATTTATGGTTTCTGCGGAAACAATTGAATTTTGTTGATTTTATGCAATTTCTAAGAGTAAGCTTATAGTTTGAGTGAAAGTAAGTGGATTTTATGTAGTCCTAGAGCAACTTTATAGTTTGAGTTGTAGTAATTGGATTTTATTGACTTTTTCCCAGCTTCTAAGTGCAAATTTTTAGTTTGAGCTGAAATAATTGGATTTATTCTATAAATTATATGTGGTTTCGAAGTGCACATGTATAGTTGGAAATGATTGGAATTTATAGATTAGATGAAGTTGTAAGTGCAAGTTTATAGTTTGAGCTAAAATAATTGGATTTTATGCATTCACTGCTTTCTGAGTGTATGTTTATGGTTTTCCTGGAAGTAATTCGACTTTTTAGATTTTATGCAGCTTCTAAGCTCACGTTTATATATTTTGAGTGGAAGATTTTATGTTGTTTCTAAGTTTTACTTATTGTGTTTCATCTTTTCTTACTATAATTTCCTTTTTCCTGAATGTCTGTTTTCTTGGTTAAGTAATTCAAGCTTTGCAATTTGTTGATTTTGTTGAGGAATAGCTATTCGTCTGATGTATAATCTTGATCTGTTCAATGCATTATCTATTATCTGCAATCAATAACTAAATCAGTGCCTTGGAGTGTCATTAACTATAAATGTTTTCCCCTTTTTTTGCTCGTAGAATTCTATTTTCTAGTTGTTACAGGTAAATATAGCTATCCTTGGAAAGTTAAACCGATTTATCATTCAATCAATGTCAATTTAATAACTCTCCATGATTACATTGTATTAGTTAATTAAATTGTAGTATAAGTAAGTATATAATCGATCACGAGTAATTTAATGCTTGTCAGAAATAAGAATGTCCACTGTAGGTCTTGAGCCCATGATCATAAGGTTAATAGCTTTGCTCTCTACACTTAATGTTGTTGGAAATGTTGTGGAGTTCCAATTATATCCACTCTTTCTCTGTTAACATTTCATTAAAATGATTTACACAGTTTAAGGCCATTGATTGCATTTATGAGAAAACCAAGGAGTTGTAGTACCTGAATGTATATGTTTACATGGACTAATTTGAACTTCTGGAATATGACTGTAGTGAAAGCTCTGAATGAAATTAAGGCATCTCTTGGATGGAGAGTGGTGTATGCATGGGTTGGAGATGATCCTTGTGGTGATGGTGATTTACCTCCATGGTCTGGTGTTACATGTTCAACGCAGGGTGATTATAGAGTAGTTACTGAATTGTAAGTTTGCTTCACCAGCTCCAAGTAGCTTTTCTCTAACTAAGCTTCAAATCTGTATGAAGTTACTCACAGAATTGAAATATCTTTACTTGTCACTTTTTCACAATGTTCTTTTCTGTTTTACCACTTGCAGGGAAGTTTATGCTGTTTCAATTGTTGGCCCATTTCCTACTGCTGTCACTAATTTATTAGATCTTACTAGACTGTAAGATTCTTCTTCTCTTGGTCTTTGTACTGGAATCTAGTTTTTACAAAACTTCAGCTACTTCTTACAGATTTTTTAACCTCATACAAATTGAGAAGTTTCCCTAAGTTCCAATATGCATGTTTTTACACCCTCCAGTCTTAATTGTTCATTAACCAACTTAAAGAATTGTTCTGCTTGGGTGTTGACTGTTACCCCAAAAAATGTGGGCGTTAGCTATAATGTGTGTAGTCATTATTTTCAAATTTGATACTACTATTTAAGGCAATAGTTAGTGGACAATTGTGCCTGTACTGTTGGATCTTTTGATTTTCATTCTGAACTTCCTAATCCATAATATTCTTTCATATGACAGGGATCTGCATAATAACAAGCTGACCGGCCCTCTTCCTTCTCAAATTGGGCGTTTGAAGCGTCTTAAAATATTGTATAATCCTTTACTTGTATACTCCTATTGAAGTATCTATATTCAGTAGCAATTATGAAGTGATATTTATGTTTAGGGCTTATGCTCTGTCTCCGAATAGCCTGTTTCTCCAGTTTCAGGATTAACATATGTTTCTCCATTAACACCTAGCTAGATGATGTTAATATGATTCAACTGTGTTGTACAAAGTGATTAAGTACATGGCTTTTCATCGAGATGAAATGTGTTAATAACAAGGAAAAGCTTACCTGATTAATGATTAATCCCAAAGAAGGGAGTGAGAGGGAGTCTGCCTGATATAATGTGCACCGAAGAAGAAAGATAAATTTTCAACTGAGAGAATATATACTCGTAGGCTTATATTCTAAAAAATAAAAACAGAGTAGGTTTATTATCAAGGAAATGTCACTTAACTAAGAAAGCCTTCTATACTCTTTCTAGGATTGAATTAGATGTATATCTCAACATCTACTTTCTGTTGCACGTATTTTACCTTTAAGTTTTCCTTTGTCCATAGGAACCTCCACGTTGTCCCAGAATGTTAACTTTAAAAGTTACTTTTATTGCTTTTTTACCCAACATACTAATATGAAATTATATCCTCTAGTTTCCTAGTGTGTTGTAATGTTGCATTATTGATTAGTTCCTTCTCCTGATTCTTTTTACCAGAAACGTAAGGTGGAATAAGCTGCAAGATGTCATTCCACCTGAAATAGGTGAACTAAAGCAGTTAACACATCTGTGAGTGCTTTTTGAGCTTCTCTAGGTGTGCTCTTTATCCATAGATCTATTTTGCTGTTATAGACTAATAACTGATATGTATATTTTTCAGCTACCTAAGTTTTAATAATTTCAAAGGTGAAATACCTAAGGAGCTTGCTAATCTTCCTGAACTTCGTTATCTCCATCTGCATGAAAATCATTTTATTGGACGAATTCCACCTGAACTGGGTTCCTTACAGTATCTTCGACACCTGTATGATACCATCTTAACTTCTTCTCTTGTTTAACATATGGAACCTGTGTCACACATGGATTTCACTTTGATCTCATGCAATTCCAGGGATGTTGGTAACAATCGTTTGGTTGGAACTATAAGAGAACTCATACGTATTGAAGGATGTTTTCCGGTTCTCCGCAACCTGTAAGTTGATTATTCTTGTTTAATTGTCATATCGCATGAAGTTATCCTGCTGCAAGTGTTATGAATTCTATGCTAATTCAAGGATAACACGTGCTGTAACTTGCAGATACTTAAACAATAATTATCTTACTGGAGGCATTCCAGCGCAACTTGCAAATCTGACGAAGTTGGAGATCTTGTAAGTTTTGGGGAGAAAGTTTCTCCGCGATTAATGCTCGTCTTTGGACAAATAACAGCTAATTTCTCCTGTACTGGTTTATAAATATTATTATGCTTAATTTGATTGCTCTGGTTATTACAGGTATCTATCCTACAATAAGATGACAGGAGTTATACCATATAGCATTGCTCATATTCCTAAGTTAACTTACTTGTAAGTCCTCCTCTATGCTCTTAGCTACTCATTTATTATACTGTATGTGGACAGTTTGGACCGAGAAAATTCAGAATTTTTGAAAGCAAATACAAAAACATCCTTAGCCTTAAGTTGAGATCTCTTTCTTTGATAGCTTTTTGGTGCAACCTGGTCTTTGTGCATGATTTAGACGCCACATTAGATATGATCAGGGACCTGCGTGTTATCATGCAGTAGTTTAGTGCTGTTTTCTGTTATCAATAGATTGCTTTATTAAATTATAAAAATATTAGCATATTGGATGCGATGTCCCAATGATTGTCTAGTGAGTGCAGTCTTTCAAGGATCTTTCTTTCTTAAAAAGATGAATTTTGAGGATTGTTCAACTTAAGGATGAACAAAAACAACATACTACTTGGTGTAATCCTACAAGTGGGGTCTGGGGAGAGTAGTGTGTACCCAAACCTTACCCCTACCTAAAAAAGGTTGAGATGTTGTTTCCAATAGACCCTCTACTCAAAACAGATGAACTCGGGACTAGTTGGCTAATGTGGTCTTGCAAGGGTCTCTTTCTTAAGGAGATAAATTTTGAGGATTGTGCAATTTATGGATGCTGTGACAAAAACATTAGATGGACTTGAAGCTTGTCAGCTGACCTTATTGGACCTTATAGTAGTAATATATAGAAATTGCCTGGAGAATTAGATGCAAGAGTGTTCTCATCAATTTTTGCACAAAAAACACAGTCCATCTCAATAGTAAAACTTCTCATGTACTCAAGATTTCAGTTGAGTTATGTAGCAACTGCACTGGTACAGAGATGCATGAATGACGTGTTAAACTTTAATTAGAAGATTGTCTGTTAGAACATAATAGTTTTATTGGTATAACGTGTTAAAAAAATTGCATGTATAATTCAAAAAAATATGAACACTGATGCATGCAAGTCTTTAAGCAATTAAACAAAGCTGATTTTGCACCTGGTTGTCAATATCCATTGCATTTCCTACCCATTTTCTTCTTTCTTCCAAAATCCATGGTCTTTCAACACTTACATTCCCCTCCATCCCCCAGATTAGAAGTCATTATTTGTACAAGTGAGAGGCTTGGATAATGCAGAAGGACCTGCATTTTGCTGGTCAAACAACCAAAGAATGTCAAGAGACAAGTCTAATTGTGTCAGATACATATCCCTGCTCAAGATATGCTGTGTTGGAAATGGGAACCTTGTTTTGAAGCTTAACCCACATATTTCCTTTGACAAAACCTTTTATCACAATATTTTCCTTTAAAGTGTTGGTGTTGGAGCTAAACCACACGAACTGAAACTAGGGCCTTCTCCATTTAAGTGGTTGTTAGTGTTGTCAAAGTGATGCTTGAGCCATGAACTGAGGCTGTGCACAAGTTTACTCCAATGAAGGTGGTAACTGGTAAGGGATATTTTCCCAAAACCAGCCAAGCAACTTCTTTTCTGAAAATATTAGATTAATTTGTGAGGTCCATGGGTTTCTTCTTGGATTGGGTTGCATTAAACAACAAATATTCAATAGATAGCATGCAAATCGATATGTCAGTTGTTAGCGAGACTGTACTGATGATCATTATTGATTACGAGCAAGAAACTGTAAATTTATTGCAAGAAAGCTAACGACGAAGTCTGAATTAGAATATTAAAGTAAGATGTTCCTTAAAGCTAGCCACTCAGATTCAAAATGCAAACTAGATGACCAATGATCCTTGCAAGTTGCAATATCCTAGATGTTCCGTTAAAGCCCAAATGGTCCTTGCAGTTGCATATGTGAGTGCTTTGTTCCTTTTATGTATTTATTTTTACGATCTTCACCCTTGACAAATCAGATGTTACACATCATTCCTTCGCTCTTCTTGATTATATACTGGTAGCTTTTGCACCAAAAAGAAAAATAGTTCTGCATGAATTTGATAGTTGATGTCTGAAGCTGAATCTGATTTCAGGTATCTCGATCATAATCAATTTTCTGGGAGGATTCCAGATGCCTTCTACAAACACCCTTTCTTGAAGGAAATGTGAGTGTTCATTTTTTGCTACTGACCATTCTTTGGCTTTGTGCAAATCCTAATAATATGATAAATTATGTCATAGGTACATTGAAGAGAACACGTTCCGGCCTGGTGTAAAACCAATTGGCCTGCACAAAGTGCTAGAACTTTCGGACGCAGATTTCCTTGTTTAGAAAAAGCCAACTCCTGTTATTTGTTCATGAGCAAAGTGTACCTAGGTTGTCGAAAGAAGCAGCCGGTCCCTGTTCTGTCAAGTATCTATTCTTTAGTTGCCAGATGAGAAAAAAGGTTCCATCTTTTTTGTGCAAAGAAGTAATCAACCAAACTTAGTTGCGGTTGCTGTAGTTAAATAGTACCACGTTATGAATCATTATTAAGTAATTGCGACATACGTCTCAATCAGTTCTCACTTGCCTTGAATTCTTGCATTTGGTTTTTGATTATACATGATGCGATTTCAGAGTTAATATGAAAGGGTAGAGAATATGCTCTGATATGACAGGTTTTGGATAGGTTTTTCATTTTTTCGTTAAATGATACAAATACATACCTAAATTATAGTATTATTTTTTTGAGTTTCACACCCTAACTATAGTTATTATCAATGTGCAAAAACACTTCGCTGAGTAGCTGATGGTATGCATGCTTTATTAGGTGCCAAAGTGGCACTCCATGTGGATAAATATTTTCACTAGAA

>M6-PGSC0003DMG400004195 M6 LRR-RLP

CTAAAACTCTTAACATAAATTGTTAAATATTTCTAACCTAAGTAGGTATTACCGAAAATCTTTAACACAGGGTGACATCGAGCTATTTAAGGAACATAAACAATTGGTAATTGCTTCACGCTATCTATTGCTCCTTTATTCATGAATTCATGTATATACAATGTAATTTTCTGTTAGAGGGGATCGTTGCACCTATGTGGTTCTGCCCTGACATCACTTATCTAGTTAAGCTCAACACTTTGAATATGTTCTTAAAAATGTGACTTTTAGAGGCTTCATAGGTCAATTTGTTGATAAAACAAAAATATTATAGAATGGGAACAGTGATATCCTCCATAAATGAAACTCAACTAAACTATTTATAAGACTATGTTTGTGACTATAATAGTGGAGTAATAATAGTAGAAAGGCTTTCCATATTATTTGTCCTGACATTTTTACTAATTGTTTTGTTGCTTAAATTGTTATGAATTGTTTTTCTTGAGCCGAATGTCTACCAGAAACAATCATCTTATCTTTTTTACACAGATAGGGATAATGTATGCATATACATTATCCTCCTCAGATCCCACTTTATGAGATTAAACTTCAGTAGTGATATTCTAAGCAATTACAAAAATATTTCCATAAAAACGAAAAATATACTAAGGAGAGACATAATATTGTGTTTTATAAAATGAAGTTGTAGTAATTTGTCTATTTGTTTACTTTCTTTAAAATTAACATGGCTTGTGAAAAAAACTTGACAACATGAGATGCTTAATGTATCGAGCTAGCGAATCTATTATTGACATAATCATCCCTTTTTTGGCAGGCCAAAATGACATCTCATCAATTTATTTGTTTCCTAGCAATGATATCTCTTATGATTATCAATATTAGTGCAAACAAGGCATGCCACCCTGATGATCTCAAGGGTCTCAATGACTTCAAAGATGGCATACATTCTGATACTTCAGGCAGACTAAGCAAATGGATTGGGCAAGATTGTTGCAATTGGCCAGGCATTTCTTGTAACTCTACAACTGACAGAGTTGTAGAAATCTATCTACCTGGTCACTATGTATCTGGTGACGATGAATCTCCAAACTTCGTGTCAAGCACAATGAGTGGTTCAATTTCTCCTTCAATTTCACTTCTCACTTCACTACAAGTTATTGATTTTAACAAACTAGTTGGACTAACAGGGCAAATTCCTGAGTCCATTGGGGTTCTCAAGAATCTCAAAGAACTCAACTTACAAACTAATCAAATCTCTAGTACAATCCCAGAGTCAGTATTCACATTGACGTCTCTAACAACTTTGAACCTTGAAAATAACCACTTGAAAGGGGAAATATCGGAGAATATTGGCAACTTACAAGCACTTCAAAAGCTTTTCTTGTCAAACAATTCGTTCACTGGAAAAATCCCAGTTTCAATAACAAAACTACATTCCATTTCCACTATCCACTTAGAAACAAATCAACTTGTTGGTGAAATACAATTGCCATTGAGTCCTGCCCAATGGCCTTTAATTAAAACTTTGGTTCTTGACAACAATAAACTTACTGGAGTTATACCTGATTCTATTGGTTATTTGACAACACTTTCATCACTCTCTATATCAAATAATCAACTCACAGGATCTATTCCTTCTTGTCTGGGAAACATAAAGCAATTGCAAATATTGAAGGTCAATAACAACAACTTCTCTGTTGAGATGTTGCCAACGTCGATTTGTGGACTTACTGAGCTAAGTGTTTTGTATGTTTCTCAAAACAAGATTCATGGACCATTACCTGGATGTCTTTCATCATTCAAGTACTTACTAGATGTCGATATACCGTTCAAACGTACAGTCACTCTTCCTATGGGAGTACACAAACAATAAGTAACGGTGACGGAGCCAGAAATTATGAAGGAAATCCAAAAATGCCTCAATTTGAAATGTTATGTATGTCTACAACAACCAAAAAAAAAAAACTTGTGTTTACTTATTATCTCTATGTTATATGAGACTCCAACTAGCCAGAATCATTGGCTAGTATGAAAAAAATCTAATACATAAAGTACTTTCTTTGTTTTATTTTTG

>M6-PGSC0003DMG400004384 M6 LRR-RLP

CAACCCTCTTTCTCAGGGATGGTCGAAGAGAGGTTCCTCTTCTGAGATCATCCTGAACACAGCAGTCAACAAAAAATATTCAAAAAAAAATAAAATATTTTTTTTGGGATTTTAATTCTACATTTCATAAAAAATCCCACAATTTCCATTGGGTTAAATTTCTTGAATTTTTTTTTTCTTTGATCTCTGTTTGCTTTTTCTGGTAAATCTTTCATCTTTACCCCCACTTTTTGAATTTTTGTTTGATTTTGCTTGAGAATATGGTTATCTTATGATGGGTTTTTTGTTTTTTCCCAAAAAAGATTCAATTTTTATTAATTCTTGTTGTTTTTATGTCTGCATTTTCAGCTAAAAATTGAATCTTGGAATTTTTGAAGACTTGGGGTTTCTGCAATGATGGAGAAAACAGAGGTAATATAGCCTATATTATAGAAATGTTAAGAAAATGAGAGACCCCATGAAGGGGTGGAGCCAGGGGTAGGGCCGAGGGGTTCAAAATTACGTTGTTTATACATAGTTAAAAGTTTTATATATATGTAGTACATGTTGAACCCCTTTGCCTTGTAGAGGTGTTTACTTCTTTATATTTTGAACGTTGTTAGTGAAAATCCTGGCTCCGCAGTTGACCTCATTGTGGCTAATATAGATAGATTTTGTTTGTAATCTTTTTTATTTTTTGTGTCTAGTGTAGAGTTGTGCAGATAAGCAACTGTGTGTTCGTGTGTCCGAGTGATTTTTGTTGCTCGGAGGAGAGTCATTTCATATCAAAAAGCTGGCGATTTCGTGTGAGAATTTGTTCAGAATCCTGGCTGTTGGCAAGAAATGAGGCGTAGATGGAACATTTATGGAATCAAGTTATTGCTTCTCACCATTTTAATTCTTCTTTTGGAGGTTCATGAATGTTGCACTCTGAATTCTGAAGGTAGTTACTATTCTCTAGTTTCTTTCTTTTCTTACTTCTGGAGGACCATAGTGTTCACAATTTGGGCCAGCTTGCATTAATTCGACTGCTCCACTGTATATCCTCTACCTCCAACCAACATAGGCATCAGTTACTATGCCCATCGAGGCTGAGGCATATGATGTGTTAGGCGAACAAAGTTCTACATTGGTAGCTAAAAAGATTGGAAATCTACAATTCTACATAAGGTGTAGGATCTCTTAACATGGTGTGAGGCCTTTTGTGAAAACCGTGCGGGCGTGCCCCTAAGCGGACAACATTGCACCATGTTAAGAGTATTTTTGGGTTGACTTAGCCCAACAAGATGAAATTACTTGCTCCCTCCATCTCATTTTATGTGGTCATGTTTTGTTGGACACAGTTAAAGAAACAAAGAATTTTGTAGCTTGTAGTCTAAAACAAGTTATAGACATTTGAGTATGGGTGTAATTTGAATATTAAGATTAAATTGTTTCTAAAGTCTAAACATGATGTTATTCTTTTTGTGACAAACTAAAAAGGAAAGTGTGCTGCGAAAATTGAGACAGAGGGAGTCGAGTTTTTGGTCTTAGCTGGGATATGAACCTTGATCTTTCATGGTTTTTTACTCACTTAGGGCTGTGTCTTCTGTCTCCCTGGGCCGTGGGCTATTCTCTAGTTTCTTTAGTCTATTCATGTTATTGGCTTATTCTTTGTTTCGCTGATTACTGTTGATTTGAGAAACAGGATTGTCGTTGTTTGAATTCAAATTGAATGTGTATTCTGATCCAAATGGAGTTCTTGAAAACTGGAATGCTGATCACTGTGACCCGTGCATGTGGTCTGGTGTCGAGTGCATGGATGGTAAAGTGTACATGTTGTAAGTCTCTTGTTATAGATACTCAAGTTTTCTAGTTAAGATTGTGTTAATAAACTTTCTCTAATGGTTAGTGTTTAGTGATTTGATTCTGGAACCAGTTTGTTGAACTAATAATGATAAACCAAATGCAACAGGGTGAAATTGTTTGTTTCGACTCTTGTGATTCTTCTTATGTCGTTAAATTAGTTGACCTATCTTCTTTTTTTCATTCTGTTCAGAAATCTCCATGGATTTTCTTTGGAAGGAACACTAGCATCAGAGCTTGGAAATCTCACTTACTTGAGATCACTGTAAGCTTTAGCTATACTAATATGCCTTTTCTTTAGGCTCCTGCTCATTAAGTCTTAAATGGTATTTTTTACCTGAACCGTCATCCGTAGAAGGTTGTCATAATCTTTGAATAGTCTTCTATCTTCTTATGTAAATTCGAAAGAACGTAAGGGATTGTTTGGTGTATGATGGACAAGCATAGATAGTCCGGGATAAAAAATTAGTAAGGGATTATCCCTTGTTTGGTTATTAATCCTGGTATAAGTTATATCAGGATTAGTAATTAGTAGCGGTATAAGTTATCCTCATCAGAGGGTGGAATAATAGTACTGGATTAGTTATCCTAGGATAAAATAGTGAAAATGACAAATATCCACGAAGTCCCTCAGACTCTTTTTCTACTAAATAAGGTGGATGGTGTTTTTGTAAAAGAACTTCTAAGAAATTTATGCAATGCATGTTATTTTTAATACAACAAACCAAACAATCACTAAGCAATAATACATTACCAACATAACTAATCCCAACATAACTTGTCTTCAAACTAAACGACCCCTAAGAGTTATGTTGGTCAAGTAAATTGGGAGGTAGAGACAAAGGTGGATAAGCTTTGGATATCTACTAATCAGCTTTCATCATCTTGTATTCTGCATTTACATTGTTCTCTGCATAAGTTGTGTTTGAATCTTAACAGTATTTCTTCTTGCTGTCTGGTTGCTTTTACAGAATGCTCTCAGGGAATCATTTTTTCGGTGCTATTCCTAAAGAATTCGGAAGACTCGATATGCTTGAAGTGCTTGACTTGAGGGATAATAATTTGAGTGGAACAATTCCAGCAGAAATAGGAGACCTGCAATCACTAAGAATCTTGTGAGTTAATGATTTTTTGTGTGCACGATAAACGGTTTGTTGTGTGGACGATAAACGGTTTGTTGTGTGCAGATGTATATATACATATGCATACACATTCATTATGCGAAGTGATTTCAAAATGCTCATCAATCGTGTTTATTTCGTTTTTCCAGGTTGATTCATAACAATAACTTTGAGGAGAGTGTTCCTTTGGAAATTGGGAAGCTTCATTTAGTCACAGAATTACAATTCGACGAAAATCTCACTTCTGCTTTTGCTTCTGGAACTCTCTGTATAAGAAAATTTCGTCATTGGTAGGATTCTTTTATTTGTTTTAGTACCCTATCTTCATTTTCAGCTGGTTTTTTGCATATCAGATGCTGGATAGTCTATCTTTGTGGCAACACACTATCTAATTGATCTAGCTCATTTCTCAATTTGATATTTTGCATGTCTTACAGCATCTGGCAAGGCAGCATGAAGCCATTCAAGACGATAACCTCCATTATTGAACCAATAAAATGTACACTTTTGCATTACCTCAGTTTCTTCCAACTGTAAGTAATTAAATAGTTTTTCACATCACGAGTCATCTAGTTAAGTTTTATTATTGAGAGACATTTTTTCTATTTCTTTAGGCCATCTCCTGGAACGGGCTTTATGGATGACCGGAAAGACTGCTCTGACAACCTTCCAAGTGAGTAAACACGTCATATTTTTTAAAATAACTCTGAAGAACCGTTTACAAATAATAATTCTTCATTGTCTAGCAGGGTTATCCAGGCCTCACATCATCCACACTATACAGCACCAAGAGGAGATTGCACGCCGTAAGTTGGCTGAACAATCCAGTAACCTTGCTGCAGCTCCTGCCAATTTCAAAGGACCGTTGGGTCCTGTCATTCAAATGCCAAGAAGCAGTGGGTCGTTTCGTGCGGTGCCAAATTCCAATGGAGCACATCCTACAAACTTACCATCACCACCTCCACAGCATGAATCTCAAGACAAATTACATCCTGTGGGACAGTCTGCAAGTGCTGTAAAACAGTCGCCTGCTGTGAAAAAGCCAACTAATAGCCAAGGCCCCCCTGCTGGAAAATCGGAAAGTACTTGGAAGTACGTAGTTGGGATTTGCATTGGAGTAATTTTGCTCATTCTTGCTGCTTCTGTGTTCTTAATCTGCCGAATCAGAGCAGCTAGAACAATTGGCCCTTGGAAGACTGGATTGAGTGGACAGCTGCAAAAAGCATTTGTTACAGGTACTTATATCATATTGTGCCAAAAAGTGATACTGCTAATATTCTGCTCCCCACCCTGATAGTAGTTGCATTGCTAACAATATTCTTAGCATAGTGTTACCTTATCAAAAAAAATATTCTTAGTGTAGTGTTGTTATAACAATTTAACTGGAGTAAAGGGAGGTACTGATATTCTTCGTCTTATGAGGAATTAGAGGACTAATATTCTCAGTGTAGCATTGTTATATACCCTTTTTCACTATATGTTTGCTAGGACGAGAAGGTACAAATGGAACTATTTCCTTCTTTCTCATGAGAAGTGAATTAGCTGAGACATCAATACTAACGCTTGAAGGCTGGAATCCTTCCCTTATACTTCTAAAATGTCGCTGATCTTTCCTCATACATTTGCTTTTGTCTAGCTTATATCCCCCCCACAAAGTTGCTTTTCAACGACAACCATAGGAACGTGCAATCCTTTATGTTAATCTATGATATTTTTCGTCTATTTTAATCCTTTTGCCCGTGAAAACATTGTTATAGGGGTTCCAAAACTGAACAGATCCGAGCTTGAAACAGCATGCGAGGATTTTAGCAACATTATTACCAGCGGCGATTCATATATAGTATACAAGGGAACACTGTCTAGCGGAGTGGAGATTGCTGTTATCTCAACTACTATAAGTTCTCTGGAAGATTGGTCCATGCACTCGGAAACAGCCTTCCGGAAGAAGGTTAATCATTTCTTTGCTTTTACAAATTAACCACCTCC

>M6-PGSC0003DMG400004529 M6 LRR-RLP

TTTTGGCCTACGTGGCTGAGTCCGTGACTTCACCTATGTTAGGCGCGTGGGTGCAATTTTTAGACAACACTGACCAATAAAAATTGGCCACGTGTTAAAATAAACTTTATTGTTGTTTATTGTTCTTCATCTTTAACTAAGTTTTCTTCTTCTTTATTGTTCTTGATCTTTAGAAAAAAAAACCTCCATTATCATTTTCATTCAAGTTAAATTTCTTCTTGCTTCGAAAAGGTAAAGTAAAATTTTAATATATTCTTTATATTTACACTTTTTTATTTCGATTGTTAGTTCTGATTTTAAATTTCTTTCACAATTTTTATGTATAAATATTAGGGTTATGACAAAATCCGAGTTGAATAAGTTGTGTATGGAGCTAGGTGAAGGATGTCCCTTGCTGAGGGATATTGTGTTGTCTCACTGTCGTCAAATAACAGATGTAGGGTTAAGTTATATTGCTAATAGGTGTACCTTGTTGGGAACTAGCCACATGGTCTATTGCCCTGGTATCACCGCTGCTGGAGTAGCCACGGTGATTACGAGCTGCACAAATTTAAAGAAAGTCCTTGTAGAGAAGTGGAAAGTGAGTCCAAGAACCAAAAGGAGAGCAGGATCCATCATCTCCTATCTCTGTGTTGATCTGTAGATCT

>M6-PGSC0003DMG400004561 M6 LRR-RLP

GTTTGATGTAAATTGTTTCTCTTGAGAAGTTGAAAGGAAGGATTTTTTTTATTTTTTTTTATTTTTTAACTACTTTGGCTTGATTAGTGAGAATAGAAAGAGGAAAAGAAAATGAAGGGCAAACAAAAGGGGTTCATTCACATAATTACATATTCTTGTAAGTAAGTTTGTTACTTTATGTAACTTAGATGGTGAATATAATCTTTATAACCTATAAGAATTGATGATTTAATCTTTCTGCTATAAGTTAAACTCTATGTACTAAATCATCTGCTATAGTTGATAAATGTAAGGGGTTAAGAACTAGGAAAGTAAGAGCTTTTGTTTAGAGAGATAGCGCGTAAACAATGAGAGAGCTTAGTGCACCAGGAGGCTGTCATTTTTTGGTTTAAGATTTGAGGTGTAACAGGACAAACATTTGGCTTCTCTATTAGGTTATTCTGGAGAAAGTCCTCTGCACTTGTGACAGTTGCTTGTTGTAAAATTAAGAGCTATCACATTTTCGAGAGTTTTACACCTCAACTATATTAGTCGTTCCCTTTTTCTATCTAAATTTGAAACAACATTTGCTCTTTTGGTAGGTGTCATCCGTTGTACTTCGTGAGGACATTGTCAATCTTCTAGATTTCATAGAGATGTTAAATAGTGGGCGTGTTCAAACTTCTTTTGACATGGATCAAATTGAAAAGTTGACATTGGAGCTGACGTTTGTGTCCGCATTTTGTTATCTTTATTACTTTCTTTTTGGAAAGTTGTAATGACGAAATGTCCTGCATATCAAATGAGATTCATGATTTGGTTCGGTCACTGTTTCATCCAAGTGGAGAAGACATGCTGGTTAATATAAAGAATCATGACCTCTTGGAGAATATCAAAAGTTATATTAACTCACATAGCTATGTTGAACCAAGTCGTGTAATCATGACCGAGGATCGTTTGGTTGAACTTTTAGATGCCATCCTCGTGTATCTCCAATGTCTACTCAACTGTAGTTCTGAATATATTTTCCCATTAGTGACTCAATGTGAGCTTCTTCAGAATGTATTTGGCAATTTAAGAGATTTCCATAGGTTGAAAGTAAATGGTTGTGTTGAGTATGAAACAATCGAATATGTCTTACCTCATCTTCAACTTATGGCTCTGAGAGTAGTAAACTTCTGATTTACTCTTTTGAATAGTCAACTGGACAAATTCGATGAATCGGATGTCTTTCTGGTTAACTCCAAGCTAGCTAATACACTTGTGGAGGTTATTCCTGTTGAACTGGAGGATATGCACATATGTTGAACAAATTTGAAAGGTTCAAAGTCGGAAGAAGTTGAACGTTTCATCAAGCAGCTCGTAGAAGCCTCTCCGGACATTCTTAGAGTATCTCTGATTCATCTACAAGAGCACATGGCTAATGCTGGTATTCCTAGCGCATCTACGTGTAACATTCATGTCATGACAGAGTTCCTATTGATTATTCTTCCTGATGTGCCCAAGGACATTATTCGTCAAGACAAATTGTTTGTTCTCTTGGCACGTGTTAGAGAACTTACCAAGGAGGTATTCGTTCTTGTTCGCAACTTAGAAGAGAATATGAATGACGCAAGTGATGCAAATCTAAACTTGCTGGAAAATATTGAACTCCTGAAGGAAGATCTCAAGAATGTTTTCTTGAAAGCCCGTGCAGACTCATCTCGTCTTTGCTTCCCCATGAGTGATGGACCGCTCTTCATGACTCTTCTGCTTACAAACTTGAAAGACTTTGTCGATTCCAATGCTTATTCAGTTACTTTGATAAAAGAAGAAATTAAGCAGGTGAAAGAAGACCTGGAAATCATAAGATCGTTCTTCGGGTATGCTGAGCAAGAGTTGAACAGAGATCTTTGGACTCATGTTCTAGATTTGGCATATGAGGCGGAACATGCCATTAACTCAATTCTTGCCCGGGATCATGGTCTATTATAGCTTATCTTCATACTTCCTGATACTGTAGAAAAGATCAAGCTTGTCAAAAAAGAGGTACAAGAGAAGATCCCCAAGAGCAGCGGTATTATTGTTGCAAATGCTCCCAACAAGCCGGTTGAAAGAAACTCATCGAGTACAGTTGGTAAAATAATCGTAGGTTTTGAGGAGGAGACAGAGTGGATAATTAGGAAGCTCACCAGTGGACCAGCAGAGATAGATGCCATTTCCATAGTCGGCATGCCAGGGCTTGGAAAAACTACTTTGGCATACAGAGTGTATAATGATAAGTTCATTGTTGATCATTTCGACGTTTGTGCTTGATGCACAGTCGACCAGGAACGTAATGCGAAAAAGTTGTGGCAGAAAATTTTCAATCAAGTTATTGGTTTGACAGAAAGATTCAGTGAGAATGACATAGATGATGACGTTGGTTATAAGCTGCGGAAACAACTATTTGCAAAACGGTACCTTATTGTCTTGGATGACATGTGGGATACTG

>M6-PGSC0003DMG400004722 M6 LRR-RLP

TTCAAGAAAATGCTTACTGTAGATACCTCATTAGTTTGGTGTAGTTACTTTTACCACACACGAGTTCATGGAACATAAGCAGAGATTGCTGCTCTTCGGATGGAGTCGTATGTGATGACATGACTGGCCATGTCACTGAACTTGACCTCAGTCTATTCAGGGACGAGAGCTTGAACTTGTCAAATTTTTGACCATTTACACCACTATTGTTCTTTCAAGTAACAGATTTGAAGGACGTATTCCAAGTATCATGGGAGAACTCATTGCTCTTCGGGTGCTGAATTTGTCTCATAATGAATTGCAAGGTAATAAACCATCATCACTTGGAAGTTTATGTGTGGTCGAATCATTGGACCTTTCATCTAACCATCTTTCAGGAGAGATACCATAACAACTGGCTTCTCTAACGTCACTTGAATTCTTAAATCTCTCCCACAATCATCTCGATGGATGCATCCCTCAAGTACCTCAATTCGCTACATTAAAGAAATATTCATATGAAGGCAATAATGGATTACGTGAATTTATGCATTGCCAAATTCATCATTGCATATGTTGGAGGAATATCAGGGGCCATAAAATTTTATGAGCTTAAGTTGGCTCGAACATCATAATTATTAAAAAAAATGTCTAAAACCCGTTGACATACTAATGACAAGGTGATATATAGAGTTGAAACAAGTTAGATATTTTTTTCCTTTTGTTTTTCAGTAATCACCTCTTATTTGTTTTAATGTCATAAATTCAACTTTCTGATGCAATATAGTTGAAAGTAGTACCTATTAATTTGAAAGTACATAGATTCTGCTTATATGGGTGCAGCTAAAGTATTTATGTACAACTATACTTCCTTTCATGATTCATAAATGTACACAACAACCTCATGACCCACTGCAATAGATTATGAAGTACCTTCACTGCAAATTGAATGAGTTGTAGCTTTCATTTCTAAGCGCGTGACTTTTCTCTGATAACGAGTGACTTTTCTGTTACATAACAAATAAGGTGACTTTCAAAGATAATTATCAATAATTGAGTGACAATATGAGAAATCAACTCTTGATGTCAGCTATATGAAATCTCAGTAGTCATATAGCTTTACCTAAACTCATCTCGATCTAGTACAAAGTATATTAGAGGTAATAATGTAGTCTAGAATTATTGTTCTATGACATAATTGTTGGATTGGTCCACTAATTTAAAGGTGAGATTGAGAACCAAACATGTCCGCATCATGGCTGATAACACAACAAAACTTTTTGGCTACTTGGTTAAGTCCTTAGCTGAATTACATATGTAACCCGTGGATAAAAATAAAAAGCAAAAAGACAAGGACAGAACAACCAAAATTTGAATGGAGAATGAGAACACTTTCTCTGATAAATATGAACAGACAAAGAAGTGTGAATAAGAGAGACAGAAGCCGAATTGGCAACATTTGGTGGCAGTCAAACATGTTTCGCTTTGATCTTTAATATTATGCACTATATAAAATCCTACGAACTCTCTACAATTAGTTCATTGAAACACAAAATATGATGATGGTTACCAAAATATCCTCTCTACTTCAGTTTTTCACTCTTTTATATCTCTTTACAGTTACTTTTGCTTCCACTGAGGAAACAACTGCCCTCTTTAAATGGAAAGCGACTTTCAAGAACAAAAATAATTCCTTATTGGCATCATGGACACCAAGTTCTGATGCATGCAGTGACTGGTATGGAGTTATATGCATTAATGGTAGGGTAAACACGTTGAATATTACAAATGCTAGTATCATTGGTACACTCTATGCTTTTCCGTTTTCATCCCTCCCTTTTCTCGAGTATCTTAATCTTAGCAGGAACAATTTATTTGGCACCATCCCACCTGAGATTGGTTATCTAACTAATCTTGTCTATCTTGACTTGCATATCAATCACATTTCAGGCTCAATCCCACCACAAATCGGCTCACTAGCCAAACTTCAGATCATCCGTTTATTTGACAATCATTTAACTGGTCCTATTCCAGAAACAATAGGGTACCTAAGGTCTCTTACTAAGCTATCATTGAGTATTAACTCTCTTACTGGTTCTATTCCTGCTTCATTGGGAAACTTGACCAACTTGTCTTTTTTGTTTCTTCATTTTAATCACCTTTCTGGATCCATTCCCGAAGAAATAGGTTACCTAAGGTCTCTTGCTCGACTAGATTTGAGGAATAACTTTCTTATTGGTTCTATTCCTACTTCCCTTGGCAACTTGAAAAATTTGCAAGTTCTGCTTCTCAATGACAACAATCTAATTGAGGAAATTCCTCCATCTATTTGCAATTTGACATCACTGGCAATCCTGTATTTGTCGAGAAACAACTTGAAAGGAAAAATTCTGCAATGCTTAGGTAATATCAGTACCCTCCAGTTTGTGATGATGTCACATAATAATCTCAGTGGAGAACTCCCTTTGTCTATTTGCAATCTAACATCACTACAAGTTCTAAATTTGGGTAGAAACAATCTGAACGGAGCAATTCCGCAATGTTTTGGCAACATGAGTGGTCATCTTGAGATTCTTGATATGCAGCACAACAATCTTTCTGGTACTTTTCAAAGCACTTTTAGCATTGGAGGAGCACTCAGAAGCTTCAATTTGCATGGCAATAAGCTAGAGGGGAAAATTCCACGATCCTTGGCAAATTGCCAACGATTGGAAGTTCTTGATCTAGGAGCTAATCTCCTCAATGACACATTCCCGATGTGGTTGGGGACTCTGCCAGAGCTACAAGTGTTAAGTTTGAGATCCAACAAATTGCATGGACCCATCAGAACTTCAAAGTTCATGAAGAACTTGTTTCCTAAGCTTCGAATCATAGACCTCGCTAGCAATGCATTTACAGCAGAGTTACCTGCAAATCTTTTTCAAAATTTTGAAGCCATGAAGAGAGTTGATCAAACTGTGGAGGATTCAAGTAATAAAGAAGGCGGTTATTACCAAGACTCCGTAACTGTCGTAACAAAGGGACTGGAGCTTGAAGTTGATTCAATCTTGTCTTTGTACATCACAATTGATTTTTCAAATAACAGATTTGAAGGACGCATTCCAAGTATTATGGGAAATCTCATTGCACTTCACATGTTGAATTTATCTCATAATGGATTGCAAGGTCATATACCGCCATCACTCGGAAAATTATCTTCAGTTGAATCATTGGACCTATCAGCTAATCATCTTGAAGGAGAGATACCTAAACAACTTACATCTCTTACATTTCTTGAATTCTTGAATCTCTCCCACAATCAGCTCCAAGGATGCATCCCAACAGGACGCCAATTTGATACATTCGAGAAGAACTCATATGAAGGTAATGATAAGTTACGTGGATTTCCTAGTTTGGAGGTTGTGGAAACAGCAGGTTTCCCAAGACAAACAACACGACACATGAGCTGGAACAAGAAAGTGATTCTACATTTCTGAATGAACTCAACTGGAAAGTGGTTCTTATGGGATATGTGTGTGGACTTATTATTGGATTCTCCATAACATATTTAATGCTTTCAGCTCGAAATCCCAATTGGCTTTCTAGGATTGCTGAAGAACTAGAATACAGAATCCTTATGTGAAGGCGAAAGAAG

>M6-PGSC0003DMG400004727 M6 LRR-RLP

TTGGTCTTTCCTTCAATAATTTTAGTGGTTTTGTGGATGTCAGCCTCTTTTCAAACCTTAAACAGCTTGGGGAACTTAGTCTTTCATATAATAATATCTCACTGACCAATGACAACAAAGTTAATGTTACTTTGCCCGAATCTCTTTATTTGTTAGAATTGGCCGCATGTGAAGTGAAAGAATTGGAGTTTCTAAGATCTGCAAACCAGCTTTTCGATTTGGATCTTTCAAATAATAAGATTCAAGGAAGAATTCCTGATTGGGCATGGTCTAACTGGATGTTTTCATTGCGAGATCTTAATATATCCCACAATATGCTAACGAGTATGGATTCAATTCCTCTTCAACATCATTTAACAATTGATTTGCGGTCAAATTTACTTCAAGGTTCACTACCTATTCCACCAAATTCCACAAGATACTTCTTCATATCCCATAATAATCTCACTGGAGAAATTCCTTCATCTATTTGCAACTTGACATCACTGATAATGCTGGATTTGGCGACAAACAATTTGCGTGGAGAAATTCTGCAGTGTTTCGGAAATATCACTTCCCTCCAGGTTTTGGATATGCGCCACAACAATCTTTCTGGGAATATTCCAACAACTTTCAGCAATGGAAGTTCACTGAGCAGCCTCAACTTGCATGGCAACAAACTAGAGGGGAAAATCCCACAATCCTTGACCAATTGCAAACAACTTGAAGTTCTTGATTTAGGAGACAATCACCTCAACGACACATTCCCAGTGTGGTTGGGGATTCTACCAAAGCTGAAAGTTCTAAGCTTGCGATTCAACAAATTGCATGGTCCCATCAGAACATCAAGGATTGAGAACATGTTTCCAGAGCTCCGAATCATAGATCTCTCTTGCAATGCCTTCTCAGGAAACTTACCTTCAAGTTTGTTTCAACATTTGAAAGCCATGAGGACAATTCATCAAACGATGAACACACCAAGTTATGAAGGAAGTCGATATTACCAAGACTCGATAGCTGTCGTATCCAAGGGATATGAGCGTGAAATTGTGAGAATCTTGTTTTTGTACACCACTATCGATCTTTCAAACAACAAATTCGAAGGACATATTCCAACTATTATGGGAGATCTCATCGCACTCCGTGTGCTGAATATATCTCATAATGGATTGGAAGGTCATATACCGTCATCACTTGGAAGTTTATCTTTGGTTGAATCACTGGACCTGTCAAGTAACCATCTTGTAGGAGATATACCAGCACAATTTGCTTCTCTTACTTCTCTTGCAGTCTTAAATCTTTCCTACAATCATCTCGAAGGGTGCATTCCTCAAGGAAATCAATTTCATACCTTTGAGAACAATTCATATGAAGGCAATGATGGATTACGTGGTTTTCCTGTTTCAATAAGTTGTGGCAATGATCATGTATCAGGTAAAAATTATGCCGTAGATGATGAAGAAAGCAATTCTGAATTTCTGAGTGACTTTCGGAAAGCTGCTCTTATGGGGTATGGAAGTGGACTATGTATTGGAATATCCATAGTATATTTCATGATGTCAACTGGAAAACCGATATGGTTTGCAAGAATCATTGTAGAGCTGGAGCACAAAATAATGATGGGAAGGAGAAAGAAGCAGCGAAGTCAAAGGAATAACAGAAGAAGAAATAATCACTAATCATATTACTGTTATTTCTTTAAGGAAAAGATGTAATAAGTTGTAGACGCATTATCGTTAAATCCTGTTTGCATAAAGTGTGTAGCATCATTTGGCCATGAAAATTGAGGGTGAG

>M6-PGSC0003DMG400005171 M6 LRR-RLP

ATGATGAAGCTAAGGCATCTGAATATAATCAATTTGTGCTTCTTGTGGGAAGATAATGATCGAGCAATTTTTGACGAATCTTCAACAACTATGCTAGAGAACTTGAAGACATTCTTCAGTATTCATATACGTGTAGATGATACAACTCCAAGATTCTGGTGGAGGTTTCCAAATATTGAATAACTTAGCTGCCTCAGTGTTGACGATTTACCTAGTTGTCCTTTGTTTCCCTCAGTGGAAGTTAATACTCGGCATCAATCTCTTGATCTGATGATAAGCTATAAGGGATTCTGGGATTCAGTTGGTTGGGAGAGCTACTTTGTCTTCCCTTCAAATCTTAGGCATCTGTCCCTCAGTGGTTGTTTTTTGACAGAAAAAATGGTTTCAAACATTGCAAGATTGAAGAAACTTGAGAAACTCTCATTAGTTGGGGGATTTCCTTTGGGGAAAATAGAGAACCGTTGCTGGGACGTCACAAATGTGGAGTTCCCTGCACTAAAATACTTGGCATTACGCTTTATGAGGATGGAAGAATGGAAAGCTTCAGAGGAATCCTTTCCTGTGCTTGAGGAGCTAACTATTAGTGTTGGTTACATCATGGAGATTCCTCCTAGTTTTGCGGATATTTTAACACTGCGACTTATTAAACTGGACAAGCTCTCCAATTCACTTGTGGTTTCAGCTTTGACTATTAAAAAGGAAATAGAAGAAAACACAGGATGTGACCGTCTCCAA

>M6-PGSC0003DMG400005172 M6 LRR-RLP

ATGGAAGTTGTTGAGACTTGTGGAAAAGGAACTCTTTCTGCTGGAGAACCTTCCACCAAACATAATATTACATCAATTTATGATGATGATGATGATGAAGTTGTGGGCTTTGAGAAAGATGCAGAAATTATAATGAAGAAAATGATACGAGGAACAAAGGAGCAGGATGTTATCTCCATCTATGGAATGCCTGGACTTGGTAAAACAACTTTGGCAAGGAAGGTGTACAACAATCCCTCTATTATTAATTACTTTGATGTTAAAGCATGGTGTGCTTTGTCGCAAGCATATAATAGGAGGACATTATTGGTTGAGATTTTCAAACAAGCTACAAATGATAAGATCAAAATCAAAGAGGACGACGATGTAGCTGACATGTTGCGCAGGGTTCTAATTGGCAAGAGATACCTCATCGTATTAGATGACATTTGGGATGTCGAGGCATGGGAAGATTTGGGAATATGTTTCCCTCAAGGTGAATATGGAAGCAGAGTAATGGTCACAACACGAATTGAACAAGTGGCTAAGCATCTTCAGCACCACAGTGATCCTTATTCTCTTAGCTTTCTAACGTCCGAAAAGAGTTGGGAATTATTGGAGAAGAAGGTATTTCGAGGAGAGAGTTGTCCTCCGGATCTACTGGAAGCAGGGTTACAAGTTGCCCTACACTGCAAGGGATTGCCTCTTGCGGTTGT

>M6-PGSC0003DMG400005226 M6 LRR-RLP

AATATTAAAAGACTTGATTTGTCTTCTAATAATTTTTCAGGATCGCTCATTTCACCTAAATTTGGTGAGTTTTCTAGTTTGACGCATCTTGATTTGTCAAATTCAGGTTTTAGATTCGGACCTCACAATTTTGAACTTCTTCTTAAGAACTTAACCCAATTAACAGAGCTTGACCTTTACGGTGTGAGCATCTCATCCACCTTTCCTTTGAATTTCTCTTCTTATTTAGCAACTCTACGACTTAGTGACACACAATTATGGGGGATACTGCCCGAAAGTGTTTTTCACCTTTCCAACCTGGAAACACTTGATTTATCATACAATTCCCAACTCACAATTAGGTTTCCCACGACCAAATGGAATAGCAGTACATCTCTCATGTACTTATCTCTACCTGAAGTGAATTTTATTGGTAAGATACCCGAATCATTTAGCTATCTAACTTCATTGCATTACTTGGATATGAGTTCTTGTAGCCTCTCAGGTCCTATTATTAAACCTCTTTGGAATCTCTCCCACATACAAAGTTTGAATCTCAGAAATAACCATCTTGAAGGACCAATTTCCCAATTCTTCAAATTTGGAAACCTCACTTGGAATTAACAACTTTGATGGCTAACTTGATTTTTTATCCTCCAACAGAAGTTGGACTCAACTTGAAGGGTTCGAAGTTTCTTCCAATTCCTTAGTTGGTCCAATTCCTTCCGATGTAAGTGGTCTGCAAAGCCTACAATCATTCTTCTTGTCATCAAACCACTTGAATGGGACTATACCATCATGGATATTCTCCCTTCCTTCACTAGATGAATTATACTTGAATGATAACTATTTCAGAGGAGAAATTCAGGAGTTCAAGTCCAAAACATTGTGGGACGTTACTCTAAAACAAAATCAGTTGCAAGGTCCTATTCCGATGTCACTCCTAAACCAACAGAGCCTATCTTATCTTCTCCTTTCGAAAAATAATCTCAGTGGACAGATTTCTTCAGCTATCTGCAGTCTAAATGCCCTGAAAGTGCTAGATCTTGGAAGTAATAATTTGAAGGGAACAATCCCACAATGTTTGGGTGAGATGAGTGGACTTTGGGTTTTTGATCTAAGCAACAACGGTCTTAGTGGGGCAATTAATACAACATTTAGTATAGGAAACACGCTCAAAGTCATTAACTTGCACGGGAATAAGCTAGAGGGGAAGTCCCACGATCTTTGGTCAATTGCAAGAGTTTGGAAATTCTAGATTTAGGTAACAATGAGTTGAGTGACACATTTCCAAAATGGTTAGGAACCCTACCGAATTTGAAGATTTTAAGCTTGAGATCAAATAAGTTCCACGGGCCCATTAAAACTTCAAGGGCTGGAAACTTGTTTGTACATCTTCGAATAATTGATCTATCATCCAATGGATTTAGTGGGAATTTACCTATGAGCCTTTTTGAGAATTTGAAAGCTATGAAGATAATTGATGAGATCACGAAAATCCCACGGTATGTGGGAGAAGACTTGTATTATTATTATAATTTTTCTGAAACAATTACAACAAAGGGGTTGGATCTTGAATTTCGTGGAGTTTTGACTACAAACATAGTTATCAATCTCTCGAAGAATAGATTTGAAGGTTGTATTCCAAGTTCCATTGGAGATCTTATTGCACTTCGTACTTTGAATTTATCTCATAATGGCTTGGAGGGTGTCATACCAACATCGCTGCAGTACTTATCTGTACTTGAATCATTAGATCTCTCATTCAACAAAATAAGCGGAGAAATTCCACAACAACTTGCATCCCTCACATTTCTTGCTGTCTTAAATCTCTCTCACAATCATTTTGTTGGATGTATCCCTAGAGGAAAACAATTTGATACGTTTGAGAATAGTTCATACCAAGGGAATGATGGGTTACGTGGATTGCCAGTGTCAAAAGATTGTGGTGGTGATGAAGGGGTACCACAAACAACAACTCCATTTGGGCTTGATCAAGGAGAAGAAGGAGACTCAACAATGATCAGTTGGCAGGCAGTTCTCATGGGTTATGGTTGCGGTCTAATTATTGGATTATCCGTAATATACATAATGTTATCAACTCAAAATCCAACATGGTTTTCGAGGATGGTTGAAGAATTGGACCACAAGATTATTACGAGAATGAAAAAATACAATAAAAGCTATTAGTGTGCAACCTAGCCAGGTATTCAACTTGATCTTTATCCTTCATAAGATTATTTTGTATATCTCAATGTATTATATATCTACCTCCTTCATCCTTAAAATTCTTGACTTCGATTCTTCATTTTTTAAAATTGCAGGATTCGAGCCTAATATGTTGTTGGTGATGCAAATATAAGCTTTTATTTTTCACATTTTGTGATGTATCTAAGTATTTATCGCCCTACGAATAAAGACATGAGCTTTCTTCCTTTGAGCTTCGGCCATAATTTTTAGAATGTTTTTAAATAGAGATTGTTAGCTTGAGTTCAGCCTCATAATTAGTTGAGTAATTAGGCTCAACTGCAAAGTCA

>M6-PGSC0003DMG400005538 M6 LRR-RLP

TTGACATTTTCCTAGAAAGACTTTGTGCATTGATTTGAGAATGACAACATTTTGTTGAATTGATTGATGATTTGATATGGAGGCAGCAGTCCATACTTGACTTTTCCCCTTTCCTCTCTTTTTGATTAGAAAATTCGCAAGTAATTAATTTCAATATGAAGATTCAAAGTTGAAGGGATTTGTTTGATAAGTAGCTTTCATTGAAATTTTTGCAGCAAGACTCATTGCAATTACTTGATAAGTCAAAGAACTGCACTAATACTTCTAGGTGCAAGAAAAAAGATAATTGGAGGGACACTCTGTGCAAAAACTTGGTGTTATTTCCCTCTTTGATAGGCCTTACACGCCGAGAATTCAGATTATTCGGGATCCTAATCTGGATATCAAAAATCGAATAGAAAATCAATTTATTTTTAGAAAAAGCCATTTCTCATTATAAATATTAGGTAAACAAGTCAATATCCTGTAATTATACCAAAAAGACACTTTGGCATATGACTTAGTCTTGTCTAATATTTGAGAATGACATTTCTCTACTTTCTTTTGAAATTTTCATAGGAAGACTTGTGCATTGACTTAAGAATAACAAACATTAATTTCCCTATGCAATATGATTCTGGATTTGTTTTTAATTTCTTTTTTTGTTTAAGAAAGCCTTTTCTTCTTCCTTTTGGAGTAGCTAGGAATGAACTTTGCCATTTAGAGCCACTTTTATAACGAGGATATATGATATATATATAAGCTCTGAATGAAAAAGTTAAGGGGTATATTAGACCCTTTTTCCTGAATAAGAAAGTAAAACAACGTACAATTATATTTTAAGAAAAAAAGATGATTCTCTACTCCTTTTTTAAGCTTGTTTCTTTAGAGAAATAAAACAACCCAAACTTGACTTGTGTTGAACAATATAAAATCCTAAGTTAGTTCATTGAAACCCAAAATAATAAAATCAAAATATGATGATGGTTCCCAGTAAAATATTTTCTCTACTTCAGTTTTTCACTCTTTTATATCTCCTTACAGTTACTTTTGCTTCGATCGAGGAAGCAACTTCCCTCCTGAAATGGAAATTAACTTTCAAGAACCAGAACAACTCCTTATTCGCTTCATGGACACCAAGTTCTAACTCATGCAGGGACTGGTACGGAGTTATATGCTTAAATGGTAGGGTAAACACGTTGAATATTACAAATTCTAGTGTCATTGGTACCCTCTATGCTTTTCCGTTTTCATCTCTCCCTTTTCTTGAATATTTTGAACTTAGCCTGAACAATTTTTCGGGAATCATTCCCCCTGAGATTGGTAATCTCACTACTCTTGTCTATCTTGACTTGTCCATGAATCAGATTTCAGGCACAATTCCGCCACAAATCGGCTCACTAGCCAAGCTTGAGACCCTCCACATCTTTGCCAACCATTTAAGTGGTTCCATTCCACAAGAAATAGGTTACCTAAGGTCTCTAACTGAGCTAGCTTTGTATACCAATTTTCTTAATGGTTCTATTCCTGCTTCATTGGGGAATTTGAACAACTTGTCCTATATGTATCTTTATGATAATCACCTTTCTGGCTCTATTCCTAAAGAAATAGGAAAACTTGTCAATCTTGTTGGAGTTTCTTTTGATACAAACCAATTAACAGGTCATATCCCTCCAGAAATAGGCAACTTGACCAATGTAAAATTGTTCTATGCTCCCTTCAATGAATTATCTGGTCCCATTCCTGCAGAAATAGGGAAGATGAAGTCACTTCAGGAATTAATCTTACATACAAACAATCTTTCTGGTCTAATTCCAAGAACTATAGGTGGCTTAACTGAGCATATCTACAACTTAATGATAATCAACTCTCTGGCCTAATTCCTAGAGAGATGGGGAATCTGAAAAACCTCAATGGTCTGGCATTATTCAATAATCAACTTTTTGGCCTAATCCCTAGTGAGTTGGGGAATCTTAGTATCCACTATCATCTAACATTATCCAATAATCAACTCACTGGTTCAATTCCTTCCTCCTTTGGTAATTTGAGAAACTTGCGAGTTCTGTATCTCCATGACAACAATCTGATTGAGGAAATTCCTTCATCTATATGCAATTTGACATTCCTCATGTTCCTGCATTTGTCGAGAAACAACTTGAAAGGAAAAATTCTGCAATGTTTGGGTAATATCAGTACTCTCCAGTTGGTGACACTGTCACATAATAATCTCAGTGGAGAGCTCCCTTTGTCTATTTGCAATTTAGCATCACTACAAATTCTAAATTTGGGCAAAAACAATATAAAGGGAGTAATTCCGCAATGTTTTGGCAACATGAGTGGTTATCTTGAGGTTCTTGATATGCAGCACAACAATCTTTCTGGGACTCTTCCAATGACTTTTAGAACTGGAACACTTAGAAGCTTCAACTTACGTGGCAATAAACTAAAGGGGAAAATTCCTCAATCTTTGGCCAATTGCAAAGAGTTGCAAGTTCTTGATTTAGGAGATAACCACCTGAATGACATCTTCCTGATGTGGTTGGGAACTCTTCCTAAGTTGAGAGTTTTAAGCTTGAGATCGAATAAGTTGCATGGACCCATTAGAACTTTAGGAAGTGAAAACATGTTTCTTGAGCTTCAAATATTGGATATCTCTTCCAATGCCTTCATGGAAAACTTACCTATGGGCCTGTTTCAACATTTGAAAGCCATGAGGACAATTGATCCATCAAAGAAGGCACCAAGTGATGGATTTTACCAAGACATGGTAGCTGTCGTAACAAAGGGATTTGAGCGTGAAATTGTGAGTATCTTGTATTTGTACACTGTTATCGACCTTTCAAGTAATAAATTTGGAGGGGAAATTCCAAGTATCATGGGGGATCTCATTGCAGTTCATACATTGAATTTATCACATAATGGATTGCAAGGTCATATACCACCATCATTTGGAGATTTATCTTCGGTTGAATCATTGGATCTATCAGGAAACCAGCTTTCGGGAGAAATACCACAACAACTGGCTTCTCTTACATCTCTTTCATTCTTAAATCTCTCCCATAATCATCTCCAAGGATGCATCCCTCAAGGACCTCAATTTCACACTTTTGAGAGCAATTCATATGAAGGCAATGATGGATTACGTGGATTCCCTGTTTCAAAAAGTTGCAGCGATGCTCGGGTATTAGATACAAATGATACTGTGTCTGGACTAGACGATGAAGAAAGCAATTCTGAATTTCTGAGTGATTTTTGGAAAGCTGCTCTTATGGGCTATGGAAGTGGACTATGTATCGGATTATCCATAATATATTTCATGTTGTCAACTGGAAAACCGATATGGCTAGCAAGAATCATTCTAGAGATGGAGCACAAAATTATGAGGGGAAGGAGAAAGAAGCAGCGAAGGCAAAGGCATTACAGAAGAAGAAGCAATCATTTCTAGACAAGTTATAAGTTTTGATTTCAGAACTTCAGGTAGTTCTAATCGATCCCCTATATACACTAATCATTTTTTTCATTTCCAATAATTTCAAAAGGCTCAGCCTCAGTTGATGTTTAACTAAGATATTTTGGATTACAGGGAGGAGCTAGTGTCTATGGAAGTGGAGAAAGCTGGGAAGAAGAAGTTATGGCAATGAATGGAAACGAATGCCAATCAGTTTTTAAGCAATAATAATTGTCTAACTGAATATTTCGTTGAACAGTCTTAGATGTGGC

>M6-PGSC0003DMG400005539 M6 LRR-RLP

ATTTGAAATGCTCCTTCAGAATTTGACCCAATTAAGAGTGCTTCATCTTAGTGGCATAAACATCTCTTCCACCATTCCTCTGAATTTTTCTTCTCATTTAACAACTCTGATCCTGTCAGAAACAGGATTGTATGGGATAATACCTGAGAGTATTTTTCACCTGCCCAACTTGGAAACACTTGACTTATTCTTCAATAATCACCTCAGTGGTTATTTTCCAAAGACCAAATGGAACAGCAGTGCATCTCTCATACAATTAGATCTCAGTGACGTGAATTTTTCTTGTAATTTGCCTGAGTCTCTTGGCTATTATCTAACTTCGGTGCGTTATTTGTCTCTTCAAAGTTGCAACCTTAGAGGGCCTATTCCTGAATCTCTTTCAAATCTCACCCGCATTGAGTATTTATACCTTAATGGTAACTCCCTGAATGGAACAATACCACCAGGGATGTTATCCCTTCCATCACTAAAGGAAATAGGCGTGAGTTTTAATCACTTATCTGGTCCGCTTGAAGATTTCAAGTCCAATTCACTAATCTGGATAAGTTTAGAGGGCAATCAGCTGCAGGGTCATCTTCCCCACTCAATTCAAAACCTTGTGAGCCTAACAGATCTTGATCTTTCTTTCAATAATTTTAGTGGTCGTGTGGATGCCAGCCTCTTTTCAAACCTTAAACAGCTTCGGTCTATTAGTCTTTCATATAATAACATCTCATTGACCAATGACAACAACGTTACTTTGCCCGAATCTCTTACTAGGTTAGAATTGGCCTCTTGTGAAGTGAAAGAGTTGGAGTTTCTAAGATCCGCAAACCAGCTTTGGCAGTTGGATCTTTCAAATAATAAGATTGAAGGAAGAATTCCTGATTGGGCATGGTCTAACTGGATGTTTTCATTGCAAGATCTTAATATATCCCACAATATGCTGACGAGTGTGGATTCAATTCCTCTTCAGTTTGCAGATACTATTAATTTGCGGTCCAATTTGCTTCAAGGTTCACTACCTATTACATCAAATTCCACAAATTACTTCTTCATATCCCATAATAATCTCACTGGAGAAATTCCTTCATCTATATGCAAGTTGACATCACTGGTAATGCTGGATTTAGCGACAAACAATTTGCGTGGAGAAATTCCGAAGTGTTTCGGAAATATCACTTCCCTCCAGGTTTTGGATATGCGCCACAACAATCTTTCTGGGAATATTCCAACAACTTTCAGCAATGGAAGTTCACTGAGCAGCCTCAACTTGCATGGCAATAAACTAGAGGGGAAAATCCCACAATCCTTGACCAATTGCAAACAACTTGAAGTTCTTGATTTAGGAGACAATCACCTCAACGACACATTCCCCGTGTGGTTGGGGACTCTGCCAAAGCTGAAAGTTCTAAGCTTGCGATTCAACAAATTGCATGGTCCCATCAGAACATCAAGGATTGAGAACATGTTTCCAGAGCTCCGAATCATAGATCTCTCTTACAATGCCTTCTCAGGAAACTTACCTTCGAGTTTGTTTCAACATTTGAAAGCCATGGGGACAATTCATCAAAAATGAACACACCAAGTTATGAAGGAAGTCGATATTACCAAGACTCGATAGCTGTCGTATCCAAGGGATATGAGCGTGAAATTGTGAGAATCTTGTTTTTGTACACCACTATCGATCTTTCAAACAACAAATTCGAAGGACATATTCCAACTATTATGGGAGATCTCATCGCACTCCGTGTGCTGAATATATCTCACAATGGATTGGAAGGTCATATACCGTCATCACTTGGAAGTTTATCTTTGGTTGAATCACTGGACCTGTCAAGTAACCATCTTGTAGGAGAGATACCAGCACAATTTGCTTCTCTTACTTCTCTTGCAGTCTTCAATCTTTCCTACAATCATCTTGAAGGGTGCATTCCTCAAGGAAATCAATTTCATACCTTTGAGAACAATTCATATGAAGGCAATGATGGATTACGTGGTTTTCCTGTTTCAAAAAGTTGTGGGACAACTTATGCCGTATCTGGACAGCTAGACGGTGAAGAAAGCAATTCTGAATTTCTGAGTGATTTTCGGAAAGCTGCTCTTATGGGGTATGGAAGTGGACTATGCATTGGAATATCCATAGTATATTTCATGATGTCAACTGGAAAACCGATATGGTTTGCAAGAATCATTGTAAAGCTGGAGCACAAAATAATGATGGGAAGGAGAAAGAAGCAACGAAGGCAAAGGAACTGCAGAAGAAGAAATAATCACTAGTAATGTTACTGTTATTTCTTTAAGGAAAATATGTAATAAGTTGTAGACGTACTATTGTTAAATCCTGTTTGCTTAAAGTGTGTAGCATTATTTGACCATGAAAATTGAGGGTGAGATAATCTTTAATGACAAGTTATATGCAATATAAAATCTTATGA

>M6-PGSC0003DMG400005590 M6 LRR-RLP

ACCTTAACACATCCTACAGAATAAAATACATCTCTACGGACTTTAGATTGAAAATTGTGGGATACAAATTAGATTTTAATAAGATCCAACTATTCATTCTTATTTTAAAATGTATATGATTTATTTATTCACCATTTAACAGGTAACAAAAGAATGAGTTCTCTTCGGAATTTACAATTGGGAAGTTCCTACTCCAACTTAAGCAGAGTCATAGAATCCCTAAAATCATTCTCTTCTCTTAAATCACTTTCTTATGAATTTAGTGATCTCACTTCTCCAACCATAATTTACGGTAAGCTCAAATTCATATATACCTTAGCTTACTTCTATTGCCACTCTTTTCTTTTGATATGTAAATTTAGTTTTCTTTTATTTTTTGGACTCTATGATATAAGCATTAAGAAATTTGAGCACGGTGGAGCACCTATACTTGGAGGGATGTTCTTTAAATGACAACTTTCTCCCAAACATTGGACAAATGACTTCTCTCTTGGTGTTCAGTATAGCATTTGGTGACAATGACGGCACCCTCCCTAATCAAGGTACCCTATTTTTCTCTTTTTTCCCTAATGCCCGCATTTTCATGTTGGAGGTCTGATTTTAGCATGTTCAGTTGATCAAACATTATAATATTTTTTGTTTATTACCAAACACAAGAAAATAAGTAACACAAGCCATCTAATTTTTCTAGAAAATATTTTCCTTCGTATCGAACACATCCATAGACATAAAAAAATATAATAAGATGTCACTCATCATAGATATTTGTTTTAGCTAAATGTTTCTTAATATATATATATAGGCTGGTGTGAACTCAAACACATTCAAGAGCTGGATTTCTCAGACAATAATTTTCGTGGAACACTGCCTTCATGTTTTGGAAACTTGACATCACTTCGACGATTGAGTTTTTTTGGGAATTACTTCATGGGAAATATAGCGTCACATTCTATTTGGAGGAGACTCACATCACTTGAGTACCTCAATATTGTGGATAACCAATTTGATGTTCCTCTGTCATTCAGCCAATTTTCCAACCATACAAAATTGATCTACTTGGATGTTAGTTCTAATACAATAATTCCAGATACCGAATTCCAAAATTGGATCCCAAATTTCCAGTTGGAGTTTTTTGCTATCGAAGGATGTAAAAACCTTCAAAAATTGCCTTCTTTCCTTCACTACCAATATGACATGAGGATTCTTGCTATAAATGGAAATCAATTGCCAAGAAACTTTCCAACATGGTTGTTAGAAAATAACACCAGACTTGCAGGGATTTATAGTAGAGATAATGCTTTCAGCGGACCATTCAAGTTGCCATCGAGTTGTCACCTCCATCTAGAGACAATTGATGTTTCTAATAACAAATTAAATGGGCATATTCCAGATAAAATGAATTCAGCATTCCCAAAGCTTACTTATTTGAACATGTCACAAAATTATCTTGAAGGTCCTATACCTTCCAAGATTAGTGGCATTCATTTACATAATCTAGACCTATCTCACAATTTCTTATTTGGAGGAGTTCCTAGTGATCTGGCAATTGGTTCTCCAGTATTAGCTTACCTTCGATGTCAAATTTTTCCAAAGGATGTCAAACCAAAGATGTTATTCTTTTTGTTCTTGAATGGAAATAACTTTGAGGGACCACTACCTTGCAACATGTTTATCACATCCCTTATTGTATTGGACGCTAGCAACAATAATTTCTCTGGAGAGATTCCGAGATGCATTAGGGATAATACAAGATTGTTACAACTTGATTTGTCCAAGAATCATCTCGATGGTTCGATTCCAGTTGAGATTTCCAATTTAAAGCTCATTCAGGTCTTAGCTATATCCGAAAACAGGCTTTCAGGCTTTATTCCTTCTTGTGTGAATTTTTTACCTCTTGAGCACATCCATCTACAGAAAAATCAATTGGGTGGTGAATTGGAACATGTACTTTTCAACTTCTCTTCTCTGATAACTTTGGATCTTGGGTACAACAATTTTACAGGAAGTATCCCACACACCATAGGCTCACTTAATACTCTCAACTACCTTCTCCTTAGCAATAATCAATTCGAAGGAGAGATTCCAACTCAGATTTGTATGCTGAACAAGTTATCTATCGTGGATTTATCTTTCAATAAGCTTTATGGTTCACTCCTTCCTTGTTTGAGTTACTTAACACAAGCCAAAAAGGATGCAGAGATAAGTAAGACCAATTATGCTACGACTTCTAGGCGTTCATGGATAACTTTTTTGGCCTGGATGCGTTTGAAAAGGCACTACCACGATAGTCATGGATTTCTAAGTGACATATTTCTGATGGATGTGGAAACTCAGGTCCAGTTTTCAACAAAAAGAAACTCATACACTTATAAGGGAAGCATTCTAAAATATATGTTGGGTATTGATCTCTCAAGTAACAGATTAACTGGGGAAATTCCCGTTGAGCTAGGGAACATGAGCAATATACATGCATTGAATCTATTGCACAACCATCTCATCGGAAGAATACCAAATACCTTCTCCAATCTACAGAAAATTGAGAGTTTAGACCTTTCCTACAACAACTTGAATGGGAGCATTCCTGTTGGTCGACTTGAGCTAAATTCTTTGGCAGTATTCAGTGTTGCATACAACAACTTATCGGGTGCAGTGCCTGATTTTAAAGCTCAATTTGGAACTTTCAACAAAAGCAGCTACGAGGGAAATCCTTTTCTTTGCGGTTATCCATTAGACAACACGTGTGGCATTCCTAAATTGTCAAATACCTCCAAGATTAATGGAGGTGAGGAATCATCAGACTTGGAGGATATTCAGTGTTTTTACATTGGCTTTGTTGTGTCTTATGGAGCGATCTTGTTGGGATTAGCTGCGGCACTCTGCTTCAATCGTTATTGGAGAAAAACATGGTTTAGGATGATAGAGGCATTGATGTTTTATTGTTACTATTCTCTCTTGGACAACGTTGTTACACCCATCAAGAATATATGGTACAGGAATGTTGGTTAATTCTAGGTTGTTGTTTTACAATATTGTTGTATGTGTATCACCATTATTAATTATATTGTCATTCTGTTTTATCAATATTCAACAAATAGGATCTGTTTGGAGTAGTTACAACTTCATAATTTAATACCCCCTGCATAGCTTCATTAGTGTTGCTATTTATGTAGCC

>M6-PGSC0003DMG400005600 M6 LRR-RLP

CCCTGAGACTTAGTTACCATCACCTTCCACTTGATTTGAGACAATGCTTTGCATATTGTGCAGTATTCCCAAAGGATACGAAATGGAAAAGAAAAATCTAATCTCTCTCTGGATGGCACATGGTTTTCTTTTATCGAAAGGAAACTTGGAGCTAGAGGATGTAGGTAATGAAGTATGGAATGAATTATACTTGAGGTCTTTCTTCCAAGAGATTGAAGTTAAATATGGTAGAACTTATTTCAAGATGCATGATCTCATTCATGATTTGGCAACATTTCTATTTTCAGCAAGCACATCAAGCAGCAATATCCGAGAAATAAATGTAAAAGGTGACCCTTACCCACATATGATGTCGATTGGTTTCTCTGAATTGGTGCCTTCTTACTCTCCTTCTCTCTTTGACAGGTTTGTCTCATTGAGGGTGCTTAATCTTAGTAAATTAGGACTTAAGCATTTACCATCTTCCATTGGAGATCTAGTACATTTAAGATACCTAAACTTGTCTGGCAATAGGAATATGCATAGTCTTCCGAAGCAGTTATGCAAAAGCTTCAAAATCTGCAGACTCTTGATCTACATGATTGCTCCTCACTTTGTTGTTTGCCAAAACAAACAAGTAAACTTGGTAGTCTCCGAAATCTTTTACTTGAAAATTGCAAAGCATTGACTTGTATGCCACCAAGGATAGCTTCTTTGACATGCCTTAAGACTCTTGGTTGCTTTGCAGTGGGAAGGAAGAAAAGTTCTCAAATTGGTGAATTACGAAACCTGAATCTCTATGGCTCAATTTCAATAACGCATCTTGAGAGAGTGAAGAATGATATGGATGCAAAAGAAGCCAATTTATCTGCAAAAGAAAATCTGTATTCTTTAAGCATGATTTGGAGTAGGATAGGACCACATAGATATGAATCAGAAGAAGTTGAAGTGCTTGAAGCCCTCAAACCACACTCCAATTTGACTTGTTTAACAATCAGTGGCTTCAGAGGAATCCGTCCCCCAGACTGGATGAATCACTCAGTTTTGAAAAATGTTGTCTCTATTGTAATTGAAGTTTGCGAAAACTGCTCATGCTTACCACCCTTTGGAGAGCTGCCTTGTCTAGAAGGTTTAGAGTTACACAAGGGGTCTGTGGAGGTGGAGTATGTTGATTCTGGATTCCCTACAAGAAGAAGGTTTCCATCTCTGAGAAAACTTATTATAGGCGACTTTGATAATCTGAAAGGATTGCTGAAAAAGGAAGGAGAAGAGCAATTCCCTGTGCTTGAAGAGATGGAGATTTACTGGTGCCCTATGTTTGTTATTCCGACCCTTTCTTCTGTCAAGAAATTGGTAGTTCATGGGGAGAAGTCAGATGCAATAGGTTTCAGTTCCATATCTAATCTCAGGCCTCTTACTTCCCTCCAAATTAGCAAAAACGAAGAAGCTACTTCGCTCCCAGAAGAGATGTTCACATGCCTTGCAAATCTCAAATACTTGAATATCAATTCCTTCACGAATCTCAACGAGCTGCCTACCAGCCTGGCTAGTCTTAATGCTTTGAAGCATCTGGATATTATTGGGTGTGATGCACTCGAGAGTCTCCCCGAGGAAGGGGTGAAAGGTTTAACTTCACTCACACAGTTATCCATAACATGCTGTGAGATGCTACAATGTTTACCGGAGGGATTGCAGCACCTAACAGCCCTCACAAATTTATCAGTTAGGGAATGTCCAACACTGGCCAAGCGATGTGAGAAGGGAATAGGAGAAGACTGGTACAAA

>M6-PGSC0003DMG400005972 M6 LRR-RLP

TAAGAAAATACTGCGTATTGCCCATTTAAGCACTTACAGAAGTTATTTAATACGAGCTTTTTCAGCAACTCTGGATGCATCATGTCTGCAGCTGATATTAAATCATTAAGCCAAGACAACACATAAGGAGGGCGAGAAGAAAGAGGATAAGCTGCTGCGAGTAATGCTCCTAAAGGGACAACAGGTACATTGAGAGCTTGGGCAGGATCAGCCTGCAGCCAAGGTAAATAAATATAAGAATATGCTGAAAGAGCTTTTTGCTGCACAATGAGAGTTGAAAGACATTTATTTTTTGCTTACAAGGGGTAGGAGCAGTCTGAGTGCAGACTTCGAAGATGTGTAATCCAGTGATGAAGCCAAAGTTGCAATAGCAGTTAATCCAGGCTCTCTTCCTTGCAATCCTTTCCAAGATAAAACCAGAAACATATTAAAATTATCTCATATCTGTATGATTCCCGGATGTAGTGAGTTTGAAAATATTTTTTTGAGACTGTGCCCACTGCACATTAACAAGGAATCAGACAATGCTCCAAGTGACTTATCTGTTAGTTGCGAAACCTTTTTACATCTAAAACCAGGTCAAATTCCTTCCAAATCAAATTGTCCCTTACACACTGTAAAAACGTATGGATAAAGTATAAAATCTTGCCTAGCTCCCTATAATCTCAATAAGGTCAAAAGATGGCTCATTTTCAGCTCTGTATGAAAGACGCTAGTAAGCAGATAAGTCACACTGTTAACCAGATCCTGTTTACCTTGGTGCACCCTATCATGGTATTTTTAAGTTAATAGGAGTACTGCACTTGGTGGCTATTACAACTATACTCCCAAAGTTGTTGTGAACTATAGCATCTTTCTTATTTTTCAGCAAAAAGCTTGTTGAAGTTGAGTCTGAAGATTTTGCTCCATGAATATTAGGAATTAAAAGAACATTAAGTCAAGAGGAGACCAGGTTAACTCAAAGTTGCACCAAGTGAAGTGGTAGCCTTTGGAATTTAATCCCAAGACTAAATCGTATGATACTCCAGCAGGTTCATGGAATAAATTTCTGGAATTATACTGCCTTCTTTATTTAAATAAAATGTTTAAATATTACTGTTTTCTTATCAACTGCAACTACAGAAAGGATTCTTTGTTACTTTTTCAAAACTCTTGCACATTAATTTACTAGTATCTTAAGTGGCTCATAAGTAGACCCGCCAGATCACGTAAGTGTAACAACGTTAATTCCATATAAACAATGAATGAATCTTACCACACTGTGATAGCCAAGCATACAGTAATATTCCTCCCATAGAATGTCCAATGGCAAGAAGTTTACCATCTTTTGGTGCAGTTTGGGCCTTAATGTATTCCATCTAGAATGAAAATAAAATAATTAGACTAGTCAGAATAATAAATAAGAGTGAAATTTCAACAAATTCTGAATTTCATTTGGTTGTGTAATCTCAAGGACAATAATATCTTTATATTCTACAACATGGGAATCTGTCACCTCACCGCAGCAGGAACATCCTCTTCCAGGTAATGATCAAAGTCCCAATCATACTTGACAATCAAGTCAAGCTGTTTCTGGAAATCTTCTATGGTAGTGGTTAGACGTTCATGCAGATCAACTAACGGAGAAACAGAACGTTGGCCTTCTTCAATGATATTAACTAGCTTTTGGCTAAGATCCTTTACTTGGTCAACAACGGCAGAGTTTTGCCCTGTCTCCAGCAGACTTAACAACTTCCCCCTTGTCTCGTTAAAGCGTTCATAAAGAAAGGAATCCTCTAATAATTTTGATATTTGATCAAATAGTTTAGCAGACATAATCCTTGATTGGCTTTCATTGAGGAAGCCAGAGACTCTTTCTGACAAGCGCATAAATGTTTCCGTAAGCTCGTTTACTACTCTTGATTCATCCCACATTGTTGCAATAGCTGTGGATTCTTCTTTGACCACAGCTGTGTCAGACTCTTCTAAGGTACTCTGGGCATCTGTTGACTGCTGTGCTGCAGAAAGAACTTTGTCAGTAGCATTATCAGCGGCAGCTTCCATCTGTTCTGAGACTGCAATTGCAGTCTTCTCAATGTTTGTGGAATCTGACTCCTGCACACTCAACCCAGCTCCACGAAGTTCAAGAACCCAAGTGTCAAATCCTTCACCAGACATGTACCGGGCGAAAGATGACTGCATAATTATTTTTAAAAGTAAAAACCGTGATTAAATAACTTTTAGAAAGAATTTAAAAAAACCAGCCATACTATGTAAAATTTCTTTGATAAGCAGGATGGACATGGTTGCCTGAAGTGATAAAGAATTCTAAACCTCCACTGAAGAGTAGGGTGCACATACTTGCCTCAAGTGATAAAAATGAACTTGAACGGATAGGAAGATGCATCAACAGAACTTTAAACAAAAAATAGACAAGAAATAATCCAAGTGAAATCAAGTTGGCATTAGTTATTGCATTAACCAGCAGACATTTAGATTTATCTCACTAAAAAAAAAAAGGATAGAAATCATCACTAGCAGCTAAGCAAAAAATCGTCATCACAATTCATATTTTAGAAATGTACAATCACAGAGAGTGCTAGCCACATGACTACCATCAACACCTGATTGATGCAGGCATGCAGCCATAAAGCCTTATAAAAAAAAGGAAGATGAAATAGATAACAAGCCAGTAATAAAAAAATCACACAAGTAAACTTTTATACGCTTGAAATGCATAAAGTATGAGAAAAGGACGAAGTTTTTAACCGACCCACCATTTTGATCACCGCACCAACATGTGTGCAGAAGGAAGTAGATGCAGATGCACTAGGAACTAGTTACCCTAGGACCTTAAACAATCCATTTTGTATTATCCAGGGTCGGGATGTGACCCAAACTACTTACAATGACCCACCCGTTGACACAGGTGGGGAAACTACTAATGCTCTTGATCCCTAATTATATAATATAATATAAGTATCCTTTTGATCAAAAAAAAAAAAGTAGTAAGGATAATACAACATCAAAACCAGGGTTAACTCTAAAAAATTTGAGCATGCCATAGAATTCTACCAGATTCAGAAACAAGAACCACTCCACGAAATAAAGAGTGAAACTTTAATGGAACCTGACATATCCACTAAGCACTGATAATATACACATTTGACATCTGTCATTATGATCTATTAAATGTGGCTATCAATGAACGATCCATGAAGATCCCCTAGTTCTGTTCACCTATACAGGCATGAATTTGCAAATAAGAGTGTCCCCGACTACTCTTTCACACAATACTGCACATACAACAAAACAGAAGAATGCACAAAGAACGGAGAGAAGGCAGACGAGTATCATACCCCAGGAGCAAGATCATATCCAATAGCATTAGTCCCCACGCCAGACAACAGAAGCAGGGGATGATTTCTCTTTGGAGCCTATCATCATCAAAAATCACACACCCTTAATGCAAAGCTCAAATCCTTGAACAAAAAAATCATGTAACAGAACTGTTAAACCTTCCTAGTATCACTCCCTGTGTTTCATTTTATATAACAGTATTTGATAGAACACAAAATTTAAAATATTCAATAGTTGAATTAGATAAAAATGGTGTCAAACCTTCTTAATGCCTTCATGTCTTATATATTGGGACCAATTCTATATGACCACATTTACTTTAAGCCGAGTATCTATCACAAACAGTCTATCTACCTTCACAAGGTACGAATAAGGTTGCATTTGCTCCACCCTCCCCGGACCTCACCTCTGAGATTACGGGTTGTATTCTATATGACCACATTTAATTGATCACAAAATTTAAAATTTGAACTTATATAGAAATGAAAGTGTCAAATATATTATTTTTTTTCAAATTGTCATACAGTTAAGAGTAACAGTTCAAATTAAAAAACATAAAAAAATTGAACCTGAGGAGGAGGAATGTAGCGCCAGAGAGCAAGTTTCCAATCGGAATTATTAACAGATACGTAGTGGAGTTCATCGGCAGTGCATAACGATGGCTTATTACAAACCTTCTGGGCTGTAGTATCGCCATCATTGGTGGTGAAAGCTCTGAGTTTAAAGGCGGAGTTCACCGCGAAGCAGTAGTTCGGGGAGTTCCGGTATGAAATGACGGCGGGAGGAAGTCGCCGGCGGTGATTTGTTAAGTGGTGGTAATGGTTGTGGTGGCGGAAGACGGCAGCGATTTGAATCGCCGATCGAATATCGGATTGAATTGGATAAACCATATTTTTTGGGATTCGTGACAATTTGGATAAAGAAAATTGGTTTGATAAATCTTTTACTCTTTTGACTTTTGAAGGTTTGAGACGTAAACTTTTTTATT

>M6-PGSC0003DMG400006027 M6 LRR-RLP

TAGAGGCGTCTACTTGCTATTTAATTCAAATCAAATTCATTCATTCATTCAACAACACATTTTTACATTGTGTGAAAAAAAACTTCATTTTCTTCTTCTTAGCTTGAAAATCTCTATCAAATTCAGCAATGCGGCCTCCTAGCCGCTGCGTATTTATCTCTTTTCTTGTTATCTCCATCTTGTCGTACCAGATCTCGAATGTGGTAGGACAAGAAGGTGATAGCGATATCGATTTAGATGATATTAAGGCTAATGCTAGTAAGTTTAGTTTCGAAAACCCAAAGATTCGCGATGCTTATATCGCGTTGCAATATTGGAAAACAGCAATGTTTTCTGATCCTTTTAACTTCACAGCTAATTGGACAGGACCTTATGTTTGTTCGTATGGTGGAGTGTTTTGTGCTCCTTCGTTAACGGATGATTCGATTAGAGTTGTAGCTGGCATTGATCTTAATCATGCTGACATTGCTGGTTCATTAGTTGCAGAACTTGGTCTTCTTACAGATCTGGTTCTTTTCCATCTTAACTCGAATCGTTTTTGTGGAGTTGTACCGAAAACTTTTAGCCATTTGAAGCTTCTTAGGGAGCTTGATTTGAGTAACAACAGGTTTGTTGGTGGGTTCCCTAAGGTAGTTCTTTCATTACCTTCTTTGAAGTTTCTGGATCTAAGGTTTAATGATTTTGAAGGACCAGTTCCTTCCGAATTGTTCGATAAAGATTTGGATGCGTTGTTTCTGAATGACAATAGATTCCGTTTTGGCATCCCGGAGAACTTGGGTAACTCCCCTGTTTCTGTTCTTGTGTTTGCTAATAACGATCTTGGAGGGTGTATTCCCGCGAGTATTGGGAAAATGGGGAAAACGTTGAATGAGTTGATATTGATGAATGATAACCTCACTGGGTGTTTGCCCATGGAGATTGGATTGTTGAATAAGTTGACAGTGTTTGATGTTAGCTTCAACAAGATTCAAGGCTCATTGCCATTAACTGTGAGCAAAATGAGGAGTGTGGAGGAACTCAATGTGGCTCACAACAAGCTTACTGGGGTTATACCAGCTAGCATTTGCCAACTTCCAAGGTTGCAGAACTTTACGTATTCGTTCAATTACTTCACTGGGGAGGCTCCGGTTTGTGCTGCGACTAGATCTGGTGATGGTCAGGAGAACTGTATCGTTGGAAAGAAGAATCAAAGATCTGCTAAAGAATGCTCTTCGGATGATGCAAAACCGTATGATTGTAAGAAATCGAAGTGTTATAGCCCATATGCTACGTCTCCTTCGACGAAACCAAAGCCAACACCAAAACCAAGAGGACCACCACCACCTACTTGGAAGTCTTCGGGTTCGCATAACAAGCGATCTCCGCCACCACCTAAATCAACTCCTCTTCCTCCACCAGCACCTTACAAAAAGTCCCCAGTGTATCGACATAGAAGTCCACCACCACCTACTCACAAAATTTCACCTGTAACTCATCATTCACCTCCTCCACCTTCACCCGTATATTACCACCCATCTCCATCGTCCCCACCACCACCAGTTTATTATTCTCCACCACCCCCGGTGTATCACGAGCCACCTACTTACAAGCCTAAATCACCACCACCACCAACACCGTCATATGAACACCCAAAAACCCCATCATCACCACCACCAACACCATCATACGAACATCCAAAAACCCCATCACCACCACCACCAACACCATCATACGAGCATCCTAAAACACCACCACCAACACCATCATACGAACATCCAAAAACACCATCACCACCACCACCAACACCATCATACGAACATCCAAAAACACCATCACCACCACCACCAACACCATCATACGAGCATCCAAAACCCCAATCACCACCACCACCAACACCGTCATACGAGCATCCAAAACCCCAATCACCACCACCACCAACACCATCATACGAACACCCAAAAACCCCATCACCACCACCACCAACACCATCATACGAACACCCAAAAACCCCATCACACCCAACTCCACCTACCCCACCTTGTAACGAACCACCACCTCCACCACCTAATTCACATTGGGAACCAAAACCATCCCCACCATACACCTACTCATCACCACCGCCTCCATCACCATCTCCTCCTCCTCCCACTTACTACTACTCATCACCACCGCCACCATCACCATCACCATCACCTCCCCCACCCACTTACTACTACTCATCACCGCCGCCACCACCACCATTTTACGAAAATATCCCTCTCCCACCAGTAATCGGAGTCTCCTACGCATCTCCACCACCACCAGTCATTCCATATTACTGAACAAGGGTTATCAATTAGCCTTTGATTTTCCATTATCAAAAAAAAAAGAATTCATTTTTTTTTATGAATTATGTATACTTTGTTATTTATTTTTTGTAATATTTTTATTTTGAGTTTGATTATATTTATGAAAATTGGGATTATGGGCGGGAGTTAGTGAGGGGTTTAACTATCAATATTTTAATATTGTTATAT

>M6-PGSC0003DMG400006076 M6 LRR-RLP

AATTCCCCTTTCAAGCCAAAAACATGGCCGCCACCGCTGACATATTCTCGGAATGTGTGATTCACAAAATACTCTGTTATCTTAGCTATGGCCAAGCATCCCGGATGAGAATTCTATCTAAAACATGGCTACAAGCTTGGTCTACTCTTCCCAACTTGGAATTCTTAGTTGAATATTTCGAAGAAACAAAAATAGTGGACGCAATTATGGAGACATACAGGGAAAACAAAATCCCCATTGACAAGTTTGGATTTTCAAATTGTTCGAAATTCAATCGTCGTCATCAAGTTTTCCCTCTGATTGATAAGTGGCTTGACACAGCTCTTCAAAATGGCGCAAGAGATTTAGTCTATGTAGATCTTCTCTACTCGGTACGATTATATCCTTTTCCTATTTCTAAAGTCTTGGCTGCAAATTCTTTAAGAGAATTAGTTCTGAAAGGTTGTAACCTTATGCATGTTTCGTCATCTAGTGGGTTCGCGAATTGTCATTCTTTGAGAAAGCTTTCTCTATCTTATATAATTTTAAGCAAAAATTTGCTTCAGACTTTACTTAATAGTTGTCCTTTCATTGTCTCTTTCAACCTTGATCATTGTTCGGGGTTGGTAAAGATTGAGCTTATGAATCTTCAAAAGATCAAGTCAGTTTCCGTATCAATAAATAGAAACCAACATGTGAAAATCCAAGCACCAACTCTTGAACACTTGTTTTATAATGGATATTTATCGGGAAATTTGGTTATCGTTGGATGTCTGAATCTAAAATCTTTAGAGCTATCACATGTGAAGATATGTATTGGATTTCTCCATAACCTAATTTCTAGATCGCAATCCCTAGAAGTATTGAAGATTCAGTATTGTGTGGGCATAAGACAGATTGAAACTTCGAATTTGGTATCACTTGAGTATATAGGATATCAAATTCCTGAAATTAAAATTGCGAAAGCTTCAAGTCAATTGAAGCATTCAAAAATCGTTCTTCATGGCAACAACAATTTAGATGCTGCATGGTTTAGTAAGTTGAGGAAGCTTCTATCGAATTCGGCCCCTTGGTCCCAAGTTCTCCTTTGTTTTCCCGAATACAATGAAATCAAGATTAAAGATTTGCAGCAGAACCAAGGAGTTGCTAGCCCACAAATGGACGTTTTAAATGTCAACATTACAAGGTCATATGGGGAGTGACCAACATTTGTGGATGCTTTGCTATGGAGTTGTAATCCTAGGAGACTCAACCTATCATCAACGGTTGAAATGATTACATGTTTCGTCAGTCGTTTGATGTATATGAAGAATCTGAGTCATTCTTCTTCGGAAAGAAGCAACCCTTGGCATAGTCAATTGAAAGAAATAAAAGTTTTTGATGGGAAAAATCAAGAGCTGCAACTCGTAAGTGGGGAACTGGAGATAAGCACCCTTTCGGAGGACAGGAAGGAAGTTTATTTTTTATTAGATTGGCAATGCAATTAATTTATCACTATTTCAAAACGAATGCACTTGTTCATTCCTAAACTTTATTAATCTGTTTTATGTCTAGCTTTAAATTCTTGCACTTTCTCACCATTGTATAAAATTGTGTTGAAGTTGAAGCATAAGATTCTTT

>M6-PGSC0003DMG400006095 M6 LRR-RLP

ATGGGGACAGCTGACATGTTGCCTATATGCCTCATTCGCAAAATACTCTGTTATCTTAGTTATGAAGAAGCATCTCGGATGAGAATTCTCTCCAAAACATGGCTACTAGCCTGGTTGGATCATCCCAACTTGGAATTCACAGTCTCGTCGGGCAAAACCATAACAATAGTGGATCATGTCATGGAGAGATATAATAACGGAAATATTCCTATTGACAAGTTTGAATTACACAACTCTACATAGTCTGATCATATTTTTCCTCTGATTGATAGGTGGCTCGGTATTGCTCTTCAAAATGGTGTAAGACATCTAGAGTACATAGATGTTGAACTTTCAACATCATATCCCTTCCCTATTTTTAAATTCTTGGAAGCAAATTTTTTACGAGAACTGGTCTTGATGGGTTGTGATCTGATGCATGTTTCGTTATCTAGTGCTAGTCAAGCGGTGACGTGTCATTCTCTGAGAAATCTTTCTCTATCTCTTGTACACTTAGACAATAACATGCTTCAGACTATACTCAGTAGTTGTCCGTTCATTGTTGATTTAATCATTGGGTATTGTAGTAGGCTGAAAAATATTGAGATCCGGAATCTTCCAAATATCAGGTCTCTTGTCATTGGTAATGACCAACTCGTTAAAATCCAAGCACCGTCTCTTGAACACTTGTATTATAGTAGTTCTTGTTTGAATAAATTGAATATTGTTGAACATAAGAATCTGAAAACTTTAGATATATCATGTACAGAGATATCTGATTTATATCTCAACCACCTTATCTCCATACCTTACTGCCTGGAGAAACTGATATTAGCAAACATTGCTTTTGGAAGATTCGACATTTGCAGGAGTCAATCCCTAAAGGTGTTAAAGATTCATAATTGCAAGATCATAGGAGCAATTGATGCTCCAAATTTGGTTTTGCTTTAGTATAAAGGAAATGATATTCCTCAGCTTAAATTTGCTCAAGAATCAAGACAATTGAGGCAATCACAAATGATTCTTTATCCCTTGTTCAATGTAGATTCGGCATGGTTTTGTCAGTTGAGGAAATCTTTATCAGATTCGACGATCTCTTGGTCTCAAGTTACCATTTACTTTCACAAATGCGAGGAGATCAACATGATAGATTTGCTACTGCAACCTAGAGTTGCTATCCCCAAAGTGGACATTTTAGATGTAAATTTTTCAAGGCCGAATTGGGAGTTTCCTACATTTGTGGATTCTTTGCTGTGGAGTTGTCATCCCAGGAAACTTAACCTACAATCAACAACTACAATTTTTACATGTTTCATGAATCGTTTCATGCAGATGAAGAATTTGCGTCGTGGTCAATTAAAAGAAGCAAATGCCTACAAATTTGATCAGAAAAGTCATAGTTGGCATCCTGTAGAACACAAAAGAGGAAGAGATTTCCCATTAAGGACCAGTAGCAAGTCGAAGAAATATTTCTTCTTATTAGATTGGTGA

>M6-PGSC0003DMG400006226 M6 LRR-RLP

ATACAAAATTTCAAGAACACAATTACTCTTGATCCTTTTGGTGTCACAAAAACTTGGACAGGCCCTGATATTTGCAATTATAAAGGCTTTTATTGTGATAATCCACCTAATAATTCATCAGCAATTTCTCTTGCTTCTATAGATTTCAATGGATTTCAACTTAGTGCACCTACATTAGATGGTTTCATTGATCAACTCCCTGACTTAGCCATATTTCATGCTAATACCAACAACTTTTCAGGCATAATTTCTTCGAAAATTTCAAATCTTCCATATCTTTAAGAACTTGATCTTAGTAACAACCAATTCATCGGTTCATTTCCCATGTCAATTCTCAATATCAAGACTCTAACTTTCTTGGATATTCGTTACAATCTCTTGACAGGAATAATCCCACCTCAAATTTTCGTACAATATCTTGATGCATTTTTTCTTAACAATAACAATTTCATTCAAAAACTTCCAGACAACATTGGATCCACCTCTGCTTTTTATTTAACCTTAGCCAACAACAAATTCGTTGGTCCAATCCCACGTAGCATTGGACAAGCTCCGAATTTATTGGAAATTTTATTCTTGAACAACTTGCTCACTGGTTGCATTCCTTATGAATTAGGCCTTTTGAAAAAAGC

>M6-PGSC0003DMG400006255 M6 LRR-RLP

ATGAATGAAGCAAGACCATACCAAAGACGAAACTTCCATAAGTCAAAGTCTTCCACAAATGAACCAAAAAGGCTAACATCAAACGAGTAAAAGGAGTTAATTCAATCTGTACAAACGCGTAGTTTTTCTTTGACATTTCTCAATCACCCTTCCCCTTATTAAGTTCCTTCTAACTGCATACTTCAGTTATATTCAAGCTAACAAGTTGCAAGAATATGACAACTCTTTACTTTCTATGGCTTTTCTTGATTCCCTTGTTTCAAATCTTAGGAAATGACATTTTCTTGGTTTCCTCTCAATGTCTTGACAATCAAAAGTCGTTGTTGCTGCAGTTGAAGGGCAGCCTTCAGTATGATTCTAGTTTGTCAAATAAATTCGCAAAATGGAACCAGAAGACAAGTGAATGTTGCAATTGGGATGGGGTTACATGTGATCTCTCTGGTCATGTGATTGCCTTGGAACTGGATAATCAGACTATTTCTAGTGGAATTGAGAATTCAACTGCTCTTTTCAGTCTTCAGTATCTTGAGAGGCTAAATTTGGCTTACAACATGTTCAATGTTGGCGTACCAGTTGGTATAGGCAACCTCACGAACTTGAAGTACCTGAATTTATCCAATGCTGGTTTTGTCGGGCAAATTCCTATGATGTTATCAAGATTAACAAAGCTAGTTACTCTCGATCTCTCAACTCTTTTCCCTTACTCTGACCAGACACTAAAACTTGAGAATCCCAATTTGAGTCATTTCATTGAGAATTCAACAGAGCTTACAGAGCTTTACCTTGATGGGGTTGATCTTTCATCTCAGAGGACTGAGTGGTGTCAATCTTTATCTTCATATTTGCCTAACTTGACCGTCTTGAGTTTGCGTGCTTGTCAAATTTCAGGCCCTATTGATGAATCACTTTCTAAGTTTAAGATTCTCTCTATCATCCGTCTTGACCAGAACAATATCTCTACCACAGTTCCAGGATATTTTGCCAGTTTCACAAACTTGACTACCTTGTCCCTCGACTCTTGTAATCTGCAAGGAGCTTTTCCTGAAAAGATCTTTCAGGTACAAGTTTTAGAGATTTTGGACTTGTCAAATAACAAGTTGCTTAGTGGTAGTATTCCGAGTTTTCCTCGAAATGGATCTTTGAGGACGATATCACTAAGCAACACCAACTTTTCGGGTTCATTACCAGAGTCCATTTCGAACCTTCAGAACCTGTCCAGGTTAGAGCTTTCCGATTGCAATTTCAATGGATCAATACCTTCCACGATAGCAAACCTTACAAATCTTGTTTATTTAGATTTTTCCTTCAACAACTTAACTGGTTCTATCCCATATTTTCAAAGATGCAAGAAACTCACCCACTTAGATCTCTCTCGTAATGGTTTAACTGGTCTCCTCTCCCCGGCTCATTTTGAAGGACTCTCAGAGCTTGTCTACATAAATTTAGGGAACAACTTACTCAACGGGATCCTTCCTGAATGTATCTTTGAGCTTCCCTCCTTGCAGACGCTTTTACTTAACAGCAATCAATTTGTTGGCCAAGTCAACCATTTTCACAATGCATCCTCCTTTCTCGATGAAATTGATTTGAGCAACAACCAACTAAATGGTTCAATTCCCAAGTCCATGTTTGACGTTGGGAGGCTTAAGGTTCTCTCACTTTCTTCCAATTTCTTTAGCGGAACAGTACCCCTTGACCTCATCGGGAAGCTGAGCAATCTTTCACGACTGGAGCTTTCTTACAATAACTTGACTGTTGATGCAAGTAGCAGTAATTCAGACGCTTTCACATTTCCCCAGTTGAACATTTTGAAACTAGCTTCGTGTCGGCTGCAAAAGTTTCCTGATCTTAAAAATCAGTCAAGGATGATCCAATTAGACCTTTCTGACAACAAAATACTGGGGGCAATACCAAATTGGATTTGGGGAATAGGTAACGGAGCTCTGACTCACCTGAATCTTTCTTTCAATCAGTTGGAGTACGTGGAACAGCCTTACAATGTTTCCAGATATCTTGTCGTCCTTGACTTGCATTCCAATAAGCTAAAAGGTGACCTACCAATTCCACCTTCCTTTGCTGCATATTTGGACTACTCGAGCAATAATTTCAGCAATTCCATCCCACTAGATATTGGCAATTATCTTGGTTTTGCCTTCTTTTTCTTGGTAGCAAACAATGGCATTACTGGAAGAATTCCCGAATCCATATGCAATGTCAGCTACCTTCAAGTTCTTGATTTCTCTAACAATTCCTTGAGTGGAACAATACCACCATGTCTACTGGAATATGGTACAAATCTTGGAGTGCTGAATCTAGGGAACAATAAACTCAATGGCGTTATACCAGATTCATTTCCAATTGGTTGTACTCTACAAACTTTAGACCTCAGTGAGAATAAGTTACAAGGTAGGCTGCCAAAATCTCTTGTGAACTGCAAGTTGTTGGAGGTCCTGAATGTTGGAAATAACAGACTTGTTGATCATTTCCCATGCATATTGAGGAACTCAAACAGTCTGAGGGTCCTAGTCTTGCGTTCCAATCAATTTAATGGAAGTGTTCAGTGTGATGCAACCAGGAATAACTGGAAAAATCTCCAGATCATAGATATAGCTTCCAACAACTTCACTGGTGTGTTAAATGCAGAATTCTTTTCAAATTGGAAAGGAATGATGGTTGCAGATGATTACGTGGAGACAGGACTCAATCATATCCAGTACAAGTTCCTCCAACTAAGTAACTTATACTATCAGGACAAAGTGACAATAACCATCAAAGGGCTAGAGCTAGAGCTTGTGAAGATTCTCAGGGTATTCACATCTATTGATTTCTCTTCAAATAGATTTCAAGGTGTGATACCAGATACGGTTGGGGATCTTAGCTCACTTTATGTTCTGAACCTATCACACAATGCCCTTGAAGGACCAATTCCAAAATCGATTGGGAAGCTACAAATGCTTGAATCACTAGACCTGTCAACAAACCACCTGTCCGGGGAGATCCCATCAGAGCTTGCAAGTCTCACATTCTTAGCAGCTTTGAACTTATCGTTCAACAAATTGTTTGGCAAAATCCCATCGACCAATCAGTTTCAAACATTCTCAGCAGATTCCTTTGGAGGAAACAGTGGCCTATGTGGGCTCCCTCTCAACAACAGTTGTCAAAGCAATGGCTCAGAGTCCCTGCCTCCACCAACTTCGCTACCAGACTCAGATGATGAATGGAAGTTCATTTTCGCAGCTGTTGGATACGTAGTAGGGGCAGCAAACACCATTTCACTTCTGTGGTTTTACGAGCCAGTGAAGAAATGGTTTGATAAGCATACGGAGAAATGCATGCTTTGGTTTTCAAGAAAGTGATCCTTAAACCCGTAAATGATGAGTTTATTTAATCCTTGCTTTGTTATAATAAACATCAAGATCCAAATTCTTATTTCCCTTGATGAATGTATAAAATTTTCTACTATCAGAAACAGCACCCACTTTTGGATTGAATAGCTTCCCACATTGCCTGCAAC

>M6-PGSC0003DMG400006268 M6 LRR-RLP

ACTTTCTATGGCTTTTCTTGATTCCCTTGTTTCAAATCTTATCAGGATATCACATTTTCTTGGTTTCCTGTCAATGTCTTCACGATCAAAAGTCGTTGTTGCTGCAGTTGAAGGGCAGCCTTCAGTATGATTCTAGTTTTTCAAATAAATTAGCAAAATGGAATCAGAAGACAAGTGAATGTTGCAATTGGGATGGGGTTACATGCGATCTCTCTGGTCATGTGATTGCCTTGGAACTGGATTATGAGACAATTTCTGGTGGAATTGAGAATTCAAGTTCTCTTTTCAGTCTACAGTATCTTGAGAAGTTAAATTTGGCTTACAACAGGTTCAGTGTTGGCATACCAGTTGGTATAGGCAACCTCACGAACTTGAAGTACCTGAATTTATCCAATGCTGGTTTTGTCGGGCAAATTCCTATGTCGTTACCAAGATTAAAAAGGCGAGTTACTCTTGATCTCTCAATCTTTTCCCTGACTTTGATCAGCCACTTAAACTTGAGAATCCCAATTTGAGACAATTCATTGAGAACTCAACAGAGCTTAGAGAGCTTTACCTTGATGGGGTTGATCTTTCATCTCAGAGGACTGAATGGTGTCAATCTTTATCTTCATATTTGCCTAAATTGACTATCTTGAGTTTGCGTGCTTGTCAAATTTCAGGCCCTATTGATGAATCACTTTCTAAGCTTCTCTTTCTCTCTTTCATCCGTCTTGAACGGAACAATCTCTCTACCACAGTTCCTGAATATTTTGCCCATTTCTCGAACTTGACTACCTTGGCCCTCGGCTCTTGTAATCTGCATGGAACATTTCCTGAAAGAATCTTTCAGGTACCAGTTTTAGAGAGTTTGGACTTGTCAAATAACGAGTTGCTTCGAGGTAGTATTCCTGTTGTTCTTCCAAATGGATCTCTGAGGAGATTATCACTAAGCTACACCAACTTTTCTGGCTCATTACCAGAGTCCATTTTGAACCTTCAGAACCTGTCCAGGTTAGAGCTCTCCAATTGCAATTTCAATGGACCAATACCTTCCACAATGGCAAACCTTACAAATCTTGTTTATTTGGACTTCTCAAACAATTTCACTGGTTCTATCCCATATTTCCGACAGTCCAAGAAACTCACCCACTTAGACCTTTCATGTAATGGTCTAACTGGTCTCTTGTCTAGAGCTCATTTCGAAGGACTCTCAGATCTTGTATACATAAATTTAGGGAACAATATACTCAACGGGAACCTTCCTGAATATATATTTGAGCTCCCCTCCTTGCAGCAGCTTTTTCTTAACAGCAATCAATTTGTTGGCCAAGTCGACGAATTCTGCAATGCATCCTCCTCTCTGTTGTATACAATTGATTTGGCAAACAACCACCTGAATGGATCGATTCCAAAGTCCATGTTTGAAATTGGGAGGCTTAAGGAACTCTTACTTTCTTCCAACTTCTTTAGTGGGACAGTGACCCTTGATCTCATTGGGAAGCTGAGAAACCTTTCAAGCCTGGAGCTATCTTACAATAACATGACTGTTGAAGTAAGTAGCAGCAATTTAACCTCTTTCACATCTCCCCAGTTGAGCATATTGAAACTAGCTTCTTGTCGGCTGCAAAAGTTTCCTGATCTTAAAAATCAGTCATGGATGATCCACTTAGACCTTTCTGACAACGAAATACTGGGGGCAATACCAAATTGGATTTGGGGAATTGGTGACGGAGCTCTGGCTCACCTGAATCTTTCCTTCAATCAGCTGGAGTACGTGGAACAGCCTTACAATGTGTCCAGCAATCTTTTTGTCCTTGACTTGCATTCCAACCGTTTAAAAGGTGACTTACCAATTCCACCTTCCTCTGCCATCTATGTGGACTATTCGAGCAATAATTTAAACAAATCCATCCCACTAGATATTGGAAATTCTCTTGCTCTTGCCTCCTTTTTCTCGGTAGCAAACAATGACATTACTGGAATAGTTCCTGAATCCATATGCAGTGTCAGCTACCTTCAAGTTCTTGATTTCTCTAACAATGCCTTGAGTGGAACAATTCCACGATGTCTACTGGAATATAGTACTACTCTTGGAGTGCTGAATCTAGGGAACAACAGACTCCATGGTGTTATACCAGATTCATTTCCAATTGGTTGTGCTCTAAAAACTCTAGACCTCAGTGAGAATAAGTTACAAGGTAGCCTCCCAAAATCGCTTGTGAACTGCAAGTTGTTGGAGGTCCTGAATGTTGGAAATTACAGACTTGTTGATCATTTCCCGTGCATGTTGAGGAACTCAAACAGTCTGAGGGTCCTAGTCTTGCGCTTCAATCAATTCAGTGGAAATCTTCAGTGTGAAGTAACCATAAATAGCTGCCCAAATCTCCAGATCATAGATATAGCTTCCAACAACTTCACTGGTGTGTTGAATGCCGAATTCTTTTCAAATTGGAGAGGAATGATGGTTGCAGATGATTACGTGGAGACAGGGCGGAATCATATCCAGTACAAGTTCCTCCAACTAAGTAACTTATACTATCAGGTCACAGTAACAATAACCAACAAAGGGATGGAGATGAAGCTTGTGAAAATTCTCAGAGTCTACACATCTATTAATTTCTCTTCAAATAGATTTCAAGGAGTGATACCAGATACTATCGGGAATCTCAGCTCACTTTATGTTCTGAACCTGTCACACAATGCCCTTGAAGGACCAATTCCAAAATCGATTGGGAAGCTACAAATGCTTGAATCACTAGACCTGTCAACAAACCACCTGTCCGGGGAGATCCCCTCAGAGCTTGCAAGTCTCACATTCTTAGCAGCTTTGAACTTATCGTTCAACAAATTGTTTGGCAAAATCCCATTAACTAATCAGTTTCAAACATTCTCAGCAGATTCCTTTGAAGGAAACAGAGGCCTATGCGGGTTCCCTCTCAACAACAATTGCGAAAGCAATGGCTTAGAGTCCCTCTCCCTGCTGCCACCAACTTTGGTACCAGACTCAGACTCAGATGATGAATGGAAGTTCATTTTCGCGGCTGTTGGATACGTAGTAGGGGCAGCAAACACTATTTCACTTCTGTGGTTTTACGAGCCAGTGAAGAAATGGTTTGATAAGCATATAGAGAAATGTTTACTTTGGTTTTCAAGA

>M6-PGSC0003DMG400006279 M6 LRR-RLP

CTTTTAGTTACGTTGATAGGTTTTTCCAAGTGTTTAAAGGTCCATCTTTGGATTTGCAAGCTAACCATGAACACGTTGATATGTGTACAGTCATATAGTCCAAGTTATTTTTATGTTATATCCAACAATGTGCATTTCATTTATTAAGTATCCCATGTTTGCTTGTTTTGACATATTTTGGTTTGTCTGTTTTTGGTTTATCAGTGCCTGTCCATCCTAGAGTTGAAAGGGTGGCGTCCTCCACCATCAAGCCGGAAATTTTTAGAACTGAACCTAGCCGTAATACAATTGGACTTCCCTGGAATTTACAGCTTTCTACAGAGCTCATCGGATCTTGAGACATTGATCATTGGTTGTTGGTACAATCACAATGAAAGAGTAAGTTTCCTTTCGACCTTTATTTATATTACTTCATGTTATTCAAATCAAACAACTTTTAGTGGAGGAATAAGCTGATATATATCAACATATTTCTAATCTTAAGTACTTCAGGAAAGAAAATTTATTGAAGTTTTTTTTAATAGCAATCTCCTCAGTCACTTTACCATCCCAATTGTGCTTCCTTCCATGAGTTCTATTTTTGTTTTTTTCCAATTCATCGTTGCCTAGGAGATATTGCAGCTTCCCTAGCTTCATTTTTAAAATATTCATCCAGTGTGGCATCCTCCTCGCTTCTCTCAAAAATGTTTGCACCTCTAGAACTTCTTGGAGAAAAATCACTAGGTCTAACATGTAGTTGCTTACTAGTGTTAATAGGAGTAAGAGAATTCACTCTTTCAATATTCGAACTCATCAGTATATCAAGAAGCGCCATGTCTATAGTATTATTGATCTTCTGAACATCAGAACTGCTGCGCAGAATTCTCAGGTGATGCATCTATCAAATTTGCAGATTTTTCAGCTACTTCAGTTTCTAGATCTGTCGGTTTCGAAATTGTATGACGTCCCTGTCTCCGGTGCATCTGTCTAGCACTTGAACTTGCAATTTCTTGTGAACTTAACGCTCCATCTTTCATGATTTCAGGCGATCGCTGCCCCCAAGTGACAACTTCATTCCCCTGAATATTGTCTTCTAATATCTTCTTGTTCCCAGCAGCCTTCTTTAACACCTGATTGCCTAAAATCGCCATGAATGGTCTTACCAACTTCAGAACCTGTAGCATCTGATTTCTCACCCAAATTTCGAACACTCTGTGCAGCTCGTCTTGCATTAATAATTTCCCTAGCATTTCCCTGATGCTTAATACCTTGCATATTAATCTGCATACTTAATCTCTGTGACCTTGATGTTTGCATGTACGTATATGAAGGTTAATAATCATATGTTATTATTTGGTACACTATCAATTCATTCGACTTTGGCTCTAATGAGCAATCTCACACGTTCAGGCAAAGCAGCCAGTAAGTTGACTTCAACTTTAACTCTTGCTGTGCTTGGCCTTGTTTGATCTGTGGTTGCTCTATCAACAACCAAAGGTCTGCCCACTGCTGATGCTAACAAATATAATGCCTCTCGAACGAAAAATTGCACCAGTAAATTCTGAAACAATATCCAAGTCAAAGCAATGGATTCTTCTTCTTTGGGATTAAACCAAGGGGTCCAACGAAAAAATTCTAAATTGATAGTCTTCCTCCA

>M6-PGSC0003DMG400006422 M6 LRR-RLP

ATGGTCAAAGGAAAAAAGGAAGTGGGAGATGATGATGAGTTAAATTATTCCTACTTTTTGGGAGTTTTGAGAGAGAGTGCCATAGTGACGAATAAAGGCAACAGGTACCAGTATGACACCATTTTGGCGTTGTTTACAAGTATGGACATGTCTAGCAATAATCTCTCTGGGGATATTCCTATAAGTTTAACACTTCTTGCAGGATTAAGATCATTTAATTTTTCTAAAAATAACCTAACTGGTAGGATTCCAAATGACATTGGTGACATGAAAGTTTTGGAATCTGTTGATCTTTCAGAAAATCAATTTTATGGTCAAATCCCACAAAGCTTTTCAAGTTTGTTCACCTTGAGCTACTTGAATCTATCTGATAACAATTTGTCAGGTATGATACCATTGAGCACTCAACTTCAAAGCTTTGATCCCACGAGTTTTCAAGGAAACAAACTCTGCGGGCTTCCACTCTCGCTGAATTGTAGTTCAAATGGTAATATTCCAAATCATGAATATGAAGATGATGAGAGTGACGAAGATGAAGTGGATTGGTTTTACATTTCAATGGCTATAGGATTTGCACTAAGCTTTTGGGGTGTTTGTGGTTCATTGCTTTTCAAGAGATCATGGAGACATGCTTATTTTTGTTTTCTAGACCGTAGCTGGGAAATGTTGCTTGCAAAGTTATCAGTATGTTGA

>M6-PGSC0003DMG400006458 M6 LRR-RLP

TTTCTCCTCTCCATTCCCTATAAGTGTTTGTGTCACCACACACAAGTAACGACCCCTCAATTTGTCAATAAACAAATAATAGGATTAGTTAATTAACAAAGGAAGAAGAAGATTATGGAATTTCAATGATGATTTGTGTTAGGATTAAGAGATCATGAAGATGAGGATGGAATTAGTTGAACCACGACGACGACAACAACAACGACAAGAAGAAAAAGGAGCCTTTGAGAGGAAGAGTAGTAGGGGAAAAAGTACAAGAAGAAAGATGACAATTAGTATTGTTGAAGAAGAAGAAGAAGAGAGCCCAAAGATTGTGGATCTAAGTGGCATGTCATTAGACTCTCTTCCTGTAAATCCTACAATCAATTTGGGAGCTATATCCAAATTACATATTTCCAACAATAATCTTCAGGTTAGTCAGTTTTCGTACACCTCAATTAATTTTTGTGAAATATTTGTTACCTTCTATCAGCAATAAGCAATATGTACTAGATGACTCTGTTCATAGAGGTTAGGATAGATGAATAAATTCACTTAGTTTTTTTTTTTTTTTGTCTTCGCTAGAATTAATTGAACAACCACTTCGATGATCACTATGTGCTATCGTGAATTATTTATTTATGAGAATTTTATTCATGATAGAATATTTTTAATCTTTGATTGGAAATTAAGTGAAAATATCTTCTTTTTGAAATAAGTGGTGGCTAAAGTATTTGATATGAAGATAAGATAATTATGAAGAAAAATGTTTTAGTCAAAATCGAGTGAAATTATTTTTTTCATTTTGACTGGAGGCGAACTAGTTGATTGCTTTGAATTGTATCAGCAACACAAGGACACTATTAAAAACACTAGTTGATTGCTTTGAATTGTATCAGCAACACAAGGACACTATTAAAAACACTATTGCATTTCATAAATGTATGATTAATTATATATTAATTTATGTCGATATTAATAGGTGGATGTGCACTACCTATATAGGTGGATCTGCCTCTAATATAATTTTGACCAAAGAATATTTTTCTAAAAATATTTTCTGACAAGCAGTCACGACTAGTATTTTGGAAAACATTAATTTTTCAATCATACCAAATACAGTCGTTGCTTCCATTTTTAGTTGCTTTTGGTTCCTTGTACTTTCACAAATATTTAGTTTTGTTTAATTTCTTCCATATTTATGGAAAGATATATTTATCATTTACCTTTCTTCTATTTTACGCAACAATAGTTATTATATGTTTCCTAGTTTTCAATTTATTTATTCAGTGTTGAGTTGATATTGAGTTTCAAAAGGTGGATAATTTTTTTATTGTTATAGTCCTAAACTAAAAATATGTAAAATGTATCAAAATATTTTTTAATTTTGTGGTTTTAAACATGTCTTGTGGAAAGTTAGAATTAAAGAGGTGTCAAAATAAGAATTTAATGTACTACTATATTTTCTTAAAATATTCCCCTTTACTTGATAATGAGTCGTTTATGTGGACTATACAGTATAATATTTTCAGGAATAAAATGCAGTACTATTTTATTCTGTGTTTAATACAAAACAAACAATATAAATTTTAACGGATCAAATTAAATCCAAATCAACCGTCCTAACGTCATATTTTGTCGTTCATCATTATTGTAAATTGACCCAATAGTAAGGATAAAGTCTTGTATAAGTTTATTTTTTACTTCATTGATATATTTAAAAAAAATATAACAACTTGAAAGGTTTTCCAATTATTCGTATTAAACGCAATAGTATATTTTTCTGGTTGTTATATGTATGTGTGAATTGTGGATTTACTTAAATATAATTGACAAAATTTGACAAAACACGATAGTACATAACGTAACAACATTTTCGTTAATTATATATCCCGTCTTTGTTTATAGTCTAATACAATGTATATGTCTCTCATTGTGTTTTACAATGTGTATGTGTCTCACTCGTCTCTCTTCTTCTATGTATTTGTATTTTATAATGTATATGTGTCTCACAATTTCAATTATAAGTACCATACAAATATCTGATACACAGATACGTCGACGGTTTATATTTTTTTTTTATATCAATGTATCTATTTCATATATGTATCGATTTTAAGTAGCATGAATTGTATCATAGATTAACACTTTATCATTTTACTTTCTCTGTCCAACAATAGTTGTCCACTATTGACTTGGCATACGGCATACCCCTTAAAAAAGAATAAATAATAGATATTACTATATTATCTTTTGACTTTATTAAATTTAGTGTTTTGAGAAATGTGTTAGATAATAAATAATATTTAACAGTAAGGGTAAAATAGATCTCTTGATTTTCTAAATTGGACAAGTATTATTGAACAACTATTTTTAGTAGGAAAAATGACAAATATACCCTTGAACTATCGTAAATGGTTTGGAGATACCCTTCGTCATACTTTTGGGACATTGGTGCCTTGTCATCCAAAAAGTAGAGCATATATACCCTTTATACTAACGGACATACACGTGTCACAACTCACAATCTTATCCACTGACCCACATTTATTAAATATCGAATCGACAGATAAGATTGCGCCACGTGTCTCTATTTAGTCTTCCGTTAGAGTGAAGGGCATATATGCTCTAGTTTTTGGATGGAAGGGGCACCAATGTCCCAAAAGTATGACGGATGATATCTGCATACCATTTACGATAGTTCGGAGTATATTTATCCTTTTTCCCTTTTTAGTATAGTGGACAACACTATTGTTGGACAGAGGAAGTACTAGACACAGGATTTTAGGGTAAGTCACATTCTTATTTGGTTGGACGATTTTCAATTATCATGTATCACTTGTGTATATAGATACATAAACACACATATGTATCTCTTCAAAGTTCCACTCATCTGCCATAGATACATAAATACATTGATACACATATATATCTCGTCAAAGCTAAGCACTCAATTATATTATATCATGGATGCACATTTACATATATACACAAATACACACACACACACATATATATATCTCTTAAAAAATTTCCACTCGTCTTTCATGGATACACAAATACACATCCATAGATACGCAATACACATATATATCTCCTCAAAACTACTCACTCATTTGCAATAGATGCACAAAAATATAGATACACTTATACATTGATATACATATGTATCTCCTCAAAGTTGCACTCTAATTTGCCATAGATGCACAAATACATATAGTCCCTCTGTAAATCGAGAGTGGAGAGAGGGAGGAGAGTGGCGAATTTTCTAGGAAAAAGGGAAATCTCATTGAGTTATGTAATTTTTTAAAACTATAAATAGGTTGAGTAGATATATATACTAGCTATGCTTGATTAATTATGTAATTTTTCATGTATAATTCTCACTTCATTGTTTTTGTTGTATGTTTGACAGAGTATACCAGAGTCTTTAACAGCAAGACTACTTAACTTGGTTGAGTTAGATATGCATTCAAATCAGCTCAACTTCATTCCTAACTCAATAGGTTGCTTATCAAAGCTCAAACTTCTCAACATTTCTGGGAATCTCCTTCTTTCGCTCCCTAAAACTATCGAGAATTGTAGGTTAGTCATCTAAACATTTCTAATCCATAACATTATCACTACAACAAAAGAGATATTTAGCAGCAATTGAGTTACTGTCATTGCATTCAAAGTCGCTAAAGCCTTTAGAACATAATAAGTATAACCTTTTCTTCCTACTTGCAGATCATTGGAAGAGTTGAATGCCAACTTCAATATGCTTACTCATTTACCGGACACCATTGGATTTGAGCTCATAAATCTCAAGAAGCTATGCATAAATTCAAATAAAATAGCATACCTTCCTTACTCCACCTCCCACTTGACCAACCTTCGCGTGTTAGATGCTAGGCTAAACTGCCTTAGATCACTCCCCGATGATCTTGAGAACCTAATCAACCTTGAAATCCTTAACGTTAGTCAAAATTTCCAGTACCTCTCAAAGTTGCCGTATTCAGTAGGATTTCTCATTTCTCTTCATGAACTAGATGTTAGTTACAACAAGATTACTGAATTACCAGATTCCATCGGATGCTTGAAGAAGCTCCAAAAGCTGAGTGTAGAAGGCAACCGCCTTGTTTCTCCCCCACCGGAGGTAGTGGAGCAAGGGATACGTGCAGTTAAGCAATACTTGTGTGAAAAGATAAATGGAATGCATGACAAGTCCCCTAAGAAGAAGTCATGGTTTGGAAAGCTAGCAAGATGTAGCACATTTAGTGGTGCTAACATACCAAGAGATGATAGAGAGGACTTTGGTGTTCCAAGCAATCGTACTATCGATGCAATTGCCTCACCACGATTTATGGGAATGCTTTCTCCTCGTCGTCTTCTATCTCCAAAAACCTACTTCTCCAAATAAGGGACATTGCAATTTGCTTGGTAGCTCTTAGCTTGTTTTCTTTTACGTTTGAAATCGTGCAATGTGTGCTAGATCAAGCTGTACATTATGTTCTGTAGATTTTGTTGTCTTTGAGATAGTACAGATTTTCATAGATGATTAGCCTCATGAGTCATGGGTATGTTTGAGAGGTAACAATGCGGTACAAATTTACCCGAACAGAGTGTTTATATCAAATCATATGAATAATTACATCGAATCAAAATTTGTTATCATAACATGTCTTGTATTATATCACTTGTACAAAAACTCATTTTGATCATTAATTATCAAAACTCAAAAAAAAAAAAATTAAAAATCTTACATAAACAGAACCATTAAAAAAGCCCCAAAAAAACAGAACAAGACTCTTAGAAAGCAAACAGAACTAATGAAAAACTTCCCAAAATCACAAATTATGAATTAGTTTCTTGCACTTGCGATTCATTTCTTTGCAAATCCGATTCACTTCTCCTATTAAACGACCATTCTCTAATGGTAGACAAAACTGGCTTCCATGATCGCGATTTATTTGACCTTTCTCCAAATTTGCACGAAGTTGTGTTAGTTGCACCACTTTTAGCTATGCAACTGAGTTTTCTAGCTATTGGAACTAAATCCATTGGAGAAACACTTGATTGAAATAAGGATTTAGGCAATAAATAGTAAATATGACCTTGTTCTAACTTAGTATCAGGGTTGAGAGGCTTTGAAGCAGCAGATAAAAGTTGGGCATGAGTGAAGACAAAGTGTTTTTGAGATTTGGCCATGAGTTCATGTACACTAATAGGATGATCAAAATCCTCTACATGACCATTCAAATGTACAACTCTAATAGTCTTTAATATATCACATGATTTGGGAGAAAAACAACAATATGTTATTTTGATGTACATATTGTTAATATTTTCCTTATTTCTCTCTAGCTAGTGTGTAAAAGGGTTCCCAAAGAAGATGTCCTTTTATATAAGGGGAAAGATGGTGCTGAAACCGGGTTGTGGTGCGGTAGTGGAATTGCTCCACCCTTAACCAAAGATCTCAGGTTCGAGCTCTGGGTATGAAGAAAATCCTATTGGGAGTGTCATCCTTGGATGGGCCTTGCAATGTGCAAATACAGATTTAGTCGGGGATGGGCCCTACAATGCACAATTCAGATTTAATCTAAGCTCGAATGCGGACTCCGGATATCGAGTAGAAACAAAAAAAATGGTGGTGGGAAAGAGTTGTGGAGATAGATGATGGGAACACGCTATTTAAGTTAGGAATTTTTTATCTACTTCACGGCTTAAGTGGCACTCTCAATAGGATTTTCTTCATACCAAGGCTCAAAGTGAGACTTGTGGTTAAGGATAGAGCAATTTCAT

>M6-PGSC0003DMG400006486 M6 LRR-RLP

CTAAGGTTACAGTGTTGCGAGGGTATAAGCAAATGTCATATTTTTGGCTTAGTACATCTCAAGTATTTCGATGCTAGTTATTGCAAACTCCATAGTCTGATAGTCCAAGCTCCATCTCTTCAATGCTTTAGATATAGAGAACCTACTGAACAGGGAAATCTTCATCCTTGCCAAATTGCTATTTTGGATGGTTACAATACTTTACAAACACTCGAGCTCGCTGGTGCCAGCATCACAGACCAACAGCTTCGAGATTTATCCTATAAGTTCCCAAACATTTCAGAATTGAATCTAACAAGATGCTATAACCTGAAAAATATAGAGATTCAGAGTGAAAAACTCAAGAAATTCACCTTATTAAAGTTGAAGAGTCTGGAAAAACTCACGATTCAAGCTCCAAATTCATTGGAATTTGATTTTCATGGTAGTAAATGGCCCTTCTCATATATGGATACTTCTTCCCTAGAAAGAGTCCGACTCAATTCTTCTTGCCCGTCATAAACTTTGGATCAGTGGACAGCAGTTGGTACACGAGCCTTCATCACTTTGTTCAAAAGTTCAACTATTCTAAAGATTTCATATTAATAATATTTTGCCGCCAGGTAGTTTACATTCTTTTTATGCTTATGTCATTTATTTATGCATTACTTGTTGAATTTGGCTTAAAATATGGCATTGACTAAATTCTTGCAGACCAAAAGCATCCTTATTTATGAAAATCCAAGAGAAATTGTTATTCCACCAAGCCACAAAATCGAGATATTCATTACATCCATGCTGCGTGTTGAATCGATCATTGGAATGTTAATGGTCTACCGTCCCTATATCATGTCCATACTTCCATGTACAGATAGCAAAGCACTACAGGTGATTTCTTGCTTTACCTCAAAAACGTCTCTTGTGCTTGTATAAACTAAGACATTAATTTGAGAAGAAAATTTGTTGTTAAATGTTGATTGGTCGTCTCATAAGCATTATTGTATAATATATCTTATGTTTTTGCTTAAATGCAGGCGATGTCTACACTAAAAGGGTGTACACAAAAGCAAAAATGTGGTAAAGAATGTCCATTCAATACCAAGTGGCATCGTAACTTAAAAGAAGTCATCAGTTGTAGTGGGACATCTGAAGAGGGAATGGCTGCCTCCATGTGGTATCTCTGGTTGAAATCCACATCTGTAATTGACCAAGTGAATAATTTCGTGTTAAAATGGAAAGATGAAGCCTAG

>M6-PGSC0003DMG400006487 M6 LRR-RLP

TATAAACCCTAACAGCAAACGGCGGCACCTACCAAAATCAGAAACCCTAATGGAAGAAAGTGTTGATGGAGAAGACAGAATATCAGAATTGCCTGTGCATATCATTCATCACATCTTATGTCGCACTAACCTCGACCTCAGAGAGGCGGCGATAACTTGTGTTTTATCCAAGAAATGGCAGTATTATTGGCTCTCAAGACCTTATTTGCTGTTCCGTCAATGTAAAGGCAACAATTGTATACCCCTGGAAAATTTTGTCAAGTTGATTGATCGACCCCTACGAATCCATGCTGAACAAAACTTACATCTTGCAAAATTCTTCCTTGCATATCGGGATCGAGAATTGGATCCTTATTTGGATCATTGGATTGAATCGGCTATTAAAGTCAATGTTTCGGTGCTGGTGATTCATGCTCCGGATATTGATTCGTCGTATTATAGCTTACCTGATGTTGTTTATGATGCGAAAAAGCTCAACACATTGTGGTTAAGCAGGTGCAAGTTTGAATTTGATATTAGCACCACTCATATAAGATTTTGTTCTCTTGTGGCTCTTTTTTTGTATAATGTTCATATTTCAGATGCCCAAATGCAAAGTGTTATAGATAAATGCCCATCCATCAGGAATATAAGAGTACTGGCTTGCAAAGGTATGAGCAAATTGTATGTTGTTGGACGGTTACATCTACACTCTTTGGCTGTACTATTATGCAAACTCGATAGTGTGATTGTCGAAGCTCCGAAACTTGAATCCTTTAGTTATACAGAATGTATTGATCAGGGAAATTTTCATCCTTGCAAAATTGATATTTTGGATGGTAACAATACTTTACAAACACTCGAGCTCACTGGCTCCAGCATCACAGACCAACAGTTTCGAGATGTATTCCACAAGTTCCCAAACATTTCAGTGTTGCATCTAACAAGATGTTATAAGCTGAAACATATAGAGATTCAGAGTGAAAAACTCAACAAAATCACCTTATCGAAGTTGGAGAGTATAGAAAAAATCATGATTCAAGCTCCAAATTTATCGCAATTTGATTTTGAGGGTGATAAAATGCCCTTCTTATCTATGGATTCTTCTTCCCTGGAAAGAGCCCGACTCAATTTCTTCTTGCCGAGGACAGTAATATCAAACTTTGGAGATGTGGATAGCAGTTGGTACACAAACCTTAATCACTTTGTTCAAAACTTCAACTATTCTAAAGGTTTGATGTTAGTAATATTTTGCCGCAAGGTACTTTCCATTCTTTTTGTGCCATTGATTTATGCATTACTTGTTGAATTTGGTTTTAAGCTATGTCATTGGCAAAATTCTTGCAGACCAAAAACATCCTTATTTATGAAAATCCAAGAGAAATTCTTATTCCACCAAGTCAAAATGTTGAGATATTCATTGTACCCATCCTGAGCATTGAATTGATCATTGGATCGATATTGCTCAACCGTCCCAGCATCATGTCCATACTTCCATGTACAGATAGCAAAGTACTCCAGGTAATTTCTTGCTTTACCTGGAAATCCTTTCAAAAAAAACGTCCCTTGTGCTTGTAGACACTAATGCAGTAGAGCTTAAGCATTATTGTATTCTCTATATCTTATGTTTTTGTTTAAATACAGGTATTGCCTGCATTAGAGCGGTGTACACAAAATCAAAATTGTGGTAAAGAATGTCCATTCAGTACCAGGTTTTTCCACAAGCATCGTATATTAGAAGAAATCATCAGTTGTACTGGGACATCTGAGGAGGAAATGGCCTCCATTTGGTATACCTGGTTGAAATCTACGTCTCTAATTGGCAAAGTGAATAATTTCATGTTCAAATGGAAAGAACAAGCTTAGACCGAAGAATCTTGGGGAATTCAGCATATAGTGTTTATTAGTGTGTTCAGAATAAAGTTCAGTTAACCCGCTTGTGGCTCTGTCAAATCCATGAAAGGTTCCTTAACCTTGCTCTAGCTCTAAATAAATTGTGCTCAAATCTGTGGTTTAATTTCATTTGTTCGATTGTACGGAGTGAAGTTTAACTGTTAACAAACAGCATAAGAGGTCTTCATTTCATTTTTTTTTTTCTGGCTCTGCCTCTTCTTCACATTCCATTCAATTTCTCTCATGCTTAGACTACTTGGAGGTGATTTAAAAGAAGTGAGCACTTGAGGTAGAGAGATGTAATGATGGATGCATATGCCTCCTACTTGTTCTGAAATCTGGTATATATTCACTGTCATACTTGAAATGTAATATGATTTTTCGTATAACTTTCTTTGTCA

>M6-PGSC0003DMG400006580 M6 LRR-RLP

TATAGAAACTTTAATTGTGAAGAATTCTCCTTACGTATGGTTACCAGGGTCTTTTTTGCAAATGGAAAAATTAAGGCATGCTCATTTTGGTAAGGCTGTTTTTGATAAGCAGGGGCTCTTTGAAGGATCCTCTAAATTGGAAAATTTGAGGATATTAAAGAAAATTGAGAAATTTCCAATTGATAGGGCGGATGTGTTATCAAGGAGGTGTCCTAATCTTCAACAACTTCATATCGAATTTGATGGACATTCTGCAGATTCTTTTTGTCTCACATTGGAGAATCTTACCCAGCTTCAAATACTTCGCGTTTCCTTTGAGCGGCCCCACATCATTGTATCTGGATTGCAATTGCCTTCAAATTTAAACAAGTTGGTACTTAAAGGAATTCATATAGAAAGTGTTATTCCCTTCATTGCGGGACTACCAAGCCTGGAGTATCTCCAATTACAAGATGTGTGTTTTCCTCAATCAAAAGAGTGGTGCCTTGGAGATATCACGTTCCATAAACTTAAGTTGTTGAAACTGGTGCAGTTAGGTATCTCAAGGTGGGATGCCTCGGAGGAATCATTTCCCTTGCTTGAAACACTAGTTATAAAAATGTGTGGTCACCTAGAGGAGATCCCAATTAGCTTTGCTGATATTCCATCATTGAAACAGATTAAATTGATTGGGCCCTGGAAAGAATCTCTGGAGGATTCAGCTGTGAGAATTAAGGAAGAAATCGAAGAGATTGAAGGATGTGATCGTATACACCTCGTCAAAGAACCGACAAATTGGGAGGCATATAGCAATTTGTCGAAGCTGTGCTTATACTGAAGTCTATTTTGTAGCTAGCCCATTCTCTTATATTGTAACACTACTACTGTTTGTTATGTTTGTTTTGATTGATCAACTCTTTTTATGTATGGTTGGAGACAAATATATATCAGATGATCTTTCACTGATTTCATGTTTTT

>M6-PGSC0003DMG400006625 M6 LRR-RLP

TATTGTTGTTCATCCTGTATTCCTATGATACATAGAACAAAAACTTTGAAGAACTTGATAAGAAATAACAAAGTTTAAAGAGTATTTTCAAAACTAGAAAATTAAAACAAGTAGAGAAAATTGAATATTAGAAACAATATACAAAATATTTTTCTTAAAGAAGATCGAGCCCACTGAATGCATAGTGTCCCCTTAAGAAAATTATTCCTCTCAAGTACCGGAGGTTAATGGAATATACCTCCCAGGATAGAACGATCTTACTCACCAGTGTATCGGTACCTAAAATCACGGTGTCAGCGAACCACTCAACGGCAGTAAAGTACACTTAGAATACTAGATTTAGTAGTACGTAGTAGAAGAAGTCCAGAAAATTAATTCTAATAAAATGAGAGAAAATCCCTCAATTTACAGAAAACAAAGGGTAGTGTGAAAACGTTCTTATTGTGCCTTACCGAAAAGGTCACAACCATTTGGAAAAGTCACAATCTTTCAGAAAGGTCACAACCTTTCATAAAAGTCACAACTTTTCATAAAAGTCACAACTCTTCACAAAAGTCGCAACTTTTCATAAAAGTCACAACTCTTCATAAAAGTCGCAACTCTTCATTTTCCATTCACACATTTTAAAACCCAACAATCCCCTGCATGAATGGGGAATGACTCCAAGACAAAGAAACGAACAAGTGTGTACTTTACAAGCAAGAATTAATTACATTTGGGTAAGTAGGTTTCCTTTTGAACTTTCTGTAGTGAACATATGTCGGATATACTCGGTCAATCGGTAGATTTGATATCTTTGAACCGTCAAGCTTTGGTGTATACCTAGACAATCATATGTCACACAATCAACCCTTTACCGTTTATGGTTCTTACGGTTATGTTCGTTTCAGCCATGAACACCTCCTGGTTTCATGAGTGTATAGAGAGATGGGCTTTTGACAATCATCTTCTTTGAAGCGGCTTACACTTCACACTCACATAGGTGATTTCTAACCGTGTTATCGCGTAGATATGCTATTTGGTCAAATCTGCCAAACTTAGCAAATCATTAAAAATATTAAGCTTCATTAACTCATTAACAAGCCTTAATGTTATATCCTTGTTCCTGAACATTGTCTTCATCATGAGAATGGATTGAGTTGGTTGACAATGCCGAAACGTCAGTCACAACTTTGTTTTTCTCCTTGAACCTAGCTCTTGGGATATCCAGTCTGCTAGGTAGAGTTACCTCCATGATGACTTGTCCTAAGCCGTAAACTTATTCCCTTAGATGATCTTTCAACTTTCTCTCTAGTTAGGTCTTTTTGTAAGTGGATCCAACACATTATCCTTTGACTTTACACAGTCAATTGTGATAATTTCACTAGAGAGTAGTTTTCTAACGATATTATGTCTATGTCTTATATGACGAGACTTTCCGTTATACATCATGCTACATACCCTACCTATTGCATCTTGAATATTAAAGTGTATGCATACTGATGCCAAATGTTTGGGCAAAAAAGAATATCTATCAAGAAATTTTGAAGTCTTTCAACTTATTCACCGGCCTTACCTAGAGCACCTAGTTCGGAGATCATTGTAGAGCTAGCGATACATGTCTTTTTGGAAGATTTCCAAGAGATTGCTCCTCCACCAAGAGTAAATACATATCCACTCGTGGATTTTACTTCATTTGCTCCCGGTGATCCAATTTGAATCACTATATCCTTTCAATACTGCTGGATATTGTTATAATGCAAGACATAGTTTTGAGTATGTTTTAAATACCTCAAAACTCTTTTCATTGTCATCCAATGAGTTCGATTGAGATCGCTCGTGAATCGAATCAGTTCATTAATAGCACATGTTATATCTGATCGCGCACACTTCGTGATATACATCATACTTCCCAATACTCTAGCACAATCCAATTGTGAGTCACTTTCGCCTTCACTCTTTTGAAATGCAAAGCTCACATTTATTAGAGTCTTGACAATAATGAATTCAAATATTTCAACTAGTCAAGTACCTTTTCAATGCAATGAGATCGTGACAATGCTAAACATTATGGAGTTCTAAGATTTTTATACCTAAGATCACATCAACAACTCCAAGATCTTTCATATCAAACTTGCTTTCTAGCATACGTTTAGTAGCATTTATGTCATTAGTGTCTCTACTGATGATCAACATATCATCAATATACAAACAAACAATGACCTTGTGATTTTAATGTAAACACATTTATCAGTTCCATCATTCTTAAATTCATTTGCCAACATGGTTTGGTCAAATTTCTTATGTCATTGTTTGGGTGCTTGTTTTAGTCCATAAAGTGACTTAACAAGTTCGCACACTTTTTTTTATTTACCAGGAACCACAAAACCCTTGGGTTGTTCCATGTAATTTTCTTCCTCCAATTCTCCATTTAAGAGAGTTGTTTTCACATTCATTTGATGAATTTAGAGATCATATACCGCAACTAATAATATTAACATCCGAATGAATGTAATTCTAGTTACTGACGAGTATGTGTCGACAATTTCAAGACCTTATTTTTGTCTAAAGCCTTTGACAAAAAGTTTTGCTTTATATTTGTCAACAATTCCATCGACTTTCATTTTCTTCTGAGGATTCACTTTGAACCCAAAAGTTTATTTCCTGGAGGAAGATCAACCAACTTCCAATAGGATTGCTCAATATTGAACCAATCTCACTATTGGCACCATCTTTCTAAAAGGATGAGTCCACATAAGACATAGAAAATGTTACAAAATCAGATCCAAAGGTAGTAGAATGTCTTTGACATTTACTACACGTTTGAATCTCTTTATTAAGTACATTTCTTTTGTTCTTTCCAAGGTCTTTTAGACCTTTTACTAGACTACTCAAGTCTAATTTTACAGTCCACAGTCTTAGGTCCTATTTTAATCCTTTTAGGAATAGGAACTTGGACTTTGACTAGACACCTCCACATTTTAAAATATTTCAAGTTGGATTTCCTTTCTTTCCATTTCTCATATGAAATAGATTGTGTCCTGAACAATCAAAACTTTTTTTGCTTTCTTTCTGAAAAGACAACTTTTTGTTGTTCCCTTTTGTTGTAAGGATAGCTTCTCCACACAAGTTTTGTGGTAAACCTGAACTTATAAGTAATGTAGCCATCATTTTCTTCAAAGTTCGATTTTTTTTCCTTCTGCAATTCCATTAGATTGAGGTGAATAGGGCAATAGTGTGATGGATAAATGCACTTTCTAAACACATTTCTGCAAAAGGAGATTTATACTCTCCATCCCTATTACTTCTTATCTTTTTTCCCAACTAATTTTCAATTTCAATATTATATT

>M6-PGSC0003DMG400006652 M6 LRR-RLP

AATAACAGCTTCATCGGTTTGTTTCCCGCTTCAATTTTAAGTTTGACACATCTTGAATACTTAGGATTGTCGAGTAATTCCCTATCTGGCCCACTGCCTAGCAACCAAAGCATGCTTCAAAAGCTAACTTCTGTGGATTTGTCATACAACTCACTGAATGGTACCGTACCATCTTGGGTGTTTAGCCTACCTTTGCTATCTTCAGTGTCGCTCCAACATAACCGATTCAGAGGACTAGCCGATGAAGTGATCAAAACAAACCCAACATTAAAACAACTGCATTTAAGCAATAATCAACTCAGTGGTTCTTTTCCTCAATCACTTGTGAATCTCACAAACCTTCAAACCCTTGGAATTTCATCAAATAACATCACCATTGATGAGGGAATGAATATCACCTTTCTTAGCCTATCATCTTTATTCTTATCATCTTGTCAACTGAAGGATTTTCCACACTTCTTGAGAAATGTAAAGAAACTTGTGTACTTGGATATTTCTAACAATAAGATTCGTGGTCAAATCCCTAACTGGTTTAGCGGCATGAGATGGGACTCGTTACAGTTCCTAAACCTTTCTCATAATTCATTGACAGGCCACCTACCACAATTTCATTACTATAACGTACAGTATCTTGATCTGAAATTTAACTCCCTTCAGGGTCCACTACCTTCATCCATTTGTAACATGAGTTCGCTTATCTTATTAGATTTATCACGCAACAACTTCAGTGACTCAGTTCCACATTGCTTGGGAAGCATGGCTAGTCTAATGGTGCTAGACTTAAGAAGGAACAATTTCACAGGGAGTCTTCCCCCATTATGTGCACACAGCACTTCGTTGAGTACCATTGTCATAAATGGTAATCAATTTGAAGGACCTGTACCTGTGTCATTGCTCAAGTGTGGTGATCTAGAAGTACTTGATGTGGGGAACAATGCTATAAATGACAGGTTTCCATCTTGGCTCGGAACTCTTGAATACTTGCAGGTCCTTATATTAAAGTCGAACAAGTTCCATGGACCTATAAGTATGTGTCAGACTAAGTTTTGCTTTCCCAAGTTGCGAATTTTTGATCTTTCTCGTAATGATTTCAGTGGCTCACTTCCTGCAAAAGTTTTTGGAAACTTCAAGGCAATGATCAAATTAGATAACGAAGACACAAGAGAGATAAAGTACATGAAATCTGTGTTGCATTGGCCGCCATTCGTCACATCGTATGAGGAGTCAGTGAGTTTGGTAATCAAAGGGCAGGATATTGAGCTACAAAGAATCAGCACAATTACGACAACCATAGATCTCTCAAGCAACCATTTCGAAGGTGTGATTCCGAAAACACTAAAGGATCTCAGATCACTTTGGCTACTCAATTTATCTCATAACAATCTCAGAGGAGATATTCCAATGAAATTAGGACAATTGAGTATGCTTGAAGCTTTAGATCTCTCTTGGAATCGGCTCACTGGAAAGATTCCACAGGAATTGACAAGAATAAACTTTCTCGCCTTCTTAAACCTCTCTCAAAATCATCTCGTCGGACCAATTCCTCAAGGTCGGCAATTCAACACATTTGAAAATGACTCATATGGTGGCAACCTTGATTTATGTGGTGTTCCTTTAACAAAGAAATGTGGAACGGGTGATTCATCGCATGTTCCTCAACAATTGGAGTCCGAAGAAGAAGGCGAGTCATATTTTTTTAGTGGATTTACGTGGGAATCAGTAGTCATAGGCTACAGTTTTGGACTAGTTGTTGGAACTGTCATGTGGAGTCTCATGTTTAAATATCGTAAGCCAAAATGGTTTGTGGAATTTTTCGATGGACTCATGCCTCACAAAAGAAGAAGGCCAAAGAAGAGAGCTCAGAGACGA

>M6-PGSC0003DMG400006655 M6 LRR-RLP

GAAATTCCTGATGTTTTCTCTAACCTCCAAGAGCTACGTACTTTACATCTTTCTTATAACAGCTTCATCGGTTCGTTTCCCGCTTCAATTTTAAGTTTGACAAATCTTGAATACTTAGACATGTCGAGTAACTCCCTATCTGGCCCACTGCCTAGCAACCCAAGCATGCTTCAAAAGCTAACTTCTGTGGATTTGTCATACAACTCACTGAATGGTACCATACCATCTTGGGTGTTTAGCCTACCTTTTCTATCTTCAGTGTTGCTCCAACATAACCGATTCAGAGGACTAGCCGATGAAGTGATCAAAATAAACCCAACATTAAAAGAACTGCATTTAAGCAATAATCAACTCAGTGGTTCTTTTCCTCAATCACTTGTGAATCTCACAAATCTTTTTACCCTTGACATTTCATCAAATAACATCACCATTGATGAGGGAATGAATATCACCTTTCTTAGCCTATCATCTTTATTCTTATCATCTTGTGAACTGAAGCATTTTCCATACTTCTTGAGAAATGTAAAGACACTTGTGTACTTGGATATTTCTAACAATAAGATTTGTGGTCAAATCCCTAACTGGTTTAGCGGCATGAGGTGGGACTCGTTGCTGTTCCTAAACCTTTCTCATAATTCATTAACAGGCCACCTACCACAATTGCATTTCCATAATCTAGGGTATCTTGATCTGAAATTTAACTCCCTTCAGGGTCCACTACCTTCATCCATTTGTAACATGAGCTCGCTTGAATTTCTTGATTTATCACGCAACAACTTCAGTGACTCAGTTTCACATTGTTTGGGAAGCATGGCTAGTCTAACGGTGCTAGACTTAAGAAGGAACAATTTCACAGGGAGTCTTCCTCCATTATGTGCACAGAGCACTTCATTGAGTACCATTGTCGTAAATGGTAATCGATTTGAAGGACCTGTTCCTGTGTCGTTGCTCAAATGTAATGGTCTACAAGTCCTTGATGTGGGGAACAATGCTATAAATGACACGTTTCCAGCTTGGCTCGGAAGTCTTCAAGAGCTGCAGGTCCTTATATTAAAGTCGAACAAGTTCCATGGACCTATAAGTACGTGTCAGACTAAGTTTTGCTTTCCCAAGTTGCGAATTTTTGATCTTTCTCGTAATGATTTCAGTGGCTCACTTCCTGCAAAAGTTTTTGGAAACTTCAAGGCAATGATCAAATTAGATGGTGAAGACACAGGAAATATCAAGTACATAAAATCTGTGTTGAATTCGTCATTTGACACATCGTATGAGGATTCAGTGAGTTTGGTAATCAAAGGGCACGATATTGAGCTACAAAGAATCAGCACAATTATGACAACCATAGATCTCTCAAGCAACCATTTTGAAGGTGTGATTCCGGAAACACTAAAGGACCTCAGCTCACTTAGGCTACTCAATTTATCCCATAACAATCTCATAGGTCATATTCCAGTGGAATTGGGGCAATTGAATAAGCTGGAAGCTTTAGATCTCTCTTGGAATCGGCTCACTGGAAAGATTCCGCAGGAATTGACAAGAATGAACTTTCTGGCAGTCTTAAACATCTCTCAAAATCATCTCATCGGACCAATTCCTCACGGTCTACAGTTCAACACATTTGAAAATGACTCGTACGGTGGTAACCTTGATTTATGTGGTCTTCCTTTAACAAAGAAATGTGGAACGAGTGATTCATCACATGTTCCTCAACAATTGGAGTCCGAAGAAGAAGGCGAGTCATATTTTTTTAGTGGATTTACATGGGAATCAGTAGTCATAGGCTACAATTTTGGACTAGTTGTTGGAACTGTCATGTGGAGTCTCATGTTTAAATATCGTAAGCCAAAATGGTTTGTGGAATTTTTCGATGGACTCATGCCTCACAAAAGAAGAAGGCCAAAGAAGAGAGCTCAGAGACGACGGACTTAA

>M6-PGSC0003DMG400006792 M6 LRR-RLP

CAACAACAAACTATCCGGAAAGTTCCCCTCGTGGATATCTGATGGTTTTCAAAATCTTAGAATCCTCAGCTTAAGGTCAAATTCATTTTCCGGAGAACTTCCTTTAGGAATGTCAAAACTGAGTTCATTGCAGGTGCTTGATCTTGCAGAAAACAACATAACTGGCGCCATTCCAACAACCATAGGAGACCTGAACGCCATGGTACAAGAGGAAAAAGTGAATGAATATCTGTTATATGGGAAGTATAGAGGGATTTACTATGAAGAAAGCTTGGTAGTGAATTTGAAAAACCAGTTCCAAAAGTACACCAAGACTCTATCACTTCTTACCTCAATAGACTTGTCGAGAAACAACCTCAACGGAGCATTTCCTGTGGAATTGACAAACTTGCATGGCCTCATAGTTTTGAACTTGTCCGGAAACCAAATCAGTGGAAAAATACCAGAAAACATTTCAAGCTTACATCAGTTAGCATCGCTTGATCTCTCGAGTAATACACTGTCTGGTGTCATCCCATCAAGCATGGAATCAATGTCATTCTTGAGCTATATTAACTTCTCAAACAACAACTTATCCGGTATGGTTCCTTATAAGGGACAGATGACAACTTTTACAGTGTCCGCGTTTGAGGGGAATCCCCATCTTTGTGGAGCTCCACTTCTAGTACTATGTCAAAATGGAAACTCGGACAATAGAACGGTACTAGAGAATGACAGCAGTGATGAATTTCCTGATAAATGGTTTTATGTTAGTGTTGCATTGGGATTTATTACAGGTATTCTTGTTCCTTACTATATCTTGCTCGTTCGAAAGCCATGGAGACGAGCATATTTTAGTTTCTTAGACACAGTCATCGACAGATTTGTCCCAGTCAGAAGCAACAGAACAACCAGCACCAAAAGATCAGATACGGGCACCAGATATGGCAAAAGAAGTGTTCAATAGGTACTAGTCCTAGT

>M6-PGSC0003DMG400006880 M6 LRR-RLP

CACAGAAGAAACACTCTCTGTAAATAGAAAAATAGAGAGTAATAGTACTGCATTTTGTAATAGAGAGAGGAGAAATGGGTCTCACCGGAGTAACATTGTTGGTCACTCTCATGGTGGTGGCCGCCGTTACTTTGCCGGCAAATGCCTGCCTGCCGGCGGACCAAGCAGCACTGATGGATATCAAAGCAGCTCTTAAGGAGCCATATTTGGGCATTTTCAACACATGGACTGGCAGCAACTGTTGCCAGGGCTGGTATGGTGTTAGCTGTGACCCGACTACGCAACGAGTTGCAGACATTGTTCTTCGGGGAGAATCCGAAGATCCGATTTACGAGAAAGCCGGCCGGTCTGGTTACATGACCGGTTCACTCTCCCCTTCTCTCTGCAAACTCGACAAGCTCACTACACTCATCGTCGCCGACTGGAAAGATATCTCCGGCGAAATTCCGGCTTGTGTCACTTCATTACCAGATCTCCGTATCCTTGAGCTCATCGGCAACAAAATCACCGGTCAAATCCCGGAAAACATCGGTCAGCTGAGTAAACTCACTGTTCTAAACCTCGCCGATAATAAAATCTGCGGTTCAATCCCCGCCTCCATTGTCAACCTCGGAAAACTAAAGCATCTCGAACTCAGCAACAATCAACTCACCGGCGAAATCCCGTCCGATATCGGAAAACTCGGCATGATGAGCCGAGCACTACTCAACAAGAACAAACTCACAGGTTCAATCCCTAACTCAATCACTCAACTTCGCCGGTTAGCCGATCTAGATCTGTCAATGAATCAAATCACCGGTTCACTCCCGGCTCAGCTCGGTTCAATGCCGGTTCTCTCAACTCTAAACCTCGATTCCAACCAAATCTCCAGTTCAATCCCCACTAATCTACTTAGCAGCAGCGGATTAAACATCTTAAACTTAAGCAGAAACTCATTAGAAGGAGAATTACCAGATGTATTTTGCTCAAAAACGTATTTCACACATCTGGATTTATCATACAATAATCTTCGAGGTACAGTTCCAAAATCATTATCATCAGCTAAGTACATTGGCCATTTGGATTTAAGCCATAATCACCTTTGTGGCCCGATTCCAAACGGGTCACCCTTCGATCATCTAGAAGCTTCTTCTTTCTCTAACAACGATTGTTTGTGTGGATCGCCATTACGTACTTGTTAATTCTTTTTTGTTTACTTACTAAGTTTGTAATTAATCGAAATTATCGGTTGATTACGCATAAATGGATGTGTAATTGTGTATTATGTTATGATTAAAAAAATATTCTTATGTGTTTAGCTTAATTATTATTATTATTGCTAAGCAATATGATGGATTATGCGGTTCTATCATGCATGCACGTGAAAATGCGTGACAGTTTACAAGAAAAATTATTGAAGAAATTGTGTGCTATTATTATTAGTAAATTGTAATATAATAAATCCCAC

>M6-PGSC0003DMG400007033 M6 LRR-RLP

ATGGCCAGATTAGTTAACTCCATTTTCTTTACCTTCTTGTACATAAGTTTCATATACACACTTTCACCTTATGTCAATGCAACAACTTCACCATCTGATATTCAAGTTCTAGTAGCCATCAAGACGTCGATAGACCCTTTTTCAATAACCCCGGATTCCTTCCTTAGTAGTTGGGATTTCAACTTAGACCCTTGTGAGACAACTGGTGCAAGTTTCTTAGGCATTCTTTGCACAGTTCCTGATCAGGTAGACACTAATGCCACGAGCCGGATAATTGAGATTGATCTAGAAGGTGATGGACTAGAGGGGTTCTTGACATCGTCTATAGGGAACTTAACAGAACTTGTTTCTCTCAATCTAGGGAGGAACAAGTTCAGAGGACCAGTGCCAGAGAGTATTACTAATTTGAGGAAACTCACAAGTCTTCAACTTTATGAGAATTTTTTCTCAGGTAGTCTTGTTGATGACATTGGTGTCCTTACAAAACTTGAAACTCTTGATGTCTCGAATAACAGGCTATCTGGCTCAATCCCTTCCTCGATTACATCACTTAGAAGTTTAACGCGATTAGACCTGTCTAACAATGAATTCACTGGAAAAATTCCTCAACTCAACGGATTATGGCAACTCTCTTCATTTGACATTTCAAATAACCAAATTTATGGGAATTTACCACAATTCCCATTGAAGATTAAAACACTATCACTAAGCCACAACTTGTTATCAGGACACATTACCCCAGTGCATAAACTTAGACATTTAAGCACATTAGACCTTAGTGATAACAGATTTTCAGGAGGAATAAACAAAGGGATCTTCATGTTACCTGAATTAAGTCACGTAAACGTTTCAGTAAACCGTTTCACAGTGTTTGAGGTGGTAGAATTCTCTGATAAAAAATCACAGCTACACACGTTAGACGTACATGCTAATCGTCTACATGGTCATTTGCCTGTGAACTTGGTTACTTACCCAAACTTAACAGAGATAAACCTTGGACATAACCTGTTTTCGGGTGAGATTCCATCAGAGTATTGGTCAAGACTCGGATTTTCATGGAGAAATCTCAACTTGGAATATAATAACCTGGAGGGATCGGTGCCTCGAGAGCTCAACAGGACTTTAGAAGGGGTTCAGGGAAGTTTTGAACATAACTGTCTCATTTGTCCAAAGGGATTGCAGCTATGTCATGGACAGAGGACAGCTTCAGAATGTCATGGAGGAGACCATAATTAG

>M6-PGSC0003DMG400007281 M6 LRR-RLP

GCTCTACGCCGCCAAATTTCTACCCTCTCTACGGTTAATTTGAACAAAACGATCAACAGCTTCATTGGAAACTGAAACCTCCAAGAAGCTCGTAAGCTCTTCGATCAAAGTTCCCACTTGACAAATGTCGTATCATGGAATTCCCTTATCGCTGGTTACTTCAAACACAACCTCATCCAACAAGCAGAGTACCTGTTTGACAAAATGCCTCACAGAGATGTCGTTTCTTGGAACACCATGTTCTCCGGATATCGCAACGTTAACAACCCAGAGAAAGTATATAGATGTTTTCTAGATATGAACAGATGTGGAGATATGAGACCCAATGAGCTCACTTTTGCTGTGTCAATTAGCTCCTTCTTGCATTTAGATTACAAGCATTTGATCCCTCAGCTTCACGGTCTCGTGCTCTGTTTGGGGATAAGTCTCAATGTCTTTGTAGGGTCAGCATTGATGAGGGGTTATGTTGATTTGGATGATTACAGAGGTTTAGCTCGAGTTTTTTATGAGATTTTGGATAAAGATGTTACGCCGTGGAATGTGTTGATTTTGGGTTATATGAAATTTGGGTGCACAAGTGAGGCGCAGAGGGCTTTTGATATGATGCCTATGAGAAATTCTTTTACTTGGAGTACTTTGATCAATGGGTATATTGAGAACAAGAAGCTTAATGAAGCTCGATTCGTTTTTGATAAGATGAGTGAGAAAGATGTGGTTTCTTGGACAGCAATGATAAGAGGGTATGTGCAATATGGAGAGTTTATGAAGGCTTTGAAATTGTTTAAGTTGATGTTGAACTCGGGTTCTCGTCCTAATCATTTTACGTTTTCAACTGTTTTAGATGCCTGTGCAGGCTACTCTGCTGTTCTGGTAGGCAATCAAGTTCATGCGTGCATCTTGAAGTCTGGTTTCCCTCTTGATGTCGTCTTGTTGACCTCCCTTCTTGACATGTATGCTAAATGTGGAGACATTGAAGTTGCTTTCTGCATTTTTGAGTCTATCCCGGCAAGAAATTTGGTAGCTTGGAATTCAATCATTGGAGGTTATGCAAGACATGGGCTTGCAGAGAGTGCAATGCAGGAGTTTGAGAGGATGCTAAAGAGTGGTATCAGGCCTGATGAAATTACTTTTATTAATTTGGTATATGCATGTGGCCATGGTGGGCTAGTCGAGGAAGGTGAGAGAATTTTCAATTCTATGGTAACAGAGTATGGATTGAAAGCAGAGATGGAACACTATGCATGTATGGTGGACTTGTATGGAAGGGCAGGTCAGCTTGAGAAAGCAGAAAAACTAATCGATGGGATGCCTTTTAAGCCTGATGTGGTCGTCTGGGGGGCATTGCTAGGAGCATGTGGCTTGCACTCATGTCTAGAGCTTGGAGAGATTGCAGCAAATGGAATCTACACATTGGAACACGACCATCCTGCAGTATTCTCGGTGCTTTCCAAAATCTATGGAGACAAAGGAGTCTGCAGTGACATAACTGGTTTGGATAAGTTGATGAAGAAATGG

>M6-PGSC0003DMG400007606 M6 LRR-RLP

ATGGACTTAACGCATGACAATTGGAAGAAGGTTGAGGAAAATCTGAACTCATTCTTTGGTACGGTATCCGAACGATGCCAATCAATTCTTTCTTTGAGCTACAACTACTTGCCCCAATATTTGAGGGCATGTTTTCTCTACATTGGAGGTTTTCCAGAAGACAGAGAAATTAATGTTTCCAAGTTGATTAGGCTATGGATTGCAGAGCAATTTGTAAAGGCAAGAAACAATAAAAGATTAGAAGTGGTGGCAGAGGAGTATCTGCAAGAGTTAATTGATAGAAGTCTAATTTTGATTGTTACGCGAAGCGATAATGGAAGGATAGAAAGTTGCAAAATTCATGATCTTCTTCGCCAACTCTGCCTAAGTGAAGGTCATACTGAAAACTATGTCATGAATGGAAAAGTTCTCGAAGCCATAGATGGTCAACGTCGAGTGATCCTTCTATCTAAATATGAAGAGAAGTATGATTATTTTCTGCGACATAGCAGTGGTATTGTCCGTACCTTTACTTCAATGGGTGTAGCTTTTCCAAAAGATATGTGTTCCATTATTTCAGAGTTCAAGTTGATTAAGGTGTTGGACGTGTTAGAAGTCCCGTACGATTTCTCTAGTGTAATACCTCAACTTGTACATTTGAGATATGTTTCTGCAAGAATTGATGAAGGTCTTTCACTAGCCAAATTGAGAAATCTACAGACCATAATTCTTCAAAGTCTTCAAAGGGAAGAATTGGAGCAGCCAGTAGATATCTGGAGTATGACAGAGATAAGACATGTGGATATTGATCGGCCACTATATATATCAAATCCTGTTGAAGCAGAACAAGCTTTGTGCCTTCTGAATAACTTGCAAACACTATATCTCCGTAACTCTCCTTTTGTTGTGGAAATCATAAGAAGAACTCCCAATCTAAAAGAGCTAAAGATTTTAGATAGTTCTGAGCGTTCTGATTGGGCTGTTATTCTTGATTCTCTCAGTATTTTACAGGATCTGGAGACACTACACATACAAAACATTGATCCGATGATTATCTCTGGAGATATTTCTCTTCCTAATCTTAAGGATTTGACATTAACAATTACTCGTATACCATGGGAAGTTGTGAATTTGCTGGCTAATTTACCCAATCTTGAGGCGCTCGTTGGGAATTGTGCATTCAATGGAACAGCTTGGAAAGTAGATGAAGATGTTGTATTTCACAAATTAAAATATCTACATCTATTTATGTGCAGAGATCTGGAAAGGTGGGAATTAGCAGCTGGTAGTGATAATTTTCCGATGCTTGAGAAACTAATGATGCTTGGGTTGGAAAAACTGGAGGAGATTCCCGAGAGTATTGGAGAAATAATGACACTAAAATCCATCCGAATAGTTAATTGCGGCTCTGGTGTAGAGACAAGTGCAAAGCAAATTCAGGAAGAGCAAGAAAGCTTGGGAAATTATGAGCTTCAACTTCAAAT

>M6-PGSC0003DMG400007684 M6 LRR-RLP

ATGAAACTCATCAACTTAGCCCTTTTTGGCTTCTTGCTCATCATTTTCTTCTCATCAAAACCCTCATTTCAAGCTTCAATCAACTATGAACCAAACCCTAGACTTTTCGATGCTTACATAGCTCTACAAGCTTGGAAACATGTCATGACTTCCGATCCAAGAAACTTCACAAAAGATTGGTATGATTACAATGTTTGTAACTACACCGGGGTTTATTGTGCTCCAGCTCCAGATAACCCTAATATTACCACTGTTGCTGGTATTGACCTCAACCATGCAAATATCTCCGGATACTTACCTGAAAAACTTGGTTTATTGACTGATTTAGCCGTCTTTCATATAAATTCCAATAGATTTCAAGGCTCAATCCCGAAATCTTTTTCTAAGCTTCGAATTTTATTCGAACTTGACGTGAGTAACAATCTTTTTTGTGGTGAGTTTCCTTCCGTTGTGTTATCTCTTCCTTCTTTGAAATTTCTTGATATTAGGTATAACCAATTCGAAGGAAAAGTCCCTTCAACCCTTTGGGATCGAACACTCGATGCTCTTTTCATAAACAACAACAAATTTCAATTTTCTTGGCCTAAAAACTTAGGTAAATCACCCGTTTCTGCTTTGGTTATGGCAAATATTAATGTAACGGGGTGTATACCATCAAGTATAGCAAAAATGTCGAAAACATTAAATGAGATTATTCTCTCGAATTCGAGTTTAAGTGGATGTTTACCACAAGAATTAGGGTTATTGAAAAATGTTAGGGTTCTTGATGTGAGTTTTAACAAGCTTGTTGGTGAGTTGCCGGAATCAATTGGTGGAATGAAGAAATTGGAACATCTTAACGTGGCACATAACAAGTTTTCCGGTGAGATTCCGATGAGTATTTGTTCATTACCAAAGTTGGAGAATTTCACTTACTCTTATAATTATTTTTGTGGTGAACCAAAGATTTGTCTCAAGTTGAAGGATAGAAATGATAAGAAAAATTGTATACCATATCGACCATCACAACGTTCAGAAAGTGAATGTAAGACATTTTATTCACGAGGTCCGGTGGATTGTAGTTCATTTGGTTGTAGATCACCGCCACCGCCACCACCGCCACCTCCACCTCCA

>M6-PGSC0003DMG400007876 M6 LRR-RLP

ATGGAAGGTGTAGCTTTAGTACTTTCACTTCTCCTTGTTGTTGTTAATTTAGATGGATTTTCTTGTGTTTATTCAGGAAGTCATTGCATTGAGAGTGAAAAAAATGCTCTTATTAAGTTCAGCCAAGGTTTCAAGAATCCGTCACGAAGCATGTTATCTTGGATGTTTGACGAAAACTGCTGCAGCTGGAAAGGTGTTGAATGTGACAACAAAACCGGGCACGTGATCACTCTTGATCTACATAAGCAATTCTTGCAAGGTGAGTTCAGGATTTCTTTACTTGGTTTGCCTTATTTAAGGCACTTAGACATGAGCCACATTGATTTCCTCGGGGAACGGATTCCTGAGTTTATTTCCAGTTTTAAGGACCTGGAGTATCTCAATTTGTCTAAAACGAATTTTAGGGGAATGATTCCTGAACATCTTGGGAATCTGTCGCGGTTGCAGTTTCTTGATCTGAGTGGTGATTTTTCACTCCGGGTTAATAACCTTGAATGGATTCACTCCCTCTTCTCTATGAAGATTCTTGATTTGAGTGGTGTTTACATGAGAAGCGCGGAAAACTGGCTACATGACATCAATTTGTTAAGTTCTCTCTCTGTATTACGTTTGTCTGCTTGTCAGCTTACTTCATTTCCTGAATTGCTTCCGACTTCTGTGAATTTCACATCCCTTCGGATTCTTGATCTTTCAATAAACTATTTTAATACTAGGATTCCTAGCTGGTTGTTCAACACAAGCCATAATCTTGTTTACCTTAATCTTTCGAGAAGTCAGCTGAGCGGTTCACTTCCAAATGCCTTTGGCAACATGAGTTCACTTAGAGTGCTTGATCTTTCTGGGAACTCTCTCCGAGGCAACCTTCCACATTCCTTTGAGAAAATGAGTTCAGTTTCCTTTCTTAATCTCTCAAGAAACTCGTTCACGGGCAATTTACCACCAACCCTGCCATCAAAGCTGAAATACTTGGATATTTCTGATAATAAAATGGAGGGTCCTTTAGCAAGAAGCATCACACAACTTAAGCAATTAGTTGTCCTCAATGTTGCTCGGAACTCTTTCAAGGACTCGATTACAGAACATTTCTTGAATTTAACTCATTTGAGAGTGTTAGACTTGTCATCCAACTCATTTATTTTCAATGTAAGCGCAACTTGGATGCCTCGTTTTCAACTTGACTCTCTAAGCCTGCAGTCTTGCGGACTTGGTGCACAGTTTCCCCAGTGGATTCAGACACAAAAAGAGTTGTCATTTATTGATATCTCTCGCACCAATATCTCAGGTGAAGTGCCTGATTGGTTCTGGAATTTTTCTGCCAAGGTTTATCACATAGACCTTTCAGGAAACAATTTCAGTGGAGAAATCCCTGAATTTACAGAAAGAGTTTGTCTTACAAAATTAGACCTGAGTGATAACAATTTTCACGGTCCTTTGCCACATTTTTCACCCAAGATGATGACATTGATTTTGGCAAGGAATTCGTTTAATGGTACCATTGCTCCTGTATGTGAATCACTTGTCATGAATAACTCTCTGAGTCTTCTAAACCTATCTTCAAATTTTCTATCAGGACAACTTTCAGATTGTTGGAGATATGGGAAGAATCTGCAAGGATTGAATTTAGGCCATAATGATCTATCAGGGGAGATACCTCATTCGATTGGAGATCTTACTAACCTTTTCTTTTTGCAATTGCAGAACAACAGTTTCTCCAAAAATCTTCCTTCATCATTGAAGAACATAACTGGACTAAAAATTCTTGATGTTTCTGAAAATAGTTTATCTGGGAATATACCATTTTGGTTAGGGGAGAGTTTAAATGCTCTTGAGATTCTTATATTAAGCAGAAACAACTATGATGGAATTATTCCCAGGGAGATTTGCCAGCTTA

>M6-PGSC0003DMG400007915 M6 LRR-RLP

ATGAGGTGGGACTCGTTGCAGTTCCTAAACCTTTCTCATAATTCATTAACAGGCCACCTACCACAATTTCATTACTATAACCTACAGTATCTTGATCTGAAATGTAACTCCCTTCAGGGTCCACTACCTTCATCCATTTGTAACATGAGTTCGCTTATCTTATTAGATTTATCACGCAACAACTTCAGTGACTCAGTTCCACATTGCTTGGGAAGCATGGCTAGTCTAACGGTGCTAGACTTAAGAAGGAACAATTTCACAGGGAGTCTTTCCCCATTATGTGCACACAGCACTTCGTTGAGTACCATTGTCATAAATGGTAATCAATTTGAAGGACCTGTACCTGTGTCGTTGCTCAAGTGTGGTGGTCTAGAAGTACTTGATGTGGGGAACAATGCTATAAATGACAGGTTTCCATCTTGGCTCGGAACTCTCGAATACCTGCAGGTCCTTATATTAAAGTCGAACAAGTTCCATGGACCTATAAGTACGTGTCTGACTAAGTTTTGCTTTCCCAAGTTGCGAATTTTTGATCTTTCTCGTAATGATTTCAGTGGCTCACTTCCTGCAAAAGTTTTTGGAAACTTCAAGGCAATGATCAAATTAGATGGCGAAGACACAGGAAATATAAAGTACATGGACTCTGTGTTGTATTCGCCATTTGTCACATCGTATGAGGATTCAGTGAGTTTGGTAATCAAAGGGCAGGATATTGAGCTATAAAGAATCAGCACAATTACGACAACCATAGATCTCTCAAGCAACCATTTTGAAGGTGTCATTCCGAAAACACTAAAGGATCTCATATCACTTTGGCTACTCAATTTATCTCATAACAATCTCAGAGGAGATATTCCAATGAAATTAGGACAATTGAGTATGCTTGAAGCTTTAGATCTCTCTTGGAATCGGCTCACTGGAAAGATTCCACAGGAATTGACAAGAATAAACTTTCTCACCTTCTTAAACCTCTCTCAAAATCATCTCGTTGGACCAATTCCTCAAGGTCGGCAATTCAACACATTTGAAAATGACTCATATGGTGGCAACCTTGATTTATGTGGTGTTCCTTTAACAAAGAAATGTGGAACGTGTGATTCATCGCATGTTCCTCAACAATTGGAGTCCGAAGAAGAAGGCGAGTCATATTTTTTTAGTGGATTTACTTGGGAATCAGTAGTCATAGGCTACAGCTGTGGACTAGTTGTTGGAACTGTCATGTGGAGTCTCATGTTTAAATATCGTAAGCCAAAATGGTTTGTGGAATTTTTTGATGGACTCATGCCTCAGAAAAGAAGAAGGCCAAAGAAGAGAGCTCAGAGACGAAGGACTT

>M6-PGSC0003DMG400007999 M6 LRR-RLP

CTCCAGTAGCATAATAACATCACTCAAAACTTCAACTTCTTCGAGCAGAATGTAAAATCAGTTTGAAGTCCCCAAACATGGCGGTGACGGACTTTTTCGTCGGCGAAATCACCACCGAACTCATAAAAAACCTTCTGCTAATAGTTAAAAAATCCACTTTATGCCGTTCAAGCGCCGAGAATCTCATCGACAGCATCAATGGTCTCCTTCCAATCATCCAAGAAATCAAACAAACCGGTGTTGAACTTCCACAGATACGTCAAACTCAGATCGACGATTTCTCCAAACTTCTCCGAGATGGTTACGAACTCGCCGGAAAAGTTATCCACTCCGGCCGTTGGAACATGTACAGGTGTCACCTTCTCATTCGACTCTAATTACAGTGAATTGAACCTCTTCCTGCTCTAATTAGGGTTATTGAATTGTACGAATCTGTTAGAAATTACTTAATTTATATTATGCCTTGATAACAGGAACCTACAGTTGGCTAGGAAAATGGAGAGGCTGGAGAAAAGAGTAGCGAGGTTCATGCAAGTTACAATGCAAGCTCATGTACTAGCGGATGTTCATCATGTTAGGTTTAGTATGGAGCAGAGATTTGACGTGCTTGAGCATAGGCTTAAAGCTATAAAAATCGGAGTTGACGATAGAAGTGGTGGTGATGGTGGAGGAGGAGGGTGTTTAGGGGAAGCTGTGAAAAGAATGGAAGAAGATGAGAAATGGTTTGAGGATAGTTTTGTAAATTTAGGTGCTGGGATTGAATTGGGGAAGAGGAAAGTGAAGGAGATGCTGATGGGTGAACAGGATAGAGGTGTGTTTGAGATTTGTGGAATTGGGGGTAGTGGCAAAACTACCTTGGCTAAAGAGATTTGTAAAGATGATCAAGTTAAAAGTAAGTAATTAATCCCAGAAAAGTTTCTCATGTAGAGTACTTTAGCTTTTTGCGGGGTTGATTTCTCGAAAGGTTAATCAACTAATAGTTATTATCTATCACTTTTCGTTTTTGATTCTGATGCAAAGAAAGTGACTTTATTAGATATTAACTATTAGTTGAGTGGACATCCGTGAAATCAACTCATATTTTGCGTTGCTGATGATCGAGTTATTAGAAGTTAATAATCACACAAAAGTTTCTCATTTCATTTTGTTTTTACTCTTCTGTTGCGGATGAAACAGTGGATAAAAGTTGAAAACTTTATTGGCTATTATCTCTATTTTATCAGTTGAATTCTAATAGAGATCAATTTTGCATGTAAAGGACTCATCAAAAGTTTTGATTGGAATACAGTTTAAATTGGTCTTAAAAACCTTTTCTTTAAATAATGGTATGGTGTCTGGGCCATGTTGTTCCAGGTTATTTCAAGGACAAGATTTTCTTTTTCACTGTTTCTCAATCTCCAAACGTGGAGCAATTAAGGAAAATGATTTGGGAAAAGATATCAGGGTGCAATCTCCATGGTTATGGATACGGGGAGATGTTACCCCAGTGGAACCTACAGTACCAATGGAATACGAAATCTGCATCCCCAGTACTCTTGATTCTTGATGATGTGTGGTCTGCATCTGTGCTAGAGCCACTAATTGTCAAGATCCCCGGCTGCAAGATCCTAGTTGTATCGCGTATCAAGTTTCCTCCATCAATCATTGACTGTATTTATGATTTAGAGTTGTTAAGGGAAGATGAAGCTATGTCCTTATTATGCCATTTTGCATTTGGACACAATTCCTTTCCGCGTGGTTTCAGCCAAAAGCTTGTCAAAGAGGTATTAGTGTATGTTTTTGGCAGCATATTAGTTTTTCTTTGCAAAAACTTCTTGTTATACTTGAGTTAAGCTAACAAAAGCTTCTTGGACTTTTTAGATTGTGGATGAATGTGAAGGGCTTCCTTTGGCTCTTAAGGTCATTGGATCTTCATTAAAGGGAAAACCTGAGATGTTCTGGATAAGTGCAAAAAACAGATTATCACGATGCCAACCTGTCTGCGAGTCTCATGAACTGCAGTTGCTTGAGCGAATGAAATTGAGTATTGACTGTTTGCCTGAGAAGGTGAGAGAGTGTTTCCTGGACTTGGGTGCTTTCCCAGAAGACAAAAGGATTCCTCTTGATGTTCTAATTAACATGTGGGTGGAGCTACATGATATTGATGAGGAGGAGGCTTTTCACATTCTTGTTGAACTTTCAGACAAAAATCTCCTAAATCTAGTCAAAGATGCACGGTAAGCGGTGGACTAAAATCTCCTCAAACAATCTTTTAATTTTTGTTAGAAGGTGATTGTTTATGGAAAAAATCAATATGTTGATAGTCTTCAATGTGGTCTTCATGTGGCAGAGCTGGAGACATGTATACAAGTTACTATGAGATATCGGTATTTCAGCATGATGTATTACGAGACCTAGCAATTCAGATGAGCAACCGTGATGATATAAATCAGAGAAAGCGATTGGTTATGCCACGAAGAGACGTAAGCTTTCCAAGAGAATGGGAAAGAAATGTGGACGAACCTTTCCATGCACGAGTTATCTCTGTGCATACAGGTATTTTTAGATCCTTGGTTACTGCTGATTGATTAATAATGAAAAGTTTATCATGTGATATAAACAGAAATATTTCTCTTTAATGCAGATGAAATGAGAGAAATAGACTGGTTCAGAATGGATTGCCCGAAAGCTGAAGTACTGATTCTCAATTTTGCCTCATCTGAGTACTTCTTGCCTCCTTTCCTGGAGAATATGCCAAAGCTAAGGGCATTGATAATCATAAACTATAGTGCTGGCAATGCAGTTCTTCATAACATGTCTGTATTCAGTCATTTAACCAACTTGAGAAGCCTTTGGTTCGAGAAGATATCCATCACTCACTTATCTGACTCCACAAATCCTCTCAATAACTTGCGGAAGATATCTCTAGTGCTTTGTGACATGAAAAACAGCTTAGATGAGTCAGATGTGGACCTCCCTGGTTTGTTCCCACAGCTTTCTGAGTTCACAATGGATCATTGCATCAACTTCAATAAGCTGCCATCAAGTATTTGCCGGTTGCATAAGCTCAACAGCCTTAGTATCACTAATTGTGATAGTCTTTATGAACTTCCATCTGATTTAGGTGAATTACAAACTCTACAAGTTTTAAGGATATATGCCTGTCCACATCTGAAAAGGCTTCCCCCTGGAATAGGTCATCTGGTGAAGCTGAAGTATCTTGACATTTCACAATGTGTTGGTTTGAGATGTCTCCCTGAAGCAATTGGTTGCTGTAGAAACTTGGAGAAGATTGATATGAGGGAGTGCCCTCAAATTGACAGTTTGCCAAGCGCTCTATCCTTTCTTGAATCATTACGTTGTGTTATTTGTGACGATGAAGTTTTTTGTCAATGGAAAGATGTTGAGAAGGCTGTACCAGGTCTATGTGTACAGGTTGCCGAGGAGTGCTATACTCTTGACTGGCTATCTCAGTAATCAAGTTTTCAATTCCCTTATCTTGGGTTTCTTTCGTGTAAATGCTCAGTCAATTTTCCAGTTCTGTATTTTCCCCTTTACATGGGGTTCATTGTATCTCTTGTAAATAATATATAGCAATATGTAAATATGTAGATGGACTACATCTCAATAGCTGATCTTTCCTTTTACTTTAAAACCATAATGCTGGAAGCAAAATTACCACAGGCAATCCAACACTAAGGCAAGTTATCTGCTATAACATGT

>M6-PGSC0003DMG400008036 M6 LRR-RLP

ATGAATAGTACTATCAAATTTCTAATATTTTTCTTCAGTGTTACAACAGCACAACCACTAAAGGGACAGAACCAATTGATTTTTGCAGATCAAAGGCTTGCTATAGTATACCCCATTATACAAAAATTCAAGAATATCATCTCTGCTGATCCTCTAGGTATTACAAAAACTTGGGTTGGTCCAAACATATGTAATTATACTGGTTTTTATTGTGAAAGTCCTCCAGACAACAATTCAGCAATTGCTCTTGCTTCTATTGATTTCAATGGTTTTCAACTTAGCGCGTTTACTCTTGATGGCTTCCTTGATCAGCTTCCTGATATAGCTATTTTCCATGCTAATTCAAACAATTTTGGTGGGACTCTGTCTCCAAAAATCGCGAATCTTCCTTATCTCTATGAACTTGATATTAGTAACAACCAATTTTTCGGCCCATTTCCTAGTCCAATTTTGGGCATGAATAGCTTAAATATATTGGATATAAGATTCAATTCTTTCACTGGATCAGTCCCACCTCAATTGTTCACTAAAGATCAATTAGACGCGCTCTTCATCAACAACAACAATTTCATGCAGATGCTTCCTGATAACATCGCGACCAGCCATGTTACATATCTTACTTTAGCTAACAACAGATTTTTCGGTCCTATCCCACATAGTATTTCCAAAATCTTGTCGAGTTTGTCTGAGATTTTGCTGCTTAACAACTTGTTAAGTGGATGTCTTCCTTACGAGATAGGATTCTTAAACGAAGCAATAGTCTTCGATGCTGGCAATAATAGACTCACCGGTCCACTTCCATTCTCATTAGGGTGTTTGGAAAATGTAGAAATGTTGAATTTTGCTTGAAATCAGATGTATGGGATAGTGCCTGATGTGATTTGTGCCCTGGGGAATTTGGCAAATTTATCTTTTTCCGATAACTATTTTACAGGATTTGGGCCAATTTGTTTGAGATTGATAGAAAATGGAGTGTTTGATTTGAGGAATAATTGTATTCCTGGTTTTCCATTTCAAAGATCAATAGCTGAATGTGTTGCATTTTTTGCTTACCCAAGGTACTGTCCTCATATGGCTACATATACTTATATTCCTTGTTGGCTTTCAAATTTCAAAACGCCTCCTCTGGATCTCCCTGAATTGGCTCCTTAGTAAGGGTAACACGGTGTACAAAGCATTCTGTGTTAGCATGGTTCGGGAAAAGATCATTAGAAAATTCTGGTATTTGATCATAAACATACATGTCCCCTTTTCGCGATCTTTTTTATTTTAATTAAATTAAGCTTGTAATTTCAATTGAATAGAGTGTGATATATCATAATAACTTGTGATCTCTGTTTATGGAGTATTGCTCAGTAGTAAATTTACGTTACTTTTATGTTATGG

>M6-PGSC0003DMG400008185 M6 LRR-RLP

TATATTTGGCTTTTTAAAAATATATAAATAAATGTGTGTAAGCACTTTTAGCTTGTCCACTTATGACCTAGTGTTTTGTTCCAATGGCTCCAACTCTATCTGCAACAATGTTTGCAGGTTTTTTGTCCCAACCTGGAAGACCTATCCATCAACAGCCTTAAAAGCATAAGTGCTCTATGCTCTCACCAACTTCCAACTGACTACTTCCCCAAACTTCAAACATTGGAAGTATCATTTTGTGGAAAATTGAGAAACTTGATGTCTCCATCAGTGGCCACAGGTCTTTTGAATCTCCGAAAATTAAGAATACAAAATTGTGATTCAATGGAAGAAGTGATCACAAAAGAGGAACAAGAAGGAGCAGGAATCATGACCTTATTTCCACTCTTAGAACAGCTGAAACTTAGTTTTCTGCCTAAGTTGAGGCATTTCTTTCTGACAGAGCATGCTCTTGAATTTCCATTTCTCACTGGCGTGTATATTGATGTCTGCCCTGAAATGAAGACGTTTGTCCAACAGGGAATATCTGTGAGTACACCCAGTCTCAAATTTATGATCTTCAATTATAGGAGGCGGAAAGTAGATGATCTGAATAAGTCGATACAGCAGATGTTTAACAAGGTATGTCTTATTCCGCTAGCTACATATTCTAATTAGGTTCTACTGCTCTCTTTCCGTTTTCGATACTTATTTAAGTTCATCAACTTTCTTTTACTTAATAATCTAGTCATCTCTCTCTTCTTTGCATTCTCTGTTTCCTGCAGAAACAAGAAGCTAGTGATGACGACGAAGCTAGTAGTGGCTATGTAATAATGATGCCGACGAATCTGAACCTAGTCATTCCAGAAGTTAGGAAGCGGGTAGGAAAAACTCAATGAAATAATGCAAGTAAAACCTCGAACATCACCTGATACCTATGCTCGTGATGGAGGTAATTAACAAGCAGAGGCGGATCTACATTATTGTAACCTATTGGTTCATGTCAACCTACTAGCTATTGGCTATAGATCTTGTCTATGTATTAGTAATATCTAAAAAATAAATGAT

>M6-PGSC0003DMG400008394 M6 LRR-RLP

CTTCCTTTGAGTACTCTACAAACAACTACCAAACGAAGACTTACTTTGAGAAGACTTTGTTGCATACAGCTCAGGAAGAAGACTTTGGTGCAGATTGCAGAGGAAGAGGAAGCAAACACAACAATGGCCGATCCTGTAATTGGTGCTACTGTTCAAGTTTTGCTTGAAAAGTTGATTTCTCTCACTATCGAGGAGGTCAACAGCTCAAGGGATTTCAACAAAGATCTCGAAATGTTGACACAAAATGTATCATTAATCCAAGCTTTCATTCATGATGTTGAAACACCACAAGAGAAACAACAGTCTGTCGAACAATGGCTCAACAGGCTTGAGAGAGTTGCTGAAGATGCTGAAAATGTGTTTGATCGATTCAGATATGAATCTCTCAAAACAAAAGTGGTGAGGAGCCCATTGAAAAAGGTCGGTGGTTTCTTTTCTCATACTGCTTTTAAGAGAAAAATGTCTCAAAAAATCAACAACATCAATAAAGAGTTGACGGCTATCAATAAGGTAGCCAAAGACCTCGGTCTACAATCACTCATGGTACCTTCTCGGAAAATACTACCAATTCGAGAAACAGATTCCTTAGTAGTTGCTTCTGATATTGTCGGTAGAGATTTGGATGTTGCTGAGATAAAGGAGAAGATTTTGAAGATGAGAGAGGAGGATATTGTTCTGTCCACCATTCCCATTGTAGGTATGGGAGGTTTAGGGAAAACAACTGTGGCTAAGAGGATTTACAATGATGAACACATGAAGCAAATCTTTAAAAAGAGAATTTGGTTGTGTCTACCTGAAATGTCTGAAACGAAGAGCTTTCTTGAACAAATCCTCGAATCATTGATAGAGAGGAAAATTGAGGTTGAAAGGAGAGATATAATAGTCAAGAAGCTACAAGATGAATTGGGAGGAAAAAAATATTTGCTAGTCCTGGATGATTTGTGGCGTGTTGACTCTACATCGTGGCATGAGTTCGTCGACACCCTGAGAAGAATAAATACATCCAGAGGAAACTGCATTCTTGTGACTACTCGTAGGAAGCAGGTGGCATCCACAGTAGCAACAGATCTTCATATCTTGGGGAAGTTAACAGAAGATCATTGTTGGTCTATTTTCAAACAAAAAGCATTTGTTGATGGCAGGGTTCCAGAGGAATTAGCGAGCATGGGCAACAAGATTGTTAAAATGTGCCAAGGTCTACCGTTGGCTGCAAGTGCATTGGGAGGGCTCTTACACAACAAAGAAAAACATGAATGGCAAGCAATTCTTGATGGCAACCTCCTTGTTACTGGTGAAGATGATAATGGAGAAAATAGCATAAAGAAAATCCTAAAACTCAGCTATGATTATCTACCATCTCCACATCTGAAAAAATGTTTTGCTTACTTTGCAATGTTTCCAAAAGATTATATGTTTGAAAAGGACCAACTAATCCAACTCTGGATGGCAGAAGGGTTTCTTCGTCCAAGTCAAGAGATCCCTGTGATGGAAGACGTTGGGCACAGGTTTTTTCAAATCTTGTTGCAGAATTCCTTGCTACAAGATGTTGTGTTAGATGAACACAACAATATAACACACTGTAAGATGCACGATCTTGTGCATGATTTGGCTGGAGATATCTTAAAATCTAGACTATTTGATCCGAAGGGCGACAATGGAGAAAAACTTTCTCAAGTTCGATACTTTGGATGTGAGTCACCAACGGATCAAATAGATAAGATATATGAGCCAGAACGTTTGTGCACACTGTTTTGGAGAAGCAATTATACATCTAAAGATATGCTGTTGAAATATAAGTTCTTGAGAGTTTTAGATTTGTCCAGTTCAGGAATCAAGGAGTTGTCAACCAAAATCGGGAAGCTGATATACTTGAGATATCTTGATCTCTCGAACACTGAGATCACAGCCTTGCCCAACTCCATTTGCGAGCTCTATAATTTGCAAACATTTAGAGTCATCAACTGCTTTTCACTCCAGGAACTTCCATATGAGATGAGAAATATGATAAGTTTGAGACACATATATTACACTTCTGTTGACGAAACAAGTGGGCATTGGGGAGGATGGTGTCTTCACAATGAACATTTTCAGATTCCACTTAATATGGGGCAATTGACTAGTCTTCAAACCCTCAAGTTTTTCAAGGTAGGTTTAGAGAAAGGTCGTCAAATAGAAGAATTAGGTCATTTGAAAAACCTAAGAGGTGAATTGACGATCAATGGTCTCCAATTGGTCTGTGATAAAGAAGAGGCTCGAACAGCATATTTACACGATAAACCAAACATCTGCAAGCTGGCATATTTATGGTCCCACGATGAATCAGAAGGCTGTGAGATCAATGATGAGCATGTGTTGGATGGTCTTCAACCGCATCCTAACTTGAAAACCATAGCAGTAGTGGACTATTTAGGGACTAAATTTCCTTCATGGTTCCGTGAAGAGTCGCTACCAAATTTGGTCAAGTTGAAATTAAGTGGAAGCAAAAGGTGCAAAGAAATTCCATCCCTTGGCCAACTGAAATTCCTTCGGCATCTTGAGCTGATAGGATTCCATGAGTTGGAATGCATTGGACCTGCTTTTTATGGTGTTGAGATTAGAAATATTGGATCAAACAGCATTATTCAAGTGTTCCCGTCATTGAAAGAACTAGTATTGGAGGATATGCGTAGCCTTATTGAGTGGAAGGGAGATGAAGTTGGAGTAAGAATGTCTCCCGGGCTTGAGAAGTTGCGGATTACAGACTGTCCATTGTTAAAAAGTATTCCGAATCAATTTGAAATCCTCCGTGAATTAGAAATTAGAGGAGTTGACAGTGAAATGCCATTGTTGAACTTGTGCAGCAACTTGACATCTCTCGTGAATCTTAGTGTCTATGATATGAAAGAGCTCACTTGTCTTCCAGATGAGATACTACGTAACAACATTTCTCTTCAACAGATAACGATTTTCGAATGTGGAGAGTTTCGTGAATTGCCACAAAGCTTGTACAATCTCCATTCTCTTTATAGATTAGACATTTACAACTGCACCAATTTCAGTTCTCTTCCTGTTCCCAAGGGAGACAACTATTTGACTTCCCTCATATTCTTTTTCTTACATAATTGTAATGGATTGATCAGTATACCAATTGGAATGCTAGATCAATGCCGGTCTCTAGAGTATTTGCATGTCAGCTGCTGTAACAACTTGGTTTCATTCCCTTTACATGTGTGGGAAATTCCTTCATTTTCAGTTTTGGAAATAACAGAATGTCCCAAATTGATTAGTGTACCCGAAGTGGGCCTTCACCATCTCACCGGGTTATTGAGATTGGGAATTGGTCCTTTCTCAGAGATGGTGGATTTTGATGCATTCCAATTGATTTTTAATGGCATTCAGAAGTTGTTGTCTCTTCGTGATCTGGAGGTGTACGGACGTGGGCACTGGGATTCTCTGCCCTATCAGCTTATGCAACTCTCTAACCTAAGAAAGATCACAATAGCTGATTTCGGAATTGAGGCTCTTCCTCCTAGACTTGACAACCTTACTTCTCTTGAAAGTTTGACGCTAGAGAGGTGCAAACGGCTACAACATCTGAACTTCTCAGATGCCATGCCCAAATTACGGCATCTGTGGATAAGTGATTGTCCATTGTTAGAAGCCCTGTCGGATGGGCTCGGCAGCATTGTTTCTTTGGAAGAATTATATTTACAGGACTGCGAAAAACTAGAGCATCTACCGTCCCGAGATGCCATGCGACGCCTCACTAAATTATGGAACATGAGAATTAAAGGATGCCCAAAGTTAGAAGAAAGTTTCACCAACTACTCCCAGTGGTCCAAAATTTCCCATATTTCAAATATTGAATTAGGTGGGTGGAGAAGGACAACCATAAGTCTCGGTTTCTCTTTCACTTTCTGAGTCGCTTGCATTTTTAATTAGAATATTATTTTCCACGAGTTCCAAAAAAGAAGTTTTGTAAGTAATGTAATAATTTTTCTTGAAATTATTTTGACATAAAAAAAAAAGCTAGATTATCAATCAATTGCATCTGTTGCTATGCTTTCAGGATTCTGCTTGTCCACTTGAACTGAATTGCTGTTATTCTAAAAATTATGTTTCTTTTCCTTATAGGTAGCCATTATAGACGTCTCTGTGGTAAATCAAGCCAAATCATATGAGCAGCAGGCAATTACGTAAGGTAAATTTTCATTTATAATTTATAGGCACTTTGTAATCTACTTGTTTCTATCTTTCTTTTTGTTGGAAACCAATTTTCGGAGGTCTATGCTGATGAATACAACATTACCTTCCTTAAAAAAAAGTTTAATCATGAAAATTTATGATTTCTGCTCATAGGTTACTAGGTACAAGGACATGGTAGAACGTCGTTGTGGTATATCTAGTCAAAATCCATGGGCAGAATCATGTTAGGCATTTTTATTTAGTTGTTAGAGGAATGCTATAACAAGTTTGTAAAGTTTCTTTTTTTTACTTTCCTTTTCTCTCATGAAATAATTTTAATGTTTAACGGGACGTCGTGTTTCACAGAGGAACCGCCACTGTTATTTTGGATGAAGAACAATCTTCAACTGTAGACTCATTATGGGACACTTTCATGAGTATTTTCGGGGAACTACAGTCTACATATGAAACATCAAAAACTCAAAAGCGTATAGGGTGCTCAATGAGAAGCTACACGGGGTCTATTCTCCATGTAAGTATTGATAACTCTCAAGGAACAATACCAAGTCTCATATATGGTAGAAATCTATATAAGTTTCTTAATAAACTCATAAAAGAATAATCTTTATGTCTTCTGCAGTTCTGGTTTTATTTTCATGCTTAAGACAGTACAGCTTTTATGAATATATGACACACTTCTCGTTCCTTTCACCTCACAGGACAATTTCTCTGTTCATTAAAACTTCCGCGAATGAAGGACTTCAACAAAGATAGTTGTGATGAATATTTTTGCAAAGGGTGACCCATTGGCTCTTAAAATGTCCTTAGCAGTATACGTTCATCTTAAACAGGTTTTCCTCTGTCCTTTTTTAGTGGTTACTTGTAATGATCCGCTTCTCCATTCTCCAAGGAGGAAACGTGTGCCAAAACTGACTTTTTACCATTTCCCGTGTTTCAGATTCATGTTTGAAACTCCAGCATGGACCTTGTCGAAATGCTACTTGTGTTAGACGTGTTTGAGTGTCATGATTGGCAAGGGAAATTGCATGTGGCGGAAATTGGAGTGGAAATACTATTTACTTGTATGACTAAGATTTGACAGCAGAGGCGCCTTTCTTTCACAGCATACGTGCCTTTTGGTACAAATAGAGTTCATTTCTCCAACAAACATTTTGAACTTTTATACTTAGCTGCTGCATCATTGCCTAGTTGTTAGCTTCTGTAATGTACATCTGCTTTTCCTCTTGAAATTTCTCTTGGCTTTTGTTCGCAAGTATAAACAAAAAAGACAGGGCCTACCATTTGCATTGTTTAAATCCAAACAATATGTATTCAAAAGTCACAAGCAATGTGTGTTATTCCAATTTCGTACAGGACTGCATTTTTGTGTCTATGTATAACATATCGCATTGCTCATTATTCTTTCAGATAGACCAGATTGCAC

>M6-PGSC0003DMG400008450 M6 LRR-RLP

TCCTTATGTTACACACCTCTTCTCTCTGAAGGTGTATGATGGGATTCATGATTGTCTTTCCTACAACAGTCACCTAAAACACTTGAGGCTTCTTAAAAGTTTGGATTTGAAGGGCCTAACATTGACAGATTCTTTGCTGAATGAAATCGGTTTGCTTGTTCATTTGAAGTACTTAATCATTCAGACGAAGGCAAAAGCTCTCCCTCCGTTATTTTCAAACCTCTGCAATCTAGAAACTCTATCGGTGTATAACATTGAGAGATCATGCATGATACTATCGCCTTGTTTTTGGAGTCTTGCAAAGCTACAAGATGTGTGGATGATGGAATGTGCTCTCTTTGAACCAACTGTGTTAGACGAGGACTCAAGGTTAGAAAATTTGAGAACATTACAGCATCTCTCCCTTCCCGGTTCAGAAGACATGGAGGATATTTTCAAAAGGTTTCCTAATCTTCAAATCCTTAGTGTCTCTATCATAGCAGAGAAGATTTGTTTTCCAAGATTGGATGTCCTTAACGAACTTGAACAACTCAATTTATCTGCTTCTTGGGATTCATTCCATGAATATACACACGGCTTCCCTTTGAGCTTGAAGAAAATGAGGTTGCAAGGGCTCGCTCTGACATCCGATTCATTGTCAAGAATTGCTAGACTACCCAACCTTCAAAAACTAAGTCTAAAAAACGCAATCATCGATGGGGGGCCAGGAGAATGGAACATGGAAGTTGTCACCTTCAAGAATCTCAAATATCTAAAATTGGATAGTGTGGAATTTTCTGAATGGCAGGTTGATGCAGAGAAATCATTTCCCGTGCTCGAGATGTTAGATATAGTTACATGTGAGAAGCTTATGGACATCCCAGATAGTTTTGGAGATATTGCGTCATTAGAACTTATCAAAGTATGGTTTAGCCCTCAGCTTAAAGAGTCAATCTTCAATATTAA

>M6-PGSC0003DMG400008492 M6 LRR-RLP

CTTTTTCTTGTTACTTGCTACTCTTCATAAAATGAAGGCCTCCTCTATAGTTTCCATTCTTTCTTTCTCAACTCTCTTGTTTCTTTCTCAACTCTCTTCATCTGAAAGATGCCATCCAAATGACAAAAAAGCCCTCCTAGAATTCAAAAAAGGTCTAGGCAATCCTGATGACTTGGGTAATTGGGATCCGAAAACTGATTGTTGTACTGATTGGTACGGACCTACTATCAAATGTGATGAAAAAACCAATCGTATTAACGTCATCGACTTCACCAAAATGGATATGGCCGGATACCTCTCCCCCGCCATCGGAGACCTTACATACCTTACAACATTCAGTATCCACAATGTACGTAATCTCACCGGTCCAATTCCATCAACAATCCTTAAACTCACTCATCTCGAATTCTTCAGAATTAGTCAAACAAACATTTCTGGACCTGTACCTGAAGTTCTTAGTAAATTGAAAAGCTTAACGTACATTAACTTATCCTATAATAAGCTTGTTGGAACTATTCCACCTTCGCTATCTCAACTTCCACACTTGGAGTTTTTACGATTAGACAGGAACAAACTTACTGGACCAATCCCGGAGTCTTTTAGCAAGTTAGCTCCAAATCTAACGTATCTTTACCTTGGACATAACCAACTTACAGGGATTGTCCCAACTTCATTTGCTGGCTGGAGTTTTGAGACAATTGACTTGTCAAGGAACATGCTTGAAGGGGATATTTCATTTTTGTTTGGTAAAGATAAGAATACATTTGAAATGCTTTTGGATCGGAATAAGTTTGAATTTGACATGTCAAAGTTGACGTTTGGGAAGAGATTATGGAGGTTGCATTTAAATCATAACAAGATTTATGGGAGTCTTCCAAAAATCATGTCCAAGAATCCGTGGCAACTCTTGAATGTGAGTTATAATAGACTTTGTGGAAAGATACCAAAGGGTGAGAATATGCAGAGATTTGAGATATATGAGTATTTCCACAACAAATGCTTGTGTGGTGCTCCACTGCCGCCCTGTAAATGATCATCACTAGTATCCTGTCTGTCGATCTATCTATGTATTTCAACTTCTCTTTGAATGTTGATGAATAATTGAGTGAGTAAACTCAAGGCATAAACATGTACTTTAATTAATTTATGAATGTTATCCTTTTTTAGTAACTTCTCGATGGAT

>M6-PGSC0003DMG400008596 M6 LRR-RLP

CTTTATTTAGGTAAAGAAGACTTTGGTACAGAGAAAGAACATTTTGTGAAGGAGCAAACACAGTAATGGCCGATCCTGTAATTGGTGCTACTGCTCAAGTTTTGCTTGACAAGTTGCTTTCTCCCACTATTGAGGAGGTCAAAAGTTTAAGGAACTGCAAGAAAGATCTCGAAATGCTGATAAAAAATGTATCCTTGATCCAAGCTTTCATTCATGATGCTGAAAGACGACAAGTCGAAGATCAGGCTGTGGAAAAATGGCTCAAGATGCTTGAGAGAGTTGCTGAAAATGCTGATAAGGTGTTTGACGAATTCAGGTATGAGTCTCTCAAAAGACAAGTGAAGATTCGAAACAATCCAATGAAAAAGGTCAGTGATTTCTTTTCTCATACAGCTTTTAAGAGCAAAATGTCTCAAAAAATCAACAACATCAATGAAGATTTGACGGCTATCAATAGGTTAGCCAAAGACCTTGGTCTCCAATCATTGATGGTTCCTCCTCGGCAAATACTACTAATTCGAGAAACAGATTCTGTAGTAGTTGCTTTGGATGTTGTTGGTAGAGACAAAGATGTTGCTGAAATAAAGGAGAAGATATTGACCATGAGAGAGGATGTTGTTCTGTCAACCATTCCAATAGTGGGTATGGGAGGTTTAGGGAAAACAACTGTCGCAAAGAGAATTTACAATGACGAACACATCAAGCAAATCTTTGAAAAGAGAGTTTGGTTGTGTCTACCTGAAATGTCGGAGACTAAGAGCTTTCTCGAACTGATCCTCGAATCATTGACAGAGAGGAAAGTTAAGGTCCAGAGCAGAGATATAATTGTCAAGAAGCTCCGAGATGAACTGACAGGAAGGAAGTATTTGCTTGTCCTGGATGATTTGTGGCGTGTTGACTCTACATTGTGGAATGAGTTTGTGGATACCTTGAGAGGAATAAATACTTCCCGGGGAAACTTCATTCTCGTGACTACTCGTATGGAACAGGTGGCGTCCACAGTAGCAGCAGTAGGTCCTCATAAGTTGGAAAAACTAGCAGAAGATCATTGTCGGTCCATTTTCAAACAAAGAGCGTTTGTTGATGGGGAAATTCCACAGGAATTAGCAAGCATGGGCAACAGGATTGTTAAAATGTGCCAAGGTCTACCGTTGGCTGCAAGTGTATTGGGAGGCCTCTTACGCAGTAAGGAAAAACATGAATGGCAGGCAATTCTTGATGGAAAGCCCCTTGTTGCAGGTGAAGATGATAATGGAGAAAATAACATAAAGAAAATCCTAAAACTCAGCTATGATTATCTACCATATCCACATCTGAAAAAATGCTTTGCTTACTTTGCAATGTTTCCAAAAGACCTTGAGTTTGAAAAGGACCAACTGATCCAACTCTGGATGGCAGAAGGCTTTCTTCATCCATGTCAAGAGACCACCGTGATGGAGGACATCGGGAACAAGTTCTTTCAAATCTTGTTGCGGAATTCCTTGCTGCAAGATGTTAAGCTAGATGAGCATAACAATATAACACACTGTAAGATGCACGATCTTGTGCATGATTTGGCTGGAGATATCTTAAAATCTAAACTAATTGATCCCAAGGGCGATGGTGGAGAAAATCTTTCTCATGCGCGATACTTTGGATGGGACTCACCAAGTGATCAAATAGATAAGATATATGAGCCAGGACGTTTGTACACACTGTTCTGGAGAAGCAATCATATATCGGAAGCTATGCTGTTGAGCTTTAAGTTCTTGAGAGTTTTAAATTTGTCCAGTTCAGGAATCAAGGAGCTGCCAGCCAAAATCAGCAAGCTAATATACTTGAGATATCTTGATCTCTCCAACACTAAGATCACAGCCTTGCCCAACTCCATTTGCAAGCTCTACAATTTGCAAACATTTAGAGTCAATGACTGCTATTCACTCAAGGAACTTCCATATGAGATGGGAAATATGATAAGTTTGAGACACATTTACTGCTATTCTCGATCTCAGTTGCCACTTAACATGGGACAATTGACTTGTCTTCAGACCCTGCAGTATTTCAATGTGGGGTTAGAGAAAGGGCGTCGAATAGGAGAATTAGGTTGTTTGAAGAACCTTAGAGGTGAATTGACGATCAATGAGCTGGAATTGGTCTGTAGTATTGAAGAAGCTCGAACAGCATATTTACAAGAGAAATCGAATATCTACAAGCTGGCATATGTGTGGTTCCACGATGAACCAGAAGGCTGTGAAACCATTGTTGAGCATGTGTTGGATGGTCTTCAACCGCATACTAACTTGAAAACCTTAGAAGTCAAGAACTATTTAGGTACTAGATTTCCTTCATGGTTCAGAGAAGAATCTCTACCAAATTTGGTCAAGTTGAAATTAAGTGGTTGCAAAAGATGCAAAGAAATTCCATGGCTTGGCCAACTGAAATTGCTTCGGCATCTTGAGCTATTAGGGTTACATAAGGTTGAATGCATCGGAACTATTTTTTATGGTATTGATGGTAACAATATTGGATCAAGCAGCAATGTCCAAGTGTTCCCATCATTGACTGAACTAGTATTGAAGGATATGTCTAGCCTTATTGAGTGGAAGGGAGATGAAATTGGAGTAAGAATGTTTCCTCTGCTTGAGAAGTTGAGGATTAGTAGCTGTCCATTGTTAAAAATTACTCCAAGTCAATTTGAAATCCTCCGTGAATTAAGAATTGAAAGAGTTGACAGTGAAATGCCATTGTTGAACTTGTGCAGTAAATTGACATCTCTCGTAAAGCTTAGTGTTTATGATGTGAAAGAGCTCACTTGTTTTCCAGATGAGATGCTACGCAACAAGGTTTCTCTTCAACACCTATTGGTCTCAGACTGTGGAGAGTTTCGTGAATTGCCACAAAGCTTATACAGTCTCCGTTCTCTTAAAAGCTTAAAGATTTACCGCTGCACCAATTTCAGTTCCTTTCCTGTTCCCAGTGGAGAGAATTATTTGACTTCCCTCCAAAGTTTTCAGTTATGGAACTGTGATGGATTGACCAGTGTACCAAGTGGGATGCTAGAGCATTGTCGGTCTCTGGAATCTTTGATGGTCTGCTACTGTAACAACTTAGTTTCCTTCCCTATATGTGTGGGGGAAATGCCTTTACTTTTTGATGTGTATCTATCAGAATGTCCCAAATTGATTAGTGTACCCGCAGGGGACCTTTACCGTCTCACTGGGTTACGGCATTTGGAAATTGGTCCTTTCTCAGAGATGGTGGATTTTGAGGCATTCCAATTGATTTTCAATGGCATTCAGCAGCTGTTGTCCCTTCGTTTTTTGTGTGTGGTAGGACATTTGCACTGGGATTCTCTGCCCTATCAGCTTATGCAACTCTCTGACCTAACAGAGATCCAAATATATCGATTCGGAATCGAGGCTCTTCCTCATAAATTTGACAACCTTGCTTCTCTTGAAAGATTAACGTTAGTGAGGTGCAAACGGCTACAACATCTGGACTTCTCAGTTGCCATGCCCAAATTACGGCATCTCCAGATACAGGATTGTCCATTGTTAGAAACACTGTCGGATGGGCTCGGCAACCTTGTTTCTTTGCAGGAGTTAGTTTTACAGAAATGCGAGAAACTAGAGCATCTGCCATCTGGAGATGCCATGCAACGCCTCACCAAATTACGTTACCTGAAAATTGAAGGCTGCCCAAAGTTAGAAGAAAGTTGCACCAATCGGATTGGCTCAAACTCCCAGTGGTCCAACATTTCCCATATTCCAAAAATTAAAGTAGGTGAAAGCATAATTCAGGACTTAAGTAAGTCTCTGTTCCTCTTTCAGTTTCTGAGTCGATTTTTTTGTTCTTTCTTTTCCACGAGTTGCTTTCCTTCTTTTTAACAAATTTTCCATTCCCCAAACTAATACATGTTATGGTACTTAAGAATTACAGAATTTTTTATCTCTCACTTGAAACTAACTGTTCTTCTTCTTATTCGTCATCAAACAAGGAATAGATACATTCCCGTCCTTTAATAATGTAAATATGTTCCTTCTCATAGGTGCCTAGATCAAATGTCATGCTGGAACGTTGCTAGCTGTGGTATATCAAGTCAAAATTTACGGTTGGCAATCAGGTTGGCTAAAATTTTCATTTAGACTTGGTAGAGGAATGCTATAACAAATTTATCGCTTCTTCCACAGGTCATTACTGTACTGTTTAATTAGGTTGGTTATGTTCTGTATGAATGCAATGTAAGCTTGGTTGCGTTTCTAGCATCATGAGGGACGTGCATCCATTTGGTTTAAGGATTCTCTACTCGCATTGGACCAAAAATCTTCATGTAAGCTTCTTCCCAGTTGTTAATCTCCTAGGTCCAATAGTTTGCTTTGTTTTAATTGTTTGTTTTGCACGATTATCGTGCAACAAAAGAGCCTATCTCTCAATCCTCAAATATTATAATGCTCTTCGTTACAGGGTATAGTGTGTTAGCACGTTGGACAAATTCCAATTAGCTGTACATATCTATGTAAGAACTCTAAATATTTATTGTAACTATGGTGTCACTTGCTTTCCACATGTAGGAGATATCAGGAATAACAGATATTTATTTAGGGGTGTAAATGACAACTTCTAATTTAATGGAAGAACTTTTAATATGAGAGCATTCATTCAGCAAATTTGACTCTGGATAAAGACAATTTCTCATTCTCTTTAGGAAAATGAAGTATTTGATTCACATAATATTGGCTTATCATGGTTGACAGTTACAATCAGTTTCTTTTGCATAAGAGAAGGTGAAACATAGAGGGGGATAAACGACCTTCTGCAAGGCGACTTTATTATCGACTATCTAGTGAAATAGTATAGGTAACAAAGCCATTTCATCATAAGCCTTGGATGCAAGACAGATTTCATTTTGGTAAAACATGTGACTTCAATTGATAGAAGAGGCTTCTCCAAATTTTGTTACCTGTCTTCACTTTTCAACAGTATTTCCTTTAATTTTCATCAGACTTATCGGAACATTGTTTACATTGGTGCCTTTGGGGACGGTGATTGCACA

>M6-PGSC0003DMG400008925 M6 LRR-RLP

TGAGGTCCTCAAACTACTTTCCCGAGCCTTTGAGGGCGTGCAATGGGACATGAAAGACCGAGAGTTCTTGAAACTAAAATTCTTGAAGTTAGACTCTTTGAACATTGAACAATGGAACATCTCATCTGATTGCTTACCTTCCCTTGGGCATCTTGTTCTGCAAAGTTGTAAGAAACTTAAGGAGTTCCCTTCATGCTTAGGCAATGTTTCAACGCTGCAGAGTGTTGAGCTACACTGGTGCAGCTGTTCTTCTGTGAACTCGATTATCTCAATTCAAGAAGATGAGGGGTTCAAGGTCCATGTATACCCTCGAGAATTCGATTCCTGA

>M6-PGSC0003DMG400009029 M6 LRR-RLP

GGTTATCCCTCGAGTAACAAGATAGTCGAATATAAAAACACTGTTCCCACACAACACACACATTGTTTACTTCACTTTTTTCAGTTGAGATTTTCTTCACTTGCACTCTCTCCGGCGGCGCGTCGAACTCACACATTCTCCCTTGTTGTTTTCCGGTGAGTTACAACAGACACTCACTCCAACGATAATCAGAAAAATCCCAAAAAGGTTTTTTCTTTTCTCTTGATTTCTAGGGTTTTTGCGCTTATGCTCTGTACGAGTCTACATTTTTGGTCATGTGATTTCACAGATCTGTGCTGGATTTTAGGTATTTTTCTTGTTTAATGTGCTTTCGTTTGTGGTTTTAGGTCGTTTTTTTTTATGTTTATTATTTTGGGGTTTGGTAGAGTAGGTTGCAATTACTGGGAGGAGGACTACAAACTTCACTATTAGGTATAAAGTTTTTTATTTTTCTCGTGAGGTTTGTTTGTTTGTAGGGATAAAAGAAGGATTTTGTTTTGTATTACAAGTTTCTGATGTTTTTAGCGGCTCTGTAAATTGTATATATGCTGCTCTGCTCTGCTCATATTATTCTTTTTGTAATGTATTGTGGGGTATTTTTTTCGTATCCAAGAAAGTTTTTATGTGCTATAGTGTGGTGAAATGGAATACATCTGTTGAGTGTACCATTTTTGGTTTTGAGTGTTTTTTACATTGTTGGAAATGAATATGGTGCATTGCAAGGTGGTTTATAAAAGAATAGAAAATTCAATCTCGTTTTCAATGATGGATGATTGTAGAAACTTTGCCATTCTTTGGACACCTGCTTGGTGTTTTGTGGTGAAACTACTGAGTTGTCAAACAATTATATCTGATTTCTATATGAGGAAGAATTTGAGCTCTTGAAATTTGGTTTCGATATATGTTAGGTTTTCTTACATGGAGTAGGTGTGCACGAACTCGTTCTTGCCTGGTACTTTGTTTTTGGTTGTTATGTTCTTAACTGCCAGAGATGAGTCTTGTAGTGGTTATATTTCTTGAAAGATAATCACTTCTATAGAAACTGAGACATGATTCACTATATTCCTTTGATTCTAGGTTTTCTTCCAATGGATGCCACAACAGCTAACTCTCAGCGCAGACCATTTTCAATTAAACTGTGGCCTCCAAGTGAGAACACAAGAAAAATGCTGGTGGAAAGAATGATGAATAATCTTTCTAGTCCAACCATTTTCACCCGCAAGTACCGCAGTCTATCCAAGGAAGAGGCTGCCAAAAATGCTGAAGAAATTGAAGATGCAGCTTTTACCATTGCTAATCAACATTATGAGAAGGAGCCTGATGGTGATGGAAGTTCTGCTGTGCAACTTTATGCCAGGGAATGCAGCAAGCTTATCCTGGAAATTCTAAAGAAGATACCCAAATCAGAGGACAAGGAAATTTCAATTTCTGAAGTGGTTCCTACTGTTCAGGAGACCTTTTTTGATATCTCAAAAGGTAAAAGGGCTTTTATCGAAGCGGAAGAGGCTCAAGAACTTTTAAAGCCATTAAAAGAGCCTGGAAATTCTTACAGCAAAATTTGTTTTAGCAATAGAAGCTTCAGTATAGATGCAGCACGTATTGCAGGCCCTATCTTGGCAGCCTTGAAGGATCAATTGAAGGAAGTTGATTTGTCAGATTTTGTTGCCGGAAGAAATGAGGCAGAAGCACTTGATGTGATGAATATATTCTCAGAAGCTCTGGAAGGTTCTAACTTGAAGTTTCTGAATCTCTCTGATAATGCTCTGGGTGAGAAGGGAGTTAGGGCATTTGGAAAGCTCCTTCAGTCTCAAACCAACTTGGAAGAATTATTTTTGATGAATGATGGGATTTCACAGGAAGCTGCAAATGCTGTTAGCGAGCTAGTTCCTTCCACCGAGAAGCTTAAGGTTCTTTATTTTCATAATAATATGACAGGTGATGAAGGGGCTGTTGCGATTGCGGAAATTGTGAAGCGTTCTCCTTTATTGGAAGATTTCAGGTGCTCTTCTACCAGGGTAGGCTCTGAAGGAGGGAGTGTCTTGTGTGAAGCACTTGGGATGTGCTCCCATTTGAAGAAGCTTGATTTGCGAGACAATATGTTTGGTCCAGAAGTTGGTCTTGTGTTGTGTAAGGCACTCAGCAAACATGAAAATCTTACGGAAATTTACCTGAGCTACCTTAATTTAGAGGATGAAGGAGCAATTGCAATAGCTAACGCTCTTAAAGATTCAGCTCCTTCACTTGCTGTCTTGGAGATGGCAGGTAATGATATAACTGCTGAAGCTGCTTCAGCAATAGCTTCTTGTATAGCTGCAAAGCAGCTTCTTGCCAAGTTAAGCTTGGGTGAGAATGAACTCAAGGATGAAGGTGCAATTCAGATTGCTAAGGCACTGGAAGGACACAGCCATCTGATTGAAGTTGATATGAGCAGCAATGCACTAAGGAGGGCTGGGGCTAGAGTGCTGGCTCAAACTGTGCTGCGCAAAGATGAGTTTAAATTACTAAATGTTAATGGGAATTTCATCTCAGAAGAAGGCGTTGACGAGTTGAAAGAGATCTTCAAGAAATCTCCTGAAATGCTTGCATCCCTGGAAGATAATGACCCGGAAGGAGAAGACGAGGATGATGAGGAGGATGAGGAGAGAGAATCTGGAGATGAGGGCAAAGACGTTGAGGATGAACTAGAATCAAAACTCAAAAACCTGGACGTCAAGCAAGAGGCAAGTCTTGACACACCAGATAGCAATTAGAATCCAGTACTTGACGACTTCTGCATGTTGCAGTTTTAGTTTTACCATTTTCTTTATGCTAAGCTGGCCATATTATGTTCCTCCAGCAAGTGCTCTAATATATAGTTCCATCAGTAATTTAAATTTGGTGGAATTGAACTGTTGCCTTAATTGGTCAAGTTAGGTCTCTGGCTGAATTATGTAGGGTACGAGAGAGAGAACTGCTATTTATGTTGAACCTTCTTTTGTTATGTAGACTTGATAATCTCCTCTTTTTTTCTTTTTCTTTTTTTTTTTATATTTTCATGATGCATTTGACTTGTTTTATT

>M6-PGSC0003DMG400009230 M6 LRR-RLP

TTCAAACATCACTACATACTACAAGGAGAAGTAGAAAGTCTACCTTATTTAAGTAATTTAGTTGAGTAATTATACATGTAACAGAAACCATCACAAACACCCAAGTTTGAAAATTTGCAGCTTGGTTTGAACCCTCCAATGTAAGCTTTATAGCTATACCTATAGCTTTGATGCTCATAACCTACAACAGAGCAACAAACAAGCTTAAAATGACTGCTACATGCAATTTTTACCAATTGTTCTTTTGAGAAATTTAGAAGGACCAGTTCCTATATTTTCTTTACTTTTCATTCGGAATAATTTTACCATGTTTCATTGGCAACACAAGCACTCTCTAAGTCAGTTCTGCAAAAAGAACTGCAATAGTAGCTCAATTAATACAAAAAATTTAAGAGAGGAACCAATTCATAAACTGTTATTTATCAAGTCTTTAAAGCAAAATACATGAGAGTATTATTCATAAGATAGTGAATTCGATTGATAAAGAATTCATAAAATAACTCGACCTCGAACTAACAAGATGTGACTAGATAAAAGCTTATCTTAACTAGCAACAAAGACTTGTTTTAAATTTATTTAAAAGACAAATTAATAAAATAAAGTATTAGCACTGCAATCTTTTATTGAGTTACACACCTACTAGTCATGGAAAAACAACAGCAACCCTGGATGCCTTATAAATTATAACAGATCAAAGAATAGTCATTTGTACAAGCATAGTGGCAGGAAAAGACATCTAGTGACATGCTAAAACGTTACAGCACCCTAAAGCTCTGGAGTAATATTTCAAGTTCCATACATCCATTTATACCTAAATGAGAAGTTCTCTTTTACTTCAGTGACAACTAGCAATGGATCGAGGACCCAGGGGCAGTTTGACTCTTAACAACTTAAAAAATATGATCCATTTTGTGGTCAAATAATGCTATTCCAATACCACAGAGAAAGGTAGAATTCGGATTAACTTTAAAAGATAAGGCGTCTTGCAAACTGACAATTTAGCTACACGGTTAACTCAGTCAGGTCAGAGCTACAGACTAAAGCACGGTTGTGACACATGTTTCTTCCCTTCCTTCCTTCTCTTTCTTAATTCTAGAAGTAGAGACTCTGTAGGGATATAGATCAATACAATAGAGGCACTAACCGTCAAGGAACCAATAATGGAGCAAACACCAATATAGACCATGATATTTGTCCGACCATAGCGAGGCTCACAATACAAGACTAGTACCAATGTTATTGCAACTGCTGATGCAGTGTACAAAAGAAAAGCTGCAAAACAAAGAGTGGAACATCAAAAAAATGATATATCAGCAAGGCAAATAAGAATGATATTCAAGGAAATACAGCACAGTACATCAGCACACCTGGTTGTGTAGCTAATGCCCATATTTCATCCACTGAATTGAGATCATGTTCACGAGGTGCATGAAGAACAATGACAACGGAACCTACTATACATAAAAGACATCCTAATACTCCCAGTTTCTTCAATTTTTCCTTCAGTATGAAATGTGCTAGAACAGCACTGCAGCAAAGCGCCAAATTATATGGAAGAAAACCGAATAAGCCATTTCCCAGAAAATAATAAAGTCAAGTGGAAATTTACCTCACAATTATACTTAAAGCTCCAAGCGGTGTCACAAGAACTGCTGGCGCATAAATGTAAGCCACAAAATTTGCAAATTCTCCAACAATCACTGCAAGGATTATGGTGAACATTGCACTCTATATTAGAATAACAAATTCAGGAACATATTCTCTGACAAATAAGAAGCAATTCAAAACTTACTAGTTACCATCCCAATCCACCAAAGAGGCTCCCGTAGATAACCATATCCACCCGAGCCTTCAGCAAGAAACAAATATCTCATTGTCAAATAGACAGAGCAACAGAGTTCATATAATCAGCTTTTAAACTAAGTTCTATTAAATGTTGATACAGAAAAGCATCTTCTATATATAAACTTTACAAGATGTGCCCGATGTATCTACTTTTAATGTGGAAACTAGCTGATAAATTACAATGTCAAGGAAATTCATTCAGCTTATTACAGTGCAGAATGATTTAAAAAATCCATTACACTGATCATTTTCAGTTCTTAAAGTCAAGATCCATTTATTATGTCTTGAGCTCATTTCTTCTCTGTAATACATGTATTAGTACTACCCAATACAGCAGTCTGTTTATTTGGAAATGATAGATCAAATAATTTGGAATTACCAACAACAATCATTTAGTGAAGCAGAATCATGAGTGCATATATTTAAATTGCTGACCCATTCTACAAGTGGTAACTATCAAAGTATCAAAGGTATTAAGTTTAAACAGGTGCTGCTGGAGTCACTCTGATTGCTAAAACAATGTCATTAATCCTAGATAACTGCCACCAAAATTCTAGGCAGCTGACTGGTCAACATTCAGAAGAATATGAATTCATTTTCAAGAACCTCTCCAAGATTATTCAAGGACTACACTTACTTGCATGATAAGTGTGAAGTGTCAATTCCAATATAAGTAACTTGATATTCATTTTCCCATAAATGTACACCCAATACTTCCCTTTCGGAGAGAAGTGTGAATTCCATTCCTTGAGTTCCAGAACAAGTTTTCTTTATTCCCACACAGATAACTGCTTTGTAAGATGCAATATCACCTTTTATAAGCAGTCATTGTGTATTAGTAATACTCACTGTGACTCTACATTTGCTGGAAGAGTTACCCCATTGCTCTTATTTTACAGTCGGGCACCTACAACATGGTGTACCAACAGAGGATAACATAGTCTTAGACTGTACTTAAGCAGAAAAAAAAATGAAGTAAGAATTTTCAGGTGTAAAACATTTGTTTCACCCGCAAAACTAAAATGGGGACAGCAGAAAAGCAGGTACCACCATACCAGTATAAAAGAAACCCAACATCATCAGTCTGTTGCCACTTGATATTTCAAAGTAAGAAGTAAAGAACAATATATTCATCAAGCATAATCTCCAATTCCATACATCTAGAAACAGCATCTGATATACTATAAAATTTGATTACTATGCCTCTTGGCTGATGGTAAATTGGTAATGCAGTAAAATGGTTAGAAATATCAATCTTCAATTGACTAATTTTGACTTGATCATATAATAGTCAGTGTCATCTATATGAGTCCACTATATCCATTCTACATATCTGGGCCTAAATCAATTTCAATACTACAAGAATTTTATATAAAGAAAAACTGGGGGTTTTTTAGAGTTGAGAGATTCTCCAAATGTCACACTTCTTTTTTGTACACTAACATACAAGGTTTTAACCTCATTCTCAATTAGCTAGCGTCATCTACAATAAATCATTTACTTTAACTGTGATTCTTACCTAAATTGCCTTAGTTGAGAGATTGTAGGGGTTTTGTGGTTATTGTTAGGATCTTCATTACCATGTGTCATGAAGAAGCTAATAAAGCAAACATTGGACTATGATAAATAATATAACAAAGAGACATATACCAATACAAAAAAATTTAACATGAGAAATTCACAAAGATAAAAAAAAAAGAGTACAAACATATAAAGAGAATAACCCCACAAAATTCACTCTTAAAGAAGTTTACTTTTGCCCTAAACAAAACTCTCTTGAGACTACATTGTGGATGCTCTAAATTGAGAAGGAAATCCTCAATTAATAGAAATTCAAAACCTGTTCTTCCAAGAAAAGGATTAGCCAAATATGGAAACTTTATATTTTCCTAATAAAATCTAGTTATCCTAAATGTTATATCCAAATATGATAAATAAATTAGGGCAAACACCTACAACTACTTGCATTTGTATAATTTTAAGCAAGGTCAACCTTCCTAACCTTAGCAAGAAGTAATGCATTGCAACACACACATATAAAGAACTGAGCATATTAAACAACTTCAGCAAAACTACAAGGTAGAAGAAAACACACAAACATCGGCAAACTGAATGGGAACACAACTAAACCTCCATGAATCCACCACTAGAAGCATTGACAAAGTTCAAAATCAACAAAAAGCAATAAATTATTGACAATTAGACCTAAAAACACTTAGCTATCAAGAAAATGAATTACTCACTGGCCCGAGTACCAGAGGCGCCTGCCCTCTGAAGACCCTTCTTCTTGATAATGAAACTGGAACCAATAAAAGCTCTAGACGCCACAGCCAATCCAAAACCAGTCAAATTAGCACTGTACATGATCCCAATATTATATCTAAACTATTGGGCAACAATTTTGGAATTTAATTCTCATCTGAACTTCCCAAGAACAAAACTTTCAACACCCAAAAGGTTTCCCTGGATATACTCAACAAGACAACACTCAAGTAAGACTAAAATAACAAGAAAAACGAAATGGGCAAGTTTGAAAAACTTGAAACCACTGTAGCTTTACTCAATTACCCCAAAATTGTAGAGCTGAGAATGGAAAAACAGCAAGAAAAACGAAATGGGTATATGAGAAAATTGGCGAAAACAGGTTTGAAATGGTGAGCTAACAGTGCAAGCTCGTTATAGACAATAATGGAGAATAGTATTACAATAAGCATAATTTTGTCATGCTTTGATTCGTTGCTTTGGGTTGATTTGGCTATGGTATGTCTTCGACAATTGGGCATAACTATGAATAAATC

>M6-PGSC0003DMG400009324 M6 LRR-RLP

AGGAGACCTAGTAAAAAAGAAAAGAAAAAAACATGGCCGACAAATTTCAGCCAGTCAACCAGAGGTACGGAGAATTGTTGGTATATGCAGTTATCATGAAATATATAAAGTCCACACACTGCTCAACTGAGCAAACCAGACACTTCGAATTGGTCAATATTTCTTTACTGGATTCAATACATGGCGGAAGCTTTTCTTTATGTTCTGCTATAAAATCTCTTTTCTTTCATTTGAGGGAAACTTGTATTGTTTTTCGGTTTCGAAAAGGAATTTGAAAAGCTGTCGTGTGTGTTTTCCACAATTCAAGCTGTGCTTCAAGATGCTCATGAGAAGCAATTGAAGGACATTGCAATTGATAATTGGTTGCAGAAACTAAATTCTGCTGCGTATGAAGTTGATGACATATTGGGCGAATGTAAAAATGAGGCAATAAGATTTGAGCAGTCTCGATTAGGGTTTTATCATCCAGGTATTATCAATTTCCGTCACAAAATTGGGAAAATGATGAAAGAGATAATGGAGAAACTAGATGCAATTGCTGAGGAAAGAAGGAAGTTTCATTTCCTTGAAAAAATTACAGAGAGACAAGCTGCCGCTGCTACGCGTGAAACAGGTGTGAGTACTGAGTAATAGTAACTTAGTTTATGTTCAATTTGTTACCACATCGTGTGTTCACCGTGATCTCTACAGTAGGATGGCAATGGGGCGGGGCGGGGTTGGAGGTGTGCAGGTGTGGGGCACAACCCCAACTTTGAGTCTGCACAAGTAGGTATTTAAATTTGTATAGAGTTAAACAAGTACACACGCGTCCTACTTGGTGTCCTACATGGCTCTTATGTGTATTATGTCATATAAGACGCATGTGTCTACTTGTTCAATTTTATACGAGTTTAAGTTATACTTGTGCACACCCAAATGAAAGTTGGAGGGCGTTGATGTCAGTTGATACCAAGTTAAAAATTCATATTAATGAATTATGCCTTTAAATTATGATTCAATTTTATATCAGTCTGTCCAAAAGATGTTCTAGTGAAAGTGTTAAACTTAGCCTGGATCTGCTATTAGAAGTGTGAATTTTTGTGGCACTAAACAATGCAATGGGTCTGGATTCATTTTTGCATTAACATTTGTTTAGACGATTTTCTTTATCGAATTTTACTGTCTAAAAAGGAAAAAACAAAGAAATAAGAAGTATACAGAGGCTGACTTCTTCATAGTATTTCTCATAAAAAAAAAACATTGATTTCTAGGATATGGGTTATTTTAAATTACAAATTTGTGAGTTAAAACAGTTCTGTTGGGAAGGATTTAGATACACGTGGATAGTATCTAGAAGTTTTTTAAATAAAAAATTAGCAAATTATGTGGGCTGGGGCGGATTGAAAACAGCAAACTTTGCAAGGCAGGGCGGGTTGAAATCTTTGCAAGTTTGTGTGGGTTTGCCCTGCACCACCCCATCTGTCATTCCTGTCTAAATGTTTGTTTTGTCTATATAATTCTTGCTGACTCATTCTAATGAGCTCAATTGTAACAAATTCTTTGTGTCCACTCTACTTGGAACAGGTTTTGTTTTAACTGAACCAAAAGTCTACGGAAGGGATAAAGAGGAGGATGAGATAGTGAAAATCTTGATAAACAATTTTAATTTTGCCGAAGAACTTCCAGTCTTCCCTATAATTGGTATGGGGGGACTAGGAAAGACGACACTTGCCCAAATGATCTTCAATGATGAGAGAGTGACTAAGCATTTCAATCCCAAAATATGGGTTTGTGTCTCAGATGATTTTGATGAGAAGAGGTTAATTAAGACAATTTTAGGAAATATTGAAAGAAGTTCTCCTCATGTCGAGGACTTGGCTTCATTTCAGAAGAAGCTCCATGAGTTATTGAATGGAAATCGATACTTGCTTGTCTTAGATGATGTTTGGAATGATGATCTAGATAAGTGGGCTAAGTTAAGAGCAGTCTTAAATGTTGGAGCAAGAGGTGCTTCTATTCTAGCTACTACTCGTCTTGAAAAGGTTGGATCAATTATGGGAACGTCGCGACCATATCATTTGTCAAATTTGTCTCCACATGATAGTTTACTGTTATTTATGCAACGCGCATTTGGGCAACAAAGAGAAGCAAATCCTAATCTAGTGGCCATTGGAAAGGAGATTGTGAAGAAATGTGGTGGTGTGCCTTTAGCGGCCAAGACTCTTGGTGGTCTTTTACGCTTCAAGAGAGAAGAGAGTGAATGGGAACATGTGAGAGATAATGAGATTTGGAGTCTGCCTCAAGATGAAAGTTCTATTTTGCCTGCTCTAAGACTGAGTTATCATCACCTTCCATTTGATTTGAGACAATGCTTTGCGTATTGTGCAGTATTCCCAAAGGACACCAAAATGATAAAGGAAAATCTCATCACTCTCTGGATGGCCCATGGTTTTCTTTTATCAAAGGGAAACTTGGAGCTAGAGTATGTGGGTAATGAAGTATGGAATGAATTATACTTGAGGTCTTTCTTCCAAGAGATTGAAGTTAAATCCGGTAATACTTATTTCAAGATACACGATCTCATCCATGATTTGGCTACATTTCTGTTTTTGGCAAGCACATCAAGCAGCAATATCCGCGAAATAAATGTCAAAGATTATAAGCATATAATGTCCATTGGTTTCGCTGAAGTGGTGTCTTCTTACTCTCCTTCGCTCTTGAAAAAGTTTGTCTCGTTAAGGGTGCTTAATCTAAGCTACTCAAAACTTGAGCAATTACCCTCTTCCATTGGAGATTTATTACATTTAAGATACCTGGACCTGTCTCGCAATAATTTCCGAAGTCTTCCAGAGAGGTTGTGCAAGCTTCAAAATCTTCAGACTCTTGATGTTCATAATTGCTACTCACTTAATTGTTTGCCAAAACAAACAAGTAAACTTAGTAGTCTCCGAAATCTTGTTCTTGATGGCTGTCCATTGACTTCTACTCCACCAAGGATAGGATTGTTGACATGCCTTAAGACTCTAGGTTTCTTTATTGTGGGAAGCAAGAAAGGTTATCAACTTGGTGAACTACGAAATTTAAACCTCCGTGGTGCAATTTCAATCACACATCTTGAGAGAGTGAAAAACGATATGGAGGCAAAAGAAGCCAATTTATCTGCAAAAGCAAATCTGCAATCTTTAAGCATGAGTTGGGATAACGATGGACCAAACAGATATGAATCTGAAGAAGTTAAAGTGCTTGAAGCCCTCAAACCACACCCCAATCTGAAATATTTAGAGATCATTGCCTTCGGAGGATTCCGTTTTCCAAGCTGGATAAATCACTCAGTTTTGGAGAAGGTCATCTCTATTAGAATTAAAAGCTGCAAAAACTGCTTGTGCTTACCACCCTTTGGGGAGCTTCCTTGTCTAGAAAGTCTAGAGTTACAAAACGGGTCTGCGGAGGTGGAGTTTGTTGAAGAGGATGATGTTCATTCTAGATTTTCCACAAGAAGAAGGTTTCCATCCCTGAAAAATCTTCGTATATGGTTCTTTCGTAATTTGAAAGGTTTGGTGAAAGAGGAAGGAGAATATAAATTTCCCATGCTTGAAGAGATGGCGATTTTACATTGCCCTCTGTTTGTTTTTCCAACCCTTTCTTCTGTCAAGAAATTAGAAGTTCACGGCAACACAAAAGCAAGAGGTTTGAGCTCCATATCTAATCTTAGCACTCTTACTTCCCTCCGCATTGGTGCTAACTACAGAGCGACTTCTCTCCCAGAAGAGATGTTCACAAGTCTTACAAATCTCGAATACTTGAGCTTCTTTGACTTCAAGAATCTCAAAGAGCTGCCTACCAGCCTGACTAGTCTCAATGCTTTGAAGCGTCTCCAAATTGAAAGTTGTGACTCACTAGAGAGCCTCCCTGAACAAGGGCTGGAAGGTTTAACTTCACTCACACAGTTATTCGTTAAATACTGTAAGATGCTAGAATGTTTACCCGAGGGATTGCAGCACCTAACTGCCCTCACAAATTTTGGAGTAACCGGTTGTCCAGAAGTGGAAAAGCGCTGTGATAAGGAATTAGGAGAAGACTGGCACAAAATTGCTCACATTCCTAATTTGAGTATATATTAGTGAGTTATTTGTAATTGTTTCTTGGGTTATTTTTTTGTCAGTCTCCCACACAGGTTATAATTCCTGCCATTGGATTATTATAGCAAAGTTTGACTGTATGAGGTAACATTTTTAAACAAAAAAGGTAAATCCGTTTAACATGACTTTTTTTTACTGTTTAATTTAATATAAAAGGATTGTACTAATTTTCTATTTGAAAAC

>M6-PGSC0003DMG400009532 M6 LRR-RLP

TGGATCGAATTTCTCAGTTACCGGCGGGAATGTTAGAGCACATCTTGTCTTTCCTTCCGTTTGAAGAAGCTGCACGAATGAGTAGAGTATCAACGACTTGGAAAAATGCATACAATTCACTCCCCTGTTTAAATTTTGAGAGGTATTTTTTAAACAAGCCAATAGAAAAACTTGTGAATGCTGTGGATCCAATTCTAACTTATCGGAAAAATCAAAAGATTTCTGTAGAAAAGTTTTCGCTACACATAAATCAACAATGGAGTAGTCTGTTTTCCAATTGGATTGATACACTCGTAGATTGTAAGATCAAAGAGTTAAGCCTAGTTTTAAGATCAACCAGAACCGGATACACGAAATTGCCTGAGGCAATTTTTAATGCCAAAGCACTAAAAGTGTTGAAATTGAGTGAATTTCATATTGAATTGCCATCTAATAGTACTATTATGTTTTCATCATCTTTGCGAGAGTTATACCTTCAATGTGTATATTTCGACGAGGAGTTTATGCAAGCTTTATGCACAAGTTGTATCAATTTAGAGGTTTTCATGGTGCGGGGCTTGAAAGGGCTTACTAGATTTCAAACTAGTTTACCTAAATTGAAAAAGCTACAGGTGACAGCGTATTATTCTAAATTACGATTTGTTGATATTAGATCACCTAATATTGAAGATCTTGACGTCTATGGCTCCAACCTCAGCTCCAACTACTTTAAGAATGAGAGTGATCTAAATGTAGTTATTATTACTAATTGTTGCAAATCCCTTAAGAGTTTGCAGCTCAATGGTGTGGCCATGACCCAAAAATGGTTTGACGAAATTTTTACCTGCCTTCAGAATATAGAGAAGTTGGATTTAGGTGATTGCGACATGTTGAAGACTGTGAATATCTCAGGCGGCTCTCTCAAACAGCTGAGAGTGTATCATTGTGATAATGTGATTGCTATTGAATTAGACACTCCTAATTTGATAGAGTTCTCGTATGATTGTTCTTCATTGCCGAATTTGAAACTAAAAGCTTCAGCTTCATTGGAAGCCAATATTGAATTCGACTCAGAAGGAGGAGAAACCCCTAACAGTGATTGGTACTCTAAGCTCGTGAAATTTCTTGATAACTTCAATCATTCCATGGCTCTTGAATTAACATGCAAAAGTGAAAAGGTATGTCCCAGAAATTCTTTAACTAGCTAGTTAATACTATATTGTTATGGATTTCTCAAAAGTTGTTTCTTTTTGCTAATACAGGGGCTAGTTATACCAAAAGATGTGAGAGAACACTTGTTTCCTCCCCTTTATGGAACTAATCGCGTGAAAGTACGCATTCATAGTCCATTGATAAGTTATTCAGTTGTAGATATTCTTGATGGTATTTTCTGGCTTTCTCCTCAATTGGACACCCTATCTTATTCTAAATTCGACTTGAACACCCTTGAGGTAAATACTTTTATCTTTCAAAGGCACCGACT

>M6-PGSC0003DMG400009755 M6 LRR-RLP

ATGGAGGAACCAAGTTATGAAGGACGTGGATATTACCAAGACTCGGTAGTTGTTGTAACAAAGGGATTGAAGCTTGAAGTTGTGAGAATTTTGTCTTTGTACACAGTTATCGATCTTTCAAGCAACAAATTCGAAGGACATATTCCTAGTGTTCTAGGAGGTCTCATTGCTCTTCGGGTGTTGAATATATCTCACAATGGATTGAAAGGCCATATACCACCATCACTTGGACGTTTATCTGTAGTGGAATCATTGGACCTTTCATTTAACTGCCTTTCGGGAGAGATACCACAACAACTTGCTTCTCTTACGTCTCTTGAATTCTTAAATCTCTCCCACAATTATCTCCAAGGATGCATCCCTCAAGGACCCCAATTTCATACCTTTGAGAACAGTTCATTTGGTGGTAATGATGGATTACGTGGATTCCCCATTTCAAAAGGTTGTGGCAATGATCTTGTGTCAGAGACAAATTATACAGTATCTGCGCTAGACGATCAAGAAAGCAATTCTGAATTTCTCAATGATTTTTGGAAAGCTGCTCTGATGGGCTATGGCAGTGGATTATGTATTGGATTATCCATAATGTATTTCATGATTTCAACTAGAAATCCGAAATGGCTTGCAAGAATCATTGAAGAAATGGAACACAAAATTAACATGCGAAGGAGAAAGAAGCAGCAAGGTCAAAGGAATTACAGAAGAAGAAATAATCGCTTATAGACAAGTTACCAATAAAGAAGAATTTGATTTCAGAACTTCAGGTATTCAAGCTAAGCTCTACACTAATCTTTTTTAGTTCATTCTACTACTAATATGCTAGGTTTTATGCTTTTATGTTATAAACAACAACTAATTAATGCTCGATACGAAGCATATTTTTCTTTCCAACAACTTTAAGAAGCTGAAGATTTGAAAAAGGATTAGCCTTAGCTAATGTTATGGATGGTTTAACAGTGAAAAACTAATGCATAATTCGGTTTTAGCTATATGATTGAAATATAATTTACAATTGTTTGACTCACTATCTTATTTTTCAGATTTTCAAATGCCAAGAATAAGAAGACGTTCGCGTAACCGATTTGGTTCTTCCTGCAGCTTATGATGTTTGATTTGATTTTTAGTTTCTTTTTATCTTTATAAGTAAGCTTTTTCTTAACTTGGAAAATGTAATATTATGAATTTTGACAATATATCAAGTTTATGTAAG

>M6-PGSC0003DMG400009899 M6 LRR-RLP

ATTCTCTAATAAATTTCTCTTTGATTATTGAGCAAAATTTTGTCACCAATGGAAGGTGTAAATTTACTACTAGTCCTTCTTCTTATAGCTAGTAGTACTATATTAGATGGATTGATATTATGCAGGGCTATGAATTCAGACATTCTTTGTGATGAAAATGAAAAAAATGCTCTTATCAAGTTCAGACAGGGTTTAAGGAATCCATCACAGTCAATGATCATGTCGTCTTGGATGCTCGAGGAAAATTGCTGCAACTGGAAAGGTGTTGAATGTGACAACACAACCGGACATGTGATCACTCTTGACCTGCACCAGCAATTCTTGCAAGGTGAGTTTGGGACTTCCTTGCTTGGCTTGTCTCACTTAAGACACTTAGACTTGAGCCAAAATGATTTACTCAAGGCCTTGCTTCCTGAGTTTATTTGCAAATTTAAGAGCCTAGAGTATCTCAATTTGCACAAAACCAACTTTAGGGGATCAATCCCTGAGTGTCTTGGAAATCGGTCGCGGTTGCAGTTACTTGATCTTGGTGATAGTTTTTCACTCCGGGTTGATAGTCTTCAATGGATTCAGCATCTTTCCAATATGAGGATTCTTGACCTGAGTGGAGTTGACCTGAGCAATGCCAAAACCTGGCTACATGACATCAATTTCCTAAGGTCTCTCGTGGAATTACGTTTGTCTTCTTGTCAACTTCCTAAATTACTTCCTGTTTTTGTAAATTTCACATCCCTTCAGGTTCTTGACCTCTCGTTAAACTATCTGGATGTTCCATTTCCTAGCTGGTTAGTCAATACAAGCTCTCAGAGTCTTGTTTATCTTAATCTCAAAAGAAGTCTGCTGCATGGTTTACTTCCAAATACCGCCTTTGGAAACATGTACTCACTTAGAGTTCTTGATCTTTCTGACAACTCTATCCGAGGCAAGCTTCCGTCTTTTGAGAACATGACTTCAGTTACTTCTCTTGATATCTCAAGGAATTCTCTTGAAGGCAAATTACCACCAACCCTGCCTTCAAAGTTGAAACATTTGGATCTTTCATCCAATAATCTGGAGGGTACTTTAGCAAGAAGCATCGCACAACTTAGGCAATTAGTTGTCCTCAATGTTGCTTGGAACTCTTTCAATGACTCGATTACTGAACATTTCTTGAATTTCAGTGATTTGAGAGTGTTAGACTTGTCATCCAACTCAATTATCCTCAATATAACTGCAACTTGGATGCCTCCCTTTCAACTTGACATTATTGGCCTGATGTCTTGCCAACTTGGTACACAGTTTCCCCAGTGGCTTCATACACAAAAGGATTTTCTCATTTATCGATATCTCATATGGTAACATCTCAGATAAAGTGCCTGATTGGTTCTGGAATCTCTCTCCAAAGGTACAACACATGAACCTTTCGCTTAACTATTTCAGAGGGGAAGTACCTCAAGTTACATCAAGACTTACTAGTCTTGAAAAACTAGACCTGAGCCAAAACAACTTTCATGGTCCTTTACCTCATTTTTCGTCCAAGATGAAGACGTTGATTTTGTCCCGGAATTCTTTTAATGGCGCCATTTCCCCTGTATGTGAATCACTTGTCATGAATAATTCTCTTATGTTTCTTGACTTATCGTTCAATAATCTATCAGGAACACTTTCAGACTGTTGGAGATATGGAAACAATCTGGCAGTATTAAAGTTAGGCAATAATAATCTGTCTGGTGAAATACCTCATTCTTTTGGATATCTTGAAACTCTCAGGTATCTGCAACTGAGAAAGAACAGGTTGTCCAAAAATCTGCCTTCATCACTGAAGAACTTACAAGAACTAAAATTACTTGATCTTAGCAGAAATTGTTTATCTGGAACGATACCATCTTTCTTAGGGGAGACTTCACAGAAGCTAATGCTTCTTTCGTTAGGAGAGAACAAGTTCGATGGAAACATTCCCCTACAGATTTGCCAGCTCAAAAACTTGATTATTTTGGACCTTAGTTCCAATGCTCTATCAGGCACCATTCCAAAGTGTGTTAAAAATTTCCTAACGATGGCCGGGGTAGAGGAGTTCCCTTCCTTGGTTTATGGTCCCTATGAAGAATACAGGAAGGATGTGATGTTGATGCTGAAAGAGAGAGGCTATGATTATAGGTTTTTCCTTGCGGCAATTTATCTATCAGATAACCATTTATCAGGGGAAATTCCTGAGGAAATCACCACACTTGTTCAATTGCAGGTATTGAATTTATCAAGGAATAATTTAACTGGAGCAATTCCTTGTGACATTAGTAAATTGCAATATTTGGAAGCACTTGATCTTTCCAGGAATTACCTGTCATCCTTTTTGCCTTCCAGCATAGTGGAGCTTTCCATGCTCGCGATCGCAAATTTTTCGTTCAATAGTTTGACAGGAAAAATTCCAATAGGTCGTCAATTTTTAACCTTCGATAATAGTTCTTATATTGGGAATCCAGAGTTGTGTGGCATGCCACTTTCCAAAAGCTGCTCGGGACATTTTCGTGAGGACGTTACTCACTGTAACACTCCCAAGAAAAAAGACGTGCAGGCCATTCATCAGCATGAGGAAGACAATTGGCTTGATGAATCATCATTCTATATCAGCATGGTAATTGGTTTCATCACAAGTTTCTGGTTATTTTGGGCTACTCTATTGGTGAAAACATCTTGGAGACATGCCTATATGAGGTATCTTAATAAAATGGGCAACAATATTTATGTGTTTGTAGCCATAAGATTACCAAATAAGCATAAACCATCAAAGATGAAAGACTGAAGTTATTTCAGGGCTTTCTGAATAAGGTAATGAAGGTCTTATTTCTGGTGCAACTTTCTTTTTTGTTGTTTTGATTTCTAGGAGCTCTTCTGGTTTTTTGTTTTTGTAAAATCTTCTAAGTACTATATTGTGTATGCTTAGTTGCATTGTTTCATCTCGTGGGATTGAACTAGGTATGTTGTTGTTGTCATAATTGTATTCTATGATATGTTTTGTGGATATCTAAGTAACTTGAGAATACTATTACTAATGTTTT

>M6-PGSC0003DMG400010183 M6 LRR-RLP

TCATCTCTTGCTCCTTGACATTTTCCAGAATGGAAATTTTCTCAATGAGTATTAATTCCCACACCTTACTGGCAACCCATGCTGTGTTTCTATTCTCATCTACTCTCAAATATGCTGCAGCAGCTACTATATACTTGGAAACGAAACTGACGAACTAGCCTTACTTGGATTCAAGTCCCAAATAACTGAGGATCCATCAAGAGTTTTCGCCTCTTGGAACCAATCTGTTCATTTATGTCAGTGGACTGGAGTAAAATGTGGCCAGACACAGGAAAGGGTCATCAGTTTGAATCTAAAAGGGCTAAGTCTGGCAGGTACAATCTCAGATATCTTTCTTTGCTCAATTCTCTTGACCTGGCAGAAAATTCATTCCATGACGAAATTCCTCAACAACTCAGCAGGCTGTCAAGGCATCAAAACCTGAATTTGAGTTTAAACTATCTAACAGGGGAAATTCCAGTTAATCTATCACATTGTGTTAACCTCAAGAGCCTTGTGCTGGACCACAACATTCTTGTGGGACAGATTCCTTACCAAGTTGGATCTTTAACAAAGTTAGTGAGGTTGTATCTCAGAAATAACAACCTGACAGGAATCTTTCCAATGTCTATTGGAAATCTTACATCTTTAGAAGAGTTGTATTTATCGTACAACAGTCTAGAGGGAGAAGTGCCAGCTTCTTTAGCTCGATTGACCAAGTTGAGACTGCTTGGATTGTCAGTAAATAGCTTCTCCGGGGAGTTCCCTCCTTCGTTGTACAATTTGTCATCCCTGGAATTAATAGCACTTTCCTGGTAATCTCAGATCCGATTTAGGCCACTACTTCCCTAATCTCCAAAGACTTTACTTGGGGAACTGTCAATTCCATGGCTCCATACAATCCTCTTTGGCCAATGCTTCAAAATTGCTACAGCTTGATTTCCCTGTAAACAAATTCACTGGAAATATACCAAAGCGTTTTGATAACTTGTGGAATTTGTTGTGGCTCAATGTTGGGAGTAACCATCTTGGATACGGTAAGAATGACGACCTTGACTTTGTAAATTCTCTTACCAACTGCAGCAGCAGTCTACAAATGCTCCATTTTGGAGACAACCAGTTTGTAGGTACATTAACTCATTCAACAGTCAACCTTTCTAGTCAGTTACAACTCCTACTCTTCTTCGGGAACAGAATTGGTGGAAGCATACCCAGAGAGATCTCAAACCTGGTGAATTTGAATCTACTCGACATGAGCAACAAAAATCTTACAGGTAGCATTCCAGATTCTATTGGAAGACTTGCAAACTTGGGAAGTCTTGACTTGCATAACAACCTCTTGACAGGGGTAATCCCTTCTTCAATAGGCAACCTCACTGAACTAGTCTATCTTTATTTAGGATTTAACAGGTTAGAGGGTAAAATACCTTTAAGGCTTTAA

>M6-PGSC0003DMG400010325 M6 LRR-RLP

ATGGTGGAACAAGAAGCAGTGTTCAATGTTCTTGAATCCATTAACTCTGATGTCTCTTGGCGTTCCCTTTTCCCTGATGATCTCTGTTCTTCTTCCCCACATGGGGTTATTTGTGATTACTTTGTTTCCACTACAAACTCATTTACTCCTCATGTTACTGAGATTAGTCTTGGTTATGTTTCAGATTATGCTCCTAATCCACCTTGTACTCCTAATAGCACACTTAATGCTTCCCTTTTTGCCCCTTTTTCTTATCTAAGAAAGATTTTTGTTTACAAATGTTTTACTGAAAATGAGGTTTCTTTGATTGACTTTGGATCTTTGCCTTCTTCTTTGGAGGAGGTGGTTTTCTTGGAGAATCCAGGTCTTGTTGGTTCTTTAAGTGATAAAGTTGGAAACTTGATGAGTTTGAGGAGGTTGGTTTTGAGTGGGAGTGGTGTTTATGGGGAAATCCCAAATGGGGTTGGTGGTTTGGTCAATTTAGAGCAGCTGACTTTATCAAGAAACAAAATTAGTGGTGAAATTCCAGTGGGGTTGATGACATTGAAGAAACTCAAGATTCTTGATTTGAGTCAAAATGAGCTTGAAAGTAGTGTTCTTGAATCTATAGGGAACTTTACTCAACTTTTGAAACTTGACTTGAGTTACAACAAGCTTACTGGGAAAATCCCTCAAAGTTTGAAGGGTTTGAAGAGTTTGGAGTTTTTGGACTTGAGTTACAATAACTTTTCCAATTCTGGGGTTCCATTGTTTTTGGCTGAGATGTCTAGTTTGAAGGAAGTCTACTTAAGTGGAAACTATCTTGGAGGGGAAATTCCAGAAATATGGGAGAATCTTGAGGGTATTGTGGGGCTAGGATTATCAAAAACAGGACTAGTTGGGAACATTCCTGTTTCTATGGGGGTGTACTTGAAAAACATATGTTATTTGGGGCTTGATAATAACAAGCTGGAAGGGGCATTGCCTGAAGAGTTTGGGGCATTGGAATATGTGAATGAGTTACATTTGGAGAACAATAATTTGAGTGGTAGACTTCCATTCTCTCCAAAATTTGTATCCAAAGTTGGAGAAAAGTTGAAGTTGGGAGGCAATCCTCAACTCTGTGTAGATGAAGGCTTGAGAATATCTAATGTTAGTGTTAGCTTGAGGCTACTAAAAATGTGCAACAATGCTATTAGTGCCAAATTTGTCCCATTCTCTGGGGCATCTTTAGTACTGGTGCCTGTTTCCCATGTTGTCTTTTTTCTTGGAATTTTGTTTCTCCTGTGCTAG

>M6-PGSC0003DMG400010328 M6 LRR-RLP

TACAAACATTATCTTATGTAGAACCCATGGAGAAAAGACTGCGACTTTCCAACAAATTTGCTTTAAAAAAGGTGTTTTTCTCAACCCCATATCTCTTTTCTTCAGTCTCTTGTCATTCTTGTGGTTTGATCAATTTTTTTGTTTCTAGTGTCAAAAGGAAGTCTGCATTGCACAAGAAGATCGTGTCAGTCAGTTACCTGATGAAATTCTGGTATATATATTGTCTTTCTTTACTGTTAAGGAAGCTGCAGATACAAGTCTCCTCTCAAGGCGTTGGCTAAACTTATGGACATATATTGATCACCTTGATTTTGATGCTACAAAATACTTGGATCAAGTGTTCGTGCAACCCAAGCTACAAAAAGGGCATATGGACAAGTATCTGACATGGGTCAACTCTGCTTTGCAAAAGTTTAAAGGCCAGAGACTTGACCAATTTCGTGTTTACTTTGATTTAATCAAATTTGCTCAGCACGACATTGACAAATGGCTTGAACTTGCCTTTTCTAAGAAAGTTCAAAGACTAGAGTTAGACTTATTAAGGTGTGGATCATTCACTCGTAATTTTGATTCCTGTTATACTTTTCCGGAGCGGCTCCTTGGTCTTGGAGATTCTTCTTCATGCTCTAATAACCTCCAAATACTCTCACTTATCAACCAGAATTTCAAGTCTCTAAAAGTGCTTCTATTGAAATCTGTAAATGTGACGGGCCAAGTTCTTGAGTTCTTTCTACGCAATTGTCCATTTCTTGAAAGAATGCAAGTTCATGGATCAGGAACATTGAAAAATTTGGAAGTTGTTGGTCCATCCCTCAAGTTGAGACACTTGGAGATATATAGCTGCTTCAATGTAAAATCACTTAAAATTTGTGATACAAATCTTATATCATTAAGGACTTCAGCAGGAAAAGTATTACTGCTTAAGAATGTCCCGATGCTGGTTGATGTAGAGATTTTGGGTATGCCATACACCCATTTATTGGATGACATATCCTCTCGTCTTTTGTGCGTTCTCTCTCAGCTGGAGGTCCTTACAGTAGATGCTTGTATATCATTGGTTAGTGAACAAAGTTTGTGTCTATCTACTCTTGTTTTACTAGCCTAGCACTAACTGGTGCCTTCTCTGTAGTTTGATTATCCGCACGAAGATTTGGAGCATTACAATTTTCCTGAACTTACGAAGCTCAAGAAATTTGTTTTTATGACATTTGGAAATTCACTAAGAGTGTCAGGCCAGAGTCTCTTAGGTTGCACGTCTATAATAAGGGCTGCCCCACAGTTGAAGGAATTTGAACTACAGGTAAGTATCATGCCTTTTCCTCTTTGGTTTACAAAAGTCTGATTTATTCATATTAGCAATTCTGGTTCATGCAGTTATTGATGCTACTTCTTTAATAAACTTAGTTATTTAAGTCGTTGAGCTTGAGTTGAAACAATTGTGAAGAGCTAAACTTGAATAAATGGTACTCTTGCATTTTAACTCATCTGACTGTGCCAATATAAAAGTCCTGCCCTTACATAGGTGTCATTTTAGATAGGTAGTAGATAGCTCTGTCGTCTCTGAGCTTTGATTTATTATCTGTAACAAAGAGCTTAAAGTCTCATCATTTCATCAAAGAGCTTAGAGTCTCATCATTCCTTCAGTTTCTTGAATATTTTTACTGAATTACTGCACTTGTTTAGCGTAATGGAATGACCTCAGAGTGTTGGGATTGTGGTTTCAGTTGTCGATTGTGTTGCTCCAAAATCTTAGGTTTCCACCTGTTTGTTGCCAGTATATTATATGATGGAATTGGGAAGTACTATACAGTGATTATTATGTGTTTAATAAAACCACTTTTTTGAGAAAAGGTAACTTGTATTATAAACACAGGAATACTCATGACGTATTCCAGTAATACTTACAATCAGAAATACAGAAGTGTTGTTGTTTTCCTATAAACTCAAGAACTCTAAGATACTAGGGATATCCTCATTTTCATGAGGATATTTTTTTATTTTTTTTTTGATAACCGAGAAATCCGTCAGTGACCCGCCCTTTGGACCAATCATAGCCTTCTAAACTCGGTGGATAATGGGCCCGCCCCTCTACCCTTCTCCACTTAAATACGGGGCTTCATGAGGATATTTATGTTTACACCAAAAACAGAAAAGAGCTAAACATTTCAGCTTCATCTTCTCCATAGAGCAGCTCTTATTCTCAAAACATCTCTGATTCCTTTCCAGCCAAATAGTCCACCAAATGCAGGCTGGGACAATCTTCCATCTCTCCCTGTGCCCTGACTGATTTCCATCCCGATTCCAACATGCCAAAGTTTCCTTTATGTTTCCTGGCATAGACCAACTGATGCCCCTTTTATTGATGAAAATTTGCCGAAGTTTCAATGTCTCCTTAAAGTGTAAAAAGAGATGATTTATTGTCTCTGTCTCACATTCACAAAAAAAACATCTAGAGCACAAATCGAGGCCCTTCTTCATACGGTTATCCTGTGTGAGAACAACCTGTTTTGCTAAGAGCCAAGTAAAACATGCCACCTTGAAGGGCACCTTAACTCTCCAGATCATTTTCCCTCCAACTGAGAATCTCTCCTGTGTATCTGGAGCCCATAACAGTTTGTCTTCACTGTTACTGAGTTCCTTGTATCGTTCCAGGATATTTAGAAATTCAACCATTCTATTCACCTCCCATTCATTTAAAGGTCTTCTGAACCTTAAACCTACTTTCTTGTCATAATTATACTGGTTGGTTTCAAATGAGAACAACCAACTAAGAAAATTGAGAAGTAAGTTGTTGCCTAGCTAGGTAGATGGGTGAAGGAGGAATGCACTGGGGTAGAGGTTTCAGAGATGAGGGATTACAAAATGTTGTTAGAACCGAGGGGAACTTCCAAATTCTATAGTAGGATTCGAAGGTGGCATCCCGAATCCCAATCAAATGACTCCAGGCTCAAAGAGTCAGAATCGAGCCTGGATGGTATTGTAGAGGAGGCTTAGGGCTTCACATGGTATCTGATGTTTTCTCTCTCTTGAATATTTATTTTTTCAAAAATCAAGTAACGATTATATAAGGGAAAAGAGAAAAAATACCCGAACTTGTATCGGATTTCTTGTTACACTTATTATTTGAAGTTGTTCTATGAATTATTTTGTAGTAGAATAATACCATAGGAAGTGTTGATGTGGCTTTTACTTTGCACCTCCATATAGTGCATAAAAAGACAAATACAAGCTAAATGATAGAGGCTCTAGAGTCCAAAAGATATAATTTTTCTTCTACTTTTTAGGCCCTTTTCTAAATCTTGAGCGGTGATCCACCACTTGTCCCACCAGGCCACCATCCATGCCCCCCACCCCCATGTCTGAGCCAGGATTTTTTTTTTTTGTAGCTGAGTAGATACACAAATTATGTTTTTCCTACTCATGAGTGGGTGCACGCACTTGGCAAAAGTTGTGTGCATAATATTTTCAATAAAAATCAAGCAATAAAAGTAAAAAAATTTATCTAGGGTGTGTTTTGTATGGAGAAAAATGTTTATCTATTTTCATGATTGGTTAGTCAATATATTTTGTGAAATATTTTTTCTAGGAAAACAAGCTCCTTAAAAATGAGGAAATTTACTTTCCTAATGGAAGTAGAAAACATAAGTTCCAAAGCAATTAAATCATATTAGCATTTATAACTATAAAAAACAATCATATAAATCACTATAGACATAGAGAAGTTAATAAAAATAATTTTAGTTACTAGTTAAATAAAATAAAAAACAATACATATCACAATAAAAGTAATAAAATAATAATTAAAATAAAAGAATTACCGCGAAACATTAGAAAAAAAACTATTGATAATCTCAACCACAAAAAAACTTTAAAGGAAAAACACGAGTACGGAGCCAAAAGGGCAAAAAAATTGACCAAAAATTCCAAAAGGGTTTGAAATCAAAAGGGTAACATTTAAACGGTAGGGGAGATGCTGTTTAAATTAGGGTAGCAAGGCGCAACTCCCTATGCGCCTTACAAGGTGCAAGAAAACAGAGCCACCATCATTGTTTATTTTTTCCATGTCAGTACTTACGGTGGTACTTTTTGCACTACAAAATAGTATAGGGGTCGTAGGGACATCACAAAGAATAAGTGTGTAACTGGAAAACTGAGATAAGTCCATAGGGTTTTTATGCCTTTTCCCACTGTGTTATTTAAAAAAGCACCAAGAGGGCTCCAACCTTTACAAGTAGAAGATTCTGTTGTTCTGTCAACCTCCCAGTCATGCAATCTCCTTCTTAATCTAATATTAAAAGATAGCCTTTCTCTGTATTCTTCAATCACATCTTTTATATGGATAGCTATGAGATCTAATTCTGAATAAGTTATTAAAGCTTCTTGATCCGGCGTACATGAAACTTTATCTTCTTCTTGCAACTAAAAGCTTTCACAAGATTAAAAGAATCAATTCCATAAGTTCTAATGTACTTTCAAACTGTGACATCATCAGATGTATTGATTAGTTTTGCTTTTGAACATCTCATATCGTTGCCATGCTCTATGTTTAACAGCTTAATCCATAAATTGTTTTGCTCTACTGTGTTTTATCTCCACAAACTAATTCGACAATAGACACTTTTTTTAATACCCAAAGTTTTTATCTATGAACTACTCTTCTTAGAGAATATAGCTCTATTTTTATGCTCTTTGAATCAACTTGCCACATCTTTCTCCTTGCATTATCGAACGGTCTTTGCACCTTAAAAAATGATAGATACAGTTTCTTGCAGGTTGCCAATCTTCTCAACCTTATGTATCACTCTCTGCCATATTATTGCACTTCTATATCTGGCATTTAGAGACTTGCTCATGTATGTCTCTGCCTGGATTCTCAATTTAAATGCAGTCAAGTGCTGTCTCAAAGTGTAATAGTAGAGATCTGTGATGCATGATCTTCCGCTCCTCAAGTCCAAATAGAAATAGGGTATAGTCTGCATATCGTGTACATGAAAAATTTCCATATTTTCATTTGTACAACTTTCCCCAACTTTTGACTCCTTTTATCCATACTATATTCATGGCATAATGTAGCATCTTACTGAATGTTTCCATTGCTTAGGAAGAGAATGGGATAATGGGGTTCTCCTATTTAACTGCATATCCTATTTATATTTTTCCATTTGTTTGCAAACCGTGTTGATTCAAAAGCTTCAGTAATATTGCCAGTTTACATGATTGTTCCACTGGGTCAAATCTTTGCTTGATCCTGTGTTGTCCCTAGTCTGAGGTGATAACTTGAGAATGAATCATTCATCGGTGCTCTCATAGTAAAGAAGGCTTCACTCAAGGAGTGACAGATTTTGACGGAGTCTTGGCTTTGATGCTAAGGAGACAAGATCCATCGATCTGAATAAGTCTAGTAACCTTTCTCAAGTCCAGCTAACAAATAATTCTTCCTATTTTAATTCTATTTAAGGCTCCAAGAATTCATTTTCTAAGAACACTGGATCTGTAGTTTATCCTTCCCTTTACAAATACATGCATTTTTTGACCTTGATATCTACTTGCCAATACTTGTTTCATCCTTGTTGTTAGGACCCCGGGAGTAATCTCATCCTTTTACCTTGTTCTTAAGAACCAATGCTGTGAATATTTCAGTCCTGCCTCTTGTAAAAATTTCTGTCTCATGAAAGAAGTGCAATGCACTGATAATGTCCCTTTCCAAAAAGTTCCTAGATTTCTGGTAAGAAGACATTATCTGCCCATCTGGCCCTAAAGTTGTTATCACCACTAGACCATCTCCATTACTTCTCCGAGAAAATTTCTTAGTAGCCAACACTCTTCCTGTGATAATAGCTCAAGGTCATTGTAACTGGAAGGCGCCATCATTGCTCGGTTTTACTAAATAACTTCTCATAGCATTACACTCATCTGATGACCTTTGGATTTGGTAGTTCGTCCATCTCTTACAAGTTAGTTTATATGGCTGTACTTTTTATTTGTATTGATCAATTTACAGTAAAACACTTAGTATTTTGTAGGGGTGTCAAAAATGAGCCCACCTATAGGTAACCCGCCCATAATTTTAAGGGTTGGGCTTAAGATAATTTGAATTAGGTTTAATCTCAACCCCCCATTTTAAGAGAATCCTCAATTGAGCCCAATTTAATTTTCAATTTCAACCCATTTTAAGACTCTTTATTTGTATTGATGTAATGTATGTTCTTAAATTGAAGGTATGAATTACTTTTGTTTTATATTTTTTAGGATTTATCTATCCATTTGTAACTTTTTTTGTAATTTTCTTAAGTTGAAATCTAAAATGTGGTTATAAAACTTAAATAAACAATATATTAAAACTATTGAGATTAAGCGGGTAAAGACCCAACCATTTTTAAGCTCATTTTAGCCCAAAGTAAACTTGGGCGGGTCAAGACTCGACCTAATTTCTATTTCAACCCATTTTAATATCTCCAATTTCAACCAACCCGCCCATTTGGCACCTCTAGTATGTTGTCTCATTCATTTAACCAGAAACACCTCAATTGTTGTTTCCAGGGTGTTTTTTAGCATTGGTTATTTGTTCCAGTTCTACTAAGTGGTTCCTTTAGGCTTTTCGTTTTTCTATATCCGTTCACCTTTTTCTGATCATACTTACCAATCCAGCAATCTGATTATGTATAATTTTTCGACATGCATTGTTGTTTGAACTCAAAAGTCTGCCTTTTAAGTTCAGTTTCCTTTCTATTAACCATAATCAGATTGCCAACACTATCATAAGATTGCTATTGCTGTTCCGCTAAATCAATGAATCCCTCTATAGCTTTTTTCATTTTCTCGAATTTGCATTAAGATTCGCCTATTCCCAAGTCTGCACTCTGATGGAATGGGTGTATGATGTTGGTAATCATCATATGCTTTGAAAACTTGAATATCACATTCCAAAGAAATTGTACACAAAAATTTCTGTGTATTCCACCTCATAAGTTTACTGTACATGCTATTGTATTTATACAAAACTAGGTCTAAATTCTGTACCAAAATAAATGAATTAAACAATGATTATTTTTACACTTGGCACCTAATTATTGGACTATTTTGTAACAAACTCTAACAGCCATTTTGCTCTTCACGTGCCTTCCTTGTGCCTAGGTTGAGAATCTTCTAGACCTTGTACAACTGGTTAGGTTGAGAATCTTCTAGACCTTCTGATTAGGTTTCCTAGAACTTTTGTTCTTTTTGGCATATGTTATTTTGGTTCTTGAAAAGAAATGTTGGAGAAATTCATTTCGGTTAGCAAGTTGGGCATGCAGCTCGAATATGTTTATATGATAAGTCACTTGTGTATTCTTTTGTTTCCTAATTTCTCCAAGTTAAGATTTTTCTCACAGAGAGTCTAGAACATGGATAATGAGTAGTGGTGATTGAATATTGTTGAATTAGTCAACTTGACTAGGAGGCATCTTACATGATAATGGGAAGTTTTCATTCAGTCGATCAAGGAACCCATATATCCCGAATCTTAAAATTTAGATCCACAACCGAGTCCGAGCAACTTAGTCTAAAATAAGTCGACCTAAACATGTTGAAAGAAGTTGCAATTCTCTCAGAAAATTGAGCACTCCACGTCTGTAATATTCTTGTCCATATTCCCCACTTTCCTTCCTCCATGTGGTAATACTTTAGAATTTGTAGAAGAACAAAGTAGAGTAGGAGAAGTCAAATGACAAGAGCGATTGATACAATCATGGTTCATCAGTGATTGGTACATAGTTTTATGTTATACTGATTCCCCAAAGTTTAATTCTTTTCATTTAACTTGTGTCCCAAGAAAAAGGCCTGTTCAAAATTTTCTTTTATTAGCTTTACTGACTTATCTTATCATTATTTTTTATTTTAATTTTAATTGGTGTTGTATATATTTACGGCAGGTGATGTCAATCCCCTTCGAGGCAAAGAAAGAGTGTAAGAAAGCAGCCATAAAATGCCCACTCCATCACCTCAAAGTGGTCAAATTTTTGGGGTTTTATACAGGCTTATTTCAACTTGAACTCGTTAGGTACTTTTTGGAAAATGGTGTTGCTCTTGAGAAAATCATTATTGATCCAGGTACTCCATTTTTTTATCGTATCCCTCCGTGCCCTAGCGAAATCAAACACACGCAAGCTTCGAGAAATCTTGCTAAGCTCCAGCTTGAATGTGAAGTGCCTCCGGCTATTGAGTTGGTCATACTTTAATGAAAATAATAATGTTATTACGGTTAAATTTGTGATACATAACGGAGGTGAATTTTTATTTCTTAGTAGATGTTCCTCATGCTTTGTTTCATTATTGAGTGATATTTTCACTTTTTGTGAGGTTTATATTGCTATATGAAATATGGGGGATTTTCAAAATGAAAATGTCGCAGTTATTCCGCCA

>M6-PGSC0003DMG400010447 M6 LRR-RLP

CAACTTGCACTTTCCAAGTCTAAAAAGCAGCCATCACATTTAACATTTATAATTTCAGAAGCCACCAAGAATTTGTCATCCAAATGTATGTATATATTATGCCATTTTTTGGCACATTCTTTTCTTATAATTTCACAAAAATGAGAGGGCTGTACTACATAGTACTTGTCCTGTTTACTATAAAGTTTTGTTCTTGCAGTGGAGTTTGCAAAGAAAATGAGAAAACAGCTTTGCTAAGGTTAAAGAAAGAAGCAAATGATCCCACAAATGTACTATCTTCTAGGGTTGATAAAGAAGATTGTTGTAACTGGGAAGGAGTTTTGTGCCACAATGTGACTGGTTCTGTCATTGAGCTATCGATTAACGGATGAGATTGGTCAGAACTTAAAGGCCTAAAGATTGATGATTTCCAGTGGTTGTCAAGTCTTGAAAATCTTGATAGTTTGGACATGTCTGGTGTGGATTTAAGTGAAGCAACAAACTGGACTGAGGTGATTAGCATGCTTCCTTTGGTTAATCTTCGTTTTTCGAATTGTAGTCTTCATTCAATACCTCCTTTGTTTGATCACAACAGTTCTGATCTTGAAAATCTTGATCTTTCTTTGAATAACTTTGGCTCTCCGATTCCCGGGTGGGTTTTCAGTTTTGGTAGTCTTGTTTCTCTTAAATTCACTGGTAGCAATTTCACTGGTTCATTTCCTGAGGGTCCTTTTAACTTGACTTCTTTCACAACATTGAGAGCTTCTAGCAACTCCTTTGGTTCTGTTTTGCCTCAATGGTTATTTGATCTTAGTAATCTTGAATATCTTGATTTGAGTTTTAGTGGTCTTGAGGGTCCAATACCAAATGGAGTTGGTGATTTTACCAAGCTTACATATCTTAGTCTTGCTTCCAATAATCTGAATTCCACTATACCAGATTGGCTCTATGGATGCAAGAATCTTGAAACACTTGCCATGAAGGATAACAGTCTTGAGGGAACAGTTTCAGACTCGATTTCAAATCTGACATCTCTAACTAGTATCGATATGTCTGAAAATTTTCTTTCTGGAAAGTTACCGAGTGTAATTGGAAAGCTATGGAAATTAGAAGATCTTGATCTCTCAGAAAATTGATTTGAAGGAGAAGTATCCGAGATTTTCAATACTATGAATGATTGCCCTCCAATAGGCTCAGGAAATTGTTCTTCTTTGAGGACCTTAAGACTAAATGACAACAAACTGATTGGAAATCTTCCAAAAAGTTTTGGGCAACTTTCTAATCTACAATTCTGTTTCATATCCAATAATAGGTTGGAAGGGGTGTTAACAGAAGAACATTTCACCAACTTGACAAATTTGAAGTTTTTCTCTGCATCTAAGAGTAATTTGACTTTAAGGGTGAGTTCAGATTGGATTCCATCATTCCAAGCTAGTGACATTGTAATGGGCGGTTGGCGCTTAGGCCCTGACTTCCCTTTGTGGATCCAGACTCAACAATCAATCATGAATTTGGACATATCAAATGCTGGTATAGTAGGTGAGGTTCCTACTTGGTTCTGAAACTTATCGTCTCAAATTCGTTTCCTCAACATGTCTCACAACCAGCTCATTGGTGAGATTCCAACTTTCTCAATTTCTGATAGCTCCATGGGAAGTGGAGGTCCTTGGTTGATATACTTGAGTTCCAACAACTTCAGTGGTTTACTCCCTCATATTCCAACCATGGTAACAGAGTTAGACCTTTCAGATAACACTTTCTCGGAAGGTTTAACAAGCTTCTTGTGTGATAAAAAGAATGAATCTTACATGTTGGAGATTCTACACCTTGGGGGAAACCGTTTCTCCGAAGAAATTCCTGATTGTTGGATGAATTGGCCAGAGTTGAGAGTTGTCAACTTGGGAGAGAATAAGCTTACCGGAGGAATCCCAAGATCCATTGGAGCTTTAAGCAATTTGAAATCTTTGGACTTGAAAAGAAACAAACTTTCTGGTCCATTGCCTTCATCCTTGAACAACTGCACAAGATTGTGGAAAATAAACATGAATGAGAACGAGTTAGATGGAAATTTACCACCTTGGTTTGGTACACTTTCAAAGCTGATAGTCCTTAGTCTTCGGTCTAATAAATTCGATGGTGAACTAGCTCCAGAACTTTGCCAGCTCAATGATCTTCAAATCTTGGATCTTGCCAACAACAATTTGTTTGGAGCTATACCTCGGTGTGTTAATAATTTCACAGCAATGGTGAATGGAAGGAAGGAAATTAGAGATGGAGACGGCGAAATGGATTATTCTTATTATGTTGGAGTTTTCAGGGAAAGTGCAAGAGTAGCAACTAAGGGCAACATGTATCAGTATGATACCATTTTGTCATTGTTTACTAGCATGGATCTTTCTAACAATAATCTTTCTGGAAACATTCCTATGAGTCTTACAAGTCTTGTTGGATTGAGATAATTAAATTTCTCCCACAATCAGCTGACTAGTAGTATTCCAAGGGACATTGACAAGATGAAGGTATTAGAATCACTTGATATCTCAAATAATCAACTCTCTGGTGAAATACCACAAAGCATTTCGAGTTTGTCCTCTTTGAGCTACTTGAATGTATCTCACAACAACTTATCAGGGCAGATTCCTGTGAGCACTCAACTTCAAAGCTTTAGCCCTTTTAGTTTTACTGGAAATAAACTGTGTGGGCTTCCACTCGTGGAGCGTTGCAGTAAAAATCCCAAGGTTGAAAATGAAGAAGAAGAAGAAGAAGAAGAAGATGATAATGAAGTGGAATGGTTCTATGTTTCCATGGCAATAGGATTCATTGTGGCCTTTTGGGGTGTAAGTGCTCCTTTGCTTTTCATAAGACCATGGAGACATGCCTATTATAGATTTTTAGATAGAACATGGAAATCATGGCACACATAATTTGTAAAGACGTGACATTGAGGAGATCTCTATTCATGTATACTGGATGAAATTTGTTTATATCTCAGGAAATCCATCACCATTCAGCGTTCTTCAAATTAAGACATTTTTATATTTGTTGTCTCCATATATATGGGAAATTTGACAATTTTGTTTTGGCAAGGTTTGAGTTATGGAATGAACAAGGAAATTAACAGTTCCT

>M6-PGSC0003DMG400010519 M6 LRR-RLP

TCAGTTCAAGTCCTCGAGAAATTTCAAATCAACATGTCTTGGATGACCGTCTGCAGCTAAACCATAACGAAAATAGTACGGTGCTCCAAACAGATTTTTGCCATTTCTATAGCCTACTTCTTCCAAAGCCTTCACAAGTGGCTCCATATATGCTGTAATTTTCCTAAACATGTTAAAAGTAAGTTACTACAAGGAATTTCTGGATGAGTTATTGGCTGAAACAAACCAAATTTCTTGAATCATACGTTACTAGTATGGCGTTGTATGATAACACCATACTGGCAACATATGACATTTTGATCAATCCAAGGCAAGTTTTTTTTTCTTTTTTGTAATGGTATCAGGCTAACTTGTATGTGCTTCAACTGGATAACAAGATTGTGCATTTTCAACACTAAGAATAATAAATAATCTAGAAGGTGATAGACAAGAGGGGTTCTTGACATCGTCTATCGGGAACGTAACAGAACTTGTCTCTCTGAATCTAGGCAGGAACAAGTTTAGAGGACCAGTGCCAGAGAGTATTACTTATTTGAGGAAACTTACAAGTCTTCAACTTTATGAGAATTTTTTCTCTAGTAGTCTTGTTAATGACATTGGTGTCCTTACAAAACTTGAGATTCTTGATGTCGCGAATAATAAGCTATCTGGTTCAATCCCTTCATCGACTACATCACTTAGAAGTTTGACGCGATTAGACCTGTCTAACAATGAATTGACAGGGAAAATTCCTCAACTCAACGGTTTATGGAAACTCCATTCATTTGACATTTCCAATAACCAAATTTAAGGGAATTTGCCACAACTTCCATCGAAGATCAAGACATTGTCACTAAGCCACAACTTGCTATCAGGACACATTACCACAGTGCATAAACTAAGACAGCTAAGAACATTAGACCTTAGTGAAAACAGATTTTCAGGAGGAATAAACAAAGGGATCTTCATGATACCTGCATCAAGTCACGTAAGCATTACAGTAAACCGGTTCACAGTGTTTGAGGTGGTAGAATTCTCTGATAAAGAATCACAGCTTCACACGTTAGACGTAAATGCCAATCGTCTACATGGTCATTTTCCTGTGAACTTGGATACTTACCCAAACTTAACAGAAATAAACCTTGGACATAACCTGTTTTCGGGTGAGATTCCATAGGAATATTGGTCAAGACTCAGGTTTTCATGGAGAAGTCTCAACTTGGAATATAATAACCTGGAAGGATCGGTTCCTCGAGAGCTCAACAAGACTTTAGAAGGGGTTCAGGGAAGTTTTGAACACAACTGTCTCATTTGTCCAAAGGGCTTTCAGCTATGTCATGGACAGAGGACAGCTTCAGAATGTCATGGAAGAGACCATAATTAGCTGAGAATGTTTTGAAAATTACATTGCATTTTTTTTAATGTTTTCCCAGTTTCCAATATAGTTTATAAGTTGCATTTTTTTTTTATGTTTTCCTAGTTTCCAATATAGTAGAAGTTCCTTAAATTTGTCAATATATTTTATTTAGAAACTCGAACTACAACATGTTTCCTATTAAGCACCTGACATGATGAACTGTGTTAGAGTTTTGCCCTGAATGTATGATATTATCAGTTATAGCAAAGTATATTTGCATAACAATATTGAACAAGAGTATCAAAATTGGTTAAGTATACGACATGAGCAAGAGTATCAAAATTAACTACTGAAGCAACATGGCAGCCAACGTTTTAAGTGCGAGCTTGTTTTAAGTTCAACTATAAGTGATCTTGTTTACTTTGTTGGTTCCAAGATTAGATAAATTCTCATAATTTATGGTGAAGCAAGGTTGCTGCAAAGAAAAGTGATTCACTCTCAACTACCTTACTAACATAAAGTCATTCTTTATTTTCTTAAACTCCATGTCAACTCAAAATACATCAAAGAAATTTTAAACGGAGGAAGCACCATACTAGTAGTAACTGGAAAAATCAAAACAAGCTATAATGAAAGTTGTCTTAATTGCTAAAGAAGCTCATGAAGACAATGACATTACAACTCATTTCTGTTGCCTAGTCTAATAAAAACAATATTTGCCACCTAAAGATATATTTTAGCAGAATTGCATGTCTAAGAACTTAAGTACCTTTTTCAGGGGAAGAAGCTGGGAACAGTCCTTTACAAGCATCTGTATTCCACAAACATGAAATCAAAAGAGCAAGTCAGGCCCGAGACTACAAGCTGCTGTATTTAAAAGCATCTCAATGGATTAGAGAAAAAACCTGACACCCACGAGTGTTAAGCTAGTAATATTATAGCTATTGACAGAAGGCTTGAAAATTGCACTACTTGTGTAATGATATGTGTAATTCATTGTTATTTTATACTCGTTCATGAGTTTTTGAAGTTGAGAAAGAGATTCTGAAAGTAACTAGTGACTGAAGATTTGAGAATAGCAATTCACACATTTTCTAAGAGTGGGAATTGTGTACAAAAATGTACTATTTCTTCTAAAGTAACGTCTGCATTTTGGAAGAAAATCTCACCTCCTATATGTGTCCGAGTGTTCCTTTGCTGTCAAATGGGGGCATCCCATACAGAAACTCAAAGCATAGGACACCAAGGCTCTAAATGTCCACACTTGCATCATGCTCCACACTTTCCACTGCAAGTTCAACAATTTACATATCAGAAAAAATGTAAAAACAGATAACAAAATGTTAAAAGCTAAACTCTCATACACGCCCATGTGAAGGTAAAAAGCAATGTATAACATTCACACATAGTCCGCCTTAAATCAAAGGTATGCACTGACCACCCAAAGTCTGCCGTTTTGAGTTCAACTTAAAAGAAGAAATATTAAAGCTGTCATTTTACTGCAATGCATTGTATATACTGTTTAGCAATGCTAAGTTTATACCTCAATATCTTCTCTGTGTATTGTGTGATTCCCATGACAGTATATTGAAAGTTCGTAAGAATTGGGAAGAAGCAATGCAGAAGAGGCAACGACTAGTTACCAGGTCATAAAAGTAACCGTAAGTATTTCAACTTCACGATGAAGCTGATGTTCAACCAGGGACTGTTTTAGCTGGCTCTTGAACAACACTTTTAATGCGACGATATGATTGCTATGTTTCCAATAAACAACTAACCTTAGTATTGGAATGTGATTGTAGAATGTTAAGTCTATCGAATAGAATTCGAAAAATCTTTTAAGCCATGTCCACAAGATCCCACCAATCCATATCAAATATAACAAATATGTTGAATGAGAGATTCTCACTCGCAATGCATAAATCCGAGGCATTGAAAAAGTAAGTAGTTTTATCAGCCATTCTTAACATGTAAAGGTTTAAAGGACTTACATACATGACCGAACTTTCCTCTTCCAAGAGGCTTCCCAATGTCGAAACCATGAAGTGTCCGCCTCTTTATATTAGCCTCTGATGAAGTCTGCAGAAGTTGAAAATTCTGTAGTCCATTTAAAAGATCAGTTAATTAAAAATAACAATAAATTCTTGTTGTTGTTGTGTACAGTATAACAGATCAGTTAATTAAAAATAACAATAACTTGTTCTTTTAATTTTCATGATAACATATGGTTATTTCACATCACAAAGTTGTACGATCCAGCTGAGCTTGATTCCATTTAGTGCGGGAAGAATACAATTTGCTTTACATGCACTGGTAGAAGATTATCTCAATTAACTGCAACTTTGTTCATACTGTCCTCTTACAGGACTTCATGAACAAGCAATGAACCACAAGTGATCGCAACTAACATAAAAGCTTAAACTCAATCAACAAATTTTAACTAACCTCAACATTCATCCTTTAGCAAGGTGGTACACTTGTGAACTATCAAGTACAACTCAACTCAATCTACATACTAATTTATTATAACTTCAAAAATAACCAAACACAGTGAGGTCAGATCATGTTACT

>M6-PGSC0003DMG400010523 M6 LRR-RLP

TTGATCTGTCAGATAATTTGCTGGATGGTGAAATTCCAG

>M6-PGSC0003DMG400010700 M6 LRR-RLP

ATGGGCATTGAAAAACTTGCATTGTTTCTGCTATACGCGTTTCTGTGTCAACTTGTTTTCTCCTCATCTTTACCTCATTTGTGCCCCAATGATCAAGCCCTTGCTCTTCTTCAATTCAAAAACACGTTTACTATAAATCTTGATGCTTCTTATAGTGGTTATTATCCTAAAACAGTTAAATGGAATAAGAGCACAGATTGTTGTTCATGGGTTGGAGTTAAGTGTTACGAGATGACAGGACAAGTGATTTAGCTTGATCTCAGTTGCAGCCAACTTCTAGGTAAGTTTCATTCCAACTGTAGTCTCTTTCAACTCTCCAATCTCAAAAGGCTTGATTTGTCTAATAATAATTTTTCAGGGTCGCTCATTTCACCTAAATTTGGTGAGCTTTCTAGTTTGACTCATCTTGATTTGTCGTATTCAAGTTTTACAGGTCTAATACCAGCAGAAATCTCTCATCTTTCTAAATTACACGTTCTTCAAATCGCGAGTGATACAGATGAGCTTAGATTCGGACCTCACAATTTTGAACTGCTCCTTAAGAACTTGACCCAACTAAGAGAGCTCAACCTTAGGTTTGTGAACATCTTTTCCACCATTCCTCTGAATTTCTCTTCTTATTTAACTACTCTGGACCTTCGGAAAACACAGTTATACGGAATATTGCCCGAAGGAGTTTTCCACCTTTCCAACTTAGAATTTCTGGATTTATCGGACAATCCCCAGCTCACTGTTAAGTTTCCCACAACCAAATGGAATAGCAGTGCATCACTCATGGAGTTAGTTCTCTATAGTGTGAATGCTGCTGGTAGGATACCTGAATCATTTGGTCATCTAACTTCACTGCGTACATTGGACATAGCTTCTTGTAATCTCTCAGGCTCTATTCCTAAACCTCTATTGAATCTCACCAACATAGAGGATTTGTACCTTGGTTATAACCATCTTGAAGGACCAATTTCTGATTTCTTTAGATTTGGAAAGCTCAGGTGGTTATCACTTAGAAATAACAACTTTGATGGCCAACTTGAGTTCTTATCCTTTAACAGATGGACGCAACTTGTTGAGCTAGATATTTCATTTAATTCCCTAACAGGTTCAATTCCTTCTAATGTAAGTGGTATGCAAAACCTACAATGTCTCAACTTGTCATCAAACCACTTGAATGGGACTATACCTTCCTGGATATTCTCCCTCCCTTCACTTGAATGGTTAGACTTGAGTGATAACCATTTCAGTGGAAACATTCAGGAGTTCAAGTCTAAAACATTGTATTCTGTTTCTCTAAAACAAAATCAGCTGCAAGGTCCTATTCCAAAGTCACTCTTAGATCTGCAGTACCTAGAAATTCTTCTCCTTTCGCACAATAATCTCAATGGAGAGATTGCTTCACCCATATGCAATCTGAAAACACTACGAGTGCTAGATTTGGGCAGCAATAATTTGGAGGGAACAATCCCACTATGTTTGGGTGAGATGAGTGGACTTTGGGTTTTGGATTTAAGCAACAATAGTCTTAGTGGGACAATTAATACTACTTTTAGTATAGGAAACTAATTTATAGTCATTAAATTTGATGGGAATAAGCTAGAGGGGAAAGTCCCGCAATCATTGATCAATTGCACATATTTGGAAGTTCTTGATTTAGGTAACAATGAGTTGAATGACACATTTCCTAAATGGTTGAGAGCCCTACCGTATCTGAAGATTCTAAGACTGAGATCAAATAAGTTCTTTGGCCCTTTAAAAGATTCATCGACTTACAAGTTTGCTCAAATTCGAATCATAGATCTCTCATCCAATGGATTTAGTGGAGATTTACCAGTGAGCCTTTTTGAGTATTTTGAAGCCATGAAAATAATTAGTGAGAACAGTGGAACCCGAGAGTATGTAGCAGATATGTCTTCATTTTATTACACAAATTCCTTTATAGTGACAACAAAGGGACTGGAGCTTGAACTTCCTCGAGTTTTGACTACAGACATGATTATCGATCTCTCAAGGAATAGATTTGAAGGTCATATCCCAAGCATTATTGGAGATCTCATTGGGCTTCGCACGTTGAACCTATCGCATAATCACTTGGAAGGTATTATACCATCATCACTGCACCAGTTATCTGTACTTGAATCATTGGATCTCTCATCCAACAAAATCAGCGGAGAAATTCCACAACAACTTGCATCCCTCAAATCACTTGAAGTCTTAAATCTCTCTCACAATCATCTTGTTGGATGCATCCCCAAAGGAAAACAATTTGATACATTTGAGAATAGTTCATACCAAGGGAATGATGGGTTACGTGGATTTCCACTCTCAAAGGATTGTGGTGGTGATGATGGGGTAGCACAAACGACGAATCCAGTTGAGCTAGATGAAGAAGGAGGAGGAGATTCATCAATGATCAGTTGGCAGGCGATTCTCATGGGTTACAGTTGTGGACTTGTTATTGGACTGTCCATAATATACATAATGTTGTCAACTCAATATCCAGCATGGTTTTCGAGGATGGATATAAAATTGGAACACAAAATTCTTACGAGAATGAGAAGGCACAAGAAAAGACATTAGTGTGTAACCTC

>M6-PGSC0003DMG400010940 M6 LRR-RLP

ATGAACTCGGTGGTGAAATGTGGAAAGGGCCATGGTATGTACCCGAACTCAGAGTCTTGGATCTCACCAACAATAACCTCACGGGTATGATCCCTCCTTCTGTTGGAAATGCCACAAAAATGATGAACTTCAGTTTGTCTGGGAATAGAGTCAGCGGCAACATTCCAAAGGAGGTCGGTAATCTGAGCCAACTTGCAGACTTGTCCTTGACTGATAATCAATTAACAGGTTCCATTCCTGTAGGACTATTTAATATCTCGTCGCTACTTGCCATAAATCTGCTATACAATAAACTTTCCGGTCCTCTCTTGCTAGATGAAGGAATTTTTTTGTCAAATCTGAAGTTTCTAAGGATATCTCGCAACCAGATTTTAGGATGCATTCCTTCCAACATATGCCAACTCAGAGAGCTCAAAGTTTTGTCCATATCTTTCAACAACATAATTGGAGACAAACCCAAAAATATTGATTGTTTATCCAATCTCCAGGAGTTTTATAATCGTCATAATTCAATAAAAGGGACTATTCCCACTTCATTGGGAAATATTTCAACTCTGCAATATCTTTATTCTGGAAACAATCTCATGGTGGGGCCAATTCCTCCAGAATTAGGGAAGCTATCAAATTTGAGGCAATTAATTTTTTCCAATAATTCTAATCTTATTGGTCAAATTCCAGAGGCTATTTTCAACATATCTTCTTTGGAATTAATCAATTTCAGTTTCAATACCCTGTCGGGTAGAATTCCAACCACTACAGGTCTTCATCTTCCGAACCTTAAAGTACTTGACTTGGGAGTCAATCAGCTTGAAGGGGAAATTCCATTGTTCATAACAAATGCTTCCAAGCTTCAGATACTGTCGCTAGAACATAACTCTCTCTCGGGAGCTATTCCTACTAATTTGGGAAATCTTCATGAGCTGCAATCACTCTTACTACATACTAATCAACTTACCAATGAACCAAGAGAGCATGAGTTGAAATTCTTCAATTCGTTGGCGAACTGTAGGATGTTGAGATATCTAGAAGTGGGTAACAATCCGTTGAGTGGCGTTCTACCAAATTCTGTTGGGAATCTCTCATCTACTATTGAAAAGATTTATATAGGAGATGCACACATCAATGGCCTCATCCCCACAAGTATAGGCAACATGAGCGGTCTTACAGCCCTAAGCCTTGGAGGAAACAACTTGGCAGGGAATATTCCTTCTGAGATTGGTAAGCTTAAACAACTCCAAGGGCTGGCTCTATATAACAATAAATTGCAGGGACGTATTCCAGAGGTGGTATGCCATTTATCGAATTTGGTTGGCTTATTACTGGATGGTAATGAGCTCTGTGGATTAATTCCAGAATGCTTTGGAAATCTTAGCATGCTACAAGCGCTTTATTTGGATTCTAATAAATTTTCATCAAAATTTCCGTTGAGCCTTTGGAAGATGAGTGGTCTTCTCTATCTAAGTGTGTCACGAAATTCAATAGAGGGAGAACTTCCATCAGATATTGGAGGACTGAAAGCCATTGTAGAACTATATCTTTACAATAACCACTTTTCAGGTGTGATACCAACCAGATTGGGGGAACTCCAAAACCTGCAGCATCTTGACCTATCCAACAATTCATTTTTTGGCCAAATTCCATTATCCTTTGCCAACTTGATAAGCTTGGAATTCTTGAATTTGTCTTTAAATGCCTTGTCCGGTACTATTCCTAAATCTTTGGAAAAACTCTCATACCTTAAAAGCATTAATGTTTCATTTAATGATTTAGAAGGTGAAATACCAAGTGGTGGCGTGTTTGCAACTTCCACTGTGCAATCATTTCTTGGGAACAAAGGTCTTTGCGGAATGCACATATTGGAGATTCCTACTTGTGCTATCACTAATCATGGAAAACAATCAAAGCTTAAGGAGGTTGTGCTAAAAATTGTTACTCCAGTGGTTATTGCATCCTTTCTGATATTCTTGTTCGTTTCAATTTGGATAATGAAACGACAGAAGAAAGGGAAGTCCAAAGATATTGAAAAGGTATTGGAGATCGGGACTCATCAATTAGTTTCTTATCACGAGATTCAACAAGCAACAAATAATTTTGATGAATCCAATTTAATTGGTGAGGGAAGCTCTGGCTCTGTGTACAAAGGCACATTGTTTGGTGGAACTGCAGTGGCCATTAAGGTTCTGGATTTGGAAAACGAGCAAGTATGCAAGAGGTTTGATACCGAATGTGAAGTGA

>M6-PGSC0003DMG400010950 M6 LRR-RLP

CATTTTTCTATTAATATTTGCACTAATGGAGAAAGCCTTCACATTTTTCCTCTCAACAATACTCCTGTTGCTTCACTATGTTATGACCCAACCGAACATTACGACTGATCAATTAGTTCTTCTCTCTTTGAAATCCCAAATTAGTTCCGATCCTTTTCATTACTTGGATGAAAGTTGGTCTCCTGCTACTTATGTTTGCCATTGGGTAGGAGTCACTTGTGACTATCGTCACCAGAGAGTGAAGTCCTTGAATCTTTTCAACATGACTCTTACCGGCAAGATTCCCAGTGATTTTGGAAACCTCACATTTCTTGTTTCTCTTGACTTGAGAAAGAATAATTTCCATGGAAATTTGCCTCAAGAAATGGTACGCTTACGTCGGCTTAAGTTTCTTAAGTTAAGTTTCAACAAATTCAGAGGGGAAATTCCTTCTTGGTTTGGGTTTTTAAAACAACTTCAAGTTTTAAATCTTAAGAATAATAGTTTCACTGGTTCCATTCCTCTGTCACTCTCAAATGCCTCAAGGTTGGACACTTTAGTAATATCTGCCAATTTACTTGAAGGAAACATCCCGGAAGAGATTGGCAATCTTCACAACCTGAAATACTTGTACATAGAAGATAATCAGCTTACGGGTTCTATACCATTCACAATTTTCAATATTTCTAGTATCCAAGTCATTGCATTTACAGACAATAGCTTATCAGGAAATCTTCCCAATGGTTTATGCAATGGTCTCCCGATACTCAAAGAGCTTTATCTATCAATAAACAAGCTTGATGGTCATATGCCTACAAGCTTGTCAAATTGTTCACAACTTCAAATGTTGTATCTATCCAGAAATGAGTTTGATGGACCAATACATAGTGAAATTGGAAGATTGAGTAACTTGCAGTTCTTGTATCTTGAGTCCAACCATTTCACAGGTATGTGTTATCTTACTAGTTTCTGGACACGTGTGTGGTCATCTAAATTGACATAAATTGTTTCAAATTATCATTGTATTAAAATAGATTTATAATTAAATTACATGAATATCG

>M6-PGSC0003DMG400010952 M6 LRR-RLP

TTCAAAGTCTCAAAACCACAGGTTACTAGTACTAATATTTGCACTAATGGAGAAAGCCTTCACTATTTTATTCTTAACAATGCTCTTGTTGCCTCACTATGTTATGGCCCAAACCAATATTAACACTGATCAATTAGCTCTTCTCTCTTTTAAATCCCAAATTAGTTCCGACCCCTTTCACTTCTTGAATGAAAGTTGGTCTCCTGCTATTTCTGTTTGCCGTTGGGTTGGAGTCATTTGTGGTTCTCGTCACCAGCGAGTGAAGTCGTTGAAGCTTTCCAACTTGGCTCTTACAGGCAGAATTCCCCGTGATTTTGGAAACCTCACATTTCTTGCTTCTCTTGACTTGGGAAGCAACCATTTCCATGGAAATTTGCCTCAAGAAATGGCACATTTGCGTCGGCTTAAGTTTCTTGATTTAAGTTTAAACAGCTTCACAGGGGAAATTCCTTCTTGGTTTGGGTTTTTACACCAACTTCAAGTTGTAAACCTTAGGAGTAACAGTTTTACTGGTTCCATTCCTCCGTCACTCTCGAATGCCTCAAGGTTGGAGATTTTAGTAATATCTACCAATTTACTTGAAGGAAACACCCCGGAAGAGTTTGGCAATCTTCACAACCTGAAATCGTTGTCCATAGAACAAAATCAGCTTACAGGTTCTATACCATTCACAATATTCAATATTCGTAGAATTGAAGTCATTTCATTTTCATATAATAGCCTATCAGGAGATCTTCCCAATGGTTTATGCAATGATCTCCCAATACTCAAAGAGCTTTATCTATCCAGAAACAAGCTTGATGGTCATATGCCTACAAGCTTGTCAAATTGTTCA

>M6-PGSC0003DMG400010964 M6 LRR-RLP

AGGACAGAATCTATTAAAGGCATAATGGTACCAATTGGCTCAGACCAACATATATGCAAGTGGAGCAAAGCTTTCAGAAACATGCCTTGTCTTAGGTTACTCATTGTCAAAGGGGAGGAGGTCTGGCATCATGACCTTATTTGTGGCCATATTGAGTATCTCCCCAGTAACTTGAAATGGCTTGATTGGTCATACTACAGTTTTGAATCTTTACCAGCATATTATGAACCAGGAAACCTTGTTGGGCTCCACATGACTTTTAGTTCTCTTGTTGAAGTTGTCAAGGAGCCGAAGGTATTATTTATCTCCTTTAGAAACAAATCATATATGAAGAAAATAAAATGATAGCTAAAGCTGAAGTTTCTATGCATTCTCTTTTTCTGAACAGGCATTTGATAAGTTGACAGTCCTCAATTTAAGTTTTTCTAGAAATTTAATCCGAACACCCAATTTTTCCGAGATTCCAAACCTGCAGAGGATTATACTGAAAAGTTGTGTAAGTTTGGTAGAAATTCATCCATCCATTGGCCAGCTCAGAAAGGTTATCTTTTTGAATATGGAAAACTGCAAAAATCTCAAATCTTTACCAAGCAGTATTCAAATGGAATCTCTTGAAATCTTCAATCTCTCAGGTTGTGAGAAGTTAGAAAAATTTCCAGAAATTCAGGGAAATATGGAATTGCTATCAGAGCTCCTTTTGGCACATACTGCAATATGGGAACTTCCCTCATCAGTTGGACTGCTCAGTGGCATTATCTTGCTTGATTTGCATTCATGTAAAAGCCTTGTAAGACTCCCAGCCAGTGTCTCTGAGATGAGAAAACTGAAAATTTTGAGTGTGAAAGGCTGCTCAAGACTTGCAAAGTTTCCTGAAAATCTAGGTGATCTAAACCAGATGGAGGAGCTCTATGCTGGTAACACTGCCATTTGGAAACTACCAGATTCTGTTAGAAACTTGAGCAAACTTAAAGTCCTATCATTAAGAAGAGGCCGGAAGGTAAAGCGTCAAACTGGTGACAGTTTGATGTTACCCTCTTCCTGGGGGTTTCGTTGTTTGAGGTTATTGAAAAGCTTAGATCTCAGTGGATGCAACTTATCTGTTAACCAAACTGCTGCTCTTATGAAATTGCCTTCTTTACTGGAACTGAATCTGAGCAGAAACAAGTTTATTTTATTACCTGATTTCATCAGTCAACTTTCTCAGCTTCGATATCTCAACATAACACAGTGTCAGGAACTTAAGGAACTTCCCAAACTTCCTTCGAGTATAGAGGAATTATATGCAGAAGATTTTTTGGCCAAGCAAAGTATCGAAAAGCTACAAATATACCCGAGGTTGAATTTGGTGTCATTCACCAATTATAGTTTTGATCAACAATCTTATACAGAGGAGAGCAATGGTAGCTCAGTTTTGGATGAGATTCTCAGCTTGTTTCTCTCAGAAGATATGGATCATGTGGTCGAGCACTCTCTTAACTCTGATCACAGGGTGACCTGTTCTATTGTCTTCCCTGAATGTGCGATTCCCAGGTGGTTTAAGCATCAGAGTGTTGATGAAAAGATATTGTTCAAACTGCCCATGAATTGGTATAATGATAAGTTCAAGGGCTTTGCTATATGTTGTGCCACTCTCATGGGAGCAGGTGTGTTGAATCCTGATTCAGGGCTATCCAAGAAGTATGATTATGCTTTCTTCAAAGCTAAATTGATATGCAAAGATGATTTAGAAGACCTTAAAGTGCTAGAGAAAGAATGTAAAGTGGGAACAGCATCTAGAACATATGGCTTGTGTGTTTGCTTTGTCTACATACCATTGTATGCTTCACTGCAGGTGTCCGGCACAGATGTTGGAAATATTAACCAGTATAGCCTATTTGAGGCATCTATCCATGGGCGCATTGTGAGACAGTGGGGAGTTCATTTGATCTATCAGGATGA

>M6-PGSC0003DMG400011045 M6 LRR-RLP

ATGGCCGATCTTGTAATTGGTGCTACTGTTAAAGTTGTGCTTGATAAGCTGCTTTCTCTCACAATTGAGGAGGCCAAAACTTTAAGGAACTGCAAGAAAAATCTAAGAATTCTGACAAAATATGTATCCATGATTCAAGCTCTCATTCATGATGCTGAAAAACGACAAGTTGATGATCAGGCTGTTGAACAATGGCTCAAGATGCTTGAGAGAGTTGCTGAAGATGCTGAAAACGTGTTTGACGAATTCAGATATGAATATCTCAAAGCACAAGTCATGAACATCCGAACCAAACTAATGGAAAAGGTCAGCAGCTTCTTTTCTCATACTGCTTTTAAGTACAAAATGTCGCGAAAAATAAACAACATCAATGAAGAGTTGAAGGCTATCAATCAGTTAGCCAATGACCTCGGTCTCAAACCACTGATGGTTCCTTCTCAGAAAATACTTCCAATTCGAGAAACAGATTCCTTAGTTGTTGCTTCGGATGTTGTTGGTAGAGACAATGATGTTGCTGAAATAAAAGAGAAGATTTTGAACATGAGAAAGGATGTTGTTCTGTGCACCATTCCCATAGTGGGTATGGGGGGTTTAGGGAAAACAACTGTGGCTAAGAGAATTTTCAATGACCAACACATCAAGCAGCAGTTCGAAAAGAGAGTTTGGTTGTGCCTACCTGAAATGGTAGAAACTAAGAGCTTTCTTGAACTGATCCTCGAATCATTGACAAAGAGGAAAGTTGAGGTCCAAAGCAGAGATATAATAGTCAAGACGCTACAAGATGCATTGGGAGAAAAAAAGTATTTGCTTGTCCTGGATGATTTGTGGCATGTTGACTCTACATCATGGCATGAGTTCATGGACACCTTGAGAGGAATAAATACATTCAGAGGAAACTGCATTCTCGTGACTACTCGTATCAAACAAGTGGCATCCACAGTAGCAGTAGATCTTCATATGTTGGGGAAATTAACAGACTATCATTGTTGGTCCATTTTCAAACAAAGAGTGTTTGTTGATGGGGAAGTTCCGGAGGAAATACTGAGCATGGAGAACAAGATAGTTGAAATGTGTCAAGGTCTACCGTTGGCTACAAGTGTGTTGGGAGGCCTCTTATGCAACAAGGAAAAACATGAATGGCGGGCAGTTCTTGATGGAAATCCCCTTGTTGCAGGGGAAGATGATAACGGGGAAAGTAGCATAAAGAAGATCCTTAAACTCAGCTATGATTATCTACCATCTCCACATCTGAAAAAGTGTTTTGCTTACTTTGCCATTTTTCCAAAAGGATTTGAGTTTGAAAAGGACCAGCTTATCCAACTCTGGATGGCAGAAGGTTTTCTCGGTCCATCTCAAGAGACCACTTTGATGGAAGACGTTGGGAACAAGGCTTTTCAACTTTTGTTGCAAAATTCCTTGCTGCAAGATGTTAAGCTAGATGATCTGAACAAAATAACATACTGTAAGATGCATGATCTTGTGCATGATTTGGCTGGAGATATTTTAAAGTCTAAACTATTTGATCAAAAGAGTATTGGAGGAGAAAATCTTTCTCAAGTCCGATACTTTGGATGGGACTCACCAAGTGATCAAATAGATAAGATAAACGAGTCAGGACGTTTGTGCACATTGTTCTGGAAAAGAATTATATCTGAAGAACTGTTGTTGAGCTTTCAGTTCTTGAGAGTTTTAAATTTGTCCAGGTCAGGCATCAAGGTATTGTCAGCCAAAATCGGCAAGCTAATATACTTGAGATATCTTGATCTCTCCAGCACTGAGATCAAAGCCTTGCCCAACTCCATTTGCAAGCTCTACAATTTGCAAACATTTAGAATCAATAGTTGTTACTCACTTAGGAAGCTCCCAGAAGAAATGGCAAATATGATAAGTTTGAGACACATATATTGCGACTTTCATTACCAAACATATCTTGATTTTGCGGGTTGGAGATCAAGGTTTATAGGTAATTATCAGTTTCAGATGCCACTTAATATGGGGCAATTGACTAGTCTTCAGACGCTACAGTTTTTCTATGTAGGTTTAAAGACAGGTCGTCGAATAGAAGAATTAGGTCATTTGAAAAACCTTAGAGGTGAATTGACGATCAAACATCTCCAATTAGTCGTAAATAAAGAAGAGGCTCGAACAGCATATTTACAGGAGAAACCAAATATCTACAAGCTGACATATTTATGGACCCATGATGAATCAGAAGGCTGTGAGATCAATGATGAGCATGTGTTGGATGGTCTTCAACCGCATCCGAACTTGAAAACCTTAGCAGTAGTGGACTATTTTGGGACTAAATTTCCTTCATGGTTCAGTGAAGGATTGCTACCAAATTTGGTCAAGTTGAAATTAAGTCGCTGCAAAAGGTGCAAAGAAATTCCATCGCTTGGCCAACTGAAGTTCCTTCGGCATCTTGAGCTGATAGGATTCCTTGAATTGGAATGCCTTGGATCTACATTTTATGGCGTTGATGTTAATGATAAAGGATCAAGCAGCAATAATGGCAATATCCAAGTGTTCCCATCACTGAAAGAACTAGTATTGCAGAATATGAATAGCCTTATTGAGTGGAAGGGTGATGAAGTTGGAGTTAGAATGTTTTCTGCGCTTGAGAAGTTGAGGATCACAAAGTGTCCACTGTTAAAAGGTACCCCACGTCAATTGGAAATCCTCCGTGAATTAAGCATTGAAGGAGTTGACAGTGAAATGCCATTGTTGACCTTGTGTAGCAACTTGACATCTCTGGTAAAGCTTATTGTGACTGATGTGGAAGAGCTCACTTGTCTTCCTGATGAGATGCTACGCAATAACGTTTCTCTTCAACAACTATTGGTCACACACTGCAGAGAGTTTCGTGAATTGCCCCAAAGCCTGTACGATCTCCATTCTCTTGAGAGCTTACAAATACTCTTCTGCACCAATTTCAGGTCTCTTCCTGTTCCCAGTGTAGAGAATCGTTTGACTTCCCTCCAAAGTCTTGATTTATCACTGTGTATTGGATTGACCAGTTTACCAAGTGGAATGCTAGAGCATTGTCGGTCTCTACAGACTTTGAAGCTCAGCTGTTGTCGCAACTTAGTTTCCTTGCCTTTACACGTATGGGAAATGCCTTCAATTTCATATTTGGGTTTATCAAATTGTCCCAAATTGGTTAGTGTACCCACAGGGGGCCTTCACAACCTCACAGGGTTAAAGGTATTGGAAATTGGTCCTTTCTCAGATACGGTGGATTTTGAGGCATTCCAATTGATTTTTAATGGCATTCAACAGCTATCATCCCTTCGTACATTGGAGGTGTATGGATGGATGCGTTGGGATTCTCTGCCTTATCAACTTATGCAACTCTCTGCCCTAACAAAGATCTGTATATATGATTTTGGAATTGAGGCTCTTCCTCATAGACTTGACAACCTTACTTCTCTTGTAACATTACATCTAATGGGGTGCCAAAGGCTACAACATGTGGACTTCTCAAATATCATGCCCAAATTACGGTATCTGGAGATCCATTATTGTCTATTGTTAGAAGGTCTGTCGGATGGGCTTGGCAACCTTGTTTCTTTGGAAGAATTAAGTTTACGTAACTGCGAGAAACTACAACATCTGCCATCCAGATATGCCATGCTGCGCCTCACTAAATTATGGAAACTTCTAATTAAAGGCTGCCCGCAGTTAGAAGAAAGTTGCACCAATCAAAGTAGCCCAAACTCCCAGTGGTCCAACATTTCCCATATTCCAAAAATTGAAGTAGGTGGGAGGATAATTCAGAAGTTACGTAAGTCTCTGTTTCTCTTTCACTTTCTGATCAGTAAAATTTTGAAATATTATTTTCCCCGAGTTCCAGAAAAGAAGTTATGTATGTAG

>M6-PGSC0003DMG400011108 M6 LRR-RLP

ATGTTTCCAAAAGATTTTGAGTTTGAGAAGGACCAACTAATCCAACTCTGGATGGCAGAAGGGTTTCTTCGTCCATGTCAAGAGACCACTGTGATGGAAGACGCCGGGAACAAGTTTTTCCAACACTTGTTGCGAAATTCCTTGCTGCAAGATGTCAAGCTAGATGAGCACAACAATATAACACACTGTAAGATGCATGATCTTGTGCATGATTTGGCCAGAGATATATTGAAATCTAAATTATTTGATCCAAAGAGAGTTGGAGGAGAAAATCTTTCTCAAGTCCGATACTTTGGATGGGACTCACCAAGTGATCAAATAGAGAGGATAAATGAGCCAGGACGTTTGTGCACATTGTTCTGGAAAAGAAATCTATCTGAAGATATGTTGTTGCGCTTTCAGTTCTTGAGAGTTTTAAATTTGTCCAGGTCAGACATCAAGGAGTTGTCAGCCTCAATCAGCAAGCTAATATACTTGAGATATCTTGATCTCTCCTACAATAAGATCAGAGCCTTGCCCAGCTCCATTTGCAAGCTCTACAATTTGCAAACTTTTAGAGTCAATAACTGTTTTTCACTCAGGAAGCTTCCAGTAGAAATGGCAAATATGATAAGTTTGAGACACATATATTGCAAGTATCATTACCAATCATGTACAGGTAAGGATCGGTTTCAGATGCCACTTAATATGGAGCAATTGACTAGTCTTCAGACGCTACAAGTTTTCTATGTAGGTTTAAAGAAAGGTAATCGAATAGAAGAATTAGGTTGTTTAAAAAAACTTAGAGGTGAATTGACGATCGAACGTCTTCAATTAGTCGGAAATAAAGAAGAGGCTCGAACAGCATATTTACAGGAGAAACCAAATATCTACAAGCTGGTGTATTCATGGTCCCGTGATGAATCAGAAGGCTGTGAGATCAATCATGAGCATGTGCTGGATGGTCTTCAACCACATCCTAACTTAAAAACCTTAGCAGTTGTGAACTATTTAGGAACTAAATTTCCTTCATGGTTCAGTGAAGAATTGCTACCAAATTTGGTCATGTTGAAATTAAGTGGTTGCAAAAGGTGCAAAGAAATTCCATCGCTTGGCCAACTGAAGTTCCTTCAGCATCTTGACCTGGTAGGATTCCTTGAATTGGAATGCATCGGACCTACATTTTATGGTGTTGAGATAATGGATAATGGATCAAGCAGCAATAATGGCAATATCCAAGTGTTCCCGTCATTGAAAGAACTAGTATTGAAGAATATGGCTAGCCTTATTGAGTGGAAGGGAGTTGAAGTTGGAGTAAGCTTGAGAAGTTGAGGATTAGTGACTGTCCCTTGTTAGAAAGTACTCCGAATCAATTTGAAATCCTATGTGAATTAGTGATTGTTGGAGTTGGCAGTGAAATGCCATTGTTGAACTTGTGCAGCAACTTGACATCTCTTGTGAATCTTGTTGTTAGTAACGTGAAAGAGCTCATTTTTCTTCCCGATGAGATGCTACGCAACAATGTTTCTCTTCAACACCTATTGGTCTCACTCTGCGGAGAGTTTCGTGAATTGCCACAAAGCTTATACAATCTCCGTTCTCTTAAGAGCTTAGAGATTATCAGCTGCACCAATTTCAGTTCTTTTCCTGTTCCCAGTGGAGAGAACTATTTGACTTCCCTCCAAAATTTTGAGTTATCCAATTGTAATGGATTGACCAGTTTACCAAGTGGAATGCTAGAGCAATGCCGGTCTCTAGAGATTTTGACGGTCAGTTACTGTAACAACTTGGTTTCCTTGCCTTTACATGTGTGGGAAATGCCTTCTCTTTCGTATTTGTGTTTATCAGAATGTCCCAAATTGGTTAGTGTACCCGCAGGGGGTCTTCACCACCTCACTGGGTTACAGAATTTGGAAATTGGTCCTTTCTCAGAGACGGTGAATTTTGAGGCATTCCAATTGATTTTTAATGGCATTCAGCAGCTATCGTCCCTTCGTACATTGGAGGTATACGGACGTACGGATTGGGATTCTCTGCCCCATCAGCTTATGCAACTCTCTGCCCTAACAGAGATCGGAATAAATGATTTCGGAATCGAGGCTCTTCCTCATAGACTTGGCAACCTTACTTCTCTTGTAACATTAGATCTGGTGAGATGCAAACGGCTACAACATGTGGACTTCTCAGATGCCATGCCCAAATTACGGTATCTGAAGATCCGTGATTGTCCATTGTTAGAAACTTTGTCGGATGGGCTCGGCAACCTTGTTTCTTTGGAAGAGATAAGTTTATGTAACTGCGAAAAACTGGAGCATCTGCCATGCAACTCCTCACTAAATTATGGAACCTGCGAATTAGTGTCTGCCCACAGTTAG

>M6-PGSC0003DMG400011116 M6 LRR-RLP

ATGTTTCCTGGGCTTGAGAAGTTGACGATTACAGAGTGTTCACTGTTAAAAAGTGCTCCAACTCAATTGGAAACCCTGCGTGAATTAGAAATTGAAAGAGTTGACAGTGAAATGCCATTGTTGAACTTGTGCAGCAACTTAACATGTCTTGTGGATCTCGATGTCAGTGATGTGAAAGAGCTCACTTGTCTACCTGATGAGATGCTACACAAGAACGTTTCTCTTCAACGCCTGTGGATCTCAGATTGCAGAGAGTTTTGTGAATTGCCACAGAGCTTGTACAATCTCCATTCCCTTAAGAGCTTACGAATATCCTTCTTCCCCAATTTCAGTTCTTTTCCTGTTCCCAGTGGAGAGAATTATTTAACTTCCCTCTGAAGGCTTCACTTACTTTATTGTGATGGATTGACCAGTTTACCAAATGGAATGCTAGAACAATGCCCGTCTCTAAAGATTTTGAGGGTCAACAACTGTAAGAACTTGGTTTCCTTCCCTTTACATGTGTGGGAAGTACCTTCACTTTCCATTTTGGATATGTCAAATTGCCCCAAATTGATTAGTCTACCCACAGGAGGCCTTCACCATCTCACTGGGTTATGACATTTGAAAATTGGTCCTTTCTCAGAGATTGTGGATTTTGAGGCATTCCAATTGATTTTTAATGGCATTCAGCCGCTATTGTCCCTTCGTACGTTGTGGGTGTACGGACATGGGCACTGGGATTCTCTGCCCTATCAGCTTATGTAACTCTCTTCCCTAATTGAGATCAGATTAAGTGATTTCGGTATCGAGGATCTTCCTCATAGTCTTTGCAATCTTACTTCTCTTGAAACATTACATCTAGTGGGGTGCAAACGGCTACAACATGTGGACTTCAGAGATGCCATGCCCAAATTACATTTTCTTGAGATCTGTGGTTGTCCATTGTTAGAAGCTCTGTCGGATGGGCTCAGCAACCTTGTTTGTTTGGAAACGTTAATTTTTTGGAACTGCGAAATACTGGAGTGTCTGCCATCCAGAGATGTCATGCGAAGCCTCACGAAATTACAGTTCCTGAAAATTAAAGACTGCCCACAGTTAGAAGAAAGTTGCACCAATTGGAGTAGCCCAAACTCCCAGTGGTCCAACATTTCCCATAATCCAAAAATTAAAGTAGGTGGGAGGATGATTCAGAACTTAA

>M6-PGSC0003DMG400011209 M6 LRR-RLP

CGACTACTAGGCGGCCACCATACTATGACTATAAATGACAAGAGATGTTCCTTCCTACTTCATTCACAAACAAGCAAATATGAAGAACAATTCACTTCAACTTATCTTTCTCATATTGTTATCTCAACTTTTCTATTTCACTTGCATAAACCATCACTTTGTTTCCTCTACTAACACCTCTTGCATTGAAACAGAGAAAAATGCACTTCTAAACCTTAAAGAAAATCTTACAGATCCTTCAGGCAAGCTATCTTCTTGGATTGATCAACAAGATTGTTGCCAATGGCTTGGTGTAACATGTAACAACAAGACCGGGCACGTCGTAAAGATTGATCTTAGGAACAAATACTCAACAGAACATGAATTGAGAGGTGTGATTAATCCATCTTTGCTCAAGTTACAACAATTGAGATACTTAGACTTAAGCATGAACAATTTTGAAGGAATCAAAGTACCTGAATTCATCGGTCGCCTCAATGAACTAAGGTATCTCAATCTATCTGGTGCATCTTTTACTGGTTTAGTCTCTTCATTCCTTGAAAACCTCTCAAATTTGCAAGTTCTTGATTTAGGAGGCTTAGATCTTAGTGAAGATGCTAATTCTTGGCTTCAAATAATCAATGATCATGTTCCTTCTCTGTTGGAGTTGCGTTTGCCTCGATGTCAACTTTTAAACATTCCTTCTTCTTTTCCATTACTAAATCTTACTTCCCTTTTGGTACTTGATCTTTCCAACAATGCTTTTAACTCATGTACATTTCCTCAATGGATATTCAAACTTAGTAGCCTTGTGTATCTGGACTTAAGCTCCAACAATATGTGTAGTGAACTTGTTGATGAGTTTGCTAAGCTCACTTTCCTTGAACATCTTGATCTGTCTTCGAACCACGGTATCAATGGGGCACTCAAGAGGAGCTTGGGGAAACTATGTAATTTGAAGAGTTTGATCCTGTCTTACAACAACATCAGTGGAGACATAATCGAGTTCATCGATGCGTTATCTGAATGCCAAAGCAATAGCTTGGAGACATTGGAGTTGAGTTTCAATAAATTGACAGGAAATCTTCCTGATGCATTAGGCAACTTGAGGAAACTGAAACATCTTCAGCTGAGGTATAATTCACTGACAGGCACAATACCTGAAAGTATAGGGAACTTATTATCTTTGGAGACATTCTACCTCACATCAAACAAAATGAGTGGGAACATCACCACGAGTATCGGACAGCTCATGTCGTTGGTCTCGTTGGATATCTCGGAGAATATGTGGGATGGTATTGTCATTGAAGCTCATTTGGTGAATCTTTCCAACTTGCAAGAGTTTTCAGTTGGGATGAAACTTAGTAAAAATATTACCTTGGCATTCGATATCAGCCCGAATTGGAATCCTTCATTCAGGCTGACATTCTTGACTATTCAGTCATGCCAATTAGGTCCTGAATTCCCACATTGGCTGAAGGATCAGAATGAGCTCACCAGCCTAATACTCAACACCGCGGGGATTTCTGATGCGGTACCTGATTGGTTTGTAGAGTTGGATTTGAAGCTTGATAATCTAGACATGGCATACAACAACTTGACAGGGAAAGTTCCAAACAAATTCAAGTTCAACTTTCAAGCCAATGTGGATTTAAGCACTAATCGCTTCGAGGGACCTCTCCCACTATGGTCTTCTAATGTTACAACATTGTATCTGAGGGATAACTTGTTTTCAGGACCTATTCCTCTCAACATTTGTGAAGCACTTCTCAATATAGAAGACTTGGATATCTCCAGGAACAATCTAAATGGCACAATTCCCTTATGCATAGGTGATATGAATCAGTTAACTACCTTAGCCCTCGATAACAACCAACTTATCGGCCAAGTTCCTGATTTCTGGGGTAAGTTACCATATCTTTACTGGATAGATATGTCAGAGAATCGCCTATCCGGACAAATTCCACACTCACTAGGCTCTCTTGCTTATCTGATGTTCTTAAGACTCTCAGGCAACAATCTCTCTGGAGAACTGCCTTTGAGTTTGAGAAATTGCACGAAAATGATCAACATTGATCTTAGTAACAATCAGTTGTCAGGGCTAATCCCAGCTTGGTTAGCGGAAACAATGAGATCACTGTTAATCCTTAGTGTAAGGAACAACAGGTTTTTCGGTCACATTCCTTTAAAAATCTGTAGCCTTTCAGGTCTTCACATACTAGACCTTTCAGGAAACAACTTATCCGGTTCTGTCCCGTCCTGTTTTGGAAACTTGGAAGGGTTCAAGGTTGAGCTAACTGATGTAGAAGCTAAACAATATCAAGGTACATTGAAGCTTGAGGCAAAAGGAAGAACACTTTCTTATGATAAAATCCTTTATCTAGTAAACAGTATCGACCTCTCGAGAAACAGTTTGTCTGGTGAAATCCCGGAAGAGTTAACAAACTTACACAAACTTGGAACCTTGAACTTGTCTATGAACCATTTGACAGGACATATACCAACAGATATCGGAAAGTTACGTTGGGTTGAAACGTTAGATCTTTCAATAAATCAGTTATCAGGTACTATCCCACCAAGCTTAGCTACATTAGACTTTTTAAGTCATTTGAACTTGTCATACAATAAGTTGACAGGAAGAATTCCTACTAGTACTCAGTTTCAAACCAAAGTTGATCCAACAATTTTCCAAGGAAATGTTGCACTTTGTGGTCCTCCTTTGGAACAATGTGTTGGTGATAGGACTACGACGACGATAACATCACAAAGTGGAGGAAATGATGAAGGGGAAACAGATGATGAAGATCAACTTGAGAAAGTATGGTTCTTTGCTGTTGTTGGTTTGGGATATATAGTTGGTTTTTGGGCATTTTTTGGCAGTTTGATCTTCAAGAAAAGATGGAGAATTGCTTATTGCAGATTCATTGAGAGGTGTATTTTGGAGTTCTATGAAATGTCTTGTAGTTCTGTGAAAAAATTGTTAGGAATCTAATAAGCTATGTTGCTCAGACTCTTCAAAAATGTCGAGGGGTGTGTGTGGGATCCTCCAAATATAGTATATTTGTGAAAGATACGACAACATTTTTAGAGAGTCTGAGAAACTTAGCTAATAAGTGGAACTATAGGAGTTTGTCTCTGCAGTTCTGTTATATGTTATGTTTTAGTTTCAATATAGCACATATTACTACTTTTTCTATATGTAAAATAATGCTACCTAAGTATTGACACTAATAGAAATGCCAGTGTTTTGGGTAGAGTTACGACATGTGAAGAGTGGTCAATTGAACACTTTTACGTAGGTCAAAATAATACTTCTTATATATGTATACATGTTCACAAGACTATTGTAGTACATATAATTAGGAAGTTGTTTAAATAGTATCTAAGACTATTCTTATACTATGCATGCATCTACTTGTATA

>M6-PGSC0003DMG400011353 M6 LRR-RLP

AATAAAACCTTATCTTCTCTGTAGAACACTCACACACACTCTCTCTCTCCTCTTTCTCTCTCTAAAACACAGCAAATCTAAGCTCGTTAATGTCGATTCAGATCTGAGAGAAGAAGAAGAAGAAGGAAGAAAAAGGACTATTGTAAATCTCGAGTTCACTGTGAAATTACACTTTTGCCCTTCCCCCTCCATTACAGTACAACAATGGCGTACGCATTCCCGGAGGAAGTACTAGAACACGTCTTCTCCTTCCTCACCTCCGACAAGGACCGTAACGCAGTATCACTAGTCTGCAAATCATGGTACGAGATCGAACGGTGGTGTAGGAAGAGAATATTCGTTGGAAACTGCTATGCTGTTAGCCCTAGGATCATGATCCGACGGTTTCCGGAAGTCAGATCTGTTGAGCTTAAAGGGAAGCCTCATTTTGCTGACTTTAATCTTGTACCGGAAGGTTGGGGAGCTTATGTGTATCCTTGGATTTTGGCTATGTCTAAATCTTATCCATGGCTTGAAGAAATTAAGCTTAAGAGGATGGTAATTACTGATGAGTCGTTGGAGTTGATTTCTAAGTCGTTTAAGAACTTTAAAGTGTTGGTTTTGTCTTCATGTGATGGGTTTACTACTGATGGACTTGCTGCTATTGCTGCTAACTGCAGGTGAAATTTTTTTTCTTGCTTGACATTTCTCTTCTGTTTTGCTCACTGTAAAGTGGCTAGTGGTGAGAGAAGTCTTAATCTGTAAAGTAATTAGATCTAATTTTGGGTATTAGTTTGGACAGTGACTTTTATTGTTGTTAAAGCTTAAGCTTTTGGATGAATGCAGTATGCATGCATTTTGGGGGTTTGTTATAATCCCTTGCTAAACACGAGAATGAAGAAGCACTTTTCTTGCAGCTGTTTGTTCTGTTTATAGGATATGTAATGTATTAGTGGAAACGAAGTATCTACTGTAGATGAATTCGTAGTGTGTGAACTTGAAGAACTGACATTTCTTCAAGTGATTATGGTATTATTGGATACGTCTTGCTGCTACATTGCATCAGTCTGGTTCATCTCACTTTAATACTTCAAAGATTAGAGCTTTTTGTTTGTTTCTTTTAGCTTCTTTTTTTTGGGGTGAACTTCTGGGAATTAATTTTTCTTGATTGAATGAGCTGATGTCAATAATTTCTTGTACTTCCTTCCCCAGATTTCTGCATTTGCATTCTATCTTTACAGATCTCTAAGGCCTTATAACAGGAAATGCTAACTTGTGGGTTTTCTTAATTGTTATAGGATTTAAACAGTTGGACTTCCTTTTAGCATTAATTTCTTGTGGTATACACTGTGTAAACATGCCCTTTAGCTTTGCTTTGTTTTGAAGTTGTTGGAGATTTGAATGTTTTCTTAATTATCTGGAGATTTCCATACGTTTGTTTCTTGGTTCTCTGGTGGGATTAATTTGTTTTATCAGGAGGGAGTTTTGTAATGGAGTATAGTTGTAGTTCAATGTTGAAATGGCAAGTGGGTTAGGAATAGGAAATGGTGATTAATTGAAATTCCAATTGGAAGATGTGTCACTCTGAGGCAAACATGATTCTAAATCTTTCTTATTTATCTTGTTCATATCGTCTGTCATTTATTATCATTTGTTGTTTTCTGTTGAGTCTGTTGATTTGCTTCAATTTATCACAGAATGATTGTGTAGATGGGTATTGTCATCTAACTCCCGTTTTGCTGCAGGAATCTGAGAAAACTAGACTTGGGAGAGAGTGAAGTGGAGGACCTTAGTGGGCATTGGTTAAGCCATTTCCCTGATAACTGCACGTCACTTGTTTCACTTAACATTGCCTGTTTGGCTTCTGAGGTCAGTCTGTTAGCTTTGGAGCGCCTGGTTACTCGCTCTCCTAATTTGAGAACTCTTCGGATAAATCGTGCTGTGCCCCTTGAGAGGCTTCCAAACTTGCTCCGCCGCACTTCCCAGTTGGTCGAATTTGGTACAGGTGTTTTCTCTGCTGATGTGCGATCTGATTTTTTCTCAAACCTGACAGAAGCTTTTTCGAGCTGCAAGCAACTTAAATGCCTGTCTGGGTTTTGGGATGTTGTACCAGCTTACCTTCCAGCTTTATATCCAGTGTGCTCAAGACTCACCTCCTTGAATTTGAGCTATGCTACCTGTCAGAACCCTGAGCTTGGCAAGCTCATAAGTCAGTGTCACAATTTGCAGCGTTTGTGGGTATGTTCTAAATGTTTTGAAGTTTTAATTGATGTTCTTGACACTGCAACTGCAAACATGTTTTTTTTTTTTTTTTTTAATAAAGAAGAAGAGGCAATGAGATTGTAACATCATACTTGCTGTGATGTCATTTAGCATATATTCTTAGTGAAGAATAAGAAAAGACCTAACTGCTACAGATCAACGAATCCACCACAACTTAATCCCAATTTAGTTGGTCTCAGCTTTATGGGTCCTCTTAAACCATTTTGCTATATTCAGGTCATTTTCATTCCGATGCTAAATTATTTGTCTTTACGCAATAATTGCCAAGGATAGGCAAAAAGTAATTCATTCTGTAGGTAAACTTCACCTATGCTGGTTAGGAAGATTCCAGGCTACATATATTTGATATTATATTGCTAAGCAGCTCACTTTTCTTGGAAGTTGAAGCCAAGTGTTACATGCTTCATATTGTGATCTGGGTTTCAGTTTTTTAATACTTGTGCCTACCCAATGCACCTAAAAATCCATCTAAATGTGATGTTTCTATGCTGAAAAATTGTATCTATTCTTCCCTAGTGGCTTCCCCAAACTGGTTCTGTAATATAAACTCCAGTATCATGTTTAAGTGCTGAATTGCTTTTCTTGCAATATATGTGACTGCAGGTGCTAGACTACATCGAAGATACTGGTCTTGAGGAACTTGCTGCCAATTGTAAGGACCTTCAAGAGCTGCGGGTGTTTCCTTCTGACCCATTTGCTGCAGAACCAAATACAACCTTGACGGAGCAAGGCCTTGTAGCTGTCTCAGATGGCTGCCCAAAGCTCCAGTCTGTTTTATACTTCTGCCGCCAAATGACTAATGCAGCTTTAGTTACTATTGCTAGGAATCGACCTAACATGATCCGTTTTCGCTTGTGTATCATTGAGCCTCGAACTCCTGATTATTTAACCCTTGGTTCATTTGATACTGGTTTTGGGGCCATAGTAGAGAACTGTAAGGAATTGCGGCGCCTTTCACTTGCTGGCCTCCTTACCGATCGTGTCTTCGAATACATTGGCGCCCATGCTAAAAAGTTGGAGATGCTTTCCATAGCTTTTGCTGGTGATAGCGATTTGGGACTCCACCATGTTCTCTCTGGTTGTGATAGCCTTCGGAAGTTGGAGATCAGAGATTGTCCGTTTGGAGACAAGGCTCTTTTGGCTAATGCTGCAAAGCTCGAGACAATGCGATCCCTTTGGATGTCTTCTTGTTCAGTAAGTTTTGAAGCATGTAAGATGCTAGCTCAGAAGATGCCTAGGCTTAATGTGGAAGTCATAGATGAGAGGGGTCCTCCAGATACAAGGCCAGAAAGCTGTCCTGTGGAGAAACTTTACATATATCGAACTGTGGCTGGACGAAGATTTGACACACCTGGTTATGTTTGGACAATGGATGAAGATGCAGCTGTAAGGTTGACCTGAGCCACACATCAAATGGAAAATTCCAGCAAGACTTCACATGGAAAGATGGACAGGTATTAGCGCTATGTTTATCCTCCTGCATTTGTTTGTCACGCCTTATGTAGTTGCAGTGCCAGTTGAGGGTTGGAACTTCACTGCTTTGAAGTAGAGGCTCCAGAGCTCATACGCCATCAATTTTATTCTTGGCTGAGAGCTACTTTACAGCAGGTAGTTTTTAATAAAGAGTTTGGACTCTGAGTTTCTGTGTATCATCTAGAGTAACAATTATTCGTTTTCTTTGGGTGGGATTTTGCTTGGGGGGTTTGGGGGCATATTTTTGCCATGTATGTGCTATTTCCAGACTTTGCTTCATTATGTAGTTCACTTTATTTATTCTATTGACTTATTCTCTATGAAATGGAGTGTCT

>M6-PGSC0003DMG400011513 M6 LRR-RLP

ATAAATAACAAAACAAACACAAAATGAAAAGATAACTCCTCTTTCTTTCCAGTAAGATCAGAACCATAAACTTCTCCGATGGCCGGCGACGACTCTCATCTGCAAAATGACGTCGTTTCAGTCCACTGTATAAACGACAGTCTCGGCGACGACGAGCTTCGGGCCGTACTGTCGAGGCTAGGAGACGACAAAGACAAGGAAGTTTTCGGGCTGGTATGTAAGCGGTGGCTTCGTATTCAAAGTACTGAAAGGAAGAAGCTTTGTGCTCGAGCCGGGCCTCACATGCTCAGAAGAATCGCGGCCCGATTCACTCGATTACATGAGCTTGATCTATCTCAATCGGTTTCTAGATCTTTCTATCCTGGTGTTACTGATTCTGATCTTTCAGTTATTGCTACTGCTTTCTCTTGCTTACGAATTCTCAATCTTCAGAACTGTAAAGGTCCGTTTTTGTATTTTTTAAAAGTTATATAGTTTGAAATTTGTTGAATTTAAATGTTCATTGCTGAAGTGAACCTGAGCTACTGCAGGAGTGGCCAAATGCACCAATGAGTTGTAGTATGTGTAGTGATATTTCACATTAATTACTGATTTAAAATCTTGAATTCGTCTTTGTGTGGGGTGGTGGAGCTGGGATTTTCACTAAGAGGGTTTTGAAATACATAAAACATAATTTTAACCATGTATAACCAGTTCGGATGAACCCCTGGGCCCTTACTAGCTCAGCCCCAAAATATGTGGGGTGATGGGTGTATTGTTACCAACTAGGACATAAAGGAAATTTTGAGATTTTGACTAAAATCTATTGTTTGAAATGTTAAAGGTTTTGACCCGATATGGAGAAGACTGGCTCTTCCTCTTTTCTATCTTGTAAATAATGGTTTTTATGATAAAAAGGGCGTAATGGGGTTTATGGAGGATTTTAGGTGAATTTATGGTTGGTTTTTTTGAAGATTTGTAGGATTTTGTGTTTCTGTGTAAAGGTTATGTCTTTTTTGCTATGTGGTTTTTGAAGTGGTAGCAAAAGGGAGGATAGTGTAGTTATATCATATGTATTCTAATTTTCAAGAGAAAGTCTAACTTTGTTGCGAATCTAAGGCTCAGATTCTTCCTTGACTTAACTTGCGCGGTTTTCTGGTCCAAATTTACAGGCATAACAGACAAAGGATTGACAGCGATTGGATCAAGTCTCTCTTCTTTACAGTCCTTAGATGTATCCTACTGCAGAAAGATCACAGACAAGGGGCTTTCAGCTGTTGCTGAGGGGTGTCATGACTTGAGAACGTTGCATCTTGCTGGCTGTAGATTTGTCTCAGATTCATTAATGAAAGCTCTCTCCAAGAATTGCCATTATGTAGAAGACCTAGGTTTGCAAGGATGCACCAACATTACCAACTCAGGTCTCTCTGTTTTGGTTGAAGGTTGCCGAAGGATAAAGTATTTGGACATTAACAAATGCAGCAATATTGGTGATATTGGGATATCTAGCGTGTCTGAAGCTTGTTCATTAACACTTAGGGCATTGAAGTTGTTGGATTGCTACAAAGTTGGTGATGACTCCATCTTATCCTTGGCCAACTAATGCAAAAATCTTGAGACACTCGTTATTGGTGGTTGCCGTAACATATCAGATGAGCCTATGAAGACACTGGCAGCTTCTTGTAGTAATAGTCTGAGGAAACTGAGGATGGATTGGTGTTTGAATATTACTGACTCTTCATTGGACTGCATCATTTCCAAATGCAGAGAGCTTGAGGTTCTTGACATTGGCTGCTGTGAGGAGTTAACTGATGCTGCATTTCAACAATTAGGAAGTGAAAACTTTATGTTGGGGATGAAGATTTTGAAGGTCAGTAACTGTCCAAAGATAACTGTTGAAGGAATTAAGAAGCTTATGAAGTCATGTGAATATCTTGAATATCTAGACGTACGGTCTTGTCCCCTAATCACTAAAGCAGGGTGTGAGGAGGCTGGAATTCAGTTTCCTGAATCCTGCAAAATAAACTTCACTGGAAGCTTAGCAGAGCCAGATGTGTTACTTTGAGCAGTACTTGATTTAGCAATGTCTTTGAAGGACTGAAGGGGATCATGCAGTGAAATCATCACATTCTTTGCCTAAGACCTAAAATATGACTTTGGAAGCTCCTGCTGTCAGTCCCTTGTTCCCATTAATTTGCCTTTTGTTAAGCATCAAGTTTTTTGTTGTAGATTATTTCCCTGCTTTTCAGTTTGTTTTTTACCCCCTGCTTTTCAGTTATTTGATATCAAACTCCTGTATATCATCCTTGTAAACTACTAGTAGGCACCATTGTTTGCTGTAGGTGTTTCCAGATTGTGAATGAAATGAGTCCATTGCTGCTAGCTTGAATGTGAACTTATACCAACAATGATGGTTTGATTGATTACATTTCTTGTACGAC

>M6-PGSC0003DMG400011708 M6 LRR-RLP

TGTTGCATGTGATTTTGTCTTTAAGTCAGTAAAGTTGGTTCTTTGTATATGTGGTTGGGAAAAATTTATGTAGGAGATATTTATCTCAAGTTCACATTTCTACTTTTTTGTGGGGCCATTGAATGGTAAGTGTTTGGATTGGAATCTCCCTAGTTTGATTTCATCTTCTCTATCATTTCAAAAAACTCAATATGGAGGATGAGACGGAAACTGAAAAATCTCCGATTACTACTCGTGTGATACTTCTTTCCCAAAAACAGTCTTATGGAATACGAGTAGAGATTGTTGTACATGGGATGGATTACAACTTGATTAGAGAACATTGGACACAACACTTCATAACTTCACAAAAACCTGTGAATTTCCCAAAAACAGTCTTATGGAATGCGAGTAGAGATTGCTGTACTTTCTCTCCCTGATGTCAACATCTCATCTCCAATACCTATGAATTTTTCTTCCTCCTTAAGGTACGTGGATCTTGAATATACTAATATACAAATTAGGTATTTGAATTTCACCGGTCATATTTCTTTCACAATCTCTAAATTGAAGCAGCTCACAAGTTTAGATCTTTCATCTAACTCCTTTTCAGGTTAAATGCTCGACGTTTTCTCTAACCTCCAAGAGCTTACATCTTTCTGACAACATGAATTATGCGGTTCCTAAACCTTTCGCATAATTCATTAACAAGCCACCTAGCTACCACAAGTTCATTACCACAATCTAGAGTATCTTGATCTTAAATTTAACTTCTTTCAGGGTCCACTACCTTCATCCATTTGTAACATGAGCTAGCTTGAATTTCTAGATTTATCACACAACAACTTCAGTAACTCGATTACAAGTTGCTTGGGAAGCATGGCTACTCTAAGGTTGTTGGACTTAAGAAGGAACAATTTCACAGGGAGTCTTCCACCATTATGTGCGCAGAGCACTTCATTGAGTACCATTTTCTTG

>M6-PGSC0003DMG400011864 M6 LRR-RLP

AATAATCGCTAGATGTGTTGTGTGATTCATTTTGAGTACTGCTATTTGATTTCTCTTTGTCATACTCTGCCAAGCTTGTTTGCCAATTGATGGCTTATCAGATTATAGTCTTTCATGAATCTTTGCTTGTGTCATGAACTCAGTTTATTGAATTGTGATGTATCATGAATGCTAAATATGCACTGTATGTTATAGAACAAATGTAGCTTGGTGATGATAATTTATGCGGCTTTGGAAGTGTTAACCAATGATTTAGAAGGTAATGTGCAGATTAAGCTGGCCAAGAAAGTATCATTATAAATAAAATGCTAAGCCGTTTAGGAATTTCTCACTGATATCTTAACGAAGTCTATGAGAATTGTGACATCCATGACAGAGGGCACACCAACAAGGGTGGGTCAAGTAATGGCAGGCATTTAGCTTGGCAAGCTTCCGAAGAAGGGGTGCACATGACACCTTAAGGGAATTCACATCAGAATGGGAAAGAAGAACTTCGTGAGAATATTTTGCACAAGATTATTTTTCACTCTCTATCCTCCGTAAGGGAGGAAAATTTCATGAGTTTGCGTTGCTCAACAAGTGCTGACATGTCTTAGGTTCAAAACTAGACGATTTCTTCCCATCTGTATAAGCTTTAGTAGGCAAAGTTACCTGGTACGTGTGCTGGTGGGAGGGAGCAGGTCATTGGTGGGTATGCACTAGCTTGCCCAGGCACCAACGTCATAAAAAGGAAAAAGCTAGTGCTGATGTGTCGTCGAACTCAATGATGATTTTTTTTGAAATTGAACGGAAAATGACATTTCACAGAACATTAAATACTTAATATTATTAAACATTTGCTAAGATTCTTACATTGTCTTAGTTAAATTTATCCCGTGTATTGTGCTTTCTCTACTTGGGAGATATACAAGTTCTGAAAAATGAGCCTGCTTATGCACTTGGCATGCTTCTGACCTTGAGTTTGTTGAAAAGAAAAGATATTGCAGAGTCCAGCTTTTGAATCCTAATGAACACCTCTGCTTGACATTCCGATGCATCTTCAGATGAATAATTGAATGTTCTCTTATCATACTCATGCTGGAATACTCTAAGTTAGTGGAGGTTTTATGCTATAGAACATCTATCCATATTCTCTTGATATCTGTGCACCACTCAAGGATAGCATGACCAAAAAAATTGTGTTTTTAAGCACCTTGGTGCTTTGTCTGATGTCCAGGTCTCAATTACATTTCATGACCATCACAACATGCTCCTTTTGTCATTGTATAATATGGGAAAGTTCTTGTTCATGTGATTCGAAGTCCTTTCTTGACTTATGAAGTTACCACGACTTGGATATGAGCTTAAATAGAGGGACATAGACAATGAAAATTCATATAGCTGGCCCCAACTGGCTTGAGATTAAGGTGTGGTAGTAGTTGTTGGTGCTGTAGTTTCCCACTGCTTTGGTACAGAACTTAAATTCCAGATGTACCAAGTTTCCAGTGACTGACTGTATTTATGAAGTTGTTACATGTAATCTGTTTCTGTGTTCTGTTATTACATTCCATCTCTTTTTCTTTCTCCGTTTTTGTTTTGTTTTGTCTAGGGTTTCTTTATCGATCATACAATAATGTGTTGATCTTCTTATTAAACAGATCACTAATGAATGCTTATCATCGATCTCGAAACTGCAATATCTTGAAGATTTGGTTCTAGAAGGCTGTTATGGAATTGATGATTACAGCCTTGCATCCCTTGAGCAAGGCTGCAAGTCACTGGAGGTATGTACTTTGGAGGACTCTGTTTAACAGACTTTGATTCTCATATTTGCCTGATTGCCTCTCCTTGAGATAAAATGAACAATAAAAAATGAAGCTAGTATTGATGTATGAAATTGCCAAAAGCTCCCAGGACGTAGCTCAATTGGCGAAGGTTGATAAACTTGTGACCTAGGTCATTAGTTCGATCCCCATGTCAAGCGAATCTGATATTTAAGTGGCGAAGGGAGAAGTGAGCCCATCATCCCCGAGTTTCGAAGGCTGCGGTTGGTCCTAAGGGTTGGCTCCTAATGGATTGCCCAAAACTTTGTCAAATTGTCCTAGCTTCAAGCACTACTTTTAGATGTACATATATATGACTTTTTTTCCCTAATTGCCTAGTCAATAATTGGAGACCTACTGTGTGAATGCCACTATAAGTACTCTGAAATAGTTAAAGAACAAGGTAACTCAGATTTTTTTTTTTGATAACCGAGAAATCCGTTTGTGACCCGCCCTTTGGACCAATCACAGCCTTCTAAACTCTGTGGATAATGGGCCCGCCCCTCTACCCTTCTCCACTTAAATACCGGGCTTCGCTTTGCATGGTGTGGGGCTTGAACCTGGGACTTAAGCCACAAATCCTCCACCTTTTGCCACTTGAGCTAGGCCTTGGGGGCAAGGTAACTCGGATTTTATGCACCCAGAATTCTACCTAATTGTCAGATGCCCCAAAGAAGTGGACTATTGTGCAAGTCAAGTGTTGCCTTGCATGTATCATGCCTTTTGGTCCTTTTTTTCCTCTTAAATAACTTTTTTCTTTACAAGGTACTATTGACTAATGTCAGTGTATGTATGTTGGCTCTTTTCAGAAACTTGATATGTCGAGCTGTCAGAATGTTAGTCATGTGGGTTTGTCATCTCTTACCAGCTGTGCTGGATGTCTGCGGCAACTTGTATTGTCGTATGGTTCTCCCGTAAGCCTTAACAAACTTGAGCATCTTTCAGTAGTATGAGTTTTAATTTGATTGTTGATTTCAAATTACTGGTATTTCAGGTTACACCTGCTGTCGCAGATAGCTTGCAGAAGCTTTCCAGGTTGCAGTGCGTCAAACTAGATGGTTGCCAAGTTACATGTTCTGGACTGAAGGCTATTGGAAATTGGTGTGTCTCACTTAAAGAACTGAGCTTAAGTAAATGTGTGGGAGTGACAGATGAAGGTCTCTGCTCCCTCGTAACAAAGCACAAAGACTTGGGGAAGCTTGACATTACTTGTTGCCGCAAGATTACTTACGTATCAATTTCCCATATAACGAATTCTTGTGCCTCCCTTACTTCTCTTAGGATGGAGTCTTGTACTCTAGTTCCAAGAGAAGCTTTTGTTCTAATTGGACAGCGTTGCCAGTACCTTGAAGAGCTTGACCTTACAGACAACGAAATTGATGATGAAGGTCACCCCTTTACCTTAAAGGTCTCAGGCTTCTGTTGTAATTTTGAAAGTTCCACCTTTATATCTGACGGTGTCCTACATATGCAGGCTTGCAGACCATTTCCAAGTGTGCCAGCCTTTCCAGCTTAAAGCTAGGGATCTGTCTGAACATAACCGATCAGGGTCTTATCCACATTGGCATGTGCTGCTCAAATCTTAAGGAGCTTGACTTATATAGGTATGTATTTCATAGTTCATATTAATTGAGATAGTGTAACTATATATCTTGTGCAGTTTTTTTTTTATATTCTTTTCTTTTTGTGATGGATTGGCTAAACTCACTTTTTAACTATGGAGTACAAGAGATTGAGTGTGCTGATTACTGGGTTATTCTTCTTGACAGGTCTGCTGGAATTTCCGACTTGGGAATATTAGCAATCTCTCGCGGTTGTAGTGGCCTTGAAATGATCAATATTGCCTATTGCAATCGTATTACTGATGGCTCATTCATATCTATATCAAAGTGTTCAAAGTTGAATACACTTGAAAGTCGAGGTTGTCCTCTTGTCACATCTTTAGGTCTTGCAGCTGTTGCTGTGGGATGTAAGCAGCTTACAACACTAGACATCAAAAATTGCCACAACATTGATGATGCCGGAATGATTCCACTTGCTCACTTCTCGACAAACCTCAAACAGGTAACCACAGTTTCTCTTTCGTCGACATAGACGAACTGGAGCATTGACATCTAATATGAGCTGGCCATCTGTCAATTACACAGATAAATTTATCCTACACTTCAGTGACGGATGTGGGGCTGCTGTCTCTTGCCAGCATCAGCGGTCTACAGAACATGACAATCCTGCACCTCAAGGGTTTATCTCCTGGTGGGCTAGGAGCTGCACTGTTGGCTTGTGGCGGACTTACTAAAGTTAAGCTGCAGACATCATTTAAATCATTGCTACCACAAGCTCTCCTTCAACATCTAGAATCCCGGGGTTGTGTTTTTCAGTGGAGAGAGAAACCATTTCAGGTATCTTTCTCATTATTTATCATTACCTACTTTTTCTCTGAAAATATGTGAGGTACTTCTATGGTTGTTGAAATATACCGTCGTATGAATCAATTAGAATTACATATTTTAGCTAAACATGACATAGTTATTACTTGATCTTGAGGGTCAATGCAGTTTCAGAGGCTTTCTTGTTTGGCATTTAATTGAGACAATAAAATTACAGTTTTGATTATCTTTGTTGTAATAATGAAAATATCCTTTTTGCTTGATCTAGTCATTAATTATCTTTCTTTTATTGGTTAAACACAGGCTGAAGTAGATCCAATATACTGGAAAATTCAGCTGGACAATTTAGACTGAAGATTTGAGCCCTCAGTTTGAGCCATTTATGCGAGTGTCCTAAGATGGATAGTCGCCCCAGAACCTCCGGAGTCCTTACTTCATCTGTTAGCACTTCATCAGGCAAGGTGCATTGTTTATGGCAACTGTTTTGCCTGTTAAGTGTCTGCAACTTGTTCGTTTGTATCCTCTAAATACAAACAGGGAAACTCAATGTCCATTCAGCATTGCCCTGGCATCAAATCAATTTTGTCAAAAATGTTATGAGATGGATTAAACAGATCCAAATCATGTTTATTGTTTAGCAATCAATATCTGGGTTTGTGAATGTTTATTCCTCTTGACTAATGTGATTTCGCATGAAGTTACTTGAAATTTACTCTCAATTAAACAGTTATTTG

>M6-PGSC0003DMG400011900 M6 LRR-RLP

TTTTAGGGAATTTGGAGAAAGAGCTATCATCTCTTACATCCATTTTAGAGAAGGAGCTATCATCCATTTTCAGATATGTCGTGCACCATGAACATAAAATTCCTAAAGATCTTCAGAGACGTACCATCAATTTGGCATATGAGGCTGAGGTAGCCATTGACTCTATTCTTGCTCAGTATAATGCTTTTTTGCATATTTTTTGCTCACTTCCTACAATTTTAAAAGAGATCATGCAAATTAATGCAGAGGTGACTGAGATGTGGTCAGATGACATTCCTCTTAATCCTCACTACGTGGCTGCTCCATTTAAACATCTGCCAACTCGACATAGCAATCCTGTGACTGATGAGGAGGTAGTGGGTTTTGAGAATAAAGCAGAAGAACTAATTGCTTATCTGATTAGAGGTACAAATGAGCTAGACGTCGTCCCAATTGTAGGCATGGGAGGACAAGGGAAAACGACAATTGCTAGAAAGTTGTACAATAATGACATCATTGTTTCTCGCTTTGATGTTCGAGCATGGTGCATCATTTCTCAAACATATAATCGGAGAGAGCTATTACAAGATATTTTTAGTCAAGTTACAGGTTCCAAAGATAAGGGAGATACGGTTGACGTTCTTGCCGATTTGTTGAGGAGAAAATTAATGGGAAAGAGATATCTCATTGTATTGGATGATATGTGGGATTGTATGGCATGGGATGACTTAAGGCTTTCTTTTCCAGATGTTGAAATGAGAAGCAGAATAGTCGTAACAACTCGACTTGAAGAAGTGGGTAAGCAAGTCAAGTACCATACTGATCCTTATTCTCTTCCATTCCTCACAACAGAAGAGAGTTGCCAATTGTTGCAGAAAAAAGTGTTTCAAAAGGAAGATTGCCCGCCTGAACTACAATATGTGAGTCAAGCAGTTGCAAAAAAATGCAAAGGACTGCCCCTAGTGGTTGTCTTGGTAGCTGGAATAATCAAAAAAAGGAAAATGGAAGAATCTTGGTGGAATGAGGTGAAAGATGCTTTATTTGACTATCTTGACAGTGAGTTCGAAGAATATAGTTTGGCTACTATGCAGTTGAGTTTTGATAACTTACCCGATTGTTTAAAGCCTTGTCTTCTTTATATGGGGATGTTTTCGGAGGACGCAAGAATTCCAGCATCTAAATTGATAAATTTATGGATTGCTGAAGGATTCGTGGAGAACACTGAATCTGGGAGATTAATGGAAGAGGAAGCTGAAGGTTACTTGATGGATCTCATTAGCAGTAACGTGGTAATAATTTCAAAAAAAAGTTATAATGGTAAAGTCAAATAATGAAAGGTTCAAGATGTTGTGCATCACTTTTGCTTGGAAAAGAGTAGAGAAGAAAAGTTTATGCTTGCAGTGAAGGGCCAATATATCCAGTTTCAACCCTTGGATTGGAAGGGAAGTCGAGTGAGCTTCAGTTTCAGTGAAGAGCTTTCCAAGTTTGCATCTTTGGTTTCCAAAACACAGAAGCCTTTCCACCAACACTTGAGGTCTCTAATAACGGCAAATGGTGGAGAATCTATTGATGTGATTCCCTTTTGTCAGATTAGTGAATTGCGACTTCTTAAAGTCTTGGATTTGATTTCTTATACTGTGGAGTTTTTGTCGTTAGCTATATTCAAACCACTAAATCAGCTGAAGTACCTCGCAGTTTGGGCAGATAAATTCTATTTTGATCCAGGAACACATCTGCCCCATATAGAAACTTTAATTGTGAAGAATTCTCCTTACGTATGGTTACCAGGGTCTTTTTTGCAAATGGAAAAATTAAGGCATGCTCATTTTGGTAAGGCTGTTTTTGATAAGCAGGGGCTCTTTGAAGGATCCTCTAAATTGGAAAATTTGAGGATATTAAAGAAAATTGAGAAATTTCCAATTGATAGGGCGGATGTGTTATCAAGGAGGTGTCCTAATCTTCAACAACTTCATATCGAATTTGATGGACATTCTGCAGATTCTTTTTGTCTCACATTGGAGAATCTTACCCAGCTTCAAATACTTCGCGTTTCCTTTGAGCGGCCCCACATCATTGTATCTGGATTGCAATTGCCTTCAAATTTAAACAAGTTGGTACTTAAAGGAATTCATATAGAAAGTGTTATTCCCTTCATTGCGGGACTACCAAGCCTGGAGTATCTCCAATTACAAGATGTGTGTTTTCCTCAATCAAAAGAGTGGTGCCTTGGAGATATCACGTTCCATAAACTTAAGTTGTTGAAACTGGTGCAGTTAGGTATCTCAAGGTGGGATGCCTCGGAGGAATCATTTCCCTTGCTTGAAACACTAGTTATAAAAATGTGTGGTCACCTAGAGGAGATCCCAATTAGCTTTGCTGATATTCCATCATTGAAACAGATTAAATTGATTGGGCCCTGGAAAGAATCTCTGGAGGATTCAGCTGTGAGAATTAAGGAAGAAATCGAAGAGATTGAAGGATGTGATCGTATACACCTCGTCAAAGAACCGACAAATTGGGAGGCATATAGCAATTTGTCGAAGCTGTGCTTATACTGAAGTCTATTTTGTAGCTAGCCCATTCTCTTATATTGTAACACTACTACTGTTTGTTATGTTTGTTTTGATTGATCAACTCTTTTTATGTATGGTTGGAGACAAATATATATCAGATGATCTTTCAC

>M6-PGSC0003DMG400013644 M6 LRR-RLP

TTTCCACATAACCGAACAAGACACATTTTCTGTCTTAGATAACTCATCCTTTCTCTTCCCTTCTTCTTCTTCTCTCTGTTCTCCACATTTCACGTACACTTTCAGAAACATTGTTGATTTACTTTGGACAGAAAAGCCACTAAATAGTGTGAGTCAATTTCTTTCCTTAAATAATTGTTTGGAGTTTTGTTTTAATGGGAAGTTTTTTTTTTTCATTTTTATGAGTGTTTTTTCCGTGTTTTTGAAAACAGTTGTTTTCCATATAGGAAAATACGGCGAAAAATAAGCTTTACTTTGCATGCCTTCATGTGGGTAAAGGTTTGAATTTGCTGTCATCGTTGATTGGCGCTTTCTAGAAACTTGTAGAAACAAAAAACTTAGCTGCGGTTTTCATTTTTCTTGATCTGCTGATTGGTTTAGATATAGCGGAATTCGTCTTTGTTTTTTGCTTTGAATAATACATACACAACGAGGAAAAGGGTAGTCTGAGTCTAATTTTTGTGGCCGGAGTTGTATTTATGTTGAATTAAATTTATCTGGGTGTTTTTTGTTTTCGAGAAATTTTCAGCGGATTATTGAAAATGGGGTCGTCAGGAATGTCGATTGATGAGATTGTGCAAGAAATTATGAGAATTCATGGATCTCTACCTACAAGGCCATCGATTGATGAGGTTGAAGCAGCAAGAGGTTTAATTGGAAATCTGGAGAAAGAAGATCAGTTGAAAATAGAAGCTATCATGAGGCAAAATAAAAGGAAAGGTGTACCAGAAGAGCTGTTCAAAATTTTGCAAGAGATGCAGAGGAACGCTGTTCTTTTTCAGAGTAAAGAACAAAAGAGGGAGGCGTTGAAACTTTTAGATCTCGACAATATTCACAGTGTGTTCGACGATTTGATTCAGAGAGCTACGAAATGTTTGCCTTCTAAATCTCACGATAATGACCCGTCTTCTTCTTCAATTGAACCGAATGATTTATCTTTGGCAAACCAAGTTTCCTTCTCCGGCGGCGGCATTAATTCTCCGACAACTACAGCTGCGGCTGCTTCTTCGTCGAGCTATTATAACTCAAAAGAGCCTATTAAAGCGTCTGAACTGTATAGTAGAGATGACAGTTACTTGAATAAGGCAAAAGCCACGTTTCTTGTGGATGGAATTGGTGTTGGGCTTCGATCTGGGGATGCTTTATCAGGGCCTAAGATTGTTGATTCCACCTTAAAGCCGGCGTCAATAAGTGGTATGTTACATCTATTTGCTGCACCGTATTTCTAGTGAATTTGATTATATAATTTGTTGTATTGTTGTGTTGACTCAAAGTTCGTATTTTTTTGATCAAGAAGACACTGGTTTCTTTGTTTGTAGGTCAAGATGGTGAGAAATTAAGTTTGATTAAGCTTGCTAGTATGATTGAAGTGTCATCCAAAAAAGGAAGTCGAGAACTTATCCTTCGGAATAAGTTGTCTGACCAAGTAGAATGGCTTCCAGATTCTATAGGGAAGCTATCAAGTTTGATCACTTTGGATCTGTCTGAGAATCGAATTACGGTCCTACCCACCACCATAGGGGGGCTTTTATCATTGCAAAAGCTAGATTTGCACTCAAACAAAATTGTTGAACTGCCTGATTGTATTGGAGATCTTCTTAACTTGGTCTATCTTGATCTCAGTGGAAATAACTTGAAGTTGTTGCCCGCGAGTTTTGCTAGGTTAGCCCACCTTCAAGAACTTGATTTGAGCTCTAACATGCTATCAGTACTCCCTGAGACAATTGGCTCCTTAGTAAGTCTTAAGAAATTGATTGTGGAAACTAATGACCTTGAGGAACTTCCTCACACTATTGGCCAATGTACTTCGCTCAAAGAGCTTCGTGTGGACTATAATCATCTTAAAGCTCTTCCTGAAGCAGTGGGACGAATTGAGTCTTTGGAGATTCTGACTGCGCGGTACAATAACATTAGGCTGTTGCCTACAACTATGTCATCCCTTACAAGTTTGAAAGAGCTGAATGTTAGTTTCAATGAAATCGAATCAGTGCCTGAGAGCTTGTGTTTTGCTACTTCACTGGTAAAGCTGAACATAAGCAACAATTTTGCAGACCTGAGATCACTACCAAGGTCCATTGGTAACCTTGAGTTGCTTGAAGAATTGGACATGAGTAACAACCAGATACGCGTTCTTCCAGATTCGTTTAGGATGCTCACAAATTTGCGGGTCCTTAAAACCGATGGAAATCCTTTGGAAGTGCCACCAAGAAGCATCGTTGAGAAGGGAGCACAGGTTTAATTTTGTTTCGTTTAACTATTATGCTTTCTGAGTTAGTGTAGTATTTAATTATCTTAGCTCTTCTCTTGGATATATCAATCACTATTGTTTTATTTGCTCGGATCCTCAGGCTGTTGTACAATATATGTCTGATCTGGTTGCCAATAGAGAGGTAAAGGCGCAACCCTTAAAGAAGAAGAAGAAATCATGGACTCATATATGCTGCTTTTCAAGTTCCAACAAACGTCAGCGCAATGGCAGTATGGACTATGTTAATGCCTGATGTGTTCCACACAAATTTGGTATGATTATATATTTCTCCTTCTTACCAAGTTAGTCATTATGGTACTGAACTTTCCATATTTCGTAAATTTCGTGTTCAGTCAAATTAGTCTAGTCTATGAGAGGTAGTTTTAAATATATATATTGTTAATAAATGGGAATGGTGCATTATACATGGTGCTGATTAGTGTTAATTAGAATAATGAAAATGAAGGGTTGTAAGGGTAACGGCTCACATGATGTCTGTGTTATCCGTTGTAGGGAAATGAGCTGTGATTTGGCTGGACGAAAGCTGCAAAATTTCAGTTGAATTCGAACATTTTTGCCAGCGAGATTCAATCAACTTTCTTCTTGTTCCAGATAAGCTCTAATCCACTGCCACGTGGTTTTTTATTCTCATTTCATGTAATAATTCTTTTAATGCTCTTTCCCTGTATTTTCCTCGATTATGATGTAAAATATCAAATGAGATGTAAAATCTTATCTTATTTTTTGACTTTCAATATATTCAATGAATTCCAGAATTTC

>M6-PGSC0003DMG400013766 M6 LRR-RLP

TACAATTAGCAATTTGCCTTGTAATGTACTGGATGCTGTTCTTGGATGCTTGCCTTGGAAAGATGCAGTGAGAACCAGTATCTTGTCAAAAGACTGGAGGTACAAATGGGTAACACGTCAAGAACTTGATTTTAATGATGAGTTTTTCAAGTCTTTCAAACAAGATGAAGAAGCCAAGAGAATCATTTACCAAGTCCTATTAGTCCATTAAGGACCAATACTGAAGTTTAGACTCTGTAGGATTACGAGTTGTCCAGGAATTCACCCTTCATGTAAGCTTAGGAAACAAATATCATTCACCTCATCATCTTTTTACATTCCAGCAACTAAGGTTTTTGGAACTTCAAGATTGCTTGTTCCACCCTCCACTCGGTTTCAAAGGGTTTGAGAAGCTTATTAATCTCGATCTTGTACGTGTCACTTTTGATCCATCAATATTTACTTATCTCATCTCTAAATGCTCGTTGCTTGAACGGTTGAGGTTGCGTTGCTGCACAAACTTTGACATCCTTGAAATTGATGCCGCTAATCTCAAATTCTATGAGTTTATTGGAAAAACAAAATCCATTTCCTTCAAGAACGCTCCTATGCTTGAAAAAGTTACTGTGGCTTTCCTTGGACGACAACTTTTGACTGATACATCTCTTGTTTGCTCTAATTTCCCAAAGTTCTTCCATTACATGCCTTCACTGCTGGATATGGATATATGCGGTACAATATTAGAGGTAAGGTACATTTGTGATATTACTATCATAACTAAATGATGAATCTGATTGGTGTCTTAAAGTAATTTTTGACTTTCTTGTAGTACTTGATCTAGGGAGGTTTACCAGAGAGTCCCCCAACTGCTCTGAACAATATCAAATCTCTCACTATTTCGTCCGTGTCTTTAAGAAACGCCGAGGTGGTTTCAAGTGCAGTTTACTTGATTACAAGCTGTCCTAAATTACAAGATTTTACACTTGAATTTGTAAGCACTCTCTTTGTCTTTTTTTTTTCATTAAGGTGATTGAATATGGTTTCTTATTCATCTGAACTCGCCTAACTAACCGTTCCTCAAATATTCTTTAGTACCAAGTGTGGGTGGGGGATATTTATGAACCTGTTGTACAGTTGCTACGAGCCCAACCATCCTCAAGCGGTGTTGTGAAGCTTCAAAGAGTTCAAGTGAACATGTTTACTGGTCTCGAGATGGAAATGGAGATTATGAAGTTTATATTGGCATCTGCACCTATACTTGAGGAGATCTTCATTTGAAATTGTATGCGTTTCCTTTTTCGTTCGTGTAAACAAATGATCGATGAGATGAAAGAATTCCGCAGAGCATCAACCAATGTTGAATTCACATTTGAAGAAAT

>M6-PGSC0003DMG400013898 M6 LRR-RLP

TATATATATATATGAGAACCCTTTTAATGTAAAAGCTCAAAAAAGTTGTATTATTGTTCTTGAAACCAAGTACTCTACCTATTAGACATGACAACTTCTTACAGTACCTTCCTTTACATTCTACTTGTGCTTCTTCTTTTTTCACTTCCACTCACAATCACATCATCAGCAACAACTGAAGCTGAAGCTCTTGTCAAATGGAAGATGAACTTACCTTCTACTTCTTTCTTGGATTCATGGTCCATTTTGAATCTCGGAAACTTGTGCAATTGGACATCTATTGTCTGCAATGCTGGTGGAACAATTTCCGAAATCAATCTTTCTGATGCAGGCCTCTCAGGCACTCTTGATCAGCTTGACTTCACTTCATTTCCGAGTCTCACAAGTTATTCTCTCAATGGCAACAACTTCAGTGGATCGATACCATCAAATATTGGTAATGCTTCTATGCTCACTTTCTTGGACCTGAGCAACAATATCTTGGAGGGTGTCATACCAGAAGAAATTGGGAAGTTAACACAACTTGAATATCTCAGTTTCTATAACAACAATATCATTGGTGTTATTCCCTACCAAATAAGTAATCTTCAGAAGGCATGGCACTTGGATCTTGGATCAAATTTTTAGAAACTCCTGATTGGTCCAAATTGAGGAATATGCCTATGTTGACATATTTGAGCTTTGGTTACAATGAACTAACATTAGAATTCCCTGAATTTGTACTCTGTTGCCATAATCTGACTTACCTCGATTTGTCGATAAACCATTTGAATGGTTCAATCCCAGAAACACTATTCACCAACTTGGATAAGCTTGAACACCTCAATCTTTCATCTAATTCATTTCAAGGTTCATTGTCACCAAATTTTACCAAGTTGACCAAGTTAAAAGAACTTCAGCTTGGTTCCAACATGTTTTCGGGCTTAATTCCTGATGAAATTTCTCTGATCAGCAGTCTTGAAGTAGTTGTACTTTTCAATAATTCACTTCAAGGAAAGATTCCATCTTCTATAGGTAGACTTAGAAATCTTCAACAACTTGATCTTCGAAAGAATCGCTTGAATTCAACCATTCCTTCTGAGCTAGGCCTTTGTACTAACCTCATTCTCTTGGCTCTATCAGGGAATGCCCTGCAAGGACCATTGCCTATATCCTTGTCTACAGTGACCAAGTTATCTGACTTGGGCTTGTCTGATAATTTTCTTTCTGGTGAGGTTCAATCATATCTCATTACCAATTGGACTGAGCTGACATCTTTGCAGCTTCAAAACAATTCATTTTCAGGGAAAATTCCACCTGAAATCAGCCAGTTGAAAAACCTTGTATATCTTTACCTCTATAGTAACAATTTCACTGGTTCCATTCCCTATCAGATTGGTAAGTTGCAAAACTTGCTGGATCTAGATGTCTCTGAGAACCAGCTTTCAGGCACAATACCTCCAACCATTGGAAACCTGACCAATCTAAGGACGTTGCAGCTTTTTCGGAACAACATCAGTGGAACCATTCCTTCTGAGATTGGCAAATTGACATCTCTTCAAACCCTTGACCTCAACACTAACCGGTTTAGTGGTGAGCTGCCCGACAGCATTTCTAACCTCGGCAGTCTGAAGCTTCTTGAGTTCCTTTCTCTTAGTAACAACCAGTTTTCAGGTGTGATCCCTTTGTCTATTTGCAATTCGACATCACTACAAACTCTAGTTTTCACAAGAAACAATTTCAGGGGAGAGATTCCACAATGTTTGGGTAATATCAGTGGCCTCTCCATTTTGGATATGCGCTACAACAATCTTTCTGGGATTCTTCCAACAAGTTTTGATATTGGATGTTCACTTAGAACCTTCAACTTTCATGGAAATGAACTACAAGGGAAAATCCCTCGATCTTTAGCCAATTGTAAAGAGTTGAAAGTTCTTGATTTAGGAGACAATCACCTCAATGATACATTCCCCATGTGGTTGGGAACTCTATCAATGTTGCAGGTTTTAAGTTTGAGATTGAATAAATTGCATGGACCCATTAGAACTTCAAGGATTGGAAACATGTTTCCTGGTCTTCGAATATTAGATCTCTCTGGCAATGCCTTCACAGGAAACTTGCCGACAAGTCTATTTCAACATTTGAAAGCCATGAAGACAACTAATCAAACAAGGAAGGCACCAATTGAAGGAAGCAACTATTACCAAGACTCGGTAGTTGTTGTAATGAAGGGACACATCTTTGAAGTTGTGAGAATCTTGTATTTATACACGAGTATGGATCTTTCAAGTAACAAATTTGAAGGACTTATTCCAAGTATTATGGGAGATCTCATTGCACTTCGGGTGTTGAATTTGTCTAACAATGGGTTGCAAGGTCATATACCGCCATCACTTGGAAATTTAGCGGTTGTAGAATCATTTGACCTTTCATTCAACCACCTTTCTGGAGAGATACCGGAACAACTTGCTTCTATAACAAGTCTTGCAGTGTTAAATATCTCCTACAATCATCTTGAAGGATGCATTCCTCAAGGACCTCAATTTGCTACATTTCAGAACAATTCATATGAAGGTAATGATAGATTACATGGATTTCCAGTTTCACAAGGTTGTGGTAATGATCAGGTATCAGAGACAAATTATACCATATCTGCACTAGACGAACAAGAAAGCAATTCGGAGTTTCTGAATGATTTTTGGAAAGCTGTTCTTATGGGCTATGGAAGTGGACTATGTATTGGATTAGCCATAATATATTTCATGATCTCAACTTGAACTCCAATATGGCTAGCAAGAATCATGGAAGAGTTGGAACACAATTATTATGCAAAGTAGAAAGAAGCAATGAGGTAAAAGGCATTACAGAAGAAGAAAATAATCGCTTCTAGACAAGTTACCAACTGAGTATTCAAGTTAAGGTCTGCTGATTTTTAGTATATAGTAATATATATTTATAGTTTCGTTACAACAACAGCTAATCGATACTTTATACTAAGTATATTTTTCTTTCCAATAATTT

>M6-PGSC0003DMG400013955 M6 LRR-RLP

TTGAATATATTATATTTTGGAATTCAAAGAAAACAAAAATAAAAATAATAAAATAGATTTAAATTTTATTTTTATTTAAAAAAAAAAGTGAAACCACACGTGACAACGTGTGTAGTATAGTGCGTGTCACGTTCGATTTTCTTCTAAAGAAAAGATTAAAATTTCTTCCATTTTGTGCTAATTAGTGAATGTGTGTTGTGTAACACAGACTTATAAGGTTGACTCCTATTGACTTGCTCTTAACAATATAATGAAAATTTAGCAACAATTATTCTTTCTATTATGACTTTTATTTCGTTTTGTGAAACACAACCCAAATAATGTTTTTTTTCTTTTTGCTACTTATTTTCCTCCTTTTAATCTGTTTTCACATTTTATTTGTTTTAGTGCTATATATATAGTGGAGTAACTTTGGTCACTGGAAATACAACACACAAGTTAATATGGAGAAGCACATCTTCTTATTGATTCTTCTCTTTCTAATTCAATTTTCTATATCACTTGCTTCCTCTAATGAGACTGACCAACAAGCTCTGTTAGCTTTCCAAAATCTTATTACAAGTCCTAATCATTTTTTGGCCAATAATTGGACCAAAAATACTTCCTTTTGCACTTGGTTTGGTGTCACTTGCACTCCAAAAAGGCAAAGGGTTATGGCCTTGGCTCTTCCTGATTTGCAACTTCAAGGCACAATTTCCCCATCTTTGGCAAATTTGTCCTTTCTCAGTGTTCTCAATCTCCAGAACAACAGCTTCCACAGTGGCATCCCTTATGGACTTGGCCACATGACTCGCTTGCGAGTGATTGACGTTAAAAACAATCAGCTCAACGGAAGTATACCAACAAGTCTATTTCAAAACCGGAGAGTTCAAGTAATTTCAATGGCTTTCAATGAACTCGGTGGTGAAATGTGGAGAGGCCCATGGTATGTACCAGAACTCAGAGTCTTAAATCTCGGGAACAATAGCCTCACGGGTATAATCCCTCCTTATGTTGGAAATGCCACAAAATTGATGAACATCAATTTGTCTGGGAATAGAATCAATGATAACATTCCAATGGAGATCAGTAATCTTAGCCAACTTGCATTTTTGTCCTTGTTCAATAATCAATTAACAGGTTCCATTCCTCCAACACTGTTTAATATCTCGTCACTACTTGGCATATATCTGTCATTCAATGGCCTTTCTGGTCCTCTCTTGCTAGATGAAGGGAATATTGTGTCAAATCTGAAGCATTTAAGTATGTCTAACAACCAAATTTCTGGTTGCATTCCTTCCAACATATGCCAACTCACAGAGCTCAAACTTTTGTCCATATCTTTCAACAATATAACTGGAGACATACCTAGAAATATTGGTTGTTTATCCAAACTCGAAAGGTTTTATATTGGTGAAAATCCAATAGAAGGGACTATTCCCACTTCATTGGGAAATATTTCTACTCTGCAATATCTTGATTGTGTAGACAATTGGATGAAGGGGCAAATTCCTCAAGAATTAGGGAAGCTATCAAATTTGAGGGAATTAAGCTTTGCCAATAATTCTAATCTTATTGGTCAAATTCCAGAGGCTATTTTCAACATATCTTCTTTGGAAGTGATCAATTTTAGTTTGAATAACCTCTCGGGGAGAATTCCAACCACTACAGGTCTTCATCTTCCGAACCTTAAAGAACTTTACTTGGGAGACAATCAGCTTGAAGGGGAAATTCCATTGTTCATAACAAATGCTTCCAAGCTTCAGACACTGGCACTAAATGATAACTCTCTCAAAGGAACTATTCCTAACAATTTGGGGAATCTTCGTGAGCTGCAAATACTGCTCCTACACACTAATCAACTTACCAATGAACCAGGAAAGCATGCGTTGAAATTCTTCAATTCTTTGGCGAACTGTAGGATGTTGAGATATCTAGAAGTGGGTAACAATCCGTTGAGTGGAGTTCTGCCAAATTCTATTGGGAATCTCTCATCTACTATTGAAAAAAATTATTTAGCAGATGCACACATCAATGGCCTCATCCCCACAAGTATAGGCAACATGACCGGTCTTACATCCTTAAGCCTTGGCGGAAACAACTTGACAGGGAATATTCCTTCTGAGATTGGTAAGCTTAAACAACTCCAAGGACTCGATCTATCTAACAATAAATTGCAAGGACATATTCCAGAAGTGGTATGCAATTTGTCTGATTTGGTTCAATTATATCTGTATGGTAATGAGCTCTCTGGATTAATTCCAGAATGCTTAGGAAATCTTACCATGCTTCAACAACTTTATTTGGCTTCTAATAAATTTTCATCAAAATTTCCATTGACCCTTTGGAAAATGAGTGGTCTTCTCTATCTAAACGTGTCACAGAATTCAATAGAGGGAGAACTTCCATCAGATATTGGAGGACTGAAAGCCATTGTAGAACTATATCTTTCTGGTAACCACTTTTCAGGCATGATATCAACCAGATTCGGGGAACTCCAAAATCTGCAGTATCTCGACCTATCGAAAAATTCATTTTTTGGCCGAATTCCATTATCCTTTGCCAACTTGATAAACTTGGAATTCTTGAATTTGTCTTTAAATGCATTGTCAGGTACTATTCCTAAGTCTTTGGAAAAACTCTCATACCTTAAAAGCATTAATGTTTCATTTAATGATTTAGAAGGTGAAATACCAAGTGGTGGTGTGTTTGCAAATTCCACTCTGCAATCATTTGTTGGGAACAAAGGTCTATGTGGAGTGCACATATTAGAGATTCCTGCTTGTGCTATCACTAATCCTGGAAAACAATCAAAGCTTAAGGAGGTTATGCTAAAAATTGTTACTCCAGTGGTTATTGCATCCTTTATGATATTCTTGTTCGTTTCAATTTGGATAATGAAACGACAGAAGAAAGGGAAGTCCAAAGATGTGGAAAAGATACCGGAGATGGGGACTCATCAATTAGTTTCTTATCACGAGATTCAACGAGCAACAAATAATTTTGATGAATCCAATTTAATTGGTGAGGGAAGCTCTAGCTCTGTGTACAAAGGCACATTGTTTGGCGGAACTGCAGTGGCCATTAAGGTTCTGGATTTGAAAAATGAGAAAGTATGCAAGAGGTTTGATACCGAATGTGAAGTGATGAGAAATGTTAGACACAGAAATCTTGTTACAGTGATTACTACATGTTCTAGTGACTATATAAGAGCCTTTGTTCTGCGATATATGCCCAATGGGAGTCTTGAGAATTGG

>M6-PGSC0003DMG400014717 M6 LRR-RLP

CGTACAAGGATATTGAGTAGACAGTTGATGTATCAGACTCGATACTTCAAAAGAATGGAGGAGAAGACGATCTGCAGCTGCAAGTGTGCATGAAGACAATATATGAAATAACTGAGCAAACATTCTTTGAAACAAACAGTTCACAATTCATATATCAGAAAAATAATTTCCCAGCACGTCTAAAACAAGCAAAAATGGAAAAGAAAAACAAAACAGTCCCAAGGATAAATCTATATTGTCACAACTTAAAAAGATTTTAGCTCCTGTCTCACCACACCAGCCCTCTGCAATGTCCACAACACACGAATCAGATCGATGCATAACCATCACAACGGTAAAAAGATTATGCAATCACACCTTGATCTTTGCCAACATGAGCTTAAACCTGTCAAAATTGTTGAGTGAGGCCCTCCTCTTCTGCACGATCAACTTCCTTTCTTAGGAACTGGACTCCCACTTGGTCTTCACATCTGCGAGGACATAAAGCAAATGAGAGTTAACCAAGTTCTTATGTAACCCTGCCGTTAGCCAAATGACTCAAATTTACCACTTCTGGCTGATGAAAACCGTTTTTCCTAATCAAAAATTCAAATTTGAATACTGCAATTCACCCAACGTACACGTATCTTGATCTCTAGCTCGGCCACTGTTTCCAGTGATATGTATTTTCTGAGGGGACTAATTAACGCATAGTCTAGTCACTTTAGTATTTAGTATTGTAACTCTAGTGTTACAAAGTGTAAGTTTAGTAGTGACGATTCATGAGATTAACCTCGAAATTTTGTGATGACTGAAAATCAGAGAAAGTAACACAAGATGCTGTTCTATATTATGAAAAAGATTAGAAAGATGAAACTCTACTTCTTTTTAAGCTTTGTATTCTAATTGATGGAATTTCTTTTCCGTATGATAGTTTTGTTTTACTTGGAAAAAGTGGGAGTAGATGTGTAACACATTGGATAAAGACAATAAAAGGACAAAAGCACAACAACCCAAATTTGACTAAAGAATGACAACATTTTCTCTTATAAAATTGAACAGACAAAGTGTGAATAAGAGTTTAGAAAAAAATGGGAGTAGATGTGTAACACATGGATAAAGACAATAAAAAGACAATAATATTTGAGTGACTTTTCTCTTACAAAACAAAGAAAAGTGACTTTCTGAGATAATTATCTATAGTTCAATGACCACATAAGAAATCAACTTTTGGCATCTACTACATATGAATTAGAGATAATGTAGTTTAGAAATTTATTAATCTAAGACTGAAAATTGTTGAAACAAATATGTCAATGTGTGATAACAAAACTATTTTTGCCACTTCGGTTAAGTCCTTGTTAGAATTAGTGTTGGTCTTTAATTATAATTTACGTAAAATATAAAATCCCATGAACCCGAAATAATAAACTCAAAATATGATGATGGTTTCCAGTAAAATATTCTCTTCACTTCAGTTTTTCACTCTTTTATATCTCCTTACAGTTACTTTTGCTTCCAATGAGGAAGCAACTGCCCTCTTGAAATGGAAAGCAACTTTCAAGAACCAGAATAATTCCTTATTGGCTTCATGGACGCCAAGTTCTAATGCATGCAGGGACTGGTATGGAGTTAAATGCTTTAATGGTAGGGTAAACACGTTGAATATTACAAGTACTAGTGTCAGTGCTACACTCTATGCTTTTCCATTTTCATCTCTCCCTTTCTTGAGAATCTTAATCTTAGCATGAACAATTTCTCTGGCACCATTCCACCTGAGATTAGTAATCTCACTAATTTTGTCTATCTTGACTTGAACTTCAATCAGATTTCAGGAACAATCCCACCACAAATCGGATCACTAGCCAAGCGTCAGATCCTCCGTATATATGGCAACCATTTAAATGGCTCTATTCCTGAAGAAATAGGTAACTTTAGGACTCTTACTAAGCTATCTTTGGGCAGGAATTTTCTTAGTGGTTCTATTCCTGCTTCATTGGGGAATTTGAACAACTTGTCTTTTTTGTTTCTTTATGAAAATCAACTTTCGGGCTCTATTCCCGAGGAAATAGGTTACCTAAGGTTTCTTAGTGTCCTAGGTTTGAGTAATAACTCTCTTAATGGTTCTATTCCTGCTTCATTCGGGAATTTGAACAACTTGTCTGATTTGTATCTTAATGATAATCACCTTTATGGATCTATTCATGAGGAAATAGGTAACCTAAGGTCTCTTATTGAGCTAGATTTGAGTACTAACTCTTTAAAAGGTTCTATTCCTGCTTCATTGGGGAATTTGAAAAACTTGTCTATTTTGTCACTTCGTGAAAATCAACTTTCTGGCTCTATTCCTCAAGAAATAGGATACCTAAGGTCTCTTACTGATCTTCGTTTGAACAATAACTTTCTTAATGGTTCAATTCCTCTTGAAATAGGTTACCTAAGGTCTCTTAGTTTCTTAGATCTGAGTATTAACTCTCTTAATGGTTCTATTCCTGCTTCATTGGGGAATTTAAGAAATGTGCAAAGAATGTTTCTTGATGAAAACAATCTCACTGAAGAAATTCCTTTATCTCTGTGCAATTTAAAATCACTTGAAATTTTGTATTTGCGGAGAAACAACTTGAAGAGAAAAGTTCCGCAATGTTTGGATAATATCAGTGGCCTCCAGGTTTTGACGATGTCACGTAATAATCTTAGTGGAGAGATCCCTTCATCCATTTCCAATTTAAACTCACTACAAATTCTAGATTTGGGAAGAAATAGTCTGGAGGGAGCAATTCCAGAATGCTTTGGCAATATTAATACCCTTCAGGCTTTTGATATGCAGAACAACAAACTTTCTGGGACTCGTCCAACAATTTTAGCATTGGAAGTTCACTTAGAAGTTTCAATTTGCATGGCAATGAACTAGAGGGGGGAAATCCCTCGATCTTTGGCCAATTGCAAAAAGTTGCAAGTTCTTGATCTAGGAAATAATCATTTCAATGACACATTCCCCATGTGGTTGGGAACTTTGCTAGAGCTGAGAGTTTTAAGATTGACATCGAATAAATTGTATGGACCCATAAGATCTTCAGGGGCTGAAATCATGTTTCCTGATCTTCGAACCATAGATCTCTCTAACAATACCTTCTCGAAAGACTTACCAACGAGTTTGTTTCAACATTTGAAAGGCATGAGGACAATTGATCAAACAATGATGGTACCAAGTTATGAAGGAGATGGAGATTACAAAGACTCGATAGTTGTTGTATCAAAGGGATTGAAGCTTGAAGTTGTGAGAATTTTGTCTTTGTACACAGTTATCGACCTTTCAAACAACAAATTTGAAGGACATATTCCTAGTGTTCTAGGAGATCTCATTGCTCTTCGGGTGTTGAATATATCTCACAATGGATTGAAAGGCCATATAACACCATCACTTGGACGTTTATCTGTAGTGGAATCATTGGACCTTTCATTTAACTGCCTTTCGGGAGAGATACCACAACAACTTGCTTCTCTTACGTCTCTTGAATTCTTAAATCTCTCCCATAATTATCTCCAAGGATGCATCCCTATGGACCCCAATTTCATACCTTTGAAAACAGTTCATTTGATGGTAATGATGGATTACGTGGATTCCCCATTTCAAAAGGTTGTGGCAATGATCCTGTGTCAGAGACAAATTATATAGTATCTGCGCTAGCTGATCAAGAAAGCAATTCTAAATTTCTCAATGATTTTTGGAAAGCTGCTCTGATGGGCTATGGAAGAGGATTATGTATTGGATTATCCATACTGTATTTCATGATCTCAACTAGAAATCCGAAATGGCTTGCAAGAATCATTGAAGAAATGGAACACAAAATTAACATGCAAAGGAGAAAGAAGCAGCGAGGTCAA

>M6-PGSC0003DMG400014718 M6 LRR-RLP

ATGATTCTCAACGATAACAATCTCATTGGGGAAATTCCTTCATCTATATGCAATTTGACATCACTTGAAATGTTTTATATGCCGAGAAACAACTTGAAGGGAAAAGTTCCGCAATGTTTGGGTAATATAAGTGGCCTCCAGGTTCTAGTGATGTCACATAATGATCTCAGTGGAGAGATGCCTCCATCTATTTCCAATTTAACGTCACTACAAATTCTAGATTTGGGCAGAAATAATCTAGAGGGAGCAATTCCACAATGTTTTGGCAATAATAGTAGCCTCCAGGCATTTGATATGCAGAATAACAAACTTTCTGGGACTCTTCCAACAAATTTTAGCATTGGAAGTTCACTTATAAGTCTCAACTTGCATGACAATGAACTAGAGGGGGAAATCCCTCGGTCTTTGGCCAATTGCAAAAAGCTGCAAGTTCTTGATCTAGGAGATAATCATCTCTATGACACATTCCCCATGTGGTTGGGAACTTTACCAGAGCTGAGAGTTCTAAGATTGACATCGAATAAACTGCATGGACCTATAAGATTTTCAGTGGCTGAAATCATGTTTCCTGATCTTCGAATCATAGATCTCTCTAGCAATGCCTTTTCGAAAGACTTACCAAAGAGTCTGTTTCAACAATTGAAAGGCATGAGGACTGTTGATCAAACAATGAAGACAACAAGTTATCTAGGAGGTAGATATAACTACCAAGACTTGGTAGTTGTTGTAACGAAGGGATTGAAGCTTGAAGTTGTGAAAATTTTGTCTTTGTACACAGTTATCGATCTTTCAAACAACAAATTTGAAGGACATATACCTAGTGTTCTAGGAGATCTTATTGCGATTCGGGTACTTAATTTATCTCATAATGGGTTGCAAGGCTATATATCACCATCACTTGGAAGTTTATCTGTACTGGAATCATTAGACCTTTCATTTAACCGCCTTTCGGGAGAGATACCACAACAACTTGCTTCTCTTACGTCTCTTGAATTCTTAAATCTCTCCCACAATTATCTCCAAGGATGCATCCCTCAAGGACCCCAATTTCATACCTTTGAGAACAGTTCATTTGATGGTAATGATGGATTACGTGGATTCCCCATTTCAAAAGGTTGTGGCAATGATCCTGTGTCAGAGACAAACTATACAGTGTCTGCGCTAGAAGATCAAGAAAGAAATTCTGAATTTTTCAATGATTTTTGGAAAGCAGCTCTGATGGGCTATGGCAGTCGACTATGTATTGGCTTATCCATAATATATTTCTTGATCTCAACTGGAAATCTAAGATGGCTTGCAAGAATCATTGAAGAACTGGAACACAAAACTATCATGCGAAGGAGAAAGAAGCAGCGAGGTCAAAAGAATTACAGAAGAAGAAATAATCGCTTCTAG

>M6-PGSC0003DMG400014719 M6 LRR-RLP

CTTCATCTATATGCAATTTGACATCACTTGAAATGTTTTATATGCCGAGAAACAACTTGAAGGGAAAAGTTCCGCAATGTTTGGGTAATATAAGTGGCCTCCAGGTTCTAGTGATGTCACATAATGATCTCAGTGGAGAGATGCCTCCATCTATTTCCAATTTAACGTCACTACAAATTCTAGATTTGGGCAGAAATAATCTAGAGGGAGCAATTCCACAATGTTTTGGCAATAATAGTAGCCTCCAGGCATTTGATATGCAGAATAACAAACTTTCTGGGACTCTTCCAACAAATTTTAGCATTGGAAGTTCACTTATAAGTCTCAACTTGCATGACAATGAACTAGAGGGGGAAATCCCTCGGTCTTTGGCCAATTGCAAAAAGCTGCAAGTTCTTGATCTAGGAGATAATCATCTCTATGACACATTCCCCATGTGGTTGGGAACTTTACCAGAGCTGAGAGTTCTAAGATTGACATCGAATAAACTGCATGGACCTATAAGATTTTCAGTGGCTGAAATCATGTTTCCTGATCTTCGAATCATAGATCTCTCTAGCAATGCCTTTTCGAAAGACTTACCAAAGAGTCTGTTTCAACAATTGAAAGGCATGAGGACTGTTGATCAAACAATGAAGACAACAAGTTATCTAGGAGGTAGATATAACTACCAAGACTTGGTAGTTGTTGTAACGAAGGGATTGAAGCTTGAAGTTGTGAAAATTTTGTCTTTGTACACAGTTATCGATCTTTCAAACAACAAATTTGAAGGACATATACCTAGTGTTCTAGGAGATCTTATTGCGATTCGGGTACTTAATTTATCTCATAATGGGTTGCAAGGCTATATATCACCATCACTTGGAAGTTTATCTGTACTGGAATCATTAGACCTTTCATTTAACCGCCTTTCGGGAGAGATACCACAACAACTTGCTTCTCTTACGTCTCTTGAATTCTTAAATCTCTCCCACAATTATCTCCAAGGATGCATCCCTCAAGGACCCCAATTTCATACCTTTGAGAACAGTTCATTTGATGGTAATGATGGATTACGTGGATTCCCCATTTCAAAAGGTTGTGGCAATGATCCTGTGTCAGAGACAAACTATACAGTGTCTGCGCTAGAAGATCAAGAAAGAAATTCTGAATTTTTCAATGATTTTTGGAAAGCAGCTCTGATGGGCTATGGCAGTCGACTATGTATTGGCTTATCCATAATATATTTCTTGATCTCAACTGGAAATCTAAGATGGCTTGCAAGAATCATTGAAGAACTGGAACACAAAACTATCATGCGAAGGAGAAAGAAGCAGCGAGGTCAAAAGAATTACAGAAGAAGAAATAATCGCTTCTAGACAAGTTACCAGTAA

>M6-PGSC0003DMG400014736 M6 LRR-RLP

GGGGATATTCCTATAAGTTTAACACTTCTTGCAGGATTAAGATCATTTAATTTTTCTAAAAATAACCTAACTGGTAGGATTCCAAATGACATTGGTGACATGAAAGTTTTGGAATCTGTTGATCTTTCAGAAAATCAATTTTATGGTCAAATCCCACAAAGCTTTTCAAGTTTGTTCACCTTGAGCTACTTGAATCTATCTGATAACAATTTGTCAGGTATGATACCATTGAGCACTCAACTTCAAAGCTTTGATCCCACGAGTTTTCAAGGAAACAAACTCTGCGGGCTTCCACTCTCGCTGAATTGTAGTTCAAATGGTAATATTCCAAATCATGAATATGAAGATGATGAGAGTGACGAAGATGAAGTGGATTGGTTTTACATTTCAATGGCTATAGGATTTGCACTAAGCTTTTGGGGTGTTTGTGGTTCATTGCTTTTCAAGAGATCATGGAGACATGCTTATTTTTGTTT

>M6-PGSC0003DMG400014738 M6 LRR-RLP

ATAGTGACGAATAAAGGCAACAGGTACCAGTATGACACCATTTTGGCGTTGTTTACAAGTATGGACATGTCTAGCAATAATCTCTCTGGGGATATTCCTATAAGTTTAACACTTCTTGCAGGATTAAGATCATTTAATTTTTCTAAAAATAACCTAACTGGTAGGATTCCAAATGACATTGGTGACATGAAAGTTTTGGAATCTGTTGATCTTTCAGAAAATCAATTTTATGGTCAAATCCCACAAAGCTTTTCAAGTTTGTTCACCTTGAGCTACTTGAATCTATCTGATAACAATTTGTCAGGTATGATACCATTGAGCACTCAACTTCAAAGCTTTGATCCCACGAGTTTTCAAGGAAACAAACTCTGCGGGCTTCCACTCTCGCTGAATTGTAGTTCAAATGGTAATATTCCAAATCATGAATATGAAGATGATGAGAGTGACGAAGATGAAGTGGATTGGTTTTACATTTCAATGGCTATAGGATTTGCACTAAGCTTTTGGGGTGTTTGTGGTTCATTGCTTTTCAAGAGATCATGGAGACATGCTTATTTTTGTTTTCTAGACCGTAGCTG

>M6-PGSC0003DMG400014739 M6 LRR-RLP

AGTCCAAAATATTACACACCATGAAAGTGTTAGCAATAACACTATGGGTTCTCTTGTTTTTTATTTTGAAAAATACCAAGTTTGTCATTTGTGTAGGGATTTGTAGAGAAAATGAGCAACGAGCATTGGAGAGCTTGAAGAAAGAAGTAGATGATCCCTTGGATATTCCCTCTTCTTGGGTTGTCGGAAAAGATTGTTGTGAATGGGATGGAGTTGTGTGCAACAATCTAACTCGTCGTGTGATTGAACTACGCATAAGTTATCTAAGGATCGATAACCTAGAGTGGTTGTCAAGTCTTTTGAGTCTTGAGAACCTGGAAATGTATCATGTGGATCTCCACAAAGCAACTAACTGGCTACAGGTTATTAACATGCTTCCTTCTTTGGTTGATCTAACGTTATCCAATTGTAGTCTTCATAATATAACAAATCTACTTCATCACAATCTTTCTTCACTTGAAACCCTCGATCTTTCTAGTACTAAGTTTAATTCACCTATTCCCAATTGGGTTTTCAACCTTTCCAATCTTGTTTCTCTTAATTTGAGTGATTGTAATTTTACTGGCCCATTTCCCGACTTTCCTGTTAACTTGACTTCTCTCACTACCTTCAAGGCTTTTGAAAACTCTTTCAATTGTCTTTTACCCGAATGGTTGTTTGATTTAAGTAATCTAGAACATCTTAACCTCCGTTCTAGTGGTATTAAAGGTGCAATACCAAATAAGAGTGGAAACACAACAACACTTAAATATCTCGATCTTTTTGATAATAACCTTAACTCCACCATACCAAATTGGTTATATCAATGCAAAGATCTAGAATCACTTTATCTTACTGAAAACCGTCTTGAGGGAACAGTTTCAAGTCTGATTTCAAATTTGAGCTCAATAACTAATATTGACCTTTCTGGTAATATGCTTACTGGAAAGTTACCAAATGTAATTGGAATTTTAGGGAAATTAGAATTTCTTTCTCTTTCAAAAAATCAATTTGAGGGACAAGTATCTGAATTATTCAAAGGTAGGAGTAATTTCTTTTCAAGGGGATTGGGAAATACTTCTTTGATGGAATTGAGACTAGATGGTAATAAACTTAATGGAACTCTTCCGAAAAGTCTTGGACAACTTTCAATGCTAGAATACTTTTTCATTTCTAACAATAAGTTGGAAGGTGTTGTAACAGAGAGTCATTTCTCTAAATTAACGCAATTACAACTCTTCTCTGCATCTAGAAATAATCTGACTTTAAAAGTGAGTAGGAATTGGATTCCACCTTTTCAAGTCACAAACATTGCGATAGGCGGGTGGAATATAGGCCCTTTGTTTCCAATGTGGCTCCGAACTCAGGAGATGATAATGAATGTTGACATATCAGATTGTGGAATACAAGGTGAGGTTCCAACTTGGTTTTGGAACTTATCTTCTCAAATTGTTTTTCTCAATCTTTCTCACAATCAATTTATTGGTGAGGTTCCAATCATCTCAACAGCTTCGTGGTTAGTTGGATTGGGAGGTCCTTGGTTAATGTATTTAGGTTCCAACAATTTTAGTGGATCACTACCACGTATTTCAACGAGTATAACTGAGTTAGGTCTCTCGAACAATTTTTTTTCAAAAGGTTTATCCAACTTCTTGTGTGAAGCAAAGAATGTATCTTATAATTTGGAGATCTTAAATCTCGGGGGAAATGATTTATCAGAAGAAATTCCTGATTGTTGGATGAATTGGCCTGAGTTGGAAGTTTTAATCTTGAGGGACAATAATTTGATTGGAAGCTTACCAAGATCTATGGAGGTTTTGAGTAATTTGCTATCATTAGACTTACGAAGAAATAGACTTAACGGTCCGTTTCCTTCATCCTTGAAAAATTGTACGAAACTGCATAAAATAGATTTAGCCGAGAATGAGTTTGCTGGGGGACTACCATCTTAGTTGGGAATGAGGTTTTCAACCTTGATAGTATTTATCCTTCGGTCCAATAAATTCAATGGTGAATTGCCTCAAGAACTTTGTCAACTCAAAGATCTTCAGATCTTAGACCTTGCAAACAATACATTTGTTGGAATCATACCAAGGTGTATTGGTAATTTCAGTGCAATGGTCAAAGGAAAAAAGGAAGTGGGAGATGATGATGAGTTAAATTATTCCTACTTTTTGGGAGTTTTGAGAGAGAGTGCCATAGTGACGAATAAAGGCAACAGGTACCAGTATGACACCATTTTGGCGTTGTTTACAAGTATGGACATGTCTAGCAATAATCTCTCTGGGGATATTCCTATAAGTTTAACACTTCTTGCAGGATTAAGATCATTTAATTTTTCTAAAAATAACCTAACTGGTAGGATTCCAAATGACATTGGTGACATGAAAGTTTTGGAATCTGTTGATCTTTCAGAAAATCAATTTTATGGTCAAATCCCACAAAGCTTTTCAAGTTTGTTCACCTTGAGCTACTTGAATCTATCTGATAACAATTTGTCAGGTATGATACCATTGAGCACTCAACTTCAAAGCTTTGATCCCACGAGTTTTCAAGGAAACAAACTCTGCGGGCTTCCACTCTCGCTGAATTGTAGTTCAAATGGTAATATTCCAAATCATGAATATGAAGATGATGAGAGTGACGAAGATGAAGTGGATTGGTTTTACATTTCAATGGCTATAGGATTTGCACTAAGCTTTTGGGGTGTTTGTGGTTCATTGCTTTTCAAGAGATCATGGAGACATGCTTATTTTTGTTTTCTAGACCGTAGCTGGGAAATGTTGCTTGCAAAGTTATCAGTATGTTGAGAGATATCATTAAGGATATTGCTATCATCAAAAGTTAGTTTGTGTGCTTACATTTCTCATGTTAGCAAACATGTTGTTTCATTTTGTATTCAAATGAATGTTTCTGAAGATGTTCAAACATTGTTGTTTCTTTCATAGTTCCAATTTTC

>M6-PGSC0003DMG400014740 M6 LRR-RLP

AGGCCCTTTGTTTCCAATGTGGCTCCGAACTCAGGAGATGATAATGAATGTTGACATATCAGATTGTGGAATACAAGGTGAGGTTCCAACTTGGTTTTGGAACTTATCTTCTCAAATTGTTTTTCTCAATCTTTCTCACAATCAATTTATTGGTGAGGTTCCAATCATCTCAACAGCTTCGTGGTTAGTTGGATTGGGAGGTCCTTGGTTAATGTATTTAGGTTCCAACAATTTTAGTGGATCACTACCACGTATTTCAACGAGTATAACTGAGTTAGGTCTCTCGAACAATTTTTTTTCAAAAGGTTTATCCAACTTCTTGTGTGAAGCAAAGAATGTATCTTATAATTTGGAGATCTTAAATCTCGGGGGAAATGATTTATCAGAAGAAATTCCTGATTGTTGGATGAATTGGCCTGAGTTGGAAGTTTTAATCTTGAGGGACAATAATTTGATTGGAAGCTTACCAAGATCTATGGAGGTTTTGAGTAATTTGCTATCATTAGACTTACGAAGAAATAGACTTAACGGTCCGTTTCCTTCATCCTTGAAAAATTGTACGAAACTGCATAAAATAGATTTAGCCGAGAATGAGTTTGCTGGGGGACTACCATCTTAGTTGGGAATGAGGTTTTCAACCTTGATAGTATTTATCCTTCGGTCCAATAAATTCAATGGTGAATTGCCTCAAGAACTTTGTCAACTCAAAGATCTTCAGATCTTAGACCTTGCAAACAATACATTTGTTGGAATCATACCAAGGTGTATTGGTAATTTCAGTGCAATGGTCAAAGGAAAAAAGGAAGTGGGAGATGATGATGAGTTAAATTATTCCTACTTTTTGGGAGTTTTGAGAGAGAGTGCCATAGTGACGAATAAAGGCAACAGGTACCAGTATGACACCATTTTGGCGTTGTTTACAAGTATGGACATGTCTAGCAATAATCTCTCTGGGGATATTCCTATAAGTTTAACACTTCTTGCAGGATTAAGATCATTTAATTTTTCTAAAAATAACCTAACTGGTAGGATTCCAAATGACATTGGTGACATGAAAGTTTTGGAATCTGTTGATCTTTCAGAAAATCAATTTTATGGTCAAATCCCACAAAGCTTTTCAAGTTTGTTCACCTTGAGCTACTTGAATCTATCTGATAACAATTTGTCAGGTATGATACCATTGAGCACTCAACTTCAAAGCTTTGATCCCACGAGTTTTCAAGGAAACAAACTCTGCGGGCTTCCACTCTCGCTGAATTGTAGTTCAAATGGTAATATTCCAAATCATGAATATGAAGATGATGAGAGTGACGAAGATGAAGTGGATTGGTTTTACATTTCAATGGCTATAGGATTTGCACTAAGCTTTTGGGGTGTTTGTGGTTCATTGCTTTTCAAGAGATCATGGAGACATGCTTATTTTTGTTTTCTAGACCGTAGCTGGGAAATGTTGCTTGCAAAGTTATCAGTATGTTGAGAGATATCATTAAGGATATTGCTATCATCAAAAGTTAGTTTGTGTGCTTACATTTCTCATGTTAGCAAACATGTTGTTTCATTTTGTATTCAAAT

>M6-PGSC0003DMG400014753 M6 LRR-RLP

AGTCCAAAATATTACACACCATGAAAGTGTTAGCAATAACACTATGGGTTCTCTTGTTTTTTATTTTGAAAAATACCAAGTTTGTCATTTGTGTAGGGATTTGTAGAGAAAATGAGCAACGAGCATTGGAGAGCTTGAAGAAAGAAGTAGATGATCCCTTGGATATTCCCTCTTCTTGGGTTGTCGGAAAAGATTGTTGTGAATGGGATGGAGTTGTGTGCAACAATCTAACTCGTCGTGTGATTGAACTACGCATAAGTTATCTAAGGATCGATAACCTAGAGTGGTTGTCAAGTCTTTTGAGTCTTGAGAACCTGGAAATGTATCATGTGGATCTCCACAAAGCAACTAACTGGCTACAGGTTATTAACATGCTTCCTTCTTTGGTTGATCTAACGTTATCCAATTGTAGTCTTCATAATATAACAAATCTACTTCATCACAATCTTTCTTCACTTGAAACCCTCGATCTTTCTAGTACTAAGTTTAATTCACCTATTCCCAATTGGGTTTTCAACCTTTCCAATCTTGTTTCTCTTAATTTGAGTGATTGTAATTTTACTGGCCCATTTCCCGACTTTCCTGTTAACTTGACTTCTCTCACTACCTTCAAGGCTTTTGAAAACTCTTTCAATTGTCTTTTACCCGAATGGTTGTTTGATTTAAGTAATCTAGAACATCTTAACCTCCGTTCTAGTGGTATTAAAGGTGCAATACCAAATAAGAGTGGAAACACAACAACACTTAAATATCTCGATCTTTTTGATAATAACCTTAACTCCACCATACCAAATTGGTTATATCAATGCAAAGATCTAGAATCACTTTATCTTACTGAAAACCGTCTTGAGGGAACAGTTTCAAGTCTGATTTCAAATTTGAGCTCAATAACTAATATTGACCTTTCTGGTAATATGCTTACTGGAAAGTTACCAAATGTAATTGGAATTTTAGGGAAATTAGAATTTCTTTCTCTTTCAAAAAATCAATTTGAGGGACAAGTATCTGAATTATTCAAAGGTAGGAGTAATTTCTTTTCAAGGGGATTGGGAAATACTTCTTTGATGGAATTGAGACTAGATGGTAATAAACTTAATGGAACTCTTCCGAAAAGTCTTGGACAACTTTCAATGCTAGAATACTTTTTCATTTCTAACAATAAGTTGGAAGGTGTTGTAACAGAGAGTCATTTCTCTAAATTAACGCAATTACAACTCTTCTCTGCATCTAGAAATAATCTGACTTTAAAAGTGAGTAGGAATTGGATTCCACCTTTTCAAGTCACAAACATTGCGATAGGCGGGTGGAATATAGGCCCTTTGTTTCCAATGTGGCTCCGAACTCAGGAGATGATAATGAATGTTGACATATCAGATTGTGGAATACAAGGTGAGGTTCCAACTTGGTTTTGGAACTTATCTTCTCAAATTGTTTTTCTCAATCTTTCTCACAATCAATTTATTGGTGAGGTTCCAATCATCTCAACAGCTTCGTGGTTAGTTGGATTGGGAGGTCCTTGGTTAATGTATTTAGGTTCCAACAATTTTAGTGGATCACTACCACGTATTTCAACGAGTATAACTGAGTTAGGTCTCTCGAACAATTTTTTTTCAAAAGGTTTATCCAACTTCTTGTGTGAAGCAAAGAATGTATCTTATAATTTGGAGATCTTAAATCTCGGGGGAAATGATTTATCAGAAGAAATTCCTGATTGTTGGATGAATTGGCCTGAGTTGGAAGTTTTAATCTTGAGGGACAATAATTTGATTGGAAGCTTACCAAGATCTATGGAGGTTTTGAGTAATTTGCTATCATTAGACTTACGAAGAAATAGACTTAACGGTCCGTTTCCTTCATCCTTGAAAAATTGTACGAAACTGCATAAAATAGATTTAGCCGAGAATGAGTTTGCTGGGGGACTACCATCTTAGTTGGGAATGAGGTTTTCAACCTTGATAGTATTTATCCTTCGGTCCAATAAATTCAATGGTGAATTGCCTCAAGAACTTTGTCAACTCAAAGATCTTCAGATCTTAGACCTTGCAAACAATACATTTGTTGGAATCATACCAAGGTGTATTGGTAATTTCAGTGCAATGGTCAAAGGAAAAAAGGAAGTGGGAGATGATGATGAGTTAAATTATTCCTACTTTTTGGGAGTTTTGAGAGAGAGTGCCATAGTGACGAATAAAGGCAACAGGTACCAGTATGACACCATTTTGGCGTTGTTTACAAGTATGGACATGTCTAGCAATAATCTCTCTGGGGATATTCCTATAAGTTTAACACTTCTTGCAGGATTAAGATCATTTAATTTTTCTAAAAATAACCTAACTGGTAGGATTCCAAATGACATTGGTGACATGAAAGTTTTGGAATCTGTTGATCTTTCAGAAAATCAATTTTATGGTCAAATCCCACAAAGCTTTTCAAGTTTGTTCACCTTGAGCTACTTGAATCTATCTGATAACAATTTGTCAGGTATGATACCATTGAGCACTCAACTTCAAAGCTTTGATCCCACGAGTTTTCAAGGAAACAAACTCTGCGGGCTTCCACTCTCGCTGAATTGTAGTTCAAATGGTAATATTCCAAATCATGAATATGAAGATGATGAGAGTGACGAAGATGAAGTGGATTGGTTTTACATTTCAATGGCTATAGGATTTGCACTAAGCTTTTGGGGTGTTTGTGGTTCATTGCTTTTCAAGAGATCATGGAGACATGCTTATTTTTGTTTTCTAGACCGTAGCTGGGAAATGTTGCTTGCAAAGTTATCAGTATGTTGAGAGATATCATTAAGGATATTGCTATCATCAAAAGTTAGTTTGTGTGCTTACATTTCTCATGTTAGCAAACATGTTGTTTCATTTTGTATTCAAAT

>M6-PGSC0003DMG400014829 M6 LRR-RLP

CAAATATCGAAATTTTCCTAAACTACTTTGCACGAAAAAACAAAGAGGAGGAGGAAGAAATAGTATTCGTAGATTACATCTGAGCTTGACAACAAATTATGCAAGATTTTGAGTGAAACAACCAGACCTTTCATGGTGTTTCTTGTTGGCTGCTAAGAAAACAATGGCGTTTTCACCTGAAAAATCAAATTTCAGGTCACCATTGAAAGGGCCACCATCTCCTAGCTGGAATGATTTGTTGCTCAAGAACAAGAAAGCACTCAATCATGTCCTTTTCACTATGCATCTTCAATCCAGGTCCAATAAGCAAACCCCATCTTCATCCCCTTCTCCACAACTTCGCAAATCCCTAGATTTCACTCTTCCCTCACTTGTATCCGATCCAATTTCTCTTCTTTCTGATCAAATACTTCTTTCCATTCTCTCTAAACTCCCTGATTCACAAAGAAATTGTAACTCTCTTGTATCTAAACGTTGGCTCAATCTCCAAGGTCGTCTGGTACGGTCGATTAAGCTTCTTGATTGGGATTTCCTTGTTTCGGGTCGGTTGTTTGTTCGGTTCCCTAATCTTATCCATGTTGATTTGGTTAATGGGTCTCTCATTTCACCGGGGAATGCAGGTGTTTTTTGCTCTAACCGGTTGCTTTCTTCTCACGTGGATTGTAGCGCTGATGCGAAAGATTGGTTTCTTAAGGAAAGATTTGTGTTGCCTTCTGATGAGATTGATAAGGGGTTGAGAATACTAGCAAGTGGGTGTCCTAATTTGCGTAGATTGTTGGTGGTAAATGCTAGTGAATTGGGATTGTTGGGGGTAGCTGAGGAATGTCCAACTTTGCAAGTGTTGGAATTGCACAGATGTAATGATCAGGTTCTTCGTGGTATAGCGGCCTGTCAGAACTTGCAGATATTGAGATTGATTGGAAATGTTGATGGGTTTTATAAATCTTTGGTTACTGATATTGGATTGACTATTTTAGCTCAGGGGTGTAAGAGGTTGGTGAAGCTGGAGCTGAGTGGTTGTGAAGGGAGCTATGAAGGGATTAAGGCAATAGCACAATGTTGTCAAATGCTCGAAGAGTTGATTCTTCGTGATCATAGGATGGAGGGTGGTTGGTTGTCTGCTCTTTCCTATTGTGAAAATTTGAACACCTTGAGGTTTATGTCTTGTAAGAGTATTGATCATTGTGGATGGTTTGATGAAGATGTAGGATCTTGCCCAACACTTGAAAGATTACATTTCGAGAAATGTCAATTAAGAAATAAGGAGAGTTTGAGAACGTTGTTTCTACTATGTCAAGAAGTTAGAGAGGTTATTTTCCACAACTGTTGGGGACTGGATAATGAAATGTTCAGTCTATCCACTGGTCTAAGGTATTGGTTATATTATTCTGTTCTCGTTTACTTTGCTTGCTGTTGCTGGAAAATTTCGTGCTGGATTAGCATTTTGTTATGCATTTGACATACATGAACTTATTGAGTTTAGGTTTTATGACCTTGAAATGTAGTTCCTTATTTTTGTTAAAAATGACTTTTTAGTGTTGCTGTAACCCCCATCCCCCACCCCTCGGGTATAATAGACGACCAAATGAAAGGATAGTTTGCTTGTAATAGTGCATTTTATTTTGGTCATAGTATACCCATTGATCTTGTTTTCTTGGTAAAAGGATAAGTGTGATAAATTTCCTTAGCTATTTGTGTTGGGGATAGATTGTTTTTTTAAGGCCAGTTTGTAGATGCATTTGTAATCATGCACCGTCTTAGTGTTTCAGTTATTAATGTTAAAATAGCCTTACCATTTCTCGAAGAGTATACACTTCCATCTTGCATTAGAGAAAACACATAATGTTGATGGCATGACAATTATGTTTCAATTCATTATTGATGCTTGATTTATCAATATCATGTGTGCTATCTTGGATTATAAAGAGTCCAGTTTTGAGTTTTATACATTGAACTTGCTTCAGTTCTGACTGGCAATTTGGTATCATATTTTGTTTTGCATTCACCTGGAGCAACAGTAAAGTTGTTTCCGTGTGACCTATAGGTCACCAATTTGAGCCGTGGAAGTAGCCACTAATGCTTGCATTAGGAGGGTATCCCCTTGCGTGAACAATGGATAATTTATGCACAAGGCTGCCCTTAAAGTTCACTTGAGTAAAGGGCCTGTAATTAACTAAATATAACATTATGTTAGGAATTTGGGTTTCACAGCTTCATTTTGGTGTTATTTTCAACAGGAGAGTGAAGTCTCTTTGCCTAGAAAGCTGTTCACTGCTCACAACTGAAGGCCTCGAATCTGCACTCCTTTCATGGAAGGAGATCCAGAGCCTCAAGGTGATATCATGTGGCAATATAAAGGATAGTGAAATCAGTCCAGCACTTTCTACCTTGTTCTCCACGCTAAAAGATTTTCAGTGGAGACCAGACACAAAATCTCTTCTTTCAGCTGGTCTTGTGGGAACTGGCATGCGAAAAAGAGGCAGTAAATTATTCAAGAAGACGTGCGACTGGAAGTCACTGCCTGGTGCATAGACTGGCTTCCTCATCATGCATTCAGACAATAAATTATTCAAGAAGACATGCGACTGGAAGTCACTGCCTGGTGTATAGACTGGCTTTCTCATCATGCATTCAGACAACTCTACTAGATTTTACCTCCTGATTTATGCATATGAGAACTCTAACACATGAAAGAGGTCAGTGCGACACTTTGCTCCAATACTGTCAGCAATGAGGCCTTGATGTCATTATCCTGTATTTGTAGTCGCTTAAGTATGTAACGTAGGAACTATAGGATCGAAGATCGGCACTTTCTCAGGGTAGAGATGGAGAAAGAGGCCTAAGTTAATGTATAGAGCTCGATGCTGTAACCTCATGAGCATTCAGCTAAAGCCATCTCGCTGTATCTTTGTCAAGAAGCTTAATGGTATTTCTGTATTTGGAACCCCAAAAACTCTTAAAAGGGGTTTTCCTATCTACGATGTTATAAAGAATTTTGGCACCTTTTATGCCATACTACTGCCTTCCACAATTGTGTGAAAGCATATTTGATATGAACTATACCAGCAAGCATGTCATTTGTGGAGGAGACGAGAACCAGAATAATTTTCTACATCAATGGCATCTATGTC

>M6-PGSC0003DMG400015121 M6 LRR-RLP

TTTAATGTTTTGCATGATTGATGTGATAGTACCTTGAGAAATCGTGATATCGCAAATACCAATTGTTCGGTCAACTCAACAACATAAGTGTTGCTATAATACATTTGAAAAACAAATCAAGCTTTGAATCAATCATGAAGAAAAACATTCTCTTATTGATTCTTCTCTTTCTAATTCAATTTTCTGTATCACTTTCTTCCTCAAATGAGACTGACCAACAAGCTCTACTGGCTTTCCAAAATCTTATTACAACTCCCAATCATTTTTTGGCCAATAATTGGACCAAAAATACTTCTTTTTGCTCTTGGTTTGGTGTCACTTGCAGTCCAAAAAGGCAAAGGGTTGTGGCCTTGGATCTTCCTAATTTGCATCTTCAAGGCACAATCTCGACGTCTTTGGCCAATTTGTCCTTCCTCAGTGTTTTCAATCTCGAGAACAACAACTTCAGCGGTGGCATCCCTTATGAACTTGGCCACTTGCCTCGCTTGCGAGTGATTGATATTCAAAACAATCAGCTCAACGGAAGTATACCAACAAGTCTATTTCAACACCGGAGAGTTCAAGTAATTTCATTGGCTTTCAATGAACTCGGTGGTGAAATGTGGAAAGGGCCATGGTATGTACCCGAACTCAGAGTCTTGGATCTCACCAACAATAACCTCACGGGTATGATCCCTCCTTCTGTTGGAAATGCCACAAAAATGATGAACTTCAGTTTGTCTGGGAATAGAGTCAGCGGCAACATTCCAAAGGAGGTCGGTAATCTGAGCCAACTTGCAGACTTGTCCTTGACTGATAATCAATTAACAGGTTCCATTCCTGTAGGACTATTTAATATCTCGTCGCTACTTGCCATAAATCTGCTATACAATAAACTTTCCGGTCCTCTCTTGCTAGATGAAGGAATTTTTTTGTCAAATCTGAAGTTTCTAAGGATATCTCGCAACCAGATTTTAGGATGCATTCCTTCCAACATATGCCAACTCAGAGAGCTCAAAGTTTTGTCCATATCTTTCAACAACATAATTGGAGACAAACCCAAAAATATTGATTGTTTATCCAATCTCCAGGAGTTTTATAATCGTCATAATTCAATAAAAGGGACTATTCCCACTTCATTGGGAAATATTTCAACTCTGCAATATCTTTATTCTGGAAACAATCTCATGGTGGGGCCAATTCCTCCAGAATTAGGGAAGCTATCAAATTTGAGGCAATTAATTTTTTCCAATAATTCTAATCTTATTGGTCAAATTCCAGAGGCTATTTTCAACATATCTTCTTTGGAATTAATCAATTTCAGTTTCAATACCCTGTCGGGTAGAATTCCAACCACTACAGGTCTTCATCTTCCGAACCTTAAAGTACTTGACTTGGGAGTCAATCAGCTTGAAGGGGAAATTCCATTGTTCATAACAAATGCTTCCAAGCTTCAGATACTGTCGCTAGAACATAACTCTCTCTCGGGAGCTATTCCTACTAATTTGGGAAATCTTCATGAGCTGCAATCACTCTTACTACATACTAATCAACTTACCAATGAACCAAGAGAGCATGAGTTGAAATTCTTCAATTCGTTGGCGAACTGTAGGATGTTGAGATATCTAGAAGTGGGTAACAATCCGTTGAGTGGCGTTCTACCAAATTCTGTTGGGAATCTCTCATCTACTATTGAAAAGATTTATATAGGAGATGCACACATCAATGGCCTCATCCCCACAAGTATAGGCAACATGAGCGGTCTTACAGCCCTAAGCCTTGGAGGAAACAACTTGGCAGGGAATATTCCTTCTGAGATTGGTAAGCTTAAACAACTCCAAGGGCTGGCTCTATATAACAATAAATTGCAGGGACGTATTCCAGAGGTGGTATGCCATTTATCGAATTTGGTTGGCTTATTACTGGATGGTAATGAGCTCTGTGGATTAATTCCAGAATGCTTTGGAAATCTTAGCATGCTACAAGCGCTTTATTTGGATTCTAATAAATTTTCATCAAAATTTCCGTTGAGCCTTTGGAAGATGAGTGGTCTTCTCTATCTAAGTGTGTCACGAAATTCAATAGAGGGAGAACTTCCATCAGATATTGGAGGACTGAAAGCCATTGTAGAACTATATCTTTACAATAACCACTTTTCAGGTGTGATACCAACCAGATTGGGGGAACTCCAAAACCTGCAGCATCTTGACCTATCCAACAATTCATTTTTTGGCCAAATTCCATTATCCTTTGCCAACTTGATAAGCTTGGAATTCTTGAATTTGTCTTTAAATGCCTTGTCCGGTACTATTCCTAAATCTTTGGAAAAACTCTCATACCTTAAAAGCATTAATGTTTCATTTAATGATTTAGAAGGTGAAATACCAAGTGGTGGCGTGTTTGCAACTTCCACTGTGCAATCATTTCTTGGGAACAAAGGTCTTTGCGGAATGCACATATTGGAGATTCCTACTTGTGCTATCACTAATCATGGAAAACAATCAAAGCTTAAGGAGGTTGTGCTAAAAATTGTTACTCCAGTGGTTATTGCATCCTTTCTGATATTCTTGTTCGTTTCAATTTGGATAATGAAACGACAGAAGAAAGGGAAGTCCAAAGATATTGAAAAGGTATTGGAGATCGGGACTCATCAATTAGTTTCTTATCACGAGATTCAACAAGCAACAAATAATTTTGATGAATCCAATTTAATTGGTGAGGGAAGCTCTGGCTCTGTGTACAAAGGCACATTGTTTGGTGGAACTGCAGTGGCCATTAAGGTTCTGGATTTGGAAAACGAGCAAGTATGCAAGAGGTTTGATACCGAATGTGAAGTGATGAGAAATGTTAGACACAGAAATCTTGTTTCAGTGATTACTACATGTTCTAGTGACTATATAAGAGCCTTTGTTCTGCAATTTATGCCCAATGGAAGTCTTGAGAATTGGCTGTACAAAGAAGATCGACACTTGAACCTTCATCAAAGAGTCACTGTAATGCTTGATGCAGCTATGGCAGTTGAATATCTACACCATGGTCATAACACTCCAATAGTTCATTGCGACCTGAAGCCAGCCAACGTTCTTTTGGATGAAGATATGATGGCTCATGTTGGTGATTTTGGAATATCTAAAATTTTAGCCATAAGCAAGTCCATGGCTTATACAGAGACATTGGGCACTCTTGGATATATTGCACCAGGTAAAAGAAAATCTACTCTCATTGATTTTCTCTTATCATAATTAAGCCTCTCCAAGTTGTAGAAGTAAAAAGAGAATTATTGTTGTATTTCAATTGAATTAACTTTTTTTTAAATCCTTTTTTAAGAATATGGCTCGGATGGAATAGTGTCTGCTAGTGGCGATGTTTATAGTTACGGCATCATGTTGATGGAGGTTTTGACGAAAAGAAGGCCAACAGATGAAGAGATATGCAATGAAAATCTTGACTTGAGGAAATGGATCACACAATCATTTTCAGGGAGTATGATGGACGTTGTGGATGCCAATCTTTTCTCCGAGGAAGAACAGATCACTTCAGAAAGTGAAATCTGCATTGCGTCCATGATAGAATTGGGTTTAGACTGCACAAAGGAAATGCCAGAATCAAGAATAAGCATGAAAGAAGTAGTGAAGAGGCTTAACAAAATCAACAACACATTTCTGGAAACATAGTAGTGATCAGTATCTATTCCGGTGGTGTTCTTTTCTGAGCTGCAAATTAATATGATGCTTTTTGTTTACTTAAT

>M6-PGSC0003DMG400015283 M6 LRR-RLP

ATGCACTCCAATACCAAAACAACATTTACGAGAGGTATCGTCTTTTGCTTCTTGACATTTTCCAAAATGGAAGTTTTCACAATGAGTTTTAATTTCCACACCTTTCTGGCTATCCGTGCTGTGTTTTTTGTGTTTCTATTGTCATTTTCTCTCCAACATGCTGCATCAGCTGCTTTTCTTGGCAATGAAACTGATAAACTTGCCTTGCTTGCATTCAAGACCCAAATAACCGAGGATCCATCAAGAGTTTTTGTCTCTTGGAATGATTCTGTTCATTTCTGTCAGTGGACTGGAGTAAAATGTGGCCTGAGACACGTAAGAGTCATCCGTTTGAATCTCAAAGGGCTGAGGCTGGCAGGTACTATCTCAGATCATCTTGGAAATCTCTCTTTTCTCAATTCTCTTGACCTTGCAGAAAATGTCTTCCATGAAAAAATTCCTCAACAACTCAGCAGGCTGCCAAGGCTTCAATACTTGAATTTGAGTTTTAACTATCTAACAGGAGAAATTCCAGTTAATCTATCACATTGTGTTAAACTCAAGAGCCTTGTCCTGGACCACAACACTCTAGTGGGAAAGATTCCTTACCAAGTTGGATCTTTAACGAAGTTAGTGAAATTGTATCTCGAAAATAACAACCTGACAGGAAACATTCCAGGTTCTATTGGAAATCTTACATCTTTAGAGGAGCTGTATTTATCGTACAACAATCTAGAGGGAGAAGTGCCAGCTTCTTTAGCTCGATTTACCAAGTTGAGGCTTCTTGGACTGTCAGTAAATAGCTTATCCGGGGATTTCCCTCCTCCGTTGTACAATTTGTCATCCCTGGAACTAATATCACTTTCCCTTAACAACTTTTCTGGTAATCTCAGATCTGATTTAGGCAACTACTTCCCTTATCTCCAAATACTTTTCTTGGCGAATTGTCAATTCATTGGTTCCATACCATCCTCTTTGGCCAATGCTTCAAAGTTGCGAGAGCTTGATTTTCCTTTAAACAATTTCACTGGAAATATACCTAAGGGTTTTGGTAACTTGTGGAATTTGTTGTGGCTCAATGTTTGGAGTAACCATCTTGGATACGGTAAGCATGATGACCTTGACTTTGTAAATTCTCTTACTAACTGCAGCAGCCTACAAATGCTCCATTTTGGAGGCAATCAGTTTGGAGGTACATTACCCCATTCAGTATGCAACCTTTCTAGTCAGTTGCAACGCCTACTCTTCTTCGAAAATAGAATTGGTGGAATCATACCAAGAGAGATCTCAAACCTGGTGAATTTGAATCTACTCTACATTGGTTCTAACAATTTTATAGGTAGTATTCCAGATTCTATTGGAAGACTTACAAACTTGGGAGCTCTCTTCTTGTTTAACAATCTCTTGACAGGTGTCATCCCTTTTTCAATAGGAAACCTCACTGTACTCTTCTATCTTTTTTTGGGACTTAACAGGTTAGAGGGTAACATACCTTCAACATTGGGAAACTGTAACCAACTGCTAAAATTGGGATTTTCTGAAAACAACCTGACAGGAACTATACCACAACAACTTTTTGGTCTCTCATCCCTCACAGATATTTATGCATCTTATAACTCTTTGACTGGTCCATTACCGGTGGATATTGGTAACTGGAATCATCTTACTTATCTTGATTTTTCTTACAACAATTTTTCGGGTATGATTCCACTAACTCTTGGAAAATGCTTGACCTTGTGGGAGATCTATATGAAGGGAAACTCCCTCCAAGGAACCATTCCCAATTTAGAAGATTTGCCGGATCTCCAATCCTTGGATCTTTCCCTAAACAACCTATCAGGGCCAATCCCTCACTTCATTGCAAATCTTACTTCCTTACACTACTTGAACTTATCTTTTAACAATCTAGAGGGTGAGGTTCCTGTCACCGGAATATTTTCTAATTTGAGTGCAGTTGTATTAAGCGGAAACTCCAAGCTTTGTGGTGGGATTCAAGAGCTTCATTTACAACCTTGTATTTATCAAAAGACACGGAAGAAGCATGTGCTTGCCCTCAAGTTTATTTTGATAATTGTTTTTGCTGCTTCATTTTCAATCCTGGCCTTGTTAGTAGTTTTCCTTTGTTGGAGAAGAAATTTGAACAATCAACCAGCACCAGAAGATAGTTCTAAATCTGCGCATTTCTACCCCAATATTTCGTATGAAGAGCTCCGCACTGCAACAGGTGGATTTTCTTCTGAAAATCTTATTGGCTCTGGCAGTTTTGGAACTGTTTATAAGGGAACTTTTCCCTCCAATGGAACGGTCGTTGCTGTGAAGGTACTCAAACTCCAGCTTGAAGGTGCTTCAAAAAGCTTCTTAGCAGAATGCCAAGCATTGAGAAACACAAGGCACCGGAATCTTGTCAAGGTCATCAGTGTATGCTCAAGTTCTGATTTCAAGGGAAATGACTTTAAAGCTTTAGTCTTTCAGTTCATGCCAAAAGGGAACTTGGACGAATGGCTACATCCAGAAAAGGAGATGCATGAGAAGAGTAGCTTGACCACACTTTAGAGAATGAATATCATAATAGATGTGGCCTCTGCACTTCATTATCTTCACCATGAATGCCAAACACCCATGATTCACTCTGATATAAAACCACAGAACATTCTTCTTGATGAAGATCTCACTGCTCATTTGGGTGATTTTGGTTTGGTGAGACTCGTTCCTGAATTCAGTAATGAATCGGATCTACATCAGTTTAGCTCACTTGGAGTTCTGGGAACCATTGGCTATGCAGCTCCAGGTAATTTTTATATATAAACAAAAACTATGAATATACCATATGGTTTCTCGTAACATTAGTGGTTTATTTATAGGCTGATCAATTTGCTTGTTGCAGAATACGGCATGGGTAGTAAGGTCTCCGTTGTAGGAGATATGTACAGTTTTGGGATCCTAATATTGGAGATATTCACTGGAAGAAGACCCATGGACTCTTTGTTCCAAGCAAGCTCTACCCTCCACCACTTTGTAGAAACAGCATTGCCAGAGAAAGTTATGGAGATTCTAGACAAAACAGCTTTTCATGGTGAGATGAGTAAAGAAACCTGTGGCTAAGAATACTGGGGCAGTATCAAGAAAGAACAGATGGAATGCTTGGTTGGCATACTTGAGATTGGTGTTGCATGTTCAGCTGAATCCCCAAGTGACAGATTGACCATGACACAAGTTTACAGTAAGTTGACACTAATAAGAGAAAAATTCCTACTAAGGGGAGATTGAGGCCACAGGCTACAAGTGAAAGCATTGGATTTAATCTAA

>M6-PGSC0003DMG400015345 M6 LRR-RLP

TATTTCTAACAATAAGATTCGTGGCCAAATCCCAAACTGGTTTAGCGGCATGAGGTGGGACAACTTGTCCTACCTAAACCTTTCGCATAACTCTTTAACAGGCCACCTACAACAATTTAATTTCCATAATCTAAGGTATCTTGATCTTAAATTTAACTCCCTTCAAGGTCCAATACCTTCATCCATTTGTAACATGAACAGGCTTGAATTTCTAGATTTATCACGCAACGACTTCAGTAACTCGATTCCAAGTTGCTTGGGAAGCATGTCTAGTCTAACAGTGTTGGACTTAAGAAGGAACAATTTCACTGGGAGTCTTCCATCGTTATGTGCGCAGAGCACTTCATTGAGAACCATTGTCTTGAATGGTAATCAATTTAAAGGAACTCTCCCTGTGTCGTTGCTCAACTGTAGTGATTTAGAAGTTCTTGATATGGGGAATAACGCTATAAATGACACATTTCCAGCTTGGCTAGGAACACTTCAACAACTGCAGGTCTTAATATTAAAGTCCAATGTGTTCCATGGACCAATTAGTACTTGTCAGACTACATTTTGCTTTCCCAAGTTGCGGATTTTTGATCTTTCGCATAACAAATTCAGTGGCTCTCTCCCTGCAAAAGTTTTTGGAAACTTCAAGGGAATGATTAGATTAGACGGTGAAGACACACGAGAGATCATGTACATGCAACCATCTATAATGCCATTTTACTCATCATCATATGAGGATTCTGTGAGTTTGGTGATCAAAGGCCTGGATATTGAGCTAGAAAGAATCAGCACGATTATGACAACCATAGACCTCTCGAGCAACCACTTTGATGGTGTCATTCCGAAAACACTAAAGGATCTCAGCTCACTTTGGCTACTCAATTTATCCCATAACAATCTCATAGGTCATATTCCAATGGAATTGGGGAAATTGAATAAGCTTGAAGCATTAGATCTCTCTTGGAATCGGCTCACTGGAAAGATTCCGCAGGAATTGACAAGGCTAACATTTCTGGAAAAGTTAAACCTTTCTCAGAATGTTCTTGTTGGACCAATTCCTCACGTTCTACAATTCAACACATTTGAAAATGACTCGTATGGCGGTAACCTTGATCTATGTGGTCCTCCTTTATCAAAGCAATGTGGAACGAGTGATCTATCCCATGTTCCTCAACCATTGGAGTCTGAAGACGAGTCATATTTTTTTAGTGGATTTACGTGGGAATCAATAGTCATTGGCTACAGCTGTGGACTAGTTGTTGGAACTGTTGTGTGGAGCCTCATGTTTAAATATCGTAAGCCAGAATGGTTTGTCGAATTTTTTGAAGGCATTTATCCCAAGAAAATGAGAAGGTCAATGAAGAGAGCTCAGAGACGACGGACTTAATAGAAGATTGCAGGAG

>M6-PGSC0003DMG400015347 M6 LRR-RLP

GTTGGTAATTAGTGAAAGCTTTTCTTCTTCCTCTGTGATGATGCATCATGTTTGCTCTCCCACTGAAGCTTTTGCTTTGCTTCACTTCAAACAATCCTTTCAAATCTCTGACCAACCTAGCTCTCGGGATGATACTGGTGTGCAATTTGTTGATGTCCCAAAAACGAAGTCTTGGAATGAGAGTAGGGATTGCTGCAGTTGGGACGGGGTAACTTGTGACATGTTAACCGGTCATGTTATCGGATTGGACCTTAGTTGCAGTCTGCTTAATGGAACTATTCATCCCAATAGCAGCCTCTTCCAACTCCATCATCTCCGGACACTAAACCTTGCTTACAATGACTTCTATTCTTCTTCAATCCCACATAACATTGGCCGATTGAGGAATTTGAGGCATCTCAACCTTTCTCATTCTTTGTTTGAAGGGGAAATCCCAACAGAAATCTCATACCTTTCCAATTTGGTTTCACTGGAACTTTTTTGTAATGGATGTGAACTTGCTGAGAGAACATTCGAAACAATTCTTCAAAACTTCACAAATCTAGAGGTACTATCTCTCTCTTATGTCAACATCTCATCTCCATTACCTGTGAATATTTCTTCTTCCTCGTTAAGGTACGTGGATCTTGAATTTACTAATCTGGGAGGTGTTCTCCCAGAGAGTCTTTTCCTTCTGCCAAACTTGGAAATGCTCTATTTGAATTACAATGATCTTCTCGAAGGAGTTTTACCAAAGATCCACCCGAGCAACACTCTGTTGGAGTTGGATATTTCCTACACAGGAATCTCCGGTGAGCTGCCTGATTCAATTGGCACCTTGAGTTCATTGAATCTCTTGAACCTCAAAAGATGTCAATTCTCTAGTTCCATTCCTGATTCCATTGGCAATCTTACACAAATGCGGGGTTTGGATTTTGGTAATAATCATTTCACCGGCTATATTCCTTCCATAATCTCTAAATTGAAGCACCTCACATCCCTAGATCTCTCGTCCAACTCCTTTTCAGGTGAAATTCCCGATATTTTTTCTAACCTCCAAGAGCTACGTTATTTATATCTTTCTGACAACAGCTTCATCGGTTTGCTTCCTTCTACAATTGTAAGCTTGACACATCTTCAATACTTAGACTTGTCAAGTAATTCCCTATCTGGCCCACTGCCTAATAATTTGAGCATGCTTCAAAAGCTAACCCAACTAGATTTGTCGCACAACTCACTGAATGGTACCATACCATCTTCGGTTTTTAGCCTCCCTTTGCTAGGTGAGTTGTGGCTCGATAATAATCGATTTAGTGGACTACCCAATGAGCTCAAAACAAACCCAACATTAGAAAGCCTGAGTTTAAGCCATAATCAACTCAGTGGTTCTTTGCTTCAATCACTTTTGAATCTCACAAACCTTTCTACCCTTGACCTTTCATCAAATAACATCACCATCGATGAGGGAATACAAATCACTTTTCCTAGCCTACAAGATTTGTTGTTATCATCTTGTGAACTGAAGGATATTCCACACTTCTTGACAAATGTAAAGAAACTTCAGGTCTTGGATATTTCTAACAATAAGATTCGTGGCCAAATCCCAAACTGGTTTAGCGGCATGAGGTGGGACAACTTGTCCTACCTAAACCTTTCGCATAACTCTTTAACAGGCCACCTACAACAATTTAATTTCCATAATCTAAGGTATCTTGATCTTAAATTTAACTCCCTTCAAGGTCCAATACCTTCATCCATTTGTAACATGAACAGGCTTGAATTTCTAGATTTATCACGCAACGACTTCAGTAACTCGATTCCAAGTTGCTTGGGAAGCATGTCTAGTCTAACAGTGTTGGACTTAAGAAGGAACAATTTCACTGGGAGTCTTCCATCGTTATGTGCGCAGAGCACTTCATTGAGAACCATTGTCTTGAATGGTAATCAATTTAAAGGAACTCTCCCTGTGTCGTTGCTCAACTGTAGTGATTTAGAAGTTCTTGATATGGGGAATAACGCTATAAATGACACATTTCCAGCTTGGCTAGGAACACTTCAACAACTGCAGGTCTTAATATTAAAGTCCAATGTGTTCCATGGACCAATTAGTACTTGTCAGACTACATTTTGCTTTCCCAAGTTGCGGATTTTTGATCTTTCGCATAACAAATTCAGTGGCTCTCTCCCTGCAAAAGTTTTTGGAAACTTCAAGGGAATGATTAGATTAGACGGTGAAGACACACGAGAGATCATGTACATGCAACCATCTATAATGCCATTTTACTCATCATCATATGAGGATTCTGTGAGTTTGGTGATCAAAGGCCTGGATATTGAGCTAGAAAGAATCAGCACGATTATGACAACCATAGACCTCTCGAGCAACCACTTTGATGGTGTCATTCCGAAAACACTAAAGGATCTCAGCTCACTTTGGCTACTCAATTTATCCCATAACAATCTCATAGGTCATATTCCAATGGAATTGGGGAAATTGAATAAGCTTGAAGCATTAGATCTCTCTTGGAATCGGCTCACTGGAAAGATTCCGCAGGAATTGACAAGGCTAACATTTCTGGAAAAGTTAAACCTTTCTCAGAATGTTCTTGTTGGACCAATTCCTCACGTTCTACAATTCAACACATTTGAAAATGACTCGTATGGCGGTAACCTTGATCTATGTGGTCCTCCTTTATCAAAGCAATGTGGAACGAGTGATCTATCCCATGTTCCTCAACCATTGGAGTCTGAAGACGAGTCATATTTTTTTAGTGGATTTACGTGGGAATCAATAGTCATTGGCTACAGCTGTGGACTAGTTGTTGGAACTGTTGTGTGGAGCCTCATGTTTAAATATCGTAAGCCAGAATGGTTTGTCGAATTTTTTGAAGGCATTTATCCCAAGAAAATGAGAAGGTCAATGAAGAGAGCTCAGAGACGACGGACTTAA

>M6-PGSC0003DMG400015348 M6 LRR-RLP

TGTCGTTGCTCAACTGTAGTGATTTAGAAGTTCTTGATATGGGGAATAACGCTATAAATGACACATTTCCAGCTTGGCTAGGAACACTTCAACAACTGCAGGTCTTAATATTAAAGTCCAATGTGTTCCATGGACCAATTAGTACTTGTCAGACTACATTTTGCTTTCCCAAGTTGCGGATTTTTGATCTTTCGCATAACAAATTCAGTGGCTCTCTCCCTGCAAAAGTTTTTGGAAACTTCAAGGGAATGATTAGATTAGACGGTGAAGACACACGAGAGATCATGTACATGCAACCATCTATAATGCCATTTTACTCATCATCATATGAGGATTCTGTGAGTTTGGTGATCAAAGGCCTGGATATTGAGCTAGAAAGAATCAGCACGATTATGACAACCATAGACCTCTCGAGCAACCACTTTGATGGTGTCATTCCGAAAACACTAAAGGATCTCAGCTCACTTTGGCTACTCAATTTATCCCATAACAATCTCATAGGTCATATTCCAATGGAATTGGGGAAATTGAATAAGCTTGAAGCATTAGATCTCTCTTGGAATCGGCTCACTGGAAAGATTCCGCAGGAATTGACAAGGCTAACATTTCTGGAAAAGTTAAACCTTTCTCAGAATGTTCTTGTTGGACCAATTCCTCACGTTCTACAATTCAACACATTTGAAAATGACTCGTATGGCGGTAACCTTGATCTATGTGGTCCTCCTTTATCAAAGCAATGTGGAACGAGTGATCTATCCCATGTTCCTCAACCATTGGAGTCTGAAGACGAGTCATATTTTTTTAGTGGATTTACGTGGGAATCAATAGTCATTGGCTACAGCTGTGGACTAGTTGTTGGAACTGTTGTGTGGAGCCTCATGTTTAAATATCGTAAGCCAGAATGGTTTGTCGAATTTTTTGAAGGCATTTATCCCAAGAAAATGAGAAGGTCAATGAAGAGAGCTCAGAGACGACGGACTTAA

>M6-PGSC0003DMG400015349 M6 LRR-RLP

ATGGTGAGTCTCTTTTTCTTTTATTCATTTCTCTGTTTGGTGTTACTGATAAGTCTATGCTTTTCTTCTTCCTTTGATCATCATATTTGTTCTTCCAGTGAAGCTTCTGCTTTGCTTAAGTTCAAACAATCCTTTGAAATCACCGACTACCTTAATTGTTGGGTTGAAGAAGACCAACATATTGATTACCCAAAAACAAAGTCTTGGAATGAGAGTACGGATTGCTGCAGTTGGGATGGAGTCACTTGTGACTTATTAAACGGTCATGTTATCGGGTTAGACCTTAGTTGTAGTCTTCTTTTTGGCACTTTTCATCCCAATAGTAGCCTCTTCCAACTTCTTCATCTCCACACACTAAACCTTTATGGCAATGCCTTCAATCATTCTTCAATCCCACATAACATTGGCCGATTGACGAATTTGAGGCATCTCAACCTCTCTGGGTCTGGCTTAAGTGGGAAAATCCCAACAGAAATCTTAGACCTTTCCAATTTGGTGTCACTTCATCTTTCTGCTGGTCTGTATGGATTACAACTTGATCAGAGAGCATTTGAAACACTGCTTCAGAACTTCACAAATCTGGAGCTACTTTCTCTCCCTGGTGTCAACATCTCATCTCCGATACCTGTGAATATTTCTTCCTCCTTAAGGTACGTGGATCTTGAATATACTAATCAGCGAGGTGTTCTCACAGAGAGCCTTTTCCTTCTGCCCAACTTGGAAACACTCTATTTGAGTAATAACTATCTTCTCAAAGGAGTTTTTCCAAAGGTCCACCCGAGCAACACTCTGTTGGAGTTGGATATTTCAGGCACAGGCATCTCTGGTGAGCTACCTGTTTCAATTGGCAACTTCAGTTCCTTGAATACCTTAATACTCGATGCATGTCAATTCTCGGGTCGCATTCCTGATTCCATTGGAAACCTGACACAAATTAGGGTGTTGTTTTTATCGCATAATCATTTCACTGGCAATATTCCTTTCACAATCTCTAAAATGAATCAGCTCTTAAGTTTAGATCTTTCTGATAATAGCTTCATCGGTCCATTTCCCTCTTCAATTCCAAGTTTCACACATCTTCAGTACTTATACTTGATGAGTAATTCCCTATCTGGACCACTTCCTATAAATGGAAGCATGCTTCAAAAGCTAACCGAACTGGATTTGTCTTACAACTCACTGAATGGTACCATACCATCTTGGGTGTTTAGCCTCCCTTTGCTATTTCAATCGGTTCTCCATAATAATCGATTTAGTGGACTACCCGATGAGCTCAAAATAAACCTACCATTAGTACACCTGGATTTAAGCCATAATCAACTCAGTGGTTCTTTTCCTCAATCACTTGCGAATCTCACAAACCTTTATTCCCTTGAACTTTCATCAAATAACATCACCGGGGATATTGCGGGAATCAATATCACTTTCCCTAGCCTAGGATTTTTGCAATTATCATCATGTGAACTGAAGGATTTTCCACACTTCTTGAGAAATGTAACGACACTTCGGTACTTGGATATTTCTAACAATAAGATTTGTGGTCAAATCCCTAACTGGTTTAGCGGCATGAAATGGAACTCGTTGCAGTCCCTAATCCTTTCTCATAATTCATTAACAGGCCATCTACCACAATTTCATTATTATAAGCTAAAGTATCTTGATCTGAAATTTAACTCCCTTCAGGGTCCACTACCTTCATCCATTTGTAACATGAGTTCACTTTCCTTATTAGATTTATCACGCAACAACTTCAGTAACTCGGTTCCAAGTTGCTTGGGAA

>M6-PGSC0003DMG400015351 M6 LRR-RLP

ATGTTAACCGGTCATGTTATCGGATTGGACCTTAGTTGCAGTCTGCTTAATGGAACTATTCATCCCAATAGCAGCCTCTTCCAACTCCATCATCTCCGGACACTAAACCTTGCTTACAATGACTTCTATTCTTCTTCAATCCCACATAACATTGGCCGATTGAGGAATTTGAGGCATCTCAACCTTTCTCATTCTTTGTTTGAAGGGGAAATCCCAACAGAAATCTCATACCTTTCCAATTTGGTTTCACTGGAACTTTTTTGTAATGGATGTGAACTTGCTGAGAGAACATTCGAAACAATTCTTCAAAACTTCACAAATCTAGAGGTACTATCTCTCTCTTATGTCAACATCTCATCTCCATTACCTGTGAATATTTCTTCTTCCTCGTTAAGGTACGTGGATCTTGAATTTACTAATCTGGGAGGTGTTCTCCCAGAGAGTCTTTTCCTTCTGCCAAACTTGGAAATGCTCTATTTGAATTACAATGATCTTCTCGAAGGAGTTTTACCAAAGATCCACCCGAGCAACACTCTGTTGGAGTTGGATATTTCCTACACAGGAATCTCCGGTGAGCTGCCTGATTCAATTGGCACCTTGAGTTCATTGAATCTCTTGAACCTCAAAAGATGTCAATTCTCTAGTTCCATTCCTGATTCCATTGGCAATCTTACACAAATGCGGGGTTTGGATTTTGGTAATAATCATTTCACCGGCTATATTCCTTCCATAATCTCTAAATTGAAGCACCTCACATCCCTAGATCTCTCGTCCAACTCCTTTTCAGGTGAAATTCCCGATATTTTTTCTAACCTCCAAGAGCTACGTTATTTATATCTTTCTGACAACAGCTTCATCGGTTTGCTTCCTTCTACAATTGTAAGCTTGACACATCTTCAATACTTAGACTTGTCAAGTAATTCCCTATCTGGCCCACTGCCTAATAATTTGAGCATGCTTCAAAAGCTAACCCAACTAGATTTGTCGCACAACTCACTGAATGGTACCATACCATCTTCGGTTTTTAGCCTCCCTTTGCTAGGTGAGTTGTGGCTCGATAATAATCGATTTAGTGGACTACCCAATGAGCTCAAAACAAACCCAACATTAGAAAGCCTGAGTTTAAGCCATAATCAACTCAGTGGTTCTTTGCTTCAATCACTTTTGAATCTCACAAACCTTTCTACCCTTGACCTTTCATCAAATAACATCACCATCGATGAGGGAATACAAATCACTTTTCCTAGCCTACAAGATTTGTTGTTATCATCTTGTGAACTGAAGGATATTCCACACTTCTTGACAAATGTAAAGAAACTTCAGGTCTTGGATATTTCTAACAATAAGATTCGTGGCCAAATCCCAAACTGGTTTAGCGGCATGAGGTGGGACAACTTGTCCTACCTAAACCTTTCGCATAACTCTTTAACAGGCCACCTACAACAATTTAATTTCCATAATCTAAGGTATCTTGATCTTAAATTTAACTCCCTTCAAGGTCCAATACCTTCATCCATTTGTAACATGAACAGGCTTGAATTTCTAGATTTATCACGCAACGACTTCAGTAACTCGATTCCAAGTTGCTTGGGAAGCATGTCTAGTCTAACAGTGTTGGACTTAAGAAGGAACAATTTCACTGGGAGTCTTCCATCGTTATGTGCGCAGAGCACTTCATTGAGAACCATTGTCTTGAATGGTAATCAATTTAAAGGAACTCTCCCTGTGTCGTTGCTCAACTGTAGTGATTTAGAAGTTCTTGATATGGGGAATAACGCTATAAATGACACATTTCCAGCTTGGCTAGGAACACTTCAACAACTGCAGGTCTTAATATTAAAGTCCAATGTGTTCCATGGACCAATTAGTACTTGTCAGACTACATTTTGCTTTCCCAAGTTGCGGATTTTTGATCTTTCGCATAACAAATTCAGTGGCTCTCTCCCTGCAAAAGTTTTTGGAAACTTCAAGGGAATGATTAGATTAGACGGTGAAGACACACGAGAGATCATGTACATGCAACCATCTATAATGCCATTTTACTCATCATCATATGAGGATTCTGTGAGTTTGGTGATCAAAGGCCTGGATATTGAGCTAGAAAGAATCAGCACGATTATGACAACCATAGACCTCTCGAGCAACCACTTTGATGGTGTCATTCCGAAAACACTAAAGGATCTCAGCTCACTTTGGCTACTCAATTTATCCCATAACAATCTCATAGGTCATATTCCAATGGAATTGGGGAAATTGAATAAGCTTGAAGCATTAGATCTCTCTTGGAATCGGCTCACTGGAAAGATTCCGCAGGAATTGACAAGGCTAACATTTCTGGAAAAGTTAAACCTTTCTCAGAATGTTCTTGTTGGACCAATTCCTCACGTTCTACAATTCAACACATTTGAAAATGACTCGTATGGCGGTAACCTTGATCTATGTGGTCCTCCTTTATCAAAGCAATGTGGAACGAGTGATCTATCCCATGTTCCTCAACCATTGGAGTCTGAAGACGAGTCATATTTTTTTAGTGGATTTACGTGGGAATCAATAGTCATTGGCTACAGCTGTGGACTAGTTGTTGGAACTGTTGTGTGGAGCCTCATGTTTAAATATCGTAAGCCAGAATGGTTTGTCGAATTTTTTGAAGGCATTTATCCCAAGAAAATGAGAAGGTCAATGAAGAGAGCTCAGAGACGACGGACTTAA

>M6-PGSC0003DMG400015352 M6 LRR-RLP

ACTTGTGACATGTTAACCGGTCATGTTATTGGATTGGACCTTAGATGCAGTCTGCTTAAAGGAACTATTCATCCCAATAGCAGCCTCTTCCAACTCCATCATCTCCATACACTAAATCTTGCTTACAATTACTTCTATTCTTCTTCAATCCCACATAACATTGGCCGATTGAGGAATTTGAGGCATCTCAACCTTTCTGGTTCATTCTTTGAAGGGGAAATCCCAACAGAAATATCATACCTTTCCAATTTGGTTTCACTTGATCTTTCTTCTTTAGTGCTGTATGGATTACAACTTGATCAGAGAACATTTGAAACAATGCTTCAGAACTTCACTAATCTGGAGGTAGTTTCTCTCCATGGTGTCGACATCTTCTCTCCGATACCTGTGAATATGTCTTTTTCCTCGTTAAGGTACGTGGATCTTGGATATACTAATCTGCGAGGTGTTATCACAGAGAGCTTTTTCCTTCTGTCAGAATTGGAAAGGCTCTATTTGAGTGACAATGATCTTCTCAAAGGAGTTTTACCAAAGATCCACCAGCAACACTCTGTTGGAGTTGGATATTTCTTACACAGGAATCTCTGGTGAGCTGCCTGATTCAGTTGGCACCTTCAGTTCCTTGAATATATTGAACCTCAAAGGATGTCAATTCTCTGGTTCCATTCCTGATTCCATTGGCAATCTTACACAAATTTGGTATTTGGATTTCAGTAATAATCATTTCACCGGCCATATTCCTTCCACAATCTCTCAATTGAAGCAGCTCACATTTCTATATCTTTCGTCCAACTCCTTTTCAGGTGAAATTCCAGATATTTTTTCTAACCTCCAAGAGCTATGTTATTTATTTCTTTATAGAAATAGTTTCATCGGTTCGTTTCCTTCTACAATTGTAAGCTTGACACATCTTCAATACTTAGACTTGTCAAGTAATTCCCTATCTGGCCCACTGCCTAATAATTTTAGCATGCTTCAAAAGCTAACCTGACTGGATTTGTGGTACAACTCACTGAATGGTATCATACCATCTTCGGTGTTTAGCCTCCCTTTGCTAGGTCAGTTGTGGCTCGATAATAATCGATTTAGTGGACTACACGATGAGCTCAAAACAAACCCAACATTAGAACGCTTGGGTTTAAGCCAAAATCAACTCAGTGGTTCTTTTCCTCATTCACTTGCTAGTCTCACAAACCTTTCTACCCTTGACCTTTCATCAAATAACATCACCGTCGATGCAGGGATACAAATCACTTTTCCTAGCCTAATAAATTTGTTGTTATCATCTTGTAAACTGAAGGATTTTCCACACTTCTTGACAAATGTAAAGAAACTTCAGGTCTTGGATATTTCTAACAATAAGATTCGTGGCCAAATCCCAAACTGGTTTAGCGGCACAAGGTGGGACAATTTGTACTACCTAAATCTTTCGCATAATTCTTTACCAGGCCACCTACAATAATTTCATTTCCATCATCTAGGGTATCTTAATCTTCAATTTAACTTCCTTCAGGGTCCAATACCTTCATCCATTTGTAACATGAACAATCTTCAATTTCTACATTTATCGCGCAACAACTTCAGTAACTCGATTCCAAGCTGCTTGGGAAGCATGTCTAGTCTAAAAGTGTTGGACTTAAGAAGGAACAATTTCACAGGGAGTCTTCCCCCCATTATGTGCGTAGAGGACTTCATTGAGAACCATTGTCTTGAATGACAATCAATTTGAAGGAACTCTCCCTATGTCGTTGCTCAACTGTATTGGTTTAGAAGTTCTTGATATGGGGAATAACACTATAAATGACACATTTCCTGCTTGGCTCGGAACACTTCAACAACTGCAGGTCTTAATATTAAAGTCCAATATGTTCCATGGACCAATTAGTACTTGTCAGACTATATTTTGCTTTCCCAAGTTGCGAATTTTTGATCTTTCCCGTAACGAATTCAGTGGCTCTCTCCCTGCAAAAGTTTTTGGAAACTTCAAGGCAATGATTAAATTAGACGATGAAGACTCAGGAGAGATCATGTACATGAAACCATCTATAATGCCATTTTACTCATCATCATATGAGGATTCTGTGAGTTTGGTGATCAAAGGCCTGGATATTGAGCTAGAAAGAATCAGCACAATTATGACAACCATAGACCTCTCGAGCAACCACTTTGATGGTGTCATTCGGAAAACACTGAAGGATCTCAGCTCACTTTGGCTACTCAATTTATCCCATAACAATCTCACAGGTGACATTCCAATGGAATTGGGGCAATTGAATACCCTTGAAGCATTAGATCTCTCTTGGAATCGGCTCACTGGAAAGATTCCGCAGGAATTGACAAGGCTAACATTTCTGGAAAAGTTAAACCTTTCTCAGAATGTCCTCGTTGGACCAATTCCTCACGGTCCACAATTCAACACATTTGAAAATGACTCCTATGGCGGAAACCTTGATTTATGTGGTCCTCCTTTATCAAAGCAATGTGGAACGAGTGATCCATCACATACTCCTCAACCATTGGACTCTGAAGAAGATGAAGATGAGTCATATTTTTTAAGTGGATTTATGTGGGAATCAGTAGTCATAGGCTACAGTTTTGGACTCGTTGTTGGAACTGTTGTGTGGAGCCTTATGTTTAAATATCATAAACCAAAATGGTTTGTGGAATTTTTTGATGGCATTTTCCCCAAGAAAATGAGAAGGTCAAAGAAGAGAAGGCCGGGACAGCGCATTTAA

>M6-PGSC0003DMG400015353 M6 LRR-RLP

TTGGATTTCGGTAATAATCATTTCACCGGCCATATTCCTTCCACAATCTCTAAATTGAAGCAGCTCACATATCTAGACATTTCGTCCAACTCCTTTACAGGTGAAATTCCAGATATTTTTTCTAACCTCCAAGAGCTACATTCTTTATATCTTTGTAACAACAGCTTCATCGGTTTGTTTCCTTCTACAATTGTAAGCTTGACACAACTTCAACGCTTAGACTTGTCCAACAACTCACTTAACGGTACCATACCATCTTCGGTGTTTAGCCTCCCTTTGCTATTTTCATTGTCGCTCCAACATAATCAATTTAGTGGAGTAGCCGATGAGTTCAAAACATGCCCAACATTAGCATACCTAGATTTAAGCCATAATCAATTCAGTGGTACTTTTCCTCAATCACTTGCGAGTCTCACAAACCTTTCTACCCTCGAGCTTTCAGCAAATAACATCACCGTCGATGAGGGAATACAAATCACTTTTCCTAGCCTAGTAGTTTTACAGTTATCATCTTGTGAACTGAAGGATTTCCCACACTTCTTGACAAATGTAAAGAAACTTCAGGTCTTGGATATTTCTAACAATAAGATTCGTGGCCAAATCCCAAAATGGTTTAGCGGCATGAGGTGGGACAGCTTGTGGTACCTAAACCTTTCGCATAATTCTTTAACAGGCCACCTACAACAATTTCATTTCCATAATCTAAGGTATCTTGATATTAAATTTAACTCCCTTCAGGGTCCACTCTCTTTGTCCATTTGTAACAGTAACACGCTTGAATTTCTAGATTTATCACGCAACAACTTCAGTAACTCGATTCCAAGCTGCTTGGGAAGCATGTCTAGTCTAAAAGTGTTGGACTTAAGAAGGAACAATTTCACAGGGAGTCTTTCTCCATTATGTGCGCAGAGCACTTCATTGAGAACCATTGTCTTGAATGACAATCAATTTGAAGGAACTCTCCCTATGTCGTTGCTCAACTGTATTGGTTTAGAAGTTCTTGATATGGGGAATAACGCTATAAATGACACATTTCCAGCTTGGCTAGGAAGACTTGAACAACTGCAGGTCCTAATATTAAAGTCCAATGTGTTCCATGGACCAATAAGTAATTGTCAGAGTACGTTTTGCTTTCCCAAGTTGCGAATTTTTGATCTTTCTAGTAATGAATTCAGTGGCTCACTTCCAGCAAATTTTCTTGGAAACTTCAAGGCAATGATTAAATTAGATGGTGAAGAGACAGGAGAGATCAAGTACATGGAACCATCTGGGAAGTCATTGTACACATCGTATGAGGATTCTGTGAGTTTGGTGATCAAAGGCCAGGACATTGAGCTAGAAAGAATCAGCACAATTATGACAACCATAGACCTCTCAAGCAACCATTTTGAAGGTGTCATTCCGAAAACACTAAAGGATCTCAGCTCACTTTGGCTACTCAATTTATCCCATAACATTTTCATAGGTCATATTCCAATGGAATTGGGACAATTGAATACGCTTGAAGCGTTAGATCTCTCTTGGAATCGGCTCACTGGAAAGATTCCGCAGGAATTGACAAGAATGAATTTTCTGGCCTTCTTAAACCTCTCTCATAATCTTCTTATTGGACCAATTCCTCAAGGTCCACAGTTCAACACATTTG

>M6-PGSC0003DMG400015449 M6 LRR-RLP

CTCATCCAACAACTATATCTCCCACAAAAAAAGTGAAAAAATCAATCTTCATTTCTTATATGATGAGCATTTCTTATCTTGAATAATTGTTATTGTGTTTAATAGATAGTTCTTTTGTATATCATCTTCCTTAGGGGCAGCCTTGGGGGTTGGTTCTTCCGAGGCAGCTTGTTAGTCGGAAATTCCCCTTAAAAAATGCAGGTGTATCGATGTTTTTTCAGTTTGTTGCTGATCTCTGCTGTTCTGTTTTCTTTCTCTTGTGCACTCTCTGATCATGAAGCATCATTGCTTGCACGGCGACAACTGTCAACACTTGCTGAAAATGGTAATTTACCCGATAACTATGAATTCGAGGTGAATGTAGAATACACATTTCCTAATTCTAGGCTTAGACGTGCTTATATTGCGTTAAAGGCATGGAAAGAAGCTATATATTCTGACCCTTCTGAATTCACTAGCAATTGGAAAGGTCCAGATGTTTGTAACTACAAAGGCGTGTTCTGTTCGCCTGCTTTAGATGATCCTAATGTCACTGTTGTTGCTGGGATTGATATGAATCATGCAGATATTGCAGGTTATTTTCCTGTAGAATTAGGATTATTGACAGATGTAGCTTTGTTTCACCTAAATTCCAATAGATTTTGTGGAATTGTTCCTGAGAGTTTTTCGAAGCTTACGTTGATGCATGAGTTTGATGTTAGTAATAATCGTCTTGTTGGCCCTTTCCCTAAGGTTGTTCTGAACATGTCTAGTCTTAAGTACTTGGATTTAAGGTTTAACAACTTTGAGGGAGAGTTGCCTCCACAGTTGTTTGACAAAGATCTTGATGCTTTGTTCTTGAATGATAATCGATTCGTGTCTACCATCCCTGAGACTCTTGGTAACTCTTCAGCTTCTGTGATTGTGTTTGCTAATAACAAGTTTCATGGTTGTATTCCCAGTAGCATTGGGAAGATGTCGAATTTGGATGAGATTGTGTTTATGAACAATGATCTTGGTGGATGTTTGCCTGTTGAAGTTGGGTTGTTAAAGAATGTAACGGTGTTCAATGCTGCTGGGAATTTGCTTTCTGGGATATTGCCAAAGACACTTGATGGTCTTTCACATGTTGAGGAATTGGACATTTCACATAATACCTTAACTGGGTTTGTTCCGGAGAATCTATGTTCATCGTTGAGTTTGAAGAAGTTTGTGTTTTCGTTTAATTACTTTAATGGAGAGGCTAATGGTTGTGAAGCGCGAACAAGGAAGGATATGAGTTTGGATGATAGGAACAATTGTTTGCCTGGTAGAGCAAAGCAAAAGTCTCAAAAAGAATGTCAACCTATTGTGAGTAAGCCAATTGATTGTAGTAAGGCTAAATGTGGAGGAACTTCAACAAAATCTCCTCCTAGTAAGAAAAATGAAGTGCCACCAAAACCTCGACCTACCCCTAAGTCACCACCCACCCCAAAGAAGGTTTCTCCTCCACCAACACAAGTCGTTTCATCGCCACCTCTAGTACATTCTCCACCACCTCTAGTACATTCTCCACCACCACGTGTCCATTCTCCTCCTGTTTCCTCTCCACCACTACCCGTCCACTCCCCTCCACCACCACATGTCCATTCTCCTCTTGTGTCCTCTCCACCTCGCGTCTACTCCCTACCACCACCTCACGTCCAATCTCCACCGCCACTCCCCGTCAACTCCCCTCCATCACCTATAGTATATTCTCCACCACCACATGTCCATTCTCCTCCTGTTTCCTCTCCACCACCACCAGTCTACTCCCCACCACCACCTCAAGTACATTCTCCACCACCACCCGTCCACTCTCCTCCTTCTCCTATTTCTTCCCCACCACCACCTATCCATTCCCCTCCACCACCTCTTATCCAATCTCCACCGCCGCCTCTAGTACATTCTCCACCACCACCTGTCCACTCTCCTCCTCCACCTGTTTCCTCTCCACCGCCACCAGTCCATTCCCCTCCACCACCACCGGTCCACTTCCCACCACCACCTCAAGTACATTCTCCACCACCACCCGTTCACTCTCCTCCACCACCACCACCACCACCCGTCCACTCTCCTCCTCCTGTTTCTTCCCCATCACCACCACCCGTCCACTCCCCACCACCACCTCTAGTACATTCTCCACCACCTCCCATCCACTCACCACCACCACCTTTCGAACATGTTGTCCTTCCACCAAATATTGGTTCAATTTACGCATCGCCACCCCCACCAATCTTCCAAGGATATTAATTAAGCAATGAATGAAGCTTCGTCTCGTTTCCACCTTTTTAGTACGTAAAATATTTAGTACATGCAGAATTACATCATCATATCTCTTAATGCAAGATTATTATCTAATATATGGTGATGTTTTTCTTTCTTTACACTGATACATATAATCTTGGAAAAG

>M6-PGSC0003DMG400015603 M6 LRR-RLP

TTTTAAGTTATTTTTATATTTGCCAAACACTCCTAGAAGTCAAAAACTGACTTAAAAGTAGGTTTGATCAACTTTTAAGTCGATTCAAATAGGCCCTAAGAATCCAAAAGATTAGGAAATAATCCTTATTATTGTAGGCTAAAACACCTATTTAGGAGTCTTCTTGTATGGTTGTAAGCATCAATTAAAAAGAAGTTGTTATTCCCTTTTCAGATGTTATAGATCTTGCTAATGACTTAAGTAACTTCTGCCTATCAGTGCACGGCATTATGCTACTACTTTTGGGTGGACTTCTCTTTCTTTTTTACGTTTTACTGATCACACTTCTTTTTTCCTTCTGGTGGCACTGCAATTCTTTTTTACTTCTTAAGTTACACTTTACCACCAATCCGAGCTATGTGGATGATGTCCAAGAGATTTAACTTAGCATATATCTTCCGGTGGTTGAATTATAGATTCCTCTACATGTGTCGAAAACACTTTCTTTCATTATTCAATTACCAGTAACCTTTTAGAAGAAAAAATTTTCAACGCTGCTTTTCTTTTCCATGATCATTCACTACTAATTAGACGATTCAGGTGACTGTTCTTGTTCTAATTACTTTTCTAGTTATTTGACCATCTTGAATATGAAGGAGCATGCTTGACGTTAAGTGAATCGTTAAAAATATTGTGTAAATTATTCATAATAATTATGTTTTATTAATTAAGAGAGCGGGAATTAGTAAACTTTCCTTCTTAATTTAGTTCAGAATTTCTATATGAGTAGCTCCTTGAATGTGGTTAAACCTTTCATTACAAGAGCTCATTTCTCTCTTCTTTTTTTACAATGTCAAATTCTAATAAACAGTTAATCAACCACTGGTATCGTTGCATCTAATTTATCTTAACATTCAAAGTTCTTACATTTCCATCTCCAAAGATGTTGCTCCATACTTCTATTCTATATGATGCGATTGAATTCCTCTTTCATTGCCAATTACCTCTACTAATAACTTCTGAGCTCTTTCTGCCCAAAAAATTGAGGAGCTACTTAACTTGCATCTGCAGAAAGAATAAATGTCTTAGTTTTACCACAGTGTTGCTCTGGTTCAGTCATTGACTATGGGCCCTGGCCATTTCTATTGCTGCACCCTTCCCAATGAACTCGTTGCAAGTTATATCCTTCCAACTTTCGCCTGGCATCCTTTCTCACGGAGGTTGAATTTTGTGAGAATGTGGCTGCTAGACTGAAGATCTTCCTTTTATGAATTGTGACTCGAAATAGAAATCTTAAGTAGAAGGAAGATTCTCTATATCGTACTAATCTATGTGTTCAGTTGACAGTTTTGTGGAAGCTGGTGGAGAAAGTTCTTCTACCCAAACCAGCCAAGACTACATCAGAAGAAAGCAAGTCTACTCAGGGATTGAAGTGGTCCATTGCCGCTGGGACAAACTTGTTGCCTGGTTTTGGTGCAAAGATTGAAAGAGAATCCAAGCTGAGGCTTAATGACTTTGCTAAAGAGCTTAGATCTTTCAGCATTGTTGACATGTCAGGTAAAATTGCAGTCAGTTCTGGCTCAGCTGCTTCAGTATTAAAGTTGGTGGTCGTCTATGCTAAATGCTGTCATAATGAATATCTTCCTTTTTTATACCTTTGATAAAAAACCTTCTATCATTTGTAGTTTTCAGAAATGTGATCACGATGTTAAAATATTTGCCTTGACTTGTTAAATTCTTAAGTCCCACTTCTTTAGAAAGTTCAACATATAAAGACCAAAGCTTTACCGTGATTTATGATTAAAAAACATTTTCTGCATACTGATTGTGCCTGAAGGAAATTTTCCAATCTTGTTTCATTTATAAGCTTCCCCCACCAAGTGCGCGACACACCAAAACCAAAGTTTTGCTGCATCTTCCTTGTATTGTGATTCTACATTCTTTGCATACTGATTAAGGATTAAAATTGCAAGTCACAAGAGGTATTGGATTGAATGTTTTAGGCTTTGGCAGGTCGTAACTTTGGAGATGAGGGGCTCTTTTTCCTAGCAGAGAGTTTAGCATATAATCAGGTAATGGCTTTTGCTCTCTCTACCCCCCTCCCTCTTCCTTTATCTGTGTAGAGTATGTCATTATTTATCATGGTCCATCTTGTTAACCCAAGAGGATAAAGCCATAATCCTGTAATGTAGACAAATAACAATACAATCCTGGCTTATTTTTTTCTGGAATAATGTTTTTGGGCAAGTTTGGTTCCTTTACCAATGTACACTGCATTTCCTTTCATGTGATTTCCAGACAGCCGAGGAAGTAAACTTTGCTGCTAATGGAATAACTGCAGATGGATTGAAGGCCTTTGATGGCATATTGCAATCAAATATTGCTCTAAAGACTCTTAACCTATCTGGAAATGCTATTGGGGATGAAGGAGCCAAGGTAATGCACAGATGATATAGGTGAAAATCCAAAGCAGTGAGTTCTTGGAATACAGATGACTAATAAGCAGATCCTAACTTTCCCTTTGCACTTACTTAACTGTTATACCAATTTAACATGTGTATTTATGGTCATGCTTTGTATTGTATTTCTCATCTCCGCCTTGGCACTTCACGATCAATCACGTCTATTTGGTGTTAATTTATGTTTGCAGTGTCTATGTGATATATTGGCAAATAACTCTGGTATTCAAAAGCTCCAGCTGAACAGCACTGGCTTAGGAGATGAGGTCTTGAACATGCCTTGAGCATTTAATTGAAATGCTGTTGCTAACTTGCGATGGTATTTTCTGCTTTCTGGTAAATTATTGCCTAAATTTCACTGCAATGGTTTGTCTGAATTTTGCAGGGAGCCAAAGCCATCGGTGAAATGTTGAAGAAAAATTCAACTTTGCGTGTTCTTGAACTAAACAATAATCTGATTGACTATACTGTATGTTATTAAGCAGAACTTGTAATTTGTATTTCCTAGTTATTAATTATCTCAATTTCTGAAGGAAAACATGTTTGATTTGGATATGGAAATAGGGATTTTCAGGTCTTGCTGGATCACTTCTTGAGAATAAAACTTTACAATCCTTGCATCTCAAGTAAGCAAAATTTATTCTGCTGTCTTTCTGCGCCCATCCTCATTTGTCGTATGTCAAATGTAGCAGTTAATTATTTTATTTTTCTTTATTGGTTTGGAGGGTGGGGGTGGGGGGAAAAGAAGTGGATTTTAATTTGTGAGGGAAAGCATATCAATTCTTATTTTCCCTTATATCTTTTAATTTATGTGCTTTAATTAACTGTTCAGCGGCAATTATGGTGGTGCTTTAGGGGCTGCAGCACTGGCTAAAGGCTTGGAGAGCAATAAGTCCTTGAGGGTACATTTCTAATCAATTCCTTCCTCCCACTGAGACCCTTCGCCTTTTGCTCTTTTTTTTGTTCTTGTAATTTGACTTGGCAAATAATTGAGCTTTTTGTCAGCCTTGATATCTTGCTTCTGCGTCATTGGACTGTTTCTTAGTATTTAACTTCTGCACTCATGTTTCCCTTTTTTTTTTTTTTTGAAAACAACCTCTCTACCTCCACAAGGTATGAGTAAGGTCTGGGTACACTCTCCCCACCCCAACCCCATTTGTGGGATTACACTGGGTATGTTGTTGTTGTTACTCATGTTTCCATTTGATTGTATAGAAGAGCTCTTCTTTTCAAAATCTTCAATTTCCTCTGTGACTGCTGACACTGGGATTAAGTGTTAAATGCACTAATGTTATTAGATAAACTTCTTCGGGGGGTTTCTTCATGTTCCTCACCAACAATCCGAGGTTGAAGCTAGATCTGCTCCACCCAAAATGGAAATGGGCTACTGTTTACATATTACTGTATATCATTAACATACACCTGTATAATCAAATGTGTTTGAAAAGATTTTTTATGTGCAGGAACTTTATTTGCAGGGGAACTCTGTTGGTGATGAAGGAGTACGGGCATTAATCTCTGGCCTATCTTTGCGTAAAGGTTTGCTTATTGGATATAATTAACCATTCTCTTTAAACATATATTGGCGTTGTTTCCTGTGCATCTTTTTTGAAAGGTCACCATCATATGTCTGCAGGGAAACTTGTCTTACTTGACCTTGCCAACAATTCAATTACTGCTAGAGGAGCATTTCATGTTGCCGAATATGTCAAAAAAAGCAAAAGCTTACTGTGGTTGAATCTCTATATGAATGACATCAAAGATGAGGTAATTGAACCACTTAGAACTTTTCACAGTTTTTTTCCTTTGTAATGTAGAAGACTCTTCTCAGCCTTGTCCTTCTCACTGCAATCAGTAGACTCTCTTGATCTATTTTATGGTTTTATTTTATGTAACAGAAGCAAAGAATAAGAAAGCTTGAGTTTTTGTCAAGTTGAAACCACTAAAATAAGATAGAAAAGATTTTTGGGTAAGAGGGTCAGGATTGATTGTCGAGAAAAAACAAAAAGAATTTGATTGGTTAGAGCTGAAGGTGATGGCTGAGTCGTTCTTGAAACCAATCATAAGTTCAGATCACATGGTTAGGAAATTTTATAATGGTACTTACCATTAAAGAATATAATTTTGAGACTAGGTTTAATCACTTTGGGGATTCTTGCCAGAAGAAAATTAGAAGGATTGCATGGTGTTGGATGAACTGAGGTCTTTTTAAATTTTGTCAGCAGTTCAGAAAGATAAAACTCAAATAATCCCAAATCTGTAGGGGTAGCTTTATTTTTTCCCTTCAAATACATATGCATATGGACAGACATATTAACTATGTAAAAATGAATCCTTTTATAGTCCTGCACAACTTTAACGTTAATATTTCAACAGTGAGAAACCTAACTGTGCTCCTCTTGGATTATAGAACTGTTTGTATCCACCATAATTAAACTGCGTTCATTGGAGTCAAAAGGAAAGCGTTGTGGTCGAAGGGGAAAGATAAAAAATTGATGATTGGAGGAAATTGGTACGGATTCAGTAACAGAAGGTTAAATCTGTTGTGAAGAGTTCAAGGGATTTGGGCTGCAAAGAAGGATTTTTTTCAGAAATTTATGGGGGTGGGTTTGCATACGACAGGGTAGTTGCTCAGAAGCACAATTAAATTTCTGCTGGAGAAGAGCTCATGTGCTGGCAACATCAGATTGAAACATAAAGTCAAACTTTAACCATATCCTTGGTAGATTAGACCAAACTAGCTAATGTTGATTACCCATTCCAGAAACTTAAGGCTATGGTTTACAATGACAGGGTACAGTGCCTACAAGCAACAAGTCCTGCACAAGAACTTGGAAAGAACTCTCTAAGTTAGCTCGCAAAAAAAATTTACTGCCTTCATTTGTGTCAATAAACTTCACATGTAATCAGTAGCCTCAAGTTTCCCACTTCTTCAATTCAACCCAGTTTCTTTTTGTTGGTATGCGATTTCCTTGGTGTTGAAGATTAAAGGCGAAGGCTTTTTGTTTTAAAAGAGAACACAAATCACATGCTTTTCACTTGAGAAACTACCGTTAATATTTGACGTCAAATTTGGGAAGGACCAAAGGTGTCCATTTTTTGCAAAGACGAAGGATTTATTTTATCCACAGTTATATTTGAAGGACAAAGTGTAACCTGTCCCCATAAATAAGGGATCATTTGAGTTTTTACATTGCAGATCCAGACTCAATCTCTTGATATGCATTGACTTTTCCTTCTTTTTTTTTCCCCCCATTTACATCAATATCCATCAGGGAGCTGAAAAGATTGCAGAAGCTTTGAAGGAGAACCGCTCGATAACTAACGTTGATCTTGTAAGAGTTGTTTCCTCTTTTCTATTTTATTCAACATTTCTTGTTTTGTTCATGGAGGAGGTTGGGTTTCTTAATCTTATCAGAGCATAATTTTCTATTTGCTCTGATGGCTGGAATTCTGCGAATTTTACAGGGGGGCAATGACATACATGCTAAGGGTATTAGTGCCATAGCAGAAGTTCTGAAAGATAATTCTGTCATTACCTCTGTAAGTTTCATCAAAATGTTAACTTTTACTGATTTATTGGACAAAATTAAATACATTATACTTCAAATTTCTCACGAACAGTTAGAACTTGGTTACAACCCCATTGGACCAGAGGGGGCAATGGCATTAGCAGAGGTACTCAAGTTTCATGGAAGTGTAAAAGATCTCATGCTTGGGTGGTGTCAGGTAATGCATGGATTGTTTCATATATGCCCTCTTTAACTATCTTTTTTAGCATGTTCACTTAATTTTTTTCCAGATCGGAGCCAAAGGTGCAGAATATTTTGCCGATATGTTGAAGTACAATAGTACCATATCAACTCTAGACTTGCGAGCCAATGGACTTAGAGACGAGGTTTCATTCAAAGTTTATAGTTTTTCTATTCTGCCACAATTTCCCTCAAAGTACATAACCATTTCCTTTTTCAGGGTGCAATCTGCCTTGCTCGGAGTTTAAAAGTGGTCAATGAATCTTTGACGACACTCAATTTAGGGTTCAATGAAATTAGAGTATGCAACTCTTTTCCAGTACAAGTACAACCACCTGTCTTGATAAAGGTATCATCTTGTATTAACTTAGTTTATGTTGCACACAGGATGAAGGAGCTTTTGCTATCGCTCAAGCACTTAAGGCAAATGAAGATGTTAGACTCACATCTTTAAATCTTGCCAGTAACTTCCTCACCAAATTAGGACAGGTAACATCAAATGATAATATTATTCTCTTATTTTAGCTGTTTTACGCGATTTTTAGTTCCTTACAATATCCGGATGGATTCCAAATTTCAGACAGCTCTCACAGATGCAAGAGACCATGTGTTTGAGATGACTGAGAAGGAGCTTGCTGTTATGTTCTAAAATATTACATTCCTTAGATCAGATATATTTTCCTCCCTAGTTGGGGCAAGGTAGATGAGTATTTTTTTTTCTATTCTGGGGGAATAGAATCGCAGCCTACCTTGATATTCTTAAGAGTCCCACCTTAGTTCCCTTTTGTGACTGAAATCATTTTGAGTGGGGAACAGTGGAAATTTTGATATCGAGATACACACTGATATTATAAAAAATGTTCCTTTTCTGAAACTTTTTTTGTGTCTGATAACGTTAGTGGCAAATACATAGAAATTACTTCTATTCACCATTTCAGATTCATTCTCTGGTTTGAAACTCTTCTGAGTTGTTGTTCTTTCCCTATGGCCTTTTAAGCTTATTTTGGTGTTGCAGGACAG

>M6-PGSC0003DMG400015672 M6 LRR-RLP

CCTTTAGTTTGTTCATTTTGTGCACAATAGGATGTGACTTAGTGGTCGATGAAGTAGTAGGAGAATTGTGAGGTTAAGAAAATGCTAGGTAATTTCTATTTATCTGCCTAGGATTTGGTGGGCAGAGACGCAGAATTACATTATACATGTGTTGGTGGTAGGTACGACATATTTAGTGTAATATTCGATGTGCACGCAAGATGGCTCGGACATCACGGTCGTTAAAGATAAGACACCGCTTAACTTAAGGGACTAATTTGATATTTTTTTTCTCCATATTCTTCTAACAGATGGAACAAGATGTGTCATTGAAGGGGATAGTGAAGATAGAATCAGTGGTTTACCAAGAAATGTTATCGATCGTATCCTTGAGCTCATACCAGTTCAAGATGCAGCAAGAACTAGTATTTTGTCCCGAAACTGGAGATATACTTGGGCAACATTTCCAAATCTGGTGCTGGATTACTGCTTCCATTGTAATCTTGAAGGAAAGTTGTTATTCAACTTCAAAGAAGCGGTAGATTGTATCCTGTTACTGCATATGGGGGACATTGTGAAGTTTCTTCTTGATACTAGAGGAATTTTTTCTGCAAATTCTTCCTATGCAATTATCGATAGATGGATTCTTTATGTCACCAGAAATGGAGTCAAGATGCTAACCCTTCGCACTTCAAATGATGATACCTATACTCTGCCCTCTTCTATCTTTAATTGCTCAACCTTGACATATTTGGAACTCTCCAACTGTGTCTTCAAACCACCCAATCCCTTTCTTGGTTTACAGAATCTTATTACCCTTCATCTGGAAGGGACAACCTTTGTGCCAGCCACATCATTTTGTGTTATCAAGGCTCCCCTTCTTGCCAAATTGGATTTGATCTCGTGTCGTGGTACGCAATACCTAAACATTGTTTCACCAGGACTGGAGTGCTTGCAAGTTCGTGACAGTCACTCATATCTTGTGCTAGATTGCTTTATGAACTGCAAGAATATAAGAATGCTAAAACTTGAATTTAACGGAGTGGTTGATGATCCAACAAACGATGAAATATCAACTTTGGAGAAGCTTCTTGTTAGCTCTCCTGCACTTGAGGTACTCCATTTAGATTCATTCTTTGTTGAGGTAAGATATTGACTACTTGTTTCTCTTGATATCTTATAGTGATGGTAATTTCTTATGATTCTAATTATTGTGTTTTCTTCAAAGCTTTTGAGTGCACGCATTGTTCCGAATGGGCAACCTTCTACACTCAACTGCTTGTGGCATCAATATTTAGGTGTAGACTTCAGCAAAATGGGTCAGATTTCTAATGTTCTGCAGTTAATTAAGAGTTCCCCCAATTTGAGTAAACTCCATATTTCGGTAAGAAAAGAAAATATATGCACGATCTGGTTCATTTTTGCACTTCATATTTGTGAACATGAGAATCTGAAATAGCTTGGTATTTTCTTCATATACATTATTAGGTCCATCGCCATGGTACCAGTGACAATGCCGAAGCAATCTTGAAGTATCT

>M6-PGSC0003DMG400015687 M6 LRR-RLP

TCCATTTATCATAAGAGTTAGTAACTATGACTATGTTCCAAAAAGGACTGAAGTAGTTATACATATCAGCAAGAATGTTAAATTCAAAATTTAATCTATTTGACTCGCGAAAACTAAACTGTTTAGCATGTCTGATTTTAAGTTTCTTCGTGATTTGTTTACAGGTTTTGAGAAACTGAGACAACTAGAAATACTTGATTTGTCATCGAACTACTTGAACCGCAGCATCTTTCAGTCTCTTAGTCAACTTTCATCTCTCAAGTCGTTGTACCTCGCGGAAAACAACATTGAATCAGGTGCTAACATTTGGTTCATCTGTTTTCCCCTTTGCACTATTAGTCTTCCTCTTTTTACAGAACTGTGCTAAAAATAATCCATCTCCTTCTAACATTTCAGGTTCTGAGAGATTATCAGGGTTAGACAAGCTGGAAATCCTGGATTTGAGCTACAATGCACTAAATGATGAAAACGTTCTCTCTGTTCTAGGTAATTAAACTTATAGTTTCTTTTGTGAAAAGCCGTTTTTCCTCAGTAGCTTCCTACATAATTATTCTGAATTCCTTGACAGAGCTAAATGTATCTAGAACGACGTTGAAGAAGCTTGATATAAGGTACAACAGATTCCGGAGTTTTATACCAAATGGAGGTATATATACAATTACTGAAGTTTTTCTTCTTTCAATTGATTTTATTCTACCTCCTTCTACATCTTGCTATGTAAGTATATCTTATTGGTACCCAATCGAAATCACAAGCTTAAAAGAACTGAAAAGTCACAAATCATTTACAAACTCCACAGGCTCCATTTGGATTTGATGAAATGTGTATAAATTCATAGACTTACATGCAAAGATGTTATTTTAATAACTCTCTGTGTCAACATTTTGGTTAAAAATATGACATTCCACCATTTTAGGTGTCATTTGCAAGTCGCTATTGCCATTTTAGTTCAAACATTTACAGTTTAAAGCGTACTTTTAATGTTTCTTTTGCATGTAAGAGTTGGGAGCTTTAAGAAATATCGAGTGGTTACTGTTGGATGGAAACACATTGGATGAGAACTTTCTCAGGAGTACTGGTGCAATGTCATCCCTTACAGTGTTATCAGTAGCTAACTGCAACCTCAGTGGAGTCCTACCTCTTCAAGGTAATTTCAGGCATACGTAAATAGAACAGTGAATAGTTGAGAATAATGAATAATTTCAGAACAATCCATGTTTCTAATATTTTCTGCATAACAACTATATAAGAACATGAATAACAATCATCTATTTGTTCATTTTATAGGCTTGTGTGATCTAAAATATCTTGAAGAACTAAGTCTCCGCCTAAACTACTTCATTGGAGAAGTCCCTGCATGTCTTGGAAACTTAACATCTCTTCGGGTAATAGATCTCAGTCAAAATTATTTTACTGGAAATATTGCCGTATCTCCACTTCCAAGTCTCTTGTCCCTTGAATATCTCTTGTTAGCAAACAACAAGTTTGAAATACCAATTTCATTCGAGCCATTTGCTAACCATTCAAAACTTAAGTTTGTCATTGCTGACCACAACTCGATAATTGTACAAGCTTATTCTAAAAGCTGGACTCCAAAGTTCCAATTGGAAGCCTTGTCTTTATCCAATTGCTCTCAAATGCCAAGTTTCCTTCAGTATCAACATCATTTACGGATTCTTTGTCTCTCGAAATGTAACATTGGTGGAAACTTCCCAAATTGGTTGTTAGAAAACAATCCCAGACTAGGAGAAGTTTATTTGGATGGAAATGCATTCAATGGATCTCTTGAATTTCTGTTCCTTCCTAATCTGAAAGCTTTTGATATCTCCAGCAACAAAATTCAAGGACAACTTCCTCCTAACATTGGGTCCATTTATCCGAATCTCATGATATTCAAAATGTCCAAAAATATGCTTGAAGGCTTGTTGCCTTCAAGTTTTGGTGACATGAAAGACCTTCAATGCTTGGATTTGTCGGGCAATAAGCTAACAGGTGATCTACCTATTGGATTGGCTCGTAAAGGATCCAAGTTAAATTTTCTGAGATTATCGAATAACATGTTGAAAGGGGAAATATTTCCAGTTTTGACTAGCATCAACAATTTTGCATATTTGTTCTTGGATGGAAATAACTTCTCAGGCCCAATTCCGCAGGCGTTATCCACTGCCTTATTATTTTCACTGGATTTAAGTTACAATAATTTGTCAGGAAACTTACCAGTGTGGCTTGGTGACATATCCTCCTTAGATTCCCTCGCACTGTCCGGAAACCATCTAAGAGGTCATATTCCGCCTGATTTTTGTAGACTTGAAGGACTTCAGGTGTTAGATCTCTCTGAAAACAACCTTGTTGGCACGATTCCATCATGTTTTAGTGCTTTACGAGCCTTAAGACGTGTGTATTTGAGCAAAAACATGTTACAAGGGGAATTTAACATGTTCTCGAATTCCCATCTACTAATGTTGGATCTTGGAGATAACAACTTCAGTGGTTCAATTCCAAAATGGTTGGACGAAACTTCTGAGTTAACTGCCTTACTCTTGAAAGGAAATCGTCTTCAAGGCACCATTCCTCCACAGTTATGTCATGCAAGTTACTTAAGAATGTTGGATCTTTCTCACAATAATATCTCAGGACCTATACCTCATTGCTTGGGAAACATAATGCAGAATCCATGGGACACAGATGTATACTACAGTCCATCATTTGGATATGGAGGATTCGAGATGTTTGGAGGAGATACCTTGATAGATGTAGATGACTCATTCGAGACAACATCAACAATTTCATTTCCAAATAGCTTTGGCACGTGGGTTGGAGTAGAGTTCACAACCAAATATAATACTTATTCATATGAAGGTAGCATTCTTAGTTACATGTTTGGTGTTGATTTATCTTGCAACCAGTTAAGTGGTGAAATACCCAATGAACTCGGTAATCTTACTGTCATTCATGCATTAAACTTGTCACATAACCACCTTACTGGAACAATACCATCAGAATTCTCAAATCTCCAGAATATTGAGAGTTTGGACATATCATACAACAACTTGACTGGGAGGCTTCCTACTCAACTTCTAAAGTTGACAGCTCTAGAAGTTTTCAGCGTGGCACACAATAATTTAACAGGTATGACTCCACAGCGTACAGCGCAGTTTGCAACTTTTGATGAAAACAGTTATGAGGGAAATCCTTCTCTTTGTGGTCCTCCATTGCACATCAGTTGCACAGAGGCTAAAGAGATTCCAACATACCTTCCTGCACCAAATTTCTCCGAAGACGATGCTTCTTTTTTAGATATGGAGTGGTTCTGTATTTCATTTCTTGTGGCATATGCAAATGTGGTGGTTGCAGTGGTTGTAGTACTTTGCGTAAACCCCTCCTGGAGAAATGTCTGGTTTTATTATATCGAGTCTTTCATGTATTCATGTTACGATTACTTTACTTCCAAAAGTTCATGACTTTCATAATTATGTAACATTTCATTATGTAAATGTATGTGTGAGTTGTTCTTGGAATGTTGGTGCCTATCAAGTTCATTAATACTAGTGAATGTTGTGTTGATTTGCTTGTGCCTTCTATAAGCATTTGCCAGATTAATTAAATTATTTCGAAATATAAAAATGTGTCATATTTTTTAGAACAGACGAATAGCCAAAATTCTTCAGTGCAATTATGGGTTTGAAAGATATGTATACTGCTCAATGAGCCATTTCTTCTATATGGTCCACTGAAATAAGTGATAAAGACTTGGTAAACTCCTAAAGTAGCTGAATTTTTTTGACATTAGCACACAAAAATTATAACTAAAAAAAAAGAAAGAAGAAGAAAATGGCTAAGGAATTAATTTTTGTTGACTAGTCAAGCACAATAATTATCAAAATTTACTCACCGTTTCAATTTGTTTGTCTTACTTTCCTTTTTAGTCTGTTTCAAAACGAATCTCTTTCCTTTTTTGCAACTCTTTACATCAAACTTTTCACGTGACATGTTTAAGATCACATGATTCAAAAGTCTTCTTTACTTTCCTAAATTCCGTGTCAAGTCAAAATCAGAAAAACAAATTGAAACGGAGGGAGTATATAGCAAGAGAAGTGTTTGCATGAAACTTCCATACAAGTGATACTAGTGGGAAATTTTTCCTCATGAGTAAGTGTCTCAAGTCAAGTCAATGATGGTGCTAGTCAACTCAAATCCAGCTCAAATTACAAGCAATGTTTTGATGAAAATAGAGTAAAATAAAGAAGTTGAAAGGGACATAACAAGAAAAGAAGTTGCACAAAATTCCACTCTAGTCTACATCAATTATGACACTATTTTCTAGGGGTACCTGCTACCTCTCACCAGCACAGATAACGAACAAGCTTCTTTTTCTTATATATAGTTTGGTTATATCTTATAGTGTTGTCAAATTGTAGGTAAAGAAACAAAATATGACTAATAGCAAGATAGTATATTGGTGGTTGTTGATTTTGTTCATGGCAAATGGATGGTTGTGTTGTTATTGTTGTTTGAATGAAGAAAAGACTGCTCTGTTGCATCTCAAGGCAAGCATAAACTACAGTATAGGCAACGATTACTTGTCATCGTGGGAGGCTAATGAAACCTCAGATTGTTGTCAATGGGAGGGAATTGTTTGCAGCAACAACACCAGAAGAGTCGTAGAGTTGTCCATTATTGTTAAGAAAATTAGTCAAGAACAACTTAGAACTCTTAATGGTACAAATACTGATGATATTTTGAATAATTGGCTTTTTAATGCATCTCTATTTATTCCCTTCAAAAATCTCAAAGCTTTGAACATACCTGGACATTCATTGGCAGGCTGGGTGAAGAATGAAGGTATGTTCTCTCCATATACGCTCCCATAATATAGAACGAAAGAGATTATACTACTCCCTCCGTTTTATATTGTATGGTGATGTTTGACTAGGCACAAAGGAATGATTTTTGGCACCTTGAAGCTTGTGGTTGTAAACATGTAATGATAATTCTATTGTGTTGTACTTTTTTTAAACGGACTAAAACAGAAAGAATGTCATATAAAATGAAACAGATAGAGTACTACATATCTGAAAAATGGTTCTAAAGATCCATTTCTAAATACAAAAAGGTGTCATTCTTTTTAAGCGGAATAAAACAGAAA

>M6-PGSC0003DMG400015706 M6 LRR-RLP

ATGAGAAATGCTTCGACATCTGGTGTAGATACAATTAGCAATTTGCCTTGTAATGTTCTCGATGTTATTCTTGGGTGCTTGCCTTGGAAAGATGCAGTAAAGACCAGTATCTTGTCAAAAGACTGGAGGTACAAATGGGTAACACGTCAAGAACTTGATTTTAATGATGAGTTTTTCAAGTCTTTGAAACAAGATGAAGAAGCCAAGATAATTGTTTACCAAGTCCTATTAGTCCATAAAGGACCAATGCTGTGTAGGATTACGAGTTGTCCTGATATTGATCATTGGATGCACTTCTTGTTGAAGAAAAATGTCCAGGAATTCACCCTTCATGTAAGATCAGAAAATCATTTACCTCATCACCTTTTTACTTTCCAGCAACTAAGGTATTTGGAACTTCAAGATTGCTTGTTCCACCCTCCACTCGGTTTCAAAGGGTTTGAGAAGCTTGTTAATCTCGACCTTATGCATGTCACTTTTGATCCATCAATATTAACAAATCTCATATCTAAAAGCCCGTTGCTTGAACAATTGAGGTTGTGTTTCATCACAAACTTTGACACCCTTGAAGTTGATGTTGCTAATCTCAAATTCTTTTGAGTTTATTAGACAAACAAAAACCATTTCCTTCAAGAATGCTCCTATGCTTGAAAAAGTTACCGTGTGTTTCATTGGACATCGACTTTTGACTGATACATCTCATTTTTTCTCTAATCTTACAAAGTTCTTTCATTACATGCCTTCACTACTGGAACTGAATCTATTCGGTTCAATATTAGGGTAACGTACATTTGTGATATTACTATCATAACCAAATGATGAATCTGATTGGTATCTTAAAGTAATTTTTAACTTGCTTCTAGTACTTGATGGAGGGAGGTGAACCAGAGAGTTTCCCAACTGCTCTAGACAATATCAGATCTCTCAATTTTTCATCCATGTCTTTAAGCAAAATCGAGGAGGTTCCAGGTGTTGTTTACTTGATTACGAGTTGTCCTAAATTACAAGATCTTACAATTGAATTTGTAAGCACTCGTTTTTTTTTCATTAAGTTGATTGAATTAATATGATTTGTTCTTCATTTGGACTCGCCAAATTAACTGTTCCTTGGAAATTCTTTTAGGACCTTGTGGTTGTGGAGGATATTGTTGAACCTGTTGTAGAATTTCTACGATGCCAATCATTCTCATTAGGTGTTGAGAAGCTTTAAAGAGTTCATGTGAACTTGTTTATGGGTCTTGAGATGGAAATGGAATTTATGAAGTTCATTTTGGCATTTGCACCTGTACTTAAGGAGATTTCCATTTTGAATTTTGCATATCTCCTTCATCGTTCGGATAAACAAATGATGGACGGGATGAAAGAAATTAGCAGAGCATCACTTAATGTTGAATTCATATTTGAATAAGTCGAGTTGGCAGATCGCAGTGAACCCCAAGAAGTCATGGAAGTGCCAT

>M6-PGSC0003DMG400015707 M6 LRR-RLP

TAGCTCCAATATTTTTTCTAATATTAATACATGGATTCTATTTTTGTTGGACAAAAATGTCCAGGAATCATCACTTGCATTTCTTAAATGCTTTCGCCTCCCTTCTCACCTATTCAAATTTCAGGATCTAAAGCACCTGGAACTTAATAGTTGTAAATTTCACCCTCCTCCCAACTTTAAAGGTTTTTGAAGTCTTGTTACCCTTAACTTTTAATTTGTAACCTTCAAGGCCATGCCATTTTAAGTCCTCCTTTAAAGTAGATCATTACTTGAGCGTGTATCGACAGTTTAGTGCAATAAGCTTAATAGCTTCATTATTGATGCCCCTAATCTAGAAATCTTTGAATTTAGCAATCGAGCAAATTCCCTTTGCTTTAGAAAAAACCCTCTACTTGAGGAAATTTTATTCGGAATGCCTTTTAACATGTTGGCGGTAATCACCATCATTGAGATATATGTTTGATGCACTACTGGATTTTGCTCGTAATTTCAGATAATTTTGTCGCAGTTTGTTTTAGTCCTTTACAGTACCATAGCCTTAGTTGATACTAGACCACATCAAAATTTTGAATTTCAAGAATTTTATCTTTTCTGAGGTTGAGTGGGTTGCATATGTGATTGACTTGATCAACCGCTACCCCAACATAGAAAGGCTTCTAATTACGGTACGTCTCATTTTTCATCTTTTGATTGTAAATATGTTGATTTTTAGCATCTTCTTGTAGTTAATCTTCTTCTTCCCTTCTCAAGTCTTATGCCTTTGAACTGTGTGCGTGTTCGTGTACACATCCCTTGCAATCCCAGGAAATGACAATGACACATATTGCAATGAAGTGGCTCAAAAGCGTGGATATGACCCTCATGTGGGTTAATAAAACTGAAATGGAATTCTTGAAGCATGTATTGGAATTTGCAATTGCACTTGAGAAAATATACATTGCAACTTCTGCAGAAATCCATAATAGGGGAATGAAAAATGATGGAGAAGATGAAGCGGTTCCCTCGAGCATCACCTAATATAGAATTTATATATCATTCAGGAGATTGACCTATCTTTGTAGCAAAAACAATATCAATTATCTGAGATTCTTATTTTCCTTTTAAATTTCATTGTGTTTTTGATCTCCTCATAAAGGAAAATAATTTATTTTTTGGATGGATGTTCCTCATAATTGAGTTTCTCAATGTAATAAAAAGGAACCTAATGGTTTTTGTATATAGTTGAAATTATTAAATTATAAGTCACGTAAGGGTATGCATACATACTTAAATATGTATTCATGTAGATTTGAATATTGTCTTTCTGTCGAAATTCTCCCAAGAATATTGAATACTTTGAAAGTTGTTAAGTGCATTTGCTTTGTACTAATGGTTCAGTTTCTTAATCTCAACTTTAAGAAGTTCTTGCAAACTCTCGTCAGTACTGGACCACGAGGAGTACTAGGAGATATGGTATAGATACAATTAG

>M6-PGSC0003DMG400015824 M6 LRR-RLP

ATGGACTATGCAATTGTGCTCTTACTTGTTTTTCTCTATCATGAAGTATTTACCGTTTCATCATCCAAATTACCTCATCTCTCTCACAAACATCAAATCGGTTCACTTCTAAAATTCAAGAACACACTTACTGTAGATACCTCAGCCTCTTATCTTTGTCAAGAGCCTTATCCAAAAATGAGTTCATGGAACATGAGCAGAGATTGTTGCTCATGGGATGGTGTCATATGCGATGAGATGACTGACCATGTAATTGAACTGGACCTCAGCTGCAGCAATCTTGCTGGGACCATCGATTCCAATAGCAGCCTCTTCCAACTCACTCATCTCCAAAGGCTTGATTTTTCTCTGAATAACTTCTCCAATTCTCACATCTCCCCTGAATTTGGCAAGTTTTCGAGCTTGACGCATCTTGATCTTTCTGACTCCAATTTCTCTGGTCAAATTCCTTCTGAAATCTCTCATCTTTCCAAGTTGCAGTCTCTTCGTCTCTATGGTCGTCTACGACTCATAGCTCACGATTTTAAATTGCTCCTTCAGAACTTGACTCAATTATTAGAGCTAGATCTTACTTCCATAAACATCTCTTCCACCATTCCTCTGAATTTTTCTTCTCATTTAACAACTCTTAGGCTGGGAGATACAACATTGTATAGGATAATACCTGAGACTATATTTCACCTGCCAAACTGAAACACTTCACTTATGGGAAAATTTTCAACTCAGTGGTTATTTTCCAAAGACCAAATAGAACAGCAGCACATCTCTCCAGAAGTTACAACTCAGTTTTGTGAATTTTCCTCATAATTTGCCCGAGTCTATTGGCTATCTAACTTCACTGCGTTATTTGTACTTCAAAATTGCAACCTTAGAGGGCCTATTCCTGAATCTCTTTCGAAGCTCACTCGCATAGAGGATTTGTACCTTCAATATAACTCCCTGAATGGAACAATACCATCAGGGATTTTCTCCCTTCCATCACTAAGTCGTTTAGTCTTGAGCAATAACCACTTTTCTGGTCAGTTTGAGGATTTCAATTCCAATTCACTAATCTTGATTGATTTAAGCAATAATCAGTTGCAAGGCCATCTTCCCAACTCAATTCAAAATCTTGTGAACCTAACAGATCTTGATCTTTCTTTCAATAATTTCCATGTGGATGTCAGCCTCTTTTCAGACCTCAAACAACTTTCGTATCTTAGTCTTTCATACAATACTGTTGATGAAAAGGAAGTTAATTCTACTTTGCCTGAGTCACTTGAGTACTTACTATTGGCCGCATGTGAAGTGAAAGAATTGGAGTTTCTAAGATTTGCAAAACAGCTTTCCAACTTGGATCTTTCAAATAATAAGCTTCAAGGAAGAATTCCTGATTGGGCATGGTCTAACTGGATGTCGTTGATAACTCTTAATATATCCCACAATATGCTGACGAGTGTGGATACAACTCCTCTTCAGATTGTATATACTATTGATTTACGGTCCAATTTGCTTCAAGGTTCACTACCTATTCCACCAAATTCCACAACATTCTTCTTCATATCCTATAATTATCTCAGTGAAGAAATTCCTTCAGATATTTGCAATTTAACATCACTTGTAATACTAGATTTGGCAAGAAACAATTTACGTGGAGAAATTCCGCCATGTTTGGGAAAAATCACTGCCCTCCAAGTTTTGGAAATGCGCCACAACAATCTTTCTGGGAACATTCCAACGACTTTCAGCAATGGAAGTTCACTGAGCAGCCTTAACTTGCATGGAAATAAACTAGAAGGGAAAATTCCCCAATCTTTGGCCAATTGCAAAAACTTGCAGCTTCTTGATTTAGGCAATAATAACCTCAACGGCACATTTCCTGTGTGGTTGGGAGCTCTGCCAGATCTGCTAGTTTTAAGCTTGAGATCAAATAAATTGCATGAACCCATCAGAACATCAAGCATTGAGAACATGTTTCCTGAGCTCCGAATCATAGATCTCTCTTATAATGCCTTTTCGGGAAACTTACCTTCGAGTTTGTTTCAACATCTGAAAGCCATGAGGACAATTGATCCATCAATGGGAACACCAAGATATCATGGAGATACATATTATGAAGATTCTATTACAGTTGCAACCAAGGGATTTTATCGTGAAATTGTGAGAATCTTGTATTTGTACACAGTTATCGATCTTTCAAGTAATAAATTTAGGGGGCAAATTCCAAGTATCATGGGGGATCTCATTGCAGTTCACATAATGAATTTATCTCATAATGGATTGCAAGGTTATATACCGTCATCATTCGGAGATTTATCTTCAGTTGAATCATTGGACCTATCAGGAAACCAGCTTTCGGGAGAGATACCACAACAACTGGCTTCTCTTACGTATCTTTCATTCTTAAATCTCTCCCACAATCATCTCCAAGGATGCATCCCTCAAGGACCTCAATTTCACACTTATGAGAGCAGCTCATATGAAGGCAATGATGGATTACGTGGATTTCCTATTTCAAAAAGTTGTGGCGATGCTCGGGTATTAGATACAAATGATACCGTATCTGGACTAGATGATGAAGAAAACAATTCTGACTTTCAGAGTGATTTTTGGAAAGGTGCTATTATGGGCTATGGAAGTGGACTATGTATTGGATTATCCATAATATATTTCATGATTTCAACTGAGACCGATATGGCTTGCAAGAATCATCATAGAGATGGAGCACAAAATAATTATGGGAAGGAGAAAGAAGCAGCGACGGCAAAGGAATTACAGAAGAAGAAACTTTAATCACTTCTAG

>M6-PGSC0003DMG400015853 M6 LRR-RLP

TATAGAAGGTGAGGGATCTCTTTTAAGGTCCCTTGTTGGTAGGGAAGCCACAGATGTTAGACTTGCTGCTATTGCTGTTGGAATCGCAAATCGTGGAGGTTTAGCAAAGCTATCTATTCGGGGAGACAATCCGTGTCGTGATGTGACAGATGCTGGTCTCAAGGCTATCGCTCGATGTTGCCCTACTCTGAGAGATCTTTCCTTGTGGAATGTGTCTTTTGTTGGTGATGAAGGCTTATCTAAGATTGCTCATGAATGCCATCTATTATAGAAGCTTGATATTTTCCAATGTCCAACAATTACCGACAATTCTTTGTTGGCCGTTGCAAAGAAATGTAATCACCTGACCTCCTTGATCATAGATTCTTGTTCAAAGATTGGGAATAAGTCTCCCAAAGTTGTGGGTCAATACTGTCTGAACCTGAAGCTCGTCATACTAAAAAATTGTCCACTCATTGGGGATAAAGGAATTGAAGATCTCTTTTATTCAGCCGGTACTGTTTTAACAGAAGTAGAGCTCCAGGCACTACACATTAGCGACATTTCTCTAAAAATTATTTCACATTATGGAACTGTACTGACAAGTCTAGTTGTTGGTGAACTTCGAAGAATAATAGCGAGGGATTTTTGGGTCATGGGCATTGGTCAATGTCTGCATAAGCTGAAGGCCCTTTCAATAAGCTCTTCCAATGGAGTCAATGATTTGGGTCTTCACGTTATCTGTAAAGCTTGCCCAAATTTAAAGTTGTTTTGTGTCCGAAAAAGTGTCATCCTATCAGATAATGGATTGGTTGCTTGTGTCAAAGCTTCAGTCTCACTTCAGAACCTTCAGTTAGAGGAATGCCACTTGATTACCCAGGCTGGATTTTTTGGTATACTTCTGAACTGTGGTAAGAAACTGAAGACTCTTTCCTTGGTGAAGTGCTTGGGTGTGATGGACTTAATTAATGTAGCTCCTTCGAAGGCTCTTTGTTGTAACTCACTGGTATCATTGACTATTCACAAATGTCTTGGAGTAGGCAATGCTACCATTGTTGTAGTAGGTAGTCTATGTCACAAACTAACTCATATAGAGCTGAGTGGGCTTCTTTGTATAACTGACGAGGGGCTCTTCCCTCTAGTTAATAACTGTGCAGCTAATTTGGTGGCGGTGAATCTAAGCGGGTGTGTCAATATTACAGACAAAGCAGTTTCAGCCATAGTTAAATTCAATGGGAGATCTCTGAAGTCTCTACTCGTAGATGGTTGCAGACATGATACTGATGCAACATTAGTAAAAATTTGGAACTCTTGCTGGATGCTTAATGTACTCGATGTTTCCAAGTGTGGAATCACCGATTCAGAGATTACAACTTTGTCCAGTGCTATACAACTTCATTTGCAGATCCTCTCACTGTCTGGTTGCCCTTTGGTGTCAGACAATAGCTTACCCCTCTTGCTCGATTTGGTTCAGAACCTTCAGGGCTTGAACATACAGCAATG

>M6-PGSC0003DMG400015878 M6 LRR-RLP

TGAATCAATCTCAGAGCTTCTGAATTTGCAAACTCTAATTTGTGATGCTTATCCATTTGATATAACTTTACCTCGGAAGATATGGATGATGAAGAACTTGAGGTATATACGTTTAAGGGAAGCCAGTTATTTACCAAGTCCTAGAATAGATAAGAATCTTGTGACAGGGATGCCAAATTTAGAGGAATTTTTTGGTCTTGGTTACTCTAGTTGTACAAATGAAGTCTTTTCTAGCATCCCCAATTTAAAGAGATTGACCATTCACGGACATTTAAGAGTAGCAGACAATATTCCATATCTTCTCTTGGATATGTCGAGCTTGACAAAACTTGAGGCATTCAAGCTTTCCTGGAGATATATTGTTGTAAAGCCCATCAAGATATTTGGTTTTCCAAAATCACTTAGAAGAGTGTCTTTAACTAGGTGTTATAAGTTTATTTGGGAAGAGATATCATCAACTTTTATCAAGTTGCCACATCTTGAAGAGCTCAAACTTAAACGTTGTGGAGCCGAAGATGATGTATGGAGATTGAGTGACAAAGACATATTCAGAAGCTTGAAGTTGTTGTTACTGAGTGAGCTAAATCTTAAGGGTTGGGTAGCAAGCAGCGATAACTTCCCAAATCTAAAACGTCTTGTTCTGAAGAGTTGCAAGGACCTGCAAGAAATTCCAACGGATTTCGTGGAAATTTGTACTTTGGAATCGATTGAGTTACATGATTGCACCACTACTGCTGAGGATTCTGTCAGAAATATCGAACAAGAACGAGAGGACATGGGATATAATATCCTTAAGGTCTACATCCATAACAGCCACAGTAAGTTATAA

>M6-PGSC0003DMG400015957 M6 LRR-RLP

ATTCTTGGAGTGCTACTGATGCCACCACTTTTAGGTGATTTGGTGAATTTGGAGCAATTGAATGTGGCACATAATATGCTTTCTAGAACAATTCCAAAAAGTATTTGTCAGCTTCCTAAGCTTCAGAATTTTACATATCCTTATAATTTCTTCACTGGTGAACCACCGGTGTATTTAGGGTTGCCGGAGTTTCATGATGAACGGAATTGTTTGCCCAAAATACCGGTGCAACGGTCTCCGGGACAGTATAAAGCGTTTTTGTCTAAAAAAATTCATTGTAGTGCTTTTAGGTGTCATAAATTTGTTCTTGTTTTGCCTTCTCCACCACCACTTTCACCGCCGTTGCCTGCCACTCCTCTGCCTGTTGTGGTGCTTATTTTTCCTCCTCCTCCACCAATTTATACTCTTCCTTCACCTTCACCACCACCTCCTCCACCTCCAGTTTATTCGCCACCACCACCCTTTCCACCTCCTCCACCACCAGTCTATTCACTGCCACTACCAACTCCATCTCCTTCACCACCAGTCTATTCACTGCCACTACCAACTCCATCTCCTCCTCCACCGCCACCACCATCACCTCCACCACCCCCACCATCTCCACCTTCTCCATCACCATTGCCTTATTGTGTACGCTCACCTCCACCTGCCTTGCAGGCCTCCTCTTCTCCTCCTTCACATTTTTTCCGTCATCTAATGAAACCAACCAATCTAATCGCTTCCTATGCTTTTTTTCTATTTTTCCTGCTGGTCCTAAACTCGACGAAGCCCCAATTCACTGTCTTCGGCCTCTCCTTCTTGATCTCACCAATCTAATTGCTTCCTA

>M6-PGSC0003DMG400016357 M6 LRR-RLP

CTGTTTCTCTAAAACAAAATCAGCTGCAAGGTCCTATTCCAAAGTCACTCTTAGATCTGCAGTACCTAGAAATTCTTCTCCTTTCGCACAATAATCTCAATGGAGAGATTGCTTCACCCATATGCAATCTGAAAACACTACGAGTGCTAGATTTGGGCAGCAATAATTTGGAGGGAACAATCCCACTATGTTTGGGTGAGATGAGTGGACTTTGGGTTTTGGATTTAAGCAACAATAGTCTTAGTGGGACAATTAATACTACTTTTAGTATAGGAAACTAATTTATAGTCATTAAATTTGATGGGAATAAGCTAGAGGGGAAAGTCCCGCAATCATTGATCAATTGCACATATTTGGAAGTTCTTGATTTAGGTAACAATGAGTTGAATGACACATTTCCTAAATGGTTGAGAGCCCTACCGTATCTGAAGATTCTAAGACTGAGATCAAATAAGTTCTTTGGCCCTTTAAAAGATTCATCGACTTACAAGTTTGCTCAAATTCGAATCATAGATCTCTCATCCAATGGATTTAGTGGAGATTTACCAGTGAGCCTTTTTGAGTATTTTGAAGCCATGAAAATAATTAGTGAGAACAGTGGAACCCGAGAGTATGTAGCAGATATGTCTTCATTTTATTACACAAATTCCTTTATAGTGACAACAAAGGGACTGGAGCTTGAACTTCCTCGAGTTTTGACTACAGACATGATTATCGATCTCTCAAGGAATAGATTTGAAGGTCATATCCCAAGCATTATTGGAGATCTCATTGGGCTTCGCACGTTGAACCTATCGCATAATCACTTGGAAGGTATTATACCATCATCACTGCACCAGTTATCTGTACTTGAATCATTGGATCTCTCATCCAACAAAATCAGCGGAGAAATTCCACAACAACTTGCATCCCTCAAATCACTTGAAGTCTTAAATCTCTCTCACAATCATCTTGTTGGATGCATCCCCAAAGGAAAACAATTTGATACATTTGAGAATAGTTCATACCAAGGGAATGATGGGTTACGTGGATTTCCACTCTCAAAGGATTGTGGTGGTGATGATGGGGTAGCACAAACGACGAATCCAGTTGAGCTAGATGAAGAAGGAGGAGGAGATTCATCAATGATCAGTTGGCAGGCGATTCTCATGGGTTACAGTTGTGGACTTGTTATTGGACTGTCCATAATATACATAATGTTGTCAACTCAATATCCAGCATGGTTTTCGAGGATGGATATAAAATTGGAACACAAAATTCTTACGAGAATGAGAAGGCACAAGAAAAGACATTAGTGTGTAACCTCGAGGTATCAACTTAATCTCTATCCTTCAGAAGATTCCATTTTATATATCCATAAATTTTCTACCTCTTTCATCGTTAAAGCTTTTAACTTTAATTCTTCATTTTTGAAAATTGCAGGATTCAAGTCTACTGCATCGTTGGGATGTAAAAATAGAAGCTTTTACTTTCATAGTTTGTTAATTCTGTTGCAACTCAACAAAATTTGATTTCATTTATCTGTAGTATGTAGAAATAAATTCAGCATTTTCAATTGTGAGGAAAAAACAAGTTCATAAAATGCAAAGGGATCAAAAGGGAAACACCAGTAGAGCAAAAAGTTTGAAATAACAGTAGAAAGCACATTCAAATTCCTAAAACAGTTCCAAATTCATCGATATATAATAACGACGACCACAATAGAGTTGGTGTAGCAGTTGAAAAGTTTGAGAAACAACCTTGGATACTAAGGTTTAAATCCTAGCAGAGATATCACATTTAGGAAGAAATCCAGCAGAACTGCTTAATTTGATGAGTCATGACTTCATATCGACAATTTTACAAATTAAACTATATTAACTAAAACTGTGGGCTGGAACCTTGTGTATTCCAGCATCTAACATTGTTCTTTCCCACCTCTCTGCGCCAATATAAATTCTTTAGAGTTGCACATAACGCCCAAAATGATTCTGTTGCGACTCAAGCAATAGTTGTGCTACTTCATTTCCCAACTCAATAGCTCCACCAAGTCTACAAGCACCCATCAGAGCCCCCCAAGACAGTAGCATCATCTTCAAAAGGCATCTTCTTTATGAAGTCCTATGCTTCGTGCAGTAGATCAACCTCACAACCATAATGTTTCATCTTAGCTGCAAGTCCAAATTCATGTGACATTGCTTCAAACAATTAGAACCCGAAATCGACAAGGTTGGCACACAGACAAGCTGATAGAACTGCCACGAAGGCAATCTCATTTGGTTGCAGCCCCTTTTACTCTCATCTTTGTTGAGAATATAATGAAATAATAGCTTAAGCTTTTAGATGAAATGGTCACACACTTCAGCAATCTTGACATACATCGTTAATGCTAGGGAAGAAATCATAGCATTCCATGTGCAAACCTGCTCCATCGCATCAAATACCTGTGTTACAGCAAATGGTTAAAAATACATGCAAGGCCTCGTACTATAAAGAGCACGCAGTTTAAAATACATGTAAGGCCTGCTAACAAGCTTTCATTAAATGATAAGCACGAGCAAAAATTAAAGACCATCGATTTGAGCGACAAAATCAAAGACCACCGAAATAGGGACAATCCGTGCAAAAACTTCTATCAATTTAGGACAGAAAATAAGACCTTTAAAATCACGAGAAGTGCAGATCCAAACCAATTTACAGGCGAATTGTTAACATAGACATCAACTGCACAAAATTCTAAACCAAGAAACAATTAATATAGTGTAATATTGTTGGCTTTGGGCAAAACATGCACAGTTTTCTTCCAAATGTCTCCAATCATATTGCAATCAGTTTCGCTACAGCACCCGCTGCTCTAGCCTAATATGAGTATCACTTCACTTTATACACATCTTCTTTTTCTTAGGAGACAAAATGCATGTGTTATTACAAGTGTTGGGACTTTCATTTAAATCTTTCTTAAATTGACTTGAGGATGTTATTTGACTTACAGTGCAATGACTTGAGAAAGATTAAACAGTCTTTACATATTGTGTGACACATTGAATGTGAATGTGTAATTGACAACTCTAAAACTGACTGAAAATGACTAGGTGACCTTCCTGGAATTGATTATTTCAAAGCAAAACTTAAATGTGCTATATAGGATTTGTTATCTGACAATTTCCACCTGCATCAAACAGTCCAGAAGCTGTTGTGGATTGTCTGTTTAAGTACTCCTTCAATTATTATTTTGTTGTCAAGGTTTATACACCAATTATTCATAGTGGTAGGAAGCTCAAAGGAAAGTGTAAAAATATCCCAATAGATAGGGAATGGAACAAGCTTTAAGGATTTTCGACTGGCACGAAGAGTTTGATTCTTAACTCTAACTCTAAGGTCTTGAATACCTGGAGTTAAGAAATTATGTTCATAACATTCAAGAACTTGAATTATTCTGTGCTCATGGGAACTAAATTAAGACATGGCATACACCTTCTTTTGCTTAAAAAAGTCGACACTACTTAAATAAAAGGACACTAATTATCAACTTGGGACAGAAAGTAAGACCTTAAAATCACAAAAAGTGCAGATCCAAACCAATTAATAAGACGAATTGTGAAAATAGACCTACATCAATTGCACAAAATTCTTAAAATGGTGTGATTTGCAGATTTTAAACTTGAGATCAAATAAGCTTCATGGCCCTATAAATGATTCAAGGACTGAGAACTTGTTTGCTAAAATTCTAGTAATAGATCTCTCATCCAATGGATTTAGTGGAGATTTACCTGTGAGCCTTTTTGAGAATTTTCAAGCCATGAAAATGATTGGTGAGAATAGTGGAACCCCAGAGTATGTAGCAGAAACATATTCCACTTTATACACAAATTCTTTGATAGTGACAACAAAGGGACTGGATCTTGAACTTCCTCAAGTTTTGACTATGGACATAATTATCGATCTCTCAAGGAATAGATTTGAAGGTTTCTATCCCAAGTATTATTGGAGATCTAATTGAACTTCGTACGTTGAACTTCCCTCATAATAACTTGAAAGGTCATATACCAGCATCAATGCAACATTTATCTGTACTTGAATCATTGGATCTCTCATTCAACAATATCGGCGGAGAAATTCCACAACAACTTGTATCCCTCACATCACTTGAAGTCTTAAATCTCTCTCA

>M6-PGSC0003DMG400016424 M6 LRR-RLP

GTTTTTTGAACATTTTAGTGCAACAATGGGCTACGTAAACCTTGTATTTTTTATGCTATTTCCCTTCCTCTGCCATCTTTCTTTCTCCTCATCCTTACCTCATTTGTGCCCCAAACATCAAGCTCTTGCTCTTCTACAATTCAAGCACATGTTTACAATAAGTTCTGTTGCTTTTGATAGACGTTGTTTCACGATAACAGGCCAGTGGATTCAGTCATATCCAAAAACTCTTTCGTGGAACAAGAGCACAGATTGTTGCTCATGGGATGGAGTTTATTGTGATGAGACGACAGGAGAAGTGATTGAGCTCAACCTCACTTGCAGCCGACTTCAAGGCAAGTTTCACTCCAACAGTAGCCTCTTTGAACTCTCCAATCTCAAAAGGCTTGATTTGTCTAGTAATAATTTCTCTGGATCACTCATTTCACCTAAATTTGGTGAGCTTTCTAGTTTGACTCATCTTGATTTGTCGTATTCAAGTTTTACAGGTCTAATACCAGCAGAAATCTCTCGTTTTTCTAAATTACAAGTTCTTAGTATCCACAGTTATCCACATGAGCTTAGATTCGTACCTCACAATTTTGAACTGCTCCTTAAGAACTTGACCCAATTAACAGAGCTCAACCTTAGGTCTGTGAACATCTCTTCCACCATTCCTCTGAATTTCTCTTCTTATTTAACTACTCTACACCTTGGGAATACACAGTTATACGGGATATTGCCGGAAAGAGTTTTCCACCTTTCCAACTTAGAATCTCTTCATTTATCAAAAAATCCCCAGCTCACTGTTAGGTTTCCCAAAACCAAATGGAACAGCAGTGCATCACTCATGGAGTTAGTTCTCAATGGTGTGAATGCTACTGGTAGGATACCTGAATCATTTGGTCATCTAACTTCACTGCGTACATTGAGAATAGCTTCTTCTAATCTCTCAGGCTCTATTCCTAAACCTCTATGGAATCTCACCAACATAGAGCTTTTGAACCTTGGTCATAACCATCTTGAAGGACCAATTTCTGATTTCTTTAGATTTGGAAAGCTCAGGGCGTTATCACTTGCAAAGAACAACTTTGA

>M6-PGSC0003DMG400016808 M6 LRR-RLP

TTAACTGAATTCATGGCATCAACTTCTTCTTCTTCTTGTTCTGGGTGCTGTCCTCGATGGAAGTACGATGTCTTCTTAAGTTTTAGAGGTGAAGATACTCGTAAAACAATTATAGGTTACTTGTATGAACGTTTGACAAGGAAGGGAATAATTGCCTTTCAAGATGATAAAAGGCTAGAGCGTGGCGATTCAATCCCAGAAGAACTCTTGAAAGCTATCCAAGACTCTCAAGTTGCCCTCATCATTTTCTCAGGGAATTATGCTACATCAAGGTGGTGCTTGGATGAACTAGTGAAGATCATGGAATGCACCAAGGAAGAAAATGAAAAAATAGTCATGCCGATATTCTATGGTGTGGAACCATCACATGTTCGAAACCAAAGTGACAGCTTTGCTGAAGCATTTGCCGAACATGAATCAAAGTATAAGGATGATGGTGAGGGGATGCAGAAGGTGAAAGGATGGAGGACTGCCCTAACTGCTGCCGCAAATCTAAAAGGATATGTATTTGGTAATGGGTGAGTTAAACTGAAATCAACGGAATAATTTATGATGACTAGCTACACATAATTATATTTTACTATAGAACAAAGAATCTCTACTTCAAAAATGATAAACAAAATCATAACTAACTTATTCACATTTTCTGAAAATAGAATGATGGTTGATTATATATGTTTCTATCTGTAAGAGTTTGGTTTCCAAGCATAGTCCTTATTCTTAAAATCATAATTATGGACTCTACAATGCGAAATACTTCTTATTCTTGACTATTTTTATATGTACATATTAGCAATATTTTTTTTTGTAGGGTTGAATCTGACTATATTGAGTGTATTGTTGATGAAATTTCGTTCAAATGCAAGAGTTCAGTATCTTATTTGCACAAAGTTGTGGGAATAGATGCTCAATTAGAGAAAGTAGAATCCCTACTAAAGATGGAAATCAATGATGTTCGGATTGTGTGGATCTGGGGAATGGGGGGAGTTGGTAAAACGACAATAGCAAAGACTATTTTTTATAACCTCTCGTCCAAATTTAAAGATGGCTGTTTCCTTGAGGATATCAAAGAAAACAAACATAGAATGCACTCTCTGCAAAATATCCTTCTCTCTAAACTATTAGGGGAAAAAGAAAATTGTGTGAACAATAAGGAGGAGGGGAAGCACCTGCTGGCTCGTAGACTTCGGTTTAAGAAGGTTTTAATTGTGCTTGATGACTTAAATCACATAGACCAGTTGGATTACCTAGCAGGGGATCTTGATTGGTTTGGCAATGGCAGTAGGATTATTGCAACCACTAGAGACAAGCGTTTGATAAGGAAGGATGTAGTACATGAAGTGGAAACACTACTTGACTGTGATGCTATTAAACTGTTCAATCAATATGCTTTCATGAAAGAAGTTCCTGATGAGTGTTTTAAGGATCTAATGTTGGAGATAGTAAGTCATGCTAAAGGCAATCCTTTAGCGTTGAAAGTGTGGGGTTCTTCATTACACGGGAAGGGTATAATTGTGTGGAGAAGTGCTCTAGATCGAATGAAGAAAAACCCTGGTTCAGAAATTATTGAAAACCTCAAAATAAGTTATGATGGGTTGGAGCCTGAAGAGCAAGAGATTTTTCTAGATATAGCATGTTTCTTCCGAGGAAATGAAAAAAAAGAAGTCATGCAAATCCTTGAGAGTTGTGATTTTCCAGCTGAATATGGATTGAGCGTCCTTATTGACAAATCTCTCGTGTTCATCTCTAGCCATAATAAAATTCAAATGCATGACTTAATACAAGATATGGGTAAATATGTAGTGAAAATGCAAAAGGATCCGGGCGAACATAGCAGATTATGGCTAGCTGAAGATTTCGAAGAAGTGATGGTCAATAATACGGTAAGTAGGCTAAACAATGCAATACTATTCAATTTTTAGTTTTTCATATTCCAAATACATATA

>M6-PGSC0003DMG400016882 M6 LRR-RLP

TGATAGAGTCCTAATTTGCTGTTCAGCAAGAAAATTGTAATCTTTTTCTTGCCCCATTGTGTTGAAGTTCCATTTTAGGTTAATTTTGGTGTGATTTGAATCATTGATAGAGTCTATAAAACAAAGGGTGTTTGTCGGTGATGTTTTTTTGAGTTCTTATTTGTGTTAATTTCATTTTCTAGGTTATTAGCCCTATAAAGTTTGTGTTTTTTGGGCTATTGATGTGTTTTGTAAGACATTGTTGAAAATTTATATGATTCTAATTTTTTTTTTCCAGAATTAATGATGGTTAATTAGAGGGTTATAGCTTCTTATGTATAACTTGTTCGTGATCCACAGCCATTTGTTCTGCCATTGCTCTATGTGAAGTGATTTAGGTGAGTCATAAGCAGATGTCTAAGTGCTACTCTTTCTTTTCATTGACTAATTGCTACTGATTTACTGTCAGTCAGAAGCTGCGAGTTGCTTCATTTGCCTTGTATCTTGCTGATTTGTTGTCATATTATTCTTGCAATCCTGCTTCGATTACATTTTTTTGAGCTGGAGGCTTGTCGAAAACAGCCTCTCTACCCTTACAAAGATAGGGGTAAGGTATGGTACATCGTACCTTCCCAGATTCGACTTGTGGTATTACAGTGGGTATGTTGTTGTTGTTGTTGTTATTTTACATAAATGATGATAAATTGAGTGGCACAATGTACTGGAAGTATTTTAGTGTCAAAATCAAATGATCCATTTGAAGATTGTGTTTCAATTAGTAGAATAACAAGCACAACACCTATTACAACTGAATAAAGAAAATTTTAAGGCGTTTTGTTTTTCTGAAGTTCTTTTGGAATCTTGCGAAGAAATTTTCCAAAAATTGTGTTTCTTTGAATCTTCTGAAACGCTTTTTTTAAAAAAATGTTTTTTTATGAAGTGTTTTTAGGGTGCTCCCACTCACAGATCTTCAATAACCAACTATCCATAAACATTTAGTCTTTGCATGTTTTAAAAAGAAAGATTTTTTGAGAAAACTCTTTCCAAGATCATCTCTAAACATGCAAGCCGAATAGCTTCTATAAATTTCTGTAATGAAAACATTAACTAATTGATGAAATTAGTAATTAGAGCATCTCTTTTCAAACCAACATTTAAAGTGTTTGCATCTGTTGATGGATGAATTCAATATGCTAATGAATTCTAACTCAGCATTTTTGGAAGATCAGCTTTCGTGGTGAAGGAATAAGCAGTTCTTTTGGAAAATAGTAAGGAGCTATGGCTTGAATGTTTATGCGAAGCGAATGTGTACCTTCTGTTTACTCTATGGATGCTAAAAAATTCTGCTAGTTGCTAAAACAGGATGAGGGCTATGGGAAATAACTTATTTTGTTTGATCCACTCATTTCTTCTACCACTGCTCTCTGTAAAGTGATTTAGGTGCACCACTTTATTATCCATGTTATAAGCATCTGTCTAGGTGCCCCCATTATGTTTCTAGGATAAGTAGAAGTTCTTGCAATTGACTAGCTGCCACTGATTGATGGTAAATACTTACCCACATCGGGTGTTTATACTTAAATAAACAGTTAACCTTTAGAATTAGCAAATTAAAGTGAAAATGCATATCACTTAACCGTGTTTAAGTGTTACTAAGTGTATATGCAAGATATATGTGTATATTCATTGTCTTGGTGTAAGCACCCATTCGGGTTAAGATTTTACCTGATTTAGTGCCTCTCGGAAGCTTTTAATTGCGATTTATGTTAGGGATTATCCATAAATGTTGATCAATTGAATGTCACAGTGGGATTTGAAGTACTTAAGTGTCAAAATCAAATGATTCACTTGAAGACCGTGCATCAATTAGGAGAAGAACAAGCATAACACACTTATTACAACTAAATAAAGAAATATCTAAGGTGTTTGTTGTGAAAGTTTCTTTGAGATCTTGGAAGAAGTTTTGAAAAATGTGTTTCTTTTGGAGTCTTCAGAAAAGCTATTTCTTGAGCAAGTTATTTTAGGGTGTTCTCCATTCACAAATCTTCAAAACCAACTATCCAAACACATTTTCTGTTTGCATGGTTTTAAAGAAAGGATTTTTCAGAAAACTTCTTTCAAAAGTTATCCGTTCAACTCTTATAAAACTTTGCGATGAATATATTAGCTAATTGTTGAAGTTTGTTATTAGACCATCTGTTTTCAAACCACATTTAAAGTGTTTGCTTCTGATGATGGGTGAGTTCAACATATTAACAAATTCTTACACAACATTTTTAGACCATACGAGTTTGTTGGTGAAGGAATGCACAGTTCTTATGGAAAATAATAAAGAGCACAGTTGAATGGATATGCAAAGTGGCGGTGTGTCTTCTGTTACTCTATGGATGCTAAAAAGTTTTTCTACTTGGTGGCATATTGTTACATTTGTCCCTTTTATATGAATCAGGTGGACCGGCAAAAATCCTCAAAAAGAGCCTTATTTGGACATTTGAGGCATGACATCTTTTACAATCTCATTACAAAAAAATGAGTGTTATGGTAAGTCACGTGCTTCTTTCATAGTAACTGTGTAATGTGATAAAATGAACCATTTAATCTACTTTCTTTCATTGTTTTCACTGATGTGTGCTTCTTTCGACATTTTCTGACGATAATTAGTCCCCTAGTTTTCAATTTATGTGCTACAGTGCTTGACAGAGTAAAGTACTGCATTTACCTTTTCAGTTAATTTACTATATTTTCTGGCCCAATTTTAGTGGTCAACATCAAACACTCGGCCTTGATTGCATGATGACGTTGTGCAATGGGAAAAATAATTGCAATAGCTAAATGGTGATGTGAAAGATCAGCCATCCCTTAACTCTGTGTTTGTGGATGAAATCCTCTCTCAGAAGAGTTCGAATGGCCATTCCTGCCCCTTGAGCATTACCAAGTAAATGACTACTTAAATCATAGACATTGCTCATAATTGGTCGAAAATTTGCCAGCATATCTGTAACTATTCTCTTAACACAAATTATAGTAGCATCATTTAAAGAAAAACAGAACTCTGAGTGCTCTTACCCTATGTCGTTCTTTGTCAGTAAACCACAGTGATAATTTCTCTCTCTTGGGTTGTCACATTGACATCTTTGCATTTTTGTAAGCACTCTTGTATGCATGTATGTATTTTAT

>M6-PGSC0003DMG400016899 M6 LRR-RLP

TTGCAATTGCAATTGCAATGCTGAGTGTATTAAATATGTAGTGTAAAACGAGAGTAGCAATAAACCTCTCATTGAAGAAGCATCAACTAAACCCACATCGTCTCTCATCAATGCTACAATCACATTTTGTGCAACCTTGCAAAAAAACGAAACGCCTCTTGTTTGAGCAAAAATGGGGGGAATTTGCTTTGTGGTTTTGCTCCTTCAAATCATAGTTACTGAAGTTGTTGGTGCTGGTGTTGGTGTTGGTGTTGGCGTTGGTGTTGGACCTGGAGGTGTTTGGTTTGGTGGAGGGATCAACAGCCCAAGCCCAGGTGTTGTCCCAAGTCAAAATTCGGCTTACACTGCTCTTCAAGCATGGAAATCTGCAATCACTGATGATCCAATGGGGATTTTGAAGAATTGGGTGGGTCCAAATGTGTGTACTTACAGAGGAATTTTCTGCACAAACTCTCAAGATTACATGGGCAACCCAACAGACCCAGTTGTTACAGGTATAGATCTCAATCATGCAAATCTTCAAGGCACTCTTGTTAAAGAACTGTCTTACTTAACAGATATTTCTCTCATTCACCTCAATACCAACAGATTTTCAGGTATAATTCCACAAACATTCAGAGATCTGTCTTCACTCGTTGAATTAGACCTTAGTAACAACCATTTCTCTGGTCCATTTCCTACTACAGTACTATTCATCCCAAATCTTCTTTATTTAGATCTCAGATTCAACAGTTTCACAGGGCCAATTCCTGAGGATCTGTTCAACAGAAAACTTGATGCTATTTTTCTTAATAATAACCAGTTTGATGGTGAACTACCTCAGAGTTTAGGGAATTCCCCTGCTTCAGTGATTAATTTTGCTAATAATAAGTTTAGTGGGAACATTCCTTTTAGTTTAGGGTACATGGGTCCGAGAATTAAGGAGATTTTGTTCTTAAACAACGACTTAAATGGTTGTATACCAGAAGGCGTAGGATTGTGGACGGATTTGCAAGTTCTAGATGTGAGTTTCAATTCACTAATGGGGCATTTGCCTGATACCCTTTCTTGTTTGAGTGGCATTGAGGTTCTCAACTTGGGACATAACAAGTTATCTGGAGATTTACCAGACTTGGTTTGCTCACTAAGAAACCTTGTGAATTTGACTCTTGCTTACAATTTCTTCTCTGAGTTGAGCCAGGATTGTGACAAACTTCCTTTGAGAAATGTGGGTATTGATTTCTCATTGAACTGTATTCCAGGAAGACAAATGCAAAGGCCTCAACCTGAATGCTCTGTCATTCCAGGTGGAAGTCTCAGTTGCCTTAGAGTACCTGCAGTAAAGCCTCTTGTTTGTGGATGAAAAAACTGCTACTGCCAGACTTAGTCGGACTCCAACGTGGGTATCAAACACCCGGTGGAAAACCAAAAAAAAAAAACTACTACTACTACTATTTTATTTATTTATGAGTGACAACAATTGAGCACTATCATCAATGCACACGCAAGTTATGACTTGTACCTTTCAAATTTATTTCTCTTCCTTATTTCTTTATGTTGCATTATGAATTGTAAATACTAGTATTAAATTAGGACACTTATGGGTTTCACCCACCTCTAGTAATCATAATTCCTCATCAGTAGTCGGGAATTCTCA

>M6-PGSC0003DMG400016982 M6 LRR-RLP

ATGGGAATGCCAAATTTGGAGTATTTGGATCTAGAGGAATGTATGAGTCTTGAAGAGGTTCACCATTCTCTGAGATGTTCCAGAAAACTCATTGAGTTAAATTTGAATGAGTGTGGAAGCCTTAATAGGTTTCCATATGTTAATGTGGAATCTCTTGAATATCTTAATCTAAGTTCCTGCTATAGTTTAGAGAAATCTCCGGAATTCCTTGGAATAATGAAGCCGGAGTTAATGATTATGATGTCATTCTTTGGAATAATGGAACTTCCATCATCTATTATTGAGCACCAAGCTGGTCCAACAAGCCTAGATTTAAGTGAAATGAATAACCTTGTAGCTCTTCCAAGCAGCATTTGTAGGATGAAAGGTTTGGTGAAGCTAATTGTGTGGGGTGCTCAAAACTTGAAAGCTTGCCAGAAGAAAAAGGTGATTTAGAAAACTTGGAGGAGCTTGATGCCAGAGATACTCTAATCTCACGACCTCCGTCTTCCATAGTCCGATTGAACAAGCTTAAATTATTAACTTTTGAAAAAGAAAAAACAGAAGACAAAGTGTACTTTGTGTTCCCTCAGGTGAATGAAGGGTTACCCTCATTGGTTGCTCTGAATCTCAGTGACTGCAATTTAATTGATGGAGGACTTCCGGAAGACATTGGATCCTTATCCTCTTTGGAAACGTTGCATCTCAATGGAAATAATTTTGAGCGTTTGCCTCGAAGCATAGCCCAGCTTGGTGCTCTTCGATCCTTGGACTTATCAGATTGCAAGAGGCTTACACAGCTGCCAGAAGACATTGGATGCTTATCCTATTTAAAAAAGTTGAATCTCAGTGGAAATAATTTTGAGTATTTGCCTCGTAGCATAGCCCAGCTTGGTGCTCTTGAATACTTGCACTTATCAGATTGCAAGAGGCTTACACAGCTGCCAGAATTTCCACATACAATAAATGCAGATTGGAGCAATTCTTCGATCTGTAATTCATTGTTTCAGAATATCTCATTATTGCACCCTGACACCTCTGATTCACATTCCTTGTCACTTAGAGTGTTTACAAGTAGGCCGAAGAATATCCCAAGTTGGTTCCACCTTCGGGGAACTGGTACAAGTGTATTAGTCAATTTGCCTATGAATTGGTATGTAACAGATAACTTCTTGGGATTTGCTGTATGTTACTCTGGCGAATTAAATGACAATACAGCTCATTTGATTCCCTTATGTGATGCTGGGATGTCGTTGATGACCCAAAAACTTGCTTTATCCAACCATGCAGAATATCTTGATGATATTAATTTTTTGTTGGTACCTCTTGGTGGCTTATGGGATGCATCTAAAGCAAATGGAAAAACACCAAATGACTGA

>M6-PGSC0003DMG400016983 M6 LRR-RLP

ATGGCATCATCTTCTTCTGCGAGTAATTCACAATACTGTCCTCCATGGAAGTACGATGTCTTTCTAAGTTTCAGAGGAGATGATACCCGGAATAACTTTACAAGTCACTTGAACAAAGGTTTGGAAAATAGGGGAATAGCCACCTTTCTAGACGATGAAAGACTAGAGGATGGCGATTCCATCTCAGAAGAACTTGTGCAAGCAATAGAAGAGTCTCAAGTTGCCGTAATCATTTTCTCAATGAATTATGCTAAGTCAAGTTGGTGCTTGAATGAAATAGTGAAGATTATGGAATGCAAGGAGAAAGAAAACAGACAAACAGTTATTCCGATCTTCTATTATGTGGATCCATCACATGTTCGATACCAAAGTGAGAGCTTTGCGGAAGCATTTGCCAAACATGAATCAAGGTATAAGGATGATGTTGAGGGGAAGCAGAAAGTGCAAGGGAGGAGAAATGCCCTAACAGCTGCCGCAAATCTAAAAGGACATGATATTCATGACCTGTGAGTTGGAAACAAATAATTGCTTTTGATGGAAAAAAAAATAACAGTATGTTTCTATATATTATCCTGAACTATAATGTTTCCAATTTAATACGGTCTTGAACATAAAAATTATGAAAATACTCTCTAATACTTGACTTCCTCACATTGTCAACAATCAATCTCTAATTTATGTAGGATTAATCAATAAATGAAAATTGATCAGATCGTCGACCACATCTCTTCCAAACTTTGCAAGAGTGCTCGTTTTTTGTCTTATTTGCAAGATGTTGTGGGAATAAATTCTCATTTAGAGGAACTAAAATCCCTACTTCAAATAGAAATCCATAATGTTCTGATTGTAGAAATCTGGGGAACAGGGGGAATTGGTAAAACGATGATAGCAAAAACCATCGTTGATACTCTATCTCATCAATTTCAAGCAGCTTGTTTTCTTGTGGATGCTAAAGAAAATGCAAGAAAGAATCAACTACATTTCTTACAAAATACCCTTCTCTCTGAATTGTTAAGAAAAAAAGATAATTACGTCAATAATAAGTATGATGGGAAGTGCATGATTCCAAACAGACTTTCTTCAATGAAGTTGTTGATTGTGCTTGATGACATTGATGAAAGAGATCATTTGGAGTATTTAGCAGGTGGTGTTGGTTGGTTTGGTAAAGGCAGCAGAGTTATTGTAACAACTAGAAACAGACAATTGATAGAGAAGGATGTTGCAATATATTAAGTGCCTACACTACCTGATGATGATACTATGCAATTGTTCAATCAACATGCTTTCAAAAAAGAAGTTCCAGATGAGTGTTTTAAGAAGTTCTCGTTGGAGGTAGTAAATCACGCTAAAGGCCTTCCTTTAGCTCTCAAGGTGTGGGGTTGTTTGCTGCATAAGAAAGGCCTAACTCAGTGGACAAGAACTGTAGACCAAATAAAGAAAAACTCTAATTCAGAAATTGTTGAAAAACTCAAAATAAGTTATGATGGGTTGGAGCCCGAAGAGCAAAAGATATTTCTAGATATCGCATGTTTCTTCCATGGACATAAAAGAAAAGAAGTCATGCAAATTGTTGATAGCTGTGATTTTGGAGCTGAATACAGATTAGATGTTCTAATTGACAAATCCCTTGGATTCATCTCTAAATATGATAGAATTGAAATGCATGACTTGATTGAAGATATGAGCAAATACATAGTGAAAATGCAAAAAGATTCCGGAAAACTTAGCAGAATATTGAAGGTTGAATATTTCGAAGATGTGATGATGGACAATATGGTAAGTAAGCTAAACAATGCAATAGTATTCATAATTCAAAGATATATATATATATATACCTTGAGTAAATTTTTATAAAATAAAATTAAGGAGGGCCAAAACTTAACCATACCAGTTCACTTGATCATTGATTAGTTTTTTTTTGGAATTAAAACTAGCATTTTATTACCAAATATCAGATCTGTACATAGTCTAGAAAAGCTGTGATGGTATGGCATTCACACAGCTGCAAACCAAACAACTAGACATCAAAAACCTATGTGCTAGTGGTAAACTATAGCTAACTATGGGTAGAAATTAAGGCTATTTAGTCTTCTCGCAAGCTTCTTCTTGAGTGATACTCCGTAGAATATGTCCTGCACAGTTCTCCTAACCACCATTTTAGGTTCACAACGTTTGGCCTGAAATATCTTCTGATTTCTCTCTATCCACACTTTGTATACTGCTGCAGCAAATGTCATCCTTAATATAGTTGTTGTTCGACCATTTCCATTGGCATAAGTGCATGCCCAGTTAAGTTCATCTTTCCAATTCATGGCCACTCTATTGATGCCCTGCCTAGACCAGATTTCACTTGAGTAGTCACAGCTGAAGAAAAGGTGTTCAATAGTCTCATTCTCCTTGTTGCATAACTCACATTGTTGTTCATCAACTTCACCCCATTGTAGCAATCTGTCTTTTATATGTAGTCTTCTAAGGGTAGCCAATCTAAGGATAAAAATCCATTTTGGAGAACCTGGATTGTTGCAAATAATTTTTCTCCACTGTACTTTTTGGAACTCACCCCTCATCAAGACATACATCTTCCTAATGGAATAACTATGAATGTTTGACAAATCACTAACTTGTAAACCAATGTCCTGTAGGCATAACTTCACCTTCAAAATCTTTTGTGTTGGGTTGGATTCCCATATTGGCCTTCCTTTTATATAGTAACAATGCACCCATCTAACCCATAATGTACCTTTTTTAGAGCACAGGTTCCAGAGCAATTTGCTAATGGCTGCTCTGTTCCACACTGTAATATCAAGAATGTGTAGGCCACCAGCAGTTCTTGGTAACCACACCTTATCCCAGGCTTGCAAAGCCTTCTTTGTTACTTCTGTACCTCCCGTCCACAAGAACCTCTTACAAGTTGATTCTATAAGTTTGATCACCTTGGCGGGTAACACAAAAATTTGCGACCAATAAATATGGATAGAGAATAATACAGATTTAATTAGCTGCTCTCTGCCTGCGTAAGATAGAAACTTTGTTGTCCAACATGTTATTCTTCCTAGCATCTTGTCAATCAAGGGTTGGCATTGGGTGATTGATAGTCGTTTAGTACTTAGAGGCCCTAAGTATCTAAAAGGTATCTCACCTCTAGTAAATCCAAGAATATTCAAGATAGCTGTCTGATTTATGACATTCACCCCTCCAAAATATACAACATTTTTGTCTTTGTTTGCAATCAGACCAGAAGCATGAGAAAATTGTTGAAAATAATCAAACAACAGCTGTACAGACCTTATGTCACCCTTGCAAAATAATAATAAGTCATCAGCAAAATTGAATAAGTTTTACCTTTGAGCACCTGGGGTGAAACTTAAATTCAGGACAGGTCATTAGGCTCTTCAATAATCGTGAGAAGTACTCCATTGCTAAAACAAAGAGAAAGGGGGACATAGGGTCCCCTTGTCTCAATCCTCTCTTAGCATCAAAAGGTACAGAAGGCCAGCCATTGATACAGATTGAGTAAGACACAGTTTGAAGGCACTCCATTATCTAGTGTATGAATTTTGTAGGGAAATTCAGATGTGTCATTACTTGCTTGACAAAGATCCATTCTAGTGAATCATATGCTTTTTGCATATCTACTTTAAGCATACATCTAGGAGAGACCCCCTTTCTTCCATAGCCCTTAACCAGTTCATGACTTAAGATAATATTGACATGCAAATTTTATACAGACATTTGAAGTTATACTTTGTAGACTAATGAACCTTTTAACAGGGAACCAAGACAATGGAAGCAATCTGGGCGCATGGTGTTCAAAAATTATGCTTTAGCAAAGAAGCCATGGAAAACATGAAAAGGCTTAGATTATTATGCATCATGAGTTTTCGTACACATGATGACTCCATTGAGTACCTGCCCAACAACTTGCGTTGGTTTGTCTGTCATCATTATCCTTGGGAGTCATTATTAGAAAATTTTGAACCTAAGAAGATTGTTTGTCTTGATCTCCAAAAGAGTTCCCTGCGTCAATTATGGACCGGAATAAAAGGTACAATTGTTGAATTCAATTTTCATTATTTTTCTTTTTGTCAAATACGTACATAGATAGACTCTCAAATTTTAAATAAATTTTGGTCTAAAGTTGTTGGGGTGGTCCAATTAGAGTCAAAACTGAGGGACCAGTTTCGTTGGCTATTTTGTATCGAGAATTTGAATTTGATGAAACGAATGAAAATTAGTTGTTGAGCTTATTAATTTTAACAACATAAAAATACTTAGCTTAGTTGAAATAAAAGATAATTTTATATAAATTGGGTTGGAGCAAATTTGATTCAATGAGAGTAAGATATTTTTTATTGGTCCACTAGCCAAATCTTAATTATCCTGGGCAGCATCCTAATTCTTCTGTGTTTTGAATAGCAGCATTTGCCGTCTCTGCGAAAGCTAGATCTTAGAGATTCAGAAAGGCTGATTCAAACACCAGATTTCACGCTAATGACAAATTTGGAGTATTTGGATCTGCGTAACTGTAGTGATCTTGAAGAGGTTCACCGTTCCTTGGAGTGTTGCAGAAAACTCATTGTGTTAAATTTGCAGAGGTGTATACGCCTAAATAAGTTTCCATGTGTTAACGTGGAATCTCTTGGATCTCTGCTTCTACTGCATTGCTCTAATTTAGAAAAATTTCCAGAAATCCTCGGAAGAATGAAGCCAGAGTTAGAGATCAAGATGAGCTGGTCTGGGCTAAGGGAAATACCATCATCTATTATTCAGCAATACGCATGTCACCTTGCGGAGTTAAGTTTGAGTGATATGAAAAAACTTGTAGCACTTCCAAGCAGCATTTGTCAGTTGAAAGGTTTGGTGAAGCAGGATGTGTCATACTGCTCAAAACTTGAAAGCTTGCTAGAAGAGATTGGTTATTTAGAAAACTTGGGAGGGGCTTCATGCAAGCTTTACTCTAATCTCACGACCTCCGTCTTCCATCATCCGCTTGAACAAGCTTAA

>M6-PGSC0003DMG400017062 M6 LRR-RLP

TATTATTTATAGTGAACAAAAAATAACTACATCATAATAATAAGATTTCAAATACAATTCTTTGGTTTTTCCTATACATGAAAGTAATAATTACTTTTTTGAACTTTATAATACAAAATGCACATAAATTTTCATAAAGTTAAATACTACTACATTAATTAAGTTATTCAAATTAATTTAGAGAAGGGGAAAAGGTTTAGTTAAGTATTTAGATACATGTTTTCATGTACCAAATAAATGCACATACTTTTATGGATTTTTTCATCTTTTAATTTTTTATTTTATACCTTTTCACTTGTAGTATTTATCATAATATTAATAATATAGTAATATGATTATTATTTAATATTTAATTAGTTTAATATTTATAATATTGGAATTAATGTAGGGGTAAAATGCTACTCTTTCTCTTTATTGTTCAAAATTCTCTTTGGTTTTTGCGGACTCCTACCGAGCTTGAATTTTATAACAATATTCTCATGTGAACATTACTTGTACCAGGTTCTGTATTAACTGAAAAACAAGTTTATGGAAGGGGCAAAGAGAAAGATGAGATAGTGAAAATCTTGATAAACAATGCCCAAGAACTTTCAGTCCTCCCAATACTTGGTTTGGGGGGACTAGGAAAGACAACTCTTGCACAAATGGTCTACAATGATCCGAGAGTTACTGAGCATTTCCATCCCAAAATATGGATTTGTGTCTCAGATGAATTTGATGAGAAGAGGTTGATAAAGGCAATCGTAGAATCTATTGAAAGAAAGTCACTCAATGACATGGACTTGGATCCACTTCAAAAGAAGCTTCAGAAGTTGCTGAATAGAGAAAGATACTTGCTTGTCTTAGATGATGTTTGGAATGAAGATCAACAGAAGTGGGATAATTTAAGAGCTGTCTTGAAGGTTGGAGCAACTGGCGCTTCTATTCTAACCACTACTCGTCTTCAAAAGGTTTGTTCAATTATGCAAACTTTGCACCCATATGAATTGTCAAATCTGTCTCAAGAAGATTGTTTGTCGTTGTTCAACCAACGTGCATTTGAGCACCTAGAAGAAATAAATCCTAACCTTGAGGCTATCGGAAAGGAGATTGTGAAAAAATGTGGTGGTGTACCCCTAGCAGCCAAGACTCTCGGAGGTATTTTGCGCTTCAAGAGAGAAGAAAGAGAATGGAAACATGTGAGAGATAGCAAGATTTGGAATTTGCCTCAAGATGAAAGTTCTATTCTGCCTGCCCTGAGACTGAGTTATCATCATCTTCCACTTGATTTGAGACAATGCTTTGCATATTGTGCGGTGTTCCCAAAGGACACCAAAATGGTAAAGGAAAATCTAATCTCTCTTTGGATAGCGCACAGTTTTCTTTCATCGAAAGGAAACTTGGAGCTAGAGGATGTGGGTAATGAAGTATGGAATGAATTATATTTGAGGTCTTTCTTCCAAGAGATTGAAGTTGAATCTGGTAAAACTTATTTCAAGATGCATGATCTCATCCATGATTTGGCTACATCTCTGTTTTCAGAAAGAGCATCAAGCAGCAATATCCGTGAAATAAATGTAAAAAAATACTCACATATAAAGTCCATTGGTTTTGTTAAAGTGGAGTCTTCCTACTCTCATTTGCTCGTGGAGAACTTTGTCTCGTTGAGGTTGCTTAATTTGAGTAACTTAGAACTCAATCAGTTACCGTCTTCCATTGGAGATCTAGTACATTTGAGATGCCTGGACTTGTCTGGCAATAGAGAAATGCGTAGTCTTCCAAAGCAGTTATGCAAGCTTCAAAATCTGCAGACTCTTGATCTACATGACTGCTGGTCACTTTCTTGTTTGCCAAAACAAACAAGTAAACTTGGTAGTCTCCGAAATCTTTTACTTGATAATTGCATTAAATTGAAGTCTACGCCACCAAGGATAGGATCTTTGACATGCCTTAAGACTCTAGGTCAATTTGTTGTTGGAAGGAAGAAAGGTTTTCAACTTGGGGAACTACGAAACCTGAATCTCTATGGCTCAATTGAAATCAGACATCTTGAGAGAGTGAAGAATGATAAGGATGCAAAAGAAGCCAATATATCTGCAAAAGAGAATCTGCAATCTTTAATCATAGAGTGGGATTGGCTTGAACGACGTAGATATGAATCAGAAGAAGTTAGAGTGATTGAAGCCCTCAAACCACACCCCAATCTGACCTCTTTAACAATCAACGACTTCAGAGGATCCCGTTTCCCTGAGTGGATGAATCACTCAGTTTTGAAAAATGTTGTCTCTATTGTAATTAAGAACTGCATAAACTGCTCGTGCTTACCACCCTTTGGTGAGCTGCCTTGTCTAGAAAGTCTAGAGTTATGGAATGGTTGTGTGGAGTATGTTGATGTTGATTCTGGGAGGTTTCCATCCTTGAGGAAACTTGTTATAGTGCACTTTAATAATCTAAAAGGATTGCTGAAAAAGGAAGGAGAAAAGCAATTCCCTGTGCTTGAAGAGATGACAATCCGCCACTGCTCTATGTTTGTTATTCCGAACCTCCCTTTGAAAGTTTATGCGACAGATGCCTCAAGTTTGGGGTCCACATCTAATCTTAGTGCTCTTACTTCCCTCAACATAAGCCATAACTATGAAGCTACTTCACTCCCAGAAGAGATGTTCAAAAGCCTTGCAAATCTCAAACACTTGAATATCTCTAGTTACAACAATCTCAAAGAGTTGCCTACCAGCCTGGCTAGTCTCAATGCTTTGAAGAGTCTGGATATTGAAAGTTGTCATGCACTAGAGAGTCTCCCCGAGGAAGGGGTGAAAGGTTTAACTTCACTCACCAAGTTGTCTGTCCGTTGCTGTAAGATGCTAAAATGTTTACCGGAGGGATTGCAGCACCTAACAGCCCTCACAACTTTAATAATTACTCGATGTCCAACACTGGAAAAGCGGTGCGAGAAGGAAATAGGAGAAGACTGGCACAAAATTGCTCACATACC

>M6-PGSC0003DMG400017072 M6 LRR-RLP

TATGCCACCAAGGATAGGATCTTTGACATGCCTTAAGACTCTTGGTTGCTTTGTAGTGGGAAGGAAGAAAAGTTCTCAACTTGGTGAATTACGAAACCTGAATCTCTACGGTTCAATTGAAATCACGCATCTTGAGAGAGTGAAGAATGATATGGATGCAAAAGAAGCCAATTTATCTGCAAAAGAAAATCTGCATTCTTTAAGCATGAGATGGGATGGCGATGAACGTCCACAAGGATATGAATCAGAAGAAGTTGAAGTGCTTGAAGCCCTCAAACCACACTCCAATCTGACTTGTTTAACAATCAGTGGCTTCAGAGGAATCCGTCTCCCAGACAGGATGAATCGCTCAGTTTTGAAAAATGTTGTCTCTATTGAAATCAGGGGTTGCAAAAACTGCTCATGCTTACCACCCTTTGGTGAGCTGCCTTGTCTAAAAAGTCTAGAGTTAAAGAACGGGTCTGCGGAATTGGAGTATGTTTATTCTGGATTCCCTACAAGAAGAAGGTTTCCATCTCTGAGAAAACTTATTATAGGCAAATTTGATAATCTGAAAGGATTGCTGAAAAAGGAAGGAGAAGAGCAATTCCCTGTGCTTGAAGAGATGTCGATTGAGTGGTGCCCTATGTTTGTTATTCCGACCTTTTCTTCTGTGAAGAAATTGGTAGCTTATGGGGACAAGTCAGATGCAATAGGTTTCAGTTCCATATCTAATCTCAGGGCTCTTACTTCCCTCTACATTCGCAATAACTATGAAGCTTCACTCCCAGAAGAGATGTTCAAAAGCCTTGCGAATCTCAAATACTTGCAAATCTCTCTCTGCTGCAATCTTAAAGAGCTGCCTATCAGCCTGGCTAGTCTCAATGCTTTGAAGCATCTGGATATTTATTTATGTGTAGCACTAGAGAGTCTCCCCAAGGAAGGGGTGAAAGGTTTAACTTCACTCACACAGTTATCCACAACAGACTGTGAGATGCTACAATGTTTACCGGAGGGATTGCAGCACCTAACAGCCCTCACAAATTTATCAGTTACGTATTGTCCAACACTGGCCAAGCGATGTGAGAAGGGAATAGGAGAAGACTGGTACAAAATTGCTCACATTCCTAATGTGTTTATCCGTTAAGTCTTATTCCTAATTGGATGTAATTTTCTGATTTTCTTTTGGAAACAAATCAACTAATTGTA

>M6-PGSC0003DMG400017081 M6 LRR-RLP

TAACGCATTATTGATCACACTGTATGTATATAAATATATGGGGTACAAGTTCCTAGCTAGTTAGATATATTCAATGGAGTACCAAAAATTGCTAATAGCTTTCTGCTTCTATTCACATTTTATACTTTTTGTCCATCAATCTCAGCTTACATATGCTGGGCAACATCTCTGTGCCCGCTATGAAGCTTTTTATTTGCTTCAATTTCAGCAAGGCCTCACAGTTGATCTTATTGGTTATGACTGTGACTATGAGGCCCTAGCTAAGACTTTGTCTTTGAATGTTACAGGAGATTGTTGTCAATGGGATGGTGTTACTTGTCATGGATTTACTGGTCACGTTATAGGACTCGATCTTTCTTGCAGTTTTCTTAACGGAACCATTAATTCTAACAGCAGCCTCACAAAACTTAGTCATCTTCAAAGACTCAACCTTGCTTTCAACAAGTTGAGTAACTTTCCACTTGTGGATAGCATTAGTGAACTCGAAAGTTTGACATATCTCAATCTTTCGCATTCAGTACATATGATAGGGGAGAAGCAGATACCACAAGTATTATCCAACTTGTCTAAATTGGTTTCACTTGACCTCTCCTTGTATTACACTGAACTCAAAGTTGATTGAAAAACCTTCAGAAGTTTGCTTCAAGACTTGACCAATTTGGAGGTACTGTTTCTTGACAACTCGTATGCATCATTTGAGTTACCTAAGAATTTTTCTTCTTCTCTTAGGTACTTAAGTCTTGAAGGTATCGGCATGTTTGGGATCATAAGTGAGTTTCAGTTATTTCATTTATCAAACTTGCAAGTGCTGAGATTTGGACAGAATCCTTTACTAACAGGCTCTTTGCCAAATTTTAATCGGAGTTCCAGTGAAAGTATTTTAGAGTTGAACTTCTTTTATACTGGTATTTCTGGGAAGGTACCCGATTCAATTGGTAATCTCCATTCTCTTTGGTATTTGAATCTCCAGATTTGTCATTTATCTGGCTCGATTCCAGAATCCATTGGCAATCTCACTGCAATCAGAGAGTTGATACTTTCAGAAAACGTCTTCATTGTTAATGTTCCCTCAACTTTCTCAAAGTTAAACAAACTCGTTCGTTTAGATCTCTCTTCAAACTATTTTCGAGGCTCAATTCTAGAATACATTGACAATCTCACTGCAATCACACAGTTGACAATTCCAGATAACAGCTTCACTGGAAATGTTCCCTCAACTATCGGAAAGTTGAACAAACTTAACTACTTATCTCTCTCTTCCAATTATTTTGAAGGCTCGATTCCAGACATCTTTGCCAACTTTTCAGATCTATTTGCTTTATTTTTTGACACTAACAATTTCACTGGTTCATTTCCCTGTTCAATTGCAACCTTGATTTGCCTTGTGAATTTAGAGTTGCAAAACAATTTACTAACTGGTTCACTTCCCTCTAATATAAGCTGATTTCAACTTCTACAATCACTTGATTTGTCCTTCAATTATTTCACTGGCACAACACCCCCTTCGTTATTCCTTCTTCCAAGATTGGTTTCTTTACATGTTCAACTCAATCAATTAACCGGGAAATTGCCAAATGTGCTCAAGATGAGTTATCTAGGATACGTACTATACCGTAGCATTGATCTTTCATACAACAAGCTGCAAGGTGAAATCCCCAATGCGATGTTATCTATGAGCATGGGTAAGCTGGATCTCTCTCATAACTTCCTCACAGGCTTTGAAAAACAAGTATGGCACTCAGAGTACTTACTATACCTTAATTTGGAGAACAATTTTCTTCAAGGGCTTTTGCATCTATCCTTTTGTGACATGTATAGTCTTGTATTCCTCATTTTAGCTCGGAACAATTTCAGTGGTTCAATCCCGGTATGTTTGGGTAACTCCAGTATTTCCATTTCTTTTTTAGACTTGCGAATGAACAATTTGCATGGAGAGATACCAAGATTCTTATCTACAGGGTTTCAATATCTTGGTTTATATGGCAATCAATTGGGGGGACAATTCCCGTGATCCTTGGTTAACTGTACAAGCTTGGAAGCTCTTGATTTGGGGAACAACAAGATAAATGACACGTTTCCGATATGGCTGCAGAAACTTCCAAACCTACAAGTTCTGATTCTGAAATCTAATCTCTTTCATGGTCCAATTGGTTATTTGGAGTCCGAGTTTCCATTTCCTGAGTTACGAATCTTTGATCTTTCCTTCAATGGGTTCACTGGAACTTTGCCATCTAATTTTTTCAAAAGTTTTAGAGGTATGATGGATGTGGATGAAGAAAAAACAGGAATTACTCGCTCCTACAAAGGGTCGACCAGGCCTTGTACAAGCATAAATCGATGTCGAAGAGATTACCTTTACCACTTTAGTTTGGTGATAAAAGGTAATGAGTTTGATATGAGAATCACGTCAATTATGACGAGTGTTGATATATCAAGCAACAGGGTTGAAGGATATATTCCAAATTCCATTGGAAATCTGAGCTCACTTGTGTTACTCAACCTGTCTCACAACAGCTTCCGTGGTCATATTCCTGCAGAATTTGCAAAGTTGCAATAGCTCGAAGCATTAGATCTCTCACGGAACAGACTCATTGGAGATATTCCAGATCCATTGTCGAGTTTGACATTTCTTGAGGTCTTAAACCTTTCCTACAATCATCTGGCTGGACGCATTCCTCTTGGGAAACAGTTCAATACATTTCCAAATGATTCTTATTGTGGAAATCCTGATTTATGTGGATTTCCACTATCAAAGGAATGTGGAAACAATAAAGAGTCACCACTTGAACACGATGATGGTGATGATGATGATTCATTTTTTGCGAGTGGGTTCACATGGCAAGCTGTGGTAATAGGTTATGGTTGTGGTCTGATATTTGGATTGTTGATGGGAGGTCTCATGTTCCTACTGGAAAAGCCAAAATGGTTTGTGAACTTTGCTAAAGATATTGCTCAACAAATTGCTGCTAAGAAGCAAACAAGGCAAAAGAAGAGACGCCAAAGACATGGTTTAAGAATGAACTAGCAATATACCAGGTATGCATTAAAAC

>M6-PGSC0003DMG400017087 M6 LRR-RLP

TCTTGTGACAGGGATGCCAAATCTAGAGGAATTTTACAGTATTTCTTACAATAGTTGTACAAATGAAGTCTTTTCTAGCATCCCCAATTTAAAGAGATTGACCATTGCGACACCTTTCAGGATAAACAATGTTCCATCTCATCCCTTGGATATGTCGAGCTTGAGAAAACTTGAGGCATTCAAGATTTCCTGGAGATATGGTTTTGTAACGCCCATCAAAAGATTTGTTTTTCCAACTTTACTTAGGAAGTTGACTTTAGCTTTCTTTGGTCATTTTATTTGGGAAGATATATCATCAACTTTTATCATGTTGCCAAATCTTGAAGAGCTCAAACTTAAACATTGTCAAGCCAAAGATAATGTATGGAGATTGAGTGACAAAGACATACTCAAAAGCTTGAAGTTGTTGTTACTGTACTACACAAATATTAAGTGTTGGGAAGCTAGCAGTGATAACTTCCCAAATCTAAAACGTCTTGTTCTGAAGAGTTGCAAGGACCTGCAAGAAATTCCAACAGATTTCGTGGAAATTTGTACTTTGGAATCGATTGAGTTACATGATTGCACCACTACTGCTGAGGATTCTGTCAGAAATATCGAACAAGAACGAGAGGATATGGGATATAATATCCTTAAGGTCTACATCCATAACAGCCACAGTAAGTTATAATTCTCTCTGAAAAGTTCTTCGATTCTTCACATTATTTAAAACCTCATTTTTTTTAAATCATACTTCTTTGAAATTAAGTTGATGACTTGTGCTTTTTATCTTGCATATGCAGGGAATTGAAGCATATTTTGAGGGATCATCTAATGCTATCTTCCGAATTCAGAACCTTGTAGTTCATATTTTTGTTTTTATATTCTGATCGTTTGTTAATTTCATGTTTTAGGAATAAGCATGTTTAATACATGTCTCGGCCAATTGTGTTTTTGTATTTTGTGAACACTAACTAGTGATGTACTCATGTAACATCTATTTTGTTTTATGTGATAGTTCGAAATAATTGAGGATGTTCAATCTAATATGCATTGCCTATGGAGTATAGTTTCACACTTTGAATCTGTTTTCCTCTGTTAATGTATAATACATGTTTTTCGTTGATGGACGGTGCTTTATGTGATTGATATCGATAACATTTCTCATGGATGGTACTTGGTAGTAACAGTATGTAATGGTGGTCGGCAGTAGTAGTTGCTGTC

>M6-PGSC0003DMG400017283 M6 LRR-RLP

ATGCAGATGACCAGTCTGAGGTTTCTCAAATTCCGGAATGCATATGTTTGTCTGGGACCTGAATTTCTTCCTGATGAGTTGAGGTGGCTTGATTGGCATGGATATCCTTCAAAAAGTCTGCCAAATAGCTTTAAGGGAGACCAACTTGTTAGTTTGAAATTGAAAAAAAGCCGCATCATACAACTTTGGAAAACCTCAAAGGATCTGGGAAAACTGAAGTACATGAACCTTAGCCATTCACAGAAGCTAATAAGGATGCCAGATTTTTCAGTTACGCCTAATCTTGAAAGGCTGGTTCTTGAAGAATGCACAAGTTTGGTAGAAATCAATTTTTCTATTGGAGATCTTGGAAAGCTAGTCTTGCTGAATCTGAAGAACTGCAGAAATTTGAAGACCCTACCAAAGAGAATTCGATTGGAAAAGCTTGAAATTCTCATTCTTTCAGGGTGCTCAAAGTTAAGAGCATTCCCAGAAATAGAAGAGAAAATGAATAGTTTAGCAGAACTGTATTTGGGCGCGACTGCTTTGAGTGAACTACCTGCATCAGTTGAGAACCTTTCAGGAGTTGGTGTATTAAATATAAGTTACTGCAAGCATCTTGAGAGTCTACCTAGCAGTATTTCTAGGTTGAAATGTCTGAAAACACTCGATGTGTCCGGCTGCTCTAAACTTAAAAATTTACCAGATAATTTGGTACTTTTAGTTGGTTTAGAGGAGCTCCACTGCACTCACACAGCCATCCAAACAATTCCGTCTTCTATATCTCTTCTTAAAAACCTTAAGCACTTATCTCTCCGTGGATGTAGTGCTATGAGTTCACAAGTAAGTAGCTCAAGTCATGGCCAGAAGTCTATGGGTGTAAATTTTCAGAATTTGTCGGGTCTTTGTTCATTGATCATGTTGGATCTTAGTGACTGCAACATTTCAGATGGAGGCATTTTAAGTAATCTTGGGTTCTTACCTTCTTTGGAGGGATTGATTCTCGATGGTAACAATTTTTCCAATATTCCAGCTGCAAGCATAAGTCGTCTCACTCGACTTAAAGCCCTTGCATTGGCTGGTTGTAGGAGGCTTGAGAGTTTACCAGAACTTCCTCCAGGTATAGAAAGGATATATGCTGATGAATGCACATCTTTGATGAGCATTGATCAGCTAACCAAATATCCAATGTTGCGTGGAGTTTCATTTACAAAATGTCATCAACTTGTAAAAAATAAACGGCATGCCTCTGTGGTTGATTCATTGTTGAAGCAGTTGCTCAAGGTAAGTATGCTTCCAAGCCACAACTCTACTATATCCACATGAAAATATAAATGAACTTCTTTTTATATAACTGGAAGGGCTTTTGTATTTGTACTCCAATTAGCTAAGGTCGCTTATGAATCTAGGTAAATAACATTCTTTTCACGGTGCATGCAGAGACTATACATGAATGGCGCATTTTATATGTATCTCCCTGGTGTGGAGATTCCTGAGTGGTTTACATACAAGAACTGGGGGACCGAATCCATCTCAGTGGCCCTGCCCCAAAATTGGTTCACACCCACATTCAGGGGGCTTACTGTTTGTGTTGTTTTTGACATGATGACTCCTTTTATTTTAGGGAAATTAATGTCGAATGATCCCTTCAATTTTCATACTACCAAATGGCTAAAAATTTTCCAACATTTTGGGATATGGTCAAGGTTCACAAGACATGATGGTTTGCAGCACGAATATAGAGTATGTGTCGGGCCAATAGGAAGTGAAAAACCTGTTGGTTTAGGTAATACTTTCCTAGCCCATGTACCATTTGGAAGTTATTGCCTGCCCAAAAATGACTTCATTCAGCTTGAGGTTGGTGTCTCTGAGCATTATCATAAGGATCTGGTGGTCAAAGGTTTGGGAGTGCATCTTGTGTATGAGAATTGA

>M6-PGSC0003DMG400017317 M6 LRR-RLP

TTTGTTGTGCATGGTTTTTATTAAACTTCATTTAGATCTAACTGGGAAAGATAAAACTCCTACTTTATTGAAATTTAGGCAAATACACACACACTTGTCTGAGATAATACTTATCCTTTGTGGTATGCAGACATGAAGCTAGAGTCATAGAGAAAATTGCGGAAGATATAATGGCTAGATTGGGTAGTCAGAGACATGCAAGCAATGCTAGAAATCTTGTTGGAATGGAGTTACACATGCACCAAGTATACAAAATGCTTGGTGTCGGGTCTGGTGGAGTCCGCTTCCTTGGAATATTGGGAATGAGCGGAGTGGGGAAGACAACTCTTGCCAGAGTTATTTATGATAACATTCGGAGTCAATTTCAAGGTGCATGTTTTCTTCACGAAGTTAGAGACCGTTCAGCAAAACAGGGACTAGAGCGCTTGCAAGAGATACTTCTTTCTAAGATCCTTGATGTAAAAAACCTAAGGATCAACGATTCTTTTGAAGGAGCTAATATGCTAAAACAAAGACTATGGTACAAAAAGGTTCTTCTTGTTCTGGATGATGTTGATCACATAGATCAGTTAGATGCATTAGCCAGGGAGCGTGAATGGTTTGGTCACGGAAGTAGAATCATCATAACAACCAAAGACAAACACTTGCTTGTTAAGTATGAGACTGAAAAGATATACAGAATGACAACTTTAAACAAATATGAAAGTTTACAACTTTTTAAACAACATGCTTTCAAGAAAAACCACCCTACCAAAGAATTTGAGGATCTATCAGCTCAAGTGATGGAGCATACTGGTGGACTTCCCCTAGCCCTGAAAGTCCTGGGAAGTTTCTTGAATGGAAGAGGTTTGGATGAATGGATAAGTGAAGTGGAGCGTCTGAAACAAATCCCAGAAAACGAAATTTTGAAAAAACTCGAACCAAGTTTCACTGATCTCCACAATATTGAGCAAAAGATATTCTTAGACATTGCATGCTTCTTTGCAGGGAAGAAGAAAGATTCAGTGACCAGAATACTTGAGAGTTTTCATTTTAGCCCTATCATTGGGATAAAAGTTCTCATGGAGAAATGTTTGATCACTATATTAAAAGGTCGCGTTATAATACACCAACTGATACAAGAAATGGGCTGGCACATTGTTCGTCGAGAAGCTTCTTACAATCCAAGAATATGTAGTAGGTTGTGGAAGCGTGAAGATATTTGTCCTGTACTAGAAAGAAATTTGGTACGTAGTTGTGAACTTATGACTTCTTACTTTATTTTGCTTTTCACATCCGCCTTTAATACTCTTGCTTATTAGGGCACTGACAAGATTGAAGGCATGTCGCTGCAGCTGACTAATGAAGAAGAAGTTAATTTTGGTGGCAAGGCCTTCATGCAGATGACCAGTCTGAGGTTTCTCAAATTCCGGAATGCATATGTTTGTCTGGGACCTGAATTTCTTCCTGATGAGTTGAGGTGGCTTGATTGGCATGGATATCCTTCAAAAAGTCTGCCAAATAGCTTTAAGGGAGACCAACTTGTTAGTTTGAAATTGAAAAAAAGCCGCATCATACAACTTTGGAAAACCTCAAAGGATCTGGGAAAACTGAAGTACATGAACCTTAGCCATTCACAGAAGCTAATAAGGATGCCAGATTTTTCAGTTACGCCTAATCTTGAAAGGCTGGTTCTTGAAGAATGCACAAGTTTGGTAGAAATCAATTTTTCTATTGGAGATCTTGGAAAGCTAGTCTTGCTGAATCTGAAGAACTGCAGAAATTTGAAGACCCTACCAAAGAGAATTCGATTGGAAAAGCTTGAAATTCTCATTCTTTCAGGGTGCTCAAAGTTAAGAGCATTCCCAGAAATAGAAGAGAAAATGAATAGTTTAGCAGAACTGTATTTGGGCGCGACTGCTTTGAGTGAACTACCTGCATCAGTTGAGAACCTTTCAGGAGTTGGTGTATTAAATATAAGTTACTGCAAGCATCTTGAGAGTCTACCTAGCAGTATTTCTAGGTTGAAATGTCTGAAAACACTCGATGTGTCCGGCTGCTCTAAACTTAAAAATTTACCAGATAATTTGGTACTTTTAGTTGGTTTAGAGGAGCTCCACTGCACTCACACAGCCATCCAAACAATTCCGTCTTCTATATCTCTTCTTAAAAACCTTAAGCACTTATCTCTCCGTGGATGTAGTGCTATGAGTTCACAAGTAAGTAGCTCAAGTCATGGCCAGAAGTCTATGGGTGTAAATTTTCAGAATTTGTCGGGTCTTTGTTCATTGATCATGTTGGATCTTAGTGACTGCAACATTTCAGATGGAGGCATTTTAAGTAATCTTGGGTTCTTACCTTCTTTGGAGGGATTGATTCTCGATGGTAACAATTTTTCCAATATTCCAGCTGCAAGCATAAGTCGTCTCACTCGACTTAAAGCCCTTGCATTGGCTGGTTGTAGGAGGCTTGAGAGTTTACCAGAACTTCCTCCAGGTATAGAAAGGATATATGCTGATGAATGCACATCTTTGATGAGCATTGATCAGCTAACCAAATATCCAATGTTGCGTGGAGTTTCATTTACAAAATGTCATCAACTTGTAAAAAATAAACGGCATGCCTCTGTGGTTGATTCATTGTTGAAGCAGTTGCTCAAGGTAAGTATGCTTCCAAGCCACAACTCTACTATATCCACATGAA

>M6-PGSC0003DMG400017337 M6 LRR-RLP

ATGACTTCACTCCAGACACTTGACTTATCTGGGAACTCTTTCCTGGAGGGTCAGTTGCATGAGCAACTGGGAAATCTGTGCAACCTAAGAGAATTGGATCTTTCTCTAAACAATTTCACAGGCGATATAACCAAATTCATCGATGGTCTTTCTAGGTGCAACGAAAGCAAGCTAGAGAGGCTGAACTTGGGATATAATGCTGGATTGGGTGGAACTCTTCCAGATTCTATCGGATTCTTGAAAAATCTCAAGTATATTGAGTTATGGAAGAATTCATTTGTTGGTATGATCCCTGATTCGATTGGAAATTTGTCATCATTGGAGTATTTGATCCTTGTATGGAACCAAATGAGTGGACAAATTCCAGTCACTTTAGGGAAACTTTCAAAGTTGGTTGTCCTAGACCTTTTTGAAAATAATTGGTGGGGTGTCATCACAGAAGCTCACTTCACAAACTTGACTAGTTTGAAGGAGTTGTATATATCCTCAAACTCATTACCTCCTAGTCCTCCTTTGGTTGCAAACGTTGACACCGAGTGGATTCCTCCTTTTAGTCTCCAGTATCTCAAGTTCAGGTCATGTCGATTTGGTCCTAAATTTCCTATGTGGATCAGACATTTTACTGAATTGCAAACTTTAACATTCAGGCGTGCTGGATTTTCAGATACTCTCCCTGATTGGTTGTGGGAAATGAACTGGGAACTTGAAGAACTGGATTTTGGTTATAATCAGCTTAAGGGCAGAGTGCCAACTACTCTAAAATTTAAGCCTGGTTCCACTGTCTATTTAATGTCTAACCTTTTTGAAGGTCCAATTCCATCTTGGTCCTCAAATGTAGCCTCGATAAGCCTGAGTAACAATTCATTTTCAGGACCTGTTCCTTTTGATATTGGCTACAGATATCCAGATTTGACTGACTTGGATCTTTCTCAAAACTCTCTCAATGGAACCATTCCCTTGTCCATTAGCAAAATCACCAACTTGGTAACTTTGGTTCTCTCGAATAATCATCTTACCGGAGAAATCCCTGATTTTTGGGACGAAGTACCAATTTTGTACACATTAGACCTGTCAAACAACAGTTTTATTGGCCGCGTCCCAAGCACCTTAAGTACTCTTAATTCCCTCAGATTCTTAATGCTAAGCAACAATCATCTTTCTGGGGAGTTCCCTCTTGGTTTCCAAAATTTCTCTTCAATGATCACTCTTGATCTTGGGAACAACCAATTTTCAGGAGAACTTCCTTCCTGGATAGGAGAAAAGATGCCAGTGTTGTTGGCCTTGAGCCTTCGATTGAACATGTTCAAAGGAGAGATTCCAAGGGAATTCTGCAACATGACATCACTTCATTTTATAGATGTTGGTTCCAACAATCTCTCTGGAAATATCCCGTCTTGTTTTGGGAATCTGAAGGGCATGACTTTGGATCCAGAATCTCAACGTTATGAAGGTCAATTGAACGTATTCTCCAAGGGGAGAGATCTCTTATACAAGGATAATCTTTATCTTGTGAATGGCATAATCCTTTCAGGCAACCATTTAACTGGTGAGATACCAGAAGAATTGACAAATCTTTCCAGGATGAATACCCTGAATTTGTCCATGAATCATTTGAGTGGAAAGATCCCTGAAGAAATTGAGAGGTTGAGACAATTGGAAACTCTTGATCTCTCTAGGAATGAGATCCATGGCACCATCCCAGCAGGCATGACAAGCTTAACATTGCTGACTCACTTGAATCTCTCGCACAATCACTTGTCAGGTAGAGTTCCCACAGCTAACCAATTTAACACCTTCAATGACCCTTCGATATACACGGGCAATCCTGATCTCTGTGGAGATCCATTGCAAAACAAATGCAAGGATGAAACAGAAGAAGAACCTGATCTTCATCAGGATCCTCCTTATCATTCTTCTCCTAGTGAATTTGGAGAAATGGGGTTCTTTGTCAGTATCGGAGTAGGATTTGTTGTGGGATTCTGGGGAGTATGCGCAATTCTAATAATAAAAAGTTCATGGAGGCATGCCTATTTACAGTTTGTTGGTTATTTGGGAGATAAGTTGTTTGTCATTTTCATTCTGCCTTGGGCTAGTTGGAGGAGATCCAGAAGCTAG

>M6-PGSC0003DMG400017666 M6 LRR-RLP

ATGTCTAAGCTCCCTATTTATAGAGAAAAAAATAGGCTTGTTGATGTCATGGATGATGTCTTTGGAAGATGCAAAATATTCAACATCTCAATGTGAATATATCAAATATGGTGGCTTCCAAATCTTTGACAAATATAGTTGCCAAATCTTTGACAAATATTGCTACCAAATTTCCACATTTGTGGCTTCCAAATCTTTCACAAAAGAAAGATAAAATAACTTTCTTTCATCAAACAAAAAGCCACATTACAACTATTATGATTTGTTCTAGCAAAATCAACTGCTACTTGGGGAAAAATTGATCATTGAACACACTTTTGTGTACATGACATTAACATCTCTCATTTTCAGATGTGAAGTTGGCATCCAAGAAGTTCCAAAAGATTCTGCATTGGCCGGGAAGATTAAGAGGAAGTGATAATGTGGTATGTATTGTTCACTTTCACCTCCCATTTTGCTGTTTCTTTGGTCCCTCTTCTTCTTGTTCCATTACTTTGTTTTCCATTGGGAATTTTCTTTTAACTGCTATGGAGGTTCAACCATATCATACGCGTCGCCAATGTTTAAAGCCCTTTTGTGATGTCATTTAGTGTTGTGATGTCATTTAGTGTAGTGATGTATTACTTCATGCTGTGATTTTAAAGAATTGCTTATCTTTTACCCATGCCTAGGCTGACGCCTTACTTTGTCTTTACTCTTTACGCATCATCTAGTGTCGCCCTGAAAAAATTTATGCAGTTAAGGTCTTTGATTCTACCTTAATCATAAATGTATCCCTTGTACTTAGGAGGTCATACAATCAGTAAGATTAATGACTTGCTTTTGTCGAAGAATGACTATGATGATTGAACGTCTGCAATTCGTTTGTCTACAGGTGGAGATATACCGCCGTTGTTACGAGAAACAACCTTTGGTTATCCAAGGTGCTGGAGCAGGAAATGATACCACAGCAGCAGGTGTCCTAGCTGATATCCTTGACATTCAAGATTTATTCCCTTGACTGAGTATTATACACTGAGATACAAACATTCGATCCTCAAAGGACGAAAAATTGACTCTTCTTGTCCCTCCTGCTTCTAGATTGGAGATTGAAATCAGGCTAAGAAATCTCACGCATTTTTTTGAAAAATTTCTCGATACACGAGAGTGCCTCGTCAGTTTACAGTAGATTATTACGCAATTTCTCTCCTGCATAGCGTTCAAATGGAACAGTTCAAACTGAGTCATTTGCCTCAGTGCTGGACTAGTTTTTAGGATTTCACTTCAGCTTATAATCAGTTTCCTTGTAATGACCATTTTCTTTATAATCTGAAAAACAGTTGATATTTCTTTTTAATATTTCTTTAGAAAACCTTATGATAGAACACCACAAAATTCACCTTTTCGGTATTTAATGTTCCTTTATGTTACATGTTCTTCTATTTTTCCCTCTTTTCTTTTATTTTTGAAGTTAATCATCAATTGGGACATGGACCCAGATAATCAAATAGGCTATGATCCCACATTCAGCAACTTGACCTATTCTTAGGACTATTAACTATACAAAAACAAGGCAAGCGTTGTAATGAATAACCTACAGTTCCCTCCAACACTATCTTCATTTATCTTTATCCCCGGCAAAATTTAGAAAGTAGAAAATGTAAAACAGAGAAAATGACATGGAAGAAAGTGTGAAGATTTACCAAAATGTAAATTCAAAATTCAAGAATGATTTCTTTTTTTTATAAAGAACAAATTTGGAGTCCCCACCCCCACCCCCTCTCTTGCTTCAATTATTGAATTGCAAAAGTAATAAACAGTCAAGAAACCAGACTCACAGACTCCATTATTTTTTTTCTATGGCTGTGAGCTGCAGACTGTAGATAAAACATAGTTGTACCAATTACAAGCAAAAGAGACATAAATCATTCTTGATTACTCATCACTCTTAGCTCTTAGCATTCAACAGGTAAAATAAAGTTGTTAGTAATTGAAGTTTGAAACATTTATCTTCTTAAAATCATTTTGTGTTTCCTTGTTTTGCTTGCACATTTCAGAAATGGAGGTTGGCTTAGCAGTTGGTGGTGCATTTCTCTCTTCAGCTTTGAATGTTTTCTTTGATAGGCTTTCTCCTCATGGTGATCTGCTCAACATGTTTTGGAAGCATAAGGATCATGTTCAGCTCTTACAAAAGTTGGAGGACATTTTGCTTGGTCTTCAGATTGTGCTAAGTGATGCAGAGAATAAGCAAACATCAAATCGACACGTGAGCCAGTGGTTTAATAAGCTTCAGAGTGCTGTGGACGGCGCTGAAAACTTGATAGAACAAGTCAATTATGAAGCTTTGAGGCTTAAGGTGGAAGGCCAGCATAAAAAATGGGCGGCCTGGGTAAGACAACACTTGCTAAAGCCGTTTACAATGATGAGAAGGTGAAGGACCATTTTGGTTTGCAAGCTTGGTTTTGTGTTTCTGACGCATATGATGCTTTCAGAATAACAAAAGGTTTACTTCAAGAAGTTGGCTCATTTGACTTGAAGGCTGATGACAATCTTAATCAGCTAAAAGTCAAATTAAAGGAAAGCTTAAAGGGGAAGAAGTTTCTTATTGTTCTGGATGATGTGTGGAATGACAACTACAATGAGTGGGATGACTTGAGAAATGTTTTTGTACAAGGAGATATAGGAAGTAAGATCATTGTGACGACACGTAAAGAGAGTGTTGCCTTGATGATGGGTAGTGAGGCAATCAACGTGGGGACTCTGTCTGATGAAGCCTCTTGGGATCTATTCAAACGACATTCACTAGAAAACAGGGATCCTAAAGAACATCCATAACTTGAAGAAATTGGAAAACAAATTGCAGACAAGTGCAAAGGGTTGCCTTTAGCTCTAAAGGCACTTGCTGGTGTTTTATGCTGCAAATCAGAGGTGGATGAGTGGAGAGACATTTTAAGAAGTGACATATGGGAGCTGCCAAGTTGTTTGAATGGTATATTACCAGCGTTGATGTTGAGCTACAATGATCTTCATGCACATTTGAAGCAATGTTTTGCTTATTGTGCAATATATCCCAAAGATTATCAATTTCGCAAAGACCAAGTTATTCACCTTTGGATTGCTAATGGTCTTGTACATCAGTTGCATTCTGGTAACCAATACTTTCTCGAGTTGAGATCAAGATCACTGTTCAAAATGGCCTCAGAGTCTTCTGAAAGAGATGTAGAGGAATTCTTAATGCATGACCTTGTCAATGATTTGGCCCAAATTGCATCTTCAAATCTTTGTATAAGTTTGGAAGAGAACAAAGGATCGCACATGTTGGAACAATGTCGGCACATGTCCTATTCAATAGGAAATAATGGTGAATTTGAGAAATTGAAATCACTCTTTAAATCAGAGCAGCTGAGGACATTAGTTCTAATCAATATCCAACTCTATTATTACAACATTAAGCTAAGCAAGAGGGTGCTGCATAACATATTGCCAAGACTAACATCCTTAAGGGCATTATCATTGTCACACTATGAGATTGTTGAGTTGCCAAATGACTTGTTCATAAAATTAAACCTCTTGAGATTTTTGGATATTTCTCGGACAAAGATTAAAAGGTTGCCAGATTCCATTTGTGTGTTGTATAACCTAGAGACACTTCTCCTATCATCTTGTTGCTGATCTGGAGGAGCTACCGCTGTAGATGGAGAAGTTGATTAACTTGCGTCATCTTGACATAAGCAACACTTGTCGCTTGAAGATGCCGCTACATCTGAGCAAGTTGAAAAGTCTCCATGTGCTAGTGGGAGCCAAGCTTCTTGTAGGTGGTCGTGGTGGTTTGAGAATGGAAGATTTGGGTGAAGCACATAACTTGTATGGATCTGTATCAGTTGTAGAGTTGCAAAATGTAGTTGATAGAAGGGAAGCTGTGAAGGCAAAGATGAGGGAGAAGAATCATGTTGACAAGTTATCATTGGAGTGGGGTGAAAGTATTAGTGCTGACAATTCACAAACAGAAAGAGACATACTTGATGAGTTACGCCCACATAAGAACATAAAAGAAGTCAAAATCACTGGATATAGAGTGACAAACTTTCCCAATTGGCTAGCTGATCCTTTGTTTCTTAAGCTAGTATTGTTGTCTATTAATAACTGCAAGAACTGCTATTCCTTGCCAGCACTAGGACAACTCCCTTGTTTAAAACTCCTTTCGATTAGAGGGATGCATGGAATAACAGAGGTAACGGAAGAATTCTATGGTAGTTTGTCCTCCAAAAAGCCTTTTAACTCTCTTGTGGAGCTTAGATTTGAAGATATGCCGGAGTGGAAGCAATGACACCTACTAGGAAGTGGAGAGTTTCCTACACTTGAGAAACTTTTCATTAAAAATTGCCCTGAACTCAGTTTGGAGATACCCATCCAACTTTCAAGTTTAAAAAGTTTTGAAGTTATTGGTTGTCCAAAGGTTGGAGTTCTTTTTTATGATGCTCAACTGTTTACCTCCCAACTTGAGGGAATGAAGGGGATTGAGGAATTAGCTATTTGGGATTGTAACTCTGTTACCTCCTTACCTTTTAGCATACTGCCCACTACCTTGAAGACAATATGCATATCTGGTTGCCAGATATTGAAATTAGATACCCCAGTTGGTGAGATGAGTATGTTTCTCGAGGTTTTGATAGTGAATGCATGTGATTAT

>M6-PGSC0003DMG400017716 M6 LRR-RLP

AATAACTCTAGGTGACTTAACTGAGCTCAAAATTTTGTATCTTTATTCTAACCAACTTTCTGGTCCCATTCCTAGAGAGTTGGGGAATTTGAAAAACCTCAATGATTTGGATCTACAAGAAAATCAACTTACTGGTCCAATTCCTGCTTCCTTTGGCAATTTGAGAAAGTTACAATTTTTGTATCTCCGTGCCAACAAACTTTCTGGTTCTATTCCTAAAGAACTTGCATATATGGATAACTTGGTAGAGCTGATACTAAGTGAAAATCAGTTTTCAGGCCATTTGCCTGAGCATCTTTGCCAAGGTGGGAAACTTGAGAACTTCACGGTGGCCAGTAACAAGTTAACAGGACCAATACCAAGAAACTTGAGCAATTGCTCCAGTTTTAAATGGGTCCGCTTTAACAACAACAGTTTCACTGGGAATTTATCTGAAGCTTTTGGTATCTATCCGAAGTTGCTGTTCATTAATTTGAGCGATAATGATTTTCATGGTGAACTCAGCAGCAAGTGGGGGAAATGCAAGAATTTGACTGATTTTCGGATTGCCAGAAATAACATTAGTGGTAGCATACCACCAGAGATTGGAAATGTCAAAGGCCTTTTAGGACTTGATCTTTCATCGAATCATTTGATTGGGCAGATTCCAAAGGAATTTGGAAAATTAACCTCTCTAGTTAATCTTTTTCTGCAAAACAACCGAATTTCTGGCAATATACCTGAGGATTTTGGGTCACTAACAAAGTTAGAGTCCCTCGATCTATCAGACAACAGATTGAATGGGTCAATCCCGATGTGTATAGTAGATTTCGTACACTTGTTTCAGTTGAACCTGAGCAACAACAAGTTTGGGCAGAAAATTCCAAAGGAGATAGGGAGGATAACTCAGCTTAATGTACTTGATTTGAGCTATAATCTTCTGGTTGGAGATATACCCCCTCAGTTAGCCAATTTGAAGGTCTTGGTAAACTTAAATCTTTCCCACAATGGCCTATCTGGGCACATTCCTACAGAACTTGATACTTTGACTGGTTTGCAGGATGTCATATTGTCATACAATGAGTTGGAAGGTCCAATCCCTAATAATAAGGCTTTTATCAATGCCTCATTGGAGGGTAATAAAGGTCTTTGCGGCAATGTAGCAGGACTCCATCCCTGCGAAATGCCGTCTTCTGTGGTGAAGAGGCACTCAATGGCGAAGGGACGTAAACTCATCCTCATCACTGTACTTCCTGTTATGGGAGCATTAGTACTGCTCTGTGTTTTCATTGGTGTTCTCATTATGTGTAATAAAAGAAGGAGAGTTAGAGATGTTGAAAGAAGGGATGGTGTTGGTTGGCTTTCGATATCCATGTTAGATGGGAAGGCATTGTACAGGGACATCTTAAATGCCACAGAAGAGTTTGATGCAAAATTTTGCATCGGGCAAGGAGGACACGGAAGTGTTTACAAGGTAAACCTTCCATCATTAGGGAATATAGCTGTGAAGAGACTTCATTCTTCATTTCAGAATACACATCCCAAAAGCTTCATGAATGAAGTAAGGGCATTGATTGGGATTAAGCACCGGAACATTGTGAACCTCTATGGCTATTGTTCGAAAGCACAACATTCACTCTTGGTTTATGAGTATGTGGAGAGGGGGAGTTTGTCTAGTATTTTGAGCAATGAAGTTGAGTCCAAGAAATTGGATTGGCTTAAAAGGGTGAATATCATCAAGGGTGTTGCTTTTGCTTTATCTTACATGCACCAAGACTGTTCACCACCGATTGTTCATCGAGACATATCAAGCAGTAATGTGTTGCTTGATTCTGAGTATGAAGCTCGTGTTGCAGATTTTGGAATAGCAAAGATTCTCAAGCCAGACTCATCCAATCGCACTGCGCTTGCAGGCACATTTGGTTATGTCGCACCTGGTAAGTTTTAA

>M6-PGSC0003DMG400017718 M6 LRR-RLP

AATAATCTTACTGGTTCAATTCCAATTACTTTAGGTGACTTAACTGAGCTCAAAGTTTTGTACCTTTATTCTAACCAACTTTCTGGTCTCATTCCTAGTGAGTTGGGGAATCTGAAAAACCTCACAGAGCTGGATATATACCATAATCAACTTAGTGGTTCAATTCCAATAACTCTAGGTGACTTAACTGAGCTCAAAATTTTGTATCTTTATTCTAACCAACTTTCTGGTCCCATTCCTAGAGAGTTGGGGAATTTGAAAAACCTCAATGATTTGGATCTACAAGAAAATCAACTTACTGGTCCAATTCCTGCTTCCTTTGGCAATTTGAGAAAGTTACAATTTTTGTATCTCCGTGCCAACAAACTTTCTGGTTCTATTCCTAAAGAACTTGCATATATGGATAACTTGGTAGAGCTGATACTAAGTGAAAATCAGTTTTCAGGCCATTTGCCTGAGCATCTTTGCCAAGGTGGGAAACTTGAGAACTTCACGGTGGCCAGTAACAAGTTAACAGGACCAATACCAAGAAACTTGAGCAATTGCTCCAGTTTTAAATGGGTCCGCTTTAACAACAACAGTTTCACTGGGAATTTATCTGAAGCTTTTGGTATCTATCCGAAGTTGCTGTTCATTAATTTGAGCGATAATGATTTTCATGGTGAACTCAGCAGCAAGTGGGGGAAATGCAAGAATTTGACTGATTTTCGGATTGCCAGAAATAACATTAGTGGTAGCATACCACCAGAGATTGGAAATGTCAAAGGCCTTTTAGGACTTGATCTTTCATCGAATCATTTGATTGGGCAGATTCCAAAGGAATTTGGAAAATTAACCTCTCTAGTTAATCTTTTTCTGCAAAACAACCGAATTTCTGGCAATATACCTGAGGATTTTGGGTCACTAACAAAGTTAGAGTCCCTCGATCTATCAGACAACAGATTGAATGGGTCAATCCCGATGTGTATAGTAGATTTCGTACACTTGTTTCAGTTGAACCTGAGCAACAACAAGTTTGGGCAGAAAATTCCAAAGGAGATAGGGAGGATAACTCAGCTTAATGTACTTGATTTGAGCTATAATCTTCTGGTTGGAGATATACCCCCTCAGTTAGCCAATTTGAAGGTCTTGGTAAACTTAAATCTTTCCCACAATGGCCTATCTGGGCACATTCCTACAGAACTTGATACTTTGACTGGTTTGCAGGATGTCATATTGTCATACAATGAGTTGGAAGGTCCAATCCCTAATAATAAGGCTTTTATCAATGCCTCATTGGAGGGTAATAAAGGTCTTTGCGGCAATGTAGCAGGACTCCATCCCTGCGAAATGCCGTCTTCTGTGGTGAAGAGGCACTCAATGGCGAAGGGACGTAAACTCATCCTCATCACTGTACTTCCTGTTATGGGAGCATTAGTACTGCTCTGTGTTTTCATTGGTGTTCTCATTATGTGTAATAAAAGAAGGAGAGTTAGAGATGTTGAAAGAAGGGATGGTGTTGGTTGGCTTTCGATATCCATGTTAGATGGGAAGGCATTGTACAGGGACATCTTAAATGCCACAGAAGAGTTTGATGCAAAATTTTGCATCGGGCAAGGAGGACACGGAAGTGTTTACAAGGTAAACCTTCCATCATTAGGGAATATAGCTGTGAAGAGACTTCATTCTTCATTTCAGAATACACATCCCAAAAGCTTCATGAATGAAGTAAGGGCATTGATTGGGATTAAGCACCGGAACATTGTGAACCTCTATGGCTATTGTTCGAAAGCACAACATTCACTCTTGGTTTATGAGTATGTGGAGAGGGGGAGTTTGTCTAGTATTTTGAGCAATGAAGTTGAGTCCAAGAAATTGGATTGGCTTAAAAGGGTGAATATCATCAAGGGTGTTGCTTTTGCTTTATCTTACATGCACCAAGACTGTTCACCACCGATTGTTCATCGAGACATATCAAGCAGTAATGTGTTGCTTGATTCTGAGTATGAAGCTCGTGTTGCAGATTTTGGAATAGCAAAGATTCTCAAGCCAGACTCATCCAATCGCACTGCGCTTGCAGGCACATTTGGTTATGTCGCACCTGGTAAGTTTTAAATAGTTCATGAACTATAACCAAACACTCCATTGTTTTTATCTCATTTACATGTTTAAATTCTTAATATTGTTCTGGAAATTGTATTGTTCAGAGCTTGCATATACAATGATGGTTACGGAAATGTGTGATGTGTATAGCTTTGGAGTATTATCATTGGAGATAGTCAAAGGAAAGCATCTTGGGGAACACATTTCTGTTCTAGCAAATTCATCGACTATAGATCCTGAGCAGCTTAGCGATTTGCTGGATGAGCGCCTTTCGTATCCTGAAGAAAGAGTAAAAGATGTTTTGGTTTTTATCATCAAGCTAGCATGCTCTTGTTTGCTTGAATCTCCAAAATCAAGGCCAACAATGCACTTCATCTCTCATAAGTTATCATCAATGGATTCACGTCCACCTATTCATCAGAGAAACCCCCATGTAAGGCGAGCAATATATAATCCTTGATATGTATGGAAATGTCACTAGTGTGAAGTAAGTATTTCAATTGCTGGGACTAGTATTTTTAGGGTGCTCATTTTGTTTGCTGTATTTAAGACCTTATATAGAAGTTTTCATTCGATTTCAAAGATTTTACCTGTATGTATAGAAGCTAGTAAATGACCCTTTCTGCTCAAGCTTTTTTCCTAACTGAATAAATGGAGTTGTT

>M6-PGSC0003DMG400017941 M6 LRR-RLP

AAGGAGACAAGATCCATCGATCTGAATAAGTCTGGTACACCTTTCTCAAGTCCAGCTAACAAATAATTCTTGCTACTTTAATTCTATTTAAGTCTCCAAGAATTCATTTTCTAATAACACTAGATCTATAGTTTGTCCTTCCCTTTACAAATACATGCATTTTTTGACCTTGATATCTACTTGCCAATACTTGTTTCATCCTTGTTGTTAGGACTCCAGTAGTAATCTCATCCTTGTACCTTGTTCTTAAGAACCAATGCTGTGAATATTGCAGTCCAGCCTCTTGTAAGAATTTCTGTCTCGTGAAAGAAGTGCAATGCACTGATAATGTCCCTTTCCAAAAAGTTCCTAGATTTCTGGTAAGAAGACATTATCTACCCATCTGGCCCTAAAGTTGTTATCACCACTAGACCATCCCGAGAAAATTTCTTAGTAGCCAACACTCTTCCCGTGATAATAGCTCAAGGTCATTGTAGCTGGAAGGCGCCTTCATGCTCAGTTTTACTAAATAACTTGTCATAGCATTACACTCATCTGATGACCTTTGGATTTGGTAGTTCGTCCATCTCTTACAAGTTAGTTTATATGGCTGTACTTTTTATTTGTATTGATCAATTTACAGTAAAACACTTAGTATTTTGTAGGGGTGTCAAAAATGAGCCCACCTATAGGTAACCCGCCCAACCTGCCCATAATTTTAAGGGTTGGGCTTAAGATAATTTGAATTAAGTTCAATCTCAACCCCCATTTTAAGAGAATCCTCAATTGAGCCCAATTTAATTTTCAATTTCAACCCATTTTAAGACTCTTTATTTGCATTGATGTAATGTATGTTCTTATATTGAAGGTATGAATTACTTTTGTTTTATATTTTTTAGGATTTATCTATCCATTTGGAACTTCTTTTGTAATTTTCTTAAGTTGAAATCTAAAATGTGGTTATAAAACTTAAATAAACAATATATTAAAATTATTGAGATTAAGTGGGTAAAGACCCAACCCATTTTTAAGCTCATTTGAGCCCAAAGTAAACTTGGGCGGGTCAAGGCTCAACCCATTTTAATATCTCCAATTTCAACCCATTTTAAGATCTCCAATTTCAACCAACCCACCCATTTGACACCTCTAGTATGTTGTCTCATTCATTTAACCAGAAACACCTCAATTGTTGTTTCCAATATGTTTTTTAGCATTGATTATTTGTTCCAGTTCTTCTACTAAGTGGTTCTTTTAGGCTTTTCGTTTTTCTATATCCATCCACCCTTTTCTGATCATACTTACCAATCCAGCAATCTGATTATGTATAATTCTTCGACATGCATTGTTGTTTGAACTCAAAGTCTGCCTTTAAGTTCAGTTTCCTTTCTATTAACCATAATCAGATTGCCTACACTATCATACGATTGCTATTGCTGTTCCGCTAAATCAATGAATCCCTCTATAGCTTTTTTCATTTTCTCGAATTTGCATTAAGATTCGCCCATTCCCAAGTCTGCACTCTGCTGGAATGGATGTATGATTGGTAATCATCATATGCTTTGAAAACTTGAATATCACATTCCAAAGAAATTGTATACACAAATTTCTGTGTATTCCACCTCATAAGTTTACTGTACATTCTATTGTATTTATACAAAACTAAGTCTAAATTCTGTACCAAAATAAATGAATTAAACAAAGATTATTTATACACTTGGCACCTAATTATTGGACTAATTTGTAACAAACACTAACAGCCATTTTGCTCTTCACGTGCCTTCCTTGTGCCTAGGTTGAGAATCTTCTAGACCTTGTACAACTGGTTAGGTTGAGAATCTTCTAGACCTTCTGATTAGGTTTCCTAGAACTTTTGTTCTTTTTGGCATATGTTATTTTGGCACAAATCTTTTGGTTCTTGAAAAGAAATGTTGGAGAAGGTTCATTTCGGTTTGCAAGTCCGGCATGCAGCTCGAATATGTCTATATGATAAGTCACTTATGTTTCTTTTGTTTCCTAATTTCTCCAAGTTAAGATTTTTCTCACAGGGAGTCTAGTCTAGATCATGGATAATGAGTAGTGGTGATTGAATATTGTTGAATTAGTCAACTTAACTAGGAGGCATCTTACATGATAATGGGAAGTTTTCATTCAGTCGATCAAGGAACCCATATCCCGAATCTTAAAATTTAGATCCACAACCGATTCCGAGCAACTTAGTCTAAATTAAGTCGACCCAAACATGTTGAAAGAAGTTGCAATTCTCTCAGAAAATTGACTTTTCATTTGCACTCCACGTCTGTAATATTCTTGTCCATATTCCCCACTTTCCTTCCTCCGTGTGGTAATACTTTAGGGAATTTGTAGAAGAACAAAGTAGTAGGAGAAGTCAAATGACTTGAGCGATTGATACAATCATGGTTCATCAGTGATTGTTACATAGTTTTATGTTTTACTGATTCCCCAAAGTTTAATTCTTTTCATTTAACCTGTGTCCTGTTCAAAATTTTCTTTTATTAACTTTACTGACTCATCTTATTATTTTTTTTATTTTAATTTTAATTTGTGCTGTATATATTTACGGCAGTTGATGTCAAGCCCCTTCAAGGCAAAGAAAGAGTGTAAGAAAGCAGCCATAAAATGCCCACTCCATCACCTCAAAGTGGTCAAATTTTCGGGGTTTTATATAGGCTTATTTCAACTTGAACTCGTTAGGTACTTTTTGGAAAATGTTGTTGCTCTTGAGAAAATCATTATTGATCCAGGTACTCCATTTTTTTATCGTACCCCTCCGTGCCCTAGCGAAATCAAACACACGCAAACTTCAAGAAATCTTGCCAAGCTCCAGGTTGAACGTGAAGTGCCTCCGGCTATTGAGTTGGTCATACTTTAATGAAAATAATAATGTTATTAGGGTTAAATTTGTGATACATATCGGAGGTGAATTTTTATTTCTTAGTAGATGTTCCTCATGCTTTGTTTCATAATTGGGTGATTTTCACTTTTTGTGAGGTTTATATTGCTATATGAAATATGGGGATTTTCAAAATGAAAATGTCGCAGTTATTCCGCCA

>M6-PGSC0003DMG400017966 M6 LRR-RLP

AATTTGACCCTATTAAGAGTGCTTGATCTTAGTGGTGTAAGCATCTCTTCCACCATTCCTATTAATTTTTCTTCTCATTTTACAACGCTGGTGCTTGGGCATTCAGAGTTGTATGGTGTAATACCGGAGAGTATTTTTCACCTTCCCAAACTAAAAATACTTGACTTATCCATCAATTATCAGCTTAATGGTTATTTCCCAAAGACCAAATGGAACAGCAGCGCATCTCTCACAGATTTAAATCTCCAAGGAGTGAATTTTTCTGGTTATTTGCCTGAGTCTCTTGGTTATCTAACGTCACTGAATAATTTGTTTCTTATTTTCTGCAATATCTGGGGGCCTATTCCTGAATCTCTTTCTAATCTCACTCACATAGAGTATATGTTCCTTGCAGGTAACTTCCTAAATGGAACAATACCATCATCGATATTCTCCCTTCCTTCACTAACTGATTTAGAATTGAGTAACAACCATTTTTCTGGTAAGCTTGAGGATATCAAGTCCAATTCACTATTATCTATTGATCTAGGCAATAATCAGCTGCAAGGCCATCTTCCCGAGTCAATTCAAAACCTTGTGAACCTCACAACTTTTGATCTTTCTTTCAATAATTTTAGTGGTAGTGTGGATGTCAGCTTCTTTTCAGACCTCAAACATCTTTCATCTCTTGGTATTTCATATAATAGTATTTCATTCACAAATGAAAACAAAGTTAATTTTACTTTGCCTGAATCACTTGTGTACTTACAGTTGGCTGCCTGTGAAGTGAAAGAATTGGAGTTTCTAAGATCCGCCAAGAAGCTTGAGCACTTGGATCTTTCTAATAATAAGCTTCAAGGAAGATTTCCAGATTGGGCATCATCTAACTGGATGTTTTCATTGAACAGTCTTAATCTATCCCACAACATGCTGACAAGTGTGGACTTAACTCCTTTTCAATCTCTAAAAACAATTAATTTGCGATCCAATTCTCTTCAAGGGTCCCTATCTATGCCACCAAATTCCACAACATACTTCTTCATATCAGAAAATAATCTTAGTGGAGAAATTCCTTCATCTATTTGCAACTTCACATCATTGGTAATGCTAGATTTGGCGAGAAACAACTTGAAGGGAGCAATTCTGCAATGTTTGGGTAATATCAGTAGCCTCCAGGTTCTGGATATGAACCACAACAGTCTTACAGGGACTCTTCCAACAACTTTTAGATTTGGAAGTTCTCTGAGAAGCCTCAACTTGCACGGCAATAAACTAGATGGGAAAATTCCCCAATCTTTGGCCAATTGCAAAGAGTTGCAAGTTCTTGATTTAGGAGATAATCATCTCATTGATATATTCCCGATGTGGTTGGGAACTCTTCCAAAGCTGCAAGTATTAAGCTTGAGATCCAATGAATTGCATGGGTCCATTAGAACTCCAACGATTGAAAACATATTTCCTAATCTTCGAATGTTAGATCTCTCTTCCAATGCCTTCACCGAAAACTTACCAACGGGTTTGTTTCAACATTTGAAAGCCATGAGGACAATTGATCAAACTATGAATGCACCACATTATGAAGGAACTGATGGATATTACCAAGACTCGGTAGCTGTCGTATCCAAGGGATTTGAGCGTGAAATAGTGAGAATCTTGTATTTGTACACCGTTATCGATCTTTCAAGTAATAAATTTGGAGGGCAGATTCCAAGTATCATGGGGGAACTCATTGCAGTTCACATATTGAATATATCTCATAATGGATTGGAAGGTCATATACCGTCATCACTTGGAAGTTTATCTTTGGTTGAATCATTGGACCTGTCAAGTAACCATCTTGTAGGAGAGATACCAGCACAATTTGCTTCTCTTACATCTCTTGAAGTCTTCAATCTTTCCTACAATCATCTTGAAGGGTGCATTCCTCAAGGAAATCAATTTCAGACCTTTGAGAACAATTCATATGAAGGTAATGATGGACTACGTGGATTCCCACTTTCAAAAGGCTGTGGTACTGATTCAGTATCAGAGACAAATGATACCGTTTCTGGACTAAATGATGAAGAAAGCAATTCTGAATTTCTGAATGATTTTTGGAAAGGTGCTCTTATGGGCTATGGGAGTGGACTATGTATTGGATTATCCATTATATATTTCATGATTTCAACTGGAAAACCGATATGGCTTGCAAGAATCATTGTAGAGCTGTAGCACAAAATAATGATGGGAAGGAGAAAGAAGCAGGGAAGGCAAAGGAATAACAGAAGAAGAAATAATCACTTCTAG

>M6-PGSC0003DMG400017967 M6 LRR-RLP

AAGGTCTCTTACTGATCTGCGGTTGTACAATAACTCCCTTAATGGTTCTATTCCAACATCATTGGGGAATTTGAACAACTTGACTATTCTGCGTCTTGATGATAATCACCTATTTGGCTTTATTCCTGAAGAAATAGGCTACCTAAGGTCTCTTACTCATCTAGATTTGTGTTCTAACACTCTTGATGGTTCTATTCCAACTTCATTGGGAAATTTGACCAGCTTGTCCATTTTGTATCTTTATGATAATCACCTTTCTGGCCCTATTCCTGAAGAAATAGGTTACCTGAGGTCTCTTACTGACCTGCGGTTGTACAATAACTCTCTTAATGGTTCTATTCCTGCTTCATTGGGGAATTTGAACAACTTGTCTGTCTTGTATCTTGATGGTAATCACCTTTCTGGCCCTATCCCTGAAGAAATAGGTTATCTAAGGTTTCTTACTGATCTACAGTTGTACAATAACTCTCTTAATGGTTCTATTCCTGCTTCATTGGGGAATTTGAGAAACTTGCAAGTTCTGTCTCTCCAACAAAACAATCTGATTGAGGAAATTCCTTCATCTATTTGCAATTTAACATCACTGACAATCCTGTATTTGTCGAGAAACAACTTGAAAGGAAAAAATCTGCAATGCTTGGGTAATATCAGTGCCCTCCAGTATGTGATAATGTCACATAATAATCTCAGTGGAGAGCTCCATCCGTCTATTTGCAATTTAACATCACTGCAAATTCTAGATTTGGGTAGAAACAATCTAAAGGGAGAAATTCCGCAATGTTTTGGCAACATGAGTGGTCATCTTGAGGTTCTGGATATGCAACACAACAATCTTTCTGGAACTATTCCAACGACTTTTAGAGCTGGAACACTTAGAAGCTTCAACTTGCATGGCAATAAACTTAAGGGGAAAATTCCCCAATCTTTGGTCAATTGCAAAGAGATGCAAGTTTTTGATTTAGGAGATAATCACCTCAACGACACATTCCCGATGTGGTTGGGAACTCTTCCTAAGTTGAGAATTTTAAGCTTGAGATCGAATAAGTTGCATGGACCGATTAGAACTTTAGGAAGTGAAAACATTTTTAATGAGCTTCGAATGTTGGATATCTCTTCCAATGCCTTCACGGAAAACTTACCGATGAGTCTGTTACAACATTTAAAAGCCATGAGGACAATTGATCAAACGATGAATGCACCAAGTGATGAAGGAGGTGAATATTACCAAGACTCGGTAGCTGTCGTAACAAAGGGATTGGAGCTTGAAGTTGTGAGAATCTTGTTTTTGTATACCACTATCGATTTTTCAGACAATAAGTTTGAAGGACATATTCCAAGTATTATGGGAGATCTCATTGCACTTCGCATGCTTAATTTATCTCATAATGGATTGCAAGGTCATATACCGCCATCACTTGGAAGATTAACTTTGGTTGAATCATTGGACCTGTCAAGTAACCACCTTGTAGGAGAGATACCAGCACAATTTGCTTCTCTAACTTCTCTTGCAGTCTTAAATCTCTCCTACAATCATCTCGAAGGGTGCATTCCTCAAGGAAATCAATTTCATACCTTTGACAACAATTCATATGTAGGTAATGATGGATTACGTGGATTCCCACATTTGAAAGGCTGTGGTACTGAAGGTAATGATTCAGAATCAGAGAAAACTGATACCGGATCTGAGCTTGATGAAGAAAGCAACTCTGAATTTCTGAATGATTTTTGGAAAGCTGCTCTTATGGGCTATGGAACTGGACTCTGTATTGGATTATCCATTATATATATCATGATTTCAACTGGAAAACTGATATGGCTTGCAAGAATCATTGTGGAGGTGGAGCATAAAATTATGATGGGAAGAAGAAAGAAGCAGCGAAAGCAAAGGAGTTACAGAAGAAGCAAATAATCGCATGTAGAAGAGTTA

>M6-PGSC0003DMG400017968 M6 LRR-RLP

ATGTTCTCCCTTCCATCACTAAGTTATTTAAGCTTGGGTTTTAACCAATTTTCTGGTCAGCTTGAGGATTTCAAGTCCAATACACTAGTCTGGATTCAGTTACAGGGCAATCGGCTGCAAGGCCATCTTCCCAAGTCAATTCAAAACCTTGTGAACCTAAAATATCTTGATCTTTCTTTCAATAATTTTAGTGGTAGTGTGGATGTCAGCCTCTTTTCAGACCTCAAACAGCTTTCGTATCTTAGTCTTTCATACAACAGTATCTCATTGACTAATGAAAAGGAAGTTAAATCTAGTTTGCCGGAGTCACTTGAGTACTTACGTTTGGCTGCATGTGATGTGAAAAGAATTGGAGTTTCTAAGATCCGCAAAGAAGCTATCTGACTTGGATCTTTCAAATAATAAGGTTCAAGGAAGAGTTCCTGATTGGGCATGGTCTACTTGGAATTTTTCAGTGTCAAATCTTAATCTATCCCACAATATGCTGACAAGCATGGACTCAATTCCTCTTCAATCTGGATATATTATTGATTTGCGGTCCAATTTTCTTAAAGGGTCACTACCTATTCCAGGGAATTCACAGCATTTTTCTTCATATCAGAAAATAATCTTAGTGGGGAAATTCCTTCATCTATTTGCAATTTGAAATCAGTGCAACTTCTAGATTTGGCGAGAAACAACTTGATGGGAGCAATTCCACAATGTTTCGGTAATATCAGTGGCCTCCTGGTTTTGGATATGTACAACAACAATCTTTCTGGGACTCTCTCAATGGCTTTTGGAATCGTAAGTGAACTTAGAAGCCTCAACTTTCATGGAAATGAACTAGAGGGGAAAATCCCTCGATCTTTGGCCAATTGTGAAGCGTTGCAAGTTCTTGATTTAGGCGATAATCACCTAAACGACACATTCCCCATGTGGTTGGGAACTCTGCCATATCTGCAAGTTTTAAGCCTGAGATCAAACAAGTTGCATGGATCTATTAGAACTTCAAGGATAGAAAA

>M6-PGSC0003DMG400017969 M6 LRR-RLP

AACTTTTTGAATTGCTTAAACAATATGGGGCTCTCAGTTGTGTTCTTAATTTTCTTTCTTTATCAGCTTTCTTGTTCTTCATCCATATTTCTGTGTCCCAAACATCAAAGCATTTCCCTTCTAAAATTCAAGGAAACATTTACTATAAATCCCTATGCTTCTTCTAGATGTTATTTCCGCGGTCAGTAGCCTTATCCAAAAACTAGTTCATGGAACATGAGCAGAGAATGTAGTTTATGGGATGGAGTCATATGCGATGAGATGACTGGCCATGTCATTGAACTTGACCTCGGCTGCAGCCGTCTTGTAGGAAAGATTGATTCCAATAGCAGCCTATTCCAACTCTCTCATCTCCAAAGACTCAACCTTTCTTGGAATGAATTCCACGGTTCTCATATCTCGCCTGAATTTGGCAAGTTCTCGAGCTTGACACATCTTGATCTTTACAACTCCAATTTCTCAGGTCAAATTCCTTCTGAAATCTCTCATCTTTTCAAGTTACACTCTCTTCGTCTCTCTGGTACTGGTCGTCTACGACTCGTAGCTCACGATTTTAAATTGCTTCTTCAGAACTTGACTCAATTAAGAGAGCTTGATCTTACTTTCATAAACATCTCTTCCACCATTCCTCTGAATTTTTCTTCTAATTTAACAACTCTGAATCTGGGATTTTCAGGATTGTATGGGATAATACCGGAGAGTATTTTTCATCTTCCTAATCTGGAAACACTTGACTTAAGTTACAACTATAAGCTCAATGGTTATTTTCCAAAAACCAAATGGAACAGCAGTGCATCCTCATATTGTTAGATCTCAGTCGCGTGAATTTTTCTGATAATTTGCCTAAGTCTATTGGCTATCTAACTTCGGTGCATTCTTTGTCTCTTCCAAATTGCAACCTTAGAGGGCCTATTCCTGAATCTCTTTCGAATCTCACCCGCATAGAGCATTTGTACCTTCAAGATAACTCCCTGAATGGAACAATACCATCAGGGATGTTCTCCCGGCTTCCATCACTAAGTCGTTTACACTTGAGTAATAACCACTTTTCTGGTGAGCTTGAGGATTTCAAGTCCAATTCACTAGAAGAGATTATTTTAGAAGGCAATCAGCTGCAAGGTCAAATTCCCAAGTCAATTCAAAACCTTGAGAACCTAAAAGGGCTTGATCTTTCCTTCAATAATTTTAGTGGCAATGTGGATATCAGCTTGTTTTCAAACCTCAAACAACTTTTGGGTCTTAGTCTTTCATATAATAAGATATCATTGATCAATGAGAACAAAATTAATTTTACATGGCCCAAATCTCTTAATGTGTTGCAATTTGCTGCATGTGAAGTGAAAGAATTGGAGTTTCTAAGATCCGCAAACCAGCTTTTGGTGTTGGATCTTTCACATAATAAGATTCAAGGAAGAATTCCTGATTGGGCATGGTCAAACTGGATGTCATTGATGACTCTTGATATATCCCACAACATGTTGACGAGTGTGGAGCCAATTCCTCTTCAGACTGTAGATACTATTGATTTACGGTCCAATTTGCTTCAAGGTTCACTACCTATTCCACCAATTTCCACAAGATACTTCTTAATATCCCAAAATAATCTCACTGAAGAAATTCCTCCATCTATTTGCAATTTGACATCACTGGTAATGCTAGATTTGGCGAGAAACAACTTGAAGGGAGAAATTCCGCAATGTTTGGGTAATATTAGTGGCCTCGAGGTTCTGGATATGCACAACAACAAACTTTCTGGGAATATTCCAACAATTTTCAGCAATGGAAGTTCACTGAGAAGCCTCAACTTGCATGGAAGTAAACTTGAGGGGAAAATTCCCCGATCTTTGGCCAATTGTGAAGACTTGCAGGTTCTTGATTTAGGAGACAATCATCTCATTGACACATTCCCCATGTGGTTGGGAACTCTTCCAAAGCTACAAGTTTTAAGCTTGAGATCATATACATTGCATGGGTCCATTAAACCTCCAAGGATTGAAACTATTTTTCCCGAGCTTCAAATCATAGATCTCTCTTACAATGCCTTCTCCGGAAACTTACCTACGAGTTTGTTTCAACATCTGAAAGCCATGAGGACAATCGATTCATCAATGGAAGCACCAAGATATCGTGGAGATACATATTACCAAGATTCTATTACAGTTACAACCAAGGGATTTATGCGTGAAATTGTGAGAATCTTGTATTTGTACACCGTTATCGATCTTTCAAGTAATAAATTTGGAGGGCAAATTCCAAGTATCATGGGAGATCTCATTGCAGTTCACACATTGAATTTATCTCATAATGGATTGCAAGGTCATATTCCACAATCATTCGGAGATTTATCTTCAGTTGAATCATTGGATCTGTCAGGAAACCAGCTTTCGGGAGATATACCACAACAACTAGTTTCTCTTACATCGCTTTCATTCTTAAATCTCTCCTACAATCATCTCCAAGGATGCATTCCTCAAGGAACTCAATTTCATACTTTTGAGAGCAATTCATGTGAAGGCAATGATGGACTACGTGGATTTCCTGTTTCAAAAAGTTGTGGCGATGCTCGGGTATTAGATACAAATGATACCGTATCTGAACTAGATGATGATGAAAACGATTCTGAATTTCTGAGTGATTTTTGGAAAGCTGCTCTTATGGGCTATGGAAGTGGACTATGTATTGGATTATCCGTAATGTATTTCATGATTTCAAGTGGAAACCCGAGATGGCTTGCAAGAATCATTGTAGAGATGGAACACAAAATAATGATGGGAAGGAAAAAGAATCAGCGAAGGAAAAAGAATTGCAGAAGAAGAAATATGTAACATTTTAATGTATTCAATGAAATGTATGCATTTCTTGTTTCAGTGAACTGTATAATTTACAAATGTATGCAGTTTATTGCATCTTCTCAACAACATCGCGTAATTTTTCCATTTATCATCAAGTGAACAAGGTAGGAACTCTTAAATCTTCTCATCAATGCATTCCTGCTGACACAACTTTGACATCTCTGCTCTCCTGGAGCTAAAAACAGGTCATTGTCTGTTCTATGAAGGGAGACGGATATGTTTTCCGTTAAACATTTAGTGACAATAAATTTTCCTTTCACAAAAACATATAGCGGTAATTGAGTTAATGTCATTGCATTCTAAATATCATAGAAACTGGTCTCTTTTGGTTGTCGCTGACTTCATTGACCATTAAACCGCACCTATAACTCTCATTATAGGCACAGTAGTATGTGAACATGTACACTTTCATTCGAAATTTCCACTACAAGACTTTGAAAAACAATACCAGTATTTAAGTTGTTACTAGA

>M6-PGSC0003DMG400018045 M6 LRR-RLP

ATCTTGATCTGAAATTTAATTCCCTTCAGGGTCCACTACCTTCGGCCATTTGTAACATGAGGAAACTTATCTTATTAGATTTATCACACAACTACTTCAGTAACTCAGTTCCACATTGCTTGGGAAGCATGGCTTTACTAACGGTGCTGGACTTAAGAAGGAACAATTTCACAGGGAGTCTTCCCCCATTATGTGCACACAGCACTTCATTGAGTACCATTGTCGTAAATGGTAATCAATTTGAAGGACATCTTCCTGTGTCGTTGCTCAATTGTAATGGTCTACAAGCCCTTGATGTGGGGAACAATGCTATAAATGACACGTTTCCAGCTTGGCTCGGAACTCTACAAGAGCTGCAGGTCCTTATATTAAAGTCGAACAAGTTTCATGGACCTATAAGTACGTGTCAAACTAAGTTTTGGTTTCCCGAGTTGCGAATTTTTGATCTTTCTCGTAATGATTTCAGTGGCTCACTTCCTGCAAAAGTTTTTCGAAACTTCAAGGCAATGATTAAATTAGATGACGAAGACACAAGAGGGATCACGTACATGGAATCTAAGTCAAATAATTTTTCATTTCAAACATCGTATGAGGATTCAGCGAGTTTGGTAATCAAAGGGCACGATATTGAGCTACAAAGAATCAGCACAATAATGACAACCATAGATCTCTCAAGCAACCATTTTGAAGGTGTCATTCCGGAAACACTAAAGGACCTCAGCTCACTTTGGCTACTCAATTTATCCCATAACAATCTCATAGGTCATATTCCAGTGGAATTGGGGCAATTGAATAAGCTGGAAGCTTTAGATCTCTCTTGGAATCGGCTCACTGGAAAGATTCCGCAAGAATTGACAAGAATAAACTTTCTGGCAGTCTTAAACATCTCTCAAAATCATCTCGTCGGACCAATTCCTCAAGGTCCACAATTCAACACATTTGGAAATGACTCGTATGGCGGCAACCTTGATTTATGTGGTCCTCCTTTATCAAAACAATGTGGAACGAGTGATTCATCACATGTTCCTCAACCAGAAGATGAAGGCGAGTCATATTTTTTAAGTGGATTTACGTGGGAATCAGTAGTCATAGGCTACAGTTTTGGTCTAGTTGTTGGAACTTTCATGTGGAGTCTCATGTTTAAATATCGTAAGCCAAAATGGTTTGTGGAATTTTTTGATGGACTCATGCCTCACAAAAGAAGAAGGTCAAAGAAGAGAGCTCAGAGACGACGATCTTAA

>M6-PGSC0003DMG400018117 M6 LRR-RLP

AGTTATAGAGATATAAAAGAGAGAGGTAGAGGAGGAGAAAAAAAAGGGAAAAAAAAAATATCGAGATGCGATCTTAACGTGTAAATCAGTTATTCATGGATTAGTTCGTCAGCCTGCCTGGCCTGCTCTTATAGTTGACGAAAAAGCTAGAGCTCACTGATCTTCACAACTCAATAACCTCCCTTCCCAATTTCATTTGATAATCTCCAGTAGCCATGGATCCAAACCCTACCAATTTCCCAATTCTCTCCTACGTCATTTCCAAGCTACCCAGTATGGGCCGTAGAACCACCGCCACCGCCGATGAATTCGACATTGAACAGCCCCAGAACCACCCTCAGCCTCCTCCAGAACCTCACTTCGACATCACCGAGCGTATGCCTCATCTCACTGACCCTAAGGTTGTCAACGCTATGCGATCAGCCGTTGCTGACGTGTCACAGACCCGGTCAATGCTCAAAACCCTGGGTGAGCGGCCTTTTCACGAGCTTGTTGATACTGCCAGGGTGAAGTTAGCTGAGATCGAGGTGGACATGTCCAAGCGTCTCGAGGAGATTGTTCTGTCTCCACGTCCGCCGGAGATGGAGAGGCAGGATTGGAGATTGGACATGGCGATTAAAGAGGATGAGTGTCGAAAGGGTGTGGAGAAGGAGAGAGAGGAGTATAAGGCGTTGATAGCTTTGGATGAATTTCACGAGGCGTATGAGAAGATGTTGAAGGATGCGGAACAGAGGTTGGAGAAGATTTATGAGACTGCTGTTGCTGGTGGTGATGTGGTATCAATTGGAGAGTCTTCTGGTGAGAAGAGTTCTGAGTTGAAAGAGGAAGTGAATGAGGAAGTGATCGGGATTTTGCAGGAGGCATCAGGTAAGAGTGTGGAAAGGGTTGATTTGTCAGGCAGGCAACTGAGGATGCTGCCTGAAGCTTTTGGGAAGATTCACTCCTTAATTGTGCTTAATTTGTCCAACAACCAGCTTATGGTATGCCCTAGTTCAGATAATCTTTCATGGTTCTTGTCTTCGACTCGTTTTACTTGGATTTGAACTTATGGTGTGTATTAATTTTCAACTGCATGATGTCTTGTTGCATTTCGTGCTACTTGATATACGGTAAATTGAACTTCTAATGATACTTTTAGCCCTATATTGTTGGCACATCCATGTCAAATGCGTAGTTTGATATTTCATTTCACATGTGGAAGTAGTAGAAGTTTGATGGAGATCATATTGTAGAGGAGCTATATAAAACTGTGGTGTTTCTTTTTCTTTTTTTTTTCCTCCTTTGGGGTGAGGGTTAGTTTCAACCTATGATTGGACTGTGAACTGCCAAACTACAGGTCCTTTGGGTTTAATTAAAAGCGTATATTTTCATCTCAATGACTTGATGCATGGTAAAAGTTTGATAACTCTTTTCTGGTTGCTGCCAGCATAATTTGAGGTACCACAAATTTACTGTCACGTGCTGGGACAAGTTGCGCTTAGAGGTTCTGAACATGGAGTTTTGCACCTTTCCCATGCTTTACTCGGAGCACTCATCTAAGTTATTTTATTTTTTCACAATCAGAAGCTTTTATCAAAACTCTGTTACATTGTTTACTGCTTCTTACTATTGTAAACTGATCATCTTTTGGCTTTTTGAGATGCCTTTTCTTTTATGAAACAAGACTCAAACTTCAAAAGAATGACACTATCAGTACTGGGGAGTCGGTAGTTCTTTCTTACTTTCAAAGTTAGGAAATCAATATCCATATCCACAAAGAATCAAGTGCTGCACTGAGCTGGGTACTTTGAACCTCACCATGCTTGAAGTTGCAATTTTGTACCTTCACTATGATTTGATCATAGAACGGAACCCCAATGCATATGATTAAAGGGGGTACTAGGTTTCTTTTGACTTGTAAGGTGCAAGAACCTTCTGAACCTACTCTAATATCTGATTATGCTAAATGTTTATCGAGCACCGGTTCCTCTTCTTGCCCTTTATCTCTTCTAGGAGAGGAACATTACTTGTCGCATTATTTTCTCATTCCTGGTTCCACTTCGTTTCTTGAATTATAACAGTGTTCTGCACTTTCGCCTCCTGATAGCTGTCTTCCAGGATCCAAAGTCTCCCAAAATATAGTATTGTACTTCCTCCATTCCAATTTATCTATCTATTTTTGCTTTCTGCTTATTGATTGACCACTTCCAAAGTTAAATCTGAAATTGATTAGTTTATATATGGAAAAATGTAAGTTAGCT

>M6-PGSC0003DMG400018163 M6 LRR-RLP

GTCAACTTAGGACACTGTTAACTTTTCTCCTATCTCTCTTAGCAAATTTTTTAGCTTCATGCTACTCATAGTTTTCCATTTTAAGTAAAAACAGCACCACAAATTTCTTTCATAAATTATGCATTTGTCAATTTAAGTTCCAAGGTTGTGTTCCATTCACAAATATTAGCATGGGAAGTGTTCTTTTTTCTTATCTGCTGTTTTTCTTGACAGTAGTTGATGTTGTTCTTGCTGTTTCTTCAGCAGATGGTTCATCTGATTCGTATCGGCTTTTGAGAATAAAGTCAGAACTTGTTGATCCATATGGAGTTCTTGAAAACTGGTCTGAAGGGACTAACATGTGTACCTGGAATGGAGTGGCATGTTCAGATGATAAATCTCATATTGTGAGGCTCAATCTTACTAGTTCAGGACTAGGAGGTCAAATATCTCAAGAATTCGCGCATCTCACTTCTCTTCGAGTGATTGATCTTTCGGATAATTTTCTTAATGGAACAATCCCTCCTGCATTAGGAGAGCTTCATAACTTGGAGGAATTGCTTCTTTTCTCTAATTTTCTCACTGGTGAGATTCCTTCGGATATCGGTCATTTGAGAAAGCTGCAAGTTCTTAGAATTGGAGCCAACATGCTGACAGGACAAGTGATACCAGAAATTGGTAAACTTTCTGAGTTGAGAGTGTTGGCTGTTGCTTATTGACAATTCAGTGGAAAGATACCAAAAGAGATTGGTAATTTGAAGCAGCTGATAAATCTAGACTTGCAGCAAAACAGTCTTAGTGGCCCCATACCAGAAGCAATTGGCGGCTGCAAAAATCTTCAAAATTTTGCAGCATCAAACAACAGGATTGATGGAAAGATTCCTGCTTCAATTGGTCAGCTCGAATCACTCGAAATCTTGAACCTGGCAAACAACAGTTTTTCTGGTTCGATTCCTGTTGAGCTGAGCCATCTTTCCAATTTGAAGTACTTGAACTTGTTTGGCAATGAATTGGAAGGCGAAATTCCTTTTGAGCTTAACAAGTTGGTTCTGGTTGAAACATTGGATTTATCAAACAACAACCTTTCAGGAACCATAAGGCTTCTCAACACTCAGCTGAAGAATCTTGAGACTTTATCACTCTCTGGCAACTTCTTAACAGGAAGCATCCCGAGTAATTTCTGTCTCCGTGATTCAAGCTTAAGCCTACTCATTCTGGCTGACAATAAGCTGTCTGGGAACTTCCCCTTGGAGTTACTGAACTGCACGTCCATACGACAACTGGATCTCTCCGGTAACAGCTTTGGAGGAATGCTGCCACGAGGCCTCGACAGGCTAGAAAGTCTCACAGACCTCTTGCTCAACAATAACAGCTTTACTGGAACTATACCCTCTGAAATAGGAAACTTGACTAACTTGGAAGATTTGTACCTGTTTCATAACATGCTCTCTGGGGGAATCCCAGTTGAAATAGGAAAGCTTCAAAGGCTGCGTGACCTATACCTCTACGAAAACCAGTTGTCAGGAAGTATTCCAAGAGAGTTGACAAACTGCTCCAGCATAACAAGCGTTGATTTCTTTGGTAACCAACTTTCAGGATCTATTCCTGATAACATTGGGAGGCTTAAGAATCTTGTTATTCTTCAGCTAAGGCAGAATGAGTTGTCAGGTCCGATTCCATCAAGCTTAGGCTACTGCAGGAAGCTTCAAAAACTGGCTTTGGCTGATAACAAACTTTCAGGATCATTGCCACCAACTTTTCGATTCCTCTCAGAGATGGACCTGATTACTCTTTACAACAATTCCCTTGAAGGTCCACTTCCTGAATCTCTTTCCCTTCTGAAAAATCTCAGCAAAATTAACTTTTCTCACAAAAAGTTCAGTGGAAAAATTTCCCCCTTAGCTGGTTTAGATTCTCTGACAGCTCTTGACTTAACAAACAACAGTTTCTCTGGTCCTATTCCCTCTGAGTTAGGTTTGTCAAAAAATCTAACCCGTCTTCGCCTTGCTAATAATTTTTTCACTGGAGAACTCCCTTCTGAATTCGGACAACTTGAAGATCTCAGGTTTCTTGATTTATCATTCAACAATTTGACCGGAAATCTAGCACCTTCCCTTGCTGGTCTAAAAAACCTTGGTCATTTTCTCCTTAGCTCTAACCAATTCTCGGGTGAAATTCCCACATGGTTAGGAGGCATAGAAGATCTTGGTGAACTCGATCTTTCGTTCAACAACTTCACTGGAACAGTTCCAACAGAACTTGGAAACTCTCCAAAATTGCTCAAACTTTCTCTCAGTTACAACAGATTATCAGGCCCCATACCACCAGAATTAGGCAATCTCACAAGTCTAAATGTCTTGAATCTTCGAAGAAACAATCTTTCTGGTTCCATCCCTTCAACTCTTCAAAAATGCCAAAAGCTCTATGAGCTAAGGCTCTCTGAAAACAACTTAACAGGCTCAATCCCTTACGAACTTGGTTCTTTATCCGAATTACAAGTAATACTAGACCTCAGCAAGAATCACATTTCAGGTGAAATCCCTTCATCACTTGGAAACCTTGTAAAGTTAGAAAGATTGAATCTTTCATTCAATCAACTTCAAGGAAAAGTCCCACCATCACTTGGAAGGTTATCAAGTTTACATAGACTCAATCTTTCATATAATCATCTCCAAGGCCAAATCCCTTCCACATTTTCAGGATTCCCACTAAGTTCTTTCATGGGAAACAATCATAACCTATGTGGCCCACCATTACTATCTTGTTCTGAATTAAAGGAACATGAGAGAATATGGGAGTTGTCAAAAGCAAGTGTTGTAGGGATAACAGTGGCTATTGTGTTAACAGCAACAGTTATATGCATGGTTCTGCTTTATATCATGCTGAGGATTTGGTGCAATTGGAGAAAAGTTACAATTTCTTGCTCAGAGAATGGTGGAATTGAGAGTAAAAGTAGACAAGGGGAAAATTGGGTATATGGAGAAGAGATCAAAAGTGGACAATATTGGAATACAACATCATTAGTTTCATCAAAAGAGAAGCAAATATCTAAAGGGACTTGCATGTTTAATCTTAGTGTAAGTTCATCAGATAGCACAGAAAAACCTTTAGTTTAAATTAAGAGGTGTTTTGTTTTAATTTGTCTCTCTACTTTGTAAATTCTTTTGTCATGTGTTGAGATTAATTTGTTTATGTGGGGGTTTACATGTAAAGAACAATCACCTTATGAAAAATCTCATCAATTCAAGGGCGGTACGCGTG

>M6-PGSC0003DMG400018308 M6 LRR-RLP

GTTGTTAGACTTTCAGATTTTTCATTTGATCTTTGTCTGTACTTTGATCAATATTCTGAAGATGAGTGTATCGAAAAAATTCATTCAACAGCTTATTGTGTTTGACATTTTCGTCGCGCTGTCAGTTACCTGTGGTGTTGTGTTCAGTTCAGGAGTTGTTGATGGGAATATCAAGTGCCTTCAAAAGGAATTAGAAGCTCTTCTCAGTTTCAAACGGGAGTTGTTAGATGTCCGTGGACGTCTTTCTTCATGGGGAAATGAAGGATTTAATCAAGATTGTTGCACATGGAGAGGTGTCCAGTGTGACAATCAAACCAGTAATGTCATTAGACTCGACCTTCGTGGTCCTTCTAGTCTCAATGCATCCGTTGCCATTGCGCCATTGATAGGTAAGATAAGTCCTGCCTTACAAGAATTGAAGCAATTAAAGTATTTGGATCTTAGTTACAATCGAATTTCAGGAGGTATACAGGACTTTCTTGGTTCTCTAAGTAAGTTGGAATACCTAAACTTATCTTGTGTTGGTGATGATTTTACGACAGTTCCTCCTAATCTTGGCGATCTTTCTAGCTTAAACACCCTTGATTTGAGCTATAATGATTTCCTATCTGTGAATAACCTTGAATGGATTTCGAGTCTTCATCAATTGAGATACCTAGCTATAGGTTATGCTAATAATCTGAATAATTCTGTATATACCTGGCTGTTTAAGTTAAGCAATCTGACTCACCTTGATCTCTCTGGTAATACTTTATATGGTCAAATTCCTGATGCTTTTTGGAATATGAAATCCTTAAAACACTTTGATCTGTCTCGGAATGCGCTTGGAGGTGGCTTCCCTAGATGTCTTGGTAACAGTAGCAATTTGAAGTTGTTGCGACTGTCCTCCAACAATCTAGATGGGCAGCTACCTGAGATAATGAATAACTTGTCATGCGTGTCACATTCTCTCGAGTACCTGAACCTAGAAGAGAATCACATTGGTGGTTCACTGACTGATGTTGTAGCTAAGTTTGCATCTCTGAGAGAGCTGAGGCTCGGGCGTAACAAGTTGAATGAATCCATCCCGGGAGCTGTTGGAAAACTACCAAGTCTTGTTCTTTTGGATTTATCTTGGAACAGAATCACGGGATCAGTCCCTGATCTTTTTCTGTTATCGTCGTTGAGGGAATTACATCTTTCTCATAATCAGTTAACTGGTGTAACGGAGAGTATTGGACGTCTTTCCAAACTTGAGAAGTTATACCTTGACTTCAATCAGCTTGAAGGCACAATATCTGAAGCTCATTTGTTCAAGCTCTCGAAATTACGTGAGCTGGACCTCTCTTATAACACTCAACTACATATTCGAGTCAGTTCAGATTGGATCCCGCCTTTTCAGTTGGATTTCATTCGTTTTACCCACTGCAAATTGGGTCCTCAATTCCCAAACTGGCTGAGGAATCAAAACAACATTTCCGAGCTTGATTTTTCAGCTTCTGGAATTTCGGGTGACGTTCCTAGTTGGTTCTGGGAGCAGTTGCCTGGATTGACCTTCTTAAACCTTTCTTACAATGATATAGGAGGAAATGTACCAGATCTCTCAAAGAAGATGAGTGATCTTCTTTGCATAGATTTGGCTACAAACAAATTCATCGGTCCAGTACCACGTTTTCCTACCAGTGTTATAACAGTTGATCTCTCCAACAACATGTTTTCCGGTACCATCTCCTTCATTTGTGACAATTTTGATTACTTGGGATATCTTAACCTCTCAGACAACAGATTGTCTGGAGAGCTTCCTCACTGCTGGACGCTTAGAAGTATCGTGCATCTTAACTTAGGGACTAATAACTTCTTCGGGGAGATACCTGATTCGATAGGCTCATTGGAGACGATGGGAATGCTGCATCTTCAAAACAATCATTTGACAGGAGAACTACCTCAGTCCCTGGCAAATTGCAAAAAGTTGAGGGTTATAGACGTTCGGTCTAATAACTTGTCTGGTGAAATTCCAGAATGGATTGGGAGTAACATATCTGATATCATTATTGTTATCTTAAAATCCAACAGGTTTAGTGGAATCATACCTTCAACCATATGCCAGCTAAAAGAACTTCAAATCTTGGACTTATCTGAAAATAAGATATCCGGGATCATACCAAAATGTATCAACAACCTCACTGCAATGACGGATGAAGAGAGCACAATGCACCAAATCAAGAGTTGGTATTTTCAGGTCGATGATCAAGGTGATGTCAAAATAAATGCCTCATACGATGAAACTGCAGTTTTGATGTGGAAGGGACGACAATTTGAATACAGCAGCATACTTGGTATGGTAAAGAGCCTCGATCTTTCAAGCAACAACATGGTGGGAGAAATCCCGGTTGAAATTACATCTCTTGTAGGTTTGCACGGTTTGAACCTGTCGAGAAACAACTTAACCGGCAGCATCCCTCTGAGGATTGGTGAAATGAGGGCACTAAATTTCCTTGACTTGAGCATAAATGATCTTACAGGGGAAATTCCTACTAGCGTTTCACAGTTAAGTCACCTGGGGGTCTTGAACTTGTCTTACAACAACTTGTCTGGTCGAATCCCTTTAGGCGAACACTTACTGACTTTCAACAACAGATCTTACATTGGAAATCATGGACTTTGCGGCTATCCACTAACAGAAGCTTGCTCTCTGGAGGAACTTCCTCGAGACTCAACGTTGAGTGATGAGGACGAGTTTATAACTCCAGAATTTTATATTAGTATGGGAATCAGCTTTATTTCTGCATTTTTGGGAGTCTCAGCTATGATGTTGAATAGAAGGTGGACTTACGGATACTAGGAAATCAATAATGTAACATCACACAGTTAAATGTAGGTTTACTTAAATACCATGATAAGGAGCTATGTAATCTCGAGTTCTTGAAACTTTGCACTTGCCTGAATTTTTATTCCATGTGCATTGTACACCACACGAATCAGCTGAT

>M6-PGSC0003DMG400018385 M6 LRR-RLP

GGCGATCTTTCTAGCTTAAACACCCTTGATTTGAGCTATAATGATTTCCTATCTGTGAATAACCTTGAATGGATTTCGAGTCTTCATCAATTGAGATACCTAGCTATAGGTTATGCTAATAATCTGAATAATTCTGTATATACCTGGCTGTTTAAGTTAAGCAATCTGACTCACCTTGATCTCTCTGGTAATACTTTATATGGTCAAATTCCTGATGCTTTTTGGAATATGAAATCCTTAAAACACTTTGATCTGTCTCGGAATGCGCTTGGAGGTGGCTTCCCTAGATGTCTTGGTAACAGTAGCAATTTGAAGTTGTTGCGACTGTCCTCCAACAATCTAGATGGGCAGCTACCTGAGATAATGAATAACTTGTCATGCGTGTCACATTCTCTCGAGTACCTGAACCTAGAAGAGAATCACATTGGTGGTTCACTGACTGATGTTGTAGCTAAGTTTGCATCTCTGAGAGAGCTGAGGCTCGGGCGTAACAAGTTGAATGAATCCATCCCGGGAGCTGTTGGAAAACTACCAAGTCTTGTTCTTTTGGATTTATCTTGGAACAGAATCACGGGATCAGTCCCTGATCTTTTTCTGTTATCGTCGTTGAGGGAATTACATCTTTCTCATAATCAGTTAACTGGTGTAACGGAGAGTATTGGACGTCTTTCCAAACTTGAGAAGTTATACCTTGACTTCAATCAGCTTGAAGGCACAATATCTGAAGCTCATTTGTTCAAGCTCTCGAAATTACGTGAGCTGGACCTCTCTTATAACACTCAACTACATATTCGAGTCAGTTCAGATTGGATCCCGCCTTTTCAGTTGGATTTCATTCGTTTTACCCACTGCAAATTGGGTCCTCAATTCCCAAACTGGCTGAGGAATCAAAACAACATTTCCGAGCTTGATTTTTCAGCTTCTGGAATTTCGGGTGACGTTCCTAGTTGGTTCTGGGAGCAGTTGCCTGGATTGACCTTCTTAAACCTTTCTTACAATGATATAGGAGGAAATGTACCAGATCTCTCAAAGAAGATGAGTGATCTTCTTTGCATAGATTTGGCTACAAACAAATTCATCGGTCCAGTACCACGTTTTCCTACCAGTGTTATAACAGTTGATCTCTCCAACAACATGTTTTCCGGTACCATCTCCTTCATTTGTGACAATTTTGATTACTTGGGATATCTTAACCTCTCAGACAACAGATTGTCTGGAGAGCTTCCTCACTGCTGGACGCTTAGAAGTATCGTGCATCTTAACTTAGGGACTAATAACTTCTTCGGGGAGATACCTGATTCGATAGGCTCATTGGAGACGATGGGAATGCTGCATCTTCAAAACAATCATTTGACAGGAGAACTACCTCAGTCCCTGGCAAATTGCAAAAAGTTGAGGGTTATAGACGTTCGGTCTAATAACTTGTCTGGTGAAATTCCAGAATGGATTGGGAGTAACATATCTGATATCATTATTGTTATCTTAAAATCCAACAGGTTTAGTGGAATCATACCTTCAACCATATGCCAGCTAAAAGAACTTCAAATCTTGGACTTATCTGAAAATAAGATATCCGGGATCATACCAAAATGTATCAACAACCTCACTGCAATGACGGATGAAGAGAGCACAATGCACCAAATCAAGAGTTGGTATTTTCAGGTCGATGATCAAGGTGATGTCAAAATAAATGCCTCATACGATGAAACTGCAGTTTTGATGTGGAAGGGACGACAATTTGAATACAGCAGCATACTTGGTATGGTAAAGAGCCTCGATCTTTCAAGCAACAACATGGTGGGAGAAATCCCGGTTGAAATTACATCTCTTGTAGGTTTGCACGGTTTGAACCTGTCGAGAAACAACTTAACCGGCAGCATCCCTCTGAGGATTGGTGAAATGAGGGCACTAAATTTCCTTGACTTGAGCATAAATGATCTTACAGGGGAAATTCCTACTAGCGTTTCACAGTTAAGTCACCTGGGGGTCTTGAACTTGTCTTACAACAACTTGTCTGGTCGAATCCCTTTAGGCGAACACTTACTGACTTTCAACAACAGATCTTACATTGGAAATCATGGACTTTGCGGCTATCCACTAACAGAAGCTTGCTCTCTGGAGGAACTTCCTCGAGACTCAACGTTGAGTGATGAGGACGAGTTTATAACTCCAGAATTTTATATTAGTATGGGAATCAGCTTTATTTCTGCATTTTTGGGAGTCTCAGCTATGATGTTGAATAGAAGGTGGACTTACGGATACTAGGAAATCAATAATGTAACATCACACAGTTAAATGTAGGTTTACTTAAATACCA

>M6-PGSC0003DMG400018574 M6 LRR-RLP

CCAACTTGAGGGAATGAAGCAGATTGTTGAATTAGATATACATGATTGTAACTCTCTTACCTTCTTTCCTTTTAGCATACTGCCCACTACCTTGAAGAGAATAGACATATCTGGTTGCCAGAAATTGAAATTGGATCCGCCAGTTGGTGAGATGAGTATGTTTCTGGAGTGTTTGAGTCTGGCAAAATGTGATTGTATAGATGATATATCACCTGAGTCGCTCCCAAGAGCACGCAGATTGTATGTATCTGATTGCCACAACCTTACTAGGTTTTTGATTCCTACTGCCACTGAAAGTCTCTGTATTTGGTATTGTGAGAATGTTGAAATACTCTCGGTGGCATGTGGGGGGACCCAGATGACGAAGTTGAATATTCAGGGCTGTAAGAAGCTGAAGTGGCTGCCAGAACGTATGCAGGAACTCCTTCCATCTCTTAAGGAACTGCATCTGATTAATTGTCCAGAAATAGAGTCCTTTCCTGAAGGAGGATTGCCCTTCAATTTACAACTCCTTATGATCAGCGAGTGCAAGAAACTGGTGAACAGCCGAAAGGAGTGGCGTTTACAGAGACTCCCCTGTCTTAGAGAGTTAACGATCGAACATGATGGCAGTGATGAAGAGATTGTTGGTGGTGAGAATTGGGAGTTGCCTTCCTCTATTCAAACACTCTATATAAAGAATCTGAAAACATTAAGCAGCCAACATCTCAAACACCTCACCTCTCTTCAATATCTAGACATTGCGGGTAATTTACCTCAAATTCAGTCAATGCTGGAACAAGGCCAGTTTTCGCATCTCACTTCGCTTCAATGTCTAGTAATCATGAATTTCCCTAATCTCCAATCACTTTCCGAATCAGCACTGCCCTCCTCCCTCTCTCAGCTGACCATCAAGGATTGCCCTAATCTCCAATCACTTCCTGAATCAGCACTGCCCTCCTCCCTCTCTCAGCTGGTGATCTCCTATTGCCCTAATCTCCAATCACTTCCTGAATCAGCACTGCCCTCCTCCCTCTCTCAGCTGACCATCTATAATTGCCCTAAGCTCCAATCTCTTCCAGTAAAAGGGATGCCCTCTACCCTCTCTAAACTAGTTATTGACAACTGCCCATTGCTCAGACCACTACTAGAATTTGATAAGGGGGAATACTGGCCAGATATTGCTCTAATTCCCGTCATATATATTGATGATAAATACCAGT

>M6-PGSC0003DMG400018577 M6 LRR-RLP

TTTCCCAATTGGCTAGCTGATCCTTTGTTTCTTAAGCTGGTGAAATTGTCTCTTAGCAACTGCAAGGACTGTTATTCTTTGCCAGCACTAGGACAACTCCCTTGTTTGAAAATCCTTTCCATTAAAGGGATGCATGGAATAACAGAGGTGACAGAAGAATTCTATGGCAGTTTGTCCTCCGAAAAGCCTTTTAACTGTCTTGAGAAGCTTGAATTTGTAGATATGCCAGAGTGGAAGCAATGGCACGTACTAGGAAATGGAGAGTTCCCTACACTTGAGAAACTTTTGATTAAAAATTGCCCTGAAGTCAGTTTGGAGACACCCATCCAACTTTCAAGTTTAAAAAGGTTTGTTGTTATTGGTTGTCCTAAGGTTGGAGATGCTCAAGTGTTTAAATCCCAACTTGAGGGAATGAAGCAGATTGTTGAATTAGGTATTAGTAACTGTAACTCTGTTACCTCCTTTCCTTTTAGCATACTTCCCACTACCTTGAAGAGAATAACCATATCTAGTTGCCAGAAATTGAAATTGGAGGCGCCAGTTGGGGAGATGAGTTATTGTAACATATTTCTGGAGTATTTGAGATTGGAAGAATGTGATTGTATAGAACATGATTTGCTCCCAAGAGCACGCTCTTTGTCTGTAGCCAATTGGCACAACCATACTAGGTTTTTGATTCCTACTGCCACTGAAACTTTCAATATTAGGAATTGTGAGAATGTTGAAATACTTTCGGTGGCATGTGGGGGGACCCAGATGACGTATTTGTATATTAGTGACTGCAAGAAGCTGAAGTGGCTGCCAGAACGTATGCAGGAACTCCTTCCATCTCTTAAGGATCTGGATCTGCGTAATTGTCCAGAAATAGAGTCCTTTCCTGAAGGAGGATTGCCCTTCAATCTACAATTCCTTTCAATCTGGAACTGCAAGAAACTGGTGAACGGACGAAAGGAGTGGCATTTACAGAGACTCCCCTGTCTCAGAGAATTAAGCATCTACCATAATGGCAGTGACGAAGAGATTGTTGGTGGTGAGAATTGGGAGTTTCCTTCCTCTATTCAAACACTTTATATAGAGAATGTGAAAACATTAAGCAGCCAACATCTCAAAAGCCTCACCTCTCTTCAACATCTTTGTATTGAGGGTAATTTACCTCAAATTCAGTCAATGCTGGAACAAGGCCAGTTTTCGCACCTCACTTCGCTTCAAAGTCTACAAATCAGGAATTTCCCTAATCTCCAATCACTACCTGAATCAGCACTGCCCTCCTCCCTCTCTCAGCTGGAGATCTCCCATTGCCCTAATCTTCAATCCCTTCCGGTAAAAGGGATGCCATCTTCCCTCTCTAAACTTTCTATTTTAGAATGTCCGTTGCTCACACCACTACTAGAATTTGACAAGGGGGAATACTGGCCAAAAATTACTAATATCTCCACCATAAAGATCGATCAGGAATGCATGTGATGATTAAA

>M6-PGSC0003DMG400018694 M6 LRR-RLP

TAAGGAGTTGCCAAATGACTTGTTTATCGAATTAAAGATCTTGAGATTTTTGGATATTTCTCAGACAGAGATTAAAAGGTTGCCAGATTCCATTTGTGTGTTGTATAACTTAGAGACACTTCTCCTTTCATCTTGTGGTTATCTCAAGGAGCTACCACTGCAGATGGAGAAGTTGATAAACCTGCGTCATCTTGACATAAGCAACACTTGGTGCTTGAAGATGCCGCTACATCTGAGCAAGTTGAAAAGCCTCCAAGTGCTAGTGGGAGCCAAGTTTCTTCTCGGTGGTTTGAGAATGGAAGATTTGGGTGAAGCACAGAACTTGTATGGATCTGTATCAGTTGTAGAGTTGCAAAATGTAGTTGATAGAAGGGAAGCTGTGAAGGCAAAGATGAAGGAGAAGAATCATGTTGAGAAGTTATATTTGGAGTGGAGTGAAAGTAGTAGTGCCGACAATTCACAAACAGAAAGAGACATACTTGATGAGCTACGCCCACATAAAAACATAAAAGAAGTCGAAATCATCGGATATAGAGGGACAACATTTCCGAATTGGCTAGCTGATCCTTTGTTTCTTAAGCTAGAACAGTTGTCTATTGCTAACTGCAAGAACTGTTTTTCCTTGCCAGCACTAGGACAGCTTCCTTGTTTGAAAATCCTTTGGATTAGAGGGATGCATGGAATAACAGGGGTGACGGAAGAATTCTATGGCAGTTTGTTCTCTAAAAAGCCTTTTAACTGTCTTGAGGAACTTATATTTGAAGATATGCCGGAGTGGAAGCAATGGCACGTACTAGGAAGTGGAGAGTTCCCTATACTTGAGAAGCTTAAAATTATAAATTGCCCTGAACTCAGTTTGGAGACAGTACCCATCCAACTTTCAAGTTTAAAAAGATTTGAAGTAATTGGTTCTCGCACGGTTGGAGTTGTTTTTGAGGGAATGAAGCAGATTGAGGAATTATATATTGGTGATTGTAACTCTCTTGCCTTCTTTCCTTTTAGCATACTGCCCACTACCTTGAAGACAATAAGGATATCTGGTTGCCAGAAATTGAAATTGGATCCGCCAGTTGGTGAGATGAGTATGTTTCTCGAGGAATTGGAACTGCATAAATGTGATTGTATAGATGATATATCACCTGAGTTACTCCCAAGAGCACGCACATTGAATGTATTTAATTGCCACAACCTTACTAGGTTTTTGATTCCTACTGCCACTGAAAGTCTCTTTATTCTGAATTGTGAGAATGTTGAAATACTTTCGGTGGCATGTGGGGGGACCCAGATGACGTATTTAAATATTCATGGCTGTGAGAAGCTGAAGTGGCTGCCAGAACGTATGCAGGAACTCCTTCCATCTCTTAAGGATCTGTATCTGATTAATTGTCCAGAAATAGAGTCCTTTCCTGAAGGAGGATTGCCCTTCAATTTACAACTCCTTTGGATCAGTAATTGCAAGAAACTGGTGAATGGCCGAAAGGAGTGGCGTTTACAGAGACTCCCCTGTCTCACACAGTTAATCATCTCTCATGATGGCAGTGACGAAGAGATTGTTGGTGGTGAGAATTGGGAGTTGCCTTCCTCTATAACAAGACTTGAGGTATACAATCTGAAAACATTAAGCAGCCAACATCTCAAAAGCCTCACCTCTCTTCAATATCTATCTATTGTGGAACAAGGCCAGTTTTCGCACCTCACTTCTCTTCAAAGTCTAGAAATCATGAATTTCCCTAATCTCCAATCACTTCCTGAATCAGCACTGCCCTCCTCCCTCTCTCAGCTGACCATCTATTTTTGCCCTAATCTCCAATCACTTTCTGAATCATCACTGCCCTCCTCCCTCTCTCAGCTGACCATCTATTATTGCCCTAAGCTCCAATCACTTTCTGAATCAGCACTGCCCTCCTCCCTCTCTCAGCTAACCATCTATAATTGCCCTAAGCTCCAATCTCTTCCAGTAAAAGGGATGCCCTCTTCCCTCTCTAAACTATCTATGTCCAAATGCCCATTGCTCACACCACTACTAGAATTTGACAAGGGGGAATACTGGCCAAATATTGCTCAATTTCCCACCATAAAGATCGATGATGAATGCATGTGATGATTAAAACGAATGGCTCTTCAACTGATGTAAGCTATTCTTTTCCCTTGGAAGCTTTTTATTTCCGTTTACTTCACTTTGTTTTTTTGTTAATTCTTTTCATTTTT

>M6-PGSC0003DMG400018722 M6 LRR-RLP

AACCAATACTTTCTTGAGTTGAGATCAAGATCATTGTTCGAAAGGGTCCCCGAGTCTGAATGGAAACCGGAGGGATTCTTAATGCATGACCTTGTCAATGATTTGGCCCAAATTGCATCTTCAAATCTTTGTATAAGGTTGGAAGAGAACAAAGCATCGCATATGTTGGAACAATGTCGGCACTTATCTTATTCAATGGGAGAACGTGACTTTGAGAAATTGAAATCACTCTTTAAATCAGAGCAGCTGAGAACATTACTTGCAATCAATATCCTTAAGCACTATTATTACAACATTAAGCTAAGCAAGAGGGTGCTGCATAACATACTGCCAAGACTAAGATCCTTAAGGGCATTATCATTGTCAGGTTATAGGATTAAGGAGTTGCCAAATGACTTGTTTATCAAATTAAAGCTCCTCAGATTTTTGGACCTTTCTTGGACAGAGATTAAAAAGTTGCCTGATTCTGTTTGTGGATTGTATAACTTGGAGACACTTCTCCTGTCATCTTGTGATTATCTTGAGGAACTACCGCTGCAGATGGAGAAGTTGATTAACTTGCGTCATCTTGACATAAGCAACACTTGTCGCTTGAAGATGCCACTACATCTGTGCAAGTTGAAAAGCCTCCAAGTGCTAATGGGAGCCAAGTTTCTTCTAGGTGGTTGGAGAATGGAAGATTTGGGTGAAGCACAGAATTTGTATGGATCTGTATCAGTTGTAGAGTTGCAAAATGTGGTTGATAGAAGGGAAGCTGTGAAGGCAAAGATGATGGAGAAGATTCATGTTGACAAGCTATCTATGGAGTGGAGTGAAAGTAGTAGTGCAGACAATTCACAAACTGAAAGAGACATACTTGATGAGCTACGCCCACATAAAAACATAAAACAACTCCAAATCACCGGATATAGAGGGACAAAATTTCCCAATTGGCTAGCTGATCCTTTGTTTCTTAAGCTGGTGAAATTGTCTCTTAGCAACTGCAAGGACTGTTATTCTTTGCCAGCACTAGGACAACTCCCTTGTTTGAAAATCCTTTCCATTAAAGGGATGCATGGAATAACAGAGGTGACAGAAGAATTCTATGGCAGTTTGTCCTCCGAAAAGCCTTTTAACTGTCTTGAGAAGCTTGAATTTGTAGATATGCCAGAGTGGAAGCAATGGCACGTACTAGGAAATGGAGAGTTCCCTACACTTGAGAAACTTTTGATTAAAAATTGCCCTGAAGTCAGTTTGGAGACACCCATCCAACTTTCAAGTTTAAAAAGGTTTGTTGTTATTGGTTGTCCTAAGGTTGGAGATGCTCAAGTGTTTAAATCCCAACTTGAGGGAATGAAGCAGATTGTTGAATTAGGTATTAGTAACTGTAACTCTGTTACCTCCTTTCCTTTTAGCATACTTCCCACTACCTTGAAGAGAATAACCATATCTAGTTGCCAGAAATTGAAATTGGAGGCGCCAGTTGGGGAGATGAGTTATTGTAACATATTTCTGGAGTATTTGAGATTGGAAGAATGTGATTGTATAGAACATGATTTGCTCCCAAGAGCACGCTCTTTGTCTGTAGCCAATTGGCACAACCATACTAGGTTTTTGATTCCTACTGCCACTGAAACTTTCAATATTAGGAATTGTGAGAATGTTGAAATACTTTCGGTGGCATGTGGGGGGACCCAGATGACGTATTTGTATATTAGTGACTGCAAGAAGCTGAAGTGGCTGCCAGAACGTATGCAGGAACTCCTTCCATCTCTTAAGGATCTGGATCTGCGTAATTGTCCAGAAATAGAGTCCTTTCCTGAAGGAGGATTGCCCTTCAATCTACAATTCCTTTCAATCTGGAACTGCAAGAAACTGGTGAACGGACGAAAGGAGTGGCATTTACAGAGACTCCCCTGTCTCAGAGAATTAAGCATCTACCATAATGGCAGTGACGAAGAGATTGTTGGTGGTGAGAATTGGGAGTTTCCTTCCTCTATTCAAACACTTTATATAGAGAATGTGAAAACATTAAGCAGCCAACATCTCAAAAGCCTCACCTCTCTTCAACATCTTTGTATTGAGGGTAATTTACCTCAAATTCAGTCAATGCTGGAACAAGGCCAGTTTTCGCACCTCACTTCGCTTCAAAGTCTACAAATCAGGAATTTCCCTAATCTCCAATCACTACCTGAATCAGCACTGCCCTCCTCCCTCTCTCAGCTGGAGATCTCCCATTGCCCTAATCTTCAATCCCTTCCGGTAAAAGGGATGCCATCTTCCCTCTCTAAACTTTCTATTTTAGAATGTCCGTTGCTCACACCACTACTAGAATTTGACAAGGGGGAATACTGGCCAAAAATTACTAATATCTCCACCATAAAGATCGATCAGGAATGCATGTGATGATTAAAACGAATGGCTCTCCAACTGATGTAAGTTATTCTTTTCCCTCGGAAGCTTTTTATTTCCGTTTACTTCACTTTGCTTTTTTGTTGATTCTTTTCATATTT

>M6-PGSC0003DMG400018806 M6 LRR-RLP

CATAGACAGACATATATAAGGTGAAAAGAAGTAGCTCCATTGTCTGTAGTATTATTTTTCCCACAAATCTCCATTTGCCGATCTTTCCAAAGCTCAATCCCAGAAAAGCGTCACCACTGCCAGCCTTCGATCCTTCGAAAGATCCGATCAGATCTCTTTGTCTTTCTCTCACTGCACTTCTCATTCTGTTTTCTTTCTCTCTACACACATACCTTTTTCTGCAACTCCATTCTCCGGTTTGTCGAAGATTCTGGAACAAGAATTTGTTAGATTCTGTGAACTATGTAATTTGTGTGCTTTCTTGTGAAAAAGGAGTCGTTTTGGTGTAGTTTGTTGTAAAGTTAGGGTTTTTTTTTGGGGAAAGTTTTAGATCGTGAGCTAGGGTTTCTGAAGATCAGAAATTGATAAAATTTGGGGTTTATTCTTTATTTTTCTTGGTGTGGTGTGAGTGAGTGTATGAGGATTTGGTGTTGTTTATGTTTTGGAGAAGAAGAGGATAGTAAGAAGGGCTACAAGAGTATGAGGGACCCGATTTTGGGGAATAACGGCGACGAATCTCCAGATGAAAATTCGGCATTTGACTGGAGGAATGTTTTTGAAGGTGTCAATGTTGCAGCAGTGGTGAGTCCACAAGCGGGCGCTGCAGGTGATCTTGGTGTTCCCAAGAATGAGGAGATTGATTTTGATAGCAACTGGTTATCCAGTGAAGTGGAGGTGAAGAATGAAAATTATAGTGGTGAGAAAATGTTGGATGTTAATTTGAATTTGGGTTTGAGTGGAGAGGCTTCATCCTCAACAGTGCTGAAAGAGGATTCTGATCGTGATACCTGTAGTAAAAGGCCCAAAGTTAACTCCTTTTCGCTGTAAGACCTGCTACTGCACCTTCATTTACAATATTTTTTTTCCAATTAGTCAAATTCTATTGAATATTAGAGTGAAGTTGGCCATAGTAAAGAACTAATACAGAGTTCATTTAAGCAACCCAAGCCTAATTACAAGGATCAGTTGTTGTTATAGGGGCGGCTCACTAATTAATTAAGGACTCTTGATTATTTCTCGTGTGTACAGTTTTACATGTGGTCAAGAGTGTTACACTACTCACTGAGTTTTGGAGAAAGTACATTCTCACATAATTCTTTTAGAGAAAAGAACATAGTATATTAACATGGAGTAGAATTATTTCTAAAGCTAGTTATATTTGGAATTTCAAAAGCTTGACCTAATATATAATGTTACTCTTTAATTTGATATAATCAAATTGTCTTTTGAGTTTTGAAGTAAGCCCTAAATGTAGGCCAACAACTCAAAAATGAAGACTGATTGATTAATATTCTTGCAGGGATTGGGACAACCACTTGTTGCAGGAAACCAGTTACCTATGTCCAATGAACGAGGGCGGTGGAGATATGAGTTTATCAAACTTACTTGGTGCAACAGATGCTGAGGGGAAAGATTCTAAAATGGAGGATTTAGATGTGCGGATGGATCTTACTGATGACTTGTTGCATATGGTATTTACTAGTTGCCACATCCGCTCCTTCTGAGTTGCTTAATCAATAAGTGTTTATACTAAGTTTCTGATTAAGGGAATATATGCCTCATCTGTTGCCAGGTGTTCTCCTTTTTGGACCATATTGATCTTTGTCGAGCTGCAAGTGTTTGTAGCCAGTGGAGAGCAGCTAGCTCTCACGAAGACTTTTGGCGGTATCTGAATTTTGAGAACAAACAGATATCCTCAAACCAATGTGAGTATCTTTTCCTCATATCCGTTCCTTCCACACTACTATGATCACTTCACATAAAATTCATGTTTCAATTAATTGCAACTTTTTGTTCATGTATGATGTAGTTGAGGACATGTGTCGGCGATACCCAAATGCAACAACAATCAACTTATATGGAACTCCTAATATTCACCCACTAGCAATGAAAGCTGTTTCTTCTTTAAGGTAATAGTTTTTTTTATGTTGTTGTCACTACTCTAGGAATGAATCACTATAAGTGACTAATTTGATACATTTCATTGTTGCCAGGAATCTTGAGACTCTGTCATTGGGCAGAGGTCAATTGGGGGAAACATTTTTTCAAGCCTTAACAGATTGCCATGTGTTGAGGAGTTTGACCATTAATGATGCCACTCTTGGAAATGGCATTCAAGAGATACCAATTTCTCATGATAGCTTGCGTCTTCTGCAGCTAGTGAAGTGTCGTGTGCTTCGAGTTTCTATCAGGTTGTATAATATTAGTCAACAATGTTAATTTGTTAAAGTAGTTAGCGTCCAATTGTGCTTAATTGTTTGTATTCTTGCAGATGTCCACAACTTGAAACGTTGTCACTTAAGCGCAGTAGCATGCCACATGCGGTGCTTAATTGCCCTCTTTTACATGACCTTGATATTGCTTCATGTCACAAACTTTCCGACGCTGCTATTCGTTCGGCTGCCACAGCATGCCCTCTTTTAGAGTCTTTGGATATGTCAAATTGTTCATGTGTCAGTGATGAAACACTGCGTGATATAGCTCAAACTTGTGGAAATCTTCGAGTTTTGGATGCTTCATACTGCCCAAACATTTCTCTTGAGGTATGTTCATTAAAATTACCTAGTACCTGAAATTGACATAGTTGGTTTATATACTAACACTTGTTACTTGTTGAAGTCTGTGAGGCTGGTTATGTTGACTGTTCTCAAGCTTCATAGTTGTGAGGGCATAACATCAGCTTCCATGGCTGCCATAGCCCATAGCTATATGTTAGAGGTTTGAGTCTTTCCTCACCCTTTTCATACTGGTAGCCATGCTGTAATCAGGAGTCCTTAAATCTCATATTGAACAGTGAATTTCACTTTTTTTCAGGTTCTGGAGCTTGATAATTGCAGTTTATTGACATCAGTGTCCTTGGATCTTCCTCGCCTTCAGAGCATCAGACTAGTACATTGTCGCAAGTATGATACAGGCTTTATAGTTTCCATATCCTGTCTCAAGGGAACTTCTTTCCATCCCTTGATTGCCGTCACTCTTTTGTGCAGTAACCTACTGTTTATATTGAAACAGGTTTATTGACCTTAACTTGCATTGTGGAATGTTATCGTCAATTACTGTTTCTAATTGCCCATTGCTTCATCGAATCAATATAACTTCAAGTGCACTCAAAGTAAGTTTATTCCTAGATGTTATATATTGTCACCTTATTTTATTGACTAGAATTATTCTATTCATTGGATAATAGTTCTCTCCTTTATTATTCCACAGAAATTAGTTTTGCAGAAGCAGGAAAGCTTGACAACAATTGCATTACAGTGTCCGAATTTGCTAGAAGTAGACCTTACTGAGTGTGAATCGCTCACCAATTCTATTTGTGAAGTTTTCAGTGATGGTGGTGGCTGTCCTGTGCTTAAATCCTTAGTTCTAGATAACTGTGAGGTTGGCAATTTTAGTTCCTCGTTGTTAACTTGTGCATTCTTGGTTGTTAACTAGAGGATGTGTTGCTGAACATGATTTTTCTAAATCTAATTATGATATATATGATCTCTTCTTAGTTTTGCTTGTCAACGGAAGGTGGTTCACTTAAAATCTTGGCAGTTGATCCGTGCAGTGTACATTTTTCATTAAGAGCTTGTTTTTGTCCTCACGTATGTCAATGTGGTTATGGACCTATGATCATAGTGCCATTTTTGCTTTTCTTGTAACAGATACTTGTGATTCTTTTTTATGCATGTTTCCAAGTAAATAAATCTAGTACTGAATTCTGACAATCAATGATCAGGTATTTTAATGAGTAATCATGAAAGATGATGAAAGGAAAAAGTATGATCCAAGTGTACTGACTGTTGTGTCATCTGGTTGTGCTTTCTTGTTTGCCGTAAGCCTGGGAAGCTGTGCTACTGGGTGCACTAAGGGCTTGTTTGATACGAGGGATAAGAGATTACTGATCCCAGGACTAGTTTTAGGATGAGTTTATCCCATGTTTGGTTGGGATAAAATCGTGGGATAACTTTTCCCTGGATTGTAGTGTCATTCTTATCCCACATTGAGGGTGGGATAACAATCCCTGGATAACTTGTTGCGAGAATCTCCTTCAATAACTCAGACAAGTTAAAAAAATATTAAAATTGTTCAAGAGGTGAAGTGGGAAAGATGAAAAGTTCTAATTCCTTTGTTTTGGCATTAAGGATAGTGATTATGTATTCCTAGATTTACCAGTAGTATTTATTTTATACTCACATTTCCAAATTAACTTGGAAGTATTTTTATCTTGTCAAAAAATAACTATTAGGGAAAACGTCTTTCTTGGTATCATAAGCTCTAAGTTTCTTTCAACATTTTGATTAGATACATGGGAGTTCTTAAGGGGACGGTGTAAGGCTTTAAGCTGAAGTAGAACATGAGAACTATTGCCCAGATGGGAGAAGAAAATAGTTCACCTTTTTTTTCCTGATTCAATGCTAGAGCTGAAGTCGCGCCATTTAAAAAAAAAATGTCAAATCGGAATTTTCTTTTGAAGCTTGACGAAAATATGTACTCCTCTCCCAATCAAGATTCCCTTTTATTGATTTGGTGATCAAGAAAAGCAAGGCTGAATTAGGTTTGGGTTTGACATTTTTGAGAGATGTACAAAATAATGAACTAAATTTGTCAATATTGTGATAGACTATTGAAGAAATTCACAACAAATCAAGTATTATATTTCGGTTACCCATTCTCCCTCCCCCGTGCCCCCTCCCTCCTCCTGGGAAAAGGGGGGAGGGACCAGGTGAGAGAAAGAGTAATATATTCAAGTTGCACCCTCCCGGGGAAAAGGGGGCTGAGAGAGGGAAAGGTTTCCACTGTTTTAGAATGGGTCCTGATTAGCATTCGCCTTCTGTGTTCTCTTTTCTGCCAGATCTGTTTTAGGCTAACACCTATTCAGAAAAGTTTTTTTTTTTTTAATATTGGAAGTGAATAATGATATATTTCTGCCTATTCAACTTGAAAAATCCGGCTGTATCTATGTAACATTTGTATGTGTTGTTATGGTCTAAATCTAATTACGGTCTTTTCTTTTTTTCAGAGCTTGACACTTGTGGCTTTCTGCAGCACTTCTTTGGTTTCTCTTTCACTTGGTGGTTGCCGTGCTTTAATTTCTCTTGCACTTAGGTGCCCATATCTTGAACAAGTCTCTTTAGATGGTTGTGATCATCTTGAAGTAGCATCATTTTGCCCAGTAAGTGATCATCGGCTCCTTCCTCTTTTCTTTACATCATCTCTGGGTTTGAGACTGGAGTTGTATGTTCACCTGCTGATTTTATGATTGTTAATTTGTTACACATGCATTTTTTTCTTTATTTATACCCAAATGGTTGATAAATGTGTTCTCTTATTCCAGGTTGGTCTTAGGTCACTTAATCTTGGTATATGCCCCAAAATGAATATGCTCCATATTGAAGCTCCACAAATTGCTTCACTTGAGCTGAAGGGCTGTGGTGTGCTATCTGAAGCATCTATTAATTGCCCCCTGTTAACCTCTTTTGATGCTTCATTCTGCAGGTTGCTTACTTTCTCTCGCTTGTCACACGTGCATATTTGTTTTAGTTCTGTCACATCTCATTAATTTCTTTCCATATATGCAGCCAACTCAAGGATGATTGTTTATCTGCCACGACTTCGTCGTGCCCCCTCATCGAGTCATTAGTATTAATGTCATGTCCATCTGTTGGCTGTGCTGGACTCCTCTCTTTGCAGTCCCTCCCAAACTTGACCTACCTTGATCTATCCTATACGTTTTTGGTTACCTTGCAACCTGTTTATGAATCATGCTTGCAGTTAAAGGTAACTTCTATCGCCTAAGCATATTATCACTTTCTCCATTACAGTGGGAAGCTAGATGAACTTGATTTAACATGCTCTATTTTGTGGAAAAAGTATGCCTATTTATCCAGTCTGGACTTGCACTCTTCTAGATATATGGGCCACTTTCCTATCTTTTTTTTTTATGGGTATCAAAGGGATGGGTTGGCTTAAGTTTGTGCTGGTCAAAATACCTAACTACAAAATTTACTCTTGTTAAGATGGGATGAACACAGGGTCATAAGCCAATTACTTTAATCTCTCCTTTTCCTTTTTCTGTTCTCCTTTTTTCCCACTTGCAGTAAAAGCTAAAACACCAGGATGCATTTAACAATTTGTGAATCAGTTCTGACTGATTGAAAGTTTACTTCAGTAGAACTGTGATTCTATTTTGACTATTACAAATTTAATAATCATTGTTTGTGGCCATATTTTTGGGGAACTAGAAGTGCTTTTTTTTTTCCTTCAGTTTTTTGGCAGTTAAGGTGTTTGACCATTCTTCTTGGAAAAGAAATGTGATCTTTAGTACCATAAGCTGTTTCTCTGTTGTTGTGGCGAAGAAGTCAAATTTGAAGCTTTTCATTAAGCAGAACCAAAAGTTTAATTTTTCAATACAAAATACCATTATAATTTTCTTACCAAGTTATACCACAGTTCTGACAAGTGACAACTGTATTAATTTAAAGAGGGAAGCCCGGTGCATTAAGCTTGCGCTATGCGCTGGGTCCGGGGAAGGGCCGGATCACTAGGGTCTATTATACGTAGCCTTACCCAGCATTTCTGCAAGAGGTTGTTTCCCATTCACATGGCAACAACGTAATCGGTTATGCCAAGGCTCCCCTTCGACAACTATATTGATTTATTATTGAAAAAGCATTTCTGTTTTCAAAGTGAATCATCAGCTCTGTACAAAGAATAAGAAGTAAATTAAGCCAATGCATTAGATCAAGTGTAAGTGATAACACAGATTTAAAGTAACATCAGTAGAATAGAAAGCTGACAATCATGGATACCAGAAAGAGCTAGGAAACAGCTAAAAGCAAGCTTAGGCTTTAGCTTTTACCTGCTCTGGAGGATCTGGAGCAACAGAAACTGTAAAGAGCAATCAGAAAACTTGTAAGAGGTTTCCATTTTGCATTGAAAAGAACTCTAGTCGAGGGCAAGGGAAAGAGAGTCAACAAAGTATAGAATAAGATTGTAGATTTGTTGGACTTCTAAAATCTACAATTATATTAGGTGTTGATGGAGTAGGATATTCTAATGCCATTCCAGGACTGATGATGTTGTCTCTGTTCTCAAATAGTTCTGTGCCACTTGGTTGGGTTGTTTATAAATGCACTTGTTAAATATTACGAAACTGTAGACTAGAATTGCCACCTTAACTCTTTCATTATTGGCCCTCCTTTTTAGACCCTTGAAGATGGCCGGCGTGGAATTGTGGAGATCCAGTTACTTTAGGCATTCTGTCATTCTTTTTTATTTTGGAGCGTAACATCTGAATATTTCTACTTTCCAACAGATGCTATAATTTTTGTTCATATCCAATGTTTGCCATTGTCTTGTGCCCAATTTGTTTCAGTGCAAGGGCATGAGTTCCTTGTATGCAGAATTTCTAGTGCTATATCTGAAATTTTCTGATGAGACTCTAGTTTAGTAACACAAGTGTGTTTCTCTTTCTCGCTGCACTAGGCACATCCTTATTGCGTTTGGGATAAACTTTTTGGATCTGCATGATACATTGATGGGCTTTTTTGGCTTACTGCCCCCCCACCCCCACCCCTCCTCCTTTCTTTTTTTTAACCCCTCTCTAAGCTTATATATATATGTTATGTTGTAGGTCCTAAAGTTGCAAGCATGCAAGTATCTCACTGATACATCTTTGGAACCCCTCTATAAAGAAAATGCTCTCCCAGCCCTTTGTGAGTTGGATTTGTCGTATGGGACCTTATGCCAATCAGCCATTGAGGAATTGCTTGCTTACTGCACACATTTGAGTCATGTCAGCTTGAATGGATGTATTAATATGCATGATTTAAATTGGGGATTTACTGGAGATCAACTGTCGCATATACCCAGTGTTAGTATACCTCATGGATCTTCCCTGGGAGAACAGCAGTTGCCAAATGAACAACCAAAGCGTTTGCTAGAGAACCTCAACTGTGTAGGCTGCCCAAATATAAAAAAGGTGTTTATTCCTATGGCACAAGGTTTCCTTTTGTCATCATTAAACCTTTCCCTGTCTGCGAATTTGAAGGAGGTTGATATAGCTTGTTACAACTTGTGCGTTCTCAATTTGAGGTATTTTCTTTAGCAATATGATTTCGTAGGTTAATGTTGTATCAATGGCTTTATTTTCATGATCATAGTTGTTACTTAATTATATTCTCTTGTTCAGCAATTGCTGTTCATTGGAATCATTGCAACTGGAATGTCCAAGATTAAGTAGCCTCTTTCTTCAGGTATTTTTGTATCTTCAATTTGGCATTTTCTTTGCTGGCTTTTTAATTATTGTAGTGGCAGCTTTGCTGCAAAAGTGTTCTTTCTAGTTGGTCTTTATCATTTCGCTTGTGCTTTTTTCCCCCTGTTGAGGTAGTATAGTTTTTTACCTTAGCACGAAGGATATGCAAACTGTACAACCATTACAAATGTCAGTCCCCTTCCTACCTTGTAGGGACTTTATGAGTTCTATCAATGATTCAACTTCATTTACAATTTCCTTTTTACATCTAAGACTCATCCATTTAGGATAGGATGATCCTTGTTTGTCTGTTTTACAAAACATTCTCGTCTGCCTCTGGTTCAGTAGGTTGGTATGGAGAATATGCATACCCTTAAGCTTGATATTCTGTAGAGAGAACTTGAGCTCCTAATCAAGTCGCTAATGTGATTGTATTGTAAGATCTAATAGCTACCTGCATATTGTCTTAGGAGTACTCAAATTAACACAATGGTCTACTAAAATAACAAGTAATCCTAATACTCTTCATGGTCTTCCTAGGCATTCATGTGGATAGATAATAAACATTTTTCAAATTTGGAAGTGTTAGAAGTCCTTAGGGATGGCATGTAGTTTGGGGGCATCGTGCCTATATTTCGTGGAGCATGATGTACTTGGGCTAAAATTACTGAAATGAGAATTTGAACTCTTAACTGGTGCTTATTTGCTTGAATATTGGAACTGCAAGTGAGAACTGAAGTCTTAGTTTTTTTTTTTTTTTTAGAAGAAGTTTTTGCTTTCTCTGAGCTGTTTTTCCGTTGTTGCTTCTATCATTACACGCCTCAGGCTTACCTCAAAGACATGCCACTTTCTAGTGTAGAGCTCGTGAAATTGATTGCCTTTCTTTCTATCATGTCTTGTAAAAAACTACTTTGATGAACTCCCTTGAAAAATCGTTGGCACACCCAGAAGGTGCTTGTGCAATTGAAATTCTTGTGACACAATTGTGATGTGGAGTTATTCTGTACTCTTGCTTTTGTTGTCCTACAGTCCTGCAACATTGATGAGGAAGCTGTTGAAGCTGCCGTATCACGATGTACGATGCTTGAGACGCTTGACGTTCGTTTTTGTCCCAAGGTCAGGAGTACTTTTGATGTTAAGAATATGCCCTGTATTGATTTTTATGTATCATTTTATTGATACAAGAGCTCTTTTGTGAATTTGCTGATAATCCTTTTTTCCACTTAACAGATTTGTCCACTGAACATGACGAGGTTGCGGGTTGCATGTCCCAGTTTGAAACGCATCTTCAGCAGCCTAGTCCCATCGTGAGGCAAAATGAAATATTTGATATGTTTGTGTAGGCTGGCTAGGGACATAATATTTGACTACAGAGGTTCATATTTTATGTATAGTTAGCTATTTATGTTGCTGTGATTGGTGAAACGTACAGCGAGCTTCATACATGTAACACCTTAAATATTTTGGTGTTGGCTTTCACATTTTATGATTTGGTTAGCGCACTTAATGCAACTTTATTTAGATCTCATGGATGAAGAAAATAAACTAGAGCCTTGTAAAGGGAAATATTAGTTAATAGTGGCCTGCTTGCCCTTTTCAGTTAATGGAAAAAAACGCAGGTTGTCAGGCTCGAGTGTGAATTTTGGTTGCCCCTAAAAATAAGGTTACAAAAAATAGCCCAAAAATAGAGTAGTCAGTCGGCTTATGTCTTTGGGAT

>M6-PGSC0003DMG400018861 M6 LRR-RLP

ATGAGCATTCTCTCGAAAACATGGCTACAAGCGTGGTTGACTCTTCCCAACTTGGAATTCTTTGTTAATTGTTGGCGAGGCTGGGAAGCGCATATAAATATAGTGGACACAACCATGGAGAGATATGGGAAGGGGAAAATTCCTATACAAAAGTTTGAATTATCAGAGATTTTTGCTGATTCTCCCCAACTCTTCCCTCTAATTGATAAGTGGCTTCTCATTGCACTTCAGAATGGTGTAAAACAACTTGTACTTCACTTTACAGCATACCCGGCGCCTATTTTAACAATCTTGACAGCAAAATCTTTAAGAGAATTGGTTCTGTACAATTGTACTCTTATGCCTGTTTCATTATCTAGTGGGGTTGTGAGTTGCAATTCTCTGAGAAAGATTTCTCTATCTTATGTAACATTAGATGAAAACATGCTTCAGACTCTACTTAATAGCTGTCCCTTTATTGTCTCTTTCATACTTGAGTATTGTTTGGGGTTGACTATGAAGCTTATAAAGATCAAGTCGGATTCCCTGAAGGTCTTGAAGATTCATCAATATTGCGGGATATGGGAGATTGATGCTCCAGATTTGGTATCACTTGACTATACAGGGAATGAAATTCCTGAACTCAACATTGTAAGAGAGTCTAGCCAATTGGAGAACTCAAAAATTATTCTTCATTGCATCTCCAGATTAAATACGTCATGGTTTTGTAAGTTGAGGAAGTTTCTATCAAATTCATCCTCTTGGTCTGAAGTTAACCTTTGTTTATTCAAATGCGATGAGATCAACATGACAGATTTGCAAATGGA

>M6-PGSC0003DMG400018865 M6 LRR-RLP

AGCTAAATATGGGCTATGCAGTTGTGCTCTCTCTTGTCTTTCTCTGTCATGAAGTATTTACTGTTTCTTCATCCATACTACCTCATCTCTCTCAAAAACATCAAAACATTTCCCTTCTAAAATTCAAGCAGTCGCTTACTATAAATACCTCAGCCTCTTATAGTTGTGAGAATCCTTATCCAAAAACGAGTTCATGAAACATGAGCAGAGATTGCTGCTCATGGGATGGAGTCGTATGCAATAACATGACTGGAAATGTCATTGAGCTTAACCTCAGTTGCAGCGGGATTGTAGGAAAGATTGATTCCAATAGCAGCCTCTTCCAACTCTCTCATCTCCAAAGGCTTGATCTCTCTTATAATAACTTCTCTAAATCTCACATCTCACCTGAATTCGGTAGATTCTCAAACTTAACAAATTTTGATCTTTCATAGGCAGATTTCTCAGGTCAAATTCCTTCTGAAATCTCTCATCTTTCCAAGTTACACTCTCTCAGTCTCTACAATTCTATTGGTCTACGACTCGTAGCCCATGATTTTAAATTGCTCCTTCAAAATTTGACTCAATTAAGAGAGCTTGATCTTAGTAACATAAACATCTCTTCCACCATTCCTCTGAATTTTTCTTCTCATTTGACAACTCTGAGACTGATAGATACAGGATTGTATGGGATAATACCTGAGAGTATATTTCACCTACCCAACCTGGAAACACTGTACTTAGGTTTCAGTGATCTCCGTGGTTATTTTCCAAAGACCAAATGGAACAGCAGTGCATCTCTAGTAGAGTTAGATCTCCTTGGAGTGAATTTTTCTGATAGTTTTCCTGAGTCTAATGGCTATCTAACTTCGGTGCGTTCTTTGTCTCTTCAAAGTTGCAACCTTAGAGGGCCTATTCCTGAATCTATTTCTAATCTCAATCATATAGAATATTTGCACAATCAAAATAACTCCCTGAATGGAACAATACCATCAGGGGTGTTCTCCCTTCCATCACTAAGATATTTACTCTTGGGTAAGAACCACTTTTCTGGTCATCTTGACGATCTCAAGTCCAATTCACTAATCTGGATTAATTTAGGTGACAACCAATTGCAAGGCTATCTTCCCAAGTCAATTCAAAACCTTGTGAACCTCACAAGTCTTTATCTTTCCTTCAATAATTTTAGTGGTCGTGTGGATGTCAGCCTCTTTTCAAACCTGAAAAGGCTTTCGGATCTTACTCTTTCAAATAATAAGGTCTCATTGACCAATGAGAACAGAGTTACTTTGCCTGAATCTCTTCAAACGTTATAATTAACCGCATGTGAAGTGAAAGAATTGGAGTTTCTAAGATCTGCAAAGCAGCTCGAGGTGTTGGATCTTTCAAATAATAAGATTCAAGGAAGAATTCCAGATTGGGCATGGTCTAACTGGATTATTTCATTGAAGAGTCTTAATATATCCCACAATATGCTGACGAGTGTGGATTCAATTCCTCTTCAGACTGTATATACTATTGATTTACGGTCCAATTTGCTTCAAGGTTCACTACCTATGCCACCAAATTCCACAAGATACTTCTTCATCTCCCAAAATAATCTCACTAAAGAAATTCCTTCATCTATTTGCAATTTGACATCACTGGTAATGCTAGATTTGGCAAGAAACAACTTGACGGGAGAAATTCCGCAATGTTTGGGTAATATTAGTGGTCTCAAGGTTCTTGATATGCAGCACAACAGGCTTATTGGTACTCTTCCAACAATTTTAAGCATTAATGGAAGTTCACTGAAAAGCCTCAACTTGCATAGCAATAAATTAGAGGGGAGAATTCCCCGATCATTGGCCAATTGCAAAGAGTTGCAGCTTCTTGATTTAGGTGATAATCATCTCACTGACACATTCCCGATGTGGTTGGGAACTCTTCCAAAGCTAAAAGTTTTAAGCTTGAGATCAAATACATTGCATGGGACCATTCAACCTCCAAGGATTGAAACTATTTTTCCCGAGCTTCAAATCTTGGATCTCTCTTACAATGCCTTCTCGGGAAACTTACCTTCGAATCTATTTCAACATCTAAAAGCTATGAGGACAAATGATCCATCAACGGAAGCACCAAGATATCCTGGAAGTACATATTACGAAGATTCGATTACAGTTACAACCAAGGGATTTGTGCGTGAAATTGTGAGAATCTTGTATCCTAGAAGTACATATTACAAAGATTCGATTACAGTTACAACCAAGGGATTTGTACGTGAAATTGTGAGAATCTTGTATCTGTACACTGTTATCGATTTTTCAAGTAATAAATTTGAAGGGCAAATTCCAAGTATCATGGGGGATCTCATTGCAGTTCACACATTAAATTTATCTCGTAATGGATTAAAAGGTCATATACCACCATCATTAGGAGATTTATCTTCAGTTGAATCATTGGATCTATCAGGAAACCAGCTTACGGGAGAGATACCAGAACAACTGGCTTCTCTTACGTCTCTTTCATTCTTAAATCTCTCCCACAATCATCTCCAAGGATGCATCCCTCAAGGATCTCAATTTCACACTTTTGAGAGCAATTCATATGAAGGGAATGATGGATTACGTGGATTTCCTGTTTCAAAAAATTGTGGAGATGCTGGGGTGTTAGATAAAAATGATACCGTATCTGGACAAGATGATGAAGAAAGCGATTCTGAATTTCAGAGTGATTTTTGGAAAGCTGCTTTTTTGGGCTATGGAAGTGGACTATGTATTGGATTATCCATAATATATTTCATGATTTCAACTGGAAACCCAAAATGGCTTGCAAGAATCATTGTAGAGCTGGAGCACAAAATAATAATGGGAAGGAGAAAGAAGCAGCGAAGGCAAAGGAACTATAGAAGAAGATAGTCAGATGAAGGTGGGGTGGAACTGAAGTTATCAAAGATGCATCTGCTGTGAAAATGATAACATTTTGTAAACTTGGAATTGATATATAATTTAATTATTTTCTAATGAGATCTGGTGGGTTAGTGCTGGTATGATCGTATCCTATTTATGAATGATTTGATATGAAGGC

>M6-PGSC0003DMG400018866 M6 LRR-RLP

ATGGCTGACATATTGCCGGGATGTCTCGTTCAAAAAATACTCAGTTTTCTTCCTAGTTATAATCAAGCATCCAAGATGAGTATTCTCTCCAAAACATGGCTGCAAGCGTGGTCGACTCTTCCCAACTTGAAATTCTTTTTATTTTGCTGGGAAGGCTGGTAAAGCCGTACAAAGATAGTGGACACAATCATGGAGAGATATGGGAAGGGGAAAATTCCTATAGAAAAGTTTGAATTATCAGAGATTTTTGCTGATTCTCTCCAACTTTTCCCTGTGGTTGATAAGTGGCTTCTGATTGCACTTCAGAATGGTGTAAAAGAACTTGTACTCCACTTTAAATCATACCCGGTGCCTATTTTAACAATCTTGGCAGCCAAATCTTTAAGAAAATTGGTTCTGCACGATTGTACTCTTATGTCTGTTTCATTATCTAGTGGGGTTGTGAATTGCAATTCTTTGAGAAAGCTTTCTCTATCTAATGTAACATTAGACGAAAACATGCTTCAGACTCTACTTAATAGCTGTCCCTTGATTGTCTCTTTCATCGTTGAGAATTGTTCGGGACTGAATGTTCTTAAGATCAAATCGGATTCACTGAAGGTCTTGAAGATTATACAATATTGCAATATATGTAACATTGATGCTCCAAATTTGGTATCCCTTGACTATAAAGGGTATGAAATTCCTGAACTTAACATTGCAAGAGAGTCAAAAATCATTCTTCATTGTCTCCTCAGTTTAAATACGGCATGGTTTTGTAAGTTGAGGAAGTTTCTATCAAATCCGTCCTCTTGGTCTGAAGTTAGCCTTTGTATATTAAAATGCGATGAGATCAACATGACAGATTTGCAAATGGACCACATAGGTTCTAGTGGAGGGGTGGACGTTTTAAATCTCAATATGCTACTGATGGATGAAATTATAGAGTGCCCAACTTTTGTGGATGCTTTGCTATGGAGTTGTCATCCTGGGAGACTTAACTTAATCTCAATTGATATTGAAACAGTTACACGTTTAATTGATCGGTTAATGTATATGAAGAATTCGAGTCATTCTACATATTCTCATGGTTGGAATAATCAATTAAAAGAAATAAAAGCTTTTGATGGAAAAAATCAGTTGCTGCAACTAGGAAGTGAGGAGCTTGCAAAAAAGATTATGGAGGGGAAGAAACCTTATTTTATATTAGATTGGTGATGGAGTTAAGTTGTCTCTATTTCGAATTGAATTCAAAAGTTTATTACTTCATTTGTTTTCCACATCTAGCTTTAAATTCTTGCAGTTCCGAAAATATGTTTCGCTGTTTTCCTTCTTCTGAGTTTGAGGGATTTTTCATTTCTCAATTATCCTACTAACAGCACAACTTATATACCAGTTTTTAAATCAGAAAGAGGGACTCTGTACTTCCCTTTCAATACCATAGAAAAACAATCCCCCTCCCTACATGTGCATTAACTTCATACTGCCTGCTGCTGGTTGATACCAAAATGAAATAAGAGTTTCCCATGAAATGGTTATTCATCTCTTACTTGGTGATGCATAACCATAGCACCTACGAATTCTGAAAAAAGTTCCAAGTTAGTTGAAGAAGCCATGATGAACAAAAATGTAACATAATGAAAGATAAAGGAAAATCAACATTAAGACTGGATAGTATCTTACTTTAACTCCTATTCATAATTCCCATACTTCCTTTTTTTCCATATCATGTTTCACTATAAAGGTTGTGGTGCACTCAGCCACTTATAATCTTCATAAACTTGATTCATCATTGGAGTAATTTTACGAGTATTACTTAAAATTAAAATACAATGTTATATTCAAGGAAAATTTATGTGTCTGCCTTCATACATATGTACTTCTCCATGATTTTGGCTAAAAACATCCATATGTTATACTCATAATCTAAAGTCTAAACTGCATCTTTAAAAACTATTCATGTGAACATAACACATCCCTATATACAAGAAAACGTTAAGCTAGGAAAAGCAGTACGGCCAAGTTAATTACATGAACAATTTTACAATATAAAATTGATGAAGAAGCAACGAATCACTTCCCAGTATCACATCACATGTGAATAGGGGTCTCAAATGGATGGTTTGGATGATGAAACTAGTCGGGTCCAAATGATCCGACTCAGTGACTCGCCCAAGAGCTACTTAGGCCAAATTGGATTGGGCTGAAATGAACTAAAAAACGAATCATATTTCAACCCACTCAATTCTTACTAAGTTTTAATTTCGCTTTATATAGAAACCATGACCCTCTCTTGCTCAAACAGATAGTCGTCTTGCACATTAGCGAGCTAATATGTCAGGGTCTCCATGATGTAGCTCAAATTCTTGATAATGTGTTGGATTGATTGTTATCCACGGACAGTACATGTGATTGTTAAGAGTCAAACAACAAACAAAGATCGTACATGTGTATCTCACAGTCTTGGTGAACAATTTGATTCAGTACGACTTCATTCGAGAGAATGA

>M6-PGSC0003DMG400018867 M6 LRR-RLP

ATGGCTGACATATTGCCGGGATGTCTCGTTCAAAAAATACTCAGTTTTCTTCCTAGTTATAATCAAGCATCCAAGATGAGTATTCTCTCCAAAACATGGCTGCAAGCGTGGTCGACTCTTCCCAACTTGAAATTCTTTTTATTTTGCTGGGAAGGCTGGTAAAGCCGTACAAAGATAGTGGACACAATCATGGAGAGATATGGGAAGGGGAAAATTCCTATAGAAAAGTTTGAATTATCAGAGATTTTTGCTGATTCTCTCCAACTTTTCCCTGTGGTTGATAAGTGGCTTCTGATTGCACTTCAGAATGGTGTAAAAGAACTTGTACTCCACTTTAAATCATACCCGGTGCCTATTTTAACAATCTTGGCAGCCAAATCTTTAAGAAAATTGGTTCTGCACGATTGTACTCTTATGTCTGTTTCATTATCTAGTGGGGTTGTGAATTGCAATTCTTTGAGAAAGCTTTCTCTATCTAATGTAACATTAGACGAAAACATGCTTCAGACTCTACTTAATAGCTGTCCCTTGATTGTCTCTTTCATCGTTGAGAATTGTTCGGGACTGAATGTTCTTAAGATCAAATCGGATTCACTGAAGGTCTTGAAGATTATACAATATTGCAATATATGTAACATTGATGCTCCAAATTTGGTATCCCTTGACTATAAAGGGTATGAAATTCCTGAACTTAACATTGCAAGAGAGTCAAAAATCATTCTTCATTGTCTCCTCAGTTTAAATACGGCATGGTTTTGTAAGTTGAGGAAGTTTCTATCAAATCCGTCCTCTTGGTCTGAAGTTAGCCTTTGTATATTAAAATGCGATGAGATCAACATGACAGATTTGCAAATGGACCACATAGGTTCTAGTGGAGGGGTGGACGTTTTAAATCTCAATATGCTACTGATGGATGAAATTATAGAGTGCCCAACTTTTGTGGATGCTTTGCTATGGAGTTGTCATCCTGGGAGACTTAACTTAATCTCAATTGATATTGAAACAGTTACACGTTTAATTGATCGGTTAATGTATATGAAGAATTCGAGTCATTCTACATATTCTCATGGTTGGAATAATCAATTAAAAGAAATAAAAGCTTTTGATGGAAAAAATCAGTTGCTGCAACTAGGAAGTGAGGAGCTTGCAAAAAAGATTATGGAGGGGAAGAAACCTTATTTTATATTAGATTGGTGA

>M6-PGSC0003DMG400018900 M6 LRR-RLP

CTATTTTAAATGAGTTGAGTACCCAACACATTTTAAAATGGGTTAATTTGTATAGGTTCTCATATAAATTATCCATTCGGTCACCCCTAGGCGAAACTGCAATCTTGTAACCCAAGGGCTCACGTGAACCCAATAGTTTTTGCCTTGAGACTTTAATTTGTATTATTAATATCTAGTGATACCTTGATTATGAACGTGATTGTTATTGGATAATATCCTGAGGATTTAGAGATATCTATAAATTTCAAATTCTGAATCTGCATCTCATCCTAAATACTTTATATTTTGCTCTTCATTAGGATTAATAAGGTAAGACATATTCCCTTGCTTGTACTTTTTCTTTTGAATTTTATTTTTATTATATATACTAGTTTATGGACACTTGCACGTGTGTCCCATAACAATGTTAAATAGATCCAGTCTATGAAAATACTTGCAATGAGTAATAATGAAACAATTACAAATAAAACTTGAAAAGTGTGACAAAAAAAACACCACATGATGATGTATTTTAATGTTTGAAATTAATTCCTTTTAAACAGTTAACTATAAATTTTGAATTTTGTTTAGTAGTTAACAAACTTTCAATAAAGTCACAAATGGCTAACTTGAACCAAATAAGAATACAGATTTACGATAGTAAGAAATTCATAGAAATTTACATTAATGAAGTCTAATAGAGAGAAAACACAAAAGGATAAACAATGAAGGAAAAAATACAACATGCACATCCTTCTTAAGACTTTAATCATAAAAATATAGATGAAAAAAATATAAAATAATTACTTAATTTTTCGAGCTCTTGTCGACACATCTTCCCTCTTAACCAATCAAAATTCAAGAGCCTTCTTCATCAAGTCAAATTAAAACATGAACCAAAATAGTACAACAATAATGTAAGTGTTTCACTACTTTAACGACATAATACATAAATGTGTCATTTAATTTGTATTCAATTGTATATATGCCCTCCAACGTTAGGTATGCACAATTAGACACTTAAATTTGAATAAAATTGAACAAGTAGACACACACATTTTATGTGGCATAATACACGTAGGATGTCATAGTATGCAAATTACCACGTAGGTGGGCCACATAAGACGTATGTATCTACTTGTTCAACTTTTGGATAGTTTAAGTGTCTACTTGTGCACACCCAAAATTGAAGGGCGTAGATGTCCAACTGAATCCAAGTTAAAGGACACATTTATGTATTATGACGAACTTTAACTATAGAATGAAATGATAAGCTTGGGAAGGGGTTTCATGAAAGTTGAAACCAAAAACAATATACGAATTCATATAGATAAAAATGGAGGAATACCAAAAGAAATCTTAAAGTTTTAAATTAAATAGTTGAAGGTTATGAATACAATAAGAATGAAAGTTACGGATATTAAGATTAGCAATATAAAAAGAAGGTATAATCCATAAATATGTCTTTTAACTTGGCTTCAACTCACATTTATGCCCTTCAACTTAGGTGTGCATAAGAAGACACTTAAACTTGTATAAAGTTGAACAAATAGACGCATACGTCCTACGTGACATCTTACATGATAATTTGTGTCCTACGTAGCATCCTACGTGTATTATGTCATGTACGAATTATGTGTCTATTTGTTCAATTTATACAAGTTTAAGTGTTTACTTGTGCACACCCAAAATTGGAGGACATAAATGTGAAATGAGACCAAGTTAAATGGCACATTTATGTATTATGCCTA

>M6-PGSC0003DMG400018901 M6 LRR-RLP

ATGTCTGCATCAGTGGCTAAAGGTTTTCTGAATCTCCGGAGACTAGACATTGAATACTGCCAATCAATGAAAGAAGTGATCACAGAAGAAGAACAACAAGGAGAAGGAATCATGACCTTATTTCCCCTGTTGGAAGAGCTGAAGCTTGATAAACTGCCTAACTTGGGGCATTTCTTTCTGACGGAGCATACTCTTCAAGCTCTTCAAATTCCATTTCTTAGAGAAGTGAAGATATCTGACTGCCCTGAAATGAAGACATTTGTCCAACAGGGAATATCTGTGAGTCTTGAAAGTGATGATGAGGTGAAAGTAATGTTCAATTCTAAGGTTTGTCTAGTAAATAGTAATAGTTATTTCTGTTCCACTTCCTGTTACCTTGACGTAAGATGATATATGGTTCATTTTTGGATTTCTACTTGAAGCTATATATATTCAATTTTCATACCTTCAACATTTGCAGGTTTCTTGTCCCAACTTGGAAGAGCTATATATCTTGGGGGCTAACAACATAAATGCTCTATGCTCTTACCAACTTCCAATTGCCTGCTTCAGCAAACTTGAAATATTACAAGTATATAATTGCGAAAAATTAAGAAACTTGATGTCTCAATCAGTGGCTAAAGGTTTTCTGAATCTCCAAAAACTAAAAATTGAAAATTGCCGATCAATGGAAGAAGTGATCACAGAAGAAGAACAAGGAGAAGGAATCATGACCTTATTTCCCCTGTTGGAAGAGCTGAAGCTTGATAGACTGCCTAACTTGGGGCATTTCTTTCTGACGGAGCATGTTCTTCAA

>M6-PGSC0003DMG400018994 M6 LRR-RLP

AGGATAAAAATTGTTTGGCTTTAATTTCAATCATGGATACTTCATGATAGTTGAAAAATCATTTCTATTTTATTATGACTTTTTGAGATAATAACTAACTATTAGTTGAGTGAACATATGAGAAATCAAAGTCGTGATTGTCTCTAGTCTCTCCAACTAAGGTATGCAGAGGACCACAATTAACGCTAATTTTTGCCTTCCATTTTCGGCTACTTAATTAGCACTATTATTCCACAGGACTTGACTTGACTTGACCTTCAAAATATTTGTCCTCATTTCAATTACGCAAACTACAAGTTACTTGTAAGACAGTAGTGAAGCTGAATCATAAGGAGAAATATATGGTCCTCTGTTATGGGTTATCTTTCTATATAAACAGGCCCAAATAACTAATGCTAGCAATATACAGTGCTTTAAACTATACCCATCGATCTCTATTAGTATAAGATCAACTAAAATGGCATGTTGGAAAATTTTAGTTGCTCTACTTGTTTTATTTCTTCTTTCCTTTCCTCTACCTGTTTCAGTCAAAGCCATAAAAGCTGTATCTTTACCTCTCACAAACACTACTGAAGCCAAAGTGAGATGTGTTAAAAGTGAGAGGAATGCTCTATTAAGATTCAAACAAGGTTTCACTGATCCCTCTGATCGACTGGCTTCTTGGGTTGGCAAAGAATGCTGCATCTGGAAAGGCATTTTTTGTGACGCTGGTTATAATGTCATCAGGCTTGATCTCAATGATAGAACAAACAATTGCTATTGGAAGAAAGTGGGATACTTGATTCCAGGCAATTCAACATGCTTGGGTGGCAACATAAGTTTTGATTTACTCGATTTGCATCACTTGGAGTATTTAGACCTTGGTGGTAATGACTTCCAAGGACTTACCACTCCCAATTTCTTGTCTTCTCTAAAGAAATTACAGTATCTTAACCTTTCTTTTTCATCTCTTATTGGTGTTCCTCCTTCACTTGGAAATCTATCAAATTTGCAGTATCTAAACTTCCGTGCGTGTTCCTATCCTTTCGATATTGTTGAAAGAAGTTGGGTCAAAGATTTTAATTGGGTTTCCGGTCTGTCTTCTTTGAAGTATCTTTATCTTTCTTATGTGAACCTTTCTTCGGCAACACATTGGCTTGATTCTATTATTAAGCTGCCATCTTTAGTGACTCTCAGTTTGCAAAACTGTAGTCTTCACTATCTTCCTTATTCCTTCCCAAACTGGAATATGACTTCTCTTTCTTATCTCTCCCTTTCACAGAATAAGTTTGTTAATTCTGAGTTGCCTAAATGGTTGTCTAATGCCACCACCATTGAGACACTTTATATGGTATACATCAACATCCAAGGCCCAATATCAAATGTTGAATGGGGTAAACTCTGTAATTTGCAGAAGTTGTATCTAGTGTAGAACAAACTTAATGGAGATATAAGTCGAGTGGTGGAAGGATTGTCCAACTGCAGCAACCCTACGATAGAGGTATTATCTCTTGGCGAAAATGTGTTGACAGGGGAATTGCCAAATTCATTAGGACATCTGAAGAACTTAAGGATTCTTAGTATTTCTTTTAATATGATCTCTGGTACTATTCCAGCATCTATTGAGCAACTTTCACAATTGGAAATTTTGTACTTGGGAGAGAATCAACTGAAGGGAGCCTTGCCTGAGAGTATTTTTAATTTTTCAGAATTAACTGAGTTGCGGCTGACAACAAATGCTTTGGAAAGCAATCTTTCACAAAATCGCTTTGAAAGACTCCATCAACTGAAAGTTTTATCCCTATCTTGTGGAAGAAGTTTTTCAGTCAACTTGAGCAATGAATGGGTTCCTCCATTTAGCCTCTATGCTATCGAGCTCAGAAGGTGCAGTTTGGGTCCCAAGTTTCCAACATGGCTAAAGATTCAAAAGCAACTTCAGTGGGTCATTCTCACCAATGGCTCCATTTCAGAACCTATACCTCCTTGGCTTTGGATGATGTGCTCACAATTTCATTTGCTCGATCTATCAAATAATTAAATTGGTGGAAGTCTACCGAGGATAGTAGATTTTCCATCGACATACCAAATCTGGACGGTGGGTTTCTATTTTAGTTACTATTATGGTGTCGTGGTGGACTTGAGTTCCAATCACTTCCATGGTTCGTTACCTCTTTGGCCAACTGTGACACATTTGAATCTTGCTAACAACTGGTTTTCAGGATACATACCCTTAAACATTGGTCATGTGATGACAAAACTTCAAGTGTTAGATCTTTCTGGAAATGCTTTTATAGGAACTATACCATATTCCATAACAAGAGTCAAGCAGCTATTGAGATTGGACCTTTCAGACAATCATTTGTCTGGGAAGATTCCTGATTGGTGGTATGATTTGCAACAACTTCAAGTCATAGATTTGTCAGGGAATAATCTTTCAGGTACTATTCCTCCTTCAGTTTGTTCACCACTTTCACTTTTTTGGTTGAGATTATGCCGCAATAACCTTTCTGGAGGGCTTCCAAAATCATTGAGGAATTGCAATAGCTTGCTTGCCCTTGATATTGGAGAGAACAAAATCACTGGCACCATACCGGAGTGGTTCGGAGAAAGTCTCTTATCTCTGCAAAAGCTCAGAATGACAGGTAACATGATTCATGGACGCATTCCACCACAACTATGCCAACTTTCTAACCTCCAAATTCTCCATCTTTCACATAACAATCTGACAGGTTCTATCCCTTCTTGCCTGGGAACCTTGAGAGGTCTCAAGTTGGTGAAATTTTACAAGTGGTTTCCCAACTACCTTTATTTCAGTTATGTTTTTACACCCAAGATGGAGTTGGTTGAGAAAGGAACAAAGCAGACTTACACTTTTACCTTAGATCAAGTGAATCTTATTGATCTTTCCTGCAACAATCTTCAGGGTGAAATCCCAAATGAGATAATAGCTCTGTCAGCCTTGAGCACTCTGAATTTATCTTGGAACCAATTATCAGGAAGAATTCCAGAGGACATAGGGAGCATGCAACAATTGGAGACACTAGACCTTTCATCTAACCATCTCTCAGGTCCTATCCCATTAAGCATGGCATCCATCACTTCATTGAGTTATCTTAACTTGTCACATAACAACCTTCATGGCCCTATCCCATCAACAAACCAATTTGGAACCTTTACTGATCCTTCGTGTTTCGAAGGTAATCCAGAACTCTGTGGGAAACCATTGACCACCAATTGCTCCTTACCCAGGAAAAGAGAAATAGAAGAAGAAGATGATGATGAAATAAGGCTTTGTTTTTTTGTGAGCGCGGGAGTGGGATTCGTTGCCATGTTTTTGGCGACATTTATAAGCTTGACGATAAAGAAGTCGTGGGGATACTGCTGCTTCCATTTCATGGAGGAAGTTGGAGAAAGAATTATCAGGTGCTGTATGGGTTTCATGAATAGGATTTTCAGCGTCACAAGGAACTAATTTCCATTGCATTATTGTTCTTTCTGTTTTGATACTGTTGCAGTGAAAATATGGCAATCTTTGAAATT

>M6-PGSC0003DMG400019071 M6 LRR-RLP

TAGGAGTAATTTCTTTTCAAGGGGATTGGGAAATACTTCTTTGATGGAATTGAGACTAGATGGTAATAAACTTAATGGAACTCTTCCGAAAAGTCTTGGACAACTTTCAATGCTAGAATACTTTTTCATTTCTAACAATAAGTTGGAAGGTGTTGTAACAGAGAGTCATTTCTCTAAATTAACGCAATTACAACTCTTCTCTGCATCTAGAAATAATCTGACTTTAAAAGTGAGTAGGAATTGGATTCCACCTTTTCAAGTCACAAACATTGCGATAGGCGGGTGGAATATAGGCCCTTTGTTTCCAATGTGGCTCCGAACTCAGGAGATGATAATGAATGTTGACATATCAGATTGTGGAATACAAGGTGAGGTTCCAACTTGGTTTTGGAACTTATCTTCTCAAATTGTTTTTCTCAATCTTTCTCACAATCAATTTATTGGTGAGGTTCCAATCATCTCAACAGCTTCGTGGTTAGTTGGATTGGGAGGTCCTTGGTTAATGTATTTAGGTTCCAACAATTTTAGTGGATCACTACCACGTATTTCAACGAGTATAACTGAGTTAGGTCTCTCGAACAATTTTTTTTCAAAAGGTTTATCCAACTTCTTGTGTGAAGCAAAGAATGTATCTTATAATTTGGAGATCTTAAATCTCGGGGGAAATGATTTATCAGAAGAAATTCCTGATTGTTGGATGAATTGGCCTGAGTTGGAAGTTTTAATCTTGAGGGACAATAATTTGATTGGAAGCTTACCAAGATCTATGGAGGTTTTGAGTAATTTGCTATCATTAGACTTACGAAGAAATAGACTTAACGGTCCGTTTCCTTCATCCTTGAAAAATTGTACGAAACTGCATAAAATAGATTTAGCCGAGAATGAGTTTGCTGGGGGACTACCATCTTAGTTGGGAATGAGGTTTTCAACCTTGATAGTATTTATCCTTCGGTCCAATAAATTCAATGGTGAATTGCCTCAAGAACTTTGTCAACTCAAAGATCTTCAGATCTTAGACCTTGCAAACAATACATTTGTTGGAATCATACCAAGGTGTATTGGTAATTTCAGTGCAATGGTCAAAGGAAAAAAGGAAGTGGGAGATGATGATGAGTTAAATTATTCCTACTTTTTGGGAGTTTTGAGAGAGAGTGC

>M6-PGSC0003DMG400019072 M6 LRR-RLP

ATGAAAGTGTTAGCAATAACACTATGGGTTCTCTTGTTTTTTATTTTGAAAAATACCAAGTTTGTCATTTGTGTAGGGATTTGTAGAGAAAATGAGCAACGAGCATTGGAGAGCTTGAAGAAAGAAGTAGATGATCCCTTGGATATTCCCTCTTCTTGGGTTGTCGGAAAAGATTGTTGTGAATGGGATGGAGTTGTGTGCAACAATCTAACTCGTCGTGTGATTGAACTACGCATAAGTTATCTAAGGATCGATAACCTAGAGTGGTTGTCAAGTCTTTTGAGTCTTGAGAACCTGGAAATGTATCATGTGGATCTCCACAAAGCAACTAACTGGCTACAGGTTATTAACATGCTTCCTTCTTTGGTTGATCTAACGTTATCCAATTGTAGTCTTCATAATATAACAAATCTACTTCATCACAATCTTTCTTCACTTGAAACCCTCGATCTTTCTAGTACTAAGTTTAATTCACCTATTCCCAATTGGGTTTTCAACCTTTCCAATCTTGTTTCTCTTAATTTGAGTGATTGTAATTTTACTGGCCCATTTCCCGACTTTCCTGTTAACTTGACTTCTCTCACTACCTTCAAGGCTTTTGAAAACTCTTTCAATTGTCTTTTACCCGAATGGTTGTTTGATTTAAGTAATCTAGAACATCTTAACCTCCGTTCTAGTGGTATTAAAGGTGCAATACCAAATAAGAGTGGAAACACAACAACACTTAAATATCTCGATCTTTTTGATAATAACCTTAACTCCACCATACCAAATTGGTTATATCAATGCAAAGATCTAGAATCACTTTATCTTACTGAAAACCGTCTTGAGGGAACAGTTTCAAGTCTGATTTCAAATTTGAGCTCAATAACTAATATTGACCTTTCTGGTAATATGCTTACTGGAAAGTTACCAAATGTAATTGGAATTTTAGGGAAATTAGAATTTCTTTCTCTTTCAAAAAATCAATTTGAGGGACAAGTATCTGAATTATTCAAAGGTAGGAGTAATTTCTTTTCAAGGGGATTGGGAAATACTTCTTTGATGGAATTGAGACTAGATGGTAATAAACTTAATGGAACTCTTCCGAAAAGTCTTGGACAACTTTCAATGCTAGAATACTTTTTCATTTCTAACAATAAGTTGGAAGGTGTTGTAACAGAGAGTCATTTCTCTAAATTAACGCAATTACAACTCTTCTCTGCATCTAGAAATAATCTGACTTTAAAAGTGAGTAGGAATTGGATTCCACCTTTTCAAGTCACAAACATTGCGATAGGCGGGTGGAATATAGGCCCTTTGTTTCCAATGTGGCTCCGAACTCAGGAGATGATAATGAATGTTGACATATCAGATTGTGGAATACAAGGTGAGGTTCCAACTTGGTTTTGGAACTTATCTTCTCAAATTGTTTTTCTCAATCTTTCTCACAATCAATTTATTGGTGAGGTTCCAATCATCTCAACAGCTTCGTGGTTAGTTGGATTGGGAGGTCCTTGGTTAATGTATTTAGGTTCCAACAATTTTAGTGGATCACTACCACGTATTTCAACGAGTATAACTGAGTTAGGTCTCTCGAACAATTTTTTTTCAAAAGGTTTATCCAACTTCTTGTGTGAAGCAAAGAATGTATCTTATAATTTGGAGATCTTAAATCTCGGGGGAAATGATTTATCAGAAGAAATTCCTGATTGTTGGATGAATTGGCCTGAGTTGGAAGTTTTAATCTTGAGGGACAATAATTTGATTGGAAGCTTACCAAGATCTATGGAGGTTTTGAGTAATTTGCTATCATTAGACTTACGAAGAAATAGACTTAACGGTCCGTTTCCTTCATCCTTGAAAAATTGTACGAAACTGCATAAAATAGATTTAGCCGAGAATGAGTTTGCTGGGGGACTACCATCTTAGTTGGGAATGAGGTTTTCAACCTTGATAGTATTTATCCTTCGGTCCAATAAATTCAATGGTGAATTGCCTCAAGAACTTTGTCAACTCAAAGATCTTCAGATCTTAGACCTTGCAAACAATACATTTGTTGGAATCATACCAAGGTGTATTGGTAATTTCAGTGCAATGGTCAAAGGAAAAAAGGAAGTGGGAGATGATGATGAGTTAAATTATTCCTACTTTTTGGGAGTTTTGAGAGAGAGTGCCATAGTGACGAATAAAGGCAACAGGTACCAGTATGACACCATTTTGGCGTTGTTTACAAGTATGGACATGTCTAGCAATAATCTCTCTGGGGATATTCCTATAAGTTTAACACTTCTTGCAGGATTAAGATCATTTAATTTTTCTAAAAATAACCTAACTGGTAGGATTCCAAATGACATTGGTGACATGAAAGTTTTGGAATCTGTTGATCTTTCAGAAAATCAATTTTATGGTCAAATCCCACAAAGCTTTTCAAGTTTGTTCACCTTGAGCTACTTGAATCTATCTGATAACAATTTGTCAGGTATGATACCATTGAGCACTCAACTTCAAAGCTTTGATCCCACGAGTTTTCAAGGAAACAAACTCTGCGGGCTTCCACTCTCGCTGAATTGTAGTTCAAATGGTAATATTCCAAATCATGAATATGAAGATGATGAGAGTGACGAAGATGAAGTGGATTGGTTTTACATTTCAATGGCTATAGGATTTGCACTAAGCTTTTGGGGTGTTTGTGGTTCATTGCTTTTCAAGAGATCATGGAGACATGCTTATTTTTGTTTTCTAGACCGTAGCTGGGAAATGTTGCTTGCAAAGTTATCAGTATGTTGA

>M6-PGSC0003DMG400019900 M6 LRR-RLP

TTAAATAAAAAATTAGCAAATTATGTGGGCTGGGGCGGATTGAAAACAGCAAACTTTGCAAGGCAGGGCGGGTTGAAATCTTTGCAAGTTTGTGTGGGTTTGCCCTGCACCACCCCATCTGTCATTCCTGTCTAAATGTTTGTTTTGTCTATATAATTCTTGCTGACTCATTCTAATGAGCTCAATTGTAACAAATTCTTTGTGTCCACTCTACTTGGAACAGGTTTTGTTTTAACTGAACCAAAAGTCTACGGAAGGGATAAAGAGGAGGATGAGATAGTGAAAATCTTGATAAACAATTTTAATTTTGCCGAAGAACTTCCAGTCTTCCCTATAATTGGTATGGGGGGACTAGGAAAGACGACACTTGCCCAAATGATCTTCAATGATGAGAGAGTGACTAAGCATTTCAATCCCAAAATATGGGTTTGTGTCTCAGATGATTTTGATGAGAAGAGGTTAATTAAGACAATTTTAGGAAATATTGAAAGAAGTTCTCCTCATGTCGAGGACTTGGCTTCATTTCAGAAGAAGCTCCATGAGTTATTGAATGGAAATCGATACTTGCTTGTCTTAGATGATGTTTGGAATGATGATCTAGATAAGTGGGCTAAGTTAAGAGCAGTCTTAAATGTTGGAGCAAGAGGTGCTTCTATTCTAGCTACTACTCGTCTTGAAAAGGTTGGATCAATTATGGGAACGTCGCGACCATATCATTTGTCAAATTTGTCTCCACATGATAGTTTACTGTTATTTATGCAACGCGCATTTGGGCAACAAAGAGAAGCAAATCCTAATCTAGTGGCCATTGGAAAGGAGATTGTGAAGAAATGTGGTGGTGTGCCTTTAGCGGCCAAGACTCTTGGTGGTCTTTTACGCTTCAAGAGAGAAGAGAGTGAATGGGAACATGTGAGAGATAATGAGATTTGGAGTCTGCCTCAAGATGAAAGTTCTATTTTGCCTGCTCTAAGACTGAGTTATCATCACCTTCCATTTGATTTGAGACAATGCTTTGCGTATTGTGCAGTATTCCCAAAGGACACCAAAATGATAAAGGAAAATCTCATCACTCTCTGGATGGCCCATGGTTTTCTTTTATCAAAGGGAAACTTGGAGCTAGAGTATGTGGGTAATGAAGTATGGAATGAATTATACTTGAGGTCTTTCTTCCAAGAGATTGAAGTTAAATCCGGTAATACTTATTTCAAGATACACGATCTCATCCATGATTTGGCTACATTTCTGTTTTTGGCAAGCACATCAAGCAGCAATATCCGCGAAATAAATGTCAAAGATTATAAGCATATAATGTCCATTGGTTTCGCTGAAGTGGTGTCTTCTTACTCTCCTTCGCTCTTGAAAAAGTTTGTCTCGTTAAGGGTGCTTAATCTAAGCTACTCAAAACTTGAGCAATTACCCTCTTCCATTGGAGATTTATTACATTTAAGATACCTGGACCTGTCTCGCAATAATTTCCGAAGTCTTCCAGAGAGGTTGTGCAAGCTTCAAAATCTTCAGACTCTTGATGTTCATAATTGCTACTCACTTAATTGTTTGCCAAAACAAACAAGTAAACTTAGTAGTCTCCGAAATCTTGTTCTTGATGGCTGTCCATTGACTTCTACTCCACCAAGGATAGGATTGTTGACATGCCTTAAGACTCTAGGTTTCTTTATTGTGGGAAGCAAGAAAGGTTATCAACTTGGTGAACTACGAAATTTAAACCTCCGTGGTGCAATTTCAATCACACATCTTGAGAGAGTGAAAAACGATATGGAGGCAAAAGAAGCCAATTTATCTGCAAAAGCAAATCTGCAATCTTTAAGCATGAGTTGGGATAACGATGGACCAAACAGATATGAATCTGAAGAAGTTAAAGTGCTTGAAGCCCTCAAACCACACCCCAATCTGAAATATTTAGAGATCATTGCCTTCGGAGGATTCCGTTTTCCAAGCTGGATAAATCACTCAGTTTTGGAGAAGGTCATCTCTATTAGAATTAAAAGCTGCAAAAACTGCTTGTGCTTACCACCCTTTGGGGAGCTTCCTTGTCTAGAAAGTCTAGAGTTACAAAACGGGTCTGCGGAGGTGGAGTTTGTTGAAGAGGATGATGTTCATTCTAGATTTTCCACAAGAAGAAGGTTTCCATCCCTGAAAAATCTTCGTATATGGTTCTTTCGTAATTTGAAAGGTTTGGTGAAAGAGGAAGGAGAATATAAATTTCCCATGCTTGAAGAGATGGCGATTTTACATTGCCCTCTGTTTGTTTTTCCAACCCTTTCTTCTGTCAAGAAATTAGAAGTTCACGGCAACACAAAAGCAAGAGGTTTGAGCTCCATATCTAATCTTAGCACTCTTACTTCCCTCCGCATTGGTGCTAACTACAGAGCGACTTCTCTCCCAGAAGAGATGTTCACAAGTCTTACAAATCTCGAATACTTGAGCTTCTTTGACTTCAAGAATCTCAAAGAGCTGCCTACCAGCCTGACTAGTCTCAATGCTTTGAAGCGTCTCCAAATTGAAAGTTGTGACTCACTAGAGAGCCTCCCTGAACAAGGGCTGGAAGGTTTAACTTCACTCACACAGTTATTCGTTAAATACTGTAAGATGCTAGAATGTTTACCCGAGGGATTGCAGCACCTAACTGCCCTCACAAATTTTGGAGTAACCGGTTGTCCAGAAGTGGAAAAGCGCTGTGATAAGGAATTAGGAGAAGACTGGCACAAAATTGCTCACATTCCTAATTTGAGTATATATTAGTGAGTTATTTGTAATTGT

>M6-PGSC0003DMG400019970 M6 LRR-RLP

GTAGATGCTGCATCTTGTGCTTGCTCTGGCAATTCCCTTCACCAGATGCGTTTTCTGAGCATGGAGGAAAGCTGGACTCTTTTATGCGATAAGGTCTTTGGGAATGGTGGTTACCCTCCGGAATTGGAGAAAATTGGGAGGTACATTGGCCATCAGTGTCAAGGGTTACCACTTGCAGTTGTTGCAATTGGTGGACTACTTTCCAAGATGAGCAAGGAAACAAGTTCTTGGGAGAACGTTGCAGAAAAAGTAGGATCACTTATGACCAGTGACACTATGGATTGTTTGAACATACTTTCTTTGAGCTACAACCACTTGCCTCAGTACTTGAAAACATGCTTCCTTTATATGGGAGTTTTTGCAGAAACTCGTGAGATTCCGGTCTGGAAGTTAATAAAATTATGGATTGCTGAGGGCTTCGTCAAGAAAGTTAATCATAAAAATCTAGAAGATGTGGCAGAGGAAAATTTGAGGGAGTTGGTTGATAGAAGTTTGGTTTTGGTGGGGAAGCATACTTCCCTAGGAAAAATCAAGACTTATAAGATGCATGATCTCATTCGAGATATATGTTTGAGAGAAGCCCAATATGAGAATTTTATACACTTCAAAACAAGGTATGACTATGATGATCTTCTAGAAAACAATAGTTGTCTACGTCGTGTAGCAGATTTCCATAAGTATACAAACAGTAGACTTTCTAAACTTAAACCACTCACACGTTCAGTTTTCTTCAAATATCCATACTTTAATTGATACCTCTCACAGTCGAGCTTTAGATTACTTAGGATATTAGATGTTTGCTTTACTACTTCTTTGTTCTGTGAACAGTCCATATCAGAACTTGTTCATTTGAGGTATCTTGCTTGTCTGTTCTTCGATGGTTTTGTGCAATCACTATGCAAGCTACAGAATATACAGAACTTAGTTATTCATGATTTAGATATGAGAACCCAGTTCCTACCATGGGAAGTTGTGAATATGCCTCAACTAAGACATATCCATACAAAAAACTTTAATCTTTTCATACCCCCACCTACATCCCTAATTGCTGAGAGGGAGAATCACTTACAAACATTAACCGGATTGATTTCCTCATCTTGTAATGAAGAGGTGTTTCTAAGAATTCCAAATTTGAAGAAGTTGGGAATTTTAATTGTTGTTGATTCAGACACTATTCAGAAGTGTTATTGTCTTGATAATCTTGTACATTTAACTCAGCTTGAGAAGCTCAAAGTTAAGTTCTGCGATTATCTGCTAGGGTTAAGTTGGTTCTTAGAAGCTCAAAGCTTAGGTTGGGTGGACCTTTCTTGGCTACACATTCCACATTGTGACAATTTCCCACCAAACCTCAAGAAGTTAACATTGTGCAGAACAAACCTACAATGGGAAGACATGAACATTCTCAGAAAATTACCTAATCTTGAGGTGCTCAAACTTAAACGTCGTGCCTTTGACGGACTAATTTGGAAACTAAGCGATGAAGACGAAGATGGATTCTTGAAGCTAAAGTTTTTACTTCTTGAGAACATGAATCTGATGCAATGGGAAGCAACAAGTTATCATTTTCCAAGCCTTGAGCACTTAGTTTTAACAGATTGCGGGTACTTGAAACAAATTCCTTTCGACTTTGCGGAGATTCAGACACTACAGCTGATTGAACTAGACAAATGTATGCGTTCTGTTCTTGTTTCAGCAGAGCAAATACAAGCGGAGCAACAAAGCTTGGGAAATGATGACCTCGTTATTCATGCGAACTATATACGTGAGTAAACATTAATAAAAAAGGTATAGACTTCTTAAAAGCATAAAGAATTTTTATTAGATAGATTCGTTCGAAATCTGAAAATGTTTTTTATATTCATCATTTTCACTTATTTATACTATAAAAGGTGCTAATTACAATTGGTTTGTGCATTGCAGACGTAACGACTAGCTGATGGAGAGTACCATATACTCAAGCAAATTTGCAGAAGTTCATGATCTTTCCTTTCTCTGTATTTCAATTTCCTTTTTTGCAAATTCAGCGAAGTTAGTTAGTGAAATAGTCATGTTTCAACTTGTTTAGGACTGAGACGTAGTTTAGTGAATCTGATATCTATATACATTCATCAGAAAGAATGCATTACCTTTGGAACGTGTATATAGTTGAATAACTTCTCTTTTTATTGAAATTGATTCTATAAATCAGTTCTCCTCAACAAAACCTTTTCGTCCATACCAAAC

>M6-PGSC0003DMG400020178 M6 LRR-RLP

AAGAAAGTGACCGTAGAACTATCTGCTACTCACTCAATTCATCCATCTTCTTCTTCTTCTCTCCTAAACCATTAATTTGTTTCAGCCATAACCAAAAAAGCTTTTGCTTTCTTGTGAAGATACCTTTTTTTTTTGCCTTTGCCTTTGCTTTTGGTTTCCATTAAAGACCCCATTTGATGACCTGTTGCTTCTCACGGAAAAGAAAACCAAAAAAGTTCCATTTTTTTTTTACCTTATATAAGCCAGCCTTTTCTATCATTCTTCTCTTCAGCTATACGGTAATTATGATACCATTCTTTGTATTCTTTTTTCTTCTTCTGCAGAATAGTTTCGCCATTTTAGACCCAATTGATTTCTTAGCTTTACAAGCGATTCGAAAAGGCTTGGATGATTTGCCCGGTTCCAACTATTTTGCTTCCTGGGATTTCACTTCTGAGCCGTGCAACTTCGCCGGAGTTTACTGCGATGGAGATAAAGTGATCGCCTTGAATCTCGGTGATCCCAGAGCTGGATCACCGGGTTTGACCGGAAACTTGGATCCTTCTATTGGGAAACTCTCTGCTCTTGCTGAGTTCACGGTGGTTCCGGGTCGGATAACCGGTGCATTGCCGGAAAGTTTTTCTCAGTTGAAGAACCTGAGATTTCTAGGAGTTAGCCGGAACTTTCTCTCCGGTGATATTCCGGCGTCATTAGGCCTGCTCCGGGAGCTACAAACTCTTGATCTTAGTTTCAATCAGCTGACTGGGAGTATCCCTTGGGCTATTGGAGCTCTTCCGGCATTGTCCAACATTATACTCTGTCACAATCATTTGTCCGGTTCGATCCCACCATTCGTATCCCAAAGATTGACCCGTTTAGACCTTAAACACAACTCACTTTCAGGTACCCTTTTGCCTAATTCACTTCCTTCATCGCTTCAATACTTGTCATTGTCATGGAACCAGTTAACCGGTCCAGTAGATTATCAGTTGAGTCGGCTGAACCAGTTGAATTACCTCGATCTTAGCTTGAACCGGTTCACCGGCTGCATTCCGGGAATCTTATTTACTTTCCCACTCACAAATCTTCAGCTACAAAGGAATCTATTTTCGGGTCCGGTTTTACCCATTTCAGAGGTGACAATTCCGACGGTGGACATTAGCTTTAACAGGTTTTTCGGGGAGATATCTCCATTGTTGTGTAATGTGGAGAATTTGTATCTGAACAATAACCGGTTCACGGGTAAGGTGCCGGCGGTGTTGGTGGACCGGTTATTATCTGCCGGAATACAAGTTTTATATTTACAGCATAATTTTCTGACGGGGATTGAGATAAATCCGAGGGTGGAAATTCCGGTGAGCAGTTCATTGTGTTTGCAGTATAATTGCATGATTCCGCCGGTTGAGACGGCTTGTCCGTTGAAAGCCGGGAAGCAGAAGAGTAGGCCTACAAATCAGTGTGTGGAGTGGAAGGGGCAAAAGAGTGAAAAAACAAACAATAATAATATTACTACTAATATGGCTTAATTTTTCTAGATTCTTTCATTTCATTTTCCTTCCTGATTATGGGCTGATGGGCTAAGTAATAGTAGTAAAAAAGAGTAAATGTGCATAATAAAGGAGAGAAAAATAGAGTTAAAAGGTGTACGAGTCATCCTCCTATTGTTTTTCTTTTGTATTAAGCCAAAACAATTTTGGCATTTCTTTTTGTAATTAGGTACTACTAGTATAATATATTCAATTACTTGGATTTACGGATCTCAAGTTGAA

>M6-PGSC0003DMG400020330 M6 LRR-RLP

ATGACAACTCTGTACTTTTTGTGGCTTTTCTTGATTCCCTTGTTTCAAATCTTATCAGGATATCACATTTTCTTGGTCTCCTCTCAATGTCATCACGATCAAAAGTCACAGTTGCTGCAGTTGAAGAGCAGCCTTCAGTATGATTCTACTTTGTCAAATATTTGGCAAGATGGAACCATAACACAAGTGAATGTTGCAATTGGGATGGGGTTACATGTAATCTCTCTGGTCATATGATTGCCTTGGAACTGGATAATCAGACTATTTCTAGTGGAATTGAGAATTCAAGTGCTCTTTTCAGTCTTCAGTATCTTGAGAGGCTAAATTTGGCTTACAACAAGTTCAATGTTGGCATACCACTTGGTATAGACAACCTCACGAATTTGAAGTACCTGAGTTTATCCAATGCTGGTTTTGTCGGGCAAATTCCTATGATGTTATCAAGATTAACAAAGCTAGTTACTCTCGATCTCTCAACTCTTTTCCCTGACTTGATAAGCCACTAAAACTTGAGAATCCCACTTTGAGTCATTTCATTGAGAACTCAACAGAGCTTAGAGAGCTTTACCTTGATGGTGTTGATCTTTCAGCTCAGGGGACGGAGTGGTGTCAATCTTTATCTTCATATTTGCCTAACTTGACCGTCTTGAGCTTGCGTGATTGTCGAATTTCAGGCCATATTCATGAATCACTTTCTAAGCTTAACTTTCTCTCTTTCATCAATCTTGGCCGGAACAATCTCTCTACCACAGTTCCTGAATATTTTGCCAATTTCTCGAGCATGACTACCTTGAACCTTGCCTCTTGTAATTTGCAAGGAACATTTCCTGAAAGAATATTTCAGGTATCAGTTTTAGAGAGTTTGAACTTGTCAACTAACGAGTTGCTTCATGGTAGTATTCCGATTTTTCTTCGAAATAGATCTTTCAGGAGGATATCACTAAGAAACACCAACTTTTCGGGTTCATTACCAGAGTCCATTTCGAACCTTCAGAACCTATCCAGGTTAGAGCTATCCAATTGCAATTTCACTGGACCAGTTCCTTCCACAATGGGAAACCTTATCAATCTTGTTCATTTGGACCTCTCTTTGAACAACTTCACTGGTTCTATCCCATATTTCCAAAGATCCAAGAAACTCGCCCACTTAGACCTCTCTCGTAATGGTTTAACTGGTCTCCTCTCCCCGACTCATTTTGAAGGACTCTCAGAGCATGTCTACATAAATTTAGGGAACAACTTACTCAACGGGATACTTCCTGAATGTATCTTTGAGCTCCCCTCCTTGCAAACGCTTTTACTTAACAGCAATCAATTTGTTGGCCAAGTCAACGATTTTCACAATGCATCCTCCTCTCTCAATGAAATTGATTTGAGCAACAACCAACTGAATGGTTCAATTCCCAAGTCCATGTTTGAAGTTGGGAGGCTTAAGGTACTCTCACTTTCTTCCAACTTCTTTAGCGGGACAGTGACCCTTGATCTCATTAGGAGGCTGAGCAACCTTTCAAGACTGGAGCTTTCTTACAATAACTTGACTGTTGATGCAAGTAGCAGTAATTCAACCTCTTTCACATTTCCCCAGTTGAACATATTGAAACTAGCTTCTTGTCGGCTGCAAAAGTTTCCTGATCTTAAAAATCAGTCAAGGATGATCCAGTTAGACCTTTCTGACAACAAAATACAGGGGGCAATACCAAATTGGATTTGGGGAATTGGTGACGGAGCTCTGACTCACCTGAATCTTTCCTTCAATCAGCTGGAGTACGTGGAACAGCCTTACAATGTTTCCAGATATCTTGTCGTCCTTGACTTGCATTCCAATAAGCTAAAAGGTGACCTACCAATTCCACCTTCCTTTGCTGCATATTTGGACTACTCGAGCAATAATTTCAGCAATTCCATCCCACTAGATATTGGCAATTATCTTGGTTTTGCCTCCTTTTTCTCGGTAGAAAACAATGGCATTACTGGAAGAATTCCCGAATCCATATGCAATGTCAGCTACCTTCAAGTTCTTGATTTCTCTAACAATTCCTCGAGTGGAACAATACCACCATGTCTACTGGAATATAGTACAAATCTTGGAGTGCTGAATCTAGGGAACAATAAACTCAATGGCGTTATACCAGATTCATTTCCAATTGGTTGTGCTCTACAAACTTTAGACCTCAGTGAGAATAAGTTACAAGGTAGGCTGCCAAAATCGCTTGTGAACTGCGAGTTGTTGGAGGTCCTGAATGTTGGAAATAACAAACTTGTTGATCATTTCCCATGCATGTTGAGGAACTCAAAGAGTCTGAGGGTCCTTGTCTTGCGCTCCAATCAATTCAATGGAAATCTTACGTGTAATGTAACCATAAATAGCTGGCAGAATCTCCAGATCATAGATATAGCTTCCAACAATTTTACTGGTATGTTGAATGCAGAATGCTTTATAACTTGGAGAGGAATGATGGTTGCAGATGACTATGTAGAGACAGGACGTAATCGCATCCAGTATAAGTTCCTCCAACTAAGTAACTTATACTATCAGGACACAGTGACAATAACCAACAAAGGGATGGAGATGAAGCTTGTGAAAATTCTCAGTCTACACAGCTATTGATTTCTCTTCAAATAGATTTCAAGGCATGATACCAGATACTGTTGGGGATGTTAGCTCCCTTTATGTTCTGAACCAGTCACACAATGCCCTTGAAGGACCAATTCCAAAATCAATTGGGATGCTACAAATGCTTGAATCACTAGACCTGTCAACAAACCACCTGTCCGGAGAGATCCCCTCAGAGCTTGCAAGTCTCACATTCTTAGCAGCTTTGAACTTATCGTTCAACAAATTGTTTGGAAAAATCCCATCGATTAATCAATTTCTAACATTCTCAGCAGATTCCTTTGAAGGAAACAGAGGCCTATGCGGGTTCCCTCTCAACAACAATTGTGAAAGCAATGGCTCAGAGTCCCTCTCCCTGCTGCCACCAACTTCGGTACCAGACTCAGACTCAGATTATGAATGGAAGTTCATTTTTGCGGCTGTTGGATACATAG

>M6-PGSC0003DMG400020520 M6 LRR-RLP

GGCTACTCCAATAACAAAAACCCCATCACCTGCCTGTACAAAAATCATAGAAGAAATCACAAGAATTTACAAATCACTCCCACCAAGACCATCCATTTTAGAAATTGAAGCTTCAATTTCTGTAATAAAATCAGTAGAAACAGAAGAAAAAATAGAACTTGATGAAATTTCCAAGAAAGTAGTGGAGATAGAAGAAAATGTTCCAGCTGAGCTTTTCTCTGTGTTACAACAAGTGAGAAAAGCTATGGTTTTGTTTCAAAGCAAAGAACAGAGGAAAGAGGCTGTTCAGTTAATTGAACTTGATAAGACTTATCAAGATTTTGATGTGTTGATTCAGGAAGCTACACAGTTGATTTCTGGAGATACCCAAATGGGAAAAATCAATTCTTTTGAGGACCCAATTGGTGAAATTGGAAAAAAAGATGAGATTTTGATGAAGGGTTTGGTTACAAGTTGTGAATTGATTACTGTTTCTTCTTCAGGTTTGGGTATTTGCTTCTCCTTTTTGCTTTTTTTGTGTCCAAGATTTCAACTTTTTGCTTGCTTGTGAGTTGTGGAGGTAGTGTGTTGTCTTTCTATCTGTCCATACTAATAGTTGTAGTATTGCTTATGTTACCATTTGTTGTTCCTTGTACTTCGATTATTTTATTATTTTGTTGTAGCTAATGTTCAGAATGCTTTGTTATGCTTTCACTTCCGTTTATTCTTTTTGTGTGTGCATCCAAGATTTCAACTTTTTGCTTGCTTGTGAGTTGTGGAGGTTGTGGAGAGCATGGATTAGGCTAGAGGTTAGTAGGTACTAGTAGTCCTAGTATTGCTTATCTGGTTTCTTGTCCTTCTGTTACTTTGTGTTGTTGTAGTATGTAGTTTGATTATCGCATTATTTTATTGGAAATCTTAGAATAAAATATGTATCCAAACACAACTTCAAAAATCGTTCAAGTTTCTAACTTCAGAATCTATGGCAAAACGCTAGCTTAGATTGATGGAAAAACATCATTTATAGTTTCTCCTTTTGGTCTTTCTTGGAACTTGAGCTCTAGTTTGCAAGATTCTTTTTCCTTTTTAGTGATGGCACTAGCTTGCTACGTATGTTGACTAATCTACTAGGTTTCCAGTTATAGGTCAGTATGTGTGGCAGTGGCGGAGCTATTGATAACCCTTTGTTTGGGATTGGGGCGTAGTTGTTGTTGGCCGTCAATGTTTAGATTGTGGAAAACATCATTTAGAGTTTTTCTTTCGGGTGTCTCTCTTGGAACTTAAATCGTAGTTAGCATTATTCGTTTTCCTTTTTAGTGATGGCACTAGCTTGCTATGTACGTCAACTAATCTACTAGGTTTCCAGCATTCGTTCAGTATGTGTGGATCCCTTCATCGGAAAATTATAATGTGCAAATAGGTCAATTTCTTTTTTATGTATAAATACTATATATGTTGTTTGGGACAGAGACATAGTTGTTGTTGTTGTATAAACTATATGTTTAATCCTGTAGCCTTCTTCGTGTGTTACTCTTTTCTATTTTAAATCCTCTTAGTAAAAGTCCTGGCTCGTTACTGATATGGCATTCAACTATACTATATCTTAATTGTTGTGTAATTGCCTTTAGGGATCAAGCACAAAGAGAAATACAGTTTAATGAAAGTAGCTGCCCTAATTGAAAATGCTGCCAAAACTCGAGCAAGAGTTGTCGATCTTCACAACAAGTTAATGGACAAGATTGAATGGCTTCCGTTGTCACTTGGGAAATTGGTGAATGTCACTGAGCTAAACGTTGCCGATAACCAAATTATGGCTCTTCCAACTACAATTGGTAGCCTTAATGCCTTAACAAAGCTCGATCTTCACTCCAACCAAATTATAAACCTTCCTGATTCCTTTGGTGAGCTAATCAATTTGACTGATCTCGACCTTCACGCGAACAGGTTGAAATCGCTGCCAGCTTCTTTCAGGAACTTAGTTAATCTGATTGATCTTGATTTGGGTTCGAATAGGTTCGCTCATTTGCCTGATTTCGTTGGGAACTTAACCTCCTTGAAGAGACTAAACATGGAAACAAATCAGCTCGAGGAACTTCCTTACACAATAGGATTTTGTTCTTCCCTTGTAGAACTGAGATTGGATTTTAATCAGCTTAAAGCTCTCCCCGAGGCAATGGGAATGCTCGAACACTTGGAGATTCTTACACTGCACATCAACAGAGTTAAAGGATTACCAACGACAATGGGGAATCTTTCTCACCTCAGGGAACTTGATGTTAGCTTCAACGAAGTTGAGAACATACCCGAAACATTCTGTTTTGCTGTCAGCTTGGAAAAGCTTAATCTTGCAAACAACTTTGCAGACTTGAAAACTTTACCAAGATCAATCGGGAACCTTGAGAATCTTGAAGAGCTTGATATTAGCAACAGTCAAATAAGAACGTTGCCCGACTCCTTCAGGCTCTTGTCAAAGTTGAAAACTTTTCGAGCTGATGAGACTCCACTAGAAGTACCACCAAGGCAAATTATAAAGCTAGGGGCTCAGGTGATTTTTATGCTTACTGTGTTTCCTGTCAATCAATCTTTAGACTTTATGGTAGTTTTTAGAGTTGATAGTTAAAAACATACTTGAACTATCGCCTTTTCCCGAGTTTCACACCCTACACTATCAACTCTTCCCTTTTCCTACATGAACTATCAATATCTATGTATTAAAGCACACCTCAACTATCACGTAGGAAAAGGAAAAAAAACAGACGGCTGAGCCGTTGAAGTGTGTTTTAATACATATATGGTGATAGTTCAAGCAGGAAAAGGGAAAAAGGGAACATCTGATAGTTGAGGTGTGAAACTCGTAAAAAACAATTGTTTGACCATAATCTCTACATTTTCTAACTTGTTTCGTTTTCTTCTTTTTTCTTGCCATCAGGTTGTTGTAGAGTACATGGCTGAATTCGTCGCTAAGAACGAGCTTCAATTGCAGCGGCCAAAGAGAAGGAGAGCTTTCTTTTCTCTGAGTTGCCTATTTCCTAACACCGAGAGGAAGAGGCGGATCTAGAGCAGGTACTGTGGGTTGAACTGAACCTAATGCTCGCGGATCGAACTACTTGTACATCCTGCAAAAAACATTTGAAAACATATACATATTATAGGATCTCACTCATTCTGAATCCACTAACTCTAGATCCTGGATTCGCCTTGGTGAGAGGTGTCGGATACCTTCAGAATCCGTCAAAGCAATTATATATATATATATATGAAAAGTTGCTCCAAAC

>M6-PGSC0003DMG400020546 M6 LRR-RLP

TTTACGTTTCACCAAGGAATGATAAATAATTTCGTCAAAAGGAATAATTGTTGTTTGAATTCAGAGATCGGCGGCGAGTAAAGAGAAATGGAGAAAGAGGGAAAAAGCGACGCCGATAACCGGAGTACAATCACAGTGAATGTTAAGTTCAGTGGTCGATCGATACCGGTGGAAATCTCCGACGAGTGCACTGTCAAACACCTCAAGTCTCTCCTTCAACCACTCACAAATGTCCTCTCTCGCGGCCAGAAACTCATCTTCAAAGGTATTTTTTCTTTCAACTCTAACCTATCAGCATTATTTCGTATACAATCGGAGCTCCAAAGAACTGAATATCAATGAATTTGATAGATGACATAATTTTTTTCATGAATAGGGAAAGTTTTGGTGGATGAAATAACATTGAAGTCATCGGAGGTTGGAAATGGTGCGAAGATCATGCTTATGGCTTCCCAGGGCTTACATCAAGGGGTAATTCTTTATTTTCCTTTACATTATACTCTATTGAGCATAACTATTTGGGATTAGTACTTATATGTGTTACTATACTTAGTTTGTTTATGGATTTGTACATAACTAGAAAATGGAGAGAACCTCTAGTATTGATAAGCTTCTCAGTTGTGTTAAAGGATAACATATTGATTACTAGAATGAATTAAGGCTACGCAGAGTAGAATGTATATTTAGATATAGGAGACTACAACTGATTTGGATTGAGGTGTAATTGATTGAGCCGTGCTTGTGACTTTCAAATTTTTGTAAAATTAAAAAACCAGAAATTCCACTGAATTTATCCCAGTTAGGTCCTGAACTTCTGCTTTCTTAAAGCTTATTTGTGTATATTGGATAGGATGGACCAATAAGAAAGGAAGCTTCCAACTTGCCCACAGCGAGGAGGATGCCTGAAGCTATGAGACCTAAAAGGGATGTTCCTCAAGTTCCTGTTGTGAAGAGCCAACTTGAACGGTGGAAGGCCACTGGGGTTGTTGCATTATCAGAATGCAATCTGAAGGTAAGACGAGAAGTAACTGACTTTGCTTTGTCCAATGATTTAGAACTTGTTCTTACTGTGAACAATCGCTATTGACGGGACAAGAGATACAATTGTTGAACTGGAACTCACTTTGATCCCAATCATATGCATATTGGAGTTGACCTAAAAATAGAATTGTTTGGTCTTTTCCTTTTTCCTTCTAATTCATTTTTTTAATTTTATATCTAATGCATTTTCTGGCTCGAGTAGGTGAGATAGCCAAGCTATTTCCTTCTCTTTCGTATACAATATGGGAAAAACCACTAAAGAAAAGAATCCATAAATTTATAAAAGTGGTCTAAGGGATTTGAACCTTTGATTGTTCTTCATTCTGGCTATAATATCAGTTTCAGTAATGGAGCTGTTAACAACTCATCCATCTGTATGGAGACTAATTTTGCAGTTTTACTCCAAATAGAACATGGCCTTTGGAGTAGATCCCCAGTATCTTTAGAAAGATCTTAGGTGTGAGTTGTAACTCCACCAGTTGTGCAGTCTAAACAAAAAACAAAATTCTATTTCCCCTCTACTATGAATAAAACTGCCAAAAGAGAGGAGTATTGTCAAATAGATTCAATACATTTGGAATAAAACAAAATCCTTCCCTGACATGTTTTATTCCAATTTAAAACATTGGCTGATAGATAGGCTGAATATTGTTCATTTTTGGGAAATTGGAGGACCAAGTGTCATGACCTTCCTCATAGGCCATAGCTTGTGACAAAATGTTGTATCTCCATTTTGTTCATCTTTTGAAATTATTTATTGCATCACCAAAACCAAAATGACCACCCCAGTGTTGTGAAAGGCGATAAGGCGAGGCGAGGCTGTTATGCCTATTTACAGTGAGGCGAGGCTAGATTAAATAGGTGAGCTATGGCGGCAATATGGCAATGGCAATGGCGAGGCGTGGCGGCGCCTTTTTTATTTTTATTTTTTTAAAAGGCCCGACTTAATAATATTAGTCAAAATTAGGGTTTCTGCAAACTTTCATTCAGTCGCGACGTGTGACTTCTTCTATCGGAATCGTCGAGCACTCGAGCTTCTTCTACTCTTCTCCTTCGCGTGAGTTGCGTCTGCCAGGTATGTCTACGAAGACCTCTTCTTCTTCAGTCAGTTCTCTTCTTCTTCTTCTCTTCTTCTCTTCTTTTCAGTTTTTCTCTACTTCAGTTTCTTCTTCTTCAGTCTGTCTGCTCACTGCTCTGTTTTCGTGGTAGTGAACTGCTCTGTTTTGCTCTGTTTTGTCGCTCACTGGCCACTGCTGAGTGCTGACTGCTCACTGCTGTCTATATTTAAAAAAAATAATTTGATCCTTCTATATTAATTTAGTTTTATACTAATTAATATAGTATTTATTTGTCAATTGTCTTTTGAATTATTGGGTCTTTAAAGTTCTAGATTTTTAGTATCTACTATTAGTTTTTGAATTGAAATATTTGCTAATATATGTTACTGCTATTGAGATTTTGGATATATGCCATTAAATGTTTTTAATTTTTGGTATTAATTAGCGCCTCAATTCAAAAAGGCGAGCGCCTCGCTGCAAGGCAAGGCAGGCCTTGTCAGCTTTTCGCCGTCCAAAACACTGGACCACCCAAAGTTTCCTCATTTGCAAGACTAAGTTTAAATTTAAGACCCAAAAAGATGCAGTCATGTAGAGAAAAGAAAAAAGAAAAAAAATACTACTGTGCAGTTCAAACTTATTGAATGAATGTTGGGTTAGTCTGAGAGCACATTTAGCCTGCATTAATCAGATAAAATCCCTCTATATATCATTTGTGCATGTACATATTGTGTATCCAGTATGAGAGTATACTATGGTCAAATGGTAAATTCATCTTAGCCAAGTAGTAGGACCAAGTTTGATTCCTTCTATCACCCAAGAGTTATTTATCTACTTGTGTTAAATCGTAAATAGCATTATGTACGTCCTGAAACTACGTTCGCAGCCGATTCTTTTTGCTTTTTCCTCCTCTTGAAGATATTGCTCTCATTACATATTGGAACTTGATAGGTCTTGCCTGATGAAGTGTTGACCTGCGGACCTTCCGCTAGAGTCCTGGATCTCAGCCACAACGCCATACAGCATCTTCCAGCCGCAATTAATAGCTTGAGTTCTCTTCAGGTCAGTAGAGTTGACTTCTTTATATTTCTTTGAAATTGCGATTTATTCACTTGTTTGAGGCAGAAATTTATTTTGGTATTGTTTCTGCTTCCCAGAAATTGATCCTCAATGGGAATGAAATTATGGATAATTCCTTAAGCTGGGAAGAGCTAGCTTCTATAAAGAGTCTTATGGTTATCTCACTGAACCAAAACCAGTAAGTCCCCCTTCATCAGCATATTACTCTCCAAGCTGAATTGGATTCACCAATATATTGCATCGTCTCTATTTAATTGAAGTAGAAACGTATTCTTTCCTGCACCATTAAATGATGGCTACACCAAACATGTATGTACATTTGAGTGTGAGGGACAATAATTGCAATGTCATAGTTCTCATTTATTGTTTTCATAAAATTAAAATATGAAAACTGAAGTATATCAATGAGTAATATAAGTTGTTGAATTTCTAGTATTCAAATGTTAAGTTGCATCATTGGAATGTTAGTCATTCCAAGTTTTCCTTTGACATACAGCTTGAAACTAAATTGTAAAATTCACTCACAGTTTTCTGTCTTATATACTTGTTTTCCTTGCACTTTTATGAGTATATATTTATTAAAAGTTACTTTCAGAATAGTCGAATACTTACAAAAACATAATTGAAAAATTGTCAATTGTCATGGTCTCTCAAATGTTTGAATAGGTTGGCTTTTACATATCAGTTTATAGTTTATTTGAACTAGTGATTGGGTATTGGCAATTTCTATGTTAACTCAATTTGTTCTTACAGAAATATTATAAAGCTTTTAATCGGTTCTTTAGTTACTCAGTTTTAACTTGTAAGTGCCAAATTGGTTATCAGACAACCCATTTGCTAAAGTATAAAAACCGTTCCAATGTATAAGATTCACATATAACTTAAGACAGTTGGACGATGGATATAAGCCCAAATACTGCTTTTGGCAGAGGGGAGAGACCAATTCTTCATGCTTGTAGTGTTTGGTGGATTGTAAATTGTACCAAGGCAAGTCTAATCATAAAACAACATTATTTATTGACTGACCCTCTCAAACATGCATGAAAACTGAAAATAAAGCTGTCTATCTTTCATTCACAAAATACATGGTATTAATTTTTTGGTCAGTGACATTCAGATGTCGTTTTGTGTTGCAATTTTATTTTCTGATACATCTTACTCTGTTTACCAACCTCAATCATTTATCATATGCCATTATTTCGTATAAAGGTTTCTTGTATGATTGTATCAAGTAGTCTTCTTTGGTATGCAGCATAGCAACGCTGCCTCCTGAGGTCGGCATTTTGACCGGTCTGAGACAACTTCATATTGCACATAACGAGTTAACAGGTCTCCCATCTGAGATAGGTCTGCTGACCAGCCTTGAGGTTCTGAAAGTTAACAACAACAGGTAATTTGAAGTACTGAAACAAAAAAAGATATTTCAATGATATGAAATCATGAGGTGGAATCAGCGTGAAATTGCATGTCCAAACGCTGATTCCATCTCACGATTCCATATCGTGATATGGTATCGCATGGCCAAACACCTACTTAGTGTTTTACGTTATTGTTCCCTCTCCTCCTTTTCTGGAGGTCATGGATGTTTCGGAGTCTGTTGATTTTCTTTTTATCTTTTATTTTTCTTTGAAGTGATAAGGACCTAAGTTAGGTTTGTAAGCTTAACCTTTTAATTGCAGGATACGCTCTATTCCGGAAAGTATAGGGGGTTGTGTTTCTCTTATTGAGGTAAAAAAGGGCATATTTTCCTCTACCCACTTGTCTTCTCTTCTCTTTCATTAACTGGAATCATGAAATCCAAAATATAGGCTGAAACCTGCAATAAGTTCCAGCGTTTATTGTTTCTTGTGATTTTATTACCTATATACAGATAACGTGATTCTATATGGTGTAGGTTGATCTTTCATCTAATCTTTTGGTAGAGCTGCCAGAAACAATTAGCAAACTAAAGGACTTAAAGGTCTGTCTTCATCTATACATTTTTCGTTTTCTCTTTTGAAGTCATTCTTCAAAATACCACTTCAGTTTACAGCTTTTTATATGCAATATAAAAAAAAAATCTTTTTTTTTTTATAAATGCCGTACAGAAAAGAACTTGCAAACGTTTTACGAAATTAGCTATATATGCTCTAGTACTGGTTGTTAGACATTGGTAAAGAATGTAACATAGTTTTCTAAGATAGGGCATAGCCTGGATGAACATTTATGTATCAGCTCTAGAACATTCAGCTATGACATTTTAAACCAAACTAGAGGATTTAGTGCCATAATCTGCTGGACCGACACCATTTGCTTGGATAAAAGCACAGCACATGATAGGTAAAATTTAATCATGGTTAAGCGAAAAACTTGGTTTTATCAATTGGCGCTAGTAAGTTTATACTGCACATTATAGGGGCATGTCACGAGTTTAAATGTTGCCACAGAAATATTCAAGTGGAGAAGGATAGATGGTAGTCCATTATCACATAGTGTCGAACTGTGTGCCAAGTGCCAACCAACTCCGAAGGATCTCTCAGTTATCAAAAGGGAAAAAATGTTGTTTTCTTTGCAAGGCACGACACAAATTTCTTTAGTCAACAATGTTTTCTGAAACATCTAGCATCATGTGATGGCCTCTTTTACCTGACAGTATGCTTTTGCCCCAATTATTTGTGTGGTTCATGTCCGATTACTGGTGCAGGCACTTTATCTTAGGAACAACGGACTCAAATCCCTTCCTAGCAGCATTTTTAAACAGTGCTGTCAACTCTCTACACTAGATCTCCATGGAACTGAAATAACTATAGATGTTATCTGCCAGGTAGTAAATCATTTACTACTTAATAACAGTAACATGCCCTAGCTAGTCTGCCTCTTACGATTTTCAAACGATCACAGATTGAAGGATGGGAGAATTTTGATGAGAGAAGACGGTCGAAGCATCAGAAACAACTGGACTTCAGAGTGAGCAGCTCTGGCAAATTTGACGAAGGAGCTGATCAAAGCTGATAAGGCCTGATACAAACTACTCGGTAACTCGGTTTATATACATCCTGAAATACATTTGCATAGTGATCCCTAGCAAACTTCACAGAAGAATCTTGTAGTACCTGTTAAGATACATCAAGAAATTATAATGAGCATACTGCTCCACTGTGCACTCCCTAAAGATGATTAAATTTTCTGAAGGCTTACTATAAAGCTTTCTGCATCTTCTCAAACTGTTACATATATAGTTGAATGAAAATTCCCTTTTAAGGATACATATTACAAATTGCTGTACCCCTA

>M6-PGSC0003DMG400020641 M6 LRR-RLP

TACTCTTATAAAACTTATTTAAATTTAGGTGATGGTTTTTATCACACAAAAACTCCGAAGACAAACCTCCCCATCATCCAAACCGCACATATCACTAAATTTGCATCACATCGAAACAAGGTGTTATAAATTCCGATGCTGATGTGAGTGATTAACAATAGTTTCAATTTCTTATCTTCGCCTCTCTCAGGTACGGCCTTCGGATTTTGATTTTTTTTTTTAAGTCTGGAAACACCTTTATCTTCCCGAATTTGATAAATTTTGTGGATTTTTTTGGTGAGATTTTGATGAAATTGTAGTTTTATAGAACCCAGATTTGTTAGATGTAGCATGTCTACAATGAGTTCAATTGAATCCTGTGTTTTTCGGCAGATAAATAGGTAAAAAAATCACTAGAAGCTCAAAAAATATTGTTTTAGAATCTATTATTTGTTTTTTCTCTCGGAGTTTTCATGTGTAAGTTGAGGGTTGAAGAATTCCAGAACAATTTTTTGTAGAGGGAAATGCTCTATATTTTCACTTTGTTCAGCTAATCATTATACTGAGTTTTATGTAGTCTGTTTGAAATTGTCAATGTGAAGAACAATTGCAAACTTTAGGTGTACTGTATTATCTATTTGATTCAAATTATCTAAAATCTACGATGTGTATTTTTAAGTGCTCACCCACGACTCAGGCCAGGGGCGGGGTCGTGGAGCTAGGTGGATTCAAGAGGTTCAGTTGAATCTCATTTGTCAGATGATTATGATAGGTAAATAGGGGGGAAACAAATGTTTTTGTGTATACGTCAAATGTTTATCTATCTCTATGAGGTAGATATAAGGTCTGCATACACTTTAACCTCCCGAGATCCCACTTGTTGGACTACACTGGGTATTATGTTGTTGTTGTTGTTCATCATATCCATTGACATAGGTCTGCATACATTCTTGCATTTTTTTTGAATACCTTTGTTAGAATCCCTTGCTCTGCCTCTGGTCATTCAAGCGTTGTTAAATACTCTTAGAATATCTAAACCCAAAATGCTGACTTATTGGAGCAAGGGGGACCTTAGAGCTATCTGTAGTTTACAAAATATTTCAAGTTAGAATGGGAATAAAGGAACTCTTTTGGGTTTGTATTTGCTGCTCCTTATGTATTGAGCTTACTTCAAAGTTGATTTCCTTTTTCGACCCCTAATCCAACCTCTTAGTGCCATTTGGTTCAACGAATTAGGTGGGCTATCCGAGAACAAATTTTGTGATTAAATTTATCCCGTGTTTGGTTGGAGGTATTAGCTAAAATTAGTATTATTTTTTACCAAAATCTTAGTATGACTTATCCCATATAGAAGGTGGATTAATTATAATCTTGAGATATTATGGTTTGAACCAATGACACACCCCTTAGGAGAGGTTTTTGTACAAGCTATTATCAAGCGTGTTTGCAGAATAACTAGTGCTTTTGAAACTCATCTTCCAAGATGAATTGTACATCTATTCCTTATGATACCATATACATATGTTTATGCCTGTGGATTATAATGTGGATCATTTGGAAAGACAGAAATAGGAGGCTTTTGAAGGGGTGGAGATGAACTTCGCTCGATTGAGTAGTAGTCTTCAATCCCTTGTAGTTTTTTTGGTGCACCCGTGTAGTTCTTGTTTGTATAGAGGATTGGGTGGCTTTTAGAGAGAACCATATATTGTAGGTTTCTCCTTTTAGGTATATTTCCTGTATATTGCCAAGTTGGCCTTGATATTACAATGAAATACTTTAACTGAGAAAAAATATTCCAAGTAACACTACTATCACAGTTGGGGATATTTTCAAGACTTCACCAAATGTAGGATACTTTTCTATATCCAGATAGTCCGGGCTCGAGATGAAGCTAAGCATACACATCAGTCTTGTAAATATTTTGTATCCTCGCTAAGATGTGCATGGCATAATTCTGAGAAAATAGCTCCCGATGGTTTTATTGTCTAGCTACACCTTGGCATGCATAAAGTACAATTAAGTAGCTTTCATTAAGGTTTTTTTTTTATTTCCTTCTGTTTTTTCTTGCATCCTTTGACTTAGGTCGCTATAAATCTATCTATCTTATGAGTCTACCTAGTATGCATTAGTAGTACTGCAGAAATTTGAGGTTATGTATTCATATTGAAGTGGACATATGCATGCACAAAGCATCTTATAATATATTGATGAGGTTACAATTTACCTTTATTAATTTTGTAATGCTTGAAAACATTAAAATAGCCTCTCTACCTTCATAAGATATAGGTAAGGGCCGTGTATCCACCCTCCCCAGATCCCACTTATGGGATTACACTGGGTATGTTGTTGTAGAGAAAATTTAATTATAAACAAAAGACAAGATATTCATGTATGGTATGCTCAATGCTGTATAATGTCAACTAATATGTGGTTTTTCTTATTCTTTGCAGTGTTGGTTTGTGTGGCTAACCTCTCAAGTTAACATCCTCCAATCATTTCTCTTTTGGTATTGGTACCAAAGACGGTAAGTATAGTTAAAACTGACCATTGGGAACATTGACATTTTACTCTGGTTGATAAGTAAACTATCTTCCTAGTTCCTACTTCTTAGGGGACTTCATAGTTCAAAAACCAGATTTCTTTGCCATAAGTCAGTGATCTCATAGGTCATGCGTTTGGAGACTTGCTCTTTCTTTTTTCGAGTCGTCTACCTCTTCTAAGACTCTAGCATGTCTTTTAATAAAATTATTGGGAAGTCCTAATAAAAAAAAAAGTTTATTGTGAACTATAGAGCTTCACACCCTTGCTGGAGTGTTAACTTGGTAGAAATGGATAATCCATCAGATGACTTCTCTGGATATATCATGCAGCTTCCTGACGATTGTTTACTCTTTATTTTCCAACACCTTGACTGTGGATCTGACCGTGAATCATTTGGTTTAACCTGTCGTCGCTGGCTTCAAATTCAAAATCTAAATAGAAGATCTTTACAGTTTCAATGCTCTTTCTCTATGCTTAATTTTTCATCGTTATCTCAAACTAACACCCGAATCAGTACATTACACTTATATAGGCTTCTGAACCGTTTCCAGAACCTAGAATCATTGTCCTTGTCTGGATGCACCGAGCTTCCAGATTTGGGGTTGTCTTTGTTACCACGTCATGGATCGAAACTGCAAAGTCTTCATCTAGACTGCTGTTTTGGGATAACTGATAGTGGTCTTTCCTGCGTGGCATCTGGCTGTTCTTTGTTGGTAATACTAAGTCTTTACCGGTGTAATGTTACAGATTATGGGTTAGAGGCCTTATCTAATTCTTGCCTAGCTTTAGAAGATCTGAATCTCTCGTATTGCTCTCGAATCTCTGATTATGGTGTAAGAGCTATTTCACAAAACTGTCGTCGTCTTAGAGCAATCAGGATATCCTACTGTAGAAATCTAACTGGTGCTGGCTTTCAAGGATGTTCTCAAACTCTGACTCATTTGGAAGCCGATTACTGTAGGCTTGAAACTAATGGAATACGGAGTATCCTAAGTGGAGGTGGTATTGAATACTTGAGTGTGTCTAACTTGATCTGGTGCACTCGTGTGAATGGTCTGGTGGCTATTAGCGTCGGATTTGCTGCAAAGTTAAGAGTGTTGAATTTTCGTTCCAGCAGAACGATTGGCGATGATTGTATCATGACAATAGCAAAGGGTTGTCCGACACTTCAGGAATGGAACTTAGCAATATGTCATGAGGTCAAGATAGCAGGTTGGGAATCTATCGCATCACATTGTCATAACTTGAAAACACTACATGTGAATCGGTGTAGAAATCTTTGTGACCGTGGACTGCAGGCATTGAGAACCGGATGCAAACGTCTCTCAATTCTTTACATCACCAGAAGTTCTCAGATCAGTGGTGTAGCGTTGGAAATTTTTAAACTTGCTAGAGGTAATGTAGAGGTCAAAGAAGAAGAAGTAATGTGTATTTGCCCTCATGGAGCCTTCAGGTTTTAATAGGTGGCAATAATCTTCAGTTCTTAAGCTGTCAGGTCTCAGTAGACATTCTGATTTTACAGACTCGTGAGGAGAAACTGTCATCCCAGTCAATCCAATTCGGATTCCCTGGAACTGACGTGTTACTCACCCGTCCACCACTGGAAACATTGTTTCCTGTCCAACTTGATGGAAATGAACGACAGTTGAGCCCACTGCACACTAGTATCAAGCTGCGTATAATTTCTTGTAATGTTTTCTTCCCATAAGAAAAAAAGTGATTCTAATAAATGCATGTACATACAATGAAACAGTTGCTTGGTGCAAATTGCAGAACCCACACTCAGAGGTGTAGCTCTTCATGTGGCATGTAGGCCACAATGCAGAACAATTGTTTGATGCATGATAAATGGAAGGATTGGTATAGAAGAATAGTAGGGAA

>M6-PGSC0003DMG400020729 M6 LRR-RLP

CGGTCTTGAGGAGATCGCGTAAGGAACTTCAAGAGCTTAGGGTGTTTCCTTCTGATCCATTTTCTCGAGGACCGAATGTATCCTTGAGAGAGCAAGGCCTTGTAGCTGTCTCAGTGGGCTGCCATAAGCTTAAGTCAGTTTTATACTTCTGCTGCATAATGACAAATGACGCCTTAGTTACTATTGGAAGGAACCGTCCTAACATGATCCAATTTCATTTGTGTATTATCAAGCCTCAAACTCCTGATTACTTAACCCTTGAACCACTTGATGCTGATCTTGGGGGCCATTGTGCAACACTGCAAGGAATTACGACGACTTTCTCTTTTCTGGCCTCCTTACAGATCGTGTGTTTGAGTACATCGGGTTCCATGCTAAGTAGTTAGAGATGATTTCCTTAGCTTTTGTGGGGGATAGCAATCTAGAAACGAAGCCTATCGCCACGCCGACCATCAAATATGCGAGTGCCTCCGTAAGTTGGAGATTGGAGACTGCCCCTTTGGTGATGAGGCTCTGTTGGCTAATGCTGCAAAGCTGGAGACAATGTGATCCCTTCAGATGTCTAATTGTTCAGTAAATTTTGATACATGTAAGTTGCCAGGGCTTAATGTTGAAGTTATAGACGAGAGGGGTCATCTGGATACGAGACCAGAAAGTTGCCCTGTTGAGAAACTTTATACATACAAGACAGTGTCAAGAAGGAGGTTCGACACTCGTGGTTTTGTTTGGATAATTGACGAAAATG

>M6-PGSC0003DMG400020918 M6 LRR-RLP

TATAATTGGCCACGGCGATGACCAGTTCTTTCTTTACTTCTTCCTACTACATATTTAAATCGTCCCCATTTCTCTCTCTCAAAGCTCCATTAGAATCAGCTAGCAGCGCCGCCCTAACATCCCTATTTTTTACGGCGGCGTCTACAATTCCGGTAGACCATTACGACGACGTTTACTAGTTTCCTTCATTCTAAATGGGCCAATCTTCCTCAGCTCACGGCTTCTCCCCCACCGATTCCAAGTATCCCCACCACCGTTGTTCCGGCACCAGCCATTCCCAATCGGAATTTTTCTACCCTACACTTTCCGAAGAAGACGATTTCCCTATTTCACAACCCTGTGAAAACCGAGATTACACTGATGGTCTTCCTGATGAATGTTTAGCCCTAATTTTCCAGTCCTTAAGCTCCGGTGATCGGAAAAGATCTTCCCTTGTCTGCCGGAGATGGCTTCTGGTTGAAGGGATTAGCCGTCACCGGCTTTCTCTCAACGCAAAAACTGAAATCATTCCGGATATACCCACCATCTTCTCTCGTTTCGATTCCGTCACCAAACTCGCTCTCCGATGTGACCGAAGATCGGTGAGCATCAACGATGAAGCTTTGATCCTCATCTCCCACCGGTGCGTTAACCTAACGGGCCTAAAGCTTCGTGGTTGCCGAGATATCACCGATTTGGGAATGGCTTGTTTGGCAAAAAATTGTGAGAAATTGAAGAAATTCTCTTGTGGGTCTTGTATGTTTGGCGCTGAAGGTATGAATGCTTTGCTTATTAACAGCATTTCACTCGAAGAACTCTCTGTTAAACGTTTACGGTGTATGAATCATACTGTATCTGTTGAACCGATTAGCATTGGATCTAATGCTGCTTGTTCGCTTAAATCTATATGTTTGAAAGAGCTTTATAATGGACAATGTTTTGGACCTTTGATCATGGGATCGAAGAATTTGAAGACGTTGAAATTGTTGCGATGTTTGGGTGATTGGGATAGAGTCTTGGAGACGATTGCAAGTAGGGAGAATCATTTAGTTGAGATTCATTTAGAGAGGGTTCAAGTGAGTGACATTGGACTCAGGGCGATATCCAAGTTTTCAGACTTGGAGATTCTTCATTTGGTGAAAGCTCCTGAATGTACAGATGCTGGAGTTGTTGATGTGGCTAGACATTGTAAGTTGTTGAGGAAGGTTCACATTGATGGATGGAGGACGAATCGAATAGGTGATGTTGGTTTGGGTGCCATCGGCGAATATAGTGTAAACCTCAAGGAATTGGTAATTATGGGTTTGAATCCTACGGTGACTAGTTTATTAGCCATTGCTTCTAATTGTCAAAAGTTAGAAAGATTAGCACTTTGTGGTAGTGAGAGTATTGGGGATGCTGAAGTAACTTGTATTGCTACAAAATGTATGGCTTTGAAGAAGCTTTGTATCAAGGGATGTGAGGTAACAGATGAAGGGATTGTGTCTTTCGCGTGGGGATGTCCTAATTTGGTGAAAATTAAGGTTAAGAAGTGCAAGAACGTAACTGGTGAAGTTGCAGATTTGTTGAGATCAAGAAGGGGATCTTTGACGGTGAATCTTGATGTTGGGGAAGTTGATGTGGAGGTTGTGGATGGTAGTGCTAGTGATGGTGGAGCATTAGAAGAAGGCATAGAGTTTCCACCGATTGCAAGAACTGTTCCCATCCTTGGTGCTGGTGGTGTCGTTGATATTGATACCCCATCGACCAGTAATTTTGGTCGATCACCAGCATCTACTTCAAGGATTGGCTTTCTTGGAGGAAGGGGCTTCGTTGCTTGCACTTTTCGTAGGTGGTCAAATGGTAACGCCAATGAGAACTTATGAAAATTGTGAACACAAGTTAGAACAGAGAAAACACTCTGTTCAACTTCATTTTTAATTGAATCAATAGAAAAGTGTGGATGTGCGACTTATCTCTCCTTCATTTGTGTGTTTGATATTTGCTTTTTGAAAATTTGATATGTAACCATAGATGCATTTAGAATCTTAGAAGTCACTTCTTGGATGATGAAATCTATCCCTGCTCTATTGACAATTGCATTATAGAGTGCTGTCTTGAACCATGTATACTGTTGCTTCTGGTATGTGAATCAAATAGATATTGTTCCATCGACTAAATGTCATCAAGGACAGTGATCATTATGCATGTTCATTGTCATTTTGCTACAAGAATGGAGTAATACATGTCAAAGCACCTTTCTATATTACCAGGAATGCACATATTACCTCGACTCCTAGACCCTCCTCTTCCCTCTTTTTTATTTTTCCGAGAAAAGATCGATCTCCAAGATTC

>M6-PGSC0003DMG400021051 M6 LRR-RLP

AAATTATTGAATAGAGACACAAAGCAAGAATTGGAAAACTTAATACTTTTGATATATATTTTGAGGCGCGAAATGGGCATTGGAAAACTTGCATTGTTTCTGCTATACGCGTTTCTCTGTCGACTTATTTTCTCCTCATCTTTACCTCATTTGTGCCCCAATGATCAAGCCCTTGCTCTTCTTCAATTCAACAACACGTTTACTATAAATCTTGATGCTTCTTCTAGTGATTACGATCCTAAAACAGTTAAATGGAATAGGAGCACAGATTGTTGTTCATGGGATGGAGTCAAGTGTGACGAGACGACAAGACAAGTGATTGAGCTTGATCTCAGTAGCAGCCAACTTCGAGGCAAGTTTCATTCCAACAGTAGCCTCTTTAAACTATCTAATCTCAAAAAGCTTGATTTGTCTTATAATGATTTTTCCGGATCGCTCATTTCACCTAAATTTGGTGAGCTTTCTAGTTTGACGCATCTTGATTTGTCGTATTCAAGTTTTACAGGCCTAATCCCAGCAGAAATCTCTCGTCTTTCTAAATTACAAGTTCTTAGTATATAGAGTAGTCTATTAAGATTCGAAACCCACAATTTTGAACTGCTCCTTAAGAACTTGACCCGATTAAGAGAGCTCCACCTTAGATCTGTGAATATCTCTTCCACCATTCCTTTGAATTTCTCTTCTTATTTAACTAATCCAGAACTTTCAAACACGCAGATATACGGGATATTGCCCGAAGGAGTTTTCCACCTTTCCAACTTGGAATCTCTTGATTTATCGGACAATCCCCAGCTCACTGTTAGGTTTCCCACATCCAAATGGAATAGCAGTGCATCACTAACGAAGTTAAATCTCAGTAGTGTGAATGCTACTGGTAGGATACTGAATCATTTGGTCATCTAACTTCACTGCGTACATTGAGAATAGCTTCTTGTAATCTCTCAGGCTCTATTCCTAAACCTCTACGGAATCTCACTAACATAGAGGAATTGTTCCTTGGAGATAACCATCTTGAAGGACCAATTTCTGATTTCTTTAGATTTGGAAAGCTCAGGTCGTTATCACTTGGAAATAACAACTTTGATGGCCAACTTGAGTTTTTATCCTTTAGCAGATGTCGCAACTGGAATCATTAGTGTTTTCATTCAATTTCCTAACTGGTCCGATTCCTTCTAATGTAAGTGGTATGCAAAACCTACAACGTCTTTACTTGTCATCAAACCTCTTGAATGGGACTATACCTTCCTGGATATTCTCCCTCCCTTCACTTAAATGGTTAGACTTGAGTGATAACCATTTCAGTGGAAACATTTTGGAGTTCAAGTCTAAAACATTAGATCGAGTTGTTGTAAGACAAAATCACCTGCAAGGTCCTATTCCAAAGTCACTCCTAAACCAGCGTGATCTATATTATCTTCTCCTTTCTCACAATAATCTCAGTGGACAGATTGCTTCAACCATCTGCAATCTGAAAACATTACGAATGCTATATTTCGGCAGCAATAATTTGGGGGGAACAATCCCACTATGTTTGGGTGAAATGAGTGAACTTTCAGTTTTAGATTTAAGCAACAATAGTCTTAGCGGGACAATTAATACAAGTTTTAGTATAGGAAACCAACTTGAGGTCATTAAACTTGACGGGAATAAGCTAGAGGGGAAATTCCCGCAATCTTTGATCAATTGCAAATATTTAGAAGTTCTTGATTTAGGTAACAATGAATTGAAAGACACATTTCCTAAATGGTTGGGAGCCCTACCTTATTTGAAGATTTTGAACTTGAGATCAAATAAGTTCTATGGCCCCATAAAAGATTCAAGGACTGACAACTTGTTTGCTCAAATTATAGTCATAGATCTCTCATCCAATAGATTTAGTGGAGATTTACCGGTGAGCCTTTGAGAATCTTGAAGCCATGAAAATAATTGGTGAGAATAGTGGAACCCGAGAGTATGTAGGTGCTATAAATTCTTATTATTACACAAATTCCTTGATAGTGACAACAAAGGGACTGGAACTTGAACTTCCTCAAGTTTTAACTACAAACATAATCATAAATCTCTCTAACAACAGATTTGAAGGTCATACCACAAGCATTATTGGAGATCTCGTTGGGCTTCGCACGCTGAACTTATCTCATAATTGCTTGGAAGGTGATATACCAGCATCACTGCACCAATTATATGTACTTGAATCATTGTATCTCTCATCAAACAAAATCAGCGGAGGAATTCCACAACAACTTGCATTCCTCACATTTCTTGCAGTCTTAAATCTCTCTCACAATCATCTTGTTGGATGCATCCCCAAAGGAAAACAATTTGATACGTTTGAGAATAGTTCATACCAAAGGAATGATGGGTTACGTGGATTGCCACTCTCAAATGATTGTGGTGGTGGTGATGGGGTACCACAAGCAACAACTCTATTTGGGCTAGATCAAGAAGAAGAAGAAGAAGGAGATTCAGCAATTATAAGTTGGAAGGCAGTTCTCATGGGTTATGGTTGTGGACTTGTTATTGGACTGTCCATAATATACATAATGTTGTCAACTCAATATCCAGAATGGTTTTCGAGGATGGATGTAAAATTGGAACACATAATTATTACAAGAATGAAAAAGCACAAGAAAAGATATTAGTTTGTTACCTTCAGGTATTCAATTTGATCTTTAATTATCAATGAATTTGCTCACCTCCTTCATCCTTAAAGCTCTTAACTTTAAATTCTTTCTTTTATGAAAATTGCAGGAT

>M6-PGSC0003DMG400021067 M6 LRR-RLP

GTTTGAAGCTAAAAATGGGGTATGCAGTTATTATTCTCTTAATTGTGTTTCTCTGTCATGAAGTTTCTTCAATATCCATATTACCTCACAAACATCAAACCATTTTGCTTCTAAAATTCAAGAACACCCTTACTATAGATCCGTCAGCGTTTTATCCAAAAACGAGTTCATGGAACATGAGCAGAGATTGTTGCTCATGGGATGGAATCATATGCGATGACATCACTGGCCATGTCATTGAACTTGACCTCAGCCGCAGCGGACTGGATGGCACCATTGATTCCAATAGCAGCTTATTCCAACTCTCTCATCTCCAAAGGCTCGATCTTTCTATGAATTACTTCTCCAATTCTCATATCTCCCCTGAATTCAGCAAGTTTTCGAGCTTGACGCATCTTGATCTTTCTGACTCCTATTTCTCAGGTCAAATTCCTTCTGAAATCTCTCATCTTTCCAAGTTACAGTCTCTTCGTATCTCCACAACTTTTTACTTTTATGGTAGTAAGAAGCTAAGACTCACAACCCATGATTTTAAATCGCTTCTTCAGAACTTGACCCAATTAAGAGAGCTTGATCTTAGTCAAGTAAACATCTCTTCCACCATTCCTCTGAATTTTTCTTCTCATTTAACAACTCTGAGTCTGACATATACAGGATTGTATGGGATAATACCTGAGAGTATATTTCACCTGCCCAACCTGGAAACACTTTACTTAGGGATCAATCATGAGCTCAATGGTTATTTTCCAAAAACCAAATGGAACAGCAGTGCATCTCTCATACAGTTAGATCTCAGTGGCATGAACTTTCCTGATAATTTGCCTGAGTCTCTTGGCTATCATCTAACTTCACTGCGTTCTTTGGACCTTGCATATTGCAACCTTAGAGGGCCTATTCCTGAATCTTTTTCAAATCTCACCCGCATTGAGTATTTGGACCTTTCATATAACTCCCTGAATGGAACAATACCACCAGGGATGTTATCCCTTCCATCACTAAATACAATAGTCTTGAGTAATAATCACTTATCTGGTCCGCTTGAAGATTTCAAGTCGAATTCACTAATCTCGATAGATTTAGAGAGCAATCAGCTGCAGGGTCATCTTCCCCACTCAATCCAAAACCTTGTGAACCTAACATCGCTTGGTCTTTCCTTCAATAATTTTAGTGGTTTTGTGGATGTCAGCCTCTTTTCAAACCTTAAACAGCTTGGGGAACTTAGTCTTTCATATAATAATATCTCACTGACCAATGACAACAAAGTTAATGTTACTTTGCCCGAATCTCTTTATTTGTTAGAATTGGCCGCATGTGAAGTGAAAGAATTGGAGTTTCTAAGATCTGCAAACCAGCTTTTCGATTTGGATCTTTCAAATAATAAGATTCAAGGAAGAATTCCTGATTGGGCATGGTCTAACTGGATGTTTTCATTGCGAGATCTTAATATATCCCACAATATGCTAACGAGTATGGATTCAATTCCTCTTCAACATCATTTAACAATTGATTTGCGGTCAAATTTACTTCAAGGTTCACTACCTATTCCACCAAATTCCACAAGATACTTCTTCATATCCCATAATAATCTCACTGGAGAAATTCCTTCATCTATTTGCAACTTGACATCACTGATAATGCTGGATTTGGCGACAAACAATTTGCGTGGAGAAATTCTGCAGTGTTTCGGAAATATCACTTCCCTCCAGGTTTTGGATATGCGCCACAACAATCTTTCTGGGAATATTCCAACAACTTTCAGCAATGGAAGTTCACTGAGCAGCCTCAACTTGCATGGCAACAAACTAGAGGGGAAAATCCCACAATCCTTGACCAATTGCAAACAACTTGAAGTTCTTGATTTAGGAGACAATCACCTCAACGACACATTCCCAGTGTGGTTGGGGATTCTACCAAAGCTGAAAGTTCTAAGCTTGCGATTCAACAAATTGCATGGTCCCATCAGAACATCAAGGATTGAGAACATGTTTCCAGAGCTCCGAATCATAGATCTCTCTTGCAATGCCTTCTCAGGAAACTTACCTTCAAGTTTGTTTCAACATTTGAAAGCCATGAGGACAATTCATCAAACGATGAACACACCAAGTTATGAAGGAAGTCGATATTAC

>M6-PGSC0003DMG400021279 M6 LRR-RLP

TGATTCTATTCTCACTAAAGATCATGTCCCCACTCTCCCCCACCCATCATATCAGTCACATCTCTAAGCCTAACCCCCCCCCCCCCTCAACCACCACTAAACCCTCCAAATAGTTCTTGCATAAAAATGATTCTTTTCTTAAAAAATCAAAATAGAACTCCAACAATTGATCAGAAGAAACCTTTACAACAAATACAATGTATCCATTCAAACTCTTTTTATTTCCTCTACTACTCATGTTTTTCCTATCTACCACCCCCCCTTCTATCTCCTCAACCAATGTAGCAAACATCAACAATCTCACACAACCAAACATGAATCCTCATGAACTAGAAACTCTTTACAAGATCATGGAATCAATTTCCAGTGACCAAAGTTGGAGAAAATCATATCCAAATCCATGCCAACAAAGTTCATCTTGGCTTGGAATTGAGTGCAAAAAAAACACCAACAACATTTACCATGTCACTAGGCTTGATTTTGGGACATATCCAAACCCCATTTGCAAGAAAACAGCAACATTCCCTTCACTCATTTTCCAACTCCCAAATCTTGAATCTCTCTTCTTTATCCAATGCTTCACACACACAAAAACAACAATCTCTATTCAAGAAAACAGAATTCTCCAACAACTTAGTATAAGATCAAATCCATCATTAATTGGCACAATCCCACCTCAAATTTCATCTCTTAAATCACTTCAAATCCTCACATTGTCACAAAACAATCTTCTTGGACAAATCCCACCACAAATCTTCACACTCAACTCACTTATTCATCTTGATTTAAGCTACAACAACCTCACAGGCCAAATCCCAAACCAAATTGGCAATCTCATCAATCTTATAGGCTTAGACTTAAGCTACAATTCATTAACAAATCCAATCCCATCCACTATTGGCCTTCTCAAATCTCTTCAAAAACTTGACTTAAGCTCAAATTCATTAACAGGCAAAATCCCTCAAACAATAGAAAAGTTACAATCTTTAAACTTCTTAGCACTTAGTAACAACAAGTTAAGTGGCAATTTCCCAAAAGGGATACATAAACTTCAAGCACTTCAATATTTTCTCATGGATGACAATCCAATGATGTTCATAACATTGCCTGATGAATTGAGTCAATTACATAAACTTCAAGAATTGAGCCTTTCAAGTTCAGGTTACTCAGGGGAAATACCATCAAGTTACTCACAACTTGTGAACTTGACTACGTTGTCTTTACAAAACAATCGATTAACAGGTGAGATACCGGTGGAATTTTCAAGATTATCACATATGTATCACTTGAATTTGAGTAGAAATTTGTTAGATGGTGTTGTGCCATTCAATTCAAGTTTTTTGAAAAGACTTGGAAGGAATTTGGACTTAAGTGGAAATCCAGGGCTATGTTTGATGAATCAACATGAAGGGGTTTATGTTGGAGTTAATGTGTGTGGGAGTAGTAGTAGTAATACTATAACTATGCCATTGAAGAAGAATAGATCATGTGAAGCTTCATGTGGAGTTGTTAGATCATTGTTTCTTGTTTGTATTTTGTGTGTTTTAGAATTGTTGTTTTGTGTTTAGGAGTTTTTAGGCTTTTGGTTGTAT

>M6-PGSC0003DMG400021415 M6 LRR-RLP

AAATGTGCCTGCATTTAAAGTCAAATTATAATTAAGATAGACTGAACAAAGTCAGAATACTAAGGGGGTAAAATGACTGACACAGTAACAAAAAGAAAAAAGGTTCAATCTAGACAGCTCTGAATCTGCAGAAGAACTTAAACAATATAGAGAAAATTGAGAGCACGATGATGCAGGTGTAAAACTTGTCTCTGTAAGATGTATTGCTCTACAGTAAAACAGACATAAAAAGGTAAAATTGAAATAGGATAGAAGGTCTACCTAATCTTGGAGCTGGATACCAAGGAAACACTTCTCTAGGCTAAAAGGCAAGTAATGGTAATAATCACTACATTAAGTAAAGCATAGAGTGGCTAGTTGAAGAGCATTCTTTGAATAAGGAGCTTTTTAGATGAGCCAAATGCACCCAAACAAATCAATAGCCAAATGGCCACTCGTTATTAGGATTGAGTGAGTGAGAAGGAGAAGGGACACTTGCATATATAGGTACTCTTTTGGTGCATAAATGAGTAATCACTCGTTGTTGATGTCAAATCCTCTGTTGTCGTTGGCATAGGCCCCCACACTACTCAAATAAGTCCCTGATTTGCTGTGACTGAAAAGGTGGTTCATATCCTCTATGTGCCTCTCACATTATGATCACCTACTACAGATGTTGACGCTCTTCTTTGGAGGTTCTCCTGGGTTAAACAAGCTCCATAAGGCACAGGTAAAGCAAAGCTCTTTAAGCATCCTACTTCCAGGGGAGCTTTGCTTTTCCAAATGCATTTCATAAGCCAAAATGTGCTACTGCCAGAAGATCTCCTAGATAGAGAATACTAGTCTTTGATCTTGACGTTACCTACTGTGTTTGCTTTCCACAAAAGAGATTCAGCTCTGGAAGATAAAGGGGGGCAGCTATGCAGCTTATTTAAAAGTGAGGACACCGTTAAATTCACAATCATTTTTGCTGCACCAGAGACCAAAAACACAGTCCTGTTCAGACCACATTTCTGAAATTGTAGCATCTTGTGTTTGTGCAATACTGTAAATATATGGATACTGCATCTTCGGTGGTGTTTGGACTAGCCAAGCATCCTCCCAGAACTTGTCTCCTTCCCATTTCCAACTTGAAGAGGTGATGGCGAAAAAATAATTTTAGTGGACTCTAATATTCCTCCATATACCATAGGGACTGTTGCTGAACCTGTTTAGTTATCCAAGGGATCAGGCATGCATTTTTACCATAATGACACCTCCAAAGTGCCTTCTCTTCTTTACCACATCTTCATAACTACTTCTGCAGTGAGCTGTTGCTCTGCATCTTCAATTTCTCTACTCTTAAGTTCTCCCTGCCTTCAAATTTGTGGCGACTGATTCCCATCTTACCAAGTTGAAGGTTCTGTGTTCTTTCTCTCTTGCATAAGCAATTCCTATGTAGTTTGTTTGTCCAACCTTTTTGTCACTGAAAACATGCAGTGGAGCAAAGAGATAGGGTATATGGGATATTATTCTGAACTTAAAACAGCTTTGGTTGGATGTTATCATCACACAGTAGCTTTGTAACTGAATTCCTTGTTAACTATGTTTGTTTTCTAATATATTATAGAAACTATTGGGAAGTTCTGAGGCTAGAAAAAGATCAAATTAAGACCATAGTTTCAAGCATTCTCTTGTCGGGTGAAGGGGTGTGAAGGTTGAAACTTTAACCCATGAGTTTGGGATATTTTTAGGCCCTATAAAATTTGATATTGTGATACTTAATGTACTTACCTTCTACTCTGCTTCTCTCTCCCATTTAGATGATTTATGTGTCGGTGTTAGAAAAGAAGAAAGATGTTCCATTGCTGATGCTGAAGTGCAAACAGTTGACGATAAATTCTTGGATTTCAAGAAATGCCATTCTCGGCATAGGTGTTCTCTTAAAGTCAACTCCTTATCTGGAGAATTTGATGATACTTCATGAAGAGGTATGTTTTCAATTGATTCTTGCTTTGAGATACCTTTTATATTCTTTTTCATTCTTTAATTTTTTCCCTTGTGTTTGTTTCATTATCCTTTACATTTTCAGTATCACATTTTGAGGCATAATATAGCCTTGGATGTCAAGCTGTTTTATGTATAGCGCCCTTTATTACTGAAATTGTTGCCCATCAGACCTACATTTTCTAATTTCTAAATATATGCGACCATAGTGTTACTGCACTGAAGACTCTTCTTCCTTTTACAATGAGTTATTAGCTTCTTGCTTTTGTTTGGCATAAAATATTACATTCAATCGTCTATTGATCTCCTACGACATCAATGGGAAGTCTTCAACATTTTAGTACTTTCCACTAAATTACTATCAGTGGTCTATCTCAATATTATACACCTTCTGACTAAAAAGTATTTTCATCTGTTTGTCTATTGTTTGACTACTATGTTGCTTATATCTCGTTTCGGATTCTTTTGTGTGTTGTTCTGCAATTTCAGTACCATAAAATTCCATTCTAAAATCCAAACATTTACTTAAGTATCTTCTAGCTGTTACACTTACACTTAACCTGATTTGTGGTTCTTCTGTATCCGTAGGGAGATTTCTTTGTCTGGGATGAAGACTTGGAAGACGAAGAGTTTGATTGGTCAGGAGACAATTCCCTCTCCTTGCATGAAAACATATTTAAAGGCTCCTTACAGAATTTGAAGCATGTTGAGGTTTCTTCAAACTGTTGTAGTTGTCAACACAAAGCTGACGCAACAGAGCGCCTTTCAGAATTTTTGAAGTCTTTGCTTGAGCATGCAAAAAATCTTGAGAAACTGGTTATAGCGCCATGGCACAATGGATGCGATATCTCTTCTGCCAATATCTCAAAAGTGATTAAGAACTTGCTAGCTTTTCCAGGAGCTTCTAATATTGCAGTTGTTTCCTTAGGATCGGTCTCCGGTGTTGACTTTTCAGGGGAGTAGAAACTTTTATATTGACCTTGTTTGTTCTAATATTTTGTTCAACTACTTTTGGAACTCTTATATTTTTGTATGTCAGTGTAAAAAAATGTTGATCATCAAGTTACTGTGTAACTGAAAGAGGAGATGTAATCTGCTATAATTGATTAAACTACTGTGGTATTATTCAAAGTTGTTTTGCATTATCAATGTATAATATGCCG

>M6-PGSC0003DMG400021962 M6 LRR-RLP

CCAATTCCATATATTATGCCTTCTTTTCAAAACCCTTACTTTGCTTGAGGAAACAATTCGAAATTAGATCCAATTCCAATGACCGGAGTTGGAGAAACCCTAGAGGACCGTATCAGTTCGTTGCCGGATGAACTTCTGATCAAAATTCTGTCTTGTTCGGCGACAAAGGATGCATACACAACATGTTTTCTCTCTAATAGGTGGCTTTATCTCTGGATTTCCATTCGTATCTTTAGTTTTGCTAATACAAATTACCGAGAAAATTTCACATCTTTTGTTAACTATATATTATCTCGTTCTGTTTCTCCCAAAATACAAAAATTCGAACTCGACTACAGTTATTGCGATGTTGAAAGCCGCTGGCAAATCACCGAATGGCTTAATTTTGCTGCTACTAGAAAAGTGGAACATCTTCTACTCTGGTCTACTGATTTTATCACTTCCCCTTTCCCATTGCCTCAATCTTTCTGCGAATGTTTGTCATTGATAACATTAGCAATAGTTTTGGGTCGTTTTAACTTTGATGACACAGTCATTGCCTGGAAGTCTTTAAAGAGCCTAAAATTGGGGAATTTTCTTATAGATGATTACAAAATTGCAAAAATATTGTCAGGCTGTCCTGTATTGGAAGATATGGAATTACATGATTTAGTTGTGCTTAATGGTCCTCATCGCTTGGAATTTATGTCTTCAAAGTTAAAGACACTGAAGCTGAATGGTCAATGGTTGATTAATGATGGAAGTCTTCGTTGTGTGGAAATTTTTGCTCCGTACATTCAGCATTTGGAAATTTCAGGAGGTCTTTATGATCTCAAGTGTATTCTCGTAGATGTGTCTGCTGTGGTTACTGCGAATCTTACTTTCAACATTATGTGTATCAAAGACATCAACGATGATTATGGTCTACATTTTGATCCTGAGGAAAACAGTTGTAGTGGCTATCATCAATTCTTTTGGTCCCTCGTCCATGGTTACCTTCTAACGTTCCGTAATGTAAAGGAGTTAACCATTGGAACTTGGTTAACAGAGGTCTGTTTTTATCTCAGAGTACTTTTCCTTGGACAGTTGAGAATTAACCTTTGTACAAATTTCACAAACTCAACTGGATGAGTTGCAGAAACTTGTGGCTTTTTTGTGTTTTTTTGTCAACTTTCGAATGTCGGGTATTTTTTTGTCATATCTGCAATACATGTATCTGAAATCACGGATTCTTCTGTGTCGATTGTTGCTTAAAGTGTTTAGATGATTAGGTTCGCTACTTGAAGATAATAGAAAAAAGTGGATATCAGGCTCTTCCTTGGGTGCGGTATATTACCATGGCTGGTGAATATGAGTTGAGACTTGCGTAAAACCACTCCGAGATGATGATCCAATCCAAATAAGTTTTGCTTGAGGCTCATATAACTCATCCAATGCTTTGAGCTCCTTTGCAGGAGATGAACTGGTTTAGCTCATAGATTTGAAGCAAATTGTTTTGTTGTGCCAATTTTTGTTATTATTGTATTTGACTTTTTTTCTTATATGTTACCCACATTTTCTTGTATAAAGATGTCTGCTGATTACTTTCGTGTTGCATATACTGTTAAAGAAATTCAATAGATTTGTCACACAACTTATGTTGCCATGGAGTTCTTTCTTTTTGTACTTATTGCTTTCTCCATTCTTTATATTACTAGTTTCATTGGGATTATTGCAACAGGTTCTGTGCATGTTGCTGTTCAAAGGGACTGTACTTCCAGAAATGAAATGCAAATATCTGATGCTGGAGTTGCATGTTAAGAGGTTTAATTTGTATGGAGTAGCTGGTCTTCTGCGAGCATCTAGTCATGTAGAGACACTCAACATAAATATATATACGGAGGCACTAGACATAGAGGGACGCGACATAGATTTATCAACAAAATCTGTAACTTCCTCTCTTCTTTTAAATCCATTCATTTTATGAGTTTAATCACCCCTCCTTTGAATTTTTCTTTTGTCATGGCAAAGCTTAACATTATTTGGTTGGTGTTGTTGCTTTCTTTGAGGCAAATGAGGTCTGAAACGGAAAGGGTTGGGTGCCATCCACTCTTTGCTTGGTTTACGTAGGCCATGAATCTTTAAGATATGAACAATCATCATAATCTGTTTAAAAAGTACCAATAAGATTTGTTGGTTCTGAAATCATTTTTTCCCCACTTTTGTGCAATATCCATGAGGCTAACTTGAATTTAGTCCTGATTAAATTCTCTTTCTCCAGCTTGTAGATACCCACTGCCACTTTGAGTTATCATATTTGGTCAAAGGAGATGATATTGATTTGCAAAATTGGATTTTATTTGGGTTTCCCAACCTCAAGAATGTTAAGATTATCAACTCCCCTGGAGAGCGGTGCTTTAAGGATCATTTCAAACAGGGCTTTGACAAGCTTTTGAAACTTTCAGAATTTCTGTTAAACAATGCAACATTTTTGGAGAAGTTCGTGATATCAAAGGGAATAATGTGCAAGATATGTTCAACCAACTGTGTGTCCACATTTTTATTTCAATTGGCTGAGAAATTGTTTTGTTGCCCAAGATCCTCTACAAATTGTGTGATTATCTACTAGTAGTGAGCTTGTAATGATTAGACTGCAAGTAGTTTTCTATCTTAGGATACACGAGATGTTGCCTTATATTGAGTTTTTGGTGAATACTCATACAATCATGACACTTTAACTTGTCTATGCAACTAATTACTTATCTTGTGGCATACCAAATGTTATGATTTCTTAGTTTCTCATTCTTGAAGCTTTCAGTTCTGATGAATCATTATATTTAAGTCATATAAGAATGTATGGTTTGATCTCTTTTTTTGATTCGTCTGAAATATCCATTCTTCAGGCAAAATTTAGATGACGATAAGGGGTTTCTAGAAATAGTTTCTGTTTTACCTTTGTTGCAATAG

>M6-PGSC0003DMG400021996 M6 LRR-RLP

CCCCTGTCTCACAAGTCTTCAAACCCACCTACCCCCATTCAACCCACCCACCCCCATCTCCTCTTCTTCAGCTCCCACACATTCCCTCTCTAGTCTCCACAGATCTGAACTTTCATTACTCAAGATTTAAAGATTCTTGAAATTCTTGAAAAAAAATGTCAACTTTAGCACCCCATTTTCGTACAGTAATGGTGAACTCACTTCCCTTAGAAGTTCTTGAACACATCTTCTCCGACATAACTTCAGATAAGGACAGGAACTCAATATCCTTGGTTTGTAAATCTTGGTATGAAGCTGAGAGGTGTTGTAGGAAAAGTGTGTTTATTGGGAACTGTTATGCTGTAAGCCCTTCAATCTTGATTAGACGTTTCCCAGACCTCAGATCTGTAACGATTAAAGGGAAGCCACATTTTGCTGATTTCGATCTTGTACCTGAAGGTTGGGGTGCATATTTCTATCCATGGGCTGTTTCAATGGCGAAAGCTTATCCATTTCTTGAGGAAATTAGGCTTAAGAGAATGGTTGTTTGTGATGAGTCATTGGAGCTGATTTCGAAGTCGTTTAAGAATTTCAGAGTCTTGGTTTTACAGTCTTGTGAAGGATTTACTACCGATGGACTTGCTGCCATTGCTGCTAACTGCAGGTTTATCTAGCATTTTTAGCTGTTTTCTTGGGTAGATTACCCTTTTCTGAGTAATGGTGAAAAAAGTCTTAATCTTTGCATCTGGGGTTAACTGTGCTTGTTTGTAGCTGAGGGGTTGAGGAAAGTTTTAATCTTTGCATCTGGGGTTTACTGTATTTGTTTGTAAGGGTGGTTGAGGAAATGGTATGTTTTTTAGAGCATTGTTGTCTGAGTACTCTAGTCGTTAATGGTGAAAAGAAGTCTTAACCTTTGCGTCTGGGGTTAACTATACTTGTTTGTAACTGAGAGGTTGAGGAAAGTTTTAATCTTTGTATCTGGGGTTAACTGTATTTGTTTGGTATTGAGGGGTTGAAGAATTGGTGTGTTTTGTAGAGTATTGTTGTCTGAGTACTTTATTTGTTAATGGTGAAAAGAAGTCTTACTCTTTGCATCTGGGGTTAGCTGTTCTTGTTTGTAACTGAGGGGTTGAGGAAAATTTTAATCTTTGCATTTGGGGTTGACTGTATTTGTTTGTAATTGAGGGTGGTTGAGGAAATGGCATGTTTTTTTTTAGAGCATTGTTGTCTGAGTACTTTAGTCGTTAATGGTGAAAAGAAGTCTTAATCTTTGCATCTGGGGTTAGCTGTACTCGTTTGTAATTGCGGGGTTGAGGAAATGGTATGTTTTGTAGAACATTGCTGTCTGAGTACTTTAGTCCTTATTGGTGGAAAAAGTCTTAATCTTTGCATCTGGGGTTTACTGTACTTGTTTGCAATTGAGGGGTTGATGAAAGTTTTAATCTTTGCATCTGGGGTTAAGTATTTGTTTGTAATTGAGGGGCTGAGGAATTTAGTCTTAAGAGAATGGTTTTTTTGTCATGAGTCATTGGAGCTGATCTCAAAGTCATTTATTCGAAGTCTTGGTTTTACAGTCTTGTGAAGGACTTACTACTGATGGACTTGTTGCCATTTCTGCTAACTGCAGGTTTATCTAGCACTTTTTAGCTGTGTTTTTCTTTGTAGATTACCCTTTTCTGGTAATGGTGAAAAAATCTTTAATCTTTGCATTTGGGGTTAGCTGTACATTTTTGAAACTGAGGGGTGAAGAAAGTTTTATTCTTTGCATCTGGGGTTGACTGTATTTGTTTGGTAGTGTGGGGTTGAGGAAATGATATGTTTTTTTGTAGAGCATTGATGTCAGAGTACTTTAGTTGTTAATGGTGAAAAGAAGTTTTAATCTTTGCATCTGGGGTTAACTGTACTTGTTTGTAACTGAGGGTTTGAGGAAAGTTTTAATCTTTGCATCTGGGGTTAATAACTGTATTTGCTTGTAATTGAGGGGTTGAGGAAATGGTAGGTTTTGTAGAGATTGTTGCTTGAGTTCTTTAGTCTTTACTGGTGAAAAAGAGTTCAAATCTTTGCATCTGTGGGGTAACTGTTTCGAGGAAATACTAGAAGGTTTTGTTGAGCACTGTTGTCTAAGTACTTTAGCCCTTACCGGTGACTTCATTAAATTATTTCTAATCTGGGATTAACTGATTTGACGGAATAGTAGGTTTTGTGGAACATTGTTGTCAGAGTACTTCAGCCCTTACTGGTTAGGATAAGTTTTAATCTATGCATCTGAGAAACTGTTCTTGTTTGTAATTGAGGCATTGGGGAAGCAGTGGGATTTTGTGGAGCATTGTTGTCTGAGTACTTTATTCTTACTGGTGAGAAAAGTTCTAATCTTTGCATCTGGGGGTTGAATGTTTCTTGTTTGTAATTGAGGGCTTGAAGAAGCAGTAGGTTTTGTGAAGCATTCTCGTCTGAGTACTTTAGCCCTTGAGATATTCCTCCCACATTGTGTGAACAATCTCTTCTCGTTGAGCATGCGTTTCCCGATCAGCTGACCTACAAGATGAAGCAACATTGTTAATCTCTCTCTCTGGTTGGTCGCTGATATTAAAGTTGGGGTTGTTATTACAAAAAAATTAGGGTAAAATTTGAAAATAAAATATAATTCCAACATTCGAAAGAGAGTTGGAAAACATGTTATAAGTTGTTTTCCAAATTTGATTTACAACTTCAAGTAGGCTTTGAACTTTTCATGACCAAACACTGGTTTTCAAATAAAGTGAAACATAATTCCGGAAAAAAGTAAATAATGTTTATGGCCAAATGGGTCCTTAGTTATGTCTGTCAGTTCATTCCTCTGTTTTGCTTGATTGGATGGTATTTCACGGTAGTTGGTGGAAAGAGTAGCAAGAATACCATAGCGAAGAAATGCCTTATTACTTTGATCCGCTTACTGGGCAAAGATTCAAGAGCATATACTTTTATAACTTGGCTACAACAAACGAAAGGGACGTAAGAGGGCACAATTTCGTCCACCTTTAGAA

>M6-PGSC0003DMG400022067 M6 LRR-RLP

ATGGCCTCGTTGCTTAAATTGAAGCATGAACTTACATTTGATTCTTCCATGCCAAAGAAAATAGTTAGGTGGAATAGAAACACTGATTGCTGCTTGTGGCCGGGTGTAAGTTGTGACGAAAAAGGTCACGTGCTTGTTCTGGAGTTGGACAACGAAGCCATTTCTGGCGGAGTTGAGAACTCAAGCAGTCTCTTCAATCTTCAACATCTTGAGAAGCTCAATCTGGCTTATAACGACCTACGTTCTCTTCCAATACCAACAGATATTTATAAGCTTGCAAACTTGACGTACCTGAATTTGTCACAAGCTGGTTTTGAAGGGCAGATTCCAGTGGAATTGTCAAGATTGACGAAGTTAGTAGTTCTTGACATTTCTACTTATAATGACATTGTAGGTTTTCCTTACTCTTTATTAAGTCTAGAGAGCCCCGATTTGGGGACACTCGTTGGAAGTCTAGCAACTCTTAGAGAACTTTACCTTGATGGCGTGAACGTTTCATTGAAGGGGAGTGAGTGGTGCTCGGCGTTATCTTCATCTCTACCTCAATTGAGAGTCTTGTCAATGAGATACTGTGAAATTTCAGGTTCAATTGATCCTGTCCTGGTAAATCTTCCTTTCCTTTCAGTTATTCATCTCGATATGAATAACTTGTCCACTATGGTTCCAGAATTTTTGGCAGATTTTACGAAATTGACAACTTTAAGTGTTCGATGGTGCAATTTGTTTGGTCCATTTCCCAGTAAAATCTTTCAGGTACCAACTCTACAACAACTTGATCTGTTAGGCAACGTAAACCTTACCGGCACTTTACCTGAATTTCCACATAAAAATGCATTTAGGGAGTTGAGGCTATCTGAGACGAGTTTCACAGGTTTGCTTCCAAATTCAATTGCTAACCTTCGAAGTTTGATTACACTAGACTTATATGATTGCAATTTCAGAGGACCTATTCCGTCAACAATGGGAAATTTAACGAACCTAGTTTCATTAGACTTGTCAAACAACAACTTTACTGGCTCAATCCCGCTGTTCCATGAGGCTAACAAGCTCAATTACATCGACATTTCTAATAACAATGGTCAACTTTCTTCTGCTCAAACTCAATTAGCCGTTCTCCTTTCCCTCCCGTCATTGCAGTACCTTTTTCTTTATACTAGTCATTTGAGTGGAGAAATCCATGAATTTCCAAATGCATCCTCCTCTGTTCTGGAAATACTTTATTTGAGCAATAACCATCTAAAAGGATCGATCCCCCGGTCTATCTTTAAACTCAATAGGCTTTCACAACTCTCACTTTCTTCCAACTCCTTCAGTGGGACCATTATTATCGAAGCGGTAAATGGACTTCCTAGGCTAACCACCCTTGACCTTTCAGACAACAAAATTGAAGGGGAAATACCAAATTGGGTTTGGACAACTGCAAAGCTGAATCTTTCTCACAATCTGCTGGAGTCTGTAGAAAAACCTTATTATATATCTACTACTTCTATTGTCATTGACTTGAGTTTCAACAGGATCAAGGGCAATCCACCATTTCTACCAGATCGCTTTGCAAACTGGAAAAGTAGTATTACTTACTTTTCAATTGCCAATAATCAATTTACAGGATCAATATCCTCTTCCATATGCAGTCTAGATCAGCTTCAATTCCTTGATATGTCAAATAACTCCATAAGCGGAAAAATACCTCCCTGTCTAATCCAAATGCTGGCTACGAGTCTCGTAGTGTTGAATATAGCGAGAAACAATCTAAGTGGTACTATTCCTGATAAATTTCCGCTGAATTGTAGCTTGGAAACTTTAGATCTCAGCAACAATATGTTAGAAGGGAAAGTTCCAAGCTCCCTGAAAAACTGTCAACCTTTGCAAGTTTTGAATATTGGAAACAACAAGATCAAAGATACATTCCCGTGCATGTTGAAGAAATTATACAGTTTTCATGTCCTAGTCATAAGGTCGAACAAGTTCTACGGGAATCTCCAGTGTTCTGTTGCTAATCAAACCTGGTCAAGGCTTCAAATTGTGGATCTTGCTTACAATAATTTCAGCGGTGCTCTGCTACCACAATACTTCTCGAGCTAGGAAGGCATGATGCAGGGCAACAATCCATACCCATGGGAGCAGTACTTAATGCTGATAATCTGTATCAAGACAAGGTGACTTTAACTATCAAAGGTTTAACAGTGGAATATGTGAAAATTCTTGTAGTTTTAACTTCCATCGATTTCTCTTGCAACAACTTTCAAGGGGAGATACCTGAGACATTGGGAGATCTCAAATCACTAATCCATCTTAACTTCTCGCACAATGCTTTGACCGGAAAAATACCAAATGCCCTTGGAAATTTGACTCAGCTTGAATCCTTAGATTTCTCAGTTAACCATTTACGTGGGAGGATTCCAGACGAGCTCGTGAGTCTCACATTTCTTGCTTTCTTGAACCTATCTTTTAATCAACTATCTGGTAGGATTCCAAGTGGTAACCAACTTCAGACGTTCTCAGCGAATTCCTTCGAAGGGAACACAGGGTTGTGCGATTTACCTTTAAAGAAAACGTGCAGTGAAACCAAAGTGAATGGATCATCACAGTTACTTAGCAGCCATTCTGAACATGAAATTGAAGGGAAATATATAAGTTTTTTCTTGGGATCTTCTGTTGGCTTTGGCATCATAATCTGGCTGCTTTTGCATAGTCAAAGATACAATGAGCTCCTTGATAGACTTCTTTTCAGAATGTTGGATCGACACAAAAAAAGGCACATGAATAAAATCTTTGAAGGTGTACAAGTATAGCTTTGATGTTTGATCTTTTACGTTCTGTTTGAAGGAGTTCATAATACTAATGTTCACTTTGTTTTTCCTTTGTTTTCTGTATGGTGAAGAGTGGTGTTTGAATTTTAATGTAAGGAATAATTTTCATGATGATCTGTTTATCATTTTTCAATTATTTGGCTCTTTTACTTATTTGCAAACATTACTGCTAGAGTCCTTATTTCTTCCTCTATTGTTTACAATCATTTTTCCAACTATCAATACTCAAAACATCAAAACAAA

>M6-PGSC0003DMG400022068 M6 LRR-RLP

ATGAAAATATCACTCCTGCTACAGTTTTCTTTCATCACCATGCAGGTGCTTTTTTTGATTAATTCTCCAGTGTATGGTCATTGTTTAGGCGATCAAAAGGCGTTGTTGCTTAAACTCAAGAATGGCCTTACATTCGATTCATCTTTGTCAACCAAACTGGTAAGATGGGATCAAAACACTGACTGTTGCCAGTGGCCAGGTGTAAGTTGTGATCTGAAAGGTCATGTGCTTGTTTTGGAACTGGACAACGAAACAATTTTCAGTGGACTTGAGAATCCAACCAGTCTATTCGATCTTCAGTATCTTGAGAAGCTGAATTTGGCTTATAATCATTTAAACTCTGTTCAAATACCGAGAGAAGTTTATAAGCTCACAAACTTAACATACCTGAATTTGTCATATGCTGGTTTTGGTGGGCAGATTCCAATGGAATTGTCAAGATTGACGAAGTTAACGTTTCTTGACCTTTCTAATGTCGCATTAAAGCTTGAGAGCGGAGATCTGAAGACACTGGTTAGAAACCTGCCAAATCTTAGAGAACTCCATCTTGATGAAGTGAACATATCATTTAAGGGGATTGAGTGGTGCTCAACTTTATCTTCATCTCTGCCTCAGTTAAGAGTGTTAAGCATGACAGATTGTGGCATTTCAAGCCCTTTTGATCCTGTCCTCTTAAATCTTCATTTCCTTTCAGTCATTCGTCTTGAGGCAACAACTTGTCCACTATAGTTCCTGAATTTCTAGCAAATTTTACAAAATTGACAACCCTCAGTCTCAGTAACTGCAACTTGAGAGGTGAATTTCCGAATAAGATCTTTCAGGTACCAACTCTACAGGAGCTTGATTTATATGGTAATGAAAAACTCACTGGCACTTTACCTGACTTTTCACAGAATGGTTCATTAAGGGAGTTAGGTCTTGGCAGCACAAATTTCACAGGTTCACTTCCGGATTCAATTGCTAACCTCACAACCTTAACCAAGATTGACTTAAGTTCATGCAATTTCTTTGGACCTATATCATCAAAAATGGAAAATCTAAGAGATTTAGTTTATTTAGACTTTTCATCCAATAGCTTTACTGGCTCAATCCCACTATTTCATAAGGCAAATAAGCTCAATTACATTGACCTTTCTCATAACAATCTTACAGGTCCACTTTCTTCTACTCATTTTGGAGGTCTTTCAGAGCTTGAGTATCTATACTTGGAAGACAATTCTATCGGTGGGACCCTTCCGACAGTTGTGTTTTCCATTCCATCATTGCAGGTGCTTGAACTTCAGAATAATCACTTCAGTGGAGAAGTCCATGAATTTGCTAGCGCGTCCTCCTCTTATTTGTATAAACTTGACTTGAGCAATAACTATCTAAATGGATCGATCCCCCGGTCTATCTTTAAACTCAATAGGCTTTTGCAACTCTTACTTTCTTCCAACTCCTTCAGTGGGACCATTAATATTGAAGCGATAAAGGGACTTCCTAGGCTCAAAAAGCTTGACCTTTCTTACAACAATTTGAGAATTGATGTACAAGGTAGTAATTCAACCTCGTTTCCCTTTCCCCAAATGAGTGAACTGAACTTGGCATCATGCCAGTTGCAAAAGTTCCCGGAACTCAAAAACCAGTCACTCATGATTGCATTAGACCTTTCATACAACAATATTTCAGGACAAGTACCCAGCTGGATTTGGAGTATTAGTTTAAGTTATCTGAATCTTTCTTGCAATTTCCTCGAGGCTCTGGAAGAACCTTACGACACATCTTCAGAATTATGGAGTGTAATTGACTTGCATAACAACAGGATCAAGGGCAATATACCCATTGTACCTACTTCTCTTATCTACCTTTCAATTGCCAATAATAAACTTACGGGATCAATACCCTCTTCCATATGCAATCTATATCAGCTTCAATTTCTTGATATGTCAAACAATTCCATAAACATCAAACTACCTCCATGTCTATTTCAAATGTTTGACTATCTCTCAGTGCTGAATCTAGGTAGAAACAGACTAAGTGGCATTATTCTTGATACATTTTTGTCAAACTGCAGTTTGAAAACTTTAGACCTCAGCAACAATAACTTAGAAGGAAAAGTACCAAGATCCCTGCAAAGATGTGCATTTATGGAGGTTTTGGACATTCGAAACAACAAGATTAGAGATACATTCCCATGTATGTTGAAAACATTACCCAGTTTGCACATCCTAGTCCTAAGGTCGAACAAGTTTTATGGAGATCTCCAGTGTCGTATAGCTAATCAAACATGGTCGAAGCTTCAAATTATAGATATTGCTTCCAACAATTTCAGGGGAGCACTGCTACCACATTACTTCTCGAGCTTGGAAGGCATGATGAAGAGCAGAAATCCAGAGCCACGACTACACTACTTGGAAGTTGAATTCATCAACTATGGCCTCTACTATCGAAACAGAGTGACTTTAACTCTCAAAGGTCAGGAGATGGAGATTGAGAATATTCTTGAAGTTTTTACATCCATCGATTTCTCCTGCAACAACTTTGAAGGAGAGATACCAGAGGTATTGGGAGATCTCAAATTGCTTTACCTTCTTAACTTATCGCACAATGCTTTGACAGGAAGAATACCAAAGGCCCTCGGAAAGTTGACTCAGCTTGGATCCTTAGATCTTTCGGTAAACCAGTTAAGTGGGAGAATTCCAGACGAGCTCGCGAGTCTCACATTCCTTACTTTCTTGAACCTATCTTTTAATCAACTTTCCGGTAGGATTCCAAGAGGAAACCAACTTCAGACATTCTCAGCAGAATCCTTTGAAGGGAACACAGGGCTGTGTGATTTTCCGCTGAAGAAACTTTGCAGTGACACCAAAATGAATGGATCGTCACAACCAAGAAGCCATTCTGACGATGAAACAGTTGATGGGAAATATATAAGTTTTGCATTGGGATCTTCATTGTGTTTTGGTATCGTTACCTGGCTGTTTTTGCATAGCACAAGATACAATGAGCTCGTTGACAGACTTCTTTTCAGAATTTTTGGTGAGCACAAAAAGGCAGGCAGAAATAGAAACAGAAGGAGGTCAAGGTAA

>M6-PGSC0003DMG400022404 M6 LRR-RLP

CTTGAGAGTTTCTTTGGCTCTTCTGGTGCACCAAAGAGTCCACATTTTTCACCATTCTCAAGACTGAGGTATGCAATAAAAATAAGAAAAAAATGCCAACAGCTTTTTCAATTCTTGTTCAATTCCAGTTTGTTTTTTTCTCTTACAACTCACGCTTTTCTCCCTGTTTTCCTTGTAATTGTTCTGAAATTTTACTGATTCTTTTCCTCTTTTTCTTTTGTTTTTTTCTGTGTGTTAAATTGATATAAGCATGGTTTTTCTGGGTCATTTTTTTTATCTGCTTTTTCAGTAGCATTTCAGAAATTTCTCAGCCTTTTCTCTCTTTAGAGTTTAGACCCATATCTCTTGCTCTGTTTTCTTGAAGCTCTAAAATGGAGTGTAGTACAACTGATTGCAGATGATGTTTTCTTTTTGAAATTCAGGTGACTCAAGATGGTTTTTTGTTTGTTTACAAATCTTGAAAGCTAGTATTTTTTTTTTTATAACCAGCAGAGTTGTTGACTATGTTTATTCAACTGAGGGGGTTAAAGCTGGAATCTTGCTGTTGAAAATTTTGTCAGCGTGTCGAAAAATAGGATAAAGATTTGAACTTTCAGTTGTTTTGATTTCAATTCAGTTTTGTTGAGATGGAAATGAATCCGAGCTTGAAAAAACCGAGGGAATCTGTAGATTTATCGAACTCATCGGAGTTAACTCCGTCAGCATTCCCAGATGAGGTGTTGGAGAAGGTGTTGTCTCTTGTTCAGTCTCATAAAGATAGAAATTCAGCATCACTGGTGTGCAAGGATTGGTATAATGCAGAGCGTTGGACAAGGACTAAGCTGTTTATTGGGAATTGCTATTCAGTTACTCCTGAGATTGTAGCAAGAAGATTCCCAAAGATTAAGAGTGTTACCCTTAAAGGGAAACCAAGATTTTCTGACTTTAATTTGGTACCAGAGAATTGGGGTGCTGATATTCAGGCTTGGCTTGATGTTTTTGCCAAAGTTTACCCCTTTCTTGAGGAGTTGAGATTGAAAAGAATGGCTGTTAGTGATGAGAGTTTGGAGTTTTTGGCAAAATCATTTCTTGGATTTAAGGCTCTGTCATTGTTGAGTTGTGATGGTTTTAGCACTGATGGGATCAGTAGCATTGCTGCTCACTGCAAGTAAGATGATCTTTTCACTCGTGTTAACTTTTGGTGGTTATTTTTAATATCAGTTAATGTTTCAGATATGGGTGTTGTTTTGATTGGATAAAAAATCTATCTTTTTTTCAAGAAATAGTTTTTATTTGATTGTAACTGGTATATTATCTAGCATCTGACTTCAAGATCTTAGCTTTCTGTTATGGTGGATTTAGAATTCTTTCAAGGCAGATTGAGGAAAAACAGTACTGTTTACAAGGTGGTCCATGCTAATTATTGTAGTAATGGAGGACAAAACAGTAGACCCATGTGTCTGGATGTGGAATGGTTAAGAAACTTGTATTTTTCATTTGAAGTGGCCATGTGGGTGAAGAAATAAAAAAGTGACATGTTCTGGGCTGGGGACATGTTAGGCTGTTTATCTTGTTATCACCACCTTGTCTTACCACACATTAAAAAGAAAGAAGAAAAAAGTATAAGTGGAAATTGAAATGGTTCCACTTTATGAATTCTTTTTCTGTTCCTTTTTTATGTTACTTTCATGATGTGACATTGACCTGTTAATGAGCATTAATTGAGGGAATTCTTTGATGTCATTGTTAGGCATAAATAACTCTTCTTATTGGTCAAGAAAGAGTTTCTACATTAACTTGGATTGTGTAACAAATATTTGATCTCTGAACCTTTTAATGTCTATTAGACCTGTTCAATGTATGGAGCTTCAACTAGTATATCACTATGTATATGTTGTCAAATTGTTGTCTAGTTTTGCAGCACCACTAGGTTATTATGGTTCATGAGGCTTTCAGGTTGGTTGTTGTCTTGGCAAATAAGGAATACCATCACAAAGGTGTATAGGTCGCATGCCAAGATTTTGCTTGTGTAGTTTTAAACGTAGAAGTTGGCATCTTTGCAATTTTAAGTTACGATAATCTGAACCAAAAAGAATGTGAGAATGTGGGATCTTTGGTTGTTCTGGTCCGCAATCGTTCCAAAGAAATGTAGTTTGAAGGATGGCTCTAATGTCATATCCACAATTCAACATACTACAGAAGTTGGACTTGTATATAACAAATGTCTTCATGTACTATCTAAATGATGCATACAAGGAGGTTCTATTTATGTTATGATAACAAGATTTCGTTTGAGTAATATTCCAAGGTATAGAAAAGCAAAACATGCTTTTCTTCATAGGCAGCAGAAATGTCGGTTTGTTTTGTGGCTCTTAATTGATGCTTATCAGCATGAAACCAAGTTATCTTAATTGCTATACCAATCTAAATGTTGGTTTCCTTCACAGGAACTTGACAGAGCTGGACATTCAGGAGAATGGCATGGATGATATTTGTGGTAGTTGGCTAAGTTGTTTTCCGGATGACTTTACATCACTGGAGGTGCTTAACTTTGCCTGTATGAACACTGAGATCAGTAAGGATGCTTTAGAGAGACTTGTCGGTAGATGCAAATCACTGAGGGTTCTGAAGGTTAACAAGAACGTCACCTTGCCTCAGTTGCAACGCCTGCTCGTCCGAGCTCCTCAGCTGATGGAGCTGGGTACAGGATGCTTTCTTCCAGATCAACTCACAAGTCGGCAGTATGCAGAACTTGAAAGTGCATTCAGCAACTGCAAGCATCTGCACTCTCTTTCAGGCTTTTGGGAAGCAAATCGTCGATATTTACCATCTCTTTATGCAGCCTGTGCTAGGCTGACTTTCCTCAACTTGAGCTATGAAGCCATTAGGAGTGGTGAATTTTCGAAGCTTCTTGCTCATTGCCCAAATTTAAGGCGCCTTTGGGTAAGAATCTACCTCAGGCTTTAACATTTACGTGATAAATCCTTTCATGATGAGGTCTTAGGTATTGATCTCCTAACATTTGACATTTTGTCGAGGAACTGAAAGTTACCACTAATTCGTCTTTTAAGTTTGAACTGCTCTATGGTTGTTTTCTCTAGGGATAAGTATTCAAGAGATGCTAATCACCTTTTTTTTCATATCATGGTAATCCATTTTTCTACTTTCAAGGTCTTCATGCCTAATCTTTCTGTTATAGGTCCTAGACACTGTCAAAGATAAGGGGTTAGAGGCTGTTGGAACCAGCTGCCCATTGCTTGAAGAATTGCGAGTCTTTCCTGCTGACCCATTTGAAGAGGATATGGACCATGGAGTGACTGAGTCAGGCTTTGTGGCTGTGTCTGCTGGTTGTCCTAAGCTTCAATATGTTCTCTATTTCTGCCGGCAAATGACTAATGCTGCTGTTGCAACAATAGTGCGTAACTGCCCTAATTTCACCCATTTTCGTCTCTGCATAATGAATCCTGGCCAGCCAGATTACTTGACAAATGAACCCATGGATGAGGCATTTGGTGCAGTGGTCAAGACTTGTAAAAAGCTCCAGAGGCTTTCTGTTTCTGGACTGTTGACTGACCTGACCTTTGAGTATATTGGAAAGTATGCCAAAAACCTTGAGACACTTTCAGTGGCTTTTGCTGGCAGCAGTGACTGGGGTATGCAGTGTGTGCTCGACGGCTGTTCTAAGCTGAGGAAGCTGGAGATAAGGGATTCTCCATTTGGAAATGCAGCACTTCTTTCTGGAATGGGGAAGTATGAATCAATGCGGTGTCTTTGGATGTCAGCTTGCAGGGTCACCATGAATGGTTGTAGAATACTTGCACAAGAGAGGCCTAGGTTGAATGTCGAGGTAATCAAAGATGAGCACAGCGATGATTATGCTGATAAATTATACGTCTATCGCTCTGTAGCAGGGCCACGAAAGGATGCCCCACCTTTTGTTGTAACTCTCTAGCACAAAGAAACAGGAACTTTGGTAAGCTCCTTCCCTTGCTCTCCTCTGTCCTATGCTTGCCAAAATAGCGTGGAGTGGAAAGATGATATGCTGATAATGCTAAGTTTTATTTTTATCTGTAGGTGTGCACTGGGATACCCTTCTATTTCAGCTAGTTTTATCTGCTACATGCATGATGTTCGTAAAGGTGAACGAGAGAAAGTAAGTAAATGAATACCTCTCCTTATGCTGCATTTACTATGTTTTACTTGGAGAAGCTTGAAATTTTCAAGGAAGATCAACAAAAGGGATGTTTGTGTGCGCTCTTACTGAACTAATGCATTATGAGAAAGAAACCATGCGAAGATTCTCTTACAGTTAGTGAGACCAAATGATCATCACTCGTGGAGAAGCAGGAACTGCAACGAATGATCACATGACGGAGTTGATTATGTCTCAGAAAAACTATCAGCAGATGAAGAGTCCTGAATCTCACAAGAACAAGCAAGATGAAGAACGCTGGTTTGTTGATTCGCGGAGTTTTTTCTTTTTCTTTTTCAAAACAATAAGTAGCATAAACATAAAGAAAGAAAAATGAAATGAGTCTCATGTAACTTGAATGATGGTGATTAGGTTTCACAGACATAAGTTATTTGGCCAAACTGGGAATGCAAAAGAAAAAAAGAAACCATGGTATTCTTTTTTTTACAATGTGATTGCTGTGCTTCATCAAAATAATTTAGTGTGACATTTACTATGTTG

>M6-PGSC0003DMG400022452 M6 LRR-RLP

TCAGTGGTACCCTTCCCGAGCTTAAGACTAATTCAATAGAAATACTTGATTTGTCTCGTAATCAATTCTCTGGTCCCATACCTCAATCGCTTAGACATCTTCTCAACCTAACTGTTGTCTTTCTTGGACAAAATAAGTTGAGCGGCGAGATTGGAGCAGAAATGTTTTCAAGCATGACAAACCTCTTAATTCTTGATCTTTCTAATAGTGGTTTATCATGGAGTGGCAATATCAACACCACTTTTCCCCTTCTTTATTATTTAGCTTTACGCTCTTGTCGTGTGAAAGATTTCCCAGATTTCATATTCAACTCTAAGAACTTATGGATTTTGGATCTCTCGGAGAATGAAATTCACGGGCAGTTACCGAAATGGTTTGGGGGTTTGGGTGCATTGGCAAACCTTAACCTCTCTCACAACTTCCTTACAAGCTTAGACCATCTACCTTGGGAGACAATCATGGTACTTGATCTTCAATCAAATTCACTCACAGGACCTTTGCCATCCCCCATTTGCACATCAACTTTCCTTTATATAATAAATTTGTCTTACAATAATTTGAGTGCCGAAATTCCCAATTGCTTGTTTACTTTTTCCTCGCTCATGGTCTTAGACTTACGAGCTAATAACTTCCAAGGACCCATCCCAAACAAATTCCAAAAGAACAGTCAGCTAATTCATATTAGTTTGAGCAAAAATCAATTGGAAGGTCCCATCCCAACATCCCTAGCCAACTGTACATCCTTAAGAGTCCTTGATTTGGGAAACAATAAAATCCATTCTACATTTCCCTCCTGGTTGGAAACATTGCAAGAGTTAGAGCTTCTCATACTGAAATCTAATAGATTTTACGGACCCATCGTTTCTTTCCAAACAAAATTGCCATTTCCAAATATGAGGATCTTTGACCTCTCCAACAATTCATTTACCGGCTCTTTACCTATGGAGGTTCTAAAAGGTTTCAAAGCTATGATGTACATGGATGCACACAAATCAGGAGTGGAGTACTATGTCGAGGAAAATTTAAGAAGTTACTATGTCGATAAGCTTTTGTATAGTGGATTGTATGTGGAATCAGTGATATTGGTGATGAAAAATCAAGAGACTAAATTCAACAAAATCTTGAAGATGTTCACTACAATCGATTTATCAAGGAACAAGTTTGAGGGAGAAATCCCGAAGTTCATTGGGAATTTTAATTCACTTCTGCTGCTGAATTTATCTCACAACAACCTCACTGGGCATATTCCTATTGAAATGAAGAACATGAGCACTCTTGAAGCTTTGGACCTTTCATTTAACCATCTTACCGGTAAAATTCCAGTGGAATTGGCAATTCTCACATTTCTTGAGGTCCTAAATCTGTCACACAACCATCTTGTTGGACCTATTCCTCAGAGTAATCAGTTCAATACATTTCCAAATGATTCCTATTTAGGAAACTCTGAATTGTGTGGATTTCCACTGTCAAATGAATGTGGGAAGCATAAGAGTGCATCAGTACCAGTTGAACAGGAAGAAGATGAACCAAGTTTTCTTAGCGAAATGACTTGGCAGTCTGTGCTAATTGGTTATGGTTGTGGCTTGTCTTTTGGTTTTGGCATAGTGTATCTCATATACCTCTTTGAATGGCCAAGATGGTTCATACACTCTTTTGAAACTATCACTCGTGACATGATATACATAGTACTGAGGGCACGTAAAAGGCGCAGGAGGAATTTTCACAGAACAAGACACTGAAAATCTATG

>M6-PGSC0003DMG400022599 M6 LRR-RLP

ATGGCTTCTCAACTTCTTTGTTTTCTTGTATTGCTAACTCTCACAACTATTTGTAAAAGTAAATCTTGTCATCCAAATGATTTAAAAGGTCTTCAAGATTTCAAGTCTGGAATTCATTCAGACACTTCTGGTAGACTAAGTAAATGGATTGGCCGCGATTGTTGTAAATGGGAAGGTATTTTTTGTAGTAATGTAACTAAAAGAGTTGTAAGTATAAGTCTACCAGGTCTCTATACTGCAGATGATGCACCAATTCAATCTGCTATGGAAGGGGAATTGTCACCTTCAATCACACTCATCAACTATCTTGAAGTTATCGATATAGGTGATCTCCTTGGTATAACGGGGAAAATCCCCTTATCGATTGGTTTACATCTACCAAATCTCAGAAAACTCAACTTTGTTAGAAACAAGTTTACTGGTGCATTACCAGAAAGCATTTGTATGTTGTCTAAACTGGAAGAATTGTATCTCCAAGAAAATGGTTTTTCGGGTTTTATTCCCTCGTGTATAGGAGGTGTCAAGAATCTAAGAAGAATGGATGTACATTCAAATAAACTTTCTGGTGTAATTCCTGAGTCTATAACAAAGTTGAAAAAATTGGAAAGTTTGTGTCTTCAAGAGAATGTCTTGACAGGGGATATACCAGAAAACATTGGTAATTTGCAAGAATTGAAAGAGGTTGATTTGTCTAATAACTTGTTAACGGGGATAATCCCTCGTTCTATTTCTAGACTAAAGTTTATGTCAGTTCTTTACCTGAACTCCAATCAGCTCGAGGGGGACATACCGTTGCCTTCAAAACCTGGGAAATTGTCTTCTCTGAGCTTTGTAAGACTGCAGAACAATCGGCTTAGTGGAACGATACCTTCATCTATTGGCTACTTGACCTCCCTTCAGAGAGTTTCACTTGCAAACAATCAGCTAAAGGGTTCTATTCCTTCGAGTATAGGTAATTTGAAGTTATTGGAAATGTTATATCTCAGTTGCAACCAATTATCTGGTCAGTTGCCAAGATCAGTAGGCGGGATTTCAAAGCTTCTTTACCTGAGTATAGCTCATAACATGATTGAAGGACCGTTGCCTCATGAAATTTCTTCCCTTTCAAATCTTCAATCGCTGGATCTTTCATTCAACCGTTTGAATATGTCAACCATTCCCAAGTGGCTCATGCAGTTGCCATCACTATCACAAATTTACTTGGCAGGATGTGAAATTCATGGTGAAATGCCAGATTACATACCCAAGTCATTGCTGGAACTAGACTTGTCCGCTAACCATCTTTCAGGAAGGATACCAGCTTGGATTGGCAGCTTTTCGCAGCTTTATTCTTTAAACCTTTCTAAGAACAAGTTTGTGTCTGAGATTCCTTCAACTGTCACAAATCTCGGCATTCTGGGAGTGTTAGACCTTCACTCAAACAAGCTAGAAGGCAGTGTGAATGCAGTTTTCCAGATAAAGAGCAGGTTTGCAGAAGGATCATTGACATATCTTGATCTGTCTGATAACAATTTCTCAACTGGTGTCGAGCAGATTGGCATAGGAGGACAAGAATATATCCAACACTTAAATTTATCGCATAACTTTCTCAAAGGTAGATTACCAACCTCTGTAGGAACATTGAAAACACTACAGACTTTGGATCTCAGTTACAATGGATTAGGAGCTAACCTGCCAACATCACTGGCCAACGTGACCATTCTGGAGAGACTGATGCTGCACAAAAATCAATTCACCGGCAGAATCCCTGAGGAGTTTTTGAAGCTGAATAAACTGAAAGAATTGAACCTTTCAGATAATCTTCTTGAAGGAAAAATACCTTATGGGAAGCCATTTCTTGATTTTCCGCAGAGCTCATTCTCTGGAAACAGAGATTTATGTGGGAAACCCCTTCTTCCCTGTAAATCTTGA

>M6-PGSC0003DMG400022785 M6 LRR-RLP

CAAAATCTCACATTCAAGAGATACTTCCTCACTGCTCCCTTTGGATCAAAAAGATATAATCGTTGGACTGGATGAAGACTTGCTGAAGATAAAAGAACGACTCACCGTGCAATCGTCAAGACTCGAAGTTGTCCCCATTATAGGGATGGGTGGTATTGGCAAGACCACACTGGCTAGAAGAGTTTATGATGATTCCTTGATTGCATACCATTTTTATGTCCGAGCATGGGCAAATGTATCTCAAGAATTTGACACTAGAGAAATTTTTCTTGGTATTCTTCGCTCCATTGGCATGGTCAACGATGAAGAAGAAAGAAACAGTACTACTGAGCAATTGGCTGAAAGAGAAAGAAACAGTACTACTGAGCAATTGGCTGAAAGAGTTTATCGAAATTTAAAAGGGAGGAAGTATCTTATTGTCCTAGATGATATGTGGTCTATCAAGGCTTGGCAGCATGTGAGGAGATCATTTCCAGATGATCGTAATGGAAGTCGAATCGTGTTGACCACAAGGCTTGTAGATGTTGCATCTTGTGTTTGCTCTGGCAATTCCCTTCACCAGATGCGTTTTCTGAGCATGGAGGAAAGCTGGGATCTTTTAAGCGGTAAGGTCTTTGGGAATGGTAGTTACCCTCCGGAATTGGAGAAAATTGGGAGGTACATTGGTCATCAGTGTCAAGGGTTGCCACTTGCAGTTGTTGCAATTGGTGGACTACTTTCCAAGACGAGCAAGGAAACAAGTTCTTGGGAGAATGTTGCAGAAAAGGTAGGATCACTTATGACAAGTGACACTATGGATTGCTTGAACATACTTTCTTTGAGCTACAACCACTTGCCTCAGTACTTGAAAACATGCTTCCTTTATATGGGAGTTTTTGCAGAAACTCGTGAGATCCCGGTCTGGAAGTTAATAAAGTTATGGATTGCTGAGGGCTTCGTCAAGAAAGTTAATCATAAAAATCTAGAAGATGTGGCAGAGGAAAATTTGAGGGAGTTGGTTGATAGAAGTTTGGTTTTGGTGGGGAAGCATACTTCACTAGGAAAAATCAAGACTTGTAAGATGCATGATCTCGTTCGAGATATGTGTTTGAGAGAAGCCAAATATGAGAATTTTATACACTTCGAAACAAGGTATAACCACGATGATCTTCTAGAAAACATTAATTGTCTACGCCGTGTAGTTGATTTCAATAAGTATAGATACAGTTTTGATATTCCTAAACTTATGCCACTCACACGTTCTCTTTTCTTGAAATATCCATACTCTGTTTGTTCCCTCTCACAGTCAAGCTTTAGATTACTTAGGATATTAGATGTTTGCTTTACATCTCGTTTGTACCGTAGTATGTCCATAACAGAACTTGTTCATTTGAGGTATCTTGCTTGTCCATCCTTCAATGGTCTTGTGCAATCACTATGCAAGCTACGGAATATACAGAACTTAGTTATTCAGGATTTACGTTCATCTGGTTTACATAGGAGAACCCAGTTCCTACCATGGGAAGTTGTGAATATGCCTAAACTAAGACATATCCATACAAAAAAGTTTAGTCTTTTCATACCCCCTACATCACTAATTGCTGAGAGGGAGAATCATTTACAAACGTTAACCGGATTGATGCCCTCATCTTGTAATGAAGAGGTGTTTCTAAGAATTCCAAATTTGAAGAAGTTGGGAATTTTAATTATTGATGATTCAGACACTATTCAGAAGTGTTATTGCCTTGATAATCTTGTACATTTAACTCAGCTTGAGAAGCTCAAAGTAGAGGTCAGAGAGAGTTGGCTAATTTTTAGGCCGACTATCAAAGGTTGGGTGGACATTCCACATTGTGACAATTTCCCACCAAACCTTAAGAAGTTAACATTGTGCACAACACACCTAGAATGGGAAGACATGAACATTCTTAGTAAGTTACCTAACCTTGAGGTGCTCAAACTTAAAGATCATGCCTTTCGCGGACATATTTGGAAACTAAGCGATGAAGACGAAGATGGATTCTTGAAGCTAAAGTTTTTACTTCTTGAGAGCTTGCATCTGATGCAATGGGAAGCAACAAGTTATCATTTTCCAAGCCTTGAGCACTTAGTCCTAACAAATTGCCGGTACTTGGAACAAATTCCTTTCGACTTTGCGGAGATTCAGACACTACAGCTGATTGAACTACACGAATGTATGCGTTCTGTTCTTGTTTCAGCAGAGCAAATACAAGAGGAGCAACAAAGCTTTGGAAATGATGACCTCATTATTCGTGTGAACTCTATCTATACTAGAGTAAACATTAATAAAAAAGGTATACACTTTTTTTAAAAGCATAAAAAAATTTGATTAGCTAGATTCCTTCGAAATCTGAAAATGATTTTTATATTCATCATTTTCACTTATTTATACTATAAAAGGTGCTAATTACAA

>M6-PGSC0003DMG400022977 M6 LRR-RLP

CTTTATTATATCTAAAGACTATTGACTTTCAAGTAATGAATCTGTTAGAGTCATTTTTGAAAAATAGCTAAGATAAGAGGATTGAAATCTTTGAATCTTTTATCTCAATGGAAGTGTCATTAAAGGAATTGGTCAAATGAAGTTGTTAGAGTC

>M6-PGSC0003DMG400022978 M6 LRR-RLP

TTGAAAGGTAATATTACTCCTTCATTGCTTGAATTGCAGCATTTGAAGTACTTGGACCTTAGTAACAATGATTTTGGTACAAGTCAAATACCGGATTTCATTGGTTCATTTCCAAGACTAGAATACCTTAATCT

>M6-PGSC0003DMG400022988 M6 LRR-RLP

CATGGGAGATGCATTAGTTCAATGGAAAAATAGTGAATTTGTGTACAATAAAACACTCGGATTGTTGAAGATCATTGATTTTTCAAGTAACAAATTAGTTGGAAGAGTTCCTGATGAAATCACACAATTAGATGGACTACTTTCACTAAACCTTTCACGAAACAACTTAACAGGACACATCATACAAGGAATTGGGATGATGGAAATGTTAGAGTCTCTTGATCTGTCTCGAAATCAGTTAACAGGACGAATTCCAGCAAGTCTTGCTCGATTAAATTTCCTAAGTGTTTTAGACTTATCGAGTAACAATTTGTCAGGGGAAATTCCTTCGAGCACTCAATTGCAAAGTTTTAATCCTTCGTCATATGCAGGCAACAATGAACTTTGTAGTCCACCACT

>M6-PGSC0003DMG400022989 M6 LRR-RLP

TTCTTGATGACATAGATCATGGTGATCATTTGGAGTACTTAGCAGGTGATGTTGGTTGGTTTGGTAATGGCAGCAGAATTGTTGTAACAACTAGAAATAGACATTTGATAGAGAAGGATAATCCAATATATGAAGTGTCTACACTGCCTAATGAAGAAGCTGTTCAATTATTCAATCAACATGCTTTCAGAAAAGAAATTCCAGATGAACGTTTTATGAAGTTCTCATTGGAGGTAGTAAATCATGCTAAAGGCCTTCCTTTAGCCCTCAAGGTGTGGGGTTCTTTGTTGCATAAGAAAGACCTAACTCAGTGGAGAAGAACTATAGACCAAATAAAGAAAAACCCTAATTCAGAAATTGTTGAAAAACTCAAAATAAGTTATGATGGGTTGGAACGTGAAGAAAAAGAGATATTTCTAGATATCGCATGTTTCTTCCGTGGATTTGAAAGAAAAGAAGTCATGCAGATCCTTGAGAGCTGTGATTTTGGAGCTGAATGCGGATTGGATGTCCTAGTTGACAAATCTCTTGTGTTCATCTCCAAATATAATGTGATTGAAATGCATGACTTGATTCAAGATATGGGTAGATATGTGGTAAAAATGCAAAAGGATTCGGGAGAACAAAGCAGACTCTGGGATGTTGAAGATTTCGAAGATGTGATGGTCAACAATACAGTAAGTAGGCTAAACAATGCTATAATATTCAGTTTTTAGTCTCCATCTTCTATTTTTTATTTAAAAGATGAAAATTGTTGGTACCCTATCAATTACAACTTCGTTCTCTTACAACATGAAAAACTTTTCGATTGGACTTTAAAGAACTAGTTGAAAATTGTATTACTAGAAATTAGAAATATAGAATTAATTGCATAAAGTATTTCTCTTTAGCGACATTTTTCTTAACTAATATTGTACCTTAATTTTATTTAATGGTCTCTTCAAATGCAATAATAAATAATACAAAACATAACGAAATTTGGCATTGCAAACAGTTAGAAAGTAGTCATATTGGCATGCAAATTTTATACAGATATTTGAAGTTATACTTTGTAAACTAATGAACCTTTTAACAGGGGACCAAGGCAATGGAAGCAATCTGGACACATGGTGTTCAAAAATTATGCTTTAGCAAAAAAGCCATGAAAAACATGAAAAGGCTTAGATTATTATCCATCATTAGTTTTAGTACACATGATGACTCCATTGAGTACCTGCCCAACAACTTGCGTTGGTTTGTCTGTCATAGTTATCCTTGGGAGTCATTATTAGAAAATTTTGAACCTAAGAAGCTTGTTTGTCTTTATCTCCAGAGGAGTTCCCTGCGTCAATTATGGACCGGAATAAAGGTACAATTGTTGAATTCAATTTTCATTATTTTTCTTTTTGTCAAATACGTACATAGATAGACTCTCAAATTTTAAATAAGTTTTGGTCTAAAATTGTTGGGGTGGTCCAATTAGAGTCAAAACTGAGGGACCAGTTTCGTTGGCTATTTTGTATCAAGAATTCGAATTTGATGAAATAGTTGTTGAGCTTTTTAATTTTAGCATAAAAATACTTAGTTTAGTTGACATAAAGGATAATTTTATATAAATTGGGTTAGAGCAAATTTGATTCAATGACTATGAGAGTAAGATATTTTTTATTGGTCTACTAGCCAAATCAAAATTATCCTGGGCAGCATCCTAATTCTCATTCTTTCTGTGTTTTGAATAGCAGCATTTGCCGTCTTTGCGAAAGCTAGATCTTAGAGATTCCGAAAGGCTGATTCAAACACCAGATTTCACGCAAATGATAAATTTGGAGTATTTGGATCTGCGTAACTGTAGTGATCTTGAAGAGGTTCACCGTTCCTTGGAGTGTTGCAGAAAACTCATTGTGTTAAATTTGCAGAGGTGTAAACGCCTAAATAAGTTTCCATGTGTTAACGTGGAATCCCTTGGATCTCTGATTCTACTACATTGCTCTAATTTGGAAAAATTTCCAGAAATCTTTGGAAGAATGAAGCCAGAGTTAGAGATCAAGATGAGCTGGTCTGGGCTAAGGGAAATACCATCATCTATTATTCAGCAATACGCATGTCATCTTACAGAGTTAAGTTTGAGTGATATGAAAAAACTTGTAGCACTTCCAAGCAGCATTTGTCAGTTGAAAGGTTTGGTGAAGCTGGATGTGTCATACTGCTCAAAACTTGAAAGCTTGCCAGAAGAGATTGGTTATTTAGAAAACTTGGAGGAGCTTCATGCTAGCTTTACTCTAATCTCACGACCTCCGTCTTCCATCATCCGCTTGAACAAGCTTAA

>M6-PGSC0003DMG400022990 M6 LRR-RLP

ATGCATTAGTTCAATGGAAAAATAGTGAATTTGTGTACAATAAAACACTCGGATTGTTGAAGATCATTGATTTTTCAAGTAACAAATTAGTTGGAAGAGTTCCTGATGAAATCACACAATTAGATGGACTACTTTCACTAAACCTTTCACGAAACAACTTAACAGGACACATCATACAAGGAATTGGGATGATGGAAATGTTAGAGTCTCTTGATCTGTCTCGAAATCAGTTAACAGGACGAATTCCAGCAAGTCTTGCTCGATTAAATTTCCTAAGTGTTTTAGACTTATCGAGTAACAATTTGTCAGGGGAAATTCCTTCGAGCACTCAATTGCAAAGTTTTAATCCTTCGTCATATGCAGGCAACAATGAACTTTGTAGTCCACCACT

>M6-PGSC0003DMG400023018 M6 LRR-RLP

ATGATCTTGCGTAACTGAAAGCATTAGTTCCTAATCTTGAACTAGATTCAAGGTGCTCTTTACTACAGTTTCTCGTACTTTCTGAATAAAAGAACTAGGTAACAAAGAGATACACACAAAATCCATTTCTACTTGTCAGTTCAATGAGGGAGAGTATGTATAAGTTGGAGTGTGCAGAGTCACAAGATATAAAACATGCCAAGTAAATCCAAAAATATTTTGTATTATACTTTCTATTCATACTAAGTAAAGGAAATTAAAGGGAGAGAGCAGTGATAAATGACAAATGGAAGAAATGAAAATGAATGAAGAGAAACGTAAGACTTTGACGATTTGGGAACTTAAAAGAGTAAAATGCGGTATAATTAATGTGAAACTCCAAGAGCTCCTTTGGTCGTTCACAAGACCTAAAGTTGTATACCAAATCAGTCAATTGCCGGATGCTCTTCTCCTTTGCTCTCTTATATCGACTGATCAAACACTCGTAATAACCGCTCTCTCAAAAACGTGGCACTATCTCTGGAATTGACTTGATACCTTTTGTTTTAGTACTTTTAATTACCATGAGCGTTATGTATCCTATGTTCATTATGTATTAGGTTATTATATTTCTTCCAAAATTAAAAAATCCTACTTGATAGCCTTGAGTCGGATGCACACGGGTGGAAAATATGAAACTCCGATTAGTTTCTGAAGAAGAGGTTTGTGCCTTGCCTGAATGTTTATGCACTTGTTCATCGTTTATAACACTAATTTTGGGATTTTGATGCTAATGTGTCCATAGCATGAAAGTCTCTAAAAACATAAAATTGAAGTGGATAGTGTTATCCAATGACCATATTGTGACTGTTGGATTGTCCTGCATTAGAAACTATGAAATTATCTTGTTTTAAAGGTTTTAGTTGCATGGAAATTAGGTCTTCAAAATTAAAGAGACTGAATTTGATATCATACAATGAAGATGAAGAAATTGATCATTCCTTGGAAATTGTTGCCCCTTATCTTCATCATTTGGGAATTTCAGGAAGTTTTCATGGTCTCAAGTGTAAGCTTGTAGACATATCCTCCATGGTTAATACTAATTAAGCTTACGTTCGACATTACATGTATCAAAGACATGTAACATGCGGTATAATATTGGATGTCTTTGATGGATGTAATGTCGAATATAAGCTTAGTACTAACCACATATGATATGGCTACAAACCTACACTTAAAAAGATGATGACTTCCTGAAATCTCTGAATAATGAATGTACTATAGGAGACAACAATTTCCACAAAATGATCAATTTCTTCATCATCATTATATGATATCAAATTCCTCTTCAATTTCGAAGACCTAATTTCGAAGCGACTAAAATCCTTAAAACGGAATGATTCCAGAGTTTCTAATGGAGAACAACCCAACAATAAATTCGCAATATGGTCCTTAGATATTACCATCCACTTCAATTTTATACTATACGGAGATTTTGAAATTATATCCACATGCACATCAAAATCTCAAAATTAGTATTATCAAATGAATGTGTATAAACATTACGGCAATGCACAAATCTCTTTTTTGAAACTGACCTGAGTTTCACATTTTCCTTTGATCAAAGTTCTAAGTTGAATTTTTTTAATTTGGGAGAGAAACAAAATTACCTAAGCCATAATCAACCTAGGATACATATCCTTTGTGATAATTAAAGTTACTAAAAGAGAAGATATCAAGTAAATTTCAAAGATAAAGCTACCTGTTTGAGAGAGCACTTGTTACGGATATCTGATTTCGTTAGAATAATAGAGAGACTGTGAAGAAAAAGAGCTTCAGGCAACTAACTGATCGGTCTACCGTTGTTGTTTCTTCTGTCTTTTCGCCGGAATCCATGGTTGGTTTTTGGTTACAGCTATTGTTTCTTCTATTTTCTCAGCTGGAATCCATGGTTTGTTGTTGGTCTACAACTGTTGTTTCTTTTATCTTCTCGCCGGAATCCAAATGTTGTTTGTCGGAATCCGGCAACTGATTGGTCGATCCAGCAACTGATCGGTCGATCCGGCAACTGATTGGTCGATCTACATCTGTTGTTTGCTGGAATCCAGAAATGCTTAGAAGGTATCGCCCCCCTTTCCCTTCAGTTTTAACTTAGGCACAAAATGAGCAAATTGAGTAATATATTGGCATAAATTTG

>M6-PGSC0003DMG400023198 M6 LRR-RLP

GCTAATAATACATCAACCTACCATTTCTAGCAACAACAGGTAACAGCTGTGGAGACTTTTTTTTTTTTTGTATAAATAAATTGTAACCCCTTCGATGGATTTATGGTTGGATTTTCCATCAAACCAAAAACTCTCCATTTTCTCTCTCACACAACTCTTCTTCTTATAGCTCCAGTTTCTTGTTAAAGAAAAAATGTCAGTGAATAGTATAACTGAGTTCTGTTCCTGAATCCTAAGTGGGGTTTGCTTTTCTCATCTTTTTTTTTTATGCTGCCCTGTTTCCTTAAACACTCGTTTTTCACTGTATAAAACACGAAAAACAAGTAAAGTTTTGTTCTTAACTACTATTATTTTGATTTGATTTGATTTGATTGGATTAACGTTTGTATTCTTGTCAATATTGATCGGCGCTTCTAGAAGTTGTACAGAAGAAGAACTCAATTATGGCTTTCTCAATATTTTTCGTTTTTTTGATCTAGGACTCTTGGAAACTAACCCCCAAACAAAGTGTATTCAAATCTGGATACTGTGGGTGGTTTCTGTGTGTAGCTGTGTGTGTGTGTGTGTTTTCCGACGAAATTGAGGGTGAAATGGGATCCTCCGCAATGTCTGTTGACGAGATTGTGGAAGAGATGATGAGGCTTAACAAATCTCTACCAATAAGGCCAGGGATAGATGAAGTGGAAGCAGCAAAAGTTTTAATTATGAATATGGAGAAGGAGGAACAAATGAAGTTAGAGACAATTGCGCGGCAAAACAAAAGAAAAGATGTACCAGAAGAGGTATTCAAAATATTGCAAGAGATGCAGAGGAATTTAGTTTGTTTTCAGAGTAAAGAACAGAAGAGAGAGGCATTAAAATTGTTAGATCTCGAAAATGTTCATTACCTGTTCGATGAATTGATTCAAAGAGCTTCGAAATGCTTGTCATCTAATCCTCAAGCTAATAATACTTCTTCAGCTTCTTCAAGGAATTCCAGCAATTTGTCTCTGGCGAATTCATCTTCCTTCAACGGTAGCAGTTTCAATTCTCCGGCAACTACTACGACTATGTCTTCGTCGAGCTTTTATAGTGAGAAAGAGCCTGTTAAAGTGTCTGAACTTGTTACAAGAGACGACAGTTACTTGAAGAAGCCGACAACTGCGTTTCAAATGGATGGAATCAGTGTTGGGCTTCGATCTGGGAACGCTTCTTCTGCTCCCCAGATTGTCGATACCACTTTGAAGCGTAGTACTGGTAAGTTTAATCTATTTCCTGTAACATATTGCTTGTGACCAATGTGTTTTTTTTATACTTTGTTAGCTTAATTGGATAAATTTTGTTTCCTATAATATGAAAATTATTAGTTTAATGGAGAAGCCATTTATCTGTAGGTCAAAATGATGAAAAAATGAGCTTGATAAAGCTTGCTAGTTTGATAGAAGTTTCAGCCAAAAAAGGAACCAAGGAGCTAATTCTTCGAAGAAAACTGTCAGACCAATTAGAATGGATTCCAGATTCACTTGGGAAGCTATCAAATTTGGTGACTTTGGATTTGTCTGAGAATCGAATTGCTGTGCTGCCAACTACCATCGGGGGTCTTTCATCGTTGCAAAAACTGGATTTGCATGGAAATAGAATTGTTGAACTTCCTGATTCAATAGGAGATCTGCTTAACTTGGTCTATCTTGATCTAAATGGCAATAACCTGAAGACATTGCCTTTGACTTTAGCGAGGTTAACTCACCTTGAGGAAGTCGATCTAAGCTCTAACATGCTATCTATACTCCCTGAGGCAGTGGGATCCCTTGTCAGTCTCAAAAAACTGATCGTTGAGACTAATGACCTAGAGGAACTTCCTCACACTATTGGTCAATGTACTTCGCTGAAAGAGCTCCGTGCTGACTATAACCGTCTTAAAGCTCTTCCTGAAGCCTTGGGGCGAATGGGTTCTTTGGAGATTCTATCCGTGCGGTACAATAACATCAGGCAATTGCCTACAACTATGGCATCCTTAACAAGTTTGAAAGAGCTCAATGTTAGTTTCAATGAGCTTGAGTCAGTGCCTGAGAGCCTGTGCTTTGCCACTACACTCATAAAACTGAACATTAGCAACAACTTTGCGGACCTGCAATCACTCCCTAGATCCATAGGAAACCTTGAGATGCTTGAGGAGCTGGATATGAGTAACAACCAGATACGAATTCTACCAGATTCATTTAGGATGCTTTCGCATTTGCGTGTACTAAAGACAGAAGGAAATCCTCTTGAAGTGCCACCAGGAAACATTATCGAGATGGGAGCACAGGTGTGTTTGAGTTTTATTTTTACTATACTATCATTATGCTCTCTGAATTTTTAATTGTTTGTTTTAGTTTTGATATTTGACTCATGAACTTTTATTTTGCTCATCAGGCTGTTGTGCAGCATATGGCTGATCTTATTGAAAAGAGGGATGCGAAGCCACAGCCCGTTAAACAGAAGAAATCATGGGCTCAGATATGCTGCTTTTCAAGATCCAACAAACGTAAGCGCAATGGTATAGACTATGTCGTCCAAGCGTGACACACTTAAAAACTCTTGACCCTATATTTATCTGGTAAGTTTGTTCTCTTTACTACCTCTGCTTCCAGCTATTAGTCTGTTGGCGAAAGAAAACAAGAAGTCTTTTATTAAGGGGAAACTGGAGGTAGTATCCAGTATCCAGCTGGTGATCCAGTATTCATGGCACTGATCCAGCAGACTGGTTGGATAGTTCTTAGCTCTTACACATTTGTTGTTTATTAATTATCCTCAATTCTTGTCCTTTGATTTTGTGAGTTAATAAATGTATATGGCGCACGAGACATGTTTGCTGCATTAATTACAATAATAAAATGTTGGGTTGTAAGGGAATGGCTCACATGCTTTTGTTATCCCTCGTTCAGTGAAATTGAGCTGTGATTGTGCTGGAAGTGAGCTCTGCCATTCTTCTTCAATTCCAACTCTGCCAACAGAATTATCAGTATTTTTTTGTTACCACATGTTTCTTTTTGCCAGTTGGTTCTATTTATCATCTTCTTTGTAATATTTGCTTGTAATTTTATAATGTATGTTCTAAAATCAGTTGTAAAAATCGATATCACACGGTTATGAAGAGGGAGGAAATATCTGTTGTTGTCGAATTCCCTCAATGTTAATGAATTATTATGAGTTGACTTCAGA

>M6-PGSC0003DMG400023274 M6 LRR-RLP

GAGGGTTTCATACCCAGAGAACAAGAAAGAACGGAGGATGTCGCTGAAGGCTTCTTGAATGAACTGACAAGACGAAGCTTGGTTCAAGTGGCTTATACATATTGGGATAGAGTTACTAAATGTAGGGTTCGTGATTTACTCCGTGATCTTGCGATAGAAAAGGCATTGGAGGTAAACTTCTTTAACATTTATGATCCAAGAAGCCACTCCATATCCTCTTTATGTATCAGACATGTGATTCATAGTCAAGGAGAAAGGTACCTCTCACTTGATCTTTCTAACTTAAAGTTGAGGTCAATTATGTTCTTCGATCCCGATTTTCGTAAGATGAGTCATATAAACTTCAGGAGTGTGTTCCAACATCTACATGTGTTGTACTTGGATATGCATGTTGGGAATGTGTCTATAGTACCTGATGCCATAGGAAGTTTGTACCACCTCAAGCTGTTAAGATTGATCCATATCCGTGATGTTCCTTCTTTCATTGGCAACCTCAAGAATCTAAATACACTTGTTGTCAATGAGGGGGCATACTCTTGCAAACTACCCCGCGAGACAGCTGACCTGATAAATCTAAGACATTTAGTTGCTCGGTATACAAAACCTCTGGTACATATAAGAAAACTCACTAGTCTTCAAGTTGTTGATGGCATTTGTTGTGATCAATGGAAAGATGTTGACCCTGTTGATTTAGTCAATCTTCAAGAATTAAGCATGGATCATATGAATAAATCTTACTCCTTAAACAACATTAGGAGCTTGAAAAACCTTAGCACTCTCAGATTGCTTTGTAAGTATGATGGATCATTTCCATCGCTTGAATTTGTTAATTGTTGTGAAAAGCTCCAGAAATTGTGGTTAAGTGGGAGAATAGAGAAACTGCCTCATCTGTTTCCAAATTCCATCACAATGATGCTTCTGAATTTCTCAGTACTGACAGAAGATCCGATGCCTATTTTGGGAATGTTGCCAAACCTAAGGAATCTCGACTTGGTTAGAGCTTATGAAGGAAAAGAAATAATGTGCAGTGATAACAGCTTCAGTCAACTGGAGTTCCTTCAACTTAGTGAACTTGAGAAGCTAGAAAGATGGCATTTAGGCACAAGTGCCATGCCTCTGATTAAAGGTCTTCATATCCATGCCTGTCCAAATTTAAAGGAGATTCCAGAGAGAATGATAGACGTGGAGCTGTTGAAGAGGAATTATATGCGGTGAGCTTTTCAAGCATATATCATGTCTTCATAATCGTAGAGGTATTGATGGGGGCATTTCTGCCAAGCACATTGGTTATTAATTTCAGGGCTGTCAATGAGAGTGGTTTGAAGATAATTGTTTTTGCCAAAAATGATTTTGTGATGTTTTAATTAATTGGTCTATTGTATGATCTCTTACCACATTTAAGACAATGCTTTATATGTTAAAATGAAATTAAAAATATTATATGGTACTCTCTC

>M6-PGSC0003DMG400023299 M6 LRR-RLP

ATGAAGCCAGAGTTAGAGATCAAGATGAGCTGGTCTGGGCTAAGGGAAATACCATCATCTATTATTCAGCAATACGCATGTCATCTTACAGAGTTAAGTTTGAGTGATATGAAAAAACTTGTAGCACTTCCAAGCAGCATTTGTCAGTTGAAAGGTTTGGTGAAGCTGGATGTGTCATACTGCTCAAAACTTGAAAGCTTGCCAGAAGAGATTGGTTATTTAGAAAACTTGGAGGAGCTTCATGCTAGCTTTACTCTAATCTCACGACCTCCGTCTTCCATCATCCGCTTGAACAAGCTTAAAATCTTGAGTTTTAGATTAGACTTGAAACCTAACTCCTTTGCCACAAAAGAATCAAAATATAGAGTTTCCTTTGTGTTCCCTAGGGTGAATGAAGGGTTACGGTTGTTGGAAATTCTGGATCTCAGCTACTGCAATCTAACAGATGGAGGACTTCCAGAAGACATTGGATGTTTATCCTTTTTGAAAAAGTTGAATCTCAGGGGAAATAATTTTGAGCGTTTGCCTCAAAGCATAGCCCAACTTGGTTCCCTTCGATCCTTGGACTTATCACAGTGCATGAGTCTTAAAGAGTTTGTAGGTGTTAACATGGCTGAAGGGTTATGGTCGTTGGAAATTCTGGATCTCAGCTACTGCAATCTAATAGATGGAGGACTTCTGGAAGCCATTGGATGTTTATCCTCTTTAAAAGAGTTGAAGCTCAATAGAAATAATTTTGAGCGTTTGCCTCAAAGCATAGCCCAACTAGGTTCTCTTCGATTGTTGGACTTATCAGAGTGCATGAGGCTTAAAGAGTTTCTAGGTGTTAAGATGGCTGAAGGGTTACGGTCGTTGGAAATTCTGAATCTCAGCTACTGCAATCTAATAGATGGAGGACTTCTGGAAGACATTGGATATTTATCCTCGTTGAAATTCTTGTATCTCATTAGAAATAATTTTGAGTGTTTGCCTCAAAGCATAGCCCAACTAGGTTCTCTTCAATTCTTGGACTTATCATATTGCAAGAGGCTCAAAGAGTTGCCAGGTTTCACGGGGATGCAATACTTGGATACTTTGGATCTCAGTTTCTGCGATCTAACAGATGTAGGACTTCTGGAAGACATTGGATCCTTATCCTCTTTGAAACACTTGAAACTCAAGGGAAATAATTTTGAGTATTTGCCTCATAGCATAGCCCAACTTGGTGCTCTTCAAACCTTGGACTTATCAGATTGCAAGAGACTTAGACAGCTGCCAGGTTTCACA

>M6-PGSC0003DMG400023396 M6 LRR-RLP

ATGTGGTTAAAGAAAGATTTCATTATATTTGCTACTCTATTTTACCATTTTTCGATCGTTTATAGCTGTCATGAAACAGAAAGATCAGCTCTGTTAAGCTTCAAATCCTTGCTGACTGATCCATCTAATCGGTTAACTTCATGGCAAGGGGGAAATTGTTGCAATTGGAAAGGAATCAAGTGCTCGAGTTCAGGTCATGTTGTAGTTGTCAACCTCCGGAATCCTAATCCCGTTGAAGTCATGATCACTATCAATAAGGAAGTTGTTTCAAATTTTAAAAACACGTCTGATTTTTCTCTTAAAGGTACTATATCTCCATTGCTCTTTACTCTTGACCATATGCGACATCTCGACCTGAGTTTCAACAACTTCATGTACTCAAAGTTACCTGTTGAGATATCGAACTTAACAAAGTTGACTTATCTCAATCTGTCGAATGCCATGTTTCAAGATTCCATAGCCACGCAATTCTCAAACCTTACATCTCTGAGGTCTCTTGATCTTTCGTGTGCTAATTTAGTACCTGATTTTTCTTCTGTTTCTGTTAGCTTGACATTTCCACTGAAGTTGGATTTTGGCTCATTGTTTTCTTTTATAAGCTATGGTTATTTGTCTAGTCCAAATCTTAGGTGGTTACAAGGACTCCGACGTCTCAGATATCTAGTATTGACTGGTGTTGATCTTTCAAAGGCGTCCGAATCATTTCATTGGGCTAAACCAATATCAAGCCTTTCAAATCTCAAGTCACTTCATTTGTCCAGCTGCAGCATTTCAGGAAGAATTCCGGTAGGGCAGTTACTTAACCTTACTAATCTTGCTACTCTAGATATGAGTTCTAACGTTCTAACATCCTCGATACCTGATCTTCTATCGAACCTCAGTACCCTCTCGGTTCTTGATTTTAGTAGCAATGATTTAGATGGTCATGTCTCTTATATTCCTCAACTCAAAGAACTTTCTGTTTCTGGCAATCCTGCTATGACCATAGATCTTGTTTCAATGTTTTCTGTTCCGTGGCCGCAGTTGACATTTCTTGACATAAGCTTCGTTCGTGTAGCTGGAATGATTCCTCCTTCTTTAAGCAACTCATCTTCATTATCCTTTTTCCGAGCTGATGGATGTTCGATCCAAGGGTCAATACCTTCTTCTGTCACAAAACTTAAGAAATTAAGTGTACTGATGCTGAATGACAACGATATCACAGGCCAACTTCCTGTATCCATGTCTAGTTTAGTAAGCCTTCAATACTTATCTATGTCTCAGAACAGATTAGAAGGTCACATACCCATTTCAATCTGTCAAATCGCCTCCCTCAAGTACCTGAATTTAGAATGGAACGATCTAACAGGACGTTTTCCGTTGTGCATACTTCAGCTTCCGAAACTTTCATCTCTTTATATTCAGAGGAATAGACTGAATGGTAATATGCCATTGTCTTTGTTCCGAAAGTCTAGATTAGATGAAATTAGTTTAGGAACTAGTGGATTGTCATTGGAAATCAATGATCAAGATCAACTCTTCGTGCAAACGTTCCAGCCCAAGATTTTGGATTTCACATCTTGCAACATGAGAGGAGAAATCCCGGATTTCTTTTCGAATTTTACAAGCCTCGTGGTTTTGAATTTCGCTAACAACAGTCTATCGGGAGCCATACCACACTGGTTGTTCAATCTCCCTTCCCTCTCTATACTATCTTTGCCTATGAATAACTTCAGGGGAATTATACCTCCAATGATTCAGCTGAAATCTTCTCATTTCCCAACAATAGTAAATTTAGCCAGAAACAAACTTCAAGGTCCTCTCCCTTCTCAGCTAGAGAATGTTAACGTCATCGATCTGTCACTTAACAATTTTGTAGGCTCGATTCCATCACGGATAGGAGAAGTTCCCGGTATCAGGTCCATATCTTTATCCGGAAACACAATTCATGGTCTAATACCTGAATCATTTTGCCAAGCGACTAACGTTCTCCAGGTTCTTGATCTCTCTAATAATAGCTTGTCAGGCACCATTCAACGTAATATGGGAAACTGCAAGTCTCTCATTTACCTTAATCTTGGACAAAACAAGCTCACTGGAAGTGTTCCAAAAGAACTTGAACGCGTTACAAGCCTTCGTTACCTTGACCTGAATGGAAATAAGCTAGAAGGATCTTTTCCAAAAATGATTGAAAACTTTCAAGATTTGGAGATTCTGATCTTGGCAGGCAATTTATTTGAAGGACCGATACCAAACTTCATCGGTGAGCTACACGGCCTCCGTATACTCGTTCTTTCATCCAACATTTTCAATGAATCTATCCCAGAAGGGCTAATGAAGTTGGAAAATCTACAATATATCGGTTTGTCCAGGAACAATCTGTCCGGTCCCATTCCTGAGAATCTCGATGGCTTGAAAATGATGACGAAAAGGAAAAATGAGGCAACCATCTTAGGTTATGTTTACTCCCTCAAGTTCACTGGTGCACAGCTAGAAATAGTCACCAAAGGACAGATACATTGGCTTGAAACAGTATATTCCTACAACACTGGATTCGACGTCTCGAGCAATGCTCTTACTGGTAAAATCCCTGAAAAAATTGGCCTTTTAAGTGGAATCCCATTCCTGAATCTCTCACATAACAATCTGTTTGGACTGATCCCAAAGACAATCGGCGATATGATTTCACTCGAGTCGTTGGATCTTAGCTACAACCATTTCAGAGGTGAAATTCCAGTGACATTGGCATTGTTAGATTTTCTCCAACATCTCAACATGTCCTACAACAATTTGAGCGGACGGGTTCCAAGGAATCCACATTTCGACACTTTGTATCAAGACGGAGCAGCATATATCGGAAACAAATACCTATGTGGTACTCCTGATGGGATGAACTGCAGTAGCAATGGTCCCTCTATCATCACTGAAACTACAGAGAACGAATATGATCAAGAAAATGTCCTGTTTGTTCTAGTTATATTTTTTGGCTATGTAACAGGCATTTCTGGATTGTTTTTGCTCTTATATTTGATGAAGGACAATTGGAGGAACAAATACTGGAGGGCAATAGACAGAATTGTGTTAAAAATAGTAAAATAG

>M6-PGSC0003DMG400023498 M6 LRR-RLP

ATGTATTCAAAGTTACCTGTTGAGATATCTAACTTAACAAAGTTGACTTATCTCAATCTGTCAAATGCTATGTTTCAAGATTCCATCACTACACAATTCTCAAACCTTACATCTCTAAGGTATCTTGATCTTTCGTGTGCTTATTTAGTACCCGATTTGTCTTCTATTACTATTAGTTTGACATTACCACCAAAGTTGGATTTTGGCTCATTGTTATCTTTTATAAGCTATGGTTATTTGTCTAGTCCAAATCTAAGGTGGTTAGAAGGACTTCGACGTCTAAGATATCTAGTATTGACAGGTGTTGATCTATCAAAGGCCTCGGAATCATTTCATTGGGCTAAACCAATATCAAGTCTTTCAAATCTCATGTCACTTGAATTGTCTAGCTGCAACATTTCAGGAAGAATTCCATTAGAACAATTGCTTAACCTTACTAGTCTTTCTACTCTAGATATGCGTTCTAACGTTCTAACATCCATGATACCTGATATGATATCGAATCTCACAACCCTCTCAGTTGTTAATTTTCGTGGCAACAATTTAGATGGTCATATCCCTTACCTTCCACAACTCGAAAGACTTTCTGTTTCTAGCAATCCTGCTATGACCATAGATCTTGTTTCAATGTTTTCAGCCCCATGGCCAAAGTTGGCATTTCTTGACATAAGCTTCACTCGGGTAGGTGGACCGATTCCTCCTTCTTTAAGCAACTCAACTTTATTATCCTTTTTCCGAGCTGATGGATGCTCGATCCAAGGGTCGATACCTTCTTCCATTACAAAACTTCAGAAATTAAGTATATTGATGCTCAATAACAATGATATCACAGGTCAACTTCCTGTTTCCATGTCCAGTTTAGTAAGCCTTCAATACTTATCTTTGTTCCAGAATAGCTTACAAGGGTATATTCCAAATTCGATTTGTCAAGTCCCCTCCCTGGAGTACCTGAATTTACAATGGAACGATCTAACAGGACGTCTTCCCTCGTGCATACTTCAGCTTCCTAAGCTTTCATTTCTTTATGTTCAGATGAATAACCTTAATGGTAATATGCCACTGTCATTGCTCCAGAAGTCTGGATTGGATCAAATCAGTTTCGGAGTTAGTGGATTGTCCGTGGAACTCGATGATCAAGATCAATCCTTTGTGCAAACTTTCCAGCCCACAGTTTTGGAATTTACATCTTGCAACATGAGAGGAGAAATCCCGGAGTTCTTTTCAAATCTGACGAGTCTCGAGATTTTGATTTTGGCTAACAACAGTCTATCAGGAGCCATACCATACTGGTTGTTCAATCTCCCATCCCTCTCTGTACTAGATTTGTCAATGAATAACTTCAAGGGAGTTATACCTCCAATGATTCAGATGAAATCTTCTGTTTTTCCAACGGTAGTAAATTTAGCCAGAAACAAACTTGAAGGTCCTATCCCTACTCAGCTCGAGAATGTTAATGTCATCAATTTGTCATTTAACAATTTTGTAGGCTCAATTCCAACACAGATAGGGGAAGTTCATGGTATCAGGTCCATATCATTATCAGGAAACAAAATCCATGGTCCAATACCTGAATCATTTTGCCAAGCGACTAACGTTCTCCAGGTTCTTGATCTCTCTAATAACAGTTTGTCAGGCACCATTCGACGTAATTTGGGAAACCGCAAGTCTCTCATTTACCTTGGTCTTGGAAAAAACAAGCTCAGTGGAAGTGTTCCAAAAGAACTTGAACGCGTTACAAGCCTTCATTACCTTGACCTGAATGGAAATGAGTTTGAAGGTTCTTTTCCAACAGTGATTGGAAAGTTTAAAGATTTGGAGATTCTGAACTTGGCAGGCAATAGATTTGAAGGAAGGATACCAAAGTTCATCGGTGAGCTACACAGCCTCCGTATACTCATGCTTGCATCAAACTCTTTCAATGAGTCTATACCGGAAGGGGTAATGAAGTTGGAAAATCTACAATATATTGGTTTATCCAGGAACAATCTGTCCGGTCCCATTCCCGAGAATCTTGATGGCTTGAAAATGATGATGAAAACACAAAATCAAACAACTATCTTAGGTTATGTTTACTCCCTCAAGTTCACTGGTGCTCAGATAGAAATTGTCACCAAAGGACAGACACATTTTCTTGTGTCAGTGTATTCCTACAACAGTGGATTCGACGTCTCAAGCAATGCTCTTACTGGTAAAATCCCTGAGAAAATTGGCCTTTTAAGTGGAATACCATTCCTGAATCTCTCACATAATAATCTTACCGGGGTTATCCCAACGACTATTGGCGAAATGATTTCCCTCGAGTCGTTGGATCTCAGCTATAACCAACTAACAGGTGAAATTCCAGTGACATTGGCACCCTTGGATTTTCTTGCATATCTCAACTTGTCTTATAACAATTTGAGCGGACGGATTCCAAGAAATCCACATTTTGACACTTTATATCAAGACGGAACAGCATATATCGGAAACAAATACTTATGTGGTACTCCTAATGGGATGAACTGCAGTAACAATGGTCCCTCTATCATCACTGAAACTACAGAGAATGGATACGATCAAGAAAATGTCCTGTTTGTTTTAGTTATATTCTTGGGATTTGTAACAGGAGTTGCTGGTGTATTTCTGCTGTTACATTTGATAGATGAGAATTGGAGGAACAGGTATTGGATTGCAGTAGACAGAATTGTGTTGAAAATAGTGAACTGTAAACTATGA

>M6-PGSC0003DMG400023549 M6 LRR-RLP

ATGGAGAAGCACATCTTCTTATTGATTCTTCTCTTTCTAATTCAATTTTCTATATCACTTGCTTCCTCTAATGAGACTGACCAACAAGCTCTGTTAGCTTTCCAAAATCTTATTACAAGTCCTAATCATTTTTTGGCCAATAATTGGACCAAAAATACTTCCTTTTGCACTTGGTTTGGTGTCACTTGCACTCCAAAAAGGCAAAGGGTTATGGCCTTGGCTCTTCCTGATTTGCAACTTCAAGGCACAATTTCCCCATCTTTGGCAAATTTGTCCTTTCTCAGTGTTCTCAATCTCCAGAACAACAGCTTCCACAGTGGCATCCCTTATGGACTTGGCCACATGACTCGCTTGCGAGTGATTGACGTTAAAAACAATCAGCTCAACGGAAGTATACCAACAAGTCTATTTCAAAACCGGAGAGTTCAAGTAATTTCAATGGCTTTCAATGAACTCGGTGGTGAAATGTGGAGAGGCCCATGGTATGTACCAGAACTCAGAGTCTTAAATCTCGGGAACAATAGCCTCACGGGTATAATCCCTCCTTATGTTGGAAATGCCACAAAATTGATGAACATCAATTTGTCTGGGAATAGAATCAATGATAACATTCCAATGGAGATCAGTAATCTTAGCCAACTTGCATTTTTGTCCTTGTTCAATAATCAATTAACAGGTTCCATTCCTCCAACACTGTTTAATATCTCGTCACTACTTGGCATATATCTGTCATTCAATGGCCTTTCTGGTCCTCTCTTGCTAGATGAAGGGAATATTGTGTCAAATCTGAAGCATTTAAGTATGTCTAACAACCAAATTTCTGGTTGCATTCCTTCCAACATATGCCAACTCACAGAGCTCAAACTTTTGTCCATATCTTTCAACAATATAACTGGAGACATACCTAGAAATATTGGTTGTTTATCCAAACTCGAAAGGTTTTATATTGGTGAAAATCCAATAGAAGGGACTATTCCCACTTCATTGGGAAATATTTCTACTCTGCAATATCTTGATTGTGTAGACAATTGGATGAAGGGGCAAATTCCTCAAGAATTAG

>M6-PGSC0003DMG400023551 M6 LRR-RLP

ATGGAGAAGCACATCTTCTTATTGATTCTTCTCTTTCTAATTCAATTTTCTATATCACTTGCTTCCTCTAATGAGACTGACCAACAAGCTCTGTTAGCTTTCCAAAATCTTATTACAAGTCCTAATCATTTTTTGGCCAATAATTGGACCAAAAATACTTCCTTTTGCACTTGGTTTGGTGTCACTTGCACTCCAAAAAGGCAAAGGGTTATGGCCTTGGCTCTTCCTGATTTGCAACTTCAAGGCACAATTTCCCCATCTTTGGCAAATTTGTCCTTTCTCAGTGTTCTCAATCTCCAGAACAACAGCTTCCACAGTGGCATCCCTTATGGACTTGGCCACATGACTCGCTTGCGAGTGATTGACGTTAAAAACAATCAGCTCAACGGAAGTATACCAACAAGTCTATTTCAAAACCGGAGAGTTCAAGTAATTTCAATGGCTTTCAATGAACTCGGTGGTGAAATGTGGAGAGGCCCATGGTATGTACCAGAACTCAGAGTCTTAAATCTCGGGAACAATAGCCTCACGGGTATAATCCCTCCTTATGTTGGAAATGCCACAAAATTGATGAACATCAATTTGTCTGGGAATAGAATCAATGATAACATTCCAATGGAGATCAGTAATCTTAGCCAACTTGCATTTTTGTCCTTGTTCAATAATCAATTAACAGGTTCCATTCCTCCAACACTGTTTAATATCTCGTCACTACTTGGCATATATCTGTCATTCAATGGCCTTTCTGGTCCTCTCTTGCTAGATGAAGGGAATATTGTGTCAAATCTGAAGCATTTAAGTATGTCTAACAACCAAATTTCTGGTTGCATTCCTTCCAACATATGCCAACTCACAGAGCTCAAACTTTTGTCCATATCTTTCAACAATATAACTGGAGACATACCTAGAAATATTGGTTGTTTATCCAAACTCGAAAGGTTTTATATTGGTGAAAATCCAATAGAAGGGACTATTCCCACTTCATTGGGAAATATTTCTACTCTGCAATATCTTGATTGTGTAGACAATTGGATGAAGGGGCAAATTCCTCAAGAATTAGGGAAGCTATCAAATTTGAGGGAATTAAGCTTTGCCAATAATTCTAATCTTATTGGTCAAATTCCAGAGGCTATTTTCAACATATCTTCTTTGGAAGTGATCAATTTTAGTTTGAATAACCTCTCGGGGAGAATTCCAACCACTACAGGTCTTCATCTTCCGAACCTTAAAGAACTTTACTTGGGAGACAATCAGCTTGAAGGGGAAATTCCATTGTTCATAA

>M6-PGSC0003DMG400023599 M6 LRR-RLP

TATTGACTTGATGGCAGCTTATACTGCTGTAATTTCTCTTCTTCAAACTCTTGATCAATCAAATATTTCAGAACTTTTTCATGATCACAGTACTGAAATGCTCGATTCTCTTCGTGCTACAGCTGAATATTTTCAAGATGTTCTTGATAACATTATCAATGAGAAAATCAAATCTTTTGAGGAAGATATTAGAGTTGTTGTTAGTGAGGCAGAGGATGTTGTTGAAATGAAGATTTCTGAAATCATCAAAGAAGAAAGCTGGACATTTGGAATTTTACAACACCATGATATGCTACCAGTTGTTGAAAAAATGGATACGACAAAGAAACAAGTGATGGAGTTTCTTTCTCATGATGCTGATCAAATTCTTGAATTAACTGCGGATTCCTTGATTGATGCTTCTTCTATGAGTGATCCAATGCTGTCAGATCACTTGAAAGATTATATTGTGCAGGGACTTGATGGTGACTTGGAGATAATAGATAAGAGATTGAGAGGACCAACGTCTTATCTAGACATTGTCACCATATCAGGTATGGGTGGCATTGGCAAAACAACACTCGCTAAAAAAGCTTATGATCATCTTCCAATCAGGTATCACTTTGACATTTTTGTTTGGGTTACAATATCTCAAGAATTTCGATATAGAAATGTATTGTTAGAAGCTTTACATTGCATTTCAAAGCAAAGAGTTATTGTGAACACAAAAGATTATGTTAATATGAATGACAGTGAGTTAGCTGATCTGGTGCAAAAGCGTCTAAAGGGTCCAAGATACCTTGTTGTTGTTGATGATATTTGGAGCAGGGATGTTTGGGATAGCATATCACAAATATTTCCTAATCGCAACAATGGGAGTCGAGTTTTATTGACTACTAGGGAAACTGATGTAGCAATATATGCAAATACTAGTAGCCCTCATAAGATGAACCTCTTGGATTTAGATAATAGTTGGAAGTTACTCCGTGATAAGGTGTTTGGACTAGAACACAATCATCCTCCTGAGTTGGAAGAAATTGGAAAAAAAATAGCAGAAAAATGTCAAGGACTTCCCTTAATAATTTCAGTGATTGCAGGGCATCTCTCTAAAACGGTGCCCAAGACATTAGAAAGTTGGAAGTATGTTGCCAAAACCTTAAGTGAAATCATTGCTAGTCATCCAAATAAATGCTTAGGAGTGGTCGGTTTGAGTTACCATCACTTGCCTAATCGACTCAAGCCGTGCTTTCTTTCTATGGGTGATTTCCCAGAGGATTTTCAAGTTGATACTTGGAGATTGATCCAATTATGGATCGCAGAAGGTTTCATAAGGATGTCTTCTGGAAGTCATGGAAGCTTGGAAGAAGTGGCTAAAGATTATTTGGAGGATCTTATCAGCAGGAACTTGATAATGGCTAGAAAAAAGAGATTCAATGGTGAGGTAAAAGCATGCGGAATACATGATCTGCTGCGTGAGTTCTGTTTGATAGAAGCTGAAATGACAAAGTTTATGCATGTTGTGAGAACTCCATTGTCACCAGCACAAAAGCCTAATTTTCGTCGCTTCAGTATTCAAGAGTATTATTCTGTTGAGGATTCTTATAAGCTCTTACCCGCTGTAGCCAGATCTATGTACTGCTTTTCTTACTTGGATCTACCTTTACAACCCCGTATTATGGTTCTCTCTATATTGCCCATCTATTGTCATGATCCTATAATACATGGATTTTTCTCTTGTTTCAACCATCTCAGGGTATTGGCCATCTTCCTTCCATTGTTCAGCTCATTTCCAGTTGTGATTACAAATTTATTTCATTTGAGATATCTTTAAGTTGGATGTTTTGACAACAATATTCCTGAATCAATCTCAGAGCTTCTGAATTTGCAAACTCTAATTTGTGATGGTCATTCTTCAAAGACAACTTTACCTAGGAAGATATGGATGATGAAGAACTTGAAGTATATATCTTTAGGGGGTATCACTTATTTACCAAGTCCTAGAATATATAAGAATCTTGTGACAGGGATGCCAAATCTAGAGAAATTTTTCAGTCTTTGTTACTCTAGTTGTACAAATGAAGTCTTTTCTAGCATTCCCAATCTAAAGATATTGACTATTAGTGCATCTTTCAAAAAAAATATTCCCAATCGTCTCTTGGATATGTCGAGCTTGAGAAAACTTCAGGCATTAAAGATTTCCTGGGACCGTTATTTGGAATACCCCTTCAAGAGATTTGTTTTTCCAACATCACTTAGGAGGTTGACTTTAACTTGGCATTCTGAGTTTATTTGGGAAGAGATATCATCAACTTTTATCATGTTGCCACATCTTGAAGAGCTCAAACTTAAATATTGTCAAGCCGAAGATGATATATGGAGATTGACTGAAAAAGACATATTCAGAAGCTTGAAGTTGTTATTACTGAGGTACCCAAATCTTAAGGGTTGGGAAGCAAGCAGCGATAACTTCCCTAATCTAGAACGCCTTGTTCTGAAGGAATGCAAGGACCTGCAAGAAATTCCAACAGATTTCGTGGAAATTTGTACTTTGGAATCGATTGAGTTACATGATTGCACCACTACTGCTGAGGATTCTGTCAGAAATATCGAACAAGAACGAGAGGACATGGGATATAATATCCTTAAGGTCTACATCCATAACAGCCACAGTAAGTTATAATTCTCTCTGAAAAGTTCTTCGATTCTTCACATTATTTAAAACCTCATTTTTATTAAATCATACTTCTTTGAGATTAAGATGATGACTTGTACTTTTTATCTTGCATATGCAGGTAATTGAAGCTTATTTTGAGGGATCATCTAATGCTATCTTCCCAATTCAGAACCTTGTACTTCATATTTTTGTTTTTATATTCTGATCGTTTGTTAATTTCATGTTTTAGGTATAAGCGTGTTTAATACA

>M6-PGSC0003DMG400023613 M6 LRR-RLP

ATTCTTCTATTTTTTATTCAATTTTCTGTATCATTTGCTACCTCAAATGAGACAGACCAACAAGCTCTACTAGCTTTCCAAAATCTTATTACAACTCCCAATCATTTTTTGGCCAATAATTGGACCAAAAATACTTCATTTTGCTTTTGGTTCGGCGTCACTTGCAGTCCAAAAAGGCAAAGGGTTGTAGCCTTGACTCTTCCTAATTTGCAACTTCAAGGCACAATTTCCCCATCTTTGGCCAATTTGTCCTTCCTCAGTGTTCTCAATCTCCAGAACAACAGCTTCCACGGTGGCATCCCTTATGGCATTGGCCACATGCCTCACTTGCGAGTGATTGATGTTAAAAACAATCAGCTCAACGGAAGTATACCAACAAGTCTATTTCAACACCGTAGAATTCAAGTAATTTCATTGGCTTTGAATGAATTCAGTGGTGAGATGTGGAGAGGGCCATGGTATGTACCCCAACTCAGAGTCTTAAATCTCAGGAATAATAGCCTCACCGGTATAATCCCTCCTTCTGTTGGAAATGCCACGAAGTTGCTGAACTTCAGTTTGTATGGGAATAAAGTTGGTGGCAACATTCCAAAGGAAATTGGTAATCTGAGCCAGCTTGCATTTTTGTTCTTGTTCAATAATCAATTAACAGGTTTCATTCCTGCAACATTGTTTAATATTTCATCACTTGTTGCCCTATCTCTTGGAACCAATAGCCTTTCCGGTCCTCTCTTGCTTGATAAAGGAAATAGTGTCTCAAATCTCAAATTTTTAAGTATATATGACAACCAAATTTCTGGTTGCATTCCTACCAACATATGCCAACTCGCAGAGCTCAAATTTTTGTCCATATCTTTCAACAACATAACCGGAGACATACCCAAAAATATTGATTGTTTATCTAAGCTCAAGGGGTTCTATATTGGCGATAATCCAATAAAAGGGACTATTCCCACTTCATTGGGAAATATTTCCACTCTGCAATATCTTTATTGTGGAAACAATCGCATAGTGGGGCAAATTCCTCCGCAATTAGGGAAGCTATCAAATTTGAGGCAATTAGGCTTTGACCATAATTTTAATCTTATTGGTCAAATTCCAGAGGCTATTTTCAACATATCTTCTTTCGAAAGGATTGATTTCAGTTTCAATAACCTCTCGGGGAGAATTCCAACCACTGCAGGTCTTCATCTTCCGAACCTTAAAGGACTTACCTTGGGATTCAATCAGCTCGAAGGAGAAATTCCATTGTTCATAACAAATGCTTCCAAGCTTGAGATACTGGGGCTAAATAATAACTTTCTCACAGGAACTATTCCTACTAATTTGGGAAATCTTCATGAGCTGCGATATCTGTTCCTACATGATAATCAACTTACCAATGAACCAAGAGAGCATGAGTTGCGATTCTTCAATTCTTTGGCAGACTGTAGAATGTTGCGATATCTACAAGTGGGTAACAATCCGTTGAATGGCACTCTGCCCAATTCAATTGGGAATTTTTCATCTACTATCGAAAACTTTAATATAGTAGATGCACACATCAATGGCCTCATCCCCACAAGTATAGGCAACATGAGTGGTCTTACAGTCCTAAGCCTTGGAGGAAACAACTTGATGGGAAATATTCCTCCTGAGATTGGTAAGCTTAAACAACTCCAAGGTCTGTATCTACGTTACAATAAATTGCAGGGACATATTTCTGAGGCAGTATGCAATTTATCTAATTTGGTTGAATTATATCTGGAACATAATGAGCTCACTGGAGTGATCCCAGAATGTATAGGAGATCTAAGCATGCTACAACTGCTTTATTTAAATTCTAATAGATTTTCATCAAAATTTCCATTGAGCCTTTGGAAAATGAGTGGTCTTCTCTTACTAGATGTGTCACTAAATTCTATAGAGGGAGAAGTTCCATCAGATATTGTAAGATTGAAAGCCATTGTAGAACTATATCTTTACGGTAACCACTTTTCAGGCATGATACCGACCAGATTGGGGGAACTCCAAAACCTGTTGTCTCTTGACCTATCAAACAATTCATTTTTTGGACAAATTCCATTATCCATTGTCAACTTGATAAGCTTGGAAT

>M6-PGSC0003DMG400023641 M6 LRR-RLP

TGCATAGCAACATCATGGAGAAGCACATTTTCTTATTGATTCTTCTCTTTTTAATTCAGTTTTCTATATCAACTGCTTTCTCAAATGAGACTGACCAACAAGCTCTACTAGCTTTCCAAAATCTTATTACAAGTCCCAATCATTTTTTGGCCAATAATTGGACCAAGACTGCGTCTTTTTGTTCTTGGTTCGGTGTCACGTGCAGTTCAAAAAGGCAAAGGGTTGTAGCCTTGACTCTTCCTAATTTGCAACTTCAAGGCACAATTTCCACGTCTTTGGCTAATTTGTCCTTTCTCAGTGTTCTCAATCTCGAGAATAACTCATTCCATGGTGGCATCCCTTATGGACTTGGCCATTTGCCTCGCTTGCGAGTGATTGATTTTCAAAACAATCAGCTCCAAGGAAGTATACCAACAAGTCTATTTCAACACCGGAGAGTTCAAATCATTTCATTGGCCTTCAATAAACTCGATGGTGAAATGTGGAAAGGCTCATGGTATGTACCCGAACTTAGAGTTTTAAATCTCAGGAACAATAGCCTCACAGGTATAATCCCTCCTTCTGTTGGAAATGCCACAAAGTTGATGAACTTTAGCTTGTCTTATAATAGAATAAACGGCAACATTCCTAAGGAGATCGGTAATCTGAGCCAACTTGCAGTTTTGTCCTTGGTTGATAATCAAGTAACAGGTTCCATTCCTACATCAATTTTTAATATCTCGTCATTACGTTCCCTAACTCTGGGAAGGAATAGCCTTTCTGGTCCTCTCTTGCTTAATGAGGGAATTTTTTTATCAAATTTGGAGCGTTTGAGTTTAACTCGCAACCAAATTTTTGGTCGCATTCCTTCCAACATCTGCCAACTCATACAACTTAAAATACTGTCCATATCTTCCAACAACATAACTGGAGAAATACCCAAAAATATTGGTTGTTTATCGAAGCTCGAGGAGTTTTATATTGGTAATAATCCAATAACAGGGACTATTCCCACTTCATTGGGAAATATTTCCACTCTGCGAAATCTTTACTGTGAAACCAATAGCTTGGAGGGGCCGATTCCTCCAGAATTGGGGAAGCTATCAAATTTGATAGAAATAGATTTTGAAGAAGTTTATAATCTCATAGGTCAGATTCCGGAGGCTATTTTTAATATATCTTCTTTGGAATACATTGCGTTAACTTCGAACAAACTCTCAGGGAGAATTCCAACCTCTACTGGTCTTCATCTTCCAAACCTCTTAGAACTTCATTTGTCTGGCAATGAGCTTGAAGGGGAAATTCCTCCGCATATCACAAATGCTTCCAAGCTTGAAAGGTTGGGGCTAGCAACCAACTTCTTCAGTGGAAGTATTCCAACTAATTTGGGAAACCTTCGTGATCTGCGACTTCTTTTCCTACATGAAAATCAACTTACCAGTGAACATGAGTTGCCATTCTTCCAATCTTTGGCAGACTGTAGGATGTTGCAATATCTAGATGTGGGTTACAATCCGTTGAATAGCATTTTGCCCAATTCAATAGGTAATCTTTCATCTACTATTGAATTCTTTGAAATGAGCAATGCTCACATCAATGGACTCATCCCCACAAGTATAGGCAACATGAGCGGTCTTATAACCTTATTCTTTCAAGATAACAACTTTATGGGAAATATTCCTCCTGAGTTTGGTAAGCTTAAACAACTCCAAGGGCTGTATCTAAATAACAATAAATTGCAGGGACATATTCCAGAGGCGGTATGCAATATATCTCATTTGGGACGATTAAATCTGGAAGGTAATAAGCTCTTTGGGTTAATTCCAGCATGTATAGGAAATCTTAGCATGCTACAACATCTTTATTTGGGTTCTAATAAATTTTCATCAAAATTTCCGTTGAGTCTTTGGAAAATGAGTGGTCTTCTCTTTCTAAACGTGTCACAAAATTCTTTAGAGGGAGAAGTTCCATCAGATATTGGAGAACTGAAAGCCATTGTAAAACTAGATCTTTCAGGTAACCACTTCTCAGGCATGATACCAAGCAGATTGGGGGAACTCCAAAACCTGCAGTCCCTTGACCTATCAAACAATTCATTTACAGGCTCAATTCCATTATCCTTTGCCAACTTGATAAGCTTAGAATTCTTGAATTTGTCTATAAATGCCTTGTCAGGTACTATTCCTAAGTCATTGGAAAAACTCACCCTTAAAAGCATTAATGTTTCATTTAATGAATTAGAAGGTGAAATACCCAATGGTGGTGTGTTTGTAAATTCAACTATGCAATCATTTCTCGGAAACAAAGGTCTATGTGGAATGAGCATATTGGAGGTTCCTGCTTGCCCTATCAACAATAATGGAAAACAATCAAAGTCTAAGGAGCTAGTGCTGAAAATTGTTATTCCACTGGTTATTTCATCCTTTCTGATATTCTTGTTGGTGTCAACTTGGATAATAAAACGAAAGAAGAAAGGGAAGTCCAGAGATGTTGAAAAGGTTCCGGAGATAAGGACATATCAATTGATTTCTTATCATGAGATTCAACGAGCAACAAATAATTTTGATGAATCAAATTTAATTGGTGTGGGAGGTTCTGGCTCTGTATACAAAGGCACATTATCTAGTGGAACTTTGGTGGCTATAAAAGTTCTGGATTTGCAAAGTGAGGAAGTATGCAAGAGGTTTGATACCGAATGTGAAGTGATGAGAAATATCAGGCACAGAAATCTTGTTCCGGTAATTACTACTTGTTCTAGTGACTATATACGAGCCTTTGTTCTGCAATATATGCCCAACAGTAGTCTTGATAATTGGTTGTACAGAGAAGATCGCCACTTGAACCTTCTTCAAAGAGTCACCATAATGCTTGATGTGGCTATGGCAATTGAATATCTGCATCATGGTCATGATACTCCGATAGTTCATTGCGACCTAAAGCCAGCCAACGTTCTTTTGGATGAAGATATGGTGGCTCATGTTGGTGATTTTGGCATCTCTAAAATTTTAGCTACAAGCAAGTTCATGGCACATACCGAGACATTAGGCACTCTTGGATATATTGCACCAGGTATAATAGTCTCTTTCTATTTCACTTATATTGACTCTTTCTTTTCTATTTTCACAAATCATAATTTAGCCCTTCCAAAATCTAGAACTCAATAAAAGCAAGTGTACATAATTATTGTTATATTTCAATTGAGTAATTTTTTTTATTTCATTTCATTCTTTTCTAAGAATATGGCTTGGATGGAATAGTGTCCACTAGTGGTGATGTTTATAGTTACGGCATCATGATGATGGAGGTTTTGTCAAAAAGAAGACCAACCGATGAGGAGATATGAAATGAAAATCTTGACTTGAGGAAATTGATAACACAGGCATTTCGGGGGACTATGATGGACATTGTGGATGCCAATCTTTTTCCTAAGAAAGAACAGATCACTTCCAAAAGTGAAATGTGCATAGCCTCCATGATAGAATTGGCTTTAGACTGCACAAATGAAAAGCCAGAATTAAGGATAACCATGAAAGAAGTAGTCAAGCGGCTTGACAAAATGAAGAACACATTTCTGAACACATAGAAGTTAGCATCTCTTCCGGTGGTGTTCTTTCTCAGCTACAATTTAACATGTTGCTCTTGAATCATATATCAT

>M6-PGSC0003DMG400023863 M6 LRR-RLP

ATGGGAAATACAGGATTGTACGGGATAATACCTGAGAGTATTTTTCACCTTCCCAATCTGGAAACACTTGTCTTACAAAACAATAATCAGCTCAGTGGTTATTTTCCGAAGACTAAATGGAACAGCAGTGCATCTCTCATAGAGTTAGATCTCAGTGGCGTGAATTTTTCTGACAATTTGCCTGAGTCTATTGGCTATCTAACTTCGGTGCATTCTTTGTCTCTTAAAAATTGCAATCTTAGAGGGACTATTCCTGAATCTCTTTTGAATCTCACTCACATAGAGGATTTGGACCTTCAATATAACTCCCTGAATGGAACAATACCATCAGGGATGTTTTCTCTTCCGTCACTAAGTCGTTTAGTCTTGAGTAATAACCACTTTTCTGGTCAGTTTGAGGATTTCAATTCCAATTCACTAATCTGGATTGATTTAAGTAATAATCAGCTGCAAGGCCATCTTCCCAAGTCGATTCAAAACCATGTGAACCTAACAGGCCTTATTCTTTCTTTCAATAATTTTGGTGGCCATGTGGATGTCAGCTTCTTTGCTGACCTCAAACAACTTTATTATCTTGATCTTTCATATAATCGTATCTCGTTGACCAATGAGAACAAACATAATGTTACTTTGCCAGGATCTCTTATGAGCTTACAATTGGCCGCATGTGAAGTGAAAGAATTGGAGTTTCTAAGATCCGCTAAGCTTCTTTGGCAGTTGGATCTTTCCAATAATAAGATTCAAGGAAGAATCCCTGATTGGGCATGGACTAACTGGATGTTCATATTGCAACGTCTTAATTTATCCCACAACATGCTGGAAAGTGTTGAGTCAATTCCTCTTCTGTTTATAGAAGCTATTGATTTGCGGTCCAATTTGCTTCAAGGTTCACTACCTATTCCACCAATTTCCACAAGATTCTTCTTCATATCCCATAATAATCTCACTGAAGAAATTCCTTCAGATATTTGCAATTTGACATCACTGGTAATGCTTGATTTGGCAAGAAACAACTTGAAGGGAGCAATTCCGCAATGTTTGGGTAATATCAGTGGCCTCGAGGTTCTGGATATGCACCACAACAGTCTTTCAGGGACTCTTCCAACAACTTTTAGAATTGGAAGTGCACTCAAAAGCTTCAACTTTCATGGCAATAAGCTGGAGGGGAAAATCCCACAATCCCTGACCAATTGCAAGCAACTCGATGTTCTTGATTTAGGAGACAATCACCTCAATGACACATTCCCCGTGTGGTTGGGGACTCTGCCAAAGCTGAAAGTTCTAAGTTTGAGATCCAATAAATTGCATGGGTCCATTAGATCTCTACCGACTGGAAACATGTTTCCTCAGCTTAGAATATTAGATCTCTCCTCCAATGCCTTCACAAAAAGCTTACCAACGAGTCTATTTCAACATTTGAAAGCCATGAGGACAGTTGATCAAACGATGAACGCACCAAGTGATGAAGGGAATCGATATTACCAAGACTCGGTAGCTCTCGTAACAAAGGGATTGGAGCTTGAAGTTGTGAGAATCTTGTTTTTGTACACCACTGTTGATCTTTCAAACAACAAGTTTGAAGGATTTATTCCAAGTATTATGGGAGATCTTATTGCACTTCGTGTGCTGAATTTATCTCATAATGGATTACAAGGTCATATACCGCCATCACTTGGAAGTTTATCTTCAGTTGAATCATTGGACCTGTCAGGTAACCATCTTGTAGGAGAGATACCAGCACGATTTGCTTCTCTTACATCTCTTGAAGTCTTAAATCTCTCCTACAATCATCTCGAAGGGTGCATTCCTCAAGGACCTCAATTCTATACATTTGAGAACAATTCATATGAAGGTAATGATAGATTACGTGGATTCCCACTTTCAAAAGGTTGTGGTAATGATGGCAATGATTCGGAGACAAATGATACCACATCTGGACTTGACGATGAAGAAAACGACTCTGAATTTCTGAATGATTTTTGGAAAGCTGCTCTTATGGGGTATGGAAGTGGACTATGTATTGGATTATCGATAATATATTTCATGTTCTCAACTAGAAATCCGAAATGGCTTGCAAGAATCATTGTAGAACTGGAACACAAAATAATGATAGGAAAGGGAAAGAAGCAGCAAAGGCAAAGGAGTTACAGAAGAAGAAATAATAGCTTAACATGA

>M6-PGSC0003DMG400023864 M6 LRR-RLP

ACAATCCTGTATTTGTCGAGAAACAACTTGAAAGGAAAAAATCTGCAATGCTTGGGTAATATCAGTGCCCTCCAGTATGTGATAATGTCACATAATAATCTCAGTGGAGAGCTCCATCCGTCTATTTGCAATTTAACATCACTGCAAATTCTAGATTTGGGTAGAAACAATCTAAAGGGAGAAATTCCGCAATGTTTTGGCAACATGAGTGGTCATCTTGAGGTTCTGGATATGCAACACAACAATCTTTCTGGAACTATTCCAACGACTTTTAGAGCTGGAACACTTAGAAGCTTCAACTTGCATGGCAATAAACTTAAGGGGAAAATTCCCCAATCTTTGGTCAATTGCAAAGAGATGCAAGTTTTTGATTTAGGAGATAATCACCTCAACGACACATTCCCGATGTGGTTGGGAACTCTTCCTAAGTTGAGAATTTTAAGCTTGAGATCGAATAAGTTGCATGGACCGATTAGAACTTTAGGAAGTGAAAACATTTTTAATGAGCTTCGAATGTTGGATATCTCTTCCAATGCCTTCACGGAAAACTTACCGATGAGTCTGTTACAACATTTAAAAGCCATGAGGACAATTGATCAAACGATGAATGCACCAAGTGATGAAGGAGGTGAATATTACCAAGACTCGGTAGCTGTCGTAACAAAGGGATTGGAGCTTGAAGTTGTGAGAATCTTGTTTTTGTATACCACTATCGATTTTTCAGACAATAAGTTTGAAGGACATATTCCAAGTATTATGGGAGATCTCATTGCACTTCGCATGCTTAATTTATCTCATAATGGATTGCAAGGTCATATACCGCCATCACTTGGAAGATTAACTTTGGTTGAATCATTGGACCTGTCAAGTAACCACCTTGTAGGAGAGATACCAGCACAATTTGCTTCTCTAACTTCTCTTGCAGTCTTAAATCTCTCCTACAATCATCTCGAAGGGTGCATTCCTCAAGGAAATCAATTTCATACCTTTGACAACAATTCATATGTAGGTAATGATGGATTACGTGGATTCCCACATTTGAAAGGCTGTGGTACTGAAGGTAATGATTCAGAATCAGAGAAAACTGATACCGGATCTGAGCTTGATGAAGAAAGCAACTCTGAATTTCTGAATGATTTTTGGAAAGCTGCTCTTATGGGCTATGGAACTGGACTCTGTATTGGATTATCCATTATATATATCATGATTTCAACTGGAAAACTGATATGGCTTGCAAGAATCATTGTGGAGGTGGAGCATAAAATTATGATGGGAAGAAGAAAGAAGCAGCGAAAGCAAAGGAGTTACAGAAGAAGCAAATAA

>M6-PGSC0003DMG400023867 M6 LRR-RLP

ATGTTTTCTCTTCCGTCACTAAGTCGTTTAGTCTTGAGTAATAACCACTTTTCTGGTCAGTTTGAGGATTTCAATTCCAATTCACTAATCTGGATTGATTTAAGTAATAATCAGCTGCAAGGCCATCTTCCCAAGTCGATTCAAAACCATGTGAACCTAACAGGCCTTATTCTTTCTTTCAATAATTTTGGTGGCCATGTGGATGTCAGCTTCTTTGCTGACCTCAAACAACTTTATTATCTTGATCTTTCATATAATCGTATCTCGTTGACCAATGAGAACAAACATAATGTTACTTTGCCAGGATCTCTTATGAGCTTACAATTGGCCGCATGTGAAGTGAAAGAATTGGAGTTTCTAAGATCCGCTAAGCTTCTTTGGCAGTTGGATCTTTCCAATAATAAGATTCAAGGAAGAATCCCTGATTGGGCATGGACTAACTGGATGTTCATATTGCAACGTCTTAATTTATCCCACAACATGCTGGAAAGTGTTGAGTCAATTCCTCTTCTGTTTATAGAAGCTATTGATTTGCGGTCCAATTTGCTTCAAGGTTCACTACCTATTCCACCAATTTCCACAAGATTCTTCTTCATATCCCATAATAATCTCACTGAAGAAATTCCTTCAGATATTTGCAATTTGACATCACTGGTAATGCTTGATTTGGCAAGAAACAACTTGAAGGGAGCAATTCCGCAATGTTTGGGTAATATCAGTGGCCTCGAGGTTCTGGATATGCACCACAACAGTCTTTCAGGGACTCTTCCAACAACTTTTAGAATTGGAAGTGCACTCAAAAGCTTCAACTTTCATGGCAATAAGCTGGAGGGGAAAATCCCACAATCCCTGACCAATTGCAAGCAACTCGATGTTCTTGATTTAGGAGACAATCACCTCAATGACACATTCCCCGTGTGGTTGGGGACTCTGCCAAAGCTGAAAGTTCTAAGTTTGAGATCCAATAAATTGCATGGGTCCATTAGATCTCTACCGACTGGAAACATGTTTCCTCAGCTTAGAATATTAGATCTCTCCTCCAATGCCTTCACAAAAAGCTTACCAACGAGTCTATTTCAACATTTGAAAGCCATGAGGACAGTTGATCAAACGATGAACGCACCAAGTGATGAAGGGAATCGATATTACCAAGACTCGGTAGCTCTCGTAACAAAGGGATTGGAGCTTGAAGTTGTGAGAATCTTGTTTTTGTACACCACTGTTGATCTTTCAAACAACAAGTTTGAAGGATTTATTCCAAGTATTATGGGAGATCTTATTGCACTTCGTGTGCTGAATTTATCTCATAATGGATTACAAGGTCATATACCGCCATCACTTGGAAGTTTATCTTCAGTTGAATCATTGGACCTGTCAGGTAACCATCTTGTAGGAGAGATACCAGCACGATTTGCTTCTCTTACATCTCTTGAAGTCTTAAATCTCTCCTACAATCATCTCGAAGGGTGCATTCCTCAAGGACCTCAATTCTATACATTTGAGAACAATTCATATGAAGGTAATGATAGATTACGTGGATTCCCACTTTCAAAAGGTTGTGGTAATGATGGCAATGATTCGGAGACAAATGATACCACATCTGGACTTGACGATGAAGAAAACGACTCTGAATTTCTGAATGATTTTTGGAAAGCTGCTCTTATGGGGTATGGAAGTGGACTATGTATTGGATTATCGATAATATATTTCATGTTCTCAACTAGAAATCCGAAATGGCTTGCAAGAATCATTGTAGAACTGGAACACAAAATAATGATAGGAAAGGGAAAGAAGCAGCAAAGGCAAAGGAGTTACAGAAGAAGAAATAATAGCTTAACATGA

>M6-PGSC0003DMG400023868 M6 LRR-RLP

ATGACTGGCTGTGTGATTGAACTTAATCTCAGTTGCAGCGGTCTTGTAGGAAAGATTGATTCCAATAGCAGTCTCTTCCAACTCTCTCATCTCCAAAGGCTCGACCTTTCTTCTAATAACTTCTCTAATTCTCACATCTCGCCTGAATTTGGAAGGTTCTCAAGCTTGACACTTCTTGATCTTTCTGACTCCTATTTCTCAGGTCATATACCTTCTGAAATCTCTCATCTTTCTCAATTACAGTCTCTTCGCCTCTCCCCGTCATTTGAAACTATTCTAAGACTCACTGCTCATGATTTGAAATTGCTCCTTCAGAATTTGACTCAATTAAGTGAGCTTGATCTTACTTCCATAAATATCTCTTCCACCATTCCTCCAAATTTTTCTTCTCATTTGACAACTATGAGGATGGGAAATACAGGATTGTACGGGATAATACCTGAGAGTATTTTTCACCTTCCCAATCTGGAAACACTTGTCTTACAAAACAATAATCAGCTCAGTGGTTATTTTCCGAAGACTAAATGGAACAGCAGTGCAT

>M6-PGSC0003DMG400023869 M6 LRR-RLP

TGCAATGCTTGGGTAATATCAGTGCCCTCCAGTATGTGATAATGTCACATAATAATCTCAGTGGAGAGCTCCATCCGTCTATTTGCAATTTAACATCACTGCAAATTCTAGATTTGGGTAGAAACAATCTAAAGGGAGAAATTCCGCAATGTTTTGGCAACATGAGTGGTCATCTTGAGGTTCTGGATATGCAACACAACAATCTTTCTGGAACTATTCCAACGACTTTTAGAGCTGGAACACTTAGAAGCTTCAACTTGCATGGCAATAAACTTAAGGGGAAAATTCCCCAATCTTTGGTCAATTGCAAAGAGATGCAAGTTTTTGATTTAGGAGATAATCACCTCAACGACACATTCCCGATGTGGTTGGGAACTCTTCCTAAGTTGAGAATTTTAAGCTTGAGATCGAATAAGTTGCATGGACCGATTAGAACTTTAGGAAGTGAAAACATTTTTAATGAGCTTCGAATGTTGGATATCTCTTCCAATGCCTTCACGGAAAACTTACCGATGAGTCTGTTACAACATTTAAAAGCCATGAGGACAATTGATCAAACGATGAATGCACCAAGTGATGAAGGAGGTGAATATTACCAAGACTCGGTAGCTGTCGTAACAAAGGGATTGGAGCTTGAAGTTGTGAGAATCTTGTTTTTGTATACCACTATCGATTTTTCAGACAATAAGTTTGAAGGACATATTCCAAGTATTATGGGAGATCTCATTGCACTTCGCATGCTTAATTTATCTCATAATGGATTGCAAGGTCATATACCGCCATCACTTGGAAGATTAACTTTGGTTGAATCATTGGACCTGTCAAGTAACCACCTTGTAGGAGAGATACCAGCACAATTTGCTTCTCTAACTTCTCTTGCAGTCTTAAATCTCTCCTACAATCATCTCGAAGGGTGCATTCCTCAAGGAAATCAATTTCATACCTTTGACAACAATTCATATGTAGGTAATGATGGATTACGTGGATTCCCACATTTGAAAGGCTGTGGTACTGAAGGTAATGATTCAGAATCAGAGAAAACTGATACCGGATCTGAGCTTGATGAAGAAAGCAACTCTGAATTTCTGAATGATTTTTGGAAAGCTGCTCTTATGGGCTATGGAACTGGACTCTGTATTGGATTATCCATTATATATATCATGATTTCAACTGGAAAACTGATATGGCTTGCAAGAATCATTGTGGAGGTGGAGCATAAAATTATGATGGGAAGAAGAAAGAAGCAGCGAAAGCAAAGGAGTTACAGAAGAAGCAAATAATCGCATGTAGAAGAGTTAAAAAT

>M6-PGSC0003DMG400024033 M6 LRR-RLP

CATCCTCTTTCTCTTCCTCTTCCAGATCTGAACAACAATGTCTTCGTCGTCATTCCCTGAACAAGTGTTGAAGCACCTCTTCTCCTTGTTAACCTCACACAAGGACCGTAACAGTATATCCTTAGTCTCTAAGTCGTGGTACGAGCTCGAGAGGCGCTCTAGGAGGAGTGTCTTCATCGGCAACTGCTACGCCGTTAGCCCTGAGATTATGATCACACGCTTCCCTCAACTCACATCTATCTATCTCAAAGGAAAGCCCCATTTTGCCGACTTTAATCTCGTTCCTGAGGGATGGGGTGCCTATCTTCAACCTTGGATTGCTGCAATGTCTCCCTCTTATCCTTGGCTTGAGGAAATTAGACTCAAGCGAATGGTTGTCACTGATGACTCCTTGGACTTGATTTCCACCTCATTTCATCATTTCAAGGTCTTGGTCTTGTTGTCTTGTGAAGGCTTTACTACCGATGGACTTGCTGCCATTGCCGCTAACTGCAGGTCGGTCCCTTTTATTGCCTTACTTATTTGATTGCTGCTTAACATAATGGGCATCTACAGAAAGAATTGAAATTGTTATTTGCACTTTTTTTCTCATGCAGAAATCTCACTCAACTAGACTTGGGAGAAAGTGAAGTGGAGGACCTAAGTAGTGATTGGCTCAGTCATTTTCCTGATGGTTGTACGTCACTGGTGTCACTTAACATTTCTTGTTTGGGTTCTGAGATCAGCTTATCACATCTGGAGAGTCTCATTGCACGTTCTCCTAATTTGAAGACACTTCGGCTCAGTCAAGCTGTCCCCCTTGAGAAACTTCCACACATATTGTCTTGTGCTTCCCAATTGGTTGAATTTGGTACAGGTGCATACTCTTCTGAGGTACAGTTACAGTCTGATGTTTTCTCAAACCTCCAAGAAGCTTTTTCAGGCTGCAAGATACTTGAAGGCTTGTCTGGATTTTGGGATGTCATGCCAGCTTACCTTCCAGCTGTATATCCAGTTTGCTCCAGACTCACCTCATTGAATTTAAGCTATGCTACTATTCAAAACCCTCATCTTAACAAGCTCATAAGTCTGTGTCATAATTTGCAGAGATTGTGGGTATGTTTACTTTACTTTACTTGCCTATGTTTTCAGTCACAGCATTCTGATAACTTAAGAGCATGATATCTTGCCTTGAAGGCAAGGGGTTGTTTATAAGAGAAGCACACAAAATTTTACCTGTTTATCGTCTTGTAAATTCTGTTGACGGCAAGGATATCTAGGATAAAAATCATGATATGCAGATACTTTCCACTCATAAGATTGGCTAAAATAATTCACCTTTACCTTAAGTTGAAGTTATAGACCTGTACATGAAGTAGGATCTAGAAGATACATTCTTTGGTTGTATAATTCTCATACGCAATGGCCATATGCAATATTTATCAATCTTAATGCTGAAGACTTTTTTACGAAATAGGTAACTTAACTTTTATGCTGAAGACTTTTTTATCTGTTCTCTTTTCCTGTGACATTCAAAGAATTTCCTGCTAATGGTTCTTTAATCTTTCTTGCTACAAGCTTGGACTAGTTGCTTAGTGTTTAAGACATTCCACTTTGAATAAATGTGACTACAGGTACTAGACTACATTGAAGATACCGGTCTCCAGGCGCTTGCCGTTACCTGTAAAGACCTGCAAGAACTTAGGGTGTTTCCTTCTGATCCCTTTGATCCAGAACCAAATGTTTCCCTGACAGAGCAGGGACTTGTAGCTGTTTCAGATGGCTGCCCTAAGCTCCAAACAGTTTTATACTTCTGCCGCCAAATGACAAATGCCGCTTTGATTACTATTGCTAGAAACAGTCCTAACATGGTTCGTTTTCGCTTGTGTATTTTGGAACCTCGAACTCCTGATTACTTGACTCTTGAACCGCTTGATGCTGGTTTTGGGGAAATTGTGGAACAATGCAAAGAGTTGCGGCGACTTTCTCTTTCTGGCCTTCTTACTGACCGTGTGTTTGAGTACATTGGGACCCATGCTAAAAAGTTAGAGATGCTTTCCGTCGCCTTTGTTGGGGATAGTGATATGGGCCTCCACCATGTACTCTCTGGATGTGAGAGCCTCAGAAAACTAGAGATCAGAGACTGCCCCTTTGGTGACGACGCTCTTTTGGCTAATGCTGCAAAACTGGAGACAATGAGATCCCTTTGGATGTCTTCTTGTTCAGTAAGTTTTGGAGCCTGTAAGCTGCTAAGTCAGAAGATGCCGAGGCTTAATATTGAAGTTATAGATGAAACGGGTCTTCTGGACACGCAGCCAAGAAGCTGCCCTGTTGATAAACTTTACATATACCGAACAGTGGCTGGACCAAGGTTTGACCTGCCTGGTTTTGTTCGGATAATGAAAGAAGATGCTGCACATTAGCAATGGAAATTCACCCTCATGCTATTAATCCTCCTTTGTGTGGCAGGTTAAGCCATAAAGTGCTATGTCGAAGACCTCACATGCCATCAACTATTACCTCTTGGCTAAGCTAATTTACTGCATCAGCTTATAAAGTGTTAAAACTCTTGTGAGGTTTGTTTGGTTTGGAATGACAGACCCAAGTTCTGTACTTATATTAGTATCCTAATCCCAGTAACCGACCCCTATGTGTTGTTATGTATCATTGGGAATAAGAATTACTACTTTGTATTTTGTTTTAGGGGGAAGGGGGTATGTTCAACAATGTACATGCTAGCGATTTGAAGACTTTGAGCATCGTTATGCAGCTAATTTCATAATTCTTTTCTATTGACTTGTTTTTCTGAAGAGATTATGCCATCATGTACTCATAAGTCAATAAAATTCATTGCATCCATGTACTAAGAATCTCTTCAGAAAATTATTTTTGGTAATCCCCATGTCAAAATTTGAACTCGAGTTCAATGGAAGGATTAATTTTGAATTCATCTTTACAAGACAATGAGAATTCAAGACTTTGAAATACTAGTGACCTACCATAGAAGAGCATCACTGGTACTCCTATCTAAGAAAGCAAATAAATGATAGAATAAGTAATGATGGAAGTTCAAACGACTTAATGAAAATATTAATTGGACAAGATGATAAACATACAGTTTTTTCTTTAAACTTTTACATCCATACACACTTGTGTCTTTTGCAACTGCCTCTTTGATGTTACTGCTCGCTCATATTCTTATACTTTTGATCCTTTAAGTGTTTCCAAAATATGTTCTATTCTGTACTTCATGACTGGTTGGCCTGATTTGGTTTTCTTGAACATTTTCTCCCTTAGTTCTTTCCTTCGAGATTCAGCTCTCTGCCTTTCTTCTTTCCTTGCTAGAATAGTGGCCTCTGTCTCCATCCTTGCCTTGTCATCCTCCTCGCGTTTTCTCTCATATATTTCTCTTAGGCTTCTAGCACTATTCTTCTTATTTCTTCTCCCTTCTTCATCCT

>M6-PGSC0003DMG400024206 M6 LRR-RLP

ATCAATCTTCTTATTCATATAGGATTGAATCAGAGAAGATTCAACAGATTGTGGACCACATTTCTACCAAATTATGCAAGACTGTGTATTCTTTATCTTCTTTGCAAGATGTTGTGGGAATAAATGCTCATTTAGAGAAACTAAAATCCCGGCTTCAGATAGAAATCAACGATGTTCGGATTGTAGGGATTTGGGGAATAGGAGGAGTCGGTAAAACAACAATAGCAAAAGCCATCTTTGATACCTTATCTAATCAATTTAAAGCTGCTTGTTTTCTTGCAGATGTTAAAGAAAATGCAAAAAAGAATCAAATGCATTCTTTACAAAATATCCTTCTATCTGAACTGTTAAGAAAAAAAGATGATTACGTTATTAATAAGTACGAAGGGAGGAGCATAATTCCGAGCAGACTTTGTTCAATGAAGGTTCTAATTGTGCTTGATGACATAGATCATAGTGATCACTTGGAGTATTTAGCAGGTGATCTTCGTTGGTTTGGTAATGGGAGCAGAGTTGTTGTAACAACTAGAAATAGACATTTGATAGAGAAGGATGATGCAATATATGAAGTGCCTACATTACCTGATCATGAAGCTTTGCAATTGTTCAATCAACATGCTTTCAAAAAAGAAGTTCCAGATGAGCGTTTTAAGAAGTTCTCGTTGGAGTTAGTAAATAATGCTAAAGGTCTTCCTTTAGCCCTCAAGGTGTGGGGTTCGTTGTTGCATAAGAAAGGCCTAACTCAGTGGGGAAGAACTGTAGACCAAATAAAGAATAACTCGATTTCAGAAATTGTTGAAAAACTCAAAATAAGTTATGATGGACTGGAGCCCGAAGAGCAAAAGATATTTCTAGATATGGCATGTTTCTTCCGAGGATATGAAAAAAAAGAAGTCATTCAAATTCTTGGAAGTTGTGATTTTGGTGTTGAATACGGATTGGATGTCCTAATCGACAAATCTCTTGTTTTCCTTTCCAAAAATGATACGATTGAAATGCATGACTTGATTCAAGATATGGGTAGATATGTGGTGAAAATGCAAAAAGATTCGGGAGAACAGAGCAGACTATGGGCCGCTGAAGATTTCGAAGAAGTGATAGTCAACAATACAGTAAGTAGGCTAAACAATGCAATAATATTAAATTTCTGGTTTTCATATTCCTAAGACATATAGGGCAATCAACTCCAGTTATTTGTTCCTCTTGCTTCATATTCTTGCAAGTACACCATTTTAGTTTTTTACTTTAGTAGGAGAAGCAAAAGTAATATAAATTTCATAACGTGGCTAGTTCATCATGTAGCCATTTTAATTAAAGCATCCTAATTCTTTTATCTTATGTTAGTAAGCCATGAATTAGTCTAGTAGCCGTTGAATATGTTTAACCCAATTTCATTCTTTGGATTAACTTCAAATTTGTATGATATTTGGATTCAAAGCTATGCTTTGTAGACTAATGATCCTTATATCAGGGGACGAAAGCAGTGGAAGCAATCTGGCTACATTACATTCAAAATCTATGCTTTAGCAAAAAAGTCATGAAAAAGATGAAAAAGCTTAGAGTACTATACATCGGTCGTTTTCATACACATGATGACACAATTGAGTACCTGCCCAACAACTTGCGTTGGTTTGAGTGTATTTGGTATCCTTGGAAATCATTGCCCGAAAATTTTGAACCCAAAAGACTTGTTCATCTTGATCTCCAGTACAGTTTGCTGCGTTATTTATGGATTGGCACAGAGGTACAATTGCTGAATTCAGTTTTAGTTTTCATT

>M6-PGSC0003DMG400024263 M6 LRR-RLP

AGATGTTATCTATAATTATATTTGAAGTGATGGATTTTAAACCTAGCACTGAGATATGTTATCTCATCCTGGTTAAAGTGTAAAAAAGACATACATTTGCTAGCTAAAGATTACGAGTGTTATCATGTCAACAGTAAATATTTGATCATCTTTGTCTGGTTGTTACATGTGAGGATGTGAGATAATCACGCGGAGAATCAGCATCATTTTCTATTGTAACTGGATAAAGAGAACTATAAATGCAATCCCTCAAATTGCAGGTATTGTTGAAGAGTTTGCAATGTTTTAAAAGAATTTCAAGTGGACAAAGAATGCGCTTTCAGGATTGAAGGTTCCTCACTTTATGCCTGTAGCTTATTTGTACATTATGGCAGCCTGTGTCTACTATAGTTTTATTACAAAAGAGTATCACTGCAGGGATCTATGATCATTTGTCACCATTTAGTCGTGATGGTTGAAATCATTGGAGGTCATTAAGCCAATGGTTTTTCATCTTAAAGTGTCCAGATTTAACTCTTTGATTGTGATGCTAAGTTGTGTAATTACTCCCCTAAAGAAGGATGGAAACTAAGGGTGAGAAGAAACTTAAATGATTGGGTGTTAACTGCACTAACCGATCTTCTTTTGTTCATGCAGGGTCATCGAGGTAACGCGGATGAAAGAGGTAAGAGTATGGTGGATTCCTAGCAGTGATGGTGAATTATCTGTGAGTTTGTAAAAAGACGATTCATAAGGGAGGTGAAATTGTTGGCGAATAGCTAGAAAAACAAATCTAGGACACTAAGGTGGCCTGTTTCACGTGGTTAGCATCCCGTCGATCTTGTTTAACACATAAGGATTCGATTACTGGTATTTCTTAGTTCATTTCGATCTAATCAATTGGAAGTTCCTTCTGTTTTATCTGTTCATTATTTTCTGAAATTTTCCATGTAGAAATTTGAATATTTGGTTGAAGCCTGATTGTATTCCCTTTTGTAAGGACTTGGACATTTCCCTTTCTTTAAGTTGTTAAGGTAAGCTGTAGAAATTTGACTGTTATGTTCTGAAATTAGTCGACTGATGCCGTTGTTTTCCCTGTAACAAAAAGGGGTTCTCTGAATAACAGGGACTTAAAACATTTTTCTCTCCTTCTGTTGGTTCAATCCAGATCAGCCTATTTGGTGTTAATTGGTTTAATTTCAACAACATTCTTGGTTTTATCCCCCACCCCCCACCTTAATCTCTTACATTTGATTTTGTAGGTTAATCTTTTTAAACCTTTTGTTAAAACTAGTTTTATGCGTTTCAGGTTGCTACTGTATTACTGGATTGTTGATAGAGAAGTTTGATCCAGATGGATAATTTACCAGCTGAATTACTTCTGGAGATTTTAGGACGGATTAAAAGAACTACTGATAGGAACTCTGTATCCCTAACTTGCAAACGCTTCCACAAATTGGATAATGGGCTGAAAATATCGATGCGCGTAGGGTGTGGTTTACATCCTGTCTGTGATGCATTAACTGCTCTATGCAACAGATTCCATAACTTGGAGAAAGTGGAAATTATTTACTCTGGTTGGATGTCTAAGTTAGGGAAGCAGTTGGATGATAACGGTCTTCTCATCCTTTCACGTTGTTGCCCTTTGCTCAAGGACCTCACATTAAGTTATTGTACTTTCATAACGGATGCTGGTCTTAGTTACTTGGCTTCTTGTTCGAAACTTACATCCTTGAAGTTGAACTTTGCTCCCAGAATTACTGGTTGTGGCATCTTGTCTCTTGTTGTGGGCTGCAAAGCCCTCAGTGTGCTTCACCTGATTAGATGTCTTAGTGTCAGTAGTATGGAGTGGCTAGAATATCTCGGAAAACATGAAATGCTTGAAGATCTCTGCATCAAGAATTGTAGGGTGATTGGAGAAGGTGATTTGATTAAGCTTGGACCTACTTGGAGAAAATTGAAGAGACTGCAATTTGAGGTTGATGCCAACTACAGATATATGAAGCTTTATGACCGTTTAGCAGTTGACCGATGGCAAAAGCAATCGATCCCATGTGAGAACATGCTTGAACTTTGTTTAGTGAACTGTGTTATCAGCCCCGGGAGAGGACTTGCATGTGTAATAGGGAAGTGCAAGAATTTGGAAAAGATTCACTTAGACATGTGTGTTGGGGTTAGGGACTGTGACATAGTATGTTTGGCCCAGAAATCAAGCAACCTTCGATCAATCTCTCTCCGAGTTCCATCAGACTTCTCCCTTCCTCTGCTACAAAACAATCCATTGCGGTTAACAGATGAAAGTCTAAAGGCAGTGGCTCAGAACTGCTCTCTGCTTGATACGGTTAGGTTGTCTTTCTCTGATGGAGAGTTCCCTTCATTTTCTTCCTTTACCTTAAATGGAATACTTATGTTAATACAAATGTGTCCAATAAGAGAACTTGTTCTTGACCATGTGTATTCTTTTGACGATACTGGAATGCAAGCTCTTCGTTCAGCACAATATCTCCAAATCTTGGAACTTGTAAGGTGCCAAGAAATCACTGATGACGGAATGCAGCTTGCTAGCCAAATTCCTCAATTATGTACATTACGGTTGAGGAAGTGCTTGGGAGTGACCGATGATGGGTTAAAACCTCTCGTTGGGTCTTACAAGTTAGACTTGTTGGTTGTAGAGGATTGTCCACAGATATCAGAAAGGGGTGTTCAAGGAGCTGCAAAGTTGGTCACCTTCAAGCAAGACTTGTCATGGATGTACTAAGCTTTCCCATTTTACAAGTATTTTTCCAGATGTTGTACAGGTTAGGTTGTTCCGTCTATACGACTTTTTGTACAGTCATTCTGTTTTTCATAACAATTCTGTAATCTGCAACACAGAATTCCCTATGTCCACATGCATGTTGCATAATAAAGGTTAATTTGAGTGTGTGTTTCCTATGCTTCGGCTACCATATTATTTGTTGTTGCTGTGTTTTTTC

>M6-PGSC0003DMG400024406 M6 LRR-RLP

AAACACAAAATATGATGATGGTTACCAAAATATCCTCTCTACTTCAGTTTTTCACTCTTTTATATCTCTTTACAGTTACTTTTGCTTCCACTGAGGAAACAACTGCCCTCTTTAAATGGAAAGCGACTTTCAAGAACAAAAATAATTCCTTATTGGCATCATGGACACCAAGTTCTGATGCATGCAGTGACTGGTATGGAGTTATATGCATTAATGGTAGGGTAAACACGTTGAATATTACAAATGCTAGTATCATTGGTACACTCTATGCTTTTCCGTTTTCATCCCTCCCTTTTCTCGAGTATCTTAATCTTAGCAGGAACAATTTATTTGGCACCATCCCACCTGAGATTGGTTATCTAACTAATCTTGTCTATCTTGACTTGCATATCAATCACATTTCAGGCTCAATCCCACCACAAATCGGCTCACTAGCCAAACTTCAGATCATCCGTTTATTTGACAATCATTTAACTGGTCCTATTCCAGAAACAATAGGGTACCTAAGGTCTCTTACTAAGCTATCATTGAGTATTAACTCTCTTACTGGTTCTATTCCTGCTTCATTGGGAAACTTGACCAACTTGTCTTTTTTGTTTCTTCATTTTAATCACCTTTCTGGATCCATTCCCGAAGAAATAGGTTACCTAAGGTCTCTTGCTCGACTAGATTTGAGGAATAACTTTCTTATTGGTTCTATTCCTACTTCCCTTGGCAACTTGAAAAATTTGCAAGTTCTGCTTCTCAATGACAACAATCTAATTGAGGAAATTCCTCCATCTATTTGCAATTTGACATCACTGGCAATCCTGTATTTGTCGAGAAACAACTTGAAAGGAAAAATTCTGCAATGCTTAGGTAATATCAGTACCCTCCAGTTTGTGATGATGTCACATAATAATCTCAGTGGAGAACTCCCTTTGTCTATTTGCAATCTAACATCACTACAAGTTCTAAATTTGGGTAGAAACAATCTGAACGGAGCAATTCCGCAATGTTTTGGCAACATGAGTGGTCATCTTGAGATTCTTGATATGCAGCACAACAATCTTTCTGGTACTTTTCAAAGCACTTTTAGCATTGGAGGAGCACTCAGAAGCTTCAATTTGCATGGCAATAAGCTAGAGGGGAAAATTCCACGATCCTTGGCAAATTGCCAACGATTGGAAGTTCTTGATCTAGGAGCTAATCTCCTCAATGACACATTCCCGATGTGGTTGGGGACTCTGCCAGAGCTACAAGTGTTAAGTTTGAGATCCAACAAA

>M6-PGSC0003DMG400024437 M6 LRR-RLP

TCTCTAACCTCCAAGAGCTACGTACTTTACATCTTTCTTATAACAGCTTCATCGGTTCATTTCCCGCTTCAATTTTAAGTTTGACACATCTTCAATACTTAGGATTGTCGAGTAATTCCCTATCTGGCCCACTGCCTAGCAACCAAAGCATGCTTCAAAAGCTAACTTTTGTGGATTTGTCATACAACTCACTGAATGGTACCATACCATCTTGGGTGTTTAGCCTACCTTTGCTATCTTCAGTGTCGCTCCAACATAACCAATTCAGAGGACTAGCCGATGAAGTGATCAAAACAAACCCAACATTAAAACAACTGCATTTAAGCAATAATCAACTCAGTGGTCCTTTTCCTCAATCACTTGTGAATCTCACAAACCTTGAAACCCTTGGAATTTCATCAAATAACATCACCATTGATGAGGGAATGAATATCACCTTTCTTAGCCTATCATCTTTATTCTTATCATCTTGTGAACTGAAGGATTTTCCACACTTCTTGAGAAATGTAAAGACACTTGTCTACTTGGATATTTCTAACAATAAGATATGTGGTCAAATCCCTAACTGGTTTAGCGGCATGAGGTGGGACTCGTTGCAGTTCCTAAACCTTTCTCATAATTCATTAACAGGCCACCTACCACAATTTCATTACTATAACCTACAGTATCTTGATCTGAAATGTAACTCCCTTCAGGGTCCACTACCTTCATCCATTTGTAACATGAGTTCGCTTATCTTATTAGATTTATCACGCAACAACTTCAGTGACTCAGTTCCACATTGCTTGGGAAGCATGGCTAGTCTAACGGTGCTAGACTTAAGAAGGAACAATTTCACAGGGAGTCTTTCCCCATTATGTGCACACAGCACTTCGTTGAGTACCATTGTCATAAATGGTAATCAATTTGAAGGACCTGTACCTGTGTCGTTGCTCAAGTGTGGTGGTCTAGAAGTACTTGATGTGGGGAACAATGCTATAAATGACAGGTTTCCATCTTGGCTCGGAACTCTCGAATACCTGCAGGTCCTTATATTAAAGTCGAACAAGTTCCATGGACCTATAAGTACGTGTCTGACTAAGTTTTGCTTTCCCAAGTTGCGAATTTTTGATCTTTCTCGTAATGATTTCAGTGGCTCACTTCCTGCAAAAGTTTTTGGAAACTTCAAGGCAATGATCAAATTAGATGGCGAAGACACAGGAAATATAAAGTACATGGACTCTGTGTTGTATTCGCCATTTGTCACATCGTATGAGGATTCAGTGAGTTTGGTAATCAAAGGGCAGGATATTGAGCTATAAAGAATCAGCACAATTACGACAACCATAGATCTCTCAAGCAACCATTTTGAAGGTGTCATTCCGAAAACACTAAAGGATCTCATATCACTTTGGCTACTCAATTTATCTCATAACAATCTCAGAGGAGATATTCCAATGAAATTAGGACAATTGAGTATGCTTGAAGCTTTAGATCTCTCTTGGAATCGGCTCACTGGAAAGATTCCACAGGAATTGACAAGAATAAACTTTCTCACCTTCTTAAACCTCTCTCAAAATCATCTCGTTGGACCAATTCCTCAAGGTCGGCAATTCAACACATTTGAAAATGACTCATATGGTGGCAACCTTGATTTATGTGGTGTTCCTTTAACAAAGAAATGTGGAACGTGTGATTCATCGCATGTTCCTCAACAATTGGAGTCCGAAGAAGAAGGCGAGTCATATTTTTTTAGTGGATTTACTTGGGAATCAGTAGTCATAGGCTACAGCTGTGGACTAGTTGTTGGAACTGTCATGTGGAGTCTCATGTTTAAATATCGTAAGCCAAAATGGTTTGTGGAATTTTTTGATGGACTCATGCCTCAGAAAAGAAGAAGGCCAAAGAAGAGAGCTCAGAGACGAAGGACTTGATGGAAGATTGAGAGGCCTAAATTGCCTTTGTTTGATTTTGTGGACTGATTAGTGAACTGTTCTGTTTTTGTCTTTTTTAATTTTCTATTTGTT

>M6-PGSC0003DMG400024438 M6 LRR-RLP

CACTACAACAAATGTGATATATTGCTACGAAATAATTAGCTACAATATAAAATTTGTCACTAATTGTTCATTTTTCATGACGAAAGTTATATTTTGTATTTAAATACATGAGGAACATACTTTTAGTGACGAAACACAAATATTTGTAGCTAAGTTTTGTCCACTATAATTTCCCTCCAAAGGATAAGTTGGCGCCATTTATTGAGTACTATTAGCTACAAATCTAAATTTATCAGTGACACATTATTTTGTCACTAAATGAAGTAGTACCCAATATATGACAAATCATTTTTTATCCTAAAATTAATTAAATTTTGGACATAAAATAGTTTCGTCACAAAAATTATGTTGATACAATAGCAACGAATTATAAATCGTAGTTAATCATTGAAGAATTTAACCATAAAATTTATGCTTTAATTGTGACCTTATTTTTTGTCACTAATAATTTGACAATTAGTGACAATTATATCATTGTCTCTATTTAACCTACAATTTAGCCAAAATAATTTTTGTCACAAAAAAATAGAATTTTCAATTGACTACAACTTTAATTTCGTAGTTGATGAATAGTGTAATTATTGGCTACGAAAAATATAGGAAAAACTGCATATCTAAGCAAACATAGCTTGTATATCTACTCAACTAACCTATAGTTTTGGAAAGGAAAAACTACCTTATTAGATATACATCCATACCGCATATACTTATTCTTCCAATACTTTCAAATAATTAGATATCTTACCACTAATTAAGAGTTTCCTACCATCTTTTGAGGAAGATATATACACCTATATATAGCGTTTCATCTACCAGATATACTAGAATAATAGTGAGTGAATCAAGCGAGATTGGAAGGGAGATACTCATGTATCCCAGATAAATAGGAGAGTGGCAAGTGAGATGGGAGGGAGACAAGCGGGATTCGTTTATGTATCCAAGATATATGTGAATCCACTTGGATACAGTGTATTTAGAATAAATTACACCAAATTGACCCAATGTATCCCAAGATCCATGTATATGGAAATATCTGGACGCACCAAAATCTTATAAGATTCGTAATATTATAAACTAGAGCGTATCTAAGTAATTAGCTCATAAACTAGTGAAATTTATGTAAGTTTTCCATTAAAATGAATAATATGTGAAGGTTCTAAATTTCATTTAATTCCGGTAAGAAATCACAATCTCATCTTATAATGAATGATTGAAATGTGTGAACTATTAAAGAAAATGGTTAGTGTAGTTTATCCTATAAGAACCCCCCGATCCAATGATAGTAACATATCTGACTAATCATCAAGATATTATTCAACAATGTTCTTCAGAAATATTCGCCACCTGATTCAAGAGATCGAATCAAAATTTTCCTCTTTTGTTATTTCATTCTTTTGATTTTATTAATTGTTTCGTTTATCCATTAAATTATTATTTCAGTTCTTCGATTACATATTCAGGATTTTGGATAATATTAGTAATTTAGTTTGTTGGTTATTTGAACTTTTACCTTGTTTGTGATAATTTGATTCTCCATCTCATAATCCCTTTCTTTACCCCATCTTGAAGAAAAAAAATTCAAAATTTTTATATCATTTGAATTAATTATGTTACTTTAAATCTCTTTCAATTAAATTCATCTGATTCTCCGAATAAATAAATTTTACGCTAAACACTAATGTCACACCTAAGATTTTGAGGGACGAGATTGACACTCATTATCTTAAACTTGGTTCATATTCGTTTTACCACACTCCAATTATTTTTCTGATTGCGCGAGTCTCACACACCTGAACTTATTTTAATATTATCATTAAAAATTATTCAATTTCAAATATAAAAACTCGTGGACACTTACAATTAATATATACAATTATTTAAATATTAAGTCCGTTCATGTAGAAAATTATTTCTCAATGTGCTTCACATTTTGACCAAATAAAATTTATGAGGTAAGAAGTGTTGAAATAATTCATAATACAAAAAAACAAAGATAAAAATATAGTTTTAATATTCAATTGAAATTCATTAAGTGAAAGAAATTATAACTTTGAGAATATATTTTATACGAAATTATTAGAGGGAAAAGAAAATAACAAATTTTACTTTCAATTAATTTTTCCCTCCACTTCAATGTGTAAATTCCCCCTCACGCCCATTTCCCATTTTCTATCCAGTAACCACGCATAACGGCACAAGGAACTGCCCACTAACTTTCTTTTCATTTTATTTTTTTAAAATAATCAAAATTTGCCCACTCTCTTTAACTATCCAAGTCCAACTCAAATGCAATACATTTTTTCAAAATAAAAGAGACTCCTCTGCATTTTCTCTCAATCTTCTTTTTTCTATCTATTTTTAAAATGATGTCCATCCAACATTTACCATGCTCTATAAGAAACAAAATATTTTCTTTGGTTAGCATAAGGAACCATTATTCACGAAGTTATTATTCAACTCCTTACAAGGATTATCTTAATGGAAGAACTCTTGATATTATCTTGGTAAGGAGATAAGTGTTTATAAATTGTGATAATCGAATAGTTAATGATTTTGTATAAAATTAACTAATTTTTTAATTCTAGTATTTGTTTCACTAATTAATGTTAATTTTAGAATTATATTTTGTTTAAATTATGATTTCAAAAAGTATATACTAAATTATATTTTTTTTCACACAGGTCCATAAAGGTGCAAAAATGACTCAATGAGTGAAGATTTTAGATAAAAGAGCACGAATAGACGAGGAAACTACATACTTTTATCTTTTTTAAATTTATCATTGTTCCCGTACCTCATCTAGGTCTTTTTTTTTTTTTGAGTGATTTCCCTTGATTTACTCAATGAATTTGTACTCAAATGTTATCCCTGAATAATTAGAAATGTAATTACTATATAGTTTCGTATGGATTTTTGGTATTATGTTCCTTTTTTTTTTTTTTTTTTTTTTATTTACAAAAGCTATGTACTTTTAAAAATTGAAGTATACATGTAATTAATGGCAAACAGTTTAACTACATATTTAAATAATACTATTATAAAACATTAAATACATTCAATTTGTTATAAATAATTTTATCCTTGGATATTTATATAATTTTAAAAATCATATTTTGGTCGCAATGCAAAATGGTATTCTTTATCTATAAAAAGTAAATCATATTACTAATGTTTTAGTTGTCACAAATTAATTTTATTCAGTGATAAAACTAATTTATTCCAAAATAGCCTTCAATATTGATAGCATAATTATATTTTTAGCAAAAAATTAACTTAAATGTATCTAATTGAGACAACATAGAGAGACAAATTATTCGTCAGTAATACCATATAGAGAGACAAATTATTCATCACTAATACCATAAAGAGAGACAAATTATTCGTCATTAATATCGTTATATACTATATCACAACTAATTTTCAGCCACAAAATTAATATCTTTTAGTGACGTAATAACACTCTTTTAACTACAAATTATGCATAGCTTTAAAGACAATTTATATCGTCACTAATTAAATTTATAATTAGGGACAAATAAAATTTGTCCGTATTAAACATATTTTTAGTGACGATATGAGTTTCTAGCTACAAAATTTTACGTTTTTTAAGACGATCAAATTCGTCATAAATGCTACATATTTGGAGACGAATTTTATTTTTGTAGTTAATAGAATCTTTTTAGCGACAAAACACTAAATTGTCTTTGTAAACTTTTAGTGACCCACGTTTAGAGACGAAAGACTGACGAAATTTTTTTCATCATAAAAATTTTTTTGTGACAAAATATTATCTTTAAGTGACAAAATAATTTGTCACTGATAATCAAACTTCTTGTAGTG

>M6-PGSC0003DMG400024768 M6 LRR-RLP

GATCACTTATCAATGAAAATGCATCTCCAGCTCCATTCCACTTATCTTCCCAATATCAACAATCCTTTCTTTCTCTCTCACACGCACATTACTTGTCTTCTTTTGCCTATGGCACACCTGTCGTCATAGCACCTAGGAATTTCCGATACCCACCTGACAAACATCGCCGGAGAATCAGGCATCTACCATGATCATCTTTGTCTGCAAGACTCGTTTTGGTCTTTCTTGTACTCTTTCTTTTTCCACCCTTCAACCCCTTCTCAACGCTCGTCTCCTCTTCCTCTTTGTCCCGACATAATACAGTTACAACCCTTTTCTTTTCTTGTAGTTTTTGGTGGGAAGAAGAAGATCCGAACAGAAAGTATTGACTAGTTGAGATTTTTTTTTTTTTTTTTTACAGAGTATTCAAGATTTGTATGTCAGCGTATTGAATGTTAAACGGATTCAAATGGGCTTAAATGAGTAGTTAGAATTCATTTAATCAACCCAAGAAACTTGGAATTGAGGTTTAGTACAAGTAGTTTTTGTTGTTGTTGTTTCTTATAGTTTTCATGGATTCTGCTCTGTTTGTTCTTAACGAGGATCTTCTTATCAGAATTCTTTCCTTCATTACCCATGATTCCGACCGCAAAGCTTTCCGTTTGGTTTGTAAAGCATTCCTCCGGGTTGACTCTTTCCACCGTACCCATCTCCGGATCCTTCGTCCTGAGTTCATTACAACCCTTTTCTCCAAATTCCCACGTATTTACTCCCTTGACCTTTCCGTTTGCCCCCAAATTAATGACGGAGCTGTTTCCATGCTGATGGGTTATGGGTTGCCGGATTGGAGTAGGAGCTTGAGGAGGCTGGTGCTGAGTAGAACCACTGGGTTGAAATCAGCCGGGTTAGAGATGCTTATGAAGTCTTGTCCTGTTTTGGAGTCTATTGACGTGTCTTACTGCTGGGGTTTTGGTGATAGGGAAGCTGCGGCTTTATCTTTTGGTGGGTCTTTAAGGGATGTAAAGCTCGACAGGTGTTTGGGTCTTACTGATGTTGGTTTGGCGAAAATAGCAATTGGGTGTCAATGCCTAGAGAAACTTAGCTTGATGTGGTGTATTGAAATTACTGATCTTGGGATTGATTTTTTGTCCAAGAAATGTACACAACTGAAGCAGCTTGACATTTCATATTTAAAGGTCTGTTTATCTTACTATCCTTCTTTTTGATCGTAGCATATCAGGTGTATGTCTTTACTGTGTTTGTGAAAATGTGGATTTGGTGTTGTTTCAGGTTACTAGTGTGTCACTGCATTCTATAAGCAGTATGGAGAAGTTGGAGCTTTTGGCTATGGTTGGGTGCGGCATTGTGGACGATGAAGGACTACATTATCTTGGAAAAGGGTGTCCTTCGCTTCAGGTGTTTAACACGTTTGTCAAACTATTGTGATCAATTGTAGAAAAATATGAGATCATAGCTGTTTTTTGGAATTTTGTTCTCCTCTCTTAAGGTGACCCTTACCATGTTGCTATCTGGTTTTTACCAATTTTAAGACTGAAAATAAACCTTGTATGCAAACACAATTGCTTCATAAATAAGCACATGTAATTTCATTTTCTTATAGCTTGTGGAATTATATTTGTTTATCAAATAGTGAAGATGGGCTTAAAGTTGTTTTTGTTAACTTAAACTCTCTCAGTTCCAATTTATATGACGTAGTTCAACTCGGGACCGAGTTAAAGAAGAAAATGGAAACTTCTAAACTTGTGGTCTTAATCATGCCATAATATTTGTATTGCTATAAAAATTTTGGAACTTGTTATCTTAAACATGCCATAACATCTATGCGGCTATAAAAGCTTCTCGTTGAGTATAGAATGAGAAGTTTTACGTTAAAGTTGGTTCAATTGGCAAAGGTTGAGGGACTTGTGTCTTTGGTCACAGATTCCCGAATGTCAAAGGCTGTGGTTGGCCCAAAGGATTGACTCCATATGGATTTCTCAGTCACTAAAAAAATTATATCCAAATATAGATATGTGTCATTTTTTTTGGAATAGACAAATACGAAAGTATTGTCACATAAAAAGGAACAGCATGAATGACTCGTTTCAGGTGTCATGGATGGAATTTGAGACAGTTAGCGCATATGACATTTTCCTGATGACCAAATTATGCAGTAAGCACATGAATCTGGAATAGAGGAAATACTTATTGGAAAATATGATTTATGGCTGGTGATATTTCCTGGTATTCTGACTGTAGGCGAACAAATGCATCACTTGATGCTGCATGATCTAGTTCTGATCCCTCTTTATCTATCTATCTATTTGTATTTTTTATGAATCCCACTTTCAGCTTTTCTAGTGATCTTTAAGCATACCAGCTGCATTCGTATTTCTTCTTTTTTCAAGTTATAGTTATAGTGGAAAACAATTTTCTATCCATGAATATCTGAAATTAGTATTCTAAAGCTCAAGCTTTAACATTTTATGTGGCTAATTTACCTCTTAGAAGAATTGGCCATGAAATTTTCGTCTGGTTAGTGAGCGTACATAAGTTTCTTGGGTTGCCATAACTATTCAGAAACTTGTATGTTTACAGGCGCTGGATGTATCAAGGTGTGACAGACTAAGCTCATCTGCCTTAGCTTTCTTGATTAATGGACATCCTTCAATGCTACAAGTCTACGCTAGTCACTGTTTCCATGTGAGTAAATTTGTTGTTAGAACCTGCATTTTGAATAATTCCTTGGGGAGTTTTTGATGTTGGTTCTGCAATTTAGATTGAGAATCTTGATGCACCAGGAGTTTCCTACTAAAGTCATCCAAGGGCTGAAGGACCTAAAGAATCTGAAAAAGCTCATCCTTGATGGAGCACTGGTTTCTGAATCATTCTTCAAGATCATTAATTTCAATTGCAAATATTTGGTCGAAATAGGGCTTGGAAAATGCAAAGGAGTGACTGACAAGGGCATTCTCCAGCTGGTATCAGGGGGTGTTAATTTAAATATCTTGAATCTCACATGCTGTAGCGAACTTACCGATAATGCAATATCAGCTATAACAGATTCTTGCCGGAGTGTTTTGTGCCTCAAGTTGGAGTGCTGCAACTTACTCACAGAGAAGAGCCTATATCACCTGGGATTACATTGTTCCTTACTTGAGGAGCTTGATCTTACTGATATTTTTGGAGTCAATGACACTGGTGAGGCTCAAGCCATTGTTCTTTTGTTTTTATTAACGTATTTATTCTGTCACCATTCAGATAAAGGTAAATTTCCATTTATTCTGAAGGACCCAGAGCAAGACATCTGATTTTCCCTCGTCTCATCTGGATTCTATTTCCATTTTTATATTCTTCAAGTTTTGGGCTTTTGAGTTGGCTTCCCATTCCTTAAGAGCAAGACATCTAATTTTCTTCTTCTTCCTTTTTCTTTTTCTTCTTGTAATGACATGTGAAGCTTTGTGCATTTGTGTCTCAAAGCTCCTCGTGTTAAGTCTTCGATGATATGGTTACTCCAAAACTTGTGCAGTTGGTGACATACCTTTAAATTAAGCATTTCAAGCAAGGAATTCAGCTATGCTGTAGGTCTGCACTTTAATGTGTTGCATCTCATCTCTTCTGCAGGACTTTATTACTTGTCTAAATGTACAAAGCTTATATGCCTAAAGTTGGGATTATGTACTAACATTACTGACAAGGGACTGTACTGTGTGGCTAGAAATTGCAGCGAGATTCGTGAACTTGATCTATACCGGTAAAGTGATTGCCCCATCTAACTTATCCTGTGAATTTGAAAATGTCTTCTTATATTGTATCTTGAAAACCATTTGCAGGTGCCAAGGAATTGGTGATGATGCACTCTATGCTTTATCAAGTGGTTGCAAAAGAATGCAGAAGTTAAACTTGTCTTATTGCAGTGAGGTTACTGATAGAGGAATTGAGTGTCTCGGCCATCTACCAGTGCTCTCTGACCTCGAGATGCGGAGTCTTTTGAATGTCACAAGCACTGGTTTAACAGCACTTGCTACGGGATGCAAGAGACTTTCTGAATTGGATGTGAAAGACTGTACTAGCATTGATGATTCAGGTTTCATGGCGCTTGCTTATTACTCTAGGAATTTGCAACAGGTATAATTTCTCAATCTTGAACTACTCTCTCTACTTCTTGTAATGTATGATGCTGTTAAATCTGCTTGGCATGTGCTTTCTGCATTACCAGTTGATTCATAATGGTTAAATAACTATACACCTTGATCAGTTTTTATCATTTCTCGAAATCCTGAGAACAGTAATTGAGTTGTAATCTTTTTTCAAAATTTTGTTCGATACAACATAGCTCTCTTCTCTTGGTCCACAATCACATTTATATAAGTGCCAGGAATATATTTTTTGTAGAAGTTATCAATAACTCTG

>M6-PGSC0003DMG400024857 M6 LRR-RLP

AATGGGGAATCTTAGTAACTCAGTTGGTTGACTACCTGAACTTTCACCTTGTTGGTGAGGGTTTGATTCCCCACATTGTAATCCCCTCTCCATTTCCCCTTCCCC

>M6-PGSC0003DMG400024958 M6 LRR-RLP

CGGCCATGGACGATATTGACGAGGACGCGCACATTTCCTCTTCTTCCTTCCTTTGATATTCCACCTTTTAACCGGTCAAATCTCTAACTCTCCTCCTTCTTCTTCCTTCTATTAATAGATTCTTCTCATTTTTCTCTTCTTTCCATAATCTCCATTTTTTCCCATTTCCAACTCTAGATCTCTATAACCCTAAAACAAAATGGGCCAATATTCATCCACCCACTGCGGCGGTGGGCCAAAACATCGCCACCGCAGCCTTCATCCTACCCCACTCCCCAATCACCACCGTTCTGCCTCCGACCATTCCCAGTCCGATTTCTTCTCCCTTATGAATTCAGATGAAGATGAGTCACCGATTCCGTTTGTTTTCGGGAACCGAGATTATACGGATGATCTTCCTGATGAATGTTTGGCTTTGATTTTCCAGTGTTTGAGTTCCGGTGACCGGAAAAAGTGTTCCCTTGTCTCCCGGAGATGGCTTTTGGTGGAGGGTCAAAGCCGTCATCGTCTTGCCCTCAACGCGAAAGTTGAGATTCTTCCGCATGTTCCCACCATCTTCACTCGTTTCGATTCTGTTACGAAACTTGCTCTCCGATGTGACCGGAAATCGGTAAGTATAAATGATGAAGCTTTGACCCTTATCTCCCTTCGTTGTTCTAAACTTATTCGCCTAAAGCTTCGCGGTTGCCGTGATGTTACTGAAGTGGGTATGTCTGCTTTTGCTCGGAATTGTAAGAGTTTGAAGAAATTCTCTTGTGGGTCTTGCATGTTTGGAGCCAAAGGTATGAATGCTTTGCTTGATAACTGTTCTTCCCTTGAAGAATTATCTGTGAAACGTCTTCGGGGTATCAATGATGGGTTTGCAGCTGACCCAATTGGGCCAGGAGCTGCTGCTTCATCGCTTAAATCTATATGTTTGAAAGAGTTATATAATGGACAATGTTTTGCACCTTTGATTACTGGGTCTAAGAATTTAAAGACTTTGAAGTTGTTGAGGTGTTTAGGTGATTGGGATAGGCTTTTTGAGACAATTGGAAGTAGGGAAAATCATGTTTCTGAGATTCATTTGGAGAGGCTTCAAGTTGGTGATACTGGACTTACAGCAATATCTAATTGTCCAAAGTTGGAGATTCTGCATTTGGTGAAGACACCAGAATGCACTGATGCTGGAGTTGTAGCAGTGGCTAATAAGTGCAAGTTATTGAGGAAGCTTCACATTGATGGATGGAGGACAAATAGGATAGGGGATGAGGGTTTAGTAGCAATTGCTGAGAATAGCTCAAATCTCAAAGAATTGGTGCTTATTGGTCTGAATCCTACGTCGACTAGTCTATTGGCAATAGCTTCTAATTGTAAAAAGTTAGAAAGATTGGCTCTTTGTGGTAGTGACACTATTGGAGATCCTGAAGTATCTTGTATTGCCACAAAATGTATGGCCTTGAAGAAGCTGTGTATCAAGGGATGTGAAGTAACGGATCAAGGGATTGAATCTTTCGCTTGGGGTTGCCCTAATTTGGTGAAAATCAAGGTGAAGAAGTGCAAGCATGTGACGGGTGATGTTGCAGATAGGTTACGAGCTAGGAGGCAATCGCTAGCAGTGAATCTTGATGGTGGGGAAATTGATATTGAACCTGTGGATAGTAGTGCAAGTGATGGTGGAGCAATAGAAGAAGCAGCAGAGTTTCAACATACTGCAGCTACACTGCCCATAATTGGTGCTAGTGATATTCCATCTACGAGCAATGTAGGACGATCATCAGCATCGAAGCCATGGTTTGGTTTTTTCGGAGGAAGGGGCCTCGTTGCTTGCACTCTCCGTAGGTGGTCAAATGGTAATGGTGAGAGCTTATGAATATTATGAAGACTAATGTTAAAATTGAGAAAACATACTGTTAAATTTGTTTAGCTGTACTTGGTCCGGTTTCAACTTTTCTCATTCATCTTTTTTAGTTTTGAATTTGTCGTCTTGAATACATTGTGCAGATATACATCTCACTTCCCAATTTTGATGAGCTTTTGTTAGATTTTGTCTTATCTTTTGTCTGTACAAGATTTGGTGTAGCAATAACCTTGTTCATGTTGGTTCTTGGATGATAAAAGTGTATCTTGCAAAGCAAATTCTCCCTCAAACATGCAT

>M6-PGSC0003DMG400025295 M6 LRR-RLP

GTGATGAATAAACAAGAAAAACCCAATCTTTCTTCTTTTTCAATTTTTTTATCATAAACTATACATACATACACCCTTTTGTTTCTTCCAAAGTCAAGATTTATTTTTTTTTCAAACCCCCATTTCCCAAAATGCATTTTCTGATGCTGGTTTTCAAGATTTTGTTAATTTCATCATTCTTTCATCTGTTCTCACTAGCCCAACAACCCCAAATTTGTGTTGAAACTGATAGTCTTGCTCTTCTTGGCTTCAAAGATAAGATTTTTAAGGATACTACTGATTTTCTGTCATCATGGACAGGTAAAGATTGTTGTGGAGGAGGTTGGGAAGGTGTAGAATGTGATGTTTCTACAGGGAGAGTGAATAGATTGGTGTTACAAACACCTTCTGAGAGAGACATTAGTGTTTACATGAAGGGTACTTTGTCATCATCTCTTGGTGATTTGGTTTTCTTGGAGACATTGATTATAAGTGGAATGAAGCATATTGCTGGTGCTATACCGGAAAGTTTTTCGAAACTTTCAAGATTGAAGCAGTTGATCTTGGATGGTAATTCACTACAAGGAGATATACCTTCTGGTTTATCTCATTTGGAATCACTTCAAACACTTTCACTTAGTGGTAATCATTTGACAGGACATGTACCTTCTGTATTAGGAAATTTCAAGAATCTGCAGCAGTTGAGTTTAACAGACAATATGCTAATTGGGGTTATCCCAATTGGTTTCAAGAATCTTGTTAACTTGCAGTCCTTGGATCTGAGCCACAATTTACTATCCGGGGTTGTTCCGAATTTTCTTGGACAGCTTCAAAATTTGACATATATTGATCTTTCAAGTAATTAATTTTCAGGGGAGTTTCCTGTTTCCTTGTGTAGTTTGCTCAAGCTGTCGTTTTTTTCGATGGATCATAATCGAATAACAGGGAGAATTCCAACTCAAATTGGGAAGATGAAAGCACTTACTTCATTGTCATTGAGTTCTAACAAGTTGACAGGTCATATCCCTGAGTCTATTGCAGGATTACCAAATCTTTGGAACCTTAGTTTGTCTAGAAATGAACTTTTAGATCCTGTGCCTATTGCTTTCTCTAAAGGGCTTCCTTCACTTTTATCAATAGATTTGTCTTATAACAAGTTCAATCTTGGAATAGTACCTGAATGGATTAGGACTAGAGAGCTTTCTGATGTCAATTTAGCAGGCTGTAAGCTTCGTGGGACGCTTCCGAATTTCACTAGGCCTGATTCATTGAACTCTATTGACTTGTCGGATAACGCCTTCACAGGTGGGATATCGAGTTTCTTTGCAAGAATGTCAAGCTTGCAAAAGGGGAAGATATCAAACAACCAGTTGAAATCTGATGTTGCTGTGATCAAATTGCCTGATGGGATTTCATATCTTGATCTTCATTCAAACCAGCTTTTCGGGTCTCTTTCTAGGATGTTAAGCAACAAGACAAGCAAGTTCTTGGAAGCAATCGACGTCTCAAACAACCAACTCTCTGGGAGCATCCCAGAGTTTGTCAGTGGCTTAAACCTGAAAGTTCTCAACATAGGAAACAACAAGATTGCAGGTCAAATCCCAACTTCAATATCGAATCTTGATAAGCTTGAGAGGTTTGATATATCAAGAAACCAGATTACTGGTACAATTCCAGTAAGTCTAGGATCACTAGTGAAGTTGCATTGGTTAGACTTATCGATCAACAGACTTTCAGGGAAAATACCAGATAGCTTGTTGGGGATTGAAGCATTGAAACATGCAAACTTCAGAGCCAACAGATTATGTGGTGAAATACCACAAGGCAGACCATTCAATATCTTCCCTGGAGCTGCCTATGCCCACAATTTATGCTTGTGTGGCAAGCCTTTGCTGCCTTGTAAGGGAACGAAACAAGATAAGTTGGGCCAGTAACTGCTGACTCCATTTATAGGAATTTATGTTTGTTAACAAACAAACAGTTTACTTTTCTTTTGTATTTTTGAATTCTAGTACTACTTGTACAACTCTATCATCTAACATTGTTAGTCAGTGTTTACTTCACATAAATTCAATATGAA

>M6-PGSC0003DMG400025383 M6 LRR-RLP

AAAAAAGAGGGAAAAGAAACATGTTGAAAGGTTATCTTTGGAGTGGAGCGGAAGTTTTGCTGACAATTCACAAACTGAAAGAGACATACTTGATGAGCTACAACCAAATACAAACATAAAAGTACTCCAAATTACCGGATATAGAGGGACAAAATTTCCAAATTGGCTAGCTGATCATTCATTTCATAAGCTAATGGAATTGTCTCTTAGCGATTGCAAGGACTGTGATTCCTTGCCAGCACTAGGACAACTTCCTTGTTTAAAATTCCTTACCATTAGAGGGATGCATCAAATAACAGAGGTGACTGAAGAGTTCTATGGTAGTTTGTCCTCCAAAAAGCCATTTAACTCCCTTGAGAAACTTGACTTTGCAGAAATGCCGGATTGGAAGCAGTGGCATGTACTGGGGAAAGGAGAATTCCCTATACTCGAGCAACTTCCAATTGAAGATTGCCCAAAGTTGATAGGAAAATTGCCTGAAAATCTTTCTTCTCTGACAAGATTGAGAATTTCGAACTGCCCTGAACTCAGTTTGGAGACACCTATCCAACTCTCAAATTCAAAAGAGTTAAAAGTTGCTGGTTGCCCTAAGGTTGGAGTTCTTTTTGAGAATGCTCAACTGTTTACATCTCAACTTGAAGGAATGAAACAAAATGTTGAATTAGTAATTACTGATTGTAAGTCTCTTACCTCCTTACCAATTAGCATTCTGCCGAGTACCTTGAAGAGAATAAGAATATCTGGTTGCGGGAAACTGAAATTGGAGGCGTCCATGAATGCTATGTTTCTCGAGGAACTGTCTTTAAAAGGATGTGATTCACCTGAGTTTTTTCCAAGAGCACCCTATTTGAATTTAAGTAGTTGCCACAACCTTAATAGGCTTTTGGTTCCTATTGCCACTGAACGACTCAGTATTAGAGGTTTTCATAATCTTGAGATACTTTCAGTGGCATGTGGGACTCAGATGACATCATTGCATATTCACGACTGCAAGATGTTGAAGTCGTTGCCAGAGCATATGCAGGAACTCCTTCCATCTCTTAAGGAACTGCAACTATGGCGTTGTCCAGAACTAGAGTCCTTTCCTGAAGGAGGATTGCCCGTCAATTTACAAGTCCTTTGGATCAACAAGTGCAAGAAACTGGTGAATGGCCGAAAGGAGTGGCATTTACAGAGACTCCTCCTTCTCGAACAGTTAGAGATCAGCCATGATGGCAGTGATCTTGCTGGTGAAAACTGGGAGTTGCCTTGCTCTATTCAAAGTCTTACCATATCCAATCTGAAAACATTAAGCAGCCAACTTCTCAAAAGCCTCACCTCTCTTAAATATCTACGTACTTATAATTTATCTCAAATTCAGTCACTGCTGGATGAAGGGCTTCCCTCATCTCTTTGTGAGCTACATTTAACTAGCCATCATCATCTCCATTCACTACCAACCGAAGGTCTTCTGCATCTCACCTGGCTTCGACATCTACAGATCTGGAATTGCCCTAATCTCCAATCTCTTCCCAAATCTGGGATGCCCTCCTCCCTCTCTGTGCTGCACATTTGGGATTGTCCTAATCTCCAATCTCTTCCTGAATTGGGGATGCCCTCCTCTCTCTCTCAGCTGTACATTTTGAATTGTCTTAATCTCCGATCTCTTCCACTAAAAGGGATGCCCCCTTCCATCTCTAAACTGTCTATTTACAAATGCCCATTGCTCAAACCACTCCTACAATTCGACAAGGGAGAATACTGGCCAAAAATTGCTCAGATTTCCACCATAATTAGTAATGGAGAATTCCAATAATGATTAAAGGAAATGGTGCTTTGATA

>M6-PGSC0003DMG400025480 M6 LRR-RLP

ATGTTGAAATTGGCATCTGTATTTGATGCTAGTAAGAATAAATTAACAGGTCCGATCCCGCAATCTTTCGGGTGCCTTGAGCAAATGGAAGTGATGGATCTGTCTTACAATCAATTGTATGGGGAAGTACCGGAAACATTGTGCAAGATCAGTAGCTTGGAGGAACTCACTTTGAAATACAACTATTTCACACAGATTGGACCGGAATGTCGGAAGTTGGTAAAGGAGAATGTACTTGATATTGGCATGAATTGCATTATTGATCTTGAGAACCAGAGAAAGCCGGAGGAATGCGAAACTTTCTTTTTACAAAAGAGAAATTGCCGGGACATGAAGTCACTAAGCTATGTACCTTGTAAGATTCATGACTTCTCTTCTAGGAACAGTTCTTCAGGTCGAGCTCATACCGGCGATGCAAGGACTCGTTCGACTACTTATGCTGCCCTCAAGAAACCTCACCACTGA

>M6-PGSC0003DMG400025546 M6 LRR-RLP

ACTACCTAGTACAGTTTGGGATATGGTTAAATTGAGATATCTGTATATTCCTAACTTCAGCCCTGAAAATAAAAAAGCATTACTCAAGAACTCTCCAAAACTTGATGATTTGGAAACCCTTTCCAATTCATATTTCGCTCGTGTTGAGGATGCAGAATTGATGCTGAGAAAAACACCTAATCTTCGAAAACTGACATGTAAAGTCGAGTGCTTAGAATACCCCCATCAGTACCATGCGTTGAATTTTCCAATACGGCTTGAAATACTAGAGCTTTATGGATCAAAAGCCTTCAAAACCATCCCCTTTTACATCTTTGCACCAAATCTCAAATACCTGATACTCTGTGGCTTTTACCTGGATTCTCAGTACTTATCAGAAACTGTTGATCATCTCAAGCACCTTGAGGTACTCGAACTGTACCGCGTTGAATTTGGTGATCATGGGGAATGGAAAGTGAGCAGTGGCAAGTTCCCTAAACTCAAAATCTTGAAACTAGAATATGTGTCCATGATGAAATGGATTGTAGCTGATGATGCCTTTCCTGACCTTGAACAATTGGTTTTGCT

>M6-PGSC0003DMG400025563 M6 LRR-RLP

TACTCCCTCTGTTTCAATTTGTTTGTTTGATTTTAACTTGACACAAAGTTTAAGAAAGTAAAGAAAACTTTTGAATCTTGGTGTCTTAAACTAAAGATATGTAGAATGTACCAAAATGCCCTTTAATCTTGTGGTCTTAAACATGTCACGTGAAAA

>M6-PGSC0003DMG400025769 M6 LRR-RLP

TGAAATTCTGCAAATTGATTAAACTGCAACACCCCACAACATTCATATCAATCAAATATAGCATTTCTCCACCATCACCTCAGCCAGCCAAAAGGGAAACCTTTTTTTTCAAAGTTACCTAAAAGAAATTTGTGTGAAAGAGCTGACCATCAACTACAAAGGTGAGATATCAACCCAAAAAAAAAAAAAGGAAAAAACGAGCTGCAGTTACTTAGGTGCTCAAAGTAAACTAGATTAGAGTACAAATACTACGAGACAAGCATAATAAAGATTCTAAATCAATAGCAAAAGAGAACTCCTAGATTATGCACTGGCTCCATTGAAATGCTTGTGGATTACCAATCAAAATGACCATGCCACTATTTCAATGCCAATCATATAAACCATTACTTCATCTGCCACAGACACTTCCACTAACAAAGGTCATCGATAACCATTCACAAAAATAAGAAAAACACTAGTAAGTTCAGCCTTCTTTAAAACTCTGCTTTGCAAGGCAATCAAACCTCATCCCAACGATGTTGGATGATGCAATCCTACAGGTCAACCTACAAATCAATAAAACAGATTTCAGAAAATTTTGAGAATGAAGTTTCGGTCATGAACACAGAATAATGCAGCTAGGCTATCTGCTTTAATCTTACTCTACTTTCAAGGGGAAACAAAATTGAAATGGAAGAAAAACAGTTGTGCAATCAAGAAAAGGAGCAAATGGGACATATTCCCTCTCATAGTTAGCAGAAGATCGAAGAAGATCGAAGGTGAGGGTGGAAGCAAAAAGACCTGTACTTCTACAAGCTGCCTTTGGATGTTGAGTGAAGGAATATTTCAAGAACTTAGTTCTCTATCTGGTTCATGTAAATCATAGCTCTAAGCTAAAATCAGGACACGAGATAAGGATCTCACTCCAAAAGTCATTAACAAGCTAATTGCCCGTTTCACGTTTCAAACCCACAGGTATCCCTTTGAATCCAAACCCAGAAACTTATATCAGCAATTATGAAGTAATCTAAAGGTTCTAAAAAATCCCAAATTAGTGTATCCTGTTAAATCTATTCTGACCAACTTGTTTAAACTTTTGAGTAATTTAAGAGAGTGAGCACACTGCAAGGGATTAACGGAAGCTTTTATGCGCACAAGGCTTACTTAGAAAATTAATCACAATTATAGAAGGAATAATTATGTATAACTAATTCTTGGCATTAAGAAAAATAAATTATATCATAAGTAAAAGCACTTTTCACCTTGAACAAGTCTCCTTTATTTCCAGTTGTTCTAAGAAAAAATTACTTTGCATTCCAAGAACCTTTTTATTTTGTCTAGACCTATCGCAGAAAACGTCTAGTAGTAGGGAACTAGTAATTTGCCTCAACAGATGAACAATTGAAAATTGGATCCAAATGGCATAGAATGGATATTGATGATTCCTACTACCTAAAAAAAACTAGTTTAGGAATGAGGTGCAGTAACAGTATAGTGGTTGTTGCATGGTTATATCTATTCTATCCGTAAGTAAGTGGCCTATTTTATTTTTTGATAATAAACAAGGTGATCTTTGACTATACCATCAGTGTATGAAGTCAGTAAACAAGAGAAAGGAACTCATAAATTTGCTAGATTTTCTAAAATAAAGTAACTAATTTCGATGGGAAAAAAATCAAAAAAACTCTTTAGGTAACTTTGAAAAGAAGGTAGCATGGGAAACGAGAAAGTGGAGTGGGGGAAAAAGTCAATAGGAGGTGGACTATAATTACACCTCTCTTTTTTGCTAACGTCCGCTACCAAAAACACAGATTCCCTCCCTTATTTTTCTAAAATATTAATAAAATATGAATACTTAACTTTTGAGTCAGGAATTATTGAAGACTATCTTTCTATTTTATAAAGATAAAGATAAAATCTATGTATTTTTTATTCTTATTCAATTTCACTTATAAAAATATATTAAATATTATATTGCTGTATAAATCTATAATAGAGGCTCATTCATGGTTTTAAATTTTCTAGGTTTAATTTTAAAGATTATGCATTGAATTCATTACATTATTAGAGTTCTTGGTTTGATAATATAGATAATTTTTTTAAGTGATCAGTATAATTTTAACTTGTTGTGAAAGGTTATTTGCCAGAGTTTTTTTGAAACCTCAGGATAATCCGCAACCGTTACAACCTCAGCCGCAATCTTCGCTCTTCGAGTGAGCACTCTGTGCGCATTGGGTAAACCCCCCTCAGGTTACTAATTTTACTTAATAATCATTAAAACTGACCTGATAGAGAAAAAAAATTTAACAATATCAGTACAAATAAGTTATATTTAGAACAAAATGAAGACAGAAACAAAGTACTTCAATATTAGCCATTCACTTTGTTTTCCCATACTTACTCAAGTACCCTTTTGCTACTACTACTATATATCAATATCAAATTATCAATATTATCATCTATGTAGCAGAGTGAACCAGAACACTTTTCTTTTTTCTTGCCTTACACTTCTACATCCCTTGCTGCCTTTTTGGTCACCAAAACAACTTCTCTAGTAGTTGGTTTACGTCTTCCTCTTTCTTCTTATGTATATAGGAGTATTATGCTGTAATCATAAATAATCAGTTCAAGAAATGATGACAGACTCAGAGCAGTACTATTCTTTCTGGTCATTTGGGGTCTTTTTCTTTCTTTCTTTGTGTCCTTCGTTTGGTCTCAACATAGATGGAACTCTCTTGTTGTCTTTCAAATACTCTGTTCTTGAAGACCCTTTATCAATTATTGATAATTGGGACTATAATGATGCAACCCCATGTTTATGGACTGGTGTAACTTGTACACCAGACATGTTTAGAGTCATTAGTTTCGTACTCCCTAATTCTAAGCTTATTGGTTCTATTCCAGAAGAAATAGGATTTATTCAACATCTTCATACTATTGATCTTTCAAATAATTTCTTGAATGGAACACTACCTGTTTCACTTCTCAATGCTTCTGAACTTCAAGTTCTTTCACTGTCAAACAATGCTATTTCTGGTGAGCTTACTGAGTCTATTGGAGGGTTGAAGAGCTTAAAGGTTCTAAATCTTTCTGTCAATGCATTTGTAGGAAATCTCCCACAAAAGCTAAGTTTTT

>M6-PGSC0003DMG400025848 M6 LRR-RLP

GTGTTGTCAAAGTTTGGTAGTTGTCTTGTTTTCAAATAGTGCTGAACAAGGCAATCAAAGAAGGTGGCCAGTCGAATCTAGTTGCTTTGAAGTATTATTTAATTGGTGTTTTCCAATTTTGGGTTTTCATTTTGTTGTTTTGGAATAGTGCAGACTAAGGCAACGAAAGAAAGTGGCCAGTCAAGTTTAGTTTCTTTAAACAAATCTGGATCACAAATTAGAAAGCCTCCTCATCGCAAAACTTCGCCTCTGAATTGGTTCCCTCGCAAGAAAGTGGATTCCTATTTGAAGAGGAAAATAAAAATGCTCCAGGTATATTTATATGCTTGAGCATTCTCTTTTTTATTGTCAATAAGTAGTAGAAGTACTGATAGTGTTTTTGGCCTGTTGTGAAATCTCAGGAAGTGGATGGTATGAACTCAACTCTTGATGAGACTTTGGGGGATGCAAATCCCCACTATTGTAGAGTTTTGAGAGAAAAAATTGCAGTAAGAGAAGCTGCTCAAAGAGCTGTGGAAGCTCGGAAGGCTGCACTTGTTGAAGCATCATGGTGCCGAATACTTCAAGCCTCAAGGTGAGCAATTTAACATTCATTTATCTGTATTTGTTGTTACATGCATTAGATCCGGTGTTTGTCTGGAGTTTTTTTTTTCTGCTCAGGAATTCATTTGCTAATTATGTGATATTGGATACATTAAATTTTAGAGATAGTTAGTTACAGTTCAATGTCTTTTGGTGATCTAATTTACAAAATAGCTAGAAAATGTATAGGTTTGGTATATCTTAGTATAAATCTACATATAATATACTCACAGCTGAGAACATGCTCTTATATATAGGAAGTTATGGAGGTGGAGAATTAGGGTAGAAGTTACTAGGTAGTCGAGTGTTGTTGTGCTGTAGTATGATTTAGGTATCGTTAGTTTCTCTAATGTATATTAATTTAATAGCTGTTGCTACACTTGTTTTGTACATTGTTATCTTGTTGTCTCTTTTTCTTACTTTCTTGCGTCAATTACATTTGTTTTGAGTCGAGAGTCTTATTGGAAACAACCTGAAGGTGGAGAGGTAAAGGTCTGCATACATTTTACTCTCTCCAAACCCCACTTGTGGGATTATACTGGGTATGTTCGTTGTATACATAATATATGCGTAATCAGTGTTATATGTTTTGTACTAGCGCCCGTAATTAATTTCGGCCGACATGCAAGAAATGGAAAAAATAGCCAATATTAGCTTGTTTATGACAGAGACATTGTTGATTGATTGAATAGAGATTTGGTCATATGTTAGTCTAGAACTTACCTTTTCTGTATCTGCCATACAGGATAGATTGTAAAGAAGCAGAACAATTGCTGATAAAGGCAGAAAAATATTCTGCAGATGCTTTTGAAGCAGCAACAGCTATTGGAGTAATCTTGTATGACATTCCTGATTGCTCACAGAAGCATTACAAAATAGAAAAGTCACCAGCAAAAGGAGGAGGACCCACCACTCATACTGTTAGAACATCTTTTGAAACTGCATTTGAGGTAGATAAACAAGTAGCAAGTGCAGTTAAGGCTGCCCTTCTTAAGCTTGCTAATTGCCCTTCTATGAACAAAGATGAAGTCAAGGAACTGCTGCACAGAATAAGCCAGAACCCTGAAACAGATGATAATCATCAAGAAATGTCTGAGTTCTCTTCAGAATGTGAATCAGATACTGCTTCCGAAGCCACAGAGTTAGAAAAAGAAAAGGAAAGGAAGTATAAAAAGAAGCAGGCGTATGAAAAGTTTAATATGCCAAATCTTGTGGAAATGATGCTGGAAAGGCTTAGATGTTTGCAAGAAGATGAACTTGCATCTCTTGCTACTATAGTTGCAACGTGTGGGTTGAACGCTGCTCTAGCTGAAGCGGAAAACAGCAAAGTGAATGTTTCAGGGTCTGCTGCTGATGACAGATCAGAATTATCTGTAGGAGACGGAACAGTTAAGGGCGCTGAGGAATTTCCAAGCCTGGATAAGTTTCTAGTTAAGCGGTTAACTAGACTTGAAAGGGAGGTATTAGAAGCCAAAAATGCTAGAAGTGAAGCTGGGGAAAGAAGTGAACAAAGTCAAAATGAATCTTATCATAAAGTGATTCATTCGAGATACCATACAAATTCAAGCCATGATCTGGCAAGTATTCTTAAGAAACCATCTTTTTCAAAATTTGAGAAGGAAATTGAAGAGGCCAAGAACAACTCCGAAACTTTGGTAAGAACTAAGTGTAAAGCTTCAGATAATTCATCTGAGGTTCCAAGCCTGGACAAGTTTTTGGTGAAGCGATTAACGAGATTTGAAAGGGAGGTTTTAGAAGCCAAAAATGCCAGGAGTGAAGCTGGGGAAAAATGTGAAAAAACACGAGATAAGTCTAGTGATAAAGTGGTTCACGCAGACTACCCTACAGATACAGGCAATGATCTGGCAAGTATTCTCAAGAAGCGTTCTTCAAAATTTGAGAAGGAAATTGAAGAGGCCAAGAACAACTCCGAGACGTTGTTAAAAAACAAGTGTAAAGCTTCAAATAGCAATGTGCATTCATTTGAGGTTCCTGACTTAGGCAGCGTGTTGGTTAAGCACTCTTCAAAGCTTGAGAAAGATATTGAAGAGGCTAAGAGGAAGAATGAAAAACTTAGTGAAATAGAGGGCAAGAATTCGAATAGATTGGGTACTGTGGCAATTGGTCGAAGAAAGAAACATGAAATGGATGTTCCTAGTTTAGAAGATTATCTTGTGAAACATATGACCAAACTTGAAAAGGAAATTCAAGAAGCAAAAAATAGGGAAAACGCTGCTGACCCTAATGCCAATGTGTCTGAGACTACCTCATTGGTAGGAAAAGAAAATGTTGACCACAATGTAAATTCTTGTAATGGAGAACAACCATCAAAGCCAGCAGACACATTTGCATTTGAGGTTGAAAACAAAGAAGCTGTGGATAGTCTGGACAAGATTCTAGTAAAGCCAGTTCATCGTCTGCAAAGACTTAAAATGCAAGAATCATCAACAAGAAGAGACTATCGTGCACAGAGGACTCAAAGAAAGATTGAAGCTAATAGTGCCACTGATAGCGAAGGTTTGGACAAGATTTTAGTGAAGCATGTTTCAAAGCTTGAAAAAGAGAGAATGTCTTTTCATGCAAAGGAAGACAATTTGCTAAATGTAAAGAAAAGGGACACCATTGGTAAACAGTTGCAGAACAATGAAGGCAGTTTGGATCAGATTTTGGTGAAACACAAGTCGAGACTTGAGAGAGAAAAGATGGCTGCTATTCAGCCAGAGGATGATGATGATCAGGTAAGACATTCAATTACTCGGAAAGCTATAAGGGAGAGGGAGTTGCAAGAAGCTTGGGGAGGCTTAAGTCTTGGGAACTCTATGCGTCCTCGTATGTCTAGACTTCAGAGAGACAAGGCAAGTGCATTAACTTCATTCTTAAGCCATTTTATATTGTTGTTCCTTTTTCTACTTCTCACGACTTTTCTCTTGTTTTCCTTGTAGTCTGCCTGGCTTAAAGCTGAAGAAGAGGAGAGGAAGGGAACCGTGGAGGAGGCGTGAAATTATGAGAACTATTAGATTCAAAATATCATAAATTCTTTGTTTTTTCCCTCCCATGCTTTCCGTTGGTCAACTTCATAACAATCAACTAAACTTTTGAATACTTCTTCACTACTCATATAGTGATCTCGTGTGAGATGGATCTTTTTTTGGTTCTTTTTTGAGGTAAATAGAATTAGAATGCTATTTTTTTTCATTTTCGAATGTGTAAACTGAAGGTTGCAATGTAAAGAAGAGTGCTGCAAATTTGTACATATTGCTTGTCTATATATTCCTTCTGTGTATATGAGTGAGCAACAAAATGTATCTGCAATATTAGAAAGAAACTATATATTGTACAAATTAAATATATAAATTAAACAAAGTACTTGAAATAATTGCAATTACCGGACATCCAAATTAAACAAAGTACTTGAAGTAATGTAAGGATAAGATGAGAGAAAATAGCGAAGATAATTTTGGCCATATTTTACCCAAGACATCCAAAATATATCATTCAATTAATACATTGTTGTTATTTACTAAAAATGAAGTAAATCTAAAATCATGTAGATGAAATTATCTCAAGCGACTTATAGCAATCTCTTGAAATCAAACTGGATTTATCCAAGAAAATATATTCAATTACATGATAAAGCCGTTAAATCCAAAAAAAAAAAAAAACACAATCCACACCCCCTCCCAACCCCCCAATTTTGTTCTTAAGCTTACATAAATTTGAATGTAATTATTGAGAGTATAGTGAAATGGTTGAGATGTCATTCAATCAACAATGTTGGATTTGAGTTCACATAAATACTTAAAAATTAGGAAATTAAATACTTAACCCCTTTTAATTTTGTGGTTTGGCATTTGGTAGACAATTATTTCAGTTGAGAGTTGAGACAGGAGGAAAGTCCTTGTGGAAAATCTCACAGTGATAGTGTGAACAACAGAACTTAGCAGTGAACACTGGTCTCATTTACTCACTTTAATAGTAATAACTTGTACAAAGTACTCAACAGTTGTACAAACACTCAACAGTATTTTACAGCTTCCAAATTTGACTCTCTCTCTCTCTCTCTCTCTCTCTCTCT

>M6-PGSC0003DMG400025864 M6 LRR-RLP

ATGATGACTATCTCAGAACATTACTATTCTTTCTGGTCATTTGGGGTCTTTTTCTTTCTTCTTTTATGTCCTTCATTTGGTCTCAACATAGATGGAACTCTCTTGTTGTCTTTCAAATACTCTGTTCTTGATGACCCTTTATCAGTTCTTGATAATTGGGACTAAAATGATGCAACCCCATGTTTATGGACTGGTGTAACTTGTGCACCAGACATGTTTAGAGTCATTAGTTTACTACTCCCAAATTCTAACCTTGTTGGTTCTGTTCCACAAGAATTAGGCTATATTCAACATCTTCATACTATTGATCTTTCAAATAATTTCTTGAATGGAACACTTCCTGTTTCACTTCTCAATGCTTCTGAACTTCAAGTTCTTTCACTGTCAAACAATGCTATTTCTGGTGAACTTACTGAATCTATTGGAGTGTTAAAGAGCTTAAAGGTTCTAAATCTTTCTGTCAATGCATTTGTAGGAAATATCCCACAAAAGCTAAGTTCTTTGCAGAATCTAAAAGTTGTTTCTTTGAGTAAAAACCTATTTTCTGGTGCAATACCAACTGGATTTCAGTTTGTTGAAGTGTTGGATCTTTCTTCTAACTTGTTGAATGGAACATTACCTGATGATTTTGGTGGTGATAGTTTGAAATACTTGAACTTTTCATCAAACAAGTTATCTGGTTTAGTGTCACCACAATTTGCCAAGAAAATACCAACAAATGCTACAATTGATCTTTCTTTCAACAGCTTTACAGGTGAAATCCCAGAGTCATTAGCTTTATCAAACCAGAAAACAGAATTCTTTGCAGGAAATACAGATCTTTGTGGCAAACCACTCAAGAAACTCTGTACAATCCCTTCAACACTTTCATCTCCACCTAATATAACCACTAATCCTCCAGCAATTGCAGCCATTCCAAA

>M6-PGSC0003DMG400025867 M6 LRR-RLP

CATCAAAAGGCCCCACACTGTGCCTAAACCACCAGAACCCTCCATTACAAAACCAAAAAATTTTGATCAAATTTGTAGTCATTTTTCAAACATGTCAAAATCAAGAACCTCTGTTTTCTTCTTTCTCTTTCTTTTTTCACTTTCATCAGTTCAATCAATTCCTCCTTCTTCAGACATAATTGCCCTTCAAGCTTTTAAAGCTGCCATTAAACCTTCCTCTATTCCTTCATATTCTTGTTTAGGTACTTGGAATTTCACTACAGACCCTTGCTCTGTTCCTCGTGTCACCCATTTCACTTGTGGCCTTTCTTGTAGCTCTGGAAACAGAGTAACTGAACTTACTCTTGATCCTGCTGGTTATACAGGGACTCTTTCTCCGTTGATTTCTAAACTCACTCAACTTGTTACTCTTGATCTTCAGAATAACAATTTTTATGGCCCAATTCCTTCTTCTCTGTCTTCTTTACCAAATCTCAAGAACCTGGTTCTTCGACTCAATTCATTTTCTGGGTCAGTCCCACCTTCCCTCACTTCTTTAAAATCTCTACTTTCTCTTGATATTTCTCATAATTTGATTTCTGGGTTACCAAATTCGATGAATGAGCTAACCAGCTTAAGAAGACTTGATCTTAGTTACAATAAGCTAACTGGGTCACTCCCAAAATTACCCCCAAATCTACTAGAACTTGCAGCGAAGGCTAATTCGTTATCTGGGTCTCTTTTAAAATCGTCGTTTTATGGGTTAAATCAACTGGAAGTTGTGGAGCTTAGTGAAAATTCGTTATTTGGAACTATTGAAACTTGGTTCTTTCAGTTACCTTCTTTACAACAAATCGACATGGCGAACAACAGCTTTACACTCGTCAAGATTCCAAAGGTTGTTGATTTGAACAGCGACCTTGTCGCCGTTGATTTGGGTTTCAACAAAATCGAAGGGTATTTACCGGTAAATTTCGCGATTTTCCCTCGATTGTCGTCGTTGTCATTTAGGTATAACAAGTTCCGGGGACCGATTCCGTTGCAGTATAGCAAAAAGGCGACGTTGAAAAGGCTGTATCTTGACGGTAATTTCTTGAATGGGTCTCCGCCGGCGGGATTTTTCGGCCGGGAAACTTCAGTTACCGGCAGTTTAGGAGATAATTGTTTACAAAAATGTCCGATTTCTTCGCAACTTTGCTTGAAATCACAGAAATCTACGTCGATTTGTCAGCAAGCTTATGGTGGGAAACCAAAGTCTTAGTTGTGTCAGTCAATGTTTCTGAGATTTTTTTTCATCTTGGTTTTCATATTGTAATTGTAGGAGTATTATATTATATTCTTTAGATAAGAAATAACCCAATATAATATTTGCTCATCTGAATTATGCCCCCTTTGTATAGAGTATAGTACTATATTCATACTAGAAAATTACGG

>M6-PGSC0003DMG400025921 M6 LRR-RLP

CTCCTACTCACTTGACTTTTTTATCCACTCTATAAAGAAAGCAAACAAAAATTCCAAATTCCCAACCTTTTTCCTCCCTTTCTTCCCACAGTCTCCGCCGTTCAATCTGCACACCTTTTTCATTGTCACACTCTGCGATTCTTCTTTATCCCCAACTTGAAGAAAACCCTAACAACCCTTTTCTTCAAGGAAAAAAGTTCAGTTTTTGAGTCATTCTGCAGAATTTTCTTGAACAAAATGAAGTTGAAGAAACCCCATCTGGGTTTTGTGTGTTTCTTGATATTCTTGTCGTTTGTTGTCTCTGTATACTCTCAGGGTAGTGCAGCAACAGATGCAGTAGTGATGCAAGAGTTGAAGAAGGGAACCAGCCCACCGAGTTCACTCAGCTGGGACGATCCAGACCCCTGTAATTGGGGTAAAGTTAAGTGTGCTAAAGATGGTCGTGTAACTAGGATTCAAGTAGGGAATCAAGGGCTAAAAGGTTCTCTACCACCAAACATGAATAACCTGACGGAACTGCAGGTGTTTGAGGTTCAACACAATGCACTTACTGGGGCGATTCCGAGTTTTGCTGGGATGAACTCGTTGCAGAGCATTCTTCTTAACAACAATGGGTTCACTTCAATTCCTAGCGACTTTTTTGAGGGTATGACTAATTTACAAAATGTTAACTTGGATTCCAATTCCTTTTCGCCATGGTCTATACCAGAGAGTTTGAAAGATGCCACTTCTCTTCAAAGTTTCTCTGCTAATTCTGCTAATATTACTGGTAAAATCCCTGATTTCTTTGGTGGAGATACCTTTGTTAGTTTAACAGATTTGCATCTAGCCTTTAATAATTTTGAGGGTCCCCTGCCCTCGAATTTTTCGGGTTCTTCTATCCAAACGTTGTGGTTAAATGGTCTCCATAGTAAGTTGAATGGTTCAATAGATGTGGTACAGAACATGACCGCATTAACCCAACTCTGGTTTTCAGGCAATAAGTTTACAGGCCCTTTGCCTGATTTTTCGGGATTGACTCAGTTGAGAGAGTGCAATTTGAGAGACAATAGCTTCACTGGTCCAGTGCCAGACTCATTGGTTAATCTTCCTTCGTTAAAGATGGTTAATTTGACAAACAACTTTTTTCAAGGGCCTACGCCAAAATTTCCATCTTCAGTGCTAGTGGATATGTTGGATAATACCAATAGTTTTTGTTTGTCACAGCCGGGTCCTTGTGACTCACAAGTTAATACGCTGTTGGCTGTGGCTAAGGATGTGGGGTATCCCACGGTGTTTGCTGAGAATTGGAAGGGGAATGATCCTTGTTCTTGGATTGGCATAACTTGTGATGGTGGGAACATTACAGTGCTGAATTTTCAGAAAATGGGACTTACTGGGACAATCTCTCCCAACTACTCGTCCATTACATCATTACAGAAGTTGATCCTAGCAAACAATTTTCTTACAGGAACTATTCCAAATGAGCTCGCATTGCTGCCAAACCTAAAGGAATTCGATGTTTCTAACAATCAAATTTACGGAAAAATCCCACCATTTAAGAGCAATGTATTGGTTAAATATGATGGCAATGTCAATATTGGGAAAGATAATCCACCTCCAGTTGCACCGTCCGGAAGAACTCCTAGTTCACCCGATGGGGGTGGAGGTGGACAAACTCATGGGAATGGTAATAAAAAATCTTCAACTGGAGTTGTTGTGGGCTCAGTGATAGGTGGTGTTTGTGGTGCGGTTGCAGTTGCTGGATTATTTGTGTTTTGTCTTTACAGGAGTAAGCGCATGCGGTCCGGTAGAGTTCAGAGTCCACACACAGTGGTTATTCATCCTCATCATTCAGGATCTGACCAAGATGTTGTTAAGATCACAGTTGCTGGTTCAAGTGTCAATGGAGGGACAACCGAAACGTACAGTTGTGGAAGCAGTGCACCTGGGGACTTGCACATTGTTGAGGCTGGTAACATGGTGATTTCCATCCAAGTTTTGAGGAATGTCACGAACAACTTCAGCGAGGAGAATATATTGGGGAGAGGTGGATTTGGAACTGTTTACAAGGGGGAGTTACATGATGGAACAAAAATTGCTGTTAAGAGGATGGAATCTGGAGTTATGAGCGAGAAGGGGTTGGATGAGTTCACGTCTGAGATTGCGGTGCTTACTAAGGTCCGCCACAGGCATTTGGTGGCACTGCTAGGATATTGCTTGGATGGAAACGAGAGGTTACTTGTATATGAATATATGCCACAAGGGACAGTCAGCAGGTATCTTTTCAACTGGAAGGAAGAAGGGATAAATCCCCTCGAATGGACAAGGAGGCTAATCATTGCACTGGATGTTGCAAGGGGTGTTGAATATCTTCATGGTTTAGCTCAACAGAGCTTCATTCATAGAGATCTGAAGCCATCTAACATTCTTTTGGGAGATGATATGAGGGCTAAAGTTGCAGATTTTGGACTTGTTCGCCTTGCTCCTGAGGGAAAAACTTCATTGGTTACTAGGTTGGCTGGAACTTTTGGCTATCTTGCACCAGAGTATGCAGGTAAACTTCATCATGTGACCTGTTAATTTTCCTTATATTTTTGCATATATTTGCTTACATAAATGGACGTATGTTAGTGTATATAAGGCCAATAATATTAGATTAATAGGCTTAACAGGTGACAGTGGAAGTAGGAACTGCTTCCTCTCTTGTTTAGTTGTTTTGGTCTGCATCTACAGACCATCAGGGAAGCCTATTCTAGCTGGATCTTTTGGAAAGTTGATAT

>M6-PGSC0003DMG400025922 M6 LRR-RLP

ATGAATTTGAAAAAACCCCATTTGAGTTTTGTGTGTTTGATGATACTCTTGTCCAATGTTGTCTCTGTATACTCTCAGGGAAGTGCTACTACAGATGCTGAAGTGATGCAAGAATTGAAGAAACAAATAATAAATCCGAATTCACTTAACTGGAACGATCCAGATCCCTGTAAATGGGATAAAGTTCAATGTACTAAGGATGGTCGGGTAATTAGGATTCAAATAGGGGATCAAGGCCTAAAAGGTTCTCTCCCATCAAACATTAATACCCTTTCTGAATTGCAAGTTTTTGAAGTTCAAAGAAATACATTCACTGGGCCGCTCCCGAGTTTTTCTGGATTGAACTCATTGCAGAGCATTCTTCTTAATGGCAATGGGTTCACTTCAATTCCTAATGACTTTTTTCAAGGTATGACTAATTTGCAAAGTGTTTACTTAGATTCCAATCCCTTTTCGTCATGGACTGTACCAGAAAGTCTGAAAAGTGCTACTTCTCTTCAAATTTTCTCCGCTTATTATGCTAATATTACTGGTAAAATCCCTGATTTCTTTGGTGGAAATACATTTTCTAGTTTAACAACTTTGCATTTGTCGTTTAATAATTTGGAGGGTCCTCTGCCCTCGAGCTTTTCGGGTTCTTCTATCCAATCGTTGTGGTTAAATGGTATTGAAGGTAAGTTGAATGGCTCAATAGATGTGATACAGAATATGACCAGATTAACACAACTCTGGTTTTCAGGCAATGAGTTTACAGGCCCTTTGCCTGATTTTTCGGGCTTAACTCAGTTGCAAGATTGTAATTTGAGAGATAATAGATTCACAGGTCCAGTGCCAGACTCTTTGGTTAATCTTCCTTCGTTAAAAGTGGTTAATTTGACAAACAACATTTTGCAAGGGCCTACGCCACAATTTCCATCTTCAGTGCTAGTGGATATGTTGGATAATACCAATAGTTTTTGTTTGTCACAGCCTGGTCCTTGTGACTCAAGGGTTAGTACACTGTTGGATGTGGTTAAGGATGTAGGGTATCCCACGGAGTTAGCTGAGAATTGGACGGGGAATGATCCTTGTTCGCGTTGGCTGGGCATAACTTGTGATGGTGGAAACATTACAGTGCTGAATTTTCAGAAAAGTGGACTTACTGGGACAATCTCTCCCAACTTGTCGTCGATTACGTCAATACAGAGGTTGATCCTAGCAGACAATTCTCTTACAGGAACTATTCCAAATGAGCTCACATTACTGCCAAACTTAACGGAATTGGATGTTTCTAACAATCAACTTTACGGAAAAATCCCACAATTTAAGAGCAGTGTAGTGGTTAAAACTGAAGGCAACGTCAATATTGGGAAAGATAAGCCACCTACATCTGCAGCTGGAAACACGTTCAAACAAATGTAG

>M6-PGSC0003DMG400025926 M6 LRR-RLP

CCATTTTGCTTTGAAGAATATATTACAATAAGCAGCCTTGGGCCTTACCCACAGGTGGACTTACAAACATGGTGGCCTTGGGTTGCATTCTCTTTTTCCTTGTATCTTTCTCTTCTTTTTCTCCATCATATTTTGCATTATCTGATATTGAAGCTGCTTCTATTGCCCGTCGCCAACTTTTAGGTAACAATGGCGAGCTTCCAAATACATACGAGTCTAAAATGACCATTAACATGAAATTCGAAAATGCCAGGTTGAAGAAAGCTTATGTTGCTCTTCAGGCATGGAAGAAAGCTATTTACTCTGATCCAACAAACTTCACAGCTAATTGGGAGGGTAATAATGTATGTGCCTATAATGGTGTCTTTTGTGACAATGCTCTTGATGATCCCAACATCTCTGTTGTTGCTGGGATTGATCTTAACCATGCAGACATAGCTGGACATCTCCCTGTGGAGCTTGGCCTCTTGGCTGATGTGAGTCTCATCCACATTAATTCTAACCGATTTTGTGGAATCATTCCCAAGAGCATTAAAAATCTTACGCTTTTGGATGAGATTGATTTCAGCAACAATCGTTTCGTGGGGCCATTCCCTGATGTTGTTCTTGAGTTGCCTAAACTCAACTATCTTGATCTTAGGTTCAACAACTTTGAAGGTCAAGTGCCCTCTGCGCTATTTGAGAAGAATCTTGATGCAATATTGATCAACAATAACCGATTCCATTCCACCATCCCTGAAAGCTTGGGCAATTCCAATGCTTCCGTGGTGGTATTGGCAAATAACAAGTTCTATGGTTGCATCCCGAGCAGTATTGGTAAAATGGGCAACTCATTAGATGAGCTTGTATTCACCAACAATGCGCTCTCTGGTTGTTTGCCTGAAGAGATTACTAAGCTCACGAGCCTAACACTGTTCGATATTAGTGGAAATAAGTTTGTTGGATCGTTACCTCAGGATTTGAAATCGATGCAGAAAGTGGAAATCTTTGACATTGCATCGAACAAGTTCATGGGAAATGTTCCAAAGAACCTTTGCAAGTTGCCTAGTTTGACGAATTTCACATTCTCTAAGAACTACTTTGAGAGCATGGATGAAACATGTACGCCATCACAGTCGAAACAAGTTAAGATAGATGGTAATGAGAATTGTTTGGGGGGAAGATCAGAGCAGAGGACGGAGAAAGAGTGTTTCCCAGTAGTGAGCAAGCCTGTTGATTGCAGTAAAGGTCATTGCGGAGTTTCAAGGGAAGGGCAGAGTCCAAAAGATCCACCTAAAACCCTGACTCCTTCAAAACCAGCAACACCAACCACTACAAAACCAAAGCCATCTCCACCACCACCAGTAGCCTCACCTCCTCCACCACCACCAGTGGCCTCACCTCCACCTCCAGTCCATTCACCACCGCCATCGGTAGCATCACCTCCACCTCCAGTCCACTCACCACCACCACCGGTCTACTCACCTCCACCTCCAGTCCAATCACCACCACCACCAGCAACACCAACCACTACAAAACCAAAGCCATCTCCACCACCACCAGTAGCCTCACCTCCTCCACCACCAAAAACCTTACCTCCACCACCACCAGTGGCCTCACCTCCACCTCCAGTCCATTCACCACAGCCACCGGTAGCATCACCTCCACCTCTAGTCCACTCACCACCACCTCCACCTCCAGTCCACTCACCACCACCACCACCACCGGTAGCATCACCTCCACCTCCAGTCCACTCACCACCACCACCAGTCCAATCACCACCACCATCGGTAGCATCTCCTCCACCTCCAGTCCACTCACCACCACCTCCAGTCCAATCACCACCACCACCGGTAGCATCACCTCCACCTCCAGTCCACTCACCACCACCACCACCGGTAGCCTCACCTCCACCTCCAGTCCAATCACCACCACCACCGGTAGCCTCACCTCCACCTCCCGTCCACTCACCACCACCACCGGTTGCCTCACCTCCACCTCCCGTCCACTCACCNCCACCACCACCGGTAGCCTCACCTCCACCTCCAGTCCAATCACCACCACCACCGGTAGCCTCACCTCCACCTCCCGTCCACTCACCACCACCACC

>M6-PGSC0003DMG400025939 M6 LRR-RLP

ACTAAGTAACAAAAAAAATTCCAATTCCATGAAGATCTTTTATTCCCTCATTATTTTTTCAATTCTCAACAATATTAATTGCCAATTACTACCACCTTTACCAGCACCAAATTTAACCTTCATAGACCAAAGTCTAGCTCTAATATTCCCAATAATACAAAATTTCAAGAACACAATTACTCTTGATCCTTTTGGTGTCACAAAAACTTGGACAGGCCCTGATATTTGCAATTATAAAGGCTTTTATTGTGATAATCCACCTAATAATTCATCAGCAATTTCTCTTGCTTCTATAGATTTCAATGGATTTCAACTTAGTGCACCTACATTAGATGGTTTCATTGATCAACTCCCTGACTTAGCCATATTTCATGCTAATACCAACAACTTTTCAGGCATAATTTCTTCGAAAATTTCAAATCTTCCATATCTTTAAGAACTTGATCTTAGTAACAACCAATTCATCGGTTCATTTCCCATGTCAATTCTCAATATCAAGACTCTAACTTTCTTGGATATTCGTTACAATCTCTTGACAGGAATAATCCCACCTCAAATTTTCGTACAATATCTTGATGCATTTTTTCTTAACAATAACAATTTCATTCAAAAACTTCCAGACAACATTGGATCCACCTCTGCTTTTTATTTAACCTTAGCCAACAACAAATTCGTTGGTCCAATCCCACGTAGCATTGGACAAGCTCCGAATTTATTGGAAATTTTATTCTTGAACAACTTGCTCACTGGTTGCATTCCTTATGAATTAGGCCTTTTGAAAAAAGCAATTGTGATCGACATTGGATTTAACACATTAACTGGTCCATTACCACGTTCATTAGGGTGTTTGGAAAGTGTGGAACAATTGAATTTTGCTGGAAATTTGTTATATGGACAAGTTACAGAAGTGATATGTTCACTAAAAAATTTAGTGAATTTTACTTTGTCATATAACTATTTTACAAAAGTAGGGCCACTATGTAGAGAATTGATCGAGAATGGAGTTCTTAATGTTGAAAAAAATTGTATTAATGGTCTTCCTAATCAGAGATCAATATTGGAGTGTGTATTATTTAGGATGCAAATTAAATTGTGTCCATTACAGAAGAGCTTGAGGATCATACCTTGTAATATTTCTAATTTTAGTCCCCATAGACCACTTAGACCAGAAAGACATTTGATTTCATACTCTGCTTTGTCAAAACATACAAAATTGTATTAATGAAACCTTTGTTTTAGTGTCTTCTTGATTTTTATTTTATTTTCTATGTTTTCTACCTGGTGTCCG

>M6-PGSC0003DMG400026064 M6 LRR-RLP

AAGCTTAGATCTCAGTGGATGTAATTTATGTGATAGCCAAGCTGCTGCTCTTATGAACTTAACCTCTCTATTGGAACTGAACCTGAGCAGAAATAAGTTTATTTATTTCCCTGATGTCTTCAGGCGACTTTCTCACCTTCGATATCTCAATATTACACACTGTCAGGAACTTAAGAAACTGCCCGTACTTCCTAAGAGTATAGAGGAATTATATGTAGAAGATTTTTTGGCAAAAGAAAGTATTGCAAAGCTACGGATGCACCCAAGGTTGAATTTGGTATCATTCACCAATTATAGTTTTGATCAACAATCTTATACAGAGGAGAGCAATGGTAGCTCAATTTCAGATGAGATTCTCAGTTTGTTTCTATCAAACAATATGGATGATGTGATCTTCCCCACTCTTAACTCTGATCATAGGGTAACCTGTTCCATTGTCTTTCCTGAACGTGCAATTCCCACATGGTTTATGCATCAGAGTGTTGAGGAAAAGATCTCGTTCAAACTGCCCATTAATTGGTATAATGATAAGTTCAAGGGCTTTGCTATATGTTGTGTGACTCTCATGGGAGCAGGTGTCTGCAGTCCTGATTCTAGGCTATCTGGGAAGTATGATTATGCTTTTATCAATGCCAAATTGATATGCAATGATCATTTGAAAGACCTTAAAGTGATAGAGAAAGAATGTAAAGTGGGCACAGCATCTAGAACATATGGCTGGTGTGTTTGCTTCGCCTACATACCATTGTATTCTTCACTGCAGGTTGGAGACATTAATCAGTATTGCCTATTTGAGGCATCTATCCATGGATGCATTGTGAGACAGTGGGGAGTTCATTTGATCTATCAGGATAAATGAATATCTTTGAAGCGCAGACTCAGAGAGATTGTATCGAGTACTCCTAAGCTGAAAAAGAGGCTAAATTATGCTGGGATATTAAGGGGATTACCAAGAAATAGTTTAAGGTAACTCGAAATACCTCTTCTTTGTTACTAGAGTTTCATAATTTGAATATGGTTAGATTAACAAGTTTAAAACTATAAGTTCGATTAGAAAAATATAAAAATCAATCAGATAACTTAAACTTAAAATTAGATAAAAATGGTTGGCATCAAGCCAAGCCCAATGTCCCTAGCATAAAGTTCTTATTTTCTGGAAAAATTACATAAATGAATCTCTTTTATTTAATAATAATTACCACTTTTAAATATACTTTTATTTTATTACTATTTTTGTCAACTTTTAGCAATACTTACTTAAACAATTCTTGAATAATTTTGTTCCTTTTATATATATAATTAAAGAATTTTCAACTTAGCAAAAATTGTTGTACCTGATCGGGAACATCGCTCATTCCCTCTTTCACTATTCTTTCTGCTCTTCTCTTTGTTTTATTTTAGGGTTTTGCCATTCTTTCTTCTTCAATTTTCAGCACAGTATTCTAATGGAAGAAGGCAAACGAAAATCGCCTAGTAGAAGTCCGCAGATACTTAGTGATGTTGATACTTCAAATCAAAAAGTTTATAGAAGGAGTCGGATGAGGGGTCAAATTTCAGCTTCTACTGCTACCGACGAAGTTAAGGATGATCCCAATTTTAATTTGGGAATTTCACAAGTATCAAGTGAGCAGAACAAGGAGGAGAGAAATGCATGTTAGGTAAATAAATGAGTTAAAGAGGTGAAAAATGTGAAAAAAGTGAAAAAAAGATGCGTTGATTTTGCATCTACATCGAAGAAAATCGTGGACAACGATGACGATTTTGAGGATTTATCTCCTAAATTTCAGTCAAAAAAATTGAAAAATAAAGTTGTGTCGGAGTAGAAGAAGCAGTGAAGGATGCAAAAACGCAGATTCGATTGGCCAAAAATGTCATTCTACCGGAGAGTAGAGATCAGGTATGTGA

>M6-PGSC0003DMG400026160 M6 LRR-RLP

ATGGAAACAAGATCCGTTAAGAAAAAAAGGCTTCTTGATGAAATCTACGCGAGTTTTTCATGTGGTGAAGACCGCATCAGTGATCTTCCTGATGCGGTCATTCACCAAATTCTTTTCTTCCTTCCAATCAAATCCATCGCCAAACTTAGTGTACTTTCTAAGCGATGGAAGCACATGTGGCACACTTTCCCAGATCTCGATTTTACAACAATCAATGTCCCATCATCATCAGGTTATAATAACTCAATCTCAAAAGTCAGCTCAAACGTTTACAAAAAATTCCATGCAAATTGTACTAGTGCTGGAGGTGCTGATTTCATAGATCAAGTGTTATCTTTTCGCGAAAAAAATTTAGGGATCAGGGTCCTCCGTTTTCGTGCAAATTTGAGTTTTTCACATCTAAACGCCCTGATTCGTGGCGCTGTTAAGCACAATGTACAAGAACTTGACGTTGAAGTAGCCACTAATGATTATTTCAATTTCCCAAGAAGTGCTATAAATAGTGACTCTTTAAGAGTTTTTAAATTAAAGTCAAAATATCCAGGATTTAGGTTACCTCCTTCATCAATTATGAAAAATGGGTTCAAAAATTTGTTCACATTGTCACTTTCGTGTCTTATTATGTACGAACAACCTTCGTTACTTGATTTATTCACGGAGACATCTTTCCCGTTGCTTAAGAAATTGAGTCTTGATGGTTGTACGGGGTTAACTTACTTACATGTAAGTTGTCGTGTACTAGAGATTTTAATTCTTGAAAATTGTTTTCAATTGGAGAATTTAGATATAACGTGTTCGAAATTGGAGAAATTGAGTGTCAACAGCTGCTTCGATTCGTATTCTAGTGGTACTGAGGTTAAGATTGTTGGTCCGAGGCTAAAAAGTATTGTTTGGTCGAATAATACTATAACGGATCAAAGTTGTTTGGAGAATTTAACTTGTTTGATTGAGGCTTTTGTTGGGTTTTTTGTTCTTCTTGAGGATCTTGGTGCAATGAAGCTTCGAAGTGTCAATGATTTCTTGTCTGGCCTCTCTCATTCACATTCTTTAGTTCTTGAAAATCAAAGTGTTGAGGTATATTTTTTATTCATTTACCAAGTTTTATTTGATGTGTGACATTTAATTTTTGCCTCTTCTATGGATCTAGCTATATGTGTGACATTTAATGTTGTCTCATAACTTGGTGTTTGTCAACGGAATTGAGTAACTCGGCCCATCAAGACTTAAGCTATTCAGATTTGAGTCTTGATTTTTCATATGTTTTTAGTTCAGTTCATTGACTACTAGATATTCTCTTGTAAATATTAGTAATTGATTATTGGACCAAAATGTTGTTGTTACATTTACCTTAGGACACGTTTTGTTCTCTACCATGATTTGAATTCTGATCTCCTTGTTATTTTGTTAGCCATTATCCTATAAATATTTATTGATTATTAGACCAAAATGTCCTTGTTACATTTACATTAGACCTCTTTTTGCTTTCTACCAAGATTATATTTGAATTCTAATCTCCCGTGATCTCCTTATTTTGCTAGATATAATCCTATAAATATTTATTGATTATTGAACCAAATTGTTCTAGTTACATTTACATTAGGCCCATTTTGCTCTCCACAAGGATTTGAATTCTAATCTCCCATGAT

>M6-PGSC0003DMG400026231 M6 LRR-RLP

TTCCTCAAATTTTCAGTACCAACTCTTTTTTTAATTCAAACATGGGCTACATGCAACTCTCAGCTGTGCTCTTAATTGTTTTTCTCTGTCAAATTGCTTTTTCTTCATCTTTGTGCCCTAAAGATCAAAGCATTTCCCTTCTAAATTTCAAGAAATCACTTACTATTGATCCCTCATATGATATTTGTCATTACAATTTTTCAAAATTGAGTTCATGGAATACGAGCAGTGATTGCTGCTTATGGGATGGGGTAACATGCGATGAGATGAGTGTCCATGTTATTGAGCTTGACATCAGTTGCAGTCAACTTGTTGGAGTGATTGATTCCAATAGTAGCTTATTCCAACTCTCTCATCTAAAAAAGCTCATTCTTTCTATGAATGACTTCTCTTCTTCACACATTTCACCTGCATTTGGAAGGTTTTCAAGGTTGACCCATCTTCATCTTTCTGACTCGCATTTCTCAGGTAAAATTCCTTCTGAAATCTTTTCCCTGTCAAATCTCACTCGCATAGAGTCCTTATACCTTAGAGGAAACTCCCTGAATGGAACAATACCATCAGGTATGTTCTCCCTTCCATCACTAATAGAATTAGACTTGAGTAATAACCAATTTTCTGGTCATCTTGAGGATTTCAAGTCCAATTCACTAAGATCTATTGATTTAAACAATAATCAGCTGCAAGGCCATCTTCCCAAGTCAATTCAAAACCTTGTGAACCTAACGTGGCTTGATCTTTCATCAAATAATTTTAGTGGCGATGTGGATGTCAGCTTATTTTCAAACATAAAAAATCTTCAGGGTCTTGATCTTTCATATAATCGTTTCTCATTAACCAATGAGAACAGGGCTAATTTTACTTTGCCAGAATCTCTTTTTAGTTTGCGATTGGCCGCATGTGAAGTAAAAGAATTGGAGTTTCTAAGACCAGTAAAGAAACTTTGGGACTTGAATCTTTCGAATAATAAGATTCAGGGAAGAATTCCAGATTGGGCATGGCCTATATGGCTTAACTTGGACCGTCTTAATTTGTCCCACAATATGTTGACAGATATGAACTCAACAAATATAAACTCAATTTCTTATCCCTCTCTATATATCATTGATTTGCGGTCCAATTTTCTTCAAGGGCCACTTCCTATTCTACCAAATTCCACACAATACTTGTTCATGTCAAACAATAATCTTAGTGAGGAAATTCCTTCATCTATTTGCAATTTGAGATCACTAAAAGTCCTAGATTTGGCGAAAAACAACTTGATGGGAGAGATACCACAATGCTTGGGTAATGTCAGTAGTAGCCTCGAGGTTTTTGATATGCATCAGAACAATCTGTCTGGGACAATTCCAACAACTTTTGGCATTGGAAGTTTACAAGTTAGGAGCCTCAACTTGCATGACAATAAGCTACAGGGGAAACTCCCACGATCTTTAGCCAATTGCAAGGAGTTGCAAGTTCTTGATCTAGGAAATAATCACCTCAACGACACATTCCCAATGTGGTTGGCAACTCTACCAAAGCTGCAAGTTTTAAGTTTGAGATCAAACAGATTGCATGGTCCCATCGGAACATCAAGGATGAGGAACTTGTTTCCTGAGCTTCGAATCTTAGATGTCGCTTACAATGCCTTAACAGAAACCTTGCCAACAAGTTTGTTTCAACATTTGAAAGCCATGAGGACAATTAATCGAACAATGAAGGCACCGGTATACCTTGGAGACGAGTACTACCAAGATTCTGTAACAATTGTATCCAAGGGAATGATGCTTGAACTTGTTAGAATTTTGACGATTTACACAGCTATCGATCTTTCAAGTAATAAATTCAGAGGACCAATTCCAAGTGTCATGGGAGATTTCATTGCACTTCACGTGTTAAACTTATCTCATAATGGATTGCAAGGTGAAATACCTCCATCACTCGGAGATTTATCTTCAGTTGAATCATTGGACCTATCAAGAAATCAACTTTCGGGAGAGATACCACAACAACTGGTTTCTCTTACGTCTCTTGCGTTCTTAAATCTCTCCCACAATCATCTCCAAGGATGCATCCCTCAAGGACCTCAAGTTCACACTTTTGAGAACAGTTCATTTGCAGGCAATGACGGATTACGTGGATTACCAATTTCGAAAGGCTGTGGTAATGATGGTGTAATAGACACAAATTACACAACATCCTCTCCAGATGAAGAAAGCAATTCCGAATTTCTGGATGATTTTTGGAAAGCTGCTCTTATGGGCTACGGAAGTGGACTATGTATTGGATTATCCATAATATATTTCATAATTTCAACTGGAAAGCCGAAATGGCTTGCAAAAATCATTTTAAAGTTGGAGCATAAACTTATGATGAGATGGAAAAAGATACAGCGATGGCATAGGAATCACAGAAGAAATAATCGCTTCTAAGATAAGTTAAGGTATTCAAGTTAAACTTTACACTGATCTTTTATTTTTTTATACTTTCGTTATATAAACTACTAATCAATATGCTCTATCCTAAGCATATTTGATTTTGAAATGGTTCTGCCTTTGTTGATGTCTCGAGCTTGA

>M6-PGSC0003DMG400026748 M6 LRR-RLP

GATTGGTAAGGTAAGGACTAGCCCCCCTTGTTTGTACTTATCCCTATTGAGATTTTTATTTCTAATAAAAGGCCGCTAGACACTTTGTCTAGTGGTAAATTTGCCAAAACAAATAAATCCTTGAAAGCATCGACTTATTAGAATGGCAACATAAAGCACATGTGTATTGGTGGATATGTTTGAAAGTTGGCTTGAATACCGTTATAACAATACATAAATGTACCATTTAACTGGCATCTATGCCCTCCAAGTGAACAAGTAAACACTTAAACTTGTGTAAAGTTGAACAAGTACATATACTGTACACATCTTATGTGGAGCTCTATGTAGCAATTTACATCCTACGTGGCATCCTACGTGTATTATACCACATAGGACACGTATGTCTACTTGTTCAATGTTATACAAGTTTATGTGTATACTTGTGCACACCCAAAATTGAAGGGCGTAGATGCCAGCTGAGCCCAAGTTAAAGAACACATTTATGTATTATGCCTTAAAAAAGTGTGATTACGTGTTCGAATAAAAGAATTGAAATTTATAATAAATATTTATGTATTTGGTAAGATGTTGTATTCGTAAATTGTTCTTTTATTAAAATAACTAAACTAAAATAACCTTAAATTGTCTTTTTTTAGAAAGTGTAAATCTAAATGTAACCTTGTAAAAAAAAAGAGGAACACTATATATCTTAGTGAAGAGGAAGTTACGAGACTTATATTGGAAAAAGTAGTTTTGAGAATTTAAAAAAAAATGAAGGACAAAATAGTAAGAGACCGGGTCAAATCAAGAGTGTTAACCCAAAAATTAATAAGTTCGGAATAATCAACTTATGACTTTTGATTGGTCTTAGTGGTTTATAAGTACTTTGAATGTTTACTAATTAATATGTACGTAAGCTAAAAATATTGTTTATAAGTTAGTTTAACGAGCCTATAAGCTTAGTCAAATACCTTCCAAATATCTCACTTAATTTTGTACTTTTTATCTCTCAAGTGAGAAGCGATAATGATCAAGAACTCACAAGTGTATTTTTTTTTTTGTAAGGTTCATACATAAATTATCAGGTGTTTGTGTTTTCTATCTGAATTATCACTATCATTAACTCTTCTTTCTTTAAAGGGAAAAGACCCAACTATGTCATTGAACTTTGAGAAAAAAACTCATAATCTATTCCATCCGTTTAGAAAATTAGTTTATCTATGTCATTTTCATTTGAGAAAATGTTCATCCATGTCATTGTTAGTTAATTAAAAATAGCTTTAATTTTGCTCAGTTATACAGCGTTTACTAGCTAGCTATTCATGTCTGAAATTAATACACAATTTGTTTTACAATTTAAAAATCAACGTAACTAATACTTGTATAAATTATGAAGAAATTTGTACATAATGTTATCTCAAATGGACTATCAATGCAATTATAGAAAATATAATAACAAAAGGATAATGACATTTGAAAAAAAGGTAATTAAGAAATGATGCATAAATCAGAGGTCAAAAATATGATAGATACTAATAATAATAATGAGATATAACTATATTAGGAAAACAAGTTTAGCAGAGGTCCTTTTCCAAGGTAGGTCCTACTTTTCATCACACAATTCTAGAGAATATGAAAGCAAGACTATTTACAAAGAATTAAGCACACAATAAATAAAGCAAAACTTGGTTTACTTGCCATGTCTTAATTGAGATGTGAAAGACTTGATTGAATGGAAAAATCTCCTAAAAAGAGATCTTATTGCTATTCAAAGTCTCAAAACCACAGGTTATTAGTACTAATCTTGCACTAATGGACAAAACCTTCACTATTTTATTCTTAACAACACTCTTGTTGCTTCACTATGTTATGGCCCAAACCAACATTACGACTGATCAATTAGCTCTTCTCTCTTTTAAATCCCAAATTAGTTCCGACCCCTTTCACTTCTTGAATGAAAGTTGGTCTCCTGCTATTTCTGTTTGCCGTTGGGTTGGAGTCATTTGTGGTTCTCGTCACCAGCGAGTGAAGTCGTTGAAGCTTTCCAACTTGGCTCTTACAGGCAGAATTCCCTGTGATTTTGGAAACCTCACATTTCTTGTTTCTCTTGACTTGGGAAGCAACCATTTCCATGGAAATTTGCCTCAAGAAATGGCACATTTGCGTAGGCTTAAGTTTCTTGATTTAAGTTTAAACAGCTTCAGAGGGGAAATTCCTTCTTGGTTTGGGTTTTTACACCAACTTCAAGTTGTAAACCTTAGGAGTAACAGTTTTACTGGTTCCATTCCTCCGTCACTCTCGAATGCCTCAAGGTTGGAGATTTTAGAAATATCTACCAATTTACTTGAGGGAAACATCCCGGAAGAGTTTGGCAATCTTCACAACCTGAAATCGTTGTCCATAGAACAAAATCAGCTTACAGGTTCTATACCATTCACAATATTCAATATTTCTAGAATTGAAGTCATTAGATTTTCAGATAATAGCCTATCAGGAGATCTTCCCAATGGTTTATGCAATGATCTCCCAATACTCAAAGAGCTTTATCTATCCGGAAACAAGCTTGATGGTCATATGCCTACAAGCTTGTCAAATTGTTCACGACTTCAAATACTGTCTCTATCCCAAAATGATTTTGATGGACCAATACATAATGAAATTGGAAGATTGAGTAACTTGCAGACATTAGAAATCGGATACAACCATTTCACAGGTATATATGTTTCATATTCTTATGAATGCTACTACTACTATATATTGTATCTAATTACCTCACATCTTTCTTTATTACACATTGAATGCAGGTATAATTCCACAAGAAATTGGAAATCTTGTTAATTTGGTGAAATTAGACATGGGGGGAAATCAGATTACTGGCTCTATCCCGATCTCTGTATTCAATAGCTCATTACTGCAATCATTGTTACTGTGGGGCAACAATCTTAGTGGATTCGTACCACAGGAGATTGGCAACTTAACCAAGATGCAGTTCTTATATCTTAACCAAAATAGGCTTACTGGTACGTATTCAAAGTTATAGTGATCTGTATTGAGCTAGTTAGTTAAACTCATTCTTTCTTCACGTGATGACAGGTGAAATACCCAAAGATACAAGAAATCTCGTTGAGTTAAAGGAATTTGATGTTGGGTTTAATAGTTTTAGTGGTTCACTTCCAACGGAGATGTTCAACATATCAGCACTGAGAATAATGTCTCCTTCATTCAATAATATATCAGCAACTCTACCACCAAACACAGGTTCTACCTTATCCAACATTGAAGAGCTTTATATGGGTCACTTAACCAATCTTGTTGGGACTATTCCTCATTCTATCTCCAATTGTTCAAAGCTTAGTAAACTAGATATTTCAGGTAACAAACTCACTGGCTTGATTCCCAATTCTCTTGGATATTTGACTCATCTACACATCCTAAACATGGAGAGAAACAATTTAACCAGCGACTCCTCGTTAAGCTTCCTCACTTCCTTAACCAATTGCAGAAGATTAACAGTTCTTTCTCTATCTTTCAACCCTCTAAATGGCATGCTGCCGGTCTCCGTAGGGAACTTCTCCAAATCTCTTGTAACCTTTTATGCCGCTGCTTGCAAGATTAACGGCCAGATTCCAAATGAAGTTGGAAACTTAACCAATTTAATAGACCTTGATCTTTCTAATAATAACTTGATTGGATCAATTCCTACCTCAACTAGTAACTTAAGAAACCTCCAACGCTTGTACTTGAACAAAAACAAACTTACAGGATTTATTGGAGATCATATATGTAAATTGCAGCATTTGGGTGTTATTTCCTTGGGTCAAAATCAACTTTCAGGATCTCTTCCTAATTGTTTAGGGAATGTTACTTCCCTTAGAGAGATACATATAAGTTCCAATAAATTGAGTTCCAATATACCACCAAGCTTAGGAAATCTTAAAGATCTAATGAAGCTTGACTTATCATCAAACAACATGGTTGGCTTTTTACTTCCGGAAATTGGAAATCTAAAAGCGGCGACATATATCGATCTGTCCATGAATCAATTCAGAAAAGGAATTCCTACAGAAATTGGAGGGTTGCAAAATCTAGTGCATCTTTCTTTGAGACACAACAAGTTGCAAGGATCTATACCTGACTCAATCAGCAACATGGTAAGTTTGGAATTTCTAGACATTTCTCATAATAATATCTTGGGAACTATTCCAATGTCTTTGGAGAAACTACAATACCTTGAGCGTTTCAATGTTTCATTCAACAAGTTGCACGGTGAAATACCCTCGGGGGGTCCTTTCAAGAACCTCTCGAGTCAATTTTTCATCAACAATGAAGCATTGTGTGGTTTGTCAAGGTTTAATGTCCCACCATGCCCCACATCATCAATGCATAGATCAAATAGGAAAAAATTGCTAGTTCTATTTCTTTTGCTAGGAATAGCACTTGTATTTGTTCCTATCGCCTTTGTGTTCCTATGGATAAGGTATAGAAGAGGTAAAAGAACTCCTCAACAAGCTGATTCATTGCCTATGGCAACAACGGAAAGAATTTCATACTATGAACAGCTCCAAGCAACTGATTCACTTAGCGAGAGTACTAATCTGATTGGTTCTGGAAGTTTTGGCTCTGTTTACAAAGGCGTTCTCAGGAGTGGGACTGTCATTGCAGTTAAAGTGTTCAATCTGCAACTGGAAGCAGCATTCAAGAGTTT

>M6-PGSC0003DMG400026750 M6 LRR-RLP

GTCTCTTTTTCACTATCTTCTGTACACAACAAACTTCAATACACCTCTGCTCTGTAATTGCAAACCATTTTCCATTTTTCTTCAACCAAAAACAACAACAATGGCTTCTTCTTCTCCAGATCCATTTTTTCTCCTCATATTTCTCACTCTTTTCCTCACCATTTCCACTTCAAACTCACTTTCTTCCTCTTCCCCATCACCCCTTTCTTCTCCATCTCCAAAACCCTCTCCTATTTCAACTCCACGAACTCATTCATCTTCATCTATTCTTGACCCAAATCAGCTCAAAGCTCTACAATCACTCAACATCCCAACAGGCAAAACCCCATGTTCCCCATCTCACAATACTACCACTATTTGTGATTCTTCAACCCCATTTAGTCACATTGTGTCATTAAACTTCATTAACTGCTCAGATGATGTTGCATTATCACTTACTGCTTTGAAATCATTGTCTACAGTGAAGGATTTGGGGTTTTACAATTGCCCCATTTCACCTATTAGGTTACCTTCTCAGCTTGCTTTGAATCTGAAATCTTTTACTTGTGTTAGTAGTTTGAAGAAATTAACTGGAGTTTGGCTTAGCAAATTGGTGAATGTTACTGATTTAACTGTTTCTCATGTTTCGATTGTTGCTAGTGGTCCATCTATAATCATGAATAGTATAAAGAATTTGCACAGTGTTACGATTTCGAATGCTAATCTTACTGGTTACTTGCCTAAACATTGGCATTCGAATCTTAGTTATGTTGATTTGTCTGGTAATAAACTGAAAGGGAGAATACCCAGCTCGTTAACTGAGCTGGAGAATCTTGTTTACCTCAATTTGTCTTCGAATTCACTTAATGGAACGATCCCCGCTTCGTTTGGTGATTTGTCTTCTTTGCAAAATGTGTCTTTGGCTTCGAATTCGTTGTCTGGATCTGTTCCGGATTCGTTTGCTGCTATTCCCGGATTAGTTCATCTTGATTTGGGGTCTAATCAGCTCAATGGGACTATTCCTAAGTTCATTTCTGATATGAAGGGATTGAAGTATTTGAATCTTGAGAGGAATAATTTTCACGGGGTTTTACCTTTTAATGCTTCGTTTATAAAGAAATTGGCTGTGTTGAAGGTGGGAGGAAATTCGAATCTTTGTTATAATCATTCGACATTGTCTTCTAAAGTGAAGCTTGGGATTGCTCCTTGTGATAAACATGGGTTGCCTGTGTCACCTCCAGCATCGAAGGATATAAGTTCGGATGATACCGATGATTCTGATGATTATGCTGATGATGAGAGTCAGCACCAGGAACATAGTCATGGACCAAGTAAGGTTGTTCTTGGTATTTCCATTGCACTTTCCTCGATTGTATTCATAATTATTTTCTTGATTCTGTTGGCTAAATGTTGTAAATGAGCATAAAAATGAAATCTTGGTTCGTTTAGTTTAGGAAATTTCTTAATTGGATTATGAAATTTGTTTAGTTAATTATACATTTGTATCATGTTAGAAGTAGAAGAGGTGGCTACTTTATTTGGTGTGATTCTTTTGAACAAGTTAAATTGCTGGAGGAAGTAGGCGGCTAAATTGATCTTTAAGTGTATGATGGGAGAGATTGCCTATTAATGAAATTTTCTTTTATGTATGTATTTTCGATTGACGGATTGAGGTACATGGTTTTGATTTCCTGATATATGGTGATATCTCAATTGCTTACGACATTTTGTTGTTTGGATTCAGCATTTGCTTCATTCATTTGTTGATTGACAATATGAAAGATTCCTTTATTTTTCATGTGTTGGTTACGAGTCATGTCTATGGATGAGATAATCTATGTCGTGTGGATCCTTCATTTGTCTTGATGTACCCCGTGTCCAATGGGAATGTGGTAAGGGTTAGGGTACTTTGCTGGAAAAGGTTGATATACCCATACCGGGTGCTTCCAAAGAAACTGTACAGAACACATATCCGTAATAGAGTCCAAGCAACATAATGGCAAGATGAAGTATCTACCGAATTAGCTCAATAGTCTATCGTCCCAGTTTTAACTGCAAAGATCAGTGTTTTCGCTGTTACTTGCGTGATTTTTCGTAGTAAGTTCTTTTGCTTTCTTTGTGCTCTACCGGACTTTCCAGTATTCCTCGTATGCCCTGTGATGCCAGTGTTTTTATGAGATCTAACATAATGCTGTGAAAACAATATTCCATTGTAGACAAAAATGTTCATTATCTGGCACTTCATTTCATTTGACAGAAGAAGCTCAATTAGTCCCCTCCTTAGCTGTTTTGCTCAGATTCTTCATAAATATGTTGGGGTTATGTGTTGGATCCTCCAAAGCTATGCTTTTTTGGAGGATCTTACACGGGTGGGGCAACACTTTTGGAGGGTCCGAGTAACAAAGCTCCTTAGTCCCCTACCTGATCACCACAAAGGATAACTAGGGGGTGCTTCCACACTGGGTTTTCCTTTCATTTTTAAATGCTTATTGTGATAATAGACTGGAGAATTCATATAGTCGATTCTAACTTGCTTGGGACAGAGGCACGAGTTATTTTTTTTATTGTCGTTTTAGTTATAACGTCTATTGTTTTGGCC

>M6-PGSC0003DMG400026840 M6 LRR-RLP

ATATACGTGTAGGGGGAGTTACTTATTTACCGAGTCCTAGAATAGATAACAATCTTGTGATAGGGATGCCAAATCTAGAGGAATTTTTTGGTCTTAGTTACTCTAGTTGTACAAATGAAGTCTTTTCTAGGATCCCCAATTTAAAGAGATTGATCATTCCTGCACATTTTAGCATACTAAACAATATTCTGTATCGTCCCTTCGATATGTCCAGCTTGACAAAACTTGAGACATTCAAGTTTTACTGGAAAGATTATTTGGAAGACCCCATCAAGAAATTTGTTTTTCCAGAATCACTTAGAAGAGTGTCTTTAACTTGGTGTTGTAAGTTTATTTGGGAAGAGATATCATCAACTTTTATCATGTTGCCACATCTTGAAGAGCTCAAACTTAAATATTGTCGAGCCGAAGATGATGTATGGAGATTGAGTGACAAAGACATATTCAGAAGCTTGAAGTTGTTGTTACTGAGCGAGCTAAATCTTAAGGGTTGGGAAGCAAGCAGCGATAACTTCCCTAATCTAAAACGCCTTGTTCTGAAGAAATGCAAGGACCTGCAAGAAATTCCAACAGATTTTGGGGAAATTTGTACTTTGGAATCGATTGAGTTACATGATTGCAGCACTACTTCTGAGGAATCTGCCAGAAATATCGAACAAGAACAAGAGGTCATGGGAAATAATATCCTTAAGGTCTACATCCATAACAGCGACAGTATCCATAACAGCGACAGTAAGTTTAATTCTCTCTGAAAAGTTCTTCGATTCTTCACATTATTTAAAACCTCGTTTTTATTAAATCATACTTCTTTGAAATTAAGATGATGACTTGTACTTTTTATCTTGCATATGCAGGTAATTGAAGCTTATTTTGAGGGATCATCTAATGCTATCTTCCGAATTCAGAACCTTGTACTTCATATTTTTGTTTTTATATTCTGATGGTTTTTTAATTTCATGTTTTAGGAATAAGCATGTTTAATACATGTATCGGCCAATTGTGTTTTTTTATTTATTTTTTAAACGGTAACTGTTGATGTA

>M6-PGSC0003DMG400027407 M6 LRR-RLP

ATGGATATTGGCTTAGCTGTTGGTGGTGCGTTTCTCTCTTCAGCTTTGAATGTTCTCTTTGATAGGCTTGCTCCTAACGGTGAGCTGATGAAGATGTTTCAGAGGGACAAACATGATGTTCGGCTTTTAAAGAAGCTGAGGATGACTTTGCTTGCTCTTCAGGCTGTGCTAAGTGATGCAGAGAACAAGCAGACATCAAATCCATATGTGAGTCAGTGGCTTAGTGAGTTTCGAGATGCTGTGGATGGTGCTGAAAACTTAATAGAAGAAGTCAACTATGAAGCTCTGAGACTTAAGGTGGAAGGTCAGCATCAAAATCTCGCAGAAACAATCACCAAGCAAGTTATTACCATAAAGGAGAAGTTGGAAGACACCATTGAAACATTGGAGGAGTTGCAAAAGCAAATTGGTCTCCTTGATCTAACAAAGTATCTTGATTCAGGTAAACAAGAAAAAAGGACATTTTCAACTTCTGTGGTTGATGAATCTGATATCTTTGGTAGGCAGAATGAAATAGAGGAATTGATTGACCGTTTATTGTCAGAGGATGCAAATGGAAAAAATCTTACTGTAGTTCCTATTGTAGGAATGGGGGGTGTTGGCAAGACAACACTTGCTAAAGCTGTTTACAATGATGAGAAAGTGAAAAACCATTTTAATTTGAAAGCTTGGTTTTGTGTATCTGAACCATTTGACGCTCTCAGAATAACAAAAGGGTTACTTCAAGAAATTGGCTCATTTGACTCGAAGGCTGATAGCAATCTTAATCAGCTACATGTCAAATTGAAGGAAATCCTAAAGGGAAAAAGGTTTCTTATTGTCCTAGATGATATGTGGAATGATAACTATAATGAGTGGGATGACTTGAGAAATCTTTTTGTAAAAGGAGATGTAGGAAGTAAGATCATTGTGACAACACGTAAAGAAAGTGTTGCCTTGGTGATGGGTAAGGAGCAAATTAGCATGGGAATTTTGTCTAGTGAAGTTTCTTGGTCTTTATTTAAAAGACATGCATTTGAATACATGGATCCTGAGGAACAACGAGAACTTAAAAAGGTCGGAAAACAAATTGTAGCTAAGTGCAAAGGACTGCCTTTAGCTCTGAAGACGCTCGCTGGCATGTTACGCTCCAAATCAGAGGTTGAAGAGTGGAAACGTATTTTGAGAAGTGAAATGTGGGAGCTGCCAGACAATGACATATTACCAGCGTTGATGTTGAGCTACAATGATCTTCCCGCACATTTAAAGCAATGTTTTTCTTATTGTGCAATCTTTCCCAAAGATTATCCATTTCGGAAAGAACAAGTCATTCAACTGTGGATTGCTAATGGTCTATTAAAGGGGTTGCAAAAAGATGAAACAATTGAAGACTTAGGCAACCTATACTTTCTCGAGTTGAGATCAAGATCACTTTTCGAAAGGGTCCGGGAGTCTTCTAAAAGGAATGAAGAGGAATTCTTAATGCATGACCTTATCAATGATCTGGCCCAAGTTGCATCTTCAAAACTTTGCATCAGGTTGGAAGATAACGAAGGATCTCATATGTTGGAAAAATGTCGACACCTATCATATTCATTGGGAGATGGTGTCTTTGAGAAGTTGAAACCACTCTACAAATCAAAGCAGCTGAGGACATTGCTTCCAATCAATATCCAGTGTGGGTACTCATTTCCTTTAAGCAAGAGGGTGTTGTATAACATATTGCCAAGACTAACATCCTTAAGGGCTTTATCATTGTCTCATTATCGGATTAAGGAGTTGCCGAATGACTTGTTTATCACATTAAAACTCCTAAGAATTTTGGACCTTTCTCAGACGGCGATAAGAAAGTTACCAGATTCCATATGTGCATTGTATAACTTGGAGATACTCCTCTTGTCTTCTTGTATATATCTTGAGGAGTTACCGCAGCATATGGAAAAGTTGATCAACTTGCGTCACCTTGACACTACTGGCACTTCTCTCTTGAAGATGCCACTACATCCAAGCAAGTTGAAAAACCTCCATGTGCTAGTAGGATTTAAGTTTATTCTAGGTGGCTGCAATGATTTGAGAATGGTAGATTTAGGTGAACTACGCAATTTGCATGGATCTATATCAGTTCTAGAGTTGCAAAATGTGGTTGATAGAAGGGAAGCTCTGAACGCAAACATGATGAAAAAGGAACATGTTGAGATGTTGTCGTTGGAGTGGAGTGAGAGTATTGCTGACAGTTCTCAAACTGAAGGAGACATACTCGATAAACTACAACCAAATACAAACATAAAGGAACTGGAAATCGCTGGATATAGAGGGACAAAATTTCCGAATTGGATGGCTGATCATTCATTTCTTAAGCTGGTGGGAGTGTCTCTTAGCAATTGCAACAACTGTGCTTCCTTGCCAGCACTAGGACAACTTCCTTCTTTGAAATTCCTTACCGTTAGAGGGATGCATCGAATAACAGAAGTGAGTGAAGAGTTCTACGGTACGTTGTCATCCAAAAAGCCATTTAACTCTCTTGAGAAGCTTGAATTTGCAGAGATGCCGGAGTGGAAGCAGTGGCATGTACTGGGGAAGGGAGAGTTCCCTGCACTTCATGACTTCTTAATTGAAGATTGCCCCAAGCTGATTGGGAAGTTGCCTGAAAAACTTTGCTCTCTGGGAGGATTGAGAATTTCAAAATGTCCTGAACTCAGTCCGGAGGCACCTATCCAACTTTCAAATTTGAAAGAGTTTAAAGTTGTAGGTTCTCCTAAAGTTGGAGTTCTTTTTGATGATGCTCAGCTGTTTACATCTCAACTTCAGGGAATGAAACAGATTGTTGAATTATGTATTCATGATTGTCACTCTCTTACCTTCTTACCAATTAGCATTCTTCCGAGTACCTTGAAGAAAATAGAGATATATCATTGCAGGAAACTGAAATTGGAGGCGTCAATGATTTCTAGAGGGGATTGTAACATGTTTCTGGAGAATTTGGTAATATATGGATGTGATTCTATAGATGATATATCACCAGAGTTTGTCCCAAGATCACACTATTTGAGTGTAAACAGTTGCCCCAACCTAACTAGGCTTTTGATTCCTACTGAGACTGAAAAACTCTATATTTGGCATTGTAAGAATCTTGAAATACTTTCAGTGGCATCTGGAACTCAGACAATGTTGCGTAATTTGAGTATTCGTGACTGTGAGAAGCTGAAGTGGCTGCCAGAATGTATGCAGGAACTCATTCCATCTCTTAAGGAACTAGAACTATGGTTTTGTACAGAAATAGTGTCCTTTCCGGAAGGAGGATTGCCCTTCAATTTACAAGTCCTTCGGATCCACTATTGTAAAAAACTGGTGAATGCGCGAAAGGGGTGGCATTTACAGAGACTCCCCTGTCTCAGAGAGTTAACCATCTTACATGATGGCAGTGACCTTGCTGGTGAAAACTGGGAGTTGCCTTGCTCTATTCGAAGGCTTACCATATCCAATCTGAAAACATTAAGCAGCCAACTTTTCAAAAGTCTTACCTCCCTTGAATATTTATCTACTGGTAACTCACTTCAAATTCAGTCACTGCTGGAAGAAGGGCTTCCCACCTCTCTTTCTCGTCTAACGTTATTTGGCAATCATGAGCTCCATTCCCTACCGATTGAAGGTCTACGACAACTCACTTCACTTCGGGATCTATTCATCTCTTCCTGTGATCAACTCCAATCTGTTCCAGAATCAGCACTGCCCTCCTCCCTCTCTGAGCTGACCATTCAGAATTGCCATAAACTCCAATACCTTCCGGTAAAAGGGATGCCCACTTCAATCTCTTCACTATCTATTTACGACTGTCCATTGCTCAAACCACTCCTTGAATTTGACAAAGGAGAATACTGGCCAAAAATTGCTCATATTTCCACCATAAATATTGATGGGGAATACCAATAA

>M6-PGSC0003DMG400027410 M6 LRR-RLP

ATGCATCGAATAACAGAAGTGAGTGAAGAGTTCTACGGTACGTTGTCATCCAAAAAGCCATTTAACTCTCTTGAGAAGCTTGAATTTGCAGAGATGCCGGAGTGGAAGCAGTGGCATGTACTGGGGAAGGGAGAGTTCCCTGCACTTCATGACTTCTTAATTGAAGATTGCCCCAAGCTGATTGGGAAGTTGCCTGAAAAACTTTGCTCTCTGGGAGGATTGAGAATTTCAAAATGTCCTGAACTCAGTCCGGAGGCACCTATCCAACTTTCAAATTTGAAAGAGTTTAAAGTTGTAGGTTCTCCTAAAGTTGGAGTTCTTTTTGATGATGCTCAGCTGTTTACATCTCAACTTCAGGGAATGAAACAGATTGTTGAATTATGTATTCATGATTGTCACTCTCTTACCTTCTTACCAATTAGCATTCTTCCGAGTACCTTGAAGAAAATAGAGATATATCATTGCAGGAAACTGAAATTGGAGGCGTCAATGATTTCTAGAGGGGATTGTAACATGTTTCTGGAGAATTTGGTAATATATGGATGTGATTCTATAGATGATATATCACCAGAGTTTGTCCCAAGATCACACTATTTGAGTGTAAACAGTTGCCCCAACCTAACTAGGCTTTTGATTCCTACTGAGACTGAAAAACTCTATATTTGGCATTGTAAGAATCTTGAAATACTTTCAGTGGCATCTGGAACTCAGACAATGTTGCGTAATTTGAGTATTCGTGACTGTGAGAAGCTGAAGTGGCTGCCAGAATGTATGCAGGAACTCATTCCATCTCTTAAGGAACTAGAACTATGGTTTTGTACAGAAATAGTGTCCTTTCCGGAAGGAGGATTGCCCTTCAATTTACAAGTCCTTCGGATCCACTATTGTAAAAAACTGGTGAATGCGCGAAAGGGGTGGCATTTACAGAGACTCCCCTGTCTCAGAGAGTTAACCATCTTACATGATGGCAGTGACCTTGCTGGTGAAAACTGGGAGTTGCCTTGCTCTATTCGAAGGCTTACCATATCCAATCTGAAAACATTAAGCAGCCAACTTTTCAAAAGTCTTACCTCCCTTGAATATTTATCTACTGGTAACTCACTTCAAATTCAGTCACTGCTGGAAGAAGGGCTTCCCACCTCTCTTTCTCGTCTAACGTTATTTGGCAATCATGAGCTCCATTCCCTACCGATTGAAGGTCTACGACAACTCACTTCACTTCGGGATCTATTCATCTCTTCCTGTGATCAACTCCAATCTGTTCCAGAATCAGCACTGCCCTCCTCCCTCTCTGAGCTGACCATTCAGAATTGCCATAAACTCCAATACCTTCCGGTAAAAGGGATGCCCACTTCAATCTCTTCACTATCTATTTACGACTGTCCATTGCTCAAACCACTCCTTGAATTTGACAAAGGAGAATACTGGCCAAAAATTGCTCATATTTCCACCATAAATATTGATGGGGAATAC

>M6-PGSC0003DMG400027469 M6 LRR-RLP

AAAGAAATTCCATTTCCACAATGTACAAATACAACATTCTTTACTTAGTACTACCATTTTTTGTAGTAGTTTGCAGTTTTTCTCTAATATGCAACGCGGTACAGAGTGACATAGATTGCTTGATGTCAATAAAAGATTCATTCGAAGATCCTTTAAATTTCTTGAATACTACGTGGAAGTTCGATGATCAGACAGAAGGTTTCATATGCAAATTTGCAGGAATACAATGCTGGCATCCTGATGAGAGCAGAGTTTTATCCATTAGTCTTCCAGACATGGGACTAAAAGGTGAGTTTCCGCGAGGCATAAAAAATTGCACTTCTATAACTTCTCTAAATCTTTCAAGCAACGAGATATATGGAACCATCCCTTCGGATATATCAAGATTAATTGGATTTATTGTAACGGTTGATCTCTCGAGCAACAATTTCTCAGGCTCGATCCCAATTAACATTGCAAATTGCTCTTTTCTTAATTCCCTCAGATTAGACAACAATAAACTACAAGGTCCAATCCCACCCGGAATTGGCTTGTTAGGTCGACTCAAGAACTTTAATGTTGCCAACAACATGTTGACAGGACCAGTGCCAAGGTTTATCAATGCCACTTTTCCACCTGAAAGTTATGCAAATAATGCAGGACTTTGTGGTAGTCCTCTGAAAGCATGTGCTACAAAACGCGTCGGGTTCCATCATATTATGTTAAGTACTGTATTGAACCGTCGTGTACAGTTTGCTTGTGGTTTTGTGACTGGTTGGTCACTTTTTACTGTACTTGGTATATATCTTTTTTTCTTTGGTTTACCTGGCGTAAAGAAGATGCTTCTATTCATCAACAAGAGAACAAAGGTAATGGTGATTGATGGAAATGAATGGCCTCAACGAGAAGAAGTCAACAACGATCCAAAGGTAAACATTACGCTTGCTTTCAACCGTTTTTTTTCCCAATAAAGTGAAACATATTATGGGTAAACGCCTCCTTAAACTTATCATTTTACTCTTAGTTGCAAGTTTTTATAGGCATACAAGTGTTATGGGACGTTTAAGACTACAACTTTAAAAATTGTCTTCTTTTGTCTTAATATTTGTGTCCAGTCAAACTATATATGTCAGATAAATTGAGACAAAAAGAGTACTACTTAATCAAACTATTGCTAAATATGGTAGCTATGTTCTCTTCCTCAGATTTTAAAGATGGAGAAGATTGTTACAAGAATGAGTTTTATGGAACTATCAAAAGCGACTTTGAATTTCAGCCAAGACAATGAAATCGGGAACGGAATGCTAGGGAAAGTGTACAAAGCACTGGTCCCAAATGGATGGACAGTTGCCATAAAGAGGTTACATGTTTCAGAGGATTTGGAGGAAGAGTTTGTTTCAGAGATCACAACTCTGGGTAGTCTGAGGCAACCAAATTTGGTGCCTATAATAGGCTTTTGTTCTGAGAGGAAAGAGCGAATTCTAATTTACAAATACATGCCTAATGGAAACTTACATGAGTGGTTGCATTCAACCGACGACAAGGCAAGACTCTTGGACTTTCCTCTAAGGGTTAAAATTGTTTTAGGAGTCGCAAAAGCCCTGACTTGGCTTCATGATGGTGGAAATTTCCATGTTGTGCATAGTAACATAAGCACACAGTGTATCTTGCTCGATGAAAATTTCAATCCTAAGTTATCCAATTTTTGGGAGGCAACACTTGCAAAAACAAATGACATAGATTCAAACTTGAGTCTATTTCCAATAGTCGAGTGTTTAGACTTTACAACATATAAAAAAGATGTTTATCGATTTGGGGTTGTGCTTCTCGAGCTTTTAACGAGGAAAGAATCTTACATATTGAGCTGCTCGAGTCAAAATTTGTCGAGCAGCTCTTTTGCCAGTCCTCTTGATGTGGATAAACTACTGTTAGGCCAAGGATATGATAATATGGTAATGCAACTACTTGAACTTGCAAGTAACTGCATGAAATTCATTCCGGATCAAAGGCCAACGATGCAACAAGTGTGTCAGACTGTTGCTGCAATTGCTCGAGTCCATGACCAAACAGGGGACTATGAAATACAAGTGCACTTGGACTAGAAGCAGGGGAGGGGCCATCAAAGGTATCAGTAGACTTTG

>M6-PGSC0003DMG400027470 M6 LRR-RLP

ATGGGACTAAAAGGTGAGTTTCCGCGAGGCATAAAAAATTGCACTTCTATAACTTCTCTAAATCTTTCAAGCAACGAGATATATGGAACCATCCCTTCGGATATATCAAGATTAATTGGATTTATTGTAACGGTTGATCTCTCGAGCAACAATTTCTCAGGCTCGATCCCAATTAACATTGCAAATTGCTCTTTTCTTAATTCCCTCAGATTAGACAACAATAAACTACAAGGTCCAATCCCACCCGGAATTGGCTTGTTAGGTCGACTCAAGAACTTTAATGTTGCCAACAACATGTTGACAGGACCAGTGCCAAGGTTTATCAATGCCACTTTTCCACCTGAAAGTTATGCAAATAATGCAGGACTTTGTGGTAGTCCTCTGAAAGCATGTGCTACAAAACGCGTCGGGTTCCATCATATTATGTTAAGTACTGTATTGAACCGTCGTGTACAGTTTGCTTGTGGTTTTGTGACTGGTTGGTCACTTTTTACTGTACTTGGTATATATCTTTTTTTCTTTGGTTTACCTGGCGTAAAGAAGATGCTTCTATTCATCAACAAGAGAACAAAGGTAATGGTGATTGATGGAAATGAATGGCCTCAACGAGAAGAAGTCAACAACGATCCAAAGGTAAACATTACGCTTGCTTTCAACCGTTTTTTTTCCCAATAA

>M6-PGSC0003DMG400027966 M6 LRR-RLP

TGGTTCAATTCTAATTACTCTGGGTGACTTAACTAAGCTCAATATTTTGTACCTTTATGAAAACCAACTTTCTGGTCCCATTCCTGCAGAAATAGGAAAGATGAAGTCACTCAAGTATTTATCCTTATGGAGTAATAATCTTTTGGGCCAATTCCAGTAACTCTAGGTGACTTAACTGAGCTCAATGTTTTGTACCTTTATTCAAACCAACTTTCTGGTCCCCTTCCCGCAGAAATAGGAAAGATGAAGTCACACAAGTATTTATCCTTATGGGATAATAAANCTCTGGGTGACTTAACTAAGCGCAATATTTTGTACCTTTATGAAAACCAACTTTCTGGTCCCCTTCCTGCAGAAATAGGAAAGATGAAGTCACTCGAGTATTTATCCTTATGGAGTAATAATCTTTCTGGTCCAATTCCAGTAACTCTAGGTGACTTAACTGAGCTCAATGTTTTGTCCCTTTATTCAAACCAACTTTCTGGTCCCATTCCTGCAGAAATAGGAAAGATGAAGTCACTTGAGTATTTATCCTTATGGAGTAATAATCTTTCCGGTCCAATTCCTAGTGAGTTGGGAAATTTGAAAAACCTCTCTAGATTGGATTTATCCTATAATCAGCTTACTGGTCTTGTTTCTGTTTCCTTTGGCAATTTGAGAAACTTGCAGACACTAAATCTCGCTGCGAACGAACTTTCGGGTTCTATTCCAAAAGAATTTGCATATTTTGATAACTTGGTTTAGTTATCAATGAGTTTCAATCACTTTTCAGGCCATTTGCCAGAGTGGCTTTGCAAAGATGGGAAACTTGAGATTTTCGTTGTGAATGGTAACAAGCTTACAGGGCAAATCCCAAGAAGCTTGAGCAAGTGCTCGAGTTTCAAAAGGGTTCGCTTCAATAACAACAGCTTTACTGGGGATTTATCTGAAACTTTTGGTATCTATCCAGAGCTTCTGTTCATTGATTTGAGCGACAATGATTTTCATGGTGAACTCAGCAGTAACTGGGGAAAATGCAAAAATTTAATTGATCTGTGTGTAGCCAGAAATAACATTAGTGGCAGCATACCACTAGAGATTGGAAATGTCAAAGGGCTTTTAGGACTTGATCTTTCATCGAATCATTTGATTGGGCAGATTCCAAAGGAATTTGGAAAATTAACTTCTTTGGTTAAGCTTTCTGTGCAGAACAACAATATCTCTGGATATATACCGGAGGAACTTGGGTCACTGACAAAGTTAGAGTCCCTCGATCTATCGTACAACAGATTGAATGGATCAATCCCGAAGTGTATAGGAGATTTCGTGCACTTGTTTCAGTTGAACCTGAGCAACAACAAGTTTGGCCAGAATATTCCAAAGGATATAAGAAGGATAACTCAGCTTAATGTACTTGATTTGAGCTATAATCTTCTCGTTGGAGATATACCCCCTCAGTTAGCCAATTTGAAGGTGTTGGTAAACTTAAATCTTTCCCACAATGGCCTATCTGGACGCATTCCTAAAGAACTTGAAAGTTCATCTGGTTTGCATGATGTTGTTTTGTCATACAATGAGTTGGAAGGTCTAATCCCTAATAATAAAGCTTTTATCAATGCCTCATTGGAGGGTAATAAAGGTCTTTGCGGAAATGTAGCAGGACTCCAGCCCTGTGAAAGGCCGTCTTCTATGGTGAAGAAGCACTCAATGGCGAAGGGACGTAAACTCATCCTCATCACTGTATTTCCTGTTATGGGAGCACTAGTACTGCTCTGCTTTTTCATTGGTGTTCTCTTTATGTGTAATAAAAGAAGGAGAGTTAGAGACTTTGAAAGAAGGGATGGTGATGATTGGCTTTCGATATCCATGTTAGATGGGAAGACATTGTACAGAGACATCTTAAACGCCACAGAAGAGTTTGATGCAAAATTTTGCATCGGGCAAGGAGGTCACGGAAGTGTTTACAAGGTGAACCTTCCATCATTAGGGAATATAGCTGTGAAGAGACTTCATTCTTCATTTCAGAATACTCATCCCAAAAGCTTCATCAATGAAGTAAGGGCATTGACTGGGCTTAAGCATCGAAACATTGTGAACCTCTACGGCTATTGTTCGAAAGCACAACACTTGTTCTTGGTTTACTAGTATGTGGAGAGGGGGAGTTTGTCTAGTATTTTGAGCAATGAAGTTGAGTCCAAGAAATTGGATTGTCTTAAAAGGGTGAATATCATCAAGGGTGTTGCTTTTGCTTTATCTTACATGCACTTAGACTGTTCACCACCGATTGTTCATCGAGACATATCAAGCAGTAATGTGTTGCTTGATTCTGAGTATGAAGCTCGTATTGCAGACTTTGGAATAGCAAAGATTCTCAAGCCGGACTCATCCAATTGCACTGCGTTTGCAGGCACATATGGTTATGTCGCACCTGGTAAGTTTTAAATAGTTTATGAACTATATCCAAACACTCTTGTTTTTATCTCATTTACATGTTTAAATTCTTAATATTGTTCTGGAAATTGTGTTGTTCAGAGCTTGCATATACAATGAAGGTTACACAAATGTGTGATGTGTATAGCTTTGGAGTATTATCCTTGGAGATAGTCAAAGGAAAGCATCTTGGGGAATACATTTCTGTGCTAGCAAATTCATCGACTATAGATCCTGAGCAGCTTAGCAATTTGCTGGATGAACGCCTTCCATATCCTGATGATAGAGTAAAAGATGTTTTGGTTTTTATCATCAATCTAGCATGCTCTTGTTTGCTTAAAACTCCAAAATCAAGGCCAACAATGCACTTCATCTCTCAGAAGTTATCATCAATGGATGCACTCCCACCTATTCATCAGAAAGCCCCCATATAAGGCAAGCTATATAATCCCCTAACATATATGGAGATGAATAACCTTTGTCGCTGAGTGGTTGTGAAGTAAGTACTGTTTCAATTGCTTGGACTAGTGGTGGTCATTTGGTTTACTCTATTAATATGTATATTGTAAGAGGTAATTATTTAATCATCTGCTTCTTTGGCGTTGAGTAAATATCAAGAATTC

>M6-PGSC0003DMG400027968 M6 LRR-RLP

ATAATAAGCTTAGTGGTTCAATTCCAATTAATCTAGGTGACTTAACAGAGCTCAAAATTTTGTACCTTTTTTCTAACCAACTTTCTGGTCCCATTCCTCGAGAATTGGGGAATTTGAATAACCTAAATGATCTGGAGCTATCCCAAAATAAGCTTACTGGTTCAATTCCAATTACTTTAGGTGACTTAACTGAGCTCAAAATTTTGTACCTTTATTCTAACCAACTTTCTGGTCTCATTCCTAGTGAGTTGGGGAATCTGAAAAACCTCAATGGACTAGATTTATCTGATAATAAGCTTAGTGGTTCAATTCCAATTAATCTAGGTGACTTAACAGAGCTCAAAATTTTGTACCTTTTTTCTAACCAACTTTCTGGTCCCATTCCTCGAGAATTGGGGAATTTGAATAACCTAAATGATCTGGAGCTATCCCAAAATAAGCTTACTGGTTCAATTCCAATTACTTTAGGTGACTTAACTGAGCTCAAAATTTTGTACCTTTATTCTAACCAACTTTCTGGTCTCATTCCTAGTGAGTTGGGGAATCTGAAAAACCTCAATGGACTAGATTTATCTGATAATAAGCTTAGTGGTTCAATTCCAATTAATCTAGGTGACTTAACAGAGCTCAAAATTTTGTACCTTTTTTCTAACCAACTTTCTGGTCCCATTCCTCGAGAATTGGGGAATTTGAATAACCTAAATGATCTGGAGCTATCCCAAAATAAGCTTACTGGTTCAATTCCAATTACTTTAGGTGACTTAACTGAGCTCAAAATTTTGTACCTTTATTCTAACCAACTTTCTGGTCTCATTCCTAGTGAGTTGGGGAATCTGAAAAACCTCAATGGACTAGATTTATCTGATAATAAGCTTAGTGGTTCAATTCCAATTAATCTAGGTGACTTAACAGAGCTCAAAATTTTGTACCTTTTTTCTAACCAACTTTCTGGTCCCATTCCTCGAGAATTGGGGAATTTGAATAACCTAAATGATCTGGAGCTATCCCAAAATAAGCTTACTGGTTCAATTCCAATTACTTTAGGTGACTTAACTGAGCTCAAAATTTTGTACCTTTATTCTAACCAACTTTCTGGTCTCATTCCTAGTGAGTTGGGGAATCTGAAAAACCTCAATGGACTAGATTTATCTGATAATAAGCTTAGTGGTTCAATTCCAATTAATCTAGGTGACTTAACAGAGCTCAAAATTTTGTACCTTTTTTCTAACCAACTTTCTGGTCCCATTCCTCGAGAATTGGGGAATTTGAATAACCTAAATGATCTGGAGCTATCCCAAAATAAGCTTACTGGTTCAATTCCAATTACTTTAGGTGACTTAACTGAGCTCAAAATTTTGTACCTTTATTCTAACCAACTTTCTGGTCTCATTCCTAGTGAGTTGGGGAATCTGAAAAACCTCAATGGACTAGATTTATCTGATAATAAGCTTAGTGGTTCAATTCCAATTAATCTAGGTGACTTAACAGAGCTCAAAATTTTGTACCTTTTTTCTAACCAACTTTCTGGTCCCATTCCTCGAGAATTGGGGAATTTGAATAACCTAAATGATCTGGAGCTATCCCAAAATAAGCTTACTGGTTCAATTCCAATTACTTTAGGTGACTTAACTGAGCTCAAAATTTTGTACCTTTATTCTAACCAACTTTCTGGTCTCATTCCTAGTGAGTTGGGGAATCTGAAAAACCTCAATGAGATAGATTTATCTGATAATAAGCTTACTGGTTCAATTCCAATTACTTTAGGTGACTTAACTGAGCTTGAAACTCTGTACCTTCATTCTAACCAACTTTCTGGTCCAATTCCTGCTTCCTTTGGCAATTTGAGAAACTTGCAATTTTTGTATCTCCACACCAACAAACTTTCTGGTTCTATTCCAAAAGAACTTGCATTTTTGGATAACTTGGTAGAGCTGATAATAAGTGAAAATCAGCTTTCAGGCCATTTGCCTGAGCATCTTTGCCAAGGTGGGAAACTTGAGAACTTCACGGTGAATAATAACAAGCTCACAGGGCCAATACCAAGCAGCTTAAGCAAGTGCTCGAGTTTCAAAAGGCTGCGCTTCAATAATAACAGTTTTACTGGGAATTTATCTGAAGCTTTTGGCATCCATCCACACCTTCATTTCATTGATTTGAGCGACAATGATTTTTATGGTGAACTCAGCAGCAACTGGGGGAAATGCGAAAATTTGATTGATCTGCGTGTAGCCAGAAATAACATTAGTGGTAGCATACCACCAGAGATTGGAAATGTCAAAGGGCTTTTAGGACTTGATCTTTCATCAAATTATTTGATTGGGCAAATTCCAAAGGAATTTGGAAAATTAACCTCTCTGGTTAATCTTCTTTTGCGACACAACCAAATTTCTGGCAATATTCCTATGGAACTTGGGTTATTGACAAAGTTAGATTACCTTGATCTATCAGATAATAGATTGAATGGTTCGATCCCACCGTTTATAGGAGATTACCAGCACCTGTTTCACTTGAACCTGAGCAACAACAAATTTGGCCAAAAAATTCCAATAGAGATTGGGAAGATAACTCAACTTAATGTACTGGATTTGAGCCGTAATCATCTTGTTGGAGAAATCCCCCCTCAGTTAGCCAATTTGAAGGTGTTGGTAAACTTAAAT

>M6-PGSC0003DMG400028001 M6 LRR-RLP

GCAAAAGAAGATTTGTGCAAAAACTTTGGTGTAGAGGAAGAAGACGACTTTGTGTGAATTAAGGAGAAAAAATAACAATGGCCGACCCTGTAATTGGTGCTACTGTTCAGGTTGTGCTTGAGAAACTACTTTCTCTCACAATTGAGGAGGCCAGAAATGTAAGGAACTGCAAGAAAAATCTCAGAATGCTGTCAAGATATGTAACCATGATCCAAGCTTTCATTCATGATGCTGAAAGACGACAAGTCGAAGATAGGGCTGTGGAAGAATGGCTCAAGATGCTTGAGAGAATTGCTGAAGATGCTGAAAATGTGTTTGACAAATTCACATATGAATCTATCAAAGCAAAAGTGATGAACAACCGAGCCAAACTAATGGAAAAGGTCAGTCACTTCTTTTCTCATACTGCTTTCAAGTACAAAATGTCTCGAAAAATCAATAAGATCAATGAAGAGTTGAGGGATATCAATCAGTTAGCCAACAACCTCGGTCTCCAATCACTGACAGATCCTTCTCGGAAAATATTACAAATCCGAGAAACAGATTCTGCAGTAGTTCCTTCAGATGTTGTTGGTAGAGACAAAGATGTTGCTGAAATAAAGGAGAAGATTTTGAACATGAGAAAGGATGCTGTTCTGTGCACCATTCCCATAGTCGGTATGGGGGGTTTAGGGAAAACTACTCTGGCTAAGAGGATTTTCAATGACCAACACATCGAGAAACACTTTGAAAAAAGAATTTGGTTGTGTCTACCTGAAATGTCTGAAATTAAGAACTTTCTTGAACTGATCCTCGAATCATTGACAGAGAGGAAAGTTGAGGTCCAGAGCAGAGATATAATAGTCAAGAAGCTCCGAGATGCATTGGGAGAAAAACAGTATTTACTTGTCCTGGATGATTTGTGGCGTGATGACTATACTTAGTGGCACGAGTTCTTGGACACCTTGAGGGGAATAAATACATCCAGAGGAAACTGCATTCTCGTGACTACTCGTTCAAAGCAGGTGGCATCCATAGTAGCAGAAGATCTTCATAAGTTGGGGAAATTAACAGATGATCAATGTTGGTCCATTTTCAAACAACGAGCATTTGTTGATGGGGAAGTTCCGGAGGAAATACTGAGTGTGGAGAACAAGATTGTTGAAATGTGTCAAGGTCTACCTTTGGCTGCTAGTGTGTTGGGAGGCCTATTTTGTAACAAGGAAAAACATGAATGGCAGGCAATTCTTGATGGCAGCTCCCTTGTTGCATCTGAAGATAGCATTAAGAATATCCTAAAACTCAGTTATGATTATCTACCATCTCCACATCTGAAGAAATGTTTTTCTTACTTTGCAATATTTCCAAAAGATTTTGAGTTTGAAAAGGACCAACTAATCCAACTCTGGATGGCAGAAGGGTTTCTTCGTCCATGTCAAGAGACCACTGTGATGGAAGACGTTGGGAACAAGTTTTTCCAACTCTTGTTACAATATTCCTTGCTGCAGGATGTTAATCTAGATAAGTACAACAATACAACACACTGTAAGATGCATGATCTTGTGCATGATTTGGCTGGAGATATTTTTAAGTCTAAACTATTTGATCAAAAGAGTGTCGGAGGAGAAAATCTTTCTCAAGTTCGATACTTTGGATGGGAATCACCAAGTGATCAAATAGATAAGATTTATGAGCCAGGACGTTTGTGCACATTGTTCTGGAAGAGCAATATATCTGATGATATGCTATTGAGCTTTCAGTTCTTGAGAGTTTTAAATTTGTCCGGGTCAGACATCAAGGAGTTGTCAGCCTCAATCGTCAAGCTAATATACTTGAGATATCTTGATCTCTCCAATACTAAGATGAACGACGGATATCTTGATCTCTCCAATACTGAGATGAACGACCTTCCCAACTCCATATGCAAGCTCTACAATTTGCAAACTTTGAGATTCTATAGCAGCTGGTATCCACTCAGGAAGCTTCCAGAAGAAATGGCAAATATGATAAGTTTGAGACACATATGCTGCTACCATTTTTTCGAATCAGATTTTCAGATGCCACTTAATATGGGGCAATTGACTAGTCTTCAGACGCTGCCGTTTTTCTATGTAGGTTCAGAGAAAGGTCGTCGAATAGAAGAACTGGGTTGTTTGAAAAATCTTAGAGGTGAATTGACGATCGAACGTCTACAATTAGTCGGAAATAAAGAAGAGGCTCGAACAGCATATTTACAGGAGAAACCAAATATCTACAAGCTGGTGTATTCATGGTCCCATGATGAACCAGAAGGCTGTGAGATCAATCATGAGCATGTGTTGGATGGTCTTCAACCACATCCGAACTTGAAAACCTTAGAAGTTGTGGACTATTTAGGGACTAAATTTGCTTCATGGTTCAGTGAAAAAATGCTAACAAATTTGGTCATGTTGAGATTAAGAGGTTGCAAAAGGTGCAAAGAAATTCCATCACTTGGCCAACTGAAATCCCTTCGGCATCTTAAGCTGATAGGATTCCTTGAATTGGAATGCATCGGACCTACATTTTATGGTGTTGATGTTAATAATAATGGATCAAGCAGCAATATCCAAGTGTTCCCGTCATTGAAAGAACTAGAATTGAACAATATGTCTAGCCTTATTGAGTGGAAAGGAGATGAAGTTGGAGTAAGAATGTTTCCTAGGCTTGAGAAGTTGACGATTAGAAACTGTCCATTGTTAAAAAGAACTCCGAATCAATTTGAAATCCTCAGTGAATTAGAAATTGTCAAAGTTGACAGTGAAATGCCATTGTTGAACTTATGCAGCAACTTGCCATCTCTCGTAGAGCTTGAAGTCGATGATATGAAAGAGCTCACTTGTTTGCCAGATGAGATGCTACGCAACAATGTTTCTCTTCAACACATATCGGTTTCCGACTGCAGAGAGTTTCATGAATTGCCACAAAGCTTGTACAATCTCCATTCTCTTAAGACCTTAAAGATAACCAGGTGCGCCAATTTCAATTCCTTTCCTGTTCCCAGTGGAGAGAATTATTTGACTTCCCTCCAAAAACTTCAGTTAAAGGATTGTGAAGGATTGAGCAGTTTACCAAGTGGAATGCTAGAGCATTGTCGGTCTCTGGAAACTTTGAGTGTCAGCTGCTGTGACAATTTAGTTTCATTCCCTTTACATGTGGGGGAAATGCCTTCACTTTCCTATTTGAATATATCACGATGTCCCAAATTGATTAGTTTACCCTCCGGGGGTATTGACCATCTCACTGAGTTAAGTGAACTGAAAATTGGTCCTTTCTCAGAGATGGTGGATTTTGAGGCATTCCAATTGATTTTTAATGGCATTCAGCAGCTGCTGTCCCTTCGTACATTGACGGTGTATGGACATGGTCACTGGGATTCTCTGCCCTATCAGTTTATGCAACTCTCTGGCTTAACAGCGATCCATATATGTGGATTCGGAATCGAGGCTCTTCCTCATAGATTTGGCAACCTTACTTCTCTTGAAACATTAATGCTACTGAGGTGCAAACGGCTACAGAACCTGGACTTCTCATATGTCATGCCCAAATTACAACATCTGTATGTTTATGAATGTCCATTGTTAGAAGCTCTGTCGGATGGGCTTGGCAACCTTGTTACTTTGGAACTGTTACATTTAGAAAACTGTGAGAAACTAGAGTATCTGCCATCTAGAGATACCATGCGACACCTCAAGAAATTACAGAGCCTGCAAATTAATGGCTGCCCAAAGTTAGAAGAAAGTTGCACCAATCGGAGTGGCCCAAACTCCCAGTGGTCCAACATTTCCCATATTCAAAGAATTGAAGTTGGTAGAATCCCCCTATATCAGTGCTTGAGTAAGTCTCTGTTTCTCTTTCAATTTCCGAGTCGCATTTTTTAATTATTATTTTCCACGAGTTAGTCTTCTCCATTCTCCAAACATTTTTCTATTCCACTTGAAACAAATTTATGAATATCTATATTACTTTTAAGATATGGTTTAATCATGAATATTTATGTATTTTTTACTCATAGGTTATTAGCTCGAAGGTCACACTAGAACGTTGTTGTGGCATTTTAAGCCAAAATCCATGGGCAGCAATTACATTAGGTGAAATTCCATTTAGAATTCTGTAACAATTTTGTCAAGTTTTTCTTTGCACTCCCCTTGTTTTTTTCCCTCATGACACAATTTTATTGTTTAACAGGGTTAGTTATGTTCGATAGAGAGGAACCAACACGTTTGTTTTGTTGAAGAACAGAATCTGCAGCCGTAGACTCTTACATGAGCATTTGGTAAAGACTACTGACTACATAATCTCTCAACATTTACTCTTCATAACATGGATATATACCAACAACTATTAAGTAAAAATAATTTAGTACACAGTCGATCATTTGCCCTAGTGAAACTATTCTTTTATGTATGGGTGTCTTTTGATAATATATAACCTTTTTCTACGAGTTAATGAAAATAAAAAGACTAACAAGTAGTTTTGGGTAAATAATGAATGAGTAGTTCAACCAACCAACAAGTACCATGTTCTAGACTTCCCCTTTTGGATGTTCACTTGAAATCTATTTTTTCTTCTTTTCTATACAATAGAAGCGTAAACTCAATATCAGGTCAGGTACTTTGGTCAGGTAGTAAAGGGTATCCGTGGACCGAGCCATTGAATTCTGTTCATGAACTATTCATAAGATGAAGTTAGGAGGTTGCTGTGCTACCAAAACTGACTTTTTACCGTCTCCTATGTTTCAGATTCTTGAATGAAACAATGGCATGAAACTTGTCGAGATGGTATACTGAAGCAGACTTCTCTTAGATGTGTTTAAGTGTCGTGATTAGACAGGGAAATTACATTAGGCGGAACTGGAGAGGAGATATTATTTAATTGTATGACTAAGATTTGACAGCAGAGGCCTTTCTTGCACAGTGTTCATTCCTTTTTGTACAAATAGTTTTTTGTTTCCAGCAAACATTTTCAACTTAATACTTAGTTGCTGCATCATTGCCTAGTTCATATACATCTTCT

>M6-PGSC0003DMG400028072 M6 LRR-RLP

ATGGACAACTATTGTAAGGACCTTACCTTGAAGAACTTAATTAGGGAATTTCCTTATGTGCAAGAACCAGATGTATCCTTGACGGAGAGAGGCCTTGTTGCTTTCTCAGCTGGCTCCACTCGGTTCTATTCTTTTGCCGCCATGGTTGTGATTGCAAGGAACTGTTCTAAGTTGATCCAATTTCGTTTTTGTTTACTCAAGCCTCCAACTTCTGATTACCTAACCCCTGAAAAGCTCAATGCTGGTTATGAGGCCATCATCCATCACTGCAAGGGATTACGGCGACTTCTCCTTTCTGGCGTCCTTATGGATCACGTGTTTCATTACATCTGGCCCATGCTAAAAAGCTAAAGATGCTCTCCAGTGGGGAATGTAATCTAGGCCTCCACTCTGGTTTTGAAAGCCATTGTAAGGTGGAGACCAGAGACAGCCCCTTCGGTGACGACAAGGCTCTGTTGGTCAGTGCTGCAAGGCTGGAGACGATGCAATATCTTTGGATGTCTTCTTGTTCAATAAGTAATGGTGCATGTGAGCTGCAAGATCAGGAGATTCCGCCTAGTTTCGGGGGTATTTGTTCATTGAAAATTATCAAACTTGTAGAGAGCCCTCAACTTGAAGATTTCTCTATGAAGATTTAG

>M6-PGSC0003DMG400028081 M6 LRR-RLP

AGCAAGCATAGTAGATTTTGGATAAATTGTTTATACATATTAAATGAATTTTTAAACACAAATACATGACAAAACCCGTAGACAGGGTTGTAGCTCTGCCCCTGATGCCCCAGTGAAGAACTTAATGAACTGTATACACAAATAGCTAAACAAACTATCTAAATCGCATTCATCATCAACAACAATTCCCCATTCTACATAAAACATAAATCATTCCCAGTTATATTATGATTTCGGATAAATAAAAAGACCTCCAGAATTCAATAAAAAAATCATGACGATATGCAGCAGCCACCACTATTCCCACCAGAATCTGTGTCCTGTCCGGGAGCAATTATCCTGGTTCCCTTTAAGGATTATGGCTTCCCATAATCAGCTCCCGGAGCTGCAATAAGCGTATTCTTGCTAATGATTTTATAAATTTCTGTTAAAACTATCATGAATGCACCCTCGATGTTGTGGATTGAAGAGCTGATGTTTCCATAAAGAAAAGATTCTCCCTTTCTGCGAATTCTTGAGATTCTTCAACTGGTACAGCTCGAAGACTTCCTAGATCACATTTGTTCATGATGAGCATAATAACTATGTTCTTGTCCGCATGACCTCTTAGTTCCTCCAGCCATCTAGCCATATGATTGAATGATTGGCGTTTTGTTAAGTCATAGACTAGCATTGCGCCAGCTGCACCTCGATAATATGCACTACTGCTTTATACCTGAGAGAGAAATGAGAGATGATTCATTGAGTTATAAAAATTACCAAATTGGCTATGCTGCTCTGACTCTTCGAAAGAGGGTCCTTTGAAAGTAGAGCGTTTTGGAGGAGCCGACACGGGTGCGGCGATATTTTTCTCGAGAGTCCAAGCAACATAGATTAACAAAAGGAAAATGTTAGTGTTACATACAAAACTGGGATCTAGGGAGGTAATATGTATGCAGACCTTACCTCTACCTTTGCGGGGTAGAGTCGTAGAGAGGTTGTTTCGATAGGCCCTCGACTCAAACAAGATGAATTGTAAGACAGGTAGATACCTTGTACAATGATAGAAACAAAATCTCAAGAAGAAACATACATATGAAAGTCTGAATCGCGATATAAATTAAAACACTCTTTAAGGAGATTTATCACAAAAACAAGAACGTCTCAAATATCTCAAACAAGTTGGGATCAACAATATGAATTCTAACTAATAACCATGTCGCTGAATTT

>M6-PGSC0003DMG400028112 M6 LRR-RLP

AAAAGCGCAAAAAAGCTCTAAGGTCCGTGGGGACTTTAAGTGCAAAGCGCAAATAAAGCGTGGGCTTTAATGAAAAATGGCGCAAAGGGAGAAAAGAATACAAATATGTATGTTTAGTCCAAGACTAATAATTATAAACATGAATGACAAATATATGACCAAAGAAATTGAAAAAAAATTATGGTTAAGTGAAATATCAATTATTTAGTTTCACTTCTTCATAAAAGTCTCATTGGCAAGGAAAAGCTTGCCTTAGAACCTTGATGACGACACTGAAGCGCACATAAAGCGAGGCGAAGCGCTCAACACGTTTTGAGCCTCGCTTCCGGGCTTAAGCACGCTTAAAGCGCGCCTTTGATAACACTGCTT

>M6-PGSC0003DMG400028156 M6 LRR-RLP

GTTCCCTCGAAACTTTGACAAAAGGGTTTCCCTGAGAAAATATGAATCAAATTTCGCAGTTACCGGCGGCAACTAATATGGATCGAATTTCGCCGGCGGCACGTAATTTGGATCGAATTTCTCAGTTACCGGCGGGAATGTTAGAGCACATCTTGTCTTTCCTTCCGTTTGAAGAAGCTGCACGAATGAGTAGAGTATCAACGACTTGGAAAAATGCATACAATTCACTCCCCTGTTTAAATTTTGAGAGGTATTTTTTAAACAAGCCAATAGAAAAACTTGTGAATGCTGTGGATCCAATTCTAACTTATCGGAAAAATCAAAAGATTTCTGTAGAAAAGTTTTCGCTACACATAAATCAACAATGGAGTAGTCTGTTTTCCAATTGGATTGATACACTCGTAGATTGTAAGATCAAAGAGTTAAGCCTAGTTTTAAGATCAACCAGAACCGGATACACGAAATTGCCTGAGGCAATTTTTAATGCCAAAGCACTAAAAGTGTTGAAATTGAGTGAATTTCATATTGAATTGCCATCTAATAGTACTATTATGTTTTCATCATCTTTGCGAGAGTTATACCTTCAATGTGTATATTTCGACGAGGAGTTTATGCAAGCTTTATGCACAAGTTGTATCAATTTAGAGGTTTTCATGGTGCGGGGCTTGAAAGGGCTTACTAGATTTCAAACTAGTTTACCTAAATTGAAAAAGCTACAGGTGACAGCGTATTATTCTAAATTACGATTTGTTGATATTAGATCACCTAATATTGAAGATCTTGACGTCTATGGCTCCAACCTCAGCTCCAACTACTTTAAGAATGAGAGTGATCTAAATGTAGTTATTATTACTAATTGTTGCAAATCCCTTAAGAGTTTGCAGCTCAATGGTGTGGCCATGACCCAAAAATGGTTTGACGAAATTTTTACCTGCCTTCAGAATATAGAGAAGTTGGATTTAGGTGATTGCGACATGTTGAAGACTGTGAATATCTCAGGCGGCTCTCTCAAACAGCTGAGAGTGTATCATTGTGATAATGTGATTGCTATTGAATTAGACACTCCTAATTTGATAGAGTTCTCGTATGATTGTTCTTCATTGCCGAATTTGAAACTAAAAGCTTCAGCTTCATTGGAAGCCAATATTGAATTCGACTCAGAAGGAGGAGAAACCCCTAACAGTGATTGGTACTCTAAGCTCGTGAAATTTCTTGATAACTTCAATCATTCCATGGCTCTTGAATTAACATGCAAAAGTGAAAAGGTATGTCCCAGAAATTCTTTAACTAGCTAGTTAATACTATATTGTTATGGATTTCTCAAAAGTTGTTTCTTTTTGCTAATACAGGGGCTAGTTATACCAAAAGATGTGAGAGAACACTTGTTTCCTCCCCTTTATGGAACTAATCGCGTGAAAGTACGCATTCATAGTCCATTGATAAGTTATTCAGTTGTAGATATTCTTGATGGTATTTTCTGGCTTTCTCCTCAATTGGACACCCTATCTTATTCTAAATTCGACTTGAACACCCTTGAGGTAAATACTTTTATCTTTCAAAGGCACCGACTCCCTTAGTGGGATTTCACTGTGTATGTTGTTGTTGATGATGATGATGATTGAAAATCATGCTTCGTAACCACTCAAACTGAACTTCCTTTTGATCTGTTCCAGCACGCATATACACTGTTGATATCGAGCATTAATTAACCCTCTCTAATATTGATTTCGTGATTATTCTTCATGAAGACAGTTTTTATCTTTATTCTTCTTCAATAGTTACTGAGTTTTCTGGTTTTACAGTTTTCATATGAAGATGCATTAGGTGAAGGTGAGAAGTTTTGTTGTACATCTCAACCTTGGAATTGTTGGAGGCATAAGTTAAAGCAAGTCAGAATACACAACTTTAAACATAAGGAAGAAAAACAGAAGTTGAAAAACTATTTTTCCACAAACACAGATATCTTAGAGATAATTGAGGTTCCCCCGACCTAGAATGCCCTGGGAACAAGTACATAACTAAATGTCCATGTCCCAATTGTTCCTCAGATTAATTTTGTAGCATTACATGCTTGGTTTGTATAGGGGATTTAATATCTAGTTCTGGCATTTAGCTGGCTTAGGTTTTTCCCCCCTTCCAGCAGAGCAATAAGCTAGGTTTTGTTTTTAGGAAGATTTTATGTTAGACATTAAACTCTTATCTCATAGCATGAAGAATTGTTTACTTCATTTGGAAGCTTTCTTATATTTCGGCGAACAGTGCCTTCTCAAACCTTAAAATGCTGTTCTTCCTACTTTTGTGGTGTTTACTTTTTTTGATATTGAAGCTCTCAGAGCAAACCAGTATTTAGAGGTTGTTTCC

>M6-PGSC0003DMG400028212 M6 LRR-RLP

CTTCTTGCTTTTATACTCCTTTCGTTTCAATTTGTCTTGCTTTCCTTTTTAGTCTATAGAAAAATGTTTCTTTACTTTTTTGGCAACTCTTCAGTTTCAACTTTACGCATGACAAGTTTAAACCATGAGATTAAACGCATTTTGGTACATTCTGCATATCCTTAGCTTAGTTAAAGACCAAAAGATTCAGAAGTCGTTCTTTACTTTCTTGAACTCCGTGCAAAGTCAAAACCAGACAAGCAAATTGAAACGGATGAAAGTACACTTTCTTGAATTGTTATCATTCTAATAACTATGGATACAGAGTTTGATGTTTGCTTACTGAAATTTCATTTCAGCTGAAGTAGTGAAGAAGTGGCAACTAGAGTGTTTACAAGAAAGCTCAAAAAAGTGTAAATTTCTCAGTTCTTTTTAGTCCTTTTTTGGTGATGGAAAAACATTTAAGACTTAAAACTCTTTTAGTTTTGCTTGTTTTATTGCTCTCAATTAGCATCTCAGAGCAACTACAATCTTCTTCACAAGCAAATATTTTATCGAGAATTCGAAATCTTCTGAACTATCCAAATGTTTTGAGTAATTGGAACAATGAAACTGATTTCTGTAACACTGAGCCAAGTTCATCTGTTACTGTTATCTGTTATGAGGATAACATAACTCAGCTTCATATAATAGGGAGCAAAGGGGCTTCTCAATTACATAATTTCTCGATCGATTCGTTTGTCACTACACTTGTTGAGCTTCCAAGTTTGAAAGTTCTAAGATTAGTTTCTCTTGGTTTATATGGACCATTGCCTAGTAAACTTTCAAGACTTTCATCATTAGAGATACTTGACCTTAGTTCAAATTTCTTTCATAGTAACATACCTCGAGAAATTTCATCGTTAACGAGCCTCCAGAGTTTAATACTAGATGGAAATAAGTTCACTGGAAGACTTCCAAATGGACTTGGTTCACTTGTTGTTTTGGCAGTTTTGAGTGTAAAGAACAATTCTTTAGATGGACATTTACCTGATATGCTCGGAAATTTGCATAATCTTCGTGTACTTTCACTTTCGAGGAATAATTTTACTGGTGATGTACCTGATCTTAGTGGTGTCGAAAACCTTCAAGTTCTAGACTTGGAAGATAATTCTCTTGGACCGAAATTTCCTCAAGTTAGCAGCAAAATCCAAAGCATTGTGTTGAGGAACAATAAGTTCACTGCAGGAATTCCGGAGAAAGTTCAATCTTACCATCAGCTTGAACATATGGATATTTCTTCAAACAGATTTATGGGGCCATTTCCTCCGTCTTTGCTGTCCTTGCCATCGATCACTTATATTAACGTTGCAGGAAATAAGTTAACAGGAATGCTTTTTGATGATAATCAATGCAATGCCGGATTAGATTTTGTGGATCTATCGACTAACCTGTTGAGTGGAAGATTACCCAATTGCCTTCTGACTGGTCCTAAGGACAGGATCGTGCGTTACTCTAACAATTGTTTAGCAACTGGAGACGGAACTCAACATCCATTTTCCTTCTGTCGGAATGAGGCGTTGGCTGTTGGTATCTTACCTCATCATCATAAACATATACCAGGTTCAAAAGTGGTTCTTGCTTTGATCATTTGTGGTAGCATCATTGGTGGAGTCGTGCTAGTTTGCGCGACTATTATAGTAGTAAGAAAATTTCTTGCAAAGATAGCTACACCGAGAAAAACAACAAGATCAATAGTAGAAAATGCAGCATCTACATATACATCAAAGTTATTCACTGATGCAAGTAAGATTCCTCTCTTCTTTACGTCTTAGCTTTGAAGTTTCAAGCATAGTACAATAATTTTGGCTTACCTGTGCACGAAATACACGATAGGAAGGCGAAAAAAGCCCAAATATACGAAGTTTCCTTTGATTCCTAATTACTACTAATTTTCCATCTTGTGTGTTTCTTCTCCGTTTACAAGACTATGTTACTCGAACAATGAAGCTGGGATCACTTAGTCTTCCATCTTATCGGACCTTTTCATCCGAAGAGCTTAAGGTGGCAACAAACAACTTTGATGCAGCAACTTTTATAGGCATGAGTTCTGATGATCAGGTGCATTTCGAGAATCACCATGTACCTTACTTTTCTTTCTTAACGTCATAATTGAGATCCTTAAACGTTCGTTTAAGAACATTTCATGGTATTATCATCCTTCATTGTTTTGACACCACAAATGAGTCAGAACCTTGCAGCCTATCAAACATATTCAGTTGCCTTAGAATTCTTAATCTATACCTTATCCCATTGAAGGTGTTTTAGAAGATCTTCTATCATGTTACTAGTCATTCATTTTCTTAGACGAGTGGATATCAAACAGGTATGGTATGATAATCCTTCTTCGTCAAGATCCATCTAGAGTTCATTAGTCCATAGAAAACAAATGTTGATACATACACTTCTGTGATACTCACGTCTGAGCGAATGTAACAACTAAAAGAGTAAACTGCATATATTATATGACAGAATTAGTAAGTATGGATGTTAACTTACTGTTTTTTTTGGATTCAGTTATACAGAGGTCAGCTGAAAGATGGTTCATACATAACCATAAGATGCCTGCAAACGAAACGAAAAAATAGCAGTCAAAATTTTATGTATCATGTAGAATTGATGTCAAAACTCAGACATCATCATCTGATCAGCACTCTTGGACACTGCTTCGAGTGTTACTTGGATGATTCGAGTGTTAGCAGGATATTTCTCGTCTTTGAGTATGTATCAAATGGTACCCTAAGAAGCTGGATCTCCGGTAATTATTCAACAAGCATTTATACTTATCGATCCCAAGATTGCAGTACAACAGTAATGTGTTATCCGATTCTGCAGATAAACATGCTAAAGGAAGACTAACGTGGACACAACGTATAGCAGCTGCAATCGGAGTAGCAAGGGGAATGCAGTTTTTGCATACTGGAAATGTACCTGGTGTTTTTTCAAATAACATAAAGATAACAGATATCTTGTTGGATCAGAACTTTGTAGCCAAAATTTGCAGCTATAATTTACCTATTCTGAATGAGAATGTAATGGAGCAGAAGAGTACAAGGTATGCCTAACTGTTCCAAATTCTAAGTTGCTCGGACTCTCCAAAAACGTTGCCACACACATGTCGGATCCTTCAAAAATGCACTTCTTTGAAGGATCCAACATGCTCCCAATGACATGATTGTTAATATATTAGTTAATCATGTGTACTATGAAGATCAAATATTATGCTAATCTCTCTATGGTAGTAGACATAATTAACTAATATATTAGGTACAAAAAAAGTTGTATTAGTTTCTTACTTCTACTAATCCACAGGGCTAATTATGAAGAAAAATTGGTTGTATATGATTTTGGAGTAATATTATTGGAGATAATTAGTGGGAAGCAGATAAATACCAAGAATGAAGTGAGAGTTACACAAAATCAGGTATGATTTAACACTTTTTTCCACTCTTCACAACAACTATGTCCAAGTCGAATGTAAGTTAGGATCAGCTATATGAATTCTCACCGACTCTTTTACTCTTCATATTATGTGCAGTTACAAGAGAGCATAATGTCTAATGCAATGTCAAGAAAGAATGTGGTTGATCCAGCAATTAGAAATTCGTGCTCGGATGAATCATTGAAGACGATGATAGAAATATGTTGCAGGTGTTTGGAACAAGATACAGAAGATATGCCCTCAGTTGAAGATGTAATATGGAATTTGCAGTTTGCTGCTCAAGTTGAGGATTCATGGAGAAAAGACTCTTCAAGTAGTGATGCCTCTCCAATTTCACATCTCTACAGCTTCAAACCCAACCTGTCGCGCAATAGCAATAGCTCGTTTGTGTGACAAGCTATTTCTAACGTGGATAACTAATCTCGATACTATTATTCCATCCTGTGACAAGAATAACTAAACTAACAATTTATGTAATAATTTTCTTTGCATAGCTTATACAAAAAAGTAAGTTTTCATACAATCTTTTTCCTAACAAGCAGATGCCATAGCAATTTCCTTGTGTATATACCAACCCAACCTTAATAGTACCTAGATATTACAAGAGCAACATATAGTGCAGAAACCTATATTAGACGTGGACGATGAAAAACAAAAACAAAAGAATGCAAAGAGGGACTAGTTAATGGAAAAGGGAAGCATTGCAAAATGATTTTTCGGACAGTAGAAACAGACTATCTATTCGTTTGCAGGAGGAGTTTCCATTTTCTGAACTTGAGGGGCTGTCACTGCCACTAAGCTAAGATCAGATGAGTTCGAATCTCCAGATCTTTCACGGATCTCCCGGTACTCCTGGCATATGGCACATAAGTGGCAGAAAAAGTGTGTGGCGAAGTCTCCACATGGAGCTTCCTGCATATATATCAGGGCACACAAAATTGTTAAATGCAGAAATACTGATGTAAGCAGTTTGCATCTTTCTATTCAGGCATGGAACCATAGTATTTCGTAGAACAGAAAAGGGGGCAGGAGCTAATGGGAGGAGAGGAAGATGAAGAATTAAGAATTCACGGTAACTGGTTATAAATGCTGCAAACTCATCCAATAACAAACTCAAAACTGAAATTTCTAACTCAGACATCATCTCAATGTAAACCAAACAGAATAGGAAACTGAAGTCAGTTCTCTTTCTAGATGGAGTCCTAATTTTCCCGATCTGAAGTTGATAGTTTATCGAACTGAAGTCCACCACTACTAATTCAAATGAAGTATAATCATCTCTCCCGTGGAGTCGATTACACACTGTAACTTTTTAATCTTTTGTAACTTCTACAGATAACATCCAATTCAAAAGAATGTATGTGGAATTTTAAGACATTTCTGATTAATTTTTCTTTCTAATGAAAAG

>M6-PGSC0003DMG400028312 M6 LRR-RLP

ATTTCTGGTGATAAACAAAAGTACATGGACATAATCAACTTAAGTTTACCTAATCATTTGACAAATGGCTTTCTTTACTTTGGAAAACATTTTGAGGATGAGATTTTGGTTCGAGAGTTGATTCAGCTTTGGATCGCGGAAGAATTTGCAGTAGGTAACCAGTTCAAGAGTGCAGAAGATGTAGCAAAGGACTATGTGAAGGAATTAATTGATAGTAATCTTGTAATGGTTGTTCGAAGGAATTTCTTTGGTGAGGTGAAAGCAATTGGTATGCATGATCTGTTGCGTGAGTTCTGCTTGACTCGATGTAAGAGTTATAGGTAAGATCTCATTTGAATCATCGATCACAAACTTTGTTACTTCCAGACTCTTTTAACTGGTTCTTACTATTTTATTTTCTTCCTTTTGGTATTTTGTTTTCTTTGCAGATCAAGTAATTCAACAGCTGCTGAGCACGAACGTGGCATTTCACTTGATTTCCGTTATAATTATAAAGAAGCTGAATATTTATATCCCCATTCTGCTATCTCCCGAATACCCTATTCTGTTGACGATATACTTGACCACAACAACTTTCTCATCTTTGACTGCAAAAGTTTGTGTTTCAAACCCAGAATTACATCACTGAGAGTGTTGGATTTGGTGTGTGTTACATCGAGCAATGCACCTTATATTCAATTCTTGATTAATTTAAGATACTTGTCCATGAAGGGGAATTTTGAGGAAGATCCATCATGGTTATCCAACCTCAAGTACTTAAAAACATTGATACTCAGAGAATCGCGCAGATTAGATCTATCTTATGCAATATGGGGTATGGTGAGCTTAAAACATTTGGAAGTGACTGGTAAACCAGCTGTCATCAGGTGCTATGTCCCAGAGTTCATGATGAACCGTCGTTTAACAAAATTACAATGTCTTTCTTTTGTACACTTACCTGAAGGCGGAGAGTTTCAAGAGTTCATGTCAAAAGTACCCAATTTAGTAAAGATGAGGTGTTTGGTTGATGAGGCTGAGCCACAAGGTGATCAGACAATATGCATTAGGCTTCCAGGTTTAGACTCTTTGAATAAACTCGAATCACTTAAATTTAAGGTCTATTTTCTAAAACATGGTGCAAAGCGACTTGAGATCAACTTTCCTAGGAACCTCAAGAAGTTGACCTTGTCATCATTTCGATTACCATGGAGTGAAATTTCAACTATAGGAAGATTAGAAAACTTGGAGGTGCTCAAGTTGGAGTCCAATGCTTTCGAGGGAGATCAGCAATGGGATGTCGATAATGAGGAGTTCCAAAACCTCAAGTTCTTGATGTTCTACAACATGAATATTCCACAATGGAACTTTTCTGATGATATGTTCTTTCCTAACCTTCAACATTTGATATTCAGAAATAGTCGACTGAAAACAATTCCTCGCGACTTTGAAAACTTACTTTTTCTGCAGATGATTGAGGTGTCTTGGTGTAAATGCAGCCGTGATGCCTTGAACACTGCTGAAGAGATTAAGAAAACTCAAACTGATATGGGAAATCATGAATTTAAGCTCATAATCAAATCTAGAGAGGATGATGATCATGAAGAAGAAGATGAAGAAGATGATCATGAGGATGCACATCAAAAGAA

>M6-PGSC0003DMG400028575 M6 LRR-RLP

AGTATTTAAAATAGGCAAAAAGAATTGATCTGTTAGCCAAAAAAATAAATACAAACGAATCCTGTTGTTTATTTGCGAGGATTTTGCTATTTTATACGTTTCAGATCTGCAGAAACTTGGAGTTTCTTTTTTCTGTTGACAGTCCAAATGAGAAGTGAAATGTTTTCCCAGCTTTTCTTGATCCTATTACTCTCCATCTGCTGTAAAATGAGTGATGCAGTGGCAGCTAATTCCAGTAAATGTCTTGAGGATCAGAAGATGTTGCTGCTGCAGCTGAGAAATAACCTTACTTATGATTCTGAAATATCTACCAAGCTAGTGAAGTGGAACCAACGGATTGACTGCTGTCAATGGGAGGGTGTCACTTGCAACGGTGAAGGCCAAGTTATTGGTCTTGACCTTAGCGCTGAATCATTCTCGGGCAGTATCACTCCGTTAGCTGATCTTAAGTTTCTCTCTAGTGTCCGTCTTGATGGGAACAATTTATCTGCACCAATTCCAGAGTTCTTTGCGGAATTCACCAATTTAACTGTCTTGAGTCTGAGTTCCTGCAACTTGATAGGAGAGGCACCTCAGAAAATATTCCAGGTACCAACTCTGCAAACAATTGACTTATCAGGAAATAGGATTCTTGGGGGTTCTTTACCTGAATTTCCTTCAAAAGGATCTCTACAAAACCTGGTTCTTAGTGATACGAAATTTTCAGGAAGTTTACCTGAGTCCATAGGGAACCTTAGGAAGTTGTCGCGTGTTGAACTTAGAGCTTGTAATTTTACAGGTCCAATTCCATCTTCTATGGAAAATCTAACCCAGCTTGTTCTTCTGGATTTCAATTTCAATAGCTTCATGGGTTCATTTCCATCTTTTAAGCTGTCCAAAAATCTCACTCATATATCCTCTGCTAGAAATCGTTTGACAGGAATATCATCAAACTGGGAAGGCTTTGAGAATCTTGAGTATCTTGACTTGAGCAGTAATTCAATTTCTGGGCTCATTCCAGAATCATTTTTTTACCTACCCTCACTTTCAGTTTTAGTTCTGTCCAACAACAAGTTTTCTGGCCAAATAACTGAATTACAAAATGTGATTTCTCCACTAAAAAGTCTTGAGTTGAGTTCCAACAAATTGGAAGGGCCAATACCTGAGTTTTTATTTGAGCTGCGTGATCTTTATGGTCTTTCACTTTCATTCAACAAGTTTAACGGTACTGTGCAGTTGAAGAAGTTTACAAACCTCAATAAACTTGTTAGTCTTGATCTGTCCCACAACAGTTTATCAGTTGACACAACTATAAGTGAATCGGACCTTGCCTTGCTACCCCAGCTGAATAGTTTCATGTTAGCATCATGCAATCTGCAGAATATTTCCTTCCTCAAGAACCAATCAAAGTTGTCGATGTTAGATCTCTCAAACAACCAGCTTACTGGTGAAATACCAAACTGGTTGGTGGAAATCAATGACGGACTTCTCCGTTTTCTGAATCTTTCTTTCAATCAATTTACGCGCCTTCAAGAACCTTATACAATTGGCTTTCTCATGAATTTCCTTGATCTGCATTCAAATCTGCTCACTGGGGTTATTCCATTACCACCACGTGCAGCTGCTTATATTGACTTCTCAGACAATAACTTTTCTTCTTTTCCACCTGACTTTGGCAAGTATCTAGTAACTGCTCGGTTCCTCTCAATTGCAAACAACAAAGTTATCAGTAGTATCCCTTCTTCGATATGCAACTCCAGCTATCTTGAAGTACTTGAGTTGTCAAACAACAGTTTGAATGGAATAATACCATCATGTTTAGCAGAAAAGAGCAGCACACTCAAAGTACTGAACCTGGGAAAGAACAATCTCATAGGAAATATACCTGAAAAGTTTTCTTATAATTGTGAGTTACAGAGCCTTGACCTCAGTCAGAATCACTTAACGGGTGTGCTTCCTCGTTCTTTGTCCAACTGCACAAAGTTAAAGGTATTAAATATTGGAAACAACAAGATCAAGGATACTTTCCCCTGCTGGTTGAAGAACATGTCAGATCTTCGTGTACTTGTATTACGCTTCAATGGTTTCCATGGGAACATTGATTGCTCACGAGTGAGTTCCAACTGGACCGCACTTCAAATCATGGATCTAGCTTCCAACAATTTAGGGGGTATTCTTCCACGAGGTTCATTCTTGGAGCTAAGAGCAATGATGGCTGACCCATCTTTAACACATTCACGCTCTGATATCTTGCACTTTGAGTCTTCTATTAGGCCGATATACTATCAGGACAGAGTCACTCTTTCTCTTAAAGGACAAGATGTTACACAGACAAAGATCTTTCTCTTCTTCACCTCCATTGACTTCTCAAGTAACAATTTTGTAGGGAACATACCAGAGATAGTTGGAGATCTCAGATCACTTTATCTTCTCAATATTTCACACAACAATCTCACGGGTCAAATCCCTCCAGCAATTGGAAATTTGAAGCAATTGGAATCGCTGGACCTCTCATTCAACAAGTTAGGTGGCAATATCCCAGAAAAGCTTGCTAGCCTCACATTTCTTTCCTTCTTAAATTTATCAGCTAATGAATTGGTTGGAATGATTCCACGAGGCAACCAAATTCAAACATTTGGAGAAATTTCTTTTGAAGGAAACAAGGGGCTATGTGGGTTTCCGCTGAACAGAACTTGCAAGAATAATTCAGCAGATGCACCTTCTGAGCCAGAAGTTGAAGTGGACGAATTCATTTCAAGGACGGAGATTTATGTAAGTGCTATATTGGGATTTGTTGTTGGTATTGGGATCATTTTCCTACCACTTCTGTTCTCAAAAAGATGGAACCAGTCGTACAACAGAATAATTGACAGATTAATTTTAAGGATATTTCAGCAGCAAGATCAAGAAAGGAGAACTAGTAGCAGTAGTGTAGCATCTTCGAAAAAGGCAGCAGGGAAATCAAGAGGCAGCCATTAATTTCACAGGATGTAACACTTCCTTGCAGCATGCACACTATTGGTACTTCTAATTGTTTCCACTATGAGTTATCGGTGTTTGTTTACCTTTCATGTTTCATTTTAGTGGCTTCTTATTAGAAAGAAGAACACAATGCAATCTTACATTCTGTTTTAGGTAACGGTCAGCGAATAGCCATAACAAA

>M6-PGSC0003DMG400028576 M6 LRR-RLP

AAAATAGGCAAAAAGAATTGATCTGTTAGCCAAAAAAATAAATACAAACGAATCCTGTTGTTTATTTGCGAGGATTTTGCTATTTTATACGTTTCAGATCTGCAGAAACTTGGAGTTTCTTTTTTCTGTTGACAGTCCAAATGAGAAGTGAAATGTTTTCCCAGCTTTTCTTGATCCTATTACTCTCCATCTGCTGTAAAATGAGTGATGCAGTGGCAGCTAATTCCAGTAAATGTCTTGAGGATCAGAAGATGTTGCTGCTGCAGCTGAGAAATAACCTTACTTATGATTCTGAAATATCTACCAAGCTAGTGAAGTGGAACCAACGGATTGACTGCTGTCAATGGGAGGGTGTCACTTGCAACGGTGAAGGCCAAGTTATTGGTCTTGACCTTAGCGCTGAATCATTCTCGGGCAGTATCACTCCGTTAGCTGATCTTAAGTTTCTCTCTAGTGTCCGTCTTGATGGGAACAATTTATCTGCACCAATTCCAGAGTTCTTTGCGGAATTCACCAATTTAACTGTCTTGAGTCTGAGTTCCTGCAACTTGATAGGAGAGGCACCTCAGAAAATATTCCAGGTACCAACTCTGCAAACAATTGACTTATCAGGAAATAGGATTCTTGGGGGTTCTTTACCTGAATTTCCTTCAAAAGGATCTCTACAAAACCTGGTTCTTAGTGATACGAAATTTTCAGGAAGTTTACCTGAGTCCATAGGGAACCTTAGGAAGTTGTCGCGTGTTGAACTTAGAGCTTGTAATTTTACAGGTCCAATTCCATCTTCTATGGAAAATCTAACCCAGCTTGTTCTTCTGGATTTCAATTTCAATAGCTTCATGGGTTCATTTCCATCTTTTAAGCTGTCCAAAAATCTCACTCATATATCCTCTGCTAGAAATCGTTTGACAGGAATATCATCAAACTGGGAAGGCTTTGAGAATCTTGAGTATCTTGACTTGAGCAGTAATTCAATTTCTGGGCTCATTCCAGAATCATTTTTTTACCTACCCTCACTTTCAGTTTTAGTTCTGTCCAACAACAAGTTTTCTGGCCAAATAACTGAATTACAAAATGTGATTTCTCCACTAAAAAGTCTTGAGTTGAGTTCCAACAAATTGGAAGGGCCAATACCTGAGTTTTTATTTGAGCTGCGTGATCTTTATGGTCTTTCACTTTCATTCAACAAGTTTAACGGTACTGTGCAGTTGAAGAAGTTTACAAACCTCAATAAACTTGTTAGTCTTGATCTGTCCCACAACAGTTTATCAGTTGACACAACTATAAGTGAATCGGACCTTGCCTTGCTACCCCAGCTGAATAGTTTCATGTTAGCATCATGCAATCTGCAGAATATTTCCTTCCTCAAGAACCAATCAAAGTTGTCGATGTTAGATCTCTCAAACAACCAGCTTACTGGTGAAATACCAAACTGGTTGGTGGAAATCAATGACGGACTTCTCCGTTTTCTGAATCTTTCTTTCAATCAATTTACGCGCCTTCAAGAACCTTATACAATTGGCTTTCTCATGAATTTCCTTGATCTGCATTCAAATCTGCTCACTGGGGTTATTCCATTACCACCACGTGCAGCTGCTTATATTGACTTCTCAGACAATAACTTTTCTTCTTTTCCACCTGACTTTGGCAAGTATCTAGTAACTGCTCGGTTCCTCTCAATTGCAAACAACAAAGTTATCAGTAGTATCCCTTCTTCGATATGCAACTCCAGCTATCTTGAAGTACTTGAGTTGTCAAACAACAGTTTGAATGGAATAATACCATCATGTTTAGCAGAAAAGAGCAGCACACTCAAAGTACTGAACCTGGGAAAGAACAATCTCATAGGAAATATACCTGAAAAGTTTTCTTATAATTGTGAGTTACAGAGCCTTGACCTCAGTCAGAATCACTTAACGGGTGTGCTTCCTCGTTCTTTGTCCAACTGCACAAAGTTAAAGGTATTAAATATTGGAAACAACAAGATCAAGGATACTTTCCCCTGCTGGTTGAAGAACATGTCAGATCTTCGTGTACTTGTATTACGCTTCAATGGTTTCCATGGGAACATTGATTGCTCACGAGTGAGTTCCAACTGGACCGCACTTCAAATCATGGATCTAGCTTCCAACAATTTAGGGGGTATTCTTCCACGAGGTTCATTCTTGGAGCTAAGAGCAATGATGGCTGACCCATCTTTAACACATTCACGCTCTGATATCTTGCACTTTGAGTCTTCTATTAGGCCGATATACTATCAGGACAGAGTCACTCTTTCTCTTAAAGGACAAGATGTTACACAGACAAAGATCTTTCTCTTCTTCACCTCCATTGACTTCTCAAGTAACAATTTTGTAGGGAACATACCAGAGATAGTTGGAGATCTCAGATCACTTTATCTTCTCAATATTTCACACAACAATCTCACGGGTCAAATCCCTCCAGCAATTGGAAATTTGAAGCAATTGGAATCGCTGGACCTCTCATTCAACAAGTTAGGTGGCAATATCCCAGAAAAGCTTGCTAGCCTCACATTTCTTTCCTTCTTAAATTTATCAGCTAATGAATTGGTTGGAATGATTCCACGAGGCAACCAAATTCAAACATTTGGAGAAATTTCTTTTGAAGGAAACAAGGGGCTATGTGGGTTTCCGCTGAACAGAACTTGCAAGAATAATTCAGCAGATGCACCTTCTGAGCCAGAAGTTGAAGTGGACGAATTCATTTCAAGGACGGAGATTTATGTAAGTGCTATATTGGGATTTGTTGTTGGTATTGGGATCATTTTCCTACCACTTCTGTTCTCAAAAAGATGGAACCAGTCGTACAACAGAATAATTGACAGATTAATTTTAAGGATATTTCAGCAGCAAGATCAAGAAAGGAGAACTAGTAGCAGTAGTGTAGCATCTTCGAAAAAGGCAGCAGGGAAATCAAGAGGCAGCCATTAATTTCACAGGATGTAACACTTCCTTGCAGCATGCACACTATTGGTACTTCTAATTGTTTCCACTATGAGTTATCGGTGTTTGTTTACCTTTCATGTTTCATTTTAGTGGCTTCTTATTAGAAAGAAGAACACAATGCAATCTTACATTCTGTTTTAGGTAACGGTCAGCGAATAGCCATAACAAACCTAATAACAACCTCTCTCTTTTTTGAAGAAACAGAAGCCAACTTTGCCATTTTCTAAGTACAAAATAGTTTATTCAAGTTCTCCCTTATTCTTCGAAGTCAGAGTCGCTCTGCGCAGACAGTAAGACTTCATCAAATTCCTTACTGAAGTCTTTATCGCCATCCTGCTCTTCAGCCACTTCTTGTTGGGAGAACTTA

>M6-PGSC0003DMG400028591 M6 LRR-RLP

TTCCAAAATTCATTTGGAACCTTTCACAAATCACCCAACTGGACCTTTCAGACAACAATTTGGATGGCAAAATTCCAGACGTTTTTTCCAACTTGCAAATGCTTACATCCTTGTCACTGCAGAACAACAACTTCACTGGCCGGTTCCCATCTTTTCTTGTGGATTTGACAAACCTTCAAGTTTTAAGGTTGGGAAACAATTCACTCAGTGGTACCCTTCCCGAGCTTAAGACTAATTCAATAGAAATACTTGATTTGTCTCGTAATCAATTCTCTGGTCCCATACCTCAATCGCTTAGACATCTTCTCAACCTAACTGTTGTCTTTCTTGGACAAAATAAGTTGAGCGGCGAGATTGGAGCAGAAATGTTTTCAAGCATGACAAACCTCTTAATTCTTGATCTTTCTAATAGTGGTTTATCATGGAGTGGCAATATCAACACCACTTTTCCCCTTCTTTATTATTTAGCTTTACGCTCTTGTCGTGTGAAAGATTTCCCAGATTTCATATTCAACTCTAAGAACTTATGGATTTTGGATCTCTCGGAGAATGAAATTCACGGGCAGTTACCGAAATGGTTTGGGGGTTTGGGTGCATTGGCAAACCTTAACCTCTCTCACAACTTCCTTACAAGCTTAGACCATCTACCTTGGGAGACAATCATGGTACTTGATCTTCAATCAAATTCACTCACAGGACCTTTGCCATCCCCCATTTGCACATCAACTTTCCTTTATATAATAAATTTGTCTTACAATAATTTGAGTGCCGAAATTCCCAATTGCTTGTTTACTTTTTCCTCGCTCATGGTCTTAGACTTACGAGCTAATAACTTCCAAGGACCCATCCCAAACAAATTCCAAAAGAACAGTCAGCTAATTCATATTAGTTTGAGCAAAAATCAATTGGAAGGTCCCATCCCAACATCCCTAGCCAACTGTACATCCTTAAGAGTCCTTGATTTGGGAAACAATAAAATCCATTCTACATTTCCCTCCTGGTTGGAAACATTGCAAGAGTTAGAGCTTCTCATACTGAAATCTAATAGATTTTACGGACCCATCGTTTCTTTCCAAACAAAATTGCCATTTCCAAATATGAGGATCTTTGACCTCTCCAACAATTCATTTACCGGCTCTTTACCTATGGAGGTTCTAAAAGGTTTCAAAGCTATGATGTACATGGATGCACACAAATCAGGAGTGGAGTACTATGTCGAGGAAAATTTAAGAAGTTACTATGTCGATAAGCTTTTGTATAGTGGATTGTATGTGGAATCAGTGATATTGGTGATGAAAAATCAAGAGACTAAATTCAACAAAATCTTGAAGATGTTCACTACAATCGATTTATCAAGGAACAAGTTTGAGGGAGAAATCCCGAAGTTCATTGGGAATTTTAATTCACTTCTGCTGCTGAATTTATCTCACAACAACCTCACTGGGCATATTCCTATTGAAATGAAGAACATGAGCACTCTTGAAGCTTTGGACCTTTCATTTAACCATCTTACCGGTAAAATTCCAGTGGAATTGGCAATTCTCACATTTCTTGAGGTCCTAAATCTGTCACACAACCATCTTGTTGGACCTATTCCTCAGAGTAATCAGTTCAATACATTTCCAAATGATTCCTATTTAGGAAACTCTGAATTGTGTGGATTTCCACTGTCAAATGAATGTGGGAAGCATAAGAGTGCATCAGTACCAGTTGAACAGGAAGAAGATGAACCAAGTTTTCTTAGCGAAATGACTTGGCAGTCTGTGCTAATTGGTTATGGTTGTGGCTTGTCTTTTGGTTTTGGCATAGTGTATCTCATATACCTCTTTGAATGGCCAAGATGGTTCATACACTCTTTTGAAACTATCACTCGTGACATGATATACATAGTACTGAGGGCACGTAAAAGGCGCAGGAGGAATTTTCACAGAACAAGACACTGAAAATCTATGAGGTAATGACTAATTAATAGTCGCGTTAATTCATTATTTTCTTGTATGCATAATCAAACATATATTTTATTAGCAATACTTAAGGTCATGAAAACTACAACATAATCCATTGATATTAATGGGAATTGGAAACTTTGTTTTTTGTACAAGCTCTAGTCCCAAAAGAGTTTCTTAGATATTCAATACTTTAAAACTTTATTTGTTTGTTTTTTTTTTTCAATAAAAAATCAAGGAAGTGTACACAATATCCTAATGTAACTCATTGTTTATCATTGCTATTCAGATGACAATCAGAAAGTATGAAGCAAGAGAGAGGTCGGAGTATGCCACTAGAGCATGGGCTTGGGAAAATAGTGATTTTATAATTTTAAAATAATGCAGATTGTTGTTTTTAGTATACTAGTTTCTGAACATGTGCGTTGTGCGTGTGTCCAATGTAATATTTTATGAATTCGTCAGGATGATTAAAATTCAATGAAAATATGTGTGTAATGAGT

>M6-PGSC0003DMG400028627 M6 LRR-RLP

GTCATTGCCATAAATATTTATATTATGTCACTTATAAAACACCCCTACAAATACAACCAAGTTTGTCACTTCATTCATAGTCCAAAATGGCCTTTTTTCTCTCAATACAAATAACACTACTACTATCTTTTTTCTTCTTTCCTCTAGGATTGTCATTCACCGTTATAATGTCGGATTCCGGTGCACCTTCAGCTCTAGTTGATGCTCCACAAACTGGTTTCTCGATGAATAATAATCGTGCACGAACCGATCCTCGTGAGCAAGAAGCCGTTTACGACATTATGAGGGCTACGGGGAACGATTGGGCCACCGATATCCCTGATGTTTGTCGTGGTCGATGGCATGGTATTGAGTGCATGCCGGATAATGACAATGTGTACCATGTTGTGTCACTTTCCTTCGGCGCATTGTCTGATGACACAGCATTTCCTACGTGTTACTCAAGTCATTCGTTCATTTCACCTTCAGTTAGGAAACTTCCACACCTTCGGACTTTATTCTTCTATCGTTGCTTTAGTAACAATCCTCAGCCCATTCCAGGATTTCTAGCCCAATTAGGGCCCACTTTGCAAACACTAGTTTTAAGGGAAAATGGGTTCACAGGGACCATTCCTAATGAATTGGGTAACCTAACTCATTTGAGAGTTCTAGATCTTCACAAAAACAATCTTAATGGTTCAATACCGATCTCTCTGGGCCGGATCATCGGTTTGAAGTCATTAGATTTGAGTGACAACAAACTAACCGGTTCAATTCCAAGTTTAACCTTCCCTCAGTTAAATATCTTGGACCTTAACCAAAATCACCTTACGGGTTCAATTCCATCCACACTCATGAGCTGTCATTCGCTTATAAAGCTAGACCTGAGCCGTAATCACCTCTCTGGTTCAATACCTGACTCGATCCACAACTTAAACAACCTCATCCTTTTGGATTTAAGCTACAATAGTCTAACAAGCCCATTCCAAACGTTACTCAAGAAATTGAATTCTCTACAAACCTTAGTCCTCAATGGAAATCCAATGGACTCAGCTACTCTATCGGATAACGTCTTCGATGGTTTCAACAACTTAATGATTTTAGGATTATCAAACATGAACCTTCAAGGTCCAATACCAAATTCACTAGGCCTATTACCTATGCTACGAGTCCTTCATCTCGATGGGAACAAGTTAAATGGTTCGATCCCATCGAGTTTTGGTAATTTAAATAACATAAGTGAGCTTAGGCTAAATAATAACATGTTGAGTGGTCCTATTCCATTTAAAAGAGACATGGTTTGGAGAATGAGAAGGAAAATGAAGCTTTCAAATAATGAAGGGCTTTGTTACAATAAAGAAAATGGGCTTGGAGATGATTTAGAGACATTGTTGGATTCAGGGATTGGACATTGTGGGGACTCTAAAACAGAGCCAATAAAAACAGTACAACACATTTCAACACTCAACAGAGCAAGTTTTGTAAGATCAAATAGCTACAAATTATTATCATCTCTTGGATGCTTGCTTTGGTCAGCTTTAGTTTTTGTTGTAGTTTGTTTGTTGTGAGTGTTTAGAGATTTTGGTACAGTTTTTTTTTCTCCTTTCTGTTTTAAATGTAAC

>M6-PGSC0003DMG400028630 M6 LRR-RLP

ATACAACACATTTAAAATCTTCAATTAGTACAAAAATACATTAAATAGTAAAAGCAGATCACTCTGGTTCTGCTCTCCGCCGTCGCCGGCAGTCTTCGTTGAGGTTTGATCGGTTCTCTTCTCTTTTATTTCATATAGAAGAACAAATAAACCAAATCAAACTAAAATTTGAGCTATTATTTATCACCGGGGTTTTTTTTTTTAGGGGAAACTGAAATTCTTCATGAAACCCTAAATTAAGAAATGAATGGGTTTTTGATTGTATGGTAGAGATGGAATTTGCAACTAGAATAAGAAAATCTCATGCTTATATTATTCATGTACAAAGAGCTTGTGATTTGATCTGTATGTAATTGTTGTTTGGTAGAGATGGACTTTGCAACTAGAATATAAAATGAATTCTATTTTTATTTTTTTACGATCTGGTGTTTGGGCCAGCTTATGCACAACTCAACTAATTGTACTGGATATGTGCTGCTTCCCACCAACAACATGTATTAGGTAACTTTTTGTACCAAGACTTGGATCGTTGGAAAGAAAGACAGACCTAGTGTTTTTTGGATTTGAAAATCATTTTCTAAAGAGCAAAATGTTTCTTAAGTTTTGAAGTGTTTTGTTTTAGTTGAAGCAAGTAATTTCTCCTACAATGAAAGGCTTCACAGAGTGCCTGAAAGAGAAGATGTTTTCTTATTTGTAATGAGCCTATTATAAGAGACATTGTCATATTATATTTGATTATGATTTCATTAGTAGAATTCTTCGCGCTGTCTTCCTTATGTTAGTTTTGATTTATATATTTTTTGTTCTATCAAGTGTTTTTATGAGTTACCTATATATTACTACATAGTGTGATTTGATGTTATCGGTTTGTTCCAATTCGTTGAATTTTTTTAAATTTCAAGCTTCAAAGCTTGAATGTCCTTGACCAAAATTCATGGCGCCGCCACTAGCTCAGGTTGTTCGGTGGTTCAGGTTGTGTGTGACAATTTCAGTCGAGGTCCTGATGCCGTAGCTCCTTTTCTGCTCTTTTGGAGAACAAAAATGTTTTTTTTTTGGTAAACCGTGAAGAATTACCATTGACAGCCAAAAATTTACACCTATAGAGCACTTTAGGTAGAGGAGCCATCTTCCAGGAAGGGTACCCTTTCCTCTAAAGCCTCAAACAAAGACAAATCTAGTTACAAAATTTCTGAACCAAATATCTACATTTACTTGCTTTCCTACTATCCTTCAGGTAATCCACTCTTTCCACTATCTCTTTCTTTATCTGTGTGATTGTAATCTATGTATTTTTTTTTTGAAACTGGTAACTATATATTTTAAATTACACCATAACAGTGCAAAAGACATTACAAAATCCCAACCAAGCTTAGGCTACTGCTTCACAAGGATCCTATCATATCAACTAGTGATTCAGCCTCCTCAGCATAGCTTTCTTTACACCAAAAATAAAACAAAAATATACAATTCATCTTGATTTTCTCCACAAAATTATATTTGTCTTCAAAAGCTCTATAGTTCCTCTCCTTCCAAATGGTCCACCATATGCAAGCAGGGACATGTTTCCACCATTTTTTCTGACTAACTGCCCCATCATTCTGGTTCCAACATTTCAGCAAATCAACACTTGTTGCTGGCATTGACCATCTGAGCCCTACCATATTCAAGAATAGTTGCCATAATTGTCCAGTGACGGGACAATGAATGAATAGATGACTATTATTTTCCCCAGCAGCTCCACACAAAAAGCATCTAGAACACAGTTGGAATCCCCTTCTTTGAAGTTTATCATGTGTGAGACAAGCATTTCTTGCTACTAGCCAAGAAAAGCATACCACCTTAAAAGGAGCCTTCATCTTCCAAATGGATCTCCATGGCCACTGATTAATTGACTGTCTGTGCCTAATCAAAATCTTATAAGCTGAGTTTACAGAATAAGTTACTTTTTTTTTTTTTTTGAAATAGGTAACTTTAGTTTACAGAATAAGTTCCTTTACTAGTGGGCTTCCATATAAGTGAATCTTCAGTGTCTTTGAGCCCTTGGAATGATTCCAAGGTCTTGAAAAATTCAACAACTCTTTCCAACTCCCAATCATGCAGTAGTCTTCTAAAAGTAATATTCCATCCATGTACTCCTTTTGCTGATTCCAGGCTAATGTTTGGCATAGTGGCAATGCTAAAGAATTCAGGGAAGAGTTCCTTTAGAGGTCCATGTCCAAGCTAGTTATCACACCAAAAAAGGATTTTCCTACCATTGCCCACTCTTAGGCTAATTTTTCCAGCCATCAAAGACTAGATGGATCTGATTGCTTTCCAGACTCCTACACCATGGGGGCTCACAACCGCCTTTGAACACCACTGGTCTTCTTGACCATACTTGTCATGAATGACATTTCTCCATAAAGCATATGTTTCAAGATTATACCTACATAACCACTTCATTAGCAAACTTTGGTTGTGAGCATTTAGGTTCCTGATGCCAAGGCCACCATCTCTTCTGGATTTGATAAGAACATCCCAATTAACCAGGTGGAGTGCTCTTTTTTCGTTGTTGCCTTGCCATATGAAATTCCTTCTGATCTTGTCAATTCTATTCCTAACCTTGCTAGGTAAAGGAAAAAGAGACATAACATAAGTAGGAAGGGCATCCAAAACACTATTGACTAGTATAACCCTACCACCTAGAGATAAGTATTGTCCCTTCCATATTGCAAGCCTCTTTTCACACCTTTCTATGACACCACTCCAAATCTGCAAGGCTTTACTTTTAGCACCCAAAGTTAACCCCAAATAGGTAGTAGGTAGGGAACCCACCACACATCCCAGAATACCTGCAAGTGCTTGAATATTGCCCACTTTATTAACCGGGAATAAACTGCTCTTATGCCAGTTGACATGAAGTCCGGAAACTGCCTCAAAAATCATGAGAATAGCTCTCAGATGTAATCTATGTATTTAGTATTGTCCCTCTGAATAGTTTCCAATTTCTTGCTCTCCAAGTGGATAGATTCTGGCTCCCCATATAGCTGTAGCTATTTCCTTCTTATATTGCTTCCAGTGTCTTCCTTTTATCCATTTTAGGCATTAGCACTTCCGTTTTTTGTATTTTAATACACGACCAAGTCTCCAGTGCTTCCTTCACTTTTGTTGTCCAGGTATATTCTGCAAATAAATGTACTTGAGTCTCCAATTTGCTTTCTCCACATAGGCAACACTTGTCATCTTCTATTGGCATTTGCAGTCTGATCATCCTCTCTTTCGTTAGCAGTTTTTCCTGATTTGCTAGCCATAAAATAAACCTGTGTCTAGGCAGCATGAGTGAGTTCCATATCAAGTCTGCTACAGGTAGTTTCAACTGCCTACCCAGTAATTGGTTGTAACTACAACTCACATTATATGCTCCATTGGTCAATAGACAGTAGGTTCCATTAGTATACCACATTCCCATGGTTAGTTTAATGGAGTTGAGCTTCCTCCAATACCAGCTGCTATCCTTAGAGGGTGTATGTGTCCAAAAATCTTGCTCGTCTTTCATATATAGTCCATGCACCCATCTTATCCATGGTGTATCTGCCTTGCTAGCAATCTGCCATATCAACTTACCAACTGATGCAATATTCCATAACTTGCATCTCTTGATGTTCACTCCACCATTCTTCTTTGGCACACAAATCTTCTCCCACGATACAAGGACTACCTTCCTCTTGTTCTCCTTATTTCCTCATAGATAATCCCTACATTTTATATCAACCTCTTTCAAAATACTTTGAGGCAAGATGAAAACATTGCTCCAAAAGTTATAGAGCGAGAAGAGTGCAGCATTTATGACCTGCAATCTGCCTGCACATGACAATGTTTTTGAATATGCACCAGTAATCCTGTTGGTTATCTTGTTGATCAGTTGATGGCAATCTATCTTTGTCCATTTCTTTGATGATAATGGCAAGCCCAAATATCTGATAGGAAGATCACCTAGTGTAAATCCTGTCAGCTGCAATATTTTTTCCTTCATAACATCATCCACCCCAGCTACAAATATATTAGACTTGTCCAGATTTGCCACAAGGCCTGTCACCCTACTGAAATGGTTCAACACCTCTATCATCCTTCTTATAGACTTAAGATCCCCTTTGCAAAATAGCATCAAGTCATCTGCAAAAATCAGATGGTTTAGCTTGGTGCTATGACACATTGGGTGAAACTTAAAGTCTGGTAGTTCTCCTACCTTCTGTAGTGCTCTAGATAGATACTCCATGACCAACACAAATAGCAATGGTGACATTGGATCCCCCTGCCTCAATCCTCTCTTCCCTTTAAAATATCCATAATTTTCGCCATTTACCTTGATGGAAAATTTTGTTGAAGTCACACAGTTCATCACTGGTCTGGTGAAGCCGTGAAGGTTTCTGGAAAACCATATCCATGAAGTGCCTCCTCAATAAACTCCCAAGATACCATATCATATGCTTTCCGAGAACAAAAATGTTGTTGTACAGGTTCTGGGGGTTCAGTAATATATTGAAGAAACTTTCCATTTAGGGTCCATTTTCATTCTTTTTTTTTTTTGTTTCTTTACTCATTTTCTTTTGTAATATTTTGGTGCAATTTGGCTTCGATTCCAAAAGTTTTAACATTTTCTATCATGGCCCCTTATGTTTTGGCTTCATAATAAGAGACAATTTTTTGACTCTTTTCGGGATACCATTGTATTTCTTAACTGTTTTGGGAGCTTAATTGAACAAAATAAAAAGACATCTACTATCATAATGTTCTCCTCTTTTCTTGTCACTGACAGATCAAGACGCCTCTTTGCACAAATTCTACACTATTGCTAGTAAGGTGGAAAAAAAATCAAATCAGAAAAAGAAAATCAAGCATGAATCAAGGCACTTTCTCTTCCCCTTAATTTGTCTGATTCTTTCTGGATTACAAACTTGAAGGCTTATTGAGATTTATCAACTTTTCTCTAACAGATAAAGGAAAAATTTAACCCTTAATAAGATCATTTAAATTCTTCTTCACAGATAGAGTAAGCATTGCGGTTGAGGGGGACATAGAAGATAGAATTAGTGTCCTACCAAGAAATGTTATTGATTGTATCCTTGAGTTCTTGCCTGTTAAAGAAGCTGCAAAAACTTGTACATTGTCCAAAAGCTGGAGATACATTTGGAGCGAGCTTCCAAAAATGGTGCTGGATTACGAGTTTTGTGATGAGTTAATAGACGAGTCTGAATCCAAATTCAGAGAAGTAGTAGATGGGATTATGTTGCTGCATATGGGCAAGATTGTGAAGTTTGATCTTGACGTTAGAGTAGAAGATTTAGCTTCATATACAGCTATTGATAGATGGGTTCTTTATGTCACCAGAAACAGTGTCAAGAAGCTACACCTTAGAATTTCTAATATTGGTGATCGTACCTATACACTGCCTCCTTGTATATTTCACTGTTCAACCCTGACACAGTTGAAACTTGCTAGATGTTCCTTTAAACCACCTGGTTCTTTTCGTGGCTTTCCCAATCTTGTAACCCTTCGGTTGGCATCAGTAACCTTTTCAGAGCATTGTGTTATCAAGGCCCCTCTTCTTGCTAATTTGACCTTGAACTGTTGTGATGGTATGCAATACCTAAACATTGTTTCACCAGTGTTGAAGTCCTTGTATGTTTGTTACCCTCACTCCTATATCGCACTAAATTGCTTCATGAACTGCAAGAATTTGAGAGATTTAGGACTTGTGTTTTCCAAAGTGGTTGATAATCCAAATCATGATCATAGATCAACGTTGGTGAAATTTCTTGGTAGCTTGCCTGCACTTGAGGTACTCCTGTTGGATTCACTTTTCATTGACGTAAGACATGAACTTGTTTTTTCATTTGAATATCTCCTATAGTGATTTCATTCAAGTGAGATTGTAATCATTGTATTTGCTCCAAAGCTTTTGAGTGCAGACGTAGTTCTATCAAGTGGTCCTCCTTTTATGCTCAACTGCTTGCGGAAGCTGTCCCTCTCTGTTGACTTTGGCAAATTGGGTCATATTTCTTGCGTTCTACAATTAATCGAGTGTTCGCCCAATTTGAATGAACTTGAGATTACGGTAAGAAATGTTAAATATATCTGGTGGTTCTTTTTTTTGCTCTTCATATATTTGAAGTTGAAGATAATGGTGTTTGCTTATACCATTAGGTCAATGGTACCGATGATAATATTGAAGCATTTGTGAAATGTCTGAAGATGGCTGACTGCTTGCAGCTACCACTCAACAAGCTAGAGTATGTGAGTTTCTATGGTTTTGCAAGTCCAATCTGTGTAGAGGCTTTTGCTATGCTATTCATTATTCAGGCACGATCTTTGCTAAGGATCTTCATTGAACAACCGCTAGGAACTAAATCCAGTAAAGATTTGAGTGGTTTACCCAGAGCTTCTCCTAAGGCTGAGCTAGTTATAAGTTAAATCGATACATTATCTATTTGTTGTCGATAAGTTTATCTAATATGTCACAAATGTTGAACCAGTTTTCTTCCATATTTACTGTTGGGAAAGTATGACTGATATCTGAATCTTCATCTTTTGACAGTTGAGCTTATTTTGGTCTCTATATCTACATGAGGCCAACTAGTAAGATTAATTTGTGAATGTTGTTGTTGGGTTTGATATCCTACTAGTTTTTGAGGATAAAAACATAAATAAAATATACTA

>M6-PGSC0003DMG400029168 M6 LRR-RLP

CCTGAAAGTTGCAGTCAAGGGAAATGCAGACTTCCAGTAGACGTCCTTCTGGTCGATTGCTGGAGTTTGTTTTTTAGAACAATGATTCTGGCCAAAGGTTCTGATATTCTGCATTGTACAGTTTGTGGTCAAAACATAGCAGATGCTGTGAGTTTTAGGGTGTGAAAGAAAGGAGAGGAAGAGAACATTTACTGGCAAAAAATATTCTTCATACAAAGCCAACCGGGAACCTTTTTTTCGAACTCGCTCAACACAAAAGAAAAGACGAAAGACGAGAATCCGCCCTTTTACCAAGTGTACCAGATAGAAGATGATAAAGGGACAACAATCGTTGCCTTGCACAAGATGATGCCCTTTCACTTCGAGGTGTCGGCTCTGAAATGATGGAAGATTGAATCATTCCTTCATTCTATTACAACTAGTGATGGCCGGGTATCATTTTTCCGGTAGGCCATGACTAAATTCGAGATTTCCTCTGTTTATTCTAAAAAATGGAGAAGACACGATAATTGCTTCTTCTTTGTTGAGATCCCCCCAGGCCTTTTCTTCTTCTTTCAGGTCCCTCCCCTCTTATTCATTTTAAATTCCATGCCACCGACTTCTTTTTCATTTTTCACAGTTAGTTCTGGTCGAAAGTTTTTGCCATGTCAGCACTTAACTTAAGCTTATATGGTGTGGTCTTCAAGTTCTTTCATTATTTATTTGGAAAATTTGAATGGTTATCAATTGGGATTGAATAAGAGTGGCAGAGTTGGCAGACCACTATTTTCAATGGAGAATTGAACGAGGAAACTACATGGAACAACCATTAGGTTTTGTTGTTAAAGACCAAGAAAAGAAAGTTTATAATTTAAAAAGATCAATTGCTGGCCTAAAGCAATCTTCAAGGCAGTGGTACCTAAAATTTCATAAGGAGGTGATTTCATTTGATTTCACCATGATCGATGAAAAACCATTGCATCTACGTGGAAAAGTCCAATGGAAAGTTTGTAATTCTTTCTTTTTATTTAGATGATATTTTATTGGTTGGAAATAATTTGGAGTATGTGAAAACTGTCAAGTCATGGCTTTCAAAGTCATTTGATATGAAAGATATGGGTGACACATACTATATATTGGGTGTTAAGATCCAGAGAAATTGTTCCAAGAAATTTTTGAGTTTATCTCAAAAAACATACATAAAGAAAATATTGAACGCTTTCGAATGAATAGTTGCAAATCCATGAAACTTTAAACCTTGAAATGTGTTCAAAGACTGAAAAAGAAATGGAAGACATGTCTCGAGTTCCATATTCTAGTGCTGTCGGGAGTCTGATGTATGCTATGATGTGCACTCGTGTACTCGCTCAGACATTTGTTATGCCGTAGGTCTTAGCAAGTATTAATCATCCTGGAAGAGATCACTTGAAAGCAGTAAAAAGAATATTTTGATACTTGAAGGGAACTGCAGATTATTCACTATGCTATAGTAGATCTAATTAATTCATAAGAGATTATACAAATGTTGATTGGGTTGGTGACCGGTGTGATAGAAAAACAGCTTATGCTTTCTTACTTATTGATAGTGCTATTTCATGGAAAAGTAAGAAACAAACTTGCACAACTCTTTCAACTATGGAATTTGAATTCGTAGCTTGTGCATCTACAGTACAAGAAACTGTTTGGTTAAAGAGATTCTTTGAGCATTTGGATATTGCAAAGAATTCTAAGGGACCTATGATTTTATATTGTGGAGAGTCATCACATATAGAAGGATCCTAAGTGTCACAGCAAGACCAAACACATTGATATCAAGTATAACTTTGTGAGAGACATAGTAGCAAGTGGAGAAATAAATTTACAATACATTCCTACACGTAAAATGATAACTAATCCTAATATCTAGAGACTTGTTTGAGAAACATGTTAATGCTCTAGGTTTGGTAGGATATGATAATATTTTGGTATTGTAAAATAACTTGATTGTTATATTGTTCTCATCAATATATGACTATTTTGAATTTATATTCATATTTCATATGCATGTAATAATGTTTTAGAAATGAAGAGTGTTGGACAAGTCGCGAGATTGGCTCTCTAACATGAACAATCGCCTCTTACGTTGGAGAACGTGAAGAGATGATTTCCGCTACCATGTTATGACTTTTTTTTATAAGCCAAACTAAGTCTCTCTAAGAAGATGATATTAATGATCAATAATGAGAGAAACCGGGTTAAACATCTAGAGGTGTTTAGACCAAGGTTTGTACGATGTTGGATAATCGAAAGAATGAGACACATGCACCAGTATAAGGTGAAAACATCATAGCACGGTTGCAACCCAAGTATTTAAAGAACCTACTTTATCTGTTGATAATATCATTTGATTAGAGCTCCGTGCATCTAAGACTTGATACTTGTGCAACATAAGATGCACTCAGAAGCTTTGTCTGCCTAAATCCAATATGATGCACAACATGCACTTCATTGACTAGAATTGCAATATGTCCTGTTTACCTTATTGCAACTTTAATAGGAGCCTCACCATGAGATACATCAAGCTAGTCGAATATGCTCTTCAAATATACATGCCAAATATACTGAGGTGTCGTTTGTGATGTTCCCCTCAAAATATTTCAAGTGACGTTCCCCTCAAAATATTTCAAATTCTTCCTTCGCTTTTCTTTTCTTCCTTTCTTTCTTTCAACTGATAAGTATAACTATCCAATTCGTATTTCGCCCTTTCTGCAGCTATGTACATAAGTTGCACTTGCTTCTCATTATGCAAATCCTCTTAGCTTCATTTGATCCATACTCGTTCTGCTGAAGATTTTTCTCATTGTATTGATATTTTATGCAGAATATTACAGACAAAAGTTTGCATCTGGTTGCTGACAATTATCAAGAATTGGAATCGTTGAACATCACTAGGTTGGATATTTTCTCTCAGTCTGCACGAGATTAGTTAAAGGAAGAGTTGTTTTTAAATGTTCTGGATAACACATAGCATCTCTTGGACATAAGGATGCCCTTTCTTGGCTTAGGCCTTTGTCTGTTGTAGGGGTGATGATAATATAAGCCCCCCTTTCTCTAAGAAGGATACTTATAATCGAGTATCTTGAGAGTATTTCTAATTGTTTGGTCTTTATAACCTTTTTGAGAGGGAACAAATATGATTATGTTTTCTTGATTGTACTGATTTTGTTTGATGTGACTAGGTGCATCAAGATGACGGATTCTGGTTTGCAGCGGATATTGCTGAACTGCTCTTCTCTACAGACTTTGAACCTTTATGCCCTTTCCACGTAACCGCTTGTTTCATTTTATAATATATCTGTCAAAAGATATGTTCCATTTCATTGCATATCGTGTGAATGCACCTTTCCTACCACTCAATGGTATACTACAAAATGCATATGTTGGAGTTTCTACTCAGATAATCTGCTTCCTTTCTTTCTTGCAGTTTAACAGATGAAGCTTATAAGATGATATCTCGTCTGCCTCATCTTAGATTTCTGGATCTCTGTGGTGCTCAGGTATAGTCTTCAGCATTTTCTTGGACTGCTTTTCATAGGTTATGCTATTGCTTGGTGCAATACCGAAACTGTCTAAGTGACATTGCTATTAACCCCAGTTGTCCTTCAGAATCTGACAGATGATGGGCTTTCTTGTATAGCGATGTGCAAAAACCTGGTGTCTCTCAATTTGACATGGTATGTATCCTAAGTATATTTCATTTTCATTGGGAGCAAAAGGAGATACAATTGTATATTATTGAACCGATATATATTCACATTTACAGGTGTGTACAAGTCAGTGATGTTGGGGTCATAGCAATAGCTCAAGGCTGCAGGTCTCTTGAATTATTAAGGTGAGTGTGGACCTAGAATTATCACTAAAATAGGAAGACCAAAATGACCTATGCTATATTACTTCAGCACCTCCACAAAGTCGACAAATATCTTATCTTGTGAACTAACTAGATAGAGATATCCTGTTTTTGAATACTCCAATGTTTGTGTAGTCATTGGGCCTCCGGCTTAAGCAATTATAAAATATAAAGTATAACCACTTCCTATTGACTACAGATTTATTTGCAATTT

>M6-PGSC0003DMG400029415 M6 LRR-RLP

TAATTTTTAAGTAGTTTACAAATATATAAATGTCATTTTAAGTCTGGATCAAATTTAGATTGTTTGACTCTTGAAAAATGAAATGTCTACATAAATTGGGACAGAGGGAGTACTTTGTACTTAACCAGGACTGGGTTCGAGTCCCCCAAGTACAAAGTTCTTTTATTAGAAAGATGTGTACACTATGTAAGACTTCTTGACACAAATTTGGATTTAGTCAACGGGATACCGGATAACGCGTGGAAAACCATAAGGCAATAAAGATACTACAATATAGTCTTTTAAGTTGTAAACATGTAATGACTTTTAGTTTTTGGTCAAGGACCAAGTTATAAGTCTTCTAATAAGGAAGATGATAAGAAGTTTCCCATAAAAGACTATGAATTGTGAAAAATGTTAAAGTCTTGCTCTTTATTATGTGTGTTTCTTGTTGTTGTTGTTGTAGTCTTTAGCAATATTTTTCGACAATGTCTCATGCTTCTTCTTCAAAAGTTTGCAAATATAGTGTTTTTTTGAGTTTTAGAGGTATAGATACACGTAAAACCTTTGTGAGTCATCTCTATAATGCTTTAGAACAAAGAGGAATTGATGTTTTTAAAGACGATGAGCGGTTGGAAACGGGAAAATCAATTCCTAATGAACTAATGAAAGCCATAGAAGAGTCTAGATTTGCCATTGTAATATTTTCCGAAAGCTATGCATCATCAAGATGGTGTTTGGAGGAGCTTGCACATATTATAAAGTGTCGAAATGATTTAGATCAGATAGTGATTCCAATTTTTTATGATGTAAGTCCATCTGATGTAAGCCATCAAAACTCACCATTCGCTGAATCGTTTTCAAAATACAAAGATGAAGAGTTGGAGAAGGCTCAAAGATGGAGAGAAGCTTGTAAGGAAGCTGGGAAAATATCAGGATACCATTTACAAGACTGCAAGTAAGATTTCTTTCTTTTGCACCATATCCTTTTTTCGTAAATACTTTGATTTTCTGTCATATTCCCTTCCCTTTTTCGTAAATGCTTTGGTTTGCTGCACTTATGTTGAGGGTCTTTCGGAAACAACCTCTCTACCTCCCCGAGGTAGGGGTAAGGTGTGCGTACACTCTAACCTCCCCAGATTGCCCCTTGTGGGATTACACTGGATATGTTGTTGTTGTTGTGAAAGAATCATTAGAAGTAATAAAATAAGATATAAATCATAAAAGAGACCACACATTTGGGTGCTAAATGTAAAGAACTCTATAATCTTATTTGATCTCTAAGTAGTTGTTCGTTTCATTGTCGTTTAGGGATGAGGCAAATTGCATCAAGAAGGTGGTTGATCACATATTACCAAACGTTATTTCAACCGTTTCGGAAAGCTTAGTGGGTACTCAAATTGAAAAAGTAATCTCATTATTACATATGAAATCTGATGACGTTTACTTCGTTGGATTATGGGGAATGAGTGGTATTGGTAAGACAGAAATCGCAAGTGTTATATATGAGAGATACCGTCATCAATTTGAAGCATATTGTTTTCTTGGTGATGTTGGAGAAATGTACCGGAAAAAGGGACTTAGATGGCTACAGAAAACTCTTATCTGTAAACTATTTGGAAAAAAGATGATTGTAACTAGTGAACATGATGGAACCATAATTATAAAGAATGGGCTTCGGTGGAAGAAAGTTTTACTCGTTCTTGACAACGTAGACCATCTAAGTCAACTAGAGCTTATAGTTGGAGGGACAGAGTGGTTTGGCAGGGGTAGTAGAATTTTGATTACTACGAGAGACAAACACCTGATAGTCAGTCATGTGAAGGAGGATAAAGTATACGAAGTCTTACTATTATCTGAGAACGAAGCTCTTGAATTGTTTTGTATGCATGCTTTCAAAAGAAAATCTCCGGAGAGAGATTTTAAGGTACTTTCAAATGAAGTAGTGAAGTATGCTGATGGACTCCCTTTAGCTCTTAAAGTTTTGGGTTCTTCTCTTTATGGACGAAACAAAGAGCAATGTAGAGATATAATTGGTAGATTGAAAAAAATCCCTAATGATGACATTCTAGGAAAACTTAAGATTGGTCTCGATGGGTTGAATAACGATGAGATGAGAACATTCCTTGATATAGCATGCTTGTACAATCATGAATATAGCTATTACGTTGAACGAATACTTAAGAGTTGTGGTATTCACTTGATAGGAATAAGCTACCTCATCGAAAAATCTCTCTTATGCATCAGATATCACTCGTTTGAGATGCATAGTTTAATAAGTCAAATGGGTCAAAATGTGTCAAGGGAAGAATACGCAAACAGCAGAGTATGGCTTATTGAGGAAGTTCATGACCTTTTTGCTGGAAAGTTGGTAAGAAATATTTTATCAAAAAAATTGATAATTTAAGAAGCTCTTAGTATTTTGTGACTTCGTTATGTTGAGTTTTTTTGACAAATAGATATATGTTCCTTTCCTTACAGAAAACAAAAAAGGTTGAAAGCCTACGGATCCCAAATGGTTACCGGTTTGATGATGATCATATCAATTATAACAAGGTATTCAAGAGAATGCAATGCTTACAGGTGTTGATACTTCATCGTAGAACTATTTGCTCGGAAAACACTGTCACTTGTTTGCCTTCTAACCTGCGGTGGATTGAATGGCCGAACTATCCTTCAAGTTCATTGCCAGAGGGGTTTGAACCATCACAACTCGTTGGGCTTTGTTTACGTGGAAGTTGTCTTGTTGAACTTTGGCCAATACCAAAGGTATAGTGATTTCTTTCTCTCCAAAACAATAGCTCCAGTTCATCACACGCTATGGTTCAAATGCTATACCAAATTAATGCTTTTGTATAAGCAATTTACATCATGTAACAGAAATAGAATAGCATATGTCTATTAGGAGTACTACTTGAGCTAACTGGTAATTCGCAGGTACGTTATGTTTTAATATTTTGTTATTTTTATGAAAACAGAAACTGTGCAACTTGAAGTATTTGAATCTAAGCGAGAGCCTTGGGTTAACGAAGACCCCTAATTTTGGTGATATGCCAAACTTGGAGACACTAAGTTTACGGGGGTGTGAGAATTTGGAAGTGGTTCATCCCTCTCTTGGACATTGCAAAATGCTTACTTATTTGAGTTTGACAGGTTGTGTCAAACTTAAGAAGCTTCCAAAATTTGTCTGCATGGAATCTCTCGAGATTCTTCTCCTTGATGAATGCACGCGCTTAAAAAGATTTCCAAAAATCTGTGGAGATATGCTGCGCTTATCAATACTGTATGTAGAATCCCCCTGGATAAGAAACTTACCCCCATCTCTCAGCGGCCTTAGTCATTTGAAACTGAAAGGTTGTGAAGTTCTTGAAAGTATTCCAGACTCTTTTCTAAATCTTGAAAGACTTCACATTTCAGGTTGCAATAAACTTACAACTTTGCCAAACAGCCTCTTTGAATCACAACAATGGATATTTCTTTATATATACCGATGTTCTGGATTGGTAGAGCTCCCCGTATCTCTTGGAGTTCAAAAGAAGCTTGGTTTTTTATCCGTAGAAGAATGTGAGAACTTAACGAAGCTTCCAAGCTCTATTCAGATGGAATCCCTTACAGAGCTCAAGATATCTAATTGCCCAAAATTATATACATTTCCAGAAATAATTGGAGATATGCATTGCTTGAAAAGTATGACTCTGGAGTCTACTGGGATTAGAGAACTGCCTTTGTCCATTGGGAATCTGAGCGGCCTCGAATATCTCAATCTGAAAAGTTGTGAAGATCTTGTAAGTCTACCAAACAGTCTCTGTAAGTTGAAGAATCTTCAACGGCTTATTCTTCGTGGCTGCAAAAAGCTTCCAGGAAACGTTGGTGATCTGCAAGAGTTTTGAGACACTTGA

>M6-PGSC0003DMG400029555 M6 LRR-RLP

ACTATAAGAATTATGAATAATATTTT

>M6-PGSC0003DMG400029556 M6 LRR-RLP

ATACTCCCGTTGTCCCAATTTAGGTGACACTTTTCATTTTTCGAGAGTCAAACAGTTTAAGTTTGACCGAGAATTTGCCCATGGTTCAATTTTTAATGAAATTATATATTTGTAAACTACGTAAAAAGTACTATAAGTCGCAATAATTGATAATTCAAAATGTTTTAAAAATATATGAAAAATTTACGATCAAAGATAGATTTGTTTGAATCTCAAAATTTGAAAAATGCCA

>M6-PGSC0003DMG400029572 M6 LRR-RLP

CAAGTTCCTAGCTAGTTAGATATATTCAATGGAGTACCAAAAATTGCTAATAGCTTTCTGCTTCTATTCACATTTTATACTTTTTGTCCATCAATCTCAGCTTACATATGCTGGGCAACATCTCTGTGCCCGCTATGAAGCTTTTTATTTGCTTCAATTTCAGCAAGGCCTCACAGTTGATCTTATTGGTTATGACTGTGACTATGAGGCCCTAGCTAAGACTTTGTCTTTGAATGTTACAGGAGATTGTTGTCAATGGGATGGTGTTACTTGTCATGGATTTACTGGTCACGTTATAGGACTCGATCTTTCTTGCAGTTTTCTTAACGGAACCATTAATTCTAACAGCAGCCTCACAAAACTTAGTCATCTTCAAAGACTCAACCTTGCTTTCAACAAGTTGAGTAACTTTCCACTTGTGGATAGCATTAGTGAACTCGAAAGTTTGACATATCTCAATCTTTCGCATTCAGTACATATGATAGGGGAGAAGCAGATACCACAAGTATTATCCAACTTGTCTAAATTGGTTTCACTTGACCTCTCCTTGTATTACACTGAACTCAAAGTTGATTGAAAAACCTTCAGAAGTTTGCTTCAAGACTTGACCAATTTGGAGGTACTGTTTCTTGACAACTCGTATGCATCATTTGAGTTACCTAAGAATTTTTCTTCTTCTCTTAGGTACTTAAGTCTTGAAGGTATCGGCATGTTTGGGATCATAAGTGAGTTTCAGTTATTTCATTTATCAAACTTGCAAGTGCTGAGATTTGGACAGAATCCTTTACTAACAGGCTCTTTGCCAAATTTTAATCGGAGTTCCAGTGAAAGTATTTTAGAGTTGAACTTCTTTTATACTGGTATTTCTGGGAAGGTACCCGATTCAATTGGTAATCTCCATTCTCTTTGGTATTTGAATCTCCAGATTTGTCATTTATCTGGCTCGATTCCAGAATCCATTGGCAATCTCACTGCAATCAGAGAGTTGATACTTTCAGAAAACGTCTTCATTGTTAATGTTCCCTCAACTTTCTCAAAGTTAAACAAACTCGTTCGTTTAGATCTCTCTTCAAACTATTTTCGAGGCTCAATTCTAGAATACATTGACAATCTCACTGCAATCACACAGTTGACAATTCCAGATAACAGCTTCACTGGAAATGTTCCCTCAACTATCGGAAAGTTGAACAAACTTAACTACTTATCTCTCTCTTCCAATTATTTTGAAGGCTCGATTCCAGACATCTTTGCCAACTTTTCAGATCTATTTGCTTTATTTTTTGACACTAACAATTTCACTGGTTCATTTCCCTGTTCAATTGCAACCTTGATTTGCCTTGTGAATTTAGAGTTGCAAAACAATTTACTAACTGGTTCACTTCCCTCTAATATAAGCTGATTTCAACTTCTACAATCACTTGATTTGTCCTTCAATTATTTCACTGGCACAACACCCCCTTCGTTATTCCTTCTTCCAAGATTGGTTTCTTTACATGTTCAACTCAATCAATTAACCGGGAAATTGCCAAATGTGCTCAAGATGAGTTATCTAGGATACGTACTATACCGTAGCATTGATCTTTCATACAACAAGCTGCAAGGTGAAATCCCCAATGCGATGTTATCTATGAGCATGGGTAAGCTGGATCTCTCTCATAACTTCCTCACAGGCTTTGAAAAACAAGTATGGCACTCAGAGTACTTACTATACCTTAATTTGGAGAACAATTTTCTTCAAGGGCTTTTGCATCTATCCTTTTGTGACATGTATAGTCTTGTATTCCTCATTTTAGCTCGGAACAATTTCAGTGGTTCAATCCCGGTATGTTTGGGTAACTCCAGTATTTCCATTTCTTTTTTAGACTTGCGAATGAACAATTTGCATGGAGAGATACCAAGATTCTTATCTACAGGGTTTCAATATCTTGGTTTATATGGCAATCAATTGGGGGGACAATTCCCGTGATCCTTGGTTAACTGTACAAGCTTGGAAGCTCTTGATTTGGGGAACAACAAGATAAATGACACGTTTCCGATATGGCTGCAGAAACTTCCAAACCTACAAGTTCTGATTCTGAAATCTAATCTCTTTCATGGTCCAATTGGTTATTTGGAGTCCGAGTTTCCATTTCCTGAGTTACGAATCTTTGATCTTTCCTTCAATGGGTTCACTGGAACTTTGCCATCTAATTTTTTCAAAAGTTTTAGAGGTATGATGGATGTGGATGAAGAAAAAACAGGAATTACTCGCTCCTACAAAGGGTCGACCAGGCCTTGTACAAGCATAAATCGATGTCGAAGAGATTACCTTTACCACTTTAGTTTGGTGATAAAAGGTAATGAGTTTGATATGAGAATCACGTCAATTATGACGAGTGTTGATATATCAAGCAACAGGGTTGAAGGATATATTCCAAATTCCATTGGAAATCTGAGCTCACTTGTGTTACTCAACCTGTCTCACAACAGCTTCCGTGGTCATATTCCTGCAGAATTTGCAAAGTTGCAATAGCTCGAAGCATTAGATCTCTCACGGAACAGACTCATTGGAGATATTCCAGATCCATTGTCGAGTTTGACATTTCTTGAGGTCTTAAACCTTTCCTACAATCATCTGGCTGGACGCATTCCTCTTGGGAAACAGTTCAATACATTTCCAAATGATTCTTATTGTGGAAATCCTGATTTATGTGGATTTCCACTATCAAAGGAATGTGGAAACAATAAAGAGTCACCACTTGAACACGATGATGGTGATGATGATGATTCATTTTTTGCGAGTGGGTTCACATGGCAAGCTGTGGTAATAGGTTATGGTTGTGGTCTGATATTTGGATTGTTGATGGGAGGTCTCATGTTCCTACTGGAAAAGCCAAAATGGTTTGTGAACTTTGCTAAAGATATTGCTCAACAAATTGCTGCTAAGAAGCAAACAAGGCAAAAGAAGAGACGCCAAAGACATGGTTTAAGAATGAACTAGCAATATACCAGGTATGCATTAAAACCTTTAGTTTTGAATCAGGAACAACACTTCAGAGGTTGTTGGATAACTATATGTCAACTGTTTTTATTTTTTCCTAAATATTCAAAACTTAATTTGTCTCTGCAGAGGTTGCTGGAGAGTTGGACAAGTTTATCAGCAAGTCAGTGCTGATGTTTGGCAAATAGCAATGGATGCATCTGTTTGTTTACTTTATGTCCATGGTTGCCAAATACGTATGTAAATGTTACAGTTTAGTTCTGTTTGTATGGTTCTTACATGTTGTTTTGTAACGTATTGTATTATATTGTATTGTTAGTACAACAATATGAATCCTGTATTATTTACTAGATGAGTCAGATCCATGCTTAGCACGAGCTCAACATTTAGGTGATTGAAATCTCTTTT

>M6-PGSC0003DMG400029597 M6 LRR-RLP

AGGGAAAATTGTATAAAATAGCAAATTATTAATTCAAATTAAATGTTATAACCACAGTTTGATTTAATTGTAACCCGTAGCAAATAGTTGTCATTCGCCTCTCTCCCTAAATTTCTCACTCGCCACTCTCGATTTCTCGCTGGGCAGTCTCTTCCTCTCCTCTCTCACTTTATACAAGCACAAATGTATAAAATGTGTTTGTGTTTGTATAAAGCGAGAGAAAACTGTATATATACATATCTTTTTGTTCGCCTCTCTCTCCAGAGAATCTCGCTCGCCACTCTCCCTCACCTCTCTCCTTTTATACAAACACAAATGTATAAATTGTGTTTGTGTTTGTATAAAGCGAAAGAAAATTGTATATACAAATACATATATTTTCATCCTATACACTTATAATTATACAAATATATATCTTCCCCTGCCCAGTTTTCTTTTGTCTTTCTCTCTTTCTCGCTTTATACAAACACATACTATACAATTGATTCTTTTGTATATATATGTATACCGAAATATACATATTAATATTCATATCAAACATAAAGTTTGTTATGGAGCGCAATTATACAAACTATAGTTATAATATACAAATATGATTTTTATGTTTGCTAAACCTATAATTTACT

>M6-PGSC0003DMG400030072 M6 LRR-RLP

CTAAGTCTCTTTTATCTACTAAAACTGCATCACTGGAAATAGTATATAATTACACTACTATCTGTGGAAGTTAAACCATATATAATATGACTTATTTGTGAACTGTGAGTTCAATTGGAAAGAAAAGTAAAGTGACCGCTTTTTCCTCACGCTCAAATATTGAGGAACCTTGGCTAATGCGAGGAGGAAAATCCATATATAAGTTTTAAATAGATTAGATGGGTGGAATATCGAGAAGAGTAATGAAACCTTTGTAGGAGTTAATGGGCTAGCTTTATCGATACTAGTGGTGTTTAAGTTCGCTGTCATCCTATCTTCCAGTGGTGGAGCCACGTTTACTGAAGAGAGGGAGGGGGGGGGGGTATAACTGAATACCATTTTGCCAAAAGATTATATTGTGTATATAGGGGAAATGTCATTTTATGTGTAGATATATACTTGTTGAATCCCCCGCACACAAGAGAACATTTTTGTGAATCCCTCAATGAAAATCTTGCCTTCACCACTATTGTCTTCAATTTATGTTCTTTCCCGTGCTCAATCATGTATCATTCTGTAATGACGTTGAACCACTAGTTATTTTCAATACATTACTGAAATGTATTGGGATATTGCCTTCATCCCTGATTTTAATTTTCAGAGTTGCTGACATGCTTGTTTTGTTAGACCATGAGGAATGCTTGGACATCTGGTGTAGATACAATTAGCAATTTGCCTTGTAATGTACTGGATGCTGTTCTTGGATGCTTGCCTTGGAAAGATGCAGTGAGAACCAGTATCTTGTCAAAAGACTGGAGGTACAAATGGGTAACACGTCAAGAACTTGATTTTAATGATGAGTTTTTCAAGTCTTTCAAACAAGATGAAGAAGCCAAGAGAATCATTTACCAAGTCCTATTAGTCCATTAAGGACCAATACTGAAGTTTAGACTCTGTAGGATTACGAGTTGTCCAGGAATTCACCCTTCATGTAAGCTTAGGAAACAAATATCATTCACCTCATCATCTTTTTACATTCCAGCAACTAAGGTTTTTGGAACTTCAAGATTGCTTGTTCCACCCTCCACTCGGTTTCAAAGGGTTTGAGAAGCTTATTAATCTCGATCTTGTACGTGTCACTTTTGATCCATCAATATTTACTTATCTCATCTCTAAATGCTCGTTGCTTGAACGGTTGAGGTTGCGTTGCTGCACAAACTTTGACATCCTTGAAATTGATGCCGCTAATCTCAAATTCTATGAGTTTATTGGAAAAACAAAATCCATTTCCTTCAAGAACGCTCCTATGCTTGAAAAAGTTACTGTGGCTTTCCTTGGACGACAACTTTTGACTGATACATCTCTTGTTTGCTCTAATTTCCCAAAGTTCTTCCATTACATGCCTTCACTGCTGGATATGGATATATGCGGTACAATATTAGAGGTAAGGTACATTTGTGATATTACTATCATAACTAAATGATGAATCTGATTGGTGTCTTAAAGTAATTTTTGACTTTCTTGTAGTACTTGATCTAGGGAGGTTTACCAGAGAGTCCCCCAACTGCTCTGAACAATATCAAATCTCTCACTATTTCGTCCGTGTCTTTAAGAAACGCCGAGGTGGTTTCAAGTGCAGTTTACTTGATTACAAGCTGTCCTAAATTACAAGATTTTACACTTGAATTTGTAAGCACTCTCTTTGTCTTTTTTTTTTCATTAAGGTGATTGAATATGGTTTCTTATTCATCTGAACTCGCCTAACTAACCGTTCCTCAAATATTCTTTAGTACCAAGTGTGGGTGGGGGATATTTATGAACCTGTTGTACAGTTGCTACGAGCCCAACCATCCTCAAGCGGTGTTGTGAAGCTTCAAAGAGTTCAAGTGAACATGTTTACTGGTCTCGAGATGGAAATGGAGATTATGAAGTTTATATTGGCATCTGCACCTATACTTGAGGAGATCTTCATTTGAAATTGTATGCGTTTCCTTTTTCGTTCGTGTAAACAAATGATCGATGAGATGAAAGAATTCCGCAGAGCATCAACCAATGTTGAATTCACATTTGAAGAAATCGATATGGAAGGTGGATATGAACATGAAGAAGTCATGGAAGTGCCATGATTTTGCTAACATATTATTTT

>M6-PGSC0003DMG400030149 M6 LRR-RLP

TGTTAGGATATACTATTAACCATGCGTTTTGATATAGTATTATTATTGAACAATTGTTTATTCATTTATTCAATTCAATAAAGTGTTACATTAAATAGATCAATTTTCATATATGTGTCCTTAACTTATATAGTAGATGATTTAGTGTATAGAGTTTAAGCTTATACACAGAAGATTAAATCATCGGTTCTTATAAGTTAAAAGTTATAGTTCATAATCGTAGATGGAAATTTTGGACAAAACCATCAGAAAGATTGTAGCACGAGATTAAATGTAATCTATCTTGATTATGAGAATGGTATAGTTCTAACTTCTCGTGCTAGTGCATTTTGTATGTATTGAACGGGACCAAGCAGAGATAGATGTTTTAGACTGACTGATAAAAACATATTCTCTAAACTGTTAAATGTACTTATATTCTTAATCTTGATATAACGATTATGATCTGTATATATTAATTATTGTTTTGATTTATTAAAAGGTGAGATTCTGAGATGGGTCAATATGCCTGGTAGATTGGATGATAATAATATATATTGGTGAAATAATTAGTTAGTTGATGGAATCCATGTCTCAGTATAGAGATTGATGATACACCTTTTTATGAGAAACTTATAAGTTTTCATGTGTAAACCCGGCCGGTAGATTTATGAATCTGGCACATGAAATAAGTTAAGTGTTGCTCTAAAGGAAATTAATCATTGAATTAAATTCGTCAGTAATTTAATTTATTGATTAGTATCTGGAATCTTAACATGGGGAATTACATAAGTGTTTAATGGTGAATTTCGAAATATAAATAGAGGAGTGCAATTACGAATTTTTAGTGGAATGATTTGTAATTTATTATGGTAAGAATAATTCAAATTAATTATTTGGAATTATTTCCATAATAGGAAGCCTCAGTAATAAATTCTGTGGTCCCTCCAGTGCCCAATTTAACTAGAACTCAGAATCCTATTTTCTAGTAGAAGAAGGAAAAGTAACGTGTATATTTTCCTCCAAGAAAGAAACGTGAGTTTTGGGGTTTTCTTGAAGAATAAGGATTTATACTTTCTTCTCCTTTTTGGACTGGGAAGACATCTCTATAAGTAGGGATTCTTCTAACCTAGAAAGAGATTTCAGTATTCTATACGATACTTGCCCACCCAAGTATTGAGAGAGTTCAATTGTTATTTGGGATCACTGTAGAAGACTGTAGAGCTGGAGTTGGATTTGCTTCAAGATATCTGCACACGCTTCAAGAGGTAATCCTGAGTTAATTTTCAGCATACCTTGTATGTGATACGTATTTGTGATTGTTGTCTATACTCATGCAAGATGTATGATTCTGGAATGCTTCCGCTGTGTATGCTATTTTCTAACAAT

>M6-PGSC0003DMG400030159 M6 LRR-RLP

ATATGGGCAACATTAAGTTTTTGTTGTTAGCTTTTTTTCTAATAGTAAATGTGGCGAATGGTTGTTGGGAGGAAGAGAGAAGTGCTCTCTTGGAGCTGCAAGCAAACATAATGAGTTTGAATGGTGAATTGGTTGTTGATTGGGCAGGATATAATGCCAATGGTTTTATAAATTGTTGTTTTTGGAAAAAAGTTAAATGCAGCTTAGCAACAGGTCGAGTGATCGAACTCGATCTTGCAGCAAGGTTTGGATCAGGGGATGATTGGAGATTTAATGCCTCTCTCTTTCTACCGTTCAAGTCCCTCCAACTTCTTTTTTTGTCTTACAGAAATATTA

>M6-PGSC0003DMG400030239 M6 LRR-RLP

TTGCAAATTGCTCTAGCATTCAACAGGTAAATTCAATCTCTTAATTCCTAGCAAAAATAGACAACCATCTTTAATTTTTTCCATTCTTGTCCATTTTTTGATTCTTTGTTTTGTTTTTCAGATTTCAGAAATGGATATTGGCTTGGCAGTTGGTGGTGCATTTCTCTCTTCAGTCCTGCAAGTCCTATTCGATAGGCTTGCTCCTCAGGGAGAGCTGCTGAATATGTTTCGAAGGGACAAGCACGATAATCGGATTTTAAAGAAGCTGAGAATGACTTTGCTCGGTCTTCAGGCTGTGCTAAGTGATGCACAGAATAAGCAGGCGTCAAATCCATATGTGAGCCAGTGGCTTAATGAGCTACAAGATGCTGTGGACAGTGCTGAGAACTTAATTGAAGAAGTCAATTATGAAGTTTTGAGACTAAAGGTGGAAGGTCAGTGTCAAAATCTAGGAGAAACAAGCAGGCCGCAGGTAAGTCACCTCAGCTTGTCCTTGAGTGATGATTTTTTTCTTAACATAAAGGATAAATTGGAAGAAAATATTGAAACACTGGAGGAGTTGCAAAAACAGATTGGTTGTCTTGACCTGAAATCGTGTCTTGATTCGGGTAAACAAGAAACTAGAAGACCTTCAACTTCTGTGGTTGATGAATCTGATATCTTTGGTAGGCAGAATGAAATAGAGGAATTGGTGGGCCGTTTATTATCTGTTGATGCAAATGGTAAAAGTTTGACTGTAATCCCTGTTGTTGGAATGGGGGGCGTGGGCAAGACAACACTTGCTAAAGCTGTTTACAATGATGAGAAGGTGAATGACCATTTTGATTTGAAAGCTTGGTTTTGTGTTTCTGAACAATATGATGCTTTCAGAATAGCAAAAGGATTGCTTCAAGAGATTGGCTTACAGGTTGATGATAATATAAATCAGATTCAGATCAAATTGAAGGAGAGCTTAAAGGGTAAAAAGTTTCTTATTGTTCTAGATGATGTTTGGAATGACCACTATAATGAGTGGGATGACTTGAGAAATCTTTTTGTGCAAGGAGATCTCGGAAGTAAGATCATTGTGACAACACGTAAGGAGAGTGTTGCCTTGATGATGGGTGGTGGGGCAATGAATGTGGGGATTCTGTCTAATGAAGTCTCTTGGGCTTTATTCAAACGACATTCATTAGAAAACAGGGATCCTGAGGAACATCTAGAACTTGAAGAGATAGGAAAGAAAATTGCAGAGAAATGCAAAGGGTTGCCTTTAGCTATAAAGACCCTTGCTGGGATGTTACGCTCAAAATCAACGATTGAAGAGTGGAAACGCATTTTGAGAAGTGAAATATGGGAGCTGCCAGGCAATGGAATGTTACCAGCATTAATGTTGAGCTACAATGAGCTTCCCCCACATTTGAAGCGATGCTTTTCCTATTGCGCAATATTTCCAAAAGATTATCAATTTTACAAAGAACAAATCATTCAACTGTGGATCGCTAATGGTCTAGTACAGAAGTTGCAAAATGATGAAACGGTTGAAGAACTAGGCAACCAGTACATTCTTGAGTTGAGATCAAGATCATTGTTAGACAGGGTCCCTGATTCTCTTAAATGGAAGGGCATGGCATTCTTAAGAGAGGACTTAGACAAATATTTTCAAATGGATGGTGAGAAATTCTTTATGCATGACCTTGTCAATGATCTTGCCCAAATTGCATCTTCAAAACACTGTACCAGGTTGGAAGACATCGAAGGATCTCATATGTTGGAACGAACTCGACACTTGTCGTATTCCATGGGTGATGGTAACTTTCTGTCACTTTCAGCAGGAGATGGTGATTTTGGTAAATTGAAAACCCTCTACAAATTGGAGCAATTAAGGACATTGCTTCCGATAAATATCCAGCTCCATTTGTTTTCAGTTAATCTAAGCAAGAGGGTGCTGCATAACATACTTCCAAGGCTAACATTTTTAAGGGCATTGTCATTTTCTCGTTATGACATCAAGGAGTTACCAATTGTCTTATTTATCAAATTAAAACTCCTAAGATTTTTGGACCTTTCTTGGACAAAGATAAAACAGTTACCAGATTCAATTTGTGTGTTGTACAACTTAGAGACACTTATCGTGTCGTCTTGTGATTATCTTGAGGAGTTACCGCTGCAGATGGGAAATTTGATCAACTTGCGTTACCTTGACATTAGGTGTTGTTCTCGGTTGAAGATGCCTCCACATCCTAGCAAGTTGAAAAGCCTCCAGGTGCTATTAGGAGTTAAATGTCTTCAAAGTGGTTTGAAACTCAAGGATTTGGGTGAACTACACAACTTGTATGGATCTCTATCAATTGTAGAGTTGCAAAATGTGGTTGATAGAAGGGAAGCTTTGAAGGCAAACATGAGGGAAAAGGAACATATTGAGAGGTTATCATTGTCGTGGGGTACAAGTATTGCTGACAATTCACAAACTGAAAGAGACATACTCGATGAACTACAACCACACACAAACATAAAAGAACTGGAAATCTCTGGATATAGAGGGACAAAATTTCCGAACTGGTTGGCTGATCATTCCTTTCTTAAGCTAGTGAAGTTGTCTCTTAGCAACTGCAACTACTGTGATTCCTAGCCAGCACTAGGACAACTTCCTTCTTTGAAATCCCTCACCATTGGGTATATGGATCGAATAACAGAGGTGACCGAAGAATTCTATGGTAGCCCATCTTCAATAAAGCCATTTAACTCTCTGGAATGGCTTGCATTTAATCAGATGACTAGGTGGAAGGAATGGCACGTACTAGGAAGTGGAGAGTTCCCTGCACTTCAGAACCTTTTAATTGAAGACTGTCCTGAGTTGATGGGGAAGTTGCCTGGAAATCTTTGTTCTTTAACAGGATTGAGAATTTCAAATTGTCCGAAATTCAATTTGGAGACACCAATTGAACTTTCAAGTCTAAAAGGGTTTGAAGTTGTTGGTTCTTTAAAGGTTGGAGTTCTTTTTGATCATGCTGAGCTATTTGCATCCCAACTTCAAGGAATGATGCAGTTAGAGTTCTTAATTATCAGAAGGTGTGACTCTCTTACCTCCTTGCATATTAGCAGTCTGCCGAAAACCTTGAAGGAAATAGTGATATCTGATTGTGATCAATTGAAATTGGAGGCGTCAGCTAGTGAGATGTTTCTCGAGAGACTGGACCTAAGTGGATGTAATTCTATAAATGAGATATCACCTGAGCTGGTCCCCAGAGCACACTATTTGAGTGTAAATAGTTGCCACAGCCTTACTAGGCTTTTGATTCCTACTGGGACTGAAGATCTCTATATTTCTGGATGTGAAAATCTTGAAATACTTTTGGTGGCATCCAGGACTCCAACGTTGTTGCGAACTATGAAAATTCATATCTGCGAGAAGCTGAAGTCGCTGCCAGAACATATGCAGGAACTTCTTCCATCTCTTAATCATCTCTACCTGTATGATTGTCCAGAAATAAAGTCCTTCCCTGAAGGAGGGTTGCCCTTCAGTCTAGTAGTCCTTGAGATCTGGTATTGCAATAAACTACAGAATGATCGAAAGGAGTGGCATTTACAGAGACTCCCTTGTCTCAGAGAACTTCAAATCTATCCCGGTTCTACTGACGAAGAGATTCAATGGGAGATGCCTTGCTCAATTCAAAAACT

>M6-PGSC0003DMG400030240 M6 LRR-RLP

GTTGTCTCTTAACCACTGCAACAACTGTGATTCCTTGCCAGCACTAGGACAACTTCCTTCTTTGAAAGTCCTCACCATTGGGTATATGGATCGAATAACTGAGGTGACCGAAGAATTCTATGGTAGCCCATCTTCAATAAAGCCATTTAACTCTCTTGAATGGCTTGAATTTAATCGGATGAACGGGTGGAAGAAATGGCACGTACTAGGAAGTGGAGAATTCCCTGCACTTCAGAACCTTTCAATTAAAGACTGTCCCAAGTTGATGGGGAAATTGCCTGGAAATCTTTGTTCTTTGACAGGATTGACAATTTCAGGTTGTCCGGAATTTATTTTGGAGACACCTGTCCAACTTTCAAGTCTAAAATGGTTTGAAGTTGTCGGTTCTTTAAAGGTTGGAGTTCTTTTTGATCATGCTGATCTATTTGCATCCCAACTTCAAGGAATGATGCAGCTAGAGTCATTAATTATCAAAAGTTGTCACTCTCTTACCTCCTTGCATATTAGCAGTCTGTCGAAAACCTTGAAGATAATAGAGATACATGATTGTGAGAAATTGAAATTGGAGCCGTCAGCTAGTGAGATGTTTCTCGAGAGACTGGTACTAACTGAATGTAATTCTATAAATGATATATCACCTGAGCTGGTCCCAAGAGCACACGATGTGAGTGTAAATAGGTGCCACAGCCTTACTAGGCTTTTGATTTCTACTGGGACTGAAGATCTCCATATTTCTGGATGTGAAAATCTTGAAATACTTTTGGTGGCATCCAGGACTCCAACGTTGTTGCGAACTATTAAAATTCGCAGCTGCAAGAAGCTGAGGTCGCTGCCAGAACATATGCAGGAACTTCTTCCATCTCTTAATCATCTCTCCCTGGAAAGTTGTCCAGAAATAAAGTCCTTCCCTGAAGGAGGGTTGCCCTTCAGTCTAGAAGTCCTTGAGATCTTGTATTGCAATAAACTGGAGAATGATCGAAAGGAGTGGCATTTACAGAGACTCCCCTGTCTCAGAGAACTTATAATCTTCCATTTGGGTAGTGACGAAGAGATTCATTGGGAGTTGCCTTGCTCAATTCAAAGACTTATCATAGACAATATGAAAACATTAAGCAGCCAACTTCTCAAAAGCCTCACCTCTCTTCAATATCTATTCACTTATACTTTACCTCAAATCCAGTCACTGATCAAAGAAGGGCTTCCCTCGTCTCTTTCTTCACTAACATTAAACAACCATCATGAGCTCCATTCACTGTCAACTGAAGGTCTTCGGGGCCTCACTTCTCTTAGGTGTCTAAAGATTGACTCCTGCAGTCAACTCCAATCTCTTCCAGAGTCAGCGCTGCCCTCCTCCCTCTCTGAGCTGACCATCCATTCTTGTCCTAAACTTCAACATCTTCCAGTAAAAGGGATGCCCTCTGCCCTCTCTAAGCTATCTATTTCAGACTGTCCAATGCTCAATCCACGTCTAGAATTTATGAAGGGGGAATACTGGCCAAATATTGCTCATATTTCCACTATACAGATCAATGGGGAATACCTGTGA

>M6-PGSC0003DMG400030496 M6 LRR-RLP

ACAAAACCTAGAGAGAGAAAATGGCCTATGCTTGTATTTCCTCACTTATGCAAACTCTGCAGCAACTCCTGCAAGCTAAATCACCTTTGATTTGTGAAAGTTCTATACAACAACATGTTGAATCTTCTTATCAAAGTCTATGTTCTCTTCAGGTTTTTCTTGAGGATACCACAAATGAGGCCAATGATATTGAAAATCTCCAGATTTTAGAAAAAAAGATCAAAGATGTAGTCTACAAAGCAGAAGATAGAGTTGATTCATGCTTAACAAACATGATTCTAGCTGATAATGAAGACGACCGAGAAAAAGCTTGTAAATTCTTCAATGAAGAATTACAACAAGTTGAAACAGAATTTGATTCTCTCAGGAAAGAGGTGATGGTGATCGAGTTCAACAAGCGTGGAAGCAAATCAACGGAGTTAGCAGCAGCAACAACTTCTTTCTCATCAATCGAGGAAACGACTTATGTTGGGATGAAAAAAGACTACAAAGCCATACTGAATTGCCTCAATGCCCAAACAAAAGAGCTAATAGTCATATCACTTGTTGGTATGGGCGGTATAGGTAAGACAACTCTTGCTAGAAAAGTTTTTGATAACTCAACTATTCGTGATCGATTTGAGAAACATGCATGGGTCACCATATCTGAACACTATAATAAGAGACAAATGCTTCTTGAAGTTGCTTCTTCAATTAGTGGAGTCAACAATCAAGAAATGAGCAATGATGAATTGATGGTGATTGTGTATAGAAGTCTCAAGGGTAGGAGATTTCTAATTGTCATAGATGATCTTTGGAGTACTGATGCTTGGGACCAAATGCGAAGAATATTTCAAAATGATAACAATAAAAGCCGAATTATACTAACCACTCGTCTCAAACGTGTTGCTGATTATGCAAGCTCCCCTGATTTTCCTCCTCATGATGTGTCTTTTCTAAGTTTTAAAGATAGTTGGAAACTATTTACCAAAAAGCTATTCAAAAAAGATTGTTGTCCTCCTCAACTACGAAAAATAGGGAAGCATATCATAGAACAATGTCAAGGATTACCTCTCTCGATTATTGTCATTGCTGGACTTCTTGGTAAGATAGGCGTGACATATGATAATTGGAAGAAAATTGAGGAAAATTTGAAGTCATTTCATGGTTCGGTTTTAGAACAGTGTCAAGCAATTCTTTCTTTGAGCTACGATTACTTGCCCCCATACTTGAAGGCGTGCTTTCTCTATATTGGAGGTTTTCCTGAAGATATGGAGTTTCGTGTATCGAAGATGATGAGCTCATGGATAGCTGAGCAATTCATAAAGGCAACAAGCGATAAAAGGTTAGAAGTTGTGGCAGAGGAGTATGTGCAAGAGTTAATGGATAGGAGTCTAATTTTGGGTAAAACACGAAAGCCTAATGGAAGATTCAAAACTTGCAAAATTCATGACCTTCTTAGGCAGTTGTGCATAAGAGAAGCTCAAATTCAAAATGTTGTGCATTTCCCTTACAGTGATCATGTTCTCACTTACTCAAATGACATAAATGATCGTAGACGTGTGATGATTCCTTTTCTGATTGAGGATTATTTTTGCGATCATCCAAGTCAAATGAGTGGTAAGATTACAACCCGCAGCTTGATCTTTATGGGAGCTGGTAAATCTTATCTAGTACCTGAAATAAATCACTGGCCTGATAGTATTTCAAACTTCAAGCTACTTAAGGTGTTGGATGCACGTGAAATTGGCTACGATTTCTCTCATATAATACCTCAACTTAACTTGTACATTTGAGGTATATTGATGCAAGAATTGAGGATCCTTCTTCACTAGCCAAATTGTTCAATCTACAAACCATAATTGTCTATTCAAGGAGAAATGTGCAGCTCCCCGCAGAGATTTGGACAATGTCACAGATAAAACATGTGGATATTGAACAAATCTATATGCCTAATCCTCTTAGTATTGAAGAGCAACAACCTTTGTTTCTGAATAACTTGCAAACACTTGCTCTCTATTCCTCGCCTTTTTTGGCAGAAATCCTAAAGAGAACACCCAATGTAGATAAGCTAAAGATTAGAGCTGCTAATACACATAATGAGTGGTCTGATTTTGTTGATTGTCTCATTAATCTACAGAGACTGGAAAAGCTAAGTATAACAACAACAGTAGACAACAGTCCGTGCATTCTGTCTAGTGCTTTTTGCGCGCCTAATCTAAAGCAAGTGAGATTAGCTTATACTTCATTGGCTTGGGAAGAAATGGATGTGCTTGCTAGTTTACCCAATCTTGAGGTGCTCAAAGCAGATAGTGCATTCAATGGAACAGATTGGACACTAAGTGAAGATGTTGTTTTTCATAAACTAAAATATCTACGAATTTGGAGCGCAGATGATCTGGAAAGATGGGAAGCAACTAGTGATAATTTTCCAATGCTTGAGCAACTAGTCCTTCAGCTTCTCGAAGAACTAGAGGAGATTCCACAAAGTTTTGGAGAAATAATGACATTAAAATTAGTCCAAGTGGATTATTGCAATCCTGCAGTCGACACTAGTGCAAGGAATATTCTAGAAGATCAACAAGGCTGGGGAAATTGTGAGCTTCAAATTCAAATTGTACCTTCGTTAAAATACTTAATTTAATAAATTCATGTGATGTTTTCAGTTTTGTGTCATTTCTTGTTTTCACCT

>M6-PGSC0003DMG400030497 M6 LRR-RLP

GAAAAGTAGCAAGTAGTAATTGATTTTATTTATGTGCTTCATTATTCATGCTTTCATCTACTTTTGACTAACAAAACCTAGAGAGAGAAAATGGCCTATGCTTGTATTTCCTCACTTATGCAAACTCTGCAGCAACTCCTGCAAGCTAAATCACCTTTGATTTGTGAAAGTTCTATACAACAACATGTTGAATCTTCTTATCAAAGTCTATGTTCTCTTCAGGTTTTTCTTGAGGATACCACAAATGAGGCCAATGATATTGAAAATCTCCAGATTTTAGAAAAAAAGATCAAAGATGTAGTCTACAAAGCAGAAGATAGAGTTGATTCATGCTTAACAAACATGATTCTAGCTGATAATGAAGACGACCGAGAAAAAGCTTGTAAATTCTTCAATGAAGAATTACAACAAGTTGAAACAGAATTTGATTCTCTCAGGAAAGAGGTGATGGTGATCGAGTTCAACAAGCGTGGAAGCAAATCAACGGAGTTAGCAGCAGCAACAACTTCTTTCTCATCAATCGAGGAAACGACTTATGTTGGGATGAAAAAAGACTACAAAGCCATACTGAATTGCCTCAATGCCCAAACAAAAGAGCTAATAGTCATATCACTTGTTGGTATGGGCGGTATAGGTAAGACAACTCTTGCTAGAAAAGTTTTTGATAACTCAACTATTCGTGATCGATTTGAGAAACATGCATGGGTCACCATATCTGAACACTATAATAAGAGACAAATGCTTCTTGAAGTTGCTTCTTCAATTAGTGGAGTCAACAATCAAGAAATGAGCAATGATGAATTGATGGTGATTGTGTATAGAAGTCTCAAGGGTAGGAGATTTCTAATTGTCATAGATGATCTTTGGAGTACTGATGCTTGGGACCAAATGCGAAGAATATTTCAAAATGATAACAATAAAAGCCGAATTATACTAACCACTCGTCTCAAACGTGTTGCTGATTATGCAAGCTCCCCTGATTTTCCTCCTCATGATGTGTCTTTTCTAAGTTTTAAAGATAGTTGGAAACTATTTACCAAAAAGCTATTCAAAAAAGATTGTTGTCCTCCTCAACTACGAAAAATAGGGAAGCATATCATAGAACAATGTCAAGGATTACCTCTCTCGATTATTGTCATTGCTGGACTTCTTGGTAAGATAGGCGTGACATATGATAATTGGAAGAAAATTGAGGAAAATTTGAAGTCATTTCATGGTTCGGTTTTAGAACAGTGTCAAGCAATTCTTTCTTTGAGCTACGATTACTTGCCCCCATACTTGAAGGCGTGCTTTCTCTATATTGGAGGTTTTCCTGAAGATATGGAGTTTCGTGTATCGAAGATGATGAGCTCATGGATAGCTGAGCAATTCATAAAGGCAACAAGCGATAAAAGGTTAGAAGTTGTGGCAGAGGAGTATGTGCAAGAGTTAATGGATAGGAGTCTAATTTTGGGTAAAACACGAAAGCCTAATGGAAGATTCAAAACTTGCAAAATTCATGACCTTCTTAGGCAGTTGTGCATAAGAGAAGCTCAAATTCAAAATGTTGTGCATTTCCCTTACAGTGATCATGTTCTCACTTACTCAAATGACATAAATGATCGTAGACGTGTGATGATTCCTTTTCTGATTGAGGATTATTTTTGCGATCATCCAAGTCAAATGAGTGGTAAGATTACAACCCGCAGCTTGATCTTTATGGGAGCTGGTAAATCTTATCTAGTACCTGAAATAAATCACTGGCCTGATAGTATTTCAAACTTCAAGCTACTTAAGGTGTTGGATGCACGTGAAATTGGCTACGATTTCTCTCATATAATACCTCAACTTAACTTGTACATTTGAGGTATATTGATGCAAGAATTGAGGATCCTTCTTCACTAGCCAAATTGTTCAATCTACAAACCATAATTGTCTATTCAAGGAGAAATGTGCAGCTCCCCGCAGAGATTTGGACAATGTCACAGATAAAACATGTGGATATTGAACAAATCTATATGCCTAATCCTCTTAGTATTGAAGAGCAACAACCTTTGTTTCTGAATAACTTGCAAACACTTGCTCTCTATTCCTCGCCTTTTTTGGCAGAAATCCTAAAGAGAACACCCAATGTAGATAAGCTAAAGATTAGAGCTGCTAATACACATAATGAGTGGTCTGATTTTGTTGATTGTCTCATTAATCTACAGAGACTGGAAAAGCTAAGTATAACAACAACAGTAGACAACAGTCCGTGCATTCTGTCTAGTGCTTTTTGCGCGCCTAATCTAAAGCAAGTGAGATTAGCTTATACTTCATTGGCTTGGGAAGAAATGGATGTGCTTGCTAGTTTACCCAATCTTGAGGTGCTCAAAGCAGATAGTGCATTCAATGGAACAGATTGGACACTAAGTGAAGATGTTGTTTTTCATAAACTAAAATATCTACGAATTTGGAGCGCAGATGATCTGGAAAGATGGGAAGCAACTAGTGATAATTTTCCAATGCTTGAGCAACTAGTCCTTCAGCTTCTCGAAGAACTAGAGGAGATTCCACAAAGTTTTGGAGAAATAATGACATTAAAATTAGTCCAAGTGGATTATTGCAATCCTGCAGTCGACACTAGTGCAAGGAATATTCTAGAAGATCAACAAGGCTGGGGAAATTGTGAGCTTCAAATTCAAATTGTACCTTCGTTAAAATAC

>M6-PGSC0003DMG400030561 M6 LRR-RLP

TGAAGTTCTTAAAGCTAAAACCCCTCAAGAAAACAGAACAATAGCACTATACTCAAATTAAAAAAAATGAACTACCATCCCTTTTTGATTCTGCTACTTCTTGTTTTTGTCACCAATCTTGAAGTTGGTTTCACTAAAACTCATCATTCCAATACAACTTGTTCTGAAAATGACCAAAAAGCTCTGATTGAATTCAAGAATGGCCTAATCGATAACTCGAATCGTTTGTCATCATGGACAGGACAAAATTGTTGCAAATGGGATGGAGTTTTTTGTGATGAAAAAACAGGAAATGTTGTCAAAATTGATTTGCATTACAAGAATTATCTTCTAGATGGTGAATATCCACAAAGGGTGTTAGATAATCACACTAAAAACTTCTTGCGAGGTGAGTTAAGTCCTTCTTTGGTTAAATTGAAACATTTGAGTTACTTAGACTTGAGTCACAACTACTTCTCAAGTATCCAAATTCCACCTTTTTTTGGATTGTTAATGAACTTGAGATTTCTTAATCTTTCCTGTGCTGGTTTTGGTGGATATATTCCCAAAAATCTTGGGAATTTGTCGAGATTAGAACATCTTTATCTTGGTGGATGTTATGGCTACGTTATCGGACCAGGAAATGATCTCAGAGTTGATAGTTTGAGTTGGATATCTAAGTTTTCTTCTTTGAAAAGTTTTGATCTTTCTCAATTGGTTATTGAAGATACTAAAGATCTTTGGAATTCAATTAACATGTTGCCCTCATTACTTTCATTGAACTTAGAAGGTTGTAACCTCAACATTTTGCCTTCTTTTAGACAAGTTAATTTCACATCTCTTAGTTCACTCAATATCCATGATAACGCGATTGGAATGGGCTTTGAGAATTCCTCCATCATCCCTCCATGGTTTTCTAATCTTACAAGGCTTATGAATCTTGATCTTGGTCAAAATAACTTTGATGATCCAACTAATGTTCTTGAGAAGCTTAGTTCACTAAGAGTATTAGATGTTTCTAGTAATGGATTTGGCGATTCGTTGCTGAGTTCATTGAGTAAGTTGAACAACTTGGTTTATTTGGATTTGTTAGGGAATGGCATCCACTTCTCTAGTCTCCATTTACTTGGAAATCTGACTTCACTTTCTGTACTTAAGTTGAGATATAACGTGTTAAAAGGGCTGATCCCGGAGGGTTTAACGAGCATTTTCTGTCAATTGAGTGTGCTTGATTTGTCAGACAATGAAATCAGTGGACTGCTTCCAACTTTTATAAGAGATCCATTGAGCTGTCTCGAGAATCATTTGAAGGAGCTATACATAGGAAATACAAAATTCAGTGACTTCTTTCCTGAAGGATTGACAATGCACAAGAATCTTGAAGTTATTGACAGTACTAATAGCTTAATATATGGTGAAATTCCATATTCAATAAGCAAGCTGTCAAATTTGAGGTTTCTAAGGATAGCAAACAACAAGTTGAATGGGAGCATTCCTTCGAGTATTGGACAGTTATCGAATCTTGAAGAGCTAGATATCTCGGAGAACTTATTCACCGGTATTGTTTCTGAACTCCACTTTGTAAAATTAAGCAAATTGATGAAATTGCACTTGTCTAAAAACGAAAACTTGGTTTTTAATGTGAGTTCCAATTGGTATCCAACTTTCCAAGTCAAAGAGATTGGGATGGCTTCAGTAAAAGTAGGGCCTGCTTTTCCTCAATGGCTACAAAAGCAGTCAATTCTTGATACTCTTATTATGTCTGATGCTAACATTTCTGATATGATCCCGAATTGGTTGAGCAATATAACGTTGTTCATGACAACTCTGGATCTCTCTGTTAACAAATTAGTCGGTGGTATAAGCGTGTTGTGTGATGCTCAACTTTTGAAATCGATTGATCTTTCAAAGAATCTGTTATCTGGACGAATCCCTTCATGTTTGAGTTACTTGGAAGAGTTGAGATCATTAAATTTGGCTGATAATAGTCTTGAAGGTGAAATCCCAAGTTCTTTAGGTGATTTGCCACTTAAGTTTTTGCATTTGCAAAAGAACAACTTACAAGGTAAAATCCCTTTGTCTTTTCAAAATTTGACATGGTTAGAAAGACTAGACTTAGGAGAAAACAAGTTGAAGATGTTTTTCCTACATGGATTGGCGAAAAGCTACAGAGTTTGGAACTCTTGAGGCTAAATTCTAATCAATTTTACGGTGTTATCCCATTAGAACTATGTCAAATCTCATCTCTATGTTGGCTAAATCTTGCTGGTAATAACTTGTATGGAACTATTCCGAGTTGTTTTGGAAACTTTTCGTGTGGTAATTTTCGTGGACCTGGCGAAGTATTTGGACAAAGAGTTGAAAGTTTTATGAAAGGAATACCACTTGAGTATGTTGGTGAACAAATTTTGTTACTTAGAATTTTGGATCTTTCTGAAAATAAACTTGTTGGAGGTATCCCCAAGGAGTTAACAAAATTGGTATACTTGCAATACTTGAATTTGTCTAGAAATAGTCTTAATGGGAGCATACCAAAAAAGATAGGTGATTTGAAGCAATTGGAGTCACTTGATTTGTCACATAACAAACTTTCTGGTTCAATTCCTCAAAGTTTAGCTTCTTTGAATTATTTGAGCTACATGAACTTGTCATATAACAGTTTTTCAGGTCCAATTCCTACTGGAAATCAGCTCCAATCTTTGGATGATCAATCATTTTATGTTGGTAATCCTAGACTTTGTGGAATATTGATCAAGAAAAGTTGCAATGGCAATGGAACATCTCATGTAAATGAACAAACTCCTCAAGGTGATGATCATGATCAAGATGTTGATGAACATAAGTGGTTTTATGCTGGAATTGCACCTGGTTTTTGTGTTGGCTTCTTGGGTGTCCTCTTCATTTTGTACAAGACAAGGAATCGTGGCGTGGTGCACGTTTTCCAATGCTTGGCTCGACGTCTATGACAAGTCGTGTATGTACTCCTATGAGAGGTTGTCTGGTTGACAGTTGACACACAAGTTTTGCAGAGATTAGAGTGTCAGAATTGTAATGCAGAACATTATAATGTAAAGATTGTAACATATATATATTTTTTTCGAACTTCATTGTACTAAAAACATAAATATTTTTTCTATAATGGCAAGCTTGTGCATGAATGTACATCTGTCTCTCATCGAGTTGCTGATCATGATCTCTCTAGACTCGAGAAGAGACTCATCATGACCTGAGATTTTGGATCTGTCACAACCCAAA

>M6-PGSC0003DMG400030790 M6 LRR-RLP

TTAGAATAGGAATAGGTTTATACAATCCTATTAGAATAGGAATAGGAATAGAAATATAAATAGGAATAGTAAATAGTATCCTACTTGGTAAAGGATTGTATTCTAGTGTCTATAAATAGGGTCTCAATGTAATAATGTAAACGACAACAATTCAATAATATTCTTTTTCAATATTTCTCATACAATCTCGAATGCGAAAGAACAAAAGAAACCCTATAATCTAAGAACAAGGAAGTAAATGATAGAAAGCTAGACACAAGAATGGAACGAATAGGCAACAAGGACTATATTAAAGTCAAAACCTCCTATTAAGAATACCAAACAAGCACCTAAACTATAGAGACACCCTCAAGAGGAATGATCCTCAAAAAAGCAATCCCATCAACATTTTCTAGCACCCTAACAAAAAAAAGTTTCACATTTAGTGGTAATATAAGGGACTCCCGTATTTGTTAGGTTCACAAGCGAATTTCCTCTGCCTTCTGTTTGTGAGACTTGGGTTATGCTTTTTTCAAGTTTGTGATGCCAAAACAGTATTTTCTAAAGGTTTGTGTGTTATATTATTTGTTAGGCTCACAAGTGAATTTCTGAGCTTGTACCGGGAAGCTACATTCATCTTGAAAGTTTAAACTTATTGTTTCTTTTGTATACCCGTTGCAGGGAAGAAGGCGTGGCACTGATCATTCCGTGATTTCAGCTCTCAAGATCCATTCCCTGGCTGAAACAAATGGCTCGGAAGACAATGAAACAGAATCTTACTGTTTGTCAGATGCTGGGTTGGCTGCTGTTGCTGCTGGCTTCAGTAAGCTTGAAAAGCTGAGCTTAATATGGTGCTCCAATGTGACACATGTGGGGCTGAGGTCTATAGCTGAAAAGTGTATATTTTTGAAGTCTTTGGATTTACAGGTAAATATCTAGTCATTTAGAACTTTGGTTTCCTCATGGTTATTTTATTGCTCTGTTTCAAGCTGTTTTGGAATTGAACTATATGATGTGCATGTTATCAAAGGTCTGAGAAGCTTTAGTTTCCTGTACCTCTTAAAGGGCTGTTATGTTGGAGATCAAGGTCTAGCAGCTGTTGGGGAGTTCTCTAAGCAACTTGAAGATTTGAATTTGAGGTTCTGTGAAGGCCTAACTGATGCAGGCTTGATTAAATTGGTAGATGGTTCTGGGAAGACGTTGAAATCTATTAGTTTGGCTGCCTGTGCAAAAGTAACTGACACCTCCTTGGAAGCTGTGGGATCTCATTGTAGATCTTTGGAGTCCTTGTCACTAGACTCAGAGTGTATCCATGATAAAGGTGTACTTGCTGTTGCCCAAGGATGCCCTCAGTTGAAGGTTCTGAAGCTACAATGCGTCAATGTTACAGATGGTGCTCTCCAAGGCGTTGGCACTTGCTGTTTGTCGCTGGAGTTGTTGGCTCTTTACAGTTTCCAAATATTTACGGACAAGTTAGTTGCTTATTCTGGTTGTCTTCCCTATTTGGATTTATTTGTTTTTAACTTACCAGCAGCATGATCGTAACTTCTATATTCATGTAAGCTTTTGTAATGTACATCTACATTTTAGGCATGAATACAATTCCTCTTAACCAATAGCTTTTCCAGTGTTCAGATTACGCAATCTAAATACCTAATCTTTAAATTTTCAACTCCATCTTTGCTAACAAAAATAATTCCTGAATATAGAGAACAGATAGTAACTTGATTCAAACTATGTGTTGCTGTCTGTCTTTTTATGGCAATAATTGCAAGCAAAAATTTAACAAAGCATGTGCTGGGATGATTGGTTTTCACAATCTGACCTATATCTTTATGCATATTTTTTCAGGAGTCTGTATGCTATTGGTAAAGGGTGCAAACGGTTGAAGTCTCTTACCTTGAATGATTGTACTTTCTTAAGTGACAAAGGCTTGGAAGCAGTTGCAGTTGGTTGCACTGGGCTCACGTATCTTGAGGTTAATGGCTGCCACAATATAGGGACATATGGACTGGAGTCTATTGCAAGATCTTGCACGTACACTTTTCTCCTCGTGCTTTATTTTGAATTTCTTTTCAACCTTTGTGTTGAATTTTAGTTATTGAAGGCAAAAAGACCTAGTGCTTACTTTTCCCCTTTCTCCTTGTTCTCTGAATTCAAGATGGATTCTTGTAAACTGCTTATATTTTTAAGGTCGTATGCATGAGTGGTGGTGCATTTGAGGAGAGTATTTAGCTTGAGAAGCACTTATGTGAGTATCTAGTAAATGTTTAGAATTACATTTAGAGGCTTCAAAAAAAAAAAAGCATTAAAAGAAGCAGCTCAAAATTGGTGCGTATAGAACTACTTTTGCACCTAAGATTTTAGGCGCCATGTTAAAGGAAAATATGCTATTCTATCAAATAGAAATGCTTGCTTGAGTTTAATTTGTATAAATTAAACAGAAAAAATGCTTCATAGAACAAGTATAGCAGATTATGTAGGGAGTGCTTATATTAGATTTTTAACGTGATTTTCTTTGAACGCATTCTCTAGAAAAGTATTGTACTCCCAAGTTAGACTAAATTGAGTTCTCTGTTATTCCAAAGCTCATGTGACAATCATGTTATTCTATTTCTGTCAGTATTACCTCATGAAACAAAATGCTATACTTTTCCAAACTCCAGGGGATGGTTTCATCCATGTTTACAGCCAGATTTTTGAATCTATTGTGTTAGATAATTAATGCGGAATGCATATAAGTTCCACCACTTTCAAAGAGGCTTACAGTAGGATGTAGGCTGATCTAGTATTATAATTTGATAGATAGAAGTGTGAAACTACCTTTGAATATAGGATAATATGTTGAATCAGAATGGTAAATTGCAGTTCTCTTATGTTTTATTTTAATTGCAGATTTTAATTTCATCATATATACTTCCTTTGAAAATGATGTTATTTCTTTCATGAAGATGTGGGCAGTAAAATTTATGTTTGTGTTCATAGACGATTGGCAGTAATTTCATTGTCTTTAATGAAATCCTGTCTTTTTTCATCGTTTAGTGTCCTCCAATTTCTTTCATCTCTTTTTCTTTAAAAATGAAGACTGAAAACCTAATCTATTATGAGTTTATGACTAGCAAACTGTTGAATGGTAGTTGATTGGTTTCACTGATAGATTGATTACTGCTATTTATGTTATGCTTGGCTCATCTTATTAATAATACTTATGAAATTCATGAATTATATCCATTGCTTGACCTGATAGTTTGTACCCCTCTAAAATTCTGTCCTGGTCCTATCTTGAGTTTTTTGTCCTTCCTTTTTCTAAGAAGTCAAAAGGTCCTAATTTGAGTTTTGGCTTCATAATGTAGGCATCTGTCTGAGTTGGCCTTGCTGTATTGTCAAAGAATTGGTAATTTCGCATTGTCCGAGATTGGCAAAGGATGCAAGTTCTTGCAAGCTTTACATTTGGTAGATTGTGCAAGTATTGGGGACGAAGCCATATGTTCTATAGCCAGAGGATGCTGCAACCTCAAAAGGCTCCACATCCGTCGATGTTATGAGGTGCTTTAATAATTTCTCTCCCTCATTACTTAGGGATTTCTCTCTTGTTCTCTAGTGATGTCAAAGTTACAGGATATCTTTACATGCGGAGGTGTTGCATATTGCTGGAGGCCTATGCATTTTTTTTATCAGTAAAGATTCAAAGATATCATTAATTACCAGCACCGAGATGGTGCTGCACATGTACGCTAGGGATCCAGAAACAACAGCAGAACTTTTCTAATACATCAGCTGACAAAAGCAAGAGCATTCCTCCAGTTCTAGTGTTTTACTATGTACACAAATCAGATACTCTCTATACCTGAAGGCACTGAATTTATCTAAGAGACTAACCTCATCTGATATGAATCATTCTCCAAACATAAAACCTCATCCGACATGAATACGAGAGTAAAAGACAAATAGTTGTATTTGATTTTGCTAACAGATTCTGCTTTATTCGTGAAGCACCTTAAGTTCCTTTCCTTCCATATTGCCCCATCTAGTGCACGCTGGTACTGTAGTTCGTACGCTTTTCTGTTTCTTATTGATGCCTCTTCCATTCCTACTGAGTGGAAGCTCCCTGGTAGTTCTTGGCATTCTCCACAATCCACATAAGCTTAGAGATGTGCAGGAACATTACTATAGTTGTGAGAGTAATATGCAATGTAGAAAAATGTGACAGCTGTCTTCCATCTCTTTCTTACAAAGTTAGCACTTGCTGCACAAGTTTATCTTCTTTTTTTAACTAGCCTCATATGCTGCCACGCGAGTCAATCTTCCTTTAACATTTGAAATCTGCTTCCAAGGCCATCTCAGGTTATGTTCACTTCCTGCCCTTTAAGTCCCTTTAATTGAACACATCACTGTTATGTGCCTTCTCTGTAGATTATGTTACCATAGCATTCAAGAGACCTGAATACTGAGTAGTCCATGGGAGGCCTAACAGCACTCCCTGTTTCCCCTTTTTCATTCTTTCTGATCTATTGTTAGTGAAATCTGATTTGTGCTGTTTCCTTTGCACTTTATGCCCAAAACTCAATTGTTTTACTTGATAATACTGCTCTTTTCATCTGATTTGAAGGTTGGAAATAAGGGAATTATAGCTGTTGGCGAGAACTGTAAATTTCTTACAGACCTTAGCCTCCGATTCTGCGATAGGTACGTGATTTTCTCTTTCTCAACTAGTATTGCCTTCTATTTTTTTTTTGTTTCTCAATCATCAAGTCTCGGCTGATAAAATTGTCAGACAGTCTCTTCCTTGCTTGCATTACTTCCTTCTATTGTCCTTTAAATTTATACTTGAATGCTCTTTGAGATGAACATTTATCTTTGCATCTCTTTTCTTTTTTGATATGGGAATTGTTGTGCTCCTTGTGGCGCCTCTGGTGCTGCAAGAGGTCGCTTGTCCATTTGAAGATGAAGATTGCCTAGGCACAATTTTCATCTGCTTCTTTAAAACTTTCTGGCCCATAGTCCGACGGAAAATGTTCTACACTGCGAGATCAACAAAAAATTGTGTCTAGATGGTTATTTGCTTTAGTAACTTCTCTTTTATGCGTATTGTTGATTGCTTATATTTTGCATAGCATATACTTTGAACTGCTTCTTTATCAAAGCTTGCTACTTAACTACCGTATTACTGGTAATTTGGCTCTAATAAGTCTTAGCCTCTATTTGATTACCAGAGTAGGGGATGAAGCACTTGTGGCTATTGGTGAAGGTTGCTCTCTGCATCATTTAAATGTCAGTGGCTGCCACCAAATTGGGGATGCTGGAATAATTGCCATAGCAAGAGGATGCCCTGAACTCAGTTACTTGGATGTGAGTGTACTCCAGGTGTGTCCACTTATACTTGCTCTTTTTATAGTGGGATACACATTGTATATGATGCATAGACGGTATGAATATGCTTACCATACCAAGGTTATTAGATAATTACATTAAGGTGATTACGTAACAACATTAAGTTATATTAGAACTACTTAGTGTAGGTAGAGCTTAAAATGGTTGCTTCCTCCTGAATCCACCGGTTAACAAGTGCATTAGGAACCCCATATTATAAAGTCCCATATGCAACACACACACACGTATTGCTCCCATATACAAGATGTTTGACAGACCAAAAAATTGTATTTTTTATCTCCATTCCGCACTGAATACTAGCACATCAATTTTTGACAGGATTTGGGAGATATGGCCATGGTGGAGCTAGGTGAAGGATGTCCCTTGCTGAGGGATATTGTGTTGTCTCACTGTCGTCAAATAACAGATGTAGGTTTAAGTTATATTGCTAATAGGTGTACCTTGTTGGAAACTAGCCACATGGTCTATTGCCCTGGTATCACCGCTGCTGGAGTAGCCACGGTGATTACGAGCTGCACAAATATAAAGAAAGTTCTTGTAGAGAAGTGGAAAGTGAGTCCAAGAACCAAAAGGAGAGCAGGATCCATCATCTCCTATCTCTGTGTTGATCTGTAGATCTCTGTACACGTGTTAGTGTTATCTATTCCTTGCAGCTCCAATCGAAAACAGCAAGGGTAAAACAGACTATCGTTGTGGATAGCCAGTTTAAGTTGCAAATTACCCCGACATAGAAAGAGGGGTGTCGTGAGAGCTTGTGTTGCACTTTTCTTTTATCTGCTATGTTCTTTCAGTGTTTTTTTTTGTCCAAATCTCACAGTAATTTACTAGTAGATCGACTTCATTTTAGAAGGTGGGGTGTCTTGTATCCCTCCCGATATGCTCGTTGATTGTCTTTATTGGGGAAAGATAGCAAATAGCCAGTTCTTTTACCATCAGTTAAAGCTTTAGCCAGCATTTCAGAATTATTCAAAGTTAACTGCTTTTTAAGGCGGAAATAAAATTTGGACAAAAACTC

>M6-PGSC0003DMG400030856 M6 LRR-RLP

AGTATATTGAGCAAAAATATTCTCTATTTTCTGCTGTCTCCTAATGAGTTTGATGCAATAATATTCTCATGTGGATGTTGCTTGCACCAGGTTCTGTATTAACCGAACCCCAAGTTTATGGAAGAGACAAAGAGGAAGATGAGATAGTGAAAATCCTGATAAACAATGTTAGTGATGCCCAACACCTTTCAGTCCTCCCAATACTTGGTATGGGGGGACTAGGAAAAACGACTCTTGCCCAAATGGTCTTCAATGACCAGAGAATTACTGAGCATTTCCATTCCAAAATATGGATTTGTGTCTCGGAAGATTTTGATGAGAAGAGGTTACTAAAGGCAATCATAGAATCTATTGAAGGAAGGCCACTACTTGGTGAGATGGACTTGGCTCCACTTCAAAAGAAGCTTCAGGAGTTGCTGAATGGAAAAAGATACTTGCTTGTCTTAGATGATGTTTGGAATGAAGATCAACAGAAGTGGGATAATTTAAGAGCAGTCCTGAAGGTTGGAGCAAGTGGTGCTTTTGTTCTAGCCACTACTCGTCTTGAAAAGGTTGGATCAATTATGGGAACTTTGCAACCATATGAATTGTCAAATCTGTCTCAAGAAGATTGTTGGTTGTTGTTCATACAACGTGCATTTGGACACCAGGAAGAAATAAATCCTAACCTTGTGGCTATTGGAAAAGAGATTGTGAAAAAAAGTGGTGGTGTGCCTCTAGCAGCCAAGACTCTTGGAGGTATTTTGCGCTTCAAGAGAGAAGAAAGAGAATGGGAACATGTGAGAGATAGTGAGATTTGGAATTTACCTCAAGAGGAAAGGTCTATTTTGCCTGCCCTAAGGCTTAGTTATCATCATCTTCCACTTGATTTGAGACAATGCTTTGCATATTGTGCAGTATTCCCAAAGGACACCAAAATGGAAAAAGAAAAGGTAATCTCTCTCTGGATGGCACACGGCTTTCTTTTATTGGAAGGAAAATTAGAGCCAGAGGATGTGGGTAATGAAGTATGGAAAGAATTATGCTTGAGCTCTTTTTTCCAAGAGATTGAAGTTAAATCTGGTAAAACTTATTTCAAGATGCATGATCTCATCCATGATTTGGCTACATCTCTGTTTTCAGCAAGCACATCAAGCAGCAATATCCGCGAAATAAATGTAAAAGGTTACCCACATAAGATGATGTCCATTGGTTTCACTGAAGTGGTGTCTTCTTACTCTCCCTCTCTGTCGCAAAAGTTTGTCTCGTTAAGAGTGCTTAATCTAAGTAACTTACACTTTGAGGAGTTATCCTCTTCCATTGGAGATCTAGTACATTTAAGATGCCTGGACTTGTCTGAGAATTCTGGAATTCATAGTCTTCCGAAGCAGTTATGCAAGCTTCAAAATCTGCAGACTCTTGATCTACATAATTGCTACTCTCTTTCTTGTTTGCCAAAACAAACAAGTAAACTTGGTAGTCTCCGAAATCTTTTACTTCATGGTTGCGATGAATTGAATTCTATGCCACCAAGGATAGGATCTTTGACATTCCTTAAGACTCTAAGTGGATTTGCTGTGGGAATTCAGAAGAAAGGTTATCAACTTGGTAAACTACGAGATGTAAATCTCTATGGCTCAATTGAAATCACACATCTTGAGAGAGTGAAGAATGTTATGGATGCAAAAGAAGCCAATTTATCAGCAAAGGGGAATCTGCACTCTTTAATCATGAATTGGAGTAGGAAAGGACCTCATAGATATGAATCAGAAGAAGTTAGAGTGATTGAAGCCCTCAAACCACACCCCAATTTGACTTGTTTAACAATCAGTGGCTTCAGAGGATTCCGTTTCCCTGAGTGGATGAATCACTCAGTTTTGAGAAATGTTGTCTCCATTACAATTAAGGGTTGCACAAACTGCTCATGCTTACCACCCTTTGGTGAGCTTCCTTGTCTAGAAAGTCTAGAGTTGTTAATGGGGTCTGCGGAGTATGTTGAAGAAGTGGATATTGATGTTCATTCTGGATTCCCTACAAGAAGAAGATTTCCATCTCTGAGAAAACTTTTTATAGGTGAATTTCATAATCTGAAAGGATTGCTGAAAAAGGAAGGAGAAGAGAAATTCCCTGTGCTTGAACGGATGACGATTTTTTATTGCCATATGTTTGTTTATACAACCCTTTCTT

>M6-PGSC0003DMG400030928 M6 LRR-RLP

ACAATACTTGTTCTTTTTTTTCTTCTCTACTTTCAACTTTCCTTCTTTTTTTTCCCTACTTAGATAGGGTTTCTGTTGAAATCTTGAACTTTTTCATGTGTAATGCAAATCTTGGTTGAAAATAAAATTTGGGGTTTGTTGTTCTTGGTGCAAAGTTTGTTTTTTTTAAATTTCCTTTGGGGGGTTTGTGTTGAAGTGAAGCAACAACCATGTCTACTCTTGTTAAATACAGTGGTGAGTTTTCTTGGTTTGTTTTCATGTATTGGATTATGGGTTTTTGAGTTTTAAGGAATTTGGCATTTGGGTGATTTAGTTTTCATAGAATAGATTCATTTTTAGTGCTCCATGAATCTATTAAGGGTTCTTGATTGTTGATATGAATATGAAGGGTGTAGGAGTAACTAGAAGAAAAAGATTGTTGGTTGGGAATTTTAGTAGCTCCATTTAGTTGGTTGGTTGGTTATCTGAACTTTCACCTTGTAAATCCTTGCCCCATTTCCCCCAATTTGAAAAAAGATTAAAAAGTTGTTGCTTTGGCTTTGAAGATGTTAATCTTGCTCCTATTTAAACTTTTGTTATGCTTTAGTAAGTTTTTGTGTGTGTGTTGTTTAGATTGATGTTAAAGATTGGCACTTTTTGATAAAAGGTGTGAATAAGAAACATTTTTCTGGTATTTGTAGCTAAGAATTTGTTTGTTGTTCAACTTTTGATCTGATTTGGTAGATCTTAGTTCTAAGTTTTTTATGTAATGAGTTCTCTGACTTGAAATGAAGTTAAAGATAGGATTTTTGTAAGAAATGGGCATGAAATTATTATCTTGGGAATTTTAGTATGTTGATTTGGTGGAAAAGATAGTTGATGAAGTTCTTGATGTTTGATCTTCAGGTGATAATGAGTTCTATAGTGGAGGATCACTTTGTTCAGCTGATTTGGGTCTTATGTTGGCTCTTGGACATGCTGATATATACTGTCCTGTTAGTAAGAGGGCACGAATTAGTGGCCCGTTTGTCGTTGAAGATAGGACTAAGAATCCATCCATTGAAGTACTTCCCAACGAATGCCTATTTGAGATCTTCAGAAGATTGGAAGGAGGCCGTGAAAGGAGTGCAGCAGCTTGCGTTTCTAAGCGTTGGCTTATGCTCTTGAGTAATATGAGGAGCTCTGATATTCGCCATACTAATCATTCAGCTGCTAATGGGGCTTCAGATGATACGAAAATGGCATCAGCTGATGAAGATCTTGAAATGGAGTGTGACGGATATCTTACTAGGTGTTTAGAAGGGAAGAAAGCTACAGACATTAGGCTCTCAGCTATAGCAGTTGGAACTTCTAGCCGTGGGGGTCTTGGAAAGCTCTCAGTTCGGGGGAGCAACTCTGTTCGGGGTATTACTAATGTTGGTCTATCAGCAATCGCCCACGGTTGTCCGTCTCTTAGGGTTCTTTCATTGTGGGATGTTCAGGATGTCGGAGATGAAGGTCTTATGGAGATTGCAAGAGGATGTCATTCATTAGAAAAGCTAGACTTAACTAAATGTCCCTCAATTTCCAACAAAGGTCTGGTTGCAATAGCAGAGAATTGCCCAAGTTTGACTTCTTTGACGATCGAATCTTGTAAAAGTATTGGAAATGAGGGCCTGCAAGCTATTGGAAGATGTTGTACCAAATTAGAGTCTCTTACTATTAAAGACTGCCCTCTTGTAGGTGATCAGGCAGTTGTTAGCCTTCTGTCGTCGGGCAATACAAGGTTGAAGAAAGTGAAACTTCATTCTCTAAACATTACGGATTTCTCCCTCGCTGTCATTGGTCACTATGGCAAGGAGATTATCGATCTTAATCTGTGTAAACTTGGTCACGTAAGTCCGAGGGGGTTTTGGGTTATGGGTGCTGCTCAGGGTTTGCAATCACTGGCTTCTTTGACTGTCACTTCGTGCATTGGACTGACGAATCCGAGTCTTGAAGCAGTGGGAAAGGGCTGCACAAATATTAAAAGCATGTGCCTTCGCGAGTGTAAACGTGTTACTGATAGTGGACTTGTTGCGTTTGCTCAAGCTGCTGGATCTCTCGAGTATCTCTTGTTGGAAGAGTGCGACAAGATCACCCAAACAGGCATACTAAATGCTGTTTCAATCTGCAGCAAGTTGAAGTCGCTTTCCCTGGTGAAGTGTTCGGGAGTCAGGGATTTGCCTCAACAAGCTTCCTTGTTGTCTCCCTGTGATTCTCTCCGATCGTTGTCCATCCGAAGCTGTGCAGGATTAGGTAGCAGAAGCTTAGCTATGGTGGGGAAGCTATGACCCCAGCTTCATCATTTGGATCTCAGAGGGCTTACTAGAATAACAGATGCTGGACTTCTCCCACTTTTGGAGAGCTCCAAGGCAGGACTAGTCAAGGTTAATCTTACTCACTGCCTGAACGTGACCGATGAAGTGGTGCTCTCATTAGCTAGGCTACACGGTGAGACTTTGGAATTGCTGAATCTTGATGGATGCAGGAGGGTTACCGATGCAAGCTTGGTGGCAATTGCTGATAATTGTCTATTACTTAACGATCTTGATGTTTCAAAGTGTTCAATAACTGATTCGGGTGTAGCTGCTTTATCCCAAGGAGTGCAATTGAATTTGCAGGTTCTTTCACTCTCAGGTTGCTCCATGGTAACAAACAAGAGCTTTTCGTCTCTTAGAACATTGGGTGAGAGCTTAATTGGTTTAAATCTCCAGCACTGCAATGCCATCGGTAGCAGCAGAGTCGAGGCTCTTGTTGAGGACTTGTGGAGGTGTGATATCCTCTCCTAAAAGCCAACTGGAGCACAAGGCTATGCATCATCCAAAGGTGTTTGAACTACGAGTCGTAGCTATTTGGTTCATTAATCTTGGTGGGATAGAAGTAAACATGATTGTGGTTTTTGGCTAACTAATGGCCTTGTGATGCTCCAGTGGCTTGGTTGTTTTCCAGGAGGCTATGACAAACAACTTTTTCCATCCATTTTTTGCAGGCTTCATCTACTTAAAAGCTTGTTCCACATCGTTCAAACTCCGGTGTCACACACGGAGGCTGCCTCTTACACAACCATGGTATCCTGAGAATACAGTTACCTAAGTCCTGGTGCTAAAGTTTTCACTAGTGTTGCTCCTTGTCTCCCCATCCGGGGATGAACTAATAATGCTTCTCTTAAACTTATATAGATATCATGCATTAGCCTTGATAGTACTACACTGCAGAATCAGACAGTTTTTTCTTAGTTCTAGTATGTTAGTTTTCGTATTAAGAAGTCGTCCTCGTTGTTATAGTTGTTACTATAAGTGTTTTACTTTTCATCTTAGTCGTCTGTAGTTGAATCAAACGATTTCTGTAGCATGGTGTCTGTTAATCTGGGGCTTGGTCATGGCAACTAGTCCATCTAGGTTGTTTATCTAGTGTCCAAGTTCTAGTTTTTGATAGGATTGTAAAAAGTCCTTGATTGTACAGGTTCCTTGTCTTTCAGATTTGTACATTGCAACTACTTTGTTATGTGTAATAAACTCTGTTTAGTTTGTCTAAATTGTGC

>M6-PGSC0003DMG400031283 M6 LRR-RLP

AACAATAGTCTTAGTGGGACAGTTAATACAACTTTTAGTATTGGAAACCAACTCCAAATCATTAACATGGACGGGAATAAGCTAGAGGGGAAGGTCCCACCATCTTTGATCAATTGCAAATATTTGGGACAGCTTGATTTAGGTAATAATGAGTTGAATGACACATTCCCAAAATGGTTGGGAGCACTATCTGATTTGCGGATTTTAAGTTTGAGATCAAATAAGTTGCATGGTCCCATAGAAGTTTCAAGGACCGAAACCATTTTTTCTCAACTTCGAATTATGGATCTCTCATCCAACAGATTTAGTGGGAAGTTACCAGCAAGTCTTTTTGATAGTTTTCAAGCCATGAAGATAATTAATGAGAACATGACAACCTCATATTTGGGAGTTCAATATGATTCTTTGACTATTACAACAAAGGGATTGGATCTTCAATTTGTTAAAGTTTTTGAAGCAAACATAGTTATCGATCTCTCTAACAATAGATTTGAAGGCTATATTCCACGTACTATTGGAGATCTCATTGGGCTTTGTACATTGAATTTGTCTCATAATAAATTGAAAGGTCATATACCAACATCACTGCACCAATTATCTGTAGTTGAGTCATTGGATCTCTCATTTAATAAAATCAGCGGAGAAATTCCACAACAACTCGCATCCCTCACATTTCTTGCAGTCTTAAATCTCTCTCACAATCATCTTGTTGGATGCATCCCCAAAGGAAAACAATTTGATACGTTTGAGAACAGTTCATACAAAGAGAATGATAGACTATGTGGATTTCCACTCTCAAAAGAATGTGTGGTGATGATGGGGTACCACAAGCGGCAACTCATTTTGGTCTTGATCAAGAAGAAGAAGGAGATTCATCAATTGTCAATTGGCAGGCAGTTCTTATGGGCTATGGTTGTGGATTAGTCATCGTATTGTTTGTAATATACATAATGTTTGCAACACGATATCCATCATGGTTTTCGAGGTTGTTTGA

>M6-PGSC0003DMG400031284 M6 LRR-RLP

AACAATAGTCTTAGTGGGACAGTTAATACAACTTTTAGTATTGGAAACCAACTCCAAATCATTAACATGGACGGGAATAAGCTAGAGGGGAAGGTCCCACCATCTTTGATCAATTGCAAATATTTGGGACAGCTTGATTTAGGTAATAATGAGTTGAATGACACATTCCCAAAATGGTTGGGAGCACTATCTGATTTGCGGATTTTAAGTTTGAGATCAAATAAGTTGCATGGTCCCATAGAAGTTTCAAGGACCGAAACCATTTTTTCTCAACTTCGAATTATGGATCTCTCATCCAACAGATTTAGTGGGAAGTTACCAGCAAGTCTTTTTGATAGTTTTCAAGCCATGAAGATAATTAATGAGAACATGACAACCTCATATTTGGGAGTTCAATATGATTCTTTGACTATTACAACAAAGGGATTGGATCTTCAATTTGTTAAAGTTTTTGAAGCAAACATAGTTATCGATCTCTCTAACAATAGATTTGAAGGCTATATTCCACGTACTATTGGAGATCTCATTGGGCTTTGTACATTGAATTTGTCTCATAATAAATTGAAAGGTCATATACCAACATCACTGCACCAATTATCTGTAGTTGAGTCATTGGATCTCTCATTTAATAAAATCAGCGGAGAAATTCCACAACAACTCGCATCCCTCACATTTCTTGCAGTCTTAAATCTCTCTCACAATCATCTTGTTGGATGCATCCCCAAAGGAAAACAATTTGATACGTTTGAGAACAGTTCATACAAAGAGAATGATAGACTATGTGGATTTCCACTCTCAAAAGAATGTGTGGTGATGATGGGGTACCACAAGCGGCAACTCATTTTGGTCTTGATCAAGAAGAAGAAGGAGATTCATCAATTGTCAATTGGCAGGCAGTTCTTATGGGCTATGGTTGTGGATTAGTCATCGTATTGTTTGTAATATACATAATGTTTGCAACACGATATCCATCATGGTTTTCGAGGTTGTTTGAAGAATTGGAACACAAAATTATTACGAG

>M6-PGSC0003DMG400031320 M6 LRR-RLP

ATGGAATCTCTTGAAATCTTCAATCTCTCAGGTTGTGAGAAGTTAGAAAAATTTCCAGAAATTCAGGGAAATATGGAATTGCTATCAGAGCTCCTTTTGGCACATACTGCAATATGGGAACTTCCCTCATCAGTTGGACTGCTCAGTGGCATTATCTTGCTTGATTTGCATTCATGTAAAAGCCTTGTAAGACTCCCAGCCAGTGTCTCTGAGATGAGAAAACTGAAAATTTTGAGTGTGAAAGGCTGCTCAAGACTTGCAAAGTTTCCTGAAAATCTAGGTGATCTAAACCAGATGGAGGAGCTCTATGCTGGTAACACTGCCATTTGGAAACTACCAGATTCTGTTAGAAACTTGAGCAAACTTAAAGTCCTATCATTAAGAAGAGGCCGGAAGGTAAAGCGTCAAACTGGTGACAGTTTGATGTTACCCTCTTCCTGGGGGTTTCGTTGTTTGAGGTTATTGAAAAGCTTAGATCTCAGTGGATGCAACTTATCTGTTAACCAAACTGCTGCTCTTATGAAATTGCCTTCTTTACTGGAACTGAATCTGAGCAGAAACAAGTTTATTTTATTACCTGATTTCATCAGTCAACTTTCTCAGCTTCGATATCTCAACATAACACAGTGTCAGGAACTTAAGGAACTTCCCAAACTTCCTTCGAGTATAGAGGAATTATATGCAGAAGATTTTTTGGCCAAGCAAAGTATCGAAAAGCTACAAATATACCCGAGGTTGAATTTGGTGTCATTCACCAATTATAGTTTTGATCAACAATCTTATACAGAGGAGAGCAATGGTAGCTCAGTTTTGGATGAGATTCTCAGCTTGTTTCTCTCAGAAGATATGGATCATGTGGTCGAGCACTCTCTTAACTCTGATCACAGGGTGACCTGTTCTATTGTCTTCCCTGAATGTGCGATTCCCAGGTGGTTTAAGCATCAGAGTGTTGATGAAAAGATATTGTTCAAACTGCCCATGAATTGGTATAATGATAAGTTCAAGGGCTTTGCTATATGTTGTGCCACTCTCATGGGAGCAGGTGTGTTGAATCCTGATTCAGGGCTATCCAAGAAGTATGATTATGCTTTCTTCAAAGCTAAATTGATATGCAAAGATGATTTAGAAGACCTTAAAGTGCTAGAGAAAGAATGTAAAGTGGGAACAGCATCTAGAACATATGGCTTGTGTGTTTGCTTTGTCTACATACCATTGTATGCTTCACTGCAGGTGTCCGGCACAGATGTTGGAAATATTAACCAGTATAGCCTATTTGAGGCATCTATCCATGGGCGCATTGTGA

>M6-PGSC0003DMG400031446 M6 LRR-RLP

ATGCCTTTTCGAAAGACTTACCAAAGAGTCTGTTTCAACAATTGAAAGGCATGAGGACTGTTGATCAAACAATGAAGACAACAAGTTATCTAGGAGGTAGATATAACTACCAAGACTTGGTAGTTGTTGTAACGAAGGGATTGAAGCTTGAAGTTGTGAAAATTTTGTCTTTGTACACAGTTATCGATCTTTCAAACAACAAATTTGAAGGACATATACCTAGTGTTCTAGGAGATCTTATTGCGATTCGGGTACTTAATTTATCTCATAATGGGTTGCAAGGCTATATATCACCATCACTTGGAAGTTTATCTGTACTGGAATCATTAGACCTTTCATTTAACCGCCTTTCGGGAGAGATACCACAACAACTTGCTTCTCTTACGTCTCTTGAATTCTTAAATCTCTCCCACAATTATCTCCAAGGATGCATCCCTCAAGGACCCCAATTTCATACCTTTGAGAACAGTTCATTTGATGGTAATGATGGATTACGTGGATTCCCCATTTCAAAAGGTTGTGGCAATGATCCTGTGTCAGAGACAAACTATACAGTGTCTGCGCTAGAAGATCAAGAAAGAAATTCTGAATTTTTCAATGATTTTTGGAAAGCAGCTCTGATGGGCTATGGCAGTCGACTATGTATTGGCTTATCCATAATATATTTCTTGATCTCAACTGGAAATCTAAGATGGCTTGCAAGAATCATTGAAGAACTGGAACACAAAACTATCATGCGAAGGAGAAAGAAGCAGCGAGGTCAAAAGAATTACAGAAGAAGAAATAATCGCTTCTAGACAAGTTACCAGTAAAGAAGAATTTGATTTCAGAACTTCAGGTATTCATGCTAAGCTTTACACTAATGTTTTTTAG

>M6-PGSC0003DMG400031451 M6 LRR-RLP

AATCTAGAGGGAGCAATTCCACAATGTTTTGGCAATAATAGTAGCCTCCAGGCATTTGATATGCAGAATAACAAACTTTCTGGGACTCTTCCAACAAATTTTAGCATTGGAAGTTCACTTATAAGTCTCAACTTGCATGACAATGAACTAGAGGGGGAAATCCCTCGGTCTTTGGCCAATTGCAAAAAGCTGCAAGTTCTTGATCTAGGAGATAATCATCTCTATGACACATTCCCCATGTGGTTGGGAACTTTACCAGAGCTGAGAGTTCTAAGATTGACATCGAATAAACTGCATGGACCTATAAGATTTTCAGTGGCTGAAATCATGTTTCCTGATCTTCGAATCATAGATCTCTCTAGCAATGCCTTTTCGAAAGACTTACCAAAGAGTCTGTTTCAACAATTGAAAGGCATGAGGACTGTTGATCAAACAATGAAGACAACAAGTTATCTAGGAGGTAGATATAACTACCAAGACTTGGTAGTTGTTGTAACGAAGGGATTGAAGCTTGAAGTTGTGAAAATTTTGTCTTTGTACACAGTTATCGATCTTTCAAACAACAAATTTGAAGGACATATACCTAGTGTTCTAGGAGATCTTATTGCGATTCGGGTACTTAATTTATCTCATAATGGGTTGCAAGGCTATATATCACCATCACTTGGAAGTTTATCTGTACTGGAATCATTAGACCTTTCATTTAACCGCCTTTCGGGAGAGATACCACAACAACTTGCTTCTCTTACGTCTCTTGAATTCTTAAATCTCTCCCACAATTATCTCCAAGGATGCATCCCTCAAGGACCCCAATTTCATACCTTTGAGAACAGTTCATTTGATGGTAATGATGGATTACGTGGATTCCCCATTTCAAAAGGTTGTGGCAATGATCCTGTGTCAGAGACAAACTATACAGTGTCTGCGCTAGAAGATCAAGAAAGAAATTCTGAATTTTTCAATGATTTTTGGAAAGCAGCTCTGATGGGCTATGGCAGTCGACTATGTATTGGCTTATCCATAATATATTTCTTGATCTCAACTGGAAATCTAAGATGGCTTGCAAGAATCATTGAAGAACTGGAACACAAAACTATCATGCGAAGGAGAAAGAAGCAGCGAGGTCAAAAGAATTACAGAAGAAGAAATAATCGCTTCTAGACAAGTTACCAGTAAAGAAGAAT

>M6-PGSC0003DMG400031478 M6 LRR-RLP

AAGGTTTGGTGAAGCTATATGTGTCAAAGTGCTCAAAACTTGAAAGCTTACCAGCAGAGATAGGTGATTTAGAAAACTTGGAGTATCTTAATGCCAGCTATACTCTAATCTCACGTCCTCCGTCTTCCATCATCCACTTGAATAAGCTTAAATCATTGACTTTTGAAAAAGATAAATCAGAAGATAGAGTGAACTTCGTGCTCCCTCCAGTGAGTGAAGGGTTACGCTCATTGGAATCTCTGTATCTCAGGTACTGCAATCTAATAGATGGAGGACTTCCAGAAGACATTGGATGTTTATCCTCTTTGAAAGATTTGAAGCTCAAGGGAAATAATTTTGAGCATTTGCCTCAAAGCATATCCAAACTTGGTGCTCTTGAATTCTTGTACTTATCAGATTGCAAGAGGCTTACACAGCTGCCAGAAGACATTGGATGTTTATCCTCTTTGAAAGATTTGAAGCTCAACGGAAATAATTTTGAGCATTTGCCTCAAAGCATATCCAAACTTGGTGCTCTTGAATTCTTGTACTTATCAGATTGCAAGAGGCTTACACAGCTGCCAGAAGACATTGGATGTTTATCCTCTTTGAAAGATTTGAAGCTCAACGGAAATAATTTTGAGCATTTGCCTCAAAGCATATCCAAACTTGGTGCTCTTGAATTCTTGTACTTATCAGATTGCAAGAGGCTTACACAGCTGCCAGAAGACATTGGATGTTTATCCTCTTTGAAAGATTTGAAGCTCAACGGAAATAATTTTGAGCATTTGCCTCAAAGCATATCCAAACTTGGTGCTCTTGAATTCTTGTACTTATCAGATTGCAAGAGGCTTACACAGCTGCCAGAAGACATTGGATGTTTATCCTCTTTGAAAGATTTGAAGCTCAACGGAAATAATTTTGAGCATTTGCCTCAAAGCATATCCAAACTTGGTGCTCTTGAATTCTTGTACTTATCAGATTGCAAGAGGCTTACACAGCTGCCAGAA

>M6-PGSC0003DMG400032130 M6 LRR-RLP

TGAAGATTTTAAGCTTGAGATCAAATAAGTTTCATGGTCCCATTAAATCTTCAGGGAATACAAACTTGTTTGCGCGGCTTCAAATTCTGGATCTATCATCTAATGGATTTAGTGGTAATTTACCCGAAAGTCTTTTTGGGAATTTACAAGCCATGAAGAAAATTGATGAGAGTACAAGAACCCCAGAATATGTTTCTGATCAATATGCTGGTTATTATGCTTATTTGACGACAATTACGACAAAGGGACAAGATTATGATTCTATTCGAATTTTGGATTCTAACATGATTATCGATCTCTCAAAGAACAGATTTGAAGGTCATATTCCAAACATTATTGGAGATCTCGTTGGACTTCGTACGTTGAACTTATCTCATAATGTCTTGGAAGGTCATATACCTGCATCATTCCAAAATTTATCTATGCTCGAATCATTGGATCTCTCATCTAATAAAATCAGCGGAGCAATTCCACCACAACTTGCATCCCTCACATTTCTTGCAGTCTTAAATCTCTCTCACAATCAACTTGTTGGATGCATCCCCAAAGGAAAACAATTTGATACGTTTGAGAACAGTTCATACCAAGGGAATGATGGGTTACGTGGACTGCCACTCTCAAAACATTGTGGTGGTGATGGTAGGGTACCACAAGCGACAATTCCAGCTGAGCTAGATCAAGAAGAAGAAGAAGAAGAAGATTCACCAATGATCTGTTGGCAGGCGGTTCTCATGGGTTACGGTTGTGGTCTAGTTATTGGACTGTCTGTAATATACATAATGTTGTCAACTCAATATCCAGCATGGTTTTCGAGGATGGTATTGAAATTGGAACACATATTTACTATGAGAATGAAAAAGCACAAGAAAAGATATTAGTGTGTAACCTCCAGGTATTCAAGTTTATCTCTATCCTTAAAAGAATTACTTTTTATATATCGATGTATTTTTTACCTATTTCATCCTCAAAGCTCTTAGCTTTAATTCTTTACTTTTGAAAATTGCAGGATTCAAGTCTACTAAGTTGCTGGGAATGCAGATAAAAGCCTTCATCTTTCATAGTTTGTGATATTTCCGTGAAGTGTTTGTTATCTTATGAATAAAGATATGACTTTTCTT

>M6-PGSC0003DMG400032286 M6 LRR-RLP

ATTTTCCATACTTCTTGAGAAATGTAAAGACACTTGTGTACTTGGATATTTCTAACAATAAGATTTGTGGTCAAATCCCTAACTGGTTTAGCGGCATGAGGTGGGACTCGTTGCTGTTCCTAAACCTTTCTCATAATTCATTAACAGGCCACCTACCACAATTGCATTTCCATAATCTAGGGTATCTTGATCTGAAATTTAACTCCCTTCAGGGTCCACTACCTTCATCCATTTGTAACATGAGCTCGCTTGAATTTCTTGATTTATCACGCAACAACTTCAGTGACTCAGTTTCACATTGTTTGGGAAGCATGGCTAGTCTAACGGTGCTAGACTTAAGAAGGAACAATTTCACAGGGAGTCTTCCTCCATTATGTGCACAGAGCACTTCATTGAGTACCATTGTCGTAAATGGTAATCGATTTGAAGGACCTGTTCCTGTGTCGTTGCTCAAATGTAATGGTCTACAAGTCCTTGATGTGGGGAACAATGCTATAAATGACACGTTTCCAGCTTGGCTCGGAAGTCTTCAAGAGCTGCAGGTCCTTATATTAAAGTCGAACAAGTTCCATGGACCTATAAGTACGTGTCAGACTAAGTTTTGCTTTCCCAAGTTGCGAATTTTTGATCTTTCTCGTAATGATTTCAGTGGCTCACTTCCTGCAAAAGTTTTTGGAAACTTCAAGGCAATGATCAAATTAGATGGTGAAGACACAGGAAATATCAAGTACATAAAATCTGTGTTGAATTCGTCATTTGACACATCGTATGAGGATTCAGTGAGTTTGGTAATCAAAGGGCACGATATTGAGCTACAAAGAATCAGCACAATTATGACAACCATAGATCTCTCAAGCAACCATTTTGAAGGTGTGATTCCGGAAACACTAAAGGACCTCAGCTCACTTAGGCTACTCAATTTATCCCATAACAATCTCATAGGTCATATTCCAGTGGAATTGGGGCAATTGAATAAGCTGGAAGCTTTAGATCTCTCTTGGAATCGGCTCACTGGAAAGATTCCGCAGGAATTGACAAGAATGAACTTTCTGGCAGTCTTAAACATCTCTCAAAATCATCTCATCGGACCAATTCCTCACGGTCTACAGTTCAACACATTTGAAAATGACTCGTACGGTGGTAACCTTGATTTATGTGGTCTTCCTTTAACAAAGAAATGTGGAACGAGTGATTCATCACATGTTCCTCAACAATTGGAGTCCGAAGAAGAAGGCGAGTCATATTTTTTTAGTGGATTTACATGGGAATCAGTAGTCATAGGCTACAATTTTGGACTAGTTGTTGGAACTGTCATGTGGAGTCTCATGTTTAAATATCGTAAGCCAAAATGGTTTGTGGAATTTTTCGATGGACTCATGCCTCACAAAAGAAGAAGGCCAAAGAAGAGAGCTCAGAGACGACGGACTTAATGGAAGATTGAGAGGGCCAA

>M6-PGSC0003DMG400032572 M6 LRR-RLP

AAAATGGCTGATGCCTTTGTGTCATTTGCAGTTCAAAAATTGGGTGATTTCCTCATACAACAAGTTTCCCTGCGTAAAAATCTGAGAAAGGAAATTGAGTGGCTGAGAAATGAGTTACTCTTCATACAGTCTTTCCTCAAAGATGCAGAACTAAAGCAATGTGTAGATCATAGAGTTCAACAATGGGTGTTTGAAATCAATTCTGTTGCGAATGATGCTGTCGCTATACTCGAGACTTACAGCTTTGAGGCTGATAAAGGTGATGACGATGAATTTGCTAGTCGTCTCAAGGCTTGCGCTTGCATATGTAGGAAAGAGAAGAAATTCTACAATGTCGCCGAGGAGATTCAATCACTCAAGCAGCGAATCATAGATATCTCTCGCAAGCGAGAAACTTATGGTATTACAAATATCAATAGTAATGCTGGAGAAGGGCCAAGTAATCAGGTTAGAACATTGAGGAGAACTACCTCATATGTGGATGACCAGGATTACATTTTTGTTGGACTTCAGGATGTTGTACAAACATTGCTATCTGAACTTCTCAAAGCAGTGCCTCGTCGAAGCGTCGTCTCCATTTATGGAATGGGCGGATTAGGCAAGACCGCTCTTGCAAGAAACCTCTACAAAAATCCTAATATAGCCTCTAGCTTCTCTACACGCGCTTGGATATGTGTTTCTCAAGAGTACAACACAATGGATCTTCTTAAGACTATCATAAAATCCATCCAAGGTCGCACCAAGGGAACTCTAGGATTGTTAGAAACTATGACAGAAGGAGATCTAGAAGTTCACCTTCGTGATCTATTGAAAGAACGCAAATACCTTGTGGTGGTTGATGATGTATGGCAGAGAGAAGCATGGGAAAGTTTGAAAAGAGCATTCCCGGATAGCAAGAATGGCAGCAGAGTTATTATTACCACGCGCAAAGAGGATGTCGCTGAAAGAGCAGACAACAGAGGTTTTGTCCATAAACTTCGTTTCCTAAGCCAAGAAGAAAGTTGGGATCTCTTTTGTAGGAAACTAGTTGATGTTCGAGCAATGGTTCCAGCAATGGAAAGTCTAGCTAAGGATATGGTGGAAAAGTGTAGAGGCTTACCTCTTGCAATTGTTGTATTGAGTGGACTACTTTCGCATAGAGGGGGGGTGGACAAATGGCAAGAAGTGAAAGATCAACTTTGGAAGAGCATTATTGAAGATAAATTTATTGAAATCTCCTGCATTCTATCATTAAGCTACAACGATTTGTCAACTGTGCTCAAGCAGTGTTTTCTATACTTTGGTATTTTTCCAGAAGATCAAGTTCTCGAGGCTGATAACATAATAAGGTTGTGGATGGCTGAGGGTTTCGTACCAAATGGAGAAGAAAGAATGGAGGATGTCGCTGAAGGCTTCTTGAATGAGCTGATAAGATGAAGCTTGGTTCAAGTGGTGCATACGTTTTGGGAAAAAGTTACTGAATGTAGGGTTCATGATTTACTCCGTGATCTTGCCATACAAAAGGCATCCGACACAAACCTCTTTGACATTTATCATCCAAGAAAGCACTCCAAATCCTCTTCATGTATCAGACTTGCCATTCATGGTCAAGGAGAAAGGTATCACTTACTTGATCTTTCTAATTTGAAGTTGAGGTCAATTATGTTCTTCGATCAGGATTTTCGTAATGTGTTCCAACATATAGATGTGTTTCGACATATATATGTGTTGTACTTGGATATTAAAGGCTGTGGTGCTATACCTGATGCCATAGGGAGTTTGTACCACCTAAAGTTGTTAAGCTTGAGAGGTATCCGTGGTCTTCCATCTTCCATTGGCAACCTCAAGAATCTACAGACACTTGCTGTCACTGACTTTGGATACAATAATTGCCAACTACCCCGCGAAACAGCTGACCTGATAAATTTAAGACATTTAGTTGTTCCGTATTCAAAACCTCTGGCACATATAAGCAAACTCACTAGTCTTCAAGTTCTTGAAAGAGTTTGTTGTGATCAATGGAAAGATGTTGATCCTGTTGATTTAGTCAATCTTCGAGAATTAAGCATGTTTCATATTAGCATATCTTACTCCCTAAACAACATTAGCAGCTTGAAAAACCTTAGCACTCTCACATTGTCTTGTCAAGATTATGATGGATCATTCCCATCCCTTGAATTTGTTAATTGTTGTGAAAAGCTCCAGAAATTGTTCTTATATGGGAGAATAGAGAAACTGCCTAATCTGTTTCCAAATTCCATCACAATGATGGCTCTGAGGGACTCAAGACTGACAGAAGATCCGATGCCTATTTTGGGAATGTTGCCAAACCTAAGGAATCTCTATTTGGTTAGTGCTTATCTTGGAAAAGAAATAATGTGCAGTGATAACAGCTTCAGTCAACTAGAGTTCCTTCATCTTTATCATCTTTGGAACCTAGAAACATGGCATTTAGGCACAAGTGCAATGCCTCTCATTAAAGGTCTTCATATCTATGAATGTCCAAACCTGAAGGAGATTCCAGAGAGAATAAAAGACGTGAAGC

>M6-PGSC0003DMG400032785 M6 LRR-RLP

TGTATAGTTCCTATATAAACATTCGGTTAGGTAGCTAGCAAATTAGTTTGCCAGAAAAGAAAGTAGAGTAATAGTGGGGATGGGAAATATTAAGTTTTTGTTGTTAGTGTTTTTTCTTATAGTAATAATGGCGAATGGTTGTTGGGAGGAAGAGAGAAGTGCTCTCTTGGATCTGCAAGCAAATATAATGAGTTCTAATGGCGAATTGTTGGTTGATTGGGAAGCTTATTATACGAATGGTTTTACAGATTGTTGTTTCTGGAAAAGGGTAAAATGCAGCTTAGCAACAGGTCGAGTGATCAAACTCGATCTCGAAACAAATTTTGGTACAGGGGATGGTTGGAGATTTAATGCCTCTCTCTTTCTTCCATTCAAGTCACTACAAGTTCTTTTATTGTCTTACGGAAATATTATAGGGTGGACAAAAAATGAAGGTATATATGGAGTATGGAGTATATTCATGACTCTTGAAAAATGAAACTATATGGAGTATATAAATTACTAACATGTGTTTTTTGGTCTAATTTTGTAATCCAGGTTTCAGTAAACTAAAGCAGCTTCCCAACCTGAAAGTACTCGATTTGCAATTTAATCATATCCATCCTAAAGATTTACTATCATCTTTACGTTGGATTTCATCTCTTGAGGTTCTAAAACTTGGTGCAGATGTTGATACCTCTTTCAACGTACCGCCGACATATAATAGCAGTATATTCTATATCCTCCTACTCTATAGTCCATAATTGGCCAATAATTTACTTAGAAGTTGTAATTTTTGGATGAAGGCACGAGTAAGAAATGTGGAGGGCTAACCAATTTAAGAGAACTCTGGTTTGAAGGTTATGCAATCAATGACATCAATAGCCTTTCTGCATTGGGTAAGCTCCCTTTAACAAAGTAGCAAAATAACAAATTTCACAAGAGAACAATTAACAAACTAACACTAAGTATGGTCTCTTACGTATTTTTTTTTTCATAAATCAAATGCAGGTGCAAATTCTGGACTGAGGAATTTAGAGAAGCTCATTTTGAATGATAATAACTTCAATTCCACCATCTTTTCATCACTCAAGATTTTTCCGTATCTCAAGCATCTCAATCTCGCCTCGAATGAAATAGATGGAAATATCGAAATGAATGGTTTGTTACGCGCCTCTTCATATCCAAGTCTTCCCCACAAAATATCTCAATTTTATTTCTTCACATATCAAATTATACTTTATTTTTTATTAACCTGTGTAGATATAATTGATTTGAGCACTTTGGAATATCTAGATCTTTCGGATAACAATATAGAGAGCTTTGCGACTACCAAAGGTACACGAATCAATGCAGTTGCTCTAATGTTGAATGTTCTATTTTAACCTTAACACATCCTACAGAACAAAATACATCTTTATGGATTTGAATAACCATGTGCCGTGGTATGTAAATTTTTTGATTGAAAACTGTGGGATACAAATTAGATTTTAATAAGATTGAACTATTCATTCTTATTTAAAAATGTATACTATTATTTATTTATTCACCATTTAACAGGTAACAAAAGAATGAGTTCTCTGCGGAATTTACTATTGGGAAGTTCCTACTCCAACTCTAGCAGAGTCATACAGTCCCTAAAACCATTCTCATCTCTTAAGTCACTTTCTTATGAAGATAGTGATCTCACTTCTCCAACCATAATTTACGGTACACTCAAATTCATATATACCTTAGCTTACTTCTATCGCCACTCTTTTCGTTTGAAATGTGAATATATTTTTCTTTTCTTTTTTTTGGACTCTATCATATAAGCATTAAGAAATCTGAGCACGATGGAGTACCTATACTTGGAGGGATGTTCTTTAAATGACAACTTTCTCCCAAACATTGGACAAATGACTTCTCTCTTGGTGTTCAGTATAGCTTTTGGTGACAATGATGGCACCCTCCCTAATCAAGGTACCCAATTTTTCTCTTTTTTCCCTAATGCCCGCATTTTCATGTTGGAGGTCTGATTTTATCATGTTCAGTTGATCAAATATTATATTATTTTTTGTTTATTACCAAACACAAGAAAATAAGTAACACAAGCCATCTAATTTTTCTAGAAAATATTTTCCTTCGTATCGAACACATCCATAGACATAAAAAAATATAATAAAATGTCACTCATCATAGATATTTGTTTTAGCTAAATGTTTCTTAATATATATATATAGGCTGGTGTGAACTCAAACACATTCAAGAGCTGGATTTCTCAGACAATAATTTTCGTGGAACACTGCCTTCATGTTTTGGAAACTTGACATCACTTCGACGATTGAGTTTTTCTGGGAATTACTTCATGCGAAATATAGCGTCACATTCTATTTGGAGGAGACTCACATCACTTGAGTACCTCGATATTGTGGATAACCAATTTGATGTTCCTCTGTCATTCAGCCAATTTTCCATCCATACAAAATTGATCTACTTGGATGTTAGTTCTAATACAATAATTCCAGATACCGAATTCCAAAATTGGATCCCAAATTTCCAGTTGGAGTTTTTTGCTATCGAAGGATGTAAAAACCTTCAAAAATTGCCTTCTTTCCTTCACTACCAATATGACTTGAGGATTCTTACTATTGAAGGAAATCAATTGTCAGGGATCTTTCCAACATGGTTGTTAGAAAATAACACCAGACTTGCAGGGATTTATAGTAGAGATAATGCTTTCAACGGACCATTCAAGTTGCCATCGAGTGTTCACCTCCATCTAGAGGCGATTGATGTATCTAATAACAAACTAAATGGATATATTCCAGAAAATATGAGTTTAGCTTTCCCAAAGCTTACTTATTTGAACATGTCACAAAATTATCTTGAAGGTCCTATACCTTCTAAGATTAGTGGCATTCATTTAGAATTTCTAGATCTATCTGTCAATAACTTGTCCGGAGAAGTTCCTGGTGATCTGGCGATTGGTTCTCCAGCATTAGTTTACCTTCGATTGTCTCACAACAAGTTAAAAGGCCAAATTTTTCCAAAGGATGTCAAACCAAGGATGTTATTGTTTTTGTACTTGAATGGAAATAACTTTGAGGGACCACTACCTAGCAACATGTTTATCACATCCCTTTTTGTACTAGACGCTAGCAACAATAATTTCTCTGGAGAGATTCCGAGATGTGTTAGGAATAATACAAGATTGTTACAGCTTGATTTGTCCAAGAATCATCTCAAAGGTTCAATTCCAGTTGAGATTTGCAATTTAAAGCTCATTCAGGTCTTAGCTATATCTGAAAACAGGCTTTCAGGCCTTATTCCTTCTTGTGTGAATTCTTTACCTGTCGAACACATCCATCTTGAGAAAAATCAATTGGGTGGTGAATTGGGACATGTACTTTTCAACTTCTCTTCTCTGATAACTTTGGATCTTGGGTACAACAATTTTACAGGAAATATCCCACACACCGTAGGTTCGCTTAATAATCTGAACTACCTTCTCCTTAGCAATAACAAATTGGAAGGGCAGATTCCAACTCAGATTTGTATGCTGAATAAGTTATCTATTGTGGATTTATCTTTCAATAAGCTTTATGGTCCACTCCTTCCTTGTTTGGGTTACTTAACACAAGCCAAAAAGGATGCAGAGATAAGTAGGACCTACTACCCTATGACTTTTGGGCATTCATGGTTAAATTTTGTGGCCTTGATACATTTGAAACAGAACTACCACGATAGTCAAGGACTTGTAATTGACACATTTTTAATGGATGTGGAAAACCAGGTCCGATTTTCAACGAAAAGAAACTCATACATTTATAAGGGAATCATTTTAAAATATATGTCGGGTATTGATCTCTCAAGTAATAGATTAACTGGTGAAATTCCCGTTGAGCTAGGGAAGATGAGTACTATACATGCACTGAATCTATCACACAACCATCTCATCGGAAGAATACCAAATAGCTTCTCCAATCTACAGAAAATTGAGAGTTTAGACCTTTCCTTCAACAACTTGAATGGGAGAATTCCTGTTGGTCTAATTGAGCTAAATTCTTTGGAAGTATTCAATGTTTCATTCAATAACTTATCCGGTGCAGTGCCTGATTTTAAAGCTCAATTTGCAACTTTCGATAAAAGCAGCTACGAGGGAAATCCTTTTCTTTGCGGTGATCCATTAGACAACAAGTGTGGCATGAGTCCTAAATTATCAAATACCTCCAAGGTTAATGGAGATGAGGAATCATCAGAATTGGAGGATATGGAGTGTTTTTACATTGGCTTTGTTGTGTCTTATGGAGCGATCTTGTTGGGATTAGCTACAGCGCTCTGCTTCAATTGTCATTGGAGAACACCATGGTTCAGGATGATAGAGTCATTGATGTTTTATTGTTACTATTTTGTGTTAGACAACATTGTTACACCCCTCAAGAGTAGATGGTGCAGGAAAGTTGGTTAATTAGGTAGTTGTTTAAAATTATTGTTGTATGTCTATCACAATTATTATGTTGTTATTC

>M6-PGSC0003DMG400032828 M6 LRR-RLP

TGCATTACTAGTCATTAACAAAAATGGCTTCATTTTCCTTTCCCATTACCACTACTGCCTTTGTTATCTTGCTTCTAAGTCTTACCTCCAATGTGTTCAACAACTCAGCAGCCAAACACAGCCATTCCTCCTCCGTTAAGGGCATAAAAAACCCGAGGCTGCAAAAAGCCTACATTGCTCTCCAAGCTTGGAAACGTGTTATCTACTCTGATCCTAACAACTTCACTTCTACTTGGGTTGGCCCTTCAGTCTGCAACTACACTGGCATATATTGCGCGCCATTCCCTAATAACACCAAACTCCAAGATGTTGCTGGCATTGATCTGAACCATGCTGATATAGCCGGTTTCCTACCTGAGGAACTCGGTCTTCTCAATGACTTAGCGTTGCTGCACCTAAACAGCAATCGCTTCTGTGGAATCCTCCCACTCTCTTTATCCAACCTCACTCTCTTGCATGAGCTTGATCTCAGTAACAACAGATTCGTAGGACCTTTTCCTAACGTTGTCCTCTCTCTACCTTCCTTGAAATATCTTGATCTTCGCTACAATGAGTTTGAAGGGCCATTGCCTCCTCCACTCTTTAGCAAAGATCTCGATGCTATTTTAGTTAACAATAACCGTTTAACTAACGTGATCCCTTCAAATTTAGGATCAAGCTCTGCTTCTGTTGTGGTTTTTGCTAATAACTACTTTGGAGGATGCTTACCACCAAGCATAGCCAACTTTGCAAACACCATAGAAGAGTTGCTCCTTACTAATACTAGCTTATCAGGGTGTTTGCCTCCTGAAGTGGGATTTTTGTACAAGCTAAAAGTTCTTGATGTGAGCAACAACAAGTTAGCAGGGACAATACCTTATACTATTGCTGGTTTAGCTCACTTGGAGCTACTAAATTTAGCACATAACATGTTTAGTGGAACTGTACCAGAGGGAATATGTGTATTACCTAAGCTCTCAAACTTCACATTCTCTTATAACTATTTATGTGAGGAGCAAGGTATTTGCAGCAACTTGACATCAAAGGGCATAGTGTACGATGATCGTCGAAATTGTTTGCCAGAAAAGCCTCTTCAGAGAAGCAAGAAGGAATGTGATGCTGTGGATGAGCATCCCATTGACTGTTTAGCATTTCATTGTGGCACTTGATACTGTTTTCATGTGTTGGTTCTACTGAAAATGTGACAAATTAGTACTAGCTTGAGTTTTCCTTGTACTTTTTTTTCAAATTTAGTAGCCAATGTGAAAAAAGAATAAAGTCAATAGTAATTCTGTAAATATGGTGTGTGATGAAGTTTTTCTTATGTTGGTTATTTAACTTTGATGCTGCTGCTATTATATGTTGTCCCAAAACATGATTCTCATTCTTGAAAAATCTTAATCAAGTTGCTACCTACATAGTGAATTCTTCTGTAATTCCATATTTTTCTTGTACTGTATATTTCATTTAGA

>M6-PGSC0003DMG400033131 M6 LRR-RLP

GTCATTTTTTTAATTTCTATTACCATCAATTTAGGGCAACCCTTTGGAGCAATTGATCTCTGTTTGCCACAATCAACAACGGAAGGTGAGATTTTTTTAAAATCTTTTTTCGCTTATAAAATCATGTAATTAATTCAAATATGTAACTAACTACTTCAAGTTGGTTTGTAAATTGTAGTACAACAAAAATTCAAATCATTTGAACTTGGTTTGGGTGTTTTTCTTCTCTAAATTATGAAATGATATTGTTTTCATTTCTTTTATTAAAGGTGAAAGAGCAATTAACTCAATCCATGGAAGTTGTTGAGACTTGTGCTAAAGGAACTCTTTCTTCTGGAGAGCCTTCAAACAAACCTGGTATACATCAATCTATGATGATGATGAAGTTGTGGGCTTTGAGAAAGAAGCAGAAATTATAATGATGAAATTGATACGAGGAACAAAGGAGCGGGATCTTATCTCCATCTATGGAATGGCAGGACTTGGTAAAAAAACTTTGGCAAGAAAGGTATACAACAATCCCTATATTGTTAATCACTTTGATGTTAAAGCATGGTGTGCTTTGTCGCAAACATATAATAGGACGACGTTATTGATTGAGATTTTCAAACAAGCTACAAATGATGAGATAGAAATCAAAGAAGACGATGATGTAGCTGACATGTTGCGCAGGGTTCTAATAGGCAAGAGATACCTCATCGTATTAGATGATATTTGGGATGTCGAGGCATGGGAAGATTTGGGTTTATGTTTCCCTCAAGGTAAATATGGAAGCAGAGTAATGGTTACAACAAGAATTGAACAAGTGGCTAAGTATCTTCAGCACCATAGTGATCCTTATTCTCTTAGCTTTATAACATCCGAACAGAGTTGGAAATTATTGGAGAAGAAGGTATTTCAAGGAGAGAGTTGTCCCCTGAATCTACTTGAAGCAGGGTTACAAGTTGCCCAACATTGCAAGGGATTGCCTCTTGTGGTTGTCCTGGTTGCTGGGTTTATTGCGAAAATGAAAAGGGAAACGTCCTTGTGGCTCGAGGTTGCAAATAACTTAAGTTCTCTTGCTTTGGGGGAGCAGAGTATGAAGGTAATACAATCAAGTTATGACCATTTAGAAGACCATTTAAAGCCTTGCCTTCTATACATGGGATTGTTTCCAGAGGACCATAAGATTCCAGTGAACGATCTGCTGAAGTTGTGGATGGATGAAGAGTTTGTATTGAATGCCGAAACAGAGAACATGGAGGAAACATCTCGAGTTTGCTTGAGTGATTTGCTTAACAGAAGCCTAGTAATGGTCTCTGGAAAGAGAGTTAATCGTGATGTAGTGTACTGCTCACTTCACGATGTAGTGCGTGAGTTCTGCTTGAGAAAACTCACAGAAGACAATTATATACAGGTTGCAGTGCCATACAATCCGTATCAACATATGCATTCCACGGAATCACGGTTATGCATTTATATTCATGATGATCTTGTTAAACAATTAGATTATTCTAAATACCAGTTGGATAAGATTCCAATGCTCGATTTCAAGGAAACAAATTCTTTGGAGTTCATTGCTCATCCAAAACTCAATACATGGAATAACCAATATTCAAATCCATTAGATTTAGTTGTTAAATTCAGATTTATTCGGGCATTGTATTTGATGGATGTGGAATTGCCAAATTCATGGGCTGCTGTACTGCAATCGCTAACTCAGTTGAGGTACCTAGCACTTTATGTCAAACAATTTCAATTGAAGTGGATATCACACCTACACCATCTCCAAACTCTGCAACTCAAAACAAGGAAGAGGATATGGCTAAGGGCTT

>M6-PGSC0003DMG400033132 M6 LRR-RLP

CCAATTGACATGGATGTTGTTGAAACTTCTGGAGAGCCTTCCACCAAACCTAGTATTACATCAATTTATGATGATGATGAAGTTGTCGGCTTTGAGAAAGATGAAGAAATTGACACGAGGAACAAAGGATCTATGGAATGGCGGGACTAGGTAAATCAACTTTGGCAAGGAAGGTGTACAACAATCCCTATATTGTTAATTACTTTGATGTTAAAATATGGTGTGCTGTGTCTCAAGCATATAGTAGGACGACTTTGTTGGTTGAGATTTTCAAACAAGCAACAAATGATAATAGCGAAATCAAAGAGGACGTTGACATAGCTGACAAGTTGCAGAAAGCTCTAAAAGGTAAGAGATACCTCATCGTATTGGATGATATATGGGAAGTCGAGGTTTATGTTTCCCTAAGGGTGAATATGGAAGTAGAGTAATGGTAACCATTCGAATTGAACAAGTGGCTACACATCTTCAACACTGCGGTGAACCTTATTCTCTTAGATTTCTAACTCCGGAAGAGAGTTGGGAATTATTGGAGAAGAAAATATTTCAAGGAGAGAGTTGTCCCTCGGACCTACTAGAAGCAGGTTTACAAGTTGCCCAACACTGCAAGGGATTGCCTCTTGTGATTGTCCTGATTGCTGGAATTATTAAGAAAATGGAAAGGCATCCGTCCATGTGGATGGAAGTTGCAAATGATTTAAGTTCTCATGTTTTAGGAGAGCAGAGTACAAAGGTGATGCAGTCAAGTTATGACCATTTAGAAGGCCACTTAAAGTCTTGCCTTCTTTACATGGCATTGTTTCCAGAAGACTATAAGATTCCAGTGTCCAATTTGATGATGTGGTGGATTGCTGAGGAGTTTGTACTGAATGTTGACAACAAGAATATGGAGGATGCATCTAGAGTTTGCTTGAATGGTCTACTTAACAGAAGCCTAGTAAAGGTCTCTCAAGGTAGTTCTAATGGTGGCGTAACTTACTGCATACTTCATGATGTAGTGCGTGAGTTTTGCTTGAGAAAACTTAAAGAGGAAAAGTTTATGCAGTTAATAGTGCCATACAATCCATATCAACAGTTTTATTCCATGGAATCGCGGATAAGTGTTTATCTTCATGATGATCTTGTTAAACAATTGGATGATTTTGAATATATCAGTTAGATAATATTCCAATGCTTGAGTCTAAGGGAAGTAGAAAGTGTTTTGGTAAATGTCCTAGGACTTTGGAGTTCATTGCTCATCCAAAATTCAACACATGGATTAGATCATCAAATCCTTTAGTTTTACTTGAAAAACTTAGACTTGTTCAGGTGTTGCATTTTCCGGATGTGAATCTGCCAAGTTGTTGGCGTAGTGCTGTGCAATCGCTAACTCACTTGAGGTACCTTGAGATTTATGTCGAAGAATTTGATTTTGAGTGGATATCTCACCTTCTCAATCTTCAAACTCTAGTAGTTCGTTCATCTAAGTCTCTGAGGATATTATCGCCTGCAGCTATGTGGAAAATGATGAAGCTAAGGCATCTGAATATAATCAATTTGTGCTTCTTGTGGGAAGATAATGATCGAGCAATTTTTGACGAATCTTCAACAACTATGCTAGAGAACTTGAAGACATTCTTCAGTATTCATATACGTGTAGATGATACAACTCCAAGATTCTGGTGGAGGTTTCCAAATATTGAATAACTTAGCTGCCTCAGTGTTGACGATTTACCTAGTTGTCCTTTGTTTCCCTCAGTGGAAGTTAATACTCGGCATCAATCTCTTGATCTGATGATAAGCTATAAGGGATTCTGGGATTCAGTTGGTTGGGAGAGCTACTTTGTCTTCCCTTCAAATCTTAGGCATCTGTCCCTCAGTGGTTGTTTTTTGACAGAAAAAATGGTTTCAAACATTGCAAGATTGAAGAAACTTGAGAAACTCTCATTAGTTGGGGGATTTCCTTTGGGGAAAATAGAGAACCGTTGCTGGGACGTCACAAATGTGGAGTTCCCTGCACTAAAATACTTGGCATTACGCTTTATGAGGATGGAAGAATGGAAAGCTTCAGAGGAATCCTTTCCTGTGCTTGAGGAGCTAACTATTAGTGTTGGTTACATCATGGAGATTCCTCCTAGTTTTGCGGATATTTTAACACTGCGACTTATTAAACTGGACAAGCTCTCCAATTCACTTGTGGTTTCAGCTTTGACTATTAAAAAGGAAATAGAAGAAAACACAGGATGTGACCGTCTCC

>M6-PGSC0003DMG400033152 M6 LRR-RLP

GTGATATACGCTTGGATAATATGAATGCAAGGTTGTGGTGGAGGTTTCCGAATCTTGAAGAACTCTGCCTCAACGTTAAAGTTGTACCTGAGTTTCCGTTGTTTCCCATAGCGGAAGTTCATAGTCACCTTCATTTTCTTACTCTGGAGTTCCCTTTAATACAATTATTTGAATTAGTTGGATGGGAGAGGTATTGTGTGTTCCCTTCAAATCTTAGGCATTTGGACCTTGCCAACTGTTTCTTGACGGAAGAAATGGTTTTAAACATTGCAAGACTGAAGAAGCTTGAGAGTCTTCAATTATCTATGGGATTTGCTTCTATGAGATATTGTTGGGATGTCACAAATGTGGAGTTTGATGCACTCAAATACTTGGGATTATTCTTTATGCAGATAGAAGAATGGAAAGCTTCAGAGGAATCCTTTCCTGTGCTTGAGAAGCTAGTTATAAGTGGATGGCCCGGTTTCAAGGATATCAAGGAGATCCCCCCAAGCTTTGCGGATATTCTAACCCTAAGACGTATTGAACTGTTTGACTGCACAGACTCTCTAGGGGTTTCAGCTTTGAATATCAAAAGAGCGATTGAAGAAAACACAGGATGTGACAGTCTCCAAGTTTTTATACAAAAGTAACTAAAT

>M6-PGSC0003DMG400033153 M6 LRR-RLP

GTGGGCTTTGAGAAACATGCAGAAATTATAATGAATAAATTGATACGAGGAACAAAGGAGCGGGATCTTATCACCATCTATGGGATGCCTGGACTAGGTAAAACAACTTTGGCAAAAAAGGTATACAACAATCCCTCTATTGTTAATTACTTTGATGTTAAAGCATGGTGTACTGTGTCACAAGCATATGATAGGCGGACGTTATTGGTTGAGATTTTCAAACAAGCTACAAATAATAAGATCATAATCGAAGAGGACGATGATGTAGCTGACATGTTGCGCAAGGTTCTAATAGGCAAGAGATACCTCATCGTATTAGATGACATTTGGGATATCAAGGCATGGGAAGATTTGGGAATATGTTTCCCTCAAGGTGAATGTGGAAGCAGAGTAATGGTCACGACACGAATTGAAGAAGTGACTAAGCATCACAGTGATCCTTATTCTCTTAGCTTTCTAACGTCCGAAGAGAGTTGGGAATTATTGGAGAAGAAGATATTTCGAGGAGAGAGTTGTCCCCCGGATCTACTGGAAGCAGGGCTACAAGTTGCCCTACACTGTAAGGGATTGCCTCTTGTGGTTGTCTTGATTGCTGGAATTATAGCAAAAACGGAAAGGGAGGCGTCCTTGTGGCTCGAGGTTGCAAATGACTTAAGTTCCCTTGCTTTAGGAGAGCAGAGTATGAAGGTGATACAATCAAGTTATGACCATTTAGAAGAACACTTAAAGCCTTGCCTTCTTTACATGGAATTGTTTCCAGAAGACCATAAGATCCCAGTGGACCATCTGCTGAAGTTGTGGATGGCTGAAGAGTTTGTATTGAATGCCGGAAAAGAGAACATGGAGGAAGCATGTCGAGTTTGCATGAGTGATTTGCTTAACAGAAGCCTAATAATGGTCTCCGGGATGAGATTTTATCGTGATGTACAATACTGCTCACTTCATGATGTAGTGCGTGAGTTCTGCTTGAGAAAACTCACAGAAGACAAGTATATGCAGGCCACAGTGCCATTCAATCCGTATCAACATTTGCATTCCACGGAATCACGGTTATGCATTTATATTCGTGATGATCTTGTTAAACAATTAGATCATTCCGACTACCAACTGGATAAGATTCCAATGCTAGATTTCAAGGAAACAAATTCTTTGGAGTTCATTGCTCATCCAAAACTCAATACATGGAATAACCAATATTCAAATCCTTTAGATTTACTTGTTAAATTAAGATTTGTTCGGGCATTGGATTTGATGGATGTGGAATTGCCAGATTCATGGGCCACTGCAATTCAATCGCTAACTGAGTTGAGATACCTTGCACTTTGTGTCAAACAATTTGAATTGAAGTGGATATCACACCTACACAATCTCCAAACTCTGCAGGTCAAGTCAAGTGAGAAGTTACACCTAAGGGCTACTACTTTATCAGAAATGACGAAGCTAAGGCATGTGAAAATAGACTGTTTTTTCATGATATGGGAAGACATTAATGACGAAGGAGGTTTTGAAACAAGTATGCTAGAGAACATGAAGAGTTTTGGAACTTGCGATATACGCTTGGATAATATGAATGCAAGGTTCTGGTGGAGGTTTCCGAATCTTGAAGAACTCGGCCTCAGCGTTAAAGATGTACCTGAGTTTCCTTTGTTTCCCATAGCGGAAGTTCATAGTCACCTTCATTTTCTTTCTCTGGAGTTTTCTATAATGGAATTATTTAATTCAGTTGGATGGGAGAGGTATTTTGTGTTGCCTTCAAGTCTTAGGCATTTGGACCTTGCCGGCTGTTTCTTGACGGAAGAAATGGTTTTAAACATTGCAAGATTGAAGAAGCTTGAGAGTCTCAAATTGCTTGTGGGATTTCCTTTTGGGAGATCAGAGAGCTGTTGCTGGGATGTCACAAATGTGGAGTTTGCTGCACTCAAATACTTGGGATTATTCTTTATGCAGATAGAAGAATGGAAAGCTTCAGAGGAATCCTTTCCTGTGCTTGAGAAGCTAGTTATAAGTGGATGTTCCAATTTCCAGGAGATCCCCCCAAGCTTTGCGGATATTCCAACACTAACACGTATTGAACTGACTAACTGCATCGACTCTCTAGGGGTTTCAGCTATGAATATCAAAAGAGAAATTGAAGAAAACACTGGATGTGACAGTCTCCAAGTTGTTATAGAAAAGTAAATAAATGTAATTAAAAGCACTTCATTGCTGCTCTGTTTCCACTTCCTACTTCTATATTGTTGCATTATTATCTGTTTTTAGACATAATACATAAACATACCATTGAACTTTGATCTTGAGCTGACAGCTATGTCCTTTGACTTTGGATGCGCACAATTAAATACTTAAGCTTGTATAAAACTTTTTATGTGACATAATACATGTAGGACACAAT

>M6-PGSC0003DMG400033154 M6 LRR-RLP

GTTGCAAATGATTTAAGTTCTCATGTTTTAGGAGAGCAGAGTACAAAGGTGATGCAGTCAAGTTATGACCATTTAGAAGGCCACTTAAAGTCTTGCCTTCTTTACATGGCATTGTTTCCAGAAGACTATAAGATTCCAGTGTCCAATTTGATGATGTGGTGGATTGCTGAGGAGTTTGTACTGAATGTTGACAACAAGAATATGGAGGATGCATCTAGAGTTTGCTTGAATGGTCTACTTAACAGAAGCCTAGTAAAGGTCTCTCAAGGTAGTTCTAATGGTGGCGTAACTTACTGCATACTTCATGATGTAGTGCGTGAGTTTTGCTTGAGAAAACTTAAAGAGGAAAAGTTTATGCAGTTAATAGTGCCATACAATCCATATCAACAGTTTTATTCCATGGAATCGCGGATAAGTGTTTATCTTCATGATGATCTTGTTAAACAATTGGATGATTTTGAATATATCAGTTAGATAATATTCCAATGCTTGAGTCTAAGGGAAGTAGAAAGTGTTTTGGTAAATGTCCTAGGACTTTGGAGTTCATTGCTCATCCAAAATTCAACACATGGATTAGATCATCAAATCCTTTAGTTTTACTTGAAAAACTTAGACTTGTTCAGGTGTTGCATTTTCCGGATGTGAATCTGCCAAGTTGTTGGCGTAGTGCTGTGCAATCGCTAACTCACTTGAGGTACCTTGAGATTTATGTCGAAGAATTTGATTTTGAGTGGATATCTCACCTTCTCAATCTTCAAACTCTAGTAGTTCGTTCATCTAAGTCTCTGAGGATATTATCGCCTGCAGCTATGTGGAAAATGATGAAGCTAAGGCATCTGAATATAATCAATTTGTGCTTCTTGTGGGAAGATAATGATCGAGCAATTTTTGACGAATCTTCAACAACTATGCTAGAGAACTTGAAGACATTCTTCAGTATTCATATACGTGTAGATGATACAACTCCAAGATTCTGGTGGAGGTTTCCAAATATTGAATAACTTAGCTGCCTCAGTGTTGACGATTTACCTAGTTGTCCTTTGTTTCCCTCAGTGGAAGTTAATACTCGGCATCAATCTCTTGATCTGATGATAAGCTATAAGGGATTCTGGGATTCAGTTGGTTGGGAGAGCTACTTTGTCTTCCCTTCAAATCTTAGGCATCTGTCCCTCAGTGGTTGTTTTTTGACAGAAAAAATGGTTTCAAACATTGCAAGATTGAAGAAACTTGAGAAACTCTCATTAGTTGGGGGATTTCCTTTGGGGAAAATAGAGAACCGTTGCTGGGACGTCACAAATGTGGAGTTCCCTGCACTAAAATACTTGGCATTACGCTTTATGAGGATGGAAGAATGGAAAGCTTCAGAGGAATCCTTTCCTGTGCTTGAGGAGCTAACTATTAGTGTTGGTTACATCATGGAGATTCCTCCTAGTTTTGCGGATATTTTAACACTGCGACTTATTAAACTGGACAAGCTCTCCAATTCACTTGTGGTTTCAGCTTTGACTATTAAAAAGGAAATAGAAGAAAACACAGGATGTGACCGTCTCC

>M6-PGSC0003DMG400033334 M6 LRR-RLP

ATCAAAGAAAACAAACACAAAATACACTCTCTCCAAAATATCCTTCTCTCTAAACTATTAGGGGAGAAAGAAAATTGTGTGGATAATAAGGAGGCGGGGAGGAGCATGTTGTCTAAGAGACTTCTGCTTAAGAAGGTTTTAATTGTGCTTGATGACATAGATCACATAGACCAGTTGGAATACCTAGCAAAGGATCTCAGTTGGTTTGGCAAGGGTAGCAGGATTATTGCAACCACTAGAGACCATTTGATAGGGGAAAATGACGTAGTACATGAAATGAATACACTACTTGACCGTGATGCTATTAAACTGTTCAATCAATATGCTTTCAAAGAAAAAGTTCCAAATGAGTGTTTTAAAGATCTAACGTCGGAAATAGTAAGTCTTGCTAAAGGCCATCCTTTAGCCCTGAAAGTATGGGGTTCTTCATTACACCGTATGGATATAACTAAGTGGGAAAGTTCCTTGGTCCGAATAAAGGAAAACTCTGGTGTAGAAATTACTAAGAACCTCAAAATAAGTTATGATGGGTTGGAGCCCAAAGATCAAAAGATATTTCTTGATATAGCATGTTTCTTCCGAGGAATTAAAAAAGAAAAAGTCATGCAAGTTCTTGAGAGCTGTTATTCTAGACCAGATTTGTGTGTTCTAGCTAATAAATCTCTTGTGTCCATTTCTGAAAATAATATGATTGAAATGCATGACTTAATTCAAGAAATGGGTAAATATATAGTGCAAATGCAAGAGCATCCGGGAGAACGTAGCAGATTATGGAATGTTGAAGATTTCAAAGAAGTTGTGGACAACAATATGTTAAGTAGGCTAAACAATGCAATAATATTCAATTTTACTTTTCATATTCCAAAGACTTATAGAATATATATATATAGTCAATTTCGTTTATTTATTCATCTTTCTTTATATTCTAGCGCGTATGTCATTTTAGTCATTTATTACTTTAAGGGAAAAGGGCTCGATATACCCCTCAACTTTGTCATGTGGAGTTGATATGCCCCTCGTTATGAAAGTGGCTCATATATACCCTTACCATTATACAAATGGCTCACATATACTCCTACCGTTACAAAATGAGCTCACATATATTCTTCATTTAATGGGAGTGAAAAATTAATTTTAAATTTATATATTTAACTTTTTTTAAAAAAAAAATCATTTAGGGTATATATGATCCTTCTAGCAAAATTCAAGGTATATTTTAATCTTTTTCATACATAAATTATTTTTTGACTTATTTGACTATCATTATTTGAGTTTCTTATTCTTATTTTATTTTTTTCTTTCATTCCTTAATTTAAAGAAAAAAATTAAAACTATTTTTTGTGGCTATATTATAATTTAAAACTATTATTTCAGCTATACTGTAATTTATATTTTGTATTTAGAAAAAATATTTGGGTCATCTATAATAAATTTTACAAGAATATTAGTGAAACATAAATAAATTTGACCATCAAAATAATAAATCTAAATTAGTCTTTGAAACAAAAAAAAAAGGTCAAAAAAAATATATGTTTGAGGATGATTATGTATATCCATATGGGATTATATTTTTATTAAAACAAAATAATTAAAAAATTAAATTAAATTAATTTTTTTACTTCCGTTAGAGGAAAAGGGTATATGTGAGCCATTTATGTAACAATAGGGGTATATATTAGCACTTTCATAACGAAGGGTATATCAACTCTAAATGTCAAAGTTGTGGGGTATATCAGACTCTTTTCCTTTACTTTATTAATAATAGAAGCAAAAATAATAGTAGTTGCCTAAGGTAGCTAATTCGTCAAGTATCGATTTTTAGTTAACGTTGCCTAATTCGTTTATGTTATG

>M6-PGSC0003DMG400033563 M6 LRR-RLP

CATCATGATTCCAGGTATACAATTCTCTATTTTCCTTTTTTTTTGTTTTTAATTTTTTCTTTTTGGATTATAAAATTTTTAAGTTTAATTAAAGCATTCAACTTTTTTTCTAATAGATTCTAAAAGAGTTGATGCTGATATTGAAGATAGATTCAGTGCTCTACCAATAAACCTTGTAGATGATATTCTAAGGCTCTTACCAGTTCACGACGCAGCAAGAACTAGTATTTTGTCAAAAAATTGGAGATATATTTGGGCCGCGCTTCCCAATTTAATGCTTGATAAGACCTTTTGCAGTAAATTAGCTGTAAAATCTCAATTTGTATTGAAAGAAACAATAGATATGATTCTCTTACAACATATTGGAGATATTGTGAAGTTTGATCTTGATGTTTCAGGAGTACAATTGTGTTCATATGCAGACATCGATAGATGGATGCTTTATGTCACGAGATATGGTGTCAAGGAATTAACCCTTTTCATGTCAAAGAATATTAACAGTACTTATAAAGTGCCTTCTTATATATTCAATTGTCAAACCCTGACACAATTGAAACTCTTCAATTGCATCTTCAAATTACCAAATTCTTTTCTTGGGTTTCAAACTCTTACCATCCTTCAGTTGAAAGAAGTAACCTTCGCGCCTGCAATAGAGTTTTATGCTATCAAAACGCCACTTCTTGTCGAATTAATCTTGACACATTGTGAGGGCACTCAATATCTTAACATTGAGTTCTCATCAGGATTGAAGTACTTGTTTGTTCGTGAGAGTCACTATAATCTTGATTTAAATAGCAATTACTTTAAGAACGTCAAAAAATTGACATACTTATACCTTGTGATTGATAATCCAATACGTGTTGATGAAAGATTAACTTTGGAAAAGCTTCTTTTTAGTTCGCCTACACTTGAGACACTTACTTTGGGTTTTTATGTACTTGAGGTAAGAAGCTAACATGTTGTTTTACTTAGATATCTACGTTATCCTTCTGTTTGTATAATTTTTCGTTAAAGCTTTTGAATGCAGGTACAGTCCCAAAGGGGCTTCATTATACACTCAACTGCTTGTGGCATCTATAGTTAGGTGTAAACTTCAACAAATTGGGTCAGACTTGTTACACTCTGGAGTTGATTAAAAGTTTCCCCAATTTGAGTAAACTTGAGATTTGGGTACGAA

>M6-PGSC0003DMG400033632 M6 LRR-RLP
