## Supplementary material for "RLP/K enrichment sequencing; a novel method to identify receptor-like protein (*RLP*) and receptor-like kinase (*RLK*) genes": Note S3

>Cf-2.1

ATGATGATGGTTTCTAGAAAAGTAGTCTCTTCACTTCAGTTTTTCACTCTTTTCTACCTCTTTACAGTTGCATTTGCTTCGACTGAGGAGGCAACTGCCCTCTTGAAATGGAAAGCAACTTTCAAGAACCAGAATAATTCCTTTTTGGCTTCATGGATTCCAAGTTCTAATGCATGCAAGGACTGGTATGGAGTTGTATGCTTTAATGGTAGGGTAAACACGTTGAATATTACAAATGCTAGTGTCATTGGTACACTCTATGCTTTTCCATTTTCATCCCTCCCTTCTCTTGAAAATCTTGATCTTAGCAAGAACAATATCTATGGTACCATTCCACCTGAGATTGGTAATCTCACAAATCTTGTCTATCTTGACTTGAACAACAATCAGATTTCAGGAACAATACCACCACAAATCGGTTTACTAGCCAAGCTTCAGATCATCCGCATATTTCACAATCAATTAAATGGATTTATTCCTAAAGAAATAGGTTACCTAAGGTCTCTTACTAAGCTATCTTTGGGTATCAACTTTCTTAGTGGTTCCATTCCTGCTTCAGTGGGGAATCTGAACAACTTGTCTTTTTTGTATCTTTACAATAATCAGCTTTCTGGCTCTATTCCTGAAGAAATAAGTTACCTAAGATCTCTTACTGAGCTAGATTTGAGTGATAATGCTCTTAATGGCTCTATTCCTGCTTCATTGGGGAATATGAACAACTTGTCTTTTTTGTTTCTTTATGGAAATCAGCTTTCTGGCTCTATTCCTGAAGAAATATGTTACCTAAGATCTCTTACTTACCTAGATTTGAGTGAGAATGCTCTTAATGGCTCTATTCCTGCTTCATTGGGGAATTTGAACAACTTGTCTTTTTTGTTTCTTTATGGAAATCAGCTTTCTGGCTCTATTCCTGAAGAAATAGGTTACCTAAGATCTCTTAATGTCCTAGGTTTGAGTGAGAATGCTCTTAATGGCTCTATTCCTGCTTCATTGGGGAATCTGAAAAACTTGTCTAGGTTGAATCTTGTTAATAATCAGCTTTCTGGCTCTATTCCTGCTTCATTGGGGAATCTGAACAACTTGTCTATGTTGTATCTTTACAATAACCAGCTTTCTGGCTCTATTCCTGCTTCATTGGGGAATCTGAACAACTTGTCTATGTTGTATCTTTACAATAATCAGCTTTCTGGCTCTATTCCTGCTTCATTGGGGAATCTGAACAACTTGTCTAGGTTGTATCTCTACAATAATCAGCTTTCTGGCTCTATTCCTGAAGAAATAGGTTACTTGAGTTCTCTTACTTATCTAGATTTGAGTAATAACTCCATTAATGGATTTATTCCTGCTTCATTTGGCAATATGAGCAACTTGGCTTTTTTGTTTCTTTATGAAAATCAGCTTGCTAGCTCTGTTCCTGAAGAAATAGGTTACCTAAGGTCTCTTAATGTCCTTGATTTGAGTGAGAATGCTCTTAATGGCTCTATTCCTGCTTCATTCGGGAATTTGAACAACTTGTCTAGGTTGAATCTTGTTAATAATCAGCTTTCTGGCTCTATTCCTGAAGAAATAGGTTACCTAAGGTCTCTTAATGTCCTTGATTTGAGTGAGAATGCTCTTAATGGCTCTATTCCTGCTTCATTCGGGAATTTGAACAACTTGTCTAGGTTGAATCTTGTTAATAATCAGCTTTCTGGCTCTATTCCTGAAGAAATAGGTTACCTAAGATCTCTTAATGACCTAGGTTTGAGTGAGAATGCTCTTAATGGCTCTATTCCTGCTTCATTGGGGAATCTGAACAACTTGTCTATGTTGTATCTTTACAATAATCAGCTTTCTGGCTCTATTCCTGAAGAAATAGGTTACTTGAGTTCTCTTACTTATCTATCTTTGGGTAATAACTCTCTTAATGGACTTATTCCTGCTTCATTTGGCAATATGAGAAATCTGCAAGCTCTGATTCTCAATGATAACAATCTCATTGGGGAAATTCCTTCATCTGTGTGCAATTTGACATCACTGGAAGTGTTGTATATGCCGAGAAACAATTTGAAGGGAAAAGTTCCGCAATGTTTGGGTAATATCAGTAACCTTCAGGTTTTGTCGATGTCATCTAATAGTTTCAGTGGAGAGCTCCCTTCATCTATTTCCAATTTAACATCACTACAAATACTTGATTTTGGCAGAAACAATCTGGAGGGAGCAATACCACAATGTTTTGGCAATATTAGTAGCCTCGAGGTTTTTGATATGCAGAACAACAAACTTTCTGGGACTCTTCCAACAAATTTTAGCATTGGATGTTCACTGATAAGTCTCAACTTGCATGGCAATGAACTAGAGGATGAAATCCCTCGGTCTTTGGACAATTGCAAAAAGCTGCAAGTTCTTGATTTAGGAGACAATCAACTCAACGACACATTTCCCATGTGGTTGGGAACTTTGCCAGAGCTGAGAGTTTTAAGGTTGACATCGAATAAATTGCATGGACCTATAAGATCATCAAGGGCTGAAATCATGTTTCCTGATCTTCGAATCATAGATCTCTCTCGCAATGCATTCTCGCAAGACTTACCAACGAGTCTATTTGAACATTTGAAAGGGATGAGGACAGTTGATAAAACAATGGAGGAACCAAGTTATGAAAGCTATTACGATGACTCGGTGGTAGTTGTGACAAAGGGATTGGAGCTTGAAATTGTGAGAATTTTGTCTTTGTACACAGTTATCGATCTTTCAAGCAACAAATTTGAAGGACATATTCCTTCTGTCCTGGGAGATCTCATTGCGATCCGTATACTTAATGTATCTCATAATGCATTGCAAGGCTATATACCATCATCACTTGGAAGTTTATCTATACTGGAATCACTAGACCTTTCGTTTAACCAACTTTCAGGAGAGATACCACAACAACTTGCTTCTCTTACGTTTCTTGAATTCTTAAATCTCTCCCACAATTATCTCCAAGGATGCATCCCTCAAGGACCTCAATTCCGTACCTTTGAGAGCAATTCATATGAAGGTAATGATGGATTACGTGGATATCCAGTTTCAAAAGGTTGTGGCAAAGATCCTGTGTCAGAGAAAAACTATACAGTGTCTGCGCTAGAAGATCAAGAAAGCAATTCTGAATTTTTCAATGATTTTTGGAAAGCAGCTCTGATGGGCTATGGAAGTGGACTGTGTATTGGCATATCCATGATATATATCTTGATCTCGACTGGAAATCTAAGATGGCTTGCAAGAATCATTGAAAAACTGGAACACAAAATTATCATGCAAAGGAGAAAGAAGCAGCGAGGTCAAAGAAATTACAGAAGAAGAAATAATCACTTCTAG

>Cf-2.2_

ATGATGATGGTTTCTAGAAAAGTAGTCTCTTCACTTCAGTTTTTCACTCTTTTCTACCTCTTTACAGTTGCATTTGCTTCGACTGAGGAGGCAACTGCCCTCTTGAAATGGAAAGCAACTTTCAAGAACCAGAATAATTCCTTTTTGGCTTCATGGATTCCAAGTTCTAATGCATGCAAGGACTGGTATGGAGTTGTATGCTTTAATGGTAGGGTAAACACGTTGAATATTACAAATGCTAGTGTCATTGGTACACTCTATGCTTTTCCATTTTCATCCCTCCCTTCTCTTGAAAATCTTGATCTTAGCAAGAACAATATCTATGGTACCATTCCACCTGAGATTGGTAATCTCACAAATCTTGTCTATCTTGACTTGAACAACAATCAGATTTCAGGAACAATACCACCACAAATCGGTTTACTAGCCAAGCTTCAGATCATCCGCATATTTCACAATCAATTAAATGGATTTATTCCTAAAGAAATAGGTTACCTAAGGTCTCTTACTAAGCTATCTTTGGGTATCAACTTTCTTAGTGGTTCCATTCCTGCTTCAGTGGGGAATCTGAACAACTTGTCTTTTTTGTATCTTTACAATAATCAGCTTTCTGGCTCTATTCCTGAAGAAATAAGTTACCTAAGATCTCTTACTGAGCTAGATTTGAGTGATAATGCTCTTAATGGCTCTATTCCTGCTTCATTGGGGAATATGAACAACTTGTCTTTTTTGTTTCTTTATGGAAATCAGCTTTCTGGCTCTATTCCTGAAGAAATATGTTACCTAAGATCTCTTACTTACCTAGATTTGAGTGAGAATGCTCTTAATGGCTCTATTCCTGCTTCATTGGGGAATTTGAACAACTTGTCTTTTTTGTTTCTTTATGGAAATCAGCTTTCTGGCTCTATTCCTGAAGAAATAGGTTACCTAAGATCTCTTAATGTCCTAGGTTTGAGTGAGAATGCTCTTAATGGCTCTATTCCTGCTTCATTGGGGAATCTGAAAAACTTGTCTAGGTTGAATCTTGTTAATAATCAGCTTTCTGGCTCTATTCCTGCTTCATTGGGGAATCTGAACAACTTGTCTATGTTGTATCTTTACAATAACCAGCTTTCTGGCTCTATTCCTGCTTCATTGGGGAATCTGAACAACTTGTCTATGTTGTATCTTTACAATAATCAGCTTTCTGGCTCTATTCCTGCTTCATTGGGGAATCTGAACAACTTGTCTAGGTTGTATCTCTACAATAATCAGCTTTCTGGCTCTATTCCTGAAGAAATAGGTTACTTGAGTTCTCTTACTTATCTAGATTTGAGTAATAACTCCATTAATGGATTTATTCCTGCTTCATTTGGCAATATGAGCAACTTGGCTTTTTTGTTTCTTTATGAAAATCAGCTTGCTAGCTCTGTTCCTGAAGAAATAGGTTACCTAAGGTCTCTTAATGTCCTTGATTTGAGTGAGAATGCTCTTAATGGCTCTATTCCTGCTTCATTCGGGAATTTGAACAACTTGTCTAGGTTGAATCTTGTTAATAATCAGCTTTCTGGCTCTATTCCTGAAGAAATAGGTTACCTAAGGTCTCTTAATGTCCTTGATTTGAGTGAGAATGCTCTTAATGGCTCTATTCCTGCTTCATTCGGGAATTTGAACAACTTGTCTAGGTTGAATCTTGTTAATAATCAGCTTTCTGGCTCTATTCCTGAAGAAATAGGTTACCTAAGATCTCTTAATGACCTAGGTTTGAGTGAGAATGCTCTTAATGGCTCTATTCCTGCTTCATTGGGGAATCTGAACAACTTGTCTATGTTGTATCTTTACAATAATCAGCTTTCTGGCTCTATTCCTGAAGAAATAGGTTACTTGAGTTCTCTTACTTATCTATCTTTGGGTAATAACTCTCTTAATGGACTTATTCCTGCTTCATTTGGCAATATGAGAAATCTGCAAGCTCTGATTCTCAATGATAACAATCTCATTGGGGAAATTCCTTCATCTGTGTGCAATTTGACATCACTGGAAGTGTTGTATATGCCGAGAAACAATTTGAAGGGAAAAGTTCCGCAATGTTTGGGTAATATCAGTAACCTTCAGGTTTTGTCGATGTCATCTAATAGTTTCAGTGGAGAGCTCCCTTCATCTATTTCCAATTTAACATCACTACAAATACTTGATTTTGGCAGAAACAATCTGGAGGGAGCAATACCACAATGTTTTGGCAATATTAGTAGCCTCGAGGTTTTTGATATGCAGAACAACAAACTTTCTGGGACTCTTCCAACAAATTTTAGCATTGGATGTTCACTGATAAGTCTCAACTTGCATGGCAATGAACTAGAGGATGAAATCCCTCGGTCTTTGGACAATTGCAAAAAGCTGCAAGTTCTTGATTTAGGAGACAATCAACTCAACGACACATTTCCCATGTGGTTGGGAACTTTGCCAGAGCTGAGAGTTTTAAGGTTGACATCGAATAAATTGCATGGACCTATAAGATCATCAAGGGCTGAAATCATGTTTCCTGATCTTCGAATCATAGATCTCTCTCGCAATGCATTCTCGCAAGACTTACCAACGAGTCTATTTGAACATTTGAAAGGGATGAGGACAGTTGATAAAACAATGGAGGAACCAAGTTATGAAAGCTATTACGATGACTCGGTGGTAGTTGTGACAAAGGGATTGGAGCTTGAAATTGTGAGAATTTTGTCTTTGTACACAGTTATCGATCTTTCAAGCAACAAATTTGAAGGACATATTCCTTCTGTCCTGGGAGATCTCATTGCGATCCGTATACTTAATGTATCTCATAATGCATTGCAAGGCTATATACCATCATCACTTGGAAGTTTATCTATACTGGAATCACTAGACCTTTCGTTTAACCAACTTTCAGGAGAGATACCACAACAACTTGCTTCTCTTACGTTTCTTGAATTCTTAAATCTCTCCCACAATTATCTCCAAGGATGCATCCCTCAAGGACCTCAATTCCGTACCTTTGAGAGCAATTCATATGAAGGTAATGATGGATTACGTGGATATCCAGTTTCAAAAGGTTGTGGCAAAGATCCTGTGTCAGAGAAAAACTATACAGTGTCTGCGCTAGAAGATCAAGAAAGCAATTCTGAATTTTTCAATGATTTTTGGAAAGCAGCTCTGATGGGCTATGGAAGTGGACTGTGTATTGGCATATCCATAATATATATCTTGATCTCGACTGGAAATCTAAGATGGCTTGCAAGAATCATTGAAGAACTGGAACACAAAATTATCATGCAAAGGAGAAAGAAGCAGCGAGGTCAAAGAAATTACAGAAGAAGAAATAATCGCTTCTAG

>Cf-4

ATGGGTTGTGTAAAACTTGTGTTTTTCATGCTATATGTCTTTCTCTTTCAACTTGTTTCCTCGTCATCCTTACCTCATTTGTGCCCCGAAGATCAAGCTCTTGCTCTTCTAGAATTCAAGAACATGTTTACCGTTAATCCTAATGCTTCTGATTATTGTTACGACAGAAGAACTCTTTCTTGGAACAAAAGCACAAGTTGCTGCTCATGGGATGGCGTTCATTGTGACGAAACGACAGGACAAGTGATTGAGCTTGACCTCCGTTGCATCCAACTTCAAGGCAAGTTTCATTCCAATAGTAGCCTCTTTCAACTCTCCAATCTCAAAAGGCTTGATTTGTCTTATAATGATTTCACTGGATCGCCCATTTCACCTAAATTTGGTGAGTTTTCAGATTTGACGCATCTCGATTTGTCGCATTCAAGTTTTAGGGGTGTAATCCCTTCTGAAATCTCTCATCTTTCTAAACTATACGTTCTTCGTATTAGTCTAAATGAGCTTACTTTTGGTCCTCACAATTTTGAATTGCTTCTTAAGAACTTGACCCAATTAAAAGTGCTCGACCTTGAATCTATCAACATCTCTTCCACTATTCCTTTGAATTTCTCTTCTCATTTAACAAATCTATGGCTTCCATACACAGAGTTACGTGGGATATTGCCCGAAAGAGTTTTCCACCTTTCCGACTTAGAATTTCTCGATTTATCAAGCAATCCCCAGCTCACGGTTAGGTTTCCCACAACCAAATGGAATAGCAGTGCATCACTCATGAAGTTATATCTCTATAATGTGAATATTGATGATAGGATACCTGAATCATTTAGCCATCTAACTTCACTTCATAAGTTGTACATGAGTCGTTCTAATCTGTCAGGGCCTATTCCTAAACCTCTATGGAATCTCACCAACATAGTGTTTTTGGACCTTAATAATAACCATCTTGAAGGACCAATTCCATCCAACGTAAGCGGACTACGTAACCTACAAATACTTTGGTTGTCATCAAACAACTTAAATGGGAGTATACCATCCTGGATATTCTCCCTTCCATCACTGATAGGGTTAGACTTGAGCAATAACACTTTCAGTGGAAAAATTCAAGAGTTCAAGTCCAAAACATTAAGTACCGTTACTCTAAAACAAAATAAGCTAAAAGGTCCTATTCCGAATTCACTCCTAAACCAGAAGAACCTACAATTCCTTCTCCTTTCACACAATAATATCAGTGGACATATTTCTTCAGCTATCTGCAATCTGAAAACATTGATATTGTTAGACTTGGGAAGTAATAATTTGGAGGGAACAATCCCGCAATGCGTGGTTGAGAGGAACGAATACCTTTCGCATTTGGATTTGAGCAACAACAGACTTAGTGGGACAATCAATACAACTTTTAGTGTTGGAAACATTTTAAGGGTCATTAGCTTGCACGGGAATAAGCTAACGGGGAAAGTCCCACGATCTATGATCAATTGCAAGTATTTGACACTACTTGATCTAGGTAACAATATGTTGAATGACACATTTCCAAACTGGTTGGGATACCTATTTCAATTGAAGATTTTAAGCTTGAGATCAAATAAGTTGCATGGTCCCATCAAATCTTCAGGGAATACAAACTTGTTTATGGGTCTTCAAATTCTTGATCTATCATCTAATGGATTTAGTGGGAATTTACCCGAAAGAATTTTGGGGAATTTGCAAACCATGAAGGAAATTGATGAGAGTACAGGATTCCCAGAGTATATTTCTGATCCATATGATATTTATTACAATTATTTGACGACAATTTCTACAAAGGGACAAGATTATGATTCTGTTCGAATTTTGGATTCTAACATGATTATCAATCTCTCAAAGAACAGATTTGAAGGTCATATTCCAAGCATTATTGGAGATCTTGTTGGACTTCGTACGTTGAACTTGTCTCACAATGTCTTGGAAGGTCATATACCGGCATCATTTCAAAATTTATCAGTACTCGAATCATTGGATCTCTCATCTAATAAAATCAGCGGAGAAATTCCGCAGCAGCTTGCATCCCTCACATTCCTTGAAGTCTTAAATCTCTCTCACAATCATCTTGTTGGATGCATCCCCAAAGGAAAACAATTTGATTCGTTCGGGAACACTTCGTACCAAGGGAATGATGGGTTACGCGGATTTCCACTCTCAAAACTTTGTGGTGGTGAAGATCAAGTGACAACTCCAGCTGAGCTAGATCAAGAAGAGGAGGAAGAAGATTCACCAATGATCAGTTGGCAGGGGGTTCTCGTGGGTTACGGTTGTGGACTTGTTATTGGACTGTCCGTAATATACATAATGTGGTCAACTCAATATCCAGCATGGTTTTCGAGGATGGATTTAAAGTTGGAACACATAATTACTACGAAAATGAAAAAGCACAAGAAAAGATATTAG

>Cf-5

ATGATGATGGTTACTAGCAAAGTATTCTCTTCACTTCAGTTTTTCACTGTTTTCTACCTCTTTACAGTTGCATTTGCTTCGACTGAGGAGGCAACTGCCCTCTTGAAATGGAAAGCAACTTTCAAGAACCAGAATAATTCCTTTTTGGCTTCATGGACGACAAGTTCTAATGCATGCAAGGACTGGTATGGAGTTGTATGCTTGAATGGTAGGGTAAACACGTTGAATATTACAAATGCCAGTGTCATTGGTACACTTTATGCTTTTCCATTTTCATCCCTCCCTTTTCTCGAGAATCTTGATCTTAGCAACAACAATATCTCTGGTACCATTCCACCTGAGATTGGTAATCTCACAAATCTTGTCTATCTTGACTTGAACACCAATCAGATTTCAGGAACAATTCCACCACAAATCGGTTCACTAGCCAAGCTTCAGATCATCCGCATATTTAACAATCATTTAAATGGCTTTATTCCTGAAGAAATAGGTTACCTAAGGTCTCTTACTAAGCTATCTTTGGGTATCAACTTTCTTAGTGGTTCTATTCCTGCTTCATTGGGCAATATGACCAACTTGTCTTTTTTATTTCTTTATGAAAATCAGCTTTCTGGCTTTATTCCTGAAGAAATAGGTTACCTAAGGTCTCTTACTAAGCTATCTTTGGATATCAACTTTCTTAGTGGTTCCATTCCTGCTTCATTGGGGAATCTGAACAACTTGTCTTTTTTGTATCTTTACAATAATCAGCTTTCTGGCTCTATTCCTGAAGAAATAGGTTACCTAAGGTCACTTACTAAGCTATCTTTGGGTATCAACTTTCTTAGTGGTTCCATTCCTGCTTCATTGGGGAATCTAAACAACTTGTCTAGGTTGGATCTTTACAATAATAAGCTTTCTGGCTCTATTCCTGAAGAAATAGGTTACCTAAGGTCTCTTACTTACCTAGATTTGGGTGAGAATGCTCTTAATGGCTCTATTCCTTCTTCATTGGGGAATCTAAACAACTTGTCTAGGTTGGATCTTTACAATAATAAGCTTTCTGGCTCTATTCCTGAAGAAATAGGTTACCTAAGGTCTCTTACTTACCTAGATTTGGGTGAGAATGCTCTTAATGGCTCTATTCCTGCTTCATTGGGGAATCTGAACAACTTGTTTATGTTGTATCTTTACAATAATCAGCTTTCTGGCTCTATTCCTGAAGAAATAGGTTACCTGAGTTCTCTTACTGAACTATATTTGGGTAATAACTCTCTTAATGGCTCTATTCCTGCTTCATTGGGGAATCTGAACAACTTGTTTATGTTGTATCTTTACAATAATCAGCTTTCTGGCTCTATTCCTGAAGAAATAGGTTACCTGAGTTCTCTTACTGAACTATTTTTGGGTAATAACTCTCTTAATGGCTCTATTCCTGCTTCATTGGGGAATCTAAACAACTTGTCTAGGTTGTATCTTTACAATAATCAGCTTTCTGGCTCTATTCCTGCTTCATTTGGCAATATGAGAAATCTGCAAACTCTGTTTCTCAGTGATAACGATCTCATTGGGGAAATTCCTTCATTTGTGTGCAATTTGACATCACTGGAAGTGTTGTATATGTCGAGAAACAATTTGAAGGGAAAAGTTCCGCAATGTTTGGGTAATATCAGTGACCTTCACATTTTGTCGATGTCATCTAATAGTTTCAGAGGAGAGCTCCCTTCATCTATTTCCAATTTAACATCACTAAAAATACTTGATTTTGGCAGAAACAATCTGGAGGGAGCAATACCACAATTTTTTGGCAATATTAGTAGCCTCCAGGTTTTTGATATGCAGAATAACAAACTTTCTGGGACTCTTCCAACAAATTTTAGCATTGGATGTTCACTGATAAGTCTCAACTTGCATGGCAATGAACTAGCAGATGAAATCCCTCGGTCTTTGGACAATTGCAAAAAGCTGCAAGTTCTTGATTTAGGAGACAATCAACTCAACGACACATTTCCCATGTGGTTGGGAACTTTGCCAGAGCTGAGAGTTTTAAGGTTGACATCGAATAAATTGCATGGACCTATAAGATCATCAGGGGCTGAAATCATGTTTCCTGATCTCCGAATCATAGATCTCTCTCGCAATGCATTCTCGCAAGACTTACCAACGAGTCTATTTGAACATTTGAAAGGGATGAGGACAGTTGATAAAACAATGGAGGAACCAAGTTATGAAAGCTATTACGATGACTCGGTGGTAGTTGTGACAAAGGGATTGGAGCTTGAAATTGTGAGAATTCTGTCTTTGTACACAATTATCGATCTTTCAAGCAACAAATTTGAAGGACATATTCCTTCTGTCCTGGGAGATCTCATTGCGATCCGTGTACTTAATGTATCTCATAATGCATTGCAAGGCTATATACCATCATCACTTGGAAGTTTATCTATACTGGAATCACTAGACCTTTCGTTTAACCAACTTTCGGGAGAGATACCACAACAACTTGCTTCTCTTACGTTTCTTGAAGTCTTAAATCTCTCCCACAATTATCTCCAAGGATGCATCCCTCAAGGACCTCAATTCCGTACCTTTGAGAGCAATTCATATGAAGGTAATGATGGATTACGTGGATATCCAGTTTCAAAAGGTTGTGGCAAAGATCCTGTGTCAGAGAAAAACTATACAGTGTCTGCGCTAGAAGATCAAGAAAGCAATTCTGAATTTTTCAATGATTTTTGGAAAGCAGCTCTGATGGGCTATGGAAGTGGACTGTGTATTGGCATATCCATAATATATATCTTGATCTCGACTGGAAATCTAAGATGGCTTGCAAGAATCATTGAAGAACTGGAACACAAAATTATCGTGCAAAGGAGAAAGAAGCAGCGAGGTCAAAGAAATTACAGAAGAAGAAATAATCGCTTCTAG

>Cf-9

ATGGATTGTGTAAAACTTGTATTCCTTATGCTATATACCTTTCTCTGTCAACTTGCTTTATCCTCATCCTTGCCTCATTTGTGCCCCGAAGATCAAGCTCTTTCTCTTCTACAATTCAAGAACATGTTTACCATTAATCCTAATGCTTCTGATTATTGTTACGACATAAGAACATACGTAGACATTCAGTCATATCCAAGAACTCTTTCTTGGAACAAAAGCACAAGTTGCTGCTCATGGGATGGCGTTCATTGTGACGAGACGACAGGACAAGTGATTGCGCTTGACCTCCGTTGCAGCCAACTTCAAGGCAAGTTTCATTCCAATAGTAGCCTCTTTCAACTCTCCAATCTCAAAAGGCTTGATTTGTCTTTTAATAATTTCACTGGATCACTCATTTCACCAAAATTTGGTGAGTTTTCAAATTTGACGCATCTCGATTTGTCGCATTCTAGTTTTACAGGTCTAATTCCTTCTGAAATCTGTCACCTTTCTAAACTACACGTTCTTCGTATATGTGATCAATATGGGCTTAGTCTTGTACCTTACAATTTTGAACTGCTCCTTAAGAACTTGACCCAATTAAGAGAGCTCAACCTTGAATCTGTAAACATCTCTTCCACTATTCCTTCAAATTTCTCTTCTCATTTAACAACTCTACAACTTTCAGGCACAGAGTTACATGGGATATTGCCCGAAAGAGTTTTTCACCTTTCCAACTTACAATCCCTTCATTTATCAGTCAATCCCCAGCTCACGGTTAGGTTTCCCACAACCAAATGGAATAGCAGTGCATCACTCATGACGTTATACGTCGATAGTGTGAATATTGCTGATAGGATACCTAAATCATTTAGCCATCTAACTTCACTTCATGAGTTGTACATGGGTCGTTGTAATCTGTCAGGGCCTATTCCTAAACCTCTATGGAATCTCACCAACATAGTGTTTTTGCACCTTGGTGATAACCATCTTGAAGGACCAATTTCCCATTTCACGATATTTGAAAAGCTCAAGAGGTTATCACTTGTAAATAACAACTTTGATGGCGGACTTGAGTTCTTATCCTTTAACACCCAACTTGAACGGCTAGATTTATCATCCAATTCCCTAACTGGTCCAATTCCATCCAACATAAGCGGACTTCAAAACCTAGAATGTCTCTACTTGTCATCAAACCACTTGAATGGGAGTATACCTTCCTGGATATTCTCCCTTCCTTCACTGGTTGAGTTAGACTTGAGCAATAACACTTTCAGTGGAAAAATTCAAGAGTTCAAGTCCAAAACATTAAGTGCCGTTACTCTAAAACAAAATAAGCTGAAAGGTCGTATTCCGAATTCACTCCTAAACCAGAAGAACCTACAATTACTTCTCCTTTCACACAATAATATCAGTGGACATATTTCTTCAGCTATCTGCAATCTGAAAACATTGATATTGTTAGACTTGGGAAGTAATAATTTGGAGGGAACAATCCCACAATGCGTGGTTGAGAGGAACGAATACCTTTCGCATTTGGATTTGAGCAAAAACAGACTTAGTGGGACAATCAATACAACTTTTAGTGTTGGAAACATTTTAAGGGTCATTAGCTTGCACGGGAATAAGCTAACGGGGAAAGTCCCACGATCTATGATCAATTGCAAGTATTTGACACTACTTGATCTAGGTAACAATATGTTGAATGACACATTTCCAAACTGGTTGGGATACCTATTTCAATTGAAGATTTTAAGCTTGAGATCAAATAAGTTGCATGGTCCCATCAAATCTTCAGGGAATACAAACTTGTTTATGGGTCTTCAAATTCTTGATCTATCATCTAATGGATTTAGTGGGAATTTACCCGAAAGAATTTTGGGGAATTTGCAAACCATGAAGGAAATTGATGAGAGTACAGGATTCCCAGAGTATATTTCTGATCCATATGATATTTATTACAATTATTTGACGACAATTTCTACAAAGGGACAAGATTATGATTCTGTTCGAATTTTGGATTCTAACATGATTATCAATCTCTCAAAGAACAGATTTGAAGGTCATATTCCAAGCATTATTGGAGATCTTGTTGGACTTCGTACGTTGAACTTGTCTCACAATGTCTTGGAAGGTCATATACCGGCATCATTTCAAAATTTATCAGTACTCGAATCTTTGGATCTCTCATCTAATAAAATCAGCGGAGAAATTCCGCAGCAGCTTGCATCCCTCACATTCCTTGAAGTCTTAAATCTCTCTCACAATCATCTTGTTGGATGCATCCCCAAAGGAAAACAATTTGATTCGTTCGGGAACACTTCGTACCAAGGGAATGATGGGTTACGCGGATTTCCACTCTCAAAACTTTGTGGTGGTGAAGATCAAGTGACAACTCCAGCTGAGCTAGATCAAGAAGAGGAGGAAGAAGATTCACCAATGATCAGTTGGCAGGGGGTTCTCGTGGGTTACGGTTGTGGACTTGTTATTGGACTGTCCGTAATATACATAATGTGGTCAACTCAATATCCAGCATGGTTTTCGAGGATGGATTTAAAGTTGGAACACATAATTACTACGAAAATGAAAAAGCACAAGAAAAGATATTAG

>LeEix1

ATGGACAAATGGAAATATGCAAGATTAGCACAGTTCCTTTTCACTTTGTCTCTACTGTTCCTAGAGACATCTTTTGGATTAGGTGGTAACAAGACCCTATGTTTAGATAAGGAGAGAGATGCTCTTCTTGAGTTCAAAAGAGGTCTTACTGATTCTTTTGATCATTTATCTACATGGGGTGATGAAGAAGATAAACAAGAATGCTGCAAATGGAAGGGTATTGAATGTGACAGAAGAACAGGTCATGTAACTGTTATTGATCTACACAATAAGTTTACTTGTTCTGCTGGTGCCAGTGCTTGTTTTGCTCCTAGATTGACAGGTAAACTTAGCCCTTCTCTGCTTGAGTTGGAGTACTTGAATTACTTGGACCTCAGTGTTAATGAATTTGAAAGAAGTGAAATACCAAGATTCATAGGCTCACTTAAGAGACTAGAGTACTTGAACCTGTCAGCTTCTTTTTTTTCTGGTGTAATTCCAATACAGTTCCAGAATCTAACTTCTTTAAGGACTCTTGATCTTGGAGAAAATAATCTTATAGTAAAGGACCTTAGATGGCTTTCTCATCTGTCTTCTCTAGAGTTTTTGAGTCTGAGTTCTAGCAACTTCCAAGTAAACAATTGGTTTCAAGAGATAACTAAGGTCCCTTCATTGAAAGAACTGGACTTGAGTGGCTGTGGACTCTCTAAGTTGGCTCCATCTCAAGCTGATTTAGCCAATTCTTCTTTTATATCTCTTTCTGTTCTTCATTTATGTTGTAATGAGTTTTCTTCTTCATCTGAATATAGCTGGGTATTCAATTTGACCACAAGCCTAACTAGCATTGACCTCCTTTATAATCAACTCAGCGGTCAAATTGATGATCGCTTTGGGACCTTGATGTATCTTGAACATCTTGATCTTGCTAATAACTTAAAGATTGAAGGTGGGGTTCCCAGTTCTTTTGGGAATTTGACACGTTTACGTCATCTGGACATGTCTAATACTCAGACAGTCCAATGGCTTCCTGAGTTGTTTCTCAGGTTATCAGGCAGTAGGAAATCACTTGAGGTTTTGGGGTTGAACGAAAACTCATTGTTTGGTTCAATTGTTAATGCAACAAGATTTTCATCCTTAAAGAAATTATACCTGCAGAAGAATATGCTGAATGGTTCTTTTATGGAAAGTGCGGGACAAGTTTCTACCCTTGAGTATCTGGATTTGTCTGAAAACCAAATGAGAGGGGCATTACCAGATTTAGCATTGTTTCCATCATTGAGAGAGTTGCATCTTGGCTCTAATCAATTTCGAGGGAGGATACCACAAGGTATCGGAAAACTTTCACAGCTTAGAATTTTGGATGTCTCGTCCAATAGACTGGAAGGACTACCAGAAAGTATGGGGCAACTATCTAATCTGGAAAGTTTTGATGCCTCTTACAATGTCCTGAAGGGAACAATTACTGAGTCCCACCTTTCAAACCTCTCCAGTTTAGTGGATTTGGACTTGTCATTCAACTCATTGGCTCTGAAGACGAGCTTCAATTGGCTTCCTCCTTTTCAGCTACAAGTTATAAGCCTGCCGTCTTGCAATTTGGGACCTTCTTTCCCAAAATGGCTTCAAAATCAGAACAACTATACTGTTCTTGATATCTCTCTTGCGAGTATTTCAGACACGCTTCCAAGTTGGTTCTCTAGCTTCCCCCCCGATCTAAAGATTTTGAATCTCTCAAACAACCAAATCAGTGGAAGAGTGTCTGACTTAATAGAGAATACATATGGGTACAGGGTTATAGATTTAAGCTATAACAACTTTTCAGGGGCTTTGCCACTAGTCCCTACCAATGTCCAAATATTTTACCTGCATAAAAATCAGTTTTTTGGATCCATCTCTTCAATTTGTCGAAGTAGAACATCTCCCACTTCTCTAGACTTGTCACACAATCAATTCTCAGGAGAACTTCCAGATTGTTGGATGAACATGACAAGTCTAGCTGTTCTTAATCTAGCTTATAACAATTTCTCTGGAGAAATTCCACATTCATTGGGTTCCTTGACAAATTTGAAGGCGTTATACATACGCCAAAACAGTTTAAGTGGAATGTTGCCTTCTTTTTCACAATGTCAGGGGTTGCAAATCTTGGATCTTGGAGGGAATAAGTTGACAGGAAGTATCCCTGGATGGATAGGGACTGATCTACTCAACTTGCGAATTCTAAGCCTGCGGTTCAACAGATTGCATGGCAGCATACCATCCATAATCTGTCAGCTTCAATTTCTTCAGATACTGGACCTTTCAGCGAATGGATTATCAGGGAAAATTCCACATTGCTTCAACAATTTCACCTTATTGTATCAAGATAATAATTCTGGTGAGCCAATGGAATTTATAGTTCAAGGTTTCTATGGTAAATTTCCTCGTCGTTACTTGTATATTGGTGACCTATTGGTTCAATGGAAAAATCAAGAGTCTGAGTACAAGAATCCTTTATTATATCTGAAGACTATTGATCTTTCAAGTAATGAATTAATTGGAGGTGTTCCTAAAGAAATAGCTGATATGAGAGGATTGAAATCTTTGAACCTCTCAAGAAATGAGCTGAATGGAACTGTCATTGAAGGAATAGGTCAAATGAGGATGTTGGAGTCTCTTGACATGTCAAGAAACCAGCTTTCTGGTGTGATACCGCAAGATCTTGCTAATTTGACTTTTCTTAGTGTGTTGGACTTGTCGAACAACCAGTTATCAGGGAGAATTCCATCAAGCACTCAACTGCAAAGTTTTGATAGATCATCCTACAGTGATAATGCTCAACTCTGTGGGCCTCCTCTTCAGGAGTGTCCTGGATATGCTCCTCCTAGCCCCCTTATTGATCATGGCAGCAACAACAATCCACAAGAACATGATGAGGAGGAGGAGTTTCCATCTCTGGAGTTTTATATATCAATGGTGCTAAGCTTTTTTGTTGCATTTTGGGGAATTTTGGGCTGTTTAATTGTCAATAGTTCTTGGAGGAATGCCTACTTCAAATTCTTGACGGACACGACAAGTTGGCTCGATATGATATCAAGAGTCTGGTTTGCAAGACTCAAGAAAAAGCTGAGGAGGGCCCGATGA

>LeEix2

ATGGGCAAAAGAACTAATCCAAGACATTTCCTTGTTACTTGGTCTTTACTGCTCCTAGAGACAGCTTTTGGATTAACTTCAAGAGAAGTTAACAAGACCCTTTGTATAGAAAAGGAGAGAGGTGCCCTTCTTGAGTTTAAAAGAGGCCTTAACGACGATTTTGGTCGTTTATCTACCTGGGGTGATGAAGAAGAATGCTGCAATTGGAAGGGTATTGAATGTGACAAAAGAACAGGTCATGTTATTGTTCTTGATCTCCACAGTGAGGTTACTTGTCCAGGCCATGCTTGTTTTGCTCCAATATTGACAGGTAAAGTTAGTCCTTCTCTACTTGAGTTGGAGTATTTGAATTTCTTGGACCTCAGTGTTAATGGATTTGAAAATAGTGAGATACCAAGATTCATAGGCTCCCTTAAGAGACTGGAGTACTTAAACCTTTCATCTTCTGATTTTTCTGGTGAAATTCCAGCACAGTTCCAGAATCTAACTTCTTTGAGGATTCTTGATCTCGGAAACAATAATCTTATAGTAAAGGACCTTGTGTGGCTTTCTCATCTCTCCTCTCTAGAATTCTTGCGCCTTGGTGGTAACGATTTCCAAGCAAGAAACTGGTTTCGAGAGATAACTAAGGTCCCTTCATTGAAAGAATTGGACTTGAGTGTTTGTGGACTCTCTAAATTCGTTCCGTCTCCAGCTGATGTAGCTAATTCATCTTTGATCTCTCTTTCTGTTCTTCATTTATGTTGTAATGAGTTTTCTACTTCATCTGAATATAGCTGGTTATTCAATTTTAGCACAAGCCTAACTAGCATAGACCTTTCTCATAATCAACTCAGTCGTCAAATTGATGATCGCTTTGGGAGCTTGATGTATCTTGAACATCTTAATCTTGCTAATAATTTTGGGGCTGAAGGTGGGGTTCCAAGTTCTTTTGGGAATTTGACACGTCTACATTATCTGGACATGTCTAACACTCAGACATACCAATGGCTTCCTGAGTTGTTTCTCAGGTTATCAGGTAGTAGGAAATCACTTGAGGTTTTGGGATTGAACGACAACTCGTTGTTTGGTTCAATTGTTAATGTGCCAAGATTTTCATCCTTGAAGAAATTATACCTGCAGAAGAATATGCTGAATGGTTTTTTCATGGAAAGAGTGGGACAAGTTTCGAGCCTCGAGTATCTAGACTTGTCTGATAACCAAATGAGAGGGCCATTACCAGATTTAGCACTTTTTCCATCATTAAGAGAGTTGCATCTTGGCTCTAATCAATTTCAAGGGAGGATACCACAAGGTATTGGAAAACTTTCACAGCTTAGAATTTTTGACGTCTCGTCCAATAGATTAGAGGGTTTACCAGAAAGTATGGGGCAACTATCAAACCTGGAAAGGTTTGATGCTTCTTATAATGTCCTGAAGGGTACAATCACAGAGTCCCACTTTTCAAACCTCTCCAGTTTAGTAGACTTAGACCTATCCTTCAACTTGTTGTCTTTGAACACGAGATTCGATTGGGTTCCTCCTTTTCAACTACAATTTATAAGGCTTCCATCTTGCAATATGGGACCTTCTTTTCCGAAATGGCTACAAACTCAGAATAACTACACTCTTCTTGATATTTCTCTTGCGAATATATCAGACATGCTACCAAGTTGGTTCTCCAATCTTCCTCCCGAGCTCAAGATTCTGAATCTCTCTAACAACCACATCAGTGGCAGAGTTTCGGAGTTCATAGTGAGTAAACAAGATTACATGATTATAGATTTAAGTTCTAACAACTTTTCAGGACATTTGCCACTAGTTCCTGCCAATATACAGATCTTTTACCTTCATAAAAATCACTTCTCTGGATCCATTTCTTCCATTTGTAGAAATACAATAGGAGCTGCCACTTCCATCGACTTGTCACGCAACCAATTTTCAGGAGAAGTTCCTGATTGTTGGATGAATATGAGTAATCTAGCTGTTCTAAATCTAGCCTACAACAATTTCTCCGGAAAAGTTCCACAATCATTAGGCTCCTTGACAAATTTGGAGGCGTTATACATACGTCAGAACAGCTTTAGAGGAATGTTGCCTTCTTTTTCACAATGTCAGCTGCTGCAAATCTTGGATATTGGAGGGAATAAGTTGACTGGAAGAATCCCAGCATGGATAGGGACTGACCTACTCCAATTGCGCATTCTAAGCCTACGTTCCAACAAATTCGATGGCAGCATTCCATCACTTATCTGCCAGCTTCAATTTCTTCAGATACTGGACCTTTCAGAAAATGGATTATCTGGGAAAATTCCACAGTGCCTCAACAACTTTACCATATTGCGTCAAGAAAATGGCTCTGGTGAGTCAATGGATTTTAAAGTCCGTTATGACTATATTCCAGGATCTTACTTGTACATAGGTGATTTACTGATTCAATGGAAAAACCAGGAGTCCGAGTACAAGAATGCTTTACTATATCTGAAGATCATTGATCTTTCAAGTAATAAATTAGTTGGAGGTATCCCTAAAGAGATAGCTGAAATGAGAGGATTGAGATCATTGAACCTCTCGAGAAATGATCTTAATGGAACTGTCGTTGAAGGAATAGGTCAAATGAAGTTGTTGGAGTCCCTCGACTTGTCAAGAAACCAACTCTCTGGCATGATCCCTCAGGGCCTTTCTAACTTGACTTTTCTTAGTGTGTTGGACTTATCGAACAACCACTTATCAGGAAGAATTCCATCAAGCACTCAACTGCAGAGTTTCGATAGATCATCCTATAGTGGCAATGCTCAACTCTGCGGTCCTCCTCTTGAAGAGTGTCCTGGATATGCTCCCCCTATCGATCGTGGAAGCAACACCAATCCACAAGAACATGATGATGATGATGAGTTCTCATCTCTGGAGTTTTATGTATCAATGGTGCTAGGTTTCTTCGTCACGTTCTGGGGAATTTTAGGCTGTTTGATTGTCAACCGTTCGTGGAGGAATGCCTACTTCACATTCTTAACAGACATGAAGAGTTGGCTCCATATGACATCAAGAGTCTGCTTTGCAAGACTGAAGGGAAAGCTAAGGAACTGA

>MCD360-1

ATGGTCATGAGTCTGTTTTTCTTTTATTCATTTCTATGTTTTGTGTTTTTAATAAGTGGATGCTTTTCTTCATCCTTTGATCATCATCTTTGCTCTCCCACTGAAGCTTCCTCTTTGCTTCAGTTTAAGCAATCCTTTCAAATATCTGATTACTCTACTCTTAAGTGTGATACTTCTTTCCCAAAAACTAAGTCTTGGAATGAGAGTAGGGATTGCTGCAGTTGGGATGGAGTCACTTGTGACTTATTAAACGGACATGTTATCGGGTTAGACCTTAGCTGCAGTCAGCTTCGTGGAAGTATTCATCCCAATAGCAGCCTCTTCCAACTTCATCATCTCCAAACAGTAAACCTTGCTTACAATAACTTTTCAACTTCTTCAATCTCACATAACATTGGCCGATGGAGAAATTTGAGGCATCTCAACCTTTCTAATTCTTTCTTTAGTGGGAAAATCCCAACAGAAATCTCATTCCTTTCCAATTTGGTTTCACTTGATCTTTCTTCTTCTTATGGATTACAACTTGATGAGAGAACATTTGAAACAATGCTTCACAACTTTACAAATCTGGAGGTACTAGCTCTCTTTCTTGGCAACATCTCATCACCGATACCTGTAAGTATTCATCCCAATAGCAGCCTCTTCCAGCTTCATCATCTCCACACACTAAACCTTGTTAACAATTTCTTTTATCCTTCTTCAATCCCAAATGGCATTGGCCGATTGAGGAATTTGAGGCATCTAAAACTCTATGGCTTTCAAGGGAAAATCCCAACAGAAATCTCATACCTTTCCAATTTGGTTTCACTTGATCTTTCTTATAGTTATGAAAGATTACAACTTGATGAGAGAACATTTGAAGCAATGTTTCAGAACTTGACAAATCTGGAGCTACTTTCTCTCTATGGTGTCAACATCTCATCTCAGATACCTGTGAATATTTCTTCTTCCTTAAGGTACCTGGATCTTGGTTATACTAATCTGCGAGGTGTTCTCACAGAGAACTTTTTCCTTCTGCCAAACTTGGAAATACTCAAATTGAGTGGGAATGATCTTGTCAAAGGAGTTTTTCCAAAGATCCACTGGAGCAACACTCTGTTAATGGAGTTGGATATTTCATCCACAGGCATCTCTGGTGAGGTGCCTGATTCAATTGGCACCTTCAGTTCCTTGAATATCTTGAACCTTGCAGGATGCCAATTCTCTGGTTCCATTCCTGATTCCATTAGCAACCTAACACAAATTAGGGAGTTGATTTTATATCATAATCATTTCACTGGCCATATTCCTTCCACAATCTCTAAATTGAAGCACCTCACACGTTTGGATCTTTCAATTAACTACTTTTCAGGTGAAATTCCTGATGTTTTCTCAAACCTCCAAGAGCTACGTACTTTACATCTTTCTTATAACAGCTTCATCGGTTCGTTTCCCGCTTCAATTTTAAGTTTGACACATCTTGAATACTTAGGATTGTCGAGTAATTCCCTATCTGGCCCACTGCCTAGCAACCAAAGCATGCTTCAAAAGCTAACCGAACTGAATTTGTCATACAACTCACTGAATGGTACCATACCATCTTGGGTGTTTAGCCTGCCTTTGCTATCTTCAGTGTCGCTCCAACATAACCGATTAAGAGGACTAGCCGATGAAGTGATCAAAACAAACCCAACATTAAAACAACTGTATTTAAGCAATAATCAACTCAGTGGTTCTTTTCCTCAATCACTTGTGAATCTCACAAACCTTGAAACCCTTGGAATTTCATCAAATAACATCACCATTGATGAGGGAATGAATATCACCTTTCTTAGCCTATCATCTTTATTCTTATCATCTTGTCAACTGAAGGATTTTCCACACTTCTTGAGAAATATAAAGACACTTCGGTACTTGGATATTTCTAACAATAAGATTTGTGGTCAAATCCCTAACTGGTTTAGCGGCATGAGGTGGGACTCGTTGCAGTTCCTAAACCTTTCTCATAATTCATTAACAGGCCACCTACCACAATTTCGTTACGATAACCTACAGTATCTTGATCTGAAATTTAACTACCTTCAGGGTCCACTAGCTTTATTCATTTGTAACATGAGGAAACTTATCTTATTAGATTTATCACACAACTACTTCAGTGACTCAGTTCCACATTGCTTGGGAAGCATGCCTAATCTAAGGGTGTTGGACTTAAGAAGGAACAATTTCACAGGGAGTCTTCCCCCATTATGTGCACAGAGCACTTCATTGAGTACCATTGTCGTAAATGGTAATCGATTTGAAGGACCTGTTCCTGTATCATTGCTCAATTGTAATGGTCTACAAGTCCTTGATGTGGGGAACAATGCTATAAATGACACGTTTCCAGCTTGGCTCGGAACTCTTCAAGAGCTGCAGGTCCTTATATTAAAGTCGAACAAGTTCCATGGACCTATAAGTACGTGTCATACTAAGTTTTGCTTTCCCAAGTTGCGAATTTTTGATCTTTCTCGTAATGATTTCAGTGGCTCACTTCCTGCAAAAGTTTTTCGAAACTTCAAGGCAATGATAAAATTAGATGGCGAAGACACAGGAAATATCAAGTACATGGAATCGATGTCGAATTTGCCATTTGTCAGATCGTATGAGGATTCAGTGAGTTTGGTAATCAAAGGGCAGGATATTGAGCTACAAAGAATCAGCACAATTATGACAACCATAGATCTCTCAAACAACCATTTTGAAGGTGTCATTCCGAAAACACTAAAGGATCTCAGCTCACTTTGGCTACTCAATTTATCCCATAACAATCTCCTAGGTCATATACCAATGGAATTGGGGCAATTGAATACGCTTGAAGCTTTAGATCTCTCCTGGAATTGGCTCACTGGAAAGATTCCGCAGGAATTGACAAGAATGAACTTTCTTGCCGTCTTAAACCTCTCTCAAAATCATCTCGTCGGACCAATTCCTCAAGGTCCACAATTCAACACATTTGAAAATGACTCGTATTGTGGCAACCTTGATTTATGTGGTCCTCCTTTATCAAAGCAATGTGAAACTAGTGATTCATCGCATGTTCCTCAACAATTGGAGTCCGAAGAAGAAGGCGAGTCATATTTTTTTAGTGGATTTACTTGGGAATCAGTAGTCATAGGCTACAGTTTTGGACTAGTTGTTGGAACTGTCATGTGGAGTCTCATGTTTAAATATCGTAAGCCAAAATGGTTTGTGGAATTTTTTGATGGACTCATGCCTCACAAAAGAAGAAGGCCAAAGAAGAGAGCTCAGAGACGAAGGACTTAA

>NbLRK1

ATGAAGTCAATTCCTACATATAGCAACATTCTTGTATGGTTCATACTATTATTCTGTTCTTGCAGTAGTAATAAGTCAATATTTGCTATAACTGAGTTTGACTTTGGAACACTAACTCTGAGTAATTTGAAGCTTCTTGGAGATGCACATTTGGGTGACAACAACAGTGTTCAGTTAACACGTGATCTTGCCGTGCCAAATTCCGGTGCCGGAAAAGCTTTATATTCAAAGCCAGTAAGATTCCGGCAGCCGGGTCTTGATTTTCCAGCGAGTTTCTCTACATTCTTTTCATTTTCAGTGACTAATTTGAACCCATCGTCTATTGGTGGCGGTCTTGCTTTTGTACTCACTCCTAATGATGAGTCAGTAGGTGATGCTGGTGGGTATATGGGAATCTTGGATTCTAAAGGCACACAAAGTGGTACAATTTTAGTTGAATTTGACACCCTTATGGATGTTGAGTTTAAAGATATTAATGGAAATCATGTTGGTTTGGATCTGAATTCAATGGTTTCAACTCAAGTTGGTGATTTAGATTCTATTGGTGTTGATCTCAAGAGTGGTGATATAGTCAATTCTTGGATTGAATATTCTGGTTCTACTGGACAGTTGAATGTGTTTGTATCATACTCTAATTTAAAGCCAAAGGAACCATTTTTATCTGTTGTTCTGAATATTGCTGAGTATGTAAATGATTTCATGTTTGTTGGGTTTTCTGGTTCAACTCAAGGGAGTACTGAGATTCATAGCATTGAGTGGTGGAGTTTTAGTTCATCATTTGATGCAAGTCCTAAGTCGGCGGCAGCGGCTGCGCCACCGCCACCAACGGCTAGTTTGATGAACCCAACGGCGGATTCCGTCATGTCGCGGCCGCCTTCTATGGCTCCTTCAGAGTCTAATAGTAGTGCAAGTATAATGCAAGATAAGAGCAGTGGGAAATGTCATAGCAATTTTTGTAAACAAGGTCCTGGAGCTGTTGTTGGGGTGGTAACTGCTAGTGCATTTTTTCTTGCATTTGCTACATTAGTACTTATTTGGTTATACACCAAAAAGTTCAAGAAAGTGAAAAATTCTGAATTTTTGGCATCTGATGTTATCAAAATGCCTAAGGAGTTTAGCTATAAAGAGCTTAAATTGGCTACAAAAGCCTTTGATTCGACGAGGATTATAGGCCACGGTGCATTTGGGACAGTTTACAAGGGCATTTTATCGGACAATGGTGGCATTGTGGCAGTGAAGAGATGTAGTCATAATGGACAGGGGAAAGCGGAGTTCTTATCTGAATTATCTATAATTGGAACACTTAGGCACAGAAATCTTGTTAGACTTCAAGGATGGTGCCATGAGAAAGGTGAAATTTTGTTAGTCTATGATCTTATGCCTAATGGGAGTCTTGATAAGGCATTATTTGAATCAAGAATGATTCTACCTTGGCTACATAGGAGGAAAATTTTGCTAGGTGTTGCTTCAGCCTTGGCATATTTACATCAAGAATGTGAAAACCAGGTGATTCACAGGGATATAAAGAGTAGTAACATTATGTTGGATGAAGGGTTCAATGCAAGATTAGGTGATTTTGGATTAGCAAGACAAGTTGAACATGACAAGTCCCCCGATGCAACGGTTGCAGCCGGGACAATGGGCTACTTGGCTCCTGAATACTTGTTAACCGGAAGAGCAACCGAAAAAACTGATGTTTTTAGCTATGGAGCAGTGGTTCTTGAAGTGGCAAGTGGAAGGAGGCCAATTGAGAGGGAAACAACAAGAGTTGAGAAAGTTGGAGTGAATAGCAACTTAGTTGAATGGGTTTGGGGATTGCATAGAGAAGGGAATTTGCTAATGGCAGCTGATTCAAGACTTTATGGTGAGTTTGATGAAGGAGAAATGAGAAGGGTACTAATGGTTGGATTAGCTTGTTCACAACCTGACCCTATGGTTAGACCAACAATGAGAAGTGTGGTCCAAATGCTAGTAGGTGAAGCTGAAGTACCTATTGTCCCAAGAACTAAGCCTTCTATGAGTTTCAGCACATCACATCTTCTAATGACTTTGCAAGATAGTGTTTCTGATTTGAATGGTATGATCACACTTTCCACTTCATCATCTGAAAACAGTTTCACTGGTGTTGCCCATGGGATGATGGACTTGGTCTAA

>NbRLK

ATGATTGCAGAACAAAGACTACATATTACTGTCCTCCTCACTTATTTCCTCTTCTGTTCGTCTCAGCTTGCTATATCCCAAATTTCCTGGAACGTAAACGCCAGTCTTTTGAATTCCACTGCCGGACTTTCCACTTTCTGGACAAACAGGCCGACACTTTTAGTTAATTCCACCACCGACTTCTCTCGTTTGGCACCCATACTCCTGCAAAGAAATGCCAGTCCACGATTCCTCTGTGGCTTCTACTGCAATTACAATGCTACAAAATGCATTTTTGGTGTCCTCTTGTTCCAGAACAGATCCGACATGCAGAATGATGTGATAAATTTCCCTCAGTTAGTTTGGTCAGCTAACAGGAACCATCCAGTGAAAACCAATGCAACCTTGCAACTAAGACAAGATGGCAACTTGATCTTGGCAGACTCTGATGGCACTCTCGTTTGGTCCACTAGTACTACTGGCAAATCCATTTCAGGCTTAAACTTGACAGAAAGAGGGAATCTCGCGCTCTTTGATAAGAGAAAACGCGTAATTTGGCAGTCTTTTGATCATCCAACGGATTCTTTGTTTCCGGGGCAGAGTTTGGTTCGCGGGCAGAAGCTCATAGCAAGTGTTTCAGCATCCAATTGGAGTGAAGGTTTGCTTTCTCTTACTGTTCTCAATGGAAGCTGGGCCACTTACATAGATTCTGACCCACCTCAGTTTTACTACACTTCAACTTATTCCTATAGTCCTTACTTCAGTTTTGACGGTCAAACCTTTGCTGCCTTACAATACCCTACTACATCGAAAGCTCAATTCATGAAGCTTGGGCCTGATGGACATTTAAGGGTATACCAATGGGACGAACCTGATTGGAAAGAGGCATCTGACATTTTGATGTCAGATGTAAGAAACTATGGGTACCCAATGGTGTGTGGTAGATATAGCATTTGCACCAATAACGGGCAATGTACTTGTCCGCCAGAGGAAAATTTATTCAGGCCATTTTCTGAGAGGAAACCAGATCTTGGATGCACAGAGCTGACATCTATTTCTTGTGATTCTCCACAGTATCATGGTCTTGTAGAGCTCAAGAATACAGCATATTTTGCATTTCAGTTCAGTCATGAACCAAGTTCGAGTATATTTTGGCCAGAGGGGAAAAAGTTGGAAGACTGCAAAATGGCCTGTCTCAGTAACTGTTCTTGCAAAGTTGCAGCTTTTCAAAATGATTTGGGTACAGATCCAAGAGGGAGTTGTTTGTTACTGAATGAAGTCTTCTCTCTCGCAGACAATGAGGACGGAATGGACAAGAGAGTATTTCTTAAGGTGCAGAATTCCTCAAAGGCACAGAATCAGTCTGCAACCATTTTTGGAGGACGGAAATCAAGACCTTATAAAGTGATAATAGGATCTTCTCTTTCAGCATTGTTCGGTATCATTTTGAGTATCACTACTTGCTTTGTTATTTTCAAAAAGAGGACGCATAAGTCCCACAAGGCTGGGGATTTTCTGGATCTAGAACCAATCTTACCTGGAATGCTAACTCGATTCTGTTACAATGAGTTGAAAATAATCACAAAAGATTTCAGCACAAAGCTTGGGGAAGGAGGATTTGGCTCTGTATATGAAGGAACACTGAGTAATGGAACCAAAATAGTTGTGAAGCATCTGGATGGTGTAGGTCAAGTAAAGGATACATTCTTAACTGAAGTAAACACAGTTGGTGGCATTCACCATGTCAATTTGGTAAAACTCATTGGATTTTGCGCCGAAAAAAGCTACAGGCTTCTAATCTACGAGTACATGGTAAATGGATCATTGGATAGGTGGATTTACCATGAAAATGGGCTTACATGGCTTACAAGGCAAGGGATAATATCGGATATTGCTAAAGGGTTAGCTTATTTACATGAGGATTGCAGCCAGAAGATAATTCATTTGGACATCAACCCACAAAATATCCTTCTGGATCAACATCTCAATGTTAAGATATCTGATTTTGGGTTGTCAAAACTAATCGAGAAAGACAAAAGCAAAGTGGTAACTAGAATGAGAGGAACACCAGGCTATCTTGCTCCTGAATGGTTGAGCTCGATAATCACTGAGAAAGTTGATGTGTATGCGTTTGGAATTGTGCTCTTGGAAATTCTCTGTGGGCGAAAGAATTTGGATTGGTCCCAGGCTGATGAAGAAGATGTCCATTTGCTTAGAGTTTTTAGGAGAAAAGCGGAAGAAGAGCAGCTCATGGATATGGTTGACAAAAACAATGAGGGTATGCAGCTCCATAAAGAAGAAGTTATGGAAATGATGAGCATTGCTGCGTGGTGTCTTCAGGGAGATTACACCAAGAGGCCTTCAATGACATGGGTGGTTAAGGCACTGGAAGGTTTGGTATCTATCGAATCCAACTTGGATTACAATTTCACAAACGTACCTCTGGTTGGGGCAGACAACCAACAGATGGAAGCCACTATCAGTACGAAATTGGCTTCAGTTTTATCTGGACCAAGGTAA

>NsSOBIR1

ATGGCCTTCACTGCTTCACAAATCCACTTCTTTTTCTTCTCCCTTTTCGCCTTTTTACTGATTGCTGTTCAAGCAAGACTGAATCTTTATCCACGAGATCATGCTGCACTTTTGCTTGTCCAAAAAGACCTGGGCATCCAAGGTCAACGCATTGCTCTACGCTGCAACTCTGCAACAATATCCTGTGAAAGGCGAAAGGCAAACAGAACACAATTGTTGAGAGTCACCCGTATTGACTTCAGATCCAATGGATTGACTGGAACTTTATCTCCTGCCATTGGAAAACTTTCTGAGCTCAAAGAACTCTCTCTTCCAAACAACCAACTCTTTGACCAAATCCCAGTTCAGATTCTTGAATGCCGTAAACTGGAGGTCCTTGACCTTGGAAACAATCTGTTTTCTGGGAAAGTTCCATCTGAATTATCATCTCTACTCCGCCTTCGAATTCTTGATCTTTCTTCAAATGAGTTGTCTGGGAATCTTAACTTCTTGAAGTATTTTCCCAACTTAGAAAAACTCTCTTTAGCTGATAACATGTTCACTGGAAAAATACCCCCTTCTTTGAAATCTTTCAGAAATCTCCGTTTCCTTAATATTTCAGGAAACAGTTTTCTTGAAGGTCCTGTGCCTGCTATGAGTCAAGTTGAGCACTTGTCAGCAGATTTGGATCGGAACACCTTCGTTCCAAAACGTTACATTCTTGCTGAAAATTCGACAAGCTCAAATCAGATATCAGCACTGGCACCTAATTCCAATTCAGGAAATGCCCCAGCTCCAGCACCGAGTCATAGTGTCGCTCCAATCCATAAAAGTAATAACAGGAAGAAAAGGAAAGTCAGAGCGTGGTTTCTTGGTTTCTTTGCTGGTTCTTTCGCAGGAGCTATATCTGCGGTACTCTTATCAGTTCTCTTTAAGCTGGTAATGTTTTTCGTCCGAAAGGGAAAGACTGATGGAACTTTAACAATATACAGTCCACTGATTAAGAAAGCCGAGGATTTGGCCTTCTTAGAGAAAGAAGATGGAGTAGCATCACTTGAAATGATTGGAAAAGGTGGATGTGGAGAAGTTTATAGAGCTGAGTTACCAGGAAGTAATGGGAAGATTATAGCTATAAAGAAGATTATACAACCCCCAATGGATGCTGCAGAACTCACCGAGGAAGATACCAAGGCTTTGAACAAGAAAATGCGACAGGTAAAATCAGAAATTCAAATTCTTGGTCAAATCAGACACAGGAATCTGCTTCCCTTACTGGCACATATGCCTAGGCCAGACTGCCATTACTTGGTATATGAATATATGAAAAATGGGAGCTTACAAGATATCCTCCAGCAAGTCACAGAAGGGACAAGGGAATTAGATTGGTTGGGACGACACCGAATTGCAGTAGGGATAGCTGCTGGACTTGAGTATCTCCATATAAACCACAGTCAATGCATAATTCACAGAGATCTAAAGCCAGCAAATGTCCTTCTTGACGATGATATGGAAGCTCGAATTGCTGATTTTGGACTTGCAAAAGCACTTCCAGATGCCCATACACATATTACGACTTCAAATGTTGCAGGAACTGTGGGATATATAGCACCAGAATACCATCAGACACTGAAGTTTACAGATAAGTGTGATATATACAGCTTTGGTGTGGTGTTGGCAGTGCTGGTTATAGGAAAGCTTCCATCAGATGAATTTTTCCAGCACACGCCTGAGATGAGTTTAGTGAAGTGGCTGAAAAATGTAATGACTTCTGAGGATCCAAAAAGGGCAATTGATCCAAAGCTGATAGGAACTGGATTTGAGGAGCAAATGCTTTTGGTTCTCAAGATAGCTTGCTTTTGTACTCTGGAAAATCCAAAGGAGAGGCCTAACAGTAAGGATGTTAGGTGTATGTTAACTCAGATCAAGCATTAA

>Sotub06g029250_(StSOBIR1)

ATGGCTTCAAATTTCCACTTTTTTCTCTTATACCTTGTGACCCTTTTCCTTTTTGCTCAAGCAAGACTGAATCTTTATCCACAAGATCATGCTGCACTTTTGCTTGTTCAAAAAGACTTAGGCATCATTGCTCTTAACAACCCCTGTACGCTCGCAGGAATATCCTGCGAGCGTAGACCGGGTAACAGAACACAAGTTCTGAGAGTTACCCGTATTGTATTCAGATCCAATGGATTGAAGGGAACTTTGTCTTCTGCTATTGGCAAACTCTCTGAGCTCAAAGAGCTTTCTCTTTCCGACAATCAACTATCTGAACAAATCCCAATTCAGATTCTTGATTGTCGGAAATTAGAGATTCTTGAACTTCAAAGAAACAGATTTTCTGGAAAGATTCCGTATGAATTGTCATCTTTAAACCGTTTAAGGGTAGTTGACTTTTCATCGAATGAGTTTTCTGGGAATCTTGATTTCTTGAAATACTTTCCTAATTTGGAAAAACTGTCTCTGGCTGATAATATGTTCACTGGAAAAATACCCTTTTCATTGAAATCTTTCAGGAATCTTCGTTTCCTCAACATTTCAGGGAATAGTTTCCTTGAAGGTCCAGTACCTGTCATGAGTCAAATTGAGCATTTATCAGCAGATTTGAATCGAAACAATGCCGTTCCCAAACGTTACATTCTTGCTGAGAATTCAACAAGGTCAAATCAGATATCTGCAATGGTGCCTGCTCCAGCTCCAGCACCGGTCAATCGTGTTGTACCGGTGGTACATAAACGTAACAAGAAGAAAAGGAAGTTAAGATCTTGGTTTCTTGGTTTCCTAGCTGGAACTTTTGCTGGGGGTATATCTGCTGTGATCTTTTCGTTGCTATTCAAGCTCGTAATGTTCTTCGTAAGAAGGGGAAAGAACGATAGTTCAAGTTTAACGATATTTAGTCCGTTGATCAAGAAAGCGGAGGACTTGGCTTTCTTAGAGATGGAAGATGGAGTGGCATCACTTGAAATGATTGGAAAAGGTGGATGTGGAGAAGTTTATAGAGCCGAGTTACCGGGGAGTAATGGGAAGATTATAGCTATAAAGAAGATTATACAATCCCCAATGGATGCTGCAGAGATCACGGAGGAAGATACTAAGGCGTTGAACAAGAAAATGCGTCAAGTTAAATCAGAAATTCAAATTGTAGGTCAAATCAGACACCGGAATCTGCTTCCATTACTGGCGCATATGCCAAGGCCAGACTGTCATTACTTGGTGTATGAATATATGAAAAATGGGAGTTTACAGGATATCCTGCAGCAAGTCACAGAAGGCACAAGAGAATTAGATTGGTTGGGACGACACAGAATTGCAGCGGGGGTAGCTGCTGGTCTTGAGTATCTCCATATAAACCATACTCAACGCATAATTCACAGAGATCTAAAGCCAGCAAATATCCTACTTGATGATGACATGGAAGCTCGAATAGCTGATTTTGGGCTTGCAAAGGCAGTTCCAGATGCTCATACACATATTACGACTTCAAATGTTGCAGGAACTATGGGATATATTGCACCAGAATATTATCAGACACTGAAGTTTACAGACAAGTGTGATATATACAGCTTCGGTGTGGTGCTAGCTGTGTTGGTTATCGGAAAGGGTCCATCGGATGAATATTTCCAACATACTTCTGAGATGAGTTTAGTTAAGTGGCTGAGAAATGTAATGACTTCTGATGATCCTAAAATAGCAATTGATCCTAAGCTGAGAGGGAATGGATATGAGGAGCAAATGCTTTTGGTTCTCAAGATAGCTTGCTTTTGTACTCTCGACAATCCAAAGGAGAGGCCTAACAGTAAGGATGTTAGGTGCATGTTAACTCAGATCAAGCATTAG

>StSERK3

ATGGATCAGTCGGTGTTGGCGATCTGGGTATTTCTCTACTTAATTGGGCTGCTTTTGAATTTGTCAATGGTCGCCGGTAACGCTGAAGGTGATGCCTTGAATGCTCTGAAGACAAATTTGGCTGATCCTAATAGTGTTCTACAGAGTTGGGATGCAACCCTTGTTAATCCTTGTACTTGGTTCCATGTAACATGCAACAATGAAAATAGTGTGACTAGAGTTGATCTAGGAAATGCAAATCTATCAGGTCAACTGGTACCACAGCTTGGCCAACTCCAGAACTTGCAGTACTTGGAACTTTATAGTAATAACATAAGCGGAAGAATTCCAAATGAACTGGGGAACTTGACAGAGTTGGTTAGTTTGGATCTTTACCTGAACAACTTAAATGGTCCTATCCCTCCCTCATTGGGCAAGCTTCAGAAACTACGCTTCCTGAGGCTCAATAATAACAGTTTGAATGAAGGTATTCCCGTGTCTCTAACCACCATTGTTGCACTTCAAGTACTTGATCTCTCAAACAACCATTTGACAGGACCAGTTCCAGTCAACGGTTCCTTTTCACTTTTTACTCCTATAAGTTTTGCTAATAATCAGTTGGAAGTTCCTCCAGTTTCTCCACCTCCTCCCCTTCCTCCTACGCCCTCATCGTCATCTTCAGTGGGCAACAGCGCAACTGGAGCTATCGCTGGAGGAGTTGCTGCAGGCGCTGCCCTTCTATTTGCAGCTCCTGCAATTTTTATTGCTTGGTGGCGTCGGAGGAAACCGCAAGACCACTTCTTTGATGTTCCTGCTGAGGAGGACCCAGAAGTTCATCTGGGACAACTCAAAAGGTTTTCCTTGCGTGAACTACAAGTTGCGTCGGATAATTTTAGCAACAGAAATATACTCGGTAGAGGCGGATTTGGTAAGGTTTATAAAGGCCGGTTAGCTGATGGCTCTTTAGTTGCAGTGGAAAGACTAAAAGAGGAACGTACTCAAGGTGGAGAGTTACAGTTTCAGACAGAAGTAGAAATGATCAGCATGGCTGTACACCGAAACCTACTTCGTTTACGGGGATTTTGCATGACACCCACTGAGCGGGTGCTTGTTTATCCGTACATGGAGAATGGAAGTGTTGCATCACGTTTAAGAGAGAGGCCTGAATCAGAGCCCCCACTTGACTGGCCAAAAAGGAAGCGTATTGCACTTGGATCTGCAAGAGGCCTTGCTTACTTGCATGATCATTGTGATCCTAAAATTATTCATCGTGACGTCAAAGCCGCAAATATCTTGTTGGATGAGGAGTTTGAAGCAGTTGTTGGGGATTTTGGGTTAGCTAAACTCATGGACTACAAGGATACTCATGTTACCACTGCTGTACGTGGTACAATTGGGCATATTGCCCCTGAATATTTATCTACTGGTAAATCTTCTGAGAAAACTGATGTGTTTGGCTATGGGGTTATGCTTCTAGAGCTCATAACTGGGCAAAGGGCTTTTGATCTTGCTCGACTTGCGAATGGTGATGATGTCATGCTGCTAGATTGGGTGAAGGGACTCCTGAAGGACAAGAAATATGAAACATTAGTTGATGCAGATCTTCAAGGTAATTACAATGAAGAAGAAGTGGAACAGCTTATTCAGGTAGCTCTACTTTGCACGCAGAGTACGCCTACGGAACGTCCAAAGATGTCAGAAGTTGTAAGAATGCTTGAAGGTGATGGCCTTGCTGAGAGGTGGGAGGAATGGCAAAAGGAGGAGATGTTCCGGCAAGATTTCAACCATGTCCACCGCCACCATACTGATTGGATAATAGCTGACTCCACTTCAAATATCCGACCGGATGAGTTGTCAGGGCCAAGATGA

>Ve1

ATGAAAATGATGGCAACTCTGTACTTCCTATGGCTTCTCTTGATTCCCTCGTTTCAAATCTTATCAGGATACCACATTTTCTTGGTTTCCTCTCAATGCCTTGACGATCAAAAGTCATTGTTGCTGCAGTTTAAGGGAAGCCTCCAATATGATTCTACTTTGTCAAAGAAATTGGCAAAATGGAACGACATGACAAGTGAATGTTGCAATTGGAATGGGGTTACATGCAATCTCTTTGGTCATGTGATCGCTTTGGAACTGGATGATGAGACTATTTCTAGTGGAATTGAGAATTCTAGTGCACTTTTCAGTCTTCAATATCTTGAGAGCCTAAATTTGGCTGACAACATGTTCAATGTTGGCATACCAGTTGGTATAGCCAACCTCACAAACTTGAAGTACCTGAATTTATCCAATGCTGGTTTTGTCGGGCAAATTCCTATAACATTATCAAGATTAACAAGGCTAGTTACTCTTGATCTCTCAACTATTCTCCCTTTTTTTGATCAGCCACTTAAACTTGAGAATCCCAATTTGAGTCATTTCATTGAGAACTCAACAGAGCTTAGAGAGCTTTACCTTGATGGGGTTGATCTTTCGTCTCAGAGGACTGAGTGGTGTCAATCTTTATCTTTACATTTGCCTAACTTGACCGTTTTGAGCTTGCGTGATTGTCAAATTTCAGGCCCTTTGGATGAATCACTTTCTAAGCTTCACTTTCTCTCTTTTGTCCAACTTGACCAGAACAATCTCTCTAGCACAGTTCCTGAATATTTTGCCAATTTCTCGAACTTGACTACATTGACCCTGGGCTCTTGTAATCTACAGGGAACATTTCCTGAAAGAATCTTTCAGGTATCAGTTTTAGAGAGTTTGGACTTGTCAATTAACAAGTTGCTTCGTGGTAGTATTCCAATTTTTTTCCGAAATGGATCTCTGAGGAGGATATCACTAAGCTACACCAACTTTTCCGGTTCATTACCAGAGTCCATTTCGAACCATCAAAATCTATCCAGGTTAGAGCTTTCTAATTGCAATTTCTATGGATCAATACCTTCCACAATGGCAAACCTTAGAAATCTTGGTTATTTGGATTTCTCCTTCAACAATTTCACTGGTTCTATCCCATATTTTCGACTGTCCAAGAAACTCACCTACTTAGACCTTTCACGTAATGGTCTAACTGGTCTCTTGTCTAGAGCTCATTTTGAAGGACTCTCAGAGCTTGTCCACATTAATTTAGGGAACAATTTACTCAGCGGGAGCCTTCCTGCATATATATTTGAGCTCCCCTCGTTGCAGCAGCTTTTTCTTTACAGAAATCAATTTGTTGGCCAAGTCGACGAATTTCGCAATGCATCCTCCTCTCCGTTGGATACAGTTGACTTGACAAACAACCACCTGAATGGATCGATTCCGAAGTCCATGTTTGAAATTGAAAGGCTTAAGGTGCTCTCACTTTCTTCCAACTTCTTTAGAGGGACAGTGCCCCTTGACCTCATTGGGAGGCTGAGCAACCTTTCAAGACTGGAGCTTTCTTACAATAACTTGACTGTTGATGCAAGTAGCAGCAATTCAACCTCTTTCACATTTCCCCAGTTGAACATATTGAAATTAGCGTCTTGTCGGCTGCAAAAGTTCCCCGATCTCAAGAATCAGTCATGGATGATGCACTTAGACCTTTCAGACAACCAAATATTGGGGGCAATACCAAATTGGATCTGGGGAATTGGTGGTGGAGGTCTCACCCACCTGAATCTTTCATTCAATCAGCTGGAGTACGTGGAACAGCCTTACACTGCTTCCAGCAATCTTGTAGTCCTTGATTTGCATTCCAACCGTTTAAAAGGTGACTTACTAATACCACCTTGCACTGCCATCTATGTGGACTACTCTAGCAATAATTTAAACAATTCCATCCCAACAGATATTGGAAAGTCTCTTGGTTTTGCCTCCTTTTTCTCGGTAGCAAACAATGGCATTACTGGAATAATTCCTGAATCCATATGCAACTGCAGCTACCTTCAAGTTCTTGATTTCTCTAACAATGCCTTGAGTGGAACAATACCACCATGTCTACTGGAATATAGTACAAAACTTGGAGTGCTGAATCTTGGGAACAATAAACTCAATGGTGTTATACCAGATTCATTTTCAATTGGTTGTGCTCTACAAACATTAGACCTCAGTGCGAATAACTTACAAGGCAGGCTGCCAAAATCGATTGTGAATTGTAAGTTGTTGGAGGTCCTGAATGTTGGAAATAACAGACTTGTTGATCATTTCCCATGCATGTTGAGGAACTCAAACAGTCTGAGGGTCCTAGTCTTGCGCTCCAATAAATTCTATGGAAATCTTATGTGTGATGTAACCAGAAATAGCTGGCAGAATCTCCAGATCATAGATATAGCTTCCAACAACTTCACTGGTGTGTTGAATGCAGAATTCTTTTCAAATTGGAGAGGAATGATGGTTGCAGATGATTACGTGGAGACAGGACGCAATCATATCCAGTATGAGTTCTTACAACTAAGTAAATTGTACTATCAGGACACAGTGACATTAACCATCAAAGGCATGGAGCTGGAGCTTGTGAAGATTCTCAGGGTCTTCACATCTATTGATTTCTCTTCCAATAGATTTCAAGGAGCGATACCAGATGCTATCGGGAATCTCAGCTCACTTTATGTTCTGAATCTGTCACACAATGCCCTTGAGGGACCAATCCCAAAATCGATTGGGAAGCTACAAATGCTTGAATCACTAGACCTGTCAACAAACCACCTGTCCGGGGAGATCCCATCAGAGCTTGCAAGTCTCACATTCTTAGCAGCTTTGAACTTATCGTTCAACAAATTGTTTGGCAAAATTCCATCAACTAATCAGTTTCAAACATTCTCAGCAGATTCCTTTGAAGGAAACAGTGGCCTATGCGGGCTCCCTCTCAACAACAGTTGTCAAAGCAATGGCTCAGCCTCAGAGTCCCTGCCTCCACCAACTCCGCTACCAGACTCAGATGATGAATGGGAGTTCATTTTTGCAGCAGTTGGATACATAGTAGGGGCAGCAAATACTATTTCAGTTGTGTGGTTTTACAAGCCAGTGAAGAAATGGTTTGATAAGCATATGGAGAAATGCTTGCTTTGGTTTTCAAGAAAGTGA

>Ve2

ATGAGATTTTTACACTTTCTATGGATCTTCTTCATCATACCCTTTTTGCAAATTTTATTAGGTAATGAGATTTTATTGGTTTCCTCTCAATGTCTTGATGATCAAAAGTCATTGTTGCTGCAGTTGAAGGGCAGCTTCCAATATGATTCTACTTTGTCAAATAAATTGGCAAGATGGAACCACAACACAAGTGAATGTTGTAACTGGAATGGGGTTACATGTGACCTCTCTGGTCATGTGATTGCCTTGGAACTGGATGATGAGAAAATTTCTAGTGGAATTGAGAATGCAAGTGCTCTTTTCAGTCTTCAGTATCTTGAGAGGCTAAATTTGGCTTACAACAAGTTCAATGTTGGCATACCAGTTGGTATAGGCAACCTCACCAACTTGACGTACCTGAATTTATCCAATGCCGGTTTTGTTGGCCAAATTCCTATGATGTTATCAAGGTTAACAAGGCTAGTTACTCTTGATCTCTCAACTCTTTTCCCTGACTTTGCCCAGCCACTAAAACTAGAGAATCCCAATTTGAGTCATTTCATTGAGAACTCAACAGAGCTTAGAGAGCTTTACCTTGATGGGGTTGATCTCTCAGCTCAGAGGACTGAGTGGTGTCAATCTTTATCTTCATATTTGCCTAACTTGACTGTCTTGAGCTTGCGTACTTGTCGAATTTCAGGCCCTATTGATGAATCACTTTCTAAGCTTCACTTTCTCTCTTTCATCCGTCTTGACCAGAACAATCTCTCTACCACAGTTCCTGAATACTTTGCCAATTTCTCAAACTTGACTACCTTGACCCTCTCCTCTTGTAATCTGCAAGGAACATTTCCTAAAAGAATCTTTCAGGTACCAGTCTTAGAGTTTTTGGACTTGTCAACTAACAAATTGCTTAGTGGTAGTATTCCGATTTTTCCTCAAATTGGATCATTGAGGACGATATCACTAAGCTACACCAAGTTTTCTGGTTCATTACCAGACACCATTTCGAACCTTCAAAACCTATCCAGGTTAGAACTCTCCAACTGCAATTTCAGTGAACCAATACCTTCCACAATGGCGAACCTTACCAATCTTGTTTATTTAGATTTCTCCTTCAACAATTTCACTGGTTCCCTCCCATATTTCCAAGGGGCCAAGAAACTCATCTACTTGGACCTTTCACGTAATGGTCTAACTGGTCTCTTGTCTAGAGCTCATTTTGAAGGACTCTCAGAACTTGTCTACATTAATTTAGGGAACAATTCACTCAACGGGAGCCTTCCTGCATATATATTTGAGCTCCCCTCGTTGAAGCAGCTTTTTCTTTACAGCAATCAATTTGTTGGCCAAGTCGACGAATTTCGCAATGCATCCTCCTCTCCGTTGGATACAGTTGACTTGAGAAACAACCACCTGAATGGATCGATTCCCAAGTCCATGTTTGAAGTTGGGAGGCTTAAGGTCCTCTCACTTTCTTCCAACTTCTTTAGAGGGACAGTTCCCCTTGACCTCATTGGGAGGCTGAGCAACCTTTCAAGACTGGAGCTTTCTTACAATAACTTGACTGTTGATGCAAGTAGCAGCAATTCAACCTCTTTCACATTTCCCCAGTTGAACATATTGAAATTAGCGTCTTGTCGGCTGCAAAAGTTCCCCGATCTCAAGAATCAGTCAAGGATGATGCACTTAGACCTTTCAGACAACCAAATATTGGGGGCAATACCAAATTGGATCTGGGGAATTGGTGGTGGAGGTCTCGCCCACCTGAATCTTTCATTCAATCAGCTGGAGTACGTGGAACAGCCTTACACTGTTTCCAGCAATCTTGCAGTCCTTGATTTGCATTCCAACCGTTTAAAAGGTGACTTACTAATACCACCTTCCACTGCCATCTATGTGGACTACTCGAGCAATAATTTAAACAATTCCATCCCAACAGATATTGGAAGATCTCTTGGTTTTGCCTCCTTTTTCTCGGTAGCAAACAATAGCATCACTGGAATAATTCCTGAATCCATATGCAACGTCAGCTACCTTCAAGTTCTTGATTTCTCTAACAATGCCTTGAGTGGAACAATACCACCATGTCTACTGGAATATAGTCCAAAACTTGGAGTGCTGAATCTAGGGAACAATAGACTCCATGGTGTTATACCAGATTCATTTCCAATTGGTTGTGCTCTAATAACTTTAGACCTCAGCAGGAATATCTTTGAAGGGAAGCTACCAAAATCGCTTGTCAACTGCACGTTGTTGGAGGTCCTGAATGTTGGAAATAACAGTCTTGTTGATCGTTTCCCATGCATGTTGAGGAACTCAACCAGCCTGAAGGTCCTAGTCTTGCGCTCCAATAAATTCAATGGAAATCTTACGTGTAATATAACCAAACATAGCTGGAAGAATCTCCAGATCATAGATATAGCTTCCAACAATTTTACTGGTATGTTGAATGCAGAATGCTTTACAAATTGGAGAGGAATGATGGTTGCAAAAGATTACGTGGAGACAGGACGCAATCATATCCAGTATGAGTTCTTACAACTAAGTAACTTGTACTATCAGGATACAGTGACATTAATCATCAAAGGCATGGAGCTGGAGCTTGTGAAGATTCTTAGGGTCTTCACATCTATTGATTTCTCTTCCAATAGATTTCAAGGAAAGATACCAGATACTGTTGGGGATCTTAGCTCACTTTATGTTTTGAACCTGTCACACAATGCCCTCGAGGGACCAATTCCAAAATCAATTGGGAAGCTACAAATGCTTGAATCACTAGACCTGTCAACAAACCACCTGTCCGGGGAGATCCCCTCAGAGCTTTCAAGTCTCACATTCTTAGCAGTTTTGAACTTATCGTTCAACAATTTGTTTGGAAAAATCCCGCAAAGTAATCAATTTGAAACATTCCCAGCAGAATCCTTTGAAGGAAACAGAGGCCTATGCGGGCTTCCTCTTAACGTCATTTGCAAAAGCGATACTTCAGAGTTGAAACCAGCACCAAGTTCTCAAGATGACTCTTATGATTGGCAGTTCATATTTACGGGTGTGGGATATGGAGTAGGGGCAGCAATCTCCATTGCACCTCTGTTGTTTTACAAGCAAGGAAACAAATACTTTGACAAACATTTGGAGAGAATGCTTAAACTGATGTTTCCTAGATACTGGTTCAGTTACACCAGATTTGACCCTGGGAAGGTTGTGGCTGTGGAACACTATGAAGATGAGACCCCAGATGACACCGAAGATGACGATGAGGGGGGAAAAGAAGCATCTCTTGGGCGTTATTGTGTCTTCTGTAGTAAACTTGATTTTCAGAAAAATGAAGCAATGCATGATCCAAAATGCACTTGTCATATGTCATCATCCCCCAATTCTTTTCCTCCTACGCCGTCCTCTTCTTCACCTTTATTAGTCATATATCACAAAAAGTTTTGA

>I3

ATGTTAGCAGAACAGAGACTGTATTTTCCTATCCTTCTCTTCATTTATCTCCTCTTTTGTTCATCTCGATTTGCTATATCGCAATCTTCATGGGACTACAACGCCACTCTTTCAAATTCTATTGCAGGGTTTGCTGGGCTTTCCTCTTTCTGGATCAACAGGCCGTCTCTTATTATTAATTCCACCACCGACAGCTTCTATGGTTCGACAACACCCGTACTTCAGCGGGGAAATGCTGGCCCACGATTCCTCTGTGGCTTCTACTGCAGGTACAATGTCACAGAATGCCTTCTTGGTATCCTTTTGTACCACAACAAATACAACGAGCAGAACGGTATGATAGATAAACCCCAGTTAGTTTGGTCTGCTAACAGGAACCGTCCAGTGAAATTCAATGCAACCTTGGAACTAGGCCAAGATGGCAACTTGGTCTTGACAGACTCTGATGGCACTCTTGTTTGGTCCACTGATACAATTGGGAAATCTGTTTCTGGCTTAAACTTAACAGAAATGGGAAATCTTGTGCTCTTTGATAAGAGAAAGCGCACAATTTGGCAGTCTTTTGATCATCCTACAGATTCTTTGCTTCCAGGGCAGAGTTTGGTTTCTGGCCGGAAGCTTGTAGCAAGCGTTTCAGCAACCAATCATAGTCAAGGTTTGCTTGCTCTTACTGTTCTCAACGGAAGTTGGGCTGCTTACATGGATACTAATCCGCCTCAATATTACTACACTTCATACTATGCTGATAGTTCTTATTACAGTTTTAATGGTCAAACCTTTACTGTTTTACACTATCCTACAAATTCGACAGCTCAATTCATGAAGATTGGGCCTGATGGACATATAAAGGTATTCCAATGGTCTGTAGTTGATTGGAATGAAGTATCTGACATTTTGTCGCCAAATGTAGATAACTGTGCGTACCCAATGGTATGTGGAAGTTATAGCATTTGTACAAATAACGGGCAATGTACTTGTCCGTCACAGGAAAACTTCTTCAGGCCATTTTCTGAGAGGAAACCAGATCTTGGATGTTCACAGCTGACTTCCGTTAACTGCAACTCTTCGCAGTATCATAGTTTCATAGAGCTCAAGAATACTACATATTTTTCATTTAACATTGATCAGGAACTAAATTCGACTAAATTGTGGCTTGGGCAGAAAAAGTTGGAAGATTGCAAAAGGGCATGTCTGAGTCACTGTTCTTGCAAAGCTGCTGTTTTTGAATATAATTGGAATGGGGATCGAAGAGGTAATTGTTTGTTACTGAATGAAGTTTTCTCTCTCAAAGACAGCGAAGAAGTAAGAGGCGAGACAGTATTTCTTAAGGTGCAGAATACCTCAAAGGCGCAGACGCAGTCTCTAATCATTCCTGGAGGAAAGAAATCAAGACCTTTCAAAGTGATAATAGGATCTACTCTTGCAGCTTTCTTTGGGATAATTTTAAGCATAGCTACTTGCTTTGTTATTTTCAGAAAGAGGACACATGAGTCCAGAAAGGCTGCGGATATTTTGGATCTAGCACCAATCTTACCGGGAATCCTAACTCGATTCTCTTACAATGAGCTGAAAATAATTACAGATGATTTCAGCAGTAAGCTTGGGGAAGGAGGATTTGGCTGTGTTTATGAAGGAACACTGAGAAATGGAACTAAAATAGCTGTGAAGAATCTGGATGGTGTAGGTCAAGTAAAGGAATCATTCTTAACAGAAGTAAAGGCGGTCGGTGGCATTCACCATATCAATCTGGTAAAACTCATTGGATTTTGTGCTGAAAAAACCCACAGGCTTCTAATCTATGAGTACATGGTGAATGGATCGCTTGATAGGTGGATTACACATGAAAACCGAGAAAATGGGCTTACATGGAGCACAAGACAGAGGATAATATCAGATATCGCCAAAGGGTTAGCGTATCTACATGAGGATTGCAGCCAAAAGATAATTCATTTGGACATCAAACCACAAAACATCCTTCTGGATCAATATCTCAATGCTAAGATATCAGATTTTGGGTTGTCGAAGCTAATTGAGAAAGATAAAAGCAAAGTCGTGACTAGAATGAGAGGAACACGGGGTTATTTAGCCCCTGAATGGTTGAGCTCGGTAATCACTGAGAAAGTTGATGTTTATGCTTTTGGAATTGTCCTCTTGGAAATTCTCTGTGGACGAAAGAATTTGGATTGGTCCCAACCTGATGAAGAAGATGTCCATTTGCTAAGTGTATTTAGGAGAAAAGCGGAACAAGAGCAGCTCATGGATATGGTTGACAAAAACAACGAAGATATGCAGCTCCACAGGGAAGCAGTGACTGAAATGATGAGCCTAGCTGCATGGTGTCTACAGGGTGATTTTTCCAAGAGGCCTTCCATGTCATTAGTGGTTAAGGCATTGGAAGGTTTGGTGACTGTTGAAACCAACTTGAATTATGATTTCACACATGTACCTGAGGTTGGGGCAGGCAACCAAGAGAGGGAAGTCATTGTCAGTTCAATATTTCCTTCAATTTTATCGGGACCAAGGTAA

>CORE

CAACTATGTTGTATTGGTTGAAGTGTTCTTCACTCGCATCCTCAGAAGATGAAGTTTAACATAAATGAGAATGCTTAAATGGATGTGTGGGCACATTAGAAGAGACAAGATTAGAAAGGAAGATATATGAGGTAAGGTGGGAGTAGCTTCATTGGTGGACAAGAAGCGGGAAACAAGGCGAAGATGGTTCGAGCATTCGAAGAGGAGGAACACAAATGCCCTAGTGAAGAGGTGCAAGAGGTTGGCAGTGATAGGTTTGAGGAAAGGTAGAGGTAGATCGAGGAAGAATTGGGGGATGTGATTATACAATAGATGAAACATCTTCAACTAATAAGGACATGACTTCAAATAGGGATCATGGAGGATCAGGGTAGACTGGTAGTAGGTGGTTGAGCATTTTCTCTCTTCCTCGTGGTTGAGCAGTGTTCCGGTCTGGAGTATCTATTGGTTTCCTTACCACTATTATCCAACCTTCCCAATTTGACACAGAGTTTAAGAAAAAGAAAAGGAAGACTTTAGAAACTTGTGGTTTAAAACAATCCTTAGATATTTATTTGATGGTAAATCAATTTGTTAGGGGTAAAAGGGAAATTTTAAAGTTGTTATTTCTAATAATACAAAAGTGACATTTTTTTTGAGGAAGCACTAAAAAGTCAAGTGTGTCACATAAATTGGGACAAAGGGAGTGTTATTTTGGTAGCATACCTTGTTCTTCATTTTGTTTCTATTGTTGTTTCTTTTACTTTTGTTATCTCTAATAATTTTATTGTCAATATTTTTCTTTTTTCAATATTTTCATAATAGCATTGCATTTCTTTCAAACCATGTTCTGAAAATAGATTTTTCTTGGGAACCATGGATTTTTATCTGTAATTTGCAACCTTTGATGCAGTTAACAAATTCTCTCCAAGAGGCCACCTCAAGAAGCAACTTAGCATCTAGACAGCTAGCTTCTTCTGCTGGATCAAACACTTCAAAGTTCTAGTCTTCGAAGACCATCTTCTTCAGGTGGCTATGGTGAAAGGGAATGAAACAGACAAAATGTCACTGCTAGCATTCAAGAATATGATCATTGATGACCCTTTCAAGATCATGGACTCCTGGAATGAAACTATACATTTCTGTGACTGGCCTGGGGTCTCATGTGGTAACCGTCATTGCCGAGTTACAGTTTTAAATCTTACTTCTCTGAAATTGAGGGGTTCTCTGTCACCAAGCATTGGTAACCTGAGCTTTCTTAATGTCCTTAAGCTTCAAAATAATAGTTTTTCAGGTGAAATCCCATCAGAGATTGGCTACTTACATAAGTTAAACGTTTTACGCCTTGATAATAACTCGTTCACTGGTCATATTCCTTCAAACATTTCTGGTTGCTTCAATCTTGTTTCTGTTGGTCTTTCGTATAATATGATGGTAGGCGAAATTCCAGCAGAATTAGGCACATTGTTGAGACTCAAACAACTTTCTCTTGTTTCTAACTCTTTGACAGGAGGAATTCCGCCTTCGTTTGGTAATCTTTCTTTACTTGATACCTTTTCTGCAAGCAAGAATAATTTGTTAGGGAAAATACCTGATGAATTATGTCAATTACTCAACTTAAAGTACTTTGTAGTGAATGAAAACAACTTGAGTAGTACCCTGCCTCCTTGTCTTTTCAATCTTTCCTCTATCGTGGCCATCGATGTTGGAACAAACCATTTAGAAGGACAATTGCCTCCATTGCTTGGTATTACTCTTCCTAAGTTAGAGTTCCTTAGCATTTATAGAAACAATGTAACTGGAAATATTCCGGGGACATTGTCAAATGCTACAAATCTTCAGTCTCTTATTGCTGGTAGAAATGGACTCACAGGAAAAGTACCACCCCTTGGAAATTTGCTGAAAATGAGGAGATTTTTAGTTGCTTTCAATGATCTTGGCAAGGAGGAGGCGGATGACCTTAGTTTTCTCTCCACATTGGTGAATGCCACCAACTTGGAGCTTGTGGAGCTCAATACAAATAATTTTGGTGGAGTCTTACCTGCATCTGTTAGTAATCTTTCGACTGAGCTTATAGAGCTTTCGTTGTCTTATAATCAAGTTTCTGGAGAAATCCCAAGAGGGATATCCAATCTCAAGAAGCTTCAGGCTTTCTTTGTAGCTTACAACAGATTTATCGGTGAAATTCCTTCTGAAATTGGTGATCTTATGTATTTGCAAGAGCTGGCTTTACTTGGAAATCAGTTCTCTGGCCAAATTCCAATCTCATTGGGAAATTTAGCTTCTTTAACCAAACTCACTTTAAGAGAAAACAATCTCCAGGGGAGAATTCCTTCCAGTTTAGGCAAATGCGACAAGTTGGAACTGTTGGATCTTGGTTCAAATAACCTTAGTGGTTTCATACCATCAGAAATTCTTGAACTCTCATCATTGTCGGAGGGCGTGGACCTATCTCAGAACCACTTAACTGGTTTCCTTCCGATGGGAATTGGGAAGTTGAGAAATCTAGGCTACTTAAACCTGTCTTATAACAAGTTGCAGGGCCAGATTCCTACCACTATAGGTACTTGTGTGAAACTTGAAGCACTAGACTTAAACAACAACAACTTTCAGGGTAGTATTCCTTCAACCATGAATAATTTGCGAGGCCTGGAATTTTTAGTCCTTTCTCACAACAATCTGTCAGGTGGAATCCCGGGATTCTTGAAGGACTTCAAATTTCTTCAGATTTTAAATCTGTCTAGCAATAATTTGGAAGGTGCTGTCCCAACTGGAGGTATCTTTAGTAATGCTACTGCGGTTTCCATCATAGGAAATAAAAATCTCTGTGGGGGTGTACCGGAGTTGGACCTTCCGGTTTGCATTGTAGGAGTAAAGAAAGAAAGGAAATCAGGTTTTCCCTTGAAAAAAGTAATTCCTGTAGTTAGTGGGCTTATAGGATTGACATTGATAGTTTGTTTTCTTGGCATACGACAGTTTAGTAGATTGAGAAAAACAACACCTACAGATATACCTGAAAACTCAACCTTAAGAATATCTTATCAGTGTCTACTTAGGGAAACTGATAGATTTTCTGCATCAAATTTGCTTGGCATGGGTGCTTTTGGGTCTGTATATAAAGGAATTTCTGAACATGATGGTACTGTTTTTGCTGTTAAGGTACTGGACCTTTCACACCATGCAGCTTCCAGGAGTTTTTTAGCTGAATGTGAGGTTCTAAAGAACATCAGACATCGTAATCTTGTAAAGGTTCTAAGTGCATGCTCAGGTATTGATTATGAAGGTAATGAGTTCAAAGCCATAGTTTATGAATACATGGATAAGGGGAATCTACAGGACTGGTTGCATTTCACTCCTCAGGAGAACTCTGAGCCACAGGAGGAACACAAGAAGCTAGGGTTTATTCAAAGATTAAACATTGCGATTGATGTTGCCTGTGCTCTGGATTATCTCCATAATGATTGCCAACCCCCTATAATTCACCGCGATCTGAAGCCAAGCAATATTCTTCTAGATGAGAACATGACTGCACATGTTGGTGATTTTGGTTTGGCAAGATTTGTTCCACCAGAAATTCCGAACTCATCAGAAAATTCAAAAAGTCTTACAGGTGTAGGGGGGACAATTGGTTATACACCTCCAGAATTAGGTATGGGAAGCGACGCCTCAACCTATGGAGATGGATACAGCTTTGGCATACTGCTCTTGGAAATGTTTACCGGGAGGAAACCAACTGATGAAATGTTCAAAGATAATTTAAACCTTCATAACTACGCGAATGCTGCTTTGCCAGACAGAGTGATGCATATTACAGATCCAATTCTTCTACAGGAACGAGATGAACTTGAAATGGAATACAAACTTCATGACAACACAAGCAGTGCTGGCGACATATTTCTGTCATTCTTAATCAACGTGATTCAAATCGGAGTTTCCTGCTCCGCTGAATCTCCAAAAGAAAGAAAGCGGATCAGTGATGTCGTTAGAGAACTCAATTCATTAAGGAAGTTGTTTTTGGAGCAAGCATACCGGAAGAAAAAGTTATAAGCTTCAACCTGCATTGACAGATGATGACAGGCATAGGGTTAGTCCTTCTCATGGTTGGTGACATGAGGTTAATATGCACTGTGAAACAAATTCATGAATTTAAGGTTTGATTCTGCTCTTATAATTTGCAGTTATAAACTTGTATGTTCCTAGTAAATGATAGCTCTTTGCACATTTCTTAGTTTAGATGCCTTATTTATTTTATTGAGATATCTATAGCTGCTAGACTAGTAGATCACTGAATTACACAATTGATCTCAAGCTGCCACCTACATGTCCAAACATCATATTTGAAGTTTTTGAGACGT

>FLS3

TCATATAAATCAAAATAGAAACACAAAACGACAACTTTTCGTCTTGTGAAATAACTCAAGTGGAGTCATTCATTCCAATATATCCTTTTTCCATTTCTCTTCATATATTTCAACTTTCAAATGCTTAGTAACATCATGGAGAAACACATTTTCTTATTGATACTTGCTATCTTAGTTCAATTTTACTTTGTTTCTTCTATATCAGCTACTATTTCCTCAAATGAGACTGATCAAGAAGCTCTACTAGCTTTTCGAAACCTTGTTACGAGTGATTCTAGTCATTTTTTAGCCAATAATTGGACAAAAAACACTTCATTTTGCTCTTGGTTTGGTGTCACTTGTAGTCCAAAAAGGCAAAGGGTTGTAGCCTTGACTCTTCCTAATTTGCAACTTCAAGGCACAATTTCGCCGTCTTTGGCCAATCTATCTTTTCTCATAGAGCTAAATCTCGCAAACAACAACTTACACAGTGAAATCCCTGATGGCATTGGCCGCTTGCCTCGTCTACGAGTGATTGATATTCAGAACAATCAGCTGCATGGAAGTATTCCAACAAGTCTATTTCAACACGGGAGTGTTCAAATCATTTCATTGGCTTTCAATAAACTCGGTGGTGAAATGTGGAACGGTACATGGTATGTACCCGAACTCAGAGTCTTAAATCTCAGGAACAATACCATTACAGGTGTGATCCCTCCTTCTATTGGAAATGCCACAAAGTTGATGAACTTCAGTTTGAATGGGAATAGAATCAACGGCAACATTCCAATGGAGATTGGTAATCTAAGCCAACTTGTTGAGTTGTCGTTGTCTCGTAATCAATTAACAGGTTCCATTCCTTCAACATTGTTTAATATCTCCTCCCTTCTCGTCGTGTCTCTGGCATACAATAGCCTTTCAGGTCCTCTGTTTCCTGATGATCGACGTAACGTTCTTTCATCAAACCTCGAGCATATAGGTGTATCATACAATCAAATCACTGGTCACATTCCTTCCAACATCTGTCAATTCACAGCTCTCAGAGTTCTGTCCATATCATACAACAACATAACTGGAGAAATACCGAGAAATATTGGTTGTTTAGCCAAGCTCGAAGAGTTTTATATCGGTTATAATGCAATAAATGGAACAATTCCTGCTTCATTAGGCAATATTTCAACTCTTCAAAATCTTCATTGCGGAAGCAATCACATGGAGGGAGAACTTCCTCCAGAATTAGGAAAGCTATCAAACTTAAGACAAATCAATTTCGAAGAAAATTATAATCTTATTGGTGAAATTCCGAATACTATTTTCAACATATCTTCTTTGGAGTTCATTGCTTTCACTTTCAACTACCTTTCAGGTAGAATTCCGAATCTTCTTCACCTTCCAAATCTTATACAACTTCTCTTAGCAAACAATCAGCTCGAAGGTGAAATTCCTCGGTACATCACAAATGCTACGAATCTTGAGCTGTTAGAGCTATCAGATAACCTTCTCACAGGTACTATTCCTAATGATTTAGGAAATCTTCGCGAGCTGCGAGATCTTTTCCTACATCATAATCAACTTACTGAGTTGGGATTCTTTGATTCTTTGGTGAAATGTAGGATGTTGAGATATGTACAAGTGGGATCGAATCCGTTGAATGATGTTCTGCCAAGTAGTATTGGCAATCTTTCATCTACTGTTGAATACTTTCATATTGGAGATGCACAAATCAATGGATTCATTCCCACTAGTACAGGCAACATGACCGGTCTTACAACGCTAGTTTTTCAAGATAACAGTTTGACAGGAAACATTCCTCGTGAGATCCGTAAGCTTAAACAACTCCAAGGTTTATTTCTAGTTAACAATGGACTACAGGGGGACATAGCAGAGGTAGTATGTGATTTATCGAATTTGGTTCGATTAGCTCTGTCTGAAAATGAGCTCTCGGGGGTGATTCCGGAATGTTTAGGAAATCTTACCATGCTACAACAACTTTTTTTAGGTTCTAACAAGTTTGAATCAAAGCTACCTTTAAGCTTTTGGAAGATGAGTAGTCTTCTATATTTAAACATGTCGCGTAATTCTATAAAGGGAGAAGTTCCATCAGATATCGGAGAACTTAAAGCTATTGTAGCAATCGATATCTCTGGTAACCATTTCTCGGGGTCGATACCAAGCAATTTGGGGGAACTTCAAACCTTGAAGTTACTTTCCTTATCGAACAATTCGTTTTCAGGTCCAATTCCATTTTCCTTTTCAAACTTGAAAAGCTTGGAATTCTTGGATTTGTCTTTGAATAACTTGTCAGGTACTATTCCTAAGTCTTTCGAAAAGCTTTTGTACCTTACAAGCATCAACGTCTCGTTTAATGTTTTAGAAGGTGAAATACCTAGTGGTGGTGTGTTTGCAAACTCCACCCTGCAATCATTTAGTGGGAACAAAGGTCTATGTGGAAGGCAAATATTGGAGGTTCCTGCTTGTGCTATCACTACTCCTGAACAACAACAATCAAAATCGAAGAAGCTTGTGCTAAAAATTGTCACTCCGATGGTTATTTCATTCTTTCTGATATTCTTGTTGGTTGTCTCGATTTGGATAATGAAACGAAAGAAGAAAGGGAAGTCCAAAGATGTTGAAAAGGTTCCGGAGATGAGGACTTATCAATTGATTTCTTATCATGAGATTCAACGAGCAACTAACAATTTTGATGAATCCAATTTGATTGGCGTGGGAGGTTCTGGCTCTGTGTACAAAGCCACATTAGCTAGTGGAATTGTGGTTGCAATTAAGGTACTGGATTTGGAAAATGAGGAAGTATGCAAAAGGTTTGATACTGAATGCGAAGTGATGAGAAATGTTAGACACAAAAACCTTGTTTCGGTGATCACTACGTGTTCTAGTGAACACATAAGAGCCTTTGTTCTGCAGTATATGCCCAACGGAAGTCTTGACAATTGGTTGTACAAAGAAGATCGCCACTTAAAACTTCGTCAAAGAGTCACCATAATGCTTGATGTAGCTATGGCAATTGAATATCTACATCATGGTAATGACACCCCAATAGTTCATTGTGACCTCAAGCCAGCCAACGTTCTTTTGGATGAAGATATGGTGGCGCGTGTTGGTGATTTTGGCATCTCAAAGATTTTAGCTGTAAGCAAATCCATGGCACATACAAAGACATTAGGCACTCTTGGATATATTGCACCAGAATATGGCTCGGAGGGAATAGTGTCCACTCGTGGTGATGTTTACAGTTATGGCATCATGCTGATGGAGGTTTTGGCAAAAAGAAGGCCAACAGGTGAAGAGATATTCAACGAAAATCTTGGCTTGAGGGAGTGGATAACGCGAGCATTTCCAAGAACTATGATGGAAGTTGTGGACGCGGATATGTTTCATGATGGAGAAAAAATTACTTCCGAAAGTGAAATATGCATACTCTCCATGATAGAACTGGCTTTAGATTGCACAAAGGCAACACCAGAATCAAGGATAACCATGAAAGATGTAGTCAAGAGGCTTAACAAAATAAAGAACACATTTGGAAACATAGAAGTAAATTAGAATCTCTTTGTGTTGTTTTTTTCTTTTAAGTTTGTAAGTTGATACTCAATCAATCTGTTTGTCATACTTTTCATTTGTCTGTTTTTTTAAAAAAATGTCTTTCTTTTTTGGCATCTCTTAAAA
