## Supplementary material for "RLP/K enrichment sequencing; a novel method to identify receptor-like protein (*RLP*) and receptor-like kinase (*RLK*) genes": Note S4

>CORE_1054

CATTCAAGAATATGATCATTGATGACCCTTTCAAGATCATGGACTCCTGGAATGAAACTATACATTTCTGTGACTGGCCTGGGGTCTCATGTGGTAACCGTCATTGCCGAGTTACAGTTT

>CORE_1116

CATTTCTGTGACTGGCCTGGGGTCTCATGTGGTAACCGTCATTGCCGAGTTACAGTTTTAAATCTTACTTCTCTGAAATTGAGGGGTTCTCTGTCACCAAGCATTGGTAACCTGAGCTTT

>CORE_1178

TCTTACTTCTCTGAAATTGAGGGGTTCTCTGTCACCAAGCATTGGTAACCTGAGCTTTCTTAATGTCCTTAAGCTTCAAAATAATAGTTTTTCAGGTGAAATCCCATCAGAGATTGGCTA

>CORE_1240

ATGTCCTTAAGCTTCAAAATAATAGTTTTTCAGGTGAAATCCCATCAGAGATTGGCTACTTACATAAGTTAAACGTTTTACGCCTTGATAATAACTCGTTCACTGGTCATATTCCTTCAA

>CORE_1302

CATAAGTTAAACGTTTTACGCCTTGATAATAACTCGTTCACTGGTCATATTCCTTCAAACATTTCTGGTTGCTTCAATCTTGTTTCTGTTGGTCTTTCGTATAATATGATGGTAGGCGAA

>CORE_1364

TTCTGGTTGCTTCAATCTTGTTTCTGTTGGTCTTTCGTATAATATGATGGTAGGCGAAATTCCAGCAGAATTAGGCACATTGTTGAGACTCAAACAACTTTCTCTTGTTTCTAACTCTTT

>CORE_1426

CAGCAGAATTAGGCACATTGTTGAGACTCAAACAACTTTCTCTTGTTTCTAACTCTTTGACAGGAGGAATTCCGCCTTCGTTTGGTAATCTTTCTTTACTTGATACCTTTTCTGCAAGCA

>CORE_1488

GGAGGAATTCCGCCTTCGTTTGGTAATCTTTCTTTACTTGATACCTTTTCTGCAAGCAAGAATAATTTGTTAGGGAAAATACCTGATGAATTATGTCAATTACTCAACTTAAAGTACTTT

>CORE_1550

TAATTTGTTAGGGAAAATACCTGATGAATTATGTCAATTACTCAACTTAAAGTACTTTGTAGTGAATGAAAACAACTTGAGTAGTACCCTGCCTCCTTGTCTTTTCAATCTTTCCTCTAT

>CORE_1674

GCCATCGATGTTGGAACAAACCATTTAGAAGGACAATTGCCTCCATTGCTTGGTATTACTCTTCCTAAGTTAGAGTTCCTTAGCATTTATAGAAACAATGTAACTGGAAATATTCCGGGG

>CORE_1736

TCCTAAGTTAGAGTTCCTTAGCATTTATAGAAACAATGTAACTGGAAATATTCCGGGGACATTGTCAAATGCTACAAATCTTCAGTCTCTTATTGCTGGTAGAAATGGACTCACAGGAAA

>CORE_1798

TGTCAAATGCTACAAATCTTCAGTCTCTTATTGCTGGTAGAAATGGACTCACAGGAAAAGTACCACCCCTTGGAAATTTGCTGAAAATGAGGAGATTTTTAGTTGCTTTCAATGATCTTG

>CORE_1860

CCACCCCTTGGAAATTTGCTGAAAATGAGGAGATTTTTAGTTGCTTTCAATGATCTTGGCAAGGAGGAGGCGGATGACCTTAGTTTTCTCTCCACATTGGTGAATGCCACCAACTTGGAG

>CORE_1922

GGAGGAGGCGGATGACCTTAGTTTTCTCTCCACATTGGTGAATGCCACCAACTTGGAGCTTGTGGAGCTCAATACAAATAATTTTGGTGGAGTCTTACCTGCATCTGTTAGTAATCTTTC

>CORE_1984

TGGAGCTCAATACAAATAATTTTGGTGGAGTCTTACCTGCATCTGTTAGTAATCTTTCGACTGAGCTTATAGAGCTTTCGTTGTCTTATAATCAAGTTTCTGGAGAAATCCCAAGAGGGA

>CORE_2046

GAGCTTATAGAGCTTTCGTTGTCTTATAATCAAGTTTCTGGAGAAATCCCAAGAGGGATATCCAATCTCAAGAAGCTTCAGGCTTTCTTTGTAGCTTACAACAGATTTATCGGTGAAATT

>CORE_2108

CAATCTCAAGAAGCTTCAGGCTTTCTTTGTAGCTTACAACAGATTTATCGGTGAAATTCCTTCTGAAATTGGTGATCTTATGTATTTGCAAGAGCTGGCTTTACTTGGAAATCAGTTCTC

>CORE_2170

CTGAAATTGGTGATCTTATGTATTTGCAAGAGCTGGCTTTACTTGGAAATCAGTTCTCTGGCCAAATTCCAATCTCATTGGGAAATTTAGCTTCTTTAACCAAACTCACTTTAAGAGAAA

>CORE_2232

CAAATTCCAATCTCATTGGGAAATTTAGCTTCTTTAACCAAACTCACTTTAAGAGAAAACAATCTCCAGGGGAGAATTCCTTCCAGTTTAGGCAAATGCGACAAGTTGGAACTGTTGGAT

>CORE_2294

TCTCCAGGGGAGAATTCCTTCCAGTTTAGGCAAATGCGACAAGTTGGAACTGTTGGATCTTGGTTCAAATAACCTTAGTGGTTTCATACCATCAGAAATTCTTGAACTCTCATCATTGTC

>CORE_2480

AAATCTAGGCTACTTAAACCTGTCTTATAACAAGTTGCAGGGCCAGATTCCTACCACTATAGGTACTTGTGTGAAACTTGAAGCACTAGACTTAAACAACAACAACTTTCAGGGTAGTAT

>CORE_2542

GTACTTGTGTGAAACTTGAAGCACTAGACTTAAACAACAACAACTTTCAGGGTAGTATTCCTTCAACCATGAATAATTTGCGAGGCCTGGAATTTTTAGTCCTTTCTCACAACAATCTGT

>CORE_2604

TCAACCATGAATAATTTGCGAGGCCTGGAATTTTTAGTCCTTTCTCACAACAATCTGTCAGGTGGAATCCCGGGATTCTTGAAGGACTTCAAATTTCTTCAGATTTTAAATCTGTCTAGC

>CORE_2666

TGGAATCCCGGGATTCTTGAAGGACTTCAAATTTCTTCAGATTTTAAATCTGTCTAGCAATAATTTGGAAGGTGCTGTCCCAACTGGAGGTATCTTTAGTAATGCTACTGCGGTTTCCAT

>CORE_2728

ATTTGGAAGGTGCTGTCCCAACTGGAGGTATCTTTAGTAATGCTACTGCGGTTTCCATCATAGGAAATAAAAATCTCTGTGGGGGTGTACCGGAGTTGGACCTTCCGGTTTGCATTGTAG

>CORE_2790

GGAAATAAAAATCTCTGTGGGGGTGTACCGGAGTTGGACCTTCCGGTTTGCATTGTAGGAGTAAAGAAAGAAAGGAAATCAGGTTTTCCCTTGAAAAAAGTAATTCCTGTAGTTAGTGGG

>CORE_2852

AAAGAAAGAAAGGAAATCAGGTTTTCCCTTGAAAAAAGTAATTCCTGTAGTTAGTGGGCTTATAGGATTGACATTGATAGTTTGTTTTCTTGGCATACGACAGTTTAGTAGATTGAGAAA

>CORE_2914

TAGGATTGACATTGATAGTTTGTTTTCTTGGCATACGACAGTTTAGTAGATTGAGAAAAACAACACCTACAGATATACCTGAAAACTCAACCTTAAGAATATCTTATCAGTGTCTACTTA

>CORE_2976

ACACCTACAGATATACCTGAAAACTCAACCTTAAGAATATCTTATCAGTGTCTACTTAGGGAAACTGATAGATTTTCTGCATCAAATTTGCTTGGCATGGGTGCTTTTGGGTCTGTATAT

>CORE_3038

AACTGATAGATTTTCTGCATCAAATTTGCTTGGCATGGGTGCTTTTGGGTCTGTATATAAAGGAATTTCTGAACATGATGGTACTGTTTTTGCTGTTAAGGTACTGGACCTTTCACACCA

>CORE_3100

GAATTTCTGAACATGATGGTACTGTTTTTGCTGTTAAGGTACTGGACCTTTCACACCATGCAGCTTCCAGGAGTTTTTTAGCTGAATGTGAGGTTCTAAAGAACATCAGACATCGTAATC

>CORE_3162

GCTTCCAGGAGTTTTTTAGCTGAATGTGAGGTTCTAAAGAACATCAGACATCGTAATCTTGTAAAGGTTCTAAGTGCATGCTCAGGTATTGATTATGAAGGTAATGAGTTCAAAGCCATA

>CORE_3224

AAAGGTTCTAAGTGCATGCTCAGGTATTGATTATGAAGGTAATGAGTTCAAAGCCATAGTTTATGAATACATGGATAAGGGGAATCTACAGGACTGGTTGCATTTCACTCCTCAGGAGAA

>CORE_3286

ATGAATACATGGATAAGGGGAATCTACAGGACTGGTTGCATTTCACTCCTCAGGAGAACTCTGAGCCACAGGAGGAACACAAGAAGCTAGGGTTTATTCAAAGATTAAACATTGCGATTG

>CORE_3348

GAGCCACAGGAGGAACACAAGAAGCTAGGGTTTATTCAAAGATTAAACATTGCGATTGATGTTGCCTGTGCTCTGGATTATCTCCATAATGATTGCCAACCCCCTATAATTCACCGCGAT

>CORE_3472

AGCCAAGCAATATTCTTCTAGATGAGAACATGACTGCACATGTTGGTGATTTTGGTTTGGCAAGATTTGTTCCACCAGAAATTCCGAACTCATCAGAAAATTCAAAAAGTCTTACAGGTG

>CORE_3534

AGATTTGTTCCACCAGAAATTCCGAACTCATCAGAAAATTCAAAAAGTCTTACAGGTGTAGGGGGGACAATTGGTTATACACCTCCAGAATTAGGTATGGGAAGCGACGCCTCAACCTAT

>CORE_3658

ATGGATACAGCTTTGGCATACTGCTCTTGGAAATGTTTACCGGGAGGAAACCAACTGATGAAATGTTCAAAGATAATTTAAACCTTCATAACTACGCGAATGCTGCTTTGCCAGACAGAG

>CORE_3720

ATGTTCAAAGATAATTTAAACCTTCATAACTACGCGAATGCTGCTTTGCCAGACAGAGTGATGCATATTACAGATCCAATTCTTCTACAGGAACGAGATGAACTTGAAATGGAATACAAA

>CORE_3782

GCATATTACAGATCCAATTCTTCTACAGGAACGAGATGAACTTGAAATGGAATACAAACTTCATGACAACACAAGCAGTGCTGGCGACATATTTCTGTCATTCTTAATCAACGTGATTCA

>CORE_3844

ATGACAACACAAGCAGTGCTGGCGACATATTTCTGTCATTCTTAATCAACGTGATTCAAATCGGAGTTTCCTGCTCCGCTGAATCTCCAAAAGAAAGAAAGCGGATCAGTGATGTCGTTA

>CORE_3906

GGAGTTTCCTGCTCCGCTGAATCTCCAAAAGAAAGAAAGCGGATCAGTGATGTCGTTAGAGAACTCAATTCATTAAGGAAGTTGTTTTTGGAGCAAGCATACCGGAAGAAAAAGTTATAA

>CORE_3968

ACTCAATTCATTAAGGAAGTTGTTTTTGGAGCAAGCATACCGGAAGAAAAAGTTATAAGCTTCAACCTGCATTGACAGATGATGACAGGCATAGGGTTAGTCCTTCTCATGGTTGGTGAC

>CORE_4092

GGTTAATATGCACTGTGAAACAAATTCATGAATTTAAGGTTTGATTCTGCTCTTATAATTTGCAGTTATAAACTTGTATGTTCCTAGTAAATGATAGCTCTTTGCACATTTCTTAGTTTA

>CORE_4154

CAGTTATAAACTTGTATGTTCCTAGTAAATGATAGCTCTTTGCACATTTCTTAGTTTAGATGCCTTATTTATTTTATTGAGATATCTATAGCTGCTAGACTAGTAGATCACTGAATTACA

>CORE_4214

TGCCTTATTTATTTTATTGAGATATCTATAGCTGCTAGACTAGTAGATCACTGAATTACACAATTGATCTCAAGCTGCCACCTACATGTCCAAACATCATATTTGAAGTTTTTGAGACGT

>CORE_682

GGAGTGTTATTTTGGTAGCATACCTTGTTCTTCATTTTGTTTCTATTGTTGTTTCTTTTACTTTTGTTATCTCTAATAATTTTATTGTCAATATTTTTCTTTTTTCAATATTTTCATAAT

>CORE_744

TTTGTTATCTCTAATAATTTTATTGTCAATATTTTTCTTTTTTCAATATTTTCATAATAGCATTGCATTTCTTTCAAACCATGTTCTGAAAATAGATTTTTCTTGGGAACCATGGATTTT

>CORE_806

TTGCATTTCTTTCAAACCATGTTCTGAAAATAGATTTTTCTTGGGAACCATGGATTTTTATCTGTAATTTGCAACCTTTGATGCAGTTAACAAATTCTCTCCAAGAGGCCACCTCAAGAA

>CORE_868

TGTAATTTGCAACCTTTGATGCAGTTAACAAATTCTCTCCAAGAGGCCACCTCAAGAAGCAACTTAGCATCTAGACAGCTAGCTTCTTCTGCTGGATCAAACACTTCAAAGTTCTAGTCT

>CORE_930

CTTAGCATCTAGACAGCTAGCTTCTTCTGCTGGATCAAACACTTCAAAGTTCTAGTCTTCGAAGACCATCTTCTTCAGGTGGCTATGGTGAAAGGGAATGAAACAGACAAAATGTCACTG

>CORE_992

AGACCATCTTCTTCAGGTGGCTATGGTGAAAGGGAATGAAACAGACAAAATGTCACTGCTAGCATTCAAGAATATGATCATTGATGACCCTTTCAAGATCATGGACTCCTGGAATGAAAC

>Cf-2.1_0

ATGATGATGGTTTCTAGAAAAGTAGTCTCTTCACTTCAGTTTTTCACTCTTTTCTACCTCTTTACAGTTGCATTTGCTTCGACTGAGGAGGCAACTGCCCTCTTGAAATGGAAAGCAACT

>Cf-2.1_122

CAAGAACCAGAATAATTCCTTTTTGGCTTCATGGATTCCAAGTTCTAATGCATGCAAGGACTGGTATGGAGTTGTATGCTTTAATGGTAGGGTAAACACGTTGAATATTACAAATGCTAG

>Cf-2.1_1281

GGTTACTTGAGTTCTCTTACTTATCTAGATTTGAGTAATAACTCCATTAATGGATTTATTCCTGCTTCATTTGGCAATATGAGCAACTTGGCTTTTTTGTTTCTTTATGAAAATCAGCTT

>Cf-2.1_1891

GTAATAACTCTCTTAATGGACTTATTCCTGCTTCATTTGGCAATATGAGAAATCTGCAAGCTCTGATTCTCAATGATAACAATCTCATTGGGGAAATTCCTTCATCTGTGTGCAATTTGA

>Cf-2.1_2013

TCACTGGAAGTGTTGTATATGCCGAGAAACAATTTGAAGGGAAAAGTTCCGCAATGTTTGGGTAATATCAGTAACCTTCAGGTTTTGTCGATGTCATCTAATAGTTTCAGTGGAGAGCTC

>Cf-2.1_2074

GTAATATCAGTAACCTTCAGGTTTTGTCGATGTCATCTAATAGTTTCAGTGGAGAGCTCCCTTCATCTATTTCCAATTTAACATCACTACAAATACTTGATTTTGGCAGAAACAATCTGG

>Cf-2.1_2196

GGAGCAATACCACAATGTTTTGGCAATATTAGTAGCCTCGAGGTTTTTGATATGCAGAACAACAAACTTTCTGGGACTCTTCCAACAAATTTTAGCATTGGATGTTCACTGATAAGTCTC

>Cf-2.1_2257

ACAAACTTTCTGGGACTCTTCCAACAAATTTTAGCATTGGATGTTCACTGATAAGTCTCAACTTGCATGGCAATGAACTAGAGGATGAAATCCCTCGGTCTTTGGACAATTGCAAAAAGC

>Cf-2.1_2440

TGCCAGAGCTGAGAGTTTTAAGGTTGACATCGAATAAATTGCATGGACCTATAAGATCATCAAGGGCTGAAATCATGTTTCCTGATCTTCGAATCATAGATCTCTCTCGCAATGCATTCT

>Cf-2.1_2501

AAGGGCTGAAATCATGTTTCCTGATCTTCGAATCATAGATCTCTCTCGCAATGCATTCTCGCAAGACTTACCAACGAGTCTATTTGAACATTTGAAAGGGATGAGGACAGTTGATAAAAC

>Cf-2.1_2562

CAAGACTTACCAACGAGTCTATTTGAACATTTGAAAGGGATGAGGACAGTTGATAAAACAATGGAGGAACCAAGTTATGAAAGCTATTACGATGACTCGGTGGTAGTTGTGACAAAGGGA

>Cf-2.1_2623

TGGAGGAACCAAGTTATGAAAGCTATTACGATGACTCGGTGGTAGTTGTGACAAAGGGATTGGAGCTTGAAATTGTGAGAATTTTGTCTTTGTACACAGTTATCGATCTTTCAAGCAACA

>Cf-2.1_2745

TTTGAAGGACATATTCCTTCTGTCCTGGGAGATCTCATTGCGATCCGTATACTTAATGTATCTCATAATGCATTGCAAGGCTATATACCATCATCACTTGGAAGTTTATCTATACTGGAA

>Cf-2.1_305

TAGCAAGAACAATATCTATGGTACCATTCCACCTGAGATTGGTAATCTCACAAATCTTGTCTATCTTGACTTGAACAACAATCAGATTTCAGGAACAATACCACCACAAATCGGTTTACT

>Cf-2.1_3050

TTCAAAAGGTTGTGGCAAAGATCCTGTGTCAGAGAAAAACTATACAGTGTCTGCGCTAGAAGATCAAGAAAGCAATTCTGAATTTTTCAATGATTTTTGGAAAGCAGCTCTGATGGGCTA

>Cf-2.1_3219

ACTGGAAATCTAAGATGGCTTGCAAGAATCATTGAAAAACTGGAACACAAAATTATCATGCAAAGGAGAAAGAAGCAGCGAGGTCAAAGAAATTACAGAAGAAGAAATAATCACTTCTAG

>Cf-2.1_366

TATCTTGACTTGAACAACAATCAGATTTCAGGAACAATACCACCACAAATCGGTTTACTAGCCAAGCTTCAGATCATCCGCATATTTCACAATCAATTAAATGGATTTATTCCTAAAGAA

>Cf-2.1_61

TTACAGTTGCATTTGCTTCGACTGAGGAGGCAACTGCCCTCTTGAAATGGAAAGCAACTTTCAAGAACCAGAATAATTCCTTTTTGGCTTCATGGATTCCAAGTTCTAATGCATGCAAGG

>Cf-4_0

ATGGGTTGTGTAAAACTTGTGTTTTTCATGCTATATGTCTTTCTCTTTCAACTTGTTTCCTCGTCATCCTTACCTCATTTGTGCCCCGAAGATCAAGCTCTTGCTCTTCTAGAATTCAAG

>Cf-4_1008

AACTTAAATGGGAGTATACCATCCTGGATATTCTCCCTTCCATCACTGATAGGGTTAGACTTGAGCAATAACACTTTCAGTGGAAAAATTCAAGAGTTCAAGTCCAAAACATTAAGTACC

>Cf-4_1071

AGCAATAACACTTTCAGTGGAAAAATTCAAGAGTTCAAGTCCAAAACATTAAGTACCGTTACTCTAAAACAAAATAAGCTAAAAGGTCCTATTCCGAATTCACTCCTAAACCAGAAGAAC

>Cf-4_1134

CTAAAACAAAATAAGCTAAAAGGTCCTATTCCGAATTCACTCCTAAACCAGAAGAACCTACAATTCCTTCTCCTTTCACACAATAATATCAGTGGACATATTTCTTCAGCTATCTGCAAT

>Cf-4_1197

TTCCTTCTCCTTTCACACAATAATATCAGTGGACATATTTCTTCAGCTATCTGCAATCTGAAAACATTGATATTGTTAGACTTGGGAAGTAATAATTTGGAGGGAACAATCCCGCAATGC

>Cf-4_126

TTTACCGTTAATCCTAATGCTTCTGATTATTGTTACGACAGAAGAACTCTTTCTTGGAACAAAAGCACAAGTTGCTGCTCATGGGATGGCGTTCATTGTGACGAAACGACAGGACAAGTG

>Cf-4_1260

ACATTGATATTGTTAGACTTGGGAAGTAATAATTTGGAGGGAACAATCCCGCAATGCGTGGTTGAGAGGAACGAATACCTTTCGCATTTGGATTTGAGCAACAACAGACTTAGTGGGACA

>Cf-4_1323

GAGAGGAACGAATACCTTTCGCATTTGGATTTGAGCAACAACAGACTTAGTGGGACAATCAATACAACTTTTAGTGTTGGAAACATTTTAAGGGTCATTAGCTTGCACGGGAATAAGCTA

>Cf-4_1386

ACAACTTTTAGTGTTGGAAACATTTTAAGGGTCATTAGCTTGCACGGGAATAAGCTAACGGGGAAAGTCCCACGATCTATGATCAATTGCAAGTATTTGACACTACTTGATCTAGGTAAC

>Cf-4_1449

AAAGTCCCACGATCTATGATCAATTGCAAGTATTTGACACTACTTGATCTAGGTAACAATATGTTGAATGACACATTTCCAAACTGGTTGGGATACCTATTTCAATTGAAGATTTTAAGC

>Cf-4_1512

TTGAATGACACATTTCCAAACTGGTTGGGATACCTATTTCAATTGAAGATTTTAAGCTTGAGATCAAATAAGTTGCATGGTCCCATCAAATCTTCAGGGAATACAAACTTGTTTATGGGT

>Cf-4_1575

TCAAATAAGTTGCATGGTCCCATCAAATCTTCAGGGAATACAAACTTGTTTATGGGTCTTCAAATTCTTGATCTATCATCTAATGGATTTAGTGGGAATTTACCCGAAAGAATTTTGGGG

>Cf-4_1638

ATTCTTGATCTATCATCTAATGGATTTAGTGGGAATTTACCCGAAAGAATTTTGGGGAATTTGCAAACCATGAAGGAAATTGATGAGAGTACAGGATTCCCAGAGTATATTTCTGATCCA

>Cf-4_1701

CAAACCATGAAGGAAATTGATGAGAGTACAGGATTCCCAGAGTATATTTCTGATCCATATGATATTTATTACAATTATTTGACGACAATTTCTACAAAGGGACAAGATTATGATTCTGTT

>Cf-4_1764

ATTTATTACAATTATTTGACGACAATTTCTACAAAGGGACAAGATTATGATTCTGTTCGAATTTTGGATTCTAACATGATTATCAATCTCTCAAAGAACAGATTTGAAGGTCATATTCCA

>Cf-4_1827

TTGGATTCTAACATGATTATCAATCTCTCAAAGAACAGATTTGAAGGTCATATTCCAAGCATTATTGGAGATCTTGTTGGACTTCGTACGTTGAACTTGTCTCACAATGTCTTGGAAGGT

>Cf-4_189

AGCACAAGTTGCTGCTCATGGGATGGCGTTCATTGTGACGAAACGACAGGACAAGTGATTGAGCTTGACCTCCGTTGCATCCAACTTCAAGGCAAGTTTCATTCCAATAGTAGCCTCTTT

>Cf-4_1890

ATTGGAGATCTTGTTGGACTTCGTACGTTGAACTTGTCTCACAATGTCTTGGAAGGTCATATACCGGCATCATTTCAAAATTTATCAGTACTCGAATCATTGGATCTCTCATCTAATAAA

>Cf-4_1953

CCGGCATCATTTCAAAATTTATCAGTACTCGAATCATTGGATCTCTCATCTAATAAAATCAGCGGAGAAATTCCGCAGCAGCTTGCATCCCTCACATTCCTTGAAGTCTTAAATCTCTCT

>Cf-4_2079

CATCTTGTTGGATGCATCCCCAAAGGAAAACAATTTGATTCGTTCGGGAACACTTCGTACCAAGGGAATGATGGGTTACGCGGATTTCCACTCTCAAAACTTTGTGGTGGTGAAGATCAA

>Cf-4_2142

GGGAATGATGGGTTACGCGGATTTCCACTCTCAAAACTTTGTGGTGGTGAAGATCAAGTGACAACTCCAGCTGAGCTAGATCAAGAAGAGGAGGAAGAAGATTCACCAATGATCAGTTGG

>Cf-4_2205

ACTCCAGCTGAGCTAGATCAAGAAGAGGAGGAAGAAGATTCACCAATGATCAGTTGGCAGGGGGTTCTCGTGGGTTACGGTTGTGGACTTGTTATTGGACTGTCCGTAATATACATAATG

>Cf-4_2268

GTTCTCGTGGGTTACGGTTGTGGACTTGTTATTGGACTGTCCGTAATATACATAATGTGGTCAACTCAATATCCAGCATGGTTTTCGAGGATGGATTTAAAGTTGGAACACATAATTACT

>Cf-4_2301

GGACTGTCCGTAATATACATAATGTGGTCAACTCAATATCCAGCATGGTTTTCGAGGATGGATTTAAAGTTGGAACACATAATTACTACGAAAATGAAAAAGCACAAGAAAAGATATTAG

>Cf-4_252

CTTGACCTCCGTTGCATCCAACTTCAAGGCAAGTTTCATTCCAATAGTAGCCTCTTTCAACTCTCCAATCTCAAAAGGCTTGATTTGTCTTATAATGATTTCACTGGATCGCCCATTTCA

>Cf-4_315

TCCAATCTCAAAAGGCTTGATTTGTCTTATAATGATTTCACTGGATCGCCCATTTCACCTAAATTTGGTGAGTTTTCAGATTTGACGCATCTCGATTTGTCGCATTCAAGTTTTAGGGGT

>Cf-4_378

TTTGGTGAGTTTTCAGATTTGACGCATCTCGATTTGTCGCATTCAAGTTTTAGGGGTGTAATCCCTTCTGAAATCTCTCATCTTTCTAAACTATACGTTCTTCGTATTAGTCTAAATGAG

>Cf-4_441

CCTTCTGAAATCTCTCATCTTTCTAAACTATACGTTCTTCGTATTAGTCTAAATGAGCTTACTTTTGGTCCTCACAATTTTGAATTGCTTCTTAAGAACTTGACCCAATTAAAAGTGCTC

>Cf-4_504

TTTGGTCCTCACAATTTTGAATTGCTTCTTAAGAACTTGACCCAATTAAAAGTGCTCGACCTTGAATCTATCAACATCTCTTCCACTATTCCTTTGAATTTCTCTTCTCATTTAACAAAT

>Cf-4_567

GAATCTATCAACATCTCTTCCACTATTCCTTTGAATTTCTCTTCTCATTTAACAAATCTATGGCTTCCATACACAGAGTTACGTGGGATATTGCCCGAAAGAGTTTTCCACCTTTCCGAC

>Cf-4_63

TCATCCTTACCTCATTTGTGCCCCGAAGATCAAGCTCTTGCTCTTCTAGAATTCAAGAACATGTTTACCGTTAATCCTAATGCTTCTGATTATTGTTACGACAGAAGAACTCTTTCTTGG

>Cf-4_630

CTTCCATACACAGAGTTACGTGGGATATTGCCCGAAAGAGTTTTCCACCTTTCCGACTTAGAATTTCTCGATTTATCAAGCAATCCCCAGCTCACGGTTAGGTTTCCCACAACCAAATGG

>Cf-4_756

AGTGCATCACTCATGAAGTTATATCTCTATAATGTGAATATTGATGATAGGATACCTGAATCATTTAGCCATCTAACTTCACTTCATAAGTTGTACATGAGTCGTTCTAATCTGTCAGGG

>Cf-4_819

TTTAGCCATCTAACTTCACTTCATAAGTTGTACATGAGTCGTTCTAATCTGTCAGGGCCTATTCCTAAACCTCTATGGAATCTCACCAACATAGTGTTTTTGGACCTTAATAATAACCAT

>Cf-4_882

CCTAAACCTCTATGGAATCTCACCAACATAGTGTTTTTGGACCTTAATAATAACCATCTTGAAGGACCAATTCCATCCAACGTAAGCGGACTACGTAACCTACAAATACTTTGGTTGTCA

>Cf-4_945

GGACCAATTCCATCCAACGTAAGCGGACTACGTAACCTACAAATACTTTGGTTGTCATCAAACAACTTAAATGGGAGTATACCATCCTGGATATTCTCCCTTCCATCACTGATAGGGTTA

>Cf-5_0

ATGATGATGGTTACTAGCAAAGTATTCTCTTCACTTCAGTTTTTCACTGTTTTCTACCTCTTTACAGTTGCATTTGCTTCGACTGAGGAGGCAACTGCCCTCTTGAAATGGAAAGCAACT

>Cf-5_124

AGAACCAGAATAATTCCTTTTTGGCTTCATGGACGACAAGTTCTAATGCATGCAAGGACTGGTATGGAGTTGTATGCTTGAATGGTAGGGTAAACACGTTGAATATTACAAATGCCAGTG

>Cf-5_1426

GGAATCTAAACAACTTGTCTAGGTTGTATCTTTACAATAATCAGCTTTCTGGCTCTATTCCTGCTTCATTTGGCAATATGAGAAATCTGCAAACTCTGTTTCTCAGTGATAACGATCTCA

>Cf-5_1488

GCTTCATTTGGCAATATGAGAAATCTGCAAACTCTGTTTCTCAGTGATAACGATCTCATTGGGGAAATTCCTTCATTTGTGTGCAATTTGACATCACTGGAAGTGTTGTATATGTCGAGA

>Cf-5_1612

ATTTGAAGGGAAAAGTTCCGCAATGTTTGGGTAATATCAGTGACCTTCACATTTTGTCGATGTCATCTAATAGTTTCAGAGGAGAGCTCCCTTCATCTATTTCCAATTTAACATCACTAA

>Cf-5_1674

TCATCTAATAGTTTCAGAGGAGAGCTCCCTTCATCTATTTCCAATTTAACATCACTAAAAATACTTGATTTTGGCAGAAACAATCTGGAGGGAGCAATACCACAATTTTTTGGCAATATT

>Cf-5_1736

ACTTGATTTTGGCAGAAACAATCTGGAGGGAGCAATACCACAATTTTTTGGCAATATTAGTAGCCTCCAGGTTTTTGATATGCAGAATAACAAACTTTCTGGGACTCTTCCAACAAATTT

>Cf-5_248

TGGTACACTTTATGCTTTTCCATTTTCATCCCTCCCTTTTCTCGAGAATCTTGATCTTAGCAACAACAATATCTCTGGTACCATTCCACCTGAGATTGGTAATCTCACAAATCTTGTCTA

>Cf-5_310

ACAACAATATCTCTGGTACCATTCCACCTGAGATTGGTAATCTCACAAATCTTGTCTATCTTGACTTGAACACCAATCAGATTTCAGGAACAATTCCACCACAAATCGGTTCACTAGCCA

>Cf-5_372

GACTTGAACACCAATCAGATTTCAGGAACAATTCCACCACAAATCGGTTCACTAGCCAAGCTTCAGATCATCCGCATATTTAACAATCATTTAAATGGCTTTATTCCTGAAGAAATAGGT

>Cf-9_0

ATGGATTGTGTAAAACTTGTATTCCTTATGCTATATACCTTTCTCTGTCAACTTGCTTTATCCTCATCCTTGCCTCATTTGTGCCCCGAAGATCAAGCTCTTTCTCTTCTACAATTCAAG

>Cf-9_1116

GGTCCAATTCCATCCAACATAAGCGGACTTCAAAACCTAGAATGTCTCTACTTGTCATCAAACCACTTGAATGGGAGTATACCTTCCTGGATATTCTCCCTTCCTTCACTGGTTGAGTTA

>Cf-9_1178

CCACTTGAATGGGAGTATACCTTCCTGGATATTCTCCCTTCCTTCACTGGTTGAGTTAGACTTGAGCAATAACACTTTCAGTGGAAAAATTCAAGAGTTCAAGTCCAAAACATTAAGTGC

>Cf-9_124

TGTTTACCATTAATCCTAATGCTTCTGATTATTGTTACGACATAAGAACATACGTAGACATTCAGTCATATCCAAGAACTCTTTCTTGGAACAAAAGCACAAGTTGCTGCTCATGGGATG

>Cf-9_186

CAGTCATATCCAAGAACTCTTTCTTGGAACAAAAGCACAAGTTGCTGCTCATGGGATGGCGTTCATTGTGACGAGACGACAGGACAAGTGATTGCGCTTGACCTCCGTTGCAGCCAACTT

>Cf-9_248

TCATTGTGACGAGACGACAGGACAAGTGATTGCGCTTGACCTCCGTTGCAGCCAACTTCAAGGCAAGTTTCATTCCAATAGTAGCCTCTTTCAACTCTCCAATCTCAAAAGGCTTGATTT

>Cf-9_310

GCAAGTTTCATTCCAATAGTAGCCTCTTTCAACTCTCCAATCTCAAAAGGCTTGATTTGTCTTTTAATAATTTCACTGGATCACTCATTTCACCAAAATTTGGTGAGTTTTCAAATTTGA

>Cf-9_434

TCTCGATTTGTCGCATTCTAGTTTTACAGGTCTAATTCCTTCTGAAATCTGTCACCTTTCTAAACTACACGTTCTTCGTATATGTGATCAATATGGGCTTAGTCTTGTACCTTACAATTT

>Cf-9_496

AACTACACGTTCTTCGTATATGTGATCAATATGGGCTTAGTCTTGTACCTTACAATTTTGAACTGCTCCTTAAGAACTTGACCCAATTAAGAGAGCTCAACCTTGAATCTGTAAACATCT

>Cf-9_62

CTCATCCTTGCCTCATTTGTGCCCCGAAGATCAAGCTCTTTCTCTTCTACAATTCAAGAACATGTTTACCATTAATCCTAATGCTTCTGATTATTGTTACGACATAAGAACATACGTAGA

>Cf-9_620

CACTATTCCTTCAAATTTCTCTTCTCATTTAACAACTCTACAACTTTCAGGCACAGAGTTACATGGGATATTGCCCGAAAGAGTTTTTCACCTTTCCAACTTACAATCCCTTCATTTATC

>Cf-9_682

ATGGGATATTGCCCGAAAGAGTTTTTCACCTTTCCAACTTACAATCCCTTCATTTATCAGTCAATCCCCAGCTCACGGTTAGGTTTCCCACAACCAAATGGAATAGCAGTGCATCACTCA

>Cf-9_806

GTTATACGTCGATAGTGTGAATATTGCTGATAGGATACCTAAATCATTTAGCCATCTAACTTCACTTCATGAGTTGTACATGGGTCGTTGTAATCTGTCAGGGCCTATTCCTAAACCTCT

>Cf-9_868

CACTTCATGAGTTGTACATGGGTCGTTGTAATCTGTCAGGGCCTATTCCTAAACCTCTATGGAATCTCACCAACATAGTGTTTTTGCACCTTGGTGATAACCATCTTGAAGGACCAATTT

>Cf-9_930

AATCTCACCAACATAGTGTTTTTGCACCTTGGTGATAACCATCTTGAAGGACCAATTTCCCATTTCACGATATTTGAAAAGCTCAAGAGGTTATCACTTGTAAATAACAACTTTGATGGC

>Cf-9_992

TTTCACGATATTTGAAAAGCTCAAGAGGTTATCACTTGTAAATAACAACTTTGATGGCGGACTTGAGTTCTTATCCTTTAACACCCAACTTGAACGGCTAGATTTATCATCCAATTCCCT

>FLS3_0

TCATATAAATCAAAATAGAAACACAAAACGACAACTTTTCGTCTTGTGAAATAACTCAAGTGGAGTCATTCATTCCAATATATCCTTTTTCCATTTCTCTTCATATATTTCAACTTTCAA

>FLS3_1054

GAGAAATACCGAGAAATATTGGTTGTTTAGCCAAGCTCGAAGAGTTTTATATCGGTTATAATGCAATAAATGGAACAATTCCTGCTTCATTAGGCAATATTTCAACTCTTCAAAATCTTC

>FLS3_1178

CGGAAGCAATCACATGGAGGGAGAACTTCCTCCAGAATTAGGAAAGCTATCAAACTTAAGACAAATCAATTTCGAAGAAAATTATAATCTTATTGGTGAAATTCCGAATACTATTTTCAA

>FLS3_124

TTAGTAACATCATGGAGAAACACATTTTCTTATTGATACTTGCTATCTTAGTTCAATTTTACTTTGTTTCTTCTATATCAGCTACTATTTCCTCAAATGAGACTGATCAAGAAGCTCTAC

>FLS3_1302

TCTTCTTTGGAGTTCATTGCTTTCACTTTCAACTACCTTTCAGGTAGAATTCCGAATCTTCTTCACCTTCCAAATCTTATACAACTTCTCTTAGCAAACAATCAGCTCGAAGGTGAAATT

>FLS3_1364

TCACCTTCCAAATCTTATACAACTTCTCTTAGCAAACAATCAGCTCGAAGGTGAAATTCCTCGGTACATCACAAATGCTACGAATCTTGAGCTGTTAGAGCTATCAGATAACCTTCTCAC

>FLS3_1426

GGTACATCACAAATGCTACGAATCTTGAGCTGTTAGAGCTATCAGATAACCTTCTCACAGGTACTATTCCTAATGATTTAGGAAATCTTCGCGAGCTGCGAGATCTTTTCCTACATCATA

>FLS3_1488

ACTATTCCTAATGATTTAGGAAATCTTCGCGAGCTGCGAGATCTTTTCCTACATCATAATCAACTTACTGAGTTGGGATTCTTTGATTCTTTGGTGAAATGTAGGATGTTGAGATATGTA

>FLS3_1550

ACTTACTGAGTTGGGATTCTTTGATTCTTTGGTGAAATGTAGGATGTTGAGATATGTACAAGTGGGATCGAATCCGTTGAATGATGTTCTGCCAAGTAGTATTGGCAATCTTTCATCTAC

>FLS3_1612

TGGGATCGAATCCGTTGAATGATGTTCTGCCAAGTAGTATTGGCAATCTTTCATCTACTGTTGAATACTTTCATATTGGAGATGCACAAATCAATGGATTCATTCCCACTAGTACAGGCA

>FLS3_1674

GAATACTTTCATATTGGAGATGCACAAATCAATGGATTCATTCCCACTAGTACAGGCAACATGACCGGTCTTACAACGCTAGTTTTTCAAGATAACAGTTTGACAGGAAACATTCCTCGT

>FLS3_1736

GACCGGTCTTACAACGCTAGTTTTTCAAGATAACAGTTTGACAGGAAACATTCCTCGTGAGATCCGTAAGCTTAAACAACTCCAAGGTTTATTTCTAGTTAACAATGGACTACAGGGGGA

>FLS3_1798

TCCGTAAGCTTAAACAACTCCAAGGTTTATTTCTAGTTAACAATGGACTACAGGGGGACATAGCAGAGGTAGTATGTGATTTATCGAATTTGGTTCGATTAGCTCTGTCTGAAAATGAGC

>FLS3_186

TTTGTTTCTTCTATATCAGCTACTATTTCCTCAAATGAGACTGATCAAGAAGCTCTACTAGCTTTTCGAAACCTTGTTACGAGTGATTCTAGTCATTTTTTAGCCAATAATTGGACAAAA

>FLS3_1860

GCAGAGGTAGTATGTGATTTATCGAATTTGGTTCGATTAGCTCTGTCTGAAAATGAGCTCTCGGGGGTGATTCCGGAATGTTTAGGAAATCTTACCATGCTACAACAACTTTTTTTAGGT

>FLS3_1922

GGGGGTGATTCCGGAATGTTTAGGAAATCTTACCATGCTACAACAACTTTTTTTAGGTTCTAACAAGTTTGAATCAAAGCTACCTTTAAGCTTTTGGAAGATGAGTAGTCTTCTATATTT

>FLS3_2046

ATGTCGCGTAATTCTATAAAGGGAGAAGTTCCATCAGATATCGGAGAACTTAAAGCTATTGTAGCAATCGATATCTCTGGTAACCATTTCTCGGGGTCGATACCAAGCAATTTGGGGGAA

>FLS3_2108

AGCAATCGATATCTCTGGTAACCATTTCTCGGGGTCGATACCAAGCAATTTGGGGGAACTTCAAACCTTGAAGTTACTTTCCTTATCGAACAATTCGTTTTCAGGTCCAATTCCATTTTC

>FLS3_2170

AAACCTTGAAGTTACTTTCCTTATCGAACAATTCGTTTTCAGGTCCAATTCCATTTTCCTTTTCAAACTTGAAAAGCTTGGAATTCTTGGATTTGTCTTTGAATAACTTGTCAGGTACTA

>FLS3_2232

TCAAACTTGAAAAGCTTGGAATTCTTGGATTTGTCTTTGAATAACTTGTCAGGTACTATTCCTAAGTCTTTCGAAAAGCTTTTGTACCTTACAAGCATCAACGTCTCGTTTAATGTTTTA

>FLS3_2294

TAAGTCTTTCGAAAAGCTTTTGTACCTTACAAGCATCAACGTCTCGTTTAATGTTTTAGAAGGTGAAATACCTAGTGGTGGTGTGTTTGCAAACTCCACCCTGCAATCATTTAGTGGGAA

>FLS3_2356

GTGAAATACCTAGTGGTGGTGTGTTTGCAAACTCCACCCTGCAATCATTTAGTGGGAACAAAGGTCTATGTGGAAGGCAAATATTGGAGGTTCCTGCTTGTGCTATCACTACTCCTGAAC

>FLS3_2418

GGTCTATGTGGAAGGCAAATATTGGAGGTTCCTGCTTGTGCTATCACTACTCCTGAACAACAACAATCAAAATCGAAGAAGCTTGTGCTAAAAATTGTCACTCCGATGGTTATTTCATTC

>FLS3_248

TTTTCGAAACCTTGTTACGAGTGATTCTAGTCATTTTTTAGCCAATAATTGGACAAAAAACACTTCATTTTGCTCTTGGTTTGGTGTCACTTGTAGTCCAAAAAGGCAAAGGGTTGTAGC

>FLS3_2480

ACAATCAAAATCGAAGAAGCTTGTGCTAAAAATTGTCACTCCGATGGTTATTTCATTCTTTCTGATATTCTTGTTGGTTGTCTCGATTTGGATAATGAAACGAAAGAAGAAAGGGAAGTC

>FLS3_2666

AGCAACTAACAATTTTGATGAATCCAATTTGATTGGCGTGGGAGGTTCTGGCTCTGTGTACAAAGCCACATTAGCTAGTGGAATTGTGGTTGCAATTAAGGTACTGGATTTGGAAAATGA

>FLS3_2728

AAGCCACATTAGCTAGTGGAATTGTGGTTGCAATTAAGGTACTGGATTTGGAAAATGAGGAAGTATGCAAAAGGTTTGATACTGAATGCGAAGTGATGAGAAATGTTAGACACAAAAACC

>FLS3_2852

TTCGGTGATCACTACGTGTTCTAGTGAACACATAAGAGCCTTTGTTCTGCAGTATATGCCCAACGGAAGTCTTGACAATTGGTTGTACAAAGAAGATCGCCACTTAAAACTTCGTCAAAG

>FLS3_2914

ACGGAAGTCTTGACAATTGGTTGTACAAAGAAGATCGCCACTTAAAACTTCGTCAAAGAGTCACCATAATGCTTGATGTAGCTATGGCAATTGAATATCTACATCATGGTAATGACACCC

>FLS3_2976

ACCATAATGCTTGATGTAGCTATGGCAATTGAATATCTACATCATGGTAATGACACCCCAATAGTTCATTGTGACCTCAAGCCAGCCAACGTTCTTTTGGATGAAGATATGGTGGCGCGT

>FLS3_3100

GTGATTTTGGCATCTCAAAGATTTTAGCTGTAAGCAAATCCATGGCACATACAAAGACATTAGGCACTCTTGGATATATTGCACCAGAATATGGCTCGGAGGGAATAGTGTCCACTCGTG

>FLS3_3286

AGATATTCAACGAAAATCTTGGCTTGAGGGAGTGGATAACGCGAGCATTTCCAAGAACTATGATGGAAGTTGTGGACGCGGATATGTTTCATGATGGAGAAAAAATTACTTCCGAAAGTG

>FLS3_3348

ATGGAAGTTGTGGACGCGGATATGTTTCATGATGGAGAAAAAATTACTTCCGAAAGTGAAATATGCATACTCTCCATGATAGAACTGGCTTTAGATTGCACAAAGGCAACACCAGAATCA

>FLS3_3534

GTAAATTAGAATCTCTTTGTGTTGTTTTTTTCTTTTAAGTTTGTAAGTTGATACTCAATCAATCTGTTTGTCATACTTTTCATTTGTCTGTTTTTTTAAAAAAATGTCTTTCTTTTTTGG

>FLS3_372

ACTCTTCCTAATTTGCAACTTCAAGGCACAATTTCGCCGTCTTTGGCCAATCTATCTTTTCTCATAGAGCTAAATCTCGCAAACAACAACTTACACAGTGAAATCCCTGATGGCATTGGC

>FLS3_434

CATAGAGCTAAATCTCGCAAACAACAACTTACACAGTGAAATCCCTGATGGCATTGGCCGCTTGCCTCGTCTACGAGTGATTGATATTCAGAACAATCAGCTGCATGGAAGTATTCCAAC

>FLS3_558

CTATTTCAACACGGGAGTGTTCAAATCATTTCATTGGCTTTCAATAAACTCGGTGGTGAAATGTGGAACGGTACATGGTATGTACCCGAACTCAGAGTCTTAAATCTCAGGAACAATACC

>FLS3_62

GAGTCATTCATTCCAATATATCCTTTTTCCATTTCTCTTCATATATTTCAACTTTCAAATGCTTAGTAACATCATGGAGAAACACATTTTCTTATTGATACTTGCTATCTTAGTTCAATT

>FLS3_620

GTGGAACGGTACATGGTATGTACCCGAACTCAGAGTCTTAAATCTCAGGAACAATACCATTACAGGTGTGATCCCTCCTTCTATTGGAAATGCCACAAAGTTGATGAACTTCAGTTTGAA

>FLS3_744

AATAGAATCAACGGCAACATTCCAATGGAGATTGGTAATCTAAGCCAACTTGTTGAGTTGTCGTTGTCTCGTAATCAATTAACAGGTTCCATTCCTTCAACATTGTTTAATATCTCCTCC

>FLS3_806

GTTGTCTCGTAATCAATTAACAGGTTCCATTCCTTCAACATTGTTTAATATCTCCTCCCTTCTCGTCGTGTCTCTGGCATACAATAGCCTTTCAGGTCCTCTGTTTCCTGATGATCGACG

>FLS3_868

TCGTCGTGTCTCTGGCATACAATAGCCTTTCAGGTCCTCTGTTTCCTGATGATCGACGTAACGTTCTTTCATCAAACCTCGAGCATATAGGTGTATCATACAATCAAATCACTGGTCACA

>FLS3_930

GTTCTTTCATCAAACCTCGAGCATATAGGTGTATCATACAATCAAATCACTGGTCACATTCCTTCCAACATCTGTCAATTCACAGCTCTCAGAGTTCTGTCCATATCATACAACAACATA

>I3_0

ATGTTAGCAGAACAGAGACTGTATTTTCCTATCCTTCTCTTCATTTATCTCCTCTTTTGTTCATCTCGATTTGCTATATCGCAATCTTCATGGGACTACAACGCCACTCTTTCAAATTCT

>I3_1054

ACTGCAACTCTTCGCAGTATCATAGTTTCATAGAGCTCAAGAATACTACATATTTTTCATTTAACATTGATCAGGAACTAAATTCGACTAAATTGTGGCTTGGGCAGAAAAAGTTGGAAG

>I3_1116

AACATTGATCAGGAACTAAATTCGACTAAATTGTGGCTTGGGCAGAAAAAGTTGGAAGATTGCAAAAGGGCATGTCTGAGTCACTGTTCTTGCAAAGCTGCTGTTTTTGAATATAATTGG

>I3_1178

CAAAAGGGCATGTCTGAGTCACTGTTCTTGCAAAGCTGCTGTTTTTGAATATAATTGGAATGGGGATCGAAGAGGTAATTGTTTGTTACTGAATGAAGTTTTCTCTCTCAAAGACAGCGA

>I3_124

CAGGGTTTGCTGGGCTTTCCTCTTTCTGGATCAACAGGCCGTCTCTTATTATTAATTCCACCACCGACAGCTTCTATGGTTCGACAACACCCGTACTTCAGCGGGGAAATGCTGGCCCAC

>I3_1240

GGGATCGAAGAGGTAATTGTTTGTTACTGAATGAAGTTTTCTCTCTCAAAGACAGCGAAGAAGTAAGAGGCGAGACAGTATTTCTTAAGGTGCAGAATACCTCAAAGGCGCAGACGCAGT

>I3_1302

GTAAGAGGCGAGACAGTATTTCTTAAGGTGCAGAATACCTCAAAGGCGCAGACGCAGTCTCTAATCATTCCTGGAGGAAAGAAATCAAGACCTTTCAAAGTGATAATAGGATCTACTCTT

>I3_1364

AATCATTCCTGGAGGAAAGAAATCAAGACCTTTCAAAGTGATAATAGGATCTACTCTTGCAGCTTTCTTTGGGATAATTTTAAGCATAGCTACTTGCTTTGTTATTTTCAGAAAGAGGAC

>I3_1426

CTTTCTTTGGGATAATTTTAAGCATAGCTACTTGCTTTGTTATTTTCAGAAAGAGGACACATGAGTCCAGAAAGGCTGCGGATATTTTGGATCTAGCACCAATCTTACCGGGAATCCTAA

>I3_1488

GAGTCCAGAAAGGCTGCGGATATTTTGGATCTAGCACCAATCTTACCGGGAATCCTAACTCGATTCTCTTACAATGAGCTGAAAATAATTACAGATGATTTCAGCAGTAAGCTTGGGGAA

>I3_1550

ATTCTCTTACAATGAGCTGAAAATAATTACAGATGATTTCAGCAGTAAGCTTGGGGAAGGAGGATTTGGCTGTGTTTATGAAGGAACACTGAGAAATGGAACTAAAATAGCTGTGAAGAA

>I3_1674

GATGGTGTAGGTCAAGTAAAGGAATCATTCTTAACAGAAGTAAAGGCGGTCGGTGGCATTCACCATATCAATCTGGTAAAACTCATTGGATTTTGTGCTGAAAAAACCCACAGGCTTCTA

>I3_1736

CCATATCAATCTGGTAAAACTCATTGGATTTTGTGCTGAAAAAACCCACAGGCTTCTAATCTATGAGTACATGGTGAATGGATCGCTTGATAGGTGGATTACACATGAAAACCGAGAAAA

>I3_1798

ATGAGTACATGGTGAATGGATCGCTTGATAGGTGGATTACACATGAAAACCGAGAAAATGGGCTTACATGGAGCACAAGACAGAGGATAATATCAGATATCGCCAAAGGGTTAGCGTATC

>I3_186

ACCGACAGCTTCTATGGTTCGACAACACCCGTACTTCAGCGGGGAAATGCTGGCCCACGATTCCTCTGTGGCTTCTACTGCAGGTACAATGTCACAGAATGCCTTCTTGGTATCCTTTTG

>I3_1860

CTTACATGGAGCACAAGACAGAGGATAATATCAGATATCGCCAAAGGGTTAGCGTATCTACATGAGGATTGCAGCCAAAAGATAATTCATTTGGACATCAAACCACAAAACATCCTTCTG

>I3_1922

TGAGGATTGCAGCCAAAAGATAATTCATTTGGACATCAAACCACAAAACATCCTTCTGGATCAATATCTCAATGCTAAGATATCAGATTTTGGGTTGTCGAAGCTAATTGAGAAAGATAA

>I3_1984

AATATCTCAATGCTAAGATATCAGATTTTGGGTTGTCGAAGCTAATTGAGAAAGATAAAAGCAAAGTCGTGACTAGAATGAGAGGAACACGGGGTTATTTAGCCCCTGAATGGTTGAGCT

>I3_2108

AATCACTGAGAAAGTTGATGTTTATGCTTTTGGAATTGTCCTCTTGGAAATTCTCTGTGGACGAAAGAATTTGGATTGGTCCCAACCTGATGAAGAAGATGTCCATTTGCTAAGTGTATT

>I3_2170

GAAAGAATTTGGATTGGTCCCAACCTGATGAAGAAGATGTCCATTTGCTAAGTGTATTTAGGAGAAAAGCGGAACAAGAGCAGCTCATGGATATGGTTGACAAAAACAACGAAGATATGC

>I3_2406

ACTGTTGAAACCAACTTGAATTATGATTTCACACATGTACCTGAGGTTGGGGCAGGCAACCAAGAGAGGGAAGTCATTGTCAGTTCAATATTTCCTTCAATTTTATCGGGACCAAGGTAA

>I3_248

CCTCTGTGGCTTCTACTGCAGGTACAATGTCACAGAATGCCTTCTTGGTATCCTTTTGTACCACAACAAATACAACGAGCAGAACGGTATGATAGATAAACCCCAGTTAGTTTGGTCTGC

>I3_310

ACAACAAATACAACGAGCAGAACGGTATGATAGATAAACCCCAGTTAGTTTGGTCTGCTAACAGGAACCGTCCAGTGAAATTCAATGCAACCTTGGAACTAGGCCAAGATGGCAACTTGG

>I3_434

GACAGACTCTGATGGCACTCTTGTTTGGTCCACTGATACAATTGGGAAATCTGTTTCTGGCTTAAACTTAACAGAAATGGGAAATCTTGTGCTCTTTGATAAGAGAAAGCGCACAATTTG

>I3_496

TAAACTTAACAGAAATGGGAAATCTTGTGCTCTTTGATAAGAGAAAGCGCACAATTTGGCAGTCTTTTGATCATCCTACAGATTCTTTGCTTCCAGGGCAGAGTTTGGTTTCTGGCCGGA

>I3_558

TCTTTTGATCATCCTACAGATTCTTTGCTTCCAGGGCAGAGTTTGGTTTCTGGCCGGAAGCTTGTAGCAAGCGTTTCAGCAACCAATCATAGTCAAGGTTTGCTTGCTCTTACTGTTCTC

>I3_62

ATCTCGATTTGCTATATCGCAATCTTCATGGGACTACAACGCCACTCTTTCAAATTCTATTGCAGGGTTTGCTGGGCTTTCCTCTTTCTGGATCAACAGGCCGTCTCTTATTATTAATTC

>I3_620

TGTAGCAAGCGTTTCAGCAACCAATCATAGTCAAGGTTTGCTTGCTCTTACTGTTCTCAACGGAAGTTGGGCTGCTTACATGGATACTAATCCGCCTCAATATTACTACACTTCATACTA

>I3_682

GAAGTTGGGCTGCTTACATGGATACTAATCCGCCTCAATATTACTACACTTCATACTATGCTGATAGTTCTTATTACAGTTTTAATGGTCAAACCTTTACTGTTTTACACTATCCTACAA

>I3_744

GATAGTTCTTATTACAGTTTTAATGGTCAAACCTTTACTGTTTTACACTATCCTACAAATTCGACAGCTCAATTCATGAAGATTGGGCCTGATGGACATATAAAGGTATTCCAATGGTCT

>I3_806

GACAGCTCAATTCATGAAGATTGGGCCTGATGGACATATAAAGGTATTCCAATGGTCTGTAGTTGATTGGAATGAAGTATCTGACATTTTGTCGCCAAATGTAGATAACTGTGCGTACCC

>I3_868

TTGATTGGAATGAAGTATCTGACATTTTGTCGCCAAATGTAGATAACTGTGCGTACCCAATGGTATGTGGAAGTTATAGCATTTGTACAAATAACGGGCAATGTACTTGTCCGTCACAGG

>I3_992

CTTCTTCAGGCCATTTTCTGAGAGGAAACCAGATCTTGGATGTTCACAGCTGACTTCCGTTAACTGCAACTCTTCGCAGTATCATAGTTTCATAGAGCTCAAGAATACTACATATTTTTC

>LeEix1_0

ATGGACAAATGGAAATATGCAAGATTAGCACAGTTCCTTTTCACTTTGTCTCTACTGTTCCTAGAGACATCTTTTGGATTAGGTGGTAACAAGACCCTATGTTTAGATAAGGAGAGAGAT

>LeEix1_1037

GAAATCACTTGAGGTTTTGGGGTTGAACGAAAACTCATTGTTTGGTTCAATTGTTAATGCAACAAGATTTTCATCCTTAAAGAAATTATACCTGCAGAAGAATATGCTGAATGGTTCTTT

>LeEix1_1098

ACAAGATTTTCATCCTTAAAGAAATTATACCTGCAGAAGAATATGCTGAATGGTTCTTTTATGGAAAGTGCGGGACAAGTTTCTACCCTTGAGTATCTGGATTTGTCTGAAAACCAAATG

>LeEix1_1159

TGGAAAGTGCGGGACAAGTTTCTACCCTTGAGTATCTGGATTTGTCTGAAAACCAAATGAGAGGGGCATTACCAGATTTAGCATTGTTTCCATCATTGAGAGAGTTGCATCTTGGCTCTA

>LeEix1_122

TCTTCTTGAGTTCAAAAGAGGTCTTACTGATTCTTTTGATCATTTATCTACATGGGGTGATGAAGAAGATAAACAAGAATGCTGCAAATGGAAGGGTATTGAATGTGACAGAAGAACAGG

>LeEix1_1220

AGGGGCATTACCAGATTTAGCATTGTTTCCATCATTGAGAGAGTTGCATCTTGGCTCTAATCAATTTCGAGGGAGGATACCACAAGGTATCGGAAAACTTTCACAGCTTAGAATTTTGGA

>LeEix1_1281

CAATTTCGAGGGAGGATACCACAAGGTATCGGAAAACTTTCACAGCTTAGAATTTTGGATGTCTCGTCCAATAGACTGGAAGGACTACCAGAAAGTATGGGGCAACTATCTAATCTGGAA

>LeEix1_1342

TCTCGTCCAATAGACTGGAAGGACTACCAGAAAGTATGGGGCAACTATCTAATCTGGAAAGTTTTGATGCCTCTTACAATGTCCTGAAGGGAACAATTACTGAGTCCCACCTTTCAAACC

>LeEix1_1403

TTTTGATGCCTCTTACAATGTCCTGAAGGGAACAATTACTGAGTCCCACCTTTCAAACCTCTCCAGTTTAGTGGATTTGGACTTGTCATTCAACTCATTGGCTCTGAAGACGAGCTTCAA

>LeEix1_1464

TCCAGTTTAGTGGATTTGGACTTGTCATTCAACTCATTGGCTCTGAAGACGAGCTTCAATTGGCTTCCTCCTTTTCAGCTACAAGTTATAAGCCTGCCGTCTTGCAATTTGGGACCTTCT

>LeEix1_1525

GGCTTCCTCCTTTTCAGCTACAAGTTATAAGCCTGCCGTCTTGCAATTTGGGACCTTCTTTCCCAAAATGGCTTCAAAATCAGAACAACTATACTGTTCTTGATATCTCTCTTGCGAGTA

>LeEix1_1586

CCCAAAATGGCTTCAAAATCAGAACAACTATACTGTTCTTGATATCTCTCTTGCGAGTATTTCAGACACGCTTCCAAGTTGGTTCTCTAGCTTCCCCCCCGATCTAAAGATTTTGAATCT

>LeEix1_1647

TCAGACACGCTTCCAAGTTGGTTCTCTAGCTTCCCCCCCGATCTAAAGATTTTGAATCTCTCAAACAACCAAATCAGTGGAAGAGTGTCTGACTTAATAGAGAATACATATGGGTACAGG

>LeEix1_1708

CAAACAACCAAATCAGTGGAAGAGTGTCTGACTTAATAGAGAATACATATGGGTACAGGGTTATAGATTTAAGCTATAACAACTTTTCAGGGGCTTTGCCACTAGTCCCTACCAATGTCC

>LeEix1_1769

TATAGATTTAAGCTATAACAACTTTTCAGGGGCTTTGCCACTAGTCCCTACCAATGTCCAAATATTTTACCTGCATAAAAATCAGTTTTTTGGATCCATCTCTTCAATTTGTCGAAGTAG

>LeEix1_183

GAAGAAGATAAACAAGAATGCTGCAAATGGAAGGGTATTGAATGTGACAGAAGAACAGGTCATGTAACTGTTATTGATCTACACAATAAGTTTACTTGTTCTGCTGGTGCCAGTGCTTGT

>LeEix1_1830

ATATTTTACCTGCATAAAAATCAGTTTTTTGGATCCATCTCTTCAATTTGTCGAAGTAGAACATCTCCCACTTCTCTAGACTTGTCACACAATCAATTCTCAGGAGAACTTCCAGATTGT

>LeEix1_1891

CATCTCCCACTTCTCTAGACTTGTCACACAATCAATTCTCAGGAGAACTTCCAGATTGTTGGATGAACATGACAAGTCTAGCTGTTCTTAATCTAGCTTATAACAATTTCTCTGGAGAAA

>LeEix1_1952

GATGAACATGACAAGTCTAGCTGTTCTTAATCTAGCTTATAACAATTTCTCTGGAGAAATTCCACATTCATTGGGTTCCTTGACAAATTTGAAGGCGTTATACATACGCCAAAACAGTTT

>LeEix1_2013

CCACATTCATTGGGTTCCTTGACAAATTTGAAGGCGTTATACATACGCCAAAACAGTTTAAGTGGAATGTTGCCTTCTTTTTCACAATGTCAGGGGTTGCAAATCTTGGATCTTGGAGGG

>LeEix1_2074

GTGGAATGTTGCCTTCTTTTTCACAATGTCAGGGGTTGCAAATCTTGGATCTTGGAGGGAATAAGTTGACAGGAAGTATCCCTGGATGGATAGGGACTGATCTACTCAACTTGCGAATTC

>LeEix1_2135

TAAGTTGACAGGAAGTATCCCTGGATGGATAGGGACTGATCTACTCAACTTGCGAATTCTAAGCCTGCGGTTCAACAGATTGCATGGCAGCATACCATCCATAATCTGTCAGCTTCAATT

>LeEix1_2196

AGCCTGCGGTTCAACAGATTGCATGGCAGCATACCATCCATAATCTGTCAGCTTCAATTTCTTCAGATACTGGACCTTTCAGCGAATGGATTATCAGGGAAAATTCCACATTGCTTCAAC

>LeEix1_2257

TTCAGATACTGGACCTTTCAGCGAATGGATTATCAGGGAAAATTCCACATTGCTTCAACAATTTCACCTTATTGTATCAAGATAATAATTCTGGTGAGCCAATGGAATTTATAGTTCAAG

>LeEix1_2318

TTTCACCTTATTGTATCAAGATAATAATTCTGGTGAGCCAATGGAATTTATAGTTCAAGGTTTCTATGGTAAATTTCCTCGTCGTTACTTGTATATTGGTGACCTATTGGTTCAATGGAA

>LeEix1_2379

TTCTATGGTAAATTTCCTCGTCGTTACTTGTATATTGGTGACCTATTGGTTCAATGGAAAAATCAAGAGTCTGAGTACAAGAATCCTTTATTATATCTGAAGACTATTGATCTTTCAAGT

>LeEix1_244

ATGTAACTGTTATTGATCTACACAATAAGTTTACTTGTTCTGCTGGTGCCAGTGCTTGTTTTGCTCCTAGATTGACAGGTAAACTTAGCCCTTCTCTGCTTGAGTTGGAGTACTTGAATT

>LeEix1_2440

ATCAAGAGTCTGAGTACAAGAATCCTTTATTATATCTGAAGACTATTGATCTTTCAAGTAATGAATTAATTGGAGGTGTTCCTAAAGAAATAGCTGATATGAGAGGATTGAAATCTTTGA

>LeEix1_2501

TGAATTAATTGGAGGTGTTCCTAAAGAAATAGCTGATATGAGAGGATTGAAATCTTTGAACCTCTCAAGAAATGAGCTGAATGGAACTGTCATTGAAGGAATAGGTCAAATGAGGATGTT

>LeEix1_2562

CTCTCAAGAAATGAGCTGAATGGAACTGTCATTGAAGGAATAGGTCAAATGAGGATGTTGGAGTCTCTTGACATGTCAAGAAACCAGCTTTCTGGTGTGATACCGCAAGATCTTGCTAAT

>LeEix1_2623

AGTCTCTTGACATGTCAAGAAACCAGCTTTCTGGTGTGATACCGCAAGATCTTGCTAATTTGACTTTTCTTAGTGTGTTGGACTTGTCGAACAACCAGTTATCAGGGAGAATTCCATCAA

>LeEix1_2684

GACTTTTCTTAGTGTGTTGGACTTGTCGAACAACCAGTTATCAGGGAGAATTCCATCAAGCACTCAACTGCAAAGTTTTGATAGATCATCCTACAGTGATAATGCTCAACTCTGTGGGCC

>LeEix1_2745

ACTCAACTGCAAAGTTTTGATAGATCATCCTACAGTGATAATGCTCAACTCTGTGGGCCTCCTCTTCAGGAGTGTCCTGGATATGCTCCTCCTAGCCCCCTTATTGATCATGGCAGCAAC

>LeEix1_2806

CTCTTCAGGAGTGTCCTGGATATGCTCCTCCTAGCCCCCTTATTGATCATGGCAGCAACAACAATCCACAAGAACATGATGAGGAGGAGGAGTTTCCATCTCTGGAGTTTTATATATCAA

>LeEix1_2867

CAATCCACAAGAACATGATGAGGAGGAGGAGTTTCCATCTCTGGAGTTTTATATATCAATGGTGCTAAGCTTTTTTGTTGCATTTTGGGGAATTTTGGGCTGTTTAATTGTCAATAGTTC

>LeEix1_2928

GTGCTAAGCTTTTTTGTTGCATTTTGGGGAATTTTGGGCTGTTTAATTGTCAATAGTTCTTGGAGGAATGCCTACTTCAAATTCTTGACGGACACGACAAGTTGGCTCGATATGATATCA

>LeEix1_2976

GTCAATAGTTCTTGGAGGAATGCCTACTTCAAATTCTTGACGGACACGACAAGTTGGCTCGATATGATATCAAGAGTCTGGTTTGCAAGACTCAAGAAAAAGCTGAGGAGGGCCCGATGA

>LeEix1_305

TGCTCCTAGATTGACAGGTAAACTTAGCCCTTCTCTGCTTGAGTTGGAGTACTTGAATTACTTGGACCTCAGTGTTAATGAATTTGAAAGAAGTGAAATACCAAGATTCATAGGCTCACT

>LeEix1_366

TTGGACCTCAGTGTTAATGAATTTGAAAGAAGTGAAATACCAAGATTCATAGGCTCACTTAAGAGACTAGAGTACTTGAACCTGTCAGCTTCTTTTTTTTCTGGTGTAATTCCAATACAG

>LeEix1_427

AGAGACTAGAGTACTTGAACCTGTCAGCTTCTTTTTTTTCTGGTGTAATTCCAATACAGTTCCAGAATCTAACTTCTTTAAGGACTCTTGATCTTGGAGAAAATAATCTTATAGTAAAGG

>LeEix1_488

CCAGAATCTAACTTCTTTAAGGACTCTTGATCTTGGAGAAAATAATCTTATAGTAAAGGACCTTAGATGGCTTTCTCATCTGTCTTCTCTAGAGTTTTTGAGTCTGAGTTCTAGCAACTT

>LeEix1_549

CTTAGATGGCTTTCTCATCTGTCTTCTCTAGAGTTTTTGAGTCTGAGTTCTAGCAACTTCCAAGTAAACAATTGGTTTCAAGAGATAACTAAGGTCCCTTCATTGAAAGAACTGGACTTG

>LeEix1_61

TAGAGACATCTTTTGGATTAGGTGGTAACAAGACCCTATGTTTAGATAAGGAGAGAGATGCTCTTCTTGAGTTCAAAAGAGGTCTTACTGATTCTTTTGATCATTTATCTACATGGGGTG

>LeEix1_610

AAGTAAACAATTGGTTTCAAGAGATAACTAAGGTCCCTTCATTGAAAGAACTGGACTTGAGTGGCTGTGGACTCTCTAAGTTGGCTCCATCTCAAGCTGATTTAGCCAATTCTTCTTTTA

>LeEix1_671

TGGCTGTGGACTCTCTAAGTTGGCTCCATCTCAAGCTGATTTAGCCAATTCTTCTTTTATATCTCTTTCTGTTCTTCATTTATGTTGTAATGAGTTTTCTTCTTCATCTGAATATAGCTG

>LeEix1_732

TCTCTTTCTGTTCTTCATTTATGTTGTAATGAGTTTTCTTCTTCATCTGAATATAGCTGGGTATTCAATTTGACCACAAGCCTAACTAGCATTGACCTCCTTTATAATCAACTCAGCGGT

>LeEix1_793

TATTCAATTTGACCACAAGCCTAACTAGCATTGACCTCCTTTATAATCAACTCAGCGGTCAAATTGATGATCGCTTTGGGACCTTGATGTATCTTGAACATCTTGATCTTGCTAATAACT

>LeEix1_854

AATTGATGATCGCTTTGGGACCTTGATGTATCTTGAACATCTTGATCTTGCTAATAACTTAAAGATTGAAGGTGGGGTTCCCAGTTCTTTTGGGAATTTGACACGTTTACGTCATCTGGA

>LeEix1_915

AAGATTGAAGGTGGGGTTCCCAGTTCTTTTGGGAATTTGACACGTTTACGTCATCTGGACATGTCTAATACTCAGACAGTCCAATGGCTTCCTGAGTTGTTTCTCAGGTTATCAGGCAGT

>LeEix1_976

TGTCTAATACTCAGACAGTCCAATGGCTTCCTGAGTTGTTTCTCAGGTTATCAGGCAGTAGGAAATCACTTGAGGTTTTGGGGTTGAACGAAAACTCATTGTTTGGTTCAATTGTTAATG

>LeEix2_0

ATGGGCAAAAGAACTAATCCAAGACATTTCCTTGTTACTTGGTCTTTACTGCTCCTAGAGACAGCTTTTGGATTAACTTCAAGAGAAGTTAACAAGACCCTTTGTATAGAAAAGGAGAGA

>LeEix2_1054

ACAACTCGTTGTTTGGTTCAATTGTTAATGTGCCAAGATTTTCATCCTTGAAGAAATTATACCTGCAGAAGAATATGCTGAATGGTTTTTTCATGGAAAGAGTGGGACAAGTTTCGAGCC

>LeEix2_1116

CTGCAGAAGAATATGCTGAATGGTTTTTTCATGGAAAGAGTGGGACAAGTTTCGAGCCTCGAGTATCTAGACTTGTCTGATAACCAAATGAGAGGGCCATTACCAGATTTAGCACTTTTT

>LeEix2_1178

GTATCTAGACTTGTCTGATAACCAAATGAGAGGGCCATTACCAGATTTAGCACTTTTTCCATCATTAAGAGAGTTGCATCTTGGCTCTAATCAATTTCAAGGGAGGATACCACAAGGTAT

>LeEix2_124

CCCTTCTTGAGTTTAAAAGAGGCCTTAACGACGATTTTGGTCGTTTATCTACCTGGGGTGATGAAGAAGAATGCTGCAATTGGAAGGGTATTGAATGTGACAAAAGAACAGGTCATGTTA

>LeEix2_1240

CATTAAGAGAGTTGCATCTTGGCTCTAATCAATTTCAAGGGAGGATACCACAAGGTATTGGAAAACTTTCACAGCTTAGAATTTTTGACGTCTCGTCCAATAGATTAGAGGGTTTACCAG

>LeEix2_1302

AAACTTTCACAGCTTAGAATTTTTGACGTCTCGTCCAATAGATTAGAGGGTTTACCAGAAAGTATGGGGCAACTATCAAACCTGGAAAGGTTTGATGCTTCTTATAATGTCCTGAAGGGT

>LeEix2_1364

TATGGGGCAACTATCAAACCTGGAAAGGTTTGATGCTTCTTATAATGTCCTGAAGGGTACAATCACAGAGTCCCACTTTTCAAACCTCTCCAGTTTAGTAGACTTAGACCTATCCTTCAA

>LeEix2_1426

TCACAGAGTCCCACTTTTCAAACCTCTCCAGTTTAGTAGACTTAGACCTATCCTTCAACTTGTTGTCTTTGAACACGAGATTCGATTGGGTTCCTCCTTTTCAACTACAATTTATAAGGC

>LeEix2_1488

TTGTCTTTGAACACGAGATTCGATTGGGTTCCTCCTTTTCAACTACAATTTATAAGGCTTCCATCTTGCAATATGGGACCTTCTTTTCCGAAATGGCTACAAACTCAGAATAACTACACT

>LeEix2_1550

ATCTTGCAATATGGGACCTTCTTTTCCGAAATGGCTACAAACTCAGAATAACTACACTCTTCTTGATATTTCTCTTGCGAATATATCAGACATGCTACCAAGTTGGTTCTCCAATCTTCC

>LeEix2_1612

TTGATATTTCTCTTGCGAATATATCAGACATGCTACCAAGTTGGTTCTCCAATCTTCCTCCCGAGCTCAAGATTCTGAATCTCTCTAACAACCACATCAGTGGCAGAGTTTCGGAGTTCA

>LeEix2_1674

GAGCTCAAGATTCTGAATCTCTCTAACAACCACATCAGTGGCAGAGTTTCGGAGTTCATAGTGAGTAAACAAGATTACATGATTATAGATTTAAGTTCTAACAACTTTTCAGGACATTTG

>LeEix2_1736

GAGTAAACAAGATTACATGATTATAGATTTAAGTTCTAACAACTTTTCAGGACATTTGCCACTAGTTCCTGCCAATATACAGATCTTTTACCTTCATAAAAATCACTTCTCTGGATCCAT

>LeEix2_1798

TAGTTCCTGCCAATATACAGATCTTTTACCTTCATAAAAATCACTTCTCTGGATCCATTTCTTCCATTTGTAGAAATACAATAGGAGCTGCCACTTCCATCGACTTGTCACGCAACCAAT

>LeEix2_186

GAAGAAGAATGCTGCAATTGGAAGGGTATTGAATGTGACAAAAGAACAGGTCATGTTATTGTTCTTGATCTCCACAGTGAGGTTACTTGTCCAGGCCATGCTTGTTTTGCTCCAATATTG

>LeEix2_1860

TCCATTTGTAGAAATACAATAGGAGCTGCCACTTCCATCGACTTGTCACGCAACCAATTTTCAGGAGAAGTTCCTGATTGTTGGATGAATATGAGTAATCTAGCTGTTCTAAATCTAGCC

>LeEix2_1922

AGGAGAAGTTCCTGATTGTTGGATGAATATGAGTAATCTAGCTGTTCTAAATCTAGCCTACAACAATTTCTCCGGAAAAGTTCCACAATCATTAGGCTCCTTGACAAATTTGGAGGCGTT

>LeEix2_1984

ACAATTTCTCCGGAAAAGTTCCACAATCATTAGGCTCCTTGACAAATTTGGAGGCGTTATACATACGTCAGAACAGCTTTAGAGGAATGTTGCCTTCTTTTTCACAATGTCAGCTGCTGC

>LeEix2_2046

ATACGTCAGAACAGCTTTAGAGGAATGTTGCCTTCTTTTTCACAATGTCAGCTGCTGCAAATCTTGGATATTGGAGGGAATAAGTTGACTGGAAGAATCCCAGCATGGATAGGGACTGAC

>LeEix2_2108

CTTGGATATTGGAGGGAATAAGTTGACTGGAAGAATCCCAGCATGGATAGGGACTGACCTACTCCAATTGCGCATTCTAAGCCTACGTTCCAACAAATTCGATGGCAGCATTCCATCACT

>LeEix2_2170

TCCAATTGCGCATTCTAAGCCTACGTTCCAACAAATTCGATGGCAGCATTCCATCACTTATCTGCCAGCTTCAATTTCTTCAGATACTGGACCTTTCAGAAAATGGATTATCTGGGAAAA

>LeEix2_2232

TGCCAGCTTCAATTTCTTCAGATACTGGACCTTTCAGAAAATGGATTATCTGGGAAAATTCCACAGTGCCTCAACAACTTTACCATATTGCGTCAAGAAAATGGCTCTGGTGAGTCAATG

>LeEix2_2294

ACAGTGCCTCAACAACTTTACCATATTGCGTCAAGAAAATGGCTCTGGTGAGTCAATGGATTTTAAAGTCCGTTATGACTATATTCCAGGATCTTACTTGTACATAGGTGATTTACTGAT

>LeEix2_2356

TTAAAGTCCGTTATGACTATATTCCAGGATCTTACTTGTACATAGGTGATTTACTGATTCAATGGAAAAACCAGGAGTCCGAGTACAAGAATGCTTTACTATATCTGAAGATCATTGATC

>LeEix2_2418

TGGAAAAACCAGGAGTCCGAGTACAAGAATGCTTTACTATATCTGAAGATCATTGATCTTTCAAGTAATAAATTAGTTGGAGGTATCCCTAAAGAGATAGCTGAAATGAGAGGATTGAGA

>LeEix2_248

TCTTGATCTCCACAGTGAGGTTACTTGTCCAGGCCATGCTTGTTTTGCTCCAATATTGACAGGTAAAGTTAGTCCTTCTCTACTTGAGTTGGAGTATTTGAATTTCTTGGACCTCAGTGT

>LeEix2_2480

AAGTAATAAATTAGTTGGAGGTATCCCTAAAGAGATAGCTGAAATGAGAGGATTGAGATCATTGAACCTCTCGAGAAATGATCTTAATGGAACTGTCGTTGAAGGAATAGGTCAAATGAA

>LeEix2_2542

TGAACCTCTCGAGAAATGATCTTAATGGAACTGTCGTTGAAGGAATAGGTCAAATGAAGTTGTTGGAGTCCCTCGACTTGTCAAGAAACCAACTCTCTGGCATGATCCCTCAGGGCCTTT

>LeEix2_2604

TTGGAGTCCCTCGACTTGTCAAGAAACCAACTCTCTGGCATGATCCCTCAGGGCCTTTCTAACTTGACTTTTCTTAGTGTGTTGGACTTATCGAACAACCACTTATCAGGAAGAATTCCA

>LeEix2_2666

CTTGACTTTTCTTAGTGTGTTGGACTTATCGAACAACCACTTATCAGGAAGAATTCCATCAAGCACTCAACTGCAGAGTTTCGATAGATCATCCTATAGTGGCAATGCTCAACTCTGCGG

>LeEix2_2728

GCACTCAACTGCAGAGTTTCGATAGATCATCCTATAGTGGCAATGCTCAACTCTGCGGTCCTCCTCTTGAAGAGTGTCCTGGATATGCTCCCCCTATCGATCGTGGAAGCAACACCAATC

>LeEix2_2790

CCTCTTGAAGAGTGTCCTGGATATGCTCCCCCTATCGATCGTGGAAGCAACACCAATCCACAAGAACATGATGATGATGATGAGTTCTCATCTCTGGAGTTTTATGTATCAATGGTGCTA

>LeEix2_2852

AGAACATGATGATGATGATGAGTTCTCATCTCTGGAGTTTTATGTATCAATGGTGCTAGGTTTCTTCGTCACGTTCTGGGGAATTTTAGGCTGTTTGATTGTCAACCGTTCGTGGAGGAA

>LeEix2_2914

TCTTCGTCACGTTCTGGGGAATTTTAGGCTGTTTGATTGTCAACCGTTCGTGGAGGAATGCCTACTTCACATTCTTAACAGACATGAAGAGTTGGCTCCATATGACATCAAGAGTCTGCT

>LeEix2_2946

TTGATTGTCAACCGTTCGTGGAGGAATGCCTACTTCACATTCTTAACAGACATGAAGAGTTGGCTCCATATGACATCAAGAGTCTGCTTTGCAAGACTGAAGGGAAAGCTAAGGAACTGA

>LeEix2_310

GTAAAGTTAGTCCTTCTCTACTTGAGTTGGAGTATTTGAATTTCTTGGACCTCAGTGTTAATGGATTTGAAAATAGTGAGATACCAAGATTCATAGGCTCCCTTAAGAGACTGGAGTACT

>LeEix2_372

GGATTTGAAAATAGTGAGATACCAAGATTCATAGGCTCCCTTAAGAGACTGGAGTACTTAAACCTTTCATCTTCTGATTTTTCTGGTGAAATTCCAGCACAGTTCCAGAATCTAACTTCT

>LeEix2_434

CCTTTCATCTTCTGATTTTTCTGGTGAAATTCCAGCACAGTTCCAGAATCTAACTTCTTTGAGGATTCTTGATCTCGGAAACAATAATCTTATAGTAAAGGACCTTGTGTGGCTTTCTCA

>LeEix2_496

GGATTCTTGATCTCGGAAACAATAATCTTATAGTAAAGGACCTTGTGTGGCTTTCTCATCTCTCCTCTCTAGAATTCTTGCGCCTTGGTGGTAACGATTTCCAAGCAAGAAACTGGTTTC

>LeEix2_558

TCCTCTCTAGAATTCTTGCGCCTTGGTGGTAACGATTTCCAAGCAAGAAACTGGTTTCGAGAGATAACTAAGGTCCCTTCATTGAAAGAATTGGACTTGAGTGTTTGTGGACTCTCTAAA

>LeEix2_62

AGCTTTTGGATTAACTTCAAGAGAAGTTAACAAGACCCTTTGTATAGAAAAGGAGAGAGGTGCCCTTCTTGAGTTTAAAAGAGGCCTTAACGACGATTTTGGTCGTTTATCTACCTGGGG

>LeEix2_620

GATAACTAAGGTCCCTTCATTGAAAGAATTGGACTTGAGTGTTTGTGGACTCTCTAAATTCGTTCCGTCTCCAGCTGATGTAGCTAATTCATCTTTGATCTCTCTTTCTGTTCTTCATTT

>LeEix2_682

TTCCGTCTCCAGCTGATGTAGCTAATTCATCTTTGATCTCTCTTTCTGTTCTTCATTTATGTTGTAATGAGTTTTCTACTTCATCTGAATATAGCTGGTTATTCAATTTTAGCACAAGCC

>LeEix2_744

TGTAATGAGTTTTCTACTTCATCTGAATATAGCTGGTTATTCAATTTTAGCACAAGCCTAACTAGCATAGACCTTTCTCATAATCAACTCAGTCGTCAAATTGATGATCGCTTTGGGAGC

>LeEix2_806

TAGCATAGACCTTTCTCATAATCAACTCAGTCGTCAAATTGATGATCGCTTTGGGAGCTTGATGTATCTTGAACATCTTAATCTTGCTAATAATTTTGGGGCTGAAGGTGGGGTTCCAAG

>LeEix2_868

TGTATCTTGAACATCTTAATCTTGCTAATAATTTTGGGGCTGAAGGTGGGGTTCCAAGTTCTTTTGGGAATTTGACACGTCTACATTATCTGGACATGTCTAACACTCAGACATACCAAT

>LeEix2_930

TTTGGGAATTTGACACGTCTACATTATCTGGACATGTCTAACACTCAGACATACCAATGGCTTCCTGAGTTGTTTCTCAGGTTATCAGGTAGTAGGAAATCACTTGAGGTTTTGGGATTG

>LeEix2_992

TCCTGAGTTGTTTCTCAGGTTATCAGGTAGTAGGAAATCACTTGAGGTTTTGGGATTGAACGACAACTCGTTGTTTGGTTCAATTGTTAATGTGCCAAGATTTTCATCCTTGAAGAAATT

>M6-PGSC0003DMG400000179-(rev)M6G-LecRK_0

CAAATATTCTCTGAGCTTTTCAATGTTTCCTTATAAACCAAAATCTTTTTTTCTTTAGCTAAAGCAAAATTCATCACCATGATTCCACAGAAAGCAATTCTCATTTTACTTTTCATCACA

>M6-PGSC0003DMG400000179-(rev)M6G-LecRK_1037

TATGTGTAGTACTTATGGTCTTTGTGGGAATTTGGGGTTCTGTAATAGTAAGTCATTGAATCCATGTAAGTGTTTACCAGGGTTCACCCCTTTGGATGGTGATTCATGGGATGCAGGGGA

>M6-PGSC0003DMG400000179-(rev)M6G-LecRK_1098

CCATGTAAGTGTTTACCAGGGTTCACCCCTTTGGATGGTGATTCATGGGATGCAGGGGATTTTTCGGGTGGCTGTCGCCATGAGAGTAATGAAGTTTGTGGTGAGAATGATAGGTTTGAA

>M6-PGSC0003DMG400000179-(rev)M6G-LecRK_1159

TTTCGGGTGGCTGTCGCCATGAGAGTAATGAAGTTTGTGGTGAGAATGATAGGTTTGAAGAGGTTGGAATGGCTAAATATGATGGAGCAAGGGTAGTTTCAGTTTCTGGAACTAGAAATG

>M6-PGSC0003DMG400000179-(rev)M6G-LecRK_122

TTCTATGGCATTTGAGATCAGTTCAGAAACAGTAAAGTCAAGAATGCAAAATGTAAACATGACCCATGTAGCTGAACTCACTAAAGAGTCTGTTTCAGCTGAAAGAAAGTTGGGATCTTT

>M6-PGSC0003DMG400000179-(rev)M6G-LecRK_1220

GGTTGGAATGGCTAAATATGATGGAGCAAGGGTAGTTTCAGTTTCTGGAACTAGAAATGAATGTGAGAGAGAATGTTTAGGTAATTGTTCATGTATTGGTTTGTATCGCAATGAGAAGAC

>M6-PGSC0003DMG400000179-(rev)M6G-LecRK_1281

TGTGAGAGAGAATGTTTAGGTAATTGTTCATGTATTGGTTTGTATCGCAATGAGAAGACTAATTTGTGCAAGAGTTTATATGGGTCATTGCTCAATCTTAGGAATCTTACTTCTGATGGC

>M6-PGSC0003DMG400000179-(rev)M6G-LecRK_1342

ATTTGTGCAAGAGTTTATATGGGTCATTGCTCAATCTTAGGAATCTTACTTCTGATGGCACTGTTGAGGATAAGCTATATGTTAGAGTTCAAGGAGGAGGGAATACTCGAAAGAATCAGA

>M6-PGSC0003DMG400000179-(rev)M6G-LecRK_1403

TGTTGAGGATAAGCTATATGTTAGAGTTCAAGGAGGAGGGAATACTCGAAAGAATCAGATTCAAGTCAGATTACTTGTGATCGAAATGATCTGTGGATTTGTGGTAATTCTGTTAGTTGG

>M6-PGSC0003DMG400000179-(rev)M6G-LecRK_1464

CAAGTCAGATTACTTGTGATCGAAATGATCTGTGGATTTGTGGTAATTCTGTTAGTTGGAATTGGGGCTTTCTTGGTAGTTAGGAGGAGAAGAGTAAGGAGGAAGAAAAAAGATGAGGAA

>M6-PGSC0003DMG400000179-(rev)M6G-LecRK_1525

TTGGGGCTTTCTTGGTAGTTAGGAGGAGAAGAGTAAGGAGGAAGAAAAAAGATGAGGAAGATGTATTTCCTATAATGAATTTGAAGGTATTTTCATACAAAGAGCTTAATGCTGCAACAA

>M6-PGSC0003DMG400000179-(rev)M6G-LecRK_1586

TGTATTTCCTATAATGAATTTGAAGGTATTTTCATACAAAGAGCTTAATGCTGCAACAAAGGGATTCTCGGAGAAGCTTGGTCATGGTGGTTTTGGAACCGTGTTCCTAGGGGAGCTATC

>M6-PGSC0003DMG400000179-(rev)M6G-LecRK_1647

GGATTCTCGGAGAAGCTTGGTCATGGTGGTTTTGGAACCGTGTTCCTAGGGGAGCTATCGGATTCTTCACTCGTAGCTGTGAAGAGACTTGAAAGACCTGGTGGTGGTGAAAAGGAATTT

>M6-PGSC0003DMG400000179-(rev)M6G-LecRK_1708

ATTCTTCACTCGTAGCTGTGAAGAGACTTGAAAGACCTGGTGGTGGTGAAAAGGAATTTCGAGCTGAGGTATGCACAATAGGGAACATTCAGCATGTTAATTTAGTGAGATTGAGGGGAT

>M6-PGSC0003DMG400000179-(rev)M6G-LecRK_1769

AGCTGAGGTATGCACAATAGGGAACATTCAGCATGTTAATTTAGTGAGATTGAGGGGATTTTGCACTGAAAATTCTCATAGGCTTCTGGTTTATGAGTACATGTCAAAGGGTCCTCTTAG

>M6-PGSC0003DMG400000179-(rev)M6G-LecRK_183

ACCCATGTAGCTGAACTCACTAAAGAGTCTGTTTCAGCTGAAAGAAAGTTGGGATCTTTAGAAGGGAGCATAAAGTTTGAATCTTTGAACAAATCTAGAGTTTTACTAAGTGGGAACTTC

>M6-PGSC0003DMG400000179-(rev)M6G-LecRK_1830

TGCACTGAAAATTCTCATAGGCTTCTGGTTTATGAGTACATGTCAAAGGGTCCTCTTAGCGCATATTTAAGGCAAGATGGTCAGAATCTAAGTTGGGATGTTAGATTCCGAATTGCTGTA

>M6-PGSC0003DMG400000179-(rev)M6G-LecRK_1891

CATATTTAAGGCAAGATGGTCAGAATCTAAGTTGGGATGTTAGATTCCGAATTGCTGTAGGGACAGCAAGAGGTATTGCTTATTTACACGAAGAGTGTCGCAATTGTATAATACATTGCG

>M6-PGSC0003DMG400000179-(rev)M6G-LecRK_1952

GACAGCAAGAGGTATTGCTTATTTACACGAAGAGTGTCGCAATTGTATAATACATTGCGACATTAAACCAGAGAATATTCTGCTACATGAAGATTTTTCAGCAAAGGTTTCTGATTTTGG

>M6-PGSC0003DMG400000179-(rev)M6G-LecRK_2013

ATTAAACCAGAGAATATTCTGCTACATGAAGATTTTTCAGCAAAGGTTTCTGATTTTGGACTTGCAAAGCTCCTTGGTAGGGACTTTAGTAGAGTGTTGGCAACCATGAGGGGTACCTGG

>M6-PGSC0003DMG400000179-(rev)M6G-LecRK_2074

TTGCAAAGCTCCTTGGTAGGGACTTTAGTAGAGTGTTGGCAACCATGAGGGGTACCTGGGGCTATGTTGCACCCGAGTGGATCTCTGGATTAGCGATCACAACGAAAGCTGATGTATATA

>M6-PGSC0003DMG400000179-(rev)M6G-LecRK_2135

CTATGTTGCACCCGAGTGGATCTCTGGATTAGCGATCACAACGAAAGCTGATGTATATAGCTATGGCATGACATTATTGGAACTGATAGGTGGTCGACGCAATGTAGAGTCACCGCCTTC

>M6-PGSC0003DMG400000179-(rev)M6G-LecRK_2196

TATGGCATGACATTATTGGAACTGATAGGTGGTCGACGCAATGTAGAGTCACCGCCTTCAGCTAAAGGTGAAGAGGGCGGAACAGAGGAGAAGTGGTTTTTCCCACCGTGGGCAGCACGT

>M6-PGSC0003DMG400000179-(rev)M6G-LecRK_2257

CTAAAGGTGAAGAGGGCGGAACAGAGGAGAAGTGGTTTTTCCCACCGTGGGCAGCACGTCAAATAGTCGAAGGAAACATTGCTGCAGTTATGGATGAAAGACTACATGGAACGTACAACC

>M6-PGSC0003DMG400000179-(rev)M6G-LecRK_2318

AATAGTCGAAGGAAACATTGCTGCAGTTATGGATGAAAGACTACATGGAACGTACAACCTCACGGAGGCAGAACGTGTGGGATTGATAGCTATTTGGTGCATTCAGGACGATGAATCAAC

>M6-PGSC0003DMG400000179-(rev)M6G-LecRK_2379

ACGGAGGCAGAACGTGTGGGATTGATAGCTATTTGGTGCATTCAGGACGATGAATCAACGAGGCCTTCCATGGGAATGGTGGTGAAAATGTTGGAAGGAGTTGTCGAGGTGACAATGCCT

>M6-PGSC0003DMG400000179-(rev)M6G-LecRK_244

AAGGGAGCATAAAGTTTGAATCTTTGAACAAATCTAGAGTTTTACTAAGTGGGAACTTCACTGTTTTGAGTGAAAACAAGACATTTGAAATGGGGTTTTTCAAGACTAATGATGAAAGTA

>M6-PGSC0003DMG400000179-(rev)M6G-LecRK_2440

GGCCTTCCATGGGAATGGTGGTGAAAATGTTGGAAGGAGTTGTCGAGGTGACAATGCCTCAACCTCCGAAGCTGCTCCAAGCATTAGTATCTGGTGAATCATTCCATGGGGTAGGAATTG

>M6-PGSC0003DMG400000179-(rev)M6G-LecRK_2501

ACCTCCGAAGCTGCTCCAAGCATTAGTATCTGGTGAATCATTCCATGGGGTAGGAATTGATTCAAGTAACGGGACATCACGAGACGGAGGTTTCAGCTCGGGTTATAATCCTCAACGTTC

>M6-PGSC0003DMG400000179-(rev)M6G-LecRK_2562

TCAAGTAACGGGACATCACGAGACGGAGGTTTCAGCTCGGGTTATAATCCTCAACGTTCAATATATTCAAAAGACTCTCAAGCTTCAAGTTAGGTCTGCTTCTCAACTAGTATGCAAGTT

>M6-PGSC0003DMG400000179-(rev)M6G-LecRK_2623

TATATTCAAAAGACTCTCAAGCTTCAAGTTAGGTCTGCTTCTCAACTAGTATGCAAGTTGATTTTTCCCAGCTTCACTGAATGTAGAGTTCTTTTTTATTGTCTTCAAAAATCCCACTGA

>M6-PGSC0003DMG400000179-(rev)M6G-LecRK_2684

TTTTTCCCAGCTTCACTGAATGTAGAGTTCTTTTTTATTGTCTTCAAAAATCCCACTGAGAATTAGTAGCAAAATGCTCATCTTGGTATTTTGTAAAATGTATCATGAATATAGAAAATC

>M6-PGSC0003DMG400000179-(rev)M6G-LecRK_2745

ATTAGTAGCAAAATGCTCATCTTGGTATTTTGTAAAATGTATCATGAATATAGAAAATCTTGTTTACTTTGCTTAAAAATTGTCACCTTTTTTTTTTTTTGTAAAGTCCAAAGAACCAAA

>M6-PGSC0003DMG400000179-(rev)M6G-LecRK_2806

GTTTACTTTGCTTAAAAATTGTCACCTTTTTTTTTTTTTGTAAAGTCCAAAGAACCAAAGCAAAACAAAACATACATTTCACTTTGGATCAGAGGGTTGGGGCGGAGCTAGTACATTACA

>M6-PGSC0003DMG400000179-(rev)M6G-LecRK_305

TGTTTTGAGTGAAAACAAGACATTTGAAATGGGGTTTTTCAAGACTAATGATGAAAGTAAATGGTATTTAGGGATTTGGTTTGCTTCAATTCCTACCCCTACTTTTGTTTGGGTTGCTAA

>M6-PGSC0003DMG400000179-(rev)M6G-LecRK_3050

ATTATAGGAGAGGGCTTCCTAACCTTAGAATGAGACAATAGTTGAAAATATCAAAAGTTATAATAGCAGAATTATTTAATGGATATATCTTTATTGGTAGATATTTCGTATAGTGAACTT

>M6-PGSC0003DMG400000179-(rev)M6G-LecRK_3111

AATAGCAGAATTATTTAATGGATATATCTTTATTGGTAGATATTTCGTATAGTGAACTTAAAATGAGATATAGGTGTATAATATAAAGCATTAAAACATGTGTTTGGATTATACCAACCA

>M6-PGSC0003DMG400000179-(rev)M6G-LecRK_3172

AATGAGATATAGGTGTATAATATAAAGCATTAAAACATGTGTTTGGATTATACCAACCAAACATTATGCAAAATTTTATATTGAGATTATTTTTGTTAATATGGTAAACCAAATGACTAA

>M6-PGSC0003DMG400000179-(rev)M6G-LecRK_3233

CATTATGCAAAATTTTATATTGAGATTATTTTTGTTAATATGGTAAACCAAATGACTAATAAGAATATACCCTCTACCTAATTTAATGGGAGAGTTAATTTTATATTTTATAAATGAGTG

>M6-PGSC0003DMG400000179-(rev)M6G-LecRK_3294

AGAATATACCCTCTACCTAATTTAATGGGAGAGTTAATTTTATATTTTATAAATGAGTGTAACGTTTGTACCTGCCCAATTATAACCACACTTTACTTTCTAAGTTTAATTTACATAACT

>M6-PGSC0003DMG400000179-(rev)M6G-LecRK_3355

ACGTTTGTACCTGCCCAATTATAACCACACTTTACTTTCTAAGTTTAATTTACATAACTGTCTTGACTACAACAATAACAACTAACCACTTGTCAATAGTTTCCTCTTTTGTAGCTAAAT

>M6-PGSC0003DMG400000179-(rev)M6G-LecRK_3416

CTTGACTACAACAATAACAACTAACCACTTGTCAATAGTTTCCTCTTTTGTAGCTAAATAATATAACAGAAATTGAAACAAGAAAGACCATATACTCTCACACAAACAAGCAAACAGAAA

>M6-PGSC0003DMG400000179-(rev)M6G-LecRK_3477

TATAACAGAAATTGAAACAAGAAAGACCATATACTCTCACACAAACAAGCAAACAGAAAAAGGAGAAATGTTTTCCTAACTTGTCAACTTTAGTAACAAAATTTTCCCAAAATTAATTTT

>M6-PGSC0003DMG400000179-(rev)M6G-LecRK_3538

GGAGAAATGTTTTCCTAACTTGTCAACTTTAGTAACAAAATTTTCCCAAAATTAATTTTTTCAAAAGAGTAAAGATTGAGGGGGTTAAAGAAAATGGTCAAAGGAGAGCAGAGGTAGACA

>M6-PGSC0003DMG400000179-(rev)M6G-LecRK_3599

CAAAAGAGTAAAGATTGAGGGGGTTAAAGAAAATGGTCAAAGGAGAGCAGAGGTAGACAACTCTGCACTCTCACCAAAGAAGTTTGTTACCTTGGAAATGGAGACACCTGGCAGTTCAGG

>M6-PGSC0003DMG400000179-(rev)M6G-LecRK_366

TGGTATTTAGGGATTTGGTTTGCTTCAATTCCTACCCCTACTTTTGTTTGGGTTGCTAATCGTGAGAAACCCATCAAGAATCCATTTTTAGCTACTTTGGAGTTTAGTGAGGATGGGAGA

>M6-PGSC0003DMG400000179-(rev)M6G-LecRK_3660

TCTGCACTCTCACCAAAGAAGTTTGTTACCTTGGAAATGGAGACACCTGGCAGTTCAGGTGGGGAGGTGCCGATATATGGGGGCTCCTCGTCTATGGAGCCGACTGAGTTACACTCGGCT

>M6-PGSC0003DMG400000179-(rev)M6G-LecRK_3721

GGGAGGTGCCGATATATGGGGGCTCCTCGTCTATGGAGCCGACTGAGTTACACTCGGCTCCATCTACCATGCTGGCGGATTATCCGCCAGTACAAGGCTTTCAAGATTTGAAGAACTTGG

>M6-PGSC0003DMG400000179-(rev)M6G-LecRK_3782

ATCTACCATGCTGGCGGATTATCCGCCAGTACAAGGCTTTCAAGATTTGAAGAACTTGGTTAGCATGGAATCTGGCAAGTTATGGACTATTGCAGGTCCCATAGCCTTTAATATTCTGTG

>M6-PGSC0003DMG400000179-(rev)M6G-LecRK_3843

AGCATGGAATCTGGCAAGTTATGGACTATTGCAGGTCCCATAGCCTTTAATATTCTGTGTAATTATGGGATTAATTCTTTCACAAGTATATTTGTTGGACATATTGGTGATGTAGAACTC

>M6-PGSC0003DMG400000179-(rev)M6G-LecRK_3904

ATTATGGGATTAATTCTTTCACAAGTATATTTGTTGGACATATTGGTGATGTAGAACTCTCTGCCGTTGCCATATCTTTATCGGTGATAGCAAATTTCTCATTCGGATTCTTGGTTAGGT

>M6-PGSC0003DMG400000179-(rev)M6G-LecRK_3965

TGCCGTTGCCATATCTTTATCGGTGATAGCAAATTTCTCATTCGGATTCTTGGTTAGGTTCTTTTTTTGTTCTTACTAATTCTCAAAGTTCTGTTCTTTTTTATATTATGTTATATGGGC

>M6-PGSC0003DMG400000179-(rev)M6G-LecRK_4026

TTTTTTTGTTCTTACTAATTCTCAAAGTTCTGTTCTTTTTTATATTATGTTATATGGGCATTGAGTATTCGTATAGCCGACCCCAACTTGCTTGAAATAGCGGGATAGTTGTTGTTATTG

>M6-PGSC0003DMG400000179-(rev)M6G-LecRK_4087

TGAGTATTCGTATAGCCGACCCCAACTTGCTTGAAATAGCGGGATAGTTGTTGTTATTGTTGTGATTGTTTGTAAAATGTAATACATCGAAGATATGTTGGATTTTGTCAGACTCTTTGA

>M6-PGSC0003DMG400000179-(rev)M6G-LecRK_4148

GTGATTGTTTGTAAAATGTAATACATCGAAGATATGTTGGATTTTGTCAGACTCTTTGATGAGACTTATGACTATAGATCACAAATAGTTTATTATGAAACAGAACTAGGATTTTGATTG

>M6-PGSC0003DMG400000179-(rev)M6G-LecRK_4209

AGACTTATGACTATAGATCACAAATAGTTTATTATGAAACAGAACTAGGATTTTGATTGTCAAATGATATAAAGTCTCAAAAAGGTTAGAATAAGCATTCTTTTAGGTTATTGGTTATGT

>M6-PGSC0003DMG400000179-(rev)M6G-LecRK_427

GTGAGAAACCCATCAAGAATCCATTTTTAGCTACTTTGGAGTTTAGTGAGGATGGGAGATTAGTTTTGAAGGAAGATTCAAGAACCATTGTATGGGAAACAACCAATTTGGAGAAAGCTA

>M6-PGSC0003DMG400000179-(rev)M6G-LecRK_4270

AAATGATATAAAGTCTCAAAAAGGTTAGAATAAGCATTCTTTTAGGTTATTGGTTATGTTAGTTTTCTAATTCTTTATCTTATTTTTGATGGATTATCAGTTAGGCATGGGTAGTGCACT

>M6-PGSC0003DMG400000179-(rev)M6G-LecRK_4331

GTTTTCTAATTCTTTATCTTATTTTTGATGGATTATCAGTTAGGCATGGGTAGTGCACTTGAGACACTTTGTGGACAAGCATTTGGTGCAGGCCAAATAGAATTGTTAGGAGTTTACTTG

>M6-PGSC0003DMG400000179-(rev)M6G-LecRK_4392

AGACACTTTGTGGACAAGCATTTGGTGCAGGCCAAATAGAATTGTTAGGAGTTTACTTGCAGAGGTCATGGATAATACTTGTTGCTTCTTGCTTCTGCATAATGCCACTCTACATCTTCT

>M6-PGSC0003DMG400000179-(rev)M6G-LecRK_4453

GAGGTCATGGATAATACTTGTTGCTTCTTGCTTCTGCATAATGCCACTCTACATCTTCTCCACGCCGATACTGAAGTTATTAGGCCAAAGAGATGACATAGCAGAGTTAGCTGGAAAATT

>M6-PGSC0003DMG400000179-(rev)M6G-LecRK_4514

ACGCCGATACTGAAGTTATTAGGCCAAAGAGATGACATAGCAGAGTTAGCTGGAAAATTTTCGATACAAATCATACCTCAAATGTTCTCCCTTGCTATCAACTTCCCAACCCAAAAGTTC

>M6-PGSC0003DMG400000179-(rev)M6G-LecRK_4575

CGATACAAATCATACCTCAAATGTTCTCCCTTGCTATCAACTTCCCAACCCAAAAGTTCTTGCAAGCACAAAGCAATGTCGCGATCCTGGCATGGGTTGGGTTCATGGCTTTAGCCATGC

>M6-PGSC0003DMG400000179-(rev)M6G-LecRK_4636

GCAAGCACAAAGCAATGTCGCGATCCTGGCATGGGTTGGGTTCATGGCTTTAGCCATGCATATCGGGGTGCTCTTTCTTTTTATCAAAGTTTTCCGATGGGGCGTTACTGGTGCTGCTGC

>M6-PGSC0003DMG400000179-(rev)M6G-LecRK_4697

ATCGGGGTGCTCTTTCTTTTTATCAAAGTTTTCCGATGGGGCGTTACTGGTGCTGCTGCAGCCTATGATGTTTCTGCTTGGGCTATTGCTCTGGCTCAAGTGGTTTATATTGTTGGCTGG

>M6-PGSC0003DMG400000179-(rev)M6G-LecRK_4758

CCTATGATGTTTCTGCTTGGGCTATTGCTCTGGCTCAAGTGGTTTATATTGTTGGCTGGTGCAAAGATAGTTGGAAGGGGTTGTCTTGGTTGGCCTTAAAGGAACTTTGGCCCTTCGTAA

>M6-PGSC0003DMG400000179-(rev)M6G-LecRK_4819

CAAAGATAGTTGGAAGGGGTTGTCTTGGTTGGCCTTAAAGGAACTTTGGCCCTTCGTAAAGCTTTCTGTTGCATCAGCAGTGATGATTTGCTTGGAGATCTGGTATTTTATGACTATAAT

>M6-PGSC0003DMG400000179-(rev)M6G-LecRK_488

AGTTTTGAAGGAAGATTCAAGAACCATTGTATGGGAAACAACCAATTTGGAGAAAGCTAGTGATGTGAAGCTTTTGGACCAAGGGAATTTGGTTCTTGTTTCTAATAAAGGTGTTTTGGT

>M6-PGSC0003DMG400000179-(rev)M6G-LecRK_4880

CTTTCTGTTGCATCAGCAGTGATGATTTGCTTGGAGATCTGGTATTTTATGACTATAATTGTTCTAACTGGTCATCTTGAGGACCCCGTCATCGCTGTTGGATCTCTTTCTATCTGGTAC

>M6-PGSC0003DMG400000179-(rev)M6G-LecRK_4941

TTCTAACTGGTCATCTTGAGGACCCCGTCATCGCTGTTGGATCTCTTTCTATCTGGTACAATCTTGCAAATATGCACTTAAAATTTATAATGTGTTCATAATTGTTCTGTTTTTCAAGCT

>M6-PGSC0003DMG400000179-(rev)M6G-LecRK_5002

TCTTGCAAATATGCACTTAAAATTTATAATGTGTTCATAATTGTTCTGTTTTTCAAGCTTAGATTCTTTCTTCTTCATTGTTTCAGTATGAACCTTAATGGATGGGAAGGCATGTTGTTC

>M6-PGSC0003DMG400000179-(rev)M6G-LecRK_5063

GATTCTTTCTTCTTCATTGTTTCAGTATGAACCTTAATGGATGGGAAGGCATGTTGTTCATTGGAATTAATGCAGCAATAAGGTATACTACTATACTAGTTATAGCAGGACATTGTCAGT

>M6-PGSC0003DMG400000179-(rev)M6G-LecRK_5124

TGGAATTAATGCAGCAATAAGGTATACTACTATACTAGTTATAGCAGGACATTGTCAGTGCACAAAAGTTAAACTCTAACGAATAGTGTAACTGCTAACGGTTGTTTTCATTTGTTGATA

>M6-PGSC0003DMG400000179-(rev)M6G-LecRK_5185

ACAAAAGTTAAACTCTAACGAATAGTGTAACTGCTAACGGTTGTTTTCATTTGTTGATATTTCAGTGTTCGCGTTTCCAACGAGCTTGGATCAGGCCATCCTAGGGCTGCAAAGTATTCC

>M6-PGSC0003DMG400000179-(rev)M6G-LecRK_5246

TCAGTGTTCGCGTTTCCAACGAGCTTGGATCAGGCCATCCTAGGGCTGCAAAGTATTCCGTCTTCGTCACAGTAGCTGAATCTCTCATGATTGGGATATTCTGCATGGTACTAATCATAT

>M6-PGSC0003DMG400000179-(rev)M6G-LecRK_5307

CTTCGTCACAGTAGCTGAATCTCTCATGATTGGGATATTCTGCATGGTACTAATCATATTAACAAAGGACCATTTTGCATTGCTTTTCACAAGCAGTGAGAAAATGCAAAAAGCTGTTTC

>M6-PGSC0003DMG400000179-(rev)M6G-LecRK_5368

ACAAAGGACCATTTTGCATTGCTTTTCACAAGCAGTGAGAAAATGCAAAAAGCTGTTTCTAAGTTGGCATATCTTCTTGCTGTTACAATGCTGCTTAATAGTGTTCAGCCAGTTATATCA

>M6-PGSC0003DMG400000179-(rev)M6G-LecRK_5429

AGTTGGCATATCTTCTTGCTGTTACAATGCTGCTTAATAGTGTTCAGCCAGTTATATCAGGTACCAAAATATGTGTGTTTCTTACTTAAAACAGTTCTCCACACACACATCAGTTGATGT

>M6-PGSC0003DMG400000179-(rev)M6G-LecRK_549

GATGTGAAGCTTTTGGACCAAGGGAATTTGGTTCTTGTTTCTAATAAAGGTGTTTTGGTATGGGAAAGTTTTGATTTTCCTACAGATACATGGCTTCCTGGGATGAACTTGACTGCAACA

>M6-PGSC0003DMG400000179-(rev)M6G-LecRK_5490

TACCAAAATATGTGTGTTTCTTACTTAAAACAGTTCTCCACACACACATCAGTTGATGTGCCAAACAAATAGTACTATTTGGTACAAATCTATGTTGCTCGGACTCGTCAAAAAATGTCA

>M6-PGSC0003DMG400000179-(rev)M6G-LecRK_5612

GGGTGTGTGTCGGATTCTTGATAAGTAGAGTATTGTTGAAGAATCCGCGCAATCAAAAAGTTAAGAGTCTGCGCAACTTTAGGTACAAACTCAAAGAATAAATTAGATTGTTCCTAATAT

>M6-PGSC0003DMG400000179-(rev)M6G-LecRK_5673

TAAGAGTCTGCGCAACTTTAGGTACAAACTCAAAGAATAAATTAGATTGTTCCTAATATTGAAGGTGTTGCTGTTGGAGGAGGATGGCAAGCTCTAGTAGCATACATTAATTTGGCTTGC

>M6-PGSC0003DMG400000179-(rev)M6G-LecRK_5734

AAGGTGTTGCTGTTGGAGGAGGATGGCAAGCTCTAGTAGCATACATTAATTTGGCTTGCTACTATGTCATTGGCCTCCCACTTGGATTCCTCTTAGGCTATAAAACAAGTTTGGGAGTTC

>M6-PGSC0003DMG400000179-(rev)M6G-LecRK_5795

CTATGTCATTGGCCTCCCACTTGGATTCCTCTTAGGCTATAAAACAAGTTTGGGAGTTCAGGTAACAACATAAATCTTTTTACTAGTATTAATTTAGCTCCGGTTTGATCATAGATTTTA

>M6-PGSC0003DMG400000179-(rev)M6G-LecRK_61

AAGCAAAATTCATCACCATGATTCCACAGAAAGCAATTCTCATTTTACTTTTCATCACAATTTCTATGGCATTTGAGATCAGTTCAGAAACAGTAAAGTCAAGAATGCAAAATGTAAACA

>M6-PGSC0003DMG400000179-(rev)M6G-LecRK_610

GGGAAAGTTTTGATTTTCCTACAGATACATGGCTTCCTGGGATGAACTTGACTGCAACAAAATGGCTGACATCTTGGAGAAGTACTAATGATCCATCACCTGGAAGGTACTCTTTAAGGC

>M6-PGSC0003DMG400000179-(rev)M6G-LecRK_6283

TCATGAAATTTGGTCAGTTTTCTAAAAATATTCAATTGAGCTAAGTTTTTGGATGAAATTTCACGTCTACTCACGAAACAAGTAAAATGCATGTCCAAACATAACTTCATATTCCAAAAA

>M6-PGSC0003DMG400000179-(rev)M6G-LecRK_6405

ATTGTTCAACACAACTTCAAAAACTCACTTTTTCAAGTTTCAACTAAATCTATGGTCAAAACTCAAACACTAGCTTGAGTTTGAGTTTTAGGAGGGAACTTTGAATTAACGATAAAGTTA

>M6-PGSC0003DMG400000179-(rev)M6G-LecRK_6466

CTCAAACACTAGCTTGAGTTTGAGTTTTAGGAGGGAACTTTGAATTAACGATAAAGTTAATAAACTATTGATGTTTGCGTCCGGATAAATTGTCTACATTACACTAATTGTCACTTCATT

>M6-PGSC0003DMG400000179-(rev)M6G-LecRK_6527

AAACTATTGATGTTTGCGTCCGGATAAATTGTCTACATTACACTAATTGTCACTTCATTTTTTTTCTCTTTTCTGTAGGGAATATGGATGGGCATGATTTTTGGAACTTTTTTGCAAACA

>M6-PGSC0003DMG400000179-(rev)M6G-LecRK_6573

TTGTCACTTCATTTTTTTTCTCTTTTCTGTAGGGAATATGGATGGGCATGATTTTTGGAACTTTTTTGCAAACAATAATTCTCTGTGTTATCGTATACAAAACCAATTGGAACGAAGAGG

>M6-PGSC0003DMG400000179-(rev)M6G-LecRK_671

ATGGCTGACATCTTGGAGAAGTACTAATGATCCATCACCTGGAAGGTACTCTTTAAGGCTCCAGCCACATAGTTATGGTGAGATAGTTCTTTTTTATAATGGTACTTATCCATATTGGTC

>M6-PGSC0003DMG400000179-(rev)M6G-LecRK_732

CAGCCACATAGTTATGGTGAGATAGTTCTTTTTTATAATGGTACTTATCCATATTGGTCCACTGGGAAATGGAGTGAAAATGCATTTGTTGATGTACCAGAAATGACTGTTCCTTACATT

>M6-PGSC0003DMG400000179-(rev)M6G-LecRK_793

CTGGGAAATGGAGTGAAAATGCATTTGTTGATGTACCAGAAATGACTGTTCCTTACATTTACAAGTTCAATTTTGAATCACCTTTTACCCCTATGGCATCATTTGGATACTCAGAGGTGA

>M6-PGSC0003DMG400000179-(rev)M6G-LecRK_854

CAAGTTCAATTTTGAATCACCTTTTACCCCTATGGCATCATTTGGATACTCAGAGGTGACATTAGAGAATGGAATGCCTCCTCCTTTAACTAGGTTCATGGTTGATTTCACTGGACAGGT

>M6-PGSC0003DMG400000179-(rev)M6G-LecRK_915

TTAGAGAATGGAATGCCTCCTCCTTTAACTAGGTTCATGGTTGATTTCACTGGACAGGTTAAACAGTTCACTTGGATCCAACAGGCGCAAAGTTGGAACATGTTTTGGTCACAGCCAGAG

>M6-PGSC0003DMG400000179-(rev)M6G-LecRK_976

AACAGTTCACTTGGATCCAACAGGCGCAAAGTTGGAACATGTTTTGGTCACAGCCAGAGAATATGTGTAGTACTTATGGTCTTTGTGGGAATTTGGGGTTCTGTAATAGTAAGTCATTGA

>M6-PGSC0003DMG400000295M6LRR-RLK_0

TTTGCCAAATGTTCATACAAACTTAGAAAAAATCACCAAAAAAAATGTCCAAAATTATTTTTTCACTTTTCATTTTTCTGTTTTTCATCATTTCTCATGGAAAATCCCAAGAATCTCCAA

>M6-PGSC0003DMG400000295M6LRR-RLK_1037

TTATTTGTTAAGTTGTTTGGGAACAAGTTATCAGGTGAAATTCCACCTGATTTTGGTCGATTTTCGAAACTTTTTGACTTCCAAGTTTCAGAAAATCAACTTGTTGGAAAGTTACCTGAG

>M6-PGSC0003DMG400000295M6LRR-RLK_1098

TTTCGAAACTTTTTGACTTCCAAGTTTCAGAAAATCAACTTGTTGGAAAGTTACCTGAGGGTATTTGCAATAACAAGGCTTTAGCTAGGATGGTTGCTTTTGGCAACAATCTAACAGGTG

>M6-PGSC0003DMG400000295M6LRR-RLK_1159

TATTTGCAATAACAAGGCTTTAGCTAGGATGGTTGCTTTTGGCAACAATCTAACAGGTGAACTGCCAAGTTCACTTGGAAGTTGTGACAGTCTGAGGTATCTTCGAGTCGAAAATAATCG

>M6-PGSC0003DMG400000295M6LRR-RLK_122

CAAGAAAAGGCCATTTTACTCCAACTAAAACAACACTGGTTTACTTCACCAAATGTCACAAAATGGATTTCATCATCAAATCATTGTTCTTGGGATGGAATAATTTGTACCCAAAATTTA

>M6-PGSC0003DMG400000295M6LRR-RLK_1220

CTGCCAAGTTCACTTGGAAGTTGTGACAGTCTGAGGTATCTTCGAGTCGAAAATAATCGACTTTCTGGTGAGGTTCCTGATGGATTATGGACAGGTAAGAGTTTGTCAATGCTTTTAATG

>M6-PGSC0003DMG400000295M6LRR-RLK_1281

TTTCTGGTGAGGTTCCTGATGGATTATGGACAGGTAAGAGTTTGTCAATGCTTTTAATGAAGAATAACTTGTTAACTGGTCAGCTTCCACATAGAGTGGCATCAAATTTGTCACAAGTTG

>M6-PGSC0003DMG400000295M6LRR-RLK_1342

GAATAACTTGTTAACTGGTCAGCTTCCACATAGAGTGGCATCAAATTTGTCACAAGTTGATATCAGCAATAACAAGTTTTCGGGGGAATTACCAGCTGGTATGGGTACTTGGCACAATCT

>M6-PGSC0003DMG400000295M6LRR-RLK_1403

ATCAGCAATAACAAGTTTTCGGGGGAATTACCAGCTGGTATGGGTACTTGGCACAATCTAAGTGAGTTCAAGGCAAGTAATAATCTCTTAAGTGGTCAAATCCCTCAAGAATTGACTCTT

>M6-PGSC0003DMG400000295M6LRR-RLK_1464

GTGAGTTCAAGGCAAGTAATAATCTCTTAAGTGGTCAAATCCCTCAAGAATTGACTCTTCTTCCAAGGATAACTAAACTTTTTCTTGATGGTAATCTACTATCTGGAAATTTTCCATCGA

>M6-PGSC0003DMG400000295M6LRR-RLK_1525

TCCAAGGATAACTAAACTTTTTCTTGATGGTAATCTACTATCTGGAAATTTTCCATCGAATATATCGTCGTGGAAGACTCTTGTCACATTAAATAGCAGAAAAAATCAGCTTTCAGGTCC

>M6-PGSC0003DMG400000295M6LRR-RLK_1586

ATATCGTCGTGGAAGACTCTTGTCACATTAAATAGCAGAAAAAATCAGCTTTCAGGTCCTATACCTTCTGCACTTGGTCTTTTACCAAATCTCATTGATTTGGACTTGTCAAGTAACCAA

>M6-PGSC0003DMG400000295M6LRR-RLK_1647

TACCTTCTGCACTTGGTCTTTTACCAAATCTCATTGATTTGGACTTGTCAAGTAACCAATTTTCAGGTGTTATTCCAACTGAATTAGGTAACTTGAGGTTAACTTCACTTAACCTGTCAT

>M6-PGSC0003DMG400000295M6LRR-RLK_1708

TTCAGGTGTTATTCCAACTGAATTAGGTAACTTGAGGTTAACTTCACTTAACCTGTCATCCAATCGCCTGTCGGGTGAAATCCCATCTCAGTTGGAAAATGCAGCTTTTGGTAAGAGCTT

>M6-PGSC0003DMG400000295M6LRR-RLK_1769

AATCGCCTGTCGGGTGAAATCCCATCTCAGTTGGAAAATGCAGCTTTTGGTAAGAGCTTCTCGGATAATCCTGGTCTTTGTGCAAGTAATCCATCAGTAGAAGTCGCGTCTTGCAAGAGG

>M6-PGSC0003DMG400000295M6LRR-RLK_183

AATGGATTTCATCATCAAATCATTGTTCTTGGGATGGAATAATTTGTACCCAAAATTTAGTTTCTGGTATTCAAATCCCTTTTGGAAATATCTCAAAACCAATTCCAAATTTCATTTGTG

>M6-PGSC0003DMG400000295M6LRR-RLK_1830

CGGATAATCCTGGTCTTTGTGCAAGTAATCCATCAGTAGAAGTCGCGTCTTGCAAGAGGGAAACTAAGTCAGATATGTTTCCTGTTGGACTTGTTGCTGCTCTTGCTAGTGTAGCAGCAG

>M6-PGSC0003DMG400000295M6LRR-RLK_1891

AACTAAGTCAGATATGTTTCCTGTTGGACTTGTTGCTGCTCTTGCTAGTGTAGCAGCAGTTACTTTCCTTGTGGCAGTTCTGTATGGTCTGTTTGTGTTGAGAAGTCATAGAAAAAGTAA

>M6-PGSC0003DMG400000295M6LRR-RLK_1952

ACTTTCCTTGTGGCAGTTCTGTATGGTCTGTTTGTGTTGAGAAGTCATAGAAAAAGTAAACAAGAATCAGTTTCGACATGGAAACAAACATCATTTCACAAATTAGACTTCACAGAATCA

>M6-PGSC0003DMG400000295M6LRR-RLK_2013

AAGAATCAGTTTCGACATGGAAACAAACATCATTTCACAAATTAGACTTCACAGAATCAGACATTGTATCCAATCTGACCGAAAACAACATAATTGGAAGTGGAGGATCTGGACAGGTTT

>M6-PGSC0003DMG400000295M6LRR-RLK_2074

CATTGTATCCAATCTGACCGAAAACAACATAATTGGAAGTGGAGGATCTGGACAGGTTTACCTCGTGCCTTTAAGCCGATCAGGTGATTATGTTGCTGTCAAGAGGATTTGGAGAAACCA

>M6-PGSC0003DMG400000295M6LRR-RLK_2135

CTCGTGCCTTTAAGCCGATCAGGTGATTATGTTGCTGTCAAGAGGATTTGGAGAAACCAAAGGTTGGATCACAAGCATGAGAAACAGTTTCTTGCTGAAGTTCAGATATTAGGCACGATT

>M6-PGSC0003DMG400000295M6LRR-RLK_2196

GGTTGGATCACAAGCATGAGAAACAGTTTCTTGCTGAAGTTCAGATATTAGGCACGATTCGACACTCCAATATAGTGAAACTCCTCTGCTGCTTCTTCAGTGAAGAGTCGAAACTTCTTG

>M6-PGSC0003DMG400000295M6LRR-RLK_2257

ACACTCCAATATAGTGAAACTCCTCTGCTGCTTCTTCAGTGAAGAGTCGAAACTTCTTGTCTACGAATATATGGAGAACAGGAGCTTGGATATATGGCTTCACTCAAAGAATAGGTTGAA

>M6-PGSC0003DMG400000295M6LRR-RLK_2318

TACGAATATATGGAGAACAGGAGCTTGGATATATGGCTTCACTCAAAGAATAGGTTGAACAATGCCTCAAGATCAACTCCACATCTGGTCTTGGAATGGCCTAGGAGGCTGCAAATTGCT

>M6-PGSC0003DMG400000295M6LRR-RLK_2379

ATGCCTCAAGATCAACTCCACATCTGGTCTTGGAATGGCCTAGGAGGCTGCAAATTGCTATAGGAGCTGCTCGTGGTTTTTGCTACATGCACCATGACTGCTCACCTCCTATTATTCATC

>M6-PGSC0003DMG400000295M6LRR-RLK_244

TTCTGGTATTCAAATCCCTTTTGGAAATATCTCAAAACCAATTCCAAATTTCATTTGTGATCTTAAAAATCTCATCTTTCTTGATTTTAACCATAATTTCATCCCTGGAAACTTCCCTGA

>M6-PGSC0003DMG400000295M6LRR-RLK_2440

AGGAGCTGCTCGTGGTTTTTGCTACATGCACCATGACTGCTCACCTCCTATTATTCATCGCGATGTGAAGTCAAGCAACATCCTATTGGATTCTCAATTCAATGCAAAAATTGCAGATTT

>M6-PGSC0003DMG400000295M6LRR-RLK_2501

GATGTGAAGTCAAGCAACATCCTATTGGATTCTCAATTCAATGCAAAAATTGCAGATTTTGGCCTTGCCAGGATGTTACTCAAGCCTGGAGATAACACAGTGACAGCAGTAGCTGGCTCT

>M6-PGSC0003DMG400000295M6LRR-RLK_2562

GCCTTGCCAGGATGTTACTCAAGCCTGGAGATAACACAGTGACAGCAGTAGCTGGCTCTTTCGGATACATTGCACCAGGTATAAGACCATCAACTTATCTATAACATCATTATAAAGTTT

>M6-PGSC0003DMG400000295M6LRR-RLK_2623

CGGATACATTGCACCAGGTATAAGACCATCAACTTATCTATAACATCATTATAAAGTTTAGAAGTTGGTTTAATCTCTTTCATGGAAAAAAGTTTCCTATGCTACACAATTCAATAACAT

>M6-PGSC0003DMG400000295M6LRR-RLK_2684

AAGTTGGTTTAATCTCTTTCATGGAAAAAAGTTTCCTATGCTACACAATTCAATAACATATCAAGGGGTCGTTTTGTTCACGTACTAGTTATGCATGAGTTGTAACACGGGTTGTTTATG

>M6-PGSC0003DMG400000295M6LRR-RLK_2745

CAAGGGGTCGTTTTGTTCACGTACTAGTTATGCATGAGTTGTAACACGGGTTGTTTATGTAGTAAATAGTAATGTAGATATTAGTACGCGAGTAGTTATTCTTAGTACACTAATTCTCGC

>M6-PGSC0003DMG400000295M6LRR-RLK_2806

GTAAATAGTAATGTAGATATTAGTACGCGAGTAGTTATTCTTAGTACACTAATTCTCGCGTAACTCATATAAATGCTAATATATTCATGATGGTATTTCAGAGTATGCGCGAAAAACTAG

>M6-PGSC0003DMG400000295M6LRR-RLK_2867

AACTCATATAAATGCTAATATATTCATGATGGTATTTCAGAGTATGCGCGAAAAACTAGAGTGACAGAGAAAATTGATGTGTATAGTTTTGGAGTCATACTATTGGAACTGGTGACTGGA

>M6-PGSC0003DMG400000295M6LRR-RLK_2928

TGACAGAGAAAATTGATGTGTATAGTTTTGGAGTCATACTATTGGAACTGGTGACTGGAAAAGAAGCCAATCTTGGAGATGAAGATTCTTGTCTTGCAGACTGGGCATGGCGCCACCTTC

>M6-PGSC0003DMG400000295M6LRR-RLK_2989

AGAAGCCAATCTTGGAGATGAAGATTCTTGTCTTGCAGACTGGGCATGGCGCCACCTTCAAAAAGGGAAACCGATGGCTGATGCATTAGATGAGGACATCAAAGAAACGCGATACTTGGA

>M6-PGSC0003DMG400000295M6LRR-RLK_305

CTTAAAAATCTCATCTTTCTTGATTTTAACCATAATTTCATCCCTGGAAACTTCCCTGATATTTACAGTTGTTCAAATCTTGAATTCTTAGATCTTTCTTATAACTACATGGTTGGTAAT

>M6-PGSC0003DMG400000295M6LRR-RLK_3050

AAAGGGAAACCGATGGCTGATGCATTAGATGAGGACATCAAAGAAACGCGATACTTGGAAGAAATCTTCGTTGTGTTTAAGCTTGGAATCTTTTGCACCAGCACATTTCCTTCATCAAGG

>M6-PGSC0003DMG400000295M6LRR-RLK_3111

AAATCTTCGTTGTGTTTAAGCTTGGAATCTTTTGCACCAGCACATTTCCTTCATCAAGGCCAACAATGAAGGAAGTTCTGCAAATCTTGATCCAATGTAGCAACAGTTCACCTACATCTG

>M6-PGSC0003DMG400000295M6LRR-RLK_3172

AACAATGAAGGAAGTTCTGCAAATCTTGATCCAATGTAGCAACAGTTCACCTACATCTGGAGAAAAGAAAAATGAAACAGAACAGGATGTTTTGCCACTCCTCAAGAACTCAAGGAGTGA

>M6-PGSC0003DMG400000295M6LRR-RLK_3233

GAAAAGAAAAATGAAACAGAACAGGATGTTTTGCCACTCCTCAAGAACTCAAGGAGTGAGAGGATTGAAGAAAATGATGATGTTGGTTTAACATCAATTATTTGATGGCATTACAATATG

>M6-PGSC0003DMG400000295M6LRR-RLK_3538

GCAAAGAATAAGACCAATAGATAATGTATCAGGTAGTGAAAACTAGCATTGTACATACACTGTATTGGATCCATAGTGTTGCTTTGCATTGTTTTGCTACATAAATTTTGCTTTTCCAAT

>M6-PGSC0003DMG400000295M6LRR-RLK_3599

GTATTGGATCCATAGTGTTGCTTTGCATTGTTTTGCTACATAAATTTTGCTTTTCCAATGTTTTCTTTACTCATAAAATAACTGTCAAAAAGGTACATTAAAACCATAAACTATTGAATG

>M6-PGSC0003DMG400000295M6LRR-RLK_366

TTTACAGTTGTTCAAATCTTGAATTCTTAGATCTTTCTTATAACTACATGGTTGGTAATCTCCCTGATGAGATTAACCGTCTTTCGAGTAATCTAAAATACCTAAACTTAACTGCCAACA

>M6-PGSC0003DMG400000295M6LRR-RLK_3660

TTTCTTTACTCATAAAATAACTGTCAAAAAGGTACATTAAAACCATAAACTATTGAATGGAGGATAGATTTCCTAAAAATCCAGAAAAATAGAAACAAGCTTGAACTACCTCAGTACTTT

>M6-PGSC0003DMG400000295M6LRR-RLK_3721

GGATAGATTTCCTAAAAATCCAGAAAAATAGAAACAAGCTTGAACTACCTCAGTACTTTAACTTTTGGCAACCATACAACAACATATCCGATGTAATCCCATGAGTCCCAAACTTTTGGC

>M6-PGSC0003DMG400000295M6LRR-RLK_3965

AATAAATCTTGTAAAGATCACACAAAAATGTGGCTACTAGAAGAACTTTATTTGTATCCTAGCTTTATTTCTGTCTAGTGAGGAAAAAATTTAAGTTTACACTGGGGATAAAGATGATCC

>M6-PGSC0003DMG400000295M6LRR-RLK_4026

GCTTTATTTCTGTCTAGTGAGGAAAAAATTTAAGTTTACACTGGGGATAAAGATGATCCTTAGAGGCACCAGGAGTGAAAAGGTCAGCATATTGATGTTTAGCTGAAGAGAGATGTGCTC

>M6-PGSC0003DMG400000295M6LRR-RLK_4087

AGAGGCACCAGGAGTGAAAAGGTCAGCATATTGATGTTTAGCTGAAGAGAGATGTGCTCATTTGGTACATTGAACTATGGCTCACTTCAAAGCTCACTCTTGCATGACGCGGTGCACAAG

>M6-PGSC0003DMG400000295M6LRR-RLK_4148

TTGGTACATTGAACTATGGCTCACTTCAAAGCTCACTCTTGCATGACGCGGTGCACAAGGAAGCAAGAAATATTTCAATCATCTTGGAGAAGAAAGTAAGAAGAGGGTTCTCTTTAGCAA

>M6-PGSC0003DMG400000295M6LRR-RLK_4209

AGCAAGAAATATTTCAATCATCTTGGAGAAGAAAGTAAGAAGAGGGTTCTCTTTAGCAAGAAGAGGGGTGATTGTAAACGTACAAACTTTGACTAGCTAGGCACTTTTGGTTGTGTCACT

>M6-PGSC0003DMG400000295M6LRR-RLK_427

CCCTGATGAGATTAACCGTCTTTCGAGTAATCTAAAATACCTAAACTTAACTGCCAACAACTTCAATGGTGATATCCCTAAAGGAATTGGTGGATTGAGTCATCTCAAAGTACTTGATCT

>M6-PGSC0003DMG400000295M6LRR-RLK_4270

AGAGGGGTGATTGTAAACGTACAAACTTTGACTAGCTAGGCACTTTTGGTTGTGTCACTCTGACCATGATCATGATTCCTCGAGATCTGTGTATTGGGAATGCGAGGCAGCCAAAAGTGC

>M6-PGSC0003DMG400000295M6LRR-RLK_4331

GACCATGATCATGATTCCTCGAGATCTGTGTATTGGGAATGCGAGGCAGCCAAAAGTGCAGCTCCAACGCCTGAACCATCATTAGAAAGCTCAATGACTATGCTGTCCGCAGCTTCTCCC

>M6-PGSC0003DMG400000295M6LRR-RLK_4392

CTCCAACGCCTGAACCATCATTAGAAAGCTCAATGACTATGCTGTCCGCAGCTTCTCCCAGTAACTCTTTCATAGTTTCCTCCAAGCAATTTCTGAACTTGGTGTAACGCTCAAACAATG

>M6-PGSC0003DMG400000295M6LRR-RLK_4453

TAACTCTTTCATAGTTTCCTCCAAGCAATTTCTGAACTTGGTGTAACGCTCAAACAATGCACCGTCCACAGCTATGACAGACCTCTGGTTCTCTCCGTCCTTCAAAGTGTCTTTTCCCAA

>M6-PGSC0003DMG400000295M6LRR-RLK_4514

CCGTCCACAGCTATGACAGACCTCTGGTTCTCTCCGTCCTTCAAAGTGTCTTTTCCCAATTTCTTGAGGATGCCCACGATTCCTGCTGCAGAAAGACGAGCTCCATGAGAGGTGATAATG

>M6-PGSC0003DMG400000295M6LRR-RLK_4575

TCTTGAGGATGCCCACGATTCCTGCTGCAGAAAGACGAGCTCCATGAGAGGTGATAATGTCGCATAGCTCCACAACTATTTTCCTCATTTTCAAGGTAGAATTAGGTACCTGTGAAAAAA

>M6-PGSC0003DMG400000295M6LRR-RLK_4636

GCATAGCTCCACAACTATTTTCCTCATTTTCAAGGTAGAATTAGGTACCTGTGAAAAAAATGTATATATAGCAAATAGGATGAGAAAGACGTTCTCCACTTAATTAAGCCCAGCCAACAT

>M6-PGSC0003DMG400000295M6LRR-RLK_4697

GTATATATAGCAAATAGGATGAGAAAGACGTTCTCCACTTAATTAAGCCCAGCCAACATTGTGATGTTCAGACAGGAGCTAGGAATTTGAGTTTATGGGTTCCGGATAAATTATTTACAC

>M6-PGSC0003DMG400000295M6LRR-RLK_4758

TGATGTTCAGACAGGAGCTAGGAATTTGAGTTTATGGGTTCCGGATAAATTATTTACACATATTAAGTGGGTTTTTTAGTACAAATACAAAGCTACTGGTTTTTTGTCCTATCATATATA

>M6-PGSC0003DMG400000295M6LRR-RLK_4819

ATTAAGTGGGTTTTTTAGTACAAATACAAAGCTACTGGTTTTTTGTCCTATCATATATAGGACAAAAAAGATGAGAACTTCAAAGACTAGTAATGTTTCACTTTAAGTCAAATGTTTGAT

>M6-PGSC0003DMG400000295M6LRR-RLK_488

TTCAATGGTGATATCCCTAAAGGAATTGGTGGATTGAGTCATCTCAAAGTACTTGATCTGATGGTGAATTTGTTTTATGGATCTTTTCCTGAAGAGATTGGTGAATTGTTGAATCTTGAA

>M6-PGSC0003DMG400000295M6LRR-RLK_4880

ACAAAAAAGATGAGAACTTCAAAGACTAGTAATGTTTCACTTTAAGTCAAATGTTTGATCTTAATACCTCTAAGATATCCTTCAGTTTATTGCCAACCACCTTGAGATCAGCCGACTCGT

>M6-PGSC0003DMG400000295M6LRR-RLK_4941

TAATACCTCTAAGATATCCTTCAGTTTATTGCCAACCACCTTGAGATCAGCCGACTCGTCGTGATGCATAGCAGCCATATCCGGAGTCCTGCATGTGTGGCAGTAAATAACCTTTTTCAC

>M6-PGSC0003DMG400000295M6LRR-RLK_5002

TGATGCATAGCAGCCATATCCGGAGTCCTGCATGTGTGGCAGTAAATAACCTTTTTCACAATATTTGACAATTCAAAGCTTACATATGCTTAATAAACGTACACGAATCTAGTCCAACGA

>M6-PGSC0003DMG400000295M6LRR-RLK_5063

TATTTGACAATTCAAAGCTTACATATGCTTAATAAACGTACACGAATCTAGTCCAACGAGAAGCATAACTTACAGTGTAAGAGATTAGTGCAATTACATAAACCTTAAGTTGCACTAATT

>M6-PGSC0003DMG400000295M6LRR-RLK_5124

AGCATAACTTACAGTGTAAGAGATTAGTGCAATTACATAAACCTTAAGTTGCACTAATTGCATGGAGAAGAATAAGGTTAAAATTGATGAATGCAAATAGGGCCTGCCAAAAAATAAATT

>M6-PGSC0003DMG400000295M6LRR-RLK_5185

ATGGAGAAGAATAAGGTTAAAATTGATGAATGCAAATAGGGCCTGCCAAAAAATAAATTACTGCCGTAATAAATGTGTTGGTCAAAGCAATACCCTACCTCAATATGAAAGGAATTTTCA

>M6-PGSC0003DMG400000295M6LRR-RLK_5246

TGCCGTAATAAATGTGTTGGTCAAAGCAATACCCTACCTCAATATGAAAGGAATTTTCAGTTTGGATGGGACATAATCACCGAATAATGAAGCTTCTTTAGCCATTCTACACAATACTCT

>M6-PGSC0003DMG400000295M6LRR-RLK_5307

TTGGATGGGACATAATCACCGAATAATGAAGCTTCTTTAGCCATTCTACACAATACTCTACGCAAAATTTCTCCAAGATACATCCCGGAGATCATCTTTTCATAAATCTGTGCAGTTTCC

>M6-PGSC0003DMG400000295M6LRR-RLK_5368

GCAAAATTTCTCCAAGATACATCCCGGAGATCATCTTTTCATAAATCTGTGCAGTTTCCGAAAGTTCAACAACAGATGTAAGATAGAAAATGTAGAAAGGAAGGGGATTGTCTTAGTTAC

>M6-PGSC0003DMG400000295M6LRR-RLK_5429

AAGTTCAACAACAGATGTAAGATAGAAAATGTAGAAAGGAAGGGGATTGTCTTAGTTACCTGCTCCCCGGGGTTTAAACTCTCAACATCAAGATTTTGGTCATATTCTGTTACTGGAAGA

>M6-PGSC0003DMG400000295M6LRR-RLK_549

TGGTGAATTTGTTTTATGGATCTTTTCCTGAAGAGATTGGTGAATTGTTGAATCTTGAAGTACTTATGTTGAGCTTGAATGGTTTTGCTCCACATGCTATACCATCACTATTCACTAAGT

>M6-PGSC0003DMG400000295M6LRR-RLK_5490

GCTCCCCGGGGTTTAAACTCTCAACATCAAGATTTTGGTCATATTCTGTTACTGGAAGATGTGATGAGCGGAAATTACCCCATTCCATGTTGATAACCTTTAAAGGCCAGTCAAAATTAC

>M6-PGSC0003DMG400000295M6LRR-RLK_5551

TGATGAGCGGAAATTACCCCATTCCATGTTGATAACCTTTAAAGGCCAGTCAAAATTACGTTTAAGTAAACAAACGTGTATGAATCTACCAAGGAAAATATTTTTAGGAAATGAATTAGG

>M6-PGSC0003DMG400000295M6LRR-RLK_5612

TTAAGTAAACAAACGTGTATGAATCTACCAAGGAAAATATTTTTAGGAAATGAATTAGGCACATATCAAGTCATGAAGGTGTTATGAAATGTCATAAGCATCTACCATCTCTCCGGATTT

>M6-PGSC0003DMG400000295M6LRR-RLK_5673

CATATCAAGTCATGAAGGTGTTATGAAATGTCATAAGCATCTACCATCTCTCCGGATTTCGGCAACAGACCATGCCATTTGGGAATCACATGAGCCCGTTCAACATATGCTGCATTGGTT

>M6-PGSC0003DMG400000295M6LRR-RLK_5734

GCAACAGACCATGCCATTTGGGAATCACATGAGCCCGTTCAACATATGCTGCATTGGTTCCAGTACCCAAAATTACTGCAGCAATGACATCAGGATTATTGTACCGACCTCCTGCTAACG

>M6-PGSC0003DMG400000295M6LRR-RLK_5795

AGTACCCAAAATTACTGCAGCAATGACATCAGGATTATTGTACCGACCTCCTGCTAACGTTCCAACAGTATCATTGACCTAAACAACCAAGTCAACTAAGAAATTCAGGCTCAGGAATTG

>M6-PGSC0003DMG400000295M6LRR-RLK_5856

CCAACAGTATCATTGACCTAAACAACCAAGTCAACTAAGAAATTCAGGCTCAGGAATTGCTTAAGATTAAAAATCAGAAATATCACACATACAAAAACCTAATACATGAAAATCATCAGC

>M6-PGSC0003DMG400000295M6LRR-RLK_5917

TAAGATTAAAAATCAGAAATATCACACATACAAAAACCTAATACATGAAAATCATCAGCTTACTTATATTTTATCGAATTTGATTGGGAAAAAAACTTACTAAAGCAGCCACACGCACAT

>M6-PGSC0003DMG400000295M6LRR-RLK_5978

ACTTATATTTTATCGAATTTGATTGGGAAAAAAACTTACTAAAGCAGCCACACGCACATCAAGGCCGGCCCTTTCCATTGCTTTTGCTAATTCTCCAACCACATCTTGGCCAACCTATGT

>M6-PGSC0003DMG400000295M6LRR-RLK_6039

AGGCCGGCCCTTTCCATTGCTTTTGCTAATTCTCCAACCACATCTTGGCCAACCTATGTTAGATTCCATGAGGAGTAGTTCAGAATTGAAATACATTAAAACATTACAAAAGGAACTCTT

>M6-PGSC0003DMG400000295M6LRR-RLK_61

TTCACTTTTCATTTTTCTGTTTTTCATCATTTCTCATGGAAAATCCCAAGAATCTCCAAATCAAGAAAAGGCCATTTTACTCCAACTAAAACAACACTGGTTTACTTCACCAAATGTCAC

>M6-PGSC0003DMG400000295M6LRR-RLK_610

ACTTATGTTGAGCTTGAATGGTTTTGCTCCACATGCTATACCATCACTATTCACTAAGTTGAAAAATTTGAAAATTTTTTGGATGACAGAAGCACATTTGATTGGAAATATCCCGGAAAA

>M6-PGSC0003DMG400000295M6LRR-RLK_6100

GATTCCATGAGGAGTAGTTCAGAATTGAAATACATTAAAACATTACAAAAGGAACTCTTTATAACGTACTTTAAATCAAATTGGTGAAAATGCAATGAAACAATGATCAACTTTGCTTGT

>M6-PGSC0003DMG400000295M6LRR-RLK_6161

TAACGTACTTTAAATCAAATTGGTGAAAATGCAATGAAACAATGATCAACTTTGCTTGTATATAGTTGTATGCATTAAATCCTCATCAATGCCAACCTATTTTAGGCCATGAGGAGTAGT

>M6-PGSC0003DMG400000295M6LRR-RLK_6222

ATAGTTGTATGCATTAAATCCTCATCAATGCCAACCTATTTTAGGCCATGAGGAGTAGTTCATCATTGAAATACAACATAACGTTGGAAAAGGAAATCCTACAACGTATTTTAAATCAAC

>M6-PGSC0003DMG400000295M6LRR-RLK_6283

ATCATTGAAATACAACATAACGTTGGAAAAGGAAATCCTACAACGTATTTTAAATCAACTCAGTGAAAAAGCAATGAAATATACGTTGTCGGACTCTCCAATAATATTGCCGCACCTTTG

>M6-PGSC0003DMG400000295M6LRR-RLK_6405

AGATCCTTCAAAATGTACTGCTTTTGAAGGATCCTACAAGCACCTGGCGAGTGGCGAAATTTTTGAAGAGTCTGAGCAACATAGATGAAATAATCATCAACTTTGCATGCATAAAGTTGT

>M6-PGSC0003DMG400000295M6LRR-RLK_6466

TTTGAAGAGTCTGAGCAACATAGATGAAATAATCATCAACTTTGCATGCATAAAGTTGTATGCATTAAATCATCATCAATGCAAAACCATACATTGAAATGAATGGACTAAATCTAAGGT

>M6-PGSC0003DMG400000295M6LRR-RLK_6527

GCATTAAATCATCATCAATGCAAAACCATACATTGAAATGAATGGACTAAATCTAAGGTTGATTGTAGGCATGATCAATCGGAAGTGAAAATCAGAAACTCTGTATGACTAACCACGTCT

>M6-PGSC0003DMG400000295M6LRR-RLK_6588

ATTGTAGGCATGATCAATCGGAAGTGAAAATCAGAAACTCTGTATGACTAACCACGTCTTCTATGGAGAAGCCCTTTGTCCATTTAATAAGAGTTCCTGATGCAATTGACAATTGTTTCA

>M6-PGSC0003DMG400000295M6LRR-RLK_6649

TATGGAGAAGCCCTTTGTCCATTTAATAAGAGTTCCTGATGCAATTGACAATTGTTTCACAGGGAAAGAGAAGGTAAAGCCTAACTCCCTTTGTCTACCAGGTGGAAGATGAAAATCATC

>M6-PGSC0003DMG400000295M6LRR-RLK_671

AAAAATTTGAAAATTTTTTGGATGACAGAAGCACATTTGATTGGAAATATCCCGGAAAATATTGGTAATATGACAAGTCTTGAATTTTTGGATTTGTCGAATAATGGATTGAGTGGTAGT

>M6-PGSC0003DMG400000295M6LRR-RLK_6710

GGGAAAGAGAAGGTAAAGCCTAACTCCCTTTGTCTACCAGGTGGAAGATGAAAATCATCACCTTCTGTAGCTACAAATTCTGCAAGTGCTGTGGCAATAAAATCAAATAACACCTGCAAT

>M6-PGSC0003DMG400000295M6LRR-RLK_6771

CTTCTGTAGCTACAAATTCTGCAAGTGCTGTGGCAATAAAATCAAATAACACCTGCAATTAACAGTTTGAAGTTTAATAATGTGAGTACAACCTTGATTGGATTAAAAAAGTTGCAGGAT

>M6-PGSC0003DMG400000295M6LRR-RLK_6832

ACAGTTTGAAGTTTAATAATGTGAGTACAACCTTGATTGGATTAAAAAAGTTGCAGGATGCATGGCTGATACAGCATAAGACTTACTTCAGATGACGATCCAGCCATCACATTCTGTGGA

>M6-PGSC0003DMG400000295M6LRR-RLK_6893

ATGGCTGATACAGCATAAGACTTACTTCAGATGACGATCCAGCCATCACATTCTGTGGAATTGAAACTTCTTTAACTTCATGTTTAACTATACGCTTTTCTTTCCCACCCAACTGCACAC

>M6-PGSC0003DMG400000295M6LRR-RLK_6954

TGAAACTTCTTTAACTTCATGTTTAACTATACGCTTTTCTTTCCCACCCAACTGCACACGCATCACACGAAAGTTTGTGCCGCCTAGATCCAATGCATAAAACAGACCTTCTTCGTCCCT

>M6-PGSC0003DMG400000295M6LRR-RLK_7015

ATCACACGAAAGTTTGTGCCGCCTAGATCCAATGCATAAAACAGACCTTCTTCGTCCCTAAGATGAAGGAATAAAAGTAATACCAGTCATTAAACTGAAGATACAATCATACATGGAAAA

>M6-PGSC0003DMG400000295M6LRR-RLK_7076

GATGAAGGAATAAAAGTAATACCAGTCATTAAACTGAAGATACAATCATACATGGAAAACATACCCCACCCCTAATGCACTACAGAGGTCTATTGGCAGCAGCTTTTGAAAATTTGTCGA

>M6-PGSC0003DMG400000295M6LRR-RLK_7137

TACCCCACCCCTAATGCACTACAGAGGTCTATTGGCAGCAGCTTTTGAAAATTTGTCGATACATGTGAATTTTTATACCCAATTTTAGTAAGTTTATCACAAGAACATGGAATTGTTACC

>M6-PGSC0003DMG400000295M6LRR-RLK_7198

CATGTGAATTTTTATACCCAATTTTAGTAAGTTTATCACAAGAACATGGAATTGTTACCAAGTCCCTACCTCCGCCTACTACTATAGGTATGCCATTTTTAAACTTTAGAGAATGTCAAA

>M6-PGSC0003DMG400000295M6LRR-RLK_7259

GTCCCTACCTCCGCCTACTACTATAGGTATGCCATTTTTAAACTTTAGAGAATGTCAAACTGATTCATATGTCATATATTCTTTTCTTATGGATCTAACTTCATTATGTTGATCACAGAG

>M6-PGSC0003DMG400000295M6LRR-RLK_732

TTGGTAATATGACAAGTCTTGAATTTTTGGATTTGTCGAATAATGGATTGAGTGGTAGTATCCCTGATGGTTTGTTTCATCTCAAGAATTTGAGTATTGTGTATTTATTTACTAACAAAT

>M6-PGSC0003DMG400000295M6LRR-RLK_7320

GATTCATATGTCATATATTCTTTTCTTATGGATCTAACTTCATTATGTTGATCACAGAGATGTCTGTGTAGACAGCACAGTAACAAATAATGAGCAGACAGAAGAAGATGGAACTGAAAA

>M6-PGSC0003DMG400000295M6LRR-RLK_7381

GTCTGTGTAGACAGCACAGTAACAAATAATGAGCAGACAGAAGAAGATGGAACTGAAAAATTTACACGCATGATAATTGTTAAAGAATGGAACTGGCAATGTAAAATTCAAAAAAGGTGG

>M6-PGSC0003DMG400000295M6LRR-RLK_7442

TTACACGCATGATAATTGTTAAAGAATGGAACTGGCAATGTAAAATTCAAAAAAGGTGGTCAAGATACAAAACAGTAGAAGAGAATATAGAATAAACATCAAACTCAGGATCCATAGATA

>M6-PGSC0003DMG400000295M6LRR-RLK_7503

AAGATACAAAACAGTAGAAGAGAATATAGAATAAACATCAAACTCAGGATCCATAGATAATGAGTGACCTGATCACTTGTCAGTTAGAAACTTCTTAGTGGACTCCAAGTTCAACAAATA

>M6-PGSC0003DMG400000295M6LRR-RLK_7564

GAGTGACCTGATCACTTGTCAGTTAGAAACTTCTTAGTGGACTCCAAGTTCAACAAATACTTCAAATAATAGAGTTTCATTTCTGCTCTTTGTGATTTATAACGATGCTCAAGGTAATCA

>M6-PGSC0003DMG400000295M6LRR-RLK_7625

TCAAATAATAGAGTTTCATTTCTGCTCTTTGTGATTTATAACGATGCTCAAGGTAATCACCTTAAGAACATCAAAAGAATCATACTTTAGGACTAAATAATAATTTGAAAAGTACTTAAT

>M6-PGSC0003DMG400000295M6LRR-RLK_7686

TTAAGAACATCAAAAGAATCATACTTTAGGACTAAATAATAATTTGAAAAGTACTTAATCAATTATCCAAACATTACGAAAGTGAAATCAATCTCCATGATCACAGCTTTATTCAAAAAA

>M6-PGSC0003DMG400000295M6LRR-RLK_7747

ATTATCCAAACATTACGAAAGTGAAATCAATCTCCATGATCACAGCTTTATTCAAAAAAGAGAAATTCTTGACACCCAAATTAGCAAAAATCAGTTTGTGCACATATACACACACAAATA

>M6-PGSC0003DMG400000295M6LRR-RLK_7808

GAAATTCTTGACACCCAAATTAGCAAAAATCAGTTTGTGCACATATACACACACAAATACTCATCCAATGTGGAGGCTATCAACTTAGCTAATAAGCATCTTTAGTTTGCCAATGAATCG

>M6-PGSC0003DMG400000295M6LRR-RLK_7869

CATCCAATGTGGAGGCTATCAACTTAGCTAATAAGCATCTTTAGTTTGCCAATGAATCGTTAGTAAAACACACAATCTGATATAACTCTCCACCCCATAAGCTAAAATCAAGAGACAACA

>M6-PGSC0003DMG400000295M6LRR-RLK_793

CCCTGATGGTTTGTTTCATCTCAAGAATTTGAGTATTGTGTATTTATTTACTAACAAATTGTCAGGGGAAATACCACAGTCAGTTTCTTCAATGAATTTAGATGTTGTTGATTTGTGTAA

>M6-PGSC0003DMG400000295M6LRR-RLK_7930

AGTAAAACACACAATCTGATATAACTCTCCACCCCATAAGCTAAAATCAAGAGACAACACTACCAAGATCACAGCTTTATTGAAAAAAGAGACAAATTCTTGACACCCAAATGAGCAAAA

>M6-PGSC0003DMG400000295M6LRR-RLK_7991

ACCAAGATCACAGCTTTATTGAAAAAAGAGACAAATTCTTGACACCCAAATGAGCAAAAATCAGTTTATACACACAAAAATACTCTCCCAGTCGGATGGTTATCAACTTAGGTAATAACC

>M6-PGSC0003DMG400000295M6LRR-RLK_8052

CAGTTTATACACACAAAAATACTCTCCCAGTCGGATGGTTATCAACTTAGGTAATAACCATCTTGAGTATGTCGATGAATCAATTGTAAAAACACAAAAAAGAACTGACATAACTCCAAC

>M6-PGSC0003DMG400000295M6LRR-RLK_8113

CTTGAGTATGTCGATGAATCAATTGTAAAAACACAAAAAAGAACTGACATAACTCCAACCCAGAAGCTAAAATAAAGAGACAACACCTCCAACCGAGTTGGAGCTAGAATTTGATGTTTA

>M6-PGSC0003DMG400000295M6LRR-RLK_8174

AGAAGCTAAAATAAAGAGACAACACCTCCAACCGAGTTGGAGCTAGAATTTGATGTTTATTGGTTCTGAACTTGTCGAGAAAAAACTATTAGGTTCATCCGAACCTGTATCTAATATGTA

>M6-PGSC0003DMG400000295M6LRR-RLK_8235

GGTTCTGAACTTGTCGAGAAAAAACTATTAGGTTCATCCGAACCTGTATCTAATATGTAGCTCCACCCCTGCCTCCAAAATCACAACTTTACTGAAAAAGAGACAAATTCTTGAACACCC

>M6-PGSC0003DMG400000295M6LRR-RLK_8296

TCCACCCCTGCCTCCAAAATCACAACTTTACTGAAAAAGAGACAAATTCTTGAACACCCAAATAAGCAAAAGACAGTTTATGCACATTCTTTTTTTGAGAAGGAGTTTATGCACATTCTT

>M6-PGSC0003DMG400000295M6LRR-RLK_8357

ATAAGCAAAAGACAGTTTATGCACATTCTTTTTTTGAGAAGGAGTTTATGCACATTCTTAAACACCCAAATTAGCAAAAAACAGTTTATCCACATTCATAAACACCCAAATAAGCATAAA

>M6-PGSC0003DMG400000295M6LRR-RLK_8418

ACACCCAAATTAGCAAAAAACAGTTTATCCACATTCATAAACACCCAAATAAGCATAAAAACAGTTTATCCACATTCTCAAACACCCATATAAGCCCAAAAAAAACAGTTTATGCACATT

>M6-PGSC0003DMG400000295M6LRR-RLK_8479

CAGTTTATCCACATTCTCAAACACCCATATAAGCCCAAAAAAAACAGTTTATGCACATTCTTGAACACCCAAATAAGCCAAAAACAGTTTATCCACACTCTTAAACACCCAAATAAGTAA

>M6-PGSC0003DMG400000295M6LRR-RLK_854

TCAGGGGAAATACCACAGTCAGTTTCTTCAATGAATTTAGATGTTGTTGATTTGTGTAACAACAGTTTGACAGGAAAAATACCAGAAGATTTCGGAAAATTGACTAAAATGACTGGATTG

>M6-PGSC0003DMG400000295M6LRR-RLK_915

ACAGTTTGACAGGAAAAATACCAGAAGATTTCGGAAAATTGACTAAAATGACTGGATTGTCTTTGTTTTTCAATCAATTATCAGGTGAAATACCATTGAGTATAGGCAAGTTATCATCAT

>M6-PGSC0003DMG400000295M6LRR-RLK_976

TTTGTTTTTCAATCAATTATCAGGTGAAATACCATTGAGTATAGGCAAGTTATCATCATTGTTATTTGTTAAGTTGTTTGGGAACAAGTTATCAGGTGAAATTCCACCTGATTTTGGTCG

>M6-PGSC0003DMG400000386M6LRR-RLP_0

ATGGTGTTTGAGAATGATAGAATCAAGAATGCTTATATTGCTTTACAAGCATGGAAACAAGCTATCATTTCAGATCCCTTTAATATCACTCTCAATTGGGTTGGTCCTAATGTATGCAGC

>M6-PGSC0003DMG400000386M6LRR-RLP_1008

CCACCTCCTCCGCCACTTTCACCCCCCTTGCCTGCCCCTCCGCCACCTATTTATTCGCCTCCGCCACCAGTTTATACTCCTCCACCTCCAGTTTATTCACCACCGCCACCACCACCCTCT

>M6-PGSC0003DMG400000386M6LRR-RLP_1260

CCCCCACCATCTCCACCTCCACCATCACCCCCACCACCAGTTTATTCACCACCACCACCGCCACCATCTCCCCCTCCACCATCACCTCCACCTCCTCCCCCACCCATTTATTCGCCACCA

>M6-PGSC0003DMG400000386M6LRR-RLP_1449

CCACCTCCTAATTCTCCACCTCCACCGCTTGCTCACTCGCCTCCACCCCCTTCACCTTACTATTACAATTCACCACCACCTCCGCCCCCAAATTCGCCGCCTCCAGCACCAAATTCGCCT

>M6-PGSC0003DMG400000386M6LRR-RLP_1512

TACAATTCACCACCACCTCCGCCCCCAAATTCGCCGCCTCCAGCACCAAATTCGCCTCCTCCTCCACCAGTATACATATACTCATCACCTCCGCCACCAAATTCACCTCCTCCTCCACCA

>M6-PGSC0003DMG400000386M6LRR-RLP_1575

CCACCAGTATACATATACTCATCACCTCCGCCACCAAATTCACCTCCTCCTCCACCACCAACATACGTATATTCTTCACCTCCTCCACCTCCTTCCCCATTGCCTTGTATAGAACCCCCT

>M6-PGSC0003DMG400000386M6LRR-RLP_189

AACCATGGTGACATTGCAGGGTACCTCCCGGAAGAGCTTGGGCTTCTTACAGATCTTGGGATTTTCCATATCAATTCAAACCGTTTCTGTGGTACAATACCAAGAAAATTCAGCAAGTTG

>M6-PGSC0003DMG400000386M6LRR-RLP_252

TTCCATATCAATTCAAACCGTTTCTGTGGTACAATACCAAGAAAATTCAGCAAGTTGAAGATACTGTTCGAGCTGGATCTGAGCAATAACAGGTTTGCTGGGAAGTTTCCTTATGTTGTT

>M6-PGSC0003DMG400000386M6LRR-RLP_315

CTGTTCGAGCTGGATCTGAGCAATAACAGGTTTGCTGGGAAGTTTCCTTATGTTGTTTTGAGTTTACCCAAGTTGATATTTTTGGATATTAGGTTCAATGAATTTGAAGGTAATGTACCT

>M6-PGSC0003DMG400000386M6LRR-RLP_378

TTACCCAAGTTGATATTTTTGGATATTAGGTTCAATGAATTTGAAGGTAATGTACCTTCACAGCTTTTTGATAAGCCATTAGACGCCATTTTTATTAATCATAATCGGTTTGCGTTTGAG

>M6-PGSC0003DMG400000386M6LRR-RLP_441

CTTTTTGATAAGCCATTAGACGCCATTTTTATTAATCATAATCGGTTTGCGTTTGAGTTGCCGGATAATTTTGGGAATTCGCCGGTTTCTGTGATAGTTCTGGCGAGTAACAGTTTCCAT

>M6-PGSC0003DMG400000386M6LRR-RLP_504

GATAATTTTGGGAATTCGCCGGTTTCTGTGATAGTTCTGGCGAGTAACAGTTTCCATGGGTGTCTTCCGGCGAGTATTGGAAATATGAGTAATCTTAATGAGGCGATTTTAATGAACAAT

>M6-PGSC0003DMG400000386M6LRR-RLP_567

CTTCCGGCGAGTATTGGAAATATGAGTAATCTTAATGAGGCGATTTTAATGAACAATGGGTTGCGTTCTTGTTTGCCGGCGGAGATTGGGTTGTTGAAAAATCTGACTGTATTTGATGTG

>M6-PGSC0003DMG400000386M6LRR-RLP_63

ATCATTTCAGATCCCTTTAATATCACTCTCAATTGGGTTGGTCCTAATGTATGCAGCTACACCGGAATCTTCTGTGCTCCGGCGTTGGATAACCCCAAAATCCGTACAGTTGCCGGCATT

>M6-PGSC0003DMG400000386M6LRR-RLP_693

AATCAGTTGATGGGTCCGTTGCCGGAGAATTTTGGTGATTTGGTGAATTTGGAGCAATTGAATGTGGCACATAATATGCTTTCTGGAACAATTCCGAAAAGAATTTGTCAGCTTCCTAAG

>M6-PGSC0003DMG400000386M6LRR-RLP_756

GTGGCACATAATATGCTTTCTGGAACAATTCCGAAAAGAATTTGTCAGCTTCCTAAGCTTGAGAATTTTACGTATTCTTATAATTTCTTCACTGGTGAACCGCCGGTGTGTTTGGGGTTG

>M6-PGSC0003DMG400000386M6LRR-RLP_882

TTTCATGATGAACGGAATTGTTTGCCCAACAGACCGGTGCAACGCTCTCAGGGACAGTGTAAGGCCTTTTTGTCTAAAAAAATTCATTGTAGTGCTTTCAGGTGTCATAAATTTGTTCCT

>M6-PGSC0003DMG400000386M6LRR-RLP_945

GCCTTTTTGTCTAAAAAAATTCATTGTAGTGCTTTCAGGTGTCATAAATTTGTTCCTGTTTTGCCACCTCCTCCGCCACTTTCACCCCCCTTGCCTGCCCCTCCGCCACCTATTTATTCG

>M6-PGSC0003DMG400000578M6L-LecRK_0

CAATCATTAAAGCATTAAAATTTACATTCCATCTGAATTACTATATTTTGTTTACTGAAACTGAAATAGGACAAAAAATTGCATTATGTACGTGAGGACTTTAAGTTCTCTTGTGTTTGG

>M6-PGSC0003DMG400000578M6L-LecRK_1008

TATATGTTTGTTGGATTTTCAGCTGCTACTGGTAACCATACACAAATCCACAACATTCTGTCATGGAATTTCACCTCAATTAGTCAAGCTTCGCTTCGAATTCCTTCAACAGAGACGTGC

>M6-PGSC0003DMG400000578M6L-LecRK_1071

TGGAATTTCACCTCAATTAGTCAAGCTTCGCTTCGAATTCCTTCAACAGAGACGTGCCAGAACAAAATCATGCTTCAAAATAGCACACAATCTGAAATTGATCATCGGAAAACTCCTAAT

>M6-PGSC0003DMG400000578M6L-LecRK_1134

AAAATCATGCTTCAAAATAGCACACAATCTGAAATTGATCATCGGAAAACTCCTAATAGTTTCTTTATTTTCCTTGCTGTTGTCATTCTTCTGGTTATCGTTCTTATTAACCTCTATTTC

>M6-PGSC0003DMG400000578M6L-LecRK_1197

TTTATTTTCCTTGCTGTTGTCATTCTTCTGGTTATCGTTCTTATTAACCTCTATTTCAGTAGCTACAAGAGAGATGACAATTCCGACGAAGCATTCACTCTGCCCCAGAAAAAGCAGAGA

>M6-PGSC0003DMG400000578M6L-LecRK_126

GTGAAATGAAGTTATCTATAGACCACATCCTTATGTACAACCTTGACAATTGATAATTTGCTACTCAAAAAATTCAAATCCCAAAAAACTTTATCACTAGCCAAGAATTAGCTCATCATG

>M6-PGSC0003DMG400000578M6L-LecRK_1260

TACAAGAGAGATGACAATTCCGACGAAGCATTCACTCTGCCCCAGAAAAAGCAGAGACCGCGGCCACCAAACAAGGCACGTCGCTTCACAATAGCTGAGATCTCCATCGCAACAAGGAAT

>M6-PGSC0003DMG400000578M6L-LecRK_1323

CCACCAAACAAGGCACGTCGCTTCACAATAGCTGAGATCTCCATCGCAACAAGGAATTTCAGCGAGTTACAAATATTAGGCAGTGATGAGAAGAGCATTACGTATAAGGCCACGATCCTG

>M6-PGSC0003DMG400000578M6L-LecRK_1386

GAGTTACAAATATTAGGCAGTGATGAGAAGAGCATTACGTATAAGGCCACGATCCTGAACGGGTGCAACGTTGTTGTAAAACGATTTTTAACTCAATTTTTCAACACACACGGGTTTGAA

>M6-PGSC0003DMG400000578M6L-LecRK_1449

TGCAACGTTGTTGTAAAACGATTTTTAACTCAATTTTTCAACACACACGGGTTTGAAAAGCATCAATTTCACAAGGAAATCAAGGCTATCACCAGAATTCGTCACCCGAATTTGGTCCCA

>M6-PGSC0003DMG400000578M6L-LecRK_1512

CAATTTCACAAGGAAATCAAGGCTATCACCAGAATTCGTCACCCGAATTTGGTCCCAATTAGAGGATGGTGCTATGACAACCAAGAAACAATAGTTGTGTACGATTTCATCCCCAATGGA

>M6-PGSC0003DMG400000578M6L-LecRK_1575

GGATGGTGCTATGACAACCAAGAAACAATAGTTGTGTACGATTTCATCCCCAATGGAAGCCTCGATAAATGGCTGTTCGGCGTTGGCGTCTTGCCTTGGACGAGACGTTTTAAGGTCGTT

>M6-PGSC0003DMG400000578M6L-LecRK_1638

GATAAATGGCTGTTCGGCGTTGGCGTCTTGCCTTGGACGAGACGTTTTAAGGTCGTTAAAGATTTAGCAGATTCTCTTGTCTACCTTCATTCAAAGCAACTCGCTCACAAAAACGTTAAA

>M6-PGSC0003DMG400000578M6L-LecRK_1701

TTAGCAGATTCTCTTGTCTACCTTCATTCAAAGCAACTCGCTCACAAAAACGTTAAAAGTAGCAGCGTGTTTCTTGACGTGAGATTCAGAGCAGTAGTGGGTGATTTCGGGTTTGTGCTT

>M6-PGSC0003DMG400000578M6L-LecRK_1764

AGCGTGTTTCTTGACGTGAGATTCAGAGCAGTAGTGGGTGATTTCGGGTTTGTGCTTACTTCAGCCGGGTCAACCCGGTTTGAGGCCATGGTGAGTCAGACAGCTGATGTGTTCGAATTC

>M6-PGSC0003DMG400000578M6L-LecRK_1827

GCCGGGTCAACCCGGTTTGAGGCCATGGTGAGTCAGACAGCTGATGTGTTCGAATTCGGAGTAGTCGTGCTAGAGATCATTGCGGGTCGGAGCAGAAAGTCCAACCCGGGAGAGCGAGAC

>M6-PGSC0003DMG400000578M6L-LecRK_189

CTCAAAAAATTCAAATCCCAAAAAACTTTATCACTAGCCAAGAATTAGCTCATCATGGATTACTTCTTCTGGTTTTTCTTCTCGTTTGCCATACTAGTTAATAGTAATTTAGGTTCGTCC

>M6-PGSC0003DMG400000578M6L-LecRK_1890

GTCGTGCTAGAGATCATTGCGGGTCGGAGCAGAAAGTCCAACCCGGGAGAGCGAGACTTGTTGGATCTTGCATGGGCAATGCATGAGGTACAACAGAAGGAAACTCTAGTGGATCGAAGA

>M6-PGSC0003DMG400000578M6L-LecRK_1953

GATCTTGCATGGGCAATGCATGAGGTACAACAGAAGGAAACTCTAGTGGATCGAAGAATGGGCGCAGTCGTGAACCTGGAGCAGGCGATTCGGGCTTTGGATATCGGGTTGCTCTGCACG

>M6-PGSC0003DMG400000578M6L-LecRK_2016

GCAGTCGTGAACCTGGAGCAGGCGATTCGGGCTTTGGATATCGGGTTGCTCTGCACGTTGAACGAGAACAAAGGAAGGCCTACTATGGAAGAAGTGGTGGAGTTCCTGAATATGGAGAAA

>M6-PGSC0003DMG400000578M6L-LecRK_2079

GAGAACAAAGGAAGGCCTACTATGGAAGAAGTGGTGGAGTTCCTGAATATGGAGAAACCGATACCGGAGTTGCCATCGGGTCGACCCGTTTGTTTGTTCCCGTACAGCAGCACCACAGGC

>M6-PGSC0003DMG400000578M6L-LecRK_2142

CCGGAGTTGCCATCGGGTCGACCCGTTTGTTTGTTCCCGTACAGCAGCACCACAGGCTTATGCAGTGGGTACGCTTGTACAGCATTCAAATGAAGTCATGTTTTTTGCTTGACCAATTGG

>M6-PGSC0003DMG400000578M6L-LecRK_2205

AGTGGGTACGCTTGTACAGCATTCAAATGAAGTCATGTTTTTTGCTTGACCAATTGGTTCTTCGGCGTTCCACGTAAGCAGCAATTATTCTTATCTGTTATACTACAAAATAATGACAGG

>M6-PGSC0003DMG400000578M6L-LecRK_2268

GGCGTTCCACGTAAGCAGCAATTATTCTTATCTGTTATACTACAAAATAATGACAGGGAAGCACATGTATATGCTATGTGACGTTATTTTCATGACATAGACAACTATTGTCTGATCTGT

>M6-PGSC0003DMG400000578M6L-LecRK_2331

CATGTATATGCTATGTGACGTTATTTTCATGACATAGACAACTATTGTCTGATCTGTGGTCGGAGTTAAATATCACAAGAACTCCTAAAACGATTTGAAATTATTGTAAATTATGTACTG

>M6-PGSC0003DMG400000578M6L-LecRK_252

TTCTTCTGGTTTTTCTTCTCGTTTGCCATACTAGTTAATAGTAATTTAGGTTCGTCCGCGGACGGACCTTCAATTAACGTGACGAAGCATATCTCTTTTCAAGATTTCAGTTCTAGTAAT

>M6-PGSC0003DMG400000578M6L-LecRK_315

GGACCTTCAATTAACGTGACGAAGCATATCTCTTTTCAAGATTTCAGTTCTAGTAATCCCAGATTGAAGCAAGATCTTACACTTCTCGGTAGTGTTATCGTCTCGGATGAAAAAAAATCC

>M6-PGSC0003DMG400000578M6L-LecRK_378

TTGAAGCAAGATCTTACACTTCTCGGTAGTGTTATCGTCTCGGATGAAAAAAAATCCGTTCAAATTCCTGATCCTGAGCGAGAAGGTGATGATCTAAAGCATCTAGCAGGACGAGCTATT

>M6-PGSC0003DMG400000578M6L-LecRK_441

ATTCCTGATCCTGAGCGAGAAGGTGATGATCTAAAGCATCTAGCAGGACGAGCTATTTATTCTTCACCTATCCGTTTCTTTGATCCTCGGACTCAAACACCGGCTTCTTTTGAAACGACT

>M6-PGSC0003DMG400000578M6L-LecRK_504

TCACCTATCCGTTTCTTTGATCCTCGGACTCAAACACCGGCTTCTTTTGAAACGACTTTCTCGTTTCAATTTCAAGTGAAGTCCAATTCAAGCAATGAGACGTCAGATCAGGGGAAATCT

>M6-PGSC0003DMG400000578M6L-LecRK_567

TTTCAATTTCAAGTGAAGTCCAATTCAAGCAATGAGACGTCAGATCAGGGGAAATCTGTTGGTGGTAGTGGCCTTACTTTTATAATCGTTCCGGATGAATTGACTGTTGGCCGTGCTGGT

>M6-PGSC0003DMG400000578M6L-LecRK_63

AAATAGGACAAAAAATTGCATTATGTACGTGAGGACTTTAAGTTCTCTTGTGTTTGGAAGAAAGTGAAATGAAGTTATCTATAGACCACATCCTTATGTACAACCTTGACAATTGATAAT

>M6-PGSC0003DMG400000578M6L-LecRK_630

GGTAGTGGCCTTACTTTTATAATCGTTCCGGATGAATTGACTGTTGGCCGTGCTGGTCCGTGGCTTGGGATGCTGAACGATTTGTGCGATGAGGATTATAAAACAGTAGCTATTGAGTTT

>M6-PGSC0003DMG400000578M6L-LecRK_693

CTTGGGATGCTGAACGATTTGTGCGATGAGGATTATAAAACAGTAGCTATTGAGTTTGATACTCGGCAGAACCCTGAATTTGGTGACCCGAATGATAATCACTTGGGCATTAATTTGGGT

>M6-PGSC0003DMG400000578M6L-LecRK_756

CGGCAGAACCCTGAATTTGGTGACCCGAATGATAATCACTTGGGCATTAATTTGGGTAGCATAGTATCAACTGCAGCGATTAATGCTTCGGATGCCGGAGTTCAATTGAATGATGGATCA

>M6-PGSC0003DMG400000578M6L-LecRK_819

GTATCAACTGCAGCGATTAATGCTTCGGATGCCGGAGTTCAATTGAATGATGGATCAATTCACAGAGTATGGATATCTTATGATGGGCGAAAGCGATTTGTTGAAATTCGTCTTGCTCCT

>M6-PGSC0003DMG400000578M6L-LecRK_882

AGAGTATGGATATCTTATGATGGGCGAAAGCGATTTGTTGAAATTCGTCTTGCTCCTGATGGCAGAGGATATCCTTCTAAACCAGTTTACTCTGGTTTACTTGATCTTTCACCTTACTTG

>M6-PGSC0003DMG400000578M6L-LecRK_945

AGAGGATATCCTTCTAAACCAGTTTACTCTGGTTTACTTGATCTTTCACCTTACTTGAATGAGTATATGTTTGTTGGATTTTCAGCTGCTACTGGTAACCATACACAAATCCACAACATT

>M6-PGSC0003DMG400000728-(rev)M6LRR-RLK_0

ATCAGTGTATTTCGTGTCTCTGAATTAGGTGGGACAAGATTCCCATAAAAGGCTCTCAAATTTTCAACCAAAAATTCAGTGCAGCACAAACACTATGCACGTTAAAGGTGCTTTGCAACT

>M6-PGSC0003DMG400000728-(rev)M6LRR-RLK_1037

ATAGACTAAAGCTTGGTATTTAAGTGGAGAAAGGTAGAAGGCTGGGCCCATTATCCACCGACTTTTGGACCGTGCACCGAAAGAGCTGGGAGGATTTCTCAGTTACAAACAAAAGATCCT

>M6-PGSC0003DMG400000728-(rev)M6LRR-RLK_1098

CTTTTGGACCGTGCACCGAAAGAGCTGGGAGGATTTCTCAGTTACAAACAAAAGATCCTTTCATGGGGCATCTAATAGACTTTCTTGAACCGAATATCCATTCCAGAAGGAGTTTCCAAA

>M6-PGSC0003DMG400000728-(rev)M6LRR-RLK_1159

CATGGGGCATCTAATAGACTTTCTTGAACCGAATATCCATTCCAGAAGGAGTTTCCAAAGGAGATGCTTCTCTAAATGTCCGCATAGTTTTTCTGCTGCAGTTCACCATGAAATTCTCAA

>M6-PGSC0003DMG400000728-(rev)M6LRR-RLK_122

ATGGATTGTGCCCATGTCTGGTTGACTAAAAATCATCTAAAGCCTATTTATAAATGATAATATTAGATGTCTGCTTCGCTCTTAGCTTCTCAGGATTTATAAAAGGAGGCAATGCATGGT

>M6-PGSC0003DMG400000728-(rev)M6LRR-RLK_1220

AGATGCTTCTCTAAATGTCCGCATAGTTTTTCTGCTGCAGTTCACCATGAAATTCTCAATGATTTAATTGCATGAATTAGTTTCTTGGATTCAACTCTTCCCTCAACAACCCACCCCCCA

>M6-PGSC0003DMG400000728-(rev)M6LRR-RLK_1281

ATTTAATTGCATGAATTAGTTTCTTGGATTCAACTCTTCCCTCAACAACCCACCCCCCACCCGCCCAAAAAAAGAACTTTATTTCTGTCAGAAGTTTGCTAGTGCATCATGTAGTGACAT

>M6-PGSC0003DMG400000728-(rev)M6LRR-RLK_1342

CGCCCAAAAAAAGAACTTTATTTCTGTCAGAAGTTTGCTAGTGCATCATGTAGTGACATCTTCTTTTCCTATGCCTACTATTCTCACAGAGATGTTCAAGCAAGCAGCATACTTCTTGAT

>M6-PGSC0003DMG400000728-(rev)M6LRR-RLK_1403

TCTTTTCCTATGCCTACTATTCTCACAGAGATGTTCAAGCAAGCAGCATACTTCTTGATGATAAGTTCGAAGTACGGCTAGGGAGTCTAAATGAAGCCTGTGCACAAGAAGGAGAGAGCC

>M6-PGSC0003DMG400000728-(rev)M6LRR-RLK_1464

TAAGTTCGAAGTACGGCTAGGGAGTCTAAATGAAGCCTGTGCACAAGAAGGAGAGAGCCATCAAAATCGAATTTCCAGGCTGCTTAGATTTCCACAGTAAGTACTTGTTGGTAAATTCTG

>M6-PGSC0003DMG400000728-(rev)M6LRR-RLK_1525

CAAAATCGAATTTCCAGGCTGCTTAGATTTCCACAGTAAGTACTTGTTGGTAAATTCTGTGTTCTAACCATCCTTTTCTTTTCGAAGGCTAACCACTATTACTACTAGGTTGTTATGTAA

>M6-PGSC0003DMG400000728-(rev)M6LRR-RLK_1586

TTCTAACCATCCTTTTCTTTTCGAAGGCTAACCACTATTACTACTAGGTTGTTATGTAATGTTCAATAATTTTGTTTCAATTTTTGTCTCCCATGATGGATCTTAAAATCAGGCAGTTTT

>M6-PGSC0003DMG400000728-(rev)M6LRR-RLK_1647

TTCAATAATTTTGTTTCAATTTTTGTCTCCCATGATGGATCTTAAAATCAGGCAGTTTTATAGTTAATATGTCTGTATCTCCAATTTGATGTTATAGAAACTACTTTGGCCTGGTTTCTA

>M6-PGSC0003DMG400000728-(rev)M6LRR-RLK_1708

AGTTAATATGTCTGTATCTCCAATTTGATGTTATAGAAACTACTTTGGCCTGGTTTCTAAATTGGATTTGAGTTCCACAGTTTTGCTGCCACTCAAGTCTTTCTGGTGTATATTTATTAG

>M6-PGSC0003DMG400000728-(rev)M6LRR-RLK_1769

TTGGATTTGAGTTCCACAGTTTTGCTGCCACTCAAGTCTTTCTGGTGTATATTTATTAGCTCTGATATAAATGAGAATCCTAACGTTCATGCAGCAACCTTACTCTTAACCTTGTCATCT

>M6-PGSC0003DMG400000728-(rev)M6LRR-RLK_183

ATTAGATGTCTGCTTCGCTCTTAGCTTCTCAGGATTTATAAAAGGAGGCAATGCATGGTTGATGGTTTTCTGATTTGCTATCAGTTGGAGATCTGTAATGTTTATTGCCTCGTGAGACTA

>M6-PGSC0003DMG400000728-(rev)M6LRR-RLK_1830

CTGATATAAATGAGAATCCTAACGTTCATGCAGCAACCTTACTCTTAACCTTGTCATCTCGTTATCTGTGCTTATACAAAAGGGCTTTAGCTTTGTTTCTAAATTAGAAAAACTTGTAAT

>M6-PGSC0003DMG400000728-(rev)M6LRR-RLK_1891

TTATCTGTGCTTATACAAAAGGGCTTTAGCTTTGTTTCTAAATTAGAAAAACTTGTAATATAGGCTCTGAATAGTTTAGTAGTCCTGATGATAGTTATCATGCAGTTGCATATAATCTTA

>M6-PGSC0003DMG400000728-(rev)M6LRR-RLK_1952

AGGCTCTGAATAGTTTAGTAGTCCTGATGATAGTTATCATGCAGTTGCATATAATCTTATTTCTTCTCAGTATTCAAGTTACAAGGAACACTGTAAAGAACAGTGGGTCTAGTCTGAAAG

>M6-PGSC0003DMG400000728-(rev)M6LRR-RLK_2013

TCTTCTCAGTATTCAAGTTACAAGGAACACTGTAAAGAACAGTGGGTCTAGTCTGAAAGACCATCACCCACTTGTTAATTACTTGTTTTATCCCTCAGTAAATTATTCCTTTTCTATGTT

>M6-PGSC0003DMG400000728-(rev)M6LRR-RLK_2074

CATCACCCACTTGTTAATTACTTGTTTTATCCCTCAGTAAATTATTCCTTTTCTATGTTGTATGAATCTTTTTGCCTTGATTTACCTATGTCTAGCGGGTGTCACATGGGTGTTGTTGTT

>M6-PGSC0003DMG400000728-(rev)M6LRR-RLK_2135

ATGAATCTTTTTGCCTTGATTTACCTATGTCTAGCGGGTGTCACATGGGTGTTGTTGTTTAAGTAGAAATTAGCAAAAAACTGTATGTACCCTTACTGGATACTTAAAACTCATGTATGG

>M6-PGSC0003DMG400000728-(rev)M6LRR-RLK_2196

AGTAGAAATTAGCAAAAAACTGTATGTACCCTTACTGGATACTTAAAACTCATGTATGGGATATGCATGCGTAGCCGAATCCATGAAACATAGTCCTTTGATGTACATGTTTTCACTTAA

>M6-PGSC0003DMG400000728-(rev)M6LRR-RLK_2257

TATGCATGCGTAGCCGAATCCATGAAACATAGTCCTTTGATGTACATGTTTTCACTTAAGGAGAATTTGTTGGTTATGCTATTTGGTAGGGAATGCATAGTTCTCATTCTTTTAGCTTTC

>M6-PGSC0003DMG400000728-(rev)M6LRR-RLK_2318

AGAATTTGTTGGTTATGCTATTTGGTAGGGAATGCATAGTTCTCATTCTTTTAGCTTTCCATCCTGTGCTGAAGTAAGTTTAACTATTTCTCACTCTCATTGTCTAGCTTTAACTATTAC

>M6-PGSC0003DMG400000728-(rev)M6LRR-RLK_2379

TCCTGTGCTGAAGTAAGTTTAACTATTTCTCACTCTCATTGTCTAGCTTTAACTATTACCTATTCTCGTTATGCCTACACATTGAATGCAGGAAATTATTAAATTTTGTATTGTGGTACT

>M6-PGSC0003DMG400000728-(rev)M6LRR-RLK_244

ATGGTTTTCTGATTTGCTATCAGTTGGAGATCTGTAATGTTTATTGCCTCGTGAGACTATCATTATACAGAGTTCATCACATAAGGAATAGCCTCCACAATATCTTGGACCTGGAGCTTG

>M6-PGSC0003DMG400000728-(rev)M6LRR-RLK_2440

ATTCTCGTTATGCCTACACATTGAATGCAGGAAATTATTAAATTTTGTATTGTGGTACTAAATACTTCTGGAACGACCTAGATATAATGGCATGTGAAAAATTGGTTTGAAGTCCCATGT

>M6-PGSC0003DMG400000728-(rev)M6LRR-RLK_2501

ATACTTCTGGAACGACCTAGATATAATGGCATGTGAAAAATTGGTTTGAAGTCCCATGTTAGATGCCAAAGATTTTAGGCCTTCTATTCTATTATTTTAATCTTCATGGCTTCTGTTAAA

>M6-PGSC0003DMG400000728-(rev)M6LRR-RLK_2562

GATGCCAAAGATTTTAGGCCTTCTATTCTATTATTTTAATCTTCATGGCTTCTGTTAAAACTTTCCAGGTTCTCTTATTTTAGAAATTGAATTTACGACCACTGATCACAGTAATCCAGT

>M6-PGSC0003DMG400000728-(rev)M6LRR-RLK_2623

TTTCCAGGTTCTCTTATTTTAGAAATTGAATTTACGACCACTGATCACAGTAATCCAGTAAGCTTTTGTTTTATAGTGAAAACTTTTCCATATGTATTTCCTGTCTTACAAATGGTTATC

>M6-PGSC0003DMG400000728-(rev)M6LRR-RLK_2684

GCTTTTGTTTTATAGTGAAAACTTTTCCATATGTATTTCCTGTCTTACAAATGGTTATCACTGTTTTTCCAGCTTTTCGTCGAAGCTTGTTGCATGATCTCAAAAGTTTAGTTATATTCT

>M6-PGSC0003DMG400000728-(rev)M6LRR-RLK_2745

TGTTTTTCCAGCTTTTCGTCGAAGCTTGTTGCATGATCTCAAAAGTTTAGTTATATTCTTTCAACCAGTAACTGTTTAATAGTTTATACTGCTTATTGACATAGTTCTCCCTTTCTTATT

>M6-PGSC0003DMG400000728-(rev)M6LRR-RLK_2806

CAACCAGTAACTGTTTAATAGTTTATACTGCTTATTGACATAGTTCTCCCTTTCTTATTTGCAGGACGTCGGAACAAGGTGCATCAGGTATGTGCAATTTTATTTAGTTCTACTTTCTTT

>M6-PGSC0003DMG400000728-(rev)M6LRR-RLK_2867

CAGGACGTCGGAACAAGGTGCATCAGGTATGTGCAATTTTATTTAGTTCTACTTTCTTTAAATAGAGTTTACTTGGATATTCTTAATATTTATTAAATTGGTGTAAGTAACTATTTAAAA

>M6-PGSC0003DMG400000728-(rev)M6LRR-RLK_2928

ATAGAGTTTACTTGGATATTCTTAATATTTATTAAATTGGTGTAAGTAACTATTTAAAATTATTTGCGTCACATTTCCAGTATTGTTTGTTGGTGGGTAAAATTTCCTTAATGTCTAGGC

>M6-PGSC0003DMG400000728-(rev)M6LRR-RLK_2989

ATTTGCGTCACATTTCCAGTATTGTTTGTTGGTGGGTAAAATTTCCTTAATGTCTAGGCAACAAATCAAAACTATAGATTCTCATTAGAAGTAAGAATAAGTGAGACTATTAGGTGGAAA

>M6-PGSC0003DMG400000728-(rev)M6LRR-RLK_305

ATTATACAGAGTTCATCACATAAGGAATAGCCTCCACAATATCTTGGACCTGGAGCTTGACTCTATCAACTTGAGGGCTTCACTTGCTATCTTTATGAGCTACACCTGCATGTAATAAAA

>M6-PGSC0003DMG400000728-(rev)M6LRR-RLK_3050

CAAATCAAAACTATAGATTCTCATTAGAAGTAAGAATAAGTGAGACTATTAGGTGGAAATTTACGATATAAAGCTGATTCCAAAGAGTGTAAAGGGCTGGACAACTGTATATTGAACTGT

>M6-PGSC0003DMG400000728-(rev)M6LRR-RLK_3111

TACGATATAAAGCTGATTCCAAAGAGTGTAAAGGGCTGGACAACTGTATATTGAACTGTACTGCCTAAAGAACTGGAGATTTTGATGGCGGTGTTTTGCTTTACTTTTTGTGGGATCTTT

>M6-PGSC0003DMG400000728-(rev)M6LRR-RLK_3172

TGCCTAAAGAACTGGAGATTTTGATGGCGGTGTTTTGCTTTACTTTTTGTGGGATCTTTGTAGCTTGAACTATCACTAGGGAATACTTCAAAGTCTACTCCTCTCGTCCTAATATTCTTT

>M6-PGSC0003DMG400000728-(rev)M6LRR-RLK_3233

AGCTTGAACTATCACTAGGGAATACTTCAAAGTCTACTCCTCTCGTCCTAATATTCTTTCTTACCTGAATGGACAGGTACACCTAGTGCCACTTGTGCGTATGATGTCTACTGTTTTGGA

>M6-PGSC0003DMG400000728-(rev)M6LRR-RLK_3294

TACCTGAATGGACAGGTACACCTAGTGCCACTTGTGCGTATGATGTCTACTGTTTTGGAAAGGTTTTGCTCGAGCTAGTAACTGGCAAGCTAGGTATCAGTGCATCCAATGACGCCAGCA

>M6-PGSC0003DMG400000728-(rev)M6LRR-RLK_3355

GGTTTTGCTCGAGCTAGTAACTGGCAAGCTAGGTATCAGTGCATCCAATGACGCCAGCATGAAGGAGTGGTTGGATGGTACACTTAAGTACATTAGTATATATGATAAAGAACTTGTCAC

>M6-PGSC0003DMG400000728-(rev)M6LRR-RLK_3416

AAGGAGTGGTTGGATGGTACACTTAAGTACATTAGTATATATGATAAAGAACTTGTCACAAACATTGTGGATCCTTCCCTAATCATAGACGAAGATTTATTAGAGGAAGTATGGGCGATG

>M6-PGSC0003DMG400000728-(rev)M6LRR-RLK_3477

ACATTGTGGATCCTTCCCTAATCATAGACGAAGATTTATTAGAGGAAGTATGGGCGATGGCTATTGTTGCCAGATCCTGCCTCAATCCCAAACCGTCAAGGAGACCCTTAATGAGATACA

>M6-PGSC0003DMG400000728-(rev)M6LRR-RLK_3538

TATTGTTGCCAGATCCTGCCTCAATCCCAAACCGTCAAGGAGACCCTTAATGAGATACATCCTTAAAGCTTTGGAAAACCCTTTGAAGGTAGTTAGGGAAGAACACACTAGCTCAGCAAG

>M6-PGSC0003DMG400000728-(rev)M6LRR-RLK_3599

CTTAAAGCTTTGGAAAACCCTTTGAAGGTAGTTAGGGAAGAACACACTAGCTCAGCAAGACTTAGAGCAACTTCATCGAGAAGTTCATGGAACGCTGCACTGTTTGGCAGCTGGCGTAGC

>M6-PGSC0003DMG400000728-(rev)M6LRR-RLK_366

TCTATCAACTTGAGGGCTTCACTTGCTATCTTTATGAGCTACACCTGCATGTAATAAAAGTAAGAGTTTAGGAAACCTAAAACCTGTCCTTTTAGGGCTTAGCAATGTAACGTCAAGAGC

>M6-PGSC0003DMG400000728-(rev)M6LRR-RLK_3660

TTAGAGCAACTTCATCGAGAAGTTCATGGAACGCTGCACTGTTTGGCAGCTGGCGTAGCTCCTCAGATGTAGCAGCCGTACCTGCTGCAGCTTCAGCACATAAGTTGGAAGGAACAAGTA

>M6-PGSC0003DMG400000728-(rev)M6LRR-RLK_3721

CTCAGATGTAGCAGCCGTACCTGCTGCAGCTTCAGCACATAAGTTGGAAGGAACAAGTAGTTTGAAGCAATCCGGAACAACAGGCTCTCAAGGAAGTGGTCCTAATGGGGACAACAACGG

>M6-PGSC0003DMG400000728-(rev)M6LRR-RLK_3782

TTGAAGCAATCCGGAACAACAGGCTCTCAAGGAAGTGGTCCTAATGGGGACAACAACGGTCATTCTTCATCAACTAGACGACAGTCCAAAGAGATTTTCCCTGAGCCATTGGAGGAGCAC

>M6-PGSC0003DMG400000728-(rev)M6LRR-RLK_3843

ATTCTTCATCAACTAGACGACAGTCCAAAGAGATTTTCCCTGAGCCATTGGAGGAGCACGATGTAGAGAGAGGACCACTGGCGGAGTAGAAGGGAGGGTGATTGTCATTAATTCTTTGTT

>M6-PGSC0003DMG400000728-(rev)M6LRR-RLK_3904

TGTAGAGAGAGGACCACTGGCGGAGTAGAAGGGAGGGTGATTGTCATTAATTCTTTGTTAATAATGCAGCAGGATCTGCATTAAGGTTGCATTTTTGTTTGGGGACACCTTGTACATTCT

>M6-PGSC0003DMG400000728-(rev)M6LRR-RLK_3965

TAATGCAGCAGGATCTGCATTAAGGTTGCATTTTTGTTTGGGGACACCTTGTACATTCTCCGTTTCCGGAATTCTTGAGTTTGCAAGCCATTGTCTTGCCCTGATCTTGAAAATGGAGAT

>M6-PGSC0003DMG400000728-(rev)M6LRR-RLK_4026

GTTTCCGGAATTCTTGAGTTTGCAAGCCATTGTCTTGCCCTGATCTTGAAAATGGAGATGCATAGTTTACCTGGAGACAGCAAACATCAAATTGTTTATTGGGGTTTTGGGATTTTAAGT

>M6-PGSC0003DMG400000728-(rev)M6LRR-RLK_4087

ATAGTTTACCTGGAGACAGCAAACATCAAATTGTTTATTGGGGTTTTGGGATTTTAAGTTCTTTTTCTTTTTCTGGGGATAGATTTGGAAAACAGGATTTAGTAGTGGTGGTGGTTCTTT

>M6-PGSC0003DMG400000728-(rev)M6LRR-RLK_4148

TTTTTCTTTTTCTGGGGATAGATTTGGAAAACAGGATTTAGTAGTGGTGGTGGTTCTTTTTGTTAATTTTGTTTTTGGGGGGTTTATGTTTGCTTCCAAATTCCAATTCTTTTCTTATAA

>M6-PGSC0003DMG400000728-(rev)M6LRR-RLK_4209

GTTAATTTTGTTTTTGGGGGGTTTATGTTTGCTTCCAAATTCCAATTCTTTTCTTATAATTCTTCTCTTTTACTAGTCTCTCCATGGAATGTAAATTATACACAATCTTTCATTTTCCAA

>M6-PGSC0003DMG400000728-(rev)M6LRR-RLK_4244

CAAATTCCAATTCTTTTCTTATAATTCTTCTCTTTTACTAGTCTCTCCATGGAATGTAAATTATACACAATCTTTCATTTTCCAATTGGAGGAGAGGGAAAAAAAGAAAAAGAGTATCCC

>M6-PGSC0003DMG400000728-(rev)M6LRR-RLK_427

AAGAGTTTAGGAAACCTAAAACCTGTCCTTTTAGGGCTTAGCAATGTAACGTCAAGAGCCATGCACTTCTAAAATTGCTGTAAGCTCAGCTTCTATAGTTCCATGCATGGCTGCGATATC

>M6-PGSC0003DMG400000728-(rev)M6LRR-RLK_488

TGCACTTCTAAAATTGCTGTAAGCTCAGCTTCTATAGTTCCATGCATGGCTGCGATATCTCTAATCTAGGCAGTTTCAATTGTTAGATTCGGTATGAGGGATGTGGCCTTAGAAGATCTT

>M6-PGSC0003DMG400000728-(rev)M6LRR-RLK_549

TAATCTAGGCAGTTTCAATTGTTAGATTCGGTATGAGGGATGTGGCCTTAGAAGATCTTGATGACATTATAGTTCTACTGTGCAGAGATATTGTGAATGCAGAAATAGAAAATTTCTCAA

>M6-PGSC0003DMG400000728-(rev)M6LRR-RLK_61

TTTCAACCAAAAATTCAGTGCAGCACAAACACTATGCACGTTAAAGGTGCTTTGCAACTTTATGGATTGTGCCCATGTCTGGTTGACTAAAAATCATCTAAAGCCTATTTATAAATGATA

>M6-PGSC0003DMG400000728-(rev)M6LRR-RLK_610

TGACATTATAGTTCTACTGTGCAGAGATATTGTGAATGCAGAAATAGAAAATTTCTCAAACTAAGAACAGTTATGTTTTTTTTTAAAATTTTTTCTTTTCTTATTATTGAACCTTCTCCA

>M6-PGSC0003DMG400000728-(rev)M6LRR-RLK_671

TAAGAACAGTTATGTTTTTTTTTAAAATTTTTTCTTTTCTTATTATTGAACCTTCTCCATACATCAATCCTTTTTTTTTTGATAACCGTGGTGCCCGGGCTAGCTTGTGCACACCTTGAC

>M6-PGSC0003DMG400000728-(rev)M6LRR-RLK_732

CATCAATCCTTTTTTTTTTGATAACCGTGGTGCCCGGGCTAGCTTGTGCACACCTTGACACAATCCCATCAATCTTAATCAACTAAAGGTAGTCTGCATATGACACTCCTCATCTTTTGT

>M6-PGSC0003DMG400000728-(rev)M6LRR-RLK_793

AATCCCATCAATCTTAATCAACTAAAGGTAGTCTGCATATGACACTCCTCATCTTTTGTACCTTTTATCTCCATGGAACCTAAGCCAGTCAGTGAACTATCTAATTTCTCTAAAACTTTT

>M6-PGSC0003DMG400000728-(rev)M6LRR-RLK_854

CTTTTATCTCCATGGAACCTAAGCCAGTCAGTGAACTATCTAATTTCTCTAAAACTTTTCATCTGCATGTTTTGGGTTCAACCAATATTCTTTCACCGGGCAACCCCTAGGGATTTGGGT

>M6-PGSC0003DMG400000728-(rev)M6LRR-RLK_915

TCTGCATGTTTTGGGTTCAACCAATATTCTTTCACCGGGCAACCCCTAGGGATTTGGGTTAGTGGCAAGAGCACAACTCATTATGTGTGGGTTAAGCTCACGTTATCAGTTTGAATCCTA

>M6-PGSC0003DMG400000728-(rev)M6LRR-RLK_976

GTGGCAAGAGCACAACTCATTATGTGTGGGTTAAGCTCACGTTATCAGTTTGAATCCTACCATAGACTAAAGCTTGGTATTTAAGTGGAGAAAGGTAGAAGGCTGGGCCCATTATCCACC

>M6-PGSC0003DMG400000789-(rev)M6L-LecRK_1054

AATCAGCTACAGAATGTTTTAATCATGAAAGGATAATCGGAGAAGGAGCATCTGCTGTTGTGTATGAAGGGGAAATTCCTTCTAGGGGATCTGTGGCTGTTAAGAGATTTGTCCAAGGGA

>M6-PGSC0003DMG400000789-(rev)M6L-LecRK_1116

TATGAAGGGGAAATTCCTTCTAGGGGATCTGTGGCTGTTAAGAGATTTGTCCAAGGGAGTAGATTGGGTCCTTCACATATTCCTTTTAATACTGAATTTGCTTCTATGGTTGGCTGTTTA

>M6-PGSC0003DMG400000789-(rev)M6L-LecRK_1178

ATTGGGTCCTTCACATATTCCTTTTAATACTGAATTTGCTTCTATGGTTGGCTGTTTAAGACACAAGAATTTGATTCAGCTTCAAGGGTGGTGTTGTGAGAGAAATGAATTGGTGTTAGT

>M6-PGSC0003DMG400000789-(rev)M6L-LecRK_124

TCACCGCAAAATCAATCACCCTCACCCAAGAACGCAACTGTTCATCATCATCTTCAACACCTCCCATTTCTGGCATTGGAAGAGCTTTCTACACATACCCAGTTCGTTTTCTTGATTCTT

>M6-PGSC0003DMG400000789-(rev)M6L-LecRK_1240

ACAAGAATTTGATTCAGCTTCAAGGGTGGTGTTGTGAGAGAAATGAATTGGTGTTAGTGTATGAATTCATGCCTAATGGTAGCCTTGACAAAATCCTCCACGAGCGATCGCATTTGACCA

>M6-PGSC0003DMG400000789-(rev)M6L-LecRK_1302

GAATTCATGCCTAATGGTAGCCTTGACAAAATCCTCCACGAGCGATCGCATTTGACCAAGTTTCTGACTTGGGAGAGAAGATTGAACATAGTTATTGGTGTGTCATCTGCTCTTATGTAT

>M6-PGSC0003DMG400000789-(rev)M6L-LecRK_1364

TCTGACTTGGGAGAGAAGATTGAACATAGTTATTGGTGTGTCATCTGCTCTTATGTATCTTCATGAAGAGTGTGAGAATCATATAATCCACAGAGATGTTAAGAGTTGCAATATAATGCT

>M6-PGSC0003DMG400000789-(rev)M6L-LecRK_1426

ATGAAGAGTGTGAGAATCATATAATCCACAGAGATGTTAAGAGTTGCAATATAATGCTTGATGCTGAGTTCAATGCCAAGCTTGGAGATTTCGGTTTAGCAGAGGTGTTTGATAATTCTA

>M6-PGSC0003DMG400000789-(rev)M6L-LecRK_1488

GCTGAGTTCAATGCCAAGCTTGGAGATTTCGGTTTAGCAGAGGTGTTTGATAATTCTAAGACAAGGGATGCTACTGTACCAGCTGGAACAATGGGATATTTTGCACCTGAGTATGTCTAT

>M6-PGSC0003DMG400000789-(rev)M6L-LecRK_1550

AAGGGATGCTACTGTACCAGCTGGAACAATGGGATATTTTGCACCTGAGTATGTCTATACTGGTATTCCAACTGTTAAAACAGATGTGTATAGCTTTGGTGTTGTGGTACTGGAAGTGGC

>M6-PGSC0003DMG400000789-(rev)M6L-LecRK_1612

GTATTCCAACTGTTAAAACAGATGTGTATAGCTTTGGTGTTGTGGTACTGGAAGTGGCATCAGGTAGAAAGCCTATCGATGAAGGTGGAGGTTTGATTACTGATTGGGTGTGGGACTTGT

>M6-PGSC0003DMG400000789-(rev)M6L-LecRK_1674

GGTAGAAAGCCTATCGATGAAGGTGGAGGTTTGATTACTGATTGGGTGTGGGACTTGTGGGAGAAAGGGAGGATAACTGAGGCTGCTGATCCTAAACTAATGGGGCGGTTTCAGAAGAAC

>M6-PGSC0003DMG400000789-(rev)M6L-LecRK_1736

GAAAGGGAGGATAACTGAGGCTGCTGATCCTAAACTAATGGGGCGGTTTCAGAAGAACGAGATGGATAGAATGCTCATTGTGGGACTTTCTTGTGTGCATCCCGATCATGAGAAGAGACC

>M6-PGSC0003DMG400000789-(rev)M6L-LecRK_1798

TGGATAGAATGCTCATTGTGGGACTTTCTTGTGTGCATCCCGATCATGAGAAGAGACCGCGAATAAGAGATGTTTTCCGTATGTTTAAAGATGAAGCTCCACTTCTCATTTTACCTCCAA

>M6-PGSC0003DMG400000789-(rev)M6L-LecRK_186

CCCATTTCTGGCATTGGAAGAGCTTTCTACACATACCCAGTTCGTTTTCTTGATTCTTTAACCAATAACACTGCTTCTTTCTTGTGTACTTTCTCTTTTACTATACTTCCAACCCCTTCT

>M6-PGSC0003DMG400000789-(rev)M6L-LecRK_1860

ATAAGAGATGTTTTCCGTATGTTTAAAGATGAAGCTCCACTTCTCATTTTACCTCCAATGAAGCCCACTGTGAGAATTCAATCTGTTTTACCAGAGAGCTGTGAAGAAATCATGAATTGG

>M6-PGSC0003DMG400000789-(rev)M6L-LecRK_1920

AAGCCCACTGTGAGAATTCAATCTGTTTTACCAGAGAGCTGTGAAGAAATCATGAATTGGGCTGCAAGAATGGATGATACACCATGGTCCACTCCTAGAACTCATTTTAGCAAGAACTAG

>M6-PGSC0003DMG400000789-(rev)M6L-LecRK_248

CAATAACACTGCTTCTTTCTTGTGTACTTTCTCTTTTACTATACTTCCAACCCCTTCTTGCCCTTTTGGTGATGGCATGGCGTTTTTGGTCACTTCTGATGTTGATTCTTTGAGCATCTC

>M6-PGSC0003DMG400000789-(rev)M6L-LecRK_310

CTTTTGGTGATGGCATGGCGTTTTTGGTCACTTCTGATGTTGATTCTTTGAGCATCTCTGATGGGTATATGGGTCTTCCGAATCCCGACTCGGAAGATTCGTTCTTGGCTGTGGAATTCA

>M6-PGSC0003DMG400000789-(rev)M6L-LecRK_372

GGGTATATGGGTCTTCCGAATCCCGACTCGGAAGATTCGTTCTTGGCTGTGGAATTCAATGCCAATGATAACCGTATTGGTGTTGATACTAAAGAAATTAGGTCTTTGGCTTCTACTAAT

>M6-PGSC0003DMG400000789-(rev)M6L-LecRK_434

CAATGATAACCGTATTGGTGTTGATACTAAAGAAATTAGGTCTTTGGCTTCTACTAATGTTTATTCAGCTGGGATTGATTTGAAAAGTGGGAAAGAAATGGCGGGTCGGATTGAGTATAA

>M6-PGSC0003DMG400000789-(rev)M6L-LecRK_496

ATTCAGCTGGGATTGATTTGAAAAGTGGGAAAGAAATGGCGGGTCGGATTGAGTATAAAGATTCAGAGAAAATGATCAGAGTTTGGATTGGGTATGAACTGCAAATTAGGCCTCCTAATC

>M6-PGSC0003DMG400000789-(rev)M6L-LecRK_558

TCAGAGAAAATGATCAGAGTTTGGATTGGGTATGAACTGCAAATTAGGCCTCCTAATCCTGTTCTTTCTACCAGAATTGATGTTTCCAATCAGTTGAAGGAGTTTATGAGGATTGGTTTC

>M6-PGSC0003DMG400000789-(rev)M6L-LecRK_62

ACCTTCCCTTTCAGCTCCTTTCTTGCCTATAAACAATGTAACTCTTTATGGTGATGCTTCATTCACCGCAAAATCAATCACCCTCACCCAAGAACGCAACTGTTCATCATCATCTTCAAC

>M6-PGSC0003DMG400000789-(rev)M6L-LecRK_620

TCTTTCTACCAGAATTGATGTTTCCAATCAGTTGAAGGAGTTTATGAGGATTGGTTTCACTGCTAAAGGGTCTGCAGTTTATAGCTTTAGTCGTTGGCGATTTAGAACGTTTGGATTGAT

>M6-PGSC0003DMG400000789-(rev)M6L-LecRK_682

CTAAAGGGTCTGCAGTTTATAGCTTTAGTCGTTGGCGATTTAGAACGTTTGGATTGATTTCGTCTCCAATATCATCGTCTTGGGATCAATCTGATGAAGGAAACTGTTTGATGTGTTTCC

>M6-PGSC0003DMG400000789-(rev)M6L-LecRK_744

TCTCCAATATCATCGTCTTGGGATCAATCTGATGAAGGAAACTGTTTGATGTGTTTCCCTGAGGAGGAAATTGGTGGGCATATTTCTGATTCTCATCATAGTAGTAGTAGTAGTGGTAGT

>M6-PGSC0003DMG400000789-(rev)M6L-LecRK_806

GGAGGAAATTGGTGGGCATATTTCTGATTCTCATCATAGTAGTAGTAGTAGTGGTAGTAAAAGTTTACTGAAATTGACTTATGGAGGGTTAGCTGCAATTGTCACACTTGTTGGTTGTGC

>M6-PGSC0003DMG400000789-(rev)M6L-LecRK_868

GTTTACTGAAATTGACTTATGGAGGGTTAGCTGCAATTGTCACACTTGTTGGTTGTGCTTTGTCTGTTGTGTTTGTTCTTGCGTTGAGAAGGAAAAAACGCGATAGAGTAGGGGAGACCA

>M6-PGSC0003DMG400000789-(rev)M6L-LecRK_930

TCTGTTGTGTTTGTTCTTGCGTTGAGAAGGAAAAAACGCGATAGAGTAGGGGAGACCAATGAACGGCAAATGTGTAGATTACAAGGAAATAGAGTGCCTCAAAGATTGTCATTATCTGAA

>M6-PGSC0003DMG400000789-(rev)M6L-LecRK_992

ACGGCAAATGTGTAGATTACAAGGAAATAGAGTGCCTCAAAGATTGTCATTATCTGAAATAAAATCAGCTACAGAATGTTTTAATCATGAAAGGATAATCGGAGAAGGAGCATCTGCTGT

>M6-PGSC0003DMG400000810M6RLKmalectin_0

GTTGCAATACCGAATTCCAGAGCATGATTAGAAGTGTAAGGAGTTGTTTGGAGCAGCTATTGCCTGAGTAAATCACCTATATCTCTGTTCTGAACAACGAAACATAGAATGAAAAATGCA

>M6-PGSC0003DMG400000810M6RLKmalectin_1037

TCTCTTACTTTTTTGCATTGCAGGAACTTAAGTTCTAATTCTTTGGGAGGTTCTCTACCGTGGGGATTGGGTCAGAAGTCTCTTGTGAAACTGTAAGTCCAGACATCAGCACTGTGTTAT

>M6-PGSC0003DMG400000810M6RLKmalectin_1098

GGGGATTGGGTCAGAAGTCTCTTGTGAAACTGTAAGTCCAGACATCAGCACTGTGTTATTGGTCATTTCTTTGTTTACTTGAAATTGTCTTCTGTTTTGCGTCTTAGAACTATGAGTTTT

>M6-PGSC0003DMG400000810M6RLKmalectin_1159

GTCATTTCTTTGTTTACTTGAAATTGTCTTCTGTTTTGCGTCTTAGAACTATGAGTTTTATGAAATCCACATACATAAATTGTTGAACGTTATTGGAGTCCTGAATATTGATCCTGCCAG

>M6-PGSC0003DMG400000810M6RLKmalectin_122

ACTTGCTGCACGAGTACCTGGGTCAAAAAGGTGTATGCTTCGCAGTTCTTTTGGTTATGACTTCTGACTAAATACTGTGGTGCTGTACGGTCTTACCACAAAAAGACGTTTATAAACATC

>M6-PGSC0003DMG400000810M6RLKmalectin_1220

GAAATCCACATACATAAATTGTTGAACGTTATTGGAGTCCTGAATATTGATCCTGCCAGATTTTTCCGATCCACTCAGTTGCTACTAGCATAACTGATAACAACTTACCTCTAGTAAATT

>M6-PGSC0003DMG400000810M6RLKmalectin_1281

TTTTCCGATCCACTCAGTTGCTACTAGCATAACTGATAACAACTTACCTCTAGTAAATTCCTTTTATTCCTAGGGCCTTTTTTTTAAAGATATTATGTCTTATTGAAGTGCTTTTTGCTC

>M6-PGSC0003DMG400000810M6RLKmalectin_1342

TTTTATTCCTAGGGCCTTTTTTTTAAAGATATTATGTCTTATTGAAGTGCTTTTTGCTCAAATTTTCCTAAAACTCTCCTCAATTATTGATATTCTTGTTAATTAATATGCGTAGAACCA

>M6-PGSC0003DMG400000810M6RLKmalectin_1403

ATTTTCCTAAAACTCTCCTCAATTATTGATATTCTTGTTAATTAATATGCGTAGAACCAAGTGCATAACTACTCTGTAGTTTAGGATATGTTGGCGGCACTTTACCGTCTTTCTCTCTTT

>M6-PGSC0003DMG400000810M6RLKmalectin_1464

TGCATAACTACTCTGTAGTTTAGGATATGTTGGCGGCACTTTACCGTCTTTCTCTCTTTTTATGGGCTAACCCCTTTTGTCAAACTGGTTTGCCGTCATGTTATTGGAAACTTGAAGCTT

>M6-PGSC0003DMG400000810M6RLKmalectin_1525

ATGGGCTAACCCCTTTTGTCAAACTGGTTTGCCGTCATGTTATTGGAAACTTGAAGCTTGCAATTCGTCATTTGAAGATCTTTTGTGCAGGGATTTATCAAATAACAAGTTTACTGGACC

>M6-PGSC0003DMG400000810M6RLKmalectin_1586

AATTCGTCATTTGAAGATCTTTTGTGCAGGGATTTATCAAATAACAAGTTTACTGGACCCCTACCAGATGGTCTAGCTTCTTCACCCTTGCAGATTGTGTAAGCTCTATTCTTGTTCTCA

>M6-PGSC0003DMG400000810M6RLKmalectin_1647

TACCAGATGGTCTAGCTTCTTCACCCTTGCAGATTGTGTAAGCTCTATTCTTGTTCTCAAAAACTTTTCATGGATTTGTCGTTGTATATTTCACTCTTTGTGAGGTACCAGTTGCTTATT

>M6-PGSC0003DMG400000810M6RLKmalectin_1708

AACTTTTCATGGATTTGTCGTTGTATATTTCACTCTTTGTGAGGTACCAGTTGCTTATTTCATGATTGTTGTAGAAAGCAGGATAAGTATTTGGGTTATAAATCACTCGGATTCTTTCAG

>M6-PGSC0003DMG400000810M6RLKmalectin_1769

ATGATTGTTGTAGAAAGCAGGATAAGTATTTGGGTTATAAATCACTCGGATTCTTTCAGTCAAAAGAAATTTATCTCGTTCTAGCTAATTTCTGCTTAGTTCTAAGAGTTACACTGTAAT

>M6-PGSC0003DMG400000810M6RLKmalectin_183

TTCTGACTAAATACTGTGGTGCTGTACGGTCTTACCACAAAAAGACGTTTATAAACATCTGTCGGAATTATGCTTAAAAACTCAATTCCAAGAAAGTTATCTTTGAAATTCCATGGCGAT

>M6-PGSC0003DMG400000810M6RLKmalectin_1830

AAAAGAAATTTATCTCGTTCTAGCTAATTTCTGCTTAGTTCTAAGAGTTACACTGTAATTTGCTCAGGTTTTTGAATGGTAATGCACTAGAAGGCCAAGTGCCCGAGGAACTTCTTTCCA

>M6-PGSC0003DMG400000810M6RLKmalectin_1891

GCTCAGGTTTTTGAATGGTAATGCACTAGAAGGCCAAGTGCCCGAGGAACTTCTTTCCATTGGGGTCCATGGTGGAGCTATTGAGTATGTCATTCTCTATTTTCTGTCTAAAGTTTTTCC

>M6-PGSC0003DMG400000810M6RLKmalectin_1952

GGGGTCCATGGTGGAGCTATTGAGTATGTCATTCTCTATTTTCTGTCTAAAGTTTTTCCTTCAGTACATTATGGTCTTCCTTTAGTAGATTTTAATTTATGAATATGTGCTTCAACTATA

>M6-PGSC0003DMG400000810M6RLKmalectin_2013

CAGTACATTATGGTCTTCCTTTAGTAGATTTTAATTTATGAATATGTGCTTCAACTATAAAAAACAAGACTGATTCGGTGAAGAGTGCCTCTGCTACTCATGTTATTAGTGTATGAATGA

>M6-PGSC0003DMG400000810M6RLKmalectin_2074

AAACAAGACTGATTCGGTGAAGAGTGCCTCTGCTACTCATGTTATTAGTGTATGAATGAATTGTGATGCTTCCCTATGTTAAAAACTAGTTTAAGTTGTTAGAAATAATGGTTTATATTG

>M6-PGSC0003DMG400000810M6RLKmalectin_2135

TGTGATGCTTCCCTATGTTAAAAACTAGTTTAAGTTGTTAGAAATAATGGTTTATATTGGAGTTTGCCCTTTGTTTCATGGATAATAAAAGTAACTTTGCTGGGTCTTCTAGACTCTAAG

>M6-PGSC0003DMG400000810M6RLKmalectin_2196

GTTTGCCCTTTGTTTCATGGATAATAAAAGTAACTTTGCTGGGTCTTCTAGACTCTAAGCATCTTTAACCTGTTCGCAAAAGCCTTTGGGATAAGTATCTTACAGAAATTAATGCAATCA

>M6-PGSC0003DMG400000810M6RLKmalectin_2257

TCTTTAACCTGTTCGCAAAAGCCTTTGGGATAAGTATCTTACAGAAATTAATGCAATCATGTGATAATGCAACTTTATAACAACCATATTATACTGAACTGGATCGCCTCTGTGCCCACA

>M6-PGSC0003DMG400000810M6RLKmalectin_2318

TGATAATGCAACTTTATAACAACCATATTATACTGAACTGGATCGCCTCTGTGCCCACAAGCTATTTCATAGCTGAAATTGTGCACACATAGTGCATCAACCACTAAGCATATGCCAATG

>M6-PGSC0003DMG400000810M6RLKmalectin_2379

CTATTTCATAGCTGAAATTGTGCACACATAGTGCATCAACCACTAAGCATATGCCAATGCTGTAACCTTTGGCACACTAACACTCGTTAAGTCGAACCCGTGATGCATTACAGTAGAGCA

>M6-PGSC0003DMG400000810M6RLKmalectin_244

TCGGAATTATGCTTAAAAACTCAATTCCAAGAAAGTTATCTTTGAAATTCCATGGCGATGCTGAGGCATTTGGGTTGGGGTGTCAATTCATCTTTCTGAAGCCATGTAAAGAAGTGTCGT

>M6-PGSC0003DMG400000810M6RLKmalectin_2440

GTAACCTTTGGCACACTAACACTCGTTAAGTCGAACCCGTGATGCATTACAGTAGAGCAGTACTACATGGCTACAACTACTGGCAAATCACATATCCGGTGATGAATCACAACTAAGCAA

>M6-PGSC0003DMG400000810M6RLKmalectin_2501

ACTACATGGCTACAACTACTGGCAAATCACATATCCGGTGATGAATCACAACTAAGCAAAATAATTGTAAGCTATCCCGTTAATGCTCAATCTGAATACCAATGCATCAAAACTGGTTTC

>M6-PGSC0003DMG400000810M6RLKmalectin_2562

TAATTGTAAGCTATCCCGTTAATGCTCAATCTGAATACCAATGCATCAAAACTGGTTTCATGCAAGTAAGTGATGTATCATACACAATGTATAGATCTAATAGAACTCAACAGATTTCAA

>M6-PGSC0003DMG400000810M6RLKmalectin_2623

GCAAGTAAGTGATGTATCATACACAATGTATAGATCTAATAGAACTCAACAGATTTCAAATTTCATTTGCAAACATGGCTGCCTGAAACCTCTTCTTTTTTTTTTGACGGTGGTGCTTGT

>M6-PGSC0003DMG400000810M6RLKmalectin_2806

CCCCACTAAGGCTTAGGCAAGGGCTGCCTAAAACCTCTTTTCAAAGCTTTATACAATTTCACTATCGTTCAACATGCCGTTTATCAAAAGCGAGTACTTCAGGGGTAAGGTCTTCACTTA

>M6-PGSC0003DMG400000810M6RLKmalectin_2867

CTATCGTTCAACATGCCGTTTATCAAAAGCGAGTACTTCAGGGGTAAGGTCTTCACTTATGTCCAATTTAAGCTAACTAGCACTGCTACAGTGTCCAGACTCAACCAAAGATTGCCAAGA

>M6-PGSC0003DMG400000810M6RLKmalectin_2928

TCCAATTTAAGCTAACTAGCACTGCTACAGTGTCCAGACTCAACCAAAGATTGCCAAGACATACAAAGAATGATTCAGCGGTGTTAAATACATCCTAACAACTCATATCCAAACTTGGAC

>M6-PGSC0003DMG400000810M6RLKmalectin_2989

TACAAAGAATGATTCAGCGGTGTTAAATACATCCTAACAACTCATATCCAAACTTGGACTTTGGTCAAATTCAAAGTCAATCTTTTGATTTAACATTGGTCAAACATTATCAAAACAATG

>M6-PGSC0003DMG400000810M6RLKmalectin_305

TGAGGCATTTGGGTTGGGGTGTCAATTCATCTTTCTGAAGCCATGTAAAGAAGTGTCGTGGAATTAATGCACATTTCAGATCTATTAGTACACGAGCTCTTATAAGCCACTGTAACTGGA

>M6-PGSC0003DMG400000810M6RLKmalectin_3050

TGGTCAAATTCAAAGTCAATCTTTTGATTTAACATTGGTCAAACATTATCAAAACAATGGGTCCAACCCAAACTTGAGAGTTCTCAATTTCAGTGTTTTTGGACCAAATCCATGCAATTC

>M6-PGSC0003DMG400000810M6RLKmalectin_3111

TCCAACCCAAACTTGAGAGTTCTCAATTTCAGTGTTTTTGGACCAAATCCATGCAATTCTATGGTTGAACCTTAATTCTGTGAAGGTAGAAACTTGATGTACTATGTCTATATTAATCCA

>M6-PGSC0003DMG400000810M6RLKmalectin_3159

CCATGCAATTCTATGGTTGAACCTTAATTCTGTGAAGGTAGAAACTTGATGTACTATGTCTATATTAATCCAATCAATCCTATTAAAATATCACCTCATACAAGTGCTAGTTTTACAATT

>M6-PGSC0003DMG400000810M6RLKmalectin_366

AATTAATGCACATTTCAGATCTATTAGTACACGAGCTCTTATAAGCCACTGTAACTGGACCATTAAAGACTAAAGTGTGAAATTTCCATTGCTCCAGCCTCTCTAAGACAGCATTCCTCT

>M6-PGSC0003DMG400000810M6RLKmalectin_427

ATTAAAGACTAAAGTGTGAAATTTCCATTGCTCCAGCCTCTCTAAGACAGCATTCCTCTTTCATATCCTACCGATCTCACTTCTAACTTGACAAATGATGTAATCTTTAATTTCAATTTG

>M6-PGSC0003DMG400000810M6RLKmalectin_488

CATATCCTACCGATCTCACTTCTAACTTGACAAATGATGTAATCTTTAATTTCAATTTGTGTGTCAAAGAAGGAAAGAACAGATCCTTTTTCTTTAACAAATATCCCTAGACTTTCTGTT

>M6-PGSC0003DMG400000810M6RLKmalectin_549

TGTCAAAGAAGGAAAGAACAGATCCTTTTTCTTTAACAAATATCCCTAGACTTTCTGTTGTTTAATTCTAGTGATATCTAATGCTGCACGTGTATATCCTTCTTGATTCTTTTAGTTCTT

>M6-PGSC0003DMG400000810M6RLKmalectin_61

GCCTGAGTAAATCACCTATATCTCTGTTCTGAACAACGAAACATAGAATGAAAAATGCACAACTTGCTGCACGAGTACCTGGGTCAAAAAGGTGTATGCTTCGCAGTTCTTTTGGTTATG

>M6-PGSC0003DMG400000810M6RLKmalectin_610

TTAATTCTAGTGATATCTAATGCTGCACGTGTATATCCTTCTTGATTCTTTTAGTTCTTATCTTTCAGAGATCTTGGAAGCCAAGGCTTAAAGGGATATATCAGTGACCAAATTGGGCTG

>M6-PGSC0003DMG400000810M6RLKmalectin_671

CTTTCAGAGATCTTGGAAGCCAAGGCTTAAAGGGATATATCAGTGACCAAATTGGGCTGTTGTCTAATTTGGTAAGCTTGTAAGTATCTCTCAACAAATTTCCTCTTAGATACTGGAATT

>M6-PGSC0003DMG400000810M6RLKmalectin_732

GTCTAATTTGGTAAGCTTGTAAGTATCTCTCAACAAATTTCCTCTTAGATACTGGAATTTCACACGTTAATCCTCCTGCATAATATCTGAATCTACTCACTTGGTAGTCCTTCAAAATAA

>M6-PGSC0003DMG400000810M6RLKmalectin_793

ACACGTTAATCCTCCTGCATAATATCTGAATCTACTCACTTGGTAGTCCTTCAAAATAATTAAAGAGGTGTTCAACAAAAAATGGTTACTGTATGAAAATATACTAGTATTTGCATATAC

>M6-PGSC0003DMG400000810M6RLKmalectin_854

AAAGAGGTGTTCAACAAAAAATGGTTACTGTATGAAAATATACTAGTATTTGCATATACTCATTATGGTGCACTGATGAATAATTCTCTTGATTGGATCATTTAAGTTGCCCCAATTTCT

>M6-PGSC0003DMG400000810M6RLKmalectin_915

ATTATGGTGCACTGATGAATAATTCTCTTGATTGGATCATTTAAGTTGCCCCAATTTCTTTTATGCACTTCTACTGTTTGTCAATCCTGCTAATTTGGCCTTCGCAAATAGTTATTCTAC

>M6-PGSC0003DMG400000810M6RLKmalectin_976

TATGCACTTCTACTGTTTGTCAATCCTGCTAATTTGGCCTTCGCAAATAGTTATTCTACCTTCTCTTACTTTTTTGCATTGCAGGAACTTAAGTTCTAATTCTTTGGGAGGTTCTCTACC

>M6-PGSC0003DMG400000819-(rev)M6LRR-RLP_0

CTCTTTTTCTACCTTAGCTTTCTATAAACATTTTTAACAAGCACCTTCCTTCTTTACTTCTTTGTTCCCTCCATTTCCCTGTTTGATTGGATACTTTTACCATTTTCCACCACTGCCCCA

>M6-PGSC0003DMG400000819-(rev)M6LRR-RLP_1054

AAGTGGAAATTTTATACTATGACAGATAATCAAAGAAATTGATAAGTTCTTTTTTCAGATTTAGTATATTTGGAATGATAGCTAATTAGCCATAGAAGTTTGATGAACCAAACTTCAATA

>M6-PGSC0003DMG400000819-(rev)M6LRR-RLP_1116

AGTATATTTGGAATGATAGCTAATTAGCCATAGAAGTTTGATGAACCAAACTTCAATAATGAATATCAGTACATTCTTGGCTTGTTTTGTTTAGCAGTTTTGACAGACCAACAGAGACAG

>M6-PGSC0003DMG400000819-(rev)M6LRR-RLP_1178

ATATCAGTACATTCTTGGCTTGTTTTGTTTAGCAGTTTTGACAGACCAACAGAGACAGAATATGTTTTAAAACACAATGATAATACAGGACAGGACAGAGCTTATACGAAATATAATCAG

>M6-PGSC0003DMG400000819-(rev)M6LRR-RLP_124

AACAAAAAAAGTACACAATTTTTTTTTTTATATTTATGAAATGAAAGCATTTTAAGGTACCCCTCTTGGAGTTTCTTCAATGTTATAAGTTCTATTTGGCTTTAATTGACAAAGTTTATT

>M6-PGSC0003DMG400000819-(rev)M6LRR-RLP_1240

TGTTTTAAAACACAATGATAATACAGGACAGGACAGAGCTTATACGAAATATAATCAGTTCGTTTCAATTTGTTTGTCTGATTTTAACTTGACACGTAGTTAAAGAATGCACCATAGTTT

>M6-PGSC0003DMG400000819-(rev)M6LRR-RLP_1426

GTATAGTTGATGCATTATACTCATATTTCTGATTTGTTTTGTTTCTATTTCCTCTGTCCAGGGTGTATCTTGTGACCTGTTTGATGGTTTTTGGTATGTTACTGACTTAAACATTGGACC

>M6-PGSC0003DMG400000819-(rev)M6LRR-RLP_1488

GTGTATCTTGTGACCTGTTTGATGGTTTTTGGTATGTTACTGACTTAAACATTGGACCTATACATGATAATTCTCTAAGTTGTGCATCAAATGTGGAATTCAGCTCCAATTTATTCACAT

>M6-PGSC0003DMG400000819-(rev)M6LRR-RLP_1550

CATGATAATTCTCTAAGTTGTGCATCAAATGTGGAATTCAGCTCCAATTTATTCACATTAAAGCACCTAAAATCTCTCACATTTTTCAATTGTTTCGTGTCGCCTCGCCATCATCCAACT

>M6-PGSC0003DMG400000819-(rev)M6LRR-RLP_1612

GCACCTAAAATCTCTCACATTTTTCAATTGTTTCGTGTCGCCTCGCCATCATCCAACTTCAATCCCAACAGAGAGGTGGGAGTTTCTTGCTAGTAGTTTGGAGTCACTCGAGTTTCGATC

>M6-PGSC0003DMG400000819-(rev)M6LRR-RLP_1674

TCCCAACAGAGAGGTGGGAGTTTCTTGCTAGTAGTTTGGAGTCACTCGAGTTTCGATCAAATCCCGGTCTCATTGGCCAAATCCCAAGTACATTTGGTAGACTCATAAAGCTCCAATCTT

>M6-PGSC0003DMG400000819-(rev)M6LRR-RLP_1736

CCCGGTCTCATTGGCCAAATCCCAAGTACATTTGGTAGACTCATAAAGCTCCAATCTTTAGTACTAGTAGAAAATGGACTCTCTGGTGAAATGCCACAAAGTCTTGGGAATTTAGTGAAC

>M6-PGSC0003DMG400000819-(rev)M6LRR-RLP_1798

ACTAGTAGAAAATGGACTCTCTGGTGAAATGCCACAAAGTCTTGGGAATTTAGTGAACTTGAAAAGACTAGTTCTTGCTGGAAACAAACTCAACGGCGAAATCCCGGATACTTTTGGAAG

>M6-PGSC0003DMG400000819-(rev)M6LRR-RLP_186

CTCTTGGAGTTTCTTCAATGTTATAAGTTCTATTTGGCTTTAATTGACAAAGTTTATTACTTTCAAGATTTTGTCAACCTAACTCATTTTCAAGCTTTTTTTGGCTTCAAAGAGTGATGG

>M6-PGSC0003DMG400000819-(rev)M6LRR-RLP_1860

AAAGACTAGTTCTTGCTGGAAACAAACTCAACGGCGAAATCCCGGATACTTTTGGAAGGTTCACTCAACTTTTGATATGTGATTTAAGCAGAAACAAACTATCTGGTGTCTTGCCTAAAT

>M6-PGSC0003DMG400000819-(rev)M6LRR-RLP_1922

ACTCAACTTTTGATATGTGATTTAAGCAGAAACAAACTATCTGGTGTCTTGCCTAAATCATTATTTGGAGGGTTGATCTCACTTTTGAAACTTGATTTGAGTTACAACAACTTACAAGGC

>M6-PGSC0003DMG400000819-(rev)M6LRR-RLP_1984

ATTTGGAGGGTTGATCTCACTTTTGAAACTTGATTTGAGTTACAACAACTTACAAGGCAAAATCCCACAAGAGATCTCACAAATGAAGAATTTAACACTTTTAGACCTTAGTAACAACAA

>M6-PGSC0003DMG400000819-(rev)M6LRR-RLP_2046

TCCCACAAGAGATCTCACAAATGAAGAATTTAACACTTTTAGACCTTAGTAACAACAAGTTATCAAGTGGATTACCTAAGTCAATTCAAGAAATGACATGTTTGGAGGAGTTAGTCTTGT

>M6-PGSC0003DMG400000819-(rev)M6LRR-RLP_2108

TCAAGTGGATTACCTAAGTCAATTCAAGAAATGACATGTTTGGAGGAGTTAGTCTTGTCAAACAACCCAATGGGTGGATTCTTGGATGTTCTTGATTGGTATAACATGATAAAGTTAACA

>M6-PGSC0003DMG400000819-(rev)M6LRR-RLP_2170

CAACCCAATGGGTGGATTCTTGGATGTTCTTGATTGGTATAACATGATAAAGTTAACAACTTTGGATTTATCAAACATGAATTTGACAGGTGAGATACCTAAGTCTATAGTAGAGTTAAA

>M6-PGSC0003DMG400000819-(rev)M6LRR-RLP_2232

TGGATTTATCAAACATGAATTTGACAGGTGAGATACCTAAGTCTATAGTAGAGTTAAAAAAGTTAAGATTTTTGGGACTTAATGACAATAAACTCATTGGTAAAATCCCAAAAAATCTTG

>M6-PGSC0003DMG400000819-(rev)M6LRR-RLP_2294

TTAAGATTTTTGGGACTTAATGACAATAAACTCATTGGTAAAATCCCAAAAAATCTTGAAAATTTGCCTAATGTAAGTGCTATTTACCTATATGGTAACAATCTGACAGGTGAACTTCAA

>M6-PGSC0003DMG400000819-(rev)M6LRR-RLP_2356

TTTGCCTAATGTAAGTGCTATTTACCTATATGGTAACAATCTGACAGGTGAACTTCAATTCTCTCAATGGTTTTATAGGAAAATGGGAAGGAGATTTGGGGCATGGGGGAATCAAAATTT

>M6-PGSC0003DMG400000819-(rev)M6LRR-RLP_2418

CTCAATGGTTTTATAGGAAAATGGGAAGGAGATTTGGGGCATGGGGGAATCAAAATTTATGTTACACTTTTGGTTTGGTGTCAAGTGATGTTCCATTTGGGGTAAAACAATGTCAACAAG

>M6-PGSC0003DMG400000819-(rev)M6LRR-RLP_248

TCAAGATTTTGTCAACCTAACTCATTTTCAAGCTTTTTTTGGCTTCAAAGAGTGATGGGGGTCTTCAAGAACATGTTTGTATGAAAAAGTTGTATTCTTTGAACACTTTGTGAACTCATT

>M6-PGSC0003DMG400000819-(rev)M6LRR-RLP_2480

TACACTTTTGGTTTGGTGTCAAGTGATGTTCCATTTGGGGTAAAACAATGTCAACAAGAACTCAAGTTTGTTATTGATGTTGAGATGAACTCAAGTTTTGTGGATTTTCAATCTTCAATG

>M6-PGSC0003DMG400000819-(rev)M6LRR-RLP_2542

CAAGTTTGTTATTGATGTTGAGATGAACTCAAGTTTTGTGGATTTTCAATCTTCAATGGGATATTTGAATGATTGTGGGATTTGGAGATGTTTTGTAGTTGAACTATTTATGGTTATTCT

>M6-PGSC0003DMG400000819-(rev)M6LRR-RLP_2604

ATTTGAATGATTGTGGGATTTGGAGATGTTTTGTAGTTGAACTATTTATGGTTATTCTACTAATGTTGAAATGAAAAGAAGGAATTTGACAATAGTGATTTACCTTTTTTTTTTGATATT

>M6-PGSC0003DMG400000819-(rev)M6LRR-RLP_2666

ATGTTGAAATGAAAAGAAGGAATTTGACAATAGTGATTTACCTTTTTTTTTTGATATTTTGTTGTTGTTTTACTTGAAATGTACCACACATTTTGGTTTGTCTTTTAACATATGAATGAA

>M6-PGSC0003DMG400000819-(rev)M6LRR-RLP_310

CTTCAAGAACATGTTTGTATGAAAAAGTTGTATTCTTTGAACACTTTGTGAACTCATTGTGTGATAAAGTGTTATCTTTGAACACTTTGTGTACTCATTTTCAAGAAAATTTAGCTTCAA

>M6-PGSC0003DMG400000819-(rev)M6LRR-RLP_372

GATAAAGTGTTATCTTTGAACACTTTGTGTACTCATTTTCAAGAAAATTTAGCTTCAAAAGAATGATGGGGTTCTTCAAGAATTCTAACTAGTTTGTTTATGTGACAGAGTTTTAAACTT

>M6-PGSC0003DMG400000819-(rev)M6LRR-RLP_434

ATGATGGGGTTCTTCAAGAATTCTAACTAGTTTGTTTATGTGACAGAGTTTTAAACTTTGAAGATTTTGTTATCTCTTTTTCAAGAAAATTTAGCTTCAAAAGAATGATGGGGTTCTTCA

>M6-PGSC0003DMG400000819-(rev)M6LRR-RLP_496

GATTTTGTTATCTCTTTTTCAAGAAAATTTAGCTTCAAAAGAATGATGGGGTTCTTCAAGAATTCTAACTAGTTTGTTTATGTGACAGAGTTTTAAACTTTGAAGATTTTGTCATCTCTT

>M6-PGSC0003DMG400000819-(rev)M6LRR-RLP_558

TTCTAACTAGTTTGTTTATGTGACAGAGTTTTAAACTTTGAAGATTTTGTCATCTCTTTTTCAAGAAACTTTTTAAGCTACAAAAAGTGATGAAGGGTTTCAAGAATTCAAGATTGTGTG

>M6-PGSC0003DMG400000819-(rev)M6LRR-RLP_62

TGTTCCCTCCATTTCCCTGTTTGATTGGATACTTTTACCATTTTCCACCACTGCCCCAAAAAAACAAAAAAAGTACACAATTTTTTTTTTTATATTTATGAAATGAAAGCATTTTAAGGT

>M6-PGSC0003DMG400000819-(rev)M6LRR-RLP_620

AAGAAACTTTTTAAGCTACAAAAAGTGATGAAGGGTTTCAAGAATTCAAGATTGTGTGTGTTAGTTATCTTTTATCTCCTCCATTTTGGTGTCTGGTGCAATGGTGATACTGAAAACATG

>M6-PGSC0003DMG400000819-(rev)M6LRR-RLP_682

AGTTATCTTTTATCTCCTCCATTTTGGTGTCTGGTGCAATGGTGATACTGAAAACATGGGAGCTCAAATGGAGAAAAATGAGATAAATTCACTTTACTCAACCATTCAAGGCTTTGTTGG

>M6-PGSC0003DMG400000819-(rev)M6LRR-RLP_744

CTCAAATGGAGAAAAATGAGATAAATTCACTTTACTCAACCATTCAAGGCTTTGTTGGTAAATGGTGGAATGGTTCAGATCTTTATCCAGATCCTTGTGGTTGGACTCCTATACAGGTAA

>M6-PGSC0003DMG400000819-(rev)M6LRR-RLP_806

TGGTGGAATGGTTCAGATCTTTATCCAGATCCTTGTGGTTGGACTCCTATACAGGTAACACAAATTTTTATATAAATTTTTCACTTTTTTAGTTACTTTTCTGCTTTATGGATTCTGTTT

>M6-PGSC0003DMG400000819-(rev)M6LRR-RLP_868

AATTTTTATATAAATTTTTCACTTTTTTAGTTACTTTTCTGCTTTATGGATTCTGTTTAGTAAAAGGTAAAAAAAAAAAAAAAACTAACTTTTTTGTGCATATGGTTATAATTGTCCTTT

>M6-PGSC0003DMG400000819-(rev)M6LRR-RLP_930

AAAGGTAAAAAAAAAAAAAAAACTAACTTTTTTGTGCATATGGTTATAATTGTCCTTTTTAATATTTTGTGGTATACATGACAAGATATGACTTCTCATTGTATATAGATTGTTTGAAAG

>M6-PGSC0003DMG400000819-(rev)M6LRR-RLP_992

TATTTTGTGGTATACATGACAAGATATGACTTCTCATTGTATATAGATTGTTTGAAAGCTTTAAGTGGAAATTTTATACTATGACAGATAATCAAAGAAATTGATAAGTTCTTTTTTCAG

>M6-PGSC0003DMG400000888M6LRR-RLP_0

TTATTGATTCAGTCTGTTGAAGACTGAGCAATATTTGTCTTGATTAATTCCATTGTGTACATATTCTGATGAAGTTTTCGTATACGGAGGGTGGAGGGGATTGATCTCTGAAGGAATTTC

>M6-PGSC0003DMG400000888M6LRR-RLP_122

AATACAGATGGTTCAAAGTAAACTCAATTACAATAATTCTAAACAATTGGAGTAGACATTAAGAAGCTTCACAGAATTTTCCTACCTATTACTTCTATCAATATATTCAATGAAAATATA

>M6-PGSC0003DMG400000888M6LRR-RLP_183

AGAAGCTTCACAGAATTTTCCTACCTATTACTTCTATCAATATATTCAATGAAAATATAAGTTGACTAGATCCAACAACAATTATTTATTCAACTGGAGTAAAGCGAATGTTCATAGAAG

>M6-PGSC0003DMG400000888M6LRR-RLP_2379

ATTTAAAAGATGAAAATTGTTGGTACCCTATCAATTACAACTTCGTTCTCTTACAACATGAAAAACTTTTCGATTGGACTTTAAAGAACTAGTTGAAAATTGTATTACTAGAAATTAGAA

>M6-PGSC0003DMG400000888M6LRR-RLP_2440

AAAACTTTTCGATTGGACTTTAAAGAACTAGTTGAAAATTGTATTACTAGAAATTAGAAATATAGAATTAATTGCATAAAGTATTTCTCTTTAGCGACATTTTTCTTAACTAATATTGTA

>M6-PGSC0003DMG400000888M6LRR-RLP_2501

ATAGAATTAATTGCATAAAGTATTTCTCTTTAGCGACATTTTTCTTAACTAATATTGTACCTTAATTTTATTTAATGGTCTCTTCAAATGCAATAATAAATAATACAAAACATAACGAAA

>M6-PGSC0003DMG400000888M6LRR-RLP_2562

TTAATTTTATTTAATGGTCTCTTCAAATGCAATAATAAATAATACAAAACATAACGAAATTTGGCATTGCAAACAGTTAGAAAGTAGTCATATTGGCATGCAAATTTTATACAGATATTT

>M6-PGSC0003DMG400000888M6LRR-RLP_61

TATTCTGATGAAGTTTTCGTATACGGAGGGTGGAGGGGATTGATCTCTGAAGGAATTTCCTAATACAGATGGTTCAAAGTAAACTCAATTACAATAATTCTAAACAATTGGAGTAGACAT

>M6-PGSC0003DMG400000888M6LRR-RLP_976

TAGGATGCTTTTGTGAAACCTAGACATATTCTACTATAGAACAAGAATCTATACCTCCAATATGAGTTAATTTAGTTGGAGAGAATACATAATAATCCTTCTAAACTATTCAAGTTTTGC

>M6-PGSC0003DMG400000915-M6G-LecRK_0

ATGGAGTTAATCTTATTCTTTCTCATTTTATCTTTGTCCGGTATTCCTGTCTTTGGTTTTTTATCAAAAGAATTTGTCTATCCCAACTTTACTGCTTCCAATTTCCAATTTGTCGATAGC

>M6-PGSC0003DMG400000915-M6G-LecRK_1037

AGGACTTGGCTTTGGTATGGACTACTTCACTACTGATTTTACCAAGCCCTTCAGGGATGGGGTAAATTTGTCCGTTTGTCAAAATCTCTGCTCAGTAGACTGTTCATGCTTGGGGATATT

>M6-PGSC0003DMG400000915-M6G-LecRK_1098

GTAAATTTGTCCGTTTGTCAAAATCTCTGCTCAGTAGACTGTTCATGCTTGGGGATATTTTATGCAAACTCGTCTGGTTCTTGTTACAAACTTGAAGATGAAATAGGGTCAATTATGTTA

>M6-PGSC0003DMG400000915-M6G-LecRK_1159

ATGCAAACTCGTCTGGTTCTTGTTACAAACTTGAAGATGAAATAGGGTCAATTATGTTAAGGACTAGTAGAAATGATGATCTGTTGGGATTTGTTAAGACCCTGGTAGGAGTTTCAACAA

>M6-PGSC0003DMG400000915-M6G-LecRK_122

CGGAAGTTTCTTGGTTTCTCGCAATGGAACATTCAAAGCTTCTATCTTCAATCCCGGTCCTGAACAAGTTAATTTCTATTTGTGTGTTATCCATGCTGAATCCAATACCATTATCTGGTC

>M6-PGSC0003DMG400000915-M6G-LecRK_1220

GACTAGTAGAAATGATGATCTGTTGGGATTTGTTAAGACCCTGGTAGGAGTTTCAACAACCTTAGGAAACAATGATAATTTTGATCAGGAAGCTGCAGACTTTCCTTTAGTTGCAACAGT

>M6-PGSC0003DMG400000915-M6G-LecRK_1281

TTAGGAAACAATGATAATTTTGATCAGGAAGCTGCAGACTTTCCTTTAGTTGCAACAGTGATCTTACCTTTCACAGGAATCTTCCTACTCATGGCGCTGGGTTTCATTATGTGGAGAAGA

>M6-PGSC0003DMG400000915-M6G-LecRK_1342

TCTTACCTTTCACAGGAATCTTCCTACTCATGGCGCTGGGTTTCATTATGTGGAGAAGATCTAGACCCCAAAGAACGGGAAAGATAGAATCCAAAAAGAGCCAACCCAATTCTCCTTTTT

>M6-PGSC0003DMG400000915-M6G-LecRK_1403

TAGACCCCAAAGAACGGGAAAGATAGAATCCAAAAAGAGCCAACCCAATTCTCCTTTTTCAGAGGACCTGGATGCCTTCTCCATCCCAGGATTACCTGTACGGTTTGAGTACAAACAGCT

>M6-PGSC0003DMG400000915-M6G-LecRK_1464

GAGGACCTGGATGCCTTCTCCATCCCAGGATTACCTGTACGGTTTGAGTACAAACAGCTTAAGGCTGCAACTGATAATTTCAAGACTCAGATCGGCACAGGAGGCTTTGGTGCTGTATAT

>M6-PGSC0003DMG400000915-M6G-LecRK_1525

AGGCTGCAACTGATAATTTCAAGACTCAGATCGGCACAGGAGGCTTTGGTGCTGTATATAAGGGTGTGCTCCCTGACAACACTCTGGTTGCAGTAAAGAAGATAATAAATGTAGGCATTC

>M6-PGSC0003DMG400000915-M6G-LecRK_1586

GGGTGTGCTCCCTGACAACACTCTGGTTGCAGTAAAGAAGATAATAAATGTAGGCATTCAAGGGCAGAGAGACTTTTGTACTGAGATTGCAATCATTGGCAATATTCACCACATAAACTT

>M6-PGSC0003DMG400000915-M6G-LecRK_1647

GGGCAGAGAGACTTTTGTACTGAGATTGCAATCATTGGCAATATTCACCACATAAACTTGGTCAAGTTAAAAGGTTTCTGTGCTCAAGAGAGACAACGCCTATTGGTTTATGAATTTATG

>M6-PGSC0003DMG400000915-M6G-LecRK_1708

TCAAGTTAAAAGGTTTCTGTGCTCAAGAGAGACAACGCCTATTGGTTTATGAATTTATGAACCGTGGATCGCTTGATCGCACTCTTTTTGGTCATGTACCTGTATTGGAATGGCAAGAAC

>M6-PGSC0003DMG400000915-M6G-LecRK_1769

CCGTGGATCGCTTGATCGCACTCTTTTTGGTCATGTACCTGTATTGGAATGGCAAGAACGGGTTGAGATAGCTCTTGGTTCTGCCCGCGGACTTGCATACTTGCACAGTGGCTGTGAACA

>M6-PGSC0003DMG400000915-M6G-LecRK_183

GAACAAGTTAATTTCTATTTGTGTGTTATCCATGCTGAATCCAATACCATTATCTGGTCTGCAAATGGTGATGCACCTATTTCAACTTCAGGAATTATGAGGTTGACCAAGAATGGTATT

>M6-PGSC0003DMG400000915-M6G-LecRK_1830

GTTGAGATAGCTCTTGGTTCTGCCCGCGGACTTGCATACTTGCACAGTGGCTGTGAACAAAAGATTGTCCACTGTGATGTGAAGCCAGAGAACATTCTCTTGCACGATAATTTTCAGGCA

>M6-PGSC0003DMG400000915-M6G-LecRK_1891

AGATTGTCCACTGTGATGTGAAGCCAGAGAACATTCTCTTGCACGATAATTTTCAGGCAAAAATATCTGATTTTGGCCTCTCCAAGCTGCTGAATCGTGAAGAATCCAGTCTATTCACAA

>M6-PGSC0003DMG400000915-M6G-LecRK_1952

AATATCTGATTTTGGCCTCTCCAAGCTGCTGAATCGTGAAGAATCCAGTCTATTCACAACTATGAGGGGGACTCGTGGCTACCTTGCTCCTGAATGGCTCACAAGTTCTGCAATCTCAGA

>M6-PGSC0003DMG400000915-M6G-LecRK_2013

ATGAGGGGGACTCGTGGCTACCTTGCTCCTGAATGGCTCACAAGTTCTGCAATCTCAGACAAAACGGATGTGTATAGCTTTGGAATGGTACTCCTTGAAATTGTAAGTGGAAGGAAAAAC

>M6-PGSC0003DMG400000915-M6G-LecRK_2074

AAACGGATGTGTATAGCTTTGGAATGGTACTCCTTGAAATTGTAAGTGGAAGGAAAAACTGCTCTTTAAGAACACAAAGTCACAGCCTTGATGATAGTGCTACTGGAGATCACTCATCAT

>M6-PGSC0003DMG400000915-M6G-LecRK_2135

CTCTTTAAGAACACAAAGTCACAGCCTTGATGATAGTGCTACTGGAGATCACTCATCATCTTCATCTGCACAGGGACTTGTTTATTTCCCTTTATTTGCATTAGAGATGCATGAGCAAGG

>M6-PGSC0003DMG400000915-M6G-LecRK_2196

TCATCTGCACAGGGACTTGTTTATTTCCCTTTATTTGCATTAGAGATGCATGAGCAAGGAAGGTACCTTGAACTTGCTGACCCAAAACTTGAAGGGCGGGTCAGTGGTGGAGATATTGAG

>M6-PGSC0003DMG400000915-M6G-LecRK_2257

GGTACCTTGAACTTGCTGACCCAAAACTTGAAGGGCGGGTCAGTGGTGGAGATATTGAGAAGTTTGTGCGGGTAGCATTGTGCTGTGTCCATGAGGAGCCATGTCTCAGGCCAACTATGG

>M6-PGSC0003DMG400000915-M6G-LecRK_2318

GTTTGTGCGGGTAGCATTGTGCTGTGTCCATGAGGAGCCATGTCTCAGGCCAACTATGGTTAGTGTTGTTAGCATGTTAGAGGGTGAGATACCACCAACTGCGCCAAGAATGGAATCTCT

>M6-PGSC0003DMG400000915-M6G-LecRK_2379

AGTGTTGTTAGCATGTTAGAGGGTGAGATACCACCAACTGCGCCAAGAATGGAATCTCTGAACTTTTTGCGCTTCTACGGGCGACGTTTTTCAGAGGCATCAAACATGGAAGAAGCTGGA

>M6-PGSC0003DMG400000915-M6G-LecRK_244

CAAATGGTGATGCACCTATTTCAACTTCAGGAATTATGAGGTTGACCAAGAATGGTATTAGTATTACTGAAAAAGATGGCAGTTTTAAGTGGTCAACTCCACCATTAAAGTTAGCAGTAT

>M6-PGSC0003DMG400000915-M6G-LecRK_2440

ACTTTTTGCGCTTCTACGGGCGACGTTTTTCAGAGGCATCAAACATGGAAGAAGCTGGAGGCCAGATTGATGCCATGTTATATCCGCAGGCAAATACTTCTCATACAACCTCTCGGAGCG

>M6-PGSC0003DMG400000915-M6G-LecRK_2495

TGGAGGCCAGATTGATGCCATGTTATATCCGCAGGCAAATACTTCTCATACAACCTCTCGGAGCGTCTCTAATGCTTACTTCTCTTATATTTCGTCCCAACAAATCTCTGGTCCAAGATA

>M6-PGSC0003DMG400000915-M6G-LecRK_305

TATTACTGAAAAAGATGGCAGTTTTAAGTGGTCAACTCCACCATTAAAGTTAGCAGTATACGCGATGCAGCTAACTGAGACTGGAAATCTCCTTCTCTTTGATCAGTTTAATGGGACACT

>M6-PGSC0003DMG400000915-M6G-LecRK_366

GCGATGCAGCTAACTGAGACTGGAAATCTCCTTCTCTTTGATCAGTTTAATGGGACACTTTGGCAAAGCTTTGATCATCCGACAGATACAATTGTAATTGGACAGAAGTTGCGTGTTGGC

>M6-PGSC0003DMG400000915-M6G-LecRK_427

GGCAAAGCTTTGATCATCCGACAGATACAATTGTAATTGGACAGAAGTTGCGTGTTGGCATGATGCTTTCAAGTGCAATAGCAGGTGATGACTTGTCAAAGGGTCATTACAGGCTTTCTT

>M6-PGSC0003DMG400000915-M6G-LecRK_488

GATGCTTTCAAGTGCAATAGCAGGTGATGACTTGTCAAAGGGTCATTACAGGCTTTCTTTGTCTGCTTCCGATGCTATGTTACAGTGGCAAGGTCTGACATACTGGAAATTGTCCATGGA

>M6-PGSC0003DMG400000915-M6G-LecRK_549

TCTGCTTCCGATGCTATGTTACAGTGGCAAGGTCTGACATACTGGAAATTGTCCATGGAAACCAAAGCTTATACTAATTCAAATTATGCAGTGGAATACATGTCTGTCAATCAAACAGGT

>M6-PGSC0003DMG400000915-M6G-LecRK_61

TATCAAAAGAATTTGTCTATCCCAACTTTACTGCTTCCAATTTCCAATTTGTCGATAGCACCGGAAGTTTCTTGGTTTCTCGCAATGGAACATTCAAAGCTTCTATCTTCAATCCCGGTC

>M6-PGSC0003DMG400000915-M6G-LecRK_610

CCAAAGCTTATACTAATTCAAATTATGCAGTGGAATACATGTCTGTCAATCAAACAGGTCTTTATCTTTTTGGTCAAGGTGGTTCTGTAGTTGTGATAATGGTGAACTTGTTACAATCCA

>M6-PGSC0003DMG400000915-M6G-LecRK_671

TTATCTTTTTGGTCAAGGTGGTTCTGTAGTTGTGATAATGGTGAACTTGTTACAATCCACTTTTAGAATTGCCAAATTGGATGGTTCAGGTCAGTTGATCGTTATTAGTTTTGTTGGTTC

>M6-PGSC0003DMG400000915-M6G-LecRK_732

TTTAGAATTGCCAAATTGGATGGTTCAGGTCAGTTGATCGTTATTAGTTTTGTTGGTTCTGACATCAAACGAGATTTTGTTGGGCCAGTGGATGGGTGTAGAGTACCTTATGTTTGCGGA

>M6-PGSC0003DMG400000915-M6G-LecRK_793

ACATCAAACGAGATTTTGTTGGGCCAGTGGATGGGTGTAGAGTACCTTATGTTTGCGGAGGGCTTGGTGTATGCACCTCTGATGTATTATCAGATAATCCTAAATGTTCCTGTCCTGCTA

>M6-PGSC0003DMG400000915-M6G-LecRK_854

GCTTGGTGTATGCACCTCTGATGTATTATCAGATAATCCTAAATGTTCCTGTCCTGCTAACTTCAATCTTGGATCACATGATTCAAGAAGCTGTGTGCCTAGTGATAGTTCCTATTCTTT

>M6-PGSC0003DMG400000915-M6G-LecRK_915

TTCAATCTTGGATCACATGATTCAAGAAGCTGTGTGCCTAGTGATAGTTCCTATTCTTTGCCAGTTTCTTGTAATTCCACCAACCACAGTAATCTGTCGAATTCTTCCTCCATGTCGTAC

>M6-PGSC0003DMG400000915-M6G-LecRK_976

CAGTTTCTTGTAATTCCACCAACCACAGTAATCTGTCGAATTCTTCCTCCATGTCGTACATAGGACTTGGCTTTGGTATGGACTACTTCACTACTGATTTTACCAAGCCCTTCAGGGATG

>M6-PGSC0003DMG400000996-(rev)M6LRR-RLK_0

GGCTGTTGTCTCTTTTTTGCCTTAAAACCTGAACTTCTCCTGAATTTTTCTACTCTACATATTACTGCAGGCATGAAGTCTTTTAATGTTTTCATTTGTGTTATGCTTTCTTGCTTATTA

>M6-PGSC0003DMG400000996-(rev)M6LRR-RLK_1037

GCATTTCAATAATCTTGGAACCGGAAGAGAAGATGACATGGATTTCTTTTCTTCATTGGTTAACATTACCAGTTTTAAAGAACTGAGTTTGAGTGTGAATAATATAGCAGGCCAGCTGCC

>M6-PGSC0003DMG400000996-(rev)M6LRR-RLK_1098

AACATTACCAGTTTTAAAGAACTGAGTTTGAGTGTGAATAATATAGCAGGCCAGCTGCCTAAGAATATTGGCAACTTAACCAATTTCCGGTCTATTGGTTTTGCTAGGAACAAACTATTC

>M6-PGSC0003DMG400000996-(rev)M6LRR-RLK_1159

AGAATATTGGCAACTTAACCAATTTCCGGTCTATTGGTTTTGCTAGGAACAAACTATTCGGACGAATACCTGATGGATTTGTTGATCTCAGCAACATGGAAGTGGTAAGTTTGGAGTATA

>M6-PGSC0003DMG400000996-(rev)M6LRR-RLK_122

TGTTCTTTCTGCAGATGCATCTTCCAGGCTTGCTGGAAATGAAACGGATCACTTAGCCCTCCTCAGTATTAAAGCACAAATAACATATGATCCTTCTGGTGTTACAAATTCTTGGAATGA

>M6-PGSC0003DMG400000996-(rev)M6LRR-RLK_1220

ACGAATACCTGATGGATTTGTTGATCTCAGCAACATGGAAGTGGTAAGTTTGGAGTATAACCAATTGACTGAAGAAATTCCAGCTAGTCTTGGTAAACTTCAAAAACTCAAGTATTTCTA

>M6-PGSC0003DMG400000996-(rev)M6LRR-RLK_1281

CAATTGACTGAAGAAATTCCAGCTAGTCTTGGTAAACTTCAAAAACTCAAGTATTTCTATGTTAACGGAAACAAACTCTCGGGGGAAATCCCTTCCTCTATTGGAAACATAACATCTTTA

>M6-PGSC0003DMG400000996-(rev)M6LRR-RLK_1342

TTAACGGAAACAAACTCTCGGGGGAAATCCCTTCCTCTATTGGAAACATAACATCTTTATATGGACTTAATCTAGCTCAAAACAACTTAGAAGGGACTATTCCTTCAGTTCTTGGAAACT

>M6-PGSC0003DMG400000996-(rev)M6LRR-RLK_1403

TGGACTTAATCTAGCTCAAAACAACTTAGAAGGGACTATTCCTTCAGTTCTTGGAAACTGTCAGTTGTTGCAAATGTTGTATCTTTCTCGAAACAGACTCAGTGGGACGATACCAAAAGA

>M6-PGSC0003DMG400000996-(rev)M6LRR-RLK_1464

CAGTTGTTGCAAATGTTGTATCTTTCTCGAAACAGACTCAGTGGGACGATACCAAAAGAGGTTCTTAGTATCTCATCCTTGTCTATCCAACTAGACCTGTCTGGAAACCAGTTAAGTGGA

>M6-PGSC0003DMG400000996-(rev)M6LRR-RLK_1525

TTCTTAGTATCTCATCCTTGTCTATCCAACTAGACCTGTCTGGAAACCAGTTAAGTGGATCACTTCCCCTTGAAGTAGGAAGTTTAGTCAATCTTGGGTTTCTTGATATCTCCGAGAACA

>M6-PGSC0003DMG400000996-(rev)M6LRR-RLK_1586

ACTTCCCCTTGAAGTAGGAAGTTTAGTCAATCTTGGGTTTCTTGATATCTCCGAGAACAAGTTATCTGGAAAACTTCCGAGCACTTTAAGCAGTTGCATCAAGTTGGAAAACCTCTACGT

>M6-PGSC0003DMG400000996-(rev)M6LRR-RLK_1647

TTATCTGGAAAACTTCCGAGCACTTTAAGCAGTTGCATCAAGTTGGAAAACCTCTACGTTCAGGAAAACATGTTTGAAGGTGTGATCCCATCATCTTTAAGTTCTTTGAGAGGGATGGAA

>M6-PGSC0003DMG400000996-(rev)M6LRR-RLK_1708

AGGAAAACATGTTTGAAGGTGTGATCCCATCATCTTTAAGTTCTTTGAGAGGGATGGAATATTTGGACTTGTCGCATAATAATTTCTCTGGTCTTATTCCAAAGTACTTTGAAACTTTCA

>M6-PGSC0003DMG400000996-(rev)M6LRR-RLK_1769

TTTGGACTTGTCGCATAATAATTTCTCTGGTCTTATTCCAAAGTACTTTGAAACTTTCATATCATTGAAAAGCTTAAACCTGTCATTTAACAATTTTGAGGGTGAAGTACCTCGAGAAGG

>M6-PGSC0003DMG400000996-(rev)M6LRR-RLK_183

CTCAGTATTAAAGCACAAATAACATATGATCCTTCTGGTGTTACAAATTCTTGGAATGATTCCTTCCATCATTGTAGTTGGCAAGGCGTTACATGTAGTGCTCGACATCAAAGGGTGACC

>M6-PGSC0003DMG400000996-(rev)M6LRR-RLK_1830

TCATTGAAAAGCTTAAACCTGTCATTTAACAATTTTGAGGGTGAAGTACCTCGAGAAGGAGTTTTCAGTAATGCAAGTGCAGCAATAGTCAACGGAAACAGAAATCTTTGTGGAGGTTCT

>M6-PGSC0003DMG400000996-(rev)M6LRR-RLK_1891

TTTTCAGTAATGCAAGTGCAGCAATAGTCAACGGAAACAGAAATCTTTGTGGAGGTTCTCAAATGCTAAAGCTACCACAATGTAAGTTTCCAATGCCGAAAAAAAGAAGATTGTCATCAA

>M6-PGSC0003DMG400000996-(rev)M6LRR-RLK_1952

AATGCTAAAGCTACCACAATGTAAGTTTCCAATGCCGAAAAAAAGAAGATTGTCATCAACCTTGAAGATAACTATATCTGTTGCTAGTGCATTATTTGGAGTAGCACTTGTTCTGATGCT

>M6-PGSC0003DMG400000996-(rev)M6LRR-RLK_2013

TTGAAGATAACTATATCTGTTGCTAGTGCATTATTTGGAGTAGCACTTGTTCTGATGCTCCTAATTTTGTGTTTCTTAAAGAGGAAACGTAGCCCTTCACTCGATTTATCAGATGATTCG

>M6-PGSC0003DMG400000996-(rev)M6LRR-RLK_2074

TAATTTTGTGTTTCTTAAAGAGGAAACGTAGCCCTTCACTCGATTTATCAGATGATTCGTTCTTGAAAATCTCCTACAGGGAACTTCTGAAAGCAACTAATGGATTCTCCTCAGATTATT

>M6-PGSC0003DMG400000996-(rev)M6LRR-RLK_2135

CTTGAAAATCTCCTACAGGGAACTTCTGAAAGCAACTAATGGATTCTCCTCAGATTATTTAATTGGTAAGGGTGGTTTTGGTTCTGTATACAAGGGAATTCTTGGTCCAGATGAAAAAAC

>M6-PGSC0003DMG400000996-(rev)M6LRR-RLK_2196

ATTGGTAAGGGTGGTTTTGGTTCTGTATACAAGGGAATTCTTGGTCCAGATGAAAAAACTGTCGCGATCAAAGTACTCAATTTGCAGCATAAAGGAGCTTTGAAAAGCTTCATTGCAGAA

>M6-PGSC0003DMG400000996-(rev)M6LRR-RLK_2257

TCGCGATCAAAGTACTCAATTTGCAGCATAAAGGAGCTTTGAAAAGCTTCATTGCAGAATGTGAAGTCATAAAGAACCTCAGGCATAGAAATCTTGTCAAGTTAGTAACTGCATGCTCAG

>M6-PGSC0003DMG400000996-(rev)M6LRR-RLK_2318

TGAAGTCATAAAGAACCTCAGGCATAGAAATCTTGTCAAGTTAGTAACTGCATGCTCAGGTACTAACTTTCAAGGCAATGATTTTAAGGCCCTCATTTACAAGTTCATGGTTAATGGTAG

>M6-PGSC0003DMG400000996-(rev)M6LRR-RLK_2379

ACTAACTTTCAAGGCAATGATTTTAAGGCCCTCATTTACAAGTTCATGGTTAATGGTAGCCTAGACGATTGGCTACATTCATTTTCTAATGATGGAAGTCTGCACGTTCGATATCTAGAT

>M6-PGSC0003DMG400000996-(rev)M6LRR-RLK_244

CCTTCCATCATTGTAGTTGGCAAGGCGTTACATGTAGTGCTCGACATCAAAGGGTGACCGTGCTGGATTTAAGCTCTAAGCAGGTTGTAGGAACATTAGTTCCTCAGATAGGAAATATGA

>M6-PGSC0003DMG400000996-(rev)M6LRR-RLK_2440

TAGACGATTGGCTACATTCATTTTCTAATGATGGAAGTCTGCACGTTCGATATCTAGATCTTTATCAGAGAGTAAACATTGCATCAGACATTGCTTTCGCGCTAGATTATCTTCATCATG

>M6-PGSC0003DMG400000996-(rev)M6LRR-RLK_2501

TTATCAGAGAGTAAACATTGCATCAGACATTGCTTTCGCGCTAGATTATCTTCATCATGGAACCCAAACACCAGTAGTTCATTGTGATTTGAAGCCAAGCAATATCCTTCTAGACAAGGA

>M6-PGSC0003DMG400000996-(rev)M6LRR-RLK_2562

ACCCAAACACCAGTAGTTCATTGTGATTTGAAGCCAAGCAATATCCTTCTAGACAAGGACATGACTGCCCGTGTTGGCGATTTTGGTTTATCAAGATTCCTCCAAGAAACATCTCAAAGG

>M6-PGSC0003DMG400000996-(rev)M6LRR-RLK_2623

TGACTGCCCGTGTTGGCGATTTTGGTTTATCAAGATTCCTCCAAGAAACATCTCAAAGGGAAACTAGCACTATTGGTAGTAAAGGATCAGTTGGCTATGCAGCTCCAGGTAAACATCTCT

>M6-PGSC0003DMG400000996-(rev)M6LRR-RLK_2867

CAGACAAATATAAAATGGAGTGATTTTGATATGGACTAACGATTTTTTTTCTGCAGAATATGGAATGGTAAGTGAAGTATCAACTTATGGTGACATATACAGCTATGGAATCATCATACT

>M6-PGSC0003DMG400000996-(rev)M6LRR-RLK_2928

GGAATGGTAAGTGAAGTATCAACTTATGGTGACATATACAGCTATGGAATCATCATACTGGAGATGTTAACAGGGAAAAAACCGACAGACGACGCCTTTTCAAATGGAATGAACCTTCAC

>M6-PGSC0003DMG400000996-(rev)M6LRR-RLK_2989

AGATGTTAACAGGGAAAAAACCGACAGACGACGCCTTTTCAAATGGAATGAACCTTCACAACTATGCTAAAATGGCTTATTCTACTGGTAGAGTCATGGAAATTGTTGATCAAATGGTTT

>M6-PGSC0003DMG400000996-(rev)M6LRR-RLK_305

GCTGGATTTAAGCTCTAAGCAGGTTGTAGGAACATTAGTTCCTCAGATAGGAAATATGAGTTTTCTTAGAGAACTAATTCTTCAAAATAACACCTTCAATAGTGAGATACCACGCGAAAT

>M6-PGSC0003DMG400000996-(rev)M6LRR-RLK_3050

CTATGCTAAAATGGCTTATTCTACTGGTAGAGTCATGGAAATTGTTGATCAAATGGTTTACCACAACTTGCAAGAAATGAAAACCAAAGATTACATTGAAGAGTGTTCAATTTGTATGTG

>M6-PGSC0003DMG400000996-(rev)M6LRR-RLK_3111

CACAACTTGCAAGAAATGAAAACCAAAGATTACATTGAAGAGTGTTCAATTTGTATGTGTAAAATTGGGATTGCTTGCACTATAGATTCACCAAAAGAAAGAATGAGAATTAGTGATGTT

>M6-PGSC0003DMG400000996-(rev)M6LRR-RLK_3172

AAATTGGGATTGCTTGCACTATAGATTCACCAAAAGAAAGAATGAGAATTAGTGATGTTATTAAAGAGTTGCAATTAGTCAAAGAAACTTTATCTTAATGTGATAGAACACTATGATATT

>M6-PGSC0003DMG400000996-(rev)M6LRR-RLK_3232

TTAAAGAGTTGCAATTAGTCAAAGAAACTTTATCTTAATGTGATAGAACACTATGATATTAGAAGTAAATCATAAAACATTATCTTTTTTGTTCTATATATTGTGAGTTGGTAAAATTTC

>M6-PGSC0003DMG400000996-(rev)M6LRR-RLK_366

TTTCTTAGAGAACTAATTCTTCAAAATAACACCTTCAATAGTGAGATACCACGCGAAATCGGAAGGCTATTCAGGTTGAAGAATTTAGTGCTAAAGGACAACTCATTTACTGGTGAAATT

>M6-PGSC0003DMG400000996-(rev)M6LRR-RLK_427

GAAGGCTATTCAGGTTGAAGAATTTAGTGCTAAAGGACAACTCATTTACTGGTGAAATTCCAGTGGAACTATCAAATTGCTCTAGACTCATTTATCTTGATTTGGATGGTAATTCACTGA

>M6-PGSC0003DMG400000996-(rev)M6LRR-RLK_488

AGTGGAACTATCAAATTGCTCTAGACTCATTTATCTTGATTTGGATGGTAATTCACTGACCGGGAAAATCCCTGTGGAGCTTGGTTTGTCTTTGAGAAATCTTCAGGTACTTTTTCTTAG

>M6-PGSC0003DMG400000996-(rev)M6LRR-RLK_549

GGGAAAATCCCTGTGGAGCTTGGTTTGTCTTTGAGAAATCTTCAGGTACTTTTTCTTAGATCAAACAATTTAACAGGTGAACTACCATACTCTTTGGGTAATCTTTCATCCCTTATTGCC

>M6-PGSC0003DMG400000996-(rev)M6LRR-RLK_61

TTACTGCAGGCATGAAGTCTTTTAATGTTTTCATTTGTGTTATGCTTTCTTGCTTATTATGTGTTCTTTCTGCAGATGCATCTTCCAGGCTTGCTGGAAATGAAACGGATCACTTAGCCC

>M6-PGSC0003DMG400000996-(rev)M6LRR-RLK_610

CAAACAATTTAACAGGTGAACTACCATACTCTTTGGGTAATCTTTCATCCCTTATTGCCTTAGCAGCAACAGAGAATAGATTAGAAGGAAGTATACCGTATTCGTTAGGCCAGTTGACTA

>M6-PGSC0003DMG400000996-(rev)M6LRR-RLK_671

AGCAGCAACAGAGAATAGATTAGAAGGAAGTATACCGTATTCGTTAGGCCAGTTGACTAACTTGAGTTATATCTCTCTTGGTGGTAATATGCTTTCTGGTTCAATTCCCATTTCAGTTTT

>M6-PGSC0003DMG400000996-(rev)M6LRR-RLK_732

TTGAGTTATATCTCTCTTGGTGGTAATATGCTTTCTGGTTCAATTCCCATTTCAGTTTTTAACTTATCCTCTCTGTATCATCTTGCTGCACCAGTTAACCAACTTAAAGGAAGTCTTCCC

>M6-PGSC0003DMG400000996-(rev)M6LRR-RLK_793

ACTTATCCTCTCTGTATCATCTTGCTGCACCAGTTAACCAACTTAAAGGAAGTCTTCCCACTGATATTGGATCCACACTTCCGAGTCTTCGCATAGTTTTTTTGTTCTCAAACTTATTAA

>M6-PGSC0003DMG400000996-(rev)M6LRR-RLK_854

TGATATTGGATCCACACTTCCGAGTCTTCGCATAGTTTTTTTGTTCTCAAACTTATTAAGTGGAGTGCTTCCAAGTTCCATTTCTAACTTAACCAATCTTGAAATTCTCTCATTATCAAG

>M6-PGSC0003DMG400000996-(rev)M6LRR-RLK_915

GGAGTGCTTCCAAGTTCCATTTCTAACTTAACCAATCTTGAAATTCTCTCATTATCAAGAAACCAGTTGTCCGGGAAAATCCCTTCTCTGGAAAAACTCCGTAATCTTCAAGGACTAGCT

>M6-PGSC0003DMG400000996-(rev)M6LRR-RLK_976

ACCAGTTGTCCGGGAAAATCCCTTCTCTGGAAAAACTCCGTAATCTTCAAGGACTAGCTATGCATTTCAATAATCTTGGAACCGGAAGAGAAGATGACATGGATTTCTTTTCTTCATTGG

>M6-PGSC0003DMG400001016-(rev)M6LRR-RLP_0

CATACATTGACAGGTGAAATACCGAAAAAATTTAGCAATCAGATTGAGTTGTTTGATCTTAGTTTCAACAGGTTTACTGGTTCACTACCAATGGAGATTTTCAACATATCGGGTATGAGA

>M6-PGSC0003DMG400001016-(rev)M6LRR-RLP_1054

TGAGCAGCATTTCAGCTTTGGAATTTCTAGACCTTTCTCATAACAATGTTTCAGGATTGATTCCCAAGTCTTTGGAGAAGCTTCTATATCTCAAGTATTTCAACGTTTCATTCAACAAGT

>M6-PGSC0003DMG400001016-(rev)M6LRR-RLP_1178

TGGTGAAATCCCCTCGAATGGTCCTTTCAAGAACCTCTCCGGTGAGTTTTTCATGTCTAATGAAGCATTGTGTGGATCTCCAAGATTTTGTGTTCCCCCATGCCACAGTAGCACATCAAA

>M6-PGSC0003DMG400001016-(rev)M6LRR-RLP_124

TTCAAGTTATTTTCAATAATGTGACTGGAACTCTCCCATTAAACATAGGTTCTACGTTACCCAACATTGAAGTTATTCATCTGAGAAACTTAAATCTTTATGGGACTATACCTCATTCTC

>M6-PGSC0003DMG400001016-(rev)M6LRR-RLP_1240

AAGCATTGTGTGGATCTCCAAGATTTTGTGTTCCCCCATGCCACAGTAGCACATCAAAGCATAAATCCAAGAGGAAAAAAGTGATTGTTCTTGTTCTACCGGGTACACTTGTATTAGTCT

>M6-PGSC0003DMG400001016-(rev)M6LRR-RLP_1302

AAATCCAAGAGGAAAAAAGTGATTGTTCTTGTTCTACCGGGTACACTTGTATTAGTCTCCATCGCCTTTGCCCTTTTATGGTTAAGGTATACAAGAGGTAAAAGAACTGATCCTCAGCAA

>M6-PGSC0003DMG400001016-(rev)M6LRR-RLP_1364

CGCCTTTGCCCTTTTATGGTTAAGGTATACAAGAGGTAAAAGAACTGATCCTCAGCAAGCAGTTGATTCATCCTTTGTCATCTCAACAAAAGGAAGAATTTCATACTATGAACTACTTCG

>M6-PGSC0003DMG400001016-(rev)M6LRR-RLP_1426

TTGATTCATCCTTTGTCATCTCAACAAAAGGAAGAATTTCATACTATGAACTACTTCGAGCAACTGATTCACTTAGTGAGATCAATTTAATTGGTTCTGGGAGTTTCGGCTCTGTTTACA

>M6-PGSC0003DMG400001016-(rev)M6LRR-RLP_1488

ACTGATTCACTTAGTGAGATCAATTTAATTGGTTCTGGGAGTTTCGGCTCTGTTTACAAAGGCATCCTCAGAGATGGGACTTTCATTGCAGCTAAAGTGTTCAACCCACAATTGCAAGTG

>M6-PGSC0003DMG400001016-(rev)M6LRR-RLP_1550

CATCCTCAGAGATGGGACTTTCATTGCAGCTAAAGTGTTCAACCCACAATTGCAAGTGGCATTTAGGAGTTTTGATACAGAATGTCAAGTTTTGCGCAACCTTCGTCATAGGAATCTCAC

>M6-PGSC0003DMG400001016-(rev)M6LRR-RLP_1674

GTCATCACCAGTTGTTCCAACCTTGAATTTAAGGCTTTAATACTCGAGTACATGCCCAATGGGAGCCTCAATAAGTGGTTGTATTCACACAACTACTTCTTAGACATCAAACATAGGCTA

>M6-PGSC0003DMG400001016-(rev)M6LRR-RLP_1736

GAGCCTCAATAAGTGGTTGTATTCACACAACTACTTCTTAGACATCAAACATAGGCTAAGTATAATGATAGATGTAGCATGTGCATTGGAATATCTACACCATGGATGCTCATTACCCGT

>M6-PGSC0003DMG400001016-(rev)M6LRR-RLP_1798

TAATGATAGATGTAGCATGTGCATTGGAATATCTACACCATGGATGCTCATTACCCGTGATTCATTGTGATTTGAAGCCTAGTAACGTCTTGTTGGATGAGGAAATGGTTGCCCACCTAA

>M6-PGSC0003DMG400001016-(rev)M6LRR-RLP_186

AACATTGAAGTTATTCATCTGAGAAACTTAAATCTTTATGGGACTATACCTCATTCTCTCTCCAATTGTTCCAAACTCACTAGTCTAGACCTTTCTCTCAATAGACTAACAGGTTTGATT

>M6-PGSC0003DMG400001016-(rev)M6LRR-RLP_1860

CATTGTGATTTGAAGCCTAGTAACGTCTTGTTGGATGAGGAAATGGTTGCCCACCTAAGTGACTTTGGAATTTCAAAGTTGCTTTCTGAAGATGAGAGTTCTTTATACACTAAAACATTA

>M6-PGSC0003DMG400001016-(rev)M6LRR-RLP_1984

CGTTTGGTTATATTGCACCAGGTACGTGATCTCTTAATCAATTGCTTTTGATTATTTCTCTTTCCTTATTTTCCACTAGTAAATGTTGAATTTCAAAATTGATTGATTGCAGAGTATGGA

>M6-PGSC0003DMG400001016-(rev)M6LRR-RLP_2046

TCCTTATTTTCCACTAGTAAATGTTGAATTTCAAAATTGATTGATTGCAGAGTATGGACTGGAGGGATTGGTGTCAATAAAATGTGATGTTTATAGTTAAGGGATCATGTTGATGGAAAC

>M6-PGSC0003DMG400001016-(rev)M6LRR-RLP_2170

ACTAGGAAAAATCCTAATGATGAAATGTTCGATGGAGATTTTAGTTTGAAGCAATGGGTGAGAAATTCACTCCCACAAGCAGCAATGGAAATTGTTGATGCCAACTTAATATCGCCACAT

>M6-PGSC0003DMG400001016-(rev)M6LRR-RLP_2232

AAATTCACTCCCACAAGCAGCAATGGAAATTGTTGATGCCAACTTAATATCGCCACATGATAATCAATTAATGAAAAAGCTAGATTGTGTGGTATCCATCATGAAAGTGGCAATAGATTG

>M6-PGSC0003DMG400001016-(rev)M6LRR-RLP_2356

GTTGAATCTCCAAAAGGAAGGATTGACATGAAAGATGTTGTGGGGAGGTTAAAGACGATCAAGATTCAACTTCTTGCATGTTGAGGAAGATCTTATTAGAAACTTTTCTTCCAAATCATC

>M6-PGSC0003DMG400001016-(rev)M6LRR-RLP_2418

GATTCAACTTCTTGCATGTTGAGGAAGATCTTATTAGAAACTTTTCTTCCAAATCATCTCATTGTAACAAACATTTGCAATAATAGTTAGCTTCAAATAAGTTATAGCATTATGTCTTTC

>M6-PGSC0003DMG400001016-(rev)M6LRR-RLP_248

CAATTGTTCCAAACTCACTAGTCTAGACCTTTCTCTCAATAGACTAACAGGTTTGATTCCCAACTCTCTTGGAAGTTTGACTCATCTACAGTCCCTAAGCTTGGCAATTAACAATTTAAT

>M6-PGSC0003DMG400001016-(rev)M6LRR-RLP_2480

TGTAACAAACATTTGCAATAATAGTTAGCTTCAAATAAGTTATAGCATTATGTCTTTCATAACATAGAGAATTGATGGGCATGTATACTTCTTAGGTCTTGAATTTGTAAAGTGGTTTTG

>M6-PGSC0003DMG400001016-(rev)M6LRR-RLP_310

ACTCTCTTGGAAGTTTGACTCATCTACAGTCCCTAAGCTTGGCAATTAACAATTTAATGGGCGATAGTAGGTTAAGCTTCTTGACTTCCTTGTCAAATTGTAGAGATTTAAAAATTATTT

>M6-PGSC0003DMG400001016-(rev)M6LRR-RLP_372

GATAGTAGGTTAAGCTTCTTGACTTCCTTGTCAAATTGTAGAGATTTAAAAATTATTTTCCTCTCTGCGAACCCTCTATATGGTGTGCTTCCAGAATCCATAGGGAACCTTTCCAGTACT

>M6-PGSC0003DMG400001016-(rev)M6LRR-RLP_434

CTCTGCGAACCCTCTATATGGTGTGCTTCCAGAATCCATAGGGAACCTTTCCAGTACTTCTCTTCTAAGGTTTAGAGCTATTGATTGTAAAATTAAGGGGGAAATTCCAAAAGGATTTGG

>M6-PGSC0003DMG400001016-(rev)M6LRR-RLP_496

TTCTAAGGTTTAGAGCTATTGATTGTAAAATTAAGGGGGAAATTCCAAAAGGATTTGGGAACTTAAGCAGCTTATTAGAACTCGATCTTTCTAGAAACGGTTTCACTGGACCAATTCCCA

>M6-PGSC0003DMG400001016-(rev)M6LRR-RLP_558

TTAAGCAGCTTATTAGAACTCGATCTTTCTAGAAACGGTTTCACTGGACCAATTCCCACAACAATCAGCAACTTGAGATTCCTTCAGAGCTTGAAGTTGAGTGTCAACAAACTTTCTGGA

>M6-PGSC0003DMG400001016-(rev)M6LRR-RLP_62

TTTCAACAGGTTTACTGGTTCACTACCAATGGAGATTTTCAACATATCGGGTATGAGAGAGATTCAAGTTATTTTCAATAATGTGACTGGAACTCTCCCATTAAACATAGGTTCTACGTT

>M6-PGSC0003DMG400001016-(rev)M6LRR-RLP_620

AATCAGCAACTTGAGATTCCTTCAGAGCTTGAAGTTGAGTGTCAACAAACTTTCTGGATATATTGGAGATGATCTATGCAAATTACAGAACTTGGGTTATTTAAACTTGACTTTGAATCA

>M6-PGSC0003DMG400001016-(rev)M6LRR-RLP_682

TTGGAGATGATCTATGCAAATTACAGAACTTGGGTTATTTAAACTTGACTTTGAATCAACTTTCAGGATCTCTTCCGAATTGTTTGGGAAACTTAACTTCCCTTAGGGAGATGATTCTAG

>M6-PGSC0003DMG400001016-(rev)M6LRR-RLP_744

TCAGGATCTCTTCCGAATTGTTTGGGAAACTTAACTTCCCTTAGGGAGATGATTCTAGGTTCCAACAAATTGCGTTCAATCATTCCAAAAAGCTTAGGAAACCTCATAAACCTCTTGAAG

>M6-PGSC0003DMG400001016-(rev)M6LRR-RLP_806

CAACAAATTGCGTTCAATCATTCCAAAAAGCTTAGGAAACCTCATAAACCTCTTGAAGCTAGACTTATCCTCAAACAACTTGGGTGGCTCTATACCTCTGGAAATTGGAAATCTAAAGGC

>M6-PGSC0003DMG400001016-(rev)M6LRR-RLP_930

ATATATATGGATCTATCAATAAATACACTCTCAAATGGAATTCCTAAAGAAATTGATAGCTTGCAAAATCTTATATACCTTTCTTTGAGAGACAACAAATTGCAAGGATCAATTCCTGGC

>M6-PGSC0003DMG400001016-(rev)M6LRR-RLP_992

GCAAAATCTTATATACCTTTCTTTGAGAGACAACAAATTGCAAGGATCAATTCCTGGCTCAATGAGCAGCATTTCAGCTTTGGAATTTCTAGACCTTTCTCATAACAATGTTTCAGGATT

>M6-PGSC0003DMG400001059M6G-LecRK_0

AAAAATTTTATACTAGAACAATAGTTCTTTAATGGCATGTCTAAAAATCCTACTATTTTTCCTCTTCATGAACATTTCTTTTGCCCAAAAAAACACAATTTTTCAAGGCCAACACCTTAG

>M6-PGSC0003DMG400001059M6G-LecRK_1037

AAAACTGTTCTTGTGCTGCTTTTGCCTCTATCACAGCTAATGGAACTGGTTGTGATATTTGGAGTAACGTATCGATTTTGAACTCACTTGAATCGGATGTTTTCATTCTTGGTGATGAAA

>M6-PGSC0003DMG400001059M6G-LecRK_1098

GAGTAACGTATCGATTTTGAACTCACTTGAATCGGATGTTTTCATTCTTGGTGATGAAAGAGGTATGTTCATATTTATCTCAAATATCCTTTTTATGTTGTTGTCGTGTGACCAGGAGGT

>M6-PGSC0003DMG400001059M6G-LecRK_122

TTGGAGAAAAATTGGAATCAGCAAACAAGGAATTCAGACTTGAGTTTTTTAGCCTTGATGCCAATAAAACACATTATATTGGCATATTCTACAATTTACCTTCTAACATGACCTTATTCC

>M6-PGSC0003DMG400001059M6G-LecRK_1281

ACAGTTGACTCTTGTGGTCCAGGGAAGTGCACTATTTCTATTGTCTAGCAAGTAACTTGAGTTAGTAAAGCGGACTCTAACGATGAATTGTGCAATGTGTATTCATAGATTTGGTGCCAA

>M6-PGSC0003DMG400001059M6G-LecRK_1342

TTAGTAAAGCGGACTCTAACGATGAATTGTGCAATGTGTATTCATAGATTTGGTGCCAATAAGAAATCGGAACAGCACAGATGTACCAAGCACAATCTCACTTCCTCCAGATTTAGCACC

>M6-PGSC0003DMG400001059M6G-LecRK_1403

AGAAATCGGAACAGCACAGATGTACCAAGCACAATCTCACTTCCTCCAGATTTAGCACCAGAACCATCTCCTTATTCAACGTCAGGATCATCTAAAGGAACTTGTATGTATTGTGTATAA

>M6-PGSC0003DMG400001059M6G-LecRK_1464

AACCATCTCCTTATTCAACGTCAGGATCATCTAAAGGAACTTGTATGTATTGTGTATAATCTTGCTCTTGAATTATTGTATATGATTGAAGGTGAAAACCTTTTTTTGATTGTTTCCTCG

>M6-PGSC0003DMG400001059M6G-LecRK_1647

ACACAATCTAGTCCTTGTTTCGAGTATATCCAGAAAAAGATCATTTACATCATCTGTATTGATGTTGTTACTGCTCCTTTCCTGAATGTTCTACACTGTTTTCCTTTTTACTTGTCATGT

>M6-PGSC0003DMG400001059M6G-LecRK_1708

ATGTTGTTACTGCTCCTTTCCTGAATGTTCTACACTGTTTTCCTTTTTACTTGTCATGTTTTCTGCACTGTCGTTTTTCTCCGTTTCCATAACTACTATGATTTGTTGCTCTTGAGCCAA

>M6-PGSC0003DMG400001059M6G-LecRK_183

CAATAAAACACATTATATTGGCATATTCTACAATTTACCTTCTAACATGACCTTATTCCCCGATGATGATCGCCCCGTTTGGGTTGCAAATCGCGATATTCCGATACAATATGCATCAGG

>M6-PGSC0003DMG400001059M6G-LecRK_1952

GAAAATGGTGGGTATGGCTTATAGCTGCAAGCGGTTTGACACCATTTGTAGGATTTTTCGCCCTATGCTACTTTCTACACGGAAAAGGTGAGCGCTCAAAGAATTTATCTAGAAGCTTGC

>M6-PGSC0003DMG400001059M6G-LecRK_2013

CCTATGCTACTTTCTACACGGAAAAGGTGAGCGCTCAAAGAATTTATCTAGAAGCTTGCACCAGTAAAACTCAACTATGTTGCTCGGACTCTGCTAAAATGCTGCCACAGTACTCCTGTC

>M6-PGSC0003DMG400001059M6G-LecRK_2074

CAGTAAAACTCAACTATGTTGCTCGGACTCTGCTAAAATGCTGCCACAGTACTCCTGTCGGATCCTCAAAAAAAAAAACGCACTACTTTTGAAGGATCTAACACACACTAAACAACATTT

>M6-PGSC0003DMG400001059M6G-LecRK_2135

ATCCTCAAAAAAAAAAACGCACTACTTTTGAAGGATCTAACACACACTAAACAACATTTTTTAAAAGTCCGAGCAATATAGAAATGCTCTATACTATGGTTTCTGATCCCACAACTTCCT

>M6-PGSC0003DMG400001059M6G-LecRK_2196

TAAAAGTCCGAGCAATATAGAAATGCTCTATACTATGGTTTCTGATCCCACAACTTCCTCTTTCACTATCAGGGAAAGCAAAAGCAACAGCTTTGCTATTGCTTAATCAAACAGCAAATA

>M6-PGSC0003DMG400001059M6G-LecRK_2257

TTCACTATCAGGGAAAGCAAAAGCAACAGCTTTGCTATTGCTTAATCAAACAGCAAATATATCTAAAAAGAGAAAAATAGACAAGAAAATGAGCCATGAAGTGCAGTTATACAGCCTCGA

>M6-PGSC0003DMG400001059M6G-LecRK_2318

TCTAAAAAGAGAAAAATAGACAAGAAAATGAGCCATGAAGTGCAGTTATACAGCCTCGAGAGCCTAGCAATCGCCACAGACAATTTTTCACCAGGAAATAAGCTCGGGGAAGGTGGATTC

>M6-PGSC0003DMG400001059M6G-LecRK_2379

GCCTAGCAATCGCCACAGACAATTTTTCACCAGGAAATAAGCTCGGGGAAGGTGGATTCGGACCAGTTTACAAGGTAAACATGAAACCTTTTATATGAATATGATTCGCGTAAGCATAAA

>M6-PGSC0003DMG400001059M6G-LecRK_244

GATGATGATCGCCCCGTTTGGGTTGCAAATCGCGATATTCCGATACAATATGCATCAGGTAACACACTCACATTGGATGATGATGGAAAGTTGAAAATTTTCTATGGAGAGGATAGTTTT

>M6-PGSC0003DMG400001059M6G-LecRK_2440

ACCAGTTTACAAGGTAAACATGAAACCTTTTATATGAATATGATTCGCGTAAGCATAAATACTTAAACGAGTCATTTTTTTTCAGGGAGAAATGCCTGATGGGCAAGAAGTTGCCATAAA

>M6-PGSC0003DMG400001059M6G-LecRK_2501

CTTAAACGAGTCATTTTTTTTCAGGGAGAAATGCCTGATGGGCAAGAAGTTGCCATAAAAAGACTTTCGACTAGTTCAGGACAAGGCCTATTGGAGTTCAAGAATGAAATTCTATTGATT

>M6-PGSC0003DMG400001059M6G-LecRK_2562

GACTTTCGACTAGTTCAGGACAAGGCCTATTGGAGTTCAAGAATGAAATTCTATTGATTGCAAAACTTCAACATACTAACCTTGTTAGGCTTCTAGGATACTGTACTCTACGCGAAGAAC

>M6-PGSC0003DMG400001059M6G-LecRK_2623

AAAACTTCAACATACTAACCTTGTTAGGCTTCTAGGATACTGTACTCTACGCGAAGAACGGATTATAGTTTATGAATACATGCACAACAAAAGCCTAGACTTCTTCCTCTTTGGTAAACA

>M6-PGSC0003DMG400001059M6G-LecRK_2684

ATTATAGTTTATGAATACATGCACAACAAAAGCCTAGACTTCTTCCTCTTTGGTAAACATCATGTTTTTCATCTTCAACAATTAGAACAACTTTGTCAAGAAATTTCATCTTTTACAATG

>M6-PGSC0003DMG400001059M6G-LecRK_2745

ATGTTTTTCATCTTCAACAATTAGAACAACTTTGTCAAGAAATTTCATCTTTTACAATGATATGAACAGATAGTAACAAAAAGGAGCTACTAAATTGGGACACACGATTCAGAATCATCG

>M6-PGSC0003DMG400001059M6G-LecRK_2806

ATGAACAGATAGTAACAAAAAGGAGCTACTAAATTGGGACACACGATTCAGAATCATCGAAGGGATTGCTCAGGGTATACTGTATCTGCATAAATATTCAAGGCTGAAAGTGATTCATAG

>M6-PGSC0003DMG400001059M6G-LecRK_2867

GGGATTGCTCAGGGTATACTGTATCTGCATAAATATTCAAGGCTGAAAGTGATTCATAGAGATTTAAAAGCAAGCAACATCTTACTTGACGCGGAAATGAATCCAAAGATATCAGACTTT

>M6-PGSC0003DMG400001059M6G-LecRK_2928

ATTTAAAAGCAAGCAACATCTTACTTGACGCGGAAATGAATCCAAAGATATCAGACTTTGGCATGGCGAGGATATTTGGAACACAAGAATCTGAAGCAAACACGAAAAGAATTGTTGGAA

>M6-PGSC0003DMG400001059M6G-LecRK_2989

CATGGCGAGGATATTTGGAACACAAGAATCTGAAGCAAACACGAAAAGAATTGTTGGAACACAGTAAGTAAATAAGCAATGTTTTCATAAACAAGTTTTTATATAATATTACTTGAAGTT

>M6-PGSC0003DMG400001059M6G-LecRK_305

ACACACTCACATTGGATGATGATGGAAAGTTGAAAATTTTCTATGGAGAGGATAGTTTTTTTATGTTAAGTTATTATAATGTAACAACAAGAAATACAAGTGCTACACTATTTGATAATG

>M6-PGSC0003DMG400001059M6G-LecRK_3050

CAGTAAGTAAATAAGCAATGTTTTCATAAACAAGTTTTTATATAATATTACTTGAAGTTCTGAATACATTTTTCGGATACTACTTCAGTGGCTATATGTCTCCAGAGTATGCCTTGAGAG

>M6-PGSC0003DMG400001059M6G-LecRK_3111

GAATACATTTTTCGGATACTACTTCAGTGGCTATATGTCTCCAGAGTATGCCTTGAGAGGCATTGTTTCAACGAAAACGGATGTATTCAGCTTTGGAGTTTTACTGTTAGAAATTGTTAG

>M6-PGSC0003DMG400001059M6G-LecRK_3172

ATTGTTTCAACGAAAACGGATGTATTCAGCTTTGGAGTTTTACTGTTAGAAATTGTTAGTGGCAAGAAAAACAACAGCTGCTATGATTCCGAACATCCACTAAACCTCATAGGACTGGTA

>M6-PGSC0003DMG400001059M6G-LecRK_3233

GCAAGAAAAACAACAGCTGCTATGATTCCGAACATCCACTAAACCTCATAGGACTGGTAAGTTTCTAGATTATATTATGAACATATCTCGCGAAATGGAGAAACATATAACTATTAGTAC

>M6-PGSC0003DMG400001059M6G-LecRK_3294

TTTCTAGATTATATTATGAACATATCTCGCGAAATGGAGAAACATATAACTATTAGTACTATGTTTTTAGGCATGGGAATTGTGGAGAGAAGAGAGAGCTTTGGAGCTAATAGATGCAAC

>M6-PGSC0003DMG400001059M6G-LecRK_3355

TGTTTTTAGGCATGGGAATTGTGGAGAGAAGAGAGAGCTTTGGAGCTAATAGATGCAACATTAATTGAATCATGCTCGCGTGATGAAGTAATGAGATGCATACATGTGGGACTCCTTTGT

>M6-PGSC0003DMG400001059M6G-LecRK_3416

TAATTGAATCATGCTCGCGTGATGAAGTAATGAGATGCATACATGTGGGACTCCTTTGTGTGCAAGATTATGCAAAAGATAGGCCTTCAATGTCAAATGTTGTTTCAATGCTTGTGAATG

>M6-PGSC0003DMG400001059M6G-LecRK_3477

GCAAGATTATGCAAAAGATAGGCCTTCAATGTCAAATGTTGTTTCAATGCTTGTGAATGATACGAAACAACCACCACCACCACCAGAACGACCAGGATTTTTCATCGAGAGAGGGGATCA

>M6-PGSC0003DMG400001059M6G-LecRK_3538

ACGAAACAACCACCACCACCACCAGAACGACCAGGATTTTTCATCGAGAGAGGGGATCAACGTGCAGAGATCTCCGAAGAAGTAGTCAGATACTCGATAAACGGGCTATCAATATCAGAG

>M6-PGSC0003DMG400001059M6G-LecRK_3599

GTGCAGAGATCTCCGAAGAAGTAGTCAGATACTCGATAAACGGGCTATCAATATCAGAGTTGAGAGCAAGATAAGATACAAGATGGTCAATTGGGAAGTAACTTATTTAACATTCTTTTG

>M6-PGSC0003DMG400001059M6G-LecRK_366

TATGTTAAGTTATTATAATGTAACAACAAGAAATACAAGTGCTACACTATTTGATAATGGAAATTTTGTGTTGGTGGAATTGAATACTAATGGTTCAGAGAATAAAACATTGTGGCAAAG

>M6-PGSC0003DMG400001059M6G-LecRK_3660

GAGAGCAAGATAAGATACAAGATGGTCAATTGGGAAGTAACTTATTTAACATTCTTTTGATGAAGTGTAAAAAGTTACCAACTTCAAGTACATACTTATAGGACTAATTTCGAGTCCTGT

>M6-PGSC0003DMG400001059M6G-LecRK_3721

GAAGTGTAAAAAGTTACCAACTTCAAGTACATACTTATAGGACTAATTTCGAGTCCTGTTTCTTTGATACTGTTGCATTATGATGCATTTTGTGAGTAGAATATTTGAGATGGATCCAGA

>M6-PGSC0003DMG400001059M6G-LecRK_3782

CTTTGATACTGTTGCATTATGATGCATTTTGTGAGTAGAATATTTGAGATGGATCCAGAATTTAAGTCAAAGATGGTAAACGAGATAAGGCAAGGCGAAGTTATGGCAATGGAAACTTCA

>M6-PGSC0003DMG400001059M6G-LecRK_3843

TTAAGTCAAAGATGGTAAACGAGATAAGGCAAGGCGAAGTTATGGCAATGGAAACTTCACCATTACCTTCCCCCTCCCGCCTGATAGCAACGTCACCAGTTACATTTAAAATGTTTTGAA

>M6-PGSC0003DMG400001059M6G-LecRK_3904

ATTACCTTCCCCCTCCCGCCTGATAGCAACGTCACCAGTTACATTTAAAATGTTTTGAATACACTTGACACTCATCGATTATATAAGTTGAAAGAGAATATGAGAGGAGTTTCTGAACTA

>M6-PGSC0003DMG400001059M6G-LecRK_3965

CACTTGACACTCATCGATTATATAAGTTGAAAGAGAATATGAGAGGAGTTTCTGAACTATCAATATTTGTATATTAGACAAAAGAGGTAACTGAATGTACATATATTTTCTCATTCTGAG

>M6-PGSC0003DMG400001059M6G-LecRK_4026

AATATTTGTATATTAGACAAAAGAGGTAACTGAATGTACATATATTTTCTCATTCTGAGCTTTACATGTTCCACAAATGGGCTGTGAATTAGTTTACACTCAATTCACTACAAAAATAAA

>M6-PGSC0003DMG400001059M6G-LecRK_4087

TTACATGTTCCACAAATGGGCTGTGAATTAGTTTACACTCAATTCACTACAAAAATAAAGAACAATTTTGGTTTTCACCTAAACTCACTACAATTCTGATGATTTAGCTTCATCAATATT

>M6-PGSC0003DMG400001059M6G-LecRK_4148

ACAATTTTGGTTTTCACCTAAACTCACTACAATTCTGATGATTTAGCTTCATCAATATTGTTATCTTTATCGCTAACAAAAACTAAACGATCACCATCTCTGATCAACTCAATTTCGTCG

>M6-PGSC0003DMG400001059M6G-LecRK_4209

TATCTTTATCGCTAACAAAAACTAAACGATCACCATCTCTGATCAACTCAATTTCGTCGATTTCAGCTCCATCTTTGCTTACAACTTTGGCCTGAGAGATACCATATCTACTAGAACCAA

>M6-PGSC0003DMG400001059M6G-LecRK_427

AATTTTGTGTTGGTGGAATTGAATACTAATGGTTCAGAGAATAAAACATTGTGGCAAAGTTTTGATTATCCTACTGATACACTATTACCTGGGATGAAACTTGGGAGAAATGTTAAAACA

>M6-PGSC0003DMG400001059M6G-LecRK_4270

TTCAGCTCCATCTTTGCTTACAACTTTGGCCTGAGAGATACCATATCTACTAGAACCAAGTTGAAGTAGCTCTTTAAAACTCTGTGGGAGTAAAATGAGTTTCCCCCCGTTGTCTCCTTT

>M6-PGSC0003DMG400001059M6G-LecRK_4331

TGAAGTAGCTCTTTAAAACTCTGTGGGAGTAAAATGAGTTTCCCCCCGTTGTCTCCTTTCTCAGGGCAACACACTGTCACTCGGGGAGCATAAGTCTTGGTTGTCATGACACTTACATTA

>M6-PGSC0003DMG400001059M6G-LecRK_4392

CAGGGCAACACACTGTCACTCGGGGAGCATAAGTCTTGGTTGTCATGACACTTACATTACTGTCGTTTGCAGATAAAAGCACGTCATGCTCATTGGTCTGCTTTGCTGACATTATCCCGA

>M6-PGSC0003DMG400001059M6G-LecRK_4453

GTCGTTTGCAGATAAAAGCACGTCATGCTCATTGGTCTGCTTTGCTGACATTATCCCGAATAATGAGTTGTGGAAGTTATTAGACCTACGTCTAGGACGCGAACGCCCTAATGATCCATC

>M6-PGSC0003DMG400001059M6G-LecRK_4514

AATGAGTTGTGGAAGTTATTAGACCTACGTCTAGGACGCGAACGCCCTAATGATCCATCTAGTCCTAGAAATGAGACTCCATGGGATGCAGGGGTGATTGTTGGTTCACTTTTGAACCTC

>M6-PGSC0003DMG400001059M6G-LecRK_4575

GTCCTAGAAATGAGACTCCATGGGATGCAGGGGTGATTGTTGGTTCACTTTTGAACCTCCCAAGAAATCTAACCCCTCGTTCCTCGGGGATAGGATCAACGGATCGAGTTCTCATCATTA

>M6-PGSC0003DMG400001059M6G-LecRK_4636

AAGAAATCTAACCCCTCGTTCCTCGGGGATAGGATCAACGGATCGAGTTCTCATCATTACCCTGGATTCGAAGAGTTCTTTGATGTCTTCATGTCCTTGTTGCTCAGCAAGGTCCCGGGG

>M6-PGSC0003DMG400001059M6G-LecRK_4697

CTGGATTCGAAGAGTTCTTTGATGTCTTCATGTCCTTGTTGCTCAGCAAGGTCCCGGGGGGTCCAATTATGTTCGTCTGGTTGATCAACGTTAGCCCCTCGATCCAAGAGGTACTTTACT

>M6-PGSC0003DMG400001059M6G-LecRK_4758

TCCAATTATGTTCGTCTGGTTGATCAACGTTAGCCCCTCGATCCAAGAGGTACTTTACTATTTCTATGTTATCTTCACAAACAGCAACATGGAGAGCTGATGAACCGTTGATCTTGGGAA

>M6-PGSC0003DMG400001059M6G-LecRK_4819

TTCTATGTTATCTTCACAAACAGCAACATGGAGAGCTGATGAACCGTTGATCTTGGGAAGCGTGACATCCCCGCCATGACGGACAATCTCCTTGAGCAAATTCAAGTTGTTTTGTTCAAC

>M6-PGSC0003DMG400001059M6G-LecRK_488

TTGATTATCCTACTGATACACTATTACCTGGGATGAAACTTGGGAGAAATGTTAAAACAGGACAGATGTGGTCACTGGTTTCTTGGATAAGCAAAGATGTACCTGCCTCGGGGTCATTTA

>M6-PGSC0003DMG400001059M6G-LecRK_4880

GTGACATCCCCGCCATGACGGACAATCTCCTTGAGCAAATTCAAGTTGTTTTGTTCAACAGCAACGCAGGCGAAATGTCCCACATCACCAGCTGATAGTTTAGCACCGTTGTCAACGAGT

>M6-PGSC0003DMG400001059M6G-LecRK_4941

CAACGCAGGCGAAATGTCCCACATCACCAGCTGATAGTTTAGCACCGTTGTCAACGAGTAACTGGATAACTGACTCGTGCTTCCCCATGATAGCCTCCCACAATGGCACATTGCCTTCTG

>M6-PGSC0003DMG400001059M6G-LecRK_5002

CTGGATAACTGACTCGTGCTTCCCCATGATAGCCTCCCACAATGGCACATTGCCTTCTGAATCTGTGTATACACAAGAAACAAAATCAAATTCTTATATCTGGCTTTGTGTCAGTGGACA

>M6-PGSC0003DMG400001059M6G-LecRK_5063

TCTGTGTATACACAAGAAACAAAATCAAATTCTTATATCTGGCTTTGTGTCAGTGGACATGGTAGGAAAAGGAGACAACTGATAGTTAGAGTGTGAAACTCGCAAAAAAAAAAACGATAG

>M6-PGSC0003DMG400001059M6G-LecRK_5124

GTAGGAAAAGGAGACAACTGATAGTTAGAGTGTGAAACTCGCAAAAAAAAAAACGATAGTTGAGGTGTGTTTTTGACTATAAACTCTCTAATGTTTGGACCAAAACAAACTTTTGGCAAC

>M6-PGSC0003DMG400001059M6G-LecRK_5185

GAGGTGTGTTTTTGACTATAAACTCTCTAATGTTTGGACCAAAACAAACTTTTGGCAACAACAGAATTTGAAAAAAGAAATGGAGCATACCTCTACTATTGGGATCAGCCCCAAAATCGA

>M6-PGSC0003DMG400001059M6G-LecRK_5229

CAAACTTTTGGCAACAACAGAATTTGAAAAAAGAAATGGAGCATACCTCTACTATTGGGATCAGCCCCAAAATCGAGCAGAAGAACCACACAGTTCTCAATGCCTTTAGACGCAGCGATA

>M6-PGSC0003DMG400001059M6G-LecRK_549

ACAGATGTGGTCACTGGTTTCTTGGATAAGCAAAGATGTACCTGCCTCGGGGTCATTTACGCTTGGTATAGACATGAATGACATTGACCAATTGATAATATGGTGGATGGGAAGTGTGTT

>M6-PGSC0003DMG400001059M6G-LecRK_61

CTCTTCATGAACATTTCTTTTGCCCAAAAAAACACAATTTTTCAAGGCCAACACCTTAGAGTTGGAGAAAAATTGGAATCAGCAAACAAGGAATTCAGACTTGAGTTTTTTAGCCTTGAT

>M6-PGSC0003DMG400001059M6G-LecRK_610

CTTGGTATAGACATGAATGACATTGACCAATTGATAATATGGTGGATGGGAAGTGTGTTTTGGACGAGTGGACGCTGGGGAAATGAGACGTTTGGCAATGTATCGCGAGTATCACACTAT

>M6-PGSC0003DMG400001059M6G-LecRK_671

GGACGAGTGGACGCTGGGGAAATGAGACGTTTGGCAATGTATCGCGAGTATCACACTATGAGTATGTTAACTTGAGTTTTGTTTCGACAGAGGATGAGAAGTATGTGACTTATTCTGTCA

>M6-PGSC0003DMG400001059M6G-LecRK_732

GTATGTTAACTTGAGTTTTGTTTCGACAGAGGATGAGAAGTATGTGACTTATTCTGTCAGTGAGACTCGAACTTTGTCTAGATATGTAATGGATACATTTGGTTTTATTAAGGAAAGAGG

>M6-PGSC0003DMG400001059M6G-LecRK_793

GAGACTCGAACTTTGTCTAGATATGTAATGGATACATTTGGTTTTATTAAGGAAAGAGGAGCAGCAGGACCTTTTGGGGTGTGTTTTTACAAGCCTAGTGCTGGTTGTGTGACAGAAGAA

>M6-PGSC0003DMG400001059M6G-LecRK_854

CAGCAGGACCTTTTGGGGTGTGTTTTTACAAGCCTAGTGCTGGTTGTGTGACAGAAGAATCGATTGGGTGCTCGATCAGAAACGATTCGTGGTTCAAGAGGAGTCCAAATAGTGTCTCTG

>M6-PGSC0003DMG400001059M6G-LecRK_915

GATTGGGTGCTCGATCAGAAACGATTCGTGGTTCAAGAGGAGTCCAAATAGTGTCTCTGGTAATAGGTTTAGGTTTGAAGACAACCACATGAGCTTATTTGATTGCAAGAGACAGTGTGA

>M6-PGSC0003DMG400001059M6G-LecRK_976

AATAGGTTTAGGTTTGAAGACAACCACATGAGCTTATTTGATTGCAAGAGACAGTGTGAGAAAAACTGTTCTTGTGCTGCTTTTGCCTCTATCACAGCTAATGGAACTGGTTGTGATATT

>M6-PGSC0003DMG400001072-(rev)M6LRR-RLP_0

ATCTCTTCTTCTCCACAAAACTTCATCTCTCTCATCTCACATTTTGCTTAAAGATTCCATTTTTAAACCAAAAATTAAAAAAAACTCCACAAACCCCATTTCTTAAATCCATTTTTCTTC

>M6-PGSC0003DMG400001072-(rev)M6LRR-RLP_1054

TTTAATGAATAATGGGTTGCGTTCTTGTTTGCCGGCGGAGATTGGGTTGTTGAAAAATCTGACTGTATTTGATGTGAGCTTTAATCAGTTGATGGGTCCGTTGCCGGAGAATTTTGGTGA

>M6-PGSC0003DMG400001072-(rev)M6LRR-RLP_124

ATCTTCACTTTTTTCATCTCACATTTGGTTAAAGATTCAATCTTTGAACAGAAAAAAGCAAAAAAAAACCAAGCATTCTTGATTTTCAGATGACTTATGTGAGATGAAGGAGAAAACTCA

>M6-PGSC0003DMG400001072-(rev)M6LRR-RLP_186

AAAAAACCAAGCATTCTTGATTTTCAGATGACTTATGTGAGATGAAGGAGAAAACTCAAACACCCATTTGCTCTCTGTTTTTTCTTGTGCTCTGTTTTTTGTGTTCAGTTTTTACTGAAC

>M6-PGSC0003DMG400001072-(rev)M6LRR-RLP_2108

CCTTCCCCATTGCCTTGTATAGAACCCCCTCCACCACCTCCTCCTTGCATTGAACCTCCCCCACCTCCCTCCCCTTCACCATCTCCTCCTCCACCACCTGTTTATCATTACAATTCTCCG

>M6-PGSC0003DMG400001072-(rev)M6LRR-RLP_2170

ACCTCCCTCCCCTTCACCATCTCCTCCTCCACCACCTGTTTATCATTACAATTCTCCGCCCCCACCATCTCTAGCACCTCCACCACCAGTTTATTTTTACAATTCTCCACCTCCACCATC

>M6-PGSC0003DMG400001072-(rev)M6LRR-RLP_2232

CACCATCTCTAGCACCTCCACCACCAGTTTATTTTTACAATTCTCCACCTCCACCATCACCACCACCTACTCCAGTATACGAAGGGCCATTACCACCAGTAATAGGAGTTCAATATGCTT

>M6-PGSC0003DMG400001072-(rev)M6LRR-RLP_2294

CCACCTACTCCAGTATACGAAGGGCCATTACCACCAGTAATAGGAGTTCAATATGCTTCACCACCTCCACCACCCTTTTATTAATTCTTTCAGAAATTTCACTAACTCTTATCCTCAGAG

>M6-PGSC0003DMG400001072-(rev)M6LRR-RLP_2356

ACCTCCACCACCCTTTTATTAATTCTTTCAGAAATTTCACTAACTCTTATCCTCAGAGCACAAAATAGTTCCTACAATTATTATTTCCTCAACTCTTCAAGCCATCAACAAAATCGAGCT

>M6-PGSC0003DMG400001072-(rev)M6LRR-RLP_2418

AAATAGTTCCTACAATTATTATTTCCTCAACTCTTCAAGCCATCAACAAAATCGAGCTAATATGTGATCACGAACCTTAAATGGTACGACAGAAAGAAAGATAAGAGGAAGGTAAAGAGG

>M6-PGSC0003DMG400001072-(rev)M6LRR-RLP_248

CCCATTTGCTCTCTGTTTTTTCTTGTGCTCTGTTTTTTGTGTTCAGTTTTTACTGAACACACAAATGCTGAGTTTACTGTCTCTGAAAATGGCCCTTTAACTGATTCTGAAGCTCAGTTC

>M6-PGSC0003DMG400001072-(rev)M6LRR-RLP_2480

TGTGATCACGAACCTTAAATGGTACGACAGAAAGAAAGATAAGAGGAAGGTAAAGAGGAGAAAGTTTCAATCTCCAATGATTTTATTTTTGAATTGTTATCAAATAGATTGAAGTTGTGG

>M6-PGSC0003DMG400001072-(rev)M6LRR-RLP_2542

AGTTTCAATCTCCAATGATTTTATTTTTGAATTGTTATCAAATAGATTGAAGTTGTGGAGAGAATAAATTGGCAAAGTAAAGATTGAAACTTTGCTGGAAAATGTATGTATAGAGAGATC

>M6-PGSC0003DMG400001072-(rev)M6LRR-RLP_2604

AATAAATTGGCAAAGTAAAGATTGAAACTTTGCTGGAAAATGTATGTATAGAGAGATCATAATCTTGAATTAGAAACCAGTTTACAGTGCTTGCTGCAACTGGTTTTATTTAATGATCAT

>M6-PGSC0003DMG400001072-(rev)M6LRR-RLP_2666

TCTTGAATTAGAAACCAGTTTACAGTGCTTGCTGCAACTGGTTTTATTTAATGATCATACTTTGATTTCTTTGTAATTTTTTCTTTTTTTTGGTTCTTCCAAACTATAGTATAAGATGAG

>M6-PGSC0003DMG400001072-(rev)M6LRR-RLP_2699

GCAACTGGTTTTATTTAATGATCATACTTTGATTTCTTTGTAATTTTTTCTTTTTTTTGGTTCTTCCAAACTATAGTATAAGATGAGCTTTTTATTTTCTTTTAAAACTTGTATTGATTC

>M6-PGSC0003DMG400001072-(rev)M6LRR-RLP_310

AAATGCTGAGTTTACTGTCTCTGAAAATGGCCCTTTAACTGATTCTGAAGCTCAGTTCATCAAACACAGGCAGCTTTTGTATTACAGAGATGAGTTTGGTGATAGAGGTGAAAATGTGAA

>M6-PGSC0003DMG400001072-(rev)M6LRR-RLP_372

AACACAGGCAGCTTTTGTATTACAGAGATGAGTTTGGTGATAGAGGTGAAAATGTGAAAATTGATCCATCTATGGTGTTTGAGAATGATAGAATCAAGAATGCTTATATTGCTTTACAAG

>M6-PGSC0003DMG400001072-(rev)M6LRR-RLP_558

GCAGCTACACTGGAATCTTCTGTGCTCCGGCGTTGGATAACCCCAAAATCCGTACAGTTGCCGGCATTGACCTTAATCATGGTGACATTGCAGGGTACCTCCCGGAAGAGCTTGGGCTTC

>M6-PGSC0003DMG400001072-(rev)M6LRR-RLP_62

TTAAACCAAAAATTAAAAAAAACTCCACAAACCCCATTTCTTAAATCCATTTTTCTTCATAGATCTTCACTTTTTTCATCTCACATTTGGTTAAAGATTCAATCTTTGAACAGAAAAAAG

>M6-PGSC0003DMG400001185M6LRR-RLK_0

TCAATGGTAGTTTCTCTTGTCTCCTTGTTTTACAACTTTTTCCTCTTTCTCATACTTTCGTCCTCCAACTTAACTGCTCAACCATCTAGCACAGACACAGACTTTTCATGCTCGTTTAAC

>M6-PGSC0003DMG400001185M6LRR-RLK_1054

CAATGAACGACCAAGTTGTAGCAGTCAAGAAAACAAAACAAGCCTCGGAGGAAATGACAATACTTCAAAAACTACATCATGCAAATCTAGTGAAACTAATGGGCTTTTCATCAGATTATC

>M6-PGSC0003DMG400001185M6LRR-RLK_1116

CTTCAAAAACTACATCATGCAAATCTAGTGAAACTAATGGGCTTTTCATCAGATTATCTTGGTAATTCCTTTTTAGTTTATGAATATGCTGAAAATGGCTCACTGGATCAATGGCTGTTT

>M6-PGSC0003DMG400001185M6LRR-RLK_1178

TAATTCCTTTTTAGTTTATGAATATGCTGAAAATGGCTCACTGGATCAATGGCTGTTTCCCGGATTATCATCATCTTCTTCTGCCTCTGTCTTAGTGGCATCACTTGGTTGGAGTCGAAG

>M6-PGSC0003DMG400001185M6LRR-RLK_124

CCTTTTCTTGTGATACTTTTATTTCATATCGTGCCCGTCCACCAAATTATTTAGATGTCGGAAGTATATCTGATCTTCTTGAAGTCAGTCGTTTAAGTATAGCTACGGCCACAGGTCTAG

>M6-PGSC0003DMG400001185M6LRR-RLK_1240

GATTATCATCATCTTCTTCTGCCTCTGTCTTAGTGGCATCACTTGGTTGGAGTCGAAGATTACACATTGCCTTGGATGTTGCAAATGCTCTGCAATACCTGCATGAACATACTCAACCTA

>M6-PGSC0003DMG400001185M6LRR-RLK_1302

CACATTGCCTTGGATGTTGCAAATGCTCTGCAATACCTGCATGAACATACTCAACCTAGTATAGTTCACGGGGACATCCTAACGTGTAGCATACTTCTTGATTCAAGGTTTAAAGCCAAA

>M6-PGSC0003DMG400001185M6LRR-RLK_1364

AGTTCACGGGGACATCCTAACGTGTAGCATACTTCTTGATTCAAGGTTTAAAGCCAAAATTGCAGGTTTCTCTACAGCCAGACATGCCACCAACTCAATGATGCTGAAAGTTGACGTGTT

>M6-PGSC0003DMG400001185M6LRR-RLK_1426

CAGGTTTCTCTACAGCCAGACATGCCACCAACTCAATGATGCTGAAAGTTGACGTGTTTGCTTTTGGGATTGTGCTGTTGGAGTTGCTTTCAGGGAAGAAAGCAATGGAATCAAAAGATT

>M6-PGSC0003DMG400001185M6LRR-RLK_1488

TTTGGGATTGTGCTGTTGGAGTTGCTTTCAGGGAAGAAAGCAATGGAATCAAAAGATTATAATGATGAGACTCTGATCATGTGCAGAGAAATTAGTGGAATCTTGGAAGTTGAAGAGAAC

>M6-PGSC0003DMG400001185M6LRR-RLK_1550

TGATGAGACTCTGATCATGTGCAGAGAAATTAGTGGAATCTTGGAAGTTGAAGAGAACAGAGAGGAGAAGTTTAGAAAATGGATGGATCCAAAGTTGAGCTTTTACCCTATTGATGATGC

>M6-PGSC0003DMG400001185M6LRR-RLK_1612

AGGAGAAGTTTAGAAAATGGATGGATCCAAAGTTGAGCTTTTACCCTATTGATGATGCTCTAAACTTGGCTGCCTTGGCGACAGCATGCACATCGGAGCAATCAGCAGAGAGGCCTAAGA

>M6-PGSC0003DMG400001185M6LRR-RLK_1674

AACTTGGCTGCCTTGGCGACAGCATGCACATCGGAGCAATCAGCAGAGAGGCCTAAGATGACAGATATTGTCTTCAACCTCTGTTTTCTTGCTCAATCATCTTTTGAAATGTATGGACCG

>M6-PGSC0003DMG400001185M6LRR-RLK_1736

AGATATTGTCTTCAACCTCTGTTTTCTTGCTCAATCATCTTTTGAAATGTATGGACCGTCTTGGACTTCAGGTGAAGCTGAGGAGATTGTCCAGATTGTTAGTCCTGTAATAGCACGTTG

>M6-PGSC0003DMG400001185M6LRR-RLK_1798

GGACTTCAGGTGAAGCTGAGGAGATTGTCCAGATTGTTAGTCCTGTAATAGCACGTTGAGTTAAAGATGGAGCGTAAATATAGTTCAAGAACTTTTGTCAGTTCGTCTCAAATTTGTAAC

>M6-PGSC0003DMG400001185M6LRR-RLK_186

AGTATATCTGATCTTCTTGAAGTCAGTCGTTTAAGTATAGCTACGGCCACAGGTCTAGCCTCAGAGGATACCGAGCTATTCCCTGACCAGCTTTTACTTGCACCTGTTAAATGCTACTGC

>M6-PGSC0003DMG400001185M6LRR-RLK_1860

AAAGATGGAGCGTAAATATAGTTCAAGAACTTTTGTCAGTTCGTCTCAAATTTGTAACATATTCTGGATTGTAAAGCCACAGGAACGATAAATAAGCTGATCTCCAGAATATATTGCAAA

>M6-PGSC0003DMG400001185M6LRR-RLK_1922

TCTGGATTGTAAAGCCACAGGAACGATAAATAAGCTGATCTCCAGAATATATTGCAAATTTGAGTTTTTTTGTCCAGTTTTTCTTTGTATTTCTCTTGCTGCTAATTTAGTTGCTATTGT

>M6-PGSC0003DMG400001185M6LRR-RLK_1984

AGTTTTTTTGTCCAGTTTTTCTTTGTATTTCTCTTGCTGCTAATTTAGTTGCTATTGTATATTCTTGTATTGGATGTTTTTCTCAACTTTGATATAAGTGGAGTGCAATTTGACCTGTAT

>M6-PGSC0003DMG400001185M6LRR-RLK_2046

TCTTGTATTGGATGTTTTTCTCAACTTTGATATAAGTGGAGTGCAATTTGACCTGTATTTCTTTCGTCCTCTCAAAAGAAAAACAGAAAAAATTAAGTTTCTGAAAAGACACGGCAAACT

>M6-PGSC0003DMG400001185M6LRR-RLK_2091

ATTTGACCTGTATTTCTTTCGTCCTCTCAAAAGAAAAACAGAAAAAATTAAGTTTCTGAAAAGACACGGCAAACTCGTTCTTATTTATATTTATTTAAGATCCAGACGAGTACATGTTTA

>M6-PGSC0003DMG400001185M6LRR-RLK_248

AGAGGATACCGAGCTATTCCCTGACCAGCTTTTACTTGCACCTGTTAAATGCTACTGCAATAGTTCTCATTATTTCTCCAATGTCACCTATCAGATCATGAAAGGCGACAGCTTTTATTC

>M6-PGSC0003DMG400001185M6LRR-RLK_310

GTTCTCATTATTTCTCCAATGTCACCTATCAGATCATGAAAGGCGACAGCTTTTATTCAGTTTCAATAGGAGCTTTTGAGAACCTTACTAATTATCACGTGGTGCAAGATATGAATCCAA

>M6-PGSC0003DMG400001185M6LRR-RLK_372

TCAATAGGAGCTTTTGAGAACCTTACTAATTATCACGTGGTGCAAGATATGAATCCAACACTAGATCCAACCAACTTGACAATTGGTGCAGAAGCTGTTTTTTCTTTGTTTTGTAAATGC

>M6-PGSC0003DMG400001185M6LRR-RLK_434

AGATCCAACCAACTTGACAATTGGTGCAGAAGCTGTTTTTTCTTTGTTTTGTAAATGCCCCACACATTCTGATCTCGAAAAGGGACTTCAGTACCTGGTCACTTATGTTTGGCAGCCTTG

>M6-PGSC0003DMG400001185M6LRR-RLK_496

CACATTCTGATCTCGAAAAGGGACTTCAGTACCTGGTCACTTATGTTTGGCAGCCTTGGGATGATGTGTTGCCTGTAAGCTATATGTTTGGAGCATCTGCAGCTGACATTCTGGCTGCCA

>M6-PGSC0003DMG400001185M6LRR-RLK_558

GATGTGTTGCCTGTAAGCTATATGTTTGGAGCATCTGCAGCTGACATTCTGGCTGCCAACAATTACAGAAATTTCACTGCTGCAATATATTCTCCTGTCTTGATACCAATAAATCTTCCT

>M6-PGSC0003DMG400001185M6LRR-RLK_62

CTCCAACTTAACTGCTCAACCATCTAGCACAGACACAGACTTTTCATGCTCGTTTAACTCTTCCTTTTCTTGTGATACTTTTATTTCATATCGTGCCCGTCCACCAAATTATTTAGATGT

>M6-PGSC0003DMG400001185M6LRR-RLK_620

TTACAGAAATTTCACTGCTGCAATATATTCTCCTGTCTTGATACCAATAAATCTTCCTATTATTCTGCAATCTTACCCCTCTTCTGCAAGCAGTAGAAAATCCAAACACGGATGGATAGT

>M6-PGSC0003DMG400001185M6LRR-RLK_682

TTCTGCAATCTTACCCCTCTTCTGCAAGCAGTAGAAAATCCAAACACGGATGGATAGTAATTACTGTTTTAAGTATTATGGGGTTGCTTGCAGTTTTTTCATTTTTCTTGATGGTATACA

>M6-PGSC0003DMG400001185M6LRR-RLK_744

ACTGTTTTAAGTATTATGGGGTTGCTTGCAGTTTTTTCATTTTTCTTGATGGTATACATGCGTCATCTAGACGAGAAGAGGAGGGCTAACTTAGCCCATAACTCTTCTACTCTTGAAACT

>M6-PGSC0003DMG400001185M6LRR-RLK_806

TCATCTAGACGAGAAGAGGAGGGCTAACTTAGCCCATAACTCTTCTACTCTTGAAACTTCTGATCTCTTTCATACCAAGAAAGCCTCAGAAGGTGAAATTACGGACCATAAGAACATTCA

>M6-PGSC0003DMG400001185M6LRR-RLK_868

ATCTCTTTCATACCAAGAAAGCCTCAGAAGGTGAAATTACGGACCATAAGAACATTCAAGATAAGCTTCTTCCTGGAGTTTCTGGCTACATTGGAAAGCCAATAATATATGACCTGAAGA

>M6-PGSC0003DMG400001185M6LRR-RLK_930

AAGCTTCTTCCTGGAGTTTCTGGCTACATTGGAAAGCCAATAATATATGACCTGAAGATCATCATGGAGGCAACTGTCGACCTCAGTGAGAGATACAGGTTAGGAGGATCAGTGTACAAG

>M6-PGSC0003DMG400001185M6LRR-RLK_992

CATGGAGGCAACTGTCGACCTCAGTGAGAGATACAGGTTAGGAGGATCAGTGTACAAGGTTACAATGAACGACCAAGTTGTAGCAGTCAAGAAAACAAAACAAGCCTCGGAGGAAATGAC

>M6-PGSC0003DMG400001252-(rev)M6LRR-RLP_0

AATTTCAGAGAGAGAAACAAAACAAAAACTCCTCAGGATGGGCAGCTTCATGTACATTGCGATCTTACTTCTTGCCATAATCTCAACCGTTCATTCCTGCCCGCCATCAGATCGGGCAGC

>M6-PGSC0003DMG400001252-(rev)M6LRR-RLP_1008

TGACGTTAGTCATAACCATCTTTGTGGCCAGATTCCGGAAGGTTCTCCCTTTGATCATCTTGAAGCTTCATCTTTTACTTACAATGATTGTCTATGTGGGAAGCCTCTTAAACCTTGTCA

>M6-PGSC0003DMG400001252-(rev)M6LRR-RLP_1071

AGCTTCATCTTTTACTTACAATGATTGTCTATGTGGGAAGCCTCTTAAACCTTGTCACTAAAGTGTGACTATTATTAGGTAAATTTAATTTTACTTTTCTTTTTTGAGATTTTTATTAAC

>M6-PGSC0003DMG400001252-(rev)M6LRR-RLP_1134

TGTGACTATTATTAGGTAAATTTAATTTTACTTTTCTTTTTTGAGATTTTTATTAACACTATATATTGGTGAATATGTTTGTATTATTATGTGATACTGAAAAGTAAAGTGTTTTTTTTT

>M6-PGSC0003DMG400001252-(rev)M6LRR-RLP_1194

ATATATTGGTGAATATGTTTGTATTATTATGTGATACTGAAAAGTAAAGTGTTTTTTTTTAGCATATATTTGTAATTTTAAATGTATAATTGTGCCTTGAGATATATGAAGGCTTTCTTA

>M6-PGSC0003DMG400001252-(rev)M6LRR-RLP_126

ATCTTTCCGAGCTGCCCTAAATGAATCTTACTTAGGAATTTTCAACTCGTGGAAAGGGTACGACTGTTGCCATGAATGGTACGGGGTGAGTTGTGATCCCACAACTCACCGCGTAGCAGA

>M6-PGSC0003DMG400001252-(rev)M6LRR-RLP_189

CTGTTGCCATGAATGGTACGGGGTGAGTTGTGATCCCACAACTCACCGCGTAGCAGACATAAACCTCCGTGGTGAATCCGAAGATCCACTCTTCGAAAAAGCTCATCGTACCGGTTACAT

>M6-PGSC0003DMG400001252-(rev)M6LRR-RLP_252

CCTCCGTGGTGAATCCGAAGATCCACTCTTCGAAAAAGCTCATCGTACCGGTTACATGACCGGTACGATCTCTCCATCTATATGCAAATTAGAAAGACTATCAAGTCTCACTATAGCTGA

>M6-PGSC0003DMG400001252-(rev)M6LRR-RLP_315

TACGATCTCTCCATCTATATGCAAATTAGAAAGACTATCAAGTCTCACTATAGCTGATTGGAAAGGCATCACTGGACCTATACCATCATGTATTACGTCCTTACCCTTCCTTCGAATCAT

>M6-PGSC0003DMG400001252-(rev)M6LRR-RLP_378

AGGCATCACTGGACCTATACCATCATGTATTACGTCCTTACCCTTCCTTCGAATCATCGACTTAGTCGGAAACAAGCTCACCGGCTCAATCCCTTCAACTATAGGTCGATTAAGTCGACT

>M6-PGSC0003DMG400001252-(rev)M6LRR-RLP_441

AGTCGGAAACAAGCTCACCGGCTCAATCCCTTCAACTATAGGTCGATTAAGTCGACTCACAGTGTTAAACGTAGCTGAAAATCAACTTACTGGCTACATACCTCGATCATTAACCAACTT

>M6-PGSC0003DMG400001252-(rev)M6LRR-RLP_504

GTTAAACGTAGCTGAAAATCAACTTACTGGCTACATACCTCGATCATTAACCAACTTATCATCTTTAATGCATTTAGATCTCCGAAACAACTTCATCTATGGAAACATCCCAACAAATTT

>M6-PGSC0003DMG400001252-(rev)M6LRR-RLP_567

TTTAATGCATTTAGATCTCCGAAACAACTTCATCTATGGAAACATCCCAACAAATTTCGGAAAATTACGGATGTTGAGTCGAGCTTTACTGAGTCGAAACAGGTTAACAGGTCATATACC

>M6-PGSC0003DMG400001252-(rev)M6LRR-RLP_63

CTTACTTCTTGCCATAATCTCAACCGTTCATTCCTGCCCGCCATCAGATCGGGCAGCATTATTATCTTTCCGAGCTGCCCTAAATGAATCTTACTTAGGAATTTTCAACTCGTGGAAAGG

>M6-PGSC0003DMG400001252-(rev)M6LRR-RLP_630

ATTACGGATGTTGAGTCGAGCTTTACTGAGTCGAAACAGGTTAACAGGTCATATACCGTATTCGATTTCGTATATTTACAGGCTTTCAGATTTGGATCTGTCATTGAACAAACTTTCTGG

>M6-PGSC0003DMG400001252-(rev)M6LRR-RLP_693

GATTTCGTATATTTACAGGCTTTCAGATTTGGATCTGTCATTGAACAAACTTTCTGGTAGCATACCTCCGTCTCTAGGTAAAATGCACGTACTTGCAACGTTAAATCTTGATGGTAACAA

>M6-PGSC0003DMG400001252-(rev)M6LRR-RLP_756

ACCTCCGTCTCTAGGTAAAATGCACGTACTTGCAACGTTAAATCTTGATGGTAACAATATTACTGGTACTATACCACCTAGTTTGATTAATTCACGTATCAACATTTTGAATTTGAGCAA

>M6-PGSC0003DMG400001252-(rev)M6LRR-RLP_819

TGGTACTATACCACCTAGTTTGATTAATTCACGTATCAACATTTTGAATTTGAGCAAAAATACTATTGAAGGTTATATTCCTGATTCATTTGATGAAAGATCTTATTTTATGGTAATGGA

>M6-PGSC0003DMG400001252-(rev)M6LRR-RLP_882

TATTGAAGGTTATATTCCTGATTCATTTGATGAAAGATCTTATTTTATGGTAATGGATTTATCTTACAACAAATTAAAAGGAAGGATTCCAAAATCGATATCGTCGGCGACGTTTATCGG

>M6-PGSC0003DMG400001252-(rev)M6LRR-RLP_945

TTACAACAAATTAAAAGGAAGGATTCCAAAATCGATATCGTCGGCGACGTTTATCGGTCACCTTGACGTTAGTCATAACCATCTTTGTGGCCAGATTCCGGAAGGTTCTCCCTTTGATCA

>M6-PGSC0003DMG400001269-(rev)M6RLKWAX-EGF_1037

ACACTTTCTTTTTCCTTGTGATGTTTTCTTTTGTCTGAAAACCTCTGATTACTTATATTTTTTTCCAGTTTTTGGTGGATTAGGATTGGTGATGATTTGTTTAGCTGTCTACTTTATCTG

>M6-PGSC0003DMG400001269-(rev)M6RLKWAX-EGF_1098

TTTCCAGTTTTTGGTGGATTAGGATTGGTGATGATTTGTTTAGCTGTCTACTTTATCTGGTGTTGCAAGAGGAGGAAATTTAATCCGCCCCACTTTCTCTCAACAAGGAAATTGTCATAT

>M6-PGSC0003DMG400001269-(rev)M6RLKWAX-EGF_1159

GTTGCAAGAGGAGGAAATTTAATCCGCCCCACTTTCTCTCAACAAGGAAATTGTCATATATATTTAAAAATGATGTTGAGGGAGGCAGTATATACTTTGGCATCCCAGTCTTCTCTTATT

>M6-PGSC0003DMG400001269-(rev)M6RLKWAX-EGF_122

AGCTGGTTGTGATGCTAAACTATATCCGAGAATCCAACTGCTTCCTCGAGGAGATTCGTACTATGCTTTAGGAAATCCGTATAATTATACAGTTGAGCTAGGGGACCCCAAGTTTCATGC

>M6-PGSC0003DMG400001269-(rev)M6RLKWAX-EGF_1220

ATTTAAAAATGATGTTGAGGGAGGCAGTATATACTTTGGCATCCCAGTCTTCTCTTATTCAGAACTTGAAGAAGCCACAAATGATTTCAATTCCTCTCGAATCCTTGGAGATGGAGGTTT

>M6-PGSC0003DMG400001269-(rev)M6RLKWAX-EGF_1342

GAACTGTTTACTATGGTGAGAGACTTCCTAACCCAACTGAGCTATGCTTCACTTCTTTCTTCATGTGCCTAATACTTGTTCATTAATGTCTAGGGAAACTTAAGGATGGACGAGAGGTTG

>M6-PGSC0003DMG400001269-(rev)M6RLKWAX-EGF_1403

CATGTGCCTAATACTTGTTCATTAATGTCTAGGGAAACTTAAGGATGGACGAGAGGTTGCTGTGAAGCGCCTTTACGAGCACAACTGCAAACGAATGGAGCAGTTCGTAAATGAAATTGA

>M6-PGSC0003DMG400001269-(rev)M6RLKWAX-EGF_1525

TCCTTACTAGGCTAAGGCACAACAATCTTGTCACTCTCTATGGCTGCACTTCAAGGCGTAGCCGTGAACTACTCCTTGTCTATGAATATATTCCTAATGGAACTCTTGCCGATCACCTCC

>M6-PGSC0003DMG400001269-(rev)M6RLKWAX-EGF_1586

CCGTGAACTACTCCTTGTCTATGAATATATTCCTAATGGAACTCTTGCCGATCACCTCCATGGTGACAGAGTGAAGGACGGATCACTCACCTGGCTAGTCCGCATGAACATTGCCATAGA

>M6-PGSC0003DMG400001269-(rev)M6RLKWAX-EGF_1708

CTGCTGGTGCATTGGCTTACCTGCATGCGTCTGACGTAATACACTGTGATGTGAAGACTAATAACATACTCCTTGATCACAACTTTAATGTTAAAGTTGCAGATTTTGGGATATCACGGC

>M6-PGSC0003DMG400001269-(rev)M6RLKWAX-EGF_1769

TAACATACTCCTTGATCACAACTTTAATGTTAAAGTTGCAGATTTTGGGATATCACGGCTTTTCCCAAATGATGTCACTCATATTTCAACTGCACCCCGGGGGACCCCTGGCTATATCGA

>M6-PGSC0003DMG400001269-(rev)M6RLKWAX-EGF_183

TATGCTTTAGGAAATCCGTATAATTATACAGTTGAGCTAGGGGACCCCAAGTTTCATGCCAAGTTGAGGCAACACAAGTGCCAGGCTTTTAACAAAAACTTATCCTTGCCAGCCTCTCCT

>M6-PGSC0003DMG400001269-(rev)M6RLKWAX-EGF_1830

TTCCCAAATGATGTCACTCATATTTCAACTGCACCCCGGGGGACCCCTGGCTATATCGATCCAAAGTATCATGAATGTTACCAGCTGACTAGTAAAAGTGATGTTTATAGCTTCGGGGTG

>M6-PGSC0003DMG400001269-(rev)M6RLKWAX-EGF_1891

CAAAGTATCATGAATGTTACCAGCTGACTAGTAAAAGTGATGTTTATAGCTTCGGGGTGGTCCTTGTCGAGCTCATTTCATCAATGCCAGCTGTGGATATGACTAGGCATAGCCAAGAGA

>M6-PGSC0003DMG400001269-(rev)M6RLKWAX-EGF_1952

CCTTGTCGAGCTCATTTCATCAATGCCAGCTGTGGATATGACTAGGCATAGCCAAGAGATTAATTTGGCTAACTTTGCAATAAACAAGATTGTAAAATGTGCATTTAATGAGTTGGTTGA

>M6-PGSC0003DMG400001269-(rev)M6RLKWAX-EGF_2013

AATTTGGCTAACTTTGCAATAAACAAGATTGTAAAATGTGCATTTAATGAGTTGGTTGATCCATCCCTGGGGTTCGACTCAGATACCAAGATTTTGGAGATGACTACTTCAGTGGCAGAG

>M6-PGSC0003DMG400001269-(rev)M6RLKWAX-EGF_2135

GGCTTTTCTATGCTTACAGACAGATAGGGACATGAGGCCTACTATGGTTGAGGTTTTGGATACTCTAAAGGAGATTCAGATTAGTGAATTTGACAATGAGAAGAGAGCTGACTCTAATCT

>M6-PGSC0003DMG400001269-(rev)M6RLKWAX-EGF_2196

ACTCTAAAGGAGATTCAGATTAGTGAATTTGACAATGAGAAGAGAGCTGACTCTAATCTCAATGGAAATGAAGCTAAAATTGTTACAGCACCCCCTTTCGCTGAATCAGAAGATAAGTTT

>M6-PGSC0003DMG400001269-(rev)M6RLKWAX-EGF_2257

ATGGAAATGAAGCTAAAATTGTTACAGCACCCCCTTTCGCTGAATCAGAAGATAAGTTTATGCTGAAACAAGTTAAATCACTACCTTCTTCAAATTCTGTTACTGATAAATGGGTTACTT

>M6-PGSC0003DMG400001269-(rev)M6RLKWAX-EGF_244

AGTTGAGGCAACACAAGTGCCAGGCTTTTAACAAAAACTTATCCTTGCCAGCCTCTCCTTCTATTTCTTTCGAAATTCTCCCAGTGAATATTCTCAACTTCATCAAATGCAACAGCACTA

>M6-PGSC0003DMG400001269-(rev)M6RLKWAX-EGF_305

TATTTCTTTCGAAATTCTCCCAGTGAATATTCTCAACTTCATCAAATGCAACAGCACTAGTAGTAGCACCCCAAACATTACCCATAAGATGAAAGATCATTTTGCTGGTTATAGAATGTA

>M6-PGSC0003DMG400001269-(rev)M6RLKWAX-EGF_366

AGTAGCACCCCAAACATTACCCATAAGATGAAAGATCATTTTGCTGGTTATAGAATGTACAATGGCTGTAAAGGTTTTAGCATATACTACAAGCTTCGCTTAGGAGATGATGAAGACGTT

>M6-PGSC0003DMG400001269-(rev)M6RLKWAX-EGF_488

AGCAGGCAATCTTCCTGCCAACTGTTCACTTATCAGATTGCCTATCCGCTTGCTATCAGATGATGGTGGTTTGTTCAATATGTTAAGCGCCGCTTTTCTAGTAGAATGGAAACTGTCTGA

>M6-PGSC0003DMG400001269-(rev)M6RLKWAX-EGF_549

GATGGTGGTTTGTTCAATATGTTAAGCGCCGCTTTTCTAGTAGAATGGAAACTGTCTGAGGACTGTTACAAATGTCACTATGGCGGAGGTCAATGCCAGACTGATATAATCAACAAATTT

>M6-PGSC0003DMG400001269-(rev)M6RLKWAX-EGF_61

TTACTGACCTGAGCTTTCCTTTCTCTCTTTCCACACAACCTGACTGTGGAATAATGTTCTTAGCTGGTTGTGATGCTAAACTATATCCGAGAATCCAACTGCTTCCTCGAGGAGATTCGT

>M6-PGSC0003DMG400001269-(rev)M6RLKWAX-EGF_610

ACTGTTACAAATGTCACTATGGCGGAGGTCAATGCCAGACTGATATAATCAACAAATTTCATTGTACTTATCCAAATAATCCACATGATCAAGGTCATCAAGAAAATAATCCACATGAAC

>M6-PGSC0003DMG400001269-(rev)M6RLKWAX-EGF_671

TTGTACTTATCCAAATAATCCACATGATCAAGGTCATCAAGAAAATAATCCACATGAACAAGGTTCAGTCACACAGTGCATCTTTCGTAGTGTCCTTTCTTGTATTCATATCACATATCT

>M6-PGSC0003DMG400001269-(rev)M6RLKWAX-EGF_732

GGTTCAGTCACACAGTGCATCTTTCGTAGTGTCCTTTCTTGTATTCATATCACATATCTATAATATTTTCCCTTTTTTTTCTTTAGATGCAAAAACAGCTAGAAGAAAGTTCGGACCGAC

>M6-PGSC0003DMG400001269-(rev)M6RLKWAX-EGF_793

AATATTTTCCCTTTTTTTTCTTTAGATGCAAAAACAGCTAGAAGAAAGTTCGGACCGACTCTGGGAGCAGGTAGTTAAATATCCTTTAACTGCAGTGTCTTTTCGATCTTTTCTATTACA

>M6-PGSC0003DMG400001269-(rev)M6RLKWAX-EGF_854

TGGGAGCAGGTAGTTAAATATCCTTTAACTGCAGTGTCTTTTCGATCTTTTCTATTACAGAATTTCCTTGCATATCGTTTGAATTACTTCTTCATTGTGGTAAAATCTCAAAACATTTCC

>M6-PGSC0003DMG400001269-(rev)M6RLKWAX-EGF_915

ATTTCCTTGCATATCGTTTGAATTACTTCTTCATTGTGGTAAAATCTCAAAACATTTCCTTAATGATAAGAGAGGTAAGTACTTTAGCTTACACAAAATGATTGGATAGTGGAGTTGAGA

>M6-PGSC0003DMG400001269-(rev)M6RLKWAX-EGF_976

AATGATAAGAGAGGTAAGTACTTTAGCTTACACAAAATGATTGGATAGTGGAGTTGAGAGGACACTTTCTTTTTCCTTGTGATGTTTTCTTTTGTCTGAAAACCTCTGATTACTTATATT

>M6-PGSC0003DMG400001271-(rev)M6RLKWAX-EGF_0

TTAGTTCAGGCAATGGGCAGAAATGATTCAACTTGTCCAAAGTCCTTTTCATGTGGAGATTTTACTGACCTGAGCTTTCCTTTCTCTCTTTCCACACAACCCGACTGTGGAATACTGCCC

>M6-PGSC0003DMG400001271-(rev)M6RLKWAX-EGF_1054

TGCTTTCCACAGGTGCAGTTTTGTGTACAGTTGGACTATTGGTTCTACTCTTCTTCTTCAGAGAAAATATTTTGCGGTATAAGTGCCTCAGGTTTTGGGAATCTATTGCTGAGGATCACC

>M6-PGSC0003DMG400001271-(rev)M6RLKWAX-EGF_1116

GAAAATATTTTGCGGTATAAGTGCCTCAGGTTTTGGGAATCTATTGCTGAGGATCACCAAAATATTGAAGCATTTTTGAAGAACTATGGGTCATATGCTCCAAAGAGATACAACTACTCA

>M6-PGSC0003DMG400001271-(rev)M6RLKWAX-EGF_1178

TATTGAAGCATTTTTGAAGAACTATGGGTCATATGCTCCAAAGAGATACAACTACTCAGAGGTCAAAAGGATGACCAGTTGCTTCAAAAACAAACTAGGTCAAGGTGGATATGGTTCTGT

>M6-PGSC0003DMG400001271-(rev)M6RLKWAX-EGF_124

CTGGTTGTGATGCTAAACCATATCCGAGAATCCAACTGCTTCCTGGAGGAGATTCGTACTATGCTTTAGGAAATCCGTATAATTATACAGTTGGGATAGGGGACCCCAAGTTTCATGCCA

>M6-PGSC0003DMG400001271-(rev)M6RLKWAX-EGF_1240

TCAAAAGGATGACCAGTTGCTTCAAAAACAAACTAGGTCAAGGTGGATATGGTTCTGTATACAAAGGAACTCTGCAGAATGGAAATCATGTGGCTGTGAAGGTCCTGAATGGACTAAAAG

>M6-PGSC0003DMG400001271-(rev)M6RLKWAX-EGF_1364

TGGTGAAGAATTCATTAACGAGGTGGCAAGTATTAGCAGGACATCACATGTTAATGTTGTGAGCCTTGAAGGATTTTGTTTTGAGGGTTGCAAAAGAGCTCTCATTTATGAATTCATGTC

>M6-PGSC0003DMG400001271-(rev)M6RLKWAX-EGF_1426

GCCTTGAAGGATTTTGTTTTGAGGGTTGCAAAAGAGCTCTCATTTATGAATTCATGTCAAATGGATCTCTTGAAAAATTTATATACGAGGAAAGATCCGACAGTGTTCGCCAATTGGATT

>M6-PGSC0003DMG400001271-(rev)M6RLKWAX-EGF_1612

CCACTCGGATTTTGCATTTTGATATAAAGCCTCATAATATCCTGCTTGATGAAGATTTTTGTCCTAAGATCTCTGACTTTGGACTAGCCAAACTGTGCATGAAAAAGGAGAGCATTGTGT

>M6-PGSC0003DMG400001271-(rev)M6RLKWAX-EGF_1860

AGAAAAAATGTTGATGTTCGAGTTGATCGAACGAGTGAAATCTACTTTCCACATTGGCTCTATCAACGAATTGAACTGGATGAAGAGCTTCAACTAATCGGGCTAATGAATGAAGAGGAG

>M6-PGSC0003DMG400001271-(rev)M6RLKWAX-EGF_1922

TCAACGAATTGAACTGGATGAAGAGCTTCAACTAATCGGGCTAATGAATGAAGAGGAGAAAGAATGTGCAAGAAAGATGGGGATGGTGAGTTTATGGTGCATACAGACTGATCCCTCAAA

>M6-PGSC0003DMG400001271-(rev)M6RLKWAX-EGF_2108

CAAGCCTTACCTATATTCTCCCTCGAGATCAGAGGTAGATTCATTGGAAGTGGGTTCATATCAATGTAGTAGATCAAACGTTTAAGTCAACTACCAGGAAAAGTTTCGTTCAATCAATGT

>M6-PGSC0003DMG400001271-(rev)M6RLKWAX-EGF_2170

AATGTAGTAGATCAAACGTTTAAGTCAACTACCAGGAAAAGTTTCGTTCAATCAATGTAGTTGTAATAATCTATCTAGATCTTCAAAGAAACGATTATATTTAACGAAGATTCAAATTTG

>M6-PGSC0003DMG400001271-(rev)M6RLKWAX-EGF_2232

GTAATAATCTATCTAGATCTTCAAAGAAACGATTATATTTAACGAAGATTCAAATTTGTGCTGCAATTATCTTTATTCATCAGGGATGTAAAATTGTTACATGTAAAATGAAAGGTTCTA

>M6-PGSC0003DMG400001271-(rev)M6RLKWAX-EGF_2263

ATTATATTTAACGAAGATTCAAATTTGTGCTGCAATTATCTTTATTCATCAGGGATGTAAAATTGTTACATGTAAAATGAAAGGTTCTAATGACTTTACATAGTCAAAATGCAAAGTTTC

>M6-PGSC0003DMG400001271-(rev)M6RLKWAX-EGF_372

ACCCCAAACATTACCCAGCAGATGAAAGATCATTTTGCTGGTTATAGAATGTACAATGGCTGTAAAGACTTTAGCGTATACTACAAGCTTCGCCTAGGCGATGATGAAGACGTGCAAGCA

>M6-PGSC0003DMG400001271-(rev)M6RLKWAX-EGF_434

TAAAGACTTTAGCGTATACTACAAGCTTCGCCTAGGCGATGATGAAGACGTGCAAGCAGGCAATCTTCCTGCAAACTGTTCACTTATCAGACTGCCGATCGGCTCGTTGTCAGATGATGG

>M6-PGSC0003DMG400001271-(rev)M6RLKWAX-EGF_496

ATCTTCCTGCAAACTGTTCACTTATCAGACTGCCGATCGGCTCGTTGTCAGATGATGGTGGTTTGTTCAATATGTTAAGTGCCACTTTTATAGTAGAATGGAAACTGTCTGAGGACTGTA

>M6-PGSC0003DMG400001271-(rev)M6RLKWAX-EGF_558

TTGTTCAATATGTTAAGTGCCACTTTTATAGTAGAATGGAAACTGTCTGAGGACTGTAACAAATGTCACTATGGCGGAGGTCAATGCCAGACTGATATAACTAACAAATTTCATTGTACA

>M6-PGSC0003DMG400001271-(rev)M6RLKWAX-EGF_620

ATGTCACTATGGCGGAGGTCAATGCCAGACTGATATAACTAACAAATTTCATTGTACATATCCAAATAATCCACATGATCAAGGTCATCAAGAAAATAACCTACTTGCTCAAGGTCTAGT

>M6-PGSC0003DMG400001271-(rev)M6RLKWAX-EGF_682

CAAATAATCCACATGATCAAGGTCATCAAGAAAATAACCTACTTGCTCAAGGTCTAGTGCATATTTCATAAAGTCCTTTCTCGTTTTCGTATCAAATATTGATAATATTTCATAGAATCT

>M6-PGSC0003DMG400001271-(rev)M6RLKWAX-EGF_744

ATTTCATAAAGTCCTTTCTCGTTTTCGTATCAAATATTGATAATATTTCATAGAATCTTTCCTCTTTTTTTAGATGCAAAAACAACTAGAAGCAAGTGGGGACCGATTCTGGGAGCAGGT

>M6-PGSC0003DMG400001271-(rev)M6RLKWAX-EGF_806

TCTTTTTTTAGATGCAAAAACAACTAGAAGCAAGTGGGGACCGATTCTGGGAGCAGGTAGTTAAATATTCATTAACTGCAAGGTTTCTTTCTTCGATCTTTAGATTTTCTTCTTCATTGT

>M6-PGSC0003DMG400001271-(rev)M6RLKWAX-EGF_868

AAATATTCATTAACTGCAAGGTTTCTTTCTTCGATCTTTAGATTTTCTTCTTCATTGTGGTAAAATCTCAAAAAATGGAAACACAAGTACTTTAGATTACATAACGTACCGATGGTGGAG

>M6-PGSC0003DMG400001271-(rev)M6RLKWAX-EGF_930

AAATCTCAAAAAATGGAAACACAAGTACTTTAGATTACATAACGTACCGATGGTGGAGTTGAGAGGACATTCTTTCTCCTGGCGATGTTTAATTTATATTCTCTGATGGTGCATGACTTC

>M6-PGSC0003DMG400001271-(rev)M6RLKWAX-EGF_992

GAGGACATTCTTTCTCCTGGCGATGTTTAATTTATATTCTCTGATGGTGCATGACTTCACCTTGCTTTCCACAGGTGCAGTTTTGTGTACAGTTGGACTATTGGTTCTACTCTTCTTCTT

>M6-PGSC0003DMG400001301-(rev)M6LRR-RLK_0

TAATTTCTTAGTAGCTAGCTTCTAAAGGCACCACTCATATTTGTAAAATAACAGACTGGGTTTTCATTTGTCAAGTCAAATGCCTCACCAGTCTTCATCTTCCTTCTCCTTGCTATTTCC

>M6-PGSC0003DMG400001301-(rev)M6LRR-RLK_1054

CCGCTCGTCAAGGTTAAATTAGCTGATTTGTTGATTGCCACAAATGGTTTCAGGACGGACAATGTTATCAACTCAACTAGAATGGGTACTACTTATAATGCTGTGTTACGTGATGGTTCT

>M6-PGSC0003DMG400001301-(rev)M6LRR-RLK_1116

TGTTATCAACTCAACTAGAATGGGTACTACTTATAATGCTGTGTTACGTGATGGTTCTGCACTTGCCATTAAGCGGCTTAATACTTGCAAACTGAGCGAGAAGCACTTTCGTGATGAGAT

>M6-PGSC0003DMG400001301-(rev)M6LRR-RLK_1178

TTGCCATTAAGCGGCTTAATACTTGCAAACTGAGCGAGAAGCACTTTCGTGATGAGATGTATAGGCTAGGACAACTTAGGCATCCTAATTTGGTGCCACTGTTGGGGTTTTGTGTTGTGG

>M6-PGSC0003DMG400001301-(rev)M6LRR-RLK_124

TATTTCTCTTGCGCATATTATCAAACACCTTGTGCCAATTACCCAAAACTGCTGTTGGATCCCTAAACCCCCCAGCTAACGATGGCTCGTTTCAGATTTCTCCTTTTTCTTCTGTTGATT

>M6-PGSC0003DMG400001301-(rev)M6LRR-RLK_1240

AGGCTAGGACAACTTAGGCATCCTAATTTGGTGCCACTGTTGGGGTTTTGTGTTGTGGAAGAGGAGAAGCTTTTGGTTTATAAACACCTATCAAATGGTACTTTGCACTCATTCTTGAAT

>M6-PGSC0003DMG400001301-(rev)M6LRR-RLK_1302

GGAGAAGCTTTTGGTTTATAAACACCTATCAAATGGTACTTTGCACTCATTCTTGAATGGGAATGCAAGTGAACTGGATTGGCCTACTCGGTTTAGAATTGGTTTCGGTGCTGCTAGGGG

>M6-PGSC0003DMG400001301-(rev)M6LRR-RLK_1364

ATGCAAGTGAACTGGATTGGCCTACTCGGTTTAGAATTGGTTTCGGTGCTGCTAGGGGCCTTGCTTGGTTACATCATGGTGTCCACCCACCTATCTTGCACCAAAACATATGTTCTAATG

>M6-PGSC0003DMG400001301-(rev)M6LRR-RLK_1426

GCTTGGTTACATCATGGTGTCCACCCACCTATCTTGCACCAAAACATATGTTCTAATGTTATTTTCCTCGATGAAGACTTTGATGCTAGAGTAATGGATTTTGGGCTGGCAAGGTTGATG

>M6-PGSC0003DMG400001301-(rev)M6LRR-RLK_1488

TTTCCTCGATGAAGACTTTGATGCTAGAGTAATGGATTTTGGGCTGGCAAGGTTGATGACATCAGATGCAAAAGAGTCTAGTTATGTGAATGGGGAGTTGGGTGAATTTGGCTATGTAGC

>M6-PGSC0003DMG400001301-(rev)M6LRR-RLK_1550

CAGATGCAAAAGAGTCTAGTTATGTGAATGGGGAGTTGGGTGAATTTGGCTATGTAGCTCCAGAGTACTCTAGCACAATGGTGCCTTCACTGAAAGGGGATGCTTATAGTTTTGGGGTAG

>M6-PGSC0003DMG400001301-(rev)M6LRR-RLK_1612

GAGTACTCTAGCACAATGGTGCCTTCACTGAAAGGGGATGCTTATAGTTTTGGGGTAGTGCTTTTGGAGTTGGCTACTGGGCAAAAACCTCTTGAAGTTATTGCTGGTGAAGAGGGATTC

>M6-PGSC0003DMG400001301-(rev)M6LRR-RLK_1674

TTTGGAGTTGGCTACTGGGCAAAAACCTCTTGAAGTTATTGCTGGTGAAGAGGGATTCAAGGGTAATTTGGTAGACTGGGTGAATCAGCTCTCTGCTTCAGGTCGCATTAAAGATGCAAT

>M6-PGSC0003DMG400001301-(rev)M6LRR-RLK_1736

GTAATTTGGTAGACTGGGTGAATCAGCTCTCTGCTTCAGGTCGCATTAAAGATGCAATTGACCAAAACATGCGTGGGAAGGGGCATGATGAAGAGATTGTGCAATTCCTAAAAGTTGCTT

>M6-PGSC0003DMG400001301-(rev)M6LRR-RLK_1798

CAAAACATGCGTGGGAAGGGGCATGATGAAGAGATTGTGCAATTCCTAAAAGTTGCTTGTAATTCTGTGGTTTCTCGGCCCAATGATAGGTGGTCTATGTATCAGGTTTATGAAGCACTT

>M6-PGSC0003DMG400001301-(rev)M6LRR-RLK_186

CTAAACCCCCCAGCTAACGATGGCTCGTTTCAGATTTCTCCTTTTTCTTCTGTTGATTCTGATTTCTCTTGTTCTTCCGTGCCTTGATGCTGCTGTAGTTGAAGATGATCTCAAGTGTTT

>M6-PGSC0003DMG400001301-(rev)M6LRR-RLK_1860

TTCTGTGGTTTCTCGGCCCAATGATAGGTGGTCTATGTATCAGGTTTATGAAGCACTTCAGAGCATGGCTGAAAAACAGGGTTTCTCCGAACAGTATGACGAATTCCCGTTACTATTTGG

>M6-PGSC0003DMG400001301-(rev)M6LRR-RLK_1922

GCATGGCTGAAAAACAGGGTTTCTCCGAACAGTATGACGAATTCCCGTTACTATTTGGCAAAGAAGGTGCTACTAGAAGTCCTGTTTGAAGCCGGAATTGCGAAATTATTACTAAATGTT

>M6-PGSC0003DMG400001301-(rev)M6LRR-RLK_1984

GAAGGTGCTACTAGAAGTCCTGTTTGAAGCCGGAATTGCGAAATTATTACTAAATGTTCCACTCCAAATGAACATGATGCATCCTGTTGTAAGTTTTCAGTTCCTATTGGTGTAAGTGGC

>M6-PGSC0003DMG400001301-(rev)M6LRR-RLK_2046

TCCAAATGAACATGATGCATCCTGTTGTAAGTTTTCAGTTCCTATTGGTGTAAGTGGCATGGCACTGTTTTGATTGGCTCAGAAGTACACCCTTTAAATATTTCTGTATTATATTTGCAA

>M6-PGSC0003DMG400001301-(rev)M6LRR-RLK_2108

CACTGTTTTGATTGGCTCAGAAGTACACCCTTTAAATATTTCTGTATTATATTTGCAAAATGCACAAATCATTCTTAAGTATTTGCTTGCTTGATATTTGTAGTATAGGAAGCAACTGAT

>M6-PGSC0003DMG400001301-(rev)M6LRR-RLK_2170

CACAAATCATTCTTAAGTATTTGCTTGCTTGATATTTGTAGTATAGGAAGCAACTGATAATTATTGTTCATGTTTGTGATTACCCATTGCCTATAGCATTAGATGTTATGTATTCAAAGG

>M6-PGSC0003DMG400001301-(rev)M6LRR-RLK_248

TTTCTCTTGTTCTTCCGTGCCTTGATGCTGCTGTAGTTGAAGATGATCTCAAGTGTTTAGAAGGTTTCAAGAAGTCTTTGGAAGATCCAGCTGGGAATCTGAATTCATGGAACTTTAAAA

>M6-PGSC0003DMG400001301-(rev)M6LRR-RLK_310

GGTTTCAAGAAGTCTTTGGAAGATCCAGCTGGGAATCTGAATTCATGGAACTTTAAAAATACTACTATCGGAGCTATCTGTAAGTTTACGGGTGTTGATTGCTGGAATGACAACGAAAAT

>M6-PGSC0003DMG400001301-(rev)M6LRR-RLK_372

TACTATCGGAGCTATCTGTAAGTTTACGGGTGTTGATTGCTGGAATGACAACGAAAATCGCATCATCAGCCTCATTCTTTCGACTATGAACCTCGGAGGGAAGGTTACAGAGCCTGTACA

>M6-PGSC0003DMG400001301-(rev)M6LRR-RLK_434

TCATCAGCCTCATTCTTTCGACTATGAACCTCGGAGGGAAGGTTACAGAGCCTGTACAGTATTGTGCCAGCTTGACAACCCTTGATCTCTCTGGTAATCGCTTTTCTGGTCCAATTCCTT

>M6-PGSC0003DMG400001301-(rev)M6LRR-RLK_496

TGTGCCAGCTTGACAACCCTTGATCTCTCTGGTAATCGCTTTTCTGGTCCAATTCCTTCTCAAATTTGTACTTGGCTACCTTTTTTGGTAACCCTAGACTTGTCCGGTAACGATTACTCC

>M6-PGSC0003DMG400001301-(rev)M6LRR-RLK_558

AATTTGTACTTGGCTACCTTTTTTGGTAACCCTAGACTTGTCCGGTAACGATTACTCCGGCCCTATCCCTGCTGATCTTGCTAAATGCACCTTTTTGAATAAATTATCGCTCAGTAATAA

>M6-PGSC0003DMG400001301-(rev)M6LRR-RLK_62

TTCATTTGTCAAGTCAAATGCCTCACCAGTCTTCATCTTCCTTCTCCTTGCTATTTCCTACCTATTTCTCTTGCGCATATTATCAAACACCTTGTGCCAATTACCCAAAACTGCTGTTGG

>M6-PGSC0003DMG400001301-(rev)M6LRR-RLK_620

CTATCCCTGCTGATCTTGCTAAATGCACCTTTTTGAATAAATTATCGCTCAGTAATAATAAGCTTACTGGAAATATTCCCCCTGAATTTTCTAGTTTAGGTAGGCTTAAAAGTTTTTCTG

>M6-PGSC0003DMG400001301-(rev)M6LRR-RLK_682

CTTACTGGAAATATTCCCCCTGAATTTTCTAGTTTAGGTAGGCTTAAAAGTTTTTCTGTGGCAAATAATCAACTCTCCGGTAGAATCCCGGCAGCTTTCGATTCTTCTAACTTTAATTTT

>M6-PGSC0003DMG400001301-(rev)M6LRR-RLK_744

AAATAATCAACTCTCCGGTAGAATCCCGGCAGCTTTCGATTCTTCTAACTTTAATTTTGAAGGAAATAGTCTCTGTGGTGGACCTTTGGGGAAATGTGGAGGACTAAGTAAGAAAAGCTT

>M6-PGSC0003DMG400001301-(rev)M6LRR-RLK_806

GAAATAGTCTCTGTGGTGGACCTTTGGGGAAATGTGGAGGACTAAGTAAGAAAAGCTTAGCTATTATTATTGCAGCAGGAGTCTTTGGTGCTGCTGCTTCTATGTTGTTGGGTTTTGGTG

>M6-PGSC0003DMG400001301-(rev)M6LRR-RLK_868

ATTATTATTGCAGCAGGAGTCTTTGGTGCTGCTGCTTCTATGTTGTTGGGTTTTGGTGCGTGGTATTGGTTTTTTACCAAGTCTGGAAAGAGGAAGAGAGGGTATGGAGTTGGGAGAGAC

>M6-PGSC0003DMG400001301-(rev)M6LRR-RLK_930

GTATTGGTTTTTTACCAAGTCTGGAAAGAGGAAGAGAGGGTATGGAGTTGGGAGAGACGACTCTGATAGTTGGGCTGATAAGCTGAGGGCTCACAAGCTCACTCAGGTTATGTTGTTCCA

>M6-PGSC0003DMG400001301-(rev)M6LRR-RLK_992

CTGATAGTTGGGCTGATAAGCTGAGGGCTCACAAGCTCACTCAGGTTATGTTGTTCCAGAAACCGCTCGTCAAGGTTAAATTAGCTGATTTGTTGATTGCCACAAATGGTTTCAGGACGG

>M6-PGSC0003DMG400001368-(rev)M6RLKmalectin_0

CCACAGATAAAAGAAGAGGAAATGCCTTGAGAGTCAACACTCAAGTCTTGCCAATGGTTGTGTATAATTGCTCCTTGGAACAACTAATATCTCCTTACACAAACGCCCATGGCAACCATT

>M6-PGSC0003DMG400001368-(rev)M6RLKmalectin_1054

AGCTGGAAGTTAAAGAAATAAACGATAGAAGTTTTTCCATTTCTATCGGAAACCAAAAAGCCGAGCGAGAGGCTGATGTAATACAATGGAGTGGAGGCTGGAGAATCCCTGTGTACAAAG

>M6-PGSC0003DMG400001368-(rev)M6RLKmalectin_1116

GAGCGAGAGGCTGATGTAATACAATGGAGTGGAGGCTGGAGAATCCCTGTGTACAAAGACTATGTGGTTAATGTTAGAAACCAAGATGGTGAGCAAAACGTGACGCTTGATTTGAGTCCT

>M6-PGSC0003DMG400001368-(rev)M6RLKmalectin_1178

TGTGGTTAATGTTAGAAACCAAGATGGTGAGCAAAACGTGACGCTTGATTTGAGTCCTAACCCAGATTCTGCGTATCAAAATGCTATCTTAAATGGCTTAGAAATTTTCAAGTTGAACGA

>M6-PGSC0003DMG400001368-(rev)M6RLKmalectin_124

TACTTTTCTTTCTCTTCTTACAACTCATAACGGCCACTATTTCTACCACGCCGCCACCGTATAACGCCACTGTCTTTGTCTTGCTTAACTGTGGCGCACAATCAGCCACCACCGATGAAA

>M6-PGSC0003DMG400001368-(rev)M6RLKmalectin_1240

CAGATTCTGCGTATCAAAATGCTATCTTAAATGGCTTAGAAATTTTCAAGTTGAACGACTCAAATGGAAACCTCTCCGTGCCGAATCCTGAATTTTTCTTTCCGAATAATTCACCACCAA

>M6-PGSC0003DMG400001368-(rev)M6RLKmalectin_1302

AATGGAAACCTCTCCGTGCCGAATCCTGAATTTTTCTTTCCGAATAATTCACCACCAAACGACAATAATAAGAGATCATCATCACACATCATTGCTGTTATCACTGCAGTAGCTGTTATT

>M6-PGSC0003DMG400001368-(rev)M6RLKmalectin_1364

CAATAATAAGAGATCATCATCACACATCATTGCTGTTATCACTGCAGTAGCTGTTATTAGCGGAATTGCTCTGCTTTCAATTCTCTGTTTCTTAATTTTCCGGCGGTGGAAAAGGGGAAA

>M6-PGSC0003DMG400001368-(rev)M6RLKmalectin_1426

GAATTGCTCTGCTTTCAATTCTCTGTTTCTTAATTTTCCGGCGGTGGAAAAGGGGAAAAGAACTTCATACAAGCGTTACAAAGTCATCATGGATCCCTTTGTCAATTACATCGAACTCAA

>M6-PGSC0003DMG400001368-(rev)M6RLKmalectin_1488

CTTCATACAAGCGTTACAAAGTCATCATGGATCCCTTTGTCAATTACATCGAACTCAACCCAAAGAACCGGCGGCTCAGGCTCATCAGCGCTACCATCCGATCTGTGCCGACACTTTTTA

>M6-PGSC0003DMG400001368-(rev)M6RLKmalectin_1550

AAGAACCGGCGGCTCAGGCTCATCAGCGCTACCATCCGATCTGTGCCGACACTTTTTACTAGAAGAGATAAAAACAGCAACATGCAATTTTGATGAAAAATTTGTGATTGGCTATGGAGG

>M6-PGSC0003DMG400001368-(rev)M6RLKmalectin_1612

AAGAGATAAAAACAGCAACATGCAATTTTGATGAAAAATTTGTGATTGGCTATGGAGGTTTTGGTAATGTATACAAAGGGTACATAGACAACGGCGCAACCATAGTTGCAGTCAAGAGAT

>M6-PGSC0003DMG400001368-(rev)M6RLKmalectin_1674

GGTAATGTATACAAAGGGTACATAGACAACGGCGCAACCATAGTTGCAGTCAAGAGATTGAATCCTTCATCAAAGCAAGGAGTCCGCGAGTTCGAAACAGAGATTCACATGTTGTCAAAG

>M6-PGSC0003DMG400001368-(rev)M6RLKmalectin_1736

TCCTTCATCAAAGCAAGGAGTCCGCGAGTTCGAAACAGAGATTCACATGTTGTCAAAGCTAAGGCATGTGCATCTGGTGTCCTTAATTGGTTATTGTGATGATAACAATGAGATGATCTT

>M6-PGSC0003DMG400001368-(rev)M6RLKmalectin_1798

GGCATGTGCATCTGGTGTCCTTAATTGGTTATTGTGATGATAACAATGAGATGATCTTGGTTTATGATTACATGTCTCATGGAACTCTTCGTGATCACTTGTACAAAACTGATAATGCTC

>M6-PGSC0003DMG400001368-(rev)M6RLKmalectin_1860

TATGATTACATGTCTCATGGAACTCTTCGTGATCACTTGTACAAAACTGATAATGCTCCACTTCCATGGAAGAAACGCCTTGAGATTTGCATAGGCGCAGCAAAAGGATTGCATTATCTC

>M6-PGSC0003DMG400001368-(rev)M6RLKmalectin_1922

TCCATGGAAGAAACGCCTTGAGATTTGCATAGGCGCAGCAAAAGGATTGCATTATCTCCATACAGGTATACGGTCACTTAATTCTATACTTCTATTTGCTGTGGTACCATATTAAAGTAT

>M6-PGSC0003DMG400001368-(rev)M6RLKmalectin_1984

CAGGTATACGGTCACTTAATTCTATACTTCTATTTGCTGTGGTACCATATTAAAGTATGTATTGAGTAAATTAAGTGATTTACTGGTAAAAAAGATAAAGATTAATTATTTTTAATATAA

>M6-PGSC0003DMG400001368-(rev)M6RLKmalectin_2108

TTATTATTATCTATAGTAGATATGATCGTTAAATTTAATTTCACAGGTAAAAAGGATATCATTATTCATCGAGATGTGAAGTCAACCAACATTTTATTGGATGATAAATGGGTTGCTAAG

>M6-PGSC0003DMG400001368-(rev)M6RLKmalectin_2170

TATTCATCGAGATGTGAAGTCAACCAACATTTTATTGGATGATAAATGGGTTGCTAAGGTTTCAGATTTTGGGTTATCCAAAATCGGTCCACTTAGTGGATCAGGAAAAACCCATGTTAG

>M6-PGSC0003DMG400001368-(rev)M6RLKmalectin_2232

CAGATTTTGGGTTATCCAAAATCGGTCCACTTAGTGGATCAGGAAAAACCCATGTTAGCACTGTTGTGAAAGGAAGCTTTGGATATCTGGATCCTGAATACTATAAGCGACAACAATTGA

>M6-PGSC0003DMG400001368-(rev)M6RLKmalectin_2294

GTTGTGAAAGGAAGCTTTGGATATCTGGATCCTGAATACTATAAGCGACAACAATTGACAGAAAAATCTGATGTGTATTCTTTTGGGGTGGTGCTATTCGAAGTGTTGTGCGCTAGACCA

>M6-PGSC0003DMG400001368-(rev)M6RLKmalectin_2356

AAAATCTGATGTGTATTCTTTTGGGGTGGTGCTATTCGAAGTGTTGTGCGCTAGACCAGCATTAATTCCTAACATGCCGAAAGGGCAAGTGAATTTGGCTGATTGGGCTTGTAGGAGCTG

>M6-PGSC0003DMG400001368-(rev)M6RLKmalectin_2418

TAATTCCTAACATGCCGAAAGGGCAAGTGAATTTGGCTGATTGGGCTTGTAGGAGCTGTAAAAAAGGAAATCTTCAACAAATAATTGATCCGAATTTGGAAGGACAAATTGCGCCAGAGT

>M6-PGSC0003DMG400001368-(rev)M6RLKmalectin_248

CCGTCGATGGGATACTGATACTCACTTTCCAAATTTCTTACCTTCCAACTTTTCAAGCATATCCACAACAGCCACAGCTTCTGAACAAGATCCTTCGGTCAATAGAGTTCCTTATACGAC

>M6-PGSC0003DMG400001368-(rev)M6RLKmalectin_2480

AAAGGAAATCTTCAACAAATAATTGATCCGAATTTGGAAGGACAAATTGCGCCAGAGTGCTTAAACAAGTTTGCAGAAGCAGCATATAATTGCTTGAAGGATCAAGGAGTCCAAAGGCCA

>M6-PGSC0003DMG400001368-(rev)M6RLKmalectin_2542

AAACAAGTTTGCAGAAGCAGCATATAATTGCTTGAAGGATCAAGGAGTCCAAAGGCCATCAATGAATGACGTTGTCTGGAATTTGGAATTTATATTAAAGCTTCAAGATGCAGCTGATAA

>M6-PGSC0003DMG400001368-(rev)M6RLKmalectin_2604

TGAATGACGTTGTCTGGAATTTGGAATTTATATTAAAGCTTCAAGATGCAGCTGATAATAGAGGTCACAAAATGGAATTGAACAGTTATCCGACTAGCCCGTCTTTCCCCATGATTATGA

>M6-PGSC0003DMG400001368-(rev)M6RLKmalectin_2666

GGTCACAAAATGGAATTGAACAGTTATCCGACTAGCCCGTCTTTCCCCATGATTATGAATGGTCATACCAATATATCTACAGACGAAGGATTCGAGGCATTTTCAGGTTCACATGAAGTT

>M6-PGSC0003DMG400001368-(rev)M6RLKmalectin_2728

TCATACCAATATATCTACAGACGAAGGATTCGAGGCATTTTCAGGTTCACATGAAGTTGGAGGAAAATACACAAGCAGTGCTACATCCATGACCAGTAATAGTGATGACAAATTGAAGAG

>M6-PGSC0003DMG400001368-(rev)M6RLKmalectin_2790

GAAAATACACAAGCAGTGCTACATCCATGACCAGTAATAGTGATGACAAATTGAAGAGTGACACTATTTTCTCAGAGATTTTGAATCCATCGGGTCGATGAGATTGTTACAACTAGTAAG

>M6-PGSC0003DMG400001368-(rev)M6RLKmalectin_2852

ACTATTTTCTCAGAGATTTTGAATCCATCGGGTCGATGAGATTGTTACAACTAGTAAGATAAATTACATATTAGTTCTTCTCTGTTTTCTTTTTCCTTTGTTGAATTTATGTCCTTATTA

>M6-PGSC0003DMG400001368-(rev)M6RLKmalectin_2914

ATTACATATTAGTTCTTCTCTGTTTTCTTTTTCCTTTGTTGAATTTATGTCCTTATTATATATGCAGATGAATTGAAGCCATTATTCAGAAGTATATAAGCTATGAAATAGTTTCACTAT

>M6-PGSC0003DMG400001368-(rev)M6RLKmalectin_310

CCACAACAGCCACAGCTTCTGAACAAGATCCTTCGGTCAATAGAGTTCCTTATACGACTGGTGCTCGTATTATGAGATCTCAATTCACCTACACTTTCCGTGTTACCCCTGGCACAATAT

>M6-PGSC0003DMG400001368-(rev)M6RLKmalectin_372

GCTCGTATTATGAGATCTCAATTCACCTACACTTTCCGTGTTACCCCTGGCACAATATTCCTCCGTCTCTATTTCTACCCTGCCAATTACTCCGGCTTCAACAAAGCTGAATCTTTTTTC

>M6-PGSC0003DMG400001368-(rev)M6RLKmalectin_434

CCGTCTCTATTTCTACCCTGCCAATTACTCCGGCTTCAACAAAGCTGAATCTTTTTTCTCCGTCACAGCTAATCATTTAACCCTCTTGAGTAACTTCAGTGCTTTCCTCACAGTCTCCGC

>M6-PGSC0003DMG400001368-(rev)M6RLKmalectin_496

TCACAGCTAATCATTTAACCCTCTTGAGTAACTTCAGTGCTTTCCTCACAGTCTCCGCAAGTAGTAGTAAAAAAGTTCAAAAAGAATATGTCATCAATGTTGATGAAACTCAGATACTCA

>M6-PGSC0003DMG400001368-(rev)M6RLKmalectin_558

AGTAGTAAAAAAGTTCAAAAAGAATATGTCATCAATGTTGATGAAACTCAGATACTCAAATTAACCTTTTCTCCTTCACCAAACTCTTATGCTTTTGTTAATGGGATTGAGATACTTTCT

>M6-PGSC0003DMG400001368-(rev)M6RLKmalectin_62

TATAATTGCTCCTTGGAACAACTAATATCTCCTTACACAAACGCCCATGGCAACCATTACTTTACTTTTCTTTCTCTTCTTACAACTCATAACGGCCACTATTTCTACCACGCCGCCACC

>M6-PGSC0003DMG400001368-(rev)M6RLKmalectin_620

AACCTTTTCTCCTTCACCAAACTCTTATGCTTTTGTTAATGGGATTGAGATACTTTCTATGCCAACTGATCTTTACATCCATGGAGACGTAAAATTGGCTGGGTATACTAATCCATATAA

>M6-PGSC0003DMG400001368-(rev)M6RLKmalectin_682

CAACTGATCTTTACATCCATGGAGACGTAAAATTGGCTGGGTATACTAATCCATATAACATCAATAACAGCACTGCTCTTGAAACTTTGTACAGGCTGAACGTGGGAGGCAATTTAGTTG

>M6-PGSC0003DMG400001368-(rev)M6RLKmalectin_806

CACAGAGGATGCAGGGATGTATCGTGTATGGGATTCAGATGATGCATTTGTTGTTGGATATGGCTATCAAACTCCTCATTTTGATGATGCCAACATCAATTACACTTCTGAAACCCCGGC

>M6-PGSC0003DMG400001368-(rev)M6RLKmalectin_868

GCTATCAAACTCCTCATTTTGATGATGCCAACATCAATTACACTTCTGAAACCCCGGCTTATTCTGCGCCAAGGATAGTTTACACAACCTCTAGGATTATGGCCAATCACAGCCAGAGCC

>M6-PGSC0003DMG400001368-(rev)M6RLKmalectin_930

TCTGCGCCAAGGATAGTTTACACAACCTCTAGGATTATGGCCAATCACAGCCAGAGCCTGGATTGGGAATTCCCTCTAAACTCTGGCTTCTATTATCTTTTCAGGCTCCATTTCTGTGAG

>M6-PGSC0003DMG400001368-(rev)M6RLKmalectin_992

TTGGGAATTCCCTCTAAACTCTGGCTTCTATTATCTTTTCAGGCTCCATTTCTGTGAGATTCAGCTGGAAGTTAAAGAAATAAACGATAGAAGTTTTTCCATTTCTATCGGAAACCAAAA

>M6-PGSC0003DMG400001369-(rev)M6RLKmalectin_0

AATTTAACAAAGAATCATCCAAATTCCAAAGAAAATAACCCACAACCTTTCCTTTCAAGCATGTCGACTATTGCTCCAGTGGGACATAACAAAGATTATTATGAAAATGAAAGGCAGCAT

>M6-PGSC0003DMG400001369-(rev)M6RLKmalectin_1037

TCCATGCCTACCAATCTCTACATCCGTGGAAACCATCGCATTGAATTGGTGGGCCAGAATGATTTTTTTTACATCAATAACAGCACTGCTCTTGAAACTTTGTACAGGCTGAACGTGGGA

>M6-PGSC0003DMG400001369-(rev)M6RLKmalectin_1098

ATTTTTTTTACATCAATAACAGCACTGCTCTTGAAACTTTGTACAGGCTGAACGTGGGAGGCAATTTAGTTGGTAGCACAGAGGATACAGGGATGTATCGTGTATGGGATTCAGATGATG

>M6-PGSC0003DMG400001369-(rev)M6RLKmalectin_122

ATGGCCTACAAGAAAAAGATGTACACCCCCAATTAACAACTTTTCCTTTACCATTTCTATTTCTCTCTGTATTTTCTTGGGTACATATATGGTCCCCTTCTTTAGTGCAGTAATCCTAAT

>M6-PGSC0003DMG400001369-(rev)M6RLKmalectin_1220

TTTGTTGTTGGATATGGCTATCAAACTCCTCATTTTGATGTCAACATCACTTACACTTCTGAAACCCCGGCTTATTCTGCGCCAACCATAATTTACACAACCTCTAGGACCATGGTCAAT

>M6-PGSC0003DMG400001369-(rev)M6RLKmalectin_1281

AAACCCCGGCTTATTCTGCGCCAACCATAATTTACACAACCTCTAGGACCATGGTCAATTACAGCAAGGGCCTAGATTGGGAATTCCCTCTAAACTCTGGTTTCTATTATCTTTTCAGGC

>M6-PGSC0003DMG400001369-(rev)M6RLKmalectin_1342

CAGCAAGGGCCTAGATTGGGAATTCCCTCTAAACTCTGGTTTCTATTATCTTTTCAGGCTCCATTTTTGTGAGATTCAGCCGGAAGTTAAGGGAGTGAACCACAGATTTTTCGACATATA

>M6-PGSC0003DMG400001369-(rev)M6RLKmalectin_1403

CATTTTTGTGAGATTCAGCCGGAAGTTAAGGGAGTGAACCACAGATTTTTCGACATATACATTGGAAACCAAATGGCAGAGCGAGCAGCGGATGTAGTTGAATGGAGTGGCGGCTGGAGA

>M6-PGSC0003DMG400001369-(rev)M6RLKmalectin_1464

TTGGAAACCAAATGGCAGAGCGAGCAGCGGATGTAGTTGAATGGAGTGGCGGCTGGAGATTCCCGGTTTACAGAGACTACGTGGTGAATGTGCCTAACCCAGACGACCGACGAAACAAGC

>M6-PGSC0003DMG400001369-(rev)M6RLKmalectin_1525

CCCGGTTTACAGAGACTACGTGGTGAATGTGCCTAACCCAGACGACCGACGAAACAAGCAAAATATTACGCTTGAATTGAGTCCTAACATGAACACAAAAGCTGTGTATGCTAATGGTAC

>M6-PGSC0003DMG400001369-(rev)M6RLKmalectin_1586

AATATTACGCTTGAATTGAGTCCTAACATGAACACAAAAGCTGTGTATGCTAATGGTACCTCAGAATTAGCGTATTCAAATGTTATCTTAAAAGGCTTAGAAATTTTCAAATTGGGTGAT

>M6-PGSC0003DMG400001369-(rev)M6RLKmalectin_1647

CAGAATTAGCGTATTCAAATGTTATCTTAAAAGGCTTAGAAATTTTCAAATTGGGTGATGCAAATGGGAGTCTGGCCCCGTCAATTCATGAATTGCCCTCAAAAGCAAACGGAAATCCGG

>M6-PGSC0003DMG400001369-(rev)M6RLKmalectin_1708

AAATGGGAGTCTGGCCCCGTCAATTCATGAATTGCCCTCAAAAGCAAACGGAAATCCGGCCGCCTTGACTCCTGAATTTGCCTCAGAAGCTCCGATCGCTTCATTAGCTTTTCCTAACAA

>M6-PGSC0003DMG400001369-(rev)M6RLKmalectin_1769

GCCTTGACTCCTGAATTTGCCTCAGAAGCTCCGATCGCTTCATTAGCTTTTCCTAACAACAAGAAGAAAAGAATATCATGGTATATCATTGTTGCTATCATCACAGGTGTGACTAGCGGT

>M6-PGSC0003DMG400001369-(rev)M6RLKmalectin_183

TCTCTCTGTATTTTCTTGGGTACATATATGGTCCCCTTCTTTAGTGCAGTAATCCTAATAGTAATTTCACTTATATTCTTCGATGAAGTCAACTCTTGCCTCACTGATTTTTACTCCATT

>M6-PGSC0003DMG400001369-(rev)M6RLKmalectin_1830

AGAAGAAAAGAATATCATGGTATATCATTGTTGCTATCATCACAGGTGTGACTAGCGGTATTGCTTTTTTTTCGATTCTGGGTTTCTTAATTTCCCGACGACGAAATGGGGGAAAAGATT

>M6-PGSC0003DMG400001369-(rev)M6RLKmalectin_1891

TGCTTTTTTTTCGATTCTGGGTTTCTTAATTTCCCGACGACGAAATGGGGGAAAAGATTCTGGTACAAGTGTTACCAAGTCATCATCACCACCATCCGATCTGTGCCGACACTTTTTAAT

>M6-PGSC0003DMG400001369-(rev)M6RLKmalectin_1952

GGTACAAGTGTTACCAAGTCATCATCACCACCATCCGATCTGTGCCGACACTTTTTAATAGCAGAGATAAAAAACGCTACCGACAATTTTGATGAAAAATTTGTGATTGGATATGGAGGT

>M6-PGSC0003DMG400001369-(rev)M6RLKmalectin_2013

CAGAGATAAAAAACGCTACCGACAATTTTGATGAAAAATTTGTGATTGGATATGGAGGTTTTGGTAATGTTTATAAAGGGTACCTTGACAATGGTACAACCACAGTTGCAGTCAAGAGAT

>M6-PGSC0003DMG400001369-(rev)M6RLKmalectin_2074

TGGTAATGTTTATAAAGGGTACCTTGACAATGGTACAACCACAGTTGCAGTCAAGAGATCGAATCCTTCATCAAAGCAAGGGGTTCGCGAGTTCCAAACAGAGATTGAGATGTTATCATC

>M6-PGSC0003DMG400001369-(rev)M6RLKmalectin_2135

AATCCTTCATCAAAGCAAGGGGTTCGCGAGTTCCAAACAGAGATTGAGATGTTATCATCAAAGCTAAGGCATCGGCATCTAGTGTCCTTAATTGGTTATTGTGATGATAAGAATGAGATG

>M6-PGSC0003DMG400001369-(rev)M6RLKmalectin_2196

AGCTAAGGCATCGGCATCTAGTGTCCTTAATTGGTTATTGTGATGATAAGAATGAGATGATCTTGGTTTATGATTACATGGCTAATGGAACTCTTCGTGATCACTTGTACAATACTCGCA

>M6-PGSC0003DMG400001369-(rev)M6RLKmalectin_2257

CTTGGTTTATGATTACATGGCTAATGGAACTCTTCGTGATCACTTGTACAATACTCGCAATGCTCCTCTTCCATGGAAGAAACGCCTTGACATTTGCATTGGTGCAGCTAAAGGATTATA

>M6-PGSC0003DMG400001369-(rev)M6RLKmalectin_2318

GCTCCTCTTCCATGGAAGAAACGCCTTGACATTTGCATTGGTGCAGCTAAAGGATTATATTACCTTCATTCAGGTAAGTAATCAATAAATTGATAAACATGTCATGATCTAACTTGCACT

>M6-PGSC0003DMG400001369-(rev)M6RLKmalectin_2379

ACCTTCATTCAGGTAAGTAATCAATAAATTGATAAACATGTCATGATCTAACTTGCACTATATTTGTTTGGTAAAAAAACTATTGGGTCAGAATAGGTCAAATAACCATCAATGTCAAGT

>M6-PGSC0003DMG400001369-(rev)M6RLKmalectin_244

TAATTTCACTTATATTCTTCGATGAAGTCAACTCTTGCCTCACTGATTTTTACTCCATTTATTTGGTGCACTTATAATAATTTACTTTTGGTATTTATTTTTTGTACAAAACTTTAACAA

>M6-PGSC0003DMG400001369-(rev)M6RLKmalectin_2440

ATTTGTTTGGTAAAAAAACTATTGGGTCAGAATAGGTCAAATAACCATCAATGTCAAGTTGCACAGCATGACTCAACCTCCCAATTTGGTTTTTGTATATTCTATTTGCATTTTGACATG

>M6-PGSC0003DMG400001369-(rev)M6RLKmalectin_2501

CACAGCATGACTCAACCTCCCAATTTGGTTTTTGTATATTCTATTTGCATTTTGACATGTTTTGATTCAATAGATTGATGGGTCATTTCGAGTTTAATCCATTAATGTGTCAAGCCAACA

>M6-PGSC0003DMG400001369-(rev)M6RLKmalectin_2562

TTGATTCAATAGATTGATGGGTCATTTCGAGTTTAATCCATTAATGTGTCAAGCCAACAAGCAATTTACAATTAAGATCGTTTAAATTTGAGTGAATTGCTGAAAAGTGAGTTCATTTTG

>M6-PGSC0003DMG400001369-(rev)M6RLKmalectin_2623

CAATTTACAATTAAGATCGTTTAAATTTGAGTGAATTGCTGAAAAGTGAGTTCATTTTGCTGTTATAGGTACCAAACATATGATCATTCATCGAGATGTCAAGTCAACTAACATCCTATT

>M6-PGSC0003DMG400001369-(rev)M6RLKmalectin_2684

GTTATAGGTACCAAACATATGATCATTCATCGAGATGTCAAGTCAACTAACATCCTATTGGATGATAAATGGGTCGCCAAGGTGTCTGATTTTGGGTTGTCCAAAGTAGGTTCAGTAGGT

>M6-PGSC0003DMG400001369-(rev)M6RLKmalectin_2745

ATGATAAATGGGTCGCCAAGGTGTCTGATTTTGGGTTGTCCAAAGTAGGTTCAGTAGGTGGGACAGATATTACCCATGTTAGCACTGCAGTGAAAGGAAGTTTTGGGTATGTAGATCCTG

>M6-PGSC0003DMG400001369-(rev)M6RLKmalectin_2806

GACAGATATTACCCATGTTAGCACTGCAGTGAAAGGAAGTTTTGGGTATGTAGATCCTGAATACTACAAGCGCCAACGATTGACAGTAAAATCAGATGTTTATTCTTTTGGGGTGGTTCT

>M6-PGSC0003DMG400001369-(rev)M6RLKmalectin_2867

TACTACAAGCGCCAACGATTGACAGTAAAATCAGATGTTTATTCTTTTGGGGTGGTTCTATTTGAAGTGTTATGTGGTAGGCCAGCAATAAATCCAAACTTGCCAAGTGGGCAAGTGAAT

>M6-PGSC0003DMG400001369-(rev)M6RLKmalectin_2928

TTGAAGTGTTATGTGGTAGGCCAGCAATAAATCCAAACTTGCCAAGTGGGCAAGTGAATTTGGCTGAATGGGCTTGTAGGAGTTGTAAAAAGAGAGAACTTGAAGAAATCATTGATCCTA

>M6-PGSC0003DMG400001369-(rev)M6RLKmalectin_2989

GGCTGAATGGGCTTGTAGGAGTTGTAAAAAGAGAGAACTTGAAGAAATCATTGATCCTAACTTGAAAGGACAAATAGCCCCAGAATGCTTAAGGAAGTTTGCAGAATTGGCATATCATTG

>M6-PGSC0003DMG400001369-(rev)M6RLKmalectin_305

TTTGGTGCACTTATAATAATTTACTTTTGGTATTTATTTTTTGTACAAAACTTTAACAAAATGATGTCGGTTTTTTATACAGAAAATTAGAAATGACTTGATCTTCGAACATCTACTGTA

>M6-PGSC0003DMG400001369-(rev)M6RLKmalectin_3050

TTGAAAGGACAAATAGCCCCAGAATGCTTAAGGAAGTTTGCAGAATTGGCATATCATTGTTTGAGAAAAGAAGGGGTTCAAAGGCCATCAATGGGTGATGTTGTAAAAACTTTAGAATTT

>M6-PGSC0003DMG400001369-(rev)M6RLKmalectin_3111

TGAGAAAAGAAGGGGTTCAAAGGCCATCAATGGGTGATGTTGTAAAAACTTTAGAATTTGCATTACAGGTTCAAGAGGGTGCAGATCATGAGGTGGAAGGATCCTCTTTTCCGTTGATTA

>M6-PGSC0003DMG400001369-(rev)M6RLKmalectin_3172

ATTACAGGTTCAAGAGGGTGCAGATCATGAGGTGGAAGGATCCTCTTTTCCGTTGATTATTAATGGTGATGCTATGAATGAAGATGAGATATTCTCAGATTTAAGTGAAGTTGAAAGAAA

>M6-PGSC0003DMG400001369-(rev)M6RLKmalectin_3233

AATGGTGATGCTATGAATGAAGATGAGATATTCTCAGATTTAAGTGAAGTTGAAAGAAATTTGACAAGCATGTCGGTCAGTAGGAGTGCCAAAGTGAACAATGTTGCCCTTTTCTCTGAG

>M6-PGSC0003DMG400001369-(rev)M6RLKmalectin_3294

TGACAAGCATGTCGGTCAGTAGGAGTGCCAAAGTGAACAATGTTGCCCTTTTCTCTGAGATTGTCAATCCATTGGGTCGATGAAATTGTTATGATCATAGTTCGATACACTGAAATTACA

>M6-PGSC0003DMG400001369-(rev)M6RLKmalectin_3331

CAATGTTGCCCTTTTCTCTGAGATTGTCAATCCATTGGGTCGATGAAATTGTTATGATCATAGTTCGATACACTGAAATTACATGTATTTTGTTTTTCAGGCTCTTTTTGTAATTTAATG

>M6-PGSC0003DMG400001369-(rev)M6RLKmalectin_366

TGATGTCGGTTTTTTATACAGAAAATTAGAAATGACTTGATCTTCGAACATCTACTGTATATATTTGCACAACAAATAGCACGTACATAGTTTTTCAAGACAACGCCATGTTAGCCATTG

>M6-PGSC0003DMG400001369-(rev)M6RLKmalectin_427

TATTTGCACAACAAATAGCACGTACATAGTTTTTCAAGACAACGCCATGTTAGCCATTGCTCTAATTTCCTTTCTCTTCTTACAACTCATAACGGCCACTATTTCTACCACGCCGCCACC

>M6-PGSC0003DMG400001369-(rev)M6RLKmalectin_61

TGTCGACTATTGCTCCAGTGGGACATAACAAAGATTATTATGAAAATGAAAGGCAGCATCCATGGCCTACAAGAAAAAGATGTACACCCCCAATTAACAACTTTTCCTTTACCATTTCTA

>M6-PGSC0003DMG400001369-(rev)M6RLKmalectin_671

ATATCTACAACAGCCACAGCTTCTGAACAAGATCCTTCGGTCGATAGAGTTCCTTATACGAATGGTGCTCGTATCATGAAATCTCGATTCACTTACACTTTCCGTGTTTCTCCTGGCGCC

>M6-PGSC0003DMG400001369-(rev)M6RLKmalectin_732

ATGGTGCTCGTATCATGAAATCTCGATTCACTTACACTTTCCGTGTTTCTCCTGGCGCCAAATTCCTCCGTCTCTATTTTTACCCCGCTAATTACTCTGGCTTCAACAAAGCTGAATCTT

>M6-PGSC0003DMG400001369-(rev)M6RLKmalectin_793

ATTCCTCCGTCTCTATTTTTACCCCGCTAATTACTCTGGCTTCAACAAAGCTGAATCTTTTTTCTCCGTCACAGCTAATCATTTAACCCTCTTGAGTAACTTCAACGCTCTCGCTGCAGT

>M6-PGSC0003DMG400001369-(rev)M6RLKmalectin_854

TTCTCCGTCACAGCTAATCATTTAACCCTCTTGAGTAACTTCAACGCTCTCGCTGCAGTTACCGATTCTAGTAAAGCTCTTCGAAAAGAATTTGTCATTAATGTTGATGAAACTCAGATT

>M6-PGSC0003DMG400001369-(rev)M6RLKmalectin_915

CCGATTCTAGTAAAGCTCTTCGAAAAGAATTTGTCATTAATGTTGATGAAACTCAGATTCTTAATGTGACCTTTTCTCCCTCGCCAAACTCTTATGCTTTTGTTAATGGGATAGAGATAC

>M6-PGSC0003DMG400001369-(rev)M6RLKmalectin_976

TAATGTGACCTTTTCTCCCTCGCCAAACTCTTATGCTTTTGTTAATGGGATAGAGATACTTTCCATGCCTACCAATCTCTACATCCGTGGAAACCATCGCATTGAATTGGTGGGCCAGAA

>M6-PGSC0003DMG400001732M6RLKmalectin_1037

TAGGTACCCTGATGACCCATTCGATAGAATATGGGCATCTGACACCATAAAAAAGGCCAATTACCTTGTTGATGTGGCTGCTGGTACTGAAAGAGTTTCAACCAAAATGCCAATTGATGT

>M6-PGSC0003DMG400001732M6RLKmalectin_1098

TACCTTGTTGATGTGGCTGCTGGTACTGAAAGAGTTTCAACCAAAATGCCAATTGATGTGAATACCTTAAATGGTGAGATGCCTCCTCAGAAAGCAATGCAGACTGCTGTAGTAGGAAGA

>M6-PGSC0003DMG400001732M6RLKmalectin_1159

ATACCTTAAATGGTGAGATGCCTCCTCAGAAAGCAATGCAGACTGCTGTAGTAGGAAGAAATGGCTCGTTAACTTATCGTCTGAATCTCGATGGTTTCCCTGGTTTTGGTTGGGCATTTA

>M6-PGSC0003DMG400001732M6RLKmalectin_122

CACTGTGTATGTTGTTGATGTTAAGAGAGATTAATAAGACATGAAGTAAAATTTCTGGGTGTTGCTAAAGGAGAAAGCAGTTCTCATTCTTTAATGAAGTTTTACCAATGGGTGTGAAAG

>M6-PGSC0003DMG400001732M6RLKmalectin_1220

TGGCTCGTTAACTTATCGTCTGAATCTCGATGGTTTCCCTGGTTTTGGTTGGGCATTTACTTATTTTGCTGAAATTGAGGATTTGAAACCTGGTGATAGTAGGAAATTCCGGTTGGTTCT

>M6-PGSC0003DMG400001732M6RLKmalectin_1281

TATTTTGCTGAAATTGAGGATTTGAAACCTGGTGATAGTAGGAAATTCCGGTTGGTTCTCCCGGGAGCTCCTGATATCAGCAAGCTTGTTGTTAATATCCAAGAGAATGCCCATGGGAAA

>M6-PGSC0003DMG400001732M6RLKmalectin_1342

CGGGAGCTCCTGATATCAGCAAGCTTGTTGTTAATATCCAAGAGAATGCCCATGGGAAATATCGTTTATATGAGCCCGGCTACTTCAACTTGTCGTTGCCCTTTGTATTATCTTTTCGTT

>M6-PGSC0003DMG400001732M6RLKmalectin_1403

TCGTTTATATGAGCCCGGCTACTTCAACTTGTCGTTGCCCTTTGTATTATCTTTTCGTTTCGGAAAGACATCCGATTCCACGATGGGGCCTCTCTTGAATGCTATGGAAGTAAACAGATA

>M6-PGSC0003DMG400001732M6RLKmalectin_1464

GGAAAGACATCCGATTCCACGATGGGGCCTCTCTTGAATGCTATGGAAGTAAACAGATATGTAAAGAGAACTGATGGCTCTTTAGACGGTAAAACCATGTCTTTTTCTTTTAACATTATT

>M6-PGSC0003DMG400001732M6RLKmalectin_1525

TAAAGAGAACTGATGGCTCTTTAGACGGTAAAACCATGTCTTTTTCTTTTAACATTATTATATTGTGGTTCGTTGAAATGATTATCTGCAGTTTATTTCCTTAGAAGGATTTTTTCTGTT

>M6-PGSC0003DMG400001732M6RLKmalectin_1586

ATTGTGGTTCGTTGAAATGATTATCTGCAGTTTATTTCCTTAGAAGGATTTTTTCTGTTGTAATGAGTATCTTTGTTCGTTAGTTTCAAAATAGTAAGGCATGTCCCTTGCTCTTTGCAT

>M6-PGSC0003DMG400001732M6RLKmalectin_1647

AATGAGTATCTTTGTTCGTTAGTTTCAAAATAGTAAGGCATGTCCCTTGCTCTTTGCATGTGTTTCAGGACCAGTTATTTCTAGTTTAGTCTCGCACTACTCTTCTGCCAATTTGATAAA

>M6-PGSC0003DMG400001732M6RLKmalectin_1708

GTTTCAGGACCAGTTATTTCTAGTTTAGTCTCGCACTACTCTTCTGCCAATTTGATAAACGAAGGTGGTGACCCATGCTTACCAGTCCCATGGTCCTGGATCCGTTGTGACTCAGATATC

>M6-PGSC0003DMG400001732M6RLKmalectin_1769

AAGGTGGTGACCCATGCTTACCAGTCCCATGGTCCTGGATCCGTTGTGACTCAGATATCAGACCAAGAATAACATCAATGTATAGTTCATCACTTTAGCTTCTTTATGCGTAGTTTCATG

>M6-PGSC0003DMG400001732M6RLKmalectin_183

TTGCTAAAGGAGAAAGCAGTTCTCATTCTTTAATGAAGTTTTACCAATGGGTGTGAAAGTTGGTGGATATATATTCATTCTTTATGCACTGCTGATAAATTTTAAACTGTCAAACAGGTT

>M6-PGSC0003DMG400001732M6RLKmalectin_1830

ACCAAGAATAACATCAATGTATAGTTCATCACTTTAGCTTCTTTATGCGTAGTTTCATGTGTTCCACTTTCTGCAATTCAGTTATGGAAGAATTAAGATTACTTTCTAGTTTTCTCTTTA

>M6-PGSC0003DMG400001732M6RLKmalectin_1891

TTCCACTTTCTGCAATTCAGTTATGGAAGAATTAAGATTACTTTCTAGTTTTCTCTTTATATTGTTCTCGCAACCTGAAGCAATGGGCTATAATTCACAACTGTTACTTTGGCAGTAAAT

>M6-PGSC0003DMG400001732M6RLKmalectin_1952

TTGTTCTCGCAACCTGAAGCAATGGGCTATAATTCACAACTGTTACTTTGGCAGTAAATTGTCTGGAAAAAATTTGACTGGAAATATTCCGTCGGAGTTGACAAAGTTGAGCGGCTTAGT

>M6-PGSC0003DMG400001732M6RLKmalectin_2013

TCTGGAAAAAATTTGACTGGAAATATTCCGTCGGAGTTGACAAAGTTGAGCGGCTTAGTTGAGCTGTAAGTCATAATATCATACTGTTAAATGTTCCATGCATATGCTGTGGTTCCCTTC

>M6-PGSC0003DMG400001732M6RLKmalectin_2074

AGCTGTAAGTCATAATATCATACTGTTAAATGTTCCATGCATATGCTGTGGTTCCCTTCTTTCTTTTCTGCTGCTGTATAATCAATATGCACCCATGTTCCTAATGCAGCTGGCTGGATG

>M6-PGSC0003DMG400001732M6RLKmalectin_2135

TCTTTTCTGCTGCTGTATAATCAATATGCACCCATGTTCCTAATGCAGCTGGCTGGATGGGAATTCACTGACTGGTCCAATACCCGATTTTAGCGGATGTCCCAACTTACAGATAATGTA

>M6-PGSC0003DMG400001732M6RLKmalectin_2196

AATTCACTGACTGGTCCAATACCCGATTTTAGCGGATGTCCCAACTTACAGATAATGTAAATTGAGCTGCCTTTTTCAATTCAACTTTTTTAGATATTCTCTTAAAACACATTAAGTGGT

>M6-PGSC0003DMG400001732M6RLKmalectin_2257

TTGAGCTGCCTTTTTCAATTCAACTTTTTTAGATATTCTCTTAAAACACATTAAGTGGTGAATCTAATCACTATGTTTCTTGCAGACATCTTGAGAACAATCAGTTGAGTGGACAGCTTC

>M6-PGSC0003DMG400001732M6RLKmalectin_2318

ATCTAATCACTATGTTTCTTGCAGACATCTTGAGAACAATCAGTTGAGTGGACAGCTTCCTTCAACCTTGGAAGACCTATCAAATTTGAAAGAACTGTAAGCAGCAGAATTATTTATTGA

>M6-PGSC0003DMG400001732M6RLKmalectin_2379

TCAACCTTGGAAGACCTATCAAATTTGAAAGAACTGTAAGCAGCAGAATTATTTATTGACTAAATGTACAATTACTTGTAGAATTTACTCAAGCACGTTAGGCTGTAGATGCTATTCAAC

>M6-PGSC0003DMG400001732M6RLKmalectin_244

GGTGGATATATATTCATTCTTTATGCACTGCTGATAAATTTTAAACTGTCAAACAGGTTTTGTAAGTTTAGATTGTGGAGGCAAAGACAACTTCACAGATGAACTTGGTCTTAAATGGAC

>M6-PGSC0003DMG400001732M6RLKmalectin_2440

AAATGTACAATTACTTGTAGAATTTACTCAAGCACGTTAGGCTGTAGATGCTATTCAACTATGAGTTCATGCATATGGCTGATAAGCTTAACTCTCATATTAAGCTTCTGCTCATCAGAA

>M6-PGSC0003DMG400001732M6RLKmalectin_2501

TGAGTTCATGCATATGGCTGATAAGCTTAACTCTCATATTAAGCTTCTGCTCATCAGAATAAGAATCTAGCATAACCCTGGTTGAGCTGCAGATCATGATCACGTCCTTGGAACTTTTTT

>M6-PGSC0003DMG400001732M6RLKmalectin_2562

AGAATCTAGCATAACCCTGGTTGAGCTGCAGATCATGATCACGTCCTTGGAACTTTTTTGGCTCCACTTGTGTCCCTTTCTCTTATTTTAGCTTGCAGTCGTATGTACTGCCTCTAGTAA

>M6-PGSC0003DMG400001732M6RLKmalectin_2623

CTCCACTTGTGTCCCTTTCTCTTATTTTAGCTTGCAGTCGTATGTACTGCCTCTAGTAAAGTTCTCTTCTCTTTCTTAATGAAAATGATGATTGCAGGTACGTGCAGAACAACATGCTAA

>M6-PGSC0003DMG400001732M6RLKmalectin_2684

TTCTCTTCTCTTTCTTAATGAAAATGATGATTGCAGGTACGTGCAGAACAACATGCTAACTGGAAGTGTGCCATCTGGTCTTCGTGATAACGGTGTGATTTTGAAGTGGGTATCATTTCT

>M6-PGSC0003DMG400001732M6RLKmalectin_2745

GGAAGTGTGCCATCTGGTCTTCGTGATAACGGTGTGATTTTGAAGTGGGTATCATTTCTGTGCTAAGTGTTTGTTTCTGAGGCCTTGTATCTTGTCATGCATTTATACTGTACTGCCATG

>M6-PGSC0003DMG400001732M6RLKmalectin_2806

GCTAAGTGTTTGTTTCTGAGGCCTTGTATCTTGTCATGCATTTATACTGTACTGCCATGGAAACTGTTGAATTTACTTTTTTAGTGCTAGAGAAACTAAATTCTTAAATGTTGGATTGAG

>M6-PGSC0003DMG400001732M6RLKmalectin_2867

AACTGTTGAATTTACTTTTTTAGTGCTAGAGAAACTAAATTCTTAAATGTTGGATTGAGGCGGGTCTAAGTAGCGTGATATGGATAGTGAGGATTCATACCGACCCCAACTAGCTTTTGG

>M6-PGSC0003DMG400001732M6RLKmalectin_2928

GGGTCTAAGTAGCGTGATATGGATAGTGAGGATTCATACCGACCCCAACTAGCTTTTGGTTAAGGTATACTTGTTGATGCCTAAACCTTTTCATTTCAGCTACCAAGGTATATCTATAGG

>M6-PGSC0003DMG400001732M6RLKmalectin_2989

AAGGTATACTTGTTGATGCCTAAACCTTTTCATTTCAGCTACCAAGGTATATCTATAGGGAGCTCATGGTAGAGACTCAATACTCCTTTCTTTATGAAAGCAGATGATCTTAACATAAGT

>M6-PGSC0003DMG400001732M6RLKmalectin_305

GTAAGTTTAGATTGTGGAGGCAAAGACAACTTCACAGATGAACTTGGTCTTAAATGGACCCCTGATACTCAAATGATTAGCGGAAATATAGCCAATATCTCTGTTGCAAATGAGACGAGG

>M6-PGSC0003DMG400001732M6RLKmalectin_3050

GCTCATGGTAGAGACTCAATACTCCTTTCTTTATGAAAGCAGATGATCTTAACATAAGTTGGCTTGTAAAGATACTCATATGTTAAACCCTCCGCGCTTGGTTGTGGTTCTTATTTCAGT

>M6-PGSC0003DMG400001732M6RLKmalectin_3111

GCTTGTAAAGATACTCATATGTTAAACCCTCCGCGCTTGGTTGTGGTTCTTATTTCAGTTACTTTAATGGTTCTTACTGCACATTCAAAGTGTCTTCAGTCTCTACTAAGCTTGACTAAT

>M6-PGSC0003DMG400001732M6RLKmalectin_3172

CTTTAATGGTTCTTACTGCACATTCAAAGTGTCTTCAGTCTCTACTAAGCTTGACTAATTGCTTTACATTTTATATGTCGAAATTACAAATCTTAAGTTTCTCTGATAGTCATGCTGCAT

>M6-PGSC0003DMG400001732M6RLKmalectin_3233

CTTTACATTTTATATGTCGAAATTACAAATCTTAAGTTTCTCTGATAGTCATGCTGCATTTCTAAATTGTTTGGCTTGTCTGTTTCCTTACTAAATCTGCTAAAACATTCTGTATTCTCC

>M6-PGSC0003DMG400001732M6RLKmalectin_3294

CTAAATTGTTTGGCTTGTCTGTTTCCTTACTAAATCTGCTAAAACATTCTGTATTCTCCTCAGTTACACTGGAAACTTTAATCTTCACGAAGGGGAAACCAGTCGCAGCCGCAAGAAAAT

>M6-PGSC0003DMG400001732M6RLKmalectin_3355

AGTTACACTGGAAACTTTAATCTTCACGAAGGGGAAACCAGTCGCAGCCGCAAGAAAATCATTATTGGATCATCAGTTGGGGCTTCCGCTCTTCTTCTTGCTACTATTGCCTCTTGCATA

>M6-PGSC0003DMG400001732M6RLKmalectin_3416

TTATTGGATCATCAGTTGGGGCTTCCGCTCTTCTTCTTGCTACTATTGCCTCTTGCATACTACTGCAAAAAGGGAAAAAAAGCTCTCCTAAGCAAGGTGTGGTGCAATACTTAATTTTGT

>M6-PGSC0003DMG400001732M6RLKmalectin_3477

ACTGCAAAAAGGGAAAAAAAGCTCTCCTAAGCAAGGTGTGGTGCAATACTTAATTTTGTCATTTTCATTATACATAATTTTTTTTGGTGGGATTCACATTATCGTTCTGCTTCTGCTTTA

>M6-PGSC0003DMG400001732M6RLKmalectin_3538

TTTTCATTATACATAATTTTTTTTGGTGGGATTCACATTATCGTTCTGCTTCTGCTTTAGGCCACCTCGAGATGAATTTACCTCCTCAAAGGTTTGTCTCTTCCTTGGGAGATGCTGCAA

>M6-PGSC0003DMG400001732M6RLKmalectin_3599

CCACCTCGAGATGAATTTACCTCCTCAAAGGTTTGTCTCTTCCTTGGGAGATGCTGCAACAGAAGCAGCACACTGCTTTACATTAGCTGAACTTGAAGAAGCGACCAAGAACTTTGAAAG

>M6-PGSC0003DMG400001732M6RLKmalectin_366

CTGATACTCAAATGATTAGCGGAAATATAGCCAATATCTCTGTTGCAAATGAGACGAGGACACAGTATATGGCCTTAAGATACTTTCCAGCCGATAACAGGAAGTATTGCTATACACTTG

>M6-PGSC0003DMG400001732M6RLKmalectin_3660

GAAGCAGCACACTGCTTTACATTAGCTGAACTTGAAGAAGCGACCAAGAACTTTGAAAGAAAGGTTGGATCAGGGGGATTTGGAGTTGTATACTATGGGAAATTGAAGGACGGGAAGGAA

>M6-PGSC0003DMG400001732M6RLKmalectin_3721

AGGTTGGATCAGGGGGATTTGGAGTTGTATACTATGGGAAATTGAAGGACGGGAAGGAAATTGCTGTCAAACTTCTTACAAATAATTCCTTTCAGGGGAAGCGAGAGTTCTCAAATGAGG

>M6-PGSC0003DMG400001732M6RLKmalectin_3782

TGCTGTCAAACTTCTTACAAATAATTCCTTTCAGGGGAAGCGAGAGTTCTCAAATGAGGTAAATAGGTGGATGCTCCTATTATATTCTAGTAGCTTGTCATGATGCATAAGATGGTCTTA

>M6-PGSC0003DMG400001732M6RLKmalectin_3843

AATAGGTGGATGCTCCTATTATATTCTAGTAGCTTGTCATGATGCATAAGATGGTCTTAAATGTCGGGACTGATGCTGTGGCTAAAAACCCATTATTCATTTGTTTAACTTTGTGAGGAA

>M6-PGSC0003DMG400001732M6RLKmalectin_3904

TGTCGGGACTGATGCTGTGGCTAAAAACCCATTATTCATTTGTTTAACTTTGTGAGGAAGATCCAGATATAAGAGGCTTCACCATTACACTTCAATGTTTTTAGATTATCGAGGAATCCA

>M6-PGSC0003DMG400001732M6RLKmalectin_3965

TCCAGATATAAGAGGCTTCACCATTACACTTCAATGTTTTTAGATTATCGAGGAATCCATTGTTTTACTTTTGGAACTCGGGGATAATGGGCCCACCCCTCTCCCTTGTTTAACTTATAT

>M6-PGSC0003DMG400001732M6RLKmalectin_4026

GTTTTACTTTTGGAACTCGGGGATAATGGGCCCACCCCTCTCCCTTGTTTAACTTATATACCAAATTTGGTGCAACAACCAGGTTTTGAAGTCATGACATGCACCTACCACACTCCTGCT

>M6-PGSC0003DMG400001732M6RLKmalectin_4087

CAAATTTGGTGCAACAACCAGGTTTTGAAGTCATGACATGCACCTACCACACTCCTGCTCTGTACTACCTCTACCTCTGATGTCGAACCAAATTCCCGGAGGCTTCCTTGACACTCACAT

>M6-PGSC0003DMG400001732M6RLKmalectin_4148

GTACTACCTCTACCTCTGATGTCGAACCAAATTCCCGGAGGCTTCCTTGACACTCACATTATTTTTGCTTTTCAAATCTGCATAACATCTCAGTATAGAAAATGAAATTTGCGTAATTTA

>M6-PGSC0003DMG400001732M6RLKmalectin_4209

TTTTTGCTTTTCAAATCTGCATAACATCTCAGTATAGAAAATGAAATTTGCGTAATTTAGAAAGGATTAATTTAACGCCAGAGGACACTTTCTCAGTGGTACTTTTTCACACTCATCAAA

>M6-PGSC0003DMG400001732M6RLKmalectin_427

ACAGTATATGGCCTTAAGATACTTTCCAGCCGATAACAGGAAGTATTGCTATACACTTGATGTGATTCCCCGAAACAGGTACCTTGTAAGAGCAACTTTTCTATATGGAAACTTTGATAA

>M6-PGSC0003DMG400001732M6RLKmalectin_4270

AAGGATTAATTTAACGCCAGAGGACACTTTCTCAGTGGTACTTTTTCACACTCATCAAATGTATATTGCATAGAAGGTCACGGGCTATAAAACCTCTAGACCAGCAATGTACTAGAACTT

>M6-PGSC0003DMG400001732M6RLKmalectin_4331

TATATTGCATAGAAGGTCACGGGCTATAAAACCTCTAGACCAGCAATGTACTAGAACTTGATTAAGTTGTTACACAGTGCTGCAACTTTTCCCTCTACATTGTGCAATTTTGGATCTGAT

>M6-PGSC0003DMG400001732M6RLKmalectin_4380

ACTAGAACTTGATTAAGTTGTTACACAGTGCTGCAACTTTTCCCTCTACATTGTGCAATTTTGGATCTGATGCAAATTAGTACAACCATATCCAGTTTATATGTGACAAATTTACAATTT

>M6-PGSC0003DMG400001732M6RLKmalectin_488

GTGATTCCCCGAAACAGGTACCTTGTAAGAGCAACTTTTCTATATGGAAACTTTGATAAGGACAATGTTTACCCAAAGTTTGATATCTCACTTGGGGCAACTCATTGGGCTACAATAGTT

>M6-PGSC0003DMG400001732M6RLKmalectin_549

ACAATGTTTACCCAAAGTTTGATATCTCACTTGGGGCAACTCATTGGGCTACAATAGTTATATCTGATGCTAATACAATGGAGTACCAAGAGGTAATATTTCTAGCCAAGGAACCTTCCG

>M6-PGSC0003DMG400001732M6RLKmalectin_610

ATCTGATGCTAATACAATGGAGTACCAAGAGGTAATATTTCTAGCCAAGGAACCTTCCGTAAGTGTCTGTTTATCCAATGCAACCACTGGGCTGCCCTTCATTTCTACACTTGAGCTCCG

>M6-PGSC0003DMG400001732M6RLKmalectin_671

AGTGTCTGTTTATCCAATGCAACCACTGGGCTGCCCTTCATTTCTACACTTGAGCTCCGACATTTTAATGGTTCAATATATATGACTGAATTTGAGAATGATTTTTTCATGAGCGTCTCT

>M6-PGSC0003DMG400001732M6RLKmalectin_732

ATTTTAATGGTTCAATATATATGACTGAATTTGAGAATGATTTTTTCATGAGCGTCTCTGCTAGGATAAATTTTGGTGCACAGAGTGATGATCCTGTCAGGTATCTTTCAGTTCTTCTGT

>M6-PGSC0003DMG400001732M6RLKmalectin_793

TAGGATAAATTTTGGTGCACAGAGTGATGATCCTGTCAGGTATCTTTCAGTTCTTCTGTTACTGTGGCAGCTTCCAAATCTATGTTCTTGCACGTTCTTGTATATTTTTAGAATATGATA

>M6-PGSC0003DMG400001732M6RLKmalectin_854

CTGTGGCAGCTTCCAAATCTATGTTCTTGCACGTTCTTGTATATTTTTAGAATATGATAGCCATTGACATTTTTGAACTGAAACAAGCAGGCTTTGGCTCTGTCCCTAGCATGGTTTTCT

>M6-PGSC0003DMG400001732M6RLKmalectin_915

CATTGACATTTTTGAACTGAAACAAGCAGGCTTTGGCTCTGTCCCTAGCATGGTTTTCTTCTGACTTACTGTTTTTGTTCGATTTCCAAAATTTTAATCTTCCCTTAATGTCCTTATTCT

>M6-PGSC0003DMG400001732M6RLKmalectin_976

TGACTTACTGTTTTTGTTCGATTTCCAAAATTTTAATCTTCCCTTAATGTCCTTATTCTCCTAGGTACCCTGATGACCCATTCGATAGAATATGGGCATCTGACACCATAAAAAAGGCCA

>M6-PGSC0003DMG400001773-(rev)M6L-LecRK_0

CCAACGGCTAGTTTGATGAACCCCACGGCGGATATAACTACATCTGCCACATCGCCCCCTCCGGAGCAACAGATTGCTCCGAAGGAGGCTAATGGTACTGCAGTACAGAAAAACAGCAGC

>M6-PGSC0003DMG400001773-(rev)M6L-LecRK_1040

ATTGAATCTTGAATTTGATGAAAAAGAAATGACAAGAGTGTTGTTAGTTGGTTTGGCTTGTTCACATCCTGACCCTATAGCTAGGCCAACAATGAGGGGGGTGGTACAAATGTTAGTAGG

>M6-PGSC0003DMG400001773-(rev)M6L-LecRK_1105

ATCCTGACCCTATAGCTAGGCCAACAATGAGGGGGGTGGTACAAATGTTAGTAGGTGAGGCTGAAGTACCAATTGTACCAAGGGCTAAACCATCAATGAGTTTTAGCACATCTCATTTGT

>M6-PGSC0003DMG400001773-(rev)M6L-LecRK_1170

GTACCAATTGTACCAAGGGCTAAACCATCAATGAGTTTTAGCACATCTCATTTGTTAATGACATTGCAAGATAGTGTTTCTGACTTGAATGGTATGATCACACTTTCAACTTCTTCATCA

>M6-PGSC0003DMG400001773-(rev)M6L-LecRK_1221

TTGTTAATGACATTGCAAGATAGTGTTTCTGACTTGAATGGTATGATCACACTTTCAACTTCTTCATCAGAAAACAGTTTCAATGGTGTTGGTGGACATGATGGCATGGACCTAGTTTAG

>M6-PGSC0003DMG400001773-(rev)M6L-LecRK_130

ATAGTTTTTGTAAGCAGGGTGCTGGTGCTGTTGTTGGGGTTGTTACTGCTGGAGCATTCTTTCTAGCTTTTGCTACATTGGTACTTATTTGGTTGTATTCGAAAAAATTCAAGAATGTGA

>M6-PGSC0003DMG400001773-(rev)M6L-LecRK_195

GCTTTTGCTACATTGGTACTTATTTGGTTGTATTCGAAAAAATTCAAGAATGTGAAGACTTCAGAGTCTATGTTGCCTTCTGATATTATCAAAATGCCAAAGGAATTTAGCTATAAGGAG

>M6-PGSC0003DMG400001773-(rev)M6L-LecRK_260

GTCTATGTTGCCTTCTGATATTATCAAAATGCCAAAGGAATTTAGCTATAAGGAGCTTAAAATCGCGACGAAAGGGTTTGATTCTACTAGGATAATAGGGCATGGTGCATTTGGTACTGT

>M6-PGSC0003DMG400001773-(rev)M6L-LecRK_325

CGACGAAAGGGTTTGATTCTACTAGGATAATAGGGCATGGTGCATTTGGTACTGTCTATAAAGGGATTTTGTCTGAGACTGGTGATATTGTTGCGGTAAAGAGGTGTAGTCATAATGGAC

>M6-PGSC0003DMG400001773-(rev)M6L-LecRK_390

ATTTTGTCTGAGACTGGTGATATTGTTGCGGTAAAGAGGTGTAGTCATAATGGACAAGGGAAAGCTGAGTTCTTTTCTGAATTGTCGATAATCGGAACACTTAGACATAGAAATCTTGTT

>M6-PGSC0003DMG400001773-(rev)M6L-LecRK_455

TGAGTTCTTTTCTGAATTGTCGATAATCGGAACACTTAGACATAGAAATCTTGTTAGGCTACAAGGATGGTGTCATGAGAAAGGTGAAATCTTGTTAGTTTATGATTTGATGCCAAATGG

>M6-PGSC0003DMG400001773-(rev)M6L-LecRK_520

GATGGTGTCATGAGAAAGGTGAAATCTTGTTAGTTTATGATTTGATGCCAAATGGTAGTCTTGACAAGGCGTTATTCGAGTCCAGAATGATCCTCTCGTGGCAACACAGGCAAAAAATCT

>M6-PGSC0003DMG400001773-(rev)M6L-LecRK_585

AAGGCGTTATTCGAGTCCAGAATGATCCTCTCGTGGCAACACAGGCAAAAAATCTTGTTAGGTGTTGCTTCTGCATTAGCATATTTACATCAAGAATGTGAAAACCAGGTGATACATAGG

>M6-PGSC0003DMG400001773-(rev)M6L-LecRK_65

GCAACAGATTGCTCCGAAGGAGGCTAATGGTACTGCAGTACAGAAAAACAGCAGCAAATGTCATAATAGTTTTTGTAAGCAGGGTGCTGGTGCTGTTGTTGGGGTTGTTACTGCTGGAGC

>M6-PGSC0003DMG400001773-(rev)M6L-LecRK_650

TGCTTCTGCATTAGCATATTTACATCAAGAATGTGAAAACCAGGTGATACATAGGGATATCAAGAGTAGTAACATTATGTTGGATGAAGGGTTCAATGCAAAGTTAGGAGATTTTGGACT

>M6-PGSC0003DMG400001773-(rev)M6L-LecRK_715

GTAGTAACATTATGTTGGATGAAGGGTTCAATGCAAAGTTAGGAGATTTTGGACTAGCACGACAAATCGAACACGACAAGTCCCCCGATGCAACGGTAGCAGCCGGTACAATGGGGTATT

>M6-PGSC0003DMG400001773-(rev)M6L-LecRK_780

ATCGAACACGACAAGTCCCCCGATGCAACGGTAGCAGCCGGTACAATGGGGTATTTGGCACCAGAATACTTGTTGACAGGTAGAGCAACTGAGAAAACTGATGTGTTTAGTTATGGGGCA

>M6-PGSC0003DMG400001773-(rev)M6L-LecRK_845

ATACTTGTTGACAGGTAGAGCAACTGAGAAAACTGATGTGTTTAGTTATGGGGCAGTGGTTCTTGAAGTGGCAAGTGGGAGGAGGCCAATTGAGAAAGAAACAGCAACTAGTGGTGTTGG

>M6-PGSC0003DMG400001773-(rev)M6L-LecRK_910

AAGTGGCAAGTGGGAGGAGGCCAATTGAGAAAGAAACAGCAACTAGTGGTGTTGGATTAAACAGCAATTTGGTTGAATGGGTGTGGGGATTACACAAAGATGGGAGTTTACTATCAGCAG

>M6-PGSC0003DMG400001773-(rev)M6L-LecRK_975

AATTTGGTTGAATGGGTGTGGGGATTACACAAAGATGGGAGTTTACTATCAGCAGTTGATTCAAAATTGAATCTTGAATTTGATGAAAAAGAAATGACAAGAGTGTTGTTAGTTGGTTTG

>M6-PGSC0003DMG400001829-(rev)M6LRR-RLP_0

TATATTTTTTTTCCAGCAATCTCAGCTCACATATGCTGGGCAACATCTCTGTGCCTGCTATGACGCTTTTTATTTGCTTCAATTTAAGAAAGGCCTCACAGTTGTTGATCAAAATATACA

>M6-PGSC0003DMG400001829-(rev)M6LRR-RLP_1054

CTTTCAGGTAACAACTTCACCGGAAATGTTCCCTCAACTTTCAAAAAGCTGAAAACACTTAACTACTTATCTCTCTCTTTCAATAATTTTGAAAGCTCAATTCCAGACAACTTCGCCTAC

>M6-PGSC0003DMG400001829-(rev)M6LRR-RLP_1116

CTACTTATCTCTCTCTTTCAATAATTTTGAAAGCTCAATTCCAGACAACTTCGCCTACTTTTCAGAGCTATCTTTTTTATACTTGCAAACTAACAATTTCACTTGCCCACTTCCCTATTT

>M6-PGSC0003DMG400001829-(rev)M6LRR-RLP_1178

CAGAGCTATCTTTTTTATACTTGCAAACTAACAATTTCACTTGCCCACTTCCCTATTTGATTGCAACCTTGACTCGCCTTCAAGAATTAAACTTGCAAAACAATTTACTAACTGGTTCAC

>M6-PGSC0003DMG400001829-(rev)M6LRR-RLP_1240

GCAACCTTGACTCGCCTTCAAGAATTAAACTTGCAAAACAATTTACTAACTGGTTCACTTCCCTCTAATTTAAGCAGACTTCAAGAGTTAGAAACACTTGATTTTTCCTTCAATTATTTC

>M6-PGSC0003DMG400001829-(rev)M6LRR-RLP_1302

CTCTAATTTAAGCAGACTTCAAGAGTTAGAAACACTTGATTTTTCCTTCAATTATTTCACTGGCACAATACCCTCTTGGTTATTCCGTCTTCCATCCTTGGTTTCTTTATCTATTCAATA

>M6-PGSC0003DMG400001829-(rev)M6LRR-RLP_1364

GCACAATACCCTCTTGGTTATTCCGTCTTCCATCCTTGGTTTCTTTATCTATTCAATACAATGAATTAACTGGGAAATTGCCAAATGAGCTCAAGAGCATTACGTTGGCAACGCTAGAAT

>M6-PGSC0003DMG400001829-(rev)M6LRR-RLP_1426

GAATTAACTGGGAAATTGCCAAATGAGCTCAAGAGCATTACGTTGGCAACGCTAGAATTGATCTTCGGAACAACAAGCTGCATGGAGAAATCCCTGATTGGATATTATCTATGAGCATGT

>M6-PGSC0003DMG400001829-(rev)M6LRR-RLP_1488

CTTCGGAACAACAAGCTGCATGGAGAAATCCCTGATTGGATATTATCTATGAGCATGTATATCCTGGATCTCTCTCATAACTTGCTCACAGGCTTTGAAAAACAAGTATGACATTCAAAG

>M6-PGSC0003DMG400001829-(rev)M6LRR-RLP_1550

CCTGGATCTCTCTCATAACTTGCTCACAGGCTTTGAAAAACAAGTATGACATTCAAAGAATTTGGTGTGGCTTAATTTGGAGAACAATTTTCTTCAAGGGACTTTGCATCAATCTATTTG

>M6-PGSC0003DMG400001829-(rev)M6LRR-RLP_1612

TGGTGTGGCTTAATTTGGAGAACAATTTTCTTCAAGGGACTTTGCATCAATCTATTTGTGACATGTTTAACCTTGAAGTCCTCATTTTGTCTCGGAACAATTTCAGTGGTTCAATCCCAG

>M6-PGSC0003DMG400001829-(rev)M6LRR-RLP_1674

ATGTTTAACCTTGAAGTCCTCATTTTGTCTCGGAACAATTTCAGTGGTTCAATCCCAGGATGTTTGGGTAACTCCATTCTTGTTTTAGACTTGCGAATGAACAATTTTTATGGAGAAATA

>M6-PGSC0003DMG400001829-(rev)M6LRR-RLP_1736

TTTGGGTAACTCCATTCTTGTTTTAGACTTGCGAATGAACAATTTTTATGGAGAAATACCAAAATTCTTACCTATAGGGTTACAATATCTTGGTGTGTATGACAATCAGTTGGGGGGGAC

>M6-PGSC0003DMG400001829-(rev)M6LRR-RLP_186

GGATGGTGTAACTTGCAATGGATTTACGGGTCATGTCATAGGCCTTGATCTTTCTTGCAGCATTCTTAACCAGACCATCAATGCTAACAGCAGCCTCACTAAACTTGGTCATCTCCAAAG

>M6-PGSC0003DMG400001829-(rev)M6LRR-RLP_1860

CCCACGATCCTTGGTTAACTGTACAAGCTTGGAAGCTCTTGATTTGGGGAACAACAATATAAATGACACGTTCCCGTATGGCTGGACAAACTTCCAAACCTACAAGTTCTGATTCTGTAA

>M6-PGSC0003DMG400001829-(rev)M6LRR-RLP_1922

ATGACACGTTCCCGTATGGCTGGACAAACTTCCAAACCTACAAGTTCTGATTCTGTAATCAAATCTCTTTCACGGTTCAGTTGGTGATTTGGATTCCGAGTTTCCATTTCCTGAGTTACG

>M6-PGSC0003DMG400001829-(rev)M6LRR-RLP_1984

ATCTCTTTCACGGTTCAGTTGGTGATTTGGATTCCGAGTTTCCATTTCCTGAGTTACGGATCTTTGACCTTTCCGTTAATGGGTTCACTGGAACTTTTCCATCTAATATTTTCAAAAGTT

>M6-PGSC0003DMG400001829-(rev)M6LRR-RLP_2046

TTTGACCTTTCCGTTAATGGGTTCACTGGAACTTTTCCATCTAATATTTTCAAAAGTTTCAGAGGTATGATGGATGTGGATGAAGGAAAAACAGGAATTACTCTAGCTATTAATAGGACT

>M6-PGSC0003DMG400001829-(rev)M6LRR-RLP_2108

AGGTATGATGGATGTGGATGAAGGAAAAACAGGAATTACTCTAGCTATTAATAGGACTCACAGAGATTACCTTTACCATGTTAGTTTGGTGATAAAAGTCAATGAATTTGATATGAGAAT

>M6-PGSC0003DMG400001829-(rev)M6LRR-RLP_2170

GAGATTACCTTTACCATGTTAGTTTGGTGATAAAAGTCAATGAATTTGATATGAGAATCACGTCAATTATGACGAGTGTTGATTTATCAAGCAACAGGTTCGAAGGAGATATTCCAAATT

>M6-PGSC0003DMG400001829-(rev)M6LRR-RLP_2294

TGGAAGTCTGAGCTCACTTGTGTTACTCAACCTGTCTCGCAACAACTTCCGTGGTCATATTCCTGCAGAAATTGCAAAGTTGCAGCAGCTCGAAGCATTATAGATCTCTCGTGGAACAGA

>M6-PGSC0003DMG400001829-(rev)M6LRR-RLP_2356

CTGCAGAAATTGCAAAGTTGCAGCAGCTCGAAGCATTATAGATCTCTCGTGGAACAGACTCATTGGAGAAGTTCCTGGTCAATTGTCGAGTTTGACATTTCTTGAGGTCTTAAACCTTTC

>M6-PGSC0003DMG400001829-(rev)M6LRR-RLP_2418

TTGGAGAAGTTCCTGGTCAATTGTCGAGTTTGACATTTCTTGAGGTCTTAAACCTTTCATACAATCATCTGGCTTGGCGCATTCCTATTGGGAAACAGTTCAATACATTTCCAGATGATT

>M6-PGSC0003DMG400001829-(rev)M6LRR-RLP_248

TTCTTAACCAGACCATCAATGCTAACAGCAGCCTCACTAAACTTGGTCATCTCCAAAGACTCAACCTTGCCTTCAGCCAGTTTGATGACGACTTTCCACTTGGATTAAGTGAACTCAGTA

>M6-PGSC0003DMG400001829-(rev)M6LRR-RLP_2542

TTGTGGAAATCTTGATTTATGTGGATTTCCACTATCAAAGAAATGTGGAAACAATAATAGGCCAGATGAGTCAACACTTGAACAAGATGATGAAGATTCATTTTTTGCGAGCGGGTTCAC

>M6-PGSC0003DMG400001829-(rev)M6LRR-RLP_2604

CAGATGAGTCAACACTTGAACAAGATGATGAAGATTCATTTTTTGCGAGCGGGTTCACATGGGAAGCTGTGGTAATAGGTTATGGTATGATATTTGGATTGTTGATGGGAGGCCTCATGT

>M6-PGSC0003DMG400001829-(rev)M6LRR-RLP_2666

GAAGCTGTGGTAATAGGTTATGGTATGATATTTGGATTGTTGATGGGAGGCCTCATGTTCCTATTGGAAAAACCAAAATGGTTTGTCAACTTTGCTGAAGATATTTCTCAACAAATTGCT

>M6-PGSC0003DMG400001829-(rev)M6LRR-RLP_2726

CTATTGGAAAAACCAAAATGGTTTGTCAACTTTGCTGAAGATATTTCTCAACAAATTGCTGCTAAGAAGCGAAGAAATCAAAAGAAGAGACGTCAAGGACGTGGTCTAAGAATGAATTAG

>M6-PGSC0003DMG400001829-(rev)M6LRR-RLP_310

AACCTTGCCTTCAGCCAGTTTGATGACGACTTTCCACTTGGATTAAGTGAACTCAGTAGTTTGACACATCTCAATCTTTCGAATACTGGATTTAATGAAGGGGAGATACAGATTCCACCA

>M6-PGSC0003DMG400001829-(rev)M6LRR-RLP_372

GACACATCTCAATCTTTCGAATACTGGATTTAATGAAGGGGAGATACAGATTCCACCAGGATTATCCAAGTTGTCTAAGTTGGTTTCACTTGACCTCTCCTCGAATGATTTCCAAGTTGG

>M6-PGSC0003DMG400001829-(rev)M6LRR-RLP_434

TATCCAAGTTGTCTAAGTTGGTTTCACTTGACCTCTCCTCGAATGATTTCCAAGTTGGCCGAACAACCTCTAGAAGTTTGCTTCAGAACTTAACTAATTTGGAGGTACTGTTGCTTGACA

>M6-PGSC0003DMG400001829-(rev)M6LRR-RLP_496

ACAACCTCTAGAAGTTTGCTTCAGAACTTAACTAATTTGGAGGTACTGTTGCTTGACAGCGTGGATGCACCATTTGAGTTACCTAAGAATTTTCCTTCTTCCCTTAGGTACTTAATTCTT

>M6-PGSC0003DMG400001829-(rev)M6LRR-RLP_558

GGATGCACCATTTGAGTTACCTAAGAATTTTCCTTCTTCCCTTAGGTACTTAATTCTTCAAGACACCAACATGTGTGGGAACATAAGTGATTCTCAGCTTTTTCATCTACCAAACTTGCA

>M6-PGSC0003DMG400001829-(rev)M6LRR-RLP_62

ACGCTTTTTATTTGCTTCAATTTAAGAAAGGCCTCACAGTTGTTGATCAAAATATACATGACTGTGACAATGAGACCCGAGCCAAGACTTTGTCCTGGAATGTTACAGGAGATTGTTGTC

>M6-PGSC0003DMG400001829-(rev)M6LRR-RLP_620

ACACCAACATGTGTGGGAACATAAGTGATTCTCAGCTTTTTCATCTACCAAACTTGCAAGTGCTGAGAATGGGATGAAATCCTTTATTAACAGGCACTTTGCCTAATTTCAATTGGAGTT

>M6-PGSC0003DMG400001829-(rev)M6LRR-RLP_682

CTGAGAATGGGATGAAATCCTTTATTAACAGGCACTTTGCCTAATTTCAATTGGAGTTTCAGTACAAGTATTTTAGAGTTGGACTTTTTCTATACTGATATTTTTGGTAAGGTACCTGAT

>M6-PGSC0003DMG400001829-(rev)M6LRR-RLP_744

TACAAGTATTTTAGAGTTGGACTTTTTCTATACTGATATTTTTGGTAAGGTACCTGATTCAATCGGCAACCTCCGTTCACTTTGGTAATTAGATCTCTCTTTAAACCGTTTATCCGGCTT

>M6-PGSC0003DMG400001829-(rev)M6LRR-RLP_806

TCGGCAACCTCCGTTCACTTTGGTAATTAGATCTCTCTTTAAACCGTTTATCCGGCTTGATTCCAGTATCCATTGGCAACCTCAGTGGAATTAGAGTGTTGAGACTTTCAAGTAACAGCT

>M6-PGSC0003DMG400001829-(rev)M6LRR-RLP_868

CCAGTATCCATTGGCAACCTCAGTGGAATTAGAGTGTTGAGACTTTCAAGTAACAGCTTCACTGGTAATGTTCCCTCATCTATCTCAAAATTAAACAAACTCTTTGATTTAGATCTCTCT

>M6-PGSC0003DMG400001829-(rev)M6LRR-RLP_930

TGGTAATGTTCCCTCATCTATCTCAAAATTAAACAAACTCTTTGATTTAGATCTCTCTTTCAACGATTTTCAAGGCTTGATTCTAGAATCCATTGGTAACCTAACTGCAATCACAAAGTT

>M6-PGSC0003DMG400001829-(rev)M6LRR-RLP_992

ACGATTTTCAAGGCTTGATTCTAGAATCCATTGGTAACCTAACTGCAATCACAAAGTTGACACTTTCAGGTAACAACTTCACCGGAAATGTTCCCTCAACTTTCAAAAAGCTGAAAACAC

>M6-PGSC0003DMG400001831-(rev)M6LRR-RLP_0

ATGGAGTACCAAAAATTGCTAATAGCTTTCTGCTTCTATTCACATTTTATACTTTTTGTCCATCAATCTCAGCTTACATATGCTGGGCAACATCTCTGTGCCCGCTATGAAGCTTTTTAT

>M6-PGSC0003DMG400001831-(rev)M6LRR-RLP_1054

GAGGCTCAATTCTAGAATACATTGACAATCTCACTGCAATCACACAGTTGACAATTCCAGATAACAGCTTCACTGGAAATGTTCCCTCAACTATCGGAAAGTTGAACAAACTTAACTACT

>M6-PGSC0003DMG400001831-(rev)M6LRR-RLP_1116

AACAGCTTCACTGGAAATGTTCCCTCAACTATCGGAAAGTTGAACAAACTTAACTACTTATCTCTCTCTTCCAATTATTTTGAAGGCTCGATTCCAGACATCTTTGCCAACTTTTCAGAT

>M6-PGSC0003DMG400001831-(rev)M6LRR-RLP_1178

TCTCTCTTCCAATTATTTTGAAGGCTCGATTCCAGACATCTTTGCCAACTTTTCAGATCTATTTGCTTTATTTTTTGACACTAACAATTTCACTGGTTCATTTCCCTGTTCAATTGCAAC

>M6-PGSC0003DMG400001831-(rev)M6LRR-RLP_124

TTCAATTTCAGCAAGGCCTCACAGTTGATCTTATTGGTTATGACTGTGACTATGAGGCCCTAGCTAAGACTTTGTCTTTGAATGTTACAGGAGATTGTTGTCAATGGGATGGTGTTACTT

>M6-PGSC0003DMG400001831-(rev)M6LRR-RLP_1240

TTGCTTTATTTTTTGACACTAACAATTTCACTGGTTCATTTCCCTGTTCAATTGCAACCTTGATTTGCCTTGTGAATTTAGAGTTGCAAAACAATTTACTAACTGGTTCACTTCCCTCTA

>M6-PGSC0003DMG400001831-(rev)M6LRR-RLP_1302

ATTTGCCTTGTGAATTTAGAGTTGCAAAACAATTTACTAACTGGTTCACTTCCCTCTAATATAAGCTGATTTCAACTTCTACAATCACTTGATTTGTCCTTCAATTATTTCACTGGCACA

>M6-PGSC0003DMG400001831-(rev)M6LRR-RLP_1364

AAGCTGATTTCAACTTCTACAATCACTTGATTTGTCCTTCAATTATTTCACTGGCACAACACCCCCTTCGTTATTCCTTCTTCCAAGATTGGTTTCTTTACATGTTCAACTCAATCAATT

>M6-PGSC0003DMG400001831-(rev)M6LRR-RLP_1426

CCCCTTCGTTATTCCTTCTTCCAAGATTGGTTTCTTTACATGTTCAACTCAATCAATTAACCGGGAAATTGCCAAATGTGCTCAAGATGAGTTATCTAGGATACGTACTATACCGTAGCA

>M6-PGSC0003DMG400001831-(rev)M6LRR-RLP_1488

GGGAAATTGCCAAATGTGCTCAAGATGAGTTATCTAGGATACGTACTATACCGTAGCATTGATCTTTCATACAACAAGCTGCAAGGTGAAATCCCCAATGCGATGTTATCTATGAGCATG

>M6-PGSC0003DMG400001831-(rev)M6LRR-RLP_1550

TCTTTCATACAACAAGCTGCAAGGTGAAATCCCCAATGCGATGTTATCTATGAGCATGGGTAAGCTGGATCTCTCTCATAACTTCCTCACAGGCTTTGAAAAACAAGTATGGCACTCAGA

>M6-PGSC0003DMG400001831-(rev)M6LRR-RLP_1612

AGCTGGATCTCTCTCATAACTTCCTCACAGGCTTTGAAAAACAAGTATGGCACTCAGAGTACTTACTATACCTTAATTTGGAGAACAATTTTCTTCAAGGGCTTTTGCATCTATCCTTTT

>M6-PGSC0003DMG400001831-(rev)M6LRR-RLP_1674

TTACTATACCTTAATTTGGAGAACAATTTTCTTCAAGGGCTTTTGCATCTATCCTTTTGTGACATGTATAGTCTTGTATTCCTCATTTTAGCTCGGAACAATTTCAGTGGTTCAATCCCG

>M6-PGSC0003DMG400001831-(rev)M6LRR-RLP_1736

CATGTATAGTCTTGTATTCCTCATTTTAGCTCGGAACAATTTCAGTGGTTCAATCCCGGTATGTTTGGGTAACTCCAGTATTTCCATTTCTTTTTTAGACTTGCGAATGAACAATTTGCA

>M6-PGSC0003DMG400001831-(rev)M6LRR-RLP_1798

GTTTGGGTAACTCCAGTATTTCCATTTCTTTTTTAGACTTGCGAATGAACAATTTGCATGGAGAGATACCAAGATTCTTATCTACAGGGTTTCAATATCTTGGTTTATATGGCAATCAAT

>M6-PGSC0003DMG400001831-(rev)M6LRR-RLP_186

GCTAAGACTTTGTCTTTGAATGTTACAGGAGATTGTTGTCAATGGGATGGTGTTACTTGTCATGGATTTACTGGTCACGTTATAGGACTCGATCTTTCTTGCAGTTTTCTTAACGGAACC

>M6-PGSC0003DMG400001831-(rev)M6LRR-RLP_1860

GAGATACCAAGATTCTTATCTACAGGGTTTCAATATCTTGGTTTATATGGCAATCAATTGGGGGGACAATTCCCGTGATCCTTGGTTAACTGTACAAGCTTGGAAGCTCTTGATTTGGGG

>M6-PGSC0003DMG400001831-(rev)M6LRR-RLP_1984

ACAAGATAAATGACACGTTTCCGATATGGCTGCAGAAACTTCCAAACCTACAAGTTCTGATTCTGAAATCTAATCTCTTTCATGGTCCAATTGGTTATTTGGAGTCCGAGTTTCCATTTC

>M6-PGSC0003DMG400001831-(rev)M6LRR-RLP_2046

CTGAAATCTAATCTCTTTCATGGTCCAATTGGTTATTTGGAGTCCGAGTTTCCATTTCCTGAGTTACGAATCTTTGATCTTTCCTTCAATGGGTTCACTGGAACTTTGCCATCTAATTTT

>M6-PGSC0003DMG400001831-(rev)M6LRR-RLP_2108

GTTACGAATCTTTGATCTTTCCTTCAATGGGTTCACTGGAACTTTGCCATCTAATTTTTTCAAAAGTTTTAGAGGTATGATGGATGTGGATGAAGAAAAAACAGGAATTACTCGCTCCTA

>M6-PGSC0003DMG400001831-(rev)M6LRR-RLP_2170

AAAGTTTTAGAGGTATGATGGATGTGGATGAAGAAAAAACAGGAATTACTCGCTCCTACAAAGGGTCGACCAGGCCTTGTACAAGCATAAATCGATGTCGAAGAGATTACCTTTACCACT

>M6-PGSC0003DMG400001831-(rev)M6LRR-RLP_2232

GGGTCGACCAGGCCTTGTACAAGCATAAATCGATGTCGAAGAGATTACCTTTACCACTTTAGTTTGGTGATAAAAGGTAATGAGTTTGATATGAGAATCACGTCAATTATGACGAGTGTT

>M6-PGSC0003DMG400001831-(rev)M6LRR-RLP_2294

TTTGGTGATAAAAGGTAATGAGTTTGATATGAGAATCACGTCAATTATGACGAGTGTTGATATATCAAGCAACAGGGTTGAAGGATATATTCCAAATTCCATTGGAAATCTGAGCTCACT

>M6-PGSC0003DMG400001831-(rev)M6LRR-RLP_2356

TATCAAGCAACAGGGTTGAAGGATATATTCCAAATTCCATTGGAAATCTGAGCTCACTTGTGTTACTCAACCTGTCTCACAACAGCTTCCGTGGTCATATTCCTGCAGAATTTGCAAAGT

>M6-PGSC0003DMG400001831-(rev)M6LRR-RLP_2418

TTACTCAACCTGTCTCACAACAGCTTCCGTGGTCATATTCCTGCAGAATTTGCAAAGTTGCAATAGCTCGAAGCATTAGATCTCTCACGGAACAGACTCATTGGAGATATTCCAGATCCA

>M6-PGSC0003DMG400001831-(rev)M6LRR-RLP_248

TGGATTTACTGGTCACGTTATAGGACTCGATCTTTCTTGCAGTTTTCTTAACGGAACCATTAATTCTAACAGCAGCCTCACAAAACTTAGTCATCTTCAAAGACTCAACCTTGCTTTCAA

>M6-PGSC0003DMG400001831-(rev)M6LRR-RLP_2542

CGAGTTTGACATTTCTTGAGGTCTTAAACCTTTCCTACAATCATCTGGCTGGACGCATTCCTCTTGGGAAACAGTTCAATACATTTCCAAATGATTCTTATTGTGGAAATCCTGATTTAT

>M6-PGSC0003DMG400001831-(rev)M6LRR-RLP_2604

CTTGGGAAACAGTTCAATACATTTCCAAATGATTCTTATTGTGGAAATCCTGATTTATGTGGATTTCCACTATCAAAGGAATGTGGAAACAATAAAGAGTCACCACTTGAACACGATGAT

>M6-PGSC0003DMG400001831-(rev)M6LRR-RLP_2666

ATTTCCACTATCAAAGGAATGTGGAAACAATAAAGAGTCACCACTTGAACACGATGATGGTGATGATGATGATTCATTTTTTGCGAGTGGGTTCACATGGCAAGCTGTGGTAATAGGTTA

>M6-PGSC0003DMG400001831-(rev)M6LRR-RLP_2728

ATGATGATGATTCATTTTTTGCGAGTGGGTTCACATGGCAAGCTGTGGTAATAGGTTATGGTTGTGGTCTGATATTTGGATTGTTGATGGGAGGTCTCATGTTCCTACTGGAAAAGCCAA

>M6-PGSC0003DMG400001831-(rev)M6LRR-RLP_310

ATTCTAACAGCAGCCTCACAAAACTTAGTCATCTTCAAAGACTCAACCTTGCTTTCAACAAGTTGAGTAACTTTCCACTTGTGGATAGCATTAGTGAACTCGAAAGTTTGACATATCTCA

>M6-PGSC0003DMG400001831-(rev)M6LRR-RLP_372

TTGAGTAACTTTCCACTTGTGGATAGCATTAGTGAACTCGAAAGTTTGACATATCTCAATCTTTCGCATTCAGTACATATGATAGGGGAGAAGCAGATACCACAAGTATTATCCAACTTG

>M6-PGSC0003DMG400001831-(rev)M6LRR-RLP_434

TTCGCATTCAGTACATATGATAGGGGAGAAGCAGATACCACAAGTATTATCCAACTTGTCTAAATTGGTTTCACTTGACCTCTCCTTGTATTACACTGAACTCAAAGTTGATTGAAAAAC

>M6-PGSC0003DMG400001831-(rev)M6LRR-RLP_496

AATTGGTTTCACTTGACCTCTCCTTGTATTACACTGAACTCAAAGTTGATTGAAAAACCTTCAGAAGTTTGCTTCAAGACTTGACCAATTTGGAGGTACTGTTTCTTGACAACTCGTATG

>M6-PGSC0003DMG400001831-(rev)M6LRR-RLP_558

AGAAGTTTGCTTCAAGACTTGACCAATTTGGAGGTACTGTTTCTTGACAACTCGTATGCATCATTTGAGTTACCTAAGAATTTTTCTTCTTCTCTTAGGTACTTAAGTCTTGAAGGTATC

>M6-PGSC0003DMG400001831-(rev)M6LRR-RLP_62

TCAATCTCAGCTTACATATGCTGGGCAACATCTCTGTGCCCGCTATGAAGCTTTTTATTTGCTTCAATTTCAGCAAGGCCTCACAGTTGATCTTATTGGTTATGACTGTGACTATGAGGC

>M6-PGSC0003DMG400001831-(rev)M6LRR-RLP_620

ATTTGAGTTACCTAAGAATTTTTCTTCTTCTCTTAGGTACTTAAGTCTTGAAGGTATCGGCATGTTTGGGATCATAAGTGAGTTTCAGTTATTTCATTTATCAAACTTGCAAGTGCTGAG

>M6-PGSC0003DMG400001831-(rev)M6LRR-RLP_682

TGTTTGGGATCATAAGTGAGTTTCAGTTATTTCATTTATCAAACTTGCAAGTGCTGAGATTTGGACAGAATCCTTTACTAACAGGCTCTTTGCCAAATTTTAATCGGAGTTCCAGTGAAA

>M6-PGSC0003DMG400001831-(rev)M6LRR-RLP_744

GGACAGAATCCTTTACTAACAGGCTCTTTGCCAAATTTTAATCGGAGTTCCAGTGAAAGTATTTTAGAGTTGAACTTCTTTTATACTGGTATTTCTGGGAAGGTACCCGATTCAATTGGT

>M6-PGSC0003DMG400001831-(rev)M6LRR-RLP_806

TTTAGAGTTGAACTTCTTTTATACTGGTATTTCTGGGAAGGTACCCGATTCAATTGGTAATCTCCATTCTCTTTGGTATTTGAATCTCCAGATTTGTCATTTATCTGGCTCGATTCCAGA

>M6-PGSC0003DMG400001831-(rev)M6LRR-RLP_868

TCCATTCTCTTTGGTATTTGAATCTCCAGATTTGTCATTTATCTGGCTCGATTCCAGAATCCATTGGCAATCTCACTGCAATCAGAGAGTTGATACTTTCAGAAAACGTCTTCATTGTTA

>M6-PGSC0003DMG400001831-(rev)M6LRR-RLP_930

ATTGGCAATCTCACTGCAATCAGAGAGTTGATACTTTCAGAAAACGTCTTCATTGTTAATGTTCCCTCAACTTTCTCAAAGTTAAACAAACTCGTTCGTTTAGATCTCTCTTCAAACTAT

>M6-PGSC0003DMG400001831-(rev)M6LRR-RLP_992

TCCCTCAACTTTCTCAAAGTTAAACAAACTCGTTCGTTTAGATCTCTCTTCAAACTATTTTCGAGGCTCAATTCTAGAATACATTGACAATCTCACTGCAATCACACAGTTGACAATTCC

>M6-PGSC0003DMG400001832-(rev)M6LRR-RLP_1550

TAGCTCGAAGCATTAGATCTCTCACGGAACAGACTCATTGGAGATATTCCAGATCCATTGTCGAGTTTGACATTTCTTGAGGTCTTAAACCTTTCCTACAATCATCTGGCTGGACGCATT

>M6-PGSC0003DMG400001832-(rev)M6LRR-RLP_1860

GTGGTCTGATATTTGGATTGTTGATGGGAGGTCTCATGTTCCTACTGGAAAAGCCAAAATGGTTTGTGAACTTTGCTAAAGATATTGCTCAACAAATTGCTGCTAAGAAGCAAACAAGGC

>M6-PGSC0003DMG400001832-(rev)M6LRR-RLP_1922

TTTGTGAACTTTGCTAAAGATATTGCTCAACAAATTGCTGCTAAGAAGCAAACAAGGCAAAAGAAGAGACGCCAAAGACATGGTTTAAGAATGAACTAGCAATATACCAGGTATGCATTA

>M6-PGSC0003DMG400001832-(rev)M6LRR-RLP_1984

GAAGAGACGCCAAAGACATGGTTTAAGAATGAACTAGCAATATACCAGGTATGCATTAAAACCTTTAGTTTTGAATCAGGAACAACACTTCAGAGGTTGTTGGATAACTATATGTCAACT

>M6-PGSC0003DMG400001832-(rev)M6LRR-RLP_2046

CTTTAGTTTTGAATCAGGAACAACACTTCAGAGGTTGTTGGATAACTATATGTCAACTGTTTTTATTTTTTCCTAAATATTCAAAACTTAATTTGTCTCTGCAGAGGTTGCTGGAGAGTT

>M6-PGSC0003DMG400001832-(rev)M6LRR-RLP_2108

TTATTTTTTCCTAAATATTCAAAACTTAATTTGTCTCTGCAGAGGTTGCTGGAGAGTTGGACAAGTTTATCAGCAAGTCAGTGCTGATGTTTGGCAAATAGCAATGGATGCATCTGTTTG

>M6-PGSC0003DMG400001832-(rev)M6LRR-RLP_2163

GTTGGACAAGTTTATCAGCAAGTCAGTGCTGATGTTTGGCAAATAGCAATGGATGCATCTGTTTGTTTACTTTATGTCCATGGTTGCCAAATACGTATGTAAATGTTACAGTTTAGTTCT

>M6-PGSC0003DMG400002012-(rev)M6LRR-RLK_0

TGTAAGTAGTTAGTCAAAATTGTGTCTAAATGAAAGGTTTTTGGTGGTCTACTGTTTCTTCATTTATATAAGGAAAGTTGGTTTGACTAAAATAACATCCCCTATTGACTTGCTTATGGT

>M6-PGSC0003DMG400002012-(rev)M6LRR-RLK_1037

AATTGAAGTATCTGGATTTGCACTCTAATGCCTTCTCAATTGATATTATGCTTCTTCTGGCCTCATTGGGGGATGTATAGTATGTAGACCTCAGTAGCAACAAGTTTGTTGGATCACTTG

>M6-PGSC0003DMG400002012-(rev)M6LRR-RLK_1098

CTCATTGGGGGATGTATAGTATGTAGACCTCAGTAGCAACAAGTTTGTTGGATCACTTGACCTGCAAGTAGGAAATTCGAGCTTTGTCTCGTCAATTCAGTATCTGAACATTAGTCATAA

>M6-PGSC0003DMG400002012-(rev)M6LRR-RLK_1159

CTGCAAGTAGGAAATTCGAGCTTTGTCTCGTCAATTCAGTATCTGAACATTAGTCATAATAACCTGGATGGAGAGCTATTTCCTCACGATGGAATGCCATATTTTGACAGTTTAGAGGCT

>M6-PGSC0003DMG400002012-(rev)M6LRR-RLK_122

AGGGAAATGTAGGGTCTCTTGCTTTTGATTTTGATGTATTGGTTATTGGTGAGTGAATTTAATGGTGTCAATTTGCATTTTGGGACATTATTGGTGTTGCTGTTGGACTTCAATGAATTT

>M6-PGSC0003DMG400002012-(rev)M6LRR-RLK_1220

ACCTGGATGGAGAGCTATTTCCTCACGATGGAATGCCATATTTTGACAGTTTAGAGGCTTTTGATGCAAGTGACAATCAGCTTACTGGTACTATTCCGTCCTTCAATTTTGTAGTCTCCC

>M6-PGSC0003DMG400002012-(rev)M6LRR-RLK_1281

TGATGCAAGTGACAATCAGCTTACTGGTACTATTCCGTCCTTCAATTTTGTAGTCTCCCTTCGAATTCTTAGGCTCGGGAACAATCAGTTGTCTGGATCACTCCCCGAAGCTCTTCTAGA

>M6-PGSC0003DMG400002012-(rev)M6LRR-RLK_1342

CGAATTCTTAGGCTCGGGAACAATCAGTTGTCTGGATCACTCCCCGAAGCTCTTCTAGAGGATAGCTCAATGATTTTATCAGAATTGGACCTAAGCCAGAATCAGCTTGCAGGTGTGTTG

>M6-PGSC0003DMG400002012-(rev)M6LRR-RLK_1403

ATAGCTCAATGATTTTATCAGAATTGGACCTAAGCCAGAATCAGCTTGCAGGTGTGTTGATCCTTTCTTCTTTAGTTAATTTGTTTGGTTGAAGTCAATCTAAGTAATTTATGGTTAATT

>M6-PGSC0003DMG400002012-(rev)M6LRR-RLK_1769

AATTGGGACGGAGGGAGTACTATCTATGATCTTAGGTTGTTGTCAGTTAGATAACAAATGAATAATATTTAGATTGCTATCTGCTGGAAAGCATGCTTGTTTTGGTTGCAATCAAATGTA

>M6-PGSC0003DMG400002012-(rev)M6LRR-RLK_183

ATGGTGTCAATTTGCATTTTGGGACATTATTGGTGTTGCTGTTGGACTTCAATGAATTTAGTGACTGGTGTTGTTGCTGCTTGTTCTTTTATCTAGGTTTATATACTCTCTTATTTCACT

>M6-PGSC0003DMG400002012-(rev)M6LRR-RLK_1830

ATAATATTTAGATTGCTATCTGCTGGAAAGCATGCTTGTTTTGGTTGCAATCAAATGTAATGTATAGAACACAATTCCTGAGCACTTCGGAATTGTGTTCTAGAGGAATGCTGTATAATT

>M6-PGSC0003DMG400002012-(rev)M6LRR-RLK_1891

GTATAGAACACAATTCCTGAGCACTTCGGAATTGTGTTCTAGAGGAATGCTGTATAATTCTCTGCTTATGTGTAATTATCATACACATTGTTCCCAATTCTTCCCTAGGTTAGAGATATT

>M6-PGSC0003DMG400002012-(rev)M6LRR-RLK_1952

CTGCTTATGTGTAATTATCATACACATTGTTCCCAATTCTTCCCTAGGTTAGAGATATTTCATACTTAGATTCAACTCTAAGGCAATTATTGCTGTAAACTTGCATTTTAATTAGTGGAA

>M6-PGSC0003DMG400002012-(rev)M6LRR-RLK_2013

ATACTTAGATTCAACTCTAAGGCAATTATTGCTGTAAACTTGCATTTTAATTAGTGGAATGTTTCAGTGTCTTTTTGATTATTTTTTTTTGCTGCATTATTAATTTCATCTCTCCTTGAG

>M6-PGSC0003DMG400002012-(rev)M6LRR-RLK_2257

CGTTATAAACAAGCTGCATAAAGATATTCTTCATGCGGTTGGAGTTCTGATGTATTTTAAGTCTTTTGTTGTTCTTTTAATTATTATATTTAGTTGTTCTTACCACAGGCCTAAACCAAG

>M6-PGSC0003DMG400002012-(rev)M6LRR-RLK_2318

TCTTTTGTTGTTCTTTTAATTATTATATTTAGTTGTTCTTACCACAGGCCTAAACCAAGCTACTTCCTTTCCAAACTATTTCATGAAGCCATTTGCTATTTTGGCAGGTCCCATTGGAGG

>M6-PGSC0003DMG400002012-(rev)M6LRR-RLK_2379

ACTTCCTTTCCAAACTATTTCATGAAGCCATTTGCTATTTTGGCAGGTCCCATTGGAGGTATAAGTGCTGTGAATCTCAAGCTTCTAAATTTATCATATAACCAGCTCTCAGGCCCCTTG

>M6-PGSC0003DMG400002012-(rev)M6LRR-RLK_244

TGACTGGTGTTGTTGCTGCTTGTTCTTTTATCTAGGTTTATATACTCTCTTATTTCACTTACATAATGATTAGTGATGATTGCAGTTTGTGTTTAAAAATTTTGCTAATTCATTTTCTGG

>M6-PGSC0003DMG400002012-(rev)M6LRR-RLK_2440

TAAGTGCTGTGAATCTCAAGCTTCTAAATTTATCATATAACCAGCTCTCAGGCCCCTTGCCTTTCAAGGTTGGGCGCTGTGCCATCATAGACCTGAGCAACAATCGGCTAACTGGAAATG

>M6-PGSC0003DMG400002012-(rev)M6LRR-RLK_2501

TTTCAAGGTTGGGCGCTGTGCCATCATAGACCTGAGCAACAATCGGCTAACTGGAAATGTGTCCCGGATCCAAGGTTGGGGAAATTATGTTGAAGTTATTGTGTTAAGCTCAAATGCATT

>M6-PGSC0003DMG400002012-(rev)M6LRR-RLK_2562

TCCCGGATCCAAGGTTGGGGAAATTATGTTGAAGTTATTGTGTTAAGCTCAAATGCATTGACAGGAACCTTTCCCAACCAAACTTCCCAGTTCTTGAGGCTTACTTCGTTAAAGATTTCA

>M6-PGSC0003DMG400002012-(rev)M6LRR-RLK_2623

CAGGAACCTTTCCCAACCAAACTTCCCAGTTCTTGAGGCTTACTTCGTTAAAGATTTCAAACAACTCACTTGAAGGTTTGTTACCAACTATGTTAGGTACATACCTCGAACTTAAAACAA

>M6-PGSC0003DMG400002012-(rev)M6LRR-RLK_2684

CAACTCACTTGAAGGTTTGTTACCAACTATGTTAGGTACATACCTCGAACTTAAAACAATTGATCTTAGCATTAACCAACTTAGTGGTACTCTCCTTCCCAGCCTTTTCAACTCTACCAA

>M6-PGSC0003DMG400002012-(rev)M6LRR-RLK_2745

GATCTTAGCATTAACCAACTTAGTGGTACTCTCCTTCCCAGCCTTTTCAACTCTACCAAATTGACTGATATTAATGTTTCTTTCAACAAATTTACCGGTAGTGTACCTATCATGGCATTT

>M6-PGSC0003DMG400002012-(rev)M6LRR-RLK_2806

TGACTGATATTAATGTTTCTTTCAACAAATTTACCGGTAGTGTACCTATCATGGCATTTAACTCAGAGAATCTAAGCTTGATTTCTCTAGATGTTTCACATAATGCATTAGCTGGTCCGT

>M6-PGSC0003DMG400002012-(rev)M6LRR-RLK_2867

CTCAGAGAATCTAAGCTTGATTTCTCTAGATGTTTCACATAATGCATTAGCTGGTCCGTTACCTCCGGGGCTGGACAAGTTCCCGGACATGGTAAATCTGGATCTCTCTGATAACAAATT

>M6-PGSC0003DMG400002012-(rev)M6LRR-RLK_2928

CCTCCGGGGCTGGACAAGTTCCCGGACATGGTAAATCTGGATCTCTCTGATAACAAATTTGAAGGTGGCCTTCCTAATGATCTCTCGGACAAATTGGAATTTTTTAATGTGGCTAATAAC

>M6-PGSC0003DMG400002012-(rev)M6LRR-RLK_2989

AAGGTGGCCTTCCTAATGATCTCTCGGACAAATTGGAATTTTTTAATGTGGCTAATAACAATTTTTCTGGACCTGTGCCACAAAATTTATGGAGGTTTCCTGACTCCTCATTCCATCCGG

>M6-PGSC0003DMG400002012-(rev)M6LRR-RLK_305

CATAATGATTAGTGATGATTGCAGTTTGTGTTTAAAAATTTTGCTAATTCATTTTCTGGGATTCTTGAAATGATTGTGCAATCTTGTCTTTCTACTTGGTGGGGTTTGTCTACTTTCTTG

>M6-PGSC0003DMG400002012-(rev)M6LRR-RLK_3050

TTTTTCTGGACCTGTGCCACAAAATTTATGGAGGTTTCCTGACTCCTCATTCCATCCGGGAAATCCCTTGCTTGTACTTCCAAAGCACGCCAAAGCTCCTTCAGAGGGAGATTCAACTTT

>M6-PGSC0003DMG400002012-(rev)M6LRR-RLK_3111

AATCCCTTGCTTGTACTTCCAAAGCACGCCAAAGCTCCTTCAGAGGGAGATTCAACTTTAAGCTTGAGAAGTCATGGTTCTCGCATGAAATCTACCATCAGGGCTGCCCTCATTGCTGGG

>M6-PGSC0003DMG400002012-(rev)M6LRR-RLK_3172

GCTTGAGAAGTCATGGTTCTCGCATGAAATCTACCATCAGGGCTGCCCTCATTGCTGGGCTGATCTGTGGTGTATCTGTTATATCTTTGTTGACATTAATAATCTACCGCAAGGCCCATC

>M6-PGSC0003DMG400002012-(rev)M6LRR-RLK_3233

GATCTGTGGTGTATCTGTTATATCTTTGTTGACATTAATAATCTACCGCAAGGCCCATCAGCGAGATGGCGGGAAGGACGATATGAAAGGAACCAAGGAAAAGAAAGGTATGGAACTATT

>M6-PGSC0003DMG400002012-(rev)M6LRR-RLK_3294

CGAGATGGCGGGAAGGACGATATGAAAGGAACCAAGGAAAAGAAAGGTATGGAACTATTCTTCTTCTAATAAGACAGGTGATCAGATCAACTGATACTAATGCCTTTTCATTTCCACAGG

>M6-PGSC0003DMG400002012-(rev)M6LRR-RLK_3355

TCTTCTAATAAGACAGGTGATCAGATCAACTGATACTAATGCCTTTTCATTTCCACAGGTTTGTCTTTATCTGATATAGAATGTGGACATGACACTAGAGAACACAGCGTGCCAGTATCA

>M6-PGSC0003DMG400002012-(rev)M6LRR-RLK_3416

TGTCTTTATCTGATATAGAATGTGGACATGACACTAGAGAACACAGCGTGCCAGTATCAACAGTTCAAAATGAGCCATTATCTTCACCTATATCTGTCATGTCATCCGCCAATCTCTCTC

>M6-PGSC0003DMG400002012-(rev)M6LRR-RLK_3477

AGTTCAAAATGAGCCATTATCTTCACCTATATCTGTCATGTCATCCGCCAATCTCTCTCCTTCTAAGGTTCAAGATCAATCCAAAAGCCCAAACTCTCTCAGGGTTTCTTCTCCTGATAA

>M6-PGSC0003DMG400002012-(rev)M6LRR-RLK_3538

TCTAAGGTTCAAGATCAATCCAAAAGCCCAAACTCTCTCAGGGTTTCTTCTCCTGATAAACTGGCTGGAGATTTACATCTACTAGATAACGCCTTGAAGGTCACTGCTGAAGAATTATCG

>M6-PGSC0003DMG400002012-(rev)M6LRR-RLK_3599

TGGCTGGAGATTTACATCTACTAGATAACGCCTTGAAGGTCACTGCTGAAGAATTATCGTGTGCTCCTGCCGAAGCTGTGGGCAGAAGTTGTCATGGAACATTGTACAAAGCTACGCTTG

>M6-PGSC0003DMG400002012-(rev)M6LRR-RLK_366

TTCTTGAAATGATTGTGCAATCTTGTCTTTCTACTTGGTGGGGTTTGTCTACTTTCTTGGCCCATTCTGAATTATTAGCTTTGTCAAAATGCTGTCTTTTTTCCTTTTTAGTTCAGTATT

>M6-PGSC0003DMG400002012-(rev)M6LRR-RLK_3660

TGCTCCTGCCGAAGCTGTGGGCAGAAGTTGTCATGGAACATTGTACAAAGCTACGCTTGGTTCAGACCAAGTTCTTGCTGTCAAATGGCTTAAGGAAGGGATAGTGAAAGGCAAGAAGGA

>M6-PGSC0003DMG400002012-(rev)M6LRR-RLK_3721

TCAGACCAAGTTCTTGCTGTCAAATGGCTTAAGGAAGGGATAGTGAAAGGCAAGAAGGAATTTGCTAGAGAAGCTAAGAAACTGGGGAGTATTAGGCATCCTAATTTAGTTTCTCTACTG

>M6-PGSC0003DMG400002012-(rev)M6LRR-RLK_3782

TTGCTAGAGAAGCTAAGAAACTGGGGAGTATTAGGCATCCTAATTTAGTTTCTCTACTGGGTTACTACTGGGGTCCCAAGGAGCATGAGCGGCTTCTCATATCAAATTACACCGATGCAC

>M6-PGSC0003DMG400002012-(rev)M6LRR-RLK_3843

TTACTACTGGGGTCCCAAGGAGCATGAGCGGCTTCTCATATCAAATTACACCGATGCACCATGTTTAGCTCTTTATCTTCTTCGTAAAGGTCAGTCTCCCCTTTTCTATCTATAGAGTAT

>M6-PGSC0003DMG400002012-(rev)M6LRR-RLK_3904

TGTTTAGCTCTTTATCTTCTTCGTAAAGGTCAGTCTCCCCTTTTCTATCTATAGAGTATATCACCTGCATTTATTTCTGTAGCTTTTCTGATTTTTGTATGGTGCTGAAGTCGTGATCCA

>M6-PGSC0003DMG400002012-(rev)M6LRR-RLK_3965

CACCTGCATTTATTTCTGTAGCTTTTCTGATTTTTGTATGGTGCTGAAGTCGTGATCCAGTTTTACCAAATAAATTAGTAGTACTCCTTCAATGAAATCAAACATAATGGACCGTGAATG

>M6-PGSC0003DMG400002012-(rev)M6LRR-RLK_4026

TTTACCAAATAAATTAGTAGTACTCCTTCAATGAAATCAAACATAATGGACCGTGAATGCGCAATACCTTGAATGTGTATATTGTTATTGTAGTTGTCAAGATTGCTCATACCTGATATG

>M6-PGSC0003DMG400002012-(rev)M6LRR-RLK_4087

CAATACCTTGAATGTGTATATTGTTATTGTAGTTGTCAAGATTGCTCATACCTGATATGCTGAAGTTTAATCACCATCTTGCCACTTGATTTCCATTCCGGTGCCTCAAGTGTACTTTTC

>M6-PGSC0003DMG400002012-(rev)M6LRR-RLK_4148

GAAGTTTAATCACCATCTTGCCACTTGATTTCCATTCCGGTGCCTCAAGTGTACTTTTCTTCCAAAAGATATTCTAGCTTGCTGGTTGAATATGTATCTGAGACTATTAATACACATTTC

>M6-PGSC0003DMG400002012-(rev)M6LRR-RLK_4209

CCAAAAGATATTCTAGCTTGCTGGTTGAATATGTATCTGAGACTATTAATACACATTTCACATAGCAACTTTCATGAAATTTATAAATGGAGCTATCTTCCAAAATCTGAACTTTTCAAT

>M6-PGSC0003DMG400002012-(rev)M6LRR-RLK_427

CCATTCTGAATTATTAGCTTTGTCAAAATGCTGTCTTTTTTCCTTTTTAGTTCAGTATTATATGTAACTCTCTGCTAGTGTCACACGGGTTTTTCATTGGGAGGAAAATAAGAGGAGGAA

>M6-PGSC0003DMG400002012-(rev)M6LRR-RLK_4270

ATAGCAACTTTCATGAAATTTATAAATGGAGCTATCTTCCAAAATCTGAACTTTTCAATTCATATATATTAGCAATGTTAGAATTACACATAGCAGAAAAGATACACACCTGTTTTTTGC

>M6-PGSC0003DMG400002012-(rev)M6LRR-RLK_4331

ATATATATTAGCAATGTTAGAATTACACATAGCAGAAAAGATACACACCTGTTTTTTGCTTTCATGGTAATACTAAAAGGTAAAACAGTTCCACCTTCGCTAATTTTGCCTTGATGGTGA

>M6-PGSC0003DMG400002012-(rev)M6LRR-RLK_4514

ATGAGAATTTTATCTCACGTGGCTCATTTTGTATTGATTACTTTGGTTGGGCTCGATTGATGTCATTTTGATAAGGGCATCCCGGTGCACTAATTAGAGACTGCTACTTTGAATTCCGTT

>M6-PGSC0003DMG400002012-(rev)M6LRR-RLK_4575

GTCATTTTGATAAGGGCATCCCGGTGCACTAATTAGAGACTGCTACTTTGAATTCCGTTTATTGCATATAGCACTGTACTTTGTTTTCTAGTTTTGAGTGATGATATGGATGCCGGTTTC

>M6-PGSC0003DMG400002012-(rev)M6LRR-RLK_4636

TTGCATATAGCACTGTACTTTGTTTTCTAGTTTTGAGTGATGATATGGATGCCGGTTTCATTCTTTGATGTATGCAGATGCAGAACCCTGTAAGTTACATCCTTTATCTCTCGATGATCG

>M6-PGSC0003DMG400002012-(rev)M6LRR-RLK_4697

TCTTTGATGTATGCAGATGCAGAACCCTGTAAGTTACATCCTTTATCTCTCGATGATCGACTCAAGATCTCTGTAGATGTTGCTCGTTGTCTAAACTACCTTCACCATGAAAGTGCCATA

>M6-PGSC0003DMG400002012-(rev)M6LRR-RLK_4758

TCAAGATCTCTGTAGATGTTGCTCGTTGTCTAAACTACCTTCACCATGAAAGTGCCATACCTCATGGCAACCTGAAATCCACAAATGTACTTATAGACACGTCTACCGTGAATGCCCTAC

>M6-PGSC0003DMG400002012-(rev)M6LRR-RLK_4819

TCATGGCAACCTGAAATCCACAAATGTACTTATAGACACGTCTACCGTGAATGCCCTACTAACTGACTATAGTCTTCATCGGTTGATGACATCTGCTGGAACAGCTGAACAGGTTCTAAA

>M6-PGSC0003DMG400002012-(rev)M6LRR-RLK_488

ATGTAACTCTCTGCTAGTGTCACACGGGTTTTTCATTGGGAGGAAAATAAGAGGAGGAAATGCAATCTACCATTTGTTTAGTGCTGCTATTTTTAGTTGAGTTAGCTAAAGGAAGCTTAG

>M6-PGSC0003DMG400002012-(rev)M6LRR-RLK_4880

ACTGACTATAGTCTTCATCGGTTGATGACATCTGCTGGAACAGCTGAACAGGTTCTAAATGCCAGTGTACTCGGTTATCGTCCACCTGAATTCGCGAGTACAAGTAAACCTTGCCCGTCA

>M6-PGSC0003DMG400002012-(rev)M6LRR-RLK_4941

CCAGTGTACTCGGTTATCGTCCACCTGAATTCGCGAGTACAAGTAAACCTTGCCCGTCATTGAAAAGTGACGTCTATGCTTTTGGTGTTATCTTGTTGGAACTTTTGACTGGAAGAAGTT

>M6-PGSC0003DMG400002012-(rev)M6LRR-RLK_5002

GAAAAGTGACGTCTATGCTTTTGGTGTTATCTTGTTGGAACTTTTGACTGGAAGAAGTTCTGCAGAAATTGTTCCTGGGAACTCAGAAGTGCTGGATTTAACCGAATGGGCGAGATTATT

>M6-PGSC0003DMG400002012-(rev)M6LRR-RLK_5063

GCAGAAATTGTTCCTGGGAACTCAGAAGTGCTGGATTTAACCGAATGGGCGAGATTATTGGCTTTTCAAGACCGTTCAATCGAGTGCTTTGATCCATTCTTATTAGGAAAGCAAAGCAAC

>M6-PGSC0003DMG400002012-(rev)M6LRR-RLK_5124

CTTTTCAAGACCGTTCAATCGAGTGCTTTGATCCATTCTTATTAGGAAAGCAAAGCAACGACGAAGACATGCATACGATTCTTGATAGCATGCTTCAGGTTGCACTAAGGTGCATTTTGC

>M6-PGSC0003DMG400002012-(rev)M6LRR-RLK_5185

CGAAGACATGCATACGATTCTTGATAGCATGCTTCAGGTTGCACTAAGGTGCATTTTGCCAGCTGATGAAAGGCCTGATATGAAATTGGTTTTCGAGCAACTTTGCTCGATAGCGCAATA

>M6-PGSC0003DMG400002012-(rev)M6LRR-RLK_5246

GCTGATGAAAGGCCTGATATGAAATTGGTTTTCGAGCAACTTTGCTCGATAGCGCAATAACACAAGTTGAGGGGAGAGGGGTAAATTTGAATACTATGTAGTTGTATATTTGTATCTTAA

>M6-PGSC0003DMG400002012-(rev)M6LRR-RLK_5307

ACAAGTTGAGGGGAGAGGGGTAAATTTGAATACTATGTAGTTGTATATTTGTATCTTAAAAGTTGATATTTCATTCATGTTCCATTTTTAATTCAATTGTTTTAGAAACAAATTCATTTG

>M6-PGSC0003DMG400002012-(rev)M6LRR-RLK_5368

GTTGATATTTCATTCATGTTCCATTTTTAATTCAATTGTTTTAGAAACAAATTCATTTGTGAATTAGCACATTAAGTGATGAATGAAGTTGTATTCTTCACATTTATTGACAAGAAATAA

>M6-PGSC0003DMG400002012-(rev)M6LRR-RLK_549

GCAATCTACCATTTGTTTAGTGCTGCTATTTTTAGTTGAGTTAGCTAAAGGAAGCTTAGATTTGGATGCTCTTTTGGAGTTTAAGAAGGGAGTTTTGAAAGACCCTTCTGGGAAAGTCCT

>M6-PGSC0003DMG400002012-(rev)M6LRR-RLK_61

ATTTATATAAGGAAAGTTGGTTTGACTAAAATAACATCCCCTATTGACTTGCTTATGGTATAGGGAAATGTAGGGTCTCTTGCTTTTGATTTTGATGTATTGGTTATTGGTGAGTGAATT

>M6-PGSC0003DMG400002012-(rev)M6LRR-RLK_610

TTGGATGCTCTTTTGGAGTTTAAGAAGGGAGTTTTGAAAGACCCTTCTGGGAAAGTCCTTTCTTCATGGGATTCTAAGTCATTGGGACCAAATGGTTGCCCCCAAAACTGGTTCGGTATC

>M6-PGSC0003DMG400002012-(rev)M6LRR-RLK_671

CTTCATGGGATTCTAAGTCATTGGGACCAAATGGTTGCCCCCAAAACTGGTTCGGTATCGGTTGCAGTGATGGCCATATCACTTCAATTGAACTAAATGATGTAGGTCTGGTTGGAGTTT

>M6-PGSC0003DMG400002012-(rev)M6LRR-RLK_732

TTGCAGTGATGGCCATATCACTTCAATTGAACTAAATGATGTAGGTCTGGTTGGAGTTTTGGATTTTGCAGCTATTAGTGGCCTAAAAATGCTGCAGAATTTGTCTGTTGCAAACAACCA

>M6-PGSC0003DMG400002012-(rev)M6LRR-RLK_793

GATTTTGCAGCTATTAGTGGCCTAAAAATGCTGCAGAATTTGTCTGTTGCAAACAACCAGTTAAGTGGGAAAATTACTGAGGAAGTTGGGTTGATTATGTCATTAGAATTTTTGGATCTT

>M6-PGSC0003DMG400002012-(rev)M6LRR-RLK_854

TAAGTGGGAAAATTACTGAGGAAGTTGGGTTGATTATGTCATTAGAATTTTTGGATCTTTCCAAGAACATGTTTAGGGGTTTTATACCCTCTAAGCTGACTAGTTTAAAGAACTTAGTAT

>M6-PGSC0003DMG400002012-(rev)M6LRR-RLK_915

CAAGAACATGTTTAGGGGTTTTATACCCTCTAAGCTGACTAGTTTAAAGAACTTAGTATCTCTTAATCTTTCTTTAAATAGTCTCGATGGAATGGTTCCCACTGGTTTTGCTAGTCTTGA

>M6-PGSC0003DMG400002012-(rev)M6LRR-RLK_976

CTTAATCTTTCTTTAAATAGTCTCGATGGAATGGTTCCCACTGGTTTTGCTAGTCTTGAGAAATTGAAGTATCTGGATTTGCACTCTAATGCCTTCTCAATTGATATTATGCTTCTTCTG

>M6-PGSC0003DMG400002017-(rev)M6G-LecRK_0

AATTCTGCTCACTAAGGCTTAGGTAGTTGTGAAAGAACACCTATGATGGTTTTGCTTCCATTGGAATCTGAACTTGAACCTCATGTTTTGGGCTTCATGTATTAGTTTTTTTCTTATCGA

>M6-PGSC0003DMG400002017-(rev)M6G-LecRK_1008

AATCTGCATCGTGATTTCTGACTAAAAATCATTCACAGAGATCTTAAGACGAGCAATATTCTGTTAGACGAAGATGTGAATCCCAAAATCTCAGATTTTGGAATGGCTAGGATTTACGGA

>M6-PGSC0003DMG400002017-(rev)M6G-LecRK_1071

TTAGACGAAGATGTGAATCCCAAAATCTCAGATTTTGGAATGGCTAGGATTTACGGAGGGAATCAAACTATAGCAAATACCAACAGGATTGCCGGTACCTAGTAGGTATTAATTGTTCAT

>M6-PGSC0003DMG400002017-(rev)M6G-LecRK_1134

CAAACTATAGCAAATACCAACAGGATTGCCGGTACCTAGTAGGTATTAATTGTTCATAGCAGTGAAATTGATCCTAACTAAATTGTATAATTGCATATCAATGCAGTGGTTACATGTCTC

>M6-PGSC0003DMG400002017-(rev)M6G-LecRK_1197

GAAATTGATCCTAACTAAATTGTATAATTGCATATCAATGCAGTGGTTACATGTCTCCTGCATATGCAATACATGGAATTTTTTCCGAGAAATTAGATGTCTTCAGTTTTGGTGTCATAT

>M6-PGSC0003DMG400002017-(rev)M6G-LecRK_126

TATTGTCATGTAGATAAGAGGAGGATGCTAACGGGAGCCCATATTGCAATTATTGTCTCCGCGACATTCAGCCTGATTGTTATTGTTTGCACTATATATGTATTCAAGATCCTGAAAAAA

>M6-PGSC0003DMG400002017-(rev)M6G-LecRK_1260

ATGCAATACATGGAATTTTTTCCGAGAAATTAGATGTCTTCAGTTTTGGTGTCATATTGTTAGAGATTATCAGCAGTAAAAGAAACAACAACTATTGTCACTCTGAAGACTCTCTCAACC

>M6-PGSC0003DMG400002017-(rev)M6G-LecRK_1323

AGATTATCAGCAGTAAAAGAAACAACAACTATTGTCACTCTGAAGACTCTCTCAACCTCCCGGACCAGGTAAAAACTGAAAAACAGAAAGAATTTCTCATTTTCTCATATATAGTCAAAC

>M6-PGSC0003DMG400002017-(rev)M6G-LecRK_1386

ACCAGGTAAAAACTGAAAAACAGAAAGAATTTCTCATTTTCTCATATATAGTCAAACCACTTTATAATAACGTTATCTATAGTGAAAATGTAGTATAGAGAACGTATAATATAACATAAT

>M6-PGSC0003DMG400002017-(rev)M6G-LecRK_1449

ATAATAACGTTATCTATAGTGAAAATGTAGTATAGAGAACGTATAATATAACATAATATGGAAGTTCAAGAAAAACTTGGTTGAGTATGATTGTTATCGATAGGTCTGAATGGAGTTGAA

>M6-PGSC0003DMG400002017-(rev)M6G-LecRK_1512

GTTCAAGAAAAACTTGGTTGAGTATGATTGTTATCGATAGGTCTGAATGGAGTTGAATATTCTTCTATCCTATACTACTACTATTTTCAAATGACAAAACTACTATAATTGGATCTTAGG

>M6-PGSC0003DMG400002017-(rev)M6G-LecRK_1575

TCTATCCTATACTACTACTATTTTCAAATGACAAAACTACTATAATTGGATCTTAGGCATGTCAACTTTGGAATGAAGGCAAGATACTCGAGTTTCCAAATCCAAATGTGTTTGTTTCAC

>M6-PGSC0003DMG400002017-(rev)M6G-LecRK_1638

AACTTTGGAATGAAGGCAAGATACTCGAGTTTCCAAATCCAAATGTGTTTGTTTCACTTGAAACACCAACACAAGTTGTTAGATGCATCATTCTGATTGGTGGGATTGCTTTGTGTGCAA

>M6-PGSC0003DMG400002017-(rev)M6G-LecRK_1701

CACCAACACAAGTTGTTAGATGCATCATTCTGATTGGTGGGATTGCTTTGTGTGCAAGAACATGCAAGTTACAGGCCAAATACAGCTACAATTGTTTCCATGCTCTGCACTGACACAGTT

>M6-PGSC0003DMG400002017-(rev)M6G-LecRK_1764

GCAAGTTACAGGCCAAATACAGCTACAATTGTTTCCATGCTCTGCACTGACACAGTTCTTCCTTCGTCTAAGCGATTGGCATATAGCAGCAAAGGCTCTTGTGATCCAAATATTTCACCT

>M6-PGSC0003DMG400002017-(rev)M6G-LecRK_1809

ACTGACACAGTTCTTCCTTCGTCTAAGCGATTGGCATATAGCAGCAAAGGCTCTTGTGATCCAAATATTTCACCTTGCTCTAGAAATTCCATTTCTATTTCAAATGTTCAAGCATGTCAA

>M6-PGSC0003DMG400002017-(rev)M6G-LecRK_189

ACATTCAGCCTGATTGTTATTGTTTGCACTATATATGTATTCAAGATCCTGAAAAAAAGGTAAAAAAAAAAAGTCAATTTTTACAATTCTTAGAGGCACAACTTTTCGCTATCTTTTGGA

>M6-PGSC0003DMG400002017-(rev)M6G-LecRK_252

AAAAAAAAAGTCAATTTTTACAATTCTTAGAGGCACAACTTTTCGCTATCTTTTGGAAAGACTTTAATTTTTACTAGATTTTATAAAGCATTGGGAGTTTAGTCCATTTTTGCATCGCTT

>M6-PGSC0003DMG400002017-(rev)M6G-LecRK_315

TTAATTTTTACTAGATTTTATAAAGCATTGGGAGTTTAGTCCATTTTTGCATCGCTTCAAAACACTCTTCCGCAACAAATTTGCTTCATAACAGGTTTACTCAAGGTAACAGTTTAGAAT

>M6-PGSC0003DMG400002017-(rev)M6G-LecRK_378

ACTCTTCCGCAACAAATTTGCTTCATAACAGGTTTACTCAAGGTAACAGTTTAGAATTACCAGTTTTCCGCTTCAAGTGCATAGAAACTGCTACTAAGAACTTCTGCTAACGAAATAAAC

>M6-PGSC0003DMG400002017-(rev)M6G-LecRK_441

TTTTCCGCTTCAAGTGCATAGAAACTGCTACTAAGAACTTCTGCTAACGAAATAAACTTGGAGAGGGGAGATTTGGACCTGTTTACAAGGTATATATATATCACTCGTGTGCTCTAATCT

>M6-PGSC0003DMG400002017-(rev)M6G-LecRK_504

AGGGGAGATTTGGACCTGTTTACAAGGTATATATATATCACTCGTGTGCTCTAATCTTCGAACTTCACTAGAAAAAGTAGAGAAAATTTCACATAGTTTATACTCATTTGATTATTAGTT

>M6-PGSC0003DMG400002017-(rev)M6G-LecRK_567

TTCACTAGAAAAAGTAGAGAAAATTTCACATAGTTTATACTCATTTGATTATTAGTTGAAGTCATGTGTAATTCATATATAGGGGATATTAGCTGATGGACAAGAGGTTGCTATTAAACA

>M6-PGSC0003DMG400002017-(rev)M6G-LecRK_63

GAATCTGAACTTGAACCTCATGTTTTGGGCTTCATGTATTAGTTTTTTTCTTATCGAGTTGCCTATTGTCATGTAGATAAGAGGAGGATGCTAACGGGAGCCCATATTGCAATTATTGTC

>M6-PGSC0003DMG400002017-(rev)M6G-LecRK_630

ATGTGTAATTCATATATAGGGGATATTAGCTGATGGACAAGAGGTTGCTATTAAACAACTTTGGAGGAGCTTAGGACAAGACATTGAAGTGTTCAAGAATGAGGTTACACTAATTTCCGA

>M6-PGSC0003DMG400002017-(rev)M6G-LecRK_693

GAGGAGCTTAGGACAAGACATTGAAGTGTTCAAGAATGAGGTTACACTAATTTCCGAACTTCAACACAGAAACTTAGTCAAACTGATGGGATTTTGCATTAACGGGGAGGAAAAGTTGAT

>M6-PGSC0003DMG400002017-(rev)M6G-LecRK_756

ACACAGAAACTTAGTCAAACTGATGGGATTTTGCATTAACGGGGAGGAAAAGTTGATGATCTATGAATACATGTCCAACAGAAGCTTAGATGTTTTCATATTGGGTGAGCTAATTAAGAA

>M6-PGSC0003DMG400002017-(rev)M6G-LecRK_819

TGAATACATGTCCAACAGAAGCTTAGATGTTTTCATATTGGGTGAGCTAATTAAGAAGAGTACTTAAACAGTACTTTTCTTGATGAATGATGGTTAAATTACTCTTTGTGCTTGGAAGGA

>M6-PGSC0003DMG400002017-(rev)M6G-LecRK_882

TTAAACAGTACTTTTCTTGATGAATGATGGTTAAATTACTCTTTGTGCTTGGAAGGAGGATCAAGATCTAATGATCAGGCGGCATTGAGTTGGAATATCGTTGAAGGGATAGCTAGAGGT

>M6-PGSC0003DMG400002017-(rev)M6G-LecRK_945

AGATCTAATGATCAGGCGGCATTGAGTTGGAATATCGTTGAAGGGATAGCTAGAGGTCTTCTTAATCTGCATCGTGATTTCTGACTAAAAATCATTCACAGAGATCTTAAGACGAGCAAT

>M6-PGSC0003DMG400002038M6RLKWAX-EGF_0

AAGAAGAATAAATTTACTATAAAAATCAATCAAATTCAACCCCCTTCTTCGTCTTCCAAAAAAAAAAAAAAAGTCATGGCTGCCCTGCTCCTCCTGCTTCTGCTAATCTGCAGCACCCCT

>M6-PGSC0003DMG400002038M6RLKWAX-EGF_1054

CCACACTGCACTCATAGCAGGTATTGTATAATGGGTCACATTTAGTTTAGTTATTACTCTGTTTTTATTTAAGTTGTTCGGATTCTTATTTTTATGTCGAGAAGATGTGGGTGTGGGATT

>M6-PGSC0003DMG400002038M6RLKWAX-EGF_1116

TTTTATTTAAGTTGTTCGGATTCTTATTTTTATGTCGAGAAGATGTGGGTGTGGGATTCATATTGGATTTGGTCAATCAAATTTGGATACTTTTATCAATATTAACGGAGAAATGCGGGA

>M6-PGSC0003DMG400002038M6RLKWAX-EGF_124

TACTTTCTGCTCGTCGTTGTGCTGATTGTGGATCTTCTCCTGTTCCTTACCCACTCAGTACCGGCCCCGATTGCGGTGACCAATCTTACAAAATACGGTGCAGTAGCAACATCCAACCCG

>M6-PGSC0003DMG400002038M6RLKWAX-EGF_1302

GTATAAAGAAATTTGTATGATAGGCCTTTTCCTTTTATCTCCTTGTAGGATTTTCTTTGTGATCAATATTTTTGCTTTGGAATAACTAAGTTTTCTACCTAATGTCTCATAAACATATAA

>M6-PGSC0003DMG400002038M6RLKWAX-EGF_1364

TCAATATTTTTGCTTTGGAATAACTAAGTTTTCTACCTAATGTCTCATAAACATATAACTCTATTTTTAAATACTTGAATTATTTTGGAGAGAAGTATCAAAACACTCCTAAACATGGCG

>M6-PGSC0003DMG400002038M6RLKWAX-EGF_1488

TTATTAGTTTTATCCCCGAACTATTGACAGCATTAAAAACACCCCTCTACTATTGCAAGATCGGGGGTGTATTTAGCTTCAATTTGTTAAGTAAATGGGTGTTTTTAAGGCTGTCAATAG

>M6-PGSC0003DMG400002038M6RLKWAX-EGF_1612

GGGATGAAACTAATAATTTGCGTCAAGTTTAGAGGTGTTTTTGTTACTTCTCTTATTAATTTTTACTGAATCTCCGTACCTGTATTTGAATCCATATCCCAGAATCTTATACTTTAGATC

>M6-PGSC0003DMG400002038M6RLKWAX-EGF_1674

TTACTGAATCTCCGTACCTGTATTTGAATCCATATCCCAGAATCTTATACTTTAGATCAGGAAGGTTAGATCTCTGCACCTGTATCGAACGCCCGCACAATTTGTTTTTTTACTCTATTT

>M6-PGSC0003DMG400002038M6RLKWAX-EGF_1736

AGGTTAGATCTCTGCACCTGTATCGAACGCCCGCACAATTTGTTTTTTTACTCTATTTTGAGACTTCAAGGAACTTATAATAATTGAAATTATCTAGTTATTAAGATGGTTGATCAGTAC

>M6-PGSC0003DMG400002038M6RLKWAX-EGF_1798

ACTTCAAGGAACTTATAATAATTGAAATTATCTAGTTATTAAGATGGTTGATCAGTACTATTTTTGGTTGTTGTGTTGATCAATACTTGAATGCTTCTTGGATTTGCAGGTTTGACATCA

>M6-PGSC0003DMG400002038M6RLKWAX-EGF_186

GGCCCCGATTGCGGTGACCAATCTTACAAAATACGGTGCAGTAGCAACATCCAACCCGAACTTTTCTTCGATACGCTGAACAATTCATATCCGATAAGTTCAATTTCTCCCGAATCACAA

>M6-PGSC0003DMG400002038M6RLKWAX-EGF_1860

TTTGGTTGTTGTGTTGATCAATACTTGAATGCTTCTTGGATTTGCAGGTTTGACATCAGGCCTGGGTGTGGCGGTGATTGCTGCTGTAATTGGTTTTTTTCTGTACAGACGTCACAAACG

>M6-PGSC0003DMG400002038M6RLKWAX-EGF_1922

TGGGTGTGGCGGTGATTGCTGCTGTAATTGGTTTTTTTCTGTACAGACGTCACAAACGAATTAAGGAAGCACGAGATCGACTAACTCGCGAACGGGAAGATATTCTAAGCTCTGGTGGTG

>M6-PGSC0003DMG400002038M6RLKWAX-EGF_1984

AAGGAAGCACGAGATCGACTAACTCGCGAACGGGAAGATATTCTAAGCTCTGGTGGTGTTAAATCTGCGAAGCTATTTACAGGCAAAGAAATCAAGAAAGCAACAAACAATTTCTCTACG

>M6-PGSC0003DMG400002038M6RLKWAX-EGF_2046

ATCTGCGAAGCTATTTACAGGCAAAGAAATCAAGAAAGCAACAAACAATTTCTCTACGGACCGCCTTCTTGGTGCTGGGGGTTATGGTGAAGTTTACTATGGTAAACTTGATGATGGTAC

>M6-PGSC0003DMG400002038M6RLKWAX-EGF_2108

GCCTTCTTGGTGCTGGGGGTTATGGTGAAGTTTACTATGGTAAACTTGATGATGGTACCGTGGTTGCTGTTAAATGTGCCAAGCTAGGGAATACTAAGGGCATTGAGCAAGTTCTCAATG

>M6-PGSC0003DMG400002038M6RLKWAX-EGF_2170

GTTGCTGTTAAATGTGCCAAGCTAGGGAATACTAAGGGCATTGAGCAAGTTCTCAATGAGGTCAGAATACTCTGTCAGGTTAACCACAAGAACCTTGTGCGCATTCTAGGTTGTTGTGTT

>M6-PGSC0003DMG400002038M6RLKWAX-EGF_2232

CAGAATACTCTGTCAGGTTAACCACAAGAACCTTGTGCGCATTCTAGGTTGTTGTGTTGAGCTTGAACAGCCATTGCTGGTTTATGAGTACGTCCCAAATGGAACTGTAAGTGATCACTT

>M6-PGSC0003DMG400002038M6RLKWAX-EGF_2294

TTGAACAGCCATTGCTGGTTTATGAGTACGTCCCAAATGGAACTGTAAGTGATCACTTGTATTTGCAACGCAAGCTTCTAACATGGGATTGCCGTCTCAGCATTGCTCATGCAACTGCAG

>M6-PGSC0003DMG400002038M6RLKWAX-EGF_2356

TTGCAACGCAAGCTTCTAACATGGGATTGCCGTCTCAGCATTGCTCATGCAACTGCAGAGGGACTTGCTTACTTGCATTTCTCTGCAGTTCCTCCTATTTACCATCGAGATGTCAAGTCT

>M6-PGSC0003DMG400002038M6RLKWAX-EGF_2418

ACTTGCTTACTTGCATTTCTCTGCAGTTCCTCCTATTTACCATCGAGATGTCAAGTCTAGCAACATTTTGCTCGATGACCAGTTGAATGCTAAGGTCGCAGACTTTGGTCTTTCACGTTT

>M6-PGSC0003DMG400002038M6RLKWAX-EGF_248

TTTCTTCGATACGCTGAACAATTCATATCCGATAAGTTCAATTTCTCCCGAATCACAAAGACTTACTATCGAACCATCACCGTTTTTATCCAACACCTGCATCACCCAGGACATTTCTAC

>M6-PGSC0003DMG400002038M6RLKWAX-EGF_2480

ACATTTTGCTCGATGACCAGTTGAATGCTAAGGTCGCAGACTTTGGTCTTTCACGTTTGGCTCATACAGACCTGAGTCATGTGTCCACCTGTGCTCAGGGTACCCTTGGTTATCTTGACC

>M6-PGSC0003DMG400002038M6RLKWAX-EGF_2542

CATACAGACCTGAGTCATGTGTCCACCTGTGCTCAGGGTACCCTTGGTTATCTTGACCCTGAGTACTACAGGAACTACCAATTGACAGACAAGAGTGATGTGTACAGTTTCGGGGTTGTG

>M6-PGSC0003DMG400002038M6RLKWAX-EGF_2604

GTACTACAGGAACTACCAATTGACAGACAAGAGTGATGTGTACAGTTTCGGGGTTGTGTTGTTGGAACTCTTGACATCCCAAAAGGCAATAGACTTTGGTCGACCACAGGATGATGTAAA

>M6-PGSC0003DMG400002038M6RLKWAX-EGF_2666

TGGAACTCTTGACATCCCAAAAGGCAATAGACTTTGGTCGACCACAGGATGATGTAAACTTGGCTATTTACGTGCAAAGGTTGGTAGAGGAAGAGAGAATAATGGATGCTGTTGATCCTG

>M6-PGSC0003DMG400002038M6RLKWAX-EGF_2728

GCTATTTACGTGCAAAGGTTGGTAGAGGAAGAGAGAATAATGGATGCTGTTGATCCTGCATTAAAGGTGGGAGCAAGCAGCGTGGAGTTGGAGACAATGAAGGCAATCGGTTTTCTGGCT

>M6-PGSC0003DMG400002038M6RLKWAX-EGF_2790

AAAGGTGGGAGCAAGCAGCGTGGAGTTGGAGACAATGAAGGCAATCGGTTTTCTGGCTGTCAGTTGCTTGGAGGAAAGAAGACAAAGCAGGCCTTCGATGAAAGAAGTAGCCGAGGAGAT

>M6-PGSC0003DMG400002038M6RLKWAX-EGF_2852

GTTGCTTGGAGGAAAGAAGACAAAGCAGGCCTTCGATGAAAGAAGTAGCCGAGGAGATTGAGTACATAATCACTATTGCTACTGCAAAGCAATAGAATGACATGTTTAAAATGGTAAATG

>M6-PGSC0003DMG400002038M6RLKWAX-EGF_2914

TACATAATCACTATTGCTACTGCAAAGCAATAGAATGACATGTTTAAAATGGTAAATGCAGTAGTTCTTATTGTGTAATTTTATAAGTTGTGAGTTAAAAGCCAGAAAGTAAGGGACATA

>M6-PGSC0003DMG400002038M6RLKWAX-EGF_2976

AGTTCTTATTGTGTAATTTTATAAGTTGTGAGTTAAAAGCCAGAAAGTAAGGGACATAAGTATTAGTATTACTAATTTGATTGGATCTTCTTCTTTTCAAAGCTTGTTGTAGATACAACT

>M6-PGSC0003DMG400002038M6RLKWAX-EGF_3038

TTAGTATTACTAATTTGATTGGATCTTCTTCTTTTCAAAGCTTGTTGTAGATACAACTGGAGAGTGAGTTTATACATACATTGATAATCTTCAAAGCTTGGCCAAACAACCTATCTCTCT

>M6-PGSC0003DMG400002038M6RLKWAX-EGF_310

TTACTATCGAACCATCACCGTTTTTATCCAACACCTGCATCACCCAGGACATTTCTACCGTAGGTGTTCAATTGAATTCTTCTCTTCCTTTCAACATTACGAGCAGCAACACAATCATTT

>M6-PGSC0003DMG400002038M6RLKWAX-EGF_3100

AGTGAGTTTATACATACATTGATAATCTTCAAAGCTTGGCCAAACAACCTATCTCTCTAGCACTTTTGGCTTCCCAAAAATGTTGCCGCACCAATATTAGATACAAAATGCATAATTTTT

>M6-PGSC0003DMG400002038M6RLKWAX-EGF_3162

CTTTTGGCTTCCCAAAAATGTTGCCGCACCAATATTAGATACAAAATGCATAATTTTTGAAGGATCCGACACACAACAGAGCGGTATTTTTGAAGATTATGAGTGCTAGTTATTAAAGTT

>M6-PGSC0003DMG400002038M6RLKWAX-EGF_3224

GATCCGACACACAACAGAGCGGTATTTTTGAAGATTATGAGTGCTAGTTATTAAAGTTGCGCTAGACACAAAGTTTGTGTCAGATCCACCCAATTCTTGTTTCATTTGATATTGGGACTC

>M6-PGSC0003DMG400002038M6RLKWAX-EGF_3286

TAGACACAAAGTTTGTGTCAGATCCACCCAATTCTTGTTTCATTTGATATTGGGACTCGTGGCGGAGGAAGAATACCAGCTACATGTTCAGTTGAATTTTGGTTCAAAAAATTTACTTGT

>M6-PGSC0003DMG400002038M6RLKWAX-EGF_3348

CGGAGGAAGAATACCAGCTACATGTTCAGTTGAATTTTGGTTCAAAAAATTTACTTGTCTTCAAGATTCAATTGACCATGTACATATTATTAACTTGGAACCAATAACTTTAAAAACTTC

>M6-PGSC0003DMG400002038M6RLKWAX-EGF_3410

AAGATTCAATTGACCATGTACATATTATTAACTTGGAACCAATAACTTTAAAAACTTCGAATCCAAAACTCACAAGCTTTAAATTATGGCTCTGCCTCTTAATTGGAATCCCACGTCCAA

>M6-PGSC0003DMG400002038M6RLKWAX-EGF_372

GGTGTTCAATTGAATTCTTCTCTTCCTTTCAACATTACGAGCAGCAACACAATCATTTACCTAAACTGTACTGGGTCTTTGCTTTCGTCTCCGCTTAATTGCTCTTCGGAGAGTTTGTGT

>M6-PGSC0003DMG400002038M6RLKWAX-EGF_434

AAACTGTACTGGGTCTTTGCTTTCGTCTCCGCTTAATTGCTCTTCGGAGAGTTTGTGTCATACTTACGTGAATGGAAGTTCTGGAAATGGAGGTGGTGTTGGCGCATGTAGGAACGCGCC

>M6-PGSC0003DMG400002038M6RLKWAX-EGF_496

CTTACGTGAATGGAAGTTCTGGAAATGGAGGTGGTGTTGGCGCATGTAGGAACGCGCCGATTTGTTGTACGTTTAGGGCTGGTGGATCGTCGACGTCGTATATGATTCGGGTTAGGGAAT

>M6-PGSC0003DMG400002038M6RLKWAX-EGF_558

TGTTGTACGTTTAGGGCTGGTGGATCGTCGACGTCGTATATGATTCGGGTTAGGGAATCGGGTTGTCGGGCTTACCGGAGTTTTGTGAATTTGGATCCGTCGTTACCGGTTAGTAGATGG

>M6-PGSC0003DMG400002038M6RLKWAX-EGF_62

AAAAAAAAAAGTCATGGCTGCCCTGCTCCTCCTGCTTCTGCTAATCTGCAGCACCCCTGCAGTACTTTCTGCTCGTCGTTGTGCTGATTGTGGATCTTCTCCTGTTCCTTACCCACTCAG

>M6-PGSC0003DMG400002038M6RLKWAX-EGF_620

TTGTCGGGCTTACCGGAGTTTTGTGAATTTGGATCCGTCGTTACCGGTTAGTAGATGGCCGGAAGCCGGGATGGAATTGCAGTGGGTTTCACCACCGGAGCCGGTTTGTACGGGTCAGTC

>M6-PGSC0003DMG400002038M6RLKWAX-EGF_682

AAGCCGGGATGGAATTGCAGTGGGTTTCACCACCGGAGCCGGTTTGTACGGGTCAGTCCGATTGTGATTCTGATTCGACCTGTGGACCCGACCCGAATTTGAATGGAAGTGTTAACAGAT

>M6-PGSC0003DMG400002038M6RLKWAX-EGF_744

TGTGATTCTGATTCGACCTGTGGACCCGACCCGAATTTGAATGGAAGTGTTAACAGATGTTTTTGTCATTCCGGGTTTCATTGGGACCCAATTGCCGCGTTATGTGCCCAAGGTAAGTTA

>M6-PGSC0003DMG400002038M6RLKWAX-EGF_806

TTGTCATTCCGGGTTTCATTGGGACCCAATTGCCGCGTTATGTGCCCAAGGTAAGTTATAAAGAGTACTTTTTATCTCGTAAAAATAAAGATCTTGGTTAAAAATGACATAAAATAGGAA

>M6-PGSC0003DMG400002038M6RLKWAX-EGF_868

GAGTACTTTTTATCTCGTAAAAATAAAGATCTTGGTTAAAAATGACATAAAATAGGAAAAAATTTAATTATAAAAAGGCTAGAAAATCATTTCTCCGATTTCGATTTGGATGTGACTTTG

>M6-PGSC0003DMG400002038M6RLKWAX-EGF_930

TTTAATTATAAAAAGGCTAGAAAATCATTTCTCCGATTTCGATTTGGATGTGACTTTGTTTTACCTATTTCTGAATTCGATACGCAGATGTGACATGCCAAGATCGTGATGGTTGTGGTA

>M6-PGSC0003DMG400002038M6RLKWAX-EGF_992

ACCTATTTCTGAATTCGATACGCAGATGTGACATGCCAAGATCGTGATGGTTGTGGTAGCGACCACACTGCACTCATAGCAGGTATTGTATAATGGGTCACATTTAGTTTAGTTATTACT

>M6-PGSC0003DMG400002045-(rev)M6RLKantifungi(GNK)_0

ATGTCTAGTTTGAATATCCCAATATACCTTGTGCTTTTCTTGATTAGCCAAAGTATTACCACAATATTAGCTCTTGATGACCTTTATTACTGCCCAAACACAACTACTTATAACAATATA

>M6-PGSC0003DMG400002045-(rev)M6RLKantifungi(GNK)_1054

GACAGGAAGAGTAATATTTTGACCAATACAATACTGACTTTAATTTGTTGTAGATTTGTGTCACCATTTGTTTACGATAAGTTTTTACTTTTAATTCCCCCCGTCCCCGCCCCCGCCCCC

>M6-PGSC0003DMG400002045-(rev)M6RLKantifungi(GNK)_1116

ACCATTTGTTTACGATAAGTTTTTACTTTTAATTCCCCCCGTCCCCGCCCCCGCCCCCACAAATATGTAAGCATGCTATCTTTTCTTTTAATTGGATAAATGACGAAATGAAAGGTATTC

>M6-PGSC0003DMG400002045-(rev)M6RLKantifungi(GNK)_1178

ATATGTAAGCATGCTATCTTTTCTTTTAATTGGATAAATGACGAAATGAAAGGTATTCAAAAAGATTTTAAGTGTGTTCTTATCTTAAATTAAATATAATTTTTATATTTGGTATAAGTC

>M6-PGSC0003DMG400002045-(rev)M6RLKantifungi(GNK)_124

ACACCTCTTACAATTCCAACCTCAATTTGCTACTTTCAAATCTTTCTTCTGCCTCTAGAGAACATCTTTTTTACAATTTCTCCATCGATGGCACCGATAATTTCGAAACAATTTACGGTC

>M6-PGSC0003DMG400002045-(rev)M6RLKantifungi(GNK)_1240

AGATTTTAAGTGTGTTCTTATCTTAAATTAAATATAATTTTTATATTTGGTATAAGTCGACATTGTGCTATTATGGATTTATGTCCCATTGATACTGACCCAAAAAAAAGTACTTACCTA

>M6-PGSC0003DMG400002045-(rev)M6RLKantifungi(GNK)_1302

TTGTGCTATTATGGATTTATGTCCCATTGATACTGACCCAAAAAAAAGTACTTACCTAAAAAAGGTATGCGTAACATTCTATTAATTACTTTAATAATATTCTTCATCTATATTCCAATC

>M6-PGSC0003DMG400002045-(rev)M6RLKantifungi(GNK)_1364

AGGTATGCGTAACATTCTATTAATTACTTTAATAATATTCTTCATCTATATTCCAATCATATGTCTTTTTTGTATCAACAAATAAAATTCAGATGAATCCGTATTTATTATTTGACCCTG

>M6-PGSC0003DMG400002045-(rev)M6RLKantifungi(GNK)_1426

GTCTTTTTTGTATCAACAAATAAAATTCAGATGAATCCGTATTTATTATTTGACCCTGATAATAATACGATGGTTGGAATATCACATCAAAAATTAAATAGAATCATTAATAGAATGAAT

>M6-PGSC0003DMG400002045-(rev)M6RLKantifungi(GNK)_1488

TAATACGATGGTTGGAATATCACATCAAAAATTAAATAGAATCATTAATAGAATGAATTTAATTCCTGAGTGGCATAGTTATAGTTGAACATTCTTCGTCGAAAAATTATATGAAGCAAC

>M6-PGSC0003DMG400002045-(rev)M6RLKantifungi(GNK)_1550

TTCCTGAGTGGCATAGTTATAGTTGAACATTCTTCGTCGAAAAATTATATGAAGCAACTAAAAAAATATTTTTTATATGTATATACTAAATCTTAAATATCATTAACGAATTTTCTGACT

>M6-PGSC0003DMG400002045-(rev)M6RLKantifungi(GNK)_1612

AAAATATTTTTTATATGTATATACTAAATCTTAAATATCATTAACGAATTTTCTGACTTGCCACTGCAGGGCATGATGGAATCCCAAAAATAGAACTACTAATTATTTGTGTAGTGGTTG

>M6-PGSC0003DMG400002045-(rev)M6RLKantifungi(GNK)_1674

ACTGCAGGGCATGATGGAATCCCAAAAATAGAACTACTAATTATTTGTGTAGTGGTTGCCATTGTAATTCTCTTGTTTCTACGATCTATTGGAGTCTTTCATTTGCTACAAAGGAGAAGA

>M6-PGSC0003DMG400002045-(rev)M6RLKantifungi(GNK)_1736

TGTAATTCTCTTGTTTCTACGATCTATTGGAGTCTTTCATTTGCTACAAAGGAGAAGAAGGATGAAAAGAAATAATAAAACTCTTAAGGAGATGAATGGTAATTAATTAAATCCCAATTA

>M6-PGSC0003DMG400002045-(rev)M6RLKantifungi(GNK)_186

CATCTTTTTTACAATTTCTCCATCGATGGCACCGATAATTTCGAAACAATTTACGGTCTTTTCATGTGTCGAGGTGATATGTTAAGTGTAAATTGTCAAGATTTTGTAACAAGAGCTGTT

>M6-PGSC0003DMG400002045-(rev)M6RLKantifungi(GNK)_1922

GTTGCAGTTGATCTTAGTGGAATTTTAACAGCTGAATCATTGCAATATGACTTTAGTACAATTGAATTTGCCACTAACTTCTTTTCTAAGGAAAACAAAATTGGTGTAGGTGGATTTGGT

>M6-PGSC0003DMG400002045-(rev)M6RLKantifungi(GNK)_1984

TGAATTTGCCACTAACTTCTTTTCTAAGGAAAACAAAATTGGTGTAGGTGGATTTGGTGATGTCTATAAGGTAACATGTTCAAATTATACCATATAATAATTTTTTAGATATAAGTTATA

>M6-PGSC0003DMG400002045-(rev)M6RLKantifungi(GNK)_2046

TCTATAAGGTAACATGTTCAAATTATACCATATAATAATTTTTTAGATATAAGTTATATATTCTGACAACGTAAATATTTTTTACACGATCATTAGGGGGAACTTGGTAATGGACAAGAG

>M6-PGSC0003DMG400002045-(rev)M6RLKantifungi(GNK)_2108

CTGACAACGTAAATATTTTTTACACGATCATTAGGGGGAACTTGGTAATGGACAAGAGATAGCTGTAAAGAGGCTATCAAGAAGGTCAAGTCAAGGAGTTGAAGAGTTTAAGAATGAAGT

>M6-PGSC0003DMG400002045-(rev)M6RLKantifungi(GNK)_2170

CTGTAAAGAGGCTATCAAGAAGGTCAAGTCAAGGAGTTGAAGAGTTTAAGAATGAAGTTGTACTTGTTGCTAAGCTTCAACATAGAAATTTAGTGAGACTCTTAGGTTTTTGCTTGGAAG

>M6-PGSC0003DMG400002045-(rev)M6RLKantifungi(GNK)_2232

CTTGTTGCTAAGCTTCAACATAGAAATTTAGTGAGACTCTTAGGTTTTTGCTTGGAAGGAGAAGAAAAAATACTCATCTACGAGTTCGTGTCCAACAAAAGCCTGGACTATTTCTTATTT

>M6-PGSC0003DMG400002045-(rev)M6RLKantifungi(GNK)_2294

AGAAAAAATACTCATCTACGAGTTCGTGTCCAACAAAAGCCTGGACTATTTCTTATTTGGTATGCTTGTTTATACGTCTCACTTTTCTTTTTGTACTCATGTTATATATAATATTTCTCT

>M6-PGSC0003DMG400002045-(rev)M6RLKantifungi(GNK)_2356

TGCTTGTTTATACGTCTCACTTTTCTTTTTGTACTCATGTTATATATAATATTTCTCTGATTTCGATGTCTCACATGATATGATTAAGATACAAAGAAACAAGCAACGTTGGATTGGTCA

>M6-PGSC0003DMG400002045-(rev)M6RLKantifungi(GNK)_2418

TCGATGTCTCACATGATATGATTAAGATACAAAGAAACAAGCAACGTTGGATTGGTCAGTACGTGATAAGATTATTAGAGGAATAGTACGAGGATTAATTTATCTTCATGAAGATTCTCG

>M6-PGSC0003DMG400002045-(rev)M6RLKantifungi(GNK)_248

CATGTGTCGAGGTGATATGTTAAGTGTAAATTGTCAAGATTTTGTAACAAGAGCTGTTAAAAAAATAATCTTGTTATGCCCCGGTAGCAAAACGGTTGTATTATGGCGGGATGATTGCAT

>M6-PGSC0003DMG400002045-(rev)M6RLKantifungi(GNK)_2480

GTGATAAGATTATTAGAGGAATAGTACGAGGATTAATTTATCTTCATGAAGATTCTCGTCCTCGAATCATACATCGTGATCTCAAAGCAAGCAATATTTTGTTAGACAAAGATATGAACC

>M6-PGSC0003DMG400002045-(rev)M6RLKantifungi(GNK)_2542

CGAATCATACATCGTGATCTCAAAGCAAGCAATATTTTGTTAGACAAAGATATGAACCCTAAAATTTCAGATTTTGGCATGGCTAGAATATTTGGAGTTGACCAAATTGAAGGAAGCACT

>M6-PGSC0003DMG400002045-(rev)M6RLKantifungi(GNK)_2604

AATTTCAGATTTTGGCATGGCTAGAATATTTGGAGTTGACCAAATTGAAGGAAGCACTAACATAATTGTTGGAACATAGTAAGTGTTGGTTTTTAAAAAAAAGAGACCTTTTTGTGAATC

>M6-PGSC0003DMG400002045-(rev)M6RLKantifungi(GNK)_2666

TAATTGTTGGAACATAGTAAGTGTTGGTTTTTAAAAAAAAGAGACCTTTTTGTGAATCGGTTTGTTATGGTATAGCAAAATGTTAATTATTGTGATTTATTGTTCTTTGTTAGCGGCTAC

>M6-PGSC0003DMG400002045-(rev)M6RLKantifungi(GNK)_2728

TGTTATGGTATAGCAAAATGTTAATTATTGTGATTTATTGTTCTTTGTTAGCGGCTACATGTCCCCGGAGTACGCAATGCATGGTCAATTCTCAGTGAAATCAGATGTATTTAGTTTTGG

>M6-PGSC0003DMG400002045-(rev)M6RLKantifungi(GNK)_2790

CCCCGGAGTACGCAATGCATGGTCAATTCTCAGTGAAATCAGATGTATTTAGTTTTGGTGTGCTTCTTTTGGAGATAATAAGTGGAAAAAGAAGAAATAATAAAGAAGATAAATTTGATG

>M6-PGSC0003DMG400002045-(rev)M6RLKantifungi(GNK)_2976

TTAACATGATTATAATTAATTGTGTTGATATTTATAGGCATGGAGGTTGTGGAAAGATAATAGAGCAATGGAAATTGTTGATCCAACTATTATTGGTGACTCAAATTCAATGTCGAAAAT

>M6-PGSC0003DMG400002045-(rev)M6RLKantifungi(GNK)_3038

GAGCAATGGAAATTGTTGATCCAACTATTATTGGTGACTCAAATTCAATGTCGAAAATCATGAGATGTATACATATTGGATTGTTATGTGTTCAAATTGATTTGGATGGAAGACCTACAA

>M6-PGSC0003DMG400002045-(rev)M6RLKantifungi(GNK)_310

AAATAATCTTGTTATGCCCCGGTAGCAAAACGGTTGTATTATGGCGGGATGATTGCATGTTACGTTACTCGAATCAATCAATATTCCCCGATCCAAACTATAGATCAATAATTAACAATC

>M6-PGSC0003DMG400002045-(rev)M6RLKantifungi(GNK)_3100

AGATGTATACATATTGGATTGTTATGTGTTCAAATTGATTTGGATGGAAGACCTACAACTACTTTGATATCTCATTTTCTCAGTACTGATTTTGCTATTCTACCAGAGCCAAATCAACCA

>M6-PGSC0003DMG400002045-(rev)M6RLKantifungi(GNK)_3162

TTTGATATCTCATTTTCTCAGTACTGATTTTGCTATTCTACCAGAGCCAAATCAACCAGCGTCTTTTAAGGATTATTCTAGATTAGGAAATATCGAGCAACTCACAAGTAATTCCATTCC

>M6-PGSC0003DMG400002045-(rev)M6RLKantifungi(GNK)_3208

CCAAATCAACCAGCGTCTTTTAAGGATTATTCTAGATTAGGAAATATCGAGCAACTCACAAGTAATTCCATTCCGTCATCTAATTTTGGGGAATTAATTACAGAAATCTACCCAAGATAG

>M6-PGSC0003DMG400002045-(rev)M6RLKantifungi(GNK)_372

CGTTACTCGAATCAATCAATATTCCCCGATCCAAACTATAGATCAATAATTAACAATCCATCTACACCTTTTGTCATGCATAACATTATTACAATTACCAATAATGAACATGAACAAAGA

>M6-PGSC0003DMG400002045-(rev)M6RLKantifungi(GNK)_434

TACACCTTTTGTCATGCATAACATTATTACAATTACCAATAATGAACATGAACAAAGAAGGTTCAAACGGAGTCTAGGGGCAATTATGGATGACGTAACGACTCGGGCTGCTAGTGATGA

>M6-PGSC0003DMG400002045-(rev)M6RLKantifungi(GNK)_496

TCAAACGGAGTCTAGGGGCAATTATGGATGACGTAACGACTCGGGCTGCTAGTGATGATCGATCGTTAGACAAGAAAATTGCAATTAGGGAAGCTAAGTTCAGTGTTGAGACAACAATTT

>M6-PGSC0003DMG400002045-(rev)M6RLKantifungi(GNK)_558

TCGTTAGACAAGAAAATTGCAATTAGGGAAGCTAAGTTCAGTGTTGAGACAACAATTTATGCCTTAGCTCAATGCATTCCTCATATGTCTTCTAAAGATTGCCTAAATTGCCTTGAAAAT

>M6-PGSC0003DMG400002045-(rev)M6RLKantifungi(GNK)_62

AATATTAGCTCTTGATGACCTTTATTACTGCCCAAACACAACTACTTATAACAATATAACAAACACCTCTTACAATTCCAACCTCAATTTGCTACTTTCAAATCTTTCTTCTGCCTCTAG

>M6-PGSC0003DMG400002045-(rev)M6RLKantifungi(GNK)_620

CTTAGCTCAATGCATTCCTCATATGTCTTCTAAAGATTGCCTAAATTGCCTTGAAAATGCCATCACAGTTTTAAACAAATGTTGTGATGTATATAGAGGTGCTAGAGTTCAATTTCCTAG

>M6-PGSC0003DMG400002045-(rev)M6RLKantifungi(GNK)_682

TCACAGTTTTAAACAAATGTTGTGATGTATATAGAGGTGCTAGAGTTCAATTTCCTAGTTGTTTTATGAGGCATGAAGTGTACCTCTTCTACAATAAAACAAACCTATCTACTTTGTCAC

>M6-PGSC0003DMG400002118M6LRR-RLP_0

CAATCCTTCAGAGATGTCTGAAGATGAGGAGCGTCCTTGTCCTTCAGATCTCACCGGTGGCGCCACTGCAAAAGCAAGGAACTGTTGCTTTAACGCCGCCGTTACCGGTGGCGGAGGGGG

>M6-PGSC0003DMG400002118M6LRR-RLP_1054

GGAGTCTTTGACCTTTGATTGTGTTGAATGCCCTATTGACTTTGAGGCATTGGAGAAGCTAGTAATCAGGTCACCTAGTTTGAAGAGTCTAAGGTTGAATCGGTTTGTTTCTATTACTCA

>M6-PGSC0003DMG400002118M6LRR-RLP_1116

TAATCAGGTCACCTAGTTTGAAGAGTCTAAGGTTGAATCGGTTTGTTTCTATTACTCAGCTGTACCGTTTGATGATTCGAGCTCCACAGCTTACCAATCTGGGAACAGGCTCTTTTGGCG

>M6-PGSC0003DMG400002118M6LRR-RLP_1178

TACCGTTTGATGATTCGAGCTCCACAGCTTACCAATCTGGGAACAGGCTCTTTTGGCGCCTCAACGGTCATTGATGAACCAGATCCTGATTATGCTTCAGCATTTGCTGCCTGCAAATCC

>M6-PGSC0003DMG400002118M6LRR-RLP_124

TTTAACTTTTCGCCACACCCAGATCAAGTTCTTGAAAATGTGCTGGAAAATGTTCTTTGCTTCTTAACTGATCGCCGTGACCGTAACGCTGCATCTCTTGTAAGCAAATCTTGGTATCGG

>M6-PGSC0003DMG400002118M6LRR-RLP_1240

AACGGTCATTGATGAACCAGATCCTGATTATGCTTCAGCATTTGCTGCCTGCAAATCCATTGTCTGTCTCTCTGGTTTCAAGGAAATTGCTCCTGAATATCTGCCTGCAATCTATCCAGT

>M6-PGSC0003DMG400002118M6LRR-RLP_1302

TCTGTCTCTCTGGTTTCAAGGAAATTGCTCCTGAATATCTGCCTGCAATCTATCCAGTTTGTGGCAATCTGACCTCTCTTAATTTAAGCTATGGTGCCAACATTAATACTGAACAATTCA

>M6-PGSC0003DMG400002118M6LRR-RLP_1364

GGCAATCTGACCTCTCTTAATTTAAGCTATGGTGCCAACATTAATACTGAACAATTCAAGTCTGTCATCAGCCGTTGCCATAAGCTCCAAGTATTGTGGGTATGTGGTTTGAATTTTCGC

>M6-PGSC0003DMG400002118M6LRR-RLP_1426

TGTCATCAGCCGTTGCCATAAGCTCCAAGTATTGTGGGTATGTGGTTTGAATTTTCGCTATATGTTCTTTTAAAGAAAATTGTACTCTGTAATAGCGTTATATTGAGTTCAATTTTGATG

>M6-PGSC0003DMG400002118M6LRR-RLP_1488

TGTTCTTTTAAAGAAAATTGTACTCTGTAATAGCGTTATATTGAGTTCAATTTTGATGTTTCAGGTATTTGATTCTGTATGTGACGAAGGACTTGAGGCAGTTGCTGCAACATGTAAGGA

>M6-PGSC0003DMG400002118M6LRR-RLP_1550

AGGTATTTGATTCTGTATGTGACGAAGGACTTGAGGCAGTTGCTGCAACATGTAAGGATCTGCGGGAGATTCGAGTTTTCCCTTTTGAAGCTCGGGAAGATGCAGATGCCCCAGTTTCTG

>M6-PGSC0003DMG400002118M6LRR-RLP_1612

CGGGAGATTCGAGTTTTCCCTTTTGAAGCTCGGGAAGATGCAGATGCCCCAGTTTCTGAAGTAGGTCTCCTTGCGATTTCTGAGGGTTGCAGTAAACTTAAGTCCATCTTATATTTCTGC

>M6-PGSC0003DMG400002118M6LRR-RLP_1674

AGGTCTCCTTGCGATTTCTGAGGGTTGCAGTAAACTTAAGTCCATCTTATATTTCTGCCAACGAATGACAAATGCGGCTGTTATAGCTATGTCAAAGAACTGCCCAGACCTTGTGGTATT

>M6-PGSC0003DMG400002118M6LRR-RLP_1736

GAATGACAAATGCGGCTGTTATAGCTATGTCAAAGAACTGCCCAGACCTTGTGGTATTCCGACTATGCATTATGGGTCGGCACTTGCCTGACCATGTTACTAATGAACCAATGGATGAAG

>M6-PGSC0003DMG400002118M6LRR-RLP_1798

CTATGCATTATGGGTCGGCACTTGCCTGACCATGTTACTAATGAACCAATGGATGAAGGCTTTGGGGCTATTGTCAAGAACTGTAAGAAGCTTACTAGACTCGCTGTATCTGGTTTACTG

>M6-PGSC0003DMG400002118M6LRR-RLP_186

CTTAACTGATCGCCGTGACCGTAACGCTGCATCTCTTGTAAGCAAATCTTGGTATCGGGCTGAGGCTTTAACCAGATCCGAAGTGTTTATTGGCAACTGTTATGCTGTTTCGCCGACCCG

>M6-PGSC0003DMG400002118M6LRR-RLP_1860

TGGGGCTATTGTCAAGAACTGTAAGAAGCTTACTAGACTCGCTGTATCTGGTTTACTGACTGATAAGGCTTTCAGTTACATTGGACAATATGGAAAACTAGTCAGAACCTTATCAGTTGC

>M6-PGSC0003DMG400002118M6LRR-RLP_1922

ATAAGGCTTTCAGTTACATTGGACAATATGGAAAACTAGTCAGAACCTTATCAGTTGCTTTTGCTGGAAACAGTGACTTGGCTCTGAAGTATGTGCTCGAGGGCTGCCCTAAGTTGCAGA

>M6-PGSC0003DMG400002118M6LRR-RLP_1984

GCTGGAAACAGTGACTTGGCTCTGAAGTATGTGCTCGAGGGCTGCCCTAAGTTGCAGAAGTTGGAGATCAGGGATTGCCCATTTGGAGATTTGTCTTTGCGTTCTGGTGTACACCACTAT

>M6-PGSC0003DMG400002118M6LRR-RLP_2046

GGAGATCAGGGATTGCCCATTTGGAGATTTGTCTTTGCGTTCTGGTGTACACCACTATTACAATATGAGGTTCCTTTGGCTGTCATCTTGTAGAGTAACTCTACAAGGTTGTCAGGAGAT

>M6-PGSC0003DMG400002118M6LRR-RLP_2108

ATATGAGGTTCCTTTGGCTGTCATCTTGTAGAGTAACTCTACAAGGTTGTCAGGAGATTGCTGGCCAATTGCCCCGTCTAGTAGTGGAAGTGATAAGTGGTGATGAGGAAGGGAGTGAGA

>M6-PGSC0003DMG400002118M6LRR-RLP_2170

GGCCAATTGCCCCGTCTAGTAGTGGAAGTGATAAGTGGTGATGAGGAAGGGAGTGAGACTGGTGAGCATGTCAATACCTTGTACATGTACCGATCTCTTGATGGACCGAGGGCTGATGTA

>M6-PGSC0003DMG400002118M6LRR-RLP_2232

TGAGCATGTCAATACCTTGTACATGTACCGATCTCTTGATGGACCGAGGGCTGATGTACCATCATTTGTGCAAATACTGTGACCGACCTTTTTAGTAGATAGGAATTTTCTGCAGTAATT

>M6-PGSC0003DMG400002118M6LRR-RLP_2294

CATTTGTGCAAATACTGTGACCGACCTTTTTAGTAGATAGGAATTTTCTGCAGTAATTGATTGGGACCTTTGAGAAATGATTAACTATTTTCTTGGGAAGAGATGCTCTCCAGAAATCTG

>M6-PGSC0003DMG400002118M6LRR-RLP_2356

GGGACCTTTGAGAAATGATTAACTATTTTCTTGGGAAGAGATGCTCTCCAGAAATCTGTTGTGACCCATATGAAGGTGCAAGTTTCCTTCTTGAGAAAACGAATGGTGCACCACTTGCAA

>M6-PGSC0003DMG400002118M6LRR-RLP_2418

GACCCATATGAAGGTGCAAGTTTCCTTCTTGAGAAAACGAATGGTGCACCACTTGCAAAGGCAGTGAGTAATCTATAATGTATTCAATGCTGGTCTCCATGGATGTCATTGTATATGGTT

>M6-PGSC0003DMG400002118M6LRR-RLP_248

AGGCTTTAACCAGATCCGAAGTGTTTATTGGCAACTGTTATGCTGTTTCGCCGACCCGGGTTACGACCCGTTTCAAATGGGTGACCTCTGTGGCCATTAAAGGGAAACCTAGGTTTGCTG

>M6-PGSC0003DMG400002118M6LRR-RLP_2480

AGTGAGTAATCTATAATGTATTCAATGCTGGTCTCCATGGATGTCATTGTATATGGTTCAGTACAGAGATTGTTTTTTTCCAGCAAAGCAATCTAGAAGACAAATGGTTCTTTGTATCAG

>M6-PGSC0003DMG400002118M6LRR-RLP_2542

ACAGAGATTGTTTTTTTCCAGCAAAGCAATCTAGAAGACAAATGGTTCTTTGTATCAGTAATCTTTTCTATTCACTTTCTTTTCTTCATATTCATGGAGCCTTCAATTAGTAAGTGGAGT

>M6-PGSC0003DMG400002118M6LRR-RLP_2604

CTTTTCTATTCACTTTCTTTTCTTCATATTCATGGAGCCTTCAATTAGTAAGTGGAGTCATTGGTAAGGCCACAGCAAGGGTAGATGTTCCAAACTGCACGGTGGCGGAGCTCAATGAGA

>M6-PGSC0003DMG400002118M6LRR-RLP_2666

GGTAAGGCCACAGCAAGGGTAGATGTTCCAAACTGCACGGTGGCGGAGCTCAATGAGATTTCTCAAAGGCTTTCGGCAGAACCAACACTTGTGGAATACACCTGCATGCTCGGTCAGGGT

>M6-PGSC0003DMG400002118M6LRR-RLP_2728

TCAAAGGCTTTCGGCAGAACCAACACTTGTGGAATACACCTGCATGCTCGGTCAGGGTACTGGCTCGTCAAGTAGTATATTTTCATACAATTGAGGCCCTTCTTATTTTTTGTTTCATTG

>M6-PGSC0003DMG400002118M6LRR-RLP_2790

GCTCGTCAAGTAGTATATTTTCATACAATTGAGGCCCTTCTTATTTTTTGTTTCATTGTTAGTTTAGAGTTGGAAGTTTTTTATTTTTAGTTGGTGATAGCCTAATGGTCAGAAGGATAC

>M6-PGSC0003DMG400002118M6LRR-RLP_2852

TTTAGAGTTGGAAGTTTTTTATTTTTAGTTGGTGATAGCCTAATGGTCAGAAGGATACCTACCGCTGTTGGATCTCTCCATCTCGCTTTCTAGAAATATTTGTTCTTTCCAGGATGTGTT

>M6-PGSC0003DMG400002118M6LRR-RLP_2914

CGCTGTTGGATCTCTCCATCTCGCTTTCTAGAAATATTTGTTCTTTCCAGGATGTGTTGTTTCGTAGCACCTGACTGTTGAATTTATTATTGTTATTTTTAGAAGAAATGTGAGATGGGT

>M6-PGSC0003DMG400002118M6LRR-RLP_2976

CGTAGCACCTGACTGTTGAATTTATTATTGTTATTTTTAGAAGAAATGTGAGATGGGTAATAAATGCCAAATATGCATGGACCAGATGATGCTCTAGTGTGTGTGTGTTTGTGTTTGTCT

>M6-PGSC0003DMG400002118M6LRR-RLP_310

ACGACCCGTTTCAAATGGGTGACCTCTGTGGCCATTAAAGGGAAACCTAGGTTTGCTGATTTCAGTTTGCTTCCTCCAGATTGGGGTGCTCACTTTACTCCTTGGGCTTCTGTTTTGGGT

>M6-PGSC0003DMG400002118M6LRR-RLP_3100

CTTTTCTTTATGTGGAATGATCAGCTTGATAGTGAAGACCCATAAATGATGTGTGGAAAGCCTTGTACAAGTACAAAAATGCAGGACCCTCCTCTCCTTCACTCACTCTCTTTGTGGGAT

>M6-PGSC0003DMG400002118M6LRR-RLP_3162

TTGTACAAGTACAAAAATGCAGGACCCTCCTCTCCTTCACTCACTCTCTTTGTGGGATCCATTTATGAGTCTTATCCTCTATTATCTTTCCAGCTGATGAATGGCCCTTCTCTGTCTCAT

>M6-PGSC0003DMG400002118M6LRR-RLP_3224

TTATGAGTCTTATCCTCTATTATCTTTCCAGCTGATGAATGGCCCTTCTCTGTCTCATCTCTTGTATCTGAAAGAAGAGCCAAGAGCAGGGAGTAGACTGTAGTTGTGGAAAGTATAGAT

>M6-PGSC0003DMG400002118M6LRR-RLP_3286

TGTATCTGAAAGAAGAGCCAAGAGCAGGGAGTAGACTGTAGTTGTGGAAAGTATAGATGGATAGTACAGCACACCCCTTATTTTTTAGCTGATGATGCCAACTTTATCATTATTCTTTTG

>M6-PGSC0003DMG400002118M6LRR-RLP_3348

AGTACAGCACACCCCTTATTTTTTAGCTGATGATGCCAACTTTATCATTATTCTTTTGGATAACAGTCTTGAGGAGCCTGCCTGTTGTGTTATCCTGTTTACTTTTCATATGCGGCTGCT

>M6-PGSC0003DMG400002118M6LRR-RLP_3388

TTTATCATTATTCTTTTGGATAACAGTCTTGAGGAGCCTGCCTGTTGTGTTATCCTGTTTACTTTTCATATGCGGCTGCTGCTGCAGCATTCTTCATCTCTATCATTGACTACAACTCAC

>M6-PGSC0003DMG400002118M6LRR-RLP_372

CAGTTTGCTTCCTCCAGATTGGGGTGCTCACTTTACTCCTTGGGCTTCTGTTTTGGGTGATTCTTATCGTGGGCTTGAGAAATTGTACCTCAAACGTATGTCCATATCTGATGATGATCT

>M6-PGSC0003DMG400002118M6LRR-RLP_434

CTTATCGTGGGCTTGAGAAATTGTACCTCAAACGTATGTCCATATCTGATGATGATCTAGGTTTGTTGGCGCGTTGTTTCCCCAGTTTCAAAGAGCTTGTTCTTGTATGCTGTGACGGTT

>M6-PGSC0003DMG400002118M6LRR-RLP_496

TTGTTGGCGCGTTGTTTCCCCAGTTTCAAAGAGCTTGTTCTTGTATGCTGTGACGGTTTTGGGACTAGTGGACTTGCTATTGTAGCTCGTGATTGCAGGTTAGTATTGTTTTGTCCGGAT

>M6-PGSC0003DMG400002118M6LRR-RLP_558

GACTAGTGGACTTGCTATTGTAGCTCGTGATTGCAGGTTAGTATTGTTTTGTCCGGATATAGATCTGTTTATGTGATAACCCTTTTGGTAAAAAGTTTAGATTTTTAATTTCTAGAAGCT

>M6-PGSC0003DMG400002118M6LRR-RLP_62

CCACTGCAAAAGCAAGGAACTGTTGCTTTAACGCCGCCGTTACCGGTGGCGGAGGGGGGATTTTTAACTTTTCGCCACACCCAGATCAAGTTCTTGAAAATGTGCTGGAAAATGTTCTTT

>M6-PGSC0003DMG400002118M6LRR-RLP_620

ATCTGTTTATGTGATAACCCTTTTGGTAAAAAGTTTAGATTTTTAATTTCTAGAAGCTGAATTCTTCTTTCTGTAATGGGAAATTAACAGCTTAAACTAGTGTAGATATATTGTGTTATT

>M6-PGSC0003DMG400002118M6LRR-RLP_682

TCTTCTTTCTGTAATGGGAAATTAACAGCTTAAACTAGTGTAGATATATTGTGTTATTTGAATTTGTGCTTCATGTGTGCTGGTGTTTGAATTAGTTATGTGATTGAGTTGTTCTTTAAC

>M6-PGSC0003DMG400002118M6LRR-RLP_744

TTTGTGCTTCATGTGTGCTGGTGTTTGAATTAGTTATGTGATTGAGTTGTTCTTTAACTGTTTTATTCTGCATTTGAAACTTAATGATACCTTTTGAGGACACTAGCATTGAAGTTGTAA

>M6-PGSC0003DMG400002118M6LRR-RLP_806

TTATTCTGCATTTGAAACTTAATGATACCTTTTGAGGACACTAGCATTGAAGTTGTAATATGTGGTGACTTGTTGTATTGGATTTGTGAGTTTGCAAGCTGTGAAATATGGTTTGGTAGT

>M6-PGSC0003DMG400002118M6LRR-RLP_868

TGGTGACTTGTTGTATTGGATTTGTGAGTTTGCAAGCTGTGAAATATGGTTTGGTAGTGTGATTTTCTTGTGCATAATAGAAGGTTTTTACTGCAGGCAAATTAGAGTGCTTGATCTGAT

>M6-PGSC0003DMG400002118M6LRR-RLP_930

TTTTCTTGTGCATAATAGAAGGTTTTTACTGCAGGCAAATTAGAGTGCTTGATCTGATTGAGTCTGAGGTATCCGACGATGAAGTGGACTGGATTTCCTACTTCCCTGAGAACAAAACGT

>M6-PGSC0003DMG400002118M6LRR-RLP_992

TCTGAGGTATCCGACGATGAAGTGGACTGGATTTCCTACTTCCCTGAGAACAAAACGTGTTTGGAGTCTTTGACCTTTGATTGTGTTGAATGCCCTATTGACTTTGAGGCATTGGAGAAG

>M6-PGSC0003DMG400002426M6LRR-RLP_1764

AATAGTATTCATATTTCAAACATATATATATATACACTCTGTTCCTCTAGCTTCATATTTCTTAAGAAATACTTAACTAATATTTGTACCTTAAATTTTGTTTAATGATGCTAACTCTTC

>M6-PGSC0003DMG400002426M6LRR-RLP_1827

AAGAAATACTTAACTAATATTTGTACCTTAAATTTTGTTTAATGATGCTAACTCTTCAAGAACTTCTAAGTATAATAATAAATAATATATAAAACATGACAAAAATAGCATTTGGGGATT

>M6-PGSC0003DMG400002426M6LRR-RLP_1890

TTCTAAGTATAATAATAAATAATATATAAAACATGACAAAAATAGCATTTGGGGATTGTACATGACATGTGAAGTTATGATTTGTTGACTCATTACCTTTTTATAGGGGACCATGACAGT

>M6-PGSC0003DMG400002478-(rev)M6LRR-RLP_0

CTTCCACTTGATTTGAGACAATGCTTTGTGTATTCTGCAGTGTTCCCAAAGGACACCAAAATGGAAAAGGAAAATCTAATCTCTCTTTGGATGGCACATGGTTTTCTTTCATCGAAAGGA

>M6-PGSC0003DMG400002478-(rev)M6LRR-RLP_1178

TTCTGGATTCCCCACAAGAATAAGGTTTCCATCCTTGAGGAAACTTGCTATATGGGACTTTCGTAATCTGAAAGGATTGCTGAAAAAGGAAGGAGAAGAGCAATTCCCTGTGCTTGAAGA

>M6-PGSC0003DMG400002478-(rev)M6LRR-RLP_124

TGGAGCTAGAGGATGTGGGTAATGAAGTATGGAATGAATTATATTTGAGGTCTTTCTTCCAAGAGATTGAAGTTAGATATGGTAAAACTTATTTCAAGATGCATGATCTCATCCATGATT

>M6-PGSC0003DMG400002478-(rev)M6LRR-RLP_1302

GAGATTTGGTGGTGCCCTATGTTTGTTATTCCGACCCTTTCTTCTGTCAAGAAATTGGAATTGGATCCAACACCTTTAAGGTCCATGTCTAATCTTAGGAATCTTACACCTTTCAGGTCC

>M6-PGSC0003DMG400002478-(rev)M6LRR-RLP_1364

GGATCCAACACCTTTAAGGTCCATGTCTAATCTTAGGAATCTTACACCTTTCAGGTCCATATCTAATCTTAGGAATCTTACTTCCCTCAACATTAGCTATAGCAATAGCAATGAAGCAAC

>M6-PGSC0003DMG400002478-(rev)M6LRR-RLP_1426

CTAATCTTAGGAATCTTACTTCCCTCAACATTAGCTATAGCAATAGCAATGAAGCAACTTCACTCCCAGAAGAGATGTTCAAAAACCTTGCAAATCTCAAATACTTGAATATCTCTAAGT

>M6-PGSC0003DMG400002478-(rev)M6LRR-RLP_1488

CTCCCAGAAGAGATGTTCAAAAACCTTGCAAATCTCAAATACTTGAATATCTCTAAGTTCGAGAATCTCAAAGAGCTGCCTACCAGCCTGGCTAGTCTCAATGCTTTGGAGAGTCTGAAG

>M6-PGSC0003DMG400002478-(rev)M6LRR-RLP_1612

AATATTGTGGTGCACTAGAGAGTCTCCCCGAGGAAGGGGTGAAAGGTTTAACTTCACTCACAAAGTTATCTGTTCATGATTGCAAGATGCTAAAATGTTTACCGGAGGGATTGCAGCACC

>M6-PGSC0003DMG400002478-(rev)M6LRR-RLP_1798

TAGGAGAAGACTGGCACAAAATTGCTCACATTCCGTATCTGGGTATTCATTACTATTAGTATTTATAATTGTTTCTTTGTTTGTGAGTCTTTTTGGTTCTTGCCATTGGATTGCATGTAA

>M6-PGSC0003DMG400002478-(rev)M6LRR-RLP_1860

TTATAATTGTTTCTTTGTTTGTGAGTCTTTTTGGTTCTTGCCATTGGATTGCATGTAATTTTTTCTAGGGTTGTTTGTTTGTAAGTCTCTCATTGGATGTAATTTTATTTTGGAAACAAA

>M6-PGSC0003DMG400002478-(rev)M6LRR-RLP_1915

TAATTTTTTCTAGGGTTGTTTGTTTGTAAGTCTCTCATTGGATGTAATTTTATTTTGGAAACAAATTAACAATCTATTTGTATTATACGTTTTGAGAATCTATTACTTATTTGTAATTGT

>M6-PGSC0003DMG400002478-(rev)M6LRR-RLP_248

TACATCTCTGTTTTCAGCAAACACATCAAACATCAATATCCGTGAAATAAATGCAGAAGTTTACTTCATTAGGGGGCACGGTAGTTTCATTGATGAGAAAGATTATATACCTAGGAAGCC

>M6-PGSC0003DMG400002478-(rev)M6LRR-RLP_310

ACTTCATTAGGGGGCACGGTAGTTTCATTGATGAGAAAGATTATATACCTAGGAAGCCCATTGGTTTTGCTAAAGTTGTGTCTTCTATCTCTCCTTCACTCTTGAAACAGTTTGTCTCGT

>M6-PGSC0003DMG400002478-(rev)M6LRR-RLP_496

ATTTAAGATATTTGAACCTGTCTCGCAATGCAATGCGTAGTCTTCCAGAAAGGTTGTGCAATCTTCAAAATCTGCTGACTCTTGATCTACGGTTGTGCTCAAACCTTTGTTGTTTGCCAA

>M6-PGSC0003DMG400002478-(rev)M6LRR-RLP_62

GGAAAAGGAAAATCTAATCTCTCTTTGGATGGCACATGGTTTTCTTTCATCGAAAGGAAACTTGGAGCTAGAGGATGTGGGTAATGAAGTATGGAATGAATTATATTTGAGGTCTTTCTT

>M6-PGSC0003DMG400002478-(rev)M6LRR-RLP_620

AACAAGTAAACTTGTTAGTCTCCGGAATCTAAAACTTGATCACTGTTCATTGAAATATATGCCACCAAGGATAGGATCTTTGACAGGCCTTAAGACTCTAAGTAACTTTGTTGTGGGCGA

>M6-PGSC0003DMG400002478-(rev)M6LRR-RLP_744

GAAGGTTGTCAACTTGGGGAACTAGGAAGCCTAAATCTCTATGGCTCATTTGAAATCACAAATCTTGAGAGAGTGAAGAATCATAAGGATGCAAAAGAAGCCAATTTATGTGCAAAAGGG

>M6-PGSC0003DMG400002478-(rev)M6LRR-RLP_806

TCTTGAGAGAGTGAAGAATCATAAGGATGCAAAAGAAGCCAATTTATGTGCAAAAGGGAATCTGCATTCATTAAGCATGAGTTGGAACATGTCCGGACCACATAGATATGAATCAGAAGA

>M6-PGSC0003DMG400002478-(rev)M6LRR-RLP_868

TGCATTCATTAAGCATGAGTTGGAACATGTCCGGACCACATAGATATGAATCAGAAGAAGATAAAGTGCTTGAAGCCCTCAAACCACACCCCAATCTGAAATATTTAGAAATCAATTGGT

>M6-PGSC0003DMG400002478-(rev)M6LRR-RLP_930

AAAGTGCTTGAAGCCCTCAAACCACACCCCAATCTGAAATATTTAGAAATCAATTGGTTCAGAGGAATCCGTCTCCCAGAGTGGATGAATCACTCAGTATTGAAAAATATTGTCTCTATT

>M6-PGSC0003DMG400002478-(rev)M6LRR-RLP_992

AGGAATCCGTCTCCCAGAGTGGATGAATCACTCAGTATTGAAAAATATTGTCTCTATTATAATTATCGAATGCACAAACTGCTCGTGCTTACCACCCTTTGGTGAGCTGCCTTGTCTAGA

>M6-PGSC0003DMG400002526M6LysM-RLK_0

TTTTCCCTTCTTGGTTGATTTTGCAGTCTCACAGAGTAATATTAGTAGTAATAAGTTCCACTAAAAGACCTTGTTTTCAGCCATTACAGGACAACTAAACCCCCCATCCCCAAACCCCAC

>M6-PGSC0003DMG400002526M6LysM-RLK_1037

TTAGTCAGTTTTTTCTTTCTCTTTTTGTTTGTGTGAGTGAGGGGATGAACCTATGTTCTAGAGGGGCACACAATCCTCTCTCATTTGTCTATGGCAGGTTAAGATACCAACTAATCCCTT

>M6-PGSC0003DMG400002526M6LysM-RLK_122

AATGTTCTGTTCTTGGTTCTCTGCTATTCCTATTTCACTTTCTTTCTCTTAAGATTCAACTCTTTTGCAATTTTTTTTCATGTTTTTCTTTAAGTTTCCATGACCCGTTTCACAAAATCT

>M6-PGSC0003DMG400002526M6LysM-RLK_1403

TACTGTGGTTGGGAGAGGATCATGACTACCTAATGCTTTGTATCCAATAGTGAACATGAGAACAATGTTCGAAAACATCTAAAAATCTCTACACTTTTTGCACTAAATAACGAAAAGTTG

>M6-PGSC0003DMG400002526M6LysM-RLK_1464

ACAATGTTCGAAAACATCTAAAAATCTCTACACTTTTTGCACTAAATAACGAAAAGTTGATTGAAGTGGATGTAATGGGAATTAGCTTGGCACCCTTGGAGCATTGCCAAAATCAAGTGC

>M6-PGSC0003DMG400002526M6LysM-RLK_1525

TGAAGTGGATGTAATGGGAATTAGCTTGGCACCCTTGGAGCATTGCCAAAATCAAGTGCCTTCGATCAGTTAGAGAGTTGGATTGCATGTATGAAGGGAAATTCGTATTTTGATGGATAA

>M6-PGSC0003DMG400002526M6LysM-RLK_1586

TCGATCAGTTAGAGAGTTGGATTGCATGTATGAAGGGAAATTCGTATTTTGATGGATAACAGGTCCTGTAGAATACATCATATAATGTATTAGTTTCAACATCAGACTATATAGCACAAT

>M6-PGSC0003DMG400002526M6LysM-RLK_1647

GGTCCTGTAGAATACATCATATAATGTATTAGTTTCAACATCAGACTATATAGCACAATAATTATATCACACAACCTGTAAGTTTTAAAGAGGACCTGGTGTGAAGATGAGCATATGGGT

>M6-PGSC0003DMG400002526M6LysM-RLK_1708

TTATATCACACAACCTGTAAGTTTTAAAGAGGACCTGGTGTGAAGATGAGCATATGGGTACTCTTATTGTGCTGTTTTATCTGAACAAGAGATCTCATGAATTCAACTAAATTATTGTAT

>M6-PGSC0003DMG400002526M6LysM-RLK_1769

TCTTATTGTGCTGTTTTATCTGAACAAGAGATCTCATGAATTCAACTAAATTATTGTATTAGCCAATCCTCAAAATTAAGTTTGCTAGGCTCAATAGTTAGCAACTTTGCCATGCTCAAT

>M6-PGSC0003DMG400002526M6LysM-RLK_183

CTTTTGCAATTTTTTTTCATGTTTTTCTTTAAGTTTCCATGACCCGTTTCACAAAATCTATCAAAGATTCAATCTTTAATACAATGGATCATATACAATCAACATTTCTTGCAATAATTT

>M6-PGSC0003DMG400002526M6LysM-RLK_1830

GCCAATCCTCAAAATTAAGTTTGCTAGGCTCAATAGTTAGCAACTTTGCCATGCTCAATAATTATCTTCATTTACTCATGGTCAATAGCTTTTCTAGTAAGCCTCTTTTGGTTAAAAGTA

>M6-PGSC0003DMG400002526M6LysM-RLK_1891

TTATCTTCATTTACTCATGGTCAATAGCTTTTCTAGTAAGCCTCTTTTGGTTAAAAGTACTTTACTTTAACGACTTTGTGTACGTCCAGCTCCTGGCGAGCCTTATCCTGTAGAAAATCA

>M6-PGSC0003DMG400002526M6LysM-RLK_1952

TTACTTTAACGACTTTGTGTACGTCCAGCTCCTGGCGAGCCTTATCCTGTAGAAAATCACACTGTTCCTGCCCCTGCTCCATCACCTTCTGTTGCTGATATTTCAGGTATTCTCAATTCT

>M6-PGSC0003DMG400002526M6LysM-RLK_2013

CTGTTCCTGCCCCTGCTCCATCACCTTCTGTTGCTGATATTTCAGGTATTCTCAATTCTCAATAGCTTACATGCATTCTTACTTTCCTGTTGTTGAGATATGTGTTTTTCTTTATTAATA

>M6-PGSC0003DMG400002526M6LysM-RLK_2074

ATAGCTTACATGCATTCTTACTTTCCTGTTGTTGAGATATGTGTTTTTCTTTATTAATATCACCCAAGAAAGACCAATAAATTTAACTTCTACTCATGGCTTGTCAAGTTTTTGTGTCCT

>M6-PGSC0003DMG400002526M6LysM-RLK_2196

TTTCTGTTAGTAGTTCCTTGCGAGGGCAGTGTCTCCATAAATTTCTATCAGTGGAACGAGAGAACTTCCTTGACAAGCTAATCCAAGAGTTTGATTATCATTCTGCAAGTTATTATCTAT

>M6-PGSC0003DMG400002526M6LysM-RLK_2257

GAACTTCCTTGACAAGCTAATCCAAGAGTTTGATTATCATTCTGCAAGTTATTATCTATTATTGCAATGAAAAATACAGACCTCTATCTGAATATATGGTTTCGTCTTAGCAGTCAACTG

>M6-PGSC0003DMG400002526M6LysM-RLK_2318

TTGCAATGAAAAATACAGACCTCTATCTGAATATATGGTTTCGTCTTAGCAGTCAACTGAATTTAACTCTATTCTATCACCAACTTTTGGCTGCGTTAAAAATCTATTAGTCATTCAGTT

>M6-PGSC0003DMG400002526M6LysM-RLK_244

CAAAGATTCAATCTTTAATACAATGGATCATATACAATCAACATTTCTTGCAATAATTTGTCTGTCAGCAATTTTCAAGATTATCCCTCTTTGTTCATCATACCCTACTCCATCAAACTG

>M6-PGSC0003DMG400002526M6LysM-RLK_2684

TTTTTGAGAAAATCTGTCAGAAGAAAATGAACAATGTAAGGAGGACAAAGTTTCCCCTTAATTAGGTATAACCTCAACTGTTTCTCTGATTTCCTCTTAATTGATAGGTTTTAATTCTTT

>M6-PGSC0003DMG400002526M6LysM-RLK_2745

TTAGGTATAACCTCAACTGTTTCTCTGATTTCCTCTTAATTGATAGGTTTTAATTCTTTCATGGCACTGAGGAGTTTATCTAACAAATCCACAATAAATTCATATATCCCTGACAGTCAT

>M6-PGSC0003DMG400002526M6LysM-RLK_2806

TGGCACTGAGGAGTTTATCTAACAAATCCACAATAAATTCATATATCCCTGACAGTCATATCCTCAATTCATCTATGCAGTAACCTTGTAATTCATTTAATAGTTCTCCTTTCAATGTTG

>M6-PGSC0003DMG400002526M6LysM-RLK_2867

CCTCAATTCATCTATGCAGTAACCTTGTAATTCATTTAATAGTTCTCCTTTCAATGTTGGTTTGTTGTTGCTACTGCTCCTTTCCCAGTGAGTTTCATACCTGATTTGATATGCTTTACT

>M6-PGSC0003DMG400002526M6LysM-RLK_305

CTGTCAGCAATTTTCAAGATTATCCCTCTTTGTTCATCATACCCTACTCCATCAAACTGTACGGACACTGGTCGCTTGTGCACTTCCTTCTTGGCTTTTAAACCCAGCCCAGAACAGACC

>M6-PGSC0003DMG400002526M6LysM-RLK_3111

GTTGTCCTCCCAATGTTGGTTTGCTAAGTTTTTTCATCCTATGGTAGTTCTCTTAATTAAAGAAAGATAATCCTCAGAAGTATTTCAGGACTCCATTTCTTAATGGAATTTTTCCCTGCA

>M6-PGSC0003DMG400002526M6LysM-RLK_3172

GAAAGATAATCCTCAGAAGTATTTCAGGACTCCATTTCTTAATGGAATTTTTCCCTGCATTAATGTGTTTTAGATAGCTACCTTTCGGTTGAGGAGTAAAAAACCTCCAATTTTCGACTA

>M6-PGSC0003DMG400002526M6LysM-RLK_3233

AATGTGTTTTAGATAGCTACCTTTCGGTTGAGGAGTAAAAAACCTCCAATTTTCGACTATATCTGCAGGAGTTGAGGAGAATCATAAGAGTCATGCTATTTACTGGTGGATAATTGGGGG

>M6-PGSC0003DMG400002526M6LysM-RLK_3294

TCTGCAGGAGTTGAGGAGAATCATAAGAGTCATGCTATTTACTGGTGGATAATTGGGGGTTTGGGTGCTGGTCTTCTCATGATTGTCGTCATTCTTGCTTTTGTTACCTGGAGGTCGTCA

>M6-PGSC0003DMG400002526M6LysM-RLK_3355

TGGGTGCTGGTCTTCTCATGATTGTCGTCATTCTTGCTTTTGTTACCTGGAGGTCGTCAAGTTGTTTTTCCAGAACTGAGAGAAGTCACACAGCAGGTTCAAATGAAAAGATTTCTCATA

>M6-PGSC0003DMG400002526M6LysM-RLK_3416

TTGTTTTTCCAGAACTGAGAGAAGTCACACAGCAGGTTCAAATGAAAAGATTTCTCATAAGTTCCAGATACTTCGAAATACAAGTTTCTGTTGTGCATCAGGAAGGTACATTTGCGGTAG

>M6-PGSC0003DMG400002526M6LysM-RLK_3477

TTCCAGATACTTCGAAATACAAGTTTCTGTTGTGCATCAGGAAGGTACATTTGCGGTAGCTCAGGGGACCAGCAAGAACCGAAAGGAGAATCTACTGACCAACAGATTAATATTCCAAAA

>M6-PGSC0003DMG400002526M6LysM-RLK_3538

CAGGGGACCAGCAAGAACCGAAAGGAGAATCTACTGACCAACAGATTAATATTCCAAAAGGTCAGAACATAGACATGCATGCTTTTTATTGATATTACTGAAGGTCCTTCCTGTTAATTT

>M6-PGSC0003DMG400002526M6LysM-RLK_3599

TCAGAACATAGACATGCATGCTTTTTATTGATATTACTGAAGGTCCTTCCTGTTAATTTCCTCCAAACTATTATGCATGAGATCAAGTCATTTGCTTATATGTATTAAAATATGCGTGTG

>M6-PGSC0003DMG400002526M6LysM-RLK_366

CGGACACTGGTCGCTTGTGCACTTCCTTCTTGGCTTTTAAACCCAGCCCAGAACAGACCCTCCCAGTGATTCAAAGCATGTTTGATGTGTTACCTAATGACATAACTGTTGAAGGAAATG

>M6-PGSC0003DMG400002526M6LysM-RLK_3660

TCCAAACTATTATGCATGAGATCAAGTCATTTGCTTATATGTATTAAAATATGCGTGTGATCTAAAATCCATCAAAGCAAATCATAAGCCACACCTGTTATAGACTTATTGAACTCCTGC

>M6-PGSC0003DMG400002526M6LysM-RLK_3721

CTAAAATCCATCAAAGCAAATCATAAGCCACACCTGTTATAGACTTATTGAACTCCTGCTAGAGAGGGGATGGGAAGTTAGGAAAAGCAAAAGGAGTTTGGTTCTATAATCATTATGTAA

>M6-PGSC0003DMG400002526M6LysM-RLK_3782

GAGAGGGGATGGGAAGTTAGGAAAAGCAAAAGGAGTTTGGTTCTATAATCATTATGTAAAAATATCCTTGAATGGAGTTTCAGATGGCTACATAAAACTAAAACTAAAACTAAGTTATTC

>M6-PGSC0003DMG400002526M6LysM-RLK_3843

ATATCCTTGAATGGAGTTTCAGATGGCTACATAAAACTAAAACTAAAACTAAGTTATTCTATAATACTTCTGAAAATGACAGGCATGGTACTGATCAAGTTATCATAGTCAAATAGAAGC

>M6-PGSC0003DMG400002526M6LysM-RLK_3904

TAATACTTCTGAAAATGACAGGCATGGTACTGATCAAGTTATCATAGTCAAATAGAAGCAAATCAGCCAAGATGTCAAATAAGCTCTCAAAATTAAATAATACTTCTTTTACCCATCTTA

>M6-PGSC0003DMG400002526M6LysM-RLK_3965

ATCAGCCAAGATGTCAAATAAGCTCTCAAAATTAAATAATACTTCTTTTACCCATCTTAAGGTATACATGTCATAAGCAAACTGTTATGTGAAGGTTACACATCAGGAACATCCCACTAA

>M6-PGSC0003DMG400002526M6LysM-RLK_4026

GTATACATGTCATAAGCAAACTGTTATGTGAAGGTTACACATCAGGAACATCCCACTAAATTTCAAGTTGTGCCCTTGAACTTGAGTTTCACGAGAATGGATGTCTAAACAGTAAACTTG

>M6-PGSC0003DMG400002526M6LysM-RLK_4087

TTCAAGTTGTGCCCTTGAACTTGAGTTTCACGAGAATGGATGTCTAAACAGTAAACTTGATGTAGTTATAGCCGTCCTATGTTGCTCAGAATCTCCAAAATGTTGCAGCACCAGTGTTGA

>M6-PGSC0003DMG400002526M6LysM-RLK_4209

CCTCAAAAAACAAACTACTTTTGAAGGATGTGACACGCACCTATCGACATTAATGAAGAGTCCGAACAAGATAATTAGCCGGGATTAGTTGAAATTGCCAAGCCTATGCTGGCATTAGTC

>M6-PGSC0003DMG400002526M6LysM-RLK_427

CCCAGTGATTCAAAGCATGTTTGATGTGTTACCTAATGACATAACTGTTGAAGGAAATGGTAAAGGTTATGTCTTTATCAGAAAGAACTGTTCTTGTGCTTATGGAATGAGAAAGTACTT

>M6-PGSC0003DMG400002526M6LysM-RLK_4270

CCGAACAAGATAATTAGCCGGGATTAGTTGAAATTGCCAAGCCTATGCTGGCATTAGTCCTTTACTCGCTTCTAATGGTGTGCCACAATGGTAAAAGGAGTATTTGAAAATGTTCATAGA

>M6-PGSC0003DMG400002526M6LysM-RLK_4331

TTACTCGCTTCTAATGGTGTGCCACAATGGTAAAAGGAGTATTTGAAAATGTTCATAGAACTTTTTCTCCTGTGCTCATCATGGGGGAAAATGTAGTTTCACTATCATGCAGGATTTCTC

>M6-PGSC0003DMG400002526M6LysM-RLK_4392

TTTTTCTCCTGTGCTCATCATGGGGGAAAATGTAGTTTCACTATCATGCAGGATTTCTCAATATAATAATTGACTATTCTTTTTTTTTAAAAAAAAAAAAAACAAACAAAAACTGACTAT

>M6-PGSC0003DMG400002526M6LysM-RLK_4453

TATAATAATTGACTATTCTTTTTTTTTAAAAAAAAAAAAAACAAACAAAAACTGACTATTTTGGGCTTCCTTGCAGTTGTAGGTACTGATGTCTTTGACATGGAAAAACCTCTTGTTTTC

>M6-PGSC0003DMG400002526M6LysM-RLK_4514

TGGGCTTCCTTGCAGTTGTAGGTACTGATGTCTTTGACATGGAAAAACCTCTTGTTTTCGCCTATGAAGACATTCTATCTTCTACCGATGGATTTTCTGACTCCAATCTTCTTGGTCATG

>M6-PGSC0003DMG400002526M6LysM-RLK_4575

CTATGAAGACATTCTATCTTCTACCGATGGATTTTCTGACTCCAATCTTCTTGGTCATGGAACCTATGGTTCTGTCTATTACGCCATCCTCCGCAACCAGGTTACTTCTCTCAACTTTAC

>M6-PGSC0003DMG400002526M6LysM-RLK_4636

ACCTATGGTTCTGTCTATTACGCCATCCTCCGCAACCAGGTTACTTCTCTCAACTTTACCTATATTGATAGATTTCCTGTCAACATGGCGTATTAACTTGCTTATGTCTTTGGAACGTTA

>M6-PGSC0003DMG400002526M6LysM-RLK_4697

ATATTGATAGATTTCCTGTCAACATGGCGTATTAACTTGCTTATGTCTTTGGAACGTTAACCTTTCAGGAAGTTGCAATCAAAAGAATGACTGCTACTAAAACAAAAGAATTTACTGCTG

>M6-PGSC0003DMG400002526M6LysM-RLK_4758

CTTTCAGGAAGTTGCAATCAAAAGAATGACTGCTACTAAAACAAAAGAATTTACTGCTGAGATGAAAGTTCTATGTAAAGTTCATCATTTGAATTTGGTAATGAATACTGAATTTCGCTG

>M6-PGSC0003DMG400002526M6LysM-RLK_4819

ATGAAAGTTCTATGTAAAGTTCATCATTTGAATTTGGTAATGAATACTGAATTTCGCTGCAGTATCCTGAGTTTTTATTTTTGGAGCTTGATGATAGTGCTACATCAATTCAAAGTTTTG

>M6-PGSC0003DMG400002526M6LysM-RLK_488

AAAGGTTATGTCTTTATCAGAAAGAACTGTTCTTGTGCTTATGGAATGAGAAAGTACTTGACAAAAACTACATATACTGTCAGAAAGAACAATGGATCTGTGTATAATATGGTTGTTGAT

>M6-PGSC0003DMG400002526M6LysM-RLK_4880

GTATCCTGAGTTTTTATTTTTGGAGCTTGATGATAGTGCTACATCAATTCAAAGTTTTGACTCCTGTCTTGGTATTATTTTGTCAAGGTAGAATTGATCGGCTATGCAGTCAGTAATGAT

>M6-PGSC0003DMG400002526M6LysM-RLK_4941

TCCTGTCTTGGTATTATTTTGTCAAGGTAGAATTGATCGGCTATGCAGTCAGTAATGATGAATTATTCCTCGTTTATGAATATGCCCAGAAGGGTTCTCTTAAGAGCCACTTGAATGATC

>M6-PGSC0003DMG400002526M6LysM-RLK_5002

ATTATTCCTCGTTTATGAATATGCCCAGAAGGGTTCTCTTAAGAGCCACTTGAATGATCCTCAGAACAAAGGTACACATCTTGAAGATAATCGTTAAGCTCAGAATTTGTGTAGTGTATA

>M6-PGSC0003DMG400002526M6LysM-RLK_5063

CAGAACAAAGGTACACATCTTGAAGATAATCGTTAAGCTCAGAATTTGTGTAGTGTATATATACATTTTTTCCAGTTGTTGATATGGAAATGACCTGTCACTGCTGTATTTTGGTGAAAA

>M6-PGSC0003DMG400002526M6LysM-RLK_5124

TACATTTTTTCCAGTTGTTGATATGGAAATGACCTGTCACTGCTGTATTTTGGTGAAAAAAACTAGATGATCTGACCTCTTATTTTGGGTAGGCCACACACCACTCTCTTGGATCATGAG

>M6-PGSC0003DMG400002526M6LysM-RLK_5185

ACTAGATGATCTGACCTCTTATTTTGGGTAGGCCACACACCACTCTCTTGGATCATGAGGGTCCAAATTGCACTTGATGCTGCTAGAGGCTTGGAATACATTCACGAGCACACCAAGCCG

>M6-PGSC0003DMG400002526M6LysM-RLK_5246

TCCAAATTGCACTTGATGCTGCTAGAGGCTTGGAATACATTCACGAGCACACCAAGCCGCATTATGTGCATCGAGATATAAAGACAAGTAACATCCTGCTTGATGATTCTTTTAGAGCAA

>M6-PGSC0003DMG400002526M6LysM-RLK_5307

TTATGTGCATCGAGATATAAAGACAAGTAACATCCTGCTTGATGATTCTTTTAGAGCAAAGGTTGTAGCTATTTTGGTGCCCAATTTTTCTCCTTTGTGGATGTCATTTTTCAAACCCAA

>M6-PGSC0003DMG400002526M6LysM-RLK_5368

GTTGTAGCTATTTTGGTGCCCAATTTTTCTCCTTTGTGGATGTCATTTTTCAAACCCAAATTCTCATTGCCTATGTTTGCAACAGATTTCAGATTTTGGTTTGTCAAAACTGATGGGGAT

>M6-PGSC0003DMG400002526M6LysM-RLK_5429

TCTCATTGCCTATGTTTGCAACAGATTTCAGATTTTGGTTTGTCAAAACTGATGGGGATAACTAATGATGCAGAAGCCTCTGCAACACGAGTTGTTGGAACCTATGGCTATGTAGCTCCA

>M6-PGSC0003DMG400002526M6LysM-RLK_549

CAAAAACTACATATACTGTCAGAAAGAACAATGGATCTGTGTATAATATGGTTGTTGATGCCTATGATGGGTTGGCTTATTTTCCAACTAATTTTACAAGGGAGGGGAAGAAAGGGGCAG

>M6-PGSC0003DMG400002526M6LysM-RLK_5490

CTAATGATGCAGAAGCCTCTGCAACACGAGTTGTTGGAACCTATGGCTATGTAGCTCCAGAGTAAGCTAACCAACAATAAGCAAGAACATAAACTGAAACATCTCTCTGATTTTGTTTGT

>M6-PGSC0003DMG400002526M6LysM-RLK_5551

GTAAGCTAACCAACAATAAGCAAGAACATAAACTGAAACATCTCTCTGATTTTGTTTGTTTCTCTTGTTCTTGCAGGTACTTGAGGGATGGCCTGGCAACTAAAAAGACTGATGTGTATG

>M6-PGSC0003DMG400002526M6LysM-RLK_5612

CTCTTGTTCTTGCAGGTACTTGAGGGATGGCCTGGCAACTAAAAAGACTGATGTGTATGCATTTGGTGTTGTTCTTTTTGAGATGCTGACAGGGAAAGAGGCCGTCACCCGGACTGAAGG

>M6-PGSC0003DMG400002526M6LysM-RLK_5673

TTTGGTGTTGTTCTTTTTGAGATGCTGACAGGGAAAGAGGCCGTCACCCGGACTGAAGGCAATGTGATGAAAACTGCAGAAAGACGCTCCTTAGTGTCCATTGTAAGTCTTATTGCTCAC

>M6-PGSC0003DMG400002526M6LysM-RLK_5734

ATGTGATGAAAACTGCAGAAAGACGCTCCTTAGTGTCCATTGTAAGTCTTATTGCTCACAAACTGTTCAAAATTGAAGGCATGCTTCCTAATGGATTCTATTTGCCTGTGTTTCCACTTC

>M6-PGSC0003DMG400002526M6LysM-RLK_5795

ACTGTTCAAAATTGAAGGCATGCTTCCTAATGGATTCTATTTGCCTGTGTTTCCACTTCTATCTGTTATGTGATTAAAAAAGATGAAAAACATCGATTTTATCATACAATTGAGTAAGAT

>M6-PGSC0003DMG400002526M6LysM-RLK_5856

TCTGTTATGTGATTAAAAAAGATGAAAAACATCGATTTTATCATACAATTGAGTAAGATCAAATTCACGAAAGCTCTGAAAAATGTTTTTGCATTCATCAGACTTGGTCAATGTATGACT

>M6-PGSC0003DMG400002526M6LysM-RLK_5917

AATTCACGAAAGCTCTGAAAAATGTTTTTGCATTCATCAGACTTGGTCAATGTATGACTAGATTCCATTGCCTATCTGGCATCTTTATCAAATTGTGAAGTTTACTGTTCTCAGGTTTAG

>M6-PGSC0003DMG400002526M6LysM-RLK_5978

ATTCCATTGCCTATCTGGCATCTTTATCAAATTGTGAAGTTTACTGTTCTCAGGTTTAGGATTAGTTCATCAACTTCCTAAATGCTTTACTTCAAAAAAAAATTGAAATTCCCTTGCCTA

>M6-PGSC0003DMG400002526M6LysM-RLK_6039

TTAGTTCATCAACTTCCTAAATGCTTTACTTCAAAAAAAAATTGAAATTCCCTTGCCTATTGTTATGTACCTACAGTGTTGTGCTGTGGCTATGTATAGGTATACTTATGCATAACATAT

>M6-PGSC0003DMG400002526M6LysM-RLK_61

TAAAAGACCTTGTTTTCAGCCATTACAGGACAACTAAACCCCCCATCCCCAAACCCCACAAAATGTTCTGTTCTTGGTTCTCTGCTATTCCTATTTCACTTTCTTTCTCTTAAGATTCAA

>M6-PGSC0003DMG400002526M6LysM-RLK_610

CTATGATGGGTTGGCTTATTTTCCAACTAATTTTACAAGGGAGGGGAAGAAAGGGGCAGTGGTTTCACTGAAGCTCATATGTGGATGCTCAAGTGGGCTGTGGAATTACCTCATGAGTTA

>M6-PGSC0003DMG400002526M6LysM-RLK_6161

ATATATATATTTACACACTTATATGCACACTATGAATCAATTTTTAGCAGTGCTTAATGCTACATAATTGCAGATGCTGGCAGCTCTGAGGAACTCACCAGACTCCACTAGTATGACAAG

>M6-PGSC0003DMG400002526M6LysM-RLK_6222

ACATAATTGCAGATGCTGGCAGCTCTGAGGAACTCACCAGACTCCACTAGTATGACAAGCTTGAAGGACCAGCTGGATCCCAGTTTGATGGATTTGTACCCTTCTGACTGCGTTTTCAAG

>M6-PGSC0003DMG400002526M6LysM-RLK_6283

TGAAGGACCAGCTGGATCCCAGTTTGATGGATTTGTACCCTTCTGACTGCGTTTTCAAGGTAAGTTGCAGCTTAACAATTATGTTTGTCAACTTTGAATCAGTTCAAGTTGTGCCTCATC

>M6-PGSC0003DMG400002526M6LysM-RLK_6344

AAGTTGCAGCTTAACAATTATGTTTGTCAACTTTGAATCAGTTCAAGTTGTGCCTCATCTATAATAACGTGCATAGAATTGGTAGGGCCACTGACTACCTCAGCTGGTTTAGTTAAAATA

>M6-PGSC0003DMG400002526M6LysM-RLK_6405

TAATAACGTGCATAGAATTGGTAGGGCCACTGACTACCTCAGCTGGTTTAGTTAAAATAATAGACAGAAGAGATAGTTTAAGATTGGAAGTTCATATATAGTTATTATCAGCTGAGTGAC

>M6-PGSC0003DMG400002526M6LysM-RLK_6466

AGACAGAAGAGATAGTTTAAGATTGGAAGTTCATATATAGTTATTATCAGCTGAGTGACATAGGTGGTTGATTTTCTAATGAAATTTCACGACTTATTTCATTCTGTCCACCATCATTGA

>M6-PGSC0003DMG400002526M6LysM-RLK_6527

AGGTGGTTGATTTTCTAATGAAATTTCACGACTTATTTCATTCTGTCCACCATCATTGATGTGTTTACTATGGTAGGTGGCTATGCTGGCTAAGCAATGTGCAGATGATGATCCAATATT

>M6-PGSC0003DMG400002526M6LysM-RLK_6588

TGTTTACTATGGTAGGTGGCTATGCTGGCTAAGCAATGTGCAGATGATGATCCAATATTGAGACCGGACATGAAGCACGTAGTGATCACACTTTCACAGATAATGTTATCCTCTGTGGAG

>M6-PGSC0003DMG400002526M6LysM-RLK_6649

GACCGGACATGAAGCACGTAGTGATCACACTTTCACAGATAATGTTATCCTCTGTGGAGTGGGAAGCAACTCTTGCTGGGAACAGTCAAGTGTTCAGTGGCATTGTGCAGGGAAGATAGG

>M6-PGSC0003DMG400002526M6LysM-RLK_671

GTTTCACTGAAGCTCATATGTGGATGCTCAAGTGGGCTGTGGAATTACCTCATGAGTTATGTGATGACAGATGATGATACTGTTGGGTCTTTGTCCAGTAGGTTTGGGGTTAGTATGGAT

>M6-PGSC0003DMG400002526M6LysM-RLK_6710

GGAAGCAACTCTTGCTGGGAACAGTCAAGTGTTCAGTGGCATTGTGCAGGGAAGATAGGTCCATATGCTAGGTGTTGCGTTGCATCCTCATATATAATGTACATGGTACTCTTGTATTTT

>M6-PGSC0003DMG400002526M6LysM-RLK_6771

CATATGCTAGGTGTTGCGTTGCATCCTCATATATAATGTACATGGTACTCTTGTATTTTTGAAGCCAATCTTCTCCATTTTACCAGTCCAGTCACATTTTTCCTATCATTCTTTATAATG

>M6-PGSC0003DMG400002526M6LysM-RLK_6832

AAGCCAATCTTCTCCATTTTACCAGTCCAGTCACATTTTTCCTATCATTCTTTATAATGTTATCTTTTTTATGCACAGAGTAATCTTCATATAGCGGATTTTTTCCTCTTTTTTCTTTTT

>M6-PGSC0003DMG400002526M6LysM-RLK_732

TGATGACAGATGATGATACTGTTGGGTCTTTGTCCAGTAGGTTTGGGGTTAGTATGGATAATATTGAGAATGTTAATGGAATTGCCAATCCTGATAATTTCACAGCAGGATCTTTATACT

>M6-PGSC0003DMG400002526M6LysM-RLK_793

TATTGAGAATGTTAATGGAATTGCCAATCCTGATAATTTCACAGCAGGATCTTTATACTATGTGCCATTAAATTCAGGCATGAGTTTCCTTTCCCCTCTTATGATTTCTGATTTGTGTTT

>M6-PGSC0003DMG400002526M6LysM-RLK_854

GTGCCATTAAATTCAGGCATGAGTTTCCTTTCCCCTCTTATGATTTCTGATTTGTGTTTTTTCCAATTCCAAATGTGTATCAGTATTTTGATTTAGGATGATTTAGGAGCTGTTTGATTA

>M6-PGSC0003DMG400002526M6LysM-RLK_915

TCCAATTCCAAATGTGTATCAGTATTTTGATTTAGGATGATTTAGGAGCTGTTTGATTAATCAAATTGTGTTAGTTGTAGTAAGTGTGATTAAATGTTGGAATTTTTAGTTGCTGAGGAA

>M6-PGSC0003DMG400002526M6LysM-RLK_976

CAAATTGTGTTAGTTGTAGTAAGTGTGATTAAATGTTGGAATTTTTAGTTGCTGAGGAAGATTAGTCAGTTTTTTCTTTCTCTTTTTGTTTGTGTGAGTGAGGGGATGAACCTATGTTCT

>M6-PGSC0003DMG400002602-(rev)M6RLKmalectin_0

AAATTAGTGGACCCAAATCATGTATTTTTGGGTCGAACAAATTTTTAATTCACCAATTTGTGAAACAAATGAAGTCTCGAGGCATAAAATAAAATTATCACTTGTATATTAAGTATAATT

>M6-PGSC0003DMG400002602-(rev)M6RLKmalectin_1054

ACTACTACTTCTTTGGTTTATAATTCGAAGGAAGAAAAACAAAAAAGGTGAGCAAATCATAAGATCGATCGATGAATTGCCAGATGATTTGTCATTGAAGAAATTAATTTCTGTAACAGG

>M6-PGSC0003DMG400002602-(rev)M6RLKmalectin_1116

GATCGATCGATGAATTGCCAGATGATTTGTCATTGAAGAAATTAATTTCTGTAACAGGGAAGATTACTGGGACAGAAAGAACAGTAGAGAGTCCAAACATTGCATTGGAGTTGAAAAACC

>M6-PGSC0003DMG400002602-(rev)M6RLKmalectin_1178

ATTACTGGGACAGAAAGAACAGTAGAGAGTCCAAACATTGCATTGGAGTTGAAAAACCGGCAGTTCACATACTCGCAGGTTTTGCACATGACAAATAACTTCCAGAGAGTCCTTGGAAAA

>M6-PGSC0003DMG400002602-(rev)M6RLKmalectin_124

TGACATTCTTGTGTTTCTGTTAAAGGAACTTGTCAAGCAATCAGCTGACGGGAGAGGTGCCTGCCAATCTCTCACAACTAGCCTTCTTACAAGAGCTGTAATTTACTCTCCTATAACTCA

>M6-PGSC0003DMG400002602-(rev)M6RLKmalectin_1240

GTTCACATACTCGCAGGTTTTGCACATGACAAATAACTTCCAGAGAGTCCTTGGAAAAGGAGGTTTTGGGACAGTTTATCTTGGCTATGTCGATAACCGCGATGTAGCTGTTAAGATGCT

>M6-PGSC0003DMG400002602-(rev)M6RLKmalectin_1302

GTTTTGGGACAGTTTATCTTGGCTATGTCGATAACCGCGATGTAGCTGTTAAGATGCTGTCCCCTTCTTCAGTTCAAGGGTTCAAGGAATTCCGGGCTGAGGTAAACTGAAGAAGTATTT

>M6-PGSC0003DMG400002602-(rev)M6RLKmalectin_1364

CCTTCTTCAGTTCAAGGGTTCAAGGAATTCCGGGCTGAGGTAAACTGAAGAAGTATTTTATAGTACATAAGACATTATATTTAGTTAACACGAATTTGACTTGACAGGCTAGTCTCCTGA

>M6-PGSC0003DMG400002602-(rev)M6RLKmalectin_1426

GTACATAAGACATTATATTTAGTTAACACGAATTTGACTTGACAGGCTAGTCTCCTGATGAGCATACATCATAAGAACTTGATTTCTATAGTTGGGTATTGTGTTGAAGGCACCCACATT

>M6-PGSC0003DMG400002602-(rev)M6RLKmalectin_1488

CATACATCATAAGAACTTGATTTCTATAGTTGGGTATTGTGTTGAAGGCACCCACATTGGTATTATTTACGAATACATGGCCAATCGAAGCTTAGACATGCAACTATCAGGTACCTGATT

>M6-PGSC0003DMG400002602-(rev)M6RLKmalectin_1550

TTATTTACGAATACATGGCCAATCGAAGCTTAGACATGCAACTATCAGGTACCTGATTTATAGTCAAACAAATGTTCTTTTAGATCACACATTTCACAAGTCTTTCATCGTTGCGTGCAG

>M6-PGSC0003DMG400002602-(rev)M6RLKmalectin_1612

GTCAAACAAATGTTCTTTTAGATCACACATTTCACAAGTCTTTCATCGTTGCGTGCAGATAGAAATCCAAATGCTTTAACATGGGAGGAAAGACTTCATATAGCATTGGATGCGGCACAA

>M6-PGSC0003DMG400002602-(rev)M6RLKmalectin_1674

AAATCCAAATGCTTTAACATGGGAGGAAAGACTTCATATAGCATTGGATGCGGCACAAGGTATATATATACACCACTAGATGAAAATAATTTCAGTCGAGTTTTAGAATGATTAATTATA

>M6-PGSC0003DMG400002602-(rev)M6RLKmalectin_1736

TATATATACACCACTAGATGAAAATAATTTCAGTCGAGTTTTAGAATGATTAATTATATATTTGACGTGTTTCTTTTAATTTTAACTGAAAAAATAATGAAGGATTGGAGTATCTACATC

>M6-PGSC0003DMG400002602-(rev)M6RLKmalectin_1798

TGACGTGTTTCTTTTAATTTTAACTGAAAAAATAATGAAGGATTGGAGTATCTACATCATGGTTGCCAGCCACCAATAATACACAGAGATATCAAGTCTAGTAATATTCTATTAGATGAT

>M6-PGSC0003DMG400002602-(rev)M6RLKmalectin_186

GCCAATCTCTCACAACTAGCCTTCTTACAAGAGCTGTAATTTACTCTCCTATAACTCATTTAGTCTCAATTTATGCGGCACAGTTGCAATTAATATGATTTTTATGACATTATTGTTGTG

>M6-PGSC0003DMG400002602-(rev)M6RLKmalectin_1860

TTGCCAGCCACCAATAATACACAGAGATATCAAGTCTAGTAATATTCTATTAGATGATAAATTCCAAGCAAATCTTGCTGATTTTGGTTTGTCAAGGACTTTACCAACTGGAGAGGGAAG

>M6-PGSC0003DMG400002602-(rev)M6RLKmalectin_1922

TCCAAGCAAATCTTGCTGATTTTGGTTTGTCAAGGACTTTACCAACTGGAGAGGGAAGTCATGTTACCACTATAATCGCGGGTAGTCCAGGCTATCTTGACCCTGAGTACGTGCTCTTCA

>M6-PGSC0003DMG400002602-(rev)M6RLKmalectin_1984

GTTACCACTATAATCGCGGGTAGTCCAGGCTATCTTGACCCTGAGTACGTGCTCTTCATTTTATTAGTCTGTCTCAAAAAGAATAACAACTATGTATATCTAATTTTAACTCTCGATTGA

>M6-PGSC0003DMG400002602-(rev)M6RLKmalectin_2046

ATTAGTCTGTCTCAAAAAGAATAACAACTATGTATATCTAATTTTAACTCTCGATTGACCTGCAGTTACTACAGGACAAATAAGCTAACGGAGAAAAGTGATGTATACAGTTTCGGTGTG

>M6-PGSC0003DMG400002602-(rev)M6RLKmalectin_2108

CAGTTACTACAGGACAAATAAGCTAACGGAGAAAAGTGATGTATACAGTTTCGGTGTGGTTCTTTTGGAAATAATAACAGGACGGCATTTACTAGGAAAACATGACAAGATTTATGTAAT

>M6-PGSC0003DMG400002602-(rev)M6RLKmalectin_2170

TTTTGGAAATAATAACAGGACGGCATTTACTAGGAAAACATGACAAGATTTATGTAATAACGTGGGTCAATGGTATGATTAATGATAACGGAGATGTTGCAAAAGTAATTGATCCAAGAT

>M6-PGSC0003DMG400002602-(rev)M6RLKmalectin_2232

TGGGTCAATGGTATGATTAATGATAACGGAGATGTTGCAAAAGTAATTGATCCAAGATTAGGAGGAGAAGTTGATGTGAATTCAGCTAAGAGAATTGTGGCATTAGCAATGGCTTGTGTT

>M6-PGSC0003DMG400002602-(rev)M6RLKmalectin_2294

AGGAGAAGTTGATGTGAATTCAGCTAAGAGAATTGTGGCATTAGCAATGGCTTGTGTTTCTCTTGAACCAACAAATAGACCAGCTATATCTGTCGTTGTCACAATAATAAAGCAATGTTT

>M6-PGSC0003DMG400002602-(rev)M6RLKmalectin_2328

GTGGCATTAGCAATGGCTTGTGTTTCTCTTGAACCAACAAATAGACCAGCTATATCTGTCGTTGTCACAATAATAAAGCAATGTTTAAGGCAGATGATTGATAATTATGACTCTAATTGA

>M6-PGSC0003DMG400002602-(rev)M6RLKmalectin_248

GTCTCAATTTATGCGGCACAGTTGCAATTAATATGATTTTTATGACATTATTGTTGTGTTTCTGTTGAAGGGACTTGTCAAACAATCAGCTGACAGGAAAGGTCCCTGCCAATCTCGCAC

>M6-PGSC0003DMG400002602-(rev)M6RLKmalectin_310

TGTTGAAGGGACTTGTCAAACAATCAGCTGACAGGAAAGGTCCCTGCCAATCTCGCACAACTACCCTTCTTAAAAAAGCTGTAATTTACGCACTCATGACCTATAAGTTTACAATTACTA

>M6-PGSC0003DMG400002602-(rev)M6RLKmalectin_372

ACCCTTCTTAAAAAAGCTGTAATTTACGCACTCATGACCTATAAGTTTACAATTACTACCTCTCTATCAATTTATGCGGCACAATTCCAAATATGATTTTAAAATACTTTAAGTTGTTAA

>M6-PGSC0003DMG400002602-(rev)M6RLKmalectin_434

TCTATCAATTTATGCGGCACAATTCCAAATATGATTTTAAAATACTTTAAGTTGTTAATTATTTTGATATATAGTAATTTTATGTTCAATTCATTATCAAAGTTTAGAATTTTGAATTGT

>M6-PGSC0003DMG400002602-(rev)M6RLKmalectin_496

TTTGATATATAGTAATTTTATGTTCAATTCATTATCAAAGTTTAGAATTTTGAATTGTGTCACATGAATTAGAACGAAGGGGTATTTATTTAATTAGAACACAGAGTAATTTGGTTATCT

>M6-PGSC0003DMG400002602-(rev)M6RLKmalectin_558

CATGAATTAGAACGAAGGGGTATTTATTTAATTAGAACACAGAGTAATTTGGTTATCTTTGTGACAGATACTTGAATGGAAACAGTTTCTCCGAAAAAATTCCTGAAGAATTGCTGGCAA

>M6-PGSC0003DMG400002602-(rev)M6RLKmalectin_62

AAACAAATGAAGTCTCGAGGCATAAAATAAAATTATCACTTGTATATTAAGTATAATTTTTATGACATTCTTGTGTTTCTGTTAAAGGAACTTGTCAAGCAATCAGCTGACGGGAGAGGT

>M6-PGSC0003DMG400002602-(rev)M6RLKmalectin_620

GACAGATACTTGAATGGAAACAGTTTCTCCGAAAAAATTCCTGAAGAATTGCTGGCAAAATCAAAAAATGAATCACTCGATTTGAGGTTTCGTTCCACTCTCAGAATCCATGAATTAATT

>M6-PGSC0003DMG400002602-(rev)M6RLKmalectin_682

AAAAAATGAATCACTCGATTTGAGGTTTCGTTCCACTCTCAGAATCCATGAATTAATTTATGAGTATACAAAAAACATCTAAGATTGATTTAGAAAATTATCTCCCCTATGATCGATATT

>M6-PGSC0003DMG400002602-(rev)M6RLKmalectin_744

AGTATACAAAAAACATCTAAGATTGATTTAGAAAATTATCTCCCCTATGATCGATATTAATTTCTCATATAATTGGTCACAGGTTAATTTGTGTAATCAACTCATCTAAAATTAAAGTCT

>M6-PGSC0003DMG400002602-(rev)M6RLKmalectin_806

TCTCATATAATTGGTCACAGGTTAATTTGTGTAATCAACTCATCTAAAATTAAAGTCTGATGTTATATTGAATTGTTATCAAGTCGTTAGATATTGAAGACACTTTTATTTACTTATAAT

>M6-PGSC0003DMG400002602-(rev)M6RLKmalectin_868

TTATATTGAATTGTTATCAAGTCGTTAGATATTGAAGACACTTTTATTTACTTATAATTACATTTCAATTAATTTTTTGTCATACAGGTATGATGAATTTCATCCTCCGAAAGGTAGTAA

>M6-PGSC0003DMG400002602-(rev)M6RLKmalectin_930

TTTCAATTAATTTTTTGTCATACAGGTATGATGAATTTCATCCTCCGAAAGGTAGTAATAACCTTAGTGCTGGTGCACTAGCAGGAATTATTGCATCCGTGGTGGTCTTGGGCATCTTGG

>M6-PGSC0003DMG400002602-(rev)M6RLKmalectin_992

CTTAGTGCTGGTGCACTAGCAGGAATTATTGCATCCGTGGTGGTCTTGGGCATCTTGGCTTTACTACTACTTCTTTGGTTTATAATTCGAAGGAAGAAAAACAAAAAAGGTGAGCAAATC

>M6-PGSC0003DMG400002618-(rev)M6LRR-RLP_0

TAAAGTAAAATAATGGGGAAAGGAGCAATTTTGAAATTGAAAAAAAAAAAAGAAAAAACAAAAGTGATCTCTTATTCCGGAAAAGCAAGGTGGGAAATCTTTACAAGACTCGTCTCTTTT

>M6-PGSC0003DMG400002618-(rev)M6LRR-RLP_1037

AGGTGTTGTATACATGGCACTTACCTGAGCCAAATGCATCACGGAATTCTGATTCTTCACCACTAAAGTCTACCACTCTGGTAGTATTGTTGGGATGGTTAGGTGCAAAGCAAAAGCATC

>M6-PGSC0003DMG400002618-(rev)M6LRR-RLP_1098

ACTAAAGTCTACCACTCTGGTAGTATTGTTGGGATGGTTAGGTGCAAAGCAAAAGCATCTAAAGAGATATGCAGAGTGGTATGCATCAAGAGGATATCATGTCATTACATTTACTTTCCC

>M6-PGSC0003DMG400002618-(rev)M6LRR-RLP_122

CTTATCAGAATCAGAAGAAGAAAGAAGAAGATTCGTCTCTCAGGTATGGTGTTCCTCTTACAATCAATCTATAATGCTTATGTTTGTTTATTTGAGAGGCCGATCCGCCCACCCTCATGA

>M6-PGSC0003DMG400002618-(rev)M6LRR-RLP_1220

TGTCTGAGATTCTAAGCTATCAAGTCGGGGGAAAGGCAGAGCAAGATATTGAAATGCTTGTGAATCATCTAGCTGATTGGTTGGAAGAAGAGCATGGAAAGAACTTGGTCTTCCACACTT

>M6-PGSC0003DMG400002618-(rev)M6LRR-RLP_1281

GAATCATCTAGCTGATTGGTTGGAAGAAGAGCATGGAAAGAACTTGGTCTTCCACACTTTCAGCAATACTGGATGGTTAACGTGAGTTCTTGATGATACTCATTAGTCTAAGATGAGAAC

>M6-PGSC0003DMG400002618-(rev)M6LRR-RLP_1342

AGCAATACTGGATGGTTAACGTGAGTTCTTGATGATACTCATTAGTCTAAGATGAGAACAGAAACATAACTATTTACATTTTCCTTAATACTAGCTTGATAATTATTGATTGACTTCTTT

>M6-PGSC0003DMG400002618-(rev)M6LRR-RLP_1403

AAACATAACTATTTACATTTTCCTTAATACTAGCTTGATAATTATTGATTGACTTCTTTTATTCTTTTGGCCTTACAGTTATGGTGTCATTTTGGAAAAGTTTCAGAAACAAGATCATGC

>M6-PGSC0003DMG400002618-(rev)M6LRR-RLP_1525

TAATGAGAAGGATTAAGGGTTGTATTGTTGATTCTGCCCCTGTCGCAGCTCCTGATCCACAGGTATTGCTGACTCAAAAATTGTGATGTTAAGATATCATTTTTATGTGCAAGTGTCAGA

>M6-PGSC0003DMG400002618-(rev)M6LRR-RLP_1586

GGTATTGCTGACTCAAAAATTGTGATGTTAAGATATCATTTTTATGTGCAAGTGTCAGAAGAGTGTTTCTTGGAAAGTAGTCTTATTTATGAAAGTTAATGCAGAGCACTTGTTTAGAGC

>M6-PGSC0003DMG400002618-(rev)M6LRR-RLP_1647

AGTGTTTCTTGGAAAGTAGTCTTATTTATGAAAGTTAATGCAGAGCACTTGTTTAGAGCTATACATTTCTATTGAAATCCAACCTTTTTCATGCTTTTATTGGACAACTATTACAAGTGT

>M6-PGSC0003DMG400002618-(rev)M6LRR-RLP_1708

TACATTTCTATTGAAATCCAACCTTTTTCATGCTTTTATTGGACAACTATTACAAGTGTCGTATATAGTCAGGTTTTAGCCATTCAGAATTGCACAGTTGTTTGGATTCTAATCAGGACC

>M6-PGSC0003DMG400002618-(rev)M6LRR-RLP_1769

TATATAGTCAGGTTTTAGCCATTCAGAATTGCACAGTTGTTTGGATTCTAATCAGGACCTGCTGTTCTCGTATTTTTATCCTGTTGCTTTTTTTTATCTTAGGGCCTTAACCATGCAGCC

>M6-PGSC0003DMG400002618-(rev)M6LRR-RLP_183

AATCAATCTATAATGCTTATGTTTGTTTATTTGAGAGGCCGATCCGCCCACCCTCATGATGGATGGGTCAAACCTGTTTTTTTTTTTTTCCATAAAAACAAACCAGATCTTTTCCGTTTA

>M6-PGSC0003DMG400002618-(rev)M6LRR-RLP_1830

CTGTTCTCGTATTTTTATCCTGTTGCTTTTTTTTATCTTAGGGCCTTAACCATGCAGCCTGCTTTTCGAAAACTATTCTTATGCTTGATAACTTGAAAGAATCTTGAGAGTTTAGCTCTC

>M6-PGSC0003DMG400002618-(rev)M6LRR-RLP_1891

CTTTTCGAAAACTATTCTTATGCTTGATAACTTGAAAGAATCTTGAGAGTTTAGCTCTCTCTGACAACTTTTTAACAGGAAGCATTCTATGCAACTTTTGTTTCACGGTTCAAGCTTTAA

>M6-PGSC0003DMG400002618-(rev)M6LRR-RLP_1952

TGACAACTTTTTAACAGGAAGCATTCTATGCAACTTTTGTTTCACGGTTCAAGCTTTAAGCCTACTTTTTCTGGCTGAGAATAAGCCATCGAGGAATTTCCCCTTGGAGCTACTAAACTG

>M6-PGSC0003DMG400002618-(rev)M6LRR-RLP_2074

CACGTCCCTGCGACTACTGGATCTCTCTAGTAACAACTTTGGAGGAATGCTTCCATGAGGCATCGACAGGCTAGAAAGTCTCACATCCTTTTCAACAATAGCAACTTTACTGGAAATATA

>M6-PGSC0003DMG400002618-(rev)M6LRR-RLP_2135

ATCGACAGGCTAGAAAGTCTCACATCCTTTTCAACAATAGCAACTTTACTGGAAATATACTGCCTCAAACAGGAAACTTGACGAACTTGGAAGATTTGTACCTAATCCATAACATGCTCT

>M6-PGSC0003DMG400002618-(rev)M6LRR-RLP_2196

GCCTCAAACAGGAAACTTGACGAACTTGGAAGATTTGTACCTAATCCATAACATGCTCTCCGGAAGAATTCCAGTGGAAATAGGAAAGCTTCAGAGGCTGTGTGAACTACCTCTATGGAA

>M6-PGSC0003DMG400002618-(rev)M6LRR-RLP_2257

GGAAGAATTCCAGTGGAAATAGGAAAGCTTCAGAGGCTGTGTGAACTACCTCTATGGAAACCAGTTGTCAGTAACTACCCCAAGAGAATTGACAAATAGCTTAGTACTCATTGATTTCTT

>M6-PGSC0003DMG400002618-(rev)M6LRR-RLP_2318

CAGTTGTCAGTAACTACCCCAAGAGAATTGACAAATAGCTTAGTACTCATTGATTTCTTCGCTAATCAATTTTCAGGCTCTATTTCTGATGCCATTGGTAGGCTTAAGAATCTTGTTATC

>M6-PGSC0003DMG400002618-(rev)M6LRR-RLP_2379

CTAATCAATTTTCAGGCTCTATTTCTGATGCCATTGGTAGGCTTAAGAATCTTGTTATCTTCAGCTAACTCAGAATGAGTTGTCTGGTCCAGTCCCATCAAGCTTAGGCTACTGCAGGAA

>M6-PGSC0003DMG400002618-(rev)M6LRR-RLP_244

GATGGGTCAAACCTGTTTTTTTTTTTTTCCATAAAAACAAACCAGATCTTTTCCGTTTACCCTTTTTGAGTTTTGACCATGAAAATCAATTTTTGAGTTTTGACCCATTGATGAAAATCA

>M6-PGSC0003DMG400002618-(rev)M6LRR-RLP_2440

CAGCTAACTCAGAATGAGTTGTCTGGTCCAGTCCCATCAAGCTTAGGCTACTGCAGGAAGCATGAAAAACTGGCCTTTGCTGATAACAAACTTTCAGGATAATTGCCACCAACTTTAAGA

>M6-PGSC0003DMG400002618-(rev)M6LRR-RLP_2562

TCTATCAGAGCTGGACCTGACTACTCTTTACAACAATTGCTTTGAAGGTCCACTTCCTGAATATCTTTCCCTTCTGAAAAATCTCAGCAAAGTTAACTTTTCTCACAATAAGTTCAGTGG

>M6-PGSC0003DMG400002618-(rev)M6LRR-RLP_2623

TATCTTTCCCTTCTGAAAAATCTCAGCAAAGTTAACTTTTCTCACAATAAGTTCAGTGGAAACATTTCTCCGCTAGCTGGTTCAAATTCTCTGACAGTTGACATGACAAACTACAGCTTC

>M6-PGSC0003DMG400002618-(rev)M6LRR-RLP_2684

ACATTTCTCCGCTAGCTGGTTCAAATTCTCTGACAGTTGACATGACAAACTACAGCTTCTCTGGTCCTATCCCATCTGAGCTAGTGGTGTCCAAAAATCTGACCCATCTTCACCTTTCTA

>M6-PGSC0003DMG400002618-(rev)M6LRR-RLP_2745

TGGTCCTATCCCATCTGAGCTAGTGGTGTCCAAAAATCTGACCCATCTTCACCTTTCTAACAATTTTTTCACTGGAAAAACCCCTTCTGAATTTGGACAACTTGAAGATCTCAGATGTCT

>M6-PGSC0003DMG400002618-(rev)M6LRR-RLP_2806

AATTTTTTCACTGGAAAAACCCCTTCTGAATTTGGACAACTTGAAGATCTCAGATGTCTTGATTTATTGTTCAAAAATTTGACTGGAGATCTAGCACCTTCCCTTGCTGGCCTGAAAACC

>M6-PGSC0003DMG400002618-(rev)M6LRR-RLP_2867

ATTTATTGTTCAAAAATTTGACTGGAGATCTAGCACCTTCCCTTGCTGGCCTGAAAACCTTGGCCGTTTTCTTCTTATATCTAATCAATTTTCGGGGAAAATTTCCACATGGTTTGTGGA

>M6-PGSC0003DMG400002618-(rev)M6LRR-RLP_2928

GGCCGTTTTCTTCTTATATCTAATCAATTTTCGGGGAAAATTTCCACATGGTTTGTGGAGGAATGGAAGGTCTTGGTGAGTTCGATCTTTCATTCAACAACTTCACTGGAACAGTTCCAG

>M6-PGSC0003DMG400002618-(rev)M6LRR-RLP_2989

AATGGAAGGTCTTGGTGAGTTCGATCTTTCATTCAACAACTTCACTGGAACAGTTCCAGCAGAACTTGGAAACTCTCCAAAATAACTCAAAACTTTTTCTCAGTTACAACAGATTATTAG

>M6-PGSC0003DMG400002618-(rev)M6LRR-RLP_305

CTTTTTGAGTTTTGACCATGAAAATCAATTTTTGAGTTTTGACCCATTGATGAAAATCAACATTAGGTTAGGGTAAGAAAAGGGAGTTTGATTGAATTATGCTGGAATATTCTGTCTTGT

>M6-PGSC0003DMG400002618-(rev)M6LRR-RLP_3050

GAACTTGGAAACTCTCCAAAATAACTCAAAACTTTTTCTCAGTTACAACAGATTATTAGGTGCCATACCACCAGAATCAGGAAATCTCACAAGTTTAAATGTCTTGAATCTTCAATGAAA

>M6-PGSC0003DMG400002618-(rev)M6LRR-RLP_3111

GCCATACCACCAGAATCAGGAAATCTCACAAGTTTAAATGTCTTGAATCTTCAATGAAACAATCTTTTTGGCTCCATTCCTTCAACTCTTCAAAAATACCAGAAGCTCTATGAGCTGAGG

>M6-PGSC0003DMG400002618-(rev)M6LRR-RLP_3172

ATCTTTTTGGCTCCATTCCTTCAACTCTTCAAAAATACCAGAAGCTCTATGAGCTGAGGCTCTCTGAGAATAATATAAGTGGCTCAATCCCTTACGAAATTGGCTCTTTATCCGAATTAC

>M6-PGSC0003DMG400002618-(rev)M6LRR-RLP_3233

CTCTGAGAATAATATAAGTGGCTCAATCCCTTACGAAATTGGCTCTTTATCCGAATTACAGGTGATATTAGACCTGAGCAAAAATCATTCTATTTTATGTGCAAGTGTCAGAATAGTGTA

>M6-PGSC0003DMG400002618-(rev)M6LRR-RLP_3294

GTGATATTAGACCTGAGCAAAAATCATTCTATTTTATGTGCAAGTGTCAGAATAGTGTATCTTCAAGCATAGTCTTATTTCCTAAGGTTAATGCAGTGCAATTGTTAAGAGCCATACATT

>M6-PGSC0003DMG400002618-(rev)M6LRR-RLP_3355

TTCAAGCATAGTCTTATTTCCTAAGGTTAATGCAGTGCAATTGTTAAGAGCCATACATTTCTATTGAAATCCAACCTTGTCATTCTTTTATTGGATAAGTATTACAAGTGTCATATATAG

>M6-PGSC0003DMG400002618-(rev)M6LRR-RLP_3416

TATTGAAATCCAACCTTGTCATTCTTTTATTGGATAAGTATTACAAGTGTCATATATAGTCCGGGCTTAGCCATTTAGGAATTGCTCAGTTGCTTGTACTCTTATCAGGACCTGCTGTTC

>M6-PGSC0003DMG400002618-(rev)M6LRR-RLP_3477

CGGGCTTAGCCATTTAGGAATTGCTCAGTTGCTTGTACTCTTATCAGGACCTGCTGTTCTCTTATTTCGATCATGTTTCTTTCTTCTATTTAGGCCTTAGACATGCAGCCTGCTTTTCAA

>M6-PGSC0003DMG400002618-(rev)M6LRR-RLP_3538

TTATTTCGATCATGTTTCTTTCTTCTATTTAGGCCTTAGACATGCAGCCTGCTTTTCAAAATATATTCTTATACTTGATAATGTGGAAGTTTTTCATTCTTGATTTAGTCTTCAAGGGAA

>M6-PGSC0003DMG400002618-(rev)M6LRR-RLP_3599

TATATTCTTATACTTGATAATGTGGAAGTTTTTCATTCTTGATTTAGTCTTCAAGGGAAGGTGGATTTCCTGGGGGTCAGCTGTGGACATTAGGAGGAGGGAGGGGAAGCTAGTACTTGT

>M6-PGSC0003DMG400002618-(rev)M6LRR-RLP_366

ATTAGGTTAGGGTAAGAAAAGGGAGTTTGATTGAATTATGCTGGAATATTCTGTCTTGTTAGCATGTGTATTTGTGAACAAGGTTCACGCACGCGTTTACTTTGACCTTTTCCGTATTAC

>M6-PGSC0003DMG400002618-(rev)M6LRR-RLP_3660

TGGATTTCCTGGGGGTCAGCTGTGGACATTAGGAGGAGGGAGGGGAAGCTAGTACTTGTTTAGTTGCATGTAATAAATGTTAATCTGTGCTTTGGTAAGGTTAAACTAATTTTGTCAAGT

>M6-PGSC0003DMG400002618-(rev)M6LRR-RLP_3721

AGTTGCATGTAATAAATGTTAATCTGTGCTTTGGTAAGGTTAAACTAATTTTGTCAAGTTCCCTTCTTTGTTTGTATCAAGGTCTGGGCTTCTGGATTCTCTGCTGCCTTTTTAAAGAAG

>M6-PGSC0003DMG400002618-(rev)M6LRR-RLP_3782

CCTTCTTTGTTTGTATCAAGGTCTGGGCTTCTGGATTCTCTGCTGCCTTTTTAAAGAAGAATAGTGTTGCAACAAAACGCATCATGACTTTAAACGACAGAGATGTGACAATAGAAGCCA

>M6-PGSC0003DMG400002618-(rev)M6LRR-RLP_3843

TAGTGTTGCAACAAAACGCATCATGACTTTAAACGACAGAGATGTGACAATAGAAGCCAAAACTTCTGTTGAAACTAAGCCTGCAGTAACTGAAGCAGCGTTGCTATTAGTACTGGAAAA

>M6-PGSC0003DMG400002618-(rev)M6LRR-RLP_3965

TCTTTGAGGTGGTTTTGAACCTTCCCACCGTAAATAGGTAAGGAGATGTCTTCCAGAATCACCTTTTCTGCAAATGTTTATCAGCCTGCAGTATTTTTAAACAGAAGTACCAATATTAAA

>M6-PGSC0003DMG400002618-(rev)M6LRR-RLP_4026

CCTTTTCTGCAAATGTTTATCAGCCTGCAGTATTTTTAAACAGAAGTACCAATATTAAATCTACTTTTTGTCTTTGTACTCTACTCAATGTGCGAAGTAGACCTTGTTGATATCATGTCC

>M6-PGSC0003DMG400002618-(rev)M6LRR-RLP_4087

TACTTTTTGTCTTTGTACTCTACTCAATGTGCGAAGTAGACCTTGTTGATATCATGTCCATTCACCAATGGATTGTAATTACATTTTCAAGTCATTAGTGTGTATAGCTAGTGCCAAAAA

>M6-PGSC0003DMG400002618-(rev)M6LRR-RLP_4148

TCACCAATGGATTGTAATTACATTTTCAAGTCATTAGTGTGTATAGCTAGTGCCAAAAATAGTTTCGTGTTTGTACTTACATTGATGGTTGGATGAAATTGCAGGAGGCTTTCTGACGTT

>M6-PGSC0003DMG400002618-(rev)M6LRR-RLP_4209

GTTTCGTGTTTGTACTTACATTGATGGTTGGATGAAATTGCAGGAGGCTTTCTGACGTTCTAGATCTATTGACATCCCGGCAACCAAGTTGCCCTCAATTATACATATACAGCAGTGCCG

>M6-PGSC0003DMG400002618-(rev)M6LRR-RLP_427

GCATGTGTATTTGTGAACAAGGTTCACGCACGCGTTTACTTTGACCTTTTCCGTATTACATTGGGTCTTGTCTTCTAATTACGTAATTAATCTCGTGCGTTTTTGTGACAGCACAAAATT

>M6-PGSC0003DMG400002618-(rev)M6LRR-RLP_4270

AGATCTATTGACATCCCGGCAACCAAGTTGCCCTCAATTATACATATACAGCAGTGCCGACAGAGTCATTCCTGCAGTTTCGGTTGAGTCCTTTGTAGAAGAGCAACGAAGAATTGGTCG

>M6-PGSC0003DMG400002618-(rev)M6LRR-RLP_4392

ATGTTAGAGCTTGCAACTTCGTTTCTACACCTCACGTTGATCATTTCAGAAATGACCCAGAATTGTATACTTCACAGCTTACCCAATTTCTAGAGGACTACGTCCTAAGCTCTTGCAAAC

>M6-PGSC0003DMG400002618-(rev)M6LRR-RLP_4453

ATTGTATACTTCACAGCTTACCCAATTTCTAGAGGACTACGTCCTAAGCTCTTGCAAACAGCCTTCTTGAATAATTCCTAATCACAACACATTGATTCTTTCTGCCCTTTTTTTCCCTCA

>M6-PGSC0003DMG400002618-(rev)M6LRR-RLP_4514

CCTTCTTGAATAATTCCTAATCACAACACATTGATTCTTTCTGCCCTTTTTTTCCCTCACTTTAACCAAATACTTGGTAAAGCTTCTGTTTTCCAACTTTAAAGCAGATATACATTGATG

>M6-PGSC0003DMG400002618-(rev)M6LRR-RLP_4575

TTAACCAAATACTTGGTAAAGCTTCTGTTTTCCAACTTTAAAGCAGATATACATTGATGTAGATTGTTACGTATACAGAGAATTAAGTTTTTTTTTCTTTTCCATTCTTATTTAATTTAG

>M6-PGSC0003DMG400002618-(rev)M6LRR-RLP_4636

GATTGTTACGTATACAGAGAATTAAGTTTTTTTTTCTTTTCCATTCTTATTTAATTTAGTTGTATCTATGGTCTCCCAGCAACAGAATTGTTGCATTGTACCCAAAATATACTCTACTTT

>M6-PGSC0003DMG400002618-(rev)M6LRR-RLP_488

TGGGTCTTGTCTTCTAATTACGTAATTAATCTCGTGCGTTTTTGTGACAGCACAAAATTTGACAGCTATATAGTCTTTGCTTGTGGGGCTGTCATTATAGTTCATCTATATTGGCAGCCC

>M6-PGSC0003DMG400002618-(rev)M6LRR-RLP_549

ACAGCTATATAGTCTTTGCTTGTGGGGCTGTCATTATAGTTCATCTATATTGGCAGCCCAATTTCTGAGGTGTGTGTGTCTTCTTTCTCTTCACCGGCAAAACAAATATATCATCATGGC

>M6-PGSC0003DMG400002618-(rev)M6LRR-RLP_61

AAGTGATCTCTTATTCCGGAAAAGCAAGGTGGGAAATCTTTACAAGACTCGTCTCTTTTTCCTTATCAGAATCAGAAGAAGAAAGAAGAAGATTCGTCTCTCAGGTATGGTGTTCCTCTT

>M6-PGSC0003DMG400002618-(rev)M6LRR-RLP_671

ACTTGTCTGGAATCCTCCAAAGACCTATAGTTGCAGCATCTGCAGTTGCTATAGCTTCTGTATCTACTGATCTTCATGAAAAATTCTGGCCATCTAAGTCATTACACAACAACAACAACA

>M6-PGSC0003DMG400002618-(rev)M6LRR-RLP_732

ATCTACTGATCTTCATGAAAAATTCTGGCCATCTAAGTCATTACACAACAACAACAACAACAATTCCCAAGAAAAGGAAAAGATAACATCTCACATATCTCTTTCTAAGCTACCCCATTT

>M6-PGSC0003DMG400002618-(rev)M6LRR-RLP_793

AATTCCCAAGAAAAGGAAAAGATAACATCTCACATATCTCTTTCTAAGCTACCCCATTTCTCCTCCTACCTCACCACAATACCTGTTCCCCTGCCTATTCTAAATAACAGTACCACTTTT

>M6-PGSC0003DMG400002618-(rev)M6LRR-RLP_854

CCTCCTACCTCACCACAATACCTGTTCCCCTGCCTATTCTAAATAACAGTACCACTTTTACTTCACTTTCAATCCTTCCTTTTCCTACTTCTTCTCTGTTGCTCAACTCTTATCAATCAG

>M6-PGSC0003DMG400002618-(rev)M6LRR-RLP_915

TTCACTTTCAATCCTTCCTTTTCCTACTTCTTCTCTGTTGCTCAACTCTTATCAATCAGCAGCATTGCCCAACTCTGCTAATCCCTCCACATATACATATACACTTCCTTCTTCACCTTC

>M6-PGSC0003DMG400002618-(rev)M6LRR-RLP_976

GCATTGCCCAACTCTGCTAATCCCTCCACATATACATATACACTTCCTTCTTCACCTTCCGAGGTGTTGTATACATGGCACTTACCTGAGCCAAATGCATCACGGAATTCTGATTCTTCA

>M6-PGSC0003DMG400002689M6LRR-RLK_0

TACACAAATTCAACTCTTTTGTTCCTTAAATTCAACTTGTGTTTTTTTGTTCTCTTCCTAACTTCCCGCGCTTTATGCCTTAAACCCAAAAAAATGAACCGTTTTTTCTCCTTCATTCTC

>M6-PGSC0003DMG400002689M6LRR-RLK_1037

TTCGAGTGAAAAACGAGTTTACGCTAATAACAATGGTGGTGGTGGTGGTGATAGTGATGGTACAACAGCTACTGATAGAAGTAAACTCGTGTTTTTCGATAGGAGGAAGCCATTTGAGCT

>M6-PGSC0003DMG400002689M6LRR-RLK_1098

ACAACAGCTACTGATAGAAGTAAACTCGTGTTTTTCGATAGGAGGAAGCCATTTGAGCTTGAGGAATTGTTAAGAGCATCAGCTGAGATGCTTGGTAAAGGGTGTTTAGGTACTGTTTAT

>M6-PGSC0003DMG400002689M6LRR-RLK_1159

AGGAATTGTTAAGAGCATCAGCTGAGATGCTTGGTAAAGGGTGTTTAGGTACTGTTTATAAGGCAGTTCTTGATGATGGGATTACAGTTGCTGTAAAAAGATTGAAAGATGCAAATCCAT

>M6-PGSC0003DMG400002689M6LRR-RLK_122

CCTCACACATCTCATCTCCGCCGTAACCGCCACCGGCGACGGCAACGACACCACCGCACTCCTCCTCTTTCAGTCACAAACTGACATCCACGGCACTCTCCTTCACAATTGGACACTTCC

>M6-PGSC0003DMG400002689M6LRR-RLK_1220

GGCAGTTCTTGATGATGGGATTACAGTTGCTGTAAAAAGATTGAAAGATGCAAATCCATGTCCAAGAAAGGAGTTTGAACAGTATATGGATGTTATTGGAAAGCTTAAGCATCCAAATAT

>M6-PGSC0003DMG400002689M6LRR-RLK_1281

CCAAGAAAGGAGTTTGAACAGTATATGGATGTTATTGGAAAGCTTAAGCATCCAAATATGGTGAAACTTAGAGCTTACTATTATGCCAAGGAAGAAAAATTGCTTGTTTATGACTACTTG

>M6-PGSC0003DMG400002689M6LRR-RLK_1342

TGAAACTTAGAGCTTACTATTATGCCAAGGAAGAAAAATTGCTTGTTTATGACTACTTGCCTAATGGGAGTTTGCATTCACTTCTTCATGGTAAACATTAGTAAATGTTCATGTCAAAGA

>M6-PGSC0003DMG400002689M6LRR-RLK_1403

TAATGGGAGTTTGCATTCACTTCTTCATGGTAAACATTAGTAAATGTTCATGTCAAAGATTAAAGATTGTTTTGTAATTCAAGATTGTCTCTTTTGGGAGTTGTTAAGGTCTATATTGCT

>M6-PGSC0003DMG400002689M6LRR-RLK_1525

GACTCTTCAAAATGTCGATGGGGTGTGTCGGATCCTCCAAAAATAGTGCATTTTTGGAGAATTCTGACGTTGTAATTCAAGATTGTCTTTTTTGGGGGTTGGTAAGGGTTATATTCCTTG

>M6-PGSC0003DMG400002689M6LRR-RLK_1586

TTCTGACGTTGTAATTCAAGATTGTCTTTTTTGGGGGTTGGTAAGGGTTATATTCCTTGGACTTTTCAAAAATATCGATGGATGTGTGTTAGATTCTCCAAAAATCTCTAGAAATAGCGC

>M6-PGSC0003DMG400002689M6LRR-RLK_1647

CTTTTCAAAAATATCGATGGATGTGTGTTAGATTCTCCAAAAATCTCTAGAAATAGCGCATTTTTGGAGAATCTTACACGGGTATGACATGTTTTGTAATTCAAGATTGTCTTTTTTGGG

>M6-PGSC0003DMG400002689M6LRR-RLK_1708

TTTTGGAGAATCTTACACGGGTATGACATGTTTTGTAATTCAAGATTGTCTTTTTTGGGGTTGGTAAGGGTTATATTGCTCGGACTTCTCATAAATGTTGGCAGGTACATGTCGGATCCT

>M6-PGSC0003DMG400002689M6LRR-RLK_1769

TGGTAAGGGTTATATTGCTCGGACTTCTCATAAATGTTGGCAGGTACATGTCGGATCCTCTCGAAATCTCCAGAAATAGTGTATCTATGGAGGATCTGATGCGGGTATGGCAACATTTTT

>M6-PGSC0003DMG400002689M6LRR-RLK_183

CTCCTCTTTCAGTCACAAACTGACATCCACGGCACTCTCCTTCACAATTGGACACTTCCGTTGAACTCCACCACCGCTTGCACAGCTCAATGGCTTGGTATAAAATGCATAAACAACCGT

>M6-PGSC0003DMG400002689M6LRR-RLK_1830

CGAAATCTCCAGAAATAGTGTATCTATGGAGGATCTGATGCGGGTATGGCAACATTTTTGGAGAGTCCGAGCAAGTTGTAGGTATGATACTAATATTGATGGGATTTTTTTTTCTTGATT

>M6-PGSC0003DMG400002689M6LRR-RLK_1891

AGAGTCCGAGCAAGTTGTAGGTATGATACTAATATTGATGGGATTTTTTTTTCTTGATTGTAATAGGAAACAGAGGACCAGGGAGAATTCCATTGGATTGGACAACAAGGATTAGCTTGG

>M6-PGSC0003DMG400002689M6LRR-RLK_1952

AATAGGAAACAGAGGACCAGGGAGAATTCCATTGGATTGGACAACAAGGATTAGCTTGGTGCTAGGGGTAGCTAGAGGGCTTGCTCACATTCATGAAGAGTATGCAGATTCAAGAATCCC

>M6-PGSC0003DMG400002689M6LRR-RLK_2013

CTAGGGGTAGCTAGAGGGCTTGCTCACATTCATGAAGAGTATGCAGATTCAAGAATCCCACATGGGAATGTGAAATCATCTAATGTGTTGCTTGACAAAAATGGTGTGGCTTGTATATCT

>M6-PGSC0003DMG400002689M6LRR-RLK_2074

ATGGGAATGTGAAATCATCTAATGTGTTGCTTGACAAAAATGGTGTGGCTTGTATATCTGATTTTGGGTTGTCATTGCTTTTGAATCCTGTTCATGCCATAGCTAGATTAGGAGGATACC

>M6-PGSC0003DMG400002689M6LRR-RLK_2135

TTTTGGGTTGTCATTGCTTTTGAATCCTGTTCATGCCATAGCTAGATTAGGAGGATACCAAGCGCCCGAACAATCTGAAATCAAGAGGCTATCTCAAAAATCTGATGTTTATAGCTTTGG

>M6-PGSC0003DMG400002689M6LRR-RLK_2196

GCGCCCGAACAATCTGAAATCAAGAGGCTATCTCAAAAATCTGATGTTTATAGCTTTGGAGTACTGTTATTAGAAGTTCTCACTGGTAGAGCTCCTTCAGAGTACCCTTCACCGACTAGG

>M6-PGSC0003DMG400002689M6LRR-RLK_2257

TACTGTTATTAGAAGTTCTCACTGGTAGAGCTCCTTCAGAGTACCCTTCACCGACTAGGCCTCGCGATGAGGAGGAAGAATTGCCAGTGGATTTGCCTAAATGGGTTCGATCAGTAGTGA

>M6-PGSC0003DMG400002689M6LRR-RLK_2318

TCGCGATGAGGAGGAAGAATTGCCAGTGGATTTGCCTAAATGGGTTCGATCAGTAGTGAGGGACGAGTGGACTGCTGAAGTGTTTGATCAAGAACTGTTGAGGTACAAAAACATTGAGGA

>M6-PGSC0003DMG400002689M6LRR-RLK_2379

GACGAGTGGACTGCTGAAGTGTTTGATCAAGAACTGTTGAGGTACAAAAACATTGAGGAGGAATTGGTGTCAATGCTACATGTAGCCATGGCATGTGTAGTGCCACTACCTGAAAAAAGG

>M6-PGSC0003DMG400002689M6LRR-RLK_244

TGAACTCCACCACCGCTTGCACAGCTCAATGGCTTGGTATAAAATGCATAAACAACCGTGTCTCCGCCGTAATCCTCCCGTCGTTCAATCTCCGGGGACCCATTACCGCTCTCTCTTCTT

>M6-PGSC0003DMG400002689M6LRR-RLK_2440

AATTGGTGTCAATGCTACATGTAGCCATGGCATGTGTAGTGCCACTACCTGAAAAAAGGCCTACTATGGTTGAAGTTGTTAAGTTGATTGAGGAGATTAGAGTTGAACAATCTCCATTAG

>M6-PGSC0003DMG400002689M6LRR-RLK_2501

TACTATGGTTGAAGTTGTTAAGTTGATTGAGGAGATTAGAGTTGAACAATCTCCATTAGGTGAAGATTATGACGAATCGCGCAATTCTCTTTCCCCCTCCCTTGCCACCACTGAAGATGG

>M6-PGSC0003DMG400002689M6LRR-RLK_2562

GAAGATTATGACGAATCGCGCAATTCTCTTTCCCCCTCCCTTGCCACCACTGAAGATGGACTCCCCGGTTACTAATTTTCGATCATCGTGGCATTGATGTTTCATAGTCCTAGTAGTATA

>M6-PGSC0003DMG400002689M6LRR-RLK_2623

TCCCCGGTTACTAATTTTCGATCATCGTGGCATTGATGTTTCATAGTCCTAGTAGTATATGTTCAAATTTGCAGCATCTTACTGCAAATTGGGAAGTTCAAAAGCTATGGACTTATGTTT

>M6-PGSC0003DMG400002689M6LRR-RLK_2684

TTCAAATTTGCAGCATCTTACTGCAAATTGGGAAGTTCAAAAGCTATGGACTTATGTTTAGTAGTTATTTGATTGCTTTGTGTTATTCCAAATTACTGCACTTTTGGAAAATCTGATGAT

>M6-PGSC0003DMG400002689M6LRR-RLK_2745

TAGTTATTTGATTGCTTTGTGTTATTCCAAATTACTGCACTTTTGGAAAATCTGATGATCCAACAAGCATATAAGAAGTGCAGCAATCAGTCCATTATGTATTTTTTTTAATGTAATGTT

>M6-PGSC0003DMG400002689M6LRR-RLK_2794

ATCTGATGATCCAACAAGCATATAAGAAGTGCAGCAATCAGTCCATTATGTATTTTTTTTAATGTAATGTTAATTCTTTGTATTTAATTCATTTTCAAGATTTGTGTTCTAAGGCAAAAC

>M6-PGSC0003DMG400002689M6LRR-RLK_305

CTCCGCCGTAATCCTCCCGTCGTTCAATCTCCGGGGACCCATTACCGCTCTCTCTTCTTTACCTTTACTCAGCCTTCTCGACCTCCGTAACAATCGCCTTAACGGAACCCTCACACCCAT

>M6-PGSC0003DMG400002689M6LRR-RLK_366

CCTTTACTCAGCCTTCTCGACCTCCGTAACAATCGCCTTAACGGAACCCTCACACCCATTACCCAATGTACTAATCTTAAGCTCATTTACCTCTCCGGCAACGATTTTTCCGGCGAAATT

>M6-PGSC0003DMG400002689M6LRR-RLK_427

CCCAATGTACTAATCTTAAGCTCATTTACCTCTCCGGCAACGATTTTTCCGGCGAAATTCCGCCGGAAATTTCTTCTCTCCGTCGTCTCCTCCGCCTTGACGTTTCAAATAACAACCTTG

>M6-PGSC0003DMG400002689M6LRR-RLK_488

GCCGGAAATTTCTTCTCTCCGTCGTCTCCTCCGCCTTGACGTTTCAAATAACAACCTTGAAGGTTCAATTCCGACCCAGATTGCGAATTTGACCCGATTGCTCACACTTCGTTTACAGAA

>M6-PGSC0003DMG400002689M6LRR-RLK_549

GGTTCAATTCCGACCCAGATTGCGAATTTGACCCGATTGCTCACACTTCGTTTACAGAACAATGAACTTTCTGGAACAATCCCAAAATCCCTAAATTCTCTTCAAAATCTTAAAGAATTG

>M6-PGSC0003DMG400002689M6LRR-RLK_61

CTTCCCGCGCTTTATGCCTTAAACCCAAAAAAATGAACCGTTTTTTCTCCTTCATTCTCCTCCTCACACATCTCATCTCCGCCGTAACCGCCACCGGCGACGGCAACGACACCACCGCAC

>M6-PGSC0003DMG400002689M6LRR-RLK_610

ATGAACTTTCTGGAACAATCCCAAAATCCCTAAATTCTCTTCAAAATCTTAAAGAATTGAACTTTTCGAATAATGAGCTTTATGGGTCTGTTCCAAATGGTTTATATAATCATTTTGGTG

>M6-PGSC0003DMG400002689M6LRR-RLK_671

CTTTTCGAATAATGAGCTTTATGGGTCTGTTCCAAATGGTTTATATAATCATTTTGGTGAAAATAGCTTTGATGGAAATGAGGGTTTATGTGGAATTGGGAATTTACCTCAATGTTCATA

>M6-PGSC0003DMG400002689M6LRR-RLK_732

AATAGCTTTGATGGAAATGAGGGTTTATGTGGAATTGGGAATTTACCTCAATGTTCATATACAGGGGAAATAGTTCCTTCAAACCCTAGCTCTTTGCCTACAACCTCCACTGCAACAATC

>M6-PGSC0003DMG400002689M6LRR-RLK_793

CAGGGGAAATAGTTCCTTCAAACCCTAGCTCTTTGCCTACAACCTCCACTGCAACAATCGAAGAACCCCACGAAAAATCGCGAAAAGGGTTGAGTAGAGGAGGGGTGGTAGCAATAGTGA

>M6-PGSC0003DMG400002689M6LRR-RLK_854

AGAACCCCACGAAAAATCGCGAAAAGGGTTGAGTAGAGGAGGGGTGGTAGCAATAGTGATGGTGAATGTGGTGGCATTGTTAGTTGTAGTGTCATTTATGGTAGCATATTACTGTGGGAA

>M6-PGSC0003DMG400002689M6LRR-RLK_915

GTGAATGTGGTGGCATTGTTAGTTGTAGTGTCATTTATGGTAGCATATTACTGTGGGAAGTATTCAAGAACTCAGAGTTTTTCAATGTCAGGAAGTGAGTGTGGGAAGAGGAGGAGTAGT

>M6-PGSC0003DMG400002689M6LRR-RLK_976

ATTCAAGAACTCAGAGTTTTTCAATGTCAGGAAGTGAGTGTGGGAAGAGGAGGAGTAGTTATTCGAGTGAAAAACGAGTTTACGCTAATAACAATGGTGGTGGTGGTGGTGATAGTGATG

>M6-PGSC0003DMG400002870-(rev)M6LRR-RLK_1054

AACTTAAGACAATTGATCTTTCTGATAACAAGTTAACCGGCGAAATCCCGTCATCTTTAGCTCAGCTGAAAAGCCTTCAATACTTGTAAGCAAAGAGAATTCTGAGATCTTATACAAACA

>M6-PGSC0003DMG400002870-(rev)M6LRR-RLK_1116

CAGCTGAAAAGCCTTCAATACTTGTAAGCAAAGAGAATTCTGAGATCTTATACAAACAAGTTTTTTTGGGGTTAAAAACTAATGATGTTTGTTCTTGAATTTTGATGATTATAGGAGATT

>M6-PGSC0003DMG400002870-(rev)M6LRR-RLK_1178

TTTTTGGGGTTAAAAACTAATGATGTTTGTTCTTGAATTTTGATGATTATAGGAGATTAAACAACAATAGTCTAACTGGAGCTATTCCTTTGGATTTGGCCAATATGACCCACCTCTCAC

>M6-PGSC0003DMG400002870-(rev)M6LRR-RLK_1240

AACAATAGTCTAACTGGAGCTATTCCTTTGGATTTGGCCAATATGACCCACCTCTCACTTATGTAATTTTCTAATAACATTTTCATGTTATGTTATGTTATATGTTCTTTATAACAACAT

>M6-PGSC0003DMG400002870-(rev)M6LRR-RLK_1302

GTAATTTTCTAATAACATTTTCATGTTATGTTATGTTATATGTTCTTTATAACAACATTTCGATATAGCAGCTAAAAAATATCCAGACAAATGAAGTACGCAGTTATAGAGAGGTTTGAC

>M6-PGSC0003DMG400002870-(rev)M6LRR-RLK_1364

ATATAGCAGCTAAAAAATATCCAGACAAATGAAGTACGCAGTTATAGAGAGGTTTGACTGTATTTTTCACTTATGGAGTTGTACTGTTTTGCATGTTTAGGGACTTGTCTTTCAACAATC

>M6-PGSC0003DMG400002870-(rev)M6LRR-RLK_1426

TTTTTCACTTATGGAGTTGTACTGTTTTGCATGTTTAGGGACTTGTCTTTCAACAATCTCAGTGGTCCTGTACCAAGGCTTCTAGCCAAAACATTCAAGTAAGGTTCTTGAAGAATTCTT

>M6-PGSC0003DMG400002870-(rev)M6LRR-RLK_1488

TGGTCCTGTACCAAGGCTTCTAGCCAAAACATTCAAGTAAGGTTCTTGAAGAATTCTTTGTGTATCACTTTTTCCTTAATCAGTAAATGAAACCAATTTTTTGTATTTATTTGTCTTTTT

>M6-PGSC0003DMG400002870-(rev)M6LRR-RLK_1550

TATCACTTTTTCCTTAATCAGTAAATGAAACCAATTTTTTGTATTTATTTGTCTTTTTTTTAACCTTATTGACACAGTATATTGGGAAATCCAATGATATGTGCAACTGGAAAAGAATCA

>M6-PGSC0003DMG400002870-(rev)M6LRR-RLK_1612

ACCTTATTGACACAGTATATTGGGAAATCCAATGATATGTGCAACTGGAAAAGAATCAGAGTGCAATGGAACAACACCAATGCCCCTCTCCTTCTCTTCAAACAATCCACAGAGTAAGAA

>M6-PGSC0003DMG400002870-(rev)M6LRR-RLK_1674

GCAATGGAACAACACCAATGCCCCTCTCCTTCTCTTCAAACAATCCACAGAGTAAGAACTAACCTTATTTTAACAGCCAAATGATAGTTTTTTTTTTTGTTTTTTTTTATGACCGTGGTG

>M6-PGSC0003DMG400002870-(rev)M6LRR-RLK_2108

TGTTTTTACTAGTCATACAGATTTATGAAGTAAATATGCTTCACTGATTTTCTCTACTGATAGATTCAATATTTATAGTTCATTCCCAGCTAAAAATATCCTGTGTTATCTCAGATGTTC

>M6-PGSC0003DMG400002870-(rev)M6LRR-RLK_2170

GATTCAATATTTATAGTTCATTCCCAGCTAAAAATATCCTGTGTTATCTCAGATGTTCAGCCTTCTGGAAAACCGAAGACTCACAAGGTTGCGTTAGCCTTTGGAACAAGTCTAGGATGC

>M6-PGSC0003DMG400002870-(rev)M6LRR-RLK_2232

TTCTGGAAAACCGAAGACTCACAAGGTTGCGTTAGCCTTTGGAACAAGTCTAGGATGCATCTTCCTACTAATTGTTGGGTTTGGATTCTTTCTATGGTGGAGACAAAAGCACAATAAGCA

>M6-PGSC0003DMG400002870-(rev)M6LRR-RLK_2294

TCCTACTAATTGTTGGGTTTGGATTCTTTCTATGGTGGAGACAAAAGCACAATAAGCAAATTTTCTTTGACAGTAATGGTGAGTATTGTTTGATCAGAACAATTTGAAGTTACGGTTTAG

>M6-PGSC0003DMG400002870-(rev)M6LRR-RLK_2356

TTCTTTGACAGTAATGGTGAGTATTGTTTGATCAGAACAATTTGAAGTTACGGTTTAGCTCATCGATCCTCTCTTATAAACGTTATTGACATAAATTGCAGAACAACATATTGAAGAAGT

>M6-PGSC0003DMG400002870-(rev)M6LRR-RLK_2418

TCGATCCTCTCTTATAAACGTTATTGACATAAATTGCAGAACAACATATTGAAGAAGTGTGCTTAGGCAACTTAAGGAGGTTCCAATTCAAGGAACTTCAGTCTGCCACAAACAATTTCA

>M6-PGSC0003DMG400002870-(rev)M6LRR-RLK_2480

TTAGGCAACTTAAGGAGGTTCCAATTCAAGGAACTTCAGTCTGCCACAAACAATTTCAGCAGCAAGAACATACTAGGGAAAGGTGGTTTTGGTAATGTTTATAAAGGACGTCTTAGTGAC

>M6-PGSC0003DMG400002870-(rev)M6LRR-RLK_2542

CAAGAACATACTAGGGAAAGGTGGTTTTGGTAATGTTTATAAAGGACGTCTTAGTGACGGGATTATTGTAGCTGTGAAAAGGCTCAAAGATGGAAATGCAGTTGGAGGCAATCAACAATT

>M6-PGSC0003DMG400002870-(rev)M6LRR-RLK_2604

TTATTGTAGCTGTGAAAAGGCTCAAAGATGGAAATGCAGTTGGAGGCAATCAACAATTTCAGACTGAAGTCGCGTTGATCAGTCTGGCAGTGCATCGGAATCTCCTCAGGTTGTATGGAT

>M6-PGSC0003DMG400002870-(rev)M6LRR-RLK_2666

ACTGAAGTCGCGTTGATCAGTCTGGCAGTGCATCGGAATCTCCTCAGGTTGTATGGATTTTGTATGACACCAACTGAAAGGTTGCTGGTGTACCCTTACATGTCTAATGGAAGTGTAGCT

>M6-PGSC0003DMG400002870-(rev)M6LRR-RLK_2728

TATGACACCAACTGAAAGGTTGCTGGTGTACCCTTACATGTCTAATGGAAGTGTAGCTTCGCGTCTCAAAGGTAAACATATCTAAAAACTGTCTCCTTTGGCCTATAATTACATTGTCCT

>M6-PGSC0003DMG400002870-(rev)M6LRR-RLK_2790

GTCTCAAAGGTAAACATATCTAAAAACTGTCTCCTTTGGCCTATAATTACATTGTCCTTATTTGCTCAGCTCATAACGTATCATTTCTTGCAGCCAAACCAACCTTGGATTGGGGTACAA

>M6-PGSC0003DMG400002870-(rev)M6LRR-RLK_2852

TGCTCAGCTCATAACGTATCATTTCTTGCAGCCAAACCAACCTTGGATTGGGGTACAAGAAAAGGAATAGCTTTGGGAGCTGCAAGGGGCTTATTATATCTACACGAACAATGTGATCCA

>M6-PGSC0003DMG400002870-(rev)M6LRR-RLK_2914

AGGAATAGCTTTGGGAGCTGCAAGGGGCTTATTATATCTACACGAACAATGTGATCCAAAGATCATTCATAGGGATGTGAAAGCTGCAAATATATTGCTTGATGATTACTGTGAAGCTGT

>M6-PGSC0003DMG400002870-(rev)M6LRR-RLK_2976

TCATTCATAGGGATGTGAAAGCTGCAAATATATTGCTTGATGATTACTGTGAAGCTGTTGTTGGAGATTTCGGATTAGCAAAGCTTTTGGATCATCACGATTCACATGTCACAACCGCGG

>M6-PGSC0003DMG400002870-(rev)M6LRR-RLK_3038

GGAGATTTCGGATTAGCAAAGCTTTTGGATCATCACGATTCACATGTCACAACCGCGGTTAGAGGAACTGTAGGACATATAGCTCCAGAATATCTCTCCACAGGACAGTCCTCTGATAAG

>M6-PGSC0003DMG400002870-(rev)M6LRR-RLK_3100

AGGAACTGTAGGACATATAGCTCCAGAATATCTCTCCACAGGACAGTCCTCTGATAAGACGGACGTCTTTGGTTTTGGTATTCTCTTGCTAGAATTAATCACTGGACAAAGAGCTCTGGA

>M6-PGSC0003DMG400002870-(rev)M6LRR-RLK_3162

ACGTCTTTGGTTTTGGTATTCTCTTGCTAGAATTAATCACTGGACAAAGAGCTCTGGAATTTGGTAAAGCAGCAAACCAGAAAGGCGCTATGCTTGATTGGGTAAGTACGACAAATCCCT

>M6-PGSC0003DMG400002870-(rev)M6LRR-RLK_3472

TAGGCGCACGTGCATTCACACTATCTAAAATATGGACTCTATAGGTATTAAGAGATGCAACGAACAAACCATCTTTAAATCCTGGATCCGTCCTTGTTATTTTTTCACTTGCAGGTAAGA

>M6-PGSC0003DMG400002870-(rev)M6LRR-RLK_3534

AACAAACCATCTTTAAATCCTGGATCCGTCCTTGTTATTTTTTCACTTGCAGGTAAGAAAGATTCAACAGGAGAAAAAGCTAGACATGTTGGTGGACAAAGACATGAAAAACGAGTACGA

>M6-PGSC0003DMG400002870-(rev)M6LRR-RLK_3596

TTCAACAGGAGAAAAAGCTAGACATGTTGGTGGACAAAGACATGAAAAACGAGTACGATAAGATAGAGTTAGAGGAAATGGTTCAAGTAGCATTACTATGCACTCAGTATCATCCAAGTC

>M6-PGSC0003DMG400002870-(rev)M6LRR-RLK_3658

ATAGAGTTAGAGGAAATGGTTCAAGTAGCATTACTATGCACTCAGTATCATCCAAGTCATAGACCAAAAATGTCTGAAGTTGTAAGAATGCTTGAAGGAGATGGACTAGCAGAGAAATGG

>M6-PGSC0003DMG400002870-(rev)M6LRR-RLK_372

CTTTTTTTCTTTGGCAGTGCAAGCTTTAATGGAGATGAAGAAGAATTTAGATGATCCACTTAATGTTTTAAACTGGGATGGAGATGCTGTTGATCCTTGTAGCTGGAATATGGTCACTTG

>M6-PGSC0003DMG400002870-(rev)M6LRR-RLK_3720

ACCAAAAATGTCTGAAGTTGTAAGAATGCTTGAAGGAGATGGACTAGCAGAGAAATGGGAAGCATCTCAAAGAGCAGAACCAACAAGATGCAGCAGAGCCAATGAATTTTCCTCTTCAGA

>M6-PGSC0003DMG400002870-(rev)M6LRR-RLK_3782

CATCTCAAAGAGCAGAACCAACAAGATGCAGCAGAGCCAATGAATTTTCCTCTTCAGAAAGATACTCTGATCTTACTGATGATTCTTCATTACTTGTACAAGCAATGGAACTATCTGGTC

>M6-PGSC0003DMG400002870-(rev)M6LRR-RLK_3844

TACTCTGATCTTACTGATGATTCTTCATTACTTGTACAAGCAATGGAACTATCTGGTCCAAGATGATAATTGAATTAAGACACAATGTCACTTTTGCTGGTGGGAGGTGACAGGTCTCTC

>M6-PGSC0003DMG400002870-(rev)M6LRR-RLK_4030

ATATTGAGCTTTGTTTATAGGTTTATACAGAAGCTGCAAATAGAATATCATACATTCTGATTGTAAAGGACAATATAATAACTTTGTAAGGCTGAGAGTTGTTTTTCTGTTTTTCCTTTT

>M6-PGSC0003DMG400002870-(rev)M6LRR-RLK_434

ATGTTTTAAACTGGGATGGAGATGCTGTTGATCCTTGTAGCTGGAATATGGTCACTTGTTCAAATGATAAATTTGTCACTAGCCTGTAAGTAGAACAATTTTCACTTATTCTAAAAATAG

>M6-PGSC0003DMG400002870-(rev)M6LRR-RLK_496

AATGATAAATTTGTCACTAGCCTGTAAGTAGAACAATTTTCACTTATTCTAAAAATAGTATTCCTAACAAGATTTCTTAAAATAGTTGTATTATTGATTATGCAGGGAGTGTCCCAGCCA

>M6-PGSC0003DMG400002870-(rev)M6LRR-RLK_558

CCTAACAAGATTTCTTAAAATAGTTGTATTATTGATTATGCAGGGAGTGTCCCAGCCAAAATTTGTCTGGCAAAATATCACCATATATCCACAACTTGACACACCTTGAACTTATGTGAG

>M6-PGSC0003DMG400002870-(rev)M6LRR-RLK_930

TATCAAGACATTTGAGTTTTGATTCTTATTCTTAGATGTTAATATGAGTGTCCTGTTTTTTACAGACTTCTACAGAGCAATAATATATCAGGATCACTTCCTATGGAGCTTGGAATGCTT

>M6-PGSC0003DMG400002870-(rev)M6LRR-RLK_992

CAGACTTCTACAGAGCAATAATATATCAGGATCACTTCCTATGGAGCTTGGAATGCTTAAAAAACTTAAGACAATTGATCTTTCTGATAACAAGTTAACCGGCGAAATCCCGTCATCTTT

>M6-PGSC0003DMG400002885M6LRR-RLP_0

ATGTGGCCTCTCTTTTTCATTTATTCACTTTTCTGTTGTGTATTTCTATGTATTTGTTCTTCCTTTGAGCATCATCTTTGCTCTCCCAGTGAAGCTTCTGCTTTGCTTCAGTTTAAGCAA

>M6-PGSC0003DMG400002885M6LRR-RLP_1054

AACTACGTTATTTAAGTCTTTCTAATTGCAGCTTCATCGGTCCGTTTCCTTCTCCACTTTTAAGCTTGACACAACTTGAAACCTTAGACTTGTCGAGAAATTCCCTATCAGGACTACTTC

>M6-PGSC0003DMG400002885M6LRR-RLP_1116

AGCTTGACACAACTTGAAACCTTAGACTTGTCGAGAAATTCCCTATCAGGACTACTTCCTAGAAATGCAAGCATGCTTCAAAAGCTAGCCGAGCTGGATTTGTCTTACAACTCACTGAAT

>M6-PGSC0003DMG400002885M6LRR-RLP_124

TTAATATATTGGACTTCTCCCAGTGTGATACGTCTTTCCCAAAAACAGTTTCATGGAATGAGAGTAACGATTGCTGCACTTGGGACGGGGTCACTTGTGACATGTTAACCGGTCATGTTA

>M6-PGSC0003DMG400002885M6LRR-RLP_1240

CCATACCATCTTGGGTGTTTAGCCTACCTTTGCTATCTTCAGTGTCGCTCCATCATAACCGGTTAAGTGGTTCTTTTCCTCAATCACCTGTGAATCTCACAAACCTTTCTACCCTTGATC

>M6-PGSC0003DMG400002885M6LRR-RLP_1302

TTAAGTGGTTCTTTTCCTCAATCACCTGTGAATCTCACAAACCTTTCTACCCTTGATCTTTCATCAAATAACATCACCCTCGATGCGGGAATACAAATCACTTTACCTAGCCTAGAAATT

>M6-PGSC0003DMG400002885M6LRR-RLP_1364

ATCAAATAACATCACCCTCGATGCGGGAATACAAATCACTTTACCTAGCCTAGAAATTATGCAGTTATCATCTTGTGAACTGAAGGATTTTCCACACTTCTTGAGAAATGTAAAGACACT

>M6-PGSC0003DMG400002885M6LRR-RLP_1426

AGTTATCATCTTGTGAACTGAAGGATTTTCCACACTTCTTGAGAAATGTAAAGACACTTATGGTGTTGGATATTTCTAACAATAAGATTCGTGGTCAAATCCCTAATTGGTTTAGCGGCA

>M6-PGSC0003DMG400002885M6LRR-RLP_1487

GGTGTTGGATATTTCTAACAATAAGATTCGTGGTCAAATCCCTAATTGGTTTAGCGGCATGAGGTGGGACTCCTTGTTGCACCTAAACCTTTCACATAACTCATTAACAGGCAACCTACA

>M6-PGSC0003DMG400002885M6LRR-RLP_186

AGTAACGATTGCTGCACTTGGGACGGGGTCACTTGTGACATGTTAACCGGTCATGTTATCGGGTTGGATCTTAGTTGTAGTCAGCTTAATGGAACTATTTATCCCAACAGCAGCCTCATC

>M6-PGSC0003DMG400002885M6LRR-RLP_248

GTTGGATCTTAGTTGTAGTCAGCTTAATGGAACTATTTATCCCAACAGCAGCCTCATCCAACTTCATCATCTTCAGACACTAAACCTTGCTAAGAACTATGTTTATCCTTCTACGATCCC

>M6-PGSC0003DMG400002885M6LRR-RLP_310

TTCATCATCTTCAGACACTAAACCTTGCTAAGAACTATGTTTATCCTTCTACGATCCCAAAAGACTTTAGCCGGTTGAGGAATTTGAAGCATCTCAATCTTTCTGATGCTTACTTTCAAG

>M6-PGSC0003DMG400002885M6LRR-RLP_372

GACTTTAGCCGGTTGAGGAATTTGAAGCATCTCAATCTTTCTGATGCTTACTTTCAAGGGGAAATCCCAACAGAAATCTCATACCTTTCCAATTTGGTTTCACTTGATCTTTCTGTTCCT

>M6-PGSC0003DMG400002885M6LRR-RLP_434

AATCCCAACAGAAATCTCATACCTTTCCAATTTGGTTTCACTTGATCTTTCTGTTCCTTCTTATGTGAACGGATTACAATTTGATCAGAGAGCATTTGAAGCAGTGCTTCAGAACTTGAC

>M6-PGSC0003DMG400002885M6LRR-RLP_558

CTAGAGGTACTTTCTCTCTTTGGTGTCAACGTTTCATCTCCGATACCTATGAATTTGTCTTCCTCCTTAAGGTATGTCGATCTTAAGGTAACCAATTTGGAAGGTGTTCTCACGGAGAGC

>M6-PGSC0003DMG400002885M6LRR-RLP_62

CTTTGAGCATCATCTTTGCTCTCCCAGTGAAGCTTCTGCTTTGCTTCAGTTTAAGCAATCCTTTAATATATTGGACTTCTCCCAGTGTGATACGTCTTTCCCAAAAACAGTTTCATGGAA

>M6-PGSC0003DMG400002885M6LRR-RLP_620

CTCCTTAAGGTATGTCGATCTTAAGGTAACCAATTTGGAAGGTGTTCTCACGGAGAGCTTATTCCTTCTGCCAAAATTGGAAACACTCTATTTGAGTAATAACTATCTTCTCAAAGGAGT

>M6-PGSC0003DMG400002885M6LRR-RLP_682

TCCTTCTGCCAAAATTGGAAACACTCTATTTGAGTAATAACTATCTTCTCAAAGGAGTTTTACCAAAGATTCATCCAAGCAGCACTCTGTTGGAGTTGGATATTTCATACACAGGAATCT

>M6-PGSC0003DMG400002885M6LRR-RLP_806

TGAGCTGCCTGATTCAATTGGCACCTTGAGTTCCTTGACTCGCTTGTACCTCTATGGATGTCAATTCTCTGGTCGCATTCCTGATTCCATTGGCAATCTTACACAAATTAGGTATTTGAA

>M6-PGSC0003DMG400002885M6LRR-RLP_868

AATTCTCTGGTCGCATTCCTGATTCCATTGGCAATCTTACACAAATTAGGTATTTGAATTTTGGTAATAATCATTTCACCGGCAATATTCCTTCCACAATCTCTAAATTGAAGCACCTCG

>M6-PGSC0003DMG400002885M6LRR-RLP_930

GGTAATAATCATTTCACCGGCAATATTCCTTCCACAATCTCTAAATTGAAGCACCTCGCAGTTTTAGCTCTTTCATCTAACTCCTTTGGAGGTGAAATTCCTAATTTTTTCTCTAACCTC

>M6-PGSC0003DMG400002886M6LRR-RLP_0

CAACTCCTTTTCAGGTGAAATTCCTGCTGTTTTCTCTAACCTCCAAGAGCTACGTACTTTACATCTTTCTTATAACAGCTTCATCGGTTCATTTCCCGCTTCAATTTTAAGTTTGACACA

>M6-PGSC0003DMG400002886M6LRR-RLP_1040

CTCTCGAATACCTGCAGGTCCTTATATTAAAGTCGAACAAGTTCCATGGACCTATAAGTACGTGTCTGACTAAGTTTTGCTTTCCCAAGTTGCGAATTTTTGATCTTTCTCGTAATGATT

>M6-PGSC0003DMG400002886M6LRR-RLP_1170

ACTTCCTGCAAAAGTTTTTGGAAACTTCAAGGCAATGATCAAATTAGATGGCGAAGACACAGGAAATATAAAGTACATGGACTCTGTGTTGTATTCGCCATTTGTCACATCGTATGAGGA

>M6-PGSC0003DMG400002886M6LRR-RLP_1235

ATATAAAGTACATGGACTCTGTGTTGTATTCGCCATTTGTCACATCGTATGAGGATTCAGTGAGTTTGGTAATCAAAGGGCAGGATATTGAGCTATAAAGAATCAGCACAATTACGACAA

>M6-PGSC0003DMG400002886M6LRR-RLP_260

CTTTGCTATCTTCAGTGTCGCTCCAACATAACCAATTCAGAGGACTAGCCGATGAAGTGATCAAAACAAACCCAACATTAAAACAACTGCATTTAAGCAATAATCAACTCAGTGGTCCTT

>M6-PGSC0003DMG400002886M6LRR-RLP_325

ACAAACCCAACATTAAAACAACTGCATTTAAGCAATAATCAACTCAGTGGTCCTTTTCCTCAATCACTTGTGAATCTCACAAACCTTGAAACCCTTGGAATTTCATCAAATAACATCACC

>M6-PGSC0003DMG400002886M6LRR-RLP_390

ACTTGTGAATCTCACAAACCTTGAAACCCTTGGAATTTCATCAAATAACATCACCATTGATGAGGGAATGAATATCACCTTTCTTAGCCTATCATCTTTATTCTTATCATCTTGTGAACT

>M6-PGSC0003DMG400002886M6LRR-RLP_455

GAATGAATATCACCTTTCTTAGCCTATCATCTTTATTCTTATCATCTTGTGAACTGAAGGATTTTCCACACTTCTTGAGAAATGTAAAGACACTTGTCTACTTGGATATTTCTAACAATA

>M6-PGSC0003DMG400002886M6LRR-RLP_65

TTTCTTATAACAGCTTCATCGGTTCATTTCCCGCTTCAATTTTAAGTTTGACACATCTTCAATACTTAGGATTGTCGAGTAATTCCCTATCTGGCCCACTGCCTAGCAACCAAAGCATGC

>M6-PGSC0003DMG400002886M6LRR-RLP_650

ATTCATTAACAGGCCACCTACCACAATTTCATTACTATAACCTACAGTATCTTGATCTGAAATGTAACTCCCTTCAGGGTCCACTACCTTCATCCATTTGTAACATGAGTTCGCTTATCT

>M6-PGSC0003DMG400002886M6LRR-RLP_845

TAGACTTAAGAAGGAACAATTTCACAGGGAGTCTTTCCCCATTATGTGCACACAGCACTTCGTTGAGTACCATTGTCATAAATGGTAATCAATTTGAAGGACCTGTACCTGTGTCGTTGC

>M6-PGSC0003DMG400002886M6LRR-RLP_910

AGTACCATTGTCATAAATGGTAATCAATTTGAAGGACCTGTACCTGTGTCGTTGCTCAAGTGTGGTGGTCTAGAAGTACTTGATGTGGGGAACAATGCTATAAATGACAGGTTTCCATCT

>M6-PGSC0003DMG400002886M6LRR-RLP_975

TGGTCTAGAAGTACTTGATGTGGGGAACAATGCTATAAATGACAGGTTTCCATCTTGGCTCGGAACTCTCGAATACCTGCAGGTCCTTATATTAAAGTCGAACAAGTTCCATGGACCTAT

>M6-PGSC0003DMG400002887M6LRR-RLP_0

AATTCCTGATGTTTTCTCTAACCTCAAAGAGCTAGTTGAATTGAATCTTTCTTTTAACAACTTCACCGGTCCATGCCCCTCTTCAATTTTAAGCTTGACAAGTCTTCAATACTTAGATTT

>M6-PGSC0003DMG400002887M6LRR-RLP_1071

CTCACTGCCTGCAGATTTTTTTAGAAACTTCAAGGCAATGACAAAAAATGGCACAGACAAAAGTGACATTAGCTATATGGAAACACCAGTTTACAGATCATTCCTTGAGTCAGTGAGCTC

>M6-PGSC0003DMG400002887M6LRR-RLP_1134

TGACATTAGCTATATGGAAACACCAGTTTACAGATCATTCCTTGAGTCAGTGAGCTCAGCCGTCATTCCTATATATAAGCTAGAAATGGCCATCAATGAAGTATACAAGGATTCAGTGAG

>M6-PGSC0003DMG400002887M6LRR-RLP_1197

CATTCCTATATATAAGCTAGAAATGGCCATCAATGAAGTATACAAGGATTCAGTGAGGTTGATGATCAAAGGAAACAATATTGAGCTAGAAAGAATCAGCACAATTGTTACAGCCATAGA

>M6-PGSC0003DMG400002887M6LRR-RLP_126

CAACTCACTGAATGGCACAATACACTCTTGGGTGTTTAGCCTCCCTTCCTTACGTGATTTGAAGCTCCATCATAACCAATTCAGTACAGTGGCTGATGAGATCAAAACAAACCCAACATT

>M6-PGSC0003DMG400002887M6LRR-RLP_1260

GATCAAAGGAAACAATATTGAGCTAGAAAGAATCAGCACAATTGTTACAGCCATAGATCTCTCAAGCAACCATTTAGAAGGTGACATTCCGAAATCACTAAAGGATCTCAGCTCACTTTG

>M6-PGSC0003DMG400002887M6LRR-RLP_1323

AAGCAACCATTTAGAAGGTGACATTCCGAAATCACTAAAGGATCTCAGCTCACTTTGGCTACTCAATTTATCCCATAACAATCTCAAAGGTGATATTCCAATGGAATTGGGGCAATTGAA

>M6-PGSC0003DMG400002887M6LRR-RLP_1386

CAATTTATCCCATAACAATCTCAAAGGTGATATTCCAATGGAATTGGGGCAATTGAATACACTTGAAGCCTTAGATCTCTCTTCGAATCGGCTCACTGGAAAGATTCCACAGGAATTGAC

>M6-PGSC0003DMG400002887M6LRR-RLP_1512

GAACTTTCTGGAGTTCTTAAACCTCTCTCAAAATCATCTCGTTGGACGCATTCCTCAAGGTTCACAATTCAGCACATTTGAAAATGACTCGTATGGTGGCAACCTTGATTTATGTGGTCC

>M6-PGSC0003DMG400002887M6LRR-RLP_1575

ACAATTCAGCACATTTGAAAATGACTCGTATGGTGGCAACCTTGATTTATGTGGTCCTCCTTTGTCAAAGCAATGTGGAACGAGTGATCGCTCGCATGTTCCTCAATCACTGGAGGAAGA

>M6-PGSC0003DMG400002887M6LRR-RLP_189

GCTCCATCATAACCAATTCAGTACAGTGGCTGATGAGATCAAAACAAACCCAACATTAGTGACTTTGTATTTAAGTCATAATCAACTCAATGGTCCTTTTCCTCTATCACTTGTGAATCT

>M6-PGSC0003DMG400002887M6LRR-RLP_252

TTTGTATTTAAGTCATAATCAACTCAATGGTCCTTTTCCTCTATCACTTGTGAATCTCACAAACCTAGCCCTCCTTGACTTTTCATCAAGTAACATCACCGGCGATGTGGGAATAAATAT

>M6-PGSC0003DMG400002887M6LRR-RLP_315

CCTAGCCCTCCTTGACTTTTCATCAAGTAACATCACCGGCGATGTGGGAATAAATATCACCTTTCCTAGACTATCAGCTTTGTTCTTATCATCTTGTGAACTGAAGGATTTTCCACACTT

>M6-PGSC0003DMG400002887M6LRR-RLP_378

TCCTAGACTATCAGCTTTGTTCTTATCATCTTGTGAACTGAAGGATTTTCCACACTTCTTGAGAAATTTAAAGACACTTCAGTTCTTGGATCTTTCTAACAATAAGATTCGTGGTCAAAT

>M6-PGSC0003DMG400002887M6LRR-RLP_441

AAATTTAAAGACACTTCAGTTCTTGGATCTTTCTAACAATAAGATTCGTGGTCAAATCCCTAACTGGTTTAGAGGCATGAGGTGGGACTCTTTGGAACAACTAAACCTTTCACGTAATTT

>M6-PGSC0003DMG400002887M6LRR-RLP_504

CTGGTTTAGAGGCATGAGGTGGGACTCTTTGGAACAACTAAACCTTTCACGTAATTTTTTGGCAGGCCATCTAGAACAATTTCATCATTATAGACTAGAGTCTCTTGATCTTAAATTTAA

>M6-PGSC0003DMG400002887M6LRR-RLP_567

AGGCCATCTAGAACAATTTCATCATTATAGACTAGAGTCTCTTGATCTTAAATTTAACTTCCTTCAGGGTTCACTACCTTCATCCATATGTAACCTAAGTAGCCTTAGAATTCTGGATTT

>M6-PGSC0003DMG400002887M6LRR-RLP_63

CACCGGTCCATGCCCCTCTTCAATTTTAAGCTTGACAAGTCTTCAATACTTAGATTTCTCTCACAACTCACTGAATGGCACAATACACTCTTGGGTGTTTAGCCTCCCTTCCTTACGTGA

>M6-PGSC0003DMG400002887M6LRR-RLP_630

TCAGGGTTCACTACCTTCATCCATATGTAACCTAAGTAGCCTTAGAATTCTGGATTTATCACGCAACAACTTCAGTAACTCGATTCCAAATTGCTTGGGAAGCATGGCTAAGCTAACGGT

>M6-PGSC0003DMG400002887M6LRR-RLP_693

CAACAACTTCAGTAACTCGATTCCAAATTGCTTGGGAAGCATGGCTAAGCTAACGGTGTTGGACTTGAGAAGGAACAATTTCAGTGGGAGTCTTCCACTGTTATGTACTCAAAGCACTTC

>M6-PGSC0003DMG400002887M6LRR-RLP_756

CTTGAGAAGGAACAATTTCAGTGGGAGTCTTCCACTGTTATGTACTCAAAGCACTTCATTGACGACCATTGTCTTGAATGGTAATCAATTTGAAGGATCTGTCCCTATGTCGTTGCTCAA

>M6-PGSC0003DMG400002887M6LRR-RLP_819

GACCATTGTCTTGAATGGTAATCAATTTGAAGGATCTGTCCCTATGTCGTTGCTCAACTGTGTTGGTTTAAAAGTTCTTGATTTAGGGAATAACGCTATAAATGACACATTTCCAGCTTG

>M6-PGSC0003DMG400002887M6LRR-RLP_882

TGGTTTAAAAGTTCTTGATTTAGGGAATAACGCTATAAATGACACATTTCCAGCTTGGCTTGGAACTCTTGAAGATCTGCAAGTCCTTATATTAAAGTCGAACAAGTTCCATGGACCTAT

>M6-PGSC0003DMG400002887M6LRR-RLP_945

AACTCTTGAAGATCTGCAAGTCCTTATATTAAAGTCGAACAAGTTCCATGGACCTATAAGTTCTAGGAAGAAGTTTTGCTTTCCTAGGTTGCGGATTTTTGATCTCTCTCATAATGCGTT

>M6-PGSC0003DMG400002902M6LRR-RLK_0

AGAAAAGAAACAGAGAAGCATTTGCAGATTGTTTCTTTAAGGGGAAGGTGGACATGTCTAGTAACAAACATGGCCGCCGTTCCCCTAATCCTTCATCTCCTTCCTCAAGCATCGTCTCCG

>M6-PGSC0003DMG400002902M6LRR-RLK_1054

GAAGGGGAGGATCACAAGGTGGTAAGGCTATGGGAGATCTTGTGTTGGTGAACAACGAAAATGGTACATTCGGATTGCCTGATTTGATGAAGGCAGCTGCTGAGGTACTCGGAAACGGAG

>M6-PGSC0003DMG400002902M6LRR-RLK_1116

GGTACATTCGGATTGCCTGATTTGATGAAGGCAGCTGCTGAGGTACTCGGAAACGGAGTGTTAGGATCAGCTTACAAGGCAAAAATGGGAAATGGAATGTCTGTTGTGGTTAAGAGATTG

>M6-PGSC0003DMG400002902M6LRR-RLK_1178

AGGATCAGCTTACAAGGCAAAAATGGGAAATGGAATGTCTGTTGTGGTTAAGAGATTGAGAGAGATGAATAAAATGAATAGAGATGTGTTTGATGCAGAAATGAGAAAACTTGGAAAGTT

>M6-PGSC0003DMG400002902M6LRR-RLK_124

AACACCATCTTTTCCTTATTCATATCTCCATTTTCTTCCTCATTTCTCTAGCTAATCCACAATTTTCTTTCTCCATTTCCGAGGACGATGCCTTGATCAAATTCAAAGAATCTTTAAAAA

>M6-PGSC0003DMG400002902M6LRR-RLK_1240

AGATGAATAAAATGAATAGAGATGTGTTTGATGCAGAAATGAGAAAACTTGGAAAGTTAAGGCACAAAAATTTACTTCAATTATTGGCATATCATTACAGAAAGGAAGAGAAATTGATGG

>M6-PGSC0003DMG400002902M6LRR-RLK_1302

CACAAAAATTTACTTCAATTATTGGCATATCATTACAGAAAGGAAGAGAAATTGATGGTATCTGAATATGTTCCCAAAGGCAGCCTATTATATCTGTTACATGGTATGACAATTTCTTTC

>M6-PGSC0003DMG400002902M6LRR-RLK_1364

TGAATATGTTCCCAAAGGCAGCCTATTATATCTGTTACATGGTATGACAATTTCTTTCATCTACAGTCAAACCTCTCTATAATAATGTCGTTTGTCTAGATAAGTTTTTGGCTGCTATAG

>M6-PGSC0003DMG400002902M6LRR-RLK_1426

ACAGTCAAACCTCTCTATAATAATGTCGTTTGTCTAGATAAGTTTTTGGCTGCTATAGCGAAATGTTGTTATCTAGAACATAAGTTTCATATCTCAACTGTTACTCCCTTTGTATTGAAA

>M6-PGSC0003DMG400002902M6LRR-RLK_1488

ATGTTGTTATCTAGAACATAAGTTTCATATCTCAACTGTTACTCCCTTTGTATTGAAACATACCTCGATGATGAGTTGGCCTACACACGCCTGTTGTTGATTGAGAACAAATATTTGACC

>M6-PGSC0003DMG400002902M6LRR-RLK_1612

CAACATCGAGGTGTGTTTTGATACAAATGGAGTGAATAGTTAAGGTAGGTTAAGAGAAAAATGGATAGTTAAAGTATGAAACTCGTGAAAAAAGGATAGTTTAGGTGTGTTTTTGACCAT

>M6-PGSC0003DMG400002902M6LRR-RLK_1674

GGATAGTTAAAGTATGAAACTCGTGAAAAAAGGATAGTTTAGGTGTGTTTTTGACCATTATGGTAAAAATCAGTTCTGAAGAAAACTTGGTCACTATTATAGTGAAATGTTATTATAAAG

>M6-PGSC0003DMG400002902M6LRR-RLK_1736

GTAAAAATCAGTTCTGAAGAAAACTTGGTCACTATTATAGTGAAATGTTATTATAAAGGATGATTGTTATAGAGAGGTTTGACTGTATATCAAATCATTAAGGTCTTCAACTTCTTTTCA

>M6-PGSC0003DMG400002902M6LRR-RLK_1798

ATTGTTATAGAGAGGTTTGACTGTATATCAAATCATTAAGGTCTTCAACTTCTTTTCAACATTTGCACGAGTAGATTTTATCCGATAATGTGATCCGATCTATACTTACCGGTCCCTTCC

>M6-PGSC0003DMG400002902M6LRR-RLK_1848

TCTTTTCAACATTTGCACGAGTAGATTTTATCCGATAATGTGATCCGATCTATACTTACCGGTCCCTTCCAAGACATCCGGTCACAATTGGGACGATACAAAGAAGATTAGCATGATCCC

>M6-PGSC0003DMG400002902M6LRR-RLK_186

TTTTCTTTCTCCATTTCCGAGGACGATGCCTTGATCAAATTCAAAGAATCTTTAAAAAACACAACATCTTTAGATTCATCTTGGCTTAAAGGTACTAGTCCATGTGACAAAAACAAGAAA

>M6-PGSC0003DMG400002902M6LRR-RLK_248

AACATCTTTAGATTCATCTTGGCTTAAAGGTACTAGTCCATGTGACAAAAACAAGAAATGGGATCGTATTCAATGCGAAGGGAACGTGGTTGAAGGCCTCCTTTTAGGTGAAGCTGGCTT

>M6-PGSC0003DMG400002902M6LRR-RLK_310

ATCGTATTCAATGCGAAGGGAACGTGGTTGAAGGCCTCCTTTTAGGTGAAGCTGGCTTATCAGGAGAACTAGATATTGATCCATTGGTTGCACTTCCAGGTCTCCGAGTTCTTGAAATTG

>M6-PGSC0003DMG400002902M6LRR-RLK_372

GGAGAACTAGATATTGATCCATTGGTTGCACTTCCAGGTCTCCGAGTTCTTGAAATTGCGAATAATTCATTTTCAGGCACAATCCCTGAATTCTTCTTGCTTGGTGCACTTAAATCTATA

>M6-PGSC0003DMG400002902M6LRR-RLK_434

TAATTCATTTTCAGGCACAATCCCTGAATTCTTCTTGCTTGGTGCACTTAAATCTATATATATAGATGGAAACCAATTCTCAGGTGAGATTCCTAAAGATTTCTTTGCCAAAATGGGGTC

>M6-PGSC0003DMG400002902M6LRR-RLK_496

TAGATGGAAACCAATTCTCAGGTGAGATTCCTAAAGATTTCTTTGCCAAAATGGGGTCTCTTAAGAAGATTTGGTTTGCACGAAACAAATTCTCCGGTTCAATTCCTGAATCTTTAACCA

>M6-PGSC0003DMG400002902M6LRR-RLK_558

AAGAAGATTTGGTTTGCACGAAACAAATTCTCCGGTTCAATTCCTGAATCTTTAACCAACTTGAAGTACTTGATGGAACTTCGTCTTGAAAGTAATGCATTTTCAGGACACGTTCCGTCA

>M6-PGSC0003DMG400002902M6LRR-RLK_62

AACAAACATGGCCGCCGTTCCCCTAATCCTTCATCTCCTTCCTCAAGCATCGTCTCCGAAAAAACACCATCTTTTCCTTATTCATATCTCCATTTTCTTCCTCATTTCTCTAGCTAATCC

>M6-PGSC0003DMG400002902M6LRR-RLK_620

GAAGTACTTGATGGAACTTCGTCTTGAAAGTAATGCATTTTCAGGACACGTTCCGTCATTATCACAAGTGAGTTTAACATCAATAGATCTTTCGAATAACAAGTTGCAAGGCGAAATCCC

>M6-PGSC0003DMG400002902M6LRR-RLK_682

CACAAGTGAGTTTAACATCAATAGATCTTTCGAATAACAAGTTGCAAGGCGAAATCCCTCAGGGGATGTCGAAATTTGGTGCTGATCCTTTTAAAGGAAATGCAGACTTATGTGGACAAC

>M6-PGSC0003DMG400002902M6LRR-RLK_744

GGGATGTCGAAATTTGGTGCTGATCCTTTTAAAGGAAATGCAGACTTATGTGGACAACAATTAGGGAAAGAATGTAATAAGGGAATCAATAAGACTCAAGGTGCACCAATGTCAAAATTG

>M6-PGSC0003DMG400002902M6LRR-RLK_806

AGGGAAAGAATGTAATAAGGGAATCAATAAGACTCAAGGTGCACCAATGTCAAAATTGAAATGGATCATTCTTGGTATAGTAGTTAGTCTTTTATTGGTGACCATTCTTTTTCGCGTCAA

>M6-PGSC0003DMG400002902M6LRR-RLK_868

GGATCATTCTTGGTATAGTAGTTAGTCTTTTATTGGTGACCATTCTTTTTCGCGTCAAGCGCAAGGAAGATCATTTCGACAAGCTTGGGAAAGAGAATCTTGATGAGGGGTTGCACGTTG

>M6-PGSC0003DMG400002902M6LRR-RLK_930

AAGGAAGATCATTTCGACAAGCTTGGGAAAGAGAATCTTGATGAGGGGTTGCACGTTGGTAGCTCGATTAGGAAGAGCACGAGCATTCATAGCAAAGGAGGTGATTCTGTTCGTGGATCA

>M6-PGSC0003DMG400002902M6LRR-RLK_992

CTCGATTAGGAAGAGCACGAGCATTCATAGCAAAGGAGGTGATTCTGTTCGTGGATCAACACGAAGGGGAGGATCACAAGGTGGTAAGGCTATGGGAGATCTTGTGTTGGTGAACAACGA

>M6-PGSC0003DMG400002906-(rev)M6LRR-RLP_0

AGTGTCCAACCTTGTCAAGCCCACTACTAAAAATGGAAAAAACAACAAATAGACCAGGCCCTTTCGTAAACCTGATATTCTAAGATAGGATTTTCGCCGCTATCTCCGCCTCTTCGCCTT

>M6-PGSC0003DMG400002906-(rev)M6LRR-RLP_1116

CTTGCTACTGATGTCTTTTTTCCTTCATGCTTCATCCTGTTTCTCTTTCTCCACAATTCCCATATTATAAAACTAGAGCTCTGTAAAAAGGTCTCATATTTGTCCCAATTTTTGTCCCCC

>M6-PGSC0003DMG400002906-(rev)M6LRR-RLP_1178

TATTATAAAACTAGAGCTCTGTAAAAAGGTCTCATATTTGTCCCAATTTTTGTCCCCCACCACAACATTATAACTTCTCTTACTTAGAGACCCTCCATGTTAAGACCTGCAAAAGAACAA

>M6-PGSC0003DMG400002906-(rev)M6LRR-RLP_124

CTCTTCACTGCTTCTGTACATGAACAACAATGTTGCCTCCATGTTACTGACCTTTATGCTTTTTTCCAGTTGTGTTCAAAAGTTTCTTCCTCTTGCTTTCTTTTCTTCTTTTCTCCCTCT

>M6-PGSC0003DMG400002906-(rev)M6LRR-RLP_1240

CAACATTATAACTTCTCTTACTTAGAGACCCTCCATGTTAAGACCTGCAAAAGAACAAAAATAGGACCAAGTCCTGTTAGCTGAATCAGATTTCAGAAACACATGGGCAAGTGTCTCATG

>M6-PGSC0003DMG400002906-(rev)M6LRR-RLP_1302

AGGACCAAGTCCTGTTAGCTGAATCAGATTTCAGAAACACATGGGCAAGTGTCTCATGTTGAGGTCTTTCACAACACCAGCATTTGGATGGGCCTTCAAGACCCCATCTTCTCATTATGT

>M6-PGSC0003DMG400002906-(rev)M6LRR-RLP_1426

AACGGGGATTTTAAATTTCCATAATCTCCACATTATGAAAGCCATTTTAAATGGTACACCACCCTTACTCCACATCCCTTTATAAATTTTGCTTTCCTCCGCCTTATGTCTGATATAATG

>M6-PGSC0003DMG400002906-(rev)M6LRR-RLP_1488

CCTTACTCCACATCCCTTTATAAATTTTGCTTTCCTCCGCCTTATGTCTGATATAATGCCAAGCAGATTTAACTGAGAAACTTCCCCTTGTCTCTGGCATCCAACAAGGTTTATCCATTG

>M6-PGSC0003DMG400002906-(rev)M6LRR-RLP_1612

TTCTTCTTTTGGAGGATAGGTTTATCCATTGTTGCTTCTTCTTTTGGAGGATGAATATTCTGCAGAATATGGTTCACCAATTCCTGTGGGAGTATCTCATGTAGTACCTCTGTATCCCAC

>M6-PGSC0003DMG400002906-(rev)M6LRR-RLP_1674

CAGAATATGGTTCACCAATTCCTGTGGGAGTATCTCATGTAGTACCTCTGTATCCCACTCCCCTTGTCTAGTTAAATCCCTAACTGTTTTATATGTATCATCACATTGAAATTCTTCTCC

>M6-PGSC0003DMG400002906-(rev)M6LRR-RLP_1736

CTTGTCTAGTTAAATCCCTAACTGTTTTATATGTATCATCACATTGAAATTCTTCTCCAGTAATTGTGTACAAATCCCTTAGGCCCGTCCAGTTATCATGCCAGACTGATACTGATCCTT

>M6-PGSC0003DMG400002906-(rev)M6LRR-RLP_1793

CAGTAATTGTGTACAAATCCCTTAGGCCCGTCCAGTTATCATGCCAGACTGATACTGATCCTTTCTCCATTTTCCATAAAATTTGGTGTTCAATGAGATCTCTAGCTTGGAGCATTTTTT

>M6-PGSC0003DMG400002906-(rev)M6LRR-RLP_186

TTTCCAGTTGTGTTCAAAAGTTTCTTCCTCTTGCTTTCTTTTCTTCTTTTCTCCCTCTCTTTGTCTCTGATTCTCATCATTTTGCAGGTGATTTTGCGATGGAGTGCAAGGGTAGTTCAA

>M6-PGSC0003DMG400002906-(rev)M6LRR-RLP_248

GTCTCTGATTCTCATCATTTTGCAGGTGATTTTGCGATGGAGTGCAAGGGTAGTTCAATTCCTCCATCTCCAGGTTAAAAGCTCCATCTTAGAATAACAATAGCATTTTCCCTATTTTGT

>M6-PGSC0003DMG400002906-(rev)M6LRR-RLP_310

TCCATCTCCAGGTTAAAAGCTCCATCTTAGAATAACAATAGCATTTTCCCTATTTTGTTGCCTTTAACTTCTATGTAATTTTGTTGTTTGTGGAAAAATATATGTTGTAGCCTTCCCTTG

>M6-PGSC0003DMG400002906-(rev)M6LRR-RLP_372

TTTAACTTCTATGTAATTTTGTTGTTTGTGGAAAAATATATGTTGTAGCCTTCCCTTGTCCCCTATCCCCTTAAGGATGATACTCTTAAATATGGGCATAGTAATTTGTCATTATTTATG

>M6-PGSC0003DMG400002906-(rev)M6LRR-RLP_434

CTATCCCCTTAAGGATGATACTCTTAAATATGGGCATAGTAATTTGTCATTATTTATGAGTCTCCTCGCAGGTGTCTTACTGTTAAAATTTGTGAATGTGCAGCTCCCTCGGTCAATATC

>M6-PGSC0003DMG400002906-(rev)M6LRR-RLP_496

TCCTCGCAGGTGTCTTACTGTTAAAATTTGTGAATGTGCAGCTCCCTCGGTCAATATCAAAGTTTGCCACATGATCTGCTAGCTTGTTTCCTCCCCTTAGAATATGTTGGAAAATAACTT

>M6-PGSC0003DMG400002906-(rev)M6LRR-RLP_558

TTTGCCACATGATCTGCTAGCTTGTTTCCTCCCCTTAGAATATGTTGGAAAATAACTTGTTTATCTTGTAGACATGATTGAATCCCTAATAATAAAAAAAAAAAAATGGTTGAATCCCTG

>M6-PGSC0003DMG400002906-(rev)M6LRR-RLP_62

TTCGTAAACCTGATATTCTAAGATAGGATTTTCGCCGCTATCTCCGCCTCTTCGCCTTAGCTCTCTTCACTGCTTCTGTACATGAACAACAATGTTGCCTCCATGTTACTGACCTTTATG

>M6-PGSC0003DMG400002906-(rev)M6LRR-RLP_620

ATCTTGTAGACATGATTGAATCCCTAATAATAAAAAAAAAAAAATGGTTGAATCCCTGTCACCATTTCTGCAATTATCCATGGACATGCCCATTTCCTTTCTAATACTTTGTACAAAAGC

>M6-PGSC0003DMG400002906-(rev)M6LRR-RLP_682

CATTTCTGCAATTATCCATGGACATGCCCATTTCCTTTCTAATACTTTGTACAAAAGCATTGAGTCAGTCTGTATGATTATTTGATTATGTTGTGACTTCTTGCAATGCTTGGACGCCTT

>M6-PGSC0003DMG400002906-(rev)M6LRR-RLP_744

AGTCAGTCTGTATGATTATTTGATTATGTTGTGACTTCTTGCAATGCTTGGACGCCTTCAGAATAGCTTTAGCTACCGCGACTGTATTAGTAGTATTTTCTATACTTGCTCCTTCCACGT

>M6-PGSC0003DMG400002906-(rev)M6LRR-RLP_806

ATAGCTTTAGCTACCGCGACTGTATTAGTAGTATTTTCTATACTTGCTCCTTCCACGTATTCTACATCTCCTTTAGCATTCCTTAAAACGAAGGCATAGGAACTAAGACCCGGATTTCCT

>M6-PGSC0003DMG400002906-(rev)M6LRR-RLP_930

AAGTTCCATTTGTATTGTATTTTACCCACCCATTCGGTGGGAATTCCCATAACCCTTTTCACTTTCATTTTAGGTTTGTAGCTTGTCCATTCTTAGAGCATATCCGGCCAGTTAACAGGA

>M6-PGSC0003DMG400002906-(rev)M6LRR-RLP_992

TTTCATTTTAGGTTTGTAGCTTGTCCATTCTTAGAGCATATCCGGCCAGTTAACAGGACATGTCATAAAGGGTTTCCTGAGTCTTGCCAGCATATACATGTTCCTGGTGATATTGTGAAT

>M6-PGSC0003DMG400002908-(rev)M6LRR-RLP_0

ATGAAATTAGGACAATTGAGTATGCTTGAAGCTTTAGATCTCTCTTGGAATCGGCTCACTGGAAAGATTCCACAGGAATTGACAAGAATAAACTTTCTCGCCTTCTTAAACCTCTCTCAA

>M6-PGSC0003DMG400002908-(rev)M6LRR-RLP_60

GGAAAGATTCCACAGGAATTGACAAGAATAAACTTTCTCGCCTTCTTAAACCTCTCTCAAAATCATCTCGTCGGACCAATTCCTCAAGGTCGGCAATTCAACACATTTGAAAATGACTCA

>M6-PGSC0003DMG400002910-(rev)M6LRR-RLP_0

AACCTACAGTATCTTGATCTGAAATGTAACTCCCTTCAGGGTCCACTACCTTCATCCATTTGTAACATGAGTTCGCTTATCTTATTAGATTTATCACGCAACAACTTCAGTGACTCAGTT

>M6-PGSC0003DMG400002910-(rev)M6LRR-RLP_1047

TCCGAAGAAGAAGGCGAGTCATATTTTTTTAGTGGATTTACTTGGGAATCAGTAGTCATAGGCTACAGCTGTGGACTAGTTGTTGGAACTGTCATGTGGAGTCTCATGTTTAAATATCGT

>M6-PGSC0003DMG400002910-(rev)M6LRR-RLP_660

ACGACAACCATAGATCTCTCAAGCAACCATTTTGAAGGTGTCATTCCGAAAACACTAAAGGATCTCATATCACTTTGGCTACTCAATTTATCTCATAACAATCTCAGAGGAGATATTCCA

>M6-PGSC0003DMG400002910-(rev)M6LRR-RLP_924

CAAGGTCGGCAATTCAACACATTTGAAAATGACTCATATGGTGGCAACCTTGATTTATGTGGTGTTCCTTTAACAAAGAAATGTGGAACGTGTGATTCATCGCATGTTCCTCAACAATTG

>M6-PGSC0003DMG400002914-(rev)M6LRR-RLP_0

AAAAAAGAACCATTCTTTGTTTCTTCTCTCTCTAACCTCAAATCTTTCTCTCTCTATTTTTGCCCTACTTTGTTGGGGTTCTTGAAAAAAACTGGATTTTTGAGGGTTCTTGAAAAAAAT

>M6-PGSC0003DMG400002914-(rev)M6LRR-RLP_1037

AAAATTCATAAGAAATGGGCATATGGATGAATACTTGTAAAAAAATTTCATTAGACAAAATTTAGTATGCTTGCTTGTTGATTTAAGGGAAAATATAGCTGACAATGTATTTGCTCCTTG

>M6-PGSC0003DMG400002914-(rev)M6LRR-RLP_1098

TTAGTATGCTTGCTTGTTGATTTAAGGGAAAATATAGCTGACAATGTATTTGCTCCTTGGATTTCAGGTGATGATGAGTTCTATTCTGGTGGATCGTTTTGTTCTGCAGATTTGGGTCTC

>M6-PGSC0003DMG400002914-(rev)M6LRR-RLP_1159

TTTCAGGTGATGATGAGTTCTATTCTGGTGGATCGTTTTGTTCTGCAGATTTGGGTCTCATGTTGTCTTTGGGTCATGCTGATGTTTACTGTCCTCCTAGGAAGAGGGCACGCATTAGTG

>M6-PGSC0003DMG400002914-(rev)M6LRR-RLP_122

GATTTTTTCATCTGGAATGCAAATCTTGGTTAAAGATATAATCTGGGTTGTGTTTTTCTTGGTTCAAAATCTGATCTTTTTTGCTTTACAGTTTGGGATTTCTTCTGAAGAACAGACATG

>M6-PGSC0003DMG400002914-(rev)M6LRR-RLP_1220

GTTGTCTTTGGGTCATGCTGATGTTTACTGTCCTCCTAGGAAGAGGGCACGCATTAGTGGCCCGTTTGTTGTCGAAGACCGGTCAAAGGATCCATCTCTTGAAATTCTTCCTGATGAATG

>M6-PGSC0003DMG400002914-(rev)M6LRR-RLP_1281

CCGTTTGTTGTCGAAGACCGGTCAAAGGATCCATCTCTTGAAATTCTTCCTGATGAATGCCTATTTGAGATCCTCAGACGATTGCCTGGTGGCCGAGAAAGGGGTGCAGCAGCTTGTGTT

>M6-PGSC0003DMG400002914-(rev)M6LRR-RLP_1342

TATTTGAGATCCTCAGACGATTGCCTGGTGGCCGAGAAAGGGGTGCAGCAGCTTGTGTTTCTAAGCGCTGGCTTACAGTGTTGAGTAGCGTGAGGAACTCTGAAATCTGCAGGAGCAAGA

>M6-PGSC0003DMG400002914-(rev)M6LRR-RLP_1403

TAAGCGCTGGCTTACAGTGTTGAGTAGCGTGAGGAACTCTGAAATCTGCAGGAGCAAGAGCTATAACAATCTAAATGACACAATAATGATCTCTAAAGATGAAGATCTTGAAGTTGAATG

>M6-PGSC0003DMG400002914-(rev)M6LRR-RLP_1464

TATAACAATCTAAATGACACAATAATGATCTCTAAAGATGAAGATCTTGAAGTTGAATGTGATGGATACCTTACTAGGTGTGTGGAAGGGAAGAAAGCTACTGATGTTAGACTTGCTGCA

>M6-PGSC0003DMG400002914-(rev)M6LRR-RLP_1525

ATGGATACCTTACTAGGTGTGTGGAAGGGAAGAAAGCTACTGATGTTAGACTTGCTGCAATTGCAGTTGGGACGTCTACCCGTGGTGGACTAGGAAAGCTTTCTATCCGAGGCAGTAACT

>M6-PGSC0003DMG400002914-(rev)M6LRR-RLP_1586

TGCAGTTGGGACGTCTACCCGTGGTGGACTAGGAAAGCTTTCTATCCGAGGCAGTAACTCAGTTCGTGGTATTACTAATGTTGGTCTATCTGCTATTGCACATGGGTGCCCTTCTCTTAG

>M6-PGSC0003DMG400002914-(rev)M6LRR-RLP_1647

GTTCGTGGTATTACTAATGTTGGTCTATCTGCTATTGCACATGGGTGCCCTTCTCTTAGGGTCCTATCATTGTGGAATGTTCCTTCTATTGGGGATGAAGGTCTCTTGGAGGTTGCAAGG

>M6-PGSC0003DMG400002914-(rev)M6LRR-RLP_1708

TCCTATCATTGTGGAATGTTCCTTCTATTGGGGATGAAGGTCTCTTGGAGGTTGCAAGGGAATGCCGTTCATTAGAAAAGCTAGATCTGAGCCACTGTCGTTCAATCTCCAACAAGGGTC

>M6-PGSC0003DMG400002914-(rev)M6LRR-RLP_1769

ATGCCGTTCATTAGAAAAGCTAGATCTGAGCCACTGTCGTTCAATCTCCAACAAGGGTCTTGTTGCAATAGCAGAGAATTGCCCGAGCTTGACTTCATTGACCATTGAATCCTGTCCAAA

>M6-PGSC0003DMG400002914-(rev)M6LRR-RLP_183

GTTCAAAATCTGATCTTTTTTGCTTTACAGTTTGGGATTTCTTCTGAAGAACAGACATGCCTACTCTTGTTAATTACAGTGGTGAGTTTCAAAAACTTCATTTCTTGGTATTTTAAGTTA

>M6-PGSC0003DMG400002914-(rev)M6LRR-RLP_1830

GTTGCAATAGCAGAGAATTGCCCGAGCTTGACTTCATTGACCATTGAATCCTGTCCAAAGATTGGAAACGAAGGCTTGCAAGCTATTGGAAGATGCTGCACCAAACTACAGTCCCTTACG

>M6-PGSC0003DMG400002914-(rev)M6LRR-RLP_1891

TTGGAAACGAAGGCTTGCAAGCTATTGGAAGATGCTGCACCAAACTACAGTCCCTTACGATTAAGGACTGCCCACTTGTTGGAGATCAGGGAGTTGCTAGTCTTCTGTCATCAGGTGCTT

>M6-PGSC0003DMG400002914-(rev)M6LRR-RLP_1952

TAAGGACTGCCCACTTGTTGGAGATCAGGGAGTTGCTAGTCTTCTGTCATCAGGTGCTTCAATGTTATCGAAAGTGAAACTACATGGTCTAAACATCACAGATTTCTCGCTTGCTGTCAT

>M6-PGSC0003DMG400002914-(rev)M6LRR-RLP_2013

ATGTTATCGAAAGTGAAACTACATGGTCTAAACATCACAGATTTCTCGCTTGCTGTCATTGGTCACTATGGCAAGCTGATTACCAGTCTTAATCTTTGTTCACTTCGTAACGTGAGTCAG

>M6-PGSC0003DMG400002914-(rev)M6LRR-RLP_2074

GTCACTATGGCAAGCTGATTACCAGTCTTAATCTTTGTTCACTTCGTAACGTGAGTCAGAAGGGGTTTTGGGTCATGGGTAATGCTCAAGGCCTCCAGTCTCTGGTTTCTTTGACAATCA

>M6-PGSC0003DMG400002914-(rev)M6LRR-RLP_2135

GGGGTTTTGGGTCATGGGTAATGCTCAAGGCCTCCAGTCTCTGGTTTCTTTGACAATCACCTTGTGCCAGGGAGCCACAGATGTCGGTCTTGAAGCAGTGGGAAAGGGATGCCCAAATCT

>M6-PGSC0003DMG400002914-(rev)M6LRR-RLP_2196

TTGTGCCAGGGAGCCACAGATGTCGGTCTTGAAGCAGTGGGAAAGGGATGCCCAAATCTCAAACATATGTGCATTCGCAAGTGTTGCTTTGTTTCTGACGGTGGACTTGTTGCTTTCGCT

>M6-PGSC0003DMG400002914-(rev)M6LRR-RLP_2257

AACATATGTGCATTCGCAAGTGTTGCTTTGTTTCTGACGGTGGACTTGTTGCTTTCGCTAAAGAGGCTGGATCTCTCGAGAGTTTAATCTTGGAGGAGTGCAACAGAATTACCCAAGTAG

>M6-PGSC0003DMG400002914-(rev)M6LRR-RLP_2318

AGAGGCTGGATCTCTCGAGAGTTTAATCTTGGAGGAGTGCAACAGAATTACCCAAGTAGGTATCCTTAATGCAGTTTCAAACTGCAGGAAGTTAAAGTCTCTTTCCCTAGTGAAGTGCAT

>M6-PGSC0003DMG400002914-(rev)M6LRR-RLP_2379

ATCCTTAATGCAGTTTCAAACTGCAGGAAGTTAAAGTCTCTTTCCCTAGTGAAGTGCATGGGAATCAAGGATCTAGCTCTACAAACTTCCATGCTGTCTCCATGTGAATCTCTTCGGTCC

>M6-PGSC0003DMG400002914-(rev)M6LRR-RLP_244

TACTCTTGTTAATTACAGTGGTGAGTTTCAAAAACTTCATTTCTTGGTATTTTAAGTTATGGGGTTGTTCATATTTCAATGAATTTTGAAGTTCAGTTATAATCTTTTGTTTTTTCATCA

>M6-PGSC0003DMG400002914-(rev)M6LRR-RLP_2440

GAATCAAGGATCTAGCTCTACAAACTTCCATGCTGTCTCCATGTGAATCTCTTCGGTCCTTGTCTATCCGAAGCTGTCCAGGATTCGGGAGCACTAGCTTAGCTATGGTCGGTAAGCTCT

>M6-PGSC0003DMG400002914-(rev)M6LRR-RLP_2501

GTCTATCCGAAGCTGTCCAGGATTCGGGAGCACTAGCTTAGCTATGGTCGGTAAGCTCTGCCCCAAGCTGCATCAATTAGATCTCAGTGGGCTTTGTGGAATAACCGATGCTGGTCTTCT

>M6-PGSC0003DMG400002914-(rev)M6LRR-RLP_2562

CCCAAGCTGCATCAATTAGATCTCAGTGGGCTTTGTGGAATAACCGATGCTGGTCTTCTCCCACTCTTGGAGAACTGTGAAGGACTTGTTAAGGTAAATCTTAGCGACTGCCTGAACTTG

>M6-PGSC0003DMG400002914-(rev)M6LRR-RLP_2623

CACTCTTGGAGAACTGTGAAGGACTTGTTAAGGTAAATCTTAGCGACTGCCTGAACTTGACAGACCAAGTGGTGCTTTCATTGGCTTCGCGACATGGAGAGACCCTTGAATTGTTGAATC

>M6-PGSC0003DMG400002914-(rev)M6LRR-RLP_2684

AGACCAAGTGGTGCTTTCATTGGCTTCGCGACATGGAGAGACCCTTGAATTGTTGAATCTCGATGGATGCAGGAAGGTTACTGACGCAAGTTTAGTGGCAATTGCAGATTATTGCCCATT

>M6-PGSC0003DMG400002914-(rev)M6LRR-RLP_2745

GATGGATGCAGGAAGGTTACTGACGCAAGTTTAGTGGCAATTGCAGATTATTGCCCATTACTTATTGATCTTGATGTTTCTAAGTGTGCAATAACTGATTCTGGTGTAGCTGCTTTATCC

>M6-PGSC0003DMG400002914-(rev)M6LRR-RLP_2806

TTATTGATCTTGATGTTTCTAAGTGTGCAATAACTGATTCTGGTGTAGCTGCTTTATCCCGTGGAGTGCAAGTGAATTTGCAGGTCCTTTCTTTATCAGGTTGCTCCATGGTATCAAACA

>M6-PGSC0003DMG400002914-(rev)M6LRR-RLP_2867

TGGAGTGCAAGTGAATTTGCAGGTCCTTTCTTTATCAGGTTGCTCCATGGTATCAAACAAGAGTGTCCCTTCTCTTAAAAAGTTGGGAGAGAATCTACTCGGCTTGAATCTCCAACATTG

>M6-PGSC0003DMG400002914-(rev)M6LRR-RLP_2928

AGTGTCCCTTCTCTTAAAAAGTTGGGAGAGAATCTACTCGGCTTGAATCTCCAACATTGCTCCATTAGTTGCAGCTCAGTCGAGCTTCTTGTGGAGGATTTGTGGAGGTGTGACATCCTC

>M6-PGSC0003DMG400002914-(rev)M6LRR-RLP_2989

CCATTAGTTGCAGCTCAGTCGAGCTTCTTGTGGAGGATTTGTGGAGGTGTGACATCCTCTCCTAAACCAAAAGGGCACAAGGAAATGCATCAAAATGGTCAAAAAGAATAAGCTGAAAGC

>M6-PGSC0003DMG400002914-(rev)M6LRR-RLP_305

GGGTTGTTCATATTTCAATGAATTTTGAAGTTCAGTTATAATCTTTTGTTTTTTCATCATGAATTACTGAGATGGAGATGGATTTGTTATGTTTAGATTCATATTAGTTGTTTATCATAA

>M6-PGSC0003DMG400002914-(rev)M6LRR-RLP_3050

CTAAACCAAAAGGGCACAAGGAAATGCATCAAAATGGTCAAAAAGAATAAGCTGAAAGCTATAATGGTTCATTCAGCCTGGCAATTACAGAAGCAAACATGAGTGTTCTTTTTTTAGCCC

>M6-PGSC0003DMG400002914-(rev)M6LRR-RLP_3111

TAATGGTTCATTCAGCCTGGCAATTACAGAAGCAAACATGAGTGTTCTTTTTTTAGCCCGTATATTCGTGGCTATCTAATACTTCAGTGTCTCGGTTGTTTTCCAGGAGGCTATGGCATC

>M6-PGSC0003DMG400002914-(rev)M6LRR-RLP_3172

ATATTCGTGGCTATCTAATACTTCAGTGTCTCGGTTGTTTTCCAGGAGGCTATGGCATCACTTTTTTCCTCCATTTTTTGCAGGCTTCGTCAAATCGAAGCCTGTTTCATCCATTTTTGG

>M6-PGSC0003DMG400002914-(rev)M6LRR-RLP_3233

TTTTTTCCTCCATTTTTTGCAGGCTTCGTCAAATCGAAGCCTGTTTCATCCATTTTTGGAGACAACTCCATCATGCCATGGTTTCCTGAGAATACAGTCACCTAGTACTGAGATTAAAAT

>M6-PGSC0003DMG400002914-(rev)M6LRR-RLP_3294

ACAACTCCATCATGCCATGGTTTCCTGAGAATACAGTCACCTAGTACTGAGATTAAAATTTTCACTCATGTTGCTCCTTGAAAGAATCAGGCCGTTTGAATCATTCTCAGCTACAACATC

>M6-PGSC0003DMG400002914-(rev)M6LRR-RLP_3355

TCACTCATGTTGCTCCTTGAAAGAATCAGGCCGTTTGAATCATTCTCAGCTACAACATCGACGGACAATATCTATCGCGCATTTGCCTCCTTGGTTAGCTACATTGCAGAAATCAGACAG

>M6-PGSC0003DMG400002914-(rev)M6LRR-RLP_3416

CGGACAATATCTATCGCGCATTTGCCTCCTTGGTTAGCTACATTGCAGAAATCAGACAGTAGTACTGTAGTTTTTCATGTTTTAGTGTTTCATTGTAAGTATTCTGTTGTGTGTCTTTTA

>M6-PGSC0003DMG400002914-(rev)M6LRR-RLP_3477

GTACTGTAGTTTTTCATGTTTTAGTGTTTCATTGTAAGTATTCTGTTGTGTGTCTTTTAGACTTTAGTTGCTTTGAGCAAATTTCTGTTGTTGTGGTGCCTGTTATTCCAGGGGCTTGGC

>M6-PGSC0003DMG400002914-(rev)M6LRR-RLP_3538

CTTTAGTTGCTTTGAGCAAATTTCTGTTGTTGTGGTGCCTGTTATTCCAGGGGCTTGGCCATTACAACCTTATGTAGGTTGTTATTTTTTAACCAGTCCCTAGTTTTTTGATAGGATTAT

>M6-PGSC0003DMG400002914-(rev)M6LRR-RLP_3599

TTACAACCTTATGTAGGTTGTTATTTTTTAACCAGTCCCTAGTTTTTTGATAGGATTATCAATGTCCTATTTTTTACCAGGTTCCACATATTCAGATTTGTACTCTTGCAACTATGTTTT

>M6-PGSC0003DMG400002914-(rev)M6LRR-RLP_3630

CCAGTCCCTAGTTTTTTGATAGGATTATCAATGTCCTATTTTTTACCAGGTTCCACATATTCAGATTTGTACTCTTGCAACTATGTTTTATATATGATGGAAGTACTAGTTTGTTTTGAA

>M6-PGSC0003DMG400002914-(rev)M6LRR-RLP_366

AATTACTGAGATGGAGATGGATTTGTTATGTTTAGATTCATATTAGTTGTTTATCATAAATAAAGTTTATAACTTTGCATAGTTACATGATTGTTGAGATGAATAGGAGCAATAATTTAT

>M6-PGSC0003DMG400002914-(rev)M6LRR-RLP_427

AAAGTTTATAACTTTGCATAGTTACATGATTGTTGAGATGAATAGGAGCAATAATTTATTGAGATGAAGATGAATTTTTTAAGGTTTAGATCCATGAATATAGTCTATAACTATGTATAG

>M6-PGSC0003DMG400002914-(rev)M6LRR-RLP_488

AGATGAAGATGAATTTTTTAAGGTTTAGATCCATGAATATAGTCTATAACTATGTATAGGTGCATGATTGTTGATAACTTGAGATAAATATGGTTTATTAATTTACTGAAATGAAGATGA

>M6-PGSC0003DMG400002914-(rev)M6LRR-RLP_549

GCATGATTGTTGATAACTTGAGATAAATATGGTTTATTAATTTACTGAAATGAAGATGATCTTTTAGGTTTAGATACAATCATGTTTTTTTTTTTTAAAGGAAGTTGGTTATGAGATAAT

>M6-PGSC0003DMG400002914-(rev)M6LRR-RLP_61

GCCCTACTTTGTTGGGGTTCTTGAAAAAAACTGGATTTTTGAGGGTTCTTGAAAAAAATTGGATTTTTTCATCTGGAATGCAAATCTTGGTTAAAGATATAATCTGGGTTGTGTTTTTCT

>M6-PGSC0003DMG400002914-(rev)M6LRR-RLP_610

TTTTAGGTTTAGATACAATCATGTTTTTTTTTTTTAAAGGAAGTTGGTTATGAGATAATAACTAAAAGTTGAGTGACTATCTAAGAAATTAACTCATATAACTTTGTATAGGTGCATGAT

>M6-PGSC0003DMG400002914-(rev)M6LRR-RLP_671

CTAAAAGTTGAGTGACTATCTAAGAAATTAACTCATATAACTTTGTATAGGTGCATGATTGTTGAGGTGAATATGAGCATTAATTTACTGAAATGGATAAAAAGGGAAAAAAGTTATGAT

>M6-PGSC0003DMG400002914-(rev)M6LRR-RLP_732

TTGAGGTGAATATGAGCATTAATTTACTGAAATGGATAAAAAGGGAAAAAAGTTATGATAAGACTAGATCCTTTGGCTTTGAAGATTAATTTCTGTGTTTGATAGATCTTTTTTTGTCTT

>M6-PGSC0003DMG400002914-(rev)M6LRR-RLP_793

GACTAGATCCTTTGGCTTTGAAGATTAATTTCTGTGTTTGATAGATCTTTTTTTGTCTTATGTTCTGAATCTATATTAGAGATTGGTAATATATTGAAGAAACTGTGGATTACTCTTTTG

>M6-PGSC0003DMG400002914-(rev)M6LRR-RLP_854

GTTCTGAATCTATATTAGAGATTGGTAATATATTGAAGAAACTGTGGATTACTCTTTTGGGTTCTGTATCTAGAAACGTGTATCTATCTGATCTGATCTGAAGATGTATGTACATAATAG

>M6-PGSC0003DMG400002914-(rev)M6LRR-RLP_915

TTCTGTATCTAGAAACGTGTATCTATCTGATCTGATCTGAAGATGTATGTACATAATAGGTAGATCTAAAGGTTCAATTCACATGTGATAGTTGCTTATCTGTTATACTTGAAGTTGAAG

>M6-PGSC0003DMG400002914-(rev)M6LRR-RLP_976

AGATCTAAAGGTTCAATTCACATGTGATAGTTGCTTATCTGTTATACTTGAAGTTGAAGTGAAAATTCATAAGAAATGGGCATATGGATGAATACTTGTAAAAAAATTTCATTAGACAAA

>M6-PGSC0003DMG400002937M6LRR-RLP_0

GATTGATTTGTAAGAAGAGAACATCCATCCTGTTTTTGGCTGAATACGAAAAAGAAAGATGAGACGTTTCTTTTTCTTTTATTCATTTCTCTGTTTTGTGTTGCTAATAAGTCGAAGTTT

>M6-PGSC0003DMG400002937M6LRR-RLP_1054

CGTTTGGATCTTTCTGACAACTCCTTTTCAGGTGAAATTCCCAACGTTTTCTCTAACCTCCAAGAGCTACGTTATTTAAATCTTTATGATAACAGCCTCATCGGTTCATTTCCTTCTACA

>M6-PGSC0003DMG400002937M6LRR-RLP_1116

AGAGCTACGTTATTTAAATCTTTATGATAACAGCCTCATCGGTTCATTTCCTTCTACAATTTCAAACTTGACACATCTTGAAAGCTTAGCCTTGTCGAGTAATTCCCTGTCTGGCCCACT

>M6-PGSC0003DMG400002937M6LRR-RLP_1240

AATAATGCGAGCATTCTTCAAAAGCTAACCCACATGGAATTGTATAACAACTCACTGAATGGTACCATACCGCCTTGGGTGTTTAACCTCCCTTTAGCATCTACATTGTGGCTCCATCAT

>M6-PGSC0003DMG400002937M6LRR-RLP_1302

TACCATACCGCCTTGGGTGTTTAACCTCCCTTTAGCATCTACATTGTGGCTCCATCATAATCAATTTAGTGGACTACCCGATGAAATCAAAACAAACCCAACATTAGAATACCTGTATTT

>M6-PGSC0003DMG400002937M6LRR-RLP_1364

AATTTAGTGGACTACCCGATGAAATCAAAACAAACCCAACATTAGAATACCTGTATTTAAGCAATAATCAACTCAGTTTTTTACACCAATCACTTGCGAATCTCACAAACCTTTCTACCC

>M6-PGSC0003DMG400002937M6LRR-RLP_1426

AATAATCAACTCAGTTTTTTACACCAATCACTTGCGAATCTCACAAACCTTTCTACCCTTGACCTTTCAACAAATAACATCACCGGTGATGCAGGAACAAATATCACCTTTCCTCGCCTA

>M6-PGSC0003DMG400002937M6LRR-RLP_1488

CCTTTCAACAAATAACATCACCGGTGATGCAGGAACAAATATCACCTTTCCTCGCCTACAAGTTTTGCAGTTATCATCTTGTGAACTGAAGGGATTTCCACACTTCTTGAGAAATGTAAA

>M6-PGSC0003DMG400002937M6LRR-RLP_1550

TTTTGCAGTTATCATCTTGTGAACTGAAGGGATTTCCACACTTCTTGAGAAATGTAAAGACACTTAAGGTGTTGGATATTTCTAACAATAAGATTCGTGGTCAAATCCCTAACTGGTTTA

>M6-PGSC0003DMG400002937M6LRR-RLP_1674

CATGAGGTGGGACTCCTTGTTGCACCTAAACCTTTCACATAACTCATTAACAGGCCACCTACAGCAACTTCATTATAATAGTCTAGAGTCTCTTGATCTTAAATTTAACTTCCTTGAGGG

>M6-PGSC0003DMG400002937M6LRR-RLP_1736

AGCAACTTCATTATAATAGTCTAGAGTCTCTTGATCTTAAATTTAACTTCCTTGAGGGAGCACTACCTTCATCCATTTGTAACATGAGCAGTCTTAGCCTTTTGGATTTGTCAAACAACT

>M6-PGSC0003DMG400002937M6LRR-RLP_1798

CTACCTTCATCCATTTGTAACATGAGCAGTCTTAGCCTTTTGGATTTGTCAAACAACTACTTCAGTGACTCGGTTCCACATTGTGTGGGCAGCATGGTTGGGCTATCTGTGCTAGACTTG

>M6-PGSC0003DMG400002937M6LRR-RLP_186

ACACTTTGAGATTATATCCGACCCTTATTCCCGATGTCCTATTCATTTCCCAAGAACAATGTCTTGGAATGAGAGTAGGGATTGCTGCATTTGGAATGGAGTCACTTGTGATATGTTAAC

>M6-PGSC0003DMG400002937M6LRR-RLP_1860

CAGTGACTCGGTTCCACATTGTGTGGGCAGCATGGTTGGGCTATCTGTGCTAGACTTGAGAAGAAATAATTTCACTGGGAGTCTTCCACCGTTTTGTGTGCAGAGCAATTCATTGAGAAC

>M6-PGSC0003DMG400002937M6LRR-RLP_1922

GAAATAATTTCACTGGGAGTCTTCCACCGTTTTGTGTGCAGAGCAATTCATTGAGAACCATTGTCTTGAATGGTAATCGTTTTGAAGGAACTGTCCCGATGTCGTTGCTCAAGTGTGATG

>M6-PGSC0003DMG400002937M6LRR-RLP_1984

GTCTTGAATGGTAATCGTTTTGAAGGAACTGTCCCGATGTCGTTGCTCAAGTGTGATGGTCTAGAAGTCCTTGATGTGGGGAACAACGCTATAAATGACACGTTTCCAGCTTGGCTCGGA

>M6-PGSC0003DMG400002937M6LRR-RLP_2108

TTCAAGAGCTGCAGGTACTTATATTAAAGTCCAACAAGTTCCATGGACCTATAAGTACTAGGCTGAAGTTTGGTTTCCCTGTGTTACGGATTTTTGATCTCTCTCATAATGAGTTTGGTG

>M6-PGSC0003DMG400002937M6LRR-RLP_2170

CTGAAGTTTGGTTTCCCTGTGTTACGGATTTTTGATCTCTCTCATAATGAGTTTGGTGGCTCACTGCCTGCAGAAGTTTTTGAAAACTTCAAAGGAATGATCAAAACAGATGACAGAGAC

>M6-PGSC0003DMG400002937M6LRR-RLP_2232

ACTGCCTGCAGAAGTTTTTGAAAACTTCAAAGGAATGATCAAAACAGATGACAGAGACAAAGGTGAGATTGAATATATGCAACCACAGTCTTACAGCGGATTTAGTAACGTAATGTATGA

>M6-PGSC0003DMG400002937M6LRR-RLP_2294

GTGAGATTGAATATATGCAACCACAGTCTTACAGCGGATTTAGTAACGTAATGTATGAGGTTTCAGTGAGGTTGGTGATCAAAAACCAGGAAATTCAGCTAGAAAAAATCACCACAATTA

>M6-PGSC0003DMG400002937M6LRR-RLP_2356

TCAGTGAGGTTGGTGATCAAAAACCAGGAAATTCAGCTAGAAAAAATCACCACAATTATGACAACCATAGATCTCTCAGGCAACCATTTTGAAGGTGTCATTCCGAAAACACTAAAGGAT

>M6-PGSC0003DMG400002937M6LRR-RLP_2418

AACCATAGATCTCTCAGGCAACCATTTTGAAGGTGTCATTCCGAAAACACTAAAGGATCTCAACTCACTTTGGCTACTCAATTTATCTCATAACAATCTTAAAGGAGATATTCCAATGGA

>M6-PGSC0003DMG400002937M6LRR-RLP_248

CTTGGAATGAGAGTAGGGATTGCTGCATTTGGAATGGAGTCACTTGTGATATGTTAACCGGTCGTGTTATCGCGTTGGACCTTAGTTGCAGTGAGCTTTGGGGAAGTATTCATCCCAACA

>M6-PGSC0003DMG400002937M6LRR-RLP_2542

GTGAAATTGAATATGCTTGAAGCGTTAGATCTCTCTTGGAATCGGCTCACTGGAAAGATTCCACAGGAATTCACAAGAATGAACTTTCTGTCATTCTTAAACCTCTCTCAAAATCTCCTT

>M6-PGSC0003DMG400002937M6LRR-RLP_2882

TTGTTGGAACTGTCATGTGGAGCCTCATGTTTAAATATCGTAAGCCAAAATGGTTTGTCAAATTTTTTGAAGGCATTTTCCCCAAGAAAATGAGAAGGTCACAGAAGAGAATGCTGAGAC

>M6-PGSC0003DMG400002937M6LRR-RLP_310

CGTGTTATCGCGTTGGACCTTAGTTGCAGTGAGCTTTGGGGAAGTATTCATCCCAACAGCAGCCTCTTCCAACTTCATCATCTCCACACACTAAACCTTGCTTACAATGACTTCAATTAT

>M6-PGSC0003DMG400002937M6LRR-RLP_372

CCTCTTCCAACTTCATCATCTCCACACACTAAACCTTGCTTACAATGACTTCAATTATTCTTCAATCCCACATAACATTGGCCGATTGAGGAATTTGAGGCATCTCAACCTTTCTTGGTC

>M6-PGSC0003DMG400002937M6LRR-RLP_434

CAATCCCACATAACATTGGCCGATTGAGGAATTTGAGGCATCTCAACCTTTCTTGGTCTCGCTTTGATGGAAAAATCCCAATAGAATTCTCATATCTTTCCAATTTGGTTTCACTTGATC

>M6-PGSC0003DMG400002937M6LRR-RLP_496

TTTGATGGAAAAATCCCAATAGAATTCTCATATCTTTCCAATTTGGTTTCACTTGATCTTTCTGCGAATGGATTACAACTTGATGAGAGAACATTTGAAACAATGCTTCAGAACTTCACA

>M6-PGSC0003DMG400002937M6LRR-RLP_558

TGCGAATGGATTACAACTTGATGAGAGAACATTTGAAACAATGCTTCAGAACTTCACAAATCTGGAGCTACTTTCTCTCTTTCAAGTCAACATCTCATCTCCGATACCTGTGAATATGTC

>M6-PGSC0003DMG400002937M6LRR-RLP_62

GACGTTTCTTTTTCTTTTATTCATTTCTCTGTTTTGTGTTGCTAATAAGTCGAAGTTTTTCTTCTTCCTTTGAGCATCATCTTTGCTCTCCCACTGAAGCTTCTGCTTTGCTTCAGTTTA

>M6-PGSC0003DMG400002937M6LRR-RLP_682

TCCTTAAGGTACGTGGATCTTGGAATCACTAATCTGCGAGGTGTTGTCCCAGAGAGTCTTTTCCTTCTGCCAAACTTGGAAACGCTCTATTTGAGTAATAACTATCTTCTCAATGGAGTT

>M6-PGSC0003DMG400002937M6LRR-RLP_744

CCTTCTGCCAAACTTGGAAACGCTCTATTTGAGTAATAACTATCTTCTCAATGGAGTTTTACCAAAGATCCACCCGAGCAACACTCTGTTAATGGAGTTGGATATTTCATACACAGGAAT

>M6-PGSC0003DMG400002937M6LRR-RLP_806

CAAAGATCCACCCGAGCAACACTCTGTTAATGGAGTTGGATATTTCATACACAGGAATCTCTGGCTCCATTCCTGATTCAATTGGCACCTTGAGTTCCTTGAATATCTTGATCCTCCAAG

>M6-PGSC0003DMG400002937M6LRR-RLP_868

GGCTCCATTCCTGATTCAATTGGCACCTTGAGTTCCTTGAATATCTTGATCCTCCAAGGATGTCAATTGTCGGGTTCCATTCCTGATTCCATAGGCAACCTAACACAAATTACGGAGTTG

>M6-PGSC0003DMG400002937M6LRR-RLP_930

TCAATTGTCGGGTTCCATTCCTGATTCCATAGGCAACCTAACACAAATTACGGAGTTGATTTTATCTTTTAATCATTTCACTGGCCATATTCCTTCAACAATCTCTAAATTGAAGCACCT

>M6-PGSC0003DMG400002937M6LRR-RLP_992

TATCTTTTAATCATTTCACTGGCCATATTCCTTCAACAATCTCTAAATTGAAGCACCTCACACGTTTGGATCTTTCTGACAACTCCTTTTCAGGTGAAATTCCCAACGTTTTCTCTAACC

>M6-PGSC0003DMG400002954M6LRR-RLP_0

TTTTCTAATAGATTCTAAAAGAGTTGATGCTGATATTGAAGATAGATTCAGTGCTCTACCAATAAACCTTGTAGATGATATTCTAAGGCTCTTACCAGTTCACGACGCAGCAAGAACTAG

>M6-PGSC0003DMG400002954M6LRR-RLP_1024

TGATTAAAAGTTTCCCCAATTTGAGTAAACTTGAGATTTGGGTACGAAATAGCAAATATATATGTATGATCTTGTTCGTTTTTGCCCTTGGCATACTTAAGTATGAGAACTAAAATGACG

>M6-PGSC0003DMG400002954M6LRR-RLP_1088

TATGATCTTGTTCGTTTTTGCCCTTGGCATACTTAAGTATGAGAACTAAAATGACGTTGTATTCTTTTAGGATAATGCTTTCGGTGACCCTGATGAAGCAGTTATGAAATATCTAGACGC

>M6-PGSC0003DMG400002954M6LRR-RLP_1152

TTTTAGGATAATGCTTTCGGTGACCCTGATGAAGCAGTTATGAAATATCTAGACGCACCATCATGTTTGGACCAAACACTCAACAAGCTTGAAGATGTTTCCATCCATATTTTTAGGCGA

>M6-PGSC0003DMG400002954M6LRR-RLP_1216

GTTTGGACCAAACACTCAACAAGCTTGAAGATGTTTCCATCCATATTTTTAGGCGATCCAAAGTTGTACTATCTTTCATAAAGTTATTGTTTGCTCATGCTCCATCTTTGTCAAGGATGA

>M6-PGSC0003DMG400002954M6LRR-RLP_128

CAAAAAATTGGAGATATATTTGGGCCGCGCTTCCCAATTTAATGCTTGATAAGACCTTTTGCAGTAAATTAGCTGTAAAATCTCAATTTGTATTGAAAGAAACAATAGATATGATTCTCT

>M6-PGSC0003DMG400002954M6LRR-RLP_1280

TGTACTATCTTTCATAAAGTTATTGTTTGCTCATGCTCCATCTTTGTCAAGGATGAGCATTAAGCCGAAAAATGCTTCTAGTTCCAAAGAAGAATTGAATGTTGCTACAGAGTTGATGTG

>M6-PGSC0003DMG400002954M6LRR-RLP_1336

GCATTAAGCCGAAAAATGCTTCTAGTTCCAAAGAAGAATTGAATGTTGCTACAGAGTTGATGTGTTTCCCCAGAGTATCTCCTAAGGCAAAGCTATTCTATCATCCAACAGTTGGATTAA

>M6-PGSC0003DMG400002954M6LRR-RLP_192

TAAATTAGCTGTAAAATCTCAATTTGTATTGAAAGAAACAATAGATATGATTCTCTTACAACATATTGGAGATATTGTGAAGTTTGATCTTGATGTTTCAGGAGTACAATTGTGTTCATA

>M6-PGSC0003DMG400002954M6LRR-RLP_256

ATTGGAGATATTGTGAAGTTTGATCTTGATGTTTCAGGAGTACAATTGTGTTCATATGCAGACATCGATAGATGGATGCTTTATGTCACGAGATATGGTGTCAAGGAATTAACCCTTTTC

>M6-PGSC0003DMG400002954M6LRR-RLP_320

TCGATAGATGGATGCTTTATGTCACGAGATATGGTGTCAAGGAATTAACCCTTTTCATGTCAAAGAATATTAACAGTACTTATAAAGTGCCTTCTTATATATTCAATTGTCAAACCCTGA

>M6-PGSC0003DMG400002954M6LRR-RLP_384

GAATATTAACAGTACTTATAAAGTGCCTTCTTATATATTCAATTGTCAAACCCTGACACAATTGAAACTCTTCAATTGCATCTTCAAATTACCAAATTCTTTTCTTGGGTTTCAAACTCT

>M6-PGSC0003DMG400002954M6LRR-RLP_448

AAACTCTTCAATTGCATCTTCAAATTACCAAATTCTTTTCTTGGGTTTCAAACTCTTACCATCCTTCAGTTGAAAGAAGTAACCTTCGCGCCTGCAATAGAGTTTTATGCTATCAAAACG

>M6-PGSC0003DMG400002954M6LRR-RLP_512

TTCAGTTGAAAGAAGTAACCTTCGCGCCTGCAATAGAGTTTTATGCTATCAAAACGCCACTTCTTGTCGAATTAATCTTGACACATTGTGAGGGCACTCAATATCTTAACATTGAGTTCT

>M6-PGSC0003DMG400002954M6LRR-RLP_576

TGTCGAATTAATCTTGACACATTGTGAGGGCACTCAATATCTTAACATTGAGTTCTCATCAGGATTGAAGTACTTGTTTGTTCGTGAGAGTCACTATAATCTTGATTTAAATAGCAATTA

>M6-PGSC0003DMG400002954M6LRR-RLP_64

AACCTTGTAGATGATATTCTAAGGCTCTTACCAGTTCACGACGCAGCAAGAACTAGTATTTTGTCAAAAAATTGGAGATATATTTGGGCCGCGCTTCCCAATTTAATGCTTGATAAGACC

>M6-PGSC0003DMG400002954M6LRR-RLP_640

TTGAAGTACTTGTTTGTTCGTGAGAGTCACTATAATCTTGATTTAAATAGCAATTACTTTAAGAACGTCAAAAAATTGACATACTTATACCTTGTGATTGATAATCCAATACGTGTTGAT

>M6-PGSC0003DMG400002954M6LRR-RLP_704

ACGTCAAAAAATTGACATACTTATACCTTGTGATTGATAATCCAATACGTGTTGATGAAAGATTAACTTTGGAAAAGCTTCTTTTTAGTTCGCCTACACTTGAGACACTTACTTTGGGTT

>M6-PGSC0003DMG400002954M6LRR-RLP_768

AACTTTGGAAAAGCTTCTTTTTAGTTCGCCTACACTTGAGACACTTACTTTGGGTTTTTATGTACTTGAGGTAAGAAGCTAACATGTTGTTTTACTTAGATATCTACGTTATCCTTCTGT

>M6-PGSC0003DMG400002954M6LRR-RLP_832

CTTGAGGTAAGAAGCTAACATGTTGTTTTACTTAGATATCTACGTTATCCTTCTGTTTGTATAATTTTTCGTTAAAGCTTTTGAATGCAGGTACAGTCCCAAAGGGGCTTCATTATACAC

>M6-PGSC0003DMG400002954M6LRR-RLP_896

TTTTTCGTTAAAGCTTTTGAATGCAGGTACAGTCCCAAAGGGGCTTCATTATACACTCAACTGCTTGTGGCATCTATAGTTAGGTGTAAACTTCAACAAATTGGGTCAGACTTGTTACAC

>M6-PGSC0003DMG400002954M6LRR-RLP_960

TTGTGGCATCTATAGTTAGGTGTAAACTTCAACAAATTGGGTCAGACTTGTTACACTCTGGAGTTGATTAAAAGTTTCCCCAATTTGAGTAAACTTGAGATTTGGGTACGAAATAGCAAA

>M6-PGSC0003DMG400002960M6LRR-RLP_0

ATACACAAGAGTAGTACATTATTTGATATCTTGAGTTACATCTAAAAATTATTTTTGTTGTAATTCCTCTCCATATTGACTTGATGGCAGCTTATACTGCTGTAATTTCTCTTCTTCAAA

>M6-PGSC0003DMG400002960M6LRR-RLP_1054

ACTAGAACACGATCACCCTCCCGAGTTGGAAGAAATTGGAAAAACAATAGCAGGAAAATGCCAAGGACTTCCCTTAATAATTTCAGTGATTGCAGGGCATCTCTCTAAAACGGTCCCCAA

>M6-PGSC0003DMG400002960M6LRR-RLP_1116

AAGGACTTCCCTTAATAATTTCAGTGATTGCAGGGCATCTCTCTAAAACGGTCCCCAAGACATTAAAAAGTTGGAAGTATGTTGCCAAAACCTTAAGTAAAATCATTGCTAGTCATCCAA

>M6-PGSC0003DMG400002960M6LRR-RLP_1178

TTAAAAAGTTGGAAGTATGTTGCCAAAACCTTAAGTAAAATCATTGCTAGTCATCCAAATAAATGCTTAGGAGTGCTCGGTTTGAGTTACCATCACTTGCCTAATCAACTCAAACCATGC

>M6-PGSC0003DMG400002960M6LRR-RLP_124

TGATCAATCAAATATTTCAGAACTCTTTCATGATCACAGTACTGAAATGCTCGATTCTCTTCGGGCTACAGCTGAATATTTTCAAGATGTTCTTGACAACAATAGCAATGAGAAAATCAA

>M6-PGSC0003DMG400002960M6LRR-RLP_1426

ACATTATTTGGAGGATCTTATCAGCAGGAACTTGATAATGGCAGTAAAAAAAAGATTCAATGGTGAGGTAAAAGTATGCGGAATACATGATCTGCTGCGTGAGTTCTGTTTGATAGAAGC

>M6-PGSC0003DMG400002960M6LRR-RLP_1550

ATGACAAAGTTTATGCATGTTGTGAGAACTCCATTGTCACCAGCACAAAAGCCTAATTTTCGTCGCTTCAGTATTCAAGAGTAATATTCTGTTGAGGATTCTTATAAGCTCTTACCCCCT

>M6-PGSC0003DMG400002960M6LRR-RLP_1612

TCGCTTCAGTATTCAAGAGTAATATTCTGTTGAGGATTCTTATAAGCTCTTACCCCCTGTAGCCAGATCTATCTACTGCTTTTCTTACTTGGATCTACCTTATGAACCCCGTATTCTTCC

>M6-PGSC0003DMG400002960M6LRR-RLP_1674

CCAGATCTATCTACTGCTTTTCTTACTTGGATCTACCTTATGAACCCCGTATTCTTCCCCCTGTATTGTCCATCTACCATCATGATCCTATAATACATGGATTTTTCTGTTGTTTCAACC

>M6-PGSC0003DMG400002960M6LRR-RLP_1736

GTATTGTCCATCTACCATCATGATCCTATAATACATGGATTTTTCTGTTGTTTCAACCATCTCAGGGTATTGGCCATCTTTCCTTTCTCGTTCAGCTCATTTCCACTTGTGATTACAAAG

>M6-PGSC0003DMG400002960M6LRR-RLP_1798

CAGGGTATTGGCCATCTTTCCTTTCTCGTTCAGCTCATTTCCACTTGTGATTACAAAGTTATTTCATTTGAGATATCTCCAAGTTCGATGTCTTGGAAATATTCCTGAATCAATCTCAAA

>M6-PGSC0003DMG400002960M6LRR-RLP_186

GGGCTACAGCTGAATATTTTCAAGATGTTCTTGACAACAATAGCAATGAGAAAATCAAATCTTTTGAGGAAGATATTAGAGTTGTTGTTAGTGAGGCAGAGGATGTTGTTGAAATGAAGA

>M6-PGSC0003DMG400002960M6LRR-RLP_1860

TTCATTTGAGATATCTCCAAGTTCGATGTCTTGGAAATATTCCTGAATCAATCTCAAAGCTTCTGAATTTGCAAACTCTAATTTGTGATGGTAATTCTTTAGAGATAACTTTAACTAGGA

>M6-PGSC0003DMG400002960M6LRR-RLP_1922

CTGAATTTGCAAACTCTAATTTGTGATGGTAATTCTTTAGAGATAACTTTAACTAGGAAGATATGGATGTTGAAGAACTTGAGGTATATACGTGTAGGGGGAGTTACTTATTTACCGAGT

>M6-PGSC0003DMG400002960M6LRR-RLP_1984

ATGGATGTTGAAGAACTTGAGGTATATACGTGTAGGGGGAGTTACTTATTTACCGAGTCCTAGAATAGATAACAATCTTGTGATAGGGATGCCAAATCTAGAGGAATTTTTTGGTCTTAG

>M6-PGSC0003DMG400002960M6LRR-RLP_2046

GAATAGATAACAATCTTGTGATAGGGATGCCAAATCTAGAGGAATTTTTTGGTCTTAGTTACTCTAGTTGTACAAATGAAGTCTTTTCTAGGATCCCCAATTTAAAGAGATTGATCATTC

>M6-PGSC0003DMG400002960M6LRR-RLP_2108

TCTAGTTGTACAAATGAAGTCTTTTCTAGGATCCCCAATTTAAAGAGATTGATCATTCCTGCACATTTTAGCATACTAAACAATATTCTGTATCGTCCCTTCGATATGTCCAGCTTGACA

>M6-PGSC0003DMG400002960M6LRR-RLP_2170

ACATTTTAGCATACTAAACAATATTCTGTATCGTCCCTTCGATATGTCCAGCTTGACAAAACTTGAGACATTCAAGTTTTACTGGAAAGATTATTTGGAAGACCCCATCAAGAAATTTGT

>M6-PGSC0003DMG400002960M6LRR-RLP_2232

TTGAGACATTCAAGTTTTACTGGAAAGATTATTTGGAAGACCCCATCAAGAAATTTGTTTTTCCAGAATCACTTAGAAGAGTGTCTTTAACTTGGTGTTGTAAGTTTATTTGGGAAGAGA

>M6-PGSC0003DMG400002960M6LRR-RLP_2294

CCAGAATCACTTAGAAGAGTGTCTTTAACTTGGTGTTGTAAGTTTATTTGGGAAGAGATATCATCAACTTTTATCATGTTGCCACATCTTGAAGAGCTCAAACTTAAATATTGTCGAGCC

>M6-PGSC0003DMG400002960M6LRR-RLP_2356

ATCAACTTTTATCATGTTGCCACATCTTGAAGAGCTCAAACTTAAATATTGTCGAGCCGAAGATGATGTATGGAGATTGAGTGACAAAGACATATTCAGAAGCTTGAAGTTGTTGTTACT

>M6-PGSC0003DMG400002960M6LRR-RLP_2418

ATGATGTATGGAGATTGAGTGACAAAGACATATTCAGAAGCTTGAAGTTGTTGTTACTGAGCGAGCTAAATCTTAAGGGTTGGGAAGCAAGCAGCGATAACTTCCCTAATCTAAAACGCC

>M6-PGSC0003DMG400002960M6LRR-RLP_2542

TCTGAAGAAATGCAAGGACCTGCAAGAAATTCCAACAGATTTTGGGGAAATTTGTACTTTGGAATCGATTGAGTTACATGATTGCAGCACTACTTCTGAGGAATCTGCCAGAAATATCGA

>M6-PGSC0003DMG400002960M6LRR-RLP_2588

GAAATTTGTACTTTGGAATCGATTGAGTTACATGATTGCAGCACTACTTCTGAGGAATCTGCCAGAAATATCGAACAAGAACAAGAGGTCATGGGAAATAATATCCTTAAGGTCTACATC

>M6-PGSC0003DMG400002960M6LRR-RLP_310

TGAAATCATCAAAGAAGAAAGCTGGACATTTGGAATTTTACAACACCATGATATGCTACCAGTTGTTGAAAAAATGGATACGACAAAGAAACAAGTGATGGAGCTTCTTTCTCATGATGC

>M6-PGSC0003DMG400002960M6LRR-RLP_372

TTGTTGAAAAAATGGATACGACAAAGAAACAAGTGATGGAGCTTCTTTCTCATGATGCTGATCAAATTCTTGAATTAACCGCGGATTCCTTGATTGACGCTTCTTCTATGAGTGATCCAA

>M6-PGSC0003DMG400002960M6LRR-RLP_434

CAAATTCTTGAATTAACCGCGGATTCCTTGATTGACGCTTCTTCTATGAGTGATCCAATGCTGTCAGATCACTTGAAAGATCATATTGTGCAGGGACTTGATGGTGACTTGGAGATAATA

>M6-PGSC0003DMG400002960M6LRR-RLP_496

GTCAGATCACTTGAAAGATCATATTGTGCAGGGACTTGATGGTGACTTGGAGATAATAGATAAGAGATTGAGAGGACCAACGCCTTATCTAGACATTGTCACCATATCAGGTATGGGTGG

>M6-PGSC0003DMG400002960M6LRR-RLP_558

AGAGATTGAGAGGACCAACGCCTTATCTAGACATTGTCACCATATCAGGTATGGGTGGCATTGGCAAAACAACACTCGCTAAAAAAGCTTATGATCATCTTCCAATTAGGTATCACTTTG

>M6-PGSC0003DMG400002960M6LRR-RLP_62

ATTCCTCTCCATATTGACTTGATGGCAGCTTATACTGCTGTAATTTCTCTTCTTCAAACTCTTGATCAATCAAATATTTCAGAACTCTTTCATGATCACAGTACTGAAATGCTCGATTCT

>M6-PGSC0003DMG400002960M6LRR-RLP_620

GGCAAAACAACACTCGCTAAAAAAGCTTATGATCATCTTCCAATTAGGTATCACTTTGACATTTTTGTTTGGGTTACAATATCTCAAGAATTTCGATATAGAAATGTATTGTTAGAAGCT

>M6-PGSC0003DMG400002960M6LRR-RLP_682

TTTTGTTTGGGTTACAATATCTCAAGAATTTCGATATAGAAATGTATTGTTAGAAGCTTTACATTGCATTTCAAAGCAAAGAGTTATTGTGAACACAAAAGATTATGATAAGATGAATGA

>M6-PGSC0003DMG400002960M6LRR-RLP_744

ATTGCATTTCAAAGCAAAGAGTTATTGTGAACACAAAAGATTATGATAAGATGAATGACAGTGAGTTAGCTGATCTGGTGCAAAAGAGTCTAAAGGGTCCTAGATACCTTGTTGTTGTTG

>M6-PGSC0003DMG400002960M6LRR-RLP_806

GAGTTAGCTGATCTGGTGCAAAAGAGTCTAAAGGGTCCTAGATACCTTGTTGTTGTTGATGATATTTGGAGTACGGATGCTTGGGATAGCATATCACAATTATTTCCTAATCGCAACAAT

>M6-PGSC0003DMG400002960M6LRR-RLP_868

TATTTGGAGTACGGATGCTTGGGATAGCATATCACAATTATTTCCTAATCGCAACAATGGGAGTCGAGTCTTATTGACTACTAGGGAAACCGATGTAGCGATGTATGCAAATACTAGTAG

>M6-PGSC0003DMG400002960M6LRR-RLP_930

GTCGAGTCTTATTGACTACTAGGGAAACCGATGTAGCGATGTATGCAAATACTAGTAGCCCTCATAAGATGAACCTCTTGGATTTAGATAACAGTTGGAAGTTACTTCGTGATAAGGTGT

>M6-PGSC0003DMG400002960M6LRR-RLP_992

CATAAGATGAACCTCTTGGATTTAGATAACAGTTGGAAGTTACTTCGTGATAAGGTGTTTGGACTAGAACACGATCACCCTCCCGAGTTGGAAGAAATTGGAAAAACAATAGCAGGAAAA

>M6-PGSC0003DMG400002961M6LRR-RLP_0

ATGACAAAGTTTATGCATGTTGTGAGAACTCCATTTTCACCAGCACAAAAGCCTAATTTTCGTCGCTTCAGTATTCAGAAGGCATTTTCAGATGTGTATGCTTATAAAGTACTACCCCCT

>M6-PGSC0003DMG400002961M6LRR-RLP_1008

TGCCAGGACCTGCAAGAAATTCCAACAGATTTTGGGGAAATTTGTACTTTGGAATCGATTGAGTTACATGATTGTAGCACTACTTCTGAGGATTCTGCCAGAAATATTGAACAAGAACAA

>M6-PGSC0003DMG400002961M6LRR-RLP_1068

GAGTTACATGATTGTAGCACTACTTCTGAGGATTCTGCCAGAAATATTGAACAAGAACAAGAGGACATGGGAAATAATATCCTTAAGGTCTACATCCATAACAGCCACAGTAAGTCATAA

>M6-PGSC0003DMG400002961M6LRR-RLP_126

AGATCTATGTACTGCTTTTCATTCTTGGATCTACCTATTGAACCCCATATTAAGCTTCTCCCTAGATTGCCCATCTACCGTCATGATCCTATAATACATGGATTTTTCTCTTGTTTCAAC

>M6-PGSC0003DMG400002961M6LRR-RLP_189

AGATTGCCCATCTACCGTCATGATCCTATAATACATGGATTTTTCTCTTGTTTCAACCATCTCAGGGTATTGGTCATCTTCCCTACCTGGTTCGGCCCATTTCCACTTGTGATTACAAAA

>M6-PGSC0003DMG400002961M6LRR-RLP_252

AGGGTATTGGTCATCTTCCCTACCTGGTTCGGCCCATTTCCACTTGTGATTACAAAATTATTTCATTTGAGATATCTCCAAGTTCGATATTTTGGAAATATTCCTGAATCAATCTCAGAG

>M6-PGSC0003DMG400002961M6LRR-RLP_315

CATTTGAGATATCTCCAAGTTCGATATTTTGGAAATATTCCTGAATCAATCTCAGAGCTTCAAAATTTGCAAACTCTAATTTATGATTGTCATTCTTCACATATGACTTTACCCGGGAAG

>M6-PGSC0003DMG400002961M6LRR-RLP_378

AATTTGCAAACTCTAATTTATGATTGTCATTCTTCACATATGACTTTACCCGGGAAGATATGGACGATGAAGAACTTGAGGTATATACATTTAGGGAAAGCTAGTAATTATATACCAAAT

>M6-PGSC0003DMG400002961M6LRR-RLP_441

ACGATGAAGAACTTGAGGTATATACATTTAGGGAAAGCTAGTAATTATATACCAAATCCTAGAATAGATAACAATCTTGTGACGGGGATGCCAAATCTAGGGGAATTTTTCAATATTTGT

>M6-PGSC0003DMG400002961M6LRR-RLP_504

ATAGATAACAATCTTGTGACGGGGATGCCAAATCTAGGGGAATTTTTCAATATTTGTTACTCTAGTTGTACAAATGAAGTTTTTTCTAGAATTCCCAATCTAGAGAGATTGAGCATTCGT

>M6-PGSC0003DMG400002961M6LRR-RLP_567

AGTTGTACAAATGAAGTTTTTTCTAGAATTCCCAATCTAGAGAGATTGAGCATTCGTGCATCTTCTAAAGTCAGAGAAAATTTTCTCTATCGTCTCTTGGATATGTCCAGCTTGAGAAAA

>M6-PGSC0003DMG400002961M6LRR-RLP_63

CGCTTCAGTATTCAGAAGGCATTTTCAGATGTGTATGCTTATAAAGTACTACCCCCTGTAGCCAGATCTATGTACTGCTTTTCATTCTTGGATCTACCTATTGAACCCCATATTAAGCTT

>M6-PGSC0003DMG400002961M6LRR-RLP_630

TCTAAAGTCAGAGAAAATTTTCTCTATCGTCTCTTGGATATGTCCAGCTTGAGAAAACTTGAGGTATTCAAGCTTTCCTGGGACAATTATTCGGAAAAGCCCATCAAGAGATTTGTTTTG

>M6-PGSC0003DMG400002961M6LRR-RLP_693

GTATTCAAGCTTTCCTGGGACAATTATTCGGAAAAGCCCATCAAGAGATTTGTTTTGCCAGCATCACTTAGGAAGTTGACTTTAGATTGCTGTAGTCATTTTATTTGGGAAGAGATATCG

>M6-PGSC0003DMG400002961M6LRR-RLP_756

TCACTTAGGAAGTTGACTTTAGATTGCTGTAGTCATTTTATTTGGGAAGAGATATCGTCAACTTTTATCATGTTGCCAAATCTTGAAGAGCTCAAACTTAAAGATTGTGAATCCGAAGAT

>M6-PGSC0003DMG400002961M6LRR-RLP_819

TTTATCATGTTGCCAAATCTTGAAGAGCTCAAACTTAAAGATTGTGAATCCGAAGATGATGTATGGAGATTGAGTGACAAAGACATATTCAAAAGCTTGAAGTTGTTGTTACTGAGCTAC

>M6-PGSC0003DMG400002961M6LRR-RLP_882

TGGAGATTGAGTGACAAAGACATATTCAAAAGCTTGAAGTTGTTGTTACTGAGCTACCCAAATCTTAAGTGTTGGGAAGCTAGCAGTGATAACTTCCCAAATCTAAAACGTCTTGTTCTG

>M6-PGSC0003DMG400002961M6LRR-RLP_945

CTTAAGTGTTGGGAAGCTAGCAGTGATAACTTCCCAAATCTAAAACGTCTTGTTCTGAAGAATTGCCAGGACCTGCAAGAAATTCCAACAGATTTTGGGGAAATTTGTACTTTGGAATCG

>M6-PGSC0003DMG400002980-(rev)M6LRR-RLP_1116

AGCTCTAAAAGCACTTGCTGGTGTTTTACGCGGCAAATCAGAGGTGGATGAGTGGAGAGACATTTTAAGAAGTGAAATATGGGAGCTGTCAATTTGTTCGAATGGTATATTACCAGCGCT

>M6-PGSC0003DMG400002980-(rev)M6LRR-RLP_1178

TTTTAAGAAGTGAAATATGGGAGCTGTCAATTTGTTCGAATGGTATATTACCAGCGCTGATGTTGAGCTACAATGATCTTCCTGCACATTTGAAGCAATGTTTTGCTTATTGTGCAGTAT

>M6-PGSC0003DMG400002980-(rev)M6LRR-RLP_124

GACATTTTGCTTGGTCTTCAAATTGTGCTAAGTGATGCAGAGAATAAGCAAGCATCAAATCAACATGTGAGCCAATGGCTTAATAAGCTCCAGAGTGCTGTGGATGGCGCTGAAAACATG

>M6-PGSC0003DMG400002980-(rev)M6LRR-RLP_1240

TTGAGCTACAATGATCTTCCTGCACATTTGAAGCAATGTTTTGCTTATTGTGCAGTATATCCCAAAGATTATCTATTTTGCAAAGACCAAGTTATTCACCTGTGGATTGCTAATGGTCTG

>M6-PGSC0003DMG400002980-(rev)M6LRR-RLP_1302

CAAAGATTATCTATTTTGCAAAGACCAAGTTATTCACCTGTGGATTGCTAATGGTCTGGTACAGCAGTTTCATTCAGGTAACCAATACTTTCTCGAGTTGAGATCAAGATCGCTGTTCGA

>M6-PGSC0003DMG400002980-(rev)M6LRR-RLP_1364

AGCAGTTTCATTCAGGTAACCAATACTTTCTCGAGTTGAGATCAAGATCGCTGTTCGAAATGGTCTCGAAGTCTTCTGAACGAGACATAGAGAAATTCTTAATGCATGACCTTGTCAATG

>M6-PGSC0003DMG400002980-(rev)M6LRR-RLP_1550

TGGAACAATGTCGGCACATGTCCTATTCAATAGGACAAGATGGTGACTTTGAGAAATTGCAATCACTCTTTAAATCAGAGCAGCTGAGGACATTACTTCCAATCAATATTCAGCTTCAGT

>M6-PGSC0003DMG400002980-(rev)M6LRR-RLP_1612

TCACTCTTTAAATCAGAGCAGCTGAGGACATTACTTCCAATCAATATTCAGCTTCAGTACCAAACTAAGCTAAGCAAGAGGGTGCTGCATAACATATTGCCAAGACTAACATCCTTAAGG

>M6-PGSC0003DMG400002980-(rev)M6LRR-RLP_1674

AACTAAGCTAAGCAAGAGGGTGCTGCATAACATATTGCCAAGACTAACATCCTTAAGGGCATTATCGTTGTTAGGCTATAAGATTAAGGAGTTGCCAGATGACTCGTTTGTCAAATTAAA

>M6-PGSC0003DMG400002980-(rev)M6LRR-RLP_186

ACATGTGAGCCAATGGCTTAATAAGCTCCAGAGTGCTGTGGATGGCGCTGAAAACATGATAGAAGAAGTCAATTATGAAGCTTTGAGGCATAAGGTGGAAGGCCAGCATCAAAATCTTGC

>M6-PGSC0003DMG400002980-(rev)M6LRR-RLP_1860

GTATAACTTAAAGGCACTTCTCCTGTCATCTTGTGCTGATCTTGAGGAGCTACCACTGCAGATGGAGAAGTTGATCAACTTACATTACCTCGACATAAGCAACACTTTTCACTTGAAGGT

>M6-PGSC0003DMG400002980-(rev)M6LRR-RLP_1922

TGGAGAAGTTGATCAACTTACATTACCTCGACATAAGCAACACTTTTCACTTGAAGGTTCCCCTACATCTGAGCAAGTTGAAAAGCCTGCAAGTGCTAATGGGAGCCAAGTTTGTTCTCG

>M6-PGSC0003DMG400002980-(rev)M6LRR-RLP_1984

CTACATCTGAGCAAGTTGAAAAGCCTGCAAGTGCTAATGGGAGCCAAGTTTGTTCTCGGTGGTTTGAGAATGGAAGATTTGGGTGAAGCACAGAACTTGTATGGATCTGTATCAGTTGTA

>M6-PGSC0003DMG400002980-(rev)M6LRR-RLP_2046

TTTGAGAATGGAAGATTTGGGTGAAGCACAGAACTTGTATGGATCTGTATCAGTTGTAGAGTTGCAAAATGTAGTTGATAGAAGGGAAGCTGTGAAGGCAAAGATGAAGGAGAAGAATCA

>M6-PGSC0003DMG400002980-(rev)M6LRR-RLP_2108

TGCAAAATGTAGTTGATAGAAGGGAAGCTGTGAAGGCAAAGATGAAGGAGAAGAATCATGTTAAGAAGTTATATTTGGAGTGGAGTGAAAGTAGTAGTGCCGACAATTCACAAACAGAAA

>M6-PGSC0003DMG400002980-(rev)M6LRR-RLP_2170

AAGAAGTTATATTTGGAGTGGAGTGAAAGTAGTAGTGCCGACAATTCACAAACAGAAAGAGACATACTTGATGAGCTATGCCCACATAAAAACATAAAAGAAGTCGAAATCATCGGATAT

>M6-PGSC0003DMG400002980-(rev)M6LRR-RLP_2232

CATACTTGATGAGCTATGCCCACATAAAAACATAAAAGAAGTCGAAATCATCGGATATAGAGGGACAACATTTCCGAATTGGTTAGCTGATCCTTTGTTTCTTAAGCTAGAACAGTTGTC

>M6-PGSC0003DMG400002980-(rev)M6LRR-RLP_2294

GGACAACATTTCCGAATTGGTTAGCTGATCCTTTGTTTCTTAAGCTAGAACAGTTGTCTATTGCTAACTGCAAGAACTGTTTTTTCTTGCCAGCACTAGGACAGCTTCCTTGTTTGAAAA

>M6-PGSC0003DMG400002980-(rev)M6LRR-RLP_2418

TTCCATTAGAGGGATGCATGGAATAACAGGGGTGACGGAAGAATTCTATGGCAGTTTGTCCTCTAAAAAGCCTTTTAATTGTCTTGAGAACCTTATATTTGAAGATATGCCGGAGTGGAA

>M6-PGSC0003DMG400002980-(rev)M6LRR-RLP_2542

TGGCACGTACTAGGAAGTGGAGAGTTCCCTATACTTGAGAAGCTTTTTATTAAAAATTGCCCTGAACTCAGTTTGGAGACACCCATCCAACTTTCAAATTTAAAAAGGTTTGAAGTTGGT

>M6-PGSC0003DMG400002980-(rev)M6LRR-RLP_2604

TGAACTCAGTTTGGAGACACCCATCCAACTTTCAAATTTAAAAAGGTTTGAAGTTGGTGGTTCTCCAAAGGTTGGAGTTGTTTTTGATGATACTGAGCTGTTTAGATCCCAACTTGAGGG

>M6-PGSC0003DMG400002980-(rev)M6LRR-RLP_2666

CTCCAAAGGTTGGAGTTGTTTTTGATGATACTGAGCTGTTTAGATCCCAACTTGAGGGAATGAAGCAGATTGTTGAATTAGATATACATGATTGTAACTCTCTTACCTTCTTTCCTTTTA

>M6-PGSC0003DMG400002980-(rev)M6LRR-RLP_2790

ACTGCCCACTACCTTGAAGAGAATAGACATATCTGGTTGCCAGAAATTGAAATTGGATCCGCCAGTTGGTGAGATGAGTATGTTTCTGGAGTGTTTGAGTCTGGCAAAATGTGATTGTAT

>M6-PGSC0003DMG400002980-(rev)M6LRR-RLP_2852

CAGTTGGTGAGATGAGTATGTTTCTGGAGTGTTTGAGTCTGGCAAAATGTGATTGTATAGATGATATATCACCTGAGTCGCTCCCAAGAGCACGCAGATTGTATGTATCTGATTGCCACA

>M6-PGSC0003DMG400002980-(rev)M6LRR-RLP_2914

GATATATCACCTGAGTCGCTCCCAAGAGCACGCAGATTGTATGTATCTGATTGCCACAACCTTACTAGGTTTTTGATTCCTACTGCCACTGAAAGTCTCTGTATTTGGTATTGTGAGAAT

>M6-PGSC0003DMG400002980-(rev)M6LRR-RLP_2976

TACTAGGTTTTTGATTCCTACTGCCACTGAAAGTCTCTGTATTTGGTATTGTGAGAATGTTGAAATACTCTCGGTGGCATGTGGGGGGACCCAGATGACGAAGTTGAATATTCAGGGCTG

>M6-PGSC0003DMG400002980-(rev)M6LRR-RLP_310

ACAAGCAACAAACAAGTAAGTGACCTCAACCTGTGCTTGACTGATGATTTTTTCCTTAACATAAAGGAAAAGTTGGAAGAAAACATTGAAACATTGGAGGTGTTGGAAAAGCAAATTGGT

>M6-PGSC0003DMG400002980-(rev)M6LRR-RLP_3162

GATTAATTGTCCAGAAATAGAGTCCTTTCCTGAAGGAGGATTGCCCTTCAATTTACAACTCCTTATGATCAGCGAGTGCAAGAAACTGGTGAACAGCCGAAAGGAGTGGCGTTTACAGAG

>M6-PGSC0003DMG400002980-(rev)M6LRR-RLP_3224

TTATGATCAGCGAGTGCAAGAAACTGGTGAACAGCCGAAAGGAGTGGCGTTTACAGAGACTCCCCTGTCTTAGAGAGTTAACGATCGAACATGATGGCAGTGATGAAGAGATTGTTGGTG

>M6-PGSC0003DMG400002980-(rev)M6LRR-RLP_3286

CCCTGTCTTAGAGAGTTAACGATCGAACATGATGGCAGTGATGAAGAGATTGTTGGTGGTGAGAATTGGGAGTTGCCTTCCTCTATTCAAACACTCTATATAAAGAATCTGAAAACATTA

>M6-PGSC0003DMG400002980-(rev)M6LRR-RLP_3410

GCCAACATCTCAAACACCTCACCTCTCTTCAATATCTAGACATTGCGGGTAATTTACCTCAAATTCAGTCAATGCTGGAACAAGGCCAGTTTTCGCATCTCACTTCGCTTCAATGTCTAG

>M6-PGSC0003DMG400002980-(rev)M6LRR-RLP_434

TTGGCTTAAAGGATCATTTTGGTTCAACTAAACAAGAAACTAGAACACCTTCAACGTCTTTGGTTGATGATGCTGGTATCTTTGGAAGGCAGAATGAAATAGAGAATTTGATTGGCCGTT

>M6-PGSC0003DMG400002980-(rev)M6LRR-RLP_496

GTTGATGATGCTGGTATCTTTGGAAGGCAGAATGAAATAGAGAATTTGATTGGCCGTTTGTTGTCTATGGATACACAGGGAAAAAATCTGGCTGTAGTTCCTATTGTTGGAATGGGCGGC

>M6-PGSC0003DMG400002980-(rev)M6LRR-RLP_62

GTGATCTGCTCAACATGTTTCGGAAGCATACGGATCATGTTAAGCTCTTAAAGAAGCTGGAGGACATTTTGCTTGGTCTTCAAATTGTGCTAAGTGATGCAGAGAATAAGCAAGCATCAA

>M6-PGSC0003DMG400002980-(rev)M6LRR-RLP_620

GTAAGACAACACTTGCTAAAGCGGTTTACAATGATGAGAGAGTGCAGAAACATTTTGGTTTGAAAGCTTGGTTTTGTGTTTCTGAGGCATATGATGCTTTCAGAATAACAAAAGGTTTAC

>M6-PGSC0003DMG400002980-(rev)M6LRR-RLP_682

AAAGCTTGGTTTTGTGTTTCTGAGGCATATGATGCTTTCAGAATAACAAAAGGTTTACTTCATGAAATTGGCTCAACTGACTTGAAGGTTGATGACAATCTTAATCAGCTACAAGTCAAA

>M6-PGSC0003DMG400002980-(rev)M6LRR-RLP_744

TGAAATTGGCTCAACTGACTTGAAGGTTGATGACAATCTTAATCAGCTACAAGTCAAATTGAAGGAAAAGCTGCATGGAAAAAAGTTTCTTGTTGTCCTTGATGACGTGTGGAATGATAA

>M6-PGSC0003DMG400002980-(rev)M6LRR-RLP_806

AGGAAAAGCTGCATGGAAAAAAGTTTCTTGTTGTCCTTGATGACGTGTGGAATGATAACTATCCTGAGTGGGATGACTTGAGAAATCTTTTTCTACAAGGAGATATAGGAAGTAAGATCA

>M6-PGSC0003DMG400002980-(rev)M6LRR-RLP_868

CCTGAGTGGGATGACTTGAGAAATCTTTTTCTACAAGGAGATATAGGAAGTAAGATCATTGTGACGACACGGAAAGAGAGTGTTGCCTTGATGATGGGTAGTGAGGCAATCAACATGGGA

>M6-PGSC0003DMG400002980-(rev)M6LRR-RLP_930

GACGACACGGAAAGAGAGTGTTGCCTTGATGATGGGTAGTGAGGCAATCAACATGGGAGTTCTGTCTAGTGAAGACTCTTGGGCTCTATTCCAACGACATTCACTAGAAAATAGAGATCC

>M6-PGSC0003DMG400002980-(rev)M6LRR-RLP_992

TGTCTAGTGAAGACTCTTGGGCTCTATTCCAACGACATTCACTAGAAAATAGAGATCCTGAGGAACATCCAGAATTTGAAGAGGTTGGAAAACAAATTGCAGACAAGTGCAAAGGGTTGC

>M6-PGSC0003DMG400003002M6LRR-RLK_0

ATGTTTGACACAATTGTATGTGATGAAACAGTGACCAGAGATGCATTGCTCAAGTTTCTTGCAGAGGTTTCTAACAATAATAGTACAATTCTTGGTACTAATTCGGGCTGGAACTCGACA

>M6-PGSC0003DMG400003002M6LRR-RLK_1054

AGGCTCCAGCAGAGTTGCTTGGAAGAGGGAAACATGGTAGTATGTACAAAGTGATGTGTGACAATCCCAAAATGACACTGGCTGTTAAGAGGATAAAAGATTGGTCAATAACTGGTAGTG

>M6-PGSC0003DMG400003002M6LRR-RLK_1116

AATCCCAAAATGACACTGGCTGTTAAGAGGATAAAAGATTGGTCAATAACTGGTAGTGAATTCAAAAAGAGAATGCAGAAACTAGACAAAATCAGGCATCCAAATGTGTTGCCTGCTGTT

>M6-PGSC0003DMG400003002M6LRR-RLK_1178

CAAAAAGAGAATGCAGAAACTAGACAAAATCAGGCATCCAAATGTGTTGCCTGCTGTTGCCTTTTATTCTTCTAGACAAGAGAAGCTTTTGGTCTATGAGTACCAAGACAATGGCAGTCT

>M6-PGSC0003DMG400003002M6LRR-RLK_124

ATCCTTGCAAGGACAAGTGGCTTGGTGTTACATGTAACGCGAAGACTCTCTCTGTGGAGAAAATCATTCTTGATGGTTATGATTTTTCATCTGGAACGTTTGATGCTACCACGATTTGTA

>M6-PGSC0003DMG400003002M6LRR-RLK_1240

TTTATTCTTCTAGACAAGAGAAGCTTTTGGTCTATGAGTACCAAGACAATGGCAGTCTTCTCCTACTTCTACAAGGTAATTGACTCTACTAGGCTATTCTTCTGTGTATTGAATTGTGCG

>M6-PGSC0003DMG400003002M6LRR-RLK_1302

CTACTTCTACAAGGTAATTGACTCTACTAGGCTATTCTTCTGTGTATTGAATTGTGCGACCATTTCATCTAGGCATATAAACAGACACTCGAACTTTGCCTTAGCTGGCAAGTAAGCACT

>M6-PGSC0003DMG400003002M6LRR-RLK_186

ATCATTCTTGATGGTTATGATTTTTCATCTGGAACGTTTGATGCTACCACGATTTGTAATGTGGAGCCTATCGCGGACACCCTTTTCCTCCTGAGTCTAAAAAATGACCTACTTCAGGGG

>M6-PGSC0003DMG400003002M6LRR-RLK_248

GGAGCCTATCGCGGACACCCTTTTCCTCCTGAGTCTAAAAAATGACCTACTTCAGGGGAAAAACTTTGATAGTATAGATAAATGTAAGCATCTAAGACATCTTGTTCTTGGTGGAAACCG

>M6-PGSC0003DMG400003002M6LRR-RLK_310

ACTTTGATAGTATAGATAAATGTAAGCATCTAAGACATCTTGTTCTTGGTGGAAACCGGTTCTCAGGTAGCATTCCACAGTCATTTTCAAGATTGAATAACTTGAAAAGCCTTGACATAT

>M6-PGSC0003DMG400003002M6LRR-RLK_372

TCAGGTAGCATTCCACAGTCATTTTCAAGATTGAATAACTTGAAAAGCCTTGACATATCCAACAATAGATTATCAGGTGTTCTCCCTGATTTATCTAGGATTTCAGGCTTGATTGAATTC

>M6-PGSC0003DMG400003002M6LRR-RLK_434

CAATAGATTATCAGGTGTTCTCCCTGATTTATCTAGGATTTCAGGCTTGATTGAATTCGAGGCTCAGAATAACCAGCTATCAGGAGAGATACCTGAATTTGATTTCTCAAACTTTGTTGC

>M6-PGSC0003DMG400003002M6LRR-RLK_496

CTCAGAATAACCAGCTATCAGGAGAGATACCTGAATTTGATTTCTCAAACTTTGTTGCATTTAATGTCTCTTTCAACAATTTCAGTGGATCGATTCCGCCTGAGAGTGATAGTTTCCCTG

>M6-PGSC0003DMG400003002M6LRR-RLK_558

AATGTCTCTTTCAACAATTTCAGTGGATCGATTCCGCCTGAGAGTGATAGTTTCCCTGTTAGTAGATACTTGGAAAATCCTCTGTTGTGTGGCTCCCCGTTGGCTGGAAAATGCTCATCA

>M6-PGSC0003DMG400003002M6LRR-RLK_62

AGAGGTTTCTAACAATAATAGTACAATTCTTGGTACTAATTCGGGCTGGAACTCGACATCTGATCCTTGCAAGGACAAGTGGCTTGGTGTTACATGTAACGCGAAGACTCTCTCTGTGGA

>M6-PGSC0003DMG400003002M6LRR-RLK_620

TAGATACTTGGAAAATCCTCTGTTGTGTGGCTCCCCGTTGGCTGGAAAATGCTCATCACTGTCCGAAGATTCAGAATCAGCTGATCATGACAAGCCAAAGAGCAGCTTGTCAAGAGATGA

>M6-PGSC0003DMG400003002M6LRR-RLK_682

CCGAAGATTCAGAATCAGCTGATCATGACAAGCCAAAGAGCAGCTTGTCAAGAGATGAAATTCTCATGTTTTCAGGTTATTTCCTCATAGGCCTGGCTCTGGTTCTTCTTATCCTACTAT

>M6-PGSC0003DMG400003002M6LRR-RLK_868

AGGTTGCTACAGTTGATCATGACAGTACTACTACATCAATGGATCAATCAAAGAAAGGCGCGAGCAATTCAGATATCTCATCAGCTGTCATTTCTTCCGATGACAGCAACACCGCGGTTT

>M6-PGSC0003DMG400003002M6LRR-RLK_930

AGCAATTCAGATATCTCATCAGCTGTCATTTCTTCCGATGACAGCAACACCGCGGTTTCACAGTCACTCGTGGTTCTTACGAGTCCTGAGGCGAATAACGGATTGAGATTTGAGGATTTG

>M6-PGSC0003DMG400003002M6LRR-RLK_992

GTCACTCGTGGTTCTTACGAGTCCTGAGGCGAATAACGGATTGAGATTTGAGGATTTGCTTAAGGCTCCAGCAGAGTTGCTTGGAAGAGGGAAACATGGTAGTATGTACAAAGTGATGTG

>M6-PGSC0003DMG400003175M6LRR-RLK_0

ATGGGTGGTAGATGGAATACTTATGGATTCAAGTTATCATATCTTGCACTTCTGATTATTCTTTATGACGTTCACGGATGTAGCTCACTCAATTCTGAAGGTAATTCTATTCTATGTTTG

>M6-PGSC0003DMG400003175M6LRR-RLK_1098

TAGTTTTTGGTAGTGAAAATTTGTGGGACAATGAAAGAGAGATTGTTGCTTGTCATGGTTGAAAATTGGTTTAAAATAGCCTTGAGCATTGAAAATAAGCACAGGTTGAAAATAACCCCT

>M6-PGSC0003DMG400003175M6LRR-RLK_1159

AAAATTGGTTTAAAATAGCCTTGAGCATTGAAAATAAGCACAGGTTGAAAATAACCCCTAATGAGTTCTAAAAAATAGATTGAAAATAGTCGTGGTGGGTTAAAAATAGCCTTGACAAGT

>M6-PGSC0003DMG400003175M6LRR-RLK_122

CTGTCCTACTGGTTTTGGTGTTTATGTCTTAGTTATATCTAGTTTGTTATGTCCATTTGAATATTGGATTGGTCTACTTTGCAAAACAAGGATTGGCATTGTTCGGACTACGATCTAAAG

>M6-PGSC0003DMG400003175M6LRR-RLK_1220

TGAGTTCTAAAAAATAGATTGAAAATAGTCGTGGTGGGTTAAAAATAGCCTTGACAAGTTAAAAATGGGTTGACAATAGTTCTTTAGAAGAACTCACATATCCCATAAGATCAATGGGGC

>M6-PGSC0003DMG400003175M6LRR-RLK_1281

AAAATGGGTTGACAATAGTTCTTTAGAAGAACTCACATATCCCATAAGATCAATGGGGCTTTGGATACTACTGTTGCTGTTTTTTTAAAGATGGTGCAGCTAAATAATGTTGAAAATGTC

>M6-PGSC0003DMG400003175M6LRR-RLK_1342

TGGATACTACTGTTGCTGTTTTTTTAAAGATGGTGCAGCTAAATAATGTTGAAAATGTCTCTCAATTTTTATTAAGCATTAGAGGGGTTTAATAAACAACTTGAAAACATAATTTGCAGA

>M6-PGSC0003DMG400003175M6LRR-RLK_1403

TCAATTTTTATTAAGCATTAGAGGGGTTTAATAAACAACTTGAAAACATAATTTGCAGAAAATCTGCATAACAAAAAGTAGTAGATTAGTAAATTGGGTGGAATAGACTAAAGATAATTA

>M6-PGSC0003DMG400003175M6LRR-RLK_1464

ATCTGCATAACAAAAAGTAGTAGATTAGTAAATTGGGTGGAATAGACTAAAGATAATTAGTTCCATGTCTAATAATCATAGTTTTCATCATCTTGCATGTTGTATCTTCCTGTGTGTTGT

>M6-PGSC0003DMG400003175M6LRR-RLK_1525

TCCATGTCTAATAATCATAGTTTTCATCATCTTGCATGTTGTATCTTCCTGTGTGTTGTGTATGACTCCTTTTTTAATCTTAACAATTTAAACGTTTTTTTCTCTATGATTGGCTCACAG

>M6-PGSC0003DMG400003175M6LRR-RLK_1586

ATGACTCCTTTTTTAATCTTAACAATTTAAACGTTTTTTTCTCTATGATTGGCTCACAGTGTGCTGTCAGAAAACAATTTTTTTGGTGCCATTCCAAAAGAATTTGGACGACTTCAAAGG

>M6-PGSC0003DMG400003175M6LRR-RLK_1647

TGCTGTCAGAAAACAATTTTTTTGGTGCCATTCCAAAAGAATTTGGACGACTTCAAAGGTTGGAAGTGCTTGACTTGACATATAATAATTTGAGTGGGAGAATTCCAGCAGAAATAGGAG

>M6-PGSC0003DMG400003175M6LRR-RLK_1708

GGAAGTGCTTGACTTGACATATAATAATTTGAGTGGGAGAATTCCAGCAGAAATAGGAGACCTGCATGCGCTAAGAAGCTTGTGAGTTCATAATTTAATTTTTGTGCATGTGAAGATGCG

>M6-PGSC0003DMG400003175M6LRR-RLK_1769

CTGCATGCGCTAAGAAGCTTGTGAGTTCATAATTTAATTTTTGTGCATGTGAAGATGCGTGCTGCCTATATTAGTTTATATATGTGAGTATTCATGAGGAAGAACCATTAAAAATATTCT

>M6-PGSC0003DMG400003175M6LRR-RLK_183

TATTGGATTGGTCTACTTTGCAAAACAAGGATTGGCATTGTTCGGACTACGATCTAAAGTGGATTGTGATCCATATGGAGTTCTTGCAAACTGGAATCTTGATCATTGTGACCCATGTAT

>M6-PGSC0003DMG400003175M6LRR-RLK_1830

CTGCCTATATTAGTTTATATATGTGAGTATTCATGAGGAAGAACCATTAAAAATATTCTTTTGAACTGTGTCTTATACCTTTTCACAGGTTGATTCGTAACAACAACTTGGAAGGAAAAA

>M6-PGSC0003DMG400003175M6LRR-RLK_1891

TGAACTGTGTCTTATACCTTTTCACAGGTTGATTCGTAACAACAACTTGGAAGGAAAAATTCCATTTGAAATTGGGAAGCTTCGTTTGCTCTCCGAATTGCAATTTGACGAAGAACTCAT

>M6-PGSC0003DMG400003175M6LRR-RLK_1952

CCATTTGAAATTGGGAAGCTTCGTTTGCTCTCCGAATTGCAATTTGACGAAGAACTCATCCCTGCTGCTGCTGGTGGAACATGCTGTTTAAATAGAAAATTTGTACACTGGTAAGGTTCT

>M6-PGSC0003DMG400003175M6LRR-RLK_2013

CTGCTGCTGCTGGTGGAACATGCTGTTTAAATAGAAAATTTGTACACTGGTAAGGTTCTTTTGGTTTTATTGGACCTTTTATATCCGTGTTCTGGTTCTTGTTACATTTATGTAACCAAA

>M6-PGSC0003DMG400003175M6LRR-RLK_2074

TGGTTTTATTGGACCTTTTATATCCGTGTTCTGGTTCTTGTTACATTTATGTAACCAAATATCAACATGCAAAGATGTCTGAAATGTATGAAATTTGTATTTAAGCCATTTCAGGCTGCC

>M6-PGSC0003DMG400003175M6LRR-RLK_2135

TCAACATGCAAAGATGTCTGAAATGTATGAAATTTGTATTTAAGCCATTTCAGGCTGCCATTATTTTCAGAATATGGTCATAATTCCTGTCATAAGGTTTTCTTGGTGGGAGTGAGGGGG

>M6-PGSC0003DMG400003175M6LRR-RLK_2196

TATTTTCAGAATATGGTCATAATTCCTGTCATAAGGTTTTCTTGGTGGGAGTGAGGGGGTGGGTGAACACAAGTTGATAAAGTTGAGTATTAATGACAACAAGGGATATTGTCGTTTATG

>M6-PGSC0003DMG400003175M6LRR-RLK_2257

GGTGAACACAAGTTGATAAAGTTGAGTATTAATGACAACAAGGGATATTGTCGTTTATGTCAAATCAAATAAAGCCTGGGAAATAGGTGATTTCATAGATGTGGCGATGCAATGACTTGT

>M6-PGSC0003DMG400003175M6LRR-RLK_2318

AAATCAAATAAAGCCTGGGAAATAGGTGATTTCATAGATGTGGCGATGCAATGACTTGTGTTGCTAATGCTTGAAATTTATTGTCTAGTTTGACATCTAGTAGTAGTACTTGACCATACA

>M6-PGSC0003DMG400003175M6LRR-RLK_2379

TGCTAATGCTTGAAATTTATTGTCTAGTTTGACATCTAGTAGTAGTACTTGACCATACATCAACTCAAACAAAAAAGCTTTGTAAACTACAAGGAGGTGATTGTCCAGTCATTGCATCCA

>M6-PGSC0003DMG400003175M6LRR-RLK_244

GATTGTGATCCATATGGAGTTCTTGCAAACTGGAATCTTGATCATTGTGACCCATGTATGTGGTTTGGAGTACAGTGTCTGGCAGGCAAAGTGCAAATGTTGTAAGTCCCTTGTTACATA

>M6-PGSC0003DMG400003175M6LRR-RLK_2440

AACTCAAACAAAAAAGCTTTGTAAACTACAAGGAGGTGATTGTCCAGTCATTGCATCCATCCAAGGCTAGGGTTGAGGATGAGGGATGCAATATATTGGAGATCTTGGGATTGATCCTGG

>M6-PGSC0003DMG400003175M6LRR-RLK_2501

CAAGGCTAGGGTTGAGGATGAGGGATGCAATATATTGGAGATCTTGGGATTGATCCTGGAATCAAGAGAGCATCAAGTTATCATATTGGTTCTAATTTATTTTAATTTTTTTATCAAGAA

>M6-PGSC0003DMG400003175M6LRR-RLK_2562

TCAAGAGAGCATCAAGTTATCATATTGGTTCTAATTTATTTTAATTTTTTTATCAAGAAACTCTTCTCAGATATCTTTAACTCTTACATAGAATACTGACTATCAAAACTAATAGCATGG

>M6-PGSC0003DMG400003175M6LRR-RLK_2623

TCTTCTCAGATATCTTTAACTCTTACATAGAATACTGACTATCAAAACTAATAGCATGGTTACTTTCAATAACACTTCTCAATCAGATATTTTCATTTGCCTGCTTCTCTCTGGTAATTT

>M6-PGSC0003DMG400003175M6LRR-RLK_2684

ACTTTCAATAACACTTCTCAATCAGATATTTTCATTTGCCTGCTTCTCTCTGGTAATTTGATGTCTGTTTCTGGCAGCATCTGTCATGGCAGTCTGATACCATTAAAGACCGTAGACTCC

>M6-PGSC0003DMG400003175M6LRR-RLK_2745

TGTCTGTTTCTGGCAGCATCTGTCATGGCAGTCTGATACCATTAAAGACCGTAGACTCCTTTATCAAACCACTTAAAGGGACACTTCTGCGTTATTTCACTATGTAAGTAACTAAATGGT

>M6-PGSC0003DMG400003175M6LRR-RLK_2806

TATCAAACCACTTAAAGGGACACTTCTGCGTTATTTCACTATGTAAGTAACTAAATGGTTTTCAATCACCATGTGTTTCTACTTACATTTTATCACTCTAGAAGCGTCTTTGTGTCAATA

>M6-PGSC0003DMG400003175M6LRR-RLK_2867

TCAATCACCATGTGTTTCTACTTACATTTTATCACTCTAGAAGCGTCTTTGTGTCAATAAAGGTTCTCCAAACAACTTGGAAGTGGCTCATTGGATAATCAGACAGGGTATTCCTCTGAC

>M6-PGSC0003DMG400003175M6LRR-RLK_2928

GGTTCTCCAAACAACTTGGAAGTGGCTCATTGGATAATCAGACAGGGTATTCCTCTGACAACCTACCAAGTGAGTGAAAAAGTTTTACCCCACATCATTAGCAGGAATTATGTTGAAAGT

>M6-PGSC0003DMG400003175M6LRR-RLK_2989

CCTACCAAGTGAGTGAAAAAGTTTTACCCCACATCATTAGCAGGAATTATGTTGAAAGTGCACAAAAGATTGTTTTCACTTTTCTTTTTTAGCAGGTTCATCTAGGCCTCACATCATCCA

>M6-PGSC0003DMG400003175M6LRR-RLK_305

GGTTTGGAGTACAGTGTCTGGCAGGCAAAGTGCAAATGTTGTAAGTCCCTTGTTACATACGATCATATCTGTTAATGAGTATTCAATTCCATAAAACTTACTTTATAATTGTAATGCTTT

>M6-PGSC0003DMG400003175M6LRR-RLK_3050

ACAAAAGATTGTTTTCACTTTTCTTTTTTAGCAGGTTCATCTAGGCCTCACATCATCCATACTATTCAGAACCATGCGAATATCGCACGCCGCAAGTTAGTAGAACAATCCAGCAACCTT

>M6-PGSC0003DMG400003175M6LRR-RLK_3111

CTATTCAGAACCATGCGAATATCGCACGCCGCAAGTTAGTAGAACAATCCAGCAACCTTGCTGCTGTTCCTGCCAATGCTGGGAAACCTTTGGGTCCTATTGTTTCAGTGCCAAGCACCA

>M6-PGSC0003DMG400003175M6LRR-RLK_3172

TGCTGTTCCTGCCAATGCTGGGAAACCTTTGGGTCCTATTGTTTCAGTGCCAAGCACCAGAAGTAGTGGGTCATTTCATGTTGTGCCAAATACCCAAGAAACACCTCCTACACCCTTACC

>M6-PGSC0003DMG400003175M6LRR-RLK_3233

AGTAGTGGGTCATTTCATGTTGTGCCAAATACCCAAGAAACACCTCCTACACCCTTACCTGCCTCGCAACGACAGCCTGAGTCTCAATCCAAACCAAATCCAATGGGACGGTCAAGTAGT

>M6-PGSC0003DMG400003175M6LRR-RLK_3294

CCTCGCAACGACAGCCTGAGTCTCAATCCAAACCAAATCCAATGGGACGGTCAAGTAGTGTTGTTAAACAGCCAACTAATAGACAAGCTCCACCTGCCCAAAAATTTGGTGATACATGGA

>M6-PGSC0003DMG400003175M6LRR-RLK_3355

TGTTAAACAGCCAACTAATAGACAAGCTCCACCTGCCCAAAAATTTGGTGATACATGGAAGTATATAGTTGGAGGTTTTGGGTTTTTTCTGCTCATTGCTGTTGTAGCCGTGTTCTTAGT

>M6-PGSC0003DMG400003175M6LRR-RLK_3416

TATATAGTTGGAGGTTTTGGGTTTTTTCTGCTCATTGCTGTTGTAGCCGTGTTCTTAGTTTACAGAAGAAGAGCTGCAAAAACAATAGGCCCTTGGAAGACTGGGTTGAGTGGACAGCTG

>M6-PGSC0003DMG400003175M6LRR-RLK_3538

AAAAGCATTTGTCACAGGTATTTCAATTAAAATTTGTGACAAGAATTACTGTTAATATTCTGTCCACATTCTGATATTCAATGACAAAATCATTGTTGGTTGATAATAGTCTTAAATCAC

>M6-PGSC0003DMG400003175M6LRR-RLK_3599

GTCCACATTCTGATATTCAATGACAAAATCATTGTTGGTTGATAATAGTCTTAAATCACAGCCATGAAAGAAGGTATAATCCTGATGTTGATTGCAAAAAATTGTTTCTCATTGCAGGGG

>M6-PGSC0003DMG400003175M6LRR-RLK_366

ATCATATCTGTTAATGAGTATTCAATTCCATAAAACTTACTTTATAATTGTAATGCTTTGTTTCTTTTTTAGTTAAGACCAAATCCGTTAGATTTATTTTAATGACAACAAATGTGCAAA

>M6-PGSC0003DMG400003175M6LRR-RLK_3660

CCATGAAAGAAGGTATAATCCTGATGTTGATTGCAAAAAATTGTTTCTCATTGCAGGGGTCCCTAAGCTTAACAGGTCTGAGCTTGAAACAGCATGTGAGGATTTTAGCAATATAATTAG

>M6-PGSC0003DMG400003175M6LRR-RLK_3721

CCTAAGCTTAACAGGTCTGAGCTTGAAACAGCATGTGAGGATTTTAGCAATATAATTAGTAGTCATGATGCATATATAATCTACAAGGGAACTTTGTCTAGTGGAGTAGAGATTGTTGTT

>M6-PGSC0003DMG400003175M6LRR-RLK_3782

GTCATGATGCATATATAATCTACAAGGGAACTTTGTCTAGTGGAGTAGAGATTGTTGTTGCCTCGACTACTCTAACTTCTCTCGAAGATTGGTCTGATAGTTCAGAAATGGCCTTCAGGA

>M6-PGSC0003DMG400003175M6LRR-RLK_3843

CTCGACTACTCTAACTTCTCTCGAAGATTGGTCTGATAGTTCAGAAATGGCCTTCAGGAAGAAGGTTTGTTGTTTCCATTTACATCAAAAATAAAATATTATCTCTCCATAGATCTCACA

>M6-PGSC0003DMG400003175M6LRR-RLK_3904

AAGGTTTGTTGTTTCCATTTACATCAAAAATAAAATATTATCTCTCCATAGATCTCACAGTCTTACACTTCACATAATGACATTGTTGTTATCTCTTCATCCTTTTCGGTTGTAAAAGAC

>M6-PGSC0003DMG400003175M6LRR-RLK_3965

CTTACACTTCACATAATGACATTGTTGTTATCTCTTCATCCTTTTCGGTTGTAAAAGACATGACATTTTGTGGTTGTGCTACTGAACAGATTGATACACTTTCAAGAGTGAACCACAAAA

>M6-PGSC0003DMG400003175M6LRR-RLK_4026

GACATTTTGTGGTTGTGCTACTGAACAGATTGATACACTTTCAAGAGTGAACCACAAAAATTTTGTTAATCTTATTGGATACTGTGAGGAGGATGAACCTTTCACTAGGATGATGGTCTT

>M6-PGSC0003DMG400003175M6LRR-RLK_4087

TTTGTTAATCTTATTGGATACTGTGAGGAGGATGAACCTTTCACTAGGATGATGGTCTTTGAGTATGCTCCAAATGGAACTCTTTCTGAGCATCTACACAGTAAGATTCTTTGTTGTCCA

>M6-PGSC0003DMG400003175M6LRR-RLK_4148

AGTATGCTCCAAATGGAACTCTTTCTGAGCATCTACACAGTAAGATTCTTTGTTGTCCACAGTTCAGCTTTAATTACTCACAATGATGGCTTGATTTAGTTCAGAATAACACTTCTTATT

>M6-PGSC0003DMG400003175M6LRR-RLK_4209

GTTCAGCTTTAATTACTCACAATGATGGCTTGATTTAGTTCAGAATAACACTTCTTATTTTATTTTGAAAAAAAAAAAAGATGTTCAGAATGACTTAAAATATAAGTTTAATTGTGTCAG

>M6-PGSC0003DMG400003175M6LRR-RLK_427

TTCTTTTTTAGTTAAGACCAAATCCGTTAGATTTATTTTAATGACAACAAATGTGCAAAAGATGAACTCATATGTTTTGACTTTTGAGCTATCTCTCATGTTGTTAATTATTTTCTTTTT

>M6-PGSC0003DMG400003175M6LRR-RLK_4270

ATTTTGAAAAAAAAAAAAGATGTTCAGAATGACTTAAAATATAAGTTTAATTGTGTCAGTTGAAGAAACCGATCCTCTTGACTGGTCTGCAAGGATGAGAGCCATAATGGGGACAGCATA

>M6-PGSC0003DMG400003175M6LRR-RLK_4331

GAAGAAACCGATCCTCTTGACTGGTCTGCAAGGATGAGAGCCATAATGGGGACAGCATATTGTCTTAAGTACATGCATGATCTGAATCCTCCAGTGCCCCATTCCAGCTTAGATTCAAAT

>M6-PGSC0003DMG400003175M6LRR-RLK_4392

GTCTTAAGTACATGCATGATCTGAATCCTCCAGTGCCCCATTCCAGCTTAGATTCAAATGCCATATTTTTGACTGACGATTATGCTGCTAAGGTCTGCTCCGTCTTCTTCAATATGTATA

>M6-PGSC0003DMG400003175M6LRR-RLK_4453

CATATTTTTGACTGACGATTATGCTGCTAAGGTCTGCTCCGTCTTCTTCAATATGTATATATAATTTATCTACCATATTGAACTAAGAAACATTTTTCATTTGTATTTCATATTGAAATA

>M6-PGSC0003DMG400003175M6LRR-RLK_4514

TAATTTATCTACCATATTGAACTAAGAAACATTTTTCATTTGTATTTCATATTGAAATAGAATTTTTTATATTTTCACAGATTACAGAGATTGGTTTCTGGTCAGAACTAATAGCCAAGT

>M6-PGSC0003DMG400003175M6LRR-RLK_4575

ATTTTTTATATTTTCACAGATTACAGAGATTGGTTTCTGGTCAGAACTAATAGCCAAGTCTAAGGACTCTAGGGATGATTTAGAAAATTCTGAGCTGCCAGCACTTGCTGATCCTGAAAC

>M6-PGSC0003DMG400003175M6LRR-RLK_4636

AAGGACTCTAGGGATGATTTAGAAAATTCTGAGCTGCCAGCACTTGCTGATCCTGAAACAAATATCTATAGTTTTGGGATTCTATTACTAGAAATAATTTCTGGAAAGTCACCTTATTCA

>M6-PGSC0003DMG400003175M6LRR-RLK_4697

ATATCTATAGTTTTGGGATTCTATTACTAGAAATAATTTCTGGAAAGTCACCTTATTCAGAAGAACAAGGATCTCTTCTGAACTGGGTAAATGTCCTTCATATGTTTCCCTTATTTTCTT

>M6-PGSC0003DMG400003175M6LRR-RLK_4758

AGAACAAGGATCTCTTCTGAACTGGGTAAATGTCCTTCATATGTTTCCCTTATTTTCTTATACTTGATCGTTACAATGCGTTCCCTAGTCTGCATGCATTTTCTTGTGTGAATGTATTTT

>M6-PGSC0003DMG400003175M6LRR-RLK_4819

ACTTGATCGTTACAATGCGTTCCCTAGTCTGCATGCATTTTCTTGTGTGAATGTATTTTCTTTCAGACTTATACCAATGCAAAAGCATATTGATAACTATTATTTTATTTAGGCGGAGCA

>M6-PGSC0003DMG400003175M6LRR-RLK_488

ATGAACTCATATGTTTTGACTTTTGAGCTATCTCTCATGTTGTTAATTATTTTCTTTTTCTTCTCTGTTTAGGGATCTTCATGGACATTCTTTAGAAGGCACATTGGCACCAGAGCTTGG

>M6-PGSC0003DMG400003175M6LRR-RLK_4880

TTCAGACTTATACCAATGCAAAAGCATATTGATAACTATTATTTTATTTAGGCGGAGCAATACCTTAAAGATAAGATGAATGTGAGCTCCTTGGTTGATCCAACACTCAAGTCCTTCAAA

>M6-PGSC0003DMG400003175M6LRR-RLK_4941

ACCTTAAAGATAAGATGAATGTGAGCTCCTTGGTTGATCCAACACTCAAGTCCTTCAAAAATAATGAGCTTATGGCTATCCGTGTGGTAATTAAAGAATGCCTTCAAGAAAATATAAGGA

>M6-PGSC0003DMG400003175M6LRR-RLK_5002

TAATGAGCTTATGGCTATCCGTGTGGTAATTAAAGAATGCCTTCAAGAAAATATAAGGAAGAGACCAACGATAACCGAAGTCATTGCAAAATTAAGGGAAGCGATTGATATATCGCCTGA

>M6-PGSC0003DMG400003175M6LRR-RLK_5063

AGACCAACGATAACCGAAGTCATTGCAAAATTAAGGGAAGCGATTGATATATCGCCTGATGCTGCAGTTCCAAGGCTTTCTCCCCTTTGGTGGGCTGAACTCGAGCTCTTATCTTCTGAG

>M6-PGSC0003DMG400003175M6LRR-RLK_549

TCTCTGTTTAGGGATCTTCATGGACATTCTTTAGAAGGCACATTGGCACCAGAGCTTGGGAATCTCACTCACTTAAGATCAATGTAAGTTTTTACCTACGACAAAATGTTTTTTGCTTTT

>M6-PGSC0003DMG400003175M6LRR-RLK_61

TTTATGACGTTCACGGATGTAGCTCACTCAATTCTGAAGGTAATTCTATTCTATGTTTGGACTGTCCTACTGGTTTTGGTGTTTATGTCTTAGTTATATCTAGTTTGTTATGTCCATTTG

>M6-PGSC0003DMG400003175M6LRR-RLK_610

ATCTCACTCACTTAAGATCAATGTAAGTTTTTACCTACGACAAAATGTTTTTTGCTTTTTTAGTTCATAAGTGAAAATTTGTGGGGAATTAAAGAGAGATTGTTGCTTGTCATGGTTGAA

>M6-PGSC0003DMG400003175M6LRR-RLK_671

AGTTCATAAGTGAAAATTTGTGGGGAATTAAAGAGAGATTGTTGCTTGTCATGGTTGAAAACTGATTAAAACTTAGTATTGGTTGAAAATAGCCTTGCGCATTGAAAATAAGCACGGGTT

>M6-PGSC0003DMG400003175M6LRR-RLK_732

CTGATTAAAACTTAGTATTGGTTGAAAATAGCCTTGCGCATTGAAAATAAGCACGGGTTAAAAATGACCCTAATGAGTTAAATGGATTGAAATAGCCTGGCAGCTTAAAAATACCTCCTT

>M6-PGSC0003DMG400003175M6LRR-RLK_793

AAATGACCCTAATGAGTTAAATGGATTGAAATAGCCTGGCAGCTTAAAAATACCTCCTTCTTTTGATTTCTTTTTTAACTTGCAATTTTTTAAAAAAAAAATTGATCCTCCTCTTCTTCT

>M6-PGSC0003DMG400003175M6LRR-RLK_854

TTTGATTTCTTTTTTAACTTGCAATTTTTTAAAAAAAAAATTGATCCTCCTCTTCTTCTTTTGGTTTTTTGGTTAAAAAAACTCCCTAAATAATCTTGTCTTGTTTGAAAGCTCTTTTGA

>M6-PGSC0003DMG400003175M6LRR-RLK_915

TGGTTTTTTGGTTAAAAAAACTCCCTAAATAATCTTGTCTTGTTTGAAAGCTCTTTTGAAAGTGCTTGCCTCTCGACAATGATAGCATTTGAAAGGAGTGTTTCTAAGGATCGTTCTTCC

>M6-PGSC0003DMG400003175M6LRR-RLK_976

GTGCTTGCCTCTCGACAATGATAGCATTTGAAAGGAGTGTTTCTAAGGATCGTTCTTCCGTTACAATATTCCATAAATCATAGGTTTCAAGATATGAATTCATTTTCACTGACCAAATCT

>M6-PGSC0003DMG400003195-(rev)M6LRR-RLP_0

GAGCTTTAAATACTTCTTAATTACAAGTTCTAGAAGGAGTCAGAGAAATTTATGAGCCATCCGGGATTGAATCCTTATCTCTCATGCACTTTATCGACCACTAGACCAAATCACTGAGGT

>M6-PGSC0003DMG400003195-(rev)M6LRR-RLP_1037

CACCCTCTTCCAACTTTCTCATCTTCAAAACCTTAATTTAGCTTACAATGATTTCTCACCTTCCCAAATTTCAAGCAAATTTGGATGGTTTCCGAGTTTGACACATCTTAATCTATCTCG

>M6-PGSC0003DMG400003195-(rev)M6LRR-RLP_1098

TCCCAAATTTCAAGCAAATTTGGATGGTTTCCGAGTTTGACACATCTTAATCTATCTCGTTCTGGTTTTTCAGGGACAGTACCTTTACAAGTCTCATATCTATCTAAGTTGATATCACTT

>M6-PGSC0003DMG400003195-(rev)M6LRR-RLP_1159

CTGGTTTTTCAGGGACAGTACCTTTACAAGTCTCATATCTATCTAAGTTGATATCACTTGATCTTTCTGCAGGTATTGAGGAGTTGAGAGTTGAACCTCACACTTTCAAAATGCTCCTTC

>M6-PGSC0003DMG400003195-(rev)M6LRR-RLP_122

CGGGTGTCTAGTCATTTATATATAAAGTCGCAACCATCTGAATTTTTCCTGTTTGAATGTCAAAGATACTCATAGCACGATCAAACTGTCAGTCAAACAAAAACAATAGAGTTGACATTT

>M6-PGSC0003DMG400003195-(rev)M6LRR-RLP_1220

TCTTTCTGCAGGTATTGAGGAGTTGAGAGTTGAACCTCACACTTTCAAAATGCTCCTTCAAAACTTAACTCAACTAAGGGAGCTTTACCTCACCTCAACATATATATCTTCATCACTTCC

>M6-PGSC0003DMG400003195-(rev)M6LRR-RLP_1281

AACTTAACTCAACTAAGGGAGCTTTACCTCACCTCAACATATATATCTTCATCACTTCCATCAAACTTGTCCTCTTCCTTGCAAGAGCTGAATCTTGTCGGAACGGAATTGTCTGGGAAA

>M6-PGSC0003DMG400003195-(rev)M6LRR-RLP_1342

CAAACTTGTCCTCTTCCTTGCAAGAGCTGAATCTTGTCGGAACGGAATTGTCTGGGAAAATCCCTGATGACATTTTCCATTTACCAAGATTGCAAATGCTTAACTTAGGGAGCAACTTGT

>M6-PGSC0003DMG400003195-(rev)M6LRR-RLP_1403

CCCTGATGACATTTTCCATTTACCAAGATTGCAAATGCTTAACTTAGGGAGCAACTTGTATCTCACAGGGCATTTGCCAACAACTCAATGGAACTGCAGCAACTCTTTAAGGGAATTAGA

>M6-PGSC0003DMG400003195-(rev)M6LRR-RLP_1464

CTCACAGGGCATTTGCCAACAACTCAATGGAACTGCAGCAACTCTTTAAGGGAATTAGATCTTTCTTCCTCAGGCTTTTCGGGGAATATACCAGATTCTATTGGTCATCTCAACTCTTTA

>M6-PGSC0003DMG400003195-(rev)M6LRR-RLP_1525

TTTCTTCCTCAGGCTTTTCGGGGAATATACCAGATTCTATTGGTCATCTCAACTCTTTAAGGAAATTAGATATATCAGGCTGCTACTTCTCTGGTAACATTCCTCCAACTATCGGTGATC

>M6-PGSC0003DMG400003195-(rev)M6LRR-RLP_1586

GAAATTAGATATATCAGGCTGCTACTTCTCTGGTAACATTCCTCCAACTATCGGTGATCTCACCGAGCTCACGAGTTTGCAACTCTTCTCCAACAACTTCAATGGTCAACTACCATCAAC

>M6-PGSC0003DMG400003195-(rev)M6LRR-RLP_1647

ACCGAGCTCACGAGTTTGCAACTCTTCTCCAACAACTTCAATGGTCAACTACCATCAACAATCTTAAAACTGGTGCAACTTGTTGAGTTTGATATCTCATCTAACAACCTTACAGGCAAC

>M6-PGSC0003DMG400003195-(rev)M6LRR-RLP_1708

TCTTAAAACTGGTGCAACTTGTTGAGTTTGATATCTCATCTAACAACCTTACAGGCAACATTCCAAATATTTTCGGTAACTTCACCAAGCTCAAATCTTTATCCTTTTCATATAACCTCT

>M6-PGSC0003DMG400003195-(rev)M6LRR-RLP_1769

TCCAAATATTTTCGGTAACTTCACCAAGCTCAAATCTTTATCCTTTTCATATAACCTCTTTACGGGCCTTTTTCCATCCTCTGTTACAAACTTAACAGAGCTAGAGAGTTTAGACCTCTC

>M6-PGSC0003DMG400003195-(rev)M6LRR-RLP_183

AAAGATACTCATAGCACGATCAAACTGTCAGTCAAACAAAAACAATAGAGTTGACATTTAACATATATGCACACAAAATTGCCTAACTATGATGCAAGCTGCATAATGATTTGACAATCA

>M6-PGSC0003DMG400003195-(rev)M6LRR-RLP_1830

ACGGGCCTTTTTCCATCCTCTGTTACAAACTTAACAGAGCTAGAGAGTTTAGACCTCTCAAATTGTTCTATAACTGGTCCTATTCCTTCTAGCGTCAGAACCGGATTCCCTAACCTCATC

>M6-PGSC0003DMG400003195-(rev)M6LRR-RLP_1891

ATTGTTCTATAACTGGTCCTATTCCTTCTAGCGTCAGAACCGGATTCCCTAACCTCATCTTGCTCTTCCTACCATATAACTCCCTTAGCGGAGAAATACCATCTTGGATATTAAATCTTC

>M6-PGSC0003DMG400003195-(rev)M6LRR-RLP_1952

GCTCTTCCTACCATATAACTCCCTTAGCGGAGAAATACCATCTTGGATATTAAATCTTCCTTCCTTAAAATACTTAGTTTTGAGAGGCAATCAACTTACTGGTCAACTTAAGGAAGTTAG

>M6-PGSC0003DMG400003195-(rev)M6LRR-RLP_2013

TCCTTAAAATACTTAGTTTTGAGAGGCAATCAACTTACTGGTCAACTTAAGGAAGTTAGATACAACTTGTTAGAAGTTGTTGATGTCGGTGACAATAAGCTGAATGGACCTATTCCAACA

>M6-PGSC0003DMG400003195-(rev)M6LRR-RLP_2074

ACAACTTGTTAGAAGTTGTTGATGTCGGTGACAATAAGCTGAATGGACCTATTCCAACATCATTCTCTAAACTAGTGAACTTGACAACACTTGATCTTTCTACAAACAACCTTTCTGGTG

>M6-PGSC0003DMG400003195-(rev)M6LRR-RLP_2135

ATTCTCTAAACTAGTGAACTTGACAACACTTGATCTTTCTACAAACAACCTTTCTGGTGGTCTAGATATTGGATTGTTTTCAAAATGCAAACAACTTCGACGTCTTGGCCTTTCTTTTAA

>M6-PGSC0003DMG400003195-(rev)M6LRR-RLP_2196

CTAGATATTGGATTGTTTTCAAAATGCAAACAACTTCGACGTCTTGGCCTTTCTTTTAACAATCTGTCTGTATTTTCCAGTCAAAAAGATGCATCCTTGCCTAGTTCTCTTGGAAGTTTG

>M6-PGSC0003DMG400003195-(rev)M6LRR-RLP_2257

ATCTGTCTGTATTTTCCAGTCAAAAAGATGCATCCTTGCCTAGTTCTCTTGGAAGTTTGTATGCATCTTCTTGCAACATAAGGGAGTTAAATTTCCTACGAGCTGCAAAATACATTGGTA

>M6-PGSC0003DMG400003195-(rev)M6LRR-RLP_2318

TGCATCTTCTTGCAACATAAGGGAGTTAAATTTCCTACGAGCTGCAAAATACATTGGTAACTTAGATCTTTCCAACAACAACATGCATGGAAAAATTCCTGATTGGGCATGGTCTAATTG

>M6-PGSC0003DMG400003195-(rev)M6LRR-RLP_2379

TTAGATCTTTCCAACAACAACATGCATGGAAAAATTCCTGATTGGGCATGGTCTAATTGGCAAGTCTCAGTGTCTTATTTGAATCTTTCTAGCAACTTTCTAACGGCTATTGACCTGCTT

>M6-PGSC0003DMG400003195-(rev)M6LRR-RLP_244

CATATATGCACACAAAATTGCCTAACTATGATGCAAGCTGCATAATGATTTGACAATCAATGCGGCTGGATTTTAATTACTTGTTCTAGAAGGAGTCAAGGAGATGTTCATGAGCAACCC

>M6-PGSC0003DMG400003195-(rev)M6LRR-RLP_2440

AAGTCTCAGTGTCTTATTTGAATCTTTCTAGCAACTTTCTAACGGCTATTGACCTGCTTCATAACTTTGAGAGTCTTTACAATCTTGATATTGGATCTAACTTGATTCAAGGAGAATTAC

>M6-PGSC0003DMG400003195-(rev)M6LRR-RLP_2501

TAACTTTGAGAGTCTTTACAATCTTGATATTGGATCTAACTTGATTCAAGGAGAATTACCTGCTCCACCTCCCTTATTGTTCCTCTTCATAGCCTCGAACAATAACTTCACCGGGAAGTT

>M6-PGSC0003DMG400003195-(rev)M6LRR-RLP_2562

GCTCCACCTCCCTTATTGTTCCTCTTCATAGCCTCGAACAATAACTTCACCGGGAAGTTACCTAATTCACCATTATGCAAAATGAGTTCCCTTTTGATTCTTGATTTGTCCAATAACAGC

>M6-PGSC0003DMG400003195-(rev)M6LRR-RLP_2623

CTAATTCACCATTATGCAAAATGAGTTCCCTTTTGATTCTTGATTTGTCCAATAACAGCCTGAGTGGAGTAATTCCAAAATGTTTAGTGAACATGAGCACAAGTCTCTCAGTGTTGGACC

>M6-PGSC0003DMG400003195-(rev)M6LRR-RLP_2684

GAGTGGAGTAATTCCAAAATGTTTAGTGAACATGAGCACAAGTCTCTCAGTGTTGGACCTGCACAACAATCATTTTAGTGGGACAATTCCAAATAACTTTGGTAAGGGAAGTAGTCTGAG

>M6-PGSC0003DMG400003195-(rev)M6LRR-RLP_2745

CACAACAATCATTTTAGTGGGACAATTCCAAATAACTTTGGTAAGGGAAGTAGTCTGAGAAGTCTCAATTTGCGGAAAAATAAACTAGAAGGAATGTTACCAAGAAAGCTTACCAATTGC

>M6-PGSC0003DMG400003195-(rev)M6LRR-RLP_2806

GTCTCAATTTGCGGAAAAATAAACTAGAAGGAATGTTACCAAGAAAGCTTACCAATTGCAGAGGGCTCGAAGTAGTTGATCTCGGAGAAAACTTGTTGAATGATACATTTCCCAAATGGT

>M6-PGSC0003DMG400003195-(rev)M6LRR-RLP_2867

AGGGCTCGAAGTAGTTGATCTCGGAGAAAACTTGTTGAATGATACATTTCCCAAATGGTTAGGATCACTTCCTAGGCTGCAAGTTCTCAGTTTGAGATCCAATAAATTACATGGCCCGAT

>M6-PGSC0003DMG400003195-(rev)M6LRR-RLP_2928

GGATCACTTCCTAGGCTGCAAGTTCTCAGTTTGAGATCCAATAAATTACATGGCCCGATTACCATTTCAAGAAACCAAGTCTTATTCTCAAAGCTGAAAATCCTTGATCTCTCTTATAAT

>M6-PGSC0003DMG400003195-(rev)M6LRR-RLP_2989

CCATTTCAAGAAACCAAGTCTTATTCTCAAAGCTGAAAATCCTTGATCTCTCTTATAATGACTTCACTGGAAACCTGTCAGAAAGGTTTTTCAACAACTTGAAATCCATGATAATAGAAG

>M6-PGSC0003DMG400003195-(rev)M6LRR-RLP_305

GCGGCTGGATTTTAATTACTTGTTCTAGAAGGAGTCAAGGAGATGTTCATGAGCAACCCAGGATTTGAATCCTGATCTGGCATGCACTTTATCGACCACTAGATCAAATCACTGGGGTGA

>M6-PGSC0003DMG400003195-(rev)M6LRR-RLP_3050

CTTCACTGGAAACCTGTCAGAAAGGTTTTTCAACAACTTGAAATCCATGATAATAGAAGATCAAACGGGAATACCTTTAACGTATATTGGAGAAGTTCGATACCATGATTCCGTGACATT

>M6-PGSC0003DMG400003195-(rev)M6LRR-RLP_3111

CAAACGGGAATACCTTTAACGTATATTGGAGAAGTTCGATACCATGATTCCGTGACATTGTCTATAAAAGGACAGCAAATTGAACTTGTGAGGATTCTTTCTATCTTCACAACCATTGAT

>M6-PGSC0003DMG400003195-(rev)M6LRR-RLP_3172

CTATAAAAGGACAGCAAATTGAACTTGTGAGGATTCTTTCTATCTTCACAACCATTGATTTCTCAAATAACAGATTTAAAGGTAATGTACCAAACTCCATTGGAAATCTTGGTTCACTTC

>M6-PGSC0003DMG400003195-(rev)M6LRR-RLP_3233

CTCAAATAACAGATTTAAAGGTAATGTACCAAACTCCATTGGAAATCTTGGTTCACTTCGCGGGTTGAATTTGTCACACAACAGCCTTATTGGACTTATACCTCAGTCTTTTGGAAATTT

>M6-PGSC0003DMG400003195-(rev)M6LRR-RLP_3294

GGGTTGAATTTGTCACACAACAGCCTTATTGGACTTATACCTCAGTCTTTTGGAAATTTATCTGTTCTTGAGTCGTTAGACCTTTCATGGAACCAACTTTCTGGGAACATTCCTCAAGAA

>M6-PGSC0003DMG400003195-(rev)M6LRR-RLP_3355

CTGTTCTTGAGTCGTTAGACCTTTCATGGAACCAACTTTCTGGGAACATTCCTCAAGAAGTTGCTTCACTGAAATCTCTTGCTGTCATGAATCTCTCACAGAACCACCTGATGGGACGCA

>M6-PGSC0003DMG400003195-(rev)M6LRR-RLP_3416

TGCTTCACTGAAATCTCTTGCTGTCATGAATCTCTCACAGAACCACCTGATGGGACGCATACCTAGAGGTCCACAGCTTGATACATTTGAAAATGACTCATATTCAGGAAATGCTGGATT

>M6-PGSC0003DMG400003195-(rev)M6LRR-RLP_3477

CCTAGAGGTCCACAGCTTGATACATTTGAAAATGACTCATATTCAGGAAATGCTGGATTATGCGGATTTCCCTTGTCAAGAAACTGCGGAGATAATGAGATGCCTCCTTCTACTGTGACT

>M6-PGSC0003DMG400003195-(rev)M6LRR-RLP_3538

GCGGATTTCCCTTGTCAAGAAACTGCGGAGATAATGAGATGCCTCCTTCTACTGTGACTGATATTGAAGACGAAGAAGATTCAGGTTTTATGGATTGGAGAGCTGCGATAATAGGATGTG

>M6-PGSC0003DMG400003195-(rev)M6LRR-RLP_3599

TATTGAAGACGAAGAAGATTCAGGTTTTATGGATTGGAGAGCTGCGATAATAGGATGTGGTTGTGGATCAGTGTTTGGGATGTTTATAGTATATATCATTTTCTTAACAGGGAGACCAAA

>M6-PGSC0003DMG400003195-(rev)M6LRR-RLP_366

GATTTGAATCCTGATCTGGCATGCACTTTATCGACCACTAGATCAAATCACTGGGGTGACGGATGTCTAGTCATTTATATATAGAGTCGCAATCATCTGAATTTATCAAGTTTGAATGTC

>M6-PGSC0003DMG400003195-(rev)M6LRR-RLP_3660

TGTGGATCAGTGTTTGGGATGTTTATAGTATATATCATTTTCTTAACAGGGAGACCAAAATGGTTTGTGAGAATTGTCAACCAACAGATCAACAGGATCCTTTTGAAAATTGAGGCAAAG

>M6-PGSC0003DMG400003195-(rev)M6LRR-RLP_3721

GGTTTGTGAGAATTGTCAACCAACAGATCAACAGGATCCTTTTGAAAATTGAGGCAAAGAGAAGAATTGAGGCAAAAAAGAGAAGACGGCAGAAAAGAAGAAGGATGAAGTTTTAGTATA

>M6-PGSC0003DMG400003195-(rev)M6LRR-RLP_3779

GAGAAGAATTGAGGCAAAAAAGAGAAGACGGCAGAAAAGAAGAAGGATGAAGTTTTAGTATATACTTCATGCATTATTGTCTTGCAACATACACAGTTTCAGCTTACACCTTGTTTTGGC

>M6-PGSC0003DMG400003195-(rev)M6LRR-RLP_427

GATGTCTAGTCATTTATATATAGAGTCGCAATCATCTGAATTTATCAAGTTTGAATGTCAAAGATACTCCTAGCGCGATCAAACCGTCAGTCAAACAGAAAAAATAGAGTTGACATTTAA

>M6-PGSC0003DMG400003195-(rev)M6LRR-RLP_488

AGATACTCCTAGCGCGATCAAACCGTCAGTCAAACAGAAAAAATAGAGTTGACATTTAACATATACAAACACAAAGTTGCCTAACTATGAATGCAAGCTGCGTAATGATTTGACGATCAA

>M6-PGSC0003DMG400003195-(rev)M6LRR-RLP_549

TATACAAACACAAAGTTGCCTAACTATGAATGCAAGCTGCGTAATGATTTGACGATCAATGATGCTGGATTTTGTTACTTGTTCTAGAAGGAGTCAAGGAAGATATTTCATACCTCAAGT

>M6-PGSC0003DMG400003195-(rev)M6LRR-RLP_61

CGGGATTGAATCCTTATCTCTCATGCACTTTATCGACCACTAGACCAAATCACTGAGGTGACGGGTGTCTAGTCATTTATATATAAAGTCGCAACCATCTGAATTTTTCCTGTTTGAATG

>M6-PGSC0003DMG400003195-(rev)M6LRR-RLP_610

ATGCTGGATTTTGTTACTTGTTCTAGAAGGAGTCAAGGAAGATATTTCATACCTCAAGTATATAAAATCTTGTGAATTATTCCTTCATTCCTCAAGTACTTATTATTTTTGTGCAGATCA

>M6-PGSC0003DMG400003195-(rev)M6LRR-RLP_671

ATAAAATCTTGTGAATTATTCCTTCATTCCTCAAGTACTTATTATTTTTGTGCAGATCATGGGCTGTCTATTTCTTCTTCTTCTTCTTCTTGTGTTGCAATATCCACTTGTTATCTTTTC

>M6-PGSC0003DMG400003195-(rev)M6LRR-RLP_732

GGCTGTCTATTTCTTCTTCTTCTTCTTCTTGTGTTGCAATATCCACTTGTTATCTTTTCAAGTCATCTATGTCGACGAGACGAGAGTTTGGTTTTGCAAAACCTGAAGCTTAAACTGTCC

>M6-PGSC0003DMG400003195-(rev)M6LRR-RLP_793

GTCATCTATGTCGACGAGACGAGAGTTTGGTTTTGCAAAACCTGAAGCTTAAACTGTCCATAGATGACTCTGCTTCTGCTGATTGTGAATCCAATGGTTACTTTCCCTATCCAAAGACAC

>M6-PGSC0003DMG400003195-(rev)M6LRR-RLP_854

AGATGACTCTGCTTCTGCTGATTGTGAATCCAATGGTTACTTTCCCTATCCAAAGACACTTTCTTGGAATAGAACCACTGATTGTTGTACTTGGGATGGTGTAACATGTGATAAAGTTAC

>M6-PGSC0003DMG400003195-(rev)M6LRR-RLP_915

TCTTGGAATAGAACCACTGATTGTTGTACTTGGGATGGTGTAACATGTGATAAAGTTACAGGTCGAATAACTCGTCTTGATCTTAGTTGTAGCCAACTTCATGGAACAATTGACTCCAAC

>M6-PGSC0003DMG400003195-(rev)M6LRR-RLP_976

GTCGAATAACTCGTCTTGATCTTAGTTGTAGCCAACTTCATGGAACAATTGACTCCAACAGCACCCTCTTCCAACTTTCTCATCTTCAAAACCTTAATTTAGCTTACAATGATTTCTCAC

>M6-PGSC0003DMG400003210-(rev)M6LRR-RLK_0

TCGCAGTATGGATCTAAAAACGTGCTGTATTATACTATTAAAAAAAACTGAGAGAGACTGGCTCCCTCATTAAGAATTATTGAACAACCAGCCAAACCAAAGCATTAAATATTCTCCTAT

>M6-PGSC0003DMG400003210-(rev)M6LRR-RLK_1054

TCAAATCTTGAATTTCTCTATTTAGATGATAACAACTTTGAAGGGGAAATTCCTGAGACCTTATGGAATGGGAGCTTACAAGAGTTAGCTCTTTCCTTGAATAAATTCAATGGTAGTATA

>M6-PGSC0003DMG400003210-(rev)M6LRR-RLK_1116

ATGGAATGGGAGCTTACAAGAGTTAGCTCTTTCCTTGAATAAATTCAATGGTAGTATATCAGAAAAGATTGGTGGTTCTCATCAGATAAATTACATTGATTTATCAGTTAACAAATTAAC

>M6-PGSC0003DMG400003210-(rev)M6LRR-RLK_1178

AAAAGATTGGTGGTTCTCATCAGATAAATTACATTGATTTATCAGTTAACAAATTAACTGGTCAGCTTCCAAGATCAGTTGGACGTCTAAAGAATCTGAACAAGCTTTTTCTGTATGATA

>M6-PGSC0003DMG400003210-(rev)M6LRR-RLK_124

GTTGTTTCCTAACTCATTCATTACCAAATACCACACTAATTAAGCATATGATTAGCTGAGGACACATATTATGAAACAACATTCTCAGGCGCCTCAGCCCAAAAATTAGTTCTTGAACTA

>M6-PGSC0003DMG400003210-(rev)M6LRR-RLK_1240

CAGCTTCCAAGATCAGTTGGACGTCTAAAGAATCTGAACAAGCTTTTTCTGTATGATAACATGTTTAGTGGTTCATTACCAGCAGAACTTGGGAACTGCACTTCTCTTGTTGCGATAAGT

>M6-PGSC0003DMG400003210-(rev)M6LRR-RLK_1302

GTTTAGTGGTTCATTACCAGCAGAACTTGGGAACTGCACTTCTCTTGTTGCGATAAGTCTCGTCACTAACTTCATTGGTGGGGAAATTCCTCTAGAGCTCTGCAACCTTCACTCCCTAAT

>M6-PGSC0003DMG400003210-(rev)M6LRR-RLK_1364

TCACTAACTTCATTGGTGGGGAAATTCCTCTAGAGCTCTGCAACCTTCACTCCCTAATAAAATTTCAAGTGTACGGAAATCAAATTCAGGGCCAAATTCCAGAATGCATTGGCAGAATAA

>M6-PGSC0003DMG400003210-(rev)M6LRR-RLK_1426

TTTCAAGTGTACGGAAATCAAATTCAGGGCCAAATTCCAGAATGCATTGGCAGAATAAGTGGACTGGAGGAGCTAGATCTTAGTGAGAACCGATTGATCGGCAAGATACCACCTGGAATC

>M6-PGSC0003DMG400003210-(rev)M6LRR-RLK_1488

ACTGGAGGAGCTAGATCTTAGTGAGAACCGATTGATCGGCAAGATACCACCTGGAATCACAAATATGACGAAACTTGTGTTACTCTCTTTGGCTCATAATAATCTTACAGGGGAGGTACC

>M6-PGSC0003DMG400003210-(rev)M6LRR-RLK_1550

ATATGACGAAACTTGTGTTACTCTCTTTGGCTCATAATAATCTTACAGGGGAGGTACCACCTAATCTTGGAAAAAACAATTTTCCTGGCTTATTTAAGGTTGATTTAGGTTATAATATCT

>M6-PGSC0003DMG400003210-(rev)M6LRR-RLK_1612

AATCTTGGAAAAAACAATTTTCCTGGCTTATTTAAGGTTGATTTAGGTTATAATATCTTCAGCGGACCTATTCCTTCTGAACTATGTAATGGTAACAGTCTTGGAGTCCTGGTTCTTGAA

>M6-PGSC0003DMG400003210-(rev)M6LRR-RLK_1674

CGGACCTATTCCTTCTGAACTATGTAATGGTAACAGTCTTGGAGTCCTGGTTCTTGAAAACAACAGTTTTAATGGCAGCTTCCCAACATATCTAGCAAAATGTGAATCCCTCTATAGGGT

>M6-PGSC0003DMG400003210-(rev)M6LRR-RLK_1736

ACAGTTTTAATGGCAGCTTCCCAACATATCTAGCAAAATGTGAATCCCTCTATAGGGTAAAACTCCCCAACAACAATCTGCAGGGGAGTATTCCTGACTACATTGAAAAGAATGAGAAGA

>M6-PGSC0003DMG400003210-(rev)M6LRR-RLK_1798

CTCCCCAACAACAATCTGCAGGGGAGTATTCCTGACTACATTGAAAAGAATGAGAAGATTTCTTACTTGAATGTTCGAGGAAATATGCTTGCAGGAAGGATTCCAGCAGCATTTGGTTAT

>M6-PGSC0003DMG400003210-(rev)M6LRR-RLK_186

CACATATTATGAAACAACATTCTCAGGCGCCTCAGCCCAAAAATTAGTTCTTGAACTAGCAGCTCAGTGTTTTAAGAGGTCAGAGATGTTCTTGTTTCTGGTTAATTGCCTTCTCTTCTT

>M6-PGSC0003DMG400003210-(rev)M6LRR-RLK_1860

TTACTTGAATGTTCGAGGAAATATGCTTGCAGGAAGGATTCCAGCAGCATTTGGTTATTGGACCAATCTTTCAACAATTGATCTTTCAGAGAATATGTTTTCTGGTTCCATACCTGCCGA

>M6-PGSC0003DMG400003210-(rev)M6LRR-RLK_1922

CCAATCTTTCAACAATTGATCTTTCAGAGAATATGTTTTCTGGTTCCATACCTGCCGAAATGGGGAAGCTGCAAAATCTTGTAAGATTGAACATTTCTTCAAACAGACTGACAGGAAAAA

>M6-PGSC0003DMG400003210-(rev)M6LRR-RLK_1984

GGGAAGCTGCAAAATCTTGTAAGATTGAACATTTCTTCAAACAGACTGACAGGAAAAATTCCTCTTCAGTTGAGTTACTCTGCAAAACTGGCAGAGTTGGATCTCAGCAACAACAATCTT

>M6-PGSC0003DMG400003210-(rev)M6LRR-RLK_2046

TCTTCAGTTGAGTTACTCTGCAAAACTGGCAGAGTTGGATCTCAGCAACAACAATCTTTCGGGTAGAATTCCGAAAGAAATTGCATCATCTTTAGTACTGACAAATCTTCTACTCCAGGA

>M6-PGSC0003DMG400003210-(rev)M6LRR-RLK_2108

GTAGAATTCCGAAAGAAATTGCATCATCTTTAGTACTGACAAATCTTCTACTCCAGGACAACAAACTTTCTGGTGCTCTTCCAGACACTTTCTCCTCCTCACAAAAGCTTGTAAAGCTGC

>M6-PGSC0003DMG400003210-(rev)M6LRR-RLK_2170

AAACTTTCTGGTGCTCTTCCAGACACTTTCTCCTCCTCACAAAAGCTTGTAAAGCTGCAGCTCGGGGACAACTTGCTTGAAGGCTCAATTCCATGCAGCCTCAGTAAACTTAGGGAACCT

>M6-PGSC0003DMG400003210-(rev)M6LRR-RLK_2232

CGGGGACAACTTGCTTGAAGGCTCAATTCCATGCAGCCTCAGTAAACTTAGGGAACCTAATGTTGCACTGAATCTGAGCATGAATAAGTTTAGTGGTCAAATTCCTAGGTGTCTGAGCAA

>M6-PGSC0003DMG400003210-(rev)M6LRR-RLK_2294

TTGCACTGAATCTGAGCATGAATAAGTTTAGTGGTCAAATTCCTAGGTGTCTGAGCAATCTAGACAATTTAGAAATCCTTGATATATCAAGCAACAACTTGTCTGGCGCAATACCTTCAG

>M6-PGSC0003DMG400003210-(rev)M6LRR-RLK_2356

GACAATTTAGAAATCCTTGATATATCAAGCAACAACTTGTCTGGCGCAATACCTTCAGAAATGGACAAGATGAGGTCCCTTTCGTTTCTCAACATATCATTTAATAACTTTTCAGGGGAG

>M6-PGSC0003DMG400003210-(rev)M6LRR-RLK_2418

GGACAAGATGAGGTCCCTTTCGTTTCTCAACATATCATTTAATAACTTTTCAGGGGAGGTACCCATTTTGTGGGGAAAACTATTAAGTTCACATCCAGGAACTTCCCAGGGAAATCCAGG

>M6-PGSC0003DMG400003210-(rev)M6LRR-RLK_248

CTCAGTGTTTTAAGAGGTCAGAGATGTTCTTGTTTCTGGTTAATTGCCTTCTCTTCTTTGCTAATTTATTTGCAGTCTTTTCGATCTCATGGCCTCAAGATTCTGTTCATCTACTTGGGT

>M6-PGSC0003DMG400003210-(rev)M6LRR-RLK_2480

CCATTTTGTGGGGAAAACTATTAAGTTCACATCCAGGAACTTCCCAGGGAAATCCAGGACTCTGCTTATCAGACACTGAAAGTAGCAGCTGCAAGCATGTGAAAAAATCACAAAGAAACT

>M6-PGSC0003DMG400003210-(rev)M6LRR-RLK_2542

TGCTTATCAGACACTGAAAGTAGCAGCTGCAAGCATGTGAAAAAATCACAAAGAAACTGGAGGACCTTGGCTGGTGTAATTAGTGGTGTGTTTTCCGTGGCAGTCATAGCTGCGGCAATA

>M6-PGSC0003DMG400003210-(rev)M6LRR-RLK_2604

GACCTTGGCTGGTGTAATTAGTGGTGTGTTTTCCGTGGCAGTCATAGCTGCGGCAATATACTTGCTAGTGACTCGGATTCAGCACCCTTCCCTGCTGAATAAGCATCGGCTCGTCAAGTA

>M6-PGSC0003DMG400003210-(rev)M6LRR-RLK_2666

TGCTAGTGACTCGGATTCAGCACCCTTCCCTGCTGAATAAGCATCGGCTCGTCAAGTATCAGGCAGAATTTGAAGATCTGCCAGATCGTATAAATTTTGAGGACATTGTACGTGCAACAG

>M6-PGSC0003DMG400003210-(rev)M6LRR-RLK_2728

GCAGAATTTGAAGATCTGCCAGATCGTATAAATTTTGAGGACATTGTACGTGCAACAGAAGGCTGGAGTGAAAAATATGTAATTGGAAGAGGCAAGCATGGAACTGTTTATAAGATGGAA

>M6-PGSC0003DMG400003210-(rev)M6LRR-RLK_2790

CTGGAGTGAAAAATATGTAATTGGAAGAGGCAAGCATGGAACTGTTTATAAGATGGAATCTGCGAAATCCAAAAAGCTTTGGGCTGTCAAGAAGGTTGACTTAGCTCAGAGAGCATTCAG

>M6-PGSC0003DMG400003210-(rev)M6LRR-RLK_2852

CGAAATCCAAAAAGCTTTGGGCTGTCAAGAAGGTTGACTTAGCTCAGAGAGCATTCAGCAATGAAATGAGAAGCCTCAATTCTGTCAGACACCGCAATTTGGTAAGATTGGGAGGACACT

>M6-PGSC0003DMG400003210-(rev)M6LRR-RLK_2914

GAAATGAGAAGCCTCAATTCTGTCAGACACCGCAATTTGGTAAGATTGGGAGGACACTGCACCAGACATGGATATGGCTTCATTCTAACAGAGTTTATCCCTGGAGGAACTCTTCATGAT

>M6-PGSC0003DMG400003210-(rev)M6LRR-RLK_3038

TTCACCAGAGAAAACCACCTGTAGTCCTGGACTGGGAGTCCCGCCATCGTATTGCTCTTGGTGTTGCTCAAGGTCTGTCTTACCTTCATCATGATTCTGTCCCTCAAATCATTCACAGAG

>M6-PGSC0003DMG400003210-(rev)M6LRR-RLK_3100

GTTGCTCAAGGTCTGTCTTACCTTCATCATGATTCTGTCCCTCAAATCATTCACAGAGATCTCAAATCAGATAATGTAATGCTGGATACTGAGATGGAGCCAAAGATTGGAGACTTCGGG

>M6-PGSC0003DMG400003210-(rev)M6LRR-RLK_3162

CAAATCAGATAATGTAATGCTGGATACTGAGATGGAGCCAAAGATTGGAGACTTCGGGATTGCTAAAACAGTATCAGACTCTGATGAAAACTCGACAAACTCCAAAATTGTTGGGACACT

>M6-PGSC0003DMG400003210-(rev)M6LRR-RLK_3224

CTAAAACAGTATCAGACTCTGATGAAAACTCGACAAACTCCAAAATTGTTGGGACACTAGGATACATTGCTCCAGGTTAGCCACACGTTACTATGCTAGTCTTCAGTAATAATTTAGTTG

>M6-PGSC0003DMG400003210-(rev)M6LRR-RLK_3286

TACATTGCTCCAGGTTAGCCACACGTTACTATGCTAGTCTTCAGTAATAATTTAGTTGGCAGATTACTTTTGATGATTTACTAAATTTTATTACTTTCTCCCCCGGTCCCCCCTCCTTCT

>M6-PGSC0003DMG400003210-(rev)M6LRR-RLK_3348

ATTACTTTTGATGATTTACTAAATTTTATTACTTTCTCCCCCGGTCCCCCCTCCTTCTCCTTGTCTGTCCATCAATTCAGAGAATGCGTACTCAGTTCAATTGACAGAGAAAAGTGATGT

>M6-PGSC0003DMG400003210-(rev)M6LRR-RLK_3410

GTCTGTCCATCAATTCAGAGAATGCGTACTCAGTTCAATTGACAGAGAAAAGTGATGTCTATAGCTATGGAGTTCTTCTACTGGAGCTCTTCTGCAGAAAAATGCCAGTAGACCCCAGCT

>M6-PGSC0003DMG400003210-(rev)M6LRR-RLK_3472

AGCTATGGAGTTCTTCTACTGGAGCTCTTCTGCAGAAAAATGCCAGTAGACCCCAGCTTTGAAGAAGGATTAGACATTGTTTCCTGGGTGAGGAAAAACTTGCACAGAAGTGACAACAAT

>M6-PGSC0003DMG400003210-(rev)M6LRR-RLK_3534

AGAAGGATTAGACATTGTTTCCTGGGTGAGGAAAAACTTGCACAGAAGTGACAACAATTTCCTTCACTTGCTTGATGAAGAAATTAGTTTTTGGTACATAGAAGAACAATGGAAGGCCCT

>M6-PGSC0003DMG400003210-(rev)M6LRR-RLK_3596

TTCACTTGCTTGATGAAGAAATTAGTTTTTGGTACATAGAAGAACAATGGAAGGCCCTGAAAATGGTGTATCTAGCGCTTCAATGTGCTGAACTCGAGGCAAGCACCAGGCCTGCAATGA

>M6-PGSC0003DMG400003210-(rev)M6LRR-RLK_372

GAGCAATCTTCCAGAGCTCTCTCAACAATTGCTCCCTTGGAATCAGTCTGTCTCACACTGCCAATGGAAAGGTGTTTCCTGCTATTCAGACATAACTTCTCATGTCAAATCATTGATTCT

>M6-PGSC0003DMG400003210-(rev)M6LRR-RLK_3720

TGTAGTAAGGTCTCTTGTCGAGTTAAATCATAGATGTAAAAGAGAAGAAATGAAACTTTGAATAGTGAAAGCCATAATCAACTTTTCCAAATTCCAAAATCTTTTTTTTCTTTGCTTACA

>M6-PGSC0003DMG400003210-(rev)M6LRR-RLK_3782

TAGTGAAAGCCATAATCAACTTTTCCAAATTCCAAAATCTTTTTTTTCTTTGCTTACATAAAAATTGAACCATTTCTCGATTTGTGTGTGTCATCCTTTCATTCAAGGGCTAAATTCTAA

>M6-PGSC0003DMG400003210-(rev)M6LRR-RLK_3844

AATTGAACCATTTCTCGATTTGTGTGTGTCATCCTTTCATTCAAGGGCTAAATTCTAACATAATCGATCCTGGGTGTTTCTGTTTAACTTTGTGACATCCACTGCAATAATATAGACTAA

>M6-PGSC0003DMG400003210-(rev)M6LRR-RLK_3906

ATCGATCCTGGGTGTTTCTGTTTAACTTTGTGACATCCACTGCAATAATATAGACTAAGAAAATGCATGGCGTTTCAAATCAATATATGGTAAAAGCAGTTTTCTCGTTCTAGCCATAGC

>M6-PGSC0003DMG400003210-(rev)M6LRR-RLK_3968

ATGCATGGCGTTTCAAATCAATATATGGTAAAAGCAGTTTTCTCGTTCTAGCCATAGCTCTTTTCATAATGGGCCACTGTCCTTATCAACTTTTGCTTGTTACTTTCTCCAGCTGCTCGA

>M6-PGSC0003DMG400003210-(rev)M6LRR-RLK_434

AATGGAAAGGTGTTTCCTGCTATTCAGACATAACTTCTCATGTCAAATCATTGATTCTAACAGACTTATTGCTACCTGGAACCTTGGACAAGGCCTTACCTAATCTCTGCCTTCTTCCTC

>M6-PGSC0003DMG400003210-(rev)M6LRR-RLK_496

GACTTATTGCTACCTGGAACCTTGGACAAGGCCTTACCTAATCTCTGCCTTCTTCCTCGCTTGGTCTCTCTTGATCTCAGTGGCAACCATTTCACTGGTGGCATCCCTGATATGCTCGCA

>M6-PGSC0003DMG400003210-(rev)M6LRR-RLK_62

TCCCTCATTAAGAATTATTGAACAACCAGCCAAACCAAAGCATTAAATATTCTCCTATTTTGGTTGTTTCCTAACTCATTCATTACCAAATACCACACTAATTAAGCATATGATTAGCTG

>M6-PGSC0003DMG400003210-(rev)M6LRR-RLK_620

GCAGCCAACTTGACACGATTCTACTCAACGACAACAGATTTTCAGGATCCATCCCACCAGAGATTTTCAAATCAAGCAAACTCATAAAAATTGATTTGGGTTTAAATCAGCTCACTGGCA

>M6-PGSC0003DMG400003210-(rev)M6LRR-RLK_682

ATTTTCAAATCAAGCAAACTCATAAAAATTGATTTGGGTTTAAATCAGCTCACTGGCACTATTCCCTCTGAGGTAATCCTTTCTACAAACCTTCAACATCTCGGGCTGTGGAACAATTTC

>M6-PGSC0003DMG400003210-(rev)M6LRR-RLK_744

TCCCTCTGAGGTAATCCTTTCTACAAACCTTCAACATCTCGGGCTGTGGAACAATTTCCTCAGTGGAAATATTCCAAAAGAGCTATTCAATTTGCCAAACTTGACACATCTCCACCTCAA

>M6-PGSC0003DMG400003210-(rev)M6LRR-RLK_806

GTGGAAATATTCCAAAAGAGCTATTCAATTTGCCAAACTTGACACATCTCCACCTCAACACAAATGAACTGACTGGACCTCTACCTGACTTCCCATCCTCGTGCTCACTTTCCGAGTTTT

>M6-PGSC0003DMG400003210-(rev)M6LRR-RLK_868

AATGAACTGACTGGACCTCTACCTGACTTCCCATCCTCGTGCTCACTTTCCGAGTTTTTTATTTATGAGAACCGTTTGTCAGGTTCTTTGCCAATTACCCTAGGCAATTGTCACAACCTT

>M6-PGSC0003DMG400003210-(rev)M6LRR-RLK_930

TTATGAGAACCGTTTGTCAGGTTCTTTGCCAATTACCCTAGGCAATTGTCACAACCTTACTTCATTCTCTGCATCATCTGCTCATCTTGGAGGAGTTATTTCCCCAGAGGTTTTAAGGGA

>M6-PGSC0003DMG400003210-(rev)M6LRR-RLK_992

CATTCTCTGCATCATCTGCTCATCTTGGAGGAGTTATTTCCCCAGAGGTTTTAAGGGACCTTTCAAATCTTGAATTTCTCTATTTAGATGATAACAACTTTGAAGGGGAAATTCCTGAGA

>M6-PGSC0003DMG400003211-(rev)M6LRR-RLK_0

TAAAATAATCTCCTATATTCATTGTTCCCTCGGACAGTCGTTGCCTACTACCCCCACACGTATCTTTAAAAAACAGGCTGACCTCAACTTAAAACCTAAATAATAGTCTCAGGTTCCAGC

>M6-PGSC0003DMG400003211-(rev)M6LRR-RLK_1037

TTATGGAATGGGAAATTACAAGAGTTGGTTCTTTCCATAAATAATTTTAATGGAAGTATATCAGAAAAGATTGGTGTTTGCCATCAGATGACTTACATTGATTTGTCAGCTAACAAATTA

>M6-PGSC0003DMG400003211-(rev)M6LRR-RLK_1098

CAGAAAAGATTGGTGTTTGCCATCAGATGACTTACATTGATTTGTCAGCTAACAAATTAAGTGGTCAGATTCCCAAGTCAGTTGGACGTCTGAAGAATTTGTACAAGCTTCTTCTGTATG

>M6-PGSC0003DMG400003211-(rev)M6LRR-RLK_1159

TGGTCAGATTCCCAAGTCAGTTGGACGTCTGAAGAATTTGTACAAGCTTCTTCTGTATGATAACATGTTTAGTGGTTCATTGCCAGCAGAAGTTGGGAACTGCACTTCTCTTGCTGAGAT

>M6-PGSC0003DMG400003211-(rev)M6LRR-RLK_122

AAAAATTAGTTTTTGAACTAGCTCCTTATTGGATGCAGCTTCTTTAAAGAAGTGTTGAAGAGTTCAGAAATGTTCTTGTTTGATCTTGTTCACTGTTTTATTTTCTTTGCCAGCATTGTA

>M6-PGSC0003DMG400003211-(rev)M6LRR-RLK_1220

AACATGTTTAGTGGTTCATTGCCAGCAGAAGTTGGGAACTGCACTTCTCTTGCTGAGATAAGTCTCGTCAGTAACTTCATCAGTGGGGAAATTCCTTCAGAAATTTGCAGCCTTCAAAAC

>M6-PGSC0003DMG400003211-(rev)M6LRR-RLK_1281

GTCTCGTCAGTAACTTCATCAGTGGGGAAATTCCTTCAGAAATTTGCAGCCTTCAAAACCTAGAAACATTTAATGCATTCTAAAATAAAATTCAGGGCCAAATTCCAGAATGCATTGGAA

>M6-PGSC0003DMG400003211-(rev)M6LRR-RLK_1342

AGAAACATTTAATGCATTCTAAAATAAAATTCAGGGCCAAATTCCAGAATGCATTGGAAGAATAAGCGGGCTGCAGGAGCTAGCTCTTTATGAGAACCGATTGACCGGCAAGTTACCGCT

>M6-PGSC0003DMG400003211-(rev)M6LRR-RLK_1403

ATAAGCGGGCTGCAGGAGCTAGCTCTTTATGAGAACCGATTGACCGGCAAGTTACCGCTTGGAATTACAAATATGACGAAACTTTCATTTCTGTCGTTGGCTCATAATAACCTTACGGGG

>M6-PGSC0003DMG400003211-(rev)M6LRR-RLK_1464

GAATTACAAATATGACGAAACTTTCATTTCTGTCGTTGGCTCATAATAACCTTACGGGGGAGGTACCACCTGATCTTGGAAAAGATAATATTCCTGGCATAGTTAAGGTTGATTTAGGTT

>M6-PGSC0003DMG400003211-(rev)M6LRR-RLK_1525

GGTACCACCTGATCTTGGAAAAGATAATATTCCTGGCATAGTTAAGGTTGATTTAGGTTATAATAACTTCAGTGGACAGATTCCTTCTAAACTGTGTAATGGCAACAGGCTTGCAGTCCT

>M6-PGSC0003DMG400003211-(rev)M6LRR-RLK_1586

AATAACTTCAGTGGACAGATTCCTTCTAAACTGTGTAATGGCAACAGGCTTGCAGTCCTGGCTCTTGAAAACAACAGGTTTAATGGCAGCTTCCCAACCTATCTAGCAAACTGCAAATCT

>M6-PGSC0003DMG400003211-(rev)M6LRR-RLK_1647

CTCTTGAAAACAACAGGTTTAATGGCAGCTTCCCAACCTATCTAGCAAACTGCAAATCTCTCTATAGGGTAAAACTTTCCAACAACAATCTGCAGGGGAGTATTCCAGACGACATTGAAA

>M6-PGSC0003DMG400003211-(rev)M6LRR-RLK_1708

CTATAGGGTAAAACTTTCCAACAACAATCTGCAGGGGAGTATTCCAGACGACATTGAAAAGAATGAGAATATTTCTTACTTGGATGTTAGACAGAATATGCTTGTAGGAAGGATTCCAGC

>M6-PGSC0003DMG400003211-(rev)M6LRR-RLK_1769

AATGAGAATATTTCTTACTTGGATGTTAGACAGAATATGCTTGTAGGAAGGATTCCAGCTGCATTTGGTTACTGGACCAATCTTTCAATGATTGATCTTTCAGAAAATATGTTTAATGGC

>M6-PGSC0003DMG400003211-(rev)M6LRR-RLK_183

GTTCAGAAATGTTCTTGTTTGATCTTGTTCACTGTTTTATTTTCTTTGCCAGCATTGTAGCGGTTTTTTCGGCATCATTGCCTCAAGATTCTGTTCATCTACTTGGGTTTCGAAGCAATC

>M6-PGSC0003DMG400003211-(rev)M6LRR-RLK_1830

CATTTGGTTACTGGACCAATCTTTCAATGATTGATCTTTCAGAAAATATGTTTAATGGCTCCATACCCACAGAACTTGGAAAGCTTCAGAATCTTGTAAAACTGAGCATTTCTTCAAACA

>M6-PGSC0003DMG400003211-(rev)M6LRR-RLK_1891

CATACCCACAGAACTTGGAAAGCTTCAGAATCTTGTAAAACTGAGCATTTCTTCAAACAGACTGACCGGACAAATTCCCTTTCAGTTGAGTCACTTTGAAAAACTGGAGGAGTTGGATCT

>M6-PGSC0003DMG400003211-(rev)M6LRR-RLK_1952

CTGACCGGACAAATTCCCTTTCAGTTGAGTCACTTTGAAAAACTGGAGGAGTTGGATCTCAGCAACAACAATCTTTCAGGAAGAATTCCTAAAGAAATTGCATCATCTTCAGTATTGACA

>M6-PGSC0003DMG400003211-(rev)M6LRR-RLK_2013

GCAACAACAATCTTTCAGGAAGAATTCCTAAAGAAATTGCATCATCTTCAGTATTGACAAATCTTCTACTCCAGGACAACAAACTTTCTGGTGCTCTTCCAGACACTTTCTCCTCTTCAC

>M6-PGSC0003DMG400003211-(rev)M6LRR-RLK_2074

TCTTCTACTCCAGGACAACAAACTTTCTGGTGCTCTTCCAGACACTTTCTCCTCTTCACAAATGCTTGTAAAGCTGCAGCTGGGAAACAACTTGCTCGAAGGCCCAATTCCATGCAGTCT

>M6-PGSC0003DMG400003211-(rev)M6LRR-RLK_2135

ATGCTTGTAAAGCTGCAGCTGGGAAACAACTTGCTCGAAGGCCCAATTCCATGCAGTCTGAGTAAGCTTATGCAACCTGTTTTTTCACTAAATCTGAGCATGAATAAGTTTACTGGTGAA

>M6-PGSC0003DMG400003211-(rev)M6LRR-RLK_2196

GTAAGCTTATGCAACCTGTTTTTTCACTAAATCTGAGCATGAATAAGTTTACTGGTGAAATTCCTAGGTGCCTAGGCAATCTTGACAAATTGGAGGTCCTTGATATATCAAGCAACAACC

>M6-PGSC0003DMG400003211-(rev)M6LRR-RLK_2257

TCCTAGGTGCCTAGGCAATCTTGACAAATTGGAGGTCCTTGATATATCAAGCAACAACCTGTCTGGTGCAATACCATCAGAAATGGAAAAGATGAGTTCCCTTTCGTTTCTCAACATATC

>M6-PGSC0003DMG400003211-(rev)M6LRR-RLK_2318

TCTGGTGCAATACCATCAGAAATGGAAAAGATGAGTTCCCTTTCGTTTCTCAACATATCATTTAATAGCTTGTCAGGGAAGGTACCCAATACATGGGAAAAACTATTAAGTTCACGTCCA

>M6-PGSC0003DMG400003211-(rev)M6LRR-RLK_2379

TTAATAGCTTGTCAGGGAAGGTACCCAATACATGGGAAAAACTATTAAGTTCACGTCCAGGATCTGCCTTGGGAAATCCAGGACTTTGCTTAATAGACACGGATGGTAGCAACTGCAAGC

>M6-PGSC0003DMG400003211-(rev)M6LRR-RLK_244

GGTTTTTTCGGCATCATTGCCTCAAGATTCTGTTCATCTACTTGGGTTTCGAAGCAATCTTCCAGAGCCCTCTCAACAATTGCTCCCTTGGAATCAATCTGTCTCACACTGTAAATGGAA

>M6-PGSC0003DMG400003211-(rev)M6LRR-RLK_2440

ATCTGCCTTGGGAAATCCAGGACTTTGCTTAATAGACACGGATGGTAGCAACTGCAAGCATGTGAAAAAATCACAAGTAAAATGGAAGACTTTGGCTGGTGTAATTAGTGGATGTGTGCT

>M6-PGSC0003DMG400003211-(rev)M6LRR-RLK_2501

GTGAAAAAATCACAAGTAAAATGGAAGACTTTGGCTGGTGTAATTAGTGGATGTGTGCTTTCCATGGCAATCATAGTTGCTGCAATGTACTTACTAGTGACTCGGATTTGGCACCCATCC

>M6-PGSC0003DMG400003211-(rev)M6LRR-RLK_2562

CCATGGCAATCATAGTTGCTGCAATGTACTTACTAGTGACTCGGATTTGGCACCCATCCCTGCTGAATAAGCATCGGCTCGTCAAGTGTCAATCAGGAATTGAAGATCTACCTGATGGTA

>M6-PGSC0003DMG400003211-(rev)M6LRR-RLK_2623

GCTGAATAAGCATCGGCTCGTCAAGTGTCAATCAGGAATTGAAGATCTACCTGATGGTATAACTTTTGAGGACATTGTACATGCAACAGAAGGCTGGAGTGAAAACTATGTTATCGGAAG

>M6-PGSC0003DMG400003211-(rev)M6LRR-RLK_2684

ACTTTTGAGGACATTGTACATGCAACAGAAGGCTGGAGTGAAAACTATGTTATCGGAAGAGGCACGCATGGAACTGTTTATAAGTTGGAATCTGCCAAATCCAATAAACTTTGGGCTGTC

>M6-PGSC0003DMG400003211-(rev)M6LRR-RLK_2745

GCACGCATGGAACTGTTTATAAGTTGGAATCTGCCAAATCCAATAAACTTTGGGCTGTCAAGAAGGTAGACTTAGCTCAGAGAGCATTCAATGATGAAATGAGAAGCCTAAATTCTGTCA

>M6-PGSC0003DMG400003211-(rev)M6LRR-RLK_2806

GAAGGTAGACTTAGCTCAGAGAGCATTCAATGATGAAATGAGAAGCCTAAATTCTGTCAGACACCGCAATTTGGTAAGATTGGGAGGATATTGCATGATACATGGATATGGCTTCATTCT

>M6-PGSC0003DMG400003211-(rev)M6LRR-RLK_2867

CACCGCAATTTGGTAAGATTGGGAGGATATTGCATGATACATGGATATGGCTTCATTCTAACAGAATTTATACCTGGAGGGACTCTTCATGATGTCCTTCACCAGAGAAAACCACCTGTA

>M6-PGSC0003DMG400003211-(rev)M6LRR-RLK_2989

CTTGGACTGGAAGTCCCGCCATCGTATTGCTCTTGGTATTGCTCAAGGTCTATCATACCTTCACCACGATTCGGTGCCTCAAATCATTCACAGAGATCTCAAATCAGATAATGTAATGCT

>M6-PGSC0003DMG400003211-(rev)M6LRR-RLK_305

CCAGAGCCCTCTCAACAATTGCTCCCTTGGAATCAATCTGTCTCACACTGTAAATGGAAAGGTGTTACCTGCTATTCAGACAAAACTTCACAAGTCAAATCATTGGATTTCAGAGACTTC

>M6-PGSC0003DMG400003211-(rev)M6LRR-RLK_3050

CACCACGATTCGGTGCCTCAAATCATTCACAGAGATCTCAAATCAGATAATGTAATGCTGGATTCTGAGATGGTGCCAAAGATTGGCGATTTCGGGATTGCTAAAATGGTATCAGATGAA

>M6-PGSC0003DMG400003211-(rev)M6LRR-RLK_3111

ATTCTGAGATGGTGCCAAAGATTGGCGATTTCGGGATTGCTAAAATGGTATCAGATGAAGAAAATTCGACGAACTCCAATATTGTTGGGACTCTAGGATACATTGCTCCAGGTTAGCCAC

>M6-PGSC0003DMG400003211-(rev)M6LRR-RLK_3172

AAATTCGACGAACTCCAATATTGTTGGGACTCTAGGATACATTGCTCCAGGTTAGCCACACGTTACTATGCTAATTTTCAGTATAATTCAGTTGGCAGATTACTTTGATAAACTAATAAC

>M6-PGSC0003DMG400003211-(rev)M6LRR-RLK_3233

GTTACTATGCTAATTTTCAGTATAATTCAGTTGGCAGATTACTTTGATAAACTAATAACTTTTACTACTTTCTTCCCTGGCCCCCTTTCTCTTTTTGTCACTCCATCTATTCAGAGAACG

>M6-PGSC0003DMG400003211-(rev)M6LRR-RLK_3294

TTACTACTTTCTTCCCTGGCCCCCTTTCTCTTTTTGTCACTCCATCTATTCAGAGAACGCATACTCAGTTCAACTGACAGAGAAAAGTGATGTCTATAGCTACGGAATTCTTCTTTTAGA

>M6-PGSC0003DMG400003211-(rev)M6LRR-RLK_3355

TACTCAGTTCAACTGACAGAGAAAAGTGATGTCTATAGCTACGGAATTCTTCTTTTAGAGCTTTTCTGCAGAAAAATGCCAGTAGATCCCAGCTTTGAAGAAGGATTAGACATTGTTTTC

>M6-PGSC0003DMG400003211-(rev)M6LRR-RLK_3416

TTTTCTGCAGAAAAATGCCAGTAGATCCCAGCTTTGAAGAAGGATTAGACATTGTTTTCTGGGTGAGGAAAAACTTGCAGAGAAGCAACAATTTCCTTAGTTTTCTTGATGAAGAAATTA

>M6-PGSC0003DMG400003211-(rev)M6LRR-RLK_3477

GGTGAGGAAAAACTTGCAGAGAAGCAACAATTTCCTTAGTTTTCTTGATGAAGAAATTAGGCTCTGGAATGTAGAAGAACAATGGAAGGCCCTGAAAATAGTGGATTTGGCGCTTCAATG

>M6-PGSC0003DMG400003211-(rev)M6LRR-RLK_366

GTGTTACCTGCTATTCAGACAAAACTTCACAAGTCAAATCATTGGATTTCAGAGACTTCCTTCTATCTGGAATCTTGGACAAGGCCTTTCCAAATCTTTGCCATCTTCCTCGTTTGATCT

>M6-PGSC0003DMG400003211-(rev)M6LRR-RLK_3782

TTACTAGACTCTATGGCCTCTTTGTAAGTTTTGTTTGAGGCTGTATCTGTAGATATAATATACCAGGTCTTTATTAGGGTGCTGATGAATATAACATGTTACCAAATTCAAAAAGAGAAT

>M6-PGSC0003DMG400003211-(rev)M6LRR-RLK_3843

ACCAGGTCTTTATTAGGGTGCTGATGAATATAACATGTTACCAAATTCAAAAAGAGAATAGATGTAAAGATAAGTAATGAAACTTTTAGTAGTGAAACCTGTGATCAACTTTCCAAATTC

>M6-PGSC0003DMG400003211-(rev)M6LRR-RLK_3904

ATGTAAAGATAAGTAATGAAACTTTTAGTAGTGAAACCTGTGATCAACTTTCCAAATTCCAACATCTCATTTTTTTCTTTTTTTGCTTACATAAAAAAATTGGACAAAGAAAATACATGA

>M6-PGSC0003DMG400003211-(rev)M6LRR-RLK_3965

ACATCTCATTTTTTTCTTTTTTTGCTTACATAAAAAAATTGGACAAAGAAAATACATGATGTTTCAGTTTAACAAAATCGTGATCTTGGGTTTTTCAGTTTAACTTTGTGACATCCAATG

>M6-PGSC0003DMG400003211-(rev)M6LRR-RLK_4026

TTTCAGTTTAACAAAATCGTGATCTTGGGTTTTTCAGTTTAACTTTGTGACATCCAATGTAGCAACATAGACTAAGAAAATGCATGATGTTTCAAATCAATTTTTGGTAAAAGCAATTTT

>M6-PGSC0003DMG400003211-(rev)M6LRR-RLK_4087

GCAACATAGACTAAGAAAATGCATGATGTTTCAAATCAATTTTTGGTAAAAGCAATTTTCACCTTCTAGCCATAGCTCTTTTCATCATGGATCACTGTCCGTAATCAGTTTTTGCTTATT

>M6-PGSC0003DMG400003211-(rev)M6LRR-RLK_4148

CCTTCTAGCCATAGCTCTTTTCATCATGGATCACTGTCCGTAATCAGTTTTTGCTTATTACTTTCTCCAGCTTCTCCAAAATCTGAACAGTTTTCAGGTATCACAATTAAAATGAGAGTG

>M6-PGSC0003DMG400003211-(rev)M6LRR-RLK_4209

TTTCTCCAGCTTCTCCAAAATCTGAACAGTTTTCAGGTATCACAATTAAAATGAGAGTGTGAAGGAACAAGGCTTTGCTACAAGCTAGTCCTTATCTTATGGACTATAACATTCCGGTTT

>M6-PGSC0003DMG400003211-(rev)M6LRR-RLK_427

TCTATCTGGAATCTTGGACAAGGCCTTTCCAAATCTTTGCCATCTTCCTCGTTTGATCTCCCTTGATCTCAGTGGCAACCATCTCACTGGTGGCATCCCTGCTATGCTTGCTAACTGCAG

>M6-PGSC0003DMG400003211-(rev)M6LRR-RLK_4270

AAGGAACAAGGCTTTGCTACAAGCTAGTCCTTATCTTATGGACTATAACATTCCGGTTTATAAAAGAAATCAAAAGAACAACCCCCACCCAAAAAGATAAGCTTTTGTTGAATTTGGTTG

>M6-PGSC0003DMG400003211-(rev)M6LRR-RLK_4331

AAAAGAAATCAAAAGAACAACCCCCACCCAAAAAGATAAGCTTTTGTTGAATTTGGTTGTACTAATGATGGTTCCTGCGAAAGAGTGCAGAAGACCTCCGCCTTACCGGTCTTTCTTTGT

>M6-PGSC0003DMG400003211-(rev)M6LRR-RLK_4392

CTAATGATGGTTCCTGCGAAAGAGTGCAGAAGACCTCCGCCTTACCGGTCTTTCTTTGTGTTCCAAACTTGCAGTAGCAGAATTATTTGCAGAAAATATAGATATGATGAATAGAAATGG

>M6-PGSC0003DMG400003211-(rev)M6LRR-RLK_4453

TCCAAACTTGCAGTAGCAGAATTATTTGCAGAAAATATAGATATGATGAATAGAAATGGCTTGTGGACAATCATTGTATTTCAAACAAATACAGATATTCAAATAAGGTAGATATAACTC

>M6-PGSC0003DMG400003211-(rev)M6LRR-RLK_4514

TGTGGACAATCATTGTATTTCAAACAAATACAGATATTCAAATAAGGTAGATATAACTCAGATATGAGTTACCAACAGGACTTCTAACAAGACTTAACAGACGAATGGCAAATACATAGG

>M6-PGSC0003DMG400003211-(rev)M6LRR-RLK_4575

ATATGAGTTACCAACAGGACTTCTAACAAGACTTAACAGACGAATGGCAAATACATAGGAATATCAGATCAGGAGAGGAAAATAATAGTCCTTGAAGAATTGATCAGAATTCAGTAGAAA

>M6-PGSC0003DMG400003211-(rev)M6LRR-RLK_4636

TATCAGATCAGGAGAGGAAAATAATAGTCCTTGAAGAATTGATCAGAATTCAGTAGAAAGGAAATATGAAAGAAGTGCAAGAATAAACTGGAAGAAATACTCTGCAGGTGACAAATAGAT

>M6-PGSC0003DMG400003211-(rev)M6LRR-RLK_4697

AAATATGAAAGAAGTGCAAGAATAAACTGGAAGAAATACTCTGCAGGTGACAAATAGATGAAAGACATACCTCAGTTACGAAAACTGCAACACCCTAGCACAATACGATGCGCACCTAAA

>M6-PGSC0003DMG400003211-(rev)M6LRR-RLK_4758

AAGACATACCTCAGTTACGAAAACTGCAACACCCTAGCACAATACGATGCGCACCTAAATAATCATCCAAGTAGGAGAAGCCCAACTTAAATGTCCCAGATTCCTTTTTCATATTTTTTA

>M6-PGSC0003DMG400003211-(rev)M6LRR-RLK_4819

ATCATCCAAGTAGGAGAAGCCCAACTTAAATGTCCCAGATTCCTTTTTCATATTTTTTATGTAGTTGACTTGGATGTGCATTACAAGACAACTGTCACAAGATTTTGAGAATTTTTCATG

>M6-PGSC0003DMG400003211-(rev)M6LRR-RLK_4880

TAGTTGACTTGGATGTGCATTACAAGACAACTGTCACAAGATTTTGAGAATTTTTCATGTTAAAATATCCACCAGAATTTGCATTGGATATATTCACATGGCACCACATATGCTGACAAA

>M6-PGSC0003DMG400003211-(rev)M6LRR-RLK_4941

AAAATATCCACCAGAATTTGCATTGGATATATTCACATGGCACCACATATGCTGACAAATCTCAGACTTCTAATAATAGAGATTCCATTCTGAGACAGATCCACTAAATTCCCAAGTTTC

>M6-PGSC0003DMG400003211-(rev)M6LRR-RLK_5002

TCAGACTTCTAATAATAGAGATTCCATTCTGAGACAGATCCACTAAATTCCCAAGTTTCTGCTCTTAGCAAGTGCAGCACTATCACATATGATGATAAGTCTAGGACCTCTAGTAATAAA

>M6-PGSC0003DMG400003211-(rev)M6LRR-RLK_5063

CTCTTAGCAAGTGCAGCACTATCACATATGATGATAAGTCTAGGACCTCTAGTAATAAAGCTTCCCTTCTGAGCCAAATCCACTAAATTCCCTAGCTTCTGCTCTTAGCAAGTGCACTTT

>M6-PGSC0003DMG400003211-(rev)M6LRR-RLK_5124

TTCCCTTCTGAGCCAAATCCACTAAATTCCCTAGCTTCTGCTCTTAGCAAGTGCACTTTTGTGCAAGACGATTAGAGACAAACTCCAGGAATTTGGTGCTTGCTCTAATACCAAAGTTAC

>M6-PGSC0003DMG400003211-(rev)M6LRR-RLK_5185

TGCAAGACGATTAGAGACAAACTCCAGGAATTTGGTGCTTGCTCTAATACCAAAGTTACAAGCAGCTCAAAAGAGAAAATATAAAACACTAAACCACATTAAAGAAAAGCTAGGAAGCTA

>M6-PGSC0003DMG400003211-(rev)M6LRR-RLK_5246

GCAGCTCAAAAGAGAAAATATAAAACACTAAACCACATTAAAGAAAAGCTAGGAAGCTAAAACCAAAATTGAAGAAGAAAAAGTTCCACACTGTTTCTCTTAACAGAACAGGTGAAATTT

>M6-PGSC0003DMG400003211-(rev)M6LRR-RLK_5307

ACCAAAATTGAAGAAGAAAAAGTTCCACACTGTTTCTCTTAACAGAACAGGTGAAATTTCTCATAAAGAGCTGATCATATCCTACATCCAATCACTTTACTGAAGTTAGAGTACAGATAA

>M6-PGSC0003DMG400003211-(rev)M6LRR-RLK_5368

CATAAAGAGCTGATCATATCCTACATCCAATCACTTTACTGAAGTTAGAGTACAGATAAATGTTTGAAAACCCTAGTGTCCAAAGATAGGGACAATAGAAATATGAAGCTGTAAAATGCT

>M6-PGSC0003DMG400003211-(rev)M6LRR-RLK_5429

GTTTGAAAACCCTAGTGTCCAAAGATAGGGACAATAGAAATATGAAGCTGTAAAATGCTCCAAGAAAAAATAATTAAATTTCTGAGAGGATTTCCAAAATAGCCTCGAGGTAAAATATTT

>M6-PGSC0003DMG400003211-(rev)M6LRR-RLK_549

AACTTGACACGATTTTCCTCAACGACAACAGGTTTTCAGGATCCATCCCACCAGAGATTTTCAAATCCAAAAGGCTTGTATACCTTGAGTTGGGTTACAATCAGCTCAATGGCACCATTC

>M6-PGSC0003DMG400003211-(rev)M6LRR-RLK_5490

AAGAAAAAATAATTAAATTTCTGAGAGGATTTCCAAAATAGCCTCGAGGTAAAATATTTATACAAGGTTAAACTTGTCTGCTTCCTCAGTCCAATAAAGCAGAGTTCAGGAGTTTTACTT

>M6-PGSC0003DMG400003211-(rev)M6LRR-RLK_5551

ACAAGGTTAAACTTGTCTGCTTCCTCAGTCCAATAAAGCAGAGTTCAGGAGTTTTACTTTGAAGAGACCGAACAAATTTAGAGAACTACTCCATTACTGGCTATGTAACCTATGGCTAAC

>M6-PGSC0003DMG400003211-(rev)M6LRR-RLK_5612

AAGAGACCGAACAAATTTAGAGAACTACTCCATTACTGGCTATGTAACCTATGGCTAACTAGATTAAATTAGAAGCTCAATGTGCATAGACGTGGTGTTCTTGGGTTATACTTAAGACCA

>M6-PGSC0003DMG400003211-(rev)M6LRR-RLK_5673

GATTAAATTAGAAGCTCAATGTGCATAGACGTGGTGTTCTTGGGTTATACTTAAGACCAGCTAACTTGAAGAAGAATTCATTAGTATACATGGTTACGGACTGAGGCACAGTGCAACAGG

>M6-PGSC0003DMG400003211-(rev)M6LRR-RLK_5734

TAACTTGAAGAAGAATTCATTAGTATACATGGTTACGGACTGAGGCACAGTGCAACAGGATAATATTTTCAACTTAAGTAAGAAATTACCATTATAAAGATAGGATGTCCTTGATCACAT

>M6-PGSC0003DMG400003211-(rev)M6LRR-RLK_5795

AATATTTTCAACTTAAGTAAGAAATTACCATTATAAAGATAGGATGTCCTTGATCACATCATACAGAATTAGATACCTAATGGAGCTTGCATGTCTGTTTAATTAACACTAAACAATGTT

>M6-PGSC0003DMG400003211-(rev)M6LRR-RLK_5856

TACAGAATTAGATACCTAATGGAGCTTGCATGTCTGTTTAATTAACACTAAACAATGTTAACTATTGCAAAATGCTCAAGTTCACAATTTTATTTATATCCTCTCCACTTGAATATTTTC

>M6-PGSC0003DMG400003211-(rev)M6LRR-RLK_5917

CTATTGCAAAATGCTCAAGTTCACAATTTTATTTATATCCTCTCCACTTGAATATTTTCATGCAACATATTTTTGTCATTTGGATATATATATGTTGAATGAAATTAAGGAACCTAAAAG

>M6-PGSC0003DMG400003211-(rev)M6LRR-RLK_5978

GCAACATATTTTTGTCATTTGGATATATATATGTTGAATGAAATTAAGGAACCTAAAAGTCAGTTCACCAGACAATCTATAATCAGTTATGTACCACATCTTTTTACCTAATACATTTTT

>M6-PGSC0003DMG400003211-(rev)M6LRR-RLK_6039

AGTTCACCAGACAATCTATAATCAGTTATGTACCACATCTTTTTACCTAATACATTTTTTGTGAATCAACACCACTATTTATACCCTATTCTACATATCAAAGGGACAAGATGAAGTTTG

>M6-PGSC0003DMG400003211-(rev)M6LRR-RLK_61

ATCTTTAAAAAACAGGCTGACCTCAACTTAAAACCTAAATAATAGTCTCAGGTTCCAGCCCAAAAATTAGTTTTTGAACTAGCTCCTTATTGGATGCAGCTTCTTTAAAGAAGTGTTGAA

>M6-PGSC0003DMG400003211-(rev)M6LRR-RLK_610

CAAATCCAAAAGGCTTGTATACCTTGAGTTGGGTTACAATCAGCTCAATGGCACCATTCCTTCTGAGGTGGGCCTTTCGACAAGCCTTCAATATCTTGGGCTGTGGAACAATTTCCTTAG

>M6-PGSC0003DMG400003211-(rev)M6LRR-RLK_6100

TGAATCAACACCACTATTTATACCCTATTCTACATATCAAAGGGACAAGATGAAGTTTGCAAGGATCAATGCCTAGTTTATACTCAGCATACAAATAAAGCTCATATAAGATACAAAACT

>M6-PGSC0003DMG400003211-(rev)M6LRR-RLK_6161

AGGATCAATGCCTAGTTTATACTCAGCATACAAATAAAGCTCATATAAGATACAAAACTAACTAAACCTCTTTTCATCTCCCAAATATAGTTTGAGCAATAAAATAGAGAGATGGAAAAG

>M6-PGSC0003DMG400003211-(rev)M6LRR-RLK_6222

CTAAACCTCTTTTCATCTCCCAAATATAGTTTGAGCAATAAAATAGAGAGATGGAAAAGTATTATTTGATCGCGAAATTATTAACCACAATAGATTGTGGCAGCAACACCAGACCTCAGT

>M6-PGSC0003DMG400003211-(rev)M6LRR-RLK_6283

TTATTTGATCGCGAAATTATTAACCACAATAGATTGTGGCAGCAACACCAGACCTCAGTCACAGTCATACTAAGTCATCTACTTTGTTGTTTTTCACATACTTTTCCGCTTCAATTTTAA

>M6-PGSC0003DMG400003211-(rev)M6LRR-RLK_6344

CAGTCATACTAAGTCATCTACTTTGTTGTTTTTCACATACTTTTCCGCTTCAATTTTAATCAAAAGCCGAATGCTGATAGCCTGATATGCATTCAACTACAGATCCTTTTTTTATTTTTG

>M6-PGSC0003DMG400003211-(rev)M6LRR-RLK_671

TCTGAGGTGGGCCTTTCGACAAGCCTTCAATATCTTGGGCTGTGGAACAATTTCCTTAGTGGAAATATTCCAAAAGAGCTATTTGGTTTGCCAAACTTGACAGATCTCTACATCCACACA

>M6-PGSC0003DMG400003211-(rev)M6LRR-RLK_6771

GGATCCTTTTCACCTAGATATTCAACATCAAGGCATCTCAGAATTCTCAAAAGCAATTTACTCCTGGCAACTTCTAATTTTGAAATCCCTAAGTTCATAAAGGGGCTGTGCCCAAGCATT

>M6-PGSC0003DMG400003211-(rev)M6LRR-RLK_6832

TCCTGGCAACTTCTAATTTTGAAATCCCTAAGTTCATAAAGGGGCTGTGCCCAAGCATTAGAAAACTAGCATTGCCTTCGAAGGTCAAATTATCGATTTAATTAATGGTACAGAGGGCTT

>M6-PGSC0003DMG400003211-(rev)M6LRR-RLK_6893

AAAACTAGCATTGCCTTCGAAGGTCAAATTATCGATTTAATTAATGGTACAGAGGGCTTTTAGAAGCGGAGAAATTGGGAACAGCATTGTACCTGTTAATTGTTGATGGAGTCATCGGAA

>M6-PGSC0003DMG400003211-(rev)M6LRR-RLK_6954

AGAAGCGGAGAAATTGGGAACAGCATTGTACCTGTTAATTGTTGATGGAGTCATCGGAATCAAAAGTAACTCAAATTAACGAAGACAAATAAATAAAAATCTCATTTCTCAACTCAAACT

>M6-PGSC0003DMG400003211-(rev)M6LRR-RLK_7008

CGGAATCAAAAGTAACTCAAATTAACGAAGACAAATAAATAAAAATCTCATTTCTCAACTCAAACTATATTGGATGGGCATAAATTTGTGGAACCCATTAAGCTTTTTTCTCCAACTTGC

>M6-PGSC0003DMG400003211-(rev)M6LRR-RLK_732

GAAATATTCCAAAAGAGCTATTTGGTTTGCCAAACTTGACAGATCTCTACATCCACACAAATAATCTAACGGGACCTCTCCCTGACTTCCCGTCCTCATGCTCACTTTCCCAGCTTCTTA

>M6-PGSC0003DMG400003211-(rev)M6LRR-RLK_793

TAATCTAACGGGACCTCTCCCTGACTTCCCGTCCTCATGCTCACTTTCCCAGCTTCTTATTCATCAGAATCGCTTCTCAGGTTCTTTGCCAATTTCCCTAGGTAATTGTCACAACCTTTC

>M6-PGSC0003DMG400003211-(rev)M6LRR-RLK_854

CATCAGAATCGCTTCTCAGGTTCTTTGCCAATTTCCCTAGGTAATTGTCACAACCTTTCTGCATTCTATGCAACATCTGCTCATCTTGGAGGAGTTATTTTGCCAGAGGTTTTTAGGGGC

>M6-PGSC0003DMG400003211-(rev)M6LRR-RLK_915

CATTCTATGCAACATCTGCTCATCTTGGAGGAGTTATTTTGCCAGAGGTTTTTAGGGGCCTTTCAAATCTCGAATTACTCTATTTAGATGACAACAACTTTGAAGGGGAAATTCCTGAGA

>M6-PGSC0003DMG400003211-(rev)M6LRR-RLK_976

TTCAAATCTCGAATTACTCTATTTAGATGACAACAACTTTGAAGGGGAAATTCCTGAGACCTTATGGAATGGGAAATTACAAGAGTTGGTTCTTTCCATAAATAATTTTAATGGAAGTAT

>M6-PGSC0003DMG400003280-(rev)M6LRR-RLK_0

TTTCAGGTGATCGAGGGATATCACACGGTGAGCTAAACTGGCCTACTCGTCTAAGGATCATCCAGGGAGTTGCTGGTGGAATGTGTTATCTTCATTCCGAGTTTGCATCGTATGCAGTGC

>M6-PGSC0003DMG400003280-(rev)M6LRR-RLK_128

AACCTCAAGTCTAGCAACATTCTACTCAATGAAAAATACGAGCCACTTCTATCAGATTACGCCTTCTATCCACTGATCAACAACACGCAAATTGTTCAATCCCTTTTCGCGTATAAAGCA

>M6-PGSC0003DMG400003280-(rev)M6LRR-RLK_192

TCTATCCACTGATCAACAACACGCAAATTGTTCAATCCCTTTTCGCGTATAAAGCACCAGAAGCCATACAACATCAACATCTCACTCCAAAATGTGACATATATTGTCTTGGAATCATAA

>M6-PGSC0003DMG400003280-(rev)M6LRR-RLK_256

CATACAACATCAACATCTCACTCCAAAATGTGACATATATTGTCTTGGAATCATAATACTTGAAATCCTTACAGGGAAATTCCCTTCTCAGTATCTCAACAATCAAAAAGGTGGAACGGA

>M6-PGSC0003DMG400003280-(rev)M6LRR-RLK_320

ATCCTTACAGGGAAATTCCCTTCTCAGTATCTCAACAATCAAAAAGGTGGAACGGACGTTGTTCAATGGGTGCAATCAGCTATAGCTGACCAAAGAGAATCCGAATTGATTGATCCAGAG

>M6-PGSC0003DMG400003280-(rev)M6LRR-RLK_384

AATGGGTGCAATCAGCTATAGCTGACCAAAGAGAATCCGAATTGATTGATCCAGAGATAGCGTCTGCAACAGATTCCGTTGAACAAATGGTGAAACTCCTTCATGTAGGAGCTGCTTGTA

>M6-PGSC0003DMG400003280-(rev)M6LRR-RLK_447

CTGCAACAGATTCCGTTGAACAAATGGTGAAACTCCTTCATGTAGGAGCTGCTTGTACTATAAGCGATCCGGATAAGAGGATTGATATGAAGGAAGCATTAAGGAGGATAGAAGAGATAA

>M6-PGSC0003DMG400003280-(rev)M6LRR-RLK_64

GGGAGTTGCTGGTGGAATGTGTTATCTTCATTCCGAGTTTGCATCGTATGCAGTGCCTCATGGAAACCTCAAGTCTAGCAACATTCTACTCAATGAAAAATACGAGCCACTTCTATCAGA

>M6-PGSC0003DMG400003313M6G-LecRK_0

TTCAAAATTACATATTCCGAAACTTTTGTACGAATTTTGATTTCATACTATAACATGAATATAAAATTTTCAATTCTTCTTTTACCTTTGCTTGTTGTTTCATCTTATTGTTCAACAATC

>M6-PGSC0003DMG400003313M6G-LecRK_1054

CAAAGAAGATTCAGCCTTGACTTAGATGGAAACCTTCGATTGTACAGCCTAGATAAGCTGAGCAACATTTGGAAAGTTTCTTGGCAGTTATTTCCAGCAGCCTGCAGGATTCATGGAGTT

>M6-PGSC0003DMG400003313M6G-LecRK_1116

CAACATTTGGAAAGTTTCTTGGCAGTTATTTCCAGCAGCCTGCAGGATTCATGGAGTTTGTGGATTGAACAGTTTATGTTCCTATGATCCTGTCTCTGGCAGAAAATGTTCTTGTGTACC

>M6-PGSC0003DMG400003313M6G-LecRK_1178

GATTGAACAGTTTATGTTCCTATGATCCTGTCTCTGGCAGAAAATGTTCTTGTGTACCAGGATACAGGATGAAAAATCCTAAAGATTGGTCCTATGGTTGTGAACCAGAATTCGATATCT

>M6-PGSC0003DMG400003313M6G-LecRK_124

GTTTATCATCTGAAGGTTCACTTTCTGCTCCACAAGATTTAATTTCGTCTTGAAATGGAAAATTCACTGCTGGTTTCTACTCTGTTGGTGATAATGCTTATATCTTTGCTATATGGTTTA

>M6-PGSC0003DMG400003313M6G-LecRK_1240

TACAGGATGAAAAATCCTAAAGATTGGTCCTATGGTTGTGAACCAGAATTCGATATCTCTTGCAACGATACAAGCTCGTTGGACTTCATTCCACTTCGCCATGTTGAGTTTTACGGGTAT

>M6-PGSC0003DMG400003313M6G-LecRK_1302

CAACGATACAAGCTCGTTGGACTTCATTCCACTTCGCCATGTTGAGTTTTACGGGTATGATATTGCATATTTTCGCAATAAAACATTGCAAGAATGTAAAGATTTGTGCTTGAAACATTG

>M6-PGSC0003DMG400003313M6G-LecRK_1364

TTGCATATTTTCGCAATAAAACATTGCAAGAATGTAAAGATTTGTGCTTGAAACATTGTGATTGTAAAGGTTTTGAATACAAGTATGTTGAGGGTAATGGCACCTATGGTTGTTACCCGA

>M6-PGSC0003DMG400003313M6G-LecRK_1426

TGTAAAGGTTTTGAATACAAGTATGTTGAGGGTAATGGCACCTATGGTTGTTACCCGAAAACACTTCTGTTCAATGGCTATGTTCAAGCTAGTTGGCCTGATTTTGTTTACCTTAAAGTG

>M6-PGSC0003DMG400003313M6G-LecRK_1488

ACTTCTGTTCAATGGCTATGTTCAAGCTAGTTGGCCTGATTTTGTTTACCTTAAAGTGCCTAAAGCACGACGTTCCTGGCAGGAGCAGTATGAAGGAAATCTTCAATGTGACGATAAAAA

>M6-PGSC0003DMG400003313M6G-LecRK_1550

AAGCACGACGTTCCTGGCAGGAGCAGTATGAAGGAAATCTTCAATGTGACGATAAAAAGGTGATGTTGGATAGAGCTTATAGAAGGAAAGAACATGGTGGATGGATGAAGCCATTCATTT

>M6-PGSC0003DMG400003313M6G-LecRK_1612

ATGTTGGATAGAGCTTATAGAAGGAAAGAACATGGTGGATGGATGAAGCCATTCATTTGGTCAGTTGTTGTAGGTGGAGTCTTTGAAATTCTCTGTGTTCTCACTTACTTTATTAAAACG

>M6-PGSC0003DMG400003313M6G-LecRK_1674

AGTTGTTGTAGGTGGAGTCTTTGAAATTCTCTGTGTTCTCACTTACTTTATTAAAACGAGAAAAAGCTCGAATGAAACCACACAAGGCTACCTTCATCTTTCGACAAGATTCAAGAAATT

>M6-PGSC0003DMG400003313M6G-LecRK_1736

AAAGCTCGAATGAAACCACACAAGGCTACCTTCATCTTTCGACAAGATTCAAGAAATTCACCTATGCTGAGCTTAAAAAGGCCACTTCCAATTTCAGTGAAGAGATAGGTCGAGGAGGTG

>M6-PGSC0003DMG400003313M6G-LecRK_1798

TATGCTGAGCTTAAAAAGGCCACTTCCAATTTCAGTGAAGAGATAGGTCGAGGAGGTGGTGGTATTGTGTACAAAGGGAAGTTGTCTGATGACAGAGTTGCAGCTATAAAGTCTCTCTAT

>M6-PGSC0003DMG400003313M6G-LecRK_186

TTCACTGCTGGTTTCTACTCTGTTGGTGATAATGCTTATATCTTTGCTATATGGTTTACTAATTCATTGGCTGATGGAACCAACACAATTGTTTGGATGGCTAATCGTGATCAACCGATA

>M6-PGSC0003DMG400003313M6G-LecRK_1860

TATTGTGTACAAAGGGAAGTTGTCTGATGACAGAGTTGCAGCTATAAAGTCTCTCTATGGAGCTAACTATCAAGGAGAAGCAGAATTTCTAGCTGAAGTGAGCACTATAGGCAACGTTAA

>M6-PGSC0003DMG400003313M6G-LecRK_1922

CTAACTATCAAGGAGAAGCAGAATTTCTAGCTGAAGTGAGCACTATAGGCAACGTTAATCACATGAATTTGATCGAAATTTGGGGATATTGTGCTGAGGGAAAGCACAGGCTTTTAGTTT

>M6-PGSC0003DMG400003313M6G-LecRK_1984

ATGAATTTGATCGAAATTTGGGGATATTGTGCTGAGGGAAAGCACAGGCTTTTAGTTTATGAGTACATGGAGTATGGTTCTTTGTCAGATAATTTGAATGCCAACAATCTTGATTGGGAA

>M6-PGSC0003DMG400003313M6G-LecRK_2046

GTACATGGAGTATGGTTCTTTGTCAGATAATTTGAATGCCAACAATCTTGATTGGGAAAAATGGTTTGAAATCGCGTTGGGAACGGCTAAAGGACTTGCATACTTGCATGAAGAATGCTT

>M6-PGSC0003DMG400003313M6G-LecRK_2108

GGTTTGAAATCGCGTTGGGAACGGCTAAAGGACTTGCATACTTGCATGAAGAATGCTTAGAATGGGTTTTGCATTGTGATGTAAAGCCTCAGAACATACTTTTAGATTCCAATTACAAGC

>M6-PGSC0003DMG400003313M6G-LecRK_2170

TGGGTTTTGCATTGTGATGTAAAGCCTCAGAACATACTTTTAGATTCCAATTACAAGCCAAAAGTTGCTGACTTTGGGCTTTCAAAGATATTTAACAGAGGTGGCTTGGATCATTCAAAC

>M6-PGSC0003DMG400003313M6G-LecRK_2232

AGTTGCTGACTTTGGGCTTTCAAAGATATTTAACAGAGGTGGCTTGGATCATTCAAACTTTTCAACGATAAGGGGGACGAGAGGATACATGGCTCCAGAATGGGTATTCAAGATGCCTAT

>M6-PGSC0003DMG400003313M6G-LecRK_2294

CAACGATAAGGGGGACGAGAGGATACATGGCTCCAGAATGGGTATTCAAGATGCCTATTACATCCAAAGTCGATGTCTATAGTTATGGCATTGTTTTATTAGAGATGATCACAGGTAAGA

>M6-PGSC0003DMG400003313M6G-LecRK_2356

TCCAAAGTCGATGTCTATAGTTATGGCATTGTTTTATTAGAGATGATCACAGGTAAGAGTCCAGAAGTTTGTGCTGATGGTGGCTCGGGGGATGATGATTCTATGGGACTAGGATTGTTG

>M6-PGSC0003DMG400003313M6G-LecRK_2418

AGAAGTTTGTGCTGATGGTGGCTCGGGGGATGATGATTCTATGGGACTAGGATTGTTGGTCACTTGGATTAGAGAAAAAATGCGCGAAAACAGTGAAAGAAAATCATGGATTCAACAAAT

>M6-PGSC0003DMG400003313M6G-LecRK_248

TTCATTGGCTGATGGAACCAACACAATTGTTTGGATGGCTAATCGTGATCAACCGATAAACGGACAAAAATCACATCTGTCGCTGTTTAAATCAGGCAATCTTGTACTGATTGACGCGAA

>M6-PGSC0003DMG400003313M6G-LecRK_2480

CTTGGATTAGAGAAAAAATGCGCGAAAACAGTGAAAGAAAATCATGGATTCAACAAATTGTGGATCCTGCATTGAATGGTAAATTCGATTTGAAAAAAATGGAGATCTTATTAGAGTTAG

>M6-PGSC0003DMG400003313M6G-LecRK_2542

GATCCTGCATTGAATGGTAAATTCGATTTGAAAAAAATGGAGATCTTATTAGAGTTAGCTTTGCAATGTTCGGAAGAAGATAGAGATGCAAGACCTACCATGTGTGAGGTGGTGGATAAA

>M6-PGSC0003DMG400003313M6G-LecRK_2604

GCAATGTTCGGAAGAAGATAGAGATGCAAGACCTACCATGTGTGAGGTGGTGGATAAAATGCTTCATCCAGAAAATTTCAAGTTAAAATTAGACATGTTAATGTAGTTTCTCAAGATGTA

>M6-PGSC0003DMG400003313M6G-LecRK_2666

TTCATCCAGAAAATTTCAAGTTAAAATTAGACATGTTAATGTAGTTTCTCAAGATGTAAAATCTTGCCAAAGGTGATATAGTACAAGCTTATATGTAATGTGTCTTGAGTAGCAAGTACT

>M6-PGSC0003DMG400003313M6G-LecRK_2728

CTTGCCAAAGGTGATATAGTACAAGCTTATATGTAATGTGTCTTGAGTAGCAAGTACTTTTGAATTCCAAAATGTAAATCCTTATAGAATGACATAGTTTCTTAACAAAATTGTGCATTA

>M6-PGSC0003DMG400003313M6G-LecRK_2785

TTTTGAATTCCAAAATGTAAATCCTTATAGAATGACATAGTTTCTTAACAAAATTGTGCATTATAGCAACTGTTATGGTCTTGTGCATCAGTGATAGATGGAAAGTGTGTATGAGTACAT

>M6-PGSC0003DMG400003313M6G-LecRK_310

GACAAAAATCACATCTGTCGCTGTTTAAATCAGGCAATCTTGTACTGATTGACGCGAATCAGATCAATGTTTGGGAATCTGGTACACAATCATCAACTTCTTCTGTTGAGTTAAGATTGT

>M6-PGSC0003DMG400003313M6G-LecRK_372

ATCAATGTTTGGGAATCTGGTACACAATCATCAACTTCTTCTGTTGAGTTAAGATTGTTGGATAATGGAAATCTTGTTCTTGTAACTTATGAAGGTCAAAGATTATGGCAAAGCTTTGAT

>M6-PGSC0003DMG400003313M6G-LecRK_434

TAATGGAAATCTTGTTCTTGTAACTTATGAAGGTCAAAGATTATGGCAAAGCTTTGATTCACCAACAGATACACTTCTTCCTGAACAACCACTTACGAAAACCTCAAAGCTTGTATCGCG

>M6-PGSC0003DMG400003313M6G-LecRK_496

CAACAGATACACTTCTTCCTGAACAACCACTTACGAAAACCTCAAAGCTTGTATCGCGTAGAAGCTTGAATAATTTCTCCTCTGGATTCTATCAGTTACATTTTAATGAAGATAATGTCC

>M6-PGSC0003DMG400003313M6G-LecRK_558

AGCTTGAATAATTTCTCCTCTGGATTCTATCAGTTACATTTTAATGAAGATAATGTCCTGCATCTTGTCTTTGATGGAATAGAGATGACAAGTGTGTTTTGGCCAAGCCCGTGGTTAATC

>M6-PGSC0003DMG400003313M6G-LecRK_62

AAAATTTTCAATTCTTCTTTTACCTTTGCTTGTTGTTTCATCTTATTGTTCAACAATCTTTAGTTTATCATCTGAAGGTTCACTTTCTGCTCCACAAGATTTAATTTCGTCTTGAAATGG

>M6-PGSC0003DMG400003313M6G-LecRK_620

TCTTGTCTTTGATGGAATAGAGATGACAAGTGTGTTTTGGCCAAGCCCGTGGTTAATCGTTTGGGATGCAGGTTAGTAGAGACGGATTCAGAATTTAAATTTGATTCGTTAAATCCTTAG

>M6-PGSC0003DMG400003313M6G-LecRK_682

GGGATGCAGGTTAGTAGAGACGGATTCAGAATTTAAATTTGATTCGTTAAATCCTTAGTTTGTTGTCACTCAACTCACATATATTATTTTTGCATGAAAAATACTAGTTTTAGTTGAACT

>M6-PGSC0003DMG400003313M6G-LecRK_744

TTGTCACTCAACTCACATATATTATTTTTGCATGAAAAATACTAGTTTTAGTTGAACTCATTATATGGTTCATTTGAGTGTATTGGTTAGGAAAGACGGATTCAGGATTTAAATTTCGTT

>M6-PGSC0003DMG400003313M6G-LecRK_806

ATATGGTTCATTTGAGTGTATTGGTTAGGAAAGACGGATTCAGGATTTAAATTTCGTTGGTTCAATATTTAGTACAACTCATTTTTGAAGTTTTCGGTTTTAACTGAACTCATTGCTGAT

>M6-PGSC0003DMG400003313M6G-LecRK_868

CAATATTTAGTACAACTCATTTTTGAAGTTTTCGGTTTTAACTGAACTCATTGCTGATATACTCCATCCATTTCTGCAGGTCGATCCACTTATAATGACAGCAAAACAGCAGTTCTTGAT

>M6-PGSC0003DMG400003313M6G-LecRK_930

TCCATCCATTTCTGCAGGTCGATCCACTTATAATGACAGCAAAACAGCAGTTCTTGATCGTTTTGGGAATTTTACATCGAGTGATCGATTTGGATTCCAATCTGCTGATTATGGTTTGGA

>M6-PGSC0003DMG400003313M6G-LecRK_992

TTGGGAATTTTACATCGAGTGATCGATTTGGATTCCAATCTGCTGATTATGGTTTGGAATTGCAAAGAAGATTCAGCCTTGACTTAGATGGAAACCTTCGATTGTACAGCCTAGATAAGC

>M6-PGSC0003DMG400003343M6RLKWAX-EGF_0

TTGGTGTGTTCTCATTACTTTATTTCTTCCCCAATCCCCTTTTTCATTCACTTCTTTATCAATTTACCAATTTTCTTTCTTTCTTTCTTTCTTCTTCCTCCTCCTCCCCTGTTCACATGT

>M6-PGSC0003DMG400003343M6RLKWAX-EGF_1054

AGGTAATTTTGCTTTAATTAGTTAACTATTTATTCAACAACTTTTTAAATAAAAACATGTTCTAGTTTAAAGAAAGAAATAATATAATGAAAATAAGTTTGACTTTTCAAAATTAGAAAT

>M6-PGSC0003DMG400003343M6RLKWAX-EGF_1116

TAGTTTAAAGAAAGAAATAATATAATGAAAATAAGTTTGACTTTTCAAAATTAGAAATTTTAGTTTGCATATACAAAAATGTGATATTTTATATATTCAACTTTCTTCAAGTCACCTAAT

>M6-PGSC0003DMG400003343M6RLKWAX-EGF_1178

GTTTGCATATACAAAAATGTGATATTTTATATATTCAACTTTCTTCAAGTCACCTAATTTGGTTATGAGTTGGGTGGTTTAACTAAATGAAATTGAAGTTTATGAGTGGGAGTTTTGACC

>M6-PGSC0003DMG400003343M6RLKWAX-EGF_124

CAAAACATGAATTTTCCTCTCCATTTCAGAACAATTCTCTGTTTGTTTATTTTCAATTTTTCATCAAACATTGTTCTTGGATCAAACTGTACCCATTCATGTGGAAACAGATTCCTTCCA

>M6-PGSC0003DMG400003343M6RLKWAX-EGF_1240

TTATGAGTTGGGTGGTTTAACTAAATGAAATTGAAGTTTATGAGTGGGAGTTTTGACCAAATTTGGTGAGAGGAGAACACCATAATTAGATTTTTTTAAAAAAAACATGGGTCCAAGTAC

>M6-PGSC0003DMG400003343M6RLKWAX-EGF_1302

TTGGTGAGAGGAGAACACCATAATTAGATTTTTTTAAAAAAAACATGGGTCCAAGTACAAATATATTATTAAAAGATGGCATGCTTAGGCGGCCTAAAAGAAAAGTAACATATAATTGAT

>M6-PGSC0003DMG400003343M6RLKWAX-EGF_1364

ATATTATTAAAAGATGGCATGCTTAGGCGGCCTAAAAGAAAAGTAACATATAATTGATGGTAAAATTTTGATTCAATTGATCGAATCTTCTTCAATAAAGTGAAAGTTTAGATAATCATT

>M6-PGSC0003DMG400003343M6RLKWAX-EGF_1426

AAATTTTGATTCAATTGATCGAATCTTCTTCAATAAAGTGAAAGTTTAGATAATCATTCAATTAGAAATAATAATAATAAAATATTTCTATACAAGTTCAATTTTCTCCGATTATAGGGT

>M6-PGSC0003DMG400003343M6RLKWAX-EGF_1488

TAGAAATAATAATAATAAAATATTTCTATACAAGTTCAATTTTCTCCGATTATAGGGTCATGGAGTGCATGTTAAGGAACTAGGAAGTCAATAGTGTATACTATTAATAAAATATAATAG

>M6-PGSC0003DMG400003343M6RLKWAX-EGF_1550

GAGTGCATGTTAAGGAACTAGGAAGTCAATAGTGTATACTATTAATAAAATATAATAGAAGAAGAATTTTTTTTTTCTCATGTTCATACTAAGTAAAGGTAAAATCAAGAAAAAAAAGAA

>M6-PGSC0003DMG400003343M6RLKWAX-EGF_1612

AGAATTTTTTTTTTCTCATGTTCATACTAAGTAAAGGTAAAATCAAGAAAAAAAAGAAATTATAATTGTTTATTAGAGCAACTTGTGATTGTCGTTTTTCATAAATTAAGGTAATAATGG

>M6-PGSC0003DMG400003343M6RLKWAX-EGF_1674

TAATTGTTTATTAGAGCAACTTGTGATTGTCGTTTTTCATAAATTAAGGTAATAATGGCAACATTAAGAAAGACAATCACGTATGCGTTTGCTTCATCAGCGGAACAACTCGTGACTATC

>M6-PGSC0003DMG400003343M6RLKWAX-EGF_1736

ATTAAGAAAGACAATCACGTATGCGTTTGCTTCATCAGCGGAACAACTCGTGACTATCTAGGTAGGACCCAATAATAGTGATGGCTGTTGCCTTTAATGTCACTTATCAATGCCAAAATT

>M6-PGSC0003DMG400003343M6RLKWAX-EGF_1798

TAGGACCCAATAATAGTGATGGCTGTTGCCTTTAATGTCACTTATCAATGCCAAAATTAAAGGCAAATCTAAAATAGTTAGATCTCGTATAAGTTGAAGTTCACGTAAAAATGACTAAAT

>M6-PGSC0003DMG400003343M6RLKWAX-EGF_186

ATCAAACATTGTTCTTGGATCAAACTGTACCCATTCATGTGGAAACAGATTCCTTCCATTTCCTTTTGGGTTTTCTCATGGCTGCCCAATTCATCTCAACTGTACCTCAAATGAAGAATT

>M6-PGSC0003DMG400003343M6RLKWAX-EGF_1860

GCAAATCTAAAATAGTTAGATCTCGTATAAGTTGAAGTTCACGTAAAAATGACTAAATGTTTGTGCATATCTAAAGTTATTCTAAGTGGGTAGTCCTGTAAATGTTCACACACTACGAGT

>M6-PGSC0003DMG400003343M6RLKWAX-EGF_1922

GTGCATATCTAAAGTTATTCTAAGTGGGTAGTCCTGTAAATGTTCACACACTACGAGTATCACTTAAATTGTGCTTCTGAAATCGAGTGTTTGTTCAATCAGACATCTAAATCACAATTA

>M6-PGSC0003DMG400003343M6RLKWAX-EGF_1984

CTTAAATTGTGCTTCTGAAATCGAGTGTTTGTTCAATCAGACATCTAAATCACAATTAATAAGTTTACGAAGACTTAACACCTTTTGCCCATACCCATACCCATACCCTATATGTGAACG

>M6-PGSC0003DMG400003343M6RLKWAX-EGF_2046

GTTTACGAAGACTTAACACCTTTTGCCCATACCCATACCCATACCCTATATGTGAACGAGTTATTATGGTATATTAACTTGATATTGTTGCAAAAACTCACAAAATTCAAATTTTGAATA

>M6-PGSC0003DMG400003343M6RLKWAX-EGF_2108

ATTATGGTATATTAACTTGATATTGTTGCAAAAACTCACAAAATTCAAATTTTGAATACTTTGATTGGTTGGTATAATGTCCTTTCATGTTATTCCATTAAGATATCCTGCCAAAATTTG

>M6-PGSC0003DMG400003343M6RLKWAX-EGF_2170

GATTGGTTGGTATAATGTCCTTTCATGTTATTCCATTAAGATATCCTGCCAAAATTTGTTTCATCATTTGGTCAAGGAAGAGGCCTTTTTTTCCTTTGTTTTCTTCATTTTCTATCAAAC

>M6-PGSC0003DMG400003343M6RLKWAX-EGF_2232

ATCATTTGGTCAAGGAAGAGGCCTTTTTTTCCTTTGTTTTCTTCATTTTCTATCAAACTCCATTTGGTTTTGAATTAACAATGGAAACAAAGTCACTATCATATTACGTGAAGATAGTTA

>M6-PGSC0003DMG400003343M6RLKWAX-EGF_2294

TTTGGTTTTGAATTAACAATGGAAACAAAGTCACTATCATATTACGTGAAGATAGTTACTCAACAATATCAATGGACCCTCCACATATAAGGTGAAAAGGAATATAAACGAATTCGTGAT

>M6-PGSC0003DMG400003343M6RLKWAX-EGF_2356

ACAATATCAATGGACCCTCCACATATAAGGTGAAAAGGAATATAAACGAATTCGTGATTACTTTTTTTTTTGTTTCCATCCAATGTTCAGTATCTGTATTGGAGTCTGAACTAGTTCAAA

>M6-PGSC0003DMG400003343M6RLKWAX-EGF_2418

TTTTTTTTTGTTTCCATCCAATGTTCAGTATCTGTATTGGAGTCTGAACTAGTTCAAATTCGCATCACCTATTAAAAAGAAACGGTCCCTATTCGGATTTTTTCCATAGCCAAGACTTGA

>M6-PGSC0003DMG400003343M6RLKWAX-EGF_248

CTTTTGGGTTTTCTCATGGCTGCCCAATTCATCTCAACTGTACCTCAAATGAAGAATTTTTCATGAACGATTTCCCAGTTCAATCAGTAACTTCAGATACCATTTTAATCAATCTTTTAG

>M6-PGSC0003DMG400003343M6RLKWAX-EGF_2480

CATCACCTATTAAAAAGAAACGGTCCCTATTCGGATTTTTTCCATAGCCAAGACTTGAACGCGATACCTCAGGTCCCATCACAACACTTGAGTTGTATGTGTCCAATTACTAATTCCATT

>M6-PGSC0003DMG400003343M6RLKWAX-EGF_2542

GATACCTCAGGTCCCATCACAACACTTGAGTTGTATGTGTCCAATTACTAATTCCATTTTTTTTTTTTTTGTGAATGCAGATGAAACAACATGCAACCCTTCAAAGTACCTATCAGGCAA

>M6-PGSC0003DMG400003343M6RLKWAX-EGF_2604

TTTTTTTTGTGAATGCAGATGAAACAACATGCAACCCTTCAAAGTACCTATCAGGCAAATGTGGAGGAACCACAAGAGTTGGTGTACTTGTTGGAGGTAATTCTACGTTGATTTAAGTTA

>M6-PGSC0003DMG400003343M6RLKWAX-EGF_2666

GGAGGAACCACAAGAGTTGGTGTACTTGTTGGAGGTAATTCTACGTTGATTTAAGTTATATACGTTGACAGTGCAGAGTCTCAACATCTAGTACTAGTTGAAATGAAATTGTTTTCGTAT

>M6-PGSC0003DMG400003343M6RLKWAX-EGF_2728

CGTTGACAGTGCAGAGTCTCAACATCTAGTACTAGTTGAAATGAAATTGTTTTCGTATTCCTTTCTTATTTCAATTTCGCTATACAAACACATTATTCTAAAGTGATCAAAACTATATAT

>M6-PGSC0003DMG400003343M6RLKWAX-EGF_2790

TTCTTATTTCAATTTCGCTATACAAACACATTATTCTAAAGTGATCAAAACTATATATATTTCTCCAGGTATTGTTGCAGGTGCTTCGTTAATGGTTAGTATAGGCCTAATATGCTGCTT

>M6-PGSC0003DMG400003343M6RLKWAX-EGF_2852

CTCCAGGTATTGTTGCAGGTGCTTCGTTAATGGTTAGTATAGGCCTAATATGCTGCTTTATTCGACGAAGATTGAATTTTCAAAACAGAACAAGAAGAAACCGCGAGTTATGTGAGACTA

>M6-PGSC0003DMG400003343M6RLKWAX-EGF_2914

CGACGAAGATTGAATTTTCAAAACAGAACAAGAAGAAACCGCGAGTTATGTGAGACTACAGGGATCAGCATCCCTGTTTATCAGTTCAAAGAAATGGAAAAGGCAACCGATTTCTTCTCT

>M6-PGSC0003DMG400003343M6RLKWAX-EGF_2976

GATCAGCATCCCTGTTTATCAGTTCAAAGAAATGGAAAAGGCAACCGATTTCTTCTCTGAGAAACGGAGACTAGGAAATGGCGCGTATGGAACAGTTTATTCTGGAAAACTTCAAAATGA

>M6-PGSC0003DMG400003343M6RLKWAX-EGF_3038

AACGGAGACTAGGAAATGGCGCGTATGGAACAGTTTATTCTGGAAAACTTCAAAATGACAATTGGGTAGCTATCAAAAGAATTAGACATAGAGATACTGATAGCATTGAACAATTCATCA

>M6-PGSC0003DMG400003343M6RLKWAX-EGF_310

ATGAACGATTTCCCAGTTCAATCAGTAACTTCAGATACCATTTTAATCAATCTTTTAGCAAAATGCAGCCGACCATTTTCAGCAATTCACAATCTTTTTAGCCCAAATTACGCACCCACT

>M6-PGSC0003DMG400003343M6RLKWAX-EGF_3100

TGGGTAGCTATCAAAAGAATTAGACATAGAGATACTGATAGCATTGAACAATTCATCAATGAGATCAAGCTCCTTTCCTCTGTCAATCATCCGAACCTGGTTCGTCTCTTGGGCTGCTCA

>M6-PGSC0003DMG400003343M6RLKWAX-EGF_3162

GATCAAGCTCCTTTCCTCTGTCAATCATCCGAACCTGGTTCGTCTCTTGGGCTGCTCAATCGAAAATGGTGAACAGATTCTTGTTTATGAGTACATGCCTAATGGAACTTTGTCACAACA

>M6-PGSC0003DMG400003343M6RLKWAX-EGF_3224

AAAATGGTGAACAGATTCTTGTTTATGAGTACATGCCTAATGGAACTTTGTCACAACATATACAAAGAGAAAAAGGGAACGGGCTTCCATGGCCCGTTCGTCTTACAATTGGTGCAGAAA

>M6-PGSC0003DMG400003343M6RLKWAX-EGF_3286

CAAAGAGAAAAAGGGAACGGGCTTCCATGGCCCGTTCGTCTTACAATTGGTGCAGAAACTGCACAAGCTATTGCTTATCTGCATAATGCAACGCACCCTCCAATATATCACAGAGATGTT

>M6-PGSC0003DMG400003343M6RLKWAX-EGF_3348

ACAAGCTATTGCTTATCTGCATAATGCAACGCACCCTCCAATATATCACAGAGATGTTAAGTCAAGTAACATACTCTTGGATTACAATTACAAGTCAAAAGTAGCAGATTTCGGACTCTC

>M6-PGSC0003DMG400003343M6RLKWAX-EGF_3410

CAAGTAACATACTCTTGGATTACAATTACAAGTCAAAAGTAGCAGATTTCGGACTCTCAAGACTTGGTATGGTTGAATCGTCTCACATTTCTACTGCACCACAAGGAACTCCAGGTTATC

>M6-PGSC0003DMG400003343M6RLKWAX-EGF_3472

CTTGGTATGGTTGAATCGTCTCACATTTCTACTGCACCACAAGGAACTCCAGGTTATCTTGATCCTCAGTATCATCAGAACTTTCAGTTGTCTGATAAGAGCGACGTTTATAGTTTTGGT

>M6-PGSC0003DMG400003343M6RLKWAX-EGF_3534

TCCTCAGTATCATCAGAACTTTCAGTTGTCTGATAAGAGCGACGTTTATAGTTTTGGTGTAGTTCTTGCTGAGATCATAACAGGACTAAAAGCAGTAGACTTTGCTCGTCCTCAAGACGA

>M6-PGSC0003DMG400003343M6RLKWAX-EGF_3596

TTCTTGCTGAGATCATAACAGGACTAAAAGCAGTAGACTTTGCTCGTCCTCAAGACGAAATAAACTTGGCTGCACTTGCTATTGACAGGATAGGAAAGGGAAACTTAGATGAAATTATCG

>M6-PGSC0003DMG400003343M6RLKWAX-EGF_3658

AACTTGGCTGCACTTGCTATTGACAGGATAGGAAAGGGAAACTTAGATGAAATTATCGATCCATTTATCGAGCTACATATGGATGCTTGGACACTTTCATCTATTCACAAGGTAGCAGAG

>M6-PGSC0003DMG400003343M6RLKWAX-EGF_372

ATGCAGCCGACCATTTTCAGCAATTCACAATCTTTTTAGCCCAAATTACGCACCCACTTCAAGAAACGGAATACTTTTCCAGAATTGCAGTTCTTCACAATCCAGCTGTTTGATTCCTTC

>M6-PGSC0003DMG400003343M6RLKWAX-EGF_3720

ATTTATCGAGCTACATATGGATGCTTGGACACTTTCATCTATTCACAAGGTAGCAGAGTTGGCATTCAGGTGTCTGGCATTTCATCGCGATATGAGGCCATCGATGATGGAAGTAGCGAT

>M6-PGSC0003DMG400003343M6RLKWAX-EGF_3782

CATTCAGGTGTCTGGCATTTCATCGCGATATGAGGCCATCGATGATGGAAGTAGCGATTGAATTGGAACAACTTAGGCTAAGTAGATGGACGAATTCGGAAGATAACATAACAGGTTCAT

>M6-PGSC0003DMG400003343M6RLKWAX-EGF_3844

TTGGAACAACTTAGGCTAAGTAGATGGACGAATTCGGAAGATAACATAACAGGTTCATCAGAGGGATCTTCAATTTCATCAACTTCAGATGCAAGTGAAAAACCATTGAATCTTTCAATT

>M6-PGSC0003DMG400003343M6RLKWAX-EGF_3906

GGGATCTTCAATTTCATCAACTTCAGATGCAAGTGAAAAACCATTGAATCTTTCAATTAAGAAGCAAGAAATGGATATATTTTCGACAGATAGTAAAATAAAGGATTATTCCCCTGTATC

>M6-PGSC0003DMG400003343M6RLKWAX-EGF_3968

AGCAAGAAATGGATATATTTTCGACAGATAGTAAAATAAAGGATTATTCCCCTGTATCAGTACAAGATCCATGGAGTAGTGAGCAAAGCTCTCCATCTTGTAGTAGTTTGCTTAATCATA

>M6-PGSC0003DMG400003343M6RLKWAX-EGF_4030

CAAGATCCATGGAGTAGTGAGCAAAGCTCTCCATCTTGTAGTAGTTTGCTTAATCATATAGTTAAGTGATTATGTAAAGGATTAAATTTCTTCATTGTAATTTACTTACAAGAACATTTA

>M6-PGSC0003DMG400003343M6RLKWAX-EGF_4092

TAAGTGATTATGTAAAGGATTAAATTTCTTCATTGTAATTTACTTACAAGAACATTTAGCAGTTGAAAATGTAGTATACTAGCAAAATATTACTACTAGCAAGAATACTACACATGCCAA

>M6-PGSC0003DMG400003343M6RLKWAX-EGF_4151

CAGTTGAAAATGTAGTATACTAGCAAAATATTACTACTAGCAAGAATACTACACATGCCAAATTTGGTCTGTACTTCTAAATGTTCTTTTTATTTCTTTTATATCAATTAGTACTTTATC

>M6-PGSC0003DMG400003343M6RLKWAX-EGF_434

GAAACGGAATACTTTTCCAGAATTGCAGTTCTTCACAATCCAGCTGTTTGATTCCTTCTACGATGATTCAAACCCATTTTGACATTCTTGATTGTGGGAAAGAAAACATGAGCTGTTATT

>M6-PGSC0003DMG400003343M6RLKWAX-EGF_496

ATGATTCAAACCCATTTTGACATTCTTGATTGTGGGAAAGAAAACATGAGCTGTTATTCTGAGCCGAATAGTGAGAACAATTTTATCAATTTGAAGAATTTGTCGAAAACAGGGTGCGGC

>M6-PGSC0003DMG400003343M6RLKWAX-EGF_558

GCCGAATAGTGAGAACAATTTTATCAATTTGAAGAATTTGTCGAAAACAGGGTGCGGCTCTCTGTTTTCTGCAATATCTGTGGAGTCAACTGTGAATAGCTCGGCGGTTTCGTTGGATGT

>M6-PGSC0003DMG400003343M6RLKWAX-EGF_62

TTTACCAATTTTCTTTCTTTCTTTCTTTCTTCTTCCTCCTCCTCCCCTGTTCACATGTCCCCCAAAACATGAATTTTCCTCTCCATTTCAGAACAATTCTCTGTTTGTTTATTTTCAATT

>M6-PGSC0003DMG400003343M6RLKWAX-EGF_620

TGTTTTCTGCAATATCTGTGGAGTCAACTGTGAATAGCTCGGCGGTTTCGTTGGATGTTCAGGTGGTTCAGTTGGGGTGGTGGGTTCATGGACATTGTCGGTGTTCTTCTAATGCGAATT

>M6-PGSC0003DMG400003343M6RLKWAX-EGF_682

GTGGTTCAGTTGGGGTGGTGGGTTCATGGACATTGTCGGTGTTCTTCTAATGCGAATTGTACTAAAATCTCGTCCCCCATTAATGGAAAACGGGGATATCGGTGTCAGTGTTTTGAAGGT

>M6-PGSC0003DMG400003343M6RLKWAX-EGF_744

TAAAATCTCGTCCCCCATTAATGGAAAACGGGGATATCGGTGTCAGTGTTTTGAAGGTTTCATCGGAGATGGCTTCTTAGCCGGCTCCGGCTGCCGGAAAGGTACGTCAGGCAGAACACA

>M6-PGSC0003DMG400003343M6RLKWAX-EGF_806

TCGGAGATGGCTTCTTAGCCGGCTCCGGCTGCCGGAAAGGTACGTCAGGCAGAACACAATTTTAATTACTTAATTAGGTGTCAAACACATTAAAATCTATCTAGTTATGAGTAACAATGA

>M6-PGSC0003DMG400003343M6RLKWAX-EGF_868

TAATTACTTAATTAGGTGTCAAACACATTAAAATCTATCTAGTTATGAGTAACAATGAGATTTGAACTCGACATATTATGTTAAAATATGTGACAATTTCTTATAAAATTTAAAATTTTA

>M6-PGSC0003DMG400003343M6RLKWAX-EGF_930

TGAACTCGACATATTATGTTAAAATATGTGACAATTTCTTATAAAATTTAAAATTTTATCCGTGGAGGGTCGTATTACCAAAAAAATTGCATTCAAATTTGAATCCTAAATCCGTCTGCA

>M6-PGSC0003DMG400003343M6RLKWAX-EGF_992

TGGAGGGTCGTATTACCAAAAAAATTGCATTCAAATTTGAATCCTAAATCCGTCTGCATTGTAGGTAATTTTGCTTTAATTAGTTAACTATTTATTCAACAACTTTTTAAATAAAAACAT

>M6-PGSC0003DMG400003380M6LRR-RLP_0

TCCATAATAGTTCTAATTCTTGTGTCACAAGGATGAATTTCAACAATGAATTGTCTGATCTGAGAGTCCCGAAATACTCGGATGTTCCAAGGGATCGAATAGATTTCTTTATATGGGAGT

>M6-PGSC0003DMG400003380M6LRR-RLP_1037

GGACTTCTTATTTATTCATTATATGATATCAAGGGGCAGAAGGAAGACACATCATTGGAGGTTATCAACCAGGCACTTGGTTTTGATCTTCCGAGAAACATTGAGTCTATTAAGGCGATG

>M6-PGSC0003DMG400003380M6LRR-RLP_1098

TTATCAACCAGGCACTTGGTTTTGATCTTCCGAGAAACATTGAGTCTATTAAGGCGATGGTCTACCTTGTCATGCAAAAGGCATTTCATTGTAACTTGCCAAGGGTTCATGGACTAGGCT

>M6-PGSC0003DMG400003380M6LRR-RLP_1159

CTACCTTGTCATGCAAAAGGCATTTCATTGTAACTTGCCAAGGGTTCATGGACTAGGCTATGTCGATTTTCTGTTGAAAAACCTGAAGGATTTCCAAGGCCGTTATTCAGATTCACTCAT

>M6-PGSC0003DMG400003380M6LRR-RLP_122

AAATTCCTTAATTGTTTTCTCCATTTGCAGAAATTCACTTTTGCAAGGGAATGTGGTATGCTAGATATCTCACAGAAAATGCTAGAAATTTGCAAGAGGTTGTGTCCATCACGTCGACAT

>M6-PGSC0003DMG400003380M6LRR-RLP_1220

GTCGATTTTCTGTTGAAAAACCTGAAGGATTTCCAAGGCCGTTATTCAGATTCACTCATTTTCCTCAAGAATCACCTTCAAGTTATTCAAACGGAATTTGAGAGCTTGCAACCTTTCTTG

>M6-PGSC0003DMG400003380M6LRR-RLP_1342

GGTTGTCGTAGAAGAGCCACACAATAAGCTCAAGACACTGAATGAAGATTGTGCTACACAGATAATTAGGAAAGCATATGAGGTGGAATATGTAGTTGATGCTTGTATTAACAAAGAAGC

>M6-PGSC0003DMG400003380M6LRR-RLP_1403

ATAATTAGGAAAGCATATGAGGTGGAATATGTAGTTGATGCTTGTATTAACAAAGAAGCTCCTCATTGGTGCCTCGAGCGTTGGCTCCTGGATATCATAGAGGAGATTACTTGTATCAAA

>M6-PGSC0003DMG400003380M6LRR-RLP_1464

CTCATTGGTGCCTCGAGCGTTGGCTCCTGGATATCATAGAGGAGATTACTTGTATCAAAGCACAGATTCAGGGAAAGATCACAGTTGAGGATTCAATGAAGAGTGTCATTGCTCGTACAT

>M6-PGSC0003DMG400003380M6LRR-RLP_1525

ACAGATTCAGGGAAAGATCACAGTTGAGGATTCAATGAAGAGTGTCATTGCTCGTACATCATCACAATTGGCAAGGACTCCAAGCATGAATGAAGAGATTGTTGGGTTTGAGGATGTCAT

>M6-PGSC0003DMG400003380M6LRR-RLP_1586

TCACAATTGGCAAGGACTCCAAGCATGAATGAAGAGATTGTTGGGTTTGAGGATGTCATAGAAAATTTAAGAAAAAAACTACTGAATGGAACCAAAGGGCAAGTTATCATTTCAATTCAC

>M6-PGSC0003DMG400003380M6LRR-RLP_1708

CATGCCAGGTTTAGGTAAGACGACTTTAGCCAACAGACTCTATTCTGACAGGTCAGTTGTTTCTCATTTTGATATTTGTGCACAATGTTGTGTGTCTCAAGTATATTCTTATAAGGACTT

>M6-PGSC0003DMG400003380M6LRR-RLP_1769

TCTCATTTTGATATTTGTGCACAATGTTGTGTGTCTCAAGTATATTCTTATAAGGACTTGTTATTGGCCTTGCTACGTGATGCTATTGGTGAGGGTTCTGTGCGTAGAGAACTTCATGCC

>M6-PGSC0003DMG400003380M6LRR-RLP_183

TAGATATCTCACAGAAAATGCTAGAAATTTGCAAGAGGTTGTGTCCATCACGTCGACATAATGCATTTGCATACTGGAAGGAGGTAATTTGCAAGAGGCTGTGCGCTATTATCATCCGGC

>M6-PGSC0003DMG400003380M6LRR-RLP_1891

TAAATTAGCTGATATGCTTCGCAAAACTTTATTGCCCCGAAGGTACCTTATCCTTGTTGATGACGTGTGGGAAAATAGTGTTTGGGATGATTTAAGAGGTTGTTTCCCAGATGTCAATAA

>M6-PGSC0003DMG400003380M6LRR-RLP_1952

GACGTGTGGGAAAATAGTGTTTGGGATGATTTAAGAGGTTGTTTCCCAGATGTCAATAACAGCAGCAGAATCATTCTGACGACAAGACATCATGAAGTTGCCAAATATGCTAGTGTTCAT

>M6-PGSC0003DMG400003380M6LRR-RLP_2074

TGATCCCCTTCATCTTCGTATGTTTGACGAAGTTGAAAGTTGGAAGTTGCTTGAAAAGAAAGTGTTTGGTGAAGAAAGTTGCTCCCCTCACCTAAAAAGTGTTGGGCAGAGAATAGCAAA

>M6-PGSC0003DMG400003380M6LRR-RLP_2135

GTGTTTGGTGAAGAAAGTTGCTCCCCTCACCTAAAAAGTGTTGGGCAGAGAATAGCAAAAATGTGTGGACAACTCCCTCTTTCAATTGTTTTGGTGGCTGGTATTCTGTCAGAGATGGAA

>M6-PGSC0003DMG400003380M6LRR-RLP_2318

AGAGCCATTGTAGACCAAAGTTATCATGTTTTACCCTGTCATCTTAAGTCTTGCTTCCTTTATTTTGGAGCATTTTTAGAGGATAGAGTGATTGACATTTCAAGGTTAATAAGGTTATGG

>M6-PGSC0003DMG400003380M6LRR-RLP_2379

ATTTTGGAGCATTTTTAGAGGATAGAGTGATTGACATTTCAAGGTTAATAAGGTTATGGATATCAGAATCATTTATAAAAAGTTGTGAAGGCAGGAGGTTGGAGGATATAGCAGAAGGTT

>M6-PGSC0003DMG400003380M6LRR-RLP_244

TGCATTTGCATACTGGAAGGAGGTAATTTGCAAGAGGCTGTGCGCTATTATCATCCGGCCAGATGATGGATTTGCATACTGGAAGAAAGTAATTTGGAAGACTAAGCAAGAATTCAGAGC

>M6-PGSC0003DMG400003380M6LRR-RLP_2440

ATCAGAATCATTTATAAAAAGTTGTGAAGGCAGGAGGTTGGAGGATATAGCAGAAGGTTACTTGGAGAATCTTATTGGAAGAAATCTAGTAATGGTTACTCAGAGGGCCGATTCAGATGG

>M6-PGSC0003DMG400003380M6LRR-RLP_2501

TTGGAGAATCTTATTGGAAGAAATCTAGTAATGGTTACTCAGAGGGCCGATTCAGATGGTAAGGTCACAGCATGTCGCCTTCATGATGTATTACTTGATTTCTGCAAGAAAAGAGCAGCT

>M6-PGSC0003DMG400003380M6LRR-RLP_2562

AGGTCACAGCATGTCGCCTTCATGATGTATTACTTGATTTCTGCAAGAAAAGAGCAGCTGAGGAGAATTTTCTACTATGGATAAATAGGTAATATGATAAGTAATGGTAATTTCAATCAA

>M6-PGSC0003DMG400003380M6LRR-RLP_2623

GGAGAATTTTCTACTATGGATAAATAGGTAATATGATAAGTAATGGTAATTTCAATCAATCAAGTATTTGAAGTTATATCTGAAAATTAATGATATGCTTTTGCTAATTGATATATTCAC

>M6-PGSC0003DMG400003380M6LRR-RLP_2684

AAGTATTTGAAGTTATATCTGAAAATTAATGATATGCTTTTGCTAATTGATATATTCACAGGGATCAGATTACCAAACCTTCTTCCTGTGTTTACTCTCACAAGCAGCATGCTCACTTGG

>M6-PGSC0003DMG400003380M6LRR-RLP_2745

GGATCAGATTACCAAACCTTCTTCCTGTGTTTACTCTCACAAGCAGCATGCTCACTTGGCCTTCACTGAAATGCATAATCTTGTAGAATGGAGTGCGTCTTGCTCATTTGTTGGCTCGGT

>M6-PGSC0003DMG400003380M6LRR-RLP_2806

TTCACTGAAATGCATAATCTTGTAGAATGGAGTGCGTCTTGCTCATTTGTTGGCTCGGTAGTACTTTCCAAGAAATATGAGCCATACTTGGCTGGTCGTCCCTTAACCTCTCGTGCTTTA

>M6-PGSC0003DMG400003380M6LRR-RLP_2867

TACTTTCCAAGAAATATGAGCCATACTTGGCTGGTCGTCCCTTAACCTCTCGTGCTTTATCAATTTCATGCATTTTATTAAATTTCAAGTTTCTAAAAGTATTGGATTTAGAGCACCGGG

>M6-PGSC0003DMG400003380M6LRR-RLP_2928

AATTTCATGCATTTTATTAAATTTCAAGTTTCTAAAAGTATTGGATTTAGAGCACCGGGTTGTTATTGATTCTATTCCAACTGAGCTCTTTTACTTGAGGTATTTATCTGCACACATTGA

>M6-PGSC0003DMG400003380M6LRR-RLP_2989

GTTATTGATTCTATTCCAACTGAGCTCTTTTACTTGAGGTATTTATCTGCACACATTGAACAGAATTCAATTCCTTCAAGCATATCCAATCTTTGGAACCTTGAAACTCTTATATTAAAG

>M6-PGSC0003DMG400003380M6LRR-RLP_305

GATGATGGATTTGCATACTGGAAGAAAGTAATTTGGAAGACTAAGCAAGAATTCAGAGCTAAATACTCCTTTCCAAAAACACTACTTGCAAACAACAAGGTAGATGATGATGATATTACT

>M6-PGSC0003DMG400003380M6LRR-RLP_3050

AGAATTCAATTCCTTCAAGCATATCCAATCTTTGGAACCTTGAAACTCTTATATTAAAGAATACATCAGTTGGGAGAGATAACACGCTATTACTACCTAGTACAATTTGGGATATGGTTA

>M6-PGSC0003DMG400003380M6LRR-RLP_3111

TACATCAGTTGGGAGAGATAACACGCTATTACTACCTAGTACAATTTGGGATATGGTTAAATTGAGACATCTGCATATTCCTAACTTCAGCCCAGAAAATGAAGAAGCATTACTGAAGAA

>M6-PGSC0003DMG400003380M6LRR-RLP_3172

TTGAGACATCTGCATATTCCTAACTTCAGCCCAGAAAATGAAGAAGCATTACTGAAGAACTCTGCAAAACTTTATGATTTGGAAACCCTTTCCTCTCCATATTTCTCTCGTGTTGAGGAT

>M6-PGSC0003DMG400003380M6LRR-RLP_3294

AGATTTGATGCTGAGAAAAACACCTAATCTTCGAAAACTGGTATGTAAAGTCCAGTGCTTAGAATACCCCCATCAGCACCATCTTTTTAATTTTCCAATACGGCTTGAAATACTAAAGCT

>M6-PGSC0003DMG400003380M6LRR-RLP_3355

GAATACCCCCATCAGCACCATCTTTTTAATTTTCCAATACGGCTTGAAATACTAAAGCTTTATGGATCAGAAGCCTTTAAAAACATCCCCTTTTGCATCTCTGCACCAAATCTCAAATGC

>M6-PGSC0003DMG400003380M6LRR-RLP_3416

ATGGATCAGAAGCCTTTAAAAACATCCCCTTTTGCATCTCTGCACCAAATCTCAAATGCTTGGAACTCTATGGCTTTTACCTGGATTCTCAGTACTTATCAGAAACTGCTGATCATCTGA

>M6-PGSC0003DMG400003380M6LRR-RLP_3477

GGAACTCTATGGCTTTTACCTGGATTCTCAGTACTTATCAGAAACTGCTGATCATCTGAAGAACCTTGAGGTACTCAAACTGTCCAGTGTTGAATTTGGTGATCATAGGGAATGGAAAGT

>M6-PGSC0003DMG400003380M6LRR-RLP_3538

AACCTTGAGGTACTCAAACTGTCCAGTGTTGAATTTGGTGATCATAGGGAATGGAAAGTGAGCAATGGCAAGTTCCCTCAACTCAAAATCTTGAAACTAGAATATTTGTCCTTGATGGAA

>M6-PGSC0003DMG400003380M6LRR-RLP_3599

GCAATGGCAAGTTCCCTCAACTCAAAATCTTGAAACTAGAATATTTGTCCTTGATGGAATGGATTGTAGCCGATGATGCCTTTCCTAACCTTGAACAATTGGTTTTGCGTGGATGTCGAG

>M6-PGSC0003DMG400003380M6LRR-RLP_366

AATACTCCTTTCCAAAAACACTACTTGCAAACAACAAGGTAGATGATGATGATATTACTAATCCCAAATTTGTGATGGAATTCATCGATGCTGTTGTGGGGAATCTCAATGTTCTAGTCA

>M6-PGSC0003DMG400003380M6LRR-RLP_3721

CTTATGGAGATCCCTTCTTGTTTCATGGACATCCTTTCTCTCAAGTACATCGAGTTAGGAAACTGCAATGAGTCGGTTGTCAAGTCAGCCATGAATATACAAGAAACACAAGTCGAAGAT

>M6-PGSC0003DMG400003380M6LRR-RLP_3782

ACTGCAATGAGTCGGTTGTCAAGTCAGCCATGAATATACAAGAAACACAAGTCGAAGATAATCAAAATACTAATTTCAAGCTCGTTCTCATCTAGGTACACTACTGAAAAAAGCTTTATT

>M6-PGSC0003DMG400003380M6LRR-RLP_3843

TCAAAATACTAATTTCAAGCTCGTTCTCATCTAGGTACACTACTGAAAAAAGCTTTATTCTGCATGATTTTGATGACTCCGAAATAATGGCCTAAATTGTATGAACTGTTTTCTCTGTTA

>M6-PGSC0003DMG400003380M6LRR-RLP_3904

GCATGATTTTGATGACTCCGAAATAATGGCCTAAATTGTATGAACTGTTTTCTCTGTTATCTTTACCTGGTGGCCTCGTTTTACATTCGGGTTTTTCTCTTGGTTTTCTTGCAGAAAAAG

>M6-PGSC0003DMG400003380M6LRR-RLP_3965

TTTACCTGGTGGCCTCGTTTTACATTCGGGTTTTTCTCTTGGTTTTCTTGCAGAAAAAGACATACCCCCAAGGCATTTAAAAGATTAGTTTTTCTTCCAGGTACTACTTATTATTTTTTC

>M6-PGSC0003DMG400003380M6LRR-RLP_4026

ATACCCCCAAGGCATTTAAAAGATTAGTTTTTCTTCCAGGTACTACTTATTATTTTTTCTTTCTTTCCTTTTTTTGAGTACACCAAATAGATAGATTCATCTTTTTTGTCTTTTCGATAT

>M6-PGSC0003DMG400003380M6LRR-RLP_4087

TCTTTCCTTTTTTTGAGTACACCAAATAGATAGATTCATCTTTTTTGTCTTTTCGATATGAAAGGGATATAATCAATTTCAACTGATATGAAAGAGAAGAAACTTACTGTGACCAGAGAT

>M6-PGSC0003DMG400003380M6LRR-RLP_4148

AAGGGATATAATCAATTTCAACTGATATGAAAGAGAAGAAACTTACTGTGACCAGAGATGTGGATGCTGATGAAGTTCAATTAGTTGTGGAGAAACTGAGAAAGTGTGGCATGCTCTAAC

>M6-PGSC0003DMG400003380M6LRR-RLP_4209

GGATGCTGATGAAGTTCAATTAGTTGTGGAGAAACTGAGAAAGTGTGGCATGCTCTAACAAAATGCTGAGCTCTGGTGAACTCAACCATGATGGATTTTCAAATAATGACAGGGTTGTAG

>M6-PGSC0003DMG400003380M6LRR-RLP_427

TCCCAAATTTGTGATGGAATTCATCGATGCTGTTGTGGGGAATCTCAATGTTCTAGTCAAGATCAATGATCCATTTTCATTGCTTTTTGTTCCAGGACCCAAGGAACAAATAGAACAAGT

>M6-PGSC0003DMG400003380M6LRR-RLP_4331

CCAACTTGTCAACACAAATGTGCTATACTCATTTTGCTTACTGTAATACCATTTCATGACACACACACAAACATTAACTGTAGTAAAGTTTTGATGGATCAGTAAATCTGAGTTCAACCC

>M6-PGSC0003DMG400003380M6LRR-RLP_4392

CACACACAAACATTAACTGTAGTAAAGTTTTGATGGATCAGTAAATCTGAGTTCAACCCATTGTAATCCGTTCAAATTCAACTCAAAGGTATCCAGAGAGTGTAGCTGGAGCAATTTGGA

>M6-PGSC0003DMG400003380M6LRR-RLP_4453

TGTAATCCGTTCAAATTCAACTCAAAGGTATCCAGAGAGTGTAGCTGGAGCAATTTGGAATATCACATGTAATTTCTTTATGAGTTATTTCGTTTAATAAAGATACTGTAAAACGTCCAA

>M6-PGSC0003DMG400003380M6LRR-RLP_4514

ATCACATGTAATTTCTTTATGAGTTATTTCGTTTAATAAAGATACTGTAAAACGTCCAACGGCTGTTGCATTCATTGTAAACTAAATACATCTCACAATGTAACTATTTGAACAATTTTT

>M6-PGSC0003DMG400003380M6LRR-RLP_4557

ACTGTAAAACGTCCAACGGCTGTTGCATTCATTGTAAACTAAATACATCTCACAATGTAACTATTTGAACAATTTTTTCATTTAACTCCTTGAGGTTTGATGTAAGTCATTAGATTTTAC

>M6-PGSC0003DMG400003380M6LRR-RLP_488

ATCAATGATCCATTTTCATTGCTTTTTGTTCCAGGACCCAAGGAACAAATAGAACAAGTGTTAAAGGAGTTGAAGTTATTGAGATTTTTTGTCTGCTTTGTTTCAAACAAATGTATAGAG

>M6-PGSC0003DMG400003380M6LRR-RLP_549

TAAAGGAGTTGAAGTTATTGAGATTTTTTGTCTGCTTTGTTTCAAACAAATGTATAGAGCCTCAATACCAACATGCTACTTTTTATACTCACGCTTTAATTGAGGCTAGCCACATCGCAA

>M6-PGSC0003DMG400003380M6LRR-RLP_61

GAGAGTCCCGAAATACTCGGATGTTCCAAGGGATCGAATAGATTTCTTTATATGGGAGTTAAAATTCCTTAATTGTTTTCTCCATTTGCAGAAATTCACTTTTGCAAGGGAATGTGGTAT

>M6-PGSC0003DMG400003380M6LRR-RLP_610

TCAATACCAACATGCTACTTTTTATACTCACGCTTTAATTGAGGCTAGCCACATCGCAATGGTTGTGTGGTTGAATTTGCCAATCTATGGAAACAGAAATCAAGACTTGGCTTCAAATGA

>M6-PGSC0003DMG400003380M6LRR-RLP_671

GTTGTGTGGTTGAATTTGCCAATCTATGGAAACAGAAATCAAGACTTGGCTTCAAATGAAGTCAGTTGTTTGCTTTCTGATTTCATGGAAATGAAGATTAAGTCCATTCAGCCAGGCATC

>M6-PGSC0003DMG400003380M6LRR-RLP_732

TCAGTTGTTTGCTTTCTGATTTCATGGAAATGAAGATTAAGTCCATTCAGCCAGGCATCAGCCGCAACAATATTTATATTGATGTCTTGAGGGCGTTGAAGTCAACCATACCACATGCTC

>M6-PGSC0003DMG400003380M6LRR-RLP_793

CCGCAACAATATTTATATTGATGTCTTGAGGGCGTTGAAGTCAACCATACCACATGCTCAAGATAAGCATGCAGCTGAGAGTGGTATTGTGGAGACTCCAACACACAATCTGATGGTTGG

>M6-PGSC0003DMG400003380M6LRR-RLP_854

GATAAGCATGCAGCTGAGAGTGGTATTGTGGAGACTCCAACACACAATCTGATGGTTGGTTTGAGTGATCAAATGGCAAACCTTCAGGAGATGCTCTGCCTTCTAAGAGACAATCTCATT

>M6-PGSC0003DMG400003380M6LRR-RLP_915

TGAGTGATCAAATGGCAAACCTTCAGGAGATGCTCTGCCTTCTAAGAGACAATCTCATTCATCTGCCAATACTAGATCTGGAATTTCATCTACAAGATATGGATTCTGTTATTGTTGATG

>M6-PGSC0003DMG400003380M6LRR-RLP_976

TCTGCCAATACTAGATCTGGAATTTCATCTACAAGATATGGATTCTGTTATTGTTGATGCAGGACTTCTTATTTATTCATTATATGATATCAAGGGGCAGAAGGAAGACACATCATTGGA

>M6-PGSC0003DMG400003406-(rev)M6RLKmalectin_1054

ATATTATTAGCAAAGCTCTTAATGAACTCTATTTCAATGTGTATATCAATGACAAGATGGCTATTTCCGGGCTAGATTTGTCATCATTGACAAATCATTTGTCTACGGCTTTTTACAAGG

>M6-PGSC0003DMG400003406-(rev)M6RLKmalectin_1116

ATTTCCGGGCTAGATTTGTCATCATTGACAAATCATTTGTCTACGGCTTTTTACAAGGATTTTGTTGTGGATTCCTCCATGGTGTCTAACCCCCTCGTTGTACGTGTTTCTCCTGTGAAT

>M6-PGSC0003DMG400003406-(rev)M6RLKmalectin_1178

TGTTGTGGATTCCTCCATGGTGTCTAACCCCCTCGTTGTACGTGTTTCTCCTGTGAATGATGCTCAAGGTTTTAGAAATGCGATTCTAAATGGCCTTGAGGTGTTTCGAATGAATAACTC

>M6-PGSC0003DMG400003406-(rev)M6RLKmalectin_124

TGATATGTTTCATTTTTTCTATCACAAATTCATTTGTCATTGCTCAAGATGATGCTGTACCTGCAAAATCATTTCCTGTATTCACTCCAGAAGATAATTTCCTCATTGATTGTGGTGCAA

>M6-PGSC0003DMG400003406-(rev)M6RLKmalectin_1240

CTCAAGGTTTTAGAAATGCGATTCTAAATGGCCTTGAGGTGTTTCGAATGAATAACTCTGTTGGTAGCTTAGATGGGGAATATGGCGTTGATGGACAAAAACAAAGTGGTCCAAGTAAAA

>M6-PGSC0003DMG400003406-(rev)M6RLKmalectin_1302

GGTAGCTTAGATGGGGAATATGGCGTTGATGGACAAAAACAAAGTGGTCCAAGTAAAACAGTGGCTTATGTTGGATTCGCGATGATGTTTGGAGCGTTTGTGGGGTTGGGTGCGATGGCA

>M6-PGSC0003DMG400003406-(rev)M6RLKmalectin_1364

GGCTTATGTTGGATTCGCGATGATGTTTGGAGCGTTTGTGGGGTTGGGTGCGATGGCAGTTAAATGGCAGAAGAGACCTCAAGATTGGCAAAAGAGGAATAGTTTTTCGTCTTGGTTACT

>M6-PGSC0003DMG400003406-(rev)M6RLKmalectin_1426

AATGGCAGAAGAGACCTCAAGATTGGCAAAAGAGGAATAGTTTTTCGTCTTGGTTACTTCCAGTTCATGCTGGTGATTCAACATTTATGTCATCATCTTCGCGAAGCAAGAGCCAATTTT

>M6-PGSC0003DMG400003406-(rev)M6RLKmalectin_1488

GTTCATGCTGGTGATTCAACATTTATGTCATCATCTTCGCGAAGCAAGAGCCAATTTTTCTCATCAAAAAACATGGGATTAGGTCAGTACTTTAGTTTTGCTGAATTGTCAGACGCCACG

>M6-PGSC0003DMG400003406-(rev)M6RLKmalectin_1550

ATCAAAAAACATGGGATTAGGTCAGTACTTTAGTTTTGCTGAATTGTCAGACGCCACGAGGAACTGGGAACCTAGTGAAATAATCGGCGTTGGTGGCTTTGGAAATGTATACTATGGAGA

>M6-PGSC0003DMG400003406-(rev)M6RLKmalectin_1612

ACTGGGAACCTAGTGAAATAATCGGCGTTGGTGGCTTTGGAAATGTATACTATGGAGAACTTGATGATGGTACAAAAGTTGCTGTCAAGAGAGGAAATCCACAATCAGAACAAGGTATAA

>M6-PGSC0003DMG400003406-(rev)M6RLKmalectin_1674

GATGATGGTACAAAAGTTGCTGTCAAGAGAGGAAATCCACAATCAGAACAAGGTATAAATGAATTCCAGACTGAAATTCAAATGTTATCTAAGTTAAGGCATCGACATTTAGTGTCGTTG

>M6-PGSC0003DMG400003406-(rev)M6RLKmalectin_1736

ATTCCAGACTGAAATTCAAATGTTATCTAAGTTAAGGCATCGACATTTAGTGTCGTTGATTGGGTATTGTGATGAAAGTGCTGAGATGATATTGGTTTATGAGTTTATGCAAAATGGTCC

>M6-PGSC0003DMG400003406-(rev)M6RLKmalectin_1798

GGTATTGTGATGAAAGTGCTGAGATGATATTGGTTTATGAGTTTATGCAAAATGGTCCATTGAGAGATCATCTCTATGGAAAGAACATGCCGCCTTTATCTTGGAAGCAACGATTAGATA

>M6-PGSC0003DMG400003406-(rev)M6RLKmalectin_186

GCAAAATCATTTCCTGTATTCACTCCAGAAGATAATTTCCTCATTGATTGTGGTGCAACTTCATCGATCACGCTTCCGGGGAATAAAGTTTTTCAACCTGATCAAAATGCCGCTAAATAT

>M6-PGSC0003DMG400003406-(rev)M6RLKmalectin_1860

AGAGATCATCTCTATGGAAAGAACATGCCGCCTTTATCTTGGAAGCAACGATTAGATATCTGTATTGGTGCTGCTCGTGGGCTTCATTACCTCCACACTGGTGCATCTACAGGGATCATA

>M6-PGSC0003DMG400003406-(rev)M6RLKmalectin_1922

TATTGGTGCTGCTCGTGGGCTTCATTACCTCCACACTGGTGCATCTACAGGGATCATACATAGGGATGTCAAGACCACCAACATTTTGCTTGATGAGAATTTTGTCGCGAAGATGGCTGA

>M6-PGSC0003DMG400003406-(rev)M6RLKmalectin_1984

GGGATGTCAAGACCACCAACATTTTGCTTGATGAGAATTTTGTCGCGAAGATGGCTGATTTTGGGCTATCTAAAGATGGACCAACAACAGAACAGACTCATGTGAGCACGGCTGTGAAGG

>M6-PGSC0003DMG400003406-(rev)M6RLKmalectin_2046

GGGCTATCTAAAGATGGACCAACAACAGAACAGACTCATGTGAGCACGGCTGTGAAGGGAAGTTTTGGCTACTTAGATCCTGAATACTTTAGGAAGCAGCAGTTGACAGACAAATCTGAT

>M6-PGSC0003DMG400003406-(rev)M6RLKmalectin_2108

TTTTGGCTACTTAGATCCTGAATACTTTAGGAAGCAGCAGTTGACAGACAAATCTGATGTCTACTCATTTGGGGTTGTACTTCTTGAAGCCTTATGCGCCCGTCCTGCTATTAATCCATC

>M6-PGSC0003DMG400003406-(rev)M6RLKmalectin_2170

ACTCATTTGGGGTTGTACTTCTTGAAGCCTTATGCGCCCGTCCTGCTATTAATCCATCATTACCAAGAGAGCAAGTGAATTTGGCAGAATGGGCAATGCAGTGGAAAAGAAAAGGGTTAT

>M6-PGSC0003DMG400003406-(rev)M6RLKmalectin_2232

CCAAGAGAGCAAGTGAATTTGGCAGAATGGGCAATGCAGTGGAAAAGAAAAGGGTTATTGGACAAGATAATTGATCCAACACTTGTAGGACATATTGATCCAGAATCAATGAAGAAATTC

>M6-PGSC0003DMG400003406-(rev)M6RLKmalectin_2294

CAAGATAATTGATCCAACACTTGTAGGACATATTGATCCAGAATCAATGAAGAAATTCGCGGAAGCAACAGAGAAGTGTTTGGCTGAATATGGTGTTGACAGACCTACTATGGGTGATGT

>M6-PGSC0003DMG400003406-(rev)M6RLKmalectin_2356

AAGCAACAGAGAAGTGTTTGGCTGAATATGGTGTTGACAGACCTACTATGGGTGATGTTTTATGGAATTTGGAGTACGCGTTGCATATGCAAGAAGCCTCGTTACAAGGAAAGACAGAAG

>M6-PGSC0003DMG400003406-(rev)M6RLKmalectin_2418

TGGAATTTGGAGTACGCGTTGCATATGCAAGAAGCCTCGTTACAAGGAAAGACAGAAGATATCGATAACAAAGCAATTGTTGCACCAATTTCTCCTGCTACAGTTGCCCATGCTAATGTC

>M6-PGSC0003DMG400003406-(rev)M6RLKmalectin_248

ATCGATCACGCTTCCGGGGAATAAAGTTTTTCAACCTGATCAAAATGCCGCTAAATATCTTTCTTATAGTGGTAAAGATGTACAAGCCTGTGCTTCTGAGAAAATCAACGTTCCTTCAAC

>M6-PGSC0003DMG400003406-(rev)M6RLKmalectin_2480

CGATAACAAAGCAATTGTTGCACCAATTTCTCCTGCTACAGTTGCCCATGCTAATGTCACAACTGATAACAAACATATTTCTTCACCAGAACAACATAAAAATCCTACTGAAGTTCAAGC

>M6-PGSC0003DMG400003406-(rev)M6RLKmalectin_2542

CTGATAACAAACATATTTCTTCACCAGAACAACATAAAAATCCTACTGAAGTTCAAGCCATAGATGACAAGTCAGTGGCTTTTGCTCAATTTGAGCCTATCAATGGACGATAAAACGAAC

>M6-PGSC0003DMG400003406-(rev)M6RLKmalectin_2604

GATGACAAGTCAGTGGCTTTTGCTCAATTTGAGCCTATCAATGGACGATAAAACGAACACTCAGCTTGTTCTATAGACAAATCAAGAATCTACAATTAAAGGTTATGAAATACATGTGTG

>M6-PGSC0003DMG400003406-(rev)M6RLKmalectin_2666

AGCTTGTTCTATAGACAAATCAAGAATCTACAATTAAAGGTTATGAAATACATGTGTGTGTATATGTATGTAGTAAAATTCAGCTTTTTATGGATGCCATTTTGCTGAAAAATGTAATTC

>M6-PGSC0003DMG400003406-(rev)M6RLKmalectin_2710

GAAATACATGTGTGTGTATATGTATGTAGTAAAATTCAGCTTTTTATGGATGCCATTTTGCTGAAAAATGTAATTCAGAACAATGTAATGTTTCACATAATATACTACTTTACTACATTG

>M6-PGSC0003DMG400003406-(rev)M6RLKmalectin_310

CTTATAGTGGTAAAGATGTACAAGCCTGTGCTTCTGAGAAAATCAACGTTCCTTCAACGTTATATGTAAACGCCAAAGTCTTCACAACTGAAGCTATTTACACGTTTCATGCCAGTACTT

>M6-PGSC0003DMG400003406-(rev)M6RLKmalectin_372

TATGTAAACGCCAAAGTCTTCACAACTGAAGCTATTTACACGTTTCATGCCAGTACTTCAGGGTTGCATTGGATCAGACTACATTTCTTTCCATTTAAATACGAAGAATATGATCTTAAA

>M6-PGSC0003DMG400003406-(rev)M6RLKmalectin_434

GTTGCATTGGATCAGACTACATTTCTTTCCATTTAAATACGAAGAATATGATCTTAAAACAGCTAAATTTTCTGTTAAAACGGATAGTTTAGTCCTTCTACGTGATTTCCAAACTGGGAA

>M6-PGSC0003DMG400003406-(rev)M6RLKmalectin_496

CTAAATTTTCTGTTAAAACGGATAGTTTAGTCCTTCTACGTGATTTCCAAACTGGGAAAGATGAGGCTGTTGTCAAGGAGTTTGTTGTTAATGTTACCTCAGAGCGTTTTGCTATAAAAT

>M6-PGSC0003DMG400003406-(rev)M6RLKmalectin_558

GAGGCTGTTGTCAAGGAGTTTGTTGTTAATGTTACCTCAGAGCGTTTTGCTATAAAATTCGAGCCAGCTCAAGGATCTGTTGCATTCGTTAATGCTATTGAGTTTGTTACTGTACCTGCA

>M6-PGSC0003DMG400003406-(rev)M6RLKmalectin_620

GCCAGCTCAAGGATCTGTTGCATTCGTTAATGCTATTGAGTTTGTTACTGTACCTGCAAAAATGCTTGATTATTCTGTTCCTGTTTTATTCCCCGTGTCACAGAGATTTGATCTTTCTAA

>M6-PGSC0003DMG400003406-(rev)M6RLKmalectin_682

TGCTTGATTATTCTGTTCCTGTTTTATTCCCCGTGTCACAGAGATTTGATCTTTCTAAAACTAATTTCCAGACAATGTATAGGCTTAATGTTGGTGGTCCTGCTTTGGATTCAACAAACG

>M6-PGSC0003DMG400003406-(rev)M6RLKmalectin_744

AATTTCCAGACAATGTATAGGCTTAATGTTGGTGGTCCTGCTTTGGATTCAACAAACGATACTCTTGGAAGAAATTGGATGTCTGATAATCAATTTCGAAACAATGCTACAGGTGCAGAG

>M6-PGSC0003DMG400003406-(rev)M6RLKmalectin_806

TCTTGGAAGAAATTGGATGTCTGATAATCAATTTCGAAACAATGCTACAGGTGCAGAGGTATCAACACAAGCCTCTGCTATTAATTATTTGATATCATCAGGAGGTTCTCCGTTGATCGC

>M6-PGSC0003DMG400003406-(rev)M6RLKmalectin_868

CAACACAAGCCTCTGCTATTAATTATTTGATATCATCAGGAGGTTCTCCGTTGATCGCGCCACCAACAGTTTATAGTTCTGCTGTAAAGATGGCGGATTCAGAAACTACAATTGCTAATT

>M6-PGSC0003DMG400003406-(rev)M6RLKmalectin_930

CCAACAGTTTATAGTTCTGCTGTAAAGATGGCGGATTCAGAAACTACAATTGCTAATTTTAACATATCATGGACTATGGATATTGATACTATATATCCTCATTTGATTCGCTTACATTTC

>M6-PGSC0003DMG400003406-(rev)M6RLKmalectin_992

CATATCATGGACTATGGATATTGATACTATATATCCTCATTTGATTCGCTTACATTTCTGCGATATTATTAGCAAAGCTCTTAATGAACTCTATTTCAATGTGTATATCAATGACAAGAT

>M6-PGSC0003DMG400003429-(rev)M6LRR-RLK_0

AAGCAACACGTCCACACTTCCAAACCACAAAAAAAAAAAAAAATGTCTTCTAGTCATTTTCTTTCATTCCTTTTCCTTTCTCTTCTAATTTCCGGGATCTTTTCGGACCTTAACGCGGAT

>M6-PGSC0003DMG400003429-(rev)M6LRR-RLK_1054

TGGAGGATTTATTGATGGCGTCCGCGGAGGTTTTGGGGAAAGGGTTAACTGGAACTACTTATAAAGCATATTTAGATAGAGATGTTGAAGTTGTTGTGAAGAGGTTGAGAAATGTTTGTG

>M6-PGSC0003DMG400003429-(rev)M6LRR-RLK_1116

AAAGCATATTTAGATAGAGATGTTGAAGTTGTTGTGAAGAGGTTGAGAAATGTTTGTGTTTCAGAAGAGGAGTTTAGGGGAAAAGTGGAGGAATTGGGAGGAATTAGTCATGGGAATTTG

>M6-PGSC0003DMG400003429-(rev)M6LRR-RLK_1178

AGAAGAGGAGTTTAGGGGAAAAGTGGAGGAATTGGGAGGAATTAGTCATGGGAATTTGGTGCCTCTTAGAGCATATTATTATGGAAGAGATGAAAAGCTTCTTGTTTATGATTCTATGCC

>M6-PGSC0003DMG400003429-(rev)M6LRR-RLK_124

CCGCTCTACTCCACCTGAGTGCGGCCTTCAAAGGCCGCACTCTTGGGTGGAACACAACCAATTCAACCCCATGTTCATGGCAAGGTGTCAAATGTGATACTACAATCAACCGTGTAATTG

>M6-PGSC0003DMG400003429-(rev)M6LRR-RLK_1240

CTCTTAGAGCATATTATTATGGAAGAGATGAAAAGCTTCTTGTTTATGATTCTATGCCTACAAGTTTGTATGCTATTTTGCATGGTAAGTATCTTATTTCTAATAAACTTTAGCTTTTGT

>M6-PGSC0003DMG400003429-(rev)M6LRR-RLK_1302

AGTTTGTATGCTATTTTGCATGGTAAGTATCTTATTTCTAATAAACTTTAGCTTTTGTTGTTTTAATAATAATATTTGCTACTTGCTTCTTTTTGGTAGTTCTAATTGGTTTGTTTTCTC

>M6-PGSC0003DMG400003429-(rev)M6LRR-RLK_1364

TTAATAATAATATTTGCTACTTGCTTCTTTTTGGTAGTTCTAATTGGTTTGTTTTCTCTGGTTTTGGTTTGGGAGTCGATCAACCTAATTAAAATTCTTAGCTTAGATTAGATTAGCTTC

>M6-PGSC0003DMG400003429-(rev)M6LRR-RLK_1426

TTTGGTTTGGGAGTCGATCAACCTAATTAAAATTCTTAGCTTAGATTAGATTAGCTTCTAGTTAATTGTTTTTTAGCTTTAGCTATGAAAACTGAAATTATGTGGAAGAGATGGTATGAA

>M6-PGSC0003DMG400003429-(rev)M6LRR-RLK_1488

TAATTGTTTTTTAGCTTTAGCTATGAAAACTGAAATTATGTGGAAGAGATGGTATGAAAACAGGGGAAAGGAAGAAGATCATTTGGTAGTTCTCTTTTACATTAGTTAATTGTCATCAGA

>M6-PGSC0003DMG400003429-(rev)M6LRR-RLK_1736

TAAAGATAATTTTTGATGTTGTGTATACTGATGTTCACATCACAACTAATGAAGGACCCATGTAATATTATTAGAGAGAGAGGGACGTACAACAAGAACTTTCTGTGTAGAGGGAAAGCC

>M6-PGSC0003DMG400003429-(rev)M6LRR-RLK_1798

TAATATTATTAGAGAGAGAGGGACGTACAACAAGAACTTTCTGTGTAGAGGGAAAGCCTCCTTAATTAGTATTTAGTTGAATAGGCCTTCTATCAACTCACATTCTATTAAATGTTATGA

>M6-PGSC0003DMG400003429-(rev)M6LRR-RLK_186

TCAACCCCATGTTCATGGCAAGGTGTCAAATGTGATACTACAATCAACCGTGTAATTGACCTCCGTCTCCCCGGATATGGTCTATCCGGTGAGATGCCCTTAAATTCCATCGGAAACTTA

>M6-PGSC0003DMG400003429-(rev)M6LRR-RLK_1860

TAATTAGTATTTAGTTGAATAGGCCTTCTATCAACTCACATTCTATTAAATGTTATGAAAACGAACTATAGAATTTTACTAGTATTAATTATTTCGTATTCTAACCAATTACTATTACTA

>M6-PGSC0003DMG400003429-(rev)M6LRR-RLK_1922

GAACTATAGAATTTTACTAGTATTAATTATTTCGTATTCTAACCAATTACTATTACTATGAGTTAGTTTTAACCTTTCTGATTTTGTTAGGCGAAGGAAAAGAAGCATTGACATGGGTAA

>M6-PGSC0003DMG400003429-(rev)M6LRR-RLK_1984

TTAGTTTTAACCTTTCTGATTTTGTTAGGCGAAGGAAAAGAAGCATTGACATGGGTAATAAGGAGCAGAATTGCATTAGGAGTTGCCAATGGTATCGAATATCTCCATTCACTAGGCGCT

>M6-PGSC0003DMG400003429-(rev)M6LRR-RLK_2046

GAGCAGAATTGCATTAGGAGTTGCCAATGGTATCGAATATCTCCATTCACTAGGCGCTAAAGTCACTCATGGCAACATCAAGTCATCTAACATCCTCCTCACACACTACTACGATGCATG

>M6-PGSC0003DMG400003429-(rev)M6LRR-RLK_2108

TCACTCATGGCAACATCAAGTCATCTAACATCCTCCTCACACACTACTACGATGCATGTCTATCTGAATTTGGCATAACACAGTTGATTTCCTCAACATTGAACTCCAAAATGACCGGAT

>M6-PGSC0003DMG400003429-(rev)M6LRR-RLK_2170

TCTGAATTTGGCATAACACAGTTGATTTCCTCAACATTGAACTCCAAAATGACCGGATACTACGCGCCTGAGGTGACAGATATTCGAAATGTTTCCCAGAAAGCTGATGTCTATAGCTTT

>M6-PGSC0003DMG400003429-(rev)M6LRR-RLK_2232

CGCGCCTGAGGTGACAGATATTCGAAATGTTTCCCAGAAAGCTGATGTCTATAGCTTTGGCACGATTCTGTTGGAACTCCTAACGGGTAAAAATCCTAGTAGTGTCATCAATGACGAGGG

>M6-PGSC0003DMG400003429-(rev)M6LRR-RLK_2294

CGATTCTGTTGGAACTCCTAACGGGTAAAAATCCTAGTAGTGTCATCAATGACGAGGGAATTGACCTGCCTAAATGGGTGAAATGTATAGTTGAAGAGAGGGGGACAACACAAGTGTTTG

>M6-PGSC0003DMG400003429-(rev)M6LRR-RLK_2356

GACCTGCCTAAATGGGTGAAATGTATAGTTGAAGAGAGGGGGACAACACAAGTGTTTGATCCTGAGTTGATTAGTTTTCAGAATTGTGATGAAGAGCAAATGGTTTCATTGCTGCATCTT

>M6-PGSC0003DMG400003429-(rev)M6LRR-RLK_2418

TGAGTTGATTAGTTTTCAGAATTGTGATGAAGAGCAAATGGTTTCATTGCTGCATCTTGCTATTTCTTGTACATCCCACCACCCCGAAAGGCGGCCGCCTATGGCTGACACTACCAGGAG

>M6-PGSC0003DMG400003429-(rev)M6LRR-RLK_248

CCGTCTCCCCGGATATGGTCTATCCGGTGAGATGCCCTTAAATTCCATCGGAAACTTAACTGAACTCCGTACTCTCAGCCTTCGGAGTAACTCTTTATCCGGTCTTCTTCCTTCTGACAT

>M6-PGSC0003DMG400003429-(rev)M6LRR-RLK_2480

TTTCTTGTACATCCCACCACCCCGAAAGGCGGCCGCCTATGGCTGACACTACCAGGAGGATCAAAGAAATTGTTATGTGAGTTTTGTTAATGTTTGATCTTTCCCCCTCATTCTTGTCCA

>M6-PGSC0003DMG400003429-(rev)M6LRR-RLK_2542

AAAGAAATTGTTATGTGAGTTTTGTTAATGTTTGATCTTTCCCCCTCATTCTTGTCCAAATGAAGGTGAAAATCATGGTTTTATTCCTTTCTTGGTTCTACAAGAGGGGTGAATTCATAC

>M6-PGSC0003DMG400003429-(rev)M6LRR-RLK_2604

AAGGTGAAAATCATGGTTTTATTCCTTTCTTGGTTCTACAAGAGGGGTGAATTCATACCTTAAAAATTGGGCACATGAATATGTGATCTCTCTATAAAACTAGATATTTTATGTATATAT

>M6-PGSC0003DMG400003429-(rev)M6LRR-RLK_2637

TTCTACAAGAGGGGTGAATTCATACCTTAAAAATTGGGCACATGAATATGTGATCTCTCTATAAAACTAGATATTTTATGTATATATATTTTGAATTAATATAATATTATTTATTGCCTC

>M6-PGSC0003DMG400003429-(rev)M6LRR-RLK_310

AACTCCGTACTCTCAGCCTTCGGAGTAACTCTTTATCCGGTCTTCTTCCTTCTGACATAGGATCATGTACAGAGCTACGCATACTTAATTTGGAAAACAACGACTTCTCCGGTTCAATTC

>M6-PGSC0003DMG400003429-(rev)M6LRR-RLK_372

TCATGTACAGAGCTACGCATACTTAATTTGGAAAACAACGACTTCTCCGGTTCAATTCCAACGACTTTTTTCAATCTGAACAATCTTATACGTGTGAGTTTTTCAGGGAACAGATTCTCC

>M6-PGSC0003DMG400003429-(rev)M6LRR-RLK_434

GACTTTTTTCAATCTGAACAATCTTATACGTGTGAGTTTTTCAGGGAACAGATTCTCCGGTGAGATATCGGACGCATTTAACAATCTGACTAGGATGAGAACTCTGTATTTGGAGAATAA

>M6-PGSC0003DMG400003429-(rev)M6LRR-RLK_496

AGATATCGGACGCATTTAACAATCTGACTAGGATGAGAACTCTGTATTTGGAGAATAACAACTTCTCTGGCTCACTTCCTGATTTGAAAAACCTCTCACAGCTTAATGAATTCAATGTGT

>M6-PGSC0003DMG400003429-(rev)M6LRR-RLK_558

TTCTCTGGCTCACTTCCTGATTTGAAAAACCTCTCACAGCTTAATGAATTCAATGTGTCTTTCAATAGATTAACCGGTTCGATCCCTTCAAGTTTAGGTCAATTCGTAGCTTCTTCATTT

>M6-PGSC0003DMG400003429-(rev)M6LRR-RLK_62

TTCATTCCTTTTCCTTTCTCTTCTAATTTCCGGGATCTTTTCGGACCTTAACGCGGATAGAGCCGCTCTACTCCACCTGAGTGCGGCCTTCAAAGGCCGCACTCTTGGGTGGAACACAAC

>M6-PGSC0003DMG400003429-(rev)M6LRR-RLK_620

CAATAGATTAACCGGTTCGATCCCTTCAAGTTTAGGTCAATTCGTAGCTTCTTCATTTCTCGGAAATTCACTCTGTGGCTCTGCTTTCTCTCTTAGTCCTTGTCCTGAAAACAATAATAT

>M6-PGSC0003DMG400003429-(rev)M6LRR-RLK_682

GAAATTCACTCTGTGGCTCTGCTTTCTCTCTTAGTCCTTGTCCTGAAAACAATAATATCACCAATCAATCGGATAAACTTTCCGGTGGAGCAATAGCTGGAATAGTAATTGGCTCGATTA

>M6-PGSC0003DMG400003429-(rev)M6LRR-RLK_744

AATCAATCGGATAAACTTTCCGGTGGAGCAATAGCTGGAATAGTAATTGGCTCGATTATCGGATTCTTCATACTACTATTAGTTTTGTTTATGTTAGTGAGATCAATTTATAGGAGTAAA

>M6-PGSC0003DMG400003429-(rev)M6LRR-RLK_806

ATTCTTCATACTACTATTAGTTTTGTTTATGTTAGTGAGATCAATTTATAGGAGTAAAAAAAGTTTCCGTCAAGTGAACGTCTCTCCTACGCCAAATCAAGTACTTTCGAGTCCTCGTGA

>M6-PGSC0003DMG400003429-(rev)M6LRR-RLK_868

GTTTCCGTCAAGTGAACGTCTCTCCTACGCCAAATCAAGTACTTTCGAGTCCTCGTGATTCAATGGTGACAGAAAATCATGATATCGAAAATGTGTATTCGGAGAGGAAAGATAGCAAGA

>M6-PGSC0003DMG400003429-(rev)M6LRR-RLK_930

ATGGTGACAGAAAATCATGATATCGAAAATGTGTATTCGGAGAGGAAAGATAGCAAGAGGAAAGTTTGTGATGATAGGACAAAAGGGATGGTGTTTTTTGGGGAAAGTTTTGAGCTTTTT

>M6-PGSC0003DMG400003429-(rev)M6LRR-RLK_992

AGTTTGTGATGATAGGACAAAAGGGATGGTGTTTTTTGGGGAAAGTTTTGAGCTTTTTAGTTTGGAGGATTTATTGATGGCGTCCGCGGAGGTTTTGGGGAAAGGGTTAACTGGAACTAC

>M6-PGSC0003DMG400003543M6LRR-RLP_0

CTTTCTTTGTTTGTTCCCTTCTTCGGACACTCAGAATATATGCTCACGCATCCATTTAAAAACCAATCTTGAAGGCTCTTCTTCTTCATCAATTCTCATTACGATTAAACAAAATAATAT

>M6-PGSC0003DMG400003543M6LRR-RLP_1008

TCAGGAACCCGGGATCTCGTTTCAGAATAGTCAGCTTATTGCAGCAGACTTAAGCAACAATGAGATCCGTGGGTTATTACCCGGGTTCCTGGGTTTAATGCCCAGATTATCATCACTTTC

>M6-PGSC0003DMG400003543M6LRR-RLP_1071

GATCCGTGGGTTATTACCCGGGTTCCTGGGTTTAATGCCCAGATTATCATCACTTTCACTAGAGAACAACAAGTTATCGGGTATGATACCGACCCAATATGCATTGAAAATGGTGTTTCC

>M6-PGSC0003DMG400003543M6LRR-RLP_1134

GAACAACAAGTTATCGGGTATGATACCGACCCAATATGCATTGAAAATGGTGTTTCCGGGTGAAGGGGTGTCTCAATTTGAGAGGCTACTGTTAGGAGGAAACTATCTATTCGGACCCAT

>M6-PGSC0003DMG400003543M6LRR-RLP_1197

AGGGGTGTCTCAATTTGAGAGGCTACTGTTAGGAGGAAACTATCTATTCGGACCCATACCGGGTCCATTGTTGGATCTGAAAGCGGGTTCAGTCACAGTGAGATTAGGAGACAACTGTTT

>M6-PGSC0003DMG400003543M6LRR-RLP_126

ACATAAAAAAATCCAGAAAATGGGTTGTTTTTCAATTTCTACTGTTCTGTTCTTTATAGTTCTCTGTGCTCTGTTCTTGCATGCGCATTCAACTACTCACTGGCAAGACATTGAAGTCTT

>M6-PGSC0003DMG400003543M6LRR-RLP_1260

TCCATTGTTGGATCTGAAAGCGGGTTCAGTCACAGTGAGATTAGGAGACAACTGTTTGTACCGGTGCCCGTTGAGATTATTTTTATGTGAAGGTGGGGAACAGAAATCATTATCGGAGTG

>M6-PGSC0003DMG400003543M6LRR-RLP_1323

GTGCCCGTTGAGATTATTTTTATGTGAAGGTGGGGAACAGAAATCATTATCGGAGTGTCAGGCTTTTGGTCCCATCATTCCTTGAGTAATGTGATAATGTCTTTACTTTGAGGTCCTCAT

>M6-PGSC0003DMG400003543M6LRR-RLP_1386

TTTTGGTCCCATCATTCCTTGAGTAATGTGATAATGTCTTTACTTTGAGGTCCTCATTTTCATTTTGTACTAGTAATTATTGTGTTGTTTCAATTTTATCTGAAGGGACTAGTAAGTATT

>M6-PGSC0003DMG400003543M6LRR-RLP_1449

TTTGTACTAGTAATTATTGTGTTGTTTCAATTTTATCTGAAGGGACTAGTAAGTATTGTATAAATATATATTTTTTCCTGCAACCTTACCCCATGTATGGGGTTTTAGATATGATCTAGA

>M6-PGSC0003DMG400003543M6LRR-RLP_1512

ATATATATTTTTTCCTGCAACCTTACCCCATGTATGGGGTTTTAGATATGATCTAGATGGATCTATGGTAGAGTTGTTATTGTTGTGAAGAGTGAGACAGAAATATAAGATTTCTGACTT

>M6-PGSC0003DMG400003543M6LRR-RLP_1564

CTAGATGGATCTATGGTAGAGTTGTTATTGTTGTGAAGAGTGAGACAGAAATATAAGATTTCTGACTTAATGTGTTGAAGTAGTAATTGTGGGACAGTTGAGGAAATTTTATTAGCGGAT

>M6-PGSC0003DMG400003543M6LRR-RLP_189

CTGTGCTCTGTTCTTGCATGCGCATTCAACTACTCACTGGCAAGACATTGAAGTCTTGAAGCAGCTGAAAAATAGCGTCGACCCCAATTCAGTGTCACCTGGCTCATGCCTGAATTCATG

>M6-PGSC0003DMG400003543M6LRR-RLP_252

GCTGAAAAATAGCGTCGACCCCAATTCAGTGTCACCTGGCTCATGCCTGAATTCATGGGATTTCTCTGTAGACCCTTGTGATAATCTCGGGGGTGAAAAATTCACTTGTGGTCTCCGTTG

>M6-PGSC0003DMG400003543M6LRR-RLP_315

CTCTGTAGACCCTTGTGATAATCTCGGGGGTGAAAAATTCACTTGTGGTCTCCGTTGCGACCTTGTCGTTTCCTCTGTTAGCCGAGTCACTGAACTCGCTCTAGACCAGTGGAATTACTC

>M6-PGSC0003DMG400003543M6LRR-RLP_378

TGTCGTTTCCTCTGTTAGCCGAGTCACTGAACTCGCTCTAGACCAGTGGAATTACTCCGGTTCAATCACTTCCGTCTCTTGGAATTTGCCTTATCTCCAAAACTTAGACATCACTAATAA

>M6-PGSC0003DMG400003543M6LRR-RLP_441

AATCACTTCCGTCTCTTGGAATTTGCCTTATCTCCAAAACTTAGACATCACTAATAATGCCTTCTCCGGTTTAGTACCTGAGTCACTCTCGAATCTAACTCGGGTTCAACGAGTCGGTTT

>M6-PGSC0003DMG400003543M6LRR-RLP_504

CTCCGGTTTAGTACCTGAGTCACTCTCGAATCTAACTCGGGTTCAACGAGTCGGTTTATCAGGTAACTCGCTATCCGGGTCAATTCCGAGTTCACTGGGTTCACTTTCAAACCTTGAAGA

>M6-PGSC0003DMG400003543M6LRR-RLP_567

TAACTCGCTATCCGGGTCAATTCCGAGTTCACTGGGTTCACTTTCAAACCTTGAAGAGCTTTACCTTGACAATAATTTCCTTGAGGGAGAAATTCCCCAAAGTTTCAACGGTCTCAAGAA

>M6-PGSC0003DMG400003543M6LRR-RLP_63

CAATCTTGAAGGCTCTTCTTCTTCATCAATTCTCATTACGATTAAACAAAATAATATAACCACACATAAAAAAATCCAGAAAATGGGTTGTTTTTCAATTTCTACTGTTCTGTTCTTTAT

>M6-PGSC0003DMG400003543M6LRR-RLP_630

CCTTGACAATAATTTCCTTGAGGGAGAAATTCCCCAAAGTTTCAACGGTCTCAAGAATTTGAAACGGCTCGAATTTCAAGGCAATCAGATTACTGGCGAGTTTCCCGAGTTGGATCAGTT

>M6-PGSC0003DMG400003543M6LRR-RLP_693

ACGGCTCGAATTTCAAGGCAATCAGATTACTGGCGAGTTTCCCGAGTTGGATCAGTTAATTAATTTGAATTTCATCGATGCTAGTGATAATGCCATCTCCGGCGAACTCCCGGCGAGTTA

>M6-PGSC0003DMG400003543M6LRR-RLP_756

TTTGAATTTCATCGATGCTAGTGATAATGCCATCTCCGGCGAACTCCCGGCGAGTTATCCGGCTTCTCTTATCGAGCTTACGATGAGAAACAACAGCATAGAAGGTAACATTCCGGCGAG

>M6-PGSC0003DMG400003543M6LRR-RLP_819

TTCTCTTATCGAGCTTACGATGAGAAACAACAGCATAGAAGGTAACATTCCGGCGAGCCTCATCGGACTGAATTTCGTACAAGTTATCGATCTGAGTCACAACAAACTCAGTGGGTCAGT

>M6-PGSC0003DMG400003543M6LRR-RLP_882

CGGACTGAATTTCGTACAAGTTATCGATCTGAGTCACAACAAACTCAGTGGGTCAGTGCCAGCGAGTCTCTTCACTCACCCATCCCTCGAACAGCTCACCCTATCATACAATCAATTCGG

>M6-PGSC0003DMG400003543M6LRR-RLP_945

GAGTCTCTTCACTCACCCATCCCTCGAACAGCTCACCCTATCATACAATCAATTCGGGTCAGTTCAGGAACCCGGGATCTCGTTTCAGAATAGTCAGCTTATTGCAGCAGACTTAAGCAA

>M6-PGSC0003DMG400003544M6LRR-RLP_0

CAAATATGGGAATTTTGGGTAGGAAAAAATCTTTCTTGTTCATCTTATTTTACATATTTTTTTCTGTTTTTGTTCCAAGGCCATTAGAATGTTGTCATTTGGTTGATAAAACATCTTTGT

>M6-PGSC0003DMG400003544M6LRR-RLP_1008

CCAATAATCTATTTACAGGGAAAATCCCATCAAGTTTTGGTAATCTAAGGAATCTACAGATACTAGATTTATCGAAAAATAAGCTCAGTGGTGAAATACCTCATCAGCTTGTGAAGTTAG

>M6-PGSC0003DMG400003544M6LRR-RLP_1071

TAGATTTATCGAAAAATAAGCTCAGTGGTGAAATACCTCATCAGCTTGTGAAGTTAGTGAGTTTACAGACATTGGATCTGTCATTTAATCCTTTGGGGCTTTCAAGAATACCAAACTGGT

>M6-PGSC0003DMG400003544M6LRR-RLP_1134

TACAGACATTGGATCTGTCATTTAATCCTTTGGGGCTTTCAAGAATACCAAACTGGTTCAAGAAACTAAGACTGTTTCGTCTCCTTTTGGCGAAGACTGGGATCAGAGGGAACCTCCCAA

>M6-PGSC0003DMG400003544M6LRR-RLP_1197

AACTAAGACTGTTTCGTCTCCTTTTGGCGAAGACTGGGATCAGAGGGAACCTCCCAAGTTGGTTGGCTTCATCGTCATTATCAACACTTGATTTATCAAACAATGGTTTGACAGGAAAAT

>M6-PGSC0003DMG400003544M6LRR-RLP_126

TTAAGCAAAAAGTCACTTCTGATCCTTCAAACTTATTGCAAACATGGACACTTTCAACTGATTGTTGCAAATCTTGGGAAGGTATTGCTTGTAATTCCAATGGAAGGGTTGTTAACGTCT

>M6-PGSC0003DMG400003544M6LRR-RLP_1260

TGGCTTCATCGTCATTATCAACACTTGATTTATCAAACAATGGTTTGACAGGAAAATTACCTACATGGATTGTTAACATGACCAACCTTTCACTCTTGAATTTATCAAACAACGCCTTGT

>M6-PGSC0003DMG400003544M6LRR-RLP_1323

CATGGATTGTTAACATGACCAACCTTTCACTCTTGAATTTATCAAACAACGCCTTGTATTCATCAATCCCTGATGAGTTCAAGAACCTGACACTTTTGATGGATCTTGATCTTCACTCAA

>M6-PGSC0003DMG400003544M6LRR-RLP_1386

CAATCCCTGATGAGTTCAAGAACCTGACACTTTTGATGGATCTTGATCTTCACTCAAACAGATTTTCAGGCAACTTGAGAGCAATTTTCTCAAAGAATTTTCAAGACCCCCTTGGTCACT

>M6-PGSC0003DMG400003544M6LRR-RLP_1449

TTTCAGGCAACTTGAGAGCAATTTTCTCAAAGAATTTTCAAGACCCCCTTGGTCACTACAATTCTATTGATCTTGCATACAACATGTTCACTGGACCATTAGACGAAAACACCGGAAATG

>M6-PGSC0003DMG400003544M6LRR-RLP_1512

CTATTGATCTTGCATACAACATGTTCACTGGACCATTAGACGAAAACACCGGAAATGAACCTGTCATGGATTCTATTTTGTCACTAAGTTTGTCCCACAATCCATTGACAGGACACATAC

>M6-PGSC0003DMG400003544M6LRR-RLP_1575

TCATGGATTCTATTTTGTCACTAAGTTTGTCCCACAATCCATTGACAGGACACATACCAAAATCACTGGGAAACTTGACAAGCTTGCAGGTAATTAAACTAGCCGAAAATGGGCTAACAG

>M6-PGSC0003DMG400003544M6LRR-RLP_1638

CACTGGGAAACTTGACAAGCTTGCAGGTAATTAAACTAGCCGAAAATGGGCTAACAGGTGGAATTCCAACAGAACTTGGGAATGCCAAGGAACTGAAAACAATTTTACTTTCAAATAATA

>M6-PGSC0003DMG400003544M6LRR-RLP_1701

TTCCAACAGAACTTGGGAATGCCAAGGAACTGAAAACAATTTTACTTTCAAATAATAATTTGGAAGGTGCAATACCAGCACAAGTTCTGAATTTGAAAGACCTTCAAGAATTTGATGTAT

>M6-PGSC0003DMG400003544M6LRR-RLP_1764

AAGGTGCAATACCAGCACAAGTTCTGAATTTGAAAGACCTTCAAGAATTTGATGTATCGGGAAATCGACTAAGCGGTAGTATCCCTCCTCATAAAGCTAATATCCCAAAGTCTGCATTTT

>M6-PGSC0003DMG400003544M6LRR-RLP_1827

ATCGACTAAGCGGTAGTATCCCTCCTCATAAAGCTAATATCCCAAAGTCTGCATTTTTGGGAAATTATGGCTTGTGTGGAGCTCCACTTCCTCCTTGTAAGCACACATAGTGGAAACCTT

>M6-PGSC0003DMG400003544M6LRR-RLP_189

GTTGCAAATCTTGGGAAGGTATTGCTTGTAATTCCAATGGAAGGGTTGTTAACGTCTCTCGCCCCGGCCTTTCTTCAGGGGACGATTTCATTATCGATACCTCAATATCTGGAACTCTTT

>M6-PGSC0003DMG400003544M6LRR-RLP_1890

ATTATGGCTTGTGTGGAGCTCCACTTCCTCCTTGTAAGCACACATAGTGGAAACCTTCAAACACAATATTTCCCTTTTCTTTATTTATTTTTCATTTTCGACATGATTGTTCGTCATAAT

>M6-PGSC0003DMG400003544M6LRR-RLP_1953

CAATATTTCCCTTTTCTTTATTTATTTTTCATTTTCGACATGATTGTTCGTCATAATGTATTGTATTTCAGTTGTAATCACTCAAATGTTATTAGTAGATACTGAACAAAGTTGATTTTT

>M6-PGSC0003DMG400003544M6LRR-RLP_2002

GTCATAATGTATTGTATTTCAGTTGTAATCACTCAAATGTTATTAGTAGATACTGAACAAAGTTGATTTTTCAAAAGCAACCTTGCCGAATTCTACATCAATATCCTCAATGGTTTAAAC

>M6-PGSC0003DMG400003544M6LRR-RLP_252

CCGGCCTTTCTTCAGGGGACGATTTCATTATCGATACCTCAATATCTGGAACTCTTTCTCCTTCTCTTGCTAATCTTTCTTTTCTTGAATTGCTTGATCTTAGTAATCTTAAGGACTTAA

>M6-PGSC0003DMG400003544M6LRR-RLP_315

CTCTTGCTAATCTTTCTTTTCTTGAATTGCTTGATCTTAGTAATCTTAAGGACTTAACAGGGCCAATTCCACCTGAATTTGGCAAGTTATCGCGCTTGAGTTACCTCTTTCTTGATACTA

>M6-PGSC0003DMG400003544M6LRR-RLP_378

CAATTCCACCTGAATTTGGCAAGTTATCGCGCTTGAGTTACCTCTTTCTTGATACTAATAAGCTCTCAGGATCTATACCTGTCACATTTAAATATCTATACCAACTTAAGAAGCTTTATC

>M6-PGSC0003DMG400003544M6LRR-RLP_441

TCTCAGGATCTATACCTGTCACATTTAAATATCTATACCAACTTAAGAAGCTTTATCTCAGTGATAATACTCTATCTGGTACTATCCCTTGGAGTATTTTCGAAAGCTTCCTTTCACTCT

>M6-PGSC0003DMG400003544M6LRR-RLP_504

ATAATACTCTATCTGGTACTATCCCTTGGAGTATTTTCGAAAGCTTCCTTTCACTCTCAGAACTTGGTTTATCAGCAAATCAACTTTCAGGTCCAATACCATCTTCAATTGGGAATTTAG

>M6-PGSC0003DMG400003544M6LRR-RLP_567

TTGGTTTATCAGCAAATCAACTTTCAGGTCCAATACCATCTTCAATTGGGAATTTAGTATCATTGACCAAGCTTGATATGAGTCGAAATAAGTTCTTTGGAAGCATTCCAGAGAGTATAG

>M6-PGSC0003DMG400003544M6LRR-RLP_63

CTGTTTTTGTTCCAAGGCCATTAGAATGTTGTCATTTGGTTGATAAAACATCTTTGTTAGACTTTAAGCAAAAAGTCACTTCTGATCCTTCAAACTTATTGCAAACATGGACACTTTCAA

>M6-PGSC0003DMG400003544M6LRR-RLP_630

TGACCAAGCTTGATATGAGTCGAAATAAGTTCTTTGGAAGCATTCCAGAGAGTATAGGAAGATTAAAAAATCTTGCATATGTTGATTTGTCTGAGAATCAGTTATCTGGAAAGATTCCAA

>M6-PGSC0003DMG400003544M6LRR-RLP_693

TAAAAAATCTTGCATATGTTGATTTGTCTGAGAATCAGTTATCTGGAAAGATTCCAAACTCTATTGGAAACCTTTCCCAAGTATCTCTAATGTATCTGAACCAAAACCAATTGAGAGGAA

>M6-PGSC0003DMG400003544M6LRR-RLP_756

TTGGAAACCTTTCCCAAGTATCTCTAATGTATCTGAACCAAAACCAATTGAGAGGAAAAATTCCTTCATCAATATCTGGTCTTAGCTCATTGGTGTTCTGCCGTTTATCCGAAAACCAGC

>M6-PGSC0003DMG400003544M6LRR-RLP_819

CTTCATCAATATCTGGTCTTAGCTCATTGGTGTTCTGCCGTTTATCCGAAAACCAGCTGAGTGGCAGTATTCCACCTTCTATTGGCAATCTTCCAAAGATCGAAAGGCTGATATTTGAGA

>M6-PGSC0003DMG400003544M6LRR-RLP_882

GCAGTATTCCACCTTCTATTGGCAATCTTCCAAAGATCGAAAGGCTGATATTTGAGAACAACAAGTTAAGTGGAAAACTGCCTGCAACACTTGGACATCTTGTAACTCTCACTGATATGT

>M6-PGSC0003DMG400003544M6LRR-RLP_945

AGTTAAGTGGAAAACTGCCTGCAACACTTGGACATCTTGTAACTCTCACTGATATGTACTTTTCCAATAATCTATTTACAGGGAAAATCCCATCAAGTTTTGGTAATCTAAGGAATCTAC

>M6-PGSC0003DMG400003551M6LRR-RLK_0

GTTAAAAAGTGCTGGCCCTTGTTTTGAAAAGTGTTCATTTGCACATTCTCCACTTCCTCCCTCACTCTTCTTCTCTGTGTCACTCTGTGCCTCTGTGCAAAGTGAGGGGGCGTCTGCCAT

>M6-PGSC0003DMG400003551M6LRR-RLK_1054

GTTCTGCAATGTTTCCATTTATCCTCCCTCACTTAGGATTGTTCAGTTGGGTTTCAATGCATTCACGAATATTGTCAAGCAAGAGTCTTCTAAATGTTTTAGTTCTTTGCAAATTTTGGA

>M6-PGSC0003DMG400003551M6LRR-RLK_1116

TCACGAATATTGTCAAGCAAGAGTCTTCTAAATGTTTTAGTTCTTTGCAAATTTTGGATCTTCAACACAACCAAATACATGGTGAATTTCCTTTGATTTTGACGAACATTTCTGCATTGA

>M6-PGSC0003DMG400003551M6LRR-RLK_1178

CAACACAACCAAATACATGGTGAATTTCCTTTGATTTTGACGAACATTTCTGCATTGACGTCGCTGGATGTGTCATGGAATTTGTTCTCTGGAAAAATCCCAAGTGCCATTGGGAACTTG

>M6-PGSC0003DMG400003551M6LRR-RLK_124

ATTAACACTCATTTTCTCCCCTTTATCATGCACCCATTTTAACCTTCACTATAAACCAAGCCAACTCCTTTTTCTTATGTTCTTGATTTCTTCTTCTTTCATGGCTGCTTATCTACTTCT

>M6-PGSC0003DMG400003551M6LRR-RLK_1240

GCTGGATGTGTCATGGAATTTGTTCTCTGGAAAAATCCCAAGTGCCATTGGGAACTTGTGGAGATTGGAGGAACTGAGAATGACAAATAACTCATTTGAAGGTGCTCTTCCGTTTGAGAT

>M6-PGSC0003DMG400003551M6LRR-RLK_1302

GATTGGAGGAACTGAGAATGACAAATAACTCATTTGAAGGTGCTCTTCCGTTTGAGATTACGAACTGCAGTGACTTGAAAGTTCTTGATCTTGAAGGGAATCGAATGACGGGTGACATTC

>M6-PGSC0003DMG400003551M6LRR-RLK_1364

AACTGCAGTGACTTGAAAGTTCTTGATCTTGAAGGGAATCGAATGACGGGTGACATTCCAATGTTTTTAGGTGATCTTAGAAGCTTGAAGACATTGTCACTGGGAAGAAATCAGTTTTCG

>M6-PGSC0003DMG400003551M6LRR-RLK_1426

GTTTTTAGGTGATCTTAGAAGCTTGAAGACATTGTCACTGGGAAGAAATCAGTTTTCGGGTTCCATTCCTTCTAGTTTTAGAAATTTGAGTAATCTCGAAAATCTCAACTTGGGAGGAAA

>M6-PGSC0003DMG400003551M6LRR-RLK_1488

CCATTCCTTCTAGTTTTAGAAATTTGAGTAATCTCGAAAATCTCAACTTGGGAGGAAATGGTCTCAATGGAAGCTTGCCCGAGGAGGTAATGGGTTTGAGCAATTTAAGCACACTGAATC

>M6-PGSC0003DMG400003551M6LRR-RLK_1550

CTCAATGGAAGCTTGCCCGAGGAGGTAATGGGTTTGAGCAATTTAAGCACACTGAATCTCAGTGGAAACAAGTTTTCTGGCAGTATGCCAGTTGATATTGGGAATCTTCAGCAGCTCTCG

>M6-PGSC0003DMG400003551M6LRR-RLK_1612

TGGAAACAAGTTTTCTGGCAGTATGCCAGTTGATATTGGGAATCTTCAGCAGCTCTCGGTTCTGAATCTGAGTAAAAATGGCTTTTCAGGCACTATTCCTTCTAGCATTGGGACTTTGTA

>M6-PGSC0003DMG400003551M6LRR-RLK_1674

TGAATCTGAGTAAAAATGGCTTTTCAGGCACTATTCCTTCTAGCATTGGGACTTTGTATAAGCTAACGGTTGTTGATTTGAGCGGACAGAATTTCTCAGGTGAAATACCATTTGATCTTG

>M6-PGSC0003DMG400003551M6LRR-RLK_1736

CTAACGGTTGTTGATTTGAGCGGACAGAATTTCTCAGGTGAAATACCATTTGATCTTGCTGGTTTGCCTAATCTACAGGTTATAGCTTTGCAAGAAAACAAGTTGTCTGGTAATGTTCCT

>M6-PGSC0003DMG400003551M6LRR-RLK_1798

TTTGCCTAATCTACAGGTTATAGCTTTGCAAGAAAACAAGTTGTCTGGTAATGTTCCTGAAGGTTTCAGTAGCTTATTAGGCATGCAATATTTGAATCTTTCTTCCAATTCCTTTTCTGG

>M6-PGSC0003DMG400003551M6LRR-RLK_186

AACTCCTTTTTCTTATGTTCTTGATTTCTTCTTCTTTCATGGCTGCTTATCTACTTCTGTTTCTTGTTTTTCTCTCTACATTATGCTCTGCTCAGAGAAACTCACAAACCCTTTTGGAAG

>M6-PGSC0003DMG400003551M6LRR-RLK_1860

GTTTCAGTAGCTTATTAGGCATGCAATATTTGAATCTTTCTTCCAATTCCTTTTCTGGCCATATACCATCTACATTTGGGTTCTTGACCTCATTAGTTGTACTTTCTTTGTCTAACAACC

>M6-PGSC0003DMG400003551M6LRR-RLK_1922

ATACCATCTACATTTGGGTTCTTGACCTCATTAGTTGTACTTTCTTTGTCTAACAACCACATAAATGGTTCGATTCCTCCCGACTTGGGCAACTGCTCTGCTCTGGAGAATTTGAATCTT

>M6-PGSC0003DMG400003551M6LRR-RLK_1984

AAATGGTTCGATTCCTCCCGACTTGGGCAACTGCTCTGCTCTGGAGAATTTGAATCTTCACTCAAATTCATTGAGTGGCCAAATACCTGCTGATCTCGGGCGTCTTTCCCACTTGAGTGT

>M6-PGSC0003DMG400003551M6LRR-RLK_2046

CAAATTCATTGAGTGGCCAAATACCTGCTGATCTCGGGCGTCTTTCCCACTTGAGTGTGCTAGATTTGGGTAGAAACAACTTAACAGGTGAAGTCCCAGTGGATATTTCCAATTGTTCAT

>M6-PGSC0003DMG400003551M6LRR-RLK_2108

GATTTGGGTAGAAACAACTTAACAGGTGAAGTCCCAGTGGATATTTCCAATTGTTCATCCTTGACATCACTCGTACTCGACTTGAACCATCTTTCAGGGAACATACCAGAGTCACTATCC

>M6-PGSC0003DMG400003551M6LRR-RLK_2170

GACATCACTCGTACTCGACTTGAACCATCTTTCAGGGAACATACCAGAGTCACTATCCAGGTTATCAAACCTAACTGTCCTTGACCTCTCTACTAACAACTTCACTGGAGAAATCCCAGC

>M6-PGSC0003DMG400003551M6LRR-RLK_2232

TATCAAACCTAACTGTCCTTGACCTCTCTACTAACAACTTCACTGGAGAAATCCCAGCTAATCTTACCATGCTCTCTAGCTTGGTGAGTTTCAATGTGTCAAACAATAATTTGGGCGGCC

>M6-PGSC0003DMG400003551M6LRR-RLK_2294

CTTACCATGCTCTCTAGCTTGGTGAGTTTCAATGTGTCAAACAATAATTTGGGCGGCCAGATTCCGGAGATGTTGGGCTCCCGTTTTAATAACTCATTGGATTATGCTGGCAACCAAGGT

>M6-PGSC0003DMG400003551M6LRR-RLK_2356

TCCGGAGATGTTGGGCTCCCGTTTTAATAACTCATTGGATTATGCTGGCAACCAAGGTTTGTGTGGGGAGCCATTGGAAAGAAGATGTGAGACATCTGGTGATGGTGGGAATAAATTGAT

>M6-PGSC0003DMG400003551M6LRR-RLK_2418

GTGGGGAGCCATTGGAAAGAAGATGTGAGACATCTGGTGATGGTGGGAATAAATTGATTATGTTCATTGCTGTGGCTGCCAGTGGAGCTCTTCTCCTTTTATCATGCTGCTGCTTATATA

>M6-PGSC0003DMG400003551M6LRR-RLK_248

CTTGTTTTTCTCTCTACATTATGCTCTGCTCAGAGAAACTCACAAACCCTTTTGGAAGTTCAAGCGCTTACTTCTTTTAAACTTAGTATTCATGATCCACTCGGTGCACTCACTGACTGG

>M6-PGSC0003DMG400003551M6LRR-RLK_2480

TTCATTGCTGTGGCTGCCAGTGGAGCTCTTCTCCTTTTATCATGCTGCTGCTTATATACTTACAACTTCTTGAGGTGGCGCAGGAAGCTCAAAGAGAAAGCAGCTGGAGAAAAGAAGCAT

>M6-PGSC0003DMG400003551M6LRR-RLK_2542

CAACTTCTTGAGGTGGCGCAGGAAGCTCAAAGAGAAAGCAGCTGGAGAAAAGAAGCATAGTCCTGCAAGGGCTAGTTCAAGGACCAGTGGTGGTCGTGGCAGTGGTGAGAATGGTGGACC

>M6-PGSC0003DMG400003551M6LRR-RLK_2604

CTGCAAGGGCTAGTTCAAGGACCAGTGGTGGTCGTGGCAGTGGTGAGAATGGTGGACCCAAACTTGTTATGTTCAATAACAAAATCACACTCGCTGAAACAATTGAAGCAACTAGAGAAT

>M6-PGSC0003DMG400003551M6LRR-RLK_2666

CTTGTTATGTTCAATAACAAAATCACACTCGCTGAAACAATTGAAGCAACTAGAGAATTTGACGAGGAACATGTGCTAAGCAGAACACATTATGGGGTGGTCTACAAGGCCTGTTACAAT

>M6-PGSC0003DMG400003551M6LRR-RLK_2728

CGAGGAACATGTGCTAAGCAGAACACATTATGGGGTGGTCTACAAGGCCTGTTACAATGATGGAATGGTGCTCTCAATTCGTAGACTTTCAAATGGAACACTAGGTGAGAACATGTTCAG

>M6-PGSC0003DMG400003551M6LRR-RLK_2790

GAATGGTGCTCTCAATTCGTAGACTTTCAAATGGAACACTAGGTGAGAACATGTTCAGAAAAGAGGCAGAATCACTTGGCAGGGTGAAACACAGGAACCTAACAGTTCTGCGTGGGTACT

>M6-PGSC0003DMG400003551M6LRR-RLK_2852

GAGGCAGAATCACTTGGCAGGGTGAAACACAGGAACCTAACAGTTCTGCGTGGGTACTATGCTGGCCCGCCTAACCTCAGACTACTTGTCCATGATTACATGCCTAACGGGAACCTTGCA

>M6-PGSC0003DMG400003551M6LRR-RLK_2914

TGGCCCGCCTAACCTCAGACTACTTGTCCATGATTACATGCCTAACGGGAACCTTGCAACACTGCTCCAAGAAGCATCCCACCAAGATGGTCATGTCCTCAACTGGCCAATGCGCCACCT

>M6-PGSC0003DMG400003551M6LRR-RLK_2976

TGCTCCAAGAAGCATCCCACCAAGATGGTCATGTCCTCAACTGGCCAATGCGCCACCTCATTGCACTTGGCATTGCTCGTGGCCTGGCTTTCCTTCACTCATCCTCAATGGTCCACGGAG

>M6-PGSC0003DMG400003551M6LRR-RLK_3038

GCACTTGGCATTGCTCGTGGCCTGGCTTTCCTTCACTCATCCTCAATGGTCCACGGAGATGTTAAGCCGCAGAATGTCCTATTTGATGCAGACTTTGAAGCCCATCTTTCAGAGTTTGGC

>M6-PGSC0003DMG400003551M6LRR-RLK_310

AGCGCTTACTTCTTTTAAACTTAGTATTCATGATCCACTCGGTGCACTCACTGACTGGGATTCTTCTTCCCCTTTTGCTCCCTGTGATTGGCGAGGTGTTTTCTGCGTCAATGGCCGGGT

>M6-PGSC0003DMG400003551M6LRR-RLK_3100

TAAGCCGCAGAATGTCCTATTTGATGCAGACTTTGAAGCCCATCTTTCAGAGTTTGGCCTAGGCAAGCTTGTAGTAGCCACACCAACCGAGCCCTCCACATCAACTTCAGTTGGTACCCT

>M6-PGSC0003DMG400003551M6LRR-RLK_3162

GCAAGCTTGTAGTAGCCACACCAACCGAGCCCTCCACATCAACTTCAGTTGGTACCCTAGGCTATATATCTCCAGAAGTAGCATTAACCGGAGAAACCACAAGAGAATCAGATGCTTATA

>M6-PGSC0003DMG400003551M6LRR-RLK_3224

TATATATCTCCAGAAGTAGCATTAACCGGAGAAACCACAAGAGAATCAGATGCTTATAGCTTTGGCATTGTTTTGCTGGAACTGCTAACAGGCAAAAGACCCTTAATGTTCACACAAGAT

>M6-PGSC0003DMG400003551M6LRR-RLK_3286

TGGCATTGTTTTGCTGGAACTGCTAACAGGCAAAAGACCCTTAATGTTCACACAAGATGAGGACATAGTGAAATGGGTGAAGAGGCAATTGCAAAGAGGGCAAATTTCCGAACTATTAGA

>M6-PGSC0003DMG400003551M6LRR-RLK_3348

ACATAGTGAAATGGGTGAAGAGGCAATTGCAAAGAGGGCAAATTTCCGAACTATTAGAGCCAGGATTGCTCGAACTGGACCCTGAATCATCAGAATGGGAAGAGTTCTTGCTAGGAATTA

>M6-PGSC0003DMG400003551M6LRR-RLK_3410

GGATTGCTCGAACTGGACCCTGAATCATCAGAATGGGAAGAGTTCTTGCTAGGAATTAAAGTAGGCTTGCTATGCACAGCACCTGACCCACTTGATCGTCCCACCATGGCCGACATTGTG

>M6-PGSC0003DMG400003551M6LRR-RLK_3534

TGCTCGAAGGCTGTCGTGTGGGACCTGATATCGCGTCTTCAGCTGATCCCACATGCCAACCTTCACCCGCCTGAATCCCCCGAGGTTCATTCTAATTCTTTTCCCTTTTTTATTATTAGC

>M6-PGSC0003DMG400003551M6LRR-RLK_3596

TCACCCGCCTGAATCCCCCGAGGTTCATTCTAATTCTTTTCCCTTTTTTATTATTAGCTCTGTATAAGTTGTGAGAGTAAATTTGCAATGCAATGCAACGCAGCAGTAGCTAGCTAGCTA

>M6-PGSC0003DMG400003551M6LRR-RLK_3658

TATAAGTTGTGAGAGTAAATTTGCAATGCAATGCAACGCAGCAGTAGCTAGCTAGCTAGCAACTTAGTTCGTTCAATTTCATGATTTTTTTTTTCTTGTAGATTTTATCAAGGTAAAAGA

>M6-PGSC0003DMG400003551M6LRR-RLK_372

CTTCTTCCCCTTTTGCTCCCTGTGATTGGCGAGGTGTTTTCTGCGTCAATGGCCGGGTCGGTGAACTTCGTCTCCCTCATTTGCAACTCAGTGGGCCACTCACTACCCAAATTGGTAACC

>M6-PGSC0003DMG400003551M6LRR-RLK_3720

CTTAGTTCGTTCAATTTCATGATTTTTTTTTTCTTGTAGATTTTATCAAGGTAAAAGAGAAACAATATTTCTCTTTTATTGATTCTTGTTTTATGCTATTATTCAAACACCTCACTCCCC

>M6-PGSC0003DMG400003551M6LRR-RLK_3782

CAATATTTCTCTTTTATTGATTCTTGTTTTATGCTATTATTCAAACACCTCACTCCCCATGGTGCCGGGATTTTTCTCCTTTGGCGTGGGGACCAAAATGTAGTGCTATCAATAAGTCAA

>M6-PGSC0003DMG400003551M6LRR-RLK_434

GAACTTCGTCTCCCTCATTTGCAACTCAGTGGGCCACTCACTACCCAAATTGGTAACCTGCGCATGCTGCGCAAGTTAAGCCTCCGGTCCAACTTCTTTAATGGCACTGTTCCAGCTTCG

>M6-PGSC0003DMG400003551M6LRR-RLK_496

CATGCTGCGCAAGTTAAGCCTCCGGTCCAACTTCTTTAATGGCACTGTTCCAGCTTCGTTATCCAAATGTACCCTTTTGCATTCTGTTTTTTTACAGGGTAACGCATTTTCCGGTAATCT

>M6-PGSC0003DMG400003551M6LRR-RLK_558

CCAAATGTACCCTTTTGCATTCTGTTTTTTTACAGGGTAACGCATTTTCCGGTAATCTTCCACCGGAAATATTCAACCTTACTGATTTACAAATCTTTAAAGTTGCCGGAAATCAGTTGT

>M6-PGSC0003DMG400003551M6LRR-RLK_62

CACTCTTCTTCTCTGTGTCACTCTGTGCCTCTGTGCAAAGTGAGGGGGCGTCTGCCATGGCCATTAACACTCATTTTCTCCCCTTTATCATGCACCCATTTTAACCTTCACTATAAACCA

>M6-PGSC0003DMG400003551M6LRR-RLK_620

CCGGAAATATTCAACCTTACTGATTTACAAATCTTTAAAGTTGCCGGAAATCAGTTGTCTGGTGAAATCCCCGGAGAGCTTCCTCAGAGTCTACGATACTTTGATCTTTCGTCAAACTTG

>M6-PGSC0003DMG400003551M6LRR-RLK_682

TGAAATCCCCGGAGAGCTTCCTCAGAGTCTACGATACTTTGATCTTTCGTCAAACTTGTTTTCAGGAGACATTCCGAGGAAATTTTCTGACCGGTCTCAGTTACTTCTGATTAATCTTTC

>M6-PGSC0003DMG400003551M6LRR-RLK_744

CAGGAGACATTCCGAGGAAATTTTCTGACCGGTCTCAGTTACTTCTGATTAATCTTTCTTATAATCGCTTTTCCGGCGAAATTCCGGCGAGTCTCGGCCGGCTTCAACAGCTTCAGTACC

>M6-PGSC0003DMG400003551M6LRR-RLK_806

AATCGCTTTTCCGGCGAAATTCCGGCGAGTCTCGGCCGGCTTCAACAGCTTCAGTACCTGTGGCTTGCGTACAACAACTTGGTAGGAACTTTGCCCTCTGCAATTGCTAACTGTTCATCG

>M6-PGSC0003DMG400003551M6LRR-RLK_868

GCTTGCGTACAACAACTTGGTAGGAACTTTGCCCTCTGCAATTGCTAACTGTTCATCGTTGGTTCACTTGAGTGCTGAAGGAAATGCTATAGGGGGTCTTATTCCGGCAGCAATTGCTGC

>M6-PGSC0003DMG400003551M6LRR-RLK_930

TTCACTTGAGTGCTGAAGGAAATGCTATAGGGGGTCTTATTCCGGCAGCAATTGCTGCTTTACCGAAGCTTCAGGTGATATCTTTATCACATAATAATCTCTCTGGTTCCTTACCAGCTT

>M6-PGSC0003DMG400003551M6LRR-RLK_992

CCGAAGCTTCAGGTGATATCTTTATCACATAATAATCTCTCTGGTTCCTTACCAGCTTCATTGTTCTGCAATGTTTCCATTTATCCTCCCTCACTTAGGATTGTTCAGTTGGGTTTCAAT

>M6-PGSC0003DMG400003659-(rev)M6OtherRLKTM_0

CAAGACCTAGCTTAGAACGAGGAAGCATCGATGAAATAATAGATGCTAATCTCTTAATGGAGGAATGCAACATGGAAATGATGCTGAAGATGGGACAACTAGGACTGAAATGTGTGGTCA

>M6-PGSC0003DMG400003659-(rev)M6OtherRLKTM_126

CCAAAGAAAGGCCTACCATGACTCAAGTGTGGCAAGAACTGGAGACTGCCTTCTTTTCTGCAGAAAATATCATAGATAAACAACCTTCAAATGGTTCCTTCAAATTGTCAAACCGATCAC

>M6-PGSC0003DMG400003659-(rev)M6OtherRLKTM_189

AAAATATCATAGATAAACAACCTTCAAATGGTTCCTTCAAATTGTCAAACCGATCACTCGTAAGGGGAAGCCATAGAATGAAACAACTTGATTATTCTCAAAGCTCAGTTAGCATAGACG

>M6-PGSC0003DMG400003659-(rev)M6OtherRLKTM_252

GGGGAAGCCATAGAATGAAACAACTTGATTATTCTCAAAGCTCAGTTAGCATAGACGGCATTAGATTGCAGAAATTTTACATGGACATAGATGGCCTATCCTTTGGGAGTGCAAAGTTGA

>M6-PGSC0003DMG400003659-(rev)M6OtherRLKTM_315

GATTGCAGAAATTTTACATGGACATAGATGGCCTATCCTTTGGGAGTGCAAAGTTGAGATGCTTAGATACCCATAGTATAAGCTTTGAAGATGACTCATTTGAACTGAGAGGAACTTCCG

>M6-PGSC0003DMG400003659-(rev)M6OtherRLKTM_378

TAGATACCCATAGTATAAGCTTTGAAGATGACTCATTTGAACTGAGAGGAACTTCCGAGGAAACAAGCACAAGTGGAAACGAGGAGCATAGGCTTAACATATAATATCAGTTAAAGTTTG

>M6-PGSC0003DMG400003659-(rev)M6OtherRLKTM_441

CAAGCACAAGTGGAAACGAGGAGCATAGGCTTAACATATAATATCAGTTAAAGTTTGATTAATGTAAGTCGGCAAAGAACATATTTTAGGTTCTCTCTATTGGGAGTTTCTCCATTCTCA

>M6-PGSC0003DMG400003659-(rev)M6OtherRLKTM_504

GTAAGTCGGCAAAGAACATATTTTAGGTTCTCTCTATTGGGAGTTTCTCCATTCTCAAGCCTCAAACCCGAGACCTTTGGTTAAAAGGTTTATAAAAACGGTACCTACCTTCTTTGTATT

>M6-PGSC0003DMG400003659-(rev)M6OtherRLKTM_63

AATGCAACATGGAAATGATGCTGAAGATGGGACAACTAGGACTGAAATGTGTGGTCAAAGTTCCCAAAGAAAGGCCTACCATGACTCAAGTGTGGCAAGAACTGGAGACTGCCTTCTTTT

>M6-PGSC0003DMG400003660-(rev)M6OtherRLKTM_0

CTTATCTACGCCTTCCACCACAGAATTCGGCCCCACCAAATGGATAGCATGTAACTAACGAAATCCACATTAGATACTGTCCTTCCCGTTTCCGTTCATCCTAAATGCCGCCGTGTCTCT

>M6-PGSC0003DMG400003660-(rev)M6OtherRLKTM_1024

TGTCAAGTTATTGGCGGGAGAAAAAACCGGTGAGGTTCGGCAACCGGTTCGGAGCACAGTATGTTTGTATTTTTCAGCTGACCAAGGCAAAGTTCCGGTGCAGGATCCGGTTCTTCTTCT

>M6-PGSC0003DMG400003660-(rev)M6OtherRLKTM_1088

TTGTATTTTTCAGCTGACCAAGGCAAAGTTCCGGTGCAGGATCCGGTTCTTCTTCTTAACGGGTCGGGTAAAGGAATCGTCAATAACATGTTCTTCGCGACCAATGTGGCTCCCTCTTAT

>M6-PGSC0003DMG400003660-(rev)M6OtherRLKTM_1152

CGGGTAAAGGAATCGTCAATAACATGTTCTTCGCGACCAATGTGGCTCCCTCTTATGCTCCGGTAGGGAAGGCACTGGTTTCTGTCACGCTTGTGGGGATTTATGGTGACGTGGCAGATG

>M6-PGSC0003DMG400003660-(rev)M6OtherRLKTM_1216

AGGGAAGGCACTGGTTTCTGTCACGCTTGTGGGGATTTATGGTGACGTGGCAGATGAGGATTTGGTGGATCAGGTCGTGAAGGAGCTATCGGGTTGGTTTGGGGAGTCGGTAGTTGGGTC

>M6-PGSC0003DMG400003660-(rev)M6OtherRLKTM_128

GCCACTCTCCCTCTCCCTTCCCAAACTCTTCTCCTCTCCACGCGCCGCCGCTCCCCACTACTAAAACCGCCGTATGCTTCCTCAAATATATCCGTTGACTCAACCTCTCATCCTAATACC

>M6-PGSC0003DMG400003660-(rev)M6OtherRLKTM_1280

GTGGATCAGGTCGTGAAGGAGCTATCGGGTTGGTTTGGGGAGTCGGTAGTTGGGTCATGGAGTTACTTGAGGATGTACAGGATCGGGTTTGCCCAACCGAACCAATGCCCACCCACTAAC

>M6-PGSC0003DMG400003660-(rev)M6OtherRLKTM_1344

ACTTGAGGATGTACAGGATCGGGTTTGCCCAACCGAACCAATGCCCACCCACTAACTTGAAGAAGAACCCGAAAATAAAACCGGGCTTGTACATTTGTGGAGATTATGTGACTAGTGCTA

>M6-PGSC0003DMG400003660-(rev)M6OtherRLKTM_1408

GAACCCGAAAATAAAACCGGGCTTGTACATTTGTGGAGATTATGTGACTAGTGCTACTTTTGATGGAGCTTTGGTTTCTGGGAAAATAGCAGCAGAAACTTTGCTACAAGATAGAGCTCT

>M6-PGSC0003DMG400003660-(rev)M6OtherRLKTM_1472

GGAGCTTTGGTTTCTGGGAAAATAGCAGCAGAAACTTTGCTACAAGATAGAGCTCTAGTTAATGTATAGCTTAATTTTGGTTGAGTTTTTTTCCAGTTGTGTAAAAGTAAAAGTTTTGAG

>M6-PGSC0003DMG400003660-(rev)M6OtherRLKTM_1536

TATAGCTTAATTTTGGTTGAGTTTTTTTCCAGTTGTGTAAAAGTAAAAGTTTTGAGTGTTCTTGATTAATTTGTATATGGATTCATAAACGTTTCGAGATCATTTTGATGCAATAATTGC

>M6-PGSC0003DMG400003660-(rev)M6OtherRLKTM_1582

AAGTTTTGAGTGTTCTTGATTAATTTGTATATGGATTCATAAACGTTTCGAGATCATTTTGATGCAATAATTGCACTATCAAGAAAGAAGACAATATGGTATGACTTGTTCTGTTGTTCC

>M6-PGSC0003DMG400003660-(rev)M6OtherRLKTM_192

AACCGCCGTATGCTTCCTCAAATATATCCGTTGACTCAACCTCTCATCCTAATACCACCGGGGTCATTGTCATTGGCGGCGGTTTAGCTGGCTTAGCAGCTTCCATTCGGCTTCAAGCCG

>M6-PGSC0003DMG400003660-(rev)M6OtherRLKTM_256

CATTGTCATTGGCGGCGGTTTAGCTGGCTTAGCAGCTTCCATTCGGCTTCAAGCCGACAACATCCCATTTCTCCTTCTCGAATCTTCTGATGCCGTCGGCGGCCGTGTTCGCTCTGATGT

>M6-PGSC0003DMG400003660-(rev)M6OtherRLKTM_320

CCATTTCTCCTTCTCGAATCTTCTGATGCCGTCGGCGGCCGTGTTCGCTCTGATGTAGTTGACGGATATACCCTTGATCGTGGTTTCCAGATTTTCATTACTGGTTACCCTGAAGCCCGA

>M6-PGSC0003DMG400003660-(rev)M6OtherRLKTM_384

GATATACCCTTGATCGTGGTTTCCAGATTTTCATTACTGGTTACCCTGAAGCCCGAAAAATCCTTGATTATGACTCTTTAGACCTCAGAAAATTCTATTCGGGGGCGAGGGTTTACTATG

>M6-PGSC0003DMG400003660-(rev)M6OtherRLKTM_448

TGATTATGACTCTTTAGACCTCAGAAAATTCTATTCGGGGGCGAGGGTTTACTATGGTGGTCGTTTCCACACCGTTGCTGATCCTCTCCGGCATTTCGCCGATGCTCTGCAATCTTTAAC

>M6-PGSC0003DMG400003660-(rev)M6OtherRLKTM_512

TTCCACACCGTTGCTGATCCTCTCCGGCATTTCGCCGATGCTCTGCAATCTTTAACCAACCCAATTGGTTCAGTTGTGGATAAATTACTTATCGGATTGACTAGATTGAAAGTTTTGACG

>M6-PGSC0003DMG400003660-(rev)M6OtherRLKTM_576

TTGGTTCAGTTGTGGATAAATTACTTATCGGATTGACTAGATTGAAAGTTTTGACGCAAGGAGATGATGAAATATTGAGTGCTGAAGAAGTTCCTACGATCGATTTATTGAAGAAGATCG

>M6-PGSC0003DMG400003660-(rev)M6OtherRLKTM_64

CCACATTAGATACTGTCCTTCCCGTTTCCGTTCATCCTAAATGCCGCCGTGTCTCTGCCTTCCCGCCACTCTCCCTCTCCCTTCCCAAACTCTTCTCCTCTCCACGCGCCGCCGCTCCCC

>M6-PGSC0003DMG400003660-(rev)M6OtherRLKTM_640

TGATGAAATATTGAGTGCTGAAGAAGTTCCTACGATCGATTTATTGAAGAAGATCGGGTTTTCTGATGCAATAGTGGAAAGGTTCTTTCGACCCTTTTTTGGTGGAATTTTCTTCGATAG

>M6-PGSC0003DMG400003660-(rev)M6OtherRLKTM_704

GATGCAATAGTGGAAAGGTTCTTTCGACCCTTTTTTGGTGGAATTTTCTTCGATAGAGAGCTGGAAACAACGTCGCGGCTGTTTAATTTCATCTTCAAATGCCTAGCTCTTGGTGACAAC

>M6-PGSC0003DMG400003660-(rev)M6OtherRLKTM_768

AAACAACGTCGCGGCTGTTTAATTTCATCTTCAAATGCCTAGCTCTTGGTGACAACACACTTCCGGCGAAGGGCATTGCCGCCATTCCGGAACAATTGGCGGCAAAATTGCCGTCGAATT

>M6-PGSC0003DMG400003660-(rev)M6OtherRLKTM_832

GGCGAAGGGCATTGCCGCCATTCCGGAACAATTGGCGGCAAAATTGCCGTCGAATTCAATATTGTTCAATACACGAGTTGTTTCCATTGATTCAGGATCGGATTCCAGCACAAAAATAAG

>M6-PGSC0003DMG400003660-(rev)M6OtherRLKTM_896

TTCAATACACGAGTTGTTTCCATTGATTCAGGATCGGATTCCAGCACAAAAATAAGAGTGACATTACAAAATGGAGAAATGTTGGAAAGTGAATTTGGGGTAATATTGGCAGTTGAAGAA

>M6-PGSC0003DMG400003660-(rev)M6OtherRLKTM_960

TACAAAATGGAGAAATGTTGGAAAGTGAATTTGGGGTAATATTGGCAGTTGAAGAACCTGAAGCTGTCAAGTTATTGGCGGGAGAAAAAACCGGTGAGGTTCGGCAACCGGTTCGGAGCA

>M6-PGSC0003DMG400003710M6LRR-RLP_0

ATGAAAGTGCAGTTTTTTCTTCATCTTTTTTCATTTTTCATCTTCTTTGTATTATTAGCACAATGCAAAACTCATCCTATTGATATACAAGGACTTCAAAATCTCAAAAATGGAATAAAC

>M6-PGSC0003DMG400003710M6LRR-RLP_1024

TAAAAGTTGTGGCTCCGAGGAATCATACGGCGTCGTTTGAGAGGTTATTGCTGGGTGGAAATTATTTGTTTGGGCCGATTCCTGGGCCTTTAATGGGCCTAAAGCCTGGTTCTGGAAATG

>M6-PGSC0003DMG400003710M6LRR-RLP_1088

TTTGTTTGGGCCGATTCCTGGGCCTTTAATGGGCCTAAAGCCTGGTTCTGGAAATGTTAGTTTGGTGGATAACTGTTTGTATATGTGTCCAGATTCGGAATACATTTGTCATGGTGGGTA

>M6-PGSC0003DMG400003710M6LRR-RLP_1137

GGAAATGTTAGTTTGGTGGATAACTGTTTGTATATGTGTCCAGATTCGGAATACATTTGTCATGGTGGGTATCAGAAGTCTTTATTGGATTGTAAAAAGTTTGGGCCTAAGATTCCTTGA

>M6-PGSC0003DMG400003710M6LRR-RLP_128

CATTAAACCCGGTTCATGCTTAAACTCATGGAACTTCTCTATAGACCCATGTGATCATATTTTCACCGACCGGTTCACCTGCGGTTTCCGATGTGATCTCATCGTCTCCGGTTTTTATCG

>M6-PGSC0003DMG400003710M6LRR-RLP_192

ACCGACCGGTTCACCTGCGGTTTCCGATGTGATCTCATCGTCTCCGGTTTTTATCGAGTCACTGAAATCAGCCTAGACCAAGCCGGTTACTCCGGTTCATTGCCTAATTCATTCAACCTC

>M6-PGSC0003DMG400003710M6LRR-RLP_256

AAATCAGCCTAGACCAAGCCGGTTACTCCGGTTCATTGCCTAATTCATTCAACCTCCCTCATCTCGAGATTTTAGATATGTCATATAATTCCCTCTCCGGTTCAATCCCGAATTCATTTT

>M6-PGSC0003DMG400003710M6LRR-RLP_320

CGAGATTTTAGATATGTCATATAATTCCCTCTCCGGTTCAATCCCGAATTCATTTTCAAATCTAACTCGGCTCCGGCGGCTCAGTCTTTCGAAAAACTCGTTCACCGGTCTAGTTACTTC

>M6-PGSC0003DMG400003710M6LRR-RLP_384

ACTCGGCTCCGGCGGCTCAGTCTTTCGAAAAACTCGTTCACCGGTCTAGTTACTTCTTCTATCGGTTCTCTATCCCGTTTACAGGAACTATTTCTCGATAACAACAAATTCACCGGTTCA

>M6-PGSC0003DMG400003710M6LRR-RLP_448

GTTCTCTATCCCGTTTACAGGAACTATTTCTCGATAACAACAAATTCACCGGTTCAATCCCGGAAAGTTTTAATGGTTTAATCAATCTGAATCGATTTGAGCTTCAACAAAATAATATTT

>M6-PGSC0003DMG400003710M6LRR-RLP_512

AAGTTTTAATGGTTTAATCAATCTGAATCGATTTGAGCTTCAACAAAATAATATTTCCGGCGAGTTACCCGATTTGAATCAGCTCATAAATCTGATCTACTTAGATTTAAGCAATAACAG

>M6-PGSC0003DMG400003710M6LRR-RLP_576

TTACCCGATTTGAATCAGCTCATAAATCTGATCTACTTAGATTTAAGCAATAACAGAATCGCCGGAAATTTCTTCACGACAAATTTACCTGATTCGATGATTGAACTCTCAATACGCAAC

>M6-PGSC0003DMG400003710M6LRR-RLP_64

GCAAAACTCATCCTATTGATATACAAGGACTTCAAAATCTCAAAAATGGAATAAACCCAAAATCCATTAAACCCGGTTCATGCTTAAACTCATGGAACTTCTCTATAGACCCATGTGATC

>M6-PGSC0003DMG400003710M6LRR-RLP_640

GAAATTTCTTCACGACAAATTTACCTGATTCGATGATTGAACTCTCAATACGCAACAATTATTTACACGGCGAGTTTCCGTTGAACTTCGAAGAGTTGAAGTTCCTACAGGTGTTAGATC

>M6-PGSC0003DMG400003710M6LRR-RLP_704

ACACGGCGAGTTTCCGTTGAACTTCGAAGAGTTGAAGTTCCTACAGGTGTTAGATCTGAGCCATAATTTGTTTTCCGGGATGATTCCGGCGGCGCTATTTCATCACTCATCTCTTCAGCA

>M6-PGSC0003DMG400003710M6LRR-RLP_768

AATTTGTTTTCCGGGATGATTCCGGCGGCGCTATTTCATCACTCATCTCTTCAGCAGCTTACACTTTCTTACAATAACTTTACGTTTTTGCAAGTGCCGGAAGATTTAGTGCTTCGAAGT

>M6-PGSC0003DMG400003710M6LRR-RLP_832

TTTCTTACAATAACTTTACGTTTTTGCAAGTGCCGGAAGATTTAGTGCTTCGAAGTAAGCTTATAGCTATTGATCTGAGTTACAACAATTTGCATGGCTTATTGCCGGCGTTTATGGCGT

>M6-PGSC0003DMG400003710M6LRR-RLP_896

AGCTATTGATCTGAGTTACAACAATTTGCATGGCTTATTGCCGGCGTTTATGGCGTCGATGCCGGAGCTGTCGGCTCTGAATTTGGAGCATAACAAGTTTACCGGCGTGATACCGATACA

>M6-PGSC0003DMG400003710M6LRR-RLP_960

GAGCTGTCGGCTCTGAATTTGGAGCATAACAAGTTTACCGGCGTGATACCGATACAGTTTGCAGTAAAAGTTGTGGCTCCGAGGAATCATACGGCGTCGTTTGAGAGGTTATTGCTGGGT

>M6-PGSC0003DMG400003714M6LRR-RLK_0

GCTCAACACATGGCGTTAGTCTTTGTCAGTGCTGCTTCTTCTTCTCCTTCTGAAACTGAAACTTTTTTTTTTTTCATCTCATACGCTCCTCTCTTCCCCACTTCCATAACAAACTCCATT

>M6-PGSC0003DMG400003714M6LRR-RLK_1037

CTGGTGTGGTCCCTGCATCGTTTTTCTGTAATGGGTCAGTAAATGCTCATGCTATTAGGATCATTGAGCTGGGTTTCAATGCAATCACGGGTTTAACTAAACCTGGGAATGCAACATGTT

>M6-PGSC0003DMG400003714M6LRR-RLK_1098

CATTGAGCTGGGTTTCAATGCAATCACGGGTTTAACTAAACCTGGGAATGCAACATGTTTGAGTGCTTTAGAGGTCTTGACCCTTCACGGGAATCGCATTAATGGTGTTTTTCCTGAATG

>M6-PGSC0003DMG400003714M6LRR-RLK_1159

AGTGCTTTAGAGGTCTTGACCCTTCACGGGAATCGCATTAATGGTGTTTTTCCTGAATGGTTGACGAATTTTTCGTCGTTGAAGGTTCTTGATATCTCCGGGAATGCAATTTCTGGAACT

>M6-PGSC0003DMG400003714M6LRR-RLK_122

CACTCACCATTCTTGATTCTTCCCCCATTTGATCTTCTTCTTCTTCTTCTTCTCTCTACCCTTACCCACCACCACTCTTTCTCATGTCAACGGCTACTTTTCTTCTCTCTCCGCCATTGG

>M6-PGSC0003DMG400003714M6LRR-RLK_1220

TGACGAATTTTTCGTCGTTGAAGGTTCTTGATATCTCCGGGAATGCAATTTCTGGAACTCTGCCAAGTGATTTAGGGAATTTAAGGTTTCTTGAGGAGCTTAGACTGGGTAACAATACAT

>M6-PGSC0003DMG400003714M6LRR-RLK_1281

GCCAAGTGATTTAGGGAATTTAAGGTTTCTTGAGGAGCTTAGACTGGGTAACAATACATTAACTGGTGATATTCCTTCTAGTATTGTGAACTTTGGTTCATTGGGGGTACTTGATTTAGG

>M6-PGSC0003DMG400003714M6LRR-RLK_1342

ACTGGTGATATTCCTTCTAGTATTGTGAACTTTGGTTCATTGGGGGTACTTGATTTAGGTGGAAATCGGTTTTCTGGATTGATTCCACAGTTCTTGGGTAAATTGACAGGGTTGAGAATG

>M6-PGSC0003DMG400003714M6LRR-RLK_1403

GAAATCGGTTTTCTGGATTGATTCCACAGTTCTTGGGTAAATTGACAGGGTTGAGAATGCTGTCTCTTATTGGGAATCGCTTTTCTGGTTCGATTCCGTATAGTTTGGGTAGTCTTTATC

>M6-PGSC0003DMG400003714M6LRR-RLK_1464

GTCTCTTATTGGGAATCGCTTTTCTGGTTCGATTCCGTATAGTTTGGGTAGTCTTTATCAGCTTGAGTTTTTTGATTTGAGCATGAATGTTTTGAATGGGAGTTTGCCTCAAAACCTTAT

>M6-PGSC0003DMG400003714M6LRR-RLK_1525

CTTGAGTTTTTTGATTTGAGCATGAATGTTTTGAATGGGAGTTTGCCTCAAAACCTTATGTTGCTTAGCAATTTGACTAGTTTGAATTTGTCCAGCAACTTGTTTTATGGTGAAATTCCA

>M6-PGSC0003DMG400003714M6LRR-RLK_1586

TGCTTAGCAATTTGACTAGTTTGAATTTGTCCAGCAACTTGTTTTATGGTGAAATTCCAAGAGGGATTGGAAGTTTGCATGGGTTAGAGGTTTTAAATGTTAGCAATTGTGGGTTCTCAG

>M6-PGSC0003DMG400003714M6LRR-RLK_1647

AGGGATTGGAAGTTTGCATGGGTTAGAGGTTTTAAATGTTAGCAATTGTGGGTTCTCAGGGAACATTCCGACTAGTATCGGAAGTTTGTTGAGGTTGACAACTCTTGATGTGAGCAAGCA

>M6-PGSC0003DMG400003714M6LRR-RLK_1708

AACATTCCGACTAGTATCGGAAGTTTGTTGAGGTTGACAACTCTTGATGTGAGCAAGCAAAACCTATCTGGTGAATTACCCTTTGACATTTTCGGGTTGCCAAGTCTACAAGTTGTGGCT

>M6-PGSC0003DMG400003714M6LRR-RLK_1769

ACCTATCTGGTGAATTACCCTTTGACATTTTCGGGTTGCCAAGTCTACAAGTTGTGGCTTTACAAGAGAACATGTTAACTGGGGATGGTCTTGAAGGTTTTAGCAGTTTATCTTGTCTGG

>M6-PGSC0003DMG400003714M6LRR-RLK_183

TTACCCACCACCACTCTTTCTCATGTCAACGGCTACTTTTCTTCTCTCTCCGCCATTGGTGGTGGTGGCACTTACATTGCTATTTATCAGTGTTTTCTCCATGGCTGAAAACGTTACAGA

>M6-PGSC0003DMG400003714M6LRR-RLK_1830

ACAAGAGAACATGTTAACTGGGGATGGTCTTGAAGGTTTTAGCAGTTTATCTTGTCTGGAATATTTGAATCTGTCATCTAATGCCTTTTATGGGCAAGTTCCGAAGACATATGGTTTCCT

>M6-PGSC0003DMG400003714M6LRR-RLK_1891

TATTTGAATCTGTCATCTAATGCCTTTTATGGGCAAGTTCCGAAGACATATGGTTTCCTTACATCCTTGAAAGTTCTTTCTATGTCTAAAAGTGGCATTAATGGCTCAATTCCAGCAGAG

>M6-PGSC0003DMG400003714M6LRR-RLK_1952

CATCCTTGAAAGTTCTTTCTATGTCTAAAAGTGGCATTAATGGCTCAATTCCAGCAGAGTTGGGTAACTGTTCCGGCCTTCAAGTGCTTGAGCTTCGAGGAAATAAGTTAACAGGTCAAA

>M6-PGSC0003DMG400003714M6LRR-RLK_2013

GGGTAACTGTTCCGGCCTTCAAGTGCTTGAGCTTCGAGGAAATAAGTTAACAGGTCAAATTCCGAAGGATTTCTCACATTTATCTCATCTAAGGATACTTGATTTGGGCCACAATGGGTT

>M6-PGSC0003DMG400003714M6LRR-RLK_2074

CCGAAGGATTTCTCACATTTATCTCATCTAAGGATACTTGATTTGGGCCACAATGGGTTAACCGGGGAAATCCCTGAAAACATCTCTAATTGTTTGTCTTTGGCTACTCTCTTGTTGGAT

>M6-PGSC0003DMG400003714M6LRR-RLK_2135

CCGGGGAAATCCCTGAAAACATCTCTAATTGTTTGTCTTTGGCTACTCTCTTGTTGGATTCTAATAATATCTCAGGTCACATACCAGAGTCGTTGTCGAAATTATCAAACTTGGAAATGC

>M6-PGSC0003DMG400003714M6LRR-RLK_2196

TAATAATATCTCAGGTCACATACCAGAGTCGTTGTCGAAATTATCAAACTTGGAAATGCTGGATCTCTCCTCAAATAATTTGAATGGATCAATTCCTTCAAGTCTGTCCCTGATTTCCAG

>M6-PGSC0003DMG400003714M6LRR-RLK_2257

GATCTCTCCTCAAATAATTTGAATGGATCAATTCCTTCAAGTCTGTCCCTGATTTCCAGTCTGAAGTATTTGAATATTTCCCATAATCATCTCGAGGGTGAGATTCCAGAGGCGTTGGGT

>M6-PGSC0003DMG400003714M6LRR-RLK_2318

TGAAGTATTTGAATATTTCCCATAATCATCTCGAGGGTGAGATTCCAGAGGCGTTGGGTTTTCGGTTCAAGGACCCTTCTTTGTTTGCTGCGAATAATGCTTTGTGTGGAAAGCCATTGA

>M6-PGSC0003DMG400003714M6LRR-RLK_2379

TCGGTTCAAGGACCCTTCTTTGTTTGCTGCGAATAATGCTTTGTGTGGAAAGCCATTGAAGGAGTGCAATGACGGTAAAAGGAAAAGAAAGAAGTTGATCTTATTCATTGTTTTGGCTGC

>M6-PGSC0003DMG400003714M6LRR-RLK_244

GTGGTGGCACTTACATTGCTATTTATCAGTGTTTTCTCCATGGCTGAAAACGTTACAGAGACGGAAATGGCAGCTTTAATGTCGTTCAAAAGGAATTTGGAAGACCCACTTGGGGTATTA

>M6-PGSC0003DMG400003714M6LRR-RLK_2440

GAGTGCAATGACGGTAAAAGGAAAAGAAAGAAGTTGATCTTATTCATTGTTTTGGCTGCTGTAGGGGCCTTTTTCGTTGCAGCGTGTTGTTGTGGGTACATCTACAGCCTTATACTTTGG

>M6-PGSC0003DMG400003714M6LRR-RLK_2501

TAGGGGCCTTTTTCGTTGCAGCGTGTTGTTGTGGGTACATCTACAGCCTTATACTTTGGCACAAAAAGTTAAGAGGTTCATCTGAAGGGAAGAAGCGAAGCCCAGGCCGTGCGAGTTCAG

>M6-PGSC0003DMG400003714M6LRR-RLK_2562

CAAAAAGTTAAGAGGTTCATCTGAAGGGAAGAAGCGAAGCCCAGGCCGTGCGAGTTCAGGAGGAGAAGGGGTTCGTGGAAGCGGAGAGAATGGAGGTCCAAAGCTTGTTATGTTTAACAA

>M6-PGSC0003DMG400003714M6LRR-RLK_2623

GGAGAAGGGGTTCGTGGAAGCGGAGAGAATGGAGGTCCAAAGCTTGTTATGTTTAACAACAAGATCACTTATGCAGATACTTTAGAAGCGACACGACAATTTGACGAGGAAAATGTGTTA

>M6-PGSC0003DMG400003714M6LRR-RLK_2684

AGATCACTTATGCAGATACTTTAGAAGCGACACGACAATTTGACGAGGAAAATGTGTTAAGCCGTGGAAAGTATGGCCTAGTATTCAAAGCCACTTTTGCAGATGGAATGGTCTTGGCTA

>M6-PGSC0003DMG400003714M6LRR-RLK_2745

CCGTGGAAAGTATGGCCTAGTATTCAAAGCCACTTTTGCAGATGGAATGGTCTTGGCTATCCGCCGCCTTCCTGATACATCAATCGAAGTGAACACATTCCGCAAGGAAGCTGAATCCCT

>M6-PGSC0003DMG400003714M6LRR-RLK_2806

CGCCGCCTTCCTGATACATCAATCGAAGTGAACACATTCCGCAAGGAAGCTGAATCCCTCGGCAAGGTGAAACATCGCAATCTGACTGTTGTTCGTGGATACTATGCAGGACCTCCACCT

>M6-PGSC0003DMG400003714M6LRR-RLK_2867

GCAAGGTGAAACATCGCAATCTGACTGTTGTTCGTGGATACTATGCAGGACCTCCACCTGATGTTCGATTTGTTGTGTACGATTACATGCCTAATGGAAATCTTGCCACACTTTTACAAG

>M6-PGSC0003DMG400003714M6LRR-RLK_2928

TGTTCGATTTGTTGTGTACGATTACATGCCTAATGGAAATCTTGCCACACTTTTACAAGAGGCATCTCATCAGGATGGCCATGTGCTCAATTGGCCAATGCGCCACCTGATTGCTCTTGG

>M6-PGSC0003DMG400003714M6LRR-RLK_2989

GCATCTCATCAGGATGGCCATGTGCTCAATTGGCCAATGCGCCACCTGATTGCTCTTGGCATTGCTCGTGGCTTAGCTTACCTTCATTCAGTTTCATTGATTCATGGGGATGTAAAGCCA

>M6-PGSC0003DMG400003714M6LRR-RLK_305

CGGAAATGGCAGCTTTAATGTCGTTCAAAAGGAATTTGGAAGACCCACTTGGGGTATTAGACGGTTGGGATTTTTCTACCACATTAGCTCCGTGTGATTGGCGTGGGGTTCTGTGTGATG

>M6-PGSC0003DMG400003714M6LRR-RLK_3050

TTGCTCGTGGCTTAGCTTACCTTCATTCAGTTTCATTGATTCATGGGGATGTAAAGCCACAAAATGTATTGTTTGATGCAGATTTTGAAGCTCATCTTTCGGATTTTGGTTTAGATAAAC

>M6-PGSC0003DMG400003714M6LRR-RLK_3111

AAATGTATTGTTTGATGCAGATTTTGAAGCTCATCTTTCGGATTTTGGTTTAGATAAACTAAGCCTAGTAGCAACCCCAGCAGAGACTTCAACATCTTCCATCCCAGTTGGAACTCTAGG

>M6-PGSC0003DMG400003714M6LRR-RLK_3172

AGCCTAGTAGCAACCCCAGCAGAGACTTCAACATCTTCCATCCCAGTTGGAACTCTAGGCTACATAGCACCTGAAGTGGCATTAACTGGACATCCCACAAAAGAAGCTGATGTATACAGT

>M6-PGSC0003DMG400003714M6LRR-RLK_3233

ACATAGCACCTGAAGTGGCATTAACTGGACATCCCACAAAAGAAGCTGATGTATACAGTTTTGGGATTGTATTGTTAGAGATACTAACAGGAAGAAAGCCAGTAATGTTCAATGGGGACG

>M6-PGSC0003DMG400003714M6LRR-RLK_3294

TGGGATTGTATTGTTAGAGATACTAACAGGAAGAAAGCCAGTAATGTTCAATGGGGACGAGGACATTGTCAAGTGGGTGAAGAGACAACTTCAAAGAGGCCAAATTTCAGAATTGCTCGA

>M6-PGSC0003DMG400003714M6LRR-RLK_3355

GACATTGTCAAGTGGGTGAAGAGACAACTTCAAAGAGGCCAAATTTCAGAATTGCTCGAGCCGGGGTTGCTTGAACTGGATCCTGAATCATCAGAATGGGAAGAGTTCTTGTTGGGAATC

>M6-PGSC0003DMG400003714M6LRR-RLK_3416

CGGGGTTGCTTGAACTGGATCCTGAATCATCAGAATGGGAAGAGTTCTTGTTGGGAATCAAAGTTGGTTTGCTTTGCACCATGCCTGATCCTCTTGAAAGACCATCGATGACCGATATTG

>M6-PGSC0003DMG400003714M6LRR-RLK_3477

AGTTGGTTTGCTTTGCACCATGCCTGATCCTCTTGAAAGACCATCGATGACCGATATTGTATTCATGCTCGAAGGCTGCAGAGTTGGTCCTGACATTCCTTCTTCAGCTGATCCTACAAC

>M6-PGSC0003DMG400003714M6LRR-RLK_3538

TTCATGCTCGAAGGCTGCAGAGTTGGTCCTGACATTCCTTCTTCAGCTGATCCTACAACTCTTCCTTCACCTATGTGATATGAAAATTCAATTCTTTGCAATGAAATCAATTTGCAAGAA

>M6-PGSC0003DMG400003714M6LRR-RLK_3599

TTCCTTCACCTATGTGATATGAAAATTCAATTCTTTGCAATGAAATCAATTTGCAAGAAAATGTCCTTCATTTTTCTCAGTTCAATGTTGTTCATGCATCTCTTCAATGTATTGTTACTC

>M6-PGSC0003DMG400003714M6LRR-RLK_3656

AAAATGTCCTTCATTTTTCTCAGTTCAATGTTGTTCATGCATCTCTTCAATGTATTGTTACTCCATTCATTATCATTTTGCTTGTACATTGTTCATTCAAGAGTTGGTTGGTTTTGTGCA

>M6-PGSC0003DMG400003714M6LRR-RLK_366

CGGTTGGGATTTTTCTACCACATTAGCTCCGTGTGATTGGCGTGGGGTTCTGTGTGATGCCGGCAGAGTTCGTGAACTCCGCCTACCTGGTTTGCAGCTGAGTGGACGGCTAACTGACCA

>M6-PGSC0003DMG400003714M6LRR-RLK_427

GGCAGAGTTCGTGAACTCCGCCTACCTGGTTTGCAGCTGAGTGGACGGCTAACTGACCAGGTTGGTAATTTACGCCAGCTACGGAGGTTAAGCTTACATTCTAACAACTTTAATGGCTCT

>M6-PGSC0003DMG400003714M6LRR-RLK_488

TTGGTAATTTACGCCAGCTACGGAGGTTAAGCTTACATTCTAACAACTTTAATGGCTCTGTCCCACGCTCTTTAGCTCAATGTGCTTTGCTACGCGCTGTTTACTTCCATTACAACTCTT

>M6-PGSC0003DMG400003714M6LRR-RLK_549

CCCACGCTCTTTAGCTCAATGTGCTTTGCTACGCGCTGTTTACTTCCATTACAACTCTTTTTCCGGTGAACTCCCGCCGGCTATCTCGAACCTCACTAATCTTCAAGTTTTAAACCTTGC

>M6-PGSC0003DMG400003714M6LRR-RLK_61

CTTTTTTTTTTTTCATCTCATACGCTCCTCTCTTCCCCACTTCCATAACAAACTCCATTGACACTCACCATTCTTGATTCTTCCCCCATTTGATCTTCTTCTTCTTCTTCTTCTCTCTAC

>M6-PGSC0003DMG400003714M6LRR-RLK_610

TCCGGTGAACTCCCGCCGGCTATCTCGAACCTCACTAATCTTCAAGTTTTAAACCTTGCTCACAACTTCCTCTCCGGTCATGTCTCTGGTAATGTTCCAGCAAGTCTTCGGTTTCTCGAT

>M6-PGSC0003DMG400003714M6LRR-RLK_671

ACAACTTCCTCTCCGGTCATGTCTCTGGTAATGTTCCAGCAAGTCTTCGGTTTCTCGATCTTTCTTCGAATCTCCTCTCCGGTGGCATTCCGGGTAACTTTTCAGCTGGTTCTCAGCTCG

>M6-PGSC0003DMG400003714M6LRR-RLK_732

TTCTTCGAATCTCCTCTCCGGTGGCATTCCGGGTAACTTTTCAGCTGGTTCTCAGCTCGAACTGTTAAACTTGTCGTTTAACCGGTTCTCCGGTGAAATTCCGGCGAGTATCGGAGCTTT

>M6-PGSC0003DMG400003714M6LRR-RLK_793

CTGTTAAACTTGTCGTTTAACCGGTTCTCCGGTGAAATTCCGGCGAGTATCGGAGCTTTACAGAAGTTAGAGTATCTCTGGCTTGATTCTAACCAACTTTACGGCACTATACCTTCAGCA

>M6-PGSC0003DMG400003714M6LRR-RLK_854

AGAAGTTAGAGTATCTCTGGCTTGATTCTAACCAACTTTACGGCACTATACCTTCAGCAATCTCTAACATTTCCACATTAATACACTTGAGCACTAGTGATAATCATCTTCAAGGACTAA

>M6-PGSC0003DMG400003714M6LRR-RLK_915

CTCTAACATTTCCACATTAATACACTTGAGCACTAGTGATAATCATCTTCAAGGACTAATCCCAGCAACAATCGGTTCACTTTCCAGTTTACAGGTAATCTCATTGTCGCAAAATCAACT

>M6-PGSC0003DMG400003714M6LRR-RLK_976

CCAGCAACAATCGGTTCACTTTCCAGTTTACAGGTAATCTCATTGTCGCAAAATCAACTATCTGGTGTGGTCCCTGCATCGTTTTTCTGTAATGGGTCAGTAAATGCTCATGCTATTAGG

>M6-PGSC0003DMG400003790M6LRR-RLK_0

TTGAAAACCAAATATTACACATGACAACATTTCATACTACCTTAGTTTACATTTTGTATGTGCTACTTCTTTTTTCACTTCCATTAAGTATCACATCTTCAGCAAGAACAGAAGCTGAAT

>M6-PGSC0003DMG400003790M6LRR-RLK_1037

ACTTTCTTAGCTCTAGCAGAAAATGACCTGCAAGGGTCATTGCCTCCATCATTCTCCTCTCTTACTAAGTTATCTGATGTGGGGTTGTCTGATAATAGTCTTTCTGGTGAAATCTCATCG

>M6-PGSC0003DMG400003790M6LRR-RLK_1098

TTACTAAGTTATCTGATGTGGGGTTGTCTGATAATAGTCTTTCTGGTGAAATCTCATCGAATTTCATCACCAATTGGACTGAGTTGACATCTTTGCAGCTTCAAAACAATATGTTTACAG

>M6-PGSC0003DMG400003790M6LRR-RLK_1159

TTTCATCACCAATTGGACTGAGTTGACATCTTTGCAGCTTCAAAACAATATGTTTACAGGGAAAATCCCACCCGAAACCAGTCAATTGACAAACCTTGTATATCTCTACCTCTTTCATAA

>M6-PGSC0003DMG400003790M6LRR-RLK_122

CTTGTCAAATGGAAGAGAAATTTACCTTCTACGTCTTTCTTGGACTCATGGTCCATTTCGAATCTCGAAAACTTGTGCAATTGGACATTTATTGTGTGTAATGATGGTGGAACAATTTCT

>M6-PGSC0003DMG400003790M6LRR-RLK_1220

AAAATCCCACCCGAAACCAGTCAATTGACAAACCTTGTATATCTCTACCTCTTTCATAACAATTTCACTGGTTCCATTCCCTATCAGATTGGGAACTTGCAAAACTTGTTGGAATTAGAC

>M6-PGSC0003DMG400003790M6LRR-RLK_1281

ATTTCACTGGTTCCATTCCCTATCAGATTGGGAACTTGCAAAACTTGTTGGAATTAGACTTTTCTGATAACCAGCTTTCAGGCATAATACCTCCAACCATTGGAAATCTGACCAATCTGA

>M6-PGSC0003DMG400003790M6LRR-RLK_1342

TTCTGATAACCAGCTTTCAGGCATAATACCTCCAACCATTGGAAATCTGACCAATCTGAAGACGTTGCAACTTTTTCGCAACAATCTCAGTGGAACCATTCCTCCTGAGATTGGAAAATT

>M6-PGSC0003DMG400003790M6LRR-RLK_1403

ACGTTGCAACTTTTTCGCAACAATCTCAGTGGAACCATTCCTCCTGAGATTGGAAAATTGATATCCCTTGTAACCATCGACATCAACACCAACCGACTAAGTGGTGAGCTGCCGGACAGC

>M6-PGSC0003DMG400003790M6LRR-RLK_1464

TATCCCTTGTAACCATCGACATCAACACCAACCGACTAAGTGGTGAGCTGCCGGACAGCATTTCTGACCTCAGTGAGCTGAAGATTCTTTCTGTATATACTAATGATTTCTCAGGCATTG

>M6-PGSC0003DMG400003790M6LRR-RLK_1525

TTCTGACCTCAGTGAGCTGAAGATTCTTTCTGTATATACTAATGATTTCTCAGGCATTGTCCCTAAGGACTTTGGAAAGAACAGTCCTCAATTGTCAAGTGCCAGCTTTGCAAATAACAG

>M6-PGSC0003DMG400003790M6LRR-RLK_1586

CCTAAGGACTTTGGAAAGAACAGTCCTCAATTGTCAAGTGCCAGCTTTGCAAATAACAGCTTCACTGGTGAACTACCTGCTGGATTGTGTAGCCCGAATCTTGAGGAGTTGACGATAAAT

>M6-PGSC0003DMG400003790M6LRR-RLK_1647

TCACTGGTGAACTACCTGCTGGATTGTGTAGCCCGAATCTTGAGGAGTTGACGATAAATGGGAACAAATTCAGTGGGAAGTTACCAGATTGCTTGAAGAACTGCACACTGCTAAGACGAG

>M6-PGSC0003DMG400003790M6LRR-RLK_1708

GAACAAATTCAGTGGGAAGTTACCAGATTGCTTGAAGAACTGCACACTGCTAAGACGAGTACGGCTTGAAGGCAACAATTTATCTGGTAATCTAGCTGATGCATTTGGAGTGCACCCGAA

>M6-PGSC0003DMG400003790M6LRR-RLK_1769

CGGCTTGAAGGCAACAATTTATCTGGTAATCTAGCTGATGCATTTGGAGTGCACCCGAATCTTGTTTACCTTTCTCTTAGTGACAACCAACTTTCAGGTGAACTTTCACCTGACTGGGGG

>M6-PGSC0003DMG400003790M6LRR-RLK_183

ATCTCGAAAACTTGTGCAATTGGACATTTATTGTGTGTAATGATGGTGGAACAATTTCTGAGATCAATCTTTCTGATGCAGCCCTCTCAGGCTCACTTGATCACCTTGATTTCACTTCAT

>M6-PGSC0003DMG400003790M6LRR-RLK_1830

TTGTTTACCTTTCTCTTAGTGACAACCAACTTTCAGGTGAACTTTCACCTGACTGGGGGAAATGTGAAAATCTCACAAGTCTAAGAATGGATGGAAACAAATTTTCTGGTGTAATCCCAA

>M6-PGSC0003DMG400003790M6LRR-RLK_1891

ATGTGAAAATCTCACAAGTCTAAGAATGGATGGAAACAAATTTTCTGGTGTAATCCCAAGTGAGCTAGGGAACCTGAGAGCGTTGCGCGTGTTAGCTTTGGAAGGAAATGAATTGACTGG

>M6-PGSC0003DMG400003790M6LRR-RLK_1952

GAGCTAGGGAACCTGAGAGCGTTGCGCGTGTTAGCTTTGGAAGGAAATGAATTGACTGGTGAAATTCCTTCTGAACTGGGGAGGTTAGACCTGCTGTACAATCTCAGCTTGAGCAAAAAC

>M6-PGSC0003DMG400003790M6LRR-RLK_2013

AAATTCCTTCTGAACTGGGGAGGTTAGACCTGCTGTACAATCTCAGCTTGAGCAAAAACAATCTTACTGGAGGCATCCCTCAGTCTGTTGGCAATTTAACTAAGCTCCAGTATCTTGATT

>M6-PGSC0003DMG400003790M6LRR-RLK_2074

TCTTACTGGAGGCATCCCTCAGTCTGTTGGCAATTTAACTAAGCTCCAGTATCTTGATTTGTCAACAAATAAGTTAAGTGGTAACATACCAGTAGATCTTGGGAAGTGTGAGAGCCTGTT

>M6-PGSC0003DMG400003790M6LRR-RLK_2135

TCAACAAATAAGTTAAGTGGTAACATACCAGTAGATCTTGGGAAGTGTGAGAGCCTGTTGAGCTTGAATCTTGGCAACAACTCATTATCAGGTGGGATTCCATCTGACCTTGGAAATTTG

>M6-PGSC0003DMG400003790M6LRR-RLK_2196

GCTTGAATCTTGGCAACAACTCATTATCAGGTGGGATTCCATCTGACCTTGGAAATTTGATGCAGTTGAGTATTCTTGTGGACCTAAGTAGCAATTCGCTAACGGGAACCATTCCACAAA

>M6-PGSC0003DMG400003790M6LRR-RLK_2257

GCAGTTGAGTATTCTTGTGGACCTAAGTAGCAATTCGCTAACGGGAACCATTCCACAAAACTTGGCCAAGCTCACCTCATTAATGCATCTCAACCTCTCACATAACAACCTCTCAGGTAG

>M6-PGSC0003DMG400003790M6LRR-RLK_2318

TTGGCCAAGCTCACCTCATTAATGCATCTCAACCTCTCACATAACAACCTCTCAGGTAGAATTCCTCCAGCATTAGCTCAGATGATCAGTCTTCAGGAGATGGATTTTTCCTACAATGAG

>M6-PGSC0003DMG400003790M6LRR-RLK_2379

TTCCTCCAGCATTAGCTCAGATGATCAGTCTTCAGGAGATGGATTTTTCCTACAATGAGTTTTCTGGTCCAATCCCAACCGACGGAGTATTTCGAGGAGCATCCGCTAGTTCTTTTCTTG

>M6-PGSC0003DMG400003790M6LRR-RLK_244

GATCAATCTTTCTGATGCAGCCCTCTCAGGCTCACTTGATCACCTTGATTTCACTTCATTTCCGAGTCTCGTGAATTTCAATCTTAATGGAAATAACTTCAGTGGATCGATACCTTCGAG

>M6-PGSC0003DMG400003790M6LRR-RLK_2440

TTCTGGTCCAATCCCAACCGACGGAGTATTTCGAGGAGCATCCGCTAGTTCTTTTCTTGGAAACTCTGGTTTGTGTGGAAATGTAGAAGGATTATCCTCATGTAATTTGGATACCCGAAA

>M6-PGSC0003DMG400003790M6LRR-RLK_2501

AACTCTGGTTTGTGTGGAAATGTAGAAGGATTATCCTCATGTAATTTGGATACCCGAAATGACAAGTCCAGAAATAATAATCAAAAGATCCTCATTGGGGTACTTGTACCGGTGGTCAGC

>M6-PGSC0003DMG400003790M6LRR-RLK_2562

ACAAGTCCAGAAATAATAATCAAAAGATCCTCATTGGGGTACTTGTACCGGTGGTCAGCCTCATACTTTTGGCAATCCTGTTTGTTGCATGTCTCGTGTCTCGAAGGAAGGCTAAGCAAT

>M6-PGSC0003DMG400003790M6LRR-RLK_2623

CATACTTTTGGCAATCCTGTTTGTTGCATGTCTCGTGTCTCGAAGGAAGGCTAAGCAATATGATGAGGAGATCAAAGCTAGCCAGATACATGAAAATACAGAATCTCTGATTTGGGAAAG

>M6-PGSC0003DMG400003790M6LRR-RLK_2684

GATGAGGAGATCAAAGCTAGCCAGATACATGAAAATACAGAATCTCTGATTTGGGAAAGAGAAGGGAAGTTTACATTTGGTGACATAGTGAAAGCTACTGAAGATTTCAGTGAAAGGAAC

>M6-PGSC0003DMG400003790M6LRR-RLK_2745

AAGGGAAGTTTACATTTGGTGACATAGTGAAAGCTACTGAAGATTTCAGTGAAAGGAACTGCATTGGAAGAGGAGGCTTTGGAAGTGTCTATAAAGCTGTTTTGCCCTCTGGGCAGATTG

>M6-PGSC0003DMG400003790M6LRR-RLK_2806

CATTGGAAGAGGAGGCTTTGGAAGTGTCTATAAAGCTGTTTTGCCCTCTGGGCAGATTGTTGCAGTCAAAAGACTCAACATGTCAGACTCAAGTGACATTCCATTGACTAATCGTCGAAG

>M6-PGSC0003DMG400003790M6LRR-RLK_2867

GCAGTCAAAAGACTCAACATGTCAGACTCAAGTGACATTCCATTGACTAATCGTCGAAGCTTTGAGAATGAGATTAGAACTTTGACAGAGGTGAGACACAGGAATATAATTAAGCTCTTT

>M6-PGSC0003DMG400003790M6LRR-RLK_2928

TTGAGAATGAGATTAGAACTTTGACAGAGGTGAGACACAGGAATATAATTAAGCTCTTTGGTTACTGTTCCAAGAATGGGTGCATGTACTTGGTTTATGAGTATATAGAAAGAGGTAGCC

>M6-PGSC0003DMG400003790M6LRR-RLK_2989

TTACTGTTCCAAGAATGGGTGCATGTACTTGGTTTATGAGTATATAGAAAGAGGTAGCCTCGGGAAAGTTCTATATGATAATGAGATGGGAATGGAACTTGGATGGGGTACAAGAGTGAA

>M6-PGSC0003DMG400003790M6LRR-RLK_305

CCGAGTCTCGTGAATTTCAATCTTAATGGAAATAACTTCAGTGGATCGATACCTTCGAGTATTGGCAATGCCTCATTGCTCACTTTCTTGGACCTGAGTAACAATATCTTGTCAGGTGTC

>M6-PGSC0003DMG400003790M6LRR-RLK_3050

GGGAAAGTTCTATATGATAATGAGATGGGAATGGAACTTGGATGGGGTACAAGAGTGAAAATTGTGCAAGGAATAGCTCATGCACTCGCTTACTTGCACCATGACTGCTCTCCACCCATT

>M6-PGSC0003DMG400003790M6LRR-RLK_3111

TTGTGCAAGGAATAGCTCATGCACTCGCTTACTTGCACCATGACTGCTCTCCACCCATTGTGCACCGTGATGTATCATTGAACAACATTTTGCTTGAATCAGAATTCGAGCCACGACTCT

>M6-PGSC0003DMG400003790M6LRR-RLK_3172

GCACCGTGATGTATCATTGAACAACATTTTGCTTGAATCAGAATTCGAGCCACGACTCTCTGACTTTGGCACAGCAAAGTTGCTAGCTTCAGACTCATCAAATTGGACCACAGTTGCTGG

>M6-PGSC0003DMG400003790M6LRR-RLK_3233

GACTTTGGCACAGCAAAGTTGCTAGCTTCAGACTCATCAAATTGGACCACAGTTGCTGGTTCTTATGGCTATATGGCACCAGGTATGTTACATACTCATTACCTAATGCATCCATATTTC

>M6-PGSC0003DMG400003790M6LRR-RLK_3294

CTTATGGCTATATGGCACCAGGTATGTTACATACTCATTACCTAATGCATCCATATTTCTGTTGATTAAGTAGTATATTCATTGGAAGGATACTTCAATAATAGTGTCTTTCTTCTTGTG

>M6-PGSC0003DMG400003790M6LRR-RLK_3355

TTGATTAAGTAGTATATTCATTGGAAGGATACTTCAATAATAGTGTCTTTCTTCTTGTGCAGAGCTTGCACTTACCATGCGTGTTACAGAAAAGTGTGATGTCTATAGTTTTGGAGTCAT

>M6-PGSC0003DMG400003790M6LRR-RLK_3416

GAGCTTGCACTTACCATGCGTGTTACAGAAAAGTGTGATGTCTATAGTTTTGGAGTCATAGCAATGGAGACTATGATGGGAAGGCATCCAGGGGAGCTTTTAACTTCATTATCAGCATCA

>M6-PGSC0003DMG400003790M6LRR-RLK_3477

CAATGGAGACTATGATGGGAAGGCATCCAGGGGAGCTTTTAACTTCATTATCAGCATCAACAACATTGTCTCCAGAAATACTTTTGAAGGATGTTCTTGACCAAAGACTTCCACCTCCCA

>M6-PGSC0003DMG400003790M6LRR-RLK_3538

AACATTGTCTCCAGAAATACTTTTGAAGGATGTTCTTGACCAAAGACTTCCACCTCCCACCAGCCACTTGGCAGAGGCAGTGGTTTTTGTGATCACAATTGCCTTGGCATGTACACGTAC

>M6-PGSC0003DMG400003790M6LRR-RLK_3599

AGCCACTTGGCAGAGGCAGTGGTTTTTGTGATCACAATTGCCTTGGCATGTACACGTACCACTCCTGAATCGCGACCAACCATGCGTTGTGTAGCACAAGAATTGTCTGTTCAGACTTTG

>M6-PGSC0003DMG400003790M6LRR-RLK_366

TTGGCAATGCCTCATTGCTCACTTTCTTGGACCTGAGTAACAATATCTTGTCAGGTGTCATACCTGAAGAGATTGGAAAATTGAATCAACTTGAGTATCTAAGTTTTTATAACAACAATA

>M6-PGSC0003DMG400003790M6LRR-RLK_3660

CTCCTGAATCGCGACCAACCATGCGTTGTGTAGCACAAGAATTGTCTGTTCAGACTTTGCCTTACCTTCCACAGCCATTGGGGACAATAGAAATGAGCAAACTAACAAGTTTTCAGAAAT

>M6-PGSC0003DMG400003790M6LRR-RLK_3721

TTACCTTCCACAGCCATTGGGGACAATAGAAATGAGCAAACTAACAAGTTTTCAGAAATAATGAGAAGATAGAACCTCATGTACAGGTGTGAAATGTACAGACTGTAGAGTTTTAGTGTT

>M6-PGSC0003DMG400003790M6LRR-RLK_3782

TGAGAAGATAGAACCTCATGTACAGGTGTGAAATGTACAGACTGTAGAGTTTTAGTGTTGTAATGCAACTTCCTCAAGATCTAAGGTCTATATAGTATCAAGGATGAAACCTTGGATCTA

>M6-PGSC0003DMG400003790M6LRR-RLK_3843

AATGCAACTTCCTCAAGATCTAAGGTCTATATAGTATCAAGGATGAAACCTTGGATCTATCCAAATTACTATGTTTCTTTTGCACTTTTCCCTAGAAAAAAAGAAATAATATCATCCTGT

>M6-PGSC0003DMG400003790M6LRR-RLK_3904

CAAATTACTATGTTTCTTTTGCACTTTTCCCTAGAAAAAAAGAAATAATATCATCCTGTTAATCATGAATGTTACAAAGTTGGTATATTCATTTGGACTAGTATGAATGTTACAAAGTTG

>M6-PGSC0003DMG400003790M6LRR-RLK_3965

ATCATGAATGTTACAAAGTTGGTATATTCATTTGGACTAGTATGAATGTTACAAAGTTGCACAATTCCCTTATCCTAGTCTTTACGTTATTGCTAGGCAAAACACTAGCAGAAAGATACA

>M6-PGSC0003DMG400003790M6LRR-RLK_4026

CAATTCCCTTATCCTAGTCTTTACGTTATTGCTAGGCAAAACACTAGCAGAAAGATACAATGTGTTCACTTGCAATAAAAATAGCAACCTAACAAAAATTTGGCAAAGAATACATCACAC

>M6-PGSC0003DMG400003790M6LRR-RLK_4087

GTGTTCACTTGCAATAAAAATAGCAACCTAACAAAAATTTGGCAAAGAATACATCACACAGATACCAGGCATGGTACTTAGCATTCCACTCCACACTGCTTGAGTATAGGTCCCTTCTGC

>M6-PGSC0003DMG400003790M6LRR-RLK_4148

ATACCAGGCATGGTACTTAGCATTCCACTCCACACTGCTTGAGTATAGGTCCCTTCTGCTTTGGTAGGAATGGGTCTATTCGCACCCAAACAAGTGAGAAGATTGAAGCAAGAAGAATTG

>M6-PGSC0003DMG400003790M6LRR-RLK_4209

TGGTAGGAATGGGTCTATTCGCACCCAAACAAGTGAGAAGATTGAAGCAAGAAGAATTGACCAGAGGACCACAATGGTGGGAGTCCTGTTCTGCCTGCCCATCAATCCCTTTAGGAAAGG

>M6-PGSC0003DMG400003790M6LRR-RLK_427

ACCTGAAGAGATTGGAAAATTGAATCAACTTGAGTATCTAAGTTTTTATAACAACAATATCGAAGGTGTGATTCCGTATCAGATTAGCAATCTGCAGAAGGTAATGCACTTGGATCTTGG

>M6-PGSC0003DMG400003790M6LRR-RLK_4270

CAGAGGACCACAATGGTGGGAGTCCTGTTCTGCCTGCCCATCAATCCCTTTAGGAAAGGATAGAGATGCACAATAACCCAAAAGGCGAAGAATAACTTTCCGAAGAGAGGACCCCATGAA

>M6-PGSC0003DMG400003790M6LRR-RLK_4331

AGAGATGCACAATAACCCAAAAGGCGAAGAATAACTTTCCGAAGAGAGGACCCCATGAACCGTAACCATTGTTGATGGCATCCGAAACACCAGCTACAACACCGACCATATTCAAGATGA

>M6-PGSC0003DMG400003790M6LRR-RLK_4392

GTAACCATTGTTGATGGCATCCGAAACACCAGCTACAACACCGACCATATTCAAGATGATCAGAGTGGTTGGTGGGATAAGAAGTGTGGTCCATTTGAAGAGGTAAAGCTCCCCAAATTC

>M6-PGSC0003DMG400003790M6LRR-RLK_4453

AGAGTGGTTGGTGGGATAAGAAGTGTGGTCCATTTGAAGAGGTAAAGCTCCCCAAATTCAGCATCATCAGCTGCTTTTGATGTTACAGTAAAGTTTGTGTCTACACCAGCAAGAACTTTA

>M6-PGSC0003DMG400003790M6LRR-RLK_4514

CATCATCAGCTGCTTTTGATGTTACAGTAAAGTTTGTGTCTACACCAGCAAGAACTTTAAGAAGGCCTTGGAAAACTGCAAACAAATGTGCTGAAACACCACCGATGACCCAAAATTGCT

>M6-PGSC0003DMG400003790M6LRR-RLK_4575

AAGGCCTTGGAAAACTGCAAACAAATGTGCTGAAACACCACCGATGACCCAAAATTGCTCGTTACGCCACCAGGCCTCAATGCTTACTCCACTCCATCGCAGCTCGAGTACACTTGTGGC

>M6-PGSC0003DMG400003790M6LRR-RLK_4636

TTACGCCACCAGGCCTCAATGCTTACTCCACTCCATCGCAGCTCGAGTACACTTGTGGCTATGATGGAGATAAAGAGAGCCATGAACCAAATGCTTGCAAAGTTGTTAAGCTACAAAACA

>M6-PGSC0003DMG400003790M6LRR-RLK_4697

TGATGGAGATAAAGAGAGCCATGAACCAAATGCTTGCAAAGTTGTTAAGCTACAAAACAAACATGGATTTTGATTTATAAAATCTAAATCAAATCTTTATATTACCGAACCAAATGACTT

>M6-PGSC0003DMG400003790M6LRR-RLK_4758

CATGGATTTTGATTTATAAAATCTAAATCAAATCTTTATATTACCGAACCAAATGACTTCAGTTCACTTACTGTGGGGACGATAAATTTTCCAGTGAGAAGGCATACAGCTGGAAGGGTG

>M6-PGSC0003DMG400003790M6LRR-RLK_4819

GTTCACTTACTGTGGGGACGATAAATTTTCCAGTGAGAAGGCATACAGCTGGAAGGGTGCAGTAAGCTAGCAGAGCAATGGAAGTGAATGGGTAAACAATGGTGTTAATGTATGCTAGCC

>M6-PGSC0003DMG400003790M6LRR-RLK_488

GAAGGTGTGATTCCGTATCAGATTAGCAATCTGCAGAAGGTAATGCACTTGGATCTTGGATCAAATTTCTTAGAAACTCCTGATTGGTTGAAGATGAAAAATATGCCTGTGTTGACGTAT

>M6-PGSC0003DMG400003790M6LRR-RLK_4880

GTAAGCTAGCAGAGCAATGGAAGTGAATGGGTAAACAATGGTGTTAATGTATGCTAGCCTTTCCAACCATTTGAGTTTTCCACCCCAGGCATACCAAAGGGGACAATGACGACTCATGAA

>M6-PGSC0003DMG400003790M6LRR-RLK_4941

TCCAACCATTTGAGTTTTCCACCCCAGGCATACCAAAGGGGACAATGACGACTCATGAAGATCTCGACAGAACCAAGAGCCCATCTCAGGACTTGGTGCAATCTATCTGATAGATTGATT

>M6-PGSC0003DMG400003790M6LRR-RLK_5002

TCTCGACAGAACCAAGAGCCCATCTCAGGACTTGGTGCAATCTATCTGATAGATTGATTGGGGCTGATCCCTTAAAAGCTGCTCTCTTCGGACAGCAATAAACTGATCTCCATCCTCTGC

>M6-PGSC0003DMG400003790M6LRR-RLK_5059

TTGGGGCTGATCCCTTAAAAGCTGCTCTCTTCGGACAGCAATAAACTGATCTCCATCCTCTGCAGTGCATCTTAAATCCAGTCAAAATGTCCTCTGTGACAGAACCATAAATCCATCCAA

>M6-PGSC0003DMG400003790M6LRR-RLK_549

CAAATTTCTTAGAAACTCCTGATTGGTTGAAGATGAAAAATATGCCTGTGTTGACGTATCTAAGCTTTGGTTATAATGAACTGAGGTTAGAATTCCCTGAATTCGTACTTCGTTGTCATA

>M6-PGSC0003DMG400003790M6LRR-RLK_61

GCTACTTCTTTTTTCACTTCCATTAAGTATCACATCTTCAGCAAGAACAGAAGCTGAATCTCTTGTCAAATGGAAGAGAAATTTACCTTCTACGTCTTTCTTGGACTCATGGTCCATTTC

>M6-PGSC0003DMG400003790M6LRR-RLK_610

AAGCTTTGGTTATAATGAACTGAGGTTAGAATTCCCTGAATTCGTACTTCGTTGTCATAATCTAACATATCTTGATTTATCGATAAATCATTTCAATGGTTCGATTCCTGAAACAGTATT

>M6-PGSC0003DMG400003790M6LRR-RLK_671

CTAACATATCTTGATTTATCGATAAATCATTTCAATGGTTCGATTCCTGAAACAGTATTTACCAACTTGATCAACCTTGAAAGACTTAATCTTTCATCTAATTCATTTCAAGGTTCATTG

>M6-PGSC0003DMG400003790M6LRR-RLK_732

CCAACTTGATCAACCTTGAAAGACTTAATCTTTCATCTAATTCATTTCAAGGTTCATTGTCACCAAATTTTACCAAGTTGTCTAAGTTAAAAGAACTTCAGCTAGGTGTTAACATGTTTT

>M6-PGSC0003DMG400003790M6LRR-RLK_793

ACCAAATTTTACCAAGTTGTCTAAGTTAAAAGAACTTCAGCTAGGTGTTAACATGTTTTCTGGTTTAATTCCTGATGAAATTGGCTTGATTACTAGTCTTGAAGTTCTTGTACTTTTCAA

>M6-PGSC0003DMG400003790M6LRR-RLK_854

GGTTTAATTCCTGATGAAATTGGCTTGATTACTAGTCTTGAAGTTCTTGTACTTTTCAACAATTCATTTGAAGGAAAGATTCCATCTTCTATAGGTAGACTCATAAATCTTCAACGGTTG

>M6-PGSC0003DMG400003790M6LRR-RLK_915

ATTCATTTGAAGGAAAGATTCCATCTTCTATAGGTAGACTCATAAATCTTCAACGGTTGGATCTTCGAAAGAACGATTTGAATTCAACTATTCCTTCTGAGCTTGGTTTTTGTACTAAAC

>M6-PGSC0003DMG400003790M6LRR-RLK_976

TCTTCGAAAGAACGATTTGAATTCAACTATTCCTTCTGAGCTTGGTTTTTGTACTAAACTCACTTTCTTAGCTCTAGCAGAAAATGACCTGCAAGGGTCATTGCCTCCATCATTCTCCTC

>M6-PGSC0003DMG400003802M6LRR-RLP_0

ATGTTTGAATTCGCCCGGAAATTGCAGAAGCGAGGCGGAGAGTTGATTAGCGAGTTTTGCAGTGAAATTGGAGTCGAAAAATGGGGTTTTAAGGTACGGTTGCACAATAGGACACCTTCA

>M6-PGSC0003DMG400003802M6LRR-RLP_1054

GGGAGAGCTCAACATTTGTGGATGCTTTGGTATGGAGTTGTCAGCCTAGGAGACTCAACGTGCATTCAACAAGAGAAAAGATCACATTTTTCATTGATCATTTGATGTATATGAAGAATT

>M6-PGSC0003DMG400003802M6LRR-RLP_1178

TCAGTCTACTTCTCATGGAAGCACGCCTCTTCTTAGTCAATTAAAAGGAGTAAAAGCCTACAAATTTGATAGGAAGAAGGAGAGTTGGCATCCTGTGGAACACAAAAGTAGTGAGCTGGT

>M6-PGSC0003DMG400003802M6LRR-RLP_1230

AAAGCCTACAAATTTGATAGGAAGAAGGAGAGTTGGCATCCTGTGGAACACAAAAGTAGTGAGCTGGTAACAACAAACTCTGAAATGTGGGATTGGTGTTATTTTTTATTAGATTGGTGA

>M6-PGSC0003DMG400003802M6LRR-RLP_186

CGGATGAGCATTCTTTCCAAAACATGGCTGCAAGCCTGGTCGACTCTTCCAAACTTGGATTTCACAGTTGATTTTTCTAAAGGCAACGATATGAAAATAGTGGATGATATAATGGAGAGA

>M6-PGSC0003DMG400003802M6LRR-RLP_248

CACAGTTGATTTTTCTAAAGGCAACGATATGAAAATAGTGGATGATATAATGGAGAGATATAGGGACGGAAAAATCCCTATAGAAAAGTTTGAATTATCAGATGATCGTTTGTTCTTTGA

>M6-PGSC0003DMG400003802M6LRR-RLP_310

GGGACGGAAAAATCCCTATAGAAAAGTTTGAATTATCAGATGATCGTTTGTTCTTTGAAAATCCTCATGGAGTTTTAGTTGTCCCTATGATTGATAAGTGGCTTGGCATTGTACTAACAA

>M6-PGSC0003DMG400003802M6LRR-RLP_372

CCTCATGGAGTTTTAGTTGTCCCTATGATTGATAAGTGGCTTGGCATTGTACTAACAAATGGTGTAAAAGACCTTACCATATCATCATACCCCTTGTTTATTTTCGCAATCTTGGCGGTA

>M6-PGSC0003DMG400003802M6LRR-RLP_434

TGTAAAAGACCTTACCATATCATCATACCCCTTGTTTATTTTCGCAATCTTGGCGGTAAAATCTGTGAGAAAATTGGTTCTGGAATCTTGTATTCTTTTGCCTAGTGTTGTTGTGAATTG

>M6-PGSC0003DMG400003802M6LRR-RLP_496

CTGTGAGAAAATTGGTTCTGGAATCTTGTATTCTTTTGCCTAGTGTTGTTGTGAATTGCAATTCTTTGAGGAAACTTTCTCTATCTTATGTAAGATTAGATGAAAACATGCTTCAGGCTC

>M6-PGSC0003DMG400003802M6LRR-RLP_558

TCTTTGAGGAAACTTTCTCTATCTTATGTAAGATTAGATGAAAACATGCTTCAGGCTCTACTTAATAGCTGTCCGTTGATTGAATCTTTCATATTTGAGTATTGCAAGGGCTTGGAAACA

>M6-PGSC0003DMG400003802M6LRR-RLP_62

TGAAATTGGAGTCGAAAAATGGGGTTTTAAGGTACGGTTGCACAATAGGACACCTTCAATTGATATATTGTCGGAATGTCTCATTCATAAAATACTCTGTTGTCTTAGTTTTAAAGAAGC

>M6-PGSC0003DMG400003802M6LRR-RLP_620

TAATAGCTGTCCGTTGATTGAATCTTTCATATTTGAGTATTGCAAGGGCTTGGAAACAATGAATCATCAAAAGATTAAGTCGGTTTCCCTGAAGTTCTTGAAGATTCGGCATTGTGGGGG

>M6-PGSC0003DMG400003802M6LRR-RLP_744

TGGGAGATTGATGCTCCAAATTTGGTATCACTTAAGTACACTTGGGAAGACATTCCTCAAATTAAAATTGTGAGAGAGTCAAGACAAATGAAGTACTCAAGAATCATTCTTGAATGCTTA

>M6-PGSC0003DMG400003802M6LRR-RLP_806

TAAAATTGTGAGAGAGTCAAGACAAATGAAGTACTCAAGAATCATTCTTGAATGCTTAGACAGTGACAATGTAAATGCTGCATGGTTTTGTAAGTTGAGGAAGTTTCTGTCAAATTCGAT

>M6-PGSC0003DMG400003802M6LRR-RLP_868

GTGACAATGTAAATGCTGCATGGTTTTGTAAGTTGAGGAAGTTTCTGTCAAATTCGATCTCTTGGTCTGAAGTTCGCATTGAATCTTATAGATGCAATGACATCAACATTCAAAATTTGC

>M6-PGSC0003DMG400003802M6LRR-RLP_930

TGGTCTGAAGTTCGCATTGAATCTTATAGATGCAATGACATCAACATTCAAAATTTGCAACTGGACCATAAAGATTCTAATCCCCACGTGGACGTTTTAAATGTCAACATTATATGGAAG

>M6-PGSC0003DMG400003802M6LRR-RLP_992

GGACCATAAAGATTCTAATCCCCACGTGGACGTTTTAAATGTCAACATTATATGGAAGAATGGGGAGAGCTCAACATTTGTGGATGCTTTGGTATGGAGTTGTCAGCCTAGGAGACTCAA

>M6-PGSC0003DMG400003814M6LRR-RLP_0

ACCCAATTGATTTCTTAGCTTTACAAGCGATTCGAAAAGGCTTGGATGATTTGCCCGGTTCCAACTATTTTGCTTCCTGGGATTTCACTTCTGAGCCGTGCAACTTCGCCGGAGTTTACT

>M6-PGSC0003DMG400003814M6LRR-RLP_1011

TGCAGTATAATTGCATGATTCCGCCGGTTGAGACGGCTTGTCCGTTGAAAGCCGGGAAGCAGAAGAGTAGGCCTACAAATCAGTGTGTGGAGTGGAAGGGGCAAAAGAGTGAAAAAACAA

>M6-PGSC0003DMG400003814M6LRR-RLP_128

GATAAAGTGATCGCCTTGAATCTCGGTGATCCCAGAGCTGGATCACCGGGTTTGACCGGAAACTTGGATCCTTCTATTGGGAAACTCTCTGCTCTTGCTGAGTTCACGGTGGTTCCGGGT

>M6-PGSC0003DMG400003814M6LRR-RLP_192

TGGATCCTTCTATTGGGAAACTCTCTGCTCTTGCTGAGTTCACGGTGGTTCCGGGTCGGATAACCGGTGCATTGCCGGAAAGTTTTTCTCAGTTGAAGAACCTGAGATTTCTAGGAGTTA

>M6-PGSC0003DMG400003814M6LRR-RLP_256

CGGTGCATTGCCGGAAAGTTTTTCTCAGTTGAAGAACCTGAGATTTCTAGGAGTTAGCCGGAACTTTCTCTCCGGTGATATTCCGGCGTCATTAGGCCTGCTCCGGGAGCTACAAACTCT

>M6-PGSC0003DMG400003814M6LRR-RLP_320

TTTCTCTCCGGTGATATTCCGGCGTCATTAGGCCTGCTCCGGGAGCTACAAACTCTTGATCTTAGTTTCAATCAGCTGACTGGGAGTATCCCTTGGGCTATTGGAGCTCTTCCGGCATTG

>M6-PGSC0003DMG400003814M6LRR-RLP_384

GTTTCAATCAGCTGACTGGGAGTATCCCTTGGGCTATTGGAGCTCTTCCGGCATTGTCCAACATTATACTCTGTCACAATCATTTGTCCGGTTCGATCCCACCATTCGTATCCCAAAGAT

>M6-PGSC0003DMG400003814M6LRR-RLP_448

TATACTCTGTCACAATCATTTGTCCGGTTCGATCCCACCATTCGTATCCCAAAGATTGACCCGTTTAGACCTTAAACACAACTCACTTTCAGGTACCCTTTTGCCTAATTCACTTCCTTC

>M6-PGSC0003DMG400003814M6LRR-RLP_512

TTAGACCTTAAACACAACTCACTTTCAGGTACCCTTTTGCCTAATTCACTTCCTTCATCGCTTCAATACTTGTCATTGTCATGGAACCAGTTAACCGGTCCAGTAGATTATCAGTTGAGT

>M6-PGSC0003DMG400003814M6LRR-RLP_576

AATACTTGTCATTGTCATGGAACCAGTTAACCGGTCCAGTAGATTATCAGTTGAGTCGGCTGAACCAGTTGAATTACCTCGATCTTAGCTTGAACCGGTTCACCGGCTGCATTCCGGGAA

>M6-PGSC0003DMG400003814M6LRR-RLP_64

CTATTTTGCTTCCTGGGATTTCACTTCTGAGCCGTGCAACTTCGCCGGAGTTTACTGCGATGGAGATAAAGTGATCGCCTTGAATCTCGGTGATCCCAGAGCTGGATCACCGGGTTTGAC

>M6-PGSC0003DMG400003814M6LRR-RLP_640

CCAGTTGAATTACCTCGATCTTAGCTTGAACCGGTTCACCGGCTGCATTCCGGGAATCTTATTTACTTTCCCACTCACAAATCTTCAGCTACAAAGGAATCTATTTTCGGGTCCGGTTTT

>M6-PGSC0003DMG400003814M6LRR-RLP_704

ACTTTCCCACTCACAAATCTTCAGCTACAAAGGAATCTATTTTCGGGTCCGGTTTTACCCATTTCAGAGGTGACAATTCCGACGGTGGACATTAGCTTTAACAGGTTTTTCGGGGAGATA

>M6-PGSC0003DMG400003814M6LRR-RLP_768

CAGAGGTGACAATTCCGACGGTGGACATTAGCTTTAACAGGTTTTTCGGGGAGATATCTCCATTGTTGTGTAATGTGGAGAATTTGTATCTGAACAATAACCGGTTCACGGGTAAGGTGC

>M6-PGSC0003DMG400003814M6LRR-RLP_832

GTTGTGTAATGTGGAGAATTTGTATCTGAACAATAACCGGTTCACGGGTAAGGTGCCGGCGGTGTTGGTGGACCGGTTATTATCTGCCGGAATACAAGTTTTATATTTACAGCATAATTT

>M6-PGSC0003DMG400003814M6LRR-RLP_896

TTGGTGGACCGGTTATTATCTGCCGGAATACAAGTTTTATATTTACAGCATAATTTTCTGACGGGGATTGAGATAAATCCGAGGGTGGAAATTCCGGTGAGCAGTTCATTGTGTTTGCAG

>M6-PGSC0003DMG400003814M6LRR-RLP_960

GGATTGAGATAAATCCGAGGGTGGAAATTCCGGTGAGCAGTTCATTGTGTTTGCAGTATAATTGCATGATTCCGCCGGTTGAGACGGCTTGTCCGTTGAAAGCCGGGAAGCAGAAGAGTA

>M6-PGSC0003DMG400003822M6LRR-RLP_5124

TGCATCTTAAATCCAGTCAAAATGTCCTCTGTGACAGAACCATAAATCCATCCAATCTGTCAAAAAAAGAAAAATGAAAACCAATTTATTAGAAGTTAGCTAAGCGAACACCCATAACTC

>M6-PGSC0003DMG400003822M6LRR-RLP_5185

AAAAAAAGAAAAATGAAAACCAATTTATTAGAAGTTAGCTAAGCGAACACCCATAACTCATAATTTATTCATGAAATATAATGCTAACCTCTTTGCCCCAGTCAGTTTTCTCTTCATACC

>M6-PGSC0003DMG400003822M6LRR-RLP_5246

AATTTATTCATGAAATATAATGCTAACCTCTTTGCCCCAGTCAGTTTTCTCTTCATACCCACAGCTAATAACATGAATAGCTTCCTTTATGAGTGTTGTCGGGTTTGTCCCTTCAGGAAG

>M6-PGSC0003DMG400003822M6LRR-RLP_5307

CAGCTAATAACATGAATAGCTTCCTTTATGAGTGTTGTCGGGTTTGTCCCTTCAGGAAGCCCACCATCTTCCTTAAGTGTGGAAGCTATGAAAACGGGGGACTGCCCGAATCGTTTCTCA

>M6-PGSC0003DMG400003822M6LRR-RLP_5368

CACCATCTTCCTTAAGTGTGGAAGCTATGAAAACGGGGGACTGCCCGAATCGTTTCTCAAAGTTCTTTTGGGACATGAGAGATGACTTCTCAAGCTCATCATATCCTTCAAGGCCTTCTT

>M6-PGSC0003DMG400003822M6LRR-RLP_5429

GTTCTTTTGGGACATGAGAGATGACTTCTCAAGCTCATCATATCCTTCAAGGCCTTCTTCAATTTCCTCAAGATCAAAGACAGGTCCAGATGGCTTCCTAGTGTAGTTCTTTCCATTCAT

>M6-PGSC0003DMG400003822M6LRR-RLP_5490

ATTTCCTCAAGATCAAAGACAGGTCCAGATGGCTTCCTAGTGTAGTTCTTTCCATTCATTGTTTTCTTTTTCTTGTTGTATAGTCCTCCAAGTCCAAGTAAGGCCTTGATGCTCTTCTTC

>M6-PGSC0003DMG400003822M6LRR-RLP_5551

TTTTCTTTTTCTTGTTGTATAGTCCTCCAAGTCCAAGTAAGGCCTTGATGCTCTTCTTCTTAGACTTGGACTTGGACTTTCTTGAACCACCACAACAGCAACTGGGCCAGCAATCACATG

>M6-PGSC0003DMG400003822M6LRR-RLP_5612

AGACTTGGACTTGGACTTTCTTGAACCACCACAACAGCAACTGGGCCAGCAATCACATGTCATCTTTTGACGCTTCTCAGAAACAGGTGGATCATAGCCGTATAACGCCTGCCTGTTAAA

>M6-PGSC0003DMG400003822M6LRR-RLP_5673

ATCTTTTGACGCTTCTCAGAAACAGGTGGATCATAGCCGTATAACGCCTGCCTGTTAAAGACAGTACCTGTTCCAACATAAACTGGTCCTTGAATTCCATCCAGGCCTTTCATGTTGATC

>M6-PGSC0003DMG400003822M6LRR-RLP_5734

CAGTACCTGTTCCAACATAAACTGGTCCTTGAATTCCATCCAGGCCTTTCATGTTGATCTGCCACCATGAAGGGAAAAAAGCATATTAATACCGGGAATAAATCCTTTGTCAACGCATTG

>M6-PGSC0003DMG400003822M6LRR-RLP_5795

CCACCATGAAGGGAAAAAAGCATATTAATACCGGGAATAAATCCTTTGTCAACGCATTGCTGTATATATAGATCAACTTACATCAAAAAAGACAACATTACGATTGGCATATCGATCGTG

>M6-PGSC0003DMG400003822M6LRR-RLP_5856

GTATATATAGATCAACTTACATCAAAAAAGACAACATTACGATTGGCATATCGATCGTGGCGATCAATACCATCAAATCTCTGGGGAAATTGGACATAGCATAGCTTCTTTCCAATTTGT

>M6-PGSC0003DMG400003822M6LRR-RLP_5917

GATCAATACCATCAAATCTCTGGGGAAATTGGACATAGCATAGCTTCTTTCCAATTTGTGGGTCCATTAAAAAGCACATGGCTTCTCTAACTGCCTTGCTGTTGTTAAGGTAGTGATCAC

>M6-PGSC0003DMG400003822M6LRR-RLP_5978

GTCCATTAAAAAGCACATGGCTTCTCTAACTGCCTTGCTGTTGTTAAGGTAGTGATCACAATCCAAGTTCAACATAAATGGTGCATTGGTCAGGACTGCAGCGACTCGAACCTATTTAGA

>M6-PGSC0003DMG400003822M6LRR-RLP_6039

TCCAAGTTCAACATAAATGGTGCATTGGTCAGGACTGCAGCGACTCGAACCTATTTAGAAAAACAGCACAAATTACATAACATTAAGAAAAGGACTTAGCAGAATGTCGAGGAAGCAAGC

>M6-PGSC0003DMG400003822M6LRR-RLP_6100

AACAGCACAAATTACATAACATTAAGAAAAGGACTTAGCAGAATGTCGAGGAAGCAAGCCAGAAACTAATATACTAATTACTAACCAAAGCATTCATGGCACCAGCTTTCTTGTGGTGTT

>M6-PGSC0003DMG400003822M6LRR-RLP_6161

GAAACTAATATACTAATTACTAACCAAAGCATTCATGGCACCAGCTTTCTTGTGGTGTTGATAGCCCGGTCGTTTCTCACGAGAAACATAGACAAGGCGGGGAAGCTCCTTACCTTCAAT

>M6-PGSC0003DMG400003822M6LRR-RLP_6222

TAGCCCGGTCGTTTCTCACGAGAAACATAGACAAGGCGGGGAAGCTCCTTACCTTCAATGTCAAGTGCACCTTCACTACCAAGATAAACTTGAATCATACCAGGATGGTCTCTGGTGTTG

>M6-PGSC0003DMG400003822M6LRR-RLP_6283

CAAGTGCACCTTCACTACCAAGATAAACTTGAATCATACCAGGATGGTCTCTGGTGTTGTTACCGGGCCATGGAGTTCCATCTTGCATTACCCATCCCTCTTCTGGTTTTTTCTGAGCTT

>M6-PGSC0003DMG400003822M6LRR-RLP_6344

ACCGGGCCATGGAGTTCCATCTTGCATTACCCATCCCTCTTCTGGTTTTTTCTGAGCTTTTGCTACTAATGCATTAATTCTTACTTTGAATTCTTCATATTCTCTCTGCCACAAGAGAGT

>M6-PGSC0003DMG400003822M6LRR-RLP_6405

GCTACTAATGCATTAATTCTTACTTTGAATTCTTCATATTCTCTCTGCCACAAGAGAGTGAGTTTTCAATTTGCAGAAAAATCTTATTTCAAGCCAAAGAAACTTAAGCCCCCGATAAAT

>M6-PGSC0003DMG400003822M6LRR-RLP_6466

GTTTTCAATTTGCAGAAAAATCTTATTTCAAGCCAAAGAAACTTAAGCCCCCGATAAATGATACTCCTTCTATCCCAACTTCTATGACATTCTTATATTTTGGGCACGTCCTAAAATGTT

>M6-PGSC0003DMG400003822M6LRR-RLP_6527

TACTCCTTCTATCCCAACTTCTATGACATTCTTATATTTTGGGCACGTCCTAAAATGTTTGACATGATTCCTTCTACAACCTTCACAATAAACTATTTGAATTTTAAACCTTGATAGGAT

>M6-PGSC0003DMG400003822M6LRR-RLP_6588

ACATGATTCCTTCTACAACCTTCACAATAAACTATTTGAATTTTAAACCTTGATAGGATTCAGTCAACATTTTAGGCACTGCATCTCCTCTGATATAGTCGTATAAAATGAGACGAAATG

>M6-PGSC0003DMG400003822M6LRR-RLP_6649

AGTCAACATTTTAGGCACTGCATCTCCTCTGATATAGTCGTATAAAATGAGACGAAATGAAGTAATAGATACATACCTTCATGGCCCTGCGATCCTTGACAAAGGTAGGCTGAACTTTAT

>M6-PGSC0003DMG400003822M6LRR-RLP_6710

GTAATAGATACATACCTTCATGGCCCTGCGATCCTTGACAAAGGTAGGCTGAACTTTATCTTTCAAGTAATCAATCTTTTCATTGAAGTAGAAATCGGGTGCTCTAGGCTCTACACTATA

>M6-PGSC0003DMG400003822M6LRR-RLP_6771

TTCAAGTAATCAATCTTTTCATTGAAGTAGAAATCGGGTGCTCTAGGCTCTACACTATATTTCTTACAAAATGGCACCCATCTCCTGGCAAATTCAGCTGTTTCTGCTAATGAATCGAAC

>M6-PGSC0003DMG400003822M6LRR-RLP_6832

TCTTACAAAATGGCACCCATCTCCTGGCAAATTCAGCTGTTTCTGCTAATGAATCGAACAACAGCATGGAAGCACCATCATCTGAAACATAACAGCTAACCTTCTCGACAGGATAATCAA

>M6-PGSC0003DMG400003822M6LRR-RLP_6893

CAGCATGGAAGCACCATCATCTGAAACATAACAGCTAACCTTCTCGACAGGATAATCAACAGCCAAGATTGACAGAACAGTGTTTGCAGTGATAATAGGAGGCTCCTTAAGAGGATCAAC

>M6-PGSC0003DMG400003822M6LRR-RLP_6954

GCCAAGATTGACAGAACAGTGTTTGCAGTGATAATAGGAGGCTCCTTAAGAGGATCAACAGTACTCACAAACACATCCACTGGGGCTAATCGATTTGGCTCGCCTTCACGTTCAAACCTC

>M6-PGSC0003DMG400003822M6LRR-RLP_7015

TACTCACAAACACATCCACTGGGGCTAATCGATTTGGCTCGCCTTCACGTTCAAACCTCAATGTCAATCGATCTAAATAAGTTTCACGGTTTATGGGTGACCATTTGGGGAACTGATCAA

>M6-PGSC0003DMG400003822M6LRR-RLP_7076

TGTCAATCGATCTAAATAAGTTTCACGGTTTATGGGTGACCATTTGGGGAACTGATCAAGTATCCATGATAATGCAAACCATATTTCACATATAACAGAGATAATCCATAATGGATAGGC

>M6-PGSC0003DMG400003822M6LRR-RLP_7137

ATCCATGATAATGCAAACCATATTTCACATATAACAGAGATAATCCATAATGGATAGGCATCATAGGCTGGTGTTAAGATCCGGAAATGGAAAAAGAATCCAAGAATTACAAGCCTGAGG

>M6-PGSC0003DMG400003822M6LRR-RLP_7198

CATAGGCTGGTGTTAAGATCCGGAAATGGAAAAAGAATCCAAGAATTACAAGCCTGAGGATAATGACTATCCGATATGGACTGATTAGACTTGATGGAAGTGGAACTTTTCGCCAGAGTG

>M6-PGSC0003DMG400003822M6LRR-RLP_7259

AATGACTATCCGATATGGACTGATTAGACTTGATGGAAGTGGAACTTTTCGCCAGAGTGGTTGCCGAGCTTCAGCCAAACTGCTTGGAAATCAGTTAAGTGGTTAGAATATTGAAATACT

>M6-PGSC0003DMG400003822M6LRR-RLP_7320

TGCCGAGCTTCAGCCAAACTGCTTGGAAATCAGTTAAGTGGTTAGAATATTGAAATACTGCAAGTTTTATATGCTTAAACTTGATCTGTTTCAATTCATACACACTCATTCCATGTTGAA

>M6-PGSC0003DMG400003822M6LRR-RLP_7381

AAGTTTTATATGCTTAAACTTGATCTGTTTCAATTCATACACACTCATTCCATGTTGAAACATTATAGTATTTGATTAATTTTGTTAATAATATTGCATGTAACTATCTAACTTTCACAA

>M6-PGSC0003DMG400003822M6LRR-RLP_7442

ATTATAGTATTTGATTAATTTTGTTAATAATATTGCATGTAACTATCTAACTTTCACAATTATAAAGTGTTCAAATTATTTAATTATACCAGTCAGTCACTGAGTGATATATAACTACAA

>M6-PGSC0003DMG400003822M6LRR-RLP_7503

ATAAAGTGTTCAAATTATTTAATTATACCAGTCAGTCACTGAGTGATATATAACTACAAGTGACAGAGACGATATATAACCGAAAATCCAATATTTAAAAGTTGTAAGGAGAATGAGGGG

>M6-PGSC0003DMG400003822M6LRR-RLP_7564

GACAGAGACGATATATAACCGAAAATCCAATATTTAAAAGTTGTAAGGAGAATGAGGGGAAGGTGGAAACAAGACCAGATATAATTTGATCGCTATGAACAAATCCAAAGACCAGGTTGA

>M6-PGSC0003DMG400003822M6LRR-RLP_7686

GATTTGTTAAAAATTTAAAGTATTTCTATAGGGGTCGTTTCCTGTTCCTTCTATGCTGTTTTCTCGATAATTCCTCACATAAAACCTTCTCGTATTAGCAACGATATAACTTCAACAAGA

>M6-PGSC0003DMG400003822M6LRR-RLP_7747

TCTCGATAATTCCTCACATAAAACCTTCTCGTATTAGCAACGATATAACTTCAACAAGATCAAACCAGAGGCGGGAATTAGCTTGAATAATCATAATTTTCAATATCTTCTTCCACTACT

>M6-PGSC0003DMG400003822M6LRR-RLP_7808

AAACCAGAGGCGGGAATTAGCTTGAATAATCATAATTTTCAATATCTTCTTCCACTACTAAGAATATATGTCAAATAACATGTTAAATTCCCCTTCTAAGTCGAAGGTCTATCGAAAAAG

>M6-PGSC0003DMG400003822M6LRR-RLP_8174

AGAGTGAAAAGAAAGAGTACATACAGATAATCATCATCTTCATCAGGTTGGTCATTTCCACCATTATTAACAAGACCTCTCTTTTCTTGACGATTCTTCCATTTCTCAACTCTTTCCTTC

>M6-PGSC0003DMG400003822M6LRR-RLP_8235

CATTATTAACAAGACCTCTCTTTTCTTGACGATTCTTCCATTTCTCAACTCTTTCCTTCCATTCTGCATCACCATAACTTTCTTTCTGCTCATCCTCCAATTCCTTCCCTGCAACTGTAA

>M6-PGSC0003DMG400003822M6LRR-RLP_8296

TTCTGCATCACCATAACTTTCTTTCTGCTCATCCTCCAATTCCTTCCCTGCAACTGTAAGTGATTTTCCATTGAAATAAAACAAAATCATGACCTTGAGTGACACATATATATAAAGTGC

>M6-PGSC0003DMG400003822M6LRR-RLP_8357

GATTTTCCATTGAAATAAAACAAAATCATGACCTTGAGTGACACATATATATAAAGTGCAACAATTAATCATTTCGAAATTAAATAATATAGCTGTACCGCTTCCAACAACTGATTGATT

>M6-PGSC0003DMG400003822M6LRR-RLP_8418

CAATTAATCATTTCGAAATTAAATAATATAGCTGTACCGCTTCCAACAACTGATTGATTAAAAGCAGCACCACTATTGGTGTTGTGATTATACTCTCCATTCTCCTGCCAAACATACAAC

>M6-PGSC0003DMG400003834-(rev)M6LRR-RLP_0

ATGAGCATTCTCTCCAAAACATGGCTACAAGAATGGTCGAATCTTCCCAACTTGGAATTCACTATTAATTGTTGGGAAGGCAATATAAATACAGCGAACACAACCATGGAGAGATATAGG

>M6-PGSC0003DMG400003834-(rev)M6LRR-RLP_1054

CCCTTATGGAGGGGGAGAAAGTTTATTTTTTATTAGATTGGTGATGTAGTTAATTTGTTTCTATTTCAAATTGAATGGACTTATTCTTTTCTGCATTGGTGTCAGAGGTATTAGATATTA

>M6-PGSC0003DMG400003834-(rev)M6LRR-RLP_1116

ATTTCAAATTGAATGGACTTATTCTTTTCTGCATTGGTGTCAGAGGTATTAGATATTAATAAAACAGAACTAAGAGCATAATTTTAAACTTGGATCTTATAGGCTTGCAAAAGCAAAAAC

>M6-PGSC0003DMG400003834-(rev)M6LRR-RLP_1178

AACAGAACTAAGAGCATAATTTTAAACTTGGATCTTATAGGCTTGCAAAAGCAAAAACAAAATTTTTAAGAAAGTAATGAGTATGGCAAATTTCTCAAAAGGAAAGAGACTTAATTTCTA

>M6-PGSC0003DMG400003834-(rev)M6LRR-RLP_124

GGAAAATCCCTATACAAAAGTTTGAATTATTAGAGACCTTTGCTAATTCTCATGAAGATTTCCCTCTCATTGATAAGTGGCTCGACATTGCACTTCAGAATGGTGTAAAACATCTTTTTC

>M6-PGSC0003DMG400003834-(rev)M6LRR-RLP_1240

TTTTTAAGAAAGTAATGAGTATGGCAAATTTCTCAAAAGGAAAGAGACTTAATTTCTACATAATTACTAGTGTTTCAACGTAACTACAAATGCCTTCGATGTCATACTTTCTCGGTAGCC

>M6-PGSC0003DMG400003834-(rev)M6LRR-RLP_1302

ATTACTAGTGTTTCAACGTAACTACAAATGCCTTCGATGTCATACTTTCTCGGTAGCCCGTTGATTTCTAATTTTCTCAAACTCTGTCTCGTGACATCATAAGAGAAACATAAAGTATCC

>M6-PGSC0003DMG400003834-(rev)M6LRR-RLP_1364

GATTTCTAATTTTCTCAAACTCTGTCTCGTGACATCATAAGAGAAACATAAAGTATCCGACATTAAGTCCAGGATGACTACAATCTCTTCATCACAGGACGTGCATGATGATATAACTAA

>M6-PGSC0003DMG400003834-(rev)M6LRR-RLP_1426

TTAAGTCCAGGATGACTACAATCTCTTCATCACAGGACGTGCATGATGATATAACTAAGGGTTTGTCCTTCCACCACGTCGAAGGGAAGTCAATGATTTGAATTTTCCATTCTTCTTCTT

>M6-PGSC0003DMG400003834-(rev)M6LRR-RLP_1488

TTGTCCTTCCACCACGTCGAAGGGAAGTCAATGATTTGAATTTTCCATTCTTCTTCTTTCTGAGTTTGTTCTAAAATCCACAAGTTGAACTATCCATGTTGCCAATATTCATAATCCATG

>M6-PGSC0003DMG400003834-(rev)M6LRR-RLP_1550

AGTTTGTTCTAAAATCCACAAGTTGAACTATCCATGTTGCCAATATTCATAATCCATGATCGCTAGTTTACCTTTCACCTCTAACAACTCGTAATACCACCACGAGTTAGATGCATTATA

>M6-PGSC0003DMG400003834-(rev)M6LRR-RLP_1612

CTAGTTTACCTTTCACCTCTAACAACTCGTAATACCACCACGAGTTAGATGCATTATATAAACCAATATTTTTTGAATTTTCGGTTTTAACATCAAATGCATCTATAGTAAGTCTGTCCT

>M6-PGSC0003DMG400003834-(rev)M6LRR-RLP_1674

CCAATATTTTTTGAATTTTCGGTTTTAACATCAAATGCATCTATAGTAAGTCTGTCCTGGAAAACAAATTTATAGATAACTCTATTGATGCAAACCCCGGGCTTGCACAGAATAGGAGGG

>M6-PGSC0003DMG400003834-(rev)M6LRR-RLP_1736

AACAAATTTATAGATAACTCTATTGATGCAAACCCCGGGCTTGCACAGAATAGGAGGGGTTATGCTTTGAGTCTCTCTCCATGATTCGTCTACTCCTAAAGTGAAAATCCAATGTTTTAT

>M6-PGSC0003DMG400003834-(rev)M6LRR-RLP_1798

TGCTTTGAGTCTCTCTCCATGATTCGTCTACTCCTAAAGTGAAAATCCAATGTTTTATGTACCCTTATATAATATGCTTTGTTGTGAAAAGAACTTTGTACTTCTTTTCTTCTGGGTCAA

>M6-PGSC0003DMG400003834-(rev)M6LRR-RLP_186

CCTCTCATTGATAAGTGGCTCGACATTGCACTTCAGAATGGTGTAAAACATCTTTTTCTTAACTTTAAATCATACCCCATGCCTATTTTAAGAATCTTGGCAGCAAAATCTTTAAGAGAA

>M6-PGSC0003DMG400003834-(rev)M6LRR-RLP_1860

CCTTATATAATATGCTTTGTTGTGAAAAGAACTTTGTACTTCTTTTCTTCTGGGTCAAAACCGAATGAACAGAACCACATGGAAGAACCATCATTTAGGTCAGGAAGAAATCTTACTTCT

>M6-PGSC0003DMG400003834-(rev)M6LRR-RLP_1920

CCGAATGAACAGAACCACATGGAAGAACCATCATTTAGGTCAGGAAGAAATCTTACTTCTCTTGTACTAGGATTGAAAATTGCAGCAAAAAATAATGGTTCCCAAATATAAAATAAACCA

>M6-PGSC0003DMG400003834-(rev)M6LRR-RLP_248

CTTTAAATCATACCCCATGCCTATTTTAAGAATCTTGGCAGCAAAATCTTTAAGAGAATTGGATGTGCAGGGTAGTATGCCTGATTCGTTATCTAGTGGTGTTGTGAATTGCAAATCATT

>M6-PGSC0003DMG400003834-(rev)M6LRR-RLP_372

AAGCTTTCTCTATCTCATCTAAGATTAGACGAAAACATGGTCCAGACTATACTTACTAGTTGTCCCTTTATTGTCTCTTTAATCCTTGAGTATTGTACTGGGAAGCTTCAAAAGATCAAG

>M6-PGSC0003DMG400003834-(rev)M6LRR-RLP_434

TCCCTTTATTGTCTCTTTAATCCTTGAGTATTGTACTGGGAAGCTTCAAAAGATCAAGTCGGATTCCTTGAAGGTCTTGAAGATACATCATCGCGGGATAGAGGAGATCGATGCTCCAAA

>M6-PGSC0003DMG400003834-(rev)M6LRR-RLP_496

ATTCCTTGAAGGTCTTGAAGATACATCATCGCGGGATAGAGGAGATCGATGCTCCAAATTTGGTATCACTTGACTATATGGGGAGTCAAATTCCCGAACTTAATAAAATTGCAAGAGACT

>M6-PGSC0003DMG400003834-(rev)M6LRR-RLP_558

GTATCACTTGACTATATGGGGAGTCAAATTCCCGAACTTAATAAAATTGCAAGAGACTCAAAAATCAGGCTTGGATGCATCAACAATTTAAATGCAGCATGGTTTTGTAAGTTGAGAAAG

>M6-PGSC0003DMG400003834-(rev)M6LRR-RLP_62

TATTAATTGTTGGGAAGGCAATATAAATACAGCGAACACAACCATGGAGAGATATAGGAAGGGGAAAATCCCTATACAAAAGTTTGAATTATTAGAGACCTTTGCTAATTCTCATGAAGA

>M6-PGSC0003DMG400003834-(rev)M6LRR-RLP_620

AATCAGGCTTGGATGCATCAACAATTTAAATGCAGCATGGTTTTGTAAGTTGAGAAAGTTTCTATCAAATTCGTCCTCGTGGTCTCATGTTTCCCTTCAATTTATGAATTGCTGTGAGAT

>M6-PGSC0003DMG400003834-(rev)M6LRR-RLP_682

TATCAAATTCGTCCTCGTGGTCTCATGTTTCCCTTCAATTTATGAATTGCTGTGAGATCAAGATGAAAGATTTGCAAATGGACCACATAGGTTCTACCCCTCGGGTGGACATTTTAAATG

>M6-PGSC0003DMG400003834-(rev)M6LRR-RLP_744

ATGAAAGATTTGCAAATGGACCACATAGGTTCTACCCCTCGGGTGGACATTTTAAATGTCGATATTCAATGGATGAATCAGAATTTTGTGGATGCTTTGCTATGGAGTTGTCATCCAAGG

>M6-PGSC0003DMG400003834-(rev)M6LRR-RLP_806

TATTCAATGGATGAATCAGAATTTTGTGGATGCTTTGCTATGGAGTTGTCATCCAAGGAGACTCAACCTACTCTCAAATATTACTCCAACGATTACACATTTCATTGATCGTTTAGTGTT

>M6-PGSC0003DMG400003834-(rev)M6LRR-RLP_868

TCAACCTACTCTCAAATATTACTCCAACGATTACACATTTCATTGATCGTTTAGTGTTTTTGACGAATTCGAGTCATTCTACTTCTCATAAAAGCTTTCCTTGGCATAGTCAATTAAAAG

>M6-PGSC0003DMG400003834-(rev)M6LRR-RLP_930

ACGAATTCGAGTCATTCTACTTCTCATAAAAGCTTTCCTTGGCATAGTCAATTAAAAGAAATAAAAGCTTTTGATGGAAAAAACCAGTCGCTACAACTCATAAATGGGGAGCTTGCAAAA

>M6-PGSC0003DMG400003834-(rev)M6LRR-RLP_992

AAAAGCTTTTGATGGAAAAAACCAGTCGCTACAACTCATAAATGGGGAGCTTGCAAAAAGGACCCTTATGGAGGGGGAGAAAGTTTATTTTTTATTAGATTGGTGATGTAGTTAATTTGT

>M6-PGSC0003DMG400003835-(rev)M6LRR-RLP_0

ATGGCTGACATGTTGCCTGAATGTTTCATTCAAAAAATACTCTCTTCTCTTAGTTTTGATGAAGCAGCGCGGATGAGCATTCTTTCCAAAACATGGCTGCAAGCCTGGTCGACTCTTCCC

>M6-PGSC0003DMG400003835-(rev)M6LRR-RLP_1056

AATTCGATTCAGTCTACTTCTCATGGAAGCAACCCTTGGTATAGTCAATTAAAAGAAGTAAAAGTCCACAAATTTGATAGGAAGAAGGAGATTTGGTATCCTGTGGAACACAAACTTGGG

>M6-PGSC0003DMG400003835-(rev)M6LRR-RLP_1107

AAAGAAGTAAAAGTCCACAAATTTGATAGGAAGAAGGAGATTTGGTATCCTGTGGAACACAAACTTGGGGAGCGGGCAACAATGAACCTTGATAGATATTATATTTTATTAGATTGGTAA

>M6-PGSC0003DMG400003835-(rev)M6LRR-RLP_132

AATGTTAAGTTTTGCAAAGGCGATATAAAGATATTGGACACTATTATGGAGAGATATGGAAACGGAAAAATCCCTATAGAAAAGTTTGATTTATCAGAGCTCTTAGGCGATTCTCATGAA

>M6-PGSC0003DMG400003835-(rev)M6LRR-RLP_198

AAAATCCCTATAGAAAAGTTTGATTTATCAGAGCTCTTAGGCGATTCTCATGAAGTTTTCCCTCTGATTGATAAGTGGCTTAACATTGCACTTCAAAATGGCGTAAACAATCTTGTCCTT

>M6-PGSC0003DMG400003835-(rev)M6LRR-RLP_264

ATTGATAAGTGGCTTAACATTGCACTTCAAAATGGCGTAAACAATCTTGTCCTTAACTATAAATCATACCCCGTGCCTTTTTTAACAATCTTGGCAGCAAAATCTTTAAGAAAATTGGTT

>M6-PGSC0003DMG400003835-(rev)M6LRR-RLP_330

TACCCCGTGCCTTTTTTAACAATCTTGGCAGCAAAATCTTTAAGAAAATTGGTTCTGAAGAGTTGTACTCTTTTGCCTATTTCTTTATCTGGTGGTGTTGTGAATCACAATTCTTTGAGT

>M6-PGSC0003DMG400003835-(rev)M6LRR-RLP_396

ACTCTTTTGCCTATTTCTTTATCTGGTGGTGTTGTGAATCACAATTCTTTGAGTAAGCTTTCTCTATCTTATGTAAAATTATACGAAAACATGCTTCAGACTCTACTTAATAGTTGTCCT

>M6-PGSC0003DMG400003835-(rev)M6LRR-RLP_462

TCTTATGTAAAATTATACGAAAACATGCTTCAGACTCTACTTAATAGTTGTCCTTTGATTGTCTCTTTCATCTTTGAGTATTGTTCGGACTTGGAAACAATTGAGCTAGTGAATCTTCAA

>M6-PGSC0003DMG400003835-(rev)M6LRR-RLP_528

TTCATCTTTGAGTATTGTTCGGACTTGGAAACAATTGAGCTAGTGAATCTTCAAAAGATCAAGTCGGTTTCCCTGAAGGTCTTGAAGATTCGGTTTAGTGGGAGCATATGGGAGATTGAT

>M6-PGSC0003DMG400003835-(rev)M6LRR-RLP_594

GTTTCCCTGAAGGTCTTGAAGATTCGGTTTAGTGGGAGCATATGGGAGATTGATGCTCCAAATTTGGTGTCATTTGAGTACACAGGGAATCAAATTCCTGAACTTCAAATTGTGAGAGAG

>M6-PGSC0003DMG400003835-(rev)M6LRR-RLP_66

GCGCGGATGAGCATTCTTTCCAAAACATGGCTGCAAGCCTGGTCGACTCTTCCCGACTTGGAATTTAATGTTAAGTTTTGCAAAGGCGATATAAAGATATTGGACACTATTATGGAGAGA

>M6-PGSC0003DMG400003835-(rev)M6LRR-RLP_660

GTGTCATTTGAGTACACAGGGAATCAAATTCCTGAACTTCAAATTGTGAGAGAGTCAAAACAACTGAAGCACTCAAAAATCATTCTTCACCGCTTAGACAATATAAATGCTGACTGGTTT

>M6-PGSC0003DMG400003835-(rev)M6LRR-RLP_792

AAGTTTCTATCAAATTCGACCTCTTGGTCTCAAGTTTCCCTTTATTTTTATGAATGCACGGAGATTAAAATGAAAGATCTGCAACAACACCACAGAGTTGCTACGCCCCAGGTGGACGTT

>M6-PGSC0003DMG400003835-(rev)M6LRR-RLP_858

AAAATGAAAGATCTGCAACAACACCACAGAGTTGCTACGCCCCAGGTGGACGTTTTAGATGTCAACATTCTGTGGCAGAATCTCGAATACCCTTCATTTGTGGATGCTTTGGTATGGAGT

>M6-PGSC0003DMG400003835-(rev)M6LRR-RLP_924

ATTCTGTGGCAGAATCTCGAATACCCTTCATTTGTGGATGCTTTGGTATGGAGTTGTCAGCCTAGGAGACTCAACTTGCAATTAACAAGAGAAATGGTTACAGTTTTCATTGATCGTTTG

>M6-PGSC0003DMG400003835-(rev)M6LRR-RLP_990

AGACTCAACTTGCAATTAACAAGAGAAATGGTTACAGTTTTCATTGATCGTTTGATGAATATGAAGAATTCGATTCAGTCTACTTCTCATGGAAGCAACCCTTGGTATAGTCAATTAAAA

>M6-PGSC0003DMG400003918-(rev)M6LRR-RLK_0

TCTATCTGGGAATCTTTTAGTAGGAGAAATACCACAAAACCTGTACAACTTGTCAAACCTTGTATATCTTGACTTACATCATAACCAACTGAATGGAAGTATTCCATCAACCATTGGAAA

>M6-PGSC0003DMG400003918-(rev)M6LRR-RLK_1037

TCCTGGATACAAACTACGAAGCCAAATTGACTGATTATGGCCTGAGTAGGTTCCTTCCCCTGCTGGATAACTATGGACTAACCAAGTTTCACAATGCAGTTGGTTATGTTGCACCAGAGT

>M6-PGSC0003DMG400003918-(rev)M6LRR-RLK_1098

GCTGGATAACTATGGACTAACCAAGTTTCACAATGCAGTTGGTTATGTTGCACCAGAGTTGGCGCAAAGTCTGAGACTAAGTGATAAATGTGACGTTTACAGTTTTGGGGTGATTCTGCT

>M6-PGSC0003DMG400003918-(rev)M6LRR-RLK_1159

GCGCAAAGTCTGAGACTAAGTGATAAATGTGACGTTTACAGTTTTGGGGTGATTCTGCTGGAGTTGGTCACTGGGAGGAAACCAGTAGAGAGCCCAGTAGCAAATGAAGTGGTGGTCTTG

>M6-PGSC0003DMG400003918-(rev)M6LRR-RLK_122

TGTCTAACTTGCACTTTTTGGATTTATCACAAAATCTGCTATCTGGGTCAATCCCTGTTGCTCTTGGGGATTTGCAAAATTTAACTCATTTTAATGTATCTTACAACCTTTTATCTGGTG

>M6-PGSC0003DMG400003918-(rev)M6LRR-RLK_1220

AGTTGGTCACTGGGAGGAAACCAGTAGAGAGCCCAGTAGCAAATGAAGTGGTGGTCTTGTGTGAATATGTTAGAGGATTGATAGAAAGGGGTGCTGCTTCAGACTGTTTCGATAGAAGTT

>M6-PGSC0003DMG400003918-(rev)M6LRR-RLK_1281

TGAATATGTTAGAGGATTGATAGAAAGGGGTGCTGCTTCAGACTGTTTCGATAGAAGTTTGAGGGGCTTTGCAGAAAATGAGCTGATTCAGGTAATGAAGTTAGGCTTGATTTGTACTTC

>M6-PGSC0003DMG400003918-(rev)M6LRR-RLK_1342

AGGGGCTTTGCAGAAAATGAGCTGATTCAGGTAATGAAGTTAGGCTTGATTTGTACTTCTGAAATATCCTCAAGGAGACCTAGCATGGCTGAGGTAGTGCAGGTTCTTGAGTCCATCAGG

>M6-PGSC0003DMG400003918-(rev)M6LRR-RLK_1403

AAATATCCTCAAGGAGACCTAGCATGGCTGAGGTAGTGCAGGTTCTTGAGTCCATCAGGAATGGCTTGGGATCATAGTAAAGAATCAGTAGAGCCATAAGACTATTTGAAACAGCAGCTG

>M6-PGSC0003DMG400003918-(rev)M6LRR-RLK_1464

TGGCTTGGGATCATAGTAAAGAATCAGTAGAGCCATAAGACTATTTGAAACAGCAGCTGTTCATAGAATGAGAAGAAAGCAGTCTAGCTGTTGTGCTAGACTCGCTTCTCATGAGATTCG

>M6-PGSC0003DMG400003918-(rev)M6LRR-RLK_1525

CATAGAATGAGAAGAAAGCAGTCTAGCTGTTGTGCTAGACTCGCTTCTCATGAGATTCGATGAGGGAATCAGAAAGGGTCTCAAATAAGAGAGACGCGCAGACCAATTAATAGTATCAGT

>M6-PGSC0003DMG400003918-(rev)M6LRR-RLK_1586

GAGGGAATCAGAAAGGGTCTCAAATAAGAGAGACGCGCAGACCAATTAATAGTATCAGTCATATTCTTGGATTTGGAATCAGAGTGATTAGGAGGTGACAGCTTTTGGTACAGAGAGAAG

>M6-PGSC0003DMG400003918-(rev)M6LRR-RLK_1647

TATTCTTGGATTTGGAATCAGAGTGATTAGGAGGTGACAGCTTTTGGTACAGAGAGAAGATTCTTTAGTATTTGATTTGTGATTCTTTTAATCAATTGGTCATTGTAAGTTTACAATAAA

>M6-PGSC0003DMG400003918-(rev)M6LRR-RLK_1708

TCTTTAGTATTTGATTTGTGATTCTTTTAATCAATTGGTCATTGTAAGTTTACAATAAAGCAATTCTTTTCCTGTAAGATGCAAAGGTATTTTGATGTCTCATATGCAGCTTGGCAACTG

>M6-PGSC0003DMG400003918-(rev)M6LRR-RLK_1769

AATTCTTTTCCTGTAAGATGCAAAGGTATTTTGATGTCTCATATGCAGCTTGGCAACTGCTTTTTATAACCATTCAATATTGCTCTTCATTTACTGTTAAGAATTACCTTTGACATGCAT

>M6-PGSC0003DMG400003918-(rev)M6LRR-RLK_183

TCTTGGGGATTTGCAAAATTTAACTCATTTTAATGTATCTTACAACCTTTTATCTGGTGCTATTCCCTCAATTGAGAGCATCAAGAAGTTTGGACCCTCTGCCTTTTTTCATAATAGTGG

>M6-PGSC0003DMG400003918-(rev)M6LRR-RLK_1830

TTTTATAACCATTCAATATTGCTCTTCATTTACTGTTAAGAATTACCTTTGACATGCATATATCAACAACCTTTGAAGGGATGAAAGAAATCGATTCATTTCTCGAATTACATATCTAGT

>M6-PGSC0003DMG400003918-(rev)M6LRR-RLK_1891

ATCAACAACCTTTGAAGGGATGAAAGAAATCGATTCATTTCTCGAATTACATATCTAGTGCTAACACAAGAATGAATGTGAATAAATAATGAGTCCATACATCTACCTGTAACTATACAA

>M6-PGSC0003DMG400003918-(rev)M6LRR-RLK_1952

TAACACAAGAATGAATGTGAATAAATAATGAGTCCATACATCTACCTGTAACTATACAACAATGAGATAATAAGAAAATATCAAGATTTAAGACCTTACTTATTGACACAAAGAATTTGG

>M6-PGSC0003DMG400003918-(rev)M6LRR-RLK_2013

ATGAGATAATAAGAAAATATCAAGATTTAAGACCTTACTTATTGACACAAAGAATTTGGAAATGGATTTCATTTGGAGCCATTTCTACATGCTATATGTTTTTCCTTGCATCAACTTTGA

>M6-PGSC0003DMG400003918-(rev)M6LRR-RLK_2074

ATGGATTTCATTTGGAGCCATTTCTACATGCTATATGTTTTTCCTTGCATCAACTTTGACTGCTAATTTAGCTATCTTCAAAATCCATAAACTGGGAGCTGGAATGCATTTGTGAAAATG

>M6-PGSC0003DMG400003918-(rev)M6LRR-RLK_2135

GCTAATTTAGCTATCTTCAAAATCCATAAACTGGGAGCTGGAATGCATTTGTGAAAATGACAGTTGGAGAGGACCAAGAACACACTTCTGGAATCATCTGCATACAATCTGGCCTTTTTT

>M6-PGSC0003DMG400003918-(rev)M6LRR-RLK_2196

AGTTGGAGAGGACCAAGAACACACTTCTGGAATCATCTGCATACAATCTGGCCTTTTTTTGTTTGTTTTCTCTGCATGTAGTTCTCAGTGCTGCTGCATTCTGATCTCATTATCCGAACA

>M6-PGSC0003DMG400003918-(rev)M6LRR-RLK_2257

TTTGTTTTCTCTGCATGTAGTTCTCAGTGCTGCTGCATTCTGATCTCATTATCCGAACATTGATAACTTATGGATCCATTAAATCACATAACACACGACCCTTAAAATCTCTGCAGAAGT

>M6-PGSC0003DMG400003918-(rev)M6LRR-RLK_2318

GATAACTTATGGATCCATTAAATCACATAACACACGACCCTTAAAATCTCTGCAGAAGTTTTGGACCTTTGGTTACGGTTTACTGTAAGGGATCATATCATTAACTCAGAGACTTACGTA

>M6-PGSC0003DMG400003918-(rev)M6LRR-RLK_2379

TGGACCTTTGGTTACGGTTTACTGTAAGGGATCATATCATTAACTCAGAGACTTACGTAACTCCAAAACTTTCTGGATACGCTGCATAACTGTGGAAATGTCCTTCTCACTTAATGTCGA

>M6-PGSC0003DMG400003918-(rev)M6LRR-RLK_244

ATTCCCTCAATTGAGAGCATCAAGAAGTTTGGACCCTCTGCCTTTTTTCATAATAGTGGTCTTTGTGGTGATCCTTTAGAAGTTTCATGCTCTGCCAGTGGCACTACTCCTGCAAAAGGG

>M6-PGSC0003DMG400003918-(rev)M6LRR-RLK_2440

TCCAAAACTTTCTGGATACGCTGCATAACTGTGGAAATGTCCTTCTCACTTAATGTCGAAAAACATAACCTGAACCACCCAGGTTCAACACAGTGGCAGGAAGAACCAGGAGTTGCATTG

>M6-PGSC0003DMG400003918-(rev)M6LRR-RLK_2501

AACATAACCTGAACCACCCAGGTTCAACACAGTGGCAGGAAGAACCAGGAGTTGCATTGATCTTAGCTACATTTAGTAGATTATCCCAAAGCTCAATCTCACCCTTCTCATTATACGACC

>M6-PGSC0003DMG400003918-(rev)M6LRR-RLK_2562

CTTAGCTACATTTAGTAGATTATCCCAAAGCTCAATCTCACCCTTCTCATTATACGACCGAATTAACCTGCTCATGTCAGCCCAACAATAGAAACCCCCACTACTTTTGGTGCACTCAAT

>M6-PGSC0003DMG400003918-(rev)M6LRR-RLK_2623

ATTAACCTGCTCATGTCAGCCCAACAATAGAAACCCCCACTACTTTTGGTGCACTCAATTCCCAGCTGCTTCAATCCACTGACAAATAGAGAAGACATCCTTCTTAGCCTCTCTCTGTTC

>M6-PGSC0003DMG400003918-(rev)M6LRR-RLK_2684

CCAGCTGCTTCAATCCACTGACAAATAGAGAAGACATCCTTCTTAGCCTCTCTCTGTTCTTTTTGATAAATTGTTGTACAAACTTTGCATCTGATAGCATCTGGATGATTAAATGCTGTG

>M6-PGSC0003DMG400003918-(rev)M6LRR-RLK_2745

TTTGATAAATTGTTGTACAAACTTTGCATCTGATAGCATCTGGATGATTAAATGCTGTGTTGGGGCTGAAATGGATGAGAATCTTGTCAACTTTTTCGCAGCAGCTAGAACATTTTCATT

>M6-PGSC0003DMG400003918-(rev)M6LRR-RLK_2806

GGGGCTGAAATGGATGAGAATCTTGTCAACTTTTTCGCAGCAGCTAGAACATTTTCATTGCAAGAATAGATGACCCCCACCCTGAAACCAGGAAGAGAAAGGTCCTTAGAGAGACCATAC

>M6-PGSC0003DMG400003918-(rev)M6LRR-RLK_2867

AAGAATAGATGACCCCCACCCTGAAACCAGGAAGAGAAAGGTCCTTAGAGAGACCATACACAATGTGTACCCTGCTCCTGTCAAAATCTTCAGAATCAATTATCTCTGCTATGCTGACAA

>M6-PGSC0003DMG400003918-(rev)M6LRR-RLK_2928

AATGTGTACCCTGCTCCTGTCAAAATCTTCAGAATCAATTATCTCTGCTATGCTGACAAATTCTTCACGTCCATAAGTTGACCCTGCCAAGATTTCATTGGATATTACATGGATGTTCTT

>M6-PGSC0003DMG400003918-(rev)M6LRR-RLK_2989

TCTTCACGTCCATAAGTTGACCCTGCCAAGATTTCATTGGATATTACATGGATGTTCTTCTCAGTTGCAAAGTCTAAGAGGTTATAAAGTGTCTCCCTAGAGAAAATATTTCCCACAGGA

>M6-PGSC0003DMG400003918-(rev)M6LRR-RLK_305

TTTGTGGTGATCCTTTAGAAGTTTCATGCTCTGCCAGTGGCACTACTCCTGCAAAAGGGAAACCAAACTTGAGTGTCTCAGCTATTGTTGCTATTGTCGCTGCTTCAATAATCCTTACAG

>M6-PGSC0003DMG400003918-(rev)M6LRR-RLK_3050

CAGTTGCAAAGTCTAAGAGGTTATAAAGTGTCTCCCTAGAGAAAATATTTCCCACAGGATTTGAGGGGTTAGAAATGATAATCCCTCGTACTTTAAGACCACGTTTCTTAGCTTGGTTAA

>M6-PGSC0003DMG400003918-(rev)M6LRR-RLK_3111

TGAGGGGTTAGAAATGATAATCCCTCGTACTTTAAGACCACGTTTCTTAGCTTGGTTAAAAGCTCGGTCAAGAGCATCAATGCTCAAGTTAAAGTTGTCTGCACTGCGGCAAGGAACAGG

>M6-PGSC0003DMG400003918-(rev)M6LRR-RLK_3172

GCTCGGTCAAGAGCATCAATGCTCAAGTTAAAGTTGTCTGCACTGCGGCAAGGAACAGGTATAATCTCCACTCCAGTTCTCCATTTGACATCCCTATCAAGACTGCAAACGCATTGAGCT

>M6-PGSC0003DMG400003918-(rev)M6LRR-RLK_3233

TAATCTCCACTCCAGTTCTCCATTTGACATCCCTATCAAGACTGCAAACGCATTGAGCTCATTTTATACACTATATACGTAAAGATGACCAAAAGATTGAAATTTATTTGGGAAGGAGAA

>M6-PGSC0003DMG400003918-(rev)M6LRR-RLK_3294

TTTTATACACTATATACGTAAAGATGACCAAAAGATTGAAATTTATTTGGGAAGGAGAATAAAACTCACTCAGGATAATATGGTGAGGGAACAAGAAAAGCATTTCCAGGATCAGCTAGG

>M6-PGSC0003DMG400003918-(rev)M6LRR-RLK_3355

AAACTCACTCAGGATAATATGGTGAGGGAACAAGAAAAGCATTTCCAGGATCAGCTAGGCAGAAGCTCAGTATCTCAAGCGCAGGAGCGGCACCACCAGTTAGAACCATCTGCGAGGGGC

>M6-PGSC0003DMG400003918-(rev)M6LRR-RLK_3416

GAAGCTCAGTATCTCAAGCGCAGGAGCGGCACCACCAGTTAGAACCATCTGCGAGGGGCTGAAGGACACTGATCTCTCCAGAGCTTGAGACATAAATTCTCCCACAGCCTGCATACAGGT

>M6-PGSC0003DMG400003918-(rev)M6LRR-RLK_3477

AAGGACACTGATCTCTCCAGAGCTTGAGACATAAATTCTCCCACAGCCTGCATACAGGTGAGGAAACATGAAAAGCAATTCAAAGAGATACCATAAATAATGTAGAATCACACAAAACAC

>M6-PGSC0003DMG400003918-(rev)M6LRR-RLK_3538

GGAAACATGAAAAGCAATTCAAAGAGATACCATAAATAATGTAGAATCACACAAAACACCTAATTGTTTTTATGAATCGATCTTCATAGTGGCTCAAGGATGATTTCATATATTGCTGTT

>M6-PGSC0003DMG400003918-(rev)M6LRR-RLK_3599

AATTGTTTTTATGAATCGATCTTCATAGTGGCTCAAGGATGATTTCATATATTGCTGTTATACAATTCTCAAACCCACAATTTAAAAAGGTAAATCCATCAGATCATCTCCCACCCTAGG

>M6-PGSC0003DMG400003918-(rev)M6LRR-RLK_366

ACCAAACTTGAGTGTCTCAGCTATTGTTGCTATTGTCGCTGCTTCAATAATCCTTACAGGCGTTTGTCTGATAACCATCATCAATATGAAGGCTCGACGGAGAAGGAGGAGAGAAGATGA

>M6-PGSC0003DMG400003918-(rev)M6LRR-RLK_3660

ACAATTCTCAAACCCACAATTTAAAAAGGTAAATCCATCAGATCATCTCCCACCCTAGGATACCCAATGACAAAGGTAGCCACTCATATTTCAAAGAGGCCAAAATTGCCTTCCATCTTC

>M6-PGSC0003DMG400003918-(rev)M6LRR-RLK_3721

ACCCAATGACAAAGGTAGCCACTCATATTTCAAAGAGGCCAAAATTGCCTTCCATCTTCAACAGATACTGCAAATCTAGCAATAGATCCTTCAACAGGTTACACAATGATGAGTATTGAG

>M6-PGSC0003DMG400003918-(rev)M6LRR-RLK_3782

CAGATACTGCAAATCTAGCAATAGATCCTTCAACAGGTTACACAATGATGAGTATTGAGGTGAGTATCTGTCAAGGAAATGATAGACCACCTCTCAGAACTACTGAGTCCATTATTCGCT

>M6-PGSC0003DMG400003918-(rev)M6LRR-RLK_3843

GAGTATCTGTCAAGGAAATGATAGACCACCTCTCAGAACTACTGAGTCCATTATTCGCTATTAACTTGCTTATAACGAAAACCATGCAAAGTAAAGCAGGTTAAACTATTCTCTTATGTG

>M6-PGSC0003DMG400003918-(rev)M6LRR-RLK_3904

TAACTTGCTTATAACGAAAACCATGCAAAGTAAAGCAGGTTAAACTATTCTCTTATGTGCTCAAACTAGATTTTTTATCATATCACACTGCAAGGGATCCTATTGTTCCGGTCAGGACAA

>M6-PGSC0003DMG400003918-(rev)M6LRR-RLK_3965

CAAACTAGATTTTTTATCATATCACACTGCAAGGGATCCTATTGTTCCGGTCAGGACAAACATCACCACAGAAAATCATCAAATGCTGAAACAGCATAAATGACTGAGCTAGATCAACAA

>M6-PGSC0003DMG400003918-(rev)M6LRR-RLK_4026

ATCACCACAGAAAATCATCAAATGCTGAAACAGCATAAATGACTGAGCTAGATCAACAATAATTATGAATTTCCAACTCAAGTTTTACCAATGTAACGCTCAAGTAGGAGAAGATATTTA

>M6-PGSC0003DMG400003918-(rev)M6LRR-RLK_4087

ATTATGAATTTCCAACTCAAGTTTTACCAATGTAACGCTCAAGTAGGAGAAGATATTTACCACAAAAAAACACCCCCATCCCTCTGGATACTGTTTCAATGCTTTGGCTGGTCAAACACA

>M6-PGSC0003DMG400003918-(rev)M6LRR-RLK_4148

ACAAAAAAACACCCCCATCCCTCTGGATACTGTTTCAATGCTTTGGCTGGTCAAACACATTTTCAGGTTTCAAAATTTTCCTATCATTTCTCTTCTTTTGTATTTTAACCTCCTTTATTA

>M6-PGSC0003DMG400003918-(rev)M6LRR-RLK_4209

TTCAGGTTTCAAAATTTTCCTATCATTTCTCTTCTTTTGTATTTTAACCTCCTTTATTAGACAGTCATGGGCTCCAAAAGCTCAAGAAAAAAATCAAAACATTCAGAACCGTGTTTATTT

>M6-PGSC0003DMG400003918-(rev)M6LRR-RLK_427

GTTTGTCTGATAACCATCATCAATATGAAGGCTCGACGGAGAAGGAGGAGAGAAGATGAAACATTCGTTGTAGAGAGCACGCCACTGGCTTCAACAGATTCAAATGTCATTATAGGAAAG

>M6-PGSC0003DMG400003918-(rev)M6LRR-RLK_4270

CAGTCATGGGCTCCAAAAGCTCAAGAAAAAAATCAAAACATTCAGAACCGTGTTTATTTTTCAAAGTGTTGCTTAGTGTCGATAGCGTAGAAGTTAGAGAATTGATGAACTTTTCAGAAG

>M6-PGSC0003DMG400003918-(rev)M6LRR-RLK_4331

CAAAGTGTTGCTTAGTGTCGATAGCGTAGAAGTTAGAGAATTGATGAACTTTTCAGAAGTTAATTGGCACTGTAAAATGAACGTGAAGCTTATTCAATGCTTTTCCTTATGTTTGTTGCA

>M6-PGSC0003DMG400003918-(rev)M6LRR-RLK_4392

AATTGGCACTGTAAAATGAACGTGAAGCTTATTCAATGCTTTTCCTTATGTTTGTTGCACTTTATAGACTTAGAAAGCTAAAGACGGGACATACTAAAACCTTGTATACAATAGGTAATT

>M6-PGSC0003DMG400003918-(rev)M6LRR-RLK_4453

TTATAGACTTAGAAAGCTAAAGACGGGACATACTAAAACCTTGTATACAATAGGTAATTTTGGGATTGTCAAATGCCATAACCAATCCCGCATCCAGGGGTTCACATTTCAAATTAAACC

>M6-PGSC0003DMG400003918-(rev)M6LRR-RLK_4514

GGGATTGTCAAATGCCATAACCAATCCCGCATCCAGGGGTTCACATTTCAAATTAAACCAGCTATAAAGCACACTGGACTAGCGGATGGGTGACTGATATGGAGTAACTGTTACACAAGC

>M6-PGSC0003DMG400003918-(rev)M6LRR-RLK_4575

CTATAAAGCACACTGGACTAGCGGATGGGTGACTGATATGGAGTAACTGTTACACAAGCATATGAACATAACATTTTTGAAATTATATGGGCATGACATGAGCCCATCCTAGGTTACAAC

>M6-PGSC0003DMG400003918-(rev)M6LRR-RLK_4636

ATGAACATAACATTTTTGAAATTATATGGGCATGACATGAGCCCATCCTAGGTTACAACTATAAACTAATACCACCCAAATAGGTATCTCGTTGATTTAATCCTATAGTCTAAACTCAAA

>M6-PGSC0003DMG400003918-(rev)M6LRR-RLK_4697

TAAACTAATACCACCCAAATAGGTATCTCGTTGATTTAATCCTATAGTCTAAACTCAAAGTAGCAACTTTAAATAATACGTACAAACTTTTTCAAGTAAATTTTTTGTTCAATTTCCTTC

>M6-PGSC0003DMG400003918-(rev)M6LRR-RLK_4758

AGCAACTTTAAATAATACGTACAAACTTTTTCAAGTAAATTTTTTGTTCAATTTCCTTCTCTAGCCTGTAAAAACTATAGCCATATGAACAACTACTAGTACTTTATTAGTTCAAGAAGA

>M6-PGSC0003DMG400003918-(rev)M6LRR-RLK_4819

TAGCCTGTAAAAACTATAGCCATATGAACAACTACTAGTACTTTATTAGTTCAAGAAGAACATATTATATCCAATACATGTCAACATTATATCAACAAACAAGTCACTAGTAGTGTGCAA

>M6-PGSC0003DMG400003918-(rev)M6LRR-RLK_488

CATTCGTTGTAGAGAGCACGCCACTGGCTTCAACAGATTCAAATGTCATTATAGGAAAGTTGGTTCTTTTCAGCAAGACTTTGCCTTCAAAATATGAGGACTGGGAGGCTGGCACAAAAG

>M6-PGSC0003DMG400003918-(rev)M6LRR-RLK_4880

ATATTATATCCAATACATGTCAACATTATATCAACAAACAAGTCACTAGTAGTGTGCAAGGAAAAATCAAAAAGCTTACCACTTTCAACTCCAACAACCCATCAAATGGCTGATAAGTAG

>M6-PGSC0003DMG400003918-(rev)M6LRR-RLK_4941

AAAAATCAAAAAGCTTACCACTTTCAACTCCAACAACCCATCAAATGGCTGATAAGTAGCTATTCCGGTTATACTCGAATCTTGAGTCATCCATCTTGACACATTTTCTGCTAGCCATTC

>M6-PGSC0003DMG400003918-(rev)M6LRR-RLK_5002

ATTCCGGTTATACTCGAATCTTGAGTCATCCATCTTGACACATTTTCTGCTAGCCATTCTTGAACTAAATCCAATGACAACTATAACCATACAAATAGAGACAAAATTCAAAATCATTGA

>M6-PGSC0003DMG400003918-(rev)M6LRR-RLK_5063

GAACTAAATCCAATGACAACTATAACCATACAAATAGAGACAAAATTCAAAATCATTGAAACAACAAACAAAAGTAGATTCCCCAACAAGAAATGTACTGCATTTGGTACATTTCTCTAG

>M6-PGSC0003DMG400003918-(rev)M6LRR-RLK_5124

CAACAAACAAAAGTAGATTCCCCAACAAGAAATGTACTGCATTTGGTACATTTCTCTAGAAAATGATTCAATATATCCAAACATTCAAAATTATTAGTACCTTGTTTTCTGCTAATCCTA

>M6-PGSC0003DMG400003918-(rev)M6LRR-RLK_5185

AATGATTCAATATATCCAAACATTCAAAATTATTAGTACCTTGTTTTCTGCTAATCCTAGCTGAATAACACCATCTGGATTACTTGACTCATCATAAGGGTCTTCAGAAGCTCTTTTCAA

>M6-PGSC0003DMG400003918-(rev)M6LRR-RLK_5246

TGAATAACACCATCTGGATTACTTGACTCATCATAAGGGTCTTCAGAAGCTCTTTTCAATCCAACATAGAAAGGGCTACTATCAATTTGTTTTGAAATAATCGAATTAGCTCTAGCAGAT

>M6-PGSC0003DMG400003918-(rev)M6LRR-RLK_5307

CAACATAGAAAGGGCTACTATCAATTTGTTTTGAAATAATCGAATTAGCTCTAGCAGATACCTGAGATTGCCCAGTAGTTCCCTTCGGTGACAGTAAAACACGTGAGTGAACCCTTTGTA

>M6-PGSC0003DMG400003918-(rev)M6LRR-RLK_5368

CTGAGATTGCCCAGTAGTTCCCTTCGGTGACAGTAAAACACGTGAGTGAACCCTTTGTAACCCAGACCCAGAAGACACTTCCGGTAAATGGGTCGACGACGTTGACCTCGCCGGAGCCGT

>M6-PGSC0003DMG400003918-(rev)M6LRR-RLK_5429

CCAGACCCAGAAGACACTTCCGGTAAATGGGTCGACGACGTTGACCTCGCCGGAGCCGTAGATTCGCCGTTATTGTCACCACCGGAGGAACGATTTCGTTTTAGGTAGAGTTGCCAAAAA

>M6-PGSC0003DMG400003918-(rev)M6LRR-RLK_549

GGTTCTTTTCAGCAAGACTTTGCCTTCAAAATATGAGGACTGGGAGGCTGGCACAAAAGCCTTGCTAGACAAGGAGTCTTTGATTGGTGGGGGGACAATTGGATCAGTTTACAGAACTTC

>M6-PGSC0003DMG400003918-(rev)M6LRR-RLK_5490

ATTCGCCGTTATTGTCACCACCGGAGGAACGATTTCGTTTTAGGTAGAGTTGCCAAAAATAGAAAAGAGCACAGGGTATAACTGAACCTAAGAAAAGACCACCCCGACCTTGAACTACAC

>M6-PGSC0003DMG400003918-(rev)M6LRR-RLK_5551

GAAAAGAGCACAGGGTATAACTGAACCTAAGAAAAGACCACCCCGACCTTGAACTACACCTTGTAAGGGTACTATCACCCTCATCGCCGTCGTGGCTCCTCCTCCGTCTCTGCCGCCGGC

>M6-PGSC0003DMG400003918-(rev)M6LRR-RLK_5612

TGTAAGGGTACTATCACCCTCATCGCCGTCGTGGCTCCTCCTCCGTCTCTGCCGCCGGCGCCGCCAGTTGAAATGGTTGTTGTTGTTGTTCTAGTTGGTGAACGGTCTCGTGACCTTGTC

>M6-PGSC0003DMG400003918-(rev)M6LRR-RLK_5673

CGCCAGTTGAAATGGTTGTTGTTGTTGTTCTAGTTGGTGAACGGTCTCGTGACCTTGTCATAACTGTGACATATTGGGGGCAGAATTAAGACGTCACCTTATCGGAGTATGACGTCAGTC

>M6-PGSC0003DMG400003918-(rev)M6LRR-RLK_5734

AACTGTGACATATTGGGGGCAGAATTAAGACGTCACCTTATCGGAGTATGACGTCAGTCAGCTATCTGGCCGGCGAGACGGTTGACGGAGAGTACGAAGGGGGGTGAGTGATGGAGAAGA

>M6-PGSC0003DMG400003918-(rev)M6LRR-RLK_5795

CTATCTGGCCGGCGAGACGGTTGACGGAGAGTACGAAGGGGGGTGAGTGATGGAGAAGAGATTTGGAGAAAATGGTGGCAAATAGTTGGTCGAATTATTATTTATCTTTTTGTCCGCCTA

>M6-PGSC0003DMG400003918-(rev)M6LRR-RLK_61

GTATATCTTGACTTACATCATAACCAACTGAATGGAAGTATTCCATCAACCATTGGAAATTTGTCTAACTTGCACTTTTTGGATTTATCACAAAATCTGCTATCTGGGTCAATCCCTGTT

>M6-PGSC0003DMG400003918-(rev)M6LRR-RLK_610

TTGCTAGACAAGGAGTCTTTGATTGGTGGGGGGACAATTGGATCAGTTTACAGAACTTCCTTTGAAGGTGGTGTATCAATTGCAGTGAAAAAGCTCGAGACATTGGGGAGAATCAGAAAT

>M6-PGSC0003DMG400003918-(rev)M6LRR-RLK_671

TTGAAGGTGGTGTATCAATTGCAGTGAAAAAGCTCGAGACATTGGGGAGAATCAGAAATCAAGATGAATTTGAGCATGAAGTTGGCCGGTTAGGGACCCTTCAGCACCCCAATCTTGTTG

>M6-PGSC0003DMG400003918-(rev)M6LRR-RLK_732

AGATGAATTTGAGCATGAAGTTGGCCGGTTAGGGACCCTTCAGCACCCCAATCTTGTTGCTTTACAAGGGTACTACTGGTCCTCTAGCATGCAGTTAATATTGTCGGAGTTTGTTCCGAA

>M6-PGSC0003DMG400003918-(rev)M6LRR-RLK_793

TTACAAGGGTACTACTGGTCCTCTAGCATGCAGTTAATATTGTCGGAGTTTGTTCCGAATGGAAACCTATACGACAATTTACATGGACTTAGTTATCCTGGAACCAGCACTGGTGCTGGT

>M6-PGSC0003DMG400003918-(rev)M6LRR-RLK_854

GAAACCTATACGACAATTTACATGGACTTAGTTATCCTGGAACCAGCACTGGTGCTGGTAACCCAGAATTGAATTGGCCAAGAAGGTTTCAGATAGCAGTTGGAACAGCTCGAGCACTCA

>M6-PGSC0003DMG400003918-(rev)M6LRR-RLK_915

CCCAGAATTGAATTGGCCAAGAAGGTTTCAGATAGCAGTTGGAACAGCTCGAGCACTCACATATCTTCACCATGATTGCAAACCTCCAGTCCTTCATCTTAATGTCAAGTCCACTAATAT

>M6-PGSC0003DMG400003918-(rev)M6LRR-RLK_976

TATCTTCACCATGATTGCAAACCTCCAGTCCTTCATCTTAATGTCAAGTCCACTAATATACTCCTGGATACAAACTACGAAGCCAAATTGACTGATTATGGCCTGAGTAGGTTCCTTCCC

>M6-PGSC0003DMG400003945-(rev)M6LRR-RLK_1037

TTTGTTAGGTGGGACAATCCCAACGGAGCTTGGTAGTATCGAATGGCTTGCAGTTTTGGACCTCCATAATCTCACCCTATTTGGTGAAATCCCATATGAGATAAGCAACTGCAAGCTTCT

>M6-PGSC0003DMG400003945-(rev)M6LRR-RLK_1098

CTCCATAATCTCACCCTATTTGGTGAAATCCCATATGAGATAAGCAACTGCAAGCTTCTTCTTCAACTGTGAGTGTATCACCATCTTAGTTTAATTTGTTTTATTTCATCATTGCTCTGA

>M6-PGSC0003DMG400003945-(rev)M6LRR-RLK_1159

TTCAACTGTGAGTGTATCACCATCTTAGTTTAATTTGTTTTATTTCATCATTGCTCTGATCTGGAATTGAACCGTCTAATATTTGCTGCAGGGATCTATCTGGGAATCTTTTAGTAGGAG

>M6-PGSC0003DMG400003945-(rev)M6LRR-RLK_122

TTCTAGTGACCCTTTTAATAGTTTAGCTTCTTGGGATCCTAATGGAACACCTTGCTTAGACTTTAGTGGTGTTTTCTGCAATTCTGTTGGAAATGTGGTGAAGATTGTGTTGTGGAATAC

>M6-PGSC0003DMG400003945-(rev)M6LRR-RLK_1220

TGGAATTGAACCGTCTAATATTTGCTGCAGGGATCTATCTGGGAATCTTTTAGTAGGAGAAATACCACAAAACCTGTACAACTTGTCAAACCTTGTATATCTTGACTTACATCATAACCA

>M6-PGSC0003DMG400003945-(rev)M6LRR-RLK_183

TTTAGTGGTGTTTTCTGCAATTCTGTTGGAAATGTGGTGAAGATTGTGTTGTGGAATACTAGTCTTGGTGGGGTTCTCTCCCCTGCCTTAGCAGAACTTAAGTCTTTGCGCATCCTTACC

>M6-PGSC0003DMG400003945-(rev)M6LRR-RLK_244

GTCTTGGTGGGGTTCTCTCCCCTGCCTTAGCAGAACTTAAGTCTTTGCGCATCCTTACCTTGTTTGGGAACCAATTTACTGGCAATATTCCTGCAGAGTATGGGGGGATTGATTCATTGT

>M6-PGSC0003DMG400003945-(rev)M6LRR-RLK_305

GTTTGGGAACCAATTTACTGGCAATATTCCTGCAGAGTATGGGGGGATTGATTCATTGTGGAAGATTAACTTCAGCTCTAATGCCTTTTTTGGTTCAATTCCTGAATTTCTTGGGGATTT

>M6-PGSC0003DMG400003945-(rev)M6LRR-RLK_366

AAGATTAACTTCAGCTCTAATGCCTTTTTTGGTTCAATTCCTGAATTTCTTGGGGATTTACCTAATATAAGGTTCCTTGATTTGTCAAGAAATGTGTATAGTGGTGAAATTCCTGTGTCT

>M6-PGSC0003DMG400003945-(rev)M6LRR-RLK_427

CTAATATAAGGTTCCTTGATTTGTCAAGAAATGTGTATAGTGGTGAAATTCCTGTGTCTTTGTTTATGAATTGTAACAAAACAAGGTTCATTTCCTTGTCCCATAACAATCTTTCAGGTC

>M6-PGSC0003DMG400003945-(rev)M6LRR-RLK_488

GTTTATGAATTGTAACAAAACAAGGTTCATTTCCTTGTCCCATAACAATCTTTCAGGTCCAATTCCAGCATCAATTGGAAACTGTCAGAGTCTTGAAGGAATAGACTTGTCTTTTAATGG

>M6-PGSC0003DMG400003945-(rev)M6LRR-RLK_549

ATTCCAGCATCAATTGGAAACTGTCAGAGTCTTGAAGGAATAGACTTGTCTTTTAATGGTCTTAGTGGAAGTTTACCTTCGCAAATTTGTGACATTCCAGGATTAGTGTACTTGTCTGTG

>M6-PGSC0003DMG400003945-(rev)M6LRR-RLK_61

AATTGGAATTGGTATCTCCCATCACTGAAAAGGAGATCCTTCTTCAATTCAAGGGTAACATTTCTAGTGACCCTTTTAATAGTTTAGCTTCTTGGGATCCTAATGGAACACCTTGCTTAG

>M6-PGSC0003DMG400003945-(rev)M6LRR-RLK_610

TTAGTGGAAGTTTACCTTCGCAAATTTGTGACATTCCAGGATTAGTGTACTTGTCTGTGAGGAGCAATGCACTATCAGGACTTGTTCAAGAACAGGTTTCTCCATGTCAAAGGTTGGAGC

>M6-PGSC0003DMG400003945-(rev)M6LRR-RLK_671

GAGCAATGCACTATCAGGACTTGTTCAAGAACAGGTTTCTCCATGTCAAAGGTTGGAGCTTTTGGATCTTGGTAGCAATGACTTCTCTGGTTTGGCTCCTTTTGGGGTTCTTGAGCTGGT

>M6-PGSC0003DMG400003945-(rev)M6LRR-RLK_732

TTGGATCTTGGTAGCAATGACTTCTCTGGTTTGGCTCCTTTTGGGGTTCTTGAGCTGGTGAATCTTACATATTTCAATGTTTCAAGTAATGGTTTTGAAGGGGAGATTGTTAACTCTGGG

>M6-PGSC0003DMG400003945-(rev)M6LRR-RLK_793

ATCTTACATATTTCAATGTTTCAAGTAATGGTTTTGAAGGGGAGATTGTTAACTCTGGGACTTGTAGTCAGCGACTGGAAGTGTTGGATGTTTCAAGGAATATTTTTTTTGGTGAAATTC

>M6-PGSC0003DMG400003945-(rev)M6LRR-RLK_854

TTGTAGTCAGCGACTGGAAGTGTTGGATGTTTCAAGGAATATTTTTTTTGGTGAAATTCCCTTGAGCATCTCGAAATGCAGCGCGTTGAAGTACTTGGACATGGCATATAACAGAATTAA

>M6-PGSC0003DMG400003945-(rev)M6LRR-RLK_915

TTGAGCATCTCGAAATGCAGCGCGTTGAAGTACTTGGACATGGCATATAACAGAATTAATGGAAGTATCCCTATTGAGCTTGCAGATTTGAATAGCCTTTCCGTTATGCGATTGGGTGAT

>M6-PGSC0003DMG400003945-(rev)M6LRR-RLK_976

GAAGTATCCCTATTGAGCTTGCAGATTTGAATAGCCTTTCCGTTATGCGATTGGGTGATAATTTGTTAGGTGGGACAATCCCAACGGAGCTTGGTAGTATCGAATGGCTTGCAGTTTTGG

>M6-PGSC0003DMG400003958-(rev)M6LRR-RLK_0

TATTTTCTAACTAACTCTCTTTTCTCCTTTGACAATTCACACCAATAAACTCCCCATTTTTCCTTCTCTCTACTGTGAAGCACCATTGAAAATCCAATGCAAAAGCTCACTAGTACTTCA

>M6-PGSC0003DMG400003958-(rev)M6LRR-RLK_1037

AAAGGTAGGACTTTGTGGGGATTGATGGTTTACTCGGTTTTTTGAATTCCTTCATCCATAAATTAACAGTGCAAGTTTGAGAGAAAATAAAGTGTTACTAGTATATATATGAATTGTTGA

>M6-PGSC0003DMG400003958-(rev)M6LRR-RLK_1098

ATTAACAGTGCAAGTTTGAGAGAAAATAAAGTGTTACTAGTATATATATGAATTGTTGAATTTCCTAACATAAGGAAAGCTTTAAGTACAACAGTAGAAGTGGCTCAAAATTGCTTTAAG

>M6-PGSC0003DMG400003958-(rev)M6LRR-RLK_1159

TTCCTAACATAAGGAAAGCTTTAAGTACAACAGTAGAAGTGGCTCAAAATTGCTTTAAGTAGCAGATTTGAATTCCAAGTATCCCGTGTTAGAATTTCTACTTCCGCCACTGTTGCACCC

>M6-PGSC0003DMG400003958-(rev)M6LRR-RLK_122

AAAAACTCAACTTTTCACTCAAATCTGAAAGTTCACAAAGTTCAGAGAAATGGGGGTTAATTATTTCCTCTTAACTTTCATTTTTGCTCTGTTTGTTCTTGAATTTTCTTGCATAGAACC

>M6-PGSC0003DMG400003958-(rev)M6LRR-RLK_1220

GCAGATTTGAATTCCAAGTATCCCGTGTTAGAATTTCTACTTCCGCCACTGTTGCACCCACACCCCCTTTTCTGGTCTTTATTCTTTCTTGTTAATTATATTTTTGGTCTGAACAGCATA

>M6-PGSC0003DMG400003958-(rev)M6LRR-RLK_1281

ACCCCCTTTTCTGGTCTTTATTCTTTCTTGTTAATTATATTTTTGGTCTGAACAGCATACTGAAATTTCTTTTCCAAATAAGGTGATGAAAGAAATGCAAGAACTTCTTTTTTTTTCCCT

>M6-PGSC0003DMG400003958-(rev)M6LRR-RLK_1342

GAAATTTCTTTTCCAAATAAGGTGATGAAAGAAATGCAAGAACTTCTTTTTTTTTCCCTTTCTACAATGATATGTGCTCTGAGATGATTTGTGCATTTTGTGGTTCCACAGTGCTGGACT

>M6-PGSC0003DMG400003958-(rev)M6LRR-RLK_1403

CTACAATGATATGTGCTCTGAGATGATTTGTGCATTTTGTGGTTCCACAGTGCTGGACTTGAGTTGTAACCAGTTGACAGGGAGCATACCAACAGAGATTTGGTTCTTGAAGAAGTTGAG

>M6-PGSC0003DMG400003958-(rev)M6LRR-RLK_1464

AGTTGTAACCAGTTGACAGGGAGCATACCAACAGAGATTTGGTTCTTGAAGAAGTTGAGTGTTCTCGAATTGCAACGTAACAGGCTGACAGGTAAAATTCCAGAGAACGTAGGAGTCCTC

>M6-PGSC0003DMG400003958-(rev)M6LRR-RLK_1525

TTCTCGAATTGCAACGTAACAGGCTGACAGGTAAAATTCCAGAGAACGTAGGAGTCCTCGGGACGTTGAAAAAGCTATATTTGGGATTCAATCAGCTATCTGGTCCGATTCCATTAAACT

>M6-PGSC0003DMG400003958-(rev)M6LRR-RLK_1586

GACGTTGAAAAAGCTATATTTGGGATTCAATCAGCTATCTGGTCCGATTCCATTAAACTTAGCTGCTGCTCCCCAGTTGGAGGTTCTAGAAGTACAAAATAACACCCTCTCTGGAGTTGT

>M6-PGSC0003DMG400003958-(rev)M6LRR-RLK_1647

GCTGCTGCTCCCCAGTTGGAGGTTCTAGAAGTACAAAATAACACCCTCTCTGGAGTTGTTCCTCCAGGTTAGTTTCCCTTTTTGAGTTCTTAACTTTTTATAAATCTTTTTTTCCTAGCT

>M6-PGSC0003DMG400003958-(rev)M6LRR-RLK_1708

CTCCAGGTTAGTTTCCCTTTTTGAGTTCTTAACTTTTTATAAATCTTTTTTTCCTAGCTTGAAGTTATATCTTTTGATTTCTGATTTTAAACCTCTTTTTTCTCCTTTTTATCTGGACTG

>M6-PGSC0003DMG400003958-(rev)M6LRR-RLK_1769

AAGTTATATCTTTTGATTTCTGATTTTAAACCTCTTTTTTCTCCTTTTTATCTGGACTGATTTGTAGCTCTAAGGAGATTAAATGAAAAATTCAATCATGAAAGCAATCCTGGTTTATGT

>M6-PGSC0003DMG400003958-(rev)M6LRR-RLK_183

TATTTCCTCTTAACTTTCATTTTTGCTCTGTTTGTTCTTGAATTTTCTTGCATAGAACCTGTTTGTGGAAATGCTGAGGTGAGAGCTTTAATGGAGATTAAAACAGCTTTGGACCCAGAA

>M6-PGSC0003DMG400003958-(rev)M6LRR-RLK_1830

TTGTAGCTCTAAGGAGATTAAATGAAAAATTCAATCATGAAAGCAATCCTGGTTTATGTGGTACTGGATTTACTTCATTGAGAGTTTGCACTGCCTGGGATCATGTCAATGTGAATCAAG

>M6-PGSC0003DMG400003958-(rev)M6LRR-RLK_1891

TACTGGATTTACTTCATTGAGAGTTTGCACTGCCTGGGATCATGTCAATGTGAATCAAGTTGGTTCTGATGTACCTGATTCCAATAATAACGGCATCCCTATAGATGTCCCAGGAGCTGC

>M6-PGSC0003DMG400003958-(rev)M6LRR-RLK_1952

GGTTCTGATGTACCTGATTCCAATAATAACGGCATCCCTATAGATGTCCCAGGAGCTGCTCATATTAGTCCGTTGCCTTGTAATCAAACTCATTGTTCAAGATCATCAAAATTTCCTCAA

>M6-PGSC0003DMG400003958-(rev)M6LRR-RLK_2013

ATATTAGTCCGTTGCCTTGTAATCAAACTCATTGTTCAAGATCATCAAAATTTCCTCAAGCAATTATCGTTGCTTCTGTGATTACAGTCACTGTAACTTTGGTAGTAGCTGTTGTTTTCG

>M6-PGSC0003DMG400003958-(rev)M6LRR-RLK_2074

AATTATCGTTGCTTCTGTGATTACAGTCACTGTAACTTTGGTAGTAGCTGTTGTTTTCGGTATATTTAGACGAAGAAGGCTAAAACAAAGAGTAGGGAACACAAGTGATGCATCTGATGA

>M6-PGSC0003DMG400003958-(rev)M6LRR-RLK_2135

ATATTTAGACGAAGAAGGCTAAAACAAAGAGTAGGGAACACAAGTGATGCATCTGATGATAGGCTTAGCACTGATCAGGCAAAGGAAATATACAAGAGAAGCCCTTCTCCACTACTTACT

>M6-PGSC0003DMG400003958-(rev)M6LRR-RLK_2196

GGCTTAGCACTGATCAGGCAAAGGAAATATACAAGAGAAGCCCTTCTCCACTACTTACTGTTGAGTACTCAAATCGTTGGGACCCTATGACCCCTGAAAAGAGTTGCAATAGCATGTTCA

>M6-PGSC0003DMG400003958-(rev)M6LRR-RLK_2257

TGAGTACTCAAATCGTTGGGACCCTATGACCCCTGAAAAGAGTTGCAATAGCATGTTCAATGAGTTCTTAAATGGATTCAAGTTCAATTTGGAAGAGGTTGAGTCTGCAACTCAGCATTT

>M6-PGSC0003DMG400003958-(rev)M6LRR-RLK_2318

GAGTTCTTAAATGGATTCAAGTTCAATTTGGAAGAGGTTGAGTCTGCAACTCAGCATTTTACAGAGACAAATCTATTGGGACGGAGTAATTTCTCTGCCGTCTATAAAGGAATTTTAAAA

>M6-PGSC0003DMG400003958-(rev)M6LRR-RLK_2379

CAGAGACAAATCTATTGGGACGGAGTAATTTCTCTGCCGTCTATAAAGGAATTTTAAAAGATGGATCTACGGTAGCTGTAAAAAGCATCAGCGTGACAAGCTGTAAGTCGGAGGAGACGG

>M6-PGSC0003DMG400003958-(rev)M6LRR-RLK_244

TTTGTGGAAATGCTGAGGTGAGAGCTTTAATGGAGATTAAAACAGCTTTGGACCCAGAAAGCAAGAAACTTTCTTCATGGACAAGTGATGGTGATCCATGTAGTGGTTCTTTTCTTGGTG

>M6-PGSC0003DMG400003958-(rev)M6LRR-RLK_2440

TGGATCTACGGTAGCTGTAAAAAGCATCAGCGTGACAAGCTGTAAGTCGGAGGAGACGGAGTTCATGGAAGGATTGAGCTTATTAACCTCCCTGAAGCATGAAAACCTTGTCAAGCTAAG

>M6-PGSC0003DMG400003958-(rev)M6LRR-RLK_2501

TTCATGGAAGGATTGAGCTTATTAACCTCCCTGAAGCATGAAAACCTTGTCAAGCTAAGAGGTTTCTGTTGTTCAAAGGGAAGAGGCGAGTGCTTTTTAATCTATGATTTGGCCTCTAAA

>M6-PGSC0003DMG400003958-(rev)M6LRR-RLK_2562

GTTTCTGTTGTTCAAAGGGAAGAGGCGAGTGCTTTTTAATCTATGATTTGGCCTCTAAAGGGAACCTTTCTCAGTATCTTGATGTTGAAGTAAACAGCAACCATGTTCTTGACTGGTCCA

>M6-PGSC0003DMG400003958-(rev)M6LRR-RLK_2623

GAACCTTTCTCAGTATCTTGATGTTGAAGTAAACAGCAACCATGTTCTTGACTGGTCCAAAAGAGTTTCGATCATCAAGGGGATTGCAAAAGGTTGGTCTTCTTTGCTTTTGTTCATCAT

>M6-PGSC0003DMG400003958-(rev)M6LRR-RLK_2684

AGAGTTTCGATCATCAAGGGGATTGCAAAAGGTTGGTCTTCTTTGCTTTTGTTCATCATTACATCTTCAACATTCTCTACAAATTTCTCAGAATATTTTAAAGTTCGGAAATATTATGGC

>M6-PGSC0003DMG400003958-(rev)M6LRR-RLK_2745

CATCTTCAACATTCTCTACAAATTTCTCAGAATATTTTAAAGTTCGGAAATATTATGGCTGTATAATATTTTCCCCATAATGTGGAGTCTTCTAGGAGTTCCATTGCATAGGCAGTCTAT

>M6-PGSC0003DMG400003958-(rev)M6LRR-RLK_2806

TATAATATTTTCCCCATAATGTGGAGTCTTCTAGGAGTTCCATTGCATAGGCAGTCTATTACCACTAAAGTGCTTCTCACAGGATTCAGGAACAGGATGTGACTGTAGATTCATTTGCAT

>M6-PGSC0003DMG400003958-(rev)M6LRR-RLK_305

CAAGAAACTTTCTTCATGGACAAGTGATGGTGATCCATGTAGTGGTTCTTTTCTTGGTGTGATTTGTAATGAACATCATAAAGTAACAAATATTACTCTGACAGATAAAGGACTTACTGG

>M6-PGSC0003DMG400003958-(rev)M6LRR-RLK_3050

AAACAACGTAATTTTCTGCCGACCACCTCGGCCTACCTAGCTCCACCCCTGTGGTTAATATATTATCAAATTACTCCTCCATCCCAAAATAAGTGTCGCTTTAGTTCATTTCACGCCCAT

>M6-PGSC0003DMG400003958-(rev)M6LRR-RLK_3111

ATTATCAAATTACTCCTCCATCCCAAAATAAGTGTCGCTTTAGTTCATTTCACGCCCATTAAGAAAGGTAATAAATGCAAGGCATAGTTTACTAAGTCACCCCTAGTTATTAGATGAAGA

>M6-PGSC0003DMG400003958-(rev)M6LRR-RLK_3172

AGAAAGGTAATAAATGCAAGGCATAGTTTACTAAGTCACCCCTAGTTATTAGATGAAGAGGAGTAGTTCTTTAATATAAAGGTATAGTTGGAACCAAGAAGTACTATTTTTTGTCTTGAC

>M6-PGSC0003DMG400003958-(rev)M6LRR-RLK_3233

AGTAGTTCTTTAATATAAAGGTATAGTTGGAACCAAGAAGTACTATTTTTTGTCTTGACTTCTAAATAGGCAATTATTTTGGGATCAACTTTTATGCTAAAGCGACACTTATTTGGGACG

>M6-PGSC0003DMG400003958-(rev)M6LRR-RLK_3273

TACTATTTTTTGTCTTGACTTCTAAATAGGCAATTATTTTGGGATCAACTTTTATGCTAAAGCGACACTTATTTGGGACGGAGGGAGTAATATGAATACAGAGAGAGACTGGAATTATTG

>M6-PGSC0003DMG400003958-(rev)M6LRR-RLK_366

ATTTGTAATGAACATCATAAAGTAACAAATATTACTCTGACAGATAAAGGACTTACTGGAAAGTTGTCACCAGCAATTGCCGAGCTGAAGTGTCTGTCTGGTCTGTTCTTGCATTATAAT

>M6-PGSC0003DMG400003958-(rev)M6LRR-RLK_427

AGTTGTCACCAGCAATTGCCGAGCTGAAGTGTCTGTCTGGTCTGTTCTTGCATTATAATTCTTTAACAGGGGAAATACCAAAGGAACTTGGGAATTTGACTGAGTTGAATGATCTTTATC

>M6-PGSC0003DMG400003958-(rev)M6LRR-RLK_488

TTTAACAGGGGAAATACCAAAGGAACTTGGGAATTTGACTGAGTTGAATGATCTTTATCTCAATGAGAACAATCTTTCTGGGATTATACCACCAGAACTTGGAAACATGGCAAGTTTACA

>M6-PGSC0003DMG400003958-(rev)M6LRR-RLK_549

AATGAGAACAATCTTTCTGGGATTATACCACCAGAACTTGGAAACATGGCAAGTTTACAAGGTTTGTAGCCTTTGTAAACATCTTTCTGTTTAATTTTGTGGGAGATTGTTTCTCTTCTT

>M6-PGSC0003DMG400003958-(rev)M6LRR-RLK_61

CCTTCTCTCTACTGTGAAGCACCATTGAAAATCCAATGCAAAAGCTCACTAGTACTTCATAAAAAACTCAACTTTTCACTCAAATCTGAAAGTTCACAAAGTTCAGAGAAATGGGGGTTA

>M6-PGSC0003DMG400003958-(rev)M6LRR-RLK_610

GTTTGTAGCCTTTGTAAACATCTTTCTGTTTAATTTTGTGGGAGATTGTTTCTCTTCTTTTCTTAGCTCCCACCTTTTTGCTGCTTTGGTGACTCAATCTTAAATTATCTGTTGTGATTT

>M6-PGSC0003DMG400003958-(rev)M6LRR-RLK_671

CTTAGCTCCCACCTTTTTGCTGCTTTGGTGACTCAATCTTAAATTATCTGTTGTGATTTCTAAGAAATAGTTGTATTAAATGATTTGTCATTTTAGAAGTTTTAAGATAAAATAATTATC

>M6-PGSC0003DMG400003958-(rev)M6LRR-RLK_732

AAGAAATAGTTGTATTAAATGATTTGTCATTTTAGAAGTTTTAAGATAAAATAATTATCCTTTTCTCATTCTACTTTTAATAGTAATTGTTCTTGAAGATCCTGTTTTGGTGACTCGAAC

>M6-PGSC0003DMG400003958-(rev)M6LRR-RLK_793

TTTCTCATTCTACTTTTAATAGTAATTGTTCTTGAAGATCCTGTTTTGGTGACTCGAACCGACAACCGTTGGAGGTAGAGAGTGCTTACAACTTGAGCAGCCCCTGCTATCAGGGAAATT

>M6-PGSC0003DMG400003958-(rev)M6LRR-RLK_854

ACAACCGTTGGAGGTAGAGAGTGCTTACAACTTGAGCAGCCCCTGCTATCAGGGAAATTGCTTGCTGTTTCATTCTACTGTTTTTTGTGTGTATTAGTGAATAATGAGTCTACGTGAATT

>M6-PGSC0003DMG400003958-(rev)M6LRR-RLK_915

TTGCTGTTTCATTCTACTGTTTTTTGTGTGTATTAGTGAATAATGAGTCTACGTGAATTTAGGTGACAGAGTAGACAAGTTGCAAAATTTCTTAGAGATCTATGCAGATTTTTTGGTTCT

>M6-PGSC0003DMG400003958-(rev)M6LRR-RLK_976

GGTGACAGAGTAGACAAGTTGCAAAATTTCTTAGAGATCTATGCAGATTTTTTGGTTCTGTAAAGGTAGGACTTTGTGGGGATTGATGGTTTACTCGGTTTTTTGAATTCCTTCATCCAT

>M6-PGSC0003DMG400003982-(rev)M6G-LecRK_0

GAGCTAAAACAGGCAACGGATGGTTTCAAAGAGGAATTAGGAAAGGGTGCTTTTGGAGCAGTATATAAAGGGAGCTTTGACAAAGGTAAAAACCTTGTGGCAGTGAAGAGGCTAGAGAAG

>M6-PGSC0003DMG400003982-(rev)M6G-LecRK_124

TGGAAGAAGGTGAAAGGGAGTTCAGAGCAGAAATGAAGGTCATTGGTAGAACTCGACACAGAAACTTGATTCGTTTGCTAGGATATTGTGCTGAAGGATCAAAAAGAGCTCTTGTTTATG

>M6-PGSC0003DMG400003982-(rev)M6G-LecRK_186

AACTTGATTCGTTTGCTAGGATATTGTGCTGAAGGATCAAAAAGAGCTCTTGTTTATGAATATATGAGCAATGGTTGCCTAGCAAATCTGTTGTTCCGAGGAGCTACTCGGCCGGATTGG

>M6-PGSC0003DMG400003982-(rev)M6G-LecRK_248

TATGAGCAATGGTTGCCTAGCAAATCTGTTGTTCCGAGGAGCTACTCGGCCGGATTGGAATCTAAGAGTAAATATTGCACTAGATGTGGCCAGGGGAATCCTCTACCTCCATGAAGAATG

>M6-PGSC0003DMG400003982-(rev)M6G-LecRK_310

TAAGAGTAAATATTGCACTAGATGTGGCCAGGGGAATCCTCTACCTCCATGAAGAATGTGAAGCTCCAATTATCCACTGTGATATAAAGCCACAAAACATCTTACTCGACGAGTTGTTGA

>M6-PGSC0003DMG400003982-(rev)M6G-LecRK_372

GCTCCAATTATCCACTGTGATATAAAGCCACAAAACATCTTACTCGACGAGTTGTTGACAGCTAAAATCTCAGATTTTGGTCTGGCCAAGCTGCTAATGCCTGATCAAACAAGAACATTC

>M6-PGSC0003DMG400003982-(rev)M6G-LecRK_434

TAAAATCTCAGATTTTGGTCTGGCCAAGCTGCTAATGCCTGATCAAACAAGAACATTCACTGGTGTAAGAGGGACTAGAGGGTATTTAGCACCAGAGTGGCAAACAAATGCCCCCATCTC

>M6-PGSC0003DMG400003982-(rev)M6G-LecRK_496

GTGTAAGAGGGACTAGAGGGTATTTAGCACCAGAGTGGCAAACAAATGCCCCCATCTCTGTCAAGGTTGATGTCTTCAGCTATGGTATTGTATTGCTAGAAATCATTTGCTGCAGAAGAA

>M6-PGSC0003DMG400003982-(rev)M6G-LecRK_558

AAGGTTGATGTCTTCAGCTATGGTATTGTATTGCTAGAAATCATTTGCTGCAGAAGAAACATTGAAGTCTGTGTGACTAAAATAGAAGAAATCCAGCTTTCTATATGGGCATATAGTTGC

>M6-PGSC0003DMG400003982-(rev)M6G-LecRK_62

ATATAAAGGGAGCTTTGACAAAGGTAAAAACCTTGTGGCAGTGAAGAGGCTAGAGAAGGTAGTGGAAGAAGGTGAAAGGGAGTTCAGAGCAGAAATGAAGGTCATTGGTAGAACTCGACA

>M6-PGSC0003DMG400003982-(rev)M6G-LecRK_620

TGAAGTCTGTGTGACTAAAATAGAAGAAATCCAGCTTTCTATATGGGCATATAGTTGCCTTGTTGAAGGTGAAGTAGACAAGCTTGTTGGAAGTGAAGAAGTAGACAAGAAAACCTTGGA

>M6-PGSC0003DMG400003982-(rev)M6G-LecRK_682

TTGAAGGTGAAGTAGACAAGCTTGTTGGAAGTGAAGAAGTAGACAAGAAAACCTTGGAAGGAATGATCTCTGTAGCGATTTGGTGCATTCAGGACGAACCAGCTCTTCGTCCTTCCATGA

>M6-PGSC0003DMG400003982-(rev)M6G-LecRK_737

GGAAGGAATGATCTCTGTAGCGATTTGGTGCATTCAGGACGAACCAGCTCTTCGTCCTTCCATGAAAAAAGTATTACACATGTTGCAGGGAATTATTGATATTCCAATTCCTCCATGTCC

>M6-PGSC0003DMG400004056M6LRR-RLP_0

ATGTACAGAGGTGTAGCAGATGCTTCTATATCTCCAATGGAGAAACATGAGAAAGAGGCATTGTACTCTGCTATTCAAGGTTTTGTTGGAAATGAGTGGAATGGTTCATTTCTTTATCCT

>M6-PGSC0003DMG400004056M6LRR-RLP_1037

CTTTATTTTGATGTAATTCATTCCTAAAACCAAGTAGTTATATGCATTGAAATTATCGTTCTTGTATGTTAACTTGACTTAACAGTGTAAAAATTCTTTACACCGTAACATAAACTCTTG

>M6-PGSC0003DMG400004056M6LRR-RLP_1098

TTGTATGTTAACTTGACTTAACAGTGTAAAAATTCTTTACACCGTAACATAAACTCTTGTCTTAACACTTTAATCAACTCTGTGAACTCTTATTTTTACTATAATTCACAGGGAGTATCA

>M6-PGSC0003DMG400004056M6LRR-RLP_1159

TTAACACTTTAATCAACTCTGTGAACTCTTATTTTTACTATAATTCACAGGGAGTATCATGTGATTTCTCCAATGGCTTCTGGCACGTTACGGATTTAACTATTGGAGACCTATATGACA

>M6-PGSC0003DMG400004056M6LRR-RLP_1220

TGATTTCTCCAATGGCTTCTGGCACGTTACGGATTTAACTATTGGAGACCTATATGACAATTCCCTTCGTTGCTCCCAAGCTGCAGAGTTCACAGAACACTTGTTTAACCTCAAGCATCT

>M6-PGSC0003DMG400004056M6LRR-RLP_1281

TCCCTTCGTTGCTCCCAAGCTGCAGAGTTCACAGAACACTTGTTTAACCTCAAGCATCTAAAAAGACTTTCCTTCTCAAATTGCTTCCTCTCAGATCAGCACAAATCAATTCCCATTTCA

>M6-PGSC0003DMG400004056M6LRR-RLP_1342

AAAGACTTTCCTTCTCAAATTGCTTCCTCTCAGATCAGCACAAATCAATTCCCATTTCAAATTGGGATAGTCTCGCAAACAGCCTTGAATCGTTAGAGTTCCGATCAAATCCTGGTCTTA

>M6-PGSC0003DMG400004056M6LRR-RLP_1403

TTGGGATAGTCTCGCAAACAGCCTTGAATCGTTAGAGTTCCGATCAAATCCTGGTCTTATTGGGACAATTCCCACAAGTTTTGGCTACCTTAAGAATCTTCAATCTTTAGTACTGCTGGA

>M6-PGSC0003DMG400004056M6LRR-RLP_1464

GGGACAATTCCCACAAGTTTTGGCTACCTTAAGAATCTTCAATCTTTAGTACTGCTGGAAAATGGACTGAAAGGAGAAATACCAGAAGCTTTAGGCAATTTAACTAAACTAAAGCGCCTC

>M6-PGSC0003DMG400004056M6LRR-RLP_1525

ATGGACTGAAAGGAGAAATACCAGAAGCTTTAGGCAATTTAACTAAACTAAAGCGCCTCGTTTTGGCAGGCAACAATTTTATTGGTCCTATTCCAACTAGTCTAGGAAGATTGACAAATC

>M6-PGSC0003DMG400004056M6LRR-RLP_1586

TTTGGCAGGCAACAATTTTATTGGTCCTATTCCAACTAGTCTAGGAAGATTGACAAATCTTTTGATACTTGACACAATCAGGAATTTTCTATCTGGTGCATTACCTGTGACTATTGGTGA

>M6-PGSC0003DMG400004056M6LRR-RLP_1647

TTGATACTTGACACAATCAGGAATTTTCTATCTGGTGCATTACCTGTGACTATTGGTGATTTATCTTCGCTCATAAAGCTTGACTTGAGCAACAATCGATTAGATGGAAAATTGCCTCGA

>M6-PGSC0003DMG400004056M6LRR-RLP_1708

TATCTTCGCTCATAAAGCTTGACTTGAGCAACAATCGATTAGATGGAAAATTGCCTCGAGAAATTGGAGGACTAAAGAGTCTAACACTACTTGATCTCAGCCACAACAATTTCTCTGATG

>M6-PGSC0003DMG400004056M6LRR-RLP_1769

AATTGGAGGACTAAAGAGTCTAACACTACTTGATCTCAGCCACAACAATTTCTCTGATGGTTTGCCAGAGACATTTCAAGAAATGGACTCCTTAGAAGAACTTGTGTTATCTGACAATCC

>M6-PGSC0003DMG400004056M6LRR-RLP_1830

TTGCCAGAGACATTTCAAGAAATGGACTCCTTAGAAGAACTTGTGTTATCTGACAATCCGATAGGCGATTATGTTACAAGAATTCAATGGAGGAACATGAAAAACTTGGAAATGTTGGAT

>M6-PGSC0003DMG400004056M6LRR-RLP_1891

TAGGCGATTATGTTACAAGAATTCAATGGAGGAACATGAAAAACTTGGAAATGTTGGATCTTTCAAACATGGGATTGAAAGGTAATATACCAAATTCCATGACAGAAATGAAGAAACTGA

>M6-PGSC0003DMG400004056M6LRR-RLP_1952

TTCAAACATGGGATTGAAAGGTAATATACCAAATTCCATGACAGAAATGAAGAAACTGAGATTCCTCAGCCTTAGTAATAACATTCTTTCAGGAATTATTCCGTCAAAATTCGAGAAAAT

>M6-PGSC0003DMG400004056M6LRR-RLP_2013

TTCCTCAGCCTTAGTAATAACATTCTTTCAGGAATTATTCCGTCAAAATTCGAGAAAATATCAACTATTAATGCTCTATACTTGGATGGAAATGATTTCACAGGGAAACTTGAATTTTCA

>M6-PGSC0003DMG400004056M6LRR-RLP_2074

CAACTATTAATGCTCTATACTTGGATGGAAATGATTTCACAGGGAAACTTGAATTTTCACAAAGGTTTTATACAAAATTAAGGAGTCGTTTTCGAGCTTCAGGTAATGTGAAACTCTGCT

>M6-PGSC0003DMG400004056M6LRR-RLP_2135

AAGGTTTTATACAAAATTAAGGAGTCGTTTTCGAGCTTCAGGTAATGTGAAACTCTGCTTGTCCCTCGAGTTAACGTCAACAAGTCATGTTCCAGAAGGTGTACAACAATGTGAACAAGA

>M6-PGSC0003DMG400004056M6LRR-RLP_2196

TCCCTCGAGTTAACGTCAACAAGTCATGTTCCAGAAGGTGTACAACAATGTGAACAAGAAAGCACAGTAGATAGTAAAGATTCAGACTCCAATTTAAAGAACAAATATTTGAGTCAGAAT

>M6-PGSC0003DMG400004056M6LRR-RLP_2257

GCACAGTAGATAGTAAAGATTCAGACTCCAATTTAAAGAACAAATATTTGAGTCAGAATTCCCACAATATTATAGCTTCTTTGGGGCATTCAGGACATGTTGTTAGTCATTTTACATTTT

>M6-PGSC0003DMG400004056M6LRR-RLP_2310

CAGAATTCCCACAATATTATAGCTTCTTTGGGGCATTCAGGACATGTTGTTAGTCATTTTACATTTTGTTATGTCGCAAGAATAATACTTATGTCTCTTCCATGGATCGTATTTTTATAA

>M6-PGSC0003DMG400004056M6LRR-RLP_366

TTCATAATTTGCTAGTTGATTCTAGTCTTATGTTTTTTTACCATTATTGACTTTTCTTGCTGTAGTTCAAAATCAAGAAAACAGTAATGTCACAATGCCATGCATGGATGTCCTTAAATT

>M6-PGSC0003DMG400004056M6LRR-RLP_427

GTAGTTCAAAATCAAGAAAACAGTAATGTCACAATGCCATGCATGGATGTCCTTAAATTTTTTATTTTATTTTAATATAAAGAAATGGATCTTGGGTCTTACTCAACCACATAAACTACA

>M6-PGSC0003DMG400004056M6LRR-RLP_488

TTATTTTATTTTAATATAAAGAAATGGATCTTGGGTCTTACTCAACCACATAAACTACATCATAGGAATTAGGATGAGTATTGTTCAAGACATTTGAGATCAATTACCCATCCCATATTC

>M6-PGSC0003DMG400004056M6LRR-RLP_549

ATAGGAATTAGGATGAGTATTGTTCAAGACATTTGAGATCAATTACCCATCCCATATTCGACTCTTGTGGGCAATATAATATGGAAGCACAACATAGGTGAACAAAGAATTCGGATGAGC

>M6-PGSC0003DMG400004056M6LRR-RLP_61

TGTACTCTGCTATTCAAGGTTTTGTTGGAAATGAGTGGAATGGTTCATTTCTTTATCCTGATCCTTGTGGCTGGACATCAATTGAGGTACTAATCTTTTGCTCTCGATGTTCTTTTGCTA

>M6-PGSC0003DMG400004056M6LRR-RLP_793

GACTCTTAATAGATCAAACTGTCCTCTCAATGGTTGTTAATTAAAGTGAAAAAGACTTTCTCAAATTCTTACTTTTTCTGTTCTGGATTTGCAAACAGTGTTATATGAGTGTGTGTAAGA

>M6-PGSC0003DMG400004056M6LRR-RLP_854

CAAATTCTTACTTTTTCTGTTCTGGATTTGCAAACAGTGTTATATGAGTGTGTGTAAGAGAGACATCACTAGTTAGGAGCTTTCCAATTTCTAACACACCTTTTCTGGTAGTACTAGTAA

>M6-PGSC0003DMG400004056M6LRR-RLP_915

GACATCACTAGTTAGGAGCTTTCCAATTTCTAACACACCTTTTCTGGTAGTACTAGTAAATATATATGTTAGTATTCTATTGGTAATTTGCAATATTTATCACTTTCTTATAGTGATTTA

>M6-PGSC0003DMG400004056M6LRR-RLP_976

ATATATGTTAGTATTCTATTGGTAATTTGCAATATTTATCACTTTCTTATAGTGATTTAGGCTTTATTTTGATGTAATTCATTCCTAAAACCAAGTAGTTATATGCATTGAAATTATCGT

>M6-PGSC0003DMG400004146-(rev)M6LRR-RLK_0

CAAACCATGAAACAAATTCATCTGTTTTGCTTGTTATTTTTGAGTGGTTTTTTCTTTGTTTCATCAAATAAGGATGAAGTTTCTATATTGTTAGCAATAAAGTCTAGTCTTGTTGATCCA

>M6-PGSC0003DMG400004146-(rev)M6LRR-RLK_1054

TGAATCTTGGGAAAAAATCTCCCTTGCAATGGTTAGACGTTTCTTCGAATATCTTGACAGGTGAGATCCCTCCTGGCTTGTGTGATTCTGGCAATCTAACAAAGCTCATTCTGTTTAATA

>M6-PGSC0003DMG400004146-(rev)M6LRR-RLK_1116

GAGATCCCTCCTGGCTTGTGTGATTCTGGCAATCTAACAAAGCTCATTCTGTTTAATAATTCCTTCTCTGGATCAATCCCATTAGGCCTCTCTAACTGTTCTTCACTTGTTCGAGTTAGA

>M6-PGSC0003DMG400004146-(rev)M6LRR-RLK_1178

CTTCTCTGGATCAATCCCATTAGGCCTCTCTAACTGTTCTTCACTTGTTCGAGTTAGAATCCAAAATAATCTGCTTTCAGGAATGATTCCGGTTGGGTTTGGAACACTTCCTATGCTCCA

>M6-PGSC0003DMG400004146-(rev)M6LRR-RLK_124

ATCATCTCAAAGATTGGAACTTTTCAAACAATGAAAGTTCTATTCATTGTAAATGGAATGGTGTGTTTTGCAACTCAAAAAGCTATGTAGAGAAGCTTGATTTGTCAAACATGAATCTAA

>M6-PGSC0003DMG400004146-(rev)M6LRR-RLK_1240

AAAATAATCTGCTTTCAGGAATGATTCCGGTTGGGTTTGGAACACTTCCTATGCTCCAAAGATTGGAATTGGCTAAAAACAATCTCACTGGTGAAATTCCTGTGGATTTCACTCTCTCCA

>M6-PGSC0003DMG400004146-(rev)M6LRR-RLK_1302

TTGGAATTGGCTAAAAACAATCTCACTGGTGAAATTCCTGTGGATTTCACTCTCTCCACCACTCTTTCTTTTATCGATGTCTCTTCAAACCATCTTGAATCATCTTTGCCTTCTAGCATT

>M6-PGSC0003DMG400004146-(rev)M6LRR-RLK_1364

TCTTTCTTTTATCGATGTCTCTTCAAACCATCTTGAATCATCTTTGCCTTCTAGCATTTTGTCCATTCCTAGCCTGCAAACCTTCATAGTCTCCAATAACAATCTTAAAGGCAACATTCC

>M6-PGSC0003DMG400004146-(rev)M6LRR-RLK_1426

CCATTCCTAGCCTGCAAACCTTCATAGTCTCCAATAACAATCTTAAAGGCAACATTCCTGACCAATTCCAAGATTGTCCTTCCCTTTCTTTGCTTGATCTTTCAAGCAACCATTTCTCGG

>M6-PGSC0003DMG400004146-(rev)M6LRR-RLK_1488

CAATTCCAAGATTGTCCTTCCCTTTCTTTGCTTGATCTTTCAAGCAACCATTTCTCGGGGAAAATTCCTCAGAGTATTGCTTCATGTGAAAAGCTGGTGAAGCTTAATCTAAGAAACAAC

>M6-PGSC0003DMG400004146-(rev)M6LRR-RLK_1550

AATTCCTCAGAGTATTGCTTCATGTGAAAAGCTGGTGAAGCTTAATCTAAGAAACAACCAATTCAGTGGTGAAATTCCAACTCACATTGCCACATTGCCAACTTTATCTATTCTTGATCT

>M6-PGSC0003DMG400004146-(rev)M6LRR-RLK_1612

TCAGTGGTGAAATTCCAACTCACATTGCCACATTGCCAACTTTATCTATTCTTGATCTATCAAACAACTCGTTAGTTGGAAAAATACCAATGGACTTTGGTAGCTCTCCAGCTCTAGAAA

>M6-PGSC0003DMG400004146-(rev)M6LRR-RLK_1674

AACAACTCGTTAGTTGGAAAAATACCAATGGACTTTGGTAGCTCTCCAGCTCTAGAAATGCTTAATCTTTCCTACAACAAGCTTGATGGTCCAGTTCCAAGAAATGGGATACTCATGACT

>M6-PGSC0003DMG400004146-(rev)M6LRR-RLK_1736

TAATCTTTCCTACAACAAGCTTGATGGTCCAGTTCCAAGAAATGGGATACTCATGACTATTAATCCCAATGATCTGATAGGAAATGCTGGTCTTTGTGGTGGCATACTTCCACCGTGTTC

>M6-PGSC0003DMG400004146-(rev)M6LRR-RLK_1798

ATCCCAATGATCTGATAGGAAATGCTGGTCTTTGTGGTGGCATACTTCCACCGTGTTCTCAAAGTCTCACCATAACTTCAAATGCGAGAAAGAATCGTGTCAATCACATAATCGTGGGAT

>M6-PGSC0003DMG400004146-(rev)M6LRR-RLK_186

GTGTTTTGCAACTCAAAAAGCTATGTAGAGAAGCTTGATTTGTCAAACATGAATCTAAGTGGCAGAGTTTCAGATCAAATTCAAGGACTTCAGAGTCTATCTTTGCTTAATCTTTGTTGC

>M6-PGSC0003DMG400004146-(rev)M6LRR-RLK_1860

AGTCTCACCATAACTTCAAATGCGAGAAAGAATCGTGTCAATCACATAATCGTGGGATTCATTGTTGGAATCTCAGTTATCTTAGCTGTTGGGATCATGGTTCTAGCAGGGAGATGGATG

>M6-PGSC0003DMG400004146-(rev)M6LRR-RLK_1922

TGTTGGAATCTCAGTTATCTTAGCTGTTGGGATCATGGTTCTAGCAGGGAGATGGATGTACAATAGATGGTACTTGTGCAACAGCTTTTTCAAGGAATTTCGGTTCAACAAGAACAACTC

>M6-PGSC0003DMG400004146-(rev)M6LRR-RLK_1984

ATAGATGGTACTTGTGCAACAGCTTTTTCAAGGAATTTCGGTTCAACAAGAACAACTCAGAATGGCCTTGGAGGCTTGTAGCATTCCAGAGGCTTAACTTCACAAGTACTGATATTCTGG

>M6-PGSC0003DMG400004146-(rev)M6LRR-RLK_2046

TGGCCTTGGAGGCTTGTAGCATTCCAGAGGCTTAACTTCACAAGTACTGATATTCTGGCTTGCCTAAAGGAGTCAAATGTGATAGGCATTGGTGGCAATGGAATTGTCTACAAGGCTGAG

>M6-PGSC0003DMG400004146-(rev)M6LRR-RLK_2108

CCTAAAGGAGTCAAATGTGATAGGCATTGGTGGCAATGGAATTGTCTACAAGGCTGAGGTTCTGAGGCCACATTCTGTTGTAGCAGTTAAAAAGTTGTGGAGATCAGATGGAGACATTGA

>M6-PGSC0003DMG400004146-(rev)M6LRR-RLK_2170

TGAGGCCACATTCTGTTGTAGCAGTTAAAAAGTTGTGGAGATCAGATGGAGACATTGAAGCAGGAGATGATCTCGTTGCAGAGGTTGATCTTTTAGGGAAGCTTCGTCACAGGAACATAG

>M6-PGSC0003DMG400004146-(rev)M6LRR-RLK_2232

GGAGATGATCTCGTTGCAGAGGTTGATCTTTTAGGGAAGCTTCGTCACAGGAACATAGTTAGGTTACTAGGCTATCTTCACAATGAGACTGATATCATGATGTTATCTGAATACATGCCT

>M6-PGSC0003DMG400004146-(rev)M6LRR-RLK_2294

GTTACTAGGCTATCTTCACAATGAGACTGATATCATGATGTTATCTGAATACATGCCTAATGGAAATCTTGGAGCTGCTTTACATGGAAAAGAAGCTGGAAAAATGCTCGTAGATTGGTT

>M6-PGSC0003DMG400004146-(rev)M6LRR-RLK_2356

GAAATCTTGGAGCTGCTTTACATGGAAAAGAAGCTGGAAAAATGCTCGTAGATTGGTTGTCGAGGTACAACGTTGCCCTTGGTATTGCTCATGGACTCGCTTATCTTCACCATGATTGTC

>M6-PGSC0003DMG400004146-(rev)M6LRR-RLK_2418

AGGTACAACGTTGCCCTTGGTATTGCTCATGGACTCGCTTATCTTCACCATGATTGTCATCCACCAGTCATACATCGCGATGTCAAGTCAAACAACATTCTCCTAGATTCAGATTTTGAG

>M6-PGSC0003DMG400004146-(rev)M6LRR-RLK_248

CAGAGTTTCAGATCAAATTCAAGGACTTCAGAGTCTATCTTTGCTTAATCTTTGTTGCAATGACTTCTCCACTTCATTGCCAAAGTCATTAGCTAATCTGACATCTTTGAAGAGCATTGA

>M6-PGSC0003DMG400004146-(rev)M6LRR-RLK_2480

ACCAGTCATACATCGCGATGTCAAGTCAAACAACATTCTCCTAGATTCAGATTTTGAGGCAAGAATTGCAGATTTTGGCTTGGCAAGGATGATGCTTCACAAGAATGAGACTGTTTCTAT

>M6-PGSC0003DMG400004146-(rev)M6LRR-RLK_2542

GAATTGCAGATTTTGGCTTGGCAAGGATGATGCTTCACAAGAATGAGACTGTTTCTATGATAGCAGGATCATATGGCTATATAGCACCAGGTTAGTTACATTTCCTGATAATGCCACTAA

>M6-PGSC0003DMG400004146-(rev)M6LRR-RLK_2604

GCAGGATCATATGGCTATATAGCACCAGGTTAGTTACATTTCCTGATAATGCCACTAACAAGAAAATTCATTTCATTTTATATAGTCTCATTTTCCTTTCTCTGTTGTTTTTGGCAGAAT

>M6-PGSC0003DMG400004146-(rev)M6LRR-RLK_2666

AAAATTCATTTCATTTTATATAGTCTCATTTTCCTTTCTCTGTTGTTTTTGGCAGAATATGGATACACATTGAAGGTTGATGAGAAGAGTGACATATACAGCTATGGGGTGGTGCTTTTG

>M6-PGSC0003DMG400004146-(rev)M6LRR-RLK_2728

ATACACATTGAAGGTTGATGAGAAGAGTGACATATACAGCTATGGGGTGGTGCTTTTGGAGCTTGTAACAGGAAAAATGCCATTAGACTCTTTGTTCGGAGAGTCGATTGACATTGTAGA

>M6-PGSC0003DMG400004146-(rev)M6LRR-RLK_2790

TTGTAACAGGAAAAATGCCATTAGACTCTTTGTTCGGAGAGTCGATTGACATTGTAGAATGGGTTAGAAGGAAAGTAAAGAACAAGGCATCAGAAGAAGCATTAGATGCTGATGTAGCTG

>M6-PGSC0003DMG400004146-(rev)M6LRR-RLK_2852

GTTAGAAGGAAAGTAAAGAACAAGGCATCAGAAGAAGCATTAGATGCTGATGTAGCTGGCCAATGTAAACATGTCCATGAAGAGATGCTTCTTGTCCTAAAAATTGCACTTCTTTGCACT

>M6-PGSC0003DMG400004146-(rev)M6LRR-RLK_2914

ATGTAAACATGTCCATGAAGAGATGCTTCTTGTCCTAAAAATTGCACTTCTTTGCACTGCCAAGCTTCCCAAAGAAAGACCATCAATGAGAGACATTATAACCATGCTTGGAGAGGCAAA

>M6-PGSC0003DMG400004146-(rev)M6LRR-RLK_2976

AGCTTCCCAAAGAAAGACCATCAATGAGAGACATTATAACCATGCTTGGAGAGGCAAAACCAAGAAGGAAAAGTATTTGCCAGAATTGGGGTTATAGTACTAGTGCTAACAAAGACAAGT

>M6-PGSC0003DMG400004146-(rev)M6LRR-RLK_3038

AGAAGGAAAAGTATTTGCCAGAATTGGGGTTATAGTACTAGTGCTAACAAAGACAAGTTAATCTTTGCTCATTCACCAGTTATAGGCCTTCTATAAGTACCAAAACCTTTTTTTTTCCAC

>M6-PGSC0003DMG400004146-(rev)M6LRR-RLK_3082

TAACAAAGACAAGTTAATCTTTGCTCATTCACCAGTTATAGGCCTTCTATAAGTACCAAAACCTTTTTTTTTCCACTTCAATATCTAGTATATCTTCATTTTCATTAGTTCATATGTAAA

>M6-PGSC0003DMG400004146-(rev)M6LRR-RLK_310

ACTTCTCCACTTCATTGCCAAAGTCATTAGCTAATCTGACATCTTTGAAGAGCATTGATGTGAGCCAGAACAACTTTGTTGGCGAATTCCCTGATGGTATTGGAATGTCGAATCCTGGTT

>M6-PGSC0003DMG400004146-(rev)M6LRR-RLK_372

AGCCAGAACAACTTTGTTGGCGAATTCCCTGATGGTATTGGAATGTCGAATCCTGGTTTGAAATATGTCAATGTTTCAAGTAATAACTTCGAGGGGTTTCTCCCTGAGGATCTTGGCAAT

>M6-PGSC0003DMG400004146-(rev)M6LRR-RLK_434

ATATGTCAATGTTTCAAGTAATAACTTCGAGGGGTTTCTCCCTGAGGATCTTGGCAATGCTATGTTGCTCGAAGTCATGGATTTTCGAGGGAGTTTCTTTGAAGGTTCAATCCCTGGATG

>M6-PGSC0003DMG400004146-(rev)M6LRR-RLK_496

TGTTGCTCGAAGTCATGGATTTTCGAGGGAGTTTCTTTGAAGGTTCAATCCCTGGATGTTACAAGAATCTGAAGAATTTGAAGTTTCTTGGCTTATCAGGGAATAATCTTTCCGGAGAGA

>M6-PGSC0003DMG400004146-(rev)M6LRR-RLK_558

AAGAATCTGAAGAATTTGAAGTTTCTTGGCTTATCAGGGAATAATCTTTCCGGAGAGATCCCTCGTGAGCTTGGTGAATTGAAGGCTATGGAGACTATGATTCTTGGATATAATCAGTTT

>M6-PGSC0003DMG400004146-(rev)M6LRR-RLK_62

ATCAAATAAGGATGAAGTTTCTATATTGTTAGCAATAAAGTCTAGTCTTGTTGATCCAATGGATCATCTCAAAGATTGGAACTTTTCAAACAATGAAAGTTCTATTCATTGTAAATGGAA

>M6-PGSC0003DMG400004146-(rev)M6LRR-RLK_620

TCGTGAGCTTGGTGAATTGAAGGCTATGGAGACTATGATTCTTGGATATAATCAGTTTGGAGGTTCAATACCAGCTGAGTTTGGTAATATGAGTAGCCTTAAGTATCTTGACTTGGCTGT

>M6-PGSC0003DMG400004146-(rev)M6LRR-RLK_682

GTTCAATACCAGCTGAGTTTGGTAATATGAGTAGCCTTAAGTATCTTGACTTGGCTGTTGGCACGTTAAGTGGACAAATTCCAGCTGAGTTAGGTAAGTTGAAGAACCTGACTACAGTGT

>M6-PGSC0003DMG400004146-(rev)M6LRR-RLK_744

ACGTTAAGTGGACAAATTCCAGCTGAGTTAGGTAAGTTGAAGAACCTGACTACAGTGTACTTGTATCAAAACAGTTTTGAAGGTAAGATTCCAGCTGAAATTGGAAATATAACATCTTTA

>M6-PGSC0003DMG400004146-(rev)M6LRR-RLK_806

GTATCAAAACAGTTTTGAAGGTAAGATTCCAGCTGAAATTGGAAATATAACATCTTTAGTTTACTTGGATCTTTCTGATAACAAGATCACTGGGGAGATTCCGAATGAGATAGCAGATTT

>M6-PGSC0003DMG400004146-(rev)M6LRR-RLK_868

ACTTGGATCTTTCTGATAACAAGATCACTGGGGAGATTCCGAATGAGATAGCAGATTTGAAGAATTTGCAGCTGCTAAATCTCATGTGCAATAACTTAACTGGTCCTATTCCTACCAAAC

>M6-PGSC0003DMG400004146-(rev)M6LRR-RLK_930

AATTTGCAGCTGCTAAATCTCATGTGCAATAACTTAACTGGTCCTATTCCTACCAAACTTGGTGAGTTAGAAAATCTTGAGATACTTGAACTATGGAAAAATTCATTAAATGGCTCTTTG

>M6-PGSC0003DMG400004146-(rev)M6LRR-RLK_992

TGAGTTAGAAAATCTTGAGATACTTGAACTATGGAAAAATTCATTAAATGGCTCTTTGCCAATGAATCTTGGGAAAAAATCTCCCTTGCAATGGTTAGACGTTTCTTCGAATATCTTGAC

>M6-PGSC0003DMG400004190M6LRR-RLP_0

AATCCACCGACGCGCAGTGCAAGCTAAATAGCGCGTTGTATGAAAAAAACTAATTTTGAGGAAAATTCCCAAAGTGAGGATTCATTTCACACACAGATCTTCTTCCTTCACTTCCATCAT

>M6-PGSC0003DMG400004190M6LRR-RLP_1037

CACATGTATAGTTGGAAATGATTGGAATTTATAGATTAGATGAAGTTGTAAGTGCAAGTTTATAGTTTGAGCTAAAATAATTGGATTTTATGCATTCACTGCTTTCTGAGTGTATGTTTA

>M6-PGSC0003DMG400004190M6LRR-RLP_1098

ATAGTTTGAGCTAAAATAATTGGATTTTATGCATTCACTGCTTTCTGAGTGTATGTTTATGGTTTTCCTGGAAGTAATTCGACTTTTTAGATTTTATGCAGCTTCTAAGCTCACGTTTAT

>M6-PGSC0003DMG400004190M6LRR-RLP_1159

GTTTTCCTGGAAGTAATTCGACTTTTTAGATTTTATGCAGCTTCTAAGCTCACGTTTATATATTTTGAGTGGAAGATTTTATGTTGTTTCTAAGTTTTACTTATTGTGTTTCATCTTTTC

>M6-PGSC0003DMG400004190M6LRR-RLP_122

ATCTCAATCTCGACTATAGTAATGTACGTATAGCTCTGCGTTTGGATTCCGGTTGATTTAATAATTCTACTCCCCTGAAGGTGATTAACCCTATTCAAATGGCTCGCCATGGGCATCTAC

>M6-PGSC0003DMG400004190M6LRR-RLP_1220

ATTTTGAGTGGAAGATTTTATGTTGTTTCTAAGTTTTACTTATTGTGTTTCATCTTTTCTTACTATAATTTCCTTTTTCCTGAATGTCTGTTTTCTTGGTTAAGTAATTCAAGCTTTGCA

>M6-PGSC0003DMG400004190M6LRR-RLP_1281

ACTATAATTTCCTTTTTCCTGAATGTCTGTTTTCTTGGTTAAGTAATTCAAGCTTTGCAATTTGTTGATTTTGTTGAGGAATAGCTATTCGTCTGATGTATAATCTTGATCTGTTCAATG

>M6-PGSC0003DMG400004190M6LRR-RLP_1342

TTGTTGATTTTGTTGAGGAATAGCTATTCGTCTGATGTATAATCTTGATCTGTTCAATGCATTATCTATTATCTGCAATCAATAACTAAATCAGTGCCTTGGAGTGTCATTAACTATAAA

>M6-PGSC0003DMG400004190M6LRR-RLP_1403

TTATCTATTATCTGCAATCAATAACTAAATCAGTGCCTTGGAGTGTCATTAACTATAAATGTTTTCCCCTTTTTTTGCTCGTAGAATTCTATTTTCTAGTTGTTACAGGTAAATATAGCT

>M6-PGSC0003DMG400004190M6LRR-RLP_1464

TTTTCCCCTTTTTTTGCTCGTAGAATTCTATTTTCTAGTTGTTACAGGTAAATATAGCTATCCTTGGAAAGTTAAACCGATTTATCATTCAATCAATGTCAATTTAATAACTCTCCATGA

>M6-PGSC0003DMG400004190M6LRR-RLP_1525

CCTTGGAAAGTTAAACCGATTTATCATTCAATCAATGTCAATTTAATAACTCTCCATGATTACATTGTATTAGTTAATTAAATTGTAGTATAAGTAAGTATATAATCGATCACGAGTAAT

>M6-PGSC0003DMG400004190M6LRR-RLP_1586

ACATTGTATTAGTTAATTAAATTGTAGTATAAGTAAGTATATAATCGATCACGAGTAATTTAATGCTTGTCAGAAATAAGAATGTCCACTGTAGGTCTTGAGCCCATGATCATAAGGTTA

>M6-PGSC0003DMG400004190M6LRR-RLP_1647

AATGCTTGTCAGAAATAAGAATGTCCACTGTAGGTCTTGAGCCCATGATCATAAGGTTAATAGCTTTGCTCTCTACACTTAATGTTGTTGGAAATGTTGTGGAGTTCCAATTATATCCAC

>M6-PGSC0003DMG400004190M6LRR-RLP_1708

AGCTTTGCTCTCTACACTTAATGTTGTTGGAAATGTTGTGGAGTTCCAATTATATCCACTCTTTCTCTGTTAACATTTCATTAAAATGATTTACACAGTTTAAGGCCATTGATTGCATTT

>M6-PGSC0003DMG400004190M6LRR-RLP_1769

TTTCTCTGTTAACATTTCATTAAAATGATTTACACAGTTTAAGGCCATTGATTGCATTTATGAGAAAACCAAGGAGTTGTAGTACCTGAATGTATATGTTTACATGGACTAATTTGAACT

>M6-PGSC0003DMG400004190M6LRR-RLP_183

TAATTCTACTCCCCTGAAGGTGATTAACCCTATTCAAATGGCTCGCCATGGGCATCTACCTTCAGTATTTCTCCTGATCTTCTCGTTATCCTTGATCCAAAGCGCTCTCGTTCACTCTAA

>M6-PGSC0003DMG400004190M6LRR-RLP_1830

GAGAAAACCAAGGAGTTGTAGTACCTGAATGTATATGTTTACATGGACTAATTTGAACTTCTGGAATATGACTGTAGTGAAAGCTCTGAATGAAATTAAGGCATCTCTTGGATGGAGAGT

>M6-PGSC0003DMG400004190M6LRR-RLP_1891

TGGAATATGACTGTAGTGAAAGCTCTGAATGAAATTAAGGCATCTCTTGGATGGAGAGTGGTGTATGCATGGGTTGGAGATGATCCTTGTGGTGATGGTGATTTACCTCCATGGTCTGGT

>M6-PGSC0003DMG400004190M6LRR-RLP_1952

TGTATGCATGGGTTGGAGATGATCCTTGTGGTGATGGTGATTTACCTCCATGGTCTGGTGTTACATGTTCAACGCAGGGTGATTATAGAGTAGTTACTGAATTGTAAGTTTGCTTCACCA

>M6-PGSC0003DMG400004190M6LRR-RLP_2013

TACATGTTCAACGCAGGGTGATTATAGAGTAGTTACTGAATTGTAAGTTTGCTTCACCAGCTCCAAGTAGCTTTTCTCTAACTAAGCTTCAAATCTGTATGAAGTTACTCACAGAATTGA

>M6-PGSC0003DMG400004190M6LRR-RLP_2074

TCCAAGTAGCTTTTCTCTAACTAAGCTTCAAATCTGTATGAAGTTACTCACAGAATTGAAATATCTTTACTTGTCACTTTTTCACAATGTTCTTTTCTGTTTTACCACTTGCAGGGAAGT

>M6-PGSC0003DMG400004190M6LRR-RLP_2135

TATCTTTACTTGTCACTTTTTCACAATGTTCTTTTCTGTTTTACCACTTGCAGGGAAGTTTATGCTGTTTCAATTGTTGGCCCATTTCCTACTGCTGTCACTAATTTATTAGATCTTACT

>M6-PGSC0003DMG400004190M6LRR-RLP_2196

ATGCTGTTTCAATTGTTGGCCCATTTCCTACTGCTGTCACTAATTTATTAGATCTTACTAGACTGTAAGATTCTTCTTCTCTTGGTCTTTGTACTGGAATCTAGTTTTTACAAAACTTCA

>M6-PGSC0003DMG400004190M6LRR-RLP_2257

ACTGTAAGATTCTTCTTCTCTTGGTCTTTGTACTGGAATCTAGTTTTTACAAAACTTCAGCTACTTCTTACAGATTTTTTAACCTCATACAAATTGAGAAGTTTCCCTAAGTTCCAATAT

>M6-PGSC0003DMG400004190M6LRR-RLP_2318

TACTTCTTACAGATTTTTTAACCTCATACAAATTGAGAAGTTTCCCTAAGTTCCAATATGCATGTTTTTACACCCTCCAGTCTTAATTGTTCATTAACCAACTTAAAGAATTGTTCTGCT

>M6-PGSC0003DMG400004190M6LRR-RLP_2379

ATGTTTTTACACCCTCCAGTCTTAATTGTTCATTAACCAACTTAAAGAATTGTTCTGCTTGGGTGTTGACTGTTACCCCAAAAAATGTGGGCGTTAGCTATAATGTGTGTAGTCATTATT

>M6-PGSC0003DMG400004190M6LRR-RLP_2440

GGTGTTGACTGTTACCCCAAAAAATGTGGGCGTTAGCTATAATGTGTGTAGTCATTATTTTCAAATTTGATACTACTATTTAAGGCAATAGTTAGTGGACAATTGTGCCTGTACTGTTGG

>M6-PGSC0003DMG400004190M6LRR-RLP_2501

CAAATTTGATACTACTATTTAAGGCAATAGTTAGTGGACAATTGTGCCTGTACTGTTGGATCTTTTGATTTTCATTCTGAACTTCCTAATCCATAATATTCTTTCATATGACAGGGATCT

>M6-PGSC0003DMG400004190M6LRR-RLP_2562

CTTTTGATTTTCATTCTGAACTTCCTAATCCATAATATTCTTTCATATGACAGGGATCTGCATAATAACAAGCTGACCGGCCCTCTTCCTTCTCAAATTGGGCGTTTGAAGCGTCTTAAA

>M6-PGSC0003DMG400004190M6LRR-RLP_2623

ATAATAACAAGCTGACCGGCCCTCTTCCTTCTCAAATTGGGCGTTTGAAGCGTCTTAAAATATTGTATAATCCTTTACTTGTATACTCCTATTGAAGTATCTATATTCAGTAGCAATTAT

>M6-PGSC0003DMG400004190M6LRR-RLP_2684

ATTGTATAATCCTTTACTTGTATACTCCTATTGAAGTATCTATATTCAGTAGCAATTATGAAGTGATATTTATGTTTAGGGCTTATGCTCTGTCTCCGAATAGCCTGTTTCTCCAGTTTC

>M6-PGSC0003DMG400004190M6LRR-RLP_2745

AGTGATATTTATGTTTAGGGCTTATGCTCTGTCTCCGAATAGCCTGTTTCTCCAGTTTCAGGATTAACATATGTTTCTCCATTAACACCTAGCTAGATGATGTTAATATGATTCAACTGT

>M6-PGSC0003DMG400004190M6LRR-RLP_2806

GATTAACATATGTTTCTCCATTAACACCTAGCTAGATGATGTTAATATGATTCAACTGTGTTGTACAAAGTGATTAAGTACATGGCTTTTCATCGAGATGAAATGTGTTAATAACAAGGA

>M6-PGSC0003DMG400004190M6LRR-RLP_2867

TGTACAAAGTGATTAAGTACATGGCTTTTCATCGAGATGAAATGTGTTAATAACAAGGAAAAGCTTACCTGATTAATGATTAATCCCAAAGAAGGGAGTGAGAGGGAGTCTGCCTGATAT

>M6-PGSC0003DMG400004190M6LRR-RLP_2928

AGCTTACCTGATTAATGATTAATCCCAAAGAAGGGAGTGAGAGGGAGTCTGCCTGATATAATGTGCACCGAAGAAGAAAGATAAATTTTCAACTGAGAGAATATATACTCGTAGGCTTAT

>M6-PGSC0003DMG400004190M6LRR-RLP_2989

TGTGCACCGAAGAAGAAAGATAAATTTTCAACTGAGAGAATATATACTCGTAGGCTTATATTCTAAAAAATAAAAACAGAGTAGGTTTATTATCAAGGAAATGTCACTTAACTAAGAAAG

>M6-PGSC0003DMG400004190M6LRR-RLP_305

CCCTCAAACGTGACGGTACACATTTCGATCCATTTACTTTTTAAACCTAATTTATTAGCAATTTAGGTGAGAAAATCAGTTGAATTTGATTGATAATAGTTTGGTTTTGTTATTTTGTAG

>M6-PGSC0003DMG400004190M6LRR-RLP_3050

TCTAAAAAATAAAAACAGAGTAGGTTTATTATCAAGGAAATGTCACTTAACTAAGAAAGCCTTCTATACTCTTTCTAGGATTGAATTAGATGTATATCTCAACATCTACTTTCTGTTGCA

>M6-PGSC0003DMG400004190M6LRR-RLP_3111

TTCTATACTCTTTCTAGGATTGAATTAGATGTATATCTCAACATCTACTTTCTGTTGCACGTATTTTACCTTTAAGTTTTCCTTTGTCCATAGGAACCTCCACGTTGTCCCAGAATGTTA

>M6-PGSC0003DMG400004190M6LRR-RLP_3172

TATTTTACCTTTAAGTTTTCCTTTGTCCATAGGAACCTCCACGTTGTCCCAGAATGTTAACTTTAAAAGTTACTTTTATTGCTTTTTTACCCAACATACTAATATGAAATTATATCCTCT

>M6-PGSC0003DMG400004190M6LRR-RLP_3233

TTTAAAAGTTACTTTTATTGCTTTTTTACCCAACATACTAATATGAAATTATATCCTCTAGTTTCCTAGTGTGTTGTAATGTTGCATTATTGATTAGTTCCTTCTCCTGATTCTTTTTAC

>M6-PGSC0003DMG400004190M6LRR-RLP_3294

TTTCCTAGTGTGTTGTAATGTTGCATTATTGATTAGTTCCTTCTCCTGATTCTTTTTACCAGAAACGTAAGGTGGAATAAGCTGCAAGATGTCATTCCACCTGAAATAGGTGAACTAAAG

>M6-PGSC0003DMG400004190M6LRR-RLP_3355

GAAACGTAAGGTGGAATAAGCTGCAAGATGTCATTCCACCTGAAATAGGTGAACTAAAGCAGTTAACACATCTGTGAGTGCTTTTTGAGCTTCTCTAGGTGTGCTCTTTATCCATAGATC

>M6-PGSC0003DMG400004190M6LRR-RLP_3416

GTTAACACATCTGTGAGTGCTTTTTGAGCTTCTCTAGGTGTGCTCTTTATCCATAGATCTATTTTGCTGTTATAGACTAATAACTGATATGTATATTTTTCAGCTACCTAAGTTTTAATA

>M6-PGSC0003DMG400004190M6LRR-RLP_3477

TTTTGCTGTTATAGACTAATAACTGATATGTATATTTTTCAGCTACCTAAGTTTTAATAATTTCAAAGGTGAAATACCTAAGGAGCTTGCTAATCTTCCTGAACTTCGTTATCTCCATCT

>M6-PGSC0003DMG400004190M6LRR-RLP_3538

TTCAAAGGTGAAATACCTAAGGAGCTTGCTAATCTTCCTGAACTTCGTTATCTCCATCTGCATGAAAATCATTTTATTGGACGAATTCCACCTGAACTGGGTTCCTTACAGTATCTTCGA

>M6-PGSC0003DMG400004190M6LRR-RLP_3599

ATGAAAATCATTTTATTGGACGAATTCCACCTGAACTGGGTTCCTTACAGTATCTTCGACACCTGTATGATACCATCTTAACTTCTTCTCTTGTTTAACATATGGAACCTGTGTCACACA

>M6-PGSC0003DMG400004190M6LRR-RLP_366

TTTAGGTGAGAAAATCAGTTGAATTTGATTGATAATAGTTTGGTTTTGTTATTTTGTAGAGTATGTATGCAGCTTCAAGTGAAATTAATTTTGTAGTTACAGAGAAAATGTAACAATGTT

>M6-PGSC0003DMG400004190M6LRR-RLP_3660

CCTGTATGATACCATCTTAACTTCTTCTCTTGTTTAACATATGGAACCTGTGTCACACATGGATTTCACTTTGATCTCATGCAATTCCAGGGATGTTGGTAACAATCGTTTGGTTGGAAC

>M6-PGSC0003DMG400004190M6LRR-RLP_3721

GATTTCACTTTGATCTCATGCAATTCCAGGGATGTTGGTAACAATCGTTTGGTTGGAACTATAAGAGAACTCATACGTATTGAAGGATGTTTTCCGGTTCTCCGCAACCTGTAAGTTGAT

>M6-PGSC0003DMG400004190M6LRR-RLP_3782

TAAGAGAACTCATACGTATTGAAGGATGTTTTCCGGTTCTCCGCAACCTGTAAGTTGATTATTCTTGTTTAATTGTCATATCGCATGAAGTTATCCTGCTGCAAGTGTTATGAATTCTAT

>M6-PGSC0003DMG400004190M6LRR-RLP_3843

TTCTTGTTTAATTGTCATATCGCATGAAGTTATCCTGCTGCAAGTGTTATGAATTCTATGCTAATTCAAGGATAACACGTGCTGTAACTTGCAGATACTTAAACAATAATTATCTTACTG

>M6-PGSC0003DMG400004190M6LRR-RLP_3904

TAATTCAAGGATAACACGTGCTGTAACTTGCAGATACTTAAACAATAATTATCTTACTGGAGGCATTCCAGCGCAACTTGCAAATCTGACGAAGTTGGAGATCTTGTAAGTTTTGGGGAG

>M6-PGSC0003DMG400004190M6LRR-RLP_3965

GGCATTCCAGCGCAACTTGCAAATCTGACGAAGTTGGAGATCTTGTAAGTTTTGGGGAGAAAGTTTCTCCGCGATTAATGCTCGTCTTTGGACAAATAACAGCTAATTTCTCCTGTACTG

>M6-PGSC0003DMG400004190M6LRR-RLP_4026

AGTTTCTCCGCGATTAATGCTCGTCTTTGGACAAATAACAGCTAATTTCTCCTGTACTGGTTTATAAATATTATTATGCTTAATTTGATTGCTCTGGTTATTACAGGTATCTATCCTACA

>M6-PGSC0003DMG400004190M6LRR-RLP_4087

TTATAAATATTATTATGCTTAATTTGATTGCTCTGGTTATTACAGGTATCTATCCTACAATAAGATGACAGGAGTTATACCATATAGCATTGCTCATATTCCTAAGTTAACTTACTTGTA

>M6-PGSC0003DMG400004190M6LRR-RLP_4148

AAGATGACAGGAGTTATACCATATAGCATTGCTCATATTCCTAAGTTAACTTACTTGTAAGTCCTCCTCTATGCTCTTAGCTACTCATTTATTATACTGTATGTGGACAGTTTGGACCGA

>M6-PGSC0003DMG400004190M6LRR-RLP_4209

TCCTCCTCTATGCTCTTAGCTACTCATTTATTATACTGTATGTGGACAGTTTGGACCGAGAAAATTCAGAATTTTTGAAAGCAAATACAAAAACATCCTTAGCCTTAAGTTGAGATCTCT

>M6-PGSC0003DMG400004190M6LRR-RLP_427

TATGTATGCAGCTTCAAGTGAAATTAATTTTGTAGTTACAGAGAAAATGTAACAATGTTGAACTTTGTAGATCTTATGTATTCTAGGTGCAAGGTTATAGTTTGAGTGGAGTTTAATTGA

>M6-PGSC0003DMG400004190M6LRR-RLP_4270

AAATTCAGAATTTTTGAAAGCAAATACAAAAACATCCTTAGCCTTAAGTTGAGATCTCTTTCTTTGATAGCTTTTTGGTGCAACCTGGTCTTTGTGCATGATTTAGACGCCACATTAGAT

>M6-PGSC0003DMG400004190M6LRR-RLP_4331

CTTTGATAGCTTTTTGGTGCAACCTGGTCTTTGTGCATGATTTAGACGCCACATTAGATATGATCAGGGACCTGCGTGTTATCATGCAGTAGTTTAGTGCTGTTTTCTGTTATCAATAGA

>M6-PGSC0003DMG400004190M6LRR-RLP_4392

GATCAGGGACCTGCGTGTTATCATGCAGTAGTTTAGTGCTGTTTTCTGTTATCAATAGATTGCTTTATTAAATTATAAAAATATTAGCATATTGGATGCGATGTCCCAATGATTGTCTAG

>M6-PGSC0003DMG400004190M6LRR-RLP_4453

GCTTTATTAAATTATAAAAATATTAGCATATTGGATGCGATGTCCCAATGATTGTCTAGTGAGTGCAGTCTTTCAAGGATCTTTCTTTCTTAAAAAGATGAATTTTGAGGATTGTTCAAC

>M6-PGSC0003DMG400004190M6LRR-RLP_4697

TCTACTCAAAACAGATGAACTCGGGACTAGTTGGCTAATGTGGTCTTGCAAGGGTCTCTTTCTTAAGGAGATAAATTTTGAGGATTGTGCAATTTATGGATGCTGTGACAAAAACATTAG

>M6-PGSC0003DMG400004190M6LRR-RLP_4758

CTTAAGGAGATAAATTTTGAGGATTGTGCAATTTATGGATGCTGTGACAAAAACATTAGATGGACTTGAAGCTTGTCAGCTGACCTTATTGGACCTTATAGTAGTAATATATAGAAATTG

>M6-PGSC0003DMG400004190M6LRR-RLP_4819

GGACTTGAAGCTTGTCAGCTGACCTTATTGGACCTTATAGTAGTAATATATAGAAATTGCCTGGAGAATTAGATGCAAGAGTGTTCTCATCAATTTTTGCACAAAAAACACAGTCCATCT

>M6-PGSC0003DMG400004190M6LRR-RLP_488

ACTTTGTAGATCTTATGTATTCTAGGTGCAAGGTTATAGTTTGAGTGGAGTTTAATTGAATTTTAAAAGTTCGAGTGAAAATAGTTAGATTTTGTAGATTATATGCAGTTTGTGGTGAAA

>M6-PGSC0003DMG400004190M6LRR-RLP_4880

TGGAGAATTAGATGCAAGAGTGTTCTCATCAATTTTTGCACAAAAAACACAGTCCATCTCAATAGTAAAACTTCTCATGTACTCAAGATTTCAGTTGAGTTATGTAGCAACTGCACTGGT

>M6-PGSC0003DMG400004190M6LRR-RLP_4941

ATAGTAAAACTTCTCATGTACTCAAGATTTCAGTTGAGTTATGTAGCAACTGCACTGGTACAGAGATGCATGAATGACGTGTTAAACTTTAATTAGAAGATTGTCTGTTAGAACATAATA

>M6-PGSC0003DMG400004190M6LRR-RLP_5002

AGAGATGCATGAATGACGTGTTAAACTTTAATTAGAAGATTGTCTGTTAGAACATAATAGTTTTATTGGTATAACGTGTTAAAAAAATTGCATGTATAATTCAAAAAAATATGAACACTG

>M6-PGSC0003DMG400004190M6LRR-RLP_5063

TTTATTGGTATAACGTGTTAAAAAAATTGCATGTATAATTCAAAAAAATATGAACACTGATGCATGCAAGTCTTTAAGCAATTAAACAAAGCTGATTTTGCACCTGGTTGTCAATATCCA

>M6-PGSC0003DMG400004190M6LRR-RLP_5124

GCATGCAAGTCTTTAAGCAATTAAACAAAGCTGATTTTGCACCTGGTTGTCAATATCCATTGCATTTCCTACCCATTTTCTTCTTTCTTCCAAAATCCATGGTCTTTCAACACTTACATT

>M6-PGSC0003DMG400004190M6LRR-RLP_5185

GCATTTCCTACCCATTTTCTTCTTTCTTCCAAAATCCATGGTCTTTCAACACTTACATTCCCCTCCATCCCCCAGATTAGAAGTCATTATTTGTACAAGTGAGAGGCTTGGATAATGCAG

>M6-PGSC0003DMG400004190M6LRR-RLP_5246

CCTCCATCCCCCAGATTAGAAGTCATTATTTGTACAAGTGAGAGGCTTGGATAATGCAGAAGGACCTGCATTTTGCTGGTCAAACAACCAAAGAATGTCAAGAGACAAGTCTAATTGTGT

>M6-PGSC0003DMG400004190M6LRR-RLP_5307

GGACCTGCATTTTGCTGGTCAAACAACCAAAGAATGTCAAGAGACAAGTCTAATTGTGTCAGATACATATCCCTGCTCAAGATATGCTGTGTTGGAAATGGGAACCTTGTTTTGAAGCTT

>M6-PGSC0003DMG400004190M6LRR-RLP_5368

GATACATATCCCTGCTCAAGATATGCTGTGTTGGAAATGGGAACCTTGTTTTGAAGCTTAACCCACATATTTCCTTTGACAAAACCTTTTATCACAATATTTTCCTTTAAAGTGTTGGTG

>M6-PGSC0003DMG400004190M6LRR-RLP_5429

CCCACATATTTCCTTTGACAAAACCTTTTATCACAATATTTTCCTTTAAAGTGTTGGTGTTGGAGCTAAACCACACGAACTGAAACTAGGGCCTTCTCCATTTAAGTGGTTGTTAGTGTT

>M6-PGSC0003DMG400004190M6LRR-RLP_549

TTTAAAAGTTCGAGTGAAAATAGTTAGATTTTGTAGATTATATGCAGTTTGTGGTGAAATTTTTGTAGTTTCAGCGAAAACAATATTGAACGTTGTACATCTTATGCAATTTCTAGGTGT

>M6-PGSC0003DMG400004190M6LRR-RLP_5490

GGAGCTAAACCACACGAACTGAAACTAGGGCCTTCTCCATTTAAGTGGTTGTTAGTGTTGTCAAAGTGATGCTTGAGCCATGAACTGAGGCTGTGCACAAGTTTACTCCAATGAAGGTGG

>M6-PGSC0003DMG400004190M6LRR-RLP_5551

CAAAGTGATGCTTGAGCCATGAACTGAGGCTGTGCACAAGTTTACTCCAATGAAGGTGGTAACTGGTAAGGGATATTTTCCCAAAACCAGCCAAGCAACTTCTTTTCTGAAAATATTAGA

>M6-PGSC0003DMG400004190M6LRR-RLP_5612

ACTGGTAAGGGATATTTTCCCAAAACCAGCCAAGCAACTTCTTTTCTGAAAATATTAGATTAATTTGTGAGGTCCATGGGTTTCTTCTTGGATTGGGTTGCATTAAACAACAAATATTCA

>M6-PGSC0003DMG400004190M6LRR-RLP_5673

AATTTGTGAGGTCCATGGGTTTCTTCTTGGATTGGGTTGCATTAAACAACAAATATTCAATAGATAGCATGCAAATCGATATGTCAGTTGTTAGCGAGACTGTACTGATGATCATTATTG

>M6-PGSC0003DMG400004190M6LRR-RLP_5734

AGATAGCATGCAAATCGATATGTCAGTTGTTAGCGAGACTGTACTGATGATCATTATTGATTACGAGCAAGAAACTGTAAATTTATTGCAAGAAAGCTAACGACGAAGTCTGAATTAGAA

>M6-PGSC0003DMG400004190M6LRR-RLP_5795

TACGAGCAAGAAACTGTAAATTTATTGCAAGAAAGCTAACGACGAAGTCTGAATTAGAATATTAAAGTAAGATGTTCCTTAAAGCTAGCCACTCAGATTCAAAATGCAAACTAGATGACC

>M6-PGSC0003DMG400004190M6LRR-RLP_5856

TTAAAGTAAGATGTTCCTTAAAGCTAGCCACTCAGATTCAAAATGCAAACTAGATGACCAATGATCCTTGCAAGTTGCAATATCCTAGATGTTCCGTTAAAGCCCAAATGGTCCTTGCAG

>M6-PGSC0003DMG400004190M6LRR-RLP_5917

TGATCCTTGCAAGTTGCAATATCCTAGATGTTCCGTTAAAGCCCAAATGGTCCTTGCAGTTGCATATGTGAGTGCTTTGTTCCTTTTATGTATTTATTTTTACGATCTTCACCCTTGACA

>M6-PGSC0003DMG400004190M6LRR-RLP_5978

GCATATGTGAGTGCTTTGTTCCTTTTATGTATTTATTTTTACGATCTTCACCCTTGACAAATCAGATGTTACACATCATTCCTTCGCTCTTCTTGATTATATACTGGTAGCTTTTGCACC

>M6-PGSC0003DMG400004190M6LRR-RLP_6039

TCAGATGTTACACATCATTCCTTCGCTCTTCTTGATTATATACTGGTAGCTTTTGCACCAAAAAGAAAAATAGTTCTGCATGAATTTGATAGTTGATGTCTGAAGCTGAATCTGATTTCA

>M6-PGSC0003DMG400004190M6LRR-RLP_61

AAAATTCCCAAAGTGAGGATTCATTTCACACACAGATCTTCTTCCTTCACTTCCATCATCTATCTCAATCTCGACTATAGTAATGTACGTATAGCTCTGCGTTTGGATTCCGGTTGATTT

>M6-PGSC0003DMG400004190M6LRR-RLP_610

TTTGTAGTTTCAGCGAAAACAATATTGAACGTTGTACATCTTATGCAATTTCTAGGTGTAAGCTTATGTGTGAGTGGAAGTAATTGAATTTTATAGTTCGAGTTGTAGTAATTGGATATT

>M6-PGSC0003DMG400004190M6LRR-RLP_6100

AAAGAAAAATAGTTCTGCATGAATTTGATAGTTGATGTCTGAAGCTGAATCTGATTTCAGGTATCTCGATCATAATCAATTTTCTGGGAGGATTCCAGATGCCTTCTACAAACACCCTTT

>M6-PGSC0003DMG400004190M6LRR-RLP_6161

TATCTCGATCATAATCAATTTTCTGGGAGGATTCCAGATGCCTTCTACAAACACCCTTTCTTGAAGGAAATGTGAGTGTTCATTTTTTGCTACTGACCATTCTTTGGCTTTGTGCAAATC

>M6-PGSC0003DMG400004190M6LRR-RLP_6222

TGAAGGAAATGTGAGTGTTCATTTTTTGCTACTGACCATTCTTTGGCTTTGTGCAAATCCTAATAATATGATAAATTATGTCATAGGTACATTGAAGAGAACACGTTCCGGCCTGGTGTA

>M6-PGSC0003DMG400004190M6LRR-RLP_6283

AATAATATGATAAATTATGTCATAGGTACATTGAAGAGAACACGTTCCGGCCTGGTGTAAAACCAATTGGCCTGCACAAAGTGCTAGAACTTTCGGACGCAGATTTCCTTGTTTAGAAAA

>M6-PGSC0003DMG400004190M6LRR-RLP_6344

ACCAATTGGCCTGCACAAAGTGCTAGAACTTTCGGACGCAGATTTCCTTGTTTAGAAAAAGCCAACTCCTGTTATTTGTTCATGAGCAAAGTGTACCTAGGTTGTCGAAAGAAGCAGCCG

>M6-PGSC0003DMG400004190M6LRR-RLP_6405

CCAACTCCTGTTATTTGTTCATGAGCAAAGTGTACCTAGGTTGTCGAAAGAAGCAGCCGGTCCCTGTTCTGTCAAGTATCTATTCTTTAGTTGCCAGATGAGAAAAAAGGTTCCATCTTT

>M6-PGSC0003DMG400004190M6LRR-RLP_6466

CCCTGTTCTGTCAAGTATCTATTCTTTAGTTGCCAGATGAGAAAAAAGGTTCCATCTTTTTTGTGCAAAGAAGTAATCAACCAAACTTAGTTGCGGTTGCTGTAGTTAAATAGTACCACG

>M6-PGSC0003DMG400004190M6LRR-RLP_6527

TGTGCAAAGAAGTAATCAACCAAACTTAGTTGCGGTTGCTGTAGTTAAATAGTACCACGTTATGAATCATTATTAAGTAATTGCGACATACGTCTCAATCAGTTCTCACTTGCCTTGAAT

>M6-PGSC0003DMG400004190M6LRR-RLP_6588

ATGAATCATTATTAAGTAATTGCGACATACGTCTCAATCAGTTCTCACTTGCCTTGAATTCTTGCATTTGGTTTTTGATTATACATGATGCGATTTCAGAGTTAATATGAAAGGGTAGAG

>M6-PGSC0003DMG400004190M6LRR-RLP_6649

TTGCATTTGGTTTTTGATTATACATGATGCGATTTCAGAGTTAATATGAAAGGGTAGAGAATATGCTCTGATATGACAGGTTTTGGATAGGTTTTTCATTTTTTCGTTAAATGATACAAA

>M6-PGSC0003DMG400004190M6LRR-RLP_671

GCTTATGTGTGAGTGGAAGTAATTGAATTTTATAGTTCGAGTTGTAGTAATTGGATATTATCCAGTTTTTAGATGAAAGCTTATAGTTTGAGTGGAAGTAGCTGGATTTAATAGGTTATG

>M6-PGSC0003DMG400004190M6LRR-RLP_6710

TATGCTCTGATATGACAGGTTTTGGATAGGTTTTTCATTTTTTCGTTAAATGATACAAATACATACCTAAATTATAGTATTATTTTTTTGAGTTTCACACCCTAACTATAGTTATTATCA

>M6-PGSC0003DMG400004190M6LRR-RLP_6771

CATACCTAAATTATAGTATTATTTTTTTGAGTTTCACACCCTAACTATAGTTATTATCAATGTGCAAAAACACTTCGCTGAGTAGCTGATGGTATGCATGCTTTATTAGGTGCCAAAGTG

>M6-PGSC0003DMG400004190M6LRR-RLP_6803

TTCACACCCTAACTATAGTTATTATCAATGTGCAAAAACACTTCGCTGAGTAGCTGATGGTATGCATGCTTTATTAGGTGCCAAAGTGGCACTCCATGTGGATAAATATTTTCACTAGAA

>M6-PGSC0003DMG400004190M6LRR-RLP_732

CCAGTTTTTAGATGAAAGCTTATAGTTTGAGTGGAAGTAGCTGGATTTAATAGGTTATGTGCAGTTTGTGGTGGATTTATGGTTTCTGCGGAAACAATTGAATTTTGTTGATTTTATGCA

>M6-PGSC0003DMG400004190M6LRR-RLP_793

CAGTTTGTGGTGGATTTATGGTTTCTGCGGAAACAATTGAATTTTGTTGATTTTATGCAATTTCTAAGAGTAAGCTTATAGTTTGAGTGAAAGTAAGTGGATTTTATGTAGTCCTAGAGC

>M6-PGSC0003DMG400004190M6LRR-RLP_854

TTCTAAGAGTAAGCTTATAGTTTGAGTGAAAGTAAGTGGATTTTATGTAGTCCTAGAGCAACTTTATAGTTTGAGTTGTAGTAATTGGATTTTATTGACTTTTTCCCAGCTTCTAAGTGC

>M6-PGSC0003DMG400004190M6LRR-RLP_915

CTTTATAGTTTGAGTTGTAGTAATTGGATTTTATTGACTTTTTCCCAGCTTCTAAGTGCAAATTTTTAGTTTGAGCTGAAATAATTGGATTTATTCTATAAATTATATGTGGTTTCGAAG

>M6-PGSC0003DMG400004190M6LRR-RLP_976

ATTTTTAGTTTGAGCTGAAATAATTGGATTTATTCTATAAATTATATGTGGTTTCGAAGTGCACATGTATAGTTGGAAATGATTGGAATTTATAGATTAGATGAAGTTGTAAGTGCAAGT

>M6-PGSC0003DMG400004195-(rev)M6LRR-RLP_0

CTAAAACTCTTAACATAAATTGTTAAATATTTCTAACCTAAGTAGGTATTACCGAAAATCTTTAACACAGGGTGACATCGAGCTATTTAAGGAACATAAACAATTGGTAATTGCTTCACG

>M6-PGSC0003DMG400004195-(rev)M6LRR-RLP_1178

GACTAACAGGGCAAATTCCTGAGTCCATTGGGGTTCTCAAGAATCTCAAAGAACTCAACTTACAAACTAATCAAATCTCTAGTACAATCCCAGAGTCAGTATTCACATTGACGTCTCTAA

>M6-PGSC0003DMG400004195-(rev)M6LRR-RLP_124

CTATTGCTCCTTTATTCATGAATTCATGTATATACAATGTAATTTTCTGTTAGAGGGGATCGTTGCACCTATGTGGTTCTGCCCTGACATCACTTATCTAGTTAAGCTCAACACTTTGAA

>M6-PGSC0003DMG400004195-(rev)M6LRR-RLP_1240

CAAACTAATCAAATCTCTAGTACAATCCCAGAGTCAGTATTCACATTGACGTCTCTAACAACTTTGAACCTTGAAAATAACCACTTGAAAGGGGAAATATCGGAGAATATTGGCAACTTA

>M6-PGSC0003DMG400004195-(rev)M6LRR-RLP_1302

TTTGAACCTTGAAAATAACCACTTGAAAGGGGAAATATCGGAGAATATTGGCAACTTACAAGCACTTCAAAAGCTTTTCTTGTCAAACAATTCGTTCACTGGAAAAATCCCAGTTTCAAT

>M6-PGSC0003DMG400004195-(rev)M6LRR-RLP_1364

CACTTCAAAAGCTTTTCTTGTCAAACAATTCGTTCACTGGAAAAATCCCAGTTTCAATAACAAAACTACATTCCATTTCCACTATCCACTTAGAAACAAATCAACTTGTTGGTGAAATAC

>M6-PGSC0003DMG400004195-(rev)M6LRR-RLP_1426

AAACTACATTCCATTTCCACTATCCACTTAGAAACAAATCAACTTGTTGGTGAAATACAATTGCCATTGAGTCCTGCCCAATGGCCTTTAATTAAAACTTTGGTTCTTGACAACAATAAA

>M6-PGSC0003DMG400004195-(rev)M6LRR-RLP_1488

GCCATTGAGTCCTGCCCAATGGCCTTTAATTAAAACTTTGGTTCTTGACAACAATAAACTTACTGGAGTTATACCTGATTCTATTGGTTATTTGACAACACTTTCATCACTCTCTATATC

>M6-PGSC0003DMG400004195-(rev)M6LRR-RLP_1550

CTGGAGTTATACCTGATTCTATTGGTTATTTGACAACACTTTCATCACTCTCTATATCAAATAATCAACTCACAGGATCTATTCCTTCTTGTCTGGGAAACATAAAGCAATTGCAAATAT

>M6-PGSC0003DMG400004195-(rev)M6LRR-RLP_1612

AATCAACTCACAGGATCTATTCCTTCTTGTCTGGGAAACATAAAGCAATTGCAAATATTGAAGGTCAATAACAACAACTTCTCTGTTGAGATGTTGCCAACGTCGATTTGTGGACTTACT

>M6-PGSC0003DMG400004195-(rev)M6LRR-RLP_1674

GGTCAATAACAACAACTTCTCTGTTGAGATGTTGCCAACGTCGATTTGTGGACTTACTGAGCTAAGTGTTTTGTATGTTTCTCAAAACAAGATTCATGGACCATTACCTGGATGTCTTTC

>M6-PGSC0003DMG400004195-(rev)M6LRR-RLP_1736

TAAGTGTTTTGTATGTTTCTCAAAACAAGATTCATGGACCATTACCTGGATGTCTTTCATCATTCAAGTACTTACTAGATGTCGATATACCGTTCAAACGTACAGTCACTCTTCCTATGG

>M6-PGSC0003DMG400004195-(rev)M6LRR-RLP_1798

TTCAAGTACTTACTAGATGTCGATATACCGTTCAAACGTACAGTCACTCTTCCTATGGGAGTACACAAACAATAAGTAACGGTGACGGAGCCAGAAATTATGAAGGAAATCCAAAAATGC

>M6-PGSC0003DMG400004195-(rev)M6LRR-RLP_186

TTGCACCTATGTGGTTCTGCCCTGACATCACTTATCTAGTTAAGCTCAACACTTTGAATATGTTCTTAAAAATGTGACTTTTAGAGGCTTCATAGGTCAATTTGTTGATAAAACAAAAAT

>M6-PGSC0003DMG400004195-(rev)M6LRR-RLP_1860

ACACAAACAATAAGTAACGGTGACGGAGCCAGAAATTATGAAGGAAATCCAAAAATGCCTCAATTTGAAATGTTATGTATGTCTACAACAACCAAAAAAAAAAAACTTGTGTTTACTTAT

>M6-PGSC0003DMG400004195-(rev)M6LRR-RLP_1922

ATTTGAAATGTTATGTATGTCTACAACAACCAAAAAAAAAAAACTTGTGTTTACTTATTATCTCTATGTTATATGAGACTCCAACTAGCCAGAATCATTGGCTAGTATGAAAAAAATCTA

>M6-PGSC0003DMG400004195-(rev)M6LRR-RLP_1954

AAAAAAAAAAACTTGTGTTTACTTATTATCTCTATGTTATATGAGACTCCAACTAGCCAGAATCATTGGCTAGTATGAAAAAAATCTAATACATAAAGTACTTTCTTTGTTTTATTTTTG

>M6-PGSC0003DMG400004195-(rev)M6LRR-RLP_248

TTCTTAAAAATGTGACTTTTAGAGGCTTCATAGGTCAATTTGTTGATAAAACAAAAATATTATAGAATGGGAACAGTGATATCCTCCATAAATGAAACTCAACTAAACTATTTATAAGAC

>M6-PGSC0003DMG400004195-(rev)M6LRR-RLP_310

TAGAATGGGAACAGTGATATCCTCCATAAATGAAACTCAACTAAACTATTTATAAGACTATGTTTGTGACTATAATAGTGGAGTAATAATAGTAGAAAGGCTTTCCATATTATTTGTCCT

>M6-PGSC0003DMG400004195-(rev)M6LRR-RLP_372

TTTGTGACTATAATAGTGGAGTAATAATAGTAGAAAGGCTTTCCATATTATTTGTCCTGACATTTTTACTAATTGTTTTGTTGCTTAAATTGTTATGAATTGTTTTTCTTGAGCCGAATG

>M6-PGSC0003DMG400004195-(rev)M6LRR-RLP_434

TTTTTACTAATTGTTTTGTTGCTTAAATTGTTATGAATTGTTTTTCTTGAGCCGAATGTCTACCAGAAACAATCATCTTATCTTTTTTACACAGATAGGGATAATGTATGCATATACATT

>M6-PGSC0003DMG400004195-(rev)M6LRR-RLP_496

CCAGAAACAATCATCTTATCTTTTTTACACAGATAGGGATAATGTATGCATATACATTATCCTCCTCAGATCCCACTTTATGAGATTAAACTTCAGTAGTGATATTCTAAGCAATTACAA

>M6-PGSC0003DMG400004195-(rev)M6LRR-RLP_558

TCCTCAGATCCCACTTTATGAGATTAAACTTCAGTAGTGATATTCTAAGCAATTACAAAAATATTTCCATAAAAACGAAAAATATACTAAGGAGAGACATAATATTGTGTTTTATAAAAT

>M6-PGSC0003DMG400004195-(rev)M6LRR-RLP_62

TAACACAGGGTGACATCGAGCTATTTAAGGAACATAAACAATTGGTAATTGCTTCACGCTATCTATTGCTCCTTTATTCATGAATTCATGTATATACAATGTAATTTTCTGTTAGAGGGG

>M6-PGSC0003DMG400004195-(rev)M6LRR-RLP_620

ATTTCCATAAAAACGAAAAATATACTAAGGAGAGACATAATATTGTGTTTTATAAAATGAAGTTGTAGTAATTTGTCTATTTGTTTACTTTCTTTAAAATTAACATGGCTTGTGAAAAAA

>M6-PGSC0003DMG400004195-(rev)M6LRR-RLP_682

TTGTAGTAATTTGTCTATTTGTTTACTTTCTTTAAAATTAACATGGCTTGTGAAAAAAACTTGACAACATGAGATGCTTAATGTATCGAGCTAGCGAATCTATTATTGACATAATCATCC

>M6-PGSC0003DMG400004195-(rev)M6LRR-RLP_744

GACAACATGAGATGCTTAATGTATCGAGCTAGCGAATCTATTATTGACATAATCATCCCTTTTTTGGCAGGCCAAAATGACATCTCATCAATTTATTTGTTTCCTAGCAATGATATCTCT

>M6-PGSC0003DMG400004195-(rev)M6LRR-RLP_806

TTTGGCAGGCCAAAATGACATCTCATCAATTTATTTGTTTCCTAGCAATGATATCTCTTATGATTATCAATATTAGTGCAAACAAGGCATGCCACCCTGATGATCTCAAGGGTCTCAATG

>M6-PGSC0003DMG400004195-(rev)M6LRR-RLP_868

ATTATCAATATTAGTGCAAACAAGGCATGCCACCCTGATGATCTCAAGGGTCTCAATGACTTCAAAGATGGCATACATTCTGATACTTCAGGCAGACTAAGCAAATGGATTGGGCAAGAT

>M6-PGSC0003DMG400004195-(rev)M6LRR-RLP_930

CAAAGATGGCATACATTCTGATACTTCAGGCAGACTAAGCAAATGGATTGGGCAAGATTGTTGCAATTGGCCAGGCATTTCTTGTAACTCTACAACTGACAGAGTTGTAGAAATCTATCT

>M6-PGSC0003DMG400004195-(rev)M6LRR-RLP_992

GCAATTGGCCAGGCATTTCTTGTAACTCTACAACTGACAGAGTTGTAGAAATCTATCTACCTGGTCACTATGTATCTGGTGACGATGAATCTCCAAACTTCGTGTCAAGCACAATGAGTG

>M6-PGSC0003DMG400004284-(rev)M6LRR-RLP_0

AACTTTTTTTCTTTCATTACCTACCATTTTTCTTGCAAAATGAGTCTCTTGTTTCTACTTGCATCTTTCTTTCTTTGTGAACTTGTTTTGTCCTCATTTGCAATATGCCCCAGAGATCAA

>M6-PGSC0003DMG400004284-(rev)M6LRR-RLP_1037

GCTATCAAATAACTCCCTGAATGGAGAAATACCGTCCTGGATATTCTCCCTTCCATTACTGAGTGAACTAGACTTGAGTAATAACCACTTTTCTGGTCAACTTAAGGAATTCAGTAACAC

>M6-PGSC0003DMG400004284-(rev)M6LRR-RLP_1098

AGTGAACTAGACTTGAGTAATAACCACTTTTCTGGTCAACTTAAGGAATTCAGTAACACTTCAGTACTAGTCGGTGTTGATATAAGCGAGAATGAGTTGCAAGGATGTCTTCCCAAATCA

>M6-PGSC0003DMG400004284-(rev)M6LRR-RLP_1159

CAGTACTAGTCGGTGTTGATATAAGCGAGAATGAGTTGCAAGGATGTCTTCCCAAATCAATTCAGAACCTTGTGAACCTAGTCTGGATTGATTTAGCAAATAACCAGTTGCAATGTCCTC

>M6-PGSC0003DMG400004284-(rev)M6LRR-RLP_122

TCTTGCCCTTGTAGAATTCAACCAGTCTTTTGTTGTTGATGCATCTGCATCTTTTATTTGTGACAAGCAGTCTTATCCTAAAACGAGTTCATGGAACATGAACAAAGATTGTTGTTCGTG

>M6-PGSC0003DMG400004284-(rev)M6LRR-RLP_1220

TCAGAACCTTGTGAACCTAGTCTGGATTGATTTAGCAAATAACCAGTTGCAATGTCCTCTTCCCAAGTCACTCCAGAACCTCAGAAATCTAAAATTACTTGATCTTTTGTCTAACAATTT

>M6-PGSC0003DMG400004284-(rev)M6LRR-RLP_1281

CCCAAGTCACTCCAGAACCTCAGAAATCTAAAATTACTTGATCTTTTGTCTAACAATTTTAGTGGGAGTGTAGATGTCAGTGTCTTTTCGAACCTCAAACAACTTTGGTATCTTGATCTG

>M6-PGSC0003DMG400004284-(rev)M6LRR-RLP_1342

GTGGGAGTGTAGATGTCAGTGTCTTTTCGAACCTCAAACAACTTTGGTATCTTGATCTGTCATACAATAGTATTTCTCTGATCAATGAGAACAAAGTCAAGTCTACATTGCCCCAATCTC

>M6-PGSC0003DMG400004284-(rev)M6LRR-RLP_1403

ATACAATAGTATTTCTCTGATCAATGAGAACAAAGTCAAGTCTACATTGCCCCAATCTCTTGAATACTTATATTTGGCTGCTTGTCAAGTAAAAGAACTGGATTTTTTGAGATCAGCAAA

>M6-PGSC0003DMG400004284-(rev)M6LRR-RLP_1464

GAATACTTATATTTGGCTGCTTGTCAAGTAAAAGAACTGGATTTTTTGAGATCAGCAAATAACTTTTATACCTTGGATCTTTCATATAACAACATTCAAGGAACAATTCCTGATGTGGTA

>M6-PGSC0003DMG400004284-(rev)M6LRR-RLP_1525

ACTTTTATACCTTGGATCTTTCATATAACAACATTCAAGGAACAATTCCTGATGTGGTATTGTCTAATTGGATGCATTCCATAAAAAATCTTAATCTAGCCCACAACATGTTGACTAGTA

>M6-PGSC0003DMG400004284-(rev)M6LRR-RLP_1586

GTCTAATTGGATGCATTCCATAAAAAATCTTAATCTAGCCCACAACATGTTGACTAGTATTGACCACATTTCATCATTTTCCCAACTGATCTCTATTAACTTATTGTCCAATTCTCTTCA

>M6-PGSC0003DMG400004284-(rev)M6LRR-RLP_1647

GACCACATTTCATCATTTTCCCAACTGATCTCTATTAACTTATTGTCCAATTCTCTTCAAGGGACACTACCAATTCCACCACCCTCTATAGAATTCTTCTTCATGTCAAATAATAATGTC

>M6-PGSC0003DMG400004284-(rev)M6LRR-RLP_1708

GGACACTACCAATTCCACCACCCTCTATAGAATTCTTCTTCATGTCAAATAATAATGTCAGTGGTAAGATCCCTTCTTCTATTTGCAATTTGACAACATTAAAAATCCTAGATTTGTCAA

>M6-PGSC0003DMG400004284-(rev)M6LRR-RLP_1769

TGGTAAGATCCCTTCTTCTATTTGCAATTTGACAACATTAAAAATCCTAGATTTGTCAAATAACAATTTGAAGGGAGAAATTCCACAATGTTTGGGTAACATGAGTGATCAACTCGAGGT

>M6-PGSC0003DMG400004284-(rev)M6LRR-RLP_183

GACAAGCAGTCTTATCCTAAAACGAGTTCATGGAACATGAACAAAGATTGTTGTTCGTGGGATGGAATAATATGTGATGAGATGACCGGTCATGTGATTGAGCTTGACCTCAGTTGCAGC

>M6-PGSC0003DMG400004284-(rev)M6LRR-RLP_1830

AACAATTTGAAGGGAGAAATTCCACAATGTTTGGGTAACATGAGTGATCAACTCGAGGTTTTGGATATGCGACGCAATAGTCTTTCTGGGAGTCTGCAAACAACTTTTAGCTTGGGAAGT

>M6-PGSC0003DMG400004284-(rev)M6LRR-RLP_1891

TGGATATGCGACGCAATAGTCTTTCTGGGAGTCTGCAAACAACTTTTAGCTTGGGAAGTAAACTCAAAAGTTTCAACTTGCATGGCAATAATCTAGAGGGAAAACTCCCAAGATCCTTGG

>M6-PGSC0003DMG400004284-(rev)M6LRR-RLP_2074

TGTGGTTGGGGACTCTTCCAAATCTACAAGTTTTAAGTTTGAGACTCAATAATTTGCATGGACCAATTAGAACTTCAACTAGTTCGAAACTGTTTCCTCAGCTTCGGATGTTAGATCTCT

>M6-PGSC0003DMG400004284-(rev)M6LRR-RLP_2135

ACCAATTAGAACTTCAACTAGTTCGAAACTGTTTCCTCAGCTTCGGATGTTAGATCTCTCTCGCAATGCCTTTACAGCAGAACTGCCAACAATTCTTTTCAGAAATCTGAAAGCAATGAG

>M6-PGSC0003DMG400004284-(rev)M6LRR-RLP_2196

CGCAATGCCTTTACAGCAGAACTGCCAACAATTCTTTTCAGAAATCTGAAAGCAATGAGGAGAATTGCCATTGATAAAACAATGAAGGCACTAGGTGATGAAGAAAAATCATATTACCAA

>M6-PGSC0003DMG400004284-(rev)M6LRR-RLP_2257

GAATTGCCATTGATAAAACAATGAAGGCACTAGGTGATGAAGAAAAATCATATTACCAAGACTCGGTAACTGTTGTAACAAAGGGAACGGAGTTTGAAATTGTGAGAATCTTGTCTTTGT

>M6-PGSC0003DMG400004284-(rev)M6LRR-RLP_2440

TCATTGCACTACGCGTGTTGAATTTATCTCACAATGAGTTGCAGGGTCATATACCACCATCATTTGGAAAGTTATCTGTAGTCGAATCATTGGACCTTTCATCCAACAGGCTTTCAGGAG

>M6-PGSC0003DMG400004284-(rev)M6LRR-RLP_2684

AAATGATGGATTGCGTGGATTCCCAGTTTCAGGAGGTTGTGGAAGTGATCGGATACCGGATACAAACAACACAACATTTGTGCCAGATGAAGAAAATGATTCAACATTTCTAAGTGAGCT

>M6-PGSC0003DMG400004284-(rev)M6LRR-RLP_2745

ACAAACAACACAACATTTGTGCCAGATGAAGAAAATGATTCAACATTTCTAAGTGAGCTCAGTTGGAAAGTGGTTCTTATGGGATATGGTACTGGATTGATTATTGGATTCTCCATAGCA

>M6-PGSC0003DMG400004284-(rev)M6LRR-RLP_2806

GTTGGAAAGTGGTTCTTATGGGATATGGTACTGGATTGATTATTGGATTCTCCATAGCATATTTGATGCTTTCTTCTCAAAATCCAAATTGGCTTTCTGGGATAGCTGAAGATTTAGAAA

>M6-PGSC0003DMG400004284-(rev)M6LRR-RLP_2867

TTTGATGCTTTCTTCTCAAAATCCAAATTGGCTTTCTGGGATAGCTGAAGATTTAGAAAACAGAATCATTATGAGAAGGCGAAAGAAGCAGCGAGGTCAAATGCATTACAGAAGAAGAAG

>M6-PGSC0003DMG400004284-(rev)M6LRR-RLP_2928

AGAATCATTATGAGAAGGCGAAAGAAGCAGCGAGGTCAAATGCATTACAGAAGAAGAAGAAGAAATAATGGCATCTAGATGTAAGTTCAAAATTTATGAGAGATGTAGTCATGTTAAGCT

>M6-PGSC0003DMG400004284-(rev)M6LRR-RLP_2989

GAAATAATGGCATCTAGATGTAAGTTCAAAATTTATGAGAGATGTAGTCATGTTAAGCTCTATACTGATTCTTTTTGTTTACTGTAACTGGTATTAAAAGAATGAGATTAGTGTTCTGTA

>M6-PGSC0003DMG400004284-(rev)M6LRR-RLP_305

ACTTGTTGGGAAGATTGATTCCAATAGCAGCCTATTCCAACTCTCTCATCTCCAAAGGCTCAATCTTTCTAATAATGACTTTGACGGTTCTCATATCTCGCCTAAATTTGGTAGGTTAGC

>M6-PGSC0003DMG400004284-(rev)M6LRR-RLP_3050

ATACTGATTCTTTTTGTTTACTGTAACTGGTATTAAAAGAATGAGATTAGTGTTCTGTATCTGACTCTTTCTTTTGTAACTTTTATGTAGGCAAAGGGTGTATCCAGTTGAAATATCAAG

>M6-PGSC0003DMG400004284-(rev)M6LRR-RLP_3111

TGACTCTTTCTTTTGTAACTTTTATGTAGGCAAAGGGTGTATCCAGTTGAAATATCAAGAAGATGATGCCTGCAACAACAAATATATCTTCTTGGAGGAAATTACTTGAGCAAGAGATGC

>M6-PGSC0003DMG400004284-(rev)M6LRR-RLP_3172

GATGATGCCTGCAACAACAAATATATCTTCTTGGAGGAAATTACTTGAGCAAGAGATGCTTTTGATATTTCCTGGAGGTTGCTACTCCTTTTTCTTCATAAAAATAAAATAAATTCTAGG

>M6-PGSC0003DMG400004284-(rev)M6LRR-RLP_3233

TTGATATTTCCTGGAGGTTGCTACTCCTTTTTCTTCATAAAAATAAAATAAATTCTAGGATTGAAGTATGTTGCTTTACTATAAACAGTTGTCACAATGAAGTGTTTTTGTTTTTCCATC

>M6-PGSC0003DMG400004284-(rev)M6LRR-RLP_3294

TGAAGTATGTTGCTTTACTATAAACAGTTGTCACAATGAAGTGTTTTTGTTTTTCCATCCAAGTGTTAGTAAAGAGAAAAAAGATATTTGTCTGCAAAATAATAGCTTTCTTTTGACATG

>M6-PGSC0003DMG400004284-(rev)M6LRR-RLP_3335

TGTTTTTGTTTTTCCATCCAAGTGTTAGTAAAGAGAAAAAAGATATTTGTCTGCAAAATAATAGCTTTCTTTTGACATGTGTATTGACTTGGAATATTACATTTACCTGGAATACAGTTT

>M6-PGSC0003DMG400004284-(rev)M6LRR-RLP_366

AATCTTTCTAATAATGACTTTGACGGTTCTCATATCTCGCCTAAATTTGGTAGGTTAGCAAGCTTGACACATCTTGATCTTTCCCAGGCAGACTTCTCAGGTCAAATTCCTTCTGAAATC

>M6-PGSC0003DMG400004284-(rev)M6LRR-RLP_427

GCTTGACACATCTTGATCTTTCCCAGGCAGACTTCTCAGGTCAAATTCCTTCTGAAATCTCTCATCTCTCCAAGTTACAGTCTGTTTTTTTATCGTCTAACAGTGAGCTTAGACTCGTAG

>M6-PGSC0003DMG400004284-(rev)M6LRR-RLP_488

TCATCTCTCCAAGTTACAGTCTGTTTTTTTATCGTCTAACAGTGAGCTTAGACTCGTAGCTTACGATTTCAAGATGTCCCTTCAAAATTTGACTCAATTAAGAGAGCTTCATCTTAGTGG

>M6-PGSC0003DMG400004284-(rev)M6LRR-RLP_61

CATCTTTCTTTCTTTGTGAACTTGTTTTGTCCTCATTTGCAATATGCCCCAGAGATCAAAGTCTTGCCCTTGTAGAATTCAACCAGTCTTTTGTTGTTGATGCATCTGCATCTTTTATTT

>M6-PGSC0003DMG400004284-(rev)M6LRR-RLP_610

TGAACATCTCTTCCACCATTCCTCTGAATTTCTCTTCTCATTTAACGACGCTGGGGCTGACGAGTACAGGAGTGTATGGGATAATACCTGAGAGTATCTTTAACCTGCCCAACCTGGAAA

>M6-PGSC0003DMG400004284-(rev)M6LRR-RLP_671

GAGTACAGGAGTGTATGGGATAATACCTGAGAGTATCTTTAACCTGCCCAACCTGGAAACACTTGACTTAAGTTCCAACGATCAGCTCAATGGTTATTTTCCAAATACCAAATGGAATAG

>M6-PGSC0003DMG400004284-(rev)M6LRR-RLP_732

CTTGACTTAAGTTCCAACGATCAGCTCAATGGTTATTTTCCAAATACCAAATGGAATAGCAGTGCATCTCTCATGGAGCTGAATCTCGCTGGGGTAAATTTTTCTGGTAATTTCCTGCCT

>M6-PGSC0003DMG400004284-(rev)M6LRR-RLP_793

GTGCATCTCTCATGGAGCTGAATCTCGCTGGGGTAAATTTTTCTGGTAATTTCCTGCCTGAATGTCTTGGCTATCTAACTTCACTGCAGAGATTGGTACTTAGTTCCTGCAATTTATCGG

>M6-PGSC0003DMG400004284-(rev)M6LRR-RLP_854

ATGTCTTGGCTATCTAACTTCACTGCAGAGATTGGTACTTAGTTCCTGCAATTTATCGGGGCAGGTTCCGAAATCTCTTTGGAACCTCACCCATCTGGAGTATATGGACCTTGAAGATAA

>M6-PGSC0003DMG400004284-(rev)M6LRR-RLP_915

CAGGTTCCGAAATCTCTTTGGAACCTCACCCATCTGGAGTATATGGACCTTGAAGATAACCGTCTTGAAGGACCGATATTTCCTCAGTTTACAAGTGGACTGCAGGATCTGAACACACTT

>M6-PGSC0003DMG400004284-(rev)M6LRR-RLP_976

GTCTTGAAGGACCGATATTTCCTCAGTTTACAAGTGGACTGCAGGATCTGAACACACTTAAGCTATCAAATAACTCCCTGAATGGAGAAATACCGTCCTGGATATTCTCCCTTCCATTAC

>M6-PGSC0003DMG400004384M6LRR-RLP_0

CAACCCTCTTTCTCAGGGATGGTCGAAGAGAGGTTCCTCTTCTGAGATCATCCTGAACACAGCAGTCAACAAAAAATATTCAAAAAAAAATAAAATATTTTTTTTGGGATTTTAATTCTA

>M6-PGSC0003DMG400004384M6LRR-RLP_1098

AAGTTCTACATTGGTAGCTAAAAAGATTGGAAATCTACAATTCTACATAAGGTGTAGGATCTCTTAACATGGTGTGAGGCCTTTTGTGAAAACCGTGCGGGCGTGCCCCTAAGCGGACAA

>M6-PGSC0003DMG400004384M6LRR-RLP_1159

TCTTAACATGGTGTGAGGCCTTTTGTGAAAACCGTGCGGGCGTGCCCCTAAGCGGACAACATTGCACCATGTTAAGAGTATTTTTGGGTTGACTTAGCCCAACAAGATGAAATTACTTGC

>M6-PGSC0003DMG400004384M6LRR-RLP_122

TTTCATAAAAAATCCCACAATTTCCATTGGGTTAAATTTCTTGAATTTTTTTTTTCTTTGATCTCTGTTTGCTTTTTCTGGTAAATCTTTCATCTTTACCCCCACTTTTTGAATTTTTGT

>M6-PGSC0003DMG400004384M6LRR-RLP_1525

CTTGATCTTTCATGGTTTTTTACTCACTTAGGGCTGTGTCTTCTGTCTCCCTGGGCCGTGGGCTATTCTCTAGTTTCTTTAGTCTATTCATGTTATTGGCTTATTCTTTGTTTCGCTGAT

>M6-PGSC0003DMG400004384M6LRR-RLP_1586

GCTATTCTCTAGTTTCTTTAGTCTATTCATGTTATTGGCTTATTCTTTGTTTCGCTGATTACTGTTGATTTGAGAAACAGGATTGTCGTTGTTTGAATTCAAATTGAATGTGTATTCTGA

>M6-PGSC0003DMG400004384M6LRR-RLP_1647

CTGTTGATTTGAGAAACAGGATTGTCGTTGTTTGAATTCAAATTGAATGTGTATTCTGATCCAAATGGAGTTCTTGAAAACTGGAATGCTGATCACTGTGACCCGTGCATGTGGTCTGGT

>M6-PGSC0003DMG400004384M6LRR-RLP_1708

CAAATGGAGTTCTTGAAAACTGGAATGCTGATCACTGTGACCCGTGCATGTGGTCTGGTGTCGAGTGCATGGATGGTAAAGTGTACATGTTGTAAGTCTCTTGTTATAGATACTCAAGTT

>M6-PGSC0003DMG400004384M6LRR-RLP_1769

CGAGTGCATGGATGGTAAAGTGTACATGTTGTAAGTCTCTTGTTATAGATACTCAAGTTTTCTAGTTAAGATTGTGTTAATAAACTTTCTCTAATGGTTAGTGTTTAGTGATTTGATTCT

>M6-PGSC0003DMG400004384M6LRR-RLP_183

TCTCTGTTTGCTTTTTCTGGTAAATCTTTCATCTTTACCCCCACTTTTTGAATTTTTGTTTGATTTTGCTTGAGAATATGGTTATCTTATGATGGGTTTTTTGTTTTTTCCCAAAAAAGA

>M6-PGSC0003DMG400004384M6LRR-RLP_1830

CTAGTTAAGATTGTGTTAATAAACTTTCTCTAATGGTTAGTGTTTAGTGATTTGATTCTGGAACCAGTTTGTTGAACTAATAATGATAAACCAAATGCAACAGGGTGAAATTGTTTGTTT

>M6-PGSC0003DMG400004384M6LRR-RLP_1891

AACCAGTTTGTTGAACTAATAATGATAAACCAAATGCAACAGGGTGAAATTGTTTGTTTCGACTCTTGTGATTCTTCTTATGTCGTTAAATTAGTTGACCTATCTTCTTTTTTTCATTCT

>M6-PGSC0003DMG400004384M6LRR-RLP_1952

ACTCTTGTGATTCTTCTTATGTCGTTAAATTAGTTGACCTATCTTCTTTTTTTCATTCTGTTCAGAAATCTCCATGGATTTTCTTTGGAAGGAACACTAGCATCAGAGCTTGGAAATCTC

>M6-PGSC0003DMG400004384M6LRR-RLP_2013

TCAGAAATCTCCATGGATTTTCTTTGGAAGGAACACTAGCATCAGAGCTTGGAAATCTCACTTACTTGAGATCACTGTAAGCTTTAGCTATACTAATATGCCTTTTCTTTAGGCTCCTGC

>M6-PGSC0003DMG400004384M6LRR-RLP_2074

TTACTTGAGATCACTGTAAGCTTTAGCTATACTAATATGCCTTTTCTTTAGGCTCCTGCTCATTAAGTCTTAAATGGTATTTTTTACCTGAACCGTCATCCGTAGAAGGTTGTCATAATC

>M6-PGSC0003DMG400004384M6LRR-RLP_2135

ATTAAGTCTTAAATGGTATTTTTTACCTGAACCGTCATCCGTAGAAGGTTGTCATAATCTTTGAATAGTCTTCTATCTTCTTATGTAAATTCGAAAGAACGTAAGGGATTGTTTGGTGTA

>M6-PGSC0003DMG400004384M6LRR-RLP_244

GATTTTGCTTGAGAATATGGTTATCTTATGATGGGTTTTTTGTTTTTTCCCAAAAAAGATTCAATTTTTATTAATTCTTGTTGTTTTTATGTCTGCATTTTCAGCTAAAAATTGAATCTT

>M6-PGSC0003DMG400004384M6LRR-RLP_2623

CCTAAGAGTTATGTTGGTCAAGTAAATTGGGAGGTAGAGACAAAGGTGGATAAGCTTTGGATATCTACTAATCAGCTTTCATCATCTTGTATTCTGCATTTACATTGTTCTCTGCATAAG

>M6-PGSC0003DMG400004384M6LRR-RLP_2684

TATCTACTAATCAGCTTTCATCATCTTGTATTCTGCATTTACATTGTTCTCTGCATAAGTTGTGTTTGAATCTTAACAGTATTTCTTCTTGCTGTCTGGTTGCTTTTACAGAATGCTCTC

>M6-PGSC0003DMG400004384M6LRR-RLP_2745

GTGTTTGAATCTTAACAGTATTTCTTCTTGCTGTCTGGTTGCTTTTACAGAATGCTCTCAGGGAATCATTTTTTCGGTGCTATTCCTAAAGAATTCGGAAGACTCGATATGCTTGAAGTG

>M6-PGSC0003DMG400004384M6LRR-RLP_2867

TGACTTGAGGGATAATAATTTGAGTGGAACAATTCCAGCAGAAATAGGAGACCTGCAATCACTAAGAATCTTGTGAGTTAATGATTTTTTGTGTGCACGATAAACGGTTTGTTGTGTGGA

>M6-PGSC0003DMG400004384M6LRR-RLP_2928

CTAAGAATCTTGTGAGTTAATGATTTTTTGTGTGCACGATAAACGGTTTGTTGTGTGGACGATAAACGGTTTGTTGTGTGCAGATGTATATATACATATGCATACACATTCATTATGCGA

>M6-PGSC0003DMG400004384M6LRR-RLP_2989

ATAAACGGTTTGTTGTGTGCAGATGTATATATACATATGCATACACATTCATTATGCGAAGTGATTTCAAAATGCTCATCAATCGTGTTTATTTCGTTTTTCCAGGTTGATTCATAACAA

>M6-PGSC0003DMG400004384M6LRR-RLP_305

CAATTTTTATTAATTCTTGTTGTTTTTATGTCTGCATTTTCAGCTAAAAATTGAATCTTGGAATTTTTGAAGACTTGGGGTTTCTGCAATGATGGAGAAAACAGAGGTAATATAGCCTAT

>M6-PGSC0003DMG400004384M6LRR-RLP_3050

TGATTTCAAAATGCTCATCAATCGTGTTTATTTCGTTTTTCCAGGTTGATTCATAACAATAACTTTGAGGAGAGTGTTCCTTTGGAAATTGGGAAGCTTCATTTAGTCACAGAATTACAA

>M6-PGSC0003DMG400004384M6LRR-RLP_3172

CGACGAAAATCTCACTTCTGCTTTTGCTTCTGGAACTCTCTGTATAAGAAAATTTCGTCATTGGTAGGATTCTTTTATTTGTTTTAGTACCCTATCTTCATTTTCAGCTGGTTTTTTGCA

>M6-PGSC0003DMG400004384M6LRR-RLP_3233

TGGTAGGATTCTTTTATTTGTTTTAGTACCCTATCTTCATTTTCAGCTGGTTTTTTGCATATCAGATGCTGGATAGTCTATCTTTGTGGCAACACACTATCTAATTGATCTAGCTCATTT

>M6-PGSC0003DMG400004384M6LRR-RLP_3294

TCAGATGCTGGATAGTCTATCTTTGTGGCAACACACTATCTAATTGATCTAGCTCATTTCTCAATTTGATATTTTGCATGTCTTACAGCATCTGGCAAGGCAGCATGAAGCCATTCAAGA

>M6-PGSC0003DMG400004384M6LRR-RLP_3416

ATAACCTCCATTATTGAACCAATAAAATGTACACTTTTGCATTACCTCAGTTTCTTCCAACTGTAAGTAATTAAATAGTTTTTCACATCACGAGTCATCTAGTTAAGTTTTATTATTGAG

>M6-PGSC0003DMG400004384M6LRR-RLP_3477

TGTAAGTAATTAAATAGTTTTTCACATCACGAGTCATCTAGTTAAGTTTTATTATTGAGAGACATTTTTTCTATTTCTTTAGGCCATCTCCTGGAACGGGCTTTATGGATGACCGGAAAG

>M6-PGSC0003DMG400004384M6LRR-RLP_3599

TGCTCTGACAACCTTCCAAGTGAGTAAACACGTCATATTTTTTAAAATAACTCTGAAGAACCGTTTACAAATAATAATTCTTCATTGTCTAGCAGGGTTATCCAGGCCTCACATCATCCA

>M6-PGSC0003DMG400004384M6LRR-RLP_366

AATTTTTGAAGACTTGGGGTTTCTGCAATGATGGAGAAAACAGAGGTAATATAGCCTATATTATAGAAATGTTAAGAAAATGAGAGACCCCATGAAGGGGTGGAGCCAGGGGTAGGGCCG

>M6-PGSC0003DMG400004384M6LRR-RLP_3660

CGTTTACAAATAATAATTCTTCATTGTCTAGCAGGGTTATCCAGGCCTCACATCATCCACACTATACAGCACCAAGAGGAGATTGCACGCCGTAAGTTGGCTGAACAATCCAGTAACCTT

>M6-PGSC0003DMG400004384M6LRR-RLP_3721

CTATACAGCACCAAGAGGAGATTGCACGCCGTAAGTTGGCTGAACAATCCAGTAACCTTGCTGCAGCTCCTGCCAATTTCAAAGGACCGTTGGGTCCTGTCATTCAAATGCCAAGAAGCA

>M6-PGSC0003DMG400004384M6LRR-RLP_3782

TGCAGCTCCTGCCAATTTCAAAGGACCGTTGGGTCCTGTCATTCAAATGCCAAGAAGCAGTGGGTCGTTTCGTGCGGTGCCAAATTCCAATGGAGCACATCCTACAAACTTACCATCACC

>M6-PGSC0003DMG400004384M6LRR-RLP_3843

GGGTCGTTTCGTGCGGTGCCAAATTCCAATGGAGCACATCCTACAAACTTACCATCACCACCTCCACAGCATGAATCTCAAGACAAATTACATCCTGTGGGACAGTCTGCAAGTGCTGTA

>M6-PGSC0003DMG400004384M6LRR-RLP_3904

CTCCACAGCATGAATCTCAAGACAAATTACATCCTGTGGGACAGTCTGCAAGTGCTGTAAAACAGTCGCCTGCTGTGAAAAAGCCAACTAATAGCCAAGGCCCCCCTGCTGGAAAATCGG

>M6-PGSC0003DMG400004384M6LRR-RLP_3965

ACAGTCGCCTGCTGTGAAAAAGCCAACTAATAGCCAAGGCCCCCCTGCTGGAAAATCGGAAAGTACTTGGAAGTACGTAGTTGGGATTTGCATTGGAGTAATTTTGCTCATTCTTGCTGC

>M6-PGSC0003DMG400004384M6LRR-RLP_4087

CTGTGTTCTTAATCTGCCGAATCAGAGCAGCTAGAACAATTGGCCCTTGGAAGACTGGATTGAGTGGACAGCTGCAAAAAGCATTTGTTACAGGTACTTATATCATATTGTGCCAAAAAG

>M6-PGSC0003DMG400004384M6LRR-RLP_4148

GAGTGGACAGCTGCAAAAAGCATTTGTTACAGGTACTTATATCATATTGTGCCAAAAAGTGATACTGCTAATATTCTGCTCCCCACCCTGATAGTAGTTGCATTGCTAACAATATTCTTA

>M6-PGSC0003DMG400004384M6LRR-RLP_4209

ATACTGCTAATATTCTGCTCCCCACCCTGATAGTAGTTGCATTGCTAACAATATTCTTAGCATAGTGTTACCTTATCAAAAAAAATATTCTTAGTGTAGTGTTGTTATAACAATTTAACT

>M6-PGSC0003DMG400004384M6LRR-RLP_4270

ATAGTGTTACCTTATCAAAAAAAATATTCTTAGTGTAGTGTTGTTATAACAATTTAACTGGAGTAAAGGGAGGTACTGATATTCTTCGTCTTATGAGGAATTAGAGGACTAATATTCTCA

>M6-PGSC0003DMG400004384M6LRR-RLP_4331

AGTAAAGGGAGGTACTGATATTCTTCGTCTTATGAGGAATTAGAGGACTAATATTCTCAGTGTAGCATTGTTATATACCCTTTTTCACTATATGTTTGCTAGGACGAGAAGGTACAAATG

>M6-PGSC0003DMG400004384M6LRR-RLP_4392

GTAGCATTGTTATATACCCTTTTTCACTATATGTTTGCTAGGACGAGAAGGTACAAATGGAACTATTTCCTTCTTTCTCATGAGAAGTGAATTAGCTGAGACATCAATACTAACGCTTGA

>M6-PGSC0003DMG400004384M6LRR-RLP_4453

ACTATTTCCTTCTTTCTCATGAGAAGTGAATTAGCTGAGACATCAATACTAACGCTTGAAGGCTGGAATCCTTCCCTTATACTTCTAAAATGTCGCTGATCTTTCCTCATACATTTGCTT

>M6-PGSC0003DMG400004384M6LRR-RLP_4575

GTCTAGCTTATATCCCCCCCACAAAGTTGCTTTTCAACGACAACCATAGGAACGTGCAATCCTTTATGTTAATCTATGATATTTTTCGTCTATTTTAATCCTTTTGCCCGTGAAAACATT

>M6-PGSC0003DMG400004384M6LRR-RLP_4636

CTTTATGTTAATCTATGATATTTTTCGTCTATTTTAATCCTTTTGCCCGTGAAAACATTGTTATAGGGGTTCCAAAACTGAACAGATCCGAGCTTGAAACAGCATGCGAGGATTTTAGCA

>M6-PGSC0003DMG400004384M6LRR-RLP_4758

ATTATTACCAGCGGCGATTCATATATAGTATACAAGGGAACACTGTCTAGCGGAGTGGAGATTGCTGTTATCTCAACTACTATAAGTTCTCTGGAAGATTGGTCCATGCACTCGGAAACA

>M6-PGSC0003DMG400004384M6LRR-RLP_4811

AGTGGAGATTGCTGTTATCTCAACTACTATAAGTTCTCTGGAAGATTGGTCCATGCACTCGGAAACAGCCTTCCGGAAGAAGGTTAATCATTTCTTTGCTTTTACAAATTAACCACCTCC

>M6-PGSC0003DMG400004384M6LRR-RLP_61

GCAGTCAACAAAAAATATTCAAAAAAAAATAAAATATTTTTTTTGGGATTTTAATTCTACATTTCATAAAAAATCCCACAATTTCCATTGGGTTAAATTTCTTGAATTTTTTTTTTCTTT

>M6-PGSC0003DMG400004384M6LRR-RLP_610

CTGGCTCCGCAGTTGACCTCATTGTGGCTAATATAGATAGATTTTGTTTGTAATCTTTTTTATTTTTTGTGTCTAGTGTAGAGTTGTGCAGATAAGCAACTGTGTGTTCGTGTGTCCGAG

>M6-PGSC0003DMG400004384M6LRR-RLP_671

ATTTTTTGTGTCTAGTGTAGAGTTGTGCAGATAAGCAACTGTGTGTTCGTGTGTCCGAGTGATTTTTGTTGCTCGGAGGAGAGTCATTTCATATCAAAAAGCTGGCGATTTCGTGTGAGA

>M6-PGSC0003DMG400004384M6LRR-RLP_732

ATTTTTGTTGCTCGGAGGAGAGTCATTTCATATCAAAAAGCTGGCGATTTCGTGTGAGAATTTGTTCAGAATCCTGGCTGTTGGCAAGAAATGAGGCGTAGATGGAACATTTATGGAATC

>M6-PGSC0003DMG400004384M6LRR-RLP_793

TTGTTCAGAATCCTGGCTGTTGGCAAGAAATGAGGCGTAGATGGAACATTTATGGAATCAAGTTATTGCTTCTCACCATTTTAATTCTTCTTTTGGAGGTTCATGAATGTTGCACTCTGA

>M6-PGSC0003DMG400004384M6LRR-RLP_854

GTTATTGCTTCTCACCATTTTAATTCTTCTTTTGGAGGTTCATGAATGTTGCACTCTGAATTCTGAAGGTAGTTACTATTCTCTAGTTTCTTTCTTTTCTTACTTCTGGAGGACCATAGT

>M6-PGSC0003DMG400004396-(rev)M6L-LecRK_0

ACATTTCTCTAACTATGGCTGTTTTTGCATATTTCACATACCTTATTTTGATAATCCCCTTTGTCACTTCATTGTCCTTCAATTTTGATAGTTTCAATCCCAATGATATGAATGTAACAT

>M6-PGSC0003DMG400004396-(rev)M6L-LecRK_1008

TGAATTTGAAAGAAACACAGGACCAAAGAGTTTTTGTACGGTGAGTTGGCTAGATGTACAAATAACTTTTCGCCGGAAGAGATGCTTGGGCATGGTGGGTTCGGAGGTGTTTATAAAGGA

>M6-PGSC0003DMG400004396-(rev)M6L-LecRK_1071

AACTTTTCGCCGGAAGAGATGCTTGGGCATGGTGGGTTCGGAGGTGTTTATAAAGGATATCTCAGGGAATATGACTCCTATATTGTTGTTAAGAGGATTTCAAGGGAGTCAAAGCAAGGA

>M6-PGSC0003DMG400004396-(rev)M6L-LecRK_1134

AGGGAATATGACTCCTATATTGTTGTTAAGAGGATTTCAAGGGAGTCAAAGCAAGGAATAAAAGAGTATGCATCAGAAGTCAGGATTATCAGCCGGTTAAAGACATAAACATTTGGTGCA

>M6-PGSC0003DMG400004396-(rev)M6L-LecRK_1197

GAGTATGCATCAGAAGTCAGGATTATCAGCCGGTTAAAGACATAAACATTTGGTGCAACTCATTGGTTGGTGCCATCAAAAAAGAGAACTTCTACTTGTCTATGAGTTTATGCCTAATGG

>M6-PGSC0003DMG400004396-(rev)M6L-LecRK_126

CAGATGCTTCTCCAGCAAACGGTGTAATTCAGCTCACCAAAAACCAGCTTGATAGTAGTGAAAGCTTTAGCGTAGGCCGAGCTACATATTCTAAAGAACTCTATCTTGGGACAAGGCCTC

>M6-PGSC0003DMG400004396-(rev)M6L-LecRK_1260

TGGTTGGTGCCATCAAAAAAGAGAACTTCTACTTGTCTATGAGTTTATGCCTAATGGAAGCTTAGATCATCATCTTTTCAAGGGAAAAAGCCATTTGACATGGCCAATAAGATTCAAGAT

>M6-PGSC0003DMG400004396-(rev)M6L-LecRK_1449

CATAAAGTCGAGCAATATTATGTTGGATTCCAATTTCAATGCTAAACTTGGGGAAAGCTTTGGCAGGCACGATGGGTTACATGGCTCCTGAATGTCTCACCACTGGCAAAGCTAGCAAGG

>M6-PGSC0003DMG400004396-(rev)M6L-LecRK_1575

ACTTCTATAGCTTTGGTGTTGTCGCGTTAGAAATAGTTTGTGGAAGGAAACCTATTGATCCTAAATCTGAGGAGCATCAAGTAAATATTATCGAATGGGCTTGGAGACTTTACGGGATGG

>M6-PGSC0003DMG400004396-(rev)M6L-LecRK_1638

AATCTGAGGAGCATCAAGTAAATATTATCGAATGGGCTTGGAGACTTTACGGGATGGGAAATCTTAGTGAAGCAGTTGATCCTAGACTCTCATCAGAGTTGAATGAACAGGAGGTGGAGC

>M6-PGSC0003DMG400004396-(rev)M6L-LecRK_1701

TTAGTGAAGCAGTTGATCCTAGACTCTCATCAGAGTTGAATGAACAGGAGGTGGAGCACTTGCTAATTGTTGGCTTATGGTGTGCTCGATCATCCAGATAACAATTGCAGACCTTCTATT

>M6-PGSC0003DMG400004396-(rev)M6L-LecRK_189

GCTTTAGCGTAGGCCGAGCTACATATTCTAAAGAACTCTATCTTGGGACAAGGCCTCTGGGAAAGTCACAGATTTCTCTACGCACTTCTCCTTTGAGATCAATTCACAGGGCAGAAATTA

>M6-PGSC0003DMG400004396-(rev)M6L-LecRK_1890

ATATATTGCAGTTATGAATCAATTACTTCACTTGCATCACCATACGACTCCAATGGTCCTCAGATTTCTGAAATAAACTCTTCAGTGACTAGAGATCATTCGACACAGCAGCATCTTCGC

>M6-PGSC0003DMG400004396-(rev)M6L-LecRK_1953

ATTTCTGAAATAAACTCTTCAGTGACTAGAGATCATTCGACACAGCAGCATCTTCGCCTTCCGCATCACTTCTGTACACACGTTGATATTGTCTTGTAAAGGAACTGGTTTTTTATTTCC

>M6-PGSC0003DMG400004396-(rev)M6L-LecRK_2003

TCTTCGCCTTCCGCATCACTTCTGTACACACGTTGATATTGTCTTGTAAAGGAACTGGTTTTTTATTTCCCTGCAGTGTCCAACTTTTGCTCTTCGTATTCTTTATTATTTATACTGTTA

>M6-PGSC0003DMG400004396-(rev)M6L-LecRK_315

TGATGGTCTTGCCTTCTTCCTCGCTCCTGCAGGTTCAGTAATTCCTGAAAAATACTCAGCTGCAGGAGAGGGCCTTGGCCTTGCAATTGATAATGGACTAAATACATCGAGAAATCAGCC

>M6-PGSC0003DMG400004396-(rev)M6L-LecRK_378

AGGAGAGGGCCTTGGCCTTGCAATTGATAATGGACTAAATACATCGAGAAATCAGCCTTTTGTTGCTGTCGAGTTTGACACCTTTTAGAACGCTTATGATCCACCAGGTGATCATGTAGG

>M6-PGSC0003DMG400004396-(rev)M6L-LecRK_441

TGCTGTCGAGTTTGACACCTTTTAGAACGCTTATGATCCACCAGGTGATCATGTAGGCGTTGATATCAACTCCATGTTATCTCTTGTTAATGTGGCCTGGCTTAGTAGAATTCCCTATGG

>M6-PGSC0003DMG400004396-(rev)M6L-LecRK_504

TATCAACTCCATGTTATCTCTTGTTAATGTGGCCTGGCTTAGTAGAATTCCCTATGGTACGAAAACTGATGCCTGGATTACTTATAATTCAACCTCAAGAAATCCTAGTGTTGTCTTCAC

>M6-PGSC0003DMG400004396-(rev)M6L-LecRK_567

AACTGATGCCTGGATTACTTATAATTCAACCTCAAGAAATCCTAGTGTTGTCTTCACCGGTTTCAGACAACAAGGGAATACCACAGTCACTGTCCTGCAGAACCTATCTTACAATCTTGA

>M6-PGSC0003DMG400004396-(rev)M6L-LecRK_63

TCACTTCATTGTCCTTCAATTTTGATAGTTTCAATCCCAATGATATGAATGTAACATATGAAGCAGATGCTTCTCCAGCAAACGGTGTAATTCAGCTCACCAAAAACCAGCTTGATAGTA

>M6-PGSC0003DMG400004396-(rev)M6L-LecRK_693

GGAATATTTGCCAGAATGGGTAACTTTTGGTTTCTCAGGTGCAACAGGAACCTTCTTTGCAATACAAAACATATATTCTTGGAATTTTACTTCTTCTTTAAAATATAGTGACAGCATAAC

>M6-PGSC0003DMG400004396-(rev)M6L-LecRK_756

ACAAAACATATATTCTTGGAATTTTACTTCTTCTTTAAAATATAGTGACAGCATAACAGATCCAGATGTACCCCAACCAAGCCCTGTGCCAGAAGATACTTCAAGCAAAAAGAAGTCAGG

>M6-PGSC0003DMG400004396-(rev)M6L-LecRK_882

GATTGGATTGATTTCTGGTGGTTGTGTTTCGGTAGCAGTATCTGTTTTTATATTGTTTGCATTTTGGAGAAAGAGGAAGGTCAGAGAAGATGAAGATGATGATATTATTGATGGTTCCAT

>M6-PGSC0003DMG400004396-(rev)M6L-LecRK_945

TTGGAGAAAGAGGAAGGTCAGAGAAGATGAAGATGATGATATTATTGATGGTTCCATGACTAATGAATTTGAAAGAAACACAGGACCAAAGAGTTTTTGTACGGTGAGTTGGCTAGATGT

>M6-PGSC0003DMG400004397-(rev)M6L-LecRK_1364

ATTGCTCAAGGCTTGGCCTCAGCGTTGTTTTATCTACACGAAGAATGGGAACAATGTGTGGTGCATAGGGACATAAAGTCGAGCAATATTATGTTGGATTCCAATTTCAATGCTAAACTT

>M6-PGSC0003DMG400004401M6L-LecRK_0

TCTCTCAAATACTTGCTTCTTATCATAATTATCCCTCTAGTCACTTCACTCTCCTTCAATTTTGATACTTTTACACCCAATGATCACAATGTAACCTATGAGAGAGATGCTTCTCCAGAA

>M6-PGSC0003DMG400004401M6L-LecRK_1054

TGCTTGGGCGGGGTGGATTCGGAGGTGTTTATAAAGGATATCTCAGTGAATCCAACCCTTATATTGCTGTTAAGAGGGTTTCAAGGGAATCACAGCAAGGAATAAAAGAGTATGTATCAG

>M6-PGSC0003DMG400004401M6L-LecRK_124

GTGCAATTCTACTCACCAAAAACCTCCTCAATCGCGATATTAATGCTAGTATAGGTCGAGCTATATATTCAAAACCACTGCATCTATGGGACAAGGCCTCTAGAAATCTCACTGATTTCA

>M6-PGSC0003DMG400004401M6L-LecRK_1302

AAGGGAAAAAGCCATTTGACATGGCCAATAAGATTCAAGATTGCTCAAGGTTTGGCCTCGGCGTTGCTATATCTACACGAAGAATGGGAACAATGTGTGGTGCATAGGGATATAAAGTCT

>M6-PGSC0003DMG400004401M6L-LecRK_1364

GTTGCTATATCTACACGAAGAATGGGAACAATGTGTGGTGCATAGGGATATAAAGTCTAGCAATATTATGTTGGATTCTAATTTCAATGCCAAGCTTGGGGATTTTGGGTTAGCTAGGCT

>M6-PGSC0003DMG400004401M6L-LecRK_1550

ATGTGTCACCACTGGCAAAGCTAACAAGGAAACAGACGTCTATAGCTTTGGTATAGTCGTGTTAGAAATAGGTTGTGGAAGGAAACCTATTGATCCTAAAGCAGAGGAGCATCGAGTAAA

>M6-PGSC0003DMG400004401M6L-LecRK_1612

TAGAAATAGGTTGTGGAAGGAAACCTATTGATCCTAAAGCAGAGGAGCATCGAGTAAACATTGTCGATTGGGTTTGGAGACTTTATGGGATGAGAAGTATTCTTGAAGCAGTTGATCCGA

>M6-PGSC0003DMG400004401M6L-LecRK_1674

GTCGATTGGGTTTGGAGACTTTATGGGATGAGAAGTATTCTTGAAGCAGTTGATCCGAAACTATCATCAGAGTTCAATGAAGAGGAGATGGAGCATTTGCTAGTTGTTGGCTTATGGTGT

>M6-PGSC0003DMG400004401M6L-LecRK_1736

ATCATCAGAGTTCAATGAAGAGGAGATGGAGCATTTGCTAGTTGTTGGCTTATGGTGTGCTCATCCAGATAATAATTGCAGGCCTTCTATTAGGCAAGCAGTTCAGGTGCTTAATTTTGA

>M6-PGSC0003DMG400004401M6L-LecRK_1798

ATCCAGATAATAATTGCAGGCCTTCTATTAGGCAAGCAGTTCAGGTGCTTAATTTTGAAGCTCCATTACCCACTCTCCCTCGGAACATGCCTGTGCCCACATATTGCAGCTTCTCACGCC

>M6-PGSC0003DMG400004401M6L-LecRK_186

ATATATTCAAAACCACTGCATCTATGGGACAAGGCCTCTAGAAATCTCACTGATTTCACTACACATTTCTCCTTTAGCATTAATTCACAGGGAATAACAAAATATGGAGATGGTTTTGCC

>M6-PGSC0003DMG400004401M6L-LecRK_1922

ACCAACTATTTCAGTTTCATCAACACATGAAACCAGCGGTACAGAGATTGCCCTTACCCAAATCCAGCATTCAGTAAACGGAAACACCACTGATGGTTCAATAAAAAGAACATCTTCTTC

>M6-PGSC0003DMG400004401M6L-LecRK_2046

TCTTCATCTTCAACATCAGTTTTGTATCCAAGTTAATATTGCCCTGTTATAACCCTGCATTTTGTCTTCCATTAGCAGTGTCCATCTTTTCCTTTCTTACTTTTTGCTGTTTCTCTTCTA

>M6-PGSC0003DMG400004401M6L-LecRK_2100

CTGCATTTTGTCTTCCATTAGCAGTGTCCATCTTTTCCTTTCTTACTTTTTGCTGTTTCTCTTCTAATGAAGAGGATTCTTTGGATTACATTTCCTGTAATTTTAACTGTTCTATTAAAC

>M6-PGSC0003DMG400004401M6L-LecRK_248

ACATTTCTCCTTTAGCATTAATTCACAGGGAATAACAAAATATGGAGATGGTTTTGCCTTCTTCCTTGCCCCTGCAGGTTCAACAATTCCTTATAATACAACAAGAGGTGGAAGTCTTGG

>M6-PGSC0003DMG400004401M6L-LecRK_310

TCCTTGCCCCTGCAGGTTCAACAATTCCTTATAATACAACAAGAGGTGGAAGTCTTGGCCTTACAAGTAATAATGAGCGACTAAATTCTTCGAGCAATCATTTTGTTGCTGTGGAGTTTG

>M6-PGSC0003DMG400004401M6L-LecRK_372

ACAAGTAATAATGAGCGACTAAATTCTTCGAGCAATCATTTTGTTGCTGTGGAGTTTGACACCTATCAGAACCCCTATGACCCAAAGGGTGATCATGTAGGTGTCGATATCAACTCTATG

>M6-PGSC0003DMG400004401M6L-LecRK_496

CTGTTGTTAATGTGACTTGGTTTAGTAGCATTCCAAATGGTACGAAAACTGATGCCTGGATCAGTTATAATTCTAACTCAAAAAATCTTACTGCTGTCTTCACTGGTTTTCAACTCCAAG

>M6-PGSC0003DMG400004401M6L-LecRK_558

AGTTATAATTCTAACTCAAAAAATCTTACTGCTGTCTTCACTGGTTTTCAACTCCAAGGGAACAATATCACAGTCACTGTCCTGCAGAACCTATCTTACAATCTTGATCTGAGGGAATAT

>M6-PGSC0003DMG400004401M6L-LecRK_62

TGATACTTTTACACCCAATGATCACAATGTAACCTATGAGAGAGATGCTTCTCCAGAAAATGGTGCAATTCTACTCACCAAAAACCTCCTCAATCGCGATATTAATGCTAGTATAGGTCG

>M6-PGSC0003DMG400004401M6L-LecRK_682

CAGAATGGGTCACTTTTGGCTTCACTGGTGGAGCGGGAAGGAGATTTGCATTTCAAAGAATCTATTCTTGGAATTTTACTTCTTCTTTAGATAATACGACAGATACAGGGGTAACCTTAA

>M6-PGSC0003DMG400004401M6L-LecRK_744

TATTCTTGGAATTTTACTTCTTCTTTAGATAATACGACAGATACAGGGGTAACCTTAACAAACACCATGCCAAAGGATGCACCAACAAAAACTAAGTTAGGACTTGTGGTTGGATTGATT

>M6-PGSC0003DMG400004401M6L-LecRK_806

CACCATGCCAAAGGATGCACCAACAAAAACTAAGTTAGGACTTGTGGTTGGATTGATTTCTGGTGGTTGTGTTTTGGTTGCAGTATGTGCTTTGGTGTTATTTGCATTTTGGAGAAAGAG

>M6-PGSC0003DMG400004401M6L-LecRK_868

GTGGTTGTGTTTTGGTTGCAGTATGTGCTTTGGTGTTATTTGCATTTTGGAGAAAGAGTAAGGTGAGAGAAGATGATTATGTCTTTGATGGTTCCATGACTAATGAATTTCAAAAAAGTG

>M6-PGSC0003DMG400004401M6L-LecRK_930

GTGAGAGAAGATGATTATGTCTTTGATGGTTCCATGACTAATGAATTTCAAAAAAGTGCAGGACCAAAGAAGTTCTTGTACAGTGAGTTGGCTAGATGTACAAATAACTTTTCACAGGAA

>M6-PGSC0003DMG400004401M6L-LecRK_992

ACCAAAGAAGTTCTTGTACAGTGAGTTGGCTAGATGTACAAATAACTTTTCACAGGAAGAGATGCTTGGGCGGGGTGGATTCGGAGGTGTTTATAAAGGATATCTCAGTGAATCCAACCC

>M6-PGSC0003DMG400004529-(rev)M6LRR-RLP_198

CATTCAAGTTAAATTTCTTCTTGCTTCGAAAAGGTAAAGTAAAATTTTAATATATTCTTTATATTTACACTTTTTTATTTCGATTGTTAGTTCTGATTTTAAATTTCTTTCACAATTTTT

>M6-PGSC0003DMG400004529-(rev)M6LRR-RLP_264

ACACTTTTTTATTTCGATTGTTAGTTCTGATTTTAAATTTCTTTCACAATTTTTATGTATAAATATTAGGGTTATGACAAAATCCGAGTTGAATAAGTTGTGTATGGAGCTAGGTGAAGG

>M6-PGSC0003DMG400004529-(rev)M6LRR-RLP_330

TAGGGTTATGACAAAATCCGAGTTGAATAAGTTGTGTATGGAGCTAGGTGAAGGATGTCCCTTGCTGAGGGATATTGTGTTGTCTCACTGTCGTCAAATAACAGATGTAGGGTTAAGTTA

>M6-PGSC0003DMG400004529-(rev)M6LRR-RLP_527

CGGTGATTACGAGCTGCACAAATTTAAAGAAAGTCCTTGTAGAGAAGTGGAAAGTGAGTCCAAGAACCAAAAGGAGAGCAGGATCCATCATCTCCTATCTCTGTGTTGATCTGTAGATCT

>M6-PGSC0003DMG400004529-(rev)M6LRR-RLP_66

ACACTGACCAATAAAAATTGGCCACGTGTTAAAATAAACTTTATTGTTGTTTATTGTTCTTCATCTTTAACTAAGTTTTCTTCTTCTTTATTGTTCTTGATCTTTAGAAAAAAAAACCTC

>M6-PGSC0003DMG400004553M6RLKWAX-EGF_0

ATCCAGATATCAGTAATTGGCTGCTTCTGTTTTTCATGTGTAAGCTCCTATATATCCCTTTAGCAATCTATATGTTATTTGGATGGACACAGATTAACATTTCATTGATCATGTGTTTTA

>M6-PGSC0003DMG400004553M6RLKWAX-EGF_1054

ATGGCTAGAAAGCTGAGTTTAGTTGCATTATGGTGTATACAAACAAATCCGATTCAACGCCCTTCAATGAGTAAAGTGGTAGAAATGCTTGAAGGTGAAGTTGGAGTGCTAGAAGTACCT

>M6-PGSC0003DMG400004553M6RLKWAX-EGF_1116

TTCAATGAGTAAAGTGGTAGAAATGCTTGAAGGTGAAGTTGGAGTGCTAGAAGTACCTCCTCAGCCTCTTCAATCTCAACCAATTGTTCATCAGATCATGGGAAGTTCTACAACATTTTC

>M6-PGSC0003DMG400004553M6RLKWAX-EGF_1178

AGCCTCTTCAATCTCAACCAATTGTTCATCAGATCATGGGAAGTTCTACAACATTTTCGTCTGATTCAATAGCTTTGTTAGATAATCCCGTTGAGGTAGACATTTATGCTGATTGATTAT

>M6-PGSC0003DMG400004553M6RLKWAX-EGF_1240

GATTCAATAGCTTTGTTAGATAATCCCGTTGAGGTAGACATTTATGCTGATTGATTATAGTCGATGTAATGTTTATGTGAAATAAACATAGCCAACCCAATAATGTTTGGGACAAAGAAT

>M6-PGSC0003DMG400004553M6RLKWAX-EGF_186

CCTTCCACTGGTAATTGCATTTCTGGTGTACAAATTCAAAAGGAGGCATTTATCAATGTACGATACGATTGAAGGCTTTTTGCAAACACAAAACAATTTCATGCCAATCAGATACAATTA

>M6-PGSC0003DMG400004553M6RLKWAX-EGF_248

ATACGATTGAAGGCTTTTTGCAAACACAAAACAATTTCATGCCAATCAGATACAATTACTCCAACATAAAGAGAATGACAAGAGGATTCAAAGAGAAATTAGGAGAGGGGGGTTACGGAA

>M6-PGSC0003DMG400004553M6RLKWAX-EGF_496

GTTGGACTCGTGCGGTATTGCGTTGAGGGAACAAAGCTTGCTCTTGTATACGACTTCATGCCCAACGGATCACTTGATAAGTACATCAGCACGAGTCAAGAAGGAAGTCCTCCGTTAAGT

>M6-PGSC0003DMG400004553M6RLKWAX-EGF_62

GCAATCTATATGTTATTTGGATGGACACAGATTAACATTTCATTGATCATGTGTTTTATCATCTCTATGGTTCGTTATAACTTCTCAAACGTGCAGTGCATTTGGGAGCAAAATTTGTAA

>M6-PGSC0003DMG400004553M6RLKWAX-EGF_868

GCAATCTCGTACAAAGCTGATGTTTACAGCTTCGGAATGCTGCTAATGGAAATGTTGGACTTGAAGAGAAATGAAGTTGCAAATGAAGAGAATTCCAGCCAATACTTTCCTTATAATATT

>M6-PGSC0003DMG400004553M6RLKWAX-EGF_930

GAAGAGAAATGAAGTTGCAAATGAAGAGAATTCCAGCCAATACTTTCCTTATAATATTTACGATAAGTTCAACCAGGGGAAAGAGATTGTGGTGGATGAAGGAGCGAATGATGATGAAAA

>M6-PGSC0003DMG400004553M6RLKWAX-EGF_992

ATAAGTTCAACCAGGGGAAAGAGATTGTGGTGGATGAAGGAGCGAATGATGATGAAAAGAAGATGGCTAGAAAGCTGAGTTTAGTTGCATTATGGTGTATACAAACAAATCCGATTCAAC

>M6-PGSC0003DMG400004561M6LRR-RLP_0

GTTTGATGTAAATTGTTTCTCTTGAGAAGTTGAAAGGAAGGATTTTTTTTATTTTTTTTTATTTTTTAACTACTTTGGCTTGATTAGTGAGAATAGAAAGAGGAAAAGAAAATGAAGGGC

>M6-PGSC0003DMG400004561M6LRR-RLP_1054

TAAGAGATTTCCATAGGTTGAAAGTAAATGGTTGTGTTGAGTATGAAACAATCGAATATGTCTTACCTCATCTTCAACTTATGGCTCTGAGAGTAGTAAACTTCTGATTTACTCTTTTGA

>M6-PGSC0003DMG400004561M6LRR-RLP_1116

TTACCTCATCTTCAACTTATGGCTCTGAGAGTAGTAAACTTCTGATTTACTCTTTTGAATAGTCAACTGGACAAATTCGATGAATCGGATGTCTTTCTGGTTAACTCCAAGCTAGCTAAT

>M6-PGSC0003DMG400004561M6LRR-RLP_1178

TCAACTGGACAAATTCGATGAATCGGATGTCTTTCTGGTTAACTCCAAGCTAGCTAATACACTTGTGGAGGTTATTCCTGTTGAACTGGAGGATATGCACATATGTTGAACAAATTTGAA

>M6-PGSC0003DMG400004561M6LRR-RLP_124

AAAAGGGGTTCATTCACATAATTACATATTCTTGTAAGTAAGTTTGTTACTTTATGTAACTTAGATGGTGAATATAATCTTTATAACCTATAAGAATTGATGATTTAATCTTTCTGCTAT

>M6-PGSC0003DMG400004561M6LRR-RLP_1364

TAGAGTATCTCTGATTCATCTACAAGAGCACATGGCTAATGCTGGTATTCCTAGCGCATCTACGTGTAACATTCATGTCATGACAGAGTTCCTATTGATTATTCTTCCTGATGTGCCCAA

>M6-PGSC0003DMG400004561M6LRR-RLP_1426

CGTGTAACATTCATGTCATGACAGAGTTCCTATTGATTATTCTTCCTGATGTGCCCAAGGACATTATTCGTCAAGACAAATTGTTTGTTCTCTTGGCACGTGTTAGAGAACTTACCAAGG

>M6-PGSC0003DMG400004561M6LRR-RLP_1488

ATTATTCGTCAAGACAAATTGTTTGTTCTCTTGGCACGTGTTAGAGAACTTACCAAGGAGGTATTCGTTCTTGTTCGCAACTTAGAAGAGAATATGAATGACGCAAGTGATGCAAATCTA

>M6-PGSC0003DMG400004561M6LRR-RLP_1550

ATTCGTTCTTGTTCGCAACTTAGAAGAGAATATGAATGACGCAAGTGATGCAAATCTAAACTTGCTGGAAAATATTGAACTCCTGAAGGAAGATCTCAAGAATGTTTTCTTGAAAGCCCG

>M6-PGSC0003DMG400004561M6LRR-RLP_1674

GACTCATCTCGTCTTTGCTTCCCCATGAGTGATGGACCGCTCTTCATGACTCTTCTGCTTACAAACTTGAAAGACTTTGTCGATTCCAATGCTTATTCAGTTACTTTGATAAAAGAAGAA

>M6-PGSC0003DMG400004561M6LRR-RLP_1736

AAACTTGAAAGACTTTGTCGATTCCAATGCTTATTCAGTTACTTTGATAAAAGAAGAAATTAAGCAGGTGAAAGAAGACCTGGAAATCATAAGATCGTTCTTCGGGTATGCTGAGCAAGA

>M6-PGSC0003DMG400004561M6LRR-RLP_1798

AGCAGGTGAAAGAAGACCTGGAAATCATAAGATCGTTCTTCGGGTATGCTGAGCAAGAGTTGAACAGAGATCTTTGGACTCATGTTCTAGATTTGGCATATGAGGCGGAACATGCCATTA

>M6-PGSC0003DMG400004561M6LRR-RLP_186

AGATGGTGAATATAATCTTTATAACCTATAAGAATTGATGATTTAATCTTTCTGCTATAAGTTAAACTCTATGTACTAAATCATCTGCTATAGTTGATAAATGTAAGGGGTTAAGAACTA

>M6-PGSC0003DMG400004561M6LRR-RLP_1860

AACAGAGATCTTTGGACTCATGTTCTAGATTTGGCATATGAGGCGGAACATGCCATTAACTCAATTCTTGCCCGGGATCATGGTCTATTATAGCTTATCTTCATACTTCCTGATACTGTA

>M6-PGSC0003DMG400004561M6LRR-RLP_1922

AATTCTTGCCCGGGATCATGGTCTATTATAGCTTATCTTCATACTTCCTGATACTGTAGAAAAGATCAAGCTTGTCAAAAAAGAGGTACAAGAGAAGATCCCCAAGAGCAGCGGTATTAT

>M6-PGSC0003DMG400004561M6LRR-RLP_2170

ATGCCATTTCCATAGTCGGCATGCCAGGGCTTGGAAAAACTACTTTGGCATACAGAGTGTATAATGATAAGTTCATTGTTGATCATTTCGACGTTTGTGCTTGATGCACAGTCGACCAGG

>M6-PGSC0003DMG400004561M6LRR-RLP_2232

AATGATAAGTTCATTGTTGATCATTTCGACGTTTGTGCTTGATGCACAGTCGACCAGGAACGTAATGCGAAAAAGTTGTGGCAGAAAATTTTCAATCAAGTTATTGGTTTGACAGAAAGA

>M6-PGSC0003DMG400004561M6LRR-RLP_2294

TAATGCGAAAAAGTTGTGGCAGAAAATTTTCAATCAAGTTATTGGTTTGACAGAAAGATTCAGTGAGAATGACATAGATGATGACGTTGGTTATAAGCTGCGGAAACAACTATTTGCAAA

>M6-PGSC0003DMG400004561M6LRR-RLP_2332

TTATTGGTTTGACAGAAAGATTCAGTGAGAATGACATAGATGATGACGTTGGTTATAAGCTGCGGAAACAACTATTTGCAAAACGGTACCTTATTGTCTTGGATGACATGTGGGATACTG

>M6-PGSC0003DMG400004561M6LRR-RLP_248

TAAACTCTATGTACTAAATCATCTGCTATAGTTGATAAATGTAAGGGGTTAAGAACTAGGAAAGTAAGAGCTTTTGTTTAGAGAGATAGCGCGTAAACAATGAGAGAGCTTAGTGCACCA

>M6-PGSC0003DMG400004561M6LRR-RLP_310

AGTAAGAGCTTTTGTTTAGAGAGATAGCGCGTAAACAATGAGAGAGCTTAGTGCACCAGGAGGCTGTCATTTTTTGGTTTAAGATTTGAGGTGTAACAGGACAAACATTTGGCTTCTCTA

>M6-PGSC0003DMG400004561M6LRR-RLP_372

GCTGTCATTTTTTGGTTTAAGATTTGAGGTGTAACAGGACAAACATTTGGCTTCTCTATTAGGTTATTCTGGAGAAAGTCCTCTGCACTTGTGACAGTTGCTTGTTGTAAAATTAAGAGC

>M6-PGSC0003DMG400004561M6LRR-RLP_434

GTTATTCTGGAGAAAGTCCTCTGCACTTGTGACAGTTGCTTGTTGTAAAATTAAGAGCTATCACATTTTCGAGAGTTTTACACCTCAACTATATTAGTCGTTCCCTTTTTCTATCTAAAT

>M6-PGSC0003DMG400004561M6LRR-RLP_496

ACATTTTCGAGAGTTTTACACCTCAACTATATTAGTCGTTCCCTTTTTCTATCTAAATTTGAAACAACATTTGCTCTTTTGGTAGGTGTCATCCGTTGTACTTCGTGAGGACATTGTCAA

>M6-PGSC0003DMG400004561M6LRR-RLP_558

AACAACATTTGCTCTTTTGGTAGGTGTCATCCGTTGTACTTCGTGAGGACATTGTCAATCTTCTAGATTTCATAGAGATGTTAAATAGTGGGCGTGTTCAAACTTCTTTTGACATGGATC

>M6-PGSC0003DMG400004561M6LRR-RLP_62

TTTTTAACTACTTTGGCTTGATTAGTGAGAATAGAAAGAGGAAAAGAAAATGAAGGGCAAACAAAAGGGGTTCATTCACATAATTACATATTCTTGTAAGTAAGTTTGTTACTTTATGTA

>M6-PGSC0003DMG400004561M6LRR-RLP_620

CTAGATTTCATAGAGATGTTAAATAGTGGGCGTGTTCAAACTTCTTTTGACATGGATCAAATTGAAAAGTTGACATTGGAGCTGACGTTTGTGTCCGCATTTTGTTATCTTTATTACTTT

>M6-PGSC0003DMG400004561M6LRR-RLP_682

TGAAAAGTTGACATTGGAGCTGACGTTTGTGTCCGCATTTTGTTATCTTTATTACTTTCTTTTTGGAAAGTTGTAATGACGAAATGTCCTGCATATCAAATGAGATTCATGATTTGGTTC

>M6-PGSC0003DMG400004561M6LRR-RLP_744

TTGGAAAGTTGTAATGACGAAATGTCCTGCATATCAAATGAGATTCATGATTTGGTTCGGTCACTGTTTCATCCAAGTGGAGAAGACATGCTGGTTAATATAAAGAATCATGACCTCTTG

>M6-PGSC0003DMG400004561M6LRR-RLP_806

ACTGTTTCATCCAAGTGGAGAAGACATGCTGGTTAATATAAAGAATCATGACCTCTTGGAGAATATCAAAAGTTATATTAACTCACATAGCTATGTTGAACCAAGTCGTGTAATCATGAC

>M6-PGSC0003DMG400004561M6LRR-RLP_868

ATATCAAAAGTTATATTAACTCACATAGCTATGTTGAACCAAGTCGTGTAATCATGACCGAGGATCGTTTGGTTGAACTTTTAGATGCCATCCTCGTGTATCTCCAATGTCTACTCAACT

>M6-PGSC0003DMG400004561M6LRR-RLP_930

GATCGTTTGGTTGAACTTTTAGATGCCATCCTCGTGTATCTCCAATGTCTACTCAACTGTAGTTCTGAATATATTTTCCCATTAGTGACTCAATGTGAGCTTCTTCAGAATGTATTTGGC

>M6-PGSC0003DMG400004561M6LRR-RLP_992

TTCTGAATATATTTTCCCATTAGTGACTCAATGTGAGCTTCTTCAGAATGTATTTGGCAATTTAAGAGATTTCCATAGGTTGAAAGTAAATGGTTGTGTTGAGTATGAAACAATCGAATA

>M6-PGSC0003DMG400004667M6LRR-RLK_0

ATCTAAAGCTGTCACTTAGCTTACTGCTGAAAAACTCATAAACCCAAAATTTTCAACAAACAATACCAAAAAAAAAAAAAAAATATCTTCAAAATTTTGAAGAATCATAGCAAAAAAAAG

>M6-PGSC0003DMG400004667M6LRR-RLK_1054

GTTGAATATATCTTTGCAAAACAATAAGTTGCAGGGTCCAATGCCACAGTTTGGACAGGGGATTAAATTGGAGATTGAGGGTACTAATAGTTTTTGTTTGGATACTCCTGGACCTTGTGA

>M6-PGSC0003DMG400004667M6LRR-RLK_1116

TTAAATTGGAGATTGAGGGTACTAATAGTTTTTGTTTGGATACTCCTGGACCTTGTGATTCGCAGGTTATGACGCTTTTAGATGTGGCGGGTGGTTTTGGATATCCGGGGTTTTTGGCTG

>M6-PGSC0003DMG400004667M6LRR-RLK_1178

CAGGTTATGACGCTTTTAGATGTGGCGGGTGGTTTTGGATATCCGGGGTTTTTGGCTGATTCTTGGAAAGGAAATGATGCTTGTAATGGATGGTCACATGTAAGTTGTGATGGATCGAAA

>M6-PGSC0003DMG400004667M6LRR-RLK_1240

TTGGAAAGGAAATGATGCTTGTAATGGATGGTCACATGTAAGTTGTGATGGATCGAAAAAGAATGTGGATGTTGTAACATTTGGTAAGCTACGTTTATCGGGTTTTATATCTCCTGCTTT

>M6-PGSC0003DMG400004667M6LRR-RLK_1302

ATGTGGATGTTGTAACATTTGGTAAGCTACGTTTATCGGGTTTTATATCTCCTGCTTTTGCTAATTTGACTAGTTTGAGGAATTTGTTATTGAATGATAACAATCTTACTGGATCTATAC

>M6-PGSC0003DMG400004667M6LRR-RLK_1364

AATTTGACTAGTTTGAGGAATTTGTTATTGAATGATAACAATCTTACTGGATCTATACCGGAAAGCTTGGCAGCATTGCCTCTGCTTCAAACACTTGATGTTTCTAATAACAATTTATCT

>M6-PGSC0003DMG400004667M6LRR-RLK_1426

AAGCTTGGCAGCATTGCCTCTGCTTCAAACACTTGATGTTTCTAATAACAATTTATCTGGTCCTGTACCTGCATTTCGATCAAATGTGAAGCTTATTAATGGTGGTAACGTGTTTCTTGG

>M6-PGSC0003DMG400004667M6LRR-RLK_1488

CTGTACCTGCATTTCGATCAAATGTGAAGCTTATTAATGGTGGTAACGTGTTTCTTGGAAAGAATATTAGTGATGGAGGGGGTAGTGGTGGATCACCGGGGTCTGGGTCGAGTAGTGATG

>M6-PGSC0003DMG400004667M6LRR-RLK_1550

AATATTAGTGATGGAGGGGGTAGTGGTGGATCACCGGGGTCTGGGTCGAGTAGTGATGGTGGAAATTCATCTGGTGGTTCGAAGGGATCCTCAGTTCAACCTGGAGTGGTTGCTGGTGTT

>M6-PGSC0003DMG400004667M6LRR-RLK_1612

AAATTCATCTGGTGGTTCGAAGGGATCCTCAGTTCAACCTGGAGTGGTTGCTGGTGTTGTTATATCTGTCGTTATATTTATCCTTGTTGTGTTGTATGTGTCTTACAAATGTTACATAAG

>M6-PGSC0003DMG400004667M6LRR-RLK_1674

TATCTGTCGTTATATTTATCCTTGTTGTGTTGTATGTGTCTTACAAATGTTACATAAGGAGACGGCATAAGAGATTTGGAAGGGTTCAGAATCCGGAAAGAAGCAATGATATGGTTAAGC

>M6-PGSC0003DMG400004667M6LRR-RLK_1736

CGGCATAAGAGATTTGGAAGGGTTCAGAATCCGGAAAGAAGCAATGATATGGTTAAGCCTGGTGTAGTGATAGGTAATGGATATGCAGGAGTCCCTAGTGAATTACAGAGCCAGAGCAGT

>M6-PGSC0003DMG400004667M6LRR-RLK_1798

TGTAGTGATAGGTAATGGATATGCAGGAGTCCCTAGTGAATTACAGAGCCAGAGCAGTGGGGATCATAGTGAGATGCCGATATTTGAAGGTGGAAATGTTGCCATTTCCTTTCAGGTGCT

>M6-PGSC0003DMG400004667M6LRR-RLK_186

TGATTGTTCTTAGATCTCTAAAACCTCCATTAATGGCACCCCACAAGCTTTTTTTTCTTGTTAACTTCTTCTTCACCTTCATTTCATTCACATTTTCCGATGACAGCACGGTCATGTCAA

>M6-PGSC0003DMG400004667M6LRR-RLK_1860

ATCATAGTGAGATGCCGATATTTGAAGGTGGAAATGTTGCCATTTCCTTTCAGGTGCTTCGACAAGTGACAAACAACTTCAGCGAAGAAAATATCTTGGGTAGAGGAGGATTTGGAGTCG

>M6-PGSC0003DMG400004667M6LRR-RLK_1922

CAAGTGACAAACAACTTCAGCGAAGAAAATATCTTGGGTAGAGGAGGATTTGGAGTCGTTTATAAGGGTGAATTACATGATGGGACTAAGATTGCTGTGAAGAGGATGGAATCTGGCGCG

>M6-PGSC0003DMG400004667M6LRR-RLK_1984

TAAGGGTGAATTACATGATGGGACTAAGATTGCTGTGAAGAGGATGGAATCTGGCGCGATGGGTACTAAAGGAATGAATGAGTTCCAAGCTGAAATCGCGGTCCTTACCAAAGTTAGGCA

>M6-PGSC0003DMG400004667M6LRR-RLK_2046

GTACTAAAGGAATGAATGAGTTCCAAGCTGAAATCGCGGTCCTTACCAAAGTTAGGCATAGGCATCTGGTCGCTCTTCTTGGTTCCTGTGTTAACGGCAATGAAAGACTTTTGGTGTATG

>M6-PGSC0003DMG400004667M6LRR-RLK_2108

CATCTGGTCGCTCTTCTTGGTTCCTGTGTTAACGGCAATGAAAGACTTTTGGTGTATGAGTACATGCCACAAGGAACTTTAAGTCAGCATTTGTTTGAATGGCAAGAACACGGGTGCCCA

>M6-PGSC0003DMG400004667M6LRR-RLK_2170

CATGCCACAAGGAACTTTAAGTCAGCATTTGTTTGAATGGCAAGAACACGGGTGCCCAATTTTGACCTGGAAGCAAAGGGTCACAATAGCATTAGATGTTGCAAGGGGAGTCGAGTATCT

>M6-PGSC0003DMG400004667M6LRR-RLK_2232

TGACCTGGAAGCAAAGGGTCACAATAGCATTAGATGTTGCAAGGGGAGTCGAGTATCTCCACAGCTTGGCACAGACAAGTTTTATCCATAGGGATTTGAAGCCCTCTAACATACTTCTTG

>M6-PGSC0003DMG400004667M6LRR-RLK_2294

AGCTTGGCACAGACAAGTTTTATCCATAGGGATTTGAAGCCCTCTAACATACTTCTTGGGGATGACATGAGAGCCAAGGTTGCAGATTTTGGATTGGTTAAGAATGCCCCCGATGGGAAA

>M6-PGSC0003DMG400004667M6LRR-RLK_2356

TGACATGAGAGCCAAGGTTGCAGATTTTGGATTGGTTAAGAATGCCCCCGATGGGAAATATTCTGTTGAGACACGGTTGGCTGGAACTTTTGGCTATCTGGCGCCTGAATATGCTGGTAA

>M6-PGSC0003DMG400004667M6LRR-RLK_2418

CTGTTGAGACACGGTTGGCTGGAACTTTTGGCTATCTGGCGCCTGAATATGCTGGTAAGTCTGTCTTTCCTCTGCTGATCTTTAATCCTTTTAGTTGTAAAACTGGTAGAAAACTAAAAC

>M6-PGSC0003DMG400004667M6LRR-RLK_248

AACTTCTTCTTCACCTTCATTTCATTCACATTTTCCGATGACAGCACGGTCATGTCAAATTTTCTGAAGGGTCTGTCTAAACCACCATCTGGTTGGGATGCTTCAAAGTCTTACTGTGAA

>M6-PGSC0003DMG400004667M6LRR-RLK_2480

GTCTTTCCTCTGCTGATCTTTAATCCTTTTAGTTGTAAAACTGGTAGAAAACTAAAACAATATCCTATCATTAGCTTTCATGTTCTTTTTAATCTTCACTATATTAGAGATAAATGGAGA

>M6-PGSC0003DMG400004667M6LRR-RLK_2542

TCCTATCATTAGCTTTCATGTTCTTTTTAATCTTCACTATATTAGAGATAAATGGAGAGGTGTTAGCTTCGTTGAAGGTCATTTCCATGTTCAAATGCTAGTTTGTTAATCTCGATCTGA

>M6-PGSC0003DMG400004667M6LRR-RLK_2604

TTAGCTTCGTTGAAGGTCATTTCCATGTTCAAATGCTAGTTTGTTAATCTCGATCTGATTCAAGGATTGATACCGCAAACGTTGTCCAGTAAATTAACTAATGTTAGCAAAAAAGAGTAC

>M6-PGSC0003DMG400004667M6LRR-RLK_2666

AGGATTGATACCGCAAACGTTGTCCAGTAAATTAACTAATGTTAGCAAAAAAGAGTACGATTTGAAAGACAAACATAAGGTAGAAATCTAGAAGCCCACTCATCCGGTTTTCAAATTGAA

>M6-PGSC0003DMG400004667M6LRR-RLK_2728

TGAAAGACAAACATAAGGTAGAAATCTAGAAGCCCACTCATCCGGTTTTCAAATTGAAATTCGGATGTTGGTACAGCATGTGTGACTGATCCCATTGTATCTAAATAGAAATTTGGCTTT

>M6-PGSC0003DMG400004667M6LRR-RLK_2790

GGATGTTGGTACAGCATGTGTGACTGATCCCATTGTATCTAAATAGAAATTTGGCTTTCAGAATATCACTCTTCTTCAAGACATTAAGTTCAACATATCTAGATTTAGTGTTGGCTGTCA

>M6-PGSC0003DMG400004667M6LRR-RLK_2852

ATATCACTCTTCTTCAAGACATTAAGTTCAACATATCTAGATTTAGTGTTGGCTGTCAATAGTTTTCTGCTCGTAACATGTTTAATTTCCTTTAGAGATCTTGTTCTGGCCTGTTTCTTG

>M6-PGSC0003DMG400004667M6LRR-RLK_2914

TTTTCTGCTCGTAACATGTTTAATTTCCTTTAGAGATCTTGTTCTGGCCTGTTTCTTGCATGTCTTAGCAAGGTTGTCTGTGATCATTATTAAATTTGGATTAAACGTGACTGCTCCTCT

>M6-PGSC0003DMG400004667M6LRR-RLK_2976

TCTTAGCAAGGTTGTCTGTGATCATTATTAAATTTGGATTAAACGTGACTGCTCCTCTTTATTTACAGCTACTGGACGAGTCACGACCAAAGTAGATGTGTATGCATTTGGAGTTGTTCT

>M6-PGSC0003DMG400004667M6LRR-RLK_3038

TTACAGCTACTGGACGAGTCACGACCAAAGTAGATGTGTATGCATTTGGAGTTGTTCTAATGGAGATCATCACTGGTAGAAAAGCATTAGATGAGACCTTGCCAGATGAGAGGTCTCATT

>M6-PGSC0003DMG400004667M6LRR-RLK_310

TCTGAAGGGTCTGTCTAAACCACCATCTGGTTGGGATGCTTCAAAGTCTTACTGTGAATGGAGTAAAGTTACTTGTGATAAATCGAGTAATACTGTTGTTTCCATTAATCTTGAATCTCA

>M6-PGSC0003DMG400004667M6LRR-RLK_3100

GAGATCATCACTGGTAGAAAAGCATTAGATGAGACCTTGCCAGATGAGAGGTCTCATTTGGTCACCTGGCTTCGCAGGGTCCTTGTCAACAAAGACAACCTCCGAAAGGCTATTGACCCA

>M6-PGSC0003DMG400004667M6LRR-RLK_3162

CACCTGGCTTCGCAGGGTCCTTGTCAACAAAGACAACCTCCGAAAGGCTATTGACCCAACTCTTGATCCTGATGAGGAAACATACGAGAGCATTTGTAAGGTCGCTGAGTTAGCAGGCCA

>M6-PGSC0003DMG400004667M6LRR-RLK_3224

TTGATCCTGATGAGGAAACATACGAGAGCATTTGTAAGGTCGCTGAGTTAGCAGGCCATTGCACTGCACGAGAACCATTCCAGCGCCCTGACATGGGACATGCAGTCAATGTCCTTGCCC

>M6-PGSC0003DMG400004667M6LRR-RLK_3286

ACTGCACGAGAACCATTCCAGCGCCCTGACATGGGACATGCAGTCAATGTCCTTGCCCCCCTCATAGAACTGTGGAAGCCTACAAGAAATGAAGACGAAGACAGTGGCATCGACCTCCAA

>M6-PGSC0003DMG400004667M6LRR-RLK_3348

CATAGAACTGTGGAAGCCTACAAGAAATGAAGACGAAGACAGTGGCATCGACCTCCAAATGAGCCTTCCTCAAATCCTTCAAAGATGGCAAGCTGATGAAGGAACTTCAAGAATGTTCGA

>M6-PGSC0003DMG400004667M6LRR-RLK_3410

GCCTTCCTCAAATCCTTCAAAGATGGCAAGCTGATGAAGGAACTTCAAGAATGTTCGATGACATCTCCTTCAGTCAAACACATTCAAGTATACCCTCGAAACCTTCTGGATTTGCAGATA

>M6-PGSC0003DMG400004667M6LRR-RLK_3472

ATCTCCTTCAGTCAAACACATTCAAGTATACCCTCGAAACCTTCTGGATTTGCAGATACTTTCAATTCAACAGATTGCAGATGACATTAAGGCAACAATCAAGAAAGGCACACAATTCAT

>M6-PGSC0003DMG400004667M6LRR-RLK_3534

CAATTCAACAGATTGCAGATGACATTAAGGCAACAATCAAGAAAGGCACACAATTCATTTGGTTAATTGGTTTCGTCTTTAACAACGCTTTTTACTTCGTCTTTTCATAGCCCTTTTGGT

>M6-PGSC0003DMG400004667M6LRR-RLK_3596

TTAATTGGTTTCGTCTTTAACAACGCTTTTTACTTCGTCTTTTCATAGCCCTTTTGGTGATAGATCATTAGACCAAAGAAGCCAAGATGGATAGCAGAGTTTATTTATGTGGTTTTGTTA

>M6-PGSC0003DMG400004667M6LRR-RLK_3658

GATCATTAGACCAAAGAAGCCAAGATGGATAGCAGAGTTTATTTATGTGGTTTTGTTAGAAGAGAAAAATTGTATTAAATTCTTTTAATCTTACATAGTTGATGTAAATAGATTTGAGTT

>M6-PGSC0003DMG400004667M6LRR-RLK_372

GTAAAGTTACTTGTGATAAATCGAGTAATACTGTTGTTTCCATTAATCTTGAATCTCAAGGTCTTTCTGGGGTTTTGCCTCCTGAGCTTAATCAACTGTCAAATCTCAAAACTCTTTCAG

>M6-PGSC0003DMG400004667M6LRR-RLK_3720

AGAAAAATTGTATTAAATTCTTTTAATCTTACATAGTTGATGTAAATAGATTTGAGTTCTTGTGTTAAACATGCAAAATATACTCTTGAAGTTTTTTACTACAAGATTTCATTATTTGTG

>M6-PGSC0003DMG400004667M6LRR-RLK_434

CTTTCTGGGGTTTTGCCTCCTGAGCTTAATCAACTGTCAAATCTCAAAACTCTTTCAGTTCAGAGGAATTCGCTTTCTGGGATTCTTCCATCTTTCGCAAACATGTCGAATTTAGCTGAG

>M6-PGSC0003DMG400004667M6LRR-RLK_496

GAGGAATTCGCTTTCTGGGATTCTTCCATCTTTCGCAAACATGTCGAATTTAGCTGAGATTTACTTGAATGACAATGGATTCACATCTGTTCCTCAAGATTTCCTTTTGGGTCTTACTAG

>M6-PGSC0003DMG400004667M6LRR-RLK_558

ACTTGAATGACAATGGATTCACATCTGTTCCTCAAGATTTCCTTTTGGGTCTTACTAGTTTGCAGACTTTTAGTATTAGTGAGAATGGGAAGTTGAGTCCTTGGCAAATTCCTAGTTATT

>M6-PGSC0003DMG400004667M6LRR-RLK_620

CAGACTTTTAGTATTAGTGAGAATGGGAAGTTGAGTCCTTGGCAAATTCCTAGTTATTTGATTGATAGCAACAATTTGGCTAATGTTTATGCTAGTAATGCTAGTTTGTTTGGTGTTATA

>M6-PGSC0003DMG400004667M6LRR-RLK_682

TGATAGCAACAATTTGGCTAATGTTTATGCTAGTAATGCTAGTTTGTTTGGTGTTATACCTGATTTTCTTGGTAAATTCCCTAATCTTCAGGATTTGAGATTGTCTTATAATAATTTTAC

>M6-PGSC0003DMG400004667M6LRR-RLK_744

ATTTTCTTGGTAAATTCCCTAATCTTCAGGATTTGAGATTGTCTTATAATAATTTTACTGGTTCTTTGCCTGCTAGTTTTGGGGATTCTGAGATTAAGAATTTATGGCTGAATAATCAGG

>M6-PGSC0003DMG400004667M6LRR-RLK_806

TCTTTGCCTGCTAGTTTTGGGGATTCTGAGATTAAGAATTTATGGCTGAATAATCAGGGTAAAGGGTTATCAGGAACTATTGATGTGATTTCAAGTATGGTTCAGTTGTCACAAGTGTGG

>M6-PGSC0003DMG400004667M6LRR-RLK_868

AGGGTTATCAGGAACTATTGATGTGATTTCAAGTATGGTTCAGTTGTCACAAGTGTGGTTACATGCTAATTCTTTTACTGGTCCTATACCCGATTTGTCGAAATGCGAAAATATATTTGA

>M6-PGSC0003DMG400004667M6LRR-RLK_930

ATGCTAATTCTTTTACTGGTCCTATACCCGATTTGTCGAAATGCGAAAATATATTTGATTTGCAGTTAAGGGATAATGATTTTACTGGTGTTATACCTGATTCGGTGATGAAACTTCCTA

>M6-PGSC0003DMG400004667M6LRR-RLK_992

CAGTTAAGGGATAATGATTTTACTGGTGTTATACCTGATTCGGTGATGAAACTTCCTAAGTTGTTGAATATATCTTTGCAAAACAATAAGTTGCAGGGTCCAATGCCACAGTTTGGACAG

>M6-PGSC0003DMG400004722M6LRR-RLP_0

TTCAAGAAAATGCTTACTGTAGATACCTCATTAGTTTGGTGTAGTTACTTTTACCACACACGAGTTCATGGAACATAAGCAGAGATTGCTGCTCTTCGGATGGAGTCGTATGTGATGACA

>M6-PGSC0003DMG400004722M6LRR-RLP_1054

AACTCTTGATGTCAGCTATATGAAATCTCAGTAGTCATATAGCTTTACCTAAACTCATCTCGATCTAGTACAAAGTATATTAGAGGTAATAATGTAGTCTAGAATTATTGTTCTATGACA

>M6-PGSC0003DMG400004722M6LRR-RLP_1116

ATCTAGTACAAAGTATATTAGAGGTAATAATGTAGTCTAGAATTATTGTTCTATGACATAATTGTTGGATTGGTCCACTAATTTAAAGGTGAGATTGAGAACCAAACATGTCCGCATCAT

>M6-PGSC0003DMG400004722M6LRR-RLP_1178

TGTTGGATTGGTCCACTAATTTAAAGGTGAGATTGAGAACCAAACATGTCCGCATCATGGCTGATAACACAACAAAACTTTTTGGCTACTTGGTTAAGTCCTTAGCTGAATTACATATGT

>M6-PGSC0003DMG400004722M6LRR-RLP_124

TGGCCATGTCACTGAACTTGACCTCAGTCTATTCAGGGACGAGAGCTTGAACTTGTCAAATTTTTGACCATTTACACCACTATTGTTCTTTCAAGTAACAGATTTGAAGGACGTATTCCA

>M6-PGSC0003DMG400004722M6LRR-RLP_1240

GATAACACAACAAAACTTTTTGGCTACTTGGTTAAGTCCTTAGCTGAATTACATATGTAACCCGTGGATAAAAATAAAAAGCAAAAAGACAAGGACAGAACAACCAAAATTTGAATGGAG

>M6-PGSC0003DMG400004722M6LRR-RLP_1302

CGTGGATAAAAATAAAAAGCAAAAAGACAAGGACAGAACAACCAAAATTTGAATGGAGAATGAGAACACTTTCTCTGATAAATATGAACAGACAAAGAAGTGTGAATAAGAGAGACAGAA

>M6-PGSC0003DMG400004722M6LRR-RLP_1364

AGAACACTTTCTCTGATAAATATGAACAGACAAAGAAGTGTGAATAAGAGAGACAGAAGCCGAATTGGCAACATTTGGTGGCAGTCAAACATGTTTCGCTTTGATCTTTAATATTATGCA

>M6-PGSC0003DMG400004722M6LRR-RLP_1426

AATTGGCAACATTTGGTGGCAGTCAAACATGTTTCGCTTTGATCTTTAATATTATGCACTATATAAAATCCTACGAACTCTCTACAATTAGTTCATTGAAACACAAAATATGATGATGGT

>M6-PGSC0003DMG400004722M6LRR-RLP_1488

ATAAAATCCTACGAACTCTCTACAATTAGTTCATTGAAACACAAAATATGATGATGGTTACCAAAATATCCTCTCTACTTCAGTTTTTCACTCTTTTATATCTCTTTACAGTTACTTTTG

>M6-PGSC0003DMG400004722M6LRR-RLP_1550

AAAATATCCTCTCTACTTCAGTTTTTCACTCTTTTATATCTCTTTACAGTTACTTTTGCTTCCACTGAGGAAACAACTGCCCTCTTTAAATGGAAAGCGACTTTCAAGAACAAAAATAAT

>M6-PGSC0003DMG400004722M6LRR-RLP_1798

TCCGTTTTCATCCCTCCCTTTTCTCGAGTATCTTAATCTTAGCAGGAACAATTTATTTGGCACCATCCCACCTGAGATTGGTTATCTAACTAATCTTGTCTATCTTGACTTGCATATCAA

>M6-PGSC0003DMG400004722M6LRR-RLP_1860

CCATCCCACCTGAGATTGGTTATCTAACTAATCTTGTCTATCTTGACTTGCATATCAATCACATTTCAGGCTCAATCCCACCACAAATCGGCTCACTAGCCAAACTTCAGATCATCCGTT

>M6-PGSC0003DMG400004722M6LRR-RLP_1922

ATTTCAGGCTCAATCCCACCACAAATCGGCTCACTAGCCAAACTTCAGATCATCCGTTTATTTGACAATCATTTAACTGGTCCTATTCCAGAAACAATAGGGTACCTAAGGTCTCTTACT

>M6-PGSC0003DMG400004722M6LRR-RLP_2170

CCTAAGGTCTCTTGCTCGACTAGATTTGAGGAATAACTTTCTTATTGGTTCTATTCCTACTTCCCTTGGCAACTTGAAAAATTTGCAAGTTCTGCTTCTCAATGACAACAATCTAATTGA

>M6-PGSC0003DMG400004722M6LRR-RLP_2232

CCCTTGGCAACTTGAAAAATTTGCAAGTTCTGCTTCTCAATGACAACAATCTAATTGAGGAAATTCCTCCATCTATTTGCAATTTGACATCACTGGCAATCCTGTATTTGTCGAGAAACA

>M6-PGSC0003DMG400004722M6LRR-RLP_2294

ATTCCTCCATCTATTTGCAATTTGACATCACTGGCAATCCTGTATTTGTCGAGAAACAACTTGAAAGGAAAAATTCTGCAATGCTTAGGTAATATCAGTACCCTCCAGTTTGTGATGATG

>M6-PGSC0003DMG400004722M6LRR-RLP_2356

GAAAGGAAAAATTCTGCAATGCTTAGGTAATATCAGTACCCTCCAGTTTGTGATGATGTCACATAATAATCTCAGTGGAGAACTCCCTTTGTCTATTTGCAATCTAACATCACTACAAGT

>M6-PGSC0003DMG400004722M6LRR-RLP_2480

AATTTGGGTAGAAACAATCTGAACGGAGCAATTCCGCAATGTTTTGGCAACATGAGTGGTCATCTTGAGATTCTTGATATGCAGCACAACAATCTTTCTGGTACTTTTCAAAGCACTTTT

>M6-PGSC0003DMG400004722M6LRR-RLP_2542

TCTTGAGATTCTTGATATGCAGCACAACAATCTTTCTGGTACTTTTCAAAGCACTTTTAGCATTGGAGGAGCACTCAGAAGCTTCAATTTGCATGGCAATAAGCTAGAGGGGAAAATTCC

>M6-PGSC0003DMG400004722M6LRR-RLP_2790

TGCATGGACCCATCAGAACTTCAAAGTTCATGAAGAACTTGTTTCCTAAGCTTCGAATCATAGACCTCGCTAGCAATGCATTTACAGCAGAGTTACCTGCAAATCTTTTTCAAAATTTTG

>M6-PGSC0003DMG400004722M6LRR-RLP_2852

GACCTCGCTAGCAATGCATTTACAGCAGAGTTACCTGCAAATCTTTTTCAAAATTTTGAAGCCATGAAGAGAGTTGATCAAACTGTGGAGGATTCAAGTAATAAAGAAGGCGGTTATTAC

>M6-PGSC0003DMG400004722M6LRR-RLP_2914

CATGAAGAGAGTTGATCAAACTGTGGAGGATTCAAGTAATAAAGAAGGCGGTTATTACCAAGACTCCGTAACTGTCGTAACAAAGGGACTGGAGCTTGAAGTTGATTCAATCTTGTCTTT

>M6-PGSC0003DMG400004722M6LRR-RLP_2976

ACTCCGTAACTGTCGTAACAAAGGGACTGGAGCTTGAAGTTGATTCAATCTTGTCTTTGTACATCACAATTGATTTTTCAAATAACAGATTTGAAGGACGCATTCCAAGTATTATGGGAA

>M6-PGSC0003DMG400004722M6LRR-RLP_3162

TCGGAAAATTATCTTCAGTTGAATCATTGGACCTATCAGCTAATCATCTTGAAGGAGAGATACCTAAACAACTTACATCTCTTACATTTCTTGAATTCTTGAATCTCTCCCACAATCAGC

>M6-PGSC0003DMG400004722M6LRR-RLP_3224

CCTAAACAACTTACATCTCTTACATTTCTTGAATTCTTGAATCTCTCCCACAATCAGCTCCAAGGATGCATCCCAACAGGACGCCAATTTGATACATTCGAGAAGAACTCATATGAAGGT

>M6-PGSC0003DMG400004722M6LRR-RLP_3286

AGGATGCATCCCAACAGGACGCCAATTTGATACATTCGAGAAGAACTCATATGAAGGTAATGATAAGTTACGTGGATTTCCTAGTTTGGAGGTTGTGGAAACAGCAGGTTTCCCAAGACA

>M6-PGSC0003DMG400004722M6LRR-RLP_3348

ATAAGTTACGTGGATTTCCTAGTTTGGAGGTTGTGGAAACAGCAGGTTTCCCAAGACAAACAACACGACACATGAGCTGGAACAAGAAAGTGATTCTACATTTCTGAATGAACTCAACTG

>M6-PGSC0003DMG400004722M6LRR-RLP_3410

ACACGACACATGAGCTGGAACAAGAAAGTGATTCTACATTTCTGAATGAACTCAACTGGAAAGTGGTTCTTATGGGATATGTGTGTGGACTTATTATTGGATTCTCCATAACATATTTAA

>M6-PGSC0003DMG400004722M6LRR-RLP_3472

GTGGTTCTTATGGGATATGTGTGTGGACTTATTATTGGATTCTCCATAACATATTTAATGCTTTCAGCTCGAAATCCCAATTGGCTTTCTAGGATTGCTGAAGAACTAGAATACAGAATC

>M6-PGSC0003DMG400004722M6LRR-RLP_434

ACAATCATCTCGATGGATGCATCCCTCAAGTACCTCAATTCGCTACATTAAAGAAATATTCATATGAAGGCAATAATGGATTACGTGAATTTATGCATTGCCAAATTCATCATTGCATAT

>M6-PGSC0003DMG400004722M6LRR-RLP_496

TATGAAGGCAATAATGGATTACGTGAATTTATGCATTGCCAAATTCATCATTGCATATGTTGGAGGAATATCAGGGGCCATAAAATTTTATGAGCTTAAGTTGGCTCGAACATCATAATT

>M6-PGSC0003DMG400004722M6LRR-RLP_558

GAGGAATATCAGGGGCCATAAAATTTTATGAGCTTAAGTTGGCTCGAACATCATAATTATTAAAAAAAATGTCTAAAACCCGTTGACATACTAATGACAAGGTGATATATAGAGTTGAAA

>M6-PGSC0003DMG400004722M6LRR-RLP_62

AGTTCATGGAACATAAGCAGAGATTGCTGCTCTTCGGATGGAGTCGTATGTGATGACATGACTGGCCATGTCACTGAACTTGACCTCAGTCTATTCAGGGACGAGAGCTTGAACTTGTCA

>M6-PGSC0003DMG400004722M6LRR-RLP_620

AAAAAAATGTCTAAAACCCGTTGACATACTAATGACAAGGTGATATATAGAGTTGAAACAAGTTAGATATTTTTTTCCTTTTGTTTTTCAGTAATCACCTCTTATTTGTTTTAATGTCAT

>M6-PGSC0003DMG400004722M6LRR-RLP_682

TTAGATATTTTTTTCCTTTTGTTTTTCAGTAATCACCTCTTATTTGTTTTAATGTCATAAATTCAACTTTCTGATGCAATATAGTTGAAAGTAGTACCTATTAATTTGAAAGTACATAGA

>M6-PGSC0003DMG400004722M6LRR-RLP_744

TCAACTTTCTGATGCAATATAGTTGAAAGTAGTACCTATTAATTTGAAAGTACATAGATTCTGCTTATATGGGTGCAGCTAAAGTATTTATGTACAACTATACTTCCTTTCATGATTCAT

>M6-PGSC0003DMG400004722M6LRR-RLP_806

GCTTATATGGGTGCAGCTAAAGTATTTATGTACAACTATACTTCCTTTCATGATTCATAAATGTACACAACAACCTCATGACCCACTGCAATAGATTATGAAGTACCTTCACTGCAAATT

>M6-PGSC0003DMG400004722M6LRR-RLP_868

GTACACAACAACCTCATGACCCACTGCAATAGATTATGAAGTACCTTCACTGCAAATTGAATGAGTTGTAGCTTTCATTTCTAAGCGCGTGACTTTTCTCTGATAACGAGTGACTTTTCT

>M6-PGSC0003DMG400004722M6LRR-RLP_930

GAGTTGTAGCTTTCATTTCTAAGCGCGTGACTTTTCTCTGATAACGAGTGACTTTTCTGTTACATAACAAATAAGGTGACTTTCAAAGATAATTATCAATAATTGAGTGACAATATGAGA

>M6-PGSC0003DMG400004722M6LRR-RLP_992

CATAACAAATAAGGTGACTTTCAAAGATAATTATCAATAATTGAGTGACAATATGAGAAATCAACTCTTGATGTCAGCTATATGAAATCTCAGTAGTCATATAGCTTTACCTAAACTCAT

>M6-PGSC0003DMG400004725M6LRR-RLP_1008

CTAACATACCTTGAAGACTATTATTACCAAGATTCTATTATAGTCATAAGCAAGGGATATGAGATTGAACTTATCAAAATTTTGACTGTTTACGCCTCTATTGATCTTTCAAGTAACAAA

>M6-PGSC0003DMG400004725M6LRR-RLP_126

AATTCCACAGCCTCTGATTCTAATGTCTGTGAAATATTTTATGGTCAGAAGCCATATCCAAAAACGAGTTCATGGAACATGAGTATAGATTGTTGTTCATGGGATGGAGTCATATGTGAT

>M6-PGSC0003DMG400004725M6LRR-RLP_1260

TCAGTTAACCAGCTTTCGGGAGAGATACCAAAACAACTTGCTTCTCTTAAGTATCTTGAATACTTCTTCACATCAGAAAATAATCTCACTGAAGAAATTCCTTCATCTATTTGCAATTTG

>M6-PGSC0003DMG400004725M6LRR-RLP_1512

TTCAGCAATGGGAGATCACTGAGGAGCCTCAACTTGCATGGCAATAAACTAGAGGGGAATATTCCCCGATCTTTGTCCAATTGCAAAGCGTTGCAGGTTCTTGATTTAGGACATAATCAT

>M6-PGSC0003DMG400004725M6LRR-RLP_1764

GATCTCTCTTACAATGCCTTCTCAGGAAACCTACCTACGAGTTTGTTTCAACATCTGAAAACCATGAGGACAATTGATCCATCCAAGACGGCACTAATTGATGGTTATTATTACCAAGAC

>M6-PGSC0003DMG400004725M6LRR-RLP_1827

ATGAGGACAATTGATCCATCCAAGACGGCACTAATTGATGGTTATTATTACCAAGACTCAGTAGTTGTAGTTACAAAGGGATTGGAGCTTGAAGTTGAGAGAATCTTGTTTTTGTACACC

>M6-PGSC0003DMG400004725M6LRR-RLP_189

ACGAGTTCATGGAACATGAGTATAGATTGTTGTTCATGGGATGGAGTCATATGTGATGAGATGACTGGCCATGTCATTGAACTTGACCTCGGCTGCAGCTGTCTCGTAGGAAAGCTTGAT

>M6-PGSC0003DMG400004725M6LRR-RLP_2205

AAAGGACCTCAATTCAATACATTTGAGATCAATTCATATGAAGGTAACGATGGATTATGTGGATACCCTTTTCCTCAAGGTTGTGGAAGTAGCAGGATGCCAGAGACAGAAAACACAGCA

>M6-PGSC0003DMG400004725M6LRR-RLP_2268

TACCCTTTTCCTCAAGGTTGTGGAAGTAGCAGGATGCCAGAGACAGAAAACACAGCATATGTTCTGGATGAAGAAAGCAATTCTACATTTCTGAGTGAATTTTGGAATGGGATTCTCATG

>M6-PGSC0003DMG400004725M6LRR-RLP_2394

GGAAGTGGACTTATTATTGGATTCTCCATAGCATCTTTCATGCTTTCGAGTCGAAATTCCAATTGGCTTTCTAGGATTGCTGAAGAACTTGAATACAGAATCGTTATGAGAAGGCGAAAG

>M6-PGSC0003DMG400004725M6LRR-RLP_252

ACTGGCCATGTCATTGAACTTGACCTCGGCTGCAGCTGTCTCGTAGGAAAGCTTGATTCCAATAGCAGCCTTTTCCAACTCTCTCATCTCCAAAGGCTCAACCTTTCTGTTAATAACTTC

>M6-PGSC0003DMG400004725M6LRR-RLP_315

AGCAGCCTTTTCCAACTCTCTCATCTCCAAAGGCTCAACCTTTCTGTTAATAACTTCTCTCGTGTGGATTCAATTCCTCTTCAGTCTCTAGATACTATTGATTTGCGGTCCAATTTGCTT

>M6-PGSC0003DMG400004725M6LRR-RLP_378

GTGGATTCAATTCCTCTTCAGTCTCTAGATACTATTGATTTGCGGTCCAATTTGCTTCAAGGTTCACTACCTATTCCACCAATTTCCACGAGATTCTTCCTCATAGCCCATAATAATCTC

>M6-PGSC0003DMG400004725M6LRR-RLP_441

TCACTACCTATTCCACCAATTTCCACGAGATTCTTCCTCATAGCCCATAATAATCTCACTGAAGAAATTCCTTCACATATTTGCAATTTGAAATCATTGGTAATGCTAGATTTGGCGAGA

>M6-PGSC0003DMG400004725M6LRR-RLP_63

CCTCATCCATGCCGCAAAGATCAAAGTATTTCCCTTCTAAAATTCAAGAAAACGTTTACTGTAAATTCCACAGCCTCTGATTCTAATGTCTGTGAAATATTTTATGGTCAGAAGCCATAT

>M6-PGSC0003DMG400004725M6LRR-RLP_945

TTACCTACGAGTCTGTTTCGACATCTGAAAACCATGAGAACAATTGATCCATCAAAGAAGAAACTAACATACCTTGAAGACTATTATTACCAAGATTCTATTATAGTCATAAGCAAGGGA

>M6-PGSC0003DMG400004727M6LRR-RLP_0

TTGGTCTTTCCTTCAATAATTTTAGTGGTTTTGTGGATGTCAGCCTCTTTTCAAACCTTAAACAGCTTGGGGAACTTAGTCTTTCATATAATAATATCTCACTGACCAATGACAACAAAG

>M6-PGSC0003DMG400004727M6LRR-RLP_1178

AGTTTATCTTTGGTTGAATCACTGGACCTGTCAAGTAACCATCTTGTAGGAGATATACCAGCACAATTTGCTTCTCTTACTTCTCTTGCAGTCTTAAATCTTTCCTACAATCATCTCGAA

>M6-PGSC0003DMG400004727M6LRR-RLP_124

TGTTACTTTGCCCGAATCTCTTTATTTGTTAGAATTGGCCGCATGTGAAGTGAAAGAATTGGAGTTTCTAAGATCTGCAAACCAGCTTTTCGATTTGGATCTTTCAAATAATAAGATTCA

>M6-PGSC0003DMG400004727M6LRR-RLP_1364

TTACGTGGTTTTCCTGTTTCAATAAGTTGTGGCAATGATCATGTATCAGGTAAAAATTATGCCGTAGATGATGAAGAAAGCAATTCTGAATTTCTGAGTGACTTTCGGAAAGCTGCTCTT

>M6-PGSC0003DMG400004727M6LRR-RLP_1550

CCGATATGGTTTGCAAGAATCATTGTAGAGCTGGAGCACAAAATAATGATGGGAAGGAGAAAGAAGCAGCGAAGTCAAAGGAATAACAGAAGAAGAAATAATCACTAATCATATTACTGT

>M6-PGSC0003DMG400004727M6LRR-RLP_1612

GAAGCAGCGAAGTCAAAGGAATAACAGAAGAAGAAATAATCACTAATCATATTACTGTTATTTCTTTAAGGAAAAGATGTAATAAGTTGTAGACGCATTATCGTTAAATCCTGTTTGCAT

>M6-PGSC0003DMG400004727M6LRR-RLP_1653

ACTAATCATATTACTGTTATTTCTTTAAGGAAAAGATGTAATAAGTTGTAGACGCATTATCGTTAAATCCTGTTTGCATAAAGTGTGTAGCATCATTTGGCCATGAAAATTGAGGGTGAG

>M6-PGSC0003DMG400004727M6LRR-RLP_186

AGTTTCTAAGATCTGCAAACCAGCTTTTCGATTTGGATCTTTCAAATAATAAGATTCAAGGAAGAATTCCTGATTGGGCATGGTCTAACTGGATGTTTTCATTGCGAGATCTTAATATAT

>M6-PGSC0003DMG400004727M6LRR-RLP_248

AGAATTCCTGATTGGGCATGGTCTAACTGGATGTTTTCATTGCGAGATCTTAATATATCCCACAATATGCTAACGAGTATGGATTCAATTCCTCTTCAACATCATTTAACAATTGATTTG

>M6-PGSC0003DMG400004727M6LRR-RLP_310

CAATATGCTAACGAGTATGGATTCAATTCCTCTTCAACATCATTTAACAATTGATTTGCGGTCAAATTTACTTCAAGGTTCACTACCTATTCCACCAAATTCCACAAGATACTTCTTCAT

>M6-PGSC0003DMG400004727M6LRR-RLP_372

CAAATTTACTTCAAGGTTCACTACCTATTCCACCAAATTCCACAAGATACTTCTTCATATCCCATAATAATCTCACTGGAGAAATTCCTTCATCTATTTGCAACTTGACATCACTGATAA

>M6-PGSC0003DMG400004727M6LRR-RLP_62

CAGCTTGGGGAACTTAGTCTTTCATATAATAATATCTCACTGACCAATGACAACAAAGTTAATGTTACTTTGCCCGAATCTCTTTATTTGTTAGAATTGGCCGCATGTGAAGTGAAAGAA

>M6-PGSC0003DMG400004727M6LRR-RLP_930

GGACAATTCATCAAACGATGAACACACCAAGTTATGAAGGAAGTCGATATTACCAAGACTCGATAGCTGTCGTATCCAAGGGATATGAGCGTGAAATTGTGAGAATCTTGTTTTTGTACA

>M6-PGSC0003DMG400004928-(rev)M6G-LecRK_0

GGATCATTCACTATGGAAATTGGAATGGTATCAACCAGCAGAAAGCAAATGAAAAATTTTCTGCTTATGTTGCACATATTGTTATGTGGTTTTGTATGCTCAATTTCTAGTGCAAAAGAT

>M6-PGSC0003DMG400004928-(rev)M6G-LecRK_1037

AAAGTGGTCCGGCGGGTGTATTAGAAGGACACCCCTTAATTGCTAGAATGGAGATGGATTTCTCAAGTACTCTGTTATAAAGCTTCCAGATACACAGTATTCACGGTTTGATGTGAGAAT

>M6-PGSC0003DMG400004928-(rev)M6G-LecRK_1098

CTCAAGTACTCTGTTATAAAGCTTCCAGATACACAGTATTCACGGTTTGATGTGAGAATGACACTCCGAGAGTGCGAGAAAGTTTGCCTAAAGAATTGTTCTTGTATGGCTTACTCAAAT

>M6-PGSC0003DMG400004928-(rev)M6G-LecRK_1159

CACTCCGAGAGTGCGAGAAAGTTTGCCTAAAGAATTGTTCTTGTATGGCTTACTCAAATCTTGATATACGCAATGGTGGCAGTGGCTGTTTGCTTTGGTATGGCGACTTAATTGACATTC

>M6-PGSC0003DMG400004928-(rev)M6G-LecRK_122

CATCACTACAACTGACTTTCTAATAGAGGGTAAGACCATTGTTTCATGTGATGAAAGCTTTGAATTGGGATTTTTTTCCCCAGGTAGTTTCGCGAATAATTGGTATATTGGAATATGGTA

>M6-PGSC0003DMG400004928-(rev)M6G-LecRK_1220

TGATATACGCAATGGTGGCAGTGGCTGTTTGCTTTGGTATGGCGACTTAATTGACATTCGAGAATTACCTGGAGGACAGGACATTTACATAAGGATAGCTAACTCAGAGTTAGGTAAAAT

>M6-PGSC0003DMG400004928-(rev)M6G-LecRK_1281

GAATTACCTGGAGGACAGGACATTTACATAAGGATAGCTAACTCAGAGTTAGGTAAAATTTATGCAACAAATTCATTACTATTTGATCAATATGAAGCATTAGCTGAATGAAGTAAACTT

>M6-PGSC0003DMG400004928-(rev)M6G-LecRK_1342

ATGCAACAAATTCATTACTATTTGATCAATATGAAGCATTAGCTGAATGAAGTAAACTTCATAATTCTTAGCTTTCGTTGTGTTTTGCTTATCTATTTGTTAAAAAGTTCACATAGCATA

>M6-PGSC0003DMG400004928-(rev)M6G-LecRK_1403

TAATTCTTAGCTTTCGTTGTGTTTTGCTTATCTATTTGTTAAAAAGTTCACATAGCATATATTTATTTTAGTTTACCCTCCATTTTTTTTCAGGCTCGAAGAAGAAAATGAAAACACTTG

>M6-PGSC0003DMG400004928-(rev)M6G-LecRK_1464

TTTATTTTAGTTTACCCTCCATTTTTTTTCAGGCTCGAAGAAGAAAATGAAAACACTTGTGCTGAGTCTATCATTGTTGAGCGGAGTGTCTGTCATTGGCCTGATCATAGGGCTGTATAT

>M6-PGSC0003DMG400004928-(rev)M6G-LecRK_1525

CTGAGTCTATCATTGTTGAGCGGAGTGTCTGTCATTGGCCTGATCATAGGGCTGTATATTTTGATAAAGGCGAGAAAGAAGAAGAGGAAGATGATTCTGAAAGATGATCTAGAACTTCCA

>M6-PGSC0003DMG400004928-(rev)M6G-LecRK_1586

TGATAAAGGCGAGAAAGAAGAAGAGGAAGATGATTCTGAAAGATGATCTAGAACTTCCACTGTTTTCCCTATCAACAGTAACTAAAGCGACCTCTAATTTCTCAGACAAGAACATGCTCG

>M6-PGSC0003DMG400004928-(rev)M6G-LecRK_1647

GTTTTCCCTATCAACAGTAACTAAAGCGACCTCTAATTTCTCAGACAAGAACATGCTCGGAGAAGGTGGATTTGGATCTGTTTACAAGGTAGCATATCTTGTAGGTTTATTATATGTACT

>M6-PGSC0003DMG400004928-(rev)M6G-LecRK_1708

GAAGGTGGATTTGGATCTGTTTACAAGGTAGCATATCTTGTAGGTTTATTATATGTACTATAATTGACAGTAAACAGATAAGTAACACTTTGAATTATCCCTGTCAAAACATCCTTAGCT

>M6-PGSC0003DMG400004928-(rev)M6G-LecRK_1769

AATTGACAGTAAACAGATAAGTAACACTTTGAATTATCCCTGTCAAAACATCCTTAGCTTCAATATGAGTCAGCTGTATGTTATGATATCGTAGGGAATCCTGGAGGAAGGACAAGAAGT

>M6-PGSC0003DMG400004928-(rev)M6G-LecRK_183

GAATTGGGATTTTTTTCCCCAGGTAGTTTCGCGAATAATTGGTATATTGGAATATGGTACAAGCACGACGATGGGCCTGACAAGTCTGTTGTATGGGTTGCCAATAGAGAAACTCCACTC

>M6-PGSC0003DMG400004928-(rev)M6G-LecRK_1830

AATATGAGTCAGCTGTATGTTATGATATCGTAGGGAATCCTGGAGGAAGGACAAGAAGTAGCTATAAAGAGGCTCTCAAAATCTTCCTTGCAAGGAGTCAACGAGTTCAAGAATGAAGTT

>M6-PGSC0003DMG400004928-(rev)M6G-LecRK_1891

CTATAAAGAGGCTCTCAAAATCTTCCTTGCAAGGAGTCAACGAGTTCAAGAATGAAGTTAATTGCATTGCCAAACTTCAGCATCGAAACCTTGTGATGCTTATAGGTTGCTGCGTTGAAG

>M6-PGSC0003DMG400004928-(rev)M6G-LecRK_1952

TTGCATTGCCAAACTTCAGCATCGAAACCTTGTGATGCTTATAGGTTGCTGCGTTGAAGGAGAAGAGAAAATTCTGGTCTATGAATACATGCCCAATAGAAGTCTTGATTTCTTCATATT

>M6-PGSC0003DMG400004928-(rev)M6G-LecRK_2013

GAAGAGAAAATTCTGGTCTATGAATACATGCCCAATAGAAGTCTTGATTTCTTCATATTTGGTATGCTTACTCTTGCTTAACACATAAAACCTCAAGTACTGTTCATAGTATAGCATTGA

>M6-PGSC0003DMG400004928-(rev)M6G-LecRK_2074

GTATGCTTACTCTTGCTTAACACATAAAACCTCAAGTACTGTTCATAGTATAGCATTGAGCTTGCAGCTTTCACCATGATTTTGAAAAAGGTTGTTAATTTCAAATGGAAGTAGATTCTG

>M6-PGSC0003DMG400004928-(rev)M6G-LecRK_2135

TTGCAGCTTTCACCATGATTTTGAAAAAGGTTGTTAATTTCAAATGGAAGTAGATTCTGTTTAACATACTAAAAGCAGTTAGAGCATCAGTTACTATACTAATTGTTGGAAGATCTGTTA

>M6-PGSC0003DMG400004928-(rev)M6G-LecRK_2196

TAACATACTAAAAGCAGTTAGAGCATCAGTTACTATACTAATTGTTGGAAGATCTGTTAACAGATGAAGGGAAGAGCTCGATACTCGATTGGCCTAAGCGCTTCAACATCATCAATGGAA

>M6-PGSC0003DMG400004928-(rev)M6G-LecRK_2257

AGATGAAGGGAAGAGCTCGATACTCGATTGGCCTAAGCGCTTCAACATCATCAATGGAATTGTTAGAGGACTTTTGTATCTCCATCAAGATTCTCGGCTAAGAATTATCCATAGAGACCT

>M6-PGSC0003DMG400004928-(rev)M6G-LecRK_2318

GTTAGAGGACTTTTGTATCTCCATCAAGATTCTCGGCTAAGAATTATCCATAGAGACCTCAAAGCTAGTAACATTTTGCTAGATGCTGACATGAACCCCAAGATCTCAGATTTTGGCATA

>M6-PGSC0003DMG400004928-(rev)M6G-LecRK_244

AGCACGACGATGGGCCTGACAAGTCTGTTGTATGGGTTGCCAATAGAGAAACTCCACTCAACAATACATCAGGTGTTATGCTGAAAATCGTTGATCCAGGACGACTTGCTCTTGTTACTT

>M6-PGSC0003DMG400004928-(rev)M6G-LecRK_2440

TAGAAGTGTGATAGGGAATGAGACGGGTGCTAATACACATCATGTGGTTGGGACACAGTAAGTGATTTCACACTGCGACCTTTTAGCATTTATCCCATGTACTACTCATATAATATCATC

>M6-PGSC0003DMG400004928-(rev)M6G-LecRK_2501

GTGATTTCACACTGCGACCTTTTAGCATTTATCCCATGTACTACTCATATAATATCATCTGTCTAACACTATTTTCTGTCTAATATGTGATGCAGTGGCTACATGTCCCCAGAATATCTA

>M6-PGSC0003DMG400004928-(rev)M6G-LecRK_2562

TCTAACACTATTTTCTGTCTAATATGTGATGCAGTGGCTACATGTCCCCAGAATATCTAGTACACGGTGTTTTCTCAATAAAATCAGATGTATTTAGCTTTGGCGTGTTAGTTCTAGAGA

>M6-PGSC0003DMG400004928-(rev)M6G-LecRK_2623

ACACGGTGTTTTCTCAATAAAATCAGATGTATTTAGCTTTGGCGTGTTAGTTCTAGAGATAATCAGTGGTCGAAGAAACAGAGGATTTGCCCGTGAAAGTCACAGTATCAACCTTCTCGG

>M6-PGSC0003DMG400004928-(rev)M6G-LecRK_2684

ATCAGTGGTCGAAGAAACAGAGGATTTGCCCGTGAAAGTCACAGTATCAACCTTCTCGGACATGTAAGTCTTGTATGTTAAAAAGTACTTACATATTCAGACCACTCGTAAGGTAATTAC

>M6-PGSC0003DMG400004928-(rev)M6G-LecRK_2745

ATGTAAGTCTTGTATGTTAAAAAGTACTTACATATTCAGACCACTCGTAAGGTAATTACAAGTGATAGTGTAACTAAACTGATAGTGCAAAAAGTCTTACTAAGTGTCGGTGTATACAAC

>M6-PGSC0003DMG400004928-(rev)M6G-LecRK_2806

GTGATAGTGTAACTAAACTGATAGTGCAAAAAGTCTTACTAAGTGTCGGTGTATACAACTTAAATCTATCAAACATGCTTCACTTACTTATACATGTTCCTTATTACTATATACTGATTG

>M6-PGSC0003DMG400004928-(rev)M6G-LecRK_2867

AAATCTATCAAACATGCTTCACTTACTTATACATGTTCCTTATTACTATATACTGATTGTGTTATTAACAGGTATGGAAACTTTATAAAGAAGGTAGACCATTGGAGCTGATTGATGTGC

>M6-PGSC0003DMG400004928-(rev)M6G-LecRK_2928

TTATTAACAGGTATGGAAACTTTATAAAGAAGGTAGACCATTGGAGCTGATTGATGTGCATATTACAGATTCATGTTATTTTACTGAAGTGCTACGGTTAATTCATGTCGCTCTATTATG

>M6-PGSC0003DMG400004928-(rev)M6G-LecRK_2989

ATTACAGATTCATGTTATTTTACTGAAGTGCTACGGTTAATTCATGTCGCTCTATTATGCGTACAACATAGTCCAGAAGACAGGCCAGACATGTCTGAAGTGGTTGTGATGTTGGCTAAT

>M6-PGSC0003DMG400004928-(rev)M6G-LecRK_305

CAATACATCAGGTGTTATGCTGAAAATCGTTGATCCAGGACGACTTGCTCTTGTTACTTCTGCTAATATGAGCATATGGTCTACTAATTCGTCAAGGATTTTGTCTGTGAAGAATCCAAT

>M6-PGSC0003DMG400004928-(rev)M6G-LecRK_3050

TACAACATAGTCCAGAAGACAGGCCAGACATGTCTGAAGTGGTTGTGATGTTGGCTAATGATACCATTTTGCCTGAAGCTAAGGAGCCTGGTTTTTTCACAGAAAGTAAATTCCCTGATT

>M6-PGSC0003DMG400004928-(rev)M6G-LecRK_3111

TACCATTTTGCCTGAAGCTAAGGAGCCTGGTTTTTTCACAGAAAGTAAATTCCCTGATTCTGAATATTCTACAAGCATGTATTCGAAAAATGAAATCACTATTACACAGTTAGATCCCCG

>M6-PGSC0003DMG400004928-(rev)M6G-LecRK_3172

GAATATTCTACAAGCATGTATTCGAAAAATGAAATCACTATTACACAGTTAGATCCCCGGTAAATTAAATTTCATTTTCTTGTAGATCTTTAAGTGTCTAGAAGCTTGAATAACAGAGGA

>M6-PGSC0003DMG400004928-(rev)M6G-LecRK_3233

AAATTAAATTTCATTTTCTTGTAGATCTTTAAGTGTCTAGAAGCTTGAATAACAGAGGAGAACATTTTATGAGTTGTATCATATTAGGAAAGAATTTACCTAACTCTTGAATTAAAAAAA

>M6-PGSC0003DMG400004928-(rev)M6G-LecRK_3416

TTGTACGACAGTTTTACAGGAACTTGAGTGTTAGTTACAAATTGCCATTGTTGTGTATGTTACATGGTTTTGTTATACTTACAGATACGTATCCACTCTGGGTGTGTGTAAGAATCTAAT

>M6-PGSC0003DMG400004928-(rev)M6G-LecRK_3458

TGCCATTGTTGTGTATGTTACATGGTTTTGTTATACTTACAGATACGTATCCACTCTGGGTGTGTGTAAGAATCTAATATGAATACTACAGTTTTTTTTGAAATTTTAATTTACTCTCCA

>M6-PGSC0003DMG400004928-(rev)M6G-LecRK_366

GCTAATATGAGCATATGGTCTACTAATTCGTCAAGGATTTTGTCTGTGAAGAATCCAATTGCGCAGTTGTTGAGTTCAGGGAATCTAATCATAAGAGATGCAAATGAAACTAAAACAGAG

>M6-PGSC0003DMG400004928-(rev)M6G-LecRK_427

CGCAGTTGTTGAGTTCAGGGAATCTAATCATAAGAGATGCAAATGAAACTAAAACAGAGAACTTCCTGTGGCAGAGTTTTGATTATCCGACTAATACGCTCTTACCAGGCATGCAACTTG

>M6-PGSC0003DMG400004928-(rev)M6G-LecRK_488

CTTCCTGTGGCAGAGTTTTGATTATCCGACTAATACGCTCTTACCAGGCATGCAACTTGGTAAGAACTTTGTGACTGGACAAGAATTTTACCTGTCCTCATGGAAGAACGAATATGATCC

>M6-PGSC0003DMG400004928-(rev)M6G-LecRK_549

AAGAACTTTGTGACTGGACAAGAATTTTACCTGTCCTCATGGAAGAACGAATATGATCCGGCTCCAGGGAAATATACGTATCATTGTGATCCTACTGGATATCCTCAGGTTGTCATGAGG

>M6-PGSC0003DMG400004928-(rev)M6G-LecRK_61

TGCTTATGTTGCACATATTGTTATGTGGTTTTGTATGCTCAATTTCTAGTGCAAAAGATACCATCACTACAACTGACTTTCTAATAGAGGGTAAGACCATTGTTTCATGTGATGAAAGCT

>M6-PGSC0003DMG400004928-(rev)M6G-LecRK_610

CTCCAGGGAAATATACGTATCATTGTGATCCTACTGGATATCCTCAGGTTGTCATGAGGAAAGGTAAAGTTAAGGTGTTCAGTGGCGGACCATGGAATGGACTCCGCTGGAGTGGTGTAC

>M6-PGSC0003DMG400004928-(rev)M6G-LecRK_671

AGGTAAAGTTAAGGTGTTCAGTGGCGGACCATGGAATGGACTCCGCTGGAGTGGTGTACTAGGATTGACAAAAAATACAATTTATACATTTAAGCTGGATTTTGATGAAAAGAAAGCAAT

>M6-PGSC0003DMG400004928-(rev)M6G-LecRK_732

GGATTGACAAAAAATACAATTTATACATTTAAGCTGGATTTTGATGAAAAGAAAGCAATTTACAGCTACGCGCTAGTGGATAATTCTGTTTTATCGAAATTAACCTTGAACAGTCATGGT

>M6-PGSC0003DMG400004928-(rev)M6G-LecRK_793

ACAGCTACGCGCTAGTGGATAATTCTGTTTTATCGAAATTAACCTTGAACAGTCATGGTATGCTTAAGCGTTGGACGTGGGATGAGAAGATAAAGGAATGGCATGTTTACCTGGCTTCAC

>M6-PGSC0003DMG400004928-(rev)M6G-LecRK_854

GCTTAAGCGTTGGACGTGGGATGAGAAGATAAAGGAATGGCATGTTTACCTGGCTTCACCAGCAGATGCCTGTGAAAATTATGGAGCATGTGGTGCCTATGGTAGTTGCAACATCATATT

>M6-PGSC0003DMG400004928-(rev)M6G-LecRK_915

GCAGATGCCTGTGAAAATTATGGAGCATGTGGTGCCTATGGTAGTTGCAACATCATATTTTCTCCTGTTTGTCGTTGTTTAGACAAGTTTGTGCCTAAAGATCTGAAAAATTGGTCAGTG

>M6-PGSC0003DMG400004928-(rev)M6G-LecRK_976

CTCCTGTTTGTCGTTGTTTAGACAAGTTTGTGCCTAAAGATCTGAAAAATTGGTCAGTGACAAAGTGGTCCGGCGGGTGTATTAGAAGGACACCCCTTAATTGCTAGAATGGAGATGGAT

>M6-PGSC0003DMG400004930-(rev)M6G-LecRK_0

ATTAAAAGGCTTAGGCATCAAATACATTGATGGAATCTAAATACTTCATACTTTTCTTACTATCTTTCTACTCTTTTATACCGAAGATTAGTTCTACTACGACAAATGTAATCAATACTA

>M6-PGSC0003DMG400004930-(rev)M6G-LecRK_1037

TGTGTTAGGAAGACACCTATTGATTGCAACAAGGAACATGGATTTCTAAAATATTCTGGAATCAAGTTGCCAGAAACTAACAACTCTCAGTACAATAAGACCATGACACTCGAAGGATGT

>M6-PGSC0003DMG400004930-(rev)M6G-LecRK_1098

TCAAGTTGCCAGAAACTAACAACTCTCAGTACAATAAGACCATGACACTCGAAGGATGTAGGCAAGTATGCTCGACGAACTGTTCTTGCACAGCTTATTCGAGCCTAGATATAAGCAATG

>M6-PGSC0003DMG400004930-(rev)M6G-LecRK_1159

GCAAGTATGCTCGACGAACTGTTCTTGCACAGCTTATTCGAGCCTAGATATAAGCAATGGAGATAAAGGTTGCTTGTTTTGGTCTGGGGAGTTGATTGATATCAGAGAGCTCTCTGGAAG

>M6-PGSC0003DMG400004930-(rev)M6G-LecRK_122

CGGTTCATAACAGATGGTGACACTATTGTTTCATCTGGTGGAACCTTTGAGCTGGGATTTTTCAGCCCCAATGGTTCCACAACGCGATATATTGGGATATGGTACAAGCAAATTCTTCCT

>M6-PGSC0003DMG400004930-(rev)M6G-LecRK_1220

GATAAAGGTTGCTTGTTTTGGTCTGGGGAGTTGATTGATATCAGAGAGCTCTCTGGAAGAGGACAAGACATTTACATACGAATGGATTCTTCAGACCTAGGTAAGAACTTATATTAACGA

>M6-PGSC0003DMG400004930-(rev)M6G-LecRK_1281

GACAAGACATTTACATACGAATGGATTCTTCAGACCTAGGTAAGAACTTATATTAACGAAATTTCTAATGATGTGTTTTTGAACTTTTAAATTTTTGAACATATGTAGACCATATTATAT

>M6-PGSC0003DMG400004930-(rev)M6G-LecRK_1342

TTTCTAATGATGTGTTTTTGAACTTTTAAATTTTTGAACATATGTAGACCATATTATATATGCAAAAGTAGAGGCCTATATCAATTCGAAGAAGCTAATTTAGAAAATGGTTGTAGTATC

>M6-PGSC0003DMG400004930-(rev)M6G-LecRK_1403

GCAAAAGTAGAGGCCTATATCAATTCGAAGAAGCTAATTTAGAAAATGGTTGTAGTATCTCAGGCAAGTTCAAACAGAAAGAAGACAGGGACAGTACTCGCGGTGAGTTTCTCGTTGTTG

>M6-PGSC0003DMG400004930-(rev)M6G-LecRK_1464

AGGCAAGTTCAAACAGAAAGAAGACAGGGACAGTACTCGCGGTGAGTTTCTCGTTGTTGGTGGCAGTGATTCTGCTAGGCCTGATTTTGTTCATGTACATACGTAAAAAGAAGAAACTAA

>M6-PGSC0003DMG400004930-(rev)M6G-LecRK_1525

GGCAGTGATTCTGCTAGGCCTGATTTTGTTCATGTACATACGTAAAAAGAAGAAACTAAAACTCAAAGAAGATTTTGAGCTGCCACAGTTTCAATTGTCGTTAATAACAAGAGCCACAGA

>M6-PGSC0003DMG400004930-(rev)M6G-LecRK_1586

CTCAAAGAAGATTTTGAGCTGCCACAGTTTCAATTGTCGTTAATAACAAGAGCCACAGATAACTTTTCGGTCAACAACCAGATTGGAGAAGGTGGATATGGACCTGTTTATAAGGTATAA

>M6-PGSC0003DMG400004930-(rev)M6G-LecRK_1647

ACTTTTCGGTCAACAACCAGATTGGAGAAGGTGGATATGGACCTGTTTATAAGGTATAAACTAAAAATTATTGGAATAGTGACCAGGCTAGAGATGACTTTCATTCTGTAATACTCGCTC

>M6-PGSC0003DMG400004930-(rev)M6G-LecRK_1708

TAAAAATTATTGGAATAGTGACCAGGCTAGAGATGACTTTCATTCTGTAATACTCGCTCGTCTTTGTAAAATATATTTCCTTCTGATATACTTTTTCCATTTCAAGGGAGTGCTCGAAGA

>M6-PGSC0003DMG400004930-(rev)M6G-LecRK_1769

CTTTGTAAAATATATTTCCTTCTGATATACTTTTTCCATTTCAAGGGAGTGCTCGAAGAGGGACAAGAAATTGCTGTGAAGAGACTTTCAAGGACTTCCATGCAAGGAATTGATGAATTC

>M6-PGSC0003DMG400004930-(rev)M6G-LecRK_183

TCAGCCCCAATGGTTCCACAACGCGATATATTGGGATATGGTACAAGCAAATTCTTCCTTATGTCCAAACAATTGTATGGGTTGCAAACAGAGAGAAACCACTCACCAATACATCTTCAG

>M6-PGSC0003DMG400004930-(rev)M6G-LecRK_1830

GACAAGAAATTGCTGTGAAGAGACTTTCAAGGACTTCCATGCAAGGAATTGATGAATTCAAGAATGAAGTTACCTATATTGCCAAGCTTCAGCATAGAAATCTTGTGAGACTTCTGGGTT

>M6-PGSC0003DMG400004930-(rev)M6G-LecRK_1891

GAATGAAGTTACCTATATTGCCAAGCTTCAGCATAGAAATCTTGTGAGACTTCTGGGTTGCTGTATTCAAGGGGAAGAAAAGATGTTGATCTATGAATACATGCCTAATAAAAGCCTGGA

>M6-PGSC0003DMG400004930-(rev)M6G-LecRK_1952

TGTATTCAAGGGGAAGAAAAGATGTTGATCTATGAATACATGCCTAATAAAAGCCTGGACTCATACATATTTGGTTTGGTTTTCTAAACTATTCTACAAGCTAAAATGACATGTTTTCAA

>M6-PGSC0003DMG400004930-(rev)M6G-LecRK_2013

CATACATATTTGGTTTGGTTTTCTAAACTATTCTACAAGCTAAAATGACATGTTTTCAAAAGTTGCTAACAAGAAAGCTTTTCTTCTTTTTTTTTTGTAGATCAAACAAAGAAAAAGTTA

>M6-PGSC0003DMG400004930-(rev)M6G-LecRK_2074

GTTGCTAACAAGAAAGCTTTTCTTCTTTTTTTTTTGTAGATCAAACAAAGAAAAAGTTACTTGATTGGTCAAGGCGTTTCGACATCATTAATGGAATTGCTCGTGGCTTATTGTATCTCC

>M6-PGSC0003DMG400004930-(rev)M6G-LecRK_2135

TGATTGGTCAAGGCGTTTCGACATCATTAATGGAATTGCTCGTGGCTTATTGTATCTCCATCAAGATTCTAGACTACGTATTATCCATAGAGACCTTAAAGCAAGCAATGTTTTGTTAGA

>M6-PGSC0003DMG400004930-(rev)M6G-LecRK_2196

CAAGATTCTAGACTACGTATTATCCATAGAGACCTTAAAGCAAGCAATGTTTTGTTAGATACAGAAATGAATCCAAAGATATCAGACTTTGGGATGGCTCGAAGTGTTGCAGGAAATGAT

>M6-PGSC0003DMG400004930-(rev)M6G-LecRK_2257

CAGAAATGAATCCAAAGATATCAGACTTTGGGATGGCTCGAAGTGTTGCAGGAAATGATATGGGAGCAAAAACATGCCATGTAGTTGGGACACAGTAAGAATCGATCATACAGAAGTACT

>M6-PGSC0003DMG400004930-(rev)M6G-LecRK_2318

GGGAGCAAAAACATGCCATGTAGTTGGGACACAGTAAGAATCGATCATACAGAAGTACTCTCATATGAAATCCCCCTTCTCATGGTGTAGTTCTTTTTTTGGAACAGCGGTTACATGTCC

>M6-PGSC0003DMG400004930-(rev)M6G-LecRK_2379

CATATGAAATCCCCCTTCTCATGGTGTAGTTCTTTTTTTGGAACAGCGGTTACATGTCCCCTGAATATGCAGTAGATGGAATCTTCTCAGTAAAATCAGATGTGTTTAGCTTTGGCGTAT

>M6-PGSC0003DMG400004930-(rev)M6G-LecRK_244

TGTCCAAACAATTGTATGGGTTGCAAACAGAGAGAAACCACTCACCAATACATCTTCAGTCGTTTTGAAGGTCAACAAGCCGGGAATACTTGCTCTTCTTAATGACAAGAATGAAACTAT

>M6-PGSC0003DMG400004930-(rev)M6G-LecRK_2440

TGAATATGCAGTAGATGGAATCTTCTCAGTAAAATCAGATGTGTTTAGCTTTGGCGTATTGGTGTTAGAGATTGTGAGTTGCAAGAAAAATAGAGGATTTGTCCATCAAGATCACAACCT

>M6-PGSC0003DMG400004930-(rev)M6G-LecRK_2501

GTGTTAGAGATTGTGAGTTGCAAGAAAAATAGAGGATTTGTCCATCAAGATCACAACCTTAACCTTCTCGGTCACGTAAGCACTCTCTCTATACATTCTTGTGATCTATATTACATATAG

>M6-PGSC0003DMG400004930-(rev)M6G-LecRK_2562

ACCTTCTCGGTCACGTAAGCACTCTCTCTATACATTCTTGTGATCTATATTACATATAGAATTAATTTGTCATGATCTAAATTACTCAAACAGGCATGGAAGCTTTACAAGGAAGATAGG

>M6-PGSC0003DMG400004930-(rev)M6G-LecRK_2623

TTAATTTGTCATGATCTAAATTACTCAAACAGGCATGGAAGCTTTACAAGGAAGATAGGTCCTTGGAACTAATTGATGAGAAGCTAGCTGAATCTTGTCATATTTCTCAAGTTTTAAGGT

>M6-PGSC0003DMG400004930-(rev)M6G-LecRK_2684

CTTGGAACTAATTGATGAGAAGCTAGCTGAATCTTGTCATATTTCTCAAGTTTTAAGGTCAATTCAAGTGGGACTATTATGTGTGCAACACTGTCCTGAAGATAGACCAAACATGTCTTC

>M6-PGSC0003DMG400004930-(rev)M6G-LecRK_2745

ATTCAAGTGGGACTATTATGTGTGCAACACTGTCCTGAAGATAGACCAAACATGTCTTCTGTGGTTCAAATGCTAGGTAATGAGAGTTTACTTGCAAAAGCTAAAGAACCAGGATTTTTC

>M6-PGSC0003DMG400004930-(rev)M6G-LecRK_2806

TGGTTCAAATGCTAGGTAATGAGAGTTTACTTGCAAAAGCTAAAGAACCAGGATTTTTCATGGAAAGAACGTTACATGATTCAGATAATTCAGGAATGCAGACAGGAAGTTCGAAAAATG

>M6-PGSC0003DMG400004930-(rev)M6G-LecRK_2867

GGAAAGAACGTTACATGATTCAGATAATTCAGGAATGCAGACAGGAAGTTCGAAAAATGAAATAACAATGACAATGTTAGATCCTCGGTAGAGTACTGTGATTGAGGCAACTACTGTTGA

>M6-PGSC0003DMG400004930-(rev)M6G-LecRK_2928

ATAACAATGACAATGTTAGATCCTCGGTAGAGTACTGTGATTGAGGCAACTACTGTTGATAAGCATGAGTTTTTAGAGGGCATGATAACAACATTGAAGCGCCTGAGAAGCGCTCAAGCA

>M6-PGSC0003DMG400004930-(rev)M6G-LecRK_2989

AGCATGAGTTTTTAGAGGGCATGATAACAACATTGAAGCGCCTGAGAAGCGCTCAAGCATCACCTCAGGGATTAAGCGTGCCTTTGAGTATTTAGAAAAATGGTTCATAATTGCCATGCT

>M6-PGSC0003DMG400004930-(rev)M6G-LecRK_305

GTTTTGAAGGTCAACAAGCCGGGAATACTTGCTCTTCTTAATGACAAGAATGAAACTATATGGTCCACGAACACCTCACGATCAGTCCAAAATCCAGTTGCAGTGTTGTTGGATTCTGGT

>M6-PGSC0003DMG400004930-(rev)M6G-LecRK_3050

ACCTCAGGGATTAAGCGTGCCTTTGAGTATTTAGAAAAATGGTTCATAATTGCCATGCTAGCTATTATAAGTAGCTATTCAACTTGAATTTTCATTTGTATCTTATCTTAGAAAGAAATG

>M6-PGSC0003DMG400004930-(rev)M6G-LecRK_3111

CTATTATAAGTAGCTATTCAACTTGAATTTTCATTTGTATCTTATCTTAGAAAGAAATGTAAATTCTAGTAAGCAAAAATGTCCAAATTATATTGCCTATGACAATACATAAAATCGAAT

>M6-PGSC0003DMG400004930-(rev)M6G-LecRK_3171

AAATTCTAGTAAGCAAAAATGTCCAAATTATATTGCCTATGACAATACATAAAATCGAATCAAACCGACAGATGCATGAGCTCTCCAATATCTGCATGGAATAGTCCAATATGTTGAAAT

>M6-PGSC0003DMG400004930-(rev)M6G-LecRK_366

GGTCCACGAACACCTCACGATCAGTCCAAAATCCAGTTGCAGTGTTGTTGGATTCTGGTAATCTTGTCCTAAAAGATGCAAATGATGACAATCCAGAAAATTTCCTATGGCAGAGTTTCA

>M6-PGSC0003DMG400004930-(rev)M6G-LecRK_427

TCTTGTCCTAAAAGATGCAAATGATGACAATCCAGAAAATTTCCTATGGCAGAGTTTCAATTTTCCAACTGATACTTTCTTGCCTGATATGAAACTCGGAAAGAATTTTAAGACTGGTAT

>M6-PGSC0003DMG400004930-(rev)M6G-LecRK_488

TTTCCAACTGATACTTTCTTGCCTGATATGAAACTCGGAAAGAATTTTAAGACTGGTATTGAAGTTTACCTTTTAGCATGGAAGAACGATAACGATCCAACTCCAGGGGAATACAGTCTC

>M6-PGSC0003DMG400004930-(rev)M6G-LecRK_549

AAGTTTACCTTTTAGCATGGAAGAACGATAACGATCCAACTCCAGGGGAATACAGTCTCCACATTGATCCTACTGGATATCCACAAGGCCTCATTAGACATGGGGCAAGGGTATCAGCTC

>M6-PGSC0003DMG400004930-(rev)M6G-LecRK_61

ATCTTTCTACTCTTTTATACCGAAGATTAGTTCTACTACGACAAATGTAATCAATACTAATCGGTTCATAACAGATGGTGACACTATTGTTTCATCTGGTGGAACCTTTGAGCTGGGATT

>M6-PGSC0003DMG400004930-(rev)M6G-LecRK_610

CATTGATCCTACTGGATATCCACAAGGCCTCATTAGACATGGGGCAAGGGTATCAGCTCGAGCAGGACCATGGAATGGTTTACGGTGGAGTGGAGCACCTGCACCACTACAAACACAAAG

>M6-PGSC0003DMG400004930-(rev)M6G-LecRK_671

GCAGGACCATGGAATGGTTTACGGTGGAGTGGAGCACCTGCACCACTACAAACACAAAGCAGCATATATACTTTTCAATTTGTTTTCAATGAAGAGGAGGTTTACTATAGTTTTTCTCTC

>M6-PGSC0003DMG400004930-(rev)M6G-LecRK_732

GCATATATACTTTTCAATTTGTTTTCAATGAAGAGGAGGTTTACTATAGTTTTTCTCTCATTAACAACTCACTGTTAACAAGGTTAGTCCTGACTAACAATGGTTTTATACAACGTTTGA

>M6-PGSC0003DMG400004930-(rev)M6G-LecRK_793

TAACAACTCACTGTTAACAAGGTTAGTCCTGACTAACAATGGTTTTATACAACGTTTGACGTGGGTGGATAGGACAAAGAGTTGGCATCTTTACCTCAATATTCCTCTGGATACTTGTGA

>M6-PGSC0003DMG400004930-(rev)M6G-LecRK_854

TGGGTGGATAGGACAAAGAGTTGGCATCTTTACCTCAATATTCCTCTGGATACTTGTGATACATATAGCTTATGTGGTGCATATGGGAACTGTGTCATAGATAGTTCTCCTGTTTGTGGA

>M6-PGSC0003DMG400004930-(rev)M6G-LecRK_915

CATATAGCTTATGTGGTGCATATGGGAACTGTGTCATAGATAGTTCTCCTGTTTGTGGATGTTTGGACAAGTTTGAACCTAAATATCCACAAAATTGGCAAACAGGAGACTGGTCACAAG

>M6-PGSC0003DMG400004930-(rev)M6G-LecRK_976

TTTGGACAAGTTTGAACCTAAATATCCACAAAATTGGCAAACAGGAGACTGGTCACAAGGGTGTGTTAGGAAGACACCTATTGATTGCAACAAGGAACATGGATTTCTAAAATATTCTGG

>M6-PGSC0003DMG400004963M6LRR-RLK_0

CTTTTATTTCCTTCCTCTCTAAGAAAAAAACAATTTAATACAAATGACTATTCAAAAAATTATCACCCTTGTGTTATTTTTTCTTACTTTGACGTCCACTTTTGTTCAAGGTGATCAATC

>M6-PGSC0003DMG400004963M6LRR-RLK_1054

AATCGATTTTCTGGAAAGCTACCGCCTTTTCTGTGTAGTGGAGGTAAATTGAGTTATATTCTTTTACTTCAAAACATGTTCTCAGGTGAACTGCCTGATGGATATGTGAAATGTCAGTCT

>M6-PGSC0003DMG400004963M6LRR-RLK_1116

TTTACTTCAAAACATGTTCTCAGGTGAACTGCCTGATGGATATGTGAAATGTCAGTCTGTTCTTCGCTTTCGAGTGAATTACAATCAGTTGGAGGGAAGAATACCACAAGAGCTTTTTAC

>M6-PGSC0003DMG400004963M6LRR-RLK_1178

TTCGCTTTCGAGTGAATTACAATCAGTTGGAGGGAAGAATACCACAAGAGCTTTTTACTCTTCCACATGTTTCAATCATTGATTTGAGTTATAATCATTTTAGTGGTCCAATTCCAACAA

>M6-PGSC0003DMG400004963M6LRR-RLK_124

TTCTTTGTACTCATGAAGAAATTTGTCACAGGAAGTTCTTTGTCTAATTGGGATATTGGAAAACCAATTTGTCAATACAAAGGTGTCGGTTGTGATGAAAGGGGAGACGTTATCAAAATT

>M6-PGSC0003DMG400004963M6LRR-RLK_1240

CCACATGTTTCAATCATTGATTTGAGTTATAATCATTTTAGTGGTCCAATTCCAACAACAATTGGAAGTGCTAGGAATTTATCAGAATTGTTTATGCAAAGTAACAAGCTTTCAGGTTTG

>M6-PGSC0003DMG400004963M6LRR-RLK_1302

TGGAAGTGCTAGGAATTTATCAGAATTGTTTATGCAAAGTAACAAGCTTTCAGGTTTGCTTCCTCATGAAATATCTACATCTTCTAATCTTGTGAAGCTTGATCTTAGCAATAACCTTTT

>M6-PGSC0003DMG400004963M6LRR-RLK_1364

CTCATGAAATATCTACATCTTCTAATCTTGTGAAGCTTGATCTTAGCAATAACCTTTTGTATGGTCCAATTCCTTCTGAAATTGGTGGTTTAAAAAGCCTCAATTTGTTACTCTTGCAAG

>M6-PGSC0003DMG400004963M6LRR-RLK_1426

GGTCCAATTCCTTCTGAAATTGGTGGTTTAAAAAGCCTCAATTTGTTACTCTTGCAAGGTAACAAGTTCAATTCTTCAATCCCCGAGTCACTTTCTTCGCTTAAATATCTCAACTATCTT

>M6-PGSC0003DMG400004963M6LRR-RLK_1488

CAAGTTCAATTCTTCAATCCCCGAGTCACTTTCTTCGCTTAAATATCTCAACTATCTTGATCTGTCTAACAATCTTTTGATAGGAAAGATCCCTGAGAGCCTAGGTGAATTGTTACCAAA

>M6-PGSC0003DMG400004963M6LRR-RLK_1550

TGTCTAACAATCTTTTGATAGGAAAGATCCCTGAGAGCCTAGGTGAATTGTTACCAAACTCCATGAACTTGTCAAACAACTTACTTTCTGGTGCTATACCTCTTTTGTTTATAAAAGGAG

>M6-PGSC0003DMG400004963M6LRR-RLK_1612

ATGAACTTGTCAAACAACTTACTTTCTGGTGCTATACCTCTTTTGTTTATAAAAGGAGGTGTTTTGGAAAGTTTTTTAGGCAATCCGGGACTTTGTGTCCCTACCTCTTTGAACTCAACA

>M6-PGSC0003DMG400004963M6LRR-RLK_1674

TTTGGAAAGTTTTTTAGGCAATCCGGGACTTTGTGTCCCTACCTCTTTGAACTCAACAAACACAAGTTTTCAAACATGTTCACATAGCTATAACCATAAGAAGAGGAACAACATTGTTTG

>M6-PGSC0003DMG400004963M6LRR-RLK_1736

CAAGTTTTCAAACATGTTCACATAGCTATAACCATAAGAAGAGGAACAACATTGTTTGGGTTATTGGGACATCGGTGGGGATAGTCATTGTTGGATTAGTCTTGTTTATCAAACGATGGT

>M6-PGSC0003DMG400004963M6LRR-RLK_1798

ATTGGGACATCGGTGGGGATAGTCATTGTTGGATTAGTCTTGTTTATCAAACGATGGTTTGGTAACAAAAAGGCGGTGATGGAACAGGATGATCATTCATTGTCATCATCATTTTTCTCC

>M6-PGSC0003DMG400004963M6LRR-RLK_186

ACCAATTTGTCAATACAAAGGTGTCGGTTGTGATGAAAGGGGAGACGTTATCAAAATTAACATATCGGCCTGGTATTTATCTGGCCAATTTCCAAGTGATGTGTGTAGTTATTTGCCAAG

>M6-PGSC0003DMG400004963M6LRR-RLK_1860

TAACAAAAAGGCGGTGATGGAACAGGATGATCATTCATTGTCATCATCATTTTTCTCCTATGATGTTAAGAGTTTCCATCGATTGAGTTTTGATCAACGTGAGATCTTTGAAGCTATGGT

>M6-PGSC0003DMG400004963M6LRR-RLK_1922

ATGTTAAGAGTTTCCATCGATTGAGTTTTGATCAACGTGAGATCTTTGAAGCTATGGTTGAGAAGAATATTGTTGGATATGGAGGGTCAGGGGCTGTCTATAAGATCGAGTTGAGTAACG

>M6-PGSC0003DMG400004963M6LRR-RLK_1984

AAGAATATTGTTGGATATGGAGGGTCAGGGGCTGTCTATAAGATCGAGTTGAGTAACGGAGGAGTAGTTGCTGCAAAGAAGCTGTGGAGTCACAAACATAAGCATTCTGTTTCTGAGGAT

>M6-PGSC0003DMG400004963M6LRR-RLK_2046

AGTAGTTGCTGCAAAGAAGCTGTGGAGTCACAAACATAAGCATTCTGTTTCTGAGGATAAATTGGTTTTGGATAAGGAACTTAAGACAGAAGTTGAGACTCTTGGTAACATAAGGCACAA

>M6-PGSC0003DMG400004963M6LRR-RLK_2108

TGGTTTTGGATAAGGAACTTAAGACAGAAGTTGAGACTCTTGGTAACATAAGGCACAAAAACATTGTGAAATTGTACTGTTATTTCTCAAGTTTGGATTGCAGCTTGTTGGTTTATGAAT

>M6-PGSC0003DMG400004963M6LRR-RLK_2170

ATTGTGAAATTGTACTGTTATTTCTCAAGTTTGGATTGCAGCTTGTTGGTTTATGAATACATGCCTAATGGAAACCTTTGGCATGCTCTTCATGGTGGGAAGTTCGTGTTGGATTGGCCT

>M6-PGSC0003DMG400004963M6LRR-RLK_2232

GCCTAATGGAAACCTTTGGCATGCTCTTCATGGTGGGAAGTTCGTGTTGGATTGGCCTATTCGTCATCAAATTGCACTTGGAATAGCTCAAGGATTGGCCTATCTTCACCATGATCTTAT

>M6-PGSC0003DMG400004963M6LRR-RLK_2294

GTCATCAAATTGCACTTGGAATAGCTCAAGGATTGGCCTATCTTCACCATGATCTTATGCCACCGATTATTCATAGAGATATCAAATCCACCAACATCCTCCTGGATATTGATTATCAAC

>M6-PGSC0003DMG400004963M6LRR-RLK_2356

CCGATTATTCATAGAGATATCAAATCCACCAACATCCTCCTGGATATTGATTATCAACCAAAAGTCGCGGATTTTGGTATAGCCAAAGTGTTGCAAGCCAGAGGAGGCAAAGATTCCAGC

>M6-PGSC0003DMG400004963M6LRR-RLK_2418

AGTCGCGGATTTTGGTATAGCCAAAGTGTTGCAAGCCAGAGGAGGCAAAGATTCCAGCACCACGGTCATCGCAGGAACTTATGGTTACTTGGCACCAGGTATAATTTTCCCAATTCATAT

>M6-PGSC0003DMG400004963M6LRR-RLK_248

TATCGGCCTGGTATTTATCTGGCCAATTTCCAAGTGATGTGTGTAGTTATTTGCCAAGATTGAAAAGTCTTCATATTGGTCATAATAATTTCCAAGGTGGTTTTCCTAAGTACCTAACAA

>M6-PGSC0003DMG400004963M6LRR-RLK_2480

CGGTCATCGCAGGAACTTATGGTTACTTGGCACCAGGTATAATTTTCCCAATTCATATTTGTTACTTCAATCTTTGAGCTTAAGAGTTATAATACATATCACAATCTTGATTGAGTAGAT

>M6-PGSC0003DMG400004963M6LRR-RLK_2542

TACTTCAATCTTTGAGCTTAAGAGTTATAATACATATCACAATCTTGATTGAGTAGATTAAATTAAATCCTTTATGTGCAAATAACATCCCATAGTTGAAGTGGCCTAATGTTACTCTAA

>M6-PGSC0003DMG400004963M6LRR-RLK_2604

TTAAATCCTTTATGTGCAAATAACATCCCATAGTTGAAGTGGCCTAATGTTACTCTAAATTGGCCATTTAATATAGTCATTGCCTAACGTTAAGAGTATATTTGACCCTTTTCCGTTCTT

>M6-PGSC0003DMG400004963M6LRR-RLK_2666

GCCATTTAATATAGTCATTGCCTAACGTTAAGAGTATATTTGACCCTTTTCCGTTCTTAATTATGGCTTTTTATGTGTAAAAACCATCACATATGCTCAAGTTTCTTGATTGAACTTTCA

>M6-PGSC0003DMG400004963M6LRR-RLK_2728

ATGGCTTTTTATGTGTAAAAACCATCACATATGCTCAAGTTTCTTGATTGAACTTTCATTTAACACATGTTTAAGTGTTCAAATTAGGCTAATGTTACTCAACTTTGGTGTAACAGAGTA

>M6-PGSC0003DMG400004963M6LRR-RLK_2790

ACACATGTTTAAGTGTTCAAATTAGGCTAATGTTACTCAACTTTGGTGTAACAGAGTATGCATATTCTTCAAAGGCAACTACAAAGTGTGATGTGTATAGTTTTGGAGTTGTTCTAATGG

>M6-PGSC0003DMG400004963M6LRR-RLK_2852

TATTCTTCAAAGGCAACTACAAAGTGTGATGTGTATAGTTTTGGAGTTGTTCTAATGGAATTGATTACTGGAAAGAAGCCAGTGGAGCCAGAATTTGGAGACAACAAGAACATTGTTTAT

>M6-PGSC0003DMG400004963M6LRR-RLK_2914

GATTACTGGAAAGAAGCCAGTGGAGCCAGAATTTGGAGACAACAAGAACATTGTTTATTGGGTATCAACAAAAGTGGAGACTAAAGAAGGTGCATTTGAAGTATTGGATAAAAAAGTCTC

>M6-PGSC0003DMG400004963M6LRR-RLK_2976

TATCAACAAAAGTGGAGACTAAAGAAGGTGCATTTGAAGTATTGGATAAAAAAGTCTCAGATTCTTTCAAAGAGGACATGATTAAGGTTCTACGTATAGCTATACGTTGTACGTATAGTA

>M6-PGSC0003DMG400004963M6LRR-RLK_3038

TCTTTCAAAGAGGACATGATTAAGGTTCTACGTATAGCTATACGTTGTACGTATAGTACACCAACACTTCGTCCTACCATGAATGAAGTTGTTCAGTTGCTAATCGAAGCGGATCCTTGC

>M6-PGSC0003DMG400004963M6LRR-RLK_310

AAAAGTCTTCATATTGGTCATAATAATTTCCAAGGTGGTTTTCCTAAGTACCTAACAAATTGTTCTTTTTTGGAAGAATTGAACATGACTAAAACATCACTTACAGGACAAATTCCTGAT

>M6-PGSC0003DMG400004963M6LRR-RLK_3100

AACACTTCGTCCTACCATGAATGAAGTTGTTCAGTTGCTAATCGAAGCGGATCCTTGCAAGTTCAATTGTTGCAACATGTCAAATAAGAAGAAGAATGACACACAAGATCAACTGATCAA

>M6-PGSC0003DMG400004963M6LRR-RLK_3162

TCAATTGTTGCAACATGTCAAATAAGAAGAAGAATGACACACAAGATCAACTGATCAATAAGCCACCAAAGAACATATATGAGTTGTAAAATATTTAGTGTATCAAATTAGTATACAAGG

>M6-PGSC0003DMG400004963M6LRR-RLK_3224

CCACCAAAGAACATATATGAGTTGTAAAATATTTAGTGTATCAAATTAGTATACAAGGTATATAGAATATGTGTACTCATGAGATATTACACCCTCAAAAGAATCAGTCTTTATGTTTTC

>M6-PGSC0003DMG400004963M6LRR-RLK_3286

TAGAATATGTGTACTCATGAGATATTACACCCTCAAAAGAATCAGTCTTTATGTTTTCTAATAAATGCTACTACTTTTTATTAGTATATAGATTCTCCAATTAGTGTTACTTTGCTTTTT

>M6-PGSC0003DMG400004963M6LRR-RLK_3329

AGTCTTTATGTTTTCTAATAAATGCTACTACTTTTTATTAGTATATAGATTCTCCAATTAGTGTTACTTTGCTTTTTTATATTTTTTTGGCCTGTTGTAAGTATTTGGGAATTTTATGAA

>M6-PGSC0003DMG400004963M6LRR-RLK_372

TTCTTTTTTGGAAGAATTGAACATGACTAAAACATCACTTACAGGACAAATTCCTGATTTGTCACCATTACAATCTTTGAAACTACTTGATCTTTCATGTAATTAATTGACAGGGGATTT

>M6-PGSC0003DMG400004963M6LRR-RLK_434

CACCATTACAATCTTTGAAACTACTTGATCTTTCATGTAATTAATTGACAGGGGATTTTCCTTTGTCAATTATTAATCTCACCAACTTAGTTATCCTGAATTTCAATGAAAATCGCCATT

>M6-PGSC0003DMG400004963M6LRR-RLK_496

TTGTCAATTATTAATCTCACCAACTTAGTTATCCTGAATTTCAATGAAAATCGCCATTTCAATCCGTGGCGATTACCAGAGGACATTTCCAGGCTTATAAATCTCAAGTGGATGATTTTA

>M6-PGSC0003DMG400004963M6LRR-RLK_558

TCCGTGGCGATTACCAGAGGACATTTCCAGGCTTATAAATCTCAAGTGGATGATTTTAACAGCATGTAATATGCATGGGACAATTCCAGTGTCAATAAGTAACATGACTTCTCTTGTTGA

>M6-PGSC0003DMG400004963M6LRR-RLK_62

TCACCCTTGTGTTATTTTTTCTTACTTTGACGTCCACTTTTGTTCAAGGTGATCAATCTCAGTTCTTTGTACTCATGAAGAAATTTGTCACAGGAAGTTCTTTGTCTAATTGGGATATTG

>M6-PGSC0003DMG400004963M6LRR-RLK_620

CATGTAATATGCATGGGACAATTCCAGTGTCAATAAGTAACATGACTTCTCTTGTTGATCTTGAATTGAGCGCGAATCGCCTTGTTGGTAAGGTGCCTAGAGAACTCGGGAAGCTGAAGA

>M6-PGSC0003DMG400004963M6LRR-RLK_682

GAATTGAGCGCGAATCGCCTTGTTGGTAAGGTGCCTAGAGAACTCGGGAAGCTGAAGAATTTGAAACTCCTTGAGCTTTTTTACAACTTACTTGATGGTGAAATCCCCGCGGAGCTAGGG

>M6-PGSC0003DMG400004963M6LRR-RLK_744

GAAACTCCTTGAGCTTTTTTACAACTTACTTGATGGTGAAATCCCCGCGGAGCTAGGGAATTTGACAGAACTTGTGGACTTAGACATGTCCGCGAATAATTTCACAGGGAGAATTCCAGA

>M6-PGSC0003DMG400004963M6LRR-RLK_806

TGACAGAACTTGTGGACTTAGACATGTCCGCGAATAATTTCACAGGGAGAATTCCAGAGTCTATAAGCCGGCTTCCTAAGCTGGAAGTGTTGCAGCTTTACCATAACGCGCTATCTGGAG

>M6-PGSC0003DMG400004963M6LRR-RLK_868

ATAAGCCGGCTTCCTAAGCTGGAAGTGTTGCAGCTTTACCATAACGCGCTATCTGGAGAGTTTCCAGCAGCACTTGCTAATTCAACAACTTTAACTATCCTGTCTCTTTATGACAATCTT

>M6-PGSC0003DMG400004963M6LRR-RLK_930

TCCAGCAGCACTTGCTAATTCAACAACTTTAACTATCCTGTCTCTTTATGACAATCTTTTTACAGGAGAAGTTCCACAACACTTTGGTCTTTCATCAGCTTTGTTGGCATTGGACTTGTC

>M6-PGSC0003DMG400004963M6LRR-RLK_992

CAGGAGAAGTTCCACAACACTTTGGTCTTTCATCAGCTTTGTTGGCATTGGACTTGTCAGAAAATCGATTTTCTGGAAAGCTACCGCCTTTTCTGTGTAGTGGAGGTAAATTGAGTTATA

>M6-PGSC0003DMG400004966M6LRR-RLK_0

GCGCTCTTCATAGTCGTCTCCTTCACACGTTTTTCTGAAGGCGGAAGAAAAAAACTTCTCTTCGTCAACTTCTTCGTCAGCTATGGCGGCGCTATTCAATTGCCGGCCGGAACTTATCCA

>M6-PGSC0003DMG400004966M6LRR-RLK_1037

TGTACACTAACAAACTCACTGGTCAGCTTCCGCAGAAGTTGGGTTCTTGGGCGAATTTTGATTTCATTGATGTATCAGAGAATAGTTTCACTGGTCCAATTCCACCGGACATGTGTAAGA

>M6-PGSC0003DMG400004966M6LRR-RLK_1098

TTTCATTGATGTATCAGAGAATAGTTTCACTGGTCCAATTCCACCGGACATGTGTAAGATGGGGACGATGAGAGGATTATTGATACTTCAGAACAATTTCACTGGCGGAATCCCTGAAAG

>M6-PGSC0003DMG400004966M6LRR-RLK_1159

GGGACGATGAGAGGATTATTGATACTTCAGAACAATTTCACTGGCGGAATCCCTGAAAGTTATGCTAATTGTACAACAATGACGCGGATTCGAGTGAGCAAAAACTCACTCTCCGGTGTG

>M6-PGSC0003DMG400004966M6LRR-RLK_122

TCTTCTCTATTTTCTCCTTCTTCCTTCTCTTCCCCGTCGCTTTTTCCGACGAGCTTCAAACACTTTTATCCATCAAATCCTCTTTATCAAACCCCATCACCAATGTATTCCAAAATTGGG

>M6-PGSC0003DMG400004966M6LRR-RLK_1220

ATGCTAATTGTACAACAATGACGCGGATTCGAGTGAGCAAAAACTCACTCTCCGGTGTGATTCCTGCAGGAATTTGGGGGCTACCAAAGCTCGAAATCCTTGATGTTGCAATGAATGAGT

>M6-PGSC0003DMG400004966M6LRR-RLK_1281

TCCTGCAGGAATTTGGGGGCTACCAAAGCTCGAAATCCTTGATGTTGCAATGAATGAGTTTGAAGGTACTATTACTTCTGTTATTGGAAATGCAAAATCTTTAGGTGAAATTGATGCGGC

>M6-PGSC0003DMG400004966M6LRR-RLK_1342

GAAGGTACTATTACTTCTGTTATTGGAAATGCAAAATCTTTAGGTGAAATTGATGCGGCTAATAACAGATTCTCTGGTGAATTGCCTTTTGATATTTCGAATGCTTCTTTCTTAGTGAAA

>M6-PGSC0003DMG400004966M6LRR-RLK_1403

ATAACAGATTCTCTGGTGAATTGCCTTTTGATATTTCGAATGCTTCTTTCTTAGTGAAAATTGATTTTAGTAACAATCAGTTTTCCGGTGAAATTCCGGGAACAATTGGGGAGCTTAAGA

>M6-PGSC0003DMG400004966M6LRR-RLK_1464

TGATTTTAGTAACAATCAGTTTTCCGGTGAAATTCCGGGAACAATTGGGGAGCTTAAGAAGATTGTTAATCTTAATTTACAGAACAACAAGTTTTCTGGTTCAATACCAGATTCTTTAGG

>M6-PGSC0003DMG400004966M6LRR-RLK_1525

ATTGTTAATCTTAATTTACAGAACAACAAGTTTTCTGGTTCAATACCAGATTCTTTAGGCTCTTGTGTTTCACTAAGCGACATTAACATGGCTAACAATTTGCTTAGTGGCTCTATTCCT

>M6-PGSC0003DMG400004966M6LRR-RLK_1586

CTTGTGTTTCACTAAGCGACATTAACATGGCTAACAATTTGCTTAGTGGCTCTATTCCTGTTTCTCTAGGATCATTACCGACTTTAACTTCATTAAACTTGTCGGAGAATCAGCTTTCAG

>M6-PGSC0003DMG400004966M6LRR-RLK_1647

TTCTCTAGGATCATTACCGACTTTAACTTCATTAAACTTGTCGGAGAATCAGCTTTCAGGGAAAATTCCGACGAGTTTATCAAATCTGAAGTTAAATCTTCTTGATTTTTCCAACAATCA

>M6-PGSC0003DMG400004966M6LRR-RLK_1708

AAAATTCCGACGAGTTTATCAAATCTGAAGTTAAATCTTCTTGATTTTTCCAACAATCAGTTGACTGGAGCAATACCAAACTCTCTATCAATTGATGCTTATAAGGGTAGCTTTGCTGGG

>M6-PGSC0003DMG400004966M6LRR-RLK_1769

TGACTGGAGCAATACCAAACTCTCTATCAATTGATGCTTATAAGGGTAGCTTTGCTGGGAATAATGGTCTTTGTAGCCAAAACATTAAGAACTTCCGTAGATGTTATGGTGAATCTGGAA

>M6-PGSC0003DMG400004966M6LRR-RLK_183

ACTTTTATCCATCAAATCCTCTTTATCAAACCCCATCACCAATGTATTCCAAAATTGGGAACCCAATACTCCTCTCTGTAAGTTCACTGGCATCACATGTAATTCCGATGGGTCAGTCAA

>M6-PGSC0003DMG400004966M6LRR-RLK_1830

TAATGGTCTTTGTAGCCAAAACATTAAGAACTTCCGTAGATGTTATGGTGAATCTGGAAAGCCTAGAGAATGGTACACTCTGTTGATCTGTTTATTGGTGGCTGTGATTGTGGTGCTCGT

>M6-PGSC0003DMG400004966M6LRR-RLK_1891

CCTAGAGAATGGTACACTCTGTTGATCTGTTTATTGGTGGCTGTGATTGTGGTGCTCGTTTCATTTGCGGGCTATTTGTTCTTGAAGAAGAAGAGTCATAAGGAACATGAGAGGTCTTTG

>M6-PGSC0003DMG400004966M6LRR-RLK_1952

CATTTGCGGGCTATTTGTTCTTGAAGAAGAAGAGTCATAAGGAACATGAGAGGTCTTTGAAGCAAAATTCTTGGAATACAAAGTCATTCCATATATTGACCTTCACAGAGGATGATATTC

>M6-PGSC0003DMG400004966M6LRR-RLK_2013

GCAAAATTCTTGGAATACAAAGTCATTCCATATATTGACCTTCACAGAGGATGATATTCTTGATGGAATCAAACACGACAATTTGATTGGAAAAGGTGGTTCTGGTAGTGTATACAGAGT

>M6-PGSC0003DMG400004966M6LRR-RLK_2074

GATGGAATCAAACACGACAATTTGATTGGAAAAGGTGGTTCTGGTAGTGTATACAGAGTGCAGCTTTCAGATGGAACAGATTTCGCGGTGAAACATATTTGGACTTCTGATTCAGGAAAC

>M6-PGSC0003DMG400004966M6LRR-RLK_2135

AGCTTTCAGATGGAACAGATTTCGCGGTGAAACATATTTGGACTTCTGATTCAGGAAACAGGAAAATATCAGGAACAACATCTCCAATGTTGGGAAAACCAGGGAAGAAATCGAAAGAAT

>M6-PGSC0003DMG400004966M6LRR-RLK_2196

GAAAATATCAGGAACAACATCTCCAATGTTGGGAAAACCAGGGAAGAAATCGAAAGAATTTGAGGCTGAGGTTGAAACACTGAGCTCAATTCGACATGTGAATGTGGTGAAATTGTACTG

>M6-PGSC0003DMG400004966M6LRR-RLK_2257

GAGGCTGAGGTTGAAACACTGAGCTCAATTCGACATGTGAATGTGGTGAAATTGTACTGTAGCATAACGAGTGATGACTCGAGCTTGTTGGTGTATGAATATATGCCAAATGGAAGTTTG

>M6-PGSC0003DMG400004966M6LRR-RLK_2318

GCATAACGAGTGATGACTCGAGCTTGTTGGTGTATGAATATATGCCAAATGGAAGTTTGTGGGATCGATTGCATACTTGCAAAAAGATGTCACTTGATTGGGAGACGAGGTACGAGATAG

>M6-PGSC0003DMG400004966M6LRR-RLK_2379

GGATCGATTGCATACTTGCAAAAAGATGTCACTTGATTGGGAGACGAGGTACGAGATAGCTCTTGGTGCAGCCAAAGGACTGGAGTATTTACACCATGGTTGTGATAAGCCAGTGATTCA

>M6-PGSC0003DMG400004966M6LRR-RLK_244

CCCAATACTCCTCTCTGTAAGTTCACTGGCATCACATGTAATTCCGATGGGTCAGTCAAAGAAATCGAGCTTTCGAGTCAAAAAATTTCTGGGTTTGTTCCTTTTGATAAAATATGTTCA

>M6-PGSC0003DMG400004966M6LRR-RLK_2440

CTTGGTGCAGCCAAAGGACTGGAGTATTTACACCATGGTTGTGATAAGCCAGTGATTCATAGAGATGTTAAATCAAGTAACATCTTGTTGGATGAGTTTTGCAAGCCAAGAATCGCCGAC

>M6-PGSC0003DMG400004966M6LRR-RLK_2501

GAGATGTTAAATCAAGTAACATCTTGTTGGATGAGTTTTGCAAGCCAAGAATCGCCGACTTTGGTCTTGCTAGGATTGCTCAAGCTGATTCAACCAAAGATACAACTCATGTCATTGCTG

>M6-PGSC0003DMG400004966M6LRR-RLK_2562

TGGTCTTGCTAGGATTGCTCAAGCTGATTCAACCAAAGATACAACTCATGTCATTGCTGGAACACATGGATATATTGCTCCTGGTAAGAAGCTCATTATATTGTAGTTGATTTTATACGT

>M6-PGSC0003DMG400004966M6LRR-RLK_2623

ACACATGGATATATTGCTCCTGGTAAGAAGCTCATTATATTGTAGTTGATTTTATACGTAGAATTTGAGTATATCTACTTAACACTAACCATGTATTCATTTTTCACCATCGGTAAAATT

>M6-PGSC0003DMG400004966M6LRR-RLK_2684

AATTTGAGTATATCTACTTAACACTAACCATGTATTCATTTTTCACCATCGGTAAAATTGACTTACGAAATAGGCAACTCCCCATATCAAGCAATATGGTATCTTGAAAGATATAGTGAA

>M6-PGSC0003DMG400004966M6LRR-RLK_2745

CTTACGAAATAGGCAACTCCCCATATCAAGCAATATGGTATCTTGAAAGATATAGTGAAAACTTGCTATAAACAGTTGGATCACTATGTTGCTCGGATCCTTCAAAAATGTTGCTGCACC

>M6-PGSC0003DMG400004966M6LRR-RLK_2806

CTTGCTATAAACAGTTGGATCACTATGTTGCTCGGATCCTTCAAAAATGTTGCTGCACCCGTGTTGGTTCCCTCAAAAATAGCGCATTCTTGGAGGATTTGACATGAGTGTGACAGCATT

>M6-PGSC0003DMG400004966M6LRR-RLK_2867

TGTTGGTTCCCTCAAAAATAGCGCATTCTTGGAGGATTTGACATGAGTGTGACAGCATTTTTGAAGGATCCGAGCAATAAGTGAATTGCACTAATAAGAGTGTAAAACAATATTAGTGTT

>M6-PGSC0003DMG400004966M6LRR-RLK_2928

TGAAGGATCCGAGCAATAAGTGAATTGCACTAATAAGAGTGTAAAACAATATTAGTGTTAATTAACTTAGATCCTATCTTGATTATCTATTCATTTGGTCATAGTGAGTGTCTCATTTCT

>M6-PGSC0003DMG400004966M6LRR-RLK_2989

TTAACTTAGATCCTATCTTGATTATCTATTCATTTGGTCATAGTGAGTGTCTCATTTCTGGCATCCTTTTTGAACTTATTTGAATTGAATTGGTGTTGCAGAATATGGATACACACACAA

>M6-PGSC0003DMG400004966M6LRR-RLK_305

AAATCGAGCTTTCGAGTCAAAAAATTTCTGGGTTTGTTCCTTTTGATAAAATATGTTCACTTAATTCATTGGAAAAGCTCTCTCTGGGTTACAATTCATTTTCTGGGGAATTTACTGATG

>M6-PGSC0003DMG400004966M6LRR-RLK_3050

CATCCTTTTTGAACTTATTTGAATTGAATTGGTGTTGCAGAATATGGATACACACACAAAGTGAACGAGAAAAGTGATGTATATAGTTTCGGAGTTGTATTGATGGAGCTTATATCTGGG

>M6-PGSC0003DMG400004966M6LRR-RLK_3111

TGAACGAGAAAAGTGATGTATATAGTTTCGGAGTTGTATTGATGGAGCTTATATCTGGGAAACGGCCAATAGAATCCGAATACGGAGAGAATGGCAACATAGTTACATGGGTGAGCAGCA

>M6-PGSC0003DMG400004966M6LRR-RLK_3172

ACGGCCAATAGAATCCGAATACGGAGAGAATGGCAACATAGTTACATGGGTGAGCAGCAAATTGAAGAGCAAGGAAAGTGTATTAAGCATAGTGGATTCAAGAATTCCTGAAGCTTTCAA

>M6-PGSC0003DMG400004966M6LRR-RLK_3233

TTGAAGAGCAAGGAAAGTGTATTAAGCATAGTGGATTCAAGAATTCCTGAAGCTTTCAAGGAAGATGCAATCAAAGTCTTGAGGATTGCTATTGTATGCACATCTAGGCTTCCCACATTA

>M6-PGSC0003DMG400004966M6LRR-RLK_3294

AAGATGCAATCAAAGTCTTGAGGATTGCTATTGTATGCACATCTAGGCTTCCCACATTAAGGCCAACAATGAGAAATGTAGTCAAAATGCTGGAAAATGCAGAACCTTGCAGATTAGTTG

>M6-PGSC0003DMG400004966M6LRR-RLK_3355

GCCAACAATGAGAAATGTAGTCAAAATGCTGGAAAATGCAGAACCTTGCAGATTAGTTGGGATAATTGTAAGCAAAGATGATGGTAGTATCAAGACAGAACAATTGAAGGATCACACCAA

>M6-PGSC0003DMG400004966M6LRR-RLK_3416

ATAATTGTAAGCAAAGATGATGGTAGTATCAAGACAGAACAATTGAAGGATCACACCAAGATGTAATTGGATCCCCCGTCTTTGCTCGTTTCACTCTTCAACCTGATCTCTGTCCTCCAT

>M6-PGSC0003DMG400004966M6LRR-RLK_3477

TGTAATTGGATCCCCCGTCTTTGCTCGTTTCACTCTTCAACCTGATCTCTGTCCTCCATGGAACATCGAATCTTGTAGCAGTATGTAAAGTTCTGAAGATGGCCTCTTGGCATTGTAACA

>M6-PGSC0003DMG400004966M6LRR-RLK_3538

AACATCGAATCTTGTAGCAGTATGTAAAGTTCTGAAGATGGCCTCTTGGCATTGTAACATATAGTAGAAGCTAGATCCATACTAGTATGTGGAATTAGATTGAATTTAGAGATGCATGGC

>M6-PGSC0003DMG400004966M6LRR-RLK_3589

TTGTAACATATAGTAGAAGCTAGATCCATACTAGTATGTGGAATTAGATTGAATTTAGAGATGCATGGCATAAGTGCTATAAGATTGGTTTAGTTATTCAAATCCCTTTTCAACCCTGCA

>M6-PGSC0003DMG400004966M6LRR-RLK_366

TAATTCATTGGAAAAGCTCTCTCTGGGTTACAATTCATTTTCTGGGGAATTTACTGATGATTTGAACAAGTGTGTTAGCTTGAATTACTTGGATGTTGGTAATAATGAATTTACTGGGTA

>M6-PGSC0003DMG400004966M6LRR-RLK_427

TTGAACAAGTGTGTTAGCTTGAATTACTTGGATGTTGGTAATAATGAATTTACTGGGTATTTCCCTGATGTATCTTCACTCAGTGAGTTAACCCATTTTTATGCAAATAACAGTGGGTTT

>M6-PGSC0003DMG400004966M6LRR-RLK_488

TCCCTGATGTATCTTCACTCAGTGAGTTAACCCATTTTTATGCAAATAACAGTGGGTTTACAGGAAAATTTCCATGGAATTCTTTTGCGAGTATGAGTAATTTGATTGTGCTTAGCCTTG

>M6-PGSC0003DMG400004966M6LRR-RLK_549

AGGAAAATTTCCATGGAATTCTTTTGCGAGTATGAGTAATTTGATTGTGCTTAGCCTTGGTGATAATCAGTTTGATCGGACACCGTTTCCTGAAGTGATTTTGAAGCTTAATAAATTGAA

>M6-PGSC0003DMG400004966M6LRR-RLK_61

TCGTCAACTTCTTCGTCAGCTATGGCGGCGCTATTCAATTGCCGGCCGGAACTTATCCACTTCTTCTCTATTTTCTCCTTCTTCCTTCTCTTCCCCGTCGCTTTTTCCGACGAGCTTCAA

>M6-PGSC0003DMG400004966M6LRR-RLK_610

GATAATCAGTTTGATCGGACACCGTTTCCTGAAGTGATTTTGAAGCTTAATAAATTGAATTGGCTCTACTTATCGAATTGTGAGCTTGAAGGGGAAATTCCTGAAGAAATTGGAAATCTT

>M6-PGSC0003DMG400004966M6LRR-RLK_671

GGCTCTACTTATCGAATTGTGAGCTTGAAGGGGAAATTCCTGAAGAAATTGGAAATCTTACAGAGCTGATTGATTTAGAGCTGTCGATGAATCACCTTACTGGTGAAATTCCTTCGGGAA

>M6-PGSC0003DMG400004966M6LRR-RLK_732

AGAGCTGATTGATTTAGAGCTGTCGATGAATCACCTTACTGGTGAAATTCCTTCGGGAATTACGAAGCTGAAGAAATTATGGCAGCTTGAGTTGTATGAAAACCAGCTTACAGGGAAGTT

>M6-PGSC0003DMG400004966M6LRR-RLK_793

ACGAAGCTGAAGAAATTATGGCAGCTTGAGTTGTATGAAAACCAGCTTACAGGGAAGTTGCCGGTTGGATTTGGAAATTTGACAAGTCTTGAATATCTTGATGCTTCGACTAATAATCTG

>M6-PGSC0003DMG400004966M6LRR-RLK_854

CGGTTGGATTTGGAAATTTGACAAGTCTTGAATATCTTGATGCTTCGACTAATAATCTGTACGGTGATTTGTCGGAAATTCGAAAGTTGAATCAATTGGTAAGTTTGCAGTTATTGCAGA

>M6-PGSC0003DMG400004966M6LRR-RLK_915

CGGTGATTTGTCGGAAATTCGAAAGTTGAATCAATTGGTAAGTTTGCAGTTATTGCAGAACCAATTTTCAGGGGAAGTACCAGCAGAATTGGGAGAGTTCAAAAAGCTGGTGAACATCTC

>M6-PGSC0003DMG400004966M6LRR-RLK_976

CAATTTTCAGGGGAAGTACCAGCAGAATTGGGAGAGTTCAAAAAGCTGGTGAACATCTCTTTGTACACTAACAAACTCACTGGTCAGCTTCCGCAGAAGTTGGGTTCTTGGGCGAATTTT

>M6-PGSC0003DMG400004972M6LRR-RLK_0

AGCTTGAAATCTTTTGTTTAAGCCGTTAACTATACCCTTACTAAATGATTCTCTATAAAATGTAATGTTTCCAGGACTTAAGAGCATTATACATTCAAAATTTTTGGAATATTTGAAGCT

>M6-PGSC0003DMG400004972M6LRR-RLK_1037

TAATTGAATTGATTATTGTAACTTTTGCATTATAGGAGGCACTCCACCTGCCAAAAGCAATCCAGCGCAGAGCAATAATAAGCACGGGACTGTTGAAGGAAGAGCCAAACATAAGTCTGC

>M6-PGSC0003DMG400004972M6LRR-RLK_1098

CCAGCGCAGAGCAATAATAAGCACGGGACTGTTGAAGGAAGAGCCAAACATAAGTCTGCTTCAAAACCTACCTGGCTTTTTGTGTTAGAAGTGGTAACTGGAGTTATTGCAGGTTCTCTC

>M6-PGSC0003DMG400004972M6LRR-RLK_1159

CAAAACCTACCTGGCTTTTTGTGTTAGAAGTGGTAACTGGAGTTATTGCAGGTTCTCTCTTCCTTGTGGCTATTTTGGCTGCCATTCAGAAGTGTAAGAACGAATCTTCTATTATCATCC

>M6-PGSC0003DMG400004972M6LRR-RLK_122

ATTAAATTTATATATCTTTTTAATCTTCTAAGGTGCCCATAGTATCTCAATTAGCATATGATTTGCTGCAACATTTTTGGCTGCAGTTTGTCTGGACATTTACAGAGTCTTAAAATTGAA

>M6-PGSC0003DMG400004972M6LRR-RLK_1220

CCTTGTGGCTATTTTGGCTGCCATTCAGAAGTGTAAGAACGAATCTTCTATTATCATCCCTTGGAAGAAATCAGCAAGTATGAAAGACCATTTGAAAGACCATATGGCAGTTTATGTAGG

>M6-PGSC0003DMG400004972M6LRR-RLK_1281

TGGAAGAAATCAGCAAGTATGAAAGACCATTTGAAAGACCATATGGCAGTTTATGTAGGTGAGTACCATGCAATTTCAGTTGGGGGATTACAGTTAAGAATTGCCTTTAAACATTTGTAT

>M6-PGSC0003DMG400004972M6LRR-RLK_1647

GGAGGGTGGGGAGGACACAATTGATGTGAAATACCACTTGTGGAACTTGTTTTCCCTACCTCCATACCGAACACACCCTTAATTCTTTCTCATATAGGATCTGATTCTTCAATTTAGTAC

>M6-PGSC0003DMG400004972M6LRR-RLK_183

TTTGCTGCAACATTTTTGGCTGCAGTTTGTCTGGACATTTACAGAGTCTTAAAATTGAAAGATTATGGCTAATGTTTAATCTTTCTGTCGTCTTGATTGATCGTGCTGGACTTCTGTTGA

>M6-PGSC0003DMG400004972M6LRR-RLK_2074

CACTTGTGGGATCACACAGGGTATAGGATCACAAGTTCAGAAGCATTAATTATAATTCACATTTCTATTAGTAACAAATGTTTCTTTGTCAGATACTGAGATGCTAAAGGACGTTGAAAG

>M6-PGSC0003DMG400004972M6LRR-RLK_2135

TTTCTATTAGTAACAAATGTTTCTTTGTCAGATACTGAGATGCTAAAGGACGTTGAAAGATATAGCAGACAAGAACTTGAGGTAGCTTGTGAAGATTTCAGCAACATTATTGGATCCTCA

>M6-PGSC0003DMG400004972M6LRR-RLK_2196

ATAGCAGACAAGAACTTGAGGTAGCTTGTGAAGATTTCAGCAACATTATTGGATCCTCACCAGACAGCTTGGTCTATAAAGGAACTGTGAAAGGCGGACCTGAATTAGCTGTAATTTCCA

>M6-PGSC0003DMG400004972M6LRR-RLK_2257

AGACAGCTTGGTCTATAAAGGAACTGTGAAAGGCGGACCTGAATTAGCTGTAATTTCCATCTGTGTCAAAGAAGAGCAGTGGACCACCTATCTAGAGCTTTATTTTCAGAAAGAGGTGAA

>M6-PGSC0003DMG400004972M6LRR-RLK_2318

TGTGTCAAAGAAGAGCAGTGGACCACCTATCTAGAGCTTTATTTTCAGAAAGAGGTGAAGCCTGTTCTCTCCTTAAAATACATGGCTACTGCTGAGCTCTCGTGCCCAATAAAAATTTCA

>M6-PGSC0003DMG400004972M6LRR-RLK_2379

CTGTTCTCTCCTTAAAATACATGGCTACTGCTGAGCTCTCGTGCCCAATAAAAATTTCATCTTTGCTGCAATGCTCTAACTTGTATGGCTTACAAACTCAGGTAGCAGAATTGGCCAGAA

>M6-PGSC0003DMG400004972M6LRR-RLK_244

ATTATGGCTAATGTTTAATCTTTCTGTCGTCTTGATTGATCGTGCTGGACTTCTGTTGACATGTATGTCTGTTGTACTTGAACTTAATTTAATACATTTGTTGTTCTCCTAGAAACTTCA

>M6-PGSC0003DMG400004972M6LRR-RLK_2440

TTTGCTGCAATGCTCTAACTTGTATGGCTTACAAACTCAGGTAGCAGAATTGGCCAGAATAAATCACGAGAACACTGCAAAACTTCTAGGTTACTGCAGAGAGAGCAGTCCCTTCACAAG

>M6-PGSC0003DMG400004972M6LRR-RLK_2501

AATCACGAGAACACTGCAAAACTTCTAGGTTACTGCAGAGAGAGCAGTCCCTTCACAAGGATGTTGGTGTTCGAGTATGCATCTAACGGGACTTTATATGAACACCTCCATTGTGAGTGG

>M6-PGSC0003DMG400004972M6LRR-RLK_2562

TGTTGGTGTTCGAGTATGCATCTAACGGGACTTTATATGAACACCTCCATTGTGAGTGGAAAAAACTATTTTTTATACATTTGCTACTGCATGTTTTAGAGGCTGAAATCCAAACTCTTT

>M6-PGSC0003DMG400004972M6LRR-RLK_2623

AAAACTATTTTTTATACATTTGCTACTGCATGTTTTAGAGGCTGAAATCCAAACTCTTTGAACTTATCACAGATGGAGAAGGATGCCAACTTTCTTGGACACGGCGAATGAAAATAGTTA

>M6-PGSC0003DMG400004972M6LRR-RLK_2684

ACTTATCACAGATGGAGAAGGATGCCAACTTTCTTGGACACGGCGAATGAAAATAGTTATTGGCATCGCTAAAGGACTGAAATATCTCCATTCAGAACTTGACCCGCCATTCACTATATC

>M6-PGSC0003DMG400004972M6LRR-RLK_2745

GGCATCGCTAAAGGACTGAAATATCTCCATTCAGAACTTGACCCGCCATTCACTATATCAGAACTGAATTCAAATGCTGTATATCTCACAGAAGATTTTTCTCCCAAGGTAAGCGAAAGA

>M6-PGSC0003DMG400004972M6LRR-RLK_2806

AACTGAATTCAAATGCTGTATATCTCACAGAAGATTTTTCTCCCAAGGTAAGCGAAAGATTTCTGTATCACTGCTGCAAAACCAATTCACTGTTAAACTTTCTGGTTCATCTTACTGTCA

>M6-PGSC0003DMG400004972M6LRR-RLK_2867

TCTGTATCACTGCTGCAAAACCAATTCACTGTTAAACTTTCTGGTTCATCTTACTGTCAATTGTGCTCATCCATAAGACAATTTTAGTTTACTCTCCACGGTTTCTGCTCATTTTGTCTT

>M6-PGSC0003DMG400004972M6LRR-RLK_2928

TGTGCTCATCCATAAGACAATTTTAGTTTACTCTCCACGGTTTCTGCTCATTTTGTCTTTGAACTCGCCGAGTTTCCTTACAAACAATTTGGGCTTACATTTTTCCATTTTGGTTTCCTT

>M6-PGSC0003DMG400004972M6LRR-RLK_2989

AACTCGCCGAGTTTCCTTACAAACAATTTGGGCTTACATTTTTCCATTTTGGTTTCCTTCAAGCTCAACCCTTTGTGCTTTTTAACCCGAGCAATAGAAAGCTGATTTTCCATACCCTTT

>M6-PGSC0003DMG400004972M6LRR-RLK_305

TGTATGTCTGTTGTACTTGAACTTAATTTAATACATTTGTTGTTCTCCTAGAAACTTCACACGATTACCGAGTGTAATGAAATCTTAAAGGGTTCAACATGTATATTCAAAAGTCATCAC

>M6-PGSC0003DMG400004972M6LRR-RLK_3050

AGCTCAACCCTTTGTGCTTTTTAACCCGAGCAATAGAAAGCTGATTTTCCATACCCTTTTCTCTCAAAGTTTAATTTATCTGGTGCAGCTTGTTGATTTTGAAAGTTGGAAAACAACATT

>M6-PGSC0003DMG400004972M6LRR-RLK_3111

TCTCAAAGTTTAATTTATCTGGTGCAGCTTGTTGATTTTGAAAGTTGGAAAACAACATTTGCCAGATCAGAAAAGAATTCCGGTGCTATCAGTAGTGAAGGAGCAATGTGTGTCCTTCCA

>M6-PGSC0003DMG400004972M6LRR-RLK_3172

CCAGATCAGAAAAGAATTCCGGTGCTATCAGTAGTGAAGGAGCAATGTGTGTCCTTCCAAATTCTCTACAGAGCCGTCATCTTGACATGCAGGGTAACATCTATGCATTTGGAGTACTAC

>M6-PGSC0003DMG400004972M6LRR-RLK_3233

TTCTCTACAGAGCCGTCATCTTGACATGCAGGGTAACATCTATGCATTTGGAGTACTACTGCTCGAAATAATCAGTGGAAGACCTGCAAATTGCAAAGATAAAGGGTGCTTAGTAGATTG

>M6-PGSC0003DMG400004972M6LRR-RLK_3294

CTCGAAATAATCAGTGGAAGACCTGCAAATTGCAAAGATAAAGGGTGCTTAGTAGATTGGGTAAGCATTTCTTTTCATGAAGTTTTTGTATAGGCATAGTACATAAATATACACTCAAAC

>M6-PGSC0003DMG400004972M6LRR-RLK_3599

CTTTATGGAATCTCAAGTTGACTCTGATATGGCAGGCCAAAGAATTCCTTGAATTGCCAGAGGTATTGCCATATGTTGTAGATCCCGAGTTGAAACATTTCAGATACGAAGACCTCAAAG

>M6-PGSC0003DMG400004972M6LRR-RLK_366

CGATTACCGAGTGTAATGAAATCTTAAAGGGTTCAACATGTATATTCAAAAGTCATCACTGACGTTATCGGCAATATTCTGTATAGTTTCTGGAAAAATCAGTAAAAAGCCTCCTATGCT

>M6-PGSC0003DMG400004972M6LRR-RLK_3660

GGTATTGCCATATGTTGTAGATCCCGAGTTGAAACATTTCAGATACGAAGACCTCAAAGTGATTTCTGAAGTGGTCAATCTCTGTATCCACCCAAGTTCGAGTAGCAGGACTTCGATGAG

>M6-PGSC0003DMG400004972M6LRR-RLK_3721

ATTTCTGAAGTGGTCAATCTCTGTATCCACCCAAGTTCGAGTAGCAGGACTTCGATGAGAGATTTGTGTGCTATGCTGGAGAGCGACATTGACACATCCATAACTGCAGAGCTCAAGACA

>M6-PGSC0003DMG400004972M6LRR-RLK_3760

AGTAGCAGGACTTCGATGAGAGATTTGTGTGCTATGCTGGAGAGCGACATTGACACATCCATAACTGCAGAGCTCAAGACATCGTCTTTAGCATGGGCTGAGCTTGCACTCTCATCCTAA

>M6-PGSC0003DMG400004972M6LRR-RLK_427

ACGTTATCGGCAATATTCTGTATAGTTTCTGGAAAAATCAGTAAAAAGCCTCCTATGCTTGCTTGAACTTGTAATTGTTCTACATAGATCCCTTTTAAGTTTTATCTCACAGTTGGTTTC

>M6-PGSC0003DMG400004972M6LRR-RLK_488

CTTGAACTTGTAATTGTTCTACATAGATCCCTTTTAAGTTTTATCTCACAGTTGGTTTCTTAGTAATCGTCTTTTATTTTTGTTTAGGCATGCCTCTGGTGCCAAGCCTACTGGTCTTTG

>M6-PGSC0003DMG400004972M6LRR-RLK_549

AGTAATCGTCTTTTATTTTTGTTTAGGCATGCCTCTGGTGCCAAGCCTACTGGTCTTTGTCGTTTATCTCAACTGAGAGTTGCTGATTTCTCCTTCAACTTCTTCTTTGGTAGCATACCG

>M6-PGSC0003DMG400004972M6LRR-RLK_61

GTAATGTTTCCAGGACTTAAGAGCATTATACATTCAAAATTTTTGGAATATTTGAAGCTTGATTAAATTTATATATCTTTTTAATCTTCTAAGGTGCCCATAGTATCTCAATTAGCATAT

>M6-PGSC0003DMG400004972M6LRR-RLK_610

GTTTATCTCAACTGAGAGTTGCTGATTTCTCCTTCAACTTCTTCTTTGGTAGCATACCGAAGTGTCTGGAGTACCTTTCAAAGTATAGTTCTTAATTCCGATCCACAAGCATTTTTTATA

>M6-PGSC0003DMG400004972M6LRR-RLK_671

GTGTCTGGAGTACCTTTCAAAGTATAGTTCTTAATTCCGATCCACAAGCATTTTTTATATTCTTTATTTCAATTTCTGAGTGCCTATTATTTTGTTCTTTCTATAGAACTAGCTTTCAAG

>M6-PGSC0003DMG400004972M6LRR-RLK_732

CTTTATTTCAATTTCTGAGTGCCTATTATTTTGTTCTTTCTATAGAACTAGCTTTCAAGGAAATTGCCTTCAAGCAAAAGATCCCAGGCAGCGTTCTACTGCTTTATGTGGTATGTGTAA

>M6-PGSC0003DMG400004972M6LRR-RLK_793

AATTGCCTTCAAGCAAAAGATCCCAGGCAGCGTTCTACTGCTTTATGTGGTATGTGTAATCTGCTAGACAACTCTTACACTTTAGCAGAATCATTAACCAAGCCTTTGAATCTTTTATGC

>M6-PGSC0003DMG400004972M6LRR-RLK_854

TGCTAGACAACTCTTACACTTTAGCAGAATCATTAACCAAGCCTTTGAATCTTTTATGCATACATTAGTGTACGAAAATGCAAGCTGGTCATGCTTACTTTTGATTGTATATCCTAAAGA

>M6-PGSC0003DMG400004972M6LRR-RLK_915

ACATTAGTGTACGAAAATGCAAGCTGGTCATGCTTACTTTTGATTGTATATCCTAAAGAGGCTTCTGCGACTAAGTTCGCCCATTTCACTAACAGTCGCAGTTGAACAGATCAACTTTAA

>M6-PGSC0003DMG400004972M6LRR-RLK_976

CTTCTGCGACTAAGTTCGCCCATTTCACTAACAGTCGCAGTTGAACAGATCAACTTTAAAATAATTGAATTGATTATTGTAACTTTTGCATTATAGGAGGCACTCCACCTGCCAAAAGCA

>M6-PGSC0003DMG400005044M6G-LecRK_0

ATGTCACCAGAGTATGCAATGGGAGGACTATTTTCTGAGAAATCTGATGTCTACAGCTTTGGTGTATTGCTACTAGAGATTGTCAGTGGAAAGAGAAACAGCGGATATTATGACCACGAA

>M6-PGSC0003DMG400005044M6G-LecRK_114

CACGAAAGGCATCTCAACCTTCTCAGTTATGTAAGTGCAAAATCAGCAGGATCTCTTGCCACAAGTTATTCACACAAACTTATAGTATAAATGATTTCAGGCATGGAAGCTACAGACCGA

>M6-PGSC0003DMG400005044M6G-LecRK_171

GCCACAAGTTATTCACACAAACTTATAGTATAAATGATTTCAGGCATGGAAGCTACAGACCGAAAGCAAAGAGTTAGACTTAATGGATAAATCAATTTTGGACTCATATTCATCTGCAAC

>M6-PGSC0003DMG400005044M6G-LecRK_228

GACCGAAAGCAAAGAGTTAGACTTAATGGATAAATCAATTTTGGACTCATATTCATCTGCAACAGTGCTGAGATGCATACATATTGGCCTGCTTTGTGTCCAGGATCATGCTGTTGATAG

>M6-PGSC0003DMG400005044M6G-LecRK_342

TGATAGGCCATCAATGCCGTCTATCGTTCTTATGCTAAGTAGTGAGATGGATCTGCCTCAACCAAAGAAACCTACATTTATATTTCAAAGATGGTTGAATTCTAATACCCAATCACAATT

>M6-PGSC0003DMG400005044M6G-LecRK_397

CCTCAACCAAAGAAACCTACATTTATATTTCAAAGATGGTTGAATTCTAATACCCAATCACAATTCAGCAAAACTCAGTCTGTTAATGATCTTACTGTCTCTGTGGCAGAAGGCCGATAA

>M6-PGSC0003DMG400005044M6G-LecRK_57

TTTGGTGTATTGCTACTAGAGATTGTCAGTGGAAAGAGAAACAGCGGATATTATGACCACGAAAGGCATCTCAACCTTCTCAGTTATGTAAGTGCAAAATCAGCAGGATCTCTTGCCACA

>M6-PGSC0003DMG400005103M6LRR-RLK_0

CTCATGGCCACCATCTTGAGAAAAAATTCTCCAATTTCTCTCGCGTGGATTTCATGTTCCCACAGAAACCAAACATCAAACATAGCCATACCAAACTTTCTTCTCTATTCTTCTTACTAG

>M6-PGSC0003DMG400005103M6LRR-RLK_1037

TTCTGGTGAGATTCCTGTTGAATTTGGAGAGTTCAAGTTTACAGAACTGTCATTGTACAGAAACATGTTTTCTGGTTCTCTTCCTCAAAATATTGGATCATGGGCAGAACTTCAGTACAT

>M6-PGSC0003DMG400005103M6LRR-RLK_1098

AACATGTTTTCTGGTTCTCTTCCTCAAAATATTGGATCATGGGCAGAACTTCAGTACATTGATGTTTCTGAGAATATGTTCACTGGTTCAATACCACCTGATATGTGCAAGAAAGGGAGT

>M6-PGSC0003DMG400005103M6LRR-RLK_1159

ATGTTTCTGAGAATATGTTCACTGGTTCAATACCACCTGATATGTGCAAGAAAGGGAGTATGACTGATCTTTTGCTTCTCCAGAACAACTTCACTGGTGGGATACCTTCGAACTATGCTA

>M6-PGSC0003DMG400005103M6LRR-RLK_122

AAAAAAAAAAATGTCAGCCAGTGCCAACTTTTTCCGGCCATGCCCAATTTTATTGATCTGTTTTTTCATATTCTTGATTTCTCCTTCTCATCAACAAGATGAGTTACAACTCCTTATGAA

>M6-PGSC0003DMG400005103M6LRR-RLK_1220

GACTGATCTTTTGCTTCTCCAGAACAACTTCACTGGTGGGATACCTTCGAACTATGCTAATTGTTTGTCGTTGCAGCGTTTAAGGGTTAGCAACAATTCACTTTCAGGAGTAGTCCCTAG

>M6-PGSC0003DMG400005103M6LRR-RLK_1281

TGTTTGTCGTTGCAGCGTTTAAGGGTTAGCAACAATTCACTTTCAGGAGTAGTCCCTAGTGGAATTTGGAGTTTGCCAGATTTGGAAATCATTGATCTCACATTGAATCTATTTGAAGGC

>M6-PGSC0003DMG400005103M6LRR-RLK_1342

GAATTTGGAGTTTGCCAGATTTGGAAATCATTGATCTCACATTGAATCTATTTGAAGGCCCAGTGACATCAAATATTGGTGAAGCAAAGTCTTTAGCACAGTTGTTCCTAGCCTACAACC

>M6-PGSC0003DMG400005103M6LRR-RLK_1403

AGTGACATCAAATATTGGTGAAGCAAAGTCTTTAGCACAGTTGTTCCTAGCCTACAACCGGTTCAATGGTCAATTACCACAGACAATATCAGAAGTTTCTTCATTAGTAGCCATTAATCT

>M6-PGSC0003DMG400005103M6LRR-RLK_1464

TTCAATGGTCAATTACCACAGACAATATCAGAAGTTTCTTCATTAGTAGCCATTAATCTGAGCGCGAATCAGTTCTCTGGTGATATTCCAGCAGCAATAGGTGAACTTAAGAAGCTTAAT

>M6-PGSC0003DMG400005103M6LRR-RLK_1525

GCGCGAATCAGTTCTCTGGTGATATTCCAGCAGCAATAGGTGAACTTAAGAAGCTTAATACTCTTCATTTAGAGTATAATTTGTTTTCTGGAAGTCTTCCAGATTCTATTGGATCATGTG

>M6-PGSC0003DMG400005103M6LRR-RLK_1586

TCTTCATTTAGAGTATAATTTGTTTTCTGGAAGTCTTCCAGATTCTATTGGATCATGTGTTTCTCTCTGTGAAATTAATCTTGCTGGTAATTCACTTTCTGGTGCAATTCCCGAAAGTCT

>M6-PGSC0003DMG400005103M6LRR-RLK_1647

TCTCTCTGTGAAATTAATCTTGCTGGTAATTCACTTTCTGGTGCAATTCCCGAAAGTCTTGGCTCTTTGCCTAGCTTGAACTCTCTCAATCTATCTGATAACAGCCTCTCAGGTCAAATT

>M6-PGSC0003DMG400005103M6LRR-RLK_1708

GCTCTTTGCCTAGCTTGAACTCTCTCAATCTATCTGATAACAGCCTCTCAGGTCAAATTCCAGCGACCCTTTCGAGTTTAAGATTAAGCCTTCTTGATTTGTCAAACAATAGGTTGAGTG

>M6-PGSC0003DMG400005103M6LRR-RLK_1769

AGCGACCCTTTCGAGTTTAAGATTAAGCCTTCTTGATTTGTCAAACAATAGGTTGAGTGGTAGCATACCGGATTCTTTATCAATAAAAGCTTTTAGTAATAGCTTTTTGGGAAATCCAGA

>M6-PGSC0003DMG400005103M6LRR-RLK_183

TTTTTCATATTCTTGATTTCTCCTTCTCATCAACAAGATGAGTTACAACTCCTTATGAATTTCAAATCCACTCTGAAAACAACACAAATTTCACATCTTTTTGACACGTGGACACCTCAG

>M6-PGSC0003DMG400005103M6LRR-RLK_1830

AGCATACCGGATTCTTTATCAATAAAAGCTTTTAGTAATAGCTTTTTGGGAAATCCAGATCTTTGTAGTGAGAATTTTGGTAGTCTCAGGCCATGTTCATCAGATCCTCACACATCTAGA

>M6-PGSC0003DMG400005103M6LRR-RLK_1891

TTTGTAGTGAGAATTTTGGTAGTCTCAGGCCATGTTCATCAGATCCTCACACATCTAGAGACCACAGGACAGTCATGTTGTGCTTGATAGCTGGGGTAGTTGTTCTTGTTCTGTCACTTA

>M6-PGSC0003DMG400005103M6LRR-RLK_1952

CCACAGGACAGTCATGTTGTGCTTGATAGCTGGGGTAGTTGTTCTTGTTCTGTCACTTACGTGTTTCGTGTATGTGAAGTTTAAGCACAATAATCAGAACACTCCCGTAAAGAGACTTGA

>M6-PGSC0003DMG400005103M6LRR-RLK_2013

TGTTTCGTGTATGTGAAGTTTAAGCACAATAATCAGAACACTCCCGTAAAGAGACTTGATTCTTGGGATATCAAACAGTTTCATGTATTGAGCTTTAGTGAAGATCAAGTCTTGAAGGCA

>M6-PGSC0003DMG400005103M6LRR-RLK_2074

CTTGGGATATCAAACAGTTTCATGTATTGAGCTTTAGTGAAGATCAAGTCTTGAAGGCATTGAAGCAAGAAAATTTGATTGGTAGAGGTGGTTCAGGGAATGTGTACAGATTAGTCTTGA

>M6-PGSC0003DMG400005103M6LRR-RLK_2135

GAAGCAAGAAAATTTGATTGGTAGAGGTGGTTCAGGGAATGTGTACAGATTAGTCTTGAATTGTGGAAAACAGTTGGCTGTGAAACACATTGTTAAATCCGATTCTGGTGACCAAAAAAG

>M6-PGSC0003DMG400005103M6LRR-RLK_2196

TGTGGAAAACAGTTGGCTGTGAAACACATTGTTAAATCCGATTCTGGTGACCAAAAAAGTTACCGGGACAGCTCAGCCATACTGGTGAAGGAGAATAGAAGATCAAAGGAGTATGATGCT

>M6-PGSC0003DMG400005103M6LRR-RLK_2257

ACCGGGACAGCTCAGCCATACTGGTGAAGGAGAATAGAAGATCAAAGGAGTATGATGCTGAGGTGACCACGTTAAGTTCCATTAGGCATGTCAATGTTGTCAAATTGTACTGTAGCATCA

>M6-PGSC0003DMG400005103M6LRR-RLK_2318

GGTGACCACGTTAAGTTCCATTAGGCATGTCAATGTTGTCAAATTGTACTGTAGCATCACAAGTGAGGACTCAAATATGCTGGTTTATGAATACTTAACTAATGGAAGCTTATGGGACCG

>M6-PGSC0003DMG400005103M6LRR-RLK_2379

AGTGAGGACTCAAATATGCTGGTTTATGAATACTTAACTAATGGAAGCTTATGGGACCGATTGCACACGTCTCAGAAAGTCAAGATGGATTGGTTGGTGAGATATGATATTGCATTAGGT

>M6-PGSC0003DMG400005103M6LRR-RLK_244

TCAAATCCACTCTGAAAACAACACAAATTTCACATCTTTTTGACACGTGGACACCTCAGAATAACATTTGCAACTTCACTGGAATTTTCTGTGATTCTGACAGCAAATTGGTTAAAGAAA

>M6-PGSC0003DMG400005103M6LRR-RLK_2440

TGCACACGTCTCAGAAAGTCAAGATGGATTGGTTGGTGAGATATGATATTGCATTAGGTGCTGCTCAAGGGCTTGAGTATCTGCATCACGGATATGACAGCCCCGTGATGCACCGGGATG

>M6-PGSC0003DMG400005103M6LRR-RLK_2501

TGCTCAAGGGCTTGAGTATCTGCATCACGGATATGACAGCCCCGTGATGCACCGGGATGTCAAGTCTAGCAACATCTTGTTGGATGAACAGATGAAGCCAAAAATTGCTGATTTTGGATT

>M6-PGSC0003DMG400005103M6LRR-RLK_2562

AAGTCTAGCAACATCTTGTTGGATGAACAGATGAAGCCAAAAATTGCTGATTTTGGATTAGCCAAAGTTTTGCATGTTAATGGTAATAAGGACTCTTCTCAGGTTGTAGCTGGGACACAT

>M6-PGSC0003DMG400005103M6LRR-RLK_2623

CCAAAGTTTTGCATGTTAATGGTAATAAGGACTCTTCTCAGGTTGTAGCTGGGACACATGGCTACATTGCTCCTGGTACATATTTTTCTCTTCCCTTTTCCATTTCATTGTCACTTAAAA

>M6-PGSC0003DMG400005103M6LRR-RLK_2684

CTACATTGCTCCTGGTACATATTTTTCTCTTCCCTTTTCCATTTCATTGTCACTTAAAAATAACTTTGGCAAATTCTAAATTTTTTCTTTCTTCATTGTTTGACACAACAGAGTATGCTT

>M6-PGSC0003DMG400005103M6LRR-RLK_2745

AACTTTGGCAAATTCTAAATTTTTTCTTTCTTCATTGTTTGACACAACAGAGTATGCTTACACAACCAAGGTGACCGAGAAGAGTGATGTCTATAGCTTCGGGGTGGTGCTAATGGAATT

>M6-PGSC0003DMG400005103M6LRR-RLK_2806

ACAACCAAGGTGACCGAGAAGAGTGATGTCTATAGCTTCGGGGTGGTGCTAATGGAATTGGTGACTGGAAAGAAGCCAGTGGATGCAGAGTACGGGGAGAACAGTGACATAGTCCAATGG

>M6-PGSC0003DMG400005103M6LRR-RLK_2867

TGACTGGAAAGAAGCCAGTGGATGCAGAGTACGGGGAGAACAGTGACATAGTCCAATGGGTTTGTAGCAAGATAAGAAACAAAACTAGCATGATCGATTTGGTGGATTCAAGCATTTTCG

>M6-PGSC0003DMG400005103M6LRR-RLK_2928

TTGTAGCAAGATAAGAAACAAAACTAGCATGATCGATTTGGTGGATTCAAGCATTTTCGAGGGGTTCAAAGAAGATGCTGTTGAGGTTCTTAAAATTGCAGTTCATTGCACATCAAGGAC

>M6-PGSC0003DMG400005103M6LRR-RLK_2989

GGGTTCAAAGAAGATGCTGTTGAGGTTCTTAAAATTGCAGTTCATTGCACATCAAGGACGCCAGCATTAAGACCTTCCATGAGGATGGTAGTTCATATGCTAGAAGAAGCCGAGCCTTGC

>M6-PGSC0003DMG400005103M6LRR-RLK_305

TAACATTTGCAACTTCACTGGAATTTTCTGTGATTCTGACAGCAAATTGGTTAAAGAAATCAATCTTTCTGAACAGAATCTATCTGGGGTTGTCTCTTTTGATTCACTATGCTCTCTGAA

>M6-PGSC0003DMG400005103M6LRR-RLK_3050

CAGCATTAAGACCTTCCATGAGGATGGTAGTTCATATGCTAGAAGAAGCCGAGCCTTGCAAGCTGACAAATGTAGTCGTAAATTCACCGAATGAGGATGGCAGGAAACAAGGACTTGCTA

>M6-PGSC0003DMG400005103M6LRR-RLK_3111

GCTGACAAATGTAGTCGTAAATTCACCGAATGAGGATGGCAGGAAACAAGGACTTGCTAACTAATGGCAAGTCATGATGTTGAACTATCTCGTTGAGAGCAGAGGAATTTTGCAGAATAT

>M6-PGSC0003DMG400005103M6LRR-RLK_3172

TAATGGCAAGTCATGATGTTGAACTATCTCGTTGAGAGCAGAGGAATTTTGCAGAATATTGAACTATGTTGTAAGTTCACAAAGAATGCTTAGTGTACATATGACACTTTTGAAGAAATA

>M6-PGSC0003DMG400005103M6LRR-RLK_3233

AACTATGTTGTAAGTTCACAAAGAATGCTTAGTGTACATATGACACTTTTGAAGAAATAGTGTATTTCACAGCTGCATTAATAAATCCAACTAGCAAGTAATGTATTTCACATCCAAATT

>M6-PGSC0003DMG400005103M6LRR-RLK_3284

AAGAAATAGTGTATTTCACAGCTGCATTAATAAATCCAACTAGCAAGTAATGTATTTCACATCCAAATTACTTATAGAAACTGAGCAGTTGATCTACCCTACAACCAATTTTATTTTTCT

>M6-PGSC0003DMG400005103M6LRR-RLK_366

AATCTTTCTGAACAGAATCTATCTGGGGTTGTCTCTTTTGATTCACTATGCTCTCTGAAATCACTAGAAAAGATTTCTCTTGGTACTAACTACTTGTATGGTAGAGTCAGCGACCATTTG

>M6-PGSC0003DMG400005103M6LRR-RLK_427

CACTAGAAAAGATTTCTCTTGGTACTAACTACTTGTATGGTAGAGTCAGCGACCATTTGAAAAACTGTACAGAGTTGCAGTATTTGGATTTGGGTAACAATTCTTTTTCAGGGGAAGTTC

>M6-PGSC0003DMG400005103M6LRR-RLK_488

AAACTGTACAGAGTTGCAGTATTTGGATTTGGGTAACAATTCTTTTTCAGGGGAAGTTCCAAATTTGTCTTCTTTAAGCCAATTGGAGTTTTTGAATCTCAATAGAAGTGGATTCTCTGG

>M6-PGSC0003DMG400005103M6LRR-RLK_549

AATTTGTCTTCTTTAAGCCAATTGGAGTTTTTGAATCTCAATAGAAGTGGATTCTCTGGTTCATTTCCATGGAGTTCACTTGGAAATCTTACTAGTTTAACTTTCTTGAGTTTGGGTGAC

>M6-PGSC0003DMG400005103M6LRR-RLK_61

ACAGAAACCAAACATCAAACATAGCCATACCAAACTTTCTTCTCTATTCTTCTTACTAGCGAAAAAAAAAAATGTCAGCCAGTGCCAACTTTTTCCGGCCATGCCCAATTTTATTGATCT

>M6-PGSC0003DMG400005103M6LRR-RLK_610

CATTTCCATGGAGTTCACTTGGAAATCTTACTAGTTTAACTTTCTTGAGTTTGGGTGACAATTCATTTAATAAAAGTCCATTCCCTCTTGAAATCTTGAATCTTGATAAGTTATATTGGG

>M6-PGSC0003DMG400005103M6LRR-RLK_671

TTCATTTAATAAAAGTCCATTCCCTCTTGAAATCTTGAATCTTGATAAGTTATATTGGGTTTACCTTACCAATAGTAGTATTGAAGGCCAAATTCCAGAAGGTATTGGAAATCTCACTCT

>M6-PGSC0003DMG400005103M6LRR-RLK_732

TACCTTACCAATAGTAGTATTGAAGGCCAAATTCCAGAAGGTATTGGAAATCTCACTCTGCTTGAAAATATTGAGCTTTCATATAATGACTTGTCAGGCAAAATCCCAGATGGAATAACC

>M6-PGSC0003DMG400005103M6LRR-RLK_793

TTGAAAATATTGAGCTTTCATATAATGACTTGTCAGGCAAAATCCCAGATGGAATAACCAAACTTTCTAAGCTTAAACAGCTTGAGATCTATTCTAATGGACTCACAGGAAAGTTCCCTG

>M6-PGSC0003DMG400005103M6LRR-RLK_854

ACTTTCTAAGCTTAAACAGCTTGAGATCTATTCTAATGGACTCACAGGAAAGTTCCCTGTGGAGTTTGGGAATCTCAGTAGCCTTGTCAATTTTGATGCTTCAAGTAACAATCTTGAAGG

>M6-PGSC0003DMG400005103M6LRR-RLK_915

GAGTTTGGGAATCTCAGTAGCCTTGTCAATTTTGATGCTTCAAGTAACAATCTTGAAGGTGATCTTTCAGAGCTCAAGTCTTTATCTCTTCTTGAATCTTTGCAGCTGTTTGAAAACCAT

>M6-PGSC0003DMG400005103M6LRR-RLK_976

ATCTTTCAGAGCTCAAGTCTTTATCTCTTCTTGAATCTTTGCAGCTGTTTGAAAACCATTTTTCTGGTGAGATTCCTGTTGAATTTGGAGAGTTCAAGTTTACAGAACTGTCATTGTACA

>M6-PGSC0003DMG400005171-(rev)M6LRR-RLP_0

ATGATGAAGCTAAGGCATCTGAATATAATCAATTTGTGCTTCTTGTGGGAAGATAATGATCGAGCAATTTTTGACGAATCTTCAACAACTATGCTAGAGAACTTGAAGACATTCTTCAGT

>M6-PGSC0003DMG400005171-(rev)M6LRR-RLP_124

ATATACGTGTAGATGATACAACTCCAAGATTCTGGTGGAGGTTTCCAAATATTGAATAACTTAGCTGCCTCAGTGTTGACGATTTACCTAGTTGTCCTTTGTTTCCCTCAGTGGAAGTTA

>M6-PGSC0003DMG400005171-(rev)M6LRR-RLP_186

AGCTGCCTCAGTGTTGACGATTTACCTAGTTGTCCTTTGTTTCCCTCAGTGGAAGTTAATACTCGGCATCAATCTCTTGATCTGATGATAAGCTATAAGGGATTCTGGGATTCAGTTGGT

>M6-PGSC0003DMG400005171-(rev)M6LRR-RLP_248

TCGGCATCAATCTCTTGATCTGATGATAAGCTATAAGGGATTCTGGGATTCAGTTGGTTGGGAGAGCTACTTTGTCTTCCCTTCAAATCTTAGGCATCTGTCCCTCAGTGGTTGTTTTTT

>M6-PGSC0003DMG400005171-(rev)M6LRR-RLP_310

AGAGCTACTTTGTCTTCCCTTCAAATCTTAGGCATCTGTCCCTCAGTGGTTGTTTTTTGACAGAAAAAATGGTTTCAAACATTGCAAGATTGAAGAAACTTGAGAAACTCTCATTAGTTG

>M6-PGSC0003DMG400005171-(rev)M6LRR-RLP_372

GAAAAAATGGTTTCAAACATTGCAAGATTGAAGAAACTTGAGAAACTCTCATTAGTTGGGGGATTTCCTTTGGGGAAAATAGAGAACCGTTGCTGGGACGTCACAAATGTGGAGTTCCCT

>M6-PGSC0003DMG400005171-(rev)M6LRR-RLP_434

ATTTCCTTTGGGGAAAATAGAGAACCGTTGCTGGGACGTCACAAATGTGGAGTTCCCTGCACTAAAATACTTGGCATTACGCTTTATGAGGATGGAAGAATGGAAAGCTTCAGAGGAATC

>M6-PGSC0003DMG400005171-(rev)M6LRR-RLP_496

TAAAATACTTGGCATTACGCTTTATGAGGATGGAAGAATGGAAAGCTTCAGAGGAATCCTTTCCTGTGCTTGAGGAGCTAACTATTAGTGTTGGTTACATCATGGAGATTCCTCCTAGTT

>M6-PGSC0003DMG400005171-(rev)M6LRR-RLP_558

CCTGTGCTTGAGGAGCTAACTATTAGTGTTGGTTACATCATGGAGATTCCTCCTAGTTTTGCGGATATTTTAACACTGCGACTTATTAAACTGGACAAGCTCTCCAATTCACTTGTGGTT

>M6-PGSC0003DMG400005171-(rev)M6LRR-RLP_615

TTTGCGGATATTTTAACACTGCGACTTATTAAACTGGACAAGCTCTCCAATTCACTTGTGGTTTCAGCTTTGACTATTAAAAAGGAAATAGAAGAAAACACAGGATGTGACCGTCTCCAA

>M6-PGSC0003DMG400005171-(rev)M6LRR-RLP_62

AGCAATTTTTGACGAATCTTCAACAACTATGCTAGAGAACTTGAAGACATTCTTCAGTATTCATATACGTGTAGATGATACAACTCCAAGATTCTGGTGGAGGTTTCCAAATATTGAATA

>M6-PGSC0003DMG400005172-(rev)M6LRR-RLP_0

ATGGAAGTTGTTGAGACTTGTGGAAAAGGAACTCTTTCTGCTGGAGAACCTTCCACCAAACATAATATTACATCAATTTATGATGATGATGATGATGAAGTTGTGGGCTTTGAGAAAGAT

>M6-PGSC0003DMG400005172-(rev)M6LRR-RLP_128

TATAATGAAGAAAATGATACGAGGAACAAAGGAGCAGGATGTTATCTCCATCTATGGAATGCCTGGACTTGGTAAAACAACTTTGGCAAGGAAGGTGTACAACAATCCCTCTATTATTAA

>M6-PGSC0003DMG400005172-(rev)M6LRR-RLP_192

GGACTTGGTAAAACAACTTTGGCAAGGAAGGTGTACAACAATCCCTCTATTATTAATTACTTTGATGTTAAAGCATGGTGTGCTTTGTCGCAAGCATATAATAGGAGGACATTATTGGTT

>M6-PGSC0003DMG400005172-(rev)M6LRR-RLP_256

ATGTTAAAGCATGGTGTGCTTTGTCGCAAGCATATAATAGGAGGACATTATTGGTTGAGATTTTCAAACAAGCTACAAATGATAAGATCAAAATCAAAGAGGACGACGATGTAGCTGACA

>M6-PGSC0003DMG400005172-(rev)M6LRR-RLP_320

CAAACAAGCTACAAATGATAAGATCAAAATCAAAGAGGACGACGATGTAGCTGACATGTTGCGCAGGGTTCTAATTGGCAAGAGATACCTCATCGTATTAGATGACATTTGGGATGTCGA

>M6-PGSC0003DMG400005172-(rev)M6LRR-RLP_384

AGGGTTCTAATTGGCAAGAGATACCTCATCGTATTAGATGACATTTGGGATGTCGAGGCATGGGAAGATTTGGGAATATGTTTCCCTCAAGGTGAATATGGAAGCAGAGTAATGGTCACA

>M6-PGSC0003DMG400005172-(rev)M6LRR-RLP_448

AAGATTTGGGAATATGTTTCCCTCAAGGTGAATATGGAAGCAGAGTAATGGTCACAACACGAATTGAACAAGTGGCTAAGCATCTTCAGCACCACAGTGATCCTTATTCTCTTAGCTTTC

>M6-PGSC0003DMG400005172-(rev)M6LRR-RLP_512

TGAACAAGTGGCTAAGCATCTTCAGCACCACAGTGATCCTTATTCTCTTAGCTTTCTAACGTCCGAAAAGAGTTGGGAATTATTGGAGAAGAAGGTATTTCGAGGAGAGAGTTGTCCTCC

>M6-PGSC0003DMG400005172-(rev)M6LRR-RLP_64

ATATTACATCAATTTATGATGATGATGATGATGAAGTTGTGGGCTTTGAGAAAGATGCAGAAATTATAATGAAGAAAATGATACGAGGAACAAAGGAGCAGGATGTTATCTCCATCTATG

>M6-PGSC0003DMG400005226M6LRR-RLP_0

AATATTAAAAGACTTGATTTGTCTTCTAATAATTTTTCAGGATCGCTCATTTCACCTAAATTTGGTGAGTTTTCTAGTTTGACGCATCTTGATTTGTCAAATTCAGGTTTTAGATTCGGA

>M6-PGSC0003DMG400005226M6LRR-RLP_1054

CCACAATGTTTGGGTGAGATGAGTGGACTTTGGGTTTTTGATCTAAGCAACAACGGTCTTAGTGGGGCAATTAATACAACATTTAGTATAGGAAACACGCTCAAAGTCATTAACTTGCAC

>M6-PGSC0003DMG400005226M6LRR-RLP_1116

TGGGGCAATTAATACAACATTTAGTATAGGAAACACGCTCAAAGTCATTAACTTGCACGGGAATAAGCTAGAGGGGAAGTCCCACGATCTTTGGTCAATTGCAAGAGTTTGGAAATTCTA

>M6-PGSC0003DMG400005226M6LRR-RLP_1178

ATAAGCTAGAGGGGAAGTCCCACGATCTTTGGTCAATTGCAAGAGTTTGGAAATTCTAGATTTAGGTAACAATGAGTTGAGTGACACATTTCCAAAATGGTTAGGAACCCTACCGAATTT

>M6-PGSC0003DMG400005226M6LRR-RLP_124

ACAATTTTGAACTTCTTCTTAAGAACTTAACCCAATTAACAGAGCTTGACCTTTACGGTGTGAGCATCTCATCCACCTTTCCTTTGAATTTCTCTTCTTATTTAGCAACTCTACGACTTA

>M6-PGSC0003DMG400005226M6LRR-RLP_1302

ATTTTAAGCTTGAGATCAAATAAGTTCCACGGGCCCATTAAAACTTCAAGGGCTGGAAACTTGTTTGTACATCTTCGAATAATTGATCTATCATCCAATGGATTTAGTGGGAATTTACCT

>M6-PGSC0003DMG400005226M6LRR-RLP_1364

GTTTGTACATCTTCGAATAATTGATCTATCATCCAATGGATTTAGTGGGAATTTACCTATGAGCCTTTTTGAGAATTTGAAAGCTATGAAGATAATTGATGAGATCACGAAAATCCCACG

>M6-PGSC0003DMG400005226M6LRR-RLP_1426

GCCTTTTTGAGAATTTGAAAGCTATGAAGATAATTGATGAGATCACGAAAATCCCACGGTATGTGGGAGAAGACTTGTATTATTATTATAATTTTTCTGAAACAATTACAACAAAGGGGT

>M6-PGSC0003DMG400005226M6LRR-RLP_1488

GTGGGAGAAGACTTGTATTATTATTATAATTTTTCTGAAACAATTACAACAAAGGGGTTGGATCTTGAATTTCGTGGAGTTTTGACTACAAACATAGTTATCAATCTCTCGAAGAATAGA

>M6-PGSC0003DMG400005226M6LRR-RLP_1550

TCTTGAATTTCGTGGAGTTTTGACTACAAACATAGTTATCAATCTCTCGAAGAATAGATTTGAAGGTTGTATTCCAAGTTCCATTGGAGATCTTATTGCACTTCGTACTTTGAATTTATC

>M6-PGSC0003DMG400005226M6LRR-RLP_1612

AAGGTTGTATTCCAAGTTCCATTGGAGATCTTATTGCACTTCGTACTTTGAATTTATCTCATAATGGCTTGGAGGGTGTCATACCAACATCGCTGCAGTACTTATCTGTACTTGAATCAT

>M6-PGSC0003DMG400005226M6LRR-RLP_1674

AATGGCTTGGAGGGTGTCATACCAACATCGCTGCAGTACTTATCTGTACTTGAATCATTAGATCTCTCATTCAACAAAATAAGCGGAGAAATTCCACAACAACTTGCATCCCTCACATTT

>M6-PGSC0003DMG400005226M6LRR-RLP_1736

TCTCTCATTCAACAAAATAAGCGGAGAAATTCCACAACAACTTGCATCCCTCACATTTCTTGCTGTCTTAAATCTCTCTCACAATCATTTTGTTGGATGTATCCCTAGAGGAAAACAATT

>M6-PGSC0003DMG400005226M6LRR-RLP_1860

ACGTTTGAGAATAGTTCATACCAAGGGAATGATGGGTTACGTGGATTGCCAGTGTCAAAAGATTGTGGTGGTGATGAAGGGGTACCACAAACAACAACTCCATTTGGGCTTGATCAAGGA

>M6-PGSC0003DMG400005226M6LRR-RLP_1922

TTGTGGTGGTGATGAAGGGGTACCACAAACAACAACTCCATTTGGGCTTGATCAAGGAGAAGAAGGAGACTCAACAATGATCAGTTGGCAGGCAGTTCTCATGGGTTATGGTTGCGGTCT

>M6-PGSC0003DMG400005226M6LRR-RLP_1984

AAGGAGACTCAACAATGATCAGTTGGCAGGCAGTTCTCATGGGTTATGGTTGCGGTCTAATTATTGGATTATCCGTAATATACATAATGTTATCAACTCAAAATCCAACATGGTTTTCGA

>M6-PGSC0003DMG400005226M6LRR-RLP_2108

GGTTGAAGAATTGGACCACAAGATTATTACGAGAATGAAAAAATACAATAAAAGCTATTAGTGTGCAACCTAGCCAGGTATTCAACTTGATCTTTATCCTTCATAAGATTATTTTGTATA

>M6-PGSC0003DMG400005226M6LRR-RLP_2170

GTGCAACCTAGCCAGGTATTCAACTTGATCTTTATCCTTCATAAGATTATTTTGTATATCTCAATGTATTATATATCTACCTCCTTCATCCTTAAAATTCTTGACTTCGATTCTTCATTT

>M6-PGSC0003DMG400005226M6LRR-RLP_2232

AATGTATTATATATCTACCTCCTTCATCCTTAAAATTCTTGACTTCGATTCTTCATTTTTTAAAATTGCAGGATTCGAGCCTAATATGTTGTTGGTGATGCAAATATAAGCTTTTATTTT

>M6-PGSC0003DMG400005226M6LRR-RLP_2294

AAATTGCAGGATTCGAGCCTAATATGTTGTTGGTGATGCAAATATAAGCTTTTATTTTTCACATTTTGTGATGTATCTAAGTATTTATCGCCCTACGAATAAAGACATGAGCTTTCTTCC

>M6-PGSC0003DMG400005226M6LRR-RLP_2356

ATTTTGTGATGTATCTAAGTATTTATCGCCCTACGAATAAAGACATGAGCTTTCTTCCTTTGAGCTTCGGCCATAATTTTTAGAATGTTTTTAAATAGAGATTGTTAGCTTGAGTTCAGC

>M6-PGSC0003DMG400005226M6LRR-RLP_2396

AGACATGAGCTTTCTTCCTTTGAGCTTCGGCCATAATTTTTAGAATGTTTTTAAATAGAGATTGTTAGCTTGAGTTCAGCCTCATAATTAGTTGAGTAATTAGGCTCAACTGCAAAGTCA

>M6-PGSC0003DMG400005226M6LRR-RLP_248

CACACAATTATGGGGGATACTGCCCGAAAGTGTTTTTCACCTTTCCAACCTGGAAACACTTGATTTATCATACAATTCCCAACTCACAATTAGGTTTCCCACGACCAAATGGAATAGCAG

>M6-PGSC0003DMG400005226M6LRR-RLP_310

ATTTATCATACAATTCCCAACTCACAATTAGGTTTCCCACGACCAAATGGAATAGCAGTACATCTCTCATGTACTTATCTCTACCTGAAGTGAATTTTATTGGTAAGATACCCGAATCAT

>M6-PGSC0003DMG400005226M6LRR-RLP_372

TCTCTCATGTACTTATCTCTACCTGAAGTGAATTTTATTGGTAAGATACCCGAATCATTTAGCTATCTAACTTCATTGCATTACTTGGATATGAGTTCTTGTAGCCTCTCAGGTCCTATT

>M6-PGSC0003DMG400005226M6LRR-RLP_434

CTATCTAACTTCATTGCATTACTTGGATATGAGTTCTTGTAGCCTCTCAGGTCCTATTATTAAACCTCTTTGGAATCTCTCCCACATACAAAGTTTGAATCTCAGAAATAACCATCTTGA

>M6-PGSC0003DMG400005226M6LRR-RLP_496

AACCTCTTTGGAATCTCTCCCACATACAAAGTTTGAATCTCAGAAATAACCATCTTGAAGGACCAATTTCCCAATTCTTCAAATTTGGAAACCTCACTTGGAATTAACAACTTTGATGGC

>M6-PGSC0003DMG400005226M6LRR-RLP_558

CCAATTTCCCAATTCTTCAAATTTGGAAACCTCACTTGGAATTAACAACTTTGATGGCTAACTTGATTTTTTATCCTCCAACAGAAGTTGGACTCAACTTGAAGGGTTCGAAGTTTCTTC

>M6-PGSC0003DMG400005226M6LRR-RLP_62

TGGTGAGTTTTCTAGTTTGACGCATCTTGATTTGTCAAATTCAGGTTTTAGATTCGGACCTCACAATTTTGAACTTCTTCTTAAGAACTTAACCCAATTAACAGAGCTTGACCTTTACGG

>M6-PGSC0003DMG400005226M6LRR-RLP_620

TTGATTTTTTATCCTCCAACAGAAGTTGGACTCAACTTGAAGGGTTCGAAGTTTCTTCCAATTCCTTAGTTGGTCCAATTCCTTCCGATGTAAGTGGTCTGCAAAGCCTACAATCATTCT

>M6-PGSC0003DMG400005226M6LRR-RLP_682

TCCTTAGTTGGTCCAATTCCTTCCGATGTAAGTGGTCTGCAAAGCCTACAATCATTCTTCTTGTCATCAAACCACTTGAATGGGACTATACCATCATGGATATTCTCCCTTCCTTCACTA

>M6-PGSC0003DMG400005226M6LRR-RLP_744

GTCATCAAACCACTTGAATGGGACTATACCATCATGGATATTCTCCCTTCCTTCACTAGATGAATTATACTTGAATGATAACTATTTCAGAGGAGAAATTCAGGAGTTCAAGTCCAAAAC

>M6-PGSC0003DMG400005226M6LRR-RLP_806

AATTATACTTGAATGATAACTATTTCAGAGGAGAAATTCAGGAGTTCAAGTCCAAAACATTGTGGGACGTTACTCTAAAACAAAATCAGTTGCAAGGTCCTATTCCGATGTCACTCCTAA

>M6-PGSC0003DMG400005226M6LRR-RLP_868

TGGGACGTTACTCTAAAACAAAATCAGTTGCAAGGTCCTATTCCGATGTCACTCCTAAACCAACAGAGCCTATCTTATCTTCTCCTTTCGAAAAATAATCTCAGTGGACAGATTTCTTCA

>M6-PGSC0003DMG400005226M6LRR-RLP_930

ACAGAGCCTATCTTATCTTCTCCTTTCGAAAAATAATCTCAGTGGACAGATTTCTTCAGCTATCTGCAGTCTAAATGCCCTGAAAGTGCTAGATCTTGGAAGTAATAATTTGAAGGGAAC

>M6-PGSC0003DMG400005226M6LRR-RLP_992

TCTGCAGTCTAAATGCCCTGAAAGTGCTAGATCTTGGAAGTAATAATTTGAAGGGAACAATCCCACAATGTTTGGGTGAGATGAGTGGACTTTGGGTTTTTGATCTAAGCAACAACGGTC

>M6-PGSC0003DMG400005254-(rev)M6LRR-RLK_0

AAAGATTTAAAAAGTAATTGGCACCTTGCTTTTTAGTGGCCTACAATGGAAAGAATGGCTTTTATTGCCAAAAGGACTTTAGGCAAAGTATGCAATAAATCCTCTTCCCACTCACTTTGA

>M6-PGSC0003DMG400005254-(rev)M6LRR-RLK_1054

TGATATTCCTGTTTCTTTATCTCCTCGTCTTGGTGTACTAGATCTCTCCTTTAACTCTTTTACTGGAGAAATCCCCCCCACAATCAAGAACTTGACGCGATTATCTGTGTTGAACCTTCA

>M6-PGSC0003DMG400005254-(rev)M6LRR-RLK_1116

CTGGAGAAATCCCCCCCACAATCAAGAACTTGACGCGATTATCTGTGTTGAACCTTCAGTTCAATTCATTGACAGGAGAAATCCCCAGCCTTGACACCCTCAGGTTAACTAACCTGAATT

>M6-PGSC0003DMG400005254-(rev)M6LRR-RLK_1178

AATTCATTGACAGGAGAAATCCCCAGCCTTGACACCCTCAGGTTAACTAACCTGAATTTGAGTTACAATATGCTAAATGGCTCCGTTCCAGACCCCCTCCAGAAGTTTCCACTTACATCA

>M6-PGSC0003DMG400005254-(rev)M6LRR-RLK_124

CTGTCTTCAGTAGTTCAATTAGTATCTGTCATAATTCTTGCCTACTGGATTGATCCATTTTCAATATGAAGAGGTAGTAAATCAAATGAAGTTTCCCTTTCATTGAACAACACAGCTCTC

>M6-PGSC0003DMG400005254-(rev)M6LRR-RLK_1240

TTACAATATGCTAAATGGCTCCGTTCCAGACCCCCTCCAGAAGTTTCCACTTACATCATTCGCAGGAAACTCTCATTTATGTGGTACACCTCTAAATAGCTGTTCCTCGACCCCGTCACC

>M6-PGSC0003DMG400005254-(rev)M6LRR-RLK_1302

CAGGAAACTCTCATTTATGTGGTACACCTCTAAATAGCTGTTCCTCGACCCCGTCACCTTCTCCAGCAGCTGATGGCTCTGCAATCCCTGAAAAACAGAAGGCTGTCCACTCTAAGAAAC

>M6-PGSC0003DMG400005254-(rev)M6LRR-RLK_1364

CCAGCAGCTGATGGCTCTGCAATCCCTGAAAAACAGAAGGCTGTCCACTCTAAGAAACTAAGTACCGGAATCATCATTGCCATTGTGGTAGTGGTTTCTTTAGTAATGTTTCTCCTAGTT

>M6-PGSC0003DMG400005254-(rev)M6LRR-RLK_1426

TACCGGAATCATCATTGCCATTGTGGTAGTGGTTTCTTTAGTAATGTTTCTCCTAGTTTTAGTAATCTCCTTCTGCTGCTTGAAAAAGAAAGTGAGTCATAGCACTAGCATAATTAAAGA

>M6-PGSC0003DMG400005254-(rev)M6LRR-RLK_1488

TAATCTCCTTCTGCTGCTTGAAAAAGAAAGTGAGTCATAGCACTAGCATAATTAAAGAGAAGGTGGCTAATGGGGGAAGAAGTGAAAAACCCGAGGACTTCGGAAGTGGTGTGCCAGATG

>M6-PGSC0003DMG400005254-(rev)M6LRR-RLK_1550

GTGGCTAATGGGGGAAGAAGTGAAAAACCCGAGGACTTCGGAAGTGGTGTGCCAGATGCTGAGAAAAACAAGTTGGTGTTTTTCGAAGGTTGCTCTTACAGCTTCAATCTGGAGGATCTT

>M6-PGSC0003DMG400005254-(rev)M6LRR-RLK_1612

GAAAAACAAGTTGGTGTTTTTCGAAGGTTGCTCTTACAGCTTCAATCTGGAGGATCTTCTTAGAGCTTCTGCTGAGGTTCTTGGAAAAGGGAGTTATGGTACAGCTTACAAAGCTGTCTT

>M6-PGSC0003DMG400005254-(rev)M6LRR-RLK_1674

GAGCTTCTGCTGAGGTTCTTGGAAAAGGGAGTTATGGTACAGCTTACAAAGCTGTCTTGGATGAGGCAACGATTGTGGTGGTGAAACGGCTGAGAGAAGTAGGGGTTGCCAAAAAGGAGT

>M6-PGSC0003DMG400005254-(rev)M6LRR-RLK_1736

GAGGCAACGATTGTGGTGGTGAAACGGCTGAGAGAAGTAGGGGTTGCCAAAAAGGAGTTTGAACAGCATATGGAGATAGTTGGAAGGGCTGGGAGGCACCCTAATATCGTTCCACTCCGC

>M6-PGSC0003DMG400005254-(rev)M6LRR-RLK_1798

ACAGCATATGGAGATAGTTGGAAGGGCTGGGAGGCACCCTAATATCGTTCCACTCCGCGCATATTATTATTCCAAGGATGAGAAACTTCTTGTTAATGAATACATGCCTGCTGGTAGCTT

>M6-PGSC0003DMG400005254-(rev)M6LRR-RLK_186

AATATGAAGAGGTAGTAAATCAAATGAAGTTTCCCTTTCATTGAACAACACAGCTCTCAGATCCCTTTCATGCATATCATATAGGTAAGTTAACTTGTGTGCCATATATTTTAGAACTAA

>M6-PGSC0003DMG400005254-(rev)M6LRR-RLK_1860

ATTATTATTCCAAGGATGAGAAACTTCTTGTTAATGAATACATGCCTGCTGGTAGCTTGTCAGCAGCCTTGCACGGTATAATTGCTAACCTTTATCTAACCTGTGTTGCTCAGACTCTTC

>M6-PGSC0003DMG400005254-(rev)M6LRR-RLK_2046

ATTTTTGAAGAGTCCGAGCAACACAACCTCTAAACTTTTTTCTTGCACATTAATTGCTTTGATCATCTGTAATGATTCATATTTTTACGCAATCTTGATAATGAATGACAGGTAATCGAG

>M6-PGSC0003DMG400005254-(rev)M6LRR-RLK_2108

TCATCTGTAATGATTCATATTTTTACGCAATCTTGATAATGAATGACAGGTAATCGAGGTATTGGAAGAACTCCACTAGATTGGGATTCAAGATTGAAGATATCTCAGGGAGCAGCAAAA

>M6-PGSC0003DMG400005254-(rev)M6LRR-RLK_2170

TGGAAGAACTCCACTAGATTGGGATTCAAGATTGAAGATATCTCAGGGAGCAGCAAAAGGAATTGCCCACATCCACACTGAAGGTGGTGTGAAATTTACTCATGGAAACATCAAGTCCTC

>M6-PGSC0003DMG400005254-(rev)M6LRR-RLK_2232

TTGCCCACATCCACACTGAAGGTGGTGTGAAATTTACTCATGGAAACATCAAGTCCTCCAATGTACTCCTGACAAGAGACCTAGATGGATGCATTTCTGATTTTGGCTTAACACCTCTGA

>M6-PGSC0003DMG400005254-(rev)M6LRR-RLK_2294

GTACTCCTGACAAGAGACCTAGATGGATGCATTTCTGATTTTGGCTTAACACCTCTGATGAACTACATATCATACAAATATAGATGTGCTGGTTACCGCGCACCAGAAGTTATTGAAACA

>M6-PGSC0003DMG400005254-(rev)M6LRR-RLK_2356

CTACATATCATACAAATATAGATGTGCTGGTTACCGCGCACCAGAAGTTATTGAAACACGGAAAGGGACACAAAAATCCGATGTTTATAGCTTTGGAGTGCTGCTCCTTGAGATGCTAAC

>M6-PGSC0003DMG400005254-(rev)M6LRR-RLK_2418

AAGGGACACAAAAATCCGATGTTTATAGCTTTGGAGTGCTGCTCCTTGAGATGCTAACAGGCAAATCCCCTCTCCCATTATCAGGGCAGGATGAGGTAGTTGATTTACCTAGGTGGGTCC

>M6-PGSC0003DMG400005254-(rev)M6LRR-RLK_248

CCCTTTCATGCATATCATATAGGTAAGTTAACTTGTGTGCCATATATTTTAGAACTAAAAGAGTAAATCTTGGCTTTCACAATAACCAAATGAATTTTTTCTATTCTGTCATTCTGAGAT

>M6-PGSC0003DMG400005254-(rev)M6LRR-RLK_2480

AAATCCCCTCTCCCATTATCAGGGCAGGATGAGGTAGTTGATTTACCTAGGTGGGTCCGGTCCGTTGTAAGAGAGGAATGGACTGCTGAAGTTTTCGACGTAGAGCTGTTGAAGTACCAG

>M6-PGSC0003DMG400005254-(rev)M6LRR-RLK_2542

CGTTGTAAGAGAGGAATGGACTGCTGAAGTTTTCGACGTAGAGCTGTTGAAGTACCAGAATATTGAAGAAGAGATGGTGCAGATGCTTCAGATTGGACTAGCTTGTGTAGCAAAGGTGCC

>M6-PGSC0003DMG400005254-(rev)M6LRR-RLK_2604

TTGAAGAAGAGATGGTGCAGATGCTTCAGATTGGACTAGCTTGTGTAGCAAAGGTGCCTGATATGAGACCTGCTATGGGAGAAGTGGTTAGAATGATTGAAGAAATTTGACAACCAGAAG

>M6-PGSC0003DMG400005254-(rev)M6LRR-RLK_2666

ATGAGACCTGCTATGGGAGAAGTGGTTAGAATGATTGAAGAAATTTGACAACCAGAAGGTGAAACGAGACCTTCCTCGGAGGACAGCAGGTCCAAGGACTCTAATGCACAAACACCTGAA

>M6-PGSC0003DMG400005254-(rev)M6LRR-RLK_2728

AACGAGACCTTCCTCGGAGGACAGCAGGTCCAAGGACTCTAATGCACAAACACCTGAATGAAAAATGCAGTTTTTAGTCATTCAGCTTTCAATCTTGATCATTTTAAGAATGCAATTTAT

>M6-PGSC0003DMG400005254-(rev)M6LRR-RLK_2790

AAATGCAGTTTTTAGTCATTCAGCTTTCAATCTTGATCATTTTAAGAATGCAATTTATCTCCATTGCCAACTTTTGCATGTTCTTGGCCATACAAGTAGTGTTATTTTTATTTTATTTTT

>M6-PGSC0003DMG400005254-(rev)M6LRR-RLK_310

GTAAATCTTGGCTTTCACAATAACCAAATGAATTTTTTCTATTCTGTCATTCTGAGATAGAGTTCTTCCTTTTTCGTTACTTCTTTAGCTGATATTATTTCTGGATTGTATAGTTTTCTT

>M6-PGSC0003DMG400005254-(rev)M6LRR-RLK_372

TTCTTCCTTTTTCGTTACTTCTTTAGCTGATATTATTTCTGGATTGTATAGTTTTCTTGCATTGTGTTAAAACTTTTTCATTGTATTTGTTAATGGTTTCTTATAGCAATCTGATTTTAT

>M6-PGSC0003DMG400005254-(rev)M6LRR-RLK_434

TGTGTTAAAACTTTTTCATTGTATTTGTTAATGGTTTCTTATAGCAATCTGATTTTATCTCTCTTTCACAGTGGCTAATGAGCTGCTTCCTCTAAGCGGAACCAACCGGCTGATTTTGTC

>M6-PGSC0003DMG400005254-(rev)M6LRR-RLK_496

CTTTCACAGTGGCTAATGAGCTGCTTCCTCTAAGCGGAACCAACCGGCTGATTTTGTCTAATGCTCTTCAAAAGTAGTCAAGGAGAGAGAGGGAAGAAGTGAAGGACTGCTGTAACTCGG

>M6-PGSC0003DMG400005254-(rev)M6LRR-RLK_558

GCTCTTCAAAAGTAGTCAAGGAGAGAGAGGGAAGAAGTGAAGGACTGCTGTAACTCGGATCTTGCTTTTGCATTTTTGTCAGCCAGATCAGTGAAAACATAATGAAGCTTCAGCCTTTAC

>M6-PGSC0003DMG400005254-(rev)M6LRR-RLK_62

TATTGCCAAAAGGACTTTAGGCAAAGTATGCAATAAATCCTCTTCCCACTCACTTTGAAAATCTGTCTTCAGTAGTTCAATTAGTATCTGTCATAATTCTTGCCTACTGGATTGATCCAT

>M6-PGSC0003DMG400005254-(rev)M6LRR-RLK_620

TGCTTTTGCATTTTTGTCAGCCAGATCAGTGAAAACATAATGAAGCTTCAGCCTTTACTGGCAGCTATTGCTTTTCTTATTCTGCTGCTATCAGTATTTCTTCATGTTATAGCTGATCTT

>M6-PGSC0003DMG400005254-(rev)M6LRR-RLK_682

AGCTATTGCTTTTCTTATTCTGCTGCTATCAGTATTTCTTCATGTTATAGCTGATCTTGATTCCGATAGACAGGCGCTACTCGACTTTGCAGAATCTGTCCCTCATATCCGAAAACTCAA

>M6-PGSC0003DMG400005254-(rev)M6LRR-RLK_744

CCGATAGACAGGCGCTACTCGACTTTGCAGAATCTGTCCCTCATATCCGAAAACTCAACTGGAATTTGGCTCTCCCTATTTGCAAATCTTGGGCTGGTATAGCTTGTAACAAAGATGGAA

>M6-PGSC0003DMG400005254-(rev)M6LRR-RLK_806

AATTTGGCTCTCCCTATTTGCAAATCTTGGGCTGGTATAGCTTGTAACAAAGATGGAACGAGAGTCATTGCAATCCATCTCCCTGCTGTAGGACTTTTTGGTCCTATCCCAGCTAATAGT

>M6-PGSC0003DMG400005254-(rev)M6LRR-RLK_868

AGTCATTGCAATCCATCTCCCTGCTGTAGGACTTTTTGGTCCTATCCCAGCTAATAGTATAGGCAAGCTAGATGCTCTTAAAGTCCTTAGTTTAAGAGCCAATTATCTTAATGGAAGTGT

>M6-PGSC0003DMG400005254-(rev)M6LRR-RLK_930

GCAAGCTAGATGCTCTTAAAGTCCTTAGTTTAAGAGCCAATTATCTTAATGGAAGTGTTCCTTCTGATCTCCTTTCGATACCGTCTCTCCAATTTGTTTACCTTCAGCATAATAGCTTCT

>M6-PGSC0003DMG400005254-(rev)M6LRR-RLK_992

TCTGATCTCCTTTCGATACCGTCTCTCCAATTTGTTTACCTTCAGCATAATAGCTTCTCTGGTGATATTCCTGTTTCTTTATCTCCTCGTCTTGGTGTACTAGATCTCTCCTTTAACTCT

>M6-PGSC0003DMG400005395-(rev)M6LRR-RLP_0

ATGGGCTGTTGTGTGAAAGCTAATGCCTCCAAGGAGATTAGAACCCGCAATCATAGACATGTAGGACTTGAAAAGATCTTTCGCCAAGTGCACGAACCTCTACTTTGCATTCTGGTGCCT

>M6-PGSC0003DMG400005395-(rev)M6LRR-RLP_1037

AAAGTTGCAACACTTGATGGGAATCGAATCAGAGCATTGCCTGATGAATGTAATTTCCCACCGTCCTAGATCATATGTGAAGTTAAGCAGGATTCTCTCTGCTATAGAGTATCACACTCT

>M6-PGSC0003DMG400005395-(rev)M6LRR-RLP_1098

CGTCCTAGATCATATGTGAAGTTAAGCAGGATTCTCTCTGCTATAGAGTATCACACTCTCTATTCTAACAACATAATGTTCTCATATTCAGTGGGCCAGTTGGTGAAGCTTGAGCGTTTG

>M6-PGSC0003DMG400005395-(rev)M6LRR-RLP_1159

ATTCTAACAACATAATGTTCTCATATTCAGTGGGCCAGTTGGTGAAGCTTGAGCGTTTGTCTGTCTCAGCAAACTTGTTAATGAGCTTGCCTGAGACTATTGGGAGATTGCAGAATGTGA

>M6-PGSC0003DMG400005395-(rev)M6LRR-RLP_122

GTGTATCTATTTCTAAGATATCAAGATAGTTACCTATATAAACCAAGTTTCCATCGAAGTTTCCGGGTGCAGGTGTATCCTCAACCTACTATAAACGGGCTTGCTAAATCCAGATTGAGG

>M6-PGSC0003DMG400005395-(rev)M6LRR-RLP_1220

TGTCTCAGCAAACTTGTTAATGAGCTTGCCTGAGACTATTGGGAGATTGCAGAATGTGAGTGATCCATTCTGATGGCTAATACGGTGCCATCTTCCGCAATTCTAAAAAGTGTGCCAGTT

>M6-PGSC0003DMG400005395-(rev)M6LRR-RLP_1281

GATCCATTCTGATGGCTAATACGGTGCCATCTTCCGCAATTCTAAAAAGTGTGCCAGTTAATATATTATTTCTTAGCTTGCTTTACTGCTTTGTTATTTTTTTTTTCCGAGTATAAAGGT

>M6-PGSC0003DMG400005395-(rev)M6LRR-RLP_1342

TATATTATTTCTTAGCTTGCTTTACTGCTTTGTTATTTTTTTTTTCCGAGTATAAAGGTGCTAAGGTGTCTAATGAAATGGTGTCAATCTTGTAGAAAATGTTCAGCATATTGATATCCA

>M6-PGSC0003DMG400005395-(rev)M6LRR-RLP_1403

TAAGGTGTCTAATGAAATGGTGTCAATCTTGTAGAAAATGTTCAGCATATTGATATCCATATTAATGGATCATTTTGCTTACAACTTTCTGTTCTTACAACTAAAAGCCCTACTTTTTTT

>M6-PGSC0003DMG400005395-(rev)M6LRR-RLP_1464

TTAATGGATCATTTTGCTTACAACTTTCTGTTCTTACAACTAAAAGCCCTACTTTTTTTTCTCTATTTCTATAGTGATGCCGCTTCTTCATGAATTTCTATTATCTTCCTGGTAGTTGAT

>M6-PGSC0003DMG400005395-(rev)M6LRR-RLP_1525

TCTATTTCTATAGTGATGCCGCTTCTTCATGAATTTCTATTATCTTCCTGGTAGTTGATGATGTCTCTTTGCTCTCTTATGTTAGTATTTTGGATTTGGAATAGGGTCCTGCAGCTCCTG

>M6-PGSC0003DMG400005395-(rev)M6LRR-RLP_1586

TGTCTCTTTGCTCTCTTATGTTAGTATTTTGGATTTGGAATAGGGTCCTGCAGCTCCTGGGGGAGGGTGGTTGAATTTGGAGAATAAGTGGCTCTATTGCAGGTTGTCTTGAGGTCCACC

>M6-PGSC0003DMG400005395-(rev)M6LRR-RLP_1647

GGAGGGTGGTTGAATTTGGAGAATAAGTGGCTCTATTGCAGGTTGTCTTGAGGTCCACCACCCTGAGTGCCGTCTGTTCTTCCTTTTTATTTCACTCTCTCCTTATATTGAATATTTTTC

>M6-PGSC0003DMG400005395-(rev)M6LRR-RLP_1708

CCTGAGTGCCGTCTGTTCTTCCTTTTTATTTCACTCTCTCCTTATATTGAATATTTTTCTTCTCAAGTTTTTTTCTTCAATGTCATTGATGAACCAAGTGTGAGAAAGAAAGACTCGGAA

>M6-PGSC0003DMG400005395-(rev)M6LRR-RLP_1769

CTCAAGTTTTTTTCTTCAATGTCATTGATGAACCAAGTGTGAGAAAGAAAGACTCGGAAAATACTTTCAATATACGTAAACTACATTATTACACAAAGCATTGTTTATAGACAATACACT

>M6-PGSC0003DMG400005395-(rev)M6LRR-RLP_183

TCCGGGTGCAGGTGTATCCTCAACCTACTATAAACGGGCTTGCTAAATCCAGATTGAGGGAAATTCATGATTAAATGTGTCTTTTAGTAAATATTAACTGTGTTACTCTGTTGGAAATGC

>M6-PGSC0003DMG400005395-(rev)M6LRR-RLP_1830

TACTTTCAATATACGTAAACTACATTATTACACAAAGCATTGTTTATAGACAATACACTACAATCCTTTTCCATGTATGGCACCATGTACAATCCTATTTGAGCACTCCCCCTCAAGCTG

>M6-PGSC0003DMG400005395-(rev)M6LRR-RLP_244

AATTCATGATTAAATGTGTCTTTTAGTAAATATTAACTGTGTTACTCTGTTGGAAATGCAAGGTTAAAGCAGTAACAATTTGTCCAAAATATAATGGCAAACACAAAATTAAAGATATGC

>M6-PGSC0003DMG400005395-(rev)M6LRR-RLP_2745

CCCCTCTGCATTCAAGAGTACAAGTTCCATTTGACTGTTTTTTTCTAAATTTTAATTCTCTGCGCATTGACAAACAAAGACTCAGAAAATATTATTAAGATACGTAAATATCACACTGAG

>M6-PGSC0003DMG400005395-(rev)M6LRR-RLP_2806

GCGCATTGACAAACAAAGACTCAGAAAATATTATTAAGATACGTAAATATCACACTGAGGTCTACTTATAGATAATACACTACAATCCTTTTCCATGTAGGACACCATGTACAACCTATT

>M6-PGSC0003DMG400005395-(rev)M6LRR-RLP_2867

CTACTTATAGATAATACACTACAATCCTTTTCCATGTAGGACACCATGTACAACCTATTGTAACACCAAGAAGTATAGTGTTCACTTTTAGGTGTATATCTGATTCTCATGGTTGCTGAC

>M6-PGSC0003DMG400005395-(rev)M6LRR-RLP_2928

AACACCAAGAAGTATAGTGTTCACTTTTAGGTGTATATCTGATTCTCATGGTTGCTGACATGGGATTATGAATTAAAGTTTTTCAATCAAGTTAACTAGCTCTATCCTCTAGTAGTGTTC

>M6-PGSC0003DMG400005395-(rev)M6LRR-RLP_2989

GGGATTATGAATTAAAGTTTTTCAATCAAGTTAACTAGCTCTATCCTCTAGTAGTGTTCATATCATTCTAAACACTTGCATATACTTTTACTCCAAATATGTTTTGATATTTGTGAGTTC

>M6-PGSC0003DMG400005395-(rev)M6LRR-RLP_305

GGTTAAAGCAGTAACAATTTGTCCAAAATATAATGGCAAACACAAAATTAAAGATATGCTGACAGCTTAACTTCAAAGGGAGATTTTAAGCTATCGTAGGATAAGTCTTAGGAGGAAATA

>M6-PGSC0003DMG400005395-(rev)M6LRR-RLP_3050

ATCATTCTAAACACTTGCATATACTTTTACTCCAAATATGTTTTGATATTTGTGAGTTCAGTTCAATATGATGATGCCGTTTTTTGTCTGTCAGAGATAGTCGTTCAGCCTATACTTAAA

>M6-PGSC0003DMG400005395-(rev)M6LRR-RLP_3111

TTCAATATGATGATGCCGTTTTTTGTCTGTCAGAGATAGTCGTTCAGCCTATACTTAAAAGTATTCTTAATTCATCTTCTTTGGAAAATGAGGCCTTTGGAATTGACAGTAATGGTTTTT

>M6-PGSC0003DMG400005395-(rev)M6LRR-RLP_3172

TATTCTTAATTCATCTTCTTTGGAAAATGAGGCCTTTGGAATTGACAGTAATGGTTTTTCACTGCAGTTGGTGCTGTTGGACATTTCAAACAATAAGTTAAAGTTTTTTCCTGAATCAAT

>M6-PGSC0003DMG400005395-(rev)M6LRR-RLP_3233

CTGCAGTTGGTGCTGTTGGACATTTCAAACAATAAGTTAAAGTTTTTTCCTGAATCAATTGGAAGTTGCTTCTCTTTGGAAGAATTGCAAGCAAATGGTACCTTCAAATTTTAGTGTCAC

>M6-PGSC0003DMG400005395-(rev)M6LRR-RLP_3294

GAAGTTGCTTCTCTTTGGAAGAATTGCAAGCAAATGGTACCTTCAAATTTTAGTGTCACTCATCCATTCTTGCTATTTCTCTATTTCCTTTTATTTTCCTGGCACTGGAACTCTGTATCA

>M6-PGSC0003DMG400005395-(rev)M6LRR-RLP_3355

ATCCATTCTTGCTATTTCTCTATTTCCTTTTATTTTCCTGGCACTGGAACTCTGTATCAAATATCCATCGAGGAAAAAAAAACATGTTACCATCAACATAAAATTAAGTGAAATTTATGG

>M6-PGSC0003DMG400005395-(rev)M6LRR-RLP_3416

TATCCATCGAGGAAAAAAAAACATGTTACCATCAACATAAAATTAAGTGAAATTTATGGCCGATGATAAAATTGGAGTGTTGGGTGACCTAGAAAGCTAAAGGAAAATCTACAAATTTTC

>M6-PGSC0003DMG400005395-(rev)M6LRR-RLP_3477

GATGATAAAATTGGAGTGTTGGGTGACCTAGAAAGCTAAAGGAAAATCTACAAATTTTCTCTGTTAAGCTTATAAGATAATATTCTTGGCACCATCCATTTGAACTGGTACATGATTTTC

>M6-PGSC0003DMG400005395-(rev)M6LRR-RLP_3538

TGTTAAGCTTATAAGATAATATTCTTGGCACCATCCATTTGAACTGGTACATGATTTTCACGTTCTGTTTCTTATTCCTCAGTGTGGCATAATGAGGTTTACTGTGTTTTTTTGGCAATC

>M6-PGSC0003DMG400005395-(rev)M6LRR-RLP_3599

GTTCTGTTTCTTATTCCTCAGTGTGGCATAATGAGGTTTACTGTGTTTTTTTGGCAATCTTGCCTCGTGATAACAATATTTCCTTTCAGATAATTCCATTGAAGAGCTTCCTGCATCAGT

>M6-PGSC0003DMG400005395-(rev)M6LRR-RLP_366

ACAGCTTAACTTCAAAGGGAGATTTTAAGCTATCGTAGGATAAGTCTTAGGAGGAAATAGCCCTGTGAAGGAGCTGTAATGAAAGTCAATTTTTTTGTCAACGAGGTAATATGTGGTGGG

>M6-PGSC0003DMG400005395-(rev)M6LRR-RLP_3660

GCCTCGTGATAACAATATTTCCTTTCAGATAATTCCATTGAAGAGCTTCCTGCATCAGTTTGCAATCTTGTTCAACTAAAATCACTCTGCCTGAACAATAACAATCTCAATCAGGTTAGT

>M6-PGSC0003DMG400005395-(rev)M6LRR-RLP_3721

GCAATCTTGTTCAACTAAAATCACTCTGCCTGAACAATAACAATCTCAATCAGGTTAGTGTCTAATTTTACCTGTGTGTATGTGTGTGTGACCCTTGTTATTTTGAGGTCATATCTAGTA

>M6-PGSC0003DMG400005395-(rev)M6LRR-RLP_3904

ACATGTGCAGTCAACTGCTGTGCATGTGTGTGTGTGTCTGTCTCTATCATCGTCACACATAAAAAGTGCAAAGAATAGTAAACCACGTGGATCAAAAGCTTACCTGTTTGTCACACCGAA

>M6-PGSC0003DMG400005395-(rev)M6LRR-RLP_3965

AAAAGTGCAAAGAATAGTAAACCACGTGGATCAAAAGCTTACCTGTTTGTCACACCGAAAAGCTGAACTTATTGTTGTATTGGTGATACTTGTATGTGCTCTCTCTCTCACTTTCTATCT

>M6-PGSC0003DMG400005395-(rev)M6LRR-RLP_4026

GCTGAACTTATTGTTGTATTGGTGATACTTGTATGTGCTCTCTCTCTCACTTTCTATCTTAATAATACTATTAAGCAGAGTTAGGAACTCTCTTCGCATGATCTGTAAGCCTAGTTCACT

>M6-PGSC0003DMG400005395-(rev)M6LRR-RLP_4087

ATAATACTATTAAGCAGAGTTAGGAACTCTCTTCGCATGATCTGTAAGCCTAGTTCACTTCCCCGTGTATGAGTGTTAAAAGGAAACTCATACATTAACCTTAAAACGTGCATCTAAAAG

>M6-PGSC0003DMG400005395-(rev)M6LRR-RLP_4148

CCCGTGTATGAGTGTTAAAAGGAAACTCATACATTAACCTTAAAACGTGCATCTAAAAGCTTTTCCTTTTAATAAGACCTCATGAAGAAACGAAGATAATCAAACAACCAGTAGCATCTA

>M6-PGSC0003DMG400005395-(rev)M6LRR-RLP_4209

TTTCCTTTTAATAAGACCTCATGAAGAAACGAAGATAATCAAACAACCAGTAGCATCTACATTGCAAACCGGCAATAAAGCCAAATACGCATAATGATCCACATTTGAGGTCATTGGATT

>M6-PGSC0003DMG400005395-(rev)M6LRR-RLP_427

CCTGTGAAGGAGCTGTAATGAAAGTCAATTTTTTTGTCAACGAGGTAATATGTGGTGGGTAAAAAGCCCAAGTGTCACACCCGAAATCCACGCTTATTGCTCAAAGTACTTCCATGCTTG

>M6-PGSC0003DMG400005395-(rev)M6LRR-RLP_4270

TTGCAAACCGGCAATAAAGCCAAATACGCATAATGATCCACATTTGAGGTCATTGGATTGAACCTCACGTATCTTCCATTTAAAGCATCATCATCCAAGTTTTTGTACTTGTGCCTATCA

>M6-PGSC0003DMG400005395-(rev)M6LRR-RLP_4331

ACCTCACGTATCTTCCATTTAAAGCATCATCATCCAAGTTTTTGTACTTGTGCCTATCAAGTATATGTTTTTACTAGATAGTCATGCTTCCTTTTGGTAACTGGATTGTTCAGATGCCTC

>M6-PGSC0003DMG400005395-(rev)M6LRR-RLP_4392

TATATGTTTTTACTAGATAGTCATGCTTCCTTTTGGTAACTGGATTGTTCAGATGCCTCTAAATTTGTTAAGAGAATGCAAGAGCTTGCAGACCATATCCCTGCATGGAAATCCGATTTC

>M6-PGSC0003DMG400005395-(rev)M6LRR-RLP_4453

AATTTGTTAAGAGAATGCAAGAGCTTGCAGACCATATCCCTGCATGGAAATCCGATTTCCATGGACCAATTTCAACAGGTTACTCACTAATATTCTACTATTTAGTTCTGCAAGTACTAA

>M6-PGSC0003DMG400005395-(rev)M6LRR-RLP_4514

TGGACCAATTTCAACAGGTTACTCACTAATATTCTACTATTTAGTTCTGCAAGTACTAAAAAAATCATTTATGAAGAATTCTATTTGCCTTTTCTTTTTCCAGATGGATGGCTTCAAGGA

>M6-PGSC0003DMG400005395-(rev)M6LRR-RLP_4575

AAATCATTTATGAAGAATTCTATTTGCCTTTTCTTTTTCCAGATGGATGGCTTCAAGGACTTTGAAGCCCGCCGTAAACAGAAGTTTGATAAACAAATTGATTCAAATGTGATAATCAGT

>M6-PGSC0003DMG400005395-(rev)M6LRR-RLP_4636

TTGAAGCCCGCCGTAAACAGAAGTTTGATAAACAAATTGATTCAAATGTGATAATCAGTTCTAAAGGGCTTGATGAGGGTGTTGACTTGTGATTCCTCTCCGCAGGCGTTTTCTATAAGG

>M6-PGSC0003DMG400005395-(rev)M6LRR-RLP_4697

TAAAGGGCTTGATGAGGGTGTTGACTTGTGATTCCTCTCCGCAGGCGTTTTCTATAAGGCTCCGCGCAATCTTGTTAATCTCAGGTTTACTATTTTTTTGCTTCAATTCCCTGATGGAAA

>M6-PGSC0003DMG400005395-(rev)M6LRR-RLP_4758

CCGCGCAATCTTGTTAATCTCAGGTTTACTATTTTTTTGCTTCAATTCCCTGATGGAAAATTTCCCTTTATTTTATGTATGACCATGTAGTAGCCACACTACACAGTTGTGCTCCTTCAG

>M6-PGSC0003DMG400005395-(rev)M6LRR-RLP_4819

TTCCCTTTATTTTATGTATGACCATGTAGTAGCCACACTACACAGTTGTGCTCCTTCAGTTGCTGTCAAGCTTTTAGTATTTCTAGTGTCTGTAATGCCTAATTTGGACAATTCATTCTT

>M6-PGSC0003DMG400005395-(rev)M6LRR-RLP_488

AAAAGCCCAAGTGTCACACCCGAAATCCACGCTTATTGCTCAAAGTACTTCCATGCTTGTTCTAGTACGTTTAATCACTTTTGGACTTTAGTATCCTTGCTGGTCCCCACACTTCCAACC

>M6-PGSC0003DMG400005395-(rev)M6LRR-RLP_4880

GCTGTCAAGCTTTTAGTATTTCTAGTGTCTGTAATGCCTAATTTGGACAATTCATTCTTGTTTACTTCATGCATGTCTCCGTTGCTCTTAGCTCCCAGCCTGGCGTGCCTGCATAGTTGC

>M6-PGSC0003DMG400005395-(rev)M6LRR-RLP_4941

TTACTTCATGCATGTCTCCGTTGCTCTTAGCTCCCAGCCTGGCGTGCCTGCATAGTTGCCTCTTCTCATTTATGTTCCAGTCTTCTAGTTTGTATTACTCTCCATTATACCCGTGCCCCT

>M6-PGSC0003DMG400005395-(rev)M6LRR-RLP_5002

CTTCTCATTTATGTTCCAGTCTTCTAGTTTGTATTACTCTCCATTATACCCGTGCCCCTAAGTGTATATAATTGTGGTAATTATTAGTAACACCAATTTCAAAAGATTGAACACGAAAAA

>M6-PGSC0003DMG400005395-(rev)M6LRR-RLP_5063

GTGTATATAATTGTGGTAATTATTAGTAACACCAATTTCAAAAGATTGAACACGAAAAAGCTTCTCATTTATGTTCCATGGAAGCAGGACCGTTTTGTCCATTTAGTTTCTGAATTTGAT

>M6-PGSC0003DMG400005395-(rev)M6LRR-RLP_5124

TTCTCATTTATGTTCCATGGAAGCAGGACCGTTTTGTCCATTTAGTTTCTGAATTTGATAGGTAGTAGCTAATCCAGTGGCCAAATTAGTTCAAAAAGAAATTGCACTCGTTTTTCTAAA

>M6-PGSC0003DMG400005395-(rev)M6LRR-RLP_5185

GTAGTAGCTAATCCAGTGGCCAAATTAGTTCAAAAAGAAATTGCACTCGTTTTTCTAAACTTCCCAGTCAACTGGTGCTATTGATGATACAATTTTTTTTGAATAAATCATTATATCTTA

>M6-PGSC0003DMG400005395-(rev)M6LRR-RLP_5246

TCCCAGTCAACTGGTGCTATTGATGATACAATTTTTTTTGAATAAATCATTATATCTTATTTACTGCAGGTTAAATTTGTTCAGGTTGGATGGAATTGCGACAGTTTTGACTTTGATGAA

>M6-PGSC0003DMG400005395-(rev)M6LRR-RLP_5307

TACTGCAGGTTAAATTTGTTCAGGTTGGATGGAATTGCGACAGTTTTGACTTTGATGAAAGGTATTGCTTCTGCTTATTTGGCATGAGATATTTCACACCTTTCTATAGTTTGTCATTGA

>M6-PGSC0003DMG400005395-(rev)M6LRR-RLP_5368

GTATTGCTTCTGCTTATTTGGCATGAGATATTTCACACCTTTCTATAGTTTGTCATTGATTGTAAGTTGTGACTTGTGCGTAATTGTATCTATGCTGACTTGCTGAGTATGTTTGTGATT

>M6-PGSC0003DMG400005395-(rev)M6LRR-RLP_5429

GTAAGTTGTGACTTGTGCGTAATTGTATCTATGCTGACTTGCTGAGTATGTTTGTGATTGTAAATGACTCAACTCTAACAAATTCTGTACATTTAGCGAGCTGTATATCACTTTAAAATC

>M6-PGSC0003DMG400005395-(rev)M6LRR-RLP_549

CTAGTACGTTTAATCACTTTTGGACTTTAGTATCCTTGCTGGTCCCCACACTTCCAACCTTGATCTGATTTTTTCAAGGTTTGAATTTTCGGGGTCTTACACCAAGTATACTAGTGGTGT

>M6-PGSC0003DMG400005395-(rev)M6LRR-RLP_5490

AAATGACTCAACTCTAACAAATTCTGTACATTTAGCGAGCTGTATATCACTTTAAAATCAAACTAACGATTTTTGTACCTAAAGATATTTGATTTTAATATGTTTGGATTGTAACGATCC

>M6-PGSC0003DMG400005395-(rev)M6LRR-RLP_5551

ACTAACGATTTTTGTACCTAAAGATATTTGATTTTAATATGTTTGGATTGTAACGATCCTTTCAACGTATCCAAGAGTAGGGCCTGGTGGCCAATAAAGTGGCTGAAAATTGTGGCCACT

>M6-PGSC0003DMG400005395-(rev)M6LRR-RLP_5612

TCAACGTATCCAAGAGTAGGGCCTGGTGGCCAATAAAGTGGCTGAAAATTGTGGCCACTTTTTATCTCCGTTCGTCGTTTTAAAATTAATTGAAACTAAGGGCGTGTTTATGATATTTTC

>M6-PGSC0003DMG400005395-(rev)M6LRR-RLP_5673

TTATCTCCGTTCGTCGTTTTAAAATTAATTGAAACTAAGGGCGTGTTTATGATATTTTCATGTTTGGTTGGTTAAATTTTTTGGAAAATATTTGCTCTAGTAAGTATTTCATACGCGAGC

>M6-PGSC0003DMG400005395-(rev)M6LRR-RLP_5734

GTTTGGTTGGTTAAATTTTTTGGAAAATATTTGCTCTAGTAAGTATTTCATACGCGAGCCATCATCTAAAATCAATTCATCGATATCAAGTCAGCAACACAAGCATTTATCTAGTGCTAT

>M6-PGSC0003DMG400005395-(rev)M6LRR-RLP_5795

TCATCTAAAATCAATTCATCGATATCAAGTCAGCAACACAAGCATTTATCTAGTGCTATATACTAGGAGTCAAGCTCACATAATCACATTAATCAATTCAGGAAGAAAATCTCCAATTTT

>M6-PGSC0003DMG400005395-(rev)M6LRR-RLP_5856

ACTAGGAGTCAAGCTCACATAATCACATTAATCAATTCAGGAAGAAAATCTCCAATTTTTCATATTCATGGCATTGGTACTCAAAGCATTTCATATTCATGGCATTGGTACTCATAGAAA

>M6-PGSC0003DMG400005395-(rev)M6LRR-RLP_5917

ATATTCATGGCATTGGTACTCAAAGCATTTCATATTCATGGCATTGGTACTCATAGAAATGCTTGTTATCTCTTTTATACCGGACCTCCATTATCCATTAACCTTTTATTCAATTAATAA

>M6-PGSC0003DMG400005395-(rev)M6LRR-RLP_5978

CTTGTTATCTCTTTTATACCGGACCTCCATTATCCATTAACCTTTTATTCAATTAATAAATACACTTTAACATTTCATAAAAAGTTTCAACTTGCCTTAATTGGAATTCAAAAAGTTAAT

>M6-PGSC0003DMG400005395-(rev)M6LRR-RLP_6039

ACACTTTAACATTTCATAAAAAGTTTCAACTTGCCTTAATTGGAATTCAAAAAGTTAATGTGGATACCTAGGCACCCAAAGACAATGAAGGGTGTAGTAATCGACAAAATGCTTTAGGGA

>M6-PGSC0003DMG400005395-(rev)M6LRR-RLP_61

TAGGACTTGAAAAGATCTTTCGCCAAGTGCACGAACCTCTACTTTGCATTCTGGTGCCTTTGTGTATCTATTTCTAAGATATCAAGATAGTTACCTATATAAACCAAGTTTCCATCGAAG

>M6-PGSC0003DMG400005395-(rev)M6LRR-RLP_610

GATCTGATTTTTTCAAGGTTTGAATTTTCGGGGTCTTACACCAAGTATACTAGTGGTGTACAAGAAGTGGACCTACACCAACACATATTTCTGGTACTCCAAAAATCTATTTAAAAAAAC

>M6-PGSC0003DMG400005395-(rev)M6LRR-RLP_6222

CCACGGGTAGGCAAGATACAACTAACGATTTGCTCTTCTTTTATGCCTTTGAACTATTTCAATTTGGTCATAAATGTAATTTAGATGAGTATACAAGTTCGTAGACACACAAATTGGCAC

>M6-PGSC0003DMG400005395-(rev)M6LRR-RLP_6283

ATTTGGTCATAAATGTAATTTAGATGAGTATACAAGTTCGTAGACACACAAATTGGCACATAATTGGCTCGAAAATTGAGTGAAAAAGTCAACCTTGAGCCTTACTGTCAAATTTGAAAA

>M6-PGSC0003DMG400005395-(rev)M6LRR-RLP_6344

AATTGGCTCGAAAATTGAGTGAAAAAGTCAACCTTGAGCCTTACTGTCAAATTTGAAAATTTGTTTACCCATAGGTGGGTCAAAAATCCGCTAAAATGGATTGTTAATCGGTCCCCAAAT

>M6-PGSC0003DMG400005395-(rev)M6LRR-RLP_6405

TGTTTACCCATAGGTGGGTCAAAAATCCGCTAAAATGGATTGTTAATCGGTCCCCAAATCGAAAGCTACATAAATCATTCTTTGGAGAATTAAAAACCAACCAAATTCCCCAATTCAAAA

>M6-PGSC0003DMG400005395-(rev)M6LRR-RLP_6466

AAAGCTACATAAATCATTCTTTGGAGAATTAAAAACCAACCAAATTCCCCAATTCAAAAACCCCCAAATTAAATATCTATGAGCTTGTATAGCAAGGTGAGACCCTCCTCTTATTGGATT

>M6-PGSC0003DMG400005395-(rev)M6LRR-RLP_6527

CCCCAAATTAAATATCTATGAGCTTGTATAGCAAGGTGAGACCCTCCTCTTATTGGATTCTAATTACTTTAAGTCACAAATAGTATTTTAGGATGAAAATTCATATCTTTCGTCGGATTG

>M6-PGSC0003DMG400005395-(rev)M6LRR-RLP_6588

AATTACTTTAAGTCACAAATAGTATTTTAGGATGAAAATTCATATCTTTCGTCGGATTGTCATAGTACTTCTTGTTATTGATTGTTGAGTGAACTCATTGTTATAGTAATGTGGGGCTGT

>M6-PGSC0003DMG400005395-(rev)M6LRR-RLP_6649

ATAGTACTTCTTGTTATTGATTGTTGAGTGAACTCATTGTTATAGTAATGTGGGGCTGTGGGAAAAAGTTATTGGATGGCAAAGAGTCAATGGAAAAAGGTATGATTAGGTTACTCTCTT

>M6-PGSC0003DMG400005395-(rev)M6LRR-RLP_671

AAGAAGTGGACCTACACCAACACATATTTCTGGTACTCCAAAAATCTATTTAAAAAAACAAAAAATCAGTTGTAATCTGTACAAGAGTCCTCAAGACTCTACTGGGATGAAAATCAGTTC

>M6-PGSC0003DMG400005395-(rev)M6LRR-RLP_6710

GAAAAAGTTATTGGATGGCAAAGAGTCAATGGAAAAAGGTATGATTAGGTTACTCTCTTCTTTTAGCATGTTTTCTTAAAAGGGTAAAGCTATATGTCAAGCGTACTTTGGATGAGCTAC

>M6-PGSC0003DMG400005395-(rev)M6LRR-RLP_6771

TTTAGCATGTTTTCTTAAAAGGGTAAAGCTATATGTCAAGCGTACTTTGGATGAGCTACTCATAGCCATAAGTTTAGTGTGTTGGCTTCCTAGAGTTGTTATGTTCGTCCCTTGTTGGGA

>M6-PGSC0003DMG400005395-(rev)M6LRR-RLP_6832

ATAGCCATAAGTTTAGTGTGTTGGCTTCCTAGAGTTGTTATGTTCGTCCCTTGTTGGGACTCCTATTATAATGATAGCGACCTCGTGCTTAGATACAACTTAGAGATTACTCATATAGGT

>M6-PGSC0003DMG400005395-(rev)M6LRR-RLP_6893

CCTATTATAATGATAGCGACCTCGTGCTTAGATACAACTTAGAGATTACTCATATAGGTCGCTTATAGTTGGTTTGCTTGCCTTAGGGCCACTTGATTGATGTTGATGCCTTTGGGCCAT

>M6-PGSC0003DMG400005395-(rev)M6LRR-RLP_6937

ATTACTCATATAGGTCGCTTATAGTTGGTTTGCTTGCCTTAGGGCCACTTGATTGATGTTGATGCCTTTGGGCCATATGTTGGTTGTTACTTCTTTCGGGGGGGGGGGGGGGGGATTGTG

>M6-PGSC0003DMG400005395-(rev)M6LRR-RLP_732

AAAATCAGTTGTAATCTGTACAAGAGTCCTCAAGACTCTACTGGGATGAAAATCAGTTCTACCTGGGAAAACTGAAAAGTGTCTGCTTCTCATTTATTCTTAAGGGACTCTTCATCTGGG

>M6-PGSC0003DMG400005395-(rev)M6LRR-RLP_793

CCTGGGAAAACTGAAAAGTGTCTGCTTCTCATTTATTCTTAAGGGACTCTTCATCTGGGTTTTCACGCTGAATACTATTGATATTGACATATTGTAGGGTGTATTCTCTTAAAAAGAAAA

>M6-PGSC0003DMG400005395-(rev)M6LRR-RLP_854

TTCACGCTGAATACTATTGATATTGACATATTGTAGGGTGTATTCTCTTAAAAAGAAAATTTTATAAAGAGTTTGTTATATCTGATTCTATCCAAAAAATTGAGTCTGATTATTTCTTTT

>M6-PGSC0003DMG400005395-(rev)M6LRR-RLP_915

TTATAAAGAGTTTGTTATATCTGATTCTATCCAAAAAATTGAGTCTGATTATTTCTTTTCAGGTTTTGGCTGATAACCTTATTGAGAATCTCCCGGTGAACCTGGGATTGCTTAAGTCTT

>M6-PGSC0003DMG400005395-(rev)M6LRR-RLP_976

GGTTTTGGCTGATAACCTTATTGAGAATCTCCCGGTGAACCTGGGATTGCTTAAGTCTTTGAAAGTTGCAACACTTGATGGGAATCGAATCAGAGCATTGCCTGATGAATGTAATTTCCC

>M6-PGSC0003DMG400005485M6LRR-RLK_0

CTAACAACGATAAATCTTAGGAATAATAGCTTCCATGGTGAAGTCCCTATGGAAATTGGCAACCTCTTGCAACTGCAACATCTTAATCTCACTTGGAATTCATTTACTGGAACCATTCCT

>M6-PGSC0003DMG400005485M6LRR-RLK_1037

GTCTACTTTACACATTGAGGACAATGAATTAGAAGGAAACATACCTCTGGAGCTCGGGCAGTGCATCCGCTTATCAAGGCTAAACCTTACAGGAAATAATCTTGTTGGTTCCATACCAAA

>M6-PGSC0003DMG400005485M6LRR-RLK_1098

TGCATCCGCTTATCAAGGCTAAACCTTACAGGAAATAATCTTGTTGGTTCCATACCAAAGGAGCTTGCAGGTCTTTCTTCTCTTTCGATTTCCTTAGCCTTAGCAAACAATTCGTTGACC

>M6-PGSC0003DMG400005485M6LRR-RLK_1159

AGCTTGCAGGTCTTTCTTCTCTTTCGATTTCCTTAGCCTTAGCAAACAATTCGTTGACCGGTTCCTTGCCAGCTGAATTTGGAAAGTTGATAAATCTCAAGGAAATGGATATTTCACATA

>M6-PGSC0003DMG400005485M6LRR-RLK_122

TAATCTGAGTTGCTGTAAAGAACTTAGATCACTTGCTCTAGAATATAATAGTCTGGTTGGTAAAGTATTACCTGATCAGTTCAGTTCACTTTCAAAGTTGAATTATTTAGGCCTTGGAAG

>M6-PGSC0003DMG400005485M6LRR-RLK_1220

TTCCTTGCCAGCTGAATTTGGAAAGTTGATAAATCTCAAGGAAATGGATATTTCACATAACAAATTATCAGGTGAAATTCCAAGTACTCTAAGTAGCTGTGTCAGCTTAGAGCGCTTCAT

>M6-PGSC0003DMG400005485M6LRR-RLK_1281

AAATTATCAGGTGAAATTCCAAGTACTCTAAGTAGCTGTGTCAGCTTAGAGCGCTTCATAGCGAATAACAACCTGTTTCGAGGAGAAATTCCTGAATCCTTGCAAGGTTTAAGAGGTTTA

>M6-PGSC0003DMG400005485M6LRR-RLK_1342

CGAATAACAACCTGTTTCGAGGAGAAATTCCTGAATCCTTGCAAGGTTTAAGAGGTTTAGAAGAAATTGATTTGTCACACAATAACATCTCAGGAGGAATACCTGAGTTTCTTGGGAAAC

>M6-PGSC0003DMG400005485M6LRR-RLK_1403

AGAAATTGATTTGTCACACAATAACATCTCAGGAGGAATACCTGAGTTTCTTGGGAAACTTCCATATCTTAGGAAGCTTGATCTTTCATTCAACGAACTTGAAGGCGAAGTACCAACTGA

>M6-PGSC0003DMG400005485M6LRR-RLK_1464

CCATATCTTAGGAAGCTTGATCTTTCATTCAACGAACTTGAAGGCGAAGTACCAACTGAAGGGATCTTTGCAAATGAAACCGCCGTCTCAATCTCAGGGAATGAAGAACTATGTGGAGGT

>M6-PGSC0003DMG400005485M6LRR-RLK_1525

GGATCTTTGCAAATGAAACCGCCGTCTCAATCTCAGGGAATGAAGAACTATGTGGAGGTCCTCCAAACTACAATTTTCCTACATGCCCTAAACAAAAAGATGCATCATCAAAGAAACACA

>M6-PGSC0003DMG400005485M6LRR-RLK_1586

TCCAAACTACAATTTTCCTACATGCCCTAAACAAAAAGATGCATCATCAAAGAAACACATTAGTTCAAGGATAAAAGTAGCAATTATTATTTCGGTTGCATTTCTATTTTTACTTTTGTG

>M6-PGSC0003DMG400005485M6LRR-RLK_1647

AGTTCAAGGATAAAAGTAGCAATTATTATTTCGGTTGCATTTCTATTTTTACTTTTGTGTTCTTTTGCAGCTTGCTATATAGTAATTAGGAAGTCAAGAAAGAGAGATCTTACTGGACGG

>M6-PGSC0003DMG400005485M6LRR-RLK_1708

CTTTTGCAGCTTGCTATATAGTAATTAGGAAGTCAAGAAAGAGAGATCTTACTGGACGGTCCTCAAGAGAAAGGCAGTCTGATCATTTTGATGATGAAGAACCTACTTTATTTAATGATC

>M6-PGSC0003DMG400005485M6LRR-RLK_1769

CTCAAGAGAAAGGCAGTCTGATCATTTTGATGATGAAGAACCTACTTTATTTAATGATCCAATTTTGACAGCAAAGATAACTTACCAAGATATATTCAAGTCAACCAATGGATTTTCTGA

>M6-PGSC0003DMG400005485M6LRR-RLK_183

AAAGTATTACCTGATCAGTTCAGTTCACTTTCAAAGTTGAATTATTTAGGCCTTGGAAGCAACAATCTAACAGGAGGTATCCCATCTTGGATAGGAAATTTTTCGACTCTTCGAGGTCTT

>M6-PGSC0003DMG400005485M6LRR-RLK_1830

ATTTTGACAGCAAAGATAACTTACCAAGATATATTCAAGTCAACCAATGGATTTTCTGAGGATAATCTAGTCGGTACAGGAAGTTTTGGTTCTGTTTACAGAGGAAAGTTTCAGGTTTTT

>M6-PGSC0003DMG400005485M6LRR-RLK_1891

ATAATCTAGTCGGTACAGGAAGTTTTGGTTCTGTTTACAGAGGAAAGTTTCAGGTTTTTGATAAAGTTATGGCAGTGAAAGTATTGAACCTTCAACAAAGAGGTGCTTTGAAAAGCTTTT

>M6-PGSC0003DMG400005485M6LRR-RLK_1952

TAAAGTTATGGCAGTGAAAGTATTGAACCTTCAACAAAGAGGTGCTTTGAAAAGCTTTTCGGATGAATGTAGAGCTTTGAAAAGCATAAGGCATCGAAATCTACTTAAGATAATAGCTGT

>M6-PGSC0003DMG400005485M6LRR-RLK_2013

GATGAATGTAGAGCTTTGAAAAGCATAAGGCATCGAAATCTACTTAAGATAATAGCTGTTTGTTCAAGCATCGATTACCAAGGTAATGATTTCAAATGCAAAGTCTTTGAGTTCATGGAA

>M6-PGSC0003DMG400005485M6LRR-RLK_2074

GTTCAAGCATCGATTACCAAGGTAATGATTTCAAATGCAAAGTCTTTGAGTTCATGGAAAATGGAAGCTTAGATGACTGGCTGCACTCGAAAGGTGATGAGCAATACCTCAATATTATCC

>M6-PGSC0003DMG400005485M6LRR-RLK_2135

TGGAAGCTTAGATGACTGGCTGCACTCGAAAGGTGATGAGCAATACCTCAATATTATCCAAAGACTAAACATCGCAATAGATGCTGCTTCAGCACTTGATTATCTCCACAACAATTGCCA

>M6-PGSC0003DMG400005485M6LRR-RLK_2196

AGACTAAACATCGCAATAGATGCTGCTTCAGCACTTGATTATCTCCACAACAATTGCCAAGTATCAATTGTTCACTGTGATTTAAAACCGAGCAACATACTCCTCGATGAAGAGATGACT

>M6-PGSC0003DMG400005485M6LRR-RLK_2257

TATCAATTGTTCACTGTGATTTAAAACCGAGCAACATACTCCTCGATGAAGAGATGACTGCCCATGTTGGTGACTTTGGATTGGCAAAATTTCTCTTCAAATCATCATGGAACAAATATA

>M6-PGSC0003DMG400005485M6LRR-RLK_2318

CCATGTTGGTGACTTTGGATTGGCAAAATTTCTCTTCAAATCATCATGGAACAAATATACATCTATTGCACTAAAGGGTTCTATTGGTTACCACCAGGTATGTCATTAATCCATGTTGAT

>M6-PGSC0003DMG400005485M6LRR-RLK_2379

TCTATTGCACTAAAGGGTTCTATTGGTTACCACCAGGTATGTCATTAATCCATGTTGATTTCTCTTGCATATCTATTTCCCTTATCAAGGGTCCTAACATGTAAGATGTTTGTGTACCTC

>M6-PGSC0003DMG400005485M6LRR-RLK_244

ACAATCTAACAGGAGGTATCCCATCTTGGATAGGAAATTTTTCGACTCTTCGAGGTCTTTCCCTTGCAATCAACAATCTACAAGGACCAATACCTCGAGATATTGGACGTTTGTCAAACT

>M6-PGSC0003DMG400005485M6LRR-RLK_2684

TTATAAATGGACTATAAAGGAAAGTAAGACAAATGAATTGAAACGAAGGGAGCTTATCTTTCTGATGCTTAAGGTGTCGAATGAACAAATCAATAACTAAGCATTCATATGAGGTTGTCA

>M6-PGSC0003DMG400005485M6LRR-RLK_2745

CTGATGCTTAAGGTGTCGAATGAACAAATCAATAACTAAGCATTCATATGAGGTTGTCATTTTCATGTGAGTGAACATATGTTCAGTGTCCCAACATAATTCATCATTATCTTGCAGAAT

>M6-PGSC0003DMG400005485M6LRR-RLK_2806

TTCATGTGAGTGAACATATGTTCAGTGTCCCAACATAATTCATCATTATCTTGCAGAATATGGATCAGGTGTAAATGTGTCCACTCTAGGAGATGTTTATAGCTTTGGTATCATGTTGCT

>M6-PGSC0003DMG400005485M6LRR-RLK_2867

GGATCAGGTGTAAATGTGTCCACTCTAGGAGATGTTTATAGCTTTGGTATCATGTTGCTAGAGTTGTGTACCGGTAGAAGGCCAACCGATGAGATCTTCAAAGATGGCCTGAATATTCAC

>M6-PGSC0003DMG400005485M6LRR-RLK_2928

AGTTGTGTACCGGTAGAAGGCCAACCGATGAGATCTTCAAAGATGGCCTGAATATTCACCAATATGTCAAATCCCATCTGCCTAGACGTGTCACGGAGATTGCTGATGCATCATTGCTTT

>M6-PGSC0003DMG400005485M6LRR-RLK_2989

ATATGTCAAATCCCATCTGCCTAGACGTGTCACGGAGATTGCTGATGCATCATTGCTTTTAGCATATGAAGAACACAATATATATGAGGACAATGCAAGTGATTTGGAAGAAAAGGCAAT

>M6-PGSC0003DMG400005485M6LRR-RLK_305

CCTTGCAATCAACAATCTACAAGGACCAATACCTCGAGATATTGGACGTTTGTCAAACTTGCAAATATTCCAAGTTTATGGAAATCAGTTGAATGGTACAATTCCTCAATCTCTTTTCAA

>M6-PGSC0003DMG400005485M6LRR-RLK_3050

GCATATGAAGAACACAATATATATGAGGACAATGCAAGTGATTTGGAAGAAAAGGCAATACTTCAAGATGATGAGTATATATCCAAGCTAAACACCAGCACTATTATACAAGAATGCTTG

>M6-PGSC0003DMG400005485M6LRR-RLK_3111

TTCAAGATGATGAGTATATATCCAAGCTAAACACCAGCACTATTATACAAGAATGCTTGATCTCAATCATGAAAATAGGACTTTTATGTAGTAGCTCATAGCCAAGAGATCGAATGCCAA

>M6-PGSC0003DMG400005485M6LRR-RLK_3172

CTCAATCATGAAAATAGGACTTTTATGTAGTAGCTCATAGCCAAGAGATCGAATGCCAATAAGCATAGCTTTGAAAGAGATTCACACAATCAAGAACTTATTTCTAGAGTCTAAGAGGAT

>M6-PGSC0003DMG400005485M6LRR-RLK_3233

AGCATAGCTTTGAAAGAGATTCACACAATCAAGAACTTATTTCTAGAGTCTAAGAGGATAAATAATAGTATAGATAGATCAGATGGTAATAAGCTAATCAAATTATCATCTTCAGAATTT

>M6-PGSC0003DMG400005485M6LRR-RLK_3294

ATAATAGTATAGATAGATCAGATGGTAATAAGCTAATCAAATTATCATCTTCAGAATTTTATTGATGAAGGCTAATATACATAGAATTACAACTTTGAAATAGATAATATTTACAATTAG

>M6-PGSC0003DMG400005485M6LRR-RLK_3355

TTGATGAAGGCTAATATACATAGAATTACAACTTTGAAATAGATAATATTTACAATTAGTACTATAGATAGCCTCACAAAATGTGAATTGTGACCAAGAAATTAAGGGATATGGTCGAGT

>M6-PGSC0003DMG400005485M6LRR-RLK_366

CAAATATTCCAAGTTTATGGAAATCAGTTGAATGGTACAATTCCTCAATCTCTTTTCAATATATCCTCTGTTTACTATTTCTCTGTTACTCAAAACTTGTTGTATGGAGAGCTTCCATCA

>M6-PGSC0003DMG400005485M6LRR-RLK_427

TATCCTCTGTTTACTATTTCTCTGTTACTCAAAACTTGTTGTATGGAGAGCTTCCATCAGATATAGGCCTTACTCTTCCGAATCTTGTGGTGTTTGCTGGTGCTGTGAACGATTTCACTG

>M6-PGSC0003DMG400005485M6LRR-RLK_488

TATAGGCCTTACTCTTCCGAATCTTGTGGTGTTTGCTGGTGCTGTGAACGATTTCACTGGACCAATTCCTGTTTCATTATCAAATGCAACAAAACTTGGTGTACTTGAATTGTCTCAAAA

>M6-PGSC0003DMG400005485M6LRR-RLK_549

CCAATTCCTGTTTCATTATCAAATGCAACAAAACTTGGTGTACTTGAATTGTCTCAAAATAAGCTCACCGGAAATGTCCCTACGAGTTTAGGACAATTGCAAGGGTTGTATAGGATGAAC

>M6-PGSC0003DMG400005485M6LRR-RLK_61

ACCTCTTGCAACTGCAACATCTTAATCTCACTTGGAATTCATTTACTGGAACCATTCCTGCTAATCTGAGTTGCTGTAAAGAACTTAGATCACTTGCTCTAGAATATAATAGTCTGGTTG

>M6-PGSC0003DMG400005485M6LRR-RLK_610

AGCTCACCGGAAATGTCCCTACGAGTTTAGGACAATTGCAAGGGTTGTATAGGATGAACTTTGAGATCAATAGTCTTGGAAGAAACACAAGTGGGGATTTGAGGTTTCTTGATTTCCTAG

>M6-PGSC0003DMG400005485M6LRR-RLK_671

TGAGATCAATAGTCTTGGAAGAAACACAAGTGGGGATTTGAGGTTTCTTGATTTCCTAGTTAACTGCACAAGTTTGCAAGTTCTTAGCTTTGAGGACAACATTTTGGGAGGTGAATTGCC

>M6-PGSC0003DMG400005485M6LRR-RLK_732

AACTGCACAAGTTTGCAAGTTCTTAGCTTTGAGGACAACATTTTGGGAGGTGAATTGCCTAAAACAATCGGTAACCTTTCCACGAGGCTAGAAATATTTGCTCTTGGTTATAATATGATA

>M6-PGSC0003DMG400005485M6LRR-RLK_793

AAACAATCGGTAACCTTTCCACGAGGCTAGAAATATTTGCTCTTGGTTATAATATGATAGTTGGTTCTCTTCCTACAGGACTAGAAAATCTTGTTAATTTGACCCTTCTATCACTGGATA

>M6-PGSC0003DMG400005485M6LRR-RLK_854

TGGTTCTCTTCCTACAGGACTAGAAAATCTTGTTAATTTGACCCTTCTATCACTGGATAATAACTACTTGAGAGGCAGTGTTCCTGAGTCTTTAGGTAAGCTTCGACGCCTGCAAGGACT

>M6-PGSC0003DMG400005485M6LRR-RLK_915

AACTACTTGAGAGGCAGTGTTCCTGAGTCTTTAGGTAAGCTTCGACGCCTGCAAGGACTGCTGTTGAATGGAAACAAGTTATCAGGAAGGATTCCATCTTCTATTGGTAACTTAACATCT

>M6-PGSC0003DMG400005485M6LRR-RLK_976

TGTTGAATGGAAACAAGTTATCAGGAAGGATTCCATCTTCTATTGGTAACTTAACATCTTTGTCTACTTTACACATTGAGGACAATGAATTAGAAGGAAACATACCTCTGGAGCTCGGGC

>M6-PGSC0003DMG400005538M6LRR-RLP_0

TTGACATTTTCCTAGAAAGACTTTGTGCATTGATTTGAGAATGACAACATTTTGTTGAATTGATTGATGATTTGATATGGAGGCAGCAGTCCATACTTGACTTTTCCCCTTTCCTCTCTT

>M6-PGSC0003DMG400005538M6LRR-RLP_1037

ATCGAGGAAGCAACTTCCCTCCTGAAATGGAAATTAACTTTCAAGAACCAGAACAACTCCTTATTCGCTTCATGGACACCAAGTTCTAACTCATGCAGGGACTGGTACGGAGTTATATGC

>M6-PGSC0003DMG400005538M6LRR-RLP_1098

TATTCGCTTCATGGACACCAAGTTCTAACTCATGCAGGGACTGGTACGGAGTTATATGCTTAAATGGTAGGGTAAACACGTTGAATATTACAAATTCTAGTGTCATTGGTACCCTCTATG

>M6-PGSC0003DMG400005538M6LRR-RLP_1159

AAATGGTAGGGTAAACACGTTGAATATTACAAATTCTAGTGTCATTGGTACCCTCTATGCTTTTCCGTTTTCATCTCTCCCTTTTCTTGAATATTTTGAACTTAGCCTGAACAATTTTTC

>M6-PGSC0003DMG400005538M6LRR-RLP_122

TGATTAGAAAATTCGCAAGTAATTAATTTCAATATGAAGATTCAAAGTTGAAGGGATTTGTTTGATAAGTAGCTTTCATTGAAATTTTTGCAGCAAGACTCATTGCAATTACTTGATAAG

>M6-PGSC0003DMG400005538M6LRR-RLP_1342

TCAGATTTCAGGCACAATTCCGCCACAAATCGGCTCACTAGCCAAGCTTGAGACCCTCCACATCTTTGCCAACCATTTAAGTGGTTCCATTCCACAAGAAATAGGTTACCTAAGGTCTCT

>M6-PGSC0003DMG400005538M6LRR-RLP_1586

ATAGGAAAACTTGTCAATCTTGTTGGAGTTTCTTTTGATACAAACCAATTAACAGGTCATATCCCTCCAGAAATAGGCAACTTGACCAATGTAAAATTGTTCTATGCTCCCTTCAATGAA

>M6-PGSC0003DMG400005538M6LRR-RLP_1647

TCCCTCCAGAAATAGGCAACTTGACCAATGTAAAATTGTTCTATGCTCCCTTCAATGAATTATCTGGTCCCATTCCTGCAGAAATAGGGAAGATGAAGTCACTTCAGGAATTAATCTTAC

>M6-PGSC0003DMG400005538M6LRR-RLP_1708

ATCTGGTCCCATTCCTGCAGAAATAGGGAAGATGAAGTCACTTCAGGAATTAATCTTACATACAAACAATCTTTCTGGTCTAATTCCAAGAACTATAGGTGGCTTAACTGAGCATATCTA

>M6-PGSC0003DMG400005538M6LRR-RLP_1769

ACAAACAATCTTTCTGGTCTAATTCCAAGAACTATAGGTGGCTTAACTGAGCATATCTACAACTTAATGATAATCAACTCTCTGGCCTAATTCCTAGAGAGATGGGGAATCTGAAAAACC

>M6-PGSC0003DMG400005538M6LRR-RLP_183

TTGATAAGTAGCTTTCATTGAAATTTTTGCAGCAAGACTCATTGCAATTACTTGATAAGTCAAAGAACTGCACTAATACTTCTAGGTGCAAGAAAAAAGATAATTGGAGGGACACTCTGT

>M6-PGSC0003DMG400005538M6LRR-RLP_1891

AATGGTCTGGCATTATTCAATAATCAACTTTTTGGCCTAATCCCTAGTGAGTTGGGGAATCTTAGTATCCACTATCATCTAACATTATCCAATAATCAACTCACTGGTTCAATTCCTTCC

>M6-PGSC0003DMG400005538M6LRR-RLP_1952

TTAGTATCCACTATCATCTAACATTATCCAATAATCAACTCACTGGTTCAATTCCTTCCTCCTTTGGTAATTTGAGAAACTTGCGAGTTCTGTATCTCCATGACAACAATCTGATTGAGG

>M6-PGSC0003DMG400005538M6LRR-RLP_2013

CTTTGGTAATTTGAGAAACTTGCGAGTTCTGTATCTCCATGACAACAATCTGATTGAGGAAATTCCTTCATCTATATGCAATTTGACATTCCTCATGTTCCTGCATTTGTCGAGAAACAA

>M6-PGSC0003DMG400005538M6LRR-RLP_2074

ATTCCTTCATCTATATGCAATTTGACATTCCTCATGTTCCTGCATTTGTCGAGAAACAACTTGAAAGGAAAAATTCTGCAATGTTTGGGTAATATCAGTACTCTCCAGTTGGTGACACTG

>M6-PGSC0003DMG400005538M6LRR-RLP_2135

TGAAAGGAAAAATTCTGCAATGTTTGGGTAATATCAGTACTCTCCAGTTGGTGACACTGTCACATAATAATCTCAGTGGAGAGCTCCCTTTGTCTATTTGCAATTTAGCATCACTACAAA

>M6-PGSC0003DMG400005538M6LRR-RLP_2257

CTAAATTTGGGCAAAAACAATATAAAGGGAGTAATTCCGCAATGTTTTGGCAACATGAGTGGTTATCTTGAGGTTCTTGATATGCAGCACAACAATCTTTCTGGGACTCTTCCAATGACT

>M6-PGSC0003DMG400005538M6LRR-RLP_244

AAAGAACTGCACTAATACTTCTAGGTGCAAGAAAAAAGATAATTGGAGGGACACTCTGTGCAAAAACTTGGTGTTATTTCCCTCTTTGATAGGCCTTACACGCCGAGAATTCAGATTATT

>M6-PGSC0003DMG400005538M6LRR-RLP_2501

ACATCTTCCTGATGTGGTTGGGAACTCTTCCTAAGTTGAGAGTTTTAAGCTTGAGATCGAATAAGTTGCATGGACCCATTAGAACTTTAGGAAGTGAAAACATGTTTCTTGAGCTTCAAA

>M6-PGSC0003DMG400005538M6LRR-RLP_2562

TAAGTTGCATGGACCCATTAGAACTTTAGGAAGTGAAAACATGTTTCTTGAGCTTCAAATATTGGATATCTCTTCCAATGCCTTCATGGAAAACTTACCTATGGGCCTGTTTCAACATTT

>M6-PGSC0003DMG400005538M6LRR-RLP_2684

AAGCCATGAGGACAATTGATCCATCAAAGAAGGCACCAAGTGATGGATTTTACCAAGACATGGTAGCTGTCGTAACAAAGGGATTTGAGCGTGAAATTGTGAGTATCTTGTATTTGTACA

>M6-PGSC0003DMG400005538M6LRR-RLP_2745

GGTAGCTGTCGTAACAAAGGGATTTGAGCGTGAAATTGTGAGTATCTTGTATTTGTACACTGTTATCGACCTTTCAAGTAATAAATTTGGAGGGGAAATTCCAAGTATCATGGGGGATCT

>M6-PGSC0003DMG400005538M6LRR-RLP_305

AAAAACTTGGTGTTATTTCCCTCTTTGATAGGCCTTACACGCCGAGAATTCAGATTATTCGGGATCCTAATCTGGATATCAAAAATCGAATAGAAAATCAATTTATTTTTAGAAAAAGCC

>M6-PGSC0003DMG400005538M6LRR-RLP_3111

TGATGGATTACGTGGATTCCCTGTTTCAAAAAGTTGCAGCGATGCTCGGGTATTAGATACAAATGATACTGTGTCTGGACTAGACGATGAAGAAAGCAATTCTGAATTTCTGAGTGATTT

>M6-PGSC0003DMG400005538M6LRR-RLP_3294

CATGTTGTCAACTGGAAAACCGATATGGCTAGCAAGAATCATTCTAGAGATGGAGCACAAAATTATGAGGGGAAGGAGAAAGAAGCAGCGAAGGCAAAGGCATTACAGAAGAAGAAGCAA

>M6-PGSC0003DMG400005538M6LRR-RLP_3355

ATTATGAGGGGAAGGAGAAAGAAGCAGCGAAGGCAAAGGCATTACAGAAGAAGAAGCAATCATTTCTAGACAAGTTATAAGTTTTGATTTCAGAACTTCAGGTAGTTCTAATCGATCCCC

>M6-PGSC0003DMG400005538M6LRR-RLP_3416

ATTTCTAGACAAGTTATAAGTTTTGATTTCAGAACTTCAGGTAGTTCTAATCGATCCCCTATATACACTAATCATTTTTTTCATTTCCAATAATTTCAAAAGGCTCAGCCTCAGTTGATG

>M6-PGSC0003DMG400005538M6LRR-RLP_3477

TATACACTAATCATTTTTTTCATTTCCAATAATTTCAAAAGGCTCAGCCTCAGTTGATGTTTAACTAAGATATTTTGGATTACAGGGAGGAGCTAGTGTCTATGGAAGTGGAGAAAGCTG

>M6-PGSC0003DMG400005538M6LRR-RLP_3538

TAACTAAGATATTTTGGATTACAGGGAGGAGCTAGTGTCTATGGAAGTGGAGAAAGCTGGGAAGAAGAAGTTATGGCAATGAATGGAAACGAATGCCAATCAGTTTTTAAGCAATAATAA

>M6-PGSC0003DMG400005538M6LRR-RLP_3579

TGGAAGTGGAGAAAGCTGGGAAGAAGAAGTTATGGCAATGAATGGAAACGAATGCCAATCAGTTTTTAAGCAATAATAATTGTCTAACTGAATATTTCGTTGAACAGTCTTAGATGTGGC

>M6-PGSC0003DMG400005538M6LRR-RLP_366

GGATCCTAATCTGGATATCAAAAATCGAATAGAAAATCAATTTATTTTTAGAAAAAGCCATTTCTCATTATAAATATTAGGTAAACAAGTCAATATCCTGTAATTATACCAAAAAGACAC

>M6-PGSC0003DMG400005538M6LRR-RLP_427

TTCTCATTATAAATATTAGGTAAACAAGTCAATATCCTGTAATTATACCAAAAAGACACTTTGGCATATGACTTAGTCTTGTCTAATATTTGAGAATGACATTTCTCTACTTTCTTTTGA

>M6-PGSC0003DMG400005538M6LRR-RLP_488

TGGCATATGACTTAGTCTTGTCTAATATTTGAGAATGACATTTCTCTACTTTCTTTTGAAATTTTCATAGGAAGACTTGTGCATTGACTTAAGAATAACAAACATTAATTTCCCTATGCA

>M6-PGSC0003DMG400005538M6LRR-RLP_549

TTTTCATAGGAAGACTTGTGCATTGACTTAAGAATAACAAACATTAATTTCCCTATGCAATATGATTCTGGATTTGTTTTTAATTTCTTTTTTTGTTTAAGAAAGCCTTTTCTTCTTCCT

>M6-PGSC0003DMG400005538M6LRR-RLP_61

GATTGATGATTTGATATGGAGGCAGCAGTCCATACTTGACTTTTCCCCTTTCCTCTCTTTTTGATTAGAAAATTCGCAAGTAATTAATTTCAATATGAAGATTCAAAGTTGAAGGGATTT

>M6-PGSC0003DMG400005538M6LRR-RLP_610

ATGATTCTGGATTTGTTTTTAATTTCTTTTTTTGTTTAAGAAAGCCTTTTCTTCTTCCTTTTGGAGTAGCTAGGAATGAACTTTGCCATTTAGAGCCACTTTTATAACGAGGATATATGA

>M6-PGSC0003DMG400005538M6LRR-RLP_793

AAGTAAAACAACGTACAATTATATTTTAAGAAAAAAAGATGATTCTCTACTCCTTTTTTAAGCTTGTTTCTTTAGAGAAATAAAACAACCCAAACTTGACTTGTGTTGAACAATATAAAA

>M6-PGSC0003DMG400005538M6LRR-RLP_854

GCTTGTTTCTTTAGAGAAATAAAACAACCCAAACTTGACTTGTGTTGAACAATATAAAATCCTAAGTTAGTTCATTGAAACCCAAAATAATAAAATCAAAATATGATGATGGTTCCCAGT

>M6-PGSC0003DMG400005538M6LRR-RLP_915

CTAAGTTAGTTCATTGAAACCCAAAATAATAAAATCAAAATATGATGATGGTTCCCAGTAAAATATTTTCTCTACTTCAGTTTTTCACTCTTTTATATCTCCTTACAGTTACTTTTGCTT

>M6-PGSC0003DMG400005538M6LRR-RLP_976

AATATTTTCTCTACTTCAGTTTTTCACTCTTTTATATCTCCTTACAGTTACTTTTGCTTCGATCGAGGAAGCAACTTCCCTCCTGAAATGGAAATTAACTTTCAAGAACCAGAACAACTC

>M6-PGSC0003DMG400005539M6LRR-RLP_1008

TACATCAAATTCCACAAATTACTTCTTCATATCCCATAATAATCTCACTGGAGAAATTCCTTCATCTATATGCAAGTTGACATCACTGGTAATGCTGGATTTAGCGACAAACAATTTGCG

>M6-PGSC0003DMG400005539M6LRR-RLP_1071

ATCTATATGCAAGTTGACATCACTGGTAATGCTGGATTTAGCGACAAACAATTTGCGTGGAGAAATTCCGAAGTGTTTCGGAAATATCACTTCCCTCCAGGTTTTGGATATGCGCCACAA

>M6-PGSC0003DMG400005539M6LRR-RLP_126

ATTGTATGGGATAATACCTGAGAGTATTTTTCACCTGCCCAACTTGGAAACACTTGACTTATTCTTCAATAATCACCTCAGTGGTTATTTTCCAAAGACCAAATGGAACAGCAGTGCATC

>M6-PGSC0003DMG400005539M6LRR-RLP_1575

GGAAGTCGATATTACCAAGACTCGATAGCTGTCGTATCCAAGGGATATGAGCGTGAAATTGTGAGAATCTTGTTTTTGTACACCACTATCGATCTTTCAAACAACAAATTCGAAGGACAT

>M6-PGSC0003DMG400005539M6LRR-RLP_1638

AGAATCTTGTTTTTGTACACCACTATCGATCTTTCAAACAACAAATTCGAAGGACATATTCCAACTATTATGGGAGATCTCATCGCACTCCGTGTGCTGAATATATCTCACAATGGATTG

>M6-PGSC0003DMG400005539M6LRR-RLP_189

CTTCAATAATCACCTCAGTGGTTATTTTCCAAAGACCAAATGGAACAGCAGTGCATCTCTCATACAATTAGATCTCAGTGACGTGAATTTTTCTTGTAATTTGCCTGAGTCTCTTGGCTA

>M6-PGSC0003DMG400005539M6LRR-RLP_1953

TATGAAGGCAATGATGGATTACGTGGTTTTCCTGTTTCAAAAAGTTGTGGGACAACTTATGCCGTATCTGGACAGCTAGACGGTGAAGAAAGCAATTCTGAATTTCTGAGTGATTTTCGG

>M6-PGSC0003DMG400005539M6LRR-RLP_2142

ACTGGAAAACCGATATGGTTTGCAAGAATCATTGTAAAGCTGGAGCACAAAATAATGATGGGAAGGAGAAAGAAGCAACGAAGGCAAAGGAACTGCAGAAGAAGAAATAATCACTAGTAA

>M6-PGSC0003DMG400005539M6LRR-RLP_2205

AGGAGAAAGAAGCAACGAAGGCAAAGGAACTGCAGAAGAAGAAATAATCACTAGTAATGTTACTGTTATTTCTTTAAGGAAAATATGTAATAAGTTGTAGACGTACTATTGTTAAATCCT

>M6-PGSC0003DMG400005539M6LRR-RLP_2268

TGTTATTTCTTTAAGGAAAATATGTAATAAGTTGTAGACGTACTATTGTTAAATCCTGTTTGCTTAAAGTGTGTAGCATTATTTGACCATGAAAATTGAGGGTGAGATAATCTTTAATGA

>M6-PGSC0003DMG400005539M6LRR-RLP_252

ACAATTAGATCTCAGTGACGTGAATTTTTCTTGTAATTTGCCTGAGTCTCTTGGCTATTATCTAACTTCGGTGCGTTATTTGTCTCTTCAAAGTTGCAACCTTAGAGGGCCTATTCCTGA

>M6-PGSC0003DMG400005539M6LRR-RLP_378

TTCAAATCTCACCCGCATTGAGTATTTATACCTTAATGGTAACTCCCTGAATGGAACAATACCACCAGGGATGTTATCCCTTCCATCACTAAAGGAAATAGGCGTGAGTTTTAATCACTT

>M6-PGSC0003DMG400005539M6LRR-RLP_441

ACCAGGGATGTTATCCCTTCCATCACTAAAGGAAATAGGCGTGAGTTTTAATCACTTATCTGGTCCGCTTGAAGATTTCAAGTCCAATTCACTAATCTGGATAAGTTTAGAGGGCAATCA

>M6-PGSC0003DMG400005539M6LRR-RLP_504

TCCGCTTGAAGATTTCAAGTCCAATTCACTAATCTGGATAAGTTTAGAGGGCAATCAGCTGCAGGGTCATCTTCCCCACTCAATTCAAAACCTTGTGAGCCTAACAGATCTTGATCTTTC

>M6-PGSC0003DMG400005539M6LRR-RLP_63

CTCTTCCACCATTCCTCTGAATTTTTCTTCTCATTTAACAACTCTGATCCTGTCAGAAACAGGATTGTATGGGATAATACCTGAGAGTATTTTTCACCTGCCCAACTTGGAAACACTTGA

>M6-PGSC0003DMG400005539M6LRR-RLP_630

TAATTTTAGTGGTCGTGTGGATGCCAGCCTCTTTTCAAACCTTAAACAGCTTCGGTCTATTAGTCTTTCATATAATAACATCTCATTGACCAATGACAACAACGTTACTTTGCCCGAATC

>M6-PGSC0003DMG400005539M6LRR-RLP_693

TCTTTCATATAATAACATCTCATTGACCAATGACAACAACGTTACTTTGCCCGAATCTCTTACTAGGTTAGAATTGGCCTCTTGTGAAGTGAAAGAGTTGGAGTTTCTAAGATCCGCAAA

>M6-PGSC0003DMG400005539M6LRR-RLP_756

TAGGTTAGAATTGGCCTCTTGTGAAGTGAAAGAGTTGGAGTTTCTAAGATCCGCAAACCAGCTTTGGCAGTTGGATCTTTCAAATAATAAGATTGAAGGAAGAATTCCTGATTGGGCATG

>M6-PGSC0003DMG400005539M6LRR-RLP_882

CTGGATGTTTTCATTGCAAGATCTTAATATATCCCACAATATGCTGACGAGTGTGGATTCAATTCCTCTTCAGTTTGCAGATACTATTAATTTGCGGTCCAATTTGCTTCAAGGTTCACT

>M6-PGSC0003DMG400005539M6LRR-RLP_945

TCCTCTTCAGTTTGCAGATACTATTAATTTGCGGTCCAATTTGCTTCAAGGTTCACTACCTATTACATCAAATTCCACAAATTACTTCTTCATATCCCATAATAATCTCACTGGAGAAAT

>M6-PGSC0003DMG400005542M6LRR-RLP_0

ATGGGGTATGCAGTTATTATGCTCTTAATTGTGTTTTTCTGTCATGAAGTTTCTTCAATATCCATATTACCTCACAAACATCAAACCATTTCACTTTTAAAATTCAAGCAGTCCTTTACT

>M6-PGSC0003DMG400005542M6LRR-RLP_1054

CCTTTGAAGATTTCAAGTCCAATTTGCTTCAAGGTTCACTACCCATTCTATCAAATTCCACAGTACTCTTCTTCATATCCCATAATAATCTCACTGGAGAAATTCCTTCATCTATTTGCA

>M6-PGSC0003DMG400005542M6LRR-RLP_1116

GTACTCTTCTTCATATCCCATAATAATCTCACTGGAGAAATTCCTTCATCTATTTGCAACTTAACATCACTGATAATGCTGGATTTGGCGACAAACAATTTGCGTGGAGAAATTCCGCAG

>M6-PGSC0003DMG400005542M6LRR-RLP_1488

TTGCGATTCAACAAATTGCATGGTCCCATCAGAACATCAAGGATTGAGAACATATTTTCAGAGCTCCGAATCATTGATCTCTCTTACAATGCCTTCTCAGGAAACTTACCTTCGAGTTTG

>M6-PGSC0003DMG400005542M6LRR-RLP_1550

GCTCCGAATCATTGATCTCTCTTACAATGCCTTCTCAGGAAACTTACCTTCGAGTTTGTTTCAACATCTGAAAGCCATGAGGACAGCTGATCCATCAAAGGAACCATCAAAATATGGTGG

>M6-PGSC0003DMG400005542M6LRR-RLP_1612

AACATCTGAAAGCCATGAGGACAGCTGATCCATCAAAGGAACCATCAAAATATGGTGGAGATTCATATTACCAAGACTCGATAGCTTTCGTATTCAAGGGATATGAGCGTGAAGTTGTGA

>M6-PGSC0003DMG400005542M6LRR-RLP_1674

TCATATTACCAAGACTCGATAGCTTTCGTATTCAAGGGATATGAGCGTGAAGTTGTGAGAATCTTGTTTTTGTATATCACTATCGATCTTTCAAACAACAAATTCGAAGGACATATTCCA

>M6-PGSC0003DMG400005542M6LRR-RLP_1798

TTATGGGAGATCTCATTGCACTCCGTGTGCTGAATATATCTCATAATGGATTGGAAGGTCCTATACCGTCATCACTTGGAAGTTTATCTTTGGTTGAATCACTGGACCTGTCAAGTAACC

>M6-PGSC0003DMG400005542M6LRR-RLP_1860

ATACCGTCATCACTTGGAAGTTTATCTTTGGTTGAATCACTGGACCTGTCAAGTAACCATCTTGTAGGAAAGATACCAGCACAATTTGCTTCTCTTACATCTCTTGAAGTCTTCAATCTT

>M6-PGSC0003DMG400005542M6LRR-RLP_1922

TGTAGGAAAGATACCAGCACAATTTGCTTCTCTTACATCTCTTGAAGTCTTCAATCTTTCCTACAATCATCTCGAAGGGTGCATTCCTCAGGGAAATCAACTTCACACCTTTGAGAACAA

>M6-PGSC0003DMG400005542M6LRR-RLP_248

ACTTGACCTCAGTTGCAGTAATCTCACTGGAACCATCGATTCCAATAGCAGCCTATTCCAACTCTCGCATCTCCAAAGGCTTGATCTTTCTTGGAATAACTTCTCCAATTCTCATATCTC

>M6-PGSC0003DMG400005542M6LRR-RLP_310

TCTCGCATCTCCAAAGGCTTGATCTTTCTTGGAATAACTTCTCCAATTCTCATATCTCCCCTGAATTCAGCAAGTTTTCGAGCTTGATGCATCTTGATCTTTCTGGGTCCTATTTCTCAG

>M6-PGSC0003DMG400005542M6LRR-RLP_372

GAATTCAGCAAGTTTTCGAGCTTGATGCATCTTGATCTTTCTGGGTCCTATTTCTCAGGTCAAATTCCTTCTGAAATCTCTCATCTTTCCAAGTTACACTCTCTTCGTATCTCCACATTT

>M6-PGSC0003DMG400005542M6LRR-RLP_434

AATTCCTTCTGAAATCTCTCATCTTTCCAAGTTACACTCTCTTCGTATCTCCACATTTTATTACTTTTACAGTAACAACCTTTACAATAAAAAGCTAAGACTCACAACCCATGGTTTTAA

>M6-PGSC0003DMG400005542M6LRR-RLP_496

ACTTTTACAGTAACAACCTTTACAATAAAAAGCTAAGACTCACAACCCATGGTTTTAAATCGCTTCTTCAAAACTTGACCCAATTAAGAGAGCTTGATCTTAGTGGCATAAACATCTCTT

>M6-PGSC0003DMG400005542M6LRR-RLP_62

CATATTACCTCACAAACATCAAACCATTTCACTTTTAAAATTCAAGCAGTCCTTTACTATACATCCCACAGTAGAATCTTATTATTGTCCGAAGCCTTATCCAAAAACGAGTTCATGGAA

>M6-PGSC0003DMG400005542M6LRR-RLP_620

CATTCCTCTTAATTTTTCTTCTCATTTCACGACTCTGAGCCTGTCAGAAACAGGATTGTAAGGGATAATACCTGAGAGTATATTTCACCTGCCCAACTTGGAAATACTTGACTTAGGGAT

>M6-PGSC0003DMG400005542M6LRR-RLP_682

GGATAATACCTGAGAGTATATTTCACCTGCCCAACTTGGAAATACTTGACTTAGGGATCAATCATGAGCTCAATGGTAATTTTCCGAAGACCAAATGGAACAGCAGTGCATCTCTCATTG

>M6-PGSC0003DMG400005542M6LRR-RLP_744

CATGAGCTCAATGGTAATTTTCCGAAGACCAAATGGAACAGCAGTGCATCTCTCATTGAGTTAGATCTCCGTGGCGTGAATTTTTCTGATAATTTGCCCGAGTCTCTTGGCTATCATCTA

>M6-PGSC0003DMG400005542M6LRR-RLP_806

AGATCTCCGTGGCGTGAATTTTTCTGATAATTTGCCCGAGTCTCTTGGCTATCATCTAACTTCGCTGCTTTATTTGTCTCTTGAAAGTTGCAACCTTAGAGGGCCTATTCCTGAATCTCT

>M6-PGSC0003DMG400005542M6LRR-RLP_868

CGCTGCTTTATTTGTCTCTTGAAAGTTGCAACCTTAGAGGGCCTATTCCTGAATCTCTTTCAAATCTCACCCGCATTGAGTATTTGCACCTTAATCGTAACTCCCTGAATGGAACAATAC

>M6-PGSC0003DMG400005542M6LRR-RLP_992

AGGGATGTTATCCTTTCCATCACTACATAGAATAGACTTGAGTAATAATCACTTATCTGGTCCCTTTGAAGATTTCAAGTCCAATTTGCTTCAAGGTTCACTACCCATTCTATCAAATTC

>M6-PGSC0003DMG400005590M6LRR-RLP_0

ACCTTAACACATCCTACAGAATAAAATACATCTCTACGGACTTTAGATTGAAAATTGTGGGATACAAATTAGATTTTAATAAGATCCAACTATTCATTCTTATTTTAAAATGTATATGAT

>M6-PGSC0003DMG400005590M6LRR-RLP_1054

ACTTGGATGTTAGTTCTAATACAATAATTCCAGATACCGAATTCCAAAATTGGATCCCAAATTTCCAGTTGGAGTTTTTTGCTATCGAAGGATGTAAAAACCTTCAAAAATTGCCTTCTT

>M6-PGSC0003DMG400005590M6LRR-RLP_1116

TTCCAGTTGGAGTTTTTTGCTATCGAAGGATGTAAAAACCTTCAAAAATTGCCTTCTTTCCTTCACTACCAATATGACATGAGGATTCTTGCTATAAATGGAAATCAATTGCCAAGAAAC

>M6-PGSC0003DMG400005590M6LRR-RLP_1178

TCACTACCAATATGACATGAGGATTCTTGCTATAAATGGAAATCAATTGCCAAGAAACTTTCCAACATGGTTGTTAGAAAATAACACCAGACTTGCAGGGATTTATAGTAGAGATAATGC

>M6-PGSC0003DMG400005590M6LRR-RLP_124

TTATTCACCATTTAACAGGTAACAAAAGAATGAGTTCTCTTCGGAATTTACAATTGGGAAGTTCCTACTCCAACTTAAGCAGAGTCATAGAATCCCTAAAATCATTCTCTTCTCTTAAAT

>M6-PGSC0003DMG400005590M6LRR-RLP_1240

CAACATGGTTGTTAGAAAATAACACCAGACTTGCAGGGATTTATAGTAGAGATAATGCTTTCAGCGGACCATTCAAGTTGCCATCGAGTTGTCACCTCCATCTAGAGACAATTGATGTTT

>M6-PGSC0003DMG400005590M6LRR-RLP_1302

AGCGGACCATTCAAGTTGCCATCGAGTTGTCACCTCCATCTAGAGACAATTGATGTTTCTAATAACAAATTAAATGGGCATATTCCAGATAAAATGAATTCAGCATTCCCAAAGCTTACT

>M6-PGSC0003DMG400005590M6LRR-RLP_1364

TAACAAATTAAATGGGCATATTCCAGATAAAATGAATTCAGCATTCCCAAAGCTTACTTATTTGAACATGTCACAAAATTATCTTGAAGGTCCTATACCTTCCAAGATTAGTGGCATTCA

>M6-PGSC0003DMG400005590M6LRR-RLP_1426

TGAACATGTCACAAAATTATCTTGAAGGTCCTATACCTTCCAAGATTAGTGGCATTCATTTACATAATCTAGACCTATCTCACAATTTCTTATTTGGAGGAGTTCCTAGTGATCTGGCAA

>M6-PGSC0003DMG400005590M6LRR-RLP_1488

CATAATCTAGACCTATCTCACAATTTCTTATTTGGAGGAGTTCCTAGTGATCTGGCAATTGGTTCTCCAGTATTAGCTTACCTTCGATGTCAAATTTTTCCAAAGGATGTCAAACCAAAG

>M6-PGSC0003DMG400005590M6LRR-RLP_1550

TTCTCCAGTATTAGCTTACCTTCGATGTCAAATTTTTCCAAAGGATGTCAAACCAAAGATGTTATTCTTTTTGTTCTTGAATGGAAATAACTTTGAGGGACCACTACCTTGCAACATGTT

>M6-PGSC0003DMG400005590M6LRR-RLP_1612

TATTCTTTTTGTTCTTGAATGGAAATAACTTTGAGGGACCACTACCTTGCAACATGTTTATCACATCCCTTATTGTATTGGACGCTAGCAACAATAATTTCTCTGGAGAGATTCCGAGAT

>M6-PGSC0003DMG400005590M6LRR-RLP_1674

ACATCCCTTATTGTATTGGACGCTAGCAACAATAATTTCTCTGGAGAGATTCCGAGATGCATTAGGGATAATACAAGATTGTTACAACTTGATTTGTCCAAGAATCATCTCGATGGTTCG

>M6-PGSC0003DMG400005590M6LRR-RLP_1736

TAGGGATAATACAAGATTGTTACAACTTGATTTGTCCAAGAATCATCTCGATGGTTCGATTCCAGTTGAGATTTCCAATTTAAAGCTCATTCAGGTCTTAGCTATATCCGAAAACAGGCT

>M6-PGSC0003DMG400005590M6LRR-RLP_1798

CAGTTGAGATTTCCAATTTAAAGCTCATTCAGGTCTTAGCTATATCCGAAAACAGGCTTTCAGGCTTTATTCCTTCTTGTGTGAATTTTTTACCTCTTGAGCACATCCATCTACAGAAAA

>M6-PGSC0003DMG400005590M6LRR-RLP_186

TCCTACTCCAACTTAAGCAGAGTCATAGAATCCCTAAAATCATTCTCTTCTCTTAAATCACTTTCTTATGAATTTAGTGATCTCACTTCTCCAACCATAATTTACGGTAAGCTCAAATTC

>M6-PGSC0003DMG400005590M6LRR-RLP_1860

GGCTTTATTCCTTCTTGTGTGAATTTTTTACCTCTTGAGCACATCCATCTACAGAAAAATCAATTGGGTGGTGAATTGGAACATGTACTTTTCAACTTCTCTTCTCTGATAACTTTGGAT

>M6-PGSC0003DMG400005590M6LRR-RLP_1922

ATTGGGTGGTGAATTGGAACATGTACTTTTCAACTTCTCTTCTCTGATAACTTTGGATCTTGGGTACAACAATTTTACAGGAAGTATCCCACACACCATAGGCTCACTTAATACTCTCAA

>M6-PGSC0003DMG400005590M6LRR-RLP_1984

GGTACAACAATTTTACAGGAAGTATCCCACACACCATAGGCTCACTTAATACTCTCAACTACCTTCTCCTTAGCAATAATCAATTCGAAGGAGAGATTCCAACTCAGATTTGTATGCTGA

>M6-PGSC0003DMG400005590M6LRR-RLP_2046

CTTCTCCTTAGCAATAATCAATTCGAAGGAGAGATTCCAACTCAGATTTGTATGCTGAACAAGTTATCTATCGTGGATTTATCTTTCAATAAGCTTTATGGTTCACTCCTTCCTTGTTTG

>M6-PGSC0003DMG400005590M6LRR-RLP_2108

GTTATCTATCGTGGATTTATCTTTCAATAAGCTTTATGGTTCACTCCTTCCTTGTTTGAGTTACTTAACACAAGCCAAAAAGGATGCAGAGATAAGTAAGACCAATTATGCTACGACTTC

>M6-PGSC0003DMG400005590M6LRR-RLP_2170

ACTTAACACAAGCCAAAAAGGATGCAGAGATAAGTAAGACCAATTATGCTACGACTTCTAGGCGTTCATGGATAACTTTTTTGGCCTGGATGCGTTTGAAAAGGCACTACCACGATAGTC

>M6-PGSC0003DMG400005590M6LRR-RLP_2232

CGTTCATGGATAACTTTTTTGGCCTGGATGCGTTTGAAAAGGCACTACCACGATAGTCATGGATTTCTAAGTGACATATTTCTGATGGATGTGGAAACTCAGGTCCAGTTTTCAACAAAA

>M6-PGSC0003DMG400005590M6LRR-RLP_2294

ATTTCTAAGTGACATATTTCTGATGGATGTGGAAACTCAGGTCCAGTTTTCAACAAAAAGAAACTCATACACTTATAAGGGAAGCATTCTAAAATATATGTTGGGTATTGATCTCTCAAG

>M6-PGSC0003DMG400005590M6LRR-RLP_2356

ACTCATACACTTATAAGGGAAGCATTCTAAAATATATGTTGGGTATTGATCTCTCAAGTAACAGATTAACTGGGGAAATTCCCGTTGAGCTAGGGAACATGAGCAATATACATGCATTGA

>M6-PGSC0003DMG400005590M6LRR-RLP_2418

AGATTAACTGGGGAAATTCCCGTTGAGCTAGGGAACATGAGCAATATACATGCATTGAATCTATTGCACAACCATCTCATCGGAAGAATACCAAATACCTTCTCCAATCTACAGAAAATT

>M6-PGSC0003DMG400005590M6LRR-RLP_248

TTCTTATGAATTTAGTGATCTCACTTCTCCAACCATAATTTACGGTAAGCTCAAATTCATATATACCTTAGCTTACTTCTATTGCCACTCTTTTCTTTTGATATGTAAATTTAGTTTTCT

>M6-PGSC0003DMG400005590M6LRR-RLP_2480

ATTGCACAACCATCTCATCGGAAGAATACCAAATACCTTCTCCAATCTACAGAAAATTGAGAGTTTAGACCTTTCCTACAACAACTTGAATGGGAGCATTCCTGTTGGTCGACTTGAGCT

>M6-PGSC0003DMG400005590M6LRR-RLP_2542

GTTTAGACCTTTCCTACAACAACTTGAATGGGAGCATTCCTGTTGGTCGACTTGAGCTAAATTCTTTGGCAGTATTCAGTGTTGCATACAACAACTTATCGGGTGCAGTGCCTGATTTTA

>M6-PGSC0003DMG400005590M6LRR-RLP_2666

TCAATTTGGAACTTTCAACAAAAGCAGCTACGAGGGAAATCCTTTTCTTTGCGGTTATCCATTAGACAACACGTGTGGCATTCCTAAATTGTCAAATACCTCCAAGATTAATGGAGGTGA

>M6-PGSC0003DMG400005590M6LRR-RLP_2728

TAGACAACACGTGTGGCATTCCTAAATTGTCAAATACCTCCAAGATTAATGGAGGTGAGGAATCATCAGACTTGGAGGATATTCAGTGTTTTTACATTGGCTTTGTTGTGTCTTATGGAG

>M6-PGSC0003DMG400005590M6LRR-RLP_2852

CTTGTTGGGATTAGCTGCGGCACTCTGCTTCAATCGTTATTGGAGAAAAACATGGTTTAGGATGATAGAGGCATTGATGTTTTATTGTTACTATTCTCTCTTGGACAACGTTGTTACACC

>M6-PGSC0003DMG400005590M6LRR-RLP_2914

TGATAGAGGCATTGATGTTTTATTGTTACTATTCTCTCTTGGACAACGTTGTTACACCCATCAAGAATATATGGTACAGGAATGTTGGTTAATTCTAGGTTGTTGTTTTACAATATTGTT

>M6-PGSC0003DMG400005590M6LRR-RLP_2976

AAGAATATATGGTACAGGAATGTTGGTTAATTCTAGGTTGTTGTTTTACAATATTGTTGTATGTGTATCACCATTATTAATTATATTGTCATTCTGTTTTATCAATATTCAACAAATAGG

>M6-PGSC0003DMG400005590M6LRR-RLP_3038

GTGTATCACCATTATTAATTATATTGTCATTCTGTTTTATCAATATTCAACAAATAGGATCTGTTTGGAGTAGTTACAACTTCATAATTTAATACCCCCTGCATAGCTTCATTAGTGTTG

>M6-PGSC0003DMG400005590M6LRR-RLP_310

ATACCTTAGCTTACTTCTATTGCCACTCTTTTCTTTTGATATGTAAATTTAGTTTTCTTTTATTTTTTGGACTCTATGATATAAGCATTAAGAAATTTGAGCACGGTGGAGCACCTATAC

>M6-PGSC0003DMG400005590M6LRR-RLP_372

TTTTTTGGACTCTATGATATAAGCATTAAGAAATTTGAGCACGGTGGAGCACCTATACTTGGAGGGATGTTCTTTAAATGACAACTTTCTCCCAAACATTGGACAAATGACTTCTCTCTT

>M6-PGSC0003DMG400005590M6LRR-RLP_434

AGGGATGTTCTTTAAATGACAACTTTCTCCCAAACATTGGACAAATGACTTCTCTCTTGGTGTTCAGTATAGCATTTGGTGACAATGACGGCACCCTCCCTAATCAAGGTACCCTATTTT

>M6-PGSC0003DMG400005590M6LRR-RLP_496

TTCAGTATAGCATTTGGTGACAATGACGGCACCCTCCCTAATCAAGGTACCCTATTTTTCTCTTTTTTCCCTAATGCCCGCATTTTCATGTTGGAGGTCTGATTTTAGCATGTTCAGTTG

>M6-PGSC0003DMG400005590M6LRR-RLP_558

TTTTTTCCCTAATGCCCGCATTTTCATGTTGGAGGTCTGATTTTAGCATGTTCAGTTGATCAAACATTATAATATTTTTTGTTTATTACCAAACACAAGAAAATAAGTAACACAAGCCAT

>M6-PGSC0003DMG400005590M6LRR-RLP_62

TACAAATTAGATTTTAATAAGATCCAACTATTCATTCTTATTTTAAAATGTATATGATTTATTTATTCACCATTTAACAGGTAACAAAAGAATGAGTTCTCTTCGGAATTTACAATTGGG

>M6-PGSC0003DMG400005590M6LRR-RLP_620

AACATTATAATATTTTTTGTTTATTACCAAACACAAGAAAATAAGTAACACAAGCCATCTAATTTTTCTAGAAAATATTTTCCTTCGTATCGAACACATCCATAGACATAAAAAAATATA

>M6-PGSC0003DMG400005590M6LRR-RLP_682

TTTTTCTAGAAAATATTTTCCTTCGTATCGAACACATCCATAGACATAAAAAAATATAATAAGATGTCACTCATCATAGATATTTGTTTTAGCTAAATGTTTCTTAATATATATATATAG

>M6-PGSC0003DMG400005590M6LRR-RLP_744

GATGTCACTCATCATAGATATTTGTTTTAGCTAAATGTTTCTTAATATATATATATAGGCTGGTGTGAACTCAAACACATTCAAGAGCTGGATTTCTCAGACAATAATTTTCGTGGAACA

>M6-PGSC0003DMG400005590M6LRR-RLP_806

GTGTGAACTCAAACACATTCAAGAGCTGGATTTCTCAGACAATAATTTTCGTGGAACACTGCCTTCATGTTTTGGAAACTTGACATCACTTCGACGATTGAGTTTTTTTGGGAATTACTT

>M6-PGSC0003DMG400005590M6LRR-RLP_868

CTTCATGTTTTGGAAACTTGACATCACTTCGACGATTGAGTTTTTTTGGGAATTACTTCATGGGAAATATAGCGTCACATTCTATTTGGAGGAGACTCACATCACTTGAGTACCTCAATA

>M6-PGSC0003DMG400005590M6LRR-RLP_930

GGAAATATAGCGTCACATTCTATTTGGAGGAGACTCACATCACTTGAGTACCTCAATATTGTGGATAACCAATTTGATGTTCCTCTGTCATTCAGCCAATTTTCCAACCATACAAAATTG

>M6-PGSC0003DMG400005590M6LRR-RLP_992

GGATAACCAATTTGATGTTCCTCTGTCATTCAGCCAATTTTCCAACCATACAAAATTGATCTACTTGGATGTTAGTTCTAATACAATAATTCCAGATACCGAATTCCAAAATTGGATCCC

>M6-PGSC0003DMG400005600M6LRR-RLP_0

CCCTGAGACTTAGTTACCATCACCTTCCACTTGATTTGAGACAATGCTTTGCATATTGTGCAGTATTCCCAAAGGATACGAAATGGAAAAGAAAAATCTAATCTCTCTCTGGATGGCACA

>M6-PGSC0003DMG400005600M6LRR-RLP_1024

TGAAAAATGTTGTCTCTATTGTAATTGAAGTTTGCGAAAACTGCTCATGCTTACCACCCTTTGGAGAGCTGCCTTGTCTAGAAGGTTTAGAGTTACACAAGGGGTCTGTGGAGGTGGAGT

>M6-PGSC0003DMG400005600M6LRR-RLP_1088

AGAGCTGCCTTGTCTAGAAGGTTTAGAGTTACACAAGGGGTCTGTGGAGGTGGAGTATGTTGATTCTGGATTCCCTACAAGAAGAAGGTTTCCATCTCTGAGAAAACTTATTATAGGCGA

>M6-PGSC0003DMG400005600M6LRR-RLP_1152

TCTGGATTCCCTACAAGAAGAAGGTTTCCATCTCTGAGAAAACTTATTATAGGCGACTTTGATAATCTGAAAGGATTGCTGAAAAAGGAAGGAGAAGAGCAATTCCCTGTGCTTGAAGAG

>M6-PGSC0003DMG400005600M6LRR-RLP_128

TTTTATCGAAAGGAAACTTGGAGCTAGAGGATGTAGGTAATGAAGTATGGAATGAATTATACTTGAGGTCTTTCTTCCAAGAGATTGAAGTTAAATATGGTAGAACTTATTTCAAGATGC

>M6-PGSC0003DMG400005600M6LRR-RLP_1280

TTACTGGTGCCCTATGTTTGTTATTCCGACCCTTTCTTCTGTCAAGAAATTGGTAGTTCATGGGGAGAAGTCAGATGCAATAGGTTTCAGTTCCATATCTAATCTCAGGCCTCTTACTTC

>M6-PGSC0003DMG400005600M6LRR-RLP_1344

GAGAAGTCAGATGCAATAGGTTTCAGTTCCATATCTAATCTCAGGCCTCTTACTTCCCTCCAAATTAGCAAAAACGAAGAAGCTACTTCGCTCCCAGAAGAGATGTTCACATGCCTTGCA

>M6-PGSC0003DMG400005600M6LRR-RLP_1408

TTAGCAAAAACGAAGAAGCTACTTCGCTCCCAGAAGAGATGTTCACATGCCTTGCAAATCTCAAATACTTGAATATCAATTCCTTCACGAATCTCAACGAGCTGCCTACCAGCCTGGCTA

>M6-PGSC0003DMG400005600M6LRR-RLP_1472

ATACTTGAATATCAATTCCTTCACGAATCTCAACGAGCTGCCTACCAGCCTGGCTAGTCTTAATGCTTTGAAGCATCTGGATATTATTGGGTGTGATGCACTCGAGAGTCTCCCCGAGGA

>M6-PGSC0003DMG400005600M6LRR-RLP_1536

GCTTTGAAGCATCTGGATATTATTGGGTGTGATGCACTCGAGAGTCTCCCCGAGGAAGGGGTGAAAGGTTTAACTTCACTCACACAGTTATCCATAACATGCTGTGAGATGCTACAATGT

>M6-PGSC0003DMG400005600M6LRR-RLP_192

GAGGTCTTTCTTCCAAGAGATTGAAGTTAAATATGGTAGAACTTATTTCAAGATGCATGATCTCATTCATGATTTGGCAACATTTCTATTTTCAGCAAGCACATCAAGCAGCAATATCCG

>M6-PGSC0003DMG400005600M6LRR-RLP_256

ATTCATGATTTGGCAACATTTCTATTTTCAGCAAGCACATCAAGCAGCAATATCCGAGAAATAAATGTAAAAGGTGACCCTTACCCACATATGATGTCGATTGGTTTCTCTGAATTGGTG

>M6-PGSC0003DMG400005600M6LRR-RLP_320

ATGTAAAAGGTGACCCTTACCCACATATGATGTCGATTGGTTTCTCTGAATTGGTGCCTTCTTACTCTCCTTCTCTCTTTGACAGGTTTGTCTCATTGAGGGTGCTTAATCTTAGTAAAT

>M6-PGSC0003DMG400005600M6LRR-RLP_576

CATGATTGCTCCTCACTTTGTTGTTTGCCAAAACAAACAAGTAAACTTGGTAGTCTCCGAAATCTTTTACTTGAAAATTGCAAAGCATTGACTTGTATGCCACCAAGGATAGCTTCTTTG

>M6-PGSC0003DMG400005600M6LRR-RLP_64

ATTCCCAAAGGATACGAAATGGAAAAGAAAAATCTAATCTCTCTCTGGATGGCACATGGTTTTCTTTTATCGAAAGGAAACTTGGAGCTAGAGGATGTAGGTAATGAAGTATGGAATGAA

>M6-PGSC0003DMG400005600M6LRR-RLP_640

TTTTACTTGAAAATTGCAAAGCATTGACTTGTATGCCACCAAGGATAGCTTCTTTGACATGCCTTAAGACTCTTGGTTGCTTTGCAGTGGGAAGGAAGAAAAGTTCTCAAATTGGTGAAT

>M6-PGSC0003DMG400005600M6LRR-RLP_704

TAAGACTCTTGGTTGCTTTGCAGTGGGAAGGAAGAAAAGTTCTCAAATTGGTGAATTACGAAACCTGAATCTCTATGGCTCAATTTCAATAACGCATCTTGAGAGAGTGAAGAATGATAT

>M6-PGSC0003DMG400005600M6LRR-RLP_768

CTGAATCTCTATGGCTCAATTTCAATAACGCATCTTGAGAGAGTGAAGAATGATATGGATGCAAAAGAAGCCAATTTATCTGCAAAAGAAAATCTGTATTCTTTAAGCATGATTTGGAGT

>M6-PGSC0003DMG400005600M6LRR-RLP_832

AAGAAGCCAATTTATCTGCAAAAGAAAATCTGTATTCTTTAAGCATGATTTGGAGTAGGATAGGACCACATAGATATGAATCAGAAGAAGTTGAAGTGCTTGAAGCCCTCAAACCACACT

>M6-PGSC0003DMG400005600M6LRR-RLP_960

ACTTGTTTAACAATCAGTGGCTTCAGAGGAATCCGTCCCCCAGACTGGATGAATCACTCAGTTTTGAAAAATGTTGTCTCTATTGTAATTGAAGTTTGCGAAAACTGCTCATGCTTACCA

>M6-PGSC0003DMG400005689M6G-LecRK_0

AAAAAAACCTGGTCAACAAAAGAAATCAAAGCCAACAGCATGCTTGTCTCTGTTTTGGTCATACTATTTTGACTGTAAACATTCATGTGTGGATTAAGGAGCTTCTCTCATATGGCTACC

>M6-PGSC0003DMG400005689M6G-LecRK_1054

TTAAATTTGATGAGAACTATACTTTGAGCCTCTCTGATTGCAAGGCCAAGTGTGAGAATGACTGCTCGTGTGTTGCCTATGCCTCAGTCGATAGTGACACCGGAATTGGCTGTCAAATTT

>M6-PGSC0003DMG400005689M6G-LecRK_1116

TGCTCGTGTGTTGCCTATGCCTCAGTCGATAGTGACACCGGAATTGGCTGTCAAATTTGGGGGAATGACACGAGTTTTGTGACAGCACAAAACAGTCTAGCCAGAGATGTTTTCTTTCTT

>M6-PGSC0003DMG400005689M6G-LecRK_1178

GAATGACACGAGTTTTGTGACAGCACAAAACAGTCTAGCCAGAGATGTTTTCTTTCTTACAAGAAAAGGTAATTCTTGAGTACAAAGTTATTCAATAGAGTGAGAGGACATGTTAATAAT

>M6-PGSC0003DMG400005689M6G-LecRK_124

GAATTCTGATTGCTTTCATCCTCGTGTTATCCTGTCACTTTTTGCGTGCAGATTTTGCTAATTTCAACTCAATATTGAAGGGAGAAAGTTTAGCTGATTACCAACAGCTGGTCTCAGCAA

>M6-PGSC0003DMG400005689M6G-LecRK_1240

GAAAAGGTAATTCTTGAGTACAAAGTTATTCAATAGAGTGAGAGGACATGTTAATAATGTTGCATCTCATATTATGATTACTTTATCCAGCTCAAGATAAATTGTTTTGTGAGGGCATCA

>M6-PGSC0003DMG400005689M6G-LecRK_1302

CATCTCATATTATGATTACTTTATCCAGCTCAAGATAAATTGTTTTGTGAGGGCATCACACTAATATCACCAATCTGCATTACTCTCATTTGCACGTCATTCAAAAAATTACTAGTACTA

>M6-PGSC0003DMG400005689M6G-LecRK_1364

AATATCACCAATCTGCATTACTCTCATTTGCACGTCATTCAAAAAATTACTAGTACTAGAACAAGCAATTTTCTCCGATCGCGATGCTGCAGGATTTGAGTTAATAAGAGTGTCATCTTC

>M6-PGSC0003DMG400005689M6G-LecRK_1426

AAGCAATTTTCTCCGATCGCGATGCTGCAGGATTTGAGTTAATAAGAGTGTCATCTTCATTTCAAAAATACACTCGAGCAAATGAACAACCATACCATAGAGAATCTTGTTTTGGAGGCT

>M6-PGSC0003DMG400005689M6G-LecRK_1488

CAAAAATACACTCGAGCAAATGAACAACCATACCATAGAGAATCTTGTTTTGGAGGCTATTAAATTATTATTTTTTGGTATTTCTAGAGTACCGACAATTATGATAGCATCATAGTGTAT

>M6-PGSC0003DMG400005689M6G-LecRK_1550

AATTATTATTTTTTGGTATTTCTAGAGTACCGACAATTATGATAGCATCATAGTGTATTCCCAAGGTAAATATGTGGAATTTTGGTCATAAACACTGACTTGCAATTTTGACAGATAGGA

>M6-PGSC0003DMG400005689M6G-LecRK_1612

AAGGTAAATATGTGGAATTTTGGTCATAAACACTGACTTGCAATTTTGACAGATAGGAAGCGGAAATGGTGGATATGGTTGACAATAGCAGTGTCACTAATGGTATTTGCGTTTATTTTC

>M6-PGSC0003DMG400005689M6G-LecRK_1674

GAAATGGTGGATATGGTTGACAATAGCAGTGTCACTAATGGTATTTGCGTTTATTTTCTCTTTATTCTGTCTGATACGAAGAAAACTCAGAGCAAGAGGTGAGCAATACATCCGCTCTAG

>M6-PGSC0003DMG400005689M6G-LecRK_1736

TATTCTGTCTGATACGAAGAAAACTCAGAGCAAGAGGTGAGCAATACATCCGCTCTAGCATTAGTCTAAGGTGACATTTTTTTTCCTTTGAAATCTTCAATTTTGCATTTGTCACCTAGG

>M6-PGSC0003DMG400005689M6G-LecRK_1798

AGTCTAAGGTGACATTTTTTTTCCTTTGAAATCTTCAATTTTGCATTTGTCACCTAGGCAAAGTCAGACAAATGGAGAAAATATTATACGAGCTTGAAGATCCTAAGACCATTTCTGGCC

>M6-PGSC0003DMG400005689M6G-LecRK_186

TTCAACTCAATATTGAAGGGAGAAAGTTTAGCTGATTACCAACAGCTGGTCTCAGCAAATGGCTTCTTCAAAATGAAATTTTTCAGTCCAGGCAAGTCAAGGAACCGCTACTTGGGTATA

>M6-PGSC0003DMG400005689M6G-LecRK_1860

GTCAGACAAATGGAGAAAATATTATACGAGCTTGAAGATCCTAAGACCATTTCTGGCCAGTACAATACTAACAAGACTGCTAGACTAAGGAAGAAGTTCAGGCATGATATACACATATTC

>M6-PGSC0003DMG400005689M6G-LecRK_1922

CAATACTAACAAGACTGCTAGACTAAGGAAGAAGTTCAGGCATGATATACACATATTCAGCCTAGAAACTATGAATATGGCCACAAACAATTTCTCGTCTTCTAATAAGTTGGGACAGGG

>M6-PGSC0003DMG400005689M6G-LecRK_1984

TAGAAACTATGAATATGGCCACAAACAATTTCTCGTCTTCTAATAAGTTGGGACAGGGCGGTTATGGACCTGTTTATAAGGTAAACGTTTAGAAGTTTGTTGTCTTAAGCTGATTTTTAG

>M6-PGSC0003DMG400005689M6G-LecRK_2046

TATGGACCTGTTTATAAGGTAAACGTTTAGAAGTTTGTTGTCTTAAGCTGATTTTTAGATTTGCTAAGTAGCAAAAATTTCAGGGGATGTTACTCGACGGGCAAGAGATAGCCATCAAGA

>M6-PGSC0003DMG400005689M6G-LecRK_2108

GCTAAGTAGCAAAAATTTCAGGGGATGTTACTCGACGGGCAAGAGATAGCCATCAAGAGACTTTCAAGAAGTTCAGGGCAAGGATTGGTTGAGTTTCAAAACGAAATTGTGCTGATTGCT

>M6-PGSC0003DMG400005689M6G-LecRK_2170

TTCAAGAAGTTCAGGGCAAGGATTGGTTGAGTTTCAAAACGAAATTGTGCTGATTGCTAAACTCCAGCACACTAATCTTGTCAGACTTTTAGGATGTTGCATTGAAGGGGAAGAGAAGAT

>M6-PGSC0003DMG400005689M6G-LecRK_2232

TCCAGCACACTAATCTTGTCAGACTTTTAGGATGTTGCATTGAAGGGGAAGAGAAGATATTAGTTTATGAATACATGATGAACAAAAGCTTAGACTTCTTCCTATTTGGTATGGATTTCC

>M6-PGSC0003DMG400005689M6G-LecRK_2294

GTTTATGAATACATGATGAACAAAAGCTTAGACTTCTTCCTATTTGGTATGGATTTCCAATTTCATGCCTAAATTGCTTAATTAATCATATTTGATAACTAATACCCCTTCCCATTTTGA

>M6-PGSC0003DMG400005689M6G-LecRK_2356

TCATGCCTAAATTGCTTAATTAATCATATTTGATAACTAATACCCCTTCCCATTTTGATTGAACAGATCCTAGCAGAAAAGATTCACTAAAATGGAATACGCGTATGAACATTATTGAAG

>M6-PGSC0003DMG400005689M6G-LecRK_2418

ACAGATCCTAGCAGAAAAGATTCACTAAAATGGAATACGCGTATGAACATTATTGAAGGAGTTGCTCAGGGACTACTTTATCTCCATAAGTATTCAAGGCTGAGGGTGATTCACAGAGAT

>M6-PGSC0003DMG400005689M6G-LecRK_248

CTTCTTCAAAATGAAATTTTTCAGTCCAGGCAAGTCAAGGAACCGCTACTTGGGTATATTTTACACTCAGCCAAGTCTTGATATGTACTCTGATGTTAATGGAGGAGGAGATGAAAAAGC

>M6-PGSC0003DMG400005689M6G-LecRK_2480

TGCTCAGGGACTACTTTATCTCCATAAGTATTCAAGGCTGAGGGTGATTCACAGAGATCTAAAAGCCAGCAATGTCTTGCTTGATGACAATATGAACCCCAAAATCTCAGATTTTGGGTT

>M6-PGSC0003DMG400005689M6G-LecRK_2542

AAGCCAGCAATGTCTTGCTTGATGACAATATGAACCCCAAAATCTCAGATTTTGGGTTGGCCAGGATATTTGGAATGCAAGAATTTGAAGCAAACACGGAGCGAATTGTTGGTACCTAGT

>M6-PGSC0003DMG400005689M6G-LecRK_2604

AGGATATTTGGAATGCAAGAATTTGAAGCAAACACGGAGCGAATTGTTGGTACCTAGTAAGTCCTTAATAGATATAATTATCTAACAATGGTGATTCACTGTGTCCTTTTTATTATGCGC

>M6-PGSC0003DMG400005689M6G-LecRK_2666

CCTTAATAGATATAATTATCTAACAATGGTGATTCACTGTGTCCTTTTTATTATGCGCTAAATTGCTACCTTTTCAGTGGTTATATGTCTCCAGAATATGCCATGAATGGCATTGTGTCC

>M6-PGSC0003DMG400005689M6G-LecRK_2728

TTGCTACCTTTTCAGTGGTTATATGTCTCCAGAATATGCCATGAATGGCATTGTGTCCATGAAGACAGATGTATTCAGTTTTGGAGTTCTAGTACTCGAGATTTTGAGTGGCAAAAGAAA

>M6-PGSC0003DMG400005689M6G-LecRK_2790

AGACAGATGTATTCAGTTTTGGAGTTCTAGTACTCGAGATTTTGAGTGGCAAAAGAAACAACAGCTGCTATCATTTAGAACGCCCACTAAACATCATCGGATATGTAAGTTCCTCTCGTA

>M6-PGSC0003DMG400005689M6G-LecRK_2852

AGCTGCTATCATTTAGAACGCCCACTAAACATCATCGGATATGTAAGTTCCTCTCGTAAAATATCAATATAGCATGATCAAGAGGCGGATGGAGTGTTTATTCAATGAGTTCAACTAAAC

>M6-PGSC0003DMG400005689M6G-LecRK_2914

ATCAATATAGCATGATCAAGAGGCGGATGGAGTGTTTATTCAATGAGTTCAACTAAACCCAATACTTTAGAAACGAAACATAAATATACAACATCAACATACCCATTGGAATCCCACAGG

>M6-PGSC0003DMG400005689M6G-LecRK_310

ACACTCAGCCAAGTCTTGATATGTACTCTGATGTTAATGGAGGAGGAGATGAAAAAGCTTTATGGATAGCTAACAGAGACGATCCGATAACGGATACATCTGGAAGTTTTATGATTGCTC

>M6-PGSC0003DMG400005689M6G-LecRK_3100

GTTTTCAATAGACCCTCGACTCAAGACACGAAACATGGATATACATGTGAAAATCACTTGAATTATAAAATTCTGAAGTATAATGAGTCTAGTTCTAAAAACTTTAAAGGTTCAACCTGT

>M6-PGSC0003DMG400005689M6G-LecRK_3162

TTATAAAATTCTGAAGTATAATGAGTCTAGTTCTAAAAACTTTAAAGGTTCAACCTGTCAACTTTACATTGTTTTGGATCCGCCCTGGCATGATCACTTAGTTGAACAAGGAAATTTCAA

>M6-PGSC0003DMG400005689M6G-LecRK_3224

TTTACATTGTTTTGGATCCGCCCTGGCATGATCACTTAGTTGAACAAGGAAATTTCAAACTTGATATGCTCAATTTGTCTCAGGCATGGGAATTGTGGAAAGCAGGCAGCGTCCTTGAAG

>M6-PGSC0003DMG400005689M6G-LecRK_3286

GATATGCTCAATTTGTCTCAGGCATGGGAATTGTGGAAAGCAGGCAGCGTCCTTGAAGAACTAACAGATCCTGTATTGACAAATGAATCAACACCAAAGAATGAAGTGATGAGATGCATT

>M6-PGSC0003DMG400005689M6G-LecRK_3348

AACAGATCCTGTATTGACAAATGAATCAACACCAAAGAATGAAGTGATGAGATGCATTCATGTAGGATTACTCTGTGTTCAAGTCAATCCTATGGACAGGCCATCCATGTCAAATGTGGT

>M6-PGSC0003DMG400005689M6G-LecRK_3410

TAGGATTACTCTGTGTTCAAGTCAATCCTATGGACAGGCCATCCATGTCAAATGTGGTTATGATGCTCACAAATGATAGCCTACAACTACCCGTCCCCAAGCAACCAGCATTTTTCATTG

>M6-PGSC0003DMG400005689M6G-LecRK_3472

ATGCTCACAAATGATAGCCTACAACTACCCGTCCCCAAGCAACCAGCATTTTTCATTGAGACAGCTATGACTGAGACTGAAACTCGCGAAGAAGTTGGACATTGTTCCACAAATGGGCTA

>M6-PGSC0003DMG400005689M6G-LecRK_3534

AGCTATGACTGAGACTGAAACTCGCGAAGAAGTTGGACATTGTTCCACAAATGGGCTATCAATTTCAGATATGGTAGCGAGATGACTTTCATCATTGCACCCCTCAAAATTGTCAAAAAT

>M6-PGSC0003DMG400005689M6G-LecRK_3596

TTTCAGATATGGTAGCGAGATGACTTTCATCATTGCACCCCTCAAAATTGTCAAAAATGGTAATATGCAATCTCCCTAATGATACAAAATAAAAACCACTGCTTGTCTCCTTCATCCAGA

>M6-PGSC0003DMG400005689M6G-LecRK_3658

ATATGCAATCTCCCTAATGATACAAAATAAAAACCACTGCTTGTCTCCTTCATCCAGATTTTACTTTATCTAGCATCGAAATTAAGTATGAGTTGTTGACAAAATAAAGTAAGCTTGTTA

>M6-PGSC0003DMG400005689M6G-LecRK_372

TGGATAGCTAACAGAGACGATCCGATAACGGATACATCTGGAAGTTTTATGATTGCTCCTGATGGCAGATTAATAATTTCTCATAAGGAAGGTAATATTACTTTGTTCAGTGCTACACCA

>M6-PGSC0003DMG400005689M6G-LecRK_3720

ACTTTATCTAGCATCGAAATTAAGTATGAGTTGTTGACAAAATAAAGTAAGCTTGTTAGAGCCTTTTGAAGATTATACTGTGTCTAAACAATTTTTACACATTTGTTTTGAATGCAAATG

>M6-PGSC0003DMG400005689M6G-LecRK_3782

CTTTTGAAGATTATACTGTGTCTAAACAATTTTTACACATTTGTTTTGAATGCAAATGAGTTTTCAGAGTATCAATTGTTACTGAAATCAGAACTCATAAATGTAGTAATTGTCACTAGT

>M6-PGSC0003DMG400005689M6G-LecRK_3844

TTCAGAGTATCAATTGTTACTGAAATCAGAACTCATAAATGTAGTAATTGTCACTAGTTTTGCAAAGATGATTAGTATCTGTACACTAGAAAATCAAATGGATGTTTCCCACGTTCCTTT

>M6-PGSC0003DMG400005689M6G-LecRK_434

TGGCAGATTAATAATTTCTCATAAGGAAGGTAATATTACTTTGTTCAGTGCTACACCAACAACAGCCACTAACTTAACTGCTATACTACTCGATAATGGGAATTTTGTGCTGAGGGAACT

>M6-PGSC0003DMG400005689M6G-LecRK_496

CAGCCACTAACTTAACTGCTATACTACTCGATAATGGGAATTTTGTGCTGAGGGAACTGAATACCAATAGCTTTGTAAATCGAACGCTATGGCAAAGCTTTGATTATCCAACTGATACAC

>M6-PGSC0003DMG400005689M6G-LecRK_558

ACCAATAGCTTTGTAAATCGAACGCTATGGCAAAGCTTTGATTATCCAACTGATACACTTCTGCCTGGAATGAAGCTTGGGATTAACTTGCGAACGGGGCATAAATGGTCATTGACTTCA

>M6-PGSC0003DMG400005689M6G-LecRK_62

ACTATTTTGACTGTAAACATTCATGTGTGGATTAAGGAGCTTCTCTCATATGGCTACCAACAGAATTCTGATTGCTTTCATCCTCGTGTTATCCTGTCACTTTTTGCGTGCAGATTTTGC

>M6-PGSC0003DMG400005689M6G-LecRK_620

GCCTGGAATGAAGCTTGGGATTAACTTGCGAACGGGGCATAAATGGTCATTGACTTCATGGGTAAATGATCAAGCACCTGGTTCAGGGTCTTTTACCTTTGGTCTTGACCCTAATGGTAC

>M6-PGSC0003DMG400005689M6G-LecRK_682

TAAATGATCAAGCACCTGGTTCAGGGTCTTTTACCTTTGGTCTTGACCCTAATGGTACAAATCAGCTGATGATCTTGTGGATGGGCAAGGTCTATTGGAAGAGTGGCCCGTGGCCTACTG

>M6-PGSC0003DMG400005689M6G-LecRK_744

CAGCTGATGATCTTGTGGATGGGCAAGGTCTATTGGAAGAGTGGCCCGTGGCCTACTGGCCATCTTGCTCTTAAATATGTCTCAAATGAGGATGAGAAATACTTTATGTATACTGTTGAA

>M6-PGSC0003DMG400005689M6G-LecRK_806

TCTTGCTCTTAAATATGTCTCAAATGAGGATGAGAAATACTTTATGTATACTGTTGAATTGGAATATCAAAGATACTTTGTGAGTCCATTTGGAATAATTCAGGATACTTTTAAAGGAAA

>M6-PGSC0003DMG400005689M6G-LecRK_868

AATATCAAAGATACTTTGTGAGTCCATTTGGAATAATTCAGGATACTTTTAAAGGAAATGCAGTGTTTGGTAATTGTAGTAATGAGACTCCCCATGCTGGCTGCGTAAAGCAGGTATTGC

>M6-PGSC0003DMG400005689M6G-LecRK_930

GTGTTTGGTAATTGTAGTAATGAGACTCCCCATGCTGGCTGCGTAAAGCAGGTATTGCCCCAGTGCAGGGCAGCTAAAAAGTACTGGTTTGAGCTTAGACAAGTTTACATGTTTGGAGAC

>M6-PGSC0003DMG400005689M6G-LecRK_992

GTGCAGGGCAGCTAAAAAGTACTGGTTTGAGCTTAGACAAGTTTACATGTTTGGAGACAGCGTTAAATTTGATGAGAACTATACTTTGAGCCTCTCTGATTGCAAGGCCAAGTGTGAGAA

>M6-PGSC0003DMG400005730M6RLKWAX-EGF_0

CCACCTAGTCTCTCTTTGATGAGTTCTCACAAATGCATGATCAAAGCTTACTTCTTTCACTCCATAATCACCATCCTATGCTTCGTGATCACGTCCTCGCCAGCCTATGGCCAAGAGGAT

>M6-PGSC0003DMG400005730M6RLKWAX-EGF_1037

AGATATCTATATTGTTAAAAAAATTTCATGTATTGATAAATTTGAAAATAATATTTTGTATAGAAAATATAATTGAGAAATTGAATATTATTTAGTTGATTGTAGTGTCTTTTCTATATT

>M6-PGSC0003DMG400005730M6RLKWAX-EGF_1098

AGAAAATATAATTGAGAAATTGAATATTATTTAGTTGATTGTAGTGTCTTTTCTATATTCTTATGTGGCTAGTTAATTGTAGTATCCTTTCTAATTTCGCATGTGGCTGTCTATTTTGTT

>M6-PGSC0003DMG400005730M6RLKWAX-EGF_1159

TATGTGGCTAGTTAATTGTAGTATCCTTTCTAATTTCGCATGTGGCTGTCTATTTTGTTGTCTAGGTCATTTATATTTTGACGATTAACATAAACCAACAATATTGTGTTCGTATACTTA

>M6-PGSC0003DMG400005730M6RLKWAX-EGF_122

ACAGTACTCCACTTGTAATAGCTCCTACACCTGTGGGAATATTCAAAATATCAGCTTCCCTTTCTGGGGTGGTGATCGACCTCAAGAATGTGGTCTTCCACAATTCGAGCTTTCATGTGA

>M6-PGSC0003DMG400005730M6RLKWAX-EGF_1220

CTAGGTCATTTATATTTTGACGATTAACATAAACCAACAATATTGTGTTCGTATACTTACTTTTCTTTATTCTCCTTTCACTATTTTTTATTCTTTCATTATTGAAAATTCTTAAGTCTC

>M6-PGSC0003DMG400005730M6RLKWAX-EGF_2257

AGCCCATATTGATAGCTCTAATCATATTTTTAAAATTTTGAGTCATGTATTATTATTCCGACTTTTTTTTTTGGACTTCAGGATAACCCGCAATCACGACAACCTCTGCAATTTGGGTGA

>M6-PGSC0003DMG400005730M6RLKWAX-EGF_244

CCAATCAGGATCCCCTTATACATATCAATGGTCATAATTTCCGCGTACTTGACATAAATGGAGACTATCAAACCATGAGAATTGCACGAAATGATCTTGAGGATGACATTTGTCCTGATA

>M6-PGSC0003DMG400005730M6RLKWAX-EGF_2440

CGCACTAGGCAAGCCATATGCAACAGACTCGACTCAAAAGGCGTAGGATTACATTATAATTATTCAATTTATATTTATCAACTAAAAAAAATAGGGTCTCTTTATAATGTTTGGCTTAAG

>M6-PGSC0003DMG400005730M6RLKWAX-EGF_2562

CTCCAAACTTGTTGAGACGACCCTGATGAAATGTGTGATCATCTCAACTTAAAACTTAATTATTAGAGAAAACACATTTTTACTAGTTAATTATATTCTCAACATTTTCTGCACTAGGGT

>M6-PGSC0003DMG400005730M6RLKWAX-EGF_2623

ATTAGAGAAAACACATTTTTACTAGTTAATTATATTCTCAACATTTTCTGCACTAGGGTCGAGATTATGCCTGAATAACAGAGGGCACTGTGGTGCGCGGTTAATGAATCCAACTCCAAA

>M6-PGSC0003DMG400005730M6RLKWAX-EGF_2684

AGATTATGCCTGAATAACAGAGGGCACTGTGGTGCGCGGTTAATGAATCCAACTCCAAAGCAGAACATGTCGTCATGGTAAATCTAGAATCTAAAAGTCTTACACGTTTGACTATAATTC

>M6-PGSC0003DMG400005730M6RLKWAX-EGF_2745

AGAACATGTCGTCATGGTAAATCTAGAATCTAAAAGTCTTACACGTTTGACTATAATTCCATTGGACTAAAACAAAATCTTCTTCCTTCGTTATAGATGTTGATGATTTGTTATACTACC

>M6-PGSC0003DMG400005730M6RLKWAX-EGF_2806

TTGGACTAAAACAAAATCTTCTTCCTTCGTTATAGATGTTGATGATTTGTTATACTACCAGAAGATCCTTTATTATGGTTTTATATCAATTCATCACTTGTTTATCTCTGCATTACAGGC

>M6-PGSC0003DMG400005730M6RLKWAX-EGF_2867

AAGATCCTTTATTATGGTTTTATATCAATTCATCACTTGTTTATCTCTGCATTACAGGCAATAATAGACATATCTCGGTGAAGATAGCCCTAGGTAATTGTCTTTTTGACCTTGTTCATG

>M6-PGSC0003DMG400005730M6RLKWAX-EGF_2928

TAATAGACATATCTCGGTGAAGATAGCCCTAGGTAATTGTCTTTTTGACCTTGTTCATGTTATATTGTCCCCACAACATAGAGATGAGACTCTTTTTACATCTGGTTAACAGTTATCTCT

>M6-PGSC0003DMG400005730M6RLKWAX-EGF_2989

ATATTGTCCCCACAACATAGAGATGAGACTCTTTTTACATCTGGTTAACAGTTATCTCTGCTGCTGGGATTGGAATAGTATTAATCTGCTACATGATCTACTGGTTTGTCTTTAAGAAAT

>M6-PGSC0003DMG400005730M6RLKWAX-EGF_305

AGACTATCAAACCATGAGAATTGCACGAAATGATCTTGAGGATGACATTTGTCCTGATAGATTCGGTAACACTAGCTTAAATGATGCCCCTTTCCGCTATGGTCCTAACTATGTGGTGCT

>M6-PGSC0003DMG400005730M6RLKWAX-EGF_3050

TGCTGGGATTGGAATAGTATTAATCTGCTACATGATCTACTGGTTTGTCTTTAAGAAATTATCTGCTACTTGCCTTTTGTTACCAAAGCAGAGGAGGGAACATCAGAATGTTGAGGCCTT

>M6-PGSC0003DMG400005730M6RLKWAX-EGF_3111

TCTGCTACTTGCCTTTTGTTACCAAAGCAGAGGAGGGAACATCAGAATGTTGAGGCCTTCCTCAAGCAATATGGATCCCTTGCCCCTAAGGTATACAGCTATTCAGACATTAGGAAAATG

>M6-PGSC0003DMG400005730M6RLKWAX-EGF_3172

TCAAGCAATATGGATCCCTTGCCCCTAAGGTATACAGCTATTCAGACATTAGGAAAATGACAAATTCACTTCAACAGAAACTTGGACAAGGGGGTTATGGAGGGGTCTATAAAGGTTACC

>M6-PGSC0003DMG400005730M6RLKWAX-EGF_3233

AAATTCACTTCAACAGAAACTTGGACAAGGGGGTTATGGAGGGGTCTATAAAGGTTACCTTAACGGCCGTCCTGTTGCTGTAAAGATTTTAAATTCTACCAAAGGGAATGGAGAAGACTT

>M6-PGSC0003DMG400005730M6RLKWAX-EGF_3294

AACGGCCGTCCTGTTGCTGTAAAGATTTTAAATTCTACCAAAGGGAATGGAGAAGACTTTATCAACGAAGTTGCAAGCATCAGTAGAACTTCACATGTGAACATAGTTACTCTGTTGGGA

>M6-PGSC0003DMG400005730M6RLKWAX-EGF_3355

TCAACGAAGTTGCAAGCATCAGTAGAACTTCACATGTGAACATAGTTACTCTGTTGGGATTCTGTTTCAATGGTCGAGAAAGAGCTCTTATTTATGAATTCATGTCGAATGGATCACTCG

>M6-PGSC0003DMG400005730M6RLKWAX-EGF_3416

CTGTTTCAATGGTCGAGAAAGAGCTCTTATTTATGAATTCATGTCGAATGGATCACTCGATCAACACATCTATGGTGAAGATTCTAAGTTGGGATGGGAAAAGTTGTACAAAATTGCACT

>M6-PGSC0003DMG400005730M6RLKWAX-EGF_3477

CAACACATCTATGGTGAAGATTCTAAGTTGGGATGGGAAAAGTTGTACAAAATTGCACTTGAGATAGGTCGAGGACTGGAGTATTTGCACCGAGGGTGCAATGCACGAATACTGCATTTT

>M6-PGSC0003DMG400005730M6RLKWAX-EGF_3538

AGATAGGTCGAGGACTGGAGTATTTGCACCGAGGGTGCAATGCACGAATACTGCATTTTGATATAAAACCTCACAACATCCTTCTTGATGAAGACTTTTGCCCCAAGATATCTGATTTTG

>M6-PGSC0003DMG400005730M6RLKWAX-EGF_3599

TATAAAACCTCACAACATCCTTCTTGATGAAGACTTTTGCCCCAAGATATCTGATTTTGGCTTAGCTAAACTATGCCTTCAGAAAGAAAGCATGATATCAATGTTGGAAGCAAGAGGAAC

>M6-PGSC0003DMG400005730M6RLKWAX-EGF_366

TTCGGTAACACTAGCTTAAATGATGCCCCTTTCCGCTATGGTCCTAACTATGTGGTGCTCATCCTGTTCTATGCTTGTCCTTTCGATATACCTTCTGAATGGAAAAATTTCACCTTTAGC

>M6-PGSC0003DMG400005730M6RLKWAX-EGF_3660

TTAGCTAAACTATGCCTTCAGAAAGAAAGCATGATATCAATGTTGGAAGCAAGAGGAACCATTGGTTACATTGCTCCGGAAGTATTCAGTAGAAATTTTGGAGGCGTCTCACACAAATCC

>M6-PGSC0003DMG400005730M6RLKWAX-EGF_3721

TTGGTTACATTGCTCCGGAAGTATTCAGTAGAAATTTTGGAGGCGTCTCACACAAATCCGATGTTTACAGCTATGGAATGATGATTCTTGAAATGGTTGGAGGAAGGAGGAACTACAGTG

>M6-PGSC0003DMG400005730M6RLKWAX-EGF_3782

TGTTTACAGCTATGGAATGATGATTCTTGAAATGGTTGGAGGAAGGAGGAACTACAGTGCTGAAAGAAGTCATCATAGTGAAATATACTTCCCGCGTTGGGCTTATCAACGCCTTGTGAC

>M6-PGSC0003DMG400005730M6RLKWAX-EGF_3843

GAAAGAAGTCATCATAGTGAAATATACTTCCCGCGTTGGGCTTATCAACGCCTTGTGACAGATGAAGACTTGAATATACAATGCATCACGACCAAAGAAGACGAAGAAATTGCTAAGAAG

>M6-PGSC0003DMG400005730M6RLKWAX-EGF_3904

ATGAAGACTTGAATATACAATGCATCACGACCAAAGAAGACGAAGAAATTGCTAAGAAGATGATACTAGTAGGCTTGTGGTGCATTCAAACTGATCCCTCACAAAGGCCACCAATGAGAA

>M6-PGSC0003DMG400005730M6RLKWAX-EGF_3965

GATACTAGTAGGCTTGTGGTGCATTCAAACTGATCCCTCACAAAGGCCACCAATGAGAAAGGCAATTGAAATGCTTGAAGGTAACTTAGAGGCCATCCAATTTCCACCAAAGCCTTTTAT

>M6-PGSC0003DMG400005730M6RLKWAX-EGF_4026

GCAATTGAAATGCTTGAAGGTAACTTAGAGGCCATCCAATTTCCACCAAAGCCTTTTATATGTTCTCCCTCGCAATCAGAAGGATCATCGATGATCCCTTTGGGGAAGCCAATAATGCCC

>M6-PGSC0003DMG400005730M6RLKWAX-EGF_4083

ATATGTTCTCCCTCGCAATCAGAAGGATCATCGATGATCCCTTTGGGGAAGCCAATAATGCCCTTTATATATTCTTCTTCGAGTTCAACACTTGATTCTACAGAACATTTGGTTATATAA

>M6-PGSC0003DMG400005730M6RLKWAX-EGF_427

TCCTGTTCTATGCTTGTCCTTTCGATATACCTTCTGAATGGAAAAATTTCACCTTTAGCTGCAATAAAACTGGAGATTCCAGTCATGGTTTCTATCCAGATCAGTCATTTATATCATTCT

>M6-PGSC0003DMG400005730M6RLKWAX-EGF_488

CAATAAAACTGGAGATTCCAGTCATGGTTTCTATCCAGATCAGTCATTTATATCATTCTGGGGACCAAAGTATGGGAATTGTGAGCGTATTGTCATAGTTCCTGTGTTGTTAACGGCATT

>M6-PGSC0003DMG400005730M6RLKWAX-EGF_61

CCATAATCACCATCCTATGCTTCGTGATCACGTCCTCGCCAGCCTATGGCCAAGAGGATAAACAGTACTCCACTTGTAATAGCTCCTACACCTGTGGGAATATTCAAAATATCAGCTTCC

>M6-PGSC0003DMG400005730M6RLKWAX-EGF_610

AGCGGTTCCAAGACCAAGGAAGTACGAAAATATTGGAGCTATTGAAGCAAGGTTTTGATGTGGTTTACAACAAAAGTGTATTATGTATGGCTTGTGAGAGGTCAGGGGGATTATGTTGGT

>M6-PGSC0003DMG400005730M6RLKWAX-EGF_671

GGTTTACAACAAAAGTGTATTATGTATGGCTTGTGAGAGGTCAGGGGGATTATGTTGGTCAGAAACAGATTTTGCGGAGTCATCTTGCCTTTGTCAGGATCGAACTTATTCTTACAACTG

>M6-PGSC0003DMG400005730M6RLKWAX-EGF_732

GAAACAGATTTTGCGGAGTCATCTTGCCTTTGTCAGGATCGAACTTATTCTTACAACTGCGGTTATACATTACTGGCTCAAGGTGTGTTTAATTAATCTCCCTATTACCTAATCTAACTT

>M6-PGSC0003DMG400005730M6RLKWAX-EGF_793

GTTATACATTACTGGCTCAAGGTGTGTTTAATTAATCTCCCTATTACCTAATCTAACTTAGTTCTGTTTACCAAATATTACTTGAATTCAATTGGCAGCTAAGAGTTTTGGTACTGAACT

>M6-PGSC0003DMG400005730M6RLKWAX-EGF_854

TTCTGTTTACCAAATATTACTTGAATTCAATTGGCAGCTAAGAGTTTTGGTACTGAACTTCTGAAACCACTATTTTTTTTCCTGTTCAAATATATACCATGATGAAATACAGGGTCGACC

>M6-PGSC0003DMG400005730M6RLKWAX-EGF_915

TGAAACCACTATTTTTTTTCCTGTTCAAATATATACCATGATGAAATACAGGGTCGACCATAAAAGCCAATAAAGCAATGTCTTGGGGCGCGAATTTTTTTGGGGCCTCTATTTTCTCTC

>M6-PGSC0003DMG400005730M6RLKWAX-EGF_976

AAAAGCCAATAAAGCAATGTCTTGGGGCGCGAATTTTTTTGGGGCCTCTATTTTCTCTCAAAGATATCTATATTGTTAAAAAAATTTCATGTATTGATAAATTTGAAAATAATATTTTGT

>M6-PGSC0003DMG400005896M6LRR-RLK_0

GAGAAATCAAAGGACAGAAGGAGACAGAAAAAAAGGTGTCATGAATTTGTACTTGTCTTTAGGGTTTGATCCACACTGCACCACAACTCTTTGCTAAACTCATCTCTCTTTTTTACAAAA

>M6-PGSC0003DMG400005896M6LRR-RLK_1037

CAAGGAAACAGTTTTAGTGGACCAATTCCTGATACAATTGGTGATTTAAATGACTTGAAGGAGCTGAATTTTAACGGAAATCAACTAGTTGGTCTGATTCCGCAAGGTTTGGCAAATCTA

>M6-PGSC0003DMG400005896M6LRR-RLK_1098

AGCTGAATTTTAACGGAAATCAACTAGTTGGTCTGATTCCGCAAGGTTTGGCAAATCTAGACTTGAGTGTGTTAGACTTGAACAATAATAAGCTCATGGGTCCTATCCCAAAGTTTAGAG

>M6-PGSC0003DMG400005896M6LRR-RLK_1159

CTTGAGTGTGTTAGACTTGAACAATAATAAGCTCATGGGTCCTATCCCAAAGTTTAGAGCTGCGAATGCTACATATTCCTCAAATTCTTTCTGTCAGCCAACTCCTGGTGTTCCTTGTAC

>M6-PGSC0003DMG400005896M6LRR-RLK_122

CACTCACATTATTAAAAAAAGGTACCATTTTTACAACAATTTATAGTTATTATGTACTTGCATCCTTATATGTTACTGTTATGTACTTTGTTCATTTGGGGTTCACAATTCTTGAGAAAA

>M6-PGSC0003DMG400005896M6LRR-RLK_1220

GCGAATGCTACATATTCCTCAAATTCTTTCTGTCAGCCAACTCCTGGTGTTCCTTGTACTCCTCAAGTTAATGCACTTTTAGATCTTCTTTGTGGTTGGAATTATCCAGCCAACCTTGCT

>M6-PGSC0003DMG400005896M6LRR-RLK_1281

CTCAAGTTAATGCACTTTTAGATCTTCTTTGTGGTTGGAATTATCCAGCCAACCTTGCTCCTGAATGGAGTGGCAATGATCCATGTACAGGTCCTTGGTTGGGAATCAGTTGCAATCCTA

>M6-PGSC0003DMG400005896M6LRR-RLK_1342

TGAATGGAGTGGCAATGATCCATGTACAGGTCCTTGGTTGGGAATCAGTTGCAATCCTAAAGGCCAGATTACAATTGTTAATCTACAGAACAAGAACCTTACGGGTACGCTTAGCCCTTC

>M6-PGSC0003DMG400005896M6LRR-RLK_1403

GGCCAGATTACAATTGTTAATCTACAGAACAAGAACCTTACGGGTACGCTTAGCCCTTCACTTGCAAACTTGGATTCACTGCTTGAAGTTCACTTAAAAAGAAATAGTTTGCACGGTCGA

>M6-PGSC0003DMG400005896M6LRR-RLK_1464

TTGCAAACTTGGATTCACTGCTTGAAGTTCACTTAAAAAGAAATAGTTTGCACGGTCGAGTTCCTGCAAACTTAACAGAGCTCAGATCCTTGAGATTGTTGGACTTAAGTGGGAACAATT

>M6-PGSC0003DMG400005896M6LRR-RLK_1525

TCCTGCAAACTTAACAGAGCTCAGATCCTTGAGATTGTTGGACTTAAGTGGGAACAATTTTGATCCACCATTACCGAAATTCCGGGATCGTGTGAAAGTCATTACAGACGGAAATGCTCA

>M6-PGSC0003DMG400005896M6LRR-RLK_1586

GATCCACCATTACCGAAATTCCGGGATCGTGTGAAAGTCATTACAGACGGAAATGCTCATCTTGTTGCTAATGTAACTGCGGCTCCTCCTCCTCTGAGTATTAGCCCATTTCCTCCATTG

>M6-PGSC0003DMG400005896M6LRR-RLK_1647

TTGTTGCTAATGTAACTGCGGCTCCTCCTCCTCTGAGTATTAGCCCATTTCCTCCATTGAGCCACAGTCCTAAATCAAGCAAGGAGTTACCATCGAAAAGCCCGTTTCCCGGTGATAATC

>M6-PGSC0003DMG400005896M6LRR-RLK_1708

CCACAGTCCTAAATCAAGCAAGGAGTTACCATCGAAAAGCCCGTTTCCCGGTGATAATCAGCCAACATTATCAGATACACCGCCCTCCCCAGAGAAAAGTTCAGGTTCAGACTCAAAGTC

>M6-PGSC0003DMG400005896M6LRR-RLK_1769

CCAACATTATCAGATACACCGCCCTCCCCAGAGAAAAGTTCAGGTTCAGACTCAAAGTCTCCCAGGGGAACCAAAGGGCAGACCACATCAGAGAACCATGATAAGACCATGATCATTGTA

>M6-PGSC0003DMG400005896M6LRR-RLK_183

ATCCTTATATGTTACTGTTATGTACTTTGTTCATTTGGGGTTCACAATTCTTGAGAAAAAAGTATGTAATTTTATGATAATCTTGAATTTTCATCATGTGTGTATATAAGTATTGTGATT

>M6-PGSC0003DMG400005896M6LRR-RLK_1830

CCAGGGGAACCAAAGGGCAGACCACATCAGAGAACCATGATAAGACCATGATCATTGTAGTTGTAAGTGCAGCTTCTTTCGTCTTCACTCTTCTTGCCGTTGTTTTGTATTTCAAGAGCA

>M6-PGSC0003DMG400005896M6LRR-RLK_1891

TGTAAGTGCAGCTTCTTTCGTCTTCACTCTTCTTGCCGTTGTTTTGTATTTCAAGAGCAAGAGAAAACGAGAAAAGGATTCAGGTACTATTGTTATCCACCCCAAAGAGCCATTCGATCA

>M6-PGSC0003DMG400005896M6LRR-RLK_1952

AGAAAACGAGAAAAGGATTCAGGTACTATTGTTATCCACCCCAAAGAGCCATTCGATCAAGATAACATTGTTAAGATCACTATCTTAGAAGATCCCATGATGTACAGCTTACAGAGTGGT

>M6-PGSC0003DMG400005896M6LRR-RLK_2013

ATAACATTGTTAAGATCACTATCTTAGAAGATCCCATGATGTACAGCTTACAGAGTGGTACTACAACAACTACTACAACTAGTGGTGGAACAAAAGGAAGTCGAGTGATTGAGATTGGAA

>M6-PGSC0003DMG400005896M6LRR-RLK_2074

TACAACAACTACTACAACTAGTGGTGGAACAAAAGGAAGTCGAGTGATTGAGATTGGAAACCTTGTCATTTCAACCCAGGACCTTCGGAGGGTAACCGATAATTTTGCACCAGAGAATGA

>M6-PGSC0003DMG400005896M6LRR-RLK_2135

CTTGTCATTTCAACCCAGGACCTTCGGAGGGTAACCGATAATTTTGCACCAGAGAATGAGCTTGGTCGTGGTGGCTTTGGAGTTGTTTACAAAGGCGTAATTGAAGATGGAATTCAAATT

>M6-PGSC0003DMG400005896M6LRR-RLK_2196

TTGGTCGTGGTGGCTTTGGAGTTGTTTACAAAGGCGTAATTGAAGATGGAATTCAAATTGCAGTGAAGAGAATGGAATCTGCCATTATCAACAGCAAGGCATTGGATGAATTTCAAGCAG

>M6-PGSC0003DMG400005896M6LRR-RLK_2257

AGTGAAGAGAATGGAATCTGCCATTATCAACAGCAAGGCATTGGATGAATTTCAAGCAGAAATAGCTGTGCTTTCTAAGGTTCGACATCGCCATTTAGTTTCACTTTTGGGTTATTCCAT

>M6-PGSC0003DMG400005896M6LRR-RLK_2318

ATAGCTGTGCTTTCTAAGGTTCGACATCGCCATTTAGTTTCACTTTTGGGTTATTCCATTGAAGGAAATGAGAGGCTTTTGGTATATGAATATATGTCAAAGGGCGCTTTGAGTAGACAT

>M6-PGSC0003DMG400005896M6LRR-RLK_2379

AAGGAAATGAGAGGCTTTTGGTATATGAATATATGTCAAAGGGCGCTTTGAGTAGACATCTTTTCCGCTGGAAGATCCTGAACTTAGAGCCTTTATCATGGACAAAGAGACTCAATATTG

>M6-PGSC0003DMG400005896M6LRR-RLK_244

GTATGTAATTTTATGATAATCTTGAATTTTCATCATGTGTGTATATAAGTATTGTGATTTCAGAGGTGTTGTTATATACTTATGTACATTAGGTGTATTTCTTGTAAATGTTGTTTCTGG

>M6-PGSC0003DMG400005896M6LRR-RLK_2440

TTTCCGCTGGAAGATCCTGAACTTAGAGCCTTTATCATGGACAAAGAGACTCAATATTGCTCTAGATGTCGCTCGAGGGATGGAGTACCTTCATAATCTGGCACACCAGAGCTTCATACA

>M6-PGSC0003DMG400005896M6LRR-RLK_2501

CTAGATGTCGCTCGAGGGATGGAGTACCTTCATAATCTGGCACACCAGAGCTTCATACATCGCGATCTCAAATCATCAAACATTCTTCTAGATGATGATTTTAGAGCAAAAGTTTCTGAT

>M6-PGSC0003DMG400005896M6LRR-RLK_2562

GCGATCTCAAATCATCAAACATTCTTCTAGATGATGATTTTAGAGCAAAAGTTTCTGATTTTGGATTAGTGAAACTCGCTCCTGACAAAGAAAGGTCCGTTGCAACAAGGCTTGCAGGAA

>M6-PGSC0003DMG400005896M6LRR-RLK_2623

TGGATTAGTGAAACTCGCTCCTGACAAAGAAAGGTCCGTTGCAACAAGGCTTGCAGGAACATTTGGATACCTTGCTCCCGAGTATGCTGGTATGTGTGCATTTTTTTCCATTGGTATCTC

>M6-PGSC0003DMG400005896M6LRR-RLK_2684

TTTGGATACCTTGCTCCCGAGTATGCTGGTATGTGTGCATTTTTTTCCATTGGTATCTCTTTCTTAGACACTCGTTTTGACTGTTCTGGTAAATATTATCATTTAAGCATCAATTGTATT

>M6-PGSC0003DMG400005896M6LRR-RLK_2745

TCTTAGACACTCGTTTTGACTGTTCTGGTAAATATTATCATTTAAGCATCAATTGTATTTTTTACGCTCAGGTAAGCTGTTACATGAACACATCTCTACCTGAACGAAACCAATACTCTT

>M6-PGSC0003DMG400005896M6LRR-RLK_2806

TTACGCTCAGGTAAGCTGTTACATGAACACATCTCTACCTGAACGAAACCAATACTCTTTTAGTCCCTCTATTTTAAGAAATGCATCAGTTCTATAATGAGAATCCAGACTGCTGAATTG

>M6-PGSC0003DMG400005896M6LRR-RLK_2867

AGTCCCTCTATTTTAAGAAATGCATCAGTTCTATAATGAGAATCCAGACTGCTGAATTGCGCTTCTGACCTTCCAAGCCCTTCCTTCTCCAGCCAAAATAGAACAGAGTAACTAATGGAG

>M6-PGSC0003DMG400005896M6LRR-RLK_2928

CTTCTGACCTTCCAAGCCCTTCCTTCTCCAGCCAAAATAGAACAGAGTAACTAATGGAGCATTTACGTTTTTCCTTTAAGTTGTCGGTGTGCCTACTAAATTGATATACCTCAGACATTG

>M6-PGSC0003DMG400005896M6LRR-RLK_2989

TTTACGTTTTTCCTTTAAGTTGTCGGTGTGCCTACTAAATTGATATACCTCAGACATTGATGTATCAATCTCGCATCATTCAGTGATATTCTCATTCTGTTCACTATCTTTGTTAAACAA

>M6-PGSC0003DMG400005896M6LRR-RLK_305

AGAGGTGTTGTTATATACTTATGTACATTAGGTGTATTTCTTGTAAATGTTGTTTCTGGTGTAACAAATTTGAATGATTTCAAGATTTTGAGTGATTTTAGAAATGGATTAGAGAATCCT

>M6-PGSC0003DMG400005896M6LRR-RLK_3050

GTATCAATCTCGCATCATTCAGTGATATTCTCATTCTGTTCACTATCTTTGTTAAACAATTCAAGTAGAAGGTTTCACATCCTCTCTCCATATGATCATTCAGAGCATAGTTATCATCTT

>M6-PGSC0003DMG400005896M6LRR-RLK_3111

CAAGTAGAAGGTTTCACATCCTCTCTCCATATGATCATTCAGAGCATAGTTATCATCTTGATTCCATAAAAAGAAACAGAAACTCTATCCCTTTAAAGATACTTTTCCCTTGAGGCATGG

>M6-PGSC0003DMG400005896M6LRR-RLK_3172

TTCCATAAAAAGAAACAGAAACTCTATCCCTTTAAAGATACTTTTCCCTTGAGGCATGGTACTGAAATATGTTTGTGTAAAGCTGTTCAACAAGCATGTCATACCTGTTAATTTTGGTGT

>M6-PGSC0003DMG400005896M6LRR-RLK_3233

CTGAAATATGTTTGTGTAAAGCTGTTCAACAAGCATGTCATACCTGTTAATTTTGGTGTAGAAACTATTGCTTACTTTATCTTTATGTTACCCGTATATTTTAGTTCTCAGTCTTGTTGA

>M6-PGSC0003DMG400005896M6LRR-RLK_3294

AAACTATTGCTTACTTTATCTTTATGTTACCCGTATATTTTAGTTCTCAGTCTTGTTGAACGAATTACCTAGTCACCTTTCATATATCCTAACTCACATTTTACCTCTCCAGTGACCGGA

>M6-PGSC0003DMG400005896M6LRR-RLK_3355

GAATTACCTAGTCACCTTTCATATATCCTAACTCACATTTTACCTCTCCAGTGACCGGAAAAGTCACAACAAAAATAGATGTCTTCAGCTTTGGGGTGGTGTTGATGGAACTTGTGACTG

>M6-PGSC0003DMG400005896M6LRR-RLK_3416

AGTCACAACAAAAATAGATGTCTTCAGCTTTGGGGTGGTGTTGATGGAACTTGTGACTGGACTAACAGCCCTTGATGAGCATCGGTCCGAGGAAACTCGATATCTAGTTGAGTGGTTCTG

>M6-PGSC0003DMG400005896M6LRR-RLK_3477

CTAACAGCCCTTGATGAGCATCGGTCCGAGGAAACTCGATATCTAGTTGAGTGGTTCTGGCAGATAAAATCGAATAAAGAAAACCTCCTTGCTTCCGTTGATCCAGCTCTTGACGTGAAG

>M6-PGSC0003DMG400005896M6LRR-RLK_3538

AGATAAAATCGAATAAAGAAAACCTCCTTGCTTCCGTTGATCCAGCTCTTGACGTGAAGGAAGATATCCACAAGAGCATTTGCACTATGGCTGAATTAGCTGGACATTGCACTGCAAGAG

>M6-PGSC0003DMG400005896M6LRR-RLK_3599

AGATATCCACAAGAGCATTTGCACTATGGCTGAATTAGCTGGACATTGCACTGCAAGAGACCCCAATCACAGGCCTGACATGTCACACGTTGTGAATGTGCTTGGCCAACTAGTTGAGTC

>M6-PGSC0003DMG400005896M6LRR-RLK_366

TAACAAATTTGAATGATTTCAAGATTTTGAGTGATTTTAGAAATGGATTAGAGAATCCTGAACTATTGAAATGGCCTTCCAAGGGGAATGATCCTTGTGGTCCTCCTGCATGGCCTCATG

>M6-PGSC0003DMG400005896M6LRR-RLK_3660

CCCAATCACAGGCCTGACATGTCACACGTTGTGAATGTGCTTGGCCAACTAGTTGAGTCTTGGAAACCCGTAGAAGAGTTAGATGAATATTCAGGAATTGACTACAGTCCACCTCTGCCA

>M6-PGSC0003DMG400005896M6LRR-RLK_3721

GGAAACCCGTAGAAGAGTTAGATGAATATTCAGGAATTGACTACAGTCCACCTCTGCCAGAAATGTTGAAGGACTGGCAGGATGAAGACACTGGAGACTTCACCAGTACTAGTCAAGATA

>M6-PGSC0003DMG400005896M6LRR-RLK_3782

AATGTTGAAGGACTGGCAGGATGAAGACACTGGAGACTTCACCAGTACTAGTCAAGATAGCAGAGGAAGTATTCCTGCTAAGCCAAATGGATTTGCGGATTCTTTTACATCCAACGATGC

>M6-PGSC0003DMG400005896M6LRR-RLK_3843

AGAGGAAGTATTCCTGCTAAGCCAAATGGATTTGCGGATTCTTTTACATCCAACGATGCTCGATAGGTGAAGTTGGTTTTGGCTACCAAAGTCCATTGCTTCCTTACTATTAGTTTTGCT

>M6-PGSC0003DMG400005896M6LRR-RLK_3904

GATAGGTGAAGTTGGTTTTGGCTACCAAAGTCCATTGCTTCCTTACTATTAGTTTTGCTAGCATTTTCGACATGAATTTTGCTTTTGGCTCGTGAAAATAATCTATGGAGGCATTGCGTC

>M6-PGSC0003DMG400005896M6LRR-RLK_3965

CATTTTCGACATGAATTTTGCTTTTGGCTCGTGAAAATAATCTATGGAGGCATTGCGTCCAGTCTATATATGGTGAATATACAGATACCGAGTTGTCATGAATGATTGATACAGAGCAAA

>M6-PGSC0003DMG400005896M6LRR-RLK_4026

GTCTATATATGGTGAATATACAGATACCGAGTTGTCATGAATGATTGATACAGAGCAAAAAACAGAAAAAACTTTTCAATTTAATGTTGTTTACACAGTTATTTGTTTTGAATGTTGCTT

>M6-PGSC0003DMG400005896M6LRR-RLK_4087

ACAGAAAAAACTTTTCAATTTAATGTTGTTTACACAGTTATTTGTTTTGAATGTTGCTTGTAAAAGTTATATTTATAATAGGTTGGTCCAATTTGCTGTATAAGTTATTTACATGACACT

>M6-PGSC0003DMG400005896M6LRR-RLK_4136

AATGTTGCTTGTAAAAGTTATATTTATAATAGGTTGGTCCAATTTGCTGTATAAGTTATTTACATGACACTCTTGAGAATCTTACGCATGAATTTGAGAGTGTTAATAATGCAAAAAAAG

>M6-PGSC0003DMG400005896M6LRR-RLK_427

ACTATTGAAATGGCCTTCCAAGGGGAATGATCCTTGTGGTCCTCCTGCATGGCCTCATGTGTTCTGTTCTAGTGACAGAGTTACCCAAATTCAGGTTCAGAGTTTGGGTTTAAAGGGTCC

>M6-PGSC0003DMG400005896M6LRR-RLK_488

TTCTGTTCTAGTGACAGAGTTACCCAAATTCAGGTTCAGAGTTTGGGTTTAAAGGGTCCTTTACCTCAGAACTTGAATCAACTTGATAAGCTTCAAAATCTTGGCCTTCAGGGGAATGGT

>M6-PGSC0003DMG400005896M6LRR-RLK_549

TACCTCAGAACTTGAATCAACTTGATAAGCTTCAAAATCTTGGCCTTCAGGGGAATGGTTTTAGTGGGAAATTACCTACTTTCAATGGATTATCTGATTTGAAATATGCATACTTGGACA

>M6-PGSC0003DMG400005896M6LRR-RLK_61

GGGTTTGATCCACACTGCACCACAACTCTTTGCTAAACTCATCTCTCTTTTTTACAAAATTCACTCACATTATTAAAAAAAGGTACCATTTTTACAACAATTTATAGTTATTATGTACTT

>M6-PGSC0003DMG400005896M6LRR-RLK_610

TAGTGGGAAATTACCTACTTTCAATGGATTATCTGATTTGAAATATGCATACTTGGACAACAATAAATTCGATACGATTCCTGGTGATTTCTTTAATGGTCTTAGCAGTGTTCAAGTTTT

>M6-PGSC0003DMG400005896M6LRR-RLK_671

AATAAATTCGATACGATTCCTGGTGATTTCTTTAATGGTCTTAGCAGTGTTCAAGTTTTGGCTTTGGATTATAATCCGTTTAATGAGAGTGGATGGTCTATTCCCATTGAGTTGCAAGAT

>M6-PGSC0003DMG400005896M6LRR-RLK_732

CTTTGGATTATAATCCGTTTAATGAGAGTGGATGGTCTATTCCCATTGAGTTGCAAGATTCAGCTCAGTTGACGAATTTTTCTTGTGTGCAATGTAATATAGTTGGACCTGTGCCTGATT

>M6-PGSC0003DMG400005896M6LRR-RLK_793

AGCTCAGTTGACGAATTTTTCTTGTGTGCAATGTAATATAGTTGGACCTGTGCCTGATTTTTTTGGGAAATTGCCTTCTCTAAGTGCGTTGAAGCTGTCGTATAATCGTTTGACCGGTGA

>M6-PGSC0003DMG400005896M6LRR-RLK_854

TTTGGGAAATTGCCTTCTCTAAGTGCGTTGAAGCTGTCGTATAATCGTTTGACCGGTGAGATACCGGACAGTTTTCGTGATTCAATGCTTCGAATTTTGTGGTTGAATAATCAAGATAGT

>M6-PGSC0003DMG400005896M6LRR-RLK_915

TACCGGACAGTTTTCGTGATTCAATGCTTCGAATTTTGTGGTTGAATAATCAAGATAGTCCTGGAATGACGGGTCCAATCGACATAATTGGTACTATGGATCAGCTTATGTTGTTATGGC

>M6-PGSC0003DMG400005896M6LRR-RLK_976

TGGAATGACGGGTCCAATCGACATAATTGGTACTATGGATCAGCTTATGTTGTTATGGCTTCAAGGAAACAGTTTTAGTGGACCAATTCCTGATACAATTGGTGATTTAAATGACTTGAA

>M6-PGSC0003DMG400005972-(rev)M6LRR-RLP_0

TAAGAAAATACTGCGTATTGCCCATTTAAGCACTTACAGAAGTTATTTAATACGAGCTTTTTCAGCAACTCTGGATGCATCATGTCTGCAGCTGATATTAAATCATTAAGCCAAGACAAC

>M6-PGSC0003DMG400005972-(rev)M6LRR-RLP_1037

GGAATAAATTTCTGGAATTATACTGCCTTCTTTATTTAAATAAAATGTTTAAATATTACTGTTTTCTTATCAACTGCAACTACAGAAAGGATTCTTTGTTACTTTTTCAAAACTCTTGCA

>M6-PGSC0003DMG400005972-(rev)M6LRR-RLP_1098

TTTTCTTATCAACTGCAACTACAGAAAGGATTCTTTGTTACTTTTTCAAAACTCTTGCACATTAATTTACTAGTATCTTAAGTGGCTCATAAGTAGACCCGCCAGATCACGTAAGTGTAA

>M6-PGSC0003DMG400005972-(rev)M6LRR-RLP_1159

TTAATTTACTAGTATCTTAAGTGGCTCATAAGTAGACCCGCCAGATCACGTAAGTGTAACAACGTTAATTCCATATAAACAATGAATGAATCTTACCACACTGTGATAGCCAAGCATACA

>M6-PGSC0003DMG400005972-(rev)M6LRR-RLP_122

ATAAGGAGGGCGAGAAGAAAGAGGATAAGCTGCTGCGAGTAATGCTCCTAAAGGGACAACAGGTACATTGAGAGCTTGGGCAGGATCAGCCTGCAGCCAAGGTAAATAAATATAAGAATA

>M6-PGSC0003DMG400005972-(rev)M6LRR-RLP_1220

ACGTTAATTCCATATAAACAATGAATGAATCTTACCACACTGTGATAGCCAAGCATACAGTAATATTCCTCCCATAGAATGTCCAATGGCAAGAAGTTTACCATCTTTTGGTGCAGTTTG

>M6-PGSC0003DMG400005972-(rev)M6LRR-RLP_1281

AATATTCCTCCCATAGAATGTCCAATGGCAAGAAGTTTACCATCTTTTGGTGCAGTTTGGGCCTTAATGTATTCCATCTAGAATGAAAATAAAATAATTAGACTAGTCAGAATAATAAAT

>M6-PGSC0003DMG400005972-(rev)M6LRR-RLP_1342

CCTTAATGTATTCCATCTAGAATGAAAATAAAATAATTAGACTAGTCAGAATAATAAATAAGAGTGAAATTTCAACAAATTCTGAATTTCATTTGGTTGTGTAATCTCAAGGACAATAAT

>M6-PGSC0003DMG400005972-(rev)M6LRR-RLP_1403

GAGTGAAATTTCAACAAATTCTGAATTTCATTTGGTTGTGTAATCTCAAGGACAATAATATCTTTATATTCTACAACATGGGAATCTGTCACCTCACCGCAGCAGGAACATCCTCTTCCA

>M6-PGSC0003DMG400005972-(rev)M6LRR-RLP_1464

CTTTATATTCTACAACATGGGAATCTGTCACCTCACCGCAGCAGGAACATCCTCTTCCAGGTAATGATCAAAGTCCCAATCATACTTGACAATCAAGTCAAGCTGTTTCTGGAAATCTTC

>M6-PGSC0003DMG400005972-(rev)M6LRR-RLP_1525

TAATGATCAAAGTCCCAATCATACTTGACAATCAAGTCAAGCTGTTTCTGGAAATCTTCTATGGTAGTGGTTAGACGTTCATGCAGATCAACTAACGGAGAAACAGAACGTTGGCCTTCT

>M6-PGSC0003DMG400005972-(rev)M6LRR-RLP_1586

TGGTAGTGGTTAGACGTTCATGCAGATCAACTAACGGAGAAACAGAACGTTGGCCTTCTTCAATGATATTAACTAGCTTTTGGCTAAGATCCTTTACTTGGTCAACAACGGCAGAGTTTT

>M6-PGSC0003DMG400005972-(rev)M6LRR-RLP_1647

AATGATATTAACTAGCTTTTGGCTAAGATCCTTTACTTGGTCAACAACGGCAGAGTTTTGCCCTGTCTCCAGCAGACTTAACAACTTCCCCCTTGTCTCGTTAAAGCGTTCATAAAGAAA

>M6-PGSC0003DMG400005972-(rev)M6LRR-RLP_1708

CCTGTCTCCAGCAGACTTAACAACTTCCCCCTTGTCTCGTTAAAGCGTTCATAAAGAAAGGAATCCTCTAATAATTTTGATATTTGATCAAATAGTTTAGCAGACATAATCCTTGATTGG

>M6-PGSC0003DMG400005972-(rev)M6LRR-RLP_1769

AATCCTCTAATAATTTTGATATTTGATCAAATAGTTTAGCAGACATAATCCTTGATTGGCTTTCATTGAGGAAGCCAGAGACTCTTTCTGACAAGCGCATAAATGTTTCCGTAAGCTCGT

>M6-PGSC0003DMG400005972-(rev)M6LRR-RLP_183

GGTACATTGAGAGCTTGGGCAGGATCAGCCTGCAGCCAAGGTAAATAAATATAAGAATATGCTGAAAGAGCTTTTTGCTGCACAATGAGAGTTGAAAGACATTTATTTTTTGCTTACAAG

>M6-PGSC0003DMG400005972-(rev)M6LRR-RLP_1830

TTCATTGAGGAAGCCAGAGACTCTTTCTGACAAGCGCATAAATGTTTCCGTAAGCTCGTTTACTACTCTTGATTCATCCCACATTGTTGCAATAGCTGTGGATTCTTCTTTGACCACAGC

>M6-PGSC0003DMG400005972-(rev)M6LRR-RLP_1891

ACTACTCTTGATTCATCCCACATTGTTGCAATAGCTGTGGATTCTTCTTTGACCACAGCTGTGTCAGACTCTTCTAAGGTACTCTGGGCATCTGTTGACTGCTGTGCTGCAGAAAGAACT

>M6-PGSC0003DMG400005972-(rev)M6LRR-RLP_1952

TGTCAGACTCTTCTAAGGTACTCTGGGCATCTGTTGACTGCTGTGCTGCAGAAAGAACTTTGTCAGTAGCATTATCAGCGGCAGCTTCCATCTGTTCTGAGACTGCAATTGCAGTCTTCT

>M6-PGSC0003DMG400005972-(rev)M6LRR-RLP_2013

GTCAGTAGCATTATCAGCGGCAGCTTCCATCTGTTCTGAGACTGCAATTGCAGTCTTCTCAATGTTTGTGGAATCTGACTCCTGCACACTCAACCCAGCTCCACGAAGTTCAAGAACCCA

>M6-PGSC0003DMG400005972-(rev)M6LRR-RLP_2074

ATGTTTGTGGAATCTGACTCCTGCACACTCAACCCAGCTCCACGAAGTTCAAGAACCCAAGTGTCAAATCCTTCACCAGACATGTACCGGGCGAAAGATGACTGCATAATTATTTTTAAA

>M6-PGSC0003DMG400005972-(rev)M6LRR-RLP_2135

TGTCAAATCCTTCACCAGACATGTACCGGGCGAAAGATGACTGCATAATTATTTTTAAAAGTAAAAACCGTGATTAAATAACTTTTAGAAAGAATTTAAAAAAACCAGCCATACTATGTA

>M6-PGSC0003DMG400005972-(rev)M6LRR-RLP_2196

TAAAAACCGTGATTAAATAACTTTTAGAAAGAATTTAAAAAAACCAGCCATACTATGTAAAATTTCTTTGATAAGCAGGATGGACATGGTTGCCTGAAGTGATAAAGAATTCTAAACCTC

>M6-PGSC0003DMG400005972-(rev)M6LRR-RLP_2257

ATTTCTTTGATAAGCAGGATGGACATGGTTGCCTGAAGTGATAAAGAATTCTAAACCTCCACTGAAGAGTAGGGTGCACATACTTGCCTCAAGTGATAAAAATGAACTTGAACGGATAGG

>M6-PGSC0003DMG400005972-(rev)M6LRR-RLP_2318

CTGAAGAGTAGGGTGCACATACTTGCCTCAAGTGATAAAAATGAACTTGAACGGATAGGAAGATGCATCAACAGAACTTTAAACAAAAAATAGACAAGAAATAATCCAAGTGAAATCAAG

>M6-PGSC0003DMG400005972-(rev)M6LRR-RLP_2379

GATGCATCAACAGAACTTTAAACAAAAAATAGACAAGAAATAATCCAAGTGAAATCAAGTTGGCATTAGTTATTGCATTAACCAGCAGACATTTAGATTTATCTCACTAAAAAAAAAAAG

>M6-PGSC0003DMG400005972-(rev)M6LRR-RLP_244

CTGAAAGAGCTTTTTGCTGCACAATGAGAGTTGAAAGACATTTATTTTTTGCTTACAAGGGGTAGGAGCAGTCTGAGTGCAGACTTCGAAGATGTGTAATCCAGTGATGAAGCCAAAGTT

>M6-PGSC0003DMG400005972-(rev)M6LRR-RLP_2440

GGCATTAGTTATTGCATTAACCAGCAGACATTTAGATTTATCTCACTAAAAAAAAAAAGGATAGAAATCATCACTAGCAGCTAAGCAAAAAATCGTCATCACAATTCATATTTTAGAAAT

>M6-PGSC0003DMG400005972-(rev)M6LRR-RLP_2501

TAGAAATCATCACTAGCAGCTAAGCAAAAAATCGTCATCACAATTCATATTTTAGAAATGTACAATCACAGAGAGTGCTAGCCACATGACTACCATCAACACCTGATTGATGCAGGCATG

>M6-PGSC0003DMG400005972-(rev)M6LRR-RLP_2562

ACAATCACAGAGAGTGCTAGCCACATGACTACCATCAACACCTGATTGATGCAGGCATGCAGCCATAAAGCCTTATAAAAAAAAGGAAGATGAAATAGATAACAAGCCAGTAATAAAAAA

>M6-PGSC0003DMG400005972-(rev)M6LRR-RLP_2623

GCCATAAAGCCTTATAAAAAAAAGGAAGATGAAATAGATAACAAGCCAGTAATAAAAAAATCACACAAGTAAACTTTTATACGCTTGAAATGCATAAAGTATGAGAAAAGGACGAAGTTT

>M6-PGSC0003DMG400005972-(rev)M6LRR-RLP_2684

CACACAAGTAAACTTTTATACGCTTGAAATGCATAAAGTATGAGAAAAGGACGAAGTTTTTAACCGACCCACCATTTTGATCACCGCACCAACATGTGTGCAGAAGGAAGTAGATGCAGA

>M6-PGSC0003DMG400005972-(rev)M6LRR-RLP_2745

AACCGACCCACCATTTTGATCACCGCACCAACATGTGTGCAGAAGGAAGTAGATGCAGATGCACTAGGAACTAGTTACCCTAGGACCTTAAACAATCCATTTTGTATTATCCAGGGTCGG

>M6-PGSC0003DMG400005972-(rev)M6LRR-RLP_2806

CACTAGGAACTAGTTACCCTAGGACCTTAAACAATCCATTTTGTATTATCCAGGGTCGGGATGTGACCCAAACTACTTACAATGACCCACCCGTTGACACAGGTGGGGAAACTACTAATG

>M6-PGSC0003DMG400005972-(rev)M6LRR-RLP_2867

TGTGACCCAAACTACTTACAATGACCCACCCGTTGACACAGGTGGGGAAACTACTAATGCTCTTGATCCCTAATTATATAATATAATATAAGTATCCTTTTGATCAAAAAAAAAAAAGTA

>M6-PGSC0003DMG400005972-(rev)M6LRR-RLP_2928

CTTGATCCCTAATTATATAATATAATATAAGTATCCTTTTGATCAAAAAAAAAAAAGTAGTAAGGATAATACAACATCAAAACCAGGGTTAACTCTAAAAAATTTGAGCATGCCATAGAA

>M6-PGSC0003DMG400005972-(rev)M6LRR-RLP_2989

AAGGATAATACAACATCAAAACCAGGGTTAACTCTAAAAAATTTGAGCATGCCATAGAATTCTACCAGATTCAGAAACAAGAACCACTCCACGAAATAAAGAGTGAAACTTTAATGGAAC

>M6-PGSC0003DMG400005972-(rev)M6LRR-RLP_305

GTAGGAGCAGTCTGAGTGCAGACTTCGAAGATGTGTAATCCAGTGATGAAGCCAAAGTTGCAATAGCAGTTAATCCAGGCTCTCTTCCTTGCAATCCTTTCCAAGATAAAACCAGAAACA

>M6-PGSC0003DMG400005972-(rev)M6LRR-RLP_3050

CTACCAGATTCAGAAACAAGAACCACTCCACGAAATAAAGAGTGAAACTTTAATGGAACCTGACATATCCACTAAGCACTGATAATATACACATTTGACATCTGTCATTATGATCTATTA

>M6-PGSC0003DMG400005972-(rev)M6LRR-RLP_3111

GACATATCCACTAAGCACTGATAATATACACATTTGACATCTGTCATTATGATCTATTAAATGTGGCTATCAATGAACGATCCATGAAGATCCCCTAGTTCTGTTCACCTATACAGGCAT

>M6-PGSC0003DMG400005972-(rev)M6LRR-RLP_3172

TGTGGCTATCAATGAACGATCCATGAAGATCCCCTAGTTCTGTTCACCTATACAGGCATGAATTTGCAAATAAGAGTGTCCCCGACTACTCTTTCACACAATACTGCACATACAACAAAA

>M6-PGSC0003DMG400005972-(rev)M6LRR-RLP_3233

ATTTGCAAATAAGAGTGTCCCCGACTACTCTTTCACACAATACTGCACATACAACAAAACAGAAGAATGCACAAAGAACGGAGAGAAGGCAGACGAGTATCATACCCCAGGAGCAAGATC

>M6-PGSC0003DMG400005972-(rev)M6LRR-RLP_3294

GAAGAATGCACAAAGAACGGAGAGAAGGCAGACGAGTATCATACCCCAGGAGCAAGATCATATCCAATAGCATTAGTCCCCACGCCAGACAACAGAAGCAGGGGATGATTTCTCTTTGGA

>M6-PGSC0003DMG400005972-(rev)M6LRR-RLP_3355

ATCCAATAGCATTAGTCCCCACGCCAGACAACAGAAGCAGGGGATGATTTCTCTTTGGAGCCTATCATCATCAAAAATCACACACCCTTAATGCAAAGCTCAAATCCTTGAACAAAAAAA

>M6-PGSC0003DMG400005972-(rev)M6LRR-RLP_3416

CTATCATCATCAAAAATCACACACCCTTAATGCAAAGCTCAAATCCTTGAACAAAAAAATCATGTAACAGAACTGTTAAACCTTCCTAGTATCACTCCCTGTGTTTCATTTTATATAACA

>M6-PGSC0003DMG400005972-(rev)M6LRR-RLP_3477

ATGTAACAGAACTGTTAAACCTTCCTAGTATCACTCCCTGTGTTTCATTTTATATAACAGTATTTGATAGAACACAAAATTTAAAATATTCAATAGTTGAATTAGATAAAAATGGTGTCA

>M6-PGSC0003DMG400005972-(rev)M6LRR-RLP_3538

ATTTGATAGAACACAAAATTTAAAATATTCAATAGTTGAATTAGATAAAAATGGTGTCAAACCTTCTTAATGCCTTCATGTCTTATATATTGGGACCAATTCTATATGACCACATTTACT

>M6-PGSC0003DMG400005972-(rev)M6LRR-RLP_366

AATAGCAGTTAATCCAGGCTCTCTTCCTTGCAATCCTTTCCAAGATAAAACCAGAAACATATTAAAATTATCTCATATCTGTATGATTCCCGGATGTAGTGAGTTTGAAAATATTTTTTT

>M6-PGSC0003DMG400005972-(rev)M6LRR-RLP_3782

TAATTGATCACAAAATTTAAAATTTGAACTTATATAGAAATGAAAGTGTCAAATATATTATTTTTTTTCAAATTGTCATACAGTTAAGAGTAACAGTTCAAATTAAAAAACATAAAAAAA

>M6-PGSC0003DMG400005972-(rev)M6LRR-RLP_3843

TTTTTTTCAAATTGTCATACAGTTAAGAGTAACAGTTCAAATTAAAAAACATAAAAAAATTGAACCTGAGGAGGAGGAATGTAGCGCCAGAGAGCAAGTTTCCAATCGGAATTATTAACA

>M6-PGSC0003DMG400005972-(rev)M6LRR-RLP_3904

GAACCTGAGGAGGAGGAATGTAGCGCCAGAGAGCAAGTTTCCAATCGGAATTATTAACAGATACGTAGTGGAGTTCATCGGCAGTGCATAACGATGGCTTATTACAAACCTTCTGGGCTG

>M6-PGSC0003DMG400005972-(rev)M6LRR-RLP_3965

TACGTAGTGGAGTTCATCGGCAGTGCATAACGATGGCTTATTACAAACCTTCTGGGCTGTAGTATCGCCATCATTGGTGGTGAAAGCTCTGAGTTTAAAGGCGGAGTTCACCGCGAAGCA

>M6-PGSC0003DMG400005972-(rev)M6LRR-RLP_4026

GTATCGCCATCATTGGTGGTGAAAGCTCTGAGTTTAAAGGCGGAGTTCACCGCGAAGCAGTAGTTCGGGGAGTTCCGGTATGAAATGACGGCGGGAGGAAGTCGCCGGCGGTGATTTGTT

>M6-PGSC0003DMG400005972-(rev)M6LRR-RLP_4087

AGTTCGGGGAGTTCCGGTATGAAATGACGGCGGGAGGAAGTCGCCGGCGGTGATTTGTTAAGTGGTGGTAATGGTTGTGGTGGCGGAAGACGGCAGCGATTTGAATCGCCGATCGAATAT

>M6-PGSC0003DMG400005972-(rev)M6LRR-RLP_4148

GTGGTGGTAATGGTTGTGGTGGCGGAAGACGGCAGCGATTTGAATCGCCGATCGAATATCGGATTGAATTGGATAAACCATATTTTTTGGGATTCGTGACAATTTGGATAAAGAAAATTG

>M6-PGSC0003DMG400005972-(rev)M6LRR-RLP_4207

CGGATTGAATTGGATAAACCATATTTTTTGGGATTCGTGACAATTTGGATAAAGAAAATTGGTTTGATAAATCTTTTACTCTTTTGACTTTTGAAGGTTTGAGACGTAAACTTTTTTATT

>M6-PGSC0003DMG400005972-(rev)M6LRR-RLP_427

TTAAAATTATCTCATATCTGTATGATTCCCGGATGTAGTGAGTTTGAAAATATTTTTTTGAGACTGTGCCCACTGCACATTAACAAGGAATCAGACAATGCTCCAAGTGACTTATCTGTT

>M6-PGSC0003DMG400005972-(rev)M6LRR-RLP_488

GACTGTGCCCACTGCACATTAACAAGGAATCAGACAATGCTCCAAGTGACTTATCTGTTAGTTGCGAAACCTTTTTACATCTAAAACCAGGTCAAATTCCTTCCAAATCAAATTGTCCCT

>M6-PGSC0003DMG400005972-(rev)M6LRR-RLP_549

TTGCGAAACCTTTTTACATCTAAAACCAGGTCAAATTCCTTCCAAATCAAATTGTCCCTTACACACTGTAAAAACGTATGGATAAAGTATAAAATCTTGCCTAGCTCCCTATAATCTCAA

>M6-PGSC0003DMG400005972-(rev)M6LRR-RLP_61

TCAGCAACTCTGGATGCATCATGTCTGCAGCTGATATTAAATCATTAAGCCAAGACAACACATAAGGAGGGCGAGAAGAAAGAGGATAAGCTGCTGCGAGTAATGCTCCTAAAGGGACAA

>M6-PGSC0003DMG400005972-(rev)M6LRR-RLP_610

CACACTGTAAAAACGTATGGATAAAGTATAAAATCTTGCCTAGCTCCCTATAATCTCAATAAGGTCAAAAGATGGCTCATTTTCAGCTCTGTATGAAAGACGCTAGTAAGCAGATAAGTC

>M6-PGSC0003DMG400005972-(rev)M6LRR-RLP_671

AGGTCAAAAGATGGCTCATTTTCAGCTCTGTATGAAAGACGCTAGTAAGCAGATAAGTCACACTGTTAACCAGATCCTGTTTACCTTGGTGCACCCTATCATGGTATTTTTAAGTTAATA

>M6-PGSC0003DMG400005972-(rev)M6LRR-RLP_732

ACTGTTAACCAGATCCTGTTTACCTTGGTGCACCCTATCATGGTATTTTTAAGTTAATAGGAGTACTGCACTTGGTGGCTATTACAACTATACTCCCAAAGTTGTTGTGAACTATAGCAT

>M6-PGSC0003DMG400005972-(rev)M6LRR-RLP_793

AGTACTGCACTTGGTGGCTATTACAACTATACTCCCAAAGTTGTTGTGAACTATAGCATCTTTCTTATTTTTCAGCAAAAAGCTTGTTGAAGTTGAGTCTGAAGATTTTGCTCCATGAAT

>M6-PGSC0003DMG400005972-(rev)M6LRR-RLP_854

TTCTTATTTTTCAGCAAAAAGCTTGTTGAAGTTGAGTCTGAAGATTTTGCTCCATGAATATTAGGAATTAAAAGAACATTAAGTCAAGAGGAGACCAGGTTAACTCAAAGTTGCACCAAG

>M6-PGSC0003DMG400005972-(rev)M6LRR-RLP_915

TAGGAATTAAAAGAACATTAAGTCAAGAGGAGACCAGGTTAACTCAAAGTTGCACCAAGTGAAGTGGTAGCCTTTGGAATTTAATCCCAAGACTAAATCGTATGATACTCCAGCAGGTTC

>M6-PGSC0003DMG400005972-(rev)M6LRR-RLP_976

AAGTGGTAGCCTTTGGAATTTAATCCCAAGACTAAATCGTATGATACTCCAGCAGGTTCATGGAATAAATTTCTGGAATTATACTGCCTTCTTTATTTAAATAAAATGTTTAAATATTAC

>M6-PGSC0003DMG400005975M6RLKWAX-EGF_0

GAAAAAACTATAGTATAGCATACAATATTTTTTTGAGTATGGTATCATTAAATTCTTTGTCAATTCTCTTGATTTCGTATCAAATGGTGATGTTTGGTTCAGTTGTAGCGCAGAATCAAA

>M6-PGSC0003DMG400005975M6RLKWAX-EGF_1159

TATTCTGCTTGTTTCCATTGTGTAACGTGGGACAAATCGCCTGCAGGATTTATGTCTGTAGCTTCTTCATAGTGATCTTGTTCTGAATTTAGTATAGCAGCATAAGCTTGTCATACTTAA

>M6-PGSC0003DMG400005975M6RLKWAX-EGF_122

CAAAATAATAGTAGTTGTGTGGATAAATTCAAATGTGGGAATCTTGGAAATGTGAGTTTTCCATTTTCTGTATCTTCACAACCTGATTGTGGATTGTACACTATAGATTGTGATGCTACT

>M6-PGSC0003DMG400005975M6RLKWAX-EGF_1220

CTTCTTCATAGTGATCTTGTTCTGAATTTAGTATAGCAGCATAAGCTTGTCATACTTAACTCTCAGTTCCTCCTCCGCGTGCAACAATTATCTAACTTTTTCGTGATTTGGTTTCGTTGC

>M6-PGSC0003DMG400005975M6RLKWAX-EGF_1281

CTCAGTTCCTCCTCCGCGTGCAACAATTATCTAACTTTTTCGTGATTTGGTTTCGTTGCAGTAACAAGAAAGTCGAGACGAGTTCTAATTGTAGGTAACAATCTATAACTTCAAATCTTT

>M6-PGSC0003DMG400005975M6RLKWAX-EGF_1342

TAACAAGAAAGTCGAGACGAGTTCTAATTGTAGGTAACAATCTATAACTTCAAATCTTTTTCATATAACCACATGTTCTCAAGTATGTTCCTTACTTTGAAATGCCAGCTTTCGTCGAAT

>M6-PGSC0003DMG400005975M6RLKWAX-EGF_1403

CATATAACCACATGTTCTCAAGTATGTTCCTTACTTTGAAATGCCAGCTTTCGTCGAATTCTGTCTACTTTTCTGTAGCCATTTTTCAAACAAATTCTGATTAGTTTCTCTGCATTTGCT

>M6-PGSC0003DMG400005975M6RLKWAX-EGF_1464

TGTCTACTTTTCTGTAGCCATTTTTCAAACAAATTCTGATTAGTTTCTCTGCATTTGCTTCAGTCTTACCTGTGGTGATTGGCTTGACCTTTTCAATCTGTCTTGGTATTGTTATCTGGC

>M6-PGSC0003DMG400005975M6RLKWAX-EGF_1525

AGTCTTACCTGTGGTGATTGGCTTGACCTTTTCAATCTGTCTTGGTATTGTTATCTGGCATTTCAAGAAAGGAAAAGGCGGTCGTTCTCACTCTTTCACAAGAAATACATTTTCTGATCC

>M6-PGSC0003DMG400005975M6RLKWAX-EGF_1586

TTCAAGAAAGGAAAAGGCGGTCGTTCTCACTCTTTCACAAGAAATACATTTTCTGATCCTTCAAGAAAAGATCTGGAAGGGGATAGCAAGTACTTTGGGGTCCCAGTCTTCTCCTACTCT

>M6-PGSC0003DMG400005975M6RLKWAX-EGF_1647

CAAGAAAAGATCTGGAAGGGGATAGCAAGTACTTTGGGGTCCCAGTCTTCTCCTACTCTACTCTCGAAGAAGCCACCAATAATTTTGATCCCTCCCAAGAGCTTGGAGATGGAGGTTTTG

>M6-PGSC0003DMG400005975M6RLKWAX-EGF_1708

TCTCGAAGAAGCCACCAATAATTTTGATCCCTCCCAAGAGCTTGGAGATGGAGGTTTTGGAACTGTATACTATGGTAATAAGCCACGTAAATCAAGTCATGACATTTCAAATCATATTTG

>M6-PGSC0003DMG400005975M6RLKWAX-EGF_1769

ACTGTATACTATGGTAATAAGCCACGTAAATCAAGTCATGACATTTCAAATCATATTTGGAAAGCATTAGGCAGAGTTTTGCATTACTGTATTAAGAATGCAATAGTTTGGTAACATGAT

>M6-PGSC0003DMG400005975M6RLKWAX-EGF_183

CATTTTCTGTATCTTCACAACCTGATTGTGGATTGTACACTATAGATTGTGATGCTACTCCCAATCCAACAATTCATCTCGGAGAAAATGTTTATAGTGTTCTGAGACAACGCTTCTCTG

>M6-PGSC0003DMG400005975M6RLKWAX-EGF_1830

AAGCATTAGGCAGAGTTTTGCATTACTGTATTAAGAATGCAATAGTTTGGTAACATGATTCAGCAGCTAACAATGCACTTCTTTTATTAGGTAAACTTCGTGATGGACGAGAAGTTGCAG

>M6-PGSC0003DMG400005975M6RLKWAX-EGF_1891

AGCAGCTAACAATGCACTTCTTTTATTAGGTAAACTTCGTGATGGACGAGAAGTTGCAGTGAAACGCCTCTATGAGCAAAGTTCCAAGAGGATGGTGCAGTTTAAGAATGAAATTGAAAT

>M6-PGSC0003DMG400005975M6RLKWAX-EGF_1952

AAACGCCTCTATGAGCAAAGTTCCAAGAGGATGGTGCAGTTTAAGAATGAAATTGAAATTCTTACTCGCTTAAGGCACCAGAATCTTGTTATGCTCTATGGTTGCACGTCAAGGCATAGT

>M6-PGSC0003DMG400005975M6RLKWAX-EGF_2013

TTACTCGCTTAAGGCACCAGAATCTTGTTATGCTCTATGGTTGCACGTCAAGGCATAGTCGTGAACTCCTCCTGGTTTATGAATACATCCCTAATGGAACAGTTGCTGATCATCTCCATG

>M6-PGSC0003DMG400005975M6RLKWAX-EGF_2074

TGAACTCCTCCTGGTTTATGAATACATCCCTAATGGAACAGTTGCTGATCATCTCCATGGAGAGAAAGCTAAGAATGGAGCCCTTATATGGCCTATACGCATGAAGATTGCCATAGAAAC

>M6-PGSC0003DMG400005975M6RLKWAX-EGF_2135

GAGAAAGCTAAGAATGGAGCCCTTATATGGCCTATACGCATGAAGATTGCCATAGAAACTGCTAGTGCTTTAGCTTATCTCCATGCTTCTGACATAATACACCGCGATGTCAAGACTTGT

>M6-PGSC0003DMG400005975M6RLKWAX-EGF_2196

CTAGTGCTTTAGCTTATCTCCATGCTTCTGACATAATACACCGCGATGTCAAGACTTGTAACATTCTACTTGACAACAATTTCTGTGTCAAAGTTGCAGATTTTGGGCTTTCGCGACTTT

>M6-PGSC0003DMG400005975M6RLKWAX-EGF_2257

CATTCTACTTGACAACAATTTCTGTGTCAAAGTTGCAGATTTTGGGCTTTCGCGACTTTTTCCAAATGATGTGACTCATATCTCAACTGCTCCACAGGGGACACCAGGATATGTTGATCC

>M6-PGSC0003DMG400005975M6RLKWAX-EGF_2318

CCAAATGATGTGACTCATATCTCAACTGCTCCACAGGGGACACCAGGATATGTTGATCCGGATTATCATGCTTGCTATCAGCTAACAAGTAAAAGTGATGTGTACAGCTTTGGAGTTGTC

>M6-PGSC0003DMG400005975M6RLKWAX-EGF_2379

ATTATCATGCTTGCTATCAGCTAACAAGTAAAAGTGATGTGTACAGCTTTGGAGTTGTCCTTATTGAGCTCATATCGTCGCTCCTTGCTGTGGATATAAGAAGGCATAGGGATGAAATTA

>M6-PGSC0003DMG400005975M6RLKWAX-EGF_244

CAATCCAACAATTCATCTCGGAGAAAATGTTTATAGTGTTCTGAGACAACGCTTCTCTGATCGTTTTCGAGTTTCAGACCATAGACTTGAGTACTTATTGGCGAGAAACAGTTGTGAAAC

>M6-PGSC0003DMG400005975M6RLKWAX-EGF_2440

TATTGAGCTCATATCGTCGCTCCTTGCTGTGGATATAAGAAGGCATAGGGATGAAATTAGTTTGGCTAACTTTGCAGTAAGTAAGATTTTGAAATGTGCATTCGAGGAGCTGATTGATCC

>M6-PGSC0003DMG400005975M6RLKWAX-EGF_2501

TTGGCTAACTTTGCAGTAAGTAAGATTTTGAAATGTGCATTCGAGGAGCTGATTGATCCATCTCTTGGTTACGAGTCCAATGCTGAAGTTAGGAGAATGACTACTTCAGTTGCAGAGCTA

>M6-PGSC0003DMG400005975M6RLKWAX-EGF_2562

CTCTTGGTTACGAGTCCAATGCTGAAGTTAGGAGAATGACTACTTCAGTTGCAGAGCTAGCTTTTCAATGTCTCCAACTCGAGAAGGAAATGAGACCAACAATGGATGAAGTGTTGGAAA

>M6-PGSC0003DMG400005975M6RLKWAX-EGF_2623

TTTTCAATGTCTCCAACTCGAGAAGGAAATGAGACCAACAATGGATGAAGTGTTGGAAATTCTAAAGGCGATTCAGAGTGGTGAGTTTGAAAACCAGAAGAAAGAAGAGATTAACGTCGT

>M6-PGSC0003DMG400005975M6RLKWAX-EGF_2684

CTAAAGGCGATTCAGAGTGGTGAGTTTGAAAACCAGAAGAAAGAAGAGATTAACGTCGTTGATAATGTAGAAGTCCCTCAGGACCCTGAATCAGAAAATGATGATGCACAATTGATGAAA

>M6-PGSC0003DMG400005975M6RLKWAX-EGF_2745

ATAATGTAGAAGTCCCTCAGGACCCTGAATCAGAAAATGATGATGCACAATTGATGAAATCTAAATTACCTGTTTCGACAAATTCTGTGAATGATAAGTGGTTTAGTTGCTCTACCACAG

>M6-PGSC0003DMG400005975M6RLKWAX-EGF_2806

TAAATTACCTGTTTCGACAAATTCTGTGAATGATAAGTGGTTTAGTTGCTCTACCACAGTTAGTATTAGCGGGTGAAATTCCTTCGATATTAAAGCTAACTTCATTACTTTTTAGCTTTC

>M6-PGSC0003DMG400005975M6RLKWAX-EGF_2867

AGTATTAGCGGGTGAAATTCCTTCGATATTAAAGCTAACTTCATTACTTTTTAGCTTTCACTCTCTTCTCATTCTTTTTTCGCTCTCTGCTTGTTTATGAGTTCAATACCTAATTGTATC

>M6-PGSC0003DMG400005975M6RLKWAX-EGF_2928

TCTCTTCTCATTCTTTTTTCGCTCTCTGCTTGTTTATGAGTTCAATACCTAATTGTATCAGAAATGTATTAAAAATAGAGATGCCTTTTTTGTAAGCAAGCCATCACCTTAACGATGTGG

>M6-PGSC0003DMG400005975M6RLKWAX-EGF_2981

TGTATCAGAAATGTATTAAAAATAGAGATGCCTTTTTTGTAAGCAAGCCATCACCTTAACGATGTGGGGGGTTAGGCTTGACCCGTACCATGTCTATAAACATCCAAAAAGTTATCAAAG

>M6-PGSC0003DMG400005975M6RLKWAX-EGF_305

CGTTTTCGAGTTTCAGACCATAGACTTGAGTACTTATTGGCGAGAAACAGTTGTGAAACTTTTGATAGAAGTATATCGTTTCCAAATTCTCCTTCTATCTCGTTTAGAATCAATGAACGA

>M6-PGSC0003DMG400005975M6RLKWAX-EGF_366

TTGATAGAAGTATATCGTTTCCAAATTCTCCTTCTATCTCGTTTAGAATCAATGAACGAAATGTTACTTTGTTCAGATGCAACAACAGTTTAGATATCAACAGGAGCATGAGTGATCATT

>M6-PGSC0003DMG400005975M6RLKWAX-EGF_427

TGTTACTTTGTTCAGATGCAACAACAGTTTAGATATCAACAGGAGCATGAGTGATCATTATTTTCGCGAATATCTGAATTATACTAAATGTAGTGGTTTTACTATATACTACAAGTATCC

>M6-PGSC0003DMG400005975M6RLKWAX-EGF_488

TTTCGCGAATATCTGAATTATACTAAATGTAGTGGTTTTACTATATACTACAAGTATCCTAGTAGAAGACGGGAAGATTCCTCCGATGCACCAGAGGGAGATATTCCTGATAACTGTTCC

>M6-PGSC0003DMG400005975M6RLKWAX-EGF_549

GTAGAAGACGGGAAGATTCCTCCGATGCACCAGAGGGAGATATTCCTGATAACTGTTCCCCGATTCAATTGCCAATTACATGGAATTCAGGATCAAACGTTTCGAATAGTAGCTTGTTTG

>M6-PGSC0003DMG400005975M6RLKWAX-EGF_61

AATTCTCTTGATTTCGTATCAAATGGTGATGTTTGGTTCAGTTGTAGCGCAGAATCAAATTCAAAATAATAGTAGTTGTGTGGATAAATTCAAATGTGGGAATCTTGGAAATGTGAGTTT

>M6-PGSC0003DMG400005975M6RLKWAX-EGF_610

GATTCAATTGCCAATTACATGGAATTCAGGATCAAACGTTTCGAATAGTAGCTTGTTTGATCTCTTAGCTGTGAATTATACAATTCAGTGGAGTCTGTCTGAAGATTGTAGTAAATGTCA

>M6-PGSC0003DMG400005975M6RLKWAX-EGF_671

CTCTTAGCTGTGAATTATACAATTCAGTGGAGTCTGTCTGAAGATTGTAGTAAATGTCACTATCAAGGAGGTCGATGTTTAACCGATAGCAACAATAGATTTCTTTGCTCCAGAGGTAAT

>M6-PGSC0003DMG400005975M6RLKWAX-EGF_732

ATCAAGGAGGTCGATGTTTAACCGATAGCAACAATAGATTTCTTTGCTCCAGAGGTAATATTTTACTTGCAATATGCTTGTCATATACAATACTCTGACCTTTGTCTAATTCTACTTCGA

>M6-PGSC0003DMG400005975M6RLKWAX-EGF_793

TTTACTTGCAATATGCTTGTCATATACAATACTCTGACCTTTGTCTAATTCTACTTCGAATAGTTGCTTCTAGTTTTTGTACTCTAGCTTCGAGATTGCCCATCTACACACCTCAACTTG

>M6-PGSC0003DMG400006027-(rev)M6LRR-RLP_0

TAGAGGCGTCTACTTGCTATTTAATTCAAATCAAATTCATTCATTCATTCAACAACACATTTTTACATTGTGTGAAAAAAAACTTCATTTTCTTCTTCTTAGCTTGAAAATCTCTATCAA

>M6-PGSC0003DMG400006027-(rev)M6LRR-RLP_1037

CTTACTGGGGTTATACCAGCTAGCATTTGCCAACTTCCAAGGTTGCAGAACTTTACGTATTCGTTCAATTACTTCACTGGGGAGGCTCCGGTTTGTGCTGCGACTAGATCTGGTGATGGT

>M6-PGSC0003DMG400006027-(rev)M6LRR-RLP_1098

CGTTCAATTACTTCACTGGGGAGGCTCCGGTTTGTGCTGCGACTAGATCTGGTGATGGTCAGGAGAACTGTATCGTTGGAAAGAAGAATCAAAGATCTGCTAAAGAATGCTCTTCGGATG

>M6-PGSC0003DMG400006027-(rev)M6LRR-RLP_1159

GGAGAACTGTATCGTTGGAAAGAAGAATCAAAGATCTGCTAAAGAATGCTCTTCGGATGATGCAAAACCGTATGATTGTAAGAAATCGAAGTGTTATAGCCCATATGCTACGTCTCCTTC

>M6-PGSC0003DMG400006027-(rev)M6LRR-RLP_122

TCAGCAATGCGGCCTCCTAGCCGCTGCGTATTTATCTCTTTTCTTGTTATCTCCATCTTGTCGTACCAGATCTCGAATGTGGTAGGACAAGAAGGTGATAGCGATATCGATTTAGATGAT

>M6-PGSC0003DMG400006027-(rev)M6LRR-RLP_1342

TTCGCATAACAAGCGATCTCCGCCACCACCTAAATCAACTCCTCTTCCTCCACCAGCACCTTACAAAAAGTCCCCAGTGTATCGACATAGAAGTCCACCACCACCTACTCACAAAATTTC

>M6-PGSC0003DMG400006027-(rev)M6LRR-RLP_1403

TACAAAAAGTCCCCAGTGTATCGACATAGAAGTCCACCACCACCTACTCACAAAATTTCACCTGTAACTCATCATTCACCTCCTCCACCTTCACCCGTATATTACCACCCATCTCCATCG

>M6-PGSC0003DMG400006027-(rev)M6LRR-RLP_1464

CTGTAACTCATCATTCACCTCCTCCACCTTCACCCGTATATTACCACCCATCTCCATCGTCCCCACCACCACCAGTTTATTATTCTCCACCACCCCCGGTGTATCACGAGCCACCTACTT

>M6-PGSC0003DMG400006027-(rev)M6LRR-RLP_1525

CCCACCACCACCAGTTTATTATTCTCCACCACCCCCGGTGTATCACGAGCCACCTACTTACAAGCCTAAATCACCACCACCACCAACACCGTCATATGAACACCCAAAAACCCCATCATC

>M6-PGSC0003DMG400006027-(rev)M6LRR-RLP_1708

ATACGAGCATCCTAAAACACCACCACCAACACCATCATACGAACATCCAAAAACACCATCACCACCACCACCAACACCATCATACGAACATCCAAAAACACCATCACCACCACCACCAAC

>M6-PGSC0003DMG400006027-(rev)M6LRR-RLP_183

CGTACCAGATCTCGAATGTGGTAGGACAAGAAGGTGATAGCGATATCGATTTAGATGATATTAAGGCTAATGCTAGTAAGTTTAGTTTCGAAAACCCAAAGATTCGCGATGCTTATATCG

>M6-PGSC0003DMG400006027-(rev)M6LRR-RLP_1952

CCACCACCAACACCATCATACGAACACCCAAAAACCCCATCACACCCAACTCCACCTACCCCACCTTGTAACGAACCACCACCTCCACCACCTAATTCACATTGGGAACCAAAACCATCC

>M6-PGSC0003DMG400006027-(rev)M6LRR-RLP_2013

CACCTTGTAACGAACCACCACCTCCACCACCTAATTCACATTGGGAACCAAAACCATCCCCACCATACACCTACTCATCACCACCGCCTCCATCACCATCTCCTCCTCCTCCCACTTACT

>M6-PGSC0003DMG400006027-(rev)M6LRR-RLP_2135

TACTCATCACCACCGCCACCATCACCATCACCATCACCTCCCCCACCCACTTACTACTACTCATCACCGCCGCCACCACCACCATTTTACGAAAATATCCCTCTCCCACCAGTAATCGGA

>M6-PGSC0003DMG400006027-(rev)M6LRR-RLP_2196

CATCACCGCCGCCACCACCACCATTTTACGAAAATATCCCTCTCCCACCAGTAATCGGAGTCTCCTACGCATCTCCACCACCACCAGTCATTCCATATTACTGAACAAGGGTTATCAATT

>M6-PGSC0003DMG400006027-(rev)M6LRR-RLP_2257

CTCCTACGCATCTCCACCACCACCAGTCATTCCATATTACTGAACAAGGGTTATCAATTAGCCTTTGATTTTCCATTATCAAAAAAAAAAGAATTCATTTTTTTTTATGAATTATGTATA

>M6-PGSC0003DMG400006027-(rev)M6LRR-RLP_2318

CCTTTGATTTTCCATTATCAAAAAAAAAAGAATTCATTTTTTTTTATGAATTATGTATACTTTGTTATTTATTTTTTGTAATATTTTTATTTTGAGTTTGATTATATTTATGAAAATTGG

>M6-PGSC0003DMG400006027-(rev)M6LRR-RLP_2375

TACTTTGTTATTTATTTTTTGTAATATTTTTATTTTGAGTTTGATTATATTTATGAAAATTGGGATTATGGGCGGGAGTTAGTGAGGGGTTTAACTATCAATATTTTAATATTGTTATAT

>M6-PGSC0003DMG400006027-(rev)M6LRR-RLP_244

TAAGGCTAATGCTAGTAAGTTTAGTTTCGAAAACCCAAAGATTCGCGATGCTTATATCGCGTTGCAATATTGGAAAACAGCAATGTTTTCTGATCCTTTTAACTTCACAGCTAATTGGAC

>M6-PGSC0003DMG400006027-(rev)M6LRR-RLP_305

TTGCAATATTGGAAAACAGCAATGTTTTCTGATCCTTTTAACTTCACAGCTAATTGGACAGGACCTTATGTTTGTTCGTATGGTGGAGTGTTTTGTGCTCCTTCGTTAACGGATGATTCG

>M6-PGSC0003DMG400006027-(rev)M6LRR-RLP_366

GACCTTATGTTTGTTCGTATGGTGGAGTGTTTTGTGCTCCTTCGTTAACGGATGATTCGATTAGAGTTGTAGCTGGCATTGATCTTAATCATGCTGACATTGCTGGTTCATTAGTTGCAG

>M6-PGSC0003DMG400006027-(rev)M6LRR-RLP_427

TAGAGTTGTAGCTGGCATTGATCTTAATCATGCTGACATTGCTGGTTCATTAGTTGCAGAACTTGGTCTTCTTACAGATCTGGTTCTTTTCCATCTTAACTCGAATCGTTTTTGTGGAGT

>M6-PGSC0003DMG400006027-(rev)M6LRR-RLP_488

CTTGGTCTTCTTACAGATCTGGTTCTTTTCCATCTTAACTCGAATCGTTTTTGTGGAGTTGTACCGAAAACTTTTAGCCATTTGAAGCTTCTTAGGGAGCTTGATTTGAGTAACAACAGG

>M6-PGSC0003DMG400006027-(rev)M6LRR-RLP_549

TACCGAAAACTTTTAGCCATTTGAAGCTTCTTAGGGAGCTTGATTTGAGTAACAACAGGTTTGTTGGTGGGTTCCCTAAGGTAGTTCTTTCATTACCTTCTTTGAAGTTTCTGGATCTAA

>M6-PGSC0003DMG400006027-(rev)M6LRR-RLP_61

TTTACATTGTGTGAAAAAAAACTTCATTTTCTTCTTCTTAGCTTGAAAATCTCTATCAAATTCAGCAATGCGGCCTCCTAGCCGCTGCGTATTTATCTCTTTTCTTGTTATCTCCATCTT

>M6-PGSC0003DMG400006027-(rev)M6LRR-RLP_610

TGTTGGTGGGTTCCCTAAGGTAGTTCTTTCATTACCTTCTTTGAAGTTTCTGGATCTAAGGTTTAATGATTTTGAAGGACCAGTTCCTTCCGAATTGTTCGATAAAGATTTGGATGCGTT

>M6-PGSC0003DMG400006027-(rev)M6LRR-RLP_671

TTTAATGATTTTGAAGGACCAGTTCCTTCCGAATTGTTCGATAAAGATTTGGATGCGTTGTTTCTGAATGACAATAGATTCCGTTTTGGCATCCCGGAGAACTTGGGTAACTCCCCTGTT

>M6-PGSC0003DMG400006027-(rev)M6LRR-RLP_732

TTCTGAATGACAATAGATTCCGTTTTGGCATCCCGGAGAACTTGGGTAACTCCCCTGTTTCTGTTCTTGTGTTTGCTAATAACGATCTTGGAGGGTGTATTCCCGCGAGTATTGGGAAAA

>M6-PGSC0003DMG400006027-(rev)M6LRR-RLP_793

TGTTCTTGTGTTTGCTAATAACGATCTTGGAGGGTGTATTCCCGCGAGTATTGGGAAAATGGGGAAAACGTTGAATGAGTTGATATTGATGAATGATAACCTCACTGGGTGTTTGCCCAT

>M6-PGSC0003DMG400006027-(rev)M6LRR-RLP_854

GGGAAAACGTTGAATGAGTTGATATTGATGAATGATAACCTCACTGGGTGTTTGCCCATGGAGATTGGATTGTTGAATAAGTTGACAGTGTTTGATGTTAGCTTCAACAAGATTCAAGGC

>M6-PGSC0003DMG400006027-(rev)M6LRR-RLP_915

AGATTGGATTGTTGAATAAGTTGACAGTGTTTGATGTTAGCTTCAACAAGATTCAAGGCTCATTGCCATTAACTGTGAGCAAAATGAGGAGTGTGGAGGAACTCAATGTGGCTCACAACA

>M6-PGSC0003DMG400006027-(rev)M6LRR-RLP_976

ATTGCCATTAACTGTGAGCAAAATGAGGAGTGTGGAGGAACTCAATGTGGCTCACAACAAGCTTACTGGGGTTATACCAGCTAGCATTTGCCAACTTCCAAGGTTGCAGAACTTTACGTA

>M6-PGSC0003DMG400006065M6LRR-RLK_0

GAAGAACATTCTCTTATTGATTCTTCTCTTTCAAATTCAATTTTCTATGTTAACTGCTTCATCAAATGAGACTGACCAACAAGCTTTACTAGCTTTCCAAAATCTTATTACAAGTCCGAG

>M6-PGSC0003DMG400006065M6LRR-RLK_1037

CTCCGGAATTAGGGAAGCTATCAAATTTGCAGCAATTATGCTTTAGGAATAATTATAATCTAATTGGTCAAATTCCAGAGGCTATTTTCAACATATCTTCTTTGGAAGTTATTGATTTCA

>M6-PGSC0003DMG400006065M6LRR-RLK_1098

AATTGGTCAAATTCCAGAGGCTATTTTCAACATATCTTCTTTGGAAGTTATTGATTTCAGTTTCAATAACCTCTCGGGGAGAATTCCAACCACTACAGGTCTTCATCTTCCAAACGTTAA

>M6-PGSC0003DMG400006065M6LRR-RLK_1159

TTCAATAACCTCTCGGGGAGAATTCCAACCACTACAGGTCTTCATCTTCCAAACGTTAAGGAACTTATCTTGGCAGTCAATCAGCTGAAAGGGGAAATTCCATTGTTCACAACAAATGCT

>M6-PGSC0003DMG400006065M6LRR-RLK_1220

AACTTATCTTGGCAGTCAATCAGCTGAAAGGGGAAATTCCATTGTTCACAACAAATGCTTCCAAGCTTGAGAGATTGGGGCTAAATAATAACTTTCTCACAGGAACTATTCCTAATAACT

>M6-PGSC0003DMG400006065M6LRR-RLK_1281

CAAGCTTGAGAGATTGGGGCTAAATAATAACTTTCTCACAGGAACTATTCCTAATAACTTGGGAAATCTTCATGAGCTGCAAATACTGCTCCTACATACTAATCAACTTGCCAATGAACC

>M6-PGSC0003DMG400006065M6LRR-RLK_1342

GGAAATCTTCATGAGCTGCAAATACTGCTCCTACATACTAATCAACTTGCCAATGAACCAAGACAGCGTGAGTTGCTATTCTTCAATTCTTTGGCGGACTGTAGGATGTTGCGATATCTA

>M6-PGSC0003DMG400006065M6LRR-RLK_1403

GACAGCGTGAGTTGCTATTCTTCAATTCTTTGGCGGACTGTAGGATGTTGCGATATCTACAAGTGGGTAACAATCAGTTGAATGGTTTTCTGCCCAATTCTATTGGGAATCTTTCATATA

>M6-PGSC0003DMG400006065M6LRR-RLK_1464

AGTGGGTAACAATCAGTTGAATGGTTTTCTGCCCAATTCTATTGGGAATCTTTCATATACTATTCAAAACTTTTATATAGGAGATGCACACATCAATGGCCTAATCCCCACAAGTATAGG

>M6-PGSC0003DMG400006065M6LRR-RLK_1586

ACATGAGCGGTCTTACATTCCTAAGCCTTGGAGGAAACAACTTGACAGGGAGTATTCCTCCTGAGATTGGTAAGCTTAAACAACTCCAAGGGTTGCATCTAACTAACAATAAATTGCAGG

>M6-PGSC0003DMG400006065M6LRR-RLK_1647

TGAGATTGGTAAGCTTAAACAACTCCAAGGGTTGCATCTAACTAACAATAAATTGCAGGGACATATTTCTGAGGCAGTATGCAATTTATCTAATTTGGTTCAATTATATCTGGAACATAA

>M6-PGSC0003DMG400006065M6LRR-RLK_1708

CATATTTCTGAGGCAGTATGCAATTTATCTAATTTGGTTCAATTATATCTGGAACATAATGAGCTCACTGGAGTGATCCAAGAATGTATAGGAGATCTAAGCATGCTACAACTGCTTTAT

>M6-PGSC0003DMG400006065M6LRR-RLK_1769

AGCTCACTGGAGTGATCCAAGAATGTATAGGAGATCTAAGCATGCTACAACTGCTTTATTTAAATTCTAATAGATTTTCATCAAAATTTCCGTTGAGCCTTTGGAAAATGAGTGGTCTTC

>M6-PGSC0003DMG400006065M6LRR-RLK_183

CAGTTCCAAAAGGCAAAGGATTGTGGCCTTGGATCTTCCTAATTTGCAACTTCAAGGACAAATTTTCCCATCTTTGGCCAATTTGTCCTTTCTCAGAGAGCTGAATCTTGGGAACAACAG

>M6-PGSC0003DMG400006065M6LRR-RLK_1830

AAATTCTAATAGATTTTCATCAAAATTTCCGTTGAGCCTTTGGAAAATGAGTGGTCTTCTCTTTCTAAACGTGTCACAAAATTCTTTAGAGGGAGAAGTTCCATCAGATATTGGAGAACT

>M6-PGSC0003DMG400006065M6LRR-RLK_1891

TTTCTAAACGTGTCACAAAATTCTTTAGAGGGAGAAGTTCCATCAGATATTGGAGAACTTAAAGTCATTATAGCGCTAGATCTTTCCAGTAATCACTTTTCAGGAATGTTACCAAGCAGA

>M6-PGSC0003DMG400006065M6LRR-RLK_1952

AAGTCATTATAGCGCTAGATCTTTCCAGTAATCACTTTTCAGGAATGTTACCAAGCAGATTGGGAGAACTCCTAAACCTGCAGTCTCTTGTCCTATCAAACAATTCATTTTTTGGACAAA

>M6-PGSC0003DMG400006065M6LRR-RLK_2013

GGGAGAACTCCTAAACCTGCAGTCTCTTGTCCTATCAAACAATTCATTTTTTGGACAAATTCCATTATCCCTTGCCAACTTGATAAGCTTAGAATTCTTGAATTTGTCTTTAAATGCATT

>M6-PGSC0003DMG400006065M6LRR-RLK_2074

CCATTATCCCTTGCCAACTTGATAAGCTTAGAATTCTTGAATTTGTCTTTAAATGCATTGTCAGGTACTATTCCGAAGTCTTTGGAAAAACTCTTATACCTTAAAAGCATTAATGTTTCA

>M6-PGSC0003DMG400006065M6LRR-RLK_2135

CAGGTACTATTCCGAAGTCTTTGGAAAAACTCTTATACCTTAAAAGCATTAATGTTTCATTTAATGATTTAGAAGGTGTAATACCGAGTGGTGGTGTGTTTGCAAATTCCACTCTGCAAT

>M6-PGSC0003DMG400006065M6LRR-RLK_2257

TTTCTTGGGAACAAAGGTCTATGTGGAGTCCACATATTGGAGATTCCTGCTTGTGCTATCACTTATCCTGGAAAACAATCAAAGCTTAAGGAGGTTGTGCTAAAAATTGCTACTCCAGTG

>M6-PGSC0003DMG400006065M6LRR-RLK_2318

CTTATCCTGGAAAACAATCAAAGCTTAAGGAGGTTGTGCTAAAAATTGCTACTCCAGTGGTTATTTCATCCTTGATGATATTCTTGTTGGTTTCAATTTGGATAATGAAACGACAGAAGA

>M6-PGSC0003DMG400006065M6LRR-RLK_2379

TATTTCATCCTTGATGATATTCTTGTTGGTTTCAATTTGGATAATGAAACGACAGAAGAAAGGAAAGTCCAAAGATGTTGAAAAGGTACCGGAGATCAGGACTCATCAATTAGTTTCTTA

>M6-PGSC0003DMG400006065M6LRR-RLK_244

ATTTTCCCATCTTTGGCCAATTTGTCCTTTCTCAGAGAGCTGAATCTTGGGAACAACAGCTTTCACGGTGGCATCCCTTATGGACTTGGCCACTTGCCTCACTTGCAAGTGATTGATATT

>M6-PGSC0003DMG400006065M6LRR-RLK_2562

TGGCTCTGTGTACAAAGGCACATTATCTGGTGGAACTTTGGTGGCCATTAAGGTTCTGGATTTGCGAAGTGAGGAAGTATGCAAAAGGTTTGACACCGAATGCGAAGTGATGAGAAATGT

>M6-PGSC0003DMG400006065M6LRR-RLK_2623

TTGCGAAGTGAGGAAGTATGCAAAAGGTTTGACACCGAATGCGAAGTGATGAGAAATGTTAGACATAGAAATCTTGTTCCGGTGATTACTACTTGTTCTAGTGACTACATAAGAGCCTTC

>M6-PGSC0003DMG400006065M6LRR-RLK_2684

GACATAGAAATCTTGTTCCGGTGATTACTACTTGTTCTAGTGACTACATAAGAGCCTTCGTTCTGCAATATATGTCAAATGGGAGTCTTGAGAATTGGTTGTACAGAGAAGAGTTCAACT

>M6-PGSC0003DMG400006065M6LRR-RLK_2745

TCTGCAATATATGTCAAATGGGAGTCTTGAGAATTGGTTGTACAGAGAAGAGTTCAACTTAAACCTTCTTCAAAGAGTCGCCATAATGCTTGATGTGGCTGTGGCAATTGAATATCTACA

>M6-PGSC0003DMG400006065M6LRR-RLK_2806

AACCTTCTTCAAAGAGTCGCCATAATGCTTGATGTGGCTGTGGCAATTGAATATCTACACCATGGGCATAACACTCCAATAGTTCATTGCGACGTAAAGCCAGCCAACGTTCTTTTGGAT

>M6-PGSC0003DMG400006065M6LRR-RLK_2989

ATGGCTTATACAGAGACATTGGGCACTCTTGGATACATTGCACCAGGTATAAAAAAAATCTTTTCTCTTTGATTTTCTCTTATCATAATTAAGCCTCTCCAAGTTCTAGAAGTAAAAAGA

>M6-PGSC0003DMG400006065M6LRR-RLK_3050

TTTCTCTTTGATTTTCTCTTATCATAATTAAGCCTCTCCAAGTTCTAGAAGTAAAAAGAGCAAGTCTTTATTTATGCAGAATTATTGTTGTATTTCAATTGAGTAACTTTTCTTCAATCC

>M6-PGSC0003DMG400006065M6LRR-RLK_3111

AAGTCTTTATTTATGCAGAATTATTGTTGTATTTCAATTGAGTAACTTTTCTTCAATCCTTTCCTAAGAATATGGCTTGGAGGGAATAGTGTCTGCTAGTGGTGATGTTTATAGCTACGG

>M6-PGSC0003DMG400006065M6LRR-RLK_3233

TCATGTTGATGGAGATTTTGACCAAAAGAAGGCCAACAGATGAAGAGATATGCAACCAAAATCTTGACTTGAGGAAATGGATCACACAATCATTTTCAGGGACTATGATGGATGTTGTGG

>M6-PGSC0003DMG400006065M6LRR-RLK_366

AAACAATCAGCTTAACGGAAGTATACCAACAAGTCTATTTCAACACCGGAGAGTTCAGAAGATTTCATTGGCTTTCAATAAACTCGGTGGTGAAATGTGGAAAGGTCCATGGTATGTACC

>M6-PGSC0003DMG400006065M6LRR-RLK_427

ATTTCATTGGCTTTCAATAAACTCGGTGGTGAAATGTGGAAAGGTCCATGGTATGTACCCAAACTCAAAGTCTTTGATCTCAGGAACAATAGCCTCACAGGTATGATCCCTCCTTCTGTT

>M6-PGSC0003DMG400006065M6LRR-RLK_488

AACTCAAAGTCTTTGATCTCAGGAACAATAGCCTCACAGGTATGATCCCTCCTTCTGTTGGAAATGCAACAAAGTTGCTGAACTTCAGTGGGAATAGAGTCTGCGGCAACATTCCAAAGG

>M6-PGSC0003DMG400006065M6LRR-RLK_549

AAATGCAACAAAGTTGCTGAACTTCAGTGGGAATAGAGTCTGCGGCAACATTCCAAAGGAAATTGGTAATCTCAGCAAACTTGCATATTTGTCCTTGGCCGATAATCAATTAACAGGTTC

>M6-PGSC0003DMG400006065M6LRR-RLK_61

TCAAATGAGACTGACCAACAAGCTTTACTAGCTTTCCAAAATCTTATTACAAGTCCGAGTCATTTTTTGGCCAGTAATTGGACCAAAAATATTTCTTTTTGCTCTTGGTTTGGTGTCACT

>M6-PGSC0003DMG400006065M6LRR-RLK_610

ATTGGTAATCTCAGCAAACTTGCATATTTGTCCTTGGCCGATAATCAATTAACAGGTTCCATTCCCGCAGTACTGTTTAATATCTCGTCGCTACTTGAAGTGCATCTGGGAATCAATAGC

>M6-PGSC0003DMG400006065M6LRR-RLK_671

TTCCCGCAGTACTGTTTAATATCTCGTCGCTACTTGAAGTGCATCTGGGAATCAATAGCCTTTCTGGTCCTCTCTTGCTAGATGAAGGGAATGCTGTGTCAAATCTGAAGTTTTTAAGTC

>M6-PGSC0003DMG400006065M6LRR-RLK_732

TTCTGGTCCTCTCTTGCTAGATGAAGGGAATGCTGTGTCAAATCTGAAGTTTTTAAGTCTATCTAACAACCAAATTTCTGGTTGCATTCCTTCCAACATATGCCAACTCACAGAGCTCGA

>M6-PGSC0003DMG400006065M6LRR-RLK_854

TTTTGTCCATATCTTACAACAACATAACTGGAGAAATACCCAGAAATATTGATTGTTTATCCAAGCTCAAAAAGTTCTTAATTGGTGATAATCCAACAAAAGGGACTATTCCCACATCAT

>M6-PGSC0003DMG400006065M6LRR-RLK_915

CAAGCTCAAAAAGTTCTTAATTGGTGATAATCCAACAAAAGGGACTATTCCCACATCATTGGGAAATATTTCCACTCTGCAATATCTTTATTGTGGAAACAATCGCATAGTGGGGCAAAT

>M6-PGSC0003DMG400006065M6LRR-RLK_976

GGAAATATTTCCACTCTGCAATATCTTTATTGTGGAAACAATCGCATAGTGGGGCAAATTCCTCCGGAATTAGGGAAGCTATCAAATTTGCAGCAATTATGCTTTAGGAATAATTATAAT

>M6-PGSC0003DMG400006076M6LRR-RLP_0

AATTCCCCTTTCAAGCCAAAAACATGGCCGCCACCGCTGACATATTCTCGGAATGTGTGATTCACAAAATACTCTGTTATCTTAGCTATGGCCAAGCATCCCGGATGAGAATTCTATCTA

>M6-PGSC0003DMG400006076M6LRR-RLP_1054

CCAAGTTCTCCTTTGTTTTCCCGAATACAATGAAATCAAGATTAAAGATTTGCAGCAGAACCAAGGAGTTGCTAGCCCACAAATGGACGTTTTAAATGTCAACATTACAAGGTCATATGG

>M6-PGSC0003DMG400006076M6LRR-RLP_1116

AAGGAGTTGCTAGCCCACAAATGGACGTTTTAAATGTCAACATTACAAGGTCATATGGGGAGTGACCAACATTTGTGGATGCTTTGCTATGGAGTTGTAATCCTAGGAGACTCAACCTAT

>M6-PGSC0003DMG400006076M6LRR-RLP_124

ATGGCTACAAGCTTGGTCTACTCTTCCCAACTTGGAATTCTTAGTTGAATATTTCGAAGAAACAAAAATAGTGGACGCAATTATGGAGACATACAGGGAAAACAAAATCCCCATTGACAA

>M6-PGSC0003DMG400006076M6LRR-RLP_1240

AACGGTTGAAATGATTACATGTTTCGTCAGTCGTTTGATGTATATGAAGAATCTGAGTCATTCTTCTTCGGAAAGAAGCAACCCTTGGCATAGTCAATTGAAAGAAATAAAAGTTTTTGA

>M6-PGSC0003DMG400006076M6LRR-RLP_1302

CTTCTTCGGAAAGAAGCAACCCTTGGCATAGTCAATTGAAAGAAATAAAAGTTTTTGATGGGAAAAATCAAGAGCTGCAACTCGTAAGTGGGGAACTGGAGATAAGCACCCTTTCGGAGG

>M6-PGSC0003DMG400006076M6LRR-RLP_1364

AAAAATCAAGAGCTGCAACTCGTAAGTGGGGAACTGGAGATAAGCACCCTTTCGGAGGACAGGAAGGAAGTTTATTTTTTATTAGATTGGCAATGCAATTAATTTATCACTATTTCAAAA

>M6-PGSC0003DMG400006076M6LRR-RLP_1426

GAAGGAAGTTTATTTTTTATTAGATTGGCAATGCAATTAATTTATCACTATTTCAAAACGAATGCACTTGTTCATTCCTAAACTTTATTAATCTGTTTTATGTCTAGCTTTAAATTCTTG

>M6-PGSC0003DMG400006076M6LRR-RLP_1479

CAAAACGAATGCACTTGTTCATTCCTAAACTTTATTAATCTGTTTTATGTCTAGCTTTAAATTCTTGCACTTTCTCACCATTGTATAAAATTGTGTTGAAGTTGAAGCATAAGATTCTTT

>M6-PGSC0003DMG400006076M6LRR-RLP_186

CAAAAATAGTGGACGCAATTATGGAGACATACAGGGAAAACAAAATCCCCATTGACAAGTTTGGATTTTCAAATTGTTCGAAATTCAATCGTCGTCATCAAGTTTTCCCTCTGATTGATA

>M6-PGSC0003DMG400006076M6LRR-RLP_248

GGATTTTCAAATTGTTCGAAATTCAATCGTCGTCATCAAGTTTTCCCTCTGATTGATAAGTGGCTTGACACAGCTCTTCAAAATGGCGCAAGAGATTTAGTCTATGTAGATCTTCTCTAC

>M6-PGSC0003DMG400006076M6LRR-RLP_310

GCTTGACACAGCTCTTCAAAATGGCGCAAGAGATTTAGTCTATGTAGATCTTCTCTACTCGGTACGATTATATCCTTTTCCTATTTCTAAAGTCTTGGCTGCAAATTCTTTAAGAGAATT

>M6-PGSC0003DMG400006076M6LRR-RLP_372

TACGATTATATCCTTTTCCTATTTCTAAAGTCTTGGCTGCAAATTCTTTAAGAGAATTAGTTCTGAAAGGTTGTAACCTTATGCATGTTTCGTCATCTAGTGGGTTCGCGAATTGTCATT

>M6-PGSC0003DMG400006076M6LRR-RLP_434

CTGAAAGGTTGTAACCTTATGCATGTTTCGTCATCTAGTGGGTTCGCGAATTGTCATTCTTTGAGAAAGCTTTCTCTATCTTATATAATTTTAAGCAAAAATTTGCTTCAGACTTTACTT

>M6-PGSC0003DMG400006076M6LRR-RLP_496

GAGAAAGCTTTCTCTATCTTATATAATTTTAAGCAAAAATTTGCTTCAGACTTTACTTAATAGTTGTCCTTTCATTGTCTCTTTCAACCTTGATCATTGTTCGGGGTTGGTAAAGATTGA

>M6-PGSC0003DMG400006076M6LRR-RLP_558

GTTGTCCTTTCATTGTCTCTTTCAACCTTGATCATTGTTCGGGGTTGGTAAAGATTGAGCTTATGAATCTTCAAAAGATCAAGTCAGTTTCCGTATCAATAAATAGAAACCAACATGTGA

>M6-PGSC0003DMG400006076M6LRR-RLP_62

CACAAAATACTCTGTTATCTTAGCTATGGCCAAGCATCCCGGATGAGAATTCTATCTAAAACATGGCTACAAGCTTGGTCTACTCTTCCCAACTTGGAATTCTTAGTTGAATATTTCGAA

>M6-PGSC0003DMG400006076M6LRR-RLP_620

ATGAATCTTCAAAAGATCAAGTCAGTTTCCGTATCAATAAATAGAAACCAACATGTGAAAATCCAAGCACCAACTCTTGAACACTTGTTTTATAATGGATATTTATCGGGAAATTTGGTT

>M6-PGSC0003DMG400006076M6LRR-RLP_682

CCAAGCACCAACTCTTGAACACTTGTTTTATAATGGATATTTATCGGGAAATTTGGTTATCGTTGGATGTCTGAATCTAAAATCTTTAGAGCTATCACATGTGAAGATATGTATTGGATT

>M6-PGSC0003DMG400006076M6LRR-RLP_744

TTGGATGTCTGAATCTAAAATCTTTAGAGCTATCACATGTGAAGATATGTATTGGATTTCTCCATAACCTAATTTCTAGATCGCAATCCCTAGAAGTATTGAAGATTCAGTATTGTGTGG

>M6-PGSC0003DMG400006076M6LRR-RLP_806

CATAACCTAATTTCTAGATCGCAATCCCTAGAAGTATTGAAGATTCAGTATTGTGTGGGCATAAGACAGATTGAAACTTCGAATTTGGTATCACTTGAGTATATAGGATATCAAATTCCT

>M6-PGSC0003DMG400006076M6LRR-RLP_868

AAGACAGATTGAAACTTCGAATTTGGTATCACTTGAGTATATAGGATATCAAATTCCTGAAATTAAAATTGCGAAAGCTTCAAGTCAATTGAAGCATTCAAAAATCGTTCTTCATGGCAA

>M6-PGSC0003DMG400006076M6LRR-RLP_930

TTAAAATTGCGAAAGCTTCAAGTCAATTGAAGCATTCAAAAATCGTTCTTCATGGCAACAACAATTTAGATGCTGCATGGTTTAGTAAGTTGAGGAAGCTTCTATCGAATTCGGCCCCTT

>M6-PGSC0003DMG400006076M6LRR-RLP_992

AATTTAGATGCTGCATGGTTTAGTAAGTTGAGGAAGCTTCTATCGAATTCGGCCCCTTGGTCCCAAGTTCTCCTTTGTTTTCCCGAATACAATGAAATCAAGATTAAAGATTTGCAGCAG

>M6-PGSC0003DMG400006086M6LysM-RLK_0

ACAAACACACATCTCTCCTCCAACAAAAAAACAAAGCAAAAAAAAATGATAACAAAACAAATATTTGGTTTTTCACTTCTTCTCTTCTTATCATTCATATTCAACCAAAACAAATGTTAT

>M6-PGSC0003DMG400006086M6LysM-RLK_1098

TCTGTGTTTAAAGGAACAATTGATGAAGAAGTGTTTGCAATCAAAAAAATGAAATGGAATGCCCGTGAAGAACTCAAAATCCTGCAAAAGGTAAAAAAAAAAAAACGTTACGTAAAATTC

>M6-PGSC0003DMG400006086M6LysM-RLK_1159

CCCGTGAAGAACTCAAAATCCTGCAAAAGGTAAAAAAAAAAAAACGTTACGTAAAATTCTACATATATACGTATCCACTCAAGAATTGCTTCTGTGTACTAGTCCTAATTTATGTGACGT

>M6-PGSC0003DMG400006086M6LysM-RLK_122

ACAAGATCCACCAAGCATAACAGGATATACTTGTAATCCCAACCAATTCAATCCATGTCAAACTTATGTGTTATACAGAGCTAGAGGACCAGAGTTTCTTGATTTAGCCTCAATAGGTGA

>M6-PGSC0003DMG400006086M6LysM-RLK_1342

ATGTCATTCTTTTCACCTTTTTTTGTTTGTGTAGGTGAACCATGGGAATTTGGTGAAGCTAGAGGGTTTTTGCATAGACCCTAAAGAAGCAAATTGCTACTTAGTCTATGAGTATGTTGA

>M6-PGSC0003DMG400006086M6LysM-RLK_1403

GAGGGTTTTTGCATAGACCCTAAAGAAGCAAATTGCTACTTAGTCTATGAGTATGTTGAAAATGGTTCACTACATTCTTGGCTTCACGGTGAAAAACCTGAAAAATTGAGCTGGAAAACA

>M6-PGSC0003DMG400006086M6LysM-RLK_1464

ATGGTTCACTACATTCTTGGCTTCACGGTGAAAAACCTGAAAAATTGAGCTGGAAAACAAGACTAAGAATTGCCACTGATGTTGCAAATGGTCTCTTATACATCCATGAACATACAAGGC

>M6-PGSC0003DMG400006086M6LysM-RLK_1525

ACTAAGAATTGCCACTGATGTTGCAAATGGTCTCTTATACATCCATGAACATACAAGGCCAAGAGTTGTCCACAAAGACATCAAGAGTAGCAACATTCTCCTAGACTCCAACATGAGAGC

>M6-PGSC0003DMG400006086M6LysM-RLK_183

ACTTATGTGTTATACAGAGCTAGAGGACCAGAGTTTCTTGATTTAGCCTCAATAGGTGATCTTTTTTCAGTAAGTAGGGTAATGATTGCAAACCCAAGTAATATTTCCAGCCCAAACACA

>M6-PGSC0003DMG400006086M6LysM-RLK_1891

GAGGAAATTGCAAGAATGGATGGATGAAAGTCTTTTGAGGGAAGAATTAACAATGGAGAGTGTTATGAATGTGATGTCTGTGGCTATTTCATGTTTGAATAAAGATCCTTCAAGAAGGCC

>M6-PGSC0003DMG400006086M6LysM-RLK_1952

GTTATGAATGTGATGTCTGTGGCTATTTCATGTTTGAATAAAGATCCTTCAAGAAGGCCAGGGATGATTGAAATTGTGTATGCCTTATCTAAAAGTATTGATCTATTTTCTGATGTTTCA

>M6-PGSC0003DMG400006086M6LysM-RLK_2013

GGATGATTGAAATTGTGTATGCCTTATCTAAAAGTATTGATCTATTTTCTGATGTTTCAGAGGAAGGACAATCTCCAAGGCAAGTCACAGCAAGATAGACTACATAATAGTACTAGCATA

>M6-PGSC0003DMG400006086M6LysM-RLK_2074

GGAAGGACAATCTCCAAGGCAAGTCACAGCAAGATAGACTACATAATAGTACTAGCATACTATGTAAAGTTGTTTTGAAGTGGAGTAGCATTTTGAGTTTTGAGGTTTCTACTTAACTTT

>M6-PGSC0003DMG400006086M6LysM-RLK_2135

ATGTAAAGTTGTTTTGAAGTGGAGTAGCATTTTGAGTTTTGAGGTTTCTACTTAACTTTAAAGTTTGCTTCTATATATGTGTAATATTACTCCTATATTATCGTTTATAGTAGTAGTGGT

>M6-PGSC0003DMG400006086M6LysM-RLK_2196

AGTTTGCTTCTATATATGTGTAATATTACTCCTATATTATCGTTTATAGTAGTAGTGGTAAATCCAATTTCGAATAAGAGCATGTTTAAGGTTGTTCTAGTGGAATATATATTAGTACCA

>M6-PGSC0003DMG400006086M6LysM-RLK_2253

GTAAATCCAATTTCGAATAAGAGCATGTTTAAGGTTGTTCTAGTGGAATATATATTAGTACCACATAAGACTCTCGTAATGAATTTATTTTTATGTTATGTTGCTTCGATTCTTCAAAAA

>M6-PGSC0003DMG400006086M6LysM-RLK_244

TTTTTTCAGTAAGTAGGGTAATGATTGCAAACCCAAGTAATATTTCCAGCCCAAACACAACTCTTGTTAATGATCAACTCCTTTTTATCCCCATCACATGTTCTTGCAATAACATCAACA

>M6-PGSC0003DMG400006086M6LysM-RLK_305

TCTTGTTAATGATCAACTCCTTTTTATCCCCATCACATGTTCTTGCAATAACATCAACACAACTTTTGGTAGCATATCTTATGCAGGCTTAAACTACAGTTTCAAACCAGGTGACACTAT

>M6-PGSC0003DMG400006086M6LysM-RLK_366

ACTTTTGGTAGCATATCTTATGCAGGCTTAAACTACAGTTTCAAACCAGGTGACACTATGTATGATATGTCAATATCCAAATACCAAAATCTTACTACTTATCAATCTGTTGAGGCTGTT

>M6-PGSC0003DMG400006086M6LysM-RLK_427

ATGATATGTCAATATCCAAATACCAAAATCTTACTACTTATCAATCTGTTGAGGCTGTTAATCCCACAGTTGTAGCAACCAACATTGATGTAGGCCAGGCCATAAAATTCCCAATTTTTT

>M6-PGSC0003DMG400006086M6LysM-RLK_549

AAGTGTCCAAACACGACACAAAATCAACCAAGACTACTCATAAGTTATGTTTTTCAACCTAATGATAACATTTCTTCTATCGCATCAAGATTCGGGATAACCCCACAATCCATAACACAA

>M6-PGSC0003DMG400006086M6LysM-RLK_61

TATTTGGTTTTTCACTTCTTCTCTTCTTATCATTCATATTCAACCAAAACAAATGTTATGCACAAGATCCACCAAGCATAACAGGATATACTTGTAATCCCAACCAATTCAATCCATGTC

>M6-PGSC0003DMG400006086M6LysM-RLK_610

ATGATAACATTTCTTCTATCGCATCAAGATTCGGGATAACCCCACAATCCATAACACAAATTAATGGAAATAATACCAATATTCTTGATACCCTTTTCATCCCTCTTTCTAATCTACCTA

>M6-PGSC0003DMG400006086M6LysM-RLK_793

AAGAAAATGACAGAAAAGGGACTGTGATAGGGTTAGCAATTGGTTTAGGGATTTGTGGGCTGCTGTTGGTTTTGGTTCTTGGTTTGTTTTACAAGGAGAAAACAGTAAAGAAAGAAAAGT

>M6-PGSC0003DMG400006086M6LysM-RLK_854

GCTGTTGGTTTTGGTTCTTGGTTTGTTTTACAAGGAGAAAACAGTAAAGAAAGAAAAGTATAGTGATGTAGAAAGGCAAAAAAGTTTGTATCTTGGATCAAAAAAGAAGTCTTTTGTTGA

>M6-PGSC0003DMG400006086M6LysM-RLK_915

AGTGATGTAGAAAGGCAAAAAAGTTTGTATCTTGGATCAAAAAAGAAGTCTTTTGTTGATAAAGATGTTGAAGTGAATTTGTTGGCTGATTTATCAGAGTGTTTGGATAAGTATAAAATG

>M6-PGSC0003DMG400006086M6LysM-RLK_976

AAGATGTTGAAGTGAATTTGTTGGCTGATTTATCAGAGTGTTTGGATAAGTATAAAATGTATAAAATGGTACAACTTTGGGAAGCAACAAATGGATTTGATGAAGGGTGTTTGATTCAAG

>M6-PGSC0003DMG400006087M6LysM-RLK_1827

AGGGAAAGTTTTGTGGGCAAAAACTAGCGATTTTTCGGAAGGGAGTGAAGAGAGGAAGGTGAGGAAATTGCAAGAATGGATGGATGAAAGTCTTTTGAGGGAAGAATTAACAATGGAGAG

>M6-PGSC0003DMG400006087M6LysM-RLK_504

ACATTGATGTAGGCCAGGCCATAAAATTCCCAATTTTTTGTAAGTGTCCAAACACGACACAAAATCAACCAAGACTACTCATAAGTTATGTTTTTCAACCTAATGATAACATTTCTTCTA

>M6-PGSC0003DMG400006088M6LysM-RLK_0

AAGGAAATGAGGGAATTCACAATTAGAGGTACTATTTTTGTAGATTGATTGGTTTTGATTTGTTTCTATAATTTGGTGTAGATTTTTTATTTTTTTTAAGATATGAAGTGTCTGTATGCA

>M6-PGSC0003DMG400006088M6LysM-RLK_1178

AATTATTGTTTAAATTGTCTATTGAAAATAGTATTTCTACCTCTAAGGTAATGATAAGTTTTGTGTATATTTTAACCTCTTCAGATCGTACTTTATGGAATTACATGAGTATGTTATTGT

>M6-PGSC0003DMG400006088M6LysM-RLK_124

TTTCTGTTGTATATTTTAGATTTAATAAGATCTTTTTAGTGTTTATGATTAAAATATTTATTATTTTTGTCAAATTTAATAAACATAAATTATTAAGTATTGACGAAGATTAAAAATGTT

>M6-PGSC0003DMG400006088M6LysM-RLK_1240

GTGTATATTTTAACCTCTTCAGATCGTACTTTATGGAATTACATGAGTATGTTATTGTTATATTGTTCCAAATTAATTATGTCTACTTAATAATTTCCCCCTTTTGTTTTGTGTAGGTGA

>M6-PGSC0003DMG400006088M6LysM-RLK_1302

TTGTTCCAAATTAATTATGTCTACTTAATAATTTCCCCCTTTTGTTTTGTGTAGGTGAACCATGGGAATTTGGTGAAGCTAGAAGGTTTTTGCATAGACCCTAAAGAAGCAAATTGCTAC

>M6-PGSC0003DMG400006088M6LysM-RLK_1364

TGGGAATTTGGTGAAGCTAGAAGGTTTTTGCATAGACCCTAAAGAAGCAAATTGCTACTTAGTCTATGAGTATGTTGAAAATGGTTCACTACATTCATGGATTCATGGTGAAAAACCTGA

>M6-PGSC0003DMG400006088M6LysM-RLK_1426

TCTATGAGTATGTTGAAAATGGTTCACTACATTCATGGATTCATGGTGAAAAACCTGAAAAATTGAGTTGGAAAACAAGACTAAGAATTGCAACTGATGTTGCAAATGGTCTATTATACA

>M6-PGSC0003DMG400006088M6LysM-RLK_1488

TTGAGTTGGAAAACAAGACTAAGAATTGCAACTGATGTTGCAAATGGTCTATTATACATTCATGAACATACAAGGCCAAGAGTTGTCCAAAAAGACATCAAAACTAGAAACATTCTCCTA

>M6-PGSC0003DMG400006088M6LysM-RLK_1550

TGAACATACAAGGCCAAGAGTTGTCCAAAAAGACATCAAAACTAGAAACATTCTCCTAGACTCCAACATGAGAGCCAAAGTTGCCAATTTTGGTCTAGCTAAATCAGGATGCAATGCTAT

>M6-PGSC0003DMG400006088M6LysM-RLK_1674

ATGCACATTGTAGGTACTCAGGGGTACATCGCCCCTGAGTACCTAACCGATGGGATTGTATCCACTAAGATGGATGTTTTCTCATTAGGAGTTGTGTTGCTCGAGCTAGTGTCAGGGAAG

>M6-PGSC0003DMG400006088M6LysM-RLK_1736

CACTAAGATGGATGTTTTCTCATTAGGAGTTGTGTTGCTCGAGCTAGTGTCAGGGAAGGAGGCTATAGAAGATGAAGGGAAAGTATTGTGGGCAAAAAATAACGATTTTTCGGAAGGGAG

>M6-PGSC0003DMG400006088M6LysM-RLK_1798

CTATAGAAGATGAAGGGAAAGTATTGTGGGCAAAAAATAACGATTTTTCGGAAGGGAGTGAAGAGAGGAAGGTGAGGAAATTGCAAGAATGGATGGATGATAGTCTTTTGAGGGAAGAGT

>M6-PGSC0003DMG400006088M6LysM-RLK_186

ATTTTTGTCAAATTTAATAAACATAAATTATTAAGTATTGACGAAGATTAAAAATGTTAAACTTTTGATAAACTTAAAGAATCAAAATTCAAAACTGTTGATTACATATTTAAGGAACTA

>M6-PGSC0003DMG400006088M6LysM-RLK_1860

GAGAGGAAGGTGAGGAAATTGCAAGAATGGATGGATGATAGTCTTTTGAGGGAAGAGTTAACAATGGAGAGTGTTGTGAATGTGATGTCTGTGGCTATTTCTTGTTTGAATAAAGATCCA

>M6-PGSC0003DMG400006088M6LysM-RLK_1922

AATGGAGAGTGTTGTGAATGTGATGTCTGTGGCTATTTCTTGTTTGAATAAAGATCCATCAAGAAGGCCAGGGATGATTGAAATTGTGTATGCATTGTCTAAAAGTATTGATCTATTTAC

>M6-PGSC0003DMG400006088M6LysM-RLK_1984

GAAGGCCAGGGATGATTGAAATTGTGTATGCATTGTCTAAAAGTATTGATCTATTTACTGATGTCTCAGAGGAAGGACTATCTCCAAGGCAAGTCACAGCAAGATAGGGGACATATCTAT

>M6-PGSC0003DMG400006088M6LysM-RLK_2046

GTCTCAGAGGAAGGACTATCTCCAAGGCAAGTCACAGCAAGATAGGGGACATATCTATTTAGCATATTATGTAAAAGTAGTTTTGAAGTGAGTAGTATTATTGTGTTTTGAGGTTTTTAC

>M6-PGSC0003DMG400006088M6LysM-RLK_2108

CATATTATGTAAAAGTAGTTTTGAAGTGAGTAGTATTATTGTGTTTTGAGGTTTTTACTTTACTTTAAAGTTTGCTTCTATATATGTGTAATATTACTCCAATATTATCTTTTCTAGTAG

>M6-PGSC0003DMG400006088M6LysM-RLK_2170

CTTTAAAGTTTGCTTCTATATATGTGTAATATTACTCCAATATTATCTTTTCTAGTAGTTGTGGAAAATCCAATTTCAAATAAGAGGATGTTTAAGGTTGGTCAAGTGGAATATATATTA

>M6-PGSC0003DMG400006088M6LysM-RLK_2227

GTTGTGGAAAATCCAATTTCAAATAAGAGGATGTTTAAGGTTGGTCAAGTGGAATATATATTATGTTGTAATGAGTGTTTTTATGTTATGTTGCTCGGATTCTTTAAAATTGTATAACTT

>M6-PGSC0003DMG400006088M6LysM-RLK_248

TTTTGATAAACTTAAAGAATCAAAATTCAAAACTGTTGATTACATATTTAAGGAACTATTTTGAACCTGTTCAGTGTTCTCCCACCTTAGAGGCTACTTTGGTCATTTTCTCAATTGATA

>M6-PGSC0003DMG400006088M6LysM-RLK_310

GAACCTGTTCAGTGTTCTCCCACCTTAGAGGCTACTTTGGTCATTTTCTCAATTGATATTTTAGTAATGTTAAAATTCCACTGCAGAATTGTTTTTCCTCCAATGGATCAGATTAAGTAT

>M6-PGSC0003DMG400006088M6LysM-RLK_372

AGTAATGTTAAAATTCCACTGCAGAATTGTTTTTCCTCCAATGGATCAGATTAAGTATTTAATTTTAATAAAATATTTAACAATTTAAATTATTAGGTCTAGTGACCTGTCGTATTCTTG

>M6-PGSC0003DMG400006088M6LysM-RLK_434

TTTTAATAAAATATTTAACAATTTAAATTATTAGGTCTAGTGACCTGTCGTATTCTTGAGACATTAGTACTTTATTTAAATATTTTAAGTGGACCATTATGGCCGTTGGGATTTGATAAT

>M6-PGSC0003DMG400006088M6LysM-RLK_496

ATTAGTACTTTATTTAAATATTTTAAGTGGACCATTATGGCCGTTGGGATTTGATAATTTATAATTTAAGCTAATCAACATCATTTAATACTCAATTTATTTGTCCTCTGATGTCTATTT

>M6-PGSC0003DMG400006088M6LysM-RLK_558

AATTTAAGCTAATCAACATCATTTAATACTCAATTTATTTGTCCTCTGATGTCTATTTTCATTAAATACTCCTCTTTTTGTTTCTCAATTATTCACTTTTTGTTTTCCTTTTTATTTTTG

>M6-PGSC0003DMG400006088M6LysM-RLK_62

TTTCTATAATTTGGTGTAGATTTTTTATTTTTTTTAAGATATGAAGTGTCTGTATGCAATGTTTTCTGTTGTATATTTTAGATTTAATAAGATCTTTTTAGTGTTTATGATTAAAATATT

>M6-PGSC0003DMG400006088M6LysM-RLK_620

TAAATACTCCTCTTTTTGTTTCTCAATTATTCACTTTTTGTTTTCCTTTTTATTTTTGTCCCTAAGGAGACCCTCCCATATGTGCCATCGGAGTCGAGTGTATATCAAAAATAATTTTTA

>M6-PGSC0003DMG400006088M6LysM-RLK_682

TAAGGAGACCCTCCCATATGTGCCATCGGAGTCGAGTGTATATCAAAAATAATTTTTATATATATTATCCTTTTCATAACTTACTTATAAAGATTTTACTAGATATATTGTTGCTGTGAT

>M6-PGSC0003DMG400006088M6LysM-RLK_744

TATTATCCTTTTCATAACTTACTTATAAAGATTTTACTAGATATATTGTTGCTGTGATAAGAATTGAGATATTAAAAATTAACAAGATGTCTAGAAAGATTTTTTTTTCTTTTTTTGCCA

>M6-PGSC0003DMG400006088M6LysM-RLK_806

ATTGAGATATTAAAAATTAACAAGATGTCTAGAAAGATTTTTTTTTCTTTTTTTGCCATAGAAAAGAGCATCTTTTATTAATAATTTATTTTATCAAGACAAAGTTCAAAATAAATTTTC

>M6-PGSC0003DMG400006088M6LysM-RLK_868

AAAGAGCATCTTTTATTAATAATTTATTTTATCAAGACAAAGTTCAAAATAAATTTTCTTCCTAAATGAAATTTCTTGTCTGTGACCTCAATTTTGTTAGTACCCTGCCTGCTTTCTCCC

>M6-PGSC0003DMG400006088M6LysM-RLK_930

TAAATGAAATTTCTTGTCTGTGACCTCAATTTTGTTAGTACCCTGCCTGCTTTCTCCCAAAAAAGCGTCATGCTTTTAGCTTGGTTTGGCTGTACTCTTTTGACTAGATTTATTAGTTGT

>M6-PGSC0003DMG400006088M6LysM-RLK_992

AAGCGTCATGCTTTTAGCTTGGTTTGGCTGTACTCTTTTGACTAGATTTATTAGTTGTCCTATTAAATAAATAAAACGTCATCAAGTCCTATTTTGCAGCCGTTACTATAATTAGTACTT

>M6-PGSC0003DMG400006095-(rev)M6LRR-RLP_0

ATGGGGACAGCTGACATGTTGCCTATATGCCTCATTCGCAAAATACTCTGTTATCTTAGTTATGAAGAAGCATCTCGGATGAGAATTCTCTCCAAAACATGGCTACTAGCCTGGTTGGAT

>M6-PGSC0003DMG400006095-(rev)M6LRR-RLP_1116

ATCAACATGATAGATTTGCTACTGCAACCTAGAGTTGCTATCCCCAAAGTGGACATTTTAGATGTAAATTTTTCAAGGCCGAATTGGGAGTTTCCTACATTTGTGGATTCTTTGCTGTGG

>M6-PGSC0003DMG400006095-(rev)M6LRR-RLP_1178

TGTAAATTTTTCAAGGCCGAATTGGGAGTTTCCTACATTTGTGGATTCTTTGCTGTGGAGTTGTCATCCCAGGAAACTTAACCTACAATCAACAACTACAATTTTTACATGTTTCATGAA

>M6-PGSC0003DMG400006095-(rev)M6LRR-RLP_124

CCAACTTGGAATTCACAGTCTCGTCGGGCAAAACCATAACAATAGTGGATCATGTCATGGAGAGATATAATAACGGAAATATTCCTATTGACAAGTTTGAATTACACAACTCTACATAGT

>M6-PGSC0003DMG400006095-(rev)M6LRR-RLP_1240

GTCATCCCAGGAAACTTAACCTACAATCAACAACTACAATTTTTACATGTTTCATGAATCGTTTCATGCAGATGAAGAATTTGCGTCGTGGTCAATTAAAAGAAGCAAATGCCTACAAAT

>M6-PGSC0003DMG400006095-(rev)M6LRR-RLP_1302

TTCATGCAGATGAAGAATTTGCGTCGTGGTCAATTAAAAGAAGCAAATGCCTACAAATTTGATCAGAAAAGTCATAGTTGGCATCCTGTAGAACACAAAAGAGGAAGAGATTTCCCATTA

>M6-PGSC0003DMG400006095-(rev)M6LRR-RLP_1350

GCCTACAAATTTGATCAGAAAAGTCATAGTTGGCATCCTGTAGAACACAAAAGAGGAAGAGATTTCCCATTAAGGACCAGTAGCAAGTCGAAGAAATATTTCTTCTTATTAGATTGGTGA

>M6-PGSC0003DMG400006095-(rev)M6LRR-RLP_186

AGATATAATAACGGAAATATTCCTATTGACAAGTTTGAATTACACAACTCTACATAGTCTGATCATATTTTTCCTCTGATTGATAGGTGGCTCGGTATTGCTCTTCAAAATGGTGTAAGA

>M6-PGSC0003DMG400006095-(rev)M6LRR-RLP_248

TCATATTTTTCCTCTGATTGATAGGTGGCTCGGTATTGCTCTTCAAAATGGTGTAAGACATCTAGAGTACATAGATGTTGAACTTTCAACATCATATCCCTTCCCTATTTTTAAATTCTT

>M6-PGSC0003DMG400006095-(rev)M6LRR-RLP_310

TAGAGTACATAGATGTTGAACTTTCAACATCATATCCCTTCCCTATTTTTAAATTCTTGGAAGCAAATTTTTTACGAGAACTGGTCTTGATGGGTTGTGATCTGATGCATGTTTCGTTAT

>M6-PGSC0003DMG400006095-(rev)M6LRR-RLP_372

GCAAATTTTTTACGAGAACTGGTCTTGATGGGTTGTGATCTGATGCATGTTTCGTTATCTAGTGCTAGTCAAGCGGTGACGTGTCATTCTCTGAGAAATCTTTCTCTATCTCTTGTACAC

>M6-PGSC0003DMG400006095-(rev)M6LRR-RLP_434

TGCTAGTCAAGCGGTGACGTGTCATTCTCTGAGAAATCTTTCTCTATCTCTTGTACACTTAGACAATAACATGCTTCAGACTATACTCAGTAGTTGTCCGTTCATTGTTGATTTAATCAT

>M6-PGSC0003DMG400006095-(rev)M6LRR-RLP_496

ACAATAACATGCTTCAGACTATACTCAGTAGTTGTCCGTTCATTGTTGATTTAATCATTGGGTATTGTAGTAGGCTGAAAAATATTGAGATCCGGAATCTTCCAAATATCAGGTCTCTTG

>M6-PGSC0003DMG400006095-(rev)M6LRR-RLP_558

TATTGTAGTAGGCTGAAAAATATTGAGATCCGGAATCTTCCAAATATCAGGTCTCTTGTCATTGGTAATGACCAACTCGTTAAAATCCAAGCACCGTCTCTTGAACACTTGTATTATAGT

>M6-PGSC0003DMG400006095-(rev)M6LRR-RLP_62

TGAAGAAGCATCTCGGATGAGAATTCTCTCCAAAACATGGCTACTAGCCTGGTTGGATCATCCCAACTTGGAATTCACAGTCTCGTCGGGCAAAACCATAACAATAGTGGATCATGTCAT

>M6-PGSC0003DMG400006095-(rev)M6LRR-RLP_620

TGGTAATGACCAACTCGTTAAAATCCAAGCACCGTCTCTTGAACACTTGTATTATAGTAGTTCTTGTTTGAATAAATTGAATATTGTTGAACATAAGAATCTGAAAACTTTAGATATATC

>M6-PGSC0003DMG400006095-(rev)M6LRR-RLP_682

CTTGTTTGAATAAATTGAATATTGTTGAACATAAGAATCTGAAAACTTTAGATATATCATGTACAGAGATATCTGATTTATATCTCAACCACCTTATCTCCATACCTTACTGCCTGGAGA

>M6-PGSC0003DMG400006095-(rev)M6LRR-RLP_744

ACAGAGATATCTGATTTATATCTCAACCACCTTATCTCCATACCTTACTGCCTGGAGAAACTGATATTAGCAAACATTGCTTTTGGAAGATTCGACATTTGCAGGAGTCAATCCCTAAAG

>M6-PGSC0003DMG400006095-(rev)M6LRR-RLP_806

GATATTAGCAAACATTGCTTTTGGAAGATTCGACATTTGCAGGAGTCAATCCCTAAAGGTGTTAAAGATTCATAATTGCAAGATCATAGGAGCAATTGATGCTCCAAATTTGGTTTTGCT

>M6-PGSC0003DMG400006095-(rev)M6LRR-RLP_868

TAAAGATTCATAATTGCAAGATCATAGGAGCAATTGATGCTCCAAATTTGGTTTTGCTTTAGTATAAAGGAAATGATATTCCTCAGCTTAAATTTGCTCAAGAATCAAGACAATTGAGGC

>M6-PGSC0003DMG400006095-(rev)M6LRR-RLP_930

TATAAAGGAAATGATATTCCTCAGCTTAAATTTGCTCAAGAATCAAGACAATTGAGGCAATCACAAATGATTCTTTATCCCTTGTTCAATGTAGATTCGGCATGGTTTTGTCAGTTGAGG

>M6-PGSC0003DMG400006095-(rev)M6LRR-RLP_992

ACAAATGATTCTTTATCCCTTGTTCAATGTAGATTCGGCATGGTTTTGTCAGTTGAGGAAATCTTTATCAGATTCGACGATCTCTTGGTCTCAAGTTACCATTTACTTTCACAAATGCGA

>M6-PGSC0003DMG400006226M6LRR-RLP_0

ATACAAAATTTCAAGAACACAATTACTCTTGATCCTTTTGGTGTCACAAAAACTTGGACAGGCCCTGATATTTGCAATTATAAAGGCTTTTATTGTGATAATCCACCTAATAATTCATCA

>M6-PGSC0003DMG400006226M6LRR-RLP_128

TCTTGCTTCTATAGATTTCAATGGATTTCAACTTAGTGCACCTACATTAGATGGTTTCATTGATCAACTCCCTGACTTAGCCATATTTCATGCTAATACCAACAACTTTTCAGGCATAAT

>M6-PGSC0003DMG400006226M6LRR-RLP_192

CAACTCCCTGACTTAGCCATATTTCATGCTAATACCAACAACTTTTCAGGCATAATTTCTTCGAAAATTTCAAATCTTCCATATCTTTAAGAACTTGATCTTAGTAACAACCAATTCATC

>M6-PGSC0003DMG400006226M6LRR-RLP_256

AAATTTCAAATCTTCCATATCTTTAAGAACTTGATCTTAGTAACAACCAATTCATCGGTTCATTTCCCATGTCAATTCTCAATATCAAGACTCTAACTTTCTTGGATATTCGTTACAATC

>M6-PGSC0003DMG400006226M6LRR-RLP_320

TCCCATGTCAATTCTCAATATCAAGACTCTAACTTTCTTGGATATTCGTTACAATCTCTTGACAGGAATAATCCCACCTCAAATTTTCGTACAATATCTTGATGCATTTTTTCTTAACAA

>M6-PGSC0003DMG400006226M6LRR-RLP_384

GGAATAATCCCACCTCAAATTTTCGTACAATATCTTGATGCATTTTTTCTTAACAATAACAATTTCATTCAAAAACTTCCAGACAACATTGGATCCACCTCTGCTTTTTATTTAACCTTA

>M6-PGSC0003DMG400006226M6LRR-RLP_448

TCATTCAAAAACTTCCAGACAACATTGGATCCACCTCTGCTTTTTATTTAACCTTAGCCAACAACAAATTCGTTGGTCCAATCCCACGTAGCATTGGACAAGCTCCGAATTTATTGGAAA

>M6-PGSC0003DMG400006226M6LRR-RLP_512

CAAATTCGTTGGTCCAATCCCACGTAGCATTGGACAAGCTCCGAATTTATTGGAAATTTTATTCTTGAACAACTTGCTCACTGGTTGCATTCCTTATGAATTAGGCCTTTTGAAAAAAGC

>M6-PGSC0003DMG400006226M6LRR-RLP_64

CTGATATTTGCAATTATAAAGGCTTTTATTGTGATAATCCACCTAATAATTCATCAGCAATTTCTCTTGCTTCTATAGATTTCAATGGATTTCAACTTAGTGCACCTACATTAGATGGTT

>M6-PGSC0003DMG400006255M6LRR-RLP_0

ATGAATGAAGCAAGACCATACCAAAGACGAAACTTCCATAAGTCAAAGTCTTCCACAAATGAACCAAAAAGGCTAACATCAAACGAGTAAAAGGAGTTAATTCAATCTGTACAAACGCGT

>M6-PGSC0003DMG400006255M6LRR-RLP_1037

TTTCCTGAAAAGATCTTTCAGGTACAAGTTTTAGAGATTTTGGACTTGTCAAATAACAAGTTGCTTAGTGGTAGTATTCCGAGTTTTCCTCGAAATGGATCTTTGAGGACGATATCACTA

>M6-PGSC0003DMG400006255M6LRR-RLP_1098

TGCTTAGTGGTAGTATTCCGAGTTTTCCTCGAAATGGATCTTTGAGGACGATATCACTAAGCAACACCAACTTTTCGGGTTCATTACCAGAGTCCATTTCGAACCTTCAGAACCTGTCCA

>M6-PGSC0003DMG400006255M6LRR-RLP_1159

CAACACCAACTTTTCGGGTTCATTACCAGAGTCCATTTCGAACCTTCAGAACCTGTCCAGGTTAGAGCTTTCCGATTGCAATTTCAATGGATCAATACCTTCCACGATAGCAAACCTTAC

>M6-PGSC0003DMG400006255M6LRR-RLP_122

TTTTTCTTTGACATTTCTCAATCACCCTTCCCCTTATTAAGTTCCTTCTAACTGCATACTTCAGTTATATTCAAGCTAACAAGTTGCAAGAATATGACAACTCTTTACTTTCTATGGCTT

>M6-PGSC0003DMG400006255M6LRR-RLP_1220

TTAGAGCTTTCCGATTGCAATTTCAATGGATCAATACCTTCCACGATAGCAAACCTTACAAATCTTGTTTATTTAGATTTTTCCTTCAACAACTTAACTGGTTCTATCCCATATTTTCAA

>M6-PGSC0003DMG400006255M6LRR-RLP_1281

ATCTTGTTTATTTAGATTTTTCCTTCAACAACTTAACTGGTTCTATCCCATATTTTCAAAGATGCAAGAAACTCACCCACTTAGATCTCTCTCGTAATGGTTTAACTGGTCTCCTCTCCC

>M6-PGSC0003DMG400006255M6LRR-RLP_1342

ATGCAAGAAACTCACCCACTTAGATCTCTCTCGTAATGGTTTAACTGGTCTCCTCTCCCCGGCTCATTTTGAAGGACTCTCAGAGCTTGTCTACATAAATTTAGGGAACAACTTACTCAA

>M6-PGSC0003DMG400006255M6LRR-RLP_1403

GCTCATTTTGAAGGACTCTCAGAGCTTGTCTACATAAATTTAGGGAACAACTTACTCAACGGGATCCTTCCTGAATGTATCTTTGAGCTTCCCTCCTTGCAGACGCTTTTACTTAACAGC

>M6-PGSC0003DMG400006255M6LRR-RLP_1464

GGATCCTTCCTGAATGTATCTTTGAGCTTCCCTCCTTGCAGACGCTTTTACTTAACAGCAATCAATTTGTTGGCCAAGTCAACCATTTTCACAATGCATCCTCCTTTCTCGATGAAATTG

>M6-PGSC0003DMG400006255M6LRR-RLP_1525

TCAATTTGTTGGCCAAGTCAACCATTTTCACAATGCATCCTCCTTTCTCGATGAAATTGATTTGAGCAACAACCAACTAAATGGTTCAATTCCCAAGTCCATGTTTGACGTTGGGAGGCT

>M6-PGSC0003DMG400006255M6LRR-RLP_1586

TTGAGCAACAACCAACTAAATGGTTCAATTCCCAAGTCCATGTTTGACGTTGGGAGGCTTAAGGTTCTCTCACTTTCTTCCAATTTCTTTAGCGGAACAGTACCCCTTGACCTCATCGGG

>M6-PGSC0003DMG400006255M6LRR-RLP_1647

AGGTTCTCTCACTTTCTTCCAATTTCTTTAGCGGAACAGTACCCCTTGACCTCATCGGGAAGCTGAGCAATCTTTCACGACTGGAGCTTTCTTACAATAACTTGACTGTTGATGCAAGTA

>M6-PGSC0003DMG400006255M6LRR-RLP_1708

GCTGAGCAATCTTTCACGACTGGAGCTTTCTTACAATAACTTGACTGTTGATGCAAGTAGCAGTAATTCAGACGCTTTCACATTTCCCCAGTTGAACATTTTGAAACTAGCTTCGTGTCG

>M6-PGSC0003DMG400006255M6LRR-RLP_1769

AGTAATTCAGACGCTTTCACATTTCCCCAGTTGAACATTTTGAAACTAGCTTCGTGTCGGCTGCAAAAGTTTCCTGATCTTAAAAATCAGTCAAGGATGATCCAATTAGACCTTTCTGAC

>M6-PGSC0003DMG400006255M6LRR-RLP_183

CAGTTATATTCAAGCTAACAAGTTGCAAGAATATGACAACTCTTTACTTTCTATGGCTTTTCTTGATTCCCTTGTTTCAAATCTTAGGAAATGACATTTTCTTGGTTTCCTCTCAATGTC

>M6-PGSC0003DMG400006255M6LRR-RLP_1830

TGCAAAAGTTTCCTGATCTTAAAAATCAGTCAAGGATGATCCAATTAGACCTTTCTGACAACAAAATACTGGGGGCAATACCAAATTGGATTTGGGGAATAGGTAACGGAGCTCTGACTC

>M6-PGSC0003DMG400006255M6LRR-RLP_1891

CAAAATACTGGGGGCAATACCAAATTGGATTTGGGGAATAGGTAACGGAGCTCTGACTCACCTGAATCTTTCTTTCAATCAGTTGGAGTACGTGGAACAGCCTTACAATGTTTCCAGATA

>M6-PGSC0003DMG400006255M6LRR-RLP_1952

CTGAATCTTTCTTTCAATCAGTTGGAGTACGTGGAACAGCCTTACAATGTTTCCAGATATCTTGTCGTCCTTGACTTGCATTCCAATAAGCTAAAAGGTGACCTACCAATTCCACCTTCC

>M6-PGSC0003DMG400006255M6LRR-RLP_2013

TTGTCGTCCTTGACTTGCATTCCAATAAGCTAAAAGGTGACCTACCAATTCCACCTTCCTTTGCTGCATATTTGGACTACTCGAGCAATAATTTCAGCAATTCCATCCCACTAGATATTG

>M6-PGSC0003DMG400006255M6LRR-RLP_2074

TGCTGCATATTTGGACTACTCGAGCAATAATTTCAGCAATTCCATCCCACTAGATATTGGCAATTATCTTGGTTTTGCCTTCTTTTTCTTGGTAGCAAACAATGGCATTACTGGAAGAAT

>M6-PGSC0003DMG400006255M6LRR-RLP_2135

AATTATCTTGGTTTTGCCTTCTTTTTCTTGGTAGCAAACAATGGCATTACTGGAAGAATTCCCGAATCCATATGCAATGTCAGCTACCTTCAAGTTCTTGATTTCTCTAACAATTCCTTG

>M6-PGSC0003DMG400006255M6LRR-RLP_2196

CCGAATCCATATGCAATGTCAGCTACCTTCAAGTTCTTGATTTCTCTAACAATTCCTTGAGTGGAACAATACCACCATGTCTACTGGAATATGGTACAAATCTTGGAGTGCTGAATCTAG

>M6-PGSC0003DMG400006255M6LRR-RLP_2257

TGGAACAATACCACCATGTCTACTGGAATATGGTACAAATCTTGGAGTGCTGAATCTAGGGAACAATAAACTCAATGGCGTTATACCAGATTCATTTCCAATTGGTTGTACTCTACAAAC

>M6-PGSC0003DMG400006255M6LRR-RLP_2318

AACAATAAACTCAATGGCGTTATACCAGATTCATTTCCAATTGGTTGTACTCTACAAACTTTAGACCTCAGTGAGAATAAGTTACAAGGTAGGCTGCCAAAATCTCTTGTGAACTGCAAG

>M6-PGSC0003DMG400006255M6LRR-RLP_2379

TAGACCTCAGTGAGAATAAGTTACAAGGTAGGCTGCCAAAATCTCTTGTGAACTGCAAGTTGTTGGAGGTCCTGAATGTTGGAAATAACAGACTTGTTGATCATTTCCCATGCATATTGA

>M6-PGSC0003DMG400006255M6LRR-RLP_244

CTTGATTCCCTTGTTTCAAATCTTAGGAAATGACATTTTCTTGGTTTCCTCTCAATGTCTTGACAATCAAAAGTCGTTGTTGCTGCAGTTGAAGGGCAGCCTTCAGTATGATTCTAGTTT

>M6-PGSC0003DMG400006255M6LRR-RLP_2440

GTTGGAGGTCCTGAATGTTGGAAATAACAGACTTGTTGATCATTTCCCATGCATATTGAGGAACTCAAACAGTCTGAGGGTCCTAGTCTTGCGTTCCAATCAATTTAATGGAAGTGTTCA

>M6-PGSC0003DMG400006255M6LRR-RLP_2501

AACTCAAACAGTCTGAGGGTCCTAGTCTTGCGTTCCAATCAATTTAATGGAAGTGTTCAGTGTGATGCAACCAGGAATAACTGGAAAAATCTCCAGATCATAGATATAGCTTCCAACAAC

>M6-PGSC0003DMG400006255M6LRR-RLP_2562

GTGATGCAACCAGGAATAACTGGAAAAATCTCCAGATCATAGATATAGCTTCCAACAACTTCACTGGTGTGTTAAATGCAGAATTCTTTTCAAATTGGAAAGGAATGATGGTTGCAGATG

>M6-PGSC0003DMG400006255M6LRR-RLP_2623

CACTGGTGTGTTAAATGCAGAATTCTTTTCAAATTGGAAAGGAATGATGGTTGCAGATGATTACGTGGAGACAGGACTCAATCATATCCAGTACAAGTTCCTCCAACTAAGTAACTTATA

>M6-PGSC0003DMG400006255M6LRR-RLP_2684

TACGTGGAGACAGGACTCAATCATATCCAGTACAAGTTCCTCCAACTAAGTAACTTATACTATCAGGACAAAGTGACAATAACCATCAAAGGGCTAGAGCTAGAGCTTGTGAAGATTCTC

>M6-PGSC0003DMG400006255M6LRR-RLP_2745

ATCAGGACAAAGTGACAATAACCATCAAAGGGCTAGAGCTAGAGCTTGTGAAGATTCTCAGGGTATTCACATCTATTGATTTCTCTTCAAATAGATTTCAAGGTGTGATACCAGATACGG

>M6-PGSC0003DMG400006255M6LRR-RLP_2806

GGTATTCACATCTATTGATTTCTCTTCAAATAGATTTCAAGGTGTGATACCAGATACGGTTGGGGATCTTAGCTCACTTTATGTTCTGAACCTATCACACAATGCCCTTGAAGGACCAAT

>M6-PGSC0003DMG400006255M6LRR-RLP_2867

GGGGATCTTAGCTCACTTTATGTTCTGAACCTATCACACAATGCCCTTGAAGGACCAATTCCAAAATCGATTGGGAAGCTACAAATGCTTGAATCACTAGACCTGTCAACAAACCACCTG

>M6-PGSC0003DMG400006255M6LRR-RLP_2928

CAAAATCGATTGGGAAGCTACAAATGCTTGAATCACTAGACCTGTCAACAAACCACCTGTCCGGGGAGATCCCATCAGAGCTTGCAAGTCTCACATTCTTAGCAGCTTTGAACTTATCGT

>M6-PGSC0003DMG400006255M6LRR-RLP_305

GACAATCAAAAGTCGTTGTTGCTGCAGTTGAAGGGCAGCCTTCAGTATGATTCTAGTTTGTCAAATAAATTCGCAAAATGGAACCAGAAGACAAGTGAATGTTGCAATTGGGATGGGGTT

>M6-PGSC0003DMG400006255M6LRR-RLP_3050

AACAAATTGTTTGGCAAAATCCCATCGACCAATCAGTTTCAAACATTCTCAGCAGATTCCTTTGGAGGAAACAGTGGCCTATGTGGGCTCCCTCTCAACAACAGTTGTCAAAGCAATGGC

>M6-PGSC0003DMG400006255M6LRR-RLP_3111

TTGGAGGAAACAGTGGCCTATGTGGGCTCCCTCTCAACAACAGTTGTCAAAGCAATGGCTCAGAGTCCCTGCCTCCACCAACTTCGCTACCAGACTCAGATGATGAATGGAAGTTCATTT

>M6-PGSC0003DMG400006255M6LRR-RLP_3172

AGAGTCCCTGCCTCCACCAACTTCGCTACCAGACTCAGATGATGAATGGAAGTTCATTTTCGCAGCTGTTGGATACGTAGTAGGGGCAGCAAACACCATTTCACTTCTGTGGTTTTACGA

>M6-PGSC0003DMG400006255M6LRR-RLP_3294

CAGTGAAGAAATGGTTTGATAAGCATACGGAGAAATGCATGCTTTGGTTTTCAAGAAAGTGATCCTTAAACCCGTAAATGATGAGTTTATTTAATCCTTGCTTTGTTATAATAAACATCA

>M6-PGSC0003DMG400006255M6LRR-RLP_3355

ATCCTTAAACCCGTAAATGATGAGTTTATTTAATCCTTGCTTTGTTATAATAAACATCAAGATCCAAATTCTTATTTCCCTTGATGAATGTATAAAATTTTCTACTATCAGAAACAGCAC

>M6-PGSC0003DMG400006255M6LRR-RLP_427

ATGTGATCTCTCTGGTCATGTGATTGCCTTGGAACTGGATAATCAGACTATTTCTAGTGGAATTGAGAATTCAACTGCTCTTTTCAGTCTTCAGTATCTTGAGAGGCTAAATTTGGCTTA

>M6-PGSC0003DMG400006255M6LRR-RLP_488

ATTGAGAATTCAACTGCTCTTTTCAGTCTTCAGTATCTTGAGAGGCTAAATTTGGCTTACAACATGTTCAATGTTGGCGTACCAGTTGGTATAGGCAACCTCACGAACTTGAAGTACCTG

>M6-PGSC0003DMG400006255M6LRR-RLP_549

ACATGTTCAATGTTGGCGTACCAGTTGGTATAGGCAACCTCACGAACTTGAAGTACCTGAATTTATCCAATGCTGGTTTTGTCGGGCAAATTCCTATGATGTTATCAAGATTAACAAAGC

>M6-PGSC0003DMG400006255M6LRR-RLP_61

AACCAAAAAGGCTAACATCAAACGAGTAAAAGGAGTTAATTCAATCTGTACAAACGCGTAGTTTTTCTTTGACATTTCTCAATCACCCTTCCCCTTATTAAGTTCCTTCTAACTGCATAC

>M6-PGSC0003DMG400006255M6LRR-RLP_610

TTTATCCAATGCTGGTTTTGTCGGGCAAATTCCTATGATGTTATCAAGATTAACAAAGCTAGTTACTCTCGATCTCTCAACTCTTTTCCCTTACTCTGACCAGACACTAAAACTTGAGAA

>M6-PGSC0003DMG400006255M6LRR-RLP_671

GTTACTCTCGATCTCTCAACTCTTTTCCCTTACTCTGACCAGACACTAAAACTTGAGAATCCCAATTTGAGTCATTTCATTGAGAATTCAACAGAGCTTACAGAGCTTTACCTTGATGGG

>M6-PGSC0003DMG400006255M6LRR-RLP_732

CCAATTTGAGTCATTTCATTGAGAATTCAACAGAGCTTACAGAGCTTTACCTTGATGGGGTTGATCTTTCATCTCAGAGGACTGAGTGGTGTCAATCTTTATCTTCATATTTGCCTAACT

>M6-PGSC0003DMG400006255M6LRR-RLP_854

ACCGTCTTGAGTTTGCGTGCTTGTCAAATTTCAGGCCCTATTGATGAATCACTTTCTAAGTTTAAGATTCTCTCTATCATCCGTCTTGACCAGAACAATATCTCTACCACAGTTCCAGGA

>M6-PGSC0003DMG400006255M6LRR-RLP_915

TTAAGATTCTCTCTATCATCCGTCTTGACCAGAACAATATCTCTACCACAGTTCCAGGATATTTTGCCAGTTTCACAAACTTGACTACCTTGTCCCTCGACTCTTGTAATCTGCAAGGAG

>M6-PGSC0003DMG400006255M6LRR-RLP_976

TTTTGCCAGTTTCACAAACTTGACTACCTTGTCCCTCGACTCTTGTAATCTGCAAGGAGCTTTTCCTGAAAAGATCTTTCAGGTACAAGTTTTAGAGATTTTGGACTTGTCAAATAACAA

>M6-PGSC0003DMG400006268-(rev)M6LRR-RLP_0

ACTTTCTATGGCTTTTCTTGATTCCCTTGTTTCAAATCTTATCAGGATATCACATTTTCTTGGTTTCCTGTCAATGTCTTCACGATCAAAAGTCGTTGTTGCTGCAGTTGAAGGGCAGCC

>M6-PGSC0003DMG400006268-(rev)M6LRR-RLP_1054

AATCTTGTTTATTTGGACTTCTCAAACAATTTCACTGGTTCTATCCCATATTTCCGACAGTCCAAGAAACTCACCCACTTAGACCTTTCATGTAATGGTCTAACTGGTCTCTTGTCTAGA

>M6-PGSC0003DMG400006268-(rev)M6LRR-RLP_1116

CAAGAAACTCACCCACTTAGACCTTTCATGTAATGGTCTAACTGGTCTCTTGTCTAGAGCTCATTTCGAAGGACTCTCAGATCTTGTATACATAAATTTAGGGAACAATATACTCAACGG

>M6-PGSC0003DMG400006268-(rev)M6LRR-RLP_1178

ATTTCGAAGGACTCTCAGATCTTGTATACATAAATTTAGGGAACAATATACTCAACGGGAACCTTCCTGAATATATATTTGAGCTCCCCTCCTTGCAGCAGCTTTTTCTTAACAGCAATC

>M6-PGSC0003DMG400006268-(rev)M6LRR-RLP_124

GTATGATTCTAGTTTTTCAAATAAATTAGCAAAATGGAATCAGAAGACAAGTGAATGTTGCAATTGGGATGGGGTTACATGCGATCTCTCTGGTCATGTGATTGCCTTGGAACTGGATTA

>M6-PGSC0003DMG400006268-(rev)M6LRR-RLP_1240

CTTCCTGAATATATATTTGAGCTCCCCTCCTTGCAGCAGCTTTTTCTTAACAGCAATCAATTTGTTGGCCAAGTCGACGAATTCTGCAATGCATCCTCCTCTCTGTTGTATACAATTGAT

>M6-PGSC0003DMG400006268-(rev)M6LRR-RLP_1302

TGTTGGCCAAGTCGACGAATTCTGCAATGCATCCTCCTCTCTGTTGTATACAATTGATTTGGCAAACAACCACCTGAATGGATCGATTCCAAAGTCCATGTTTGAAATTGGGAGGCTTAA

>M6-PGSC0003DMG400006268-(rev)M6LRR-RLP_1364

CAAACAACCACCTGAATGGATCGATTCCAAAGTCCATGTTTGAAATTGGGAGGCTTAAGGAACTCTTACTTTCTTCCAACTTCTTTAGTGGGACAGTGACCCTTGATCTCATTGGGAAGC

>M6-PGSC0003DMG400006268-(rev)M6LRR-RLP_1426

CTCTTACTTTCTTCCAACTTCTTTAGTGGGACAGTGACCCTTGATCTCATTGGGAAGCTGAGAAACCTTTCAAGCCTGGAGCTATCTTACAATAACATGACTGTTGAAGTAAGTAGCAGC

>M6-PGSC0003DMG400006268-(rev)M6LRR-RLP_1488

AAACCTTTCAAGCCTGGAGCTATCTTACAATAACATGACTGTTGAAGTAAGTAGCAGCAATTTAACCTCTTTCACATCTCCCCAGTTGAGCATATTGAAACTAGCTTCTTGTCGGCTGCA

>M6-PGSC0003DMG400006268-(rev)M6LRR-RLP_1550

TAACCTCTTTCACATCTCCCCAGTTGAGCATATTGAAACTAGCTTCTTGTCGGCTGCAAAAGTTTCCTGATCTTAAAAATCAGTCATGGATGATCCACTTAGACCTTTCTGACAACGAAA

>M6-PGSC0003DMG400006268-(rev)M6LRR-RLP_1612

TTTCCTGATCTTAAAAATCAGTCATGGATGATCCACTTAGACCTTTCTGACAACGAAATACTGGGGGCAATACCAAATTGGATTTGGGGAATTGGTGACGGAGCTCTGGCTCACCTGAAT

>M6-PGSC0003DMG400006268-(rev)M6LRR-RLP_1674

GGGGGCAATACCAAATTGGATTTGGGGAATTGGTGACGGAGCTCTGGCTCACCTGAATCTTTCCTTCAATCAGCTGGAGTACGTGGAACAGCCTTACAATGTGTCCAGCAATCTTTTTGT

>M6-PGSC0003DMG400006268-(rev)M6LRR-RLP_1736

CCTTCAATCAGCTGGAGTACGTGGAACAGCCTTACAATGTGTCCAGCAATCTTTTTGTCCTTGACTTGCATTCCAACCGTTTAAAAGGTGACTTACCAATTCCACCTTCCTCTGCCATCT

>M6-PGSC0003DMG400006268-(rev)M6LRR-RLP_1798

GACTTGCATTCCAACCGTTTAAAAGGTGACTTACCAATTCCACCTTCCTCTGCCATCTATGTGGACTATTCGAGCAATAATTTAAACAAATCCATCCCACTAGATATTGGAAATTCTCTT

>M6-PGSC0003DMG400006268-(rev)M6LRR-RLP_186

ATTGGGATGGGGTTACATGCGATCTCTCTGGTCATGTGATTGCCTTGGAACTGGATTATGAGACAATTTCTGGTGGAATTGAGAATTCAAGTTCTCTTTTCAGTCTACAGTATCTTGAGA

>M6-PGSC0003DMG400006268-(rev)M6LRR-RLP_1860

GGACTATTCGAGCAATAATTTAAACAAATCCATCCCACTAGATATTGGAAATTCTCTTGCTCTTGCCTCCTTTTTCTCGGTAGCAAACAATGACATTACTGGAATAGTTCCTGAATCCAT

>M6-PGSC0003DMG400006268-(rev)M6LRR-RLP_1922

TTGCCTCCTTTTTCTCGGTAGCAAACAATGACATTACTGGAATAGTTCCTGAATCCATATGCAGTGTCAGCTACCTTCAAGTTCTTGATTTCTCTAACAATGCCTTGAGTGGAACAATTC

>M6-PGSC0003DMG400006268-(rev)M6LRR-RLP_1984

AGTGTCAGCTACCTTCAAGTTCTTGATTTCTCTAACAATGCCTTGAGTGGAACAATTCCACGATGTCTACTGGAATATAGTACTACTCTTGGAGTGCTGAATCTAGGGAACAACAGACTC

>M6-PGSC0003DMG400006268-(rev)M6LRR-RLP_2046

ATGTCTACTGGAATATAGTACTACTCTTGGAGTGCTGAATCTAGGGAACAACAGACTCCATGGTGTTATACCAGATTCATTTCCAATTGGTTGTGCTCTAAAAACTCTAGACCTCAGTGA

>M6-PGSC0003DMG400006268-(rev)M6LRR-RLP_2108

GTGTTATACCAGATTCATTTCCAATTGGTTGTGCTCTAAAAACTCTAGACCTCAGTGAGAATAAGTTACAAGGTAGCCTCCCAAAATCGCTTGTGAACTGCAAGTTGTTGGAGGTCCTGA

>M6-PGSC0003DMG400006268-(rev)M6LRR-RLP_2170

AAGTTACAAGGTAGCCTCCCAAAATCGCTTGTGAACTGCAAGTTGTTGGAGGTCCTGAATGTTGGAAATTACAGACTTGTTGATCATTTCCCGTGCATGTTGAGGAACTCAAACAGTCTG

>M6-PGSC0003DMG400006268-(rev)M6LRR-RLP_2232

TGGAAATTACAGACTTGTTGATCATTTCCCGTGCATGTTGAGGAACTCAAACAGTCTGAGGGTCCTAGTCTTGCGCTTCAATCAATTCAGTGGAAATCTTCAGTGTGAAGTAACCATAAA

>M6-PGSC0003DMG400006268-(rev)M6LRR-RLP_2294

TCCTAGTCTTGCGCTTCAATCAATTCAGTGGAAATCTTCAGTGTGAAGTAACCATAAATAGCTGCCCAAATCTCCAGATCATAGATATAGCTTCCAACAACTTCACTGGTGTGTTGAATG

>M6-PGSC0003DMG400006268-(rev)M6LRR-RLP_2356

TGCCCAAATCTCCAGATCATAGATATAGCTTCCAACAACTTCACTGGTGTGTTGAATGCCGAATTCTTTTCAAATTGGAGAGGAATGATGGTTGCAGATGATTACGTGGAGACAGGGCGG

>M6-PGSC0003DMG400006268-(rev)M6LRR-RLP_2418

ATTCTTTTCAAATTGGAGAGGAATGATGGTTGCAGATGATTACGTGGAGACAGGGCGGAATCATATCCAGTACAAGTTCCTCCAACTAAGTAACTTATACTATCAGGTCACAGTAACAAT

>M6-PGSC0003DMG400006268-(rev)M6LRR-RLP_248

ACAATTTCTGGTGGAATTGAGAATTCAAGTTCTCTTTTCAGTCTACAGTATCTTGAGAAGTTAAATTTGGCTTACAACAGGTTCAGTGTTGGCATACCAGTTGGTATAGGCAACCTCACG

>M6-PGSC0003DMG400006268-(rev)M6LRR-RLP_2480

ATATCCAGTACAAGTTCCTCCAACTAAGTAACTTATACTATCAGGTCACAGTAACAATAACCAACAAAGGGATGGAGATGAAGCTTGTGAAAATTCTCAGAGTCTACACATCTATTAATT

>M6-PGSC0003DMG400006268-(rev)M6LRR-RLP_2542

AACAAAGGGATGGAGATGAAGCTTGTGAAAATTCTCAGAGTCTACACATCTATTAATTTCTCTTCAAATAGATTTCAAGGAGTGATACCAGATACTATCGGGAATCTCAGCTCACTTTAT

>M6-PGSC0003DMG400006268-(rev)M6LRR-RLP_2604

TTCAAATAGATTTCAAGGAGTGATACCAGATACTATCGGGAATCTCAGCTCACTTTATGTTCTGAACCTGTCACACAATGCCCTTGAAGGACCAATTCCAAAATCGATTGGGAAGCTACA

>M6-PGSC0003DMG400006268-(rev)M6LRR-RLP_2790

TCTCACATTCTTAGCAGCTTTGAACTTATCGTTCAACAAATTGTTTGGCAAAATCCCATTAACTAATCAGTTTCAAACATTCTCAGCAGATTCCTTTGAAGGAAACAGAGGCCTATGCGG

>M6-PGSC0003DMG400006268-(rev)M6LRR-RLP_2852

CTAATCAGTTTCAAACATTCTCAGCAGATTCCTTTGAAGGAAACAGAGGCCTATGCGGGTTCCCTCTCAACAACAATTGCGAAAGCAATGGCTTAGAGTCCCTCTCCCTGCTGCCACCAA

>M6-PGSC0003DMG400006268-(rev)M6LRR-RLP_2914

CCTCTCAACAACAATTGCGAAAGCAATGGCTTAGAGTCCCTCTCCCTGCTGCCACCAACTTTGGTACCAGACTCAGACTCAGATGATGAATGGAAGTTCATTTTCGCGGCTGTTGGATAC

>M6-PGSC0003DMG400006268-(rev)M6LRR-RLP_2976

GGTACCAGACTCAGACTCAGATGATGAATGGAAGTTCATTTTCGCGGCTGTTGGATACGTAGTAGGGGCAGCAAACACTATTTCACTTCTGTGGTTTTACGAGCCAGTGAAGAAATGGTT

>M6-PGSC0003DMG400006268-(rev)M6LRR-RLP_3016

TTCGCGGCTGTTGGATACGTAGTAGGGGCAGCAAACACTATTTCACTTCTGTGGTTTTACGAGCCAGTGAAGAAATGGTTTGATAAGCATATAGAGAAATGTTTACTTTGGTTTTCAAGA

>M6-PGSC0003DMG400006268-(rev)M6LRR-RLP_372

TGAAGTACCTGAATTTATCCAATGCTGGTTTTGTCGGGCAAATTCCTATGTCGTTACCAAGATTAAAAAGGCGAGTTACTCTTGATCTCTCAATCTTTTCCCTGACTTTGATCAGCCACT

>M6-PGSC0003DMG400006268-(rev)M6LRR-RLP_434

TTAAAAAGGCGAGTTACTCTTGATCTCTCAATCTTTTCCCTGACTTTGATCAGCCACTTAAACTTGAGAATCCCAATTTGAGACAATTCATTGAGAACTCAACAGAGCTTAGAGAGCTTT

>M6-PGSC0003DMG400006268-(rev)M6LRR-RLP_496

CTTGAGAATCCCAATTTGAGACAATTCATTGAGAACTCAACAGAGCTTAGAGAGCTTTACCTTGATGGGGTTGATCTTTCATCTCAGAGGACTGAATGGTGTCAATCTTTATCTTCATAT

>M6-PGSC0003DMG400006268-(rev)M6LRR-RLP_558

TGATGGGGTTGATCTTTCATCTCAGAGGACTGAATGGTGTCAATCTTTATCTTCATATTTGCCTAAATTGACTATCTTGAGTTTGCGTGCTTGTCAAATTTCAGGCCCTATTGATGAATC

>M6-PGSC0003DMG400006268-(rev)M6LRR-RLP_62

GTTTCCTGTCAATGTCTTCACGATCAAAAGTCGTTGTTGCTGCAGTTGAAGGGCAGCCTTCAGTATGATTCTAGTTTTTCAAATAAATTAGCAAAATGGAATCAGAAGACAAGTGAATGT

>M6-PGSC0003DMG400006268-(rev)M6LRR-RLP_620

CTAAATTGACTATCTTGAGTTTGCGTGCTTGTCAAATTTCAGGCCCTATTGATGAATCACTTTCTAAGCTTCTCTTTCTCTCTTTCATCCGTCTTGAACGGAACAATCTCTCTACCACAG

>M6-PGSC0003DMG400006268-(rev)M6LRR-RLP_682

TCTAAGCTTCTCTTTCTCTCTTTCATCCGTCTTGAACGGAACAATCTCTCTACCACAGTTCCTGAATATTTTGCCCATTTCTCGAACTTGACTACCTTGGCCCTCGGCTCTTGTAATCTG

>M6-PGSC0003DMG400006268-(rev)M6LRR-RLP_744

TGAATATTTTGCCCATTTCTCGAACTTGACTACCTTGGCCCTCGGCTCTTGTAATCTGCATGGAACATTTCCTGAAAGAATCTTTCAGGTACCAGTTTTAGAGAGTTTGGACTTGTCAAA

>M6-PGSC0003DMG400006268-(rev)M6LRR-RLP_806

GAACATTTCCTGAAAGAATCTTTCAGGTACCAGTTTTAGAGAGTTTGGACTTGTCAAATAACGAGTTGCTTCGAGGTAGTATTCCTGTTGTTCTTCCAAATGGATCTCTGAGGAGATTAT

>M6-PGSC0003DMG400006268-(rev)M6LRR-RLP_868

GAGTTGCTTCGAGGTAGTATTCCTGTTGTTCTTCCAAATGGATCTCTGAGGAGATTATCACTAAGCTACACCAACTTTTCTGGCTCATTACCAGAGTCCATTTTGAACCTTCAGAACCTG

>M6-PGSC0003DMG400006268-(rev)M6LRR-RLP_930

AAGCTACACCAACTTTTCTGGCTCATTACCAGAGTCCATTTTGAACCTTCAGAACCTGTCCAGGTTAGAGCTCTCCAATTGCAATTTCAATGGACCAATACCTTCCACAATGGCAAACCT

>M6-PGSC0003DMG400006268-(rev)M6LRR-RLP_992

GGTTAGAGCTCTCCAATTGCAATTTCAATGGACCAATACCTTCCACAATGGCAAACCTTACAAATCTTGTTTATTTGGACTTCTCAAACAATTTCACTGGTTCTATCCCATATTTCCGAC

>M6-PGSC0003DMG400006279M6LRR-RLP_0

CTTTTAGTTACGTTGATAGGTTTTTCCAAGTGTTTAAAGGTCCATCTTTGGATTTGCAAGCTAACCATGAACACGTTGATATGTGTACAGTCATATAGTCCAAGTTATTTTTATGTTATA

>M6-PGSC0003DMG400006279M6LRR-RLP_1024

GCGATCGCTGCCCCCAAGTGACAACTTCATTCCCCTGAATATTGTCTTCTAATATCTTCTTGTTCCCAGCAGCCTTCTTTAACACCTGATTGCCTAAAATCGCCATGAATGGTCTTACCA

>M6-PGSC0003DMG400006279M6LRR-RLP_1088

CCCAGCAGCCTTCTTTAACACCTGATTGCCTAAAATCGCCATGAATGGTCTTACCAACTTCAGAACCTGTAGCATCTGATTTCTCACCCAAATTTCGAACACTCTGTGCAGCTCGTCTTG

>M6-PGSC0003DMG400006279M6LRR-RLP_1152

ACCTGTAGCATCTGATTTCTCACCCAAATTTCGAACACTCTGTGCAGCTCGTCTTGCATTAATAATTTCCCTAGCATTTCCCTGATGCTTAATACCTTGCATATTAATCTGCATACTTAA

>M6-PGSC0003DMG400006279M6LRR-RLP_1216

ATTTCCCTAGCATTTCCCTGATGCTTAATACCTTGCATATTAATCTGCATACTTAATCTCTGTGACCTTGATGTTTGCATGTACGTATATGAAGGTTAATAATCATATGTTATTATTTGG

>M6-PGSC0003DMG400006279M6LRR-RLP_128

TGTGCATTTCATTTATTAAGTATCCCATGTTTGCTTGTTTTGACATATTTTGGTTTGTCTGTTTTTGGTTTATCAGTGCCTGTCCATCCTAGAGTTGAAAGGGTGGCGTCCTCCACCATC

>M6-PGSC0003DMG400006279M6LRR-RLP_1280

ACCTTGATGTTTGCATGTACGTATATGAAGGTTAATAATCATATGTTATTATTTGGTACACTATCAATTCATTCGACTTTGGCTCTAATGAGCAATCTCACACGTTCAGGCAAAGCAGCC

>M6-PGSC0003DMG400006279M6LRR-RLP_1344

CAATTCATTCGACTTTGGCTCTAATGAGCAATCTCACACGTTCAGGCAAAGCAGCCAGTAAGTTGACTTCAACTTTAACTCTTGCTGTGCTTGGCCTTGTTTGATCTGTGGTTGCTCTAT

>M6-PGSC0003DMG400006279M6LRR-RLP_1408

GACTTCAACTTTAACTCTTGCTGTGCTTGGCCTTGTTTGATCTGTGGTTGCTCTATCAACAACCAAAGGTCTGCCCACTGCTGATGCTAACAAATATAATGCCTCTCGAACGAAAAATTG

>M6-PGSC0003DMG400006279M6LRR-RLP_1472

AAAGGTCTGCCCACTGCTGATGCTAACAAATATAATGCCTCTCGAACGAAAAATTGCACCAGTAAATTCTGAAACAATATCCAAGTCAAAGCAATGGATTCTTCTTCTTTGGGATTAAAC

>M6-PGSC0003DMG400006279M6LRR-RLP_1516

AACGAAAAATTGCACCAGTAAATTCTGAAACAATATCCAAGTCAAAGCAATGGATTCTTCTTCTTTGGGATTAAACCAAGGGGTCCAACGAAAAAATTCTAAATTGATAGTCTTCCTCCA

>M6-PGSC0003DMG400006279M6LRR-RLP_192

TTGGTTTATCAGTGCCTGTCCATCCTAGAGTTGAAAGGGTGGCGTCCTCCACCATCAAGCCGGAAATTTTTAGAACTGAACCTAGCCGTAATACAATTGGACTTCCCTGGAATTTACAGC

>M6-PGSC0003DMG400006279M6LRR-RLP_256

AATTTTTAGAACTGAACCTAGCCGTAATACAATTGGACTTCCCTGGAATTTACAGCTTTCTACAGAGCTCATCGGATCTTGAGACATTGATCATTGGTTGTTGGTACAATCACAATGAAA

>M6-PGSC0003DMG400006279M6LRR-RLP_320

GAGCTCATCGGATCTTGAGACATTGATCATTGGTTGTTGGTACAATCACAATGAAAGAGTAAGTTTCCTTTCGACCTTTATTTATATTACTTCATGTTATTCAAATCAAACAACTTTTAG

>M6-PGSC0003DMG400006279M6LRR-RLP_384

TTCCTTTCGACCTTTATTTATATTACTTCATGTTATTCAAATCAAACAACTTTTAGTGGAGGAATAAGCTGATATATATCAACATATTTCTAATCTTAAGTACTTCAGGAAAGAAAATTT

>M6-PGSC0003DMG400006279M6LRR-RLP_448

TAAGCTGATATATATCAACATATTTCTAATCTTAAGTACTTCAGGAAAGAAAATTTATTGAAGTTTTTTTTAATAGCAATCTCCTCAGTCACTTTACCATCCCAATTGTGCTTCCTTCCA

>M6-PGSC0003DMG400006279M6LRR-RLP_512

TTTTTTTAATAGCAATCTCCTCAGTCACTTTACCATCCCAATTGTGCTTCCTTCCATGAGTTCTATTTTTGTTTTTTTCCAATTCATCGTTGCCTAGGAGATATTGCAGCTTCCCTAGCT

>M6-PGSC0003DMG400006279M6LRR-RLP_576

ATTTTTGTTTTTTTCCAATTCATCGTTGCCTAGGAGATATTGCAGCTTCCCTAGCTTCATTTTTAAAATATTCATCCAGTGTGGCATCCTCCTCGCTTCTCTCAAAAATGTTTGCACCTC

>M6-PGSC0003DMG400006279M6LRR-RLP_64

CCATGAACACGTTGATATGTGTACAGTCATATAGTCCAAGTTATTTTTATGTTATATCCAACAATGTGCATTTCATTTATTAAGTATCCCATGTTTGCTTGTTTTGACATATTTTGGTTT

>M6-PGSC0003DMG400006279M6LRR-RLP_640

AAAATATTCATCCAGTGTGGCATCCTCCTCGCTTCTCTCAAAAATGTTTGCACCTCTAGAACTTCTTGGAGAAAAATCACTAGGTCTAACATGTAGTTGCTTACTAGTGTTAATAGGAGT

>M6-PGSC0003DMG400006279M6LRR-RLP_704

CTTGGAGAAAAATCACTAGGTCTAACATGTAGTTGCTTACTAGTGTTAATAGGAGTAAGAGAATTCACTCTTTCAATATTCGAACTCATCAGTATATCAAGAAGCGCCATGTCTATAGTA

>M6-PGSC0003DMG400006279M6LRR-RLP_768

TCACTCTTTCAATATTCGAACTCATCAGTATATCAAGAAGCGCCATGTCTATAGTATTATTGATCTTCTGAACATCAGAACTGCTGCGCAGAATTCTCAGGTGATGCATCTATCAAATTT

>M6-PGSC0003DMG400006279M6LRR-RLP_832

CTTCTGAACATCAGAACTGCTGCGCAGAATTCTCAGGTGATGCATCTATCAAATTTGCAGATTTTTCAGCTACTTCAGTTTCTAGATCTGTCGGTTTCGAAATTGTATGACGTCCCTGTC

>M6-PGSC0003DMG400006279M6LRR-RLP_896

TTCAGCTACTTCAGTTTCTAGATCTGTCGGTTTCGAAATTGTATGACGTCCCTGTCTCCGGTGCATCTGTCTAGCACTTGAACTTGCAATTTCTTGTGAACTTAACGCTCCATCTTTCAT

>M6-PGSC0003DMG400006279M6LRR-RLP_960

ATCTGTCTAGCACTTGAACTTGCAATTTCTTGTGAACTTAACGCTCCATCTTTCATGATTTCAGGCGATCGCTGCCCCCAAGTGACAACTTCATTCCCCTGAATATTGTCTTCTAATATC

>M6-PGSC0003DMG400006280M6LRR-RLK_0

CCAAAGCCAAAAATTACTATCATATTTTATCAGTATCAATGGAATCTTTCAAAATACTTGTTGCATCTTCTTCTTCTTCTTCTTACCTTTGATGTTTTCTAAAGTATTTGAACTTCAATT

>M6-PGSC0003DMG400006280M6LRR-RLK_1054

GAGATTCCTGGATGTATCGCGGAGCTGCCAAAGTTGGAAATGTTGAATCTTTGGAGGAATAATTTCACTGGATCGATTCCTTCTAAGCTTGGGATGAACGGTAAACTAGTTGAAATTGAT

>M6-PGSC0003DMG400006280M6LRR-RLK_1116

TTTCACTGGATCGATTCCTTCTAAGCTTGGGATGAACGGTAAACTAGTTGAAATTGATCTTTCTAGTAATAGACTCACTGGCTTGATACCAAAATCTCTATGCTTTGGGAGGAATTTGAA

>M6-PGSC0003DMG400006280M6LRR-RLK_1178

CTAGTAATAGACTCACTGGCTTGATACCAAAATCTCTATGCTTTGGGAGGAATTTGAAAATCTTGATTCTGCTTGATAATTTTCTGTTTGGACCTTTACCTGATGATTTTGGGCAGTGTC

>M6-PGSC0003DMG400006280M6LRR-RLK_124

GTTATCTAAATGGCTACTTCTAACACAAGTCTCTTCTTTTTCGCGTATTTCCTACTCGTGTTCCTCATTACTCCATCTCAATCGCATAACCTGTCTCTGAGACGACAGGCTAAAACTCTA

>M6-PGSC0003DMG400006280M6LRR-RLK_1240

TTGATTCTGCTTGATAATTTTCTGTTTGGACCTTTACCTGATGATTTTGGGCAGTGTCGAACCTTGTCCAGAGTCAGAATGGGACAGAATTACTTGAGTGGATCAATACCAACAGGGTTT

>M6-PGSC0003DMG400006280M6LRR-RLK_1302

CTTGTCCAGAGTCAGAATGGGACAGAATTACTTGAGTGGATCAATACCAACAGGGTTTCTTTATTTGCCTGAGTTGTCACTGGTGGAACTGCAGAACAACTACATCAGTGGACAACTCTC

>M6-PGSC0003DMG400006280M6LRR-RLK_1364

ATTTGCCTGAGTTGTCACTGGTGGAACTGCAGAACAACTACATCAGTGGACAACTCTCGAACGAGAAAATCTCAGCGTCTTCTAAACTTGAAGGGCTGAACCTGTCGAACAATCGCTTGT

>M6-PGSC0003DMG400006280M6LRR-RLK_1426

GAGAAAATCTCAGCGTCTTCTAAACTTGAAGGGCTGAACCTGTCGAACAATCGCTTGTCTGGTGCACTTCCCAGTGCTATTGGAAACTATTCAGGGCTGAAGAATCTTGTGTTAACTGGA

>M6-PGSC0003DMG400006280M6LRR-RLK_1488

TGCACTTCCCAGTGCTATTGGAAACTATTCAGGGCTGAAGAATCTTGTGTTAACTGGAAATGGTTTCTCAGGAGATATCCCTTCTGATATTGGCAGACTAAAAAGCATCTTAAAGCTGGA

>M6-PGSC0003DMG400006280M6LRR-RLK_1550

GTTTCTCAGGAGATATCCCTTCTGATATTGGCAGACTAAAAAGCATCTTAAAGCTGGACCTGAGTAGGAACAACTTCTCTGGCACAATCCCCCCTCAGATTAGTAACTGTCTTTCCTTAA

>M6-PGSC0003DMG400006280M6LRR-RLK_1612

AGTAGGAACAACTTCTCTGGCACAATCCCCCCTCAGATTAGTAACTGTCTTTCCTTAACTTACTTGGATTTGAGCCAAAATCAACTTTCTGGTCCTATCCCAGTTCAAATTGCTCAAATT

>M6-PGSC0003DMG400006280M6LRR-RLK_1674

CTTGGATTTGAGCCAAAATCAACTTTCTGGTCCTATCCCAGTTCAAATTGCTCAAATTCACATCTTAAATTACATCAATATTTCCTGGAATCACTTCAACGATAGCCTTCCCGCGGAGAT

>M6-PGSC0003DMG400006280M6LRR-RLK_1736

TCTTAAATTACATCAATATTTCCTGGAATCACTTCAACGATAGCCTTCCCGCGGAGATTGGCTCGATGAAGAGCTTAACTTCAGCAGATTTTTCCCACAATAACTTATCTGGATCAATAC

>M6-PGSC0003DMG400006280M6LRR-RLK_1798

TCGATGAAGAGCTTAACTTCAGCAGATTTTTCCCACAATAACTTATCTGGATCAATACCTGAAACAGGACAGTATTTATATTTCAACTCAACTTCCTTCATTGGCAACCCTTATCTCTCT

>M6-PGSC0003DMG400006280M6LRR-RLK_186

CCTCATTACTCCATCTCAATCGCATAACCTGTCTCTGAGACGACAGGCTAAAACTCTAGTTTCATTGAAACAAGCATTTGTAGTATCCTCTGTTCCTAGTACTCTGTCTACTTGGAACAT

>M6-PGSC0003DMG400006280M6LRR-RLK_1860

AACAGGACAGTATTTATATTTCAACTCAACTTCCTTCATTGGCAACCCTTATCTCTCTGGATCCGACTCGACTCCTAGCAACATTACATCCAACTCACCGTCAAAACTTGGAGACGGAAG

>M6-PGSC0003DMG400006280M6LRR-RLK_1922

CCGACTCGACTCCTAGCAACATTACATCCAACTCACCGTCAAAACTTGGAGACGGAAGTGACAACAGAACTAAGGTTCCAACAATATACAAGTTCATATTTGCATTTGGGCTCTTATTCT

>M6-PGSC0003DMG400006280M6LRR-RLK_1984

AACAGAACTAAGGTTCCAACAATATACAAGTTCATATTTGCATTTGGGCTCTTATTCTGCTCCCTCATTTTTGTTGTCTTAGCAATAATCAAGACAAGAAAGGGGAGTAAGAATTCAAAT

>M6-PGSC0003DMG400006280M6LRR-RLK_2046

CCTCATTTTTGTTGTCTTAGCAATAATCAAGACAAGAAAGGGGAGTAAGAATTCAAATTTGTGGAAGCTGACAGCATTTCAGAAGCTTGAATTCGGAAGTGAAGACGTCTTGCAGTGCTT

>M6-PGSC0003DMG400006280M6LRR-RLK_2108

GGAAGCTGACAGCATTTCAGAAGCTTGAATTCGGAAGTGAAGACGTCTTGCAGTGCTTGAAAGACAACAACGTCATAGGGAGAGGTGGAGCAGGGATAGTGTATAAAGGAACTATGCCAA

>M6-PGSC0003DMG400006280M6LRR-RLK_2170

GACAACAACGTCATAGGGAGAGGTGGAGCAGGGATAGTGTATAAAGGAACTATGCCAAATGGTGATCATGTCGCGGTGAAGAAGTTGGGAATAAGCAAAGGCTCACATGATAACGGCCTA

>M6-PGSC0003DMG400006280M6LRR-RLK_2232

TGATCATGTCGCGGTGAAGAAGTTGGGAATAAGCAAAGGCTCACATGATAACGGCCTATCTGCTGAACTTAAAACATTAGGAAAGATCAGGCATAGGTACATTGTGAGACTGCTCGCGTT

>M6-PGSC0003DMG400006280M6LRR-RLK_2294

CTGAACTTAAAACATTAGGAAAGATCAGGCATAGGTACATTGTGAGACTGCTCGCGTTTTGTTCAAACAAGGAAATCAACTTGCTAGTTTATGAGTACATGCTAAATGGAAGCTTAGGTG

>M6-PGSC0003DMG400006280M6LRR-RLK_2356

TCAAACAAGGAAATCAACTTGCTAGTTTATGAGTACATGCTAAATGGAAGCTTAGGTGAAGTGCTTCATGGGAAGAACGGCGGGCAACTCCAATGGGACACTAGGCTAAAAATTGCCATA

>M6-PGSC0003DMG400006280M6LRR-RLK_2418

GCTTCATGGGAAGAACGGCGGGCAACTCCAATGGGACACTAGGCTAAAAATTGCCATAGAAGCTGCCAAGGGCCTTTCTTATTTGCACCATGATTGCTCCCCTATGATAATCCACCGCGA

>M6-PGSC0003DMG400006280M6LRR-RLK_248

CATTGAAACAAGCATTTGTAGTATCCTCTGTTCCTAGTACTCTGTCTACTTGGAACATGTCAAATTATATGTCTATATGTTGTTGGACCGGTATCACGTGTGATGATACCAAATCAGTAA

>M6-PGSC0003DMG400006280M6LRR-RLK_2480

CTGCCAAGGGCCTTTCTTATTTGCACCATGATTGCTCCCCTATGATAATCCACCGCGATGTCAAGTCCAACAATATATTGTTGAACTCTGAACTTGAAGCTCATGTTGCAGATTTTGGCT

>M6-PGSC0003DMG400006280M6LRR-RLK_2542

AAGTCCAACAATATATTGTTGAACTCTGAACTTGAAGCTCATGTTGCAGATTTTGGCTTAGCCAAGTACTTTCATAATAATGGTACTTCTGAGTGCATGTCTGCAATTGCAGGGTCTTAT

>M6-PGSC0003DMG400006280M6LRR-RLK_2604

CAAGTACTTTCATAATAATGGTACTTCTGAGTGCATGTCTGCAATTGCAGGGTCTTATGGCTACATTGCTCCAGGTAACTAGTTTGTTCATACATTAGTTGATTTAATTTCTTCTCGACT

>M6-PGSC0003DMG400006280M6LRR-RLK_2666

ACATTGCTCCAGGTAACTAGTTTGTTCATACATTAGTTGATTTAATTTCTTCTCGACTGGCCTATTATGCACTCGGACCAATCTACAACCTACTAATAATGGTCTTTACATGTTATCTTT

>M6-PGSC0003DMG400006280M6LRR-RLK_2728

TATTATGCACTCGGACCAATCTACAACCTACTAATAATGGTCTTTACATGTTATCTTTTTGTATAGAATATGCATACACGCTGAAAATTGATGAGAAAAGCGATGTGTATAGCTTTGGAG

>M6-PGSC0003DMG400006280M6LRR-RLK_2790

ATAGAATATGCATACACGCTGAAAATTGATGAGAAAAGCGATGTGTATAGCTTTGGAGTGGTGTTGTTGGAGCTTATAACAGGACGAAAGCCAGTTGGAAATTTTGGAGAAGAAGGAATG

>M6-PGSC0003DMG400006280M6LRR-RLK_2852

GTTGTTGGAGCTTATAACAGGACGAAAGCCAGTTGGAAATTTTGGAGAAGAAGGAATGGACATTGTCCAATGGGCGAAAACGGAGACAAATTGGAGCAAAGAAGAGGTGGTGAAAATCTT

>M6-PGSC0003DMG400006280M6LRR-RLK_2914

TTGTCCAATGGGCGAAAACGGAGACAAATTGGAGCAAAGAAGAGGTGGTGAAAATCTTGGATGAGAGGCTAAAAAATGTTGCAATTGTAGAAGCTATGCAAGTATTTTTTGTAGCAATGC

>M6-PGSC0003DMG400006280M6LRR-RLK_2976

GAGAGGCTAAAAAATGTTGCAATTGTAGAAGCTATGCAAGTATTTTTTGTAGCAATGCTTTGTGTTGAAGAGTACAGCATTGAGAGGCCTACAATGAGGGAAGTAGTCCAAATGCTTTCT

>M6-PGSC0003DMG400006280M6LRR-RLK_3038

TGTTGAAGAGTACAGCATTGAGAGGCCTACAATGAGGGAAGTAGTCCAAATGCTTTCTCAAGCTAAACAACCAAATACTTTCCAAATCCAATGATCTAATTGTGGCTCTACTAATTCTTA

>M6-PGSC0003DMG400006280M6LRR-RLK_310

AATTATATGTCTATATGTTGTTGGACCGGTATCACGTGTGATGATACCAAATCAGTAACTACCATTGATATATCCAATCTAAACATTTCTGGCTCTTTATCACCTGATATTCATGAGCTC

>M6-PGSC0003DMG400006280M6LRR-RLK_3100

CTAAACAACCAAATACTTTCCAAATCCAATGATCTAATTGTGGCTCTACTAATTCTTATAGTATATCTTTTTTTGTTTCTTGTGTGCTTGGGAAGACCCCTTTTGTTAGCTTTGCAAACA

>M6-PGSC0003DMG400006280M6LRR-RLK_3162

ATATCTTTTTTTGTTTCTTGTGTGCTTGGGAAGACCCCTTTTGTTAGCTTTGCAAACAAATTACAATTATGTTAGATCTTAGGGCATGATATCTAACTATCCTCTCAACTTGGAATAGTG

>M6-PGSC0003DMG400006280M6LRR-RLK_3224

ACAATTATGTTAGATCTTAGGGCATGATATCTAACTATCCTCTCAACTTGGAATAGTGTATTGGATGTGTAGAACTAGTATTAGTATCCGCGTGATGTGTGGTGAATATCAAAAGAAATT

>M6-PGSC0003DMG400006280M6LRR-RLK_372

CATTGATATATCCAATCTAAACATTTCTGGCTCTTTATCACCTGATATTCATGAGCTCACCAGACTTCGCATCCTGAATATTTCTAACAACTTGTTAGGTGGGAACTTGAGCTGGGAGTA

>M6-PGSC0003DMG400006280M6LRR-RLK_434

GACTTCGCATCCTGAATATTTCTAACAACTTGTTAGGTGGGAACTTGAGCTGGGAGTATCGAAAGTTTAACGTACTTCAAGTGTTGGATGCTTATAACAACAATTTCACTGGTCCACTCC

>M6-PGSC0003DMG400006280M6LRR-RLK_496

AAGTTTAACGTACTTCAAGTGTTGGATGCTTATAACAACAATTTCACTGGTCCACTCCCCTTGGGAGTTACTCAACTTCTGCAGCTGAAGTATTTGAATTTCGGGGGTAACTACTTTTCG

>M6-PGSC0003DMG400006280M6LRR-RLK_558

GGGAGTTACTCAACTTCTGCAGCTGAAGTATTTGAATTTCGGGGGTAACTACTTTTCGGGGAAAATTCCTCTGAGTTATGGTAGTTTTAATCAGCTTGAGTTCCTGTCTCTTGCTGGGAA

>M6-PGSC0003DMG400006280M6LRR-RLK_62

GCATCTTCTTCTTCTTCTTCTTACCTTTGATGTTTTCTAAAGTATTTGAACTTCAATTACTAGTTATCTAAATGGCTACTTCTAACACAAGTCTCTTCTTTTTCGCGTATTTCCTACTCG

>M6-PGSC0003DMG400006280M6LRR-RLK_620

AAATTCCTCTGAGTTATGGTAGTTTTAATCAGCTTGAGTTCCTGTCTCTTGCTGGGAATGACTTGCACGGTCCTATACCAAGGGAGCTGGGCAACGTTACGAGCCTCAGGTGGTTACAAT

>M6-PGSC0003DMG400006280M6LRR-RLK_682

TTGCACGGTCCTATACCAAGGGAGCTGGGCAACGTTACGAGCCTCAGGTGGTTACAATTGGGTTATTATAATCAATTTGATGAGGGGATTCCGCCAGAGTTGGGGAAACTTGTAAATTTG

>M6-PGSC0003DMG400006280M6LRR-RLK_744

TTATTATAATCAATTTGATGAGGGGATTCCGCCAGAGTTGGGGAAACTTGTAAATTTGGTTCATCTAGATCTTTCTAGCTGTAACTTGACGGGTTCGATTCCAGCAGAATTGGGCAATCT

>M6-PGSC0003DMG400006280M6LRR-RLK_806

ATCTAGATCTTTCTAGCTGTAACTTGACGGGTTCGATTCCAGCAGAATTGGGCAATCTTAATATGTTGGATACTCTTTTCTTGCAAAAAAATCAACTTACTGGTGTATTTCCTCCTCAGC

>M6-PGSC0003DMG400006280M6LRR-RLK_868

ATGTTGGATACTCTTTTCTTGCAAAAAAATCAACTTACTGGTGTATTTCCTCCTCAGCTAGGGAATTTGACAAGGTTAAAATCTCTTGATATATCCGTCAATGAACTCACAGGAGAGATA

>M6-PGSC0003DMG400006280M6LRR-RLK_930

GAATTTGACAAGGTTAAAATCTCTTGATATATCCGTCAATGAACTCACAGGAGAGATACCAGTTGACTTGTCAGGACTCAAGGAGCTCACATTGTTGAACCTCTTCATCAACAATTTGCA

>M6-PGSC0003DMG400006280M6LRR-RLK_992

TTGACTTGTCAGGACTCAAGGAGCTCACATTGTTGAACCTCTTCATCAACAATTTGCACGGTGAGATTCCTGGATGTATCGCGGAGCTGCCAAAGTTGGAAATGTTGAATCTTTGGAGGA

>M6-PGSC0003DMG400006364-(rev)M6LRR-RLK_0

ATGTATCTTGCAAATTATGGGATTAGAGGTAACCTTCACCATCTAAATTTCACTTCTTTTCCCCATCTCCAATTGATTAATTTTTCTAATAACTCACTCAATGGTACACTTCCAAAAAAC

>M6-PGSC0003DMG400006364-(rev)M6LRR-RLK_1054

ATTTAGGTAACTTGCACTCTCTAGCGTATCTTGATCTCAGTTATAACATGCTCAGTGGTGAGATACCGTGGCACATAGGGAGTTTGAGAAGTTTGGAAAGAATGAATCTTTCGCGTAATA

>M6-PGSC0003DMG400006364-(rev)M6LRR-RLK_1116

ATACCGTGGCACATAGGGAGTTTGAGAAGTTTGGAAAGAATGAATCTTTCGCGTAATAACTTGTCTGGTTCAATCCCGCCTAGTTTCAATGAACGTGTTAGTTTGAGGTCAATTGATATA

>M6-PGSC0003DMG400006364-(rev)M6LRR-RLK_1178

GTCTGGTTCAATCCCGCCTAGTTTCAATGAACGTGTTAGTTTGAGGTCAATTGATATATCTTGCAATCAATTGGTTGGTCCCCTTCCCAAGATCGTGGCATTTCAAAATGCATCGCGTGA

>M6-PGSC0003DMG400006364-(rev)M6LRR-RLK_124

TCAATCTTTCAAGATTAAGTTTCCTATATTTGAGTTACAATCATTTTGATGGTGTGATTCCACATGAAATAGGGCTATTGAAGAATCTCAAGCACTTAGACTTGTCGGTAAATCTTTTTA

>M6-PGSC0003DMG400006364-(rev)M6LRR-RLK_1240

GCAATCAATTGGTTGGTCCCCTTCCCAAGATCGTGGCATTTCAAAATGCATCGCGTGAAGAATTAAGAGATAACAAGGATTTATGCAGCAATGATCATATTGGTATGAGACCTTGCTCTT

>M6-PGSC0003DMG400006364-(rev)M6LRR-RLK_1302

TTAAGAGATAACAAGGATTTATGCAGCAATGATCATATTGGTATGAGACCTTGCTCTTCATTGAGGAAAAAAGAGAGAACAAGTAGAAAGCTTATACTGACCTTAACTCTGTCTCTAATT

>M6-PGSC0003DMG400006364-(rev)M6LRR-RLK_1364

GAGGAAAAAAGAGAGAACAAGTAGAAAGCTTATACTGACCTTAACTCTGTCTCTAATTTTGGTTGCCTTGCTTCTACTTGTGATTACTTGTATACTTTTTCGAATGAAAAGGAAAAGAAA

>M6-PGSC0003DMG400006364-(rev)M6LRR-RLK_1426

TTGCCTTGCTTCTACTTGTGATTACTTGTATACTTTTTCGAATGAAAAGGAAAAGAAATAGAAGTATTCAGCCTAGAGAACCATCCAGCAATTCTTTTTCAGTTAGGGACTTTGATGGGA

>M6-PGSC0003DMG400006364-(rev)M6LRR-RLK_1488

AGTATTCAGCCTAGAGAACCATCCAGCAATTCTTTTTCAGTTAGGGACTTTGATGGGAAAATAGCACATGAAAACATAATTGCAGCAACAGAAAACTTTGATAGCAAATATTGCATTGGG

>M6-PGSC0003DMG400006364-(rev)M6LRR-RLK_1550

AGCACATGAAAACATAATTGCAGCAACAGAAAACTTTGATAGCAAATATTGCATTGGGAAGGGAGGACATGGAAGTGTCTATAAGGTTGAGTTACCATGTGGTCAAGTTGTTGCAGTGAA

>M6-PGSC0003DMG400006364-(rev)M6LRR-RLK_1612

GAGGACATGGAAGTGTCTATAAGGTTGAGTTACCATGTGGTCAAGTTGTTGCAGTGAAAAAAGTTCATGCTTTAGAAGATGAAGAATCTGATGACAACTTGATCAAAAGCTTTTCAAAAG

>M6-PGSC0003DMG400006364-(rev)M6LRR-RLK_1674

GTTCATGCTTTAGAAGATGAAGAATCTGATGACAACTTGATCAAAAGCTTTTCAAAAGAGATACAAGCTCTAGTAAACATCCGCCATCGGAATATTGTGAAGCTTTATGGGTTCTGTTCA

>M6-PGSC0003DMG400006364-(rev)M6LRR-RLK_1736

ACAAGCTCTAGTAAACATCCGCCATCGGAATATTGTGAAGCTTTATGGGTTCTGTTCACACGCGCGCCATTCATTCTTGGTTTATGAGTTGTTGGAAGGAGGAAACTTGTCTCAAAACCT

>M6-PGSC0003DMG400006364-(rev)M6LRR-RLK_1798

CGCGCCATTCATTCTTGGTTTATGAGTTGTTGGAAGGAGGAAACTTGTCTCAAAACCTTAGCAATGAGGGTAGAGCAAGAGATTTAAATTGGCTCAAGAGGGTTGATATTGTCAAGGGGA

>M6-PGSC0003DMG400006364-(rev)M6LRR-RLK_186

CATGAAATAGGGCTATTGAAGAATCTCAAGCACTTAGACTTGTCGGTAAATCTTTTTATTGGCTTTATCCCTTCATCTATTGGTAACTTGATTAAACTTGAGTTGTTATTTTTATCAGTT

>M6-PGSC0003DMG400006364-(rev)M6LRR-RLK_1860

AATGAGGGTAGAGCAAGAGATTTAAATTGGCTCAAGAGGGTTGATATTGTCAAGGGGATAGCAAACGCGTTGTGTTACTTGCATCACGCGTGTTCACCTCCTATAGTTCATAGAGATATA

>M6-PGSC0003DMG400006364-(rev)M6LRR-RLK_1922

AAACGCGTTGTGTTACTTGCATCACGCGTGTTCACCTCCTATAGTTCATAGAGATATATCAAGTAACAATGTTCTCTTAGATGATGAAGGCAATCCCCATGTTTCTGATTTTGGTACTGC

>M6-PGSC0003DMG400006364-(rev)M6LRR-RLK_1984

GTAACAATGTTCTCTTAGATGATGAAGGCAATCCCCATGTTTCTGATTTTGGTACTGCTAAACTCTTAAGGCCTAACTCCACCAACTGGACATCGTTTGCTGGAACTCTTGGCTATGTAG

>M6-PGSC0003DMG400006364-(rev)M6LRR-RLK_2170

GAAACAAATGGAGTATATTTTTAACTAGGAATTCATTGCTTCAAAATTGTTCACTCTTTCACACAATTTTCTAATTTGTTTGTAGAGCTTGCTTACACAATGAAGGTAAATGAGAAGAGT

>M6-PGSC0003DMG400006364-(rev)M6LRR-RLK_2232

ACAATTTTCTAATTTGTTTGTAGAGCTTGCTTACACAATGAAGGTAAATGAGAAGAGTGATGTTTACAGCTTCGGGGTGATATCACTAGAAGTTATTATAGGTCATCATCCAGGTGATAT

>M6-PGSC0003DMG400006364-(rev)M6LRR-RLK_2294

TTTACAGCTTCGGGGTGATATCACTAGAAGTTATTATAGGTCATCATCCAGGTGATATTACTCAAGCTATATCATCATTTTCGCCAGCATCAGGAGCGAATGGAACGCTGTTAAAGGAAT

>M6-PGSC0003DMG400006364-(rev)M6LRR-RLK_2356

CAAGCTATATCATCATTTTCGCCAGCATCAGGAGCGAATGGAACGCTGTTAAAGGAATTGATTGACAAACGGACACTAGCTCCTGGAAAACAAGAAGCAGAAGAATTGATCAAGATCACA

>M6-PGSC0003DMG400006364-(rev)M6LRR-RLK_2418

TGACAAACGGACACTAGCTCCTGGAAAACAAGAAGCAGAAGAATTGATCAAGATCACAAAACTAGCATTCGCGTGTCTTCATCAGAGTCCTCTAGCTCGGCCAAGTATGAAGCAAGTTTG

>M6-PGSC0003DMG400006364-(rev)M6LRR-RLK_248

CTTTATCCCTTCATCTATTGGTAACTTGATTAAACTTGAGTTGTTATTTTTATCAGTTAATGAACTCTATGGCCCTGTCCCTTCAAGTTTTGGTAACTTAAAATCCCTTGTGCATTTGTG

>M6-PGSC0003DMG400006364-(rev)M6LRR-RLK_2480

TAGCATTCGCGTGTCTTCATCAGAGTCCTCTAGCTCGGCCAAGTATGAAGCAAGTTTGTGCATCTTTATCAAAAGAAAATTGGCCTTCAAAAGGCTTGTTCTCCACTGTTACATTAGGCC

>M6-PGSC0003DMG400006364-(rev)M6LRR-RLK_2512

GCTCGGCCAAGTATGAAGCAAGTTTGTGCATCTTTATCAAAAGAAAATTGGCCTTCAAAAGGCTTGTTCTCCACTGTTACATTAGGCCAGTTGATTGAACCAACACTACTAAGATGTTGA

>M6-PGSC0003DMG400006364-(rev)M6LRR-RLK_310

AACTCTATGGCCCTGTCCCTTCAAGTTTTGGTAACTTAAAATCCCTTGTGCATTTGTGTTTGCTTAGAAATAAACTCAATGGATCACTTCCAAAAGAGTTTGAAAATCTTACACATATGC

>M6-PGSC0003DMG400006364-(rev)M6LRR-RLK_372

CTTAGAAATAAACTCAATGGATCACTTCCAAAAGAGTTTGAAAATCTTACACATATGCAAATTTTTCAAGTAGCAGAGAACAATTTTTCTGGTCATTTGCCACAAAATGTGTGCCTTGGT

>M6-PGSC0003DMG400006364-(rev)M6LRR-RLK_434

TTTTCAAGTAGCAGAGAACAATTTTTCTGGTCATTTGCCACAAAATGTGTGCCTTGGTGGATCACTTGCTAAATTCATAGCATATGACAATAAGTTTATTGGTCATGTCCCAAAAACCTT

>M6-PGSC0003DMG400006364-(rev)M6LRR-RLK_496

CACTTGCTAAATTCATAGCATATGACAATAAGTTTATTGGTCATGTCCCAAAAACCTTGAAAAATTGCTCTACGCTATCAAGAGTTAGGCTTGATGGTAACCAATTGTCTAGCAATATAT

>M6-PGSC0003DMG400006364-(rev)M6LRR-RLK_558

AATTGCTCTACGCTATCAAGAGTTAGGCTTGATGGTAACCAATTGTCTAGCAATATATCAGAAGCTTTTGGTGTCTATCCAAGTCTAGTTTACATGGATTTGAGTCACAATAAACTATAT

>M6-PGSC0003DMG400006364-(rev)M6LRR-RLK_62

CCATCTCCAATTGATTAATTTTTCTAATAACTCACTCAATGGTACACTTCCAAAAAACATGTTCAATCTTTCAAGATTAAGTTTCCTATATTTGAGTTACAATCATTTTGATGGTGTGAT

>M6-PGSC0003DMG400006364-(rev)M6LRR-RLK_620

AGCTTTTGGTGTCTATCCAAGTCTAGTTTACATGGATTTGAGTCACAATAAACTATATGGCGAGTTATCGTCACAATGGGGCTTTTCTCATAATCTAACAAGTTTGAAGATCTCCAAGAA

>M6-PGSC0003DMG400006364-(rev)M6LRR-RLK_682

AGTTATCGTCACAATGGGGCTTTTCTCATAATCTAACAAGTTTGAAGATCTCCAAGAACAACTTGTTTGGTGCCATACCAGTTGAAATAGGAAATTTGATCAAACTAGAAATGCTTGATC

>M6-PGSC0003DMG400006364-(rev)M6LRR-RLK_744

TTGTTTGGTGCCATACCAGTTGAAATAGGAAATTTGATCAAACTAGAAATGCTTGATCTCTCTTCAAATCACTTGACCGGCGAAATTCCTAAAAGTTTAGAGAGGCTAACTCTCTTGCTG

>M6-PGSC0003DMG400006364-(rev)M6LRR-RLK_806

TTCAAATCACTTGACCGGCGAAATTCCTAAAAGTTTAGAGAGGCTAACTCTCTTGCTGGTACTTGATTTACATGAGAATATAATTTCAGGAAAAATACCTATAGAAGTGGGGAAGTTGTC

>M6-PGSC0003DMG400006364-(rev)M6LRR-RLK_868

TTGATTTACATGAGAATATAATTTCAGGAAAAATACCTATAGAAGTGGGGAAGTTGTCTAAACTTACAAGACTTAACTTAGGTGCGAATAATATGAGTGGAACGATTCCAGGAGAAATAG

>M6-PGSC0003DMG400006364-(rev)M6LRR-RLK_930

CTTACAAGACTTAACTTAGGTGCGAATAATATGAGTGGAACGATTCCAGGAGAAATAGGAGATTGTAGGCAGCTTTGGTACTTGAATTTAAGCAAGAACATGCTGAACACAACTATTCCT

>M6-PGSC0003DMG400006364-(rev)M6LRR-RLK_992

TTGTAGGCAGCTTTGGTACTTGAATTTAAGCAAGAACATGCTGAACACAACTATTCCTTCTAATTTAGGTAACTTGCACTCTCTAGCGTATCTTGATCTCAGTTATAACATGCTCAGTGG

>M6-PGSC0003DMG400006422M6LRR-RLP_0

ATGGTCAAAGGAAAAAAGGAAGTGGGAGATGATGATGAGTTAAATTATTCCTACTTTTTGGGAGTTTTGAGAGAGAGTGCCATAGTGACGAATAAAGGCAACAGGTACCAGTATGACACC

>M6-PGSC0003DMG400006422M6LRR-RLP_128

GTTGTTTACAAGTATGGACATGTCTAGCAATAATCTCTCTGGGGATATTCCTATAAGTTTAACACTTCTTGCAGGATTAAGATCATTTAATTTTTCTAAAAATAACCTAACTGGTAGGAT

>M6-PGSC0003DMG400006422M6LRR-RLP_192

CTTCTTGCAGGATTAAGATCATTTAATTTTTCTAAAAATAACCTAACTGGTAGGATTCCAAATGACATTGGTGACATGAAAGTTTTGGAATCTGTTGATCTTTCAGAAAATCAATTTTAT

>M6-PGSC0003DMG400006422M6LRR-RLP_256

ACATTGGTGACATGAAAGTTTTGGAATCTGTTGATCTTTCAGAAAATCAATTTTATGGTCAAATCCCACAAAGCTTTTCAAGTTTGTTCACCTTGAGCTACTTGAATCTATCTGATAACA

>M6-PGSC0003DMG400006422M6LRR-RLP_320

CCCACAAAGCTTTTCAAGTTTGTTCACCTTGAGCTACTTGAATCTATCTGATAACAATTTGTCAGGTATGATACCATTGAGCACTCAACTTCAAAGCTTTGATCCCACGAGTTTTCAAGG

>M6-PGSC0003DMG400006422M6LRR-RLP_384

GGTATGATACCATTGAGCACTCAACTTCAAAGCTTTGATCCCACGAGTTTTCAAGGAAACAAACTCTGCGGGCTTCCACTCTCGCTGAATTGTAGTTCAAATGGTAATATTCCAAATCAT

>M6-PGSC0003DMG400006422M6LRR-RLP_573

GCACTAAGCTTTTGGGGTGTTTGTGGTTCATTGCTTTTCAAGAGATCATGGAGACATGCTTATTTTTGTTTTCTAGACCGTAGCTGGGAAATGTTGCTTGCAAAGTTATCAGTATGTTGA

>M6-PGSC0003DMG400006422M6LRR-RLP_64

TTTTGAGAGAGAGTGCCATAGTGACGAATAAAGGCAACAGGTACCAGTATGACACCATTTTGGCGTTGTTTACAAGTATGGACATGTCTAGCAATAATCTCTCTGGGGATATTCCTATAA

>M6-PGSC0003DMG400006458M6LRR-RLP_0

TTTCTCCTCTCCATTCCCTATAAGTGTTTGTGTCACCACACACAAGTAACGACCCCTCAATTTGTCAATAAACAAATAATAGGATTAGTTAATTAACAAAGGAAGAAGAAGATTATGGAA

>M6-PGSC0003DMG400006458M6LRR-RLP_1037

TTCTGACAAGCAGTCACGACTAGTATTTTGGAAAACATTAATTTTTCAATCATACCAAATACAGTCGTTGCTTCCATTTTTAGTTGCTTTTGGTTCCTTGTACTTTCACAAATATTTAGT

>M6-PGSC0003DMG400006458M6LRR-RLP_1098

CAGTCGTTGCTTCCATTTTTAGTTGCTTTTGGTTCCTTGTACTTTCACAAATATTTAGTTTTGTTTAATTTCTTCCATATTTATGGAAAGATATATTTATCATTTACCTTTCTTCTATTT

>M6-PGSC0003DMG400006458M6LRR-RLP_1403

AAAGAGGTGTCAAAATAAGAATTTAATGTACTACTATATTTTCTTAAAATATTCCCCTTTACTTGATAATGAGTCGTTTATGTGGACTATACAGTATAATATTTTCAGGAATAAAATGCA

>M6-PGSC0003DMG400006458M6LRR-RLP_1586

TCCAAATCAACCGTCCTAACGTCATATTTTGTCGTTCATCATTATTGTAAATTGACCCAATAGTAAGGATAAAGTCTTGTATAAGTTTATTTTTTACTTCATTGATATATTTAAAAAAAA

>M6-PGSC0003DMG400006458M6LRR-RLP_1647

AGTAAGGATAAAGTCTTGTATAAGTTTATTTTTTACTTCATTGATATATTTAAAAAAAATATAACAACTTGAAAGGTTTTCCAATTATTCGTATTAAACGCAATAGTATATTTTTCTGGT

>M6-PGSC0003DMG400006458M6LRR-RLP_1708

TAACAACTTGAAAGGTTTTCCAATTATTCGTATTAAACGCAATAGTATATTTTTCTGGTTGTTATATGTATGTGTGAATTGTGGATTTACTTAAATATAATTGACAAAATTTGACAAAAC

>M6-PGSC0003DMG400006458M6LRR-RLP_1769

TTATATGTATGTGTGAATTGTGGATTTACTTAAATATAATTGACAAAATTTGACAAAACACGATAGTACATAACGTAACAACATTTTCGTTAATTATATATCCCGTCTTTGTTTATAGTC

>M6-PGSC0003DMG400006458M6LRR-RLP_1830

GATAGTACATAACGTAACAACATTTTCGTTAATTATATATCCCGTCTTTGTTTATAGTCTAATACAATGTATATGTCTCTCATTGTGTTTTACAATGTGTATGTGTCTCACTCGTCTCTC

>M6-PGSC0003DMG400006458M6LRR-RLP_1891

ATACAATGTATATGTCTCTCATTGTGTTTTACAATGTGTATGTGTCTCACTCGTCTCTCTTCTTCTATGTATTTGTATTTTATAATGTATATGTGTCTCACAATTTCAATTATAAGTACC

>M6-PGSC0003DMG400006458M6LRR-RLP_1952

CTTCTATGTATTTGTATTTTATAATGTATATGTGTCTCACAATTTCAATTATAAGTACCATACAAATATCTGATACACAGATACGTCGACGGTTTATATTTTTTTTTTATATCAATGTAT

>M6-PGSC0003DMG400006458M6LRR-RLP_2013

ACAAATATCTGATACACAGATACGTCGACGGTTTATATTTTTTTTTTATATCAATGTATCTATTTCATATATGTATCGATTTTAAGTAGCATGAATTGTATCATAGATTAACACTTTATC

>M6-PGSC0003DMG400006458M6LRR-RLP_2074

ATTTCATATATGTATCGATTTTAAGTAGCATGAATTGTATCATAGATTAACACTTTATCATTTTACTTTCTCTGTCCAACAATAGTTGTCCACTATTGACTTGGCATACGGCATACCCCT

>M6-PGSC0003DMG400006458M6LRR-RLP_244

GGAAAAAGTACAAGAAGAAAGATGACAATTAGTATTGTTGAAGAAGAAGAAGAAGAGAGCCCAAAGATTGTGGATCTAAGTGGCATGTCATTAGACTCTCTTCCTGTAAATCCTACAATC

>M6-PGSC0003DMG400006458M6LRR-RLP_2684

GAGTATATTTATCCTTTTTCCCTTTTTAGTATAGTGGACAACACTATTGTTGGACAGAGGAAGTACTAGACACAGGATTTTAGGGTAAGTCACATTCTTATTTGGTTGGACGATTTTCAA

>M6-PGSC0003DMG400006458M6LRR-RLP_2745

AGTACTAGACACAGGATTTTAGGGTAAGTCACATTCTTATTTGGTTGGACGATTTTCAATTATCATGTATCACTTGTGTATATAGATACATAAACACACATATGTATCTCTTCAAAGTTC

>M6-PGSC0003DMG400006458M6LRR-RLP_2989

ATATATATATCTCTTAAAAAATTTCCACTCGTCTTTCATGGATACACAAATACACATCCATAGATACGCAATACACATATATATCTCCTCAAAACTACTCACTCATTTGCAATAGATGCA

>M6-PGSC0003DMG400006458M6LRR-RLP_305

CAAAGATTGTGGATCTAAGTGGCATGTCATTAGACTCTCTTCCTGTAAATCCTACAATCAATTTGGGAGCTATATCCAAATTACATATTTCCAACAATAATCTTCAGGTTAGTCAGTTTT

>M6-PGSC0003DMG400006458M6LRR-RLP_3050

AGATACGCAATACACATATATATCTCCTCAAAACTACTCACTCATTTGCAATAGATGCACAAAAATATAGATACACTTATACATTGATATACATATGTATCTCCTCAAAGTTGCACTCTA

>M6-PGSC0003DMG400006458M6LRR-RLP_3111

AAAATATAGATACACTTATACATTGATATACATATGTATCTCCTCAAAGTTGCACTCTAATTTGCCATAGATGCACAAATACATATAGTCCCTCTGTAAATCGAGAGTGGAGAGAGGGAG

>M6-PGSC0003DMG400006458M6LRR-RLP_3172

TTGCCATAGATGCACAAATACATATAGTCCCTCTGTAAATCGAGAGTGGAGAGAGGGAGGAGAGTGGCGAATTTTCTAGGAAAAAGGGAAATCTCATTGAGTTATGTAATTTTTTAAAAC

>M6-PGSC0003DMG400006458M6LRR-RLP_3233

GAGTGGCGAATTTTCTAGGAAAAAGGGAAATCTCATTGAGTTATGTAATTTTTTAAAACTATAAATAGGTTGAGTAGATATATATACTAGCTATGCTTGATTAATTATGTAATTTTTCAT

>M6-PGSC0003DMG400006458M6LRR-RLP_3294

TAAATAGGTTGAGTAGATATATATACTAGCTATGCTTGATTAATTATGTAATTTTTCATGTATAATTCTCACTTCATTGTTTTTGTTGTATGTTTGACAGAGTATACCAGAGTCTTTAAC

>M6-PGSC0003DMG400006458M6LRR-RLP_3355

ATAATTCTCACTTCATTGTTTTTGTTGTATGTTTGACAGAGTATACCAGAGTCTTTAACAGCAAGACTACTTAACTTGGTTGAGTTAGATATGCATTCAAATCAGCTCAACTTCATTCCT

>M6-PGSC0003DMG400006458M6LRR-RLP_3416

CAAGACTACTTAACTTGGTTGAGTTAGATATGCATTCAAATCAGCTCAACTTCATTCCTAACTCAATAGGTTGCTTATCAAAGCTCAAACTTCTCAACATTTCTGGGAATCTCCTTCTTT

>M6-PGSC0003DMG400006458M6LRR-RLP_3477

CTCAATAGGTTGCTTATCAAAGCTCAAACTTCTCAACATTTCTGGGAATCTCCTTCTTTCGCTCCCTAAAACTATCGAGAATTGTAGGTTAGTCATCTAAACATTTCTAATCCATAACAT

>M6-PGSC0003DMG400006458M6LRR-RLP_3538

CTCCCTAAAACTATCGAGAATTGTAGGTTAGTCATCTAAACATTTCTAATCCATAACATTATCACTACAACAAAAGAGATATTTAGCAGCAATTGAGTTACTGTCATTGCATTCAAAGTC

>M6-PGSC0003DMG400006458M6LRR-RLP_3599

TCACTACAACAAAAGAGATATTTAGCAGCAATTGAGTTACTGTCATTGCATTCAAAGTCGCTAAAGCCTTTAGAACATAATAAGTATAACCTTTTCTTCCTACTTGCAGATCATTGGAAG

>M6-PGSC0003DMG400006458M6LRR-RLP_366

TTTGGGAGCTATATCCAAATTACATATTTCCAACAATAATCTTCAGGTTAGTCAGTTTTCGTACACCTCAATTAATTTTTGTGAAATATTTGTTACCTTCTATCAGCAATAAGCAATATG

>M6-PGSC0003DMG400006458M6LRR-RLP_3660

TAAAGCCTTTAGAACATAATAAGTATAACCTTTTCTTCCTACTTGCAGATCATTGGAAGAGTTGAATGCCAACTTCAATATGCTTACTCATTTACCGGACACCATTGGATTTGAGCTCAT

>M6-PGSC0003DMG400006458M6LRR-RLP_3721

TTGAATGCCAACTTCAATATGCTTACTCATTTACCGGACACCATTGGATTTGAGCTCATAAATCTCAAGAAGCTATGCATAAATTCAAATAAAATAGCATACCTTCCTTACTCCACCTCC

>M6-PGSC0003DMG400006458M6LRR-RLP_3782

ATCTCAAGAAGCTATGCATAAATTCAAATAAAATAGCATACCTTCCTTACTCCACCTCCCACTTGACCAACCTTCGCGTGTTAGATGCTAGGCTAAACTGCCTTAGATCACTCCCCGATG

>M6-PGSC0003DMG400006458M6LRR-RLP_3843

CTTGACCAACCTTCGCGTGTTAGATGCTAGGCTAAACTGCCTTAGATCACTCCCCGATGATCTTGAGAACCTAATCAACCTTGAAATCCTTAACGTTAGTCAAAATTTCCAGTACCTCTC

>M6-PGSC0003DMG400006458M6LRR-RLP_3904

CTTGAGAACCTAATCAACCTTGAAATCCTTAACGTTAGTCAAAATTTCCAGTACCTCTCAAAGTTGCCGTATTCAGTAGGATTTCTCATTTCTCTTCATGAACTAGATGTTAGTTACAAC

>M6-PGSC0003DMG400006458M6LRR-RLP_3965

AGTTGCCGTATTCAGTAGGATTTCTCATTTCTCTTCATGAACTAGATGTTAGTTACAACAAGATTACTGAATTACCAGATTCCATCGGATGCTTGAAGAAGCTCCAAAAGCTGAGTGTAG

>M6-PGSC0003DMG400006458M6LRR-RLP_4026

GATTACTGAATTACCAGATTCCATCGGATGCTTGAAGAAGCTCCAAAAGCTGAGTGTAGAAGGCAACCGCCTTGTTTCTCCCCCACCGGAGGTAGTGGAGCAAGGGATACGTGCAGTTAA

>M6-PGSC0003DMG400006458M6LRR-RLP_4148

AATACTTGTGTGAAAAGATAAATGGAATGCATGACAAGTCCCCTAAGAAGAAGTCATGGTTTGGAAAGCTAGCAAGATGTAGCACATTTAGTGGTGCTAACATACCAAGAGATGATAGAG

>M6-PGSC0003DMG400006458M6LRR-RLP_4209

TGGAAAGCTAGCAAGATGTAGCACATTTAGTGGTGCTAACATACCAAGAGATGATAGAGAGGACTTTGGTGTTCCAAGCAATCGTACTATCGATGCAATTGCCTCACCACGATTTATGGG

>M6-PGSC0003DMG400006458M6LRR-RLP_427

TACACCTCAATTAATTTTTGTGAAATATTTGTTACCTTCTATCAGCAATAAGCAATATGTACTAGATGACTCTGTTCATAGAGGTTAGGATAGATGAATAAATTCACTTAGTTTTTTTTT

>M6-PGSC0003DMG400006458M6LRR-RLP_4270

GACTTTGGTGTTCCAAGCAATCGTACTATCGATGCAATTGCCTCACCACGATTTATGGGAATGCTTTCTCCTCGTCGTCTTCTATCTCCAAAAACCTACTTCTCCAAATAAGGGACATTG

>M6-PGSC0003DMG400006458M6LRR-RLP_4331

TGCTTTCTCCTCGTCGTCTTCTATCTCCAAAAACCTACTTCTCCAAATAAGGGACATTGCAATTTGCTTGGTAGCTCTTAGCTTGTTTTCTTTTACGTTTGAAATCGTGCAATGTGTGCT

>M6-PGSC0003DMG400006458M6LRR-RLP_4392

ATTTGCTTGGTAGCTCTTAGCTTGTTTTCTTTTACGTTTGAAATCGTGCAATGTGTGCTAGATCAAGCTGTACATTATGTTCTGTAGATTTTGTTGTCTTTGAGATAGTACAGATTTTCA

>M6-PGSC0003DMG400006458M6LRR-RLP_4453

ATCAAGCTGTACATTATGTTCTGTAGATTTTGTTGTCTTTGAGATAGTACAGATTTTCATAGATGATTAGCCTCATGAGTCATGGGTATGTTTGAGAGGTAACAATGCGGTACAAATTTA

>M6-PGSC0003DMG400006458M6LRR-RLP_4514

GATGATTAGCCTCATGAGTCATGGGTATGTTTGAGAGGTAACAATGCGGTACAAATTTACCCGAACAGAGTGTTTATATCAAATCATATGAATAATTACATCGAATCAAAATTTGTTATC

>M6-PGSC0003DMG400006458M6LRR-RLP_4575

CGAACAGAGTGTTTATATCAAATCATATGAATAATTACATCGAATCAAAATTTGTTATCATAACATGTCTTGTATTATATCACTTGTACAAAAACTCATTTTGATCATTAATTATCAAAA

>M6-PGSC0003DMG400006458M6LRR-RLP_4636

AACATGTCTTGTATTATATCACTTGTACAAAAACTCATTTTGATCATTAATTATCAAAACTCAAAAAAAAAAAAATTAAAAATCTTACATAAACAGAACCATTAAAAAAGCCCCAAAAAA

>M6-PGSC0003DMG400006458M6LRR-RLP_4697

CAAAAAAAAAAAAATTAAAAATCTTACATAAACAGAACCATTAAAAAAGCCCCAAAAAAACAGAACAAGACTCTTAGAAAGCAAACAGAACTAATGAAAAACTTCCCAAAATCACAAATT

>M6-PGSC0003DMG400006458M6LRR-RLP_4758

AGAACAAGACTCTTAGAAAGCAAACAGAACTAATGAAAAACTTCCCAAAATCACAAATTATGAATTAGTTTCTTGCACTTGCGATTCATTTCTTTGCAAATCCGATTCACTTCTCCTATT

>M6-PGSC0003DMG400006458M6LRR-RLP_4819

GAATTAGTTTCTTGCACTTGCGATTCATTTCTTTGCAAATCCGATTCACTTCTCCTATTAAACGACCATTCTCTAATGGTAGACAAAACTGGCTTCCATGATCGCGATTTATTTGACCTT

>M6-PGSC0003DMG400006458M6LRR-RLP_488

CTAGATGACTCTGTTCATAGAGGTTAGGATAGATGAATAAATTCACTTAGTTTTTTTTTTTTTTTGTCTTCGCTAGAATTAATTGAACAACCACTTCGATGATCACTATGTGCTATCGTG

>M6-PGSC0003DMG400006458M6LRR-RLP_4880

ACGACCATTCTCTAATGGTAGACAAAACTGGCTTCCATGATCGCGATTTATTTGACCTTTCTCCAAATTTGCACGAAGTTGTGTTAGTTGCACCACTTTTAGCTATGCAACTGAGTTTTC

>M6-PGSC0003DMG400006458M6LRR-RLP_4941

TCCAAATTTGCACGAAGTTGTGTTAGTTGCACCACTTTTAGCTATGCAACTGAGTTTTCTAGCTATTGGAACTAAATCCATTGGAGAAACACTTGATTGAAATAAGGATTTAGGCAATAA

>M6-PGSC0003DMG400006458M6LRR-RLP_5002

GCTATTGGAACTAAATCCATTGGAGAAACACTTGATTGAAATAAGGATTTAGGCAATAAATAGTAAATATGACCTTGTTCTAACTTAGTATCAGGGTTGAGAGGCTTTGAAGCAGCAGAT

>M6-PGSC0003DMG400006458M6LRR-RLP_5063

AGTAAATATGACCTTGTTCTAACTTAGTATCAGGGTTGAGAGGCTTTGAAGCAGCAGATAAAAGTTGGGCATGAGTGAAGACAAAGTGTTTTTGAGATTTGGCCATGAGTTCATGTACAC

>M6-PGSC0003DMG400006458M6LRR-RLP_5124

AAGTTGGGCATGAGTGAAGACAAAGTGTTTTTGAGATTTGGCCATGAGTTCATGTACACTAATAGGATGATCAAAATCCTCTACATGACCATTCAAATGTACAACTCTAATAGTCTTTAA

>M6-PGSC0003DMG400006458M6LRR-RLP_5185

ATAGGATGATCAAAATCCTCTACATGACCATTCAAATGTACAACTCTAATAGTCTTTAATATATCACATGATTTGGGAGAAAAACAACAATATGTTATTTTGATGTACATATTGTTAATA

>M6-PGSC0003DMG400006458M6LRR-RLP_5246

TATCACATGATTTGGGAGAAAAACAACAATATGTTATTTTGATGTACATATTGTTAATATTTTCCTTATTTCTCTCTAGCTAGTGTGTAAAAGGGTTCCCAAAGAAGATGTCCTTTTATA

>M6-PGSC0003DMG400006458M6LRR-RLP_5307

TTCCTTATTTCTCTCTAGCTAGTGTGTAAAAGGGTTCCCAAAGAAGATGTCCTTTTATATAAGGGGAAAGATGGTGCTGAAACCGGGTTGTGGTGCGGTAGTGGAATTGCTCCACCCTTA

>M6-PGSC0003DMG400006458M6LRR-RLP_549

TTTTGTCTTCGCTAGAATTAATTGAACAACCACTTCGATGATCACTATGTGCTATCGTGAATTATTTATTTATGAGAATTTTATTCATGATAGAATATTTTTAATCTTTGATTGGAAATT

>M6-PGSC0003DMG400006458M6LRR-RLP_5490

ATGGGCCTTGCAATGTGCAAATACAGATTTAGTCGGGGATGGGCCCTACAATGCACAATTCAGATTTAATCTAAGCTCGAATGCGGACTCCGGATATCGAGTAGAAACAAAAAAAATGGT

>M6-PGSC0003DMG400006458M6LRR-RLP_5551

AGATTTAATCTAAGCTCGAATGCGGACTCCGGATATCGAGTAGAAACAAAAAAAATGGTGGTGGGAAAGAGTTGTGGAGATAGATGATGGGAACACGCTATTTAAGTTAGGAATTTTTTA

>M6-PGSC0003DMG400006458M6LRR-RLP_5612

TGGGAAAGAGTTGTGGAGATAGATGATGGGAACACGCTATTTAAGTTAGGAATTTTTTATCTACTTCACGGCTTAAGTGGCACTCTCAATAGGATTTTCTTCATACCAAGGCTCAAAGTG

>M6-PGSC0003DMG400006458M6LRR-RLP_5643

ACACGCTATTTAAGTTAGGAATTTTTTATCTACTTCACGGCTTAAGTGGCACTCTCAATAGGATTTTCTTCATACCAAGGCTCAAAGTGAGACTTGTGGTTAAGGATAGAGCAATTTCAT

>M6-PGSC0003DMG400006458M6LRR-RLP_61

TTGTCAATAAACAAATAATAGGATTAGTTAATTAACAAAGGAAGAAGAAGATTATGGAATTTCAATGATGATTTGTGTTAGGATTAAGAGATCATGAAGATGAGGATGGAATTAGTTGAA

>M6-PGSC0003DMG400006458M6LRR-RLP_610

TTATTTATTTATGAGAATTTTATTCATGATAGAATATTTTTAATCTTTGATTGGAAATTAAGTGAAAATATCTTCTTTTTGAAATAAGTGGTGGCTAAAGTATTTGATATGAAGATAAGA

>M6-PGSC0003DMG400006458M6LRR-RLP_671

GTGAAAATATCTTCTTTTTGAAATAAGTGGTGGCTAAAGTATTTGATATGAAGATAAGATAATTATGAAGAAAAATGTTTTAGTCAAAATCGAGTGAAATTATTTTTTTCATTTTGACTG

>M6-PGSC0003DMG400006458M6LRR-RLP_732

ATTATGAAGAAAAATGTTTTAGTCAAAATCGAGTGAAATTATTTTTTTCATTTTGACTGGAGGCGAACTAGTTGATTGCTTTGAATTGTATCAGCAACACAAGGACACTATTAAAAACAC

>M6-PGSC0003DMG400006458M6LRR-RLP_793

GGCGAACTAGTTGATTGCTTTGAATTGTATCAGCAACACAAGGACACTATTAAAAACACTAGTTGATTGCTTTGAATTGTATCAGCAACACAAGGACACTATTAAAAACACTATTGCATT

>M6-PGSC0003DMG400006458M6LRR-RLP_854

GTTGATTGCTTTGAATTGTATCAGCAACACAAGGACACTATTAAAAACACTATTGCATTTCATAAATGTATGATTAATTATATATTAATTTATGTCGATATTAATAGGTGGATGTGCACT

>M6-PGSC0003DMG400006458M6LRR-RLP_915

ATAAATGTATGATTAATTATATATTAATTTATGTCGATATTAATAGGTGGATGTGCACTACCTATATAGGTGGATCTGCCTCTAATATAATTTTGACCAAAGAATATTTTTCTAAAAATA

>M6-PGSC0003DMG400006458M6LRR-RLP_976

CTATATAGGTGGATCTGCCTCTAATATAATTTTGACCAAAGAATATTTTTCTAAAAATATTTTCTGACAAGCAGTCACGACTAGTATTTTGGAAAACATTAATTTTTCAATCATACCAAA

>M6-PGSC0003DMG400006486M6LRR-RLP_0

CTAAGGTTACAGTGTTGCGAGGGTATAAGCAAATGTCATATTTTTGGCTTAGTACATCTCAAGTATTTCGATGCTAGTTATTGCAAACTCCATAGTCTGATAGTCCAAGCTCCATCTCTT

>M6-PGSC0003DMG400006486M6LRR-RLP_1056

GTAAAGAATGTCCATTCAATACCAAGTGGCATCGTAACTTAAAAGAAGTCATCAGTTGTAGTGGGACATCTGAAGAGGGAATGGCTGCCTCCATGTGGTATCTCTGGTTGAAATCCACAT

>M6-PGSC0003DMG400006486M6LRR-RLP_1109

AGTTGTAGTGGGACATCTGAAGAGGGAATGGCTGCCTCCATGTGGTATCTCTGGTTGAAATCCACATCTGTAATTGACCAAGTGAATAATTTCGTGTTAAAATGGAAAGATGAAGCCTAG

>M6-PGSC0003DMG400006486M6LRR-RLP_198

ACTTTACAAACACTCGAGCTCGCTGGTGCCAGCATCACAGACCAACAGCTTCGAGATTTATCCTATAAGTTCCCAAACATTTCAGAATTGAATCTAACAAGATGCTATAACCTGAAAAAT

>M6-PGSC0003DMG400006486M6LRR-RLP_264

AAGTTCCCAAACATTTCAGAATTGAATCTAACAAGATGCTATAACCTGAAAAATATAGAGATTCAGAGTGAAAAACTCAAGAAATTCACCTTATTAAAGTTGAAGAGTCTGGAAAAACTC

>M6-PGSC0003DMG400006486M6LRR-RLP_330

AGTGAAAAACTCAAGAAATTCACCTTATTAAAGTTGAAGAGTCTGGAAAAACTCACGATTCAAGCTCCAAATTCATTGGAATTTGATTTTCATGGTAGTAAATGGCCCTTCTCATATATG

>M6-PGSC0003DMG400006486M6LRR-RLP_396

CCAAATTCATTGGAATTTGATTTTCATGGTAGTAAATGGCCCTTCTCATATATGGATACTTCTTCCCTAGAAAGAGTCCGACTCAATTCTTCTTGCCCGTCATAAACTTTGGATCAGTGG

>M6-PGSC0003DMG400006486M6LRR-RLP_462

CTAGAAAGAGTCCGACTCAATTCTTCTTGCCCGTCATAAACTTTGGATCAGTGGACAGCAGTTGGTACACGAGCCTTCATCACTTTGTTCAAAAGTTCAACTATTCTAAAGATTTCATAT

>M6-PGSC0003DMG400006486M6LRR-RLP_528

ACACGAGCCTTCATCACTTTGTTCAAAAGTTCAACTATTCTAAAGATTTCATATTAATAATATTTTGCCGCCAGGTAGTTTACATTCTTTTTATGCTTATGTCATTTATTTATGCATTAC

>M6-PGSC0003DMG400006486M6LRR-RLP_594

GCCGCCAGGTAGTTTACATTCTTTTTATGCTTATGTCATTTATTTATGCATTACTTGTTGAATTTGGCTTAAAATATGGCATTGACTAAATTCTTGCAGACCAAAAGCATCCTTATTTAT

>M6-PGSC0003DMG400006486M6LRR-RLP_66

TTCGATGCTAGTTATTGCAAACTCCATAGTCTGATAGTCCAAGCTCCATCTCTTCAATGCTTTAGATATAGAGAACCTACTGAACAGGGAAATCTTCATCCTTGCCAAATTGCTATTTTG

>M6-PGSC0003DMG400006486M6LRR-RLP_660

GCTTAAAATATGGCATTGACTAAATTCTTGCAGACCAAAAGCATCCTTATTTATGAAAATCCAAGAGAAATTGTTATTCCACCAAGCCACAAAATCGAGATATTCATTACATCCATGCTG

>M6-PGSC0003DMG400006486M6LRR-RLP_726

GAAATTGTTATTCCACCAAGCCACAAAATCGAGATATTCATTACATCCATGCTGCGTGTTGAATCGATCATTGGAATGTTAATGGTCTACCGTCCCTATATCATGTCCATACTTCCATGT

>M6-PGSC0003DMG400006486M6LRR-RLP_792

ATCATTGGAATGTTAATGGTCTACCGTCCCTATATCATGTCCATACTTCCATGTACAGATAGCAAAGCACTACAGGTGATTTCTTGCTTTACCTCAAAAACGTCTCTTGTGCTTGTATAA

>M6-PGSC0003DMG400006486M6LRR-RLP_858

GCACTACAGGTGATTTCTTGCTTTACCTCAAAAACGTCTCTTGTGCTTGTATAAACTAAGACATTAATTTGAGAAGAAAATTTGTTGTTAAATGTTGATTGGTCGTCTCATAAGCATTAT

>M6-PGSC0003DMG400006486M6LRR-RLP_924

ATTTGAGAAGAAAATTTGTTGTTAAATGTTGATTGGTCGTCTCATAAGCATTATTGTATAATATATCTTATGTTTTTGCTTAAATGCAGGCGATGTCTACACTAAAAGGGTGTACACAAA

>M6-PGSC0003DMG400006486M6LRR-RLP_990

CTTATGTTTTTGCTTAAATGCAGGCGATGTCTACACTAAAAGGGTGTACACAAAAGCAAAAATGTGGTAAAGAATGTCCATTCAATACCAAGTGGCATCGTAACTTAAAAGAAGTCATCA

>M6-PGSC0003DMG400006487M6LRR-RLP_0

TATAAACCCTAACAGCAAACGGCGGCACCTACCAAAATCAGAAACCCTAATGGAAGAAAGTGTTGATGGAGAAGACAGAATATCAGAATTGCCTGTGCATATCATTCATCACATCTTATG

>M6-PGSC0003DMG400006487M6LRR-RLP_1054

GATAAAATGCCCTTCTTATCTATGGATTCTTCTTCCCTGGAAAGAGCCCGACTCAATTTCTTCTTGCCGAGGACAGTAATATCAAACTTTGGAGATGTGGATAGCAGTTGGTACACAAAC

>M6-PGSC0003DMG400006487M6LRR-RLP_1116

CTTGCCGAGGACAGTAATATCAAACTTTGGAGATGTGGATAGCAGTTGGTACACAAACCTTAATCACTTTGTTCAAAACTTCAACTATTCTAAAGGTTTGATGTTAGTAATATTTTGCCG

>M6-PGSC0003DMG400006487M6LRR-RLP_1178

ATCACTTTGTTCAAAACTTCAACTATTCTAAAGGTTTGATGTTAGTAATATTTTGCCGCAAGGTACTTTCCATTCTTTTTGTGCCATTGATTTATGCATTACTTGTTGAATTTGGTTTTA

>M6-PGSC0003DMG400006487M6LRR-RLP_124

ACTAACCTCGACCTCAGAGAGGCGGCGATAACTTGTGTTTTATCCAAGAAATGGCAGTATTATTGGCTCTCAAGACCTTATTTGCTGTTCCGTCAATGTAAAGGCAACAATTGTATACCC

>M6-PGSC0003DMG400006487M6LRR-RLP_1240

GTACTTTCCATTCTTTTTGTGCCATTGATTTATGCATTACTTGTTGAATTTGGTTTTAAGCTATGTCATTGGCAAAATTCTTGCAGACCAAAAACATCCTTATTTATGAAAATCCAAGAG

>M6-PGSC0003DMG400006487M6LRR-RLP_1302

ATGTCATTGGCAAAATTCTTGCAGACCAAAAACATCCTTATTTATGAAAATCCAAGAGAAATTCTTATTCCACCAAGTCAAAATGTTGAGATATTCATTGTACCCATCCTGAGCATTGAA

>M6-PGSC0003DMG400006487M6LRR-RLP_1364

TCTTATTCCACCAAGTCAAAATGTTGAGATATTCATTGTACCCATCCTGAGCATTGAATTGATCATTGGATCGATATTGCTCAACCGTCCCAGCATCATGTCCATACTTCCATGTACAGA

>M6-PGSC0003DMG400006487M6LRR-RLP_1426

TCATTGGATCGATATTGCTCAACCGTCCCAGCATCATGTCCATACTTCCATGTACAGATAGCAAAGTACTCCAGGTAATTTCTTGCTTTACCTGGAAATCCTTTCAAAAAAAACGTCCCT

>M6-PGSC0003DMG400006487M6LRR-RLP_1488

AAAGTACTCCAGGTAATTTCTTGCTTTACCTGGAAATCCTTTCAAAAAAAACGTCCCTTGTGCTTGTAGACACTAATGCAGTAGAGCTTAAGCATTATTGTATTCTCTATATCTTATGTT

>M6-PGSC0003DMG400006487M6LRR-RLP_1550

CTTGTAGACACTAATGCAGTAGAGCTTAAGCATTATTGTATTCTCTATATCTTATGTTTTTGTTTAAATACAGGTATTGCCTGCATTAGAGCGGTGTACACAAAATCAAAATTGTGGTAA

>M6-PGSC0003DMG400006487M6LRR-RLP_1612

TTTAAATACAGGTATTGCCTGCATTAGAGCGGTGTACACAAAATCAAAATTGTGGTAAAGAATGTCCATTCAGTACCAGGTTTTTCCACAAGCATCGTATATTAGAAGAAATCATCAGTT

>M6-PGSC0003DMG400006487M6LRR-RLP_1674

TGTCCATTCAGTACCAGGTTTTTCCACAAGCATCGTATATTAGAAGAAATCATCAGTTGTACTGGGACATCTGAGGAGGAAATGGCCTCCATTTGGTATACCTGGTTGAAATCTACGTCT

>M6-PGSC0003DMG400006487M6LRR-RLP_1736

TGGGACATCTGAGGAGGAAATGGCCTCCATTTGGTATACCTGGTTGAAATCTACGTCTCTAATTGGCAAAGTGAATAATTTCATGTTCAAATGGAAAGAACAAGCTTAGACCGAAGAATC

>M6-PGSC0003DMG400006487M6LRR-RLP_1798

TTGGCAAAGTGAATAATTTCATGTTCAAATGGAAAGAACAAGCTTAGACCGAAGAATCTTGGGGAATTCAGCATATAGTGTTTATTAGTGTGTTCAGAATAAAGTTCAGTTAACCCGCTT

>M6-PGSC0003DMG400006487M6LRR-RLP_186

TTGGCTCTCAAGACCTTATTTGCTGTTCCGTCAATGTAAAGGCAACAATTGTATACCCCTGGAAAATTTTGTCAAGTTGATTGATCGACCCCTACGAATCCATGCTGAACAAAACTTACA

>M6-PGSC0003DMG400006487M6LRR-RLP_1860

GGAATTCAGCATATAGTGTTTATTAGTGTGTTCAGAATAAAGTTCAGTTAACCCGCTTGTGGCTCTGTCAAATCCATGAAAGGTTCCTTAACCTTGCTCTAGCTCTAAATAAATTGTGCT

>M6-PGSC0003DMG400006487M6LRR-RLP_1922

CTCTGTCAAATCCATGAAAGGTTCCTTAACCTTGCTCTAGCTCTAAATAAATTGTGCTCAAATCTGTGGTTTAATTTCATTTGTTCGATTGTACGGAGTGAAGTTTAACTGTTAACAAAC

>M6-PGSC0003DMG400006487M6LRR-RLP_1984

TCTGTGGTTTAATTTCATTTGTTCGATTGTACGGAGTGAAGTTTAACTGTTAACAAACAGCATAAGAGGTCTTCATTTCATTTTTTTTTTTCTGGCTCTGCCTCTTCTTCACATTCCATT

>M6-PGSC0003DMG400006487M6LRR-RLP_2046

TAAGAGGTCTTCATTTCATTTTTTTTTTTCTGGCTCTGCCTCTTCTTCACATTCCATTCAATTTCTCTCATGCTTAGACTACTTGGAGGTGATTTAAAAGAAGTGAGCACTTGAGGTAGA

>M6-PGSC0003DMG400006487M6LRR-RLP_2108

TTCTCTCATGCTTAGACTACTTGGAGGTGATTTAAAAGAAGTGAGCACTTGAGGTAGAGAGATGTAATGATGGATGCATATGCCTCCTACTTGTTCTGAAATCTGGTATATATTCACTGT

>M6-PGSC0003DMG400006487M6LRR-RLP_2153

CACTTGAGGTAGAGAGATGTAATGATGGATGCATATGCCTCCTACTTGTTCTGAAATCTGGTATATATTCACTGTCATACTTGAAATGTAATATGATTTTTCGTATAACTTTCTTTGTCA

>M6-PGSC0003DMG400006487M6LRR-RLP_248

AAAATTTTGTCAAGTTGATTGATCGACCCCTACGAATCCATGCTGAACAAAACTTACATCTTGCAAAATTCTTCCTTGCATATCGGGATCGAGAATTGGATCCTTATTTGGATCATTGGA

>M6-PGSC0003DMG400006487M6LRR-RLP_310

GCAAAATTCTTCCTTGCATATCGGGATCGAGAATTGGATCCTTATTTGGATCATTGGATTGAATCGGCTATTAAAGTCAATGTTTCGGTGCTGGTGATTCATGCTCCGGATATTGATTCG

>M6-PGSC0003DMG400006487M6LRR-RLP_372

ATCGGCTATTAAAGTCAATGTTTCGGTGCTGGTGATTCATGCTCCGGATATTGATTCGTCGTATTATAGCTTACCTGATGTTGTTTATGATGCGAAAAAGCTCAACACATTGTGGTTAAG

>M6-PGSC0003DMG400006487M6LRR-RLP_434

ATTATAGCTTACCTGATGTTGTTTATGATGCGAAAAAGCTCAACACATTGTGGTTAAGCAGGTGCAAGTTTGAATTTGATATTAGCACCACTCATATAAGATTTTGTTCTCTTGTGGCTC

>M6-PGSC0003DMG400006487M6LRR-RLP_496

TGCAAGTTTGAATTTGATATTAGCACCACTCATATAAGATTTTGTTCTCTTGTGGCTCTTTTTTTGTATAATGTTCATATTTCAGATGCCCAAATGCAAAGTGTTATAGATAAATGCCCA

>M6-PGSC0003DMG400006487M6LRR-RLP_558

TTTGTATAATGTTCATATTTCAGATGCCCAAATGCAAAGTGTTATAGATAAATGCCCATCCATCAGGAATATAAGAGTACTGGCTTGCAAAGGTATGAGCAAATTGTATGTTGTTGGACG

>M6-PGSC0003DMG400006487M6LRR-RLP_62

TTGATGGAGAAGACAGAATATCAGAATTGCCTGTGCATATCATTCATCACATCTTATGTCGCACTAACCTCGACCTCAGAGAGGCGGCGATAACTTGTGTTTTATCCAAGAAATGGCAGT

>M6-PGSC0003DMG400006487M6LRR-RLP_620

TCAGGAATATAAGAGTACTGGCTTGCAAAGGTATGAGCAAATTGTATGTTGTTGGACGGTTACATCTACACTCTTTGGCTGTACTATTATGCAAACTCGATAGTGTGATTGTCGAAGCTC

>M6-PGSC0003DMG400006487M6LRR-RLP_682

CATCTACACTCTTTGGCTGTACTATTATGCAAACTCGATAGTGTGATTGTCGAAGCTCCGAAACTTGAATCCTTTAGTTATACAGAATGTATTGATCAGGGAAATTTTCATCCTTGCAAA

>M6-PGSC0003DMG400006487M6LRR-RLP_744

ACTTGAATCCTTTAGTTATACAGAATGTATTGATCAGGGAAATTTTCATCCTTGCAAAATTGATATTTTGGATGGTAACAATACTTTACAAACACTCGAGCTCACTGGCTCCAGCATCAC

>M6-PGSC0003DMG400006487M6LRR-RLP_806

ATATTTTGGATGGTAACAATACTTTACAAACACTCGAGCTCACTGGCTCCAGCATCACAGACCAACAGTTTCGAGATGTATTCCACAAGTTCCCAAACATTTCAGTGTTGCATCTAACAA

>M6-PGSC0003DMG400006487M6LRR-RLP_868

CAACAGTTTCGAGATGTATTCCACAAGTTCCCAAACATTTCAGTGTTGCATCTAACAAGATGTTATAAGCTGAAACATATAGAGATTCAGAGTGAAAAACTCAACAAAATCACCTTATCG

>M6-PGSC0003DMG400006487M6LRR-RLP_930

TTATAAGCTGAAACATATAGAGATTCAGAGTGAAAAACTCAACAAAATCACCTTATCGAAGTTGGAGAGTATAGAAAAAATCATGATTCAAGCTCCAAATTTATCGCAATTTGATTTTGA

>M6-PGSC0003DMG400006487M6LRR-RLP_992

TGGAGAGTATAGAAAAAATCATGATTCAAGCTCCAAATTTATCGCAATTTGATTTTGAGGGTGATAAAATGCCCTTCTTATCTATGGATTCTTCTTCCCTGGAAAGAGCCCGACTCAATT

>M6-PGSC0003DMG400006498-(rev)M6RLKWAX-EGF_2583

CTAGCTTGAAAGGGGTATTCAACATGTGAATGTGATGCAGGGTGACCTGTCATTTGCTGATTGGACTAGAAGTTAGTTAATTAGATATTTTATCATTTTCAGTTAGTTAGGTTGTTAGAA

>M6-PGSC0003DMG400006502M6LRR-RLK_0

CCATGGCCTATTATCTTGACAAAATCTTCCTTCTCTTATTTCTCTTCACACTTAGTACTCCCTCAAAATCAGATAATATGACAGATATTGAAGCACTCTTTGCTATCAAGAATGAAATAA

>M6-PGSC0003DMG400006502M6LRR-RLK_1054

TGAGTTTTTCAGCAACCTTTCAAGTAATCTGGCGTTTGTAGACTTGTCAAGCAATCATTTTTCAGGACATATCCCGATAGACATTGGGAAATTCGTCAACTTGGAAGAGCTTAGGCTGAG

>M6-PGSC0003DMG400006502M6LRR-RLK_1178

AACCAACTCACAGGTATCATCCCTTCGACTATTGGGAAGCTTTATAAGCTACGATATTTACACCTGTCCCAAAACAGAATTTCGGGTAGTATACCCTTTTCCATTGCAAATCTGACCCTT

>M6-PGSC0003DMG400006502M6LRR-RLK_124

TCCCTTTGAATCATTAAGTTCATGGAATGAGTCTGTTCCTTTATGTCAATGGCGTGGCGTTGTATGTGGAACGCGAAATCAACGAGTCATTGAACTAAACCTTCTTGACCATAAGCTAGC

>M6-PGSC0003DMG400006502M6LRR-RLK_1240

CCTGTCCCAAAACAGAATTTCGGGTAGTATACCCTTTTCCATTGCAAATCTGACCCTTCTAACAGAGTTAAATCTTGATCATAACAATCTTAGAGGTAGAATACCTTCAAGTGTTGCAAA

>M6-PGSC0003DMG400006502M6LRR-RLK_1302

CAGAGTTAAATCTTGATCATAACAATCTTAGAGGTAGAATACCTTCAAGTGTTGCAAAATGTCAAAGTTTGTTGTACCTTAACCTGAACCAAAACAATTTAATCGGTTTGATACCTAAAG

>M6-PGSC0003DMG400006502M6LRR-RLK_1364

CAAAGTTTGTTGTACCTTAACCTGAACCAAAACAATTTAATCGGTTTGATACCTAAAGAAATTTGTCTAATCTCTTCCTTAGTTGTTCTAAACCTTTCCAGTAACAACTTGACTGGCGCC

>M6-PGSC0003DMG400006502M6LRR-RLK_1426

TTGTCTAATCTCTTCCTTAGTTGTTCTAAACCTTTCCAGTAACAACTTGACTGGCGCCCTTCCCAAAGAAATCGAAAGCCTGAAAAATCTCTTATCATTAGATGTCTCTTTGAATAATCT

>M6-PGSC0003DMG400006502M6LRR-RLK_1488

CCAAAGAAATCGAAAGCCTGAAAAATCTCTTATCATTAGATGTCTCTTTGAATAATCTCAATGGTGAAATTCCTAGTACACTTGGGAGTTGTGTGACAATTGTTAACTTAACAATGAAAA

>M6-PGSC0003DMG400006502M6LRR-RLK_1550

GGTGAAATTCCTAGTACACTTGGGAGTTGTGTGACAATTGTTAACTTAACAATGAAAAGGAATTTATTCCGCGGGATTATTCCTTCCACTTTCAGTTCACTAAAGAGTCTTCAAGTTTTA

>M6-PGSC0003DMG400006502M6LRR-RLK_1612

TTTATTCCGCGGGATTATTCCTTCCACTTTCAGTTCACTAAAGAGTCTTCAAGTTTTAGATCTTTCGCGGAATAAGCTATCAGGTATGATTCCAAAGTACTTGGAAGGATTTGCCTTGCA

>M6-PGSC0003DMG400006502M6LRR-RLK_1674

TTTCGCGGAATAAGCTATCAGGTATGATTCCAAAGTACTTGGAAGGATTTGCCTTGCATTTCTTAAACTTATCGTTTAATGATTTTGAAGGTGGCCTTCCACAACGAGGCATCTTTGAGA

>M6-PGSC0003DMG400006502M6LRR-RLK_1798

GAGTGTAGACTCTTTCGTTGGAAATCTTAGAATCTGTGGTGGTGTACCTGGCTTGAAACTTCCCAATTGCAACTTCAGTCATTCAAAGAAAATGAACTTTAAGCTTGTTATTTTAGTAAT

>M6-PGSC0003DMG400006502M6LRR-RLK_186

TATGTGGAACGCGAAATCAACGAGTCATTGAACTAAACCTTCTTGACCATAAGCTAGCCGGTGTGTTATCACCCTTTGTTGGAAATTTGAGCTTCCTCATACGACTAGACATCGGAAACA

>M6-PGSC0003DMG400006502M6LRR-RLK_1860

CCAATTGCAACTTCAGTCATTCAAAGAAAATGAACTTTAAGCTTGTTATTTTAGTAATATTAGGAATTTTGGGGCTAGTTGTTATGGTTTGTGCTTTTTTCTTCTATCGATTTGGAAGGT

>M6-PGSC0003DMG400006502M6LRR-RLK_1922

GGAATTTTGGGGCTAGTTGTTATGGTTTGTGCTTTTTTCTTCTATCGATTTGGAAGGTCCAAAAGAACAATTCCTTCATTAGATAACAATCTAAACCAACTCATAGCTATGTCCTATCAA

>M6-PGSC0003DMG400006502M6LRR-RLK_1984

AAGAACAATTCCTTCATTAGATAACAATCTAAACCAACTCATAGCTATGTCCTATCAAAGTATTCTCAAAGTTACCAACGGGTTTTCTACGAGCAACTTGATTGGTGTTGGGAGTCATGG

>M6-PGSC0003DMG400006502M6LRR-RLK_2046

TTCTCAAAGTTACCAACGGGTTTTCTACGAGCAACTTGATTGGTGTTGGGAGTCATGGATATGTTTACAAGGGAATTTTAGAAACGGATGGTAAACACGTTGCTATAAAAGTGTTGAATC

>M6-PGSC0003DMG400006502M6LRR-RLK_2108

GTTTACAAGGGAATTTTAGAAACGGATGGTAAACACGTTGCTATAAAAGTGTTGAATCTTTTACAATATGGAGCTATCAAAAGTTTCATAGCAGAATGTGAGGCCTTGAGAAATGTTAGA

>M6-PGSC0003DMG400006502M6LRR-RLK_2170

ACAATATGGAGCTATCAAAAGTTTCATAGCAGAATGTGAGGCCTTGAGAAATGTTAGACATCGGAATTTAGTTAAGCTACTTACTGCATGTTCTGGTGTTGATTATCATGGCAACGAATT

>M6-PGSC0003DMG400006502M6LRR-RLK_2232

GGAATTTAGTTAAGCTACTTACTGCATGTTCTGGTGTTGATTATCATGGCAACGAATTCAAGGCACTGATTTACGAGTTTATGGCCAACGGAAGTTTGGAGGACTGGTTGCATCCAGATA

>M6-PGSC0003DMG400006502M6LRR-RLK_2294

GCACTGATTTACGAGTTTATGGCCAACGGAAGTTTGGAGGACTGGTTGCATCCAGATAATTCAAGACCGAATGTTCAGCCAAGAAGGTTAGGATTTCTACAAAGGCTGAATATTGCTATT

>M6-PGSC0003DMG400006502M6LRR-RLK_2356

AAGACCGAATGTTCAGCCAAGAAGGTTAGGATTTCTACAAAGGCTGAATATTGCTATTGATGTTGCTAGTGCAATTCATTATCTTCATAATGATTGTCAAATATCAATAGTTCATTGTGA

>M6-PGSC0003DMG400006502M6LRR-RLK_2418

TTGCTAGTGCAATTCATTATCTTCATAATGATTGTCAAATATCAATAGTTCATTGTGATCTCAAACCAAGCAATATTCTCCTTGACAATGAATTGGTTGCCCATGTTGGGGATTTTGGAT

>M6-PGSC0003DMG400006502M6LRR-RLK_248

GTGTTATCACCCTTTGTTGGAAATTTGAGCTTCCTCATACGACTAGACATCGGAAACAATACCATTTCTGGCAATATCCCTCCTGAATTAGGCCGTTTGACAAGCTTACGCCATCTATAT

>M6-PGSC0003DMG400006502M6LRR-RLK_2480

AAACCAAGCAATATTCTCCTTGACAATGAATTGGTTGCCCATGTTGGGGATTTTGGATTGTCAAGATTCCTACATCTTACTGACGAAACCACTTGTAGAATTCAGACAAGCTCTTCAACT

>M6-PGSC0003DMG400006502M6LRR-RLK_2542

AAGATTCCTACATCTTACTGACGAAACCACTTGTAGAATTCAGACAAGCTCTTCAACTTTTAAAGGATCTATTGGCTACATAGCTCCAGGTAAATTTCACTCTATATAAATGCCTCTAAA

>M6-PGSC0003DMG400006502M6LRR-RLK_2604

AAGGATCTATTGGCTACATAGCTCCAGGTAAATTTCACTCTATATAAATGCCTCTAAAGACTTTAGTGACTCTGGATGCAATGACATTAACTCAATTGCCTTTATATTTTCATTGCAGAG

>M6-PGSC0003DMG400006502M6LRR-RLK_2666

TTAGTGACTCTGGATGCAATGACATTAACTCAATTGCCTTTATATTTTCATTGCAGAGTATGGTATGGGAAGCGAGGCATCAACGCAAGGTGATGTGTATAGTTTTGGTATCGTCTTACT

>M6-PGSC0003DMG400006502M6LRR-RLK_2728

GTATGGGAAGCGAGGCATCAACGCAAGGTGATGTGTATAGTTTTGGTATCGTCTTACTTGAAATGTTAACTGGGAAAAGACCAACAGATGACACGTTTGGCAGCGATTTAAGCCTTCATG

>M6-PGSC0003DMG400006502M6LRR-RLK_2790

ATGTTAACTGGGAAAAGACCAACAGATGACACGTTTGGCAGCGATTTAAGCCTTCATGATTTTGTTAGGAATGCCATGCCTGATGGAGCACTTGAGATAGTAGATCCAGTTCTTAATTTT

>M6-PGSC0003DMG400006502M6LRR-RLK_2852

TGTTAGGAATGCCATGCCTGATGGAGCACTTGAGATAGTAGATCCAGTTCTTAATTTTGAAGAGGAAGAGATAAGCACAGAGAGGTCTCAAATTCCAAGGTTCATGAGAAGGCAAAAAAT

>M6-PGSC0003DMG400006502M6LRR-RLK_2914

AGGAAGAGATAAGCACAGAGAGGTCTCAAATTCCAAGGTTCATGAGAAGGCAAAAAATGGTAGAGGGTTTAATTTCCTTATTTGGAGTTGGTATTGATTGTTCAATGTATGATTCAAGTA

>M6-PGSC0003DMG400006502M6LRR-RLK_2976

GAGGGTTTAATTTCCTTATTTGGAGTTGGTATTGATTGTTCAATGTATGATTCAAGTAAAAGAAAAAACATGAAGGAGGTTGTTAGAGAGTTGTGTTCTATAAGAGATTCTATTGTTGGT

>M6-PGSC0003DMG400006502M6LRR-RLK_3009

GATTGTTCAATGTATGATTCAAGTAAAAGAAAAAACATGAAGGAGGTTGTTAGAGAGTTGTGTTCTATAAGAGATTCTATTGTTGGTTGTACAATATGAAACACTTTTCTTGTTTTGTTT

>M6-PGSC0003DMG400006502M6LRR-RLK_310

CATTTCTGGCAATATCCCTCCTGAATTAGGCCGTTTGACAAGCTTACGCCATCTATATTTGCAAAACAATTCATTTTATGGTGAAATTCCTGTCAATCTATCGTTTTGTAGTAACCTCGT

>M6-PGSC0003DMG400006502M6LRR-RLK_372

AAAACAATTCATTTTATGGTGAAATTCCTGTCAATCTATCGTTTTGTAGTAACCTCGTTGAGTTATGGGCATACAAGAACAATTTTGTAGGTGTTCTTCCAAGGGAGCTTGGTTTACTTA

>M6-PGSC0003DMG400006502M6LRR-RLK_434

TTATGGGCATACAAGAACAATTTTGTAGGTGTTCTTCCAAGGGAGCTTGGTTTACTTACCAAACTCAAATATTTAGATTTCGGTTACAACAAGCTCATTGGTGAAATCCCAAAATCTTAT

>M6-PGSC0003DMG400006502M6LRR-RLK_496

ACTCAAATATTTAGATTTCGGTTACAACAAGCTCATTGGTGAAATCCCAAAATCTTATGGTAACTTTTCAAGTCTATTGGAAATGTACCTTTTGGATAATGATCTCGAGGGTAAAATCCC

>M6-PGSC0003DMG400006502M6LRR-RLK_558

ACTTTTCAAGTCTATTGGAAATGTACCTTTTGGATAATGATCTCGAGGGTAAAATCCCCGATGAGTTTGGGAAACTGAAGAGTTTGGAAATATTTGACGCGGACTTTAATAGGTTATCTG

>M6-PGSC0003DMG400006502M6LRR-RLK_62

TCAAAATCAGATAATATGACAGATATTGAAGCACTCTTTGCTATCAAGAATGAAATAATTGATCCCTTTGAATCATTAAGTTCATGGAATGAGTCTGTTCCTTTATGTCAATGGCGTGGC

>M6-PGSC0003DMG400006502M6LRR-RLK_620

GAGTTTGGGAAACTGAAGAGTTTGGAAATATTTGACGCGGACTTTAATAGGTTATCTGGTAGAATTCCGAGCTCATTGTACAATCTTACCTCATTGAAAGTCATTGATGTGTCATTTAAC

>M6-PGSC0003DMG400006502M6LRR-RLK_682

AATTCCGAGCTCATTGTACAATCTTACCTCATTGAAAGTCATTGATGTGTCATTTAACCAACTCGAAGGTACTCTGCCTCGGGATTTAGGGATCAACCTTCCTAACCTGGAGTGTCTCAG

>M6-PGSC0003DMG400006502M6LRR-RLK_806

AGTGAGAATCAATTCACTGGATCGCTTCCTCATTCACTTTTCAACCTCACAGGACTCGCGTACCTTTATGTTGGAAAAAATGACTTAGGAGGACGTGTTCCTAGGTTCGATAAGTTGCAT

>M6-PGSC0003DMG400006502M6LRR-RLK_868

CCTTTATGTTGGAAAAAATGACTTAGGAGGACGTGTTCCTAGGTTCGATAAGTTGCATAATTTGTATATTTTAAGCATGGAAAATAACCAATTTGATAATGACTTGACCCTCATGTCCTC

>M6-PGSC0003DMG400006502M6LRR-RLK_930

TGTATATTTTAAGCATGGAAAATAACCAATTTGATAATGACTTGACCCTCATGTCCTCCTTGACAAATGCAACTAACCTCAACCGTCTGTATATGCAAAACAACGCGTTTGGAGGGACAT

>M6-PGSC0003DMG400006502M6LRR-RLK_992

ACAAATGCAACTAACCTCAACCGTCTGTATATGCAAAACAACGCGTTTGGAGGGACATTTCATGAGTTTTTCAGCAACCTTTCAAGTAATCTGGCGTTTGTAGACTTGTCAAGCAATCAT

>M6-PGSC0003DMG400006563-(rev)M6LRR-RLK_0

AGTAAACATACAGAGCTAAGGAGATCAAGTGTCATTACATTTCTGTATATTTAGGTCTCTGAAATGCATAGCATAGTTATTTAAAATTTATGTTTACTTGTGAGTAAGATTTTGGAACTT

>M6-PGSC0003DMG400006563-(rev)M6LRR-RLK_1037

GTAAGTGCGAGTTTGTTATTGTTTTGTTCTTCTTTTGTTAAGCTTTATCTGTATATTTTTACTTCAAGATTGTTCAACTAGTAGATCTCTGAAAATGTCTAGCTTCTGCTTGCTATACAG

>M6-PGSC0003DMG400006563-(rev)M6LRR-RLK_1098

CTTCAAGATTGTTCAACTAGTAGATCTCTGAAAATGTCTAGCTTCTGCTTGCTATACAGTGATCTCAGCTGGAATCAACTTTCTGGATCTATACCACAGAATAAGCTTTCCAACAATATG

>M6-PGSC0003DMG400006563-(rev)M6LRR-RLK_1159

ATCTCAGCTGGAATCAACTTTCTGGATCTATACCACAGAATAAGCTTTCCAACAATATGACAACTATGTAAGCCTTTTGATTTGGCCTCGATAGTTTCTTCATATAGTTATATAAATTGA

>M6-PGSC0003DMG400006563-(rev)M6LRR-RLK_122

GTTGTATTTTTAAAGGTGTGAAATTATGAACCAATGCAGGTATTCTTTAAAGCCAATAATTGAAAAGTTGGATAAACATATGTTTTTTGCTCATGGAGCTGTTAATTGTATAACGCCTGG

>M6-PGSC0003DMG400006563-(rev)M6LRR-RLK_1220

AACTATGTAAGCCTTTTGATTTGGCCTCGATAGTTTCTTCATATAGTTATATAAATTGACAGATGTTGGCTTTTGTTTCCACAGCATTTGAACTGGATTGCCTGGGTTCCTTTTACTTAT

>M6-PGSC0003DMG400006563-(rev)M6LRR-RLK_1281

GATGTTGGCTTTTGTTTCCACAGCATTTGAACTGGATTGCCTGGGTTCCTTTTACTTATTAGCATATTCTAATGTATATTTCAAATGCATGATGCAGTATTCTATCACATAACCGTCTTA

>M6-PGSC0003DMG400006563-(rev)M6LRR-RLK_1342

GCATATTCTAATGTATATTTCAAATGCATGATGCAGTATTCTATCACATAACCGTCTTAATGGATCTATACCCAAGAGCTTTTCTCTCCTTCCTTCCCTGCAGAAACTGTGAGAATCTGG

>M6-PGSC0003DMG400006563-(rev)M6LRR-RLK_1403

GGATCTATACCCAAGAGCTTTTCTCTCCTTCCTTCCCTGCAGAAACTGTGAGAATCTGGACTTAGTCTTGCAATATCTACTATTTCACTGAAGGAAATGTACCATTAACGAATTGGTGGG

>M6-PGSC0003DMG400006563-(rev)M6LRR-RLK_1464

TTAGTCTTGCAATATCTACTATTTCACTGAAGGAAATGTACCATTAACGAATTGGTGGGACAAATTTGTCACAGGTCTCTTGAGAACAACTTTCTGAATGGTTCCATTTCCACTGACATT

>M6-PGSC0003DMG400006563-(rev)M6LRR-RLK_1525

AAATTTGTCACAGGTCTCTTGAGAACAACTTTCTGAATGGTTCCATTTCCACTGACATTTGGCAAAACAAGATCTTTAATACAACTTCCAGACTTGTGATGTAAGATGTGAAGCAGAAAA

>M6-PGSC0003DMG400006563-(rev)M6LRR-RLK_1586

GCAAAACAAGATCTTTAATACAACTTCCAGACTTGTGATGTAAGATGTGAAGCAGAAAAATAACAGATCTTATTTCAATAGAAGTATTTACTTCCAGAGTCAAACCTTTGAATGAGGATT

>M6-PGSC0003DMG400006563-(rev)M6LRR-RLK_1647

AACAGATCTTATTTCAATAGAAGTATTTACTTCCAGAGTCAAACCTTTGAATGAGGATTTCTTTCTAACAACTCTTTCATTGATTTTGCAAACCTAGTGATCTGCGGAATAATTCAGTTT

>M6-PGSC0003DMG400006563-(rev)M6LRR-RLK_1708

TTTCTAACAACTCTTTCATTGATTTTGCAAACCTAGTGATCTGCGGAATAATTCAGTTTCAAACATTTCGGGTGCTTTTGAACCTCATGTAAATGTCACACTAAGGTATGATTTCTTCTT

>M6-PGSC0003DMG400006563-(rev)M6LRR-RLK_1769

AACATTTCGGGTGCTTTTGAACCTCATGTAAATGTCACACTAAGGTATGATTTCTTCTTTCAAATTCAAAAATAGAACAATTTGAATGTCGGTTTATGTCCTCATGGGCCACATAAATAG

>M6-PGSC0003DMG400006563-(rev)M6LRR-RLK_183

GAAAAGTTGGATAAACATATGTTTTTTGCTCATGGAGCTGTTAATTGTATAACGCCTGGCCATTTCTTTCGTGATTAAGGTGGTGTCTTGTGAGGACCTTGATTGGTCCAACATATACCT

>M6-PGSC0003DMG400006563-(rev)M6LRR-RLK_1830

AAATTCAAAAATAGAACAATTTGAATGTCGGTTTATGTCCTCATGGGCCACATAAATAGGACAAACAAATGGGTAAAGAAACTAGAATAACAGAAGCAATATGCCATACCAAAAAGGGAA

>M6-PGSC0003DMG400006563-(rev)M6LRR-RLK_1891

CAAACAAATGGGTAAAGAAACTAGAATAACAGAAGCAATATGCCATACCAAAAAGGGAAAGTACGTCGATTTTCCTCGTGTGTGTGACAACAATTACTTAAATTACTTGAAGAAGCACAA

>M6-PGSC0003DMG400006563-(rev)M6LRR-RLK_1952

TACGTCGATTTTCCTCGTGTGTGTGACAACAATTACTTAAATTACTTGAAGAAGCACAAATGCATGGAATCACTTCTGTTATGTTTTATCCACACACACCGATACACATGCTTTTAGCAA

>M6-PGSC0003DMG400006563-(rev)M6LRR-RLK_2013

GCATGGAATCACTTCTGTTATGTTTTATCCACACACACCGATACACATGCTTTTAGCAAATTCATATATGGTGCTTTATAAAATGTAAACAAAATGGAAAGAACAAGGAGTTGAGAGAAC

>M6-PGSC0003DMG400006563-(rev)M6LRR-RLK_2074

TCATATATGGTGCTTTATAAAATGTAAACAAAATGGAAAGAACAAGGAGTTGAGAGAACCTTTGTGGTTATCAGTTTTTTCATTGTTTGAAAAATTTAAGTATGCTTAAAAAGGAGGTAT

>M6-PGSC0003DMG400006563-(rev)M6LRR-RLK_2135

TTGTGGTTATCAGTTTTTTCATTGTTTGAAAAATTTAAGTATGCTTAAAAAGGAGGTATAGATAGAAAAGGGAATTCTGAAATTCTTAGTCACATATTTTCATCTAGTTGCAGTGGAAAA

>M6-PGSC0003DMG400006563-(rev)M6LRR-RLK_2196

ATAGAAAAGGGAATTCTGAAATTCTTAGTCACATATTTTCATCTAGTTGCAGTGGAAAAGGATGAAATGATGAAGGGCCCTATAATATTGGTTATGGTCCTGATCAGCGTAAATCTTGAA

>M6-PGSC0003DMG400006563-(rev)M6LRR-RLK_2257

ATGAAATGATGAAGGGCCCTATAATATTGGTTATGGTCCTGATCAGCGTAAATCTTGAAAAAGTACAAAGTTTTACCCAATAATATAAATGAGAAGTGTGTGTGTGTGGGGGTGGGGGGT

>M6-PGSC0003DMG400006563-(rev)M6LRR-RLK_2318

AGTACAAAGTTTTACCCAATAATATAAATGAGAAGTGTGTGTGTGTGGGGGTGGGGGGTAAAAAGATATAAATAGAGGGAAACTATGAGCATAAAAGGAGTTTTTGTCTTTTGTTGAGTC

>M6-PGSC0003DMG400006563-(rev)M6LRR-RLK_2379

AAAGATATAAATAGAGGGAAACTATGAGCATAAAAGGAGTTTTTGTCTTTTGTTGAGTCTCTCATTTATTCGTTGAAAGGAGGCCTGAACAAATGGAGAGGAAGGAAGATTCTGTAAAAT

>M6-PGSC0003DMG400006563-(rev)M6LRR-RLK_2440

TCATTTATTCGTTGAAAGGAGGCCTGAACAAATGGAGAGGAAGGAAGATTCTGTAAAATTTGAGTCCACTACATGCCTTATGATGATTTTTTAATGTGACACAATGTAATTCTAATCTAA

>M6-PGSC0003DMG400006563-(rev)M6LRR-RLK_2989

GCAACTTAGATTCTAATAGTTTGTGAAACTGAGGGAGATTTTCTGCAGTATATCACATCAAGACAAGTGATATAAACTGGTATCATTAAATTCTTGAAGAAGGAAAACATCACATTTAGG

>M6-PGSC0003DMG400006563-(rev)M6LRR-RLK_3050

GACAAGTGATATAAACTGGTATCATTAAATTCTTGAAGAAGGAAAACATCACATTTAGGAAAGTGGAGAATGGAAATTGTGCCAAACAAAAAGAGTTATTGGTCTAAAATTACTTGTACA

>M6-PGSC0003DMG400006563-(rev)M6LRR-RLK_3111

AGTGGAGAATGGAAATTGTGCCAAACAAAAAGAGTTATTGGTCTAAAATTACTTGTACATCTTTGTTACTGTCTCTGGAGTTCATGTTTTCAGTTATGCACTCATTCTTGTTGTCCAATT

>M6-PGSC0003DMG400006563-(rev)M6LRR-RLK_3172

TTTGTTACTGTCTCTGGAGTTCATGTTTTCAGTTATGCACTCATTCTTGTTGTCCAATTATTCATAAATATCTGATTTTGACAGTGAATTGAGGTCATCTTTCAAAAATTACTTTCTGAT

>M6-PGSC0003DMG400006563-(rev)M6LRR-RLK_3233

TCATAAATATCTGATTTTGACAGTGAATTGAGGTCATCTTTCAAAAATTACTTTCTGATCTCTTTCTATCTCATGAGGTCAATTCATCTTGAATTCTTAATTGAAAATCTGCTCAATGTT

>M6-PGSC0003DMG400006563-(rev)M6LRR-RLK_3294

CTTTCTATCTCATGAGGTCAATTCATCTTGAATTCTTAATTGAAAATCTGCTCAATGTTTCTCAAAAGAAACATGTCAATAAAAATGTTTTAGATTTAGTACTATGTTAAAATTCATTTA

>M6-PGSC0003DMG400006563-(rev)M6LRR-RLK_3355

TCAAAAGAAACATGTCAATAAAAATGTTTTAGATTTAGTACTATGTTAAAATTCATTTATTATTTTCATAGTCTTCCCCTAACTTTAGCAGACGATTTCCAGGTTCCAAAGCAATCCCAT

>M6-PGSC0003DMG400006563-(rev)M6LRR-RLK_3416

ATTTTCATAGTCTTCCCCTAACTTTAGCAGACGATTTCCAGGTTCCAAAGCAATCCCATCTGCAGCAGTACAAGTATCAGAAACATTGGTAAATACTGTGGCCCTGACTTTGGAGCAGAT

>M6-PGSC0003DMG400006563-(rev)M6LRR-RLK_3477

GCAGCAGTACAAGTATCAGAAACATTGGTAAATACTGTGGCCCTGACTTTGGAGCAGATGACGACGAACCTAGTAATTCAACAAATCCAAGTGGTGTTTGTCCTATTCATGCTTGTCCAA

>M6-PGSC0003DMG400006563-(rev)M6LRR-RLK_3538

CGACGAACCTAGTAATTCAACAAATCCAAGTGGTGTTTGTCCTATTCATGCTTGTCCAACTGACAACTATTATGAATATGTTCCTGCATCACCTAAACCATGCTTTTGTGCCTCACCTCT

>M6-PGSC0003DMG400006563-(rev)M6LRR-RLK_3599

GACAACTATTATGAATATGTTCCTGCATCACCTAAACCATGCTTTTGTGCCTCACCTCTTAGAATTGGGTACAGGTTGAAAAGTCCTAGTATTTCTTATTTTGATCCGTACAAGCAACTA

>M6-PGSC0003DMG400006563-(rev)M6LRR-RLK_3660

GAATTGGGTACAGGTTGAAAAGTCCTAGTATTTCTTATTTTGATCCGTACAAGCAACTATTCGAATCTTATGTGACTAGTTCCCTTCAGTTGGACCTTTACCAATTGTGGATCGATTCAT

>M6-PGSC0003DMG400006563-(rev)M6LRR-RLK_3721

CGAATCTTATGTGACTAGTTCCCTTCAGTTGGACCTTTACCAATTGTGGATCGATTCATATTTTTGGGAAAAAGGACCTCGTCTGAGGATGTATTTGAAGCTTTTCCCGGTGGTTGGGGA

>M6-PGSC0003DMG400006563-(rev)M6LRR-RLK_3782

TTTTGGGAAAAAGGACCTCGTCTGAGGATGTATTTGAAGCTTTTCCCGGTGGTTGGGGACAATACATTTAATGAAAGTGAGATTCTTCGTATCAGCGAGATTTTTGCCTCTTGGGAATTT

>M6-PGSC0003DMG400006563-(rev)M6LRR-RLK_3843

ATACATTTAATGAAAGTGAGATTCTTCGTATCAGCGAGATTTTTGCCTCTTGGGAATTTCGAGGAAGTCACCTATTTGGACCATATGAGCTCCTGAACTTCACACTCCTCGGACCTTATT

>M6-PGSC0003DMG400006563-(rev)M6LRR-RLK_3904

AGGAAGTCACCTATTTGGACCATATGAGCTCCTGAACTTCACACTCCTCGGACCTTATTCACACTGTAAGTACTCACAGCTTAAATTGATTCTAATTCAACTTGTAACAACTTAGGCTGT

>M6-PGSC0003DMG400006563-(rev)M6LRR-RLK_3965

CACTGTAAGTACTCACAGCTTAAATTGATTCTAATTCAACTTGTAACAACTTAGGCTGTGTTACTACAATTGCATGACAATTGAATCATTGACATGCAGCATGTGGATAAGATTGAAATA

>M6-PGSC0003DMG400006563-(rev)M6LRR-RLK_4026

TACTACAATTGCATGACAATTGAATCATTGACATGCAGCATGTGGATAAGATTGAAATATTTGACTATCATTGCCACTTCTAGCCAACAAGTATAAGGATCTTAATTCATATAATCAAAT

>M6-PGSC0003DMG400006563-(rev)M6LRR-RLK_4087

TGACTATCATTGCCACTTCTAGCCAACAAGTATAAGGATCTTAATTCATATAATCAAATAGTTTAGATAAGCAACAAAAGACTATGATCCATGATCTTCTAAGGACTTCATATATATTGG

>M6-PGSC0003DMG400006563-(rev)M6LRR-RLK_4148

TTTAGATAAGCAACAAAAGACTATGATCCATGATCTTCTAAGGACTTCATATATATTGGGTTGAAAAAGACAATGAAGTCTTTGTTTATATCCAAAGATTTTTTAAGACTTCAGATATTT

>M6-PGSC0003DMG400006563-(rev)M6LRR-RLK_4209

TGAAAAAGACAATGAAGTCTTTGTTTATATCCAAAGATTTTTTAAGACTTCAGATATTTTACGTTGAGGAACCTTACTTGATTAAAAGGTTGAAACTTCAAGGGGTTCAAACTTAGATAG

>M6-PGSC0003DMG400006563-(rev)M6LRR-RLK_427

TTGTATCCCTTTTCGACTTTCAGCTTGTGCATTTCTTCATCTGTTGTATTCCCTTTCTCTCTTTTGATAATGTTTGACCCTTTATTTGCATTGTGGTTTCTCTTTCCTATTCAGTCTATC

>M6-PGSC0003DMG400006563-(rev)M6LRR-RLK_4270

CGTTGAGGAACCTTACTTGATTAAAAGGTTGAAACTTCAAGGGGTTCAAACTTAGATAGTTTGGTCCAGTCTCTCTTCAAGAGCTTCAATTCTGGAGGTCTCAGCCCTTACAGTTTTTGT

>M6-PGSC0003DMG400006563-(rev)M6LRR-RLK_4331

TGGTCCAGTCTCTCTTCAAGAGCTTCAATTCTGGAGGTCTCAGCCCTTACAGTTTTTGTTGGGATGAAAGTGGATAAAAAAAAATGGACCTTAGGCCTCCCAAAAAGCTAGCTCATGAGG

>M6-PGSC0003DMG400006563-(rev)M6LRR-RLK_4453

GGGATTGTCCAAAATCATATAAGGAGAATAACCCTTGTCACTCCAACACCTTGCTGCACTCCCAAGAATTGACATTGGAGCGTGGACAATATAATAAGGGGGGGTTCAACACCGGAAACA

>M6-PGSC0003DMG400006563-(rev)M6LRR-RLK_4514

CCAAGAATTGACATTGGAGCGTGGACAATATAATAAGGGGGGGTTCAACACCGGAAACAATAAATTGGGATCAGTGTGGATCTGAGACCATGTGGGAAATAAAGAAGAAAAATATTACTC

>M6-PGSC0003DMG400006563-(rev)M6LRR-RLK_488

TTTTGATAATGTTTGACCCTTTATTTGCATTGTGGTTTCTCTTTCCTATTCAGTCTATCAGGAAAATGGTGTGAGGGTTCAATAAGTTCACTAAACATCTGAAGACTAAGATGCTAATAT

>M6-PGSC0003DMG400006563-(rev)M6LRR-RLK_5246

CTTTGTACAAGCTTTGATTTCTTATATTATCTGCAGTGAATCCTGAAATGAAAGGTAAAAAACAAAGCAAAGGCGTTGTGATAGCTTGCATTGTAGCAGCAGGTGCTTTTGCTGCCTTTG

>M6-PGSC0003DMG400006563-(rev)M6LRR-RLK_5307

ACAAAGCAAAGGCGTTGTGATAGCTTGCATTGTAGCAGCAGGTGCTTTTGCTGCCTTTGTTTCCTCAATTGCCACTATATTAATCACAAGACGACGTGCTAAGTATCAAAAGATCTTGTC

>M6-PGSC0003DMG400006563-(rev)M6LRR-RLK_5368

TCCTCAATTGCCACTATATTAATCACAAGACGACGTGCTAAGTATCAAAAGATCTTGTCAAGAAAACGTTTGTGTAAGTCCTCTTATTAGATTAGCTATAATTTTACTTTATCAAGTCTG

>M6-PGSC0003DMG400006563-(rev)M6LRR-RLK_5429

GAAAACGTTTGTGTAAGTCCTCTTATTAGATTAGCTATAATTTTACTTTATCAAGTCTGCTATTTTTAATTCTTTCTGCAAAATTGATGACTATGGGTTTTTGTTTGCAGCTTCAAGCCT

>M6-PGSC0003DMG400006563-(rev)M6LRR-RLK_549

GAAAATGGTGTGAGGGTTCAATAAGTTCACTAAACATCTGAAGACTAAGATGCTAATATTGATAGTGATCAGTTCAATTTCCACCTTTAACTTTTTTAAATTGCATCGATCCTAGCAGAA

>M6-PGSC0003DMG400006563-(rev)M6LRR-RLK_5490

ATTTTTAATTCTTTCTGCAAAATTGATGACTATGGGTTTTTGTTTGCAGCTTCAAGCCTCTCTATAAAAGTAGATGGCGTTAAAAGTTTCACCTTCAGAGAAATGACTTCGGCCACTAAC

>M6-PGSC0003DMG400006563-(rev)M6LRR-RLK_5551

CTATAAAAGTAGATGGCGTTAAAAGTTTCACCTTCAGAGAAATGACTTCGGCCACTAACAACTTTGATACCTCAACTCAAGTTGGTGAAGGAGGATATGGTTCCGTCTTTAGGGGAATTT

>M6-PGSC0003DMG400006563-(rev)M6LRR-RLK_5612

CTTTGATACCTCAACTCAAGTTGGTGAAGGAGGATATGGTTCCGTCTTTAGGGGAATTTTAGCTGACAAAACAGTTGTTGCTATCAAGCGAGCAAAAGTAGGATCCCTGCAAGGGCAAAA

>M6-PGSC0003DMG400006563-(rev)M6LRR-RLK_5673

GCTGACAAAACAGTTGTTGCTATCAAGCGAGCAAAAGTAGGATCCCTGCAAGGGCAAAAGGAATTCTTGACCGAAATAGAACTATTATCAAGGTTACATCACCGTAATCTGGTTGTGTTG

>M6-PGSC0003DMG400006563-(rev)M6LRR-RLK_5734

AATTCTTGACCGAAATAGAACTATTATCAAGGTTACATCACCGTAATCTGGTTGTGTTGCTTGGATACTGTGACGAAGAAGGAGAGCAGGTATGACAAAAGCTGAAAACCTTTTCAATTC

>M6-PGSC0003DMG400006563-(rev)M6LRR-RLK_5795

TGGATACTGTGACGAAGAAGGAGAGCAGGTATGACAAAAGCTGAAAACCTTTTCAATTCCAACTGTTTTAGTTACTAATAATTCACACCATTCATCATTTTTAATCATAGTTAGTCATAT

>M6-PGSC0003DMG400006563-(rev)M6LRR-RLK_5856

ACTGTTTTAGTTACTAATAATTCACACCATTCATCATTTTTAATCATAGTTAGTCATATTTATTTAGTGTAGCATCATAAACCTTCTTGTATTCTTTCTCACTTCAGTTGCTAATGATCA

>M6-PGSC0003DMG400006563-(rev)M6LRR-RLK_5917

ATTTAGTGTAGCATCATAAACCTTCTTGTATTCTTTCTCACTTCAGTTGCTAATGATCAGACTGAATGTGTAGCAGACTAAATGCTTTCTTGAGCTACTGAAGTCTTCTTGTTTAGCACA

>M6-PGSC0003DMG400006563-(rev)M6LRR-RLK_5978

CTGAATGTGTAGCAGACTAAATGCTTTCTTGAGCTACTGAAGTCTTCTTGTTTAGCACAAAGAGTGATATAGTCGAGCAATTATGAGTCTTCTGTATACATTTGCAGATGCTGATTTATG

>M6-PGSC0003DMG400006563-(rev)M6LRR-RLK_6039

GAGTGATATAGTCGAGCAATTATGAGTCTTCTGTATACATTTGCAGATGCTGATTTATGAGTTCATGCCCAATGGAACTTTGCGAGACTGGCTTTCTGGTAAGAACTGTTTCCTGGTTGT

>M6-PGSC0003DMG400006563-(rev)M6LRR-RLK_61

AAATGCATAGCATAGTTATTTAAAATTTATGTTTACTTGTGAGTAAGATTTTGGAACTTGCGTTGTATTTTTAAAGGTGTGAAATTATGAACCAATGCAGGTATTCTTTAAAGCCAATAA

>M6-PGSC0003DMG400006563-(rev)M6LRR-RLK_610

ATAGTGATCAGTTCAATTTCCACCTTTAACTTTTTTAAATTGCATCGATCCTAGCAGAACTCTTATTCTATTTTTACTTTTCTTTACATGTTAGATGATGATGGAGAGAAAGGATATGAA

>M6-PGSC0003DMG400006563-(rev)M6LRR-RLK_6100

TTCATGCCCAATGGAACTTTGCGAGACTGGCTTTCTGGTAAGAACTGTTTCCTGGTTGTTTCTAGCTTTCATATGATTAATCCTCCTCAATTTCTTTCTTTTAGTTTAGTAGTGAAGATC

>M6-PGSC0003DMG400006563-(rev)M6LRR-RLK_6161

CTAGCTTTCATATGATTAATCCTCCTCAATTTCTTTCTTTTAGTTTAGTAGTGAAGATCCAATGCGGAGTTGCAATGTTTATAGATGTTCCTTGCACATGTTGTGTCATTAGTAGTACTA

>M6-PGSC0003DMG400006563-(rev)M6LRR-RLK_6222

ATGCGGAGTTGCAATGTTTATAGATGTTCCTTGCACATGTTGTGTCATTAGTAGTACTATGGTGAATTGATAATACAGGCTTCAGTATTTTATTGTGTTTGTGGAGAAACCATTGTCAAG

>M6-PGSC0003DMG400006563-(rev)M6LRR-RLK_6283

GTGAATTGATAATACAGGCTTCAGTATTTTATTGTGTTTGTGGAGAAACCATTGTCAAGGTGGTGATCGATCTAAAAGTTGCATAATGGAAAAGATATCGCAGGAAATCTAATAGTCAGA

>M6-PGSC0003DMG400006563-(rev)M6LRR-RLK_6344

GGTGATCGATCTAAAAGTTGCATAATGGAAAAGATATCGCAGGAAATCTAATAGTCAGAGAAACATAGATTCCATCTTCTTTTTCCTTATCAGAACAAAAAAATTTGTTAGTTTCTTTAT

>M6-PGSC0003DMG400006563-(rev)M6LRR-RLK_6405

AACATAGATTCCATCTTCTTTTTCCTTATCAGAACAAAAAAATTTGTTAGTTTCTTTATGCGGCTGTGATAAACTTCAAGAAACTTTTGTATTGCTATCTAAGTAGGTGATTTCCATCAT

>M6-PGSC0003DMG400006563-(rev)M6LRR-RLK_6466

GGCTGTGATAAACTTCAAGAAACTTTTGTATTGCTATCTAAGTAGGTGATTTCCATCATTTGACAGCCAAATGTAAAAAAATTTTGAAGTTTGGAGCAAGGTTGGGGATCGCACTGGGGG

>M6-PGSC0003DMG400006563-(rev)M6LRR-RLK_6527

GACAGCCAAATGTAAAAAAATTTTGAAGTTTGGAGCAAGGTTGGGGATCGCACTGGGGGCATCTAAGGGCATCCTTTACCTCCATACTGAAGCTGATCCACCCATATTCCAAAGGGATAT

>M6-PGSC0003DMG400006563-(rev)M6LRR-RLK_6588

TCTAAGGGCATCCTTTACCTCCATACTGAAGCTGATCCACCCATATTCCAAAGGGATATTAAAGCCAGCAACATTCTTTTAGACTCTAAACTCACTGCCAAAGTTGCTGATTTTGGATTG

>M6-PGSC0003DMG400006563-(rev)M6LRR-RLK_6649

AAGCCAGCAACATTCTTTTAGACTCTAAACTCACTGCCAAAGTTGCTGATTTTGGATTGTCTCGGCTTGCACCTGTCCAGGATGATGAAGGTCTTCTGCCTAATCATGTGTCAACAATGG

>M6-PGSC0003DMG400006563-(rev)M6LRR-RLK_671

CTTATTCTATTTTTACTTTTCTTTACATGTTAGATGATGATGGAGAGAAAGGATATGAAGGTATAACAGATATAATGAATTCATGATACATGTTGTGAAAAGACAACTTACCAAAAAGTT

>M6-PGSC0003DMG400006563-(rev)M6LRR-RLK_6710

TCGGCTTGCACCTGTCCAGGATGATGAAGGTCTTCTGCCTAATCATGTGTCAACAATGGTGAAGGGGACACCTGTAAGTCTGTCTATGCTCTAGATTTAAGGAAATAATTGTGCAATGAT

>M6-PGSC0003DMG400006563-(rev)M6LRR-RLK_6771

AAGGGGACACCTGTAAGTCTGTCTATGCTCTAGATTTAAGGAAATAATTGTGCAATGATCTTACTGATTTTCTTATGTATATGCACAGGGTTACCTTGATCCCGAATATTTCTTGACCCG

>M6-PGSC0003DMG400006563-(rev)M6LRR-RLK_6832

TACTGATTTTCTTATGTATATGCACAGGGTTACCTTGATCCCGAATATTTCTTGACCCGTAAAATGACAGATAAAAGTGATGTCTACAGCATTGGGGTTGTATTTCTGGAGATCTTAACT

>M6-PGSC0003DMG400006563-(rev)M6LRR-RLK_6893

AAATGACAGATAAAAGTGATGTCTACAGCATTGGGGTTGTATTTCTGGAGATCTTAACTGGCATGCATCCTATCTCACATGGAAAAAACATTGTCCGTGAGGTATTTCTGTCGGCTGAAC

>M6-PGSC0003DMG400006563-(rev)M6LRR-RLK_6954

CATGCATCCTATCTCACATGGAAAAAACATTGTCCGTGAGGTATTTCTGTCGGCTGAACACCTATTGGAATACAAAAACAGTGAATATTTGTCTAGCATTTTAGCAATAAAAGCATAGGT

>M6-PGSC0003DMG400006563-(rev)M6LRR-RLK_7015

CTATTGGAATACAAAAACAGTGAATATTTGTCTAGCATTTTAGCAATAAAAGCATAGGTGTATGTATATAACCATTAACTGTCCATGACAGGTAAAAATGGCTCATAAATCGGGAGTAAT

>M6-PGSC0003DMG400006563-(rev)M6LRR-RLK_7076

ATGTATATAACCATTAACTGTCCATGACAGGTAAAAATGGCTCATAAATCGGGAGTAATGTTCTCCATTATGGATAAGAACATGGGATCTTACCCCTCCGAATGTGCGGAGAAGATGGTG

>M6-PGSC0003DMG400006563-(rev)M6LRR-RLK_7137

TCTCCATTATGGATAAGAACATGGGATCTTACCCCTCCGAATGTGCGGAGAAGATGGTGGAGTTGGCCCTGAAGTGTTGCCAGGACAAACCAGAAGACAGACCCTCAATGCTGGAGGTGG

>M6-PGSC0003DMG400006563-(rev)M6LRR-RLK_7198

GTTGGCCCTGAAGTGTTGCCAGGACAAACCAGAAGACAGACCCTCAATGCTGGAGGTGGTTAGGACGCTAGAAACCATACTCCAAATGATGCCATATACTGACGCCGATCCTTTGGATGA

>M6-PGSC0003DMG400006563-(rev)M6LRR-RLK_7259

AGGACGCTAGAAACCATACTCCAAATGATGCCATATACTGACGCCGATCCTTTGGATGATAAAGCCAGTTTCAGTGAAACTACATCATCAGCTTCATTTTCTAATACTAGAAGTGGAGAC

>M6-PGSC0003DMG400006563-(rev)M6LRR-RLK_732

TATAACAGATATAATGAATTCATGATACATGTTGTGAAAAGACAACTTACCAAAAAGTTATATATGAGAAAGGAATAAATCGAAACATATTTGGACTCCCACATTATGTTTGAATTTTTG

>M6-PGSC0003DMG400006563-(rev)M6LRR-RLK_7320

AAGCCAGTTTCAGTGAAACTACATCATCAGCTTCATTTTCTAATACTAGAAGTGGAGACCTATTTATATCATCATCCAATGTATCAGGTGGTGATCTTATTAGTGGTGTTACTCTTAATA

>M6-PGSC0003DMG400006563-(rev)M6LRR-RLK_7381

ATTTATATCATCATCCAATGTATCAGGTGGTGATCTTATTAGTGGTGTTACTCTTAATATTACACCTCGTTAACATTAGCTTGTCTTCCACTGCATAAACATTGGCTTGTACACTATGAC

>M6-PGSC0003DMG400006563-(rev)M6LRR-RLK_7442

ACACCTCGTTAACATTAGCTTGTCTTCCACTGCATAAACATTGGCTTGTACACTATGACTAGTAGGAAGTCTATACGACTTGTCTTTTATGATGTATACGAGTGTACAATAATACCATCT

>M6-PGSC0003DMG400006563-(rev)M6LRR-RLK_7498

GACTAGTAGGAAGTCTATACGACTTGTCTTTTATGATGTATACGAGTGTACAATAATACCATCTTACTGCTTCTAGATTTATGTATATTCCTCAAGAATCTGTTGTGACAGACAACCCTT

>M6-PGSC0003DMG400006563-(rev)M6LRR-RLK_793

ATATGAGAAAGGAATAAATCGAAACATATTTGGACTCCCACATTATGTTTGAATTTTTGAAAGACAGATTATGGAATATTTTATCATGATAGAACATGTTGGTAGAAATAAACATGAAGT

>M6-PGSC0003DMG400006563-(rev)M6LRR-RLK_854

AGACAGATTATGGAATATTTTATCATGATAGAACATGTTGGTAGAAATAAACATGAAGTTAGGTCCTCATCTTGCACACTTATTGAAGATCAAATGCTGATTCTGATATTGCAGAAGTGT

>M6-PGSC0003DMG400006563-(rev)M6LRR-RLK_915

GGTCCTCATCTTGCACACTTATTGAAGATCAAATGCTGATTCTGATATTGCAGAAGTGTCAGAAATTGCTCTTTGCAAGGATCCATTCCTGATTTCAGCAGAATTGCAAACCTAAGCTAT

>M6-PGSC0003DMG400006563-(rev)M6LRR-RLK_976

GAAATTGCTCTTTGCAAGGATCCATTCCTGATTTCAGCAGAATTGCAAACCTAAGCTATCTGTAAGTGCGAGTTTGTTATTGTTTTGTTCTTCTTTTGTTAAGCTTTATCTGTATATTTT

>M6-PGSC0003DMG400006580-(rev)M6LRR-RLP_0

TATAGAAACTTTAATTGTGAAGAATTCTCCTTACGTATGGTTACCAGGGTCTTTTTTGCAAATGGAAAAATTAAGGCATGCTCATTTTGGTAAGGCTGTTTTTGATAAGCAGGGGCTCTT

>M6-PGSC0003DMG400006580-(rev)M6LRR-RLP_130

TCTAAATTGGAAAATTTGAGGATATTAAAGAAAATTGAGAAATTTCCAATTGATAGGGCGGATGTGTTATCAAGGAGGTGTCCTAATCTTCAACAACTTCATATCGAATTTGATGGACAT

>M6-PGSC0003DMG400006580-(rev)M6LRR-RLP_260

CTTTTTGTCTCACATTGGAGAATCTTACCCAGCTTCAAATACTTCGCGTTTCCTTTGAGCGGCCCCACATCATTGTATCTGGATTGCAATTGCCTTCAAATTTAAACAAGTTGGTACTTA

>M6-PGSC0003DMG400006580-(rev)M6LRR-RLP_325

CACATCATTGTATCTGGATTGCAATTGCCTTCAAATTTAAACAAGTTGGTACTTAAAGGAATTCATATAGAAAGTGTTATTCCCTTCATTGCGGGACTACCAAGCCTGGAGTATCTCCAA

>M6-PGSC0003DMG400006580-(rev)M6LRR-RLP_390

TATAGAAAGTGTTATTCCCTTCATTGCGGGACTACCAAGCCTGGAGTATCTCCAATTACAAGATGTGTGTTTTCCTCAATCAAAAGAGTGGTGCCTTGGAGATATCACGTTCCATAAACT

>M6-PGSC0003DMG400006580-(rev)M6LRR-RLP_520

AAACTGGTGCAGTTAGGTATCTCAAGGTGGGATGCCTCGGAGGAATCATTTCCCTTGCTTGAAACACTAGTTATAAAAATGTGTGGTCACCTAGAGGAGATCCCAATTAGCTTTGCTGAT

>M6-PGSC0003DMG400006580-(rev)M6LRR-RLP_585

ACTAGTTATAAAAATGTGTGGTCACCTAGAGGAGATCCCAATTAGCTTTGCTGATATTCCATCATTGAAACAGATTAAATTGATTGGGCCCTGGAAAGAATCTCTGGAGGATTCAGCTGT

>M6-PGSC0003DMG400006580-(rev)M6LRR-RLP_65

AAAAATTAAGGCATGCTCATTTTGGTAAGGCTGTTTTTGATAAGCAGGGGCTCTTTGAAGGATCCTCTAAATTGGAAAATTTGAGGATATTAAAGAAAATTGAGAAATTTCCAATTGATA

>M6-PGSC0003DMG400006580-(rev)M6LRR-RLP_715

GAAGAAATCGAAGAGATTGAAGGATGTGATCGTATACACCTCGTCAAAGAACCGACAAATTGGGAGGCATATAGCAATTTGTCGAAGCTGTGCTTATACTGAAGTCTATTTTGTAGCTAG

>M6-PGSC0003DMG400006580-(rev)M6LRR-RLP_780

GGCATATAGCAATTTGTCGAAGCTGTGCTTATACTGAAGTCTATTTTGTAGCTAGCCCATTCTCTTATATTGTAACACTACTACTGTTTGTTATGTTTGTTTTGATTGATCAACTCTTTT

>M6-PGSC0003DMG400006580-(rev)M6LRR-RLP_837

CATTCTCTTATATTGTAACACTACTACTGTTTGTTATGTTTGTTTTGATTGATCAACTCTTTTTATGTATGGTTGGAGACAAATATATATCAGATGATCTTTCACTGATTTCATGTTTTT

>M6-PGSC0003DMG400006652M6LRR-RLP_1071

GATCTTTCTCGTAATGATTTCAGTGGCTCACTTCCTGCAAAAGTTTTTGGAAACTTCAAGGCAATGATCAAATTAGATAACGAAGACACAAGAGAGATAAAGTACATGAAATCTGTGTTG

>M6-PGSC0003DMG400006652M6LRR-RLP_1134

ATGATCAAATTAGATAACGAAGACACAAGAGAGATAAAGTACATGAAATCTGTGTTGCATTGGCCGCCATTCGTCACATCGTATGAGGAGTCAGTGAGTTTGGTAATCAAAGGGCAGGAT

>M6-PGSC0003DMG400006652M6LRR-RLP_1260

CTACAAAGAATCAGCACAATTACGACAACCATAGATCTCTCAAGCAACCATTTCGAAGGTGTGATTCCGAAAACACTAAAGGATCTCAGATCACTTTGGCTACTCAATTTATCTCATAAC

>M6-PGSC0003DMG400006652M6LRR-RLP_1323

ATTCCGAAAACACTAAAGGATCTCAGATCACTTTGGCTACTCAATTTATCTCATAACAATCTCAGAGGAGATATTCCAATGAAATTAGGACAATTGAGTATGCTTGAAGCTTTAGATCTC

>M6-PGSC0003DMG400006652M6LRR-RLP_441

AAGGATTTTCCACACTTCTTGAGAAATGTAAAGAAACTTGTGTACTTGGATATTTCTAACAATAAGATTCGTGGTCAAATCCCTAACTGGTTTAGCGGCATGAGATGGGACTCGTTACAG

>M6-PGSC0003DMG400006652M6LRR-RLP_882

CCTGTGTCATTGCTCAAGTGTGGTGATCTAGAAGTACTTGATGTGGGGAACAATGCTATAAATGACAGGTTTCCATCTTGGCTCGGAACTCTTGAATACTTGCAGGTCCTTATATTAAAG

>M6-PGSC0003DMG400006652M6LRR-RLP_945

GACAGGTTTCCATCTTGGCTCGGAACTCTTGAATACTTGCAGGTCCTTATATTAAAGTCGAACAAGTTCCATGGACCTATAAGTATGTGTCAGACTAAGTTTTGCTTTCCCAAGTTGCGA

>M6-PGSC0003DMG400006655M6LRR-RLP_1197

TTAGATGGTGAAGACACAGGAAATATCAAGTACATAAAATCTGTGTTGAATTCGTCATTTGACACATCGTATGAGGATTCAGTGAGTTTGGTAATCAAAGGGCACGATATTGAGCTACAA

>M6-PGSC0003DMG400006655M6LRR-RLP_1386

CTAAAGGACCTCAGCTCACTTAGGCTACTCAATTTATCCCATAACAATCTCATAGGTCATATTCCAGTGGAATTGGGGCAATTGAATAAGCTGGAAGCTTTAGATCTCTCTTGGAATCGG

>M6-PGSC0003DMG400006655M6LRR-RLP_1512

GGAAAGATTCCGCAGGAATTGACAAGAATGAACTTTCTGGCAGTCTTAAACATCTCTCAAAATCATCTCATCGGACCAATTCCTCACGGTCTACAGTTCAACACATTTGAAAATGACTCG

>M6-PGSC0003DMG400006655M6LRR-RLP_1638

GGTAACCTTGATTTATGTGGTCTTCCTTTAACAAAGAAATGTGGAACGAGTGATTCATCACATGTTCCTCAACAATTGGAGTCCGAAGAAGAAGGCGAGTCATATTTTTTTAGTGGATTT

>M6-PGSC0003DMG400006655M6LRR-RLP_315

CCAACATTAAAAGAACTGCATTTAAGCAATAATCAACTCAGTGGTTCTTTTCCTCAATCACTTGTGAATCTCACAAATCTTTTTACCCTTGACATTTCATCAAATAACATCACCATTGAT

>M6-PGSC0003DMG400006655M6LRR-RLP_378

GTGAATCTCACAAATCTTTTTACCCTTGACATTTCATCAAATAACATCACCATTGATGAGGGAATGAATATCACCTTTCTTAGCCTATCATCTTTATTCTTATCATCTTGTGAACTGAAG

>M6-PGSC0003DMG400006655M6LRR-RLP_630

CATAATTCATTAACAGGCCACCTACCACAATTGCATTTCCATAATCTAGGGTATCTTGATCTGAAATTTAACTCCCTTCAGGGTCCACTACCTTCATCCATTTGTAACATGAGCTCGCTT

>M6-PGSC0003DMG400006655M6LRR-RLP_693

AAATTTAACTCCCTTCAGGGTCCACTACCTTCATCCATTTGTAACATGAGCTCGCTTGAATTTCTTGATTTATCACGCAACAACTTCAGTGACTCAGTTTCACATTGTTTGGGAAGCATG

>M6-PGSC0003DMG400006667M6OtherRLKTM_0

CTTTCACCTACACTGCTCTTATTTTGAGAAGATGTAGTTTTGAAGCATTTAATGCTCACTTTTCTCTCTCTATTTCTCCTATGCTGTCTTTCACCACTTCATTCTTGAGGGCACCGATAA

>M6-PGSC0003DMG400006667M6OtherRLKTM_1054

GCACCAAAAGATTTGAGATTTATTAAGATTGCAGCTGGGGGATATCATGTTTGTGGGATCCTAGAAGGGGTGAATTCTCAAGTGTATTGCTGGGGAAGGAGCATGAACCTTGAAGAAGAA

>M6-PGSC0003DMG400006667M6OtherRLKTM_1116

AGAAGGGGTGAATTCTCAAGTGTATTGCTGGGGAAGGAGCATGAACCTTGAAGAAGAATTCTCTGTTGCTCAACTCAATGTTGAATTGGCAGCCCCTAGTGATCCAATTATATCTGTTGT

>M6-PGSC0003DMG400006667M6OtherRLKTM_1178

CTGTTGCTCAACTCAATGTTGAATTGGCAGCCCCTAGTGATCCAATTATATCTGTTGTTGGTGGTAAGTTTCATGCTTGTGGGATTAGGAGCTATGACCGTCATGTCGTTTGCTGGGGTT

>M6-PGSC0003DMG400006667M6OtherRLKTM_124

GACACAAAAGGGGCTAAAAAAAATGTCATTATCTTCTCTTTTTTGTGTTTCTTGAACTGAAATATGCAGCTTCTTGGCTACAAATTTTGTTTAATTGTATTGATAACGAGGGTATTATAA

>M6-PGSC0003DMG400006667M6OtherRLKTM_1240

GGTAAGTTTCATGCTTGTGGGATTAGGAGCTATGACCGTCATGTCGTTTGCTGGGGTTACAGAGTTGAGAAAAGCACACCACCTCCTAGTGGAGTTAGGCTTTATGAGATAGCAGCTGGT

>M6-PGSC0003DMG400006667M6OtherRLKTM_1302

AGTTGAGAAAAGCACACCACCTCCTAGTGGAGTTAGGCTTTATGAGATAGCAGCTGGTGACTACTTCACTTGTGGTATCCTTGCGGAAATTTCAATTTTGCCTGTTTGTTGGGGGTTTGG

>M6-PGSC0003DMG400006667M6OtherRLKTM_1364

ACTTCACTTGTGGTATCCTTGCGGAAATTTCAATTTTGCCTGTTTGTTGGGGGTTTGGTTTTCCCTCATCGCTACCACTCGCTGTTTCTCCTGGAGTCTGCAAGCCTAGACCCTGTGCAT

>M6-PGSC0003DMG400006667M6OtherRLKTM_1426

CCCTCATCGCTACCACTCGCTGTTTCTCCTGGAGTCTGCAAGCCTAGACCCTGTGCATCTGGCTTCTATGAGTTTAACAACGGAAGTGCAACTTGCAAGTCTCCTGATTCTCGCATTTGC

>M6-PGSC0003DMG400006667M6OtherRLKTM_1488

CTTCTATGAGTTTAACAACGGAAGTGCAACTTGCAAGTCTCCTGATTCTCGCATTTGCCTTCCCTGCACCAATGGCTGCCCTGCTGAAATGTATCAACAGGTTGAATGCACTTCATCTAC

>M6-PGSC0003DMG400006667M6OtherRLKTM_1550

CCTGCACCAATGGCTGCCCTGCTGAAATGTATCAACAGGTTGAATGCACTTCATCTACGGACAGTCAGTGCACGTATAATTGTTCAAGTTGTATCTCTGTTGACTGCATAAACAGCTGTT

>M6-PGSC0003DMG400006667M6OtherRLKTM_1612

AGTCAGTGCACGTATAATTGTTCAAGTTGTATCTCTGTTGACTGCATAAACAGCTGTTCTACTGCTATTTCTGGAAAGAAGAACGCTAAATTTTGGTCACTCCAGTTACCAGTAATTGTT

>M6-PGSC0003DMG400006667M6OtherRLKTM_1674

TGCTATTTCTGGAAAGAAGAACGCTAAATTTTGGTCACTCCAGTTACCAGTAATTGTTGCTGAGGTTGCATTTGCAGTATTCTTGGTGAGTGTTGTATCTCTAACTTCGATCGTATATGT

>M6-PGSC0003DMG400006667M6OtherRLKTM_1736

AGGTTGCATTTGCAGTATTCTTGGTGAGTGTTGTATCTCTAACTTCGATCGTATATGTTCGCTACAAATTAAGGAACTGTAGATGTTCAGGGAAAGGTCCTAGTCCTAGGAAGAATGGTA

>M6-PGSC0003DMG400006667M6OtherRLKTM_1798

TACAAATTAAGGAACTGTAGATGTTCAGGGAAAGGTCCTAGTCCTAGGAAGAATGGTACTTTCCCAAAGGAAATTGCTAAAGATAGGGCTGATTTGGATGATCTTAAAATAAGGAGAGCT

>M6-PGSC0003DMG400006667M6OtherRLKTM_186

TATGCAGCTTCTTGGCTACAAATTTTGTTTAATTGTATTGATAACGAGGGTATTATAATTATTTGGGATGAAGGAAAGTTGTGAATTTTGATCCCACCTACCCACCCATGGTATGTTTGA

>M6-PGSC0003DMG400006667M6OtherRLKTM_1860

CCCAAAGGAAATTGCTAAAGATAGGGCTGATTTGGATGATCTTAAAATAAGGAGAGCTCAGATGTTTACTTATGAAGATCTTGAGAGAGCAACTGAGGGATTCAAAGAAGAATCACAAGT

>M6-PGSC0003DMG400006667M6OtherRLKTM_1922

TGTTTACTTATGAAGATCTTGAGAGAGCAACTGAGGGATTCAAAGAAGAATCACAAGTTGGAAAGGGTAGCTTTTCGTGTGTGTTCAAGGGCGTTTTGAAGGACGGTACTGTGGTTGCTG

>M6-PGSC0003DMG400006667M6OtherRLKTM_1984

AAGGGTAGCTTTTCGTGTGTGTTCAAGGGCGTTTTGAAGGACGGTACTGTGGTTGCTGTCAAGAGGGCTATAATGTCATCTGACATGAAGAAGAATTCAAAGGAGTTCCACAATGAGCTA

>M6-PGSC0003DMG400006667M6OtherRLKTM_2046

GAGGGCTATAATGTCATCTGACATGAAGAAGAATTCAAAGGAGTTCCACAATGAGCTAGACTTGCTGTCCAGGTTGAATCATGCTCATTTGCTCAATTTGCTAGGTTATTGTGAAGAAGG

>M6-PGSC0003DMG400006667M6OtherRLKTM_2108

TGCTGTCCAGGTTGAATCATGCTCATTTGCTCAATTTGCTAGGTTATTGTGAAGAAGGTGGAGAGAGACTTCTAGTTTATGAGTACATGGCTAATGGCACGTTGCATGAACATCTACACG

>M6-PGSC0003DMG400006667M6OtherRLKTM_2170

GAGAGACTTCTAGTTTATGAGTACATGGCTAATGGCACGTTGCATGAACATCTACACGGGAAAAAGAAGGAGCAATTGGATTGGATAAGAAGGGTAACCATTGCAGTCCAAGCTGCTCGG

>M6-PGSC0003DMG400006667M6OtherRLKTM_2232

AAAGAAGGAGCAATTGGATTGGATAAGAAGGGTAACCATTGCAGTCCAAGCTGCTCGGGGAATCGAATATTTGCATGGTTATGCATGTCCACCTGTGATTCACAGAGACATCAAGTCCTC

>M6-PGSC0003DMG400006667M6OtherRLKTM_2294

TCGAATATTTGCATGGTTATGCATGTCCACCTGTGATTCACAGAGACATCAAGTCCTCAAACATCCTTATAGATGAAGAACACAATGCTCGAGTAGCTGATTTTGGGCTTTCCTTGCTTG

>M6-PGSC0003DMG400006667M6OtherRLKTM_2356

ATCCTTATAGATGAAGAACACAATGCTCGAGTAGCTGATTTTGGGCTTTCCTTGCTTGGACCTGCTAATAGCAGTTCCCCATTAGCTGAGTTACCAGCAGGGACACTTGGGTACCTTGAT

>M6-PGSC0003DMG400006667M6OtherRLKTM_2418

TGCTAATAGCAGTTCCCCATTAGCTGAGTTACCAGCAGGGACACTTGGGTACCTTGATCCCGAGTACTACAGACTACATTATCTTACAACCAAATCTGATGTCTATAGCTTTGGTGTTTT

>M6-PGSC0003DMG400006667M6OtherRLKTM_248

TTGGGATGAAGGAAAGTTGTGAATTTTGATCCCACCTACCCACCCATGGTATGTTTGAACTATTTTTTTTTCCGTGTGTTTCATAAATCAACTCAGCTAATATGGAGAAGGAGAGGTGTT

>M6-PGSC0003DMG400006667M6OtherRLKTM_2480

AGTACTACAGACTACATTATCTTACAACCAAATCTGATGTCTATAGCTTTGGTGTTTTGCTTTTGGAAATTCTCAGTGGTCGGAAAGCTATTGACATGCAATACGATGAAGGGAACATAG

>M6-PGSC0003DMG400006667M6OtherRLKTM_2542

TTGGAAATTCTCAGTGGTCGGAAAGCTATTGACATGCAATACGATGAAGGGAACATAGTGGAATGGGCAGTCCCATTAATCAAAGCTGGTGAAATAGAAGCAATACTGGATCCAGTTTTG

>M6-PGSC0003DMG400006667M6OtherRLKTM_2604

ATGGGCAGTCCCATTAATCAAAGCTGGTGAAATAGAAGCAATACTGGATCCAGTTTTGAAATCACCTTCTGATGGTGAAGCTCTTAGAAGAATCGCTAATATAGCCAGCAAATGCGTGAG

>M6-PGSC0003DMG400006667M6OtherRLKTM_2666

CACCTTCTGATGGTGAAGCTCTTAGAAGAATCGCTAATATAGCCAGCAAATGCGTGAGGATGAGAGGGAAAGAGAGGCCGTCAATGGATAAAGTAACAACAGCTTTGGAGAGAGCACTTG

>M6-PGSC0003DMG400006667M6OtherRLKTM_2728

AGAGGGAAAGAGAGGCCGTCAATGGATAAAGTAACAACAGCTTTGGAGAGAGCACTTGCTCAATTGATGGGTAGTCCAAGCAATGACCAGCCTATCTTGCCAACAGAGGTTGTTCTAGGA

>M6-PGSC0003DMG400006667M6OtherRLKTM_2790

ATTGATGGGTAGTCCAAGCAATGACCAGCCTATCTTGCCAACAGAGGTTGTTCTAGGAAGCAGCAGAATGCACAAGAAGTCCTCATCAAATCGATCAACGTCAGAAACAACAGATGTTGC

>M6-PGSC0003DMG400006667M6OtherRLKTM_2852

GCAGAATGCACAAGAAGTCCTCATCAAATCGATCAACGTCAGAAACAACAGATGTTGCAGAAACTGAGGATCAGAGGTATGTCGAATTCAGAGCTCCTTCGTGGATTACATTCCCAAGTG

>M6-PGSC0003DMG400006667M6OtherRLKTM_2914

ACTGAGGATCAGAGGTATGTCGAATTCAGAGCTCCTTCGTGGATTACATTCCCAAGTGTAGCATCATCTCAGAGAAGAAAGTCTTCAGTATCGGACGCAGATGTTGAAGCAAAGAATTTA

>M6-PGSC0003DMG400006667M6OtherRLKTM_2976

ATCATCTCAGAGAAGAAAGTCTTCAGTATCGGACGCAGATGTTGAAGCAAAGAATTTAGAAAGTAGGAACTGTGGAAATGGAACTGATGGATTGAGAAGTTTGGAAGAAGAAATTGGACC

>M6-PGSC0003DMG400006667M6OtherRLKTM_3038

GTAGGAACTGTGGAAATGGAACTGATGGATTGAGAAGTTTGGAAGAAGAAATTGGACCAGCTTCTCCTCATGAACATTTGTTCTTGAAACACAACTTCTAATTAACATAACACTAAAAAG

>M6-PGSC0003DMG400006667M6OtherRLKTM_310

TTTTTTTTTCCGTGTGTTTCATAAATCAACTCAGCTAATATGGAGAAGGAGAGGTGTTATTTTGGTTTTATCAGAAGGATAAAGGTGAAAGAAAGCACTGACTTTTTGTTTGTGTACTTT

>M6-PGSC0003DMG400006667M6OtherRLKTM_3100

TCTCCTCATGAACATTTGTTCTTGAAACACAACTTCTAATTAACATAACACTAAAAAGGAAATGCTCTGCAGGTTGATGCTTAATTATTTTGTTAGTGGAAGAAATGTATACAAGTAAAA

>M6-PGSC0003DMG400006667M6OtherRLKTM_3162

TGCTCTGCAGGTTGATGCTTAATTATTTTGTTAGTGGAAGAAATGTATACAAGTAAAAGGTAAATGTTGTGAATTCTTGATAAATGTGATCAAGAATTCCCCACTCAACCATTGGCGATG

>M6-PGSC0003DMG400006667M6OtherRLKTM_3224

AATGTTGTGAATTCTTGATAAATGTGATCAAGAATTCCCCACTCAACCATTGGCGATGAGCTTTAGCTAGATGTTTGATCCTGTAATTGTAACTTCAATCTTTCTTCTTCTTTTTATGCA

>M6-PGSC0003DMG400006667M6OtherRLKTM_372

TGGTTTTATCAGAAGGATAAAGGTGAAAGAAAGCACTGACTTTTTGTTTGTGTACTTTGTTTAATTTTTAATTTGTGTAATGGACAGTTTAAATAAGTGGTGTGTGGTGATGTGGAAATG

>M6-PGSC0003DMG400006667M6OtherRLKTM_434

AATTTTTAATTTGTGTAATGGACAGTTTAAATAAGTGGTGTGTGGTGATGTGGAAATGTAGATACTTTGTAAGAGCTTTTATGCTCCTTGTGGTATTTTCAAAAGTATCAGGGTTTGGAT

>M6-PGSC0003DMG400006667M6OtherRLKTM_496

TACTTTGTAAGAGCTTTTATGCTCCTTGTGGTATTTTCAAAAGTATCAGGGTTTGGATCAATGTCTTCAATTGCTATTTCATATGGTGAATATGGTTCTGTTTTTTGTGGGTTGAAGTCA

>M6-PGSC0003DMG400006667M6OtherRLKTM_558

GTCTTCAATTGCTATTTCATATGGTGAATATGGTTCTGTTTTTTGTGGGTTGAAGTCAGATGGATCTCATTTGGTCAGTTGCTATGGCTCTACTTCTTCTATAATATATTCAACTCCAGC

>M6-PGSC0003DMG400006667M6OtherRLKTM_62

TCTCTCTCTATTTCTCCTATGCTGTCTTTCACCACTTCATTCTTGAGGGCACCGATAACTTTGACACAAAAGGGGCTAAAAAAAATGTCATTATCTTCTCTTTTTTGTGTTTCTTGAACT

>M6-PGSC0003DMG400006667M6OtherRLKTM_620

GATCTCATTTGGTCAGTTGCTATGGCTCTACTTCTTCTATAATATATTCAACTCCAGCTCATTTCCCTTTTATTGGTCTTACTGCTGGAAATGGCTTTGTATGTGGACTTTTGATGGATT

>M6-PGSC0003DMG400006667M6OtherRLKTM_682

TTCCCTTTTATTGGTCTTACTGCTGGAAATGGCTTTGTATGTGGACTTTTGATGGATTCTTACCAGCCTTATTGTTGGGGGAAAAGTAATTTTGTACAAATGGGAGTGCCTCAGCCTATG

>M6-PGSC0003DMG400006667M6OtherRLKTM_744

CCAGCCTTATTGTTGGGGGAAAAGTAATTTTGTACAAATGGGAGTGCCTCAGCCTATGATCAAAGGGTCTCAATACTTGGAAATATCTGCAGGTGAAAATCATTTGTGTGGACTAAGGCA

>M6-PGSC0003DMG400006667M6OtherRLKTM_806

AAGGGTCTCAATACTTGGAAATATCTGCAGGTGAAAATCATTTGTGTGGACTAAGGCAACCTTTAATGGGGAAGCATAGGAACACTTCACTTGTTGATTGCTGGGGTTATAACATGACTA

>M6-PGSC0003DMG400006667M6OtherRLKTM_868

TTAATGGGGAAGCATAGGAACACTTCACTTGTTGATTGCTGGGGTTATAACATGACTACAAACAATGAGTTTGAAGGTCAGATCCACTCTATTTCAGCTGGTTCTGAGTTTAATTGTGCT

>M6-PGSC0003DMG400006667M6OtherRLKTM_930

CAATGAGTTTGAAGGTCAGATCCACTCTATTTCAGCTGGTTCTGAGTTTAATTGTGCTTTGTTTTCTGTCAATAAAAGTGTTTTATGTTGGGGGGATGAAACTAGTAGCCAGGTTATTAC

>M6-PGSC0003DMG400006667M6OtherRLKTM_992

TTTCTGTCAATAAAAGTGTTTTATGTTGGGGGGATGAAACTAGTAGCCAGGTTATTACCCTAGCACCAAAAGATTTGAGATTTATTAAGATTGCAGCTGGGGGATATCATGTTTGTGGGA

>M6-PGSC0003DMG400006729-(rev)M6RLKWAX-EGF_0

TTGTAACTACCCTTTTCAACCTCCTCAGCTTTATCTCGGTGACACAACTATTCAAATCACACGCATATCGCTCGCTGGGCAACTAACAGTCTTACAATTTATATCGAGAGATTGTTAAAA

>M6-PGSC0003DMG400006729-(rev)M6RLKWAX-EGF_124

GGGATAAACACGGAGAATAACACACCATGGATAGCATTACCACTGTCGTTTTCTGTGAATGACACCACAAATAAGTTCATCGCAATTGGCTGTGACACATATGTTATAGTTCAGGGTTAT

>M6-PGSC0003DMG400006729-(rev)M6RLKWAX-EGF_186

CACCACAAATAAGTTCATCGCAATTGGCTGTGACACATATGTTATAGTTCAGGGTTATTATTCCAAGTTTGAAAGTAGTAAAATCGAGGATTATTATTCCTACATAACAGGATGATGCAT

>M6-PGSC0003DMG400006729-(rev)M6RLKWAX-EGF_248

CCAAGTTTGAAAGTAGTAAAATCGAGGATTATTATTCCTACATAACAGGATGATGCATGTCCATGTGCAATAGTATTGATGATGATAAATGTTCTGGTGTTGGTTGTTGCATGACATCCA

>M6-PGSC0003DMG400006729-(rev)M6RLKWAX-EGF_310

ATGTGCAATAGTATTGATGATGATAAATGTTCTGGTGTTGGTTGTTGCATGACATCCATCCCTAATGGGGCCTGGAATGTTACTATAACACTAAGGAGCTATTATAATCATACTTATGTG

>M6-PGSC0003DMG400006729-(rev)M6RLKWAX-EGF_372

TAATGGGGCCTGGAATGTTACTATAACACTAAGGAGCTATTATAATCATACTTATGTGAAAGATAACCCTAGTTACGTTGTAGCTATGGTTTTGTAGTTGAAGAAGCTAATTTTAGTTAT

>M6-PGSC0003DMG400006729-(rev)M6RLKWAX-EGF_434

ATAACCCTAGTTACGTTGTAGCTATGGTTTTGTAGTTGAAGAAGCTAATTTTAGTTATTCTAAGAATTATCTAAGAAGTTTGCAGAATATGGAGGAGCTTCCTCTTGTGCTAGATTGGGT

>M6-PGSC0003DMG400006729-(rev)M6RLKWAX-EGF_496

AGAATTATCTAAGAAGTTTGCAGAATATGGAGGAGCTTCCTCTTGTGCTAGATTGGGTAATTGGTGAAGAAACATGTGAAATAGCTAAGAGAAACTCGACTACATATGTATGTAAGAGTA

>M6-PGSC0003DMG400006729-(rev)M6RLKWAX-EGF_62

GCATATCGCTCGCTGGGCAACTAACAGTCTTACAATTTATATCGAGAGATTGTTAAAAGTCCGGGATAAACACGGAGAATAACACACCATGGATAGCATTACCACTGTCGTTTTCTGTGA

>M6-PGSC0003DMG400006792M6LRR-RLP_0

CAACAACAAACTATCCGGAAAGTTCCCCTCGTGGATATCTGATGGTTTTCAAAATCTTAGAATCCTCAGCTTAAGGTCAAATTCATTTTCCGGAGAACTTCCTTTAGGAATGTCAAAACT

>M6-PGSC0003DMG400006792M6LRR-RLP_130

CAGGTGCTTGATCTTGCAGAAAACAACATAACTGGCGCCATTCCAACAACCATAGGAGACCTGAACGCCATGGTACAAGAGGAAAAAGTGAATGAATATCTGTTATATGGGAAGTATAGA

>M6-PGSC0003DMG400006792M6LRR-RLP_195

CGCCATGGTACAAGAGGAAAAAGTGAATGAATATCTGTTATATGGGAAGTATAGAGGGATTTACTATGAAGAAAGCTTGGTAGTGAATTTGAAAAACCAGTTCCAAAAGTACACCAAGAC

>M6-PGSC0003DMG400006792M6LRR-RLP_260

ATGAAGAAAGCTTGGTAGTGAATTTGAAAAACCAGTTCCAAAAGTACACCAAGACTCTATCACTTCTTACCTCAATAGACTTGTCGAGAAACAACCTCAACGGAGCATTTCCTGTGGAAT

>M6-PGSC0003DMG400006792M6LRR-RLP_325

CTTACCTCAATAGACTTGTCGAGAAACAACCTCAACGGAGCATTTCCTGTGGAATTGACAAACTTGCATGGCCTCATAGTTTTGAACTTGTCCGGAAACCAAATCAGTGGAAAAATACCA

>M6-PGSC0003DMG400006792M6LRR-RLP_390

GCATGGCCTCATAGTTTTGAACTTGTCCGGAAACCAAATCAGTGGAAAAATACCAGAAAACATTTCAAGCTTACATCAGTTAGCATCGCTTGATCTCTCGAGTAATACACTGTCTGGTGT

>M6-PGSC0003DMG400006792M6LRR-RLP_455

CAAGCTTACATCAGTTAGCATCGCTTGATCTCTCGAGTAATACACTGTCTGGTGTCATCCCATCAAGCATGGAATCAATGTCATTCTTGAGCTATATTAACTTCTCAAACAACAACTTAT

>M6-PGSC0003DMG400006792M6LRR-RLP_520

AGCATGGAATCAATGTCATTCTTGAGCTATATTAACTTCTCAAACAACAACTTATCCGGTATGGTTCCTTATAAGGGACAGATGACAACTTTTACAGTGTCCGCGTTTGAGGGGAATCCC

>M6-PGSC0003DMG400006792M6LRR-RLP_585

TCCTTATAAGGGACAGATGACAACTTTTACAGTGTCCGCGTTTGAGGGGAATCCCCATCTTTGTGGAGCTCCACTTCTAGTACTATGTCAAAATGGAAACTCGGACAATAGAACGGTACT

>M6-PGSC0003DMG400006792M6LRR-RLP_65

TCAGCTTAAGGTCAAATTCATTTTCCGGAGAACTTCCTTTAGGAATGTCAAAACTGAGTTCATTGCAGGTGCTTGATCTTGCAGAAAACAACATAACTGGCGCCATTCCAACAACCATAG

>M6-PGSC0003DMG400006792M6LRR-RLP_650

GAGCTCCACTTCTAGTACTATGTCAAAATGGAAACTCGGACAATAGAACGGTACTAGAGAATGACAGCAGTGATGAATTTCCTGATAAATGGTTTTATGTTAGTGTTGCATTGGGATTTA

>M6-PGSC0003DMG400006792M6LRR-RLP_715

AGCAGTGATGAATTTCCTGATAAATGGTTTTATGTTAGTGTTGCATTGGGATTTATTACAGGTATTCTTGTTCCTTACTATATCTTGCTCGTTCGAAAGCCATGGAGACGAGCATATTTT

>M6-PGSC0003DMG400006792M6LRR-RLP_780

TCTTGTTCCTTACTATATCTTGCTCGTTCGAAAGCCATGGAGACGAGCATATTTTAGTTTCTTAGACACAGTCATCGACAGATTTGTCCCAGTCAGAAGCAACAGAACAACCAGCACCAA

>M6-PGSC0003DMG400006792M6LRR-RLP_840

CTTAGACACAGTCATCGACAGATTTGTCCCAGTCAGAAGCAACAGAACAACCAGCACCAAAAGATCAGATACGGGCACCAGATATGGCAAAAGAAGTGTTCAATAGGTACTAGTCCTAGT

>M6-PGSC0003DMG400006880-(rev)M6LRR-RLP_0

CACAGAAGAAACACTCTCTGTAAATAGAAAAATAGAGAGTAATAGTACTGCATTTTGTAATAGAGAGAGGAGAAATGGGTCTCACCGGAGTAACATTGTTGGTCACTCTCATGGTGGTGG

>M6-PGSC0003DMG400006880-(rev)M6LRR-RLP_1024

AAAATCATTATCATCAGCTAAGTACATTGGCCATTTGGATTTAAGCCATAATCACCTTTGTGGCCCGATTCCAAACGGGTCACCCTTCGATCATCTAGAAGCTTCTTCTTTCTCTAACAA

>M6-PGSC0003DMG400006880-(rev)M6LRR-RLP_1088

CCGATTCCAAACGGGTCACCCTTCGATCATCTAGAAGCTTCTTCTTTCTCTAACAACGATTGTTTGTGTGGATCGCCATTACGTACTTGTTAATTCTTTTTTGTTTACTTACTAAGTTTG

>M6-PGSC0003DMG400006880-(rev)M6LRR-RLP_1152

TGTGTGGATCGCCATTACGTACTTGTTAATTCTTTTTTGTTTACTTACTAAGTTTGTAATTAATCGAAATTATCGGTTGATTACGCATAAATGGATGTGTAATTGTGTATTATGTTATGA

>M6-PGSC0003DMG400006880-(rev)M6LRR-RLP_1216

CGAAATTATCGGTTGATTACGCATAAATGGATGTGTAATTGTGTATTATGTTATGATTAAAAAAATATTCTTATGTGTTTAGCTTAATTATTATTATTATTGCTAAGCAATATGATGGAT

>M6-PGSC0003DMG400006880-(rev)M6LRR-RLP_128

ACTTTGCCGGCAAATGCCTGCCTGCCGGCGGACCAAGCAGCACTGATGGATATCAAAGCAGCTCTTAAGGAGCCATATTTGGGCATTTTCAACACATGGACTGGCAGCAACTGTTGCCAG

>M6-PGSC0003DMG400006880-(rev)M6LRR-RLP_1280

ATATTCTTATGTGTTTAGCTTAATTATTATTATTATTGCTAAGCAATATGATGGATTATGCGGTTCTATCATGCATGCACGTGAAAATGCGTGACAGTTTACAAGAAAAATTATTGAAGA

>M6-PGSC0003DMG400006880-(rev)M6LRR-RLP_1327

ATGATGGATTATGCGGTTCTATCATGCATGCACGTGAAAATGCGTGACAGTTTACAAGAAAAATTATTGAAGAAATTGTGTGCTATTATTATTAGTAAATTGTAATATAATAAATCCCAC

>M6-PGSC0003DMG400006880-(rev)M6LRR-RLP_192

TTAAGGAGCCATATTTGGGCATTTTCAACACATGGACTGGCAGCAACTGTTGCCAGGGCTGGTATGGTGTTAGCTGTGACCCGACTACGCAACGAGTTGCAGACATTGTTCTTCGGGGAG

>M6-PGSC0003DMG400006880-(rev)M6LRR-RLP_256

TGGTGTTAGCTGTGACCCGACTACGCAACGAGTTGCAGACATTGTTCTTCGGGGAGAATCCGAAGATCCGATTTACGAGAAAGCCGGCCGGTCTGGTTACATGACCGGTTCACTCTCCCC

>M6-PGSC0003DMG400006880-(rev)M6LRR-RLP_320

GATCCGATTTACGAGAAAGCCGGCCGGTCTGGTTACATGACCGGTTCACTCTCCCCTTCTCTCTGCAAACTCGACAAGCTCACTACACTCATCGTCGCCGACTGGAAAGATATCTCCGGC

>M6-PGSC0003DMG400006880-(rev)M6LRR-RLP_384

GCAAACTCGACAAGCTCACTACACTCATCGTCGCCGACTGGAAAGATATCTCCGGCGAAATTCCGGCTTGTGTCACTTCATTACCAGATCTCCGTATCCTTGAGCTCATCGGCAACAAAA

>M6-PGSC0003DMG400006880-(rev)M6LRR-RLP_448

GGCTTGTGTCACTTCATTACCAGATCTCCGTATCCTTGAGCTCATCGGCAACAAAATCACCGGTCAAATCCCGGAAAACATCGGTCAGCTGAGTAAACTCACTGTTCTAAACCTCGCCGA

>M6-PGSC0003DMG400006880-(rev)M6LRR-RLP_512

CAAATCCCGGAAAACATCGGTCAGCTGAGTAAACTCACTGTTCTAAACCTCGCCGATAATAAAATCTGCGGTTCAATCCCCGCCTCCATTGTCAACCTCGGAAAACTAAAGCATCTCGAA

>M6-PGSC0003DMG400006880-(rev)M6LRR-RLP_576

TCTGCGGTTCAATCCCCGCCTCCATTGTCAACCTCGGAAAACTAAAGCATCTCGAACTCAGCAACAATCAACTCACCGGCGAAATCCCGTCCGATATCGGAAAACTCGGCATGATGAGCC

>M6-PGSC0003DMG400006880-(rev)M6LRR-RLP_64

GAGAGGAGAAATGGGTCTCACCGGAGTAACATTGTTGGTCACTCTCATGGTGGTGGCCGCCGTTACTTTGCCGGCAAATGCCTGCCTGCCGGCGGACCAAGCAGCACTGATGGATATCAA

>M6-PGSC0003DMG400006880-(rev)M6LRR-RLP_640

CAATCAACTCACCGGCGAAATCCCGTCCGATATCGGAAAACTCGGCATGATGAGCCGAGCACTACTCAACAAGAACAAACTCACAGGTTCAATCCCTAACTCAATCACTCAACTTCGCCG

>M6-PGSC0003DMG400006880-(rev)M6LRR-RLP_704

CTCAACAAGAACAAACTCACAGGTTCAATCCCTAACTCAATCACTCAACTTCGCCGGTTAGCCGATCTAGATCTGTCAATGAATCAAATCACCGGTTCACTCCCGGCTCAGCTCGGTTCA

>M6-PGSC0003DMG400006880-(rev)M6LRR-RLP_768

ATCTAGATCTGTCAATGAATCAAATCACCGGTTCACTCCCGGCTCAGCTCGGTTCAATGCCGGTTCTCTCAACTCTAAACCTCGATTCCAACCAAATCTCCAGTTCAATCCCCACTAATC

>M6-PGSC0003DMG400006880-(rev)M6LRR-RLP_832

TCTCTCAACTCTAAACCTCGATTCCAACCAAATCTCCAGTTCAATCCCCACTAATCTACTTAGCAGCAGCGGATTAAACATCTTAAACTTAAGCAGAAACTCATTAGAAGGAGAATTACC

>M6-PGSC0003DMG400006880-(rev)M6LRR-RLP_896

AGCAGCGGATTAAACATCTTAAACTTAAGCAGAAACTCATTAGAAGGAGAATTACCAGATGTATTTTGCTCAAAAACGTATTTCACACATCTGGATTTATCATACAATAATCTTCGAGGT

>M6-PGSC0003DMG400006880-(rev)M6LRR-RLP_960

TTTGCTCAAAAACGTATTTCACACATCTGGATTTATCATACAATAATCTTCGAGGTACAGTTCCAAAATCATTATCATCAGCTAAGTACATTGGCCATTTGGATTTAAGCCATAATCACC

>M6-PGSC0003DMG400006912M6RLKantifungi(GNK)_0

AATTTAACGTGCAGACTTGGAAACATTTCAGGGAGAAAAGGGCATAGCGGATAATAGAAGATGCGAATGAAGTGCTAAGGGTGGTTCAAATTGTGATGCTATGTACTCAAGAGTCGCCTA

>M6-PGSC0003DMG400006912M6RLKantifungi(GNK)_147

TTAAACTACTTACACAAGAAAAATTATATATATAGAAGTGACTGTTCCTTCAAAGCCTCCTTTCATTGATATGGTTCCGAACAACATGGTTCTAGTCATCATCATCACCCATTTCATCAT

>M6-PGSC0003DMG400006912M6RLKantifungi(GNK)_49

GATAATAGAAGATGCGAATGAAGTGCTAAGGGTGGTTCAAATTGTGATGCTATGTACTCAAGAGTCGCCTATTATGCGTTCAGATATTTCAACAATTATTAAACTACTTACACAAGAAAA

>M6-PGSC0003DMG400006912M6RLKantifungi(GNK)_98

CTATGTACTCAAGAGTCGCCTATTATGCGTTCAGATATTTCAACAATTATTAAACTACTTACACAAGAAAAATTATATATATAGAAGTGACTGTTCCTTCAAAGCCTCCTTTCATTGATA

>M6-PGSC0003DMG400006923M6LRR-RLK_0

ATTTTGAAGTCACTGTTAAATGGGTTTCTTCAAGATTTTTGTGATAGTGTTGGTTATTTGCGCTTTCTTTATAGCAAATACTTATCAGTTACAGTCCTATGAAATTCAAGCTCTTCTTCA

>M6-PGSC0003DMG400006923M6LRR-RLK_1008

TGGTTTTGTTGATATTTCTAATAATAGATTAGTTGGTGGGCTTCCTTCTTGCTTGAACACTATTTCAGATAAGCGAATTGTTAAGGTTGGTGGAAATTGCTTGTCTCTTGACACTCAATA

>M6-PGSC0003DMG400006923M6LRR-RLK_1071

TTCAGATAAGCGAATTGTTAAGGTTGGTGGAAATTGCTTGTCTCTTGACACTCAATATCAGCATTCAGAGGGCTATTGTAAACAAGCTAATTCGGATAAGAAACAGATCACAGGAAAACA

>M6-PGSC0003DMG400006923M6LRR-RLK_1134

TTCAGAGGGCTATTGTAAACAAGCTAATTCGGATAAGAAACAGATCACAGGAAAACAAATAGCATTGTTGGCAGGTGTTATTGGGGGAATTGTAATTGTTGTGGTGTTTCTGTTAGTTGT

>M6-PGSC0003DMG400006923M6LRR-RLK_1197

ATTGTTGGCAGGTGTTATTGGGGGAATTGTAATTGTTGTGGTGTTTCTGTTAGTTGTCCTTTTCATCTTCCGTAGAAAACGACATGCGCGTAACATGGTGGATAAGCACATTCCTCCGAA

>M6-PGSC0003DMG400006923M6LRR-RLK_126

GAAGCACCTGGAATATCCAGTTCAGTTGGATGTTTGGGAGAATTACCATGGAGATTTTTGCAGCTTGACTTCAACACTGCATATGAGCATTACCTGTCAAGATAACTCTGTTACTGAACT

>M6-PGSC0003DMG400006923M6LRR-RLK_1260

CATCTTCCGTAGAAAACGACATGCGCGTAACATGGTGGATAAGCACATTCCTCCGAAAGTTGTGCAAAATACACAGCCCGGGGTTCCCTCTGAACTCCTAGCAAACGCCAGTGAGTTGAT

>M6-PGSC0003DMG400006923M6LRR-RLK_1323

GCAAAATACACAGCCCGGGGTTCCCTCTGAACTCCTAGCAAACGCCAGTGAGTTGATGTTTTTCTGATTTCATTTTTAATCTGTTTAATTTATATGAAGGAAATGTTCTCCAGAAAAGCT

>M6-PGSC0003DMG400006923M6LRR-RLK_1386

CTGATTTCATTTTTAATCTGTTTAATTTATATGAAGGAAATGTTCTCCAGAAAAGCTTGATCTCAATCTTCGAATTCTCAGGAGTCATTTCTCAAGTGGCAAACACAGGATCACAAGGTG

>M6-PGSC0003DMG400006923M6LRR-RLK_1449

CAATCTTCGAATTCTCAGGAGTCATTTCTCAAGTGGCAAACACAGGATCACAAGGTGCCCCATCATATCGAGTGTTCTCCATGGAAGAATTGCTAGAAGCAACAGAAAATTTTGATCAGT

>M6-PGSC0003DMG400006923M6LRR-RLK_1506

CCCCATCATATCGAGTGTTCTCCATGGAAGAATTGCTAGAAGCAACAGAAAATTTTGATCAGTCAGCATTACTTGGTGAAGGGTCAGTTGGAAAAGTATGGCTTCAAAACAATACTTATG

>M6-PGSC0003DMG400006923M6LRR-RLK_189

CTTGACTTCAACACTGCATATGAGCATTACCTGTCAAGATAACTCTGTTACTGAACTCAAAATCAAAGGAGATAAGTTAGTTAAGTTGAATGAATTTCATGGATTTGCAATTCCAAACCA

>M6-PGSC0003DMG400006923M6LRR-RLK_252

CAAAGGAGATAAGTTAGTTAAGTTGAATGAATTTCATGGATTTGCAATTCCAAACCAAACATTATCTGAGGGTTTCTCCATTGATTCTTTTGTTACCACATTGACAAGGCTAAGTAGCTT

>M6-PGSC0003DMG400006923M6LRR-RLK_315

ATCTGAGGGTTTCTCCATTGATTCTTTTGTTACCACATTGACAAGGCTAAGTAGCTTAAAGGTTCTTACTTTAGTTTCTCTTGGTATTTGGGGACCCCTTCCTGATAAAATTCATAGGTT

>M6-PGSC0003DMG400006923M6LRR-RLK_378

TCTTACTTTAGTTTCTCTTGGTATTTGGGGACCCCTTCCTGATAAAATTCATAGGTTAGCTTCACTTGAACTTTTGGATATGAGTTCAAATTTTCTATTTGGTTCTGTTCCTTTTCAGAT

>M6-PGSC0003DMG400006923M6LRR-RLK_441

ACTTGAACTTTTGGATATGAGTTCAAATTTTCTATTTGGTTCTGTTCCTTTTCAGATGTCAAGATTGATTAAGCTTCATACTTTGACATTAGATGCCAATTTTTTCAATGAAACTTTCCC

>M6-PGSC0003DMG400006923M6LRR-RLK_504

ATTGATTAAGCTTCATACTTTGACATTAGATGCCAATTTTTTCAATGAAACTTTCCCTGATTGGTTTGATTCCTTACACAATCTTACTATATTAAGCTTCAAGAACAACAGGTTGAAGGG

>M6-PGSC0003DMG400006923M6LRR-RLK_567

GTTTGATTCCTTACACAATCTTACTATATTAAGCTTCAAGAACAACAGGTTGAAGGGTCAGTTTCCTTCTTCAATGTCTAAAATCACAACCCTCACTGACATTGTTCTGTCACACAATGT

>M6-PGSC0003DMG400006923M6LRR-RLK_63

TTTCTTTATAGCAAATACTTATCAGTTACAGTCCTATGAAATTCAAGCTCTTCTTCAGTTGAGGAAGCACCTGGAATATCCAGTTCAGTTGGATGTTTGGGAGAATTACCATGGAGATTT

>M6-PGSC0003DMG400006923M6LRR-RLK_630

TCCTTCTTCAATGTCTAAAATCACAACCCTCACTGACATTGTTCTGTCACACAATGTGCTTTCTGGTAAATTGCCCGATTTAACGGCCTTGTCTAAGTTGCTTTTGTTGGATTTAAGGGA

>M6-PGSC0003DMG400006923M6LRR-RLK_693

TGGTAAATTGCCCGATTTAACGGCCTTGTCTAAGTTGCTTTTGTTGGATTTAAGGGAAAATCATTTGGATTCTGAATTGCCACCCTTGCCAAAAGGACTTACCACAGTTCTTCTTAGCAG

>M6-PGSC0003DMG400006923M6LRR-RLK_756

TTTGGATTCTGAATTGCCACCCTTGCCAAAAGGACTTACCACAGTTCTTCTTAGCAGTAACGTTTTTACGGGTGAAATCCCAGTAGAATTTGGCACACTAAATCAACTTCAACACCTTGA

>M6-PGSC0003DMG400006923M6LRR-RLK_819

TTTTACGGGTGAAATCCCAGTAGAATTTGGCACACTAAATCAACTTCAACACCTTGATCTGTCGAACAATTCTCTCAGTGGAATGCTGCCTGCTGGCTTATTCTCTTTGTCTAGCATTAG

>M6-PGSC0003DMG400006923M6LRR-RLK_882

GAACAATTCTCTCAGTGGAATGCTGCCTGCTGGCTTATTCTCTTTGTCTAGCATTAGTTACTTGAATTTAGCGTCTAATGTTCTTAGTGGTTCACTTCCTAGCCATCTCAATTGTGGGGG

>M6-PGSC0003DMG400006923M6LRR-RLK_945

GAATTTAGCGTCTAATGTTCTTAGTGGTTCACTTCCTAGCCATCTCAATTGTGGGGGTGAACTTGGTTTTGTTGATATTTCTAATAATAGATTAGTTGGTGGGCTTCCTTCTTGCTTGAA

>M6-PGSC0003DMG400006942-(rev)M6LRR-RLK_0

GGTGCTTACTCAGGGAGTCCACTCCATTAACAGGATTAACAATAAATTAAACTTTACAAATTCTCTAAATCCATAATTCCAAGGTTCCACTTCATCATTCAAAGTCAAATGTCATCTTCT

>M6-PGSC0003DMG400006942-(rev)M6LRR-RLK_1054

GGCTCATAGGCTTACTCAGGTTACATTGTTTAAGAAACCGCTTGTGAAGGTTAAGTTGGCGGATCTGTTGGCTGCCACGAACAATTTTAGTACGTCGAGTGTGATAAACTCGACTAGGAC

>M6-PGSC0003DMG400006942-(rev)M6LRR-RLK_1116

ATCTGTTGGCTGCCACGAACAATTTTAGTACGTCGAGTGTGATAAACTCGACTAGGACGGGGACTACATTTAGAGCTGTTCTACGTGATGGTTCTACGCTTGCTATTAAACGGCTTAAAG

>M6-PGSC0003DMG400006942-(rev)M6LRR-RLK_1178

ACTACATTTAGAGCTGTTCTACGTGATGGTTCTACGCTTGCTATTAAACGGCTTAAAGCTTGCAAGCTGAGTGAGAAGTTGTTTCGAATGGAAATGAATGGACTAGGACAAGTTAGGCAT

>M6-PGSC0003DMG400006942-(rev)M6LRR-RLK_124

TTTTGCTTTTTTCATACACTTATATTTCTCATGTAAACAATTCATCAAACACCTTCTATGCATCCCTACTCCCTAAAGCTCAAAGATGAAACTTTTCAGATTTTTCAATCTTCCCCTTAT

>M6-PGSC0003DMG400006942-(rev)M6LRR-RLK_1240

CAAGCTGAGTGAGAAGTTGTTTCGAATGGAAATGAATGGACTAGGACAAGTTAGGCATCCTAATTTGGTACCACTTTTGGGCTTTTGTGTTGTTGAGGAGGAAAAACTTTTGGTTTATAA

>M6-PGSC0003DMG400006942-(rev)M6LRR-RLK_1302

ATTTGGTACCACTTTTGGGCTTTTGTGTTGTTGAGGAGGAAAAACTTTTGGTTTATAAGCACCTGTCAAATGGTACTTTGTATTCATTATTGAAAGGGAGCGCAAGTGTGTTGGATTGGC

>M6-PGSC0003DMG400006942-(rev)M6LRR-RLK_1364

CTGTCAAATGGTACTTTGTATTCATTATTGAAAGGGAGCGCAAGTGTGTTGGATTGGCCTACTAGGTTTAGGATTGGCTTGGGTGCTGCAAGGGGCCTTGCTTGGCTTCATCATGGTTGT

>M6-PGSC0003DMG400006942-(rev)M6LRR-RLK_1426

TAGGTTTAGGATTGGCTTGGGTGCTGCAAGGGGCCTTGCTTGGCTTCATCATGGTTGTCAACCACCAATATTGCACCAAAACATATGTTCTAACGTTATTTTCCTTGATGAGGATTTTGA

>M6-PGSC0003DMG400006942-(rev)M6LRR-RLK_1488

CACCAATATTGCACCAAAACATATGTTCTAACGTTATTTTCCTTGATGAGGATTTTGATGCTAGAATAATGGATTTTGGGTTGGCAAGGCTGGTGACACCTCCAGATGCAAAAGAGACTA

>M6-PGSC0003DMG400006942-(rev)M6LRR-RLK_1550

AGAATAATGGATTTTGGGTTGGCAAGGCTGGTGACACCTCCAGATGCAAAAGAGACTAGTTTTGTGAATGGGGAGTTGGGTGAATTTGGCTATGTTGCTCCGGAGTACTCAAGCACAATT

>M6-PGSC0003DMG400006942-(rev)M6LRR-RLK_1674

CTTCACTGAAAGGGGATGCTTACAGCTTTGGGGTTGTGCTTTTGGAGTTGGCTACTGGGCAAAGACCTCTAGAAATCACTGCTGCTGATGAAGGTTTTAAGGGAAATTTGGTGGACTGGG

>M6-PGSC0003DMG400006942-(rev)M6LRR-RLK_1736

AGACCTCTAGAAATCACTGCTGCTGATGAAGGTTTTAAGGGAAATTTGGTGGACTGGGTAAATCAGCTCTCTGTTTCTGGTCGGATAAAAGATGCAATTGATAAGCATATTTGTAGAAAG

>M6-PGSC0003DMG400006942-(rev)M6LRR-RLK_1798

TCAGCTCTCTGTTTCTGGTCGGATAAAAGATGCAATTGATAAGCATATTTGTAGAAAGGGCCATGATGAAGAGATTGTAAAATTCCTCAAAATTGCTTGCAATTGTTTAATTTCTCGGCC

>M6-PGSC0003DMG400006942-(rev)M6LRR-RLK_186

TCCCTACTCCCTAAAGCTCAAAGATGAAACTTTTCAGATTTTTCAATCTTCCCCTTATAGTATTCATCTTATTGATTCTTGTATTTCAACCCCTTCTTCATTGTACTGCTGTTGCTGGTG

>M6-PGSC0003DMG400006942-(rev)M6LRR-RLK_1860

ATGATGAAGAGATTGTAAAATTCCTCAAAATTGCTTGCAATTGTTTAATTTCTCGGCCCAAGGAGAGGTGGTCTATGTATCAGGTATATGAAGCACTGAAGAGCATGGCTGAGAAACATG

>M6-PGSC0003DMG400006942-(rev)M6LRR-RLK_1922

GAGAGGTGGTCTATGTATCAGGTATATGAAGCACTGAAGAGCATGGCTGAGAAACATGGTTTCTCTGAACATTATGATGAATTTCCCTTACTATTTAACAAACAAGAGACCAGCAGTCCC

>M6-PGSC0003DMG400006942-(rev)M6LRR-RLK_1984

CTCTGAACATTATGATGAATTTCCCTTACTATTTAACAAACAAGAGACCAGCAGTCCCATTTGATACATGAATGGGAAAGCATTACGAGATGCTTCACATCTGGTGAACACAAATGCATT

>M6-PGSC0003DMG400006942-(rev)M6LRR-RLK_2046

GATACATGAATGGGAAAGCATTACGAGATGCTTCACATCTGGTGAACACAAATGCATTATGGTACGTTCTTCACTTCTCTTAGGTGCAAGTGGAATCAGGCACGATACATATGGTTCTGT

>M6-PGSC0003DMG400006942-(rev)M6LRR-RLK_2108

TACGTTCTTCACTTCTCTTAGGTGCAAGTGGAATCAGGCACGATACATATGGTTCTGTGAAGTACTACCACAGATATTGCTGTATTGTACTGTGAAGTGTCAAATCATTCTGAGGTGTTC

>M6-PGSC0003DMG400006942-(rev)M6LRR-RLK_2170

TACTACCACAGATATTGCTGTATTGTACTGTGAAGTGTCAAATCATTCTGAGGTGTTCTGCTCGAAATTTATGTTACAGATAGCTAATATTATTGGTATCTTAGGTTTATGTTTCTCTTT

>M6-PGSC0003DMG400006942-(rev)M6LRR-RLK_2232

CGAAATTTATGTTACAGATAGCTAATATTATTGGTATCTTAGGTTTATGTTTCTCTTTGCTCATACTTATGCATTAAACAAGGCAGTAAGCAAAAGCTGTCACAGTTTATTTGAAAGCTG

>M6-PGSC0003DMG400006942-(rev)M6LRR-RLK_2294

ATACTTATGCATTAAACAAGGCAGTAAGCAAAAGCTGTCACAGTTTATTTGAAAGCTGTAATACCACTAATTTAGCTGTATATAACTGACCTGGGCATTGGCCTTTTTAGGGTCGAGATA

>M6-PGSC0003DMG400006942-(rev)M6LRR-RLK_2356

ACCACTAATTTAGCTGTATATAACTGACCTGGGCATTGGCCTTTTTAGGGTCGAGATATCTTCAGTATGTCCCTTACAGTCATTTATGGTTTGTAAGATCAAATCACACTAGATTTCCCT

>M6-PGSC0003DMG400006942-(rev)M6LRR-RLK_2418

CAGTATGTCCCTTACAGTCATTTATGGTTTGTAAGATCAAATCACACTAGATTTCCCTGATAGATTGGTTCGGTAGAATATCTGCCTCCACACAGATATTAATTCATCTAACCGGAATGC

>M6-PGSC0003DMG400006942-(rev)M6LRR-RLK_248

TTCATCTTATTGATTCTTGTATTTCAACCCCTTCTTCATTGTACTGCTGTTGCTGGTGCTGTAGCTGAAGATGATATAAAGTGTTTAAAAGGAGTCAAGAACTCCTTAACAGATCCAAAA

>M6-PGSC0003DMG400006942-(rev)M6LRR-RLK_2480

GATTGGTTCGGTAGAATATCTGCCTCCACACAGATATTAATTCATCTAACCGGAATGCTATCAAGCAGAGTTATTAGTATTTACTCCTTCTGATTAATTTTGGATGATGTTGATCAGTAA

>M6-PGSC0003DMG400006942-(rev)M6LRR-RLK_2542

AAGCAGAGTTATTAGTATTTACTCCTTCTGATTAATTTTGGATGATGTTGATCAGTAATTGTAGGTCCTTTTGCAAAATTTGTGTCTGAATGGGAATAATTTTCACTAAAATGGAAGTTG

>M6-PGSC0003DMG400006942-(rev)M6LRR-RLK_2604

AGGTCCTTTTGCAAAATTTGTGTCTGAATGGGAATAATTTTCACTAAAATGGAAGTTGAATGGGCTTTCTAAAAGTTATGTGCCCTCTTGAATTTGATTTGGCAGCTGAAATTTTCATTC

>M6-PGSC0003DMG400006942-(rev)M6LRR-RLK_2666

GGCTTTCTAAAAGTTATGTGCCCTCTTGAATTTGATTTGGCAGCTGAAATTTTCATTCACTTGATTCTAATCTGCTTCTTTACAATCTCAGAAACATAATGCACAATTAAAGTTTCCAAG

>M6-PGSC0003DMG400006942-(rev)M6LRR-RLK_2728

GATTCTAATCTGCTTCTTTACAATCTCAGAAACATAATGCACAATTAAAGTTTCCAAGTAGGAAAAAAAATAATTCTTGTTGACAGTTGAAACTCTTTGCAATAGTAGTTATCAAACCTC

>M6-PGSC0003DMG400006942-(rev)M6LRR-RLK_2790

AAAAAAAATAATTCTTGTTGACAGTTGAAACTCTTTGCAATAGTAGTTATCAAACCTCTTTACTTTTTCCCTCAAGCTCAAGCTCGGATGCGTGTTTCCATGTGCTCTCCTGCTTCCCTG

>M6-PGSC0003DMG400006942-(rev)M6LRR-RLK_2852

CTTTTTCCCTCAAGCTCAAGCTCGGATGCGTGTTTCCATGTGCTCTCCTGCTTCCCTGAAAAGTCATTTTGTAATATTTCAAAACAATAACTTGTAGTCTAATAGTAAATGAAGGGAAAA

>M6-PGSC0003DMG400006942-(rev)M6LRR-RLK_310

AGCTGAAGATGATATAAAGTGTTTAAAAGGAGTCAAGAACTCCTTAACAGATCCAAAAGGGAACTTGAATTCATGGAATTTTGCTAATTCTACAGTGGGGTTTATATGTAAATTTGTTGG

>M6-PGSC0003DMG400006942-(rev)M6LRR-RLK_372

ACTTGAATTCATGGAATTTTGCTAATTCTACAGTGGGGTTTATATGTAAATTTGTTGGTGCTTCTTGCTGGAATGACCGTGAAAATCGTCTCATAAATCTTGAACTTCGGGATATGAATC

>M6-PGSC0003DMG400006942-(rev)M6LRR-RLK_434

TCTTGCTGGAATGACCGTGAAAATCGTCTCATAAATCTTGAACTTCGGGATATGAATCTTGGAGGTAATGTTCCAGATTCGTTAAAGTATTGTCGAAGCTTGCAATATCTTGATCTTTCT

>M6-PGSC0003DMG400006942-(rev)M6LRR-RLK_496

AGGTAATGTTCCAGATTCGTTAAAGTATTGTCGAAGCTTGCAATATCTTGATCTTTCTGGTAATCGGATATCTGGTTCAATTCCTTCTGATATTTGTACTTGGCTACCTTACTTGGTAAC

>M6-PGSC0003DMG400006942-(rev)M6LRR-RLK_558

ATCGGATATCTGGTTCAATTCCTTCTGATATTTGTACTTGGCTACCTTACTTGGTAACCCTTGATTTGTCGAACAATGAATTTACCGGTTCTATTCCGTCTGATCTTGTTAGTTGTTCCT

>M6-PGSC0003DMG400006942-(rev)M6LRR-RLK_62

CTCTAAATCCATAATTCCAAGGTTCCACTTCATCATTCAAAGTCAAATGTCATCTTCTATTCTTTTGCTTTTTTCATACACTTATATTTCTCATGTAAACAATTCATCAAACACCTTCTA

>M6-PGSC0003DMG400006942-(rev)M6LRR-RLK_620

GATTTGTCGAACAATGAATTTACCGGTTCTATTCCGTCTGATCTTGTTAGTTGTTCCTATTTGAATAAATTGATGCTTAATGATAATAGGCTTTCTGGGAATATACCACCTCAGTTTTCT

>M6-PGSC0003DMG400006942-(rev)M6LRR-RLK_682

GAATAAATTGATGCTTAATGATAATAGGCTTTCTGGGAATATACCACCTCAGTTTTCTAGTTTGGGGAGGCTTAAAATATTTTCTGTGGCAAACAATGATCTTTCTGGTAGGATTCCGGA

>M6-PGSC0003DMG400006942-(rev)M6LRR-RLK_744

TGGGGAGGCTTAAAATATTTTCTGTGGCAAACAATGATCTTTCTGGTAGGATTCCGGAAGCTTTTGATTCGGCTGAGTCGTTTGATTTTGGAGGAAATGATGGGCTTTGTGGTGGTCCGT

>M6-PGSC0003DMG400006942-(rev)M6LRR-RLK_806

TTTGATTCGGCTGAGTCGTTTGATTTTGGAGGAAATGATGGGCTTTGTGGTGGTCCGTTGGGGAAATGTGGGAGGCTTAGTAAGAAAAGTTTAGCTATTATTATTGCTGCCGGTGTGTTT

>M6-PGSC0003DMG400006942-(rev)M6LRR-RLK_868

GAAATGTGGGAGGCTTAGTAAGAAAAGTTTAGCTATTATTATTGCTGCCGGTGTGTTTGGTGCTGCTGCTTCTCTGTTGTTGGGTTTTGGGGCTTGGTATTGGTATTTTACGAAGGCGGG

>M6-PGSC0003DMG400006942-(rev)M6LRR-RLK_930

CTGCTGCTTCTCTGTTGTTGGGTTTTGGGGCTTGGTATTGGTATTTTACGAAGGCGGGGACAAGGAGGAAGATGGGGTATGGATTAGGGAGAGTTGACTCGGAAAGGTGGGCGGATAAGT

>M6-PGSC0003DMG400006942-(rev)M6LRR-RLK_992

AGGAGGAAGATGGGGTATGGATTAGGGAGAGTTGACTCGGAAAGGTGGGCGGATAAGTTGAAGGCTCATAGGCTTACTCAGGTTACATTGTTTAAGAAACCGCTTGTGAAGGTTAAGTTG

>M6-PGSC0003DMG400007033-(rev)M6LRR-RLP_0

ATGGCCAGATTAGTTAACTCCATTTTCTTTACCTTCTTGTACATAAGTTTCATATACACACTTTCACCTTATGTCAATGCAACAACTTCACCATCTGATATTCAAGTTCTAGTAGCCATC

>M6-PGSC0003DMG400007033-(rev)M6LRR-RLP_1054

CATGGAGAAATCTCAACTTGGAATATAATAACCTGGAGGGATCGGTGCCTCGAGAGCTCAACAGGACTTTAGAAGGGGTTCAGGGAAGTTTTGAACATAACTGTCTCATTTGTCCAAAGG

>M6-PGSC0003DMG400007033-(rev)M6LRR-RLP_124

CGTCGATAGACCCTTTTTCAATAACCCCGGATTCCTTCCTTAGTAGTTGGGATTTCAACTTAGACCCTTGTGAGACAACTGGTGCAAGTTTCTTAGGCATTCTTTGCACAGTTCCTGATC

>M6-PGSC0003DMG400007033-(rev)M6LRR-RLP_186

GACCCTTGTGAGACAACTGGTGCAAGTTTCTTAGGCATTCTTTGCACAGTTCCTGATCAGGTAGACACTAATGCCACGAGCCGGATAATTGAGATTGATCTAGAAGGTGATGGACTAGAG

>M6-PGSC0003DMG400007033-(rev)M6LRR-RLP_248

AGACACTAATGCCACGAGCCGGATAATTGAGATTGATCTAGAAGGTGATGGACTAGAGGGGTTCTTGACATCGTCTATAGGGAACTTAACAGAACTTGTTTCTCTCAATCTAGGGAGGAA

>M6-PGSC0003DMG400007033-(rev)M6LRR-RLP_310

TCTTGACATCGTCTATAGGGAACTTAACAGAACTTGTTTCTCTCAATCTAGGGAGGAACAAGTTCAGAGGACCAGTGCCAGAGAGTATTACTAATTTGAGGAAACTCACAAGTCTTCAAC

>M6-PGSC0003DMG400007033-(rev)M6LRR-RLP_372

TTCAGAGGACCAGTGCCAGAGAGTATTACTAATTTGAGGAAACTCACAAGTCTTCAACTTTATGAGAATTTTTTCTCAGGTAGTCTTGTTGATGACATTGGTGTCCTTACAAAACTTGAA

>M6-PGSC0003DMG400007033-(rev)M6LRR-RLP_434

TGAGAATTTTTTCTCAGGTAGTCTTGTTGATGACATTGGTGTCCTTACAAAACTTGAAACTCTTGATGTCTCGAATAACAGGCTATCTGGCTCAATCCCTTCCTCGATTACATCACTTAG

>M6-PGSC0003DMG400007033-(rev)M6LRR-RLP_496

TTGATGTCTCGAATAACAGGCTATCTGGCTCAATCCCTTCCTCGATTACATCACTTAGAAGTTTAACGCGATTAGACCTGTCTAACAATGAATTCACTGGAAAAATTCCTCAACTCAACG

>M6-PGSC0003DMG400007033-(rev)M6LRR-RLP_558

TTAACGCGATTAGACCTGTCTAACAATGAATTCACTGGAAAAATTCCTCAACTCAACGGATTATGGCAACTCTCTTCATTTGACATTTCAAATAACCAAATTTATGGGAATTTACCACAA

>M6-PGSC0003DMG400007033-(rev)M6LRR-RLP_62

TTCACCTTATGTCAATGCAACAACTTCACCATCTGATATTCAAGTTCTAGTAGCCATCAAGACGTCGATAGACCCTTTTTCAATAACCCCGGATTCCTTCCTTAGTAGTTGGGATTTCAA

>M6-PGSC0003DMG400007033-(rev)M6LRR-RLP_620

ATGGCAACTCTCTTCATTTGACATTTCAAATAACCAAATTTATGGGAATTTACCACAATTCCCATTGAAGATTAAAACACTATCACTAAGCCACAACTTGTTATCAGGACACATTACCCC

>M6-PGSC0003DMG400007033-(rev)M6LRR-RLP_682

CATTGAAGATTAAAACACTATCACTAAGCCACAACTTGTTATCAGGACACATTACCCCAGTGCATAAACTTAGACATTTAAGCACATTAGACCTTAGTGATAACAGATTTTCAGGAGGAA

>M6-PGSC0003DMG400007033-(rev)M6LRR-RLP_744

CATAAACTTAGACATTTAAGCACATTAGACCTTAGTGATAACAGATTTTCAGGAGGAATAAACAAAGGGATCTTCATGTTACCTGAATTAAGTCACGTAAACGTTTCAGTAAACCGTTTC

>M6-PGSC0003DMG400007033-(rev)M6LRR-RLP_806

CAAAGGGATCTTCATGTTACCTGAATTAAGTCACGTAAACGTTTCAGTAAACCGTTTCACAGTGTTTGAGGTGGTAGAATTCTCTGATAAAAAATCACAGCTACACACGTTAGACGTACA

>M6-PGSC0003DMG400007033-(rev)M6LRR-RLP_868

TGTTTGAGGTGGTAGAATTCTCTGATAAAAAATCACAGCTACACACGTTAGACGTACATGCTAATCGTCTACATGGTCATTTGCCTGTGAACTTGGTTACTTACCCAAACTTAACAGAGA

>M6-PGSC0003DMG400007033-(rev)M6LRR-RLP_930

AATCGTCTACATGGTCATTTGCCTGTGAACTTGGTTACTTACCCAAACTTAACAGAGATAAACCTTGGACATAACCTGTTTTCGGGTGAGATTCCATCAGAGTATTGGTCAAGACTCGGA

>M6-PGSC0003DMG400007033-(rev)M6LRR-RLP_992

CCTTGGACATAACCTGTTTTCGGGTGAGATTCCATCAGAGTATTGGTCAAGACTCGGATTTTCATGGAGAAATCTCAACTTGGAATATAATAACCTGGAGGGATCGGTGCCTCGAGAGCT

>M6-PGSC0003DMG400007073-(rev)M6G-LecRK_0

CCAATCTCAAGAAGCAAATGTATCATATAACAAACAAGCTAAAACAACAAACTAAACATGTCTCCTTCATGGCCTACTACACTTCTTGCCTCTCTCTTTCTTTTCTCCCAAATATTTTCT

>M6-PGSC0003DMG400007073-(rev)M6G-LecRK_1024

TTATTGTTGTTGTTATATTGTTGTAAACGTAGCTCTATAAGATAATGATTAGTGTGTCCAGTACAATAAGTGGGTCCACAACTAGTAGTCCAATAAATGAGTAGTACTTCCTATTTAATG

>M6-PGSC0003DMG400007073-(rev)M6G-LecRK_1083

AGTACAATAAGTGGGTCCACAACTAGTAGTCCAATAAATGAGTAGTACTTCCTATTTAATGTCAATTATATTTTTCTTTTTCCCTTTTTGAAAAAATATTATGTTTTGGTTGAATTAAGA

>M6-PGSC0003DMG400007073-(rev)M6G-LecRK_128

TCAAGTTCCAGCTGAAAACACCTTTAAATTCGTCGATGAAGGCGAATTAGGAGATTATATTGTAGAATACAGAGCAAATTATCGCGTTATTAGTATTTTCAATAACCCATTTCAGTTCTG

>M6-PGSC0003DMG400007073-(rev)M6G-LecRK_192

GAATACAGAGCAAATTATCGCGTTATTAGTATTTTCAATAACCCATTTCAGTTCTGTTTCTACAACACAACTCCTAATGCATGGACATTAGCATTGCGAATGGGCACTGTCCGTTCTGAA

>M6-PGSC0003DMG400007073-(rev)M6G-LecRK_256

ACACAACTCCTAATGCATGGACATTAGCATTGCGAATGGGCACTGTCCGTTCTGAATCACTCATGCGTTGGGTGTGGGAAGCAAATAGGGGAAACCCCGTTAAAGAAAACGCGACATTCA

>M6-PGSC0003DMG400007073-(rev)M6G-LecRK_384

ACTAACAGAAATCTTGTATTGGCGGATGCAGACGGTTGAATCGCTTGGCAGACCAACACGGCCAATAAGGGGGTCACCGGGTTCAAGTTGTTACCTAACGGTAATGTTGTGCTACATGAC

>M6-PGSC0003DMG400007073-(rev)M6G-LecRK_448

ATAAGGGGGTCACCGGGTTCAAGTTGTTACCTAACGGTAATGTTGTGCTACATGACTCCAAGGGTAAATTTGTCTGGCAGAGTTTCAACTATCCCACTGACACATTACTAGTGGGCCAAA

>M6-PGSC0003DMG400007073-(rev)M6G-LecRK_576

TTGTCCGGCCCGAATAAGCTTGTGAGCTGGGCCTCGGTGAAAAAGAATGCGAACGGGCCTTATAGCTTGGTGGTCCAGCCGAGATTGTTCGCTGTCTACAAGGGAAACAAGGTAGACGTG

>M6-PGSC0003DMG400007073-(rev)M6G-LecRK_64

CTTCATGGCCTACTACACTTCTTGCCTCTCTCTTTCTTTTCTCCCAAATATTTTCTTGCATTGCTCAAGTTCCAGCTGAAAACACCTTTAAATTCGTCGATGAAGGCGAATTAGGAGATT

>M6-PGSC0003DMG400007073-(rev)M6G-LecRK_640

GCTTGGTGGTCCAGCCGAGATTGTTCGCTGTCTACAAGGGAAACAAGGTAGACGTGGAATTAGCTTGGTTCGATTATGGGAATAACACGTTGGAATCGATAAAATTGAATAGTGGGAATC

>M6-PGSC0003DMG400007073-(rev)M6G-LecRK_704

TTGGTTCGATTATGGGAATAACACGTTGGAATCGATAAAATTGAATAGTGGGAATCAAAGATTGAAATTGGATTATAGATTTGCAAAATCAACACAGAGAAGTTCACATGTGATGGCATT

>M6-PGSC0003DMG400007073-(rev)M6G-LecRK_768

AAATTGGATTATAGATTTGCAAAATCAACACAGAGAAGTTCACATGTGATGGCATTTACTAAATACAACACAACATTGACATATCTTAGACTTGAAATTGATGGAAATCTAAAGGCTTAT

>M6-PGSC0003DMG400007073-(rev)M6G-LecRK_832

ACAACACAACATTGACATATCTTAGACTTGAAATTGATGGAAATCTAAAGGCTTATACTTTCGTTAGAGATGAAGAAGCAGATGAATTCCGTTGGAAAGTAACTTACAGTAGGATTTAAA

>M6-PGSC0003DMG400007073-(rev)M6G-LecRK_896

TAGAGATGAAGAAGCAGATGAATTCCGTTGGAAAGTAACTTACAGTAGGATTTAAAACGTTTCATATTTGAAACGTCTAAATCCATGTGCACATTATCCTTTACGAGCAAATTTCACGGG

>M6-PGSC0003DMG400007073-(rev)M6G-LecRK_960

TATTTGAAACGTCTAAATCCATGTGCACATTATCCTTTACGAGCAAATTTCACGGGTTATATTATTATTGTTGTTGTTATATTGTTGTAAACGTAGCTCTATAAGATAATGATTAGTGTG

>M6-PGSC0003DMG400007234-(rev)M6LRR-RLK_0

CTCATCTTTACTTGCCTTATTTTCCTCAACCAATTTTTTCTTACAATCTCATAACCATGAACCATTTCTATCTCACTACAATTTTCTTCTTTCTCATACAATTTTCACTTGTTTTCTCTA

>M6-PGSC0003DMG400007234-(rev)M6LRR-RLK_1054

AATCCCTGAAGGGATTGGTGAGTTACCAAATCTCGAATTGTTAGCTCTTTGGAACAATTCACTCACTGGAATTTTACCACGAATGTTAGGATCAAATGCTAAGTTACAAAAACTTGATGT

>M6-PGSC0003DMG400007234-(rev)M6LRR-RLK_1116

TCACTGGAATTTTACCACGAATGTTAGGATCAAATGCTAAGTTACAAAAACTTGATGTGTCCTCAAACAATTTGACTGGTCCAATTCCCCAAAATCTCTGCCTCAGCAACAACTTAGTTA

>M6-PGSC0003DMG400007234-(rev)M6LRR-RLK_1178

TCAAACAATTTGACTGGTCCAATTCCCCAAAATCTCTGCCTCAGCAACAACTTAGTTAAACTCATTCTGTTTTCGAACGAGTTTACTGGTGAGATTCCTTCATCCTTAACTAACTGCAAC

>M6-PGSC0003DMG400007234-(rev)M6LRR-RLK_124

AAATTTTCCTCTTCAACTCATTTCACTTTTATCGTTAAAATCTTCATTTCAAGATCCAAACAATACATTTCAAGATTGGAATCCTACATCAACTTTCTCAAATTTTGGTTCACAACCCTT

>M6-PGSC0003DMG400007234-(rev)M6LRR-RLK_1240

CATTCTGTTTTCGAACGAGTTTACTGGTGAGATTCCTTCATCCTTAACTAACTGCAACGCTTTATCCAGGTTGAGAATCCAAGACAACAAACTCAATGGCACGATTCCATCAGGATTCGG

>M6-PGSC0003DMG400007234-(rev)M6LRR-RLK_1302

TATCCAGGTTGAGAATCCAAGACAACAAACTCAATGGCACGATTCCATCAGGATTCGGTTTTTTGCCTGATTTCACTTACATGGACATCAGCAAGAACAAATTTTCGGGGCAAATCCCAA

>M6-PGSC0003DMG400007234-(rev)M6LRR-RLK_1364

TTGCCTGATTTCACTTACATGGACATCAGCAAGAACAAATTTTCGGGGCAAATCCCAAAAGATTTTGGAAATGCCCCGAAAATGATGTACTTGAACATTTCTGAGAATTTTTTCGGTAGT

>M6-PGSC0003DMG400007234-(rev)M6LRR-RLK_1426

TTTTGGAAATGCCCCGAAAATGATGTACTTGAACATTTCTGAGAATTTTTTCGGTAGTAACTTGCCTGATAATATCTGGAACGCCCCGAATTTACAGATATTCTCAGCAAGTAAGAGTGG

>M6-PGSC0003DMG400007234-(rev)M6LRR-RLK_1488

TGCCTGATAATATCTGGAACGCCCCGAATTTACAGATATTCTCAGCAAGTAAGAGTGGCCTTGTTGGAAATCTCCCTGATTTCAAAGGATGTCAAAGTCTTTACAAAATCGAACTCGAAG

>M6-PGSC0003DMG400007234-(rev)M6LRR-RLK_1550

GTTGGAAATCTCCCTGATTTCAAAGGATGTCAAAGTCTTTACAAAATCGAACTCGAAGGAAACAATCTCAATGGTAGCATTCCATGGGACATTGAACATTGTGAAAAGCTTATTTCTCTA

>M6-PGSC0003DMG400007234-(rev)M6LRR-RLK_1612

CAATCTCAATGGTAGCATTCCATGGGACATTGAACATTGTGAAAAGCTTATTTCTCTAAATCTTCGACGAAATTCATTTACAGGAATCATTCCTTGGGAAATATCTGGTATTCCTTCGAT

>M6-PGSC0003DMG400007234-(rev)M6LRR-RLK_1674

TTCGACGAAATTCATTTACAGGAATCATTCCTTGGGAAATATCTGGTATTCCTTCGATAACAGACGTTGATTTGTCACATAATTTTCTCACAGGAACAATCCCTTCCAATTTCGAAAAGT

>M6-PGSC0003DMG400007234-(rev)M6LRR-RLK_1736

GACGTTGATTTGTCACATAATTTTCTCACAGGAACAATCCCTTCCAATTTCGAAAAGTCTAGCACGTTGGAACATTTCAACGTGTCGTATAATCAGCTCACTGGTCCATTGCCTTCTTCA

>M6-PGSC0003DMG400007234-(rev)M6LRR-RLK_1798

CACGTTGGAACATTTCAACGTGTCGTATAATCAGCTCACTGGTCCATTGCCTTCTTCAGGGTCTATGTTCTCAACGTTCCATCCTTCGTCCTTCGTAGGAAACGAAGGTCTTTGTGGTAC

>M6-PGSC0003DMG400007234-(rev)M6LRR-RLK_186

ATACATTTCAAGATTGGAATCCTACATCAACTTTCTCAAATTTTGGTTCACAACCCTTTTGGTGTTCATGGTCAGGCATCAAATGTGACACAAAAACAAGTCACATTACATCACTTAATC

>M6-PGSC0003DMG400007234-(rev)M6LRR-RLK_1860

CTATGTTCTCAACGTTCCATCCTTCGTCCTTCGTAGGAAACGAAGGTCTTTGTGGTACTATCATACATAATCCGTGCAGGACTGATGAACTCAAAGATGGAAAAACAGATTTTTCGAACC

>M6-PGSC0003DMG400007234-(rev)M6LRR-RLK_1922

ATACATAATCCGTGCAGGACTGATGAACTCAAAGATGGAAAAACAGATTTTTCGAACCATTCAAAGAAAACAGATGGTGCATTGATAGTATGGATTGCGTCTGCAGCATTTGGAGTAGTA

>M6-PGSC0003DMG400007234-(rev)M6LRR-RLK_1984

AAAGAAAACAGATGGTGCATTGATAGTATGGATTGCGTCTGCAGCATTTGGAGTAGTAATTATCATTCTTATTGTGGCAATTCGGTGTTTTCATTCAAATTACAATCGAAGATTTCCATG

>M6-PGSC0003DMG400007234-(rev)M6LRR-RLK_2046

TCATTCTTATTGTGGCAATTCGGTGTTTTCATTCAAATTACAATCGAAGATTTCCATGTGACAAGGAAATCGGACCGTGGAAGTTAACAGCATTCAAAAGGTTGAATTTCACAGCAGATG

>M6-PGSC0003DMG400007234-(rev)M6LRR-RLK_2108

AAGGAAATCGGACCGTGGAAGTTAACAGCATTCAAAAGGTTGAATTTCACAGCAGATGATGTACTCGAAAGCATAGCAATCACAGATAAGATCATCGGTATGGGGTCTGCAGGTACTTAA

>M6-PGSC0003DMG400007234-(rev)M6LRR-RLK_2170

ACTCGAAAGCATAGCAATCACAGATAAGATCATCGGTATGGGGTCTGCAGGTACTTAACGGATCTATATTGTTAGTTATTATGAATTCAATTGAACCCGCAATTTTTCATTAAGTATATG

>M6-PGSC0003DMG400007234-(rev)M6LRR-RLK_2232

TCTATATTGTTAGTTATTATGAATTCAATTGAACCCGCAATTTTTCATTAAGTATATGTGGGGGGGAAATCATACAAAATTAGTGTACATATGTTTTTTAAAAAATGCTAACATCACCTA

>M6-PGSC0003DMG400007234-(rev)M6LRR-RLK_2270

AATTTTTCATTAAGTATATGTGGGGGGGAAATCATACAAAATTAGTGTACATATGTTTTTTAAAAAATGCTAACATCACCTAGACAAGTTGGCAATGTGGAATTTATTAATAAAAATCTC

>M6-PGSC0003DMG400007234-(rev)M6LRR-RLK_248

TGTTCATGGTCAGGCATCAAATGTGACACAAAAACAAGTCACATTACATCACTTAATCTCTCAAAGAGGAATCTTTCTGGTAAAATACCTCAAAATATTGCATCCTTGATACACTTACAT

>M6-PGSC0003DMG400007234-(rev)M6LRR-RLK_310

AAAGAGGAATCTTTCTGGTAAAATACCTCAAAATATTGCATCCTTGATACACTTACATCACTTGAATTTAAGTGGAAATTCATTTGATGGACCTTTACCATCCATCATTTTTGAATTTCC

>M6-PGSC0003DMG400007234-(rev)M6LRR-RLK_372

TGAATTTAAGTGGAAATTCATTTGATGGACCTTTACCATCCATCATTTTTGAATTTCCATTTCTTAGAACACTTGATATTAGTCATAATTATTTCAATTCAGCATTTCCTCTTGACGGGC

>M6-PGSC0003DMG400007234-(rev)M6LRR-RLK_434

CTTAGAACACTTGATATTAGTCATAATTATTTCAATTCAGCATTTCCTCTTGACGGGCTAACGAATCTCAAGTCCCTAGTACACTTGAATGCCTATAGCAACAGCTTTACAGGCGCGTTG

>M6-PGSC0003DMG400007234-(rev)M6LRR-RLK_496

GAATCTCAAGTCCCTAGTACACTTGAATGCCTATAGCAACAGCTTTACAGGCGCGTTGCCTATAGAGCTTGTTGGGCTACAAAATCTTGAGTATTTGAATCTTGGTGGCAGTTACTTCAC

>M6-PGSC0003DMG400007234-(rev)M6LRR-RLK_558

TAGAGCTTGTTGGGCTACAAAATCTTGAGTATTTGAATCTTGGTGGCAGTTACTTCACTGGTGAAATTCCGGCAAGTTATGGGAATTTTAACAAGTTGAAATTCCTACATTTAGCCGGAA

>M6-PGSC0003DMG400007234-(rev)M6LRR-RLK_62

CATTTCTATCTCACTACAATTTTCTTCTTTCTCATACAATTTTCACTTGTTTTCTCTACTTCAAATTTTCCTCTTCAACTCATTTCACTTTTATCGTTAAAATCTTCATTTCAAGATCCA

>M6-PGSC0003DMG400007234-(rev)M6LRR-RLK_620

GAAATTCCGGCAAGTTATGGGAATTTTAACAAGTTGAAATTCCTACATTTAGCCGGAAATTCACTGAGTGGGAAAATTCCCGAGGAATTGAGATTGTTGAACCAACTCGAACATCTTGAA

>M6-PGSC0003DMG400007234-(rev)M6LRR-RLK_682

ACTGAGTGGGAAAATTCCCGAGGAATTGAGATTGTTGAACCAACTCGAACATCTTGAAATTGGATACAACAATTATACTGGAAATGTTCCTCCTGGATTTTCCTCGTTATCGAATTTAGC

>M6-PGSC0003DMG400007234-(rev)M6LRR-RLK_744

GATACAACAATTATACTGGAAATGTTCCTCCTGGATTTTCCTCGTTATCGAATTTAGCCTATCTTGACATCTCACAAGCTAATCTTTCAGGTGAAATTCCAATTCAGCTTGGAAATTTAA

>M6-PGSC0003DMG400007234-(rev)M6LRR-RLK_806

CTTGACATCTCACAAGCTAATCTTTCAGGTGAAATTCCAATTCAGCTTGGAAATTTAACAAAGATTGAATCCTTATTTCTTTTCAAGAACCATTTTATCGGTACAATTCCCTCGAGTTTT

>M6-PGSC0003DMG400007234-(rev)M6LRR-RLK_868

GATTGAATCCTTATTTCTTTTCAAGAACCATTTTATCGGTACAATTCCCTCGAGTTTTTCCAAACTCACATTGCTGAAATCATTGGATTTATCGGATAATCGTCTCTCAGGCAACATTCC

>M6-PGSC0003DMG400007234-(rev)M6LRR-RLK_930

AACTCACATTGCTGAAATCATTGGATTTATCGGATAATCGTCTCTCAGGCAACATTCCTTTTGGATTTTCAGAGTTGAAAGAGCTAATCATGTTGTATTTGATGAACAATAACTTATCAG

>M6-PGSC0003DMG400007234-(rev)M6LRR-RLK_992

GGATTTTCAGAGTTGAAAGAGCTAATCATGTTGTATTTGATGAACAATAACTTATCAGGTGAAATCCCTGAAGGGATTGGTGAGTTACCAAATCTCGAATTGTTAGCTCTTTGGAACAAT

>M6-PGSC0003DMG400007281M6LRR-RLP_0

GCTCTACGCCGCCAAATTTCTACCCTCTCTACGGTTAATTTGAACAAAACGATCAACAGCTTCATTGGAAACTGAAACCTCCAAGAAGCTCGTAAGCTCTTCGATCAAAGTTCCCACTTG

>M6-PGSC0003DMG400007281M6LRR-RLP_1088

GAGGATGCTAAAGAGTGGTATCAGGCCTGATGAAATTACTTTTATTAATTTGGTATATGCATGTGGCCATGGTGGGCTAGTCGAGGAAGGTGAGAGAATTTTCAATTCTATGGTAACAGA

>M6-PGSC0003DMG400007281M6LRR-RLP_1395

AATGGAATCTACACATTGGAACACGACCATCCTGCAGTATTCTCGGTGCTTTCCAAAATCTATGGAGACAAAGGAGTCTGCAGTGACATAACTGGTTTGGATAAGTTGATGAAGAAATGG

>M6-PGSC0003DMG400007281M6LRR-RLP_192

CTGTTTGACAAAATGCCTCACAGAGATGTCGTTTCTTGGAACACCATGTTCTCCGGATATCGCAACGTTAACAACCCAGAGAAAGTATATAGATGTTTTCTAGATATGAACAGATGTGGA

>M6-PGSC0003DMG400007281M6LRR-RLP_256

ACGTTAACAACCCAGAGAAAGTATATAGATGTTTTCTAGATATGAACAGATGTGGAGATATGAGACCCAATGAGCTCACTTTTGCTGTGTCAATTAGCTCCTTCTTGCATTTAGATTACA

>M6-PGSC0003DMG400007281M6LRR-RLP_320

ACCCAATGAGCTCACTTTTGCTGTGTCAATTAGCTCCTTCTTGCATTTAGATTACAAGCATTTGATCCCTCAGCTTCACGGTCTCGTGCTCTGTTTGGGGATAAGTCTCAATGTCTTTGT

>M6-PGSC0003DMG400007281M6LRR-RLP_384

ATCCCTCAGCTTCACGGTCTCGTGCTCTGTTTGGGGATAAGTCTCAATGTCTTTGTAGGGTCAGCATTGATGAGGGGTTATGTTGATTTGGATGATTACAGAGGTTTAGCTCGAGTTTTT

>M6-PGSC0003DMG400007281M6LRR-RLP_448

CATTGATGAGGGGTTATGTTGATTTGGATGATTACAGAGGTTTAGCTCGAGTTTTTTATGAGATTTTGGATAAAGATGTTACGCCGTGGAATGTGTTGATTTTGGGTTATATGAAATTTG

>M6-PGSC0003DMG400007281M6LRR-RLP_576

AGTGAGGCGCAGAGGGCTTTTGATATGATGCCTATGAGAAATTCTTTTACTTGGAGTACTTTGATCAATGGGTATATTGAGAACAAGAAGCTTAATGAAGCTCGATTCGTTTTTGATAAG

>M6-PGSC0003DMG400007281M6LRR-RLP_64

TTGGAAACTGAAACCTCCAAGAAGCTCGTAAGCTCTTCGATCAAAGTTCCCACTTGACAAATGTCGTATCATGGAATTCCCTTATCGCTGGTTACTTCAAACACAACCTCATCCAACAAG

>M6-PGSC0003DMG400007281M6LRR-RLP_640

TCAATGGGTATATTGAGAACAAGAAGCTTAATGAAGCTCGATTCGTTTTTGATAAGATGAGTGAGAAAGATGTGGTTTCTTGGACAGCAATGATAAGAGGGTATGTGCAATATGGAGAGT

>M6-PGSC0003DMG400007281M6LRR-RLP_704

GAAAGATGTGGTTTCTTGGACAGCAATGATAAGAGGGTATGTGCAATATGGAGAGTTTATGAAGGCTTTGAAATTGTTTAAGTTGATGTTGAACTCGGGTTCTCGTCCTAATCATTTTAC

>M6-PGSC0003DMG400007281M6LRR-RLP_768

GCTTTGAAATTGTTTAAGTTGATGTTGAACTCGGGTTCTCGTCCTAATCATTTTACGTTTTCAACTGTTTTAGATGCCTGTGCAGGCTACTCTGCTGTTCTGGTAGGCAATCAAGTTCAT

>M6-PGSC0003DMG400007281M6LRR-RLP_896

CTTGAAGTCTGGTTTCCCTCTTGATGTCGTCTTGTTGACCTCCCTTCTTGACATGTATGCTAAATGTGGAGACATTGAAGTTGCTTTCTGCATTTTTGAGTCTATCCCGGCAAGAAATTT

>M6-PGSC0003DMG400007281M6LRR-RLP_960

TGTGGAGACATTGAAGTTGCTTTCTGCATTTTTGAGTCTATCCCGGCAAGAAATTTGGTAGCTTGGAATTCAATCATTGGAGGTTATGCAAGACATGGGCTTGCAGAGAGTGCAATGCAG

>M6-PGSC0003DMG400007299-(rev)M6RLKmalectin_0

CTTCATTGTAGGTTGTTTCCCAGCATCCTATAGTTGATTAGCTAAACACTTGTTTCATGGATGCCAAGAAATGACCTGGTCTTTGTACCAAAAGCAAAGAAACCTCCAAGTTTCTCTTTG

>M6-PGSC0003DMG400007299-(rev)M6RLKmalectin_1024

CATACGTTGCACCACTTGCTGTCTACGAGAGCTGTAGATCGATGGGTTATGATTCAAACCTCAACAAGAACTATAACCTCACTTGGGTTTTCAATGTTGATCCCAATTTTATATATCTTG

>M6-PGSC0003DMG400007299-(rev)M6RLKmalectin_1088

CAAGAACTATAACCTCACTTGGGTTTTCAATGTTGATCCCAATTTTATATATCTTGTGAGGCTTCACTGGTGTGATTTCAGCATCGATGCCACGAAATACAAATCCATGGTCTTTAATGT

>M6-PGSC0003DMG400007299-(rev)M6RLKmalectin_1152

CACTGGTGTGATTTCAGCATCGATGCCACGAAATACAAATCCATGGTCTTTAATGTATATATTAATGGACAAATTGCAGAAACAGAAGTCGATTTAGTTGCAATGAAAAGTGGAAAAAAG

>M6-PGSC0003DMG400007299-(rev)M6RLKmalectin_1216

ATGGACAAATTGCAGAAACAGAAGTCGATTTAGTTGCAATGAAAAGTGGAAAAAAGGAAGTTGCAATCGTAAAAGATTATGTGACTCGTGTTAATGATAAAGAAGGAGGTCAGCTTTGGG

>M6-PGSC0003DMG400007299-(rev)M6RLKmalectin_128

AGGCGTACAGGATTGTTTATTGTCCTGTACGCCTTTTTTCTTTCGTTTGTCAACCATGTTAACAAAATTATCCCACATTTTGTTACTAGTGATCTTATGTTACATCTCTTCTATTCCTCG

>M6-PGSC0003DMG400007299-(rev)M6RLKmalectin_1280

AATCGTAAAAGATTATGTGACTCGTGTTAATGATAAAGAAGGAGGTCAGCTTTGGGTAGCAGTGCATCCATTAGTGCATGCATTAGGAGTTGGAAGTGAGATAGCTAATGCTATTGTAAA

>M6-PGSC0003DMG400007299-(rev)M6RLKmalectin_1344

CATCCATTAGTGCATGCATTAGGAGTTGGAAGTGAGATAGCTAATGCTATTGTAAATGGAATAGAGATATTTAAGATTGGCGGTGGAGATACGAGCTTGGCTGGAAGGAATCCTACAATG

>M6-PGSC0003DMG400007299-(rev)M6RLKmalectin_1408

AGATATTTAAGATTGGCGGTGGAGATACGAGCTTGGCTGGAAGGAATCCTACAATGTCAAAATTAATGCAAAAATATCAGGAAGAACATAGTAAAGAAGCTTCGGGGGCATTCTCTAATC

>M6-PGSC0003DMG400007299-(rev)M6RLKmalectin_1472

AATGCAAAAATATCAGGAAGAACATAGTAAAGAAGCTTCGGGGGCATTCTCTAATCAGGACAAATCACATAGTAAAACTGTTATAATCAGTGGTGCTGCTGGTGGAGTTGCTGCTGCATT

>M6-PGSC0003DMG400007299-(rev)M6RLKmalectin_1600

CTGCTGTTTTGGTCTTTGTTGCTTTGAAGAGGAAAAAAAGATCCCCGGGAGCTGAACCAGGCATAACTAGTTGGCTGCCGATTCATGGGAACTCTCATTCTTCTGGTAGCAGCAAGTCAG

>M6-PGSC0003DMG400007299-(rev)M6RLKmalectin_1664

AACTAGTTGGCTGCCGATTCATGGGAACTCTCATTCTTCTGGTAGCAGCAAGTCAGCTTCTCGAAGAAGTGATGGTAGCACAGCCATTTCATCAGATGCTGCTTCTAACTGCCGATATTT

>M6-PGSC0003DMG400007299-(rev)M6RLKmalectin_1728

AGAAGTGATGGTAGCACAGCCATTTCATCAGATGCTGCTTCTAACTGCCGATATTTTTCACTAGCTGAGATCAAACAGGCCACTAAAAACTTCGACGAATCATATGTCATTGGAGTTGGT

>M6-PGSC0003DMG400007299-(rev)M6RLKmalectin_1769

TAACTGCCGATATTTTTCACTAGCTGAGATCAAACAGGCCACTAAAAACTTCGACGAATCATATGTCATTGGAGTTGGTGGATTTGGAAAAGTGTACAAAGGAGTTATCGATGGTGACAC

>M6-PGSC0003DMG400007299-(rev)M6RLKmalectin_192

AAATTATCCCACATTTTGTTACTAGTGATCTTATGTTACATCTCTTCTATTCCTCGCTCGCTTCATGCTGCATTAATCAATGATCAAAACTCATTGTTCTTAGCATGTGGTTCTCAAAAT

>M6-PGSC0003DMG400007299-(rev)M6RLKmalectin_256

ATGCTGCATTAATCAATGATCAAAACTCATTGTTCTTAGCATGTGGTTCTCAAAATGGTGGAGTTGATGAAGATATGATAAAATGGGAATCAGACACGAAGTATCTCGTGTCTGGTGATA

>M6-PGSC0003DMG400007299-(rev)M6RLKmalectin_320

TGATGAAGATATGATAAAATGGGAATCAGACACGAAGTATCTCGTGTCTGGTGATAAATCAGTCATGACACAACCTGAATCCCCTGACCCTTTGGCTCCATCTATTGTACCATATACAAA

>M6-PGSC0003DMG400007299-(rev)M6RLKmalectin_384

ATGACACAACCTGAATCCCCTGACCCTTTGGCTCCATCTATTGTACCATATACAAATGCAAGAGTTTTTCAATCAGAATCTTCATATAATCTTCCTGTTACAAACCAAACAACACATGTT

>M6-PGSC0003DMG400007299-(rev)M6RLKmalectin_448

TTTTTCAATCAGAATCTTCATATAATCTTCCTGTTACAAACCAAACAACACATGTTATGCTTAGGCTTCGTTTCAACCCAACGCCTTATCCTAATTTCAATATTTCTAAATCATTTATGA

>M6-PGSC0003DMG400007299-(rev)M6RLKmalectin_512

GCTTCGTTTCAACCCAACGCCTTATCCTAATTTCAATATTTCTAAATCATTTATGACTGTTTTTGCTGGTAAAGTGACATTGTTAAATAATTTAAGCGCGTATATAACTGCAGAAGCCTT

>M6-PGSC0003DMG400007299-(rev)M6RLKmalectin_576

GCTGGTAAAGTGACATTGTTAAATAATTTAAGCGCGTATATAACTGCAGAAGCCTTGTCACAAAGTTGCATTACCAAAGAGTACTTATTGTCTCCTGTGGGTTCTCCAACTTTGGAGATC

>M6-PGSC0003DMG400007299-(rev)M6RLKmalectin_64

CAAGAAATGACCTGGTCTTTGTACCAAAAGCAAAGAAACCTCCAAGTTTCTCTTTGTATTGCGTAGGCGTACAGGATTGTTTATTGTCCTGTACGCCTTTTTTCTTTCGTTTGTCAACCA

>M6-PGSC0003DMG400007299-(rev)M6RLKmalectin_640

GTTGCATTACCAAAGAGTACTTATTGTCTCCTGTGGGTTCTCCAACTTTGGAGATCAAGATTAAGCCTTTTGGTGATTCATCATTTGCATTCATAAATGGAATTGAAATGATCACCACGC

>M6-PGSC0003DMG400007299-(rev)M6RLKmalectin_704

GCCTTTTGGTGATTCATCATTTGCATTCATAAATGGAATTGAAATGATCACCACGCCAGATATATTTTATGGTCCGGATCCTTTGATGGTTGGATTCAGTACAAGTATTCAAAATTCTAT

>M6-PGSC0003DMG400007299-(rev)M6RLKmalectin_768

TTTTATGGTCCGGATCCTTTGATGGTTGGATTCAGTACAAGTATTCAAAATTCTATTACTCAGCAAAGTATTTCTGTAAAGGAAAACAATATGGAGTTAGCATATAGGCTCAATGTTGGT

>M6-PGSC0003DMG400007299-(rev)M6RLKmalectin_832

AAAGTATTTCTGTAAAGGAAAACAATATGGAGTTAGCATATAGGCTCAATGTTGGTGGACAATACATTTCTCCAAAAAACGATTCAGGTGGGTTAATAAGAAGTTGGTATGAAGATACTC

>M6-PGSC0003DMG400007299-(rev)M6RLKmalectin_896

CATTTCTCCAAAAAACGATTCAGGTGGGTTAATAAGAAGTTGGTATGAAGATACTCCCTACTTATATGGTGCTTCAATAGGCATTACTATGAAGAGTAACAGGACAATTCAATACGATAA

>M6-PGSC0003DMG400007299-(rev)M6RLKmalectin_960

TATGGTGCTTCAATAGGCATTACTATGAAGAGTAACAGGACAATTCAATACGATAACATACCAACATACGTTGCACCACTTGCTGTCTACGAGAGCTGTAGATCGATGGGTTATGATTCA

>M6-PGSC0003DMG400007307-(rev)M6RLKmalectin_0

TTGATTAAGCTAAAACCCTTTCAACAATGGATGCCTAGAAGGCAAGAAATGATCTCGTCTTCGCCATAAATCAAACACAAAAGATCAATCTCTCATTTTTATTGTTTCATAGGCGTACAG

>M6-PGSC0003DMG400007307-(rev)M6RLKmalectin_1054

TAGATCGATGGGTTTTGAGTCGAATATCAACATGAATTATAACCTTACTTGGGTTTTCAGAGTTGATGCAAACTTTTCCTACCTTGTGAGGTTTCACTGGTGTGATTTCATTGTCGATTC

>M6-PGSC0003DMG400007307-(rev)M6RLKmalectin_1116

TTGATGCAAACTTTTCCTACCTTGTGAGGTTTCACTGGTGTGATTTCATTGTCGATTCCACTAAGGTCAATTCGTTAGTCTTTAATGTATATATAAACGGACAAATCGAAGAAAGTGGAG

>M6-PGSC0003DMG400007307-(rev)M6RLKmalectin_1178

AAGGTCAATTCGTTAGTCTTTAATGTATATATAAACGGACAAATCGAAGAAAGTGGAGTCGATTTAGTTGCAATGCAACATGGGGAAAAGGCAATTCCTTTAGTGAAAGACTATGTGACT

>M6-PGSC0003DMG400007307-(rev)M6RLKmalectin_124

CATTGCTTTCCTGTACGCCTTTTTTTTCTTTTTTGTTAACACCATGTTAACAAAAAATCGTATTTTGCTAATATTCATATGCCTTTCTTTATTTTCACGATCATTACATGCTGCAGTAGG

>M6-PGSC0003DMG400007307-(rev)M6RLKmalectin_1240

TTTAGTTGCAATGCAACATGGGGAAAAGGCAATTCCTTTAGTGAAAGACTATGTGACTCATGTTGAAGATAAAGAAGGAGATGATCTGCTTTGGATAGCAGTGCATCCATCAGGAGATAA

>M6-PGSC0003DMG400007307-(rev)M6RLKmalectin_1302

TTGAAGATAAAGAAGGAGATGATCTGCTTTGGATAGCAGTGCATCCATCAGGAGATAATGCTGATTTAGCTAATGCTATTGTAAATGGACTAGAGATATTTAAGATCAACGGTGGCCACA

>M6-PGSC0003DMG400007307-(rev)M6RLKmalectin_1364

GATTTAGCTAATGCTATTGTAAATGGACTAGAGATATTTAAGATCAACGGTGGCCACACGAGCTTGGCAGGTCAGAATCCTGCAATGTCAGATTTAATGATGAAGTATCAGGAAGAACAT

>M6-PGSC0003DMG400007307-(rev)M6RLKmalectin_1426

CTTGGCAGGTCAGAATCCTGCAATGTCAGATTTAATGATGAAGTATCAGGAAGAACATAGTAAAGAAGAAGCTCAACGGGGATTCACTGATCAGGACAAATCACATAGTCATACTGTTAT

>M6-PGSC0003DMG400007307-(rev)M6RLKmalectin_1488

AAGAAGAAGCTCAACGGGGATTCACTGATCAGGACAAATCACATAGTCATACTGTTATAATCAGTGGTGCTGCTGGTGGAGTTGCTGCTGCATTTGGTCTAGCTGCTGTTTTAGTTTTTG

>M6-PGSC0003DMG400007307-(rev)M6RLKmalectin_1550

AGTGGTGCTGCTGGTGGAGTTGCTGCTGCATTTGGTCTAGCTGCTGTTTTAGTTTTTGTTGCTTTGAAGAGGAAAAGAAAATCCCTAGGAGGTGAATCAGGCACGACTAGTTGGCTACCG

>M6-PGSC0003DMG400007307-(rev)M6RLKmalectin_1674

ATGGCAACTCTCATTCTTCCTCTAGCAAATCAGCTTCCGGAAGAAGTGGTAGTGGTAGCACAATCATTTCATCAGATGTTGCTAGTAACTGTCGATACTTTTCACTTGCTGAGATCAAAC

>M6-PGSC0003DMG400007307-(rev)M6RLKmalectin_1736

ATCATTTCATCAGATGTTGCTAGTAACTGTCGATACTTTTCACTTGCTGAGATCAAACAAGCTACTAAAAACTTTGACGAATCATATGTCATTGGAGTTGGTGGATTTGGAAAGGTATAC

>M6-PGSC0003DMG400007307-(rev)M6RLKmalectin_1798

TACTAAAAACTTTGACGAATCATATGTCATTGGAGTTGGTGGATTTGGAAAGGTATACAAAGGAGTTATCGATGGTGACACGAAAGTTGCAATCAAAAGATCAAATCCATCTTCAGAACA

>M6-PGSC0003DMG400007307-(rev)M6RLKmalectin_186

TTTTGCTAATATTCATATGCCTTTCTTTATTTTCACGATCATTACATGCTGCAGTAGGTGATCATACCTCATTTTTCATAGCATGTGGTTCTGAAAATGGTGGAACAGATGAAGATGCTA

>M6-PGSC0003DMG400007307-(rev)M6RLKmalectin_1860

GAGTTATCGATGGTGACACGAAAGTTGCAATCAAAAGATCAAATCCATCTTCAGAACAAGGAGTCAATGAATTCCAGACAGAAATAGAGATGCTCTCAAAGCTAAGGCATCGACATTTAG

>M6-PGSC0003DMG400007307-(rev)M6RLKmalectin_1922

GTCAATGAATTCCAGACAGAAATAGAGATGCTCTCAAAGCTAAGGCATCGACATTTAGTGTCCTTGATCGGATTTTGTGAAGAGAATAATGAAATGGTCCTTGTTTATGACCACATGGCA

>M6-PGSC0003DMG400007307-(rev)M6RLKmalectin_1984

CTTGATCGGATTTTGTGAAGAGAATAATGAAATGGTCCTTGTTTATGACCACATGGCACATGGAACATTAAGGGAGCATCTTTATAAAGGTAACAAAACTATACTATCTTGGAAGCAAAG

>M6-PGSC0003DMG400007307-(rev)M6RLKmalectin_2046

GAACATTAAGGGAGCATCTTTATAAAGGTAACAAAACTATACTATCTTGGAAGCAAAGGTTGGAGATTTGTATTGGATCAGCCAGAGGACTTCATTACCTTCACACAGGAGCTAAATACA

>M6-PGSC0003DMG400007307-(rev)M6RLKmalectin_2108

GAGATTTGTATTGGATCAGCCAGAGGACTTCATTACCTTCACACAGGAGCTAAATACACTATTATACATAGAGATGTGAAAAGTACAAACATTTTGTTAAATGACAAATGGGTTGCTAAG

>M6-PGSC0003DMG400007307-(rev)M6RLKmalectin_2170

TATACATAGAGATGTGAAAAGTACAAACATTTTGTTAAATGACAAATGGGTTGCTAAGGTTTCAGATTTTGGACTTTCAAAAACAGGTCCAAACATGAACCAAGGACATGTTTCAACAGT

>M6-PGSC0003DMG400007307-(rev)M6RLKmalectin_2232

CAGATTTTGGACTTTCAAAAACAGGTCCAAACATGAACCAAGGACATGTTTCAACAGTCGTAAAGGGTAGCTTCGGGTACTTAGATCCTGAATACTTCAGAAGACAACAATTGACTGAAA

>M6-PGSC0003DMG400007307-(rev)M6RLKmalectin_2294

AAGGGTAGCTTCGGGTACTTAGATCCTGAATACTTCAGAAGACAACAATTGACTGAAAAATCAGATGTGTACTCATATGGTGTTGTCCTGTTTGAAGTACTGTGTGCAAGGCCAGCACTT

>M6-PGSC0003DMG400007307-(rev)M6RLKmalectin_2418

CAAATCTTCCGAAAGAACAAGTTAGTCTTGCAGATTGGGCATTAGCTTGTCAAAGAAAAGGGAATTTAGAAGATATTATTGATCCACATTTGAAAGGCAAGATCAATCCTGAATGCTTGA

>M6-PGSC0003DMG400007307-(rev)M6RLKmalectin_248

CATACCTCATTTTTCATAGCATGTGGTTCTGAAAATGGTGGAACAGATGAAGATGCTAGAAAATGGGAATCGGACGCGAAGTATCTCATATCAGGTGATAAATCAGTCATGACACAACCT

>M6-PGSC0003DMG400007307-(rev)M6RLKmalectin_2480

AATTTAGAAGATATTATTGATCCACATTTGAAAGGCAAGATCAATCCTGAATGCTTGAAAAAATTTGCAGAAACAGCTGAGAAATGCTTAGCTGACAATGGTATAAATCGTCCGTCCATG

>M6-PGSC0003DMG400007307-(rev)M6RLKmalectin_2542

ATTTGCAGAAACAGCTGAGAAATGCTTAGCTGACAATGGTATAAATCGTCCGTCCATGGGAGATGTGCTATGGAATCTTGAATATGCCCTTCAGATACAAGAAAATCCAGATGGAGTTTT

>M6-PGSC0003DMG400007307-(rev)M6RLKmalectin_2604

ATGTGCTATGGAATCTTGAATATGCCCTTCAGATACAAGAAAATCCAGATGGAGTTTTATCAACTTCCGAGGCAACGAAACAAGTTGATCATGACAATGCTGATCAACACAGTTTGATAG

>M6-PGSC0003DMG400007307-(rev)M6RLKmalectin_2790

TCTTTTCACAAATTGTGAATCCAACAGGAAGGTAATAGAAAATATACACACATATATATAAGTAGTTGTTTCTTATGAATGGTTATAATAAGTATACAAAACTGTACATTTCACTATATA

>M6-PGSC0003DMG400007307-(rev)M6RLKmalectin_310

ATGGGAATCGGACGCGAAGTATCTCATATCAGGTGATAAATCAGTCATGACACAACCTCAATCTCCTGACCCCTCTGTTCCATCTGTTGTACCATATGTAAATGCAAGATTTTTTCAATC

>M6-PGSC0003DMG400007307-(rev)M6RLKmalectin_372

CTCCTGACCCCTCTGTTCCATCTGTTGTACCATATGTAAATGCAAGATTTTTTCAATCAGAATCTTCATATAACCTTCCTGTTACAAATAGATCAGCTCATATTTTGCTTAGGCTTCATT

>M6-PGSC0003DMG400007307-(rev)M6RLKmalectin_434

TCTTCATATAACCTTCCTGTTACAAATAGATCAGCTCATATTTTGCTTAGGCTTCATTTCTACCCGACTTCTTATCCTAATTTCAACGTTTCCAATTCATACATCTCTCTTTCTGCTGGT

>M6-PGSC0003DMG400007307-(rev)M6RLKmalectin_496

CCCGACTTCTTATCCTAATTTCAACGTTTCCAATTCATACATCTCTCTTTCTGCTGGTAAAGTTACGTTGTTAAATAATTTCAGTGCATATATAACAGCACAAGCATGGTCACAAAATTA

>M6-PGSC0003DMG400007307-(rev)M6RLKmalectin_558

TTACGTTGTTAAATAATTTCAGTGCATATATAACAGCACAAGCATGGTCACAAAATTACATTATTAAAGAATACATTTTGTATCCTATGGATTCTCCAACTTTGGAGATCAAAATTAGTC

>M6-PGSC0003DMG400007307-(rev)M6RLKmalectin_62

GCCATAAATCAAACACAAAAGATCAATCTCTCATTTTTATTGTTTCATAGGCGTACAGAAAACATTGCTTTCCTGTACGCCTTTTTTTTCTTTTTTGTTAACACCATGTTAACAAAAAAT

>M6-PGSC0003DMG400007307-(rev)M6RLKmalectin_620

ATTAAAGAATACATTTTGTATCCTATGGATTCTCCAACTTTGGAGATCAAAATTAGTCCTGTTGGTAACTCATCACTTGCATTCATCAATGGAATTGAAATGATCACCTCACCTGATTTG

>M6-PGSC0003DMG400007307-(rev)M6RLKmalectin_682

TGGTAACTCATCACTTGCATTCATCAATGGAATTGAAATGATCACCTCACCTGATTTGTTTTACGGCCCTGATCCTGTGATGGCTGGGTTTAGTACTGATGAAGAACCGGATTCTTCTGG

>M6-PGSC0003DMG400007307-(rev)M6RLKmalectin_744

ACGGCCCTGATCCTGTGATGGCTGGGTTTAGTACTGATGAAGAACCGGATTCTTCTGGTGACGGAGATGTTGTTTCTCAGCAAACTATATCTGTAAGGGAAAATAATATGGAGCTGATGT

>M6-PGSC0003DMG400007307-(rev)M6RLKmalectin_806

GGAGATGTTGTTTCTCAGCAAACTATATCTGTAAGGGAAAATAATATGGAGCTGATGTTTAGGGTCAATGTAGGTGGACAATTTATTCCTCCGAAAAATGATTCAGGAGGGTTAATGAGG

>M6-PGSC0003DMG400007307-(rev)M6RLKmalectin_868

GGTCAATGTAGGTGGACAATTTATTCCTCCGAAAAATGATTCAGGAGGGTTAATGAGGAGTTGGTATGAAGATACTCCCTACGTATATGGCGGTTCAGGTGTTGGTATTGCTATGAGCAC

>M6-PGSC0003DMG400007307-(rev)M6RLKmalectin_930

GGTATGAAGATACTCCCTACGTATATGGCGGTTCAGGTGTTGGTATTGCTATGAGCACCAATATGACAATTCGTTATGATGACTTACCATCATATGTTGCACCCCTTGATGTGTATGATA

>M6-PGSC0003DMG400007307-(rev)M6RLKmalectin_992

ATGACAATTCGTTATGATGACTTACCATCATATGTTGCACCCCTTGATGTGTATGATAGTTGTAGATCGATGGGTTTTGAGTCGAATATCAACATGAATTATAACCTTACTTGGGTTTTC

>M6-PGSC0003DMG400007389M6OtherRLKTM_0

TTGTTACATCCTCTGTTCTTCTCAAGAAGTTATAAATCTTTCATTCTTTTCGAGTGTGCACCTTTGATATGGAGGAGAGACAATTGTTGTTTGAGAAGTATGAAATTGGGACATTATTAG

>M6-PGSC0003DMG400007389M6OtherRLKTM_1008

CTGCATTTTTCAATGCATTCGAGTTGATTTCATCGATGTCATCTGGTTTCGACTTGTCTAGCTTGTTTGAGATCAAAGGGAAGGCATCATCCATGTTCACTTCAAGGAGCACTGCCCGGG

>M6-PGSC0003DMG400007389M6OtherRLKTM_1071

TGTTTGAGATCAAAGGGAAGGCATCATCCATGTTCACTTCAAGGAGCACTGCCCGGGATGTCATCCGTAAAATGGAGAAAATGGCTAAAGTTATGAGGTATAAGGTTTATAGGGTAAAAC

>M6-PGSC0003DMG400007389M6OtherRLKTM_1134

TCCGTAAAATGGAGAAAATGGCTAAAGTTATGAGGTATAAGGTTTATAGGGTAAAACCATTCAAGTTGAAGATGCAATGCCCCGAGGAAGGACGAAAAGGGCGGTTGTTGGTCACCGCTG

>M6-PGSC0003DMG400007389M6OtherRLKTM_1197

AGTTGAAGATGCAATGCCCCGAGGAAGGACGAAAAGGGCGGTTGTTGGTCACCGCTGAGGTGTTTAAGGTGGCCCCGGAAGTCACCGTGGTCGAGCTCTCTAAGTCCTCCGGTGACACGT

>M6-PGSC0003DMG400007389M6OtherRLKTM_126

GAACTTTTGCGAAAGTGTACTATGGTAAGGAACTCGAAACAGGGGAAAGTGTAGCCATTAAAGTGATAAAAAAAGATCAAGTTCAGAAAGAAGGAATGATGGAGCAAATCACAAGAGAAA

>M6-PGSC0003DMG400007389M6OtherRLKTM_1260

TTAAGGTGGCCCCGGAAGTCACCGTGGTCGAGCTCTCTAAGTCCTCCGGTGACACGTTGGAGTATAACAAGTTTTGTGAAGAGGAAGTTAGGCCTGCATTGAAGGACATTGTTTGGACAT

>M6-PGSC0003DMG400007389M6OtherRLKTM_1323

ATAACAAGTTTTGTGAAGAGGAAGTTAGGCCTGCATTGAAGGACATTGTTTGGACATGGCAAGGGACAGGTACTGGAAATGCTGATATTGATAACAAAAAGAACTTGGAGCAATAAAATT

>M6-PGSC0003DMG400007389M6OtherRLKTM_1386

GGACAGGTACTGGAAATGCTGATATTGATAACAAAAAGAACTTGGAGCAATAAAATTTGGTACATGTGCATCTTTCTCCCTCTTCTTTTGTGTTTGTGTTTCGCTCCCCTGATCGGCTGA

>M6-PGSC0003DMG400007389M6OtherRLKTM_1449

ATGTGCATCTTTCTCCCTCTTCTTTTGTGTTTGTGTTTCGCTCCCCTGATCGGCTGATAGAAATAAATCAACCATCGGAGATATTCGTGTTACTTTGTCAATATTGATTTGTACATAAAT

>M6-PGSC0003DMG400007389M6OtherRLKTM_1512

TAAATCAACCATCGGAGATATTCGTGTTACTTTGTCAATATTGATTTGTACATAAATTGTGTGTTATACTGACCACTTGGGATTTGTATTGTTTGCATGTTGTGATTCACTTAATTTCCT

>M6-PGSC0003DMG400007389M6OtherRLKTM_1575

TTATACTGACCACTTGGGATTTGTATTGTTTGCATGTTGTGATTCACTTAATTTCCTCTAGTTATTCAAGCACGGGGATCTTGTTCAAAAAGATATATTTCACCTCCCGGCACCCTTTAT

>M6-PGSC0003DMG400007389M6OtherRLKTM_1624

AATTTCCTCTAGTTATTCAAGCACGGGGATCTTGTTCAAAAAGATATATTTCACCTCCCGGCACCCTTTATCTATTTGTAAGTTGCAAGTTGAAGTTGCAATATGGACATTTTGTTCAGA

>M6-PGSC0003DMG400007389M6OtherRLKTM_189

TGATAAAAAAAGATCAAGTTCAGAAAGAAGGAATGATGGAGCAAATCACAAGAGAAATCTCTGTGATGGGCCTTGTTCGTCATCCAAATATTGTTGAACTCAAAGAAGTCATGGCTACAA

>M6-PGSC0003DMG400007389M6OtherRLKTM_252

TGATGGGCCTTGTTCGTCATCCAAATATTGTTGAACTCAAAGAAGTCATGGCTACAAAGTCTAAAATCTTCTTTATAATGGAGTATGTTAAAGGGGGTGAGCTATTCGCAAAATTAGTAT

>M6-PGSC0003DMG400007389M6OtherRLKTM_315

AAATCTTCTTTATAATGGAGTATGTTAAAGGGGGTGAGCTATTCGCAAAATTAGTATCCAAAGGCAAATTTAAAGAAGATACAGCAAGAAAGTACTTTCAACAATTGATAAGTGCTGTTG

>M6-PGSC0003DMG400007389M6OtherRLKTM_378

GCAAATTTAAAGAAGATACAGCAAGAAAGTACTTTCAACAATTGATAAGTGCTGTTGATTACTGTCATAGTCGAGGTGTTTATCATCGCGATTTGAAGCCTGAGAACCTACTCCTTGATG

>M6-PGSC0003DMG400007389M6OtherRLKTM_441

GTCATAGTCGAGGTGTTTATCATCGCGATTTGAAGCCTGAGAACCTACTCCTTGATGAGAATGAGGACTTGAAGATATCTGATTTTGGGTTATCGGCTTTGCCTGATGAGCAGTTGAGTA

>M6-PGSC0003DMG400007389M6OtherRLKTM_504

AGGACTTGAAGATATCTGATTTTGGGTTATCGGCTTTGCCTGATGAGCAGTTGAGTAGAAATGATGGTTTGCTTCATACACAGTGTGGGACACCAGCATATGTTGCTCCTGAGGTTTTAA

>M6-PGSC0003DMG400007389M6OtherRLKTM_567

ATGGTTTGCTTCATACACAGTGTGGGACACCAGCATATGTTGCTCCTGAGGTTTTAAGGAGGAAAGGTTATGATGGTGCAAAAGCTGATATTTGGTCATGTGGGGTTATTTTGTATGTGC

>M6-PGSC0003DMG400007389M6OtherRLKTM_63

TTGATATGGAGGAGAGACAATTGTTGTTTGAGAAGTATGAAATTGGGACATTATTAGGCAAAGGAACTTTTGCGAAAGTGTACTATGGTAAGGAACTCGAAACAGGGGAAAGTGTAGCCA

>M6-PGSC0003DMG400007389M6OtherRLKTM_630

AAGGTTATGATGGTGCAAAAGCTGATATTTGGTCATGTGGGGTTATTTTGTATGTGCTTCTAGCTGGTTACCTACCATTTCAAGATGAGAATGTGATGAACATGTACAAGAAGATTTTCA

>M6-PGSC0003DMG400007389M6OtherRLKTM_693

CTGGTTACCTACCATTTCAAGATGAGAATGTGATGAACATGTACAAGAAGATTTTCAAGGCGTATTTCGAGTTCCCTCCATGGTTTTCAATGGATTCGAGGCGTTTGATATCTAAACTTT

>M6-PGSC0003DMG400007389M6OtherRLKTM_756

ATTTCGAGTTCCCTCCATGGTTTTCAATGGATTCGAGGCGTTTGATATCTAAACTTTTAATGGCTGATCCGGACAGGAGGATCACTATTCAAGGTATCATGAGGGTTCCATGGTATAGGA

>M6-PGSC0003DMG400007389M6OtherRLKTM_819

CTGATCCGGACAGGAGGATCACTATTCAAGGTATCATGAGGGTTCCATGGTATAGGAAGGATTTCGCGATGCCACGGGCTTTTTCAATCCAAGATTTTCAAAAGTTAGAAAAGGAACAAG

>M6-PGSC0003DMG400007389M6OtherRLKTM_882

TCGCGATGCCACGGGCTTTTTCAATCCAAGATTTTCAAAAGTTAGAAAAGGAACAAGAAAATGATCATGTTGAAGAAGGAATGAGCAGCAAACAAGGAGGGATGATTAGGAAATCCCCTT

>M6-PGSC0003DMG400007389M6OtherRLKTM_945

ATCATGTTGAAGAAGGAATGAGCAGCAAACAAGGAGGGATGATTAGGAAATCCCCTTCATCCCCTGCATTTTTCAATGCATTCGAGTTGATTTCATCGATGTCATCTGGTTTCGACTTGT

>M6-PGSC0003DMG400007455M6G-LecRK_0

ATTTTTTAGAGGAATGAGGTGGAAAAAGGAAAAAAAAGAAATTGGGTCCTTTGTGATTTCTTGCATTGGCAGTAACCAACGTCCGAAGGCCAAATCTCATCGATTGTGACATTGTAACAA

>M6-PGSC0003DMG400007455M6G-LecRK_1037

GCCTCACTACCAATCTTACAGACTTATAATGGATTCTTCACCATCTCATGAAGAATCTGCTCCTCCAACTGATGTGGCACCTCCAAAGTTTAGTCCTCCGCCTACACCTTCTAGTCCATC

>M6-PGSC0003DMG400007455M6G-LecRK_1098

CCTCCAACTGATGTGGCACCTCCAAAGTTTAGTCCTCCGCCTACACCTTCTAGTCCATCTGATAGCTCACCACCACCTCCATCTTCTAGTCCACCTCCTAGTTCACCGCCACCTGTATCT

>M6-PGSC0003DMG400007455M6G-LecRK_122

TATACATAATTCATCAATCTTCATAAACCTATCATCATAATCATAATCAATACAACATTATTGTAAACAAAACAAATATTCCTTCTTGTATATACACACAATATTAAAAAATTATAAACT

>M6-PGSC0003DMG400007455M6G-LecRK_1220

TAGTCCACCTCCTAGTTCACCTCCACCTGAATCAACTAGTCCGCCACCTAGTTCACCACCACCTCAATCTTCTAGTCCACCACCAGCCTCACCTCCACCTGAACCAACTCCTTCTTCTCC

>M6-PGSC0003DMG400007455M6G-LecRK_1281

CCTCAATCTTCTAGTCCACCACCAGCCTCACCTCCACCTGAACCAACTCCTTCTTCTCCTGTCAGTTCACCTCCACCAACTTCATCACCTCCCCCTGCATTGTCACCTCCGCCACACGTA

>M6-PGSC0003DMG400007455M6G-LecRK_1342

TCAGTTCACCTCCACCAACTTCATCACCTCCCCCTGCATTGTCACCTCCGCCACACGTACAGAATCCAGAACCTCCTCCAGCACCACAATCACATGGAAATTCATCCCCACCACAATCTC

>M6-PGSC0003DMG400007455M6G-LecRK_1403

GAATCCAGAACCTCCTCCAGCACCACAATCACATGGAAATTCATCCCCACCACAATCTCCTGAACCTAAAAAAGGTTCATCATCATCTTCACCACCTTCACTACCTTCATCCAACTCACC

>M6-PGSC0003DMG400007455M6G-LecRK_1464

GAACCTAAAAAAGGTTCATCATCATCTTCACCACCTTCACTACCTTCATCCAACTCACCTGGGGATGGAGGTGGAGGTGGAAATTCACCACCATCCCCAATATCTAAGTCATCAGATAAT

>M6-PGSC0003DMG400007455M6G-LecRK_1525

GGGATGGAGGTGGAGGTGGAAATTCACCACCATCCCCAATATCTAAGTCATCAGATAATTCGCCTCCTTCACCAGCCTCAGGCCCTCCTAAAGATTCGCCACCTTCACCAGCTGCATTAT

>M6-PGSC0003DMG400007455M6G-LecRK_1586

GCCTCCTTCACCAGCCTCAGGCCCTCCTAAAGATTCGCCACCTTCACCAGCTGCATTATCTCCAAGTAACTCACCATCTTCTGATTCACCCTCAAACTCATCAACACCAACATCTGTAAT

>M6-PGSC0003DMG400007455M6G-LecRK_1647

CCAAGTAACTCACCATCTTCTGATTCACCCTCAAACTCATCAACACCAACATCTGTAATACAATGGTCACCACCACCTCCATCCTCTAATGGTGCACAATCTTCTCTTAATCCTCCTTCA

>M6-PGSC0003DMG400007455M6G-LecRK_1708

AATGGTCACCACCACCTCCATCCTCTAATGGTGCACAATCTTCTCTTAATCCTCCTTCATCACAAGTTCCTAGTAACCATAGTTCCGGAAACCATTCAACAGGAAGTCCAAAGCCTGCTG

>M6-PGSC0003DMG400007455M6G-LecRK_1769

ACAAGTTCCTAGTAACCATAGTTCCGGAAACCATTCAACAGGAAGTCCAAAGCCTGCTGGTGGCAACAGCAATGGTACTGCAGAGACAGTTTCCATTGGCGTTGTAATTGTTCTTTTTTT

>M6-PGSC0003DMG400007455M6G-LecRK_183

TGTAAACAAAACAAATATTCCTTCTTGTATATACACACAATATTAAAAAATTATAAACTTTTTTACATTCTTGATTGTGTATGTGGGAAATTGGGCAGCAAATTTGCAATTTTTGATCCT

>M6-PGSC0003DMG400007455M6G-LecRK_1830

GGCAACAGCAATGGTACTGCAGAGACAGTTTCCATTGGCGTTGTAATTGTTCTTTTTTTGGTTGGTATCATCGGAGCAGTTGCATGGTGCATACGGAAGCGAAAGAAGAAGAATTCTGTC

>M6-PGSC0003DMG400007455M6G-LecRK_1891

TTGGTATCATCGGAGCAGTTGCATGGTGCATACGGAAGCGAAAGAAGAAGAATTCTGTCCATTGTAGTGGTTATGTCAAGCCAATCTCTGCGTGCTCCTCTCCAAAATCAGGTAAATTTT

>M6-PGSC0003DMG400007455M6G-LecRK_1952

TTGTAGTGGTTATGTCAAGCCAATCTCTGCGTGCTCCTCTCCAAAATCAGGTAAATTTTCAGAAACTATAAAATCATTCCTTTTAGTTTGTTATTCAGTGTTCAAGATGCTTGCAAGAAA

>M6-PGSC0003DMG400007455M6G-LecRK_2013

GAAACTATAAAATCATTCCTTTTAGTTTGTTATTCAGTGTTCAAGATGCTTGCAAGAAAAATCAATGTAATATTTGTAGTTGAGACCTTTCAAGTGAAATCTGAGAAGATGTGAAAGTTG

>M6-PGSC0003DMG400007455M6G-LecRK_2074

TCAATGTAATATTTGTAGTTGAGACCTTTCAAGTGAAATCTGAGAAGATGTGAAAGTTGTGTACAGAGTCTGCTTTATTGAAGGTCAATGAATCAGCACCTGAGAGAAATGGAACGAGCT

>M6-PGSC0003DMG400007455M6G-LecRK_2135

TACAGAGTCTGCTTTATTGAAGGTCAATGAATCAGCACCTGAGAGAAATGGAACGAGCTCTAATTTCCTCAATTCTCCAGCAGACCCTGGCGGATTCAACAGTTCAAAGGCATGGTTTAC

>M6-PGSC0003DMG400007455M6G-LecRK_2196

AATTTCCTCAATTCTCCAGCAGACCCTGGCGGATTCAACAGTTCAAAGGCATGGTTTACTTATCAAGAACTTGTTGAGGCCACTAATGATTTTTCAGCGAATAATGAATTAGGGAAGGGT

>M6-PGSC0003DMG400007455M6G-LecRK_2257

ATCAAGAACTTGTTGAGGCCACTAATGATTTTTCAGCGAATAATGAATTAGGGAAGGGTGGATTTGGTTCTGTATACAAAGGCTATCTAGCAGATGGAAGATATGTCGCGGTAAAACAGC

>M6-PGSC0003DMG400007455M6G-LecRK_2318

ATTTGGTTCTGTATACAAAGGCTATCTAGCAGATGGAAGATATGTCGCGGTAAAACAGCTAAATATTGGTGGGAGCCAGGGCGAACGGGAATTTAGAGCTGAAGTTGAAATCATTAGCCG

>M6-PGSC0003DMG400007455M6G-LecRK_2379

AATATTGGTGGGAGCCAGGGCGAACGGGAATTTAGAGCTGAAGTTGAAATCATTAGCCGGATACACCATCGTCATTTGGTTTCACTTGTAGGTTATTGCATTTCAGAGAACAAAAGGCTC

>M6-PGSC0003DMG400007455M6G-LecRK_244

TTTACATTCTTGATTGTGTATGTGGGAAATTGGGCAGCAAATTTGCAATTTTTGATCCTGTATCTAATGCTTTGTGGATGTTTTCACAATTTGATTCACCTTTTTGGAGTGGGGTTTCCT

>M6-PGSC0003DMG400007455M6G-LecRK_2440

TACACCATCGTCATTTGGTTTCACTTGTAGGTTATTGCATTTCAGAGAACAAAAGGCTCCTCGTTTATGACTATGTATCCAATAATACCCTTTACTTCCATCTTCATGGTAAGCATAACA

>M6-PGSC0003DMG400007455M6G-LecRK_2501

CGTTTATGACTATGTATCCAATAATACCCTTTACTTCCATCTTCATGGTAAGCATAACATTTGTAGATTTAATTATTTCATTTGTTATGTTTGGTCACTAAAGCGAGCTGTTGATACTTG

>M6-PGSC0003DMG400007455M6G-LecRK_2562

TGTAGATTTAATTATTTCATTTGTTATGTTTGGTCACTAAAGCGAGCTGTTGATACTTGCAAGTTACAGCACAAGGGAGGCCTGTCATGAATTGGCCAACACGTGTTAAAATTGCTGTTG

>M6-PGSC0003DMG400007455M6G-LecRK_2623

AGTTACAGCACAAGGGAGGCCTGTCATGAATTGGCCAACACGTGTTAAAATTGCTGTTGGTGCAGCTCGTGGAATAGCTTATCTGCATGAAGATTGTAATACTGCTCTCTTCCTCAATTC

>M6-PGSC0003DMG400007455M6G-LecRK_2684

GCAGCTCGTGGAATAGCTTATCTGCATGAAGATTGTAATACTGCTCTCTTCCTCAATTCTTATGAGTTATCCATGTTTTATTCATGTGATATCTTTCATTGGTTGAACAAACGTCCCCCT

>M6-PGSC0003DMG400007455M6G-LecRK_2745

ATGAGTTATCCATGTTTTATTCATGTGATATCTTTCATTGGTTGAACAAACGTCCCCCTTATAGTTAAAAGAGAATGTACAGTTAAAAACTTTACCACAACTTGGAATTAGTTTTAATGA

>M6-PGSC0003DMG400007455M6G-LecRK_2806

TAGTTAAAAGAGAATGTACAGTTAAAAACTTTACCACAACTTGGAATTAGTTTTAATGATCCCTAAAGCTGTGGCGGGATCATGGATGGAAATTTAAGCTCCACAACCACCTTCTTCCTA

>M6-PGSC0003DMG400007455M6G-LecRK_2867

CCTAAAGCTGTGGCGGGATCATGGATGGAAATTTAAGCTCCACAACCACCTTCTTCCTACTCTTCCTAAAAGCATACCTTAACGAAAGGAAAATAAAAAAGAGAGAGAAAACAAAACATG

>M6-PGSC0003DMG400007455M6G-LecRK_2928

CTTCCTAAAAGCATACCTTAACGAAAGGAAAATAAAAAAGAGAGAGAAAACAAAACATGTAGTAATAATTTTCAGCTCATAGAAGTAGAAGGCCTTCCAGTTGGCATTTCCTTTTTGTAA

>M6-PGSC0003DMG400007455M6G-LecRK_2989

GTAATAATTTTCAGCTCATAGAAGTAGAAGGCCTTCCAGTTGGCATTTCCTTTTTGTAATGTAATGATATTACTTATGCCATTTAGAATTGAGTTCTAAGGTCCTTTTTCATGCCATAAA

>M6-PGSC0003DMG400007455M6G-LecRK_305

ATCTAATGCTTTGTGGATGTTTTCACAATTTGATTCACCTTTTTGGAGTGGGGTTTCCTTAGAAAGGGTAAATTCTACATAATCTTGAATTTCTATATCTTTTTCTTGATTTGGGTGTCT

>M6-PGSC0003DMG400007455M6G-LecRK_3050

TAATGATATTACTTATGCCATTTAGAATTGAGTTCTAAGGTCCTTTTTCATGCCATAAATTACCTCGTTATATAACTTTTTGGCATGAGTGTGGATTATCATATCTGCATTAAGATTGTA

>M6-PGSC0003DMG400007455M6G-LecRK_3111

ACCTCGTTATATAACTTTTTGGCATGAGTGTGGATTATCATATCTGCATTAAGATTGTAAGTAGTGTTCTGTTCCTTAATCCTTTTATGTTATTCATGTTTATTTTTATTTCAGTTCGTT

>M6-PGSC0003DMG400007455M6G-LecRK_3172

TAGTGTTCTGTTCCTTAATCCTTTTATGTTATTCATGTTTATTTTTATTTCAGTTCGTTTGTATGGCTGAAAAAAAATGTTTCTCCTCTTTCCTGCCCCAAAAGTTAAGAGAGAAAGCAG

>M6-PGSC0003DMG400007455M6G-LecRK_3233

TATGGCTGAAAAAAAATGTTTCTCCTCTTTCCTGCCCCAAAAGTTAAGAGAGAAAGCAGGTGGTAAAACTTTTTACCATATGACGAATCGGATTGTGATATTAGATTATGTGTTCCTATC

>M6-PGSC0003DMG400007455M6G-LecRK_3294

GGTAAAACTTTTTACCATATGACGAATCGGATTGTGATATTAGATTATGTGTTCCTATCATATCACTTACATTCTTTTTATTGATCCCTACTCCACTAGCTGAACAGCTGTAGTAAATTA

>M6-PGSC0003DMG400007455M6G-LecRK_3355

ATCACTTACATTCTTTTTATTGATCCCTACTCCACTAGCTGAACAGCTGTAGTAAATTAAATGATGCAATCTGAGCTACTTCCAGAACCAACTTTCTCCTATTATCCCTATAAGAGGACT

>M6-PGSC0003DMG400007455M6G-LecRK_3416

TGATGCAATCTGAGCTACTTCCAGAACCAACTTTCTCCTATTATCCCTATAAGAGGACTTGAAGTAAAAAGAGAAAATGGAACGTATAAATAGTTTAAGCTCAATGGAAGTAGCAAGATC

>M6-PGSC0003DMG400007455M6G-LecRK_3477

AAGTAAAAAGAGAAAATGGAACGTATAAATAGTTTAAGCTCAATGGAAGTAGCAAGATCTTGATATTGGATTTCCCTTTGGTAATGTTATAATATAACTTCCGCAATTTAGAATTGTGTC

>M6-PGSC0003DMG400007455M6G-LecRK_3538

GATATTGGATTTCCCTTTGGTAATGTTATAATATAACTTCCGCAATTTAGAATTGTGTCATCAATTTTTATGCATCTCGTAAATTATCTCGATTATATGACCTTTTGGCAGTATAAATAA

>M6-PGSC0003DMG400007455M6G-LecRK_3599

CAATTTTTATGCATCTCGTAAATTATCTCGATTATATGACCTTTTGGCAGTATAAATAATGAATTCTAATCACTTCTCTAATTAGTTACTTAAAAGATGTTTTTGTAGCTGAATGTACCA

>M6-PGSC0003DMG400007455M6G-LecRK_366

GAAAGGGTAAATTCTACATAATCTTGAATTTCTATATCTTTTTCTTGATTTGGGTGTCTGGGTATTCGAATAAATCTGTTTGGTGTTGTGGATGTTTCTTGAAACTGCATTTGGTGATCT

>M6-PGSC0003DMG400007455M6G-LecRK_3660

AATTCTAATCACTTCTCTAATTAGTTACTTAAAAGATGTTTTTGTAGCTGAATGTACCAAATGAGGCTCAAAATCGAACTAAGACTGAACAAAACATGATTGGTTCCATTGAGAGAACAG

>M6-PGSC0003DMG400007455M6G-LecRK_3721

TGAGGCTCAAAATCGAACTAAGACTGAACAAAACATGATTGGTTCCATTGAGAGAACAGAAAAAATTGTATGACTAAGTTCTTTATACACTCTTGTATTGCACCATGTAGTTACATTCCC

>M6-PGSC0003DMG400007455M6G-LecRK_3782

AAAATTGTATGACTAAGTTCTTTATACACTCTTGTATTGCACCATGTAGTTACATTCCCGAGAAAGAAATAATTTAATCCATGAGCAGGCATAGTCTCCTTGAACATGAGTATATCTTCC

>M6-PGSC0003DMG400007455M6G-LecRK_3843

GAAAGAAATAATTTAATCCATGAGCAGGCATAGTCTCCTTGAACATGAGTATATCTTCCACACTGAAGTGAGCGCATCAGGCTCTGCATTTGTAGTGCATTATATGCTTTTCGCTTGCCA

>M6-PGSC0003DMG400007455M6G-LecRK_3904

ACTGAAGTGAGCGCATCAGGCTCTGCATTTGTAGTGCATTATATGCTTTTCGCTTGCCAATATTCACTCTACGACAAGCATTTGAATAAATCTTCACTGATCGAGCCAGGTTGAAAAGTG

>M6-PGSC0003DMG400007455M6G-LecRK_3965

ATTCACTCTACGACAAGCATTTGAATAAATCTTCACTGATCGAGCCAGGTTGAAAAGTGAGAGAAACATGTTCAACTGCTCTTGTGAAGCATATAGTGAAAAGCAAGTGACAAACTGGTT

>M6-PGSC0003DMG400007455M6G-LecRK_4026

AGAAACATGTTCAACTGCTCTTGTGAAGCATATAGTGAAAAGCAAGTGACAAACTGGTTTCTCCATATAACAAAGCTGAATGTGTTAATCATCACTGAGTTATTATGCTATACTGTTGAA

>M6-PGSC0003DMG400007455M6G-LecRK_4087

TCCATATAACAAAGCTGAATGTGTTAATCATCACTGAGTTATTATGCTATACTGTTGAATAAGTAATGAAAAGTTAAGAAAAGTAGACCAGGCGAAGAGATTGTGATTTTCTATGCTCAT

>M6-PGSC0003DMG400007455M6G-LecRK_4148

AGTAATGAAAAGTTAAGAAAAGTAGACCAGGCGAAGAGATTGTGATTTTCTATGCTCATTACCTTCATTCTTGTTACTTGAGCTACCTTTAATTATATAGTTAAAGTATGATACTCCTTT

>M6-PGSC0003DMG400007455M6G-LecRK_4209

CCTTCATTCTTGTTACTTGAGCTACCTTTAATTATATAGTTAAAGTATGATACTCCTTTTTTATTTGGTTCTTAATTAATTCTTTTGGTGTGGATGACAGGTTATCCTCGTATTATTCAC

>M6-PGSC0003DMG400007455M6G-LecRK_427

GTATTCGAATAAATCTGTTTGGTGTTGTGGATGTTTCTTGAAACTGCATTTGGTGATCTGGGGTACTTTTATTTTGGTCATAGTTGCTGGTGATAATCATTATTTGGCTGCTTTGTTTAG

>M6-PGSC0003DMG400007455M6G-LecRK_4270

TATTTGGTTCTTAATTAATTCTTTTGGTGTGGATGACAGGTTATCCTCGTATTATTCACAGGGACATTAAGTCATCAAACATTCTTTTAGATGATAACTTTGAGGCCCATGTATGTCTAA

>M6-PGSC0003DMG400007455M6G-LecRK_4331

GGACATTAAGTCATCAAACATTCTTTTAGATGATAACTTTGAGGCCCATGTATGTCTAAGTCTTCGAACTCCTATATCCACATTCTTTGTCTCACGAGCCGCCTTTTACAATTAATTTAC

>M6-PGSC0003DMG400007455M6G-LecRK_4392

CTTCGAACTCCTATATCCACATTCTTTGTCTCACGAGCCGCCTTTTACAATTAATTTACTTTCAACAGGTTGCTGATTTTGGACTAGCTAAATTAGCTCAGGATGCAGAAAGTCATATCA

>M6-PGSC0003DMG400007455M6G-LecRK_4453

TCAACAGGTTGCTGATTTTGGACTAGCTAAATTAGCTCAGGATGCAGAAAGTCATATCACAACACGTGTCGTGGGAACATTTGGGTATGTTTCAGTGCCAACTTTTATATTAAGTTGACA

>M6-PGSC0003DMG400007455M6G-LecRK_4514

ACACGTGTCGTGGGAACATTTGGGTATGTTTCAGTGCCAACTTTTATATTAAGTTGACAGCATAAAATAAATATTTTATTCCATCTTTTTGAATTTCACATATTTATTCTAATTCCTTAG

>M6-PGSC0003DMG400007455M6G-LecRK_4575

ATAAAATAAATATTTTATTCCATCTTTTTGAATTTCACATATTTATTCTAATTCCTTAGATCTGACTCAGATACATGGCTCCTGAATATGCATCAACTGGCAAGCTGACTGAAAAATCTG

>M6-PGSC0003DMG400007455M6G-LecRK_4636

CTGACTCAGATACATGGCTCCTGAATATGCATCAACTGGCAAGCTGACTGAAAAATCTGATGTATATTCCTTTGGCGTGGTGCTTCTGGAGCTAATTACTGGAAGGAAGTCAGTGGATAC

>M6-PGSC0003DMG400007455M6G-LecRK_4697

GTATATTCCTTTGGCGTGGTGCTTCTGGAGCTAATTACTGGAAGGAAGTCAGTGGATACATCTCAGCCTTCAGGGCAAGAGAATCTAGTTGAGTGGGTATGTTTGTAGCCAATCTATTAT

>M6-PGSC0003DMG400007455M6G-LecRK_4758

CTCAGCCTTCAGGGCAAGAGAATCTAGTTGAGTGGGTATGTTTGTAGCCAATCTATTATTTGTCTTTCTCGAAATCCATATTTTCATCAGGTCTCCTTTATTACCTCTTTGTTAGAAATT

>M6-PGSC0003DMG400007455M6G-LecRK_4819

GTCTTTCTCGAAATCCATATTTTCATCAGGTCTCCTTTATTACCTCTTTGTTAGAAATTTTGACATTAATATCCGACCTAAGGCAATGAAAGTGCATATCAGCATGCAGAGAGAACTTAA

>M6-PGSC0003DMG400007455M6G-LecRK_488

GGTACTTTTATTTTGGTCATAGTTGCTGGTGATAATCATTATTTGGCTGCTTTGTTTAGAAAGATTGAATCTTTTTTACTCTGATAAGGTTCTGCAGTGTGTGGAGGGATTGTTCTTATG

>M6-PGSC0003DMG400007455M6G-LecRK_4880

GACATTAATATCCGACCTAAGGCAATGAAAGTGCATATCAGCATGCAGAGAGAACTTAATTTTTAGTCCTTTTAGCTAGGATAACCTACAAAATTTCATTTGAGTCAGGATTTGGTTGTT

>M6-PGSC0003DMG400007455M6G-LecRK_4941

TTTAGTCCTTTTAGCTAGGATAACCTACAAAATTTCATTTGAGTCAGGATTTGGTTGTTCCTCCGTCTTTTCCCATACTGCACAAGAACTTGGTAGCTGATATGTGAAGAAACCGCCATT

>M6-PGSC0003DMG400007455M6G-LecRK_5002

TCCGTCTTTTCCCATACTGCACAAGAACTTGGTAGCTGATATGTGAAGAAACCGCCATTTTGCTACTGTACAGTAGGCCTCTAGTTGGTCTGCTAGTGAAGTTCCTCTTCTTTATTTTTC

>M6-PGSC0003DMG400007455M6G-LecRK_5063

GCTACTGTACAGTAGGCCTCTAGTTGGTCTGCTAGTGAAGTTCCTCTTCTTTATTTTTCCCTTTTTTACCATGAAATGTGCCTTATCCTGGGAAGAATGTGCTTGCAATGCTGTAAGCCT

>M6-PGSC0003DMG400007455M6G-LecRK_5124

TTTTTTACCATGAAATGTGCCTTATCCTGGGAAGAATGTGCTTGCAATGCTGTAAGCCTGAAACATTGTGTCTACATTTTTATTTTTATTTTACAACAATCAAACAGCTTGATCAAGTTA

>M6-PGSC0003DMG400007455M6G-LecRK_5185

AACATTGTGTCTACATTTTTATTTTTATTTTACAACAATCAAACAGCTTGATCAAGTTATCCGCTTTAATTTCATTGATCCTTTAATGTTAGAAAACTGAAAAACCATTAACAGTAGCAG

>M6-PGSC0003DMG400007455M6G-LecRK_5246

CGCTTTAATTTCATTGATCCTTTAATGTTAGAAAACTGAAAAACCATTAACAGTAGCAGCCGAAAACTTGTATCTTTTATCATGCTGAATGTGTGAATTCCATTTACTATAGTTATTTTA

>M6-PGSC0003DMG400007455M6G-LecRK_5307

GAAAACTTGTATCTTTTATCATGCTGAATGTGTGAATTCCATTTACTATAGTTATTTTATTTTCGAACTCTTTTGTTTTCTTATCGTCAGTGCATAGTAGCAGTAGCCAGTTTACAAACT

>M6-PGSC0003DMG400007455M6G-LecRK_5368

TTCGAACTCTTTTGTTTTCTTATCGTCAGTGCATAGTAGCAGTAGCCAGTTTACAAACTTTTAATGCGTTGTATTCTGGTACTAGTTTCCACTGCTAAATTTGAACCAGGCTTAAAAATA

>M6-PGSC0003DMG400007455M6G-LecRK_5429

TAATGCGTTGTATTCTGGTACTAGTTTCCACTGCTAAATTTGAACCAGGCTTAAAAATAACTACTCCATTATATTGTACTGTAACTTGATTTTTTTGTTGTCCAGGCACGACCTTTGCTT

>M6-PGSC0003DMG400007455M6G-LecRK_549

AGATTGAATCTTTTTTACTCTGATAAGGTTCTGCAGTGTGTGGAGGGATTGTTCTTATGAATTATAGCTAAAAAGGAGATATTGTTCTGCTTTAGACCTGTCACTAAATTCTGTGTCCTG

>M6-PGSC0003DMG400007455M6G-LecRK_5490

TACTCCATTATATTGTACTGTAACTTGATTTTTTTGTTGTCCAGGCACGACCTTTGCTTAGTCGAGCACTTCAAAAGGAGGAATTTGATCTGTTAGCAGATCCGTGCCTTGAAAAGAACT

>M6-PGSC0003DMG400007455M6G-LecRK_5551

TCGAGCACTTCAAAAGGAGGAATTTGATCTGTTAGCAGATCCGTGCCTTGAAAAGAACTACGTTGGCACGGAGATGTTCCGAATGATTGAGGCTGCTGCATCTTGCGTGCGTCACTCATC

>M6-PGSC0003DMG400007455M6G-LecRK_5612

GTTGGCACGGAGATGTTCCGAATGATTGAGGCTGCTGCATCTTGCGTGCGTCACTCATCTGCAAAGAGACCAGCAATGGGACAGGTAACTCTTGTTCTGTTATATTGAAAACCTATGCAA

>M6-PGSC0003DMG400007455M6G-LecRK_5673

CAAAGAGACCAGCAATGGGACAGGTAACTCTTGTTCTGTTATATTGAAAACCTATGCAAAAGACACCTTCAAGACTCTTCTTGTGAAGGGCATTTACAACATTCTGAATTCTGATCATGG

>M6-PGSC0003DMG400007455M6G-LecRK_5734

GACACCTTCAAGACTCTTCTTGTGAAGGGCATTTACAACATTCTGAATTCTGATCATGGTTATGCTTCGTAGATCATGAGAGCTTTCGATGGTATGGCTATACAAGATCTGTCAAACGGG

>M6-PGSC0003DMG400007455M6G-LecRK_5795

ATGCTTCGTAGATCATGAGAGCTTTCGATGGTATGGCTATACAAGATCTGTCAAACGGGATGAAAGTTGGTGAAAGTGCAATACATAGCGCTGCACTACAATCTGCAGAAATAAGTTGGT

>M6-PGSC0003DMG400007455M6G-LecRK_5856

GAAAGTTGGTGAAAGTGCAATACATAGCGCTGCACTACAATCTGCAGAAATAAGTTGGTTTAGGAAAATGGCATTTGGTAAGCAAGATTTTAGTTCGGATTTTTTCAGCGAAAGTAATCA

>M6-PGSC0003DMG400007455M6G-LecRK_5917

AGGAAAATGGCATTTGGTAAGCAAGATTTTAGTTCGGATTTTTTCAGCGAAAGTAATCAGAATAGTAGAGAGTCAAATGAACATCCTTAGAGGATTTTATAGAACAAAAAGGCTTATTGA

>M6-PGSC0003DMG400007455M6G-LecRK_5978

ATAGTAGAGAGTCAAATGAACATCCTTAGAGGATTTTATAGAACAAAAAGGCTTATTGAAGCTATGTATGCATCTGTTGATACATCAAAGGTACCTTGTTAGCTTAGAGATGCCTTGGTT

>M6-PGSC0003DMG400007455M6G-LecRK_6023

AAAAGGCTTATTGAAGCTATGTATGCATCTGTTGATACATCAAAGGTACCTTGTTAGCTTAGAGATGCCTTGGTTTAGCTTTCATATGTATAAGTTCCCTTTTGGCTTTGAGTAATTTTC

>M6-PGSC0003DMG400007455M6G-LecRK_61

TGCATTGGCAGTAACCAACGTCCGAAGGCCAAATCTCATCGATTGTGACATTGTAACAAAATATACATAATTCATCAATCTTCATAAACCTATCATCATAATCATAATCAATACAACATT

>M6-PGSC0003DMG400007455M6G-LecRK_610

TTATAGCTAAAAAGGAGATATTGTTCTGCTTTAGACCTGTCACTAAATTCTGTGTCCTGTGGCAACATTTTGATTTCGCTTAAAATCTGATATGTTTAAGGTTGAGGTTGAGGTTCTTGT

>M6-PGSC0003DMG400007455M6G-LecRK_732

TGTTGAAAATCCTAGCTGTTTAACTTGGGGGTGGTGTGTGTGTATACTATGAGTGAATTGGTTGGTGAGTTTGGATAAATTTTGGAATAAGAGGAAGTAATTTACTGCATATTAGGGATA

>M6-PGSC0003DMG400007455M6G-LecRK_793

TTGGTGAGTTTGGATAAATTTTGGAATAAGAGGAAGTAATTTACTGCATATTAGGGATATTAGTTTTATGTCAGCAAGTCTGCATTGTATAAGAAATGTTGTAAATGTAGCTAGCTTTGG

>M6-PGSC0003DMG400007455M6G-LecRK_854

AGTTTTATGTCAGCAAGTCTGCATTGTATAAGAAATGTTGTAAATGTAGCTAGCTTTGGTGGTGGTGAGATGTTAAAACTTTAATCATCCACCTCCAAGTTATAGAATTTGAAGTTCCAT

>M6-PGSC0003DMG400007455M6G-LecRK_915

GTGGTGAGATGTTAAAACTTTAATCATCCACCTCCAAGTTATAGAATTTGAAGTTCCATTATCTGTGACTTCTCTCCGGGATCGTTACTCTCTTTCGACAAATGCAAATATAGTCTGTCC

>M6-PGSC0003DMG400007455M6G-LecRK_976

TCTGTGACTTCTCTCCGGGATCGTTACTCTCTTTCGACAAATGCAAATATAGTCTGTCCTAGCCTCACTACCAATCTTACAGACTTATAATGGATTCTTCACCATCTCATGAAGAATCTG

>M6-PGSC0003DMG400007517-(rev)M6LRR-RLK_0

TTACTCCTGGTGGTGTTAATTATGAAGGTTAGTTCTTGTTATTTTTAATAAAGATTGAAACTTTGAGACATTATTGGAATGAGATTGATCATATTAGAAAATATCAAGAATTTAAAAGTA

>M6-PGSC0003DMG400007517-(rev)M6LRR-RLK_1054

ATTACTTTTTAACTTTTTTTTTTTTTTTAAACAGGCTTCTGCAGAGCAATAATATAACAGGACCAATTCCAACTGAACTTGGAATGCTTCAGAAACTTAATACAATTGATCTTTCTGATA

>M6-PGSC0003DMG400007517-(rev)M6LRR-RLK_1116

CCAATTCCAACTGAACTTGGAATGCTTCAGAAACTTAATACAATTGATCTTTCTGATAATCAACTCACTGGGAAAATCCCTTCATCTTTAGCTCAATTAAAAAGCCTTCAATACTTGTAA

>M6-PGSC0003DMG400007517-(rev)M6LRR-RLK_1178

ACTCACTGGGAAAATCCCTTCATCTTTAGCTCAATTAAAAAGCCTTCAATACTTGTAAGCAAAGAAAATTTTAAAGATTTTGATATAAAAATGAGTTCTTGAATTATAAAAAGCTAATGA

>M6-PGSC0003DMG400007517-(rev)M6LRR-RLK_124

TTACTCCTGTTGGTGTTAATTATGAAGGTTGTTTCTTGTGGTTTTTCTCATTAAGATTGAAACTTTATGACTTTTTTGTTGAAAAAATCATATGAGGTGTTTGAAAATTTTGAAAGGAAT

>M6-PGSC0003DMG400007517-(rev)M6LRR-RLK_1240

AGAAAATTTTAAAGATTTTGATATAAAAATGAGTTCTTGAATTATAAAAAGCTAATGATTATGGTTTTTTTTTCTTTTTCAATTGTGTTATAGGAGACTAAACAACAATAGTCTTAGTGG

>M6-PGSC0003DMG400007517-(rev)M6LRR-RLK_1302

GGTTTTTTTTTCTTTTTCAATTGTGTTATAGGAGACTAAACAACAATAGTCTTAGTGGAGCTGTTCCTTTGGAATTGTCTAATATGACTCAACTTACTCTTCTGTAAGTTCCTTAGCTTC

>M6-PGSC0003DMG400007517-(rev)M6LRR-RLK_1364

GTTCCTTTGGAATTGTCTAATATGACTCAACTTACTCTTCTGTAAGTTCCTTAGCTTCATTTAGTTTTGTTTATTTTCTCATACTTGAAGGGGAGCCTCGGAGCAATAGTAAACTTGTCT

>M6-PGSC0003DMG400007517-(rev)M6LRR-RLK_1426

AGTTTTGTTTATTTTCTCATACTTGAAGGGGAGCCTCGGAGCAATAGTAAACTTGTCTCCTTGTTATCTATATAGGTCACGCGTTCGAGTTAGGGGTAGGCTGCCTATAGTATAGCCTTT

>M6-PGSC0003DMG400007517-(rev)M6LRR-RLK_1488

GTTATCTATATAGGTCACGCGTTCGAGTTAGGGGTAGGCTGCCTATAGTATAGCCTTTCCCCGGACCTTGCGTGAACGTGGAATGCTTTGTGTACCTTTTTCCTAATTGCTTTCTGTTTT

>M6-PGSC0003DMG400007517-(rev)M6LRR-RLK_1550

GGACCTTGCGTGAACGTGGAATGCTTTGTGTACCTTTTTCCTAATTGCTTTCTGTTTTCATGAGTATATTTTGATGGGATTTTTGTTGTTTTTTTCTAGAGACTTGTCTTTTAATAATTT

>M6-PGSC0003DMG400007517-(rev)M6LRR-RLK_1612

AGTATATTTTGATGGGATTTTTGTTGTTTTTTTCTAGAGACTTGTCTTTTAATAATTTGAGTGGTCCTGTGCCAAGGCTTCTAGCTAAAACATTCAAGTAAGTTTCTTGAACAATTCTTT

>M6-PGSC0003DMG400007517-(rev)M6LRR-RLK_1674

GGTCCTGTGCCAAGGCTTCTAGCTAAAACATTCAAGTAAGTTTCTTGAACAATTCTTTGAATTTCATTTTTTTTCCCTTTGCTAAAGATTAGAGAAAATAAATCATTTTTTCTCTCATTG

>M6-PGSC0003DMG400007517-(rev)M6LRR-RLK_1736

TTCATTTTTTTTCCCTTTGCTAAAGATTAGAGAAAATAAATCATTTTTTCTCTCATTGATGCAGCCTTTTGGGAAATCCTATTATATGTGCAACTGGAAAAGAACCAGAATGCAATGGAA

>M6-PGSC0003DMG400007517-(rev)M6LRR-RLK_1798

AGCCTTTTGGGAAATCCTATTATATGTGCAACTGGAAAAGAACCAGAATGCAATGGAACAACAACATCCATGCCTTTGCCCTTCACTTTGAACAACTCACAAAGTAAGAGTCTTGAATCA

>M6-PGSC0003DMG400007517-(rev)M6LRR-RLK_186

CTTTATGACTTTTTTGTTGAAAAAATCATATGAGGTGTTTGAAAATTTTGAAAGGAATGAAATTTTGATCAAATTTTAGAAATATCAAGCATTTTGAAATATGCTTACCCTTGTTGGAGT

>M6-PGSC0003DMG400007517-(rev)M6LRR-RLK_1860

AACATCCATGCCTTTGCCCTTCACTTTGAACAACTCACAAAGTAAGAGTCTTGAATCATCCTCCTAAGTATTTTTTCGAGCTAGCTTGTGTGCCTCTCGACTATTCCACAAATCTTGGTT

>M6-PGSC0003DMG400007517-(rev)M6LRR-RLK_1922

CCTAAGTATTTTTTCGAGCTAGCTTGTGTGCCTCTCGACTATTCCACAAATCTTGGTTGTTGTCAAGTCTCCTGCTTTCAATGTCGTTATGATTAACTACAACTCTATTATAAGGATTCA

>M6-PGSC0003DMG400007517-(rev)M6LRR-RLK_1984

TCAAGTCTCCTGCTTTCAATGTCGTTATGATTAACTACAACTCTATTATAAGGATTCATCATGAAGTCTTGGATAGTTTCTGTGTTGTACTATTCACAACTGAAAAGTCTCTTGTTATAT

>M6-PGSC0003DMG400007517-(rev)M6LRR-RLK_2046

GAAGTCTTGGATAGTTTCTGTGTTGTACTATTCACAACTGAAAAGTCTCTTGTTATATCTCAGATGCTCAGCCTTTTAGAAGATCTAAAACTCACAAAATTGCCTTAGCCTTTGGGACAA

>M6-PGSC0003DMG400007517-(rev)M6LRR-RLK_2108

GATGCTCAGCCTTTTAGAAGATCTAAAACTCACAAAATTGCCTTAGCCTTTGGGACAAGTCTTGGATGCATCTGCCTACTAATTATTGGGTTTGGAATATTTCTGTGGTGGAGACAAAAG

>M6-PGSC0003DMG400007517-(rev)M6LRR-RLK_2170

TGGATGCATCTGCCTACTAATTATTGGGTTTGGAATATTTCTGTGGTGGAGACAAAAGCACAACAAGCAAATTTTCTTTGACATCAATGGTGAGTATGATATGCAGCTAAAACCTATATT

>M6-PGSC0003DMG400007517-(rev)M6LRR-RLK_2232

ACAAGCAAATTTTCTTTGACATCAATGGTGAGTATGATATGCAGCTAAAACCTATATTGCTCGAATCTCGAGATGCTAGTAATGTCTTGCTTTAACCAAGAGGATCATGCTTGTTGATAA

>M6-PGSC0003DMG400007517-(rev)M6LRR-RLK_2294

GAATCTCGAGATGCTAGTAATGTCTTGCTTTAACCAAGAGGATCATGCTTGTTGATAATTTCAGAACAACATCATGAACAAGTGTGTTTGGGACACTTGACGAGGTTCCAATTTAGGGAA

>M6-PGSC0003DMG400007517-(rev)M6LRR-RLK_2356

AGAACAACATCATGAACAAGTGTGTTTGGGACACTTGACGAGGTTCCAATTTAGGGAACTTCAAGTTGCCACAAACAACTTCAGCAACAAGAAGATACTGGGAAAAGGCGGTTTTGGAAA

>M6-PGSC0003DMG400007517-(rev)M6LRR-RLK_2418

AAGTTGCCACAAACAACTTCAGCAACAAGAAGATACTGGGAAAAGGCGGTTTTGGAAACGTGTACAAAGGTTGTCTTAGTGATGGCACGATTGTTGCTGTAAAAAGGCTCAAAGATGGAA

>M6-PGSC0003DMG400007517-(rev)M6LRR-RLK_248

TTTTGATCAAATTTTAGAAATATCAAGCATTTTGAAATATGCTTACCCTTGTTGGAGTTGATTATGAAGGTTAGTTCTTGTGGTTTTTCTCATGAAGATTGAAACTTTATGACTTTTTAT

>M6-PGSC0003DMG400007517-(rev)M6LRR-RLK_2480

TACAAAGGTTGTCTTAGTGATGGCACGATTGTTGCTGTAAAAAGGCTCAAAGATGGAAATGCAATCGGAGGAGACAAACAATTCCAAACAGAAGTTGAAATGATCAGCTTGGCAGTGCAT

>M6-PGSC0003DMG400007517-(rev)M6LRR-RLK_2542

AATCGGAGGAGACAAACAATTCCAAACAGAAGTTGAAATGATCAGCTTGGCAGTGCATCGCAATCTCCTACGTTTGTATGGGTTCTCCATGACACCAACTGAAAGGCTTCTGGTTTACCC

>M6-PGSC0003DMG400007517-(rev)M6LRR-RLK_2604

ATCTCCTACGTTTGTATGGGTTCTCCATGACACCAACTGAAAGGCTTCTGGTTTACCCTTACATGTCTAATGGAAGCGTTGCTTCTCGTCTCAAAGGTAAAAAGAAACAACCTTAGAGAA

>M6-PGSC0003DMG400007517-(rev)M6LRR-RLK_2666

ATGTCTAATGGAAGCGTTGCTTCTCGTCTCAAAGGTAAAAAGAAACAACCTTAGAGAACTAAATATTCTGCAAGTATTCTTAAATATGGGGGATGGTTGAGATCCCGTCATCCTTAACCA

>M6-PGSC0003DMG400007517-(rev)M6LRR-RLK_2728

ATATTCTGCAAGTATTCTTAAATATGGGGGATGGTTGAGATCCCGTCATCCTTAACCATAGTAGGCTTACCTGTGTGAACCGGAATAGTCGAACTAGTGGACTTCAGATACCTGATGGTA

>M6-PGSC0003DMG400007517-(rev)M6LRR-RLK_2790

AGGCTTACCTGTGTGAACCGGAATAGTCGAACTAGTGGACTTCAGATACCTGATGGTAGATTTCTCTATTTCTGTTGTTTCTTGCAGCCAAGCCACCATTGGACTGGACTACAAGAAAAA

>M6-PGSC0003DMG400007517-(rev)M6LRR-RLK_2852

TCTCTATTTCTGTTGTTTCTTGCAGCCAAGCCACCATTGGACTGGACTACAAGAAAAAGAATTGCATTAGGAGCTGCAAGAGGTTTACTATATCTACATGAGCAGTGTGATCCAAAGATT

>M6-PGSC0003DMG400007517-(rev)M6LRR-RLK_2914

TGCATTAGGAGCTGCAAGAGGTTTACTATATCTACATGAGCAGTGTGATCCAAAGATTATCCACAGGGATGTGAAGGCAGCAAACATATTGCTCGACGATGACTGTGAGGCTGTCGTGGG

>M6-PGSC0003DMG400007517-(rev)M6LRR-RLK_2976

ACAGGGATGTGAAGGCAGCAAACATATTGCTCGACGATGACTGTGAGGCTGTCGTGGGAGATTTTGGATTGGCAAAGCTATTGGATCATCGCGATTCACATGTCACAACTGCAGTTAGGG

>M6-PGSC0003DMG400007517-(rev)M6LRR-RLK_3038

TTTGGATTGGCAAAGCTATTGGATCATCGCGATTCACATGTCACAACTGCAGTTAGGGGAACTGTAGGACATATAGCTCCTGAGTATCTCTCTACAGGACAGTCCTCTGATAAAACAGAT

>M6-PGSC0003DMG400007517-(rev)M6LRR-RLK_310

TATGAAGGTTAGTTCTTGTGGTTTTTCTCATGAAGATTGAAACTTTATGACTTTTTATTTGAAAAAATCATATGGGGTGTTTGAAATTTTTGAAAGGAATGAGATTTTGATCAAATTTTA

>M6-PGSC0003DMG400007517-(rev)M6LRR-RLK_3100

TGTAGGACATATAGCTCCTGAGTATCTCTCTACAGGACAGTCCTCTGATAAAACAGATGTTTTCGGGTTTGGAATTCTCCTGCTAGAGTTGATATCTGGCCAGAAAGCTCTAGAATTTGG

>M6-PGSC0003DMG400007517-(rev)M6LRR-RLK_3162

TCGGGTTTGGAATTCTCCTGCTAGAGTTGATATCTGGCCAGAAAGCTCTAGAATTTGGTAAAGCAGCAAATCAGAAAGGCATTATGCTTGATTGGGTAAGTATTACACATTCCAAGAAGT

>M6-PGSC0003DMG400007517-(rev)M6LRR-RLK_372

AAAAATCATATGGGGTGTTTGAAATTTTTGAAAGGAATGAGATTTTGATCAAATTTTAGAAATATTAAGAATTTTGAAATATGTTTACCCCTGTTGGAGTTGATTATGAAGGCTAGTTCT

>M6-PGSC0003DMG400007517-(rev)M6LRR-RLK_434

TATTAAGAATTTTGAAATATGTTTACCCCTGTTGGAGTTGATTATGAAGGCTAGTTCTCATAAAGATTGAAACTTTGATACTTTTTTGTGAAAAATTCATAAGGGGTTTCTGATTTTTTT

>M6-PGSC0003DMG400007517-(rev)M6LRR-RLK_496

AAGATTGAAACTTTGATACTTTTTTGTGAAAAATTCATAAGGGGTTTCTGATTTTTTTTGAAAGGAATGAGATTTGAACAAATTTAAAATATTTCAAGAATTTCATCTATTTTAACTTTC

>M6-PGSC0003DMG400007517-(rev)M6LRR-RLK_558

AGGAATGAGATTTGAACAAATTTAAAATATTTCAAGAATTTCATCTATTTTAACTTTCTTGTTATGTTGATTTTGGACAGTGCAAGCTTTAATGGAGATCAAGAACTCATTACATGATCC

>M6-PGSC0003DMG400007517-(rev)M6LRR-RLK_62

TTGAGACATTATTGGAATGAGATTGATCATATTAGAAAATATCAAGAATTTAAAAGTATGATTTACTCCTGTTGGTGTTAATTATGAAGGTTGTTTCTTGTGGTTTTTCTCATTAAGATT

>M6-PGSC0003DMG400007517-(rev)M6LRR-RLK_620

TATGTTGATTTTGGACAGTGCAAGCTTTAATGGAGATCAAGAACTCATTACATGATCCACTTGGTGTTTTGAATTGGGATGAAAGTTCTGTTGATCCATGTAGCTGGAATATGATCACTT

>M6-PGSC0003DMG400007517-(rev)M6LRR-RLK_682

GGTGTTTTGAATTGGGATGAAAGTTCTGTTGATCCATGTAGCTGGAATATGATCACTTGTTCAAATGATAAATTTGTCACTGGCTTGTAAGTATACCCGATTGTCTGGTATCTGCATTGG

>M6-PGSC0003DMG400007517-(rev)M6LRR-RLK_744

AAATGATAAATTTGTCACTGGCTTGTAAGTATACCCGATTGTCTGGTATCTGCATTGGAGCCCGACTAACAAAACCTTTTCTATTCCTAATACTTGAACCGGAGACCTCTAGTTAAGGAT

>M6-PGSC0003DMG400007517-(rev)M6LRR-RLK_806

CGACTAACAAAACCTTTTCTATTCCTAATACTTGAACCGGAGACCTCTAGTTAAGGATGAAAGTGCTTACAAGTTTTCTTGAATTGTGCAGGGCTTGTTCAAGCCAAAGTTTGTCTGGGA

>M6-PGSC0003DMG400007517-(rev)M6LRR-RLK_868

GTGCTTACAAGTTTTCTTGAATTGTGCAGGGCTTGTTCAAGCCAAAGTTTGTCTGGGAAAATATCACCATACATCCACAACTTGACTCACTTGGATCTACTGTGAGTACTTTATATGAAG

>M6-PGSC0003DMG400007517-(rev)M6LRR-RLK_930

ATCACCATACATCCACAACTTGACTCACTTGGATCTACTGTGAGTACTTTATATGAAGAAAAAATAATGAAATTTCAAATGATTGATTATTTGAATCAAGAATATAGAAAATAGTAGTAC

>M6-PGSC0003DMG400007517-(rev)M6LRR-RLK_992

AATAATGAAATTTCAAATGATTGATTATTTGAATCAAGAATATAGAAAATAGTAGTACTAATATTACTTTTTAACTTTTTTTTTTTTTTTAAACAGGCTTCTGCAGAGCAATAATATAAC

>M6-PGSC0003DMG400007606M6LRR-RLP_0

ATGGACTTAACGCATGACAATTGGAAGAAGGTTGAGGAAAATCTGAACTCATTCTTTGGTACGGTATCCGAACGATGCCAATCAATTCTTTCTTTGAGCTACAACTACTTGCCCCAATAT

>M6-PGSC0003DMG400007606M6LRR-RLP_1024

CGATGATTATCTCTGGAGATATTTCTCTTCCTAATCTTAAGGATTTGACATTAACAATTACTCGTATACCATGGGAAGTTGTGAATTTGCTGGCTAATTTACCCAATCTTGAGGCGCTCG

>M6-PGSC0003DMG400007606M6LRR-RLP_1088

TATACCATGGGAAGTTGTGAATTTGCTGGCTAATTTACCCAATCTTGAGGCGCTCGTTGGGAATTGTGCATTCAATGGAACAGCTTGGAAAGTAGATGAAGATGTTGTATTTCACAAATT

>M6-PGSC0003DMG400007606M6LRR-RLP_1152

TGTGCATTCAATGGAACAGCTTGGAAAGTAGATGAAGATGTTGTATTTCACAAATTAAAATATCTACATCTATTTATGTGCAGAGATCTGGAAAGGTGGGAATTAGCAGCTGGTAGTGAT

>M6-PGSC0003DMG400007606M6LRR-RLP_128

ATGTTTTCTCTACATTGGAGGTTTTCCAGAAGACAGAGAAATTAATGTTTCCAAGTTGATTAGGCTATGGATTGCAGAGCAATTTGTAAAGGCAAGAAACAATAAAAGATTAGAAGTGGT

>M6-PGSC0003DMG400007606M6LRR-RLP_1280

GATGCTTGAGAAACTAATGATGCTTGGGTTGGAAAAACTGGAGGAGATTCCCGAGAGTATTGGAGAAATAATGACACTAAAATCCATCCGAATAGTTAATTGCGGCTCTGGTGTAGAGAC

>M6-PGSC0003DMG400007606M6LRR-RLP_1343

AGAAATAATGACACTAAAATCCATCCGAATAGTTAATTGCGGCTCTGGTGTAGAGACAAGTGCAAAGCAAATTCAGGAAGAGCAAGAAAGCTTGGGAAATTATGAGCTTCAACTTCAAAT

>M6-PGSC0003DMG400007606M6LRR-RLP_192

CTATGGATTGCAGAGCAATTTGTAAAGGCAAGAAACAATAAAAGATTAGAAGTGGTGGCAGAGGAGTATCTGCAAGAGTTAATTGATAGAAGTCTAATTTTGATTGTTACGCGAAGCGAT

>M6-PGSC0003DMG400007606M6LRR-RLP_320

GATAGAAAGTTGCAAAATTCATGATCTTCTTCGCCAACTCTGCCTAAGTGAAGGTCATACTGAAAACTATGTCATGAATGGAAAAGTTCTCGAAGCCATAGATGGTCAACGTCGAGTGAT

>M6-PGSC0003DMG400007606M6LRR-RLP_384

AACTATGTCATGAATGGAAAAGTTCTCGAAGCCATAGATGGTCAACGTCGAGTGATCCTTCTATCTAAATATGAAGAGAAGTATGATTATTTTCTGCGACATAGCAGTGGTATTGTCCGT

>M6-PGSC0003DMG400007606M6LRR-RLP_448

CTAAATATGAAGAGAAGTATGATTATTTTCTGCGACATAGCAGTGGTATTGTCCGTACCTTTACTTCAATGGGTGTAGCTTTTCCAAAAGATATGTGTTCCATTATTTCAGAGTTCAAGT

>M6-PGSC0003DMG400007606M6LRR-RLP_512

TTCAATGGGTGTAGCTTTTCCAAAAGATATGTGTTCCATTATTTCAGAGTTCAAGTTGATTAAGGTGTTGGACGTGTTAGAAGTCCCGTACGATTTCTCTAGTGTAATACCTCAACTTGT

>M6-PGSC0003DMG400007606M6LRR-RLP_64

TATCCGAACGATGCCAATCAATTCTTTCTTTGAGCTACAACTACTTGCCCCAATATTTGAGGGCATGTTTTCTCTACATTGGAGGTTTTCCAGAAGACAGAGAAATTAATGTTTCCAAGT

>M6-PGSC0003DMG400007606M6LRR-RLP_640

GATATGTTTCTGCAAGAATTGATGAAGGTCTTTCACTAGCCAAATTGAGAAATCTACAGACCATAATTCTTCAAAGTCTTCAAAGGGAAGAATTGGAGCAGCCAGTAGATATCTGGAGTA

>M6-PGSC0003DMG400007606M6LRR-RLP_704

AATTCTTCAAAGTCTTCAAAGGGAAGAATTGGAGCAGCCAGTAGATATCTGGAGTATGACAGAGATAAGACATGTGGATATTGATCGGCCACTATATATATCAAATCCTGTTGAAGCAGA

>M6-PGSC0003DMG400007606M6LRR-RLP_768

ATAAGACATGTGGATATTGATCGGCCACTATATATATCAAATCCTGTTGAAGCAGAACAAGCTTTGTGCCTTCTGAATAACTTGCAAACACTATATCTCCGTAACTCTCCTTTTGTTGTG

>M6-PGSC0003DMG400007606M6LRR-RLP_832

TGTGCCTTCTGAATAACTTGCAAACACTATATCTCCGTAACTCTCCTTTTGTTGTGGAAATCATAAGAAGAACTCCCAATCTAAAAGAGCTAAAGATTTTAGATAGTTCTGAGCGTTCTG

>M6-PGSC0003DMG400007606M6LRR-RLP_896

AAGAAGAACTCCCAATCTAAAAGAGCTAAAGATTTTAGATAGTTCTGAGCGTTCTGATTGGGCTGTTATTCTTGATTCTCTCAGTATTTTACAGGATCTGGAGACACTACACATACAAAA

>M6-PGSC0003DMG400007606M6LRR-RLP_960

GTTATTCTTGATTCTCTCAGTATTTTACAGGATCTGGAGACACTACACATACAAAACATTGATCCGATGATTATCTCTGGAGATATTTCTCTTCCTAATCTTAAGGATTTGACATTAACA

>M6-PGSC0003DMG400007634-(rev)M6LRR-RLK_0

TATATATTCATTTTTATCAATTTGGCATTTTCTTCACACCTCCAATGTTGACCAAAATCAACTCATAATTCTAACCCCACCTTTTTTTTTTTTTCATGGAGTTTTCCCATTTTCCGACCG

>M6-PGSC0003DMG400007634-(rev)M6LRR-RLK_1037

GGGTTTAAACATTTGGAAATTATGAATCTGTTTGGTAACCAATTGTATGGAGCAATTCCAGAGAGCATAGCCAAGATTCCATCTTTGAAGGTATTCAAAGTGTTTAGGAATAAACTGAAT

>M6-PGSC0003DMG400007634-(rev)M6LRR-RLK_1098

AGAGCATAGCCAAGATTCCATCTTTGAAGGTATTCAAAGTGTTTAGGAATAAACTGAATGGGAGCTTACCATCTGAAATGGGACTTCATTCGAAACTCGAATCTTTTGAAGTTTCACTGA

>M6-PGSC0003DMG400007634-(rev)M6LRR-RLK_1159

GAGCTTACCATCTGAAATGGGACTTCATTCGAAACTCGAATCTTTTGAAGTTTCACTGAATTCCTTCACTGGAAACTTGCCAGAGCATTTGTGTGCTGGAGGAACTCTTTTTGGTGCTGT

>M6-PGSC0003DMG400007634-(rev)M6LRR-RLK_122

ATCTTCGCCGGAAAACAGTCCCTTTATCTCATTCTTCTACTAAACTTCATACCCATTTTCGTAACTTCACAATCTCCGGCCACTACTGAACGTGACACTCTACTGAAGATTAAGCGTCAA

>M6-PGSC0003DMG400007634-(rev)M6LRR-RLK_1220

TCCTTCACTGGAAACTTGCCAGAGCATTTGTGTGCTGGAGGAACTCTTTTTGGTGCTGTAGCTTATGTTAACAATTTGTCTGGTGAAATACCAAAATCACTCGGAACTTGTTCTAGCTTA

>M6-PGSC0003DMG400007634-(rev)M6LRR-RLK_1281

CTTATGTTAACAATTTGTCTGGTGAAATACCAAAATCACTCGGAACTTGTTCTAGCTTACGTTCGATTCAGCTCTACAAGAATCAGTTTTCGGGTGAAATTCCATCTGGGGTTTGGACTT

>M6-PGSC0003DMG400007634-(rev)M6LRR-RLK_1403

GTTGACATGACAAGTTTGTTGTTGAATGATAACTCATTTTCCGGGGAGCTACCGAGCAAAATTGCATACAATTTTACGCGGTTAGAGATCAGTAACAACAAGTTTTCTGGTGAAATTCCT

>M6-PGSC0003DMG400007634-(rev)M6LRR-RLK_1464

TTGCATACAATTTTACGCGGTTAGAGATCAGTAACAACAAGTTTTCTGGTGAAATTCCTGTGGGGATATCTTCTTGGAGGAGTTTAGTGGTGCTTTTGGCAAGTAACAATTCATTTTCAG

>M6-PGSC0003DMG400007634-(rev)M6LRR-RLK_1525

GGGGATATCTTCTTGGAGGAGTTTAGTGGTGCTTTTGGCAAGTAACAATTCATTTTCAGGTCGGATTCCAGTGGAATTGACTAGTCTTTCTCAGATAACTCAGCTAGAGCTTGATGGTAA

>M6-PGSC0003DMG400007634-(rev)M6LRR-RLK_1586

CGGATTCCAGTGGAATTGACTAGTCTTTCTCAGATAACTCAGCTAGAGCTTGATGGTAATTCGTTATCAGGAGAACTTCCAGCCGATATAATATCGTGGAAGTCGTTGACAATATTGGAT

>M6-PGSC0003DMG400007634-(rev)M6LRR-RLK_1647

CGTTATCAGGAGAACTTCCAGCCGATATAATATCGTGGAAGTCGTTGACAATATTGGATCTTGCCAGGAACAAACTCTCTGGCAAGATTCCAGCAGCTATAGGTTTAATCCCTGATCTTG

>M6-PGSC0003DMG400007634-(rev)M6LRR-RLK_1769

GCTCTAGATTTGTCTCAAAATCAATTTTCAGGTCCAATTCCACCTCAACTAGGTGTAAAAAGGATCACTAGCCTTAACGTGTCGTCCAATCAACTTATCGGGAATATCCCTGATGCATTT

>M6-PGSC0003DMG400007634-(rev)M6LRR-RLK_183

TAACTTCACAATCTCCGGCCACTACTGAACGTGACACTCTACTGAAGATTAAGCGTCAATGGGGAAACCCAGTAGCACTTGACTCATGGAACTCTACTTCTTCTCCTTGTAGCTGGCCGG

>M6-PGSC0003DMG400007634-(rev)M6LRR-RLK_1830

GGATCACTAGCCTTAACGTGTCGTCCAATCAACTTATCGGGAATATCCCTGATGCATTTGCTAACCTAGCCTTTGAAAACAGTTTCTTGAACAATCCCAGTCTTTGTACCACCAACTCAC

>M6-PGSC0003DMG400007634-(rev)M6LRR-RLK_1952

CCATATCTTCCAAGTTGTAATAATGCGAAAGTCGCCAATTCCAAAAGATTGTCCCATAGAGTCCTAGCCCTGATTCTAGTCCTTGCATTCGCTGTTTTCCTATTTAGTGTTATATCGACC

>M6-PGSC0003DMG400007634-(rev)M6LRR-RLK_2013

TCCTAGCCCTGATTCTAGTCCTTGCATTCGCTGTTTTCCTATTTAGTGTTATATCGACCTTGTTCATGGTTAGAGACTACAGGAGGAAGAAGCATAAGCGCGATGTTGCAAGCTGGAAGT

>M6-PGSC0003DMG400007634-(rev)M6LRR-RLK_2074

GTTCATGGTTAGAGACTACAGGAGGAAGAAGCATAAGCGCGATGTTGCAAGCTGGAAGTTGACTTCATTCCAGAGGTTGGATTTCACAGAGGCGAACATTTTGTCAAGTTTGACGGAAAA

>M6-PGSC0003DMG400007634-(rev)M6LRR-RLK_2196

ACATGATTGGAAGTGGTGGATCGGGTAAGGTGTACAGAATTTCCGTTGGCAGACCAAATGAGTATGTTGCTGTGAAGAGGATTTGGAGTGATAGGAAAGTAGACTACATTCTGGAGAGGG

>M6-PGSC0003DMG400007634-(rev)M6LRR-RLK_2257

GTATGTTGCTGTGAAGAGGATTTGGAGTGATAGGAAAGTAGACTACATTCTGGAGAGGGAATTCTTGGCTGAGGTTCAGATATTGGGATCAATTAGACATTCCAATATCGTGAAGTTATT

>M6-PGSC0003DMG400007634-(rev)M6LRR-RLK_2318

TTCTTGGCTGAGGTTCAGATATTGGGATCAATTAGACATTCCAATATCGTGAAGTTATTGTGTTGTATATCAAGCGAAGATTCAAAGTTGCTCGTTTACGAGTACATGGTAAATCACAGT

>M6-PGSC0003DMG400007634-(rev)M6LRR-RLK_2379

GTTGTATATCAAGCGAAGATTCAAAGTTGCTCGTTTACGAGTACATGGTAAATCACAGTCTTGATGGATGGCTTCATGGGAAGAAGAGAGTATCCTTGAGTAATAAAGTCATGGATTGGC

>M6-PGSC0003DMG400007634-(rev)M6LRR-RLK_2440

TGATGGATGGCTTCATGGGAAGAAGAGAGTATCCTTGAGTAATAAAGTCATGGATTGGCCAAAAAGATTGGAGGTTGCCATTGGAGCTGCTCAAGGACTTTGTTATATGCACCATGACTG

>M6-PGSC0003DMG400007634-(rev)M6LRR-RLK_2501

AAAAGATTGGAGGTTGCCATTGGAGCTGCTCAAGGACTTTGTTATATGCACCATGACTGCACTCCACCAATCATTCATCGCGATGTAAAAACAAGCAATATCTTGTTGGATTCTGATTTC

>M6-PGSC0003DMG400007634-(rev)M6LRR-RLK_2562

CTCCACCAATCATTCATCGCGATGTAAAAACAAGCAATATCTTGTTGGATTCTGATTTCACAGCCAAAATAGCTGATTTTGGGTTGGCCAAGATTTTAGAGAAGAAAGGAGAGCTTAACA

>M6-PGSC0003DMG400007634-(rev)M6LRR-RLK_2684

ATGTCTGCTGTTGCGGGTTCCTTTGGTTACATAGCTCCAGGTAAAAGCAAAATCCAAGCTTGAACTTCGTATTACATCAATTTCGTTAACATTATATCAAAAGTATGACTAACTAATAAT

>M6-PGSC0003DMG400007634-(rev)M6LRR-RLK_2745

GAACTTCGTATTACATCAATTTCGTTAACATTATATCAAAAGTATGACTAACTAATAATTTACTGTTTGGTTACAGAGTATGCTTATACAACTAAAGTGAACGAGAAGATTGATATCTAT

>M6-PGSC0003DMG400007634-(rev)M6LRR-RLK_2806

ACTGTTTGGTTACAGAGTATGCTTATACAACTAAAGTGAACGAGAAGATTGATATCTATAGCTTCGGGGTGGTGCTATTGGAGCTAGTGACTGGAAGACAACCAAATTTTGGGGACGAGC

>M6-PGSC0003DMG400007634-(rev)M6LRR-RLK_2867

CTTCGGGGTGGTGCTATTGGAGCTAGTGACTGGAAGACAACCAAATTTTGGGGACGAGCATACAAGCCTGGCAGAATGGGCATGGAAGCAGCACGGAGAAGGGAACACTGCCATCGATAA

>M6-PGSC0003DMG400007634-(rev)M6LRR-RLK_2928

ACAAGCCTGGCAGAATGGGCATGGAAGCAGCACGGAGAAGGGAACACTGCCATCGATAATATGTTAGATACAGATATCAAAGAAACATGTTACTTGGAAGAAATGAAGACTGTGTTCAGA

>M6-PGSC0003DMG400007634-(rev)M6LRR-RLK_2989

TGTTAGATACAGATATCAAAGAAACATGTTACTTGGAAGAAATGAAGACTGTGTTCAGACTCGGGCTTATATGCACGAGCAACTTACCTGCAAGCAGGCCTTCCATGAAAGAGATTCTGC

>M6-PGSC0003DMG400007634-(rev)M6LRR-RLK_305

ATTGAGTGTGATGATGGGAAGGTTACCGGAATAATAATTCAAGAAAAAGATATTACTGTTGAAATTCCAAGTTCTATTTGTGAACTTAAAAATCTTACTTTTCTTAATCTCCGGTTGAAT

>M6-PGSC0003DMG400007634-(rev)M6LRR-RLK_3050

CGGGCTTATATGCACGAGCAACTTACCTGCAAGCAGGCCTTCCATGAAAGAGATTCTGCAGATACTTCATCGATGCAAGTCCTTTAGGTATTCTGGAGGAAAATCACCAGACACAGAGTA

>M6-PGSC0003DMG400007634-(rev)M6LRR-RLK_3172

ATGTTGCTCCTCTCCTTAGCGGAAACAACAGTGAAAAATACATTGCTAGTTACAAACGTATTAATTCCAATAAGGTAATAGATGATAGTAGTGATGATGGTCTAATAATTTCCAGTGTGT

>M6-PGSC0003DMG400007634-(rev)M6LRR-RLK_3233

TAATTCCAATAAGGTAATAGATGATAGTAGTGATGATGGTCTAATAATTTCCAGTGTGTAATATTGGAGATGCCGAAGTAAACTGAGTTGTTGTAAAATGATCATAGAATAGTGCCACTT

>M6-PGSC0003DMG400007634-(rev)M6LRR-RLK_3294

TATTGGAGATGCCGAAGTAAACTGAGTTGTTGTAAAATGATCATAGAATAGTGCCACTTAAGTAAAGTACAAGTTTGAATTTCTAGTGTACTCTTGAATTTGTTCACTGCTTGTAGATTG

>M6-PGSC0003DMG400007634-(rev)M6LRR-RLK_3338

AGAATAGTGCCACTTAAGTAAAGTACAAGTTTGAATTTCTAGTGTACTCTTGAATTTGTTCACTGCTTGTAGATTGAGTGAAATGCATCATTTAACTTTTGCTAGTTATGTAGATTGTAA

>M6-PGSC0003DMG400007634-(rev)M6LRR-RLK_366

AAATTCCAAGTTCTATTTGTGAACTTAAAAATCTTACTTTTCTTAATCTCCGGTTGAATTATCTTCCCGGAAAATTCCCAACTTTTCTTTACAAATGTTCCAATTTGCAGCATTTGGATC

>M6-PGSC0003DMG400007634-(rev)M6LRR-RLK_427

TCTTCCCGGAAAATTCCCAACTTTTCTTTACAAATGTTCCAATTTGCAGCATTTGGATCTTTCTCAGAATTACTTTGTGGGTTCCATTCCGGAGGATATTCACCGGCTTGGGAAACTCAA

>M6-PGSC0003DMG400007634-(rev)M6LRR-RLK_488

TCTCAGAATTACTTTGTGGGTTCCATTCCGGAGGATATTCACCGGCTTGGGAAACTCAAGTATCTTAATCTGGGTGGTAATAATTTCACCGGAGATATCCCGTCGTCAGTCGGGAATTTG

>M6-PGSC0003DMG400007634-(rev)M6LRR-RLK_549

ATCTTAATCTGGGTGGTAATAATTTCACCGGAGATATCCCGTCGTCAGTCGGGAATTTGACGGAGCTGGAGACTTTGTGCATGCATTCGAACTTGTTTGATGGGAGTTTTCCGGCGGAAA

>M6-PGSC0003DMG400007634-(rev)M6LRR-RLK_61

CTCATAATTCTAACCCCACCTTTTTTTTTTTTTCATGGAGTTTTCCCATTTTCCGACCGGCATCTTCGCCGGAAAACAGTCCCTTTATCTCATTCTTCTACTAAACTTCATACCCATTTT

>M6-PGSC0003DMG400007634-(rev)M6LRR-RLK_610

GGAGCTGGAGACTTTGTGCATGCATTCGAACTTGTTTGATGGGAGTTTTCCGGCGGAAATCGGAAACTTAACCAATCTTGAAAGTTTGGGGTTGGAGTTTAATGGGTTTTCTCCAATGGC

>M6-PGSC0003DMG400007634-(rev)M6LRR-RLK_732

TACCGCCGGAGTTTGGGAAGCTGAAGAAGATAAAGTATATTTGGATGAGAGAGACGAAATTAATCGGTGAAATCCCTGAAAGTTTTGGGGAATTTCAAAATCTTGAGCTTATTGATTTTG

>M6-PGSC0003DMG400007634-(rev)M6LRR-RLK_793

AATCGGTGAAATCCCTGAAAGTTTTGGGGAATTTCAAAATCTTGAGCTTATTGATTTTGCTTACAACAATCTGGAAGGAAAAATCCCTTCTGGGTTGTTTCTGTTGAAGAATTTGACGAT

>M6-PGSC0003DMG400007634-(rev)M6LRR-RLK_854

TACAACAATCTGGAAGGAAAAATCCCTTCTGGGTTGTTTCTGTTGAAGAATTTGACGATGATGTACTTGTTCAGTAATCGGCTTTCGGGTCGAATCCCTGAAACTTTTGATTCATCAAAG

>M6-PGSC0003DMG400007634-(rev)M6LRR-RLK_915

TGTACTTGTTCAGTAATCGGCTTTCGGGTCGAATCCCTGAAACTTTTGATTCATCAAAGTTGATGGAACTTGATGTTTCCAACAACAATAATTTAACTGGAACAATCCCAGAAAGTTTTG

>M6-PGSC0003DMG400007634-(rev)M6LRR-RLK_976

GATGGAACTTGATGTTTCCAACAACAATAATTTAACTGGAACAATCCCAGAAAGTTTTGGAGGGTTTAAACATTTGGAAATTATGAATCTGTTTGGTAACCAATTGTATGGAGCAATTCC

>M6-PGSC0003DMG400007684M6LRR-RLP_0

ATGAAACTCATCAACTTAGCCCTTTTTGGCTTCTTGCTCATCATTTTCTTCTCATCAAAACCCTCATTTCAAGCTTCAATCAACTATGAACCAAACCCTAGACTTTTCGATGCTTACATA

>M6-PGSC0003DMG400007684M6LRR-RLP_126

CAAGCTTGGAAACATGTCATGACTTCCGATCCAAGAAACTTCACAAAAGATTGGTATGATTACAATGTTTGTAACTACACCGGGGTTTATTGTGCTCCAGCTCCAGATAACCCTAATATT

>M6-PGSC0003DMG400007684M6LRR-RLP_189

AATGTTTGTAACTACACCGGGGTTTATTGTGCTCCAGCTCCAGATAACCCTAATATTACCACTGTTGCTGGTATTGACCTCAACCATGCAAATATCTCCGGATACTTACCTGAAAAACTT

>M6-PGSC0003DMG400007684M6LRR-RLP_252

GTTGCTGGTATTGACCTCAACCATGCAAATATCTCCGGATACTTACCTGAAAAACTTGGTTTATTGACTGATTTAGCCGTCTTTCATATAAATTCCAATAGATTTCAAGGCTCAATCCCG

>M6-PGSC0003DMG400007684M6LRR-RLP_315

TTGACTGATTTAGCCGTCTTTCATATAAATTCCAATAGATTTCAAGGCTCAATCCCGAAATCTTTTTCTAAGCTTCGAATTTTATTCGAACTTGACGTGAGTAACAATCTTTTTTGTGGT

>M6-PGSC0003DMG400007684M6LRR-RLP_378

TTTTCTAAGCTTCGAATTTTATTCGAACTTGACGTGAGTAACAATCTTTTTTGTGGTGAGTTTCCTTCCGTTGTGTTATCTCTTCCTTCTTTGAAATTTCTTGATATTAGGTATAACCAA

>M6-PGSC0003DMG400007684M6LRR-RLP_441

CCTTCCGTTGTGTTATCTCTTCCTTCTTTGAAATTTCTTGATATTAGGTATAACCAATTCGAAGGAAAAGTCCCTTCAACCCTTTGGGATCGAACACTCGATGCTCTTTTCATAAACAAC

>M6-PGSC0003DMG400007684M6LRR-RLP_504

GGAAAAGTCCCTTCAACCCTTTGGGATCGAACACTCGATGCTCTTTTCATAAACAACAACAAATTTCAATTTTCTTGGCCTAAAAACTTAGGTAAATCACCCGTTTCTGCTTTGGTTATG

>M6-PGSC0003DMG400007684M6LRR-RLP_567

TTTCAATTTTCTTGGCCTAAAAACTTAGGTAAATCACCCGTTTCTGCTTTGGTTATGGCAAATATTAATGTAACGGGGTGTATACCATCAAGTATAGCAAAAATGTCGAAAACATTAAAT

>M6-PGSC0003DMG400007684M6LRR-RLP_63

TCATTTCAAGCTTCAATCAACTATGAACCAAACCCTAGACTTTTCGATGCTTACATAGCTCTACAAGCTTGGAAACATGTCATGACTTCCGATCCAAGAAACTTCACAAAAGATTGGTAT

>M6-PGSC0003DMG400007684M6LRR-RLP_630

ATTAATGTAACGGGGTGTATACCATCAAGTATAGCAAAAATGTCGAAAACATTAAATGAGATTATTCTCTCGAATTCGAGTTTAAGTGGATGTTTACCACAAGAATTAGGGTTATTGAAA

>M6-PGSC0003DMG400007684M6LRR-RLP_693

ATTCTCTCGAATTCGAGTTTAAGTGGATGTTTACCACAAGAATTAGGGTTATTGAAAAATGTTAGGGTTCTTGATGTGAGTTTTAACAAGCTTGTTGGTGAGTTGCCGGAATCAATTGGT

>M6-PGSC0003DMG400007684M6LRR-RLP_756

AGGGTTCTTGATGTGAGTTTTAACAAGCTTGTTGGTGAGTTGCCGGAATCAATTGGTGGAATGAAGAAATTGGAACATCTTAACGTGGCACATAACAAGTTTTCCGGTGAGATTCCGATG

>M6-PGSC0003DMG400007684M6LRR-RLP_819

AAGAAATTGGAACATCTTAACGTGGCACATAACAAGTTTTCCGGTGAGATTCCGATGAGTATTTGTTCATTACCAAAGTTGGAGAATTTCACTTACTCTTATAATTATTTTTGTGGTGAA

>M6-PGSC0003DMG400007684M6LRR-RLP_882

TGTTCATTACCAAAGTTGGAGAATTTCACTTACTCTTATAATTATTTTTGTGGTGAACCAAAGATTTGTCTCAAGTTGAAGGATAGAAATGATAAGAAAAATTGTATACCATATCGACCA

>M6-PGSC0003DMG400007684M6LRR-RLP_945

ATTTGTCTCAAGTTGAAGGATAGAAATGATAAGAAAAATTGTATACCATATCGACCATCACAACGTTCAGAAAGTGAATGTAAGACATTTTATTCACGAGGTCCGGTGGATTGTAGTTCA

>M6-PGSC0003DMG400007709M6LRR-RLK_0

TTTTCTCCATCTGTCTCTAACTATTCCTCAAAGTTTTATCTGAGTAACGACTTCTCAAATGCAGATCATTGTTATCCAGAGGTATGCTTGGGTCACCTGAAAAGGTACACCCTTAAGGAA

>M6-PGSC0003DMG400007709M6LRR-RLK_124

GGACTGCCACTGACCATTTCAGCTCAAAACGTATTTTGGGAAAGGGAGGATTTGGGGTTGTGTACAAGGCCAGCTTAAACAATGGTACTGTTGTGGCCGTCAAACGATTGAACGACTACA

>M6-PGSC0003DMG400007709M6LRR-RLK_186

TACAAGGCCAGCTTAAACAATGGTACTGTTGTGGCCGTCAAACGATTGAACGACTACAATGCTGTTGGTGGTGAAAATCAATTTCAGACTGAAGTAGAATTGATTAGTTTGGCTGTCCAT

>M6-PGSC0003DMG400007709M6LRR-RLK_248

TGTTGGTGGTGAAAATCAATTTCAGACTGAAGTAGAATTGATTAGTTTGGCTGTCCATCGGAATCTTCTCCATCTTCTGGGGTTTTGTTCAACTGAAAGTGAACGGCTTCTTGTTTACCC

>M6-PGSC0003DMG400007709M6LRR-RLK_309

AATCTTCTCCATCTTCTGGGGTTTTGTTCAACTGAAAGTGAACGGCTTCTTGTTTACCCCTATATGCCAAATGGAAGTGTAGCATCACGATTAAAAGGTTAGCTCTTGCCTCCATGTGTT

>M6-PGSC0003DMG400007709M6LRR-RLK_62

AGATCATTGTTATCCAGAGGTATGCTTGGGTCACCTGAAAAGGTACACCCTTAAGGAACTGCGGACTGCCACTGACCATTTCAGCTCAAAACGTATTTTGGGAAAGGGAGGATTTGGGGT

>M6-PGSC0003DMG400007862-(rev)M6LRR-RLK_0

AGTTGAGGCCACTGTTATGATATGTGAATACTAATTGTGGTCATCTCCTGTGGCAGAGATCTGAGCAACAACCATATTGGCGGAAGCATTCCGTCTAATCTGCCAGTTACCTTGCGGAAT

>M6-PGSC0003DMG400007862-(rev)M6LRR-RLK_1037

TAATATCACTGTATCCCTAGTCGTTTGTTGGAGATATCATTTGACATGTAACATATCCCACAAAAAAGATATATAATCATAGAAGACAATAGGCATGTATGTCGAAACCGCGGAAACAAA

>M6-PGSC0003DMG400007862-(rev)M6LRR-RLK_1098

AAAAAAGATATATAATCATAGAAGACAATAGGCATGTATGTCGAAACCGCGGAAACAAATATCTTTTCAAGTTCAATGTTACTTAATACTGCTCCTTATTTTCCAATAATGCCTCACTTT

>M6-PGSC0003DMG400007862-(rev)M6LRR-RLK_1159

TCTTTTCAAGTTCAATGTTACTTAATACTGCTCCTTATTTTCCAATAATGCCTCACTTTTGTACTATCTACCAGTGGTTATAACCAAACATATAACTTTTCTAGCTCCCAATACCCCCCT

>M6-PGSC0003DMG400007862-(rev)M6LRR-RLK_122

GTAAGTTCTACCATTTGACTTATGAAGGATCCCCAGAAACACGGGAAAAAGAAAGAAAAGAAAGAGGACATATTGCTAAACCTAGCTTTACTTATTGACTAAAACAGTTGCTATTCTTCA

>M6-PGSC0003DMG400007862-(rev)M6LRR-RLK_1220

TACTATCTACCAGTGGTTATAACCAAACATATAACTTTTCTAGCTCCCAATACCCCCCTTTAAAGCTTGAGAACTTGAATCGTCTGTGCTATGCTTCATACACACAAGTAAACCTTTCAG

>M6-PGSC0003DMG400007862-(rev)M6LRR-RLK_1281

AAAGCTTGAGAACTTGAATCGTCTGTGCTATGCTTCATACACACAAGTAAACCTTTCAGAAACTTCTTGTTTACAAGTTCCTGAAATCTTACTTAGTAAAAAATGGATGTACTTGTATGC

>M6-PGSC0003DMG400007862-(rev)M6LRR-RLK_1342

ACTTCTTGTTTACAAGTTCCTGAAATCTTACTTAGTAAAAAATGGATGTACTTGTATGCTATGTTGCTTGGACTCGTATTGGATCCTCCCAAAGTAATCCGACACTGGTGCAACAACATA

>M6-PGSC0003DMG400007862-(rev)M6LRR-RLK_1403

TGTTGCTTGGACTCGTATTGGATCCTCCCAAAGTAATCCGACACTGGTGCAACAACATATTTGGAAAGTCCGAGCAACATAGCTTGTATGCTGCATGACCAGAAATATAGACTTAACGTC

>M6-PGSC0003DMG400007862-(rev)M6LRR-RLK_1464

TGGAAAGTCCGAGCAACATAGCTTGTATGCTGCATGACCAGAAATATAGACTTAACGTCTGACATGGATATGAATCCTAAACTTGCCCTTAGCATAAATGAAAGCTCAATGAGTAAAAGT

>M6-PGSC0003DMG400007862-(rev)M6LRR-RLK_1525

ACATGGATATGAATCCTAAACTTGCCCTTAGCATAAATGAAAGCTCAATGAGTAAAAGTAAGAATAAAACAAATAAGAAATTGACCTGAAAAAAGAAAAATAGAATCCTGACAAGCTTGT

>M6-PGSC0003DMG400007862-(rev)M6LRR-RLK_1586

GAATAAAACAAATAAGAAATTGACCTGAAAAAAGAAAAATAGAATCCTGACAAGCTTGTCATTTAAATACATGAATTCTATCATATGGCCAGTCCCTCAAAACTTGGAAGTAAACAACTC

>M6-PGSC0003DMG400007862-(rev)M6LRR-RLK_1647

TTTAAATACATGAATTCTATCATATGGCCAGTCCCTCAAAACTTGGAAGTAAACAACTCATATAGACACAATAAATAGACTGCAGATATAGGTATAGATTTACAAACCGTAGTACAATAT

>M6-PGSC0003DMG400007862-(rev)M6LRR-RLK_1708

ATAGACACAATAAATAGACTGCAGATATAGGTATAGATTTACAAACCGTAGTACAATATGCATATGAAAACATACAAACAGTTAAGTTATGATATTCATGACCTGTAAAATAGACAAAAT

>M6-PGSC0003DMG400007862-(rev)M6LRR-RLK_1769

ATATGAAAACATACAAACAGTTAAGTTATGATATTCATGACCTGTAAAATAGACAAAATTGCAAGTTATTTAGTTTAGCAAGCTGGCTCCTGGTCTAAACTTGATCATAATTTAAGTTCA

>M6-PGSC0003DMG400007862-(rev)M6LRR-RLK_183

AAGAGGACATATTGCTAAACCTAGCTTTACTTATTGACTAAAACAGTTGCTATTCTTCACCTTGGACAGTTTTCTTTCAGCTAACGCATTCACTGGAAGTATCCCTACTTCTTTGTCTTC

>M6-PGSC0003DMG400007862-(rev)M6LRR-RLK_1830

CAAGTTATTTAGTTTAGCAAGCTGGCTCCTGGTCTAAACTTGATCATAATTTAAGTTCACATTCCATAGTGGCTATACTCTAGTTAGGTCACCCACCTTAATCATATATCGAGCCCTTAA

>M6-PGSC0003DMG400007862-(rev)M6LRR-RLK_1891

TTCCATAGTGGCTATACTCTAGTTAGGTCACCCACCTTAATCATATATCGAGCCCTTAACTTTCAGGAGGCAAATAACATGGAAGTCAGGCAGTCTTATTTATTAAGCTGACTTAAATGG

>M6-PGSC0003DMG400007862-(rev)M6LRR-RLK_1952

TTCAGGAGGCAAATAACATGGAAGTCAGGCAGTCTTATTTATTAAGCTGACTTAAATGGTACAAGTTGGTAGATATATTTGATATATGCAAAGATCAATGTATCAGTTGTTGGATCATTT

>M6-PGSC0003DMG400007862-(rev)M6LRR-RLK_2013

CAAGTTGGTAGATATATTTGATATATGCAAAGATCAATGTATCAGTTGTTGGATCATTTTGTTACCATCTGATACATGCCTTGTCGATTATGATCAGTGATGGTTTGTACTGTCTTAGTT

>M6-PGSC0003DMG400007862-(rev)M6LRR-RLK_2074

TTACCATCTGATACATGCCTTGTCGATTATGATCAGTGATGGTTTGTACTGTCTTAGTTACCGGTTTTGTTTTTCGTGCAGAAAAGATGGAAATCTTTTCAACAGTAGTACTGCTCCTTT

>M6-PGSC0003DMG400007862-(rev)M6LRR-RLK_2135

CGGTTTTGTTTTTCGTGCAGAAAAGATGGAAATCTTTTCAACAGTAGTACTGCTCCTTTACCTCCACCCACATCATCAACAGCACCTCCCATTTCTGGGTCTCCAACTTCAGCACAGATG

>M6-PGSC0003DMG400007862-(rev)M6LRR-RLK_2196

CTCCACCCACATCATCAACAGCACCTCCCATTTCTGGGTCTCCAACTTCAGCACAGATGCCATCTTCTCAAAGCAAACCTAAAACACAGGCCGATGGACCATCAGCATCTGCAGAATCAA

>M6-PGSC0003DMG400007862-(rev)M6LRR-RLK_2257

ATCTTCTCAAAGCAAACCTAAAACACAGGCCGATGGACCATCAGCATCTGCAGAATCAAGTTCAACAGGAAAAAAAAAGTCTTTATCTGTTAAAAGGGTGGTCTGGATATCAATTGCAGC

>M6-PGSC0003DMG400007862-(rev)M6LRR-RLK_2318

TCAACAGGAAAAAAAAAGTCTTTATCTGTTAAAAGGGTGGTCTGGATATCAATTGCAGCAGTTTTGTCATTTATCATATTGGTTTTAGCAATCCTTCTCTGTCTGCCAAAGTGCCTCAGG

>M6-PGSC0003DMG400007862-(rev)M6LRR-RLK_2379

TTTTGTCATTTATCATATTGGTTTTAGCAATCCTTCTCTGTCTGCCAAAGTGCCTCAGGAAATGGCACGAGACTCAAAGAACCCCCAAACACCATGAAATAGCCCCTTATATGGGTGCTA

>M6-PGSC0003DMG400007862-(rev)M6LRR-RLK_244

TTGGACAGTTTTCTTTCAGCTAACGCATTCACTGGAAGTATCCCTACTTCTTTGTCTTCCCTGAGTCAACTTTCAGCTATGTAAGTATCATTTATAGTGTAGATTAAAAGTTGCTTTCTT

>M6-PGSC0003DMG400007862-(rev)M6LRR-RLK_2440

ATGGCACGAGACTCAAAGAACCCCCAAACACCATGAAATAGCCCCTTATATGGGTGCTAGAGAGAATCTTGGAGATAGTGGTTCATTAGTCCAACCTGGCCATGATATAGAAAAAGGTGC

>M6-PGSC0003DMG400007862-(rev)M6LRR-RLK_2501

GAGAATCTTGGAGATAGTGGTTCATTAGTCCAACCTGGCCATGATATAGAAAAAGGTGCTACAAAATACTCAATATCTATGTTATTTTATGATATCTTTTATCACTGTTTCTTAAAGTTC

>M6-PGSC0003DMG400007862-(rev)M6LRR-RLK_305

TGAGTCAACTTTCAGCTATGTAAGTATCATTTATAGTGTAGATTAAAAGTTGCTTTCTTCATGAAGAATCAAACTCAGGTATTTCGTTGTGTGTCAGGTCTCTGAATGGCAACCATTTGA

>M6-PGSC0003DMG400007862-(rev)M6LRR-RLK_3172

TTAGAACTGAAAAGTTAATGAATCTCAAAGTTTAACCTTCTACTCTTTAGTCTTTAGTGATAAGTGGCTTTCTTATGTGTTAAACAGCTCCCCTAGTTGTGACGCCAAAAGAAAAGCAAC

>M6-PGSC0003DMG400007862-(rev)M6LRR-RLK_3233

AAGTGGCTTTCTTATGTGTTAAACAGCTCCCCTAGTTGTGACGCCAAAAGAAAAGCAACCAAGAAGACCGGCTCCTATTCCAAAGCCACAAGATGAGCAGCAGGTAAATGTGCAAACCAT

>M6-PGSC0003DMG400007862-(rev)M6LRR-RLK_3294

AGAAGACCGGCTCCTATTCCAAAGCCACAAGATGAGCAGCAGGTAAATGTGCAAACCATGAGTGCTGTGCCAAAGAAGGATGATAGTGAGATAAACCTTGGCAGATTGGATATTGATTTG

>M6-PGSC0003DMG400007862-(rev)M6LRR-RLK_3477

GTCGAGCCGATTGCGCCTGCTGAAGATACAGCAACGAGGCTTCCAACCAGGCCACTTCCACTAACTTCTGTGAAATCTTACACAATTGCATCACTTCAGCAATACACTAATAGCTTCTCT

>M6-PGSC0003DMG400007862-(rev)M6LRR-RLK_3538

TAACTTCTGTGAAATCTTACACAATTGCATCACTTCAGCAATACACTAATAGCTTCTCTCAAGATAATCTGTTGGGATCAGGAATGCTGGGTAGTGTATACAGGGTGGAGCTTCCTAATG

>M6-PGSC0003DMG400007862-(rev)M6LRR-RLK_3599

AGATAATCTGTTGGGATCAGGAATGCTGGGTAGTGTATACAGGGTGGAGCTTCCTAATGGAAAGGTATAACTTTCAATCGGCATGAATATATTATGATGTACCAATTTATTAACCGTGGT

>M6-PGSC0003DMG400007862-(rev)M6LRR-RLK_366

TGAAGAATCAAACTCAGGTATTTCGTTGTGTGTCAGGTCTCTGAATGGCAACCATTTGACTGGAGAAATTCCTGATTCCTTTCAAGGCCTTACCGCTTTGGTCAATCTGTATGAAAGCAA

>M6-PGSC0003DMG400007862-(rev)M6LRR-RLK_3660

AAGGTATAACTTTCAATCGGCATGAATATATTATGATGTACCAATTTATTAACCGTGGTTTCCTACTTCCACTAGATTGCGAGAGTCTTGACTTACCACTTTTTCTCTCCTTGCTTTTAT

>M6-PGSC0003DMG400007862-(rev)M6LRR-RLK_3721

CCTACTTCCACTAGATTGCGAGAGTCTTGACTTACCACTTTTTCTCTCCTTGCTTTTATTTCTCCATCATGTTTGGTATTCTGGAATGCTAAATTATCAGCAAGTAGCATCAAAATTGCC

>M6-PGSC0003DMG400007862-(rev)M6LRR-RLK_3782

CTCCATCATGTTTGGTATTCTGGAATGCTAAATTATCAGCAAGTAGCATCAAAATTGCCTCTTTATCTCTAGTATAAATTGTTACATTGCTTAGATTTATCAAGAATGGTTGTGATACAA

>M6-PGSC0003DMG400007862-(rev)M6LRR-RLK_3843

TTTATCTCTAGTATAAATTGTTACATTGCTTAGATTTATCAAGAATGGTTGTGATACAAGCACAATTATATGCAAAAAGGCTGTTTTTAATGGAAAATTCACCTTCATATATCCTTAATT

>M6-PGSC0003DMG400007862-(rev)M6LRR-RLK_3904

ACAATTATATGCAAAAAGGCTGTTTTTAATGGAAAATTCACCTTCATATATCCTTAATTAGAGCACTCCTTAGCAAATCTATTAAATGAGTTCTACCTATTGTATCTGATATAATGTTTC

>M6-PGSC0003DMG400007862-(rev)M6LRR-RLK_3965

AGCACTCCTTAGCAAATCTATTAAATGAGTTCTACCTATTGTATCTGATATAATGTTTCTTGAATTACTTTAGCTAGTTGAGACCTGCTTAAGGACAAAAGTTGTAATTGCAATATTTTT

>M6-PGSC0003DMG400007862-(rev)M6LRR-RLK_4026

GAATTACTTTAGCTAGTTGAGACCTGCTTAAGGACAAAAGTTGTAATTGCAATATTTTTCCTCTAGGTTTCTTCTAGTTCTTTTCTTTTCCTCCTTGTTCCAAATTAATCTTCAACTAGT

>M6-PGSC0003DMG400007862-(rev)M6LRR-RLK_4087

TCTAGGTTTCTTCTAGTTCTTTTCTTTTCCTCCTTGTTCCAAATTAATCTTCAACTAGTATTTTTTTTTTCCAGTAACCAACATCATCGACGAAATCTTGTCCTAGAAGTGTATTACAAA

>M6-PGSC0003DMG400007862-(rev)M6LRR-RLK_4148

TTTTTTTTTCCAGTAACCAACATCATCGACGAAATCTTGTCCTAGAAGTGTATTACAAATTTGTTAAAATGAAAAACAGCCTTCTAGCTATATGATATTACTATTTTTCTGCCTTGACTT

>M6-PGSC0003DMG400007862-(rev)M6LRR-RLK_4209

TGTTAAAATGAAAAACAGCCTTCTAGCTATATGATATTACTATTTTTCTGCCTTGACTTTTGTCTCACTTTATAAGAGGAACCGCCAGATACTGCTGCATCTTTTTTCTGGTCATGATTT

>M6-PGSC0003DMG400007862-(rev)M6LRR-RLK_427

GGAGAAATTCCTGATTCCTTTCAAGGCCTTACCGCTTTGGTCAATCTGTATGAAAGCAAACAAAAGCATTTACATACTTCCATTTAGTTTTTTTCATATGAATAACTGGTTTGGGTGTTT

>M6-PGSC0003DMG400007862-(rev)M6LRR-RLK_4270

GTCTCACTTTATAAGAGGAACCGCCAGATACTGCTGCATCTTTTTTCTGGTCATGATTTTAACTATTAAAGTAAGTATATGCATGCTGTTTCATTTATATAAATTCCTTGGGGATGTATA

>M6-PGSC0003DMG400007862-(rev)M6LRR-RLK_4331

ACTATTAAAGTAAGTATATGCATGCTGTTTCATTTATATAAATTCCTTGGGGATGTATATGCATTTCTCTGGCATGTAAATTTTACCGCTCTTCCTTCTCTTGCACTATGCACCACATCA

>M6-PGSC0003DMG400007862-(rev)M6LRR-RLK_4392

CATTTCTCTGGCATGTAAATTTTACCGCTCTTCCTTCTCTTGCACTATGCACCACATCATTCAAGTGGTTGCAACATTTCACTTCTGACACATGCTCCTTTTGGATGCTTCTGCAGTTGC

>M6-PGSC0003DMG400007862-(rev)M6LRR-RLK_4453

CAAGTGGTTGCAACATTTCACTTCTGACACATGCTCCTTTTGGATGCTTCTGCAGTTGCTTGCTGTCAAAAAACTGGACAAAAGAGTTTGTGATCAACAAAAGGATGATGAGTTTCTTGA

>M6-PGSC0003DMG400007862-(rev)M6LRR-RLK_4514

GCTGTCAAAAAACTGGACAAAAGAGTTTGTGATCAACAAAAGGATGATGAGTTTCTTGACCTAGTAAACAATATTGACAGAATTCGTCATGCTAATGTTGTTGAGCTGATGGGCTATTGT

>M6-PGSC0003DMG400007862-(rev)M6LRR-RLK_4575

TAGTAAACAATATTGACAGAATTCGTCATGCTAATGTTGTTGAGCTGATGGGCTATTGTGCAGAGCATGGTCAAAGGCTTCTGGTCTATGAGTATTGCTGTAACGGGACACTACAAGATG

>M6-PGSC0003DMG400007862-(rev)M6LRR-RLK_4636

AGAGCATGGTCAAAGGCTTCTGGTCTATGAGTATTGCTGTAACGGGACACTACAAGATGCACTGCATTTCGACGATGAATTCAAGAGACAACTTTCATGGAATACTCGCATGAGGATGGC

>M6-PGSC0003DMG400007862-(rev)M6LRR-RLK_4697

CTGCATTTCGACGATGAATTCAAGAGACAACTTTCATGGAATACTCGCATGAGGATGGCTCTTGGAGCTGCAAGAGCTTTAGAGTATGTCAAATAACATTACATACATTTAATTATGCAC

>M6-PGSC0003DMG400007862-(rev)M6LRR-RLK_4758

TTGGAGCTGCAAGAGCTTTAGAGTATGTCAAATAACATTACATACATTTAATTATGCACATCATGTTTGTACACGCTCGCGTTCATGCGTGTGTGTATTAGTGTTTTGGGACCTATAATC

>M6-PGSC0003DMG400007862-(rev)M6LRR-RLK_4819

CATGTTTGTACACGCTCGCGTTCATGCGTGTGTGTATTAGTGTTTTGGGACCTATAATCAGTGTTTCTGCACATACCATATGATAAATCAGTGCTATTAAACACTAAACAATAAAATGGT

>M6-PGSC0003DMG400007862-(rev)M6LRR-RLK_488

AAAAGCATTTACATACTTCCATTTAGTTTTTTTCATATGAATAACTGGTTTGGGTGTTTATCTTTCAACAGTGATTTGTCAAGTAACAATTTAAGTGGATCATTGCCACCATCTGTAGGA

>M6-PGSC0003DMG400007862-(rev)M6LRR-RLK_4880

TGTTTCTGCACATACCATATGATAAATCAGTGCTATTAAACACTAAACAATAAAATGGTTCTGATATAGTGGCAAATTCAATAACAAGCTCATTTTAGTCCTAACTTTCTTGTGTTGTGT

>M6-PGSC0003DMG400007862-(rev)M6LRR-RLK_4941

TGATATAGTGGCAAATTCAATAACAAGCTCATTTTAGTCCTAACTTTCTTGTGTTGTGTATATTACATGATAGAAAGCCACACAAACGACTTGGAATGGTCTTAACTATATGTAAAATGT

>M6-PGSC0003DMG400007862-(rev)M6LRR-RLK_5002

ATTACATGATAGAAAGCCACACAAACGACTTGGAATGGTCTTAACTATATGTAAAATGTTAGATGTAGCTGTCTTTTGCTTCCTCTCCTTTCTTTGCTTCTGAATGTTCAAGAAAAAGAT

>M6-PGSC0003DMG400007862-(rev)M6LRR-RLK_5063

GATGTAGCTGTCTTTTGCTTCCTCTCCTTTCTTTGCTTCTGAATGTTCAAGAAAAAGATGAAGAAAATTAAATTAATTGGTTTGCCACGCTATGCTGTAAAAGGGAAGCCCTATGCACAA

>M6-PGSC0003DMG400007862-(rev)M6LRR-RLK_5307

GAGACAACTTCACCGTTGCTAAGCAGTCTGATTCTCATTTCAAGAATTGCATGAAAAAGTGAGGTAGACTGCTAGAGCATGTTTGTAATTAATAATTATTCTCGTTTGATGCAATGCAGG

>M6-PGSC0003DMG400007862-(rev)M6LRR-RLK_5368

AGGTAGACTGCTAGAGCATGTTTGTAATTAATAATTATTCTCGTTTGATGCAATGCAGGTATCTCCATGAAGTCTGTGAGCCACCTATTATTCATAGAAACTTCAAGTCTGCCAATCTTC

>M6-PGSC0003DMG400007862-(rev)M6LRR-RLK_5429

TCTCCATGAAGTCTGTGAGCCACCTATTATTCATAGAAACTTCAAGTCTGCCAATCTTCTGCTTGATGAGGAGTTGGCTGTACATGTTTCAGATTGTGGTTTGGCTCCACTTATGTCATC

>M6-PGSC0003DMG400007862-(rev)M6LRR-RLK_549

CTTTCAACAGTGATTTGTCAAGTAACAATTTAAGTGGATCATTGCCACCATCTGTAGGAAATTTGTCTTCATTGACCACTCTGTGAGTAATTCAACATATTATACAGTTTGTTTGTAACT

>M6-PGSC0003DMG400007862-(rev)M6LRR-RLK_5490

CTTGATGAGGAGTTGGCTGTACATGTTTCAGATTGTGGTTTGGCTCCACTTATGTCATCTGGTGCTGTGAGTCAGGTAACTTGTGACCCTAATGTTATGGAGTAGAAAGAATTGAAATGC

>M6-PGSC0003DMG400007862-(rev)M6LRR-RLK_5551

GTGCTGTGAGTCAGGTAACTTGTGACCCTAATGTTATGGAGTAGAAAGAATTGAAATGCTTCCTTGGATTATTAGTTTCCTCTATGAGCTGCAAGTGGAAAGAAAAATGGATTCCTAGTG

>M6-PGSC0003DMG400007862-(rev)M6LRR-RLK_5612

CCTTGGATTATTAGTTTCCTCTATGAGCTGCAAGTGGAAAGAAAAATGGATTCCTAGTGTTTGATAGAAGTGGAATGGGAGCTGAATAAATAATTTAAAAAAGGAACAGTAGTGATGGTC

>M6-PGSC0003DMG400007862-(rev)M6LRR-RLK_5673

TGATAGAAGTGGAATGGGAGCTGAATAAATAATTTAAAAAAGGAACAGTAGTGATGGTCAAAGTGTTTTCAACAAGATGTTGGAATAAGGAGCAGTAATAATGAATTCCTGTAAATGTGT

>M6-PGSC0003DMG400007862-(rev)M6LRR-RLK_5734

AGTGTTTTCAACAAGATGTTGGAATAAGGAGCAGTAATAATGAATTCCTGTAAATGTGTAAAGTATATTCTTTGGGAAATGGTTGGTTCTCACCCTGTAGGATAGGGTGAAGTGTTGTCA

>M6-PGSC0003DMG400007862-(rev)M6LRR-RLK_5795

AGTATATTCTTTGGGAAATGGTTGGTTCTCACCCTGTAGGATAGGGTGAAGTGTTGTCAAAGGTGCGCTTAAAGCACGCTTGAACCCTGAAGTGAGGCTCAAAATGTGTTGAGCATTTCA

>M6-PGSC0003DMG400007862-(rev)M6LRR-RLK_6039

AATGGTGAAGGTGCTCTTCTCATGATGTGTTTTATATGAACAGAAAATTTCTCTAGGATGCATGTGGATCATCTATCTTGCTGTTATTTGTTGTTCTTTTGATAAAGTAGTCTTGGGATT

>M6-PGSC0003DMG400007862-(rev)M6LRR-RLK_61

TGAGCAACAACCATATTGGCGGAAGCATTCCGTCTAATCTGCCAGTTACCTTGCGGAATATGTAAGTTCTACCATTTGACTTATGAAGGATCCCCAGAAACACGGGAAAAAGAAAGAAAA

>M6-PGSC0003DMG400007862-(rev)M6LRR-RLK_610

TTTGTCTTCATTGACCACTCTGTGAGTAATTCAACATATTATACAGTTTGTTTGTAACTTTATAGTTTGATCAATATGATAAAACTAACTTCTTGATTTCCAGATATTTGCAGAATAATC

>M6-PGSC0003DMG400007862-(rev)M6LRR-RLK_6100

ATGTGGATCATCTATCTTGCTGTTATTTGTTGTTCTTTTGATAAAGTAGTCTTGGGATTGATTACTGTTACGTTCAATTTAAGGTATGTTAAAAACTATATCAAGTTGACATTACACTGA

>M6-PGSC0003DMG400007862-(rev)M6LRR-RLK_6161

TTACTGTTACGTTCAATTTAAGGTATGTTAAAAACTATATCAAGTTGACATTACACTGATTTTTCAGTTGTCGGGACAACTTCTCACAACATATGGCTACGGTGCTCCCGAATTTGAGTC

>M6-PGSC0003DMG400007862-(rev)M6LRR-RLK_6222

TTTCAGTTGTCGGGACAACTTCTCACAACATATGGCTACGGTGCTCCCGAATTTGAGTCAGGGATTTACACTTCCCAAAGTGATGTTTACAGTTTTGGTGTGGTGATGCTGGAACTAATA

>M6-PGSC0003DMG400007862-(rev)M6LRR-RLK_6283

GGATTTACACTTCCCAAAGTGATGTTTACAGTTTTGGTGTGGTGATGCTGGAACTAATAACAGGCAGAATGTCATATGACCGGTAAAAAAGCCTTCTGCATCTTTTTGAGCTAATATAGT

>M6-PGSC0003DMG400007862-(rev)M6LRR-RLK_6344

AGGCAGAATGTCATATGACCGGTAAAAAAGCCTTCTGCATCTTTTTGAGCTAATATAGTTGTGCCTCATTTCTCAGTTAACACACCTGCATATCATATTCAGGACACGAAGTCGAGGAGA

>M6-PGSC0003DMG400007862-(rev)M6LRR-RLK_6405

TGCCTCATTTCTCAGTTAACACACCTGCATATCATATTCAGGACACGAAGTCGAGGAGAGCAGTTCTTGGTCAGGTGGGCAATCCCTCAGCTACATGATATTGATCTCTTGACAAGGATG

>M6-PGSC0003DMG400007862-(rev)M6LRR-RLK_6466

AGTTCTTGGTCAGGTGGGCAATCCCTCAGCTACATGATATTGATCTCTTGACAAGGATGGTTGATCCTGCTCTCGATGGAAAATACCCCATTAAATCACTATCACACTTTGCTGACATAA

>M6-PGSC0003DMG400007862-(rev)M6LRR-RLK_6527

TGATCCTGCTCTCGATGGAAAATACCCCATTAAATCACTATCACACTTTGCTGACATAATTTCCCGCTGTGTTCAGGTGAGTTATTCTGTTGTTCCTTCATGTGAAACACATAAATGCTT

>M6-PGSC0003DMG400007862-(rev)M6LRR-RLK_6588

TCCCGCTGTGTTCAGGTGAGTTATTCTGTTGTTCCTTCATGTGAAACACATAAATGCTTACTAATATGCATTGTCTAATCGCTGAGTGTACCGAATAGTCTTAGATTGTTCGTCATCTCA

>M6-PGSC0003DMG400007862-(rev)M6LRR-RLK_6649

TAATATGCATTGTCTAATCGCTGAGTGTACCGAATAGTCTTAGATTGTTCGTCATCTCAATCGGTTGGAAGGATATTTCTAGGATCTAGTTGTGGAATAGGTTTGCCTTCAAATGTCACT

>M6-PGSC0003DMG400007862-(rev)M6LRR-RLK_671

ATAGTTTGATCAATATGATAAAACTAACTTCTTGATTTCCAGATATTTGCAGAATAATCAGCTTTCTGGAACTCTTGATGTTTTGCAGGATCTTCCCCTTGCAAATTTGTACTCTTCCAA

>M6-PGSC0003DMG400007862-(rev)M6LRR-RLK_6710

CGGTTGGAAGGATATTTCTAGGATCTAGTTGTGGAATAGGTTTGCCTTCAAATGTCACTTAACTTTTAAACTCTTATACAGCATGAGCGAGAATTCAGGCCACCAATGTCTGAAGTTGTT

>M6-PGSC0003DMG400007862-(rev)M6LRR-RLK_6771

ACTTTTAAACTCTTATACAGCATGAGCGAGAATTCAGGCCACCAATGTCTGAAGTTGTTCAGGACCTCATACAAATGATAAGAAGAGAGTCTCACAAAAGATCTGATGGTGAATGATTTG

>M6-PGSC0003DMG400007862-(rev)M6LRR-RLK_6832

GGACCTCATACAAATGATAAGAAGAGAGTCTCACAAAAGATCTGATGGTGAATGATTTGAGTTAGACGCGCATACTGGGAATTCAGCCGATGAGAAACTAAGATTATTTCTTCTTCGAAG

>M6-PGSC0003DMG400007862-(rev)M6LRR-RLK_6893

TTAGACGCGCATACTGGGAATTCAGCCGATGAGAAACTAAGATTATTTCTTCTTCGAAGGAGCTGAGATACTCGGAAAAAGAAAGGCGGTTTGCAGGCCAAGCACCGTGCTAAAGACAAC

>M6-PGSC0003DMG400007862-(rev)M6LRR-RLK_6954

GCTGAGATACTCGGAAAAAGAAAGGCGGTTTGCAGGCCAAGCACCGTGCTAAAGACAACTTCAAACTGAAGCAAAATGCACCTCAAAGAAGCGCGATTCATGAAAATACCAACACATGCA

>M6-PGSC0003DMG400007862-(rev)M6LRR-RLK_7015

CAAACTGAAGCAAAATGCACCTCAAAGAAGCGCGATTCATGAAAATACCAACACATGCAGGCGACTTAATGACCTGTAAAAACAGCTTGCTCATCTGATTCTTGTCAAGCTGCATCATTT

>M6-PGSC0003DMG400007862-(rev)M6LRR-RLK_7076

CGACTTAATGACCTGTAAAAACAGCTTGCTCATCTGATTCTTGTCAAGCTGCATCATTTTAATGTTCATTTAGGCTTACATGTGAGCACCAATGCATATAATTCATTGATCTTGACAAAA

>M6-PGSC0003DMG400007862-(rev)M6LRR-RLK_7137

ATGTTCATTTAGGCTTACATGTGAGCACCAATGCATATAATTCATTGATCTTGACAAAATGGTATATTTTCTTGAAGTTGTATATCTTTACTTGCTGCAACATGTACAAACTTCATTTGG

>M6-PGSC0003DMG400007862-(rev)M6LRR-RLK_7198

GTATATTTTCTTGAAGTTGTATATCTTTACTTGCTGCAACATGTACAAACTTCATTTGGTTGTACAAGGATACCTTCACTCTTTCTCATGTACTTGTTGCACATGCCAAAGTTATTTACT

>M6-PGSC0003DMG400007862-(rev)M6LRR-RLK_7259

GTACAAGGATACCTTCACTCTTTCTCATGTACTTGTTGCACATGCCAAAGTTATTTACTACTAGTCATTACCTAAGGAGTTGGAGACCTACAACCACTCAACCATGTAACAATATGTAAA

>M6-PGSC0003DMG400007862-(rev)M6LRR-RLK_732

CTTTCTGGAACTCTTGATGTTTTGCAGGATCTTCCCCTTGCAAATTTGTACTCTTCCAAACTAGATTTTACTCAAATATAATTCAGTTCACTAGAATGATAGAATCTTACTAACATCTCT

>M6-PGSC0003DMG400007862-(rev)M6LRR-RLK_793

TAGATTTTACTCAAATATAATTCAGTTCACTAGAATGATAGAATCTTACTAACATCTCTCCACCCTCCCTGTGTCCTTTCTAGGAATGTGGAAAACAACTTGTTTTCTGGGCCCATACCT

>M6-PGSC0003DMG400007862-(rev)M6LRR-RLK_854

ACCCTCCCTGTGTCCTTTCTAGGAATGTGGAAAACAACTTGTTTTCTGGGCCCATACCTCAAAAGTTGTTCTCTATTCCCAATTTCAAGTAAGTACATTCATTTTATGCTTTCATATACC

>M6-PGSC0003DMG400007862-(rev)M6LRR-RLK_915

AAAGTTGTTCTCTATTCCCAATTTCAAGTAAGTACATTCATTTTATGCTTTCATATACCCGTTTCATTTTTGTTTATGTTGAGTCAACATCTTGTATCTTAAAATGTAATCTTGGTCACA

>M6-PGSC0003DMG400007862-(rev)M6LRR-RLK_976

TTTCATTTTTGTTTATGTTGAGTCAACATCTTGTATCTTAAAATGTAATCTTGGTCACATTTAATATCACTGTATCCCTAGTCGTTTGTTGGAGATATCATTTGACATGTAACATATCCC

>M6-PGSC0003DMG400007876-(rev)M6LRR-RLP_0

ATGGAAGGTGTAGCTTTAGTACTTTCACTTCTCCTTGTTGTTGTTAATTTAGATGGATTTTCTTGTGTTTATTCAGGAAGTCATTGCATTGAGAGTGAAAAAAATGCTCTTATTAAGTTC

>M6-PGSC0003DMG400007876-(rev)M6LRR-RLP_1054

ACTCTTTCAAGGACTCGATTACAGAACATTTCTTGAATTTAACTCATTTGAGAGTGTTAGACTTGTCATCCAACTCATTTATTTTCAATGTAAGCGCAACTTGGATGCCTCGTTTTCAAC

>M6-PGSC0003DMG400007876-(rev)M6LRR-RLP_1116

TTGTCATCCAACTCATTTATTTTCAATGTAAGCGCAACTTGGATGCCTCGTTTTCAACTTGACTCTCTAAGCCTGCAGTCTTGCGGACTTGGTGCACAGTTTCCCCAGTGGATTCAGACA

>M6-PGSC0003DMG400007876-(rev)M6LRR-RLP_1178

CTCTCTAAGCCTGCAGTCTTGCGGACTTGGTGCACAGTTTCCCCAGTGGATTCAGACACAAAAAGAGTTGTCATTTATTGATATCTCTCGCACCAATATCTCAGGTGAAGTGCCTGATTG

>M6-PGSC0003DMG400007876-(rev)M6LRR-RLP_124

AAGGTTTCAAGAATCCGTCACGAAGCATGTTATCTTGGATGTTTGACGAAAACTGCTGCAGCTGGAAAGGTGTTGAATGTGACAACAAAACCGGGCACGTGATCACTCTTGATCTACATA

>M6-PGSC0003DMG400007876-(rev)M6LRR-RLP_1240

AAGAGTTGTCATTTATTGATATCTCTCGCACCAATATCTCAGGTGAAGTGCCTGATTGGTTCTGGAATTTTTCTGCCAAGGTTTATCACATAGACCTTTCAGGAAACAATTTCAGTGGAG

>M6-PGSC0003DMG400007876-(rev)M6LRR-RLP_1302

TGGAATTTTTCTGCCAAGGTTTATCACATAGACCTTTCAGGAAACAATTTCAGTGGAGAAATCCCTGAATTTACAGAAAGAGTTTGTCTTACAAAATTAGACCTGAGTGATAACAATTTT

>M6-PGSC0003DMG400007876-(rev)M6LRR-RLP_1364

CCCTGAATTTACAGAAAGAGTTTGTCTTACAAAATTAGACCTGAGTGATAACAATTTTCACGGTCCTTTGCCACATTTTTCACCCAAGATGATGACATTGATTTTGGCAAGGAATTCGTT

>M6-PGSC0003DMG400007876-(rev)M6LRR-RLP_1426

GTCCTTTGCCACATTTTTCACCCAAGATGATGACATTGATTTTGGCAAGGAATTCGTTTAATGGTACCATTGCTCCTGTATGTGAATCACTTGTCATGAATAACTCTCTGAGTCTTCTAA

>M6-PGSC0003DMG400007876-(rev)M6LRR-RLP_1488

GGTACCATTGCTCCTGTATGTGAATCACTTGTCATGAATAACTCTCTGAGTCTTCTAAACCTATCTTCAAATTTTCTATCAGGACAACTTTCAGATTGTTGGAGATATGGGAAGAATCTG

>M6-PGSC0003DMG400007876-(rev)M6LRR-RLP_1550

ATCTTCAAATTTTCTATCAGGACAACTTTCAGATTGTTGGAGATATGGGAAGAATCTGCAAGGATTGAATTTAGGCCATAATGATCTATCAGGGGAGATACCTCATTCGATTGGAGATCT

>M6-PGSC0003DMG400007876-(rev)M6LRR-RLP_1612

GATTGAATTTAGGCCATAATGATCTATCAGGGGAGATACCTCATTCGATTGGAGATCTTACTAACCTTTTCTTTTTGCAATTGCAGAACAACAGTTTCTCCAAAAATCTTCCTTCATCAT

>M6-PGSC0003DMG400007876-(rev)M6LRR-RLP_1674

AACCTTTTCTTTTTGCAATTGCAGAACAACAGTTTCTCCAAAAATCTTCCTTCATCATTGAAGAACATAACTGGACTAAAAATTCTTGATGTTTCTGAAAATAGTTTATCTGGGAATATA

>M6-PGSC0003DMG400007876-(rev)M6LRR-RLP_1736

GAACATAACTGGACTAAAAATTCTTGATGTTTCTGAAAATAGTTTATCTGGGAATATACCATTTTGGTTAGGGGAGAGTTTAAATGCTCTTGAGATTCTTATATTAAGCAGAAACAACTA

>M6-PGSC0003DMG400007876-(rev)M6LRR-RLP_1771

AAAATAGTTTATCTGGGAATATACCATTTTGGTTAGGGGAGAGTTTAAATGCTCTTGAGATTCTTATATTAAGCAGAAACAACTATGATGGAATTATTCCCAGGGAGATTTGCCAGCTTA

>M6-PGSC0003DMG400007876-(rev)M6LRR-RLP_186

TGGAAAGGTGTTGAATGTGACAACAAAACCGGGCACGTGATCACTCTTGATCTACATAAGCAATTCTTGCAAGGTGAGTTCAGGATTTCTTTACTTGGTTTGCCTTATTTAAGGCACTTA

>M6-PGSC0003DMG400007876-(rev)M6LRR-RLP_248

ATTCTTGCAAGGTGAGTTCAGGATTTCTTTACTTGGTTTGCCTTATTTAAGGCACTTAGACATGAGCCACATTGATTTCCTCGGGGAACGGATTCCTGAGTTTATTTCCAGTTTTAAGGA

>M6-PGSC0003DMG400007876-(rev)M6LRR-RLP_310

TGAGCCACATTGATTTCCTCGGGGAACGGATTCCTGAGTTTATTTCCAGTTTTAAGGACCTGGAGTATCTCAATTTGTCTAAAACGAATTTTAGGGGAATGATTCCTGAACATCTTGGGA

>M6-PGSC0003DMG400007876-(rev)M6LRR-RLP_372

GAGTATCTCAATTTGTCTAAAACGAATTTTAGGGGAATGATTCCTGAACATCTTGGGAATCTGTCGCGGTTGCAGTTTCTTGATCTGAGTGGTGATTTTTCACTCCGGGTTAATAACCTT

>M6-PGSC0003DMG400007876-(rev)M6LRR-RLP_434

GTCGCGGTTGCAGTTTCTTGATCTGAGTGGTGATTTTTCACTCCGGGTTAATAACCTTGAATGGATTCACTCCCTCTTCTCTATGAAGATTCTTGATTTGAGTGGTGTTTACATGAGAAG

>M6-PGSC0003DMG400007876-(rev)M6LRR-RLP_496

GGATTCACTCCCTCTTCTCTATGAAGATTCTTGATTTGAGTGGTGTTTACATGAGAAGCGCGGAAAACTGGCTACATGACATCAATTTGTTAAGTTCTCTCTCTGTATTACGTTTGTCTG

>M6-PGSC0003DMG400007876-(rev)M6LRR-RLP_558

GAAAACTGGCTACATGACATCAATTTGTTAAGTTCTCTCTCTGTATTACGTTTGTCTGCTTGTCAGCTTACTTCATTTCCTGAATTGCTTCCGACTTCTGTGAATTTCACATCCCTTCGG

>M6-PGSC0003DMG400007876-(rev)M6LRR-RLP_62

TTGTGTTTATTCAGGAAGTCATTGCATTGAGAGTGAAAAAAATGCTCTTATTAAGTTCAGCCAAGGTTTCAAGAATCCGTCACGAAGCATGTTATCTTGGATGTTTGACGAAAACTGCTG

>M6-PGSC0003DMG400007876-(rev)M6LRR-RLP_620

TCAGCTTACTTCATTTCCTGAATTGCTTCCGACTTCTGTGAATTTCACATCCCTTCGGATTCTTGATCTTTCAATAAACTATTTTAATACTAGGATTCCTAGCTGGTTGTTCAACACAAG

>M6-PGSC0003DMG400007876-(rev)M6LRR-RLP_682

TTGATCTTTCAATAAACTATTTTAATACTAGGATTCCTAGCTGGTTGTTCAACACAAGCCATAATCTTGTTTACCTTAATCTTTCGAGAAGTCAGCTGAGCGGTTCACTTCCAAATGCCT

>M6-PGSC0003DMG400007876-(rev)M6LRR-RLP_744

AATCTTGTTTACCTTAATCTTTCGAGAAGTCAGCTGAGCGGTTCACTTCCAAATGCCTTTGGCAACATGAGTTCACTTAGAGTGCTTGATCTTTCTGGGAACTCTCTCCGAGGCAACCTT

>M6-PGSC0003DMG400007876-(rev)M6LRR-RLP_806

CAACATGAGTTCACTTAGAGTGCTTGATCTTTCTGGGAACTCTCTCCGAGGCAACCTTCCACATTCCTTTGAGAAAATGAGTTCAGTTTCCTTTCTTAATCTCTCAAGAAACTCGTTCAC

>M6-PGSC0003DMG400007876-(rev)M6LRR-RLP_868

ATTCCTTTGAGAAAATGAGTTCAGTTTCCTTTCTTAATCTCTCAAGAAACTCGTTCACGGGCAATTTACCACCAACCCTGCCATCAAAGCTGAAATACTTGGATATTTCTGATAATAAAA

>M6-PGSC0003DMG400007876-(rev)M6LRR-RLP_930

AATTTACCACCAACCCTGCCATCAAAGCTGAAATACTTGGATATTTCTGATAATAAAATGGAGGGTCCTTTAGCAAGAAGCATCACACAACTTAAGCAATTAGTTGTCCTCAATGTTGCT

>M6-PGSC0003DMG400007876-(rev)M6LRR-RLP_992

GGGTCCTTTAGCAAGAAGCATCACACAACTTAAGCAATTAGTTGTCCTCAATGTTGCTCGGAACTCTTTCAAGGACTCGATTACAGAACATTTCTTGAATTTAACTCATTTGAGAGTGTT

>M6-PGSC0003DMG400007899-(rev)M6OtherRLKTM_0

TCGCCGGCGGCAGGAATTGCCACCGGAAACACGTAGAAGACGGGTGAATCAATCACCCGTTTTTGTCTTCTCCGTTCACCAAATCTCTTTTTATTAACAGATTCAAACACCCTTTTCAAT

>M6-PGSC0003DMG400007899-(rev)M6OtherRLKTM_1054

ACGCGTTTCATATCATTTCGTTATCTGAAGGGTTTGATTTGTCACCATTGTTCGAAGAGAATAAGAGGAATGAGAAAGAACAGATGAGATTCGCAACTACTAGGTCAGCAAGTAGTGTGA

>M6-PGSC0003DMG400007899-(rev)M6OtherRLKTM_1116

AAGAGGAATGAGAAAGAACAGATGAGATTCGCAACTACTAGGTCAGCAAGTAGTGTGATATCTAAGCTTGAGGAGGTAGCAAAGACATCGAATTTCAGCGTAAAAAAGAGTGATTCATGT

>M6-PGSC0003DMG400007899-(rev)M6OtherRLKTM_1178

TAAGCTTGAGGAGGTAGCAAAGACATCGAATTTCAGCGTAAAAAAGAGTGATTCATGTGTAAGGTTACAAGGACAAGTGACTGGGAGAAAAGGGAAATTGGGAATTGCTGCTGATATATT

>M6-PGSC0003DMG400007899-(rev)M6OtherRLKTM_124

AGTCTCTTTTTTCAATCAATTTGGAACAAAAACAAATGGCTCCGGAAGAGAAATGTGGAGCTTTGAACGGAAAATATGAACTTGGTCGGCTTTTAGGCCATGGAACATTCGCGAAAGTTT

>M6-PGSC0003DMG400007899-(rev)M6OtherRLKTM_1240

GGTTACAAGGACAAGTGACTGGGAGAAAAGGGAAATTGGGAATTGCTGCTGATATATTTGCTGTGACAAATTCATTTCTAGTTGTGGAAGTGAATAAAGCTAGTGGTGATACATTGGAGT

>M6-PGSC0003DMG400007899-(rev)M6OtherRLKTM_1299

GCTGTGACAAATTCATTTCTAGTTGTGGAAGTGAATAAAGCTAGTGGTGATACATTGGAGTATAATCAATTCTGCAGAAAAGAGCTTAGACCAGCACTTAAGGATATTGTTTGGACATCA

>M6-PGSC0003DMG400007899-(rev)M6OtherRLKTM_186

TTGAACGGAAAATATGAACTTGGTCGGCTTTTAGGCCATGGAACATTCGCGAAAGTTTATCACGCACGAAACGTGCAGAATGGTAAAAATGTTGCCATGAAAGTAGTGGGGAAAGAGAAA

>M6-PGSC0003DMG400007899-(rev)M6OtherRLKTM_248

CGCACGAAACGTGCAGAATGGTAAAAATGTTGCCATGAAAGTAGTGGGGAAAGAGAAAGTCATTAAAGTCGGTATGATGGATCAGATCAAGCGAGAAATATCCGTAATGAAAATGGTGAA

>M6-PGSC0003DMG400007899-(rev)M6OtherRLKTM_310

TTAAAGTCGGTATGATGGATCAGATCAAGCGAGAAATATCCGTAATGAAAATGGTGAAAAACCCGAATATAGTTGAGCTCCACGAAGTCATGGCGAGTAAAACTAAGATTTACTTCGCCA

>M6-PGSC0003DMG400007899-(rev)M6OtherRLKTM_372

CCGAATATAGTTGAGCTCCACGAAGTCATGGCGAGTAAAACTAAGATTTACTTCGCCATGGAGTTAGTTAGAGGAGGTGAGCTTTTCGCGAAGATCGCGAAAGGCAAGCTGAGAGAGGAT

>M6-PGSC0003DMG400007899-(rev)M6OtherRLKTM_434

GTTAGTTAGAGGAGGTGAGCTTTTCGCGAAGATCGCGAAAGGCAAGCTGAGAGAGGATGTTGCTAGAGGTTACTTCCAGCAACTGATCTCAGCTATCGAATTCTGTCATAGCAGAGGAGT

>M6-PGSC0003DMG400007899-(rev)M6OtherRLKTM_496

CTAGAGGTTACTTCCAGCAACTGATCTCAGCTATCGAATTCTGTCATAGCAGAGGAGTATTTCATCGCGATTTGAAACCTGAGAACTTGTTATTAGATGAAGAAGGAAATCTCAAAGTGA

>M6-PGSC0003DMG400007899-(rev)M6OtherRLKTM_558

CATCGCGATTTGAAACCTGAGAACTTGTTATTAGATGAAGAAGGAAATCTCAAAGTGACTGATTTCGGGCTAAGTGCATTTACAGAACATTTAAGACAGGACGGATTGCTTCACACAACT

>M6-PGSC0003DMG400007899-(rev)M6OtherRLKTM_62

TTGTCTTCTCCGTTCACCAAATCTCTTTTTATTAACAGATTCAAACACCCTTTTCAATCTTTAGTCTCTTTTTTCAATCAATTTGGAACAAAAACAAATGGCTCCGGAAGAGAAATGTGG

>M6-PGSC0003DMG400007899-(rev)M6OtherRLKTM_620

TTTCGGGCTAAGTGCATTTACAGAACATTTAAGACAGGACGGATTGCTTCACACAACTTGTGGAACTCCGGCTTATGTTGCCCCTGAAGTTATTGGTAAAAAAGGCTAAGATGGTTCAAA

>M6-PGSC0003DMG400007899-(rev)M6OtherRLKTM_682

GAACTCCGGCTTATGTTGCCCCTGAAGTTATTGGTAAAAAAGGCTAAGATGGTTCAAAAGCGGATATTTGGTCCTGTGGAGTGATCCTCTATGTTTTATTAGCCGGTTTTTTACCATTTC

>M6-PGSC0003DMG400007899-(rev)M6OtherRLKTM_744

GATATTTGGTCCTGTGGAGTGATCCTCTATGTTTTATTAGCCGGTTTTTTACCATTTCAAGAAGAAAACATCATGGCGATGTATAAGAAGATTTATAGGGGTGATTTCAAATGTCCACCT

>M6-PGSC0003DMG400007899-(rev)M6OtherRLKTM_806

AGAAAACATCATGGCGATGTATAAGAAGATTTATAGGGGTGATTTCAAATGTCCACCTTGGTTTTCTTCAGAAGCTAGGAGGTTGATCACAAAGATGTTGGATCCGAATCCCAATTCAAG

>M6-PGSC0003DMG400007899-(rev)M6OtherRLKTM_868

TTTCTTCAGAAGCTAGGAGGTTGATCACAAAGATGTTGGATCCGAATCCCAATTCAAGAATCACCACCTCGAAGATCATGGATTCGTCATGGTTCAAGAAATCAATCCCAAGGACGTTAA

>M6-PGSC0003DMG400007899-(rev)M6OtherRLKTM_930

ACCACCTCGAAGATCATGGATTCGTCATGGTTCAAGAAATCAATCCCAAGGACGTTAAGGAACAAAGATGAGGAAGAATTTGCATTTGCCTCAGATAAGGCTTGTAAAGAAGTTGAGACA

>M6-PGSC0003DMG400007899-(rev)M6OtherRLKTM_992

CAAAGATGAGGAAGAATTTGCATTTGCCTCAGATAAGGCTTGTAAAGAAGTTGAGACAATGAACGCGTTTCATATCATTTCGTTATCTGAAGGGTTTGATTTGTCACCATTGTTCGAAGA

>M6-PGSC0003DMG400007915-(rev)M6LRR-RLP_128

TTCATCCATTTGTAACATGAGTTCGCTTATCTTATTAGATTTATCACGCAACAACTTCAGTGACTCAGTTCCACATTGCTTGGGAAGCATGGCTAGTCTAACGGTGCTAGACTTAAGAAG

>M6-PGSC0003DMG400007915-(rev)M6LRR-RLP_704

GCAGGATATTGAGCTATAAAGAATCAGCACAATTACGACAACCATAGATCTCTCAAGCAACCATTTTGAAGGTGTCATTCCGAAAACACTAAAGGATCTCATATCACTTTGGCTACTCAA

>M6-PGSC0003DMG400007915-(rev)M6LRR-RLP_960

TTCTTAAACCTCTCTCAAAATCATCTCGTTGGACCAATTCCTCAAGGTCGGCAATTCAACACATTTGAAAATGACTCATATGGTGGCAACCTTGATTTATGTGGTGTTCCTTTAACAAAG

>M6-PGSC0003DMG400007948M6LRR-RLK_0

TTAATAATTAATTCTAAAATAACTTATTATTTTCGACCGTGAAACATTATGGCCATGGCTGACACAGAAACAGAGAGAAAAATAGCCAGACTAAATATTGGAAGTCCCCACTTCCCTTTT

>M6-PGSC0003DMG400007948M6LRR-RLK_1037

CTCCGGCAGAACTCGGCCAAAACGAAGAGTTACGACAGGGTAGCCCGTTAAACCGTAAAGAAAACAAATTACATAAAAGATCCTTACTCATTATTGGAGTTTCAACAGCTTGTTTAGTCC

>M6-PGSC0003DMG400007948M6LRR-RLK_1098

AAACAAATTACATAAAAGATCCTTACTCATTATTGGAGTTTCAACAGCTTGTTTAGTCCTTTTATGTTCAGTTATACTATTAGCATTAGCGTCGAAGAAGCACAGGAACTCGAAGAAATT

>M6-PGSC0003DMG400007948M6LRR-RLK_1159

TTATGTTCAGTTATACTATTAGCATTAGCGTCGAAGAAGCACAGGAACTCGAAGAAATTAGGAGAAACTAAGAAATCCGCATTCGATCCAAGTGTTAGCGGCAATGCAGAAGCAGTAATA

>M6-PGSC0003DMG400007948M6LRR-RLK_122

CTTTCTCAAAAATCTTCACGTGAATCTGACTCTGCCTTTTGCCATATCTTCTTAGACTGCTGCTTTTCTCTTCACTCTTCCACCTCTTTTGTCACGTATTGCACCTCTACACATATTCCA

>M6-PGSC0003DMG400007948M6LRR-RLK_1220

GAGAAACTAAGAAATCCGCATTCGATCCAAGTGTTAGCGGCAATGCAGAAGCAGTAATAAGAATTGAAGAAGACAACAATGAACTGGAAGAGAAGGTGAAGAGAGTACAACAAGGAATGC

>M6-PGSC0003DMG400007948M6LRR-RLK_1281

AATTGAAGAAGACAACAATGAACTGGAAGAGAAGGTGAAGAGAGTACAACAAGGAATGCAGCAAGTGATGGGTATAAGTGGGAGCTTAGTTTTCTGTGCAGGTGAGGTGCAGGTGTATAC

>M6-PGSC0003DMG400007948M6LRR-RLK_1342

CAAGTGATGGGTATAAGTGGGAGCTTAGTTTTCTGTGCAGGTGAGGTGCAGGTGTATACCCTGGAGCAGCTGATGAGAGCTTCTGCTGAGCTTTTAGGTAGAGGAACAATGGGGACTACT

>M6-PGSC0003DMG400007948M6LRR-RLK_1403

TGGAGCAGCTGATGAGAGCTTCTGCTGAGCTTTTAGGTAGAGGAACAATGGGGACTACTTACAAAGCAGTGCTTGATAACCGTCTGATTGTATGTGTTAAGAGATTGGACGGTGGGAGAT

>M6-PGSC0003DMG400007948M6LRR-RLK_1464

CAAAGCAGTGCTTGATAACCGTCTGATTGTATGTGTTAAGAGATTGGACGGTGGGAGATTGGCAGGTACAAGTCAAGAGGAGTTTGAGCAGCATATGGAATCAGTGGGCGGGCTTAGGCA

>M6-PGSC0003DMG400007948M6LRR-RLK_1525

GCAGGTACAAGTCAAGAGGAGTTTGAGCAGCATATGGAATCAGTGGGCGGGCTTAGGCACCCGAATTTGGTTCCTTTTCGGGCTTATTTTCAGGCCCGACAAGAAAGGCTTTTGGTTTAT

>M6-PGSC0003DMG400007948M6LRR-RLK_1586

CGAATTTGGTTCCTTTTCGGGCTTATTTTCAGGCCCGACAAGAAAGGCTTTTGGTTTATGATTATCAACCCAATGGCAGCCTGTCTTCTCTTATTCATGGTACGTTCTTCTCTGTATATC

>M6-PGSC0003DMG400007948M6LRR-RLK_1647

TTATCAACCCAATGGCAGCCTGTCTTCTCTTATTCATGGTACGTTCTTCTCTGTATATCTTGTACTTATTATTTATTAAGATGCGCATAAACTATACGAAAAATATTATTAATGCAATTA

>M6-PGSC0003DMG400007948M6LRR-RLK_1708

GTACTTATTATTTATTAAGATGCGCATAAACTATACGAAAAATATTATTAATGCAATTATTTTGACCTTTGATAAGTAAAATATAATATAATGAACGGAGTATTTTGAAGGTGTTGTCCA

>M6-PGSC0003DMG400007948M6LRR-RLK_1769

TTGACCTTTGATAAGTAAAATATAATATAATGAACGGAGTATTTTGAAGGTGTTGTCCAATCAAATAGGGTCCATACACCATACCCTATTGTTACAGTTTCTCAACCCCAATGTTCTCAC

>M6-PGSC0003DMG400007948M6LRR-RLK_183

GCTTTTCTCTTCACTCTTCCACCTCTTTTGTCACGTATTGCACCTCTACACATATTCCACAACAATTCAATCTGGAAATTTTTTCTTAAACAAGATGAAAATACAACAATTCTTACTACT

>M6-PGSC0003DMG400007948M6LRR-RLK_1830

CAAATAGGGTCCATACACCATACCCTATTGTTACAGTTTCTCAACCCCAATGTTCTCACAAACACAGAGAATCTTGCAGCTGACTCTGATGACATTTTTGGACACATATTTCTGTCATCG

>M6-PGSC0003DMG400007948M6LRR-RLK_1891

ACACAGAGAATCTTGCAGCTGACTCTGATGACATTTTTGGACACATATTTCTGTCATCGTAAAATACTTTTATGTTTGTTTAGTGTCCTAATCATAGATTATAGTATTACTTTATTGAAG

>M6-PGSC0003DMG400007948M6LRR-RLK_1952

AAATACTTTTATGTTTGTTTAGTGTCCTAATCATAGATTATAGTATTACTTTATTGAAGCTAGTTCTGTCTGCATTAAGAATGTTATTAATTTTGTTTCACGTGTGGTTGCGTAATGTGA

>M6-PGSC0003DMG400007948M6LRR-RLK_2013

AGTTCTGTCTGCATTAAGAATGTTATTAATTTTGTTTCACGTGTGGTTGCGTAATGTGAAGTTAATAAGAGTACGGTGGTTAGCATTAATAATTCATAAAGTAATATAAATAATAGCTTT

>M6-PGSC0003DMG400007948M6LRR-RLK_2074

TTAATAAGAGTACGGTGGTTAGCATTAATAATTCATAAAGTAATATAAATAATAGCTTTATTTCATATTTAATGCCACGACGGTACTGAATTTTTTGGTCAATCTGACGTGGCATATGGA

>M6-PGSC0003DMG400007948M6LRR-RLK_2135

TTCATATTTAATGCCACGACGGTACTGAATTTTTTGGTCAATCTGACGTGGCATATGGACTGGTCAACCTGCCACGTGTTCAGGTTCCAAGTCATCAAGAGCAAAGCCACTTCACTGGAC

>M6-PGSC0003DMG400007948M6LRR-RLK_2196

GGTCAACCTGCCACGTGTTCAGGTTCCAAGTCATCAAGAGCAAAGCCACTTCACTGGACTTCATGTTTGAAAATAGCAGAGGATGTAACTCAAGGACTTTCCTACATCCACCAAGCATGG

>M6-PGSC0003DMG400007948M6LRR-RLK_2257

CATGTTTGAAAATAGCAGAGGATGTAACTCAAGGACTTTCCTACATCCACCAAGCATGGAGGCTTGTCCATGGCAATCTCAAGTCATCTAACGTCCTCCTCGGCTCCGATTTCGAGGCGT

>M6-PGSC0003DMG400007948M6LRR-RLK_2318

GCTTGTCCATGGCAATCTCAAGTCATCTAACGTCCTCCTCGGCTCCGATTTCGAGGCGTGCATAACTGACTACTGTCTCTCCATCCTAGCTGTCCCCTCAGACGATGACAATCCTGATTC

>M6-PGSC0003DMG400007948M6LRR-RLK_2379

ATAACTGACTACTGTCTCTCCATCCTAGCTGTCCCCTCAGACGATGACAATCCTGATTCTGTAGCCTATCAGGCTCCAGAAATTCGGAAACTGAATCACAACAACCACAACCACCACCGC

>M6-PGSC0003DMG400007948M6LRR-RLK_244

ACAATTCAATCTGGAAATTTTTTCTTAAACAAGATGAAAATACAACAATTCTTACTACTAACCATTCTTCTGTTTTTATCTGTTTCTTTTCATTCCACTGTTTCGTTGATTCCTTCAGAT

>M6-PGSC0003DMG400007948M6LRR-RLK_2440

TAGCCTATCAGGCTCCAGAAATTCGGAAACTGAATCACAACAACCACAACCACCACCGCCAAGCTAGTGCCAAGGCTGATGTCTATTCGTTTGGCGTTCTTTTGCTTGAACTTCTTACGG

>M6-PGSC0003DMG400007948M6LRR-RLK_2501

AGCTAGTGCCAAGGCTGATGTCTATTCGTTTGGCGTTCTTTTGCTTGAACTTCTTACGGGGAGGCATCCTTCGGAACATCCATATCTAATGCCAGATGATATGATACATTGGGTGAAGTC

>M6-PGSC0003DMG400007948M6LRR-RLK_2562

AGGCATCCTTCGGAACATCCATATCTAATGCCAGATGATATGATACATTGGGTGAAGTCTACTAGAGAGGATCATGATGGTAGTATTGGAGAAGATAGTAAGTTGGAGATGCTTCTTGAA

>M6-PGSC0003DMG400007948M6LRR-RLK_2623

CTAGAGAGGATCATGATGGTAGTATTGGAGAAGATAGTAAGTTGGAGATGCTTCTTGAAGTTGCTATGGCTTGTAGAGTGAGCTCACCAGAGCAAAGACCAACAATGTGGCAAGTGTTAA

>M6-PGSC0003DMG400007948M6LRR-RLK_2684

TGCTATGGCTTGTAGAGTGAGCTCACCAGAGCAAAGACCAACAATGTGGCAAGTGTTAAAGATGATACAAGAGATTAAAGAAGCTGTTGTTATGGAAGATAGTCATGAAATGGACCTCCT

>M6-PGSC0003DMG400007948M6LRR-RLK_2745

ATGATACAAGAGATTAAAGAAGCTGTTGTTATGGAAGATAGTCATGAAATGGACCTCCTCACTGGCCCTCCTAAATCCTAATTCTTAGTTATTATAAAATTGAATTTGTATTATATTCAC

>M6-PGSC0003DMG400007948M6LRR-RLK_2804

CACTGGCCCTCCTAAATCCTAATTCTTAGTTATTATAAAATTGAATTTGTATTATATTCACTACATGAGACTTTTCAATATTAGTAATTGCAAAAGAAAGGAACAAAAGGGGAAAGACTT

>M6-PGSC0003DMG400007948M6LRR-RLK_305

CCATTCTTCTGTTTTTATCTGTTTCTTTTCATTCCACTGTTTCGTTGATTCCTTCAGATGCATCAGCATTGTTAGCATTCAAATATAAAGCTGATTTGGACAACAAACTTGCATTTTCTG

>M6-PGSC0003DMG400007948M6LRR-RLK_366

ATCAGCATTGTTAGCATTCAAATATAAAGCTGATTTGGACAACAAACTTGCATTTTCTGCTAATACAAGTTTCAGATTCTGCAAATGGAAAGGGATACAATGCTCGGAGAAGAAAGTTAT

>M6-PGSC0003DMG400007948M6LRR-RLK_427

AATACAAGTTTCAGATTCTGCAAATGGAAAGGGATACAATGCTCGGAGAAGAAAGTTATTCGTATTGTTATCGAAAGTTTTAGCTTACGAGGTACGTTTCCAGCTAATACATTGTCTATG

>M6-PGSC0003DMG400007948M6LRR-RLK_488

GTATTGTTATCGAAAGTTTTAGCTTACGAGGTACGTTTCCAGCTAATACATTGTCTATGCTTGATCAACTGAGAGTATTGAGTTTACATAACAACTCACTTACCGGTCCGATTCCGGACC

>M6-PGSC0003DMG400007948M6LRR-RLK_549

TGATCAACTGAGAGTATTGAGTTTACATAACAACTCACTTACCGGTCCGATTCCGGACCTTTCTCCACTTATTAATCTCAAAGTTCTTTTCCTGGACCATAATTTGTTCACCGGTTCAAT

>M6-PGSC0003DMG400007948M6LRR-RLK_61

ACACAGAAACAGAGAGAAAAATAGCCAGACTAAATATTGGAAGTCCCCACTTCCCTTTTTTCTTTCTCAAAAATCTTCACGTGAATCTGACTCTGCCTTTTGCCATATCTTCTTAGACTG

>M6-PGSC0003DMG400007948M6LRR-RLK_610

TCTCCACTTATTAATCTCAAAGTTCTTTTCCTGGACCATAATTTGTTCACCGGTTCAATACCATCTTCCATTTTCACTCTTCATAGGCTTAAAACCCTTGATCTCTCTTACAATAACCTC

>M6-PGSC0003DMG400007948M6LRR-RLK_671

CATCTTCCATTTTCACTCTTCATAGGCTTAAAACCCTTGATCTCTCTTACAATAACCTCACCGGTTCAATATCCGTTGCAATCAACGGGTTGAACCGGTTGTACTATCTCCGGCTTGACT

>M6-PGSC0003DMG400007948M6LRR-RLK_732

CGGTTCAATATCCGTTGCAATCAACGGGTTGAACCGGTTGTACTATCTCCGGCTTGACTCGAACCGGTTAAACGGATCAATCCCACCGCTGAACCAATCTACCCTCCATATCTTCAACAT

>M6-PGSC0003DMG400007948M6LRR-RLK_793

AACCGGTTAAACGGATCAATCCCACCGCTGAACCAATCTACCCTCCATATCTTCAACATTTCTCACAATGCCCTTTCTGGTCCTATTCCGGTAACGAAAACGTTATCCCGGTTTAAAACG

>M6-PGSC0003DMG400007948M6LRR-RLK_854

CTCACAATGCCCTTTCTGGTCCTATTCCGGTAACGAAAACGTTATCCCGGTTTAAAACGGCGTCGTTTTCAGATAATAAAGGACTTTGTGGTGAGATCGTACATAAAGAATGCCGTCCAA

>M6-PGSC0003DMG400007948M6LRR-RLK_915

GTCGTTTTCAGATAATAAAGGACTTTGTGGTGAGATCGTACATAAAGAATGCCGTCCAATACAACCGTTTTTCAGTCCATCCACCGCTGCCGCCACTAAAATTACGCCGCCGCCGTCTAA

>M6-PGSC0003DMG400007948M6LRR-RLK_976

CAACCGTTTTTCAGTCCATCCACCGCTGCCGCCACTAAAATTACGCCGCCGCCGTCTAAAACTCCGGCAGAACTCGGCCAAAACGAAGAGTTACGACAGGGTAGCCCGTTAAACCGTAAA

>M6-PGSC0003DMG400007999M6LRR-RLP_0

CTCCAGTAGCATAATAACATCACTCAAAACTTCAACTTCTTCGAGCAGAATGTAAAATCAGTTTGAAGTCCCCAAACATGGCGGTGACGGACTTTTTCGTCGGCGAAATCACCACCGAAC

>M6-PGSC0003DMG400007999M6LRR-RLP_1037

TATTAACTATTAGTTGAGTGGACATCCGTGAAATCAACTCATATTTTGCGTTGCTGATGATCGAGTTATTAGAAGTTAATAATCACACAAAAGTTTCTCATTTCATTTTGTTTTTACTCT

>M6-PGSC0003DMG400007999M6LRR-RLP_1098

CGAGTTATTAGAAGTTAATAATCACACAAAAGTTTCTCATTTCATTTTGTTTTTACTCTTCTGTTGCGGATGAAACAGTGGATAAAAGTTGAAAACTTTATTGGCTATTATCTCTATTTT

>M6-PGSC0003DMG400007999M6LRR-RLP_1159

TGTTGCGGATGAAACAGTGGATAAAAGTTGAAAACTTTATTGGCTATTATCTCTATTTTATCAGTTGAATTCTAATAGAGATCAATTTTGCATGTAAAGGACTCATCAAAAGTTTTGATT

>M6-PGSC0003DMG400007999M6LRR-RLP_122

ATAAAAAACCTTCTGCTAATAGTTAAAAAATCCACTTTATGCCGTTCAAGCGCCGAGAATCTCATCGACAGCATCAATGGTCTCCTTCCAATCATCCAAGAAATCAAACAAACCGGTGTT

>M6-PGSC0003DMG400007999M6LRR-RLP_1220

CAGTTGAATTCTAATAGAGATCAATTTTGCATGTAAAGGACTCATCAAAAGTTTTGATTGGAATACAGTTTAAATTGGTCTTAAAAACCTTTTCTTTAAATAATGGTATGGTGTCTGGGC

>M6-PGSC0003DMG400007999M6LRR-RLP_1281

AATACAGTTTAAATTGGTCTTAAAAACCTTTTCTTTAAATAATGGTATGGTGTCTGGGCCATGTTGTTCCAGGTTATTTCAAGGACAAGATTTTCTTTTTCACTGTTTCTCAATCTCCAA

>M6-PGSC0003DMG400007999M6LRR-RLP_1342

TGTTGTTCCAGGTTATTTCAAGGACAAGATTTTCTTTTTCACTGTTTCTCAATCTCCAAACGTGGAGCAATTAAGGAAAATGATTTGGGAAAAGATATCAGGGTGCAATCTCCATGGTTA

>M6-PGSC0003DMG400007999M6LRR-RLP_1403

GTGGAGCAATTAAGGAAAATGATTTGGGAAAAGATATCAGGGTGCAATCTCCATGGTTATGGATACGGGGAGATGTTACCCCAGTGGAACCTACAGTACCAATGGAATACGAAATCTGCA

>M6-PGSC0003DMG400007999M6LRR-RLP_1464

GATACGGGGAGATGTTACCCCAGTGGAACCTACAGTACCAATGGAATACGAAATCTGCATCCCCAGTACTCTTGATTCTTGATGATGTGTGGTCTGCATCTGTGCTAGAGCCACTAATTG

>M6-PGSC0003DMG400007999M6LRR-RLP_1525

CCCAGTACTCTTGATTCTTGATGATGTGTGGTCTGCATCTGTGCTAGAGCCACTAATTGTCAAGATCCCCGGCTGCAAGATCCTAGTTGTATCGCGTATCAAGTTTCCTCCATCAATCAT

>M6-PGSC0003DMG400007999M6LRR-RLP_1586

AAGATCCCCGGCTGCAAGATCCTAGTTGTATCGCGTATCAAGTTTCCTCCATCAATCATTGACTGTATTTATGATTTAGAGTTGTTAAGGGAAGATGAAGCTATGTCCTTATTATGCCAT

>M6-PGSC0003DMG400007999M6LRR-RLP_1647

ACTGTATTTATGATTTAGAGTTGTTAAGGGAAGATGAAGCTATGTCCTTATTATGCCATTTTGCATTTGGACACAATTCCTTTCCGCGTGGTTTCAGCCAAAAGCTTGTCAAAGAGGTAT

>M6-PGSC0003DMG400007999M6LRR-RLP_1708

TGCATTTGGACACAATTCCTTTCCGCGTGGTTTCAGCCAAAAGCTTGTCAAAGAGGTATTAGTGTATGTTTTTGGCAGCATATTAGTTTTTCTTTGCAAAAACTTCTTGTTATACTTGAG

>M6-PGSC0003DMG400007999M6LRR-RLP_1769

GTGTATGTTTTTGGCAGCATATTAGTTTTTCTTTGCAAAAACTTCTTGTTATACTTGAGTTAAGCTAACAAAAGCTTCTTGGACTTTTTAGATTGTGGATGAATGTGAAGGGCTTCCTTT

>M6-PGSC0003DMG400007999M6LRR-RLP_183

TCATCGACAGCATCAATGGTCTCCTTCCAATCATCCAAGAAATCAAACAAACCGGTGTTGAACTTCCACAGATACGTCAAACTCAGATCGACGATTTCTCCAAACTTCTCCGAGATGGTT

>M6-PGSC0003DMG400007999M6LRR-RLP_1830

AAGCTAACAAAAGCTTCTTGGACTTTTTAGATTGTGGATGAATGTGAAGGGCTTCCTTTGGCTCTTAAGGTCATTGGATCTTCATTAAAGGGAAAACCTGAGATGTTCTGGATAAGTGCA

>M6-PGSC0003DMG400007999M6LRR-RLP_1891

CTCTTAAGGTCATTGGATCTTCATTAAAGGGAAAACCTGAGATGTTCTGGATAAGTGCAAAAAACAGATTATCACGATGCCAACCTGTCTGCGAGTCTCATGAACTGCAGTTGCTTGAGC

>M6-PGSC0003DMG400007999M6LRR-RLP_1952

AAACAGATTATCACGATGCCAACCTGTCTGCGAGTCTCATGAACTGCAGTTGCTTGAGCGAATGAAATTGAGTATTGACTGTTTGCCTGAGAAGGTGAGAGAGTGTTTCCTGGACTTGGG

>M6-PGSC0003DMG400007999M6LRR-RLP_2013

ATGAAATTGAGTATTGACTGTTTGCCTGAGAAGGTGAGAGAGTGTTTCCTGGACTTGGGTGCTTTCCCAGAAGACAAAAGGATTCCTCTTGATGTTCTAATTAACATGTGGGTGGAGCTA

>M6-PGSC0003DMG400007999M6LRR-RLP_2074

CTTTCCCAGAAGACAAAAGGATTCCTCTTGATGTTCTAATTAACATGTGGGTGGAGCTACATGATATTGATGAGGAGGAGGCTTTTCACATTCTTGTTGAACTTTCAGACAAAAATCTCC

>M6-PGSC0003DMG400007999M6LRR-RLP_2135

TGATATTGATGAGGAGGAGGCTTTTCACATTCTTGTTGAACTTTCAGACAAAAATCTCCTAAATCTAGTCAAAGATGCACGGTAAGCGGTGGACTAAAATCTCCTCAAACAATCTTTTAA

>M6-PGSC0003DMG400007999M6LRR-RLP_2196

AATCTAGTCAAAGATGCACGGTAAGCGGTGGACTAAAATCTCCTCAAACAATCTTTTAATTTTTGTTAGAAGGTGATTGTTTATGGAAAAAATCAATATGTTGATAGTCTTCAATGTGGT

>M6-PGSC0003DMG400007999M6LRR-RLP_2257

TTTGTTAGAAGGTGATTGTTTATGGAAAAAATCAATATGTTGATAGTCTTCAATGTGGTCTTCATGTGGCAGAGCTGGAGACATGTATACAAGTTACTATGAGATATCGGTATTTCAGCA

>M6-PGSC0003DMG400007999M6LRR-RLP_2318

TCATGTGGCAGAGCTGGAGACATGTATACAAGTTACTATGAGATATCGGTATTTCAGCATGATGTATTACGAGACCTAGCAATTCAGATGAGCAACCGTGATGATATAAATCAGAGAAAG

>M6-PGSC0003DMG400007999M6LRR-RLP_2379

ATGTATTACGAGACCTAGCAATTCAGATGAGCAACCGTGATGATATAAATCAGAGAAAGCGATTGGTTATGCCACGAAGAGACGTAAGCTTTCCAAGAGAATGGGAAAGAAATGTGGACG

>M6-PGSC0003DMG400007999M6LRR-RLP_244

ACTTCCACAGATACGTCAAACTCAGATCGACGATTTCTCCAAACTTCTCCGAGATGGTTACGAACTCGCCGGAAAAGTTATCCACTCCGGCCGTTGGAACATGTACAGGTGTCACCTTCT

>M6-PGSC0003DMG400007999M6LRR-RLP_2440

ATTGGTTATGCCACGAAGAGACGTAAGCTTTCCAAGAGAATGGGAAAGAAATGTGGACGAACCTTTCCATGCACGAGTTATCTCTGTGCATACAGGTATTTTTAGATCCTTGGTTACTGC

>M6-PGSC0003DMG400007999M6LRR-RLP_2501

CCTTTCCATGCACGAGTTATCTCTGTGCATACAGGTATTTTTAGATCCTTGGTTACTGCTGATTGATTAATAATGAAAAGTTTATCATGTGATATAAACAGAAATATTTCTCTTTAATGC

>M6-PGSC0003DMG400007999M6LRR-RLP_2562

ATTGATTAATAATGAAAAGTTTATCATGTGATATAAACAGAAATATTTCTCTTTAATGCAGATGAAATGAGAGAAATAGACTGGTTCAGAATGGATTGCCCGAAAGCTGAAGTACTGATT

>M6-PGSC0003DMG400007999M6LRR-RLP_2623

ATGAAATGAGAGAAATAGACTGGTTCAGAATGGATTGCCCGAAAGCTGAAGTACTGATTCTCAATTTTGCCTCATCTGAGTACTTCTTGCCTCCTTTCCTGGAGAATATGCCAAAGCTAA

>M6-PGSC0003DMG400007999M6LRR-RLP_2684

CAATTTTGCCTCATCTGAGTACTTCTTGCCTCCTTTCCTGGAGAATATGCCAAAGCTAAGGGCATTGATAATCATAAACTATAGTGCTGGCAATGCAGTTCTTCATAACATGTCTGTATT

>M6-PGSC0003DMG400007999M6LRR-RLP_2745

GCATTGATAATCATAAACTATAGTGCTGGCAATGCAGTTCTTCATAACATGTCTGTATTCAGTCATTTAACCAACTTGAGAAGCCTTTGGTTCGAGAAGATATCCATCACTCACTTATCT

>M6-PGSC0003DMG400007999M6LRR-RLP_2806

GTCATTTAACCAACTTGAGAAGCCTTTGGTTCGAGAAGATATCCATCACTCACTTATCTGACTCCACAAATCCTCTCAATAACTTGCGGAAGATATCTCTAGTGCTTTGTGACATGAAAA

>M6-PGSC0003DMG400007999M6LRR-RLP_2867

CTCCACAAATCCTCTCAATAACTTGCGGAAGATATCTCTAGTGCTTTGTGACATGAAAAACAGCTTAGATGAGTCAGATGTGGACCTCCCTGGTTTGTTCCCACAGCTTTCTGAGTTCAC

>M6-PGSC0003DMG400007999M6LRR-RLP_2928

AGCTTAGATGAGTCAGATGTGGACCTCCCTGGTTTGTTCCCACAGCTTTCTGAGTTCACAATGGATCATTGCATCAACTTCAATAAGCTGCCATCAAGTATTTGCCGGTTGCATAAGCTC

>M6-PGSC0003DMG400007999M6LRR-RLP_305

GAACTCGCCGGAAAAGTTATCCACTCCGGCCGTTGGAACATGTACAGGTGTCACCTTCTCATTCGACTCTAATTACAGTGAATTGAACCTCTTCCTGCTCTAATTAGGGTTATTGAATTG

>M6-PGSC0003DMG400007999M6LRR-RLP_3050

CAGCCTTAGTATCACTAATTGTGATAGTCTTTATGAACTTCCATCTGATTTAGGTGAATTACAAACTCTACAAGTTTTAAGGATATATGCCTGTCCACATCTGAAAAGGCTTCCCCCTGG

>M6-PGSC0003DMG400007999M6LRR-RLP_3111

CAAACTCTACAAGTTTTAAGGATATATGCCTGTCCACATCTGAAAAGGCTTCCCCCTGGAATAGGTCATCTGGTGAAGCTGAAGTATCTTGACATTTCACAATGTGTTGGTTTGAGATGT

>M6-PGSC0003DMG400007999M6LRR-RLP_3172

TAGGTCATCTGGTGAAGCTGAAGTATCTTGACATTTCACAATGTGTTGGTTTGAGATGTCTCCCTGAAGCAATTGGTTGCTGTAGAAACTTGGAGAAGATTGATATGAGGGAGTGCCCTC

>M6-PGSC0003DMG400007999M6LRR-RLP_3233

CCCTGAAGCAATTGGTTGCTGTAGAAACTTGGAGAAGATTGATATGAGGGAGTGCCCTCAAATTGACAGTTTGCCAAGCGCTCTATCCTTTCTTGAATCATTACGTTGTGTTATTTGTGA

>M6-PGSC0003DMG400007999M6LRR-RLP_3294

ATTGACAGTTTGCCAAGCGCTCTATCCTTTCTTGAATCATTACGTTGTGTTATTTGTGACGATGAAGTTTTTTGTCAATGGAAAGATGTTGAGAAGGCTGTACCAGGTCTATGTGTACAG

>M6-PGSC0003DMG400007999M6LRR-RLP_3355

ATGAAGTTTTTTGTCAATGGAAAGATGTTGAGAAGGCTGTACCAGGTCTATGTGTACAGGTTGCCGAGGAGTGCTATACTCTTGACTGGCTATCTCAGTAATCAAGTTTTCAATTCCCTT

>M6-PGSC0003DMG400007999M6LRR-RLP_3416

TGCCGAGGAGTGCTATACTCTTGACTGGCTATCTCAGTAATCAAGTTTTCAATTCCCTTATCTTGGGTTTCTTTCGTGTAAATGCTCAGTCAATTTTCCAGTTCTGTATTTTCCCCTTTA

>M6-PGSC0003DMG400007999M6LRR-RLP_3477

CTTGGGTTTCTTTCGTGTAAATGCTCAGTCAATTTTCCAGTTCTGTATTTTCCCCTTTACATGGGGTTCATTGTATCTCTTGTAAATAATATATAGCAATATGTAAATATGTAGATGGAC

>M6-PGSC0003DMG400007999M6LRR-RLP_3538

TGGGGTTCATTGTATCTCTTGTAAATAATATATAGCAATATGTAAATATGTAGATGGACTACATCTCAATAGCTGATCTTTCCTTTTACTTTAAAACCATAATGCTGGAAGCAAAATTAC

>M6-PGSC0003DMG400007999M6LRR-RLP_3582

AATATGTAGATGGACTACATCTCAATAGCTGATCTTTCCTTTTACTTTAAAACCATAATGCTGGAAGCAAAATTACCACAGGCAATCCAACACTAAGGCAAGTTATCTGCTATAACATGT

>M6-PGSC0003DMG400007999M6LRR-RLP_366

TTCGACTCTAATTACAGTGAATTGAACCTCTTCCTGCTCTAATTAGGGTTATTGAATTGTACGAATCTGTTAGAAATTACTTAATTTATATTATGCCTTGATAACAGGAACCTACAGTTG

>M6-PGSC0003DMG400007999M6LRR-RLP_427

CGAATCTGTTAGAAATTACTTAATTTATATTATGCCTTGATAACAGGAACCTACAGTTGGCTAGGAAAATGGAGAGGCTGGAGAAAAGAGTAGCGAGGTTCATGCAAGTTACAATGCAAG

>M6-PGSC0003DMG400007999M6LRR-RLP_488

TAGGAAAATGGAGAGGCTGGAGAAAAGAGTAGCGAGGTTCATGCAAGTTACAATGCAAGCTCATGTACTAGCGGATGTTCATCATGTTAGGTTTAGTATGGAGCAGAGATTTGACGTGCT

>M6-PGSC0003DMG400007999M6LRR-RLP_549

CATGTACTAGCGGATGTTCATCATGTTAGGTTTAGTATGGAGCAGAGATTTGACGTGCTTGAGCATAGGCTTAAAGCTATAAAAATCGGAGTTGACGATAGAAGTGGTGGTGATGGTGGA

>M6-PGSC0003DMG400007999M6LRR-RLP_61

TTTGAAGTCCCCAAACATGGCGGTGACGGACTTTTTCGTCGGCGAAATCACCACCGAACTCATAAAAAACCTTCTGCTAATAGTTAAAAAATCCACTTTATGCCGTTCAAGCGCCGAGAA

>M6-PGSC0003DMG400007999M6LRR-RLP_610

AGCATAGGCTTAAAGCTATAAAAATCGGAGTTGACGATAGAAGTGGTGGTGATGGTGGAGGAGGAGGGTGTTTAGGGGAAGCTGTGAAAAGAATGGAAGAAGATGAGAAATGGTTTGAGG

>M6-PGSC0003DMG400007999M6LRR-RLP_671

AGGAGGGTGTTTAGGGGAAGCTGTGAAAAGAATGGAAGAAGATGAGAAATGGTTTGAGGATAGTTTTGTAAATTTAGGTGCTGGGATTGAATTGGGGAAGAGGAAAGTGAAGGAGATGCT

>M6-PGSC0003DMG400007999M6LRR-RLP_732

AGTTTTGTAAATTTAGGTGCTGGGATTGAATTGGGGAAGAGGAAAGTGAAGGAGATGCTGATGGGTGAACAGGATAGAGGTGTGTTTGAGATTTGTGGAATTGGGGGTAGTGGCAAAACT

>M6-PGSC0003DMG400007999M6LRR-RLP_793

TGGGTGAACAGGATAGAGGTGTGTTTGAGATTTGTGGAATTGGGGGTAGTGGCAAAACTACCTTGGCTAAAGAGATTTGTAAAGATGATCAAGTTAAAAGTAAGTAATTAATCCCAGAAA

>M6-PGSC0003DMG400007999M6LRR-RLP_854

CTTGGCTAAAGAGATTTGTAAAGATGATCAAGTTAAAAGTAAGTAATTAATCCCAGAAAAGTTTCTCATGTAGAGTACTTTAGCTTTTTGCGGGGTTGATTTCTCGAAAGGTTAATCAAC

>M6-PGSC0003DMG400007999M6LRR-RLP_915

TTTCTCATGTAGAGTACTTTAGCTTTTTGCGGGGTTGATTTCTCGAAAGGTTAATCAACTAATAGTTATTATCTATCACTTTTCGTTTTTGATTCTGATGCAAAGAAAGTGACTTTATTA

>M6-PGSC0003DMG400007999M6LRR-RLP_976

ATAGTTATTATCTATCACTTTTCGTTTTTGATTCTGATGCAAAGAAAGTGACTTTATTAGATATTAACTATTAGTTGAGTGGACATCCGTGAAATCAACTCATATTTTGCGTTGCTGATG

>M6-PGSC0003DMG400008036M6LRR-RLP_0

ATGAATAGTACTATCAAATTTCTAATATTTTTCTTCAGTGTTACAACAGCACAACCACTAAAGGGACAGAACCAATTGATTTTTGCAGATCAAAGGCTTGCTATAGTATACCCCATTATA

>M6-PGSC0003DMG400008036M6LRR-RLP_1024

TAGCTGAATGTGTTGCATTTTTTGCTTACCCAAGGTACTGTCCTCATATGGCTACATATACTTATATTCCTTGTTGGCTTTCAAATTTCAAAACGCCTCCTCTGGATCTCCCTGAATTGG

>M6-PGSC0003DMG400008036M6LRR-RLP_1088

TATTCCTTGTTGGCTTTCAAATTTCAAAACGCCTCCTCTGGATCTCCCTGAATTGGCTCCTTAGTAAGGGTAACACGGTGTACAAAGCATTCTGTGTTAGCATGGTTCGGGAAAAGATCA

>M6-PGSC0003DMG400008036M6LRR-RLP_1152

TAAGGGTAACACGGTGTACAAAGCATTCTGTGTTAGCATGGTTCGGGAAAAGATCATTAGAAAATTCTGGTATTTGATCATAAACATACATGTCCCCTTTTCGCGATCTTTTTTATTTTA

>M6-PGSC0003DMG400008036M6LRR-RLP_1216

TTCTGGTATTTGATCATAAACATACATGTCCCCTTTTCGCGATCTTTTTTATTTTAATTAAATTAAGCTTGTAATTTCAATTGAATAGAGTGTGATATATCATAATAACTTGTGATCTCT

>M6-PGSC0003DMG400008036M6LRR-RLP_1266

ATTTTAATTAAATTAAGCTTGTAATTTCAATTGAATAGAGTGTGATATATCATAATAACTTGTGATCTCTGTTTATGGAGTATTGCTCAGTAGTAAATTTACGTTACTTTTATGTTATGG

>M6-PGSC0003DMG400008036M6LRR-RLP_128

CAAGAATATCATCTCTGCTGATCCTCTAGGTATTACAAAAACTTGGGTTGGTCCAAACATATGTAATTATACTGGTTTTTATTGTGAAAGTCCTCCAGACAACAATTCAGCAATTGCTCT

>M6-PGSC0003DMG400008036M6LRR-RLP_192

AATTATACTGGTTTTTATTGTGAAAGTCCTCCAGACAACAATTCAGCAATTGCTCTTGCTTCTATTGATTTCAATGGTTTTCAACTTAGCGCGTTTACTCTTGATGGCTTCCTTGATCAG

>M6-PGSC0003DMG400008036M6LRR-RLP_256

TTGATTTCAATGGTTTTCAACTTAGCGCGTTTACTCTTGATGGCTTCCTTGATCAGCTTCCTGATATAGCTATTTTCCATGCTAATTCAAACAATTTTGGTGGGACTCTGTCTCCAAAAA

>M6-PGSC0003DMG400008036M6LRR-RLP_320

TATAGCTATTTTCCATGCTAATTCAAACAATTTTGGTGGGACTCTGTCTCCAAAAATCGCGAATCTTCCTTATCTCTATGAACTTGATATTAGTAACAACCAATTTTTCGGCCCATTTCC

>M6-PGSC0003DMG400008036M6LRR-RLP_384

CTTCCTTATCTCTATGAACTTGATATTAGTAACAACCAATTTTTCGGCCCATTTCCTAGTCCAATTTTGGGCATGAATAGCTTAAATATATTGGATATAAGATTCAATTCTTTCACTGGA

>M6-PGSC0003DMG400008036M6LRR-RLP_448

TTTTGGGCATGAATAGCTTAAATATATTGGATATAAGATTCAATTCTTTCACTGGATCAGTCCCACCTCAATTGTTCACTAAAGATCAATTAGACGCGCTCTTCATCAACAACAACAATT

>M6-PGSC0003DMG400008036M6LRR-RLP_512

ACCTCAATTGTTCACTAAAGATCAATTAGACGCGCTCTTCATCAACAACAACAATTTCATGCAGATGCTTCCTGATAACATCGCGACCAGCCATGTTACATATCTTACTTTAGCTAACAA

>M6-PGSC0003DMG400008036M6LRR-RLP_576

ATGCTTCCTGATAACATCGCGACCAGCCATGTTACATATCTTACTTTAGCTAACAACAGATTTTTCGGTCCTATCCCACATAGTATTTCCAAAATCTTGTCGAGTTTGTCTGAGATTTTG

>M6-PGSC0003DMG400008036M6LRR-RLP_64

GACAGAACCAATTGATTTTTGCAGATCAAAGGCTTGCTATAGTATACCCCATTATACAAAAATTCAAGAATATCATCTCTGCTGATCCTCTAGGTATTACAAAAACTTGGGTTGGTCCAA

>M6-PGSC0003DMG400008036M6LRR-RLP_640

TCGGTCCTATCCCACATAGTATTTCCAAAATCTTGTCGAGTTTGTCTGAGATTTTGCTGCTTAACAACTTGTTAAGTGGATGTCTTCCTTACGAGATAGGATTCTTAAACGAAGCAATAG

>M6-PGSC0003DMG400008036M6LRR-RLP_704

CAACTTGTTAAGTGGATGTCTTCCTTACGAGATAGGATTCTTAAACGAAGCAATAGTCTTCGATGCTGGCAATAATAGACTCACCGGTCCACTTCCATTCTCATTAGGGTGTTTGGAAAA

>M6-PGSC0003DMG400008036M6LRR-RLP_768

GCTGGCAATAATAGACTCACCGGTCCACTTCCATTCTCATTAGGGTGTTTGGAAAATGTAGAAATGTTGAATTTTGCTTGAAATCAGATGTATGGGATAGTGCCTGATGTGATTTGTGCC

>M6-PGSC0003DMG400008036M6LRR-RLP_832

TGTTGAATTTTGCTTGAAATCAGATGTATGGGATAGTGCCTGATGTGATTTGTGCCCTGGGGAATTTGGCAAATTTATCTTTTTCCGATAACTATTTTACAGGATTTGGGCCAATTTGTT

>M6-PGSC0003DMG400008036M6LRR-RLP_896

TTTGGCAAATTTATCTTTTTCCGATAACTATTTTACAGGATTTGGGCCAATTTGTTTGAGATTGATAGAAAATGGAGTGTTTGATTTGAGGAATAATTGTATTCCTGGTTTTCCATTTCA

>M6-PGSC0003DMG400008036M6LRR-RLP_960

ATAGAAAATGGAGTGTTTGATTTGAGGAATAATTGTATTCCTGGTTTTCCATTTCAAAGATCAATAGCTGAATGTGTTGCATTTTTTGCTTACCCAAGGTACTGTCCTCATATGGCTACA

>M6-PGSC0003DMG400008146-(rev)M6LRR-RLK_0

ATGGATTCTTGATTGTTCTATCTGGTTGTTTTGGAGGGAAAAGGAGGAAGAAATTGAAGAAAAGGTGTTTTCTTGAAAAAAAAAAAAACTCAAGAATGGCAATTTTGAGGTTTCTGTTAA

>M6-PGSC0003DMG400008146-(rev)M6LRR-RLK_1054

TGGAACTATTCCTCCTGAGTTTGGTCAGCTTAACAGTTTGCAAGAATTGATTGTTTCTGGTAATAACCTTTATGGAGAGATTCCAATATCAGTTCTTAAGTGCAAGAATCTCAATAAGCT

>M6-PGSC0003DMG400008146-(rev)M6LRR-RLK_1116

ATAACCTTTATGGAGAGATTCCAATATCAGTTCTTAAGTGCAAGAATCTCAATAAGCTTGACTTGAGCAACAACAAGTTCAATGGTACCATACCGGCAGATATCTGCAATACGTCCAGAT

>M6-PGSC0003DMG400008146-(rev)M6LRR-RLK_1178

TTGAGCAACAACAAGTTCAATGGTACCATACCGGCAGATATCTGCAATACGTCCAGATTGCAATATTTGCTGTTGGGGGAGAACTCGATTCGAGGGGAGATACCTCATGAGATAGGGAAT

>M6-PGSC0003DMG400008146-(rev)M6LRR-RLK_124

GTTTTTTCTAGTTGGTTTGTTATCAAGATTCCAACTTGTGAATTCCCAGTTGGTATTTCATGATCAAGATGTAGTGGAAGCAATTGGGAAAGAGCTATAAGTACCTGGATGGGGTGTGAT

>M6-PGSC0003DMG400008146-(rev)M6LRR-RLK_1240

ATATTTGCTGTTGGGGGAGAACTCGATTCGAGGGGAGATACCTCATGAGATAGGGAATTGTGTAAAGTTGCTTGAGTTGGAAATGGGGAGTAATGAACTTACTGGAAGTATCCCTCCTGA

>M6-PGSC0003DMG400008146-(rev)M6LRR-RLK_1302

TAAAGTTGCTTGAGTTGGAAATGGGGAGTAATGAACTTACTGGAAGTATCCCTCCTGAGATTGGTCATATGAAGAACTTGCAGATTTCGTTGAATTTGAGTCATAATCGTCTCCACGGAA

>M6-PGSC0003DMG400008146-(rev)M6LRR-RLK_1364

GGTCATATGAAGAACTTGCAGATTTCGTTGAATTTGAGTCATAATCGTCTCCACGGAAAATTGCCTCAGGATTTAGGAAAACTTGACAAGTTGGTTTCTTTAGATGTATCTGATAATCAG

>M6-PGSC0003DMG400008146-(rev)M6LRR-RLK_1426

GCCTCAGGATTTAGGAAAACTTGACAAGTTGGTTTCTTTAGATGTATCTGATAATCAGCTCTCGGGGAATATTCCGCCGGCATTGAAGGGCATGCAGAGTTTGATAGAGGTTGATTTTTC

>M6-PGSC0003DMG400008146-(rev)M6LRR-RLK_1488

CGGGGAATATTCCGCCGGCATTGAAGGGCATGCAGAGTTTGATAGAGGTTGATTTTTCGAGCAACCAATTCACTGGTCCAATACCTACTTTTGCACCATTTCAGAAAAGTCCTAATTCAA

>M6-PGSC0003DMG400008146-(rev)M6LRR-RLK_1550

AACCAATTCACTGGTCCAATACCTACTTTTGCACCATTTCAGAAAAGTCCTAATTCAAGTTTTCTTGGAAACAAAGATCTCTGTGGTGATCCTTTGAGTGTTGGTTGTGGGGATTTAAAT

>M6-PGSC0003DMG400008146-(rev)M6LRR-RLK_1612

TCTTGGAAACAAAGATCTCTGTGGTGATCCTTTGAGTGTTGGTTGTGGGGATTTAAATGGTTATGACCACAGCGTTTACGGTCACAAGGTCTCTTACCGAATCGTTTTGGCTGTTATCGG

>M6-PGSC0003DMG400008146-(rev)M6LRR-RLK_1674

ATGACCACAGCGTTTACGGTCACAAGGTCTCTTACCGAATCGTTTTGGCTGTTATCGGTTCTGGTTTGGCAGTTTTTGTATGTGTCACAGTGGTTGTTTTGCTGTTCATGATGAGGGAGA

>M6-PGSC0003DMG400008146-(rev)M6LRR-RLK_1736

GGTTTGGCAGTTTTTGTATGTGTCACAGTGGTTGTTTTGCTGTTCATGATGAGGGAGAAACAAGAGAAAGCTGCCAAGGAAGCAGCTGGAAAAACCATTGATGAAATTTCTAGCAAACCA

>M6-PGSC0003DMG400008146-(rev)M6LRR-RLK_1798

AGAGAAAGCTGCCAAGGAAGCAGCTGGAAAAACCATTGATGAAATTTCTAGCAAACCAGTGATACTAGCGGGAAACGTCTTTGTTGACAATCTAAAGCAAGCAATTGATTTTGATGCAGT

>M6-PGSC0003DMG400008146-(rev)M6LRR-RLK_186

ATCAAGATGTAGTGGAAGCAATTGGGAAAGAGCTATAAGTACCTGGATGGGGTGTGATGAACACAACTGATTTCTGCTCTTGGCATGGAGTTATTTGCAGCTCAAACAGTTCAATGGTGG

>M6-PGSC0003DMG400008146-(rev)M6LRR-RLK_1860

TACTAGCGGGAAACGTCTTTGTTGACAATCTAAAGCAAGCAATTGATTTTGATGCAGTTGTAAAGGCAGTCATGAAAGACTCGAATAAGATTTGTGTTGGTACTTTCAGCACTGTCTATA

>M6-PGSC0003DMG400008146-(rev)M6LRR-RLK_1922

AAGGCAGTCATGAAAGACTCGAATAAGATTTGTGTTGGTACTTTCAGCACTGTCTATAAAGCAGACATGCCTTCCGGGATGATTTTGTCGGTGAAAAAGCTAAAGTCAATGGACAAAACT

>M6-PGSC0003DMG400008146-(rev)M6LRR-RLK_1984

AGACATGCCTTCCGGGATGATTTTGTCGGTGAAAAAGCTAAAGTCAATGGACAAAACTATAATTCATCACCAGAGTAAGATGATCAGAGAGCTTGAAAAACTTAGTAAGCTCGTTCATGA

>M6-PGSC0003DMG400008146-(rev)M6LRR-RLK_2046

TTCATCACCAGAGTAAGATGATCAGAGAGCTTGAAAAACTTAGTAAGCTCGTTCATGATAATCTTGCTCGTCCTATCGGATTTGGAATCTATGAAGATGTCGTTCTTCTGTTGCATGAAT

>M6-PGSC0003DMG400008146-(rev)M6LRR-RLK_2108

CTTGCTCGTCCTATCGGATTTGGAATCTATGAAGATGTCGTTCTTCTGTTGCATGAATATTATGCCAAAGGAACGTTGACTCAATATCTTCATAACTCTAGCCAGAAACCTGAATATAAG

>M6-PGSC0003DMG400008146-(rev)M6LRR-RLK_2170

TGCCAAAGGAACGTTGACTCAATATCTTCATAACTCTAGCCAGAAACCTGAATATAAGCCCGACTGGCCAACAAGACTTGCCATTGCCACTGGAGTTGCAGAAGGATTGGCATTCCTTCA

>M6-PGSC0003DMG400008146-(rev)M6LRR-RLK_2232

ACTGGCCAACAAGACTTGCCATTGCCACTGGAGTTGCAGAAGGATTGGCATTCCTTCATCATGTTGCTATCATCCATCTTGACATTTCTTCCGGGAATGTGCTTCTTGATTCTAATTTCA

>M6-PGSC0003DMG400008146-(rev)M6LRR-RLK_2294

GTTGCTATCATCCATCTTGACATTTCTTCCGGGAATGTGCTTCTTGATTCTAATTTCAGGCCTTTGGTTAGTGAAGTTGAAATCTCTAGGCTTCTCGATCCATCTAGAGGGACTGCAAGT

>M6-PGSC0003DMG400008146-(rev)M6LRR-RLK_2340

ATTCTAATTTCAGGCCTTTGGTTAGTGAAGTTGAAATCTCTAGGCTTCTCGATCCATCTAGAGGGACTGCAAGTATTAGTGCTGTTGCTGGTTCATTCGGATATATTCCTCCAGGTACAT

>M6-PGSC0003DMG400008146-(rev)M6LRR-RLK_248

ACAACTGATTTCTGCTCTTGGCATGGAGTTATTTGCAGCTCAAACAGTTCAATGGTGGAAAGACTTGATCTTTCTGGTTTTAGATTACAAGGTAATGTAACTCTAATATCTGAGCTCAAA

>M6-PGSC0003DMG400008146-(rev)M6LRR-RLK_310

ACTTGATCTTTCTGGTTTTAGATTACAAGGTAATGTAACTCTAATATCTGAGCTCAAAGGTTTAAAGTGGCTGGACCTTTCTAATAATAACTTCCAAGGTTCTATTCCATTTGCATTTGG

>M6-PGSC0003DMG400008146-(rev)M6LRR-RLK_372

TAAAGTGGCTGGACCTTTCTAATAATAACTTCCAAGGTTCTATTCCATTTGCATTTGGGAATTTATCTGAGCTTCAATATCTTGATTTGTCTTTTAACAAATTTGGAAACTCAATACCTA

>M6-PGSC0003DMG400008146-(rev)M6LRR-RLK_434

TTATCTGAGCTTCAATATCTTGATTTGTCTTTTAACAAATTTGGAAACTCAATACCTAGTGAATTAGGTAAGCTCAGAAGCCTAAAGGCATTGAACCTTTCGAAAAACTGGCTTACAGGA

>M6-PGSC0003DMG400008146-(rev)M6LRR-RLK_496

ATTAGGTAAGCTCAGAAGCCTAAAGGCATTGAACCTTTCGAAAAACTGGCTTACAGGAGCTATACCTGATGAGCTTGAGGGGTTAGAGAATTTGCAAGATTTTCAAATATTTACTAACAA

>M6-PGSC0003DMG400008146-(rev)M6LRR-RLK_558

TACCTGATGAGCTTGAGGGGTTAGAGAATTTGCAAGATTTTCAAATATTTACTAACAACTTGAATGGTTCTATCCCAATGTGGATTGGTAATTTGACCAATCTTAAGGTTTTTGCTGCTT

>M6-PGSC0003DMG400008146-(rev)M6LRR-RLK_62

AGGTGTTTTCTTGAAAAAAAAAAAAACTCAAGAATGGCAATTTTGAGGTTTCTGTTAACTGTGTTTTTTCTAGTTGGTTTGTTATCAAGATTCCAACTTGTGAATTCCCAGTTGGTATTT

>M6-PGSC0003DMG400008146-(rev)M6LRR-RLK_620

AATGGTTCTATCCCAATGTGGATTGGTAATTTGACCAATCTTAAGGTTTTTGCTGCTTATGAGAATGAATTTAGTGGTGGTGTTCCTGTTAACTTAGGATTGTACTCTGATCTTTCTTTG

>M6-PGSC0003DMG400008146-(rev)M6LRR-RLK_682

GAATGAATTTAGTGGTGGTGTTCCTGTTAACTTAGGATTGTACTCTGATCTTTCTTTGTTGAACCTTCACTCAAACCAGCTTGAAGGTACTATTCCTGAGAGCATTTGTGCTAATGGGAA

>M6-PGSC0003DMG400008146-(rev)M6LRR-RLK_744

ACCTTCACTCAAACCAGCTTGAAGGTACTATTCCTGAGAGCATTTGTGCTAATGGGAATCTTGAATTTCTTGTTCTAACTCAGAATAAGTTGACCGGAAAGATTCCTGATTCGATTGGGA

>M6-PGSC0003DMG400008146-(rev)M6LRR-RLK_806

GAATTTCTTGTTCTAACTCAGAATAAGTTGACCGGAAAGATTCCTGATTCGATTGGGAATTGTAAAGGACTTTCTAGTATTAGAATTGGGAATAACAAGTTGATTGGAGGTATCCCTAAA

>M6-PGSC0003DMG400008146-(rev)M6LRR-RLK_868

TAAAGGACTTTCTAGTATTAGAATTGGGAATAACAAGTTGATTGGAGGTATCCCTAAATCAATTGGGAATATCAGAAGTCTTACTTATTTTGAAGCAGATAGTAATAGTTTGTCTGGTGA

>M6-PGSC0003DMG400008146-(rev)M6LRR-RLK_930

TTGGGAATATCAGAAGTCTTACTTATTTTGAAGCAGATAGTAATAGTTTGTCTGGTGAAATTGTCACCGAGTTCGCAAAATGCTCAAATCTCACTCTACTTAATTTAGCTTCAAATAAGT

>M6-PGSC0003DMG400008146-(rev)M6LRR-RLK_992

GTCACCGAGTTCGCAAAATGCTCAAATCTCACTCTACTTAATTTAGCTTCAAATAAGTTTAATGGAACTATTCCTCCTGAGTTTGGTCAGCTTAACAGTTTGCAAGAATTGATTGTTTCT

>M6-PGSC0003DMG400008185M6LRR-RLP_0

TATATTTGGCTTTTTAAAAATATATAAATAAATGTGTGTAAGCACTTTTAGCTTGTCCACTTATGACCTAGTGTTTTGTTCCAATGGCTCCAACTCTATCTGCAACAATGTTTGCAGGTT

>M6-PGSC0003DMG400008185M6LRR-RLP_496

AAATGAAGACGTTTGTCCAACAGGGAATATCTGTGAGTACACCCAGTCTCAAATTTATGATCTTCAATTATAGGAGGCGGAAAGTAGATGATCTGAATAAGTCGATACAGCAGATGTTTA

>M6-PGSC0003DMG400008185M6LRR-RLP_558

TTCAATTATAGGAGGCGGAAAGTAGATGATCTGAATAAGTCGATACAGCAGATGTTTAACAAGGTATGTCTTATTCCGCTAGCTACATATTCTAATTAGGTTCTACTGCTCTCTTTCCGT

>M6-PGSC0003DMG400008185M6LRR-RLP_62

ATGACCTAGTGTTTTGTTCCAATGGCTCCAACTCTATCTGCAACAATGTTTGCAGGTTTTTTGTCCCAACCTGGAAGACCTATCCATCAACAGCCTTAAAAGCATAAGTGCTCTATGCTC

>M6-PGSC0003DMG400008185M6LRR-RLP_620

GGTATGTCTTATTCCGCTAGCTACATATTCTAATTAGGTTCTACTGCTCTCTTTCCGTTTTCGATACTTATTTAAGTTCATCAACTTTCTTTTACTTAATAATCTAGTCATCTCTCTCTT

>M6-PGSC0003DMG400008185M6LRR-RLP_682

GATACTTATTTAAGTTCATCAACTTTCTTTTACTTAATAATCTAGTCATCTCTCTCTTCTTTGCATTCTCTGTTTCCTGCAGAAACAAGAAGCTAGTGATGACGACGAAGCTAGTAGTGG

>M6-PGSC0003DMG400008185M6LRR-RLP_744

GCATTCTCTGTTTCCTGCAGAAACAAGAAGCTAGTGATGACGACGAAGCTAGTAGTGGCTATGTAATAATGATGCCGACGAATCTGAACCTAGTCATTCCAGAAGTTAGGAAGCGGGTAG

>M6-PGSC0003DMG400008185M6LRR-RLP_806

GTAATAATGATGCCGACGAATCTGAACCTAGTCATTCCAGAAGTTAGGAAGCGGGTAGGAAAAACTCAATGAAATAATGCAAGTAAAACCTCGAACATCACCTGATACCTATGCTCGTGA

>M6-PGSC0003DMG400008185M6LRR-RLP_868

AACTCAATGAAATAATGCAAGTAAAACCTCGAACATCACCTGATACCTATGCTCGTGATGGAGGTAATTAACAAGCAGAGGCGGATCTACATTATTGTAACCTATTGGTTCATGTCAACC

>M6-PGSC0003DMG400008185M6LRR-RLP_927

GGAGGTAATTAACAAGCAGAGGCGGATCTACATTATTGTAACCTATTGGTTCATGTCAACCTACTAGCTATTGGCTATAGATCTTGTCTATGTATTAGTAATATCTAAAAAATAAATGAT

>M6-PGSC0003DMG400008296M6LRR-RLK_0

TTATGTACTCCTAGGTTTCGAGTAGGTGACGAAGTTCTTGTTGTCTTTTCTTGTGTGGGGGTACATACAACTTGTTCTTTAACACGGAGCATCCTAGATTAGAGGCATGCTTGGTATTTT

>M6-PGSC0003DMG400008296M6LRR-RLK_1037

ACTCTCTTTCTGGGAAAATCCCATCTGAACTTGGCCTCTGTATCAATCTTTTTACCTTGAACATGTATACTAATCAATTTACTGGAAGCATACCTCCTGAGCTTGGAAATTTAGAAAACT

>M6-PGSC0003DMG400008296M6LRR-RLK_1098

CATGTATACTAATCAATTTACTGGAAGCATACCTCCTGAGCTTGGAAATTTAGAAAACTTGCAAGTGCTCAGATTGCATAACAATAAGTTGAACTCCAGTATACCTGCCTCAATATTCCA

>M6-PGSC0003DMG400008296M6LRR-RLK_1159

CAAGTGCTCAGATTGCATAACAATAAGTTGAACTCCAGTATACCTGCCTCAATATTCCACCTGAAGTCATTAACTCATTTAGGACTCTCACAGAATGAACTAACTGGTAATATTCCTCCC

>M6-PGSC0003DMG400008296M6LRR-RLK_122

AACAAATTCATAGAATGCAAGTCATTTTCCTTAGTGTGAATGGATGACTAGTTGACTGCTACTGAAAACGGTGATTCACTGTACGATTCACTAAGAAATGTGTATAAAGGCTTCATTGTC

>M6-PGSC0003DMG400008296M6LRR-RLK_1220

TGAAGTCATTAACTCATTTAGGACTCTCACAGAATGAACTAACTGGTAATATTCCTCCCGAGTTGGGATCTTTAACGTCACTAGAAGTGCTTACCCTTCACTCAAATAGGTTATCGGGGG

>M6-PGSC0003DMG400008296M6LRR-RLK_1281

GTTGGGATCTTTAACGTCACTAGAAGTGCTTACCCTTCACTCAAATAGGTTATCGGGGGAGATCCCATCAACTATAACAAACCTGGCAAACTTGACATATTTGTCTTTGGGCTTTAATTT

>M6-PGSC0003DMG400008296M6LRR-RLK_1342

ATCCCATCAACTATAACAAACCTGGCAAACTTGACATATTTGTCTTTGGGCTTTAATTTATTGACTGGATCACTTCCATCGGAATTTGGGTTACTCTATAATCTGAAGAATCTAACTGCA

>M6-PGSC0003DMG400008296M6LRR-RLK_1403

TGACTGGATCACTTCCATCGGAATTTGGGTTACTCTATAATCTGAAGAATCTAACTGCAAATGATAACCTCCTGGAAGGATCTATTCCATCGAGCATAACAAATTGTTCTCATCTTCTTG

>M6-PGSC0003DMG400008296M6LRR-RLK_1464

TGATAACCTCCTGGAAGGATCTATTCCATCGAGCATAACAAATTGTTCTCATCTTCTTGTTTTAACCCTTGCTTATAATAGAATAACGGGGAAAATACCAAATGGGTTGGGGCAATTGTC

>M6-PGSC0003DMG400008296M6LRR-RLK_1525

TTAACCCTTGCTTATAATAGAATAACGGGGAAAATACCAAATGGGTTGGGGCAATTGTCCAATCTTACATTTTTGTCTTTGGGATCAAACAAAATGATGGGGGAGATTCCTGAAGATCTC

>M6-PGSC0003DMG400008296M6LRR-RLK_1586

ATCTTACATTTTTGTCTTTGGGATCAAACAAAATGATGGGGGAGATTCCTGAAGATCTCTTCAACTGTTCAATGCTTGAAATTCTAGATCTGAGTGACAACAATTTCAGCGGGAAACTCA

>M6-PGSC0003DMG400008296M6LRR-RLK_1647

CAACTGTTCAATGCTTGAAATTCTAGATCTGAGTGACAACAATTTCAGCGGGAAACTCAAATCAATGATTGGCAGACTCTCTAAACTTCGAGTTTTAAGAGCTCGTACTAATTCATTTCT

>M6-PGSC0003DMG400008296M6LRR-RLK_1708

TCAATGATTGGCAGACTCTCTAAACTTCGAGTTTTAAGAGCTCGTACTAATTCATTTCTTGGGCCAATCCCACCAGAGATTGGTAAACTGAGTCAACTCATGGATATAGTGCTTCACGAA

>M6-PGSC0003DMG400008296M6LRR-RLK_1769

GGCCAATCCCACCAGAGATTGGTAAACTGAGTCAACTCATGGATATAGTGCTTCACGAAAACAGTTTCTCAGGTGCGATACCACCAGAAATTTCAATGCTTTCAAACCTTCAGGGTCTTT

>M6-PGSC0003DMG400008296M6LRR-RLK_183

CTGAAAACGGTGATTCACTGTACGATTCACTAAGAAATGTGTATAAAGGCTTCATTGTCAGCATAAGTATTTTCAGTGAGTGAAGCTTTTTTCTTTTGAAACAATGATGTCAAAGACAGT

>M6-PGSC0003DMG400008296M6LRR-RLK_1830

CAGTTTCTCAGGTGCGATACCACCAGAAATTTCAATGCTTTCAAACCTTCAGGGTCTTTTGCTATCTGAAAACAAGCTGGAAGGTGAACTTCCTGTGCAACTTTTTGAGCTCAAACAGCT

>M6-PGSC0003DMG400008296M6LRR-RLK_1891

CTATCTGAAAACAAGCTGGAAGGTGAACTTCCTGTGCAACTTTTTGAGCTCAAACAGCTCAATGAACTCCGGCTGAAGAACAATAATTTCTTTGGTCCACTACCGCAACATATATCCAAA

>M6-PGSC0003DMG400008296M6LRR-RLK_1952

ATGAACTCCGGCTGAAGAACAATAATTTCTTTGGTCCACTACCGCAACATATATCCAAACTAGAATCACTGTCGCTCATGGATCTGAGTGGAAATAAGCTTAATGGCACAATCCCAGAGA

>M6-PGSC0003DMG400008296M6LRR-RLK_2013

AGAATCACTGTCGCTCATGGATCTGAGTGGAAATAAGCTTAATGGCACAATCCCAGAGAGCATGGCGAGCCTCCGCAGATTGATGACCGTAGATCTTTCACACAACCTTCTTACCGGAAC

>M6-PGSC0003DMG400008296M6LRR-RLK_2074

ATGGCGAGCCTCCGCAGATTGATGACCGTAGATCTTTCACACAACCTTCTTACCGGAACTATTCCAAGAGCAGTACTTGCCAGCATGAGAAGAATGCAACTTTACTTGAACGTTTCCAGC

>M6-PGSC0003DMG400008296M6LRR-RLK_2135

TTCCAAGAGCAGTACTTGCCAGCATGAGAAGAATGCAACTTTACTTGAACGTTTCCAGCAATTTGTTGCACGGAGAGATCCCAGATGAGATAGGCGTGTTAGAAATGGTTCAAGAGATTG

>M6-PGSC0003DMG400008296M6LRR-RLK_2196

TTTGTTGCACGGAGAGATCCCAGATGAGATAGGCGTGTTAGAAATGGTTCAAGAGATTGACATGTCAAATAATAAACTGTCAGGCAGCATTCCCAAATCCCTAGAACGCTGCAAAAACTT

>M6-PGSC0003DMG400008296M6LRR-RLK_2257

ATGTCAAATAATAAACTGTCAGGCAGCATTCCCAAATCCCTAGAACGCTGCAAAAACTTATTTTCACTGGACCTATCAGGAAACATGCTCTCTGGTCCTGCTCCAGGTGAAATTCTCACC

>M6-PGSC0003DMG400008296M6LRR-RLK_2318

TTTCACTGGACCTATCAGGAAACATGCTCTCTGGTCCTGCTCCAGGTGAAATTCTCACCAAATTAAGAGAGCTTGTGTTCTTGAACCTCTCAAGGAACAGATTAGAAGGCAGTCTTCCTG

>M6-PGSC0003DMG400008296M6LRR-RLK_2379

ATTAAGAGAGCTTGTGTTCTTGAACCTCTCAAGGAACAGATTAGAAGGCAGTCTTCCTGAAATAGCAGGATTATCACATCTTCGCTCTCTTGATGTTTCACAAAACAAGTTCAATGGAAT

>M6-PGSC0003DMG400008296M6LRR-RLK_244

CATAAGTATTTTCAGTGAGTGAAGCTTTTTTCTTTTGAAACAATGATGTCAAAGACAGTTGTATATGCATTAGCCATATTTTCTATTACCTTTCTCATACCTCTGTCATCTGGGCAAAAT

>M6-PGSC0003DMG400008296M6LRR-RLK_2440

ATAGCAGGATTATCACATCTTCGCTCTCTTGATGTTTCACAAAACAAGTTCAATGGAATCATTCCCGAGAGATTTGCCAATATGACTGCATTGAAATATCTCAACCTTTCTTTCAACCAA

>M6-PGSC0003DMG400008296M6LRR-RLK_2501

TTCCCGAGAGATTTGCCAATATGACTGCATTGAAATATCTCAACCTTTCTTTCAACCAACTTGAAGGTCACATTCCAAAGGGGGGTGTATTCAACAATATAAGGTCGGAGGATTTACTGG

>M6-PGSC0003DMG400008296M6LRR-RLK_2562

TGAAGGTCACATTCCAAAGGGGGGTGTATTCAACAATATAAGGTCGGAGGATTTACTGGGAAATCCATCTCTAAGTGGAACAAAGTTTCTCAGTCCATGCAACATAAAAAGAAACGGAAC

>M6-PGSC0003DMG400008296M6LRR-RLK_2623

AATCCATCTCTAAGTGGAACAAAGTTTCTCAGTCCATGCAACATAAAAAGAAACGGAACAAGTTCTCATGGTTTTTCCAAGAAAACCTGGATCATACTTGCAGCACTTGGATCTGTTTTC

>M6-PGSC0003DMG400008296M6LRR-RLK_2684

GTTCTCATGGTTTTTCCAAGAAAACCTGGATCATACTTGCAGCACTTGGATCTGTTTTCAGTCTCATTCTTCTTGTCCTGGGAATATTTTTATTTCATCGGTACATGAAGAAGAAGAAAA

>M6-PGSC0003DMG400008296M6LRR-RLK_2745

TCTCATTCTTCTTGTCCTGGGAATATTTTTATTTCATCGGTACATGAAGAAGAAGAAAAATGTGAATGAAGCGGAGGATACAAATCCAAAGTATACCGCAGCACTGAGCCTCCAGAGATT

>M6-PGSC0003DMG400008296M6LRR-RLK_2806

GTGAATGAAGCGGAGGATACAAATCCAAAGTATACCGCAGCACTGAGCCTCCAGAGATTTTATCAAAAGGATTTGGAACATGCTACCAATAATTTCAGTCCGGAAAACATTATTGGAGCC

>M6-PGSC0003DMG400008296M6LRR-RLK_2867

ATCAAAAGGATTTGGAACATGCTACCAATAATTTCAGTCCGGAAAACATTATTGGAGCCAGCAGTTTAAGTACTGTGTACAAAGGTACACTGGAAGATGGGAAGATTGTAGCAGTTAAGA

>M6-PGSC0003DMG400008296M6LRR-RLK_2928

CAGTTTAAGTACTGTGTACAAAGGTACACTGGAAGATGGGAAGATTGTAGCAGTTAAGAAGCTGAATCACCAGTTCTCAGCAGAATCTGGTAAATGCTTTGATAGGGAAGTCAAGACTCT

>M6-PGSC0003DMG400008296M6LRR-RLK_2989

CTGAATCACCAGTTCTCAGCAGAATCTGGTAAATGCTTTGATAGGGAAGTCAAGACTCTGAGCCAACTCAGACACAGGAACCTAGTTAAGGTGCTAGGTTACGCTTGGGAAAGCAAGAAG

>M6-PGSC0003DMG400008296M6LRR-RLK_305

TATATGCATTAGCCATATTTTCTATTACCTTTCTCATACCTCTGTCATCTGGGCAAAATCCAAGCTTTGAAGTTGAAGTTGCTGCTTTGAAAGCTTTCAAGAGCTCAATCTCTGACGACC

>M6-PGSC0003DMG400008296M6LRR-RLK_3050

GCCAACTCAGACACAGGAACCTAGTTAAGGTGCTAGGTTACGCTTGGGAAAGCAAGAAGCTAAGGGCTTTAGTTTTAGAATACATGGAGAATGGGAACTTGGACAACATGATTTATGGTC

>M6-PGSC0003DMG400008296M6LRR-RLK_3111

AAGGGCTTTAGTTTTAGAATACATGGAGAATGGGAACTTGGACAACATGATTTATGGTCAAGTGGAGGATAACTGGACGTTGTCCAATAGGATTGACATTTTAGTTTCAGTTGCAAGTGG

>M6-PGSC0003DMG400008296M6LRR-RLK_3172

GTGGAGGATAACTGGACGTTGTCCAATAGGATTGACATTTTAGTTTCAGTTGCAAGTGGACTGTCATACCTGCATTCAGGCTATGATTTTCCAATAGTGCACTGCGACATGAAGCCTTCA

>M6-PGSC0003DMG400008296M6LRR-RLK_3233

TGTCATACCTGCATTCAGGCTATGATTTTCCAATAGTGCACTGCGACATGAAGCCTTCAAATATTCTTCTGGATAAAAATATGGAGGCACATGTGAGTGACTTTGGGACAGCTAGGATGT

>M6-PGSC0003DMG400008296M6LRR-RLK_3355

GGTATTCATCTCCAGGATGGGAGCAGCATATCATCAGCATCCGCATTTGAAGGAACGATTGGTTACATGGCGCCAGGTAATAATTCTTACAACTTCTTGACCCACACTTTACTAACTAGA

>M6-PGSC0003DMG400008296M6LRR-RLK_3416

GTTACATGGCGCCAGGTAATAATTCTTACAACTTCTTGACCCACACTTTACTAACTAGATTATGTGTTACAATATCATGAATACAAGATGTCTACAATGTAAACGAGCTTATATGGGTAC

>M6-PGSC0003DMG400008296M6LRR-RLK_3477

ATGTGTTACAATATCATGAATACAAGATGTCTACAATGTAAACGAGCTTATATGGGTACCACAAATAATATTTTGTCCATGATGAATGTTTCAAATACATTCCATTTTAGCATCATTCAG

>M6-PGSC0003DMG400008296M6LRR-RLK_3538

CAAATAATATTTTGTCCATGATGAATGTTTCAAATACATTCCATTTTAGCATCATTCAGTAGTAACATAACTTCTGTCAATCACAATACGCAGCACAGCAGATTTTTTTTTCCTTTCAAT

>M6-PGSC0003DMG400008296M6LRR-RLK_3599

GTAACATAACTTCTGTCAATCACAATACGCAGCACAGCAGATTTTTTTTTCCTTTCAATCACAAAATACTGATTTAACTTTTATGTGGAAGAATAGACAAGCAAGTAAATGATGCAATGT

>M6-PGSC0003DMG400008296M6LRR-RLK_366

AAGCTTTGAAGTTGAAGTTGCTGCTTTGAAAGCTTTCAAGAGCTCAATCTCTGACGACCCTTTCAGTGCACTTGTGGATTGGACTGATGCAAATCACCATTGCAACTGGTCTGGAATCAT

>M6-PGSC0003DMG400008296M6LRR-RLK_3660

CAAAATACTGATTTAACTTTTATGTGGAAGAATAGACAAGCAAGTAAATGATGCAATGTTTAGAACATATAATCTGTAAAGTATTTTTTTAAATGATAATGTCAGTTTTAATCTATCCAA

>M6-PGSC0003DMG400008296M6LRR-RLK_3721

AGAACATATAATCTGTAAAGTATTTTTTTAAATGATAATGTCAGTTTTAATCTATCCAAAAATCTAGCGCAAGCTTTGAAAATTTGAATTTCTTGGACATTTTACATGCTCCTAATTTAA

>M6-PGSC0003DMG400008296M6LRR-RLK_3782

ATCTAGCGCAAGCTTTGAAAATTTGAATTTCTTGGACATTTTACATGCTCCTAATTTAATTTGAGACCTCACTTACATGAAATGTTATCTATGAAACAGAGTTTGCATATATGAGGAAAG

>M6-PGSC0003DMG400008296M6LRR-RLK_3843

TGAGACCTCACTTACATGAAATGTTATCTATGAAACAGAGTTTGCATATATGAGGAAAGTGACCACAAAAGTAGATGTATTCAGCTTTGGTGTCATTGTGATGGAGATCATTACAAAAAG

>M6-PGSC0003DMG400008296M6LRR-RLK_3904

ACCACAAAAGTAGATGTATTCAGCTTTGGTGTCATTGTGATGGAGATCATTACAAAAAGAAGGCCAACAAGTCTTACCGGAGCAGATGAATTACCAATGACTTTGCATCAAATTGTTCAG

>M6-PGSC0003DMG400008296M6LRR-RLK_3965

GGCCAACAAGTCTTACCGGAGCAGATGAATTACCAATGACTTTGCATCAAATTGTTCAGAATGCCCTTGCGAATGGCATAAACAAGCTAGTTCAGATTGTGGATCCTAATCTAGCTTCGT

>M6-PGSC0003DMG400008296M6LRR-RLK_4026

TGCCCTTGCGAATGGCATAAACAAGCTAGTTCAGATTGTGGATCCTAATCTAGCTTCGTATGTCTCCAAGAAACAGGATGTAGTAGAGGGACTTCTTAACTTGGCTTTGTCCTGCACCTG

>M6-PGSC0003DMG400008296M6LRR-RLK_4087

GTCTCCAAGAAACAGGATGTAGTAGAGGGACTTCTTAACTTGGCTTTGTCCTGCACCTGTCCTGATCCTGAAGATAGACCTGACATGGAACAAATTCTGTCTTCTCTTTCAAAGCTAAGT

>M6-PGSC0003DMG400008296M6LRR-RLK_4148

CTGATCCTGAAGATAGACCTGACATGGAACAAATTCTGTCTTCTCTTTCAAAGCTAAGTAAGATGGACTGCATGCCTTCGCATTTGGTAAAAGATTAAATTTGATGTAGAAGAAGCTGGA

>M6-PGSC0003DMG400008296M6LRR-RLK_4209

GATGGACTGCATGCCTTCGCATTTGGTAAAAGATTAAATTTGATGTAGAAGAAGCTGGAATATCATCACTCTTAATGTGAACCACCACCAAGTTTCCCATGGTGGCTAATGGAACTAGTT

>M6-PGSC0003DMG400008296M6LRR-RLK_427

TTCAGTGCACTTGTGGATTGGACTGATGCAAATCACCATTGCAACTGGTCTGGAATCATATGTGATCCTTCTTCAAATCATGTCATCAACATTTCGCTTATTGAGACACAGCTCAAAGGC

>M6-PGSC0003DMG400008296M6LRR-RLK_4270

ATCATCACTCTTAATGTGAACCACCACCAAGTTTCCCATGGTGGCTAATGGAACTAGTTCATAACTTAAAATGTATTTACTATTTAAATAAATTTTCCATATTTCAAGCCACAAATGAAC

>M6-PGSC0003DMG400008296M6LRR-RLK_4331

TAACTTAAAATGTATTTACTATTTAAATAAATTTTCCATATTTCAAGCCACAAATGAACTCTCGAGTGTTCCTGCTTTGCTAGAGAGATATAATGGAACTGTCCCATCAGATTATGTTGA

>M6-PGSC0003DMG400008296M6LRR-RLK_488

GTGATCCTTCTTCAAATCATGTCATCAACATTTCGCTTATTGAGACACAGCTCAAAGGCGAAATTTCTCCGTTTCTAGGAAACCTCTCCAAACTCCAGGTTCTTGATCTAACTTTGAATT

>M6-PGSC0003DMG400008296M6LRR-RLK_549

AATTTCTCCGTTTCTAGGAAACCTCTCCAAACTCCAGGTTCTTGATCTAACTTTGAATTCATTTACTGGAAATATCCCACCCCAGCTGGGGCATTGCACAGACCTGGTTGAGCTTGTTTT

>M6-PGSC0003DMG400008296M6LRR-RLK_61

TACATACAACTTGTTCTTTAACACGGAGCATCCTAGATTAGAGGCATGCTTGGTATTTTGTAACAAATTCATAGAATGCAAGTCATTTTCCTTAGTGTGAATGGATGACTAGTTGACTGC

>M6-PGSC0003DMG400008296M6LRR-RLK_610

TTTACTGGAAATATCCCACCCCAGCTGGGGCATTGCACAGACCTGGTTGAGCTTGTTTTTAACCAAAACTCTCTTTTTGGTGAAATTCCTGCCGAGCTGGAAAACCTAAAAAAACTGCAG

>M6-PGSC0003DMG400008296M6LRR-RLK_671

ACCAAAACTCTCTTTTTGGTGAAATTCCTGCCGAGCTGGAAAACCTAAAAAAACTGCAGTTAATAGATTTTGGAAATAACTTTCTAAATGGGAGTATACCTGATAGTATATGCAACTGCA

>M6-PGSC0003DMG400008296M6LRR-RLK_732

AATAGATTTTGGAAATAACTTTCTAAATGGGAGTATACCTGATAGTATATGCAACTGCACTGAATTGTTGTTAGTAGGCCTCAACAACAACAATCTCACAGGTAAATTACCATCTGAAAT

>M6-PGSC0003DMG400008296M6LRR-RLK_793

GAATTGTTGTTAGTAGGCCTCAACAACAACAATCTCACAGGTAAATTACCATCTGAAATTGGCAATTTGGCTAATCTTCAGTTGTTTGTAGCTTATACAAACAATTTGGTTGGTTCTATT

>M6-PGSC0003DMG400008296M6LRR-RLK_854

GCAATTTGGCTAATCTTCAGTTGTTTGTAGCTTATACAAACAATTTGGTTGGTTCTATTCCTACCTCCATTGGAATGCTGACAGCATTGCAAACCCTTGATCTGAGCGAAAACCAGTTAT

>M6-PGSC0003DMG400008296M6LRR-RLK_915

TACCTCCATTGGAATGCTGACAGCATTGCAAACCCTTGATCTGAGCGAAAACCAGTTATCTGGACCTATACCACCAGCAATTGGCAACTTGTCAAGTTTAGGAATTCTTCAGTTACATCT

>M6-PGSC0003DMG400008296M6LRR-RLK_976

GGACCTATACCACCAGCAATTGGCAACTTGTCAAGTTTAGGAATTCTTCAGTTACATCTCAACTCTCTTTCTGGGAAAATCCCATCTGAACTTGGCCTCTGTATCAATCTTTTTACCTTG

>M6-PGSC0003DMG400008394-(rev)M6LRR-RLP_0

CTTCCTTTGAGTACTCTACAAACAACTACCAAACGAAGACTTACTTTGAGAAGACTTTGTTGCATACAGCTCAGGAAGAAGACTTTGGTGCAGATTGCAGAGGAAGAGGAAGCAAACACA

>M6-PGSC0003DMG400008394-(rev)M6LRR-RLP_1037

GAAGCAGGTGGCATCCACAGTAGCAACAGATCTTCATATCTTGGGGAAGTTAACAGAAGATCATTGTTGGTCTATTTTCAAACAAAAAGCATTTGTTGATGGCAGGGTTCCAGAGGAATT

>M6-PGSC0003DMG400008394-(rev)M6LRR-RLP_1159

CGAGCATGGGCAACAAGATTGTTAAAATGTGCCAAGGTCTACCGTTGGCTGCAAGTGCATTGGGAGGGCTCTTACACAACAAAGAAAAACATGAATGGCAAGCAATTCTTGATGGCAACC

>M6-PGSC0003DMG400008394-(rev)M6LRR-RLP_1220

GGGAGGGCTCTTACACAACAAAGAAAAACATGAATGGCAAGCAATTCTTGATGGCAACCTCCTTGTTACTGGTGAAGATGATAATGGAGAAAATAGCATAAAGAAAATCCTAAAACTCAG

>M6-PGSC0003DMG400008394-(rev)M6LRR-RLP_1281

CTTGTTACTGGTGAAGATGATAATGGAGAAAATAGCATAAAGAAAATCCTAAAACTCAGCTATGATTATCTACCATCTCCACATCTGAAAAAATGTTTTGCTTACTTTGCAATGTTTCCA

>M6-PGSC0003DMG400008394-(rev)M6LRR-RLP_1342

ATGATTATCTACCATCTCCACATCTGAAAAAATGTTTTGCTTACTTTGCAATGTTTCCAAAAGATTATATGTTTGAAAAGGACCAACTAATCCAACTCTGGATGGCAGAAGGGTTTCTTC

>M6-PGSC0003DMG400008394-(rev)M6LRR-RLP_1464

CCAAGTCAAGAGATCCCTGTGATGGAAGACGTTGGGCACAGGTTTTTTCAAATCTTGTTGCAGAATTCCTTGCTACAAGATGTTGTGTTAGATGAACACAACAATATAACACACTGTAAG

>M6-PGSC0003DMG400008394-(rev)M6LRR-RLP_1525

AGAATTCCTTGCTACAAGATGTTGTGTTAGATGAACACAACAATATAACACACTGTAAGATGCACGATCTTGTGCATGATTTGGCTGGAGATATCTTAAAATCTAGACTATTTGATCCGA

>M6-PGSC0003DMG400008394-(rev)M6LRR-RLP_1586

GCACGATCTTGTGCATGATTTGGCTGGAGATATCTTAAAATCTAGACTATTTGATCCGAAGGGCGACAATGGAGAAAAACTTTCTCAAGTTCGATACTTTGGATGTGAGTCACCAACGGA

>M6-PGSC0003DMG400008394-(rev)M6LRR-RLP_1647

GGCGACAATGGAGAAAAACTTTCTCAAGTTCGATACTTTGGATGTGAGTCACCAACGGATCAAATAGATAAGATATATGAGCCAGAACGTTTGTGCACACTGTTTTGGAGAAGCAATTAT

>M6-PGSC0003DMG400008394-(rev)M6LRR-RLP_1708

AAATAGATAAGATATATGAGCCAGAACGTTTGTGCACACTGTTTTGGAGAAGCAATTATACATCTAAAGATATGCTGTTGAAATATAAGTTCTTGAGAGTTTTAGATTTGTCCAGTTCAG

>M6-PGSC0003DMG400008394-(rev)M6LRR-RLP_1769

ATCTAAAGATATGCTGTTGAAATATAAGTTCTTGAGAGTTTTAGATTTGTCCAGTTCAGGAATCAAGGAGTTGTCAACCAAAATCGGGAAGCTGATATACTTGAGATATCTTGATCTCTC

>M6-PGSC0003DMG400008394-(rev)M6LRR-RLP_183

ACTATCGAGGAGGTCAACAGCTCAAGGGATTTCAACAAAGATCTCGAAATGTTGACACAAAATGTATCATTAATCCAAGCTTTCATTCATGATGTTGAAACACCACAAGAGAAACAACAG

>M6-PGSC0003DMG400008394-(rev)M6LRR-RLP_1830

ATCAAGGAGTTGTCAACCAAAATCGGGAAGCTGATATACTTGAGATATCTTGATCTCTCGAACACTGAGATCACAGCCTTGCCCAACTCCATTTGCGAGCTCTATAATTTGCAAACATTT

>M6-PGSC0003DMG400008394-(rev)M6LRR-RLP_1952

AGTCATCAACTGCTTTTCACTCCAGGAACTTCCATATGAGATGAGAAATATGATAAGTTTGAGACACATATATTACACTTCTGTTGACGAAACAAGTGGGCATTGGGGAGGATGGTGTCT

>M6-PGSC0003DMG400008394-(rev)M6LRR-RLP_2013

AGACACATATATTACACTTCTGTTGACGAAACAAGTGGGCATTGGGGAGGATGGTGTCTTCACAATGAACATTTTCAGATTCCACTTAATATGGGGCAATTGACTAGTCTTCAAACCCTC

>M6-PGSC0003DMG400008394-(rev)M6LRR-RLP_2074

ACAATGAACATTTTCAGATTCCACTTAATATGGGGCAATTGACTAGTCTTCAAACCCTCAAGTTTTTCAAGGTAGGTTTAGAGAAAGGTCGTCAAATAGAAGAATTAGGTCATTTGAAAA

>M6-PGSC0003DMG400008394-(rev)M6LRR-RLP_2135

GTTTTTCAAGGTAGGTTTAGAGAAAGGTCGTCAAATAGAAGAATTAGGTCATTTGAAAAACCTAAGAGGTGAATTGACGATCAATGGTCTCCAATTGGTCTGTGATAAAGAAGAGGCTCG

>M6-PGSC0003DMG400008394-(rev)M6LRR-RLP_2196

CTAAGAGGTGAATTGACGATCAATGGTCTCCAATTGGTCTGTGATAAAGAAGAGGCTCGAACAGCATATTTACACGATAAACCAAACATCTGCAAGCTGGCATATTTATGGTCCCACGAT

>M6-PGSC0003DMG400008394-(rev)M6LRR-RLP_2257

CAGCATATTTACACGATAAACCAAACATCTGCAAGCTGGCATATTTATGGTCCCACGATGAATCAGAAGGCTGTGAGATCAATGATGAGCATGTGTTGGATGGTCTTCAACCGCATCCTA

>M6-PGSC0003DMG400008394-(rev)M6LRR-RLP_2379

TTGAAAACCATAGCAGTAGTGGACTATTTAGGGACTAAATTTCCTTCATGGTTCCGTGAAGAGTCGCTACCAAATTTGGTCAAGTTGAAATTAAGTGGAAGCAAAAGGTGCAAAGAAATT

>M6-PGSC0003DMG400008394-(rev)M6LRR-RLP_244

ATGTATCATTAATCCAAGCTTTCATTCATGATGTTGAAACACCACAAGAGAAACAACAGTCTGTCGAACAATGGCTCAACAGGCTTGAGAGAGTTGCTGAAGATGCTGAAAATGTGTTTG

>M6-PGSC0003DMG400008394-(rev)M6LRR-RLP_2501

ATCCCTTGGCCAACTGAAATTCCTTCGGCATCTTGAGCTGATAGGATTCCATGAGTTGGAATGCATTGGACCTGCTTTTTATGGTGTTGAGATTAGAAATATTGGATCAAACAGCATTAT

>M6-PGSC0003DMG400008394-(rev)M6LRR-RLP_2562

TGCATTGGACCTGCTTTTTATGGTGTTGAGATTAGAAATATTGGATCAAACAGCATTATTCAAGTGTTCCCGTCATTGAAAGAACTAGTATTGGAGGATATGCGTAGCCTTATTGAGTGG

>M6-PGSC0003DMG400008394-(rev)M6LRR-RLP_2623

AAGTGTTCCCGTCATTGAAAGAACTAGTATTGGAGGATATGCGTAGCCTTATTGAGTGGAAGGGAGATGAAGTTGGAGTAAGAATGTCTCCCGGGCTTGAGAAGTTGCGGATTACAGACT

>M6-PGSC0003DMG400008394-(rev)M6LRR-RLP_2684

GGGAGATGAAGTTGGAGTAAGAATGTCTCCCGGGCTTGAGAAGTTGCGGATTACAGACTGTCCATTGTTAAAAAGTATTCCGAATCAATTTGAAATCCTCCGTGAATTAGAAATTAGAGG

>M6-PGSC0003DMG400008394-(rev)M6LRR-RLP_2745

CCATTGTTAAAAAGTATTCCGAATCAATTTGAAATCCTCCGTGAATTAGAAATTAGAGGAGTTGACAGTGAAATGCCATTGTTGAACTTGTGCAGCAACTTGACATCTCTCGTGAATCTT

>M6-PGSC0003DMG400008394-(rev)M6LRR-RLP_2806

TTGACAGTGAAATGCCATTGTTGAACTTGTGCAGCAACTTGACATCTCTCGTGAATCTTAGTGTCTATGATATGAAAGAGCTCACTTGTCTTCCAGATGAGATACTACGTAACAACATTT

>M6-PGSC0003DMG400008394-(rev)M6LRR-RLP_2928

CTTCAACAGATAACGATTTTCGAATGTGGAGAGTTTCGTGAATTGCCACAAAGCTTGTACAATCTCCATTCTCTTTATAGATTAGACATTTACAACTGCACCAATTTCAGTTCTCTTCCT

>M6-PGSC0003DMG400008394-(rev)M6LRR-RLP_2989

ATCTCCATTCTCTTTATAGATTAGACATTTACAACTGCACCAATTTCAGTTCTCTTCCTGTTCCCAAGGGAGACAACTATTTGACTTCCCTCATATTCTTTTTCTTACATAATTGTAATG

>M6-PGSC0003DMG400008394-(rev)M6LRR-RLP_305

TGTCGAACAATGGCTCAACAGGCTTGAGAGAGTTGCTGAAGATGCTGAAAATGTGTTTGATCGATTCAGATATGAATCTCTCAAAACAAAAGTGGTGAGGAGCCCATTGAAAAAGGTCGG

>M6-PGSC0003DMG400008394-(rev)M6LRR-RLP_3050

TCCCAAGGGAGACAACTATTTGACTTCCCTCATATTCTTTTTCTTACATAATTGTAATGGATTGATCAGTATACCAATTGGAATGCTAGATCAATGCCGGTCTCTAGAGTATTTGCATGT

>M6-PGSC0003DMG400008394-(rev)M6LRR-RLP_3111

TTGATCAGTATACCAATTGGAATGCTAGATCAATGCCGGTCTCTAGAGTATTTGCATGTCAGCTGCTGTAACAACTTGGTTTCATTCCCTTTACATGTGTGGGAAATTCCTTCATTTTCA

>M6-PGSC0003DMG400008394-(rev)M6LRR-RLP_3172

GCTGCTGTAACAACTTGGTTTCATTCCCTTTACATGTGTGGGAAATTCCTTCATTTTCAGTTTTGGAAATAACAGAATGTCCCAAATTGATTAGTGTACCCGAAGTGGGCCTTCACCATC

>M6-PGSC0003DMG400008394-(rev)M6LRR-RLP_3233

TTTGGAAATAACAGAATGTCCCAAATTGATTAGTGTACCCGAAGTGGGCCTTCACCATCTCACCGGGTTATTGAGATTGGGAATTGGTCCTTTCTCAGAGATGGTGGATTTTGATGCATT

>M6-PGSC0003DMG400008394-(rev)M6LRR-RLP_3355

AATTGATTTTTAATGGCATTCAGAAGTTGTTGTCTCTTCGTGATCTGGAGGTGTACGGACGTGGGCACTGGGATTCTCTGCCCTATCAGCTTATGCAACTCTCTAACCTAAGAAAGATCA

>M6-PGSC0003DMG400008394-(rev)M6LRR-RLP_3416

TGGGCACTGGGATTCTCTGCCCTATCAGCTTATGCAACTCTCTAACCTAAGAAAGATCACAATAGCTGATTTCGGAATTGAGGCTCTTCCTCCTAGACTTGACAACCTTACTTCTCTTGA

>M6-PGSC0003DMG400008394-(rev)M6LRR-RLP_3477

ATAGCTGATTTCGGAATTGAGGCTCTTCCTCCTAGACTTGACAACCTTACTTCTCTTGAAAGTTTGACGCTAGAGAGGTGCAAACGGCTACAACATCTGAACTTCTCAGATGCCATGCCC

>M6-PGSC0003DMG400008394-(rev)M6LRR-RLP_3538

GTTTGACGCTAGAGAGGTGCAAACGGCTACAACATCTGAACTTCTCAGATGCCATGCCCAAATTACGGCATCTGTGGATAAGTGATTGTCCATTGTTAGAAGCCCTGTCGGATGGGCTCG

>M6-PGSC0003DMG400008394-(rev)M6LRR-RLP_3599

ATTACGGCATCTGTGGATAAGTGATTGTCCATTGTTAGAAGCCCTGTCGGATGGGCTCGGCAGCATTGTTTCTTTGGAAGAATTATATTTACAGGACTGCGAAAAACTAGAGCATCTACC

>M6-PGSC0003DMG400008394-(rev)M6LRR-RLP_366

CGATTCAGATATGAATCTCTCAAAACAAAAGTGGTGAGGAGCCCATTGAAAAAGGTCGGTGGTTTCTTTTCTCATACTGCTTTTAAGAGAAAAATGTCTCAAAAAATCAACAACATCAAT

>M6-PGSC0003DMG400008394-(rev)M6LRR-RLP_3660

AGCATTGTTTCTTTGGAAGAATTATATTTACAGGACTGCGAAAAACTAGAGCATCTACCGTCCCGAGATGCCATGCGACGCCTCACTAAATTATGGAACATGAGAATTAAAGGATGCCCA

>M6-PGSC0003DMG400008394-(rev)M6LRR-RLP_3782

GTTAGAAGAAAGTTTCACCAACTACTCCCAGTGGTCCAAAATTTCCCATATTTCAAATATTGAATTAGGTGGGTGGAGAAGGACAACCATAAGTCTCGGTTTCTCTTTCACTTTCTGAGT

>M6-PGSC0003DMG400008394-(rev)M6LRR-RLP_3843

GAATTAGGTGGGTGGAGAAGGACAACCATAAGTCTCGGTTTCTCTTTCACTTTCTGAGTCGCTTGCATTTTTAATTAGAATATTATTTTCCACGAGTTCCAAAAAAGAAGTTTTGTAAGT

>M6-PGSC0003DMG400008394-(rev)M6LRR-RLP_3904

CTTGCATTTTTAATTAGAATATTATTTTCCACGAGTTCCAAAAAAGAAGTTTTGTAAGTAATGTAATAATTTTTCTTGAAATTATTTTGACATAAAAAAAAAAGCTAGATTATCAATCAA

>M6-PGSC0003DMG400008394-(rev)M6LRR-RLP_3965

TGTAATAATTTTTCTTGAAATTATTTTGACATAAAAAAAAAAGCTAGATTATCAATCAATTGCATCTGTTGCTATGCTTTCAGGATTCTGCTTGTCCACTTGAACTGAATTGCTGTTATT

>M6-PGSC0003DMG400008394-(rev)M6LRR-RLP_4026

GCATCTGTTGCTATGCTTTCAGGATTCTGCTTGTCCACTTGAACTGAATTGCTGTTATTCTAAAAATTATGTTTCTTTTCCTTATAGGTAGCCATTATAGACGTCTCTGTGGTAAATCAA

>M6-PGSC0003DMG400008394-(rev)M6LRR-RLP_4087

AAAAATTATGTTTCTTTTCCTTATAGGTAGCCATTATAGACGTCTCTGTGGTAAATCAAGCCAAATCATATGAGCAGCAGGCAATTACGTAAGGTAAATTTTCATTTATAATTTATAGGC

>M6-PGSC0003DMG400008394-(rev)M6LRR-RLP_4148

CAAATCATATGAGCAGCAGGCAATTACGTAAGGTAAATTTTCATTTATAATTTATAGGCACTTTGTAATCTACTTGTTTCTATCTTTCTTTTTGTTGGAAACCAATTTTCGGAGGTCTAT

>M6-PGSC0003DMG400008394-(rev)M6LRR-RLP_4209

TTTGTAATCTACTTGTTTCTATCTTTCTTTTTGTTGGAAACCAATTTTCGGAGGTCTATGCTGATGAATACAACATTACCTTCCTTAAAAAAAAGTTTAATCATGAAAATTTATGATTTC

>M6-PGSC0003DMG400008394-(rev)M6LRR-RLP_427

GTTTCTTTTCTCATACTGCTTTTAAGAGAAAAATGTCTCAAAAAATCAACAACATCAATAAAGAGTTGACGGCTATCAATAAGGTAGCCAAAGACCTCGGTCTACAATCACTCATGGTAC

>M6-PGSC0003DMG400008394-(rev)M6LRR-RLP_4270

TGATGAATACAACATTACCTTCCTTAAAAAAAAGTTTAATCATGAAAATTTATGATTTCTGCTCATAGGTTACTAGGTACAAGGACATGGTAGAACGTCGTTGTGGTATATCTAGTCAAA

>M6-PGSC0003DMG400008394-(rev)M6LRR-RLP_4331

CTCATAGGTTACTAGGTACAAGGACATGGTAGAACGTCGTTGTGGTATATCTAGTCAAAATCCATGGGCAGAATCATGTTAGGCATTTTTATTTAGTTGTTAGAGGAATGCTATAACAAG

>M6-PGSC0003DMG400008394-(rev)M6LRR-RLP_4392

CCATGGGCAGAATCATGTTAGGCATTTTTATTTAGTTGTTAGAGGAATGCTATAACAAGTTTGTAAAGTTTCTTTTTTTTACTTTCCTTTTCTCTCATGAAATAATTTTAATGTTTAACG

>M6-PGSC0003DMG400008394-(rev)M6LRR-RLP_4453

TGTAAAGTTTCTTTTTTTTACTTTCCTTTTCTCTCATGAAATAATTTTAATGTTTAACGGGACGTCGTGTTTCACAGAGGAACCGCCACTGTTATTTTGGATGAAGAACAATCTTCAACT

>M6-PGSC0003DMG400008394-(rev)M6LRR-RLP_4514

ACGTCGTGTTTCACAGAGGAACCGCCACTGTTATTTTGGATGAAGAACAATCTTCAACTGTAGACTCATTATGGGACACTTTCATGAGTATTTTCGGGGAACTACAGTCTACATATGAAA

>M6-PGSC0003DMG400008394-(rev)M6LRR-RLP_4575

AGACTCATTATGGGACACTTTCATGAGTATTTTCGGGGAACTACAGTCTACATATGAAACATCAAAAACTCAAAAGCGTATAGGGTGCTCAATGAGAAGCTACACGGGGTCTATTCTCCA

>M6-PGSC0003DMG400008394-(rev)M6LRR-RLP_4636

TCAAAAACTCAAAAGCGTATAGGGTGCTCAATGAGAAGCTACACGGGGTCTATTCTCCATGTAAGTATTGATAACTCTCAAGGAACAATACCAAGTCTCATATATGGTAGAAATCTATAT

>M6-PGSC0003DMG400008394-(rev)M6LRR-RLP_4697

TAAGTATTGATAACTCTCAAGGAACAATACCAAGTCTCATATATGGTAGAAATCTATATAAGTTTCTTAATAAACTCATAAAAGAATAATCTTTATGTCTTCTGCAGTTCTGGTTTTATT

>M6-PGSC0003DMG400008394-(rev)M6LRR-RLP_4758

GTTTCTTAATAAACTCATAAAAGAATAATCTTTATGTCTTCTGCAGTTCTGGTTTTATTTTCATGCTTAAGACAGTACAGCTTTTATGAATATATGACACACTTCTCGTTCCTTTCACCT

>M6-PGSC0003DMG400008394-(rev)M6LRR-RLP_4819

CATGCTTAAGACAGTACAGCTTTTATGAATATATGACACACTTCTCGTTCCTTTCACCTCACAGGACAATTTCTCTGTTCATTAAAACTTCCGCGAATGAAGGACTTCAACAAAGATAGT

>M6-PGSC0003DMG400008394-(rev)M6LRR-RLP_488

AGAGTTGACGGCTATCAATAAGGTAGCCAAAGACCTCGGTCTACAATCACTCATGGTACCTTCTCGGAAAATACTACCAATTCGAGAAACAGATTCCTTAGTAGTTGCTTCTGATATTGT

>M6-PGSC0003DMG400008394-(rev)M6LRR-RLP_4880

CAGGACAATTTCTCTGTTCATTAAAACTTCCGCGAATGAAGGACTTCAACAAAGATAGTTGTGATGAATATTTTTGCAAAGGGTGACCCATTGGCTCTTAAAATGTCCTTAGCAGTATAC

>M6-PGSC0003DMG400008394-(rev)M6LRR-RLP_4941

TGATGAATATTTTTGCAAAGGGTGACCCATTGGCTCTTAAAATGTCCTTAGCAGTATACGTTCATCTTAAACAGGTTTTCCTCTGTCCTTTTTTAGTGGTTACTTGTAATGATCCGCTTC

>M6-PGSC0003DMG400008394-(rev)M6LRR-RLP_5002

TCATCTTAAACAGGTTTTCCTCTGTCCTTTTTTAGTGGTTACTTGTAATGATCCGCTTCTCCATTCTCCAAGGAGGAAACGTGTGCCAAAACTGACTTTTTACCATTTCCCGTGTTTCAG

>M6-PGSC0003DMG400008394-(rev)M6LRR-RLP_5063

CATTCTCCAAGGAGGAAACGTGTGCCAAAACTGACTTTTTACCATTTCCCGTGTTTCAGATTCATGTTTGAAACTCCAGCATGGACCTTGTCGAAATGCTACTTGTGTTAGACGTGTTTG

>M6-PGSC0003DMG400008394-(rev)M6LRR-RLP_5185

TGTCATGATTGGCAAGGGAAATTGCATGTGGCGGAAATTGGAGTGGAAATACTATTTACTTGTATGACTAAGATTTGACAGCAGAGGCGCCTTTCTTTCACAGCATACGTGCCTTTTGGT

>M6-PGSC0003DMG400008394-(rev)M6LRR-RLP_5246

GTATGACTAAGATTTGACAGCAGAGGCGCCTTTCTTTCACAGCATACGTGCCTTTTGGTACAAATAGAGTTCATTTCTCCAACAAACATTTTGAACTTTTATACTTAGCTGCTGCATCAT

>M6-PGSC0003DMG400008394-(rev)M6LRR-RLP_5307

AAATAGAGTTCATTTCTCCAACAAACATTTTGAACTTTTATACTTAGCTGCTGCATCATTGCCTAGTTGTTAGCTTCTGTAATGTACATCTGCTTTTCCTCTTGAAATTTCTCTTGGCTT

>M6-PGSC0003DMG400008394-(rev)M6LRR-RLP_5368

CCTAGTTGTTAGCTTCTGTAATGTACATCTGCTTTTCCTCTTGAAATTTCTCTTGGCTTTTGTTCGCAAGTATAAACAAAAAAGACAGGGCCTACCATTTGCATTGTTTAAATCCAAACA

>M6-PGSC0003DMG400008394-(rev)M6LRR-RLP_5429

GTTCGCAAGTATAAACAAAAAAGACAGGGCCTACCATTTGCATTGTTTAAATCCAAACAATATGTATTCAAAAGTCACAAGCAATGTGTGTTATTCCAATTTCGTACAGGACTGCATTTT

>M6-PGSC0003DMG400008394-(rev)M6LRR-RLP_5488

ATATGTATTCAAAAGTCACAAGCAATGTGTGTTATTCCAATTTCGTACAGGACTGCATTTTTGTGTCTATGTATAACATATCGCATTGCTCATTATTCTTTCAGATAGACCAGATTGCAC

>M6-PGSC0003DMG400008394-(rev)M6LRR-RLP_549

TCTCGGAAAATACTACCAATTCGAGAAACAGATTCCTTAGTAGTTGCTTCTGATATTGTCGGTAGAGATTTGGATGTTGCTGAGATAAAGGAGAAGATTTTGAAGATGAGAGAGGAGGAT

>M6-PGSC0003DMG400008394-(rev)M6LRR-RLP_61

GCATACAGCTCAGGAAGAAGACTTTGGTGCAGATTGCAGAGGAAGAGGAAGCAAACACAACAATGGCCGATCCTGTAATTGGTGCTACTGTTCAAGTTTTGCTTGAAAAGTTGATTTCTC

>M6-PGSC0003DMG400008394-(rev)M6LRR-RLP_610

GTAGAGATTTGGATGTTGCTGAGATAAAGGAGAAGATTTTGAAGATGAGAGAGGAGGATATTGTTCTGTCCACCATTCCCATTGTAGGTATGGGAGGTTTAGGGAAAACAACTGTGGCTA

>M6-PGSC0003DMG400008394-(rev)M6LRR-RLP_671

TGTTCTGTCCACCATTCCCATTGTAGGTATGGGAGGTTTAGGGAAAACAACTGTGGCTAAGAGGATTTACAATGATGAACACATGAAGCAAATCTTTAAAAAGAGAATTTGGTTGTGTCT

>M6-PGSC0003DMG400008394-(rev)M6LRR-RLP_732

AGGATTTACAATGATGAACACATGAAGCAAATCTTTAAAAAGAGAATTTGGTTGTGTCTACCTGAAATGTCTGAAACGAAGAGCTTTCTTGAACAAATCCTCGAATCATTGATAGAGAGG

>M6-PGSC0003DMG400008394-(rev)M6LRR-RLP_854

AATTGAGGTTGAAAGGAGAGATATAATAGTCAAGAAGCTACAAGATGAATTGGGAGGAAAAAAATATTTGCTAGTCCTGGATGATTTGTGGCGTGTTGACTCTACATCGTGGCATGAGTT

>M6-PGSC0003DMG400008394-(rev)M6LRR-RLP_915

AAATATTTGCTAGTCCTGGATGATTTGTGGCGTGTTGACTCTACATCGTGGCATGAGTTCGTCGACACCCTGAGAAGAATAAATACATCCAGAGGAAACTGCATTCTTGTGACTACTCGT

>M6-PGSC0003DMG400008394-(rev)M6LRR-RLP_976

TCGACACCCTGAGAAGAATAAATACATCCAGAGGAAACTGCATTCTTGTGACTACTCGTAGGAAGCAGGTGGCATCCACAGTAGCAACAGATCTTCATATCTTGGGGAAGTTAACAGAAG

>M6-PGSC0003DMG400008403M6LRR-RLK_0

AGTGACCATATGCATGAAAAATCAAATTAGTTATTTGCTGATCTGAAGCAAATTTTTGAACAATGGCAGCCACTCGGCTGTTACTCTGCTTTATCGTCTTCTCGTCTGGCATTGATTTCA

>M6-PGSC0003DMG400008403M6LRR-RLK_122

TACACTGTTACAGATCCCAGAGATGGTAAGAACTTTGTATAAAATTGTCGGATCCAATATTTAAACTTTATGAGTTTTACTATTTGTTGCAGTTTTTGTGAACTTTTACTTATAAATTTA

>M6-PGSC0003DMG400008403M6LRR-RLK_1281

ATTTACGTAAGTTTCTCACTTTTTTATACTTCAAACCAAGCCACAACTTAGTTTTTGAAATTGTTGTTTCACCAACGAAGTAGTTGTTAACAATTGATTGTGTTTATTGTTTCTTTCAGG

>M6-PGSC0003DMG400008403M6LRR-RLK_1342

TGTTGTTTCACCAACGAAGTAGTTGTTAACAATTGATTGTGTTTATTGTTTCTTTCAGGGGATTATCAACAATGGGACTGAGAGGAAAATTAAGTGGTGACATTGGAGGGCTCACTGAAT

>M6-PGSC0003DMG400008403M6LRR-RLK_1403

ATTATCAACAATGGGACTGAGAGGAAAATTAAGTGGTGACATTGGAGGGCTCACTGAATTAATTTCCTTGTGAGTTCGGTTTTTAAATCCAATAAACTTCTGTATATGACACCAACAATT

>M6-PGSC0003DMG400008403M6LRR-RLK_1464

ATTTCCTTGTGAGTTCGGTTTTTAAATCCAATAAACTTCTGTATATGACACCAACAATTATTAGCAATAAGCTTGTATTTGTCACCATAATTTGCTCACTTCAGTTGCTAAACTTGCACT

>M6-PGSC0003DMG400008403M6LRR-RLK_1525

TAGCAATAAGCTTGTATTTGTCACCATAATTTGCTCACTTCAGTTGCTAAACTTGCACTGCAGAGACCTGTCGTTCAACCGAGGACTTACAGGTTCCCTTTCTCCGCGGATAGGAGATCT

>M6-PGSC0003DMG400008403M6LRR-RLK_1586

AGAGACCTGTCGTTCAACCGAGGACTTACAGGTTCCCTTTCTCCGCGGATAGGAGATCTGCAGAAACTAAACATATTGTAAGACTACGAAAACTTGATCATTTTGATTGTATAGTGTATA

>M6-PGSC0003DMG400008403M6LRR-RLK_1769

TTTCTTCTTCGTAATTTGTATCATGATTCAATGACATACATAACTTACTTGTACGCGAATGAATTTATTTACTTTTCAGAATCCTCGCGGGCTGTAGCTTCAGTGGCAGTATACCGAGGG

>M6-PGSC0003DMG400008403M6LRR-RLK_183

TAAACTTTATGAGTTTTACTATTTGTTGCAGTTTTTGTGAACTTTTACTTATAAATTTATGTTCTGCGTTGAAAGTACTGAGTTCAGATAATAATGTTTTGTTATGAATAAGTGTGAACT

>M6-PGSC0003DMG400008403M6LRR-RLK_1830

AATTTATTTACTTTTCAGAATCCTCGCGGGCTGTAGCTTCAGTGGCAGTATACCGAGGGAGTTGGGAAGACTTGCTGAGTTATCCTTCTTGTAAGAATTTCTCAGCAACATCAACTTGTG

>M6-PGSC0003DMG400008403M6LRR-RLK_1891

TTGGGAAGACTTGCTGAGTTATCCTTCTTGTAAGAATTTCTCAGCAACATCAACTTGTGATCGTTTTTGGTGTATCATCAACTCATTTAGTTCCATGTAAATTTCTTTGACAGGGCACTT

>M6-PGSC0003DMG400008403M6LRR-RLK_1952

CGTTTTTGGTGTATCATCAACTCATTTAGTTCCATGTAAATTTCTTTGACAGGGCACTTAACTCAAATAATTTCACCGGTGAGATTCCACGAACTCTTGGCAATCTGTCAAAACTGTACT

>M6-PGSC0003DMG400008403M6LRR-RLK_2013

CTCAAATAATTTCACCGGTGAGATTCCACGAACTCTTGGCAATCTGTCAAAACTGTACTGGCTTGATCTAGCAGATAACCTGTTAACAGGACCAATTCCTGTTTCAACCTCCTCAAGTCC

>M6-PGSC0003DMG400008403M6LRR-RLK_2074

CTTGATCTAGCAGATAACCTGTTAACAGGACCAATTCCTGTTTCAACCTCCTCAAGTCCGGGGTTAGACCTTCTAAAAAAAGCAAAACACTTGTGAGTAAACTTTATATGTCCATATTAC

>M6-PGSC0003DMG400008403M6LRR-RLK_2135

GGTTAGACCTTCTAAAAAAAGCAAAACACTTGTGAGTAAACTTTATATGTCCATATTACATGCTGTTCTTAAGTTGGTATATATACCTCTTAAGGTCTGACTTCTACGTTATGTCATTAT

>M6-PGSC0003DMG400008403M6LRR-RLK_2196

GCTGTTCTTAAGTTGGTATATATACCTCTTAAGGTCTGACTTCTACGTTATGTCATTATCGCAGCCATTTTAACAAAAACCAACTTTCAGGCTCCATTCCGGACATTCTTTTTAGTGCTG

>M6-PGSC0003DMG400008403M6LRR-RLK_2257

CAGCCATTTTAACAAAAACCAACTTTCAGGCTCCATTCCGGACATTCTTTTTAGTGCTGACATGGTGCTAATACATGTGTAAGTCAAGAAACTAAAGGGTGATTCGCGCTATTCTTCCTG

>M6-PGSC0003DMG400008403M6LRR-RLK_2318

ATGGTGCTAATACATGTGTAAGTCAAGAAACTAAAGGGTGATTCGCGCTATTCTTCCTGCCCTCCTTCTCACTTTGTGAACTTCTGATATGGCAGACTGTTTGATGGAAATCAACTTAGC

>M6-PGSC0003DMG400008403M6LRR-RLK_2379

CTCCTTCTCACTTTGTGAACTTCTGATATGGCAGACTGTTTGATGGAAATCAACTTAGCGGGAGCATTCCATTTACACTAGGACTTGTTCAGACACTTGAGGTTCTGTGAGTAGAATTGA

>M6-PGSC0003DMG400008403M6LRR-RLK_244

TTCTGCGTTGAAAGTACTGAGTTCAGATAATAATGTTTTGTTATGAATAAGTGTGAACTTGGTTGATTGCAGTAACGATACTTCGATCACTGAAAGATCAGTGGGAAAATACGCCGCCTA

>M6-PGSC0003DMG400008403M6LRR-RLK_2440

GAGCATTCCATTTACACTAGGACTTGTTCAGACACTTGAGGTTCTGTGAGTAGAATTGACCGTCTTATTTATCCTATCTTTCGTGCTACTTCACCTTCTGTTTTGGATTATATGCTTGTA

>M6-PGSC0003DMG400008403M6LRR-RLK_2501

GTCTTATTTATCCTATCTTTCGTGCTACTTCACCTTCTGTTTTGGATTATATGCTTGTACTCTTTTATCTTTTGCTGGTATTCATGTAACCTAACATATACAGTCGGCTCGATAGAAATG

>M6-PGSC0003DMG400008403M6LRR-RLK_2562

CTTTTATCTTTTGCTGGTATTCATGTAACCTAACATATACAGTCGGCTCGATAGAAATGCCTTAAATGGTAGCGTCCCATCAAACCTCAACAATCTCACAAGCGTTGTAGAGTTGTAAGT

>M6-PGSC0003DMG400008403M6LRR-RLK_2623

TTAAATGGTAGCGTCCCATCAAACCTCAACAATCTCACAAGCGTTGTAGAGTTGTAAGTATACTTTTTCCCATCATTGCTCACTCAATTGTCTGCTTGAAATCAATAACTTGTAGATTGA

>M6-PGSC0003DMG400008403M6LRR-RLK_2684

ACTTTTTCCCATCATTGCTCACTCAATTGTCTGCTTGAAATCAATAACTTGTAGATTGAAGTAGTCCTTTTGTGCTCTTATTTCTGTCTTCAGGAACTTAGCTCACAACGAACTGTCCGG

>M6-PGSC0003DMG400008403M6LRR-RLK_2745

TAGTCCTTTTGTGCTCTTATTTCTGTCTTCAGGAACTTAGCTCACAACGAACTGTCCGGTCCACTACCAGATTTAACTGGAATGAATTCCCTTAATTATCTGTAAGGAAAATATCCAGAA

>M6-PGSC0003DMG400008403M6LRR-RLK_2806

CACTACCAGATTTAACTGGAATGAATTCCCTTAATTATCTGTAAGGAAAATATCCAGAATCCTGATTTTCCCTAGTTGTTTCTTAATGTTGAAGTGATACATCATATTTTGTTTCTCTTC

>M6-PGSC0003DMG400008403M6LRR-RLK_2867

CTGATTTTCCCTAGTTGTTTCTTAATGTTGAAGTGATACATCATATTTTGTTTCTCTTCCAGGGACTTGAGCAATAATTCCTTCCATAAATCGGAGGCGCCAATATGGTTCTCGACTCTA

>M6-PGSC0003DMG400008403M6LRR-RLK_2928

GGGACTTGAGCAATAATTCCTTCCATAAATCGGAGGCGCCAATATGGTTCTCGACTCTAGAGTCTCTCACCACTCTGTGAGTATTAATCAGTTATGAATGCAACTATGAATTTCCCAAAA

>M6-PGSC0003DMG400008403M6LRR-RLK_2989

GTCTCTCACCACTCTGTGAGTATTAATCAGTTATGAATGCAACTATGAATTTCCCAAAAAGTTTGATGAATCTAAGTAGGAAGAACTGATGTAATGCTCAGAGTCATCGAGTATGGCTCA

>M6-PGSC0003DMG400008403M6LRR-RLK_305

GTTGATTGCAGTAACGATACTTCGATCACTGAAAGATCAGTGGGAAAATACGCCGCCTAGTTGGCAGAGATCGGATGATCCTTGTGGAACATCTTGGGAAGGTGTAACCTGCAACAACTC

>M6-PGSC0003DMG400008403M6LRR-RLK_3050

TTTGATGAATCTAAGTAGGAAGAACTGATGTAATGCTCAGAGTCATCGAGTATGGCTCACTCCATGGATCAGTGCCTCAGAAACTTTTCGCTTTACCTCAATTACAACAAGTGTGAGTCT

>M6-PGSC0003DMG400008403M6LRR-RLK_3111

CCATGGATCAGTGCCTCAGAAACTTTTCGCTTTACCTCAATTACAACAAGTGTGAGTCTCGCGATCCTAACTGGATATTCCATGTCTATTACCTGCAATGAAGAGGTCGTAATGATCATA

>M6-PGSC0003DMG400008403M6LRR-RLK_3172

CGATCCTAACTGGATATTCCATGTCTATTACCTGCAATGAAGAGGTCGTAATGATCATAATAACTTATGCAGGAAACTGAGGAACAACGCATTGAATGACACATTGAACATGGGTGGCAT

>M6-PGSC0003DMG400008403M6LRR-RLK_3233

AACTTATGCAGGAAACTGAGGAACAACGCATTGAATGACACATTGAACATGGGTGGCATCAGTGGCAGGCAACTCACACTTGTTGATTTACAGAACAACGAGATTTCCTCAATAACACTT

>M6-PGSC0003DMG400008403M6LRR-RLK_3294

GTGGCAGGCAACTCACACTTGTTGATTTACAGAACAACGAGATTTCCTCAATAACACTTGGCTCTGGATACAAAAATACCTTGATGTAAGATGGTTCATTTCTCCTTTTCGTCGATTAGA

>M6-PGSC0003DMG400008403M6LRR-RLK_3355

CTCTGGATACAAAAATACCTTGATGTAAGATGGTTCATTTCTCCTTTTCGTCGATTAGATGATTGTTTTTTTTCTTTCTCCTTGATTGCTTTGTCGTTGTCCTCAGACTGATCGGAAACC

>M6-PGSC0003DMG400008403M6LRR-RLK_3416

ATTGTTTTTTTTCTTTCTCCTTGATTGCTTTGTCGTTGTCCTCAGACTGATCGGAAACCCTGTGTGTGACACGGCCCTTGGAAACACGAACTACTGTCAGCTTCAGCAGCAATCTGCAAA

>M6-PGSC0003DMG400008403M6LRR-RLK_3477

GTGTGTGACACGGCCCTTGGAAACACGAACTACTGTCAGCTTCAGCAGCAATCTGCAAAGCCTTACTCAACCAGCCTAGCTAACTGTGGGAGAAAATCATGTCCTGCTGATCAAAAAGTT

>M6-PGSC0003DMG400008403M6LRR-RLK_3538

CTTACTCAACCAGCCTAGCTAACTGTGGGAGAAAATCATGTCCTGCTGATCAAAAAGTTAGCCCACAAAGTTGTGATTGTGCTTATCCGTACCAGGGAACATTCTACTTTAGAGGACCCT

>M6-PGSC0003DMG400008403M6LRR-RLK_3599

CCCACAAAGTTGTGATTGTGCTTATCCGTACCAGGGAACATTCTACTTTAGAGGACCCTCTTTTAGAGAGTTGTCCAATGATAATACGTTCCATTCACTCGAAATGAGCCTGTGGGTGAA

>M6-PGSC0003DMG400008403M6LRR-RLK_366

TGGCAGAGATCGGATGATCCTTGTGGAACATCTTGGGAAGGTGTAACCTGCAACAACTCAAGGGTAACTGCATTGTGAGTATGCTGTTTTCTTCCAATATATCTAGGGATCGTTTGATTT

>M6-PGSC0003DMG400008403M6LRR-RLK_3660

TTTAGAGAGTTGTCCAATGATAATACGTTCCATTCACTCGAAATGAGCCTGTGGGTGAAGTTGGATCTCACGCCTGGTTCGGTTTCTCTGCAAAACCCCTTCTTCAACATTGATGATTAT

>M6-PGSC0003DMG400008403M6LRR-RLK_3721

TGGATCTCACGCCTGGTTCGGTTTCTCTGCAAAACCCCTTCTTCAACATTGATGATTATCTTCAGGTGCAGTTAGAATTATTTCCACCAACAGGAAAGTATTTTAATAGATCAGAGGTTG

>M6-PGSC0003DMG400008403M6LRR-RLK_3782

TCAGGTGCAGTTAGAATTATTTCCACCAACAGGAAAGTATTTTAATAGATCAGAGGTTGAGAGGATTGGATTCTCCTTGAGTAACCAAACTTATAAGCCGCCTCATGAGTTTGGACCGTA

>M6-PGSC0003DMG400008403M6LRR-RLK_3843

AGGATTGGATTCTCCTTGAGTAACCAAACTTATAAGCCGCCTCATGAGTTTGGACCGTACTATTTTATAGCATCTCCTTACACTTTCCAAGGTATAATTACTGAGTTTCTTTACAACTGT

>M6-PGSC0003DMG400008403M6LRR-RLK_3904

ATTTTATAGCATCTCCTTACACTTTCCAAGGTATAATTACTGAGTTTCTTTACAACTGTAACTTCTGGACCAATACTGACTTAAAACTAAAACTGTTTCAGCCGAACATGGAGAAACTTC

>M6-PGSC0003DMG400008403M6LRR-RLK_3965

CTTCTGGACCAATACTGACTTAAAACTAAAACTGTTTCAGCCGAACATGGAGAAACTTCTATTAGCTCAAAACAAGTTATAGGAATAGCTACCGGGTGTACCATCCTGGTCCTCTTGCTC

>M6-PGSC0003DMG400008403M6LRR-RLK_4026

TTAGCTCAAAACAAGTTATAGGAATAGCTACCGGGTGTACCATCCTGGTCCTCTTGCTCGTTGCTCTTGCAATATATGCTATTCAACAAAAGAAACTTGCTGAAAGAGCTATTGGATTGA

>M6-PGSC0003DMG400008403M6LRR-RLK_4087

TGCTCTTGCAATATATGCTATTCAACAAAAGAAACTTGCTGAAAGAGCTATTGGATTGAGTAGACCATTTGGTAATGTCATCTTTTACTGAAATAAACCGATGAGTTATAACTGCATATA

>M6-PGSC0003DMG400008403M6LRR-RLK_4148

AGACCATTTGGTAATGTCATCTTTTACTGAAATAAACCGATGAGTTATAACTGCATATACATTCATGCCTCTCTATAACAGTTATCCTCTATGATTACATTTCACTATAACGACCAAGTT

>M6-PGSC0003DMG400008403M6LRR-RLK_4209

TTCATGCCTCTCTATAACAGTTATCCTCTATGATTACATTTCACTATAACGACCAAGTTTTCTTTGGAACTGATTTTTCATGTTGTATTGAATTATATGTTTTCTATAACAACATCTCGC

>M6-PGSC0003DMG400008403M6LRR-RLK_4270

CTTTGGAACTGATTTTTCATGTTGTATTGAATTATATGTTTTCTATAACAACATCTCGCTATAGCAGCCAAAGAAATATCAGAACAAATGACGTCTTTATAGAGAGGTTTGACTATATGT

>M6-PGSC0003DMG400008403M6LRR-RLK_4331

TAGCAGCCAAAGAAATATCAGAACAAATGACGTCTTTATAGAGAGGTTTGACTATATGTCCATAATGCAACTAGGATTAAACTAAAATGTCTGATTACTTTGACCTTCTTTGTCCAGCAT

>M6-PGSC0003DMG400008403M6LRR-RLK_4392

ATAATGCAACTAGGATTAAACTAAAATGTCTGATTACTTTGACCTTCTTTGTCCAGCATCGTGGGCTCCAAGTGGAAACGACAGTGAAGGTGCACCACAACTAAAGGGAGCCAGATGGTT

>M6-PGSC0003DMG400008403M6LRR-RLK_4453

TGGGCTCCAAGTGGAAACGACAGTGAAGGTGCACCACAACTAAAGGGAGCCAGATGGTTTTCTTATGATGAACTTAAGAAATGCACGGGAAACTTCTCGGAGAGAAATGAGATTGGTTCC

>M6-PGSC0003DMG400008403M6LRR-RLK_4514

CTTATGATGAACTTAAGAAATGCACGGGAAACTTCTCGGAGAGAAATGAGATTGGTTCCGGTGGCTATGGGAAGGTTTGCTACCAAAATCCACAGTAGATATAGAATAAAGGGATAAACT

>M6-PGSC0003DMG400008403M6LRR-RLK_4575

TGGCTATGGGAAGGTTTGCTACCAAAATCCACAGTAGATATAGAATAAAGGGATAAACTCTTAGTTACTTTGCTTATTTGCGTCTGTTCTGTTTCCTAGGTATACAGAGGAACACTCGCA

>M6-PGSC0003DMG400008403M6LRR-RLK_4636

TAGTTACTTTGCTTATTTGCGTCTGTTCTGTTTCCTAGGTATACAGAGGAACACTCGCAAATGGGCAAGTAATTGCTATCAAAAGGGCTCAACATGGATCCATGCAAGGTGGACAAGAGT

>M6-PGSC0003DMG400008403M6LRR-RLK_4697

TGGGCAAGTAATTGCTATCAAAAGGGCTCAACATGGATCCATGCAAGGTGGACAAGAGTTCAAGACAGAAATTGAATTGCTTTCACGGGTTCATCACAAGAACCTAGTTGGTCTGGTTGG

>M6-PGSC0003DMG400008403M6LRR-RLK_4758

AAGACAGAAATTGAATTGCTTTCACGGGTTCATCACAAGAACCTAGTTGGTCTGGTTGGATTCTGTTTTGAGCAAGGGGAACAAGTACTGGTATACGAGTATATGCCAAATGGTTCACTC

>M6-PGSC0003DMG400008403M6LRR-RLK_4819

TCTGTTTTGAGCAAGGGGAACAAGTACTGGTATACGAGTATATGCCAAATGGTTCACTCAGGGAGAATTTGTCAGGTAAGTTACATCCTCCTATACATTAAAGACTGCTTCCTATTTATA

>M6-PGSC0003DMG400008403M6LRR-RLK_4880

GGAGAATTTGTCAGGTAAGTTACATCCTCCTATACATTAAAGACTGCTTCCTATTTATAGTAATTTACACAAAAAAGTTCCTAATATCTCCATTTCGCTGTAGAAGTTAATTTCCACTAT

>M6-PGSC0003DMG400008403M6LRR-RLK_4941

AATTTACACAAAAAAGTTCCTAATATCTCCATTTCGCTGTAGAAGTTAATTTCCACTATGCTCTACAGGGAAAACGAGCATTTATCTAGATTGGAAGAGGAGACTTCGTGTTGCGCTTGG

>M6-PGSC0003DMG400008403M6LRR-RLK_5002

TCTACAGGGAAAACGAGCATTTATCTAGATTGGAAGAGGAGACTTCGTGTTGCGCTTGGTTCAGCCAGAGGATTAGCTTATCTGCATGAGCTTGCCAATCCTCCTATAATTCACAGAGAT

>M6-PGSC0003DMG400008403M6LRR-RLK_5063

CAGCCAGAGGATTAGCTTATCTGCATGAGCTTGCCAATCCTCCTATAATTCACAGAGATGTGAAATCAACTAACATTTTGTTGGATCAGAACTTAACAGCAAAGGTCGGGGATTTTGGCT

>M6-PGSC0003DMG400008403M6LRR-RLK_5124

GAAATCAACTAACATTTTGTTGGATCAGAACTTAACAGCAAAGGTCGGGGATTTTGGCTTGTCCAAACTCGTATCAGATAGCTCAAAGGGTCATGTTTCCACTCAAGTTAAAGGCACATT

>M6-PGSC0003DMG400008403M6LRR-RLK_5185

TCCAAACTCGTATCAGATAGCTCAAAGGGTCATGTTTCCACTCAAGTTAAAGGCACATTGGTCAGTACCACATCCTGCACATTTTCGCGTTCTATTTCAAGAAAAAGATTGAATTTTGTT

>M6-PGSC0003DMG400008403M6LRR-RLK_5246

TCAGTACCACATCCTGCACATTTTCGCGTTCTATTTCAAGAAAAAGATTGAATTTTGTTGAACTCTTGAAGCATGAAATTGAATTTAAGACTATTGATTACATTGCAGGGTTATCTAGAT

>M6-PGSC0003DMG400008403M6LRR-RLK_5307

ACTCTTGAAGCATGAAATTGAATTTAAGACTATTGATTACATTGCAGGGTTATCTAGATCCTGAATACTATATGACTCAACAACTAACCGAGAAGAGCGATGTGTATAGCTTTGGTGTTG

>M6-PGSC0003DMG400008403M6LRR-RLK_5368

TGAATACTATATGACTCAACAACTAACCGAGAAGAGCGATGTGTATAGCTTTGGTGTTGTAATGCTAGAACTCATAACTGCAAAACAACCAATAGAGAAGGGGAAATACGTTGTTCGAGA

>M6-PGSC0003DMG400008403M6LRR-RLK_5429

ATGCTAGAACTCATAACTGCAAAACAACCAATAGAGAAGGGGAAATACGTTGTTCGAGAGATGAGAACAGCAATAAATAAGAACGACGAGGAGCATTATGGCTTGAGCAATATGATCGAT

>M6-PGSC0003DMG400008403M6LRR-RLK_5490

TGAGAACAGCAATAAATAAGAACGACGAGGAGCATTATGGCTTGAGCAATATGATCGATCCAGTCATTAGAAACATGCCAAATCTAATTGGTTTTACAAGGTTTGTGGACGTGGCGATGC

>M6-PGSC0003DMG400008403M6LRR-RLK_5551

AGTCATTAGAAACATGCCAAATCTAATTGGTTTTACAAGGTTTGTGGACGTGGCGATGCAATGTGTTGAAGAGGCTGCTGCTGATCGTCCAACGATGAGTGAAGTGGTAAAAATGCTTGA

>M6-PGSC0003DMG400008403M6LRR-RLK_5612

TGTGTTGAAGAGGCTGCTGCTGATCGTCCAACGATGAGTGAAGTGGTAAAAATGCTTGAAAGTATTCTTCAAAATGATGGTCTGGAAACGAATTCTACGTCTGCATCATCATCGATAACG

>M6-PGSC0003DMG400008403M6LRR-RLK_5673

GTATTCTTCAAAATGATGGTCTGGAAACGAATTCTACGTCTGCATCATCATCGATAACGGATTTTGGGACAGCAATAGCTGCTTCTAGACATCCTTATAACAAGGAGGCTCTACAGAGAA

>M6-PGSC0003DMG400008403M6LRR-RLK_5734

TTTTGGGACAGCAATAGCTGCTTCTAGACATCCTTATAACAAGGAGGCTCTACAGAGAAAGGAGATCAATGATACTCATGCGTTTGATTACAGTGGTGGATACACTCTTCCAACAAATGT

>M6-PGSC0003DMG400008403M6LRR-RLK_5795

GAGATCAATGATACTCATGCGTTTGATTACAGTGGTGGATACACTCTTCCAACAAATGTGGAACCAAAATAGTGACAACCAAAAAAAAAAGGACTTGTCTCTACTGTCTCTCATTCTTTA

>M6-PGSC0003DMG400008403M6LRR-RLK_5856

AACCAAAATAGTGACAACCAAAAAAAAAAGGACTTGTCTCTACTGTCTCTCATTCTTTAATTTATTCTCTAGAAAAAAAATTGGAAAATTGGTGAAAGGGATAAGAGAATTTAGTTGTTG

>M6-PGSC0003DMG400008403M6LRR-RLK_5917

TTATTCTCTAGAAAAAAAATTGGAAAATTGGTGAAAGGGATAAGAGAATTTAGTTGTTGTATTGTCCCTATATTGGCTACTTTTCGCGAAACAATCACGTATAGTATCAGGATCGAAGTT

>M6-PGSC0003DMG400008403M6LRR-RLK_5978

TTGTCCCTATATTGGCTACTTTTCGCGAAACAATCACGTATAGTATCAGGATCGAAGTTCAGATTTTCTATTGCCAAGACTTACTGTCTAGTGGTACTAGTGAGAGTCTCCTATACACTA

>M6-PGSC0003DMG400008403M6LRR-RLK_6039

GATTTTCTATTGCCAAGACTTACTGTCTAGTGGTACTAGTGAGAGTCTCCTATACACTAAGTGTGAGTTCGATTTTTACCACCCCCGATCCTCAAATGTAATAGAAAAAAGTTAAAAGGA

>M6-PGSC0003DMG400008403M6LRR-RLK_61

AATGGCAGCCACTCGGCTGTTACTCTGCTTTATCGTCTTCTCGTCTGGCATTGATTTCATATACACTGTTACAGATCCCAGAGATGGTAAGAACTTTGTATAAAATTGTCGGATCCAATA

>M6-PGSC0003DMG400008403M6LRR-RLK_6100

TGTGAGTTCGATTTTTACCACCCCCGATCCTCAAATGTAATAGAAAAAAGTTAAAAGGAAAAAATTTAGATTTTGTACCATCAAGGCTGCAGTCTAGTGGTATACCTAAGAGGTCTTCCA

>M6-PGSC0003DMG400008403M6LRR-RLK_6161

AAATTTAGATTTTGTACCATCAAGGCTGCAGTCTAGTGGTATACCTAAGAGGTCTTCCATACACTAAGTGTGAGTTCGATTTTTTACTATCTCCTTTCCCATTCCCTGATTCTACAATGT

>M6-PGSC0003DMG400008403M6LRR-RLK_6198

GGTATACCTAAGAGGTCTTCCATACACTAAGTGTGAGTTCGATTTTTTACTATCTCCTTTCCCATTCCCTGATTCTACAATGTAATAGAAAAAAATCAAAATTAAAAAGTTCAGATTTTC

>M6-PGSC0003DMG400008403M6LRR-RLK_671

TATTAAACTCCATATGACTAGTGCAGAGTTTAATACCGGAAACAAAACATCATGTTAGTTATATGGAAACTATATTAGCTTACACGTTTTTTACATTGAAAACCAAATGTTGTATAAGTA

>M6-PGSC0003DMG400008403M6LRR-RLK_732

TATGGAAACTATATTAGCTTACACGTTTTTTACATTGAAAACCAAATGTTGTATAAGTAAAATGAAAATTACTTCTTTTTTTTTTAGGAAAAACTACATAATTAGGACATAGGTCGCTAT

>M6-PGSC0003DMG400008403M6LRR-RLK_793

ATGAAAATTACTTCTTTTTTTTTTAGGAAAAACTACATAATTAGGACATAGGTCGCTATCGTAAATACTATTATTATCTATACTTTCAAAATACTATGTATCTTACCACTAATTAAGAAT

>M6-PGSC0003DMG400008450M6LRR-RLP_0

TCCTTATGTTACACACCTCTTCTCTCTGAAGGTGTATGATGGGATTCATGATTGTCTTTCCTACAACAGTCACCTAAAACACTTGAGGCTTCTTAAAAGTTTGGATTTGAAGGGCCTAAC

>M6-PGSC0003DMG400008450M6LRR-RLP_128

ATTCTTTGCTGAATGAAATCGGTTTGCTTGTTCATTTGAAGTACTTAATCATTCAGACGAAGGCAAAAGCTCTCCCTCCGTTATTTTCAAACCTCTGCAATCTAGAAACTCTATCGGTGT

>M6-PGSC0003DMG400008450M6LRR-RLP_192

AAAAGCTCTCCCTCCGTTATTTTCAAACCTCTGCAATCTAGAAACTCTATCGGTGTATAACATTGAGAGATCATGCATGATACTATCGCCTTGTTTTTGGAGTCTTGCAAAGCTACAAGA

>M6-PGSC0003DMG400008450M6LRR-RLP_256

GAGAGATCATGCATGATACTATCGCCTTGTTTTTGGAGTCTTGCAAAGCTACAAGATGTGTGGATGATGGAATGTGCTCTCTTTGAACCAACTGTGTTAGACGAGGACTCAAGGTTAGAA

>M6-PGSC0003DMG400008450M6LRR-RLP_320

TGATGGAATGTGCTCTCTTTGAACCAACTGTGTTAGACGAGGACTCAAGGTTAGAAAATTTGAGAACATTACAGCATCTCTCCCTTCCCGGTTCAGAAGACATGGAGGATATTTTCAAAA

>M6-PGSC0003DMG400008450M6LRR-RLP_384

AACATTACAGCATCTCTCCCTTCCCGGTTCAGAAGACATGGAGGATATTTTCAAAAGGTTTCCTAATCTTCAAATCCTTAGTGTCTCTATCATAGCAGAGAAGATTTGTTTTCCAAGATT

>M6-PGSC0003DMG400008450M6LRR-RLP_448

AATCTTCAAATCCTTAGTGTCTCTATCATAGCAGAGAAGATTTGTTTTCCAAGATTGGATGTCCTTAACGAACTTGAACAACTCAATTTATCTGCTTCTTGGGATTCATTCCATGAATAT

>M6-PGSC0003DMG400008450M6LRR-RLP_512

TTAACGAACTTGAACAACTCAATTTATCTGCTTCTTGGGATTCATTCCATGAATATACACACGGCTTCCCTTTGAGCTTGAAGAAAATGAGGTTGCAAGGGCTCGCTCTGACATCCGATT

>M6-PGSC0003DMG400008450M6LRR-RLP_64

AACAGTCACCTAAAACACTTGAGGCTTCTTAAAAGTTTGGATTTGAAGGGCCTAACATTGACAGATTCTTTGCTGAATGAAATCGGTTTGCTTGTTCATTTGAAGTACTTAATCATTCAG

>M6-PGSC0003DMG400008450M6LRR-RLP_640

AGAATTGCTAGACTACCCAACCTTCAAAAACTAAGTCTAAAAAACGCAATCATCGATGGGGGGCCAGGAGAATGGAACATGGAAGTTGTCACCTTCAAGAATCTCAAATATCTAAAATTG

>M6-PGSC0003DMG400008450M6LRR-RLP_704

CAGGAGAATGGAACATGGAAGTTGTCACCTTCAAGAATCTCAAATATCTAAAATTGGATAGTGTGGAATTTTCTGAATGGCAGGTTGATGCAGAGAAATCATTTCCCGTGCTCGAGATGT

>M6-PGSC0003DMG400008450M6LRR-RLP_768

GGAATTTTCTGAATGGCAGGTTGATGCAGAGAAATCATTTCCCGTGCTCGAGATGTTAGATATAGTTACATGTGAGAAGCTTATGGACATCCCAGATAGTTTTGGAGATATTGCGTCATT

>M6-PGSC0003DMG400008450M6LRR-RLP_825

AGATATAGTTACATGTGAGAAGCTTATGGACATCCCAGATAGTTTTGGAGATATTGCGTCATTAGAACTTATCAAAGTATGGTTTAGCCCTCAGCTTAAAGAGTCAATCTTCAATATTAA

>M6-PGSC0003DMG400008482M6RLKWAX-EGF_1426

GGCCGTGCCTTGGGGCATATCGATTGAAGTAGATTCTGATTCATATGTTATGCTTAATTAGATACTCATAGTTGCCATTAATGACAAAGTAATTAATCCTGGAATTTTCGTCGGGTGTAA

>M6-PGSC0003DMG400008482M6RLKWAX-EGF_1488

TACTCATAGTTGCCATTAATGACAAAGTAATTAATCCTGGAATTTTCGTCGGGTGTAATCACAACTTGTGCTAGTTGTAAAACAAAGAAGATATAAAGACATAGCCAAAGGGATTAAATT

>M6-PGSC0003DMG400008482M6RLKWAX-EGF_1550

AACTTGTGCTAGTTGTAAAACAAAGAAGATATAAAGACATAGCCAAAGGGATTAAATTAATGTGGTATGTTTCTCTGTTTGGCATGTTAGATATTGATGAATGTCAAAATTCAACTTCCA

>M6-PGSC0003DMG400008482M6RLKWAX-EGF_1612

TGGTATGTTTCTCTGTTTGGCATGTTAGATATTGATGAATGTCAAAATTCAACTTCCAATAAGTGTGGACACAACAATTGTATAAACACGTTGGGGAGTTACAAATGTTCTTGTCGAAAA

>M6-PGSC0003DMG400008482M6RLKWAX-EGF_1674

GTGTGGACACAACAATTGTATAAACACGTTGGGGAGTTACAAATGTTCTTGTCGAAAAGGATACACTGATGATGGTGAAAACAATGGTCTTGTCTGTATTGCAACAAACTCCGAGTTCCC

>M6-PGSC0003DMG400008482M6RLKWAX-EGF_1736

ACACTGATGATGGTGAAAACAATGGTCTTGTCTGTATTGCAACAAACTCCGAGTTCCCATGGATCAAGTTTTCCGTTGGTAAGTTTCTCAGATATATATATATATATATATATATATATA

>M6-PGSC0003DMG400008482M6RLKWAX-EGF_1860

TATATTGAAGTGATCATAGGAATTCACTTCCTGTTATTATAAACACAGGTATGGGAGTTGGCTTTATATCCCTAGTGGTTGGGACAATCTTGCTCTATTTTTGTATCAACAAACGAAGAC

>M6-PGSC0003DMG400008482M6RLKWAX-EGF_1922

TTTATATCCCTAGTGGTTGGGACAATCTTGCTCTATTTTTGTATCAACAAACGAAGACTCATTAAAAGTAGGGAGAAGTTCTTCCAACAAAATGGTGGTTTACTCTTGAAACAACAAATC

>M6-PGSC0003DMG400008482M6RLKWAX-EGF_1984

TAAAAGTAGGGAGAAGTTCTTCCAACAAAATGGTGGTTTACTCTTGAAACAACAAATCTCCTCTAAAAAGGGTGGTGTGGAAGCAACCAAAATTTTTACAGCTGAGGAGTTGAAGAAGGC

>M6-PGSC0003DMG400008482M6RLKWAX-EGF_2046

CTAAAAAGGGTGGTGTGGAAGCAACCAAAATTTTTACAGCTGAGGAGTTGAAGAAGGCGACAAATAACTATGCTAGTGATAGAATTCTTGGTCGCGGTGGAACTGGAATTGTCTATAAAG

>M6-PGSC0003DMG400008482M6RLKWAX-EGF_2108

AATAACTATGCTAGTGATAGAATTCTTGGTCGCGGTGGAACTGGAATTGTCTATAAAGGTATTCTACCTGATAACCGTATAGTTGCTATTAAGAAATCTAAGATTGTGGATGAGAATCAA

>M6-PGSC0003DMG400008482M6RLKWAX-EGF_2170

TCTACCTGATAACCGTATAGTTGCTATTAAGAAATCTAAGATTGTGGATGAGAATCAAATTGAACAGTTCATCAATGAGGTACTTATTCTTACTCAAGTAAACCATAGAAACGTCGTGAG

>M6-PGSC0003DMG400008482M6RLKWAX-EGF_2232

AACAGTTCATCAATGAGGTACTTATTCTTACTCAAGTAAACCATAGAAACGTCGTGAGACTCTTTGGATGTTGTTTGGAAGCTGAAGTGCCTTTACTGGTTTACGAATACGTTTCTCATG

>M6-PGSC0003DMG400008482M6RLKWAX-EGF_2294

TTTGGATGTTGTTTGGAAGCTGAAGTGCCTTTACTGGTTTACGAATACGTTTCTCATGGAACTCTTTACGAGCATATCCACAATCAAAATGGGTTATCTTTGCAAAATCGGTTGAGAATT

>M6-PGSC0003DMG400008482M6RLKWAX-EGF_2356

TCTTTACGAGCATATCCACAATCAAAATGGGTTATCTTTGCAAAATCGGTTGAGAATTGCTTCAGAGACAGCAAGTTCACTTGCTTACCTTCATTCATCAGCATCGATGCCAATAATTCA

>M6-PGSC0003DMG400008482M6RLKWAX-EGF_2418

CAGAGACAGCAAGTTCACTTGCTTACCTTCATTCATCAGCATCGATGCCAATAATTCATAGAGACGTTAAATCTACTAATATATTATTGGATGATGGTTACACAGCAAAAGTGGCAGATT

>M6-PGSC0003DMG400008482M6RLKWAX-EGF_2480

GACGTTAAATCTACTAATATATTATTGGATGATGGTTACACAGCAAAAGTGGCAGATTTTGGAGCTTCAAGATTAGTCCCTCTTGACCAAACACGTGTTGCTACATTGGTTCAAGGAACG

>M6-PGSC0003DMG400008482M6RLKWAX-EGF_2542

AGCTTCAAGATTAGTCCCTCTTGACCAAACACGTGTTGCTACATTGGTTCAAGGAACGTTAGGGTACTTAGATCCTGAATATTTCCACACAGGTCAATTGACAGACAAAAGTGACGTTTA

>M6-PGSC0003DMG400008482M6RLKWAX-EGF_2604

GGTACTTAGATCCTGAATATTTCCACACAGGTCAATTGACAGACAAAAGTGACGTTTATAGTTTTGGAGTAGTTCTTGCAGAACTTTTAACAGGGATGAAACCTATTTTGAGGGACACAA

>M6-PGSC0003DMG400008482M6RLKWAX-EGF_2666

TTTGGAGTAGTTCTTGCAGAACTTTTAACAGGGATGAAACCTATTTTGAGGGACACAATTAGTGACAAGGACAAATGTTTGGTGGAGTACTTTGTCTCATCGATGAACAAGAATAGCTTA

>M6-PGSC0003DMG400008482M6RLKWAX-EGF_2728

TGACAAGGACAAATGTTTGGTGGAGTACTTTGTCTCATCGATGAACAAGAATAGCTTATTTCAAATTGTTGATTGTAGAGTTGTAAGAGAAGGAGGTCTTGAGCAACTTCAAAAGATAGC

>M6-PGSC0003DMG400008482M6RLKWAX-EGF_2790

AAATTGTTGATTGTAGAGTTGTAAGAGAAGGAGGTCTTGAGCAACTTCAAAAGATAGCTGAGCTCATTAAGAGTTGCCTTCACTTGCACGGAGAAGATAGGCCTACGATGAAGGAAGTGG

>M6-PGSC0003DMG400008482M6RLKWAX-EGF_2914

GGAACTTGAGAGTTTAAGAAAGTTCACTAGCCTTTGGACTAATGGAAAGGAACATGAAGATGAGAAGGAAGTTGAATTAACAGATCTTTATACAATTCCAATTGATTCTAATATAGGTAT

>M6-PGSC0003DMG400008482M6RLKWAX-EGF_2976

AGAAGGAAGTTGAATTAACAGATCTTTATACAATTCCAATTGATTCTAATATAGGTATGGACAACTTCTCTGGAAAATATCCTAGCTCCTACACAAATAGCAGCACCATCTTTTCTGGAC

>M6-PGSC0003DMG400008482M6RLKWAX-EGF_3038

AACTTCTCTGGAAAATATCCTAGCTCCTACACAAATAGCAGCACCATCTTTTCTGGACAATATAGTTCGGATAGTTCTTCATTACTATACAATACAAATTTACCGCGGTGATCGATCCAG

>M6-PGSC0003DMG400008482M6RLKWAX-EGF_3100

TAGTTCGGATAGTTCTTCATTACTATACAATACAAATTTACCGCGGTGATCGATCCAGAATAAAAGGTCAATATATGAAGATTTTTCTCTGTTTTGTAAGTTCCTTTTTAGCCATCTTTC

>M6-PGSC0003DMG400008482M6RLKWAX-EGF_3162

AAAGGTCAATATATGAAGATTTTTCTCTGTTTTGTAAGTTCCTTTTTAGCCATCTTTCTTTCTTGTTGAATGTTGTTATGTATTTTTATTTGCTTCTGAATTTAATCATCAATATATTGT

>M6-PGSC0003DMG400008482M6RLKWAX-EGF_3224

TTGTTGAATGTTGTTATGTATTTTTATTTGCTTCTGAATTTAATCATCAATATATTGTTGATTGTATGATTGCCAAGAACTTTTAGATTGTATGATTGACAAGAACTTTTAGTACATCGA

>M6-PGSC0003DMG400008482M6RLKWAX-EGF_496

AGCACAAATGAACGTTTTTTTAAAGTTTGATACATATTTGAGTCTTTTCCCTATAAACCAATTGAAATGGAGGAATTAAAAAAAGTAAGAGGTATAAATTGAAAATAAGAAAAGAAATAC

>M6-PGSC0003DMG400008482M6RLKWAX-EGF_558

TGAAATGGAGGAATTAAAAAAAGTAAGAGGTATAAATTGAAAATAAGAAAAGAAATACTTTAGTAGGTTGGGTGTCAAGAAGGGTAGGATATCAAAATCATAGGGACATTTATATTTATT

>M6-PGSC0003DMG400008482M6RLKWAX-EGF_620

GTAGGTTGGGTGTCAAGAAGGGTAGGATATCAAAATCATAGGGACATTTATATTTATTGTAGGATAGTGAGTGCCTTTTTAACACCACACACTAGAAATTAATGAATCGAAGTACCTAAT

>M6-PGSC0003DMG400008482M6RLKWAX-EGF_682

GATAGTGAGTGCCTTTTTAACACCACACACTAGAAATTAATGAATCGAAGTACCTAATTATCTGCCTGATTGAAGCGTGGCCATAACGATAGAGGTGAAGTTGACTCCTCACTAAGGTTT

>M6-PGSC0003DMG400008482M6RLKWAX-EGF_744

TGCCTGATTGAAGCGTGGCCATAACGATAGAGGTGAAGTTGACTCCTCACTAAGGTTTTGCTCTCATCAAAATTTGACTAATTCAAACACTTACCAATATATAATAGTCACCCATCAACC

>M6-PGSC0003DMG400008482M6RLKWAX-EGF_806

CTCATCAAAATTTGACTAATTCAAACACTTACCAATATATAATAGTCACCCATCAACCTGACTTGTGGTATAGTGCGGAGGTACTATTCCAATTTTAATTAGAGATTTTAAGTTTCGAAC

>M6-PGSC0003DMG400008482M6RLKWAX-EGF_868

TTGTGGTATAGTGCGGAGGTACTATTCCAATTTTAATTAGAGATTTTAAGTTTCGAACTCTGAGTATAGAGAAAATTTTGTTAGGAGCGTCACCTCAAAATGGGCCTTGCAGTGCATGAT

>M6-PGSC0003DMG400008482M6RLKWAX-EGF_930

AGTATAGAGAAAATTTTGTTAGGAGCGTCACCTCAAAATGGGCCTTGCAGTGCATGATTTAGATTTAATCAGAGCTCCAATATAGAAACCGAATAACAAATTTAAGAAGATATTTTCTTT

>M6-PGSC0003DMG400008482M6RLKWAX-EGF_992

ATTTAATCAGAGCTCCAATATAGAAACCGAATAACAAATTTAAGAAGATATTTTCTTTATCTTGTATTTAAACTAAGTTAATACATGTAATTTTACTAAAAATATTTTGCCTTATATTTA

>M6-PGSC0003DMG400008492M6LRR-RLP_1008

TGCTCCACTGCCGCCCTGTAAATGATCATCACTAGTATCCTGTCTGTCGATCTATCTATGTATTTCAACTTCTCTTTGAATGTTGATGAATAATTGAGTGAGTAAACTCAAGGCATAAAC

>M6-PGSC0003DMG400008492M6LRR-RLP_1063

CTATGTATTTCAACTTCTCTTTGAATGTTGATGAATAATTGAGTGAGTAAACTCAAGGCATAAACATGTACTTTAATTAATTTATGAATGTTATCCTTTTTTAGTAACTTCTCGATGGAT

>M6-PGSC0003DMG400008492M6LRR-RLP_126

AAAAGCCCTCCTAGAATTCAAAAAAGGTCTAGGCAATCCTGATGACTTGGGTAATTGGGATCCGAAAACTGATTGTTGTACTGATTGGTACGGACCTACTATCAAATGTGATGAAAAAAC

>M6-PGSC0003DMG400008492M6LRR-RLP_189

GAAAACTGATTGTTGTACTGATTGGTACGGACCTACTATCAAATGTGATGAAAAAACCAATCGTATTAACGTCATCGACTTCACCAAAATGGATATGGCCGGATACCTCTCCCCCGCCAT

>M6-PGSC0003DMG400008492M6LRR-RLP_252

TATTAACGTCATCGACTTCACCAAAATGGATATGGCCGGATACCTCTCCCCCGCCATCGGAGACCTTACATACCTTACAACATTCAGTATCCACAATGTACGTAATCTCACCGGTCCAAT

>M6-PGSC0003DMG400008492M6LRR-RLP_315

CCTTACATACCTTACAACATTCAGTATCCACAATGTACGTAATCTCACCGGTCCAATTCCATCAACAATCCTTAAACTCACTCATCTCGAATTCTTCAGAATTAGTCAAACAAACATTTC

>M6-PGSC0003DMG400008492M6LRR-RLP_378

AACAATCCTTAAACTCACTCATCTCGAATTCTTCAGAATTAGTCAAACAAACATTTCTGGACCTGTACCTGAAGTTCTTAGTAAATTGAAAAGCTTAACGTACATTAACTTATCCTATAA

>M6-PGSC0003DMG400008492M6LRR-RLP_441

TGTACCTGAAGTTCTTAGTAAATTGAAAAGCTTAACGTACATTAACTTATCCTATAATAAGCTTGTTGGAACTATTCCACCTTCGCTATCTCAACTTCCACACTTGGAGTTTTTACGATT

>M6-PGSC0003DMG400008492M6LRR-RLP_504

TGTTGGAACTATTCCACCTTCGCTATCTCAACTTCCACACTTGGAGTTTTTACGATTAGACAGGAACAAACTTACTGGACCAATCCCGGAGTCTTTTAGCAAGTTAGCTCCAAATCTAAC

>M6-PGSC0003DMG400008492M6LRR-RLP_567

GAACAAACTTACTGGACCAATCCCGGAGTCTTTTAGCAAGTTAGCTCCAAATCTAACGTATCTTTACCTTGGACATAACCAACTTACAGGGATTGTCCCAACTTCATTTGCTGGCTGGAG

>M6-PGSC0003DMG400008492M6LRR-RLP_630

TTACCTTGGACATAACCAACTTACAGGGATTGTCCCAACTTCATTTGCTGGCTGGAGTTTTGAGACAATTGACTTGTCAAGGAACATGCTTGAAGGGGATATTTCATTTTTGTTTGGTAA

>M6-PGSC0003DMG400008492M6LRR-RLP_693

GACAATTGACTTGTCAAGGAACATGCTTGAAGGGGATATTTCATTTTTGTTTGGTAAAGATAAGAATACATTTGAAATGCTTTTGGATCGGAATAAGTTTGAATTTGACATGTCAAAGTT

>M6-PGSC0003DMG400008492M6LRR-RLP_756

GAATACATTTGAAATGCTTTTGGATCGGAATAAGTTTGAATTTGACATGTCAAAGTTGACGTTTGGGAAGAGATTATGGAGGTTGCATTTAAATCATAACAAGATTTATGGGAGTCTTCC

>M6-PGSC0003DMG400008492M6LRR-RLP_819

TGGGAAGAGATTATGGAGGTTGCATTTAAATCATAACAAGATTTATGGGAGTCTTCCAAAAATCATGTCCAAGAATCCGTGGCAACTCTTGAATGTGAGTTATAATAGACTTTGTGGAAA

>M6-PGSC0003DMG400008492M6LRR-RLP_882

CATGTCCAAGAATCCGTGGCAACTCTTGAATGTGAGTTATAATAGACTTTGTGGAAAGATACCAAAGGGTGAGAATATGCAGAGATTTGAGATATATGAGTATTTCCACAACAAATGCTT

>M6-PGSC0003DMG400008492M6LRR-RLP_945

AAAGGGTGAGAATATGCAGAGATTTGAGATATATGAGTATTTCCACAACAAATGCTTGTGTGGTGCTCCACTGCCGCCCTGTAAATGATCATCACTAGTATCCTGTCTGTCGATCTATCT

>M6-PGSC0003DMG400008493M6G-LecRK_0

TAAAAAGTTGTATATAGCTAGTGTCCGGCATGCTTTTATTTCAACTATAATGGCTTTTGATTTTCAATCACAACAAGAAATCATGTACGTATAGATTATTATCTAAATTTTCATGTTTTT

>M6-PGSC0003DMG400008493M6G-LecRK_1054

TTTGTACTTATGCAACTTTGTTCTTTAAATGTAGGGAGTTTTGAGAGATGGAACTTGTGTTGCCATTAAATGTCTCTCTGCTGAGTCAAAACAAGGGACTAAAGAATTCTTAACAGAGAT

>M6-PGSC0003DMG400008493M6G-LecRK_1116

CCATTAAATGTCTCTCTGCTGAGTCAAAACAAGGGACTAAAGAATTCTTAACAGAGATTAATATGATTTCAAATACCCAACATCCAAATCTTGTTCAGTTGATTGGTTGTTGTGTTGAGA

>M6-PGSC0003DMG400008493M6G-LecRK_1178

ATGATTTCAAATACCCAACATCCAAATCTTGTTCAGTTGATTGGTTGTTGTGTTGAGAGTGGCAATAGAATGTTGATATATGAATACTTGAAGAATAACAGCCTTGCCAGTGCTTTACTA

>M6-PGSC0003DMG400008493M6G-LecRK_124

TTTGTCTAAGTAGATTTTATAGTAGTTGATTATTTTTTTTTCGTATTTCCTTTTCATTAATTTGCAGAAATATTAGGTGCTACAAATTTAAAATTTTGAATATTTGAGCAATGGGTTGGA

>M6-PGSC0003DMG400008493M6G-LecRK_1240

CAATAGAATGTTGATATATGAATACTTGAAGAATAACAGCCTTGCCAGTGCTTTACTAGGTAATGTTCTCACAGATCAGTTTGAAAAAAAAAAACTTATTTTGAACCAAATGAACATGTC

>M6-PGSC0003DMG400008493M6G-LecRK_1302

ATGTTCTCACAGATCAGTTTGAAAAAAAAAAACTTATTTTGAACCAAATGAACATGTCATAGTTGAAGTTAGGAATGTTAAATGAGCAGATTGGACTGAATCTGGACGGGTTAAAATTGT

>M6-PGSC0003DMG400008493M6G-LecRK_1364

TTGAAGTTAGGAATGTTAAATGAGCAGATTGGACTGAATCTGGACGGGTTAAAATTGTTATTTCAGGTTCTAATGGTAAACGCGTTGCCCTGAATTGGCCTCAGAGAGTTGCTATATGTT

>M6-PGSC0003DMG400008493M6G-LecRK_1426

TCAGGTTCTAATGGTAAACGCGTTGCCCTGAATTGGCCTCAGAGAGTTGCTATATGTTTAGGTACAGCATCTGGTTTAGCATTTCTTCATGAGGAAGCCTCACCACCGATTGTTCATCGA

>M6-PGSC0003DMG400008493M6G-LecRK_1488

TACAGCATCTGGTTTAGCATTTCTTCATGAGGAAGCCTCACCACCGATTGTTCATCGAGATATTAAGGCTAGTAATATCCTTATAGATGAAAACCTTCATCCTAAAATTGGAGATTTTGG

>M6-PGSC0003DMG400008493M6G-LecRK_1550

TTAAGGCTAGTAATATCCTTATAGATGAAAACCTTCATCCTAAAATTGGAGATTTTGGTCTGGCTAAGCTTTTTCCTGAAAATGTCACTCATCTAAGTACGCGAGTAGCAGGAACAATGT

>M6-PGSC0003DMG400008493M6G-LecRK_1612

GCTAAGCTTTTTCCTGAAAATGTCACTCATCTAAGTACGCGAGTAGCAGGAACAATGTGAGTAGTTACATTGAATTTCAGTTTCATAACATTCGATATTCCTCTCCTGTCTCGTCTCAGT

>M6-PGSC0003DMG400008493M6G-LecRK_1674

AGTTACATTGAATTTCAGTTTCATAACATTCGATATTCCTCTCCTGTCTCGTCTCAGTAAGGTATTCTGATGTGCAGAGGTTACCTTGCTCCGGAGTATGCCTTGTTTGGACAACTCACG

>M6-PGSC0003DMG400008493M6G-LecRK_1736

TATTCTGATGTGCAGAGGTTACCTTGCTCCGGAGTATGCCTTGTTTGGACAACTCACGAAAAAGGCTGATGTCTATAGCTTTGGAGTTCTTGTACTTGAAATCATAAGTGGAAGAAGCAG

>M6-PGSC0003DMG400008493M6G-LecRK_1798

AGGCTGATGTCTATAGCTTTGGAGTTCTTGTACTTGAAATCATAAGTGGAAGAAGCAGTAGTAAATCAGCATTTGGAGTGGATCTCTTGGTTCTGGTGGAATGGGTAGGATTATACTTTC

>M6-PGSC0003DMG400008493M6G-LecRK_186

TGCAGAAATATTAGGTGCTACAAATTTAAAATTTTGAATATTTGAGCAATGGGTTGGAATTGCTTTGGAGTTTTGCAATCATACAAAAACAATAGAGAGACTCAAATCCAAGGTAAGTAC

>M6-PGSC0003DMG400008493M6G-LecRK_1860

AAATCAGCATTTGGAGTGGATCTCTTGGTTCTGGTGGAATGGGTAGGATTATACTTTCAGCTTTATGATCTTCAAGATTCATCAGTAATCAGCATTTTGAAAGTTTACGCGTTGCAGGTA

>M6-PGSC0003DMG400008493M6G-LecRK_1922

TTATGATCTTCAAGATTCATCAGTAATCAGCATTTTGAAAGTTTACGCGTTGCAGGTATGGAAGCTAAGAGAAGAAGCACGGCTTCTGGAAATCATTGATCCAGAGCTAACAGAGTATCC

>M6-PGSC0003DMG400008493M6G-LecRK_1984

AGCTAAGAGAAGAAGCACGGCTTCTGGAAATCATTGATCCAGAGCTAACAGAGTATCCAGAAACTGAATTGCTGCGGTTCATCAAAGTTGCTCTCTTTTGCACACAATCAGCACCTACCC

>M6-PGSC0003DMG400008493M6G-LecRK_2046

ACTGAATTGCTGCGGTTCATCAAAGTTGCTCTCTTTTGCACACAATCAGCACCTACCCAAAGACCTAATATGAAACAAGTGATTGAGATGTTGTCTAAGGAAGTCAACCTCAACGAGAAG

>M6-PGSC0003DMG400008493M6G-LecRK_2108

ACCTAATATGAAACAAGTGATTGAGATGTTGTCTAAGGAAGTCAACCTCAACGAGAAGTCGTTAACTGAACCAGGAGTTTATAGACCACATAGCTCGAAACGATCAAGTTATGGTAGTCT

>M6-PGSC0003DMG400008493M6G-LecRK_2170

TAACTGAACCAGGAGTTTATAGACCACATAGCTCGAAACGATCAAGTTATGGTAGTCTGCAGAAATCATCCTCTGCGGGGAAGAATGGGGTGCAATCTGTAAATCCTTCTGTAACATCAA

>M6-PGSC0003DMG400008493M6G-LecRK_2232

AAATCATCCTCTGCGGGGAAGAATGGGGTGCAATCTGTAAATCCTTCTGTAACATCAACAAAGTTTGAGAGCTTTCAGAGTGTCACCCAAATGCTTCCTAGATAATGTTAACTGGTGTTC

>M6-PGSC0003DMG400008493M6G-LecRK_2294

GTTTGAGAGCTTTCAGAGTGTCACCCAAATGCTTCCTAGATAATGTTAACTGGTGTTCTTATTCTTCATACTTGTATATTGTCGTTCTTCTAACGTGGGCTATCGAGGAGTAAATGTATT

>M6-PGSC0003DMG400008493M6G-LecRK_2356

TCTTCATACTTGTATATTGTCGTTCTTCTAACGTGGGCTATCGAGGAGTAAATGTATTGTCTCTCATTAGACTTACCTTTTTCATACATTTTCTCATGTAATGTAAAGTTGGCAGTAGCT

>M6-PGSC0003DMG400008493M6G-LecRK_2418

CTCATTAGACTTACCTTTTTCATACATTTTCTCATGTAATGTAAAGTTGGCAGTAGCTGATAAGGGAGAAAGAGAGACAAATCACACAGAATGAGCTACTATGTAACTACAAATTTCAGA

>M6-PGSC0003DMG400008493M6G-LecRK_248

CTTTGGAGTTTTGCAATCATACAAAAACAATAGAGAGACTCAAATCCAAGGTAAGTACACTCATAACTCTTATTTCTGTTTTCTCAATTACATGCTTAAAACAAATCGTTCTTATAAAAT

>M6-PGSC0003DMG400008493M6G-LecRK_2480

AGGGAGAAAGAGAGACAAATCACACAGAATGAGCTACTATGTAACTACAAATTTCAGATATGAAGATGAAGTTTACAGCAGAGACTATGAACACAGTAAAGGATAGAAACTTTTCTCTAT

>M6-PGSC0003DMG400008493M6G-LecRK_2542

AAGATGAAGTTTACAGCAGAGACTATGAACACAGTAAAGGATAGAAACTTTTCTCTATGCACATATTAGCTCCTAAAACTAGAAATGAAGGTAAAAACTTCCTGATAGAAGTTATTCTCA

>M6-PGSC0003DMG400008493M6G-LecRK_2604

ATATTAGCTCCTAAAACTAGAAATGAAGGTAAAAACTTCCTGATAGAAGTTATTCTCAAACTCATAACATTTAAGTTAACGTATAAAGGATGACTGCTTTACAAGATCAACACAGTAGCA

>M6-PGSC0003DMG400008493M6G-LecRK_2666

CATAACATTTAAGTTAACGTATAAAGGATGACTGCTTTACAAGATCAACACAGTAGCATTTTCATTGTGCAGCAGAGAATAAACCTTCAAGATGCAGCAGTAGTAGCTTGAGGCTTTGCA

>M6-PGSC0003DMG400008493M6G-LecRK_2728

CATTGTGCAGCAGAGAATAAACCTTCAAGATGCAGCAGTAGTAGCTTGAGGCTTTGCAGAGGCAGCAACAGAAGTTTCTATTCCTGTCTTTGCATCTTTTCCTGGTTCCGAAGATGGTTT

>M6-PGSC0003DMG400008493M6G-LecRK_2790

CAGCAACAGAAGTTTCTATTCCTGTCTTTGCATCTTTTCCTGGTTCCGAAGATGGTTTTGCAGAAGGCTTCATATGCTCAGGGGGGTTGTGTTTGAGTTTCACAAGTATTTGACGCATTC

>M6-PGSC0003DMG400008493M6G-LecRK_2852

GAAGGCTTCATATGCTCAGGGGGGTTGTGTTTGAGTTTCACAAGTATTTGACGCATTCTAGAGAGTTCAGTTGGCCTGCCAGGTTTGGGTCCTTGGATGACATTGATGGATGGCTGCTTC

>M6-PGSC0003DMG400008493M6G-LecRK_2914

GAGTTCAGTTGGCCTGCCAGGTTTGGGTCCTTGGATGACATTGATGGATGGCTGCTTCTCCTTATCCTTCTTTCCTTTCTTCTCTATGGTCGGGACCTCCTTTGGGATAAGCTCTCCAAT

>M6-PGSC0003DMG400008493M6G-LecRK_2976

TATCCTTCTTTCCTTTCTTCTCTATGGTCGGGACCTCCTTTGGGATAAGCTCTCCAATTGCCCTCCAATAAGCCTTATTAGCTTCAGTATGGAACTTCTCTTGGTTGGCCAAGAATAACT

>M6-PGSC0003DMG400008493M6G-LecRK_3038

CTCCAATAAGCCTTATTAGCTTCAGTATGGAACTTCTCTTGGTTGGCCAAGAATAACTGGAAAAAAAGAAAAAGATGACACATGAACTGCAACACATTGATTGACTAAGGAGGAACTTGC

>M6-PGSC0003DMG400008493M6G-LecRK_310

ATAACTCTTATTTCTGTTTTCTCAATTACATGCTTAAAACAAATCGTTCTTATAAAATTCAGAATCCAAAAAATTTAAAAATTTGACGCATCTGCTCTACTTTTTGTAGTCCATCCAAAT

>M6-PGSC0003DMG400008493M6G-LecRK_3100

AAAAAGAAAAAGATGACACATGAACTGCAACACATTGATTGACTAAGGAGGAACTTGCCGTACTGTGGAACTATGTTTAGAGATTTGACAAACCATAAAACCAAAGACATAGTTCTGTGG

>M6-PGSC0003DMG400008493M6G-LecRK_3162

CTGTGGAACTATGTTTAGAGATTTGACAAACCATAAAACCAAAGACATAGTTCTGTGGAGATTAGCAAATAAAACTCACCAAAGCACAAAAACTCAGCATATATTGTTCTAAAGGTTATC

>M6-PGSC0003DMG400008493M6G-LecRK_3224

TAGCAAATAAAACTCACCAAAGCACAAAAACTCAGCATATATTGTTCTAAAGGTTATCAGCACATTTTGATCTCTTGTGGAAAACTTCCACTTTCAAAAATTTTAATTTACACCTCTTAC

>M6-PGSC0003DMG400008493M6G-LecRK_3286

CATTTTGATCTCTTGTGGAAAACTTCCACTTTCAAAAATTTTAATTTACACCTCTTACGCAGTATTAATGCTAAATCATATGAAATGTAAACAGACAAAATGTAGAAACTGCTTTTCTCT

>M6-PGSC0003DMG400008493M6G-LecRK_3348

TATTAATGCTAAATCATATGAAATGTAAACAGACAAAATGTAGAAACTGCTTTTCTCTGTGTCCCTACCTTGGTTGAGTCGAAAGTACCAATACAATGATTTTTTCAAAACTAATAACTT

>M6-PGSC0003DMG400008493M6G-LecRK_3410

CCCTACCTTGGTTGAGTCGAAAGTACCAATACAATGATTTTTTCAAAACTAATAACTTAAAAAGGAGAGTAACAAGTACTATGATTAAAAATTGGATGGTACTTTTACTAGGTTAACTAG

>M6-PGSC0003DMG400008493M6G-LecRK_3472

AGGAGAGTAACAAGTACTATGATTAAAAATTGGATGGTACTTTTACTAGGTTAACTAGGTTAATCTCCCTGTCTCAACACTCATATCCTATTCCGAAGAGTTGAAAGCAATGATTAGGAA

>M6-PGSC0003DMG400008493M6G-LecRK_3534

ATCTCCCTGTCTCAACACTCATATCCTATTCCGAAGAGTTGAAAGCAATGATTAGGAAGAGAAGGTAACTAGATGCATGACATGTCAGACCTTCCAGCTAAAGCAACTGTTTGGAAAACA

>M6-PGSC0003DMG400008493M6G-LecRK_3596

AGGTAACTAGATGCATGACATGTCAGACCTTCCAGCTAAAGCAACTGTTTGGAAAACAAACATCCAGCACAAGCAAATTTATGCCAAACTAGTTAACAAACCGTGACTTTAAGCAAAATG

>M6-PGSC0003DMG400008493M6G-LecRK_3658

TCCAGCACAAGCAAATTTATGCCAAACTAGTTAACAAACCGTGACTTTAAGCAAAATGCCTCGGAAAGACTTTTAATTTATATAGCTATAGGTCTCACCTTCTCCTTCTCCCGGTTAGCA

>M6-PGSC0003DMG400008493M6G-LecRK_372

AATCCAAAAAATTTAAAAATTTGACGCATCTGCTCTACTTTTTGTAGTCCATCCAAATTTGAGAAGTACTTCACATAATTTTCATTTTCTTTTTGTGTTTTTCATTGTGTAGAGTTCACG

>M6-PGSC0003DMG400008493M6G-LecRK_3720

GGAAAGACTTTTAATTTATATAGCTATAGGTCTCACCTTCTCCTTCTCCCGGTTAGCAGCTATGTTGTTCTCACATCTAAGTTTCCATTTCCTATAGTACTCTGCTTTGTATTCCTCTGC

>M6-PGSC0003DMG400008493M6G-LecRK_3782

TGTTGTTCTCACATCTAAGTTTCCATTTCCTATAGTACTCTGCTTTGTATTCCTCTGCTTCTCTGATTATTTCCCTAACAATTTCTTTCTCCCTTTTCTCCTTCTCCTCCAGCCGTATGG

>M6-PGSC0003DMG400008493M6G-LecRK_434

GAAGTACTTCACATAATTTTCATTTTCTTTTTGTGTTTTTCATTGTGTAGAGTTCACGACCAACAATGTGAGGGTTTTTTCCTATAACTCATTGAGATCAGCAACAAGTCACTTCCATCC

>M6-PGSC0003DMG400008493M6G-LecRK_496

ACAATGTGAGGGTTTTTTCCTATAACTCATTGAGATCAGCAACAAGTCACTTCCATCCATCTAACAAAATAGGAGGTGGTGGTTTTGGAGTTGTTTATAAGGTAATCATCTCTTGTTTAT

>M6-PGSC0003DMG400008493M6G-LecRK_558

AACAAAATAGGAGGTGGTGGTTTTGGAGTTGTTTATAAGGTAATCATCTCTTGTTTATGGTGTCATGCCTTTTAATGGTCCTTGTGGATGCTAGTTATTAAGTACTGAGTTTAACATCTA

>M6-PGSC0003DMG400008493M6G-LecRK_62

TTCAATCACAACAAGAAATCATGTACGTATAGATTATTATCTAAATTTTCATGTTTTTCCCATTTGTCTAAGTAGATTTTATAGTAGTTGATTATTTTTTTTTCGTATTTCCTTTTCATT

>M6-PGSC0003DMG400008493M6G-LecRK_620

TCATGCCTTTTAATGGTCCTTGTGGATGCTAGTTATTAAGTACTGAGTTTAACATCTATACGTTCAAAATTTAGAAGAAGTCACCTAAAAGATAACTATAATAATTTAAATCATTTGAAG

>M6-PGSC0003DMG400008493M6G-LecRK_682

TTCAAAATTTAGAAGAAGTCACCTAAAAGATAACTATAATAATTTAAATCATTTGAAGTGGGATTTGCTAGTGAGTAGAAAAATGAGAGCATGTAAGGATAGATGCAGCCTAATGATCAA

>M6-PGSC0003DMG400008493M6G-LecRK_930

CTGATACAAACACCACCATCATAAAAATTAAATTAAATAAGGTAGATTACCTGCTACATTAATAATGTAAAAATATTTTACACTATCAATATATATAGGTTAAATCTTTTATGTATTTCC

>M6-PGSC0003DMG400008493M6G-LecRK_992

TAATGTAAAAATATTTTACACTATCAATATATATAGGTTAAATCTTTTATGTATTTCCACTCTTTGTACTTATGCAACTTTGTTCTTTAAATGTAGGGAGTTTTGAGAGATGGAACTTGT

>M6-PGSC0003DMG400008499-(rev)M6RLKWAX-EGF_0

ATGCTAATTTTGACATTAGCCACTCCCCAAATTATCCCCTCTAACACCACTTCTCCTCCGCCCACAAACACCACATCTCCGCCAACCACGACGATCACTAAAGCTGCGATCATTACAAAA

>M6-PGSC0003DMG400008499-(rev)M6RLKWAX-EGF_1054

AATTATATGATTAAAATGTAATTCTTCTCAAGTCAATACGATAAACAAGTTAAATTTAAAATGTTTGTCAAACTTCGCATGCATAATCTTAGTTATAAGACAAGAGAGAGTTATTCGAAT

>M6-PGSC0003DMG400008499-(rev)M6RLKWAX-EGF_124

GATGTCCGAAAAAGTGTGGGAACCTAACAGTTCCATACCCTTTTGGGATTGGCTTAGGATCTGGTTGTGCCTTGGACCGGAATTTTGAAATCAATTGTGATACTAATACAACAGATTCTC

>M6-PGSC0003DMG400008499-(rev)M6RLKWAX-EGF_1240

GCTCCTGGGGGAGGGAGAGAAAAATAAAATTAAAAAAATTATTATAAATTGAAACAAAGAGAATAAAACTATTTTATTGGGGGAAAAAATAACAATTTGAGTTGAGACTTTTTTGAGGTG

>M6-PGSC0003DMG400008499-(rev)M6RLKWAX-EGF_1302

ATAAAACTATTTTATTGGGGGAAAAAATAACAATTTGAGTTGAGACTTTTTTGAGGTGAATCAGTCTAGCAGTGAGGTCGCTATTAACTTATCATTTTTTTTTAATATAGAAATGATTAA

>M6-PGSC0003DMG400008499-(rev)M6RLKWAX-EGF_1364

AGTCTAGCAGTGAGGTCGCTATTAACTTATCATTTTTTTTTAATATAGAAATGATTAATATTCAAAGTTGTACAAATATTAGTTAGCTTTTAAAATATTTAACATATGTGATAATGAATA

>M6-PGSC0003DMG400008499-(rev)M6RLKWAX-EGF_1426

CAAAGTTGTACAAATATTAGTTAGCTTTTAAAATATTTAACATATGTGATAATGAATATGGTTTAAATACTTGGAAAAAGAGCGACGCTAATTAAATACTCCTACTTGCCATTAGTTAAC

>M6-PGSC0003DMG400008499-(rev)M6RLKWAX-EGF_1488

TTAAATACTTGGAAAAAGAGCGACGCTAATTAAATACTCCTACTTGCCATTAGTTAACAAAACAATGTTGGACCGTACAACTTCACAAGTAAATACGTAAAAAATAAAGATAAAAAAACA

>M6-PGSC0003DMG400008499-(rev)M6RLKWAX-EGF_1550

CAATGTTGGACCGTACAACTTCACAAGTAAATACGTAAAAAATAAAGATAAAAAAACAAAGTGTCACAAATTAATTTATTTGATCCACCAGCCAACGTAATATATATTTATGTACGCTAT

>M6-PGSC0003DMG400008499-(rev)M6RLKWAX-EGF_1612

GTCACAAATTAATTTATTTGATCCACCAGCCAACGTAATATATATTTATGTACGCTATGATTTTGACAGATAAAAAGAAATTACTATCTAATGTCATGATTTTCATTCACCATATTGATC

>M6-PGSC0003DMG400008499-(rev)M6RLKWAX-EGF_1674

TTGACAGATAAAAAGAAATTACTATCTAATGTCATGATTTTCATTCACCATATTGATCGATTAATCTTTCTTTAATTTTGGGTGACGCAACGTTTGGTTAATTGTACCAGATATTGATGA

>M6-PGSC0003DMG400008499-(rev)M6RLKWAX-EGF_1736

AATCTTTCTTTAATTTTGGGTGACGCAACGTTTGGTTAATTGTACCAGATATTGATGAATGTGCTAATCCAAATACCAATTTGTGTGAACAAAATTGTATAAACATACCGGGGAGTTACA

>M6-PGSC0003DMG400008499-(rev)M6RLKWAX-EGF_1798

GCTAATCCAAATACCAATTTGTGTGAACAAAATTGTATAAACATACCGGGGAGTTACAATTGTTCATGTCCAAAAGGGTACACTGGTGATGGAAGAAAGAATGGTCGTGGTTGTATTGCA

>M6-PGSC0003DMG400008499-(rev)M6RLKWAX-EGF_186

GGTTGTGCCTTGGACCGGAATTTTGAAATCAATTGTGATACTAATACAACAGATTCTCCAACACCATACATAGGGAATATTCCAGTGTATGACATCTCTGATGCTGAAATGCGCGTATCG

>M6-PGSC0003DMG400008499-(rev)M6RLKWAX-EGF_1860

TTCATGTCCAAAAGGGTACACTGGTGATGGAAGAAAGAATGGTCGTGGTTGTATTGCACCAAACTCGAACTCAGAGTTCCCATGGATTAAGTTTTCCGTTGGTAAGTTTCTCAGAAATTC

>M6-PGSC0003DMG400008499-(rev)M6RLKWAX-EGF_1922

ACTCGAACTCAGAGTTCCCATGGATTAAGTTTTCCGTTGGTAAGTTTCTCAGAAATTCGACTTGACTCTACTTGCATATAGATAAACGTACTGATTTGTATTTCAATTTATCTGTCTTAG

>M6-PGSC0003DMG400008499-(rev)M6RLKWAX-EGF_1984

TGACTCTACTTGCATATAGATAAACGTACTGATTTGTATTTCAATTTATCTGTCTTAGTTTGATTGGACACGAAATTTAAGAAAATAAATTTGTTGGATAAACACAGGTATGGGAGTTGG

>M6-PGSC0003DMG400008499-(rev)M6RLKWAX-EGF_2046

ATTGGACACGAAATTTAAGAAAATAAATTTGTTGGATAAACACAGGTATGGGAGTTGGTTTTATGTCCCTAGTAGTTGGGACAACTTGGCTCTATTTCAGCATCAAGAAAAGAAAACTCA

>M6-PGSC0003DMG400008499-(rev)M6RLKWAX-EGF_2108

ATGTCCCTAGTAGTTGGGACAACTTGGCTCTATTTCAGCATCAAGAAAAGAAAACTCATTAAACTTCGCGAGAAATTCTTCCAACAAAATGGTGGTTTGCTGTTGAAACAACGATTATCT

>M6-PGSC0003DMG400008499-(rev)M6RLKWAX-EGF_2170

ACTTCGCGAGAAATTCTTCCAACAAAATGGTGGTTTGCTGTTGAAACAACGATTATCTTCTAACGAGGGTGGTGTGGAAGCAACCAAAGTTTTTACAGCAGAAGAGTTGAAGAAGGCTAC

>M6-PGSC0003DMG400008499-(rev)M6RLKWAX-EGF_2232

ACGAGGGTGGTGTGGAAGCAACCAAAGTTTTTACAGCAGAAGAGTTGAAGAAGGCTACAAACAACTATGCTAGTGATAGAATTCTTGGTCGTGGTGGAAATGGAATTGTCTATAAAGGTA

>M6-PGSC0003DMG400008499-(rev)M6RLKWAX-EGF_2294

AACTATGCTAGTGATAGAATTCTTGGTCGTGGTGGAAATGGAATTGTCTATAAAGGTATACTACCCGATAATCTCATAGTTGCTATTAAAAAGTCTAAGTTTGTGGACGAGGATCAAATT

>M6-PGSC0003DMG400008499-(rev)M6RLKWAX-EGF_2356

ACCCGATAATCTCATAGTTGCTATTAAAAAGTCTAAGTTTGTGGACGAGGATCAAATTGAACAGTTCATCAATGAGGTTCTTATTCTTACTCAAGTCAACCATAGAAACGTGGTGAGACT

>M6-PGSC0003DMG400008499-(rev)M6RLKWAX-EGF_2418

AGTTCATCAATGAGGTTCTTATTCTTACTCAAGTCAACCATAGAAACGTGGTGAGACTCTTTGGATGTTGTTTGGAAGCCGAAGTTCCTTTACTAGTTTACGAATACATTTCTAATGGTA

>M6-PGSC0003DMG400008499-(rev)M6RLKWAX-EGF_248

ACCATACATAGGGAATATTCCAGTGTATGACATCTCTGATGCTGAAATGCGCGTATCGACTAACATAAATCGGAGGTGTTATTCATCGACGGGCATGTTGCTTCGAAATGATCCTTCTTG

>M6-PGSC0003DMG400008499-(rev)M6RLKWAX-EGF_2480

GGATGTTGTTTGGAAGCCGAAGTTCCTTTACTAGTTTACGAATACATTTCTAATGGTACTCTTTACGAGCATATCCACAACCAAAATGGAGCACCTTGGTTATCATGGAAGAATCGGCTA

>M6-PGSC0003DMG400008499-(rev)M6RLKWAX-EGF_2542

TTACGAGCATATCCACAACCAAAATGGAGCACCTTGGTTATCATGGAAGAATCGGCTAAGAGTTGCTAGTGAGACAGCAAGTGCACTTGCTTACCTTCATTCATCCGCGTCAATGCCTAT

>M6-PGSC0003DMG400008499-(rev)M6RLKWAX-EGF_2604

TTGCTAGTGAGACAGCAAGTGCACTTGCTTACCTTCATTCATCCGCGTCAATGCCTATAATTCATAGAGATGTCAAGTCTGCCAATTTATTGTTAGATGATGTTTACACCGCGAAAGTGG

>M6-PGSC0003DMG400008499-(rev)M6RLKWAX-EGF_2666

CATAGAGATGTCAAGTCTGCCAATTTATTGTTAGATGATGTTTACACCGCGAAAGTGGCAGATTTTGGAGCTTCAAGACTAATTCCTCTTGATCAAACACATATTGCTACATTGGTTCAA

>M6-PGSC0003DMG400008499-(rev)M6RLKWAX-EGF_2728

TTTTGGAGCTTCAAGACTAATTCCTCTTGATCAAACACATATTGCTACATTGGTTCAAGGGACATTAGGATACCTAGATCCTGAATATTTTCGTTTAAGTAAATTGACCGAGAAAAGTGA

>M6-PGSC0003DMG400008499-(rev)M6RLKWAX-EGF_2790

CATTAGGATACCTAGATCCTGAATATTTTCGTTTAAGTAAATTGACCGAGAAAAGTGATGTTTATAGTTTTGGAGTAGTCCTTGCAGAACTTTTAACAGGGATAAAACCTATTTCGAGGG

>M6-PGSC0003DMG400008499-(rev)M6RLKWAX-EGF_2852

TATAGTTTTGGAGTAGTCCTTGCAGAACTTTTAACAGGGATAAAACCTATTTCGAGGGATAAAAATAATGAGGACAAGAATTTGGCAGAGTACTTCATTTTGTCCATGAGAAAGAATCAA

>M6-PGSC0003DMG400008499-(rev)M6RLKWAX-EGF_2914

AAATAATGAGGACAAGAATTTGGCAGAGTACTTCATTTTGTCCATGAGAAAGAATCAATTGTTTCAAATTCTTGATCGTAGAGTGGTAAGAGAAGGAAGCCTTGAACAACTCCAAAAGGT

>M6-PGSC0003DMG400008499-(rev)M6RLKWAX-EGF_2976

TTCAAATTCTTGATCGTAGAGTGGTAAGAGAAGGAAGCCTTGAACAACTCCAAAAGGTGGCTGAGCTAGTGAAAAATTGTCTTAGATTGCACGGTGAACATAGGCCTACGATGAAGGAAG

>M6-PGSC0003DMG400008499-(rev)M6RLKWAX-EGF_3038

GAGCTAGTGAAAAATTGTCTTAGATTGCACGGTGAACATAGGCCTACGATGAAGGAAGTAGCAATGGAACTTGAGAGTTTGCGAAAGTTCACAAAAAATAACCTTTGGGCTGATGGACAC

>M6-PGSC0003DMG400008499-(rev)M6RLKWAX-EGF_310

ACATAAATCGGAGGTGTTATTCATCGACGGGCATGTTGCTTCGAAATGATCCTTCTTGGATGAATTTAGGAACCTCGAGTCCTTACAGTTTCTCTACCCTCAACAGGTACGTACCTAATT

>M6-PGSC0003DMG400008499-(rev)M6RLKWAX-EGF_3100

AATGGAACTTGAGAGTTTGCGAAAGTTCACAAAAAATAACCTTTGGGCTGATGGACACGGACATGAAGAGAACGAAGATGAATCAACAGATCTTTATACAATTCCAATTGACTCGAATAC

>M6-PGSC0003DMG400008499-(rev)M6RLKWAX-EGF_3162

ATGAAGAGAACGAAGATGAATCAACAGATCTTTATACAATTCCAATTGACTCGAATACAGGTATCAACAAATTCTCTGGACAATACAGTTCCAACTCCAACACAAATAGCAGCATGTTTT

>M6-PGSC0003DMG400008499-(rev)M6RLKWAX-EGF_3200

ATTCCAATTGACTCGAATACAGGTATCAACAAATTCTCTGGACAATACAGTTCCAACTCCAACACAAATAGCAGCATGTTTTCTTCATTGATATATACTAGTACAAATATTCCGCGGTGA

>M6-PGSC0003DMG400008499-(rev)M6RLKWAX-EGF_372

AATTTAGGAACCTCGAGTCCTTACAGTTTCTCTACCCTCAACAGGTACGTACCTAATTATAAATTTTCTCTTGAAACCTTATAAACTTTAATTTTGAATTCACATATATATATATACAGG

>M6-PGSC0003DMG400008499-(rev)M6RLKWAX-EGF_434

ATTTTCTCTTGAAACCTTATAAACTTTAATTTTGAATTCACATATATATATATACAGGTTCACCGTTGTTGGTTGTGATGAAGCTGCTATAATATTTGGAGGTGACTTTGCTAATGGCTG

>M6-PGSC0003DMG400008499-(rev)M6RLKWAX-EGF_496

CCGTTGTTGGTTGTGATGAAGCTGCTATAATATTTGGAGGTGACTTTGCTAATGGCTGCCCCACGGTCTGTATTAGTACGAGTCAGGTAACTGAAGGAAGGTGTATGGGTAATGGTTGCT

>M6-PGSC0003DMG400008499-(rev)M6RLKWAX-EGF_558

ACGGTCTGTATTAGTACGAGTCAGGTAACTGAAGGAAGGTGTATGGGTAATGGTTGCTGCCAAATAACAATACCAAAAGGCTTGAAAATTTTCAACACAACGATGCAAAGTTTGAAAAAT

>M6-PGSC0003DMG400008499-(rev)M6RLKWAX-EGF_62

CACAAACACCACATCTCCGCCAACCACGACGATCACTAAAGCTGCGATCATTACAAAAGCAGGATGTCCGAAAAAGTGTGGGAACCTAACAGTTCCATACCCTTTTGGGATTGGCTTAGG

>M6-PGSC0003DMG400008499-(rev)M6RLKWAX-EGF_620

AATAACAATACCAAAAGGCTTGAAAATTTTCAACACAACGATGCAAAGTTTGAAAAATCACACTGGAGTTTGGTCTTTCAATCCATGTGGCTATTCATTTCTTGGTGAAGCTACTCGATT

>M6-PGSC0003DMG400008499-(rev)M6RLKWAX-EGF_682

CTGGAGTTTGGTCTTTCAATCCATGTGGCTATTCATTTCTTGGTGAAGCTACTCGATTCGAATTCATGGGTTTACAAGATCTACGCGATCTTAATTTTAAAAAGAGGTCTTTGGATAATG

>M6-PGSC0003DMG400008499-(rev)M6RLKWAX-EGF_744

TTCATGGGTTTACAAGATCTACGCGATCTTAATTTTAAAAAGAGGTCTTTGGATAATGTCCCTATTGTGCTAGATTGGGCTATTGGAACTCTTAGTTGTGTTGAAGCACAGAAGAGTAGT

>M6-PGSC0003DMG400008499-(rev)M6RLKWAX-EGF_806

TATTGTGCTAGATTGGGCTATTGGAACTCTTAGTTGTGTTGAAGCACAGAAGAGTAGTAACGATTATGCTTGTTTGAATAATAGCCAATGTGTTGATTCCAACACTGGCCTTGGTGGTTA

>M6-PGSC0003DMG400008499-(rev)M6RLKWAX-EGF_868

ATTATGCTTGTTTGAATAATAGCCAATGTGTTGATTCCAACACTGGCCTTGGTGGTTACAGGTGTGCCTGTAACCCGGGATATGAAGGCAATCCATATATCGGCCCAGGTTGTCAAGGTA

>M6-PGSC0003DMG400008499-(rev)M6RLKWAX-EGF_930

TGTGCCTGTAACCCGGGATATGAAGGCAATCCATATATCGGCCCAGGTTGTCAAGGTAACTTCATTTCTCTACTTCCCTCACATGTTTCAACTTCCTTTTATTTGTAAAATTTAGATATT

>M6-PGSC0003DMG400008499-(rev)M6RLKWAX-EGF_992

CATTTCTCTACTTCCCTCACATGTTTCAACTTCCTTTTATTTGTAAAATTTAGATATTTAAAAATTATATGATTAAAATGTAATTCTTCTCAAGTCAATACGATAAACAAGTTAAATTTA

>M6-PGSC0003DMG400008508-(rev)M6G-LecRK_0

CATGGAAATGAAGAACAATCATTACTTTCTGCTACTCTTTTTATGCTTCAGTCTCAACAGTAATCTCTGTATCGGAAGGGACACCATTTCTGCGACTGAATCTCTTTCTTTTGGTGAAAC

>M6-PGSC0003DMG400008508-(rev)M6G-LecRK_1054

TTATGGATAACTTCAATTGTGAGAGTACCTTCTTCTCAAAATACTAATATCACAGTTGGGGAAGCCTCACAGTGTAGATCTGCATGTTTCAACGATTGCTCTTGCACTGCTTATACTTAT

>M6-PGSC0003DMG400008508-(rev)M6G-LecRK_1178

GTTCCGGTACCTGTTCTATCTGGACAGGGGATCTGTTCAATCTGCAACAACTCAGCACAACTGAAACAGAAAGGACTATATTTGTCAAACGCAGCTCACCTGAAGGTTAGAATAGCATAG

>M6-PGSC0003DMG400008508-(rev)M6G-LecRK_124

GTCTCTTCAGGTGAGATATTTGAGCTTGGTTTTTTCAAACCAGGTAACTCTCTTAAATATTATCTAGGCATATGGTACAAGAACGTTATTTTATCGCAAACAGTAATTTGGGTAGCAAAT

>M6-PGSC0003DMG400008508-(rev)M6G-LecRK_1240

GAAACAGAAAGGACTATATTTGTCAAACGCAGCTCACCTGAAGGTTAGAATAGCATAGTATAACATGCTTTGATTCTTTATTTCATATTAGTACAATGTTATGAATAGAATTCCTTATTA

>M6-PGSC0003DMG400008508-(rev)M6G-LecRK_1302

ACATGCTTTGATTCTTTATTTCATATTAGTACAATGTTATGAATAGAATTCCTTATTAACGTTAGCACTTAACAACAATGATTCTCCTTTCAGCTCAAACTAATGCTAAGAAATCAAAGA

>M6-PGSC0003DMG400008508-(rev)M6G-LecRK_1364

TAGCACTTAACAACAATGATTCTCCTTTCAGCTCAAACTAATGCTAAGAAATCAAAGAAACTAAAAGCAATCATTTCGTCAATATCAACTTTGATGTTTCTACTCATAGGCAGCATTAGC

>M6-PGSC0003DMG400008508-(rev)M6G-LecRK_1426

AAAAGCAATCATTTCGTCAATATCAACTTTGATGTTTCTACTCATAGGCAGCATTAGCTACATTTACTATAGAAGAAGAATGGCAAAAAGAGCAGGTATTTGCTGAATGAATCTAACTTT

>M6-PGSC0003DMG400008508-(rev)M6G-LecRK_1488

TTTACTATAGAAGAAGAATGGCAAAAAGAGCAGGTATTTGCTGAATGAATCTAACTTTGGAAGATATTTTCTCTTATACCATTTTGGACTTAATTCCAAAGGTCATCCTCAATCCAATTG

>M6-PGSC0003DMG400008508-(rev)M6G-LecRK_1550

GATATTTTCTCTTATACCATTTTGGACTTAATTCCAAAGGTCATCCTCAATCCAATTGAGGTAATGAAATCCCTCCCACTCCCCTTTTCTATTTCAGATAGTAGTAAAGGTACTCAAGGG

>M6-PGSC0003DMG400008508-(rev)M6G-LecRK_1612

AATGAAATCCCTCCCACTCCCCTTTTCTATTTCAGATAGTAGTAAAGGTACTCAAGGGGCACGTAAATCTCACTGGCACAAAGCTGAAGGAGAAGCAAAAGTATTAATGAATGAAAACAG

>M6-PGSC0003DMG400008508-(rev)M6G-LecRK_1674

GTAAATCTCACTGGCACAAAGCTGAAGGAGAAGCAAAAGTATTAATGAATGAAAACAGTGATGAAGCGATTGATGTTCCATACTTTCACTTGGAGACTATTTTGGAAGCTACAGACAACT

>M6-PGSC0003DMG400008508-(rev)M6G-LecRK_1798

AAATGCGAATAAGCTCGGACAAGGAGGATTTGGTCCTGTTTACAAGGTTAAAACTTCATTCCTACATCTATGTGGACTGGCATTGACCCATAGTAATGAGATGAATTTTTTTTGCAGGGT

>M6-PGSC0003DMG400008508-(rev)M6G-LecRK_186

TCTAGGCATATGGTACAAGAACGTTATTTTATCGCAAACAGTAATTTGGGTAGCAAATAGGGATAAACCACTTGATTATGGTGCTGCTGGGATGATAATTAATCAAGGAAACCTGGTGCT

>M6-PGSC0003DMG400008508-(rev)M6G-LecRK_1860

TACATCTATGTGGACTGGCATTGACCCATAGTAATGAGATGAATTTTTTTTGCAGGGTATCTTTCCGGGTGAAAAAGAAATTGCGGTAAAAACGTTATCTAGTCAATCAGGACAAGGCAT

>M6-PGSC0003DMG400008508-(rev)M6G-LecRK_1984

GAATTTAAGAATGAAGTGACTCTAATTGCAAAGCTTCAACATCGAAATCTGGTGAGGCTATTGGGCTATTGCATCAATGCAACCGAACAAATATTACTTTATGAATATATGCCGAATAAA

>M6-PGSC0003DMG400008508-(rev)M6G-LecRK_2046

GGGCTATTGCATCAATGCAACCGAACAAATATTACTTTATGAATATATGCCGAATAAAAGTTTAGATACATTCATATTTGGTGAGTTAGCCTTAATAAATTCTTTATTCTCAAAGTCATA

>M6-PGSC0003DMG400008508-(rev)M6G-LecRK_2108

TAGATACATTCATATTTGGTGAGTTAGCCTTAATAAATTCTTTATTCTCAAAGTCATACATTTCTTATGATGATCTAATTTTATTGACGTACCAAACTTTGTTATGGTAACAATGCAAGA

>M6-PGSC0003DMG400008508-(rev)M6G-LecRK_2170

TCTTATGATGATCTAATTTTATTGACGTACCAAACTTTGTTATGGTAACAATGCAAGATGGAACACTTTGTCAATTATTGGATTGGAAGAAACGTTATGACATCATATTGGGCATTGCAC

>M6-PGSC0003DMG400008508-(rev)M6G-LecRK_2232

ACACTTTGTCAATTATTGGATTGGAAGAAACGTTATGACATCATATTGGGCATTGCACGGGGTCTATCTTATCTTCATCATGATTCACGACTAAGGATCATTCATAGAGATTTAAAAACC

>M6-PGSC0003DMG400008508-(rev)M6G-LecRK_2294

TCTATCTTATCTTCATCATGATTCACGACTAAGGATCATTCATAGAGATTTAAAAACCAGTAACATTTTATTAGATGAGGAGATGAACCCGAAAATTTCAGATTTTGGCTTGGCAAGGAT

>M6-PGSC0003DMG400008508-(rev)M6G-LecRK_2418

GAAGGAAAAGTAACAGAAGCAAATACAAAGAAAATAGTGGGGACCTAGTAAGTATAATATTCCTTCCTGTCATTAATTTCTTACAAAAGGAGCTTGGGAGAAAATCTTATACATAATGTT

>M6-PGSC0003DMG400008508-(rev)M6G-LecRK_248

ATAAACCACTTGATTATGGTGCTGCTGGGATGATAATTAATCAAGGAAACCTGGTGCTTCTTGATAGATTTCAAGGCGAAGTTTGGTCGGCACTTGCTGGAAACATCAATCCAGATATTT

>M6-PGSC0003DMG400008508-(rev)M6G-LecRK_2480

CTTCCTGTCATTAATTTCTTACAAAAGGAGCTTGGGAGAAAATCTTATACATAATGTTTTGTTGACCTGGTGAAATCTAGTGGCTATATGTCTCCTGAATATGCACTAGATGGATTGTTT

>M6-PGSC0003DMG400008508-(rev)M6G-LecRK_2542

TGACCTGGTGAAATCTAGTGGCTATATGTCTCCTGAATATGCACTAGATGGATTGTTTTCCATTAAATCAGATGTATTCAGTTTTGGAGTAGTCGTACTTGAGATAATCAGTGGAAGAAG

>M6-PGSC0003DMG400008508-(rev)M6G-LecRK_2604

TTAAATCAGATGTATTCAGTTTTGGAGTAGTCGTACTTGAGATAATCAGTGGAAGAAGGAATACAGGATTTTATCAGTCAGAAGAAGCCTTAAACTTGTTGGGCTATGTAAGTGAAGAAA

>M6-PGSC0003DMG400008508-(rev)M6G-LecRK_2666

ACAGGATTTTATCAGTCAGAAGAAGCCTTAAACTTGTTGGGCTATGTAAGTGAAGAAACTTCCCATGACTTAATTATTATATCTTCTTCTTAGCTTTGTCTCTCTGATATAAAAATTATC

>M6-PGSC0003DMG400008508-(rev)M6G-LecRK_2728

CCATGACTTAATTATTATATCTTCTTCTTAGCTTTGTCTCTCTGATATAAAAATTATCACAGGCATGGAAATTGTGGACAGAAAAAGCTGAGACACAATTAATAGAGAAGTCATTACTTG

>M6-PGSC0003DMG400008508-(rev)M6G-LecRK_2790

GCATGGAAATTGTGGACAGAAAAAGCTGAGACACAATTAATAGAGAAGTCATTACTTGAATCATGCAACAGAAGTGAAGCAATAAAGTGCATAAATATTGCGCTCCTGTGTGTACAAGAA

>M6-PGSC0003DMG400008508-(rev)M6G-LecRK_2852

ATGCAACAGAAGTGAAGCAATAAAGTGCATAAATATTGCGCTCCTGTGTGTACAAGAAGATCCAAATCATCGTCCTAATATGTCAGATGTCATTTTGATGCTGGGAGGGGAAGGTACTAA

>M6-PGSC0003DMG400008508-(rev)M6G-LecRK_2976

CCAACACCTAATAGACCAGCTTTTGTAATAAGGACGCATGCTTCTAGTACATCTTCTTCTTCCTCCGATAAGAAATACATCGTATCCAACAATCAGGTGACGATTACAGTTGAAGAAGGA

>M6-PGSC0003DMG400008508-(rev)M6G-LecRK_3038

CTCCGATAAGAAATACATCGTATCCAACAATCAGGTGACGATTACAGTTGAAGAAGGACGTTAGCAGGAATAGTGAAGGCATTTAATTCGTATGGAAAGGGCTAGGGTGGCACTAGCTAG

>M6-PGSC0003DMG400008508-(rev)M6G-LecRK_310

GATAGATTTCAAGGCGAAGTTTGGTCGGCACTTGCTGGAAACATCAATCCAGATATTTCAGTTACTGCACTTCTTCGTGATGATGGGAATTTGATTTTGAGTGTTGTGTCAAATTCATCA

>M6-PGSC0003DMG400008508-(rev)M6G-LecRK_3100

AGCAGGAATAGTGAAGGCATTTAATTCGTATGGAAAGGGCTAGGGTGGCACTAGCTAGGACTTATAGATGAGCCATATTATACTTTTTTAACCTCTACTTTGGTCCTTAAAAAGTCCAGG

>M6-PGSC0003DMG400008508-(rev)M6G-LecRK_3162

TATAGATGAGCCATATTATACTTTTTTAACCTCTACTTTGGTCCTTAAAAAGTCCAGGGATACTTGACCAAATGTCGTCTTCTCTATATATAATAGTTAGGACAAAAGTGGAAATGTTGG

>M6-PGSC0003DMG400008508-(rev)M6G-LecRK_3224

CTTGACCAAATGTCGTCTTCTCTATATATAATAGTTAGGACAAAAGTGGAAATGTTGGGAAAACTTTCAAGTGTTTGAACTCAAACATATACTATATTTTTGGTTTAGAATATAGTAGGC

>M6-PGSC0003DMG400008508-(rev)M6G-LecRK_3286

ACTTTCAAGTGTTTGAACTCAAACATATACTATATTTTTGGTTTAGAATATAGTAGGCCTTAGCAATCAAGTACAAGAATAATTCGATTATTCCTGATTCCTTCAAGCTTCAAATATATA

>M6-PGSC0003DMG400008508-(rev)M6G-LecRK_372

TACTGCACTTCTTCGTGATGATGGGAATTTGATTTTGAGTGTTGTGTCAAATTCATCAACACCCTTACTACTTTGGCAAAGTTTTGATCACCCAACACACACTTTTATGCCTGGTGCTAA

>M6-PGSC0003DMG400008508-(rev)M6G-LecRK_434

CCTTACTACTTTGGCAAAGTTTTGATCACCCAACACACACTTTTATGCCTGGTGCTAAGCTTGGATATGACAAACGTACACAAAGAAAACAGGTTCTGGTTTCATGGAAGAATTCGAACG

>M6-PGSC0003DMG400008508-(rev)M6G-LecRK_496

GGATATGACAAACGTACACAAAGAAAACAGGTTCTGGTTTCATGGAAGAATTCGAACGATCCAGCACCAGGATTGTATTCTATGGAAATGGACCCAAAGCATGCCCAGTTTGTTTTAAAA

>M6-PGSC0003DMG400008508-(rev)M6G-LecRK_558

AGCACCAGGATTGTATTCTATGGAAATGGACCCAAAGCATGCCCAGTTTGTTTTAAAATGGAATAGGACTACAGAGTATTGGGCAAGTGGTTCATGGGATGGCCAAAGATTCAGCTCAGT

>M6-PGSC0003DMG400008508-(rev)M6G-LecRK_62

ATCTCTGTATCGGAAGGGACACCATTTCTGCGACTGAATCTCTTTCTTTTGGTGAAACTCTAGTCTCTTCAGGTGAGATATTTGAGCTTGGTTTTTTCAAACCAGGTAACTCTCTTAAAT

>M6-PGSC0003DMG400008508-(rev)M6G-LecRK_620

ATAGGACTACAGAGTATTGGGCAAGTGGTTCATGGGATGGCCAAAGATTCAGCTCAGTGCCTGAGATGAGGTTAAACTACATATATAATTACAGCTATATCGACAATGAGAATGAGTCAT

>M6-PGSC0003DMG400008508-(rev)M6G-LecRK_682

GAGATGAGGTTAAACTACATATATAATTACAGCTATATCGACAATGAGAATGAGTCATATTTTACATATTCCCTTTACAATAGTTCAATCTTATCAATATTCATTATGGATGTCTCGGGA

>M6-PGSC0003DMG400008508-(rev)M6G-LecRK_744

TACATATTCCCTTTACAATAGTTCAATCTTATCAATATTCATTATGGATGTCTCGGGACAAATTAAGCAACTAACATGGTTGGATGATAGCAGTGTTTGGAATCTATTCTGGTCTCAACC

>M6-PGSC0003DMG400008508-(rev)M6G-LecRK_806

TTAAGCAACTAACATGGTTGGATGATAGCAGTGTTTGGAATCTATTCTGGTCTCAACCAAGACCAGATTGTGAGGTTTATGCTAAATGTGGGGCATTTGGAGTCTGTGATGAAGCTAATG

>M6-PGSC0003DMG400008508-(rev)M6G-LecRK_868

CCAGATTGTGAGGTTTATGCTAAATGTGGGGCATTTGGAGTCTGTGATGAAGCTAATGCAACCTGCAATTGTTTGAGTGGTTTCAAGCAAAGATCAGATACAGAATGGAATTCAGATGAT

>M6-PGSC0003DMG400008508-(rev)M6G-LecRK_930

CTGCAATTGTTTGAGTGGTTTCAAGCAAAGATCAGATACAGAATGGAATTCAGATGATTATTCTGGTGGCTGTGTAAGAGACCAAAAGGTGCAGTGTAATGCAATTACTGAAGACACAGA

>M6-PGSC0003DMG400008508-(rev)M6G-LecRK_992

CTGGTGGCTGTGTAAGAGACCAAAAGGTGCAGTGTAATGCAATTACTGAAGACACAGATAGTTTATGGATAACTTCAATTGTGAGAGTACCTTCTTCTCAAAATACTAATATCACAGTTG

>M6-PGSC0003DMG400008523M6G-LecRK_0

AAACAAGAAACTTGTTTTGACTAGTTTCATGGACTCCAAAGAAGAAAAATCTCCTTCCGTAAACACTTGCACTCCTTATTTTCTCTCATTCTTTCAGTCATCAATTTCTCTATGTACGTA

>M6-PGSC0003DMG400008523M6G-LecRK_1830

AAAGTATTAATGAATGAAAACAGTGATGAAGCGATTGATGTTCCATACTTTCACTTGGAGACTATTTTGGAAGCTACAGACAACTTCTCAAATGCGAATAAGCTCGGACAAGGAGGATTT

>M6-PGSC0003DMG400008523M6G-LecRK_2013

AATGAGATGAATTTTTTTTGCAGGGTATCTTTCCGGGTGAAAAAGAAATTGCGGTAAAAACGTTATCTAGTCAATCAGGACAAGGCATAGATGAATTTAAGAATGAAGTGACTCTAATTG

>M6-PGSC0003DMG400008523M6G-LecRK_2501

ACCCGAAAATTTCAGATTTTGGCTTGGCAAGGATTGTTGAAGGAAAAGTAACAGAAGCAAATACAAAGAAAATAGTGGGGACCTAGTAAGTATAATATTCCTTCCTGTCATTAATTTCTT

>M6-PGSC0003DMG400008523M6G-LecRK_3050

ATATGTCAGATGTCATTTTGATGCTGGGAGGGGAAGGTACTAACCTTCCAACACCTAATAGACCAGCTTTTGTAATAAGGACGCATGCTTCTAGTACATCTTCTTCTTCCTCCGATAAGA

>M6-PGSC0003DMG400008523M6G-LecRK_61

AACACTTGCACTCCTTATTTTCTCTCATTCTTTCAGTCATCAATTTCTCTATGTACGTAGCATGGAAATGAAGAACAATCATTACTTTCTGCTACTCTTTTTATGCTTCAGTCTCAACAG

>M6-PGSC0003DMG400008524M6G-LecRK_0

GTGAAGTTTCTCCACTTGCCATAATCTTGTCTCCAACCCCATTAAGAGAAAAAAAAAATAAAATTAAACAACAGAGCAAGAAAAGCTGGAGCTCAAGAAGGTGAATGATACTCAAAGAAG

>M6-PGSC0003DMG400008524M6G-LecRK_1037

TTTAGCTAGTTTTCTCAAAATGCATGAAAAGGAAGAGATGCTCTATTTCATCTTTTAAGTTTAATTCTTGAGGCTATTTCTTGAATTCATTTTTTTTTTTAAAAAATAAAATAAAATTGA

>M6-PGSC0003DMG400008524M6G-LecRK_1098

TAATTCTTGAGGCTATTTCTTGAATTCATTTTTTTTTTTAAAAAATAAAATAAAATTGAGGGCCTAAGAAAGGGATTACAATGTGGGGAATCGAATACTCACACACGTAGACAACTAACT

>M6-PGSC0003DMG400008524M6G-LecRK_1159

GCCTAAGAAAGGGATTACAATGTGGGGAATCGAATACTCACACACGTAGACAACTAACTGAGTTATTGAGATTCTCCTTTATCTTGATTTCACATGATTTGCTTATATTTTGTTGTGATC

>M6-PGSC0003DMG400008524M6G-LecRK_122

AAATACAGGAAGAAAAATTGTTTGGATTTTCTAAAATCCAATTTAATTTAAGGTAAATTAGTTTTAGGTACTTTTAAACTTTATTTTGGAGTAAATATAATGTGTATGTTATTCATTTTC

>M6-PGSC0003DMG400008524M6G-LecRK_1220

GTTATTGAGATTCTCCTTTATCTTGATTTCACATGATTTGCTTATATTTTGTTGTGATCCAAGTTGTCTATCTATTTTAGCTTTAGGTTATCAGTTCTGTTCCTATTTCAGGCTAATTGC

>M6-PGSC0003DMG400008524M6G-LecRK_1281

AGTTGTCTATCTATTTTAGCTTTAGGTTATCAGTTCTGTTCCTATTTCAGGCTAATTGCAAATGATTTGAGGCTTTGAAGGTTTAAGAGGATGTTGGACTTAAACAAAATAGAACCTGAA

>M6-PGSC0003DMG400008524M6G-LecRK_1342

ATGATTTGAGGCTTTGAAGGTTTAAGAGGATGTTGGACTTAAACAAAATAGAACCTGAAGAAGGGAAGCTAGACTTGGATCTTCGTCGCGTTAACTATGATCTTGATGACAAATCTCATC

>M6-PGSC0003DMG400008524M6G-LecRK_1403

AGGGAAGCTAGACTTGGATCTTCGTCGCGTTAACTATGATCTTGATGACAAATCTCATCTTAGATATGGTGATTGGGTCACTTTCAAGAGCCATTCATCTAAAAAAGTTAGAATCTCTCG

>M6-PGSC0003DMG400008524M6G-LecRK_1464

AGATATGGTGATTGGGTCACTTTCAAGAGCCATTCATCTAAAAAAGTTAGAATCTCTCGACCTGAATCTCCGAGAGGTGAAAATTCAGATAGACAACCTCAGGATGCAGATTATCCCTCA

>M6-PGSC0003DMG400008524M6G-LecRK_1525

CTGAATCTCCGAGAGGTGAAAATTCAGATAGACAACCTCAGGATGCAGATTATCCCTCACTTAGCTATGAACTCGAAAGTGAGATTCTAGCCAGGTTTCCTAGGTGGGAGTATTGGAAAC

>M6-PGSC0003DMG400008524M6G-LecRK_1586

TAGCTATGAACTCGAAAGTGAGATTCTAGCCAGGTTTCCTAGGTGGGAGTATTGGAAACTTAGCCTCGTAAACAAGAGATGTTCAATGCTTCTAAAGAGCGGTGAGATATTTGAGATTCG

>M6-PGSC0003DMG400008524M6G-LecRK_1647

AGCCTCGTAAACAAGAGATGTTCAATGCTTCTAAAGAGCGGTGAGATATTTGAGATTCGTAAAGAGAATGGATTTAAAGAACCTTCGGTATATATGTTAGCAAGTGGGGAGACTAACTGG

>M6-PGSC0003DMG400008524M6G-LecRK_1708

AAGAGAATGGATTTAAAGAACCTTCGGTATATATGTTAGCAAGTGGGGAGACTAACTGGTGGACATTTGATCGAGAGTTTAAGTCCCGGAGGAAAATCCCTGACTTACCATCAGATGTGT

>M6-PGSC0003DMG400008524M6G-LecRK_1769

GACATTTGATCGAGAGTTTAAGTCCCGGAGGAAAATCCCTGACTTACCATCAGATGTGTGCTTCACATTTGGCGATAAGGAATCGCTTTGTGCAGGCACACATTTACTTGTTTCGGGTAG

>M6-PGSC0003DMG400008524M6G-LecRK_183

TTTTAGGTACTTTTAAACTTTATTTTGGAGTAAATATAATGTGTATGTTATTCATTTTCTTTTTTATGGCTTTTATGGTTCTTGAGTGATTGATTTTTGGATTTTCATCTTGGTTGTTGC

>M6-PGSC0003DMG400008524M6G-LecRK_1830

TTCACATTTGGCGATAAGGAATCGCTTTGTGCAGGCACACATTTACTTGTTTCGGGTAGGGAAATCGATGGCCTAGTTATATGGAGGTTCGAATTAGCAACGAATTGTTGGTACAAAGGT

>M6-PGSC0003DMG400008524M6G-LecRK_1891

AAATCGATGGCCTAGTTATATGGAGGTTCGAATTAGCAACGAATTGTTGGTACAAAGGTCCCTCTATGGTTAATCCAAGATGTTTATTTGCATCAGCTACTTGTGGCACCTCTGTTTTTG

>M6-PGSC0003DMG400008524M6G-LecRK_1952

CTCTATGGTTAATCCAAGATGTTTATTTGCATCAGCTACTTGTGGCACCTCTGTTTTTGTTGCTGGTGGTGTTGGTATTATGGCGAATAGTGAGGTATACGACACAGCTGAGAAATACAA

>M6-PGSC0003DMG400008524M6G-LecRK_2013

GCTGGTGGTGTTGGTATTATGGCGAATAGTGAGGTATACGACACAGCTGAGAAATACAACCCCGATAGCAGATCGTGGGACCTACTACCGAGGATGAAGAGGAAGAGGAAGCTTTGTTCA

>M6-PGSC0003DMG400008524M6G-LecRK_2074

CCGATAGCAGATCGTGGGACCTACTACCGAGGATGAAGAGGAAGAGGAAGCTTTGTTCAGGATGTTACATGGACAAGAAATTCTACGTGATCGGGGGGAGGAATGAGAACGGAGAGCTTA

>M6-PGSC0003DMG400008524M6G-LecRK_2135

ATGTTACATGGACAAGAAATTCTACGTGATCGGGGGGAGGAATGAGAACGGAGAGCTTACCTGTGGAGAATTCTTCGATGAGGGTATGAACAAGTGGGAGCTGATTCCGGACATGTTGAA

>M6-PGSC0003DMG400008524M6G-LecRK_2196

TGTGGAGAATTCTTCGATGAGGGTATGAACAAGTGGGAGCTGATTCCGGACATGTTGAAGGATGATCCAGTCCTAACATGCCACTCACCACCCCTCATTGCTGTCGTGAACAACGAGCTG

>M6-PGSC0003DMG400008524M6G-LecRK_2257

ATGATCCAGTCCTAACATGCCACTCACCACCCCTCATTGCTGTCGTGAACAACGAGCTGTACTCACTTGAGGCATCTTCAAACCAGCTGAAGCTTTACCTGAAAAAAACCAACACTTGGA

>M6-PGSC0003DMG400008524M6G-LecRK_2318

CTCACTTGAGGCATCTTCAAACCAGCTGAAGCTTTACCTGAAAAAAACCAACACTTGGAAAATGTTAGGACAAGTGCCAGTGAGAGCTGATTCAAACAGAGGATGGGGCATTGCTTTCAA

>M6-PGSC0003DMG400008524M6G-LecRK_2379

ATGTTAGGACAAGTGCCAGTGAGAGCTGATTCAAACAGAGGATGGGGCATTGCTTTCAAGTCATTGGGAAATGAACTTCTTGTAATCGGAGCTGCTTCGTCTTCAGCCTCTTATTCGGGT

>M6-PGSC0003DMG400008524M6G-LecRK_244

TTTTATGGCTTTTATGGTTCTTGAGTGATTGATTTTTGGATTTTCATCTTGGTTGTTGCATGTTATTTGCTTTTTGCGTGTGTGTGTGTGGATTTCATGGTTCTTGAGTTTATTCTGATT

>M6-PGSC0003DMG400008524M6G-LecRK_2440

CATTGGGAAATGAACTTCTTGTAATCGGAGCTGCTTCGTCTTCAGCCTCTTATTCGGGTAATTCCATGGCTATATATACTTGCTGTCCAGATCCTGATGTAATTGATTTGCAGTGGAAAC

>M6-PGSC0003DMG400008524M6G-LecRK_2501

TTCCATGGCTATATATACTTGCTGTCCAGATCCTGATGTAATTGATTTGCAGTGGAAACCTCTAGACAGTGGTCGAAATCGGCTTAGTAGTTTCATCTTAAACTGTTCTGTTATGGTAGC

>M6-PGSC0003DMG400008524M6G-LecRK_2562

CTAGACAGTGGTCGAAATCGGCTTAGTAGTTTCATCTTAAACTGTTCTGTTATGGTAGCTTGAATAATGTCATTTATCTAGCAAACAAATTATGTATTGTGATCAAAATATATAAGTGGC

>M6-PGSC0003DMG400008524M6G-LecRK_2623

GAATAATGTCATTTATCTAGCAAACAAATTATGTATTGTGATCAAAATATATAAGTGGCATATGGCCACTGGTTTCTGATATTTTCTTGCATATATTCACTACTGTTTGCTCTGCTATGT

>M6-PGSC0003DMG400008524M6G-LecRK_2682

ATATGGCCACTGGTTTCTGATATTTTCTTGCATATATTCACTACTGTTTGCTCTGCTATGTTTGATGTAATTTGCATTTATTGTTGCATTATTATGATTTATATTGAAGGTGTTATTCTA

>M6-PGSC0003DMG400008524M6G-LecRK_305

GTTATTTGCTTTTTGCGTGTGTGTGTGTGGATTTCATGGTTCTTGAGTTTATTCTGATTCAATAATGAATTCTTGAAAGTTAGTATAGTTTTTCATATTGAATTTACATGTGGAAACTAC

>M6-PGSC0003DMG400008524M6G-LecRK_366

ATAATGAATTCTTGAAAGTTAGTATAGTTTTTCATATTGAATTTACATGTGGAAACTACTTTTGATTTTGACTTTTATGTTATAAGCATTCTAGGTGGAAAAGTTATTTAAAAATTGTTA

>M6-PGSC0003DMG400008524M6G-LecRK_427

TTGATTTTGACTTTTATGTTATAAGCATTCTAGGTGGAAAAGTTATTTAAAAATTGTTATATATAACTTAAATCCTTTCGACTTTTGTCATTTCAGGCATTTATGTTTGTTATTGGGTAT

>M6-PGSC0003DMG400008524M6G-LecRK_488

TATAACTTAAATCCTTTCGACTTTTGTCATTTCAGGCATTTATGTTTGTTATTGGGTATATGTGTTATCAATTACTACAATTACCTAAAAAGAAAAATTGTGTCTTATTACTCTCAACAT

>M6-PGSC0003DMG400008524M6G-LecRK_549

GTGTTATCAATTACTACAATTACCTAAAAAGAAAAATTGTGTCTTATTACTCTCAACATAATTTCGTTGTCTAGCAAGATACGTCTAGTTCTCCCTCTACTTACCTATCCATTAGGAAAG

>M6-PGSC0003DMG400008524M6G-LecRK_61

AATTAAACAACAGAGCAAGAAAAGCTGGAGCTCAAGAAGGTGAATGATACTCAAAGAAGTGAAATACAGGAAGAAAAATTGTTTGGATTTTCTAAAATCCAATTTAATTTAAGGTAAATT

>M6-PGSC0003DMG400008524M6G-LecRK_610

TTTCGTTGTCTAGCAAGATACGTCTAGTTCTCCCTCTACTTACCTATCCATTAGGAAAGTGGAACTTTCTTGACCTTCGAAAAGGTATAAAATGGTTTTATAAATATGGTATGTTATATG

>M6-PGSC0003DMG400008524M6G-LecRK_671

GAACTTTCTTGACCTTCGAAAAGGTATAAAATGGTTTTATAAATATGGTATGTTATATGGACTAATTTGTTAGGTGCTACTTGCTTGCTTCAAGCACAATACTTTGATTTCTTTTTAAAG

>M6-PGSC0003DMG400008524M6G-LecRK_732

CTAATTTGTTAGGTGCTACTTGCTTGCTTCAAGCACAATACTTTGATTTCTTTTTAAAGTTTTAGACAAAGGCGGAGCCAAGATTTAAAATTTATGGGTTTTAGATTCTAATCTTTTATG

>M6-PGSC0003DMG400008524M6G-LecRK_793

TTAGACAAAGGCGGAGCCAAGATTTAAAATTTATGGGTTTTAGATTCTAATCTTTTATGTTACTGGGTTCTAAATTGATAAAATGTATAAATTCAAAAGATTTTAAAGACAATTACAAGA

>M6-PGSC0003DMG400008524M6G-LecRK_854

ACTGGGTTCTAAATTGATAAAATGTATAAATTCAAAAGATTTTAAAGACAATTACAAGATTTAAACTAAAACTACTTTATTCAGACTTAACCCTTAGCTGATATTCTAGCTCCGCCCCTA

>M6-PGSC0003DMG400008524M6G-LecRK_915

TAAACTAAAACTACTTTATTCAGACTTAACCCTTAGCTGATATTCTAGCTCCGCCCCTAGACTTTTTTATTAGCATCCTCAAATACTAAAGTGGTGTTATTGCCCCTTCTTTTGTCACAC

>M6-PGSC0003DMG400008524M6G-LecRK_976

CTTTTTTATTAGCATCCTCAAATACTAAAGTGGTGTTATTGCCCCTTCTTTTGTCACACTTTTTAGCTAGTTTTCTCAAAATGCATGAAAAGGAAGAGATGCTCTATTTCATCTTTTAAG

>M6-PGSC0003DMG400008581-(rev)M6LRR-RLK_0

ATGAATTTATTAACCATTTTCATACTCTTCTTGTTCTACTATTACAGTGTTTTTGCTAACGGAGTGGTTGCGAAAGTTGATTTGAATGATGAGGTGTCTATTCTACTTTCTATTAAAGAA

>M6-PGSC0003DMG400008581-(rev)M6LRR-RLK_1054

TGTGGAATAACAGCTTTTCAGGTCCTTTGCCAAGTGATCTTGGAAGAAATTCGCCATTGCAATGGGTGGATATTTCATCCAACTCATTTACCGGTCCAATTCCAGCAGGATTATGCGCCA

>M6-PGSC0003DMG400008581-(rev)M6LRR-RLK_1116

TGGGTGGATATTTCATCCAACTCATTTACCGGTCCAATTCCAGCAGGATTATGCGCCAAAGGTGACCTCACTAAGCTTATCATGTTTAACAATGCTTTCTCAGGTCCAATTCCAACTGGT

>M6-PGSC0003DMG400008581-(rev)M6LRR-RLK_1178

TGACCTCACTAAGCTTATCATGTTTAACAATGCTTTCTCAGGTCCAATTCCAACTGGTTTATCAACCTGCACGCCCCTTGTTCGTGTAAGAATGCAGAACAATCTTCTTTCGGGGACGAT

>M6-PGSC0003DMG400008581-(rev)M6LRR-RLK_124

TCGTTGATCCATTGGATCATCTTCATGATTGGACAGTGCCCAATAATGCAGCTGCAGGGAACAATAGAAGTATTATTGTTCCCTGCAGCTGGACGGGAGTTGAATGCAACTCCCATGGGG

>M6-PGSC0003DMG400008581-(rev)M6LRR-RLK_1240

CAACCTGCACGCCCCTTGTTCGTGTAAGAATGCAGAACAATCTTCTTTCGGGGACGATCCCAGCAGGTTTTGGTAAACTTGGGAAACTACAGAGGCTGGAACTGGCAAACAATAGTCTGA

>M6-PGSC0003DMG400008581-(rev)M6LRR-RLK_1302

GCAGGTTTTGGTAAACTTGGGAAACTACAGAGGCTGGAACTGGCAAACAATAGTCTGACAGGACAAATCCCAAGTGATCTTGCTTCTTCTACCTCTCTTTCTTTTATTGATTTCTCTAGA

>M6-PGSC0003DMG400008581-(rev)M6LRR-RLK_1488

TCCGACAATAAGATGACAGGCGAAATCCCGGATCAATTCCAGGATTGTCCTTCTCTTACTGTTCTGGATCTATCCACAAACCATTTCACTGGAGATCTTCCAGCTAGCATTGCTTCCTGT

>M6-PGSC0003DMG400008581-(rev)M6LRR-RLK_1550

TCTGGATCTATCCACAAACCATTTCACTGGAGATCTTCCAGCTAGCATTGCTTCCTGTGAGAAGTTAGTAACTTTGAACCTTCGAAATAATCAATTAAACGGTCCAATTCCAAGAGCGAT

>M6-PGSC0003DMG400008581-(rev)M6LRR-RLK_1612

AGTTAGTAACTTTGAACCTTCGAAATAATCAATTAAACGGTCCAATTCCAAGAGCGATTTCCATGATGCCAACATTAGCTATTCTAGATCTATCCAATAATTCTTTAACTGGTGGGATAC

>M6-PGSC0003DMG400008581-(rev)M6LRR-RLK_1674

ATGATGCCAACATTAGCTATTCTAGATCTATCCAATAATTCTTTAACTGGTGGGATACCTGAGAACTTTGGCAATTCACCAGCACTAGAAATGCTGAATGTTTCTCATAACAAACTGGAG

>M6-PGSC0003DMG400008581-(rev)M6LRR-RLK_1736

GAACTTTGGCAATTCACCAGCACTAGAAATGCTGAATGTTTCTCATAACAAACTGGAGGGTCCTGTACCGGAAAATGGCATGTTGAGAACAATCAATCCGGATGACCTGATTGGAAATGC

>M6-PGSC0003DMG400008581-(rev)M6LRR-RLK_1798

CTGTACCGGAAAATGGCATGTTGAGAACAATCAATCCGGATGACCTGATTGGAAATGCCGGTCTATGTGGTGGAGTGCTTCCTCCTTGTTCTCATAATGCAGCATACACATCAAAGCAGA

>M6-PGSC0003DMG400008581-(rev)M6LRR-RLK_186

AATAGAAGTATTATTGTTCCCTGCAGCTGGACGGGAGTTGAATGCAACTCCCATGGGGCAGTCGAGAAGCTTGATCTGTCCCACATGAATCTCACTGGCACAGTGTCTAATGATATCCAA

>M6-PGSC0003DMG400008581-(rev)M6LRR-RLK_1860

CTATGTGGTGGAGTGCTTCCTCCTTGTTCTCATAATGCAGCATACACATCAAAGCAGAAGAGCTTGCACGTGAAGCACATCATTACAGGATGGCTAATTGGTGTGGCAGCTGTTCTGCTT

>M6-PGSC0003DMG400008581-(rev)M6LRR-RLK_1922

CTTGCACGTGAAGCACATCATTACAGGATGGCTAATTGGTGTGGCAGCTGTTCTGCTTTTTGTGACAGCAGGTCTTGTAGCTAGATCTCTATACAAGAGATGGCATGAAAATGGGAGCTG

>M6-PGSC0003DMG400008581-(rev)M6LRR-RLK_1984

TGACAGCAGGTCTTGTAGCTAGATCTCTATACAAGAGATGGCATGAAAATGGGAGCTGCTTTGAACCAAGCTTTGAGATGAGCCGTGGTGAATGGCCATGGAGGTTGATGGCATTCCAGA

>M6-PGSC0003DMG400008581-(rev)M6LRR-RLK_2046

GAACCAAGCTTTGAGATGAGCCGTGGTGAATGGCCATGGAGGTTGATGGCATTCCAGAGGCTAGGATTCACTAGTAATGACATCTTGGCTTGCCTTAAAGAGTCAAATGTCATTGGAATG

>M6-PGSC0003DMG400008581-(rev)M6LRR-RLK_2108

AGGATTCACTAGTAATGACATCTTGGCTTGCCTTAAAGAGTCAAATGTCATTGGAATGGGAGCAACAGGGGTTGTATACAAAGCTGAAATGCAACGCGAGAATATGGTTGTTGCAGTAAA

>M6-PGSC0003DMG400008581-(rev)M6LRR-RLK_2170

CAACAGGGGTTGTATACAAAGCTGAAATGCAACGCGAGAATATGGTTGTTGCAGTAAAGAAGCTGTGGAAATCTGGAACTGATATTGAGATGGGGCACAGTGATGATCTCGTAGGTGAGG

>M6-PGSC0003DMG400008581-(rev)M6LRR-RLK_2232

CTGTGGAAATCTGGAACTGATATTGAGATGGGGCACAGTGATGATCTCGTAGGTGAGGTGAAAGTTCTTGGAAAATTGAGGCATCGAAATATTGTCAGGCTTCTGGGGTTCCTTCACAAC

>M6-PGSC0003DMG400008581-(rev)M6LRR-RLK_2294

AGTTCTTGGAAAATTGAGGCATCGAAATATTGTCAGGCTTCTGGGGTTCCTTCACAACAAACGTGATGCAATGATAATATACGAGTATATGCAAAATGGTAGCCTTGGAGAGGTTTTACA

>M6-PGSC0003DMG400008581-(rev)M6LRR-RLK_2356

GTGATGCAATGATAATATACGAGTATATGCAAAATGGTAGCCTTGGAGAGGTTTTACATGGTAAACAAGCAGCAGGGAGACTGCTTGTAGATTGGGTCACTCGATATAACATAGCCCTTG

>M6-PGSC0003DMG400008581-(rev)M6LRR-RLK_2418

AAACAAGCAGCAGGGAGACTGCTTGTAGATTGGGTCACTCGATATAACATAGCCCTTGGAGTGGCACAAGGCCTTGCTTATCTCCATCATTACTGCCATCCACCAGTTATTCACCGGGAT

>M6-PGSC0003DMG400008581-(rev)M6LRR-RLK_248

CGAGAAGCTTGATCTGTCCCACATGAATCTCACTGGCACAGTGTCTAATGATATCCAAAAGTTGAAGAGTTTGACTTCTCTAAATCTATGCTGCAATGAGTTTTCATCGCCATTGCCGAA

>M6-PGSC0003DMG400008581-(rev)M6LRR-RLK_2480

GGCACAAGGCCTTGCTTATCTCCATCATTACTGCCATCCACCAGTTATTCACCGGGATGTCAAGTCAAATAACATACTTCTTGATGCAAATCTAGAGGCAAGAATTGCAGATTTTGGTTT

>M6-PGSC0003DMG400008581-(rev)M6LRR-RLK_2542

AGTCAAATAACATACTTCTTGATGCAAATCTAGAGGCAAGAATTGCAGATTTTGGTTTGGCAAGAACGATGCTCAAGAAGAATGAAACAGTTTCCATGGTTGCTGGATCTTATGGATACA

>M6-PGSC0003DMG400008581-(rev)M6LRR-RLK_2604

AGAACGATGCTCAAGAAGAATGAAACAGTTTCCATGGTTGCTGGATCTTATGGATACATAGCCCCCGGTAAGTCCTTCAAATTTTTCATCAATGTTTACTCTAGAGTGATTTTTGTTACT

>M6-PGSC0003DMG400008581-(rev)M6LRR-RLK_2666

CCCCGGTAAGTCCTTCAAATTTTTCATCAATGTTTACTCTAGAGTGATTTTTGTTACTAATATTTAACAATCATTTATCTGGATTTTGACTTTGCAGAGTACGGATACACGTTGAAAGTG

>M6-PGSC0003DMG400008581-(rev)M6LRR-RLK_2728

TTTAACAATCATTTATCTGGATTTTGACTTTGCAGAGTACGGATACACGTTGAAAGTGGATGAGAAGAGTGATATTTACAGCTATGGAGTTGTTCTCATGGAGCTCCTAACGGGTAAACG

>M6-PGSC0003DMG400008581-(rev)M6LRR-RLK_2790

AGAAGAGTGATATTTACAGCTATGGAGTTGTTCTCATGGAGCTCCTAACGGGTAAACGACCTCTAGATCCTGAATTTGGGGAATCTGTTGACATTGTAGAGTGGTTTCGGATGAAGATCC

>M6-PGSC0003DMG400008581-(rev)M6LRR-RLK_2852

CTAGATCCTGAATTTGGGGAATCTGTTGACATTGTAGAGTGGTTTCGGATGAAGATCCGAGACAACAAGTCTTTAGAAGAAGCACTTGACCCAAATGTAGGAGCAACACAACATGTTCAA

>M6-PGSC0003DMG400008581-(rev)M6LRR-RLK_2914

CAACAAGTCTTTAGAAGAAGCACTTGACCCAAATGTAGGAGCAACACAACATGTTCAAGAAGAAATGTTGTTAGTTTTAAGGATAGCAATTCTATGTATAGCCAAACTCCCCAAGGACAG

>M6-PGSC0003DMG400008581-(rev)M6LRR-RLK_2976

AAATGTTGTTAGTTTTAAGGATAGCAATTCTATGTATAGCCAAACTCCCCAAGGACAGACCATCCATGAGGGATGTTTTGACGATGCTCGAAGAGGCAAAGCCTCGGAGAAAAAGCAATA

>M6-PGSC0003DMG400008581-(rev)M6LRR-RLK_3038

TCCATGAGGGATGTTTTGACGATGCTCGAAGAGGCAAAGCCTCGGAGAAAAAGCAATAGCAACAGTGGAGGTAGCCATGATGCTACTACTAACAAAGATAAGCCAATATTCAGTACATCA

>M6-PGSC0003DMG400008581-(rev)M6LRR-RLK_310

TGAAGAGTTTGACTTCTCTAAATCTATGCTGCAATGAGTTTTCATCGCCATTGCCGAAAAGTTTATCAAATCTAACTGCATTGAGAAGCATTGATGTGAGCCAGAACTATTTTGTCTATG

>M6-PGSC0003DMG400008581-(rev)M6LRR-RLK_372

TTATCAAATCTAACTGCATTGAGAAGCATTGATGTGAGCCAGAACTATTTTGTCTATGACTTCCCTGTAGGTCTTGGAATGTCAGAAGCACTTATGTATTTGAATGCTTCGAGTAATAAC

>M6-PGSC0003DMG400008581-(rev)M6LRR-RLK_434

CCCTGTAGGTCTTGGAATGTCAGAAGCACTTATGTATTTGAATGCTTCGAGTAATAACTTCTCCGGTTATCTTCCTGAGGATATTGGAAATGCTACATTGCTCGAGACATTGGATTTTCG

>M6-PGSC0003DMG400008581-(rev)M6LRR-RLK_496

CCGGTTATCTTCCTGAGGATATTGGAAATGCTACATTGCTCGAGACATTGGATTTTCGAGGGAATTTCTTCGAAGGTTCAATTCCTAAATCTTATAGGAATTTGGGGAAATTGAAGTTTT

>M6-PGSC0003DMG400008581-(rev)M6LRR-RLK_558

AATTTCTTCGAAGGTTCAATTCCTAAATCTTATAGGAATTTGGGGAAATTGAAGTTTTTAGGTCTTTCAGGGAACAATTTGACTGGACACATTCCAGGAGAGTTGGGGCAGCTTTCGTCG

>M6-PGSC0003DMG400008581-(rev)M6LRR-RLK_62

AGTGGTTGCGAAAGTTGATTTGAATGATGAGGTGTCTATTCTACTTTCTATTAAAGAAAGTCTCGTTGATCCATTGGATCATCTTCATGATTGGACAGTGCCCAATAATGCAGCTGCAGG

>M6-PGSC0003DMG400008581-(rev)M6LRR-RLK_620

TCTTTCAGGGAACAATTTGACTGGACACATTCCAGGAGAGTTGGGGCAGCTTTCGTCGCTTGAGACTGTAGTACTGGGATACAATGTTTTCGAAGGTGGAATTCCAGCTGAATTTGGAAA

>M6-PGSC0003DMG400008581-(rev)M6LRR-RLK_682

AGACTGTAGTACTGGGATACAATGTTTTCGAAGGTGGAATTCCAGCTGAATTTGGAAATCTGACAAATCTCAAGTATCTTGATCTTGCTATTGGAAATCTTGGAGGTTCAATTCCTTCTG

>M6-PGSC0003DMG400008581-(rev)M6LRR-RLK_744

ACAAATCTCAAGTATCTTGATCTTGCTATTGGAAATCTTGGAGGTTCAATTCCTTCTGAGCTAGGAAAGCTGAAGCTGCTCGATACGATTTTCTTGTACAAGAACAAGCTTGAAGGCAAG

>M6-PGSC0003DMG400008581-(rev)M6LRR-RLK_806

AGGAAAGCTGAAGCTGCTCGATACGATTTTCTTGTACAAGAACAAGCTTGAAGGCAAGATTCCGCCAGAGATTGGCAATATGACTTCATTGCAGTTGCTTGATCTGTCTGATAACATGCT

>M6-PGSC0003DMG400008581-(rev)M6LRR-RLK_868

CGCCAGAGATTGGCAATATGACTTCATTGCAGTTGCTTGATCTGTCTGATAACATGCTTACGGGTGAAATTCCAGCTAAGATTGCTGAACTTAAGAATTTGCAGCTTTTGAATATAATGT

>M6-PGSC0003DMG400008581-(rev)M6LRR-RLK_930

GGTGAAATTCCAGCTAAGATTGCTGAACTTAAGAATTTGCAGCTTTTGAATATAATGTCAAACAAATTATCAGGTTCAGTTCCCCCTGGGATTGGAGGTTTGACTCAGCTGGAAGTAGTT

>M6-PGSC0003DMG400008581-(rev)M6LRR-RLK_992

CAAATTATCAGGTTCAGTTCCCCCTGGGATTGGAGGTTTGACTCAGCTGGAAGTAGTTGAGCTGTGGAATAACAGCTTTTCAGGTCCTTTGCCAAGTGATCTTGGAAGAAATTCGCCATT

>M6-PGSC0003DMG400008596-(rev)M6LRR-RLP_0

CTTTATTTAGGTAAAGAAGACTTTGGTACAGAGAAAGAACATTTTGTGAAGGAGCAAACACAGTAATGGCCGATCCTGTAATTGGTGCTACTGCTCAAGTTTTGCTTGACAAGTTGCTTT

>M6-PGSC0003DMG400008596-(rev)M6LRR-RLP_1037

GCAGAAGATCATTGTCGGTCCATTTTCAAACAAAGAGCGTTTGTTGATGGGGAAATTCCACAGGAATTAGCAAGCATGGGCAACAGGATTGTTAAAATGTGCCAAGGTCTACCGTTGGCT

>M6-PGSC0003DMG400008596-(rev)M6LRR-RLP_1098

AGGAATTAGCAAGCATGGGCAACAGGATTGTTAAAATGTGCCAAGGTCTACCGTTGGCTGCAAGTGTATTGGGAGGCCTCTTACGCAGTAAGGAAAAACATGAATGGCAGGCAATTCTTG

>M6-PGSC0003DMG400008596-(rev)M6LRR-RLP_122

CCCACTATTGAGGAGGTCAAAAGTTTAAGGAACTGCAAGAAAGATCTCGAAATGCTGATAAAAAATGTATCCTTGATCCAAGCTTTCATTCATGATGCTGAAAGACGACAAGTCGAAGAT

>M6-PGSC0003DMG400008596-(rev)M6LRR-RLP_1220

GGAAAGCCCCTTGTTGCAGGTGAAGATGATAATGGAGAAAATAACATAAAGAAAATCCTAAAACTCAGCTATGATTATCTACCATATCCACATCTGAAAAAATGCTTTGCTTACTTTGCA

>M6-PGSC0003DMG400008596-(rev)M6LRR-RLP_1281

AACTCAGCTATGATTATCTACCATATCCACATCTGAAAAAATGCTTTGCTTACTTTGCAATGTTTCCAAAAGACCTTGAGTTTGAAAAGGACCAACTGATCCAACTCTGGATGGCAGAAG

>M6-PGSC0003DMG400008596-(rev)M6LRR-RLP_1342

GTTTCCAAAAGACCTTGAGTTTGAAAAGGACCAACTGATCCAACTCTGGATGGCAGAAGGCTTTCTTCATCCATGTCAAGAGACCACCGTGATGGAGGACATCGGGAACAAGTTCTTTCA

>M6-PGSC0003DMG400008596-(rev)M6LRR-RLP_1403

TTTCTTCATCCATGTCAAGAGACCACCGTGATGGAGGACATCGGGAACAAGTTCTTTCAAATCTTGTTGCGGAATTCCTTGCTGCAAGATGTTAAGCTAGATGAGCATAACAATATAACA

>M6-PGSC0003DMG400008596-(rev)M6LRR-RLP_1464

TCTTGTTGCGGAATTCCTTGCTGCAAGATGTTAAGCTAGATGAGCATAACAATATAACACACTGTAAGATGCACGATCTTGTGCATGATTTGGCTGGAGATATCTTAAAATCTAAACTAA

>M6-PGSC0003DMG400008596-(rev)M6LRR-RLP_1525

CTGTAAGATGCACGATCTTGTGCATGATTTGGCTGGAGATATCTTAAAATCTAAACTAATTGATCCCAAGGGCGATGGTGGAGAAAATCTTTCTCATGCGCGATACTTTGGATGGGACTC

>M6-PGSC0003DMG400008596-(rev)M6LRR-RLP_1586

GATCCCAAGGGCGATGGTGGAGAAAATCTTTCTCATGCGCGATACTTTGGATGGGACTCACCAAGTGATCAAATAGATAAGATATATGAGCCAGGACGTTTGTACACACTGTTCTGGAGA

>M6-PGSC0003DMG400008596-(rev)M6LRR-RLP_1647

CAAGTGATCAAATAGATAAGATATATGAGCCAGGACGTTTGTACACACTGTTCTGGAGAAGCAATCATATATCGGAAGCTATGCTGTTGAGCTTTAAGTTCTTGAGAGTTTTAAATTTGT

>M6-PGSC0003DMG400008596-(rev)M6LRR-RLP_1769

AGTTCAGGAATCAAGGAGCTGCCAGCCAAAATCAGCAAGCTAATATACTTGAGATATCTTGATCTCTCCAACACTAAGATCACAGCCTTGCCCAACTCCATTTGCAAGCTCTACAATTTG

>M6-PGSC0003DMG400008596-(rev)M6LRR-RLP_1830

ATCTCTCCAACACTAAGATCACAGCCTTGCCCAACTCCATTTGCAAGCTCTACAATTTGCAAACATTTAGAGTCAATGACTGCTATTCACTCAAGGAACTTCCATATGAGATGGGAAATA

>M6-PGSC0003DMG400008596-(rev)M6LRR-RLP_1891

AACATTTAGAGTCAATGACTGCTATTCACTCAAGGAACTTCCATATGAGATGGGAAATATGATAAGTTTGAGACACATTTACTGCTATTCTCGATCTCAGTTGCCACTTAACATGGGACA

>M6-PGSC0003DMG400008596-(rev)M6LRR-RLP_1952

ATAAGTTTGAGACACATTTACTGCTATTCTCGATCTCAGTTGCCACTTAACATGGGACAATTGACTTGTCTTCAGACCCTGCAGTATTTCAATGTGGGGTTAGAGAAAGGGCGTCGAATA

>M6-PGSC0003DMG400008596-(rev)M6LRR-RLP_2013

TGACTTGTCTTCAGACCCTGCAGTATTTCAATGTGGGGTTAGAGAAAGGGCGTCGAATAGGAGAATTAGGTTGTTTGAAGAACCTTAGAGGTGAATTGACGATCAATGAGCTGGAATTGG

>M6-PGSC0003DMG400008596-(rev)M6LRR-RLP_2135

TGTAGTATTGAAGAAGCTCGAACAGCATATTTACAAGAGAAATCGAATATCTACAAGCTGGCATATGTGTGGTTCCACGATGAACCAGAAGGCTGTGAAACCATTGTTGAGCATGTGTTG

>M6-PGSC0003DMG400008596-(rev)M6LRR-RLP_2196

CATATGTGTGGTTCCACGATGAACCAGAAGGCTGTGAAACCATTGTTGAGCATGTGTTGGATGGTCTTCAACCGCATACTAACTTGAAAACCTTAGAAGTCAAGAACTATTTAGGTACTA

>M6-PGSC0003DMG400008596-(rev)M6LRR-RLP_2257

TGGTCTTCAACCGCATACTAACTTGAAAACCTTAGAAGTCAAGAACTATTTAGGTACTAGATTTCCTTCATGGTTCAGAGAAGAATCTCTACCAAATTTGGTCAAGTTGAAATTAAGTGG

>M6-PGSC0003DMG400008596-(rev)M6LRR-RLP_2318

TTTCCTTCATGGTTCAGAGAAGAATCTCTACCAAATTTGGTCAAGTTGAAATTAAGTGGTTGCAAAAGATGCAAAGAAATTCCATGGCTTGGCCAACTGAAATTGCTTCGGCATCTTGAG

>M6-PGSC0003DMG400008596-(rev)M6LRR-RLP_2379

GCAAAAGATGCAAAGAAATTCCATGGCTTGGCCAACTGAAATTGCTTCGGCATCTTGAGCTATTAGGGTTACATAAGGTTGAATGCATCGGAACTATTTTTTATGGTATTGATGGTAACA

>M6-PGSC0003DMG400008596-(rev)M6LRR-RLP_244

GGCTGTGGAAAAATGGCTCAAGATGCTTGAGAGAGTTGCTGAAAATGCTGATAAGGTGTTTGACGAATTCAGGTATGAGTCTCTCAAAAGACAAGTGAAGATTCGAAACAATCCAATGAA

>M6-PGSC0003DMG400008596-(rev)M6LRR-RLP_2440

ATTAGGGTTACATAAGGTTGAATGCATCGGAACTATTTTTTATGGTATTGATGGTAACAATATTGGATCAAGCAGCAATGTCCAAGTGTTCCCATCATTGACTGAACTAGTATTGAAGGA

>M6-PGSC0003DMG400008596-(rev)M6LRR-RLP_2501

ATTGGATCAAGCAGCAATGTCCAAGTGTTCCCATCATTGACTGAACTAGTATTGAAGGATATGTCTAGCCTTATTGAGTGGAAGGGAGATGAAATTGGAGTAAGAATGTTTCCTCTGCTT

>M6-PGSC0003DMG400008596-(rev)M6LRR-RLP_2562

TGTCTAGCCTTATTGAGTGGAAGGGAGATGAAATTGGAGTAAGAATGTTTCCTCTGCTTGAGAAGTTGAGGATTAGTAGCTGTCCATTGTTAAAAATTACTCCAAGTCAATTTGAAATCC

>M6-PGSC0003DMG400008596-(rev)M6LRR-RLP_2684

CGTGAATTAAGAATTGAAAGAGTTGACAGTGAAATGCCATTGTTGAACTTGTGCAGTAAATTGACATCTCTCGTAAAGCTTAGTGTTTATGATGTGAAAGAGCTCACTTGTTTTCCAGAT

>M6-PGSC0003DMG400008596-(rev)M6LRR-RLP_2745

TGACATCTCTCGTAAAGCTTAGTGTTTATGATGTGAAAGAGCTCACTTGTTTTCCAGATGAGATGCTACGCAACAAGGTTTCTCTTCAACACCTATTGGTCTCAGACTGTGGAGAGTTTC

>M6-PGSC0003DMG400008596-(rev)M6LRR-RLP_2806

GATGCTACGCAACAAGGTTTCTCTTCAACACCTATTGGTCTCAGACTGTGGAGAGTTTCGTGAATTGCCACAAAGCTTATACAGTCTCCGTTCTCTTAAAAGCTTAAAGATTTACCGCTG

>M6-PGSC0003DMG400008596-(rev)M6LRR-RLP_2867

GAATTGCCACAAAGCTTATACAGTCTCCGTTCTCTTAAAAGCTTAAAGATTTACCGCTGCACCAATTTCAGTTCCTTTCCTGTTCCCAGTGGAGAGAATTATTTGACTTCCCTCCAAAGT

>M6-PGSC0003DMG400008596-(rev)M6LRR-RLP_2989

TCAGTTATGGAACTGTGATGGATTGACCAGTGTACCAAGTGGGATGCTAGAGCATTGTCGGTCTCTGGAATCTTTGATGGTCTGCTACTGTAACAACTTAGTTTCCTTCCCTATATGTGT

>M6-PGSC0003DMG400008596-(rev)M6LRR-RLP_305

GACGAATTCAGGTATGAGTCTCTCAAAAGACAAGTGAAGATTCGAAACAATCCAATGAAAAAGGTCAGTGATTTCTTTTCTCATACAGCTTTTAAGAGCAAAATGTCTCAAAAAATCAAC

>M6-PGSC0003DMG400008596-(rev)M6LRR-RLP_3050

TCTCTGGAATCTTTGATGGTCTGCTACTGTAACAACTTAGTTTCCTTCCCTATATGTGTGGGGGAAATGCCTTTACTTTTTGATGTGTATCTATCAGAATGTCCCAAATTGATTAGTGTA

>M6-PGSC0003DMG400008596-(rev)M6LRR-RLP_3111

GGGAAATGCCTTTACTTTTTGATGTGTATCTATCAGAATGTCCCAAATTGATTAGTGTACCCGCAGGGGACCTTTACCGTCTCACTGGGTTACGGCATTTGGAAATTGGTCCTTTCTCAG

>M6-PGSC0003DMG400008596-(rev)M6LRR-RLP_3233

ATGGTGGATTTTGAGGCATTCCAATTGATTTTCAATGGCATTCAGCAGCTGTTGTCCCTTCGTTTTTTGTGTGTGGTAGGACATTTGCACTGGGATTCTCTGCCCTATCAGCTTATGCAA

>M6-PGSC0003DMG400008596-(rev)M6LRR-RLP_3294

GTTTTTTGTGTGTGGTAGGACATTTGCACTGGGATTCTCTGCCCTATCAGCTTATGCAACTCTCTGACCTAACAGAGATCCAAATATATCGATTCGGAATCGAGGCTCTTCCTCATAAAT

>M6-PGSC0003DMG400008596-(rev)M6LRR-RLP_3355

CTCTGACCTAACAGAGATCCAAATATATCGATTCGGAATCGAGGCTCTTCCTCATAAATTTGACAACCTTGCTTCTCTTGAAAGATTAACGTTAGTGAGGTGCAAACGGCTACAACATCT

>M6-PGSC0003DMG400008596-(rev)M6LRR-RLP_3416

GACAACCTTGCTTCTCTTGAAAGATTAACGTTAGTGAGGTGCAAACGGCTACAACATCTGGACTTCTCAGTTGCCATGCCCAAATTACGGCATCTCCAGATACAGGATTGTCCATTGTTA

>M6-PGSC0003DMG400008596-(rev)M6LRR-RLP_3477

ACTTCTCAGTTGCCATGCCCAAATTACGGCATCTCCAGATACAGGATTGTCCATTGTTAGAAACACTGTCGGATGGGCTCGGCAACCTTGTTTCTTTGCAGGAGTTAGTTTTACAGAAAT

>M6-PGSC0003DMG400008596-(rev)M6LRR-RLP_3538

AACACTGTCGGATGGGCTCGGCAACCTTGTTTCTTTGCAGGAGTTAGTTTTACAGAAATGCGAGAAACTAGAGCATCTGCCATCTGGAGATGCCATGCAACGCCTCACCAAATTACGTTA

>M6-PGSC0003DMG400008596-(rev)M6LRR-RLP_3599

GAGAAACTAGAGCATCTGCCATCTGGAGATGCCATGCAACGCCTCACCAAATTACGTTACCTGAAAATTGAAGGCTGCCCAAAGTTAGAAGAAAGTTGCACCAATCGGATTGGCTCAAAC

>M6-PGSC0003DMG400008596-(rev)M6LRR-RLP_366

AGGTCAGTGATTTCTTTTCTCATACAGCTTTTAAGAGCAAAATGTCTCAAAAAATCAACAACATCAATGAAGATTTGACGGCTATCAATAGGTTAGCCAAAGACCTTGGTCTCCAATCAT

>M6-PGSC0003DMG400008596-(rev)M6LRR-RLP_3721

CCAGTGGTCCAACATTTCCCATATTCCAAAAATTAAAGTAGGTGAAAGCATAATTCAGGACTTAAGTAAGTCTCTGTTCCTCTTTCAGTTTCTGAGTCGATTTTTTTGTTCTTTCTTTTC

>M6-PGSC0003DMG400008596-(rev)M6LRR-RLP_3782

TTAAGTAAGTCTCTGTTCCTCTTTCAGTTTCTGAGTCGATTTTTTTGTTCTTTCTTTTCCACGAGTTGCTTTCCTTCTTTTTAACAAATTTTCCATTCCCCAAACTAATACATGTTATGG

>M6-PGSC0003DMG400008596-(rev)M6LRR-RLP_3843

CGAGTTGCTTTCCTTCTTTTTAACAAATTTTCCATTCCCCAAACTAATACATGTTATGGTACTTAAGAATTACAGAATTTTTTATCTCTCACTTGAAACTAACTGTTCTTCTTCTTATTC

>M6-PGSC0003DMG400008596-(rev)M6LRR-RLP_3904

CTTAAGAATTACAGAATTTTTTATCTCTCACTTGAAACTAACTGTTCTTCTTCTTATTCGTCATCAAACAAGGAATAGATACATTCCCGTCCTTTAATAATGTAAATATGTTCCTTCTCA

>M6-PGSC0003DMG400008596-(rev)M6LRR-RLP_3965

CATCAAACAAGGAATAGATACATTCCCGTCCTTTAATAATGTAAATATGTTCCTTCTCATAGGTGCCTAGATCAAATGTCATGCTGGAACGTTGCTAGCTGTGGTATATCAAGTCAAAAT

>M6-PGSC0003DMG400008596-(rev)M6LRR-RLP_4026

GGTGCCTAGATCAAATGTCATGCTGGAACGTTGCTAGCTGTGGTATATCAAGTCAAAATTTACGGTTGGCAATCAGGTTGGCTAAAATTTTCATTTAGACTTGGTAGAGGAATGCTATAA

>M6-PGSC0003DMG400008596-(rev)M6LRR-RLP_4087

ACGGTTGGCAATCAGGTTGGCTAAAATTTTCATTTAGACTTGGTAGAGGAATGCTATAACAAATTTATCGCTTCTTCCACAGGTCATTACTGTACTGTTTAATTAGGTTGGTTATGTTCT

>M6-PGSC0003DMG400008596-(rev)M6LRR-RLP_4148

AATTTATCGCTTCTTCCACAGGTCATTACTGTACTGTTTAATTAGGTTGGTTATGTTCTGTATGAATGCAATGTAAGCTTGGTTGCGTTTCTAGCATCATGAGGGACGTGCATCCATTTG

>M6-PGSC0003DMG400008596-(rev)M6LRR-RLP_4209

ATGAATGCAATGTAAGCTTGGTTGCGTTTCTAGCATCATGAGGGACGTGCATCCATTTGGTTTAAGGATTCTCTACTCGCATTGGACCAAAAATCTTCATGTAAGCTTCTTCCCAGTTGT

>M6-PGSC0003DMG400008596-(rev)M6LRR-RLP_427

CATCAATGAAGATTTGACGGCTATCAATAGGTTAGCCAAAGACCTTGGTCTCCAATCATTGATGGTTCCTCCTCGGCAAATACTACTAATTCGAGAAACAGATTCTGTAGTAGTTGCTTT

>M6-PGSC0003DMG400008596-(rev)M6LRR-RLP_4270

TTAAGGATTCTCTACTCGCATTGGACCAAAAATCTTCATGTAAGCTTCTTCCCAGTTGTTAATCTCCTAGGTCCAATAGTTTGCTTTGTTTTAATTGTTTGTTTTGCACGATTATCGTGC

>M6-PGSC0003DMG400008596-(rev)M6LRR-RLP_4331

ATCTCCTAGGTCCAATAGTTTGCTTTGTTTTAATTGTTTGTTTTGCACGATTATCGTGCAACAAAAGAGCCTATCTCTCAATCCTCAAATATTATAATGCTCTTCGTTACAGGGTATAGT

>M6-PGSC0003DMG400008596-(rev)M6LRR-RLP_4392

CAAAAGAGCCTATCTCTCAATCCTCAAATATTATAATGCTCTTCGTTACAGGGTATAGTGTGTTAGCACGTTGGACAAATTCCAATTAGCTGTACATATCTATGTAAGAACTCTAAATAT

>M6-PGSC0003DMG400008596-(rev)M6LRR-RLP_4453

GTTAGCACGTTGGACAAATTCCAATTAGCTGTACATATCTATGTAAGAACTCTAAATATTTATTGTAACTATGGTGTCACTTGCTTTCCACATGTAGGAGATATCAGGAATAACAGATAT

>M6-PGSC0003DMG400008596-(rev)M6LRR-RLP_4514

ATTGTAACTATGGTGTCACTTGCTTTCCACATGTAGGAGATATCAGGAATAACAGATATTTATTTAGGGGTGTAAATGACAACTTCTAATTTAATGGAAGAACTTTTAATATGAGAGCAT

>M6-PGSC0003DMG400008596-(rev)M6LRR-RLP_4575

ATTTAGGGGTGTAAATGACAACTTCTAATTTAATGGAAGAACTTTTAATATGAGAGCATTCATTCAGCAAATTTGACTCTGGATAAAGACAATTTCTCATTCTCTTTAGGAAAATGAAGT

>M6-PGSC0003DMG400008596-(rev)M6LRR-RLP_4636

ATTCAGCAAATTTGACTCTGGATAAAGACAATTTCTCATTCTCTTTAGGAAAATGAAGTATTTGATTCACATAATATTGGCTTATCATGGTTGACAGTTACAATCAGTTTCTTTTGCATA

>M6-PGSC0003DMG400008596-(rev)M6LRR-RLP_4697

TTGATTCACATAATATTGGCTTATCATGGTTGACAGTTACAATCAGTTTCTTTTGCATAAGAGAAGGTGAAACATAGAGGGGGATAAACGACCTTCTGCAAGGCGACTTTATTATCGACT

>M6-PGSC0003DMG400008596-(rev)M6LRR-RLP_4758

AGAAGGTGAAACATAGAGGGGGATAAACGACCTTCTGCAAGGCGACTTTATTATCGACTATCTAGTGAAATAGTATAGGTAACAAAGCCATTTCATCATAAGCCTTGGATGCAAGACAGA

>M6-PGSC0003DMG400008596-(rev)M6LRR-RLP_4819

CTAGTGAAATAGTATAGGTAACAAAGCCATTTCATCATAAGCCTTGGATGCAAGACAGATTTCATTTTGGTAAAACATGTGACTTCAATTGATAGAAGAGGCTTCTCCAAATTTTGTTAC

>M6-PGSC0003DMG400008596-(rev)M6LRR-RLP_488

ATGGTTCCTCCTCGGCAAATACTACTAATTCGAGAAACAGATTCTGTAGTAGTTGCTTTGGATGTTGTTGGTAGAGACAAAGATGTTGCTGAAATAAAGGAGAAGATATTGACCATGAGA

>M6-PGSC0003DMG400008596-(rev)M6LRR-RLP_4880

TCATTTTGGTAAAACATGTGACTTCAATTGATAGAAGAGGCTTCTCCAAATTTTGTTACCTGTCTTCACTTTTCAACAGTATTTCCTTTAATTTTCATCAGACTTATCGGAACATTGTTT

>M6-PGSC0003DMG400008596-(rev)M6LRR-RLP_4912

AGAAGAGGCTTCTCCAAATTTTGTTACCTGTCTTCACTTTTCAACAGTATTTCCTTTAATTTTCATCAGACTTATCGGAACATTGTTTACATTGGTGCCTTTGGGGACGGTGATTGCACA

>M6-PGSC0003DMG400008596-(rev)M6LRR-RLP_549

ATGTTGTTGGTAGAGACAAAGATGTTGCTGAAATAAAGGAGAAGATATTGACCATGAGAGAGGATGTTGTTCTGTCAACCATTCCAATAGTGGGTATGGGAGGTTTAGGGAAAACAACTG

>M6-PGSC0003DMG400008596-(rev)M6LRR-RLP_61

AGTAATGGCCGATCCTGTAATTGGTGCTACTGCTCAAGTTTTGCTTGACAAGTTGCTTTCTCCCACTATTGAGGAGGTCAAAAGTTTAAGGAACTGCAAGAAAGATCTCGAAATGCTGAT

>M6-PGSC0003DMG400008596-(rev)M6LRR-RLP_610

GGATGTTGTTCTGTCAACCATTCCAATAGTGGGTATGGGAGGTTTAGGGAAAACAACTGTCGCAAAGAGAATTTACAATGACGAACACATCAAGCAAATCTTTGAAAAGAGAGTTTGGTT

>M6-PGSC0003DMG400008596-(rev)M6LRR-RLP_671

GCAAAGAGAATTTACAATGACGAACACATCAAGCAAATCTTTGAAAAGAGAGTTTGGTTGTGTCTACCTGAAATGTCGGAGACTAAGAGCTTTCTCGAACTGATCCTCGAATCATTGACA

>M6-PGSC0003DMG400008596-(rev)M6LRR-RLP_793

GAGGAAAGTTAAGGTCCAGAGCAGAGATATAATTGTCAAGAAGCTCCGAGATGAACTGACAGGAAGGAAGTATTTGCTTGTCCTGGATGATTTGTGGCGTGTTGACTCTACATTGTGGAA

>M6-PGSC0003DMG400008596-(rev)M6LRR-RLP_854

GGAAGGAAGTATTTGCTTGTCCTGGATGATTTGTGGCGTGTTGACTCTACATTGTGGAATGAGTTTGTGGATACCTTGAGAGGAATAAATACTTCCCGGGGAAACTTCATTCTCGTGACT

>M6-PGSC0003DMG400008596-(rev)M6LRR-RLP_915

AGTTTGTGGATACCTTGAGAGGAATAAATACTTCCCGGGGAAACTTCATTCTCGTGACTACTCGTATGGAACAGGTGGCGTCCACAGTAGCAGCAGTAGGTCCTCATAAGTTGGAAAAAC

>M6-PGSC0003DMG400008596-(rev)M6LRR-RLP_976

TCGTATGGAACAGGTGGCGTCCACAGTAGCAGCAGTAGGTCCTCATAAGTTGGAAAAACTAGCAGAAGATCATTGTCGGTCCATTTTCAAACAAAGAGCGTTTGTTGATGGGGAAATTCC

>M6-PGSC0003DMG400008822M6L-LecRK_0

ATGTTCTTCAATCTTTTCATACTAGTAGCACTCTTTTTAGTTGATTTTTCACCAGCTTCTTGTGAAGTTGATGCATTCATTTACAATGGATTCCAATCAGCTAATCTTAGCTTGGATGGC

>M6-PGSC0003DMG400008822M6L-LecRK_1008

GCTACTAAGGGATTCAGAGAAAAGGAGGTATTGGGAGCTGGAGGATTTGGTAAAGTTTATAAAGGAGTGATGCCTATCACTAAACTTGAGATAGCTGTAAAGAAGATATCTCATGAATCA

>M6-PGSC0003DMG400008822M6L-LecRK_1071

GGAGTGATGCCTATCACTAAACTTGAGATAGCTGTAAAGAAGATATCTCATGAATCAAGACAAGGTATGAAGGAATTTGTTTCGGAGATTGTAAGTATTGGTCGCATGCAACACAGGAAT

>M6-PGSC0003DMG400008822M6L-LecRK_1134

GGTATGAAGGAATTTGTTTCGGAGATTGTAAGTATTGGTCGCATGCAACACAGGAATGTAGTACAACTTTTGGGTTATTGCAGGCGAAAAGGGGAGTTGATTTTGATATATGAATACATG

>M6-PGSC0003DMG400008822M6L-LecRK_1197

CAACTTTTGGGTTATTGCAGGCGAAAAGGGGAGTTGATTTTGATATATGAATACATGTCTAATGGAAGTCTAGACAAGTATTTATACGATCAACCAAGATACACCCTCGATTGGAACCAA

>M6-PGSC0003DMG400008822M6L-LecRK_126

CAGTTCACATCTGATGATCTTTTGTTGTTAACAAATTCAGGAACACAAAATCAGGGCCATGCTTTCTATCCAAATCCAATTCATTTCAAGAATTCATCAAATGGTACTGTATTTTCTTTC

>M6-PGSC0003DMG400008822M6L-LecRK_1260

GGAAGTCTAGACAAGTATTTATACGATCAACCAAGATACACCCTCGATTGGAACCAAAGATTCAAAGTCATTAGAGGTGTAGCCTCAGGACTATTTTTCCTACACGAAGAATGTGATCAC

>M6-PGSC0003DMG400008822M6L-LecRK_1323

AAAGTCATTAGAGGTGTAGCCTCAGGACTATTTTTCCTACACGAAGAATGTGATCACGTAGTGGTTCATAGAGATGTTAAGGCCAGTAATGTCTTGTTAGATGGTGAACTAAATGGCAGG

>M6-PGSC0003DMG400008822M6L-LecRK_1386

GTTCATAGAGATGTTAAGGCCAGTAATGTCTTGTTAGATGGTGAACTAAATGGCAGGTTAGGTGATTTTGGACTCGCAAGGCTGTATGGTCATGGGACCGATCCTCAGTCTACTCGTGTT

>M6-PGSC0003DMG400008822M6L-LecRK_1449

GATTTTGGACTCGCAAGGCTGTATGGTCATGGGACCGATCCTCAGTCTACTCGTGTTGTTGGTACTCTTGGTTACCTTGCACCAGAGCATACTAGAACTGGCAGGGCAACACCGAGTAGT

>M6-PGSC0003DMG400008822M6L-LecRK_1575

TTTTCATTTGGGGCATTTCTGCTTGAAGTTGCATGTGGTAGGAGGCCGATACAGCCAAGACAAGACGGTGATGATCTGATTTTGGTCGATTGGGTGTTCTCGTGCTGGAATAGGGGTAAT

>M6-PGSC0003DMG400008822M6L-LecRK_1638

GACGGTGATGATCTGATTTTGGTCGATTGGGTGTTCTCGTGCTGGAATAGGGGTAATATACTTGATGCTGCTGATCCAAACATAGGCATTGATTTTGTTCCAGGGCAAGTGGAGTTGGTC

>M6-PGSC0003DMG400008822M6L-LecRK_1701

GATGCTGCTGATCCAAACATAGGCATTGATTTTGTTCCAGGGCAAGTGGAGTTGGTCTTAAAGTTAGGCTTGTTCTGTTCTCATTCAGATCCTTCGTGTAGGCCAACCATGCGACAAATC

>M6-PGSC0003DMG400008822M6L-LecRK_1764

TTAGGCTTGTTCTGTTCTCATTCAGATCCTTCGTGTAGGCCAACCATGCGACAAATCTTGTTGTTCTTGGATGGTGTTGTGGCCTTACCAGAGTTATCAGAACTCGGTGTTTCATCAGCT

>M6-PGSC0003DMG400008822M6L-LecRK_1827

TTCTTGGATGGTGTTGTGGCCTTACCAGAGTTATCAGAACTCGGTGTTTCATCAGCTAACCTAACATTTGAACATCGCGGAGGTTTTGATGATTTTGTCAAGTCATATCCAGCTTCTTTG

>M6-PGSC0003DMG400008822M6L-LecRK_1887

CTAACATTTGAACATCGCGGAGGTTTTGATGATTTTGTCAAGTCATATCCAGCTTCTTTGGGCTATGCATATTCCTGCTCTCCATCTGTAACAGACTCCTTTCTCTCTGGTGGCCGTTGA

>M6-PGSC0003DMG400008822M6L-LecRK_189

TTCTATCCAAATCCAATTCATTTCAAGAATTCATCAAATGGTACTGTATTTTCTTTCTCCACAACTTTCGTGTTCGCGATAAGGTCTGATTATGGAAATTTGAGTGGTCATGGACTGGCT

>M6-PGSC0003DMG400008822M6L-LecRK_252

ACTTTCGTGTTCGCGATAAGGTCTGATTATGGAAATTTGAGTGGTCATGGACTGGCTTTCGTTATCGCGCCACATAAAGGACTTCAGGGGTCTTTGGCAAATCACTATCTTGGTCTTTTT

>M6-PGSC0003DMG400008822M6L-LecRK_315

ATCGCGCCACATAAAGGACTTCAGGGGTCTTTGGCAAATCACTATCTTGGTCTTTTTAACTCTAATAATAATGGGAATAGATCAAACCATGTCGTTGGAGTTGAGCTCGATACAATCATT

>M6-PGSC0003DMG400008822M6L-LecRK_378

AATAATAATGGGAATAGATCAAACCATGTCGTTGGAGTTGAGCTCGATACAATCATTAGCGAGGATTTTGGTGATATCAATGACAATCATGTTGGAATTGATATAAATGGATTGAGGTCT

>M6-PGSC0003DMG400008822M6L-LecRK_441

GATTTTGGTGATATCAATGACAATCATGTTGGAATTGATATAAATGGATTGAGGTCTGTAGCATATCATACAGCAGGTTATTTTGATGATACTGGTTTTTTTCATAACTTGACTTTAATT

>M6-PGSC0003DMG400008822M6L-LecRK_504

TATCATACAGCAGGTTATTTTGATGATACTGGTTTTTTTCATAACTTGACTTTAATTAGTGGCCAGGAAATGCAAGTTTGGATTGATTATGATGGGAGGACTAAGCAAATGAATGTAACA

>M6-PGSC0003DMG400008822M6L-LecRK_567

CAGGAAATGCAAGTTTGGATTGATTATGATGGGAGGACTAAGCAAATGAATGTAACAGTAGCTCAATTACATATGGAAAAACCAGTTAGACCACTTTTGGCTTTGAAATATGATCTTTCG

>M6-PGSC0003DMG400008822M6L-LecRK_63

GAAGTTGATGCATTCATTTACAATGGATTCCAATCAGCTAATCTTAGCTTGGATGGCATAGCACAGTTCACATCTGATGATCTTTTGTTGTTAACAAATTCAGGAACACAAAATCAGGGC

>M6-PGSC0003DMG400008822M6L-LecRK_630

CAATTACATATGGAAAAACCAGTTAGACCACTTTTGGCTTTGAAATATGATCTTTCGTCTATTCTTGAGCAGAGTATGTATGTTGGTTTTTCATCGTCAACAGGTTCAGTCCCGACACAT

>M6-PGSC0003DMG400008822M6L-LecRK_693

CTTGAGCAGAGTATGTATGTTGGTTTTTCATCGTCAACAGGTTCAGTCCCGACACATCATTATATCTTGGGATGGAGCTTCAAGACAAATGGGAAAGCTCAAGAACTCTCGCAACTTCCT

>M6-PGSC0003DMG400008822M6L-LecRK_756

ATCTTGGGATGGAGCTTCAAGACAAATGGGAAAGCTCAAGAACTCTCGCAACTTCCTAAGCTTCCTCCTCTTGGGAGTAAAGGGACGTCAAGATTTGTAACGATTGGTTTGCCTATAATC

>M6-PGSC0003DMG400008822M6L-LecRK_819

CCTCCTCTTGGGAGTAAAGGGACGTCAAGATTTGTAACGATTGGTTTGCCTATAATCTCATTGGTTTCACTTGTTGCAGCTGTGTTAGCGGTGGTTTTTTATGTAAGAAGGAAGAAGTAT

>M6-PGSC0003DMG400008822M6L-LecRK_882

GTTTCACTTGTTGCAGCTGTGTTAGCGGTGGTTTTTTATGTAAGAAGGAAGAAGTATGAAGAAATTCATGAAGATTGGGAACGCGAGTATAGACTACAAAGGTTTAAGTATAAAGAATTG

>M6-PGSC0003DMG400008822M6L-LecRK_945

ATTCATGAAGATTGGGAACGCGAGTATAGACTACAAAGGTTTAAGTATAAAGAATTGTACATTGCTACTAAGGGATTCAGAGAAAAGGAGGTATTGGGAGCTGGAGGATTTGGTAAAGTT

>M6-PGSC0003DMG400008897-(rev)M6L-LecRK_0

ACTAGATTTGACTATATTGTGTGATTATTTTTCATATATACATATTGAAGTTTGACTGGCCTCAAGAAAATGACATTACAGAGTTTACATTGTAGACTTGAGCATTAGGAGATAGATAGT

>M6-PGSC0003DMG400008897-(rev)M6L-LecRK_1054

GCTTGGCTTGCCCGTGATTTCTTCAGTTTCATTTGTGATTGTAGTCCTGGTAATAATTTATTGCATAGGAAAGAAGATGAAGTTTGCTGAATTGCAAGAAGAGTGGGAGCTTGATTATGG

>M6-PGSC0003DMG400008897-(rev)M6L-LecRK_1116

GCATAGGAAAGAAGATGAAGTTTGCTGAATTGCAAGAAGAGTGGGAGCTTGATTATGGACCACACAGGTTCAAGTACAAGGATTTGTACACTGCCACAAAGGGCTTTAGCAGGAAGGATT

>M6-PGSC0003DMG400008897-(rev)M6L-LecRK_1178

CACAGGTTCAAGTACAAGGATTTGTACACTGCCACAAAGGGCTTTAGCAGGAAGGATTTGTTGGGATCCGGCGGATTTGGTAGAGTCTACAGAGGGGTATTGCCCATATCCAACATTCAG

>M6-PGSC0003DMG400008897-(rev)M6L-LecRK_124

TTTGTGATTGTTATCAGCACTAATCCTAGGAGAGTTCTGAGCAGACATGGAGAAATTGTTTTTAGCCATGATTTTTCAGTTTCTTGGTTTGCTTTTTCATGCTCCATTAGGCTATGCAGC

>M6-PGSC0003DMG400008897-(rev)M6L-LecRK_1240

GGGATCCGGCGGATTTGGTAGAGTCTACAGAGGGGTATTGCCCATATCCAACATTCAGATAGCAGTCAAGAGGGTCTCTCATGAATCAAGACAAGGGATGAGGGAATTTGTTTCAGAAAT

>M6-PGSC0003DMG400008897-(rev)M6L-LecRK_1302

CAGTCAAGAGGGTCTCTCATGAATCAAGACAAGGGATGAGGGAATTTGTTTCAGAAATTGTTAGTGTTGGCCGGCTACGTCACAGGAATTTAGTACCACTTTTGGGTTATTGCAGGCGTA

>M6-PGSC0003DMG400008897-(rev)M6L-LecRK_1364

AGTGTTGGCCGGCTACGTCACAGGAATTTAGTACCACTTTTGGGTTATTGCAGGCGTAAAGGAGAGTTACTTCTGGTTTACGAATGCATGCCTAACGGAAGCCTAGACAAGTATTTGTTT

>M6-PGSC0003DMG400008897-(rev)M6L-LecRK_1426

AGAGTTACTTCTGGTTTACGAATGCATGCCTAACGGAAGCCTAGACAAGTATTTGTTTGACAAACCAAGCTGCACTCTTGATTGGAACCAAAGATTTCGAGTCATCAAAGGTGTAGCATC

>M6-PGSC0003DMG400008897-(rev)M6L-LecRK_1488

AACCAAGCTGCACTCTTGATTGGAACCAAAGATTTCGAGTCATCAAAGGTGTAGCATCAGGACTAGTCTATCTTCATGAAGAGTGGGAACAAGTAGTGATTCATAGAGATATAAAAGCGA

>M6-PGSC0003DMG400008897-(rev)M6L-LecRK_1550

CTAGTCTATCTTCATGAAGAGTGGGAACAAGTAGTGATTCATAGAGATATAAAAGCGAGTAACGTGTTGTTAGATGTTGAATTAAATGGAAGATTAGGAGATTTTGGGCTAGCAAGATTG

>M6-PGSC0003DMG400008897-(rev)M6L-LecRK_1612

CGTGTTGTTAGATGTTGAATTAAATGGAAGATTAGGAGATTTTGGGCTAGCAAGATTGTATGATCATGGGACAGATCCTCAGACTACGCATATCGTTGGAACTCTCGGATACCTTGCCCC

>M6-PGSC0003DMG400008897-(rev)M6L-LecRK_1674

ATCATGGGACAGATCCTCAGACTACGCATATCGTTGGAACTCTCGGATACCTTGCCCCAGAGCATACAAGAACAGGCAAAGCCACGACTAGAACTGATGTATATTCATTTGGAGCATTTT

>M6-PGSC0003DMG400008897-(rev)M6L-LecRK_1736

CATACAAGAACAGGCAAAGCCACGACTAGAACTGATGTATATTCATTTGGAGCATTTTTACTTGAGGTGGCTTGTGGGAGAAGGCCAATAGATCAAAGGGTATCGGACGAGGATCTCGTT

>M6-PGSC0003DMG400008897-(rev)M6L-LecRK_1798

TGAGGTGGCTTGTGGGAGAAGGCCAATAGATCAAAGGGTATCGGACGAGGATCTCGTTTTGGTTGACTATGTATTCTCTTGTTGGACTAGAGGTGATATTCTTGAAGCCATTGATCCAAA

>M6-PGSC0003DMG400008897-(rev)M6L-LecRK_186

TAGCCATGATTTTTCAGTTTCTTGGTTTGCTTTTTCATGCTCCATTAGGCTATGCAGCTGCAGAAGATCTCGATTTCACCTACAACGGGTTCCACTCAACAAACTTGAGTCTTGATGGAA

>M6-PGSC0003DMG400008897-(rev)M6L-LecRK_1860

TTGACTATGTATTCTCTTGTTGGACTAGAGGTGATATTCTTGAAGCCATTGATCCAAACTTGGGCAATAACTATGTAAAAGAAGAGATTGAGTTGGTACTAAAACTTGGACTATTGTGCT

>M6-PGSC0003DMG400008897-(rev)M6L-LecRK_1922

GGCAATAACTATGTAAAAGAAGAGATTGAGTTGGTACTAAAACTTGGACTATTGTGCTCTCAGACAGAACCAGCAGCTAGGCCAAATATGAGGCAAGTGTTGATGTACTTGGAAAATGCC

>M6-PGSC0003DMG400008897-(rev)M6L-LecRK_1984

GACAGAACCAGCAGCTAGGCCAAATATGAGGCAAGTGTTGATGTACTTGGAAAATGCCCTGCCTTTGCCAGAGTTATCACTGATGCAAACTTCAAATTTTAGCTTAACTTTTGCAGGCTT

>M6-PGSC0003DMG400008897-(rev)M6L-LecRK_2046

CTTTGCCAGAGTTATCACTGATGCAAACTTCAAATTTTAGCTTAACTTTTGCAGGCTTTGATCATTTTGCAATGTCATGGTCTTCTTCTGTTACTGACTCCCTTCTCTCCGGTGGTCGAT

>M6-PGSC0003DMG400008897-(rev)M6L-LecRK_2108

CATTTTGCAATGTCATGGTCTTCTTCTGTTACTGACTCCCTTCTCTCCGGTGGTCGATGACTCGGGCGAAATTTTAATATATTCCTAGTAAAAATAAAACTTGTTATATAAGCTATATTT

>M6-PGSC0003DMG400008897-(rev)M6L-LecRK_2153

TCCGGTGGTCGATGACTCGGGCGAAATTTTAATATATTCCTAGTAAAAATAAAACTTGTTATATAAGCTATATTTCTTCCACCTAAGGTTGTATACTTGGTTCATTTAGGCATCTTGGAG

>M6-PGSC0003DMG400008897-(rev)M6L-LecRK_248

GAAGATCTCGATTTCACCTACAACGGGTTCCACTCAACAAACTTGAGTCTTGATGGAATAGCAGAGTTCACATCAGATGGCTTGTTAAGGCTAACTAATGCTTCTCAATTGCAGGAGGGT

>M6-PGSC0003DMG400008897-(rev)M6L-LecRK_310

AGAGTTCACATCAGATGGCTTGTTAAGGCTAACTAATGCTTCTCAATTGCAGGAGGGTCATGCTTTTTATCCTAGACCTATCAATTTCATGAATTTAGCATCCAATGGTTCAGATTTCTC

>M6-PGSC0003DMG400008897-(rev)M6L-LecRK_372

CTTTTTATCCTAGACCTATCAATTTCATGAATTTAGCATCCAATGGTTCAGATTTCTCTTTCTCAACCACATTTGTCTTTGCCATAGTGCCTCAGTACTCGATTTTAGCTGGTCATGGTA

>M6-PGSC0003DMG400008897-(rev)M6L-LecRK_434

TCAACCACATTTGTCTTTGCCATAGTGCCTCAGTACTCGATTTTAGCTGGTCATGGTATGTCTTTTGTGATTGCACCAGCTAGAAGGCTTCCAGAAGCACTTCCAAGCTCTTATCTTGGC

>M6-PGSC0003DMG400008897-(rev)M6L-LecRK_496

TTTTGTGATTGCACCAGCTAGAAGGCTTCCAGAAGCACTTCCAAGCTCTTATCTTGGCTTGTTTAATGATTCTAGTAATGGAAATGCCACCAGCCATGTTGTTGCGGTGGAGTTTGACAC

>M6-PGSC0003DMG400008897-(rev)M6L-LecRK_558

TTAATGATTCTAGTAATGGAAATGCCACCAGCCATGTTGTTGCGGTGGAGTTTGACACGCTACAGAACCGTGAGTTTGATGATATTGACAGCAACCATGTTGGCATTGACATCAATGACC

>M6-PGSC0003DMG400008897-(rev)M6L-LecRK_62

CAAGAAAATGACATTACAGAGTTTACATTGTAGACTTGAGCATTAGGAGATAGATAGTCATATTTGTGATTGTTATCAGCACTAATCCTAGGAGAGTTCTGAGCAGACATGGAGAAATTG

>M6-PGSC0003DMG400008897-(rev)M6L-LecRK_620

CAGAACCGTGAGTTTGATGATATTGACAGCAACCATGTTGGCATTGACATCAATGACCTAAAATCTGTAATAGCCAAGCCGGCCAGTTATTATAACAGTAATGAGCAATCCTTCAATAAC

>M6-PGSC0003DMG400008897-(rev)M6L-LecRK_682

ATCTGTAATAGCCAAGCCGGCCAGTTATTATAACAGTAATGAGCAATCCTTCAATAACTTAACTCTAGCTAGTGGCCAGCCAATGCAAGTTTGGGTGGAATATGATTCTGTGGATAAGCA

>M6-PGSC0003DMG400008897-(rev)M6L-LecRK_744

CTCTAGCTAGTGGCCAGCCAATGCAAGTTTGGGTGGAATATGATTCTGTGGATAAGCATATGAATGTCACAATGGCTCCACTGCACGTTGCAAAACCAGACCGTCCTCTCCTGTCCTTAG

>M6-PGSC0003DMG400008897-(rev)M6L-LecRK_806

AATGTCACAATGGCTCCACTGCACGTTGCAAAACCAGACCGTCCTCTCCTGTCCTTAGTCTACGATCTTTCATCTGCCACGGATGAAAACGTGTCTGTTGGCTTCTCAGCATCCACCGGT

>M6-PGSC0003DMG400008897-(rev)M6L-LecRK_868

CGATCTTTCATCTGCCACGGATGAAAACGTGTCTGTTGGCTTCTCAGCATCCACCGGTGCAGTTGTTTCAACACATTATATCCTTGGATGGAGCTTCAAATTGAATGGAATGGCTCGAGG

>M6-PGSC0003DMG400008897-(rev)M6L-LecRK_930

TTGTTTCAACACATTATATCCTTGGATGGAGCTTCAAATTGAATGGAATGGCTCGAGGGCTTGATCTCTCTCAACTTCCTAAGCTTCCTCGAGTAGTACCTAAGAAAGAATCCAAGCTTT

>M6-PGSC0003DMG400008897-(rev)M6L-LecRK_992

GATCTCTCTCAACTTCCTAAGCTTCCTCGAGTAGTACCTAAGAAAGAATCCAAGCTTTTATTGCTTGGCTTGCCCGTGATTTCTTCAGTTTCATTTGTGATTGTAGTCCTGGTAATAATT

>M6-PGSC0003DMG400008925-(rev)M6LRR-RLP_0

TGAGGTCCTCAAACTACTTTCCCGAGCCTTTGAGGGCGTGCAATGGGACATGAAAGACCGAGAGTTCTTGAAACTAAAATTCTTGAAGTTAGACTCTTTGAACATTGAACAATGGAACAT

>M6-PGSC0003DMG400008925-(rev)M6LRR-RLP_104

TTGAACAATGGAACATCTCATCTGATTGCTTACCTTCCCTTGGGCATCTTGTTCTGCAAAGTTGTAAGAAACTTAAGGAGTTCCCTTCATGCTTAGGCAATGTTTCAACGCTGCAGAGTG

>M6-PGSC0003DMG400008925-(rev)M6LRR-RLP_156

TCTGCAAAGTTGTAAGAAACTTAAGGAGTTCCCTTCATGCTTAGGCAATGTTTCAACGCTGCAGAGTGTTGAGCTACACTGGTGCAGCTGTTCTTCTGTGAACTCGATTATCTCAATTCA

>M6-PGSC0003DMG400008925-(rev)M6LRR-RLP_208

TCAACGCTGCAGAGTGTTGAGCTACACTGGTGCAGCTGTTCTTCTGTGAACTCGATTATCTCAATTCAAGAAGATGAGGGGTTCAAGGTCCATGTATACCCTCGAGAATTCGATTCCTGA

>M6-PGSC0003DMG400008925-(rev)M6LRR-RLP_52

AAAGACCGAGAGTTCTTGAAACTAAAATTCTTGAAGTTAGACTCTTTGAACATTGAACAATGGAACATCTCATCTGATTGCTTACCTTCCCTTGGGCATCTTGTTCTGCAAAGTTGTAAG

>M6-PGSC0003DMG400009029M6LRR-RLP_0

GGTTATCCCTCGAGTAACAAGATAGTCGAATATAAAAACACTGTTCCCACACAACACACACATTGTTTACTTCACTTTTTTCAGTTGAGATTTTCTTCACTTGCACTCTCTCCGGCGGCG

>M6-PGSC0003DMG400009029M6LRR-RLP_1054

TCACTATATTCCTTTGATTCTAGGTTTTCTTCCAATGGATGCCACAACAGCTAACTCTCAGCGCAGACCATTTTCAATTAAACTGTGGCCTCCAAGTGAGAACACAAGAAAAATGCTGGT

>M6-PGSC0003DMG400009029M6LRR-RLP_1116

GCAGACCATTTTCAATTAAACTGTGGCCTCCAAGTGAGAACACAAGAAAAATGCTGGTGGAAAGAATGATGAATAATCTTTCTAGTCCAACCATTTTCACCCGCAAGTACCGCAGTCTAT

>M6-PGSC0003DMG400009029M6LRR-RLP_1178

AGAATGATGAATAATCTTTCTAGTCCAACCATTTTCACCCGCAAGTACCGCAGTCTATCCAAGGAAGAGGCTGCCAAAAATGCTGAAGAAATTGAAGATGCAGCTTTTACCATTGCTAAT

>M6-PGSC0003DMG400009029M6LRR-RLP_124

GAACTCACACATTCTCCCTTGTTGTTTTCCGGTGAGTTACAACAGACACTCACTCCAACGATAATCAGAAAAATCCCAAAAAGGTTTTTTCTTTTCTCTTGATTTCTAGGGTTTTTGCGC

>M6-PGSC0003DMG400009029M6LRR-RLP_1240

GGAAGAGGCTGCCAAAAATGCTGAAGAAATTGAAGATGCAGCTTTTACCATTGCTAATCAACATTATGAGAAGGAGCCTGATGGTGATGGAAGTTCTGCTGTGCAACTTTATGCCAGGGA

>M6-PGSC0003DMG400009029M6LRR-RLP_1302

ATTATGAGAAGGAGCCTGATGGTGATGGAAGTTCTGCTGTGCAACTTTATGCCAGGGAATGCAGCAAGCTTATCCTGGAAATTCTAAAGAAGATACCCAAATCAGAGGACAAGGAAATTT

>M6-PGSC0003DMG400009029M6LRR-RLP_1364

AGCAAGCTTATCCTGGAAATTCTAAAGAAGATACCCAAATCAGAGGACAAGGAAATTTCAATTTCTGAAGTGGTTCCTACTGTTCAGGAGACCTTTTTTGATATCTCAAAAGGTAAAAGG

>M6-PGSC0003DMG400009029M6LRR-RLP_1426

TTCTGAAGTGGTTCCTACTGTTCAGGAGACCTTTTTTGATATCTCAAAAGGTAAAAGGGCTTTTATCGAAGCGGAAGAGGCTCAAGAACTTTTAAAGCCATTAAAAGAGCCTGGAAATTC

>M6-PGSC0003DMG400009029M6LRR-RLP_1488

TTATCGAAGCGGAAGAGGCTCAAGAACTTTTAAAGCCATTAAAAGAGCCTGGAAATTCTTACAGCAAAATTTGTTTTAGCAATAGAAGCTTCAGTATAGATGCAGCACGTATTGCAGGCC

>M6-PGSC0003DMG400009029M6LRR-RLP_1550

AGCAAAATTTGTTTTAGCAATAGAAGCTTCAGTATAGATGCAGCACGTATTGCAGGCCCTATCTTGGCAGCCTTGAAGGATCAATTGAAGGAAGTTGATTTGTCAGATTTTGTTGCCGGA

>M6-PGSC0003DMG400009029M6LRR-RLP_1612

CTTGGCAGCCTTGAAGGATCAATTGAAGGAAGTTGATTTGTCAGATTTTGTTGCCGGAAGAAATGAGGCAGAAGCACTTGATGTGATGAATATATTCTCAGAAGCTCTGGAAGGTTCTAA

>M6-PGSC0003DMG400009029M6LRR-RLP_1674

ATGAGGCAGAAGCACTTGATGTGATGAATATATTCTCAGAAGCTCTGGAAGGTTCTAACTTGAAGTTTCTGAATCTCTCTGATAATGCTCTGGGTGAGAAGGGAGTTAGGGCATTTGGAA

>M6-PGSC0003DMG400009029M6LRR-RLP_1736

AAGTTTCTGAATCTCTCTGATAATGCTCTGGGTGAGAAGGGAGTTAGGGCATTTGGAAAGCTCCTTCAGTCTCAAACCAACTTGGAAGAATTATTTTTGATGAATGATGGGATTTCACAG

>M6-PGSC0003DMG400009029M6LRR-RLP_1798

CCTTCAGTCTCAAACCAACTTGGAAGAATTATTTTTGATGAATGATGGGATTTCACAGGAAGCTGCAAATGCTGTTAGCGAGCTAGTTCCTTCCACCGAGAAGCTTAAGGTTCTTTATTT

>M6-PGSC0003DMG400009029M6LRR-RLP_186

AATCAGAAAAATCCCAAAAAGGTTTTTTCTTTTCTCTTGATTTCTAGGGTTTTTGCGCTTATGCTCTGTACGAGTCTACATTTTTGGTCATGTGATTTCACAGATCTGTGCTGGATTTTA

>M6-PGSC0003DMG400009029M6LRR-RLP_1860

CTGCAAATGCTGTTAGCGAGCTAGTTCCTTCCACCGAGAAGCTTAAGGTTCTTTATTTTCATAATAATATGACAGGTGATGAAGGGGCTGTTGCGATTGCGGAAATTGTGAAGCGTTCTC

>M6-PGSC0003DMG400009029M6LRR-RLP_1922

AATAATATGACAGGTGATGAAGGGGCTGTTGCGATTGCGGAAATTGTGAAGCGTTCTCCTTTATTGGAAGATTTCAGGTGCTCTTCTACCAGGGTAGGCTCTGAAGGAGGGAGTGTCTTG

>M6-PGSC0003DMG400009029M6LRR-RLP_1984

ATTGGAAGATTTCAGGTGCTCTTCTACCAGGGTAGGCTCTGAAGGAGGGAGTGTCTTGTGTGAAGCACTTGGGATGTGCTCCCATTTGAAGAAGCTTGATTTGCGAGACAATATGTTTGG

>M6-PGSC0003DMG400009029M6LRR-RLP_2046

AAGCACTTGGGATGTGCTCCCATTTGAAGAAGCTTGATTTGCGAGACAATATGTTTGGTCCAGAAGTTGGTCTTGTGTTGTGTAAGGCACTCAGCAAACATGAAAATCTTACGGAAATTT

>M6-PGSC0003DMG400009029M6LRR-RLP_2108

GAAGTTGGTCTTGTGTTGTGTAAGGCACTCAGCAAACATGAAAATCTTACGGAAATTTACCTGAGCTACCTTAATTTAGAGGATGAAGGAGCAATTGCAATAGCTAACGCTCTTAAAGAT

>M6-PGSC0003DMG400009029M6LRR-RLP_2170

GAGCTACCTTAATTTAGAGGATGAAGGAGCAATTGCAATAGCTAACGCTCTTAAAGATTCAGCTCCTTCACTTGCTGTCTTGGAGATGGCAGGTAATGATATAACTGCTGAAGCTGCTTC

>M6-PGSC0003DMG400009029M6LRR-RLP_2232

CTCCTTCACTTGCTGTCTTGGAGATGGCAGGTAATGATATAACTGCTGAAGCTGCTTCAGCAATAGCTTCTTGTATAGCTGCAAAGCAGCTTCTTGCCAAGTTAAGCTTGGGTGAGAATG

>M6-PGSC0003DMG400009029M6LRR-RLP_2294

ATAGCTTCTTGTATAGCTGCAAAGCAGCTTCTTGCCAAGTTAAGCTTGGGTGAGAATGAACTCAAGGATGAAGGTGCAATTCAGATTGCTAAGGCACTGGAAGGACACAGCCATCTGATT

>M6-PGSC0003DMG400009029M6LRR-RLP_2356

CAAGGATGAAGGTGCAATTCAGATTGCTAAGGCACTGGAAGGACACAGCCATCTGATTGAAGTTGATATGAGCAGCAATGCACTAAGGAGGGCTGGGGCTAGAGTGCTGGCTCAAACTGT

>M6-PGSC0003DMG400009029M6LRR-RLP_2418

TTGATATGAGCAGCAATGCACTAAGGAGGGCTGGGGCTAGAGTGCTGGCTCAAACTGTGCTGCGCAAAGATGAGTTTAAATTACTAAATGTTAATGGGAATTTCATCTCAGAAGAAGGCG

>M6-PGSC0003DMG400009029M6LRR-RLP_248

GCTCTGTACGAGTCTACATTTTTGGTCATGTGATTTCACAGATCTGTGCTGGATTTTAGGTATTTTTCTTGTTTAATGTGCTTTCGTTTGTGGTTTTAGGTCGTTTTTTTTTATGTTTAT

>M6-PGSC0003DMG400009029M6LRR-RLP_2480

CGCAAAGATGAGTTTAAATTACTAAATGTTAATGGGAATTTCATCTCAGAAGAAGGCGTTGACGAGTTGAAAGAGATCTTCAAGAAATCTCCTGAAATGCTTGCATCCCTGGAAGATAAT

>M6-PGSC0003DMG400009029M6LRR-RLP_2666

GACGTTGAGGATGAACTAGAATCAAAACTCAAAAACCTGGACGTCAAGCAAGAGGCAAGTCTTGACACACCAGATAGCAATTAGAATCCAGTACTTGACGACTTCTGCATGTTGCAGTTT

>M6-PGSC0003DMG400009029M6LRR-RLP_2728

TGACACACCAGATAGCAATTAGAATCCAGTACTTGACGACTTCTGCATGTTGCAGTTTTAGTTTTACCATTTTCTTTATGCTAAGCTGGCCATATTATGTTCCTCCAGCAAGTGCTCTAA

>M6-PGSC0003DMG400009029M6LRR-RLP_2790

TTTACCATTTTCTTTATGCTAAGCTGGCCATATTATGTTCCTCCAGCAAGTGCTCTAATATATAGTTCCATCAGTAATTTAAATTTGGTGGAATTGAACTGTTGCCTTAATTGGTCAAGT

>M6-PGSC0003DMG400009029M6LRR-RLP_2852

TAGTTCCATCAGTAATTTAAATTTGGTGGAATTGAACTGTTGCCTTAATTGGTCAAGTTAGGTCTCTGGCTGAATTATGTAGGGTACGAGAGAGAGAACTGCTATTTATGTTGAACCTTC

>M6-PGSC0003DMG400009029M6LRR-RLP_310

TTTTTCTTGTTTAATGTGCTTTCGTTTGTGGTTTTAGGTCGTTTTTTTTTATGTTTATTATTTTGGGGTTTGGTAGAGTAGGTTGCAATTACTGGGAGGAGGACTACAAACTTCACTATT

>M6-PGSC0003DMG400009029M6LRR-RLP_372

TTGGGGTTTGGTAGAGTAGGTTGCAATTACTGGGAGGAGGACTACAAACTTCACTATTAGGTATAAAGTTTTTTATTTTTCTCGTGAGGTTTGTTTGTTTGTAGGGATAAAAGAAGGATT

>M6-PGSC0003DMG400009029M6LRR-RLP_434

ATAAAGTTTTTTATTTTTCTCGTGAGGTTTGTTTGTTTGTAGGGATAAAAGAAGGATTTTGTTTTGTATTACAAGTTTCTGATGTTTTTAGCGGCTCTGTAAATTGTATATATGCTGCTC

>M6-PGSC0003DMG400009029M6LRR-RLP_496

TTTGTATTACAAGTTTCTGATGTTTTTAGCGGCTCTGTAAATTGTATATATGCTGCTCTGCTCTGCTCATATTATTCTTTTTGTAATGTATTGTGGGGTATTTTTTTCGTATCCAAGAAA

>M6-PGSC0003DMG400009029M6LRR-RLP_558

CTGCTCATATTATTCTTTTTGTAATGTATTGTGGGGTATTTTTTTCGTATCCAAGAAAGTTTTTATGTGCTATAGTGTGGTGAAATGGAATACATCTGTTGAGTGTACCATTTTTGGTTT

>M6-PGSC0003DMG400009029M6LRR-RLP_62

TTGTTTACTTCACTTTTTTCAGTTGAGATTTTCTTCACTTGCACTCTCTCCGGCGGCGCGTCGAACTCACACATTCTCCCTTGTTGTTTTCCGGTGAGTTACAACAGACACTCACTCCAA

>M6-PGSC0003DMG400009029M6LRR-RLP_620

TTATGTGCTATAGTGTGGTGAAATGGAATACATCTGTTGAGTGTACCATTTTTGGTTTTGAGTGTTTTTTACATTGTTGGAAATGAATATGGTGCATTGCAAGGTGGTTTATAAAAGAAT

>M6-PGSC0003DMG400009029M6LRR-RLP_682

TGTTTTTTACATTGTTGGAAATGAATATGGTGCATTGCAAGGTGGTTTATAAAAGAATAGAAAATTCAATCTCGTTTTCAATGATGGATGATTGTAGAAACTTTGCCATTCTTTGGACAC

>M6-PGSC0003DMG400009029M6LRR-RLP_744

AATTCAATCTCGTTTTCAATGATGGATGATTGTAGAAACTTTGCCATTCTTTGGACACCTGCTTGGTGTTTTGTGGTGAAACTACTGAGTTGTCAAACAATTATATCTGATTTCTATATG

>M6-PGSC0003DMG400009029M6LRR-RLP_806

TTGGTGTTTTGTGGTGAAACTACTGAGTTGTCAAACAATTATATCTGATTTCTATATGAGGAAGAATTTGAGCTCTTGAAATTTGGTTTCGATATATGTTAGGTTTTCTTACATGGAGTA

>M6-PGSC0003DMG400009029M6LRR-RLP_868

AGAATTTGAGCTCTTGAAATTTGGTTTCGATATATGTTAGGTTTTCTTACATGGAGTAGGTGTGCACGAACTCGTTCTTGCCTGGTACTTTGTTTTTGGTTGTTATGTTCTTAACTGCCA

>M6-PGSC0003DMG400009029M6LRR-RLP_930

TGCACGAACTCGTTCTTGCCTGGTACTTTGTTTTTGGTTGTTATGTTCTTAACTGCCAGAGATGAGTCTTGTAGTGGTTATATTTCTTGAAAGATAATCACTTCTATAGAAACTGAGACA

>M6-PGSC0003DMG400009029M6LRR-RLP_992

TGAGTCTTGTAGTGGTTATATTTCTTGAAAGATAATCACTTCTATAGAAACTGAGACATGATTCACTATATTCCTTTGATTCTAGGTTTTCTTCCAATGGATGCCACAACAGCTAACTCT

>M6-PGSC0003DMG400009068-(rev)M6LRR-RLK_0

CGTGATTAAGAGTGGTTAATTTCCTTTCCCTTTTTCGTAATTGCAATCTGTGTCTTCCTTTAGTGAAGAGCTTCAAAAAATGGCGGAAATGGTGGATAATGCTCTTCTGGTGCTCCCTTG

>M6-PGSC0003DMG400009068-(rev)M6LRR-RLK_1054

GTAGAGATGAAGGGTGCTGTTACTAGAAGTTGATGCCTTTGTAGTTTAGCTCAGAGGAGGAGAAGGATGGCTACAAAGTTCCTGTTTTTTTCAGTTTTATTCTGCACAGCTCTGTTCTTG

>M6-PGSC0003DMG400009068-(rev)M6LRR-RLK_1116

AAGGATGGCTACAAAGTTCCTGTTTTTTTCAGTTTTATTCTGCACAGCTCTGTTCTTGCTAGCAAGTAGTGAACCATATGAGGACAAAGAAGCGCTTCTTGATTTCCTTAATAATGTTAA

>M6-PGSC0003DMG400009068-(rev)M6LRR-RLK_1178

CAAGTAGTGAACCATATGAGGACAAAGAAGCGCTTCTTGATTTCCTTAATAATGTTAATCATACCCGATACCTGAATTGGGATGAGAGAACTTCTGCTTGCAGTTCTTGGACTGGAGTCA

>M6-PGSC0003DMG400009068-(rev)M6LRR-RLK_124

ACTGGCACCACGTGCACCCACTTCTCCTCTTCTTCTAAATCCAAGCAAATTGTTAAAGAATTGATTTTGTGTATTACTGTTAACTTCTTCATAGCATAGTAGTAGTAGTTCCACTTTTGT

>M6-PGSC0003DMG400009068-(rev)M6LRR-RLK_1240

ACCCGATACCTGAATTGGGATGAGAGAACTTCTGCTTGCAGTTCTTGGACTGGAGTCACTTGTAATCATGACAAATCAAGAATTATAGCAATTAGGTTGCCTGGAGTTGGATTCCGTGGT

>M6-PGSC0003DMG400009068-(rev)M6LRR-RLK_1302

TAATCATGACAAATCAAGAATTATAGCAATTAGGTTGCCTGGAGTTGGATTCCGTGGTTCGATTCCTGGAAACACTCTTAGCCGCTTGTCTGATCTTCAGATCTTGAGTTTAAGGTCTAA

>M6-PGSC0003DMG400009068-(rev)M6LRR-RLK_1364

TTCCTGGAAACACTCTTAGCCGCTTGTCTGATCTTCAGATCTTGAGTTTAAGGTCTAATAGTTTTAGTGGCAGTTTACCTTCTGATTTTGCAAACCTTGGAAATCTTACTTCTATTTATC

>M6-PGSC0003DMG400009068-(rev)M6LRR-RLK_1426

TTTAGTGGCAGTTTACCTTCTGATTTTGCAAACCTTGGAAATCTTACTTCTATTTATCTTCAATCCAATAACTTTCAAGGTCCATTGCCTGCTGATTTTTCTGCTTGGAAAAGCCTTTCG

>M6-PGSC0003DMG400009068-(rev)M6LRR-RLK_1488

ATCCAATAACTTTCAAGGTCCATTGCCTGCTGATTTTTCTGCTTGGAAAAGCCTTTCGGTTTTGAATTTGTCAAATAATGATTTTAGTGGGAGTATACCTTCTTCAATATCGAACTTGAC

>M6-PGSC0003DMG400009068-(rev)M6LRR-RLK_1550

TGAATTTGTCAAATAATGATTTTAGTGGGAGTATACCTTCTTCAATATCGAACTTGACTCATTTGACTGCTCTGGTTCTTGCTAATAACTCGCTTTCTGGCAATATTCCTGATCTCAATC

>M6-PGSC0003DMG400009068-(rev)M6LRR-RLK_1612

TTGACTGCTCTGGTTCTTGCTAATAACTCGCTTTCTGGCAATATTCCTGATCTCAATCTCCCTAGTTTGCAAATATTAGATCTATCAAACAATAACTTCACTGGAAGTATCCCCAATTCT

>M6-PGSC0003DMG400009068-(rev)M6LRR-RLK_1674

TAGTTTGCAAATATTAGATCTATCAAACAATAACTTCACTGGAAGTATCCCCAATTCTCTTCAAAGATTTCCTGGTTCAGCTTTTGCTGGTAACCGGCTTTCTCCTGCAAATTTTTCACC

>M6-PGSC0003DMG400009068-(rev)M6LRR-RLK_1736

AAAGATTTCCTGGTTCAGCTTTTGCTGGTAACCGGCTTTCTCCTGCAAATTTTTCACCTTCATTTCCTCCAGTTCCTCCTCCAAGTGTCCTACCGAAAAAGAAATCGTTTAAACTCCGTG

>M6-PGSC0003DMG400009068-(rev)M6LRR-RLK_1798

TTTCCTCCAGTTCCTCCTCCAAGTGTCCTACCGAAAAAGAAATCGTTTAAACTCCGTGAACCTGCAATACTTGGAATTGTAATTGGAGGGTGTGTTTTAGGTTTCTTGGTAGTAGCTGCA

>M6-PGSC0003DMG400009068-(rev)M6LRR-RLK_186

GATTTTGTGTATTACTGTTAACTTCTTCATAGCATAGTAGTAGTAGTTCCACTTTTGTCTCCACGTGCTTTTCCCACTCTTTTCGCTTCTTTGTTAACCAAATACATACATATACCCCAT

>M6-PGSC0003DMG400009068-(rev)M6LRR-RLK_1860

TGCAATACTTGGAATTGTAATTGGAGGGTGTGTTTTAGGTTTCTTGGTAGTAGCTGCAGTGCTGATTATGTGCTTTTCGAAGAAAGAGGGTAAAAGTGGGGCCACAGAGAAATCAATCAA

>M6-PGSC0003DMG400009068-(rev)M6LRR-RLK_1922

TGATTATGTGCTTTTCGAAGAAAGAGGGTAAAAGTGGGGCCACAGAGAAATCAATCAAGAAAGAAGATTTAGTCAGGAAGGGAGTTTCCAGCAGTCAGCATGGAGTAGGGAATCTCGCAT

>M6-PGSC0003DMG400009068-(rev)M6LRR-RLK_1984

GAAGATTTAGTCAGGAAGGGAGTTTCCAGCAGTCAGCATGGAGTAGGGAATCTCGCATTCTTTGAGGGTTGTAATCTTGCATTTGACCTTGAAGATCTGTTGAGAGCTTCAGCGGAAGTG

>M6-PGSC0003DMG400009068-(rev)M6LRR-RLK_2046

TGAGGGTTGTAATCTTGCATTTGACCTTGAAGATCTGTTGAGAGCTTCAGCGGAAGTGCTTGGAAAGGGAACATTTGGCACTACATATAAGGCAGCTTTGGAGGATTCTACCACAGTTGT

>M6-PGSC0003DMG400009068-(rev)M6LRR-RLK_2108

GAAAGGGAACATTTGGCACTACATATAAGGCAGCTTTGGAGGATTCTACCACAGTTGTGGTGAAGAGATTGAAGGAGAGTGTGGGAAGAAAAGATTTTGAGCAACAGATGGAGGTTGTTG

>M6-PGSC0003DMG400009068-(rev)M6LRR-RLK_2170

AAGAGATTGAAGGAGAGTGTGGGAAGAAAAGATTTTGAGCAACAGATGGAGGTTGTTGGCAATATCAGGCATGAGAATGTGGCTCCACTGAGGGCATACTACTATTCCAAGGATGAGAAG

>M6-PGSC0003DMG400009068-(rev)M6LRR-RLK_2232

TATCAGGCATGAGAATGTGGCTCCACTGAGGGCATACTACTATTCCAAGGATGAGAAGCTCATGGTGTATGATTTCTACAGCCAAGGGAGTGCATCTTTGATGCTGCATGGTATGTTAAG

>M6-PGSC0003DMG400009068-(rev)M6LRR-RLK_2294

TGGTGTATGATTTCTACAGCCAAGGGAGTGCATCTTTGATGCTGCATGGTATGTTAAGCACCCAATTTCTACTAATCTTGCTGTTTTATTCTTGACCGATCCATAGAGAACTGGTTATCA

>M6-PGSC0003DMG400009068-(rev)M6LRR-RLK_2356

CAATTTCTACTAATCTTGCTGTTTTATTCTTGACCGATCCATAGAGAACTGGTTATCAGGACCAAAAAGATAATGCAGATCATTTTTAGCTATGATGTGTCTTGTGCAGTAGTATTTGAA

>M6-PGSC0003DMG400009068-(rev)M6LRR-RLK_2418

CAAAAAGATAATGCAGATCATTTTTAGCTATGATGTGTCTTGTGCAGTAGTATTTGAAATTGCTTGTAGGCTAATTCCAACATAACCAATTACCTTTGAATTAATAGAGTATCTGCTGTT

>M6-PGSC0003DMG400009068-(rev)M6LRR-RLK_248

ACGTGCTTTTCCCACTCTTTTCGCTTCTTTGTTAACCAAATACATACATATACCCCATTTCTTTATTTGCTTTCTTCTTCTTCAAATTAACTGCCTTTCAATCATTTATTGCCTCTCACC

>M6-PGSC0003DMG400009068-(rev)M6LRR-RLK_2480

CTTGTAGGCTAATTCCAACATAACCAATTACCTTTGAATTAATAGAGTATCTGCTGTTGTTGTCCTCTTGATAGTTTTTTTTTATCATCAATAGCTTATAGGACAGCATAAGAGTCTCGG

>M6-PGSC0003DMG400009068-(rev)M6LRR-RLK_2542

TCCTCTTGATAGTTTTTTTTTATCATCAATAGCTTATAGGACAGCATAAGAGTCTCGGAGTCATGTCTAAGTGCCATTTTTGTTTTAATTTGTCAAAATTCAATAACATCATGTTGCTCG

>M6-PGSC0003DMG400009068-(rev)M6LRR-RLK_2604

ATGTCTAAGTGCCATTTTTGTTTTAATTTGTCAAAATTCAATAACATCATGTTGCTCGCGAACTTTGCTTTGGCGCTATAGTCAACTTTCATCATTGTTAGGTGGATAGTCTCTTAACTG

>M6-PGSC0003DMG400009068-(rev)M6LRR-RLK_2666

CTTTGCTTTGGCGCTATAGTCAACTTTCATCATTGTTAGGTGGATAGTCTCTTAACTGCACCTTCAAGTATTCCTATCATTATATCTGATGCACTCTGTAATTGCAGCAAAGAGGTCTGC

>M6-PGSC0003DMG400009068-(rev)M6LRR-RLK_2728

TTCAAGTATTCCTATCATTATATCTGATGCACTCTGTAATTGCAGCAAAGAGGTCTGCTGATCGAATTCCTCTGGACTGGGAAACCCGACTACGCATTGCCATTGGTGCAGCTAGAGGCA

>M6-PGSC0003DMG400009068-(rev)M6LRR-RLK_2790

CGAATTCCTCTGGACTGGGAAACCCGACTACGCATTGCCATTGGTGCAGCTAGAGGCATTGCTCAAATACATGGACAGTCTGGGGGGAGGCTTGTCCACGGGAACATAAAATCCTCAAAC

>M6-PGSC0003DMG400009068-(rev)M6LRR-RLK_2852

TCAAATACATGGACAGTCTGGGGGGAGGCTTGTCCACGGGAACATAAAATCCTCAAACGTATTCCTCAACTCCCAAGGTTTCGGTTGTATTTCTGATCTTGGCTTGGCAACTATAATGGG

>M6-PGSC0003DMG400009068-(rev)M6LRR-RLK_2914

TCCTCAACTCCCAAGGTTTCGGTTGTATTTCTGATCTTGGCTTGGCAACTATAATGGGTCCAATTGCTACACCAATTGTGCGAGCTGCTGGTTATCAACCTCCAGAAGTTACAGATTCAA

>M6-PGSC0003DMG400009068-(rev)M6LRR-RLK_2976

ATTGCTACACCAATTGTGCGAGCTGCTGGTTATCAACCTCCAGAAGTTACAGATTCAAGAAAAGTATCTCAAGCCACTGATGTCTACAGCTTTGGTGTCCTAATACTTGAACTTCTCACT

>M6-PGSC0003DMG400009068-(rev)M6LRR-RLK_3038

AGTATCTCAAGCCACTGATGTCTACAGCTTTGGTGTCCTAATACTTGAACTTCTCACTGGAAAGTCACCTACACATGCCACAGGTACAAGCGACATTGTCCACTTAGTCAGGTGGGTACA

>M6-PGSC0003DMG400009068-(rev)M6LRR-RLK_310

TTATTTGCTTTCTTCTTCTTCAAATTAACTGCCTTTCAATCATTTATTGCCTCTCACCCACTTCCCCCCCTTCTTCTCATTTAATCATTGTTTCTCTCTAGCTACTTACTTAATTCTGCA

>M6-PGSC0003DMG400009068-(rev)M6LRR-RLK_3100

AGTCACCTACACATGCCACAGGTACAAGCGACATTGTCCACTTAGTCAGGTGGGTACATTCTGTCGTTCGTGAGGAGTGGACTGCTGAAGTATTCGACGTGGAGCTTTTGAGGTATCCAA

>M6-PGSC0003DMG400009068-(rev)M6LRR-RLK_3162

GTCGTTCGTGAGGAGTGGACTGCTGAAGTATTCGACGTGGAGCTTTTGAGGTATCCAAATATAGAAGAAGAAATGGTAGAGATGTTACAAATAGGTTTGACTTGTGTATCCAGAATGCCA

>M6-PGSC0003DMG400009068-(rev)M6LRR-RLK_3224

AGAAGAAGAAATGGTAGAGATGTTACAAATAGGTTTGACTTGTGTATCCAGAATGCCAGAGCAGAGACCAAAAATGACTGAAGTAGTAAAGATGGTAGAGGGTGTTAGGAGAGTGAATAC

>M6-PGSC0003DMG400009068-(rev)M6LRR-RLK_3286

AGAGACCAAAAATGACTGAAGTAGTAAAGATGGTAGAGGGTGTTAGGAGAGTGAATACGGGGACTCGAACATCCACTGAAGCTTCTACACCGAACCTGACACCACCGATGACTGAAATAG

>M6-PGSC0003DMG400009068-(rev)M6LRR-RLK_3348

ACTCGAACATCCACTGAAGCTTCTACACCGAACCTGACACCACCGATGACTGAAATAGGGTCGTCTTCCATTACACACTGATATTTCTTTGTTATTTAGTGATTAGTTACTGTTCTATAT

>M6-PGSC0003DMG400009068-(rev)M6LRR-RLK_3410

GTCTTCCATTACACACTGATATTTCTTTGTTATTTAGTGATTAGTTACTGTTCTATATATGCTTAATTGTTTCATTGTTATAAGTTTTTTCCTCAGACTTCTGTCAAACATTATTTCAGC

>M6-PGSC0003DMG400009068-(rev)M6LRR-RLK_3441

ATTTAGTGATTAGTTACTGTTCTATATATGCTTAATTGTTTCATTGTTATAAGTTTTTTCCTCAGACTTCTGTCAAACATTATTTCAGCTACAAAATTGAATAGAAATATTTACTTGATC

>M6-PGSC0003DMG400009068-(rev)M6LRR-RLK_372

TCCCCCCCTTCTTCTCATTTAATCATTGTTTCTCTCTAGCTACTTACTTAATTCTGCATTTACTTCCCATCATTTTCCTCTTCATCTACTCCTTCTTTTCAGTTTACTCTGCCAAGTGTT

>M6-PGSC0003DMG400009068-(rev)M6LRR-RLK_434

CTTCCCATCATTTTCCTCTTCATCTACTCCTTCTTTTCAGTTTACTCTGCCAAGTGTTTGATTTTTTGCCTCACTGGTACTTGCTTGTATACTATACAGTTTCTGAATTTATGGGGTTTT

>M6-PGSC0003DMG400009068-(rev)M6LRR-RLK_496

TTTTTGCCTCACTGGTACTTGCTTGTATACTATACAGTTTCTGAATTTATGGGGTTTTCCGTATTCTCCATTTTGATGTGATTTGGCTATCTGGGTATTGCTATTCAGGTTCATTTCAAT

>M6-PGSC0003DMG400009068-(rev)M6LRR-RLK_558

ATTCTCCATTTTGATGTGATTTGGCTATCTGGGTATTGCTATTCAGGTTCATTTCAATGTTCTTTTTGTTATTGCCGACAGTTTTCAAATTATGCATACTTTGCTTTAAGGTCAAGAAAT

>M6-PGSC0003DMG400009068-(rev)M6LRR-RLK_62

GTGAAGAGCTTCAAAAAATGGCGGAAATGGTGGATAATGCTCTTCTGGTGCTCCCTTGCCTTACTGGCACCACGTGCACCCACTTCTCCTCTTCTTCTAAATCCAAGCAAATTGTTAAAG

>M6-PGSC0003DMG400009068-(rev)M6LRR-RLK_620

TTTTTGTTATTGCCGACAGTTTTCAAATTATGCATACTTTGCTTTAAGGTCAAGAAATGTGTTAAAAATTCAATCTTTGTATGTGTGTGTGCTTGGAAATAGGTGAAGAACGTGAAATGC

>M6-PGSC0003DMG400009068-(rev)M6LRR-RLK_806

ATTTGTAGCTGGTTCTTGTTTTTACTTTAGCTGTTCTGTTCATTGAAACCTAACTTTTACTTAACTTGCGTGGTTGACCCAAAAACAAAATGTGTTCTGAGTGTCTACGTAACTTTAGTT

>M6-PGSC0003DMG400009068-(rev)M6LRR-RLK_868

AACTTGCGTGGTTGACCCAAAAACAAAATGTGTTCTGAGTGTCTACGTAACTTTAGTTGCCCTATAAATTGTATATTTATGGTTAAGAAACGGTTCTCTAGTTTAAATGGTAATTATTTA

>M6-PGSC0003DMG400009068-(rev)M6LRR-RLK_930

TATAAATTGTATATTTATGGTTAAGAAACGGTTCTCTAGTTTAAATGGTAATTATTTATAGTGAATAATGAACATCATGCCGTTTCAATGTCTGATTCCATTTTTATTTTTTTTTGAAAT

>M6-PGSC0003DMG400009068-(rev)M6LRR-RLK_992

GAATAATGAACATCATGCCGTTTCAATGTCTGATTCCATTTTTATTTTTTTTTGAAATTCTTGTAGAGATGAAGGGTGCTGTTACTAGAAGTTGATGCCTTTGTAGTTTAGCTCAGAGGA

>M6-PGSC0003DMG400009114M6RLKWAX-EGF_0

CCACCTTTCCTCAGACTCCACTCGTGGGACTACACTGAGTATGTTGTTGCACTATGTTCAACTAGGAATACCTTTTCGAATTATCTGTTTTAAGATGTGGTTTATTCAGATTGTAAATGG

>M6-PGSC0003DMG400009114M6RLKWAX-EGF_1037

GATGACTAACAAATTCAAGGAAAAACTAGGCGAAGGAGCCTATGGAACTGTCTTTAAAGGAAAACTGCGTAGTGGTCCTTTTGTTGCAGTGAAGATGATGCACCAATCTATGGCGAGTGG

>M6-PGSC0003DMG400009114M6RLKWAX-EGF_1098

AAACTGCGTAGTGGTCCTTTTGTTGCAGTGAAGATGATGCACCAATCTATGGCGAGTGGGCAAGAATTTATCAGTGAAGTTTCCACAATTGGAAGGATTCACCATGTTAACATTGTGCAA

>M6-PGSC0003DMG400009114M6RLKWAX-EGF_1159

AAGAATTTATCAGTGAAGTTTCCACAATTGGAAGGATTCACCATGTTAACATTGTGCAACTCGTTGGATTCTGTGTAGAGGGATCAAAACGTGCTTTGGTTTATGAGTTCATGCTCAATG

>M6-PGSC0003DMG400009114M6RLKWAX-EGF_122

GTAAAATTATTGAAATATGGTATTTTGATAACAGAAATATCTCCTTTCTGGCATCCGAGCAATTTAACTGTACTTCCAAGTTTCAAAGGTGTCATGATTTACAACAGATTAGGTTGTCAA

>M6-PGSC0003DMG400009114M6RLKWAX-EGF_1220

CGTTGGATTCTGTGTAGAGGGATCAAAACGTGCTTTGGTTTATGAGTTCATGCTCAATGGTTCTTTAGACAAGTACATATCTCCCCGAGAAGGAACTGTTTCCTTGAGTTACAAGCAAAT

>M6-PGSC0003DMG400009114M6RLKWAX-EGF_1281

TCTTTAGACAAGTACATATCTCCCCGAGAAGGAACTGTTTCCTTGAGTTACAAGCAAATGTTCGACATATCTCTTGGAGTGGCCTCTGGCATAGACTACCTACATCGGGGCTGTGACATG

>M6-PGSC0003DMG400009114M6RLKWAX-EGF_1342

TCGACATATCTCTTGGAGTGGCCTCTGGCATAGACTACCTACATCGGGGCTGTGACATGCAGATCTTACATTTTGATATTAAGCCTCACAACATTCTTCTTGATGAAAATTTCAATCCCA

>M6-PGSC0003DMG400009114M6RLKWAX-EGF_1464

ATATCTGACTTTGGGCTTGCGAAACTGTACTCCTCAGATGATAGTATTGTGAGTCTAACTGCAGCTAGAGGGACAATGGGTTACATGGCACCAGAGCTGTTTTATAAAAATATTGGAGGA

>M6-PGSC0003DMG400009114M6RLKWAX-EGF_1525

CAGCTAGAGGGACAATGGGTTACATGGCACCAGAGCTGTTTTATAAAAATATTGGAGGAGTGTCACACAAAGCCGATGTATATAGCTATGGCATGTTATTGATGGAAATGGCAGGCAGAA

>M6-PGSC0003DMG400009114M6RLKWAX-EGF_1586

GTCACACAAAGCCGATGTATATAGCTATGGCATGTTATTGATGGAAATGGCAGGCAGAAGGAAGAACATGAACCCATTTGAGGACCACTTAAGCCAAATCTACTTTCCTACATGGATTTA

>M6-PGSC0003DMG400009114M6RLKWAX-EGF_1647

AAGAACATGAACCCATTTGAGGACCACTTAAGCCAAATCTACTTTCCTACATGGATTTACGACCAATTTAATGAAGGAAATGATATAGAGATACAGGATGCAAGCGAGGAAGATAGGAGA

>M6-PGSC0003DMG400009114M6RLKWAX-EGF_1708

ACCAATTTAATGAAGGAAATGATATAGAGATACAGGATGCAAGCGAGGAAGATAGGAGACTTGTCAAGAAGATGATGATTGTGGCTTTGTGGTGCATACAAATGAAGCCTGCTGATCGCC

>M6-PGSC0003DMG400009114M6RLKWAX-EGF_1769

TGTCAAGAAGATGATGATTGTGGCTTTGTGGTGCATACAAATGAAGCCTGCTGATCGCCCTGCCATGAACAAAGTAGTTGAAATGCTTGAAGGAAATGTTGAACTCCTACAGATGCCCCC

>M6-PGSC0003DMG400009114M6RLKWAX-EGF_183

ATTTAACTGTACTTCCAAGTTTCAAAGGTGTCATGATTTACAACAGATTAGGTTGTCAATCAAAGCAGGTCGAAGATGTGTCATGGTTCTTTCATTTATTATATCCATTCTCTATGGGGT

>M6-PGSC0003DMG400009114M6RLKWAX-EGF_1830

GCCATGAACAAAGTAGTTGAAATGCTTGAAGGAAATGTTGAACTCCTACAGATGCCCCCAAGACCTTTTATAGCTCCTCGTGAGATTGCTGGAGACAGTATTGAGACAATACAAATTTCC

>M6-PGSC0003DMG400009114M6RLKWAX-EGF_1891

GACCTTTTATAGCTCCTCGTGAGATTGCTGGAGACAGTATTGAGACAATACAAATTTCCAGTGACGTATAGGTGCCGGTGGTTGGCTGCTTACTGTGCTGTGCAAACTACAGTCAAAATT

>M6-PGSC0003DMG400009114M6RLKWAX-EGF_1952

TGACGTATAGGTGCCGGTGGTTGGCTGCTTACTGTGCTGTGCAAACTACAGTCAAAATTCACTTAATTTAGTTCACGGCATCCAATACTATCAGAAGAGCTCAATCTCAGGCCAACATTC

>M6-PGSC0003DMG400009114M6RLKWAX-EGF_2013

CTTAATTTAGTTCACGGCATCCAATACTATCAGAAGAGCTCAATCTCAGGCCAACATTCACTTGAGATGTCACTGTGGATTAAAAGAGTTGAACGAAAAGATGCCTATATCTTCGAGCAA

>M6-PGSC0003DMG400009114M6RLKWAX-EGF_2074

TTGAGATGTCACTGTGGATTAAAAGAGTTGAACGAAAAGATGCCTATATCTTCGAGCAAGTAGTCATTTCTGTCCCATGGTGGTGACACTAGTTCCTCCTAGTTTTTTAGGTAAGAATTT

>M6-PGSC0003DMG400009114M6RLKWAX-EGF_2135

AGTCATTTCTGTCCCATGGTGGTGACACTAGTTCCTCCTAGTTTTTTAGGTAAGAATTTGAATGTAATATGATAATTTAGTAAAGTGTAAATTAGTTTGCATGTTGGTGTTGTAAATTCT

>M6-PGSC0003DMG400009114M6RLKWAX-EGF_2196

ATGTAATATGATAATTTAGTAAAGTGTAAATTAGTTTGCATGTTGGTGTTGTAAATTCTTGGAAAAGGTAACTATCAAGCATAGATGTATCAAACAAGGAAATGATTACACCTTCAATTT

>M6-PGSC0003DMG400009114M6RLKWAX-EGF_2252

TCTTGGAAAAGGTAACTATCAAGCATAGATGTATCAAACAAGGAAATGATTACACCTTCAATTTTTTGTGTTTAAACTACCATTGATCGTCTTTATGGATTGAGAGCATCTTCTGTTTCT

>M6-PGSC0003DMG400009114M6RLKWAX-EGF_244

AAAGCAGGTCGAAGATGTGTCATGGTTCTTTCATTTATTATATCCATTCTCTATGGGGTCTGTTCTTTGAAGTTATAAGAAAATTTGGATCTCTTTTGTATTCAACTCTCCCCTGGGCCC

>M6-PGSC0003DMG400009114M6RLKWAX-EGF_305

GTTCTTTGAAGTTATAAGAAAATTTGGATCTCTTTTGTATTCAACTCTCCCCTGGGCCCTCTCTTTATGCGTATCAGTTTCAATTCTCAGCAACTTATTTTGAGATGTAGCATCTGTATT

>M6-PGSC0003DMG400009114M6RLKWAX-EGF_366

TCTTTATGCGTATCAGTTTCAATTCTCAGCAACTTATTTTGAGATGTAGCATCTGTATTTTACTCATATTCCTATATTTGGACAAATGCAGGTAGCATCCACTACTATGGCGGTACGTAT

>M6-PGSC0003DMG400009114M6RLKWAX-EGF_427

ACTCATATTCCTATATTTGGACAAATGCAGGTAGCATCCACTACTATGGCGGTACGTATTTGAGATACTGGAGTGCAATAACTTCATTACACTAAGCATTTCATTGAAAGGAGATCGTTA

>M6-PGSC0003DMG400009114M6RLKWAX-EGF_488

GAGATACTGGAGTGCAATAACTTCATTACACTAAGCATTTCATTGAAAGGAGATCGTTACAGCAATATGTTTACCATATCTAGAATCTGCGTTCATTGCACTATATAGCTGTAAGCTTTT

>M6-PGSC0003DMG400009114M6RLKWAX-EGF_549

GCAATATGTTTACCATATCTAGAATCTGCGTTCATTGCACTATATAGCTGTAAGCTTTTTCTTCAAAACTTTAATGATTTTACATCCTCTAACTTTATGTTTCCTTTGTGATTTGCCTGT

>M6-PGSC0003DMG400009114M6RLKWAX-EGF_61

CTAGGAATACCTTTTCGAATTATCTGTTTTAAGATGTGGTTTATTCAGATTGTAAATGGTGGTAAAATTATTGAAATATGGTATTTTGATAACAGAAATATCTCCTTTCTGGCATCCGAG

>M6-PGSC0003DMG400009114M6RLKWAX-EGF_610

TTCAAAACTTTAATGATTTTACATCCTCTAACTTTATGTTTCCTTTGTGATTTGCCTGTTTTGACAGTTCTTACTATATTGTACGGTGGAATAGTTATAGGTGAGTCTTCTTATCTTACT

>M6-PGSC0003DMG400009114M6RLKWAX-EGF_671

TGACAGTTCTTACTATATTGTACGGTGGAATAGTTATAGGTGAGTCTTCTTATCTTACTCAAGGATAAGCATTCTTCGCACACTCAGAATTCTGAAAAATAATTAGCTTTTGAAGCTTTT

>M6-PGSC0003DMG400009114M6RLKWAX-EGF_732

AGGATAAGCATTCTTCGCACACTCAGAATTCTGAAAAATAATTAGCTTTTGAAGCTTTTCTCAACCCATTTACCCCTATGAAAAAGATGTACTGGTTTTCTTGTCGATGACTTCCATTTT

>M6-PGSC0003DMG400009114M6RLKWAX-EGF_793

CAACCCATTTACCCCTATGAAAAAGATGTACTGGTTTTCTTGTCGATGACTTCCATTTTTATTTTTGAAATCATGCAGTATCAATTCTATCGTTACGACTTCTCTTTGGATTGGTGTTCC

>M6-PGSC0003DMG400009114M6RLKWAX-EGF_854

TTTTTGAAATCATGCAGTATCAATTCTATCGTTACGACTTCTCTTTGGATTGGTGTTCCTGACTGCATTAATAGTGTATAAATGGAGAAGACGACACTTATCTATGTATCAATCAATTGA

>M6-PGSC0003DMG400009114M6RLKWAX-EGF_915

ACTGCATTAATAGTGTATAAATGGAGAAGACGACACTTATCTATGTATCAATCAATTGAAGACTTCCTGCAAGTCCAGAGCAACCTCTTGCCGATTAAATACTCCTACTCTGAGATTAAA

>M6-PGSC0003DMG400009114M6RLKWAX-EGF_976

ACTTCCTGCAAGTCCAGAGCAACCTCTTGCCGATTAAATACTCCTACTCTGAGATTAAAAAGATGACTAACAAATTCAAGGAAAAACTAGGCGAAGGAGCCTATGGAACTGTCTTTAAAG

>M6-PGSC0003DMG400009180M6LRR-RLK_0

ATTAATCAAAATAAAAATAGGAAAGTTCTGACAAAGCCTTTCTTCTGTTCCCGAGCTGAAGCTTGGGGTTTTGTGTTTTGTGGTGCCCGGTACTGGTGGTGAAAGAAAACTCCGACGACT

>M6-PGSC0003DMG400009180M6LRR-RLK_1037

TGGACAGCTAACAAAATGGAATGCCAATGGTGGTGATCCTTGTGGAGAATCCTGGACAGGTATTACTTGCTCGGGCAATAGAGTTACTGAAATGTGAGTTCCTCTCTTCCTTACTGTTGT

>M6-PGSC0003DMG400009180M6LRR-RLK_1098

ATTACTTGCTCGGGCAATAGAGTTACTGAAATGTGAGTTCCTCTCTTCCTTACTGTTGTAAGTAATTATTGTTGATGTTTTCCTTCTTTATCTTCTTCATCATGTTTCAATACTTGGAAT

>M6-PGSC0003DMG400009180M6LRR-RLK_1159

GTAATTATTGTTGATGTTTTCCTTCTTTATCTTCTTCATCATGTTTCAATACTTGGAATTGTTGACTTGGTGATTTCTGATGCTTTCTTCCCTTTTCATTTTCCTCCAGCAAGATATCAC

>M6-PGSC0003DMG400009180M6LRR-RLK_122

GACTGAGGTGGAAGAGGAGAAAACGACGCCGTGTAGATGAGGTTGCTGCTGCTGTTGCCGATACTCATCGGCATTTTGCTTTATTGTGCTAATTATGCCATTGCCGACACAGATCCAAAT

>M6-PGSC0003DMG400009180M6LRR-RLK_1220

TTGACTTGGTGATTTCTGATGCTTTCTTCCCTTTTCATTTTCCTCCAGCAAGATATCACGTCTTGGACTCTCTGGTTCACTGGGCTACCAATTGGCAAGTCTTACATCGGTGACTAATTT

>M6-PGSC0003DMG400009180M6LRR-RLK_1281

CTTGGACTCTCTGGTTCACTGGGCTACCAATTGGCAAGTCTTACATCGGTGACTAATTTGTAAGATAAATGGCCTTCAATAATGTCCTCATAAATGTTTCTTCGATCTATATCTATATCA

>M6-PGSC0003DMG400009180M6LRR-RLK_1342

AAGATAAATGGCCTTCAATAATGTCCTCATAAATGTTTCTTCGATCTATATCTATATCAAATTCTTACACACTTCTAATTTTGCAGTGACATAAGTAATAACAATTTGGGAAACCAGTTA

>M6-PGSC0003DMG400009180M6LRR-RLK_1403

TTCTTACACACTTCTAATTTTGCAGTGACATAAGTAATAACAATTTGGGAAACCAGTTACCTTATCAGCTTCCTCCAAATGTGCAAAGATTGTATGTCTCGTAGTTAGTATATTTTCATC

>M6-PGSC0003DMG400009180M6LRR-RLK_1464

TTATCAGCTTCCTCCAAATGTGCAAAGATTGTATGTCTCGTAGTTAGTATATTTTCATCATTCCAGATGCTATTTTTCCACCTATCTTTCTTAAACCTTGCCTTGCAATGCAGAAACCTT

>M6-PGSC0003DMG400009180M6LRR-RLK_1525

TCCAGATGCTATTTTTCCACCTATCTTTCTTAAACCTTGCCTTGCAATGCAGAAACCTTGCTGCTAATGGTTTTAATGGTGCCCTTCCATATTCCGTTTCGCAGATGACTTCCCTCCGAT

>M6-PGSC0003DMG400009180M6LRR-RLK_1586

TGCTAATGGTTTTAATGGTGCCCTTCCATATTCCGTTTCGCAGATGACTTCCCTCCGATACTTGTATGAAAAATCTTTGAATTTGACTGCAATTTGCTTTCCAGCAATAAGCTTTTGGTT

>M6-PGSC0003DMG400009180M6LRR-RLK_1647

TTGTATGAAAAATCTTTGAATTTGACTGCAATTTGCTTTCCAGCAATAAGCTTTTGGTTTAAGGTTGCGCTTTTCAATGACTTTAATATTTTTATCTGGCTGTAGAAATGTCAGTCATAA

>M6-PGSC0003DMG400009180M6LRR-RLK_1708

AGGTTGCGCTTTTCAATGACTTTAATATTTTTATCTGGCTGTAGAAATGTCAGTCATAATCAAATTCAAGGACAAGTGACTGTCGCGTTCGACTCTCTGTCTTCTCTCGATACATTGTAA

>M6-PGSC0003DMG400009180M6LRR-RLK_1769

AAATTCAAGGACAAGTGACTGTCGCGTTCGACTCTCTGTCTTCTCTCGATACATTGTAAGCCCTATCTTCTCTCATGCCATTTCTTGAGTGTTTTAATATTACAAAGTTGTGGTCATAAC

>M6-PGSC0003DMG400009180M6LRR-RLK_183

TACTCATCGGCATTTTGCTTTATTGTGCTAATTATGCCATTGCCGACACAGATCCAAATGACGGTCAGCTATTGCTTTTAATCATTCTCATTTATTGTTTCAATCTGTTTGTCTGTATGC

>M6-PGSC0003DMG400009180M6LRR-RLK_1830

CCTATCTTCTCTCATGCCATTTCTTGAGTGTTTTAATATTACAAAGTTGTGGTCATAACTCATATATGAGGTGTTGTAGATAAAACATGTCATTCTTACTTCATTAGTATAAACTTTTCT

>M6-PGSC0003DMG400009180M6LRR-RLK_1891

ATATATGAGGTGTTGTAGATAAAACATGTCATTCTTACTTCATTAGTATAAACTTTTCTTCTCGCATCCGAGTTGTTGACTTGTTGCTTTGGGAAAGAAAGTCAATATCATGAAAGACAG

>M6-PGSC0003DMG400009180M6LRR-RLK_1952

TCGCATCCGAGTTGTTGACTTGTTGCTTTGGGAAAGAAAGTCAATATCATGAAAGACAGTATCTGATCCTTGTTGTAATCTGTAAACTTATGTGATTGTCCCAGTTAAGTCAACTAAGAA

>M6-PGSC0003DMG400009180M6LRR-RLK_2013

TCTGATCCTTGTTGTAATCTGTAAACTTATGTGATTGTCCCAGTTAAGTCAACTAAGAAGCTGTATACATTTCGAGATATTCTTAAGTCAAGTCTTAGAAACCTCTAAATGACTAGTTTA

>M6-PGSC0003DMG400009180M6LRR-RLK_2074

TGTATACATTTCGAGATATTCTTAAGTCAAGTCTTAGAAACCTCTAAATGACTAGTTTATCATTCTGTCTCATAGTTTTTGATTGATTGATCTGTACGCTTCAGGGATTTCTCATTCAAT

>M6-PGSC0003DMG400009180M6LRR-RLK_2135

ATTCTGTCTCATAGTTTTTGATTGATTGATCTGTACGCTTCAGGGATTTCTCATTCAATGCTATGACGGGTGATCTTCCTCAAAGCTTCAAGACACTGACTAGTATGAATAAAATGTGAG

>M6-PGSC0003DMG400009180M6LRR-RLK_2196

TATGACGGGTGATCTTCCTCAAAGCTTCAAGACACTGACTAGTATGAATAAAATGTGAGCTAATAAATTTTCTGCTTCGTCCTTGTAGACAATTGGTTGAGTCCTTCAGTGGTGAAGTTT

>M6-PGSC0003DMG400009180M6LRR-RLK_2257

AATAAATTTTCTGCTTCGTCCTTGTAGACAATTGGTTGAGTCCTTCAGTGGTGAAGTTTACATGTTGAAGTATGCAGGTATTTGCAGAACAACCAGTTTACAGGAACTATTGATATCTTA

>M6-PGSC0003DMG400009180M6LRR-RLK_2318

ATGTTGAAGTATGCAGGTATTTGCAGAACAACCAGTTTACAGGAACTATTGATATCTTAGCCAATCTTCCTCTTGATGATTTGTGAGTTTTCAGTCTATGCTTCTTTTTAGATGTTATCC

>M6-PGSC0003DMG400009180M6LRR-RLK_2379

CAATCTTCCTCTTGATGATTTGTGAGTTTTCAGTCTATGCTTCTTTTTAGATGTTATCCCTGATGTTTTATCTTATTTATATAGACTCGATTGAAAGCATGGTTGAAAGTCATGTGTATT

>M6-PGSC0003DMG400009180M6LRR-RLK_244

CGGTCAGCTATTGCTTTTAATCATTCTCATTTATTGTTTCAATCTGTTTGTCTGTATGCTTACTGATCATTAATGATTTAATGGAGGTTGATTGATTTGGAAAGTCAATGAGTATCAATC

>M6-PGSC0003DMG400009180M6LRR-RLK_2440

GATGTTTTATCTTATTTATATAGACTCGATTGAAAGCATGGTTGAAAGTCATGTGTATTCCTTTATGCTTTAGGAATGTTGAGAACAACCGTTTTACTGGCTGGGTTCCTGATCACTTGA

>M6-PGSC0003DMG400009180M6LRR-RLK_2501

TTTATGCTTTAGGAATGTTGAGAACAACCGTTTTACTGGCTGGGTTCCTGATCACTTGAAAGGGATCACGAAGTAAGTATATCATGTTAGAGTAATAAATTTTGAAACTTATATCAGAAG

>M6-PGSC0003DMG400009180M6LRR-RLK_2562

GGGATCACGAAGTAAGTATATCATGTTAGAGTAATAAATTTTGAAACTTATATCAGAAGTATTTTGAATTTATTCAGGGTTTGTGACATTTCTGTTACAGATCAAATGGTAACTCATGGA

>M6-PGSC0003DMG400009180M6LRR-RLK_2623

TTTTGAATTTATTCAGGGTTTGTGACATTTCTGTTACAGATCAAATGGTAACTCATGGAACTCAGGATCTGCACCCCCTCCTCCACCTGGCACACCCCCTGCAAGCAGGCCGCACCACAA

>M6-PGSC0003DMG400009180M6LRR-RLK_2684

TCAGGATCTGCACCCCCTCCTCCACCTGGCACACCCCCTGCAAGCAGGCCGCACCACAAATCTGGAGGCAATAATAGCCCTACTGATGGTGCAAGCAGTGGTGATGGTGGTGGTAAATCG

>M6-PGSC0003DMG400009180M6LRR-RLK_2745

CTGGAGGCAATAATAGCCCTACTGATGGTGCAAGCAGTGGTGATGGTGGTGGTAAATCGGGGATAGGGGGTGGAGCCATTGCAGGCATAGTGATATCTGTTCTAGTTGTTGGGGCAATTG

>M6-PGSC0003DMG400009180M6LRR-RLK_2806

GATAGGGGGTGGAGCCATTGCAGGCATAGTGATATCTGTTCTAGTTGTTGGGGCAATTGTAGCATTCTTTGTTATCAAGAAGAGATTGAAGCGGTCATCGACGGACATTGAGAAACATGA

>M6-PGSC0003DMG400009180M6LRR-RLK_2867

GCATTCTTTGTTATCAAGAAGAGATTGAAGCGGTCATCGACGGACATTGAGAAACATGATAATCAGCCATTTGCTCCTCTTGCTCCCCCCCAGGAAGTACATGGTACTTCTTTTCATAAT

>M6-PGSC0003DMG400009180M6LRR-RLK_2928

ATCAGCCATTTGCTCCTCTTGCTCCCCCCCAGGAAGTACATGGTACTTCTTTTCATAATTTCTTGGTTTTATCAGTTAAGATGTCTTTTAGACCCTTTCATGGCCTCTGTGATTGTTATT

>M6-PGSC0003DMG400009180M6LRR-RLK_2989

CTTGGTTTTATCAGTTAAGATGTCTTTTAGACCCTTTCATGGCCTCTGTGATTGTTATTTAACTGTGGAACATATCAATCAACCACACTATCATCAACCTCATTCCTGTATCCTGCTTAA

>M6-PGSC0003DMG400009180M6LRR-RLK_305

ACTGATCATTAATGATTTAATGGAGGTTGATTGATTTGGAAAGTCAATGAGTATCAATCTACCTGACAATGAACTCGGCAACCGTCACAAATTTGTTGATATTTTCCTTACCTGGATAGT

>M6-PGSC0003DMG400009180M6LRR-RLK_3050

ACTGTGGAACATATCAATCAACCACACTATCATCAACCTCATTCCTGTATCCTGCTTAAAAAGGTGTAAAATAAACCTGCTGCATAAAATGGAAATTTTATGAAAGTGAATTTGGAGAAT

>M6-PGSC0003DMG400009180M6LRR-RLK_3111

AGGTGTAAAATAAACCTGCTGCATAAAATGGAAATTTTATGAAAGTGAATTTGGAGAATCAATCTTATCCTTTGTGCTTTATCACTGCCTGTGGTGTATGCCTAATAGTGGATCAACACA

>M6-PGSC0003DMG400009180M6LRR-RLK_3172

ATCTTATCCTTTGTGCTTTATCACTGCCTGTGGTGTATGCCTAATAGTGGATCAACACAGCCACTGTCTTCGCTATAAAAGAAAAACACCAAAGTAAAAACAAACTTTATATCGAAATGT

>M6-PGSC0003DMG400009180M6LRR-RLK_3233

CACTGTCTTCGCTATAAAAGAAAAACACCAAAGTAAAAACAAACTTTATATCGAAATGTAAAACTGCTAAGGCTGATAGGTCCTATCTTCCATGGAAGCAAAACTTCTTGCTTACTGAAA

>M6-PGSC0003DMG400009180M6LRR-RLK_3294

AACTGCTAAGGCTGATAGGTCCTATCTTCCATGGAAGCAAAACTTCTTGCTTACTGAAAAGTTCCAAGAACTTGATTGTAGAAATCTGAACTGATGTTTATACTCTGTATTGCCTGTAGA

>M6-PGSC0003DMG400009180M6LRR-RLK_3355

TTCCAAGAACTTGATTGTAGAAATCTGAACTGATGTTTATACTCTGTATTGCCTGTAGATGCTGATAGTGGGATAAAGAGAACTTCAGATTTTTTGATAATGAAGAAAACTTCAGCTTTA

>M6-PGSC0003DMG400009180M6LRR-RLK_3416

CTGATAGTGGGATAAAGAGAACTTCAGATTTTTTGATAATGAAGAAAACTTCAGCTTTATGATCTCTTTCAGTGGATCTGGTCCTATGCTTGTTATAAGTTAGCTCCACTTGATGGTTTT

>M6-PGSC0003DMG400009180M6LRR-RLK_3477

ATCTCTTTCAGTGGATCTGGTCCTATGCTTGTTATAAGTTAGCTCCACTTGATGGTTTTCCTTCCTATGAAACTACTATTCATAAATCTTATTGACCTTTATAGAACAACTATTGCTTCA

>M6-PGSC0003DMG400009180M6LRR-RLK_3538

TTCCTATGAAACTACTATTCATAAATCTTATTGACCTTTATAGAACAACTATTGCTTCAAAATGCAACTTTCAGCTAAGTAAAAGTTTCTCCCATTCTAAACCATTTGTGCTCACCACCA

>M6-PGSC0003DMG400009180M6LRR-RLK_3599

ATGCAACTTTCAGCTAAGTAAAAGTTTCTCCCATTCTAAACCATTTGTGCTCACCACCAAATTCACCTCTTTGCAAATTCTAATTCTTTGCTTTTCTCAACTCCATCACTCGTGGTGCAT

>M6-PGSC0003DMG400009180M6LRR-RLK_366

CCTGACAATGAACTCGGCAACCGTCACAAATTTGTTGATATTTTCCTTACCTGGATAGTATATTGTCTTATTGTATATGTAAAAATGCGACAAGTTCCTTTCTAATCAAGGCATGAATCA

>M6-PGSC0003DMG400009180M6LRR-RLK_3660

TTCACCTCTTTGCAAATTCTAATTCTTTGCTTTTCTCAACTCCATCACTCGTGGTGCATAGTTATGCCTTTTAAGCCTCATCTCACTTCCGTATCTCATTAATCCGAACCTGAGTATGCA

>M6-PGSC0003DMG400009180M6LRR-RLK_3721

TTATGCCTTTTAAGCCTCATCTCACTTCCGTATCTCATTAATCCGAACCTGAGTATGCATTTGCCTGTTTGTAACACTTACAGTCTTGTGCTTCTTCTCTGTCTGGATTTTCTGTTCATT

>M6-PGSC0003DMG400009180M6LRR-RLK_3782

TGCCTGTTTGTAACACTTACAGTCTTGTGCTTCTTCTCTGTCTGGATTTTCTGTTCATTCATTGTCCTTTTTTTTTACTGCAGAATTGAAAACTAATGAAGCTTCCTCTGCGCCAATAGT

>M6-PGSC0003DMG400009180M6LRR-RLK_3843

TTGTCCTTTTTTTTTACTGCAGAATTGAAAACTAATGAAGCTTCCTCTGCGCCAATAGTGAAAGTATTTGAAGCTCCTGCAGTAGTAAATCTAAGACCTCCACCTATTGAACGCCACAAG

>M6-PGSC0003DMG400009180M6LRR-RLK_3904

AAGTATTTGAAGCTCCTGCAGTAGTAAATCTAAGACCTCCACCTATTGAACGCCACAAGTCATTTGATGAAGTTGACATAGCAGCAAAACCAATTGTCCCACCCAAGAAAGTTAATACAG

>M6-PGSC0003DMG400009180M6LRR-RLK_3965

ATTTGATGAAGTTGACATAGCAGCAAAACCAATTGTCCCACCCAAGAAAGTTAATACAGCTAAAATAGATGCAAGACAATATTCAATCGCAGACCTACAGATGGCGACTGATAGCTTCAA

>M6-PGSC0003DMG400009180M6LRR-RLK_4026

AAAATAGATGCAAGACAATATTCAATCGCAGACCTACAGATGGCGACTGATAGCTTCAATGTTGATAACCTTATTGGCGAGGGATCATTTGGGCGTGTTTATAGGGCTCAGTTCGAAGAT

>M6-PGSC0003DMG400009180M6LRR-RLK_4087

TTGATAACCTTATTGGCGAGGGATCATTTGGGCGTGTTTATAGGGCTCAGTTCGAAGATGGAAAGGTATGATTTCTCTCTTATGACTGACTGACTATCTATGTGAAGAAAATGGCTCTCT

>M6-PGSC0003DMG400009180M6LRR-RLK_4148

AAAGGTATGATTTCTCTCTTATGACTGACTGACTATCTATGTGAAGAAAATGGCTCTCTCTAATTGGATAAAAAAATCATGAATTTTTTATGTGCTGAGAATGAATGTTTAAGCAAGATA

>M6-PGSC0003DMG400009180M6LRR-RLK_4209

AATTGGATAAAAAAATCATGAATTTTTTATGTGCTGAGAATGAATGTTTAAGCAAGATAGTTGAAATGACAGATAAACGTTCAATATGAAAAAGTACCTATCTAAAGAAAATGAAAGAAG

>M6-PGSC0003DMG400009180M6LRR-RLK_427

ATTGTCTTATTGTATATGTAAAAATGCGACAAGTTCCTTTCTAATCAAGGCATGAATCATATAGAGGGAGCTCTCTTGGATTCTTATATATGTTATGTATGTATTGGCCATGTCGGTGGA

>M6-PGSC0003DMG400009180M6LRR-RLK_4270

TGAAATGACAGATAAACGTTCAATATGAAAAAGTACCTATCTAAAGAAAATGAAAGAAGTTTGATCATTAAAAATTAGTTGAAAAATTATGTTTGTTCAAAACTCAGTTCGGAGCTACTA

>M6-PGSC0003DMG400009180M6LRR-RLK_4331

TGATCATTAAAAATTAGTTGAAAAATTATGTTTGTTCAAAACTCAGTTCGGAGCTACTATCATCTTGTCCTGTGTAACTGAACCATCCTCATAAGTTCCTTTCTAATTTTTGAAATGCCA

>M6-PGSC0003DMG400009180M6LRR-RLK_4392

ATCTTGTCCTGTGTAACTGAACCATCCTCATAAGTTCCTTTCTAATTTTTGAAATGCCATAGGTTCTTGCAGTCAAGAAAATCAACTCTTCTGCACTCCAGAATCCTGAAGATTTTCTCG

>M6-PGSC0003DMG400009180M6LRR-RLK_4453

GGTTCTTGCAGTCAAGAAAATCAACTCTTCTGCACTCCAGAATCCTGAAGATTTTCTCGATATTGTTTCTGAGATATCACGGTTGCACCATCCAAATGTTACTGAGCTGGTTGGATATTG

>M6-PGSC0003DMG400009180M6LRR-RLK_4514

ATTGTTTCTGAGATATCACGGTTGCACCATCCAAATGTTACTGAGCTGGTTGGATATTGTTCAGAGCACGGGCAGCATCTGCTGGTTTATGAATTCCACAAAAATGGTTCTCTGCATGAT

>M6-PGSC0003DMG400009180M6LRR-RLK_4575

CAGAGCACGGGCAGCATCTGCTGGTTTATGAATTCCACAAAAATGGTTCTCTGCATGATTTCCTGCATCTGTCAGATGAGGAAAGCAAGCCTCTAACATGGAATAGCCGAGTTAAGATTG

>M6-PGSC0003DMG400009180M6LRR-RLK_4636

CCTGCATCTGTCAGATGAGGAAAGCAAGCCTCTAACATGGAATAGCCGAGTTAAGATTGCACTAGGCACAGCAAGAGCACTAGAGTGAGTTTCCATGATCTACAAAGATATTTGATGCCA

>M6-PGSC0003DMG400009180M6LRR-RLK_4697

CTAGGCACAGCAAGAGCACTAGAGTGAGTTTCCATGATCTACAAAGATATTTGATGCCATTTTATAATGTTCACGGAGCTCATAACAAGGATTTATAATTTCCTTGTACCATTGGTTGAA

>M6-PGSC0003DMG400009180M6LRR-RLK_4758

TTATAATGTTCACGGAGCTCATAACAAGGATTTATAATTTCCTTGTACCATTGGTTGAAATACTAGTCTCTGTCATATATTTTCTGGTATGTGTAGTCAAACAATAATGAAGTTTCTTTT

>M6-PGSC0003DMG400009180M6LRR-RLK_488

TAGAGGGAGCTCTCTTGGATTCTTATATATGTTATGTATGTATTGGCCATGTCGGTGGACTTTTAGTACTGTTGAATCTTTGCATTATTACTTCCATTCATTCTGAGAACTGACATATTG

>M6-PGSC0003DMG400009180M6LRR-RLK_549

TTTAGTACTGTTGAATCTTTGCATTATTACTTCCATTCATTCTGAGAACTGACATATTGAAACTTCGTCAGACATATTTGACTTAGTTACCATTTTCAGCCAAGTTGCTCCAGAAGATTT

>M6-PGSC0003DMG400009180M6LRR-RLK_61

CTTGGGGTTTTGTGTTTTGTGGTGCCCGGTACTGGTGGTGAAAGAAAACTCCGACGACTGTGACTGAGGTGGAAGAGGAGAAAACGACGCCGTGTAGATGAGGTTGCTGCTGCTGTTGCC

>M6-PGSC0003DMG400009180M6LRR-RLK_610

ACTTCGTCAGACATATTTGACTTAGTTACCATTTTCAGCCAAGTTGCTCCAGAAGATTTTGGCTTTTCTTCTTTCTGCTGCTTGGAAGCGGGACAGGGTAGTGGGGGTGGGACGCCAAAA

>M6-PGSC0003DMG400009180M6LRR-RLK_671

GCTTTTCTTCTTTCTGCTGCTTGGAAGCGGGACAGGGTAGTGGGGGTGGGACGCCAAAACAAGTGTGATCAAGGGCATTATAACCTAAAAGACTCTTCTCCCAGTGATAATACACAAATA

>M6-PGSC0003DMG400009180M6LRR-RLK_732

AGTGTGATCAAGGGCATTATAACCTAAAAGACTCTTCTCCCAGTGATAATACACAAATAGGTTCATGGCTTCATCAACGCACTTGTTGCGTTTCACCATGTTTATTGATGTCTTGTAGAT

>M6-PGSC0003DMG400009180M6LRR-RLK_793

TTCATGGCTTCATCAACGCACTTGTTGCGTTTCACCATGTTTATTGATGTCTTGTAGATTTACCACTCGTTAACAGTTTGTCCACCGTTTGATTACTCTTCAAGCTATTATTAATGAAAA

>M6-PGSC0003DMG400009180M6LRR-RLK_854

ACCACTCGTTAACAGTTTGTCCACCGTTTGATTACTCTTCAAGCTATTATTAATGAAAAGAAACTTTTTGTTTGAACAGTATAAGAATATGTTGACCTGTAAGTATGCCTAGCTGATTTT

>M6-PGSC0003DMG400009180M6LRR-RLK_915

AACTTTTTGTTTGAACAGTATAAGAATATGTTGACCTGTAAGTATGCCTAGCTGATTTTTTTGGTGGCGCTTTGATTTTCAGCTTCTGCTCTGAGAGTTCTGTATAGCAGCCTAAATTCA

>M6-PGSC0003DMG400009180M6LRR-RLK_976

TGGTGGCGCTTTGATTTTCAGCTTCTGCTCTGAGAGTTCTGTATAGCAGCCTAAATTCACCTGGACAGCTAACAAAATGGAATGCCAATGGTGGTGATCCTTGTGGAGAATCCTGGACAG

>M6-PGSC0003DMG400009181M6LRR-RLK_0

GAGGAGAGGAGAATAGTATGGTAAAGTCTCTCTCTCTCTCCCACGTATATTACACGCTTTCAAATATCCATAGAATAAACAAAGTCCACCAGCCATCTATCTATCTAGCCGTCCCAAAAG

>M6-PGSC0003DMG400009181M6LRR-RLK_1054

GACTTCAGGGGCCAGATGCCCCAAGGGTTTGGGAGTTGCTCCAGTTTGGTTACTTTGAGAGCAGGATTCAACCATCTTTCGGGATTCATTCCTGATGATATTTATAGTGTGTCAACACTG

>M6-PGSC0003DMG400009181M6LRR-RLK_1116

AGGATTCAACCATCTTTCGGGATTCATTCCTGATGATATTTATAGTGTGTCAACACTGCAAGAAATCTCTTTACCTGGAAATAAATTTTCTGGACCCATCCCAGAAAGCATTGTCAACCT

>M6-PGSC0003DMG400009181M6LRR-RLK_1178

AAATCTCTTTACCTGGAAATAAATTTTCTGGACCCATCCCAGAAAGCATTGTCAACCTTGTCAACCTCAGAATCCTCGCACTCTATGGCAATGAATTGACAGGTTTGATCCCTCAAGATA

>M6-PGSC0003DMG400009181M6LRR-RLK_124

CGCTTTATTTCAAAAACTCATACCTCACCTTTTTGCTACATAAAAATTGCACCTTGTCAACATCAACTACACTCTCTCTCTTCTCTTCATGGGGTTAGACTAAAGTCTGCACGATCAACA

>M6-PGSC0003DMG400009181M6LRR-RLK_1240

AACCTCAGAATCCTCGCACTCTATGGCAATGAATTGACAGGTTTGATCCCTCAAGATATTGGAAGGCTCTCTAGATTGGAACAGTTGCTCCTCCATATAAACAATCTAAATGGCACTGTG

>M6-PGSC0003DMG400009181M6LRR-RLK_1302

AAGGCTCTCTAGATTGGAACAGTTGCTCCTCCATATAAACAATCTAAATGGCACTGTGCCGCCATCGCTGATGACTTGCACCCGTCTCACTGTGCTCAATTTAAGGGTCAACTTCTTGGA

>M6-PGSC0003DMG400009181M6LRR-RLK_1364

CATCGCTGATGACTTGCACCCGTCTCACTGTGCTCAATTTAAGGGTCAACTTCTTGGAAGGTGAACTCTCAGGTCTTGATTTCTCCAACCTTAGTCGACTTGGCATAATCGACCTTGGAA

>M6-PGSC0003DMG400009181M6LRR-RLK_1426

GAACTCTCAGGTCTTGATTTCTCCAACCTTAGTCGACTTGGCATAATCGACCTTGGAAATAACTTCTTCACTGGAAGCATTCCACAAAGCCTTTTTTCATGCAGGTCATTAACTGCAATC

>M6-PGSC0003DMG400009181M6LRR-RLK_1488

CTTCTTCACTGGAAGCATTCCACAAAGCCTTTTTTCATGCAGGTCATTAACTGCAATCCGTCTGGCTACTAACTATCTGACAGGAGATATCTTGCCTGGCATAACGTCTTTACAAGCTCT

>M6-PGSC0003DMG400009181M6LRR-RLK_1550

TGGCTACTAACTATCTGACAGGAGATATCTTGCCTGGCATAACGTCTTTACAAGCTCTGTCCTTCCTCTCGGTTTCCAATAAAAGCCTAACCAATTTTGCAGGGGCAATAGAAGTCCTTA

>M6-PGSC0003DMG400009181M6LRR-RLK_1612

TTCCTCTCGGTTTCCAATAAAAGCCTAACCAATTTTGCAGGGGCAATAGAAGTCCTTAAGGGTTGTAAGAATCTTACCACACTAATCCTCACCAAAAACTTTTATAATGAAACGTTGCCT

>M6-PGSC0003DMG400009181M6LRR-RLK_1674

TTGTAAGAATCTTACCACACTAATCCTCACCAAAAACTTTTATAATGAAACGTTGCCTGATAATGAGAACTTGATTGGGTCTGAAGATTTTCAAAATCTCCAAATTCTGGGTTTAGGTGG

>M6-PGSC0003DMG400009181M6LRR-RLK_1736

ATGAGAACTTGATTGGGTCTGAAGATTTTCAAAATCTCCAAATTCTGGGTTTAGGTGGTTGTAATTTCACCGGACAGATACCCACATGGCTGGTTAAACTTGGGAGGGTAGAGGTTCTAG

>M6-PGSC0003DMG400009181M6LRR-RLK_1798

AATTTCACCGGACAGATACCCACATGGCTGGTTAAACTTGGGAGGGTAGAGGTTCTAGACCTTTCTATGAATCAAATCACAGGCAAAATTCCAGGTTGGTTGGGGACTTTACAGAATCTA

>M6-PGSC0003DMG400009181M6LRR-RLK_186

TCAACTACACTCTCTCTCTTCTCTTCATGGGGTTAGACTAAAGTCTGCACGATCAACAGGACAAAGAATTGCCAAAAAAATACCCACATGGTTGATTGGAGTATCTTGGCATCTTCAACT

>M6-PGSC0003DMG400009181M6LRR-RLK_1860

TTCTATGAATCAAATCACAGGCAAAATTCCAGGTTGGTTGGGGACTTTACAGAATCTATTTTACCTGGATTTGTCTCAGAATTTTCTATATGGAGGCTTCCCAGGTGAACTTACTCAACT

>M6-PGSC0003DMG400009181M6LRR-RLK_1922

ACCTGGATTTGTCTCAGAATTTTCTATATGGAGGCTTCCCAGGTGAACTTACTCAACTGCAAAGACTCGCATCACAAGAGGCTGCTGATCAGGTCGACAGAAGTGCTTTAGAATTGCCTG

>M6-PGSC0003DMG400009181M6LRR-RLK_1984

AGACTCGCATCACAAGAGGCTGCTGATCAGGTCGACAGAAGTGCTTTAGAATTGCCTGTGTTTGTTCAGCCAAACAATGCCTCAAATCAGCAATATAATCTACTGTCAAACCTGCCACCA

>M6-PGSC0003DMG400009181M6LRR-RLK_2046

TGTTCAGCCAAACAATGCCTCAAATCAGCAATATAATCTACTGTCAAACCTGCCACCAGCTATTTACCTTGGAAACAACAACCTTGATGGCAATATTCCAACTAAGATTGGCCAGCTGAA

>M6-PGSC0003DMG400009181M6LRR-RLK_2108

TTTACCTTGGAAACAACAACCTTGATGGCAATATTCCAACTAAGATTGGCCAGCTGAAGTACATTCATGTTCTTGATCTGAGCAAGAACAACTTTACAGGAAACATTCCAGAAACAATAT

>M6-PGSC0003DMG400009181M6LRR-RLK_2170

ATTCATGTTCTTGATCTGAGCAAGAACAACTTTACAGGAAACATTCCAGAAACAATATCCAACCTCACTAATCTGGAGAAACTAGACCTCTCTGCCAACAACCTCTCAGGTGAAATTCCT

>M6-PGSC0003DMG400009181M6LRR-RLK_2232

CCTCACTAATCTGGAGAAACTAGACCTCTCTGCCAACAACCTCTCAGGTGAAATTCCTTCTTCACTCAAGGGTCTTCATTTTTTGTCTTCTTTCAGTGTTGCTCACAACAATCTTGAGGG

>M6-PGSC0003DMG400009181M6LRR-RLK_2294

CACTCAAGGGTCTTCATTTTTTGTCTTCTTTCAGTGTTGCTCACAACAATCTTGAGGGACCTATTCCAACAGGAGGTCAGTTTGACACATTTCCTATCACTAGTTTCTTAGGTAATCCAG

>M6-PGSC0003DMG400009181M6LRR-RLK_2356

ATTCCAACAGGAGGTCAGTTTGACACATTTCCTATCACTAGTTTCTTAGGTAATCCAGGACTGTGTGGTCAAATCCTGCAGCACCCTTGCCCTGATCGATCAGGAACTACACAACCTTCT

>M6-PGSC0003DMG400009181M6LRR-RLK_2418

GTGTGGTCAAATCCTGCAGCACCCTTGCCCTGATCGATCAGGAACTACACAACCTTCTGCAGTTAGAAAAACCGCAAAAAGGAAAATCTTAATTGGACTCATCCTAGGGATCTCCTTTGG

>M6-PGSC0003DMG400009181M6LRR-RLK_248

AAAGAATTGCCAAAAAAATACCCACATGGTTGATTGGAGTATCTTGGCATCTTCAACTGTCTTTCTTCTTCATCTTTAGTTGTCTAAAAATAATAATGATCATGAATAATAAAGACAGCC

>M6-PGSC0003DMG400009181M6LRR-RLK_2480

TTAGAAAAACCGCAAAAAGGAAAATCTTAATTGGACTCATCCTAGGGATCTCCTTTGGCATTGCCTTCACAGTCATCATCATAGCATTCTGGATATTCTCCAAGAGGAGGATCCTTCCAA

>M6-PGSC0003DMG400009181M6LRR-RLK_2542

GCCTTCACAGTCATCATCATAGCATTCTGGATATTCTCCAAGAGGAGGATCCTTCCAAGAGGTGACGCCGAGAAAAATGACATGGAAATTGTGTCCTACAACTCCACTTCTGGATTGTCT

>M6-PGSC0003DMG400009181M6LRR-RLK_2604

TGACGCCGAGAAAAATGACATGGAAATTGTGTCCTACAACTCCACTTCTGGATTGTCTGCTGAGATTGGAAAGGATAACAGCATGCTTGTTATGTTTCCAACCAATAAGGATCAGATTAA

>M6-PGSC0003DMG400009181M6LRR-RLK_2666

AGATTGGAAAGGATAACAGCATGCTTGTTATGTTTCCAACCAATAAGGATCAGATTAATGATCTCACAATTTTTGACATTCTGAGAGCCACGAATAACTTCAACCAAGCAAACATAGTAG

>M6-PGSC0003DMG400009181M6LRR-RLK_2728

CTCACAATTTTTGACATTCTGAGAGCCACGAATAACTTCAACCAAGCAAACATAGTAGGTTGTGGAGGCTTTGGCCTGGTCTATAAAGCAACTTTAGCAGATGGAACTACACTGGCTGTT

>M6-PGSC0003DMG400009181M6LRR-RLK_2790

TGGAGGCTTTGGCCTGGTCTATAAAGCAACTTTAGCAGATGGAACTACACTGGCTGTTAAGAAACTTTCAGGAGATATGGGTTTAATTGAAAGAGAATTTAAAGCAGAGGTGGAAGTTTT

>M6-PGSC0003DMG400009181M6LRR-RLK_2852

AACTTTCAGGAGATATGGGTTTAATTGAAAGAGAATTTAAAGCAGAGGTGGAAGTTTTATCCACTGCCCAACATGAGAATTTGGTCTCTCTTCAAGGTTACTGTGTGCATGATGGCTGTA

>M6-PGSC0003DMG400009181M6LRR-RLK_2914

ACTGCCCAACATGAGAATTTGGTCTCTCTTCAAGGTTACTGTGTGCATGATGGCTGTAGGTTGCTGATCTATTCCTATATGCAAAACGGAAGTCTGGACTACTGGTTGCACGAGAAAACT

>M6-PGSC0003DMG400009181M6LRR-RLK_2976

GCTGATCTATTCCTATATGCAAAACGGAAGTCTGGACTACTGGTTGCACGAGAAAACTGATGGAGCATTCCTACTTGATTGGCCAACTCGACTGAAGATTGCACAGGGGGCAAGTTGTGG

>M6-PGSC0003DMG400009181M6LRR-RLK_3038

GAGCATTCCTACTTGATTGGCCAACTCGACTGAAGATTGCACAGGGGGCAAGTTGTGGGCTGGCTTACATGCATCAGATATGTGAACCACATATTGTTCATCGTGACATAAAGTCCAGTA

>M6-PGSC0003DMG400009181M6LRR-RLK_3100

GCTTACATGCATCAGATATGTGAACCACATATTGTTCATCGTGACATAAAGTCCAGTAACATCCTCCTTGATGAAAAGTTTAAAGCACATGTGGCAGATTTCGGATTGTCCAGAATGATT

>M6-PGSC0003DMG400009181M6LRR-RLK_3162

CCTCCTTGATGAAAAGTTTAAAGCACATGTGGCAGATTTCGGATTGTCCAGAATGATTCTTCCTTACCAAACTCACGTCACCACTGAACTGGTTGGTACCCTCGGCTACATCCCACCAGA

>M6-PGSC0003DMG400009181M6LRR-RLK_3224

CTTACCAAACTCACGTCACCACTGAACTGGTTGGTACCCTCGGCTACATCCCACCAGAGTATAGTCAATCATGGATAGCCACTTTGAGAGGAGATGTCTATAGTTTTGGAGTTGTGATGT

>M6-PGSC0003DMG400009181M6LRR-RLK_3286

AGTCAATCATGGATAGCCACTTTGAGAGGAGATGTCTATAGTTTTGGAGTTGTGATGTTAGAACTCCTGGCTGGCAGAAGACCTGTGGACATGAGCAAACCAAAGATGTCAAGGGAATTG

>M6-PGSC0003DMG400009181M6LRR-RLK_3348

ACTCCTGGCTGGCAGAAGACCTGTGGACATGAGCAAACCAAAGATGTCAAGGGAATTGGTTGTATGGGTGCACCTAATGAGAAATGAGGGTAAACAAGAAGAAATATTTGATCCTATACT

>M6-PGSC0003DMG400009181M6LRR-RLK_3410

TATGGGTGCACCTAATGAGAAATGAGGGTAAACAAGAAGAAATATTTGATCCTATACTGAGAGACAAGGGTTTCGAAGAAGATATGCTTCAAGTGCTTGATGTTGCATGTATGTGTGTCA

>M6-PGSC0003DMG400009181M6LRR-RLK_3472

GACAAGGGTTTCGAAGAAGATATGCTTCAAGTGCTTGATGTTGCATGTATGTGTGTCAGCCAGAATCCTTTCAAAAGGCCAACTATAGCAGAAGTGGTTGAGTGGCTCAACAGAGTAGTA

>M6-PGSC0003DMG400009181M6LRR-RLK_3534

GAATCCTTTCAAAAGGCCAACTATAGCAGAAGTGGTTGAGTGGCTCAACAGAGTAGTATCCAACCAGGGAGCACCTAAATAAGGTGGTGGTACACATAAATTTGATGTAGATGTGAGTGT

>M6-PGSC0003DMG400009181M6LRR-RLK_3596

ACCAGGGAGCACCTAAATAAGGTGGTGGTACACATAAATTTGATGTAGATGTGAGTGTCTGTATAGATGAACTTTTGCTTTTTACACGTAGGAAGTTAACATACAGGAATAAATGTATAA

>M6-PGSC0003DMG400009181M6LRR-RLK_3658

ATAGATGAACTTTTGCTTTTTACACGTAGGAAGTTAACATACAGGAATAAATGTATAAAGTGAACAATTTCCTCAATAAATTACATTTACGATTTTGCGTCATGGCTAACATTGGATGAC

>M6-PGSC0003DMG400009181M6LRR-RLK_434

CTTCTTCTTTACCCCACCATAAATCTTTCAATCCAATAATATTGATAGTTGTTCTACTTCTTTCTTCAATTGCTACCATCTGTCATGCTTCTTGCAATCAGCTTGATCGTGACTCTTTGT

>M6-PGSC0003DMG400009181M6LRR-RLK_496

TCTTCAATTGCTACCATCTGTCATGCTTCTTGCAATCAGCTTGATCGTGACTCTTTGTTATCATTCTCTGTTGGTATCTCTTCACCTTCTCCTTTGAATTGGTCCTCTTCCGTGGATTGC

>M6-PGSC0003DMG400009181M6LRR-RLK_558

ATTCTCTGTTGGTATCTCTTCACCTTCTCCTTTGAATTGGTCCTCTTCCGTGGATTGCTGCACCTTGTGGGAAGGTGTTGGTTGTGATGATAATGGCCGAGTAACCACTCTCTGGCTTCC

>M6-PGSC0003DMG400009181M6LRR-RLK_62

AATATCCATAGAATAAACAAAGTCCACCAGCCATCTATCTATCTAGCCGTCCCAAAAGTAAACGCTTTATTTCAAAAACTCATACCTCACCTTTTTGCTACATAAAAATTGCACCTTGTC

>M6-PGSC0003DMG400009181M6LRR-RLK_620

CCTTGTGGGAAGGTGTTGGTTGTGATGATAATGGCCGAGTAACCACTCTCTGGCTTCCCTCAAGAAGCCTTTTTGGAAACATAACCCCTGCTATTGCAAATTTGAGCAAACTCTCTCAAC

>M6-PGSC0003DMG400009181M6LRR-RLK_682

AGAAGCCTTTTTGGAAACATAACCCCTGCTATTGCAAATTTGAGCAAACTCTCTCAACTCAGCCTGTCAAATAATCGATTTATTGGTCCCCTCCCAGATGGATTTTTCAAGTCATTTAGT

>M6-PGSC0003DMG400009181M6LRR-RLK_744

CCTGTCAAATAATCGATTTATTGGTCCCCTCCCAGATGGATTTTTCAAGTCATTTAGTAGCTTGCAGATCATTGACTTGAGTTATAACCGTTTATCAGGACGATTGCCGCTATCAGATAG

>M6-PGSC0003DMG400009181M6LRR-RLK_806

TGCAGATCATTGACTTGAGTTATAACCGTTTATCAGGACGATTGCCGCTATCAGATAGATTAACATCACCCATCAAGACAGTAAATCTCTCTAGCAACCACTTCAATGGGACAATTCTAT

>M6-PGSC0003DMG400009181M6LRR-RLK_868

ACATCACCCATCAAGACAGTAAATCTCTCTAGCAACCACTTCAATGGGACAATTCTATCATCATTTCTCGAGCCAGCAATTAATTTAGAGAGTTTCGATATTAGCAACAATAGCTTCTCT

>M6-PGSC0003DMG400009181M6LRR-RLK_930

ATTTCTCGAGCCAGCAATTAATTTAGAGAGTTTCGATATTAGCAACAATAGCTTCTCTGGTCCAATACCCTCCTTTATTTGCAGCTACTCAGCCGCTGTCAGAGTTCTTGATTTCACTTC

>M6-PGSC0003DMG400009181M6LRR-RLK_992

CAATACCCTCCTTTATTTGCAGCTACTCAGCCGCTGTCAGAGTTCTTGATTTCACTTCCAATGACTTCAGGGGCCAGATGCCCCAAGGGTTTGGGAGTTGCTCCAGTTTGGTTACTTTGA

>M6-PGSC0003DMG400009196M6LRR-RLK_0

TTCTTGAAAGATGACATTGAGAAATTTGACTTACCTGATTTGTTGAAAGCCTCTGCTGAAGTATTGGGAAGTGGAGTGTTTGGTTCAACTTATAAAGCTGCACTTAGTACTGGTCCTGTC

>M6-PGSC0003DMG400009196M6LRR-RLK_128

TAAGAGGTTTAGACATATGAATAAAGTTGGTAAAGAAGATTTCCATGAACATATGAGAAGGCTTGGAAGATTGAGTCATAAGAACTTGCTTCCTGTTATTGCTTTCTATTATAGGAAAGA

>M6-PGSC0003DMG400009196M6LRR-RLK_190

TTGGAAGATTGAGTCATAAGAACTTGCTTCCTGTTATTGCTTTCTATTATAGGAAAGAGGAGAAGCTTCTTGTCTTTGAATATGTCAACAATGTCAGCCTTGCTGTTTATCTTCATGGTA

>M6-PGSC0003DMG400009196M6LRR-RLK_64

TGGGAAGTGGAGTGTTTGGTTCAACTTATAAAGCTGCACTTAGTACTGGTCCTGTCATGGTTGTTAAGAGGTTTAGACATATGAATAAAGTTGGTAAAGAAGATTTCCATGAACATATGA

>M6-PGSC0003DMG400009230-(rev)M6LRR-RLP_0

TTCAAACATCACTACATACTACAAGGAGAAGTAGAAAGTCTACCTTATTTAAGTAATTTAGTTGAGTAATTATACATGTAACAGAAACCATCACAAACACCCAAGTTTGAAAATTTGCAG

>M6-PGSC0003DMG400009230-(rev)M6LRR-RLP_1054

CACATGTTTCTTCCCTTCCTTCCTTCTCTTTCTTAATTCTAGAAGTAGAGACTCTGTAGGGATATAGATCAATACAATAGAGGCACTAACCGTCAAGGAACCAATAATGGAGCAAACACC

>M6-PGSC0003DMG400009230-(rev)M6LRR-RLP_1116

TATAGATCAATACAATAGAGGCACTAACCGTCAAGGAACCAATAATGGAGCAAACACCAATATAGACCATGATATTTGTCCGACCATAGCGAGGCTCACAATACAAGACTAGTACCAATG

>M6-PGSC0003DMG400009230-(rev)M6LRR-RLP_1178

TAGACCATGATATTTGTCCGACCATAGCGAGGCTCACAATACAAGACTAGTACCAATGTTATTGCAACTGCTGATGCAGTGTACAAAAGAAAAGCTGCAAAACAAAGAGTGGAACATCAA

>M6-PGSC0003DMG400009230-(rev)M6LRR-RLP_124

GTTTGAACCCTCCAATGTAAGCTTTATAGCTATACCTATAGCTTTGATGCTCATAACCTACAACAGAGCAACAAACAAGCTTAAAATGACTGCTACATGCAATTTTTACCAATTGTTCTT

>M6-PGSC0003DMG400009230-(rev)M6LRR-RLP_1240

TGCAACTGCTGATGCAGTGTACAAAAGAAAAGCTGCAAAACAAAGAGTGGAACATCAAAAAAATGATATATCAGCAAGGCAAATAAGAATGATATTCAAGGAAATACAGCACAGTACATC

>M6-PGSC0003DMG400009230-(rev)M6LRR-RLP_1302

ATGATATATCAGCAAGGCAAATAAGAATGATATTCAAGGAAATACAGCACAGTACATCAGCACACCTGGTTGTGTAGCTAATGCCCATATTTCATCCACTGAATTGAGATCATGTTCACG

>M6-PGSC0003DMG400009230-(rev)M6LRR-RLP_1364

CACCTGGTTGTGTAGCTAATGCCCATATTTCATCCACTGAATTGAGATCATGTTCACGAGGTGCATGAAGAACAATGACAACGGAACCTACTATACATAAAAGACATCCTAATACTCCCA

>M6-PGSC0003DMG400009230-(rev)M6LRR-RLP_1426

GCATGAAGAACAATGACAACGGAACCTACTATACATAAAAGACATCCTAATACTCCCAGTTTCTTCAATTTTTCCTTCAGTATGAAATGTGCTAGAACAGCACTGCAGCAAAGCGCCAAA

>M6-PGSC0003DMG400009230-(rev)M6LRR-RLP_1488

CTTCAATTTTTCCTTCAGTATGAAATGTGCTAGAACAGCACTGCAGCAAAGCGCCAAATTATATGGAAGAAAACCGAATAAGCCATTTCCCAGAAAATAATAAAGTCAAGTGGAAATTTA

>M6-PGSC0003DMG400009230-(rev)M6LRR-RLP_1550

ATGGAAGAAAACCGAATAAGCCATTTCCCAGAAAATAATAAAGTCAAGTGGAAATTTACCTCACAATTATACTTAAAGCTCCAAGCGGTGTCACAAGAACTGCTGGCGCATAAATGTAAG

>M6-PGSC0003DMG400009230-(rev)M6LRR-RLP_1612

ACAATTATACTTAAAGCTCCAAGCGGTGTCACAAGAACTGCTGGCGCATAAATGTAAGCCACAAAATTTGCAAATTCTCCAACAATCACTGCAAGGATTATGGTGAACATTGCACTCTAT

>M6-PGSC0003DMG400009230-(rev)M6LRR-RLP_1674

AAAATTTGCAAATTCTCCAACAATCACTGCAAGGATTATGGTGAACATTGCACTCTATATTAGAATAACAAATTCAGGAACATATTCTCTGACAAATAAGAAGCAATTCAAAACTTACTA

>M6-PGSC0003DMG400009230-(rev)M6LRR-RLP_1736

GAATAACAAATTCAGGAACATATTCTCTGACAAATAAGAAGCAATTCAAAACTTACTAGTTACCATCCCAATCCACCAAAGAGGCTCCCGTAGATAACCATATCCACCCGAGCCTTCAGC

>M6-PGSC0003DMG400009230-(rev)M6LRR-RLP_1798

CCATCCCAATCCACCAAAGAGGCTCCCGTAGATAACCATATCCACCCGAGCCTTCAGCAAGAAACAAATATCTCATTGTCAAATAGACAGAGCAACAGAGTTCATATAATCAGCTTTTAA

>M6-PGSC0003DMG400009230-(rev)M6LRR-RLP_186

ACAGAGCAACAAACAAGCTTAAAATGACTGCTACATGCAATTTTTACCAATTGTTCTTTTGAGAAATTTAGAAGGACCAGTTCCTATATTTTCTTTACTTTTCATTCGGAATAATTTTAC

>M6-PGSC0003DMG400009230-(rev)M6LRR-RLP_1860

AACAAATATCTCATTGTCAAATAGACAGAGCAACAGAGTTCATATAATCAGCTTTTAAACTAAGTTCTATTAAATGTTGATACAGAAAAGCATCTTCTATATATAAACTTTACAAGATGT

>M6-PGSC0003DMG400009230-(rev)M6LRR-RLP_1922

AGTTCTATTAAATGTTGATACAGAAAAGCATCTTCTATATATAAACTTTACAAGATGTGCCCGATGTATCTACTTTTAATGTGGAAACTAGCTGATAAATTACAATGTCAAGGAAATTCA

>M6-PGSC0003DMG400009230-(rev)M6LRR-RLP_1984

GATGTATCTACTTTTAATGTGGAAACTAGCTGATAAATTACAATGTCAAGGAAATTCATTCAGCTTATTACAGTGCAGAATGATTTAAAAAATCCATTACACTGATCATTTTCAGTTCTT

>M6-PGSC0003DMG400009230-(rev)M6LRR-RLP_2046

GCTTATTACAGTGCAGAATGATTTAAAAAATCCATTACACTGATCATTTTCAGTTCTTAAAGTCAAGATCCATTTATTATGTCTTGAGCTCATTTCTTCTCTGTAATACATGTATTAGTA

>M6-PGSC0003DMG400009230-(rev)M6LRR-RLP_2108

TCAAGATCCATTTATTATGTCTTGAGCTCATTTCTTCTCTGTAATACATGTATTAGTACTACCCAATACAGCAGTCTGTTTATTTGGAAATGATAGATCAAATAATTTGGAATTACCAAC

>M6-PGSC0003DMG400009230-(rev)M6LRR-RLP_2170

CCAATACAGCAGTCTGTTTATTTGGAAATGATAGATCAAATAATTTGGAATTACCAACAACAATCATTTAGTGAAGCAGAATCATGAGTGCATATATTTAAATTGCTGACCCATTCTACA

>M6-PGSC0003DMG400009230-(rev)M6LRR-RLP_2232

ATCATTTAGTGAAGCAGAATCATGAGTGCATATATTTAAATTGCTGACCCATTCTACAAGTGGTAACTATCAAAGTATCAAAGGTATTAAGTTTAAACAGGTGCTGCTGGAGTCACTCTG

>M6-PGSC0003DMG400009230-(rev)M6LRR-RLP_2294

GTAACTATCAAAGTATCAAAGGTATTAAGTTTAAACAGGTGCTGCTGGAGTCACTCTGATTGCTAAAACAATGTCATTAATCCTAGATAACTGCCACCAAAATTCTAGGCAGCTGACTGG

>M6-PGSC0003DMG400009230-(rev)M6LRR-RLP_2356

CTAAAACAATGTCATTAATCCTAGATAACTGCCACCAAAATTCTAGGCAGCTGACTGGTCAACATTCAGAAGAATATGAATTCATTTTCAAGAACCTCTCCAAGATTATTCAAGGACTAC

>M6-PGSC0003DMG400009230-(rev)M6LRR-RLP_2418

CATTCAGAAGAATATGAATTCATTTTCAAGAACCTCTCCAAGATTATTCAAGGACTACACTTACTTGCATGATAAGTGTGAAGTGTCAATTCCAATATAAGTAACTTGATATTCATTTTC

>M6-PGSC0003DMG400009230-(rev)M6LRR-RLP_248

GAAATTTAGAAGGACCAGTTCCTATATTTTCTTTACTTTTCATTCGGAATAATTTTACCATGTTTCATTGGCAACACAAGCACTCTCTAAGTCAGTTCTGCAAAAAGAACTGCAATAGTA

>M6-PGSC0003DMG400009230-(rev)M6LRR-RLP_2480

ACTTGCATGATAAGTGTGAAGTGTCAATTCCAATATAAGTAACTTGATATTCATTTTCCCATAAATGTACACCCAATACTTCCCTTTCGGAGAGAAGTGTGAATTCCATTCCTTGAGTTC

>M6-PGSC0003DMG400009230-(rev)M6LRR-RLP_2542

AAATGTACACCCAATACTTCCCTTTCGGAGAGAAGTGTGAATTCCATTCCTTGAGTTCCAGAACAAGTTTTCTTTATTCCCACACAGATAACTGCTTTGTAAGATGCAATATCACCTTTT

>M6-PGSC0003DMG400009230-(rev)M6LRR-RLP_2604

ACAAGTTTTCTTTATTCCCACACAGATAACTGCTTTGTAAGATGCAATATCACCTTTTATAAGCAGTCATTGTGTATTAGTAATACTCACTGTGACTCTACATTTGCTGGAAGAGTTACC

>M6-PGSC0003DMG400009230-(rev)M6LRR-RLP_2666

GCAGTCATTGTGTATTAGTAATACTCACTGTGACTCTACATTTGCTGGAAGAGTTACCCCATTGCTCTTATTTTACAGTCGGGCACCTACAACATGGTGTACCAACAGAGGATAACATAG

>M6-PGSC0003DMG400009230-(rev)M6LRR-RLP_2728

TGCTCTTATTTTACAGTCGGGCACCTACAACATGGTGTACCAACAGAGGATAACATAGTCTTAGACTGTACTTAAGCAGAAAAAAAAATGAAGTAAGAATTTTCAGGTGTAAAACATTTG

>M6-PGSC0003DMG400009230-(rev)M6LRR-RLP_2790

AGACTGTACTTAAGCAGAAAAAAAAATGAAGTAAGAATTTTCAGGTGTAAAACATTTGTTTCACCCGCAAAACTAAAATGGGGACAGCAGAAAAGCAGGTACCACCATACCAGTATAAAA

>M6-PGSC0003DMG400009230-(rev)M6LRR-RLP_2852

ACCCGCAAAACTAAAATGGGGACAGCAGAAAAGCAGGTACCACCATACCAGTATAAAAGAAACCCAACATCATCAGTCTGTTGCCACTTGATATTTCAAAGTAAGAAGTAAAGAACAATA

>M6-PGSC0003DMG400009230-(rev)M6LRR-RLP_2914

CCCAACATCATCAGTCTGTTGCCACTTGATATTTCAAAGTAAGAAGTAAAGAACAATATATTCATCAAGCATAATCTCCAATTCCATACATCTAGAAACAGCATCTGATATACTATAAAA

>M6-PGSC0003DMG400009230-(rev)M6LRR-RLP_2976

CATCAAGCATAATCTCCAATTCCATACATCTAGAAACAGCATCTGATATACTATAAAATTTGATTACTATGCCTCTTGGCTGATGGTAAATTGGTAATGCAGTAAAATGGTTAGAAATAT

>M6-PGSC0003DMG400009230-(rev)M6LRR-RLP_3038

ATTACTATGCCTCTTGGCTGATGGTAAATTGGTAATGCAGTAAAATGGTTAGAAATATCAATCTTCAATTGACTAATTTTGACTTGATCATATAATAGTCAGTGTCATCTATATGAGTCC

>M6-PGSC0003DMG400009230-(rev)M6LRR-RLP_310

TTTCATTGGCAACACAAGCACTCTCTAAGTCAGTTCTGCAAAAAGAACTGCAATAGTAGCTCAATTAATACAAAAAATTTAAGAGAGGAACCAATTCATAAACTGTTATTTATCAAGTCT

>M6-PGSC0003DMG400009230-(rev)M6LRR-RLP_3100

CTTCAATTGACTAATTTTGACTTGATCATATAATAGTCAGTGTCATCTATATGAGTCCACTATATCCATTCTACATATCTGGGCCTAAATCAATTTCAATACTACAAGAATTTTATATAA

>M6-PGSC0003DMG400009230-(rev)M6LRR-RLP_3162

TATCCATTCTACATATCTGGGCCTAAATCAATTTCAATACTACAAGAATTTTATATAAAGAAAAACTGGGGGTTTTTTAGAGTTGAGAGATTCTCCAAATGTCACACTTCTTTTTTGTAC

>M6-PGSC0003DMG400009230-(rev)M6LRR-RLP_3224

AAACTGGGGGTTTTTTAGAGTTGAGAGATTCTCCAAATGTCACACTTCTTTTTTGTACACTAACATACAAGGTTTTAACCTCATTCTCAATTAGCTAGCGTCATCTACAATAAATCATTT

>M6-PGSC0003DMG400009230-(rev)M6LRR-RLP_3286

ACATACAAGGTTTTAACCTCATTCTCAATTAGCTAGCGTCATCTACAATAAATCATTTACTTTAACTGTGATTCTTACCTAAATTGCCTTAGTTGAGAGATTGTAGGGGTTTTGTGGTTA

>M6-PGSC0003DMG400009230-(rev)M6LRR-RLP_3348

TAACTGTGATTCTTACCTAAATTGCCTTAGTTGAGAGATTGTAGGGGTTTTGTGGTTATTGTTAGGATCTTCATTACCATGTGTCATGAAGAAGCTAATAAAGCAAACATTGGACTATGA

>M6-PGSC0003DMG400009230-(rev)M6LRR-RLP_3410

TAGGATCTTCATTACCATGTGTCATGAAGAAGCTAATAAAGCAAACATTGGACTATGATAAATAATATAACAAAGAGACATATACCAATACAAAAAAATTTAACATGAGAAATTCACAAA

>M6-PGSC0003DMG400009230-(rev)M6LRR-RLP_3472

TAATATAACAAAGAGACATATACCAATACAAAAAAATTTAACATGAGAAATTCACAAAGATAAAAAAAAAAGAGTACAAACATATAAAGAGAATAACCCCACAAAATTCACTCTTAAAGA

>M6-PGSC0003DMG400009230-(rev)M6LRR-RLP_372

AATTAATACAAAAAATTTAAGAGAGGAACCAATTCATAAACTGTTATTTATCAAGTCTTTAAAGCAAAATACATGAGAGTATTATTCATAAGATAGTGAATTCGATTGATAAAGAATTCA

>M6-PGSC0003DMG400009230-(rev)M6LRR-RLP_3782

AATTAGGGCAAACACCTACAACTACTTGCATTTGTATAATTTTAAGCAAGGTCAACCTTCCTAACCTTAGCAAGAAGTAATGCATTGCAACACACACATATAAAGAACTGAGCATATTAA

>M6-PGSC0003DMG400009230-(rev)M6LRR-RLP_3844

AACCTTAGCAAGAAGTAATGCATTGCAACACACACATATAAAGAACTGAGCATATTAAACAACTTCAGCAAAACTACAAGGTAGAAGAAAACACACAAACATCGGCAAACTGAATGGGAA

>M6-PGSC0003DMG400009230-(rev)M6LRR-RLP_3906

CTTCAGCAAAACTACAAGGTAGAAGAAAACACACAAACATCGGCAAACTGAATGGGAACACAACTAAACCTCCATGAATCCACCACTAGAAGCATTGACAAAGTTCAAAATCAACAAAAA

>M6-PGSC0003DMG400009230-(rev)M6LRR-RLP_3968

ACTAAACCTCCATGAATCCACCACTAGAAGCATTGACAAAGTTCAAAATCAACAAAAAGCAATAAATTATTGACAATTAGACCTAAAAACACTTAGCTATCAAGAAAATGAATTACTCAC

>M6-PGSC0003DMG400009230-(rev)M6LRR-RLP_4030

TAAATTATTGACAATTAGACCTAAAAACACTTAGCTATCAAGAAAATGAATTACTCACTGGCCCGAGTACCAGAGGCGCCTGCCCTCTGAAGACCCTTCTTCTTGATAATGAAACTGGAA

>M6-PGSC0003DMG400009230-(rev)M6LRR-RLP_4092

CCGAGTACCAGAGGCGCCTGCCCTCTGAAGACCCTTCTTCTTGATAATGAAACTGGAACCAATAAAAGCTCTAGACGCCACAGCCAATCCAAAACCAGTCAAATTAGCACTGTACATGAT

>M6-PGSC0003DMG400009230-(rev)M6LRR-RLP_4154

TAAAAGCTCTAGACGCCACAGCCAATCCAAAACCAGTCAAATTAGCACTGTACATGATCCCAATATTATATCTAAACTATTGGGCAACAATTTTGGAATTTAATTCTCATCTGAACTTCC

>M6-PGSC0003DMG400009230-(rev)M6LRR-RLP_4216

ATATTATATCTAAACTATTGGGCAACAATTTTGGAATTTAATTCTCATCTGAACTTCCCAAGAACAAAACTTTCAACACCCAAAAGGTTTCCCTGGATATACTCAACAAGACAACACTCA

>M6-PGSC0003DMG400009230-(rev)M6LRR-RLP_4278

AACAAAACTTTCAACACCCAAAAGGTTTCCCTGGATATACTCAACAAGACAACACTCAAGTAAGACTAAAATAACAAGAAAAACGAAATGGGCAAGTTTGAAAAACTTGAAACCACTGTA

>M6-PGSC0003DMG400009230-(rev)M6LRR-RLP_434

AGCAAAATACATGAGAGTATTATTCATAAGATAGTGAATTCGATTGATAAAGAATTCATAAAATAACTCGACCTCGAACTAACAAGATGTGACTAGATAAAAGCTTATCTTAACTAGCAA

>M6-PGSC0003DMG400009230-(rev)M6LRR-RLP_4340

AGACTAAAATAACAAGAAAAACGAAATGGGCAAGTTTGAAAAACTTGAAACCACTGTAGCTTTACTCAATTACCCCAAAATTGTAGAGCTGAGAATGGAAAAACAGCAAGAAAAACGAAA

>M6-PGSC0003DMG400009230-(rev)M6LRR-RLP_4402

TACTCAATTACCCCAAAATTGTAGAGCTGAGAATGGAAAAACAGCAAGAAAAACGAAATGGGTATATGAGAAAATTGGCGAAAACAGGTTTGAAATGGTGAGCTAACAGTGCAAGCTCGT

>M6-PGSC0003DMG400009230-(rev)M6LRR-RLP_4464

TATATGAGAAAATTGGCGAAAACAGGTTTGAAATGGTGAGCTAACAGTGCAAGCTCGTTATAGACAATAATGGAGAATAGTATTACAATAAGCATAATTTTGTCATGCTTTGATTCGTTG

>M6-PGSC0003DMG400009230-(rev)M6LRR-RLP_4523

ATAGACAATAATGGAGAATAGTATTACAATAAGCATAATTTTGTCATGCTTTGATTCGTTGCTTTGGGTTGATTTGGCTATGGTATGTCTTCGACAATTGGGCATAACTATGAATAAATC

>M6-PGSC0003DMG400009230-(rev)M6LRR-RLP_496

ATAACTCGACCTCGAACTAACAAGATGTGACTAGATAAAAGCTTATCTTAACTAGCAACAAAGACTTGTTTTAAATTTATTTAAAAGACAAATTAATAAAATAAAGTATTAGCACTGCAA

>M6-PGSC0003DMG400009230-(rev)M6LRR-RLP_558

GACTTGTTTTAAATTTATTTAAAAGACAAATTAATAAAATAAAGTATTAGCACTGCAATCTTTTATTGAGTTACACACCTACTAGTCATGGAAAAACAACAGCAACCCTGGATGCCTTAT

>M6-PGSC0003DMG400009230-(rev)M6LRR-RLP_62

TGAGTAATTATACATGTAACAGAAACCATCACAAACACCCAAGTTTGAAAATTTGCAGCTTGGTTTGAACCCTCCAATGTAAGCTTTATAGCTATACCTATAGCTTTGATGCTCATAACC

>M6-PGSC0003DMG400009230-(rev)M6LRR-RLP_620

TTATTGAGTTACACACCTACTAGTCATGGAAAAACAACAGCAACCCTGGATGCCTTATAAATTATAACAGATCAAAGAATAGTCATTTGTACAAGCATAGTGGCAGGAAAAGACATCTAG

>M6-PGSC0003DMG400009230-(rev)M6LRR-RLP_682

TATAACAGATCAAAGAATAGTCATTTGTACAAGCATAGTGGCAGGAAAAGACATCTAGTGACATGCTAAAACGTTACAGCACCCTAAAGCTCTGGAGTAATATTTCAAGTTCCATACATC

>M6-PGSC0003DMG400009230-(rev)M6LRR-RLP_744

ATGCTAAAACGTTACAGCACCCTAAAGCTCTGGAGTAATATTTCAAGTTCCATACATCCATTTATACCTAAATGAGAAGTTCTCTTTTACTTCAGTGACAACTAGCAATGGATCGAGGAC

>M6-PGSC0003DMG400009230-(rev)M6LRR-RLP_806

TATACCTAAATGAGAAGTTCTCTTTTACTTCAGTGACAACTAGCAATGGATCGAGGACCCAGGGGCAGTTTGACTCTTAACAACTTAAAAAATATGATCCATTTTGTGGTCAAATAATGC

>M6-PGSC0003DMG400009230-(rev)M6LRR-RLP_868

GGGCAGTTTGACTCTTAACAACTTAAAAAATATGATCCATTTTGTGGTCAAATAATGCTATTCCAATACCACAGAGAAAGGTAGAATTCGGATTAACTTTAAAAGATAAGGCGTCTTGCA

>M6-PGSC0003DMG400009230-(rev)M6LRR-RLP_930

CCAATACCACAGAGAAAGGTAGAATTCGGATTAACTTTAAAAGATAAGGCGTCTTGCAAACTGACAATTTAGCTACACGGTTAACTCAGTCAGGTCAGAGCTACAGACTAAAGCACGGTT

>M6-PGSC0003DMG400009230-(rev)M6LRR-RLP_992

GACAATTTAGCTACACGGTTAACTCAGTCAGGTCAGAGCTACAGACTAAAGCACGGTTGTGACACATGTTTCTTCCCTTCCTTCCTTCTCTTTCTTAATTCTAGAAGTAGAGACTCTGTA

>M6-PGSC0003DMG400009238M6LRR-RLK_0

ATGAAACACAAAACATTTGTTCCTTTGTTCCCTCCCCTTATTTCACCTGAGCTCAGAGAGTATATGCAAATCTTGTACATAGATTTTCTTGAAACCCCTTATCATTTTCTTGAAGTTTTC

>M6-PGSC0003DMG400009238M6LRR-RLK_1054

ACAGATTAAATGGTACAATTCCTCAGGACCTCTTATTCCCAATGAAATTGCAGTTTGTTGATCTCACTTCCAATAAATTAGAGGGACCTGTTCCTTCAAACATGTCGATAAATTTGATCA

>M6-PGSC0003DMG400009238M6LRR-RLK_1116

CTCACTTCCAATAAATTAGAGGGACCTGTTCCTTCAAACATGTCGATAAATTTGATCAGGTTGAGATTGGGGCAAAATGCTTTGAATGGATCATTTCCATCTGCTTCATTTGAAAGCCTT

>M6-PGSC0003DMG400009238M6LRR-RLK_1178

GAGATTGGGGCAAAATGCTTTGAATGGATCATTTCCATCTGCTTCATTTGAAAGCCTTCAAAGTTTGACCTACTTGGAACTGGACAACAACCAGTTAACTGGACCAATTCCTTCTGAATT

>M6-PGSC0003DMG400009238M6LRR-RLK_124

TTCTGACTGTCAAAGCCAAAAACAAAAAAAAAATGAACTTAAGGTATTCTCCCCATATAGTTATTACTATATTCTTGTCTTTTTCTCTTCAAATTGTGTTTTCTCAATTGCCCCCAAATC

>M6-PGSC0003DMG400009238M6LRR-RLK_1240

GTTTGACCTACTTGGAACTGGACAACAACCAGTTAACTGGACCAATTCCTTCTGAATTGGGAAAGTGCCAAAAGATGGCCCTTTTGAACTTAGCCCAGAATAAGCTGAGTGGTGTTATTC

>M6-PGSC0003DMG400009238M6LRR-RLK_1364

TGAGTTGGGTGATATTTCTAATCTTCAGGTACTGAGTCTTCAGTCCAATAACCTAGTTGGAGAAATTCCAAGTAACATTTCACAGTTGAACAGATTGCAGAGGATCAATTTCAGCTAGAA

>M6-PGSC0003DMG400009238M6LRR-RLK_1426

AAATTCCAAGTAACATTTCACAGTTGAACAGATTGCAGAGGATCAATTTCAGCTAGAATTAGTTGACTGGTTCCATTCCAAGTTCATTATCAAGTTTGAGAAATCTCACAAACTTGAATT

>M6-PGSC0003DMG400009238M6LRR-RLK_1488

TTGACTGGTTCCATTCCAAGTTCATTATCAAGTTTGAGAAATCTCACAAACTTGAATTTGCAGGGGAATAATCTAAGTGGTCGAATTCCGGTTGACATTAGTAACTTAAATATGCTATTA

>M6-PGSC0003DMG400009238M6LRR-RLK_1550

GGGGAATAATCTAAGTGGTCGAATTCCGGTTGACATTAGTAACTTAAATATGCTATTAGAACTCCAACTTGGAGGGAACCAACTCGGTGGGCCTATTCCTGATATGCCTTTGAGTTTACA

>M6-PGSC0003DMG400009238M6LRR-RLK_1612

TCCAACTTGGAGGGAACCAACTCGGTGGGCCTATTCCTGATATGCCTTTGAGTTTACAGATTGCTTTGAATCTGAGTGGCAATCTTTTTCAAGGACCTATACCAAGTAGTTTTTCAAGAT

>M6-PGSC0003DMG400009238M6LRR-RLK_1674

GCTTTGAATCTGAGTGGCAATCTTTTTCAAGGACCTATACCAAGTAGTTTTTCAAGATTGACTTCATTGGAAGTTTTGGATCTCTCTTACAACAGGTTCTCGGGTCAGATTCCAGACTAC

>M6-PGSC0003DMG400009238M6LRR-RLK_1798

CGGGAATGAGAGGATTGACTAGGTTGGTGCTTTCAAACAATCAACTGTCTGGAGTTATTCCAAAGTTTGAAAGATTTGTCAGTGTTGAGACAGATGGAAATGGGGATCTGATATATCCAT

>M6-PGSC0003DMG400009238M6LRR-RLK_186

ATTACTATATTCTTGTCTTTTTCTCTTCAAATTGTGTTTTCTCAATTGCCCCCAAATCAAATTAATGACATGAGAAGTGTCTATGATCTGTTTCAGAATGAAACTGGTGCTTCTTTTGTA

>M6-PGSC0003DMG400009238M6LRR-RLK_1860

AAGTTTGAAAGATTTGTCAGTGTTGAGACAGATGGAAATGGGGATCTGATATATCCATACACAGTTAATCCACCACAAGCTGCTGCAAAAAAGAGAAAATCAATAGTTGTTGCAGTTGTT

>M6-PGSC0003DMG400009238M6LRR-RLK_1922

AGTTAATCCACCACAAGCTGCTGCAAAAAAGAGAAAATCAATAGTTGTTGCAGTTGTTGTCCCAATTGCATGTGTAGCAACAATTGCTGTATTAGTGGTGATTGCTATTTCAATCTCTAG

>M6-PGSC0003DMG400009238M6LRR-RLK_1984

CAATTGCATGTGTAGCAACAATTGCTGTATTAGTGGTGATTGCTATTTCAATCTCTAGAAGATATTACAGGATCAATGATGAACATTTTCACTCAGGATTAGAAATTTCTCAATCCCCTG

>M6-PGSC0003DMG400009238M6LRR-RLK_2046

TATTACAGGATCAATGATGAACATTTTCACTCAGGATTAGAAATTTCTCAATCCCCTGTCGTTCAGGGAAAGGTTTTAACTGCAAACAGCATTCACAAGTCAAATATAGACTTCACAAAG

>M6-PGSC0003DMG400009238M6LRR-RLK_2108

TCAGGGAAAGGTTTTAACTGCAAACAGCATTCACAAGTCAAATATAGACTTCACAAAGGCCATGGTAGCAGTGTCTGAGCCCTCAAATGTTGTGTTCAAGACAAGATTTTCAACCTACTA

>M6-PGSC0003DMG400009238M6LRR-RLK_2170

TGGTAGCAGTGTCTGAGCCCTCAAATGTTGTGTTCAAGACAAGATTTTCAACCTACTATAAAGCTGTTATGCCGTCGGGTACAACCTATTTTGTCAAGAAGCTTAATTGGAGTGACAAAA

>M6-PGSC0003DMG400009238M6LRR-RLK_2232

GCTGTTATGCCGTCGGGTACAACCTATTTTGTCAAGAAGCTTAATTGGAGTGACAAAATATTTCAGTTGGGAAGCCATGAACTATTTGGGGAAGAGCTAAAAAATATTGGAAAGCTGAAT

>M6-PGSC0003DMG400009238M6LRR-RLK_2294

TCAGTTGGGAAGCCATGAACTATTTGGGGAAGAGCTAAAAAATATTGGAAAGCTGAATAATTCAAATGTTATGATTCCTCTAGGCTATCTTTTGGCTGCTGATAGTGCTTATCTCTTCTA

>M6-PGSC0003DMG400009238M6LRR-RLK_2356

CAAATGTTATGATTCCTCTAGGCTATCTTTTGGCTGCTGATAGTGCTTATCTCTTCTACGAATTTGCCCCCATTGGCTCGTTATATGATGTTCTTCGTGGTAGCTTGGGCTACTCGTTAG

>M6-PGSC0003DMG400009238M6LRR-RLK_2418

TTTGCCCCCATTGGCTCGTTATATGATGTTCTTCGTGGTAGCTTGGGCTACTCGTTAGATTGGGCAAGTCGCTACAGCATAGCTATTGGAGTAGCTCAGGGTTTAGCTTTTCTTCATGGA

>M6-PGSC0003DMG400009238M6LRR-RLK_248

TAATGACATGAGAAGTGTCTATGATCTGTTTCAGAATGAAACTGGTGCTTCTTTTGTATGGAATGGAACTGATAAAGCTTCAACCCCATGTTCTTGGAAAGGGGTTTCTTGCAATTCTGA

>M6-PGSC0003DMG400009238M6LRR-RLK_2480

GGCAAGTCGCTACAGCATAGCTATTGGAGTAGCTCAGGGTTTAGCTTTTCTTCATGGATGTGAAAAGGGTCCAATCCTCCTCCTCGACCTTTCCAGCAAGAGCATTCTCTTGAAGTCCCA

>M6-PGSC0003DMG400009238M6LRR-RLK_2542

AAAAGGGTCCAATCCTCCTCCTCGACCTTTCCAGCAAGAGCATTCTCTTGAAGTCCCAGAATGAGGCTCAGATAGGAGACATTGAACTCTACAAGGTGATGGATCCTTCGAAGAGCACAG

>M6-PGSC0003DMG400009238M6LRR-RLK_2604

GAGGCTCAGATAGGAGACATTGAACTCTACAAGGTGATGGATCCTTCGAAGAGCACAGGCAGCTTTTCTGCTGTTGCTGGTTCTGTCGGTTATATTCCTCCAGGTAATGAACTATTACTT

>M6-PGSC0003DMG400009238M6LRR-RLK_2666

CTTTTCTGCTGTTGCTGGTTCTGTCGGTTATATTCCTCCAGGTAATGAACTATTACTTGGTTAATGGTAAAAGTTTAAGATCCTTCAACATTGATTTATACTCCTTAAGGGTAATGTGAC

>M6-PGSC0003DMG400009238M6LRR-RLK_2728

AATGGTAAAAGTTTAAGATCCTTCAACATTGATTTATACTCCTTAAGGGTAATGTGACGTTTTTTGTCCTAACTGTCGTACCATGCTTTAAACTCATCTTTTTGTGTTTAATCTTCATAT

>M6-PGSC0003DMG400009238M6LRR-RLK_2790

TTTGTCCTAACTGTCGTACCATGCTTTAAACTCATCTTTTTGTGTTTAATCTTCATATAGTTATTGTTGTTTTCAAACTTAAGACTTCATTATATCCTCCCACCATGAAACGGTGTTGAA

>M6-PGSC0003DMG400009238M6LRR-RLK_2852

ATTGTTGTTTTCAAACTTAAGACTTCATTATATCCTCCCACCATGAAACGGTGTTGAACTATCTTTCGATATCTAAAGTGAAACGGTGTGTCAGTATAGCTTAGAGAGGGTGGGGTGCAT

>M6-PGSC0003DMG400009238M6LRR-RLK_2914

CTTTCGATATCTAAAGTGAAACGGTGTGTCAGTATAGCTTAGAGAGGGTGGGGTGCATTAAAATGTAACATGAGAAGCCTCAAATTTGAGCATATATTTAGTGATGCATTTCCTTTTTGT

>M6-PGSC0003DMG400009238M6LRR-RLK_2976

ATGTAACATGAGAAGCCTCAAATTTGAGCATATATTTAGTGATGCATTTCCTTTTTGTGTTACAGAATATGCATACACAATGAGAGTCACAATGGCTGGAAATGTATATAGCTTTGGGGT

>M6-PGSC0003DMG400009238M6LRR-RLK_3038

CAGAATATGCATACACAATGAGAGTCACAATGGCTGGAAATGTATATAGCTTTGGGGTTGTCCTGCTGGAATTGTTGACAGGAAGGCCTGCAGTTAGTCAGGGAACCGAGTTAGCCAAGT

>M6-PGSC0003DMG400009238M6LRR-RLK_310

ATGGAACTGATAAAGCTTCAACCCCATGTTCTTGGAAAGGGGTTTCTTGCAATTCTGATAATTCTTCCCTTACCAAAGTTACCTTTTCATTGTTCACTATTTCTAGCTCTGAATTCTTGC

>M6-PGSC0003DMG400009238M6LRR-RLK_3100

CTGCTGGAATTGTTGACAGGAAGGCCTGCAGTTAGTCAGGGAACCGAGTTAGCCAAGTCAGTGCTGAGCAACTCTGAGAAGCACAGCAAGTGGGATCACATTCTGGATTCTAGCATTTCT

>M6-PGSC0003DMG400009238M6LRR-RLK_3162

GCTGAGCAACTCTGAGAAGCACAGCAAGTGGGATCACATTCTGGATTCTAGCATTTCTAAAACATCGCTCAACGTTAGGAGCCAGATGTTAGCAGTCCTCAAATTGGCTCTAGCTTGTGT

>M6-PGSC0003DMG400009238M6LRR-RLK_3224

CATCGCTCAACGTTAGGAGCCAGATGTTAGCAGTCCTCAAATTGGCTCTAGCTTGTGTCAGTGTATCACCAGAAGGCCGGCCAAAGATGAAGAGTGTGCTTCGAGTGCTTCTCCATGCAA

>M6-PGSC0003DMG400009238M6LRR-RLK_3286

GTATCACCAGAAGGCCGGCCAAAGATGAAGAGTGTGCTTCGAGTGCTTCTCCATGCAAGGTAATCGGACGTCAAAGGAAAGTTAAGGAGAAAACTGCCAAGGTGGTTTTCTTCTTTTTCT

>M6-PGSC0003DMG400009238M6LRR-RLK_3348

ATCGGACGTCAAAGGAAAGTTAAGGAGAAAACTGCCAAGGTGGTTTTCTTCTTTTTCTTTTGGGGAGGAGCTGCAAGTGGTGAAGGTGTTGTGAGCTATGCAGAATATGCCAGTTTTGCC

>M6-PGSC0003DMG400009238M6LRR-RLK_3410

GGGAGGAGCTGCAAGTGGTGAAGGTGTTGTGAGCTATGCAGAATATGCCAGTTTTGCCAGTTTATACATATTATATTAATGTAAACATATAGATATATGATGATAGTAGTTACATAACTT

>M6-PGSC0003DMG400009238M6LRR-RLK_3472

TATACATATTATATTAATGTAAACATATAGATATATGATGATAGTAGTTACATAACTTTGAATATGATGTTAGTTCTCAGTGTAACATAGTCCATTTTTCTTGTTCATGTGTAATATATA

>M6-PGSC0003DMG400009238M6LRR-RLK_3508

GATGATAGTAGTTACATAACTTTGAATATGATGTTAGTTCTCAGTGTAACATAGTCCATTTTTCTTGTTCATGTGTAATATATATAATATATAAAAGGTCTTTGGGAGTGAATCCAATTT

>M6-PGSC0003DMG400009238M6LRR-RLK_372

TCTTCCCTTACCAAAGTTACCTTTTCATTGTTCACTATTTCTAGCTCTGAATTCTTGCCTGTTATTTGTCAAATTGATACTTTGGAGTCTATTGATGTTTCCCAAAACTTTTTGAGCTCT

>M6-PGSC0003DMG400009238M6LRR-RLK_434

TATTTGTCAAATTGATACTTTGGAGTCTATTGATGTTTCCCAAAACTTTTTGAGCTCTATCCCAAATGAGTTCATCACTGTTTGTGGGGGAATTAGTGGGTTGAAATTGTTGAACTTTAG

>M6-PGSC0003DMG400009238M6LRR-RLK_558

AATAAATTGGAGGGTTTTTTGCCTACTTTCACTGGTTTTGGAAAGTTGGAGTCTTTGGACTTTTCTGACAATAAGATGAATGGGAAAGTTGACTTGCAGTTGGATGGATTAGACTCACTC

>M6-PGSC0003DMG400009238M6LRR-RLK_62

TATGCAAATCTTGTACATAGATTTTCTTGAAACCCCTTATCATTTTCTTGAAGTTTTCAGCTTTCTGACTGTCAAAGCCAAAAACAAAAAAAAAATGAACTTAAGGTATTCTCCCCATAT

>M6-PGSC0003DMG400009238M6LRR-RLK_620

TTCTGACAATAAGATGAATGGGAAAGTTGACTTGCAGTTGGATGGATTAGACTCACTCAAGAGTTTGGACCTTAGCTTTAACATGTTTAATGGTTCAGTTCCTACCAGTCTTGGAAAATT

>M6-PGSC0003DMG400009238M6LRR-RLK_682

GTTTGGACCTTAGCTTTAACATGTTTAATGGTTCAGTTCCTACCAGTCTTGGAAAATTTAATCTTCTGGAGGAGCTTCATCTTTCTGCAAATGCTTTTGGAGGTGAATTCCCCACTCAAA

>M6-PGSC0003DMG400009238M6LRR-RLK_744

CTTCTGGAGGAGCTTCATCTTTCTGCAAATGCTTTTGGAGGTGAATTCCCCACTCAAATTGTGAACTTTGGCAACTTAACTTTGATTGACCTTTCTCTAAACAACCTATCTGGTGTAATT

>M6-PGSC0003DMG400009238M6LRR-RLK_806

GAACTTTGGCAACTTAACTTTGATTGACCTTTCTCTAAACAACCTATCTGGTGTAATTCCTGATAGATTAGGTGAACTTTCAAAATTGCAAGTCTTGATTTTGTCAACCAATAGATTGAG

>M6-PGSC0003DMG400009238M6LRR-RLK_868

ATAGATTAGGTGAACTTTCAAAATTGCAAGTCTTGATTTTGTCAACCAATAGATTGAGTGGCACAATCCCACAATCCCTTAGGAATATCACAACGCTGACGCGTTTTGCTGCAAATCAGA

>M6-PGSC0003DMG400009238M6LRR-RLK_930

ACAATCCCACAATCCCTTAGGAATATCACAACGCTGACGCGTTTTGCTGCAAATCAGAATAATTTTGTTGGAAATATACCCTTTGGTATAACCACATAACTGAGGAATTTGGACCTCAGT

>M6-PGSC0003DMG400009238M6LRR-RLK_992

TTTTGTTGGAAATATACCCTTTGGTATAACCACATAACTGAGGAATTTGGACCTCAGTTTTAACAGATTAAATGGTACAATTCCTCAGGACCTCTTATTCCCAATGAAATTGCAGTTTGT

>M6-PGSC0003DMG400009324-(rev)M6LRR-RLP_0

AGGAGACCTAGTAAAAAAGAAAAGAAAAAAACATGGCCGACAAATTTCAGCCAGTCAACCAGAGGTACGGAGAATTGTTGGTATATGCAGTTATCATGAAATATATAAAGTCCACACACT

>M6-PGSC0003DMG400009324-(rev)M6LRR-RLP_1037

GTTAAACTTAGCCTGGATCTGCTATTAGAAGTGTGAATTTTTGTGGCACTAAACAATGCAATGGGTCTGGATTCATTTTTGCATTAACATTTGTTTAGACGATTTTCTTTATCGAATTTT

>M6-PGSC0003DMG400009324-(rev)M6LRR-RLP_1098

TGGGTCTGGATTCATTTTTGCATTAACATTTGTTTAGACGATTTTCTTTATCGAATTTTACTGTCTAAAAAGGAAAAAACAAAGAAATAAGAAGTATACAGAGGCTGACTTCTTCATAGT

>M6-PGSC0003DMG400009324-(rev)M6LRR-RLP_1159

TGTCTAAAAAGGAAAAAACAAAGAAATAAGAAGTATACAGAGGCTGACTTCTTCATAGTATTTCTCATAAAAAAAAAACATTGATTTCTAGGATATGGGTTATTTTAAATTACAAATTTG

>M6-PGSC0003DMG400009324-(rev)M6LRR-RLP_122

TCAACTGAGCAAACCAGACACTTCGAATTGGTCAATATTTCTTTACTGGATTCAATACATGGCGGAAGCTTTTCTTTATGTTCTGCTATAAAATCTCTTTTCTTTCATTTGAGGGAAACT

>M6-PGSC0003DMG400009324-(rev)M6LRR-RLP_1220

TTCTCATAAAAAAAAAACATTGATTTCTAGGATATGGGTTATTTTAAATTACAAATTTGTGAGTTAAAACAGTTCTGTTGGGAAGGATTTAGATACACGTGGATAGTATCTAGAAGTTTT

>M6-PGSC0003DMG400009324-(rev)M6LRR-RLP_1281

AGTTAAAACAGTTCTGTTGGGAAGGATTTAGATACACGTGGATAGTATCTAGAAGTTTTTTAAATAAAAAATTAGCAAATTATGTGGGCTGGGGCGGATTGAAAACAGCAAACTTTGCAA

>M6-PGSC0003DMG400009324-(rev)M6LRR-RLP_1342

AAATAAAAAATTAGCAAATTATGTGGGCTGGGGCGGATTGAAAACAGCAAACTTTGCAAGGCAGGGCGGGTTGAAATCTTTGCAAGTTTGTGTGGGTTTGCCCTGCACCACCCCATCTGT

>M6-PGSC0003DMG400009324-(rev)M6LRR-RLP_1403

CAGGGCGGGTTGAAATCTTTGCAAGTTTGTGTGGGTTTGCCCTGCACCACCCCATCTGTCATTCCTGTCTAAATGTTTGTTTTGTCTATATAATTCTTGCTGACTCATTCTAATGAGCTC

>M6-PGSC0003DMG400009324-(rev)M6LRR-RLP_1464

TTCCTGTCTAAATGTTTGTTTTGTCTATATAATTCTTGCTGACTCATTCTAATGAGCTCAATTGTAACAAATTCTTTGTGTCCACTCTACTTGGAACAGGTTTTGTTTTAACTGAACCAA

>M6-PGSC0003DMG400009324-(rev)M6LRR-RLP_1525

TTGTAACAAATTCTTTGTGTCCACTCTACTTGGAACAGGTTTTGTTTTAACTGAACCAAAAGTCTACGGAAGGGATAAAGAGGAGGATGAGATAGTGAAAATCTTGATAAACAATTTTAA

>M6-PGSC0003DMG400009324-(rev)M6LRR-RLP_1586

GTCTACGGAAGGGATAAAGAGGAGGATGAGATAGTGAAAATCTTGATAAACAATTTTAATTTTGCCGAAGAACTTCCAGTCTTCCCTATAATTGGTATGGGGGGACTAGGAAAGACGACA

>M6-PGSC0003DMG400009324-(rev)M6LRR-RLP_1647

TTGCCGAAGAACTTCCAGTCTTCCCTATAATTGGTATGGGGGGACTAGGAAAGACGACACTTGCCCAAATGATCTTCAATGATGAGAGAGTGACTAAGCATTTCAATCCCAAAATATGGG

>M6-PGSC0003DMG400009324-(rev)M6LRR-RLP_1708

TGCCCAAATGATCTTCAATGATGAGAGAGTGACTAAGCATTTCAATCCCAAAATATGGGTTTGTGTCTCAGATGATTTTGATGAGAAGAGGTTAATTAAGACAATTTTAGGAAATATTGA

>M6-PGSC0003DMG400009324-(rev)M6LRR-RLP_1769

TGTGTCTCAGATGATTTTGATGAGAAGAGGTTAATTAAGACAATTTTAGGAAATATTGAAAGAAGTTCTCCTCATGTCGAGGACTTGGCTTCATTTCAGAAGAAGCTCCATGAGTTATTG

>M6-PGSC0003DMG400009324-(rev)M6LRR-RLP_183

GCGGAAGCTTTTCTTTATGTTCTGCTATAAAATCTCTTTTCTTTCATTTGAGGGAAACTTGTATTGTTTTTCGGTTTCGAAAAGGAATTTGAAAAGCTGTCGTGTGTGTTTTCCACAATT

>M6-PGSC0003DMG400009324-(rev)M6LRR-RLP_1830

GAAGTTCTCCTCATGTCGAGGACTTGGCTTCATTTCAGAAGAAGCTCCATGAGTTATTGAATGGAAATCGATACTTGCTTGTCTTAGATGATGTTTGGAATGATGATCTAGATAAGTGGG

>M6-PGSC0003DMG400009324-(rev)M6LRR-RLP_1891

TGGAAATCGATACTTGCTTGTCTTAGATGATGTTTGGAATGATGATCTAGATAAGTGGGCTAAGTTAAGAGCAGTCTTAAATGTTGGAGCAAGAGGTGCTTCTATTCTAGCTACTACTCG

>M6-PGSC0003DMG400009324-(rev)M6LRR-RLP_1952

AAGTTAAGAGCAGTCTTAAATGTTGGAGCAAGAGGTGCTTCTATTCTAGCTACTACTCGTCTTGAAAAGGTTGGATCAATTATGGGAACGTCGCGACCATATCATTTGTCAAATTTGTCT

>M6-PGSC0003DMG400009324-(rev)M6LRR-RLP_2013

TTGAAAAGGTTGGATCAATTATGGGAACGTCGCGACCATATCATTTGTCAAATTTGTCTCCACATGATAGTTTACTGTTATTTATGCAACGCGCATTTGGGCAACAAAGAGAAGCAAATC

>M6-PGSC0003DMG400009324-(rev)M6LRR-RLP_2074

ACATGATAGTTTACTGTTATTTATGCAACGCGCATTTGGGCAACAAAGAGAAGCAAATCCTAATCTAGTGGCCATTGGAAAGGAGATTGTGAAGAAATGTGGTGGTGTGCCTTTAGCGGC

>M6-PGSC0003DMG400009324-(rev)M6LRR-RLP_2135

AATCTAGTGGCCATTGGAAAGGAGATTGTGAAGAAATGTGGTGGTGTGCCTTTAGCGGCCAAGACTCTTGGTGGTCTTTTACGCTTCAAGAGAGAAGAGAGTGAATGGGAACATGTGAGA

>M6-PGSC0003DMG400009324-(rev)M6LRR-RLP_2196

AGACTCTTGGTGGTCTTTTACGCTTCAAGAGAGAAGAGAGTGAATGGGAACATGTGAGAGATAATGAGATTTGGAGTCTGCCTCAAGATGAAAGTTCTATTTTGCCTGCTCTAAGACTGA

>M6-PGSC0003DMG400009324-(rev)M6LRR-RLP_2257

TAATGAGATTTGGAGTCTGCCTCAAGATGAAAGTTCTATTTTGCCTGCTCTAAGACTGAGTTATCATCACCTTCCATTTGATTTGAGACAATGCTTTGCGTATTGTGCAGTATTCCCAAA

>M6-PGSC0003DMG400009324-(rev)M6LRR-RLP_2318

TATCATCACCTTCCATTTGATTTGAGACAATGCTTTGCGTATTGTGCAGTATTCCCAAAGGACACCAAAATGATAAAGGAAAATCTCATCACTCTCTGGATGGCCCATGGTTTTCTTTTA

>M6-PGSC0003DMG400009324-(rev)M6LRR-RLP_2379

ACACCAAAATGATAAAGGAAAATCTCATCACTCTCTGGATGGCCCATGGTTTTCTTTTATCAAAGGGAAACTTGGAGCTAGAGTATGTGGGTAATGAAGTATGGAATGAATTATACTTGA

>M6-PGSC0003DMG400009324-(rev)M6LRR-RLP_244

TATTGTTTTTCGGTTTCGAAAAGGAATTTGAAAAGCTGTCGTGTGTGTTTTCCACAATTCAAGCTGTGCTTCAAGATGCTCATGAGAAGCAATTGAAGGACATTGCAATTGATAATTGGT

>M6-PGSC0003DMG400009324-(rev)M6LRR-RLP_2440

AAAGGGAAACTTGGAGCTAGAGTATGTGGGTAATGAAGTATGGAATGAATTATACTTGAGGTCTTTCTTCCAAGAGATTGAAGTTAAATCCGGTAATACTTATTTCAAGATACACGATCT

>M6-PGSC0003DMG400009324-(rev)M6LRR-RLP_2501

TCTTTCTTCCAAGAGATTGAAGTTAAATCCGGTAATACTTATTTCAAGATACACGATCTCATCCATGATTTGGCTACATTTCTGTTTTTGGCAAGCACATCAAGCAGCAATATCCGCGAA

>M6-PGSC0003DMG400009324-(rev)M6LRR-RLP_2562

TCCATGATTTGGCTACATTTCTGTTTTTGGCAAGCACATCAAGCAGCAATATCCGCGAAATAAATGTCAAAGATTATAAGCATATAATGTCCATTGGTTTCGCTGAAGTGGTGTCTTCTT

>M6-PGSC0003DMG400009324-(rev)M6LRR-RLP_2623

AAATGTCAAAGATTATAAGCATATAATGTCCATTGGTTTCGCTGAAGTGGTGTCTTCTTACTCTCCTTCGCTCTTGAAAAAGTTTGTCTCGTTAAGGGTGCTTAATCTAAGCTACTCAAA

>M6-PGSC0003DMG400009324-(rev)M6LRR-RLP_2928

GAAATCTTGTTCTTGATGGCTGTCCATTGACTTCTACTCCACCAAGGATAGGATTGTTGACATGCCTTAAGACTCTAGGTTTCTTTATTGTGGGAAGCAAGAAAGGTTATCAACTTGGTG

>M6-PGSC0003DMG400009324-(rev)M6LRR-RLP_2989

ATGCCTTAAGACTCTAGGTTTCTTTATTGTGGGAAGCAAGAAAGGTTATCAACTTGGTGAACTACGAAATTTAAACCTCCGTGGTGCAATTTCAATCACACATCTTGAGAGAGTGAAAAA

>M6-PGSC0003DMG400009324-(rev)M6LRR-RLP_305

AGCTGTGCTTCAAGATGCTCATGAGAAGCAATTGAAGGACATTGCAATTGATAATTGGTTGCAGAAACTAAATTCTGCTGCGTATGAAGTTGATGACATATTGGGCGAATGTAAAAATGA

>M6-PGSC0003DMG400009324-(rev)M6LRR-RLP_3050

CTACGAAATTTAAACCTCCGTGGTGCAATTTCAATCACACATCTTGAGAGAGTGAAAAACGATATGGAGGCAAAAGAAGCCAATTTATCTGCAAAAGCAAATCTGCAATCTTTAAGCATG

>M6-PGSC0003DMG400009324-(rev)M6LRR-RLP_3111

ATATGGAGGCAAAAGAAGCCAATTTATCTGCAAAAGCAAATCTGCAATCTTTAAGCATGAGTTGGGATAACGATGGACCAAACAGATATGAATCTGAAGAAGTTAAAGTGCTTGAAGCCC

>M6-PGSC0003DMG400009324-(rev)M6LRR-RLP_3172

TTGGGATAACGATGGACCAAACAGATATGAATCTGAAGAAGTTAAAGTGCTTGAAGCCCTCAAACCACACCCCAATCTGAAATATTTAGAGATCATTGCCTTCGGAGGATTCCGTTTTCC

>M6-PGSC0003DMG400009324-(rev)M6LRR-RLP_3233

AAACCACACCCCAATCTGAAATATTTAGAGATCATTGCCTTCGGAGGATTCCGTTTTCCAAGCTGGATAAATCACTCAGTTTTGGAGAAGGTCATCTCTATTAGAATTAAAAGCTGCAAA

>M6-PGSC0003DMG400009324-(rev)M6LRR-RLP_3294

GCTGGATAAATCACTCAGTTTTGGAGAAGGTCATCTCTATTAGAATTAAAAGCTGCAAAAACTGCTTGTGCTTACCACCCTTTGGGGAGCTTCCTTGTCTAGAAAGTCTAGAGTTACAAA

>M6-PGSC0003DMG400009324-(rev)M6LRR-RLP_3416

GGGTCTGCGGAGGTGGAGTTTGTTGAAGAGGATGATGTTCATTCTAGATTTTCCACAAGAAGAAGGTTTCCATCCCTGAAAAATCTTCGTATATGGTTCTTTCGTAATTTGAAAGGTTTG

>M6-PGSC0003DMG400009324-(rev)M6LRR-RLP_3477

GAAGGTTTCCATCCCTGAAAAATCTTCGTATATGGTTCTTTCGTAATTTGAAAGGTTTGGTGAAAGAGGAAGGAGAATATAAATTTCCCATGCTTGAAGAGATGGCGATTTTACATTGCC

>M6-PGSC0003DMG400009324-(rev)M6LRR-RLP_3538

GAAAGAGGAAGGAGAATATAAATTTCCCATGCTTGAAGAGATGGCGATTTTACATTGCCCTCTGTTTGTTTTTCCAACCCTTTCTTCTGTCAAGAAATTAGAAGTTCACGGCAACACAAA

>M6-PGSC0003DMG400009324-(rev)M6LRR-RLP_3599

CTGTTTGTTTTTCCAACCCTTTCTTCTGTCAAGAAATTAGAAGTTCACGGCAACACAAAAGCAAGAGGTTTGAGCTCCATATCTAATCTTAGCACTCTTACTTCCCTCCGCATTGGTGCT

>M6-PGSC0003DMG400009324-(rev)M6LRR-RLP_366

CAGAAACTAAATTCTGCTGCGTATGAAGTTGATGACATATTGGGCGAATGTAAAAATGAGGCAATAAGATTTGAGCAGTCTCGATTAGGGTTTTATCATCCAGGTATTATCAATTTCCGT

>M6-PGSC0003DMG400009324-(rev)M6LRR-RLP_3660

CAAGAGGTTTGAGCTCCATATCTAATCTTAGCACTCTTACTTCCCTCCGCATTGGTGCTAACTACAGAGCGACTTCTCTCCCAGAAGAGATGTTCACAAGTCTTACAAATCTCGAATACT

>M6-PGSC0003DMG400009324-(rev)M6LRR-RLP_3721

CTACAGAGCGACTTCTCTCCCAGAAGAGATGTTCACAAGTCTTACAAATCTCGAATACTTGAGCTTCTTTGACTTCAAGAATCTCAAAGAGCTGCCTACCAGCCTGACTAGTCTCAATGC

>M6-PGSC0003DMG400009324-(rev)M6LRR-RLP_3782

AGCTTCTTTGACTTCAAGAATCTCAAAGAGCTGCCTACCAGCCTGACTAGTCTCAATGCTTTGAAGCGTCTCCAAATTGAAAGTTGTGACTCACTAGAGAGCCTCCCTGAACAAGGGCTG

>M6-PGSC0003DMG400009324-(rev)M6LRR-RLP_3843

TGAAGCGTCTCCAAATTGAAAGTTGTGACTCACTAGAGAGCCTCCCTGAACAAGGGCTGGAAGGTTTAACTTCACTCACACAGTTATTCGTTAAATACTGTAAGATGCTAGAATGTTTAC

>M6-PGSC0003DMG400009324-(rev)M6LRR-RLP_3904

AGGTTTAACTTCACTCACACAGTTATTCGTTAAATACTGTAAGATGCTAGAATGTTTACCCGAGGGATTGCAGCACCTAACTGCCCTCACAAATTTTGGAGTAACCGGTTGTCCAGAAGT

>M6-PGSC0003DMG400009324-(rev)M6LRR-RLP_3965

GAGGGATTGCAGCACCTAACTGCCCTCACAAATTTTGGAGTAACCGGTTGTCCAGAAGTGGAAAAGCGCTGTGATAAGGAATTAGGAGAAGACTGGCACAAAATTGCTCACATTCCTAAT

>M6-PGSC0003DMG400009324-(rev)M6LRR-RLP_4087

GAGTATATATTAGTGAGTTATTTGTAATTGTTTCTTGGGTTATTTTTTTGTCAGTCTCCCACACAGGTTATAATTCCTGCCATTGGATTATTATAGCAAAGTTTGACTGTATGAGGTAAC

>M6-PGSC0003DMG400009324-(rev)M6LRR-RLP_4148

CACAGGTTATAATTCCTGCCATTGGATTATTATAGCAAAGTTTGACTGTATGAGGTAACATTTTTAAACAAAAAAGGTAAATCCGTTTAACATGACTTTTTTTTACTGTTTAATTTAATA

>M6-PGSC0003DMG400009324-(rev)M6LRR-RLP_4181

AGCAAAGTTTGACTGTATGAGGTAACATTTTTAAACAAAAAAGGTAAATCCGTTTAACATGACTTTTTTTTACTGTTTAATTTAATATAAAAGGATTGTACTAATTTTCTATTTGAAAAC

>M6-PGSC0003DMG400009324-(rev)M6LRR-RLP_427

CAATAAGATTTGAGCAGTCTCGATTAGGGTTTTATCATCCAGGTATTATCAATTTCCGTCACAAAATTGGGAAAATGATGAAAGAGATAATGGAGAAACTAGATGCAATTGCTGAGGAAA

>M6-PGSC0003DMG400009324-(rev)M6LRR-RLP_549

AGGAAGTTTCATTTCCTTGAAAAAATTACAGAGAGACAAGCTGCCGCTGCTACGCGTGAAACAGGTGTGAGTACTGAGTAATAGTAACTTAGTTTATGTTCAATTTGTTACCACATCGTG

>M6-PGSC0003DMG400009324-(rev)M6LRR-RLP_61

GAGGTACGGAGAATTGTTGGTATATGCAGTTATCATGAAATATATAAAGTCCACACACTGCTCAACTGAGCAAACCAGACACTTCGAATTGGTCAATATTTCTTTACTGGATTCAATACA

>M6-PGSC0003DMG400009324-(rev)M6LRR-RLP_610

CAGGTGTGAGTACTGAGTAATAGTAACTTAGTTTATGTTCAATTTGTTACCACATCGTGTGTTCACCGTGATCTCTACAGTAGGATGGCAATGGGGCGGGGCGGGGTTGGAGGTGTGCAG

>M6-PGSC0003DMG400009324-(rev)M6LRR-RLP_976

ATGCCTTTAAATTATGATTCAATTTTATATCAGTCTGTCCAAAAGATGTTCTAGTGAAAGTGTTAAACTTAGCCTGGATCTGCTATTAGAAGTGTGAATTTTTGTGGCACTAAACAATGC

>M6-PGSC0003DMG400009380M6LRR-RLK_0

AAAGCTTACACCATTTTAACAAGCTCAGCAAACACCCATTTCACCATTATTACAGAATAATCTCTGTATCTTCTTCCTGTGCATGAAAAACCATGCAAATGAAACTATTACTCTTCTTTC

>M6-PGSC0003DMG400009380M6LRR-RLK_1037

CAAGACACTTGAACTTGCTTACAACCCTTTTGCATCGAGTCAGTTTCCTCCTGAACTCGGTAACTTGACTAATCTTGAGACATTATGGCTAAGCATGTGTAGTCTTGTTGGTTCAATTCC

>M6-PGSC0003DMG400009380M6LRR-RLK_1098

AACTTGACTAATCTTGAGACATTATGGCTAAGCATGTGTAGTCTTGTTGGTTCAATTCCAAAAAGTATTGAGAAATTGAGTCGATTGACTAATTTTGATGTGTCCAATAATGGACTGGTT

>M6-PGSC0003DMG400009380M6LRR-RLK_1159

AAAGTATTGAGAAATTGAGTCGATTGACTAATTTTGATGTGTCCAATAATGGACTGGTTGGGTCAATACCAAGTGCAATTTTCCAGCTTAATAGTATTGTTCAAATTGAGCTCTACAATA

>M6-PGSC0003DMG400009380M6LRR-RLK_122

AGTACATTCCCTTTGATTTTTGCTTTAAATCAAGATGGGTTGTATCTACAAAAACTGAAACACTCTTTGTCAACCTCAGATCAAGGGGTATTTTCTACTTGGTATGAAAATGATCCTACC

>M6-PGSC0003DMG400009380M6LRR-RLK_1220

GTCAATACCAAGTGCAATTTTCCAGCTTAATAGTATTGTTCAAATTGAGCTCTACAATAATTCCCTCACCGGAGAATTGCCTTCGGGATGGTCTAACTTGACCAGGTTGAGAAGATTCGA

>M6-PGSC0003DMG400009380M6LRR-RLK_1281

TCCCTCACCGGAGAATTGCCTTCGGGATGGTCTAACTTGACCAGGTTGAGAAGATTCGATGTGTCGACTAACAAGTTAAATGGGACTATTCCTGATGAGTTGTGCGAGTTGCCACTTGAG

>M6-PGSC0003DMG400009380M6LRR-RLK_1342

TGTCGACTAACAAGTTAAATGGGACTATTCCTGATGAGTTGTGCGAGTTGCCACTTGAGTCACTCAATTTATTTGAGAATCAATTTGAGGGGTTTCTTCCAGAAAGTATAGCTAACTCGC

>M6-PGSC0003DMG400009380M6LRR-RLK_1403

ACTCAATTTATTTGAGAATCAATTTGAGGGGTTTCTTCCAGAAAGTATAGCTAACTCGCCGAATCTGTATGAGCTTAAGTTATTCTCAAACAGATTTTCTGGTTCATTGCCTAGTGAACT

>M6-PGSC0003DMG400009380M6LRR-RLK_1464

AATCTGTATGAGCTTAAGTTATTCTCAAACAGATTTTCTGGTTCATTGCCTAGTGAACTGGGGAAGAACTCGGCTTTACAGTATCTTGATGTTTCATACAATACATTTTCTGGTAAAATT

>M6-PGSC0003DMG400009380M6LRR-RLK_1525

GGAAGAACTCGGCTTTACAGTATCTTGATGTTTCATACAATACATTTTCTGGTAAAATTCCTGAAAGTTTGTGTGAGATGGGAGCTTTAGAGGATATTATAGTGATATATAATTCGTTCT

>M6-PGSC0003DMG400009380M6LRR-RLK_1586

TGAAAGTTTGTGTGAGATGGGAGCTTTAGAGGATATTATAGTGATATATAATTCGTTCTCTGGGAGTATTCCGGCTAGTATTGGCAACTGCCGGAGTTTACTTAGGGTCAGGTTTCGGGC

>M6-PGSC0003DMG400009380M6LRR-RLK_1647

GGGAGTATTCCGGCTAGTATTGGCAACTGCCGGAGTTTACTTAGGGTCAGGTTTCGGGCTAATAAGCTATGCGGGGAAGTCCCAACTGAGTTTTGGAGTTTGCCTCAGGTTTATCTCTTA

>M6-PGSC0003DMG400009380M6LRR-RLK_1708

ATAAGCTATGCGGGGAAGTCCCAACTGAGTTTTGGAGTTTGCCTCAGGTTTATCTCTTAGACCTTTTTGGCAATGCATTTTCAGGAAATATATCACACATGATTTCTGGTGCCAAAAATT

>M6-PGSC0003DMG400009380M6LRR-RLK_1769

CCTTTTTGGCAATGCATTTTCAGGAAATATATCACACATGATTTCTGGTGCCAAAAATTTGTCTAACCTACAGATATCAAGAAACAAATTCTCAGGGGTAATACCTAGTGAAGTAGGAAA

>M6-PGSC0003DMG400009380M6LRR-RLK_183

ACTCTTTGTCAACCTCAGATCAAGGGGTATTTTCTACTTGGTATGAAAATGATCCTACCCCATGTAACTGGACAGGTGTCACCTGTAATGACGCCGGAGATTCTCCCTCCGTCGTCGCTG

>M6-PGSC0003DMG400009380M6LRR-RLK_1830

TCTAACCTACAGATATCAAGAAACAAATTCTCAGGGGTAATACCTAGTGAAGTAGGAAAGTTGAAGAACTTGGTTGAGTTTTCCGCGAGTCATAATGAGCTAACGGGAGAACTTCCAGAC

>M6-PGSC0003DMG400009380M6LRR-RLK_1891

TGAAGAACTTGGTTGAGTTTTCCGCGAGTCATAATGAGCTAACGGGAGAACTTCCAGACACATTAGTGCAGCTAGGGCAGTTAGGAACCCTTGATCTTAGTTTCAATGAGCTATCAGGGA

>M6-PGSC0003DMG400009380M6LRR-RLK_1952

ATTAGTGCAGCTAGGGCAGTTAGGAACCCTTGATCTTAGTTTCAATGAGCTATCAGGGAAAATCCCCTTCGGAATTCACACAATGAAGCAACTCAGTGAGCTTGACTTGGCAAACAATGG

>M6-PGSC0003DMG400009380M6LRR-RLK_2013

ATCCCCTTCGGAATTCACACAATGAAGCAACTCAGTGAGCTTGACTTGGCAAACAATGGATTTTCTGGGGAAATTCCGGAGGAAATTGGGACTTTGCCCGTGCTTAATTATCTTGATCTT

>M6-PGSC0003DMG400009380M6LRR-RLK_2074

TTTCTGGGGAAATTCCGGAGGAAATTGGGACTTTGCCCGTGCTTAATTATCTTGATCTTTCTGGGAATTACTTCTCAGGGGAAATCCCACTCAGTCTGCAAAGCTTGAAGCTTAATAAGC

>M6-PGSC0003DMG400009380M6LRR-RLK_2135

TGGGAATTACTTCTCAGGGGAAATCCCACTCAGTCTGCAAAGCTTGAAGCTTAATAAGCTAAATTTGTCTAATAATCAGCTGTCAGGGATGATTCCTGCAGTTTTTGATAAGGGTGTTTA

>M6-PGSC0003DMG400009380M6LRR-RLK_2196

AATTTGTCTAATAATCAGCTGTCAGGGATGATTCCTGCAGTTTTTGATAAGGGTGTTTATAGAGACAGCTTTCGAGGTAATCCAGGTTTGTGTCAAGGTGTTGCTGGTCTTTGTCCTACG

>M6-PGSC0003DMG400009380M6LRR-RLK_2257

GAGACAGCTTTCGAGGTAATCCAGGTTTGTGTCAAGGTGTTGCTGGTCTTTGTCCTACGAAAGGTAGAGGACAGCATGAAGGATACTTATGGACTTTGAGAGCTATCTACACAGTTGCTG

>M6-PGSC0003DMG400009380M6LRR-RLK_2318

AGGTAGAGGACAGCATGAAGGATACTTATGGACTTTGAGAGCTATCTACACAGTTGCTGGCTTCGTTTTTCTTGTCGGGATTGCTATGTTCATTTGGAAGTATCAGAAATTCAAGAAGAT

>M6-PGSC0003DMG400009380M6LRR-RLK_2379

TTCGTTTTTCTTGTCGGGATTGCTATGTTCATTTGGAAGTATCAGAAATTCAAGAAGATTAAGAAAGGAAACACTATGACAAAGTGGACATCATTCCATAAGCTTGGATTTAGTGAATTT

>M6-PGSC0003DMG400009380M6LRR-RLK_244

ATGTAACTGGACAGGTGTCACCTGTAATGACGCCGGAGATTCTCCCTCCGTCGTCGCTGTTAACCTCTCCGGTGCTTCTCTTGCCGGACCCTTTCCGGTTTTCCTCTGCCATCTTACTTC

>M6-PGSC0003DMG400009380M6LRR-RLK_2440

AGAAAGGAAACACTATGACAAAGTGGACATCATTCCATAAGCTTGGATTTAGTGAATTTGAAATACCCGTTGGCCTAGATGAAGCTAATGTAATTGGCAATGGAGCTTCAGGAAGAGTGT

>M6-PGSC0003DMG400009380M6LRR-RLK_2501

AATACCCGTTGGCCTAGATGAAGCTAATGTAATTGGCAATGGAGCTTCAGGAAGAGTGTACAAAGCTGTCCTAAGCAATGGTGAGGCAGTAGCTGTCAAGAAGCTATGGGAGAGAACAGT

>M6-PGSC0003DMG400009380M6LRR-RLK_2562

AAAGCTGTCCTAAGCAATGGTGAGGCAGTAGCTGTCAAGAAGCTATGGGAGAGAACAGTTAAAGATGAAACCCCTTATGGTGCTCTTGAGTCTGATAAAGACGAGTTTGAAATTGAAGTT

>M6-PGSC0003DMG400009380M6LRR-RLK_2623

AAGATGAAACCCCTTATGGTGCTCTTGAGTCTGATAAAGACGAGTTTGAAATTGAAGTTGAAACTCTGGGTAAAATTAGGCACAAGAATATTGTGAGATTGTGGTGCTGTTGCGATACTG

>M6-PGSC0003DMG400009380M6LRR-RLK_2684

AACTCTGGGTAAAATTAGGCACAAGAATATTGTGAGATTGTGGTGCTGTTGCGATACTGGGGATAGCAAGCTCTTGGTATATGAGTACATGCCAAATGGAAGTTTGGGCGATTTGCTGCA

>M6-PGSC0003DMG400009380M6LRR-RLK_2745

GATAGCAAGCTCTTGGTATATGAGTACATGCCAAATGGAAGTTTGGGCGATTTGCTGCACAGTTGCAAGGCCAAATTGTTGGATTGGCCGTTGAGGTTCAAGATAGCTTTAGATGCTGCT

>M6-PGSC0003DMG400009380M6LRR-RLK_2806

GTTGCAAGGCCAAATTGTTGGATTGGCCGTTGAGGTTCAAGATAGCTTTAGATGCTGCTGAGGGGCTCTCTTATTTGCACCATGGTTGTGTTCCTCCAATTGTTCACCGTGATGTTAAGT

>M6-PGSC0003DMG400009380M6LRR-RLK_2867

GGGGCTCTCTTATTTGCACCATGGTTGTGTTCCTCCAATTGTTCACCGTGATGTTAAGTCAAACAACATATTGCTGGATGATGAGTTTCGAGCCAAAATTTCAGATTTTGGTGTGGCAAA

>M6-PGSC0003DMG400009380M6LRR-RLK_2928

AACAACATATTGCTGGATGATGAGTTTCGAGCCAAAATTTCAGATTTTGGTGTGGCAAAAATTGTTAAAGCAGGCAGCAAAGGTGACGTTGAATCCATGTCTGTAATTGCTGGTTCCTGT

>M6-PGSC0003DMG400009380M6LRR-RLK_2989

TTGTTAAAGCAGGCAGCAAAGGTGACGTTGAATCCATGTCTGTAATTGCTGGTTCCTGTGGTTACATTGCACCAGGTATTGATTTGTTCAAAACTATATCACTGGCTTTAGTCATTTTTA

>M6-PGSC0003DMG400009380M6LRR-RLK_305

AACCTCTCCGGTGCTTCTCTTGCCGGACCCTTTCCGGTTTTCCTCTGCCATCTTACTTCGCTTTCGTCACTCTCGCTTTCGAATAATCTTATTAATTCTACTCTTCCGGTTTCTATTTCT

>M6-PGSC0003DMG400009380M6LRR-RLK_3050

TTACATTGCACCAGGTATTGATTTGTTCAAAACTATATCACTGGCTTTAGTCATTTTTACCGATATATTGGCTCATCTAGCTTATATTTTCGGAAAGATTGCAATTTCATTTTAAAGTAG

>M6-PGSC0003DMG400009380M6LRR-RLK_3111

GATATATTGGCTCATCTAGCTTATATTTTCGGAAAGATTGCAATTTCATTTTAAAGTAGCCAAAACTAACATTAGTGAAATTCATTGATTGCAGAGTATGCATATACTCTTCATGTGAAT

>M6-PGSC0003DMG400009380M6LRR-RLK_3172

AAAACTAACATTAGTGAAATTCATTGATTGCAGAGTATGCATATACTCTTCATGTGAATGAAAAAAGTGACATATATAGCTTTGGAGTGGTCATTTTGGAGCTGGTGACAGGCAAAAGAC

>M6-PGSC0003DMG400009380M6LRR-RLK_3233

AAAAAGTGACATATATAGCTTTGGAGTGGTCATTTTGGAGCTGGTGACAGGCAAAAGACCAGTCAGTCCAGAATTCGGGGAGAAAGATCTAGCTACTTGGGTACACACGACGTTGAACGA

>M6-PGSC0003DMG400009380M6LRR-RLK_3294

GTCAGTCCAGAATTCGGGGAGAAAGATCTAGCTACTTGGGTACACACGACGTTGAACGAGAAAGGAGTTGATCAGTTGCTCGACCCAAATCTAAACTCCAGCTTCAAAGAACATATATGC

>M6-PGSC0003DMG400009380M6LRR-RLK_3355

AAGGAGTTGATCAGTTGCTCGACCCAAATCTAAACTCCAGCTTCAAAGAACATATATGCAAGGTTCTTGATGTTGGTCTACGTTGTCTTAACCAAACTCCAGCTAATCGCCCCTCAATGC

>M6-PGSC0003DMG400009380M6LRR-RLK_3416

GGTTCTTGATGTTGGTCTACGTTGTCTTAACCAAACTCCAGCTAATCGCCCCTCAATGCACAGAGTGGTGAAAATGATCCAAGAATCAGTTCCTTATAATGTGCCAGAAATGGAAAACAA

>M6-PGSC0003DMG400009380M6LRR-RLK_3477

AGAGTGGTGAAAATGATCCAAGAATCAGTTCCTTATAATGTGCCAGAAATGGAAAACAAGAATGGTAAACTTTCCCCTCACTTTCCAAAGTCTGTCTAGTTAATATATTAGATCAAGTAA

>M6-PGSC0003DMG400009380M6LRR-RLK_3538

ATGGTAAACTTTCCCCTCACTTTCCAAAGTCTGTCTAGTTAATATATTAGATCAAGTAATAGAGCAGGAGTGTTTCTTGTCAAGAATTGGCAAGGAAGAAGCTGCATAATGAAAAATTAG

>M6-PGSC0003DMG400009380M6LRR-RLK_3599

GAGCAGGAGTGTTTCTTGTCAAGAATTGGCAAGGAAGAAGCTGCATAATGAAAAATTAGTAGTCTTTTTCAAGAATTAAGCATATTGTACTCTTTGAATTCTACAACCCTTTATGAATCT

>M6-PGSC0003DMG400009380M6LRR-RLK_366

TTTCGTCACTCTCGCTTTCGAATAATCTTATTAATTCTACTCTTCCGGTTTCTATTTCTGAATGCCGTAGCCTCACGTACCTTGACCTTTCTCAGAATCTCATCGGTGGAACTATCCCCG

>M6-PGSC0003DMG400009380M6LRR-RLK_3660

GTCTTTTTCAAGAATTAAGCATATTGTACTCTTTGAATTCTACAACCCTTTATGAATCTATATTACGAGGATTCATCATGAATTAACAAATTTTACGCTAGCTGTCAGCCTACCGCAACA

>M6-PGSC0003DMG400009380M6LRR-RLK_3721

ATTACGAGGATTCATCATGAATTAACAAATTTTACGCTAGCTGTCAGCCTACCGCAACACAGATGGGGTACTGAAAGTTGTAAGTATTCTTAACTAATGCCTTTGATTAAGTTTGTTAAA

>M6-PGSC0003DMG400009380M6LRR-RLK_3776

AACACAGATGGGGTACTGAAAGTTGTAAGTATTCTTAACTAATGCCTTTGATTAAGTTTGTTAAACTTAGGTCCTACTTTGCTGTCCATCATTCACTGAGCTGGGGCCTTTAGAAGTAAA

>M6-PGSC0003DMG400009380M6LRR-RLK_427

ATGCCGTAGCCTCACGTACCTTGACCTTTCTCAGAATCTCATCGGTGGAACTATCCCCGACACTATTTCCGATCTTCCTTACCTCAGGTAACTCAGTTTGAACCTCAATTTTGCATTTCC

>M6-PGSC0003DMG400009380M6LRR-RLK_488

ACTATTTCCGATCTTCCTTACCTCAGGTAACTCAGTTTGAACCTCAATTTTGCATTTCCCCTGTTTTTTTTTTCTTTCAATTTTTCGAAAAATATCAACTAATTGTGACTTACATTTAGC

>M6-PGSC0003DMG400009380M6LRR-RLK_549

TGTTTTTTTTTTCTTTCAATTTTTCGAAAAATATCAACTAATTGTGACTTACATTTAGCCATATATGCGTATTTGGGTCTGTTTTAAGGTGAATATAAGATCTATAGTTAGTGTATATAA

>M6-PGSC0003DMG400009380M6LRR-RLK_61

CTCTGTATCTTCTTCCTGTGCATGAAAAACCATGCAAATGAAACTATTACTCTTCTTTCTGAGTACATTCCCTTTGATTTTTGCTTTAAATCAAGATGGGTTGTATCTACAAAAACTGAA

>M6-PGSC0003DMG400009380M6LRR-RLK_610

TATATGCGTATTTGGGTCTGTTTTAAGGTGAATATAAGATCTATAGTTAGTGTATATAACATAAACATCTCGCATTAGCAAGGTCCGGGGAAGGGGCTGTAAGCAATGATTCAGCTGAAC

>M6-PGSC0003DMG400009380M6LRR-RLK_671

TAAACATCTCGCATTAGCAAGGTCCGGGGAAGGGGCTGTAAGCAATGATTCAGCTGAACCCACATAACCTGCCCGGATTTGCTTGCTGGTGTTATAGTAACTCTAAATTACTATTCATTG

>M6-PGSC0003DMG400009380M6LRR-RLK_732

ACATAACCTGCCCGGATTTGCTTGCTGGTGTTATAGTAACTCTAAATTACTATTCATTGTTTGAATAAAAATGCAGATGGAACGAAATGACTATTATGATCTTTTATTGCAGATTTTGTT

>M6-PGSC0003DMG400009380M6LRR-RLK_793

TGAATAAAAATGCAGATGGAACGAAATGACTATTATGATCTTTTATTGCAGATTTTGTTATTCTTCTTGTTTCATTAGTATATATTATGATCTTTTATTGCAGGTATCTTGATCTTAGCG

>M6-PGSC0003DMG400009380M6LRR-RLK_854

TCTTCTTGTTTCATTAGTATATATTATGATCTTTTATTGCAGGTATCTTGATCTTAGCGGATGCTATTTTTCGGGGAATATTCCGGCAAGTTTCGGAAGATTCAGGCAACTGGAGACTCT

>M6-PGSC0003DMG400009380M6LRR-RLK_915

TGCTATTTTTCGGGGAATATTCCGGCAAGTTTCGGAAGATTCAGGCAACTGGAGACTCTTATTCTGACTGAGAACATACTTACTGGTGAAGTTCCTGCTGCATTAGGTAATGTAACGAGT

>M6-PGSC0003DMG400009380M6LRR-RLK_976

TTCTGACTGAGAACATACTTACTGGTGAAGTTCCTGCTGCATTAGGTAATGTAACGAGTCTCAAGACACTTGAACTTGCTTACAACCCTTTTGCATCGAGTCAGTTTCCTCCTGAACTCG

>M6-PGSC0003DMG400009417-(rev)M6C-LecRK_0

AACACAATTAGGAGTCCAAGCTGAATGAAAAGAGAAGAGAACACTGGAGCTTATTTTTTTATAGATTTTTTTTTTAAAGTTTCAGTGTTCTCTTCTCTTTTCGTTCAGCTTGGACAAGCT

>M6-PGSC0003DMG400009417-(rev)M6C-LecRK_1037

CCCTTTTCCTAATTTCTTTTCTTCTTCTTAGATAAATGAAGAAGTTAGATCAAGTATTATTAAAACACTAGTGCAACTGAAACTTATTTGGACCTCTCAACATGGTTTTTCTTAGTGTAT

>M6-PGSC0003DMG400009417-(rev)M6C-LecRK_1098

AAAACACTAGTGCAACTGAAACTTATTTGGACCTCTCAACATGGTTTTTCTTAGTGTATGTTTTTTATTTTTAATTGGTCAAGTTATTATGTCTCATCTCTTTGGCACTGTTGTTTATTG

>M6-PGSC0003DMG400009417-(rev)M6C-LecRK_1159

TTTTTATTTTTAATTGGTCAAGTTATTATGTCTCATCTCTTTGGCACTGTTGTTTATTGTTTCTCAAGCAGGAAGATGTGATCCTGGTTGGTTTAGCAGCCCAAATGGGAAAAAGTGTTT

>M6-PGSC0003DMG400009417-(rev)M6C-LecRK_122

GTTTAATGCAAGTTTGGCTTTGTTTGTCAGAACTTGGTACTTTCATTTATCCTTCATACTTGAATCTAGCACACCTTCCCTCTCAACAAAGGAAAAGTGAAAACTGAGAACACACTGTCG

>M6-PGSC0003DMG400009417-(rev)M6C-LecRK_1220

TCTCAAGCAGGAAGATGTGATCCTGGTTGGTTTAGCAGCCCAAATGGGAAAAAGTGTTTCAAATATAATGCGACCTCTTTGCAATGGAATGAATCAGAGACTTACTGCAATGGTTTTGGA

>M6-PGSC0003DMG400009417-(rev)M6C-LecRK_1281

AATATAATGCGACCTCTTTGCAATGGAATGAATCAGAGACTTACTGCAATGGTTTTGGAGGAAACCTGGCTGCCATAACATCTTCAGAAGAACTGAACTTTGCCCAAAAGCTCTGTGGCA

>M6-PGSC0003DMG400009417-(rev)M6C-LecRK_1342

AAACCTGGCTGCCATAACATCTTCAGAAGAACTGAACTTTGCCCAAAAGCTCTGTGGCAATGATAGTAACGGATGCTGGGTTGGCGGAAGGAGCATCAATACCACAGCTGGGTTGGGTTG

>M6-PGSC0003DMG400009417-(rev)M6C-LecRK_1403

GATAGTAACGGATGCTGGGTTGGCGGAAGGAGCATCAATACCACAGCTGGGTTGGGTTGGAAATGGTCTGATAATGCTTCTAGCTGGAATGAGAGTATTGCTCCTAAGACAACTTTGGAC

>M6-PGSC0003DMG400009417-(rev)M6C-LecRK_1464

AATGGTCTGATAATGCTTCTAGCTGGAATGAGAGTATTGCTCCTAAGACAACTTTGGACTCAAGTTGCAAAAATTTCTCATGCTATGGCTTTAAGTCGGTGGATTTGTGTACATCGATAA

>M6-PGSC0003DMG400009417-(rev)M6C-LecRK_1525

AAGTTGCAAAAATTTCTCATGCTATGGCTTTAAGTCGGTGGATTTGTGTACATCGATAACCAATGGTGCTATTATAGCTGAGAGATGCAATGTAACTCATGCTTCTATATGCACACTTGA

>M6-PGSC0003DMG400009417-(rev)M6C-LecRK_1586

AATGGTGCTATTATAGCTGAGAGATGCAATGTAACTCATGCTTCTATATGCACACTTGATGCTGGTAAGTGATTGCAGACCAATTTATTTCTAATTGTTTCTTTCACCCAGTGTAATTTT

>M6-PGSC0003DMG400009417-(rev)M6C-LecRK_1647

CTGGTAAGTGATTGCAGACCAATTTATTTCTAATTGTTTCTTTCACCCAGTGTAATTTTGAAAAGTAGGAGCAAAACTCTTTCCCTGATATAGTGTGGTTATAATATGTGATTAAAAGCC

>M6-PGSC0003DMG400009417-(rev)M6C-LecRK_1708

AAAGTAGGAGCAAAACTCTTTCCCTGATATAGTGTGGTTATAATATGTGATTAAAAGCCTCATAGTCCCCTTTTGTTTTTTCCATGAGAAGTGCATTTTAGCTAACCATTATACAGGCCT

>M6-PGSC0003DMG400009417-(rev)M6C-LecRK_1769

ATAGTCCCCTTTTGTTTTTTCCATGAGAAGTGCATTTTAGCTAACCATTATACAGGCCTACATTGTTATCTGGATAAATCTATTACATTCCTTAATTGGTTTTTAAATAATCCTCTAAAT

>M6-PGSC0003DMG400009417-(rev)M6C-LecRK_183

GAATCTAGCACACCTTCCCTCTCAACAAAGGAAAAGTGAAAACTGAGAACACACTGTCGATGTCAAACCTCCACTAATTTGAGTGATATCACACAATCCATAACACGGAGGCCAAGCAAC

>M6-PGSC0003DMG400009417-(rev)M6C-LecRK_1830

ATTGTTATCTGGATAAATCTATTACATTCCTTAATTGGTTTTTAAATAATCCTCTAAATTTTATATTAGGGAGCAAGTGTCGTCATATGCGTTGCCACAGGGAGTATCTCATTATTCTTG

>M6-PGSC0003DMG400009417-(rev)M6C-LecRK_1891

TATATTAGGGAGCAAGTGTCGTCATATGCGTTGCCACAGGGAGTATCTCATTATTCTTGCAGTTGTGAGTGGGTTGATCCTTTGCACTACTTTGGCTATTGTGATATGGCTTCTTGTATT

>M6-PGSC0003DMG400009417-(rev)M6C-LecRK_1952

GTTGTGAGTGGGTTGATCCTTTGCACTACTTTGGCTATTGTGATATGGCTTCTTGTATTTCGGCGAAGTAAAAGACGCAAAAGATCATCGCGTGCATCATTAGCATTAGTTCCTCCGTCT

>M6-PGSC0003DMG400009417-(rev)M6C-LecRK_2013

GGCGAAGTAAAAGACGCAAAAGATCATCGCGTGCATCATTAGCATTAGTTCCTCCGTCTTGGAAAATTTTCAATCGTGATGAAGTAAAGTCCATGACAAAGAATTTCAGTGAAGCAAATC

>M6-PGSC0003DMG400009417-(rev)M6C-LecRK_2074

GAAAATTTTCAATCGTGATGAAGTAAAGTCCATGACAAAGAATTTCAGTGAAGCAAATCGTCTCCTTGGGGATGCCAAAACAGGTGGTACCTATAGTGGGCTTTTACCTGATGGTTCAAG

>M6-PGSC0003DMG400009417-(rev)M6C-LecRK_2135

CTCCTTGGGGATGCCAAAACAGGTGGTACCTATAGTGGGCTTTTACCTGATGGTTCAAGGGTGGCTGTGAAAAGATTGAAGAGGTGTGGTATTCAGAGGAAGAAAGAGTTCTACTCTGAG

>M6-PGSC0003DMG400009417-(rev)M6C-LecRK_2196

TGGCTGTGAAAAGATTGAAGAGGTGTGGTATTCAGAGGAAGAAAGAGTTCTACTCTGAGATTGGAAGAGTTGCGAGACTTCATCATTCAAATCTGGTGTCTATCAAAGGATGTTGCTATC

>M6-PGSC0003DMG400009417-(rev)M6C-LecRK_2257

TGGAAGAGTTGCGAGACTTCATCATTCAAATCTGGTGTCTATCAAAGGATGTTGCTATCATCATGGTGACCGCTATATTCTGTATGAGTTTATTATCAATGGACCCCTGGATAGGTGGCT

>M6-PGSC0003DMG400009417-(rev)M6C-LecRK_2318

CATGGTGACCGCTATATTCTGTATGAGTTTATTATCAATGGACCCCTGGATAGGTGGCTACACCATATTCCTAGAGGAGGTAGGAGTTTGGACTGGAGCATGAGAATGAAGGTCGCCACC

>M6-PGSC0003DMG400009417-(rev)M6C-LecRK_2379

ACCATATTCCTAGAGGAGGTAGGAGTTTGGACTGGAGCATGAGAATGAAGGTCGCCACCACCCTTGCTCAAGGGATTGCGTAAGTAGTGTTCCAACATTATCTTCATGTGCATTGTAGAT

>M6-PGSC0003DMG400009417-(rev)M6C-LecRK_244

GTCAAACCTCCACTAATTTGAGTGATATCACACAATCCATAACACGGAGGCCAAGCAACGACTACTCCTATCAGGTTCATAAGACACAGGGGTAGAGGGCAATGACCTGGTGTCAACATC

>M6-PGSC0003DMG400009417-(rev)M6C-LecRK_2440

CCTTGCTCAAGGGATTGCGTAAGTAGTGTTCCAACATTATCTTCATGTGCATTGTAGATTAGGAATAAATATTTTAACTTATCCGTGAGGTGGATTGATTTTAGTCAGAAATGTGAGTTT

>M6-PGSC0003DMG400009417-(rev)M6C-LecRK_2501

GGAATAAATATTTTAACTTATCCGTGAGGTGGATTGATTTTAGTCAGAAATGTGAGTTTATTCTTTAAATTTTATCATACTTCCCTTCTTTGTCATGAGTATGAACTCATATATACATGT

>M6-PGSC0003DMG400009417-(rev)M6C-LecRK_2562

TCTTTAAATTTTATCATACTTCCCTTCTTTGTCATGAGTATGAACTCATATATACATGTTTGGGTTGATGTCCAATGAAGATGTTATCGTGGAAGTCAAACAAGGGTGAAAATAGATGAT

>M6-PGSC0003DMG400009417-(rev)M6C-LecRK_2623

GGGTTGATGTCCAATGAAGATGTTATCGTGGAAGTCAAACAAGGGTGAAAATAGATGATAAGAATGCATTATCTATATTGTTAAGAAAAAAGGGTGTGTTATCTGTCTTGCAATATTCCT

>M6-PGSC0003DMG400009417-(rev)M6C-LecRK_2684

GAATGCATTATCTATATTGTTAAGAAAAAAGGGTGTGTTATCTGTCTTGCAATATTCCTTTTCCTATTTTCTTTTTCCTTCCCTAGATTTTTCATTCAGAAGAAAGTCAAATTCTGCACA

>M6-PGSC0003DMG400009417-(rev)M6C-LecRK_2745

TCCTATTTTCTTTTTCCTTCCCTAGATTTTTCATTCAGAAGAAAGTCAAATTCTGCACATCTTGACCAAGATACTTCTTGTTAACATATAATGCAATTAATAAATGAAAACCTTTTTGGT

>M6-PGSC0003DMG400009417-(rev)M6C-LecRK_2806

TTGACCAAGATACTTCTTGTTAACATATAATGCAATTAATAAATGAAAACCTTTTTGGTTTACAACATCCGTTCCAGTACTAAGGAGAATTCACTTCCTAGGGATCTACCTATCTGCTTT

>M6-PGSC0003DMG400009417-(rev)M6C-LecRK_2867

ACAACATCCGTTCCAGTACTAAGGAGAATTCACTTCCTAGGGATCTACCTATCTGCTTTTCCTACTCCAGCTAGATCTTACCATTGCCAAACGTGTTTTTGCAGCACTGGCCTTCTGGCT

>M6-PGSC0003DMG400009417-(rev)M6C-LecRK_2928

CTACTCCAGCTAGATCTTACCATTGCCAAACGTGTTTTTGCAGCACTGGCCTTCTGGCTTCTCATGTTTCCTTTCAGGTTTGGGGTGGATCTTTCCTTTCTCAATTATCACATACTCGGT

>M6-PGSC0003DMG400009417-(rev)M6C-LecRK_2989

TCATGTTTCCTTTCAGGTTTGGGGTGGATCTTTCCTTTCTCAATTATCACATACTCGGTTAGGTCGCCCGTGTGTGTGTTTTTCAATTTAAGTGTTTGAACAGGAACTCTCTTCATGAAT

>M6-PGSC0003DMG400009417-(rev)M6C-LecRK_305

CTACTCCTATCAGGTTCATAAGACACAGGGGTAGAGGGCAATGACCTGGTGTCAACATCGTTCTTGGGCCTTCTTATTTCGTAAGCTTAGTTTGAGATGGCAGTGGAGGCCAATTGTGGT

>M6-PGSC0003DMG400009417-(rev)M6C-LecRK_3050

GGTCGCCCGTGTGTGTGTTTTTCAATTTAAGTGTTTGAACAGGAACTCTCTTCATGAATTTCACTAAAATATTGTAGGAAAGCTATGATTGATTTTCTATTCGGGGATTCATTATTAACT

>M6-PGSC0003DMG400009417-(rev)M6C-LecRK_3111

CACTAAAATATTGTAGGAAAGCTATGATTGATTTTCTATTCGGGGATTCATTATTAACTATTTATGACCCTATTTATTGGTGACAGAATTGTTGAACTTTTATCTAGGGAATATTTCCTT

>M6-PGSC0003DMG400009417-(rev)M6C-LecRK_3172

TTATGACCCTATTTATTGGTGACAGAATTGTTGAACTTTTATCTAGGGAATATTTCCTTGACATGGAAATTATGGCAAAAATTTTTTTCAGGGGTTTGTGAGACTACTCACCACCTAACC

>M6-PGSC0003DMG400009417-(rev)M6C-LecRK_3233

CATGGAAATTATGGCAAAAATTTTTTTCAGGGGTTTGTGAGACTACTCACCACCTAACCCCCAACAACAAATATACATGGAAGAAATATTGAAGTGTACAAAAAGAAGCCAATATATTAT

>M6-PGSC0003DMG400009417-(rev)M6C-LecRK_3294

CAACAACAAATATACATGGAAGAAATATTGAAGTGTACAAAAAGAAGCCAATATATTATTCCAATTATTCACGATGATATAGTGAAATAATTAGCATAAGATAGACCAAACCTTAGGCTT

>M6-PGSC0003DMG400009417-(rev)M6C-LecRK_3355

CAATTATTCACGATGATATAGTGAAATAATTAGCATAAGATAGACCAAACCTTAGGCTTATCTACATCACCATATTTAATATATGCAATGTATTATGTATGGTCCCTGGTAGTTTTAACT

>M6-PGSC0003DMG400009417-(rev)M6C-LecRK_3416

CTACATCACCATATTTAATATATGCAATGTATTATGTATGGTCCCTGGTAGTTTTAACTTTTCATGGACTTCTTTGTTAAGAGGTATTATTTTCCATAATTTATTCAGTGTAAGTTGAAA

>M6-PGSC0003DMG400009417-(rev)M6C-LecRK_3477

TCATGGACTTCTTTGTTAAGAGGTATTATTTTCCATAATTTATTCAGTGTAAGTTGAAAAATATAGGCCACTTGAGTAAGCTGGGTTCATTTCATAATGTACATACTTTTAAAGATTTCG

>M6-PGSC0003DMG400009417-(rev)M6C-LecRK_3538

TATAGGCCACTTGAGTAAGCTGGGTTCATTTCATAATGTACATACTTTTAAAGATTTCGATAGTTTGCAAGTAGGTCTTTGTAATTTTTATATGCTACAAGGAATCCAGAGAGAAGCGAG

>M6-PGSC0003DMG400009417-(rev)M6C-LecRK_3599

AGTTTGCAAGTAGGTCTTTGTAATTTTTATATGCTACAAGGAATCCAGAGAGAAGCGAGTGAGAAAAGATGAAATGCTTAAGGAAATGGATACCAGCGACATCTTAGTAAGAAAATAAAA

>M6-PGSC0003DMG400009417-(rev)M6C-LecRK_366

TCTTGGGCCTTCTTATTTCGTAAGCTTAGTTTGAGATGGCAGTGGAGGCCAATTGTGGTGAACCGTGAATCTTTCAATCTTCAAATCTTGGAGACCTTATCTGAAATGAATTCTTATTTT

>M6-PGSC0003DMG400009417-(rev)M6C-LecRK_3660

AGAAAAGATGAAATGCTTAAGGAAATGGATACCAGCGACATCTTAGTAAGAAAATAAAAACTGCATATTACTTTCCTGTTGTCTTCATGAGCTTTTGTTCCAAGTGTCTTACCTTGTTCT

>M6-PGSC0003DMG400009417-(rev)M6C-LecRK_3721

TGCATATTACTTTCCTGTTGTCTTCATGAGCTTTTGTTCCAAGTGTCTTACCTTGTTCTAGGGCGGATGTCTATGGAAAGTGAAGTCATAGAAAATAGGATGTCTATGGAGGTCACTGTT

>M6-PGSC0003DMG400009417-(rev)M6C-LecRK_3782

GGCGGATGTCTATGGAAAGTGAAGTCATAGAAAATAGGATGTCTATGGAGGTCACTGTTATAAGCGAAGCAACCTAGATGCCTAATATATCGATGGATAGATACTTTAAGACCAGTCATG

>M6-PGSC0003DMG400009417-(rev)M6C-LecRK_3843

AAGCGAAGCAACCTAGATGCCTAATATATCGATGGATAGATACTTTAAGACCAGTCATGCCTATGGCATATTTGTTAAAAAAGGGAAAATAAACAGAGACAATTCAGGGTCTATTACTAG

>M6-PGSC0003DMG400009417-(rev)M6C-LecRK_3904

TATGGCATATTTGTTAAAAAAGGGAAAATAAACAGAGACAATTCAGGGTCTATTACTAGTTTATTACCTTTGTAATTTATGGAATTGAAGGATCTAGCATGTTAATCAACTGCATAAGGT

>M6-PGSC0003DMG400009417-(rev)M6C-LecRK_3965

TATTACCTTTGTAATTTATGGAATTGAAGGATCTAGCATGTTAATCAACTGCATAAGGTGAAAATGAGTGGTCTGGTTGTGTTATGGTTGTATTTTTAACCATGAAAGGTGGTGATAGTT

>M6-PGSC0003DMG400009417-(rev)M6C-LecRK_4026

AAATGAGTGGTCTGGTTGTGTTATGGTTGTATTTTTAACCATGAAAGGTGGTGATAGTTGTTAAAACTCTGTTTTGTGTCACTGTGTCAGCTAGTGCACAAGAGGAAGTGCGAGTGGACA

>M6-PGSC0003DMG400009417-(rev)M6C-LecRK_4087

TAAAACTCTGTTTTGTGTCACTGTGTCAGCTAGTGCACAAGAGGAAGTGCGAGTGGACAAAAGCATGACATTGTTTGTTTGCTAATCTCTGTGATGTCAGTTTCTTTGCTTATATTGTTT

>M6-PGSC0003DMG400009417-(rev)M6C-LecRK_4148

AGCATGACATTGTTTGTTTGCTAATCTCTGTGATGTCAGTTTCTTTGCTTATATTGTTTAATGTTTACTTTCATCATATGTGACAGGTTTCTTCACGACAAAATGAAGCCACAAGTTGTG

>M6-PGSC0003DMG400009417-(rev)M6C-LecRK_4209

TGTTTACTTTCATCATATGTGACAGGTTTCTTCACGACAAAATGAAGCCACAAGTTGTGCATAGGGATATCCGTGCAAGCAATGTTCTTCTTGACGAAGACTTTGGCGCCCATCTAATGG

>M6-PGSC0003DMG400009417-(rev)M6C-LecRK_427

ACCGTGAATCTTTCAATCTTCAAATCTTGGAGACCTTATCTGAAATGAATTCTTATTTTGAAGCTGATCCCTCAATTGTTAGCATAATCAAGTAAATAAATCCATTAATGAAGATAAAGA

>M6-PGSC0003DMG400009417-(rev)M6C-LecRK_4270

TAGGGATATCCGTGCAAGCAATGTTCTTCTTGACGAAGACTTTGGCGCCCATCTAATGGGAGTTGGTCTGTCTAAGTTTGTACCATGGGAAATTATGCATGAGAGGAGAGTCATGGCAGG

>M6-PGSC0003DMG400009417-(rev)M6C-LecRK_4331

GTTGGTCTGTCTAAGTTTGTACCATGGGAAATTATGCATGAGAGGAGAGTCATGGCAGGTGGTACTCATGGGTATCTTGCTCCGGAGTTTGTTTACAGAAATGAGCTTACTACAAAGAGT

>M6-PGSC0003DMG400009417-(rev)M6C-LecRK_4392

GTACTCATGGGTATCTTGCTCCGGAGTTTGTTTACAGAAATGAGCTTACTACAAAGAGTGACGTCTACAGCTTTGGAGTTCTCTTGCTTGAAATAATTAGTGGACGAAGACCAGCACAGG

>M6-PGSC0003DMG400009417-(rev)M6C-LecRK_4453

CGTCTACAGCTTTGGAGTTCTCTTGCTTGAAATAATTAGTGGACGAAGACCAGCACAGGCAGTTGATTCTGTTGGATGGCAAAGTATATTTGAATGGGCCACTCCTCTGGTGCAGGCTCA

>M6-PGSC0003DMG400009417-(rev)M6C-LecRK_4514

GTTGATTCTGTTGGATGGCAAAGTATATTTGAATGGGCCACTCCTCTGGTGCAGGCTCATCGATATGTTGAACTGTTGGATCCTCTCATATCATCTTCTTCCTCTTCTCAGATTCCTGAA

>M6-PGSC0003DMG400009417-(rev)M6C-LecRK_4575

GATATGTTGAACTGTTGGATCCTCTCATATCATCTTCTTCCTCTTCTCAGATTCCTGAAGTTGGAGTCATCCAAAAAGTTGTTGATCTTGTTTATTCTTGTACACAACATGTACCATCAA

>M6-PGSC0003DMG400009417-(rev)M6C-LecRK_4636

TGGAGTCATCCAAAAAGTTGTTGATCTTGTTTATTCTTGTACACAACATGTACCATCAATGCGTCCGAGGATGTCACACGTCGTCCATCAGCTACAACAATTAGCTCAGCCCCCTATTGT

>M6-PGSC0003DMG400009417-(rev)M6C-LecRK_4697

CGTCCGAGGATGTCACACGTCGTCCATCAGCTACAACAATTAGCTCAGCCCCCTATTGTAAAGTAAGTGGATAAGAATTATAGTTCAGCATTTCTTTTGACTGAAATCAGCCTGCAATCT

>M6-PGSC0003DMG400009417-(rev)M6C-LecRK_4758

AGTAAGTGGATAAGAATTATAGTTCAGCATTTCTTTTGACTGAAATCAGCCTGCAATCTTTTATCCTAGTTGCAAGGGCTAATTGTATTAGGCATTATTTTTGAACTTGACATGCAAGGC

>M6-PGSC0003DMG400009417-(rev)M6C-LecRK_488

AGCTGATCCCTCAATTGTTAGCATAATCAAGTAAATAAATCCATTAATGAAGATAAAGATACTTTTAGGGTGAAAAGTTATCTTTAGTTGTGGTATTAGAAATAACTAGTTCAGAATACC

>M6-PGSC0003DMG400009417-(rev)M6C-LecRK_4880

TGAGGCTTGAACCAGTCCATGATGTTTGAGATGTAATACAACAACTAATCTGTATCACTAAAGCCAGTATGCTTTTCATCTGCTGCTGCTTTGCATTGTTTACTTGTAACACCTGTATTT

>M6-PGSC0003DMG400009417-(rev)M6C-LecRK_4941

AGCCAGTATGCTTTTCATCTGCTGCTGCTTTGCATTGTTTACTTGTAACACCTGTATTTTTAAAATTTGTTTCTGTTTCCTGTGTTAAAATATCACTATGCATAACAACCAATTCAATAA

>M6-PGSC0003DMG400009417-(rev)M6C-LecRK_549

CTTTTAGGGTGAAAAGTTATCTTTAGTTGTGGTATTAGAAATAACTAGTTCAGAATACCCGAGCTTTATTGTCCATTATGAGGTATTTATTAGTTATAAACAAGTTCATGATACCTGAGT

>M6-PGSC0003DMG400009417-(rev)M6C-LecRK_61

TAGATTTTTTTTTTAAAGTTTCAGTGTTCTCTTCTCTTTTCGTTCAGCTTGGACAAGCTTAGTTTAATGCAAGTTTGGCTTTGTTTGTCAGAACTTGGTACTTTCATTTATCCTTCATAC

>M6-PGSC0003DMG400009417-(rev)M6C-LecRK_610

AGCTTTATTGTCCATTATGAGGTATTTATTAGTTATAAACAAGTTCATGATACCTGAGTTTTACCTACCTAGATACAAACAACTATTCCTACCTAGCACTTTCCAGTCCCCGCAATCTTA

>M6-PGSC0003DMG400009417-(rev)M6C-LecRK_671

TACCTACCTAGATACAAACAACTATTCCTACCTAGCACTTTCCAGTCCCCGCAATCTTACTAGTATTATTTACAACTGAAGTCGATGTTCAAACTTAAGGAAACAACAGAAAGAAAAAAC

>M6-PGSC0003DMG400009417-(rev)M6C-LecRK_732

AGTATTATTTACAACTGAAGTCGATGTTCAAACTTAAGGAAACAACAGAAAGAAAAAACATTAAAACTACAAATATAGAAAGATGGTGAAAAGAATCAACAAATGTGTACAAGAGTTGGA

>M6-PGSC0003DMG400009417-(rev)M6C-LecRK_793

TAAAACTACAAATATAGAAAGATGGTGAAAAGAATCAACAAATGTGTACAAGAGTTGGAGATCATGTGGCTAAATGGATGGGGTTAGTGCAGGAATGTCAAGTTTGAGCCACTTTTATAA

>M6-PGSC0003DMG400009417-(rev)M6C-LecRK_854

TCATGTGGCTAAATGGATGGGGTTAGTGCAGGAATGTCAAGTTTGAGCCACTTTTATAACATTAGGAACACATATAGACTGATAGAATAACATTAGGGGTATATATGTACTGATAGTTTA

>M6-PGSC0003DMG400009417-(rev)M6C-LecRK_915

TTAGGAACACATATAGACTGATAGAATAACATTAGGGGTATATATGTACTGATAGTTTAATGGAGGGTAAAATTAAACCTTTTCAATAGTACAGCTGTACAGGAAGTGAGGGACAAATAT

>M6-PGSC0003DMG400009417-(rev)M6C-LecRK_976

GGAGGGTAAAATTAAACCTTTTCAATAGTACAGCTGTACAGGAAGTGAGGGACAAATATGACCCTTTTCCTAATTTCTTTTCTTCTTCTTAGATAAATGAAGAAGTTAGATCAAGTATTA

>M6-PGSC0003DMG400009532-(rev)M6LRR-RLP_0

TGGATCGAATTTCTCAGTTACCGGCGGGAATGTTAGAGCACATCTTGTCTTTCCTTCCGTTTGAAGAAGCTGCACGAATGAGTAGAGTATCAACGACTTGGAAAAATGCATACAATTCAC

>M6-PGSC0003DMG400009532-(rev)M6LRR-RLP_1024

ATTGGAAGCCAATATTGAATTCGACTCAGAAGGAGGAGAAACCCCTAACAGTGATTGGTACTCTAAGCTCGTGAAATTTCTTGATAACTTCAATCATTCCATGGCTCTTGAATTAACATG

>M6-PGSC0003DMG400009532-(rev)M6LRR-RLP_1088

AAGCTCGTGAAATTTCTTGATAACTTCAATCATTCCATGGCTCTTGAATTAACATGCAAAAGTGAAAAGGTATGTCCCAGAAATTCTTTAACTAGCTAGTTAATACTATATTGTTATGGA

>M6-PGSC0003DMG400009532-(rev)M6LRR-RLP_1152

AAAAGGTATGTCCCAGAAATTCTTTAACTAGCTAGTTAATACTATATTGTTATGGATTTCTCAAAAGTTGTTTCTTTTTGCTAATACAGGGGCTAGTTATACCAAAAGATGTGAGAGAAC

>M6-PGSC0003DMG400009532-(rev)M6LRR-RLP_1216

AAGTTGTTTCTTTTTGCTAATACAGGGGCTAGTTATACCAAAAGATGTGAGAGAACACTTGTTTCCTCCCCTTTATGGAACTAATCGCGTGAAAGTACGCATTCATAGTCCATTGATAAG

>M6-PGSC0003DMG400009532-(rev)M6LRR-RLP_128

TTAAATTTTGAGAGGTATTTTTTAAACAAGCCAATAGAAAAACTTGTGAATGCTGTGGATCCAATTCTAACTTATCGGAAAAATCAAAAGATTTCTGTAGAAAAGTTTTCGCTACACATA

>M6-PGSC0003DMG400009532-(rev)M6LRR-RLP_1280

CCTCCCCTTTATGGAACTAATCGCGTGAAAGTACGCATTCATAGTCCATTGATAAGTTATTCAGTTGTAGATATTCTTGATGGTATTTTCTGGCTTTCTCCTCAATTGGACACCCTATCT

>M6-PGSC0003DMG400009532-(rev)M6LRR-RLP_1342

AGTTGTAGATATTCTTGATGGTATTTTCTGGCTTTCTCCTCAATTGGACACCCTATCTTATTCTAAATTCGACTTGAACACCCTTGAGGTAAATACTTTTATCTTTCAAAGGCACCGACT

>M6-PGSC0003DMG400009532-(rev)M6LRR-RLP_192

TTCTAACTTATCGGAAAAATCAAAAGATTTCTGTAGAAAAGTTTTCGCTACACATAAATCAACAATGGAGTAGTCTGTTTTCCAATTGGATTGATACACTCGTAGATTGTAAGATCAAAG

>M6-PGSC0003DMG400009532-(rev)M6LRR-RLP_256

ATGGAGTAGTCTGTTTTCCAATTGGATTGATACACTCGTAGATTGTAAGATCAAAGAGTTAAGCCTAGTTTTAAGATCAACCAGAACCGGATACACGAAATTGCCTGAGGCAATTTTTAA

>M6-PGSC0003DMG400009532-(rev)M6LRR-RLP_320

CTAGTTTTAAGATCAACCAGAACCGGATACACGAAATTGCCTGAGGCAATTTTTAATGCCAAAGCACTAAAAGTGTTGAAATTGAGTGAATTTCATATTGAATTGCCATCTAATAGTACT

>M6-PGSC0003DMG400009532-(rev)M6LRR-RLP_384

CACTAAAAGTGTTGAAATTGAGTGAATTTCATATTGAATTGCCATCTAATAGTACTATTATGTTTTCATCATCTTTGCGAGAGTTATACCTTCAATGTGTATATTTCGACGAGGAGTTTA

>M6-PGSC0003DMG400009532-(rev)M6LRR-RLP_448

TTCATCATCTTTGCGAGAGTTATACCTTCAATGTGTATATTTCGACGAGGAGTTTATGCAAGCTTTATGCACAAGTTGTATCAATTTAGAGGTTTTCATGGTGCGGGGCTTGAAAGGGCT

>M6-PGSC0003DMG400009532-(rev)M6LRR-RLP_512

TTATGCACAAGTTGTATCAATTTAGAGGTTTTCATGGTGCGGGGCTTGAAAGGGCTTACTAGATTTCAAACTAGTTTACCTAAATTGAAAAAGCTACAGGTGACAGCGTATTATTCTAAA

>M6-PGSC0003DMG400009532-(rev)M6LRR-RLP_576

TTCAAACTAGTTTACCTAAATTGAAAAAGCTACAGGTGACAGCGTATTATTCTAAATTACGATTTGTTGATATTAGATCACCTAATATTGAAGATCTTGACGTCTATGGCTCCAACCTCA

>M6-PGSC0003DMG400009532-(rev)M6LRR-RLP_64

AGAAGCTGCACGAATGAGTAGAGTATCAACGACTTGGAAAAATGCATACAATTCACTCCCCTGTTTAAATTTTGAGAGGTATTTTTTAAACAAGCCAATAGAAAAACTTGTGAATGCTGT

>M6-PGSC0003DMG400009532-(rev)M6LRR-RLP_640

TGTTGATATTAGATCACCTAATATTGAAGATCTTGACGTCTATGGCTCCAACCTCAGCTCCAACTACTTTAAGAATGAGAGTGATCTAAATGTAGTTATTATTACTAATTGTTGCAAATC

>M6-PGSC0003DMG400009532-(rev)M6LRR-RLP_704

TACTTTAAGAATGAGAGTGATCTAAATGTAGTTATTATTACTAATTGTTGCAAATCCCTTAAGAGTTTGCAGCTCAATGGTGTGGCCATGACCCAAAAATGGTTTGACGAAATTTTTACC

>M6-PGSC0003DMG400009532-(rev)M6LRR-RLP_768

GTTTGCAGCTCAATGGTGTGGCCATGACCCAAAAATGGTTTGACGAAATTTTTACCTGCCTTCAGAATATAGAGAAGTTGGATTTAGGTGATTGCGACATGTTGAAGACTGTGAATATCT

>M6-PGSC0003DMG400009532-(rev)M6LRR-RLP_832

GAATATAGAGAAGTTGGATTTAGGTGATTGCGACATGTTGAAGACTGTGAATATCTCAGGCGGCTCTCTCAAACAGCTGAGAGTGTATCATTGTGATAATGTGATTGCTATTGAATTAGA

>M6-PGSC0003DMG400009532-(rev)M6LRR-RLP_896

TCTCTCAAACAGCTGAGAGTGTATCATTGTGATAATGTGATTGCTATTGAATTAGACACTCCTAATTTGATAGAGTTCTCGTATGATTGTTCTTCATTGCCGAATTTGAAACTAAAAGCT

>M6-PGSC0003DMG400009532-(rev)M6LRR-RLP_960

ATTTGATAGAGTTCTCGTATGATTGTTCTTCATTGCCGAATTTGAAACTAAAAGCTTCAGCTTCATTGGAAGCCAATATTGAATTCGACTCAGAAGGAGGAGAAACCCCTAACAGTGATT

>M6-PGSC0003DMG400009561M6LRR-RLK_0

AAAACTACAAAAACAAAAACAAAAATCTTTCATTTGTAGGAGCCCTTTGAGAGGGCGCGCACCTGCAAGTGCGCGCACCAGAAAAAAAAAAAAGTTTAAAAAAATAAAGAAGACGGTCTC

>M6-PGSC0003DMG400009561M6LRR-RLK_1037

ACCAACAACACTTTTTCTTTTTTTTCTTAAAAAACAATCCTCTCTTCTTCTTTCAAGTTCTTGTGAAAGGATTTCATAGAACAAACTTCTGTAACAGGTACGCGATTTTAGACTATGGCT

>M6-PGSC0003DMG400009561M6LRR-RLK_1098

TGTGAAAGGATTTCATAGAACAAACTTCTGTAACAGGTACGCGATTTTAGACTATGGCTTAATATGTAGTTCGTCTTTATATGTGATTACATCTAGTGAAACCAAATTCATTATTGATTT

>M6-PGSC0003DMG400009561M6LRR-RLK_1159

ATATGTAGTTCGTCTTTATATGTGATTACATCTAGTGAAACCAAATTCATTATTGATTTATTAACTAACTAAACATAAGTATTTTTCTTCTTATAAAATAGGCAATGATGGTCTTTGTGG

>M6-PGSC0003DMG400009561M6LRR-RLK_122

TTCTCTATGCAATCATGGCATCACAAAAAAACTACAAAAACAAAAACAAAAACAAAAACAAACATGTCCTCCTCCTTGCTATCATCATGTGTTCATTAGCATTTGTGACAGAGGGAAAAT

>M6-PGSC0003DMG400009561M6LRR-RLK_1220

TAACTAACTAAACATAAGTATTTTTCTTCTTATAAAATAGGCAATGATGGTCTTTGTGGTCCTCCTTTAGCAAAATCATGCAATAAGGAAGATGAACAAAAGAAGCAAGAATCATCATCA

>M6-PGSC0003DMG400009561M6LRR-RLK_1281

CTCCTTTAGCAAAATCATGCAATAAGGAAGATGAACAAAAGAAGCAAGAATCATCATCATCGTCATCGTCATCAGGTTGGAAGATTGCCCTTATTGTGGTTATAGTAGTTGTAGTGATAG

>M6-PGSC0003DMG400009561M6LRR-RLK_1342

GTCATCGTCATCAGGTTGGAAGATTGCCCTTATTGTGGTTATAGTAGTTGTAGTGATAGGCATTATTGTGTTTGTACTAATCAATCGTCGCAAGAAGAATCACCAACAAGAAGTAGTACT

>M6-PGSC0003DMG400009561M6LRR-RLK_1403

ATTATTGTGTTTGTACTAATCAATCGTCGCAAGAAGAATCACCAACAAGAAGTAGTACTTGGAGGATCATCATTAAGTTCTTCCTCACCAACAAGTCAAGACCAAAAACTAATCCCTCAA

>M6-PGSC0003DMG400009561M6LRR-RLK_1464

GAGGATCATCATTAAGTTCTTCCTCACCAACAAGTCAAGACCAAAAACTAATCCCTCAATCTCATGATCATCTAAACAAAATGGAACAAGGCCAATCATCAGCTGCATCTACTCCTGATC

>M6-PGSC0003DMG400009561M6LRR-RLK_1525

TCATGATCATCTAAACAAAATGGAACAAGGCCAATCATCAGCTGCATCTACTCCTGATCGCGCGTCTAATGATGGCGGTAAAAGAGCTGAAGTTGCTGGACAAAAACTTCTATTCTTGAA

>M6-PGSC0003DMG400009561M6LRR-RLK_1586

GCGTCTAATGATGGCGGTAAAAGAGCTGAAGTTGCTGGACAAAAACTTCTATTCTTGAAAGATGACATTGAGAAATTTGACTTACCTGATTTGTTGAAAGCCTCTGCTGAAGTATTGGGA

>M6-PGSC0003DMG400009561M6LRR-RLK_183

ACATGTCCTCCTCCTTGCTATCATCATGTGTTCATTAGCATTTGTGACAGAGGGAAAATTATCAGAGCCTGAGGTTCTTCTCAAGTTTCGTCAATCATTAAAGTTCGATGGAGATCCCTT

>M6-PGSC0003DMG400009561M6LRR-RLK_2196

TTCTGTCTCAATTTAAATGTCTTACTTTCACTCTTGTTCTTTTCTAAAAAGTGTGTCGGTTTCTATATTTAGTAAGTTTTCAAATTGTGACATTCTACCTGCTTAGTTTAAGACCACAAA

>M6-PGSC0003DMG400009561M6LRR-RLK_2257

TCTATATTTAGTAAGTTTTCAAATTGTGACATTCTACCTGCTTAGTTTAAGACCACAAAATTAAAAAAAATTCTTTATTTTTTTTAAAACTATTTACTGACTTAGATTAAGACACTTAAA

>M6-PGSC0003DMG400009561M6LRR-RLK_2318

TAAAAAAAATTCTTTATTTTTTTTAAAACTATTTACTGACTTAGATTAAGACACTTAAATTGAAAGGAAGGATGTACTCCTGCATCTATCTTTACTTGTCTAGTATATGGGATGTCCAAT

>M6-PGSC0003DMG400009561M6LRR-RLK_2379

GAAAGGAAGGATGTACTCCTGCATCTATCTTTACTTGTCTAGTATATGGGATGTCCAATAATACTTGTCCAGTTTATAAAACTAATGAATAATTTTTCATTTTATGTCTATTTTACTTTT

>M6-PGSC0003DMG400009561M6LRR-RLK_244

TCAGAGCCTGAGGTTCTTCTCAAGTTTCGTCAATCATTAAAGTTCGATGGAGATCCCTTCTCGACGTGGGATATTAATGTACCACCTTGCATCAAAGAGAACAATAAGCCTAAATGGAAC

>M6-PGSC0003DMG400009561M6LRR-RLK_2501

TATTAAATACTATTCATTTTTTAATGTTTAAATGACTTATATTTAAATAAAAGTGATTTAATAAAACTACCCCTATTATTTACTGCTTCATAATCCATGTGCCAAGTCAGTAGTGTACGA

>M6-PGSC0003DMG400009561M6LRR-RLK_2562

TAAAACTACCCCTATTATTTACTGCTTCATAATCCATGTGCCAAGTCAGTAGTGTACGACGGACAGAGGGAGAGGGAGTAATATTTTTTTCATAGTTTTGACCACCTTTTCTTGATTTTT

>M6-PGSC0003DMG400009561M6LRR-RLK_2623

GACAGAGGGAGAGGGAGTAATATTTTTTTCATAGTTTTGACCACCTTTTCTTGATTTTTCAGGTAATAGCAAATCACGTGGTAACCAAAGTCTTGATTGGCCAACTCGTTTGAAAATTGT

>M6-PGSC0003DMG400009561M6LRR-RLK_2684

GGTAATAGCAAATCACGTGGTAACCAAAGTCTTGATTGGCCAACTCGTTTGAAAATTGTGAAAGGAGTAAGCAAAGGAATTTTATACCTTTACAATGAGCTACCAAGCTTAACATCACCT

>M6-PGSC0003DMG400009561M6LRR-RLK_2745

AAGGAGTAAGCAAAGGAATTTTATACCTTTACAATGAGCTACCAAGCTTAACATCACCTCATGGTCACCTCAAGTCTTCAAATGTTCTTCTAACTGAAAACTTTGAGGCAGTTCTCACAG

>M6-PGSC0003DMG400009561M6LRR-RLK_2806

TGGTCACCTCAAGTCTTCAAATGTTCTTCTAACTGAAAACTTTGAGGCAGTTCTCACAGACTATGCCTTACTACCAGTAGTAAATGCTGAACATGCACATGAACACATGATCTCATACAA

>M6-PGSC0003DMG400009561M6LRR-RLK_2867

TATGCCTTACTACCAGTAGTAAATGCTGAACATGCACATGAACACATGATCTCATACAAAGCACCAGAGTTAAAACAATCAGGCAAAATCAATAGAAAAACTGATGTTTGGACACTTGGT

>M6-PGSC0003DMG400009561M6LRR-RLK_2928

CACCAGAGTTAAAACAATCAGGCAAAATCAATAGAAAAACTGATGTTTGGACACTTGGTATGTTAATCCTTGAGATCTTGACTGGAAAATTCCCTTCAAATCTTTTAGGCAAAGGAACAC

>M6-PGSC0003DMG400009561M6LRR-RLK_2989

GTTAATCCTTGAGATCTTGACTGGAAAATTCCCTTCAAATCTTTTAGGCAAAGGAACACAAGATAGCGATGATTTAGCGACATGGGTTAATACTACTCTTGGAGATGAATCATCTGAGAA

>M6-PGSC0003DMG400009561M6LRR-RLK_305

CGACGTGGGATATTAATGTACCACCTTGCATCAAAGAGAACAATAAGCCTAAATGGAACAATCTATTTTGCGAAAGTGGAAAGGTTTATGGTTTGAATCTTGAAAATCTTGGGTTAAGTG

>M6-PGSC0003DMG400009561M6LRR-RLK_3050

GATAGCGATGATTTAGCGACATGGGTTAATACTACTCTTGGAGATGAATCATCTGAGAAAGAAGTGTTTGATAAAGAAATGAAAGGAACAAAAGATTGTGAGAGTGAAATGATGAAGTTA

>M6-PGSC0003DMG400009561M6LRR-RLK_3111

AAGTGTTTGATAAAGAAATGAAAGGAACAAAAGATTGTGAGAGTGAAATGATGAAGTTATTGAAGATTGGACTAAGTTGTTGTGAGGTTGATGTGGAGAAGAGATGGGATATAAAAGAGG

>M6-PGSC0003DMG400009561M6LRR-RLK_3172

GAAGATTGGACTAAGTTGTTGTGAGGTTGATGTGGAGAAGAGATGGGATATAAAAGAGGCAGTTGAAAGGATTGATGAAGTAAAAGAGAAAGGTGATTTCAGCTCTAATGTTGCAACTAA

>M6-PGSC0003DMG400009561M6LRR-RLK_3233

GTTGAAAGGATTGATGAAGTAAAAGAGAAAGGTGATTTCAGCTCTAATGTTGCAACTAATGAAGTTGACAATATGCATACTTCAAGATAAAAAAATATTTTTGATTGATAAGCTCATGAT

>M6-PGSC0003DMG400009561M6LRR-RLK_3294

AAGTTGACAATATGCATACTTCAAGATAAAAAAATATTTTTGATTGATAAGCTCATGATCATAATATGGAAATTTTGATTATACTGGATGCATATTGCATGACTATATATACTTTGGGTA

>M6-PGSC0003DMG400009561M6LRR-RLK_3355

TAATATGGAAATTTTGATTATACTGGATGCATATTGCATGACTATATATACTTTGGGTATATGATTTTGTTTTGTCATGGTGGAGGATAAATTGAACCATGATGTGGGGTTACTACAAAA

>M6-PGSC0003DMG400009561M6LRR-RLK_3532

AATTTTTGTTTGCATATTGTAAATTTGTCCTTTATGAATCTATAGTCTAGGTGTGTTTCTTTTATTTTTTTTATTATTCATTCATCAAAGCTTATTTGGGACTTGCTTTGCAAAATTGGC

>M6-PGSC0003DMG400009561M6LRR-RLK_366

TCTATTTTGCGAAAGTGGAAAGGTTTATGGTTTGAATCTTGAAAATCTTGGGTTAAGTGGTACCATTGATTTGGACATACTTAAAGACTTGCCAAATTTGAGAACCATAAGTGTTTTCAA

>M6-PGSC0003DMG400009561M6LRR-RLK_427

ACCATTGATTTGGACATACTTAAAGACTTGCCAAATTTGAGAACCATAAGTGTTTTCAAAAATAAATTTGAAGGTTCCTTACCAATATTAAATAAATTGCCTGCATTAAAGAGTGCTTAT

>M6-PGSC0003DMG400009561M6LRR-RLK_488

ATAAATTTGAAGGTTCCTTACCAATATTAAATAAATTGCCTGCATTAAAGAGTGCTTATTTCTCAAATAACAAGTTTTCTGGTCCAATTGATCAAAAGATATTTGAAGGGATGAATTCAT

>M6-PGSC0003DMG400009561M6LRR-RLK_549

CTCAAATAACAAGTTTTCTGGTCCAATTGATCAAAAGATATTTGAAGGGATGAATTCATTAAAGAAACTTCACCTAGCAAATAATGAATTTACAGGACCACTCCCTCCTATTTTTGGTGA

>M6-PGSC0003DMG400009561M6LRR-RLK_61

CCTGCAAGTGCGCGCACCAGAAAAAAAAAAAAGTTTAAAAAAATAAAGAAGACGGTCTCTCTTCTCTATGCAATCATGGCATCACAAAAAAACTACAAAAACAAAAACAAAAACAAAAAC

>M6-PGSC0003DMG400009561M6LRR-RLK_610

AAGAAACTTCACCTAGCAAATAATGAATTTACAGGACCACTCCCTCCTATTTTTGGTGATATGCCAAATCTAAGGGAGTTGAATATTCAAAACAATAAGTTTGAGGGTCCAATTCCTCCT

>M6-PGSC0003DMG400009561M6LRR-RLK_671

TGCCAAATCTAAGGGAGTTGAATATTCAAAACAATAAGTTTGAGGGTCCAATTCCTCCTTCCTATTCACATCTATATTTTCCTGCTTATGATGGTAATTCCTACCTCGATCTTTTCCTTT

>M6-PGSC0003DMG400009561M6LRR-RLK_732

CTATTCACATCTATATTTTCCTGCTTATGATGGTAATTCCTACCTCGATCTTTTCCTTTTTTATATTTCATTAATTATTGTGTCCAATGAATTAAACACTTTTTTTGTTTATCATTATGC

>M6-PGSC0003DMG400009561M6LRR-RLK_793

TATATTTCATTAATTATTGTGTCCAATGAATTAAACACTTTTTTTGTTTATCATTATGCCATGTTTGTCTTCCAATCCTAGATCATTAAAAAGATCATATCATTGGTTACATAATAAACA

>M6-PGSC0003DMG400009561M6LRR-RLK_854

TGTTTGTCTTCCAATCCTAGATCATTAAAAAGATCATATCATTGGTTACATAATAAACAAAAGATCATTAGATTCGGGTCATCCATGTAAATAATAAGGGAAGAGACACAACTATTCAGA

>M6-PGSC0003DMG400009561M6LRR-RLK_915

AGATCATTAGATTCGGGTCATCCATGTAAATAATAAGGGAAGAGACACAACTATTCAGATTATTTCTGAAAAGTAGTAGTACTCCATCTCTTTAGAATTATAAAAGAATTGTCTTAATAG

>M6-PGSC0003DMG400009561M6LRR-RLK_976

ATTTCTGAAAAGTAGTAGTACTCCATCTCTTTAGAATTATAAAAGAATTGTCTTAATAGAAACCAACAACACTTTTTCTTTTTTTTCTTAAAAAACAATCCTCTCTTCTTCTTTCAAGTT

>M6-PGSC0003DMG400009689M6RLKmalectin_1054

GCTGCTGGTAATGATGCTAGTGCTAATTTTAATGTGACGTGGGTGTTCAATGTGGATGTTGGTTTCCAGTATTTCGTTCGATTACATTTGTGTGATATTGTGAGTACAGCTGCTAATCAG

>M6-PGSC0003DMG400009689M6RLKmalectin_1116

TTTCCAGTATTTCGTTCGATTACATTTGTGTGATATTGTGAGTACAGCTGCTAATCAGCTGTTGTTTAATATCTATGTTGATTCGTCGAATATAGCTTCTGATTTTGATCTGAGTTCTAA

>M6-PGSC0003DMG400009689M6RLKmalectin_1178

TGTTTAATATCTATGTTGATTCGTCGAATATAGCTTCTGATTTTGATCTGAGTTCTAAAGTTCAGAGCAGATTGGCTACCGCGTATTACATGGATTTTGTGACACCACAAGCTAATAGCA

>M6-PGSC0003DMG400009689M6RLKmalectin_124

TCCAAATCTGGGCAATCTCATTCTTGGAGCATATTTAATCAAAAAGGTTCAATCTTTTTCTTCTTCAGCTGTAGTTATAATTGAAATTTTTTCCAATTCTTGGATTGTTGGGGTTTTGAA

>M6-PGSC0003DMG400009689M6RLKmalectin_1240

CAGAGCAGATTGGCTACCGCGTATTACATGGATTTTGTGACACCACAAGCTAATAGCAACAACATTAGCATTAGTATTGGCCCATCTTCAAAGAGTGGTTTTCCAGATGCCTTTCTAAAT

>M6-PGSC0003DMG400009689M6RLKmalectin_1302

CATTAGCATTAGTATTGGCCCATCTTCAAAGAGTGGTTTTCCAGATGCCTTTCTAAATGGAGTGGAACTGCTGAAGTTGAATAACTCTCAAGGCAGCCTTGCTGTTGTTTCCTCTGTTCC

>M6-PGSC0003DMG400009689M6RLKmalectin_1364

TGGAACTGCTGAAGTTGAATAACTCTCAAGGCAGCCTTGCTGTTGTTTCCTCTGTTCCTCCTTCCTCGAGCTCGGGCTCGAAAAAGAATATCGGAGTCATTGTGGGCGCGATTGTAGGTG

>M6-PGSC0003DMG400009689M6RLKmalectin_1426

TCCTCGAGCTCGGGCTCGAAAAAGAATATCGGAGTCATTGTGGGCGCGATTGTAGGTGTGACACTTGTGTTGCTCGTTGTTGGTTTTCTGTTTTGTATGCGCAGAAGAAGAAAGCAGGAA

>M6-PGSC0003DMG400009689M6RLKmalectin_1488

ACTTGTGTTGCTCGTTGTTGGTTTTCTGTTTTGTATGCGCAGAAGAAGAAAGCAGGAACAACTCGGCCTGTCGAAAACATGGATTCCTTTGTCTATTGGTGGTGGTGGCTTGTCTCATAC

>M6-PGSC0003DMG400009689M6RLKmalectin_1550

TCGGCCTGTCGAAAACATGGATTCCTTTGTCTATTGGTGGTGGTGGCTTGTCTCATACTATGGGAAGCAAATACTCGAACGGGACGACCTTAAGTGCTGCTTCTAACTTGAGTTATCGCA

>M6-PGSC0003DMG400009689M6RLKmalectin_1612

GGAAGCAAATACTCGAACGGGACGACCTTAAGTGCTGCTTCTAACTTGAGTTATCGCATCCCTTTTGCAGCCATGCTAGCAGCAACCAAAAAGTTTGATGAGAGTTTGGTCATTGGGATA

>M6-PGSC0003DMG400009689M6RLKmalectin_1674

TTTTGCAGCCATGCTAGCAGCAACCAAAAAGTTTGATGAGAGTTTGGTCATTGGGATAGGTGGCTTTGGTAAGGTATTCAAGGGCGTTTTAAACGATGGGACGAACGTAGCTATTAAGAG

>M6-PGSC0003DMG400009689M6RLKmalectin_1736

GCTTTGGTAAGGTATTCAAGGGCGTTTTAAACGATGGGACGAACGTAGCTATTAAGAGGGGTAATCATAGCTCCCAACAAGGTCTTAGAGAGTTCCAAACTGAAATAGAGATGCTCTCTC

>M6-PGSC0003DMG400009689M6RLKmalectin_1798

AATCATAGCTCCCAACAAGGTCTTAGAGAGTTCCAAACTGAAATAGAGATGCTCTCTCAGTTCCGTCATCGTCATCTTGTTTCATTGATTGGATATTGTGATGAAAAGAATGAGATGATT

>M6-PGSC0003DMG400009689M6RLKmalectin_186

CTTCAGCTGTAGTTATAATTGAAATTTTTTCCAATTCTTGGATTGTTGGGGTTTTGAAATGTTGTCATTACATAGTTTTTTGTTTGTATTGTTGTTGGGGTTTTTCTGTGTTTGTTCTGG

>M6-PGSC0003DMG400009689M6RLKmalectin_1860

CCGTCATCGTCATCTTGTTTCATTGATTGGATATTGTGATGAAAAGAATGAGATGATTCTTGTGTACGAGTACATGGAAAATGGGACCCTCAAGAGCCATCTGTATGGTTCGGATATGCC

>M6-PGSC0003DMG400009689M6RLKmalectin_1922

TGTACGAGTACATGGAAAATGGGACCCTCAAGAGCCATCTGTATGGTTCGGATATGCCTAGTATGAGCTGGAAACAGAGGCTGGAGATTTGCATTGGGGCAGCCAGAGGTCTACACTACC

>M6-PGSC0003DMG400009689M6RLKmalectin_1984

ATGAGCTGGAAACAGAGGCTGGAGATTTGCATTGGGGCAGCCAGAGGTCTACACTACCTTCATACTAGCTATGCCAAAGCAGTTATACATCGTGATGTCAAATCAGCAAACATATTGCTT

>M6-PGSC0003DMG400009689M6RLKmalectin_2046

TACTAGCTATGCCAAAGCAGTTATACATCGTGATGTCAAATCAGCAAACATATTGCTTGATGAGAATATGATGGCAAAAGTAGCTGATTTCGGACTTTCTAAGGCTGGACCTGAGCTCGA

>M6-PGSC0003DMG400009689M6RLKmalectin_2108

AGAATATGATGGCAAAAGTAGCTGATTTCGGACTTTCTAAGGCTGGACCTGAGCTCGATCAAACTCATGTTAGCACGGCAGTGAAAGGAAGCTTCGGGTACCTAGATCCTGAATATTTCA

>M6-PGSC0003DMG400009689M6RLKmalectin_2170

ACTCATGTTAGCACGGCAGTGAAAGGAAGCTTCGGGTACCTAGATCCTGAATATTTCAGGAGGCAGCAACTGACTGAAAAAAGTGATGTTTACTCTTTTGGTGTTGTTCTGTTTGAAGTT

>M6-PGSC0003DMG400009689M6RLKmalectin_2232

GCAGCAACTGACTGAAAAAAGTGATGTTTACTCTTTTGGTGTTGTTCTGTTTGAAGTTCTTTGTGCTAGGCCTGTAATCGATCCGTCTCTTCCAAGGGAGATGGTTAATTTAGCTGAATG

>M6-PGSC0003DMG400009689M6RLKmalectin_2294

GTGCTAGGCCTGTAATCGATCCGTCTCTTCCAAGGGAGATGGTTAATTTAGCTGAATGGGCGATGAAGTGGCAGAAGAAGGGACAACTAGAGCAAATCATAGATCCCAATCTTGCTGGAA

>M6-PGSC0003DMG400009689M6RLKmalectin_2356

ATGAAGTGGCAGAAGAAGGGACAACTAGAGCAAATCATAGATCCCAATCTTGCTGGAAAAATAAGACCGGATTCCCTAAGGAAGTTCGGTGAAACAGCAGAGAAATGCTTGGCTGATTTT

>M6-PGSC0003DMG400009689M6RLKmalectin_2418

AAGACCGGATTCCCTAAGGAAGTTCGGTGAAACAGCAGAGAAATGCTTGGCTGATTTTGGTGTGGACAGACCTTCAATGGGCGACGTGCTGTGGAACCTCGAGTATGCACTTCAACTACA

>M6-PGSC0003DMG400009689M6RLKmalectin_248

TGTCATTACATAGTTTTTTGTTTGTATTGTTGTTGGGGTTTTTCTGTGTTTGTTCTGGATTTGATCCTTCTGATGAGTACTTAATAGACTGTGGATCTTCAAAAGATACTAATTTTGGTA

>M6-PGSC0003DMG400009689M6RLKmalectin_2480

TGGACAGACCTTCAATGGGCGACGTGCTGTGGAACCTCGAGTATGCACTTCAACTACAAGAGGCAGTTATTCAAGTCGATCCTGACGAGAACAGTAGCAATCTCATCGGAGAACTCTCTC

>M6-PGSC0003DMG400009689M6RLKmalectin_2542

GCAGTTATTCAAGTCGATCCTGACGAGAACAGTAGCAATCTCATCGGAGAACTCTCTCCTCAGGTAAACGATTTCAGTCACGTCGATGCCACTACTTCAGCAATGCAATTCGAAGGGTCG

>M6-PGSC0003DMG400009689M6RLKmalectin_2604

GGTAAACGATTTCAGTCACGTCGATGCCACTACTTCAGCAATGCAATTCGAAGGGTCGAATCTAGATGATCTATCTGGAGTTTCCATGAGTAAAGTTTTCTCACAATTGGTTAAGTCTGA

>M6-PGSC0003DMG400009689M6RLKmalectin_2666

TAGATGATCTATCTGGAGTTTCCATGAGTAAAGTTTTCTCACAATTGGTTAAGTCTGAAGGAAGATAAATTTGGTGAAAGATTGTATACAATTGTACTCTTTTTTTTCTTAGCTCCTCTT

>M6-PGSC0003DMG400009689M6RLKmalectin_2728

AGATAAATTTGGTGAAAGATTGTATACAATTGTACTCTTTTTTTTCTTAGCTCCTCTTTTAATTCCACTTTTTTGTTCATTAATCATTTGAAGAATGTACTTTTTATTTTTTTAAACTAT

>M6-PGSC0003DMG400009689M6RLKmalectin_2776

AGCTCCTCTTTTAATTCCACTTTTTTGTTCATTAATCATTTGAAGAATGTACTTTTTATTTTTTTAAACTATAGTTTTACATGTACTACTTTTGTTATGAATATAAAAATTATTCCAGCA

>M6-PGSC0003DMG400009689M6RLKmalectin_310

GATCCTTCTGATGAGTACTTAATAGACTGTGGATCTTCAAAAGATACTAATTTTGGTAATAGGGTTTTTGTGTCTGATAAATCTGGTTCGAAGTTTCTGTCAGTTTCTCAAGATATATTA

>M6-PGSC0003DMG400009689M6RLKmalectin_372

GGTTTTTGTGTCTGATAAATCTGGTTCGAAGTTTCTGTCAGTTTCTCAAGATATATTAGCTGATAATACAAGGTCTAGTGATTCACCCCTTTATCAAACAGCTAGAGTATTTACTCAAAC

>M6-PGSC0003DMG400009689M6RLKmalectin_434

ATAATACAAGGTCTAGTGATTCACCCCTTTATCAAACAGCTAGAGTATTTACTCAAACTTCTAGTTATAAATTCCCTATTACTAAAGTTGGAAGGCATTGGATCCGTCTTCATTTTTCCC

>M6-PGSC0003DMG400009689M6RLKmalectin_496

AGTTATAAATTCCCTATTACTAAAGTTGGAAGGCATTGGATCCGTCTTCATTTTTCCCCATTTGTGTACCAATCTTATGATATGAGTTCAGCAACTTTCTCTGTTTCAACTCAACAAAAT

>M6-PGSC0003DMG400009689M6RLKmalectin_558

TGTGTACCAATCTTATGATATGAGTTCAGCAACTTTCTCTGTTTCAACTCAACAAAATGTTCTCCTTGGAAATTTTAGTCCTAAAAAAGATGTATCTGTTAAGGAGTTTTCTGTTAACGT

>M6-PGSC0003DMG400009689M6RLKmalectin_62

CTTTACTACCCCTTCCCAAAAAAAAAAAAAAAAAACTTGTGATTTGGCTTCTATTCACCCATTCCAAATCTGGGCAATCTCATTCTTGGAGCATATTTAATCAAAAAGGTTCAATCTTTT

>M6-PGSC0003DMG400009689M6RLKmalectin_620

TCCTTGGAAATTTTAGTCCTAAAAAAGATGTATCTGTTAAGGAGTTTTCTGTTAACGTAACGACGGGTGATCTTGTTATCACTTTTTCCCCTTTGAGTGATTCATTTGCTTATATAAATG

>M6-PGSC0003DMG400009689M6RLKmalectin_682

ACGGGTGATCTTGTTATCACTTTTTCCCCTTTGAGTGATTCATTTGCTTATATAAATGCTTTGGAAGTTGTCTCTGTACCTGATAGTCTTATAACTAATGATGCCTCGACGTTTGGTCCA

>M6-PGSC0003DMG400009689M6RLKmalectin_744

GGAAGTTGTCTCTGTACCTGATAGTCTTATAACTAATGATGCCTCGACGTTTGGTCCATCTGGAGTATTTAGTGGTATGTATGCACAAGCACTTGAAACAGTTGCTAGAGTAAATATGGG

>M6-PGSC0003DMG400009689M6RLKmalectin_806

GAGTATTTAGTGGTATGTATGCACAAGCACTTGAAACAGTTGCTAGAGTAAATATGGGTGGACCAGCTGTATCGTCTGAAAACGATACATTATGGAGGAGTTGGGTGCCTGATCGGGCTT

>M6-PGSC0003DMG400009689M6RLKmalectin_868

CCAGCTGTATCGTCTGAAAACGATACATTATGGAGGAGTTGGGTGCCTGATCGGGCTTTCTTGACACAGCCGGATTTTGCAAAGTCCTTGTCTAAGATTGGATCTGTAAAGTATTTAGCG

>M6-PGSC0003DMG400009689M6RLKmalectin_930

GACACAGCCGGATTTTGCAAAGTCCTTGTCTAAGATTGGATCTGTAAAGTATTTAGCGGGTGGAGCAACATCGGATATAGCACCACCGAGTGTTTATGGTACTTGTACTAAGATGAATGT

>M6-PGSC0003DMG400009689M6RLKmalectin_992

GAGCAACATCGGATATAGCACCACCGAGTGTTTATGGTACTTGTACTAAGATGAATGTGGATGCTGCTGGTAATGATGCTAGTGCTAATTTTAATGTGACGTGGGTGTTCAATGTGGATG

>M6-PGSC0003DMG400009755M6LRR-RLP_0

ATGGAGGAACCAAGTTATGAAGGACGTGGATATTACCAAGACTCGGTAGTTGTTGTAACAAAGGGATTGAAGCTTGAAGTTGTGAGAATTTTGTCTTTGTACACAGTTATCGATCTTTCA

>M6-PGSC0003DMG400009755M6LRR-RLP_1040

ATTTTCAAATGCCAAGAATAAGAAGACGTTCGCGTAACCGATTTGGTTCTTCCTGCAGCTTATGATGTTTGATTTGATTTTTAGTTTCTTTTTATCTTTATAAGTAAGCTTTTTCTTAAC

>M6-PGSC0003DMG400009755M6LRR-RLP_1090

TCCTGCAGCTTATGATGTTTGATTTGATTTTTAGTTTCTTTTTATCTTTATAAGTAAGCTTTTTCTTAACTTGGAAAATGTAATATTATGAATTTTGACAATATATCAAGTTTATGTAAG

>M6-PGSC0003DMG400009755M6LRR-RLP_130

TCGAAGGACATATTCCTAGTGTTCTAGGAGGTCTCATTGCTCTTCGGGTGTTGAATATATCTCACAATGGATTGAAAGGCCATATACCACCATCACTTGGACGTTTATCTGTAGTGGAAT

>M6-PGSC0003DMG400009755M6LRR-RLP_195

AATGGATTGAAAGGCCATATACCACCATCACTTGGACGTTTATCTGTAGTGGAATCATTGGACCTTTCATTTAACTGCCTTTCGGGAGAGATACCACAACAACTTGCTTCTCTTACGTCT

>M6-PGSC0003DMG400009755M6LRR-RLP_390

AACAGTTCATTTGGTGGTAATGATGGATTACGTGGATTCCCCATTTCAAAAGGTTGTGGCAATGATCTTGTGTCAGAGACAAATTATACAGTATCTGCGCTAGACGATCAAGAAAGCAAT

>M6-PGSC0003DMG400009755M6LRR-RLP_455

TCTTGTGTCAGAGACAAATTATACAGTATCTGCGCTAGACGATCAAGAAAGCAATTCTGAATTTCTCAATGATTTTTGGAAAGCTGCTCTGATGGGCTATGGCAGTGGATTATGTATTGG

>M6-PGSC0003DMG400009755M6LRR-RLP_585

ATGTATTTCATGATTTCAACTAGAAATCCGAAATGGCTTGCAAGAATCATTGAAGAAATGGAACACAAAATTAACATGCGAAGGAGAAAGAAGCAGCAAGGTCAAAGGAATTACAGAAGA

>M6-PGSC0003DMG400009755M6LRR-RLP_650

CAAAATTAACATGCGAAGGAGAAAGAAGCAGCAAGGTCAAAGGAATTACAGAAGAAGAAATAATCGCTTATAGACAAGTTACCAATAAAGAAGAATTTGATTTCAGAACTTCAGGTATTC

>M6-PGSC0003DMG400009755M6LRR-RLP_715

GCTTATAGACAAGTTACCAATAAAGAAGAATTTGATTTCAGAACTTCAGGTATTCAAGCTAAGCTCTACACTAATCTTTTTTAGTTCATTCTACTACTAATATGCTAGGTTTTATGCTTT

>M6-PGSC0003DMG400009755M6LRR-RLP_780

CTACACTAATCTTTTTTAGTTCATTCTACTACTAATATGCTAGGTTTTATGCTTTTATGTTATAAACAACAACTAATTAATGCTCGATACGAAGCATATTTTTCTTTCCAACAACTTTAA

>M6-PGSC0003DMG400009755M6LRR-RLP_845

ACAACAACTAATTAATGCTCGATACGAAGCATATTTTTCTTTCCAACAACTTTAAGAAGCTGAAGATTTGAAAAAGGATTAGCCTTAGCTAATGTTATGGATGGTTTAACAGTGAAAAAC

>M6-PGSC0003DMG400009755M6LRR-RLP_910

ATTTGAAAAAGGATTAGCCTTAGCTAATGTTATGGATGGTTTAACAGTGAAAAACTAATGCATAATTCGGTTTTAGCTATATGATTGAAATATAATTTACAATTGTTTGACTCACTATCT

>M6-PGSC0003DMG400009755M6LRR-RLP_975

TTCGGTTTTAGCTATATGATTGAAATATAATTTACAATTGTTTGACTCACTATCTTATTTTTCAGATTTTCAAATGCCAAGAATAAGAAGACGTTCGCGTAACCGATTTGGTTCTTCCTG

>M6-PGSC0003DMG400009843M6LRR-RLK_0

TTAGAGACTATTTTGACCCTACGGCCAAAATTAATGGACCATATTTGTCATTTTCTTTCTTCTTCACCCTACCCCTCGTTTTCCCACTTCACTCTTCTCCATAGTTCCTCACTGTCTCCG

>M6-PGSC0003DMG400009843M6LRR-RLK_1037

CCAAAACGCACAGTCGCAAGGTATTCTTATTACTCCTCCACCTCAGCACAAACACAAAAAAGTTGGTGTTGTTTTGGGTTTTGTAGTTGGAACATTGATCTTGATTGCAGCTGTTCTCTG

>M6-PGSC0003DMG400009843M6LRR-RLK_1098

GTTGGTGTTGTTTTGGGTTTTGTAGTTGGAACATTGATCTTGATTGCAGCTGTTCTCTGTCTTTTTGCTTTTGTGAAAAAGAGAAGAGAATAAACTGAAACTGAATCAAAAGCAACAAAA

>M6-PGSC0003DMG400009843M6LRR-RLK_1159

TTTTTGCTTTTGTGAAAAAGAGAAGAGAATAAACTGAAACTGAATCAAAAGCAACAAAATGTACCATTGAGACCATCACTAATAGTGCGGCGAACGCAACGGTTTCTGAGCCAGATGATA

>M6-PGSC0003DMG400009843M6LRR-RLK_122

AATCCAGTCCATTTTTCAGACCTCCTTTAACGCACAACATGACAACAACAAACACTAAACTTGGCCTCTAGTCACTAAACCAAATCACTCGACATTTTTAGTTGCATAACTTGCAATGTC

>M6-PGSC0003DMG400009843M6LRR-RLK_1220

TACCATTGAGACCATCACTAATAGTGCGGCGAACGCAACGGTTTCTGAGCCAGATGATAGTAGCCAAGAGATTAAATTAGAAAAGGAAATGAAAGTGCTTCAAGCCCCCAAACAACAGAT

>M6-PGSC0003DMG400009843M6LRR-RLK_1281

AGCCAAGAGATTAAATTAGAAAAGGAAATGAAAGTGCTTCAAGCCCCCAAACAACAGATGAAGAGTGGAAATCTGATATTTTGTTCAGGGGAAACAGAGTTGTATTCTTTAGAACAACTA

>M6-PGSC0003DMG400009843M6LRR-RLK_1342

AGAGTGGAAATCTGATATTTTGTTCAGGGGAAACAGAGTTGTATTCTTTAGAACAACTAATGAGAGCATCAGCAGAGTTGCTTGGAAGAGGCACAATTGGAACAACGTATAAAGCTCTGA

>M6-PGSC0003DMG400009843M6LRR-RLK_1403

GAGAGCATCAGCAGAGTTGCTTGGAAGAGGCACAATTGGAACAACGTATAAAGCTCTGATGGCTAGTCAGTTAATCGTTTCAGTAAAGCGATTGGATGCTGGTAAAACTTCGATTACGAG

>M6-PGSC0003DMG400009843M6LRR-RLK_1464

GCTAGTCAGTTAATCGTTTCAGTAAAGCGATTGGATGCTGGTAAAACTTCGATTACGAGTGCAGAAGCGTTTGAGCAACACATGGAGTCAGTTGGGATGTTGAGGCACCCAAATTTGGTT

>M6-PGSC0003DMG400009843M6LRR-RLK_1525

CAGAAGCGTTTGAGCAACACATGGAGTCAGTTGGGATGTTGAGGCACCCAAATTTGGTTGCAGTCAGAGCATACTTTCAGGCTAAACAAGAAAGTCTTGTTATCTATGATTATCAGCCTA

>M6-PGSC0003DMG400009843M6LRR-RLK_1586

AGTCAGAGCATACTTTCAGGCTAAACAAGAAAGTCTTGTTATCTATGATTATCAGCCTAATGGCAGTCTCTTCAATCTAATTCATGGTAAAGTTTCTAACTTTTTAATATATCATTACTA

>M6-PGSC0003DMG400009843M6LRR-RLK_1647

GGCAGTCTCTTCAATCTAATTCATGGTAAAGTTTCTAACTTTTTAATATATCATTACTAGTTCATCTGAAATGGATCAATGTTTATTTTTCATTTTTGGTAATTCTGCTAATTGGTCAAT

>M6-PGSC0003DMG400009843M6LRR-RLK_1708

TCATCTGAAATGGATCAATGTTTATTTTTCATTTTTGGTAATTCTGCTAATTGGTCAATGTGAAGTCAATTTGGTGGACCTGTGATGGAGAAAAAAGTGGAAGAGTGAGTTGTCTGTGGC

>M6-PGSC0003DMG400009843M6LRR-RLK_1769

GAAGTCAATTTGGTGGACCTGTGATGGAGAAAAAAGTGGAAGAGTGAGTTGTCTGTGGCTACTGGCTAATCACTATCTAGCTTTTTTTTTTTCTTTACAGAAATTAGTTGTAGTGTTTTG

>M6-PGSC0003DMG400009843M6LRR-RLK_1830

CTGGCTAATCACTATCTAGCTTTTTTTTTTTCTTTACAGAAATTAGTTGTAGTGTTTTGGATCTTTCAGTGGTTGCTATGGTTGTAGTAGCAGGGGACTTTTTTAGCTTTTAGTGTGGAT

>M6-PGSC0003DMG400009843M6LRR-RLK_1891

TCTTTCAGTGGTTGCTATGGTTGTAGTAGCAGGGGACTTTTTTAGCTTTTAGTGTGGATGCCTAACATGGCTCCCATTGAGTTCACTACCGACAATTGCAAAGTTTGTGTTTACAGGGCA

>M6-PGSC0003DMG400009843M6LRR-RLK_1952

CTAACATGGCTCCCATTGAGTTCACTACCGACAATTGCAAAGTTTGTGTTTACAGGGCAGCCAAATAGTAATGTGTTTTAGTTTCAGGCTGTCACTGACTGACCGGTATGGGTTGATAGC

>M6-PGSC0003DMG400009843M6LRR-RLK_2013

CAAATAGTAATGTGTTTTAGTTTCAGGCTGTCACTGACTGACCGGTATGGGTTGATAGCATTATATTCCTCAAATATTTTTATCATGCATTGAGGCAAATGGTCCTTGGCTCTTGCGTCT

>M6-PGSC0003DMG400009843M6LRR-RLK_2074

TATATTCCTCAAATATTTTTATCATGCATTGAGGCAAATGGTCCTTGGCTCTTGCGTCTGTCTTTTTGGTATTTATGAGTATGTTTGCTTCTTCTTTAAGTTGAATAGGATGTTGACAGT

>M6-PGSC0003DMG400009843M6LRR-RLK_2135

CTTTTTGGTATTTATGAGTATGTTTGCTTCTTCTTTAAGTTGAATAGGATGTTGACAGTAACTTTTTGGTGTTTTCCTTTTACTTGTCCAAATGTTCTCTGTTTCTTAAATATTAAATTT

>M6-PGSC0003DMG400009843M6LRR-RLK_2196

CTTTTTGGTGTTTTCCTTTTACTTGTCCAAATGTTCTCTGTTTCTTAAATATTAAATTTACTTTAAGTGAACGTTAACGTTGTTAGATAGAAGGATTAAGGGAAAGATCTAGAACATCCT

>M6-PGSC0003DMG400009843M6LRR-RLK_2257

TTTAAGTGAACGTTAACGTTGTTAGATAGAAGGATTAAGGGAAAGATCTAGAACATCCTTGGGAACTCTTAGTGTAGTGACAGAAATGGTATGAATTTGCTTAATTTACATTTTTTTGAA

>M6-PGSC0003DMG400009843M6LRR-RLK_2318

GGAACTCTTAGTGTAGTGACAGAAATGGTATGAATTTGCTTAATTTACATTTTTTTGAATCTTAGGTATTGCATTTTTTGTATCCCCTTGTGATACGCTATCAAGAAATTGGGATTTCAT

>M6-PGSC0003DMG400009843M6LRR-RLK_2379

TTAGGTATTGCATTTTTTGTATCCCCTTGTGATACGCTATCAAGAAATTGGGATTTCATTTTCATAGATTACTCTGCTCAAGAAGATTGATTCCTCTATATAATTGTGTGTGTTTTTTAA

>M6-PGSC0003DMG400009843M6LRR-RLK_2440

TCATAGATTACTCTGCTCAAGAAGATTGATTCCTCTATATAATTGTGTGTGTTTTTTAAGTGGTTGGTCTTTTCACCTAAAACCTGATAATCTGTTGTTCGTTTACTTATATATTGTGTT

>M6-PGSC0003DMG400009843M6LRR-RLK_2501

GGTTGGTCTTTTCACCTAAAACCTGATAATCTGTTGTTCGTTTACTTATATATTGTGTTCAATTTTCTATTGTTCGTGGCAGGTTCAAGATCAACAAGGGCAATGCCTCTTCACTGGACA

>M6-PGSC0003DMG400009843M6LRR-RLK_2562

ATTTTCTATTGTTCGTGGCAGGTTCAAGATCAACAAGGGCAATGCCTCTTCACTGGACATCGTGTCTAAAAATAGCAGAAGATGTGGCTCAGGGCCTTGCTTACATCCATCAAGCGTCAA

>M6-PGSC0003DMG400009843M6LRR-RLK_2623

GTGTCTAAAAATAGCAGAAGATGTGGCTCAGGGCCTTGCTTACATCCATCAAGCGTCAAATCTCACTCATGGCAACTTGAAATCCTCCAACGTCCTTCTAGGGTCTGATTTTGAGGCCTG

>M6-PGSC0003DMG400009843M6LRR-RLK_2684

CTCACTCATGGCAACTTGAAATCCTCCAACGTCCTTCTAGGGTCTGATTTTGAGGCCTGTCTAACAGACTACTCCATCATCGCGCTTGCAGATATTTCTTCGGAGGATGATCCTGATTCA

>M6-PGSC0003DMG400009843M6LRR-RLK_2745

TAACAGACTACTCCATCATCGCGCTTGCAGATATTTCTTCGGAGGATGATCCTGATTCAGCATGCTATAAAGCACCAGAAGTGCGCAAATCAGCACGAAGAGCCACACCAGGGTCTGACG

>M6-PGSC0003DMG400009843M6LRR-RLK_2806

ATGCTATAAAGCACCAGAAGTGCGCAAATCAGCACGAAGAGCCACACCAGGGTCTGACGTATATGCCTATGGCATTCTGTTATTAGAGCTTTTAACGGGTAAACCTCCATCTCAGCATCC

>M6-PGSC0003DMG400009843M6LRR-RLK_2867

TATGCCTATGGCATTCTGTTATTAGAGCTTTTAACGGGTAAACCTCCATCTCAGCATCCCCATCTCTCTCCCCCTGATGTGCCAGATTGGGTTAGAGCTATGAGGGAGGATGATAATGAG

>M6-PGSC0003DMG400009843M6LRR-RLK_2928

ATCTCTCTCCCCCTGATGTGCCAGATTGGGTTAGAGCTATGAGGGAGGATGATAATGAGGAGGATAGATGGCTTGCAATGCTAGTTGACCTTGCTAGTATATGTAGCTTGACATCACCAG

>M6-PGSC0003DMG400009843M6LRR-RLK_2989

GGATAGATGGCTTGCAATGCTAGTTGACCTTGCTAGTATATGTAGCTTGACATCACCAGAACAGAGACCAACAATGAGGCAAATCTTGAAAATGATACAAGATATAAAGGATAGTGCAAT

>M6-PGSC0003DMG400009843M6LRR-RLK_3050

CAGAGACCAACAATGAGGCAAATCTTGAAAATGATACAAGATATAAAGGATAGTGCAATGGTGGAGAATAACAAAAGAGATGCACATAATGGATACTCATAGCTTCTTCTGAATTGCAGT

>M6-PGSC0003DMG400009843M6LRR-RLK_3111

TGGAGAATAACAAAAGAGATGCACATAATGGATACTCATAGCTTCTTCTGAATTGCAGTTACACCATTAACCAGCGAGGGTACATGGTCAATTGTAAGTAAGTGTCATCTCCCCCTTGGC

>M6-PGSC0003DMG400009843M6LRR-RLK_3172

CACCATTAACCAGCGAGGGTACATGGTCAATTGTAAGTAAGTGTCATCTCCCCCTTGGCAAACTGCCATTGATCGACAGCATATCTGTAATTGAGGTTTGCTCAGATAGACCAAATGACT

>M6-PGSC0003DMG400009843M6LRR-RLK_3233

ACTGCCATTGATCGACAGCATATCTGTAATTGAGGTTTGCTCAGATAGACCAAATGACTCATCAACTATAATGGTGCTTGTACCTTTACTATTCAGCAGTAAACAACCTAAAACTGACAG

>M6-PGSC0003DMG400009843M6LRR-RLK_3294

TCAACTATAATGGTGCTTGTACCTTTACTATTCAGCAGTAAACAACCTAAAACTGACAGGTTAAGGTCTATCTGCTAGAACATGGTCAACTTAGTCACTGAAGATTACGACACATTTTGT

>M6-PGSC0003DMG400009843M6LRR-RLK_3355

TAAGGTCTATCTGCTAGAACATGGTCAACTTAGTCACTGAAGATTACGACACATTTTGTTCGAAATCCCTCTCTTTTTTGGTCACACAAACATATGGTGCCTATATTTTTAGGGCATTTT

>M6-PGSC0003DMG400009843M6LRR-RLK_3416

GAAATCCCTCTCTTTTTTGGTCACACAAACATATGGTGCCTATATTTTTAGGGCATTTTGTAGTAGGTGGTTTTGGTTGATTCTTTTTCTAAAGAACTTCAAAATGTATTCGCTTTACCT

>M6-PGSC0003DMG400009843M6LRR-RLK_3477

AGTAGGTGGTTTTGGTTGATTCTTTTTCTAAAGAACTTCAAAATGTATTCGCTTTACCTTAGTGGCAGAAGAATATTGTATAATCTTGAATGGTAAAGCTAACCCAAATTCTGAACTTTC

>M6-PGSC0003DMG400009843M6LRR-RLK_3531

TACCTTAGTGGCAGAAGAATATTGTATAATCTTGAATGGTAAAGCTAACCCAAATTCTGAACTTTCCCTTCTCTGCATGGTTGGTTAGTATCAACTTAGCAGGAGAAGAGTTGTTTAATG

>M6-PGSC0003DMG400009843M6LRR-RLK_366

CCCTCCGACGCTGTTTCTTTATTGTCTTTCAAGTCAAAAGCTGACCTTGACAACAAGCTTCATTACACACTTAACGAGCGTTTTGATTACTGTCAATGGCGAGGAGTGAAATGTGTACAG

>M6-PGSC0003DMG400009843M6LRR-RLK_427

ATTACACACTTAACGAGCGTTTTGATTACTGTCAATGGCGAGGAGTGAAATGTGTACAGGGTCGTGTTGTGCGTTTTGTTCTTCAAGGGTTTAGTCTCCGGGGTACTTTTCCGGCGAATA

>M6-PGSC0003DMG400009843M6LRR-RLK_488

TCGTGTTGTGCGTTTTGTTCTTCAAGGGTTTAGTCTCCGGGGTACTTTTCCGGCGAATAGTTTAACTCATCTTGACCAACTCCGAATCCTTAATCTTAGGAATAACTCACTTTCCGGTCC

>M6-PGSC0003DMG400009843M6LRR-RLK_549

TTAACTCATCTTGACCAACTCCGAATCCTTAATCTTAGGAATAACTCACTTTCCGGTCCTATTCCTGATCTTTCTGGTCTTCCCAACCTCAAAACTCTGTTTCTTGATCATAATTTTTTC

>M6-PGSC0003DMG400009843M6LRR-RLK_61

CTTCACCCTACCCCTCGTTTTCCCACTTCACTCTTCTCCATAGTTCCTCACTGTCTCCGAAAATCCAGTCCATTTTTCAGACCTCCTTTAACGCACAACATGACAACAACAAACACTAAA

>M6-PGSC0003DMG400009843M6LRR-RLK_610

TTCCTGATCTTTCTGGTCTTCCCAACCTCAAAACTCTGTTTCTTGATCATAATTTTTTCTCTGGAACTTTTCCCCTTTCTGTTCTTTCTATTCGCCGCCTTGTTATTCTTGATCTTTCTC

>M6-PGSC0003DMG400009843M6LRR-RLK_671

TGGAACTTTTCCCCTTTCTGTTCTTTCTATTCGCCGCCTTGTTATTCTTGATCTTTCTCGGAACAATCTCACTGGTTCACTCCCTGTTAGGCTTACTGTTCTTGACCGGTTGAACTATCT

>M6-PGSC0003DMG400009843M6LRR-RLK_732

AACAATCTCACTGGTTCACTCCCTGTTAGGCTTACTGTTCTTGACCGGTTGAACTATCTCCGGCTTGATTCCAACTGGTTTAACGGTTCGATTCCGCCGTTGAACCAGACCCAACTCCAA

>M6-PGSC0003DMG400009843M6LRR-RLK_793

GGCTTGATTCCAACTGGTTTAACGGTTCGATTCCGCCGTTGAACCAGACCCAACTCCAAATATTCAATGTGTCTAATAATAACCTTACTGGTCCAGTACCTGTTACTCCAACTCTGAAGA

>M6-PGSC0003DMG400009843M6LRR-RLK_854

ATTCAATGTGTCTAATAATAACCTTACTGGTCCAGTACCTGTTACTCCAACTCTGAAGAAATTTAACATACGCTCGTTCTTACGGAACCCTAGTCTTTGTGGTGAGGTTGTCGACAAGCC

>M6-PGSC0003DMG400009843M6LRR-RLK_915

TTTAACATACGCTCGTTCTTACGGAACCCTAGTCTTTGTGGTGAGGTTGTCGACAAGCCGTGCCGTTCGGCTCCATTCTTTGATTCGCCTTCCTCTGCTGCCTCTCCGCCCACACCGTTA

>M6-PGSC0003DMG400009843M6LRR-RLK_976

GCCGTTCGGCTCCATTCTTTGATTCGCCTTCCTCTGCTGCCTCTCCGCCCACACCGTTATACCAAAACGCACAGTCGCAAGGTATTCTTATTACTCCTCCACCTCAGCACAAACACAAAA

>M6-PGSC0003DMG400009899M6LRR-RLP_0

ATTCTCTAATAAATTTCTCTTTGATTATTGAGCAAAATTTTGTCACCAATGGAAGGTGTAAATTTACTACTAGTCCTTCTTCTTATAGCTAGTAGTACTATATTAGATGGATTGATATTA

>M6-PGSC0003DMG400009899M6LRR-RLP_1037

TTCATCCAATAATCTGGAGGGTACTTTAGCAAGAAGCATCGCACAACTTAGGCAATTAGTTGTCCTCAATGTTGCTTGGAACTCTTTCAATGACTCGATTACTGAACATTTCTTGAATTT

>M6-PGSC0003DMG400009899M6LRR-RLP_1098

GTCCTCAATGTTGCTTGGAACTCTTTCAATGACTCGATTACTGAACATTTCTTGAATTTCAGTGATTTGAGAGTGTTAGACTTGTCATCCAACTCAATTATCCTCAATATAACTGCAACT

>M6-PGSC0003DMG400009899M6LRR-RLP_1159

GTGATTTGAGAGTGTTAGACTTGTCATCCAACTCAATTATCCTCAATATAACTGCAACTTGGATGCCTCCCTTTCAACTTGACATTATTGGCCTGATGTCTTGCCAACTTGGTACACAGT

>M6-PGSC0003DMG400009899M6LRR-RLP_122

CAGGGCTATGAATTCAGACATTCTTTGTGATGAAAATGAAAAAAATGCTCTTATCAAGTTCAGACAGGGTTTAAGGAATCCATCACAGTCAATGATCATGTCGTCTTGGATGCTCGAGGA

>M6-PGSC0003DMG400009899M6LRR-RLP_1220

GATGCCTCCCTTTCAACTTGACATTATTGGCCTGATGTCTTGCCAACTTGGTACACAGTTTCCCCAGTGGCTTCATACACAAAAGGATTTTCTCATTTATCGATATCTCATATGGTAACA

>M6-PGSC0003DMG400009899M6LRR-RLP_1281

CCCCAGTGGCTTCATACACAAAAGGATTTTCTCATTTATCGATATCTCATATGGTAACATCTCAGATAAAGTGCCTGATTGGTTCTGGAATCTCTCTCCAAAGGTACAACACATGAACCT

>M6-PGSC0003DMG400009899M6LRR-RLP_1342

TCAGATAAAGTGCCTGATTGGTTCTGGAATCTCTCTCCAAAGGTACAACACATGAACCTTTCGCTTAACTATTTCAGAGGGGAAGTACCTCAAGTTACATCAAGACTTACTAGTCTTGAA

>M6-PGSC0003DMG400009899M6LRR-RLP_1403

CGCTTAACTATTTCAGAGGGGAAGTACCTCAAGTTACATCAAGACTTACTAGTCTTGAAAAACTAGACCTGAGCCAAAACAACTTTCATGGTCCTTTACCTCATTTTTCGTCCAAGATGA

>M6-PGSC0003DMG400009899M6LRR-RLP_1464

ACTAGACCTGAGCCAAAACAACTTTCATGGTCCTTTACCTCATTTTTCGTCCAAGATGAAGACGTTGATTTTGTCCCGGAATTCTTTTAATGGCGCCATTTCCCCTGTATGTGAATCACT

>M6-PGSC0003DMG400009899M6LRR-RLP_1525

ACGTTGATTTTGTCCCGGAATTCTTTTAATGGCGCCATTTCCCCTGTATGTGAATCACTTGTCATGAATAATTCTCTTATGTTTCTTGACTTATCGTTCAATAATCTATCAGGAACACTT

>M6-PGSC0003DMG400009899M6LRR-RLP_1586

TCATGAATAATTCTCTTATGTTTCTTGACTTATCGTTCAATAATCTATCAGGAACACTTTCAGACTGTTGGAGATATGGAAACAATCTGGCAGTATTAAAGTTAGGCAATAATAATCTGT

>M6-PGSC0003DMG400009899M6LRR-RLP_1647

AGACTGTTGGAGATATGGAAACAATCTGGCAGTATTAAAGTTAGGCAATAATAATCTGTCTGGTGAAATACCTCATTCTTTTGGATATCTTGAAACTCTCAGGTATCTGCAACTGAGAAA

>M6-PGSC0003DMG400009899M6LRR-RLP_1708

GGTGAAATACCTCATTCTTTTGGATATCTTGAAACTCTCAGGTATCTGCAACTGAGAAAGAACAGGTTGTCCAAAAATCTGCCTTCATCACTGAAGAACTTACAAGAACTAAAATTACTT

>M6-PGSC0003DMG400009899M6LRR-RLP_1769

ACAGGTTGTCCAAAAATCTGCCTTCATCACTGAAGAACTTACAAGAACTAAAATTACTTGATCTTAGCAGAAATTGTTTATCTGGAACGATACCATCTTTCTTAGGGGAGACTTCACAGA

>M6-PGSC0003DMG400009899M6LRR-RLP_183

AGACAGGGTTTAAGGAATCCATCACAGTCAATGATCATGTCGTCTTGGATGCTCGAGGAAAATTGCTGCAACTGGAAAGGTGTTGAATGTGACAACACAACCGGACATGTGATCACTCTT

>M6-PGSC0003DMG400009899M6LRR-RLP_1830

TCTTAGCAGAAATTGTTTATCTGGAACGATACCATCTTTCTTAGGGGAGACTTCACAGAAGCTAATGCTTCTTTCGTTAGGAGAGAACAAGTTCGATGGAAACATTCCCCTACAGATTTG

>M6-PGSC0003DMG400009899M6LRR-RLP_1891

CTAATGCTTCTTTCGTTAGGAGAGAACAAGTTCGATGGAAACATTCCCCTACAGATTTGCCAGCTCAAAAACTTGATTATTTTGGACCTTAGTTCCAATGCTCTATCAGGCACCATTCCA

>M6-PGSC0003DMG400009899M6LRR-RLP_1952

AGCTCAAAAACTTGATTATTTTGGACCTTAGTTCCAATGCTCTATCAGGCACCATTCCAAAGTGTGTTAAAAATTTCCTAACGATGGCCGGGGTAGAGGAGTTCCCTTCCTTGGTTTATG

>M6-PGSC0003DMG400009899M6LRR-RLP_2013

GTGTGTTAAAAATTTCCTAACGATGGCCGGGGTAGAGGAGTTCCCTTCCTTGGTTTATGGTCCCTATGAAGAATACAGGAAGGATGTGATGTTGATGCTGAAAGAGAGAGGCTATGATTA

>M6-PGSC0003DMG400009899M6LRR-RLP_2074

CCCTATGAAGAATACAGGAAGGATGTGATGTTGATGCTGAAAGAGAGAGGCTATGATTATAGGTTTTTCCTTGCGGCAATTTATCTATCAGATAACCATTTATCAGGGGAAATTCCTGAG

>M6-PGSC0003DMG400009899M6LRR-RLP_2135

GGTTTTTCCTTGCGGCAATTTATCTATCAGATAACCATTTATCAGGGGAAATTCCTGAGGAAATCACCACACTTGTTCAATTGCAGGTATTGAATTTATCAAGGAATAATTTAACTGGAG

>M6-PGSC0003DMG400009899M6LRR-RLP_2196

AATCACCACACTTGTTCAATTGCAGGTATTGAATTTATCAAGGAATAATTTAACTGGAGCAATTCCTTGTGACATTAGTAAATTGCAATATTTGGAAGCACTTGATCTTTCCAGGAATTA

>M6-PGSC0003DMG400009899M6LRR-RLP_2257

ATTCCTTGTGACATTAGTAAATTGCAATATTTGGAAGCACTTGATCTTTCCAGGAATTACCTGTCATCCTTTTTGCCTTCCAGCATAGTGGAGCTTTCCATGCTCGCGATCGCAAATTTT

>M6-PGSC0003DMG400009899M6LRR-RLP_2318

TGTCATCCTTTTTGCCTTCCAGCATAGTGGAGCTTTCCATGCTCGCGATCGCAAATTTTTCGTTCAATAGTTTGACAGGAAAAATTCCAATAGGTCGTCAATTTTTAACCTTCGATAATA

>M6-PGSC0003DMG400009899M6LRR-RLP_2379

GTTCAATAGTTTGACAGGAAAAATTCCAATAGGTCGTCAATTTTTAACCTTCGATAATAGTTCTTATATTGGGAATCCAGAGTTGTGTGGCATGCCACTTTCCAAAAGCTGCTCGGGACA

>M6-PGSC0003DMG400009899M6LRR-RLP_244

ATTGCTGCAACTGGAAAGGTGTTGAATGTGACAACACAACCGGACATGTGATCACTCTTGACCTGCACCAGCAATTCTTGCAAGGTGAGTTTGGGACTTCCTTGCTTGGCTTGTCTCACT

>M6-PGSC0003DMG400009899M6LRR-RLP_2440

TCTTATATTGGGAATCCAGAGTTGTGTGGCATGCCACTTTCCAAAAGCTGCTCGGGACATTTTCGTGAGGACGTTACTCACTGTAACACTCCCAAGAAAAAAGACGTGCAGGCCATTCAT

>M6-PGSC0003DMG400009899M6LRR-RLP_2501

TTCGTGAGGACGTTACTCACTGTAACACTCCCAAGAAAAAAGACGTGCAGGCCATTCATCAGCATGAGGAAGACAATTGGCTTGATGAATCATCATTCTATATCAGCATGGTAATTGGTT

>M6-PGSC0003DMG400009899M6LRR-RLP_2562

GCATGAGGAAGACAATTGGCTTGATGAATCATCATTCTATATCAGCATGGTAATTGGTTTCATCACAAGTTTCTGGTTATTTTGGGCTACTCTATTGGTGAAAACATCTTGGAGACATGC

>M6-PGSC0003DMG400009899M6LRR-RLP_2684

ATATGAGGTATCTTAATAAAATGGGCAACAATATTTATGTGTTTGTAGCCATAAGATTACCAAATAAGCATAAACCATCAAAGATGAAAGACTGAAGTTATTTCAGGGCTTTCTGAATAA

>M6-PGSC0003DMG400009899M6LRR-RLP_2745

AAATAAGCATAAACCATCAAAGATGAAAGACTGAAGTTATTTCAGGGCTTTCTGAATAAGGTAATGAAGGTCTTATTTCTGGTGCAACTTTCTTTTTTGTTGTTTTGATTTCTAGGAGCT

>M6-PGSC0003DMG400009899M6LRR-RLP_2806

TAATGAAGGTCTTATTTCTGGTGCAACTTTCTTTTTTGTTGTTTTGATTTCTAGGAGCTCTTCTGGTTTTTTGTTTTTGTAAAATCTTCTAAGTACTATATTGTGTATGCTTAGTTGCAT

>M6-PGSC0003DMG400009899M6LRR-RLP_2867

TCTGGTTTTTTGTTTTTGTAAAATCTTCTAAGTACTATATTGTGTATGCTTAGTTGCATTGTTTCATCTCGTGGGATTGAACTAGGTATGTTGTTGTTGTCATAATTGTATTCTATGATA

>M6-PGSC0003DMG400009899M6LRR-RLP_2915

CTTAGTTGCATTGTTTCATCTCGTGGGATTGAACTAGGTATGTTGTTGTTGTCATAATTGTATTCTATGATATGTTTTGTGGATATCTAAGTAACTTGAGAATACTATTACTAATGTTTT

>M6-PGSC0003DMG400009899M6LRR-RLP_305

CCTGCACCAGCAATTCTTGCAAGGTGAGTTTGGGACTTCCTTGCTTGGCTTGTCTCACTTAAGACACTTAGACTTGAGCCAAAATGATTTACTCAAGGCCTTGCTTCCTGAGTTTATTTG

>M6-PGSC0003DMG400009899M6LRR-RLP_366

AGACACTTAGACTTGAGCCAAAATGATTTACTCAAGGCCTTGCTTCCTGAGTTTATTTGCAAATTTAAGAGCCTAGAGTATCTCAATTTGCACAAAACCAACTTTAGGGGATCAATCCCT

>M6-PGSC0003DMG400009899M6LRR-RLP_427

AATTTAAGAGCCTAGAGTATCTCAATTTGCACAAAACCAACTTTAGGGGATCAATCCCTGAGTGTCTTGGAAATCGGTCGCGGTTGCAGTTACTTGATCTTGGTGATAGTTTTTCACTCC

>M6-PGSC0003DMG400009899M6LRR-RLP_488

GTGTCTTGGAAATCGGTCGCGGTTGCAGTTACTTGATCTTGGTGATAGTTTTTCACTCCGGGTTGATAGTCTTCAATGGATTCAGCATCTTTCCAATATGAGGATTCTTGACCTGAGTGG

>M6-PGSC0003DMG400009899M6LRR-RLP_549

GTTGATAGTCTTCAATGGATTCAGCATCTTTCCAATATGAGGATTCTTGACCTGAGTGGAGTTGACCTGAGCAATGCCAAAACCTGGCTACATGACATCAATTTCCTAAGGTCTCTCGTG

>M6-PGSC0003DMG400009899M6LRR-RLP_61

ATTTACTACTAGTCCTTCTTCTTATAGCTAGTAGTACTATATTAGATGGATTGATATTATGCAGGGCTATGAATTCAGACATTCTTTGTGATGAAAATGAAAAAAATGCTCTTATCAAGT

>M6-PGSC0003DMG400009899M6LRR-RLP_610

TTGACCTGAGCAATGCCAAAACCTGGCTACATGACATCAATTTCCTAAGGTCTCTCGTGGAATTACGTTTGTCTTCTTGTCAACTTCCTAAATTACTTCCTGTTTTTGTAAATTTCACAT

>M6-PGSC0003DMG400009899M6LRR-RLP_671

ATTACGTTTGTCTTCTTGTCAACTTCCTAAATTACTTCCTGTTTTTGTAAATTTCACATCCCTTCAGGTTCTTGACCTCTCGTTAAACTATCTGGATGTTCCATTTCCTAGCTGGTTAGT

>M6-PGSC0003DMG400009899M6LRR-RLP_732

CTTCAGGTTCTTGACCTCTCGTTAAACTATCTGGATGTTCCATTTCCTAGCTGGTTAGTCAATACAAGCTCTCAGAGTCTTGTTTATCTTAATCTCAAAAGAAGTCTGCTGCATGGTTTA

>M6-PGSC0003DMG400009899M6LRR-RLP_793

ATACAAGCTCTCAGAGTCTTGTTTATCTTAATCTCAAAAGAAGTCTGCTGCATGGTTTACTTCCAAATACCGCCTTTGGAAACATGTACTCACTTAGAGTTCTTGATCTTTCTGACAACT

>M6-PGSC0003DMG400009899M6LRR-RLP_854

TCCAAATACCGCCTTTGGAAACATGTACTCACTTAGAGTTCTTGATCTTTCTGACAACTCTATCCGAGGCAAGCTTCCGTCTTTTGAGAACATGACTTCAGTTACTTCTCTTGATATCTC

>M6-PGSC0003DMG400009899M6LRR-RLP_915

ATCCGAGGCAAGCTTCCGTCTTTTGAGAACATGACTTCAGTTACTTCTCTTGATATCTCAAGGAATTCTCTTGAAGGCAAATTACCACCAACCCTGCCTTCAAAGTTGAAACATTTGGAT

>M6-PGSC0003DMG400009899M6LRR-RLP_976

GGAATTCTCTTGAAGGCAAATTACCACCAACCCTGCCTTCAAAGTTGAAACATTTGGATCTTTCATCCAATAATCTGGAGGGTACTTTAGCAAGAAGCATCGCACAACTTAGGCAATTAG

>M6-PGSC0003DMG400009941-(rev)M6LRR-RLK_0

CTTCAAATACCCTTAAACCTATCCTAAAATTTGTACAAACCAGACTGAACTAACAGTGTAACAATGTCACTCCCCAAAAAAATATCCCTTTTCCTCCAAATTTTCATCTTTTTGGTTTTC

>M6-PGSC0003DMG400009941-(rev)M6LRR-RLK_1037

ACTGAACATCAAATTGTTTTAAAATACTCAATAGTAATAGTTGTTTCTTGAGTGACGTGCAAAGTCTAGAGTTGATAAGTAAAAGTGAATGGAGTGAGAATATCATAAAAAGTCTCTTTG

>M6-PGSC0003DMG400009941-(rev)M6LRR-RLK_1098

AAGTCTAGAGTTGATAAGTAAAAGTGAATGGAGTGAGAATATCATAAAAAGTCTCTTTGAGAAAAACTGTGTGTTTATAGTGAAATGATGTTAGAGTATGAGTGTTAATAGAGAGGTTCT

>M6-PGSC0003DMG400009941-(rev)M6LRR-RLK_1159

AAAAACTGTGTGTTTATAGTGAAATGATGTTAGAGTATGAGTGTTAATAGAGAGGTTCTTGTGTAACATATATGTATATGTGCAAAATATTATGCATATAAAGTTTTGAAGTCATAATTC

>M6-PGSC0003DMG400009941-(rev)M6LRR-RLK_122

CATTAATGCAAACTCTGATCTTGAAACCCTTTTGAAGCTCAAAGAATCCATGGTTGCTCCTGGAACTTCTGCACTTCTTGATTGGAACAACAACACAAATTACCCTTTTTCCCATTGTTC

>M6-PGSC0003DMG400009941-(rev)M6LRR-RLK_1342

CGCATACCATCTGAACTATCTGGTTTAGAGAGTTTGATGTCGTTTGATTTGTCTTTAAACCAACTGACGGGAGAAATACCAGAGAGTTTTGTGAAGTTGCAGAATTTGACATTGATTAAC

>M6-PGSC0003DMG400009941-(rev)M6LRR-RLK_1403

AACTGACGGGAGAAATACCAGAGAGTTTTGTGAAGTTGCAGAATTTGACATTGATTAACTTGTTTAGAAACAACTTGCATGGTCCAATTCCCCCTTTTATTGGTGACCTTCCAAATCTTG

>M6-PGSC0003DMG400009941-(rev)M6LRR-RLK_1464

GTTTAGAAACAACTTGCATGGTCCAATTCCCCCTTTTATTGGTGACCTTCCAAATCTTGAAGTGTTGCAGATTTGGGGAAACAATTTTACTCTCGAATTACCGGAAAATCTTGGGCGCAA

>M6-PGSC0003DMG400009941-(rev)M6LRR-RLK_1525

GTGTTGCAGATTTGGGGAAACAATTTTACTCTCGAATTACCGGAAAATCTTGGGCGCAACGGGAGGTTTTTGTTTCTTGATATTTCTGTTAATCATTTTACTGGAAGGATACCACCTGAT

>M6-PGSC0003DMG400009941-(rev)M6LRR-RLK_1586

GGAGGTTTTTGTTTCTTGATATTTCTGTTAATCATTTTACTGGAAGGATACCACCTGATTTGTGTAAAGGGGGGAAGTTAAAGACACTTATTCTAATGGAGAATTACTTCTTTGGTCCTA

>M6-PGSC0003DMG400009941-(rev)M6LRR-RLK_1647

GTGTAAAGGGGGGAAGTTAAAGACACTTATTCTAATGGAGAATTACTTCTTTGGTCCTATTCCTGAACAACTTGGTGAGTGCAAATCGCTTACTCGAATTCGCGTTAGGAAGAATTACTT

>M6-PGSC0003DMG400009941-(rev)M6LRR-RLK_1708

CCTGAACAACTTGGTGAGTGCAAATCGCTTACTCGAATTCGCGTTAGGAAGAATTACTTAAATGGTACTATTCCAGTTGGTTTTTTCAAGTTACCTGCATTGGATATGCTTGAACTTGAC

>M6-PGSC0003DMG400009941-(rev)M6LRR-RLK_1769

ATGGTACTATTCCAGTTGGTTTTTTCAAGTTACCTGCATTGGATATGCTTGAACTTGACAACAACTATTTCACTGGTGAGCTGCCAACGGAGATAAACGCGAATAATCTCACTAAACTTG

>M6-PGSC0003DMG400009941-(rev)M6LRR-RLK_183

GGAACTTCTGCACTTCTTGATTGGAACAACAACACAAATTACCCTTTTTCCCATTGTTCTTTTTCTGGTGTTACATGTAACAATAACTCTCATGTTATATCTATAAACATCACTAATGTT

>M6-PGSC0003DMG400009941-(rev)M6LRR-RLK_1830

CAACTATTTCACTGGTGAGCTGCCAACGGAGATAAACGCGAATAATCTCACTAAACTTGTACTTTCCAACAACTGGATCACGGGGAACATTCCTCCGTCATTAGGGAACTTGAAGAATCT

>M6-PGSC0003DMG400009941-(rev)M6LRR-RLK_1891

CTTTCCAACAACTGGATCACGGGGAACATTCCTCCGTCATTAGGGAACTTGAAGAATCTAGTAACTCTATCACTTGATATGAACAGATTGTCCGGTGAAATTCCTCAAGAAATTGCGAGT

>M6-PGSC0003DMG400009941-(rev)M6LRR-RLK_1952

TAACTCTATCACTTGATATGAACAGATTGTCCGGTGAAATTCCTCAAGAAATTGCGAGTTTGAATAAACTCGTGACCATCAACTTGAGCGGCAACAATTTAACAGGTGAAATCCCGAGTT

>M6-PGSC0003DMG400009941-(rev)M6LRR-RLK_2013

GAATAAACTCGTGACCATCAACTTGAGCGGCAACAATTTAACAGGTGAAATCCCGAGTTCAATTGCGCTTTGTTCAGAGCTAACATTGGTTGACTTGAGCAGAAATCAACTGGTTGGTGA

>M6-PGSC0003DMG400009941-(rev)M6LRR-RLK_2074

ATTGCGCTTTGTTCAGAGCTAACATTGGTTGACTTGAGCAGAAATCAACTGGTTGGTGAAGTGCCAAAAGAAATCACCAAGTTAAATAGCTTGAATGCGCTGAACTTGTCAAGAAACCAA

>M6-PGSC0003DMG400009941-(rev)M6LRR-RLK_2135

TGCCAAAAGAAATCACCAAGTTAAATAGCTTGAATGCGCTGAACTTGTCAAGAAACCAACTGAGTGGCGCCATTCCTGGAGAAGTAGGAGTGATGAATGGCTTGACAGTTTTGGATCTTT

>M6-PGSC0003DMG400009941-(rev)M6LRR-RLK_2196

GAGTGGCGCCATTCCTGGAGAAGTAGGAGTGATGAATGGCTTGACAGTTTTGGATCTTTCTTACAATGATCTTTCTGGAAGGAGACCGACCAACGGACAACTAAAGTTCTTCAATGACAC

>M6-PGSC0003DMG400009941-(rev)M6LRR-RLK_2257

TACAATGATCTTTCTGGAAGGAGACCGACCAACGGACAACTAAAGTTCTTCAATGACACTTATTTTGTAGGAAATCCAAAACTCTGTTCGCCTCATGCTACTATTTGCCCGTCAGCTTCC

>M6-PGSC0003DMG400009941-(rev)M6LRR-RLK_2318

ATTTTGTAGGAAATCCAAAACTCTGTTCGCCTCATGCTACTATTTGCCCGTCAGCTTCCAATTCACCACAAAATGCGCTCAAAATCCATGCTGGGAAGTTCACAACTACCCAATTGGTGA

>M6-PGSC0003DMG400009941-(rev)M6LRR-RLK_2379

TTCACCACAAAATGCGCTCAAAATCCATGCTGGGAAGTTCACAACTACCCAATTGGTGATTACAATAATCATCTTAGTCACTGTTGCATTGCTGTTGGCAGTTACCGTGTTGTTCATCAA

>M6-PGSC0003DMG400009941-(rev)M6LRR-RLK_244

TTTCTGGTGTTACATGTAACAATAACTCTCATGTTATATCTATAAACATCACTAATGTTCCTCTATTTGGTACTATTCCACCTGAAATTGGTCTTTTACAAAATCTTGAAAATCTTATTA

>M6-PGSC0003DMG400009941-(rev)M6LRR-RLK_2440

ACAATAATCATCTTAGTCACTGTTGCATTGCTGTTGGCAGTTACCGTGTTGTTCATCAAGAAGGAAAAGTTCAAGAATTCGAAACTTTGGAAGTTAACAGCATTCCAGAAACTTGATTTC

>M6-PGSC0003DMG400009941-(rev)M6LRR-RLK_2501

AGGAAAAGTTCAAGAATTCGAAACTTTGGAAGTTAACAGCATTCCAGAAACTTGATTTCAGAGCTGAGGATGTTTTGGAGTGTTTAAAAGAGGAGAACATAATTGGGAAAGGTGGAGCTG

>M6-PGSC0003DMG400009941-(rev)M6LRR-RLK_2562

AGCTGAGGATGTTTTGGAGTGTTTAAAAGAGGAGAACATAATTGGGAAAGGTGGAGCTGGCGTTGTGTACCGAGGGTCTATGTCAAATGGCATCGACGTTGCAATTAAGAAACTTGTAGG

>M6-PGSC0003DMG400009941-(rev)M6LRR-RLK_2623

GTTGTGTACCGAGGGTCTATGTCAAATGGCATCGACGTTGCAATTAAGAAACTTGTAGGCCGAGGAACCGGACACCATGATCACGGATTCTCAGCTGAAATCCAAACACTAGGAAGGATC

>M6-PGSC0003DMG400009941-(rev)M6LRR-RLK_2684

GAGGAACCGGACACCATGATCACGGATTCTCAGCTGAAATCCAAACACTAGGAAGGATCAGGCACAGAAACATCGTACGATTACTAGGATATGTCTCAAACAAAGACACAAACTTGTTGT

>M6-PGSC0003DMG400009941-(rev)M6LRR-RLK_2745

GCACAGAAACATCGTACGATTACTAGGATATGTCTCAAACAAAGACACAAACTTGTTGTTGTACGAATACATGTCGAATGGGAGCTTAGGTGAAATGTTACATGGTGCCAAAGGAGCACA

>M6-PGSC0003DMG400009941-(rev)M6LRR-RLK_2806

TACGAATACATGTCGAATGGGAGCTTAGGTGAAATGTTACATGGTGCCAAAGGAGCACATTTGAAATGGGAGACGAGGTACCGTATTGCTGTGGAAGCTGCAAAGGGATTGTGTTATTTG

>M6-PGSC0003DMG400009941-(rev)M6LRR-RLK_2867

TGAAATGGGAGACGAGGTACCGTATTGCTGTGGAAGCTGCAAAGGGATTGTGTTATTTGCACCATGATTGTTCGCCTTCGATTATTCATAGAGATGTCAAGTCCAATAATATTCTGCTGG

>M6-PGSC0003DMG400009941-(rev)M6LRR-RLK_2928

CCATGATTGTTCGCCTTCGATTATTCATAGAGATGTCAAGTCCAATAATATTCTGCTGGATTCTGATTACGAGGCTCATGTTGCTGATTTTGGTCTAGCCAAATTCTTGCAGGATGCTGG

>M6-PGSC0003DMG400009941-(rev)M6LRR-RLK_2989

TCTGATTACGAGGCTCATGTTGCTGATTTTGGTCTAGCCAAATTCTTGCAGGATGCTGGTGCATCAGAGTGCATGTCCTCTATTGCTGGCTCATATGGTTACATTGCTCCAGGTAAATTA

>M6-PGSC0003DMG400009941-(rev)M6LRR-RLK_305

TCTATTTGGTACTATTCCACCTGAAATTGGTCTTTTACAAAATCTTGAAAATCTTATTATTTTTGGTGATAATATTACTGGTACACTCCCTTTAGAAATGTCACAACTTTCTTCTATTAA

>M6-PGSC0003DMG400009941-(rev)M6LRR-RLK_3050

CATCAGAGTGCATGTCCTCTATTGCTGGCTCATATGGTTACATTGCTCCAGGTAAATTAATACAGTTACAGTATTTTTCATATTTATACAATTATGAATTTTCCGTATTAGTATTCTTTT

>M6-PGSC0003DMG400009941-(rev)M6LRR-RLK_3111

ACAGTTACAGTATTTTTCATATTTATACAATTATGAATTTTCCGTATTAGTATTCTTTTATTGCTGGCTCATATGGTAATCATATTTATACAATTAGGTGGCATATAATGTAGTTACTGG

>M6-PGSC0003DMG400009941-(rev)M6LRR-RLK_3172

TGCTGGCTCATATGGTAATCATATTTATACAATTAGGTGGCATATAATGTAGTTACTGGTAAAATTTAATGATAATCTTAAAATGTGATACTAATTAACTTTCTATAATAGATTAAATTA

>M6-PGSC0003DMG400009941-(rev)M6LRR-RLK_3233

AAATTTAATGATAATCTTAAAATGTGATACTAATTAACTTTCTATAATAGATTAAATTATGATGATACTAATTAACCTTAAGCTATGTTGCTCAAACTCTTAAAAAATGTCAGTGGGGTA

>M6-PGSC0003DMG400009941-(rev)M6LRR-RLK_3416

TGAAGAATCAGAGAAAAATAAACCCTTAAGACCTTTTATATACCTGTGTTATTGGTCAATATTTTCCTATTTCATGAAAGATGCACTTCACCAACTTTTAGTACCAATTACCAAGACCTA

>M6-PGSC0003DMG400009941-(rev)M6LRR-RLK_3477

TTTTCCTATTTCATGAAAGATGCACTTCACCAACTTTTAGTACCAATTACCAAGACCTATGAACTTTTAGAATTTACTGAACTGGGGTTTAAAGTAGTAGGTATTTAAAAGCTATGCTCC

>M6-PGSC0003DMG400009941-(rev)M6LRR-RLK_3538

AACTTTTAGAATTTACTGAACTGGGGTTTAAAGTAGTAGGTATTTAAAAGCTATGCTCCAGGACAAAATCTATTTAAAACATTTGTCCAAAACATTAAATTGTGCAATTAATGTTTTTCT

>M6-PGSC0003DMG400009941-(rev)M6LRR-RLK_3599

GACAAAATCTATTTAAAACATTTGTCCAAAACATTAAATTGTGCAATTAATGTTTTTCTGAGGTGCATTTTTGTTTGGATATGAAACAGAGTATGCATACACATTGAAAGTTGACCAAAA

>M6-PGSC0003DMG400009941-(rev)M6LRR-RLK_366

TTTGGTGATAATATTACTGGTACACTCCCTTTAGAAATGTCACAACTTTCTTCTATTAAACATGTTAATCTTTCTTACAACAATTTTTCTGGTCCTTTTCCTAGAGAAATCTTGTTGGGG

>M6-PGSC0003DMG400009941-(rev)M6LRR-RLK_3660

GGTGCATTTTTGTTTGGATATGAAACAGAGTATGCATACACATTGAAAGTTGACCAAAAGAGTGATGTATACAGTTTTGGAGTAGTACTGTTGGAACTTATCACAGGTCACAAGCCAGTT

>M6-PGSC0003DMG400009941-(rev)M6LRR-RLK_3721

GTGATGTATACAGTTTTGGAGTAGTACTGTTGGAACTTATCACAGGTCACAAGCCAGTTGGTGAATTCGGAGACGGTGTAGATATAGTCAGATGGGTAAATAAAACAATGTCCGAATTAT

>M6-PGSC0003DMG400009941-(rev)M6LRR-RLK_3782

TGAATTCGGAGACGGTGTAGATATAGTCAGATGGGTAAATAAAACAATGTCCGAATTATCTCAGCCATCTGATGCAGCCTCAGTTTTAGCAGTCGTTGACTCGAGGCTACATAGTTACCC

>M6-PGSC0003DMG400009941-(rev)M6LRR-RLK_3843

CAGCCATCTGATGCAGCCTCAGTTTTAGCAGTCGTTGACTCGAGGCTACATAGTTACCCTCTTGCAAGTGTTATCAATTTGTTCAAGATTGCTATAATGTGTGTTGAAGAAGAGAGTTGT

>M6-PGSC0003DMG400009941-(rev)M6LRR-RLK_3904

TTGCAAGTGTTATCAATTTGTTCAAGATTGCTATAATGTGTGTTGAAGAAGAGAGTTGTGCTAGGCCTACTATGAGGGAAGTTGTTCACATGCTTACAAATCTTCCTCAGTCTACTACTA

>M6-PGSC0003DMG400009941-(rev)M6LRR-RLK_3965

TAGGCCTACTATGAGGGAAGTTGTTCACATGCTTACAAATCTTCCTCAGTCTACTACTACTACTCCTACTCTCCTTGCCCTTTAAAATTGTGCGGATATTAAGTGTCTGGTTGAAAACTC

>M6-PGSC0003DMG400009941-(rev)M6LRR-RLK_4026

ACTCCTACTCTCCTTGCCCTTTAAAATTGTGCGGATATTAAGTGTCTGGTTGAAAACTCGTGGAGTTTGAGGCCGGGAACACGAGCCTCATGAGTCTATTTGGGTATGGGGAACAAAAAA

>M6-PGSC0003DMG400009941-(rev)M6LRR-RLK_4087

GGAGTTTGAGGCCGGGAACACGAGCCTCATGAGTCTATTTGGGTATGGGGAACAAAAAAATTGTGCAAGTAGAGATATGTATTTGAAATGTATTGCTTGTTTTTATATTTTTGAAGTAGT

>M6-PGSC0003DMG400009941-(rev)M6LRR-RLK_4148

TGTGCAAGTAGAGATATGTATTTGAAATGTATTGCTTGTTTTTATATTTTTGAAGTAGTTGTATAAAAAGGGGCATGAAAAAAAGAAGGAAAAAGACTCCTTTAATTACTTATTATGGGT

>M6-PGSC0003DMG400009941-(rev)M6LRR-RLK_4209

TATAAAAAGGGGCATGAAAAAAAGAAGGAAAAAGACTCCTTTAATTACTTATTATGGGTATCGAGTATAATTATGTGGCCTTAGGAAATAAATATCAATATGGCAATTAGTCAATATTTG

>M6-PGSC0003DMG400009941-(rev)M6LRR-RLK_4264

GGGTATCGAGTATAATTATGTGGCCTTAGGAAATAAATATCAATATGGCAATTAGTCAATATTTGCTTTTAATAACATGGCCTATGTAATGTTTTCAAGAGTTTGCGAAGTTAAAATTTG

>M6-PGSC0003DMG400009941-(rev)M6LRR-RLK_427

ATGTTAATCTTTCTTACAACAATTTTTCTGGTCCTTTTCCTAGAGAAATCTTGTTGGGGTTAATAAAGCTTGAATCTTTTGACATTTATAACAACAATTTCACTGGTGAACTTCCTAATG

>M6-PGSC0003DMG400009941-(rev)M6LRR-RLK_488

AATAAAGCTTGAATCTTTTGACATTTATAACAACAATTTCACTGGTGAACTTCCTAATGAGTTTGTAAAGTTGAAAAAGTTGGAAACTTTACATCTTGGTGGAAACTATTTTCATGGTGA

>M6-PGSC0003DMG400009941-(rev)M6LRR-RLK_549

TTTGTAAAGTTGAAAAAGTTGGAAACTTTACATCTTGGTGGAAACTATTTTCATGGTGAAATACCAGAAGTTTATTCTCATATTGTAAGTTTAAAGTGGTTGGGTTTAGAGGGAAATTCA

>M6-PGSC0003DMG400009941-(rev)M6LRR-RLK_61

CAATGTCACTCCCCAAAAAAATATCCCTTTTCCTCCAAATTTTCATCTTTTTGGTTTTCACCATTAATGCAAACTCTGATCTTGAAACCCTTTTGAAGCTCAAAGAATCCATGGTTGCTC

>M6-PGSC0003DMG400009941-(rev)M6LRR-RLK_610

TACCAGAAGTTTATTCTCATATTGTAAGTTTAAAGTGGTTGGGTTTAGAGGGAAATTCACTTACTGGGAAAATACCAAAGAGTTTGGCTTTGTTACCAAATCTTGAAGAACTTAGATTGG

>M6-PGSC0003DMG400009941-(rev)M6LRR-RLK_671

TACTGGGAAAATACCAAAGAGTTTGGCTTTGTTACCAAATCTTGAAGAACTTAGATTGGGTTATTATAATAGTTATGAAGGGGGTATTCCATCTGAGTTTGGTAATATTAGTACACTTAA

>M6-PGSC0003DMG400009941-(rev)M6LRR-RLK_732

TATTATAATAGTTATGAAGGGGGTATTCCATCTGAGTTTGGTAATATTAGTACACTTAAACTTCTTGATCTTGGAAATTGTAATCTTGATGGTGAAGTTCCTCCAAGTCTTGGAAATTTG

>M6-PGSC0003DMG400009941-(rev)M6LRR-RLK_793

TTCTTGATCTTGGAAATTGTAATCTTGATGGTGAAGTTCCTCCAAGTCTTGGAAATTTGAAGAAGTTGCATACTTTGTTTCTACAAGTGAACAGACTTACAGGTATTCATGAACTCACTA

>M6-PGSC0003DMG400009941-(rev)M6LRR-RLK_854

GAAGTTGCATACTTTGTTTCTACAAGTGAACAGACTTACAGGTATTCATGAACTCACTATTTGTTTAGTTATACAATCAAATCTCTCTCTAATAACGTCGTTTGTCTGAATATATATTAG

>M6-PGSC0003DMG400009941-(rev)M6LRR-RLK_915

TGTTTAGTTATACAATCAAATCTCTCTCTAATAACGTCGTTTGTCTGAATATATATTAGTTGACATGGTTAAATATTGTTATAGGAAACATATTTGTTACAAATCATTTTCCAAGATGTA

>M6-PGSC0003DMG400009941-(rev)M6LRR-RLK_976

GACATGGTTAAATATTGTTATAGGAAACATATTTGTTACAAATCATTTTCCAAGATGTAAGACTGAACATCAAATTGTTTTAAAATACTCAATAGTAATAGTTGTTTCTTGAGTGACGTG

>M6-PGSC0003DMG400010032M6RLKWAX-EGF_0

TGGATATGCGCTGCGTGTGCGTAAAAGAAGCTTCTTCCCCGGTGTATTTTAAAGTCTACCTTCCTTGTTTGTATCTCCGATTACTATTTGATAGCTTCATTGTTAGGAAATCCAGTTTTT

>M6-PGSC0003DMG400010032M6RLKWAX-EGF_1037

ATCTTGATTGGTATGGATAATGATACAATGAAGAGTTCTGCCATGTTCCTGGAGTTTCAGACGATGGAATTAGCGTGGGGATTGAAAGGGGATTGTGCATGTCATGACGATGCAAATTGT

>M6-PGSC0003DMG400010032M6RLKWAX-EGF_1098

CGATGGAATTAGCGTGGGGATTGAAAGGGGATTGTGCATGTCATGACGATGCAAATTGTACCAACGTTTCTCTTCCGGGGAACCGAAAAGGTTTCCGATGCCGGTGTAAAGATGGATTCG

>M6-PGSC0003DMG400010032M6RLKWAX-EGF_122

AGAGCTGTTTTTGGAGTTGAGATATTTCAAATTTTGGACTCTTTAATTGCTACTATTTGCTTCCAATCAAAGAAAGGAAAATTTGCAGACGAAGGAAGAGAGAGAAGTAAAAGCTAAGGT

>M6-PGSC0003DMG400010032M6RLKWAX-EGF_1220

GGTGATGGCTTCCGCGACGGTGATGGGTGCCGTAAAGGTTAGTTCTTTTTTTTTTCAAGTGTTAAAATAGTTAATTTGCAACATTATTCACCTCCTTGTTGAATTGTGTGAATCATCTTA

>M6-PGSC0003DMG400010032M6RLKWAX-EGF_1403

TATAAAGATATGGGAAGTTCAAAATGGAGATGTTAAATTTGACGTTCAAAATTCAATAAAATCTTAGTTTTATTTTACATAACATATATTTCGTTTCAATGATATTAATTTCAATTGAAC

>M6-PGSC0003DMG400010032M6RLKWAX-EGF_1464

TCTTAGTTTTATTTTACATAACATATATTTCGTTTCAATGATATTAATTTCAATTGAACTCAAGAGTTCCATCTACACGTATTGGTAGATATTTTAAATTTTAGTACTTGGAGATATATC

>M6-PGSC0003DMG400010032M6RLKWAX-EGF_1525

AAGAGTTCCATCTACACGTATTGGTAGATATTTTAAATTTTAGTACTTGGAGATATATCATTAATGGAAATGCCGTTGAACTCAAAATGTCTACTTAATTTGGTATATGGCTTTTCAATA

>M6-PGSC0003DMG400010032M6RLKWAX-EGF_1586

TAATGGAAATGCCGTTGAACTCAAAATGTCTACTTAATTTGGTATATGGCTTTTCAATAGAGACAACGTTTATATTATATTAGGGGTAAAATTCAAAGTTAATTGCTAGATGGCCTAATT

>M6-PGSC0003DMG400010032M6RLKWAX-EGF_1647

GACAACGTTTATATTATATTAGGGGTAAAATTCAAAGTTAATTGCTAGATGGCCTAATTATTGTTTGACTATGGGCAAAAAGTGACGTGTGAGATGATAAGATGACCAGTAGTTTTTATA

>M6-PGSC0003DMG400010032M6RLKWAX-EGF_1708

TGTTTGACTATGGGCAAAAAGTGACGTGTGAGATGATAAGATGACCAGTAGTTTTTATATTTGGTTTGACCAGTAGCAACACGCCACGAAGAAGGGCTGTTGTTAATTTCAATGGAGATT

>M6-PGSC0003DMG400010032M6RLKWAX-EGF_1769

TGGTTTGACCAGTAGCAACACGCCACGAAGAAGGGCTGTTGTTAATTTCAATGGAGATTTACATGAATTGACTATGATTAAACTAATGATATTTTAGTGTGATTAAGTTATTTAGTAAAT

>M6-PGSC0003DMG400010032M6RLKWAX-EGF_183

TCCAATCAAAGAAAGGAAAATTTGCAGACGAAGGAAGAGAGAGAAGTAAAAGCTAAGGTAGACATTAATACATGAAACCAGTCCACGGTACTCTTTACTCTCTACTTTCCCTTCTATGGA

>M6-PGSC0003DMG400010032M6RLKWAX-EGF_1830

CATGAATTGACTATGATTAAACTAATGATATTTTAGTGTGATTAAGTTATTTAGTAAATCCAAATATTCCACAAGTTGTGTTAAAACTTAAAAGAGCACTGCATATGCGGCTGGTGTAGT

>M6-PGSC0003DMG400010032M6RLKWAX-EGF_1891

AAATATTCCACAAGTTGTGTTAAAACTTAAAAGAGCACTGCATATGCGGCTGGTGTAGTCAATAAAGCCTTCTCTTGATTTTGATATGGTCGAATTTGGGTTCTAAGCTGTCGACTGTTG

>M6-PGSC0003DMG400010032M6RLKWAX-EGF_1952

ATAAAGCCTTCTCTTGATTTTGATATGGTCGAATTTGGGTTCTAAGCTGTCGACTGTTGAAAGTCTCCTATATGACCTCCTCATGCGACTTGGGTTGTGTTGATATTCTTTTATTACTCG

>M6-PGSC0003DMG400010032M6RLKWAX-EGF_2013

AGTCTCCTATATGACCTCCTCATGCGACTTGGGTTGTGTTGATATTCTTTTATTACTCGAATTTATAGTCTTTAGCTAGTTTTTATTAAAAGATAAAAAATTCTATTTACGAACAATCAT

>M6-PGSC0003DMG400010032M6RLKWAX-EGF_2074

TTTATAGTCTTTAGCTAGTTTTTATTAAAAGATAAAAAATTCTATTTACGAACAATCATGGGGGTTCAGCTTTATCTTATCATTCCTAATAAAGTAATGAAATTCTAGTAATTTATAAGT

>M6-PGSC0003DMG400010032M6RLKWAX-EGF_2135

GGGTTCAGCTTTATCTTATCATTCCTAATAAAGTAATGAAATTCTAGTAATTTATAAGTATGAAGAAAATAGGTATTTTATGATCATTACCAATGAGAATGTTTATATGATGCAAAAATA

>M6-PGSC0003DMG400010032M6RLKWAX-EGF_2196

GAAGAAAATAGGTATTTTATGATCATTACCAATGAGAATGTTTATATGATGCAAAAATATAAAAACTAAAAAAAAATTGAGTTTTCAAACGCAAACTAACTTTAACAAAGGTATATTTAA

>M6-PGSC0003DMG400010032M6RLKWAX-EGF_2257

AAAACTAAAAAAAAATTGAGTTTTCAAACGCAAACTAACTTTAACAAAGGTATATTTAACCTTTTCACATTCCATCAAAAAAATTGGAAAAAATAAATTTCTTTAAAAATGATAGTCTAG

>M6-PGSC0003DMG400010032M6RLKWAX-EGF_2318

TTTTCACATTCCATCAAAAAAATTGGAAAAAATAAATTTCTTTAAAAATGATAGTCTAGTCTACTGTGTTCTCCTCTCCCTTTCATATTCACCCATAGTTTCCATTCCCATTACCCTACA

>M6-PGSC0003DMG400010032M6RLKWAX-EGF_2379

TACTGTGTTCTCCTCTCCCTTTCATATTCACCCATAGTTTCCATTCCCATTACCCTACATATTAGTTTAGATTATACGTCCAACAATAATTGTTCACTGTTGATTTGGCACACTCCTTAA

>M6-PGSC0003DMG400010032M6RLKWAX-EGF_244

ACATTAATACATGAAACCAGTCCACGGTACTCTTTACTCTCTACTTTCCCTTCTATGGAATTCGGAAAATAGCATTTATTGACAAACAATGAACTCCCCACCAACATCAACTGCACCCCC

>M6-PGSC0003DMG400010032M6RLKWAX-EGF_2623

TAAATTATCTCTTAATTTTCTAAATTGGATAAGTATTGTTAGACAACAATTTTTAGAGTAGTAAAGAACTATTGTTGGAAGAATGGAGTACATAAATACTTCGATGGAAGTTTAATTTTC

>M6-PGSC0003DMG400010032M6RLKWAX-EGF_2806

GATAAAAAGATATAATATGAAAAATAATTTGCTTGGTACTAAACATACCCTTGATCACTTTTTTTTGGTCATGTGGGACATTGGGACTTGGTCTCACTAACTTTTTCCCCACTAGGTTCC

>M6-PGSC0003DMG400010032M6RLKWAX-EGF_2867

TTTTTGGTCATGTGGGACATTGGGACTTGGTCTCACTAACTTTTTCCCCACTAGGTTCCTAACTTCAGCCTGCCACCATTTCCGCGTATTAGGCTTTTAGGTATTACTCACAGTACCTTC

>M6-PGSC0003DMG400010032M6RLKWAX-EGF_2928

ACTTCAGCCTGCCACCATTTCCGCGTATTAGGCTTTTAGGTATTACTCACAGTACCTTCCACTTTCCTTTCTTCTTCCTAGCTCCTGCACATGGACCCAAGACAAATAATATTATTCTTG

>M6-PGSC0003DMG400010032M6RLKWAX-EGF_2989

CTTTCCTTTCTTCTTCCTAGCTCCTGCACATGGACCCAAGACAAATAATATTATTCTTGTGAAGATATTTGATTCTATGAGAAGCATTTTAAAAAGTTTGTGATTTTAAATGCACATTTT

>M6-PGSC0003DMG400010032M6RLKWAX-EGF_305

TCGGAAAATAGCATTTATTGACAAACAATGAACTCCCCACCAACATCAACTGCACCCCCTTAATTTCCTCAATAAATGGTCTTCAACAAGTCAAACTCTTCAACTTTTTTCTTCCTTTTT

>M6-PGSC0003DMG400010032M6RLKWAX-EGF_3050

AAGATATTTGATTCTATGAGAAGCATTTTAAAAAGTTTGTGATTTTAAATGCACATTTTTGTGGATATAAAATAATGTTGTTAGCATTAATATGGGGATTCTAATATCAGTTCTTTTTTT

>M6-PGSC0003DMG400010032M6RLKWAX-EGF_3111

TGGATATAAAATAATGTTGTTAGCATTAATATGGGGATTCTAATATCAGTTCTTTTTTTCAAGATGTGTCATTCTTTCTTCCATTGACAAGAAAGGAAGAGTGTTGACTTTGTAACATTG

>M6-PGSC0003DMG400010032M6RLKWAX-EGF_3172

AGATGTGTCATTCTTTCTTCCATTGACAAGAAAGGAAGAGTGTTGACTTTGTAACATTGTTATAAAGTAGTTGTCGGTACAATGATATTTCCTATCACCAGTCACCAATGTGCCATCTCT

>M6-PGSC0003DMG400010032M6RLKWAX-EGF_3233

ATAAAGTAGTTGTCGGTACAATGATATTTCCTATCACCAGTCACCAATGTGCCATCTCTGGTCTTTCTTAATAAGATGAAGAAGGATCTGAGTCTTTATGAGACTGCCGATTCGCGGTGG

>M6-PGSC0003DMG400010032M6RLKWAX-EGF_3294

TCTTTCTTAATAAGATGAAGAAGGATCTGAGTCTTTATGAGACTGCCGATTCGCGGTGGTAATGTAACTTATCAAAATGACAAGTTTGAGCTCGATAACCGGAATTTTGTTGAATTGAGC

>M6-PGSC0003DMG400010032M6RLKWAX-EGF_3355

ATGTAACTTATCAAAATGACAAGTTTGAGCTCGATAACCGGAATTTTGTTGAATTGAGCTCTCAAATGAATTTTCTTGTCTTGAAGAGTTGAACACTTCAATTAGTTATCTTACCCAATT

>M6-PGSC0003DMG400010032M6RLKWAX-EGF_3416

TCAAATGAATTTTCTTGTCTTGAAGAGTTGAACACTTCAATTAGTTATCTTACCCAATTGCATATTACATGACACGTTCAAAGATCTTACTTTAAAATAAAATAAAATAAAGAAGCATTT

>M6-PGSC0003DMG400010032M6RLKWAX-EGF_3477

ATATTACATGACACGTTCAAAGATCTTACTTTAAAATAAAATAAAATAAAGAAGCATTTACTTGTTTGTTCTATGACTCGAAGATGGTGAATGTGATTGATGATGAGGTTTGAATTACTC

>M6-PGSC0003DMG400010032M6RLKWAX-EGF_3538

TTGTTTGTTCTATGACTCGAAGATGGTGAATGTGATTGATGATGAGGTTTGAATTACTCTATTCCGTAGATTATCAATTGGACAATCCCCAGGTCAATGATTTTCTTGCCTTTTTAAGTT

>M6-PGSC0003DMG400010032M6RLKWAX-EGF_3599

TTCCGTAGATTATCAATTGGACAATCCCCAGGTCAATGATTTTCTTGCCTTTTTAAGTTTATTTTGAGGGGGACAGCACATGGTCACTTGATGTCCACATGTTTGAACGGGTGAAAGTCA

>M6-PGSC0003DMG400010032M6RLKWAX-EGF_366

AATTTCCTCAATAAATGGTCTTCAACAAGTCAAACTCTTCAACTTTTTTCTTCCTTTTTCATGGGTAAATTAGCTATCACTCATTTTCATTTTCTCTTTGCAGAAGCAGATGATTGTTCT

>M6-PGSC0003DMG400010032M6RLKWAX-EGF_3660

TTTTGAGGGGGACAGCACATGGTCACTTGATGTCCACATGTTTGAACGGGTGAAAGTCAGAGGATGAACAAGACCTGGATTGTTGAATAATTTCTTATTTTATTATTGAGTAAGCGTTAT

>M6-PGSC0003DMG400010032M6RLKWAX-EGF_3721

GGATGAACAAGACCTGGATTGTTGAATAATTTCTTATTTTATTATTGAGTAAGCGTTATGATAAAAGGATCTGGAATCAATGCCCCGTCTTAAACCAAACGTGTAATGAGACGGCTTATT

>M6-PGSC0003DMG400010032M6RLKWAX-EGF_3782

TAAAAGGATCTGGAATCAATGCCCCGTCTTAAACCAAACGTGTAATGAGACGGCTTATTATTATGCCAGTTTAATTTTAAACAACTTAGTACTTCACCTGAATCAATGGCATAATTTGAA

>M6-PGSC0003DMG400010032M6RLKWAX-EGF_3843

TATGCCAGTTTAATTTTAAACAACTTAGTACTTCACCTGAATCAATGGCATAATTTGAAGGGATGAAAGTTGAAACTGATTCATCTTCATTTTTACCAGTACATTCCATGTCCAGGAAAC

>M6-PGSC0003DMG400010032M6RLKWAX-EGF_3904

GATGAAAGTTGAAACTGATTCATCTTCATTTTTACCAGTACATTCCATGTCCAGGAAACTATCTGTTAAGACAACATCTTGACTTTGGAGATTTAAAACTTAACTGACTAGCACGTTTCC

>M6-PGSC0003DMG400010032M6RLKWAX-EGF_3965

TCTGTTAAGACAACATCTTGACTTTGGAGATTTAAAACTTAACTGACTAGCACGTTTCCAGTGAATTGATTTCATTTGTTGATCATGTGCTTGCTTCATGACTCTGGGCGGGCAAACATT

>M6-PGSC0003DMG400010032M6RLKWAX-EGF_4026

TGAATTGATTTCATTTGTTGATCATGTGCTTGCTTCATGACTCTGGGCGGGCAAACATTTTCATTCTGACGATGTTCTAAGATACTTTATCTATATATTCAATCAACTCTGGGAGGGCAT

>M6-PGSC0003DMG400010032M6RLKWAX-EGF_4087

CATTCTGACGATGTTCTAAGATACTTTATCTATATATTCAATCAACTCTGGGAGGGCATACATTTTCATTCTAACGATGTTCTAAGATACTTTGCCATATGCTCCTTCCTTCTACATCTT

>M6-PGSC0003DMG400010032M6RLKWAX-EGF_4148

ATTTTCATTCTAACGATGTTCTAAGATACTTTGCCATATGCTCCTTCCTTCTACATCTTTCTCTATGATTATCTTCAGTATGTCGTCTTTATGCTAACTTCTTTATATTTGTCTCTCAAA

>M6-PGSC0003DMG400010032M6RLKWAX-EGF_4209

TCTATGATTATCTTCAGTATGTCGTCTTTATGCTAACTTCTTTATATTTGTCTCTCAAATCAGTTTCAAGATGCAATCCTTCCAGATACCTCTCTGGCCGATGTGGTGGAACTACCAGAA

>M6-PGSC0003DMG400010032M6RLKWAX-EGF_427

TGGGTAAATTAGCTATCACTCATTTTCATTTTCTCTTTGCAGAAGCAGATGATTGTTCTCTTTGATTCAATTTCGAGCCATGGGTTTTCATCAGATCGTTATTATTAGATTCGTTTTCTC

>M6-PGSC0003DMG400010032M6RLKWAX-EGF_4270

AGTTTCAAGATGCAATCCTTCCAGATACCTCTCTGGCCGATGTGGTGGAACTACCAGAATTGGTGTGCTCGTTGGAGGTATATTTCTTTCTTTTTTTGGTTAACTTGTTCAGAAAATGTT

>M6-PGSC0003DMG400010032M6RLKWAX-EGF_4331

GGTGTGCTCGTTGGAGGTATATTTCTTTCTTTTTTTGGTTAACTTGTTCAGAAAATGTTGGTAAATAGTAACCATTACTTCTCAACTCGTTTAATGCACTAGTGTTCAATTCTGCATCTC

>M6-PGSC0003DMG400010032M6RLKWAX-EGF_4392

TAAATAGTAACCATTACTTCTCAACTCGTTTAATGCACTAGTGTTCAATTCTGCATCTCCTGCTCCTCTGTTTCAATTTGGCAACACCTCAATTACCAAATGCGCACTCTTTAGTTTTCT

>M6-PGSC0003DMG400010032M6RLKWAX-EGF_4453

GCTCCTCTGTTTCAATTTGGCAACACCTCAATTACCAAATGCGCACTCTTTAGTTTTCTATAAACGAGCTTGCAAGCCATCACAGATTGGTTTTGTGTTCTTCATATGTACGAGATTTTG

>M6-PGSC0003DMG400010032M6RLKWAX-EGF_4514

AAACGAGCTTGCAAGCCATCACAGATTGGTTTTGTGTTCTTCATATGTACGAGATTTTGCAAAAGAGTTGCTGTAGTTGCTAATTGATCAGTCATCAGTAAACTTCCGAGACTGCTTACT

>M6-PGSC0003DMG400010032M6RLKWAX-EGF_4575

AAAGAGTTGCTGTAGTTGCTAATTGATCAGTCATCAGTAAACTTCCGAGACTGCTTACTTTTACTGCCGTTTGATTTAATATAACTAGTGAGTTTGATCTCTTCCTGGTCTAATGAAATT

>M6-PGSC0003DMG400010032M6RLKWAX-EGF_4636

TACTGCCGTTTGATTTAATATAACTAGTGAGTTTGATCTCTTCCTGGTCTAATGAAATTTCAACCACCTACAGGGATTATTGCTGGAGCTGGTTTAATGGCTGCTTTAGCTGTTCTTTGC

>M6-PGSC0003DMG400010032M6RLKWAX-EGF_4697

AACCACCTACAGGGATTATTGCTGGAGCTGGTTTAATGGCTGCTTTAGCTGTTCTTTGCTACTGCATTCGAAGACGTTCTTCATCTCTCAAGAAAAGAATGAGCGCAAGACGCCTTCTCT

>M6-PGSC0003DMG400010032M6RLKWAX-EGF_4758

CTGCATTCGAAGACGTTCTTCATCTCTCAAGAAAAGAATGAGCGCAAGACGCCTTCTCTCTGAAGCTGCAGGCAGCAACAGTGTTCATGTATTTCAATACAAAGAAATTGAGAGGGCAAC

>M6-PGSC0003DMG400010032M6RLKWAX-EGF_4819

GAAGCTGCAGGCAGCAACAGTGTTCATGTATTTCAATACAAAGAAATTGAGAGGGCAACAAATAGTTTCTCTGAGAAACAGAGGCTTGGAATAGGGGCTTATGGCACAGTTTATGCTGGA

>M6-PGSC0003DMG400010032M6RLKWAX-EGF_488

TTGATTCAATTTCGAGCCATGGGTTTTCATCAGATCGTTATTATTAGATTCGTTTTCTCAATTTTATTGGGTTGGTTATCATCGTCAGTTGAGGCTACTAATTCAACTAAGTGCAATCAG

>M6-PGSC0003DMG400010032M6RLKWAX-EGF_4880

ATAGTTTCTCTGAGAAACAGAGGCTTGGAATAGGGGCTTATGGCACAGTTTATGCTGGAAAGCTCCACAGTGATGAATGGGTTGCAATTAAGAAACTAAGACACCGAGATCCGGATGGTG

>M6-PGSC0003DMG400010032M6RLKWAX-EGF_4941

GCTCCACAGTGATGAATGGGTTGCAATTAAGAAACTAAGACACCGAGATCCGGATGGTGTTGAACAAGTAATGAATGAGGTTAAGCTTTTGTCTTCTGTAAGTCATCCAAACTTAGTCCG

>M6-PGSC0003DMG400010032M6RLKWAX-EGF_5002

GAACAAGTAATGAATGAGGTTAAGCTTTTGTCTTCTGTAAGTCATCCAAACTTAGTCCGCCTCTTAGGTTGTTGTATAGAAAATGGTGAACAGATTCTTGTTTATGAATTTATGCCTAAC

>M6-PGSC0003DMG400010032M6RLKWAX-EGF_5063

TCTTAGGTTGTTGTATAGAAAATGGTGAACAGATTCTTGTTTATGAATTTATGCCTAACGGAACTGTAGCTCAGCATCTACAGAGAGAAAGGGGTTCAGGACTTCCTTGGACAATACGTC

>M6-PGSC0003DMG400010032M6RLKWAX-EGF_5124

AACTGTAGCTCAGCATCTACAGAGAGAAAGGGGTTCAGGACTTCCTTGGACAATACGTCTCACTATCGCAACAGAAACTGCTCATGCAATTGCTCACCTTCACTCAGCAATGAATCCACC

>M6-PGSC0003DMG400010032M6RLKWAX-EGF_5185

ACTATCGCAACAGAAACTGCTCATGCAATTGCTCACCTTCACTCAGCAATGAATCCACCAATATACCACAGGGATATCAAATCAAGTAACATACTATTGGATTACAACTTCAACTCAAAG

>M6-PGSC0003DMG400010032M6RLKWAX-EGF_5246

TATACCACAGGGATATCAAATCAAGTAACATACTATTGGATTACAACTTCAACTCAAAGGTTGCAGACTTTGGTCTTTCCAGATTTGGCATGACAGATGATTCCCACATTTCTACAGCAC

>M6-PGSC0003DMG400010032M6RLKWAX-EGF_5307

TGCAGACTTTGGTCTTTCCAGATTTGGCATGACAGATGATTCCCACATTTCTACAGCACCACAAGGTACTCCAGGTTATGTGGATCCTCAGTACCATCAGAATTACCATCTTTCCGATAA

>M6-PGSC0003DMG400010032M6RLKWAX-EGF_5368

CAAGGTACTCCAGGTTATGTGGATCCTCAGTACCATCAGAATTACCATCTTTCCGATAAGAGTGATGTGTACAGCTTCGGGGTGGTTCTTGTAGAAATCATCACAGCAATGAAAGTGGTT

>M6-PGSC0003DMG400010032M6RLKWAX-EGF_5429

GTGATGTGTACAGCTTCGGGGTGGTTCTTGTAGAAATCATCACAGCAATGAAAGTGGTTGATTTCTCTAGATCACACAGTGAGATTAATTTGGCTGCACTGGCAATAGACAGAATAGGTA

>M6-PGSC0003DMG400010032M6RLKWAX-EGF_549

TTTTATTGGGTTGGTTATCATCGTCAGTTGAGGCTACTAATTCAACTAAGTGCAATCAGTACTGTGGAGCTGCTGGTTCTAATGGTCTGCGTGTTTCTTATCCATTTGGTTTCTCAGAGG

>M6-PGSC0003DMG400010032M6RLKWAX-EGF_5490

TTTCTCTAGATCACACAGTGAGATTAATTTGGCTGCACTGGCAATAGACAGAATAGGTAAGGGTCGTGTGGATGAGATTATCGATCCATTTCTCGAGCCACACAGAGATGCATGGACTCT

>M6-PGSC0003DMG400010032M6RLKWAX-EGF_5551

GGTCGTGTGGATGAGATTATCGATCCATTTCTCGAGCCACACAGAGATGCATGGACTCTATCATCAGTTCATAGAGTTGCAGAGCTTGCTTTCAGATGCCTTGCATTTCACAAGGATATG

>M6-PGSC0003DMG400010032M6RLKWAX-EGF_5612

CATCAGTTCATAGAGTTGCAGAGCTTGCTTTCAGATGCCTTGCATTTCACAAGGATATGAGGCCTTCAATGACAGAAGTGGCAGATGAGCTAGAACAGGTCAGGCTCAGCAGTTGGGCAT

>M6-PGSC0003DMG400010032M6RLKWAX-EGF_5673

GCCTTCAATGACAGAAGTGGCAGATGAGCTAGAACAGGTCAGGCTCAGCAGTTGGGCATCATTGGAGGACAATGTATGCATGACATCATCAGTGAATTCCTCTTGCTCATCTCCCCGTAG

>M6-PGSC0003DMG400010032M6RLKWAX-EGF_5734

TTGGAGGACAATGTATGCATGACATCATCAGTGAATTCCTCTTGCTCATCTCCCCGTAGTATGAGTGAGACATCTTTTCGTAGTACAACAACTAAGAAAGGAGTAGGGAGTAGGAGGTTG

>M6-PGSC0003DMG400010032M6RLKWAX-EGF_5795

TGAGTGAGACATCTTTTCGTAGTACAACAACTAAGAAAGGAGTAGGGAGTAGGAGGTTGATTGTTCCGCAGAAGATAGCGAATTCACTGGCAATCATGGAGGAAATAAAAGACAGTTCCC

>M6-PGSC0003DMG400010032M6RLKWAX-EGF_5856

TGTTCCGCAGAAGATAGCGAATTCACTGGCAATCATGGAGGAAATAAAAGACAGTTCCCCTGTTTCTGTGCAGGATCCTTGGTTTAGTGAAGAGAGTCCCCCTTCAACTAACCGATTATT

>M6-PGSC0003DMG400010032M6RLKWAX-EGF_5917

GTTTCTGTGCAGGATCCTTGGTTTAGTGAAGAGAGTCCCCCTTCAACTAACCGATTATTAGGTAATTCTGGTCGATGATGACTTGAAATAAAAGAAGTTTGCTCTAGGTAGAAAATTGAG

>M6-PGSC0003DMG400010032M6RLKWAX-EGF_5978

GTAATTCTGGTCGATGATGACTTGAAATAAAAGAAGTTTGCTCTAGGTAGAAAATTGAGGGGGATCACCTGTAATCTATTACCACCTTTTCTTGCCTTCTAATCTAGTGATAATATCAAA

>M6-PGSC0003DMG400010032M6RLKWAX-EGF_6039

GGATCACCTGTAATCTATTACCACCTTTTCTTGCCTTCTAATCTAGTGATAATATCAAAGAGTATTAGAGCACTTCTCTTTGTATCGAAATTTTATTGCTTCATACGCAGAATTATACAT

>M6-PGSC0003DMG400010032M6RLKWAX-EGF_61

TCCTTGTTTGTATCTCCGATTACTATTTGATAGCTTCATTGTTAGGAAATCCAGTTTTTGCAGAGCTGTTTTTGGAGTTGAGATATTTCAAATTTTGGACTCTTTAATTGCTACTATTTG

>M6-PGSC0003DMG400010032M6RLKWAX-EGF_610

CTGTGGAGCTGCTGGTTCTAATGGTCTGCGTGTTTCTTATCCATTTGGTTTCTCAGAGGGTTGTGGGATTCGTTTAGATTGTACTGAAAGTAGTGGAGAAATCAGAATTGGGGAATATAT

>M6-PGSC0003DMG400010032M6RLKWAX-EGF_6100

GTATTAGAGCACTTCTCTTTGTATCGAAATTTTATTGCTTCATACGCAGAATTATACATCATGAAGAATGTGAAATAGTATTTGCAATGCCATGAATTTGAATCGTCCTTGCGCAGTTGT

>M6-PGSC0003DMG400010032M6RLKWAX-EGF_6161

TGAAGAATGTGAAATAGTATTTGCAATGCCATGAATTTGAATCGTCCTTGCGCAGTTGTGTTCGTTATTTTTCTGTTATTTACAACAAGTGTAATCCTACATCATACATCATCTTCAAGA

>M6-PGSC0003DMG400010032M6RLKWAX-EGF_6222

TCGTTATTTTTCTGTTATTTACAACAAGTGTAATCCTACATCATACATCATCTTCAAGAATTTGAGTTAAATAATTTATAAAGTCATATTATATAACAGAATTAATTGTTACACTAATAG

>M6-PGSC0003DMG400010032M6RLKWAX-EGF_6283

TTGAGTTAAATAATTTATAAAGTCATATTATATAACAGAATTAATTGTTACACTAATAGAAATGCCCAAGCTTTGAATTTAGCAAATTCAATTTGCAAATAACAACAATGAAAAGCAAGA

>M6-PGSC0003DMG400010032M6RLKWAX-EGF_6344

ATGCCCAAGCTTTGAATTTAGCAAATTCAATTTGCAAATAACAACAATGAAAAGCAAGAATCTTTTCAAGATTTTTGAGGTTCAGGTCCTAAAATACCATTAACAACAGCTCCGGCAACC

>M6-PGSC0003DMG400010032M6RLKWAX-EGF_6405

CTTTTCAAGATTTTTGAGGTTCAGGTCCTAAAATACCATTAACAACAGCTCCGGCAACCAAAACTCCAACAAATTCAGCTCCGGCCACCGCTCCAGCTTGCGCCAACCCACCGAAACTCA

>M6-PGSC0003DMG400010032M6RLKWAX-EGF_6466

AACTCCAACAAATTCAGCTCCGGCCACCGCTCCAGCTTGCGCCAACCCACCGAAACTCACTGTACTCTCGATCTTCTCCTCCGCTGCTTCAATCCCCTTCTCAGCCGTCACTAACCCTTT

>M6-PGSC0003DMG400010032M6RLKWAX-EGF_6588

CCGCCGCTTCAATTCCCTTCTCTGCCGCTGCGATTCCCTTGTCTGCCGCCTCAAATCCCTCCTTCACTAATTCTTCTGCCTCCTCTTCTACTTTCTTAAGCTTCTTTCTCAGTCCGAAGA

>M6-PGSC0003DMG400010032M6RLKWAX-EGF_6649

CTTCACTAATTCTTCTGCCTCCTCTTCTACTTTCTTAAGCTTCTTTCTCAGTCCGAAGAGGCCAATGTCCGCAGCCTTGGACGGCATATCCAGCACTTTCAGTACCGTTGTCGCCGTCAC

>M6-PGSC0003DMG400010032M6RLKWAX-EGF_671

TGTGGGATTCGTTTAGATTGTACTGAAAGTAGTGGAGAAATCAGAATTGGGGAATATATAATACAGAATGTAACTTCAGAGACTCTGATGGTGAACTTCCCGATGAATTGCAGCCGTCCT

>M6-PGSC0003DMG400010032M6RLKWAX-EGF_6710

CCAATGTCCGCAGCCTTGGACGGCATATCCAGCACTTTCAGTACCGTTGTCGCCGTCACCAGAGAGATCAACTGCCTCCGCCCCGGGAGAGCCGCCGCCGCCTGTTTGGTAGGAGGATTT

>M6-PGSC0003DMG400010032M6RLKWAX-EGF_6771

GAGAGATCAACTGCCTCCGCCCCGGGAGAGCCGCCGCCGCCTGTTTGGTAGGAGGATTTGTGTTTGCATTTAGCACACAACGGGATGTGAAAATCCTTACTGTAAGTGAAGTAGTGGTGG

>M6-PGSC0003DMG400010032M6RLKWAX-EGF_6832

GTTTGCATTTAGCACACAACGGGATGTGAAAATCCTTACTGTAAGTGAAGTAGTGGTGGAAGCCATGGATTTGCTCTCTGTGCAATCAAAATGAGGTTTGCAGTTTGCACAACTCCCTTG

>M6-PGSC0003DMG400010032M6RLKWAX-EGF_6893

GCCATGGATTTGCTCTCTGTGCAATCAAAATGAGGTTTGCAGTTTGCACAACTCCCTTGACACCACAATCACAAACAAGTGTGTGGACTGATATTTTTTGAACGGACAAGAGAAAATACA

>M6-PGSC0003DMG400010032M6RLKWAX-EGF_6924

GAGGTTTGCAGTTTGCACAACTCCCTTGACACCACAATCACAAACAAGTGTGTGGACTGATATTTTTTGAACGGACAAGAGAAAATACAAAAAAATTACTCCTAAAACTTGAACCAGAAA

>M6-PGSC0003DMG400010032M6RLKWAX-EGF_732

TACAGAATGTAACTTCAGAGACTCTGATGGTGAACTTCCCGATGAATTGCAGCCGTCCTATTGAGGATCTACAGGAATTTGATCGAACCAACTTTGGTATGACTTGGAGAAATGGACTGC

>M6-PGSC0003DMG400010032M6RLKWAX-EGF_793

TGAGGATCTACAGGAATTTGATCGAACCAACTTTGGTATGACTTGGAGAAATGGACTGCTTCTACACAATTGTAAAGTCCCGAGGAGTGAGTGTACCATACCATCGGAAATTTTAAGCAC

>M6-PGSC0003DMG400010032M6RLKWAX-EGF_854

CTACACAATTGTAAAGTCCCGAGGAGTGAGTGTACCATACCATCGGAAATTTTAAGCACGCGTTTGAATATACAGTCATGTGATTCGAAGAAAGAGAATGTCAGTTGTTATTCAGAGGCG

>M6-PGSC0003DMG400010032M6RLKWAX-EGF_915

GTTTGAATATACAGTCATGTGATTCGAAGAAAGAGAATGTCAGTTGTTATTCAGAGGCGAGGGCGGATTATTTAGATTACCAGAAACTGAAGAATACAGGGTGTGGAACTGTGGTTTCGT

>M6-PGSC0003DMG400010032M6RLKWAX-EGF_976

GGCGGATTATTTAGATTACCAGAAACTGAAGAATACAGGGTGTGGAACTGTGGTTTCGTCCATCTTGATTGGTATGGATAATGATACAATGAAGAGTTCTGCCATGTTCCTGGAGTTTCA

>M6-PGSC0003DMG400010120M6RLKmalectin_0

TATGCGGTTAACTTGAGACTTCGCGCTCTTATGCAAAATATGATATTCAAGTTCCTAATGTACCTACATTTTTTCACTCTTATTCCTTCTAGAAATTCGATGAACATTAACATAGGTATT

>M6-PGSC0003DMG400010120M6RLKmalectin_130

ATCTCATTTGCACACTGTACAACTGTATTTCATACAATGAGCAATTGTGAATAGAGGATATATATTCCCAGAAGGACTAAGTATCACTTTGCTTGTAATCTTCTACACTTCTGTACGTTC

>M6-PGSC0003DMG400010120M6RLKmalectin_195

TCCCAGAAGGACTAAGTATCACTTTGCTTGTAATCTTCTACACTTCTGTACGTTCTATTGATGTTACTTTGGACTTAGAAGACCAAAATACTGGGCATGAGTAACTTGTATGTATCTCAT

>M6-PGSC0003DMG400010120M6RLKmalectin_260

ACTTTGGACTTAGAAGACCAAAATACTGGGCATGAGTAACTTGTATGTATCTCATATTATTATGGCTTGTATTTGTTTTTGTATTCCTAGTTCAATCTCGTTATGTGTCAATTGTTAATG

>M6-PGSC0003DMG400010120M6RLKmalectin_325

CTTGTATTTGTTTTTGTATTCCTAGTTCAATCTCGTTATGTGTCAATTGTTAATGTAATACCATTTTGTGTGGGTTCATCATAAATCAATTAACTTCATGTTGGTTGTCAATCCATTGGA

>M6-PGSC0003DMG400010120M6RLKmalectin_390

TTGTGTGGGTTCATCATAAATCAATTAACTTCATGTTGGTTGTCAATCCATTGGATCCCTCATTCTTTATCCAAGCTATTTTCTGTTATTCTTAGTGTAAATGTCTTATTTCTTTATCAA

>M6-PGSC0003DMG400010120M6RLKmalectin_455

TTTATCCAAGCTATTTTCTGTTATTCTTAGTGTAAATGTCTTATTTCTTTATCAACCAAAAGGGCTCTTAATAGAAGTCAAGTATGTTAAACGAGTGATTCTGTTTATTGAAAGAATTGG

>M6-PGSC0003DMG400010120M6RLKmalectin_520

TCTTAATAGAAGTCAAGTATGTTAAACGAGTGATTCTGTTTATTGAAAGAATTGGATTATAAAGGATCCTAATGTAGGAATAATGTAATTTTATTGTTTCACTAGGTTAAGTGTTTTTAA

>M6-PGSC0003DMG400010120M6RLKmalectin_585

ATCCTAATGTAGGAATAATGTAATTTTATTGTTTCACTAGGTTAAGTGTTTTTAAGAGATCAAGATGGAGGGGTGTAAATGTTAATGTCTTGATGGAGTATGTTGAATTTCAGATTTTTT

>M6-PGSC0003DMG400010120M6RLKmalectin_65

ACATTTTTTCACTCTTATTCCTTCTAGAAATTCGATGAACATTAACATAGGTATTGGTCTGCTCCATCTCATTTGCACACTGTACAACTGTATTTCATACAATGAGCAATTGTGAATAGA

>M6-PGSC0003DMG400010120M6RLKmalectin_650

TGGAGGGGTGTAAATGTTAATGTCTTGATGGAGTATGTTGAATTTCAGATTTTTTCGAGTCAGGGCAAGTGAAACTAACTAAAACTACCTGCTCTCTTTGTATATAGGTAGTTCTTTGAT

>M6-PGSC0003DMG400010120M6RLKmalectin_715

CAAGTGAAACTAACTAAAACTACCTGCTCTCTTTGTATATAGGTAGTTCTTTGATTGAAGGAATGAAGAGCAGACATGAGGATACATCACCATGATGAAGAACTTCCTTCAGGTTCATTC

>M6-PGSC0003DMG400010120M6RLKmalectin_845

AGTGATCTTCCAAATGTGTCAATTCAGAGCTCAATCTTAATATTCTCCTATACTGTGTTAATTACAGCATCACAAGTTGTTGTATCCGAGTTTGTATGATAGTGGGTGAGGTTGTATCTA

>M6-PGSC0003DMG400010183-(rev)M6LRR-RLP_0

TCATCTCTTGCTCCTTGACATTTTCCAGAATGGAAATTTTCTCAATGAGTATTAATTCCCACACCTTACTGGCAACCCATGCTGTGTTTCTATTCTCATCTACTCTCAAATATGCTGCAG

>M6-PGSC0003DMG400010183-(rev)M6LRR-RLP_1054

AACTGCAGCAGCAGTCTACAAATGCTCCATTTTGGAGACAACCAGTTTGTAGGTACATTAACTCATTCAACAGTCAACCTTTCTAGTCAGTTACAACTCCTACTCTTCTTCGGGAACAGA

>M6-PGSC0003DMG400010183-(rev)M6LRR-RLP_1116

TCATTCAACAGTCAACCTTTCTAGTCAGTTACAACTCCTACTCTTCTTCGGGAACAGAATTGGTGGAAGCATACCCAGAGAGATCTCAAACCTGGTGAATTTGAATCTACTCGACATGAG

>M6-PGSC0003DMG400010183-(rev)M6LRR-RLP_1178

GTGGAAGCATACCCAGAGAGATCTCAAACCTGGTGAATTTGAATCTACTCGACATGAGCAACAAAAATCTTACAGGTAGCATTCCAGATTCTATTGGAAGACTTGCAAACTTGGGAAGTC

>M6-PGSC0003DMG400010183-(rev)M6LRR-RLP_124

TACTATATACTTGGAAACGAAACTGACGAACTAGCCTTACTTGGATTCAAGTCCCAAATAACTGAGGATCCATCAAGAGTTTTCGCCTCTTGGAACCAATCTGTTCATTTATGTCAGTGG

>M6-PGSC0003DMG400010183-(rev)M6LRR-RLP_1240

AAAAATCTTACAGGTAGCATTCCAGATTCTATTGGAAGACTTGCAAACTTGGGAAGTCTTGACTTGCATAACAACCTCTTGACAGGGGTAATCCCTTCTTCAATAGGCAACCTCACTGAA

>M6-PGSC0003DMG400010183-(rev)M6LRR-RLP_1300

GACTTGCATAACAACCTCTTGACAGGGGTAATCCCTTCTTCAATAGGCAACCTCACTGAACTAGTCTATCTTTATTTAGGATTTAACAGGTTAGAGGGTAAAATACCTTTAAGGCTTTAA

>M6-PGSC0003DMG400010183-(rev)M6LRR-RLP_186

TGAGGATCCATCAAGAGTTTTCGCCTCTTGGAACCAATCTGTTCATTTATGTCAGTGGACTGGAGTAAAATGTGGCCAGACACAGGAAAGGGTCATCAGTTTGAATCTAAAAGGGCTAAG

>M6-PGSC0003DMG400010183-(rev)M6LRR-RLP_248

GAGTAAAATGTGGCCAGACACAGGAAAGGGTCATCAGTTTGAATCTAAAAGGGCTAAGTCTGGCAGGTACAATCTCAGATATCTTTCTTTGCTCAATTCTCTTGACCTGGCAGAAAATTC

>M6-PGSC0003DMG400010183-(rev)M6LRR-RLP_310

GCAGGTACAATCTCAGATATCTTTCTTTGCTCAATTCTCTTGACCTGGCAGAAAATTCATTCCATGACGAAATTCCTCAACAACTCAGCAGGCTGTCAAGGCATCAAAACCTGAATTTGA

>M6-PGSC0003DMG400010183-(rev)M6LRR-RLP_372

CATGACGAAATTCCTCAACAACTCAGCAGGCTGTCAAGGCATCAAAACCTGAATTTGAGTTTAAACTATCTAACAGGGGAAATTCCAGTTAATCTATCACATTGTGTTAACCTCAAGAGC

>M6-PGSC0003DMG400010183-(rev)M6LRR-RLP_434

AAACTATCTAACAGGGGAAATTCCAGTTAATCTATCACATTGTGTTAACCTCAAGAGCCTTGTGCTGGACCACAACATTCTTGTGGGACAGATTCCTTACCAAGTTGGATCTTTAACAAA

>M6-PGSC0003DMG400010183-(rev)M6LRR-RLP_496

TGCTGGACCACAACATTCTTGTGGGACAGATTCCTTACCAAGTTGGATCTTTAACAAAGTTAGTGAGGTTGTATCTCAGAAATAACAACCTGACAGGAATCTTTCCAATGTCTATTGGAA

>M6-PGSC0003DMG400010183-(rev)M6LRR-RLP_558

GTGAGGTTGTATCTCAGAAATAACAACCTGACAGGAATCTTTCCAATGTCTATTGGAAATCTTACATCTTTAGAAGAGTTGTATTTATCGTACAACAGTCTAGAGGGAGAAGTGCCAGCT

>M6-PGSC0003DMG400010183-(rev)M6LRR-RLP_62

ACCTTACTGGCAACCCATGCTGTGTTTCTATTCTCATCTACTCTCAAATATGCTGCAGCAGCTACTATATACTTGGAAACGAAACTGACGAACTAGCCTTACTTGGATTCAAGTCCCAAA

>M6-PGSC0003DMG400010183-(rev)M6LRR-RLP_620

TACATCTTTAGAAGAGTTGTATTTATCGTACAACAGTCTAGAGGGAGAAGTGCCAGCTTCTTTAGCTCGATTGACCAAGTTGAGACTGCTTGGATTGTCAGTAAATAGCTTCTCCGGGGA

>M6-PGSC0003DMG400010183-(rev)M6LRR-RLP_682

TAGCTCGATTGACCAAGTTGAGACTGCTTGGATTGTCAGTAAATAGCTTCTCCGGGGAGTTCCCTCCTTCGTTGTACAATTTGTCATCCCTGGAATTAATAGCACTTTCCTGGTAATCTC

>M6-PGSC0003DMG400010183-(rev)M6LRR-RLP_744

CCTCCTTCGTTGTACAATTTGTCATCCCTGGAATTAATAGCACTTTCCTGGTAATCTCAGATCCGATTTAGGCCACTACTTCCCTAATCTCCAAAGACTTTACTTGGGGAACTGTCAATT

>M6-PGSC0003DMG400010183-(rev)M6LRR-RLP_806

CCGATTTAGGCCACTACTTCCCTAATCTCCAAAGACTTTACTTGGGGAACTGTCAATTCCATGGCTCCATACAATCCTCTTTGGCCAATGCTTCAAAATTGCTACAGCTTGATTTCCCTG

>M6-PGSC0003DMG400010183-(rev)M6LRR-RLP_868

GGCTCCATACAATCCTCTTTGGCCAATGCTTCAAAATTGCTACAGCTTGATTTCCCTGTAAACAAATTCACTGGAAATATACCAAAGCGTTTTGATAACTTGTGGAATTTGTTGTGGCTC

>M6-PGSC0003DMG400010183-(rev)M6LRR-RLP_930

CAAATTCACTGGAAATATACCAAAGCGTTTTGATAACTTGTGGAATTTGTTGTGGCTCAATGTTGGGAGTAACCATCTTGGATACGGTAAGAATGACGACCTTGACTTTGTAAATTCTCT

>M6-PGSC0003DMG400010183-(rev)M6LRR-RLP_992

TTGGGAGTAACCATCTTGGATACGGTAAGAATGACGACCTTGACTTTGTAAATTCTCTTACCAACTGCAGCAGCAGTCTACAAATGCTCCATTTTGGAGACAACCAGTTTGTAGGTACAT

>M6-PGSC0003DMG400010230-(rev)M6LRR-RLK_0

AAAAAAGAGAAAAAATCTTGTAGAGAGAGAAACAAGAGAGAGGAAAAAACTTTTTTTTTCCCACCATTTTCCCATATTTTCCCTCTTGTTCCTCCTCAACCATTTTTCTGAGAAAAAAAA

>M6-PGSC0003DMG400010230-(rev)M6LRR-RLK_1037

AAGCTATGGGTGGTAGATGGAACACTTATGGAATCAAATTGTCGTATCTCGCTCTTTTGATTATCATTTTGGATATTCGCGGTTGTTGTTCTCTTAATTCGGAAGGTAATTCTTATGCTT

>M6-PGSC0003DMG400010230-(rev)M6LRR-RLK_1098

TATCATTTTGGATATTCGCGGTTGTTGTTCTCTTAATTCGGAAGGTAATTCTTATGCTTGAGTACAGTTCTATTGGTTTTGTAGTTCCGTCTTAGGCTAGTTTTAAGTGTTTGGTATGTT

>M6-PGSC0003DMG400010230-(rev)M6LRR-RLK_1159

GTACAGTTCTATTGGTTTTGTAGTTCCGTCTTAGGCTAGTTTTAAGTGTTTGGTATGTTTTCTTGTTTGCTAATTATTGTTGGTGTGAGAAACAGGATTGGCACTATTGAGATTCCGCGT

>M6-PGSC0003DMG400010230-(rev)M6LRR-RLK_122

TAGTTAAAATGAAGCCTCTGTTTTGAAATCACCCTGCAAACTGCAGTCAAGAAAAAAAAGAAAAGATATTACACTATTTTTTACAAAAAAAAGAAAAAAAAGATTGATTTTTTCTTGTGT

>M6-PGSC0003DMG400010230-(rev)M6LRR-RLK_1220

CTTGTTTGCTAATTATTGTTGGTGTGAGAAACAGGATTGGCACTATTGAGATTCCGCGTGAAAGTGGATTCTGATCCGTATGGCATTCTTGAAAACTGGAATTCAGATCATTATGACCCG

>M6-PGSC0003DMG400010230-(rev)M6LRR-RLK_1281

AAGTGGATTCTGATCCGTATGGCATTCTTGAAAACTGGAATTCAGATCATTATGACCCGTGCTTGTGGTCTGGTGTCCAGTGTATGGACGGGAGAGTGCAAATGTTGTAAGTCTCTTGTT

>M6-PGSC0003DMG400010230-(rev)M6LRR-RLK_1342

CTTGTGGTCTGGTGTCCAGTGTATGGACGGGAGAGTGCAAATGTTGTAAGTCTCTTGTTAAATATACCCAAAAATTTCCTAGTGTGAATCCTTTTCTTATAAATTTATATGTGTTGATTT

>M6-PGSC0003DMG400010230-(rev)M6LRR-RLK_1403

ATATACCCAAAAATTTCCTAGTGTGAATCCTTTTCTTATAAATTTATATGTGTTGATTTGCTATCGACAAGCCGGTTGAAATGATAATGACAGACAAAGTGCAAAAACTGAGTTGTGAGT

>M6-PGSC0003DMG400010230-(rev)M6LRR-RLK_1464

TATCGACAAGCCGGTTGAAATGATAATGACAGACAAAGTGCAAAAACTGAGTTGTGAGTTTTTCTGTTTGTGGTTCTTCGGATATTGTTAAATCATTTGTCTGTTCTCTATTTAGGGACC

>M6-PGSC0003DMG400010230-(rev)M6LRR-RLK_1525

TTCTGTTTGTGGTTCTTCGGATATTGTTAAATCATTTGTCTGTTCTCTATTTAGGGACCTTCATGGATATTCGTTGAAAGGAAAACTCGCAGCAGAACTTGGAAATCTTACTCACTTAAA

>M6-PGSC0003DMG400010230-(rev)M6LRR-RLK_1586

CATGGATATTCGTTGAAAGGAAAACTCGCAGCAGAACTTGGAAATCTTACTCACTTAAAATCACTGTAAGCTTCAGCTATACTTATATACTTCTCATTTGTTTCTTTAGCTCATATTCCT

>M6-PGSC0003DMG400010230-(rev)M6LRR-RLK_1647

CACTGTAAGCTTCAGCTATACTTATATACTTCTCATTTGTTTCTTTAGCTCATATTCCTGTCCTCACCATCTGAAAAAAGATTGTTCCAATCATTGATGTTTTTTTCTACTTACAAAATG

>M6-PGSC0003DMG400010230-(rev)M6LRR-RLK_1708

CCTCACCATCTGAAAAAAGATTGTTCCAATCATTGATGTTTTTTTCTACTTACAAAATGCTTTCATGATTCTGGTCTTAGTCTAGAAACAATATTTAGTGTGCTATGTCATGGAACTAAT

>M6-PGSC0003DMG400010230-(rev)M6LRR-RLK_1769

TTCATGATTCTGGTCTTAGTCTAGAAACAATATTTAGTGTGCTATGTCATGGAACTAATACTTCCCTATCGTGGTAGACACTATCTTATCACTAAACAATTTGAAATTGCACTATACTTA

>M6-PGSC0003DMG400010230-(rev)M6LRR-RLK_183

AAAGATATTACACTATTTTTTACAAAAAAAAGAAAAAAAAGATTGATTTTTTCTTGTGTGATTTCAATTTTCTTCTTCAATCTCTGTTATTTCTTTAATTTTTTTCAAAAAAGAAAAAAA

>M6-PGSC0003DMG400010230-(rev)M6LRR-RLK_1830

TTCCCTATCGTGGTAGACACTATCTTATCACTAAACAATTTGAAATTGCACTATACTTACATGTCTGGTCAGTACAGGTTTCCATTTTTCGTTCCCATTTGTTACTTAGGAGATGTCTCA

>M6-PGSC0003DMG400010230-(rev)M6LRR-RLK_1891

TGTCTGGTCAGTACAGGTTTCCATTTTTCGTTCCCATTTGTTACTTAGGAGATGTCTCAAAATATAAAGAGCGTCATATACATTAGGATATTTAGCATATTTGGTCAGGCAAATTGCACC

>M6-PGSC0003DMG400010230-(rev)M6LRR-RLK_1952

ATATAAAGAGCGTCATATACATTAGGATATTTAGCATATTTGGTCAGGCAAATTGCACCAAAGAAAAACTTTACCTTTTTTTTTTCCAAAGAAAAATGTATAAAGTGAATTGGGTAGATT

>M6-PGSC0003DMG400010230-(rev)M6LRR-RLK_2013

AGAAAAACTTTACCTTTTTTTTTTCCAAAGAAAAATGTATAAAGTGAATTGGGTAGATTTATGGTAATCAATGTTTCACCTTCTGTTCTGCATCTATGTACTATGTATTAATAAATTTTC

>M6-PGSC0003DMG400010230-(rev)M6LRR-RLK_2074

TGGTAATCAATGTTTCACCTTCTGTTCTGCATCTATGTACTATGTATTAATAAATTTTCTTCCTCTTTGAATGCTTTTCTTCCACAGTGTGCTTTCTGAAAACCGTCTTTTTGGTGCTAT

>M6-PGSC0003DMG400010230-(rev)M6LRR-RLK_2135

CCTCTTTGAATGCTTTTCTTCCACAGTGTGCTTTCTGAAAACCGTCTTTTTGGTGCTATTCCTAAAGAATTTGGACGACTCAGAATGCTGGAAGTGCTTGACATTAAGAACAACAATTTG

>M6-PGSC0003DMG400010230-(rev)M6LRR-RLK_2196

CTAAAGAATTTGGACGACTCAGAATGCTGGAAGTGCTTGACATTAAGAACAACAATTTGAGTGGAAGAATTCCAGCAGTAATAGGAGATTTGCAATCACTTAGAAGCTTGTAAGTTGATA

>M6-PGSC0003DMG400010230-(rev)M6LRR-RLK_2257

TGGAAGAATTCCAGCAGTAATAGGAGATTTGCAATCACTTAGAAGCTTGTAAGTTGATAATTTGTTGTGTTTCCGTCTTATAATATATTTGTATGTATTTATGCTACAACATTGCTAACA

>M6-PGSC0003DMG400010230-(rev)M6LRR-RLK_2318

TTGTTGTGTTTCCGTCTTATAATATATTTGTATGTATTTATGCTACAACATTGCTAACAAAATGTTCTTGAATTGTGCTTGATACTTCTTTTCCAGGTTGATTAGGGACAATAACTTCAA

>M6-PGSC0003DMG400010230-(rev)M6LRR-RLK_2379

ATGTTCTTGAATTGTGCTTGATACTTCTTTTCCAGGTTGATTAGGGACAATAACTTCAAAGGGAAGATCCCTTTGGAGATCAGAAAGCTTCATCTGCTCTCGGAATTGCAATTTGATGAT

>M6-PGSC0003DMG400010230-(rev)M6LRR-RLK_244

TTTCAATTTTCTTCTTCAATCTCTGTTATTTCTTTAATTTTTTTCAAAAAAGAAAAAAATAAATTCTGTCCCACCGCCTGCATTTTCCTATTGCAGTTATTAATTTTTGGCTTTTTTTTC

>M6-PGSC0003DMG400010230-(rev)M6LRR-RLK_2440

GGAAGATCCCTTTGGAGATCAGAAAGCTTCATCTGCTCTCGGAATTGCAATTTGATGATTACCTCACCTCTGGTGTTGCTGTCGGAACTGGTTGCATAAATAGAAAATTTGGGCGTTGGT

>M6-PGSC0003DMG400010230-(rev)M6LRR-RLK_2501

CCTCACCTCTGGTGTTGCTGTCGGAACTGGTTGCATAAATAGAAAATTTGGGCGTTGGTAAGACTTCCGTAGTTGTTTGGTCCTTTTATCTTCATGTTTCAGCTGTGCTGCATTTTAAAG

>M6-PGSC0003DMG400010230-(rev)M6LRR-RLK_2562

GACTTCCGTAGTTGTTTGGTCCTTTTATCTTCATGTTTCAGCTGTGCTGCATTTTAAAGTCGCAGACTTGTTTCTATGTTCAATTATATTGAAAATGCATAAAATCCATCATTTGAGGAA

>M6-PGSC0003DMG400010230-(rev)M6LRR-RLK_2623

GCAGACTTGTTTCTATGTTCAATTATATTGAAAATGCATAAAATCCATCATTTGAGGAACATATACAAATATACTACTTTCATACGAACTACCTCCTGCTATAAGTTTGGTATTAACATT

>M6-PGSC0003DMG400010230-(rev)M6LRR-RLK_2684

TATACAAATATACTACTTTCATACGAACTACCTCCTGCTATAAGTTTGGTATTAACATTAAATTAAGTGCAAACAATTATCCCATGGAGTTGCACTGAAGATCTTTGAGACATTTTCTAT

>M6-PGSC0003DMG400010230-(rev)M6LRR-RLK_2745

ATTAAGTGCAAACAATTATCCCATGGAGTTGCACTGAAGATCTTTGAGACATTTTCTATGCATAGTGTGTCCCCACTAAACTCTCATACTTCCCTTACAATAGTGAAGCAACTATGATGG

>M6-PGSC0003DMG400010230-(rev)M6LRR-RLK_2806

ATAGTGTGTCCCCACTAAACTCTCATACTTCCCTTACAATAGTGAAGCAACTATGATGGTGATTGTCGAAAGCATTGCCATGGGGCACTTGATACTTTCCTGTCTCCAACCCTAATATTC

>M6-PGSC0003DMG400010230-(rev)M6LRR-RLK_2867

ATTGTCGAAAGCATTGCCATGGGGCACTTGATACTTTCCTGTCTCCAACCCTAATATTCAGAGGGGATTAGACTTTAGTGTTTCTAAAGTTGTGATTCATCTTCTTATTCATCAAGACCT

>M6-PGSC0003DMG400010230-(rev)M6LRR-RLK_2928

AGGGGATTAGACTTTAGTGTTTCTAAAGTTGTGATTCATCTTCTTATTCATCAAGACCTGTTTATGTGGAAATTTGACTGATCTATTGCATAATATTTGTTTATTTTGTCGACTTTTCTG

>M6-PGSC0003DMG400010230-(rev)M6LRR-RLK_2989

TTATGTGGAAATTTGACTGATCTATTGCATAATATTTGTTTATTTTGTCGACTTTTCTGTACATTAACCAATAAAATTGCTGGAAATGTGGTTATTCTTTCCCATGAGTAGAAGAACACT

>M6-PGSC0003DMG400010230-(rev)M6LRR-RLK_305

AATTCTGTCCCACCGCCTGCATTTTCCTATTGCAGTTATTAATTTTTGGCTTTTTTTTCAAGAAATGTGACGAATTGGCATTTCCATTTCTGCAATAAAAAGCAGAAAAAGATTTCATTC

>M6-PGSC0003DMG400010230-(rev)M6LRR-RLK_3050

CATTAACCAATAAAATTGCTGGAAATGTGGTTATTCTTTCCCATGAGTAGAAGAACACTTGGATATTATCGTATTCCTGTTTTGTAGCATGAAACTTCAGCCAGCTATATGAATAAATTT

>M6-PGSC0003DMG400010230-(rev)M6LRR-RLK_3111

GATATTATCGTATTCCTGTTTTGTAGCATGAAACTTCAGCCAGCTATATGAATAAATTTATTCTATTCATCCCGTTGACTATGTTTGAATTCTGGATCGAGATACTCATCTTGGACAAGT

>M6-PGSC0003DMG400010230-(rev)M6LRR-RLK_3172

TCTATTCATCCCGTTGACTATGTTTGAATTCTGGATCGAGATACTCATCTTGGACAAGTACCTTCAGCTTTACCCTTATAACTGATGACAGATTACTGACTACCTCTTAATATGCTAATG

>M6-PGSC0003DMG400010230-(rev)M6LRR-RLK_3233

CTTCAGCTTTACCCTTATAACTGATGACAGATTACTGACTACCTCTTAATATGCTAATGTTTTATTTCTCAATCACGCACTTTTAAGTTTCATACTCCGCTCTGAAAATTTGGTGACTTT

>M6-PGSC0003DMG400010230-(rev)M6LRR-RLK_3294

TTATTTCTCAATCACGCACTTTTAAGTTTCATACTCCGCTCTGAAAATTTGGTGACTTTTTCTTGCAGCATCTGGCATGGCAGTTTGAGGCCATATAAGACTATAGGCTCCTTTCTCAGA

>M6-PGSC0003DMG400010230-(rev)M6LRR-RLK_3355

CTTGCAGCATCTGGCATGGCAGTTTGAGGCCATATAAGACTATAGGCTCCTTTCTCAGACCAATTAAAGGGATGCTAGCATGGTATTTTAGTTTTTTCACACTGTAAGTAACTAAATACG

>M6-PGSC0003DMG400010230-(rev)M6LRR-RLK_3416

AATTAAAGGGATGCTAGCATGGTATTTTAGTTTTTTCACACTGTAAGTAACTAAATACGTTTTTTTCACCTTATGCTCTCTAAATAAGTTTAATGGTTCAGAAGCAACTTTTTTGTTTCC

>M6-PGSC0003DMG400010230-(rev)M6LRR-RLK_3477

TTTTTCACCTTATGCTCTCTAAATAAGTTTAATGGTTCAGAAGCAACTTTTTTGTTTCCTAAAAGGTTCCCACGTTTTTTGGATGCTCAGGCAGACTTCTGCTCAGACAACCTACCAAGT

>M6-PGSC0003DMG400010230-(rev)M6LRR-RLK_3538

AAAGGTTCCCACGTTTTTTGGATGCTCAGGCAGACTTCTGCTCAGACAACCTACCAAGTGAGTGAAAATATTTTGTGACATATCTTTTATTCGATTTCAAAGAATTGTACACAAAAAAAA

>M6-PGSC0003DMG400010230-(rev)M6LRR-RLK_3599

GTGAAAATATTTTGTGACATATCTTTTATTCGATTTCAAAGAATTGTACACAAAAAAAAGAAAGTATTTTATTCTCTTTTGTTTTTGCAGGTTCAACTAGGCCTCACAATATCCACGCTG

>M6-PGSC0003DMG400010230-(rev)M6LRR-RLK_366

GAAATGTGACGAATTGGCATTTCCATTTCTGCAATAAAAAGCAGAAAAAGATTTCATTCTTGTCTGGTAAGTTTTACTCTATACATTTTACCATCTTTTGGGTTTTTGGATTTTTATTTT

>M6-PGSC0003DMG400010230-(rev)M6LRR-RLK_3660

AAGTATTTTATTCTCTTTTGTTTTTGCAGGTTCAACTAGGCCTCACAATATCCACGCTGTAGAGAATCAAGCAAACATTGAACGCCGTAAGTTAGCTGAACAATCCAGTAACTTTGCTGC

>M6-PGSC0003DMG400010230-(rev)M6LRR-RLK_3782

CTCCTGCTAACGGAGGGAAACCATTGGGACCTGTTGATCCTATTCCAAGCAGCAGAAGTAGTGGTTCATTTCGTGCTGTGCCAAGTACTGAAGGAGCTCCTCCACCTCCTCTGACTACAC

>M6-PGSC0003DMG400010230-(rev)M6LRR-RLK_3843

TGGTTCATTTCGTGCTGTGCCAAGTACTGAAGGAGCTCCTCCACCTCCTCTGACTACACCATCACGGACCACACATGATCATCATCCGCCTCCAAATCCTGGGGGACACTCTAATGGTGC

>M6-PGSC0003DMG400010230-(rev)M6LRR-RLK_3904

TCACGGACCACACATGATCATCATCCGCCTCCAAATCCTGGGGGACACTCTAATGGTGCTGTTAAACAGCCAATTGCTAGTCAAGCTCCACCTGGTGGAAAGTCTGGAAGTACATGGAAG

>M6-PGSC0003DMG400010230-(rev)M6LRR-RLK_3965

TTAAACAGCCAATTGCTAGTCAAGCTCCACCTGGTGGAAAGTCTGGAAGTACATGGAAGTATATAGGAATTGGCATTGGGGCTTTTCTGGTTGCTGTTTTTGTGTGCCTGATAATCATCT

>M6-PGSC0003DMG400010230-(rev)M6LRR-RLK_4026

TATAGGAATTGGCATTGGGGCTTTTCTGGTTGCTGTTTTTGTGTGCCTGATAATCATCTGCAAGAGCAAAGCTGCACAAACAATAGGCCCTTGGAAGACTGGATTGAGTGGACAGCTGCA

>M6-PGSC0003DMG400010230-(rev)M6LRR-RLK_4148

AAGCATTTATTACAGGTACTTTTGATTTAATATTGTGTCAAAGCTGATGAATCCATATGTTCTAGTTCAACTGTTGGATTCATTTTTCTCTGGTGGATATAATCCGAATCTTCAACTCCT

>M6-PGSC0003DMG400010230-(rev)M6LRR-RLK_4209

CTAGTTCAACTGTTGGATTCATTTTTCTCTGGTGGATATAATCCGAATCTTCAACTCCTACAATAGTTTTTGTTTCATACTCCAAGATGATGTTAAAAGCATGTATTAATACTCCTGTGC

>M6-PGSC0003DMG400010230-(rev)M6LRR-RLK_427

GTCTGGTAAGTTTTACTCTATACATTTTACCATCTTTTGGGTTTTTGGATTTTTATTTTTCTTTTGATTGCTTGCAAATTCAGTTATTCCCTCATCATCATCTTTTTCTTTTTCTTTTTC

>M6-PGSC0003DMG400010230-(rev)M6LRR-RLK_4270

AATAGTTTTTGTTTCATACTCCAAGATGATGTTAAAAGCATGTATTAATACTCCTGTGCTATATTGTTTCTCATGTCTTATGCATGAAATCAATTTTGCAGGGGTCCCAAAACTAAACAG

>M6-PGSC0003DMG400010230-(rev)M6LRR-RLK_4331

TATTGTTTCTCATGTCTTATGCATGAAATCAATTTTGCAGGGGTCCCAAAACTAAACAGGCCGGAGCTTGAAAATGCATGCGAGGATTTTAGCAACATAGTTTGCCATCAAGATACATTT

>M6-PGSC0003DMG400010230-(rev)M6LRR-RLK_4392

CGGAGCTTGAAAATGCATGCGAGGATTTTAGCAACATAGTTTGCCATCAAGATACATTTACAGTCTACAAGGGAACTATATCCAGTGGAGTAGAGATTGCTGTTGTGTCAACTGCTGTAA

>M6-PGSC0003DMG400010230-(rev)M6LRR-RLK_4453

AGTCTACAAGGGAACTATATCCAGTGGAGTAGAGATTGCTGTTGTGTCAACTGCTGTAAATTCTTTGAAAGATTGGTCCAAGCATTCTGAATTGGCCTTCAGGAAGAAGGTTAGTAGTTG

>M6-PGSC0003DMG400010230-(rev)M6LRR-RLK_4514

TCTTTGAAAGATTGGTCCAAGCATTCTGAATTGGCCTTCAGGAAGAAGGTTAGTAGTTGTCATCTCCTTCACAAATAAAAAAACTGCTTCATAACTTATTACCATCTCACTCTTTCTGTT

>M6-PGSC0003DMG400010230-(rev)M6LRR-RLK_4575

ATCTCCTTCACAAATAAAAAAACTGCTTCATAACTTATTACCATCTCACTCTTTCTGTTGAGATATCCTAGTCGTCTCGGTTTTTTTACTTATAAGATTTCTACTTATTGTGCACTTGGG

>M6-PGSC0003DMG400010230-(rev)M6LRR-RLK_4636

GATATCCTAGTCGTCTCGGTTTTTTTACTTATAAGATTTCTACTTATTGTGCACTTGGGTAGATCGATTCACTCTCAAGGATAAACCACAAGAATTTTGTTAATCTAATTGGATATTGCG

>M6-PGSC0003DMG400010230-(rev)M6LRR-RLK_4697

GATCGATTCACTCTCAAGGATAAACCACAAGAATTTTGTTAATCTAATTGGATATTGCGAGGAAGATGAACCTTTCACTAGGATGATGGTCTTTGAGTATGCAACAAATGGAACACTTTT

>M6-PGSC0003DMG400010230-(rev)M6LRR-RLK_4758

GAAGATGAACCTTTCACTAGGATGATGGTCTTTGAGTATGCAACAAATGGAACACTTTTCGAGCATCTGCATGGTAAACTGGTTAATTGTTCAAGTTTCAGTCTAATCATTTTTGTGATT

>M6-PGSC0003DMG400010230-(rev)M6LRR-RLK_4819

AGCATCTGCATGGTAAACTGGTTAATTGTTCAAGTTTCAGTCTAATCATTTTTGTGATTTCGTTTAGATTAATGCAGTCTACATGTTTTGTTGTTTCAGATGAAGAACTTGAACCTCTTG

>M6-PGSC0003DMG400010230-(rev)M6LRR-RLK_4880

GTTTAGATTAATGCAGTCTACATGTTTTGTTGTTTCAGATGAAGAACTTGAACCTCTTGACTGGCCTGCAAGGGTACGAGCTATAATGGGAACAGCATATTGTCTCGAGTACATGCACAA

>M6-PGSC0003DMG400010230-(rev)M6LRR-RLK_4941

TGGCCTGCAAGGGTACGAGCTATAATGGGAACAGCATATTGTCTCGAGTACATGCACAATTTGAATCCTCCTCTGTCGCATTCTGATGTGAATTCACACTCCATATTTTTTACGGACGAT

>M6-PGSC0003DMG400010230-(rev)M6LRR-RLK_5002

TGAATCCTCCTCTGTCGCATTCTGATGTGAATTCACACTCCATATTTTTTACGGACGATTATGCTGCTAAGGTATGCTTCATTTTGTATACCACATTGAATTTAGAAACGTTTTACTGTC

>M6-PGSC0003DMG400010230-(rev)M6LRR-RLK_5063

TGCTGCTAAGGTATGCTTCATTTTGTATACCACATTGAATTTAGAAACGTTTTACTGTCTTGAATTAAACGAATCGTTTTTTATTTACACAGATTACAGAGCTTGCTTTCTGGTCAGATA

>M6-PGSC0003DMG400010230-(rev)M6LRR-RLK_5124

GAATTAAACGAATCGTTTTTTATTTACACAGATTACAGAGCTTGCTTTCTGGTCAGATATAATGGTGAAGTCAAAGTCCTCTAGTGGTGGTGATGTAGAGAACTCTGAGCTACCACCACT

>M6-PGSC0003DMG400010230-(rev)M6LRR-RLK_5185

ATGGTGAAGTCAAAGTCCTCTAGTGGTGGTGATGTAGAGAACTCTGAGCTACCACCACTTTCTGATCCCGAAACAAATGTCTACAATTTTGGGATTTTGTTGCTGGAAGTAATTTCTGGG

>M6-PGSC0003DMG400010230-(rev)M6LRR-RLK_5246

CTGATCCCGAAACAAATGTCTACAATTTTGGGATTTTGTTGCTGGAAGTAATTTCTGGGAAGTCACCTTACTCGGAACGAGATTCTCTTCTGAGCTGGGTAAAGTCTCAATCTATGTAAC

>M6-PGSC0003DMG400010230-(rev)M6LRR-RLK_5307

GTCACCTTACTCGGAACGAGATTCTCTTCTGAGCTGGGTAAAGTCTCAATCTATGTAACTCTAATTTTCTTCGGTGTTTAATATAGACTAGTGAAACGTTCTTCTCTGCTTTTGTTTAGG

>M6-PGSC0003DMG400010230-(rev)M6LRR-RLK_5368

TAATTTTCTTCGGTGTTTAATATAGACTAGTGAAACGTTCTTCTCTGCTTTTGTTTAGGCGGAGCAATGCGTTAACGATAGACAGTTGGTTGATCCAAAATTGAAGTCGTTCAAGAATAA

>M6-PGSC0003DMG400010230-(rev)M6LRR-RLK_5429

GAGCAATGCGTTAACGATAGACAGTTGGTTGATCCAAAATTGAAGTCGTTCAAGAATAATGAGCTTATGGTTATGTGTGAGGTAATGAGAGAATGTATTGGAGAAGATTCAAGGAAGAGA

>M6-PGSC0003DMG400010230-(rev)M6LRR-RLK_549

TTTCTTTTTCTCTCTCTCTCTTTTTCTTTTCATGGTTTGGCTTGTGAAGTGTTCTTAGGTTTTTCACCAACCCCTTTTCAAGAAACATATTTTTACTAATTTTTTTTCTTTTGTGTTTAT

>M6-PGSC0003DMG400010230-(rev)M6LRR-RLK_5490

AGCTTATGGTTATGTGTGAGGTAATGAGAGAATGTATTGGAGAAGATTCAAGGAAGAGACCAACAATAAAAGAAGTGATTAAAAAGCTAAGGGAAGCGATTGATATCTCCCCTGATGCTG

>M6-PGSC0003DMG400010230-(rev)M6LRR-RLK_5551

AACAATAAAAGAAGTGATTAAAAAGCTAAGGGAAGCGATTGATATCTCCCCTGATGCTGCAGTTCCAAGGCTATCTCCCCTGTGGTGGGCTGAACTTGAGATCCTATCCTCTGAGGCAGC

>M6-PGSC0003DMG400010230-(rev)M6LRR-RLK_5612

GTTCCAAGGCTATCTCCCCTGTGGTGGGCTGAACTTGAGATCCTATCCTCTGAGGCAGCCTAATATACTGACTAATGTGTCTTCTACTTTTGTAAGTAGATAGTTGCTTTGCTATGCATA

>M6-PGSC0003DMG400010230-(rev)M6LRR-RLK_5673

AATATACTGACTAATGTGTCTTCTACTTTTGTAAGTAGATAGTTGCTTTGCTATGCATAAATGTTCTGAAATATACTTCACTTAATGTCAACTGTGTACCACATTTTTCTGTTCTAAATA

>M6-PGSC0003DMG400010230-(rev)M6LRR-RLK_61

CACCATTTTCCCATATTTTCCCTCTTGTTCCTCCTCAACCATTTTTCTGAGAAAAAAAAAATAGTTAAAATGAAGCCTCTGTTTTGAAATCACCCTGCAAACTGCAGTCAAGAAAAAAAA

>M6-PGSC0003DMG400010230-(rev)M6LRR-RLK_610

TTTCACCAACCCCTTTTCAAGAAACATATTTTTACTAATTTTTTTTCTTTTGTGTTTATTCATTTTCCAGATAAAAATTGAATCTTGGAGATTAATCAATCGAGTTTGTGGAAGTGTTTG

>M6-PGSC0003DMG400010230-(rev)M6LRR-RLK_671

ATTTTCCAGATAAAAATTGAATCTTGGAGATTAATCAATCGAGTTTGTGGAAGTGTTTGGTGAGGGAGATGTTGTTATAGTCTAAATTTTAGAAACTGTGAAAAATGAGAGACCCCAATT

>M6-PGSC0003DMG400010230-(rev)M6LRR-RLK_732

GAGGGAGATGTTGTTATAGTCTAAATTTTAGAAACTGTGAAAAATGAGAGACCCCAATTTTGTCCATTGTGTCAATAGGGTTAATAGTTTTTCATTTAACTTGTAATATAGTTTTACATT

>M6-PGSC0003DMG400010230-(rev)M6LRR-RLK_793

GTCCATTGTGTCAATAGGGTTAATAGTTTTTCATTTAACTTGTAATATAGTTTTACATTGTAAAAATTGAATGATAAGCAAGCAGTGGTGTATTTGCTCTCCTTTTTTTCAACTTGTTTA

>M6-PGSC0003DMG400010230-(rev)M6LRR-RLK_854

AAAAATTGAATGATAAGCAAGCAGTGGTGTATTTGCTCTCCTTTTTTTCAACTTGTTTATATTTACGAGTGAATTCAATTGTTTGGAGAGTAATTTTGTACAAGAAAGCTGGCGATTTTG

>M6-PGSC0003DMG400010230-(rev)M6LRR-RLK_915

TTTACGAGTGAATTCAATTGTTTGGAGAGTAATTTTGTACAAGAAAGCTGGCGATTTTGTATAGAGTCCTGGCTGTTGACAGATATTGTTACTGCCTCAACAACTGATTCATATGGTTGT

>M6-PGSC0003DMG400010230-(rev)M6LRR-RLK_976

TAGAGTCCTGGCTGTTGACAGATATTGTTACTGCCTCAACAACTGATTCATATGGTTGTGCAAGCTATGGGTGGTAGATGGAACACTTATGGAATCAAATTGTCGTATCTCGCTCTTTTG

>M6-PGSC0003DMG400010240-(rev)M6G-LecRK_0

TCTTCTTTGCTTGTAATAACACTTTGTTCCTTATTCATAGTAACATTAGCTAATGTTCCTCCTTCCAAAACCTTCAAGTACATAAATGAAGGCACATTTGGCGAATACTCCGTTGAATAT

>M6-PGSC0003DMG400010240-(rev)M6G-LecRK_1040

AAAGTTGCCAGTATGCAGGGGAGGATCAAATGTGGACTATTATAAGGTGGAAGGAGTTGAACATTTTACTAGTGATTATAATGAAGGAGTTGGACCTATAAAAATAAGTGAATGCAAAAA

>M6-PGSC0003DMG400010240-(rev)M6G-LecRK_1105

TTACTAGTGATTATAATGAAGGAGTTGGACCTATAAAAATAAGTGAATGCAAAAATAGGTGCAGCAAAGATTGCAAATGTGTGGGCTTTTTCTACAAACAAGAGTCATCAAAATGTTTGG

>M6-PGSC0003DMG400010240-(rev)M6G-LecRK_1170

AAAGATTGCAAATGTGTGGGCTTTTTCTACAAACAAGAGTCATCAAAATGTTTGGTTGTACCTGAGCTTGGAGCTTTGGTTAAAGTGGACAATTCCTCCCATGTTGGCTACATCAAGATC

>M6-PGSC0003DMG400010240-(rev)M6G-LecRK_1235

GCTTGGAGCTTTGGTTAAAGTGGACAATTCCTCCCATGTTGGCTACATCAAGATCTCAAAGTAGATTAAGTTATTACTGATTTGTATTTATATCAAGCAAGCATTTTATATTTTTTTAAA

>M6-PGSC0003DMG400010240-(rev)M6G-LecRK_1288

TCTCAAAGTAGATTAAGTTATTACTGATTTGTATTTATATCAAGCAAGCATTTTATATTTTTTTAAAATTTCAGTAATAATAATGTAACTTTTGTCAAAATATAATGATAAAAAATGCAC

>M6-PGSC0003DMG400010240-(rev)M6G-LecRK_130

ATCGTGTGTTGGACATTGTCAACTTCCCTTTCACCCTTTGTTTCTTCAATTCTACTCCAAACACCTTCATACTTGGGCAACGAATGGGTCATCGTAGATCTGAGTCCATTATGCGATGGG

>M6-PGSC0003DMG400010240-(rev)M6G-LecRK_195

TTCATACTTGGGCAACGAATGGGTCATCGTAGATCTGAGTCCATTATGCGATGGGTCTGGGAGGCTAATCGTGGTAAGCCTGTTCGCGAAAACGCCACTCTAACGTTTGGTACTAATGGG

>M6-PGSC0003DMG400010240-(rev)M6G-LecRK_260

TAATCGTGGTAAGCCTGTTCGCGAAAACGCCACTCTAACGTTTGGTACTAATGGGAATTTGGTACTAACTGATGTCAATGGCACGGTAGCTTGGCATACCAATACTGTCAACAAAGGTGT

>M6-PGSC0003DMG400010240-(rev)M6G-LecRK_325

TAACTGATGTCAATGGCACGGTAGCTTGGCATACCAATACTGTCAACAAAGGTGTAACTGGGCTAAAACTTTTGCCAAATGGCAACTTGGTTTTACATGACAAAAATGATAGATTTGTTT

>M6-PGSC0003DMG400010240-(rev)M6G-LecRK_390

AAACTTTTGCCAAATGGCAACTTGGTTTTACATGACAAAAATGATAGATTTGTTTGGCAAAGTTTTGATCACCCAACGGATACACTTTTGGTGGGCCAAAGTTTAAGGGCTAAAGGCCCA

>M6-PGSC0003DMG400010240-(rev)M6G-LecRK_455

TGATCACCCAACGGATACACTTTTGGTGGGCCAAAGTTTAAGGGCTAAAGGCCCATCTAAGATTGTTAGTAGAAAGTCCAGTAAAGAACCTGTTGATGGGCCTTATAGTTTTATCCTAGA

>M6-PGSC0003DMG400010240-(rev)M6G-LecRK_520

TTAGTAGAAAGTCCAGTAAAGAACCTGTTGATGGGCCTTATAGTTTTATCCTAGAAGGGAGACATTGGGCCATGTATTATACAAGTGTAAATTCACAAAAGCCTTTACCATATTTCAAAT

>M6-PGSC0003DMG400010240-(rev)M6G-LecRK_585

TGGGCCATGTATTATACAAGTGTAAATTCACAAAAGCCTTTACCATATTTCAAATCAGATGATTTTGGTAATGGGAAAGGCTCTTTAGCCAATTTTGATTTCTACTGTGAGCCAGAATAT

>M6-PGSC0003DMG400010240-(rev)M6G-LecRK_65

CAAAACCTTCAAGTACATAAATGAAGGCACATTTGGCGAATACTCCGTTGAATATTTTGCAGATTATCGTGTGTTGGACATTGTCAACTTCCCTTTCACCCTTTGTTTCTTCAATTCTAC

>M6-PGSC0003DMG400010240-(rev)M6G-LecRK_650

TGGTAATGGGAAAGGCTCTTTAGCCAATTTTGATTTCTACTGTGAGCCAGAATATGGTGAAAATTATGCTTTTGAGCTAGGATTTAGGATCAACATGACCAATTCTCCTAGCTCTGGCAC

>M6-PGSC0003DMG400010240-(rev)M6G-LecRK_715

ATGCTTTTGAGCTAGGATTTAGGATCAACATGACCAATTCTCCTAGCTCTGGCACATCTATATTTACTAGGCCCAAATACAATAGTACATACACTATGTTTAGAGTTGATATGGATGGTG

>M6-PGSC0003DMG400010240-(rev)M6G-LecRK_780

ACTAGGCCCAAATACAATAGTACATACACTATGTTTAGAGTTGATATGGATGGTGATTTAAGAATGTACACTTATGAACCAAATGTTGATTGGGGAGCATGGGAAGTCACATATCGACTT

>M6-PGSC0003DMG400010240-(rev)M6G-LecRK_845

GTACACTTATGAACCAAATGTTGATTGGGGAGCATGGGAAGTCACATATCGACTTCTCGATAGGGACAACGATGGAAAAAGTGAATGCAAGCTACCGAGGAGGTGTGGATCATTAGGCGT

>M6-PGSC0003DMG400010240-(rev)M6G-LecRK_910

ACAACGATGGAAAAAGTGAATGCAAGCTACCGAGGAGGTGTGGATCATTAGGCGTTTGTGAGGACAGCCAGTGTGTGGCGTGTCCGAGCCCTCAAGGCTTGATTGGGTGGAGTAAGGGAT

>M6-PGSC0003DMG400010240-(rev)M6G-LecRK_975

AGCCAGTGTGTGGCGTGTCCGAGCCCTCAAGGCTTGATTGGGTGGAGTAAGGGATGTGCACCACCAAAGTTGCCAGTATGCAGGGGAGGATCAAATGTGGACTATTATAAGGTGGAAGGA

>M6-PGSC0003DMG400010325-(rev)M6LRR-RLP_0

ATGGTGGAACAAGAAGCAGTGTTCAATGTTCTTGAATCCATTAACTCTGATGTCTCTTGGCGTTCCCTTTTCCCTGATGATCTCTGTTCTTCTTCCCCACATGGGGTTATTTGTGATTAC

>M6-PGSC0003DMG400010325-(rev)M6LRR-RLP_1024

GTGGTAGACTTCCATTCTCTCCAAAATTTGTATCCAAAGTTGGAGAAAAGTTGAAGTTGGGAGGCAATCCTCAACTCTGTGTAGATGAAGGCTTGAGAATATCTAATGTTAGTGTTAGCT

>M6-PGSC0003DMG400010325-(rev)M6LRR-RLP_1088

CAATCCTCAACTCTGTGTAGATGAAGGCTTGAGAATATCTAATGTTAGTGTTAGCTTGAGGCTACTAAAAATGTGCAACAATGCTATTAGTGCCAAATTTGTCCCATTCTCTGGGGCATC

>M6-PGSC0003DMG400010325-(rev)M6LRR-RLP_1152

CTAAAAATGTGCAACAATGCTATTAGTGCCAAATTTGTCCCATTCTCTGGGGCATCTTTAGTACTGGTGCCTGTTTCCCATGTTGTCTTTTTTCTTGGAATTTTGTTTCTCCTGTGCTAG

>M6-PGSC0003DMG400010325-(rev)M6LRR-RLP_128

CACTACAAACTCATTTACTCCTCATGTTACTGAGATTAGTCTTGGTTATGTTTCAGATTATGCTCCTAATCCACCTTGTACTCCTAATAGCACACTTAATGCTTCCCTTTTTGCCCCTTT

>M6-PGSC0003DMG400010325-(rev)M6LRR-RLP_192

CCTAATCCACCTTGTACTCCTAATAGCACACTTAATGCTTCCCTTTTTGCCCCTTTTTCTTATCTAAGAAAGATTTTTGTTTACAAATGTTTTACTGAAAATGAGGTTTCTTTGATTGAC

>M6-PGSC0003DMG400010325-(rev)M6LRR-RLP_256

TAAGAAAGATTTTTGTTTACAAATGTTTTACTGAAAATGAGGTTTCTTTGATTGACTTTGGATCTTTGCCTTCTTCTTTGGAGGAGGTGGTTTTCTTGGAGAATCCAGGTCTTGTTGGTT

>M6-PGSC0003DMG400010325-(rev)M6LRR-RLP_320

TTTGCCTTCTTCTTTGGAGGAGGTGGTTTTCTTGGAGAATCCAGGTCTTGTTGGTTCTTTAAGTGATAAAGTTGGAAACTTGATGAGTTTGAGGAGGTTGGTTTTGAGTGGGAGTGGTGT

>M6-PGSC0003DMG400010325-(rev)M6LRR-RLP_384

GATAAAGTTGGAAACTTGATGAGTTTGAGGAGGTTGGTTTTGAGTGGGAGTGGTGTTTATGGGGAAATCCCAAATGGGGTTGGTGGTTTGGTCAATTTAGAGCAGCTGACTTTATCAAGA

>M6-PGSC0003DMG400010325-(rev)M6LRR-RLP_448

AAATCCCAAATGGGGTTGGTGGTTTGGTCAATTTAGAGCAGCTGACTTTATCAAGAAACAAAATTAGTGGTGAAATTCCAGTGGGGTTGATGACATTGAAGAAACTCAAGATTCTTGATT

>M6-PGSC0003DMG400010325-(rev)M6LRR-RLP_512

TAGTGGTGAAATTCCAGTGGGGTTGATGACATTGAAGAAACTCAAGATTCTTGATTTGAGTCAAAATGAGCTTGAAAGTAGTGTTCTTGAATCTATAGGGAACTTTACTCAACTTTTGAA

>M6-PGSC0003DMG400010325-(rev)M6LRR-RLP_576

AATGAGCTTGAAAGTAGTGTTCTTGAATCTATAGGGAACTTTACTCAACTTTTGAAACTTGACTTGAGTTACAACAAGCTTACTGGGAAAATCCCTCAAAGTTTGAAGGGTTTGAAGAGT

>M6-PGSC0003DMG400010325-(rev)M6LRR-RLP_64

CCCTTTTCCCTGATGATCTCTGTTCTTCTTCCCCACATGGGGTTATTTGTGATTACTTTGTTTCCACTACAAACTCATTTACTCCTCATGTTACTGAGATTAGTCTTGGTTATGTTTCAG

>M6-PGSC0003DMG400010325-(rev)M6LRR-RLP_640

TGAGTTACAACAAGCTTACTGGGAAAATCCCTCAAAGTTTGAAGGGTTTGAAGAGTTTGGAGTTTTTGGACTTGAGTTACAATAACTTTTCCAATTCTGGGGTTCCATTGTTTTTGGCTG

>M6-PGSC0003DMG400010325-(rev)M6LRR-RLP_704

TTTGGACTTGAGTTACAATAACTTTTCCAATTCTGGGGTTCCATTGTTTTTGGCTGAGATGTCTAGTTTGAAGGAAGTCTACTTAAGTGGAAACTATCTTGGAGGGGAAATTCCAGAAAT

>M6-PGSC0003DMG400010325-(rev)M6LRR-RLP_768

AGTTTGAAGGAAGTCTACTTAAGTGGAAACTATCTTGGAGGGGAAATTCCAGAAATATGGGAGAATCTTGAGGGTATTGTGGGGCTAGGATTATCAAAAACAGGACTAGTTGGGAACATT

>M6-PGSC0003DMG400010325-(rev)M6LRR-RLP_832

ATCTTGAGGGTATTGTGGGGCTAGGATTATCAAAAACAGGACTAGTTGGGAACATTCCTGTTTCTATGGGGGTGTACTTGAAAAACATATGTTATTTGGGGCTTGATAATAACAAGCTGG

>M6-PGSC0003DMG400010325-(rev)M6LRR-RLP_896

TATGGGGGTGTACTTGAAAAACATATGTTATTTGGGGCTTGATAATAACAAGCTGGAAGGGGCATTGCCTGAAGAGTTTGGGGCATTGGAATATGTGAATGAGTTACATTTGGAGAACAA

>M6-PGSC0003DMG400010325-(rev)M6LRR-RLP_960

TTGCCTGAAGAGTTTGGGGCATTGGAATATGTGAATGAGTTACATTTGGAGAACAATAATTTGAGTGGTAGACTTCCATTCTCTCCAAAATTTGTATCCAAAGTTGGAGAAAAGTTGAAG

>M6-PGSC0003DMG400010328-(rev)M6LRR-RLP_0

TACAAACATTATCTTATGTAGAACCCATGGAGAAAAGACTGCGACTTTCCAACAAATTTGCTTTAAAAAAGGTGTTTTTCTCAACCCCATATCTCTTTTCTTCAGTCTCTTGTCATTCTT

>M6-PGSC0003DMG400010328-(rev)M6LRR-RLP_1037

TAGATGCTTGTATATCATTGGTTAGTGAACAAAGTTTGTGTCTATCTACTCTTGTTTTACTAGCCTAGCACTAACTGGTGCCTTCTCTGTAGTTTGATTATCCGCACGAAGATTTGGAGC

>M6-PGSC0003DMG400010328-(rev)M6LRR-RLP_1098

AGCCTAGCACTAACTGGTGCCTTCTCTGTAGTTTGATTATCCGCACGAAGATTTGGAGCATTACAATTTTCCTGAACTTACGAAGCTCAAGAAATTTGTTTTTATGACATTTGGAAATTC

>M6-PGSC0003DMG400010328-(rev)M6LRR-RLP_1159

TACAATTTTCCTGAACTTACGAAGCTCAAGAAATTTGTTTTTATGACATTTGGAAATTCACTAAGAGTGTCAGGCCAGAGTCTCTTAGGTTGCACGTCTATAATAAGGGCTGCCCCACAG

>M6-PGSC0003DMG400010328-(rev)M6LRR-RLP_122

GGTTTGATCAATTTTTTTGTTTCTAGTGTCAAAAGGAAGTCTGCATTGCACAAGAAGATCGTGTCAGTCAGTTACCTGATGAAATTCTGGTATATATATTGTCTTTCTTTACTGTTAAGG

>M6-PGSC0003DMG400010328-(rev)M6LRR-RLP_1281

GAAGGAATTTGAACTACAGGTAAGTATCATGCCTTTTCCTCTTTGGTTTACAAAAGTCTGATTTATTCATATTAGCAATTCTGGTTCATGCAGTTATTGATGCTACTTCTTTAATAAACT

>M6-PGSC0003DMG400010328-(rev)M6LRR-RLP_1342

TTTATTCATATTAGCAATTCTGGTTCATGCAGTTATTGATGCTACTTCTTTAATAAACTTAGTTATTTAAGTCGTTGAGCTTGAGTTGAAACAATTGTGAAGAGCTAAACTTGAATAAAT

>M6-PGSC0003DMG400010328-(rev)M6LRR-RLP_1464

TACTCTTGCATTTTAACTCATCTGACTGTGCCAATATAAAAGTCCTGCCCTTACATAGGTGTCATTTTAGATAGGTAGTAGATAGCTCTGTCGTCTCTGAGCTTTGATTTATTATCTGTA

>M6-PGSC0003DMG400010328-(rev)M6LRR-RLP_1525

TCATTTTAGATAGGTAGTAGATAGCTCTGTCGTCTCTGAGCTTTGATTTATTATCTGTAACAAAGAGCTTAAAGTCTCATCATTTCATCAAAGAGCTTAGAGTCTCATCATTCCTTCAGT

>M6-PGSC0003DMG400010328-(rev)M6LRR-RLP_1586

AAAGAGCTTAAAGTCTCATCATTTCATCAAAGAGCTTAGAGTCTCATCATTCCTTCAGTTTCTTGAATATTTTTACTGAATTACTGCACTTGTTTAGCGTAATGGAATGACCTCAGAGTG

>M6-PGSC0003DMG400010328-(rev)M6LRR-RLP_1708

GGGATTGTGGTTTCAGTTGTCGATTGTGTTGCTCCAAAATCTTAGGTTTCCACCTGTTTGTTGCCAGTATATTATATGATGGAATTGGGAAGTACTATACAGTGATTATTATGTGTTTAA

>M6-PGSC0003DMG400010328-(rev)M6LRR-RLP_183

TGTCAGTCAGTTACCTGATGAAATTCTGGTATATATATTGTCTTTCTTTACTGTTAAGGAAGCTGCAGATACAAGTCTCCTCTCAAGGCGTTGGCTAAACTTATGGACATATATTGATCA

>M6-PGSC0003DMG400010328-(rev)M6LRR-RLP_244

GCTGCAGATACAAGTCTCCTCTCAAGGCGTTGGCTAAACTTATGGACATATATTGATCACCTTGATTTTGATGCTACAAAATACTTGGATCAAGTGTTCGTGCAACCCAAGCTACAAAAA

>M6-PGSC0003DMG400010328-(rev)M6LRR-RLP_2745

GGTTTCAAATGAGAACAACCAACTAAGAAAATTGAGAAGTAAGTTGTTGCCTAGCTAGGTAGATGGGTGAAGGAGGAATGCACTGGGGTAGAGGTTTCAGAGATGAGGGATTACAAAATG

>M6-PGSC0003DMG400010328-(rev)M6LRR-RLP_2806

GATGGGTGAAGGAGGAATGCACTGGGGTAGAGGTTTCAGAGATGAGGGATTACAAAATGTTGTTAGAACCGAGGGGAACTTCCAAATTCTATAGTAGGATTCGAAGGTGGCATCCCGAAT

>M6-PGSC0003DMG400010328-(rev)M6LRR-RLP_2867

GTTAGAACCGAGGGGAACTTCCAAATTCTATAGTAGGATTCGAAGGTGGCATCCCGAATCCCAATCAAATGACTCCAGGCTCAAAGAGTCAGAATCGAGCCTGGATGGTATTGTAGAGGA

>M6-PGSC0003DMG400010328-(rev)M6LRR-RLP_2989

CTTAGGGCTTCACATGGTATCTGATGTTTTCTCTCTCTTGAATATTTATTTTTTCAAAAATCAAGTAACGATTATATAAGGGAAAAGAGAAAAAATACCCGAACTTGTATCGGATTTCTT

>M6-PGSC0003DMG400010328-(rev)M6LRR-RLP_3050

CAAGTAACGATTATATAAGGGAAAAGAGAAAAAATACCCGAACTTGTATCGGATTTCTTGTTACACTTATTATTTGAAGTTGTTCTATGAATTATTTTGTAGTAGAATAATACCATAGGA

>M6-PGSC0003DMG400010328-(rev)M6LRR-RLP_3111

TACACTTATTATTTGAAGTTGTTCTATGAATTATTTTGTAGTAGAATAATACCATAGGAAGTGTTGATGTGGCTTTTACTTTGCACCTCCATATAGTGCATAAAAAGACAAATACAAGCT

>M6-PGSC0003DMG400010328-(rev)M6LRR-RLP_3172

TGTTGATGTGGCTTTTACTTTGCACCTCCATATAGTGCATAAAAAGACAAATACAAGCTAAATGATAGAGGCTCTAGAGTCCAAAAGATATAATTTTTCTTCTACTTTTTAGGCCCTTTT

>M6-PGSC0003DMG400010328-(rev)M6LRR-RLP_3233

ATGATAGAGGCTCTAGAGTCCAAAAGATATAATTTTTCTTCTACTTTTTAGGCCCTTTTCTAAATCTTGAGCGGTGATCCACCACTTGTCCCACCAGGCCACCATCCATGCCCCCCACCC

>M6-PGSC0003DMG400010328-(rev)M6LRR-RLP_3355

ATGTCTGAGCCAGGATTTTTTTTTTTTGTAGCTGAGTAGATACACAAATTATGTTTTTCCTACTCATGAGTGGGTGCACGCACTTGGCAAAAGTTGTGTGCATAATATTTTCAATAAAAA

>M6-PGSC0003DMG400010328-(rev)M6LRR-RLP_3416

ACTCATGAGTGGGTGCACGCACTTGGCAAAAGTTGTGTGCATAATATTTTCAATAAAAATCAAGCAATAAAAGTAAAAAAATTTATCTAGGGTGTGTTTTGTATGGAGAAAAATGTTTAT

>M6-PGSC0003DMG400010328-(rev)M6LRR-RLP_3477

AAGCAATAAAAGTAAAAAAATTTATCTAGGGTGTGTTTTGTATGGAGAAAAATGTTTATCTATTTTCATGATTGGTTAGTCAATATATTTTGTGAAATATTTTTTCTAGGAAAACAAGCT

>M6-PGSC0003DMG400010328-(rev)M6LRR-RLP_3599

TTAAAAATGAGGAAATTTACTTTCCTAATGGAAGTAGAAAACATAAGTTCCAAAGCAATTAAATCATATTAGCATTTATAACTATAAAAAACAATCATATAAATCACTATAGACATAGAG

>M6-PGSC0003DMG400010328-(rev)M6LRR-RLP_366

GCATATGGACAAGTATCTGACATGGGTCAACTCTGCTTTGCAAAAGTTTAAAGGCCAGAGACTTGACCAATTTCGTGTTTACTTTGATTTAATCAAATTTGCTCAGCACGACATTGACAA

>M6-PGSC0003DMG400010328-(rev)M6LRR-RLP_3660

AATCATATTAGCATTTATAACTATAAAAAACAATCATATAAATCACTATAGACATAGAGAAGTTAATAAAAATAATTTTAGTTACTAGTTAAATAAAATAAAAAACAATACATATCACAA

>M6-PGSC0003DMG400010328-(rev)M6LRR-RLP_3721

GTTAATAAAAATAATTTTAGTTACTAGTTAAATAAAATAAAAAACAATACATATCACAATAAAAGTAATAAAATAATAATTAAAATAAAAGAATTACCGCGAAACATTAGAAAAAAAACT

>M6-PGSC0003DMG400010328-(rev)M6LRR-RLP_3782

AAAGTAATAAAATAATAATTAAAATAAAAGAATTACCGCGAAACATTAGAAAAAAAACTATTGATAATCTCAACCACAAAAAAACTTTAAAGGAAAAACACGAGTACGGAGCCAAAAGGG

>M6-PGSC0003DMG400010328-(rev)M6LRR-RLP_3843

TGATAATCTCAACCACAAAAAAACTTTAAAGGAAAAACACGAGTACGGAGCCAAAAGGGCAAAAAAATTGACCAAAAATTCCAAAAGGGTTTGAAATCAAAAGGGTAACATTTAAACGGT

>M6-PGSC0003DMG400010328-(rev)M6LRR-RLP_3904

AAAAAATTGACCAAAAATTCCAAAAGGGTTTGAAATCAAAAGGGTAACATTTAAACGGTAGGGGAGATGCTGTTTAAATTAGGGTAGCAAGGCGCAACTCCCTATGCGCCTTACAAGGTG

>M6-PGSC0003DMG400010328-(rev)M6LRR-RLP_4026

AGAAAACAGAGCCACCATCATTGTTTATTTTTTCCATGTCAGTACTTACGGTGGTACTTTTTGCACTACAAAATAGTATAGGGGTCGTAGGGACATCACAAAGAATAAGTGTGTAACTGG

>M6-PGSC0003DMG400010328-(rev)M6LRR-RLP_4087

TGCACTACAAAATAGTATAGGGGTCGTAGGGACATCACAAAGAATAAGTGTGTAACTGGAAAACTGAGATAAGTCCATAGGGTTTTTATGCCTTTTCCCACTGTGTTATTTAAAAAAGCA

>M6-PGSC0003DMG400010328-(rev)M6LRR-RLP_4148

AACTGAGATAAGTCCATAGGGTTTTTATGCCTTTTCCCACTGTGTTATTTAAAAAAGCACCAAGAGGGCTCCAACCTTTACAAGTAGAAGATTCTGTTGTTCTGTCAACCTCCCAGTCAT

>M6-PGSC0003DMG400010328-(rev)M6LRR-RLP_427

CTTGACCAATTTCGTGTTTACTTTGATTTAATCAAATTTGCTCAGCACGACATTGACAAATGGCTTGAACTTGCCTTTTCTAAGAAAGTTCAAAGACTAGAGTTAGACTTATTAAGGTGT

>M6-PGSC0003DMG400010328-(rev)M6LRR-RLP_4270

AATCTCCTTCTTAATCTAATATTAAAAGATAGCCTTTCTCTGTATTCTTCAATCACATCTTTTATATGGATAGCTATGAGATCTAATTCTGAATAAGTTATTAAAGCTTCTTGATCCGGC

>M6-PGSC0003DMG400010328-(rev)M6LRR-RLP_4331

TTATATGGATAGCTATGAGATCTAATTCTGAATAAGTTATTAAAGCTTCTTGATCCGGCGTACATGAAACTTTATCTTCTTCTTGCAACTAAAAGCTTTCACAAGATTAAAAGAATCAAT

>M6-PGSC0003DMG400010328-(rev)M6LRR-RLP_4392

ACATGAAACTTTATCTTCTTCTTGCAACTAAAAGCTTTCACAAGATTAAAAGAATCAATTCCATAAGTTCTAATGTACTTTCAAACTGTGACATCATCAGATGTATTGATTAGTTTTGCT

>M6-PGSC0003DMG400010328-(rev)M6LRR-RLP_4453

CATAAGTTCTAATGTACTTTCAAACTGTGACATCATCAGATGTATTGATTAGTTTTGCTTTTGAACATCTCATATCGTTGCCATGCTCTATGTTTAACAGCTTAATCCATAAATTGTTTT

>M6-PGSC0003DMG400010328-(rev)M6LRR-RLP_4514

TGAACATCTCATATCGTTGCCATGCTCTATGTTTAACAGCTTAATCCATAAATTGTTTTGCTCTACTGTGTTTTATCTCCACAAACTAATTCGACAATAGACACTTTTTTTAATACCCAA

>M6-PGSC0003DMG400010328-(rev)M6LRR-RLP_4575

TCTACTGTGTTTTATCTCCACAAACTAATTCGACAATAGACACTTTTTTTAATACCCAAAGTTTTTATCTATGAACTACTCTTCTTAGAGAATATAGCTCTATTTTTATGCTCTTTGAAT

>M6-PGSC0003DMG400010328-(rev)M6LRR-RLP_4636

TTTTTATCTATGAACTACTCTTCTTAGAGAATATAGCTCTATTTTTATGCTCTTTGAATCAACTTGCCACATCTTTCTCCTTGCATTATCGAACGGTCTTTGCACCTTAAAAAATGATAG

>M6-PGSC0003DMG400010328-(rev)M6LRR-RLP_4697

ACTTGCCACATCTTTCTCCTTGCATTATCGAACGGTCTTTGCACCTTAAAAAATGATAGATACAGTTTCTTGCAGGTTGCCAATCTTCTCAACCTTATGTATCACTCTCTGCCATATTAT

>M6-PGSC0003DMG400010328-(rev)M6LRR-RLP_4819

CACTTCTATATCTGGCATTTAGAGACTTGCTCATGTATGTCTCTGCCTGGATTCTCAATTTAAATGCAGTCAAGTGCTGTCTCAAAGTGTAATAGTAGAGATCTGTGATGCATGATCTTC

>M6-PGSC0003DMG400010328-(rev)M6LRR-RLP_488

GGCTTGAACTTGCCTTTTCTAAGAAAGTTCAAAGACTAGAGTTAGACTTATTAAGGTGTGGATCATTCACTCGTAATTTTGATTCCTGTTATACTTTTCCGGAGCGGCTCCTTGGTCTTG

>M6-PGSC0003DMG400010328-(rev)M6LRR-RLP_4880

AAATGCAGTCAAGTGCTGTCTCAAAGTGTAATAGTAGAGATCTGTGATGCATGATCTTCCGCTCCTCAAGTCCAAATAGAAATAGGGTATAGTCTGCATATCGTGTACATGAAAAATTTC

>M6-PGSC0003DMG400010328-(rev)M6LRR-RLP_4941

CTCCTCAAGTCCAAATAGAAATAGGGTATAGTCTGCATATCGTGTACATGAAAAATTTCCATATTTTCATTTGTACAACTTTCCCCAACTTTTGACTCCTTTTATCCATACTATATTCAT

>M6-PGSC0003DMG400010328-(rev)M6LRR-RLP_5002

TATTTTCATTTGTACAACTTTCCCCAACTTTTGACTCCTTTTATCCATACTATATTCATGGCATAATGTAGCATCTTACTGAATGTTTCCATTGCTTAGGAAGAGAATGGGATAATGGGG

>M6-PGSC0003DMG400010328-(rev)M6LRR-RLP_5063

CATAATGTAGCATCTTACTGAATGTTTCCATTGCTTAGGAAGAGAATGGGATAATGGGGTTCTCCTATTTAACTGCATATCCTATTTATATTTTTCCATTTGTTTGCAAACCGTGTTGAT

>M6-PGSC0003DMG400010328-(rev)M6LRR-RLP_5124

CTCCTATTTAACTGCATATCCTATTTATATTTTTCCATTTGTTTGCAAACCGTGTTGATTCAAAAGCTTCAGTAATATTGCCAGTTTACATGATTGTTCCACTGGGTCAAATCTTTGCTT

>M6-PGSC0003DMG400010328-(rev)M6LRR-RLP_5185

AAAAGCTTCAGTAATATTGCCAGTTTACATGATTGTTCCACTGGGTCAAATCTTTGCTTGATCCTGTGTTGTCCCTAGTCTGAGGTGATAACTTGAGAATGAATCATTCATCGGTGCTCT

>M6-PGSC0003DMG400010328-(rev)M6LRR-RLP_5307

TAGTAAAGAAGGCTTCACTCAAGGAGTGACAGATTTTGACGGAGTCTTGGCTTTGATGCTAAGGAGACAAGATCCATCGATCTGAATAAGTCTAGTAACCTTTCTCAAGTCCAGCTAACA

>M6-PGSC0003DMG400010328-(rev)M6LRR-RLP_5368

AGGAGACAAGATCCATCGATCTGAATAAGTCTAGTAACCTTTCTCAAGTCCAGCTAACAAATAATTCTTCCTATTTTAATTCTATTTAAGGCTCCAAGAATTCATTTTCTAAGAACACTG

>M6-PGSC0003DMG400010328-(rev)M6LRR-RLP_5429

TAATTCTTCCTATTTTAATTCTATTTAAGGCTCCAAGAATTCATTTTCTAAGAACACTGGATCTGTAGTTTATCCTTCCCTTTACAAATACATGCATTTTTTGACCTTGATATCTACTTG

>M6-PGSC0003DMG400010328-(rev)M6LRR-RLP_549

ATCATTCACTCGTAATTTTGATTCCTGTTATACTTTTCCGGAGCGGCTCCTTGGTCTTGGAGATTCTTCTTCATGCTCTAATAACCTCCAAATACTCTCACTTATCAACCAGAATTTCAA

>M6-PGSC0003DMG400010328-(rev)M6LRR-RLP_5490

TCTGTAGTTTATCCTTCCCTTTACAAATACATGCATTTTTTGACCTTGATATCTACTTGCCAATACTTGTTTCATCCTTGTTGTTAGGACCCCGGGAGTAATCTCATCCTTTTACCTTGT

>M6-PGSC0003DMG400010328-(rev)M6LRR-RLP_5551

AATACTTGTTTCATCCTTGTTGTTAGGACCCCGGGAGTAATCTCATCCTTTTACCTTGTTCTTAAGAACCAATGCTGTGAATATTTCAGTCCTGCCTCTTGTAAAAATTTCTGTCTCATG

>M6-PGSC0003DMG400010328-(rev)M6LRR-RLP_5612

TTAAGAACCAATGCTGTGAATATTTCAGTCCTGCCTCTTGTAAAAATTTCTGTCTCATGAAAGAAGTGCAATGCACTGATAATGTCCCTTTCCAAAAAGTTCCTAGATTTCTGGTAAGAA

>M6-PGSC0003DMG400010328-(rev)M6LRR-RLP_5734

CATTATCTGCCCATCTGGCCCTAAAGTTGTTATCACCACTAGACCATCTCCATTACTTCTCCGAGAAAATTTCTTAGTAGCCAACACTCTTCCTGTGATAATAGCTCAAGGTCATTGTAA

>M6-PGSC0003DMG400010328-(rev)M6LRR-RLP_5795

CGAGAAAATTTCTTAGTAGCCAACACTCTTCCTGTGATAATAGCTCAAGGTCATTGTAACTGGAAGGCGCCATCATTGCTCGGTTTTACTAAATAACTTCTCATAGCATTACACTCATCT

>M6-PGSC0003DMG400010328-(rev)M6LRR-RLP_5856

GGAAGGCGCCATCATTGCTCGGTTTTACTAAATAACTTCTCATAGCATTACACTCATCTGATGACCTTTGGATTTGGTAGTTCGTCCATCTCTTACAAGTTAGTTTATATGGCTGTACTT

>M6-PGSC0003DMG400010328-(rev)M6LRR-RLP_5917

TGACCTTTGGATTTGGTAGTTCGTCCATCTCTTACAAGTTAGTTTATATGGCTGTACTTTTTATTTGTATTGATCAATTTACAGTAAAACACTTAGTATTTTGTAGGGGTGTCAAAAATG

>M6-PGSC0003DMG400010328-(rev)M6LRR-RLP_61

TTTAAAAAAGGTGTTTTTCTCAACCCCATATCTCTTTTCTTCAGTCTCTTGTCATTCTTGTGGTTTGATCAATTTTTTTGTTTCTAGTGTCAAAAGGAAGTCTGCATTGCACAAGAAGAT

>M6-PGSC0003DMG400010328-(rev)M6LRR-RLP_610

GATTCTTCTTCATGCTCTAATAACCTCCAAATACTCTCACTTATCAACCAGAATTTCAAGTCTCTAAAAGTGCTTCTATTGAAATCTGTAAATGTGACGGGCCAAGTTCTTGAGTTCTTT

>M6-PGSC0003DMG400010328-(rev)M6LRR-RLP_6466

CAACCCGCCCATTTGGCACCTCTAGTATGTTGTCTCATTCATTTAACCAGAAACACCTCAATTGTTGTTTCCAGGGTGTTTTTTAGCATTGGTTATTTGTTCCAGTTCTACTAAGTGGTT

>M6-PGSC0003DMG400010328-(rev)M6LRR-RLP_6527

TTGTTGTTTCCAGGGTGTTTTTTAGCATTGGTTATTTGTTCCAGTTCTACTAAGTGGTTCCTTTAGGCTTTTCGTTTTTCTATATCCGTTCACCTTTTTCTGATCATACTTACCAATCCA

>M6-PGSC0003DMG400010328-(rev)M6LRR-RLP_6588

TTTAGGCTTTTCGTTTTTCTATATCCGTTCACCTTTTTCTGATCATACTTACCAATCCAGCAATCTGATTATGTATAATTTTTCGACATGCATTGTTGTTTGAACTCAAAAGTCTGCCTT

>M6-PGSC0003DMG400010328-(rev)M6LRR-RLP_6649

AATCTGATTATGTATAATTTTTCGACATGCATTGTTGTTTGAACTCAAAAGTCTGCCTTTTAAGTTCAGTTTCCTTTCTATTAACCATAATCAGATTGCCAACACTATCATAAGATTGCT

>M6-PGSC0003DMG400010328-(rev)M6LRR-RLP_671

CTCTAAAAGTGCTTCTATTGAAATCTGTAAATGTGACGGGCCAAGTTCTTGAGTTCTTTCTACGCAATTGTCCATTTCTTGAAAGAATGCAAGTTCATGGATCAGGAACATTGAAAAATT

>M6-PGSC0003DMG400010328-(rev)M6LRR-RLP_6710

AAGTTCAGTTTCCTTTCTATTAACCATAATCAGATTGCCAACACTATCATAAGATTGCTATTGCTGTTCCGCTAAATCAATGAATCCCTCTATAGCTTTTTTCATTTTCTCGAATTTGCA

>M6-PGSC0003DMG400010328-(rev)M6LRR-RLP_6832

AAGATTCGCCTATTCCCAAGTCTGCACTCTGATGGAATGGGTGTATGATGTTGGTAATCATCATATGCTTTGAAAACTTGAATATCACATTCCAAAGAAATTGTACACAAAAATTTCTGT

>M6-PGSC0003DMG400010328-(rev)M6LRR-RLP_6893

CATATGCTTTGAAAACTTGAATATCACATTCCAAAGAAATTGTACACAAAAATTTCTGTGTATTCCACCTCATAAGTTTACTGTACATGCTATTGTATTTATACAAAACTAGGTCTAAAT

>M6-PGSC0003DMG400010328-(rev)M6LRR-RLP_6954

ATTCCACCTCATAAGTTTACTGTACATGCTATTGTATTTATACAAAACTAGGTCTAAATTCTGTACCAAAATAAATGAATTAAACAATGATTATTTTTACACTTGGCACCTAATTATTGG

>M6-PGSC0003DMG400010328-(rev)M6LRR-RLP_7015

TGTACCAAAATAAATGAATTAAACAATGATTATTTTTACACTTGGCACCTAATTATTGGACTATTTTGTAACAAACTCTAACAGCCATTTTGCTCTTCACGTGCCTTCCTTGTGCCTAGG

>M6-PGSC0003DMG400010328-(rev)M6LRR-RLP_7137

GAGAATCTTCTAGACCTTGTACAACTGGTTAGGTTGAGAATCTTCTAGACCTTCTGATTAGGTTTCCTAGAACTTTTGTTCTTTTTGGCATATGTTATTTTGGTTCTTGAAAAGAAATGT

>M6-PGSC0003DMG400010328-(rev)M6LRR-RLP_7198

GTTTCCTAGAACTTTTGTTCTTTTTGGCATATGTTATTTTGGTTCTTGAAAAGAAATGTTGGAGAAATTCATTTCGGTTAGCAAGTTGGGCATGCAGCTCGAATATGTTTATATGATAAG

>M6-PGSC0003DMG400010328-(rev)M6LRR-RLP_7259

GAGAAATTCATTTCGGTTAGCAAGTTGGGCATGCAGCTCGAATATGTTTATATGATAAGTCACTTGTGTATTCTTTTGTTTCCTAATTTCTCCAAGTTAAGATTTTTCTCACAGAGAGTC

>M6-PGSC0003DMG400010328-(rev)M6LRR-RLP_7320

ACTTGTGTATTCTTTTGTTTCCTAATTTCTCCAAGTTAAGATTTTTCTCACAGAGAGTCTAGAACATGGATAATGAGTAGTGGTGATTGAATATTGTTGAATTAGTCAACTTGACTAGGA

>M6-PGSC0003DMG400010328-(rev)M6LRR-RLP_7381

GAACATGGATAATGAGTAGTGGTGATTGAATATTGTTGAATTAGTCAACTTGACTAGGAGGCATCTTACATGATAATGGGAAGTTTTCATTCAGTCGATCAAGGAACCCATATATCCCGA

>M6-PGSC0003DMG400010328-(rev)M6LRR-RLP_7442

CATCTTACATGATAATGGGAAGTTTTCATTCAGTCGATCAAGGAACCCATATATCCCGAATCTTAAAATTTAGATCCACAACCGAGTCCGAGCAACTTAGTCTAAAATAAGTCGACCTAA

>M6-PGSC0003DMG400010328-(rev)M6LRR-RLP_7503

CTTAAAATTTAGATCCACAACCGAGTCCGAGCAACTTAGTCTAAAATAAGTCGACCTAAACATGTTGAAAGAAGTTGCAATTCTCTCAGAAAATTGAGCACTCCACGTCTGTAATATTCT

>M6-PGSC0003DMG400010328-(rev)M6LRR-RLP_7564

ATGTTGAAAGAAGTTGCAATTCTCTCAGAAAATTGAGCACTCCACGTCTGTAATATTCTTGTCCATATTCCCCACTTTCCTTCCTCCATGTGGTAATACTTTAGAATTTGTAGAAGAACA

>M6-PGSC0003DMG400010328-(rev)M6LRR-RLP_7625

TCCATATTCCCCACTTTCCTTCCTCCATGTGGTAATACTTTAGAATTTGTAGAAGAACAAAGTAGAGTAGGAGAAGTCAAATGACAAGAGCGATTGATACAATCATGGTTCATCAGTGAT

>M6-PGSC0003DMG400010328-(rev)M6LRR-RLP_7686

GTAGAGTAGGAGAAGTCAAATGACAAGAGCGATTGATACAATCATGGTTCATCAGTGATTGGTACATAGTTTTATGTTATACTGATTCCCCAAAGTTTAATTCTTTTCATTTAACTTGTG

>M6-PGSC0003DMG400010328-(rev)M6LRR-RLP_7747

GTACATAGTTTTATGTTATACTGATTCCCCAAAGTTTAATTCTTTTCATTTAACTTGTGTCCCAAGAAAAAGGCCTGTTCAAAATTTTCTTTTATTAGCTTTACTGACTTATCTTATCAT

>M6-PGSC0003DMG400010328-(rev)M6LRR-RLP_793

GAAGTTGTTGGTCCATCCCTCAAGTTGAGACACTTGGAGATATATAGCTGCTTCAATGTAAAATCACTTAAAATTTGTGATACAAATCTTATATCATTAAGGACTTCAGCAGGAAAAGTA

>M6-PGSC0003DMG400010328-(rev)M6LRR-RLP_7930

CGAGGCAAAGAAAGAGTGTAAGAAAGCAGCCATAAAATGCCCACTCCATCACCTCAAAGTGGTCAAATTTTTGGGGTTTTATACAGGCTTATTTCAACTTGAACTCGTTAGGTACTTTTT

>M6-PGSC0003DMG400010328-(rev)M6LRR-RLP_7991

GTCAAATTTTTGGGGTTTTATACAGGCTTATTTCAACTTGAACTCGTTAGGTACTTTTTGGAAAATGGTGTTGCTCTTGAGAAAATCATTATTGATCCAGGTACTCCATTTTTTTATCGT

>M6-PGSC0003DMG400010328-(rev)M6LRR-RLP_8052

AAAATGGTGTTGCTCTTGAGAAAATCATTATTGATCCAGGTACTCCATTTTTTTATCGTATCCCTCCGTGCCCTAGCGAAATCAAACACACGCAAGCTTCGAGAAATCTTGCTAAGCTCC

>M6-PGSC0003DMG400010328-(rev)M6LRR-RLP_8174

CTTGAATGTGAAGTGCCTCCGGCTATTGAGTTGGTCATACTTTAATGAAAATAATAATGTTATTACGGTTAAATTTGTGATACATAACGGAGGTGAATTTTTATTTCTTAGTAGATGTTC

>M6-PGSC0003DMG400010328-(rev)M6LRR-RLP_8235

ATTACGGTTAAATTTGTGATACATAACGGAGGTGAATTTTTATTTCTTAGTAGATGTTCCTCATGCTTTGTTTCATTATTGAGTGATATTTTCACTTTTTGTGAGGTTTATATTGCTATA

>M6-PGSC0003DMG400010328-(rev)M6LRR-RLP_8283

AGTAGATGTTCCTCATGCTTTGTTTCATTATTGAGTGATATTTTCACTTTTTGTGAGGTTTATATTGCTATATGAAATATGGGGGATTTTCAAAATGAAAATGTCGCAGTTATTCCGCCA

>M6-PGSC0003DMG400010328-(rev)M6LRR-RLP_854

AATCACTTAAAATTTGTGATACAAATCTTATATCATTAAGGACTTCAGCAGGAAAAGTATTACTGCTTAAGAATGTCCCGATGCTGGTTGATGTAGAGATTTTGGGTATGCCATACACCC

>M6-PGSC0003DMG400010328-(rev)M6LRR-RLP_915

ACTGCTTAAGAATGTCCCGATGCTGGTTGATGTAGAGATTTTGGGTATGCCATACACCCATTTATTGGATGACATATCCTCTCGTCTTTTGTGCGTTCTCTCTCAGCTGGAGGTCCTTAC

>M6-PGSC0003DMG400010348-(rev)M6LRR-RLK_0

CTCTGTCTTCATCTCTTCGTCATTAAATGCAACGTCGCCACAACTAACCAGAAGTGATATGACATTTAGGTTGACATTTTTGGCAAACATCTCATAAAGGCAATATTAGCACTGAGCCAC

>M6-PGSC0003DMG400010348-(rev)M6LRR-RLK_1037

TTCATCCACCTTTACCCTTCTTTCAGTAATAACTTATGCTCTTCTGTGAATATTTCTCTAGGGGTTTTCCCATCTTTGTTTTGTTAGTGCAAAGTTTTAATGATAAACAACTTATTCTTG

>M6-PGSC0003DMG400010348-(rev)M6LRR-RLK_1098

GGGTTTTCCCATCTTTGTTTTGTTAGTGCAAAGTTTTAATGATAAACAACTTATTCTTGCATCAAAAAGGAAAGAATCAAGAATGTGTACATCCTTGACTCTGTAAAAAAACAATGGAGG

>M6-PGSC0003DMG400010348-(rev)M6LRR-RLK_122

ATGAAGCAAGAAATGTGATAACTTCTCGTTGTTGGTTAGGATGTTGCATAATTTGTCACCTCTATATTGATTCTACAGTTACTTCGCAGGGTGAATACTACTGGATATATGGATCAGAAG

>M6-PGSC0003DMG400010348-(rev)M6LRR-RLK_1403

TAAATATGAACAATTAACTCTCTTGAACTTATCACTTAGAAAAGGTAAATGAAAGACAAAAATGGAAGAAATTTTTAACAACTCTCTTGAACATTGAATAATTGAAAAAATGTCAAAAGT

>M6-PGSC0003DMG400010348-(rev)M6LRR-RLK_1464

ATGGAAGAAATTTTTAACAACTCTCTTGAACATTGAATAATTGAAAAAATGTCAAAAGTTCAATTAGTATGGACTAGATGGAGTAAAATATATAGTTTATATTTTATTAGTCCATATATC

>M6-PGSC0003DMG400010348-(rev)M6LRR-RLK_1525

AATTAGTATGGACTAGATGGAGTAAAATATATAGTTTATATTTTATTAGTCCATATATCATAAAACAATTTTTTTTTTGTTTCTAGTCAATAGTAATATGAATTTTACACATCATATTAG

>M6-PGSC0003DMG400010348-(rev)M6LRR-RLK_1586

AAAACAATTTTTTTTTTGTTTCTAGTCAATAGTAATATGAATTTTACACATCATATTAGGCCTCTTATTGTAATTTGAAGTAATTCCATTGACCCATCCCTGTGTAACACACTTGTGGAG

>M6-PGSC0003DMG400010348-(rev)M6LRR-RLK_1647

CTCTTATTGTAATTTGAAGTAATTCCATTGACCCATCCCTGTGTAACACACTTGTGGAGATGTGTAACACATTATGAAGGGTAGAGAGGACTATTGATAGACACAAAAATCACGCCAAAA

>M6-PGSC0003DMG400010348-(rev)M6LRR-RLK_1708

GTGTAACACATTATGAAGGGTAGAGAGGACTATTGATAGACACAAAAATCACGCCAAAACTAATGGCAGTATTCAAAATGCAAGGCCACAAGTAGTGATCAGAAAGACCAAGATTCTTGC

>M6-PGSC0003DMG400010348-(rev)M6LRR-RLK_1769

AATGGCAGTATTCAAAATGCAAGGCCACAAGTAGTGATCAGAAAGACCAAGATTCTTGCACATTTCATTGAAGGCAAAAAACATTTTCCCTGTAGGTCACTAATGCCATTTTCTAGTCTT

>M6-PGSC0003DMG400010348-(rev)M6LRR-RLK_183

CTATATTGATTCTACAGTTACTTCGCAGGGTGAATACTACTGGATATATGGATCAGAAGACAATTGTGTTTTACGGTTTAGTGTCGAGGCGTGTGTCATGATGTCAGATTTTGTCAGTGG

>M6-PGSC0003DMG400010348-(rev)M6LRR-RLK_1830

ATTTCATTGAAGGCAAAAAACATTTTCCCTGTAGGTCACTAATGCCATTTTCTAGTCTTCACTAAAAGAAGTTGTTGAACTCCAAATTGTCACTCTAAACCAAAAAATACTCCAGTCATC

>M6-PGSC0003DMG400010348-(rev)M6LRR-RLK_1891

CTAAAAGAAGTTGTTGAACTCCAAATTGTCACTCTAAACCAAAAAATACTCCAGTCATCAATTCTTGTTTTAGGAAAAAGAGAAACCTGTCAGTTATAGTTATGGGAAGAAGTTGCGATC

>M6-PGSC0003DMG400010348-(rev)M6LRR-RLK_1952

TTCTTGTTTTAGGAAAAAGAGAAACCTGTCAGTTATAGTTATGGGAAGAAGTTGCGATCTTCTCTTTGCTCTTTCGGTTTTCATTCTGCTCCATACTTCACTATCTATTAGTACTGATGA

>M6-PGSC0003DMG400010348-(rev)M6LRR-RLK_2013

CTCTTTGCTCTTTCGGTTTTCATTCTGCTCCATACTTCACTATCTATTAGTACTGATGAAGCTGCTCTTCTTGCCTTCAAATCTCACATTTCTTCTTCTGATCCTAACAACATCTTAGCA

>M6-PGSC0003DMG400010348-(rev)M6LRR-RLK_2074

CTGCTCTTCTTGCCTTCAAATCTCACATTTCTTCTTCTGATCCTAACAACATCTTAGCAACCAACTGGTCTTCTTCCACCCCTGTTTGCACCTGGATTGGAATCACTTGCACCTCCCCAC

>M6-PGSC0003DMG400010348-(rev)M6LRR-RLK_2196

CATCGAGTCGCAGCTTTAGACATTTCTAGTATGCAACTTCATGGTACCATTTCTCCACACCTTGGAAACCTCTCATTTCTTGTTTCCCTCAACATTAGTAACAACTCTTTCCGTGGAGAT

>M6-PGSC0003DMG400010348-(rev)M6LRR-RLK_2257

TTGGAAACCTCTCATTTCTTGTTTCCCTCAACATTAGTAACAACTCTTTCCGTGGAGATTTGCCCAAAGAGTTGGCTCATTTGCAGAGGTTGAAATTGATTGATGTCTCAAGCAATAACT

>M6-PGSC0003DMG400010348-(rev)M6LRR-RLK_2318

GCCCAAAGAGTTGGCTCATTTGCAGAGGTTGAAATTGATTGATGTCTCAAGCAATAACTTCACTGGTGCCATTCCATCATTTTTAAGTTTGCTAGTAGACCTTTGCATTTTGCGCCTGTC

>M6-PGSC0003DMG400010348-(rev)M6LRR-RLK_2379

ACTGGTGCCATTCCATCATTTTTAAGTTTGCTAGTAGACCTTTGCATTTTGCGCCTGTCGAGCAACCAATTTTCAGGAAAAATCCCATCTTCACTTTCCAATCTAACAAAGCTGGAGGTG

>M6-PGSC0003DMG400010348-(rev)M6LRR-RLK_244

AATTGTGTTTTACGGTTTAGTGTCGAGGCGTGTGTCATGATGTCAGATTTTGTCAGTGGCTTACACTTACATACATACGATTTTTTACCCTGCAGAGATACTTGTTTTAGAGCATAAGCT

>M6-PGSC0003DMG400010348-(rev)M6LRR-RLK_2440

GCAACCAATTTTCAGGAAAAATCCCATCTTCACTTTCCAATCTAACAAAGCTGGAGGTGTTGAAAATAAAGGGAAATCTTCTCGAAGGAGAGATCCCTCGAGAACTTGGTGATCTTCATT

>M6-PGSC0003DMG400010348-(rev)M6LRR-RLK_2501

GAAAATAAAGGGAAATCTTCTCGAAGGAGAGATCCCTCGAGAACTTGGTGATCTTCATTACATGACTGCTTTAAACCTGGGAAGTAACCACCTTACTGGCTCTATACCACCATCAATTTA

>M6-PGSC0003DMG400010348-(rev)M6LRR-RLK_2562

ATGACTGCTTTAAACCTGGGAAGTAACCACCTTACTGGCTCTATACCACCATCAATTTATAACATTACAACCATGCGAATCATTGCTCTTACCAACAATAATCTTAGTGGTGAGCTTCCA

>M6-PGSC0003DMG400010348-(rev)M6LRR-RLK_2623

ACATTACAACCATGCGAATCATTGCTCTTACCAACAATAATCTTAGTGGTGAGCTTCCAACAACTATATGTGACCATCTTCCAAACTTGGAAGGGCTTTACCTCTCAGAAAACATCCTAG

>M6-PGSC0003DMG400010348-(rev)M6LRR-RLK_2684

AACTATATGTGACCATCTTCCAAACTTGGAAGGGCTTTACCTCTCAGAAAACATCCTAGGTGGCGTTATTCCACCAAACTTGGAAAAATGCAAAAACCTCAAAATTTTGTCATTGTCTGT

>M6-PGSC0003DMG400010348-(rev)M6LRR-RLK_2745

GGCGTTATTCCACCAAACTTGGAAAAATGCAAAAACCTCAAAATTTTGTCATTGTCTGTAAACGAGTTCACTGGAACCATACCAAGAGAGTTAGGCAACATAACAGATCTTACAGATATA

>M6-PGSC0003DMG400010348-(rev)M6LRR-RLK_2806

ACGAGTTCACTGGAACCATACCAAGAGAGTTAGGCAACATAACAGATCTTACAGATATAGCTCTTGGAATACAGCACTTGGGAGGTAGTATTGCATCTCCTAGCAAGAAATTTTGAATTT

>M6-PGSC0003DMG400010348-(rev)M6LRR-RLK_2867

TCTTGGAATACAGCACTTGGGAGGTAGTATTGCATCTCCTAGCAAGAAATTTTGAATTTTCTTCTTTTTCACTCTTCCACAAATTTGGTACTCATATATATATATATATATATATATTTC

>M6-PGSC0003DMG400010348-(rev)M6LRR-RLK_2928

TTCTTTTTCACTCTTCCACAAATTTGGTACTCATATATATATATATATATATATATTTCCCCACTTGACAGGAAAGATACCAGTGGAGCTAGGTAATCTTAAGAAACTACAGATTCTGTC

>M6-PGSC0003DMG400010348-(rev)M6LRR-RLK_2989

CACTTGACAGGAAAGATACCAGTGGAGCTAGGTAATCTTAAGAAACTACAGATTCTGTCATTAAACGACTGTAAGCTTACTGGTTCCATCCCCGCAAGCATTTTCAACATGTCAGCACTG

>M6-PGSC0003DMG400010348-(rev)M6LRR-RLK_305

TACACTTACATACATACGATTTTTTACCCTGCAGAGATACTTGTTTTAGAGCATAAGCTACTATACAATGAAGTAAATGATGTGCTAAACCTATTAATTATTACCAATCTCAAATAGCAT

>M6-PGSC0003DMG400010348-(rev)M6LRR-RLK_3050

TAAACGACTGTAAGCTTACTGGTTCCATCCCCGCAAGCATTTTCAACATGTCAGCACTGCAGCTCCTAGGAATATCAGAAAACAGGCTTTCAGGTACTCTACCTTCAGATTTAGGTCGTG

>M6-PGSC0003DMG400010348-(rev)M6LRR-RLK_3111

GCTCCTAGGAATATCAGAAAACAGGCTTTCAGGTACTCTACCTTCAGATTTAGGTCGTGGAATGCCCAACCTAGAAGGATTCTCTTGTTTTCAGAATAGTTTGAGTGGTTTTCTCCCTGC

>M6-PGSC0003DMG400010348-(rev)M6LRR-RLK_3172

ATGCCCAACCTAGAAGGATTCTCTTGTTTTCAGAATAGTTTGAGTGGTTTTCTCCCTGCTTCTATCTCAAACTCTTCTAGACTCAGAGTACTTGAACTCTCATATAACAGTTTTACAGGC

>M6-PGSC0003DMG400010348-(rev)M6LRR-RLK_3233

CTATCTCAAACTCTTCTAGACTCAGAGTACTTGAACTCTCATATAACAGTTTTACAGGCCCAATTCCTGAATCAGTTAGTGACTTAGAATACATTGAGATTTTGAACTTGGGGGGGAATA

>M6-PGSC0003DMG400010348-(rev)M6LRR-RLK_3294

AATTCCTGAATCAGTTAGTGACTTAGAATACATTGAGATTTTGAACTTGGGGGGGAATAATTTTTTCAGCAATTCAGCATTGAGCTTCTTGACATCTTTGACAAACTGTAGGAAACTGAA

>M6-PGSC0003DMG400010348-(rev)M6LRR-RLK_3477

TGATTCTCTGCAAATTTTCCAAGGATGGTATTGTAAACTGAAAGGCTTCATTCCTGGAGAAATTGGCAATCTTACCGGTGTGATAAAGATGGATTTGTCACAAAATGAGTTGACTGGATA

>M6-PGSC0003DMG400010348-(rev)M6LRR-RLK_3538

ATTGGCAATCTTACCGGTGTGATAAAGATGGATTTGTCACAAAATGAGTTGACTGGATATATTCCAAAAACCATCCAAGGCTTGTCAAACCTTCAAGAACTTTACTTAGGTGGCAACATG

>M6-PGSC0003DMG400010348-(rev)M6LRR-RLK_3599

TTCCAAAAACCATCCAAGGCTTGTCAAACCTTCAAGAACTTTACTTAGGTGGCAACATGATAAAAGGAACCATACCAGATGTTATGTGCAATTTAAATAAACTTGGAGCATTAGATTTGT

>M6-PGSC0003DMG400010348-(rev)M6LRR-RLK_366

TATACAATGAAGTAAATGATGTGCTAAACCTATTAATTATTACCAATCTCAAATAGCATTGATGAAAAAATTCTCTCAAGTTTCATAAACAAACATTACAATGTTTTTGCACTGTATTTT

>M6-PGSC0003DMG400010348-(rev)M6LRR-RLK_3660

AAAAGGAACCATACCAGATGTTATGTGCAATTTAAATAAACTTGGAGCATTAGATTTGTCAAAAAATCAAGCTTCTGGTTCCATACCACCATGCTTAGGGAACGTTACCAATTTGAGGTA

>M6-PGSC0003DMG400010348-(rev)M6LRR-RLK_3782

TTTATCTAGCTAACAACAGGCTGAACTGGACCTTACCTTCAAGCTTGGGGAGCCTTCAAGATCTCCTAGAATTCAATGTTTCGTCCAATTTATTAACTGGGGAAGTTCCTATAGAGATTG

>M6-PGSC0003DMG400010348-(rev)M6LRR-RLK_3843

TCTCCTAGAATTCAATGTTTCGTCCAATTTATTAACTGGGGAAGTTCCTATAGAGATTGGAAATTTAAAGGTTGCAACACTAGTTGATCTGTCAAAAAATGATTTTAATGGTAAGATCCC

>M6-PGSC0003DMG400010348-(rev)M6LRR-RLK_3904

AATTTAAAGGTTGCAACACTAGTTGATCTGTCAAAAAATGATTTTAATGGTAAGATCCCTAGCACACTAGGGGGTCTGGATAGAATGCTTAGTCTTTCTCTAGCACATAATAAATTATAT

>M6-PGSC0003DMG400010348-(rev)M6LRR-RLK_3965

GCACACTAGGGGGTCTGGATAGAATGCTTAGTCTTTCTCTAGCACATAATAAATTATATGGGCCTATTCCAGATTCATTTGGCAAAATGCTGGCCTTGGAATTCTTGGATTTGAGCAATA

>M6-PGSC0003DMG400010348-(rev)M6LRR-RLK_4026

GCCTATTCCAGATTCATTTGGCAAAATGCTGGCCTTGGAATTCTTGGATTTGAGCAATAACAAACTTAGTGGTGAAATTCCAAAGTCATTAGAAGCTCTCGTGTATCTCAAATACTTGAA

>M6-PGSC0003DMG400010348-(rev)M6LRR-RLK_4148

TCTCATTTAATGAACTCAGTGGAGAGATTCCCACGGGTGGTCCTTTTGCAAATGCCACAAGTCAATCTTTCTTGAAAAATTATGCCCTTTGTGGCGACTCTAAGTTTCATGTTTCACCAT

>M6-PGSC0003DMG400010348-(rev)M6LRR-RLK_4209

TCAATCTTTCTTGAAAAATTATGCCCTTTGTGGCGACTCTAAGTTTCATGTTTCACCATGCATCATCAAATCTCCCAAGAGGTCAAAGAAAAAAAAGGCAATATTGGTTTTGGGAGTGGG

>M6-PGSC0003DMG400010348-(rev)M6LRR-RLK_427

ATGAAAAAATTCTCTCAAGTTTCATAAACAAACATTACAATGTTTTTGCACTGTATTTTACTGACAATAAGAAGATACTCCTATGTGAAACCGTCATCAGCAATGAAGGCTCTGTTGTTA

>M6-PGSC0003DMG400010348-(rev)M6LRR-RLK_4270

ATCATCAAATCTCCCAAGAGGTCAAAGAAAAAAAAGGCAATATTGGTTTTGGGAGTGGGTATGCTTTTTCTTGCATTAGCCCTCACATATGTATTTTTGAGATTGAGAAAGAAAAAAAAG

>M6-PGSC0003DMG400010348-(rev)M6LRR-RLK_4331

TGCTTTTTCTTGCATTAGCCCTCACATATGTATTTTTGAGATTGAGAAAGAAAAAAAAGAATGCAGGTCAATCTGATATGTCTCTGATAAAATGTCATGAAAGAATTTCCTATTATGAAC

>M6-PGSC0003DMG400010348-(rev)M6LRR-RLK_4392

TGCAGGTCAATCTGATATGTCTCTGATAAAATGTCATGAAAGAATTTCCTATTATGAACTTTAACAAGCAACCGAAAGATTCAGTGAAAGCAACTTGCTTGGTAACGGGAACTTCTGCAA

>M6-PGSC0003DMG400010348-(rev)M6LRR-RLK_4453

TAACAAGCAACCGAAAGATTCAGTGAAAGCAACTTGCTTGGTAACGGGAACTTCTGCAAGGTCTACAAAGGGAGACTAAAGGATGGTACTCTTTTGGCAGCAAAGGTATTCAATGTGCAA

>M6-PGSC0003DMG400010348-(rev)M6LRR-RLK_4575

GGAGGGTGCATTCAAAAGTTTTGATACAGAATGTGAGATGTTGCGCAACCTCCGCCACCGAAATCTGACCAAAGTCATCACCAGTTGCTCCAACCTTGATTTCAAAGCCTTAGTGTTGGA

>M6-PGSC0003DMG400010348-(rev)M6LRR-RLK_4636

AATCTGACCAAAGTCATCACCAGTTGCTCCAACCTTGATTTCAAAGCCTTAGTGTTGGAATACATGTCCAATGGGACACTTGATAAATGGCTATACTCTCACAACTTGTTCTTGGATTTA

>M6-PGSC0003DMG400010348-(rev)M6LRR-RLK_4697

ACATGTCCAATGGGACACTTGATAAATGGCTATACTCTCACAACTTGTTCTTGGATTTATTTCAGAGATTAGATATAATGATAGATGTTGCATCTGCAATGGTATATCTCCACAGTGGTT

>M6-PGSC0003DMG400010348-(rev)M6LRR-RLK_4758

TCAGAGATTAGATATAATGATAGATGTTGCATCTGCAATGGTATATCTCCACAGTGGTTGTTCAAATCCGGTGGTGCATTGTGACTTAAAGCCAAGCAATGTCTTGCTAGATCAAGACAT

>M6-PGSC0003DMG400010348-(rev)M6LRR-RLK_488

TGACAATAAGAAGATACTCCTATGTGAAACCGTCATCAGCAATGAAGGCTCTGTTGTTAAGAGAATTTGTGCTTTCCCGATCTAAAGAGAAACTTGGACCAATATGTTGATTTTGCCATA

>M6-PGSC0003DMG400010348-(rev)M6LRR-RLK_4880

TTGGCCATGTCAGTGATTTTGGAATTGCAAAATTGTTAGGTGCAGGGGAGACTTTTGTCCAAACAAGGACAATAGCAACCATTGGATATATTGCTCCAGGTATTAAAACTTTTACATCTT

>M6-PGSC0003DMG400010348-(rev)M6LRR-RLK_549

AGAATTTGTGCTTTCCCGATCTAAAGAGAAACTTGGACCAATATGTTGATTTTGCCATATCAAGCCATAATTTATACTGTAAACACCCAAATAAAGCCACAGGAACACCAACACAAGCAG

>M6-PGSC0003DMG400010348-(rev)M6LRR-RLK_61

ACATTTAGGTTGACATTTTTGGCAAACATCTCATAAAGGCAATATTAGCACTGAGCCACTGATGAAGCAAGAAATGTGATAACTTCTCGTTGTTGGTTAGGATGTTGCATAATTTGTCAC

>M6-PGSC0003DMG400010348-(rev)M6LRR-RLK_610

AAGCCATAATTTATACTGTAAACACCCAAATAAAGCCACAGGAACACCAACACAAGCAGCAATGAGCTTTGGCTCCCAACCAATATGATTACTATAGACCAAAAAGAATGTTGCTGCAAA

>M6-PGSC0003DMG400010348-(rev)M6LRR-RLK_671

ATGAGCTTTGGCTCCCAACCAATATGATTACTATAGACCAAAAAGAATGTTGCTGCAAAAGCCAAGAGCATGCTAACTATTGAGACAAACAGCATTGTGAGTCCAAACAATAACTTGGCT

>M6-PGSC0003DMG400010348-(rev)M6LRR-RLK_732

CCAAGAGCATGCTAACTATTGAGACAAACAGCATTGTGAGTCCAAACAATAACTTGGCTGGCAACGACACAAGAAAATCACCTTCAGCATAGTGGGATGTCACAATGGACAAGAACATAA

>M6-PGSC0003DMG400010348-(rev)M6LRR-RLK_793

CAACGACACAAGAAAATCACCTTCAGCATAGTGGGATGTCACAATGGACAAGAACATAATGATGGAAACTATTGAGGTGAATAGTGCCACTGCATCTGAATTGACAAAGATCCTAAATCC

>M6-PGSC0003DMG400010348-(rev)M6LRR-RLK_854

ATGGAAACTATTGAGGTGAATAGTGCCACTGCATCTGAATTGACAAAGATCCTAAATCCATACAATTTTAGCAATATCAGAATGCAATTATCACTGTTATTACCGCCTGGCACAGTGAAT

>M6-PGSC0003DMG400010348-(rev)M6LRR-RLK_915

ACAATTTTAGCAATATCAGAATGCAATTATCACTGTTATTACCGCCTGGCACAGTGAATCCAGCAGCAAAGACCATGGTGGTAATCAAAGTAGCGATAATCATACAAGAATTTGCAGTGT

>M6-PGSC0003DMG400010348-(rev)M6LRR-RLK_976

AGCAGCAAAGACCATGGTGGTAATCAAAGTAGCGATAATCATACAAGAATTTGCAGTGTCTTTCATCCACCTTTACCCTTCTTTCAGTAATAACTTATGCTCTTCTGTGAATATTTCTCT

>M6-PGSC0003DMG400010350-(rev)M6LRR-RLK_0

TTTCTTTCATTTTTATCTTCCGCCAAATTGGTACTAATCATGTATTTTCCTCACTTAACAGGAGAGATTCCAGCAAGCATTTCTAACATGTCAGAACTGCAGAACCTAGGATTTGCAAGA

>M6-PGSC0003DMG400010350-(rev)M6LRR-RLK_1116

TGCCGAGAATCCCTTGGACGGTTTTTTGCCTGCCTCCATTGAAAATTTCTCTGATTCTCTGCAAATTTTCCAAGGATGGTATTGTAAACTGAAAGGCTTCATTCCCGGAGAAATTGGCAA

>M6-PGSC0003DMG400010350-(rev)M6LRR-RLK_1302

CCAAGGCTTGTCGAACCTTCAAGAACTTAACTTAGGTGGCAACATGATAAAAGGAATCATACCAGATGTTATCTGCAATTTATATAATCTTGGAGCATTAGATTTGTCAGGAAATCAAGT

>M6-PGSC0003DMG400010350-(rev)M6LRR-RLK_1364

CAGATGTTATCTGCAATTTATATAATCTTGGAGCATTAGATTTGTCAGGAAATCAAGTTTCTGGTTCCATACCGCCATGCTTAGGGAACACTACCAGTTTGAGGTATCTTTATCTAGCTA

>M6-PGSC0003DMG400010350-(rev)M6LRR-RLK_1426

GGTTCCATACCGCCATGCTTAGGGAACACTACCAGTTTGAGGTATCTTTATCTAGCTAACAACAGGCTGAATTTGAGCTTACCTTCAAGCTTGTGGACTCTTCAAGATCTCATAAAATTC

>M6-PGSC0003DMG400010350-(rev)M6LRR-RLK_1488

CAGGCTGAATTTGAGCTTACCTTCAAGCTTGTGGACTCTTCAAGATCTCATAAAATTCAATGTTTCGTCCAATTTATTAAGTGGGAAAGTTCCTATTGAGATTGGAAATTTAAAGGTTGC

>M6-PGSC0003DMG400010350-(rev)M6LRR-RLK_1550

TTTCGTCCAATTTATTAAGTGGGAAAGTTCCTATTGAGATTGGAAATTTAAAGGTTGCAACACTAGTTGATCTGTCAAAAAATGATTTTAGTGGTAAGATCCCTAGCACGCGGGGGTCTG

>M6-PGSC0003DMG400010350-(rev)M6LRR-RLK_1612

CTAGTTGATCTGTCAAAAAATGATTTTAGTGGTAAGATCCCTAGCACGCGGGGGTCTGGATAGATTGATGAAACTTTCTCTAGCACATAATAAATTAGATGGGCCTATTCCAGATTCATT

>M6-PGSC0003DMG400010350-(rev)M6LRR-RLK_1674

GATTGATGAAACTTTCTCTAGCACATAATAAATTAGATGGGCCTATTCCAGATTCATTTGGCAAAATGCTGGCCTTAGAATTCTTGGATTTGAGCAATAACAATCTAACCGGTGAAATTC

>M6-PGSC0003DMG400010350-(rev)M6LRR-RLK_1736

AAAATGCTGGCCTTAGAATTCTTGGATTTGAGCAATAACAATCTAACCGGTGAAATTCCAAAGTCATTAGAAGCTCTTGTGTATCTCAAATACTTGAACTTCTCATTTAATGAACTTAGT

>M6-PGSC0003DMG400010350-(rev)M6LRR-RLK_1798

GTCATTAGAAGCTCTTGTGTATCTCAAATACTTGAACTTCTCATTTAATGAACTTAGTGGAGAGATTCCCACGGGTGGTCCTTTTGCAAATGCCACAAGTCGATCTTTCTTGAAAAATTA

>M6-PGSC0003DMG400010350-(rev)M6LRR-RLK_1922

CTTTGTGGTGACTCTAAGTTTCATGTTTCACCATGCGTCATCAAATCTCCGAAGAAGTCAAAGAGAAAAAAGATAATCTTGGTTTTGTACATCCTTTTGGGAGTGGGTATGCTATTTCTT

>M6-PGSC0003DMG400010350-(rev)M6LRR-RLK_1984

GAGAAAAAAGATAATCTTGGTTTTGTACATCCTTTTGGGAGTGGGTATGCTATTTCTTGCATTAGCCCTCGCATATGTATTTTTAAGATGGAGAAAGATAAAGAAGAATGCAGATCAAGC

>M6-PGSC0003DMG400010350-(rev)M6LRR-RLK_2046

TAGCCCTCGCATATGTATTTTTAAGATGGAGAAAGATAAAGAAGAATGCAGATCAAGCAGATGTCTTTCCCGTAAAAGGGAAACGTGAAAGAATTTCTTACTATGAACTTGAACAAGCAA

>M6-PGSC0003DMG400010350-(rev)M6LRR-RLK_2108

GTCTTTCCCGTAAAAGGGAAACGTGAAAGAATTTCTTACTATGAACTTGAACAAGCAACAGAAGGATTCGATGAAAGCAACTTGCTTGGTAGTGGGAGTTTCAGCAAGGTTTTCAAAGGG

>M6-PGSC0003DMG400010350-(rev)M6LRR-RLK_2161

AGCAACAGAAGGATTCGATGAAAGCAACTTGCTTGGTAGTGGGAGTTTCAGCAAGGTTTTCAAAGGGATATTTAAGGATGGTACTCTTTTGGCAGCAAAGGTATTCAATGTGCAATTGGA

>M6-PGSC0003DMG400010350-(rev)M6LRR-RLK_62

AGAGATTCCAGCAAGCATTTCTAACATGTCAGAACTGCAGAACCTAGGATTTGCAAGAAACAGGCTTTCAGGTAATTGTTGTGTAACAAAAAATCTGAAGTGTTTATGTAAATATATGAC

>M6-PGSC0003DMG400010350-(rev)M6LRR-RLK_620

GCCCACATTTCATATTTTTACATTCTCAAGTACTAACATGTCAACTCATATATTTCCCCCTCTTGACAGGAGAGATACCAATGGAACTAGGTTATCTTCAGAAACTACTGTTTCTGGCAT

>M6-PGSC0003DMG400010350-(rev)M6LRR-RLK_806

TAGGAATTGCAGAAAACAGGCTTTCAGGTACTCTACCTTCAGATTTAGGTCGTGGAATGCCCAACCTAGACGGATTCTATTGTTATCAGAATAGTTTGAGTGGTTTTCTCCCTGCTTCTA

>M6-PGSC0003DMG400010350-(rev)M6LRR-RLK_868

AACCTAGACGGATTCTATTGTTATCAGAATAGTTTGAGTGGTTTTCTCCCTGCTTCTATCTCAAATTCTTCAAGACTCAGAGTACTTGAACTCTCATATAACAGTTTCACAGGCCCAATT

>M6-PGSC0003DMG400010350-(rev)M6LRR-RLK_930

AAATTCTTCAAGACTCAGAGTACTTGAACTCTCATATAACAGTTTCACAGGCCCAATTCCTGAATCAGTTAGTGACTTAGAATACATTGAGGTTTTGAACTTAGGGGCGAATAATTTTGT

>M6-PGSC0003DMG400010350-(rev)M6LRR-RLK_992

AATCAGTTAGTGACTTAGAATACATTGAGGTTTTGAACTTAGGGGCGAATAATTTTGTCAGCAATTCAGCATTGAGCTTCTTGACATCTTTGGCAAACTGTAGGAAACTGAAAGAAATCA

>M6-PGSC0003DMG400010351-(rev)M6LRR-RLK_2542

CTTGGTAACGGGAACTTCTGCAAGGTCTACAAAGGGAGACTAAAGGATGGTACTCTTTTGGCAGCAAAGGTATTCAATGTGCAATTGGAGGGTGCATTCAAAAGTTTTGATACAGAATGT

>M6-PGSC0003DMG400010351-(rev)M6LRR-RLK_2852

TATATCTCCACAGTGGTTGTTCAAATCCGGTGGTGCATTGTGACTTAAAGCCAAGCAATGTCTTGCTAGATCAAGACATGGTTGGCCATGTCAGTGATTTTGGAATTGCAAAATTGTTAG

>M6-PGSC0003DMG400010400M6LRR-RLK_0

GGCATTTAAATCGAGCCTTCGATTGGAACCTCACCACATTTTGTCCAGAAATTGGAGCTCAGACACACACTTCTGCAGCTGGTATGGAGTTTCATGCTTGCACCAGAGAGTTGTAGCTTT

>M6-PGSC0003DMG400010400M6LRR-RLK_1037

GCAATGCTTCAAAGCTTGAAAATCTTGAAGTTGCAAACAATAGTTTTACCGGACCGATTCCCATGATGTTAGGTAACCTGGTGGACCTTATAACTCTAAATCTGCAAAATAATCTGTTGG

>M6-PGSC0003DMG400010400M6LRR-RLK_1098

CATGATGTTAGGTAACCTGGTGGACCTTATAACTCTAAATCTGCAAAATAATCTGTTGGAGAACAATCCTGGAGCCACACAACTTGACTTCTTGAATTCATTGGTAACCAGCAGGAATCT

>M6-PGSC0003DMG400010400M6LRR-RLK_1159

AACAATCCTGGAGCCACACAACTTGACTTCTTGAATTCATTGGTAACCAGCAGGAATCTGGTGTACCTGATTCTTGCATCTAATCCTTTGAATGGAATTTTTACGGAATCAATAGGAAAT

>M6-PGSC0003DMG400010400M6LRR-RLK_1220

TGTACCTGATTCTTGCATCTAATCCTTTGAATGGAATTTTTACGGAATCAATAGGAAATTTATCAAGCAATATCAGGGTCCTTGATGCATTCGACTGTGGAATTAGAGGCAGTATGCCAA

>M6-PGSC0003DMG400010400M6LRR-RLK_1281

ATCAAGCAATATCAGGGTCCTTGATGCATTCGACTGTGGAATTAGAGGCAGTATGCCAACAAACATTGGAAACTTAAGTGGCTTGATTTTCCTGGGATTAGCAAGTAATAACATTGTCGG

>M6-PGSC0003DMG400010400M6LRR-RLK_1342

AACATTGGAAACTTAAGTGGCTTGATTTTCCTGGGATTAGCAAGTAATAACATTGTCGGTAATGTGCCCCCCTCGTTTGTTGGCCTGCAAAACCTGGAAAGGCTGTATCTTACTAGAAAT

>M6-PGSC0003DMG400010400M6LRR-RLK_1403

ATGTGCCCCCCTCGTTTGTTGGCCTGCAAAACCTGGAAAGGCTGTATCTTACTAGAAATAAATTAGAAGGGACATTTCCAGCTGAACTGTGCAGCATAGGAAGATTAGGTCTACTGCATT

>M6-PGSC0003DMG400010400M6LRR-RLK_1464

ATTAGAAGGGACATTTCCAGCTGAACTGTGCAGCATAGGAAGATTAGGTCTACTGCATTTGGGAGAAAACAGACTGTCAGGAGAAGTACCCTCGTGTATCGGAAATCTGACTGAATTAAG

>M6-PGSC0003DMG400010400M6LRR-RLK_1525

GGAGAAAACAGACTGTCAGGAGAAGTACCCTCGTGTATCGGAAATCTGACTGAATTAAGAGTAATGTCAATTGCTGCCAACAATTTCAGCTCAAAGTTACCTTCAAGTTTCTGGAGATTA

>M6-PGSC0003DMG400010400M6LRR-RLK_1586

TAATGTCAATTGCTGCCAACAATTTCAGCTCAAAGTTACCTTCAAGTTTCTGGAGATTAGTAAAGCTTGATGATCTGAACTTATCGAGAAACCTGCTGCATGGTTTCTTCCCTTCTGATG

>M6-PGSC0003DMG400010400M6LRR-RLK_1647

AAAGCTTGATGATCTGAACTTATCGAGAAACCTGCTGCATGGTTTCTTCCCTTCTGATGTGGGAACTCTGAACGCAATGAGGATCATGGACCTCTCTTTCAATAGGTATTTTGGTGAAAT

>M6-PGSC0003DMG400010400M6LRR-RLK_1708

GGAACTCTGAACGCAATGAGGATCATGGACCTCTCTTTCAATAGGTATTTTGGTGAAATACCCAATTCCATAGGATCTCTTCAATATCTAGTTTCACTAGATATGTCAAGAAATGCATTA

>M6-PGSC0003DMG400010400M6LRR-RLK_1769

CCAATTCCATAGGATCTCTTCAATATCTAGTTTCACTAGATATGTCAAGAAATGCATTACAAGGTCCCATACCGGCCACATTTAGCAATCTAATAGTTCTTGAGGCTCTGGATTTGTCTA

>M6-PGSC0003DMG400010400M6LRR-RLK_183

TGCTCTTCTTGACCTTAGTAGCAATGATTTGAATGGCAATTTACCAAGTGAGTTGGGCTTCTTGCAACAACTGAGACTTTTTAATGTGACTGGGAATTCTCTTGATGGAACTATTCCTCT

>M6-PGSC0003DMG400010400M6LRR-RLK_1830

AGGTCCCATACCGGCCACATTTAGCAATCTAATAGTTCTTGAGGCTCTGGATTTGTCTACTAATGCTCTATCAGGTGTGATTCCCAAATCTCTTGAGCTATTGCGAGATCTCAAGTATCT

>M6-PGSC0003DMG400010400M6LRR-RLK_1891

AATGCTCTATCAGGTGTGATTCCCAAATCTCTTGAGCTATTGCGAGATCTCAAGTATCTCAATGTTTCTTTCAATAACTTGCAAGGAGAAGTTCCAAAGAAAGGGGTATTTGCAAATGTT

>M6-PGSC0003DMG400010400M6LRR-RLK_1952

ATGTTTCTTTCAATAACTTGCAAGGAGAAGTTCCAAAGAAAGGGGTATTTGCAAATGTTAATTACCGGTTCTTGATGGGAAATCCAAGACTTTGTGGAGCACCTGACTTATATATTCCTA

>M6-PGSC0003DMG400010400M6LRR-RLK_2013

TTACCGGTTCTTGATGGGAAATCCAAGACTTTGTGGAGCACCTGACTTATATATTCCTATATGTCCAGCTCAAGGTAGAAAGACAACAAGGAAAAAGTCCATTATTCTTAGAACAGCAGT

>M6-PGSC0003DMG400010400M6LRR-RLK_2074

TGTCCAGCTCAAGGTAGAAAGACAACAAGGAAAAAGTCCATTATTCTTAGAACAGCAGTACCTGTTGCTGTAACATTTCTGATATTGGTCGCGTTGTTTTTCACATGGATGATTTGGTCA

>M6-PGSC0003DMG400010400M6LRR-RLK_2135

CTGTTGCTGTAACATTTCTGATATTGGTCGCGTTGTTTTTCACATGGATGATTTGGTCAAGAAAAAAACATGCGATTGGAAACAATGAATCGGATTACCCTCCTAGAATTGCACATCAGA

>M6-PGSC0003DMG400010400M6LRR-RLK_2196

AAAAAAACATGCGATTGGAAACAATGAATCGGATTACCCTCCTAGAATTGCACATCAGAAAATCACTTATTATGAGCTTCTCCAAGCTACAGAAAATTTCAGTCAGTCCAAGCTAATAGG

>M6-PGSC0003DMG400010400M6LRR-RLK_2257

ATCACTTATTATGAGCTTCTCCAAGCTACAGAAAATTTCAGTCAGTCCAAGCTAATAGGAAGTGGAAGTTCCGGTACAGTCTACCAAGGTACATTTTCTGGTGGTGCTGTCTTTGCAATC

>M6-PGSC0003DMG400010400M6LRR-RLK_2318

GTGGAAGTTCCGGTACAGTCTACCAAGGTACATTTTCTGGTGGTGCTGTCTTTGCAATCAAAGTTTTTGATATGCAATGGCAACGGGGTTTAAGAAATTTTGATTCAGAATGTGAAATTT

>M6-PGSC0003DMG400010400M6LRR-RLK_2379

AGTTTTTGATATGCAATGGCAACGGGGTTTAAGAAATTTTGATTCAGAATGTGAAATTTTGAGCAATGTAAGACACAGAAACTTGGTCAAGATAGTGTCGACTTGTTCAAATTTGGATTT

>M6-PGSC0003DMG400010400M6LRR-RLK_244

TTGCAACAACTGAGACTTTTTAATGTGACTGGGAATTCTCTTGATGGAACTATTCCTCTGAATATAAGCCGTTGTCGTGTACTGCAAGAACTTCATTTATCAGATAACATGATCAAAGGA

>M6-PGSC0003DMG400010400M6LRR-RLK_2440

AGCAATGTAAGACACAGAAACTTGGTCAAGATAGTGTCGACTTGTTCAAATTTGGATTTCACAGCAATTGTGCTGGAGTACATGCCTAATGGAAGCTTAGACAAAAGATTATACTCTGAT

>M6-PGSC0003DMG400010400M6LRR-RLK_2501

CAGCAATTGTGCTGGAGTACATGCCTAATGGAAGCTTAGACAAAAGATTATACTCTGATGAAAATTGCTTGAGCCTGGTTGAAAGACTCAATATAATGATCGATGTAGCACTAGCCATGG

>M6-PGSC0003DMG400010400M6LRR-RLK_2562

AAATTGCTTGAGCCTGGTTGAAAGACTCAATATAATGATCGATGTAGCACTAGCCATGGAGTATCTTCATCATGATTATACAGTGCCCATTGTGCACTGTGATCTGAAGCCAGGTAACGT

>M6-PGSC0003DMG400010400M6LRR-RLK_2623

TATCTTCATCATGATTATACAGTGCCCATTGTGCACTGTGATCTGAAGCCAGGTAACGTGCTTCTTGATCAGGATCTGACTGCTCATGTTGCAGATTTTGGAATTGCAAAAATGTTGGCA

>M6-PGSC0003DMG400010400M6LRR-RLK_2684

TTCTTGATCAGGATCTGACTGCTCATGTTGCAGATTTTGGAATTGCAAAAATGTTGGCACAAGAAGGAAACATGGCTCAAACCAAGACTCTCGGAACTATTGGTTACATTGCTCCAGGTA

>M6-PGSC0003DMG400010400M6LRR-RLK_2745

AGAAGGAAACATGGCTCAAACCAAGACTCTCGGAACTATTGGTTACATTGCTCCAGGTATGATAATGTCTTGAAATGAAGGTGTCCTATGCTGGTTACTTTTTTCCTTTATGTCTACCAT

>M6-PGSC0003DMG400010400M6LRR-RLK_2806

ATAATGTCTTGAAATGAAGGTGTCCTATGCTGGTTACTTTTTTCCTTTATGTCTACCATAGCTTTATACTGTAACATGGATAACTACACATTTTAACTGGGAATTTGTTGAAACAGAATA

>M6-PGSC0003DMG400010400M6LRR-RLK_2867

CTTTATACTGTAACATGGATAACTACACATTTTAACTGGGAATTTGTTGAAACAGAATATGGTTTAGACGGGCAAATTTCCACTAGCAGTGATGTTTATAGCTTTGGGATTTTGTTGCTT

>M6-PGSC0003DMG400010400M6LRR-RLK_2928

GTTTAGACGGGCAAATTTCCACTAGCAGTGATGTTTATAGCTTTGGGATTTTGTTGCTTGAGACATTCACAAGAAAGAAGCCTACAGATGACTTGTTTAGCGGAGACTTAAGCTTGCATA

>M6-PGSC0003DMG400010400M6LRR-RLK_2989

GACATTCACAAGAAAGAAGCCTACAGATGACTTGTTTAGCGGAGACTTAAGCTTGCATAAATGGACATCTCTCTCCTTTCCTGATGCTGTACTGGATATTCTGGATGCTGACCTTGTTAG

>M6-PGSC0003DMG400010400M6LRR-RLK_305

ATATAAGCCGTTGTCGTGTACTGCAAGAACTTCATTTATCAGATAACATGATCAAAGGAAGCATACCTCAAGAACTTGGTCGTTTAAGTCAACTTAGAATTTTGCGTCTCAACGACAATA

>M6-PGSC0003DMG400010400M6LRR-RLK_3050

TGGACATCTCTCTCCTTTCCTGATGCTGTACTGGATATTCTGGATGCTGACCTTGTTAGCGACATTGGCTTTACAATTGATGAAAACCAAAGCCAAATCAAACAATTGCTGGTTTTGATA

>M6-PGSC0003DMG400010400M6LRR-RLK_3111

ACATTGGCTTTACAATTGATGAAAACCAAAGCCAAATCAAACAATTGCTGGTTTTGATAATAAATGTTGCATTCCTGTGTTTGAAAGAGTTGCCAGAGGAAAGGATAAACATGAGAGAGG

>M6-PGSC0003DMG400010400M6LRR-RLK_3172

AAATGTTGCATTCCTGTGTTTGAAAGAGTTGCCAGAGGAAAGGATAAACATGAGAGAGGTTGTGGTGCAGCTAAACAAGATAAGAGCTGAATTAACCAAGCTTTTAACTCAATCCCATTC

>M6-PGSC0003DMG400010400M6LRR-RLK_3233

GTGGTGCAGCTAAACAAGATAAGAGCTGAATTAACCAAGCTTTTAACTCAATCCCATTCAAAATTTGAGAAAGAGGATTGATGCAGCAACAACCAAATTCAGTTTTCCACCTTCTATGGC

>M6-PGSC0003DMG400010400M6LRR-RLK_3270

AGCTTTTAACTCAATCCCATTCAAAATTTGAGAAAGAGGATTGATGCAGCAACAACCAAATTCAGTTTTCCACCTTCTATGGCCTGTCAAAACATCTGTTATTTCTTAACATTTTCTTTC

>M6-PGSC0003DMG400010400M6LRR-RLK_366

CATACCTCAAGAACTTGGTCGTTTAAGTCAACTTAGAATTTTGCGTCTCAACGACAATAATCTGACAGGAAAGATTCCACCTTTATTAGGAAACTTATCAAAGCTGGAATATTTTTATCT

>M6-PGSC0003DMG400010400M6LRR-RLK_427

CTGACAGGAAAGATTCCACCTTTATTAGGAAACTTATCAAAGCTGGAATATTTTTATCTGCATGAAAATGATTTGGAAGGTGAAATACCTGATGAAATTGGTGATCTTACCAATTTGAGG

>M6-PGSC0003DMG400010400M6LRR-RLK_488

ATGAAAATGATTTGGAAGGTGAAATACCTGATGAAATTGGTGATCTTACCAATTTGAGGCTCCTTTCCTTTCGTGGCAATGACTTCACTGGATCGATACCTGCTTCAATTTTCAATATTT

>M6-PGSC0003DMG400010400M6LRR-RLK_549

CCTTTCCTTTCGTGGCAATGACTTCACTGGATCGATACCTGCTTCAATTTTCAATATTTCAAGATTACAAATTGTGGATCTCTCGATAAATGCCTTATCAGGGGAACTTCCAAGTGATTT

>M6-PGSC0003DMG400010400M6LRR-RLK_61

GACACACACTTCTGCAGCTGGTATGGAGTTTCATGCTTGCACCAGAGAGTTGTAGCTTTGCAACTTCCTAATTTGGCCATTCAAGGCAGAGTTTCTTCAGAAATTGCCAACTTGACTCAA

>M6-PGSC0003DMG400010400M6LRR-RLK_610

AGATTACAAATTGTGGATCTCTCGATAAATGCCTTATCAGGGGAACTTCCAAGTGATTTGGGTAATTATCTTCCTATCCTTGAACATCTTTTTCTGGACAGCAACAGGATCACGGGGAAA

>M6-PGSC0003DMG400010400M6LRR-RLK_671

GTAATTATCTTCCTATCCTTGAACATCTTTTTCTGGACAGCAACAGGATCACGGGGAAAATCCCGGTTTCTTTATCAAATGCATCAAAGATTACTCATCTCTTCTTTACTGAAAATGATG

>M6-PGSC0003DMG400010400M6LRR-RLK_732

CCCGGTTTCTTTATCAAATGCATCAAAGATTACTCATCTCTTCTTTACTGAAAATGATGTGCAAGGAAATATTCCAAATGAGTTTGGTAAGTTGCATGAACTAGTCTGGTTTGAATTTGA

>M6-PGSC0003DMG400010400M6LRR-RLK_793

CAAGGAAATATTCCAAATGAGTTTGGTAAGTTGCATGAACTAGTCTGGTTTGAATTTGAATATAATCGAATTTCTGGTACCATTCCATCCAGCTTATTCAATATTTCGTCTCTTGAAATC

>M6-PGSC0003DMG400010400M6LRR-RLK_854

ATAATCGAATTTCTGGTACCATTCCATCCAGCTTATTCAATATTTCGTCTCTTGAAATCCTCAAAGCCAGGCATAATTACTTAAATGGGCATCTTCCATATGATCTTGGTACTTGGCTAC

>M6-PGSC0003DMG400010400M6LRR-RLK_915

CAAAGCCAGGCATAATTACTTAAATGGGCATCTTCCATATGATCTTGGTACTTGGCTACCTAACTTGCAGGAGATTTTCCTTTCACACAACCAATTCAGTGGAGATCTACCCGCTGCAAT

>M6-PGSC0003DMG400010400M6LRR-RLK_976

AACTTGCAGGAGATTTTCCTTTCACACAACCAATTCAGTGGAGATCTACCCGCTGCAATATGCAATGCTTCAAAGCTTGAAAATCTTGAAGTTGCAAACAATAGTTTTACCGGACCGATT

>M6-PGSC0003DMG400010427M6RLKWAX-EGF_0

TTCTAGCTTATTGGCTAACTCTTGTCACTCCTATCTCCACAGGTGTATTCATAAGTATCGGAGTACTGCTACTACTGTTTGGGAGCTATTTATTGTACAAAACAATCAAGAAGAGAAGGA

>M6-PGSC0003DMG400010427M6RLKWAX-EGF_1008

GTTGCATGAACTTGAATGGTAAGAAAAGGCCTACTATGAAGGAAGTAACTACAGCATTGGAAGCCATCAGATCACAAATGCAATCTACAGCAGTAACAAATTTTCAACTGGATAAGGGTG

>M6-PGSC0003DMG400010427M6RLKWAX-EGF_1071

CCATCAGATCACAAATGCAATCTACAGCAGTAACAAATTTTCAACTGGATAAGGGTGTCACAGCAGAAAGAACTGTGATATCAGAAGTTAACTACACATGGACAAACACAAGTACCTCAT

>M6-PGSC0003DMG400010427M6RLKWAX-EGF_1134

CAGAAAGAACTGTGATATCAGAAGTTAACTACACATGGACAAACACAAGTACCTCATCATCAGATATCCATCCATTGTTGTTTGAAACTAGTTAATTATTACTCCCTCTATCGGAGAAAG

>M6-PGSC0003DMG400010427M6RLKWAX-EGF_1188

CATCATCAGATATCCATCCATTGTTGTTTGAAACTAGTTAATTATTACTCCCTCTATCGGAGAAAGAATGTGTGTGTCTGTTTGTTCGACTTTACAATAGTTTAAATGTCTACTTCTACA

>M6-PGSC0003DMG400010427M6RLKWAX-EGF_126

GACAGAAACAAAAGTTTTTCAAGCGAAATGGCGGCCTCTTATTACAACAGCAGCTATCATCCAATAATGAAGGTACTATTGACAAAGGAAAACTTTTCACTGCTAAAGAGCTGGAGAAGG

>M6-PGSC0003DMG400010427M6RLKWAX-EGF_189

ATAATGAAGGTACTATTGACAAAGGAAAACTTTTCACTGCTAAAGAGCTGGAGAAGGCAACAGACCACTTCAATGAAAATCGCATACTTGGTCGTGGAGGTCAAGGTACGGTATACAAGG

>M6-PGSC0003DMG400010427M6RLKWAX-EGF_252

ACCACTTCAATGAAAATCGCATACTTGGTCGTGGAGGTCAAGGTACGGTATACAAGGGCATGTTACTTGATGGAAAGATTGTAGCAGTGAAAAAGTCTAAGCTGGTAGACGAAAACCAGT

>M6-PGSC0003DMG400010427M6RLKWAX-EGF_315

TACTTGATGGAAAGATTGTAGCAGTGAAAAAGTCTAAGCTGGTAGACGAAAACCAGTTGGATCAGTTTATCAACGAGGTGGCTGTTCTTTCACAAATCAATCACAGGAATGTGGTGAAGC

>M6-PGSC0003DMG400010427M6RLKWAX-EGF_378

AGTTTATCAACGAGGTGGCTGTTCTTTCACAAATCAATCACAGGAATGTGGTGAAGCTACTTGGCTGTTGTTTGGAAACAGAAGTTCCTCTATTAGTATACGAGTTCATAATTAATGGCA

>M6-PGSC0003DMG400010427M6RLKWAX-EGF_441

GCTGTTGTTTGGAAACAGAAGTTCCTCTATTAGTATACGAGTTCATAATTAATGGCACACTTTACAGCCTCATACATAATGAGAACAATGAGTTTCCCTTTACTTGGAGCACGCGTTTGA

>M6-PGSC0003DMG400010427M6RLKWAX-EGF_504

ACAGCCTCATACATAATGAGAACAATGAGTTTCCCTTTACTTGGAGCACGCGTTTGAGAATTGCCACAGAGGTAGCTGGAGCATTGGCTTATCTACACTCAGCAACTTCTGTTCCAGTCT

>M6-PGSC0003DMG400010427M6RLKWAX-EGF_567

CCACAGAGGTAGCTGGAGCATTGGCTTATCTACACTCAGCAACTTCTGTTCCAGTCTATCATTGGGACATCAAATCAAGCAATATACTTTTAGACGAAAAATACCGTGCCAAGGTATCAG

>M6-PGSC0003DMG400010427M6RLKWAX-EGF_63

TACTGCTACTACTGTTTGGGAGCTATTTATTGTACAAAACAATCAAGAAGAGAAGGAACAAACGACAGAAACAAAAGTTTTTCAAGCGAAATGGCGGCCTCTTATTACAACAGCAGCTAT

>M6-PGSC0003DMG400010427M6RLKWAX-EGF_630

GGGACATCAAATCAAGCAATATACTTTTAGACGAAAAATACCGTGCCAAGGTATCAGATTTCGGCACCTCAAGGTCTATTGCAATTGATCAAACTCACCTAACGACCGGAGTCCAAGGGA

>M6-PGSC0003DMG400010427M6RLKWAX-EGF_693

GCACCTCAAGGTCTATTGCAATTGATCAAACTCACCTAACGACCGGAGTCCAAGGGACATTTGGCTATTTAGATCCTGAGTACTTCCAATCCAGCCAATATACTGAAAAAAGTGATGTCT

>M6-PGSC0003DMG400010427M6RLKWAX-EGF_756

GCTATTTAGATCCTGAGTACTTCCAATCCAGCCAATATACTGAAAAAAGTGATGTCTATAGCTTCGGAGTTGTTCTTGCTGAACTCTTAACAGGAAAGAAAGCAATTTCAACCACAACAA

>M6-PGSC0003DMG400010427M6RLKWAX-EGF_819

TCGGAGTTGTTCTTGCTGAACTCTTAACAGGAAAGAAAGCAATTTCAACCACAACAAACCAAGACAGAAGTTTAGCAACAAACTTCCTTTTGGCAACAGATAGAAATCTCCTTGATACGA

>M6-PGSC0003DMG400010427M6RLKWAX-EGF_882

ACAGAAGTTTAGCAACAAACTTCCTTTTGGCAACAGATAGAAATCTCCTTGATACGATTATCTATACAAAAATTTCACAAGAAGGTAGAAAAGAGGATATAATGGCGGCTGCTAATATAG

>M6-PGSC0003DMG400010427M6RLKWAX-EGF_945

ATACAAAAATTTCACAAGAAGGTAGAAAAGAGGATATAATGGCGGCTGCTAATATAGCGTATCGTTGCATGAACTTGAATGGTAAGAAAAGGCCTACTATGAAGGAAGTAACTACAGCAT

>M6-PGSC0003DMG400010447-(rev)M6LRR-RLP_0

CAACTTGCACTTTCCAAGTCTAAAAAGCAGCCATCACATTTAACATTTATAATTTCAGAAGCCACCAAGAATTTGTCATCCAAATGTATGTATATATTATGCCATTTTTTGGCACATTCT

>M6-PGSC0003DMG400010447-(rev)M6LRR-RLP_1037

AATTTTCTTTCTGGAAAGTTACCGAGTGTAATTGGAAAGCTATGGAAATTAGAAGATCTTGATCTCTCAGAAAATTGATTTGAAGGAGAAGTATCCGAGATTTTCAATACTATGAATGAT

>M6-PGSC0003DMG400010447-(rev)M6LRR-RLP_1098

ATCTCTCAGAAAATTGATTTGAAGGAGAAGTATCCGAGATTTTCAATACTATGAATGATTGCCCTCCAATAGGCTCAGGAAATTGTTCTTCTTTGAGGACCTTAAGACTAAATGACAACA

>M6-PGSC0003DMG400010447-(rev)M6LRR-RLP_1159

CCCTCCAATAGGCTCAGGAAATTGTTCTTCTTTGAGGACCTTAAGACTAAATGACAACAAACTGATTGGAAATCTTCCAAAAAGTTTTGGGCAACTTTCTAATCTACAATTCTGTTTCAT

>M6-PGSC0003DMG400010447-(rev)M6LRR-RLP_122

TCTTATAATTTCACAAAAATGAGAGGGCTGTACTACATAGTACTTGTCCTGTTTACTATAAAGTTTTGTTCTTGCAGTGGAGTTTGCAAAGAAAATGAGAAAACAGCTTTGCTAAGGTTA

>M6-PGSC0003DMG400010447-(rev)M6LRR-RLP_1220

CTGATTGGAAATCTTCCAAAAAGTTTTGGGCAACTTTCTAATCTACAATTCTGTTTCATATCCAATAATAGGTTGGAAGGGGTGTTAACAGAAGAACATTTCACCAACTTGACAAATTTG

>M6-PGSC0003DMG400010447-(rev)M6LRR-RLP_1281

CCAATAATAGGTTGGAAGGGGTGTTAACAGAAGAACATTTCACCAACTTGACAAATTTGAAGTTTTTCTCTGCATCTAAGAGTAATTTGACTTTAAGGGTGAGTTCAGATTGGATTCCAT

>M6-PGSC0003DMG400010447-(rev)M6LRR-RLP_1342

GTTTTTCTCTGCATCTAAGAGTAATTTGACTTTAAGGGTGAGTTCAGATTGGATTCCATCATTCCAAGCTAGTGACATTGTAATGGGCGGTTGGCGCTTAGGCCCTGACTTCCCTTTGTG

>M6-PGSC0003DMG400010447-(rev)M6LRR-RLP_1403

TTCCAAGCTAGTGACATTGTAATGGGCGGTTGGCGCTTAGGCCCTGACTTCCCTTTGTGGATCCAGACTCAACAATCAATCATGAATTTGGACATATCAAATGCTGGTATAGTAGGTGAG

>M6-PGSC0003DMG400010447-(rev)M6LRR-RLP_1464

TCCAGACTCAACAATCAATCATGAATTTGGACATATCAAATGCTGGTATAGTAGGTGAGGTTCCTACTTGGTTCTGAAACTTATCGTCTCAAATTCGTTTCCTCAACATGTCTCACAACC

>M6-PGSC0003DMG400010447-(rev)M6LRR-RLP_1525

TCCTACTTGGTTCTGAAACTTATCGTCTCAAATTCGTTTCCTCAACATGTCTCACAACCAGCTCATTGGTGAGATTCCAACTTTCTCAATTTCTGATAGCTCCATGGGAAGTGGAGGTCC

>M6-PGSC0003DMG400010447-(rev)M6LRR-RLP_1586

CTCATTGGTGAGATTCCAACTTTCTCAATTTCTGATAGCTCCATGGGAAGTGGAGGTCCTTGGTTGATATACTTGAGTTCCAACAACTTCAGTGGTTTACTCCCTCATATTCCAACCATG

>M6-PGSC0003DMG400010447-(rev)M6LRR-RLP_1647

GGTTGATATACTTGAGTTCCAACAACTTCAGTGGTTTACTCCCTCATATTCCAACCATGGTAACAGAGTTAGACCTTTCAGATAACACTTTCTCGGAAGGTTTAACAAGCTTCTTGTGTG

>M6-PGSC0003DMG400010447-(rev)M6LRR-RLP_1708

AACAGAGTTAGACCTTTCAGATAACACTTTCTCGGAAGGTTTAACAAGCTTCTTGTGTGATAAAAAGAATGAATCTTACATGTTGGAGATTCTACACCTTGGGGGAAACCGTTTCTCCGA

>M6-PGSC0003DMG400010447-(rev)M6LRR-RLP_1769

AAAAAGAATGAATCTTACATGTTGGAGATTCTACACCTTGGGGGAAACCGTTTCTCCGAAGAAATTCCTGATTGTTGGATGAATTGGCCAGAGTTGAGAGTTGTCAACTTGGGAGAGAAT

>M6-PGSC0003DMG400010447-(rev)M6LRR-RLP_183

AGTTTTGTTCTTGCAGTGGAGTTTGCAAAGAAAATGAGAAAACAGCTTTGCTAAGGTTAAAGAAAGAAGCAAATGATCCCACAAATGTACTATCTTCTAGGGTTGATAAAGAAGATTGTT

>M6-PGSC0003DMG400010447-(rev)M6LRR-RLP_1830

AAATTCCTGATTGTTGGATGAATTGGCCAGAGTTGAGAGTTGTCAACTTGGGAGAGAATAAGCTTACCGGAGGAATCCCAAGATCCATTGGAGCTTTAAGCAATTTGAAATCTTTGGACT

>M6-PGSC0003DMG400010447-(rev)M6LRR-RLP_1891

GCTTACCGGAGGAATCCCAAGATCCATTGGAGCTTTAAGCAATTTGAAATCTTTGGACTTGAAAAGAAACAAACTTTCTGGTCCATTGCCTTCATCCTTGAACAACTGCACAAGATTGTG

>M6-PGSC0003DMG400010447-(rev)M6LRR-RLP_1952

AAAAGAAACAAACTTTCTGGTCCATTGCCTTCATCCTTGAACAACTGCACAAGATTGTGGAAAATAAACATGAATGAGAACGAGTTAGATGGAAATTTACCACCTTGGTTTGGTACACTT

>M6-PGSC0003DMG400010447-(rev)M6LRR-RLP_2013

AAATAAACATGAATGAGAACGAGTTAGATGGAAATTTACCACCTTGGTTTGGTACACTTTCAAAGCTGATAGTCCTTAGTCTTCGGTCTAATAAATTCGATGGTGAACTAGCTCCAGAAC

>M6-PGSC0003DMG400010447-(rev)M6LRR-RLP_2074

AAAGCTGATAGTCCTTAGTCTTCGGTCTAATAAATTCGATGGTGAACTAGCTCCAGAACTTTGCCAGCTCAATGATCTTCAAATCTTGGATCTTGCCAACAACAATTTGTTTGGAGCTAT

>M6-PGSC0003DMG400010447-(rev)M6LRR-RLP_2135

TGCCAGCTCAATGATCTTCAAATCTTGGATCTTGCCAACAACAATTTGTTTGGAGCTATACCTCGGTGTGTTAATAATTTCACAGCAATGGTGAATGGAAGGAAGGAAATTAGAGATGGA

>M6-PGSC0003DMG400010447-(rev)M6LRR-RLP_2196

CTCGGTGTGTTAATAATTTCACAGCAATGGTGAATGGAAGGAAGGAAATTAGAGATGGAGACGGCGAAATGGATTATTCTTATTATGTTGGAGTTTTCAGGGAAAGTGCAAGAGTAGCAA

>M6-PGSC0003DMG400010447-(rev)M6LRR-RLP_2257

CGGCGAAATGGATTATTCTTATTATGTTGGAGTTTTCAGGGAAAGTGCAAGAGTAGCAACTAAGGGCAACATGTATCAGTATGATACCATTTTGTCATTGTTTACTAGCATGGATCTTTC

>M6-PGSC0003DMG400010447-(rev)M6LRR-RLP_2318

AAGGGCAACATGTATCAGTATGATACCATTTTGTCATTGTTTACTAGCATGGATCTTTCTAACAATAATCTTTCTGGAAACATTCCTATGAGTCTTACAAGTCTTGTTGGATTGAGATAA

>M6-PGSC0003DMG400010447-(rev)M6LRR-RLP_2379

ACAATAATCTTTCTGGAAACATTCCTATGAGTCTTACAAGTCTTGTTGGATTGAGATAATTAAATTTCTCCCACAATCAGCTGACTAGTAGTATTCCAAGGGACATTGACAAGATGAAGG

>M6-PGSC0003DMG400010447-(rev)M6LRR-RLP_244

GAAAGAAGCAAATGATCCCACAAATGTACTATCTTCTAGGGTTGATAAAGAAGATTGTTGTAACTGGGAAGGAGTTTTGTGCCACAATGTGACTGGTTCTGTCATTGAGCTATCGATTAA

>M6-PGSC0003DMG400010447-(rev)M6LRR-RLP_2440

AAATTTCTCCCACAATCAGCTGACTAGTAGTATTCCAAGGGACATTGACAAGATGAAGGTATTAGAATCACTTGATATCTCAAATAATCAACTCTCTGGTGAAATACCACAAAGCATTTC

>M6-PGSC0003DMG400010447-(rev)M6LRR-RLP_2501

TTAGAATCACTTGATATCTCAAATAATCAACTCTCTGGTGAAATACCACAAAGCATTTCGAGTTTGTCCTCTTTGAGCTACTTGAATGTATCTCACAACAACTTATCAGGGCAGATTCCT

>M6-PGSC0003DMG400010447-(rev)M6LRR-RLP_2562

GTTTGTCCTCTTTGAGCTACTTGAATGTATCTCACAACAACTTATCAGGGCAGATTCCTGTGAGCACTCAACTTCAAAGCTTTAGCCCTTTTAGTTTTACTGGAAATAAACTGTGTGGGC

>M6-PGSC0003DMG400010447-(rev)M6LRR-RLP_2623

GAGCACTCAACTTCAAAGCTTTAGCCCTTTTAGTTTTACTGGAAATAAACTGTGTGGGCTTCCACTCGTGGAGCGTTGCAGTAAAAATCCCAAGGTTGAAAATGAAGAAGAAGAAGAAGA

>M6-PGSC0003DMG400010447-(rev)M6LRR-RLP_2684

CCACTCGTGGAGCGTTGCAGTAAAAATCCCAAGGTTGAAAATGAAGAAGAAGAAGAAGAAGAAGAAGATGATAATGAAGTGGAATGGTTCTATGTTTCCATGGCAATAGGATTCATTGTG

>M6-PGSC0003DMG400010447-(rev)M6LRR-RLP_2745

AAGAAGATGATAATGAAGTGGAATGGTTCTATGTTTCCATGGCAATAGGATTCATTGTGGCCTTTTGGGGTGTAAGTGCTCCTTTGCTTTTCATAAGACCATGGAGACATGCCTATTATA

>M6-PGSC0003DMG400010447-(rev)M6LRR-RLP_2806

CTTTTGGGGTGTAAGTGCTCCTTTGCTTTTCATAAGACCATGGAGACATGCCTATTATAGATTTTTAGATAGAACATGGAAATCATGGCACACATAATTTGTAAAGACGTGACATTGAGG

>M6-PGSC0003DMG400010447-(rev)M6LRR-RLP_2867

TTTTTAGATAGAACATGGAAATCATGGCACACATAATTTGTAAAGACGTGACATTGAGGAGATCTCTATTCATGTATACTGGATGAAATTTGTTTATATCTCAGGAAATCCATCACCATT

>M6-PGSC0003DMG400010447-(rev)M6LRR-RLP_2928

ATCTCTATTCATGTATACTGGATGAAATTTGTTTATATCTCAGGAAATCCATCACCATTCAGCGTTCTTCAAATTAAGACATTTTTATATTTGTTGTCTCCATATATATGGGAAATTTGA

>M6-PGSC0003DMG400010447-(rev)M6LRR-RLP_2985

TTCAGCGTTCTTCAAATTAAGACATTTTTATATTTGTTGTCTCCATATATATGGGAAATTTGACAATTTTGTTTTGGCAAGGTTTGAGTTATGGAATGAACAAGGAAATTAACAGTTCCT

>M6-PGSC0003DMG400010447-(rev)M6LRR-RLP_305

AACTGGGAAGGAGTTTTGTGCCACAATGTGACTGGTTCTGTCATTGAGCTATCGATTAACGGATGAGATTGGTCAGAACTTAAAGGCCTAAAGATTGATGATTTCCAGTGGTTGTCAAGT

>M6-PGSC0003DMG400010447-(rev)M6LRR-RLP_366

GATGAGATTGGTCAGAACTTAAAGGCCTAAAGATTGATGATTTCCAGTGGTTGTCAAGTCTTGAAAATCTTGATAGTTTGGACATGTCTGGTGTGGATTTAAGTGAAGCAACAAACTGGA

>M6-PGSC0003DMG400010447-(rev)M6LRR-RLP_427

TGAAAATCTTGATAGTTTGGACATGTCTGGTGTGGATTTAAGTGAAGCAACAAACTGGACTGAGGTGATTAGCATGCTTCCTTTGGTTAATCTTCGTTTTTCGAATTGTAGTCTTCATTC

>M6-PGSC0003DMG400010447-(rev)M6LRR-RLP_488

GAGGTGATTAGCATGCTTCCTTTGGTTAATCTTCGTTTTTCGAATTGTAGTCTTCATTCAATACCTCCTTTGTTTGATCACAACAGTTCTGATCTTGAAAATCTTGATCTTTCTTTGAAT

>M6-PGSC0003DMG400010447-(rev)M6LRR-RLP_549

TACCTCCTTTGTTTGATCACAACAGTTCTGATCTTGAAAATCTTGATCTTTCTTTGAATAACTTTGGCTCTCCGATTCCCGGGTGGGTTTTCAGTTTTGGTAGTCTTGTTTCTCTTAAAT

>M6-PGSC0003DMG400010447-(rev)M6LRR-RLP_61

CCACCAAGAATTTGTCATCCAAATGTATGTATATATTATGCCATTTTTTGGCACATTCTTTTCTTATAATTTCACAAAAATGAGAGGGCTGTACTACATAGTACTTGTCCTGTTTACTAT

>M6-PGSC0003DMG400010447-(rev)M6LRR-RLP_610

CTTTGGCTCTCCGATTCCCGGGTGGGTTTTCAGTTTTGGTAGTCTTGTTTCTCTTAAATTCACTGGTAGCAATTTCACTGGTTCATTTCCTGAGGGTCCTTTTAACTTGACTTCTTTCAC

>M6-PGSC0003DMG400010447-(rev)M6LRR-RLP_671

ACTGGTAGCAATTTCACTGGTTCATTTCCTGAGGGTCCTTTTAACTTGACTTCTTTCACAACATTGAGAGCTTCTAGCAACTCCTTTGGTTCTGTTTTGCCTCAATGGTTATTTGATCTT

>M6-PGSC0003DMG400010447-(rev)M6LRR-RLP_732

CATTGAGAGCTTCTAGCAACTCCTTTGGTTCTGTTTTGCCTCAATGGTTATTTGATCTTAGTAATCTTGAATATCTTGATTTGAGTTTTAGTGGTCTTGAGGGTCCAATACCAAATGGAG

>M6-PGSC0003DMG400010447-(rev)M6LRR-RLP_793

TAATCTTGAATATCTTGATTTGAGTTTTAGTGGTCTTGAGGGTCCAATACCAAATGGAGTTGGTGATTTTACCAAGCTTACATATCTTAGTCTTGCTTCCAATAATCTGAATTCCACTAT

>M6-PGSC0003DMG400010447-(rev)M6LRR-RLP_854

GGTGATTTTACCAAGCTTACATATCTTAGTCTTGCTTCCAATAATCTGAATTCCACTATACCAGATTGGCTCTATGGATGCAAGAATCTTGAAACACTTGCCATGAAGGATAACAGTCTT

>M6-PGSC0003DMG400010447-(rev)M6LRR-RLP_915

CAGATTGGCTCTATGGATGCAAGAATCTTGAAACACTTGCCATGAAGGATAACAGTCTTGAGGGAACAGTTTCAGACTCGATTTCAAATCTGACATCTCTAACTAGTATCGATATGTCTG

>M6-PGSC0003DMG400010447-(rev)M6LRR-RLP_976

GGGAACAGTTTCAGACTCGATTTCAAATCTGACATCTCTAACTAGTATCGATATGTCTGAAAATTTTCTTTCTGGAAAGTTACCGAGTGTAATTGGAAAGCTATGGAAATTAGAAGATCT

>M6-PGSC0003DMG400010466-(rev)M6L-LecRK_0

CGTGTTTCCTTTGGATTCTCTGCTTCATTCACACCCCTCTTCTTCACCCCCTGTTTCCAGTTTTCTTCAAAAAAAATCACTATTTTCTAAGCTCAAAAAATAGCTAAAAATGAAATCTTT

>M6-PGSC0003DMG400010466-(rev)M6L-LecRK_1037

TGTTGAATTCAAAGATATTAATGGTAATCATGTTGGTTTGGATCTAAATTCAATGATTTCAACTGAAATTGGTAATTTGGATTCCATTGATATTGATCTCAAGAGTGGTGATTTGGTTAA

>M6-PGSC0003DMG400010466-(rev)M6L-LecRK_1098

ACTGAAATTGGTAATTTGGATTCCATTGATATTGATCTCAAGAGTGGTGATTTGGTTAATTGTTGGATTGAATATTCTGGTTCTAATGGAGAGATGGATATATTTGTGTCATATACTAAT

>M6-PGSC0003DMG400010466-(rev)M6L-LecRK_1159

GTTGGATTGAATATTCTGGTTCTAATGGAGAGATGGATATATTTGTGTCATATACTAATCTCAAGCCAAAAGAATCATTTTTATCAGTTAACATCAATCTTGCTGAGTATGTAAATGATT

>M6-PGSC0003DMG400010466-(rev)M6L-LecRK_122

GCAATTTCATCTTCATTGTGTAATGGGGTTTTGCTAAAATCTAGCAATGAAGTCAATTCTTGAAGAACTTTCCAGCAACATATTCCTGTAAAAGTCATCCTGTTGTAGTGTTGTTCTCTC

>M6-PGSC0003DMG400010466-(rev)M6L-LecRK_1220

CAAGCCAAAAGAATCATTTTTATCAGTTAACATCAATCTTGCTGAGTATGTAAATGATTTCATGTTTGTGGGGTTTTCTGGTTCAACTCAAGGGAGTACTGAGATTCATAATATTGTGTG

>M6-PGSC0003DMG400010466-(rev)M6L-LecRK_1281

ATGTTTGTGGGGTTTTCTGGTTCAACTCAAGGGAGTACTGAGATTCATAATATTGTGTGGTGGAGTTTCAGTTCATCATTTGATGCAACTCCAAACTCGGGGGAGGTGGTGGCTGAGCCA

>M6-PGSC0003DMG400010466-(rev)M6L-LecRK_1342

GGAGTTTCAGTTCATCATTTGATGCAACTCCAAACTCGGGGGAGGTGGTGGCTGAGCCACCACCGCCAACGACTAGTTTGATGAACCCAACGGCAGATTCTGTAACGTTGCCAACACCTT

>M6-PGSC0003DMG400010466-(rev)M6L-LecRK_1403

ACCGCCAACGACTAGTTTGATGAACCCAACGGCAGATTCTGTAACGTTGCCAACACCTTCTATGGCTCCATCAGAGTCTAATAGTAGCACAGGTGTATTGCAAGAAAAGAGTAGTAGGAA

>M6-PGSC0003DMG400010466-(rev)M6L-LecRK_1464

ATGGCTCCATCAGAGTCTAATAGTAGCACAGGTGTATTGCAAGAAAAGAGTAGTAGGAAATGTCATAGCAATTTTTGTAGACAAGGTCCTGGAGCTGTTGTTGGTGTGGTGACTGCTGGT

>M6-PGSC0003DMG400010466-(rev)M6L-LecRK_1525

GTCATAGCAATTTTTGTAGACAAGGTCCTGGAGCTGTTGTTGGTGTGGTGACTGCTGGTGCATTTTTTCTTGCATTTGCTACATTGGTACTTATTTGGTTATACTCCAAAAGATTCAAGA

>M6-PGSC0003DMG400010466-(rev)M6L-LecRK_1586

ATTTTTTCTTGCATTTGCTACATTGGTACTTATTTGGTTATACTCCAAAAGATTCAAGAGAGTGAAAAATTCTGAAATTGTGGCATCTGATATCATCAAAATGCCTAAGGAGTTCAGCTA

>M6-PGSC0003DMG400010466-(rev)M6L-LecRK_1647

GTGAAAAATTCTGAAATTGTGGCATCTGATATCATCAAAATGCCTAAGGAGTTCAGCTATAAAGAGCTTAAATTGGCCACAAAAGGTTTCATTCCAACGAGAATTATAGGCCGTGGTGCA

>M6-PGSC0003DMG400010466-(rev)M6L-LecRK_1708

AAGAGCTTAAATTGGCCACAAAAGGTTTCATTCCAACGAGAATTATAGGCCGTGGTGCATTTGGGACTGTTTACAAGGGCATTTTATCGGATACCGGTGGCATTGTTGCAGTCAAGAGAT

>M6-PGSC0003DMG400010466-(rev)M6L-LecRK_1769

TGGGACTGTTTACAAGGGCATTTTATCGGATACCGGTGGCATTGTTGCAGTCAAGAGATGTAGTCATAATGGACAAGGGAAAGCAGAGTTCTTATCTGAATTATCTATAATTGGAACCCT

>M6-PGSC0003DMG400010466-(rev)M6L-LecRK_183

GAAGAACTTTCCAGCAACATATTCCTGTAAAAGTCATCCTGTTGTAGTGTTGTTCTCTCCTGTCTACTTTCTTGCGTTTTCACTGTTTCTAAGCTAAAAAAAAAGCTAAAAAACAAATCT

>M6-PGSC0003DMG400010466-(rev)M6L-LecRK_1830

AGTCATAATGGACAAGGGAAAGCAGAGTTCTTATCTGAATTATCTATAATTGGAACCCTTAGGCATAGAAATCTTGTTAGGCTTCAAGGATGGTGCCATGAGAAAGGTGAAATTTTGTTG

>M6-PGSC0003DMG400010466-(rev)M6L-LecRK_1891

GGCATAGAAATCTTGTTAGGCTTCAAGGATGGTGCCATGAGAAAGGTGAAATTTTGTTGGTTTACGATCTTATGCCTAATGGGAGTCTTGATAAGGTACTATTTGAATCAAGAATGGTTC

>M6-PGSC0003DMG400010466-(rev)M6L-LecRK_1952

TTACGATCTTATGCCTAATGGGAGTCTTGATAAGGTACTATTTGAATCAAGAATGGTTCTTTCATGGTCACATAGGCGAAAAATCTTGATAGGTGTTGCTTCTGCCTTGGCATATTTGCA

>M6-PGSC0003DMG400010466-(rev)M6L-LecRK_2013

TCATGGTCACATAGGCGAAAAATCTTGATAGGTGTTGCTTCTGCCTTGGCATATTTGCATCAAGAATGCGAAAACCAGGTGATTCATAGGGACATTAAGAGTAGTAACATCATGTTGGAT

>M6-PGSC0003DMG400010466-(rev)M6L-LecRK_2074

AAGAATGCGAAAACCAGGTGATTCATAGGGACATTAAGAGTAGTAACATCATGTTGGATGAAGGGTTCAATGCAAGATTAGGAGATTTTGGATTAGCAAGGCAAGTTGAGCATGACAAGT

>M6-PGSC0003DMG400010466-(rev)M6L-LecRK_2135

AGGGTTCAATGCAAGATTAGGAGATTTTGGATTAGCAAGGCAAGTTGAGCATGACAAGTCCCCCGATGCAACGGTAGCAGCCGGGACAATGGGCTACTTGGCTCCAGAGTACTTGTTAAC

>M6-PGSC0003DMG400010466-(rev)M6L-LecRK_2196

CCCGATGCAACGGTAGCAGCCGGGACAATGGGCTACTTGGCTCCAGAGTACTTGTTAACCGGTAGGGCAACCGAAAAAACTGATGTTTTCAGCTATGGAGCAGTGGTTCTTGAAGTGGCT

>M6-PGSC0003DMG400010466-(rev)M6L-LecRK_2257

GTAGGGCAACCGAAAAAACTGATGTTTTCAGCTATGGAGCAGTGGTTCTTGAAGTGGCTTGTGGAAGGAGGCCAATTGAGAGGGAAACAACTAAAGTTGAGAAAGTTAGAGTGAATAGCA

>M6-PGSC0003DMG400010466-(rev)M6L-LecRK_2318

TGGAAGGAGGCCAATTGAGAGGGAAACAACTAAAGTTGAGAAAGTTAGAGTGAATAGCAACTTGGTTGAATGGGTATGGGGGGTGCATAGAGAAGGGAATTTACTAAATGCAGCTGATTC

>M6-PGSC0003DMG400010466-(rev)M6L-LecRK_2379

TTGGTTGAATGGGTATGGGGGGTGCATAGAGAAGGGAATTTACTAAATGCAGCTGATTCAAGACTTTGTGGTGAGTTTGATGAACAAGAAATGAGAAGGGTTCTAATGATTGGACTAGCT

>M6-PGSC0003DMG400010466-(rev)M6L-LecRK_244

GTCTACTTTCTTGCGTTTTCACTGTTTCTAAGCTAAAAAAAAAGCTAAAAAACAAATCTTTGTGCAATTTGATTTTCATTGTGTAATGGGGTTTTGCTAAATCAATTCTTGAAGAAATTT

>M6-PGSC0003DMG400010466-(rev)M6L-LecRK_2440

GACTTTGTGGTGAGTTTGATGAACAAGAAATGAGAAGGGTTCTAATGATTGGACTAGCTTGTTCACATCCCGACCCTACGGCTAGACCAACAATGAGAGGTGTAGTTCAAATGCTGGAAG

>M6-PGSC0003DMG400010466-(rev)M6L-LecRK_2501

TTCACATCCCGACCCTACGGCTAGACCAACAATGAGAGGTGTAGTTCAAATGCTGGAAGGTGAATCTGAAATCCCAATTGTCCCAAGAACTAGACCTTCTATGAGTTTTAGCACATCACA

>M6-PGSC0003DMG400010466-(rev)M6L-LecRK_2562

GAATCTGAAATCCCAATTGTCCCAAGAACTAGACCTTCTATGAGTTTTAGCACATCACATCTTCTAATGAGTTTGCAAGATAGTGTTTCTGACTTGAATGGTTTGATCACACTTTCCCCT

>M6-PGSC0003DMG400010466-(rev)M6L-LecRK_2623

TTCTAATGAGTTTGCAAGATAGTGTTTCTGACTTGAATGGTTTGATCACACTTTCCCCTTCATCATCCGAAAGTAGCTTCACCGGTGGTCGCAATGGCAATGGCATGGAGTTGGTCTAAA

>M6-PGSC0003DMG400010466-(rev)M6L-LecRK_2684

ATCATCCGAAAGTAGCTTCACCGGTGGTCGCAATGGCAATGGCATGGAGTTGGTCTAAATGCAAGACTAAAAAAGACTTGACTTTTCGAGTTAATGCACCTATAAGAGCCCAACAATCCC

>M6-PGSC0003DMG400010466-(rev)M6L-LecRK_2745

CAAGACTAAAAAAGACTTGACTTTTCGAGTTAATGCACCTATAAGAGCCCAACAATCCCACATATGGGGCCACCATCAAGGCGGAACAGACACAATGACTACTCTGATTGGGCGTGGACA

>M6-PGSC0003DMG400010466-(rev)M6L-LecRK_2806

ATATGGGGCCACCATCAAGGCGGAACAGACACAATGACTACTCTGATTGGGCGTGGACAAGGAAAAAGTGGCGTCTTTTTTGTTTTGTTCATAAATTCATTAATTTTATTCATAATATAT

>M6-PGSC0003DMG400010466-(rev)M6L-LecRK_2867

GAAAAAGTGGCGTCTTTTTTGTTTTGTTCATAAATTCATTAATTTTATTCATAATATATATGTACCAAATAAGCTTTAAGATTTAGTAATGTAATTTTCTCCTTTTCCTTTCATATACTA

>M6-PGSC0003DMG400010466-(rev)M6L-LecRK_2915

TCATAATATATATGTACCAAATAAGCTTTAAGATTTAGTAATGTAATTTTCTCCTTTTCCTTTCATATACTATAGTCCATTGAGTGCAAGATTAGAGATATTTAGTGGGGGAAAAATAAA

>M6-PGSC0003DMG400010466-(rev)M6L-LecRK_305

GTGCAATTTGATTTTCATTGTGTAATGGGGTTTTGCTAAATCAATTCTTGAAGAAATTTCCTTCTGTAAAAGCCATCATGTTGTAGTGTTGTTCTCTCCTTAATCCTCACTCTACTGTGT

>M6-PGSC0003DMG400010466-(rev)M6L-LecRK_366

TTCTGTAAAAGCCATCATGTTGTAGTGTTGTTCTCTCCTTAATCCTCACTCTACTGTGTCAGTTTTCTACTTTTTTGCATTTTCACTGTTTCTAATCCCAAAAAAAAGCCTACCCTTTTT

>M6-PGSC0003DMG400010466-(rev)M6L-LecRK_427

GTTTTCTACTTTTTTGCATTTTCACTGTTTCTAATCCCAAAAAAAAGCCTACCCTTTTTGTATCTTCACCAAATTCTTGTGATTTGGTGTGAAAAATCTTACCTTTTGTACACTTTGATC

>M6-PGSC0003DMG400010466-(rev)M6L-LecRK_488

ATCTTCACCAAATTCTTGTGATTTGGTGTGAAAAATCTTACCTTTTGTACACTTTGATCTTCATTCTCTGATGGGTTTTTTCTAGAATCTAGCAATGACATCAAATCTTGAAAAAAAATT

>M6-PGSC0003DMG400010466-(rev)M6L-LecRK_549

CATTCTCTGATGGGTTTTTTCTAGAATCTAGCAATGACATCAAATCTTGAAAAAAAATTCAGGGGCTTTCTTGTATGGCTCATCTTATTGTTCTGTTTCTGTAGTATTAGTTCAACATTT

>M6-PGSC0003DMG400010466-(rev)M6L-LecRK_61

TTTCTTCAAAAAAAATCACTATTTTCTAAGCTCAAAAAATAGCTAAAAATGAAATCTTTGTGCAATTTCATCTTCATTGTGTAATGGGGTTTTGCTAAAATCTAGCAATGAAGTCAATTC

>M6-PGSC0003DMG400010466-(rev)M6L-LecRK_610

GGGGCTTTCTTGTATGGCTCATCTTATTGTTCTGTTTCTGTAGTATTAGTTCAACATTTGGTAAAACTGATTTCGATTTTGAAAATTTAACACTAAGTAGTTTAAAACTTCTTGGTGATG

>M6-PGSC0003DMG400010466-(rev)M6L-LecRK_671

TAAAACTGATTTCGATTTTGAAAATTTAACACTAAGTAGTTTAAAACTTCTTGGTGATGCACACTTGAGTAACAACACTGTAAGGTTGACAGGTGATCTTGCTGTACCAAATTCCGGTGC

>M6-PGSC0003DMG400010466-(rev)M6L-LecRK_732

CACTTGAGTAACAACACTGTAAGGTTGACAGGTGATCTTGCTGTACCAAATTCCGGTGCCGGAAAAGTTCTGTATTCAAAACCAGTAAGATTCCGGCAACCGGGGTTTGATTTTCCGGCG

>M6-PGSC0003DMG400010466-(rev)M6L-LecRK_793

GAAAAGTTCTGTATTCAAAACCAGTAAGATTCCGGCAACCGGGGTTTGATTTTCCGGCGAGTTTCTCGACATTCTTTACATTTTCAGTTACTAATTTGAACCCATCATCGATTGGTGGTG

>M6-PGSC0003DMG400010466-(rev)M6L-LecRK_854

TTTCTCGACATTCTTTACATTTTCAGTTACTAATTTGAACCCATCATCGATTGGTGGTGGTCTTGCTTTTGTGCTTACGCCGGGTGATGAGTTAGTAGGTGATGCTGGTGGGTATATGGG

>M6-PGSC0003DMG400010466-(rev)M6L-LecRK_915

CTTGCTTTTGTGCTTACGCCGGGTGATGAGTTAGTAGGTGATGCTGGTGGGTATATGGGAATCTTGGATGCTAAAGGGACACAAATTGGTACAATTGCTGTGGAATTTGACACCCTTATG

>M6-PGSC0003DMG400010466-(rev)M6L-LecRK_976

TCTTGGATGCTAAAGGGACACAAATTGGTACAATTGCTGTGGAATTTGACACCCTTATGGATGTTGAATTCAAAGATATTAATGGTAATCATGTTGGTTTGGATCTAAATTCAATGATTT

>M6-PGSC0003DMG400010484-(rev)M6RLKWAX-EGF_0

GATTCATTCTATTTCCATGCATAGTTGTTGTTCTAATGACTGAGTTACAAAGAAGGCATTTGTCGATTTATGATGCAATTGAAAATTTCCTTCACGGTGAGAACAATTTCGCGCCCATTA

>M6-PGSC0003DMG400010484-(rev)M6RLKWAX-EGF_1053

TATCTAATTAGTTCTTCAATAAGCTCAATAGAGTCTGTGAAGTTGCTTGAGAGTTGTTCTGATTCTGCAAAAATAGATGTAGTTGTTGCTTGACATCACATATTGTGTACGTAATTCGTG

>M6-PGSC0003DMG400010484-(rev)M6RLKWAX-EGF_124

TTCCAACTCCAACATAAGAAAGATGACTAGGAACTTCAAAGAGAAACTAGGCCAAGGAGGTTATGGTTCAATATATAAAGGTAAGCTTCAAAGTGGACCTAATGTAGCAGTGAAGATATT

>M6-PGSC0003DMG400010484-(rev)M6RLKWAX-EGF_372

ACTTCATGCCTAACAGATCACTTGAAAAGTACATCACACCTCAAGATGGAGGAGCTGATCAACTGTTAAGTTGGGACAGAAAGTTTGAAATTGCTCTTGGAGTTGCTCGAGGTATTGAAT

>M6-PGSC0003DMG400010484-(rev)M6RLKWAX-EGF_558

ATGAGAATTTCATTCCAAAACTATCTGATTTTGGTCTTGCAAAATCATACCCGGCTGATAAGAGCATTGTTACTCTAACACCAGCCAGGGGAACAATTGGTTATGCAGCTCCAGAGCTGA

>M6-PGSC0003DMG400010484-(rev)M6RLKWAX-EGF_62

TCGATTTATGATGCAATTGAAAATTTCCTTCACGGTGAGAACAATTTCGCGCCCATTAGATATTCCAACTCCAACATAAGAAAGATGACTAGGAACTTCAAAGAGAAACTAGGCCAAGGA

>M6-PGSC0003DMG400010484-(rev)M6RLKWAX-EGF_620

AGCATTGTTACTCTAACACCAGCCAGGGGAACAATTGGTTATGCAGCTCCAGAGCTGATTAACAGAAGCATTGGTCCAATATATTATAAGGCATATGTCTATAGCTTATAAGGCATATGT

>M6-PGSC0003DMG400010484-(rev)M6RLKWAX-EGF_682

CAGAAGCATTGGTCCAATATATTATAAGGCATATGTCTATAGCTTATAAGGCATATGTCTATAGCTTTGGAATGTTACTCATTGATATCGTAGGCTTGAAAACAAACTCGGCAGCCAAAG

>M6-PGSC0003DMG400010484-(rev)M6RLKWAX-EGF_744

AGCTTTGGAATGTTACTCATTGATATCGTAGGCTTGAAAACAAACTCGGCAGCCAAAGAGGACATATCAAGCCAGTATTTCCCATATTGGATTTATGATTAACTCGACAAGGGAAAGGAG

>M6-PGSC0003DMG400010484-(rev)M6RLKWAX-EGF_806

CATATCAAGCCAGTATTTCCCATATTGGATTTATGATTAACTCGACAAGGGAAAGGAGATTGAAGTACTAGACGAGACACATGAAGACGTAAAGAAGCTGACTTTAGTTGCTTTGTGGTG

>M6-PGSC0003DMG400010484-(rev)M6RLKWAX-EGF_992

ACAAGCACTGCAAATGCCTCCCAGGCCCTCTGAATTGCAGGAACCATGTTCGTTGGAATTCTATCTAATTAGTTCTTCAATAAGCTCAATAGAGTCTGTGAAGTTGCTTGAGAGTTGTTC

>M6-PGSC0003DMG400010503-(rev)M6OtherRLKTM_0

ATGGTCCCATCATTCATTTCCCATTCAGACTCAGTCATCAACCAAAACATTGTGGCTATGATCCTGGCTTTGAACTACACTGTAACAATAAAAAAGACACCATCCTTGAGCTTCCCTCCT

>M6-PGSC0003DMG400010503-(rev)M6OtherRLKTM_65

GGCTTTGAACTACACTGTAACAATAAAAAAGACACCATCCTTGAGCTTCCCTCCTCTGTTCACTTAGTTGTTGAAAAGATTGATTATGTCTCACAGCAAATTCACCTTCATGATCCTGAA

>M6-PGSC0003DMG400010504-(rev)M6OtherRLKTM_0

CAAGTAGAAATGAAAGAGAGAGTCGAGTTAGACTTGAAAAGTTTTTGGAAGACTACAAGGCCATCAGACCGACACGATATGCTTATGCTGATATTAAGAAGATAACAGATGATTTTAATG

>M6-PGSC0003DMG400010504-(rev)M6OtherRLKTM_128

GGAGAAGGGAGTTATGGAACTGTATACAAAGGCAGACTTTCCAGTGAAATCTATGTTGCAGTAAAAGTCCTACACGATTCCAAGGGTAAAGGGGAAGAATTTATCAATGAAATAGGTACA

>M6-PGSC0003DMG400010504-(rev)M6OtherRLKTM_448

TGAGTATCTTCACCAGGGATGTGATCAGCAAATCCTACATTTCGATATCAAACCACAAAACATCCTTTTAGACCAGAACTTGAACCCCAAAATATGTGATTTCGGCCTAGCCAAACTATG

>M6-PGSC0003DMG400010504-(rev)M6OtherRLKTM_576

AAAAAAGTGCGGTCACTATGACAGCTGCTAGGGGAACCATTGGTTACATTGCACCAGAAGTGTTATCAAGAAACTTTGGTAAAGTTTCTCATAAGTCTGATATTTATAGCTTTGGGATGG

>M6-PGSC0003DMG400010504-(rev)M6OtherRLKTM_64

CAGACCGACACGATATGCTTATGCTGATATTAAGAAGATAACAGATGATTTTAATGAAAAGTTAGGAGAAGGGAGTTATGGAACTGTATACAAAGGCAGACTTTCCAGTGAAATCTATGT

>M6-PGSC0003DMG400010504-(rev)M6OtherRLKTM_640

ATCAAGAAACTTTGGTAAAGTTTCTCATAAGTCTGATATTTATAGCTTTGGGATGGTCTTGTTAGAAATGGTTGGTGGGAGGATAAAGATGGATCCTAAGATGAATAACCACAGCAAAGT

>M6-PGSC0003DMG400010504-(rev)M6OtherRLKTM_704

GAAATGGTTGGTGGGAGGATAAAGATGGATCCTAAGATGAATAACCACAGCAAAGTGAATTCTTTGGAATGGATATATAGACATTTGGAGAAAGGGGAAGAATTGAAGATTCGGATAGAG

>M6-PGSC0003DMG400010504-(rev)M6OtherRLKTM_768

TGGAATGGATATATAGACATTTGGAGAAAGGGGAAGAATTGAAGATTCGGATAGAGGAAGACGGAGATGGTACAATTGTGAGAAAGTTAGCTATTGTTGGACTTTGGTGCATACAGTGGC

>M6-PGSC0003DMG400010504-(rev)M6OtherRLKTM_822

AGGAAGACGGAGATGGTACAATTGTGAGAAAGTTAGCTATTGTTGGACTTTGGTGCATACAGTGGCATCCCAATGATAGGCCTTCAATAAAAGAAGTTATTCAGATGATGGAAGGAGATG

>M6-PGSC0003DMG400010519M6LRR-RLP_0

TCAGTTCAAGTCCTCGAGAAATTTCAAATCAACATGTCTTGGATGACCGTCTGCAGCTAAACCATAACGAAAATAGTACGGTGCTCCAAACAGATTTTTGCCATTTCTATAGCCTACTTC

>M6-PGSC0003DMG400010519M6LRR-RLP_1054

GCCAATCGTCTACATGGTCATTTTCCTGTGAACTTGGATACTTACCCAAACTTAACAGAAATAAACCTTGGACATAACCTGTTTTCGGGTGAGATTCCATAGGAATATTGGTCAAGACTC

>M6-PGSC0003DMG400010519M6LRR-RLP_1116

AAACCTTGGACATAACCTGTTTTCGGGTGAGATTCCATAGGAATATTGGTCAAGACTCAGGTTTTCATGGAGAAGTCTCAACTTGGAATATAATAACCTGGAAGGATCGGTTCCTCGAGA

>M6-PGSC0003DMG400010519M6LRR-RLP_1178

TTTCATGGAGAAGTCTCAACTTGGAATATAATAACCTGGAAGGATCGGTTCCTCGAGAGCTCAACAAGACTTTAGAAGGGGTTCAGGGAAGTTTTGAACACAACTGTCTCATTTGTCCAA

>M6-PGSC0003DMG400010519M6LRR-RLP_124

AAAGCCTTCACAAGTGGCTCCATATATGCTGTAATTTTCCTAAACATGTTAAAAGTAAGTTACTACAAGGAATTTCTGGATGAGTTATTGGCTGAAACAAACCAAATTTCTTGAATCATA

>M6-PGSC0003DMG400010519M6LRR-RLP_1302

CTTTCAGCTATGTCATGGACAGAGGACAGCTTCAGAATGTCATGGAAGAGACCATAATTAGCTGAGAATGTTTTGAAAATTACATTGCATTTTTTTTAATGTTTTCCCAGTTTCCAATAT

>M6-PGSC0003DMG400010519M6LRR-RLP_1364

TGAGAATGTTTTGAAAATTACATTGCATTTTTTTTAATGTTTTCCCAGTTTCCAATATAGTTTATAAGTTGCATTTTTTTTTTATGTTTTCCTAGTTTCCAATATAGTAGAAGTTCCTTA

>M6-PGSC0003DMG400010519M6LRR-RLP_1426

TATAAGTTGCATTTTTTTTTTATGTTTTCCTAGTTTCCAATATAGTAGAAGTTCCTTAAATTTGTCAATATATTTTATTTAGAAACTCGAACTACAACATGTTTCCTATTAAGCACCTGA

>M6-PGSC0003DMG400010519M6LRR-RLP_1488

TGTCAATATATTTTATTTAGAAACTCGAACTACAACATGTTTCCTATTAAGCACCTGACATGATGAACTGTGTTAGAGTTTTGCCCTGAATGTATGATATTATCAGTTATAGCAAAGTAT

>M6-PGSC0003DMG400010519M6LRR-RLP_1550

ATGAACTGTGTTAGAGTTTTGCCCTGAATGTATGATATTATCAGTTATAGCAAAGTATATTTGCATAACAATATTGAACAAGAGTATCAAAATTGGTTAAGTATACGACATGAGCAAGAG

>M6-PGSC0003DMG400010519M6LRR-RLP_1612

GCATAACAATATTGAACAAGAGTATCAAAATTGGTTAAGTATACGACATGAGCAAGAGTATCAAAATTAACTACTGAAGCAACATGGCAGCCAACGTTTTAAGTGCGAGCTTGTTTTAAG

>M6-PGSC0003DMG400010519M6LRR-RLP_1674

AAAATTAACTACTGAAGCAACATGGCAGCCAACGTTTTAAGTGCGAGCTTGTTTTAAGTTCAACTATAAGTGATCTTGTTTACTTTGTTGGTTCCAAGATTAGATAAATTCTCATAATTT

>M6-PGSC0003DMG400010519M6LRR-RLP_1736

ACTATAAGTGATCTTGTTTACTTTGTTGGTTCCAAGATTAGATAAATTCTCATAATTTATGGTGAAGCAAGGTTGCTGCAAAGAAAAGTGATTCACTCTCAACTACCTTACTAACATAAA

>M6-PGSC0003DMG400010519M6LRR-RLP_186

CTACAAGGAATTTCTGGATGAGTTATTGGCTGAAACAAACCAAATTTCTTGAATCATACGTTACTAGTATGGCGTTGTATGATAACACCATACTGGCAACATATGACATTTTGATCAATC

>M6-PGSC0003DMG400010519M6LRR-RLP_1922

AGCACCATACTAGTAGTAACTGGAAAAATCAAAACAAGCTATAATGAAAGTTGTCTTAATTGCTAAAGAAGCTCATGAAGACAATGACATTACAACTCATTTCTGTTGCCTAGTCTAATA

>M6-PGSC0003DMG400010519M6LRR-RLP_1984

CTAAAGAAGCTCATGAAGACAATGACATTACAACTCATTTCTGTTGCCTAGTCTAATAAAAACAATATTTGCCACCTAAAGATATATTTTAGCAGAATTGCATGTCTAAGAACTTAAGTA

>M6-PGSC0003DMG400010519M6LRR-RLP_2046

CAATATTTGCCACCTAAAGATATATTTTAGCAGAATTGCATGTCTAAGAACTTAAGTACCTTTTTCAGGGGAAGAAGCTGGGAACAGTCCTTTACAAGCATCTGTATTCCACAAACATGA

>M6-PGSC0003DMG400010519M6LRR-RLP_2108

TTTCAGGGGAAGAAGCTGGGAACAGTCCTTTACAAGCATCTGTATTCCACAAACATGAAATCAAAAGAGCAAGTCAGGCCCGAGACTACAAGCTGCTGTATTTAAAAGCATCTCAATGGA

>M6-PGSC0003DMG400010519M6LRR-RLP_2170

AAAAGAGCAAGTCAGGCCCGAGACTACAAGCTGCTGTATTTAAAAGCATCTCAATGGATTAGAGAAAAAACCTGACACCCACGAGTGTTAAGCTAGTAATATTATAGCTATTGACAGAAG

>M6-PGSC0003DMG400010519M6LRR-RLP_2232

AGAAAAAACCTGACACCCACGAGTGTTAAGCTAGTAATATTATAGCTATTGACAGAAGGCTTGAAAATTGCACTACTTGTGTAATGATATGTGTAATTCATTGTTATTTTATACTCGTTC

>M6-PGSC0003DMG400010519M6LRR-RLP_2294

GAAAATTGCACTACTTGTGTAATGATATGTGTAATTCATTGTTATTTTATACTCGTTCATGAGTTTTTGAAGTTGAGAAAGAGATTCTGAAAGTAACTAGTGACTGAAGATTTGAGAATA

>M6-PGSC0003DMG400010519M6LRR-RLP_2356

GTTTTTGAAGTTGAGAAAGAGATTCTGAAAGTAACTAGTGACTGAAGATTTGAGAATAGCAATTCACACATTTTCTAAGAGTGGGAATTGTGTACAAAAATGTACTATTTCTTCTAAAGT

>M6-PGSC0003DMG400010519M6LRR-RLP_2418

TTCACACATTTTCTAAGAGTGGGAATTGTGTACAAAAATGTACTATTTCTTCTAAAGTAACGTCTGCATTTTGGAAGAAAATCTCACCTCCTATATGTGTCCGAGTGTTCCTTTGCTGTC

>M6-PGSC0003DMG400010519M6LRR-RLP_248

ACTAGTATGGCGTTGTATGATAACACCATACTGGCAACATATGACATTTTGATCAATCCAAGGCAAGTTTTTTTTTCTTTTTTGTAATGGTATCAGGCTAACTTGTATGTGCTTCAACTG

>M6-PGSC0003DMG400010519M6LRR-RLP_2480

TCTGCATTTTGGAAGAAAATCTCACCTCCTATATGTGTCCGAGTGTTCCTTTGCTGTCAAATGGGGGCATCCCATACAGAAACTCAAAGCATAGGACACCAAGGCTCTAAATGTCCACAC

>M6-PGSC0003DMG400010519M6LRR-RLP_2604

ATCATGCTCCACACTTTCCACTGCAAGTTCAACAATTTACATATCAGAAAAAATGTAAAAACAGATAACAAAATGTTAAAAGCTAAACTCTCATACACGCCCATGTGAAGGTAAAAAGCA

>M6-PGSC0003DMG400010519M6LRR-RLP_2666

AGATAACAAAATGTTAAAAGCTAAACTCTCATACACGCCCATGTGAAGGTAAAAAGCAATGTATAACATTCACACATAGTCCGCCTTAAATCAAAGGTATGCACTGACCACCCAAAGTCT

>M6-PGSC0003DMG400010519M6LRR-RLP_2728

ATAACATTCACACATAGTCCGCCTTAAATCAAAGGTATGCACTGACCACCCAAAGTCTGCCGTTTTGAGTTCAACTTAAAAGAAGAAATATTAAAGCTGTCATTTTACTGCAATGCATTG

>M6-PGSC0003DMG400010519M6LRR-RLP_2790

TTTTGAGTTCAACTTAAAAGAAGAAATATTAAAGCTGTCATTTTACTGCAATGCATTGTATATACTGTTTAGCAATGCTAAGTTTATACCTCAATATCTTCTCTGTGTATTGTGTGATTC

>M6-PGSC0003DMG400010519M6LRR-RLP_2852

TACTGTTTAGCAATGCTAAGTTTATACCTCAATATCTTCTCTGTGTATTGTGTGATTCCCATGACAGTATATTGAAAGTTCGTAAGAATTGGGAAGAAGCAATGCAGAAGAGGCAACGAC

>M6-PGSC0003DMG400010519M6LRR-RLP_2914

GACAGTATATTGAAAGTTCGTAAGAATTGGGAAGAAGCAATGCAGAAGAGGCAACGACTAGTTACCAGGTCATAAAAGTAACCGTAAGTATTTCAACTTCACGATGAAGCTGATGTTCAA

>M6-PGSC0003DMG400010519M6LRR-RLP_2976

TACCAGGTCATAAAAGTAACCGTAAGTATTTCAACTTCACGATGAAGCTGATGTTCAACCAGGGACTGTTTTAGCTGGCTCTTGAACAACACTTTTAATGCGACGATATGATTGCTATGT

>M6-PGSC0003DMG400010519M6LRR-RLP_3038

GGACTGTTTTAGCTGGCTCTTGAACAACACTTTTAATGCGACGATATGATTGCTATGTTTCCAATAAACAACTAACCTTAGTATTGGAATGTGATTGTAGAATGTTAAGTCTATCGAATA

>M6-PGSC0003DMG400010519M6LRR-RLP_310

GCAAGTTTTTTTTTCTTTTTTGTAATGGTATCAGGCTAACTTGTATGTGCTTCAACTGGATAACAAGATTGTGCATTTTCAACACTAAGAATAATAAATAATCTAGAAGGTGATAGACAA

>M6-PGSC0003DMG400010519M6LRR-RLP_3100

AATAAACAACTAACCTTAGTATTGGAATGTGATTGTAGAATGTTAAGTCTATCGAATAGAATTCGAAAAATCTTTTAAGCCATGTCCACAAGATCCCACCAATCCATATCAAATATAACA

>M6-PGSC0003DMG400010519M6LRR-RLP_3162

TCGAAAAATCTTTTAAGCCATGTCCACAAGATCCCACCAATCCATATCAAATATAACAAATATGTTGAATGAGAGATTCTCACTCGCAATGCATAAATCCGAGGCATTGAAAAAGTAAGT

>M6-PGSC0003DMG400010519M6LRR-RLP_3224

TGTTGAATGAGAGATTCTCACTCGCAATGCATAAATCCGAGGCATTGAAAAAGTAAGTAGTTTTATCAGCCATTCTTAACATGTAAAGGTTTAAAGGACTTACATACATGACCGAACTTT

>M6-PGSC0003DMG400010519M6LRR-RLP_3286

TTATCAGCCATTCTTAACATGTAAAGGTTTAAAGGACTTACATACATGACCGAACTTTCCTCTTCCAAGAGGCTTCCCAATGTCGAAACCATGAAGTGTCCGCCTCTTTATATTAGCCTC

>M6-PGSC0003DMG400010519M6LRR-RLP_3348

TTCCAAGAGGCTTCCCAATGTCGAAACCATGAAGTGTCCGCCTCTTTATATTAGCCTCTGATGAAGTCTGCAGAAGTTGAAAATTCTGTAGTCCATTTAAAAGATCAGTTAATTAAAAAT

>M6-PGSC0003DMG400010519M6LRR-RLP_3410

GAAGTCTGCAGAAGTTGAAAATTCTGTAGTCCATTTAAAAGATCAGTTAATTAAAAATAACAATAAATTCTTGTTGTTGTTGTGTACAGTATAACAGATCAGTTAATTAAAAATAACAAT

>M6-PGSC0003DMG400010519M6LRR-RLP_3472

ATAAATTCTTGTTGTTGTTGTGTACAGTATAACAGATCAGTTAATTAAAAATAACAATAACTTGTTCTTTTAATTTTCATGATAACATATGGTTATTTCACATCACAAAGTTGTACGATC

>M6-PGSC0003DMG400010519M6LRR-RLP_3534

TGTTCTTTTAATTTTCATGATAACATATGGTTATTTCACATCACAAAGTTGTACGATCCAGCTGAGCTTGATTCCATTTAGTGCGGGAAGAATACAATTTGCTTTACATGCACTGGTAGA

>M6-PGSC0003DMG400010519M6LRR-RLP_3596

TGAGCTTGATTCCATTTAGTGCGGGAAGAATACAATTTGCTTTACATGCACTGGTAGAAGATTATCTCAATTAACTGCAACTTTGTTCATACTGTCCTCTTACAGGACTTCATGAACAAG

>M6-PGSC0003DMG400010519M6LRR-RLP_3658

TATCTCAATTAACTGCAACTTTGTTCATACTGTCCTCTTACAGGACTTCATGAACAAGCAATGAACCACAAGTGATCGCAACTAACATAAAAGCTTAAACTCAATCAACAAATTTTAACT

>M6-PGSC0003DMG400010519M6LRR-RLP_372

ACAAGATTGTGCATTTTCAACACTAAGAATAATAAATAATCTAGAAGGTGATAGACAAGAGGGGTTCTTGACATCGTCTATCGGGAACGTAACAGAACTTGTCTCTCTGAATCTAGGCAG

>M6-PGSC0003DMG400010519M6LRR-RLP_3720

GAACCACAAGTGATCGCAACTAACATAAAAGCTTAAACTCAATCAACAAATTTTAACTAACCTCAACATTCATCCTTTAGCAAGGTGGTACACTTGTGAACTATCAAGTACAACTCAACT

>M6-PGSC0003DMG400010519M6LRR-RLP_3782

TCAACATTCATCCTTTAGCAAGGTGGTACACTTGTGAACTATCAAGTACAACTCAACTCAATCTACATACTAATTTATTATAACTTCAAAAATAACCAAACACAGTGAGGTCAGATCATG

>M6-PGSC0003DMG400010519M6LRR-RLP_434

GGTTCTTGACATCGTCTATCGGGAACGTAACAGAACTTGTCTCTCTGAATCTAGGCAGGAACAAGTTTAGAGGACCAGTGCCAGAGAGTATTACTTATTTGAGGAAACTTACAAGTCTTC

>M6-PGSC0003DMG400010519M6LRR-RLP_496

AAGTTTAGAGGACCAGTGCCAGAGAGTATTACTTATTTGAGGAAACTTACAAGTCTTCAACTTTATGAGAATTTTTTCTCTAGTAGTCTTGTTAATGACATTGGTGTCCTTACAAAACTT

>M6-PGSC0003DMG400010519M6LRR-RLP_558

TTATGAGAATTTTTTCTCTAGTAGTCTTGTTAATGACATTGGTGTCCTTACAAAACTTGAGATTCTTGATGTCGCGAATAATAAGCTATCTGGTTCAATCCCTTCATCGACTACATCACT

>M6-PGSC0003DMG400010519M6LRR-RLP_62

CATAACGAAAATAGTACGGTGCTCCAAACAGATTTTTGCCATTTCTATAGCCTACTTCTTCCAAAGCCTTCACAAGTGGCTCCATATATGCTGTAATTTTCCTAAACATGTTAAAAGTAA

>M6-PGSC0003DMG400010519M6LRR-RLP_620

TTCTTGATGTCGCGAATAATAAGCTATCTGGTTCAATCCCTTCATCGACTACATCACTTAGAAGTTTGACGCGATTAGACCTGTCTAACAATGAATTGACAGGGAAAATTCCTCAACTCA

>M6-PGSC0003DMG400010519M6LRR-RLP_682

AGTTTGACGCGATTAGACCTGTCTAACAATGAATTGACAGGGAAAATTCCTCAACTCAACGGTTTATGGAAACTCCATTCATTTGACATTTCCAATAACCAAATTTAAGGGAATTTGCCA

>M6-PGSC0003DMG400010519M6LRR-RLP_744

TTTATGGAAACTCCATTCATTTGACATTTCCAATAACCAAATTTAAGGGAATTTGCCACAACTTCCATCGAAGATCAAGACATTGTCACTAAGCCACAACTTGCTATCAGGACACATTAC

>M6-PGSC0003DMG400010519M6LRR-RLP_806

TTCCATCGAAGATCAAGACATTGTCACTAAGCCACAACTTGCTATCAGGACACATTACCACAGTGCATAAACTAAGACAGCTAAGAACATTAGACCTTAGTGAAAACAGATTTTCAGGAG

>M6-PGSC0003DMG400010519M6LRR-RLP_868

GTGCATAAACTAAGACAGCTAAGAACATTAGACCTTAGTGAAAACAGATTTTCAGGAGGAATAAACAAAGGGATCTTCATGATACCTGCATCAAGTCACGTAAGCATTACAGTAAACCGG

>M6-PGSC0003DMG400010519M6LRR-RLP_930

AAACAAAGGGATCTTCATGATACCTGCATCAAGTCACGTAAGCATTACAGTAAACCGGTTCACAGTGTTTGAGGTGGTAGAATTCTCTGATAAAGAATCACAGCTTCACACGTTAGACGT

>M6-PGSC0003DMG400010519M6LRR-RLP_992

CAGTGTTTGAGGTGGTAGAATTCTCTGATAAAGAATCACAGCTTCACACGTTAGACGTAAATGCCAATCGTCTACATGGTCATTTTCCTGTGAACTTGGATACTTACCCAAACTTAACAG

>M6-PGSC0003DMG400010577-(rev)M6RLKmalectin_0

TTTGATGGTAGGCACACTATTCCGCTTCAAAATGTTTGTTTATCGCACTTGTAATTCTAACAATCAATTGAACATGTTTCTCTCTCAGTTATTCATTCTCTTATTGTGTATTCAAAAAAT

>M6-PGSC0003DMG400010577-(rev)M6RLKmalectin_1037

TGATAAATGAAGGCCAAAGGAATTTCACTATTGTCATAAACAATCAGAATGCTGAAGATGATGCTGAAGGGATCAAATGGAGTGGGGGGCATGGAATTTCTGTATACAGGGACTATGTTG

>M6-PGSC0003DMG400010577-(rev)M6RLKmalectin_1159

ATTATGGAGGGTGATAGAAGGGAAGGGAAGCGTAACCTTACTATAGTTTTGCAGCCAAACTTTGCTTCAATCGGTAAACATGCTAATGCCATCTTAAATGGGCTAGAAGTCTTCAAGATA

>M6-PGSC0003DMG400010577-(rev)M6RLKmalectin_122

TCATATCTACTGCAACTAACATTCATGAAAACGTTGCCATATCATGTGGTGCCTCTGCTAATTCCCCTGCACCAGATGGACGACTATGGATTGGAGATTCTACTTTCACTTCCTCATTTC

>M6-PGSC0003DMG400010577-(rev)M6RLKmalectin_1220

TTGCTTCAATCGGTAAACATGCTAATGCCATCTTAAATGGGCTAGAAGTCTTCAAGATAAGCAACCCTGACAACAATCTTGGTAGTGTGAGCCCAGTGCATCCTGTGACAAGTTCAACAC

>M6-PGSC0003DMG400010577-(rev)M6RLKmalectin_1281

CAACCCTGACAACAATCTTGGTAGTGTGAGCCCAGTGCATCCTGTGACAAGTTCAACACCAGAAAAATCAGAGGAAACTGTGCTGTTCTATACAAAGAATCAAATTGCAACTGCACTGAT

>M6-PGSC0003DMG400010577-(rev)M6RLKmalectin_1342

GAAAAATCAGAGGAAACTGTGCTGTTCTATACAAAGAATCAAATTGCAACTGCACTGATATTTATAGTCACTTTGATTAACGTTGCTGTTTATTACATTAGGCGTAATGCAGAAATAAAG

>M6-PGSC0003DMG400010577-(rev)M6RLKmalectin_1403

TTATAGTCACTTTGATTAACGTTGCTGTTTATTACATTAGGCGTAATGCAGAAATAAAGTCTGGCAAGACATACAATGGAATATCTTCAGGAGTGCATCAGTGTCGTCAATTTTCACTGG

>M6-PGSC0003DMG400010577-(rev)M6RLKmalectin_1464

TGGCAAGACATACAATGGAATATCTTCAGGAGTGCATCAGTGTCGTCAATTTTCACTGGATGAGATGGAAAGATCAACGAACAATTTTGATCCTCAGCTTGTCATTGGTAGCGGTGGATA

>M6-PGSC0003DMG400010577-(rev)M6RLKmalectin_1586

GTACAGTATACAAAGGGGATATTGATGGTGGAGAGACCACTGTAGCAGTTAAGCGATCGAAACCAGGATCTAGCCAGGGGGAGAAAGAGTTTTGGACGGAAATCAATATGTTATCTACAC

>M6-PGSC0003DMG400010577-(rev)M6RLKmalectin_1647

ACCAGGATCTAGCCAGGGGGAGAAAGAGTTTTGGACGGAAATCAATATGTTATCTACACACCGCCATGAGAACCTTCTCTCCCTAATTGGTTACTGCAATGAAGGTCAAGAGATGTTATT

>M6-PGSC0003DMG400010577-(rev)M6RLKmalectin_1708

CGCCATGAGAACCTTCTCTCCCTAATTGGTTACTGCAATGAAGGTCAAGAGATGTTATTAGTTTATGATTATATGCCTCGAGGATCACTTGCTGACAAACTGTACAAAATGGACAAAAAT

>M6-PGSC0003DMG400010577-(rev)M6RLKmalectin_1769

TTTATGATTATATGCCTCGAGGATCACTTGCTGACAAACTGTACAAAATGGACAAAAATAGCTCATCTCTCTCTTGGGAACGGAGACTCAAGATTGCTATGGGTGCTGCACGTGGACTGG

>M6-PGSC0003DMG400010577-(rev)M6RLKmalectin_183

TTCCCCTGCACCAGATGGACGACTATGGATTGGAGATTCTACTTTCACTTCCTCATTTCTACACTTAACTGGAAAATCTATTAATTCAAGAGTACCTCACCAAGCTGCCTTGTCCGATCC

>M6-PGSC0003DMG400010577-(rev)M6RLKmalectin_1830

CTCATCTCTCTCTTGGGAACGGAGACTCAAGATTGCTATGGGTGCTGCACGTGGACTGGATTTCCTTCATACATCTCAAAATAGGGTGATACATCGTGATATCAAAAGCTCAAACATTCT

>M6-PGSC0003DMG400010577-(rev)M6RLKmalectin_1891

TTCCTTCATACATCTCAAAATAGGGTGATACATCGTGATATCAAAAGCTCAAACATTCTATTGGATGAAAACTGGGAAAGTAAGATCTCAGATTTTGGGTTGTCCAAAATGGGGCCTGGA

>M6-PGSC0003DMG400010577-(rev)M6RLKmalectin_2013

TGAATCAGCTACTCATGTTAGTACACAAGTCAAAGGCACATTTGGGTACCTCGATCCTGAGTACTTCCTGACTAATAGATTGACATGGAAAACTGATGTATATGCTTTTGGAGTAGTACT

>M6-PGSC0003DMG400010577-(rev)M6RLKmalectin_2074

TACTTCCTGACTAATAGATTGACATGGAAAACTGATGTATATGCTTTTGGAGTAGTACTATTTGAATTGCTCTCAGGAAGACCTGCAGTCGATATGAGCCTGACTGAGGAACAACATGGA

>M6-PGSC0003DMG400010577-(rev)M6RLKmalectin_2135

TTGAATTGCTCTCAGGAAGACCTGCAGTCGATATGAGCCTGACTGAGGAACAACATGGACTTGTAGCATGGGCCAAACAATGCATCAAGGAAGGAGAAATTAATAAGCTTATTGATCAGA

>M6-PGSC0003DMG400010577-(rev)M6RLKmalectin_2196

TGTAGCATGGGCCAAACAATGCATCAAGGAAGGAGAAATTAATAAGCTTATTGATCAGAATCTGTTGGGGTCCATCTCATCAACTTGTCTAAAGGCGTTTATAGGAATTTCTGCTAAATG

>M6-PGSC0003DMG400010577-(rev)M6RLKmalectin_2257

CTGTTGGGGTCCATCTCATCAACTTGTCTAAAGGCGTTTATAGGAATTTCTGCTAAATGCTTCGATGGTCGTCCGCAAGAACGGCCTGCGATGTCTGAAGTGGTGAAAAGCCTGGAATTA

>M6-PGSC0003DMG400010577-(rev)M6RLKmalectin_2318

TCGATGGTCGTCCGCAAGAACGGCCTGCGATGTCTGAAGTGGTGAAAAGCCTGGAATTAGCATTAGTATTTCAGCAGAACGAAGGTGAAGGTATCATTTCATTTGATGACACATCAACGT

>M6-PGSC0003DMG400010577-(rev)M6RLKmalectin_2379

ATTAGTATTTCAGCAGAACGAAGGTGAAGGTATCATTTCATTTGATGACACATCAACGTCATCTCAGTTACAGATTGAAGGAGAGAGGGCATCCATTAAAGAGGATTGTAATGGCGGTGA

>M6-PGSC0003DMG400010577-(rev)M6RLKmalectin_244

CACTTAACTGGAAAATCTATTAATTCAAGAGTACCTCACCAAGCTGCCTTGTCCGATCCAGTTCCTTATAACTCTGCTCGAACGTCTCGCCATCAGTTTACCTATCAATTTTCAATGAAA

>M6-PGSC0003DMG400010577-(rev)M6RLKmalectin_2440

TCTCAGTTACAGATTGAAGGAGAGAGGGCATCCATTAAAGAGGATTGTAATGGCGGTGATACAACAGAAAGGAGTGCCATCTCAAGGGAAAAGGTAAAATCTGAGGATAAAAGTCCCTAC

>M6-PGSC0003DMG400010577-(rev)M6RLKmalectin_2501

CAACAGAAAGGAGTGCCATCTCAAGGGAAAAGGTAAAATCTGAGGATAAAAGTCCCTACACTGCTTCACCAAGATGGTGGGATGTTCGTAGCCATTTCAGAAAAGCACCACCAAAGCCAG

>M6-PGSC0003DMG400010577-(rev)M6RLKmalectin_2562

TGCTTCACCAAGATGGTGGGATGTTCGTAGCCATTTCAGAAAAGCACCACCAAAGCCAGGGAATTTTGTTTATCCAGATAGCCAAATATCACAGCACCCAAATTTAAGGATATTTTCTTT

>M6-PGSC0003DMG400010577-(rev)M6RLKmalectin_2623

AATTTTGTTTATCCAGATAGCCAAATATCACAGCACCCAAATTTAAGGATATTTTCTTTCTCTGAACTCAAGGCCGCCACCAGAAAATTTAGCAAGGACACGGTGCTGGGAGAAGGTGGT

>M6-PGSC0003DMG400010577-(rev)M6RLKmalectin_2684

CTGAACTCAAGGCCGCCACCAGAAAATTTAGCAAGGACACGGTGCTGGGAGAAGGTGGTTTTGGTAAAGTTTACAAGGGTTACCTTGCTGAGTCGCTTTCTTCCAAGAGTGGCCGGACAG

>M6-PGSC0003DMG400010577-(rev)M6RLKmalectin_2745

TGGTAAAGTTTACAAGGGTTACCTTGCTGAGTCGCTTTCTTCCAAGAGTGGCCGGACAGTAATTGCTGTCAAGCAATTGAGTTCTGAAAGATTTCAAGAGTGGCAGGTGATTCCTCTGTG

>M6-PGSC0003DMG400010577-(rev)M6RLKmalectin_2806

ATTGCTGTCAAGCAATTGAGTTCTGAAAGATTTCAAGAGTGGCAGGTGATTCCTCTGTGCTAAATATGTGAACTTCTAGTTTAAGTTTTCTAATTGAAGCTATTTCTAGTTTATGCCTGG

>M6-PGSC0003DMG400010577-(rev)M6RLKmalectin_2867

AAATATGTGAACTTCTAGTTTAAGTTTTCTAATTGAAGCTATTTCTAGTTTATGCCTGGCACTAGCTTGCTTCATTTTAATGATGCTGCAATTGCTCTCTTCTTTATTGTCTGTCTGCGA

>M6-PGSC0003DMG400010577-(rev)M6RLKmalectin_2928

CTAGCTTGCTTCATTTTAATGATGCTGCAATTGCTCTCTTCTTTATTGTCTGTCTGCGAGAAATATAGATTTACGTAGAAGCACTTGAAATTGATTTATTACTTAGTAGTAACACATTCT

>M6-PGSC0003DMG400010577-(rev)M6RLKmalectin_2989

AATATAGATTTACGTAGAAGCACTTGAAATTGATTTATTACTTAGTAGTAACACATTCTAGGTAAATGTTGGTTATATATCTCTTGCAAATTTCATTTCATTTGTTAATTATGCAGTCTG

>M6-PGSC0003DMG400010577-(rev)M6RLKmalectin_3050

GTAAATGTTGGTTATATATCTCTTGCAAATTTCATTTCATTTGTTAATTATGCAGTCTGAGGTGAGCATTCTTGGAAGACTTTCTCATCCTAACCTCATCAAACTCTTGGGTTACTGCCA

>M6-PGSC0003DMG400010577-(rev)M6RLKmalectin_3111

GTGAGCATTCTTGGAAGACTTTCTCATCCTAACCTCATCAAACTCTTGGGTTACTGCCAGGAAGATAAAAAACTACTGCTTGTCTATGAGTTTATGCCAAAGGGCAGCTTGAACAACCAT

>M6-PGSC0003DMG400010577-(rev)M6RLKmalectin_3172

AAGATAAAAAACTACTGCTTGTCTATGAGTTTATGCCAAAGGGCAGCTTGAACAACCATCTTTTTGGAAGTAAGAGATTATCTATACGTTCCGTCAATAATATATAACATATTGTAGATC

>M6-PGSC0003DMG400010577-(rev)M6RLKmalectin_3233

TTTTGGAAGTAAGAGATTATCTATACGTTCCGTCAATAATATATAACATATTGTAGATCAAAGCAGCAAGGAATGTGCCTAGCTACTTCATGCTCCACCTAAACATATTTGTTATCTTTG

>M6-PGSC0003DMG400010577-(rev)M6RLKmalectin_3294

AGCAGCAAGGAATGTGCCTAGCTACTTCATGCTCCACCTAAACATATTTGTTATCTTTGTTTGCTTAGGGCACTCTGTTGCTTTGTCACTTACATGGAATGTAAGGGTTCAAATTATGAT

>M6-PGSC0003DMG400010577-(rev)M6RLKmalectin_3355

TGCTTAGGGCACTCTGTTGCTTTGTCACTTACATGGAATGTAAGGGTTCAAATTATGATTGGCGCAGCTCGAGGCTTGGCATTCCTACATGCATCAGAAAAGCAAGTCATTTACAGAGAC

>M6-PGSC0003DMG400010577-(rev)M6RLKmalectin_3416

GCGCAGCTCGAGGCTTGGCATTCCTACATGCATCAGAAAAGCAAGTCATTTACAGAGACTTCAAAGCCTCAAATATATTACTTGATGGCGTATGTTCTCTTTACTTCTCCCTAGGAATAT

>M6-PGSC0003DMG400010577-(rev)M6RLKmalectin_3477

CAAAGCCTCAAATATATTACTTGATGGCGTATGTTCTCTTTACTTCTCCCTAGGAATATACCTTTATTAGCGCAAGTAATAAAGGTATCACAAGCCCTTTTCTTACTGGGCCCCTTTTTG

>M6-PGSC0003DMG400010577-(rev)M6RLKmalectin_3538

CTTTATTAGCGCAAGTAATAAAGGTATCACAAGCCCTTTTCTTACTGGGCCCCTTTTTGGTTTACACACACTGTTTTGCCTTAAGTTTTGGACCTATTTATCACAGAGAGACAACACCAG

>M6-PGSC0003DMG400010577-(rev)M6RLKmalectin_3599

TTACACACACTGTTTTGCCTTAAGTTTTGGACCTATTTATCACAGAGAGACAACACCAGACGTATAACAGTGAAAACATTAGAGAAAGGTTGTTATTAATATAACTGTTCCATGACATTC

>M6-PGSC0003DMG400010577-(rev)M6RLKmalectin_366

GGGACACAAGTTCATACGCCTGCACTTCAAACCAGCTTCATACAAAGGTTTCATCAAGTCCAAACCCATTTTTACTGTCAAAACTGATCAGCATACCCTGCTCAGTGACTTCATGCCTAC

>M6-PGSC0003DMG400010577-(rev)M6RLKmalectin_3660

GTATAACAGTGAAAACATTAGAGAAAGGTTGTTATTAATATAACTGTTCCATGACATTCTTTATTTCTTTAATGTTTCAGTCTTACAATGCGAAGATAGCAGATTTTGGCATTGCAAAAC

>M6-PGSC0003DMG400010577-(rev)M6RLKmalectin_3721

TATTTCTTTAATGTTTCAGTCTTACAATGCGAAGATAGCAGATTTTGGCATTGCAAAACAAGGTATTTCGGCTAGTCAATCACATGTGACAACACAGATACTTGGAACGTACGGTTATGC

>M6-PGSC0003DMG400010577-(rev)M6RLKmalectin_3782

GGTATTTCGGCTAGTCAATCACATGTGACAACACAGATACTTGGAACGTACGGTTATGCTGCTCCTGAGTATATTGCAACAGGTAAACTTGGAGTCTGTTGGGCTTCTGCACTTGGAAAT

>M6-PGSC0003DMG400010577-(rev)M6RLKmalectin_3843

CTCCTGAGTATATTGCAACAGGTAAACTTGGAGTCTGTTGGGCTTCTGCACTTGGAAATTCTACTTCTCTTGAATCACATGCATCTTGTGCTTACAAGTTTAAACATATAGTCAAGTGTA

>M6-PGSC0003DMG400010577-(rev)M6RLKmalectin_3904

TACTTCTCTTGAATCACATGCATCTTGTGCTTACAAGTTTAAACATATAGTCAAGTGTATTATTTGTTTTCTACAGGATTTTTGTACTTGAAGAGCGATGTCTATGCTTTTGGTGTCTTT

>M6-PGSC0003DMG400010577-(rev)M6RLKmalectin_3965

ATTTGTTTTCTACAGGATTTTTGTACTTGAAGAGCGATGTCTATGCTTTTGGTGTCTTTTTAGTGGAAATGCTAACAGGTCTACGAGTAATAGACTTAAACCGTCCAAGCAACCAGCATA

>M6-PGSC0003DMG400010577-(rev)M6RLKmalectin_4026

AGTGGAAATGCTAACAGGTCTACGAGTAATAGACTTAAACCGTCCAAGCAACCAGCATAATCTAATTGACTGGATTAAGCCTCATTTATCTGACAGAAGGAAGTTGAAGGACAGAATCGA

>M6-PGSC0003DMG400010577-(rev)M6RLKmalectin_4087

CTAATTGACTGGATTAAGCCTCATTTATCTGACAGAAGGAAGTTGAAGGACAGAATCGATTCACGGCTGGGAGGAAAATATCCATCCAGAGCTGCAGTACAAATAGCACAACTGGCACTA

>M6-PGSC0003DMG400010577-(rev)M6RLKmalectin_4148

CACGGCTGGGAGGAAAATATCCATCCAGAGCTGCAGTACAAATAGCACAACTGGCACTATCATGTCTTGGAAATGAACCTAAAAGCAGACCATCGATGAAGGAAATAGTGGAGAAACTAG

>M6-PGSC0003DMG400010577-(rev)M6RLKmalectin_4209

ATGTCTTGGAAATGAACCTAAAAGCAGACCATCGATGAAGGAAATAGTGGAGAAACTAGAACAGATTGAAGTTACCAATGAAAGATCCAAGGACCATGCTTCTCCTTCTAATCCAAGGAA

>M6-PGSC0003DMG400010577-(rev)M6RLKmalectin_427

AAACCCATTTTTACTGTCAAAACTGATCAGCATACCCTGCTCAGTGACTTCATGCCTACCCTTGCTGCAGGCATCAACTATTTCAAAAAGGAATTTTGCATCAATGTAAAAGAAAGTGAA

>M6-PGSC0003DMG400010577-(rev)M6RLKmalectin_4270

CAGATTGAAGTTACCAATGAAAGATCCAAGGACCATGCTTCTCCTTCTAATCCAAGGAACCGGGCCTATCCACTCCTTAGAAGAGCAGAGTGAATATGATACTACTTTTTACAATATGAA

>M6-PGSC0003DMG400010577-(rev)M6RLKmalectin_4331

GGGCCTATCCACTCCTTAGAAGAGCAGAGTGAATATGATACTACTTTTTACAATATGAAGAAGCTACTTGTTAAAATTTTCAACTGTGAGTACTGTCACTTTAGGCGAGATCGTGCCAGT

>M6-PGSC0003DMG400010577-(rev)M6RLKmalectin_4697

TCATTCCTTTGTCTAGAAAATCCAAATCTTGGCAGAATTGTTTATCGGAGAATATTTGTGTCCCCATTTCCTTCTATATTACTATAGCTCCTGTGGATCCTGGAAAGGTTTCATGACTGC

>M6-PGSC0003DMG400010577-(rev)M6RLKmalectin_4758

CCCCATTTCCTTCTATATTACTATAGCTCCTGTGGATCCTGGAAAGGTTTCATGACTGCTATAAATCAGTAATGATGCCTTCTATGTAGTTCTATTAGTCTCTCAAAAGAATTCTGAGAG

>M6-PGSC0003DMG400010577-(rev)M6RLKmalectin_4819

TAAATCAGTAATGATGCCTTCTATGTAGTTCTATTAGTCTCTCAAAAGAATTCTGAGAGTTAACAGACATAAAATTCTAGAATATGATTTTTTTTTGCAATCATTGATCAATGGATCGGG

>M6-PGSC0003DMG400010577-(rev)M6RLKmalectin_488

TTGCTGCAGGCATCAACTATTTCAAAAAGGAATTTTGCATCAATGTAAAAGAAAGTGAAACATTAAGCATAACATTCATACCATCTCGAAAATCAGGCTTTTCAGAAGACACTTATGCTT

>M6-PGSC0003DMG400010577-(rev)M6RLKmalectin_4880

AACAGACATAAAATTCTAGAATATGATTTTTTTTTGCAATCATTGATCAATGGATCGGGTCTTATCTTTACTGCTCCTTGTTCTAACCTGAAGAGGGATCACAGATTTGCCTTTTCTGAT

>M6-PGSC0003DMG400010577-(rev)M6RLKmalectin_4941

TTATCTTTACTGCTCCTTGTTCTAACCTGAAGAGGGATCACAGATTTGCCTTTTCTGATCCTACTCTTCAGGTTATGTATGTGTCTAGGGCACAAAGTCCCCATCTTTGTTGATTTGATG

>M6-PGSC0003DMG400010577-(rev)M6RLKmalectin_5002

TACTCTTCAGGTTATGTATGTGTCTAGGGCACAAAGTCCCCATCTTTGTTGATTTGATGAATATTCTCCTCCACACTATCATCCTTCTCCACATCCTTTCTCATTTGTATATGTTTCTCT

>M6-PGSC0003DMG400010577-(rev)M6RLKmalectin_5063

TATTCTCCTCCACACTATCATCCTTCTCCACATCCTTTCTCATTTGTATATGTTTCTCTGTTGCTACTACATTGTTTTTGTGGTATCTCCTCTTGAAGTTTTGTTCTTCCGTGTCCTTCG

>M6-PGSC0003DMG400010577-(rev)M6RLKmalectin_5185

TATAGCTGTGGGTGTTCAACATAACATTCCTCCTAGTTATGTCTCTAAATCATACACGTTTCACAATATTTGGGACATTGTCAACTTGATTCAGATACACTTCTCAACTATCTCTCCATT

>M6-PGSC0003DMG400010577-(rev)M6RLKmalectin_5246

CACAATATTTGGGACATTGTCAACTTGATTCAGATACACTTCTCAACTATCTCTCCATTTTGTTGTTTCATCCTATGTTGATGCGCATTGGAAACTCGCCCACCAGATCTACTTCAACTT

>M6-PGSC0003DMG400010577-(rev)M6RLKmalectin_5307

GTTGTTTCATCCTATGTTGATGCGCATTGGAAACTCGCCCACCAGATCTACTTCAACTTCCAATCTTGCATAGCTTGGTCTGTTCTGATTCTTGATGGTCATATTATCCTGTATGTAATG

>M6-PGSC0003DMG400010577-(rev)M6RLKmalectin_5368

AATCTTGCATAGCTTGGTCTGTTCTGATTCTTGATGGTCATATTATCCTGTATGTAATGAATTATAAATCATTGTTGCTAGAGAAAACATTGTTTCCTTCCCAAAAAATATTTGGTGGAA

>M6-PGSC0003DMG400010577-(rev)M6RLKmalectin_549

ATTAAGCATAACATTCATACCATCTCGAAAATCAGGCTTTTCAGAAGACACTTATGCTTTTGTTAATGCAATCGAGATTGTTTCCATATCGTCTGGTCTGTATTTCACACCAGATGGCGA

>M6-PGSC0003DMG400010577-(rev)M6RLKmalectin_61

AATCAATTGAACATGTTTCTCTCTCAGTTATTCATTCTCTTATTGTGTATTCAAAAAATCATCATATCTACTGCAACTAACATTCATGAAAACGTTGCCATATCATGTGGTGCCTCTGCT

>M6-PGSC0003DMG400010577-(rev)M6RLKmalectin_610

GTTAATGCAATCGAGATTGTTTCCATATCGTCTGGTCTGTATTTCACACCAGATGGCGATCAGGGAGTCCCTGTTGTTGGACGAAATTACAGATTCTACATTGACAATAGCACAGCACTG

>M6-PGSC0003DMG400010577-(rev)M6RLKmalectin_732

GACAATTCAGCGAATAAATGTTGGCGGAAACTCTATTTCATCCTTGGAAGATGCAACCATGTTTCGGGACTGGGAAGATGACACTAATTACCTGATTCAAGTTGGTGCCTTTTCGATCAA

>M6-PGSC0003DMG400010577-(rev)M6RLKmalectin_793

TTTCGGGACTGGGAAGATGACACTAATTACCTGATTCAAGTTGGTGCCTTTTCGATCAACAGAGCCGTTGATATCAGATATGCATCCTCAGCAACTCACATTGCACCAAAAGAGGTTTAC

>M6-PGSC0003DMG400010577-(rev)M6RLKmalectin_854

GAGCCGTTGATATCAGATATGCATCCTCAGCAACTCACATTGCACCAAAAGAGGTTTACCAAACGGCCAGGTCAGTGGGTGTTCACTGCCATTCGAATTTCTGTAATCTCACATGGAACA

>M6-PGSC0003DMG400010577-(rev)M6RLKmalectin_976

CCACTTGATTTGGGATTCAGATACCTTGTTAGGCTTCACTTTTGTGAAATTGAACCTACAATGATAAATGAAGGCCAAAGGAATTTCACTATTGTCATAAACAATCAGAATGCTGAAGAT

>M6-PGSC0003DMG400010700-(rev)M6LRR-RLP_0

ATGGGCATTGAAAAACTTGCATTGTTTCTGCTATACGCGTTTCTGTGTCAACTTGTTTTCTCCTCATCTTTACCTCATTTGTGCCCCAATGATCAAGCCCTTGCTCTTCTTCAATTCAAA

>M6-PGSC0003DMG400010700-(rev)M6LRR-RLP_1054

CGCAACTTGTTGAGCTAGATATTTCATTTAATTCCCTAACAGGTTCAATTCCTTCTAATGTAAGTGGTATGCAAAACCTACAATGTCTCAACTTGTCATCAAACCACTTGAATGGGACTA

>M6-PGSC0003DMG400010700-(rev)M6LRR-RLP_1116

AGTGGTATGCAAAACCTACAATGTCTCAACTTGTCATCAAACCACTTGAATGGGACTATACCTTCCTGGATATTCTCCCTCCCTTCACTTGAATGGTTAGACTTGAGTGATAACCATTTC

>M6-PGSC0003DMG400010700-(rev)M6LRR-RLP_1178

TTCCTGGATATTCTCCCTCCCTTCACTTGAATGGTTAGACTTGAGTGATAACCATTTCAGTGGAAACATTCAGGAGTTCAAGTCTAAAACATTGTATTCTGTTTCTCTAAAACAAAATCA

>M6-PGSC0003DMG400010700-(rev)M6LRR-RLP_124

CGTTTACTATAAATCTTGATGCTTCTTATAGTGGTTATTATCCTAAAACAGTTAAATGGAATAAGAGCACAGATTGTTGTTCATGGGTTGGAGTTAAGTGTTACGAGATGACAGGACAAG

>M6-PGSC0003DMG400010700-(rev)M6LRR-RLP_1240

GAAACATTCAGGAGTTCAAGTCTAAAACATTGTATTCTGTTTCTCTAAAACAAAATCAGCTGCAAGGTCCTATTCCAAAGTCACTCTTAGATCTGCAGTACCTAGAAATTCTTCTCCTTT

>M6-PGSC0003DMG400010700-(rev)M6LRR-RLP_1302

CAAGGTCCTATTCCAAAGTCACTCTTAGATCTGCAGTACCTAGAAATTCTTCTCCTTTCGCACAATAATCTCAATGGAGAGATTGCTTCACCCATATGCAATCTGAAAACACTACGAGTG

>M6-PGSC0003DMG400010700-(rev)M6LRR-RLP_1488

TGGGTTTTGGATTTAAGCAACAATAGTCTTAGTGGGACAATTAATACTACTTTTAGTATAGGAAACTAATTTATAGTCATTAAATTTGATGGGAATAAGCTAGAGGGGAAAGTCCCGCAA

>M6-PGSC0003DMG400010700-(rev)M6LRR-RLP_1550

AAACTAATTTATAGTCATTAAATTTGATGGGAATAAGCTAGAGGGGAAAGTCCCGCAATCATTGATCAATTGCACATATTTGGAAGTTCTTGATTTAGGTAACAATGAGTTGAATGACAC

>M6-PGSC0003DMG400010700-(rev)M6LRR-RLP_1612

TGATCAATTGCACATATTTGGAAGTTCTTGATTTAGGTAACAATGAGTTGAATGACACATTTCCTAAATGGTTGAGAGCCCTACCGTATCTGAAGATTCTAAGACTGAGATCAAATAAGT

>M6-PGSC0003DMG400010700-(rev)M6LRR-RLP_1674

CCTAAATGGTTGAGAGCCCTACCGTATCTGAAGATTCTAAGACTGAGATCAAATAAGTTCTTTGGCCCTTTAAAAGATTCATCGACTTACAAGTTTGCTCAAATTCGAATCATAGATCTC

>M6-PGSC0003DMG400010700-(rev)M6LRR-RLP_1736

TGGCCCTTTAAAAGATTCATCGACTTACAAGTTTGCTCAAATTCGAATCATAGATCTCTCATCCAATGGATTTAGTGGAGATTTACCAGTGAGCCTTTTTGAGTATTTTGAAGCCATGAA

>M6-PGSC0003DMG400010700-(rev)M6LRR-RLP_1798

CCAATGGATTTAGTGGAGATTTACCAGTGAGCCTTTTTGAGTATTTTGAAGCCATGAAAATAATTAGTGAGAACAGTGGAACCCGAGAGTATGTAGCAGATATGTCTTCATTTTATTACA

>M6-PGSC0003DMG400010700-(rev)M6LRR-RLP_186

AAGAGCACAGATTGTTGTTCATGGGTTGGAGTTAAGTGTTACGAGATGACAGGACAAGTGATTTAGCTTGATCTCAGTTGCAGCCAACTTCTAGGTAAGTTTCATTCCAACTGTAGTCTC

>M6-PGSC0003DMG400010700-(rev)M6LRR-RLP_1860

ATTAGTGAGAACAGTGGAACCCGAGAGTATGTAGCAGATATGTCTTCATTTTATTACACAAATTCCTTTATAGTGACAACAAAGGGACTGGAGCTTGAACTTCCTCGAGTTTTGACTACA

>M6-PGSC0003DMG400010700-(rev)M6LRR-RLP_1922

TTCCTTTATAGTGACAACAAAGGGACTGGAGCTTGAACTTCCTCGAGTTTTGACTACAGACATGATTATCGATCTCTCAAGGAATAGATTTGAAGGTCATATCCCAAGCATTATTGGAGA

>M6-PGSC0003DMG400010700-(rev)M6LRR-RLP_1984

TGATTATCGATCTCTCAAGGAATAGATTTGAAGGTCATATCCCAAGCATTATTGGAGATCTCATTGGGCTTCGCACGTTGAACCTATCGCATAATCACTTGGAAGGTATTATACCATCAT

>M6-PGSC0003DMG400010700-(rev)M6LRR-RLP_2046

ATTGGGCTTCGCACGTTGAACCTATCGCATAATCACTTGGAAGGTATTATACCATCATCACTGCACCAGTTATCTGTACTTGAATCATTGGATCTCTCATCCAACAAAATCAGCGGAGAA

>M6-PGSC0003DMG400010700-(rev)M6LRR-RLP_2356

CGAATCCAGTTGAGCTAGATGAAGAAGGAGGAGGAGATTCATCAATGATCAGTTGGCAGGCGATTCTCATGGGTTACAGTTGTGGACTTGTTATTGGACTGTCCATAATATACATAATGT

>M6-PGSC0003DMG400010700-(rev)M6LRR-RLP_2418

ATTCTCATGGGTTACAGTTGTGGACTTGTTATTGGACTGTCCATAATATACATAATGTTGTCAACTCAATATCCAGCATGGTTTTCGAGGATGGATATAAAATTGGAACACAAAATTCTT

>M6-PGSC0003DMG400010700-(rev)M6LRR-RLP_2462

AATATACATAATGTTGTCAACTCAATATCCAGCATGGTTTTCGAGGATGGATATAAAATTGGAACACAAAATTCTTACGAGAATGAGAAGGCACAAGAAAAGACATTAGTGTGTAACCTC

>M6-PGSC0003DMG400010700-(rev)M6LRR-RLP_248

TTAGCTTGATCTCAGTTGCAGCCAACTTCTAGGTAAGTTTCATTCCAACTGTAGTCTCTTTCAACTCTCCAATCTCAAAAGGCTTGATTTGTCTAATAATAATTTTTCAGGGTCGCTCAT

>M6-PGSC0003DMG400010700-(rev)M6LRR-RLP_372

CCTAAATTTGGTGAGCTTTCTAGTTTGACTCATCTTGATTTGTCGTATTCAAGTTTTACAGGTCTAATACCAGCAGAAATCTCTCATCTTTCTAAATTACACGTTCTTCAAATCGCGAGT

>M6-PGSC0003DMG400010700-(rev)M6LRR-RLP_434

TCTAATACCAGCAGAAATCTCTCATCTTTCTAAATTACACGTTCTTCAAATCGCGAGTGATACAGATGAGCTTAGATTCGGACCTCACAATTTTGAACTGCTCCTTAAGAACTTGACCCA

>M6-PGSC0003DMG400010700-(rev)M6LRR-RLP_496

CAGATGAGCTTAGATTCGGACCTCACAATTTTGAACTGCTCCTTAAGAACTTGACCCAACTAAGAGAGCTCAACCTTAGGTTTGTGAACATCTTTTCCACCATTCCTCTGAATTTCTCTT

>M6-PGSC0003DMG400010700-(rev)M6LRR-RLP_62

CTCATCTTTACCTCATTTGTGCCCCAATGATCAAGCCCTTGCTCTTCTTCAATTCAAAAACACGTTTACTATAAATCTTGATGCTTCTTATAGTGGTTATTATCCTAAAACAGTTAAATG

>M6-PGSC0003DMG400010700-(rev)M6LRR-RLP_620

TTTAACTACTCTGGACCTTCGGAAAACACAGTTATACGGAATATTGCCCGAAGGAGTTTTCCACCTTTCCAACTTAGAATTTCTGGATTTATCGGACAATCCCCAGCTCACTGTTAAGTT

>M6-PGSC0003DMG400010700-(rev)M6LRR-RLP_682

ACCTTTCCAACTTAGAATTTCTGGATTTATCGGACAATCCCCAGCTCACTGTTAAGTTTCCCACAACCAAATGGAATAGCAGTGCATCACTCATGGAGTTAGTTCTCTATAGTGTGAATG

>M6-PGSC0003DMG400010700-(rev)M6LRR-RLP_744

ACAACCAAATGGAATAGCAGTGCATCACTCATGGAGTTAGTTCTCTATAGTGTGAATGCTGCTGGTAGGATACCTGAATCATTTGGTCATCTAACTTCACTGCGTACATTGGACATAGCT

>M6-PGSC0003DMG400010700-(rev)M6LRR-RLP_806

TGGTAGGATACCTGAATCATTTGGTCATCTAACTTCACTGCGTACATTGGACATAGCTTCTTGTAATCTCTCAGGCTCTATTCCTAAACCTCTATTGAATCTCACCAACATAGAGGATTT

>M6-PGSC0003DMG400010700-(rev)M6LRR-RLP_868

GTAATCTCTCAGGCTCTATTCCTAAACCTCTATTGAATCTCACCAACATAGAGGATTTGTACCTTGGTTATAACCATCTTGAAGGACCAATTTCTGATTTCTTTAGATTTGGAAAGCTCA

>M6-PGSC0003DMG400010700-(rev)M6LRR-RLP_930

CTTGGTTATAACCATCTTGAAGGACCAATTTCTGATTTCTTTAGATTTGGAAAGCTCAGGTGGTTATCACTTAGAAATAACAACTTTGATGGCCAACTTGAGTTCTTATCCTTTAACAGA

>M6-PGSC0003DMG400010701M6RLKWAX-EGF_1089

GTGAAGTTGCTTGAGAGTTGTTCTGATTCTGCAAAAATAGATGTAGTTGTTGCTTGACATCACATATTGTGTACGTAATTCGTGTTGTTCTGAATAAAATATCAAGTTCATAATAAATTT

>M6-PGSC0003DMG400010701M6RLKWAX-EGF_520

TCTACATTTTGACATCAAGCCACACAACATACTTTTGGATGAGAATTTCATTCCAAAACTATCTGATTTTGGTCTTGCAAAATCATACCCGGCTGATAAGAGCATTGTTACTCTAACACC

>M6-PGSC0003DMG400010746M6RLKmalectin_0

CATGACTCAAAATTCTTTCCCAAAGCAAAGTGAAAAAGTTGAAGATGAAAGATGAAGTTGAATTCCATCTACCTTTGGAAAAAGCAAATTTCAAAATATTCAAAAGAAAAAACAACATAG

>M6-PGSC0003DMG400010746M6RLKmalectin_1037

GAGCTTAAACGGAATGTTATGGTCAGTGAGGATTCATGTAGCTGATAGCCGATCCAACTCGCTTGGGATTGAGGCATAGTTACAAGTAACTTGATTTTTATTAGGACAAGATAAATGAGT

>M6-PGSC0003DMG400010746M6RLKmalectin_1098

CTTGGGATTGAGGCATAGTTACAAGTAACTTGATTTTTATTAGGACAAGATAAATGAGTGGCAATCTTGTTCATGAACTAAAGGTTTAAATATGCAGAGTTTGTAAGTATAGACTGTGGA

>M6-PGSC0003DMG400010746M6RLKmalectin_1159

CAATCTTGTTCATGAACTAAAGGTTTAAATATGCAGAGTTTGTAAGTATAGACTGTGGAAGTGCAAGCAATTATACGGACACGAGTACAGGATTAGCATGGACTTCAGATGCTGGAATGA

>M6-PGSC0003DMG400010746M6RLKmalectin_122

ATGTGAGTGAACATTACATAAAAACACCTCAATGACAAAAAAGTTAAAGGTAAACCAACAAAAGATCAATTAAGATCCACTTTTAGAGAGTCTTGAACTTCACTAAATACTAAAATATTA

>M6-PGSC0003DMG400010746M6RLKmalectin_1220

TGCAAGCAATTATACGGACACGAGTACAGGATTAGCATGGACTTCAGATGCTGGAATGATGGGGCAAGGAAAACCAGTTGTGGTAGTGAATGCAAATGTAAACTCACAACAATATCAAAG

>M6-PGSC0003DMG400010746M6RLKmalectin_1281

GGGCAAGGAAAACCAGTTGTGGTAGTGAATGCAAATGTAAACTCACAACAATATCAAAGGCGCAGAGACTTTCCAGCAGACAGCAACAAATACTGTTATACTTTAAATACCAAGGAGAGA

>M6-PGSC0003DMG400010746M6RLKmalectin_1342

GCAGAGACTTTCCAGCAGACAGCAACAAATACTGTTATACTTTAAATACCAAGGAGAGAAGGCGATATCTTGTCCGAGCAACATTTCTATATGGAAGTCCTGCAGCAGAGGGAACATACC

>M6-PGSC0003DMG400010746M6RLKmalectin_1403

GCGATATCTTGTCCGAGCAACATTTCTATATGGAAGTCCTGCAGCAGAGGGAACATACCCCAAATTTGAGCTCTACTTGGATGCAACTAAATGGGGAACTATAACCATTTCAGAATCTTC

>M6-PGSC0003DMG400010746M6RLKmalectin_1464

AAATTTGAGCTCTACTTGGATGCAACTAAATGGGGAACTATAACCATTTCAGAATCTTCAAGGATATATGTGAACGAAATGATCATAAGAGCACCTTCAAACTCTATCGATGTGTGCTTG

>M6-PGSC0003DMG400010746M6RLKmalectin_1525

GGATATATGTGAACGAAATGATCATAAGAGCACCTTCAAACTCTATCGATGTGTGCTTGTGTTGTGCAACAACAGAGTCCCCCTTCATATCTACTCTTGAGCTGAGACCTCTAAACTTGT

>M6-PGSC0003DMG400010746M6RLKmalectin_1586

TTGTGCAACAACAGAGTCCCCCTTCATATCTACTCTTGAGCTGAGACCTCTAAACTTGTCAATGTATGCCACAGACTATGAAGATAACTTTTATTTAAAAGTGGCAGCAAGAGTAGATTT

>M6-PGSC0003DMG400010746M6RLKmalectin_1647

ATGTATGCCACAGACTATGAAGATAACTTTTATTTAAAAGTGGCAGCAAGAGTAGATTTTGGAGCTCAAAGCAAGGAACCTATAAGGTATAGTAAATTTTGAGAATATGCTTTAGTCATG

>M6-PGSC0003DMG400010746M6RLKmalectin_1708

GAGCTCAAAGCAAGGAACCTATAAGGTATAGTAAATTTTGAGAATATGCTTTAGTCATGCTTGTACGTTCTCTTTGGATCCTGCGTTTTTTAAGAGTCTTTTAGTCCTGTTAGTAATTTA

>M6-PGSC0003DMG400010746M6RLKmalectin_1769

TGTACGTTCTCTTTGGATCCTGCGTTTTTTAAGAGTCTTTTAGTCCTGTTAGTAATTTATATGCCAATAGAAAATATAGTGAACTTATAGCATCTTTTTTTTTTTAATTGTAATATTGGT

>M6-PGSC0003DMG400010746M6RLKmalectin_183

AAGATCAATTAAGATCCACTTTTAGAGAGTCTTGAACTTCACTAAATACTAAAATATTATACTTCAAGTTTATCAAACAAAACACAACTTATCTTCTTCTTCTTCTTTCATCTTCCATGG

>M6-PGSC0003DMG400010746M6RLKmalectin_1830

TGCCAATAGAAAATATAGTGAACTTATAGCATCTTTTTTTTTTTAATTGTAATATTGGTCTAATATGCACTTAAATGGAGCGACAATGATCAGTGAGGATTCATATAGCGTAACTTGCTT

>M6-PGSC0003DMG400010746M6RLKmalectin_1891

AATATGCACTTAAATGGAGCGACAATGATCAGTGAGGATTCATATAGCGTAACTTGCTTGGGAATGAGGCATAGTTGTTGCTGTTTGTGAGGTGATACAAAAGCAGATAAACTATATGTT

>M6-PGSC0003DMG400010746M6RLKmalectin_1952

GAATGAGGCATAGTTGTTGCTGTTTGTGAGGTGATACAAAAGCAGATAAACTATATGTTTTGACTTTTTCAGCTCTCACAAGATCATAACATGAATAGATCTGTCTCACAGGTATCCTGA

>M6-PGSC0003DMG400010746M6RLKmalectin_2013

GACTTTTTCAGCTCTCACAAGATCATAACATGAATAGATCTGTCTCACAGGTATCCTGATGATCCCTATGACCGTATATGGGATTCGGATCTTAAAAAAAGGCCAAATTTTCTAGTAGAT

>M6-PGSC0003DMG400010746M6RLKmalectin_2074

ATCCCTATGACCGTATATGGGATTCGGATCTTAAAAAAAGGCCAAATTTTCTAGTAGATGTGGCGGCAGGCACAGAAAGAATCAACACAACAAAGTACATAGATACAAACACTAGAGAAT

>M6-PGSC0003DMG400010746M6RLKmalectin_2135

GGCGGCAGGCACAGAAAGAATCAACACAACAAAGTACATAGATACAAACACTAGAGAATACCCGCCCGTTAAAGTAATGCAGACTGCAGTAGTTGGGACTAAAGGAAAGCTCAGCTATAG

>M6-PGSC0003DMG400010746M6RLKmalectin_2196

CCGCCCGTTAAAGTAATGCAGACTGCAGTAGTTGGGACTAAAGGAAAGCTCAGCTATAGGCTGAATCTTGATGACTTTCCAGCGAATGCACGAGCTTTTGCATACTTTGCTGAGATTGAA

>M6-PGSC0003DMG400010746M6RLKmalectin_2257

TGAATCTTGATGACTTTCCAGCGAATGCACGAGCTTTTGCATACTTTGCTGAGATTGAAGATTTGGGGATGAATGAGACCAGGAAGTTTAAAATGGAGCACCCTTACGTTCCGGACTACA

>M6-PGSC0003DMG400010746M6RLKmalectin_2318

TTTGGGGATGAATGAGACCAGGAAGTTTAAAATGGAGCACCCTTACGTTCCGGACTACAGCAATGCTGTGGTGAATATAGCAGAGAATGCAAATGGAAGTTATACTTTGTATGAACCTAG

>M6-PGSC0003DMG400010746M6RLKmalectin_2379

AATGCTGTGGTGAATATAGCAGAGAATGCAAATGGAAGTTATACTTTGTATGAACCTAGCTACATGAACATAACATTGGATTTTATACTGTCATTCTCCTTTGTGAAGACTCCTGACTCA

>M6-PGSC0003DMG400010746M6RLKmalectin_244

CTTCAAGTTTATCAAACAAAACACAACTTATCTTCTTCTTCTTCTTTCATCTTCCATGGGTTTTGTTTTACAATCCTTTGTTGTACTTCTTTGCTTCACTCCAGCAATTGTGTGTCAAGT

>M6-PGSC0003DMG400010746M6RLKmalectin_2440

ACATGAACATAACATTGGATTTTATACTGTCATTCTCCTTTGTGAAGACTCCTGACTCAACTCGAGGTCCACTACTAAGTGCAATGGAAATATGTAGATACGTGCAAATTGCTACAAAGA

>M6-PGSC0003DMG400010746M6RLKmalectin_2501

TCGAGGTCCACTACTAAGTGCAATGGAAATATGTAGATACGTGCAAATTGCTACAAAGACCGATGAACAAGATGGTACAATGACCATTCCCTGTTTAGTCTAGTTATTTCTCTCTATCCA

>M6-PGSC0003DMG400010746M6RLKmalectin_2562

GATGAACAAGATGGTACAATGACCATTCCCTGTTTAGTCTAGTTATTTCTCTCTATCCATATATGGTAATTGTGCTACTTCTACGCATATTTTTACCAAAAATAGCTAATTTATTTAAAG

>M6-PGSC0003DMG400010746M6RLKmalectin_2623

TATGGTAATTGTGCTACTTCTACGCATATTTTTACCAAAAATAGCTAATTTATTTAAAGCTTCTACTTGGAAGTGAGTACTCTCAACGCCTTCCGCTCTATGTCATTGGGAAGTGATTGG

>M6-PGSC0003DMG400010746M6RLKmalectin_2684

TCTACTTGGAAGTGAGTACTCTCAACGCCTTCCGCTCTATGTCATTGGGAAGTGATTGGTCAGATGAAGACGGTGATCCTTGTGTACCTACACAATGGGAATGGGTGACTTGCAGCACAA

>M6-PGSC0003DMG400010746M6RLKmalectin_2745

AGATGAAGACGGTGATCCTTGTGTACCTACACAATGGGAATGGGTGACTTGCAGCACAACTAGTCCACCAAGAATCACAAAAATGTGAGTTATATTGCTACTATATGATATATGGATATA

>M6-PGSC0003DMG400010746M6RLKmalectin_2806

AGTCCACCAAGAATCACAAAAATGTGAGTTATATTGCTACTATATGATATATGGATATAATGCTAGTCACAGGAAGAAAACAAAAGGACATTATTTGCAACTACTACTACCCACTGTCCT

>M6-PGSC0003DMG400010746M6RLKmalectin_2867

GCTAGTCACAGGAAGAAAACAAAAGGACATTATTTGCAACTACTACTACCCACTGTCCTGCTATAACCCGACAGTCAGTTACAGATAAGAGATGATAAAACTTTAACACAGCATTAGACA

>M6-PGSC0003DMG400010746M6RLKmalectin_2928

TATAACCCGACAGTCAGTTACAGATAAGAGATGATAAAACTTTAACACAGCATTAGACAGAAATGGAAACTCAGGTCAGTGCTTAAAGTATAGTCTTTCCCAGTCAATTTGATATTTTAC

>M6-PGSC0003DMG400010746M6RLKmalectin_2989

AATGGAAACTCAGGTCAGTGCTTAAAGTATAGTCTTTCCCAGTCAATTTGATATTTTACTTTAATTTCATCAGAACACTGTCAGGAAAGAATGTTACGGGTGAAATTCCTCGCGAGCTAC

>M6-PGSC0003DMG400010746M6RLKmalectin_305

TTTGTTTTACAATCCTTTGTTGTACTTCTTTGCTTCACTCCAGCAATTGTGTGTCAAGTAACAGGTATGTGTTCTTTACATCTTTACCCTTTACTATACCACATAATCTTGAAATAAAAA

>M6-PGSC0003DMG400010746M6RLKmalectin_3050

TAATTTCATCAGAACACTGTCAGGAAAGAATGTTACGGGTGAAATTCCTCGCGAGCTACAACATATGGAAGGACTGACAGAGTTGTAAGTTTGCTTATAAAACAAGTCAAATTTTAATCT

>M6-PGSC0003DMG400010746M6RLKmalectin_3111

CATATGGAAGGACTGACAGAGTTGTAAGTTTGCTTATAAAACAAGTCAAATTTTAATCTTACTATTTCAGTTCCTCATATTTTCTACTCTTTAGGTGGTTGGATGGAAACTCCCTGACCG

>M6-PGSC0003DMG400010746M6RLKmalectin_3172

CTATTTCAGTTCCTCATATTTTCTACTCTTTAGGTGGTTGGATGGAAACTCCCTGACCGGACCAATCCCTGATATGAGCAATCTTGTTAATTTGAGAATTATGTAAGATCTTTTCTAGTG

>M6-PGSC0003DMG400010746M6RLKmalectin_3233

CCAATCCCTGATATGAGCAATCTTGTTAATTTGAGAATTATGTAAGATCTTTTCTAGTGGATCACTATTACTACCACTTCAACAAGTAAATCTTTCTCTTTTCTGTTTTTCCTTTTTAAC

>M6-PGSC0003DMG400010746M6RLKmalectin_3294

TCACTATTACTACCACTTCAACAAGTAAATCTTTCTCTTTTCTGTTTTTCCTTTTTAACTAGATTGTTGTATTTTGTCAACATGCAGACATCTTGAGAACAACAAGTTGACTGGTCCAAT

>M6-PGSC0003DMG400010746M6RLKmalectin_3355

GATTGTTGTATTTTGTCAACATGCAGACATCTTGAGAACAACAAGTTGACTGGTCCAATACCTTCATACCTTGGAGGTTTACCAAGCTTATTAGAACTGTATGTTGTGATCTGGATAACA

>M6-PGSC0003DMG400010746M6RLKmalectin_3416

CTTCATACCTTGGAGGTTTACCAAGCTTATTAGAACTGTATGTTGTGATCTGGATAACAAAATCTTAACTTTCTTTAGCGTATAATGCTTCATAATTTGGCTACTCCAAATTTTGAGTAT

>M6-PGSC0003DMG400010746M6RLKmalectin_3477

ATCTTAACTTTCTTTAGCGTATAATGCTTCATAATTTGGCTACTCCAAATTTTGAGTATCCAACATGGATAGTTGGCAGTCTTCTTTATATTTTTACTTGTCATAAAACACCTTGTATCA

>M6-PGSC0003DMG400010746M6RLKmalectin_3538

AACATGGATAGTTGGCAGTCTTCTTTATATTTTTACTTGTCATAAAACACCTTGTATCAATTTTGGAAATTTTGTTTCTTTGTGGCAGAGATGTACAGAATAACAGCCTAACCGGAGAAA

>M6-PGSC0003DMG400010746M6RLKmalectin_3599

TTTGGAAATTTTGTTTCTTTGTGGCAGAGATGTACAGAATAACAGCCTAACCGGAGAAATACCGTCATCTTTATTAACAGGAAAAGTAACTTTCAAGTAAGTATAACTTTAATTCTGTAC

>M6-PGSC0003DMG400010746M6RLKmalectin_366

CAGGTATGTGTTCTTTACATCTTTACCCTTTACTATACCACATAATCTTGAAATAAAAAATCAATCTTGGCTTCAAATAACCTGAATTTCAATAGAATAAAAAAAAATTGCCACAGAGAG

>M6-PGSC0003DMG400010746M6RLKmalectin_3660

CCGTCATCTTTATTAACAGGAAAAGTAACTTTCAAGTAAGTATAACTTTAATTCTGTACTGGATTGTTGTACCCATAATACCATGAGAATATGTTAGCAACTAGATCAAAGTAGCTATTT

>M6-PGSC0003DMG400010746M6RLKmalectin_3721

GATTGTTGTACCCATAATACCATGAGAATATGTTAGCAACTAGATCAAAGTAGCTATTTTACTATAGACTGATTATATTCCTTAGTTTGGATTATCTATAGCATAAACAAAGTTTCATTT

>M6-PGSC0003DMG400010746M6RLKmalectin_3782

CTATAGACTGATTATATTCCTTAGTTTGGATTATCTATAGCATAAACAAAGTTTCATTTTTTATGTCACATTATGCTAGGAAGAGAGCAAATGAAATTCATGAATTTTCATAAATTTCTT

>M6-PGSC0003DMG400010746M6RLKmalectin_3843

TATGTCACATTATGCTAGGAAGAGAGCAAATGAAATTCATGAATTTTCATAAATTTCTTGTAAGTGAGAACTAGGAAGTAACATGTACATTTATCTTTGAACATAGTAAAAGAAAGAGAA

>M6-PGSC0003DMG400010746M6RLKmalectin_3904

AAGTGAGAACTAGGAAGTAACATGTACATTTATCTTTGAACATAGTAAAAGAAAGAGAAATTTTCTTGCAACTTTCGGAGTACCACATGGATGTGTGTGCACTGATGGAAGATCTGTTAC

>M6-PGSC0003DMG400010746M6RLKmalectin_3965

TTTCTTGCAACTTTCGGAGTACCACATGGATGTGTGTGCACTGATGGAAGATCTGTTACAATCAGGAAATGAACGTCACTTAAAAATGTGTTCCATGTCTCACAGGCATGAAGGAAATCC

>M6-PGSC0003DMG400010746M6RLKmalectin_4026

TCAGGAAATGAACGTCACTTAAAAATGTGTTCCATGTCTCACAGGCATGAAGGAAATCCTGATCTAAGACCCGAGTCAAAGCATAGTACACGTTACAAAGTGATACTAGGAGCTTCAATT

>M6-PGSC0003DMG400010746M6RLKmalectin_4087

ATCTAAGACCCGAGTCAAAGCATAGTACACGTTACAAAGTGATACTAGGAGCTTCAATTGGAGGACTTGTGATTATATTGGTTCTATTTGCTGCCAGTATATTCTTTTTATGCCACTTCA

>M6-PGSC0003DMG400010746M6RLKmalectin_4148

AGGACTTGTGATTATATTGGTTCTATTTGCTGCCAGTATATTCTTTTTATGCCACTTCAGAACAAAAGTATCTCATCAAAAAGGTGATACAAATCCTTAAGTGTCTGATCTTAACTTTTA

>M6-PGSC0003DMG400010746M6RLKmalectin_4209

ACAAAAGTATCTCATCAAAAAGGTGATACAAATCCTTAAGTGTCTGATCTTAACTTTTAATTCTATTGCTCCGACACCTTTTCATATAAATCATTTCATCTGTACCTCCCAGGTGAATCT

>M6-PGSC0003DMG400010746M6RLKmalectin_427

CAATCTTGGCTTCAAATAACCTGAATTTCAATAGAATAAAAAAAAATTGCCACAGAGAGACATTTTCTACTTGATTTTCATTAGGTCAAGATAGAGGGTTTTTTTATTTATTTTATATAT

>M6-PGSC0003DMG400010746M6RLKmalectin_4270

TCTATTGCTCCGACACCTTTTCATATAAATCATTTCATCTGTACCTCCCAGGTGAATCTATGCGCACAAACACTAAACCGTCAACAACTTGTTCAATGGCAAGAGGTGGATCCTTAATGG

>M6-PGSC0003DMG400010746M6RLKmalectin_4331

GCGCACAAACACTAAACCGTCAACAACTTGTTCAATGGCAAGAGGTGGATCCTTAATGGATGAAGGTGTGGCATACTATATCCCACAATCCGAGATAGAAGAAGCAACTGAAAATTTTTC

>M6-PGSC0003DMG400010746M6RLKmalectin_4392

GAAGGTGTGGCATACTATATCCCACAATCCGAGATAGAAGAAGCAACTGAAAATTTTTCAAAGAAAATTGGAAAGGGAAGTTTTGGACCTGTTTACTATGGAAGACTTAGGGATGGGAAA

>M6-PGSC0003DMG400010746M6RLKmalectin_4453

AGAAAATTGGAAAGGGAAGTTTTGGACCTGTTTACTATGGAAGACTTAGGGATGGGAAAGAAGTTGCTGTCAAAACTATGGCAGATTCATCAAGCCATGGAACCAAACAGTTCGCTACAG

>M6-PGSC0003DMG400010746M6RLKmalectin_4514

AGTTGCTGTCAAAACTATGGCAGATTCATCAAGCCATGGAACCAAACAGTTCGCTACAGAGGTATATTAAGGTTGCTATACTTTTGGTATCATCTGAATAAAACATATTAATTTTCTATT

>M6-PGSC0003DMG400010746M6RLKmalectin_4575

GTATATTAAGGTTGCTATACTTTTGGTATCATCTGAATAAAACATATTAATTTTCTATTTGGAGTAGCTTATTTAAGCTGCTAATTGGCTTTAAGCCCAAATAATAGCAGCTGGCACAGC

>M6-PGSC0003DMG400010746M6RLKmalectin_4636

GAGTAGCTTATTTAAGCTGCTAATTGGCTTTAAGCCCAAATAATAGCAGCTGGCACAGCTATGGTAATAAGGACTATGAAATTGTCTAAGCTAACACATTTGACACGGCGTAAAACTGAT

>M6-PGSC0003DMG400010746M6RLKmalectin_4697

TGGTAATAAGGACTATGAAATTGTCTAAGCTAACACATTTGACACGGCGTAAAACTGATCATAAGTCAGACTTCATACATTATAAGACATGACTAAAGAAATTGTTCCTGTTGCTTATTT

>M6-PGSC0003DMG400010746M6RLKmalectin_4758

TAAGTCAGACTTCATACATTATAAGACATGACTAAAGAAATTGTTCCTGTTGCTTATTTCATTCCTGTACAAATTAGAAAATTCACAACTTGATATGATATTACTTTTGGTTGAAAATTT

>M6-PGSC0003DMG400010746M6RLKmalectin_4819

TTCCTGTACAAATTAGAAAATTCACAACTTGATATGATATTACTTTTGGTTGAAAATTTATACCGTTTGTGAAATAACATACTTGCAGGTGGCTCTCTTGTCAAGGATCCATCACAGAAA

>M6-PGSC0003DMG400010746M6RLKmalectin_4880

ACCGTTTGTGAAATAACATACTTGCAGGTGGCTCTCTTGTCAAGGATCCATCACAGAAACTTGGTTCCATTGATCGGATACTATGAGGATGACCATCAACGTATGCTTGTCTATGAATAC

>M6-PGSC0003DMG400010746M6RLKmalectin_4941

TGGTTCCATTGATCGGATACTATGAGGATGACCATCAACGTATGCTTGTCTATGAATACATGCATAATGGAACTTTGAGAGATCATATAACTGGTCTGCCCCTTTATGTGTACTATAATT

>M6-PGSC0003DMG400010746M6RLKmalectin_5002

GCATAATGGAACTTTGAGAGATCATATAACTGGTCTGCCCCTTTATGTGTACTATAATTGCTTTACATTTTTCTGCATTATTTCCTTCTTGAACTGAACATCACACCACATATTTGTGAA

>M6-PGSC0003DMG400010746M6RLKmalectin_5063

TTTACATTTTTCTGCATTATTTCCTTCTTGAACTGAACATCACACCACATATTTGTGAACAGAATCCACTGATAAAAAGCATTTGGACTGGCTAGCACGTCTTAATATTGCAGAAGATGC

>M6-PGSC0003DMG400010746M6RLKmalectin_5124

GAATCCACTGATAAAAAGCATTTGGACTGGCTAGCACGTCTTAATATTGCAGAAGATGCAGCCAAAGGTTACCAGAAATTTTTCATTGTCAATAACTTAGAAGTATGTATTCTCCTAGTT

>M6-PGSC0003DMG400010746M6RLKmalectin_5185

CCAAAGGTTACCAGAAATTTTTCATTGTCAATAACTTAGAAGTATGTATTCTCCTAGTTAGTCCGGAATCACAAGATAATATTTTCAGCCTCCTTGTAGGCCTTGAGTACTTACATACTG

>M6-PGSC0003DMG400010746M6RLKmalectin_5246

TCCGGAATCACAAGATAATATTTTCAGCCTCCTTGTAGGCCTTGAGTACTTACATACTGGATGCAACCCCTCTATCATTCATCGAGATGTGAAAACTAGCAACATTCTCTTAGATATAAA

>M6-PGSC0003DMG400010746M6RLKmalectin_5307

TGCAACCCCTCTATCATTCATCGAGATGTGAAAACTAGCAACATTCTCTTAGATATAAATATGCGAGCAAAAGTATCTGATTTTGGACTATCAAGGCAAGCTGAAGAAGATTTAACACAT

>M6-PGSC0003DMG400010746M6RLKmalectin_5368

TGCGAGCAAAAGTATCTGATTTTGGACTATCAAGGCAAGCTGAAGAAGATTTAACACATGTCTCAAGCGTGGCACGAGGAACTGTCGGCTACCTTGATCCTGAGTAGGCCATAATCTATC

>M6-PGSC0003DMG400010746M6RLKmalectin_5429

CTCAAGCGTGGCACGAGGAACTGTCGGCTACCTTGATCCTGAGTAGGCCATAATCTATCATGAGACTGAATATTTTGCTAGTGATTTTCATTCTTTGTCCTCATGTTTCTGCATTTCCAA

>M6-PGSC0003DMG400010746M6RLKmalectin_549

ATATATAGTGAACTTCTAGTGTTTCTTCTTTTGGTATAATATTAGTGGGAGCTAGATGAGCTTAAATGACTAGACAGAGTTGCTCGGATGAAAAGCACCCTCATAATCCTTAGTGGTTGT

>M6-PGSC0003DMG400010746M6RLKmalectin_5490

GAGACTGAATATTTTGCTAGTGATTTTCATTCTTTGTCCTCATGTTTCTGCATTTCCAACAGGTACTATGCAAATCAGCAATTGACAGAGAAAAGTGATGTATATAGCTTTGGGGTTGTC

>M6-PGSC0003DMG400010746M6RLKmalectin_5551

GGTACTATGCAAATCAGCAATTGACAGAGAAAAGTGATGTATATAGCTTTGGGGTTGTCCTTTTGGAATTGATCTCTGGAAGAAAGCCTTTTTCTTCTGAGGAATATGGTGCTGATTGGA

>M6-PGSC0003DMG400010746M6RLKmalectin_5612

TTTGGAATTGATCTCTGGAAGAAAGCCTTTTTCTTCTGAGGAATATGGTGCTGATTGGAGCATTGTTCACTGGGTAAAGTTTATCATACTTTCTATATATTTCAGGGATGCTATGCCATC

>M6-PGSC0003DMG400010746M6RLKmalectin_5673

ATTGTTCACTGGGTAAAGTTTATCATACTTTCTATATATTTCAGGGATGCTATGCCATCCACTGTGATAATAACATATAGTTGAGCTTCAGCAGGGAAACATGGCCAGTGAGGATTCATA

>M6-PGSC0003DMG400010746M6RLKmalectin_5734

CTGTGATAATAACATATAGTTGAGCTTCAGCAGGGAAACATGGCCAGTGAGGATTCATATAGCGAACCCCAACTTGTTTGAAATTGAGGAATAGTTGTTGTAAATATCACCCACCGGCAT

>M6-PGSC0003DMG400010746M6RLKmalectin_5795

GCGAACCCCAACTTGTTTGAAATTGAGGAATAGTTGTTGTAAATATCACCCACCGGCATCACGTTTTCATAATGCATATAAAAAAATTCAGATATCTCTTCTACAAAAATCACAATAACT

>M6-PGSC0003DMG400010746M6RLKmalectin_5856

CGTTTTCATAATGCATATAAAAAAATTCAGATATCTCTTCTACAAAAATCACAATAACTATCAAAGCCAAAGCCAGCTCAGTAGAAAGTGCAGCAAAACACAAGAAATCAATAAGAAGCT

>M6-PGSC0003DMG400010746M6RLKmalectin_5917

CAAAGCCAAAGCCAGCTCAGTAGAAAGTGCAGCAAAACACAAGAAATCAATAAGAAGCTGATCAGGGGAATATAAATTTTCAGGGTGGATAGTTATTGTAGTGATATATAAAGGATTCAG

>M6-PGSC0003DMG400010746M6RLKmalectin_5978

TCAGGGGAATATAAATTTTCAGGGTGGATAGTTATTGTAGTGATATATAAAGGATTCAGGAAATGAACATAAGAAAAGTTAATGATTTGAAATGCTGGTATAGTGGTGAATCACCACACA

>M6-PGSC0003DMG400010746M6RLKmalectin_6039

AATGAACATAAGAAAAGTTAATGATTTGAAATGCTGGTATAGTGGTGAATCACCACACATAAAGTTCTGTCTGTCTGAGAAAAAATCGAAAGACGAAAAACATGGTTCTTTTCTCTGGAT

>M6-PGSC0003DMG400010746M6RLKmalectin_61

ATTCCATCTACCTTTGGAAAAAGCAAATTTCAAAATATTCAAAAGAAAAAACAACATAGGAATGTGAGTGAACATTACATAAAAACACCTCAATGACAAAAAAGTTAAAGGTAAACCAAC

>M6-PGSC0003DMG400010746M6RLKmalectin_6100

AAGTTCTGTCTGTCTGAGAAAAAATCGAAAGACGAAAAACATGGTTCTTTTCTCTGGATAGAAGCAGAAGATACTCTATCAAGTTCTGGATGTGCTGTAGTATGCATGCAAATAAAATGT

>M6-PGSC0003DMG400010746M6RLKmalectin_6161

AAGCAGAAGATACTCTATCAAGTTCTGGATGTGCTGTAGTATGCATGCAAATAAAATGTTGTGGAGTATTATGAAGTTGCTAATTAGAAAAACTGAAATAAATTGAATACAATCTGCTGA

>M6-PGSC0003DMG400010746M6RLKmalectin_6222

TGGAGTATTATGAAGTTGCTAATTAGAAAAACTGAAATAAATTGAATACAATCTGCTGAAATAAATTCATTGTTAAGTTGTTAGATAGAACACACTTTATTAAGTTAATTATATTATGCT

>M6-PGSC0003DMG400010746M6RLKmalectin_6283

TAAATTCATTGTTAAGTTGTTAGATAGAACACACTTTATTAAGTTAATTATATTATGCTTCAACAGATCTGACTAACAATATCTTGTAGGCAAGATCATTGATTCGCAAAGGAGATGTGA

>M6-PGSC0003DMG400010746M6RLKmalectin_6344

AACAGATCTGACTAACAATATCTTGTAGGCAAGATCATTGATTCGCAAAGGAGATGTGATAAGTATCATGGATCCTGCACTAGTAGGACATGTCAAAGTTGAGTCAGTATGGAGAATAGC

>M6-PGSC0003DMG400010746M6RLKmalectin_6405

AGTATCATGGATCCTGCACTAGTAGGACATGTCAAAGTTGAGTCAGTATGGAGAATAGCAGAAGTTGCAATACAATGTGTGGAAAGGCATGGCACATCCAGGCCAAGAATGCATGAAATT

>M6-PGSC0003DMG400010746M6RLKmalectin_6466

AAGTTGCAATACAATGTGTGGAAAGGCATGGCACATCCAGGCCAAGAATGCATGAAATTCTATCAGCCATACAGGATGCAATCAAGATCGAAAAGGGGATTGATAAACTGTCTTCATCAG

>M6-PGSC0003DMG400010746M6RLKmalectin_6527

ATCAGCCATACAGGATGCAATCAAGATCGAAAAGGGGATTGATAAACTGTCTTCATCAGGAAGTTCAAAGGCTCAATCTTCCAGAAAGACACTCTTAACAAGCTTCCTCGACGTAGAAAG

>M6-PGSC0003DMG400010746M6RLKmalectin_6588

AGTTCAAAGGCTCAATCTTCCAGAAAGACACTCTTAACAAGCTTCCTCGACGTAGAAAGCCCTGACATATCCAATAGCTCACTTACCCCTTCTGCTAGGTGAACAAGTAACTAACAAGAA

>M6-PGSC0003DMG400010746M6RLKmalectin_6771

TTGGATGTACATTAGTACATAATGTCTATTATTGTAACTGTACATTAACACAGACATTAATAGGATGTTACACACTAACAAGTGAATCATATTTGGGGGGCAATAGTCAGTACATTATGT

>M6-PGSC0003DMG400010746M6RLKmalectin_6832

AGGATGTTACACACTAACAAGTGAATCATATTTGGGGGGCAATAGTCAGTACATTATGTCTCAACTCAGAAACTACAGTATCTAACCATATACCTCTAAGTTGCATGAAAGTAGCATCAT

>M6-PGSC0003DMG400010746M6RLKmalectin_6893

CAACTCAGAAACTACAGTATCTAACCATATACCTCTAAGTTGCATGAAAGTAGCATCATTGAGCAATGTTTCATCCTCTATCTCTACTTCTTGTTGTAATTTGGACTAGTTTGGAGTTAA

>M6-PGSC0003DMG400010746M6RLKmalectin_976

GTCACCAAAGGGAGCAAAAAAGGGTGGGAGCTCCTAGGGAGGGGTAAAAAAGAGAAGAGATGAGCTTAAACGGAATGTTATGGTCAGTGAGGATTCATGTAGCTGATAGCCGATCCAACT

>M6-PGSC0003DMG400010940-(rev)M6LRR-RLP_1008

GGATGTTGAGATATCTAGAAGTGGGTAACAATCCGTTGAGTGGCGTTCTACCAAATTCTGTTGGGAATCTCTCATCTACTATTGAAAAGATTTATATAGGAGATGCACACATCAATGGCC

>M6-PGSC0003DMG400010940-(rev)M6LRR-RLP_1197

GGAATATTCCTTCTGAGATTGGTAAGCTTAAACAACTCCAAGGGCTGGCTCTATATAACAATAAATTGCAGGGACGTATTCCAGAGGTGGTATGCCATTTATCGAATTTGGTTGGCTTAT

>M6-PGSC0003DMG400010940-(rev)M6LRR-RLP_1260

AATTGCAGGGACGTATTCCAGAGGTGGTATGCCATTTATCGAATTTGGTTGGCTTATTACTGGATGGTAATGAGCTCTGTGGATTAATTCCAGAATGCTTTGGAAATCTTAGCATGCTAC

>M6-PGSC0003DMG400010940-(rev)M6LRR-RLP_1323

ATGGTAATGAGCTCTGTGGATTAATTCCAGAATGCTTTGGAAATCTTAGCATGCTACAAGCGCTTTATTTGGATTCTAATAAATTTTCATCAAAATTTCCGTTGAGCCTTTGGAAGATGA

>M6-PGSC0003DMG400010940-(rev)M6LRR-RLP_1386

TTTATTTGGATTCTAATAAATTTTCATCAAAATTTCCGTTGAGCCTTTGGAAGATGAGTGGTCTTCTCTATCTAAGTGTGTCACGAAATTCAATAGAGGGAGAACTTCCATCAGATATTG

>M6-PGSC0003DMG400010940-(rev)M6LRR-RLP_1449

TTCTCTATCTAAGTGTGTCACGAAATTCAATAGAGGGAGAACTTCCATCAGATATTGGAGGACTGAAAGCCATTGTAGAACTATATCTTTACAATAACCACTTTTCAGGTGTGATACCAA

>M6-PGSC0003DMG400010940-(rev)M6LRR-RLP_1512

TGAAAGCCATTGTAGAACTATATCTTTACAATAACCACTTTTCAGGTGTGATACCAACCAGATTGGGGGAACTCCAAAACCTGCAGCATCTTGACCTATCCAACAATTCATTTTTTGGCC

>M6-PGSC0003DMG400010940-(rev)M6LRR-RLP_1575

TGGGGGAACTCCAAAACCTGCAGCATCTTGACCTATCCAACAATTCATTTTTTGGCCAAATTCCATTATCCTTTGCCAACTTGATAAGCTTGGAATTCTTGAATTTGTCTTTAAATGCCT

>M6-PGSC0003DMG400010940-(rev)M6LRR-RLP_1638

CATTATCCTTTGCCAACTTGATAAGCTTGGAATTCTTGAATTTGTCTTTAAATGCCTTGTCCGGTACTATTCCTAAATCTTTGGAAAAACTCTCATACCTTAAAAGCATTAATGTTTCAT

>M6-PGSC0003DMG400010940-(rev)M6LRR-RLP_1701

GTACTATTCCTAAATCTTTGGAAAAACTCTCATACCTTAAAAGCATTAATGTTTCATTTAATGATTTAGAAGGTGAAATACCAAGTGGTGGCGTGTTTGCAACTTCCACTGTGCAATCAT

>M6-PGSC0003DMG400010940-(rev)M6LRR-RLP_1827

GGAACAAAGGTCTTTGCGGAATGCACATATTGGAGATTCCTACTTGTGCTATCACTAATCATGGAAAACAATCAAAGCTTAAGGAGGTTGTGCTAAAAATTGTTACTCCAGTGGTTATTG

>M6-PGSC0003DMG400010940-(rev)M6LRR-RLP_189

AACTTGCAGACTTGTCCTTGACTGATAATCAATTAACAGGTTCCATTCCTGTAGGACTATTTAATATCTCGTCGCTACTTGCCATAAATCTGCTATACAATAAACTTTCCGGTCCTCTCT

>M6-PGSC0003DMG400010940-(rev)M6LRR-RLP_1890

GAAAACAATCAAAGCTTAAGGAGGTTGTGCTAAAAATTGTTACTCCAGTGGTTATTGCATCCTTTCTGATATTCTTGTTCGTTTCAATTTGGATAATGAAACGACAGAAGAAAGGGAAGT

>M6-PGSC0003DMG400010940-(rev)M6LRR-RLP_1953

TTCTGATATTCTTGTTCGTTTCAATTTGGATAATGAAACGACAGAAGAAAGGGAAGTCCAAAGATATTGAAAAGGTATTGGAGATCGGGACTCATCAATTAGTTTCTTATCACGAGATTC

>M6-PGSC0003DMG400010940-(rev)M6LRR-RLP_252

ATATCTCGTCGCTACTTGCCATAAATCTGCTATACAATAAACTTTCCGGTCCTCTCTTGCTAGATGAAGGAATTTTTTTGTCAAATCTGAAGTTTCTAAGGATATCTCGCAACCAGATTT

>M6-PGSC0003DMG400010940-(rev)M6LRR-RLP_315

ATGAAGGAATTTTTTTGTCAAATCTGAAGTTTCTAAGGATATCTCGCAACCAGATTTTAGGATGCATTCCTTCCAACATATGCCAACTCAGAGAGCTCAAAGTTTTGTCCATATCTTTCA

>M6-PGSC0003DMG400010940-(rev)M6LRR-RLP_378

GCATTCCTTCCAACATATGCCAACTCAGAGAGCTCAAAGTTTTGTCCATATCTTTCAACAACATAATTGGAGACAAACCCAAAAATATTGATTGTTTATCCAATCTCCAGGAGTTTTATA

>M6-PGSC0003DMG400010940-(rev)M6LRR-RLP_441

TAATTGGAGACAAACCCAAAAATATTGATTGTTTATCCAATCTCCAGGAGTTTTATAATCGTCATAATTCAATAAAAGGGACTATTCCCACTTCATTGGGAAATATTTCAACTCTGCAAT

>M6-PGSC0003DMG400010940-(rev)M6LRR-RLP_504

ATAATTCAATAAAAGGGACTATTCCCACTTCATTGGGAAATATTTCAACTCTGCAATATCTTTATTCTGGAAACAATCTCATGGTGGGGCCAATTCCTCCAGAATTAGGGAAGCTATCAA

>M6-PGSC0003DMG400010940-(rev)M6LRR-RLP_567

ATTCTGGAAACAATCTCATGGTGGGGCCAATTCCTCCAGAATTAGGGAAGCTATCAAATTTGAGGCAATTAATTTTTTCCAATAATTCTAATCTTATTGGTCAAATTCCAGAGGCTATTT

>M6-PGSC0003DMG400010940-(rev)M6LRR-RLP_630

GGCAATTAATTTTTTCCAATAATTCTAATCTTATTGGTCAAATTCCAGAGGCTATTTTCAACATATCTTCTTTGGAATTAATCAATTTCAGTTTCAATACCCTGTCGGGTAGAATTCCAA

>M6-PGSC0003DMG400010940-(rev)M6LRR-RLP_693

TATCTTCTTTGGAATTAATCAATTTCAGTTTCAATACCCTGTCGGGTAGAATTCCAACCACTACAGGTCTTCATCTTCCGAACCTTAAAGTACTTGACTTGGGAGTCAATCAGCTTGAAG

>M6-PGSC0003DMG400010940-(rev)M6LRR-RLP_819

TTCCATTGTTCATAACAAATGCTTCCAAGCTTCAGATACTGTCGCTAGAACATAACTCTCTCTCGGGAGCTATTCCTACTAATTTGGGAAATCTTCATGAGCTGCAATCACTCTTACTAC

>M6-PGSC0003DMG400010940-(rev)M6LRR-RLP_882

CGGGAGCTATTCCTACTAATTTGGGAAATCTTCATGAGCTGCAATCACTCTTACTACATACTAATCAACTTACCAATGAACCAAGAGAGCATGAGTTGAAATTCTTCAATTCGTTGGCGA

>M6-PGSC0003DMG400010940-(rev)M6LRR-RLP_945

ATCAACTTACCAATGAACCAAGAGAGCATGAGTTGAAATTCTTCAATTCGTTGGCGAACTGTAGGATGTTGAGATATCTAGAAGTGGGTAACAATCCGTTGAGTGGCGTTCTACCAAATT

>M6-PGSC0003DMG400010950-(rev)M6LRR-RLP_0

CATTTTTCTATTAATATTTGCACTAATGGAGAAAGCCTTCACATTTTTCCTCTCAACAATACTCCTGTTGCTTCACTATGTTATGACCCAACCGAACATTACGACTGATCAATTAGTTCT

>M6-PGSC0003DMG400010950-(rev)M6LRR-RLP_132

ATCCCAAATTAGTTCCGATCCTTTTCATTACTTGGATGAAAGTTGGTCTCCTGCTACTTATGTTTGCCATTGGGTAGGAGTCACTTGTGACTATCGTCACCAGAGAGTGAAGTCCTTGAA

>M6-PGSC0003DMG400010950-(rev)M6LRR-RLP_198

CCATTGGGTAGGAGTCACTTGTGACTATCGTCACCAGAGAGTGAAGTCCTTGAATCTTTTCAACATGACTCTTACCGGCAAGATTCCCAGTGATTTTGGAAACCTCACATTTCTTGTTTC

>M6-PGSC0003DMG400010950-(rev)M6LRR-RLP_264

GACTCTTACCGGCAAGATTCCCAGTGATTTTGGAAACCTCACATTTCTTGTTTCTCTTGACTTGAGAAAGAATAATTTCCATGGAAATTTGCCTCAAGAAATGGTACGCTTACGTCGGCT

>M6-PGSC0003DMG400010950-(rev)M6LRR-RLP_396

GTTAAGTTTCAACAAATTCAGAGGGGAAATTCCTTCTTGGTTTGGGTTTTTAAAACAACTTCAAGTTTTAAATCTTAAGAATAATAGTTTCACTGGTTCCATTCCTCTGTCACTCTCAAA

>M6-PGSC0003DMG400010950-(rev)M6LRR-RLP_462

TTTAAATCTTAAGAATAATAGTTTCACTGGTTCCATTCCTCTGTCACTCTCAAATGCCTCAAGGTTGGACACTTTAGTAATATCTGCCAATTTACTTGAAGGAAACATCCCGGAAGAGAT

>M6-PGSC0003DMG400010950-(rev)M6LRR-RLP_528

GGACACTTTAGTAATATCTGCCAATTTACTTGAAGGAAACATCCCGGAAGAGATTGGCAATCTTCACAACCTGAAATACTTGTACATAGAAGATAATCAGCTTACGGGTTCTATACCATT

>M6-PGSC0003DMG400010950-(rev)M6LRR-RLP_594

CAACCTGAAATACTTGTACATAGAAGATAATCAGCTTACGGGTTCTATACCATTCACAATTTTCAATATTTCTAGTATCCAAGTCATTGCATTTACAGACAATAGCTTATCAGGAAATCT

>M6-PGSC0003DMG400010950-(rev)M6LRR-RLP_66

GTTGCTTCACTATGTTATGACCCAACCGAACATTACGACTGATCAATTAGTTCTTCTCTCTTTGAAATCCCAAATTAGTTCCGATCCTTTTCATTACTTGGATGAAAGTTGGTCTCCTGC

>M6-PGSC0003DMG400010950-(rev)M6LRR-RLP_660

TATTTCTAGTATCCAAGTCATTGCATTTACAGACAATAGCTTATCAGGAAATCTTCCCAATGGTTTATGCAATGGTCTCCCGATACTCAAAGAGCTTTATCTATCAATAAACAAGCTTGA

>M6-PGSC0003DMG400010950-(rev)M6LRR-RLP_726

ATGCAATGGTCTCCCGATACTCAAAGAGCTTTATCTATCAATAAACAAGCTTGATGGTCATATGCCTACAAGCTTGTCAAATTGTTCACAACTTCAAATGTTGTATCTATCCAGAAATGA

>M6-PGSC0003DMG400010950-(rev)M6LRR-RLP_792

TACAAGCTTGTCAAATTGTTCACAACTTCAAATGTTGTATCTATCCAGAAATGAGTTTGATGGACCAATACATAGTGAAATTGGAAGATTGAGTAACTTGCAGTTCTTGTATCTTGAGTC

>M6-PGSC0003DMG400010950-(rev)M6LRR-RLP_858

AATACATAGTGAAATTGGAAGATTGAGTAACTTGCAGTTCTTGTATCTTGAGTCCAACCATTTCACAGGTATGTGTTATCTTACTAGTTTCTGGACACGTGTGTGGTCATCTAAATTGAC

>M6-PGSC0003DMG400010950-(rev)M6LRR-RLP_920

TCACAGGTATGTGTTATCTTACTAGTTTCTGGACACGTGTGTGGTCATCTAAATTGACATAAATTGTTTCAAATTATCATTGTATTAAAATAGATTTATAATTAAATTACATGAATATCG

>M6-PGSC0003DMG400010952-(rev)M6LRR-RLP_0

TTCAAAGTCTCAAAACCACAGGTTACTAGTACTAATATTTGCACTAATGGAGAAAGCCTTCACTATTTTATTCTTAACAATGCTCTTGTTGCCTCACTATGTTATGGCCCAAACCAATAT

>M6-PGSC0003DMG400010952-(rev)M6LRR-RLP_130

CAATTAGCTCTTCTCTCTTTTAAATCCCAAATTAGTTCCGACCCCTTTCACTTCTTGAATGAAAGTTGGTCTCCTGCTATTTCTGTTTGCCGTTGGGTTGGAGTCATTTGTGGTTCTCGT

>M6-PGSC0003DMG400010952-(rev)M6LRR-RLP_195

TTGGTCTCCTGCTATTTCTGTTTGCCGTTGGGTTGGAGTCATTTGTGGTTCTCGTCACCAGCGAGTGAAGTCGTTGAAGCTTTCCAACTTGGCTCTTACAGGCAGAATTCCCCGTGATTT

>M6-PGSC0003DMG400010952-(rev)M6LRR-RLP_260

TGAAGTCGTTGAAGCTTTCCAACTTGGCTCTTACAGGCAGAATTCCCCGTGATTTTGGAAACCTCACATTTCTTGCTTCTCTTGACTTGGGAAGCAACCATTTCCATGGAAATTTGCCTC

>M6-PGSC0003DMG400010952-(rev)M6LRR-RLP_325

ACATTTCTTGCTTCTCTTGACTTGGGAAGCAACCATTTCCATGGAAATTTGCCTCAAGAAATGGCACATTTGCGTCGGCTTAAGTTTCTTGATTTAAGTTTAAACAGCTTCACAGGGGAA

>M6-PGSC0003DMG400010952-(rev)M6LRR-RLP_390

ACATTTGCGTCGGCTTAAGTTTCTTGATTTAAGTTTAAACAGCTTCACAGGGGAAATTCCTTCTTGGTTTGGGTTTTTACACCAACTTCAAGTTGTAAACCTTAGGAGTAACAGTTTTAC

>M6-PGSC0003DMG400010952-(rev)M6LRR-RLP_520

CCTCCGTCACTCTCGAATGCCTCAAGGTTGGAGATTTTAGTAATATCTACCAATTTACTTGAAGGAAACACCCCGGAAGAGTTTGGCAATCTTCACAACCTGAAATCGTTGTCCATAGAA

>M6-PGSC0003DMG400010952-(rev)M6LRR-RLP_585

AAACACCCCGGAAGAGTTTGGCAATCTTCACAACCTGAAATCGTTGTCCATAGAACAAAATCAGCTTACAGGTTCTATACCATTCACAATATTCAATATTCGTAGAATTGAAGTCATTTC

>M6-PGSC0003DMG400010952-(rev)M6LRR-RLP_65

TTTTATTCTTAACAATGCTCTTGTTGCCTCACTATGTTATGGCCCAAACCAATATTAACACTGATCAATTAGCTCTTCTCTCTTTTAAATCCCAAATTAGTTCCGACCCCTTTCACTTCT

>M6-PGSC0003DMG400010952-(rev)M6LRR-RLP_650

TTACAGGTTCTATACCATTCACAATATTCAATATTCGTAGAATTGAAGTCATTTCATTTTCATATAATAGCCTATCAGGAGATCTTCCCAATGGTTTATGCAATGATCTCCCAATACTCA

>M6-PGSC0003DMG400010952-(rev)M6LRR-RLP_715

AATAGCCTATCAGGAGATCTTCCCAATGGTTTATGCAATGATCTCCCAATACTCAAAGAGCTTTATCTATCCAGAAACAAGCTTGATGGTCATATGCCTACAAGCTTGTCAAATTGTTCA

>M6-PGSC0003DMG400010963-(rev)M6RLKWAX-EGF_0

ATTCTCATATGTTACGCTTAATAAGACTCCCTTATAGGAATGACATGGAATCTCCAAGATCACAGGATTTAAAGTACTTTTAGGGATGTTCAGTACAAAGATTAATAAAGCAGAGATTAG

>M6-PGSC0003DMG400010963-(rev)M6RLKWAX-EGF_1037

ACAATCTACCAAACGACCCCTTAGTGTGTTATAGGCACCTTTTGTTTAATATCACAAGATCCAAATGTATTTTTTATGTTTTTAAATTTCGTGCAAAATCAAACTAAGACATATAAAATG

>M6-PGSC0003DMG400010963-(rev)M6RLKWAX-EGF_1098

CAAATGTATTTTTTATGTTTTTAAATTTCGTGCAAAATCAAACTAAGACATATAAAATGAAACAAAGAGAGTAACAATATAACAATATCCCACTGTGCCCCTTTATTTTAACTTGTTTTT

>M6-PGSC0003DMG400010963-(rev)M6RLKWAX-EGF_1159

ACAAAGAGAGTAACAATATAACAATATCCCACTGTGCCCCTTTATTTTAACTTGTTTTTAGCATGTAACATGTCTTATTTTCTAAGTTACTAATCTCAGTTTTTATGATGATTTCTTGTT

>M6-PGSC0003DMG400010963-(rev)M6RLKWAX-EGF_1220

CATGTAACATGTCTTATTTTCTAAGTTACTAATCTCAGTTTTTATGATGATTTCTTGTTAATTTCCACAGACATTGATGAGTGTGCAAGTCCAAAAAGGAATCGTTGTCCCAACAATACT

>M6-PGSC0003DMG400010963-(rev)M6RLKWAX-EGF_1281

TTTCCACAGACATTGATGAGTGTGCAAGTCCAAAAAGGAATCGTTGTCCCAACAATACTCGCTGTGTTAATACTCCTGGCACTTACCGTTGTGACTCAAATAATGTGAGACATATGCTAG

>M6-PGSC0003DMG400010963-(rev)M6RLKWAX-EGF_1342

CTGTGTTAATACTCCTGGCACTTACCGTTGTGACTCAAATAATGTGAGACATATGCTAGCTAAACAACTCTCCATCGGTATTGTATAACTCTCTTCTTTGTCTTCTTTTTTCTCCTCTTG

>M6-PGSC0003DMG400010963-(rev)M6RLKWAX-EGF_1403

AAACAACTCTCCATCGGTATTGTATAACTCTCTTCTTTGTCTTCTTTTTTCTCCTCTTGTTTCTTTGTTAGATTAACTTAAGCTTTCTATAGATTTTTATTTGAGACAAATTACTCCTTT

>M6-PGSC0003DMG400010963-(rev)M6RLKWAX-EGF_1464

TCTTTGTTAGATTAACTTAAGCTTTCTATAGATTTTTATTTGAGACAAATTACTCCTTTGAACTGTCCCAACAAATGATTCATAATCCTATTCATGTTTATTCTAGCTAACTGCTCTTTA

>M6-PGSC0003DMG400010963-(rev)M6RLKWAX-EGF_1525

ACTGTCCCAACAAATGATTCATAATCCTATTCATGTTTATTCTAGCTAACTGCTCTTTAGAAAATTAGAAGCTCAACTTAATATATGCTTTAAAATTACAAAGGGGCCTGGATAAGCATT

>M6-PGSC0003DMG400010963-(rev)M6RLKWAX-EGF_1586

AAATTAGAAGCTCAACTTAATATATGCTTTAAAATTACAAAGGGGCCTGGATAAGCATTTAAAGCTCTTTCTCTTAAATTCTCAAAATGAATTTATGCTTGTTTTAGATGAAATTGGTTT

>M6-PGSC0003DMG400010963-(rev)M6RLKWAX-EGF_1647

AAGCTCTTTCTCTTAAATTCTCAAAATGAATTTATGCTTGTTTTAGATGAAATTGGTTTGCTCACAAGCAGGTATTGGAGCAGCAATTACTTTTGTGATCCTGGTAGCAGTTTGTCTTTG

>M6-PGSC0003DMG400010963-(rev)M6RLKWAX-EGF_1708

TCACAAGCAGGTATTGGAGCAGCAATTACTTTTGTGATCCTGGTAGCAGTTTGTCTTTGGCTACGCAAATGGCTCCGGAAGAGGGAAGAAAACAAAGCAAAGCAAAAGTATTTCAAGAGG

>M6-PGSC0003DMG400010963-(rev)M6RLKWAX-EGF_1769

TACGCAAATGGCTCCGGAAGAGGGAAGAAAACAAAGCAAAGCAAAAGTATTTCAAGAGGAATGGTGGATTGCTTTTGCGACAACGTATTTCCTTAATTGGAGAAAGTAGTGGTGGCTCTT

>M6-PGSC0003DMG400010963-(rev)M6RLKWAX-EGF_1830

TGGTGGATTGCTTTTGCGACAACGTATTTCCTTAATTGGAGAAAGTAGTGGTGGCTCTTTGCCAAAGCTCTTTTTGAAAGAGGAGTTGGAGAAAGCAACAGACAATTTCAATGAAATTCG

>M6-PGSC0003DMG400010963-(rev)M6RLKWAX-EGF_1891

CCAAAGCTCTTTTTGAAAGAGGAGTTGGAGAAAGCAACAGACAATTTCAATGAAATTCGAATTCTTGGTAAAGGAGGGGCTGGGACTGTTTACAAAGGAATGTTATCAGATGGGAGCATT

>M6-PGSC0003DMG400010963-(rev)M6RLKWAX-EGF_2013

AGCTGTGAAGAAGTCCAATGCAGTGGATAAAGATCAAATTGAACAGTTTATAAATGAGATACTCATTCTCTCCCAAATAAATCACAGACACATTGTCAAGGTACTGTTGATTTTTGTACT

>M6-PGSC0003DMG400010963-(rev)M6RLKWAX-EGF_2074

CTCATTCTCTCCCAAATAAATCACAGACACATTGTCAAGGTACTGTTGATTTTTGTACTAGGCTCCTGCTACCATTTTATTATGAATGGAAGAAGAGAGTAACACAATTTGACGATAGAG

>M6-PGSC0003DMG400010963-(rev)M6RLKWAX-EGF_2135

GCTCCTGCTACCATTTTATTATGAATGGAAGAAGAGAGTAACACAATTTGACGATAGAGAAATTTATAATGCCTAAAGAGGATTGAAGAGAGAACTTATAATATTAGAAAGACTCCTATA

>M6-PGSC0003DMG400010963-(rev)M6RLKWAX-EGF_2196

ATTTATAATGCCTAAAGAGGATTGAAGAGAGAACTTATAATATTAGAAAGACTCCTATACTCTTACAATATAACACACTAGACTTACACACACGCGCAGTCGCACACACAGAGCTGCTAC

>M6-PGSC0003DMG400010963-(rev)M6RLKWAX-EGF_2257

CTTACAATATAACACACTAGACTTACACACACGCGCAGTCGCACACACAGAGCTGCTACATCAACTAGTGCTCCTATTCACTTCAACCATAATAAACGAACAGTTTAGGATAACTCATTA

>M6-PGSC0003DMG400010963-(rev)M6RLKWAX-EGF_2318

CAACTAGTGCTCCTATTCACTTCAACCATAATAAACGAACAGTTTAGGATAACTCATTAGTCACTCATGAACCAAAGATAACTCAAAATTACAATTGAACCTAGCACTTGATCACACTTT

>M6-PGSC0003DMG400010963-(rev)M6RLKWAX-EGF_2379

CACTCATGAACCAAAGATAACTCAAAATTACAATTGAACCTAGCACTTGATCACACTTTTAACATTTAAGTTTACTGTTTACAACAGGTACTTGGTTGTTGCTTAGAAACTCAAGTCCCA

>M6-PGSC0003DMG400010963-(rev)M6RLKWAX-EGF_2440

ACATTTAAGTTTACTGTTTACAACAGGTACTTGGTTGTTGCTTAGAAACTCAAGTCCCATTATTAGTGTATGAGTACATCTCGAATGGAACCTTGTCATCCCATATTCATGGAAAGCTCA

>M6-PGSC0003DMG400010963-(rev)M6RLKWAX-EGF_2501

ATTAGTGTATGAGTACATCTCGAATGGAACCTTGTCATCCCATATTCATGGAAAGCTCAGTCACAGTTCTAACCCTACTGTCTCAAAGTCTGATTTGGATGATCAGATTCTTCCGCACCC

>M6-PGSC0003DMG400010963-(rev)M6RLKWAX-EGF_2562

CACAGTTCTAACCCTACTGTCTCAAAGTCTGATTTGGATGATCAGATTCTTCCGCACCCAGCAATAATATTATCGTGGGATCATCGGGTGCGAATTGCTGCAGAAATAGCAGGGGCACTA

>M6-PGSC0003DMG400010963-(rev)M6RLKWAX-EGF_2684

TTACATGCACTCATGTGCTTCTACTCCCATTCTTCACAGAGATATAAAATCAAGCAACATATTACTAGACGATAATTTTAGAGCAGTGGTTTCTGATTTTGGACTATCGAGATTGTTGTC

>M6-PGSC0003DMG400010963-(rev)M6RLKWAX-EGF_2745

TTACTAGACGATAATTTTAGAGCAGTGGTTTCTGATTTTGGACTATCGAGATTGTTGTCTGTTGACAAGACTCATTTAACAACAATGGTAGGGGGTACATTTGGTTACATAGATCCAGAA

>M6-PGSC0003DMG400010963-(rev)M6RLKWAX-EGF_2806

TTGACAAGACTCATTTAACAACAATGGTAGGGGGTACATTTGGTTACATAGATCCAGAATACTTCCGCTCAGGTCAACTTACAGACAAATGTGATGTCTATGCCTTCGGTGTAATTCTTG

>M6-PGSC0003DMG400010963-(rev)M6RLKWAX-EGF_2928

GAGCTTCTAACAAGTCAAAGAGTTGTCACTTCTAACCAACCAGAAGATCCAGGTTTGGCTTTCCGTTTCACATTAGCATTGAAAGAAAATCGTATAATTGAGATTGTAGACCCTGAAATT

>M6-PGSC0003DMG400010963-(rev)M6RLKWAX-EGF_2989

TCCGTTTCACATTAGCATTGAAAGAAAATCGTATAATTGAGATTGTAGACCCTGAAATTGTAAAAGAAGTTGAGGATGAACATGTGATTCCTGCTGTGGCCAAACTTGCCAAGAGATGCT

>M6-PGSC0003DMG400010963-(rev)M6RLKWAX-EGF_3172

GAAAAATGAGGCAAGATATGCCTCATAATGAGAGTTTTCAAGATAACATTTCACCAAAAAGTGAAAGTTCATGTAGTCATACTTCTGATTGTACCGAAGAAGATTATCAGAACTCAGTGT

>M6-PGSC0003DMG400010963-(rev)M6RLKWAX-EGF_3233

TGAAAGTTCATGTAGTCATACTTCTGATTGTACCGAAGAAGATTATCAGAACTCAGTGTCTCACAATGAAATGCATTATGACAGAAAGGGAAGTTTGTAACTTTTATAACTTCAACTTTT

>M6-PGSC0003DMG400010963-(rev)M6RLKWAX-EGF_3294

CACAATGAAATGCATTATGACAGAAAGGGAAGTTTGTAACTTTTATAACTTCAACTTTTATCTTTTGTAATTGTAATGTCGAAGTAATTGCATGTATGGACACATAATCAAATGAAGATT

>M6-PGSC0003DMG400010963-(rev)M6RLKWAX-EGF_3332

ACTTTTATAACTTCAACTTTTATCTTTTGTAATTGTAATGTCGAAGTAATTGCATGTATGGACACATAATCAAATGAAGATTAGCATGATTTGCATTAATATTTAAGTCGTATATATGAT

>M6-PGSC0003DMG400010964M6LRR-RLP_1008

AACTGGTGACAGTTTGATGTTACCCTCTTCCTGGGGGTTTCGTTGTTTGAGGTTATTGAAAAGCTTAGATCTCAGTGGATGCAACTTATCTGTTAACCAAACTGCTGCTCTTATGAAATT

>M6-PGSC0003DMG400010964M6LRR-RLP_1071

CTTAGATCTCAGTGGATGCAACTTATCTGTTAACCAAACTGCTGCTCTTATGAAATTGCCTTCTTTACTGGAACTGAATCTGAGCAGAAACAAGTTTATTTTATTACCTGATTTCATCAG

>M6-PGSC0003DMG400010964M6LRR-RLP_1134

TTTACTGGAACTGAATCTGAGCAGAAACAAGTTTATTTTATTACCTGATTTCATCAGTCAACTTTCTCAGCTTCGATATCTCAACATAACACAGTGTCAGGAACTTAAGGAACTTCCCAA

>M6-PGSC0003DMG400010964M6LRR-RLP_1197

TTCTCAGCTTCGATATCTCAACATAACACAGTGTCAGGAACTTAAGGAACTTCCCAAACTTCCTTCGAGTATAGAGGAATTATATGCAGAAGATTTTTTGGCCAAGCAAAGTATCGAAAA

>M6-PGSC0003DMG400010964M6LRR-RLP_1260

TTCGAGTATAGAGGAATTATATGCAGAAGATTTTTTGGCCAAGCAAAGTATCGAAAAGCTACAAATATACCCGAGGTTGAATTTGGTGTCATTCACCAATTATAGTTTTGATCAACAATC

>M6-PGSC0003DMG400010964M6LRR-RLP_1323

AATATACCCGAGGTTGAATTTGGTGTCATTCACCAATTATAGTTTTGATCAACAATCTTATACAGAGGAGAGCAATGGTAGCTCAGTTTTGGATGAGATTCTCAGCTTGTTTCTCTCAGA

>M6-PGSC0003DMG400010964M6LRR-RLP_1386

AGAGGAGAGCAATGGTAGCTCAGTTTTGGATGAGATTCTCAGCTTGTTTCTCTCAGAAGATATGGATCATGTGGTCGAGCACTCTCTTAACTCTGATCACAGGGTGACCTGTTCTATTGT

>M6-PGSC0003DMG400010964M6LRR-RLP_1449

GGATCATGTGGTCGAGCACTCTCTTAACTCTGATCACAGGGTGACCTGTTCTATTGTCTTCCCTGAATGTGCGATTCCCAGGTGGTTTAAGCATCAGAGTGTTGATGAAAAGATATTGTT

>M6-PGSC0003DMG400010964M6LRR-RLP_1575

GCCCATGAATTGGTATAATGATAAGTTCAAGGGCTTTGCTATATGTTGTGCCACTCTCATGGGAGCAGGTGTGTTGAATCCTGATTCAGGGCTATCCAAGAAGTATGATTATGCTTTCTT

>M6-PGSC0003DMG400010964M6LRR-RLP_1638

AGCAGGTGTGTTGAATCCTGATTCAGGGCTATCCAAGAAGTATGATTATGCTTTCTTCAAAGCTAAATTGATATGCAAAGATGATTTAGAAGACCTTAAAGTGCTAGAGAAAGAATGTAA

>M6-PGSC0003DMG400010964M6LRR-RLP_1701

TAAATTGATATGCAAAGATGATTTAGAAGACCTTAAAGTGCTAGAGAAAGAATGTAAAGTGGGAACAGCATCTAGAACATATGGCTTGTGTGTTTGCTTTGTCTACATACCATTGTATGC

>M6-PGSC0003DMG400010964M6LRR-RLP_1764

AACAGCATCTAGAACATATGGCTTGTGTGTTTGCTTTGTCTACATACCATTGTATGCTTCACTGCAGGTGTCCGGCACAGATGTTGGAAATATTAACCAGTATAGCCTATTTGAGGCATC

>M6-PGSC0003DMG400010964M6LRR-RLP_1818

TGCTTCACTGCAGGTGTCCGGCACAGATGTTGGAAATATTAACCAGTATAGCCTATTTGAGGCATCTATCCATGGGCGCATTGTGAGACAGTGGGGAGTTCATTTGATCTATCAGGATGA

>M6-PGSC0003DMG400010964M6LRR-RLP_252

ATGACTTTTAGTTCTCTTGTTGAAGTTGTCAAGGAGCCGAAGGTATTATTTATCTCCTTTAGAAACAAATCATATATGAAGAAAATAAAATGATAGCTAAAGCTGAAGTTTCTATGCATT

>M6-PGSC0003DMG400010964M6LRR-RLP_315

AACAAATCATATATGAAGAAAATAAAATGATAGCTAAAGCTGAAGTTTCTATGCATTCTCTTTTTCTGAACAGGCATTTGATAAGTTGACAGTCCTCAATTTAAGTTTTTCTAGAAATTT

>M6-PGSC0003DMG400010964M6LRR-RLP_945

TGTTAGAAACTTGAGCAAACTTAAAGTCCTATCATTAAGAAGAGGCCGGAAGGTAAAGCGTCAAACTGGTGACAGTTTGATGTTACCCTCTTCCTGGGGGTTTCGTTGTTTGAGGTTATT

>M6-PGSC0003DMG400011045-(rev)M6LRR-RLP_0

ATGGCCGATCTTGTAATTGGTGCTACTGTTAAAGTTGTGCTTGATAAGCTGCTTTCTCTCACAATTGAGGAGGCCAAAACTTTAAGGAACTGCAAGAAAAATCTAAGAATTCTGACAAAA

>M6-PGSC0003DMG400011045-(rev)M6LRR-RLP_1098

GTGTTGGGAGGCCTCTTATGCAACAAGGAAAAACATGAATGGCGGGCAGTTCTTGATGGAAATCCCCTTGTTGCAGGGGAAGATGATAACGGGGAAAGTAGCATAAAGAAGATCCTTAAA

>M6-PGSC0003DMG400011045-(rev)M6LRR-RLP_1159

ATCCCCTTGTTGCAGGGGAAGATGATAACGGGGAAAGTAGCATAAAGAAGATCCTTAAACTCAGCTATGATTATCTACCATCTCCACATCTGAAAAAGTGTTTTGCTTACTTTGCCATTT

>M6-PGSC0003DMG400011045-(rev)M6LRR-RLP_122

TGTATCCATGATTCAAGCTCTCATTCATGATGCTGAAAAACGACAAGTTGATGATCAGGCTGTTGAACAATGGCTCAAGATGCTTGAGAGAGTTGCTGAAGATGCTGAAAACGTGTTTGA

>M6-PGSC0003DMG400011045-(rev)M6LRR-RLP_1220

CAGCTATGATTATCTACCATCTCCACATCTGAAAAAGTGTTTTGCTTACTTTGCCATTTTTCCAAAAGGATTTGAGTTTGAAAAGGACCAGCTTATCCAACTCTGGATGGCAGAAGGTTT

>M6-PGSC0003DMG400011045-(rev)M6LRR-RLP_1281

CCAAAAGGATTTGAGTTTGAAAAGGACCAGCTTATCCAACTCTGGATGGCAGAAGGTTTTCTCGGTCCATCTCAAGAGACCACTTTGATGGAAGACGTTGGGAACAAGGCTTTTCAACTT

>M6-PGSC0003DMG400011045-(rev)M6LRR-RLP_1342

TCGGTCCATCTCAAGAGACCACTTTGATGGAAGACGTTGGGAACAAGGCTTTTCAACTTTTGTTGCAAAATTCCTTGCTGCAAGATGTTAAGCTAGATGATCTGAACAAAATAACATACT

>M6-PGSC0003DMG400011045-(rev)M6LRR-RLP_1403

GTTGCAAAATTCCTTGCTGCAAGATGTTAAGCTAGATGATCTGAACAAAATAACATACTGTAAGATGCATGATCTTGTGCATGATTTGGCTGGAGATATTTTAAAGTCTAAACTATTTGA

>M6-PGSC0003DMG400011045-(rev)M6LRR-RLP_1464

AAGATGCATGATCTTGTGCATGATTTGGCTGGAGATATTTTAAAGTCTAAACTATTTGATCAAAAGAGTATTGGAGGAGAAAATCTTTCTCAAGTCCGATACTTTGGATGGGACTCACCA

>M6-PGSC0003DMG400011045-(rev)M6LRR-RLP_1525

AAAAGAGTATTGGAGGAGAAAATCTTTCTCAAGTCCGATACTTTGGATGGGACTCACCAAGTGATCAAATAGATAAGATAAACGAGTCAGGACGTTTGTGCACATTGTTCTGGAAAAGAA

>M6-PGSC0003DMG400011045-(rev)M6LRR-RLP_1586

TGATCAAATAGATAAGATAAACGAGTCAGGACGTTTGTGCACATTGTTCTGGAAAAGAATTATATCTGAAGAACTGTTGTTGAGCTTTCAGTTCTTGAGAGTTTTAAATTTGTCCAGGTC

>M6-PGSC0003DMG400011045-(rev)M6LRR-RLP_1708

GCATCAAGGTATTGTCAGCCAAAATCGGCAAGCTAATATACTTGAGATATCTTGATCTCTCCAGCACTGAGATCAAAGCCTTGCCCAACTCCATTTGCAAGCTCTACAATTTGCAAACAT

>M6-PGSC0003DMG400011045-(rev)M6LRR-RLP_1769

CAGCACTGAGATCAAAGCCTTGCCCAACTCCATTTGCAAGCTCTACAATTTGCAAACATTTAGAATCAATAGTTGTTACTCACTTAGGAAGCTCCCAGAAGAAATGGCAAATATGATAAG

>M6-PGSC0003DMG400011045-(rev)M6LRR-RLP_1830

AGAATCAATAGTTGTTACTCACTTAGGAAGCTCCCAGAAGAAATGGCAAATATGATAAGTTTGAGACACATATATTGCGACTTTCATTACCAAACATATCTTGATTTTGCGGGTTGGAGA

>M6-PGSC0003DMG400011045-(rev)M6LRR-RLP_1891

TGAGACACATATATTGCGACTTTCATTACCAAACATATCTTGATTTTGCGGGTTGGAGATCAAGGTTTATAGGTAATTATCAGTTTCAGATGCCACTTAATATGGGGCAATTGACTAGTC

>M6-PGSC0003DMG400011045-(rev)M6LRR-RLP_1952

AAGGTTTATAGGTAATTATCAGTTTCAGATGCCACTTAATATGGGGCAATTGACTAGTCTTCAGACGCTACAGTTTTTCTATGTAGGTTTAAAGACAGGTCGTCGAATAGAAGAATTAGG

>M6-PGSC0003DMG400011045-(rev)M6LRR-RLP_2013

CAGACGCTACAGTTTTTCTATGTAGGTTTAAAGACAGGTCGTCGAATAGAAGAATTAGGTCATTTGAAAAACCTTAGAGGTGAATTGACGATCAAACATCTCCAATTAGTCGTAAATAAA

>M6-PGSC0003DMG400011045-(rev)M6LRR-RLP_2074

ATTTGAAAAACCTTAGAGGTGAATTGACGATCAAACATCTCCAATTAGTCGTAAATAAAGAAGAGGCTCGAACAGCATATTTACAGGAGAAACCAAATATCTACAAGCTGACATATTTAT

>M6-PGSC0003DMG400011045-(rev)M6LRR-RLP_2135

AGAGGCTCGAACAGCATATTTACAGGAGAAACCAAATATCTACAAGCTGACATATTTATGGACCCATGATGAATCAGAAGGCTGTGAGATCAATGATGAGCATGTGTTGGATGGTCTTCA

>M6-PGSC0003DMG400011045-(rev)M6LRR-RLP_2257

CGCATCCGAACTTGAAAACCTTAGCAGTAGTGGACTATTTTGGGACTAAATTTCCTTCATGGTTCAGTGAAGGATTGCTACCAAATTTGGTCAAGTTGAAATTAAGTCGCTGCAAAAGGT

>M6-PGSC0003DMG400011045-(rev)M6LRR-RLP_2318

GTTCAGTGAAGGATTGCTACCAAATTTGGTCAAGTTGAAATTAAGTCGCTGCAAAAGGTGCAAAGAAATTCCATCGCTTGGCCAACTGAAGTTCCTTCGGCATCTTGAGCTGATAGGATT

>M6-PGSC0003DMG400011045-(rev)M6LRR-RLP_244

AATTCAGATATGAATATCTCAAAGCACAAGTCATGAACATCCGAACCAAACTAATGGAAAAGGTCAGCAGCTTCTTTTCTCATACTGCTTTTAAGTACAAAATGTCGCGAAAAATAAACA

>M6-PGSC0003DMG400011045-(rev)M6LRR-RLP_2440

TTGAATTGGAATGCCTTGGATCTACATTTTATGGCGTTGATGTTAATGATAAAGGATCAAGCAGCAATAATGGCAATATCCAAGTGTTCCCATCACTGAAAGAACTAGTATTGCAGAATA

>M6-PGSC0003DMG400011045-(rev)M6LRR-RLP_2501

CAGCAATAATGGCAATATCCAAGTGTTCCCATCACTGAAAGAACTAGTATTGCAGAATATGAATAGCCTTATTGAGTGGAAGGGTGATGAAGTTGGAGTTAGAATGTTTTCTGCGCTTGA

>M6-PGSC0003DMG400011045-(rev)M6LRR-RLP_2562

AATAGCCTTATTGAGTGGAAGGGTGATGAAGTTGGAGTTAGAATGTTTTCTGCGCTTGAGAAGTTGAGGATCACAAAGTGTCCACTGTTAAAAGGTACCCCACGTCAATTGGAAATCCTC

>M6-PGSC0003DMG400011045-(rev)M6LRR-RLP_2623

AGTTGAGGATCACAAAGTGTCCACTGTTAAAAGGTACCCCACGTCAATTGGAAATCCTCCGTGAATTAAGCATTGAAGGAGTTGACAGTGAAATGCCATTGTTGACCTTGTGTAGCAACT

>M6-PGSC0003DMG400011045-(rev)M6LRR-RLP_2684

TGAATTAAGCATTGAAGGAGTTGACAGTGAAATGCCATTGTTGACCTTGTGTAGCAACTTGACATCTCTGGTAAAGCTTATTGTGACTGATGTGGAAGAGCTCACTTGTCTTCCTGATGA

>M6-PGSC0003DMG400011045-(rev)M6LRR-RLP_2745

ACATCTCTGGTAAAGCTTATTGTGACTGATGTGGAAGAGCTCACTTGTCTTCCTGATGAGATGCTACGCAATAACGTTTCTCTTCAACAACTATTGGTCACACACTGCAGAGAGTTTCGT

>M6-PGSC0003DMG400011045-(rev)M6LRR-RLP_2806

TGCTACGCAATAACGTTTCTCTTCAACAACTATTGGTCACACACTGCAGAGAGTTTCGTGAATTGCCCCAAAGCCTGTACGATCTCCATTCTCTTGAGAGCTTACAAATACTCTTCTGCA

>M6-PGSC0003DMG400011045-(rev)M6LRR-RLP_2867

ATTGCCCCAAAGCCTGTACGATCTCCATTCTCTTGAGAGCTTACAAATACTCTTCTGCACCAATTTCAGGTCTCTTCCTGTTCCCAGTGTAGAGAATCGTTTGACTTCCCTCCAAAGTCT

>M6-PGSC0003DMG400011045-(rev)M6LRR-RLP_2928

AATTTCAGGTCTCTTCCTGTTCCCAGTGTAGAGAATCGTTTGACTTCCCTCCAAAGTCTTGATTTATCACTGTGTATTGGATTGACCAGTTTACCAAGTGGAATGCTAGAGCATTGTCGG

>M6-PGSC0003DMG400011045-(rev)M6LRR-RLP_2989

ATTTATCACTGTGTATTGGATTGACCAGTTTACCAAGTGGAATGCTAGAGCATTGTCGGTCTCTACAGACTTTGAAGCTCAGCTGTTGTCGCAACTTAGTTTCCTTGCCTTTACACGTAT

>M6-PGSC0003DMG400011045-(rev)M6LRR-RLP_305

GGTCAGCAGCTTCTTTTCTCATACTGCTTTTAAGTACAAAATGTCGCGAAAAATAAACAACATCAATGAAGAGTTGAAGGCTATCAATCAGTTAGCCAATGACCTCGGTCTCAAACCACT

>M6-PGSC0003DMG400011045-(rev)M6LRR-RLP_3050

TCTACAGACTTTGAAGCTCAGCTGTTGTCGCAACTTAGTTTCCTTGCCTTTACACGTATGGGAAATGCCTTCAATTTCATATTTGGGTTTATCAAATTGTCCCAAATTGGTTAGTGTACC

>M6-PGSC0003DMG400011045-(rev)M6LRR-RLP_3111

GAAATGCCTTCAATTTCATATTTGGGTTTATCAAATTGTCCCAAATTGGTTAGTGTACCCACAGGGGGCCTTCACAACCTCACAGGGTTAAAGGTATTGGAAATTGGTCCTTTCTCAGAT

>M6-PGSC0003DMG400011045-(rev)M6LRR-RLP_3172

CAGGGGGCCTTCACAACCTCACAGGGTTAAAGGTATTGGAAATTGGTCCTTTCTCAGATACGGTGGATTTTGAGGCATTCCAATTGATTTTTAATGGCATTCAACAGCTATCATCCCTTC

>M6-PGSC0003DMG400011045-(rev)M6LRR-RLP_3233

GGTGGATTTTGAGGCATTCCAATTGATTTTTAATGGCATTCAACAGCTATCATCCCTTCGTACATTGGAGGTGTATGGATGGATGCGTTGGGATTCTCTGCCTTATCAACTTATGCAACT

>M6-PGSC0003DMG400011045-(rev)M6LRR-RLP_3294

ACATTGGAGGTGTATGGATGGATGCGTTGGGATTCTCTGCCTTATCAACTTATGCAACTCTCTGCCCTAACAAAGATCTGTATATATGATTTTGGAATTGAGGCTCTTCCTCATAGACTT

>M6-PGSC0003DMG400011045-(rev)M6LRR-RLP_3355

CTGCCCTAACAAAGATCTGTATATATGATTTTGGAATTGAGGCTCTTCCTCATAGACTTGACAACCTTACTTCTCTTGTAACATTACATCTAATGGGGTGCCAAAGGCTACAACATGTGG

>M6-PGSC0003DMG400011045-(rev)M6LRR-RLP_3416

CAACCTTACTTCTCTTGTAACATTACATCTAATGGGGTGCCAAAGGCTACAACATGTGGACTTCTCAAATATCATGCCCAAATTACGGTATCTGGAGATCCATTATTGTCTATTGTTAGA

>M6-PGSC0003DMG400011045-(rev)M6LRR-RLP_3477

TTCTCAAATATCATGCCCAAATTACGGTATCTGGAGATCCATTATTGTCTATTGTTAGAAGGTCTGTCGGATGGGCTTGGCAACCTTGTTTCTTTGGAAGAATTAAGTTTACGTAACTGC

>M6-PGSC0003DMG400011045-(rev)M6LRR-RLP_3538

GTCTGTCGGATGGGCTTGGCAACCTTGTTTCTTTGGAAGAATTAAGTTTACGTAACTGCGAGAAACTACAACATCTGCCATCCAGATATGCCATGCTGCGCCTCACTAAATTATGGAAAC

>M6-PGSC0003DMG400011045-(rev)M6LRR-RLP_3599

GAAACTACAACATCTGCCATCCAGATATGCCATGCTGCGCCTCACTAAATTATGGAAACTTCTAATTAAAGGCTGCCCGCAGTTAGAAGAAAGTTGCACCAATCAAAGTAGCCCAAACTC

>M6-PGSC0003DMG400011045-(rev)M6LRR-RLP_366

ATCAATGAAGAGTTGAAGGCTATCAATCAGTTAGCCAATGACCTCGGTCTCAAACCACTGATGGTTCCTTCTCAGAAAATACTTCCAATTCGAGAAACAGATTCCTTAGTTGTTGCTTCG

>M6-PGSC0003DMG400011045-(rev)M6LRR-RLP_3660

CTAATTAAAGGCTGCCCGCAGTTAGAAGAAAGTTGCACCAATCAAAGTAGCCCAAACTCCCAGTGGTCCAACATTTCCCATATTCCAAAAATTGAAGTAGGTGGGAGGATAATTCAGAAG

>M6-PGSC0003DMG400011045-(rev)M6LRR-RLP_3721

AGTGGTCCAACATTTCCCATATTCCAAAAATTGAAGTAGGTGGGAGGATAATTCAGAAGTTACGTAAGTCTCTGTTTCTCTTTCACTTTCTGATCAGTAAAATTTTGAAATATTATTTTC

>M6-PGSC0003DMG400011045-(rev)M6LRR-RLP_3753

GAAGTAGGTGGGAGGATAATTCAGAAGTTACGTAAGTCTCTGTTTCTCTTTCACTTTCTGATCAGTAAAATTTTGAAATATTATTTTCCCCGAGTTCCAGAAAAGAAGTTATGTATGTAG

>M6-PGSC0003DMG400011045-(rev)M6LRR-RLP_427

TGGTTCCTTCTCAGAAAATACTTCCAATTCGAGAAACAGATTCCTTAGTTGTTGCTTCGGATGTTGTTGGTAGAGACAATGATGTTGCTGAAATAAAAGAGAAGATTTTGAACATGAGAA

>M6-PGSC0003DMG400011045-(rev)M6LRR-RLP_488

TGTTGTTGGTAGAGACAATGATGTTGCTGAAATAAAAGAGAAGATTTTGAACATGAGAAAGGATGTTGTTCTGTGCACCATTCCCATAGTGGGTATGGGGGGTTTAGGGAAAACAACTGT

>M6-PGSC0003DMG400011045-(rev)M6LRR-RLP_61

CAATTGAGGAGGCCAAAACTTTAAGGAACTGCAAGAAAAATCTAAGAATTCTGACAAAATATGTATCCATGATTCAAGCTCTCATTCATGATGCTGAAAAACGACAAGTTGATGATCAGG

>M6-PGSC0003DMG400011045-(rev)M6LRR-RLP_610

CTAAGAGAATTTTCAATGACCAACACATCAAGCAGCAGTTCGAAAAGAGAGTTTGGTTGTGCCTACCTGAAATGGTAGAAACTAAGAGCTTTCTTGAACTGATCCTCGAATCATTGACAA

>M6-PGSC0003DMG400011045-(rev)M6LRR-RLP_671

CCTACCTGAAATGGTAGAAACTAAGAGCTTTCTTGAACTGATCCTCGAATCATTGACAAAGAGGAAAGTTGAGGTCCAAAGCAGAGATATAATAGTCAAGACGCTACAAGATGCATTGGG

>M6-PGSC0003DMG400011045-(rev)M6LRR-RLP_732

AGGAAAGTTGAGGTCCAAAGCAGAGATATAATAGTCAAGACGCTACAAGATGCATTGGGAGAAAAAAAGTATTTGCTTGTCCTGGATGATTTGTGGCATGTTGACTCTACATCATGGCAT

>M6-PGSC0003DMG400011045-(rev)M6LRR-RLP_793

AAAAAAAGTATTTGCTTGTCCTGGATGATTTGTGGCATGTTGACTCTACATCATGGCATGAGTTCATGGACACCTTGAGAGGAATAAATACATTCAGAGGAAACTGCATTCTCGTGACTA

>M6-PGSC0003DMG400011045-(rev)M6LRR-RLP_854

GTTCATGGACACCTTGAGAGGAATAAATACATTCAGAGGAAACTGCATTCTCGTGACTACTCGTATCAAACAAGTGGCATCCACAGTAGCAGTAGATCTTCATATGTTGGGGAAATTAAC

>M6-PGSC0003DMG400011045-(rev)M6LRR-RLP_915

CGTATCAAACAAGTGGCATCCACAGTAGCAGTAGATCTTCATATGTTGGGGAAATTAACAGACTATCATTGTTGGTCCATTTTCAAACAAAGAGTGTTTGTTGATGGGGAAGTTCCGGAG

>M6-PGSC0003DMG400011045-(rev)M6LRR-RLP_976

ACTATCATTGTTGGTCCATTTTCAAACAAAGAGTGTTTGTTGATGGGGAAGTTCCGGAGGAAATACTGAGCATGGAGAACAAGATAGTTGAAATGTGTCAAGGTCTACCGTTGGCTACAA

>M6-PGSC0003DMG400011071M6L-LecRK_1054

CAATACGGAGATTCAAGAATGCTGACGTGATTGAGCAGTGGGAGCTTGAGATTGGTCCTCATAGATATTCGTATCATGAACTCAAGAAAGGTACTAGAGGTTTTAAGGAGAGTGAACTAC

>M6-PGSC0003DMG400011071M6L-LecRK_1116

AGATATTCGTATCATGAACTCAAGAAAGGTACTAGAGGTTTTAAGGAGAGTGAACTACTCGGGCGAGGTGGATTTGGGAAAGTTTACAAAGGTGTTTTGAGAAATTCAAAAGCAGAAATT

>M6-PGSC0003DMG400011071M6L-LecRK_1178

GCGAGGTGGATTTGGGAAAGTTTACAAAGGTGTTTTGAGAAATTCAAAAGCAGAAATTGCTGTGAAACGAATTTCTCACGAATCTAAACAAGGGTTGAGGGAATTTGTGTCTGAGATTGC

>M6-PGSC0003DMG400011071M6L-LecRK_124

CTTGCTTTAAAATGTTCAATTTTCTCACATTCCTCTGCTTTGTCATTTCCTTCTCAATTCTTGCTTCATCTCAACTTGATGAATTCATTTTCACAAAATTCAATCAACCAAACAACAACA

>M6-PGSC0003DMG400011071M6L-LecRK_1240

TGAAACGAATTTCTCACGAATCTAAACAAGGGTTGAGGGAATTTGTGTCTGAGATTGCGAGCATTGGAAGACTCCGTCATAGGAATTTGGTTCAATTGTTAGGATGGTGTAGACGTCGTG

>M6-PGSC0003DMG400011071M6L-LecRK_1302

ATTGGAAGACTCCGTCATAGGAATTTGGTTCAATTGTTAGGATGGTGTAGACGTCGTGGTGATTTGTTGCTTGTGTATGAGTTTATGCCTAATGGAAGTTTGGATAACTTCTTGTTTGAT

>M6-PGSC0003DMG400011071M6L-LecRK_1364

TTTGTTGCTTGTGTATGAGTTTATGCCTAATGGAAGTTTGGATAACTTCTTGTTTGATGAGCCAAAAATGGTGTTGACATGGGAACAAAGGTTCAGGATTATTAAAGGGGTTGCGTCTGG

>M6-PGSC0003DMG400011071M6L-LecRK_1426

CAAAAATGGTGTTGACATGGGAACAAAGGTTCAGGATTATTAAAGGGGTTGCGTCTGGTTTACTTTACTTACATGAAGGCTATGAACAAGTCGTGCTACATCGAGATATTAAGGCAAGTA

>M6-PGSC0003DMG400011071M6L-LecRK_1488

CTTTACTTACATGAAGGCTATGAACAAGTCGTGCTACATCGAGATATTAAGGCAAGTAATGTGTTACTAGATGGTGAATTAAATGGTAAGTTAGGGGATTTTGGACTTGCTAGATTGTAT

>M6-PGSC0003DMG400011071M6L-LecRK_1550

GTTACTAGATGGTGAATTAAATGGTAAGTTAGGGGATTTTGGACTTGCTAGATTGTATGAGCATGGGTCTAATCCGTCCACGACTAAAGTTGTAGGTACATTAGGGTACCTTGCACCAGA

>M6-PGSC0003DMG400011071M6L-LecRK_1612

ATGGGTCTAATCCGTCCACGACTAAAGTTGTAGGTACATTAGGGTACCTTGCACCAGAATTACCAAGAATAGGACGAGCCACTACAAAGACTGATGTGTTTGCCTTTGGTGCTTTGTTAC

>M6-PGSC0003DMG400011071M6L-LecRK_1674

CCAAGAATAGGACGAGCCACTACAAAGACTGATGTGTTTGCCTTTGGTGCTTTGTTACTCGAGGTGGTTTGTGGACGTAGGCCAATTGAACACAAAGCAGGACCCGAGGAGTTTGTTCTC

>M6-PGSC0003DMG400011071M6L-LecRK_1736

GGTGGTTTGTGGACGTAGGCCAATTGAACACAAAGCAGGACCCGAGGAGTTTGTTCTCGTGGACTTGGTGTGGGATAAATGGAGAGAAGGTAAAATACTTGACATTGTGGACTATAGATT

>M6-PGSC0003DMG400011071M6L-LecRK_1798

ACTTGGTGTGGGATAAATGGAGAGAAGGTAAAATACTTGACATTGTGGACTATAGATTGAAAGGCGAATACAATGAGAGTGAGGTTTTGATGGTGTTAAAATTGGGACTAATGTGCTCAA

>M6-PGSC0003DMG400011071M6L-LecRK_186

GCTTCATCTCAACTTGATGAATTCATTTTCACAAAATTCAATCAACCAAACAACAACATAACCTTAAGTGGAGTAGCAAAAATAAGCCAAAATGGGTTTATTCAATTAACCAATGACACA

>M6-PGSC0003DMG400011071M6L-LecRK_1860

GGCGAATACAATGAGAGTGAGGTTTTGATGGTGTTAAAATTGGGACTAATGTGCTCAAATAACAAGCCATTGGCGCGACCAAGTATTAGGCAAGTTGTAAGATACTTTGAAGGTGAAATT

>M6-PGSC0003DMG400011071M6L-LecRK_1922

CAAGCCATTGGCGCGACCAAGTATTAGGCAAGTTGTAAGATACTTTGAAGGTGAAATTGGGATGCCTGAGGCTCCAACAACACAAGGTTCTGATGATCTTGAAGGATTTGGATTCGATGA

>M6-PGSC0003DMG400011071M6L-LecRK_1984

TGCCTGAGGCTCCAACAACACAAGGTTCTGATGATCTTGAAGGATTTGGATTCGATGAAAGTGAACGTTGGAATTCCTTAGCATCGACTAGTATTATGATCAAGTCAAGTTCATTTACAT

>M6-PGSC0003DMG400011071M6L-LecRK_2046

GAACGTTGGAATTCCTTAGCATCGACTAGTATTATGATCAAGTCAAGTTCATTTACATGTTTGTCTAATGGAGATGAAAATAGTACTAATTATGTTTCCTCTTCTACTTCACCACTTCCA

>M6-PGSC0003DMG400011071M6L-LecRK_2108

GTCTAATGGAGATGAAAATAGTACTAATTATGTTTCCTCTTCTACTTCACCACTTCCACTTCTATATAGTAGTGAATGTACCTAGATACTAATGTGAACTAGCATTGCAAACTTTTCTAT

>M6-PGSC0003DMG400011071M6L-LecRK_2170

TATATAGTAGTGAATGTACCTAGATACTAATGTGAACTAGCATTGCAAACTTTTCTATATTTTCCTGAAGAACACTAATGTACATTGTTTTTTCCTTTTCTTTGTGTTACCTCTTTGTAG

>M6-PGSC0003DMG400011071M6L-LecRK_2232

TCCTGAAGAACACTAATGTACATTGTTTTTTCCTTTTCTTTGTGTTACCTCTTTGTAGTAGTTTTTTTCCCACAGAAAACTTAGGAAGCTTCCCGGTATTTCAGTGAGAACCTGTTTTCC

>M6-PGSC0003DMG400011071M6L-LecRK_2294

TTTTTTCCCACAGAAAACTTAGGAAGCTTCCCGGTATTTCAGTGAGAACCTGTTTTCCACCATGTGTTTTCCCAGTTACCTGGTTTGATGGGAAATGGGTGGAAAAATCATGTTTTATAA

>M6-PGSC0003DMG400011071M6L-LecRK_248

CTTAAGTGGAGTAGCAAAAATAAGCCAAAATGGGTTTATTCAATTAACCAATGACACAAGTAGACTAATAGGACATGCTTTTTACTCTTCACCTTTCCACTTCAAGACTACTAATGCCTC

>M6-PGSC0003DMG400011071M6L-LecRK_310

GACTAATAGGACATGCTTTTTACTCTTCACCTTTCCACTTCAAGACTACTAATGCCTCTACCTTCTCTTTCTCAACGTGTTTCGCGTTGGCCATAGTCCCCGAGTATCCAAAACTCGGGG

>M6-PGSC0003DMG400011071M6L-LecRK_372

TTCTCTTTCTCAACGTGTTTCGCGTTGGCCATAGTCCCCGAGTATCCAAAACTCGGGGGCCATGGCCTAGCGTTCACAATTTCTCCGTCAAAAGACTTTAGTACAGCTCTTCCGAGCCAG

>M6-PGSC0003DMG400011071M6L-LecRK_434

TGGCCTAGCGTTCACAATTTCTCCGTCAAAAGACTTTAGTACAGCTCTTCCGAGCCAGTACTTAGGTCTGCTCAATGCTAGTGATATTGGTAACTTCTCTAACCATATTTTTGCTGTTGA

>M6-PGSC0003DMG400011071M6L-LecRK_496

TAGGTCTGCTCAATGCTAGTGATATTGGTAACTTCTCTAACCATATTTTTGCTGTTGAATTCGATACGGTACGTGATTTTGAGTTTGGTGATATTGATGATAACCATGTTGGTGTTAATA

>M6-PGSC0003DMG400011071M6L-LecRK_558

GATACGGTACGTGATTTTGAGTTTGGTGATATTGATGATAACCATGTTGGTGTTAATATTAACAGCTTAGAATCAAATAAATCAGCTGCAGCTGGCTATTTTATTAACGATCAAAATGTG

>M6-PGSC0003DMG400011071M6L-LecRK_620

CAGCTTAGAATCAAATAAATCAGCTGCAGCTGGCTATTTTATTAACGATCAAAATGTGAAACAAGATTTGAATCTTAAAAGTGGGAAAGTCATTCTTGTTTGGGTTGAATATGATTCTGT

>M6-PGSC0003DMG400011071M6L-LecRK_682

AAGATTTGAATCTTAAAAGTGGGAAAGTCATTCTTGTTTGGGTTGAATATGATTCTGTTACAAAATTAATTAATGTAACTCTTTCACCAAATTCTTTAAAACCAAAAATTCCTCTTTTAT

>M6-PGSC0003DMG400011071M6L-LecRK_744

AAATTAATTAATGTAACTCTTTCACCAAATTCTTTAAAACCAAAAATTCCTCTTTTATCTTATAATTTTGATGTCTCTACAATTTTTAAAGAAAATATGTATGTTGGCTTCTCTGCTTCT

>M6-PGSC0003DMG400011071M6L-LecRK_806

TAATTTTGATGTCTCTACAATTTTTAAAGAAAATATGTATGTTGGCTTCTCTGCTTCTACTGGTTTGCTTGCTAGTTCACATTTCATTTTGGGTTGGAGTTTTAAGTTAAATGGAGAAGC

>M6-PGSC0003DMG400011071M6L-LecRK_868

GTTTGCTTGCTAGTTCACATTTCATTTTGGGTTGGAGTTTTAAGTTAAATGGAGAAGCAAAATCACTTGACTTGGATTCTCTACCATCACTTCCTGGACCTAAGAAGAAGCACAGAAGAC

>M6-PGSC0003DMG400011071M6L-LecRK_930

TCACTTGACTTGGATTCTCTACCATCACTTCCTGGACCTAAGAAGAAGCACAGAAGACTAATTATTGTCATCTCGATTATAGTGTTTGTGTTCGCGTTAATTTCAATCTTGATTGCTTTT

>M6-PGSC0003DMG400011071M6L-LecRK_992

TATTGTCATCTCGATTATAGTGTTTGTGTTCGCGTTAATTTCAATCTTGATTGCTTTTTATACAATACGGAGATTCAAGAATGCTGACGTGATTGAGCAGTGGGAGCTTGAGATTGGTCC

>M6-PGSC0003DMG400011108M6LRR-RLP_1071

TGCAAAGAAATTCCATCGCTTGGCCAACTGAAGTTCCTTCAGCATCTTGACCTGGTAGGATTCCTTGAATTGGAATGCATCGGACCTACATTTTATGGTGTTGAGATAATGGATAATGGA

>M6-PGSC0003DMG400011108M6LRR-RLP_1134

CTTGAATTGGAATGCATCGGACCTACATTTTATGGTGTTGAGATAATGGATAATGGATCAAGCAGCAATAATGGCAATATCCAAGTGTTCCCGTCATTGAAAGAACTAGTATTGAAGAAT

>M6-PGSC0003DMG400011108M6LRR-RLP_1197

AGCAATAATGGCAATATCCAAGTGTTCCCGTCATTGAAAGAACTAGTATTGAAGAATATGGCTAGCCTTATTGAGTGGAAGGGAGTTGAAGTTGGAGTAAGCTTGAGAAGTTGAGGATTA

>M6-PGSC0003DMG400011108M6LRR-RLP_126

TTGTTGCGAAATTCCTTGCTGCAAGATGTCAAGCTAGATGAGCACAACAATATAACACACTGTAAGATGCATGATCTTGTGCATGATTTGGCCAGAGATATATTGAAATCTAAATTATTT

>M6-PGSC0003DMG400011108M6LRR-RLP_1260

AGCCTTATTGAGTGGAAGGGAGTTGAAGTTGGAGTAAGCTTGAGAAGTTGAGGATTAGTGACTGTCCCTTGTTAGAAAGTACTCCGAATCAATTTGAAATCCTATGTGAATTAGTGATTG

>M6-PGSC0003DMG400011108M6LRR-RLP_1323

GTCCCTTGTTAGAAAGTACTCCGAATCAATTTGAAATCCTATGTGAATTAGTGATTGTTGGAGTTGGCAGTGAAATGCCATTGTTGAACTTGTGCAGCAACTTGACATCTCTTGTGAATC

>M6-PGSC0003DMG400011108M6LRR-RLP_1386

TTGGCAGTGAAATGCCATTGTTGAACTTGTGCAGCAACTTGACATCTCTTGTGAATCTTGTTGTTAGTAACGTGAAAGAGCTCATTTTTCTTCCCGATGAGATGCTACGCAACAATGTTT

>M6-PGSC0003DMG400011108M6LRR-RLP_1512

AACACCTATTGGTCTCACTCTGCGGAGAGTTTCGTGAATTGCCACAAAGCTTATACAATCTCCGTTCTCTTAAGAGCTTAGAGATTATCAGCTGCACCAATTTCAGTTCTTTTCCTGTTC

>M6-PGSC0003DMG400011108M6LRR-RLP_1575

GTTCTCTTAAGAGCTTAGAGATTATCAGCTGCACCAATTTCAGTTCTTTTCCTGTTCCCAGTGGAGAGAACTATTTGACTTCCCTCCAAAATTTTGAGTTATCCAATTGTAATGGATTGA

>M6-PGSC0003DMG400011108M6LRR-RLP_1638

GAGAGAACTATTTGACTTCCCTCCAAAATTTTGAGTTATCCAATTGTAATGGATTGACCAGTTTACCAAGTGGAATGCTAGAGCAATGCCGGTCTCTAGAGATTTTGACGGTCAGTTACT

>M6-PGSC0003DMG400011108M6LRR-RLP_1701

TACCAAGTGGAATGCTAGAGCAATGCCGGTCTCTAGAGATTTTGACGGTCAGTTACTGTAACAACTTGGTTTCCTTGCCTTTACATGTGTGGGAAATGCCTTCTCTTTCGTATTTGTGTT

>M6-PGSC0003DMG400011108M6LRR-RLP_1764

ACTTGGTTTCCTTGCCTTTACATGTGTGGGAAATGCCTTCTCTTTCGTATTTGTGTTTATCAGAATGTCCCAAATTGGTTAGTGTACCCGCAGGGGGTCTTCACCACCTCACTGGGTTAC

>M6-PGSC0003DMG400011108M6LRR-RLP_1827

AATGTCCCAAATTGGTTAGTGTACCCGCAGGGGGTCTTCACCACCTCACTGGGTTACAGAATTTGGAAATTGGTCCTTTCTCAGAGACGGTGAATTTTGAGGCATTCCAATTGATTTTTA

>M6-PGSC0003DMG400011108M6LRR-RLP_189

AAGATGCATGATCTTGTGCATGATTTGGCCAGAGATATATTGAAATCTAAATTATTTGATCCAAAGAGAGTTGGAGGAGAAAATCTTTCTCAAGTCCGATACTTTGGATGGGACTCACCA

>M6-PGSC0003DMG400011108M6LRR-RLP_1890

TGGAAATTGGTCCTTTCTCAGAGACGGTGAATTTTGAGGCATTCCAATTGATTTTTAATGGCATTCAGCAGCTATCGTCCCTTCGTACATTGGAGGTATACGGACGTACGGATTGGGATT

>M6-PGSC0003DMG400011108M6LRR-RLP_1953

TTCAGCAGCTATCGTCCCTTCGTACATTGGAGGTATACGGACGTACGGATTGGGATTCTCTGCCCCATCAGCTTATGCAACTCTCTGCCCTAACAGAGATCGGAATAAATGATTTCGGAA

>M6-PGSC0003DMG400011108M6LRR-RLP_2016

CCCATCAGCTTATGCAACTCTCTGCCCTAACAGAGATCGGAATAAATGATTTCGGAATCGAGGCTCTTCCTCATAGACTTGGCAACCTTACTTCTCTTGTAACATTAGATCTGGTGAGAT

>M6-PGSC0003DMG400011108M6LRR-RLP_2079

CTCTTCCTCATAGACTTGGCAACCTTACTTCTCTTGTAACATTAGATCTGGTGAGATGCAAACGGCTACAACATGTGGACTTCTCAGATGCCATGCCCAAATTACGGTATCTGAAGATCC

>M6-PGSC0003DMG400011108M6LRR-RLP_2142

GGCTACAACATGTGGACTTCTCAGATGCCATGCCCAAATTACGGTATCTGAAGATCCGTGATTGTCCATTGTTAGAAACTTTGTCGGATGGGCTCGGCAACCTTGTTTCTTTGGAAGAGA

>M6-PGSC0003DMG400011108M6LRR-RLP_2205

GTCCATTGTTAGAAACTTTGTCGGATGGGCTCGGCAACCTTGTTTCTTTGGAAGAGATAAGTTTATGTAACTGCGAAAAACTGGAGCATCTGCCATGCAACTCCTCACTAAATTATGGAA

>M6-PGSC0003DMG400011108M6LRR-RLP_252

AAGAGAGTTGGAGGAGAAAATCTTTCTCAAGTCCGATACTTTGGATGGGACTCACCAAGTGATCAAATAGAGAGGATAAATGAGCCAGGACGTTTGTGCACATTGTTCTGGAAAAGAAAT

>M6-PGSC0003DMG400011108M6LRR-RLP_315

CAAATAGAGAGGATAAATGAGCCAGGACGTTTGTGCACATTGTTCTGGAAAAGAAATCTATCTGAAGATATGTTGTTGCGCTTTCAGTTCTTGAGAGTTTTAAATTTGTCCAGGTCAGAC

>M6-PGSC0003DMG400011108M6LRR-RLP_378

GAAGATATGTTGTTGCGCTTTCAGTTCTTGAGAGTTTTAAATTTGTCCAGGTCAGACATCAAGGAGTTGTCAGCCTCAATCAGCAAGCTAATATACTTGAGATATCTTGATCTCTCCTAC

>M6-PGSC0003DMG400011108M6LRR-RLP_504

ATCAGAGCCTTGCCCAGCTCCATTTGCAAGCTCTACAATTTGCAAACTTTTAGAGTCAATAACTGTTTTTCACTCAGGAAGCTTCCAGTAGAAATGGCAAATATGATAAGTTTGAGACAC

>M6-PGSC0003DMG400011108M6LRR-RLP_567

TGTTTTTCACTCAGGAAGCTTCCAGTAGAAATGGCAAATATGATAAGTTTGAGACACATATATTGCAAGTATCATTACCAATCATGTACAGGTAAGGATCGGTTTCAGATGCCACTTAAT

>M6-PGSC0003DMG400011108M6LRR-RLP_63

TTTCTTCGTCCATGTCAAGAGACCACTGTGATGGAAGACGCCGGGAACAAGTTTTTCCAACACTTGTTGCGAAATTCCTTGCTGCAAGATGTCAAGCTAGATGAGCACAACAATATAACA

>M6-PGSC0003DMG400011108M6LRR-RLP_630

TGCAAGTATCATTACCAATCATGTACAGGTAAGGATCGGTTTCAGATGCCACTTAATATGGAGCAATTGACTAGTCTTCAGACGCTACAAGTTTTCTATGTAGGTTTAAAGAAAGGTAAT

>M6-PGSC0003DMG400011108M6LRR-RLP_693

CAATTGACTAGTCTTCAGACGCTACAAGTTTTCTATGTAGGTTTAAAGAAAGGTAATCGAATAGAAGAATTAGGTTGTTTAAAAAAACTTAGAGGTGAATTGACGATCGAACGTCTTCAA

>M6-PGSC0003DMG400011108M6LRR-RLP_756

GAAGAATTAGGTTGTTTAAAAAAACTTAGAGGTGAATTGACGATCGAACGTCTTCAATTAGTCGGAAATAAAGAAGAGGCTCGAACAGCATATTTACAGGAGAAACCAAATATCTACAAG

>M6-PGSC0003DMG400011108M6LRR-RLP_819

GGAAATAAAGAAGAGGCTCGAACAGCATATTTACAGGAGAAACCAAATATCTACAAGCTGGTGTATTCATGGTCCCGTGATGAATCAGAAGGCTGTGAGATCAATCATGAGCATGTGCTG

>M6-PGSC0003DMG400011108M6LRR-RLP_882

TATTCATGGTCCCGTGATGAATCAGAAGGCTGTGAGATCAATCATGAGCATGTGCTGGATGGTCTTCAACCACATCCTAACTTAAAAACCTTAGCAGTTGTGAACTATTTAGGAACTAAA

>M6-PGSC0003DMG400011108M6LRR-RLP_945

CTTCAACCACATCCTAACTTAAAAACCTTAGCAGTTGTGAACTATTTAGGAACTAAATTTCCTTCATGGTTCAGTGAAGAATTGCTACCAAATTTGGTCATGTTGAAATTAAGTGGTTGC

>M6-PGSC0003DMG400011116M6LRR-RLP_0

ATGTTTCCTGGGCTTGAGAAGTTGACGATTACAGAGTGTTCACTGTTAAAAAGTGCTCCAACTCAATTGGAAACCCTGCGTGAATTAGAAATTGAAAGAGTTGACAGTGAAATGCCATTG

>M6-PGSC0003DMG400011116M6LRR-RLP_1008

CTGCCATCCAGAGATGTCATGCGAAGCCTCACGAAATTACAGTTCCTGAAAATTAAAGACTGCCCACAGTTAGAAGAAAGTTGCACCAATTGGAGTAGCCCAAACTCCCAGTGGTCCAAC

>M6-PGSC0003DMG400011116M6LRR-RLP_1060

TTAAAGACTGCCCACAGTTAGAAGAAAGTTGCACCAATTGGAGTAGCCCAAACTCCCAGTGGTCCAACATTTCCCATAATCCAAAAATTAAAGTAGGTGGGAGGATGATTCAGAACTTAA

>M6-PGSC0003DMG400011116M6LRR-RLP_126

TTGTGCAGCAACTTAACATGTCTTGTGGATCTCGATGTCAGTGATGTGAAAGAGCTCACTTGTCTACCTGATGAGATGCTACACAAGAACGTTTCTCTTCAACGCCTGTGGATCTCAGAT

>M6-PGSC0003DMG400011116M6LRR-RLP_189

CTACCTGATGAGATGCTACACAAGAACGTTTCTCTTCAACGCCTGTGGATCTCAGATTGCAGAGAGTTTTGTGAATTGCCACAGAGCTTGTACAATCTCCATTCCCTTAAGAGCTTACGA

>M6-PGSC0003DMG400011116M6LRR-RLP_252

GAGTTTTGTGAATTGCCACAGAGCTTGTACAATCTCCATTCCCTTAAGAGCTTACGAATATCCTTCTTCCCCAATTTCAGTTCTTTTCCTGTTCCCAGTGGAGAGAATTATTTAACTTCC

>M6-PGSC0003DMG400011116M6LRR-RLP_378

AGGCTTCACTTACTTTATTGTGATGGATTGACCAGTTTACCAAATGGAATGCTAGAACAATGCCCGTCTCTAAAGATTTTGAGGGTCAACAACTGTAAGAACTTGGTTTCCTTCCCTTTA

>M6-PGSC0003DMG400011116M6LRR-RLP_441

CCGTCTCTAAAGATTTTGAGGGTCAACAACTGTAAGAACTTGGTTTCCTTCCCTTTACATGTGTGGGAAGTACCTTCACTTTCCATTTTGGATATGTCAAATTGCCCCAAATTGATTAGT

>M6-PGSC0003DMG400011116M6LRR-RLP_504

TGGGAAGTACCTTCACTTTCCATTTTGGATATGTCAAATTGCCCCAAATTGATTAGTCTACCCACAGGAGGCCTTCACCATCTCACTGGGTTATGACATTTGAAAATTGGTCCTTTCTCA

>M6-PGSC0003DMG400011116M6LRR-RLP_63

CAATTGGAAACCCTGCGTGAATTAGAAATTGAAAGAGTTGACAGTGAAATGCCATTGTTGAACTTGTGCAGCAACTTAACATGTCTTGTGGATCTCGATGTCAGTGATGTGAAAGAGCTC

>M6-PGSC0003DMG400011116M6LRR-RLP_630

GTGGATTTTGAGGCATTCCAATTGATTTTTAATGGCATTCAGCCGCTATTGTCCCTTCGTACGTTGTGGGTGTACGGACATGGGCACTGGGATTCTCTGCCCTATCAGCTTATGTAACTC

>M6-PGSC0003DMG400011116M6LRR-RLP_693

TTGTGGGTGTACGGACATGGGCACTGGGATTCTCTGCCCTATCAGCTTATGTAACTCTCTTCCCTAATTGAGATCAGATTAAGTGATTTCGGTATCGAGGATCTTCCTCATAGTCTTTGC

>M6-PGSC0003DMG400011116M6LRR-RLP_756

CTAATTGAGATCAGATTAAGTGATTTCGGTATCGAGGATCTTCCTCATAGTCTTTGCAATCTTACTTCTCTTGAAACATTACATCTAGTGGGGTGCAAACGGCTACAACATGTGGACTTC

>M6-PGSC0003DMG400011116M6LRR-RLP_819

ACTTCTCTTGAAACATTACATCTAGTGGGGTGCAAACGGCTACAACATGTGGACTTCAGAGATGCCATGCCCAAATTACATTTTCTTGAGATCTGTGGTTGTCCATTGTTAGAAGCTCTG

>M6-PGSC0003DMG400011116M6LRR-RLP_882

GCCATGCCCAAATTACATTTTCTTGAGATCTGTGGTTGTCCATTGTTAGAAGCTCTGTCGGATGGGCTCAGCAACCTTGTTTGTTTGGAAACGTTAATTTTTTGGAACTGCGAAATACTG

>M6-PGSC0003DMG400011116M6LRR-RLP_945

GGGCTCAGCAACCTTGTTTGTTTGGAAACGTTAATTTTTTGGAACTGCGAAATACTGGAGTGTCTGCCATCCAGAGATGTCATGCGAAGCCTCACGAAATTACAGTTCCTGAAAATTAAA

>M6-PGSC0003DMG400011160-(rev)M6LRR-RLK_1054

TCTAATAACCATTTCTCTGGTCATGTTCCTGATTCTTTGGGTCTGTTAAACAGTCTTCAGTATCTGTAAGTGTATTAAACTTTCAAAATTTTCATTTTTGTGTGATTATCAAGATTATGT

>M6-PGSC0003DMG400011160-(rev)M6LRR-RLK_1116

TCTGTAAGTGTATTAAACTTTCAAAATTTTCATTTTTGTGTGATTATCAAGATTATGTTTTTAGAGTTGAATGGTCTAAAACACACCTGAAGTATCACGTTTTTGCGAGTTTTCTACGAA

>M6-PGSC0003DMG400011160-(rev)M6LRR-RLK_1178

AGAGTTGAATGGTCTAAAACACACCTGAAGTATCACGTTTTTGCGAGTTTTCTACGAATTATCACCTAAAACATATCTCACCGCCATCTCATAGTCGATGTGTGTTTTGATAAATAGTTG

>M6-PGSC0003DMG400011160-(rev)M6LRR-RLK_1240

CACCTAAAACATATCTCACCGCCATCTCATAGTCGATGTGTGTTTTGATAAATAGTTGGTGATGGTTTAAGAAGGAAAAGTGAACACATGATAGTTTAGGTAGAAAATGTGCGAAATCGT

>M6-PGSC0003DMG400011160-(rev)M6LRR-RLK_1302

TGGTTTAAGAAGGAAAAGTGAACACATGATAGTTTAGGTAGAAAATGTGCGAAATCGTGATACTTCAAAGTGTGGTTTTTACCATTATCTCTTGTTTTTCTGAGTACCCTTTTTCCTTTA

>M6-PGSC0003DMG400011160-(rev)M6LRR-RLK_1364

CTTCAAAGTGTGGTTTTTACCATTATCTCTTGTTTTTCTGAGTACCCTTTTTCCTTTAATTGTTGTTTTCTCATGTAGACAATGTTTAATTGCTATCTTATTCTTTGACTATATGTTTTT

>M6-PGSC0003DMG400011160-(rev)M6LRR-RLK_1550

TTGTCTGGTGCTATTCCTCTTTCTTTAGCCAGGGTACCTCAGCTTGCTTTCTTGTGAGTATTGCTTCTTTAAATCTATGACTTTTGTTTGATTTTGTTTTATTCTTTTGTCTGATTGTCC

>M6-PGSC0003DMG400011160-(rev)M6LRR-RLK_1612

GCTTCTTTAAATCTATGACTTTTGTTTGATTTTGTTTTATTCTTTTGTCTGATTGTCCTGAGAATTGGAATTATCTAGTCACTTTCTTGAATCAGCCTTATGTTAAAGCTTTTTTGGTTT

>M6-PGSC0003DMG400011160-(rev)M6LRR-RLK_1674

AATTGGAATTATCTAGTCACTTTCTTGAATCAGCCTTATGTTAAAGCTTTTTTGGTTTAAAGGAAATTGAATAGTTTGTTCTGTTTAGGGGTATTATTATCTTTTAACTTATTATTCATT

>M6-PGSC0003DMG400011160-(rev)M6LRR-RLK_186

ACAAAATAGACAGACTGATAAAGAACCTGTTTCTTTTCCTTTACACTCTCTCACTTTTTTTTTTTTAGTTTTTGACTGAAAAAAGACACAGTTTTTACTCATTTATATGAACCAAAATGT

>M6-PGSC0003DMG400011160-(rev)M6LRR-RLK_1860

ACCCCAACTTGTTTGGGACTGAGGCGTTGTTGTCGTTGTCGGATAGTGTTGTCTTTTTTTTGTATAAAGAAATGAATAGTGTGTTCTGATTAGGGGTATTATTATATTTTAACTTATTTT

>M6-PGSC0003DMG400011160-(rev)M6LRR-RLK_1984

TTTTATTATATTTAGTATGACGATGATATGCAATGACCTTAAATGAATGTGGGGATTCATTATGCGGACCCCAACTTGTTTGTAACTGAGGCGTTGTTGTTTTTGTTGTCATCGAATAGT

>M6-PGSC0003DMG400011160-(rev)M6LRR-RLK_2046

TGCGGACCCCAACTTGTTTGTAACTGAGGCGTTGTTGTTTTTGTTGTCATCGAATAGTGTGTTCTGTTTTTGTGTAAAGGAAATGAAGAGTGTGTTCTGATCAGGGGTATTATTGTATTT

>M6-PGSC0003DMG400011160-(rev)M6LRR-RLK_2170

TTATTTTTTACTTTTATTTTATTTAGTATAATGATGATATGCAGTGAACTTAAATGAAAGTGGGGATTCATATTGAACTTGTTTGGGATTGAGACGTTGTTGTTGTTGTCGAATAGTGTG

>M6-PGSC0003DMG400011160-(rev)M6LRR-RLK_2232

GGGATTCATATTGAACTTGTTTGGGATTGAGACGTTGTTGTTGTTGTCGAATAGTGTGTCTGCTTTGTGTAAAGGAAATGAATAGTGTGTTCTGTTCAGGGTTATTATTATATTCTATAC

>M6-PGSC0003DMG400011160-(rev)M6LRR-RLK_2294

CTTTGTGTAAAGGAAATGAATAGTGTGTTCTGTTCAGGGTTATTATTATATTCTATACTTATATTTCACTTTTATTTAATCTAGAATGACGATGATATGCAACGAGCTTAAATGAAAGTG

>M6-PGSC0003DMG400011160-(rev)M6LRR-RLK_2356

ATTTCACTTTTATTTAATCTAGAATGACGATGATATGCAACGAGCTTAAATGAAAGTGGCTATTCATATTGCCGACCCCAACTTTTTTTCTTGGTTGAGACTGAGGCGTTGTTCGCGTCG

>M6-PGSC0003DMG400011160-(rev)M6LRR-RLK_248

TTTTAGTTTTTGACTGAAAAAAGACACAGTTTTTACTCATTTATATGAACCAAAATGTTAAATAAATTCATGGGGTTTGCTTAATATATGTGTTGAATCATTTGATTTTACTTTCTTGGT

>M6-PGSC0003DMG400011160-(rev)M6LRR-RLK_310

TAAATTCATGGGGTTTGCTTAATATATGTGTTGAATCATTTGATTTTACTTTCTTGGTTTTTTTTTTTTATCATCAATGGCTTCCATTTTCTTGATTTTTGTTCTCTCTTTGCTTCTGTT

>M6-PGSC0003DMG400011160-(rev)M6LRR-RLK_372

TTTTTTTATCATCAATGGCTTCCATTTTCTTGATTTTTGTTCTCTCTTTGCTTCTGTTTTACCCTTTTCTTTGTTTTTCATATGAAGCTAGGAATCATGAAGGTAAACAAAAAAAAAAAA

>M6-PGSC0003DMG400011160-(rev)M6LRR-RLK_434

CCTTTTCTTTGTTTTTCATATGAAGCTAGGAATCATGAAGGTAAACAAAAAAAAAAAAGAACCCCATTTGCTTATTTTTCAGCTTTTGATTTTCAAAATTTGATCTTTTGTTTTGATGTA

>M6-PGSC0003DMG400011160-(rev)M6LRR-RLK_496

CCCATTTGCTTATTTTTCAGCTTTTGATTTTCAAAATTTGATCTTTTGTTTTGATGTATAGTGGAAGCTTTGATTGCTATAAGGAGAAGTTTGGATGACCCAAAAGGAGCTTTAAGTAAT

>M6-PGSC0003DMG400011160-(rev)M6LRR-RLK_558

GGAAGCTTTGATTGCTATAAGGAGAAGTTTGGATGACCCAAAAGGAGCTTTAAGTAATTGGGATGAAGATTCTGTTGATCCTTGTAGTTGGGCTATGATCAGTTGTTCTTCTGAAAATCT

>M6-PGSC0003DMG400011160-(rev)M6LRR-RLK_620

ATGAAGATTCTGTTGATCCTTGTAGTTGGGCTATGATCAGTTGTTCTTCTGAAAATCTTGTTATTGCACTGTATGTTTAATAGTGTGTTTTTTTTTGGAATTTTATGGTACTATATGATA

>M6-PGSC0003DMG400011160-(rev)M6LRR-RLK_682

ATTGCACTGTATGTTTAATAGTGTGTTTTTTTTTGGAATTTTATGGTACTATATGATAATATTGGTTGTTAAAGATTGAAACTTTTACTGTTTTATCAACTAAGAAAGTATTTTTTTTTT

>M6-PGSC0003DMG400011160-(rev)M6LRR-RLK_744

TGGTTGTTAAAGATTGAAACTTTTACTGTTTTATCAACTAAGAAAGTATTTTTTTTTTGTTGTAGAGGAGCTCCAAGTCAAGATTTATCTGGGGTTTTGTCTGGGATGATATCAAATCTT

>M6-PGSC0003DMG400011160-(rev)M6LRR-RLK_806

TAGAGGAGCTCCAAGTCAAGATTTATCTGGGGTTTTGTCTGGGATGATATCAAATCTTACAAACCTTAAACAAGTGTAAGAGACAATTTTTTAATGTTTTTGTTGTATGTAATTTCAAAT

>M6-PGSC0003DMG400011160-(rev)M6LRR-RLK_868

ACCTTAAACAAGTGTAAGAGACAATTTTTTAATGTTTTTGTTGTATGTAATTTCAAATTTTGATGACCACATTTGGTTAATTTTCTTGTATGTTGAAAATTGTGTAGTTTATTGCAAAAC

>M6-PGSC0003DMG400011160-(rev)M6LRR-RLK_930

ATGACCACATTTGGTTAATTTTCTTGTATGTTGAAAATTGTGTAGTTTATTGCAAAACAACAACATTTCTGGGCATATTCCAAGAGAGCTAGGGAAACTTCCTAATCTTGAAACATTGGA

>M6-PGSC0003DMG400011160-(rev)M6LRR-RLK_992

ACATTTCTGGGCATATTCCAAGAGAGCTAGGGAAACTTCCTAATCTTGAAACATTGGATCTGTCTAATAACCATTTCTCTGGTCATGTTCCTGATTCTTTGGGTCTGTTAAACAGTCTTC

>M6-PGSC0003DMG400011205M6LRR-RLK_0

ATGTCTATGATTGTTTCTGATCCATTTCATGTACTTGAAAGTTGGAATTCTTCCAACGTTCATGTATGTAATTGGACAGGAATTGTATGTGACAAGAAAATGCATAGAGTTGTTGAGCTT

>M6-PGSC0003DMG400011205M6LRR-RLK_1054

TCCCAAATACCTTTGCTAATCTTCCGCAACTGAGGAGGCTTTTGTTGCACGATAATCACCTCTCTGGAACGATTCCCCCGAGCTTAGGTGATTGTGTTAACTTGGAAATACTTGATCTTT

>M6-PGSC0003DMG400011205M6LRR-RLK_1116

TCTGGAACGATTCCCCCGAGCTTAGGTGATTGTGTTAACTTGGAAATACTTGATCTTTCTCACAACAGAATCACGGGGACGATTCCAAGTGCAGTTGCTGGATTGAGTAGTTTGAAGCTC

>M6-PGSC0003DMG400011205M6LRR-RLK_1178

CAACAGAATCACGGGGACGATTCCAAGTGCAGTTGCTGGATTGAGTAGTTTGAAGCTCTATCTGAACTTGTCTAGTAATTACCTACATGGTCCAATTCCTTTAGAGCTCAGCAAAATGGA

>M6-PGSC0003DMG400011205M6LRR-RLK_124

TTAGTCATCATTCGTTACGTGGAACGATTTCTCCTTCTCTCTCTGGGCTCTCTTTCTTGCAGATTCTTGATTTGTCTGGTAACTTGTTAGAAGGACAAATTCCAGCTGAATTAGGCAATC

>M6-PGSC0003DMG400011205M6LRR-RLK_1240

TGAACTTGTCTAGTAATTACCTACATGGTCCAATTCCTTTAGAGCTCAGCAAAATGGATATGGTGTTGGCAATTGACTTGTCTTTGAATAATCTCTCGTCTAATGTCCCTTCACAGCTTG

>M6-PGSC0003DMG400011205M6LRR-RLK_1302

GTGTTGGCAATTGACTTGTCTTTGAATAATCTCTCGTCTAATGTCCCTTCACAGCTTGGCAGCTGCATTGCTCTCGAATACTTGAATCTATCGCGTAATTCCTTAGAAGGACCTCTTCCA

>M6-PGSC0003DMG400011205M6LRR-RLK_1364

CTGCATTGCTCTCGAATACTTGAATCTATCGCGTAATTCCTTAGAAGGACCTCTTCCATCTTCGATAGGAAGATTGCCTTACCTTAAGGAAATCGACGTTTCATTCAATGTGTTGAGTGG

>M6-PGSC0003DMG400011205M6LRR-RLK_1426

CGATAGGAAGATTGCCTTACCTTAAGGAAATCGACGTTTCATTCAATGTGTTGAGTGGGGAAATACCACAGACATTTCAAGCATCGTCGACTTTGCAGAACCTCAACTTCTCCTATAACC

>M6-PGSC0003DMG400011205M6LRR-RLK_1488

ATACCACAGACATTTCAAGCATCGTCGACTTTGCAGAACCTCAACTTCTCCTATAACCGTTTCTCGGGGAACGTGACAGATACTGGTGCATTTTCATCACTAACCATCAGCTCATTCATG

>M6-PGSC0003DMG400011205M6LRR-RLK_1550

CTCGGGGAACGTGACAGATACTGGTGCATTTTCATCACTAACCATCAGCTCATTCATGGGAAATTTCAAGCTTTGTGGCTCAGTACAAGGGATGAGAAGTTGCCATAAAAAGAGGGGTAG

>M6-PGSC0003DMG400011205M6LRR-RLK_1612

ATTTCAAGCTTTGTGGCTCAGTACAAGGGATGAGAAGTTGCCATAAAAAGAGGGGTAGACATTTCATTATCACAATCCTCCTTTCATTGCTTATAACTCCTATATTTTGCATAGTCGGGT

>M6-PGSC0003DMG400011205M6LRR-RLK_1674

TTCATTATCACAATCCTCCTTTCATTGCTTATAACTCCTATATTTTGCATAGTCGGGTACCCTCTCCTCCTAAGATCAAGATTCAGGAACCAATTCGCGAGTTTTAATAAAAGCAAGGTA

>M6-PGSC0003DMG400011205M6LRR-RLK_1736

TCTCCTCCTAAGATCAAGATTCAGGAACCAATTCGCGAGTTTTAATAAAAGCAAGGTAATTGAAGATGTGGAGCAAGGTGGTGGGAGGAAAGAGCAAAAGTACCCGAGGATATCTCGTAT

>M6-PGSC0003DMG400011205M6LRR-RLK_1798

AAGATGTGGAGCAAGGTGGTGGGAGGAAAGAGCAAAAGTACCCGAGGATATCTCGTATGCAGCTGATTGAAGCAACGGGAGGTTTTAGCAGCTCGAGTTTAATAGGAGCAGGAAGATATG

>M6-PGSC0003DMG400011205M6LRR-RLK_186

ATTCTTGATTTGTCTGGTAACTTGTTAGAAGGACAAATTCCAGCTGAATTAGGCAATCTTGTTAAGCTAAATCAGCTTAGTTTGTCTTCTAATCTCCTCGAAGGGGAAATCCCAAACGAG

>M6-PGSC0003DMG400011205M6LRR-RLK_1860

CTGATTGAAGCAACGGGAGGTTTTAGCAGCTCGAGTTTAATAGGAGCAGGAAGATATGGGCGTGTTTATAAAGGAGTCCTGAAGAACAACATGAGCATAGCTGTTAAAGTAATTGACACA

>M6-PGSC0003DMG400011205M6LRR-RLK_1922

TGTTTATAAAGGAGTCCTGAAGAACAACATGAGCATAGCTGTTAAAGTAATTGACACAAAAGGAGGTGGAGAGATATCAGGTAGCTTCAAAAGGGAATGTCAAATCTTGAAAAGGACTAG

>M6-PGSC0003DMG400011205M6LRR-RLK_1984

GAGGTGGAGAGATATCAGGTAGCTTCAAAAGGGAATGTCAAATCTTGAAAAGGACTAGACATAGGAACTTGATAAGGATCATAACGACATGTAGCAGACCGGACTTCAAAGCTCTCGTGT

>M6-PGSC0003DMG400011205M6LRR-RLK_2046

AGGAACTTGATAAGGATCATAACGACATGTAGCAGACCGGACTTCAAAGCTCTCGTGTTCCCTCTGATGTCAAACGGAAGCCTAGAGAACCATCTGTACCCAAGCCACAGGCTGAGACAT

>M6-PGSC0003DMG400011205M6LRR-RLK_2108

TCTGATGTCAAACGGAAGCCTAGAGAACCATCTGTACCCAAGCCACAGGCTGAGACATCAACTTGACTTAGCTCAATTGGTTAACATTTGCAGTGATGTAGCAGAAGGAGTGTCCTATCT

>M6-PGSC0003DMG400011205M6LRR-RLK_2170

TTGACTTAGCTCAATTGGTTAACATTTGCAGTGATGTAGCAGAAGGAGTGTCCTATCTCCACCACTATTCGCCTGTTAAAGTCGTGCATTGTGATCTAAAACCAAGCAATATCCTTCTTG

>M6-PGSC0003DMG400011205M6LRR-RLK_2232

CACTATTCGCCTGTTAAAGTCGTGCATTGTGATCTAAAACCAAGCAATATCCTTCTTGATTACAACATGACAGCTTTGGTAACAGATTTTGGGATTTCAACATTAGTAAAAGCTGTTGAG

>M6-PGSC0003DMG400011205M6LRR-RLK_2294

CAACATGACAGCTTTGGTAACAGATTTTGGGATTTCAACATTAGTAAAAGCTGTTGAGGAAAATACTAATCCTATCGACGAATCAGTTAATTACAACTCAACAGATGGATTACTTTGTGG

>M6-PGSC0003DMG400011205M6LRR-RLK_2356

ATACTAATCCTATCGACGAATCAGTTAATTACAACTCAACAGATGGATTACTTTGTGGATCAATTGGTTACATTGCTCCTGGTTAGTAACAAATAATAGTTCTATTCTATCTAATTCATT

>M6-PGSC0003DMG400011205M6LRR-RLK_2418

ATTGGTTACATTGCTCCTGGTTAGTAACAAATAATAGTTCTATTCTATCTAATTCATTTCAATTTATATGACTAAAGTTTCTGTTTCTTTGATTGTAGAATATGGAATGGGAAGGAGGGC

>M6-PGSC0003DMG400011205M6LRR-RLK_248

TAAGCTAAATCAGCTTAGTTTGTCTTCTAATCTCCTCGAAGGGGAAATCCCAAACGAGTTAGGATTCCTTCATGAATTGAAGTATCTTGATTTGGGAACAAACAATCTTTCTGGTGCAAT

>M6-PGSC0003DMG400011205M6LRR-RLK_2480

TTTATATGACTAAAGTTTCTGTTTCTTTGATTGTAGAATATGGAATGGGAAGGAGGGCTTCAACAAAAGGAGATGTATTTAGCTATGGAGTTTTGTTACTTGAGATTGTGACAGGTAAAC

>M6-PGSC0003DMG400011205M6LRR-RLK_2542

ACAAAAGGAGATGTATTTAGCTATGGAGTTTTGTTACTTGAGATTGTGACAGGTAAACGTCCCACAGACGTTCTATTTCAACAAGGCTCGAGCTTACACGAATGGGTGAAGAGTCAGTAT

>M6-PGSC0003DMG400011205M6LRR-RLK_2604

CACAGACGTTCTATTTCAACAAGGCTCGAGCTTACACGAATGGGTGAAGAGTCAGTATTCACATAAACTCGAGGCAATGATTGATGAAGCATTACAAAGATGTACAACAGTTCAACGCGA

>M6-PGSC0003DMG400011205M6LRR-RLK_2666

ATAAACTCGAGGCAATGATTGATGAAGCATTACAAAGATGTACAACAGTTCAACGCGATAGCAATATATGGCACGATATCGTGTTAGAGTTGATTGAACTTGGCTTAATGTGCACTCAGT

>M6-PGSC0003DMG400011205M6LRR-RLK_2728

AATATATGGCACGATATCGTGTTAGAGTTGATTGAACTTGGCTTAATGTGCACTCAGTACAATCCATCAACTAGGCCAACAATGTTGGATGTTGCACTTGAAATTGGAAGGTTAAAACAG

>M6-PGSC0003DMG400011205M6LRR-RLK_2790

TCCATCAACTAGGCCAACAATGTTGGATGTTGCACTTGAAATTGGAAGGTTAAAACAGTATGTTTCAACTTCTTCAAACACTGTTACTGAAGATGTCACAACAAAAAGTTGAACAGTTTT

>M6-PGSC0003DMG400011205M6LRR-RLK_2852

TTTCAACTTCTTCAAACACTGTTACTGAAGATGTCACAACAAAAAGTTGAACAGTTTTTTTTTTTATATATCTAACAATGTACCTTTTAACTTTATTCATTGTACTTCTATGTTCTTTTT

>M6-PGSC0003DMG400011205M6LRR-RLK_2914

TTTATATATCTAACAATGTACCTTTTAACTTTATTCATTGTACTTCTATGTTCTTTTTTTTTTTTACCATTTTAATTGTTCCTTCTTGGGGTGCGGACCTTCTCTGGATTCTGCGTGAAT

>M6-PGSC0003DMG400011205M6LRR-RLK_2946

ATTCATTGTACTTCTATGTTCTTTTTTTTTTTTACCATTTTAATTGTTCCTTCTTGGGGTGCGGACCTTCTCTGGATTCTGCGTGAATGTGAGATGGTTCGTGCACCAAGACTACCCTTC

>M6-PGSC0003DMG400011205M6LRR-RLK_310

GATTCCTTCATGAATTGAAGTATCTTGATTTGGGAACAAACAATCTTTCTGGTGCAATTCCTCTGTCTTTGTTCTGTAATTGCTCTGCTTCATTACAATATATGGACCTTTCTAACAATT

>M6-PGSC0003DMG400011205M6LRR-RLK_372

CTGTCTTTGTTCTGTAATTGCTCTGCTTCATTACAATATATGGACCTTTCTAACAATTCGCTTCGCGGTGAGATCCCTATGAATGATCATTGTGAGTTGAGTGGTTTGAAGTTTCTTCTT

>M6-PGSC0003DMG400011205M6LRR-RLK_434

TCGCGGTGAGATCCCTATGAATGATCATTGTGAGTTGAGTGGTTTGAAGTTTCTTCTTTTATGGTCTAATGAACTTGTTGGTGAAGTTCCTATTGCTCTTTCGAAATCTACAAAGCTAGA

>M6-PGSC0003DMG400011205M6LRR-RLK_496

GGTCTAATGAACTTGTTGGTGAAGTTCCTATTGCTCTTTCGAAATCTACAAAGCTAGAATGGCTTGATCTTGAATCGAATTCGTTGAGTGGTGAACTTCCTTCTGATATTGTAAGTAAAA

>M6-PGSC0003DMG400011205M6LRR-RLK_558

CTTGATCTTGAATCGAATTCGTTGAGTGGTGAACTTCCTTCTGATATTGTAAGTAAAATGCCAAAGTTACAGTTTTTGTATTTGTCTTACAACAACTTTGATAGTCATAGTGGTAATACT

>M6-PGSC0003DMG400011205M6LRR-RLK_62

TGTATGTAATTGGACAGGAATTGTATGTGACAAGAAAATGCATAGAGTTGTTGAGCTTGATCTTAGTCATCATTCGTTACGTGGAACGATTTCTCCTTCTCTCTCTGGGCTCTCTTTCTT

>M6-PGSC0003DMG400011205M6LRR-RLK_620

AAAGTTACAGTTTTTGTATTTGTCTTACAACAACTTTGATAGTCATAGTGGTAATACTGACCTGACGCCTTTTTTCGCGTCTTTAGTTAATTCTACTAACTTGCAAGAACTTGAGTTAGC

>M6-PGSC0003DMG400011205M6LRR-RLK_682

TGACGCCTTTTTTCGCGTCTTTAGTTAATTCTACTAACTTGCAAGAACTTGAGTTAGCTGGGAATAATCTTGGTGGGGAGTTACCTCCTATTATCGGTAACATCTCCAAGAACCTTGCAC

>M6-PGSC0003DMG400011205M6LRR-RLK_744

AATAATCTTGGTGGGGAGTTACCTCCTATTATCGGTAACATCTCCAAGAACCTTGCACAGATTCATCTTGACGATAATCTTATCTATGGTCCAATCCCAACACAAATTTCGAGTCTTGTA

>M6-PGSC0003DMG400011205M6LRR-RLK_806

TCATCTTGACGATAATCTTATCTATGGTCCAATCCCAACACAAATTTCGAGTCTTGTAAATATTACTCTATTGAATTTGTCTAGTAATCATCTGAATGGTACAATTCCTCCTGAGTTGTG

>M6-PGSC0003DMG400011205M6LRR-RLK_868

TTACTCTATTGAATTTGTCTAGTAATCATCTGAATGGTACAATTCCTCCTGAGTTGTGTCGAATGGGGAAACTTGAGAGGCTTTATCTGTCGAATAATTCACTCTCTGGTGTCATTCCTT

>M6-PGSC0003DMG400011205M6LRR-RLK_930

ATGGGGAAACTTGAGAGGCTTTATCTGTCGAATAATTCACTCTCTGGTGTCATTCCTTCTGCTTTTGGAAATGTTTCTCATTTAGGCCTTCTTGATCTGTCGAAAAACAACCTCTCTGGA

>M6-PGSC0003DMG400011205M6LRR-RLK_992

TTTTGGAAATGTTTCTCATTTAGGCCTTCTTGATCTGTCGAAAAACAACCTCTCTGGATTAATCCCAAATACCTTTGCTAATCTTCCGCAACTGAGGAGGCTTTTGTTGCACGATAATCA

>M6-PGSC0003DMG400011209M6LRR-RLP_0

CGACTACTAGGCGGCCACCATACTATGACTATAAATGACAAGAGATGTTCCTTCCTACTTCATTCACAAACAAGCAAATATGAAGAACAATTCACTTCAACTTATCTTTCTCATATTGTT

>M6-PGSC0003DMG400011209M6LRR-RLP_1037

TGAGTTTCAATAAATTGACAGGAAATCTTCCTGATGCATTAGGCAACTTGAGGAAACTGAAACATCTTCAGCTGAGGTATAATTCACTGACAGGCACAATACCTGAAAGTATAGGGAACT

>M6-PGSC0003DMG400011209M6LRR-RLP_1098

ACATCTTCAGCTGAGGTATAATTCACTGACAGGCACAATACCTGAAAGTATAGGGAACTTATTATCTTTGGAGACATTCTACCTCACATCAAACAAAATGAGTGGGAACATCACCACGAG

>M6-PGSC0003DMG400011209M6LRR-RLP_1159

TTATCTTTGGAGACATTCTACCTCACATCAAACAAAATGAGTGGGAACATCACCACGAGTATCGGACAGCTCATGTCGTTGGTCTCGTTGGATATCTCGGAGAATATGTGGGATGGTATT

>M6-PGSC0003DMG400011209M6LRR-RLP_122

CTCAACTTTTCTATTTCACTTGCATAAACCATCACTTTGTTTCCTCTACTAACACCTCTTGCATTGAAACAGAGAAAAATGCACTTCTAAACCTTAAAGAAAATCTTACAGATCCTTCAG

>M6-PGSC0003DMG400011209M6LRR-RLP_1220

TCGGACAGCTCATGTCGTTGGTCTCGTTGGATATCTCGGAGAATATGTGGGATGGTATTGTCATTGAAGCTCATTTGGTGAATCTTTCCAACTTGCAAGAGTTTTCAGTTGGGATGAAAC

>M6-PGSC0003DMG400011209M6LRR-RLP_1281

CATTGAAGCTCATTTGGTGAATCTTTCCAACTTGCAAGAGTTTTCAGTTGGGATGAAACTTAGTAAAAATATTACCTTGGCATTCGATATCAGCCCGAATTGGAATCCTTCATTCAGGCT

>M6-PGSC0003DMG400011209M6LRR-RLP_1342

AGTAAAAATATTACCTTGGCATTCGATATCAGCCCGAATTGGAATCCTTCATTCAGGCTGACATTCTTGACTATTCAGTCATGCCAATTAGGTCCTGAATTCCCACATTGGCTGAAGGAT

>M6-PGSC0003DMG400011209M6LRR-RLP_1403

CATTCTTGACTATTCAGTCATGCCAATTAGGTCCTGAATTCCCACATTGGCTGAAGGATCAGAATGAGCTCACCAGCCTAATACTCAACACCGCGGGGATTTCTGATGCGGTACCTGATT

>M6-PGSC0003DMG400011209M6LRR-RLP_1464

GAATGAGCTCACCAGCCTAATACTCAACACCGCGGGGATTTCTGATGCGGTACCTGATTGGTTTGTAGAGTTGGATTTGAAGCTTGATAATCTAGACATGGCATACAACAACTTGACAGG

>M6-PGSC0003DMG400011209M6LRR-RLP_1525

TTTGTAGAGTTGGATTTGAAGCTTGATAATCTAGACATGGCATACAACAACTTGACAGGGAAAGTTCCAAACAAATTCAAGTTCAACTTTCAAGCCAATGTGGATTTAAGCACTAATCGC

>M6-PGSC0003DMG400011209M6LRR-RLP_1586

AAGTTCCAAACAAATTCAAGTTCAACTTTCAAGCCAATGTGGATTTAAGCACTAATCGCTTCGAGGGACCTCTCCCACTATGGTCTTCTAATGTTACAACATTGTATCTGAGGGATAACT

>M6-PGSC0003DMG400011209M6LRR-RLP_1647

CGAGGGACCTCTCCCACTATGGTCTTCTAATGTTACAACATTGTATCTGAGGGATAACTTGTTTTCAGGACCTATTCCTCTCAACATTTGTGAAGCACTTCTCAATATAGAAGACTTGGA

>M6-PGSC0003DMG400011209M6LRR-RLP_1708

TTTTCAGGACCTATTCCTCTCAACATTTGTGAAGCACTTCTCAATATAGAAGACTTGGATATCTCCAGGAACAATCTAAATGGCACAATTCCCTTATGCATAGGTGATATGAATCAGTTA

>M6-PGSC0003DMG400011209M6LRR-RLP_1769

TCTCCAGGAACAATCTAAATGGCACAATTCCCTTATGCATAGGTGATATGAATCAGTTAACTACCTTAGCCCTCGATAACAACCAACTTATCGGCCAAGTTCCTGATTTCTGGGGTAAGT

>M6-PGSC0003DMG400011209M6LRR-RLP_183

CATTGAAACAGAGAAAAATGCACTTCTAAACCTTAAAGAAAATCTTACAGATCCTTCAGGCAAGCTATCTTCTTGGATTGATCAACAAGATTGTTGCCAATGGCTTGGTGTAACATGTAA

>M6-PGSC0003DMG400011209M6LRR-RLP_1830

TACCTTAGCCCTCGATAACAACCAACTTATCGGCCAAGTTCCTGATTTCTGGGGTAAGTTACCATATCTTTACTGGATAGATATGTCAGAGAATCGCCTATCCGGACAAATTCCACACTC

>M6-PGSC0003DMG400011209M6LRR-RLP_1891

CCATATCTTTACTGGATAGATATGTCAGAGAATCGCCTATCCGGACAAATTCCACACTCACTAGGCTCTCTTGCTTATCTGATGTTCTTAAGACTCTCAGGCAACAATCTCTCTGGAGAA

>M6-PGSC0003DMG400011209M6LRR-RLP_1952

TAGGCTCTCTTGCTTATCTGATGTTCTTAAGACTCTCAGGCAACAATCTCTCTGGAGAACTGCCTTTGAGTTTGAGAAATTGCACGAAAATGATCAACATTGATCTTAGTAACAATCAGT

>M6-PGSC0003DMG400011209M6LRR-RLP_2013

GCCTTTGAGTTTGAGAAATTGCACGAAAATGATCAACATTGATCTTAGTAACAATCAGTTGTCAGGGCTAATCCCAGCTTGGTTAGCGGAAACAATGAGATCACTGTTAATCCTTAGTGT

>M6-PGSC0003DMG400011209M6LRR-RLP_2074

TCAGGGCTAATCCCAGCTTGGTTAGCGGAAACAATGAGATCACTGTTAATCCTTAGTGTAAGGAACAACAGGTTTTTCGGTCACATTCCTTTAAAAATCTGTAGCCTTTCAGGTCTTCAC

>M6-PGSC0003DMG400011209M6LRR-RLP_2135

GGAACAACAGGTTTTTCGGTCACATTCCTTTAAAAATCTGTAGCCTTTCAGGTCTTCACATACTAGACCTTTCAGGAAACAACTTATCCGGTTCTGTCCCGTCCTGTTTTGGAAACTTGG

>M6-PGSC0003DMG400011209M6LRR-RLP_2196

ACTAGACCTTTCAGGAAACAACTTATCCGGTTCTGTCCCGTCCTGTTTTGGAAACTTGGAAGGGTTCAAGGTTGAGCTAACTGATGTAGAAGCTAAACAATATCAAGGTACATTGAAGCT

>M6-PGSC0003DMG400011209M6LRR-RLP_2257

GGGTTCAAGGTTGAGCTAACTGATGTAGAAGCTAAACAATATCAAGGTACATTGAAGCTTGAGGCAAAAGGAAGAACACTTTCTTATGATAAAATCCTTTATCTAGTAAACAGTATCGAC

>M6-PGSC0003DMG400011209M6LRR-RLP_2318

AGGCAAAAGGAAGAACACTTTCTTATGATAAAATCCTTTATCTAGTAAACAGTATCGACCTCTCGAGAAACAGTTTGTCTGGTGAAATCCCGGAAGAGTTAACAAACTTACACAAACTTG

>M6-PGSC0003DMG400011209M6LRR-RLP_2379

CTCGAGAAACAGTTTGTCTGGTGAAATCCCGGAAGAGTTAACAAACTTACACAAACTTGGAACCTTGAACTTGTCTATGAACCATTTGACAGGACATATACCAACAGATATCGGAAAGTT

>M6-PGSC0003DMG400011209M6LRR-RLP_244

AAGCTATCTTCTTGGATTGATCAACAAGATTGTTGCCAATGGCTTGGTGTAACATGTAACAACAAGACCGGGCACGTCGTAAAGATTGATCTTAGGAACAAATACTCAACAGAACATGAA

>M6-PGSC0003DMG400011209M6LRR-RLP_2440

ACCTTGAACTTGTCTATGAACCATTTGACAGGACATATACCAACAGATATCGGAAAGTTACGTTGGGTTGAAACGTTAGATCTTTCAATAAATCAGTTATCAGGTACTATCCCACCAAGC

>M6-PGSC0003DMG400011209M6LRR-RLP_2501

GTTGGGTTGAAACGTTAGATCTTTCAATAAATCAGTTATCAGGTACTATCCCACCAAGCTTAGCTACATTAGACTTTTTAAGTCATTTGAACTTGTCATACAATAAGTTGACAGGAAGAA

>M6-PGSC0003DMG400011209M6LRR-RLP_2562

AGCTACATTAGACTTTTTAAGTCATTTGAACTTGTCATACAATAAGTTGACAGGAAGAATTCCTACTAGTACTCAGTTTCAAACCAAAGTTGATCCAACAATTTTCCAAGGAAATGTTGC

>M6-PGSC0003DMG400011209M6LRR-RLP_2623

CCTACTAGTACTCAGTTTCAAACCAAAGTTGATCCAACAATTTTCCAAGGAAATGTTGCACTTTGTGGTCCTCCTTTGGAACAATGTGTTGGTGATAGGACTACGACGACGATAACATCA

>M6-PGSC0003DMG400011209M6LRR-RLP_2684

TTTGTGGTCCTCCTTTGGAACAATGTGTTGGTGATAGGACTACGACGACGATAACATCACAAAGTGGAGGAAATGATGAAGGGGAAACAGATGATGAAGATCAACTTGAGAAAGTATGGT

>M6-PGSC0003DMG400011209M6LRR-RLP_2745

AAGTGGAGGAAATGATGAAGGGGAAACAGATGATGAAGATCAACTTGAGAAAGTATGGTTCTTTGCTGTTGTTGGTTTGGGATATATAGTTGGTTTTTGGGCATTTTTTGGCAGTTTGAT

>M6-PGSC0003DMG400011209M6LRR-RLP_2806

TTTGCTGTTGTTGGTTTGGGATATATAGTTGGTTTTTGGGCATTTTTTGGCAGTTTGATCTTCAAGAAAAGATGGAGAATTGCTTATTGCAGATTCATTGAGAGGTGTATTTTGGAGTTC

>M6-PGSC0003DMG400011209M6LRR-RLP_2867

TCAAGAAAAGATGGAGAATTGCTTATTGCAGATTCATTGAGAGGTGTATTTTGGAGTTCTATGAAATGTCTTGTAGTTCTGTGAAAAAATTGTTAGGAATCTAATAAGCTATGTTGCTCA

>M6-PGSC0003DMG400011209M6LRR-RLP_305

ACAAGACCGGGCACGTCGTAAAGATTGATCTTAGGAACAAATACTCAACAGAACATGAATTGAGAGGTGTGATTAATCCATCTTTGCTCAAGTTACAACAATTGAGATACTTAGACTTAA

>M6-PGSC0003DMG400011209M6LRR-RLP_3050

TACGACAACATTTTTAGAGAGTCTGAGAAACTTAGCTAATAAGTGGAACTATAGGAGTTTGTCTCTGCAGTTCTGTTATATGTTATGTTTTAGTTTCAATATAGCACATATTACTACTTT

>M6-PGSC0003DMG400011209M6LRR-RLP_3111

TCTCTGCAGTTCTGTTATATGTTATGTTTTAGTTTCAATATAGCACATATTACTACTTTTTCTATATGTAAAATAATGCTACCTAAGTATTGACACTAATAGAAATGCCAGTGTTTTGGG

>M6-PGSC0003DMG400011209M6LRR-RLP_3172

CTATATGTAAAATAATGCTACCTAAGTATTGACACTAATAGAAATGCCAGTGTTTTGGGTAGAGTTACGACATGTGAAGAGTGGTCAATTGAACACTTTTACGTAGGTCAAAATAATACT

>M6-PGSC0003DMG400011209M6LRR-RLP_3233

GAGTTACGACATGTGAAGAGTGGTCAATTGAACACTTTTACGTAGGTCAAAATAATACTTCTTATATATGTATACATGTTCACAAGACTATTGTAGTACATATAATTAGGAAGTTGTTTA

>M6-PGSC0003DMG400011209M6LRR-RLP_3281

AAAATAATACTTCTTATATATGTATACATGTTCACAAGACTATTGTAGTACATATAATTAGGAAGTTGTTTAAATAGTATCTAAGACTATTCTTATACTATGCATGCATCTACTTGTATA

>M6-PGSC0003DMG400011209M6LRR-RLP_366

GAGAGGTGTGATTAATCCATCTTTGCTCAAGTTACAACAATTGAGATACTTAGACTTAAGCATGAACAATTTTGAAGGAATCAAAGTACCTGAATTCATCGGTCGCCTCAATGAACTAAG

>M6-PGSC0003DMG400011209M6LRR-RLP_427

ATGAACAATTTTGAAGGAATCAAAGTACCTGAATTCATCGGTCGCCTCAATGAACTAAGGTATCTCAATCTATCTGGTGCATCTTTTACTGGTTTAGTCTCTTCATTCCTTGAAAACCTC

>M6-PGSC0003DMG400011209M6LRR-RLP_488

ATCTCAATCTATCTGGTGCATCTTTTACTGGTTTAGTCTCTTCATTCCTTGAAAACCTCTCAAATTTGCAAGTTCTTGATTTAGGAGGCTTAGATCTTAGTGAAGATGCTAATTCTTGGC

>M6-PGSC0003DMG400011209M6LRR-RLP_549

AAATTTGCAAGTTCTTGATTTAGGAGGCTTAGATCTTAGTGAAGATGCTAATTCTTGGCTTCAAATAATCAATGATCATGTTCCTTCTCTGTTGGAGTTGCGTTTGCCTCGATGTCAACT

>M6-PGSC0003DMG400011209M6LRR-RLP_61

ATTCACAAACAAGCAAATATGAAGAACAATTCACTTCAACTTATCTTTCTCATATTGTTATCTCAACTTTTCTATTTCACTTGCATAAACCATCACTTTGTTTCCTCTACTAACACCTCT

>M6-PGSC0003DMG400011209M6LRR-RLP_610

CAAATAATCAATGATCATGTTCCTTCTCTGTTGGAGTTGCGTTTGCCTCGATGTCAACTTTTAAACATTCCTTCTTCTTTTCCATTACTAAATCTTACTTCCCTTTTGGTACTTGATCTT

>M6-PGSC0003DMG400011209M6LRR-RLP_671

TAAACATTCCTTCTTCTTTTCCATTACTAAATCTTACTTCCCTTTTGGTACTTGATCTTTCCAACAATGCTTTTAACTCATGTACATTTCCTCAATGGATATTCAAACTTAGTAGCCTTG

>M6-PGSC0003DMG400011209M6LRR-RLP_732

CAACAATGCTTTTAACTCATGTACATTTCCTCAATGGATATTCAAACTTAGTAGCCTTGTGTATCTGGACTTAAGCTCCAACAATATGTGTAGTGAACTTGTTGATGAGTTTGCTAAGCT

>M6-PGSC0003DMG400011209M6LRR-RLP_793

TATCTGGACTTAAGCTCCAACAATATGTGTAGTGAACTTGTTGATGAGTTTGCTAAGCTCACTTTCCTTGAACATCTTGATCTGTCTTCGAACCACGGTATCAATGGGGCACTCAAGAGG

>M6-PGSC0003DMG400011209M6LRR-RLP_854

CTTTCCTTGAACATCTTGATCTGTCTTCGAACCACGGTATCAATGGGGCACTCAAGAGGAGCTTGGGGAAACTATGTAATTTGAAGAGTTTGATCCTGTCTTACAACAACATCAGTGGAG

>M6-PGSC0003DMG400011209M6LRR-RLP_915

CTTGGGGAAACTATGTAATTTGAAGAGTTTGATCCTGTCTTACAACAACATCAGTGGAGACATAATCGAGTTCATCGATGCGTTATCTGAATGCCAAAGCAATAGCTTGGAGACATTGGA

>M6-PGSC0003DMG400011209M6LRR-RLP_976

ATAATCGAGTTCATCGATGCGTTATCTGAATGCCAAAGCAATAGCTTGGAGACATTGGAGTTGAGTTTCAATAAATTGACAGGAAATCTTCCTGATGCATTAGGCAACTTGAGGAAACTG

>M6-PGSC0003DMG400011251M6G-LecRK_0

ATGTTAATTTCTCCAAAAAGGAAATTTATTTTGGGATTTTTTAGTCCAAATGTTTCTAATAATCAAAGGTTTGTTGGTATATGGTATGTTGATGGTCCAAAAAATTCATTTGTTTGGGTT

>M6-PGSC0003DMG400011251M6G-LecRK_1054

CATTTGTGACATCTTATCTAAAAGATTAAGTTGATAACATGTGTTTCATAACTTACTTTTGTTTTTAACACAACCTCTCATGTGCTTCTTGAATCCTTGATATTGGATGGCAGTGAGATT

>M6-PGSC0003DMG400011251M6G-LecRK_1116

TTTTAACACAACCTCTCATGTGCTTCTTGAATCCTTGATATTGGATGGCAGTGAGATTCGAACCTAAGAACTATGCCTGTTCGGATACAATGTTGCATTTGTGACATCCCAAAGTCAAAG

>M6-PGSC0003DMG400011251M6G-LecRK_1178

CCTAAGAACTATGCCTGTTCGGATACAATGTTGCATTTGTGACATCCCAAAGTCAAAGAGCATATTTTTATTTACTTAATTACGTTTCCAACAGGTACTCATTAGGAGTAAATACTCGCG

>M6-PGSC0003DMG400011251M6G-LecRK_124

ACAGAGATAAACCAATTTTTGATAAAAATGGTGTTTTTACTATTGAGAAAAATGGGAATTTAGTGGTGAAAAATGGTCGTGGCGATTTAATGTGGACGAATAATGTTACTGCTATGAATA

>M6-PGSC0003DMG400011251M6G-LecRK_1240

TATTTTTATTTACTTAATTACGTTTCCAACAGGTACTCATTAGGAGTAAATACTCGCGGATCACCACAAATTGTGATTTTGGATGGAGAGAAGAGACGTTGGCGGAGTGGATATTGGGAT

>M6-PGSC0003DMG400011251M6G-LecRK_1302

ACCACAAATTGTGATTTTGGATGGAGAGAAGAGACGTTGGCGGAGTGGATATTGGGATGGACGCATATTCACAGGAGTTACAGATATGAAGCCCGAGTATCTACATGGTTTCAAGCTTTA

>M6-PGSC0003DMG400011251M6G-LecRK_1364

GCATATTCACAGGAGTTACAGATATGAAGCCCGAGTATCTACATGGTTTCAAGCTTTACAATGAAGGAGATATGCTATACTTCACTTACACTGTTGCAGATTCTTCTGATTTGGTGAGGT

>M6-PGSC0003DMG400011251M6G-LecRK_1426

GAAGGAGATATGCTATACTTCACTTACACTGTTGCAGATTCTTCTGATTTGGTGAGGTTCCATATTAGTCCAACCGGATACGAGCTGGAACAAAGGTGGGATAAAGATAACGAAAACTGG

>M6-PGSC0003DMG400011251M6G-LecRK_1488

TATTAGTCCAACCGGATACGAGCTGGAACAAAGGTGGGATAAAGATAACGAAAACTGGAGTATAATACAGTCTCATCCATCAGGTGATTGTGACTTGTATAACTTATGTGGGAATTTCGC

>M6-PGSC0003DMG400011251M6G-LecRK_1550

TAATACAGTCTCATCCATCAGGTGATTGTGACTTGTATAACTTATGTGGGAATTTCGCGAAATGTGATGTCACATATTTGAAAAAATGCATTTGTTTAGTTGGATTTGTACCGAAAGATT

>M6-PGSC0003DMG400011251M6G-LecRK_1612

TGTGATGTCACATATTTGAAAAAATGCATTTGTTTAGTTGGATTTGTACCGAAAGATTTGGGGCAGTGGAATGCGAGGAACTGGTCAGAAGGGTGTGTTAGGAGGAAAGAACTGGAATGC

>M6-PGSC0003DMG400011251M6G-LecRK_1674

GCAGTGGAATGCGAGGAACTGGTCAGAAGGGTGTGTTAGGAGGAAAGAACTGGAATGCAGAGGAAATAACAGTGCGTTGAAGAGTGTTAGTGGAAAGAAGGACGGATTCTTTGAGATTGA

>M6-PGSC0003DMG400011251M6G-LecRK_1736

GAAATAACAGTGCGTTGAAGAGTGTTAGTGGAAAGAAGGACGGATTCTTTGAGATTGAGAAAATCAAGTTGCCGGATTTTGCTGATACTGCATATTTACAAAATATCGATGAGTGTCGAA

>M6-PGSC0003DMG400011251M6G-LecRK_1798

ATCAAGTTGCCGGATTTTGCTGATACTGCATATTTACAAAATATCGATGAGTGTCGAAGCATGTGCCTTGAAAATTGTTCATGTACTGCTTATGCTTTTGTTAGCGGAATCAATTGCATG

>M6-PGSC0003DMG400011251M6G-LecRK_186

GTGGTGAAAAATGGTCGTGGCGATTTAATGTGGACGAATAATGTTACTGCTATGAATAATAATTGTACAGCACATCTTTCTGATAATGGAAATCTTAGGATGTTTAATGGCAATAACAAG

>M6-PGSC0003DMG400011251M6G-LecRK_1860

GTGCCTTGAAAATTGTTCATGTACTGCTTATGCTTTTGTTAGCGGAATCAATTGCATGATGTGGAGTGGAGATTTAGTCGATATGCAGCAGTTTCAGGAAGGCGGAAACACTCTCTATGT

>M6-PGSC0003DMG400011251M6G-LecRK_1922

GGAGTGGAGATTTAGTCGATATGCAGCAGTTTCAGGAAGGCGGAAACACTCTCTATGTTCGCCTTGCTGATTCTGAATTTGGTAAAAATGTGTACTAACTTTTCTGTTGGTATCATTTTA

>M6-PGSC0003DMG400011251M6G-LecRK_1984

CTTGCTGATTCTGAATTTGGTAAAAATGTGTACTAACTTTTCTGTTGGTATCATTTTAGATGTTCTTATTATAAACTTTCTTGAAACAGCTGGAAGCAACAGGACAGTCAAAATTGTGGT

>M6-PGSC0003DMG400011251M6G-LecRK_2046

TTCTTATTATAAACTTTCTTGAAACAGCTGGAAGCAACAGGACAGTCAAAATTGTGGTAATACCCGTTATGGTAGCCGGGGCTTTACTTGTTTGTATGGTAGTTTTTCTACTATGCAAGT

>M6-PGSC0003DMG400011251M6G-LecRK_2108

CCCGTTATGGTAGCCGGGGCTTTACTTGTTTGTATGGTAGTTTTTCTACTATGCAAGTACAAAGCCAAAACTCGAGGTAAGATTTCAGAAAGGCTGGTGTTTTACGACAGTCATTTGTTG

>M6-PGSC0003DMG400011251M6G-LecRK_2170

AGCCAAAACTCGAGGTAAGATTTCAGAAAGGCTGGTGTTTTACGACAGTCATTTGTTGTTATGCAATATACCTATATCTTCTTCAATGTCCCATTTTCAGTGTCCAATAGGATCAACCAA

>M6-PGSC0003DMG400011251M6G-LecRK_2232

GCAATATACCTATATCTTCTTCAATGTCCCATTTTCAGTGTCCAATAGGATCAACCAAATGGAGACAAGCGATCCAACTAGGAGCGGAGAGTTTTCTATGAACACATCAGGAGCAGGAGA

>M6-PGSC0003DMG400011251M6G-LecRK_2294

AGACAAGCGATCCAACTAGGAGCGGAGAGTTTTCTATGAACACATCAGGAGCAGGAGACTTGAGTATTGAAGGGCATCAAGGGAGTGGTTCAGAACTTATATTCTTCAGTTTCAGCGGTG

>M6-PGSC0003DMG400011251M6G-LecRK_2356

AGTATTGAAGGGCATCAAGGGAGTGGTTCAGAACTTATATTCTTCAGTTTCAGCGGTGTAGCAGCAGCTACCGACAACTTTTCTAACGAAAACAAGCTGGGACAAGGTGGATTTGGCCCT

>M6-PGSC0003DMG400011251M6G-LecRK_2418

AGCAGCTACCGACAACTTTTCTAACGAAAACAAGCTGGGACAAGGTGGATTTGGCCCTGTCTACAAGGTTGGTTTTCATTCAATTGACAAGTGTGTCTTACTTTTCTAGAGAAGTATGAC

>M6-PGSC0003DMG400011251M6G-LecRK_248

TTGTACAGCACATCTTTCTGATAATGGAAATCTTAGGATGTTTAATGGCAATAACAAGGAGTTGTGGCAGAGTTTTCAGCATCCTACAGATACATTTTTACCAGAAATGAGATTTTATTT

>M6-PGSC0003DMG400011251M6G-LecRK_2480

ACAAGGTTGGTTTTCATTCAATTGACAAGTGTGTCTTACTTTTCTAGAGAAGTATGACATTATTGTGTTAAACTAACGTCGATGTTTGTTTATTTAGGGAAAACTACTATGTGGCTTAGA

>M6-PGSC0003DMG400011251M6G-LecRK_2542

TTGTGTTAAACTAACGTCGATGTTTGTTTATTTAGGGAAAACTACTATGTGGCTTAGAAATTGCAGTGAAGAGACTTTCAAGAAAGTCAGGACAAGGTGTGGAGGAATTCAAGAATGAGA

>M6-PGSC0003DMG400011251M6G-LecRK_2604

GCAGTGAAGAGACTTTCAAGAAAGTCAGGACAAGGTGTGGAGGAATTCAAGAATGAGATCAAGCTAATCGCCAAATTACAACATAGAAATCTTGTTAGACTTATGGGATGCTGCATTGAA

>M6-PGSC0003DMG400011251M6G-LecRK_2666

GCTAATCGCCAAATTACAACATAGAAATCTTGTTAGACTTATGGGATGCTGCATTGAAGGAGAAGAAAAGATGCTTCTTTACGAGTACATGGCAAACAGAAGTTTAGATTCATTTCTATT

>M6-PGSC0003DMG400011251M6G-LecRK_2728

AAGAAAAGATGCTTCTTTACGAGTACATGGCAAACAGAAGTTTAGATTCATTTCTATTTGGTTAGTACTCTAGATATGCTATAAATTATTTCATCGATAGAGAGATCGATCACTTACAAT

>M6-PGSC0003DMG400011251M6G-LecRK_2790

TAGTACTCTAGATATGCTATAAATTATTTCATCGATAGAGAGATCGATCACTTACAATTATTTTGGTTGGTAAATTTTTTGACAGATCCTGTTAAGCAAGCTCAATTAGACTGGAGGAAA

>M6-PGSC0003DMG400011251M6G-LecRK_2852

TTGGTTGGTAAATTTTTTGACAGATCCTGTTAAGCAAGCTCAATTAGACTGGAGGAAACGCTTCAACATTATTGAAGGGATTGCACGAGGGCTATTGTATCTCCATAGAGATTCGCGACT

>M6-PGSC0003DMG400011251M6G-LecRK_2914

TCAACATTATTGAAGGGATTGCACGAGGGCTATTGTATCTCCATAGAGATTCGCGACTCAGAATAATCCATAGGGATTTAAAGGCTAGTAACATTTTGCTAGATGAAGATATGAACCCGA

>M6-PGSC0003DMG400011251M6G-LecRK_2976

ATAATCCATAGGGATTTAAAGGCTAGTAACATTTTGCTAGATGAAGATATGAACCCGAAAATTTCAGACTTTGGTATGGCTAGGATATTTGGTGGAAATGAAAATGAAGCAAATACAAAT

>M6-PGSC0003DMG400011251M6G-LecRK_3038

TTCAGACTTTGGTATGGCTAGGATATTTGGTGGAAATGAAAATGAAGCAAATACAAATAGAGTAGTTGGGACATAGTGAGTATCATTTGATTTTTATACCTTTAAGAGTGAGTACCTTTT

>M6-PGSC0003DMG400011251M6G-LecRK_310

TGTGGCAGAGTTTTCAGCATCCTACAGATACATTTTTACCAGAAATGAGATTTTATTTGGATGAAGTTTTGAGGTCATGGACTAGTGATAGTGATCCTTCACCTGGACGGTACGTAATCT

>M6-PGSC0003DMG400011251M6G-LecRK_3100

TAGTTGGGACATAGTGAGTATCATTTGATTTTTATACCTTTAAGAGTGAGTACCTTTTAACTTGTTCAAAGAACTAACTCCATCAACGCGTTTCTTTTTTTGTTTTTACTTAAATCAGCG

>M6-PGSC0003DMG400011251M6G-LecRK_3162

TGTTCAAAGAACTAACTCCATCAACGCGTTTCTTTTTTTGTTTTTACTTAAATCAGCGGATATATGGCTCCTGAATATGCAATGGAAGGCTTGTTCTCCGGAAAATCTGATGTTTATAGC

>M6-PGSC0003DMG400011251M6G-LecRK_3224

TATGGCTCCTGAATATGCAATGGAAGGCTTGTTCTCCGGAAAATCTGATGTTTATAGCTTTGGTGTACTATTACTGGAGATCATCTGTGGACGGAGGAACACAAGCTTTCGTTCAAATGA

>M6-PGSC0003DMG400011251M6G-LecRK_3286

GTGTACTATTACTGGAGATCATCTGTGGACGGAGGAACACAAGCTTTCGTTCAAATGAACACTCTGGCATTATTGGTTATGTATGTCCAACAACTACTTTTCAATATCTGTTGTTAAGTT

>M6-PGSC0003DMG400011251M6G-LecRK_3348

TCTGGCATTATTGGTTATGTATGTCCAACAACTACTTTTCAATATCTGTTGTTAAGTTAAATCATTTAACAACAATTCTACGTTTTTCTTTTTAGGCATGGCAGAAGTGGGATGAAGGTA

>M6-PGSC0003DMG400011251M6G-LecRK_3410

CATTTAACAACAATTCTACGTTTTTCTTTTTAGGCATGGCAGAAGTGGGATGAAGGTACACCAATGGACTTAGTAGACCGTTCTATTTGGGACGAGTGTCAATATGATGAAGCATTGAGA

>M6-PGSC0003DMG400011251M6G-LecRK_3472

AATGGACTTAGTAGACCGTTCTATTTGGGACGAGTGTCAATATGATGAAGCATTGAGATGTATACAGTTAGCACTAATTTGTGTACAAGACATGGCAGTTCACAGACCGAGCATATCATC

>M6-PGSC0003DMG400011251M6G-LecRK_3534

TACAGTTAGCACTAATTTGTGTACAAGACATGGCAGTTCACAGACCGAGCATATCATCCATCGTGTTAATGTTGGAGACGGACAATATACGATTGCCCTTGCCTAGGCAACCTACATATA

>M6-PGSC0003DMG400011251M6G-LecRK_3596

GTGTTAATGTTGGAGACGGACAATATACGATTGCCCTTGCCTAGGCAACCTACATATACCTCAATGAGAAAACATGAAAATGCAGAAACATGGAATGAGAAGCAGGATTTTTCTGCAAAC

>M6-PGSC0003DMG400011251M6G-LecRK_3629

CCCTTGCCTAGGCAACCTACATATACCTCAATGAGAAAACATGAAAATGCAGAAACATGGAATGAGAAGCAGGATTTTTCTGCAAACAATGTCACAATTAGTGTTATATTTGGTAGATGA

>M6-PGSC0003DMG400011251M6G-LecRK_372

GAAGTTTTGAGGTCATGGACTAGTGATAGTGATCCTTCACCTGGACGGTACGTAATCTTGTATCGTTTGATCATCTTATCTGAAATCTTTTATTTACTTAATTTTCTTTTGTACATGCCC

>M6-PGSC0003DMG400011251M6G-LecRK_434

TCGTTTGATCATCTTATCTGAAATCTTTTATTTACTTAATTTTCTTTTGTACATGCCCCCTCACGTTTGGTATGAATTTTTTACATAATGTGTGGTGGTGAGACTCTCTGATAGCATGTT

>M6-PGSC0003DMG400011251M6G-LecRK_496

ACGTTTGGTATGAATTTTTTACATAATGTGTGGTGGTGAGACTCTCTGATAGCATGTTGCGTTTGTGACATCTCATTTAAAAGCTTAATTGAGAATGTGCTTTTTTATTTTACGTAATTT

>M6-PGSC0003DMG400011251M6G-LecRK_558

TTGTGACATCTCATTTAAAAGCTTAATTGAGAATGTGCTTTTTTATTTTACGTAATTTTGTTTTGAACACAACCCCTCACGTGTTTGACTCATGGAAGTTGAAAGACAGAAGGGTTGTTG

>M6-PGSC0003DMG400011251M6G-LecRK_62

TCAAAGGTTTGTTGGTATATGGTATGTTGATGGTCCAAAAAATTCATTTGTTTGGGTTGCAAACAGAGATAAACCAATTTTTGATAAAAATGGTGTTTTTACTATTGAGAAAAATGGGAA

>M6-PGSC0003DMG400011251M6G-LecRK_620

TTGAACACAACCCCTCACGTGTTTGACTCATGGAAGTTGAAAGACAGAAGGGTTGTTGGTCAGTCACGGACTATTTGATTTCATTATGACAATTCAATGTTAATGACCTCATTCAATATA

>M6-PGSC0003DMG400011251M6G-LecRK_682

GTCACGGACTATTTGATTTCATTATGACAATTCAATGTTAATGACCTCATTCAATATATATATATTTTTTTGATCCTTCACTATTGTGGGCTGGATGGATAGCTAGTAGTTGTGAGTAGC

>M6-PGSC0003DMG400011251M6G-LecRK_744

TATTTTTTTGATCCTTCACTATTGTGGGCTGGATGGATAGCTAGTAGTTGTGAGTAGCAGTCCCCTTTTTATAGATTAAAGGTGATGATTTTTTGATATGGGGTGGTAGGGAGACTCGAA

>M6-PGSC0003DMG400011251M6G-LecRK_806

CCCTTTTTATAGATTAAAGGTGATGATTTTTTGATATGGGGTGGTAGGGAGACTCGAACCTACAAACTTTGCTTGCCCTGATATGGTAATGCATTTGTGACATCTTATCGAAAAGATTAA

>M6-PGSC0003DMG400011251M6G-LecRK_868

CAAACTTTGCTTGCCCTGATATGGTAATGCATTTGTGACATCTTATCGAAAAGATTAAGTTGAAAACATGTTTTTCTTTACTTAATTTTGTTTTTAACACGACCCCTCACGGTTTCTTGA

>M6-PGSC0003DMG400011251M6G-LecRK_930

AAAACATGTTTTTCTTTACTTAATTTTGTTTTTAACACGACCCCTCACGGTTTCTTGAATCTTTGATATTGGGTGATAGTGAGACTCGAAACCTAAGAACTCTACTTGCACTGATATGGT

>M6-PGSC0003DMG400011251M6G-LecRK_992

TTGATATTGGGTGATAGTGAGACTCGAAACCTAAGAACTCTACTTGCACTGATATGGTAATGCATTTGTGACATCTTATCTAAAAGATTAAGTTGATAACATGTGTTTCATAACTTACTT

>M6-PGSC0003DMG400011304-(rev)M6RLKWAX-EGF_0

CCAGTTGCATCTTCCTAGATGAGGACATGACCGTAAAACTTTTTGGGTTAGAGCTCCCCACGAATGCAAACACCGATAACAAAAAGCATAGCGATGTTTACAACTTGGGGTTGGTTCTTT

>M6-PGSC0003DMG400011304-(rev)M6RLKWAX-EGF_124

GGTTATTACAGGTAGCAGCTCAGATCATCTTCCATCAAAGATAGCTTTACAGAAGATAACAAGTGGGAAGCTAGAAGAGATAGTCGATCCACGTCTTTACTATCACGAGCAGCCTATTTT

>M6-PGSC0003DMG400011304-(rev)M6RLKWAX-EGF_186

GTGGGAAGCTAGAAGAGATAGTCGATCCACGTCTTTACTATCACGAGCAGCCTATTTTCCGGAGGGAACAAATAGAGATAGTTGCAGACCTTGCAACAAGATGCATAATATTTGGTTGCC

>M6-PGSC0003DMG400011304-(rev)M6RLKWAX-EGF_248

AGGGAACAAATAGAGATAGTTGCAGACCTTGCAACAAGATGCATAATATTTGGTTGCCAAAATGGGAAATTTCACATTGGAGATGTTTCAAGAGAACTTGTTAACATAACAAAAGATGGA

>M6-PGSC0003DMG400011304-(rev)M6RLKWAX-EGF_310

TGGGAAATTTCACATTGGAGATGTTTCAAGAGAACTTGTTAACATAACAAAAGATGGAATTGATGGTAGAAGTAGGAGATGTCCATCAACTCATAATCTTGAAGAAACTTTTTCCAATTC

>M6-PGSC0003DMG400011304-(rev)M6RLKWAX-EGF_372

ATGGTAGAAGTAGGAGATGTCCATCAACTCATAATCTTGAAGAAACTTTTTCCAATTCAAGCCTTCTTCAGATGATATCTATGTCTCCTGATTCAATACATGTCCCTGCAGCCAGAGGGT

>M6-PGSC0003DMG400011304-(rev)M6RLKWAX-EGF_434

CTTCTTCAGATGATATCTATGTCTCCTGATTCAATACATGTCCCTGCAGCCAGAGGGTTCTCTTAAGCTATGCTATGTTGCCAATGTCAACGGCCGCCTGTCAGTGCATTAAATTAGTTG

>M6-PGSC0003DMG400011304-(rev)M6RLKWAX-EGF_496

TTAAGCTATGCTATGTTGCCAATGTCAACGGCCGCCTGTCAGTGCATTAAATTAGTTGGAAAAAGAGGAAGGCTCTCACACTATTGACATGCTGTAATTATGTACAGTAATGATTAACCT

>M6-PGSC0003DMG400011304-(rev)M6RLKWAX-EGF_558

AAGAGGAAGGCTCTCACACTATTGACATGCTGTAATTATGTACAGTAATGATTAACCTACCTGCTGCTGTTATTTCCATCTTCTTATGCATAAGCAATCCTATTTTTGTTTCAATAAACT

>M6-PGSC0003DMG400011304-(rev)M6RLKWAX-EGF_611

AACCTACCTGCTGCTGTTATTTCCATCTTCTTATGCATAAGCAATCCTATTTTTGTTTCAATAAACTTCCAAATCACATTGAAAAATGGTAATATATCCTGATAGTATTGCAGTTTGTAT

>M6-PGSC0003DMG400011304-(rev)M6RLKWAX-EGF_62

AATGCAAACACCGATAACAAAAAGCATAGCGATGTTTACAACTTGGGGTTGGTTCTTTTGGAGGTTATTACAGGTAGCAGCTCAGATCATCTTCCATCAAAGATAGCTTTACAGAAGATA

>M6-PGSC0003DMG400011353-(rev)M6LRR-RLP_1037

TGCATCAGTCTGGTTCATCTCACTTTAATACTTCAAAGATTAGAGCTTTTTGTTTGTTTCTTTTAGCTTCTTTTTTTTGGGGTGAACTTCTGGGAATTAATTTTTCTTGATTGAATGAGC

>M6-PGSC0003DMG400011353-(rev)M6LRR-RLP_1098

TTTAGCTTCTTTTTTTTGGGGTGAACTTCTGGGAATTAATTTTTCTTGATTGAATGAGCTGATGTCAATAATTTCTTGTACTTCCTTCCCCAGATTTCTGCATTTGCATTCTATCTTTAC

>M6-PGSC0003DMG400011353-(rev)M6LRR-RLP_1159

ATGTCAATAATTTCTTGTACTTCCTTCCCCAGATTTCTGCATTTGCATTCTATCTTTACAGATCTCTAAGGCCTTATAACAGGAAATGCTAACTTGTGGGTTTTCTTAATTGTTATAGGA

>M6-PGSC0003DMG400011353-(rev)M6LRR-RLP_122

AAGGAAGAAAAAGGACTATTGTAAATCTCGAGTTCACTGTGAAATTACACTTTTGCCCTTCCCCCTCCATTACAGTACAACAATGGCGTACGCATTCCCGGAGGAAGTACTAGAACACGT

>M6-PGSC0003DMG400011353-(rev)M6LRR-RLP_1220

ATCTCTAAGGCCTTATAACAGGAAATGCTAACTTGTGGGTTTTCTTAATTGTTATAGGATTTAAACAGTTGGACTTCCTTTTAGCATTAATTTCTTGTGGTATACACTGTGTAAACATGC

>M6-PGSC0003DMG400011353-(rev)M6LRR-RLP_1281

TAAACAGTTGGACTTCCTTTTAGCATTAATTTCTTGTGGTATACACTGTGTAAACATGCCCTTTAGCTTTGCTTTGTTTTGAAGTTGTTGGAGATTTGAATGTTTTCTTAATTATCTGGA

>M6-PGSC0003DMG400011353-(rev)M6LRR-RLP_1342

TTTAGCTTTGCTTTGTTTTGAAGTTGTTGGAGATTTGAATGTTTTCTTAATTATCTGGAGATTTCCATACGTTTGTTTCTTGGTTCTCTGGTGGGATTAATTTGTTTTATCAGGAGGGAG

>M6-PGSC0003DMG400011353-(rev)M6LRR-RLP_1403

TTTCCATACGTTTGTTTCTTGGTTCTCTGGTGGGATTAATTTGTTTTATCAGGAGGGAGTTTTGTAATGGAGTATAGTTGTAGTTCAATGTTGAAATGGCAAGTGGGTTAGGAATAGGAA

>M6-PGSC0003DMG400011353-(rev)M6LRR-RLP_1464

TTGTAATGGAGTATAGTTGTAGTTCAATGTTGAAATGGCAAGTGGGTTAGGAATAGGAAATGGTGATTAATTGAAATTCCAATTGGAAGATGTGTCACTCTGAGGCAAACATGATTCTAA

>M6-PGSC0003DMG400011353-(rev)M6LRR-RLP_1525

GGTGATTAATTGAAATTCCAATTGGAAGATGTGTCACTCTGAGGCAAACATGATTCTAAATCTTTCTTATTTATCTTGTTCATATCGTCTGTCATTTATTATCATTTGTTGTTTTCTGTT

>M6-PGSC0003DMG400011353-(rev)M6LRR-RLP_1586

CTTTCTTATTTATCTTGTTCATATCGTCTGTCATTTATTATCATTTGTTGTTTTCTGTTGAGTCTGTTGATTTGCTTCAATTTATCACAGAATGATTGTGTAGATGGGTATTGTCATCTA

>M6-PGSC0003DMG400011353-(rev)M6LRR-RLP_1647

GTCTGTTGATTTGCTTCAATTTATCACAGAATGATTGTGTAGATGGGTATTGTCATCTAACTCCCGTTTTGCTGCAGGAATCTGAGAAAACTAGACTTGGGAGAGAGTGAAGTGGAGGAC

>M6-PGSC0003DMG400011353-(rev)M6LRR-RLP_1708

TCCCGTTTTGCTGCAGGAATCTGAGAAAACTAGACTTGGGAGAGAGTGAAGTGGAGGACCTTAGTGGGCATTGGTTAAGCCATTTCCCTGATAACTGCACGTCACTTGTTTCACTTAACA

>M6-PGSC0003DMG400011353-(rev)M6LRR-RLP_1769

TAGTGGGCATTGGTTAAGCCATTTCCCTGATAACTGCACGTCACTTGTTTCACTTAACATTGCCTGTTTGGCTTCTGAGGTCAGTCTGTTAGCTTTGGAGCGCCTGGTTACTCGCTCTCC

>M6-PGSC0003DMG400011353-(rev)M6LRR-RLP_183

CCCCTCCATTACAGTACAACAATGGCGTACGCATTCCCGGAGGAAGTACTAGAACACGTCTTCTCCTTCCTCACCTCCGACAAGGACCGTAACGCAGTATCACTAGTCTGCAAATCATGG

>M6-PGSC0003DMG400011353-(rev)M6LRR-RLP_1830

GCCTGTTTGGCTTCTGAGGTCAGTCTGTTAGCTTTGGAGCGCCTGGTTACTCGCTCTCCTAATTTGAGAACTCTTCGGATAAATCGTGCTGTGCCCCTTGAGAGGCTTCCAAACTTGCTC

>M6-PGSC0003DMG400011353-(rev)M6LRR-RLP_1891

ATTTGAGAACTCTTCGGATAAATCGTGCTGTGCCCCTTGAGAGGCTTCCAAACTTGCTCCGCCGCACTTCCCAGTTGGTCGAATTTGGTACAGGTGTTTTCTCTGCTGATGTGCGATCTG

>M6-PGSC0003DMG400011353-(rev)M6LRR-RLP_1952

CCGCACTTCCCAGTTGGTCGAATTTGGTACAGGTGTTTTCTCTGCTGATGTGCGATCTGATTTTTTCTCAAACCTGACAGAAGCTTTTTCGAGCTGCAAGCAACTTAAATGCCTGTCTGG

>M6-PGSC0003DMG400011353-(rev)M6LRR-RLP_2013

TTTTTCTCAAACCTGACAGAAGCTTTTTCGAGCTGCAAGCAACTTAAATGCCTGTCTGGGTTTTGGGATGTTGTACCAGCTTACCTTCCAGCTTTATATCCAGTGTGCTCAAGACTCACC

>M6-PGSC0003DMG400011353-(rev)M6LRR-RLP_2074

TTTGGGATGTTGTACCAGCTTACCTTCCAGCTTTATATCCAGTGTGCTCAAGACTCACCTCCTTGAATTTGAGCTATGCTACCTGTCAGAACCCTGAGCTTGGCAAGCTCATAAGTCAGT

>M6-PGSC0003DMG400011353-(rev)M6LRR-RLP_2135

CTTGAATTTGAGCTATGCTACCTGTCAGAACCCTGAGCTTGGCAAGCTCATAAGTCAGTGTCACAATTTGCAGCGTTTGTGGGTATGTTCTAAATGTTTTGAAGTTTTAATTGATGTTCT

>M6-PGSC0003DMG400011353-(rev)M6LRR-RLP_2196

CACAATTTGCAGCGTTTGTGGGTATGTTCTAAATGTTTTGAAGTTTTAATTGATGTTCTTGACACTGCAACTGCAAACATGTTTTTTTTTTTTTTTTTTAATAAAGAAGAAGAGGCAATG

>M6-PGSC0003DMG400011353-(rev)M6LRR-RLP_2257

ACACTGCAACTGCAAACATGTTTTTTTTTTTTTTTTTTAATAAAGAAGAAGAGGCAATGAGATTGTAACATCATACTTGCTGTGATGTCATTTAGCATATATTCTTAGTGAAGAATAAGA

>M6-PGSC0003DMG400011353-(rev)M6LRR-RLP_2318

ATTGTAACATCATACTTGCTGTGATGTCATTTAGCATATATTCTTAGTGAAGAATAAGAAAAGACCTAACTGCTACAGATCAACGAATCCACCACAACTTAATCCCAATTTAGTTGGTCT

>M6-PGSC0003DMG400011353-(rev)M6LRR-RLP_2379

AGACCTAACTGCTACAGATCAACGAATCCACCACAACTTAATCCCAATTTAGTTGGTCTCAGCTTTATGGGTCCTCTTAAACCATTTTGCTATATTCAGGTCATTTTCATTCCGATGCTA

>M6-PGSC0003DMG400011353-(rev)M6LRR-RLP_244

TCTCCTTCCTCACCTCCGACAAGGACCGTAACGCAGTATCACTAGTCTGCAAATCATGGTACGAGATCGAACGGTGGTGTAGGAAGAGAATATTCGTTGGAAACTGCTATGCTGTTAGCC

>M6-PGSC0003DMG400011353-(rev)M6LRR-RLP_2440

GCTTTATGGGTCCTCTTAAACCATTTTGCTATATTCAGGTCATTTTCATTCCGATGCTAAATTATTTGTCTTTACGCAATAATTGCCAAGGATAGGCAAAAAGTAATTCATTCTGTAGGT

>M6-PGSC0003DMG400011353-(rev)M6LRR-RLP_2501

TTATTTGTCTTTACGCAATAATTGCCAAGGATAGGCAAAAAGTAATTCATTCTGTAGGTAAACTTCACCTATGCTGGTTAGGAAGATTCCAGGCTACATATATTTGATATTATATTGCTA

>M6-PGSC0003DMG400011353-(rev)M6LRR-RLP_2562

ACTTCACCTATGCTGGTTAGGAAGATTCCAGGCTACATATATTTGATATTATATTGCTAAGCAGCTCACTTTTCTTGGAAGTTGAAGCCAAGTGTTACATGCTTCATATTGTGATCTGGG

>M6-PGSC0003DMG400011353-(rev)M6LRR-RLP_2623

CAGCTCACTTTTCTTGGAAGTTGAAGCCAAGTGTTACATGCTTCATATTGTGATCTGGGTTTCAGTTTTTTAATACTTGTGCCTACCCAATGCACCTAAAAATCCATCTAAATGTGATGT

>M6-PGSC0003DMG400011353-(rev)M6LRR-RLP_2684

TCAGTTTTTTAATACTTGTGCCTACCCAATGCACCTAAAAATCCATCTAAATGTGATGTTTCTATGCTGAAAAATTGTATCTATTCTTCCCTAGTGGCTTCCCCAAACTGGTTCTGTAAT

>M6-PGSC0003DMG400011353-(rev)M6LRR-RLP_2745

CTATGCTGAAAAATTGTATCTATTCTTCCCTAGTGGCTTCCCCAAACTGGTTCTGTAATATAAACTCCAGTATCATGTTTAAGTGCTGAATTGCTTTTCTTGCAATATATGTGACTGCAG

>M6-PGSC0003DMG400011353-(rev)M6LRR-RLP_2806

AAACTCCAGTATCATGTTTAAGTGCTGAATTGCTTTTCTTGCAATATATGTGACTGCAGGTGCTAGACTACATCGAAGATACTGGTCTTGAGGAACTTGCTGCCAATTGTAAGGACCTTC

>M6-PGSC0003DMG400011353-(rev)M6LRR-RLP_2867

GCTAGACTACATCGAAGATACTGGTCTTGAGGAACTTGCTGCCAATTGTAAGGACCTTCAAGAGCTGCGGGTGTTTCCTTCTGACCCATTTGCTGCAGAACCAAATACAACCTTGACGGA

>M6-PGSC0003DMG400011353-(rev)M6LRR-RLP_2928

GAGCTGCGGGTGTTTCCTTCTGACCCATTTGCTGCAGAACCAAATACAACCTTGACGGAGCAAGGCCTTGTAGCTGTCTCAGATGGCTGCCCAAAGCTCCAGTCTGTTTTATACTTCTGC

>M6-PGSC0003DMG400011353-(rev)M6LRR-RLP_2989

AAGGCCTTGTAGCTGTCTCAGATGGCTGCCCAAAGCTCCAGTCTGTTTTATACTTCTGCCGCCAAATGACTAATGCAGCTTTAGTTACTATTGCTAGGAATCGACCTAACATGATCCGTT

>M6-PGSC0003DMG400011353-(rev)M6LRR-RLP_305

CGAGATCGAACGGTGGTGTAGGAAGAGAATATTCGTTGGAAACTGCTATGCTGTTAGCCCTAGGATCATGATCCGACGGTTTCCGGAAGTCAGATCTGTTGAGCTTAAAGGGAAGCCTCA

>M6-PGSC0003DMG400011353-(rev)M6LRR-RLP_3050

CCAAATGACTAATGCAGCTTTAGTTACTATTGCTAGGAATCGACCTAACATGATCCGTTTTCGCTTGTGTATCATTGAGCCTCGAACTCCTGATTATTTAACCCTTGGTTCATTTGATAC

>M6-PGSC0003DMG400011353-(rev)M6LRR-RLP_3111

CGCTTGTGTATCATTGAGCCTCGAACTCCTGATTATTTAACCCTTGGTTCATTTGATACTGGTTTTGGGGCCATAGTAGAGAACTGTAAGGAATTGCGGCGCCTTTCACTTGCTGGCCTC

>M6-PGSC0003DMG400011353-(rev)M6LRR-RLP_3172

GTTTTGGGGCCATAGTAGAGAACTGTAAGGAATTGCGGCGCCTTTCACTTGCTGGCCTCCTTACCGATCGTGTCTTCGAATACATTGGCGCCCATGCTAAAAAGTTGGAGATGCTTTCCA

>M6-PGSC0003DMG400011353-(rev)M6LRR-RLP_3233

TACCGATCGTGTCTTCGAATACATTGGCGCCCATGCTAAAAAGTTGGAGATGCTTTCCATAGCTTTTGCTGGTGATAGCGATTTGGGACTCCACCATGTTCTCTCTGGTTGTGATAGCCT

>M6-PGSC0003DMG400011353-(rev)M6LRR-RLP_3294

GCTTTTGCTGGTGATAGCGATTTGGGACTCCACCATGTTCTCTCTGGTTGTGATAGCCTTCGGAAGTTGGAGATCAGAGATTGTCCGTTTGGAGACAAGGCTCTTTTGGCTAATGCTGCA

>M6-PGSC0003DMG400011353-(rev)M6LRR-RLP_3355

GGAAGTTGGAGATCAGAGATTGTCCGTTTGGAGACAAGGCTCTTTTGGCTAATGCTGCAAAGCTCGAGACAATGCGATCCCTTTGGATGTCTTCTTGTTCAGTAAGTTTTGAAGCATGTA

>M6-PGSC0003DMG400011353-(rev)M6LRR-RLP_3416

GCTCGAGACAATGCGATCCCTTTGGATGTCTTCTTGTTCAGTAAGTTTTGAAGCATGTAAGATGCTAGCTCAGAAGATGCCTAGGCTTAATGTGGAAGTCATAGATGAGAGGGGTCCTCC

>M6-PGSC0003DMG400011353-(rev)M6LRR-RLP_3477

ATGCTAGCTCAGAAGATGCCTAGGCTTAATGTGGAAGTCATAGATGAGAGGGGTCCTCCAGATACAAGGCCAGAAAGCTGTCCTGTGGAGAAACTTTACATATATCGAACTGTGGCTGGA

>M6-PGSC0003DMG400011353-(rev)M6LRR-RLP_3538

ATACAAGGCCAGAAAGCTGTCCTGTGGAGAAACTTTACATATATCGAACTGTGGCTGGACGAAGATTTGACACACCTGGTTATGTTTGGACAATGGATGAAGATGCAGCTGTAAGGTTGA

>M6-PGSC0003DMG400011353-(rev)M6LRR-RLP_3599

AAGATTTGACACACCTGGTTATGTTTGGACAATGGATGAAGATGCAGCTGTAAGGTTGACCTGAGCCACACATCAAATGGAAAATTCCAGCAAGACTTCACATGGAAAGATGGACAGGTA

>M6-PGSC0003DMG400011353-(rev)M6LRR-RLP_366

AGGATCATGATCCGACGGTTTCCGGAAGTCAGATCTGTTGAGCTTAAAGGGAAGCCTCATTTTGCTGACTTTAATCTTGTACCGGAAGGTTGGGGAGCTTATGTGTATCCTTGGATTTTG

>M6-PGSC0003DMG400011353-(rev)M6LRR-RLP_3660

TGAGCCACACATCAAATGGAAAATTCCAGCAAGACTTCACATGGAAAGATGGACAGGTATTAGCGCTATGTTTATCCTCCTGCATTTGTTTGTCACGCCTTATGTAGTTGCAGTGCCAGT

>M6-PGSC0003DMG400011353-(rev)M6LRR-RLP_3721

AGCGCTATGTTTATCCTCCTGCATTTGTTTGTCACGCCTTATGTAGTTGCAGTGCCAGTTGAGGGTTGGAACTTCACTGCTTTGAAGTAGAGGCTCCAGAGCTCATACGCCATCAATTTT

>M6-PGSC0003DMG400011353-(rev)M6LRR-RLP_3782

AGGGTTGGAACTTCACTGCTTTGAAGTAGAGGCTCCAGAGCTCATACGCCATCAATTTTATTCTTGGCTGAGAGCTACTTTACAGCAGGTAGTTTTTAATAAAGAGTTTGGACTCTGAGT

>M6-PGSC0003DMG400011353-(rev)M6LRR-RLP_3843

TCTTGGCTGAGAGCTACTTTACAGCAGGTAGTTTTTAATAAAGAGTTTGGACTCTGAGTTTCTGTGTATCATCTAGAGTAACAATTATTCGTTTTCTTTGGGTGGGATTTTGCTTGGGGG

>M6-PGSC0003DMG400011353-(rev)M6LRR-RLP_3904

CTGTGTATCATCTAGAGTAACAATTATTCGTTTTCTTTGGGTGGGATTTTGCTTGGGGGGTTTGGGGGCATATTTTTGCCATGTATGTGCTATTTCCAGACTTTGCTTCATTATGTAGTT

>M6-PGSC0003DMG400011353-(rev)M6LRR-RLP_3950

TTTTGCTTGGGGGGTTTGGGGGCATATTTTTGCCATGTATGTGCTATTTCCAGACTTTGCTTCATTATGTAGTTCACTTTATTTATTCTATTGACTTATTCTCTATGAAATGGAGTGTCT

>M6-PGSC0003DMG400011353-(rev)M6LRR-RLP_427

TTGCTGACTTTAATCTTGTACCGGAAGGTTGGGGAGCTTATGTGTATCCTTGGATTTTGGCTATGTCTAAATCTTATCCATGGCTTGAAGAAATTAAGCTTAAGAGGATGGTAATTACTG

>M6-PGSC0003DMG400011353-(rev)M6LRR-RLP_488

TATGTCTAAATCTTATCCATGGCTTGAAGAAATTAAGCTTAAGAGGATGGTAATTACTGATGAGTCGTTGGAGTTGATTTCTAAGTCGTTTAAGAACTTTAAAGTGTTGGTTTTGTCTTC

>M6-PGSC0003DMG400011353-(rev)M6LRR-RLP_549

GAGTCGTTGGAGTTGATTTCTAAGTCGTTTAAGAACTTTAAAGTGTTGGTTTTGTCTTCATGTGATGGGTTTACTACTGATGGACTTGCTGCTATTGCTGCTAACTGCAGGTGAAATTTT

>M6-PGSC0003DMG400011353-(rev)M6LRR-RLP_61

CTAAAACACAGCAAATCTAAGCTCGTTAATGTCGATTCAGATCTGAGAGAAGAAGAAGAAGAAGGAAGAAAAAGGACTATTGTAAATCTCGAGTTCACTGTGAAATTACACTTTTGCCCT

>M6-PGSC0003DMG400011353-(rev)M6LRR-RLP_610

GTGATGGGTTTACTACTGATGGACTTGCTGCTATTGCTGCTAACTGCAGGTGAAATTTTTTTTCTTGCTTGACATTTCTCTTCTGTTTTGCTCACTGTAAAGTGGCTAGTGGTGAGAGAA

>M6-PGSC0003DMG400011353-(rev)M6LRR-RLP_671

TTCTTGCTTGACATTTCTCTTCTGTTTTGCTCACTGTAAAGTGGCTAGTGGTGAGAGAAGTCTTAATCTGTAAAGTAATTAGATCTAATTTTGGGTATTAGTTTGGACAGTGACTTTTAT

>M6-PGSC0003DMG400011353-(rev)M6LRR-RLP_732

CTTAATCTGTAAAGTAATTAGATCTAATTTTGGGTATTAGTTTGGACAGTGACTTTTATTGTTGTTAAAGCTTAAGCTTTTGGATGAATGCAGTATGCATGCATTTTGGGGGTTTGTTAT

>M6-PGSC0003DMG400011353-(rev)M6LRR-RLP_793

TTGTTAAAGCTTAAGCTTTTGGATGAATGCAGTATGCATGCATTTTGGGGGTTTGTTATAATCCCTTGCTAAACACGAGAATGAAGAAGCACTTTTCTTGCAGCTGTTTGTTCTGTTTAT

>M6-PGSC0003DMG400011353-(rev)M6LRR-RLP_854

TCCCTTGCTAAACACGAGAATGAAGAAGCACTTTTCTTGCAGCTGTTTGTTCTGTTTATAGGATATGTAATGTATTAGTGGAAACGAAGTATCTACTGTAGATGAATTCGTAGTGTGTGA

>M6-PGSC0003DMG400011353-(rev)M6LRR-RLP_915

GATATGTAATGTATTAGTGGAAACGAAGTATCTACTGTAGATGAATTCGTAGTGTGTGAACTTGAAGAACTGACATTTCTTCAAGTGATTATGGTATTATTGGATACGTCTTGCTGCTAC

>M6-PGSC0003DMG400011353-(rev)M6LRR-RLP_976

TTGAAGAACTGACATTTCTTCAAGTGATTATGGTATTATTGGATACGTCTTGCTGCTACATTGCATCAGTCTGGTTCATCTCACTTTAATACTTCAAAGATTAGAGCTTTTTGTTTGTTT

>M6-PGSC0003DMG400011509-(rev)M6LRR-RLK_0

ATGAAAACATGGAATGCATCCACACACTTTTGCCATTGGTCTGGAGTAACTTGTGGCTGTAAACATGTTAGAGTCATTCAACTAAACGTTGCGAACCAAAAACTAGATGGCCCCTTGTCA

>M6-PGSC0003DMG400011509-(rev)M6LRR-RLK_1054

ACTTACTGGTACTATTCCAATCAATTTTGGTAAAGTTAAGAAACTGCAAATTTTGGATTTGTCTCAGAATAGATTTTCAGGGAAAATTCCGTCATCTCTAGGAAATCTTAGTGTAGTATC

>M6-PGSC0003DMG400011509-(rev)M6LRR-RLK_1116

CTCAGAATAGATTTTCAGGGAAAATTCCGTCATCTCTAGGAAATCTTAGTGTAGTATCCATCCTTCTTCTGCACGATAAAAATCTCACCGGTGAGATCCCAGCAAGTTTAGGAAATTGCA

>M6-PGSC0003DMG400011509-(rev)M6LRR-RLK_1178

CTTCTTCTGCACGATAAAAATCTCACCGGTGAGATCCCAGCAAGTTTAGGAAATTGCAATTACGTGATTGAAATATATGTTGCTAAAAATAACCTGTTAGGACAGATACCTAAGGACCTT

>M6-PGSC0003DMG400011509-(rev)M6LRR-RLK_124

TCATTGGGAACATGAGTTTCCTCAGGTCTCTATACCTGTCAAACAATAGCTTCCGTGGTGAAATCCCTTCTGAGATTGGTCGTTTGAGAAGGCTACACCGATTGTATCTTGGTAACAACT

>M6-PGSC0003DMG400011509-(rev)M6LRR-RLK_1240

CGTGATTGAAATATATGTTGCTAAAAATAACCTGTTAGGACAGATACCTAAGGACCTTTTTGCTCTCTTCCTTAGTTGCTGTGGACATTTCTGAGAACCATTTAGATGGTTTCATTCCTC

>M6-PGSC0003DMG400011509-(rev)M6LRR-RLK_1302

CTCTCTTCCTTAGTTGCTGTGGACATTTCTGAGAACCATTTAGATGGTTTCATTCCTCTTGAAGTTGGAAATATGATAAATCTTGAATATCTTAATGTTTCCGTAAATAACTTGACCGGT

>M6-PGSC0003DMG400011509-(rev)M6LRR-RLK_1364

AGTTGGAAATATGATAAATCTTGAATATCTTAATGTTTCCGTAAATAACTTGACCGGTAAAATCCCAAGTACTATTGGTAGCTGTGTGACACTAGAAGCCTTAGACATGAAAGGGAACTT

>M6-PGSC0003DMG400011509-(rev)M6LRR-RLK_1426

TCCCAAGTACTATTGGTAGCTGTGTGACACTAGAAGCCTTAGACATGAAAGGGAACTTCTTTCAAGGGATTATACCTCCTTCTTTCAGCTCTTTACGAGGTCTCCGTGTTTTAGACCTTT

>M6-PGSC0003DMG400011509-(rev)M6LRR-RLK_1488

CAAGGGATTATACCTCCTTCTTTCAGCTCTTTACGAGGTCTCCGTGTTTTAGACCTTTCTAGAAATAATTTGTCTGGCCAAGTACCAAAATACTTGGAAGATTTCAAGTTTCAACTCTTG

>M6-PGSC0003DMG400011509-(rev)M6LRR-RLK_1550

AAATAATTTGTCTGGCCAAGTACCAAAATACTTGGAAGATTTCAAGTTTCAACTCTTGAATTTAGCGTTCAATGATTTCGAAGGAGCGTTACCAAATGAAGGCGTCATTAAGAATGCAAG

>M6-PGSC0003DMG400011509-(rev)M6LRR-RLK_1612

TAGCGTTCAATGATTTCGAAGGAGCGTTACCAAATGAAGGCGTCATTAAGAATGCAAGTGCTATATCAGAAATTGGAAATCCAAAACTTTGTGGGGGTGTGCCTGACATACATCTTCCGG

>M6-PGSC0003DMG400011509-(rev)M6LRR-RLK_1674

ATATCAGAAATTGGAAATCCAAAACTTTGTGGGGGTGTGCCTGACATACATCTTCCGGGATGTGACATCAAAAGATCGAAAAAATTTGGTTCGAGGTTTATCCTCAAGATTGTCATCTCT

>M6-PGSC0003DMG400011509-(rev)M6LRR-RLK_1736

TGACATCAAAAGATCGAAAAAATTTGGTTCGAGGTTTATCCTCAAGATTGTCATCTCTGTTGTCTTTGGTATACTAGGCCTTGGTATGCTTATCACTTTACTATTTTGTTTCTTGTTAAA

>M6-PGSC0003DMG400011509-(rev)M6LRR-RLK_1798

TCTTTGGTATACTAGGCCTTGGTATGCTTATCACTTTACTATTTTGTTTCTTGTTAAAGAAACCGAAAAGGCCGCCTGTTTCAAGTTCATTGGGTGAGTCTCTCATAAATGTGTCTTACC

>M6-PGSC0003DMG400011509-(rev)M6LRR-RLK_186

ATCCCTTCTGAGATTGGTCGTTTGAGAAGGCTACACCGATTGTATCTTGGTAACAACTCATTTCATGGAGAGATCCCCTCAAATTTATCAAGGTGTTTGAATCTTGTATCTCTTGTCCTT

>M6-PGSC0003DMG400011509-(rev)M6LRR-RLK_1860

CCGAAAAGGCCGCCTGTTTCAAGTTCATTGGGTGAGTCTCTCATAAATGTGTCTTACCGAAGTTTGCTTCAAGCTACTACTGGATTCTCTGAAGACAACTTGATTGGTGCTGGGAGCTAA

>M6-PGSC0003DMG400011509-(rev)M6LRR-RLK_1922

TTTGCTTCAAGCTACTACTGGATTCTCTGAAGACAACTTGATTGGTGCTGGGAGCTAAGGATCTGTTTATAAAGGAACTCTCGATGGAGGAATGGTTGTCGCAGTTAAAGTGCTAAACCT

>M6-PGSC0003DMG400011509-(rev)M6LRR-RLK_1984

CTGTTTATAAAGGAACTCTCGATGGAGGAATGGTTGTCGCAGTTAAAGTGCTAAACCTCTCGCGACATGGAGCTTCCAAAAGCTTTATGGCTGAGTGTGAAGTCTTGAGAAACATTAGAC

>M6-PGSC0003DMG400011509-(rev)M6LRR-RLK_2046

CGACATGGAGCTTCCAAAAGCTTTATGGCTGAGTGTGAAGTCTTGAGAAACATTAGACATCGGAATTTGGTCAAGGTTCTCACAGCATGTTCAGGAGTTGACTATCGAGGGAACGATGTC

>M6-PGSC0003DMG400011509-(rev)M6LRR-RLK_2108

GAATTTGGTCAAGGTTCTCACAGCATGTTCAGGAGTTGACTATCGAGGGAACGATGTCAAGGCTTTGGTTTACGAGTTTATGGTTAATGGAAGCCTCGAAGACTGGTTGCATCCTAGTCC

>M6-PGSC0003DMG400011509-(rev)M6LRR-RLK_2170

CTTTGGTTTACGAGTTTATGGTTAATGGAAGCCTCGAAGACTGGTTGCATCCTAGTCCAAGTGAAGATACAAGTCAGGCAGCTGAAACAAAAAAGTTGAATATTCTTCAAAGGCTCAACA

>M6-PGSC0003DMG400011509-(rev)M6LRR-RLK_2232

GAAGATACAAGTCAGGCAGCTGAAACAAAAAAGTTGAATATTCTTCAAAGGCTCAACATTGTTATAGATGTTGCTTCTGCAATCAATTATCTTCATCTCCACTGCGAGACTCCGATAGTT

>M6-PGSC0003DMG400011509-(rev)M6LRR-RLK_2294

TATAGATGTTGCTTCTGCAATCAATTATCTTCATCTCCACTGCGAGACTCCGATAGTTCACTGTGATCTCAAGCCAAGCAACATTCTTCTAGACAACCAATTGGTCGGGCACATTGGTGA

>M6-PGSC0003DMG400011509-(rev)M6LRR-RLK_2356

GTGATCTCAAGCCAAGCAACATTCTTCTAGACAACCAATTGGTCGGGCACATTGGTGATTTTGGTTTGGCAAAGTTCCTTCAACCAACTGCACAAAATTCTTCTATTTCAGAAGGAAGCT

>M6-PGSC0003DMG400011509-(rev)M6LRR-RLK_2418

GGTTTGGCAAAGTTCCTTCAACCAACTGCACAAAATTCTTCTATTTCAGAAGGAAGCTCTGGTCTTGTTAGAGGAACAATTGGTTACACTGCTCCAGGTAACAATTAATACATAAACAAA

>M6-PGSC0003DMG400011509-(rev)M6LRR-RLK_248

TCATGGAGAGATCCCCTCAAATTTATCAAGGTGTTTGAATCTTGTATCTCTTGTCCTTGAAGGTAACAAGTTAGTAGGAAGCCTTCCGCAGGAGCTAGGTGCCTTGTCCAAACTCGAGTA

>M6-PGSC0003DMG400011509-(rev)M6LRR-RLK_2480

TCTTGTTAGAGGAACAATTGGTTACACTGCTCCAGGTAACAATTAATACATAAACAAACACTCGAACTTGGTATATGCCCTTTTTTTCTCTCTCATTTCTTCTATTTTTTCTTGAAAATG

>M6-PGSC0003DMG400011509-(rev)M6LRR-RLK_2542

CGAACTTGGTATATGCCCTTTTTTTCTCTCTCATTTCTTCTATTTTTTCTTGAAAATGATATATTCATAGCAAAACCACACTGCTTAACCATAACTTGGTTCGACAGAGTTAATTTATCA

>M6-PGSC0003DMG400011509-(rev)M6LRR-RLK_2604

ATTCATAGCAAAACCACACTGCTTAACCATAACTTGGTTCGACAGAGTTAATTTATCAATATCATAAACTATCTCTTAATTGTCACACACATACTTCCATTACTTTTATAAAATCAAGTC

>M6-PGSC0003DMG400011509-(rev)M6LRR-RLK_2666

CATAAACTATCTCTTAATTGTCACACACATACTTCCATTACTTTTATAAAATCAAGTCATACTAAGTTATGTCCCTATTAGGAAAGTTATTGTTCCTCGTCTTTTTTTTTGTTTTGTGAA

>M6-PGSC0003DMG400011509-(rev)M6LRR-RLK_2728

TAAGTTATGTCCCTATTAGGAAAGTTATTGTTCCTCGTCTTTTTTTTTGTTTTGTGAATTTTAGAATATGGGATGGGAAGCGAGCTATCGACTTGTGGTGATGTGTACAGCTTCGGGATC

>M6-PGSC0003DMG400011509-(rev)M6LRR-RLK_2790

AGAATATGGGATGGGAAGCGAGCTATCGACTTGTGGTGATGTGTACAGCTTCGGGATCCTTTTACTAGAAATGTTCACCGGAAAAAGGCCTACTGATGGAATGTTTAGGGATGGGTTAGA

>M6-PGSC0003DMG400011509-(rev)M6LRR-RLK_2852

TACTAGAAATGTTCACCGGAAAAAGGCCTACTGATGGAATGTTTAGGGATGGGTTAGACCTTCCAAGCTTCGCTAAACATGCATTGCTTAATGGAGCTATGGAAGTCATAGATCCATCCC

>M6-PGSC0003DMG400011509-(rev)M6LRR-RLK_2914

CCAAGCTTCGCTAAACATGCATTGCTTAATGGAGCTATGGAAGTCATAGATCCATCCCTTATTTATGGAAGTGAAGAAGATGAGAAAGGTAAAAACACAAATATCTACCAGAATAAGGTG

>M6-PGSC0003DMG400011509-(rev)M6LRR-RLK_2976

TTATGGAAGTGAAGAAGATGAGAAAGGTAAAAACACAAATATCTACCAGAATAAGGTGTATCTGGTTTCTGTTCTTCGAGTCGGGGTTGCTTGTTCTGCATATTCTGGAGCTGAAAGAAT

>M6-PGSC0003DMG400011509-(rev)M6LRR-RLK_3038

TGGTTTCTGTTCTTCGAGTCGGGGTTGCTTGTTCTGCATATTCTGGAGCTGAAAGAATGAACATCACTGAAACTGTTTCCCAATTGTACTCTATCAAGGAAGCTCTTCTGAAATCAGAAA

>M6-PGSC0003DMG400011509-(rev)M6LRR-RLK_310

GTAACAAGTTAGTAGGAAGCCTTCCGCAGGAGCTAGGTGCCTTGTCCAAACTCGAGTACTTGTTACTCACTCGAAACAACTTGACAGGTGAAATCCCTTCTTCTTTTGGAAACTTAACAT

>M6-PGSC0003DMG400011509-(rev)M6LRR-RLK_372

TTACTCACTCGAAACAACTTGACAGGTGAAATCCCTTCTTCTTTTGGAAACTTAACATCATTGATAGGATTCTATGCACCATTGAATAATCTTCAGGGGAAAATCCCTGACTCTTTTGGG

>M6-PGSC0003DMG400011509-(rev)M6LRR-RLK_434

GATAGGATTCTATGCACCATTGAATAATCTTCAGGGGAAAATCCCTGACTCTTTTGGGCAGCTGAAAAACTTAGAAATTATCGGAGTAGCTGCAAACCATTTGTCAGGTACTATCCCTTC

>M6-PGSC0003DMG400011509-(rev)M6LRR-RLK_496

TGAAAAACTTAGAAATTATCGGAGTAGCTGCAAACCATTTGTCAGGTACTATCCCTTCTGAAATCTTCAACATATCTTCCGTCACAACATTTGATGTTGGAATGAACCAAATCCAAGGTA

>M6-PGSC0003DMG400011509-(rev)M6LRR-RLK_558

ATCTTCAACATATCTTCCGTCACAACATTTGATGTTGGAATGAACCAAATCCAAGGTACACTTCCATCAAGTTTGGGAATCACCCTCCCGAATCTTGAATTATTTATCATCGGTGGGAAC

>M6-PGSC0003DMG400011509-(rev)M6LRR-RLK_62

ACATGTTAGAGTCATTCAACTAAACGTTGCGAACCAAAAACTAGATGGCCCCTTGTCATCCTTCATTGGGAACATGAGTTTCCTCAGGTCTCTATACCTGTCAAACAATAGCTTCCGTGG

>M6-PGSC0003DMG400011509-(rev)M6LRR-RLK_620

TCCATCAAGTTTGGGAATCACCCTCCCGAATCTTGAATTATTTATCATCGGTGGGAACAATATTTCTGGATCGATCCCATCTACATTGTCCAATTCATCAAAGCTAGTGTATTTTCTTGC

>M6-PGSC0003DMG400011509-(rev)M6LRR-RLK_682

TTTCTGGATCGATCCCATCTACATTGTCCAATTCATCAAAGCTAGTGTATTTTCTTGCTGAAAGAAATCAACTTACTGGAAGTGTACCAAGTCTTGAGAACCTGAATGGGCTTCAACAGT

>M6-PGSC0003DMG400011509-(rev)M6LRR-RLK_744

AGAAATCAACTTACTGGAAGTGTACCAAGTCTTGAGAACCTGAATGGGCTTCAACAGTTGACAATTCCTGGGAATTATCTTGGAACTGGAGAACCAGATGATCTAAGCTTCATAGCCTCT

>M6-PGSC0003DMG400011509-(rev)M6LRR-RLK_806

AATTCCTGGGAATTATCTTGGAACTGGAGAACCAGATGATCTAAGCTTCATAGCCTCTCTGACTAACGTGTCCCGTTTCAGAATTTTGGAGATTCAGTTCAATAGTTTTGGAGGGGTTCT

>M6-PGSC0003DMG400011509-(rev)M6LRR-RLK_868

CTAACGTGTCCCGTTTCAGAATTTTGGAGATTCAGTTCAATAGTTTTGGAGGGGTTCTGCCTGCATCCTTTAGAAATCTTTCAACTGAGCTTCAGGTTGTGCAGTTGTCATACAACAGAA

>M6-PGSC0003DMG400011509-(rev)M6LRR-RLK_930

GCATCCTTTAGAAATCTTTCAACTGAGCTTCAGGTTGTGCAGTTGTCATACAACAGAATACGTGGTAACATACCGTCTGAGATTGAAAATTTCGTCAACGTGGAAGAATTTCAGGTGAGG

>M6-PGSC0003DMG400011509-(rev)M6LRR-RLK_992

TGGTAACATACCGTCTGAGATTGAAAATTTCGTCAACGTGGAAGAATTTCAGGTGAGGGAAAACTTACTGGTACTATTCCAATCAATTTTGGTAAAGTTAAGAAACTGCAAATTTTGGAT

>M6-PGSC0003DMG400011513-(rev)M6LRR-RLP_0

ATAAATAACAAAACAAACACAAAATGAAAAGATAACTCCTCTTTCTTTCCAGTAAGATCAGAACCATAAACTTCTCCGATGGCCGGCGACGACTCTCATCTGCAAAATGACGTCGTTTCA

>M6-PGSC0003DMG400011513-(rev)M6LRR-RLP_1008

TTTGAAGTGGTAGCAAAAGGGAGGATAGTGTAGTTATATCATATGTATTCTAATTTTCAAGAGAAAGTCTAACTTTGTTGCGAATCTAAGGCTCAGATTCTTCCTTGACTTAACTTGCGC

>M6-PGSC0003DMG400011513-(rev)M6LRR-RLP_1071

AAAGTCTAACTTTGTTGCGAATCTAAGGCTCAGATTCTTCCTTGACTTAACTTGCGCGGTTTTCTGGTCCAAATTTACAGGCATAACAGACAAAGGATTGACAGCGATTGGATCAAGTCT

>M6-PGSC0003DMG400011513-(rev)M6LRR-RLP_1134

CTGGTCCAAATTTACAGGCATAACAGACAAAGGATTGACAGCGATTGGATCAAGTCTCTCTTCTTTACAGTCCTTAGATGTATCCTACTGCAGAAAGATCACAGACAAGGGGCTTTCAGC

>M6-PGSC0003DMG400011513-(rev)M6LRR-RLP_1197

TTTACAGTCCTTAGATGTATCCTACTGCAGAAAGATCACAGACAAGGGGCTTTCAGCTGTTGCTGAGGGGTGTCATGACTTGAGAACGTTGCATCTTGCTGGCTGTAGATTTGTCTCAGA

>M6-PGSC0003DMG400011513-(rev)M6LRR-RLP_126

TGTATAAACGACAGTCTCGGCGACGACGAGCTTCGGGCCGTACTGTCGAGGCTAGGAGACGACAAAGACAAGGAAGTTTTCGGGCTGGTATGTAAGCGGTGGCTTCGTATTCAAAGTACT

>M6-PGSC0003DMG400011513-(rev)M6LRR-RLP_1260

TGAGGGGTGTCATGACTTGAGAACGTTGCATCTTGCTGGCTGTAGATTTGTCTCAGATTCATTAATGAAAGCTCTCTCCAAGAATTGCCATTATGTAGAAGACCTAGGTTTGCAAGGATG

>M6-PGSC0003DMG400011513-(rev)M6LRR-RLP_1323

AATGAAAGCTCTCTCCAAGAATTGCCATTATGTAGAAGACCTAGGTTTGCAAGGATGCACCAACATTACCAACTCAGGTCTCTCTGTTTTGGTTGAAGGTTGCCGAAGGATAAAGTATTT

>M6-PGSC0003DMG400011513-(rev)M6LRR-RLP_1386

CATTACCAACTCAGGTCTCTCTGTTTTGGTTGAAGGTTGCCGAAGGATAAAGTATTTGGACATTAACAAATGCAGCAATATTGGTGATATTGGGATATCTAGCGTGTCTGAAGCTTGTTC

>M6-PGSC0003DMG400011513-(rev)M6LRR-RLP_1449

TAACAAATGCAGCAATATTGGTGATATTGGGATATCTAGCGTGTCTGAAGCTTGTTCATTAACACTTAGGGCATTGAAGTTGTTGGATTGCTACAAAGTTGGTGATGACTCCATCTTATC

>M6-PGSC0003DMG400011513-(rev)M6LRR-RLP_1512

ACTTAGGGCATTGAAGTTGTTGGATTGCTACAAAGTTGGTGATGACTCCATCTTATCCTTGGCCAACTAATGCAAAAATCTTGAGACACTCGTTATTGGTGGTTGCCGTAACATATCAGA

>M6-PGSC0003DMG400011513-(rev)M6LRR-RLP_1575

CAACTAATGCAAAAATCTTGAGACACTCGTTATTGGTGGTTGCCGTAACATATCAGATGAGCCTATGAAGACACTGGCAGCTTCTTGTAGTAATAGTCTGAGGAAACTGAGGATGGATTG

>M6-PGSC0003DMG400011513-(rev)M6LRR-RLP_1638

TATGAAGACACTGGCAGCTTCTTGTAGTAATAGTCTGAGGAAACTGAGGATGGATTGGTGTTTGAATATTACTGACTCTTCATTGGACTGCATCATTTCCAAATGCAGAGAGCTTGAGGT

>M6-PGSC0003DMG400011513-(rev)M6LRR-RLP_1701

GAATATTACTGACTCTTCATTGGACTGCATCATTTCCAAATGCAGAGAGCTTGAGGTTCTTGACATTGGCTGCTGTGAGGAGTTAACTGATGCTGCATTTCAACAATTAGGAAGTGAAAA

>M6-PGSC0003DMG400011513-(rev)M6LRR-RLP_1764

CATTGGCTGCTGTGAGGAGTTAACTGATGCTGCATTTCAACAATTAGGAAGTGAAAACTTTATGTTGGGGATGAAGATTTTGAAGGTCAGTAACTGTCCAAAGATAACTGTTGAAGGAAT

>M6-PGSC0003DMG400011513-(rev)M6LRR-RLP_1827

GTTGGGGATGAAGATTTTGAAGGTCAGTAACTGTCCAAAGATAACTGTTGAAGGAATTAAGAAGCTTATGAAGTCATGTGAATATCTTGAATATCTAGACGTACGGTCTTGTCCCCTAAT

>M6-PGSC0003DMG400011513-(rev)M6LRR-RLP_189

AAAGACAAGGAAGTTTTCGGGCTGGTATGTAAGCGGTGGCTTCGTATTCAAAGTACTGAAAGGAAGAAGCTTTGTGCTCGAGCCGGGCCTCACATGCTCAGAAGAATCGCGGCCCGATTC

>M6-PGSC0003DMG400011513-(rev)M6LRR-RLP_1890

GCTTATGAAGTCATGTGAATATCTTGAATATCTAGACGTACGGTCTTGTCCCCTAATCACTAAAGCAGGGTGTGAGGAGGCTGGAATTCAGTTTCCTGAATCCTGCAAAATAAACTTCAC

>M6-PGSC0003DMG400011513-(rev)M6LRR-RLP_1953

AGCAGGGTGTGAGGAGGCTGGAATTCAGTTTCCTGAATCCTGCAAAATAAACTTCACTGGAAGCTTAGCAGAGCCAGATGTGTTACTTTGAGCAGTACTTGATTTAGCAATGTCTTTGAA

>M6-PGSC0003DMG400011513-(rev)M6LRR-RLP_2016

CTTAGCAGAGCCAGATGTGTTACTTTGAGCAGTACTTGATTTAGCAATGTCTTTGAAGGACTGAAGGGGATCATGCAGTGAAATCATCACATTCTTTGCCTAAGACCTAAAATATGACTT

>M6-PGSC0003DMG400011513-(rev)M6LRR-RLP_2079

AAGGGGATCATGCAGTGAAATCATCACATTCTTTGCCTAAGACCTAAAATATGACTTTGGAAGCTCCTGCTGTCAGTCCCTTGTTCCCATTAATTTGCCTTTTGTTAAGCATCAAGTTTT

>M6-PGSC0003DMG400011513-(rev)M6LRR-RLP_2142

CTCCTGCTGTCAGTCCCTTGTTCCCATTAATTTGCCTTTTGTTAAGCATCAAGTTTTTTGTTGTAGATTATTTCCCTGCTTTTCAGTTTGTTTTTTACCCCCTGCTTTTCAGTTATTTGA

>M6-PGSC0003DMG400011513-(rev)M6LRR-RLP_2205

TAGATTATTTCCCTGCTTTTCAGTTTGTTTTTTACCCCCTGCTTTTCAGTTATTTGATATCAAACTCCTGTATATCATCCTTGTAAACTACTAGTAGGCACCATTGTTTGCTGTAGGTGT

>M6-PGSC0003DMG400011513-(rev)M6LRR-RLP_2268

ACTCCTGTATATCATCCTTGTAAACTACTAGTAGGCACCATTGTTTGCTGTAGGTGTTTCCAGATTGTGAATGAAATGAGTCCATTGCTGCTAGCTTGAATGTGAACTTATACCAACAAT

>M6-PGSC0003DMG400011513-(rev)M6LRR-RLP_2300

AGGCACCATTGTTTGCTGTAGGTGTTTCCAGATTGTGAATGAAATGAGTCCATTGCTGCTAGCTTGAATGTGAACTTATACCAACAATGATGGTTTGATTGATTACATTTCTTGTACGAC

>M6-PGSC0003DMG400011513-(rev)M6LRR-RLP_252

AAGAAGCTTTGTGCTCGAGCCGGGCCTCACATGCTCAGAAGAATCGCGGCCCGATTCACTCGATTACATGAGCTTGATCTATCTCAATCGGTTTCTAGATCTTTCTATCCTGGTGTTACT

>M6-PGSC0003DMG400011513-(rev)M6LRR-RLP_315

TTACATGAGCTTGATCTATCTCAATCGGTTTCTAGATCTTTCTATCCTGGTGTTACTGATTCTGATCTTTCAGTTATTGCTACTGCTTTCTCTTGCTTACGAATTCTCAATCTTCAGAAC

>M6-PGSC0003DMG400011513-(rev)M6LRR-RLP_378

GATCTTTCAGTTATTGCTACTGCTTTCTCTTGCTTACGAATTCTCAATCTTCAGAACTGTAAAGGTCCGTTTTTGTATTTTTTAAAAGTTATATAGTTTGAAATTTGTTGAATTTAAATG

>M6-PGSC0003DMG400011513-(rev)M6LRR-RLP_441

GGTCCGTTTTTGTATTTTTTAAAAGTTATATAGTTTGAAATTTGTTGAATTTAAATGTTCATTGCTGAAGTGAACCTGAGCTACTGCAGGAGTGGCCAAATGCACCAATGAGTTGTAGTA

>M6-PGSC0003DMG400011513-(rev)M6LRR-RLP_504

GCTGAAGTGAACCTGAGCTACTGCAGGAGTGGCCAAATGCACCAATGAGTTGTAGTATGTGTAGTGATATTTCACATTAATTACTGATTTAAAATCTTGAATTCGTCTTTGTGTGGGGTG

>M6-PGSC0003DMG400011513-(rev)M6LRR-RLP_567

GTGATATTTCACATTAATTACTGATTTAAAATCTTGAATTCGTCTTTGTGTGGGGTGGTGGAGCTGGGATTTTCACTAAGAGGGTTTTGAAATACATAAAACATAATTTTAACCATGTAT

>M6-PGSC0003DMG400011513-(rev)M6LRR-RLP_63

CCATAAACTTCTCCGATGGCCGGCGACGACTCTCATCTGCAAAATGACGTCGTTTCAGTCCACTGTATAAACGACAGTCTCGGCGACGACGAGCTTCGGGCCGTACTGTCGAGGCTAGGA

>M6-PGSC0003DMG400011513-(rev)M6LRR-RLP_756

GTTACCAACTAGGACATAAAGGAAATTTTGAGATTTTGACTAAAATCTATTGTTTGAAATGTTAAAGGTTTTGACCCGATATGGAGAAGACTGGCTCTTCCTCTTTTCTATCTTGTAAAT

>M6-PGSC0003DMG400011513-(rev)M6LRR-RLP_819

AAAGGTTTTGACCCGATATGGAGAAGACTGGCTCTTCCTCTTTTCTATCTTGTAAATAATGGTTTTTATGATAAAAAGGGCGTAATGGGGTTTATGGAGGATTTTAGGTGAATTTATGGT

>M6-PGSC0003DMG400011513-(rev)M6LRR-RLP_882

TTTTATGATAAAAAGGGCGTAATGGGGTTTATGGAGGATTTTAGGTGAATTTATGGTTGGTTTTTTTGAAGATTTGTAGGATTTTGTGTTTCTGTGTAAAGGTTATGTCTTTTTTGCTAT

>M6-PGSC0003DMG400011513-(rev)M6LRR-RLP_945

TTTTGAAGATTTGTAGGATTTTGTGTTTCTGTGTAAAGGTTATGTCTTTTTTGCTATGTGGTTTTTGAAGTGGTAGCAAAAGGGAGGATAGTGTAGTTATATCATATGTATTCTAATTTT

>M6-PGSC0003DMG400011580M6LRR-RLK_0

AATAAAGGTTGAATCTTTCTCATTTTTATCCAACCCCTTTTCAAAATCTTGCATGTATATAGGAAGATCCCCACCATTATTGGAAAAATGTTATAGGTTCTTGGTTTTAGATATAGTATA

>M6-PGSC0003DMG400011580M6LRR-RLK_1037

TTCCAGGATTCGTTTAACCATTTGGGGAATTGGAATCCAAATGATGAAACTCCTTGTGGCTGGGTAGGTGTGAATTGCACTTCTGATTATAATCCAGTTGTGCAATCTCTTTATCTGAGC

>M6-PGSC0003DMG400011580M6LRR-RLK_1098

GGGTAGGTGTGAATTGCACTTCTGATTATAATCCAGTTGTGCAATCTCTTTATCTGAGCTATATGAATCTTTCAGGCACGTTGAGTTCTAGCATTGGTGGTCTTGAATATTTAGCTTATC

>M6-PGSC0003DMG400011580M6LRR-RLK_1159

TATGAATCTTTCAGGCACGTTGAGTTCTAGCATTGGTGGTCTTGAATATTTAGCTTATCTTGACCTGTCTTACAATCAATTCACCGGGAACATTCCCAAAGAGATTGGAAATTGCTCAAA

>M6-PGSC0003DMG400011580M6LRR-RLK_122

CTTTGATTTTATCTTTTCCTCAATTCATGAGTTTTATGGTCTTTGCTCTGTCATCATCTAAAGGAATTACTTGAGAGTTTTTTGCTTTTTCTGGGTTTTTGGTGGAATTCAGCTCTTTTT

>M6-PGSC0003DMG400011580M6LRR-RLK_1220

GACCTGTCTTACAATCAATTCACCGGGAACATTCCCAAAGAGATTGGAAATTGCTCAAAATTGCAGTCTCTCCAGCTCCATTTTAATACTTTTTACGGTCCGATTCCTGCTGAACTGTAT

>M6-PGSC0003DMG400011580M6LRR-RLK_1281

TGCAGTCTCTCCAGCTCCATTTTAATACTTTTTACGGTCCGATTCCTGCTGAACTGTATAACCTGTCTAATTTGAAGGATGTGAACATGAGCAGCAACATGATCTCCGGTCCTATTGCTG

>M6-PGSC0003DMG400011580M6LRR-RLK_1342

CCTGTCTAATTTGAAGGATGTGAACATGAGCAGCAACATGATCTCCGGTCCTATTGCTGAGGAGTTTGGGAAACTCTCGTCTCTAGTTACTTTCGTTGCATACACCAATAATCTTACCGG

>M6-PGSC0003DMG400011580M6LRR-RLK_1403

GAGTTTGGGAAACTCTCGTCTCTAGTTACTTTCGTTGCATACACCAATAATCTTACCGGTCCAGTGCCTCGATCTATTGGAAGTTTGAAAAATTTGACAATATTTCGAGTGGGGCAGAAT

>M6-PGSC0003DMG400011580M6LRR-RLK_1464

CAGTGCCTCGATCTATTGGAAGTTTGAAAAATTTGACAATATTTCGAGTGGGGCAGAATGCACTTTCTGGAAGTTTACCTGCAGAGATAGGTGGTTGTGAGAGCTTGGAATCTCTTGGTC

>M6-PGSC0003DMG400011580M6LRR-RLK_1525

ACTTTCTGGAAGTTTACCTGCAGAGATAGGTGGTTGTGAGAGCTTGGAATCTCTTGGTCTAACACAGAATTGCCTGGAAGGAAACATACCAAAAGAACTTGGCATGCTTAGTAAGTTGAA

>M6-PGSC0003DMG400011580M6LRR-RLK_1586

ACACAGAATTGCCTGGAAGGAAACATACCAAAAGAACTTGGCATGCTTAGTAAGTTGAAAGAGCTTGTGCTTTGGGGAAATCAGTTCTCTGGTTATATTCCAAAGGAACTTGGAAATTTA

>M6-PGSC0003DMG400011580M6LRR-RLK_1647

AGCTTGTGCTTTGGGGAAATCAGTTCTCTGGTTATATTCCAAAGGAACTTGGAAATTTAACACAAATTCAATTGCTTGCTTTATATCAGAATAACCTGATTGGAGATATTCCTGCTGAAA

>M6-PGSC0003DMG400011580M6LRR-RLK_1708

ACAAATTCAATTGCTTGCTTTATATCAGAATAACCTGATTGGAGATATTCCTGCTGAAATAGGTAAACTCAAAACTTTGACGAAGTTATACCTATACCGGAATGGATTAAATGGTAGCAT

>M6-PGSC0003DMG400011580M6LRR-RLK_1769

GGTAAACTCAAAACTTTGACGAAGTTATACCTATACCGGAATGGATTAAATGGTAGCATTCCAAGGGAGATTGGGAATCTTTCTATGGCAACTGAAATTGATTTCTCGGAAAACTTTCTG

>M6-PGSC0003DMG400011580M6LRR-RLK_183

AGGAATTACTTGAGAGTTTTTTGCTTTTTCTGGGTTTTTGGTGGAATTCAGCTCTTTTTGACAAGGTGATGGACTCAATAGCCTAGAGGTATGCTAAAGCTTAGATCTTTAATGAGTTTC

>M6-PGSC0003DMG400011580M6LRR-RLK_1830

CAAGGGAGATTGGGAATCTTTCTATGGCAACTGAAATTGATTTCTCGGAAAACTTTCTGAAAGGAGAAATTCCAGTTGAGTTTGGCCAAATAAAAAGTCTGAAATTGTTGTTTCTTTTCC

>M6-PGSC0003DMG400011580M6LRR-RLK_1891

AGGAGAAATTCCAGTTGAGTTTGGCCAAATAAAAAGTCTGAAATTGTTGTTTCTTTTCCAGAATCAGCTCGAAGGCGTTATACCAGATGAACTTACTACTCTAAAGAACTTGATTAGTCT

>M6-PGSC0003DMG400011580M6LRR-RLK_1952

AATCAGCTCGAAGGCGTTATACCAGATGAACTTACTACTCTAAAGAACTTGATTAGTCTTGATTTGTCAATTAATCACCTAACAGGTCCTATTCCATTTGGTTTTCAGTATCAAAAGGAA

>M6-PGSC0003DMG400011580M6LRR-RLK_2013

ATTTGTCAATTAATCACCTAACAGGTCCTATTCCATTTGGTTTTCAGTATCAAAAGGAATTGATTCAGTTACAGCTTTTTGAGAATTCTTTAACTGGTACTATTCCTCAACGTCTTGGCA

>M6-PGSC0003DMG400011580M6LRR-RLK_2074

GATTCAGTTACAGCTTTTTGAGAATTCTTTAACTGGTACTATTCCTCAACGTCTTGGCATTTATAGTCGACTTTGGGTGCTTGATTTAAATAACAATCAACTCACAGGAAGAATTCCTCC

>M6-PGSC0003DMG400011580M6LRR-RLK_2135

TATAGTCGACTTTGGGTGCTTGATTTAAATAACAATCAACTCACAGGAAGAATTCCTCCATTCGTTTGTCAGAATTCCAATTTGATTTTGCTGAATCTTGCATCAAATAAATTGCATGGA

>M6-PGSC0003DMG400011580M6LRR-RLK_2196

TCGTTTGTCAGAATTCCAATTTGATTTTGCTGAATCTTGCATCAAATAAATTGCATGGATATATTCCATCTGGTGTTCTTAAATGTGATTCATTGGTGCAACTTCGTCTGAACGATAACA

>M6-PGSC0003DMG400011580M6LRR-RLK_2257

TATTCCATCTGGTGTTCTTAAATGTGATTCATTGGTGCAACTTCGTCTGAACGATAACAGGCTAACAGGGACATTTCCTTCTGAGTTGTGCAAATTGATAAACCTGTCTGCTGTTGAACT

>M6-PGSC0003DMG400011580M6LRR-RLK_2318

CTAACAGGGACATTTCCTTCTGAGTTGTGCAAATTGATAAACCTGTCTGCTGTTGAACTGGGGCAGAACAAGTTCACTGGTCCAATACCTCCTGATATTGGATATTGCCAGAAGTTGCAG

>M6-PGSC0003DMG400011580M6LRR-RLK_2379

GGCAGAACAAGTTCACTGGTCCAATACCTCCTGATATTGGATATTGCCAGAAGTTGCAGAGGCTTGATTTCTCAGGCAATAGTTTCAACCAGTTGCCAAGGGAGATAGGAAATCTTACCA

>M6-PGSC0003DMG400011580M6LRR-RLK_244

CAAGGTGATGGACTCAATAGCCTAGAGGTATGCTAAAGCTTAGATCTTTAATGAGTTTCTTATTTATTCCTGAATTATTCCTTTTATATAGTTGAGTTTCTGGGCATTTCTGAGTTGATT

>M6-PGSC0003DMG400011580M6LRR-RLK_2440

GCTTGATTTCTCAGGCAATAGTTTCAACCAGTTGCCAAGGGAGATAGGAAATCTTACCAGGCTTGTGACATTCAATGTCTCAGCAAATTCGCTCACTGGACCGATACCTCCAGAAATTCG

>M6-PGSC0003DMG400011580M6LRR-RLK_2501

CTTGTGACATTCAATGTCTCAGCAAATTCGCTCACTGGACCGATACCTCCAGAAATTCGGAACTGCAAGGCACTACAAAGACTGGATCTCAGCAAGAACAGATTTACTGATGTTATACCT

>M6-PGSC0003DMG400011580M6LRR-RLK_2562

ACTGCAAGGCACTACAAAGACTGGATCTCAGCAAGAACAGATTTACTGATGTTATACCTGATGACATTGGTAGCTTGTCGCAGCTAGAACGTCTTTTGCTTTCAGAGAATAAGTTATCTG

>M6-PGSC0003DMG400011580M6LRR-RLK_2623

TGACATTGGTAGCTTGTCGCAGCTAGAACGTCTTTTGCTTTCAGAGAATAAGTTATCTGGAAAGATACCAGCAGCATTGGGGAGCCTCTCTCATTTGACTGAGCTGCAGATGGGCAGTAA

>M6-PGSC0003DMG400011580M6LRR-RLK_2684

AAGATACCAGCAGCATTGGGGAGCCTCTCTCATTTGACTGAGCTGCAGATGGGCAGTAATCTATTGTCAGGTGAAATACCATCAGAGTTGGGTAATCTTAGTGGCTTACAAATTGCAATG

>M6-PGSC0003DMG400011580M6LRR-RLK_2745

TATTGTCAGGTGAAATACCATCAGAGTTGGGTAATCTTAGTGGCTTACAAATTGCAATGGATCTTAGCAACAACAAGCTCTCTGGCAGTATACCACCCAACCTTGGTAATCTTATCCTAT

>M6-PGSC0003DMG400011580M6LRR-RLK_2806

TCTTAGCAACAACAAGCTCTCTGGCAGTATACCACCCAACCTTGGTAATCTTATCCTATTAGAGTACCTATATCTCAACAACAATCATTTGAGTGGTGAAATACCAAGCACATTTGGAAA

>M6-PGSC0003DMG400011580M6LRR-RLK_2867

GAGTACCTATATCTCAACAACAATCATTTGAGTGGTGAAATACCAAGCACATTTGGAAACCTGACAAGTCTCTTGGGCATTGACTTCTCATACAATGACCTGACTGGACCACTGCCGGAT

>M6-PGSC0003DMG400011580M6LRR-RLK_2928

TGACAAGTCTCTTGGGCATTGACTTCTCATACAATGACCTGACTGGACCACTGCCGGATATACCACTCTTTCGGAACATGGATATCAGCAGCTTCATTGGAAACAAAGGTCTTTGCGGTG

>M6-PGSC0003DMG400011580M6LRR-RLK_2989

ACCACTCTTTCGGAACATGGATATCAGCAGCTTCATTGGAAACAAAGGTCTTTGCGGTGGGCCTCTAGGGGAATGTAATGCATCTCCAGCTTATGATGCCAATAATCCTCCTAGAGTCGA

>M6-PGSC0003DMG400011580M6LRR-RLK_305

ATTTATTCCTGAATTATTCCTTTTATATAGTTGAGTTTCTGGGCATTTCTGAGTTGATTGGATTCTTTGAAGTTAGTTGATTAATTGGTATGCATAGAACTTTTTCTGGTTTGTCTAGTT

>M6-PGSC0003DMG400011580M6LRR-RLK_3050

CCTCTAGGGGAATGTAATGCATCTCCAGCTTATGATGCCAATAATCCTCCTAGAGTCGAGAGTGCAGATTCTCCTCGAGCAAAGATTATTACTGCAGTTGCCGGTGTAATTGGTGGAGTT

>M6-PGSC0003DMG400011580M6LRR-RLK_3111

GTGCAGATTCTCCTCGAGCAAAGATTATTACTGCAGTTGCCGGTGTAATTGGTGGAGTTTCTCTTGTTTTGATTGTGGTAGTTTTGTACTATATGAAGCAGCATCCAGTTGAGATGGTTG

>M6-PGSC0003DMG400011580M6LRR-RLK_3172

TCTTGTTTTGATTGTGGTAGTTTTGTACTATATGAAGCAGCATCCAGTTGAGATGGTTGCAACCCAGGATAAAGACATGGCATCTTCAGATCCAGACATTTATTTCCGTCCCAAAGAGGG

>M6-PGSC0003DMG400011580M6LRR-RLK_3233

ACCCAGGATAAAGACATGGCATCTTCAGATCCAGACATTTATTTCCGTCCCAAAGAGGGGTTCACTTTCCAGGACTTGGTTGAGGCTACAAACAATTTTCAAGATTGTTATGTTCTTGGA

>M6-PGSC0003DMG400011580M6LRR-RLK_3294

TCACTTTCCAGGACTTGGTTGAGGCTACAAACAATTTTCAAGATTGTTATGTTCTTGGAAGAGGAGCCGTCGGAACAGTTTACAAAGCAGTCATGCAGTCTGGACAAACGATTGCAGTCA

>M6-PGSC0003DMG400011580M6LRR-RLK_3355

AGGAGCCGTCGGAACAGTTTACAAAGCAGTCATGCAGTCTGGACAAACGATTGCAGTCAAGAAGCTTGCTTCTAACAGAGAGGGTAATAACATCGACAATAGCTTCCGTGCAGAGATCTT

>M6-PGSC0003DMG400011580M6LRR-RLK_3416

AAGCTTGCTTCTAACAGAGAGGGTAATAACATCGACAATAGCTTCCGTGCAGAGATCTTAACTCTAGGAAAGATTAGGCATCGTAACATCGTAAAGCTATATGGCTTTTGCTATCATCAG

>M6-PGSC0003DMG400011580M6LRR-RLK_3477

CTCTAGGAAAGATTAGGCATCGTAACATCGTAAAGCTATATGGCTTTTGCTATCATCAGGGTTCTAATTTACTTCTCTACGAGTATATGGCCAGAGGTAGCTTGGGTGAACTGCTTCACA

>M6-PGSC0003DMG400011580M6LRR-RLK_3538

TTCTAATTTACTTCTCTACGAGTATATGGCCAGAGGTAGCTTGGGTGAACTGCTTCACAGTACATCTTGTAGATTGGATTGGCCAACTCGCTTTATGGTGGCTGTTGGAGCTGCTCAAGG

>M6-PGSC0003DMG400011580M6LRR-RLK_3599

ACATCTTGTAGATTGGATTGGCCAACTCGCTTTATGGTGGCTGTTGGAGCTGCTCAAGGACTTTCTTACTTACATCATGATTGCAAGCCCCGGATCATCCACCGTGATATAAAGTCCAAT

>M6-PGSC0003DMG400011580M6LRR-RLK_366

ATTCTTTGAAGTTAGTTGATTAATTGGTATGCATAGAACTTTTTCTGGTTTGTCTAGTTTATCTAGTTAGTATTGTCTCTTCTTGATGATTGACCTTAAGGGTGTGGAGGGTAGGACTTG

>M6-PGSC0003DMG400011580M6LRR-RLK_3660

TTTCTTACTTACATCATGATTGCAAGCCCCGGATCATCCACCGTGATATAAAGTCCAATAATATTCTGATTGATGAGAAGTTTGAAGCTCATGTTGGTGATTTTGGTTTGGCAAAAGTTG

>M6-PGSC0003DMG400011580M6LRR-RLK_3721

TATTCTGATTGATGAGAAGTTTGAAGCTCATGTTGGTGATTTTGGTTTGGCAAAAGTTGTTGATATGCCTCAATCTAAGTCGATGTCTGCAGTTGCTGGCTCATATGGCTACATAGCACC

>M6-PGSC0003DMG400011580M6LRR-RLK_3782

GATATGCCTCAATCTAAGTCGATGTCTGCAGTTGCTGGCTCATATGGCTACATAGCACCTGGTATGTCTTTCTGTTTTCTCCTGGACAACTTCTTATGTTCCCTTGTTATCTATAGAAAT

>M6-PGSC0003DMG400011580M6LRR-RLK_3843

GTATGTCTTTCTGTTTTCTCCTGGACAACTTCTTATGTTCCCTTGTTATCTATAGAAATATGTCGAGAAATGTACTGCCATTTGATCCTTATTCCCACATATTTATCTGTATCTTCACTC

>M6-PGSC0003DMG400011580M6LRR-RLK_3904

GTCGAGAAATGTACTGCCATTTGATCCTTATTCCCACATATTTATCTGTATCTTCACTCTGAAAGTGTTACCTTTTTCTGTTGATATGAATAGGATTTTCTTGTTGAGATTAGATAGTTA

>M6-PGSC0003DMG400011580M6LRR-RLK_3965

AAAGTGTTACCTTTTTCTGTTGATATGAATAGGATTTTCTTGTTGAGATTAGATAGTTAGAATTGTATGCAATGTCTATTTCTTCGATTATCTGAAGACAACAATGAAATGCTTTTAACT

>M6-PGSC0003DMG400011580M6LRR-RLK_4026

ATTGTATGCAATGTCTATTTCTTCGATTATCTGAAGACAACAATGAAATGCTTTTAACTTTTATCTGTGTATACTTCTCATTTTATATGTGTTCTATGCCAGGAGAAACCATACAAACTG

>M6-PGSC0003DMG400011580M6LRR-RLK_4087

TATCTGTGTATACTTCTCATTTTATATGTGTTCTATGCCAGGAGAAACCATACAAACTGCTCTATGTATCTTAGGCAAACTTTGGGGACTCTTCCTGATGCATATGGAGCATTCTTGTTC

>M6-PGSC0003DMG400011580M6LRR-RLK_4148

CTATGTATCTTAGGCAAACTTTGGGGACTCTTCCTGATGCATATGGAGCATTCTTGTTCCCACATATTTATGTTCAGTTTAGATATTCGACCATGTGTTGTTCACTTTGAAAGTCTAATT

>M6-PGSC0003DMG400011580M6LRR-RLK_4209

ACATATTTATGTTCAGTTTAGATATTCGACCATGTGTTGTTCACTTTGAAAGTCTAATTCTTCTCTTTGTGATATGAAATGCATATCTTGTTGAAGTTTGATAGTTAAAATTGTTTACGA

>M6-PGSC0003DMG400011580M6LRR-RLK_427

TCTAGTTAGTATTGTCTCTTCTTGATGATTGACCTTAAGGGTGTGGAGGGTAGGACTTGGATTAGGGTAGTAGGTGGTAATGCGTTGTCTTGTTGTCCGTTCATGTTAGTAGTCATACTA

>M6-PGSC0003DMG400011580M6LRR-RLK_4270

TCTCTTTGTGATATGAAATGCATATCTTGTTGAAGTTTGATAGTTAAAATTGTTTACGAACAAGTTAATTTCTTTGATGAAATGACAGAATACAAGAAAAAATGCTTTTAACTTTTCTCT

>M6-PGSC0003DMG400011580M6LRR-RLK_4331

AAGTTAATTTCTTTGATGAAATGACAGAATACAAGAAAAAATGCTTTTAACTTTTCTCTGTCTATGTAGCTTTTAATATCCCTCTCGTAATTTTCATTCTGTGTTATAAAACCATTTTTT

>M6-PGSC0003DMG400011580M6LRR-RLK_4392

CTATGTAGCTTTTAATATCCCTCTCGTAATTTTCATTCTGTGTTATAAAACCATTTTTTCCTTTTCAATTGATGTTTTCTACCTTCTCTGTTGCAGAATATGCTTACACCATGAAAGTTA

>M6-PGSC0003DMG400011580M6LRR-RLK_4453

TTTTCAATTGATGTTTTCTACCTTCTCTGTTGCAGAATATGCTTACACCATGAAAGTTACGGAAAAATGTGATATCTATAGTTATGGGGTAGTCCTTTTGGAGTTGCTAACTGGAAAAGC

>M6-PGSC0003DMG400011580M6LRR-RLK_4514

GAAAAATGTGATATCTATAGTTATGGGGTAGTCCTTTTGGAGTTGCTAACTGGAAAAGCACCCGTACAGCCCCTAGAACAAGGTGGTGATCTTGTCTCATGGGTGAAGCATTATGTTAGG

>M6-PGSC0003DMG400011580M6LRR-RLK_4575

CCGTACAGCCCCTAGAACAAGGTGGTGATCTTGTCTCATGGGTGAAGCATTATGTTAGGAACCATTCATTGACACCAGGGGTACTCGATAGCCGATTGGATTTGGAAGATGTAATTACTG

>M6-PGSC0003DMG400011580M6LRR-RLK_4636

CCATTCATTGACACCAGGGGTACTCGATAGCCGATTGGATTTGGAAGATGTAATTACTGTTAGTCATATGCTTACAGTTCTTAAAATTGCTCTAATGTGCACTAGCATGTCCCCTTATGA

>M6-PGSC0003DMG400011580M6LRR-RLK_4697

AGTCATATGCTTACAGTTCTTAAAATTGCTCTAATGTGCACTAGCATGTCCCCTTATGATCGTCCTTCAATGCGCGAGGTTGTTCTGATGCTGATTGAGTCTGAAGAGCAAGAAGGGAAC

>M6-PGSC0003DMG400011580M6LRR-RLK_4758

GTCCTTCAATGCGCGAGGTTGTTCTGATGCTGATTGAGTCTGAAGAGCAAGAAGGGAACTTTCTCTCATCTCCAGTTTATGATCCCCCTCTGAAAGATAATTCCTTCTAGATGCATGTAA

>M6-PGSC0003DMG400011580M6LRR-RLK_4819

TCTCTCATCTCCAGTTTATGATCCCCCTCTGAAAGATAATTCCTTCTAGATGCATGTAAGAAGCACATCTCAATTAACTTTGATAATTTTTTAGTACTAATTCTCCTTGTATTTTTTTTC

>M6-PGSC0003DMG400011580M6LRR-RLK_488

TTAGGGTAGTAGGTGGTAATGCGTTGTCTTGTTGTCCGTTCATGTTAGTAGTCATACTATTGCTCATTCTGTTACTATATTTTGGTTCCTTATACCTCGATTGTGTACTGTTTTGTTGTA

>M6-PGSC0003DMG400011580M6LRR-RLK_4880

AGCACATCTCAATTAACTTTGATAATTTTTTAGTACTAATTCTCCTTGTATTTTTTTTCTTAATGGCTAAATTATAGATATTCCCATGTATATTTCTCATAGTGAAAATTTCATCAATCA

>M6-PGSC0003DMG400011580M6LRR-RLK_4941

AATGGCTAAATTATAGATATTCCCATGTATATTTCTCATAGTGAAAATTTCATCAATCACTTTCTAGAACTAGTTTGGAGGTGATTGATGTTATTTCAATAGTTCTCCATCTGATGGTTA

>M6-PGSC0003DMG400011580M6LRR-RLK_4995

AATCACTTTCTAGAACTAGTTTGGAGGTGATTGATGTTATTTCAATAGTTCTCCATCTGATGGTTATGGAAAATAATATGCTTTATGCTTTTCTCACTATATATTGAGGGCTACCTAGAT

>M6-PGSC0003DMG400011580M6LRR-RLK_61

GGAAGATCCCCACCATTATTGGAAAAATGTTATAGGTTCTTGGTTTTAGATATAGTATAATCTTTGATTTTATCTTTTCCTCAATTCATGAGTTTTATGGTCTTTGCTCTGTCATCATCT

>M6-PGSC0003DMG400011580M6LRR-RLK_793

GACTACACTGGTTATGTTGTTGTTGTTGTTGTATTTTATAAGGATTGACACATAGTTGGATTAACAGGTCAAACTGAGACTGTCTTATTGGAAGTTTGTTATGATGTCTGGAGATTTTGA

>M6-PGSC0003DMG400011580M6LRR-RLK_854

TAACAGGTCAAACTGAGACTGTCTTATTGGAAGTTTGTTATGATGTCTGGAGATTTTGAATCAAGAAGTGGTTTGGTTCTTCTAATTTGGATAAGTGCTTTGGTGGCTGCTGTACTATTG

>M6-PGSC0003DMG400011580M6LRR-RLK_915

CAAGAAGTGGTTTGGTTCTTCTAATTTGGATAAGTGCTTTGGTGGCTGCTGTACTATTGCTGGTTTCTCCTGCCGAAGGATTGAATCAGGAAGGAATGTACCTTCTTGAATTGAAGAAGA

>M6-PGSC0003DMG400011580M6LRR-RLK_976

GGTTTCTCCTGCCGAAGGATTGAATCAGGAAGGAATGTACCTTCTTGAATTGAAGAAGAATTTCCAGGATTCGTTTAACCATTTGGGGAATTGGAATCCAAATGATGAAACTCCTTGTGG

>M6-PGSC0003DMG400011610-(rev)M6LRR-RLP_0

ATGGTTAATAGTATATCCCAACAAATTGATCAAACAGCGAAGGTACCGACATATCTTGGTAAATCTGGAGAGAGAGATTACAATGATTCTATAACAATTTCAACAAAGGGAATGGAGTAT

>M6-PGSC0003DMG400011610-(rev)M6LRR-RLP_130

GAATTTTGACAATTCTCAAGACTATTGATCTTTCAAGTAACAAATTTGAAGGACATATTCCAAGTAATCTCGGAGATCTCATTGCGCTCCGGGTGTTAAATTTGTCTCATAATAGATTGC

>M6-PGSC0003DMG400011610-(rev)M6LRR-RLP_455

TGATGGATTACAAGGATTCCCTATTTCGAAAGGTTGTGGAAGTAATTGGACACCGGAGACAAATAATAGGGATTATGAGTCAGATGATGAAGAAAGCAATTCTGAATTTCTAAATGATTT

>M6-PGSC0003DMG400011610-(rev)M6LRR-RLP_65

TGGAGAGAGAGATTACAATGATTCTATAACAATTTCAACAAAGGGAATGGAGTATGAACTGGATAGAATTTTGACAATTCTCAAGACTATTGATCTTTCAAGTAACAAATTTGAAGGACA

>M6-PGSC0003DMG400011611-(rev)M6LRR-RLP_0

GCAATTTGATGAACTTGCAATTGATGTATTTGGAGAGAAACAACCTCAGTGGAAAAATTCCTTAATGTCTAGTTAATATCAGTGGACTCGAGGTATTGAAAATAGAAGATAATAATCTCA

>M6-PGSC0003DMG400011611-(rev)M6LRR-RLP_1208

GTATCCATTATATATATCATGATCTCAACTGGAAATCTAAAATGGCTTGCAAATATTTTTGAAAAGTTAGAAGAAAAAATTATTATGAGACAGATAAGGAAGAAGCGAAGTCAAAGGTAA

>M6-PGSC0003DMG400011611-(rev)M6LRR-RLP_192

AGGGAGCAATTCCACAATGTTTTGGCAACATGGGTGGTCATCTTGAGGTTTTGGATATCCACCAAAACAATCTTTATGGGACTCTTCCAGAAACTTTTAGCAATGGAAGTGTACTCAGAA

>M6-PGSC0003DMG400011611-(rev)M6LRR-RLP_256

AAACAATCTTTATGGGACTCTTCCAGAAACTTTTAGCAATGGAAGTGTACTCAGAATCTTAGTCTTGCATTACAATGAATTAGAAGGAAAAATCCCCCGATCTTTAGTCAATTGTAAAAA

>M6-PGSC0003DMG400011611-(rev)M6LRR-RLP_320

TTGCATTACAATGAATTAGAAGGAAAAATCCCCCGATCTTTAGTCAATTGTAAAAACTTGGAAATTCTTGATTTAGGAGACAATCACTTCAACGACACATTTCCAATATGGTTTGAAACT

>M6-PGSC0003DMG400011611-(rev)M6LRR-RLP_576

CTAGTCTATTTCAAAAATTGAAAGCCATGAGGACAATTGATCAAACAGCAAAGGTACCAACATATCTTGGTAAATCTGGAGAGAGAGATTACAATGATTCTGTAACAGTTTCAACGAAGG

>M6-PGSC0003DMG400011611-(rev)M6LRR-RLP_64

ATGTCTAGTTAATATCAGTGGACTCGAGGTATTGAAAATAGAAGATAATAATCTCAGTGAAGATATCCCTTCGTCTATTTGCAATCTAACATCACTAAGGATTTTAGATTTGGGTAGAAA

>M6-PGSC0003DMG400011611-(rev)M6LRR-RLP_640

TCTTGGTAAATCTGGAGAGAGAGATTACAATGATTCTGTAACAGTTTCAACGAAGGGAATGGAGTATGAACTGGATAGAATTTTGACACTTTTCAAGACTATCGATCTTTCAAGTAACAA

>M6-PGSC0003DMG400011611-(rev)M6LRR-RLP_704

TATGAACTGGATAGAATTTTGACACTTTTCAAGACTATCGATCTTTCAAGTAACAAATTTGAAGGACATATTCCAAGTAGTTTGGGAGATCTCATTGCACTTAAGGTGCTAAATTTGTCT

>M6-PGSC0003DMG400011611-(rev)M6LRR-RLP_768

GACATATTCCAAGTAGTTTGGGAGATCTCATTGCACTTAAGGTGCTAAATTTGTCTCATAACAGATTGCAAGGTAATATACCATCATCACTTGAAAGTTTATATTTAGTCGAATCATTGG

>M6-PGSC0003DMG400011708-(rev)M6LRR-RLP_0

TGTTGCATGTGATTTTGTCTTTAAGTCAGTAAAGTTGGTTCTTTGTATATGTGGTTGGGAAAAATTTATGTAGGAGATATTTATCTCAAGTTCACATTTCTACTTTTTTGTGGGGCCATT

>M6-PGSC0003DMG400011708-(rev)M6LRR-RLP_130

TGTTTGGATTGGAATCTCCCTAGTTTGATTTCATCTTCTCTATCATTTCAAAAAACTCAATATGGAGGATGAGACGGAAACTGAAAAATCTCCGATTACTACTCGTGTGATACTTCTTTC

>M6-PGSC0003DMG400011708-(rev)M6LRR-RLP_195

AGGATGAGACGGAAACTGAAAAATCTCCGATTACTACTCGTGTGATACTTCTTTCCCAAAAACAGTCTTATGGAATACGAGTAGAGATTGTTGTACATGGGATGGATTACAACTTGATTA

>M6-PGSC0003DMG400011708-(rev)M6LRR-RLP_260

TCTTATGGAATACGAGTAGAGATTGTTGTACATGGGATGGATTACAACTTGATTAGAGAACATTGGACACAACACTTCATAACTTCACAAAAACCTGTGAATTTCCCAAAAACAGTCTTA

>M6-PGSC0003DMG400011708-(rev)M6LRR-RLP_325

GACACAACACTTCATAACTTCACAAAAACCTGTGAATTTCCCAAAAACAGTCTTATGGAATGCGAGTAGAGATTGCTGTACTTTCTCTCCCTGATGTCAACATCTCATCTCCAATACCTA

>M6-PGSC0003DMG400011708-(rev)M6LRR-RLP_390

GTAGAGATTGCTGTACTTTCTCTCCCTGATGTCAACATCTCATCTCCAATACCTATGAATTTTTCTTCCTCCTTAAGGTACGTGGATCTTGAATATACTAATATACAAATTAGGTATTTG

>M6-PGSC0003DMG400011708-(rev)M6LRR-RLP_455

TTCCTCCTTAAGGTACGTGGATCTTGAATATACTAATATACAAATTAGGTATTTGAATTTCACCGGTCATATTTCTTTCACAATCTCTAAATTGAAGCAGCTCACAAGTTTAGATCTTTC

>M6-PGSC0003DMG400011708-(rev)M6LRR-RLP_520

GTCATATTTCTTTCACAATCTCTAAATTGAAGCAGCTCACAAGTTTAGATCTTTCATCTAACTCCTTTTCAGGTTAAATGCTCGACGTTTTCTCTAACCTCCAAGAGCTTACATCTTTCT

>M6-PGSC0003DMG400011708-(rev)M6LRR-RLP_585

TTTTCAGGTTAAATGCTCGACGTTTTCTCTAACCTCCAAGAGCTTACATCTTTCTGACAACATGAATTATGCGGTTCCTAAACCTTTCGCATAATTCATTAACAAGCCACCTAGCTACCA

>M6-PGSC0003DMG400011708-(rev)M6LRR-RLP_65

TTATGTAGGAGATATTTATCTCAAGTTCACATTTCTACTTTTTTGTGGGGCCATTGAATGGTAAGTGTTTGGATTGGAATCTCCCTAGTTTGATTTCATCTTCTCTATCATTTCAAAAAA

>M6-PGSC0003DMG400011708-(rev)M6LRR-RLP_650

ATTATGCGGTTCCTAAACCTTTCGCATAATTCATTAACAAGCCACCTAGCTACCACAAGTTCATTACCACAATCTAGAGTATCTTGATCTTAAATTTAACTTCTTTCAGGGTCCACTACC

>M6-PGSC0003DMG400011708-(rev)M6LRR-RLP_780

TGTAACATGAGCTAGCTTGAATTTCTAGATTTATCACACAACAACTTCAGTAACTCGATTACAAGTTGCTTGGGAAGCATGGCTACTCTAAGGTTGTTGGACTTAAGAAGGAACAATTTC

>M6-PGSC0003DMG400011708-(rev)M6LRR-RLP_837

ATTACAAGTTGCTTGGGAAGCATGGCTACTCTAAGGTTGTTGGACTTAAGAAGGAACAATTTCACAGGGAGTCTTCCACCATTATGTGCGCAGAGCACTTCATTGAGTACCATTTTCTTG

>M6-PGSC0003DMG400011792M6RLKWAX-EGF_0

TTTGTTAAATTATTTCGATTTGTTAGATAAGTTTTGAAAATTGAGTTTATTGAAAAGAAAATATTTTTTAAAATGCAAAGGGTATAGTTGGGTCTATTGTCAAGTTTAACCTTGAATTCC

>M6-PGSC0003DMG400011792M6RLKWAX-EGF_1054

GTATCAGATATTAATGAATGTGCTCATTCAAGTACCAATATGTGTGAACACAATTGCATAGACACGCCAGGGAGTTACACTTGTACTTGTCTAGAAGGATTCAGTGGTGATGGCAGAAAG

>M6-PGSC0003DMG400011792M6RLKWAX-EGF_1116

CACGCCAGGGAGTTACACTTGTACTTGTCTAGAAGGATTCAGTGGTGATGGCAGAAAGGATGATGGTGGTTGTATTGCACCAAATTTGAACTATGAGTTCCCATGGATCAAGTTTTCTAT

>M6-PGSC0003DMG400011792M6RLKWAX-EGF_1178

ATGGTGGTTGTATTGCACCAAATTTGAACTATGAGTTCCCATGGATCAAGTTTTCTATTGGTATGTTTCTCAAAAATTGAACTTAGCTTGTTTATTGAAGGAATTCATCTCCTTTAAATT

>M6-PGSC0003DMG400011792M6RLKWAX-EGF_124

ATGACAATTATTTTTAGCATACTCCCTCCGTCTAATAGTATTCTTGGTTTCAGCATGCTCTTTAAGAAATTACTCACCTCTATAAAGTAAGAAATGTTTTCACTAAATTACCGTTAGTTT

>M6-PGSC0003DMG400011792M6RLKWAX-EGF_1240

ATGTTTCTCAAAAATTGAACTTAGCTTGTTTATTGAAGGAATTCATCTCCTTTAAATTTTTGTTGTTATAAACACAGGTATGGGAGTCGGCTTTATATCCCTCGTGGTTGGGACAATTTT

>M6-PGSC0003DMG400011792M6RLKWAX-EGF_1302

TTGTTATAAACACAGGTATGGGAGTCGGCTTTATATCCCTCGTGGTTGGGACAATTTTGCTCTATTTTTGTATCAAGAAACGAAAACTCATTAAAAATAGGGAAAAGTTCTTCCAACAAA

>M6-PGSC0003DMG400011792M6RLKWAX-EGF_1364

TATTTTTGTATCAAGAAACGAAAACTCATTAAAAATAGGGAAAAGTTCTTCCAACAAAATGGTGGTTTACTCTTGAAACAACAAATCTCTTCTATAAAGGGTGGTGTGGAAGCAACCAAA

>M6-PGSC0003DMG400011792M6RLKWAX-EGF_1426

TGGTTTACTCTTGAAACAACAAATCTCTTCTATAAAGGGTGGTGTGGAAGCAACCAAAATATTTACAGCTGACGAGTTGAAGAAGGCTACGAATAACTATGCCAGTGATAGAATTCTTGG

>M6-PGSC0003DMG400011792M6RLKWAX-EGF_1488

TTACAGCTGACGAGTTGAAGAAGGCTACGAATAACTATGCCAGTGATAGAATTCTTGGTCGTGGTGGAAATGGAATTGTCTATAAAGGTATTCTACCTGATAATCGCATAGTTGCAATTA

>M6-PGSC0003DMG400011792M6RLKWAX-EGF_1550

GGTGGAAATGGAATTGTCTATAAAGGTATTCTACCTGATAATCGCATAGTTGCAATTAAAAAATCTAAGTTTGTGGATGAGAATCAAGTTGAACAGTTCATCAATGAGGTACTTATTCTT

>M6-PGSC0003DMG400011792M6RLKWAX-EGF_1612

ATCTAAGTTTGTGGATGAGAATCAAGTTGAACAGTTCATCAATGAGGTACTTATTCTTACTCAAGTTAACCATAGAAATGTAGTGAGACTCTTCGGGTGTTGTTTGGAAGATGAAGTGCC

>M6-PGSC0003DMG400011792M6RLKWAX-EGF_1674

AAGTTAACCATAGAAATGTAGTGAGACTCTTCGGGTGTTGTTTGGAAGATGAAGTGCCTTTACTGGTTTATGAATACGTTTCTCATGGAACTCTTTACGAGCATATCCACAACCAAAATG

>M6-PGSC0003DMG400011792M6RLKWAX-EGF_1736

CTGGTTTATGAATACGTTTCTCATGGAACTCTTTACGAGCATATCCACAACCAAAATGGAGTCCCATGGTTATCTTTGCAAAATCGATTGAGAATTGCTTCAGAGACAGCAAGTTCACTT

>M6-PGSC0003DMG400011792M6RLKWAX-EGF_1798

CCCATGGTTATCTTTGCAAAATCGATTGAGAATTGCTTCAGAGACAGCAAGTTCACTTGCTTACCTTCATTCATCTGCATCCATGCCAATAATTCATAGAGACGTCAAATCCACTAATAT

>M6-PGSC0003DMG400011792M6RLKWAX-EGF_186

TAAGAAATTACTCACCTCTATAAAGTAAGAAATGTTTTCACTAAATTACCGTTAGTTTAATATTAGTATCAATTTAATATAGTTTTTTGGTTATGTAAAAGTTCATTGATCTAATAAAGA

>M6-PGSC0003DMG400011792M6RLKWAX-EGF_1860

ACCTTCATTCATCTGCATCCATGCCAATAATTCATAGAGACGTCAAATCCACTAATATATTATTGGATGATGTTTATACAGCAAAAGTGGCGGATTTTGGAGCTTCAAGATTAGTCCCTC

>M6-PGSC0003DMG400011792M6RLKWAX-EGF_1922

TTGGATGATGTTTATACAGCAAAAGTGGCGGATTTTGGAGCTTCAAGATTAGTCCCTCTTGACCAAACACATGTTGCTACATTGGTTCAAGGAACATTAGGGTACTTAGATCCTGAATAT

>M6-PGSC0003DMG400011792M6RLKWAX-EGF_1984

CCAAACACATGTTGCTACATTGGTTCAAGGAACATTAGGGTACTTAGATCCTGAATATTTCCACACAGGTGAATTGACAAATAAAAGTGACGTTTATAGTTTTGGAGTAGTTCTTGCCGA

>M6-PGSC0003DMG400011792M6RLKWAX-EGF_2046

ACACAGGTGAATTGACAAATAAAAGTGACGTTTATAGTTTTGGAGTAGTTCTTGCCGAACTTTTAACAGGGATGACACCTATTTTTAGGGATACAAGTGACAAGTACAAATGTTTGGTGG

>M6-PGSC0003DMG400011792M6RLKWAX-EGF_2108

TTAACAGGGATGACACCTATTTTTAGGGATACAAGTGACAAGTACAAATGTTTGGTGGAGCATTTTATCTCGTCGATGAACAAGAATAACATATTTGAAATTCTTGATTATAGAGTTGTA

>M6-PGSC0003DMG400011792M6RLKWAX-EGF_2170

TTTTATCTCGTCGATGAACAAGAATAACATATTTGAAATTCTTGATTATAGAGTTGTAAGAGAAGGAGGTGTTGAGCAACTTCAAAAGATAGCTGAGCTCATTAAGGGTTGCCTTCAGTT

>M6-PGSC0003DMG400011792M6RLKWAX-EGF_2294

GGGGAAGATAGGCCTATGATGAAGGAAGTGGCTATGGAACTTGAGAGTTTAAGAAAGTTCACTAGCCCATGGACTAATGAACACGGACATGAAGATCATAATGAAGTTGAATTAAAAGAT

>M6-PGSC0003DMG400011792M6RLKWAX-EGF_2356

TAGCCCATGGACTAATGAACACGGACATGAAGATCATAATGAAGTTGAATTAAAAGATCGGTATATAATTCCAATTGATTCTAATACAGGTATCGACAACTTCTCTGGACAATATCCCGG

>M6-PGSC0003DMG400011792M6RLKWAX-EGF_2418

ATATAATTCCAATTGATTCTAATACAGGTATCGACAACTTCTCTGGACAATATCCCGGCTCCTACACAAATAACAACATCTTTTCTGGACAATATAGTTCGGATAAATCTTCATTACTAC

>M6-PGSC0003DMG400011792M6RLKWAX-EGF_248

TTAGTATCAATTTAATATAGTTTTTTGGTTATGTAAAAGTTCATTGATCTAATAAAGATAAATTTGAAAAGAAAAAAATGATAATTTTTTCCTTATTTATGGAATTATATACTTTGGATT

>M6-PGSC0003DMG400011792M6RLKWAX-EGF_2480

TACACAAATAACAACATCTTTTCTGGACAATATAGTTCGGATAAATCTTCATTACTACCCAATAGAAATATCCCGCGGTGATCCAAAATAAAATATTAATATGATAATTAGTGACAATAT

>M6-PGSC0003DMG400011792M6RLKWAX-EGF_2542

TAGAAATATCCCGCGGTGATCCAAAATAAAATATTAATATGATAATTAGTGACAATATAATTTTCTGTGTTTTGTAAGTTTCTTTTCTGGCTATCTTTCTTTCATATTCAATGTATGTTG

>M6-PGSC0003DMG400011792M6RLKWAX-EGF_2588

TAGTGACAATATAATTTTCTGTGTTTTGTAAGTTTCTTTTCTGGCTATCTTTCTTTCATATTCAATGTATGTTGTTATGTATTTTACTTGCTTTTGAATTTAATCATCAATATATTGTTC

>M6-PGSC0003DMG400011792M6RLKWAX-EGF_310

TTTGAAAAGAAAAAAATGATAATTTTTTCCTTATTTATGGAATTATATACTTTGGATTGAATGTAAAGCTATAAACACTGTATAATTGAATGTAAAGCTAAAAACACTACTCCCTCTGTC

>M6-PGSC0003DMG400011792M6RLKWAX-EGF_62

ATTTTTTAAAATGCAAAGGGTATAGTTGGGTCTATTGTCAAGTTTAACCTTGAATTCCTAAAATGACAATTATTTTTAGCATACTCCCTCCGTCTAATAGTATTCTTGGTTTCAGCATGC

>M6-PGSC0003DMG400011792M6RLKWAX-EGF_682

ACATGAAAGTTCATCTTAGATCATTTTTCAAAAGAGCTCTAACTTTGATTAGGCAAAGAAACAACTTCAGTGAAAAACATGTTTCTCTCAGTGAGTTCACCCCAATTAGCTAACTTGTTA

>M6-PGSC0003DMG400011792M6RLKWAX-EGF_744

AACTTCAGTGAAAAACATGTTTCTCTCAGTGAGTTCACCCCAATTAGCTAACTTGTTAGCCATCATATTGCCTTCCTTAAGGCAATGTTGGAATAATTTTCACCACTTGCCCAGAGGGAT

>M6-PGSC0003DMG400011792M6RLKWAX-EGF_806

TCATATTGCCTTCCTTAAGGCAATGTTGGAATAATTTTCACCACTTGCCCAGAGGGATTGAGTTCCTTGTTGATCATGTTGATGATGATTTGAGAATCAGATTCAATAATGACTTGATGG

>M6-PGSC0003DMG400011792M6RLKWAX-EGF_868

TTCCTTGTTGATCATGTTGATGATGATTTGAGAATCAGATTCAATAATGACTTGATGGAAGTTATTGTACAAGCACCATGACATCCGTTATTGTTCTGAGATAATGATCATTAACTAGAT

>M6-PGSC0003DMG400011792M6RLKWAX-EGF_930

TATTGTACAAGCACCATGACATCCGTTATTGTTCTGAGATAATGATCATTAACTAGATTAGCTAGTGAACCTATATAAAGGCAAAGCGCGAGGGATTAACGTGGTATATTTCTCTGTTTT

>M6-PGSC0003DMG400011792M6RLKWAX-EGF_992

TAGTGAACCTATATAAAGGCAAAGCGCGAGGGATTAACGTGGTATATTTCTCTGTTTTGCATGTATCAGATATTAATGAATGTGCTCATTCAAGTACCAATATGTGTGAACACAATTGCA

>M6-PGSC0003DMG400011793M6RLKWAX-EGF_1054

TTTGTTGGTAAAAATTGAAGGGAAAGTAAATTTTAAAAAAAGATTATCTAAGTTTAAGGCTGATGTGGTGTTTTTTTAACCTATCACTCATCTAAAAGAATTGAGATCACTTGCTATGAT

>M6-PGSC0003DMG400011793M6RLKWAX-EGF_1116

ATGTGGTGTTTTTTTAACCTATCACTCATCTAAAAGAATTGAGATCACTTGCTATGATATGTGGCAAGCGAGGGGTGTTTTAAAATTCAGTTAGTCAAGTAGAAAAATGTTTTTAAGAAT

>M6-PGSC0003DMG400011793M6RLKWAX-EGF_124

ATATTTACTTTGGATTGAATGTAAAGCTAAAAATACTATATAATTGAATGTAAAGCTAAAAACACTACATAATTGAATGTAAAGCTAAAAATACTATTCCCTCACTCCCAGTTTATATCG

>M6-PGSC0003DMG400011793M6RLKWAX-EGF_1240

ATAGTTTATGGACAAAATTAATAATTCTCGCTAAGTTTGAAGGTGTTTTCAATACTTCTCTTCTTCAAAAGGACTAACTTTAGTGAAGAACATGTTTCTCCCAGTGAGTTCACCCCAATT

>M6-PGSC0003DMG400011793M6RLKWAX-EGF_1302

CTTCAAAAGGACTAACTTTAGTGAAGAACATGTTTCTCCCAGTGAGTTCACCCCAATTAGCTAACTTGTCAACCACCATATTTCCTTCTTTATGGCGATGTAGGAATATATAGTTCCCAC

>M6-PGSC0003DMG400011793M6RLKWAX-EGF_1364

AACTTGTCAACCACCATATTTCCTTCTTTATGGCGATGTAGGAATATATAGTTCCCACTAGAAGTGAAGTTCCATGTTGGTAGAAGAATTTTCATCACTTGCCAAAAGGGGTTGAGTTCC

>M6-PGSC0003DMG400011793M6RLKWAX-EGF_186

CACTACATAATTGAATGTAAAGCTAAAAATACTATTCCCTCACTCCCAGTTTATATCGCTTGATCTTCTTGCTTTTACTACTAGCTGGAAAGTGATCCTTCTTTTTTGTTTTAATAGTAA

>M6-PGSC0003DMG400011793M6RLKWAX-EGF_248

ATCTTCTTGCTTTTACTACTAGCTGGAAAGTGATCCTTCTTTTTTGTTTTAATAGTAAAAAACAAATGCTTTAATAGGTAGGGTCACAATAATTGGTAGGAATCAAAATTGTGATAAACA

>M6-PGSC0003DMG400011793M6RLKWAX-EGF_2852

ATCCTGGGATAAGTTTTAGTACCCTTCAAATCTTAGCTCTATCCAAGATCGATTTATATGCCAATAAACAATATACATAAATTCTAGTACTTAAAACTTTTGGTAATAATGTCAACTAGT

>M6-PGSC0003DMG400011793M6RLKWAX-EGF_2914

AATAAACAATATACATAAATTCTAGTACTTAAAACTTTTGGTAATAATGTCAACTAGTGAGTTAAAACTGAGAAATATTGCTACTGATGATTCATATGACCGATCGTGATTCGTAAGTCT

>M6-PGSC0003DMG400011793M6RLKWAX-EGF_2976

TAAAACTGAGAAATATTGCTACTGATGATTCATATGACCGATCGTGATTCGTAAGTCTAATGATTTAGTAGTTATAATGCTTAATTAAATACTCTTTGCTAGTTTCCATTAACAACTTCT

>M6-PGSC0003DMG400011793M6RLKWAX-EGF_3038

ATTTAGTAGTTATAATGCTTAATTAAATACTCTTTGCTAGTTTCCATTAACAACTTCTACAAATAAATATGTTAAAAGAAAAGATATAAAGGCAACGCGCGAGGGATTAATGTGGTATAT

>M6-PGSC0003DMG400011793M6RLKWAX-EGF_310

CAAATGCTTTAATAGGTAGGGTCACAATAATTGGTAGGAATCAAAATTGTGATAAACATCATTGGGACACTAATCACTAGAAATTAACTCAATAGTGCGTAGTGCTTTATTTGCCTTATT

>M6-PGSC0003DMG400011793M6RLKWAX-EGF_3100

ATAAATATGTTAAAAGAAAAGATATAAAGGCAACGCGCGAGGGATTAATGTGGTATATTTCTCTGTTTTGCATGTCAGATATTGATGAATGTGCTGATTCAAGTTCCAATATGTGTGAAC

>M6-PGSC0003DMG400011793M6RLKWAX-EGF_3162

CTGTTTTGCATGTCAGATATTGATGAATGTGCTGATTCAAGTTCCAATATGTGTGAACACAATTGCATAAACACGCCCGGAAATTTCACTTGTTCCTGCTTAAAAGGCTACACTGGCGAT

>M6-PGSC0003DMG400011793M6RLKWAX-EGF_3224

TTGCATAAACACGCCCGGAAATTTCACTTGTTCCTGCTTAAAAGGCTACACTGGCGATGGTAAAAAGAATGGTCGTGGTTGTATTGCACCAAACTCGAACTATGAGTTCCCATGGATCAA

>M6-PGSC0003DMG400011793M6RLKWAX-EGF_3286

AAAAGAATGGTCGTGGTTGTATTGCACCAAACTCGAACTATGAGTTCCCATGGATCAAGTTTTCTGTTGGTAAGTTTCTCAGAAATTCAACTTGTTTCTACTTGCATTCTTCTTCCCTAG

>M6-PGSC0003DMG400011793M6RLKWAX-EGF_372

TGGGACACTAATCACTAGAAATTAACTCAATAGTGCGTAGTGCTTTATTTGCCTTATTGGAGCATGGAGCCTAACACTCGAGGTGAAGTTGACTCATCACCTAAACAGGTGGATCGAGTT

>M6-PGSC0003DMG400011793M6RLKWAX-EGF_434

CATGGAGCCTAACACTCGAGGTGAAGTTGACTCATCACCTAAACAGGTGGATCGAGTTAGATATGGATTGATCGAAAATAAATCAGATGAAAATAAATAAAATATTTGACTCAATCTATA

>M6-PGSC0003DMG400011793M6RLKWAX-EGF_496

ATGGATTGATCGAAAATAAATCAGATGAAAATAAATAAAATATTTGACTCAATCTATATTTAAGACAAGTAAAAATGAATAAATCCAAAAGATAATATAGATATTCATATTATCCATAAA

>M6-PGSC0003DMG400011793M6RLKWAX-EGF_558

AGACAAGTAAAAATGAATAAATCCAAAAGATAATATAGATATTCATATTATCCATAAATTATCTGACCAACCAAATTATTATTTTTTCTTCTTTTAGGATGTGTTTGGTATGAAAAAAAT

>M6-PGSC0003DMG400011793M6RLKWAX-EGF_62

TTGATCTAAATAAGGATAAATTTGAAAAGAAAAAAATTAATATTTTCTTCACTATTTATGGAATATTTACTTTGGATTGAATGTAAAGCTAAAAATACTATATAATTGAATGTAAAGCTA

>M6-PGSC0003DMG400011793M6RLKWAX-EGF_620

CTGACCAACCAAATTATTATTTTTTCTTCTTTTAGGATGTGTTTGGTATGAAAAAAATATTTTAACGTTTTCAGAGAAAATATTTATCTAATTAAAGTAAAGTTTTGAAAAGAAATATTT

>M6-PGSC0003DMG400011793M6RLKWAX-EGF_682

TAACGTTTTCAGAGAAAATATTTATCTAATTAAAGTAAAGTTTTGAAAAGAAATATTTTTTTATGAGGAAATAATTTTTTCTTGAATTAGAGGAAAATAAGTCCAATTAAAATTTTTTTT

>M6-PGSC0003DMG400011793M6RLKWAX-EGF_744

ATGAGGAAATAATTTTTTCTTGAATTAGAGGAAAATAAGTCCAATTAAAATTTTTTTTTTCAAAATATTTAAATCAACTGGCAACTATTTTTTAAAAACATTTTTCTTCACACCAAACAC

>M6-PGSC0003DMG400011793M6RLKWAX-EGF_806

AAATATTTAAATCAACTGGCAACTATTTTTTAAAAACATTTTTCTTCACACCAAACACATCCTTAATAAGTTACTTTTTTTATAGGATGTTACACCCTTACGTACTTTAAGGGGAGTGCA

>M6-PGSC0003DMG400011793M6RLKWAX-EGF_868

TTAATAAGTTACTTTTTTTATAGGATGTTACACCCTTACGTACTTTAAGGGGAGTGCAATTGATAGATACTCATAATTTATTCATATTTTACGTGGATAGTATGTTGATCCATATTTGAC

>M6-PGSC0003DMG400011793M6RLKWAX-EGF_930

ATAGATACTCATAATTTATTCATATTTTACGTGGATAGTATGTTGATCCATATTTGACAAATTTTTAAATGATTAAATAACCAACCCATTTAAAATGAAAGAAGAAAGTTGACGTTAAAG

>M6-PGSC0003DMG400011793M6RLKWAX-EGF_992

TTTTAAATGATTAAATAACCAACCCATTTAAAATGAAAGAAGAAAGTTGACGTTAAAGATTATTTGTTGGTAAAAATTGAAGGGAAAGTAAATTTTAAAAAAAGATTATCTAAGTTTAAG

>M6-PGSC0003DMG400011864M6LRR-RLP_0

AATAATCGCTAGATGTGTTGTGTGATTCATTTTGAGTACTGCTATTTGATTTCTCTTTGTCATACTCTGCCAAGCTTGTTTGCCAATTGATGGCTTATCAGATTATAGTCTTTCATGAAT

>M6-PGSC0003DMG400011864M6LRR-RLP_1037

CTTCAGATGAATAATTGAATGTTCTCTTATCATACTCATGCTGGAATACTCTAAGTTAGTGGAGGTTTTATGCTATAGAACATCTATCCATATTCTCTTGATATCTGTGCACCACTCAAG

>M6-PGSC0003DMG400011864M6LRR-RLP_1098

GAGGTTTTATGCTATAGAACATCTATCCATATTCTCTTGATATCTGTGCACCACTCAAGGATAGCATGACCAAAAAAATTGTGTTTTTAAGCACCTTGGTGCTTTGTCTGATGTCCAGGT

>M6-PGSC0003DMG400011864M6LRR-RLP_1159

TAGCATGACCAAAAAAATTGTGTTTTTAAGCACCTTGGTGCTTTGTCTGATGTCCAGGTCTCAATTACATTTCATGACCATCACAACATGCTCCTTTTGTCATTGTATAATATGGGAAAG

>M6-PGSC0003DMG400011864M6LRR-RLP_122

TTGCTTGTGTCATGAACTCAGTTTATTGAATTGTGATGTATCATGAATGCTAAATATGCACTGTATGTTATAGAACAAATGTAGCTTGGTGATGATAATTTATGCGGCTTTGGAAGTGTT

>M6-PGSC0003DMG400011864M6LRR-RLP_1220

CAATTACATTTCATGACCATCACAACATGCTCCTTTTGTCATTGTATAATATGGGAAAGTTCTTGTTCATGTGATTCGAAGTCCTTTCTTGACTTATGAAGTTACCACGACTTGGATATG

>M6-PGSC0003DMG400011864M6LRR-RLP_1281

CTTGTTCATGTGATTCGAAGTCCTTTCTTGACTTATGAAGTTACCACGACTTGGATATGAGCTTAAATAGAGGGACATAGACAATGAAAATTCATATAGCTGGCCCCAACTGGCTTGAGA

>M6-PGSC0003DMG400011864M6LRR-RLP_1342

CTTAAATAGAGGGACATAGACAATGAAAATTCATATAGCTGGCCCCAACTGGCTTGAGATTAAGGTGTGGTAGTAGTTGTTGGTGCTGTAGTTTCCCACTGCTTTGGTACAGAACTTAAA

>M6-PGSC0003DMG400011864M6LRR-RLP_1403

AAGGTGTGGTAGTAGTTGTTGGTGCTGTAGTTTCCCACTGCTTTGGTACAGAACTTAAATTCCAGATGTACCAAGTTTCCAGTGACTGACTGTATTTATGAAGTTGTTACATGTAATCTG

>M6-PGSC0003DMG400011864M6LRR-RLP_1464

CCAGATGTACCAAGTTTCCAGTGACTGACTGTATTTATGAAGTTGTTACATGTAATCTGTTTCTGTGTTCTGTTATTACATTCCATCTCTTTTTCTTTCTCCGTTTTTGTTTTGTTTTGT

>M6-PGSC0003DMG400011864M6LRR-RLP_1525

TCTGTGTTCTGTTATTACATTCCATCTCTTTTTCTTTCTCCGTTTTTGTTTTGTTTTGTCTAGGGTTTCTTTATCGATCATACAATAATGTGTTGATCTTCTTATTAAACAGATCACTAA

>M6-PGSC0003DMG400011864M6LRR-RLP_1586

AGGGTTTCTTTATCGATCATACAATAATGTGTTGATCTTCTTATTAAACAGATCACTAATGAATGCTTATCATCGATCTCGAAACTGCAATATCTTGAAGATTTGGTTCTAGAAGGCTGT

>M6-PGSC0003DMG400011864M6LRR-RLP_1647

AATGCTTATCATCGATCTCGAAACTGCAATATCTTGAAGATTTGGTTCTAGAAGGCTGTTATGGAATTGATGATTACAGCCTTGCATCCCTTGAGCAAGGCTGCAAGTCACTGGAGGTAT

>M6-PGSC0003DMG400011864M6LRR-RLP_1708

TGGAATTGATGATTACAGCCTTGCATCCCTTGAGCAAGGCTGCAAGTCACTGGAGGTATGTACTTTGGAGGACTCTGTTTAACAGACTTTGATTCTCATATTTGCCTGATTGCCTCTCCT

>M6-PGSC0003DMG400011864M6LRR-RLP_1769

ACTTTGGAGGACTCTGTTTAACAGACTTTGATTCTCATATTTGCCTGATTGCCTCTCCTTGAGATAAAATGAACAATAAAAAATGAAGCTAGTATTGATGTATGAAATTGCCAAAAGCTC

>M6-PGSC0003DMG400011864M6LRR-RLP_183

TGTATGTTATAGAACAAATGTAGCTTGGTGATGATAATTTATGCGGCTTTGGAAGTGTTAACCAATGATTTAGAAGGTAATGTGCAGATTAAGCTGGCCAAGAAAGTATCATTATAAATA

>M6-PGSC0003DMG400011864M6LRR-RLP_1830

AGATAAAATGAACAATAAAAAATGAAGCTAGTATTGATGTATGAAATTGCCAAAAGCTCCCAGGACGTAGCTCAATTGGCGAAGGTTGATAAACTTGTGACCTAGGTCATTAGTTCGATC

>M6-PGSC0003DMG400011864M6LRR-RLP_1891

AGGACGTAGCTCAATTGGCGAAGGTTGATAAACTTGTGACCTAGGTCATTAGTTCGATCCCCATGTCAAGCGAATCTGATATTTAAGTGGCGAAGGGAGAAGTGAGCCCATCATCCCCGA

>M6-PGSC0003DMG400011864M6LRR-RLP_2013

TTCGAAGGCTGCGGTTGGTCCTAAGGGTTGGCTCCTAATGGATTGCCCAAAACTTTGTCAAATTGTCCTAGCTTCAAGCACTACTTTTAGATGTACATATATATGACTTTTTTTCCCTAA

>M6-PGSC0003DMG400011864M6LRR-RLP_2074

ATTGTCCTAGCTTCAAGCACTACTTTTAGATGTACATATATATGACTTTTTTTCCCTAATTGCCTAGTCAATAATTGGAGACCTACTGTGTGAATGCCACTATAAGTACTCTGAAATAGT

>M6-PGSC0003DMG400011864M6LRR-RLP_2135

GCCTAGTCAATAATTGGAGACCTACTGTGTGAATGCCACTATAAGTACTCTGAAATAGTTAAAGAACAAGGTAACTCAGATTTTTTTTTTTGATAACCGAGAAATCCGTTTGTGACCCGC

>M6-PGSC0003DMG400011864M6LRR-RLP_244

CCAATGATTTAGAAGGTAATGTGCAGATTAAGCTGGCCAAGAAAGTATCATTATAAATAAAATGCTAAGCCGTTTAGGAATTTCTCACTGATATCTTAACGAAGTCTATGAGAATTGTGA

>M6-PGSC0003DMG400011864M6LRR-RLP_2440

ACCCAGAATTCTACCTAATTGTCAGATGCCCCAAAGAAGTGGACTATTGTGCAAGTCAAGTGTTGCCTTGCATGTATCATGCCTTTTGGTCCTTTTTTTCCTCTTAAATAACTTTTTTCT

>M6-PGSC0003DMG400011864M6LRR-RLP_2501

GTTGCCTTGCATGTATCATGCCTTTTGGTCCTTTTTTTCCTCTTAAATAACTTTTTTCTTTACAAGGTACTATTGACTAATGTCAGTGTATGTATGTTGGCTCTTTTCAGAAACTTGATA

>M6-PGSC0003DMG400011864M6LRR-RLP_2562

ACAAGGTACTATTGACTAATGTCAGTGTATGTATGTTGGCTCTTTTCAGAAACTTGATATGTCGAGCTGTCAGAATGTTAGTCATGTGGGTTTGTCATCTCTTACCAGCTGTGCTGGATG

>M6-PGSC0003DMG400011864M6LRR-RLP_2623

TCGAGCTGTCAGAATGTTAGTCATGTGGGTTTGTCATCTCTTACCAGCTGTGCTGGATGTCTGCGGCAACTTGTATTGTCGTATGGTTCTCCCGTAAGCCTTAACAAACTTGAGCATCTT

>M6-PGSC0003DMG400011864M6LRR-RLP_2684

TGCGGCAACTTGTATTGTCGTATGGTTCTCCCGTAAGCCTTAACAAACTTGAGCATCTTTCAGTAGTATGAGTTTTAATTTGATTGTTGATTTCAAATTACTGGTATTTCAGGTTACACC

>M6-PGSC0003DMG400011864M6LRR-RLP_2745

AGTAGTATGAGTTTTAATTTGATTGTTGATTTCAAATTACTGGTATTTCAGGTTACACCTGCTGTCGCAGATAGCTTGCAGAAGCTTTCCAGGTTGCAGTGCGTCAAACTAGATGGTTGC

>M6-PGSC0003DMG400011864M6LRR-RLP_2806

CTGTCGCAGATAGCTTGCAGAAGCTTTCCAGGTTGCAGTGCGTCAAACTAGATGGTTGCCAAGTTACATGTTCTGGACTGAAGGCTATTGGAAATTGGTGTGTCTCACTTAAAGAACTGA

>M6-PGSC0003DMG400011864M6LRR-RLP_2867

AGTTACATGTTCTGGACTGAAGGCTATTGGAAATTGGTGTGTCTCACTTAAAGAACTGAGCTTAAGTAAATGTGTGGGAGTGACAGATGAAGGTCTCTGCTCCCTCGTAACAAAGCACAA

>M6-PGSC0003DMG400011864M6LRR-RLP_2928

TTAAGTAAATGTGTGGGAGTGACAGATGAAGGTCTCTGCTCCCTCGTAACAAAGCACAAAGACTTGGGGAAGCTTGACATTACTTGTTGCCGCAAGATTACTTACGTATCAATTTCCCAT

>M6-PGSC0003DMG400011864M6LRR-RLP_2989

ACTTGGGGAAGCTTGACATTACTTGTTGCCGCAAGATTACTTACGTATCAATTTCCCATATAACGAATTCTTGTGCCTCCCTTACTTCTCTTAGGATGGAGTCTTGTACTCTAGTTCCAA

>M6-PGSC0003DMG400011864M6LRR-RLP_305

ATGCTAAGCCGTTTAGGAATTTCTCACTGATATCTTAACGAAGTCTATGAGAATTGTGACATCCATGACAGAGGGCACACCAACAAGGGTGGGTCAAGTAATGGCAGGCATTTAGCTTGG

>M6-PGSC0003DMG400011864M6LRR-RLP_3050

AACGAATTCTTGTGCCTCCCTTACTTCTCTTAGGATGGAGTCTTGTACTCTAGTTCCAAGAGAAGCTTTTGTTCTAATTGGACAGCGTTGCCAGTACCTTGAAGAGCTTGACCTTACAGA

>M6-PGSC0003DMG400011864M6LRR-RLP_3111

GAAGCTTTTGTTCTAATTGGACAGCGTTGCCAGTACCTTGAAGAGCTTGACCTTACAGACAACGAAATTGATGATGAAGGTCACCCCTTTACCTTAAAGGTCTCAGGCTTCTGTTGTAAT

>M6-PGSC0003DMG400011864M6LRR-RLP_3172

ACGAAATTGATGATGAAGGTCACCCCTTTACCTTAAAGGTCTCAGGCTTCTGTTGTAATTTTGAAAGTTCCACCTTTATATCTGACGGTGTCCTACATATGCAGGCTTGCAGACCATTTC

>M6-PGSC0003DMG400011864M6LRR-RLP_3233

TGAAAGTTCCACCTTTATATCTGACGGTGTCCTACATATGCAGGCTTGCAGACCATTTCCAAGTGTGCCAGCCTTTCCAGCTTAAAGCTAGGGATCTGTCTGAACATAACCGATCAGGGT

>M6-PGSC0003DMG400011864M6LRR-RLP_3294

AGTGTGCCAGCCTTTCCAGCTTAAAGCTAGGGATCTGTCTGAACATAACCGATCAGGGTCTTATCCACATTGGCATGTGCTGCTCAAATCTTAAGGAGCTTGACTTATATAGGTATGTAT

>M6-PGSC0003DMG400011864M6LRR-RLP_3355

TATCCACATTGGCATGTGCTGCTCAAATCTTAAGGAGCTTGACTTATATAGGTATGTATTTCATAGTTCATATTAATTGAGATAGTGTAACTATATATCTTGTGCAGTTTTTTTTTTATA

>M6-PGSC0003DMG400011864M6LRR-RLP_3416

CATAGTTCATATTAATTGAGATAGTGTAACTATATATCTTGTGCAGTTTTTTTTTTATATTCTTTTCTTTTTGTGATGGATTGGCTAAACTCACTTTTTAACTATGGAGTACAAGAGATT

>M6-PGSC0003DMG400011864M6LRR-RLP_3477

CTTTTCTTTTTGTGATGGATTGGCTAAACTCACTTTTTAACTATGGAGTACAAGAGATTGAGTGTGCTGATTACTGGGTTATTCTTCTTGACAGGTCTGCTGGAATTTCCGACTTGGGAA

>M6-PGSC0003DMG400011864M6LRR-RLP_3538

GTGTGCTGATTACTGGGTTATTCTTCTTGACAGGTCTGCTGGAATTTCCGACTTGGGAATATTAGCAATCTCTCGCGGTTGTAGTGGCCTTGAAATGATCAATATTGCCTATTGCAATCG

>M6-PGSC0003DMG400011864M6LRR-RLP_3599

TTAGCAATCTCTCGCGGTTGTAGTGGCCTTGAAATGATCAATATTGCCTATTGCAATCGTATTACTGATGGCTCATTCATATCTATATCAAAGTGTTCAAAGTTGAATACACTTGAAAGT

>M6-PGSC0003DMG400011864M6LRR-RLP_366

TCCATGACAGAGGGCACACCAACAAGGGTGGGTCAAGTAATGGCAGGCATTTAGCTTGGCAAGCTTCCGAAGAAGGGGTGCACATGACACCTTAAGGGAATTCACATCAGAATGGGAAAG

>M6-PGSC0003DMG400011864M6LRR-RLP_3660

TTACTGATGGCTCATTCATATCTATATCAAAGTGTTCAAAGTTGAATACACTTGAAAGTCGAGGTTGTCCTCTTGTCACATCTTTAGGTCTTGCAGCTGTTGCTGTGGGATGTAAGCAGC

>M6-PGSC0003DMG400011864M6LRR-RLP_3721

AGGTTGTCCTCTTGTCACATCTTTAGGTCTTGCAGCTGTTGCTGTGGGATGTAAGCAGCTTACAACACTAGACATCAAAAATTGCCACAACATTGATGATGCCGGAATGATTCCACTTGC

>M6-PGSC0003DMG400011864M6LRR-RLP_3782

ACAACACTAGACATCAAAAATTGCCACAACATTGATGATGCCGGAATGATTCCACTTGCTCACTTCTCGACAAACCTCAAACAGGTAACCACAGTTTCTCTTTCGTCGACATAGACGAAC

>M6-PGSC0003DMG400011864M6LRR-RLP_3843

ACTTCTCGACAAACCTCAAACAGGTAACCACAGTTTCTCTTTCGTCGACATAGACGAACTGGAGCATTGACATCTAATATGAGCTGGCCATCTGTCAATTACACAGATAAATTTATCCTA

>M6-PGSC0003DMG400011864M6LRR-RLP_3904

GAGCATTGACATCTAATATGAGCTGGCCATCTGTCAATTACACAGATAAATTTATCCTACACTTCAGTGACGGATGTGGGGCTGCTGTCTCTTGCCAGCATCAGCGGTCTACAGAACATG

>M6-PGSC0003DMG400011864M6LRR-RLP_3965

CTTCAGTGACGGATGTGGGGCTGCTGTCTCTTGCCAGCATCAGCGGTCTACAGAACATGACAATCCTGCACCTCAAGGGTTTATCTCCTGGTGGGCTAGGAGCTGCACTGTTGGCTTGTG

>M6-PGSC0003DMG400011864M6LRR-RLP_4026

AATCCTGCACCTCAAGGGTTTATCTCCTGGTGGGCTAGGAGCTGCACTGTTGGCTTGTGGCGGACTTACTAAAGTTAAGCTGCAGACATCATTTAAATCATTGCTACCACAAGCTCTCCT

>M6-PGSC0003DMG400011864M6LRR-RLP_4087

GGACTTACTAAAGTTAAGCTGCAGACATCATTTAAATCATTGCTACCACAAGCTCTCCTTCAACATCTAGAATCCCGGGGTTGTGTTTTTCAGTGGAGAGAGAAACCATTTCAGGTATCT

>M6-PGSC0003DMG400011864M6LRR-RLP_4148

AACATCTAGAATCCCGGGGTTGTGTTTTTCAGTGGAGAGAGAAACCATTTCAGGTATCTTTCTCATTATTTATCATTACCTACTTTTTCTCTGAAAATATGTGAGGTACTTCTATGGTTG

>M6-PGSC0003DMG400011864M6LRR-RLP_4209

CTCATTATTTATCATTACCTACTTTTTCTCTGAAAATATGTGAGGTACTTCTATGGTTGTTGAAATATACCGTCGTATGAATCAATTAGAATTACATATTTTAGCTAAACATGACATAGT

>M6-PGSC0003DMG400011864M6LRR-RLP_427

AGCTTCCGAAGAAGGGGTGCACATGACACCTTAAGGGAATTCACATCAGAATGGGAAAGAAGAACTTCGTGAGAATATTTTGCACAAGATTATTTTTCACTCTCTATCCTCCGTAAGGGA

>M6-PGSC0003DMG400011864M6LRR-RLP_4270

GAAATATACCGTCGTATGAATCAATTAGAATTACATATTTTAGCTAAACATGACATAGTTATTACTTGATCTTGAGGGTCAATGCAGTTTCAGAGGCTTTCTTGTTTGGCATTTAATTGA

>M6-PGSC0003DMG400011864M6LRR-RLP_4331

TTACTTGATCTTGAGGGTCAATGCAGTTTCAGAGGCTTTCTTGTTTGGCATTTAATTGAGACAATAAAATTACAGTTTTGATTATCTTTGTTGTAATAATGAAAATATCCTTTTTGCTTG

>M6-PGSC0003DMG400011864M6LRR-RLP_4392

CAATAAAATTACAGTTTTGATTATCTTTGTTGTAATAATGAAAATATCCTTTTTGCTTGATCTAGTCATTAATTATCTTTCTTTTATTGGTTAAACACAGGCTGAAGTAGATCCAATATA

>M6-PGSC0003DMG400011864M6LRR-RLP_4453

CTAGTCATTAATTATCTTTCTTTTATTGGTTAAACACAGGCTGAAGTAGATCCAATATACTGGAAAATTCAGCTGGACAATTTAGACTGAAGATTTGAGCCCTCAGTTTGAGCCATTTAT

>M6-PGSC0003DMG400011864M6LRR-RLP_4514

GGAAAATTCAGCTGGACAATTTAGACTGAAGATTTGAGCCCTCAGTTTGAGCCATTTATGCGAGTGTCCTAAGATGGATAGTCGCCCCAGAACCTCCGGAGTCCTTACTTCATCTGTTAG

>M6-PGSC0003DMG400011864M6LRR-RLP_4575

GAGTGTCCTAAGATGGATAGTCGCCCCAGAACCTCCGGAGTCCTTACTTCATCTGTTAGCACTTCATCAGGCAAGGTGCATTGTTTATGGCAACTGTTTTGCCTGTTAAGTGTCTGCAAC

>M6-PGSC0003DMG400011864M6LRR-RLP_4636

CTTCATCAGGCAAGGTGCATTGTTTATGGCAACTGTTTTGCCTGTTAAGTGTCTGCAACTTGTTCGTTTGTATCCTCTAAATACAAACAGGGAAACTCAATGTCCATTCAGCATTGCCCT

>M6-PGSC0003DMG400011864M6LRR-RLP_4697

GTTCGTTTGTATCCTCTAAATACAAACAGGGAAACTCAATGTCCATTCAGCATTGCCCTGGCATCAAATCAATTTTGTCAAAAATGTTATGAGATGGATTAAACAGATCCAAATCATGTT

>M6-PGSC0003DMG400011864M6LRR-RLP_4758

CATCAAATCAATTTTGTCAAAAATGTTATGAGATGGATTAAACAGATCCAAATCATGTTTATTGTTTAGCAATCAATATCTGGGTTTGTGAATGTTTATTCCTCTTGACTAATGTGATTT

>M6-PGSC0003DMG400011864M6LRR-RLP_4803

ATCCAAATCATGTTTATTGTTTAGCAATCAATATCTGGGTTTGTGAATGTTTATTCCTCTTGACTAATGTGATTTCGCATGAAGTTACTTGAAATTTACTCTCAATTAAACAGTTATTTG

>M6-PGSC0003DMG400011864M6LRR-RLP_488

GAACTTCGTGAGAATATTTTGCACAAGATTATTTTTCACTCTCTATCCTCCGTAAGGGAGGAAAATTTCATGAGTTTGCGTTGCTCAACAAGTGCTGACATGTCTTAGGTTCAAAACTAG

>M6-PGSC0003DMG400011864M6LRR-RLP_61

ATACTCTGCCAAGCTTGTTTGCCAATTGATGGCTTATCAGATTATAGTCTTTCATGAATCTTTGCTTGTGTCATGAACTCAGTTTATTGAATTGTGATGTATCATGAATGCTAAATATGC

>M6-PGSC0003DMG400011864M6LRR-RLP_732

AAAAGCTAGTGCTGATGTGTCGTCGAACTCAATGATGATTTTTTTTGAAATTGAACGGAAAATGACATTTCACAGAACATTAAATACTTAATATTATTAAACATTTGCTAAGATTCTTAC

>M6-PGSC0003DMG400011864M6LRR-RLP_793

ATGACATTTCACAGAACATTAAATACTTAATATTATTAAACATTTGCTAAGATTCTTACATTGTCTTAGTTAAATTTATCCCGTGTATTGTGCTTTCTCTACTTGGGAGATATACAAGTT

>M6-PGSC0003DMG400011864M6LRR-RLP_854

TGTCTTAGTTAAATTTATCCCGTGTATTGTGCTTTCTCTACTTGGGAGATATACAAGTTCTGAAAAATGAGCCTGCTTATGCACTTGGCATGCTTCTGACCTTGAGTTTGTTGAAAAGAA

>M6-PGSC0003DMG400011864M6LRR-RLP_915

GAAAAATGAGCCTGCTTATGCACTTGGCATGCTTCTGACCTTGAGTTTGTTGAAAAGAAAAGATATTGCAGAGTCCAGCTTTTGAATCCTAATGAACACCTCTGCTTGACATTCCGATGC

>M6-PGSC0003DMG400011864M6LRR-RLP_976

GATATTGCAGAGTCCAGCTTTTGAATCCTAATGAACACCTCTGCTTGACATTCCGATGCATCTTCAGATGAATAATTGAATGTTCTCTTATCATACTCATGCTGGAATACTCTAAGTTAG

>M6-PGSC0003DMG400011900-(rev)M6LRR-RLP_0

TTTTAGGGAATTTGGAGAAAGAGCTATCATCTCTTACATCCATTTTAGAGAAGGAGCTATCATCCATTTTCAGATATGTCGTGCACCATGAACATAAAATTCCTAAAGATCTTCAGAGAC

>M6-PGSC0003DMG400011900-(rev)M6LRR-RLP_124

CATCAATTTGGCATATGAGGCTGAGGTAGCCATTGACTCTATTCTTGCTCAGTATAATGCTTTTTTGCATATTTTTTGCTCACTTCCTACAATTTTAAAAGAGATCATGCAAATTAATGC

>M6-PGSC0003DMG400011900-(rev)M6LRR-RLP_1240

CTTGATGGATCTCATTAGCAGTAACGTGGTAATAATTTCAAAAAAAAGTTATAATGGTAAAGTCAAATAATGAAAGGTTCAAGATGTTGTGCATCACTTTTGCTTGGAAAAGAGTAGAGA

>M6-PGSC0003DMG400011900-(rev)M6LRR-RLP_1302

TCAAATAATGAAAGGTTCAAGATGTTGTGCATCACTTTTGCTTGGAAAAGAGTAGAGAAGAAAAGTTTATGCTTGCAGTGAAGGGCCAATATATCCAGTTTCAACCCTTGGATTGGAAGG

>M6-PGSC0003DMG400011900-(rev)M6LRR-RLP_1488

AGAAGCCTTTCCACCAACACTTGAGGTCTCTAATAACGGCAAATGGTGGAGAATCTATTGATGTGATTCCCTTTTGTCAGATTAGTGAATTGCGACTTCTTAAAGTCTTGGATTTGATTT

>M6-PGSC0003DMG400011900-(rev)M6LRR-RLP_1550

GTGATTCCCTTTTGTCAGATTAGTGAATTGCGACTTCTTAAAGTCTTGGATTTGATTTCTTATACTGTGGAGTTTTTGTCGTTAGCTATATTCAAACCACTAAATCAGCTGAAGTACCTC

>M6-PGSC0003DMG400011900-(rev)M6LRR-RLP_1612

TACTGTGGAGTTTTTGTCGTTAGCTATATTCAAACCACTAAATCAGCTGAAGTACCTCGCAGTTTGGGCAGATAAATTCTATTTTGATCCAGGAACACATCTGCCCCATATAGAAACTTT

>M6-PGSC0003DMG400011900-(rev)M6LRR-RLP_1674

TTTGGGCAGATAAATTCTATTTTGATCCAGGAACACATCTGCCCCATATAGAAACTTTAATTGTGAAGAATTCTCCTTACGTATGGTTACCAGGGTCTTTTTTGCAAATGGAAAAATTAA

>M6-PGSC0003DMG400011900-(rev)M6LRR-RLP_186

TTTTGCATATTTTTTGCTCACTTCCTACAATTTTAAAAGAGATCATGCAAATTAATGCAGAGGTGACTGAGATGTGGTCAGATGACATTCCTCTTAATCCTCACTACGTGGCTGCTCCAT

>M6-PGSC0003DMG400011900-(rev)M6LRR-RLP_1922

AGGAGGTGTCCTAATCTTCAACAACTTCATATCGAATTTGATGGACATTCTGCAGATTCTTTTTGTCTCACATTGGAGAATCTTACCCAGCTTCAAATACTTCGCGTTTCCTTTGAGCGG

>M6-PGSC0003DMG400011900-(rev)M6LRR-RLP_310

ACATCTGCCAACTCGACATAGCAATCCTGTGACTGATGAGGAGGTAGTGGGTTTTGAGAATAAAGCAGAAGAACTAATTGCTTATCTGATTAGAGGTACAAATGAGCTAGACGTCGTCCC

>M6-PGSC0003DMG400011900-(rev)M6LRR-RLP_496

TGTTTCTCGCTTTGATGTTCGAGCATGGTGCATCATTTCTCAAACATATAATCGGAGAGAGCTATTACAAGATATTTTTAGTCAAGTTACAGGTTCCAAAGATAAGGGAGATACGGTTGA

>M6-PGSC0003DMG400011900-(rev)M6LRR-RLP_558

TATTACAAGATATTTTTAGTCAAGTTACAGGTTCCAAAGATAAGGGAGATACGGTTGACGTTCTTGCCGATTTGTTGAGGAGAAAATTAATGGGAAAGAGATATCTCATTGTATTGGATG

>M6-PGSC0003DMG400011900-(rev)M6LRR-RLP_62

TCCATTTTCAGATATGTCGTGCACCATGAACATAAAATTCCTAAAGATCTTCAGAGACGTACCATCAATTTGGCATATGAGGCTGAGGTAGCCATTGACTCTATTCTTGCTCAGTATAAT

>M6-PGSC0003DMG400011900-(rev)M6LRR-RLP_992

TGGTGGAATGAGGTGAAAGATGCTTTATTTGACTATCTTGACAGTGAGTTCGAAGAATATAGTTTGGCTACTATGCAGTTGAGTTTTGATAACTTACCCGATTGTTTAAAGCCTTGTCTT

>M6-PGSC0003DMG400011932-(rev)M6LRR-RLK_0

ATGTCACTACTAGCATTCAAGAATATGATCATTGATGACCCTTTCAAGATCATGGACTCCTGGAATGAAACTATACATTTTTGTGACTGGCCTGGGGTTTCATGTGGTCGACGTCATCCC

>M6-PGSC0003DMG400011932-(rev)M6LRR-RLK_1054

CAAGATGGATATCCAATCTCAAGAAGCTTCAGGCTTTCTTTGTAGCTTACAACAGATTTCACGGTGAAATTCCGTCTGAAATTGGTGATCTGATATATTTGCAAGAGCTGGCTTTACTTG

>M6-PGSC0003DMG400011932-(rev)M6LRR-RLK_1178

TCAGTTCTCTGGCAAAATTCCAATCTCATTGGGAAATTTAGCTTCTTTAACCAAACTCAGTTTAAGAGAAAACAATCTCCAGGGGAGAATTCCTTCCAGTTTAGGCAAATGCCACAATTT

>M6-PGSC0003DMG400011932-(rev)M6LRR-RLK_124

TTACAGTTTTAAATCTTACTTCTCTGAAATTGAGGGGTTCTCTGTCACCAAGCATTGGTAATCTGAGCTTTCTTTATGTCCTTAAGCTCCAAAATAATAGTTTTTCAGGTGAAATTCCAT

>M6-PGSC0003DMG400011932-(rev)M6LRR-RLK_1550

GGGTAGTATTCCTTCAACCATGAATAGTTTGCAAGGCCTGGAACTTTTAGTCCTTTCTCACAACAATCTGTCAGGTGGAATCCCGGGATTCTTGAAGGACTTCAAATTTCTGCAGATTTT

>M6-PGSC0003DMG400011932-(rev)M6LRR-RLK_1612

ACAATCTGTCAGGTGGAATCCCGGGATTCTTGAAGGACTTCAAATTTCTGCAGATTTTAAATCTGTCTAGCAATACTTTGGAAGGTGCTGTCCCAACAGGAGGCATCTTTAGTAATGCTA

>M6-PGSC0003DMG400011932-(rev)M6LRR-RLK_1674

CTGTCTAGCAATACTTTGGAAGGTGCTGTCCCAACAGGAGGCATCTTTAGTAATGCTACTGTGGTTTCCATCATAGGAAATAGAAATCTCTGTGGGGGTGTACCAGAGTTGGACCTTCCG

>M6-PGSC0003DMG400011932-(rev)M6LRR-RLK_186

CTGAGCTTTCTTTATGTCCTTAAGCTCCAAAATAATAGTTTTTCAGGTGAAATTCCATCAGAGATTGGCTACTTACATAAGTTAAAGTTTTTACGCCTTGATAACAACTCGTTCACTGGT

>M6-PGSC0003DMG400011932-(rev)M6LRR-RLK_1860

GTTAGTGGGCTTATAGGATTGACATTCATAGTTTGTTTTCTTGGCATACGACAGTTTAGTAGATCGAGAAAACCAACACCTACATATCTACCTGAAAACTCAACCTTAAGAGTATCTTAT

>M6-PGSC0003DMG400011932-(rev)M6LRR-RLK_1922

ATCGAGAAAACCAACACCTACATATCTACCTGAAAACTCAACCTTAAGAGTATCTTATCAGTGTCTACTTAGGGAAACTGATAGATTTTCTGCATCAAATTTGCTTGGAATGGGTGCTTT

>M6-PGSC0003DMG400011932-(rev)M6LRR-RLK_1984

GTCTACTTAGGGAAACTGATAGATTTTCTGCATCAAATTTGCTTGGAATGGGTGCTTTTGGGTCTGTATATAAAGGAATTTCTGAACACGATGGTACGTTTTTTGCTGTTAAGGTACTGG

>M6-PGSC0003DMG400011932-(rev)M6LRR-RLK_2046

TCTGTATATAAAGGAATTTCTGAACACGATGGTACGTTTTTTGCTGTTAAGGTACTGGACCTTTCACACCATGCAGCTTCCAGGAGTTTTTTAGCTGAATGTGAGGTTCTAAAGAACATT

>M6-PGSC0003DMG400011932-(rev)M6LRR-RLK_2170

ATCGTAATCTTGTAAAGGTTCTAAGTGCATGCTCAGGTATTGATTATGAAGGTAATGAGTTCAAGGCCATAGTTTATGAATACATGGATAAGGGAAGTCTACATGACTGGTTACATTTCA

>M6-PGSC0003DMG400011932-(rev)M6LRR-RLK_2232

AAGGCCATAGTTTATGAATACATGGATAAGGGAAGTCTACATGACTGGTTACATTTCACTCCTCAGGAGAAATCTGAGTCACAGGAAGAGCACAAGAAGCTCGGGTTTATTCAAAGATTA

>M6-PGSC0003DMG400011932-(rev)M6LRR-RLK_248

GATTGGCTACTTACATAAGTTAAAGTTTTTACGCCTTGATAACAACTCGTTCACTGGTCATATTCCATCAAACATTTCAGGATGCTTCAATCTTGTTTCTGTTGTTCTTTCGTATAATAT

>M6-PGSC0003DMG400011932-(rev)M6LRR-RLK_2480

TTTTGGTTTGGCAAGATTTGTTCCACCAGCAATTCCAAACTCATCAGCAAATTCTAAAAGCTCTACAGGTGTAGGGGGAACTATTGGTTATACACCTCCAGGTAATAATAAATCAAGATT

>M6-PGSC0003DMG400011932-(rev)M6LRR-RLK_2542

CTACAGGTGTAGGGGGAACTATTGGTTATACACCTCCAGGTAATAATAAATCAAGATTTCTGCTGTCATATATGTCTTTTGATAAGAACATCTCTAATTCTGTAGTATTTCTTGTTGACA

>M6-PGSC0003DMG400011932-(rev)M6LRR-RLK_2604

CTGTCATATATGTCTTTTGATAAGAACATCTCTAATTCTGTAGTATTTCTTGTTGACAGAATTAGGTATGGGAAGCGACGCCTCAATCTATGGAGATGTGTACAGCTTTGGCATCCTACT

>M6-PGSC0003DMG400011932-(rev)M6LRR-RLK_2728

GAAATGTTTACCGGGAGGAAACCAACTGATGAAATGTTCAAAGATAATTTAAACCTTCATAAATATGCCAATGCTGCTTTGCCAGACAGAGTGATGCATATTACAGATCCCATTCTTCTC

>M6-PGSC0003DMG400011932-(rev)M6LRR-RLK_2790

ATATGCCAATGCTGCTTTGCCAGACAGAGTGATGCATATTACAGATCCCATTCTTCTCCAGGAAAGAGATGAACTTGGAATGAAATACAATGTTGATGACAATACAAGCAGTGCTGGCGA

>M6-PGSC0003DMG400011932-(rev)M6LRR-RLK_2852

AAAGAGATGAACTTGGAATGAAATACAATGTTGATGACAATACAAGCAGTGCTGGCGACATATTTCTGTCATTCTTAGTTAAGGTGATTCAAATCGGAGTTTCCTGCTCCGTTGAATCTC

>M6-PGSC0003DMG400011932-(rev)M6LRR-RLK_2965

GAATCTCCAAAAGAAAGAAAGCAGATCAGTGATGTCGTTGGAGAACTCAATTCATTAAGGAAGTTGTTCTTGGAGCAAGCATACCGGAAAGAAAAGGTTATAAGCTTCAGCCTGCATTGA

>M6-PGSC0003DMG400011932-(rev)M6LRR-RLK_310

TTCCATCAAACATTTCAGGATGCTTCAATCTTGTTTCTGTTGTTCTTTCGTATAATATGATGGTAGGCAAAATTCCAGCAGAATTAGGCACATTGTTGAAACTCAAACAACTTTCTCTTG

>M6-PGSC0003DMG400011932-(rev)M6LRR-RLK_372

GTAGGCAAAATTCCAGCAGAATTAGGCACATTGTTGAAACTCAAACAACTTTCTCTTGTTTCTAACTATTTTACAGGAGGAATTCCGCCTTCATTTGGTAATCTTTCCTTACTTGATACC

>M6-PGSC0003DMG400011932-(rev)M6LRR-RLK_434

TAACTATTTTACAGGAGGAATTCCGCCTTCATTTGGTAATCTTTCCTTACTTGATACCTTTTCCGCAAGCAAGAATAATTTGTTAGGGAAAATACCTGATGAATTATGTCAATTGCTCAA

>M6-PGSC0003DMG400011932-(rev)M6LRR-RLK_62

GAATGAAACTATACATTTTTGTGACTGGCCTGGGGTTTCATGTGGTCGACGTCATCCCCGAGTTACAGTTTTAAATCTTACTTCTCTGAAATTGAGGGGTTCTCTGTCACCAAGCATTGG

>M6-PGSC0003DMG400011932-(rev)M6LRR-RLK_682

TTGGTATTACTCTTCCTAATTTAGAGTTCCTTAGCATTTATAGAAACAATGTAACTGGAAATATTCCGGTAACATTGTCAAATGCTACGAATCTTCAGTCTCTTATTGCTAGTAGAAATG

>M6-PGSC0003DMG400011989-(rev)M6LRR-RLK_0

CACAACTTCATCACAAAACCAAAAACAATCAATCATTTAGCCTTGATGATGGTTCCCACAATATTTTATAACCTTCCCTTCTTATATCTTTTCATAGTTATAACTTTTGCTTCCACTGAG

>M6-PGSC0003DMG400011989-(rev)M6LRR-RLK_1054

CTATTCCTGGTTCATTTGGCAATTTGAGAAACTTGCAATTTCTGTATCTCCGTGCCAACAAACTTTCTGGTTCTATTCCAAAAGAACTTGCATATTTGGATAACTTGGTTATAATGGAAA

>M6-PGSC0003DMG400011989-(rev)M6LRR-RLK_1116

CTTTCTGGTTCTATTCCAAAAGAACTTGCATATTTGGATAACTTGGTTATAATGGAAATGGATGAAAATCAGTTTTCAGGCAATTTGCCTGAGCATCTTTGTCAAGGTGGAAAACTTGAG

>M6-PGSC0003DMG400011989-(rev)M6LRR-RLK_1178

TGAAAATCAGTTTTCAGGCAATTTGCCTGAGCATCTTTGTCAAGGTGGAAAACTTGAGAACTTCACGGTGAATAGTAACAAGCTGACAGGGCCAATACCAAGAAGCTTTAGCAAGTGCTC

>M6-PGSC0003DMG400011989-(rev)M6LRR-RLK_124

CAACTGCTCTTCTTAAATGGAAAGCAACATTCCTAAACCAGAACAATTCCTTACAGGCTTCATGGACACTAAGTAGTCCTGCTGATACCAAGAATTCTTCCAGAATTGGAGCGACGAGTT

>M6-PGSC0003DMG400011989-(rev)M6LRR-RLK_1302

TTCAAAAGAGTTCGCTTGGATAACAACAGTTTTACTGGGAATTTATCTGAAGCTTTTGGCGTCCATCCACACCTTTACTTCATTGATTTGAGTGATAATGATTTTCATGGTGAACTCAGC

>M6-PGSC0003DMG400011989-(rev)M6LRR-RLK_1364

CCATCCACACCTTTACTTCATTGATTTGAGTGATAATGATTTTCATGGTGAACTCAGCAGAAACTGGGGGAAATGCAAGAAATTGACTGATTTTCAGATAGCCAGAAATAACATTAGTGG

>M6-PGSC0003DMG400011989-(rev)M6LRR-RLK_1426

ACTGGGGGAAATGCAAGAAATTGACTGATTTTCAGATAGCCAGAAATAACATTAGTGGTAGCATACCACCAGAGATTGGAAACATCAAAGGTCTTTTGGGACTTGATCTTTCAGCAAATC

>M6-PGSC0003DMG400011989-(rev)M6LRR-RLK_1488

ATACCACCAGAGATTGGAAACATCAAAGGTCTTTTGGGACTTGATCTTTCAGCAAATCATTTGATTGGGCGGATACCAAAGGAATTTGGAAAATTAACCTCTCTGGTTAAGCTTTCTGTG

>M6-PGSC0003DMG400011989-(rev)M6LRR-RLK_1550

GATTGGGCGGATACCAAAGGAATTTGGAAAATTAACCTCTCTGGTTAAGCTTTCTGTGCAAAATAACTATATTTCTGGCAATATTCCTATAGAACTTGGGGCACTGACAAAGTTAGATTC

>M6-PGSC0003DMG400011989-(rev)M6LRR-RLK_1612

ATAACTATATTTCTGGCAATATTCCTATAGAACTTGGGGCACTGACAAAGTTAGATTCCCTAGATCTGTCAAACAATAGATTGAATGGGTCGATCCCAACGTTTATAGGAGATTACCAGC

>M6-PGSC0003DMG400011989-(rev)M6LRR-RLK_1674

GATCTGTCAAACAATAGATTGAATGGGTCGATCCCAACGTTTATAGGAGATTACCAGCACCTATTTCACTTGAACCTGAGTTGTAACAAGTTTGGGCAGAAAATTCCAAAGGAGATAGGG

>M6-PGSC0003DMG400011989-(rev)M6LRR-RLK_1736

ATTTCACTTGAACCTGAGTTGTAACAAGTTTGGGCAGAAAATTCCAAAGGAGATAGGGAGGATAACTCATCTTAGTGTACTTGATTTGAGCTATAATCTTCTTGATGGAGAAATACCTGC

>M6-PGSC0003DMG400011989-(rev)M6LRR-RLK_1798

TAACTCATCTTAGTGTACTTGATTTGAGCTATAATCTTCTTGATGGAGAAATACCTGCTCAGTTAGCTAGTTTGCTGGACTTGGCAAACTTGAATCTTTCCCACAACAGCCTTGCTGGGC

>M6-PGSC0003DMG400011989-(rev)M6LRR-RLK_186

TGGACACTAAGTAGTCCTGCTGATACCAAGAATTCTTCCAGAATTGGAGCGACGAGTTCTGATGCATGCAAGGGCTGGTATGGAGTTGTATGAGTTAATGGTCGGGTTAACAGGTTGAAT

>M6-PGSC0003DMG400011989-(rev)M6LRR-RLK_1860

TTAGCTAGTTTGCTGGACTTGGCAAACTTGAATCTTTCCCACAACAGCCTTGCTGGGCGTATTCCTGAAGAATTTGAAAGTTTGACCAGTTTGCAGGATGTTATCTTGTCATACAATGAG

>M6-PGSC0003DMG400011989-(rev)M6LRR-RLK_1922

TCCTGAAGAATTTGAAAGTTTGACCAGTTTGCAGGATGTTATCTTGTCATACAATGAGTTGGAAGGTCCAATCCCTAATAATAAAGCTTTCATGAATGCCTCATTAGAAGGTAATAAAGA

>M6-PGSC0003DMG400011989-(rev)M6LRR-RLK_1984

AAGGTCCAATCCCTAATAATAAAGCTTTCATGAATGCCTCATTAGAAGGTAATAAAGATCTTTGTGGCAATGTGATTGGATTCCAACCTTGCGAAAGGCCATCTTCTATTGTGAAGAAGC

>M6-PGSC0003DMG400011989-(rev)M6LRR-RLK_2046

TGTGGCAATGTGATTGGATTCCAACCTTGCGAAAGGCCATCTTCTATTGTGAAGAAGCACTCAATGGCGAAGGGACATAAACTCATCCTCATCACTGTACTTCCTATTATGGGAGCACTA

>M6-PGSC0003DMG400011989-(rev)M6LRR-RLK_2108

AATGGCGAAGGGACATAAACTCATCCTCATCACTGTACTTCCTATTATGGGAGCACTAGTACTACTCTGTGCTTTCATTGGTGTTCTCTTTATGTGTAATAAAAGAAGGAGAGTTAGAGA

>M6-PGSC0003DMG400011989-(rev)M6LRR-RLK_2170

TACTCTGTGCTTTCATTGGTGTTCTCTTTATGTGTAATAAAAGAAGGAGAGTTAGAGACGTTGAAAGAAGGGATTCCAGTGATGATGTTGATGGTTTGATTTACATATCCACATTAAATG

>M6-PGSC0003DMG400011989-(rev)M6LRR-RLK_2232

GAAAGAAGGGATTCCAGTGATGATGTTGATGGTTTGATTTACATATCCACATTAAATGGAAATGCATTGTACTGGGATATCTTAAAAGCCACAAAGGAGTTTGATGCAATGTTTTGCATC

>M6-PGSC0003DMG400011989-(rev)M6LRR-RLK_2294

TGCATTGTACTGGGATATCTTAAAAGCCACAAAGGAGTTTGATGCAATGTTTTGCATCGGAAAAGGAGGATCTGGAAGCGTTTACAAGGTAAAACTTCCAACAATAGAAAATGTAGCTGT

>M6-PGSC0003DMG400011989-(rev)M6LRR-RLK_2356

AAGGAGGATCTGGAAGCGTTTACAAGGTAAAACTTCCAACAATAGAAAATGTAGCTGTGAAGAGACTTCATTCTTCATTTGAGGTTACCCATCGCAAAAGCTTCATGAATGAGATAAGGT

>M6-PGSC0003DMG400011989-(rev)M6LRR-RLK_2418

AGACTTCATTCTTCATTTGAGGTTACCCATCGCAAAAGCTTCATGAATGAGATAAGGTCATTGACAAGTATCAAGCATCGCAACATTGTGAAACTCTATGCCTTCTGTACAAATGCACAA

>M6-PGSC0003DMG400011989-(rev)M6LRR-RLK_2542

CATTCTTGGTTTACGAGTACATGGAGAAGGGGAGCTTGTCTAGTATTTTGAGCAATGAAGTAGAGTCCAAGAAATTGGATTGGCTTACAAGGGTGAATATCATCAAAGGTGTTGCTTATG

>M6-PGSC0003DMG400011989-(rev)M6LRR-RLK_2604

GAGTCCAAGAAATTGGATTGGCTTACAAGGGTGAATATCATCAAAGGTGTTGCTTATGCTTTATCTTACATGCACCATGATTGCTCACCCCCGATTGTTCATCGAGACATATCAAGCAGT

>M6-PGSC0003DMG400011989-(rev)M6LRR-RLK_2666

ATCTTACATGCACCATGATTGCTCACCCCCGATTGTTCATCGAGACATATCAAGCAGTAATGTTTTGCTTGATTCCGAGTATGAAGCTCGTGTTTCAGATTTTGGCATAGCTAAGATTCT

>M6-PGSC0003DMG400011989-(rev)M6LRR-RLK_2790

CCAGACTCATCCAATTGTACTGCACTTGCAGGCACATATGGCTATGTAGCACCAGGTAAGATCTAACTTCTTATCCTTATATTAGTTCATGAAATATATCTGCAAAATCTATATTGTTTC

>M6-PGSC0003DMG400011989-(rev)M6LRR-RLK_2852

CTAACTTCTTATCCTTATATTAGTTCATGAAATATATCTGCAAAATCTATATTGTTTCTATCTCATTTGAAACTTTATTGTTTCAGAGCTTGCCTATACAATGAAGGTGACGGAAATGTG

>M6-PGSC0003DMG400011989-(rev)M6LRR-RLK_2976

GTCTATAGCTTTGGAGTATTAGCATTGGAGGTAATCAAAGGAAAGCATCTTGGGGAATACATTACTGTGCTAGCAAATTCATCGACAAGAGATCATGTGCAGCATAGCGATTTGCTAGAT

>M6-PGSC0003DMG400011989-(rev)M6LRR-RLK_3038

TACTGTGCTAGCAAATTCATCGACAAGAGATCATGTGCAGCATAGCGATTTGCTAGATGAACGCCTTCCATATCCTGAAGATAGAGTAAAAGAGGTTTTGGTTTTTATCATCAAGCTAGC

>M6-PGSC0003DMG400011989-(rev)M6LRR-RLK_3100

GCCTTCCATATCCTGAAGATAGAGTAAAAGAGGTTTTGGTTTTTATCATCAAGCTAGCAAGCTCATGTTTGCTTGAAACTCCAAAATCAAGGCCAACAATGCACTTCATCTCCAGTATGT

>M6-PGSC0003DMG400011989-(rev)M6LRR-RLK_3224

AATGGATCCACCTGTTTCTCAAATAACTTCTATGAAGGCGAAAAGAGTCCCCGAGATGTAAGGTGTCAGTGTATGAAAAAAAAAGAAGCAAGTTTAGACACCAAGTGGTTGTGAAGTAAG

>M6-PGSC0003DMG400011989-(rev)M6LRR-RLK_3286

GTGTCAGTGTATGAAAAAAAAAGAAGCAAGTTTAGACACCAAGTGGTTGTGAAGTAAGAATAAGAAGCTATTATTACTGGTACTAGTATTTGTGTATTTTATTGTTAGTTATGCTTATCT

>M6-PGSC0003DMG400011989-(rev)M6LRR-RLK_3348

AGAAGCTATTATTACTGGTACTAGTATTTGTGTATTTTATTGTTAGTTATGCTTATCTAGTTTAGTGTTGTACTAGAGTGGATACAATATCAAGTAGAAGGTAATATTTTAATTTTCTTG

>M6-PGSC0003DMG400011989-(rev)M6LRR-RLK_372

TATGTTGATCTTAGTATGAACCAGCTCTCTGGCACCATCCCACCTGAAATTGGTAAGCTCGCTAATCTTGTCTACCTTGACTTGTCTATCAATCATATTTCGGGCACAATCCCACAACAA

>M6-PGSC0003DMG400011989-(rev)M6LRR-RLK_434

TAATCTTGTCTACCTTGACTTGTCTATCAATCATATTTCGGGCACAATCCCACAACAAATCAGCTCACTAATAAAGCTTGAGACCCTCCACATATTTGACAACCAATTAAATGGTTCCAT

>M6-PGSC0003DMG400011989-(rev)M6LRR-RLK_496

GCTCACTAATAAAGCTTGAGACCCTCCACATATTTGACAACCAATTAAATGGTTCCATTCCGAGGGAAATAGGTCGCCTAAGGTCTCTTACTGATCTAGCTTTGAGTTCTAACTTTCTTA

>M6-PGSC0003DMG400011989-(rev)M6LRR-RLK_62

ATTTTATAACCTTCCCTTCTTATATCTTTTCATAGTTATAACTTTTGCTTCCACTGAGGAAGCAACTGCTCTTCTTAAATGGAAAGCAACATTCCTAAACCAGAACAATTCCTTACAGGC

>M6-PGSC0003DMG400011989-(rev)M6LRR-RLK_682

AGCTTTCTGGCTCCATTCCTTCAGAAATAGGAAAACTTGTCAATCTTGTTGAAGCTTATATTAGTAGAAATCAATTAACAGGTCATATTCCTCCAGAAATAGGTAATTTGATCAATGCAA

>M6-PGSC0003DMG400011989-(rev)M6LRR-RLK_744

AGTAGAAATCAATTAACAGGTCATATTCCTCCAGAAATAGGTAATTTGATCAATGCAAAAGTGTTGTATGCTTTTTCCAATAACTTGTCTGGTCCCATTCCTGATGAAATTGGGAAAATG

>M6-PGSC0003DMG400011989-(rev)M6LRR-RLK_806

GTTGTATGCTTTTTCCAATAACTTGTCTGGTCCCATTCCTGATGAAATTGGGAAAATGAAGTCACTTGAGAACTTAAGCTTCCAGACAAACAATCTTTCTGGTCCAATTCCCAAAACAAT

>M6-PGSC0003DMG400011989-(rev)M6LRR-RLK_992

TGATTTTGGGAATCTGAAAAAATTGAATGATCTGCAATTATCCAATAATCAACTTACTGGTCCTATTCCTGGTTCATTTGGCAATTTGAGAAACTTGCAATTTCTGTATCTCCGTGCCAA

>M6-PGSC0003DMG400012218M6LRR-RLK_0

TAGCTATTGTGTATTTATGAGAAAATATTCGAAGTTTGATCAATCATAGCGACGCTGGTAAAGGCCCACACTTCCTTTACTCCGCCGCTCCTTCTATTATCTCCTTCTCCATTTTCTGCT

>M6-PGSC0003DMG400012218M6LRR-RLK_1037

TTATGCTCTCAATACCTTCAAGGAAGCTATATATGAAGACCCTCTTCTGGTTTTGTCAAATTGGAATGGTTTAGATTCAGATCCTTGTGGTTGGGCTGGAGTCTTTTGTTCTATGGCTCA

>M6-PGSC0003DMG400012218M6LRR-RLK_1098

TGGAATGGTTTAGATTCAGATCCTTGTGGTTGGGCTGGAGTCTTTTGTTCTATGGCTCAAGATCATGTCATGAAAATGTAAGTTAATTGTCATCAATTTTATGTGCAATACATGGATAGG

>M6-PGSC0003DMG400012218M6LRR-RLK_1159

ATCATGTCATGAAAATGTAAGTTAATTGTCATCAATTTTATGTGCAATACATGGATAGGTTGTTTGGTTAATGATTTTTCTCAACTAAAATGCAATCTAGTAAATCTTTTATGGACTCAA

>M6-PGSC0003DMG400012218M6LRR-RLK_122

ATTTTTTGTTCCCTTATAATTTTAACATATAGTTTTTATAACCCCAAAATTCTTCTTCTTCAGCTCATAAATTCTCTGTTTCTAGACACCAAATCTCTCTTTGAACCCAAGTTTTATTTT

>M6-PGSC0003DMG400012218M6LRR-RLK_1220

GTTTGGTTAATGATTTTTCTCAACTAAAATGCAATCTAGTAAATCTTTTATGGACTCAAGTTTAGTGCAAATGAGTCTCTAATACGCATTTCCACTGGATTGTGTGACAGAAACATTTCT

>M6-PGSC0003DMG400012218M6LRR-RLK_1281

TTAGTGCAAATGAGTCTCTAATACGCATTTCCACTGGATTGTGTGACAGAAACATTTCTGGCGCTTCGTTGAAGGGGTTTCTTTCACCCAACTTGCATTTACTCTCCTCTTTGCAAGAAC

>M6-PGSC0003DMG400012218M6LRR-RLK_1342

CGCTTCGTTGAAGGGGTTTCTTTCACCCAACTTGCATTTACTCTCCTCTTTGCAAGAACTGTAAGTTATTTCCGAATAGATTTGTAAGGATTACATAGTTCTGGTTGTTCTTTTATGTGG

>M6-PGSC0003DMG400012218M6LRR-RLK_1403

TAAGTTATTTCCGAATAGATTTGTAAGGATTACATAGTTCTGGTTGTTCTTTTATGTGGTTTTGTTAAAGCTTATGTGATTCCCATGCAATATGCAGAATTTTGCATGGAAATGTGCTGA

>M6-PGSC0003DMG400012218M6LRR-RLK_1464

TTGTTAAAGCTTATGTGATTCCCATGCAATATGCAGAATTTTGCATGGAAATGTGCTGATTGGTACAATTCCTAAGGAGATTGGTCTATTGAAAAACCTGAAGGCCTTGGATTTGGGTGC

>M6-PGSC0003DMG400012218M6LRR-RLK_1525

GGTACAATTCCTAAGGAGATTGGTCTATTGAAAAACCTGAAGGCCTTGGATTTGGGTGCTAACCAGCTAACAGGACCAATTCCTTCCGAGCTTGGCAATTTGACAAACATTATGAAAATG

>M6-PGSC0003DMG400012218M6LRR-RLK_1586

ACCAGCTAACAGGACCAATTCCTTCCGAGCTTGGCAATTTGACAAACATTATGAAAATGTTAGTCTAACTTGAATTTTGTAGGTTCATTTGTACTTGCCTTTACCTATCCTTGTTGAATC

>M6-PGSC0003DMG400012218M6LRR-RLK_1647

AGTCTAACTTGAATTTTGTAGGTTCATTTGTACTTGCCTTTACCTATCCTTGTTGAATCTCCCTTACTTTTCTAGCTTCCATGTGTCTCTGCAGAAACCTACAATCTAATGGATTAACAG

>M6-PGSC0003DMG400012218M6LRR-RLK_1708

CCTTACTTTTCTAGCTTCCATGTGTCTCTGCAGAAACCTACAATCTAATGGATTAACAGGGAAATTGCCTGCTGAGCTTGGTAATTTGAAATACCTTGAGGAACTTCGGTTGGACAGGAA

>M6-PGSC0003DMG400012218M6LRR-RLK_1769

AAATTGCCTGCTGAGCTTGGTAATTTGAAATACCTTGAGGAACTTCGGTTGGACAGGAACAAACTCCAAGGACTAGTGCCTGCTAATACTGGATCAGATTTTACATCCGGTGTACGTGGA

>M6-PGSC0003DMG400012218M6LRR-RLK_183

AGCTCATAAATTCTCTGTTTCTAGACACCAAATCTCTCTTTGAACCCAAGTTTTATTTTTTTTAACTAAGTTGGTGACACTTAATAGTAGATTCCACTCAAAAGGGTTGTTACAGATTTA

>M6-PGSC0003DMG400012218M6LRR-RLK_1830

AACTCCAAGGACTAGTGCCTGCTAATACTGGATCAGATTTTACATCCGGTGTACGTGGAATGTGAGTTGATTAGCTACGTCTTGTACTTTATCGTTTTTCTTTTGAATTTTCTGGTATGT

>M6-PGSC0003DMG400012218M6LRR-RLK_1891

GTGAGTTGATTAGCTACGTCTTGTACTTTATCGTTTTTCTTTTGAATTTTCTGGTATGTGGATAAGGATTAATGCACGATTCTCTGTTTACTTGCAAGATATTTTTGGTTGTCCTTAGAG

>M6-PGSC0003DMG400012218M6LRR-RLK_1952

ATAAGGATTAATGCACGATTCTCTGTTTACTTGCAAGATATTTTTGGTTGTCCTTAGAGCGTTTTCCGTCTTGGCCCTCTAGGATTCTGTCAGATATTTGAAGGATTGAGTGACCAAAGA

>M6-PGSC0003DMG400012218M6LRR-RLK_2013

TTTTCCGTCTTGGCCCTCTAGGATTCTGTCAGATATTTGAAGGATTGAGTGACCAAAGAATCTAATTTTCTGATCTTTATGACAGTCATTGTCTGGAATGATGATGACAGAATCCAGGAA

>M6-PGSC0003DMG400012218M6LRR-RLK_2074

CTAATTTTCTGATCTTTATGACAGTCATTGTCTGGAATGATGATGACAGAATCCAGGAAAGAGATGCTTATGGGATAATGTCTGTTCTTTCTGTCAGCATTGTTTGATATGTACTGCCTG

>M6-PGSC0003DMG400012218M6LRR-RLK_2135

AGATGCTTATGGGATAATGTCTGTTCTTTCTGTCAGCATTGTTTGATATGTACTGCCTGCAGATTGACACTGATGGCTGTTTTACTTGAGTCTAGTGTAGTACATTCATTGTCTGTTGTT

>M6-PGSC0003DMG400012218M6LRR-RLK_2196

GATTGACACTGATGGCTGTTTTACTTGAGTCTAGTGTAGTACATTCATTGTCTGTTGTTTGATGGGAGGATGGTGAATTGGGGGACCCCAATGGAGTGTGTTTTTGCTATAGCCTTTAAT

>M6-PGSC0003DMG400012218M6LRR-RLK_2257

ATGGGAGGATGGTGAATTGGGGGACCCCAATGGAGTGTGTTTTTGCTATAGCCTTTAATATTTGACCAAATGAGAGCAATGTTACAGAGTTCAGAATGTATGTTTCAAAAGTTTAACGAT

>M6-PGSC0003DMG400012218M6LRR-RLK_2318

TTGACCAAATGAGAGCAATGTTACAGAGTTCAGAATGTATGTTTCAAAAGTTTAACGATGATGTTCTTGGAATTGATCTGTATTTCTGTGAAGAAAAATAAGTAAAAACTGTTTGATGTC

>M6-PGSC0003DMG400012218M6LRR-RLK_2379

TGTTCTTGGAATTGATCTGTATTTCTGTGAAGAAAAATAAGTAAAAACTGTTTGATGTCCTGTATATTTGACCCAACTGATTTTGCTATAGCCTTACGTCCATCACATGTATCTTTATCT

>M6-PGSC0003DMG400012218M6LRR-RLK_244

TTAACTAAGTTGGTGACACTTAATAGTAGATTCCACTCAAAAGGGTTGTTACAGATTTAGATTCTTGAATTCACTCTTAAGCTTTGTTAAGTTTTCTTTAGTGTTTATCTTTCTCCCCTC

>M6-PGSC0003DMG400012218M6LRR-RLK_2440

GTATATTTGACCCAACTGATTTTGCTATAGCCTTACGTCCATCACATGTATCTTTATCTGATTGTTTTTTTTCTTTTCTGTTGTACCCATTTAGGTATGCCTCTGGTGCCAGCGCTACAG

>M6-PGSC0003DMG400012218M6LRR-RLK_2501

TTGTTTTTTTTCTTTTCTGTTGTACCCATTTAGGTATGCCTCTGGTGCCAGCGCTACAGGTTTTTGTCGCACATCCCAGCTCAAAGTGGCTGATTTCTCATTCAACTTTCTTATTGGAAG

>M6-PGSC0003DMG400012218M6LRR-RLK_2562

TTTTGTCGCACATCCCAGCTCAAAGTGGCTGATTTCTCATTCAACTTTCTTATTGGAAGCATACCAAAGTGTTTGGGTTACCTTCCAAAGTCAGTTCTTAATTCTACGCCATAATTCTTG

>M6-PGSC0003DMG400012218M6LRR-RLK_2623

TACCAAAGTGTTTGGGTTACCTTCCAAAGTCAGTTCTTAATTCTACGCCATAATTCTTGTTTTACCTCATCTTCTCTATTCCTGAATGCCTATTAGTTTTGTTGCTGTAAATCTATATCC

>M6-PGSC0003DMG400012218M6LRR-RLK_2684

TTACCTCATCTTCTCTATTCCTGAATGCCTATTAGTTTTGTTGCTGTAAATCTATATCCAAAGTAGCACTCTGATGCTCTACCAGTGTTGTTTCTCTTACCTTTTCTGTTTCTTAATGCC

>M6-PGSC0003DMG400012218M6LRR-RLK_2745

AGTAGCACTCTGATGCTCTACCAGTGTTGTTTCTCTTACCTTTTCTGTTTCTTAATGCCTGTTATTTTGTGATTCTATAGATCTAGTTTTCAAGGAAATTGCCTTCAGCACAAAGACCCG

>M6-PGSC0003DMG400012218M6LRR-RLK_2806

TTATTTTGTGATTCTATAGATCTAGTTTTCAAGGAAATTGCCTTCAGCACAAAGACCCGAAGCAACGGTCTGCTGCTTTATGTGGTAAGTAAACAATCATAGTTACAATTTTCAACACTA

>M6-PGSC0003DMG400012218M6LRR-RLK_2867

GCAACGGTCTGCTGCTTTATGTGGTAAGTAAACAATCATAGTTACAATTTTCAACACTATGCATAGGAGAATTGATGTCTCAACCTTTTTTCTTTAATAGTTGCAGTCAAATAGATTTAT

>M6-PGSC0003DMG400012218M6LRR-RLK_2928

CATAGGAGAATTGATGTCTCAACCTTTTTTCTTTAATAGTTGCAGTCAAATAGATTTATTTAAGAAACTGAACTGGTTGTTTTGACATTGCATTATAGGTAGCACTCCGCCTCCCACTAG

>M6-PGSC0003DMG400012218M6LRR-RLK_2989

AAGAAACTGAACTGGTTGTTTTGACATTGCATTATAGGTAGCACTCCGCCTCCCACTAGCCATGCATCGACACTCAGCAATAAGCACCAGCCTGTTGAAGAACCGAGACATCAGACTTCT

>M6-PGSC0003DMG400012218M6LRR-RLK_305

TTCTTGAATTCACTCTTAAGCTTTGTTAAGTTTTCTTTAGTGTTTATCTTTCTCCCCTCACTTGTAAAAAAGACAACTTTATCTTTTTCCAGCTTTTTTTTTTGTTTGTTTTCATCTACT

>M6-PGSC0003DMG400012218M6LRR-RLK_3050

ATGCATCGACACTCAGCAATAAGCACCAGCCTGTTGAAGAACCGAGACATCAGACTTCTTCCAAGCCCACTTGGCTTTTGGCTTTAGAAGTGGTAACAGGAGTTATGGTGGGTTCTCTCT

>M6-PGSC0003DMG400012218M6LRR-RLK_3111

CAAGCCCACTTGGCTTTTGGCTTTAGAAGTGGTAACAGGAGTTATGGTGGGTTCTCTCTTCATTGTAGGTCTTGTGACTGGCCTTCAGAGGCTTAAGATCAAATCCTCAATTATACCTTG

>M6-PGSC0003DMG400012218M6LRR-RLK_3172

ATTGTAGGTCTTGTGACTGGCCTTCAGAGGCTTAAGATCAAATCCTCAATTATACCTTGGAAGAAATCAGGAAGCGAGAAAGATCATATGACAATTTATGTAGGTTAGTACTATCATGCA

>M6-PGSC0003DMG400012218M6LRR-RLK_3233

AGAAATCAGGAAGCGAGAAAGATCATATGACAATTTATGTAGGTTAGTACTATCATGCACTTGAACTTGAGCACTTAGTACTTTCAAAAGCATTGAGCCTAATCTTCGTTTCTGTTTACC

>M6-PGSC0003DMG400012218M6LRR-RLK_3294

TGAACTTGAGCACTTAGTACTTTCAAAAGCATTGAGCCTAATCTTCGTTTCTGTTTACCGCTGATGATCTTTGTAGATACTGGTATGCTGAAAGATGTTGTAAGATACAGCAGACAAGAA

>M6-PGSC0003DMG400012218M6LRR-RLK_3355

TGATGATCTTTGTAGATACTGGTATGCTGAAAGATGTTGTAAGATACAGCAGACAAGAACTTGAAGTGGCTTGTGAAGATTTCAGCAACATCATTGGATCCTCACCAGATAGCATAGTTT

>M6-PGSC0003DMG400012218M6LRR-RLK_3416

TGAAGTGGCTTGTGAAGATTTCAGCAACATCATTGGATCCTCACCAGATAGCATAGTTTACAAAGGAACCATGAAAGGTGGACCTGAAATCGCTGTAATATCCCTTTGCGTCAAAGAAGA

>M6-PGSC0003DMG400012218M6LRR-RLK_3477

AAAGGAACCATGAAAGGTGGACCTGAAATCGCTGTAATATCCCTTTGCGTCAAAGAAGAGCACTGGACAGCCTATCTAGAACTTTATTTTCAGAAAGAGGCAAGACTCAATTCTTTCTTC

>M6-PGSC0003DMG400012218M6LRR-RLK_3538

ACTGGACAGCCTATCTAGAACTTTATTTTCAGAAAGAGGCAAGACTCAATTCTTTCTTCATCATCACTATTTATCTCTTCCAAAAGTTTTCATCTTTAGTTCCATGCTCTAACTTCTGTA

>M6-PGSC0003DMG400012218M6LRR-RLK_3599

CATCACTATTTATCTCTTCCAAAAGTTTTCATCTTTAGTTCCATGCTCTAACTTCTGTAAATTTACATTTAGGTGGCAGAATTGGCCAGAATAAATCACGAGAACGCTGGGAAACTTCTA

>M6-PGSC0003DMG400012218M6LRR-RLK_366

TTGTAAAAAAGACAACTTTATCTTTTTCCAGCTTTTTTTTTTGTTTGTTTTCATCTACTGTCTTGGGTCTTTTCTTGTTTTCTCTTTCAATTTTGTTTGGATTATTGCTTTTTGAATCTT

>M6-PGSC0003DMG400012218M6LRR-RLK_3660

TTTACATTTAGGTGGCAGAATTGGCCAGAATAAATCACGAGAACGCTGGGAAACTTCTAGGCTACTGCAGAGAGAGTAGTCCCTTTACGAGGATGTTGGTGTTTGAATATGCATCTAATG

>M6-PGSC0003DMG400012218M6LRR-RLK_3721

CTACTGCAGAGAGAGTAGTCCCTTTACGAGGATGTTGGTGTTTGAATATGCATCTAATGGGACATTATATGAACACCTTCATTGTGAGTAAAGTGAATTAATTTTCTCATGCACTTCATT

>M6-PGSC0003DMG400012218M6LRR-RLK_3782

ACATTATATGAACACCTTCATTGTGAGTAAAGTGAATTAATTTTCTCATGCACTTCATTTTATGTTTTCCAAAGCAAACTCCTAACTCTTTGAATGTATCATAGATGGAGAAGGATGCCA

>M6-PGSC0003DMG400012218M6LRR-RLK_3843

ATGTTTTCCAAAGCAAACTCCTAACTCTTTGAATGTATCATAGATGGAGAAGGATGCCAACTATCTTGGACACGCCGGATGAAAATAGTTATTGGCATCGCTAAGGGACTGACATATTTC

>M6-PGSC0003DMG400012218M6LRR-RLK_3904

TATCTTGGACACGCCGGATGAAAATAGTTATTGGCATCGCTAAGGGACTGACATATTTCCATTCAGAACTTGACCCTCCATTTACTATATCAGAACTGAATTCAAATTCTGTATATCTCA

>M6-PGSC0003DMG400012218M6LRR-RLK_3941

CGCTAAGGGACTGACATATTTCCATTCAGAACTTGACCCTCCATTTACTATATCAGAACTGAATTCAAATTCTGTATATCTCACCGAAGATTTTTCTCCCAAGGTAAACGAAAGACCATT

>M6-PGSC0003DMG400012218M6LRR-RLK_427

CTTGGGTCTTTTCTTGTTTTCTCTTTCAATTTTGTTTGGATTATTGCTTTTTGAATCTTTTGCTTTCTGGGTTTTTCTTCTTTTTGTTGATTGCTTAAGTTCTTGGAGACTGTTTCTTTA

>M6-PGSC0003DMG400012218M6LRR-RLK_488

GCTTTCTGGGTTTTTCTTCTTTTTGTTGATTGCTTAAGTTCTTGGAGACTGTTTCTTTAGATGGGTTCTTTCAGTTCTGGATTTTTGTAGAAACTTAGTGTGTTCTTGTGGTGTGGAATT

>M6-PGSC0003DMG400012218M6LRR-RLK_549

TGGGTTCTTTCAGTTCTGGATTTTTGTAGAAACTTAGTGTGTTCTTGTGGTGTGGAATTATTTGGGGGAAAAGATGAGACTTTGTGTTGTATTTCAAGTTTTAGTTACTGTTGGACTAAT

>M6-PGSC0003DMG400012218M6LRR-RLK_61

AGGCCCACACTTCCTTTACTCCGCCGCTCCTTCTATTATCTCCTTCTCCATTTTCTGCTTGATTTTTTGTTCCCTTATAATTTTAACATATAGTTTTTATAACCCCAAAATTCTTCTTCT

>M6-PGSC0003DMG400012218M6LRR-RLK_610

TTGGGGGAAAAGATGAGACTTTGTGTTGTATTTCAAGTTTTAGTTACTGTTGGACTAATTCTTGGGATTTGTGATGCTTTTCCATCAAATGAAGGTGTGTATTATTAACTTAAGTTCAGA

>M6-PGSC0003DMG400012218M6LRR-RLK_671

TTGGGATTTGTGATGCTTTTCCATCAAATGAAGGTGTGTATTATTAACTTAAGTTCAGATCTTGATGTAGTTACTGCTGCAAAGATTCAATTTTTGTCTTTTAATTGAGCATTTAGTGTT

>M6-PGSC0003DMG400012218M6LRR-RLK_732

TTGATGTAGTTACTGCTGCAAAGATTCAATTTTTGTCTTTTAATTGAGCATTTAGTGTTGTAATGTAATGTTGTTTAGGGAAGATCTTGTATTTTCTTGTTTTGAGATACTTGCTATATT

>M6-PGSC0003DMG400012218M6LRR-RLK_976

ATTCAAAGAGTAGGTAGTTTATTCTGAGTGAATCTGATTGGGGACTCTTTGTTGATACAGTTTATGCTCTCAATACCTTCAAGGAAGCTATATATGAAGACCCTCTTCTGGTTTTGTCAA

>M6-PGSC0003DMG400012307-(rev)M6LRR-RLK_0

GTTTTTTAGAAGTTTTGAACTTTAGATTCTTGATGCTCTTTTTGTTCTTCTTATGTGACTACAAAAGCTAAGTTTTTGAGGGGTTGAGGGTATTCATAATCTTTGAAATGGTTAAAGTTT

>M6-PGSC0003DMG400012307-(rev)M6LRR-RLK_128

AGGGAGTGGTGGGGGATTGTTTCTTTGGGTTTCTTGGTGGGGTTGTTTCTGTTGTCACAAGACACTGTTGCTTTTCTTGAAAATGATAAGCAAGCATTGTTGGATTTTGTTAATCAGTTG

>M6-PGSC0003DMG400012307-(rev)M6LRR-RLK_192

CTGTTGCTTTTCTTGAAAATGATAAGCAAGCATTGTTGGATTTTGTTAATCAGTTGCCTCATTTGCACACTTTAAACTGGGATGCAAATTCTTCCGTTTGCAAGAACTGGACTGGAGTGG

>M6-PGSC0003DMG400012307-(rev)M6LRR-RLK_256

GCACACTTTAAACTGGGATGCAAATTCTTCCGTTTGCAAGAACTGGACTGGAGTGGGGTGTAGTGAAGATGGTTCTAGAGTGATAGCACTGAGGTTACCTGGGGTTGGGTTCAATGGTCC

>M6-PGSC0003DMG400012307-(rev)M6LRR-RLK_320

GAAGATGGTTCTAGAGTGATAGCACTGAGGTTACCTGGGGTTGGGTTCAATGGTCCAATTCCTAATAATACATTGAGTCGTTTGACTGCATTGCAGATCTTGAGCCTTAGATCTAATGGT

>M6-PGSC0003DMG400012307-(rev)M6LRR-RLK_384

ATAATACATTGAGTCGTTTGACTGCATTGCAGATCTTGAGCCTTAGATCTAATGGTATCAATGGCACTTTCCCTATGGACTTTGATAACCTGAAAAACTTGTCTTATCTTTACCTTCATT

>M6-PGSC0003DMG400012307-(rev)M6LRR-RLK_448

CACTTTCCCTATGGACTTTGATAACCTGAAAAACTTGTCTTATCTTTACCTTCATTACAACAATTTCTCAGGTCCTCTGCCTTTTGATTGCTCTGTTTGGCAAAATCTCACCAATTTGAA

>M6-PGSC0003DMG400012307-(rev)M6LRR-RLK_512

TTCTCAGGTCCTCTGCCTTTTGATTGCTCTGTTTGGCAAAATCTCACCAATTTGAACTTGTCAAACAACAGATTCAATGGCACCGTTCCTAGCTCGATATCGGGTTTAAGTCATCTTACT

>M6-PGSC0003DMG400012307-(rev)M6LRR-RLK_576

ACAACAGATTCAATGGCACCGTTCCTAGCTCGATATCGGGTTTAAGTCATCTTACTCCCCTTAATCTTGCTAACAACTCACTTTCTGGTACCGTTCTTGATTTTCATTTTCCCAACTTGC

>M6-PGSC0003DMG400012307-(rev)M6LRR-RLK_64

AAGCTAAGTTTTTGAGGGGTTGAGGGTATTCATAATCTTTGAAATGGTTAAAGTTTTGAGCTTTAGGGAGTGGTGGGGGATTGTTTCTTTGGGTTTCTTGGTGGGGTTGTTTCTGTTGTC

>M6-PGSC0003DMG400012307-(rev)M6LRR-RLK_640

TCTTGCTAACAACTCACTTTCTGGTACCGTTCTTGATTTTCATTTTCCCAACTTGCAGCTACTGAATTTGCCAAACAATAACCTTATAGGTATTGTGCCTAAATCACTTCAAAAATTCCC

>M6-PGSC0003DMG400012307-(rev)M6LRR-RLK_704

AATTTGCCAAACAATAACCTTATAGGTATTGTGCCTAAATCACTTCAAAAATTCCCAAAGAATGTGTTCATTAGTAATGATATATCTTTACTTGACTATCCTGTTTCGAATTCTTCTATT

>M6-PGSC0003DMG400012307-(rev)M6LRR-RLK_764

AATGTGTTCATTAGTAATGATATATCTTTACTTGACTATCCTGTTTCGAATTCTTCTATTGTTTCCCTGCCTCAGCAACCAAATCCAAAACTCAAGAATGATGGTAAATTGAGTGAGCGG

>M6-PGSC0003DMG400012443M6LRR-RLK_0

CTTTGTCTAATCTTAATATAACTGTTCCTTTGATTTGAAATCTCAATCTTTATCAGCTTAAATAACAATTGTATAATCTTTTTTTTTCTTTATTAGATTCTGTCCCTCTTTCTTGAAATT

>M6-PGSC0003DMG400012443M6LRR-RLK_1037

GTTCTTGATTTGAGTCATAATGTGTTGCTTGGACCAGTGTTGAGGGTGTTTGATGGATTGGAATCAATCCATTCTCTCAATATATCAAGCAATTTGTTCACTGGAAATTTCAGTGAGTTT

>M6-PGSC0003DMG400012443M6LRR-RLK_1098

AATCAATCCATTCTCTCAATATATCAAGCAATTTGTTCACTGGAAATTTCAGTGAGTTTGGTGAATTCCCTAACCTTGTTGCATTTAACACAAGCAACAATTCGTTTACTGGTAGTTTCA

>M6-PGSC0003DMG400012443M6LRR-RLK_1159

TGAATTCCCTAACCTTGTTGCATTTAACACAAGCAACAATTCGTTTACTGGTAGTTTCAAGTTTGAAATTTGCAGTTTTTCCAAGAAGCTTAAGGTTCTGGATATATCACTTAATCATCT

>M6-PGSC0003DMG400012443M6LRR-RLK_122

CTAAAAGAGTTTGCTTGAGTTGAAAGTTGGGTTCTTGCTTCTCTTACAAGTGATGGGGTGTTGAAAAATCTTATCTTTTTCATTTTTTCTTCCTAGTTTTAGGCTTTGAGTTTTGTAATT

>M6-PGSC0003DMG400012443M6LRR-RLK_1220

TTTGAAATTTGCAGTTTTTCCAAGAAGCTTAAGGTTCTGGATATATCACTTAATCATCTCACTGGTGATCTTGGAGGACTGGATAATTGCAGTTCATTGCTCCAGCAGCTACATGTGGAT

>M6-PGSC0003DMG400012443M6LRR-RLK_1281

CTGGTGATCTTGGAGGACTGGATAATTGCAGTTCATTGCTCCAGCAGCTACATGTGGATTCTAATGATCTTGGGGGTCACCTTCCGGACTCGTTGTATTCAATGACATCTTTGGAGCAAC

>M6-PGSC0003DMG400012443M6LRR-RLK_1342

TAATGATCTTGGGGGTCACCTTCCGGACTCGTTGTATTCAATGACATCTTTGGAGCAACTTTCACTGTCTGCCAATAATTTCTCAGGCCAGCTAAGCCCACAGCTTAGTAAGCTTTCCAA

>M6-PGSC0003DMG400012443M6LRR-RLK_1403

TCACTGTCTGCCAATAATTTCTCAGGCCAGCTAAGCCCACAGCTTAGTAAGCTTTCCAAACTGAAATCCTTAGTTTTATCAGGAAATCGCTTTCATGGTTTGCTTCCTAATGTGTTTGGT

>M6-PGSC0003DMG400012443M6LRR-RLK_1464

TGAAATCCTTAGTTTTATCAGGAAATCGCTTTCATGGTTTGCTTCCTAATGTGTTTGGTAATTTGACATTGTTAGAACAGCTAGCTGCACATTCTAATAGATTTTCAGGACCATTGCCCT

>M6-PGSC0003DMG400012443M6LRR-RLK_1525

TTTGACATTGTTAGAACAGCTAGCTGCACATTCTAATAGATTTTCAGGACCATTGCCCTCTACGATTTCGTATCTTTCTGTGCTTAGGGTGCTTGATCTTAGGAATAATTCTTTGTCTGG

>M6-PGSC0003DMG400012443M6LRR-RLK_1586

ACGATTTCGTATCTTTCTGTGCTTAGGGTGCTTGATCTTAGGAATAATTCTTTGTCTGGTCCTGTTGATCTTGATTTTACTAAATTGACAAGTTTGTGCACTCTTGACCTTGCAACTAAC

>M6-PGSC0003DMG400012443M6LRR-RLK_1647

CTGTTGATCTTGATTTTACTAAATTGACAAGTTTGTGCACTCTTGACCTTGCAACTAACCATTTCAAAGGTAACCTTCCTGTATCGCTCTCTAGTCGGGAGTTGAAAATCATGAGTCTTG

>M6-PGSC0003DMG400012443M6LRR-RLK_1708

TTTCAAAGGTAACCTTCCTGTATCGCTCTCTAGTCGGGAGTTGAAAATCATGAGTCTTGCCAAAAATGAATTCACCGGGCCAATTCCTGAGAACTATGCAAACCTTTCATCCCTTGTGTT

>M6-PGSC0003DMG400012443M6LRR-RLK_1769

AAAAATGAATTCACCGGGCCAATTCCTGAGAACTATGCAAACCTTTCATCCCTTGTGTTCCTCTCATTGTCCAATAATTCCCTTTCGAATTTATCTGGAGCTTTATCTGTTCTGCAGCAC

>M6-PGSC0003DMG400012443M6LRR-RLK_1830

TCTCATTGTCCAATAATTCCCTTTCGAATTTATCTGGAGCTTTATCTGTTCTGCAGCACTGCAGAAATCTTTCAACTCTTATTCTCACCAGGAACTTCCGTGGTGAAGAGATTCCGAAAA

>M6-PGSC0003DMG400012443M6LRR-RLK_1891

CAGAAATCTTTCAACTCTTATTCTCACCAGGAACTTCCGTGGTGAAGAGATTCCGAAAAATGTGAGTGGATTTGAGAACCTGATGATATTTGCATTGGGGAACTGTGGTTTGGAGGGGCG

>M6-PGSC0003DMG400012443M6LRR-RLK_1952

GTGAGTGGATTTGAGAACCTGATGATATTTGCATTGGGGAACTGTGGTTTGGAGGGGCGAATTCCGATATGGTTATACAATTGCAGTAAATTGCAAGTGCTTGACCTGTCATGGAATCAT

>M6-PGSC0003DMG400012443M6LRR-RLK_2013

TTCCGATATGGTTATACAATTGCAGTAAATTGCAAGTGCTTGACCTGTCATGGAATCATTTGGATGGCGAAATTCCTCCTTGGATTGGTGAAATGGAAAAATTGTTCTACTTGGATTTCT

>M6-PGSC0003DMG400012443M6LRR-RLK_2074

GGATGGCGAAATTCCTCCTTGGATTGGTGAAATGGAAAAATTGTTCTACTTGGATTTCTCGAATAATTCACTCACAGGTGAAATCCCAAAAAATTTAACTGATCTAAAGAGCCTCATTTC

>M6-PGSC0003DMG400012443M6LRR-RLK_2135

AATAATTCACTCACAGGTGAAATCCCAAAAAATTTAACTGATCTAAAGAGCCTCATTTCCCCACACAACTACGCATCTAGTCTGAATTCTCCCACTGGTATACCGTTGTTTGTCAAGAGA

>M6-PGSC0003DMG400012443M6LRR-RLK_2196

CACACAACTACGCATCTAGTCTGAATTCTCCCACTGGTATACCGTTGTTTGTCAAGAGAAATCAGAGCGGTAGTGGTTTGCAGTACAATCAGGCTTCAAGCTTCCCTCCGTCTATCTTAT

>M6-PGSC0003DMG400012443M6LRR-RLK_2257

TCAGAGCGGTAGTGGTTTGCAGTACAATCAGGCTTCAAGCTTCCCTCCGTCTATCTTATTGAGTAATAACAGACTAAACGGGACAATCTGGCCCGAAATTGGTCGTCTGAAACAACTTCA

>M6-PGSC0003DMG400012443M6LRR-RLK_2318

AGTAATAACAGACTAAACGGGACAATCTGGCCCGAAATTGGTCGTCTGAAACAACTTCATGTCTTGGATCTCAGTAAGAACAACATTACGGGGACTATTCCTAGCTCGATTTCAAATATG

>M6-PGSC0003DMG400012443M6LRR-RLK_2379

TCTTGGATCTCAGTAAGAACAACATTACGGGGACTATTCCTAGCTCGATTTCAAATATGGGGAACCTGGAAGTTTTGGATCTTTCATGTAATGATCTCAATGGATCAATTCCTGCTTCAT

>M6-PGSC0003DMG400012443M6LRR-RLK_2440

GAACCTGGAAGTTTTGGATCTTTCATGTAATGATCTCAATGGATCAATTCCTGCTTCATTCAATAAGCTCACATTCCTTTCAAAGTTCAATGTAGCTAATAATCACTTGCAGGGAGCGAT

>M6-PGSC0003DMG400012443M6LRR-RLK_2501

AATAAGCTCACATTCCTTTCAAAGTTCAATGTAGCTAATAATCACTTGCAGGGAGCGATTCCAACAGGGGGGCAGTTCTTGAGCTTTCCCAACTCGAGCTTTGAGGGTAATCCTGGACTT

>M6-PGSC0003DMG400012443M6LRR-RLK_2562

CAACAGGGGGGCAGTTCTTGAGCTTTCCCAACTCGAGCTTTGAGGGTAATCCTGGACTTTGTGGAAAAATCATTTCTCCTTGTGCTGCCAGCAATTTGGACCTCCGACCGGCTAGTCCTC

>M6-PGSC0003DMG400012443M6LRR-RLK_2623

TGGAAAAATCATTTCTCCTTGTGCTGCCAGCAATTTGGACCTCCGACCGGCTAGTCCTCTTGCTTCTAGTAGTAGTAGGCTTGGCCGAGGCGGAATTATTGGAATTACAATCAGCATAGG

>M6-PGSC0003DMG400012443M6LRR-RLK_2684

GCTTCTAGTAGTAGTAGGCTTGGCCGAGGCGGAATTATTGGAATTACAATCAGCATAGGGGTAGGAATTGCACTTCTTCTTGCAATAGTGCTGCTTAGAGTGTCTAGAAGAGATGCTGGT

>M6-PGSC0003DMG400012443M6LRR-RLK_2745

TAGGAATTGCACTTCTTCTTGCAATAGTGCTGCTTAGAGTGTCTAGAAGAGATGCTGGTCACCAGATTGGGGACTTCGAGGAAGATTTCAGCAGACCACCTCGATCGTCTGATACTTTTG

>M6-PGSC0003DMG400012443M6LRR-RLK_2806

CCAGATTGGGGACTTCGAGGAAGATTTCAGCAGACCACCTCGATCGTCTGATACTTTTGTTCCTTCTAAGTTGGTACTTTTTCAGAATTCTGATTGCAAGGAACTGACTGTTGCAGACTT

>M6-PGSC0003DMG400012443M6LRR-RLK_2867

CCTTCTAAGTTGGTACTTTTTCAGAATTCTGATTGCAAGGAACTGACTGTTGCAGACTTGCTTAAATCTACAAACAACTTCAACCAGTCGAACATTGTTGGATGTGGAGGATTTGGTCTT

>M6-PGSC0003DMG400012443M6LRR-RLK_2928

TTAAATCTACAAACAACTTCAACCAGTCGAACATTGTTGGATGTGGAGGATTTGGTCTTGTTTACAAGGCGGAACTTCCTAATGGCATAAAGACTGCGATCAAGAGGCTTTCGGGGGATT

>M6-PGSC0003DMG400012443M6LRR-RLK_2989

TTACAAGGCGGAACTTCCTAATGGCATAAAGACTGCGATCAAGAGGCTTTCGGGGGATTGTGGTCAGATGGAGCGTGAATTCCAAGCTGAAGTGGAAGCCCTCTCGAGAGCTCAGCACAA

>M6-PGSC0003DMG400012443M6LRR-RLK_305

GCTTTAATTACAAGTAGTTGGTAGGCTTTTTTCTTTGAGATTTTTGAGGTGTTTTCTTGGGATTTGTGTTATTTGAAGTTTCTGCAACTGGGGTTCTCTTCTTCCTTTACTATAATAATA

>M6-PGSC0003DMG400012443M6LRR-RLK_3050

GGTCAGATGGAGCGTGAATTCCAAGCTGAAGTGGAAGCCCTCTCGAGAGCTCAGCACAAAAACCTTGTATCCCTTCAAGGTTACTGTCAGCATGGGAGTGATAGATTGCTGATATATTCT

>M6-PGSC0003DMG400012443M6LRR-RLK_3111

ACCTTGTATCCCTTCAAGGTTACTGTCAGCATGGGAGTGATAGATTGCTGATATATTCTTACATGGAAAATGGAAGCTTGGACTATTGGCTACATGAAAGAGTCGACGGGAGCTCATTAA

>M6-PGSC0003DMG400012443M6LRR-RLK_3172

CATGGAAAATGGAAGCTTGGACTATTGGCTACATGAAAGAGTCGACGGGAGCTCATTAACATGGGACATTAGGTTAAAGATTGCACAAGGAGCTGCTCGCGGATTAGCCTATTTGCATAA

>M6-PGSC0003DMG400012443M6LRR-RLK_3233

TGGGACATTAGGTTAAAGATTGCACAAGGAGCTGCTCGCGGATTAGCCTATTTGCATAAGGAACCAAATATAGTTCATCGCGACATTAAAACCAGCAACATTCTTTTGAATGAGAGATTT

>M6-PGSC0003DMG400012443M6LRR-RLK_3294

AACCAAATATAGTTCATCGCGACATTAAAACCAGCAACATTCTTTTGAATGAGAGATTTGAAGCTCATCTAGCTGATTTTGGACTATCAAGGCTGTTGCATCCCTATGATACTCATGTCA

>M6-PGSC0003DMG400012443M6LRR-RLK_3355

AGCTCATCTAGCTGATTTTGGACTATCAAGGCTGTTGCATCCCTATGATACTCATGTCACGACAGATCTCGTTGGAACCTTAGGATACATTCCTCCTGAATATAGTCAAACATTAACTGC

>M6-PGSC0003DMG400012443M6LRR-RLK_3416

ACAGATCTCGTTGGAACCTTAGGATACATTCCTCCTGAATATAGTCAAACATTAACTGCTACTTTTCGGGGGGATGTTTACAGCTTTGGTGTTGTTCTACTTGAGCTATTGACAGGCAAG

>M6-PGSC0003DMG400012443M6LRR-RLK_3477

CTTTTCGGGGGGATGTTTACAGCTTTGGTGTTGTTCTACTTGAGCTATTGACAGGCAAGCGTCCCGTGGAAGTATGCAGGGGGAAAAACTGCAGGGACTTGGTATCATGGGTTTTTCAAC

>M6-PGSC0003DMG400012443M6LRR-RLK_3538

TCCCGTGGAAGTATGCAGGGGGAAAAACTGCAGGGACTTGGTATCATGGGTTTTTCAACTGAAATCTGAGAACAGAGTGGAGGAAATATTCGATACATCGATATGGGATACAAGTTACGA

>M6-PGSC0003DMG400012443M6LRR-RLK_3599

AAATCTGAGAACAGAGTGGAGGAAATATTCGATACATCGATATGGGATACAAGTTACGAGAGGCAGCTTTTGGAGGTGTTAACTATAGCTTGTCAATGCATAGTGCAAGATCCACGCCAG

>M6-PGSC0003DMG400012443M6LRR-RLK_366

ATTTGTGTTATTTGAAGTTTCTGCAACTGGGGTTCTCTTCTTCCTTTACTATAATAATAAGGGTGGTGGTGCCCTGCTGCTACCGTTAAGTAGTCTTGATTTTTTCAGTCTTGGGATCTG

>M6-PGSC0003DMG400012443M6LRR-RLK_3660

GGCAGCTTTTGGAGGTGTTAACTATAGCTTGTCAATGCATAGTGCAAGATCCACGCCAGAGGCCCTCGATCGATCAAGTTGTCTTATGGCTCGAGGCAATCGGAAGTGTAAAAGAAAGAT

>M6-PGSC0003DMG400012443M6LRR-RLK_3721

GCCCTCGATCGATCAAGTTGTCTTATGGCTCGAGGCAATCGGAAGTGTAAAAGAAAGATGAGATTCAACTAGTTTTTCCTTGTAGACTTGGCTTCTTTTGCTTACAAAGTGAAAAATATC

>M6-PGSC0003DMG400012443M6LRR-RLK_3782

GATTCAACTAGTTTTTCCTTGTAGACTTGGCTTCTTTTGCTTACAAAGTGAAAAATATCAGTGTAAAGCAACAGAAAATTTCAATTTTATAGAGTTAGTTATAAAACTTATGAACAGCTT

>M6-PGSC0003DMG400012443M6LRR-RLK_3843

TGTAAAGCAACAGAAAATTTCAATTTTATAGAGTTAGTTATAAAACTTATGAACAGCTTTGTTTGGTTCACTTTGTTTCATACACAAAAAATAGTGATAGTTGTGTTTGTATACATATGG

>M6-PGSC0003DMG400012443M6LRR-RLK_3889

TTATGAACAGCTTTGTTTGGTTCACTTTGTTTCATACACAAAAAATAGTGATAGTTGTGTTTGTATACATATGGTTAACTTGGGGACAAGTCCTTTACCAATATACAACTAATTACTCCC

>M6-PGSC0003DMG400012443M6LRR-RLK_427

GGTGGTGGTGCCCTGCTGCTACCGTTAAGTAGTCTTGATTTTTTCAGTCTTGGGATCTGATTTGATTTTTGGTACTAGGGGATCTGAGGAAGGTTCTTTCTTGGTGCTAATTTGCATCAA

>M6-PGSC0003DMG400012443M6LRR-RLK_488

TTGATTTTTGGTACTAGGGGATCTGAGGAAGGTTCTTTCTTGGTGCTAATTTGCATCAAAGGTGGAAACTTTTCTTCATTTGGAACTGTGAATTTGGAATACATGGTGGTGGAATCTTGA

>M6-PGSC0003DMG400012443M6LRR-RLK_549

GTGGAAACTTTTCTTCATTTGGAACTGTGAATTTGGAATACATGGTGGTGGAATCTTGATTTTGAGCTGTTTATATAAAGATTTGTGTGTTTTCATCAATGGTGATTTGGGAGTTTCTGC

>M6-PGSC0003DMG400012443M6LRR-RLK_61

ATAACAATTGTATAATCTTTTTTTTTCTTTATTAGATTCTGTCCCTCTTTCTTGAAATTAACTAAAAGAGTTTGCTTGAGTTGAAAGTTGGGTTCTTGCTTCTCTTACAAGTGATGGGGT

>M6-PGSC0003DMG400012443M6LRR-RLK_610

TTGAGCTGTTTATATAAAGATTTGTGTGTTTTCATCAATGGTGATTTGGGAGTTTCTGCCAATGAGTTTTCTGTGTTGGGTGTTTTTGGCTTATCTTTTTTGTACATCTTTGAGTCTTGA

>M6-PGSC0003DMG400012443M6LRR-RLK_671

ATGAGTTTTCTGTGTTGGGTGTTTTTGGCTTATCTTTTTTGTACATCTTTGAGTCTTGAAACCCCAGTTCAAACCTGTCATCCATATGATTTGTTGGCATTGAAAGAAATTGCTGGCAAT

>M6-PGSC0003DMG400012443M6LRR-RLK_732

CCCCAGTTCAAACCTGTCATCCATATGATTTGTTGGCATTGAAAGAAATTGCTGGCAATCTAACAGATGGGGTTATTCTATCAGCTTGGTCTAATGAACCTAATTGCTGTAAATGGGATG

>M6-PGSC0003DMG400012443M6LRR-RLK_793

AACAGATGGGGTTATTCTATCAGCTTGGTCTAATGAACCTAATTGCTGTAAATGGGATGGGGTTGTCTGTGGTAATGTTTCTGCTCAAAGTAGAGTGATCAGGCTAAATTTGTCAAGAAA

>M6-PGSC0003DMG400012443M6LRR-RLK_854

GTTGTCTGTGGTAATGTTTCTGCTCAAAGTAGAGTGATCAGGCTAAATTTGTCAAGAAAAGGTTTGAGAGGTGTGGTTTCACAGTCCTTGGAGAAATTGGATCAGTTGAAATTGCTCGAT

>M6-PGSC0003DMG400012443M6LRR-RLK_915

GTTTGAGAGGTGTGGTTTCACAGTCCTTGGAGAAATTGGATCAGTTGAAATTGCTCGATCTTTCGCACAATCACTTGGAAGGTGGATTGCCTTTGGACTTGTCCAAATGGAAGCAGTTGG

>M6-PGSC0003DMG400012443M6LRR-RLK_976

TTCGCACAATCACTTGGAAGGTGGATTGCCTTTGGACTTGTCCAAATGGAAGCAGTTGGAAGTTCTTGATTTGAGTCATAATGTGTTGCTTGGACCAGTGTTGAGGGTGTTTGATGGATT

>M6-PGSC0003DMG400012514-(rev)M6LRR-RLK_0

GAAATCGGGATTCTTAATAATTTGGTGATTTTAGCACTGGATACTAACCATATTTCTGGTAATATCCCAATCTCCACATTCAATATCTCCTCATTACAGGTTTTGGTACTGAATCACAAC

>M6-PGSC0003DMG400012514-(rev)M6LRR-RLK_126

ATTGGATCCTTTCCAAGGGAGGTTGGCAATTTGACAGAATTGCAAATTCTGGATCTTGCCGATAATAATCTTACAGGTATTCAAAAGGGGTATGCAAATCATTACTAGCAAGATTTATAA

>M6-PGSC0003DMG400012514-(rev)M6LRR-RLK_1386

TGTTTCAGGATTGATTCCCAAGTCTTTGGAGAAGCTTCTATATCTCAAGTATTTCAACGTTTCATTCAACAAGTTGGTTGGTGAAATCCCCTCGAATGGTCCTTTCAAGAACCTCTCCGG

>M6-PGSC0003DMG400012514-(rev)M6LRR-RLK_189

AATAATCTTACAGGTATTCAAAAGGGGTATGCAAATCATTACTAGCAAGATTTATAATTTCTTAAATCTTAGTAATTACTAACTATACATCTACATGCATACATTGACAGGTGAAATACC

>M6-PGSC0003DMG400012514-(rev)M6LRR-RLK_1890

AGTGGCATTTAGGAGTTTTGATACAGAATGTCAAGTTTTGCGCAACCTTCGTCATAGGAATCTCACCAAAGTCATCACCAGTTGTTCCAACCTTGAATTTAAGGCTTTAATACTCGAGTA

>M6-PGSC0003DMG400012514-(rev)M6LRR-RLK_252

AAATCTTAGTAATTACTAACTATACATCTACATGCATACATTGACAGGTGAAATACCGAAAAAATTTAGCAATCAGATTGAGTTGTTTGATCTTAGTTTCAACAGGTTTACTGGTTCACT

>M6-PGSC0003DMG400012514-(rev)M6LRR-RLK_2583

AATTAATGAAAAAGCTAGATTGTGTGGTATCCATCATGAAAGTGGCAATAGATTGTTGTGTTGAATCTCCAAAAGGAAGGATTGACATGAAAGATGTTGTGGGGAGGTTAAAGACGATCA

>M6-PGSC0003DMG400012514-(rev)M6LRR-RLK_441

TCTCCCATTAAACATAGGTTCTACGTTACCCAACATTGAAGTTATTCATCTGAGAAACTTAAATCTTTATGGGACTATACCTCATTCTCTCTCCAATTGTTCCAAACTCACTAGTCTAGA

>M6-PGSC0003DMG400012514-(rev)M6LRR-RLK_63

ATCCCAATCTCCACATTCAATATCTCCTCATTACAGGTTTTGGTACTGAATCACAACAATCTAATTGGATCCTTTCCAAGGGAGGTTGGCAATTTGACAGAATTGCAAATTCTGGATCTT

>M6-PGSC0003DMG400012594-(rev)M6LRR-RLK_0

ATGCTGTAAGCAGTAACGAAATACTATTTCCATGGACAGAATCCCTCTGCCGCAGATGTTCCCATTGACAATTTAAACACATTTTGAACCTCCTTTTCTTTTCTCTCAAAAGGTGTAGCT

>M6-PGSC0003DMG400012594-(rev)M6LRR-RLK_1037

TTTACATGCTATTCTTGTTATTTCCCGAAGCTCTGAAAACTAGATACTTCTGATGATCGTATATATGAGCATAAATCTCAACCTGCTAACAGTGTGAAGAAGCTTTCTTAGATATTGGGC

>M6-PGSC0003DMG400012594-(rev)M6LRR-RLK_1098

TATATGAGCATAAATCTCAACCTGCTAACAGTGTGAAGAAGCTTTCTTAGATATTGGGCATGAATTTTGTAGCATCTTCTAATACTACTATTCTTCTTCATTAGTCAAGTATTTACTACA

>M6-PGSC0003DMG400012594-(rev)M6LRR-RLK_1159

GAATTTTGTAGCATCTTCTAATACTACTATTCTTCTTCATTAGTCAAGTATTTACTACAATGTACACATGGACTGATTAATTAGCCGAGTTCTGTCTGTTTAAGATTGATCAAACTTTTT

>M6-PGSC0003DMG400012594-(rev)M6LRR-RLK_122

TGGATGCACTTATTAATTTTTAATGTGCTACATAGCTATGTGATGACTAATGTATAGCCCCTTAATTTTTACAGATTGTGTGGTCCCTAAGTTTTAGAACTGATTTGCTTACGACCTCTT

>M6-PGSC0003DMG400012594-(rev)M6LRR-RLK_1281

TTCTTATTGTTATTGTTTTCTTTTTGCTAAATTGAAGGAGGCTTAACAATAACAGTTTGACTGGGCAAATTCCGATTCTTTTAACCACAGTCACTTCACTTCAAGTGCTGTAAGTATAAA

>M6-PGSC0003DMG400012594-(rev)M6LRR-RLK_1342

TGGGCAAATTCCGATTCTTTTAACCACAGTCACTTCACTTCAAGTGCTGTAAGTATAAAGCTGGCACTTTTTTATCATGTGGTTTTTTTCGCCATCAGTTTTTTCTAATTCTTAGGTACT

>M6-PGSC0003DMG400012594-(rev)M6LRR-RLK_1403

TGGCACTTTTTTATCATGTGGTTTTTTTCGCCATCAGTTTTTTCTAATTCTTAGGTACTAATGTTCATTCCTCGCAATGCAGTGATTTGTCAAACAATAATTTAACAGGACCTATTCCAG

>M6-PGSC0003DMG400012594-(rev)M6LRR-RLK_1464

TGTTCATTCCTCGCAATGCAGTGATTTGTCAAACAATAATTTAACAGGACCTATTCCAGTCAATGGTTCCTTCTCGCTTTTTACTCCTATCAGGTCTGTGAAAAGCACTAGTTTATATTT

>M6-PGSC0003DMG400012594-(rev)M6LRR-RLK_1525

AATGGTTCCTTCTCGCTTTTTACTCCTATCAGGTCTGTGAAAAGCACTAGTTTATATTTTAAGTTCTGTCACTGGAATCATGATAATGAAATTCTTACATCTGAATGTCTATATTTCAGT

>M6-PGSC0003DMG400012594-(rev)M6LRR-RLK_1586

AGTTCTGTCACTGGAATCATGATAATGAAATTCTTACATCTGAATGTCTATATTTCAGTTTTGCTAATAATCCATTGGATACTCCTCCACCTTCTCCACCTCCACCTATTTCTCCAACAC

>M6-PGSC0003DMG400012594-(rev)M6LRR-RLK_1647

TGCTAATAATCCATTGGATACTCCTCCACCTTCTCCACCTCCACCTATTTCTCCAACACCCACTTCTCCAGGTTTGTTAAAATAATTTATGTTTTATGGGTTAAGACACTGCCCCCCCCC

>M6-PGSC0003DMG400012594-(rev)M6LRR-RLK_1708

ACTTCTCCAGGTTTGTTAAAATAATTTATGTTTTATGGGTTAAGACACTGCCCCCCCCCCCCCCCCCCCCCCCCCAAAGGAAGAAGGCAAAAGGGCTGACTGTTTTAGGGGAAGAGATTT

>M6-PGSC0003DMG400012594-(rev)M6LRR-RLK_1769

CCCCCCCCCCCCCCAAAGGAAGAAGGCAAAAGGGCTGACTGTTTTAGGGGAAGAGATTTGCACATTATAATGAGTTTTAGTAGTGTATGTGAGATAAATTTTACAGGTGATTTTGGGGTT

>M6-PGSC0003DMG400012594-(rev)M6LRR-RLK_183

TTAATTTTTACAGATTGTGTGGTCCCTAAGTTTTAGAACTGATTTGCTTACGACCTCTTTTTCCTAATTGCAGGGAACTTTATAGTAATAATATAAGTGGAAGAATTCCTTATGAACTGG

>M6-PGSC0003DMG400012594-(rev)M6LRR-RLK_1830

ACATTATAATGAGTTTTAGTAGTGTATGTGAGATAAATTTTACAGGTGATTTTGGGGTTATCTACTGTTTGATAGGTTTGAATAAATAGATTAAACTTACAAGTGATATTGAAGAGTGGA

>M6-PGSC0003DMG400012594-(rev)M6LRR-RLK_1891

CTACTGTTTGATAGGTTTGAATAAATAGATTAAACTTACAAGTGATATTGAAGAGTGGATAGAAGTGGCCTACATCAGGAAATGCAACTGTATGCATTGGATATTGTGACTATAAAGGAC

>M6-PGSC0003DMG400012594-(rev)M6LRR-RLK_1952

GAAGTGGCCTACATCAGGAAATGCAACTGTATGCATTGGATATTGTGACTATAAAGGACCCAAACTTTCAGAATTTATTGTCATACATTCTCTCAATTCTTTCCCCATGTGTCTTATTTC

>M6-PGSC0003DMG400012594-(rev)M6LRR-RLK_2013

AAACTTTCAGAATTTATTGTCATACATTCTCTCAATTCTTTCCCCATGTGTCTTATTTCTCACATGGTATCGGAGCTTTGGTGAAAAATTAGTTTCAGTGCATTTACCAGCAACACGCGG

>M6-PGSC0003DMG400012594-(rev)M6LRR-RLK_2135

GAAAAATCTCCTTGCTGTGCAATTCTCATTGACCATGAATGCTGTCAAAATGTATGTATCCTTGCCGTCTCTGACTACTGTACATAAATTAACACCACCAAAAGGCCGAATAACAAACAA

>M6-PGSC0003DMG400012594-(rev)M6LRR-RLK_2196

TTGCCGTCTCTGACTACTGTACATAAATTAACACCACCAAAAGGCCGAATAACAAACAAGTACCTGTTGTTATGAATAAACTGAAGGATCTTTACCTATTTCTGCATCTTTTCTATTTTG

>M6-PGSC0003DMG400012594-(rev)M6LRR-RLK_2257

ACCTGTTGTTATGAATAAACTGAAGGATCTTTACCTATTTCTGCATCTTTTCTATTTTGTTAGCACTTTAGCTTTACAACATAGTCCTCGGTGCGAGTGTAAATGGTGAAACTACCATAT

>M6-PGSC0003DMG400012594-(rev)M6LRR-RLK_2318

AGCACTTTAGCTTTACAACATAGTCCTCGGTGCGAGTGTAAATGGTGAAACTACCATATCCCTTTTCTCAATTTTTTTTAACTACTTGTGTTAAGTTTTTATCATCCATCAAAAAAAGGG

>M6-PGSC0003DMG400012594-(rev)M6LRR-RLK_2379

CTTTTCTCAATTTTTTTTAACTACTTGTGTTAAGTTTTTATCATCCATCAAAAAAAGGGAATTAAAACTTAAATTTTGAAATTCCTAGGTATGGGAAACAGTGCAACCGGGGCGATTGCT

>M6-PGSC0003DMG400012594-(rev)M6LRR-RLK_244

TCCTAATTGCAGGGAACTTTATAGTAATAATATAAGTGGAAGAATTCCTTATGAACTGGGGAATTTGACGAATTTGGTGAGCTTGGATCTTTACCTGAACAAGTTAGTTGGTCCAATCCC

>M6-PGSC0003DMG400012594-(rev)M6LRR-RLK_2440

TTAAAACTTAAATTTTGAAATTCCTAGGTATGGGAAACAGTGCAACCGGGGCGATTGCTGGCGGTGTTGCTGCAGGGGCTGCTCTTCTGTTTGCAGCTCCTGCGATTTTGCTTGCCTGGT

>M6-PGSC0003DMG400012594-(rev)M6LRR-RLK_2501

CGGTGTTGCTGCAGGGGCTGCTCTTCTGTTTGCAGCTCCTGCGATTTTGCTTGCCTGGTGGCGCCGAAGGAAACCGGAAGACCACTTCTTTGATGTTCCAGGTGGGAAGCCTCAATCAAA

>M6-PGSC0003DMG400012594-(rev)M6LRR-RLK_2562

CGCCGAAGGAAACCGGAAGACCACTTCTTTGATGTTCCAGGTGGGAAGCCTCAATCAAACACCTGAATTAGTTTTTTGTTTTCATTTAATTTTAATTTAATGAGTTCTTGCAATTTTACA

>M6-PGSC0003DMG400012594-(rev)M6LRR-RLK_2623

CCTGAATTAGTTTTTTGTTTTCATTTAATTTTAATTTAATGAGTTCTTGCAATTTTACATTCTTTTGCAGCTGAGGAGGATCCAGAAGTTCATCTTGGACAACTCAAGAGGTTCTCCTTG

>M6-PGSC0003DMG400012594-(rev)M6LRR-RLK_2684

CTTTTGCAGCTGAGGAGGATCCAGAAGTTCATCTTGGACAACTCAAGAGGTTCTCCTTGCGCGAATTACAAGTTGCAACTGACAGTTTTAGCAACAAAAATATACTTGGAAGAGGAGGAT

>M6-PGSC0003DMG400012594-(rev)M6LRR-RLK_2745

CGAATTACAAGTTGCAACTGACAGTTTTAGCAACAAAAATATACTTGGAAGAGGAGGATTTGGCAAGGTTTACAAGGGCAGGTTGGCTGATGGCTCTTTAGTTGCTGTTAAAAGATTAAA

>M6-PGSC0003DMG400012594-(rev)M6LRR-RLK_2806

GGCAAGGTTTACAAGGGCAGGTTGGCTGATGGCTCTTTAGTTGCTGTTAAAAGATTAAAAGAGGAGCGTACTCAAGGCGGGGAGTTACAGTTTCAGACAGAAGTTGAAATGATTAGTATG

>M6-PGSC0003DMG400012594-(rev)M6LRR-RLK_2867

AGGAGCGTACTCAAGGCGGGGAGTTACAGTTTCAGACAGAAGTTGAAATGATTAGTATGGCCGTGCATCGAAATCTACTTCGTTTATGGGGCTTTTGCATGACTGCAACTGAACGGTTGC

>M6-PGSC0003DMG400012594-(rev)M6LRR-RLK_2928

CGTGCATCGAAATCTACTTCGTTTATGGGGCTTTTGCATGACTGCAACTGAACGGTTGCTTGTTTATCCATACATGGCGAATGGAAGTGTTGCTTCACGATTAAGAGGTATCTCTTAGTC

>M6-PGSC0003DMG400012594-(rev)M6LRR-RLK_2989

GTTTATCCATACATGGCGAATGGAAGTGTTGCTTCACGATTAAGAGGTATCTCTTAGTCTTGAATTTTCCATCCTTCTTTTTCTTTGCTCCATTGAGTCGAAATACATCAGATCAAGCAC

>M6-PGSC0003DMG400012594-(rev)M6LRR-RLK_305

AATTTGACGAATTTGGTGAGCTTGGATCTTTACCTGAACAAGTTAGTTGGTCCAATCCCCGACACTTTGGGCAAGCTTCAGAAACTACGTTTCCTGTATGGGTTACACATCCTTTTTCCT

>M6-PGSC0003DMG400012594-(rev)M6LRR-RLK_3050

GAATTTTCCATCCTTCTTTTTCTTTGCTCCATTGAGTCGAAATACATCAGATCAAGCACTTGAAAAAGCTGAAAAATGATAACAACGTTTTTTTTGAACTTTTAAGTTATGGTCTGGTAA

>M6-PGSC0003DMG400012594-(rev)M6LRR-RLK_3111

GAAAAAGCTGAAAAATGATAACAACGTTTTTTTTGAACTTTTAAGTTATGGTCTGGTAATATAAGTTATCTGGAAAAGTTACTGCAGATGTGTACATCATTGTGTAAACTTGGCATTATT

>M6-PGSC0003DMG400012594-(rev)M6LRR-RLK_3172

TAAGTTATCTGGAAAAGTTACTGCAGATGTGTACATCATTGTGTAAACTTGGCATTATTGGATATTGTCGTGGTTGTATTAGGAGTTAGTTTGTTATAGTTAGAATATTCAATCCCGTAG

>M6-PGSC0003DMG400012594-(rev)M6LRR-RLK_3233

ATATTGTCGTGGTTGTATTAGGAGTTAGTTTGTTATAGTTAGAATATTCAATCCCGTAGTTAGAAGCTGTTGTTCTTTGTGATTTTACCCGTGGTCTATCAAGCAATTTCAAGAGCGTTG

>M6-PGSC0003DMG400012594-(rev)M6LRR-RLK_3294

AGAAGCTGTTGTTCTTTGTGATTTTACCCGTGGTCTATCAAGCAATTTCAAGAGCGTTGCTATCCTTGATGTCCTACCAAATATTAAGATTTCCGTTCAAGTTTTATGTTCTGAATTCAT

>M6-PGSC0003DMG400012594-(rev)M6LRR-RLK_3355

ATCCTTGATGTCCTACCAAATATTAAGATTTCCGTTCAAGTTTTATGTTCTGAATTCATAGTGTCAGATATGGAAAGTAATCTTCATTTATGCATGAACCGCATATATCAATTAATTAAA

>M6-PGSC0003DMG400012594-(rev)M6LRR-RLK_3416

TGTCAGATATGGAAAGTAATCTTCATTTATGCATGAACCGCATATATCAATTAATTAAATTGCGTTCAAGCTAAAATGGGAACTTGAACCAGTTTACTTGACATCTTGAACTTGTCCTTA

>M6-PGSC0003DMG400012594-(rev)M6LRR-RLK_3477

GCGTTCAAGCTAAAATGGGAACTTGAACCAGTTTACTTGACATCTTGAACTTGTCCTTACAGTAATTTATTTTTCTTGTAGAGCGGCCAGAGTCAGATCCACCACTTGGCTGGCCAATAA

>M6-PGSC0003DMG400012594-(rev)M6LRR-RLK_3538

GTAATTTATTTTTCTTGTAGAGCGGCCAGAGTCAGATCCACCACTTGGCTGGCCAATAAGGAAGTGTATAGCACTTGGATCTGCAAGAGGCCTTGCTTATTTGCATGATCATTGCGACCC

>M6-PGSC0003DMG400012594-(rev)M6LRR-RLK_3599

AAGTGTATAGCACTTGGATCTGCAAGAGGCCTTGCTTATTTGCATGATCATTGCGACCCTAAGATCATCCACCGTGATGTCAAAGCTGCAAATATATTGTTGGATGAGGAGTATGAAGCA

>M6-PGSC0003DMG400012594-(rev)M6LRR-RLK_3660

AGATCATCCACCGTGATGTCAAAGCTGCAAATATATTGTTGGATGAGGAGTATGAAGCAGTTGTTGGGGATTTTGGGTTAGCCAAACTCATGGACTACAAGGATACTCATGTTACTACTG

>M6-PGSC0003DMG400012594-(rev)M6LRR-RLK_3721

TGTTGGGGATTTTGGGTTAGCCAAACTCATGGACTACAAGGATACTCATGTTACTACTGCTGTACGTGGTACGATTGGGCATATCGCCCCGGAGTATTTATCCACGGGTAAATCTTCAGA

>M6-PGSC0003DMG400012594-(rev)M6LRR-RLK_3782

GTACGTGGTACGATTGGGCATATCGCCCCGGAGTATTTATCCACGGGTAAATCTTCAGAGAAAACTGATGTGTTTGGCTACGGAGTTATGCTTCTAGAGCTCATAACTGGGCAAAGGGCA

>M6-PGSC0003DMG400012594-(rev)M6LRR-RLK_3843

AAACTGATGTGTTTGGCTACGGAGTTATGCTTCTAGAGCTCATAACTGGGCAAAGGGCATTTGATCTTGCTCGACTTGCAAATGATGATGATGTCATGTTGCTAGATTGGGTAAGTATTT

>M6-PGSC0003DMG400012594-(rev)M6LRR-RLK_3904

TGATCTTGCTCGACTTGCAAATGATGATGATGTCATGTTGCTAGATTGGGTAAGTATTTACTAGCCTACATGCTCTAGTTATTCTACTAGCTTATGTCTTTTTCTCTTTTCTTTATTCGT

>M6-PGSC0003DMG400012594-(rev)M6LRR-RLK_4026

AAACTTTATACTACTTACAATTGCTAGTTTTTGTCGTGCAACTGACTTCAAACATGGAAGTGAGCACGGGTGAGAAGATCCCACTTTCCTGCATGCATCTCCCTATAAGCAGAACTCTAT

>M6-PGSC0003DMG400012594-(rev)M6LRR-RLK_4087

GAGCACGGGTGAGAAGATCCCACTTTCCTGCATGCATCTCCCTATAAGCAGAACTCTATCATTGCTGTACCCAGCATGATGAAAAGCTGATTTGTAGACCCACTACAAATTAGGACTGGC

>M6-PGSC0003DMG400012594-(rev)M6LRR-RLK_4148

TTGCTGTACCCAGCATGATGAAAAGCTGATTTGTAGACCCACTACAAATTAGGACTGGCGAATAAGACACTGTCATGTCCCTTCTCTTCCCCTGCCTTTTCCTTTAATATCCCCAATCCC

>M6-PGSC0003DMG400012594-(rev)M6LRR-RLK_4209

ATAAGACACTGTCATGTCCCTTCTCTTCCCCTGCCTTTTCCTTTAATATCCCCAATCCCGTTCCTGTTCTGTTATCGCTTCCCTACCCCCGTATGTAACCCACATCCCATATTCTCGTTT

>M6-PGSC0003DMG400012594-(rev)M6LRR-RLK_4270

TCCTGTTCTGTTATCGCTTCCCTACCCCCGTATGTAACCCACATCCCATATTCTCGTTTTATATCCTCATCCATCAATGAGAAATCTCAGCTTCATCTCATCTCCCTCACTCTTTTGCAC

>M6-PGSC0003DMG400012594-(rev)M6LRR-RLK_4331

TATCCTCATCCATCAATGAGAAATCTCAGCTTCATCTCATCTCCCTCACTCTTTTGCACAGATTACACACACTCATAGCTCTCTTTGTTCTTTACTCAAGTCTCAGGGCTGCAGTTCTGT

>M6-PGSC0003DMG400012594-(rev)M6LRR-RLK_4392

ATTACACACACTCATAGCTCTCTTTGTTCTTTACTCAAGTCTCAGGGCTGCAGTTCTGTTTAAAGTAAAAAGTTTTATATTGACCCCCCCCAAAAAAAAAATAATAATAATAAAAAAATT

>M6-PGSC0003DMG400012594-(rev)M6LRR-RLK_4453

AAAGTAAAAAGTTTTATATTGACCCCCCCCAAAAAAAAAATAATAATAATAAAAAAATTGTTAGCTCTAGTAGGTTTTGCATTGCAGTTTGGCATGTATGATTTCTGTAATCTTTGATAC

>M6-PGSC0003DMG400012594-(rev)M6LRR-RLK_4514

TAGCTCTAGTAGGTTTTGCATTGCAGTTTGGCATGTATGATTTCTGTAATCTTTGATACTACATCATACGAAACATCCAAATTTCCAAAATAATTTGATTCCTGGGACTTGTATTTGTGC

>M6-PGSC0003DMG400012594-(rev)M6LRR-RLK_4575

CATCATACGAAACATCCAAATTTCCAAAATAATTTGATTCCTGGGACTTGTATTTGTGCACATTCTTCATGTAATCTTTTTGTTTCTCTTTCTAGAACTTAACCATCTTGCGAAAACATT

>M6-PGSC0003DMG400012594-(rev)M6LRR-RLK_4636

ATTCTTCATGTAATCTTTTTGTTTCTCTTTCTAGAACTTAACCATCTTGCGAAAACATTTGTTGCATTAGCTGTATGGTCAGGGTGGATCACTTTCAATATCGTTTAGTAGCAGATTGTT

>M6-PGSC0003DMG400012594-(rev)M6LRR-RLK_4697

TTGCATTAGCTGTATGGTCAGGGTGGATCACTTTCAATATCGTTTAGTAGCAGATTGTTCTGGAATTGCTTTTTAGTGTTAAAATCTTTTTCCCTTTGGTTAGCAAAAGCACTCATATAG

>M6-PGSC0003DMG400012594-(rev)M6LRR-RLK_4758

GGAATTGCTTTTTAGTGTTAAAATCTTTTTCCCTTTGGTTAGCAAAAGCACTCATATAGCATATTGCTATATCAGAAGATTTAGATACTTTTTCCCTGCCATATTGACTGTCGCCTTCCG

>M6-PGSC0003DMG400012594-(rev)M6LRR-RLK_4819

TATTGCTATATCAGAAGATTTAGATACTTTTTCCCTGCCATATTGACTGTCGCCTTCCGTGCAGATAAAACTACATCTCTTTTTTTTTCCTTTGTATGGAATTGCTAGGCTATTTCTCTG

>M6-PGSC0003DMG400012594-(rev)M6LRR-RLK_488

TTATTTTCTGAACATTTTGCTTAGTCAAGAAAGGCCACCTTCTATGTTAGTGGCTCTGTACTCTGTAGTACTATTTCTCTTGTTTAGTTATTTGGAAATAGGGATAGACTTTGCTCCTTA

>M6-PGSC0003DMG400012594-(rev)M6LRR-RLK_4880

CAGATAAAACTACATCTCTTTTTTTTTCCTTTGTATGGAATTGCTAGGCTATTTCTCTGTATTTGCATCACAGAGTAGGTGAGATAGAGAATACACTTTACGAGAGGGGACTTCTTTGGG

>M6-PGSC0003DMG400012594-(rev)M6LRR-RLK_4941

TTTGCATCACAGAGTAGGTGAGATAGAGAATACACTTTACGAGAGGGGACTTCTTTGGGAGGATCCTCTAGAAAGGGAACCCACCCTTTAGGACTTGGGAGTGCAGAAGATGGTCAATTC

>M6-PGSC0003DMG400012594-(rev)M6LRR-RLK_5002

GATCCTCTAGAAAGGGAACCCACCCTTTAGGACTTGGGAGTGCAGAAGATGGTCAATTCTTTCACTATGGTTTAAGTATAGAAAATTGAAAGACAAGTGGCTATCCGAACTTGGAAAGTA

>M6-PGSC0003DMG400012594-(rev)M6LRR-RLK_5063

TCACTATGGTTTAAGTATAGAAAATTGAAAGACAAGTGGCTATCCGAACTTGGAAAGTAAATAAAGTGGCTATCTGAACTTGGAAATTCAGTATCTCCCTATAGGGCTGCTTATCAAAAA

>M6-PGSC0003DMG400012594-(rev)M6LRR-RLK_5124

TAAAGTGGCTATCTGAACTTGGAAATTCAGTATCTCCCTATAGGGCTGCTTATCAAAAATCTCAACAACTTCTATGGTGAATGCTTTGTTATTTTAGTGATGAAATTAATTATTTGCTGA

>M6-PGSC0003DMG400012594-(rev)M6LRR-RLK_5185

TCAACAACTTCTATGGTGAATGCTTTGTTATTTTAGTGATGAAATTAATTATTTGCTGACATGATGAGGAAAGTCCGCATGCACTGTGATTTCTTTTTAAAAAATGGCACTCAATTGAAA

>M6-PGSC0003DMG400012594-(rev)M6LRR-RLK_5246

TGATGAGGAAAGTCCGCATGCACTGTGATTTCTTTTTAAAAAATGGCACTCAATTGAAAAAAATGGTTATGTACCTTATTTGACCTCAAAAATTGTGTTAAAATTTCAGGTAAAGGGACT

>M6-PGSC0003DMG400012594-(rev)M6LRR-RLK_5307

AATGGTTATGTACCTTATTTGACCTCAAAAATTGTGTTAAAATTTCAGGTAAAGGGACTTCTTAAGGACGAGAAGTATGAAACATTAGTAGATGCAGATCTTCAGGGTAATTACAATGAA

>M6-PGSC0003DMG400012594-(rev)M6LRR-RLK_5368

TTAAGGACGAGAAGTATGAAACATTAGTAGATGCAGATCTTCAGGGTAATTACAATGAAGAAGAGGTGAAACAGCTTATTCAGGTAGCTCTACTCTGCACACAGAGCTCTCCTATGGAAC

>M6-PGSC0003DMG400012594-(rev)M6LRR-RLK_5429

AGAGGTGAAACAGCTTATTCAGGTAGCTCTACTCTGCACACAGAGCTCTCCTATGGAACGTCCAAAGATGTCGGAAGTGGTCAGAATGCTTGAAGGTGATGGCCTTGCTGAGAGGTGGGA

>M6-PGSC0003DMG400012594-(rev)M6LRR-RLK_549

TCTGTAGTACTATTTCTCTTGTTTAGTTATTTGGAAATAGGGATAGACTTTGCTCCTTATTGGCCAAGTGGCAGACATGAAGATAAAAACATTTCAATCACCTGGAGTATCATTACCAAT

>M6-PGSC0003DMG400012594-(rev)M6LRR-RLK_5490

CCAAAGATGTCGGAAGTGGTCAGAATGCTTGAAGGTGATGGCCTTGCTGAGAGGTGGGAAGAATGGCAGAAGGAAGAGATGTTCCGTCAAGACTTCAACCATGCTCACCATCCTCACACT

>M6-PGSC0003DMG400012594-(rev)M6LRR-RLK_5551

AATGGCAGAAGGAAGAGATGTTCCGTCAAGACTTCAACCATGCTCACCATCCTCACACTGATTGGATTATTGCAGACTCCACTTATAATCTACGACCCGATGAGTTATCAGGGCCAAGAT

>M6-PGSC0003DMG400012594-(rev)M6LRR-RLK_61

CCATTGACAATTTAAACACATTTTGAACCTCCTTTTCTTTTCTCTCAAAAGGTGTAGCTTGTGGATGCACTTATTAATTTTTAATGTGCTACATAGCTATGTGATGACTAATGTATAGCC

>M6-PGSC0003DMG400012594-(rev)M6LRR-RLK_610

GGCCAAGTGGCAGACATGAAGATAAAAACATTTCAATCACCTGGAGTATCATTACCAATGAGACAGCATCATAATTTTCTATGTGGCTTAAATTATTCATAAAGAGAGAGAGAGGAAGAG

>M6-PGSC0003DMG400012594-(rev)M6LRR-RLK_732

GGCGGAAGAAGAAAACCTGAAGCAGCTCATAGAAGTACCCAAAGTTGGTGTCTTAGGGTGTCCAAAGTGCTGGACATGCATGTCTGAAAATTCAAAAAATATACAGGAAGTTAGACACAC

>M6-PGSC0003DMG400012594-(rev)M6LRR-RLK_793

CCAAAGTGCTGGACATGCATGTCTGAAAATTCAAAAAATATACAGGAAGTTAGACACACACATTCTCAATGTGTGTCCAACACCAGTACTTGACAATGAAATTTGTCCATCATCCTTGGC

>M6-PGSC0003DMG400012594-(rev)M6LRR-RLK_854

ATTCTCAATGTGTGTCCAACACCAGTACTTGACAATGAAATTTGTCCATCATCCTTGGCAGTCATCGTTCCTATTTCCCTTATATAGGCTTGCAAAAGGCTAGGTGGATTTGAGTGGTTG

>M6-PGSC0003DMG400012594-(rev)M6LRR-RLK_915

TCATCGTTCCTATTTCCCTTATATAGGCTTGCAAAAGGCTAGGTGGATTTGAGTGGTTGATAAAATTGTGGTTGCTGCTCTCTATAAAGGTTCATTGGCGTGCTGCTGAACAGGATAAAA

>M6-PGSC0003DMG400012594-(rev)M6LRR-RLK_976

AAAATTGTGGTTGCTGCTCTCTATAAAGGTTCATTGGCGTGCTGCTGAACAGGATAAAAAATTTACATGCTATTCTTGTTATTTCCCGAAGCTCTGAAAACTAGATACTTCTGATGATCG

>M6-PGSC0003DMG400012816-(rev)M6RLKmalectin_0

GCAAAAGGGAAATGTATTTCGAGAAATTATAAGCAAATTGAAGAAAATTCAGAGCAGAAATTGGACTTAAAAATGCTCAAATTTCACTTTTTGTTGTTCTTCTTTGCTTTCAAGATTGCT

>M6-PGSC0003DMG400012816-(rev)M6RLKmalectin_1037

GGTTGTGGTTAATTCTATGAGTATGTTATTGGATGTAAGATAGGATTCCTAATTATTCGTAACGTAATTGCACGTGCTCGTCAGTATAACCCTGTATATTTGCAAAATTTCTAATCGCTT

>M6-PGSC0003DMG400012816-(rev)M6RLKmalectin_1098

ACGTAATTGCACGTGCTCGTCAGTATAACCCTGTATATTTGCAAAATTTCTAATCGCTTCACTCCATGATATGATGAAGTATCTTGAGTCTTGTGTTAAAGAGGAAAAAGAAATAAGTGA

>M6-PGSC0003DMG400012816-(rev)M6RLKmalectin_1159

CTCCATGATATGATGAAGTATCTTGAGTCTTGTGTTAAAGAGGAAAAAGAAATAAGTGAGGTGTCTTCACAAACTTAATGAGCAGCTAGAAGTCTGGTTTTTTCATAAGAGATAAATTTT

>M6-PGSC0003DMG400012816-(rev)M6RLKmalectin_122

TTGTCTACCTGGTTAGCTCCTTGTTATGGTTCTCTTTTTCTTTTTGTATAATGTGAATTTGATTAAGGGGTTTAAAGGGTGTCTATATTATGTAGTTATAATGTGTTTGAGTAACTTTTT

>M6-PGSC0003DMG400012816-(rev)M6RLKmalectin_1220

TGTCTTCACAAACTTAATGAGCAGCTAGAAGTCTGGTTTTTTCATAAGAGATAAATTTTTTTGGTGCAAACATACTAGGTGATTCTTCTCATATGTCCCCTGGAATTAGTCGAGGTGCAC

>M6-PGSC0003DMG400012816-(rev)M6RLKmalectin_1403

TTTATAAGACGTTATTGCTTCCCTATATGTACTTTTTCTGCTGTTCTTTCCATTGCTAGCACTGTTACATGAAAAGAAAAAGGCTGCTCCTTTATTATTGATTTGCCTCTTTCAAGTTGC

>M6-PGSC0003DMG400012816-(rev)M6RLKmalectin_1464

CTGTTACATGAAAAGAAAAAGGCTGCTCCTTTATTATTGATTTGCCTCTTTCAAGTTGCTCTTTTGTTTACTAATCTAATTATCTATCCTGTTTATTCACTGCAGCACCATATGTAACGC

>M6-PGSC0003DMG400012816-(rev)M6RLKmalectin_1525

TTTTGTTTACTAATCTAATTATCTATCCTGTTTATTCACTGCAGCACCATATGTAACGCGAATAAGCTGTGGAGCTCAAAATGATGTTCGTAGTTCCCCAACAAATACCCTCTGGCATAA

>M6-PGSC0003DMG400012816-(rev)M6RLKmalectin_1586

ATAAGCTGTGGAGCTCAAAATGATGTTCGTAGTTCCCCAACAAATACCCTCTGGCATAAAGATTTTGGATATACAGGAGGCATACGTACGAAAGCAAACCGCCCAAGTTACATAACACCT

>M6-PGSC0003DMG400012816-(rev)M6RLKmalectin_1647

ATTTTGGATATACAGGAGGCATACGTACGAAAGCAAACCGCCCAAGTTACATAACACCTCCACTCGACACATTACGGTATTTCCCTCTATCAGAGGGTCCTGATAATTGTTACAATATCG

>M6-PGSC0003DMG400012816-(rev)M6RLKmalectin_1708

ACTCGACACATTACGGTATTTCCCTCTATCAGAGGGTCCTGATAATTGTTACAATATCGATAAAGTGCCACATGGCCACTATTCAGTGAGGATCTACTTTGGATTGGTTAAAGAACCAAG

>M6-PGSC0003DMG400012816-(rev)M6RLKmalectin_1769

AAAGTGCCACATGGCCACTATTCAGTGAGGATCTACTTTGGATTGGTTAAAGAACCAAGTTTTGATAACGAACCTCTATTTGATGTTTCTGTTGAAGGGACCTTGGTTTATTCTTTGCCG

>M6-PGSC0003DMG400012816-(rev)M6RLKmalectin_183

ATTAAGGGGTTTAAAGGGTGTCTATATTATGTAGTTATAATGTGTTTGAGTAACTTTTTTTTTTTTGATAACTGTGATGTTTGGGTCAGTTTGTGTGCAGCTTGACTAATGGATGATGTA

>M6-PGSC0003DMG400012816-(rev)M6RLKmalectin_1830

TTGATAACGAACCTCTATTTGATGTTTCTGTTGAAGGGACCTTGGTTTATTCTTTGCCGTCTGGTTGGAGTAATCACGATGATGAACAAGCATTTGTTGAAACTCTTGTATTTCTTGAGA

>M6-PGSC0003DMG400012816-(rev)M6RLKmalectin_1891

TGGTTGGAGTAATCACGATGATGAACAAGCATTTGTTGAAACTCTTGTATTTCTTGAGAATGGTACTACCTCACTTTGCTTCCACAGCACTGGTCATGGAGATCCTGCTATACTCGCAAT

>M6-PGSC0003DMG400012816-(rev)M6RLKmalectin_1952

GGTACTACCTCACTTTGCTTCCACAGCACTGGTCATGGAGATCCTGCTATACTCGCAATCGAGATTCAAGAAGTAGATGAGAAAGCATACTATTCTGGTCCTGGATTTGGTCGAGGGACA

>M6-PGSC0003DMG400012816-(rev)M6RLKmalectin_2013

AGATTCAAGAAGTAGATGAGAAAGCATACTATTCTGGTCCTGGATTTGGTCGAGGGACAATCTTTAGAACTAGCAAAAGATTGAGCTGTGGAGATCAGAATCCAAAGTTTGACGTTGACT

>M6-PGSC0003DMG400012816-(rev)M6RLKmalectin_2074

CTTTAGAACTAGCAAAAGATTGAGCTGTGGAGATCAGAATCCAAAGTTTGACGTTGACTACGATGGGGACCGTTGGGGTGGGGATCGGTTCTGGAGCTCAATTATGACCTTTGGTCAGAA

>M6-PGSC0003DMG400012816-(rev)M6RLKmalectin_2135

GATGGGGACCGTTGGGGTGGGGATCGGTTCTGGAGCTCAATTATGACCTTTGGTCAGAATTCTGATAAGCCTATATCTACTAAAAGCACCATAAAGTCAGCGTCTGTAGCTCCAAACTTC

>M6-PGSC0003DMG400012816-(rev)M6RLKmalectin_2196

CTGATAAGCCTATATCTACTAAAAGCACCATAAAGTCAGCGTCTGTAGCTCCAAACTTCTATCCACAAGCTCTTTACCAAACTGCTCTTGTTAGTACCGACAACGATCCGGATTTAACGC

>M6-PGSC0003DMG400012816-(rev)M6RLKmalectin_2257

TCCACAAGCTCTTTACCAAACTGCTCTTGTTAGTACCGACAACGATCCGGATTTAACGCACACAATAGAAGTCGATCCCAACAAAAACTACTCTGTCTGGTTACACTTTGCCGAAATTGA

>M6-PGSC0003DMG400012816-(rev)M6RLKmalectin_2318

ACAATAGAAGTCGATCCCAACAAAAACTACTCTGTCTGGTTACACTTTGCCGAAATTGATCCTTCAGTTACAGATGTGGGACAAAGGGTTATTGACATTTCGATTAATGGTAACATTGTG

>M6-PGSC0003DMG400012816-(rev)M6RLKmalectin_2379

CTTCAGTTACAGATGTGGGACAAAGGGTTATTGACATTTCGATTAATGGTAACATTGTGTATCGAGATATTGACATTGTGAGAATAGCGGGGGGTGTTAACAGTGCTCTTGTACTGAACA

>M6-PGSC0003DMG400012816-(rev)M6RLKmalectin_244

TTTTTGATAACTGTGATGTTTGGGTCAGTTTGTGTGCAGCTTGACTAATGGATGATGTAAAATTTGTGTTTGAATTCAATTTGTTTTGGTTTGATTGGTTAATGTTAACTCAGAGCAGCA

>M6-PGSC0003DMG400012816-(rev)M6RLKmalectin_2440

TCGAGATATTGACATTGTGAGAATAGCGGGGGGTGTTAACAGTGCTCTTGTACTGAACACAACCATTCCAGTCAGTGGCAGGATTTTGACAATCACATTACATCCAACAACGGGGACTCA

>M6-PGSC0003DMG400012816-(rev)M6RLKmalectin_2501

ACCATTCCAGTCAGTGGCAGGATTTTGACAATCACATTACATCCAACAACGGGGACTCATGCCATCATTAGTGCCATTGAAATTTTTGAGGTTATACCAGTTGAATCTAAAACATTACCA

>M6-PGSC0003DMG400012816-(rev)M6RLKmalectin_2562

CCATCATTAGTGCCATTGAAATTTTTGAGGTTATACCAGTTGAATCTAAAACATTACCAGAGGAAGGTAAGTTAAATGTTTCACTTGTACTTCTTGGATGCTACATGTTTTTTCATCTCA

>M6-PGSC0003DMG400012816-(rev)M6RLKmalectin_2623

GGAAGGTAAGTTAAATGTTTCACTTGTACTTCTTGGATGCTACATGTTTTTTCATCTCATACACTTTACACACTGGAAAGGGGCCAAATGTAGTATGTTATTTTTTCCTTTGTTCCTTGG

>M6-PGSC0003DMG400012816-(rev)M6RLKmalectin_2684

CACTTTACACACTGGAAAGGGGCCAAATGTAGTATGTTATTTTTTCCTTTGTTCCTTGGTCAAGCTTGTGCCTTTCTAATTAAGATAATCTTTATCTTCTTTTCTTTTTCAGTAAAGGCT

>M6-PGSC0003DMG400012816-(rev)M6RLKmalectin_2745

AAGCTTGTGCCTTTCTAATTAAGATAATCTTTATCTTCTTTTCTTTTTCAGTAAAGGCTTTACAGTCACTGAAACTTGCACTTGGACTTCCCCTTCGTTTTGGGTGGAATGGTGATCCAT

>M6-PGSC0003DMG400012816-(rev)M6RLKmalectin_2806

ACAGTCACTGAAACTTGCACTTGGACTTCCCCTTCGTTTTGGGTGGAATGGTGATCCATGTTTTCCGCAACATCCGTGGAGTGGAGTAGATTGTCAGTTTGACAACTCCAGCAATAAATG

>M6-PGSC0003DMG400012816-(rev)M6RLKmalectin_2867

TTTCCGCAACATCCGTGGAGTGGAGTAGATTGTCAGTTTGACAACTCCAGCAATAAATGGGTCATCGATGGGCTGTACGTTCTTGACCTTGACCTTAGATAGAAAAGTTCCTCCCATTCC

>M6-PGSC0003DMG400012816-(rev)M6RLKmalectin_2928

TCATCGATGGGCTGTACGTTCTTGACCTTGACCTTAGATAGAAAAGTTCCTCCCATTCCACACACGTGCACAAGCACACACCTGTAACACTCATTGACATATATTCCTTCTATTGATGTT

>M6-PGSC0003DMG400012816-(rev)M6RLKmalectin_2989

ACACGTGCACAAGCACACACCTGTAACACTCATTGACATATATTCCTTCTATTGATGTTCTGCACCCCAATTTAAAAGTTTTGCTGGACTATACTTGGCTTTCTTTCTATGTTATGTGAT

>M6-PGSC0003DMG400012816-(rev)M6RLKmalectin_305

ATTTGTGTTTGAATTCAATTTGTTTTGGTTTGATTGGTTAATGTTAACTCAGAGCAGCATTTGGACTCAAAATGCTCAAATTCCACTTTTTGCTCTTCTTCTTTGCTTTCAGGATTGATT

>M6-PGSC0003DMG400012816-(rev)M6RLKmalectin_3050

GCACCCCAATTTAAAAGTTTTGCTGGACTATACTTGGCTTTCTTTCTATGTTATGTGATTTTATTATTTTATCTTAATGTGCGTCCTCTTTATCTGTTTTTCCTGTAAGCACAGTGACCT

>M6-PGSC0003DMG400012816-(rev)M6RLKmalectin_3111

TATTATTTTATCTTAATGTGCGTCCTCTTTATCTGTTTTTCCTGTAAGCACAGTGACCTGGACAACCAAGGTTTACGAGGTTTCTTACCAGATGATGTTTCTCAATTGCGTCATCTTCAG

>M6-PGSC0003DMG400012816-(rev)M6RLKmalectin_3172

ACAACCAAGGTTTACGAGGTTTCTTACCAGATGATGTTTCTCAATTGCGTCATCTTCAGAGCATGTAAGTTTAAATGCTGCTTTTTTTGAATATACTATTTGGATGAAAGATGTGTATCT

>M6-PGSC0003DMG400012816-(rev)M6RLKmalectin_3233

CATGTAAGTTTAAATGCTGCTTTTTTTGAATATACTATTTGGATGAAAGATGTGTATCTTCCAACGGTCTTCTACTGCTTAAAATGGAGTTAATGATCAATATGAGTTTTAGCCAATATA

>M6-PGSC0003DMG400012816-(rev)M6RLKmalectin_3294

CAACGGTCTTCTACTGCTTAAAATGGAGTTAATGATCAATATGAGTTTTAGCCAATATATTGACACTTTTTTCAAAATTATTTGAACATAATTTGTCTGAAGGAATTTCATACTAAACGG

>M6-PGSC0003DMG400012816-(rev)M6RLKmalectin_3355

GACACTTTTTTCAAAATTATTTGAACATAATTTGTCTGAAGGAATTTCATACTAAACGGTTAAATAACTTCCAGTTTAAGAGGTTTTCAATATTTGAGCATCTTTTGGAATGTGTTTTCG

>M6-PGSC0003DMG400012816-(rev)M6RLKmalectin_3416

AAATAACTTCCAGTTTAAGAGGTTTTCAATATTTGAGCATCTTTTGGAATGTGTTTTCGTTCATGTATATTGCAATGCTGGCGTCCACTTCTCAAACATGATCTTCTCTGACTTATGCTT

>M6-PGSC0003DMG400012816-(rev)M6RLKmalectin_3477

CATGTATATTGCAATGCTGGCGTCCACTTCTCAAACATGATCTTCTCTGACTTATGCTTTGCTTGTGTGCATTTTACAGAAATTTGAGTGGAAACAACATTCACGGGCCTATTCCCTCAG

>M6-PGSC0003DMG400012816-(rev)M6RLKmalectin_3538

CTTGTGTGCATTTTACAGAAATTTGAGTGGAAACAACATTCACGGGCCTATTCCCTCAGCACTTGGAACAATAACTAGTCTGGAGAAATTGTGAGTACATTTGTGGAGAGTTGTGCTTTT

>M6-PGSC0003DMG400012816-(rev)M6RLKmalectin_3599

CTTGGAACAATAACTAGTCTGGAGAAATTGTGAGTACATTTGTGGAGAGTTGTGCTTTTCTGTACTTGATATTAGATTTCTTAGCTGCCAGAGTAATAGTATATGAAAATATGGAAACAG

>M6-PGSC0003DMG400012816-(rev)M6RLKmalectin_366

TGGACTCAAAATGCTCAAATTCCACTTTTTGCTCTTCTTCTTTGCTTTCAGGATTGATTTTTGTCTACCTGGTTAGGTTCTTTTATGTTTATTTTTTCCTTTTGTATGATGTGAATTTGA

>M6-PGSC0003DMG400012816-(rev)M6RLKmalectin_3660

GTACTTGATATTAGATTTCTTAGCTGCCAGAGTAATAGTATATGAAAATATGGAAACAGAATAGTTTTGTTAACTACTTTGTGAGTATGTTAGGCATTCAATATATAATTATTGTTCATT

>M6-PGSC0003DMG400012816-(rev)M6RLKmalectin_3721

TAGTTTTGTTAACTACTTTGTGAGTATGTTAGGCATTCAATATATAATTATTGTTCATTATCATCTATATGCGCCGCGAGATTAGTGAATGTGCATAATGTTCGTTGGTTGTTTATGTGA

>M6-PGSC0003DMG400012816-(rev)M6RLKmalectin_3782

CATCTATATGCGCCGCGAGATTAGTGAATGTGCATAATGTTCGTTGGTTGTTTATGTGATTAGATCCTCACTGTGTGATTACCTAACAAAAGTTGCTCAACAGGGATCTTTCATACAATT

>M6-PGSC0003DMG400012816-(rev)M6RLKmalectin_3843

AGATCCTCACTGTGTGATTACCTAACAAAAGTTGCTCAACAGGGATCTTTCATACAATTCCTTCAACAGGTCAATTCCTGAAAGCCTTGGGGAATTGACGTCGTTACTGACACTGTAAGA

>M6-PGSC0003DMG400012816-(rev)M6RLKmalectin_3904

TTCAACAGGTCAATTCCTGAAAGCCTTGGGGAATTGACGTCGTTACTGACACTGTAAGATACTTCAACAAGCTCAGTTGAGGACATTTGATGGATCAATAAAGAGCTCTAATTTACTACT

>M6-PGSC0003DMG400012816-(rev)M6RLKmalectin_3965

CTTCAACAAGCTCAGTTGAGGACATTTGATGGATCAATAAAGAGCTCTAATTTACTACTTCATTTATTACAGGAATCTCAACGGCAATTTGCTCTCCGGAAAAATTCCTGCAGCGTTAGG

>M6-PGSC0003DMG400012816-(rev)M6RLKmalectin_4026

ATTTATTACAGGAATCTCAACGGCAATTTGCTCTCCGGAAAAATTCCTGCAGCGTTAGGCGGAAGGCTCTTGCACAGAGCTAGCTTCAAGTATGTTCTCCTAATTGCACGTTCTTAGCAT

>M6-PGSC0003DMG400012816-(rev)M6RLKmalectin_4087

GAAGGCTCTTGCACAGAGCTAGCTTCAAGTATGTTCTCCTAATTGCACGTTCTTAGCATAAGGCTCCGTCTTAGTGAAGCACCCTTCAGAAATTGCATCGTAACTATAGATACATACAAT

>M6-PGSC0003DMG400012816-(rev)M6RLKmalectin_427

TGTCTACCTGGTTAGGTTCTTTTATGTTTATTTTTTCCTTTTGTATGATGTGAATTTGATGAAGGACTTTAAAGGGTGTCTAAATTATGTAGTTTTAACGTGTTTTGAGTAACTTTCTTC

>M6-PGSC0003DMG400012816-(rev)M6RLKmalectin_4453

ATTGAAACGGATAGAACATCACTTTCACGTAGTTTTTCTCGTGTCCAACTTTTTATCGTCAACAATACATACTACTTATGGTCGATATTCAATTTCTAGATGCGTCGATGAAGGTCTCCT

>M6-PGSC0003DMG400012816-(rev)M6RLKmalectin_4514

ACAATACATACTACTTATGGTCGATATTCAATTTCTAGATGCGTCGATGAAGGTCTCCTTGTCATCATTTTGTGTGGATGCTAACTGTGGAAAGAACAAAGAAGTTTATAATCATAGCAT

>M6-PGSC0003DMG400012816-(rev)M6RLKmalectin_4575

TCATCATTTTGTGTGGATGCTAACTGTGGAAAGAACAAAGAAGTTTATAATCATAGCATTTGTTAAAACACACATACAATTATAAGTTAAGGTTAAACAAGTGTACTGTTAAACTCCATT

>M6-PGSC0003DMG400012816-(rev)M6RLKmalectin_4636

GTTAAAACACACATACAATTATAAGTTAAGGTTAAACAAGTGTACTGTTAAACTCCATTGATTTAGCATCCACTATTGGTGGTTAGGCCTAACACCAACTTCCGCAATCCACTAGCTACT

>M6-PGSC0003DMG400012816-(rev)M6RLKmalectin_4697

TTTAGCATCCACTATTGGTGGTTAGGCCTAACACCAACTTCCGCAATCCACTAGCTACTTCAATGGTTGGATGTTAACTATTACCACATAATCAACAAATGAAATATACTAGGGTTCTTT

>M6-PGSC0003DMG400012816-(rev)M6RLKmalectin_4758

AATGGTTGGATGTTAACTATTACCACATAATCAACAAATGAAATATACTAGGGTTCTTTCTGATAGCTAGGTAAACAGGGGTTAGCTAGGGTTCTTTCTGGGCCACATCACTATTCATTC

>M6-PGSC0003DMG400012816-(rev)M6RLKmalectin_4819

GATAGCTAGGTAAACAGGGGTTAGCTAGGGTTCTTTCTGGGCCACATCACTATTCATTCAGGCTATAACCAATTGACGTAAGTCCAAATTCTAACAGTAACATTTAGCATATTTCCTCAA

>M6-PGSC0003DMG400012816-(rev)M6RLKmalectin_4880

GCTATAACCAATTGACGTAAGTCCAAATTCTAACAGTAACATTTAGCATATTTCCTCAAAAGAAATGCTGCGGTAAATAGGTGTCAATGTCATTTGTGTTAAGGCAGAGATGCTAATTAA

>M6-PGSC0003DMG400012816-(rev)M6RLKmalectin_4941

GAAATGCTGCGGTAAATAGGTGTCAATGTCATTTGTGTTAAGGCAGAGATGCTAATTAAACCTTACCATTCATGTGTTGAAACTCATATGACAATTCATCGGTGTATTAACTCGAGGATC

>M6-PGSC0003DMG400012816-(rev)M6RLKmalectin_5002

CTTACCATTCATGTGTTGAAACTCATATGACAATTCATCGGTGTATTAACTCGAGGATCAGATAAAAAACCTGTGACTTTTTTTGTTCATTAGTACTTGTGCTAAATACTTACTTTGGAT

>M6-PGSC0003DMG400012816-(rev)M6RLKmalectin_5063

ATAAAAAACCTGTGACTTTTTTTGTTCATTAGTACTTGTGCTAAATACTTACTTTGGATGCTTTCAGTTTTACTGATAATGCAGGCCTTTGTGGTATTCCTGGATTACCAACATGTGGAC

>M6-PGSC0003DMG400012816-(rev)M6RLKmalectin_5124

TTTCAGTTTTACTGATAATGCAGGCCTTTGTGGTATTCCTGGATTACCAACATGTGGACCACATCTTACTGTTGGTGCCAAAATCGGCATTGGCTTAGGGGCGTGTGTCGCTGTTGTAGT

>M6-PGSC0003DMG400012816-(rev)M6RLKmalectin_5185

CATCTTACTGTTGGTGCCAAAATCGGCATTGGCTTAGGGGCGTGTGTCGCTGTTGTAGTTATCGCTACATGCATAACATGTTGGTGGAAAAGGCGGCAAAATATCCTCCGCGCTAAAAGA

>M6-PGSC0003DMG400012816-(rev)M6RLKmalectin_5246

TCGCTACATGCATAACATGTTGGTGGAAAAGGCGGCAAAATATCCTCCGCGCTAAAAGAATGGCTGGTAAGAGTTTTTCAACATAACATGCTCTATATCCATAGTCAAATGTATTTCCCA

>M6-PGSC0003DMG400012816-(rev)M6RLKmalectin_5307

GGCTGGTAAGAGTTTTTCAACATAACATGCTCTATATCCATAGTCAAATGTATTTCCCATCTCTTTCATGTGTACTGTTGTCAACATGGTGTTTTTGAGAAATGAATTTCGAGTGAAGAT

>M6-PGSC0003DMG400012816-(rev)M6RLKmalectin_5368

TCTTTCATGTGTACTGTTGTCAACATGGTGTTTTTGAGAAATGAATTTCGAGTGAAGATGTATCTCTGCACGATGAAATTAAACGAGTTTCATGCTTTCTCATAGATGAATCACTGTTTT

>M6-PGSC0003DMG400012816-(rev)M6RLKmalectin_5429

ATCTCTGCACGATGAAATTAAACGAGTTTCATGCTTTCTCATAGATGAATCACTGTTTTTTTTTCCTCTTCATTTTTTATACAGTGAGAGACGCTCCTTATGCGAAGAAAAGAACACATT

>M6-PGSC0003DMG400012816-(rev)M6RLKmalectin_5490

TTTCCTCTTCATTTTTTATACAGTGAGAGACGCTCCTTATGCGAAGAAAAGAACACATTTAAACCACATCCAAATGACAAGACACCATCACAATGGTCATCATCACGAACATACTCGGAC

>M6-PGSC0003DMG400012816-(rev)M6RLKmalectin_5551

AACCACATCCAAATGACAAGACACCATCACAATGGTCATCATCACGAACATACTCGGACAGCTGTTGAGAATGGACCTCCTTTGCTTGTCTGAGTTCGAACAATTTTTGAGTCAACTCAC

>M6-PGSC0003DMG400012816-(rev)M6RLKmalectin_5612

CTGTTGAGAATGGACCTCCTTTGCTTGTCTGAGTTCGAACAATTTTTGAGTCAACTCACACTAGAAACTTCTTTCACGTCAAGAGGATTCAACAATTTATACATATACCAACAAAAACAT

>M6-PGSC0003DMG400012816-(rev)M6RLKmalectin_5673

TAGAAACTTCTTTCACGTCAAGAGGATTCAACAATTTATACATATACCAACAAAAACATTGTTCGTATCCTACTTACACAGTATAATTTTTTGACAAAGGAAGGATCATATGAATCTCAT

>M6-PGSC0003DMG400012816-(rev)M6RLKmalectin_5734

TTCGTATCCTACTTACACAGTATAATTTTTTGACAAAGGAAGGATCATATGAATCTCATCCGCAGACCTTGTAACTCCATCGCAAGCTTAGTGAGAAGATTTCCAGATACAATGCTGCTG

>M6-PGSC0003DMG400012816-(rev)M6RLKmalectin_5795

GCAGACCTTGTAACTCCATCGCAAGCTTAGTGAGAAGATTTCCAGATACAATGCTGCTGTTTTGTGGCTAGAAAATGTGTATAGTGGTTATACAGATCAGGAATCAATTTTAAATGCCAT

>M6-PGSC0003DMG400012816-(rev)M6RLKmalectin_5856

TTGTGGCTAGAAAATGTGTATAGTGGTTATACAGATCAGGAATCAATTTTAAATGCCATAGATGTAATTCAGTGCAAGAGACCTATGTTGTTCGGACTCTTCAAAAATGTCGACGGGTGT

>M6-PGSC0003DMG400012816-(rev)M6RLKmalectin_6039

GGAGAGTCCGAGCAACTTAGCAAGAGACAGTGTGTTGTATATTTTAACTATATAAATGTATTCTTGTCATTAGTTTTGGAGCTTTTGGTTAAAGACACTCGTGTAAACTGTGAAATGGTG

>M6-PGSC0003DMG400012816-(rev)M6RLKmalectin_6090

ATAAATGTATTCTTGTCATTAGTTTTGGAGCTTTTGGTTAAAGACACTCGTGTAAACTGTGAAATGGTGAATTACAATAGAACCACAGAATGTGATCGCATTATACGGTCTCAATTGAAC

>M6-PGSC0003DMG400012816-(rev)M6RLKmalectin_61

TGGACTTAAAAATGCTCAAATTTCACTTTTTGTTGTTCTTCTTTGCTTTCAAGATTGCTTTTTGTCTACCTGGTTAGCTCCTTGTTATGGTTCTCTTTTTCTTTTTGTATAATGTGAATT

>M6-PGSC0003DMG400012816-(rev)M6RLKmalectin_610

TGTTTGAATTCGATTTGTTTGGTTTGATTGTCTAATGTTAACTAATTGCAAGTACTTAGCTGTTTGTGGTTGGTTCTATGATTATGTTGTTGGATTTAGTCATGGACCTTATGGTGCCTA

>M6-PGSC0003DMG400012816-(rev)M6RLKmalectin_671

GTTTGTGGTTGGTTCTATGATTATGTTGTTGGATTTAGTCATGGACCTTATGGTGCCTAAATTTTCAATATTGGAATTGAAGCAAATTGAAGAAAACTCATAGCAGCATTCTCAAAAAAA

>M6-PGSC0003DMG400012816-(rev)M6RLKmalectin_732

TTTTCAATATTGGAATTGAAGCAAATTGAAGAAAACTCATAGCAGCATTCTCAAAAAAAAATCCAACTTTTGTTGTTCTTCTTTAACTCTCCCCGTCTCAGCTCCTTTTATGTTTTTTAT

>M6-PGSC0003DMG400012816-(rev)M6RLKmalectin_793

TCCAACTTTTGTTGTTCTTCTTTAACTCTCCCCGTCTCAGCTCCTTTTATGTTTTTTATGCAATGTGAATTTAGTTGTAATTATGTAGCTGTAATTGAGTCTGCATTAAAATGTTTTAGT

>M6-PGSC0003DMG400012816-(rev)M6RLKmalectin_854

AATGTGAATTTAGTTGTAATTATGTAGCTGTAATTGAGTCTGCATTAAAATGTTTTAGTTTTGATTCACTAATCTTAACTTGTGTTGGAAGTGCAACAAGCATGTGAGATGTTGCGATTA

>M6-PGSC0003DMG400012816-(rev)M6RLKmalectin_915

TGATTCACTAATCTTAACTTGTGTTGGAAGTGCAACAAGCATGTGAGATGTTGCGATTAGACTCTTTATATGGTCTTGGAAAATAGTTAGCTCATGTGCTAGATTTTTGCAAGTACTTGG

>M6-PGSC0003DMG400012816-(rev)M6RLKmalectin_976

CTCTTTATATGGTCTTGGAAAATAGTTAGCTCATGTGCTAGATTTTTGCAAGTACTTGGCTGGTTGTGGTTAATTCTATGAGTATGTTATTGGATGTAAGATAGGATTCCTAATTATTCG

>M6-PGSC0003DMG400012817-(rev)M6LRR-RLK_0

GGCTTAATCACTTTTCCAGAAAAAAATCCACTAATCCATTCATCTCCTTCAACAATGCATCTTCAAATCTTGCTTATACTTCTATTACCTACATTGATTCTCTCAATAAACCAAGAATCC

>M6-PGSC0003DMG400012817-(rev)M6LRR-RLK_1037

TTCACGCAATTTATACGAATTAAAGCTCTTTGGTAATAGTTTAAATGGAACTTTACCTAAAGATCTTGGTAAATTTTCTCCATTGGTATGGATTGATGTTTCAAACAATGAGTTTTCGGG

>M6-PGSC0003DMG400012817-(rev)M6LRR-RLK_1098

GATCTTGGTAAATTTTCTCCATTGGTATGGATTGATGTTTCAAACAATGAGTTTTCGGGTGAAATTCCGGTGAATTTGTGCGGAAATGGGGTTTTAGAGGAGGTTTTGATGATAGATAAC

>M6-PGSC0003DMG400012817-(rev)M6LRR-RLK_1159

AAATTCCGGTGAATTTGTGCGGAAATGGGGTTTTAGAGGAGGTTTTGATGATAGATAACTCGTTTTCCGGTGGAATTCCGGTGAGTTTAAGCCAATGCCGGAGCTTATTACGTGTGAGAT

>M6-PGSC0003DMG400012817-(rev)M6LRR-RLK_122

TTATTTACACACCATAAAGCTTGGATTTGATGACCCAAATGGTGTTTTTTCAAACTGGAATCTCCATGATAACTCCTCACCCTGTAACTGGTTTGGAGTAAAATGCGACTCTTTAACTCG

>M6-PGSC0003DMG400012817-(rev)M6LRR-RLK_1220

GTTTTCCGGTGGAATTCCGGTGAGTTTAAGCCAATGCCGGAGCTTATTACGTGTGAGATTAGCTCATAATAAGTTCTCAGGTGATGTCCCTGTGGAATTTTGGGGGCTGCCACGCCTTTC

>M6-PGSC0003DMG400012817-(rev)M6LRR-RLK_1281

GCTCATAATAAGTTCTCAGGTGATGTCCCTGTGGAATTTTGGGGGCTGCCACGCCTTTCGCTGCTTGAGTTAACGGACAATTCATTTTCTGGTGGAATCGTGAAAACTATAGCTGGTGCA

>M6-PGSC0003DMG400012817-(rev)M6LRR-RLK_1342

TGCTTGAGTTAACGGACAATTCATTTTCTGGTGGAATCGTGAAAACTATAGCTGGTGCATCGAATTTATCCGCTTTGATTTTGTCAAAGAACGAATTTTCGGGTACTATTCCTGAAGAGA

>M6-PGSC0003DMG400012817-(rev)M6LRR-RLK_1403

GAATTTATCCGCTTTGATTTTGTCAAAGAACGAATTTTCGGGTACTATTCCTGAAGAGATTGGCTTTTTGGAAAGTTTGGTTGATTTTGTGGGAAATGATAACAAGTTTTCAGGGTCATT

>M6-PGSC0003DMG400012817-(rev)M6LRR-RLK_1464

GGCTTTTTGGAAAGTTTGGTTGATTTTGTGGGAAATGATAACAAGTTTTCAGGGTCATTACCGGTTAGTATAGTGAATCTTGAGCAATTGGGAAGAATGGATTTTCACAACAATGAATTA

>M6-PGSC0003DMG400012817-(rev)M6LRR-RLK_1525

CGGTTAGTATAGTGAATCTTGAGCAATTGGGAAGAATGGATTTTCACAACAATGAATTAAGTGGTAAGTTTCCTAGTGGGGTTCATTCTTTGAAGAAATTGAATGAATTGAACTTGGCAA

>M6-PGSC0003DMG400012817-(rev)M6LRR-RLK_1586

TGGTAAGTTTCCTAGTGGGGTTCATTCTTTGAAGAAATTGAATGAATTGAACTTGGCAAACAATGATCTTTCTGGAGAAATTCCCCGGGAAATCGGGAGCTTGTCTGTTTTGAACTATCT

>M6-PGSC0003DMG400012817-(rev)M6LRR-RLK_1647

AATGATCTTTCTGGAGAAATTCCCCGGGAAATCGGGAGCTTGTCTGTTTTGAACTATCTTGACCTATCAGGAAACAAGTTTTCGGGGGAAATTCCAGTTGCGTTGCAGAATTTGAAGCTC

>M6-PGSC0003DMG400012817-(rev)M6LRR-RLK_1708

ACCTATCAGGAAACAAGTTTTCGGGGGAAATTCCAGTTGCGTTGCAGAATTTGAAGCTCAATCAGCTGAATTTATCGAATAATGGCCTTTCGGGTGGTATTCCTCCTTCATATGCAAAGG

>M6-PGSC0003DMG400012817-(rev)M6LRR-RLK_1769

TCAGCTGAATTTATCGAATAATGGCCTTTCGGGTGGTATTCCTCCTTCATATGCAAAGGGAATGTACAAGAATAGCTTTCTGGGGAATCCCGGTTTATGTGGAGATATTGGAGGCTTATG

>M6-PGSC0003DMG400012817-(rev)M6LRR-RLK_183

CTCCATGATAACTCCTCACCCTGTAACTGGTTTGGAGTAAAATGCGACTCTTTAACTCGTTCTGTTACATCTATTGACCTCTCCAACACCAATATCGCCGGCCCATTTCCGGCTTCTCTT

>M6-PGSC0003DMG400012817-(rev)M6LRR-RLK_1830

ATGTACAAGAATAGCTTTCTGGGGAATCCCGGTTTATGTGGAGATATTGGAGGCTTATGTGATGGAAAAGATGAAGGTAAAACTGCTGGTTATGTATGGTTACTGAGATTGCTTTTCGTA

>M6-PGSC0003DMG400012817-(rev)M6LRR-RLK_1891

ATGGAAAAGATGAAGGTAAAACTGCTGGTTATGTATGGTTACTGAGATTGCTTTTCGTACTTGCTGTTTTGGTGTTTGTAGTTGGGGTAGTTTCGTTCTATTGGAAGTATAGGAATTACA

>M6-PGSC0003DMG400012817-(rev)M6LRR-RLK_1952

TGCTGTTTTGGTGTTTGTAGTTGGGGTAGTTTCGTTCTATTGGAAGTATAGGAATTACAAGAAAGCAAAAAGGTTGGATAGATCGAAATGGACCTTGACGTCGTTTCATAAGTTAGGTTT

>M6-PGSC0003DMG400012817-(rev)M6LRR-RLK_2013

AAAGCAAAAAGGTTGGATAGATCGAAATGGACCTTGACGTCGTTTCATAAGTTAGGTTTCGATGAGTATGAAGTACTGGAAGCTCTAGATGAAGACAACTTGATTGGTAGTGGTTCTTCC

>M6-PGSC0003DMG400012817-(rev)M6LRR-RLK_2074

ATGAGTATGAAGTACTGGAAGCTCTAGATGAAGACAACTTGATTGGTAGTGGTTCTTCCGGGAAGGTTTACAAGGTCGTTTTGAGTAATGGTGAGGCTGCTGCTGTGAAAAAACTTTCAA

>M6-PGSC0003DMG400012817-(rev)M6LRR-RLK_2135

GAAGGTTTACAAGGTCGTTTTGAGTAATGGTGAGGCTGCTGCTGTGAAAAAACTTTCAAGAAGTTTGAAAAAAACAGATGAGAGTTGTGACATTGAGAAAGGTAACTATCAGGATGATAT

>M6-PGSC0003DMG400012817-(rev)M6LRR-RLK_2196

AGTTTGAAAAAAACAGATGAGAGTTGTGACATTGAGAAAGGTAACTATCAGGATGATATATTTGAAGCGGAGGTTGAGACATTGGGCAAGATTCGACACAAGAACATCGTTAGGCTATGG

>M6-PGSC0003DMG400012817-(rev)M6LRR-RLK_2257

TTGAAGCGGAGGTTGAGACATTGGGCAAGATTCGACACAAGAACATCGTTAGGCTATGGTGTTGTTGTACAACAAGGGGTTGCAAACTTTTGGTTTACGAGTACATGCCTAATGGAAGCT

>M6-PGSC0003DMG400012817-(rev)M6LRR-RLK_2318

TTGTTGTACAACAAGGGGTTGCAAACTTTTGGTTTACGAGTACATGCCTAATGGAAGCTTGGGCGATTTGCTACACAGCAGCAAAAGTGGGTTGTTGGATTGGCCTAAGAGATTTAAGAT

>M6-PGSC0003DMG400012817-(rev)M6LRR-RLK_2379

GGCGATTTGCTACACAGCAGCAAAAGTGGGTTGTTGGATTGGCCTAAGAGATTTAAGATAGCTATGGATGCTGCAGAGGGACTCTCATATTTGCATCATGATTGTGCTCCTCCGATTGTT

>M6-PGSC0003DMG400012817-(rev)M6LRR-RLK_244

CTGTTACATCTATTGACCTCTCCAACACCAATATCGCCGGCCCATTTCCGGCTTCTCTTCTTTGCCGGCTCAAGTATATCAAGTACATTTCATTCTACAATAACTCCATTAACTCGACAC

>M6-PGSC0003DMG400012817-(rev)M6LRR-RLK_2440

CTATGGATGCTGCAGAGGGACTCTCATATTTGCATCATGATTGTGCTCCTCCGATTGTTCACAGAGACTTGAAGTCGAACAATATCTTGTTGGACGGGGAGTTTGGAGCTCGAGTTGCTG

>M6-PGSC0003DMG400012817-(rev)M6LRR-RLK_2501

CAGAGACTTGAAGTCGAACAATATCTTGTTGGACGGGGAGTTTGGAGCTCGAGTTGCTGATTTTGGTGTGGCAAAGGCGATGGATGTTGATGACAAGGGAACCACGTCTATGTCAGTCAT

>M6-PGSC0003DMG400012817-(rev)M6LRR-RLK_2562

TTTGGTGTGGCAAAGGCGATGGATGTTGATGACAAGGGAACCACGTCTATGTCAGTCATTGCAGGGTCTTGCGGTTATATTGCTCCAGGTTAGAGGATTAATCCTTCACTTTGGGAGTTC

>M6-PGSC0003DMG400012817-(rev)M6LRR-RLK_2623

CAGGGTCTTGCGGTTATATTGCTCCAGGTTAGAGGATTAATCCTTCACTTTGGGAGTTCTACTAACATGATTTTGTTGAATTTGGTTTATGCCTTCCATTGCAAATTGTAGTTAAGATGT

>M6-PGSC0003DMG400012817-(rev)M6LRR-RLK_2684

CTAACATGATTTTGTTGAATTTGGTTTATGCCTTCCATTGCAAATTGTAGTTAAGATGTATGCCTAAATATAACATATTTCATATATTGTTATATTATAATTAAGCGTCGAATAAGGTCT

>M6-PGSC0003DMG400012817-(rev)M6LRR-RLK_2867

GATTGTTAAACGTCCAATATCTATAATCAGGCAGACAGCTATATATGAGTAATTTTGGTCCATTCTTGAGCTATGTTGCTCAAATCCTTCAAAAAATGTGGCATGTCTTTGTCGGATTCT

>M6-PGSC0003DMG400012817-(rev)M6LRR-RLK_2928

ATTCTTGAGCTATGTTGCTCAAATCCTTCAAAAAATGTGGCATGTCTTTGTCGGATTCTCCCAAAGTTATTCAATTTTGGAGGATCCAATACAGGCGCAACAACATTTTTGGAGGATCCG

>M6-PGSC0003DMG400012817-(rev)M6LRR-RLK_2989

CAAAGTTATTCAATTTTGGAGGATCCAATACAGGCGCAACAACATTTTTGGAGGATCCGAGCAACATAGTACTTGAGTGCCCTTGTACATTAACACTAAGGTTTGCTCCAAGAACAGGGC

>M6-PGSC0003DMG400012817-(rev)M6LRR-RLK_305

TTGCCGGCTCAAGTATATCAAGTACATTTCATTCTACAATAACTCCATTAACTCGACACTTCCGGTGGAGGAGTTATCCGCTTGTAAATCTCTTGTCCATCTAGATTTAGCTCAAAATTT

>M6-PGSC0003DMG400012817-(rev)M6LRR-RLK_3050

CAACATAGTACTTGAGTGCCCTTGTACATTAACACTAAGGTTTGCTCCAAGAACAGGGCATTCTTTGTTTACATTTGCTTTCTGGTTCTGGTAGTGCTCCTCCGGTTGAACTAAATTCTG

>M6-PGSC0003DMG400012817-(rev)M6LRR-RLK_3111

TCTTTGTTTACATTTGCTTTCTGGTTCTGGTAGTGCTCCTCCGGTTGAACTAAATTCTGTTCGTCTTGCAGAATATGCTTACACACTTCAGGTGAACGAGAAGAGTGATATATACAGTTT

>M6-PGSC0003DMG400012817-(rev)M6LRR-RLK_3172

CGTCTTGCAGAATATGCTTACACACTTCAGGTGAACGAGAAGAGTGATATATACAGTTTTGGCGTGGTAATCCTCGAGCTAGTGACAGGGAAACTCCCTGTAGGTCCTGAATACGGGGAA

>M6-PGSC0003DMG400012817-(rev)M6LRR-RLK_3233

GCGTGGTAATCCTCGAGCTAGTGACAGGGAAACTCCCTGTAGGTCCTGAATACGGGGAAAAGGATTTGGTGAAGTGGGTTTGTGCTACTCTAGACCAGAAGGGTAGAGATCATGTTATTG

>M6-PGSC0003DMG400012817-(rev)M6LRR-RLK_3294

GGATTTGGTGAAGTGGGTTTGTGCTACTCTAGACCAGAAGGGTAGAGATCATGTTATTGACCCGAAACTCGACTCTTGTTTCAAGGAGGACATAAGCAAAGTCCTAAAAATTGGCCTCCT

>M6-PGSC0003DMG400012817-(rev)M6LRR-RLK_3355

CCGAAACTCGACTCTTGTTTCAAGGAGGACATAAGCAAAGTCCTAAAAATTGGCCTCCTCTGCACTAGCCCCCTCCCAATTAACCGACCCTCGATGAGAAAAGTCGTAAAAATGCTGCAG

>M6-PGSC0003DMG400012817-(rev)M6LRR-RLK_3416

GCACTAGCCCCCTCCCAATTAACCGACCCTCGATGAGAAAAGTCGTAAAAATGCTGCAGGAAGTTGGTGGCGGAGACCAGCTCAAGACAGCATTAACAGATGGCAAGTTGACCCCTTATT

>M6-PGSC0003DMG400012817-(rev)M6LRR-RLK_3477

AGTTGGTGGCGGAGACCAGCTCAAGACAGCATTAACAGATGGCAAGTTGACCCCTTATTACCACGAAGACGCATCAGATCAAGGAAATGTAGCTTAAAGATAGTCAACATATCTTTGCTT

>M6-PGSC0003DMG400012817-(rev)M6LRR-RLK_3538

CACGAAGACGCATCAGATCAAGGAAATGTAGCTTAAAGATAGTCAACATATCTTTGCTTAGATGTAGCGACATGGACAGTGAAGATTCATAGGCATAGTCGTTGTATCGTGAAAGTGAAG

>M6-PGSC0003DMG400012817-(rev)M6LRR-RLK_3599

ATGTAGCGACATGGACAGTGAAGATTCATAGGCATAGTCGTTGTATCGTGAAAGTGAAGATGTGTGAATTTTTTTATTTGCTAATTGCTTAATACTTTGTCTATCAATACGTGATTTTTG

>M6-PGSC0003DMG400012817-(rev)M6LRR-RLK_3651

AGTGAAGATGTGTGAATTTTTTTATTTGCTAATTGCTTAATACTTTGTCTATCAATACGTGATTTTTGGTTTGATTTTGGAGAGAAGGGGAAAACAGAATTGAGGTGAAATGTAAAAGGA

>M6-PGSC0003DMG400012817-(rev)M6LRR-RLK_366

CCGGTGGAGGAGTTATCCGCTTGTAAATCTCTTGTCCATCTAGATTTAGCTCAAAATTTGTTAGTGGGTAGTCTTCCATCGAGTTTGGCGGAGCTTCCTGAGCTGAAATACCTTGATTTG

>M6-PGSC0003DMG400012817-(rev)M6LRR-RLK_427

TAGTGGGTAGTCTTCCATCGAGTTTGGCGGAGCTTCCTGAGCTGAAATACCTTGATTTGACCGGAAATAACTTTACCGGCGAAATTCCGGCGAGTTTTGGGGCATTCCGGCGACTTGAAG

>M6-PGSC0003DMG400012817-(rev)M6LRR-RLK_488

CGGAAATAACTTTACCGGCGAAATTCCGGCGAGTTTTGGGGCATTCCGGCGACTTGAAGTTCTGGGTCTGGTTGAAAATTTGTTAACTGGGACTATCCCGCTGGAGATTGGAAACATTTC

>M6-PGSC0003DMG400012817-(rev)M6LRR-RLK_549

CTGGGTCTGGTTGAAAATTTGTTAACTGGGACTATCCCGCTGGAGATTGGAAACATTTCGAGTTTGAAGCAGCTGAATTTGTCGTACAACCCGTTTTCTCCGGGTCGGATCCCGCCGGAG

>M6-PGSC0003DMG400012817-(rev)M6LRR-RLK_61

TTCAAATCTTGCTTATACTTCTATTACCTACATTGATTCTCTCAATAAACCAAGAATCCCTTTATTTACACACCATAAAGCTTGGATTTGATGACCCAAATGGTGTTTTTTCAAACTGGA

>M6-PGSC0003DMG400012817-(rev)M6LRR-RLK_610

GTTTGAAGCAGCTGAATTTGTCGTACAACCCGTTTTCTCCGGGTCGGATCCCGCCGGAGATTGGGAACCTCACGAATCTAGAGGTGCTATGGTTAACTGACTGTGGGTTAATCGGTGAGG

>M6-PGSC0003DMG400012817-(rev)M6LRR-RLK_671

TGGGAACCTCACGAATCTAGAGGTGCTATGGTTAACTGACTGTGGGTTAATCGGTGAGGTTCCGGGTACGTTAAGGGGATTAAATAAGCTTGTTAACTTGGACCTTGCGTTAAACAACTT

>M6-PGSC0003DMG400012817-(rev)M6LRR-RLK_732

CCGGGTACGTTAAGGGGATTAAATAAGCTTGTTAACTTGGACCTTGCGTTAAACAACTTGTACGGTCCTATTCCGAGCTGGCTCACTGAGTTAACTAGTGTTGAGCAAATTGAGCTGTAC

>M6-PGSC0003DMG400012817-(rev)M6LRR-RLK_793

ACGGTCCTATTCCGAGCTGGCTCACTGAGTTAACTAGTGTTGAGCAAATTGAGCTGTACAATAACTCGTTCTCTGGCGAGTTTCCGGTGAATGGGTGGTCGAATATGACCTCGTTGAGGC

>M6-PGSC0003DMG400012817-(rev)M6LRR-RLK_854

TAACTCGTTCTCTGGCGAGTTTCCGGTGAATGGGTGGTCGAATATGACCTCGTTGAGGCGGGTCGACCTGTCGATGAATCGGGTTACCGGGTCGATCCCGAGCGGGTTGTGTGAGTTGCC

>M6-PGSC0003DMG400012817-(rev)M6LRR-RLK_915

GTCGACCTGTCGATGAATCGGGTTACCGGGTCGATCCCGAGCGGGTTGTGTGAGTTGCCACTTGAGTCGCTCAATCTTTATGAGAATCAATTGTATGGTGAGTTACCTATAACCATTGCA

>M6-PGSC0003DMG400012817-(rev)M6LRR-RLK_976

TTGAGTCGCTCAATCTTTATGAGAATCAATTGTATGGTGAGTTACCTATAACCATTGCAAATTCACGCAATTTATACGAATTAAAGCTCTTTGGTAATAGTTTAAATGGAACTTTACCTA

>M6-PGSC0003DMG400012929M6LRR-RLK_0

GCTGCCTGCAGTAACACCTTGAAAACCATATCTCATAGAAGCAAAAAACAAGAGTCTGGTATTGTTGCTGGTTAAATGATGAATGTTGTTGTAGTTGTGATCCTTAGTTTTTTCACATTT

>M6-PGSC0003DMG400012929M6LRR-RLK_1037

TGGGATGGAACTAGAGGATTTGTTAAGGGCTTCAGCTTATGTAGTAGGTAAGAGTAGGAGTGGGATAGTCTACAAGGTGGTGGCTGGCGCTACGGCTGTTGCTGTTCGACGATTAAGCGA

>M6-PGSC0003DMG400012929M6LRR-RLK_1098

GGGATAGTCTACAAGGTGGTGGCTGGCGCTACGGCTGTTGCTGTTCGACGATTAAGCGAGGGGGATGTCACGTGGAAGTTCAAGGAGTTTGAGACGGAGGTGGAAGCCATTGGGAGAGTA

>M6-PGSC0003DMG400012929M6LRR-RLK_1159

GGGATGTCACGTGGAAGTTCAAGGAGTTTGAGACGGAGGTGGAAGCCATTGGGAGAGTACAACATCCAAATGTGGTGAGGCTTAGGGCTTATTACTATGCAAGTGATGAGAAATTGCTGG

>M6-PGSC0003DMG400012929M6LRR-RLK_122

AATCTCTGTTATATTTTCATTGAATTCAGATGGTGTTTCTCTGTTAGCTTTGAAATCTGCAATTTCTAATGACCCATCTGGGTTTCTTTCTTCTTGGTCGGAGTTAGACTCATCAGCATG

>M6-PGSC0003DMG400012929M6LRR-RLK_1220

ACATCCAAATGTGGTGAGGCTTAGGGCTTATTACTATGCAAGTGATGAGAAATTGCTGGTAACTGATTTCATCCGCAATGGCACCTTGCACAATGCTTTACATGGTAATCTCACTTCATT

>M6-PGSC0003DMG400012929M6LRR-RLK_1281

ACTGATTTCATCCGCAATGGCACCTTGCACAATGCTTTACATGGTAATCTCACTTCATTCTTACTTGGAATTTTCTTTACTGTTAAAAGAAAATTCACTCTAAACATACTACTTACGGAG

>M6-PGSC0003DMG400012929M6LRR-RLK_1342

TACTTGGAATTTTCTTTACTGTTAAAAGAAAATTCACTCTAAACATACTACTTACGGAGTATTATTCTAAATCACTATAAGGATAAACTTTTAGTAGTGCTGGAGTAAGTTATCCTGCCT

>M6-PGSC0003DMG400012929M6LRR-RLK_1403

TTATTCTAAATCACTATAAGGATAAACTTTTAGTAGTGCTGGAGTAAGTTATCCTGCCTATGCGAGTGAGAAAGGGTGTACTTTTTTGGAGAAAATTAGATCAAATCCTCATAAGCAGTA

>M6-PGSC0003DMG400012929M6LRR-RLK_1464

GCGAGTGAGAAAGGGTGTACTTTTTTGGAGAAAATTAGATCAAATCCTCATAAGCAGTATTATCTCTGTGTCCATGGTACTTCAGGGGTTTTATTAATATATCAGACAACCTTGAAGGTG

>M6-PGSC0003DMG400012929M6LRR-RLK_1525

ATCTCTGTGTCCATGGTACTTCAGGGGTTTTATTAATATATCAGACAACCTTGAAGGTGGCATTGTATTTGATATTTATGCAAAAAAAGAAGTTTAATGTCTACTCCATTGAAGTAAGTG

>M6-PGSC0003DMG400012929M6LRR-RLK_1586

ATTGTATTTGATATTTATGCAAAAAAAGAAGTTTAATGTCTACTCCATTGAAGTAAGTGTTTACAACCTAAGATTCCTTAAAGCTGCTCATGATTATTCGTTGAAGTATTCTCTAGGCAC

>M6-PGSC0003DMG400012929M6LRR-RLK_1647

TACAACCTAAGATTCCTTAAAGCTGCTCATGATTATTCGTTGAAGTATTCTCTAGGCACATTTAAGAATTTCTGTCATGGTTTCAAGCATTCTCTGGTAAATATTAATGATCCCAAATTA

>M6-PGSC0003DMG400012929M6LRR-RLK_1708

TTAAGAATTTCTGTCATGGTTTCAAGCATTCTCTGGTAAATATTAATGATCCCAAATTATATGCTTTTAAGTCGTTAGCTGCCTACTGTCCTTATCTTCTTTTGATCAAGAATAGGCAAC

>M6-PGSC0003DMG400012929M6LRR-RLK_1769

TGCTTTTAAGTCGTTAGCTGCCTACTGTCCTTATCTTCTTTTGATCAAGAATAGGCAACCAAATGTAATTATGATGAACAAGAACTAGTTTGGGGTAGTGGGGTGGAGAGGGGAATAATA

>M6-PGSC0003DMG400012929M6LRR-RLK_183

ATTTCTAATGACCCATCTGGGTTTCTTTCTTCTTGGTCGGAGTTAGACTCATCAGCATGCCATTGGAATGGAATCACATGCGATGAAAATCAGAAGGTGACTTCTATTTCTCTGTCGTCG

>M6-PGSC0003DMG400012929M6LRR-RLK_1830

AATGTAATTATGATGAACAAGAACTAGTTTGGGGTAGTGGGGTGGAGAGGGGAATAATACGTGCATGGTTCATTTGTTTTTATCTGAGCAATTGGCATTTGTGTTTTCTGCTTCCTGGAA

>M6-PGSC0003DMG400012929M6LRR-RLK_1891

TGCATGGTTCATTTGTTTTTATCTGAGCAATTGGCATTTGTGTTTTCTGCTTCCTGGAAAACATCTATGCAAGTGTTAAGCTATTGAGACCTTTTGGGCTCCACAATTCAACATCTGTTG

>M6-PGSC0003DMG400012929M6LRR-RLK_1952

CATCTATGCAAGTGTTAAGCTATTGAGACCTTTTGGGCTCCACAATTCAACATCTGTTGGTTTATTGTAATTCACTAGTTTCTGGTCCCCATTATGGCATTTTCCCCTTGTTTTCTGCTT

>M6-PGSC0003DMG400012929M6LRR-RLK_2013

TTATTGTAATTCACTAGTTTCTGGTCCCCATTATGGCATTTTCCCCTTGTTTTCTGCTTCCTAGGGAGTACATTTTATGTGGCCTAAGATACATTGAGAGAGGCTTTAAAGTTGAAGTAA

>M6-PGSC0003DMG400012929M6LRR-RLK_2074

TAGGGAGTACATTTTATGTGGCCTAAGATACATTGAGAGAGGCTTTAAAGTTGAAGTAAACATTCGGAAGTTACACTAAGGAAAAAAAACATTAGATCTATCAATTGGATGTTAAGCTAT

>M6-PGSC0003DMG400012929M6LRR-RLK_2135

ATTCGGAAGTTACACTAAGGAAAAAAAACATTAGATCTATCAATTGGATGTTAAGCTATTCGTTCGATTCACGTTCATTAGGAATTGAACTAAGCATGGTTACACATAAAATGTGAATCC

>M6-PGSC0003DMG400012929M6LRR-RLK_2196

GTTCGATTCACGTTCATTAGGAATTGAACTAAGCATGGTTACACATAAAATGTGAATCCAAAGTGTGAAATTTTTGCAGAACAAAATATCGACTAATCATAAAATGTTGTAGTTCCTCCT

>M6-PGSC0003DMG400012929M6LRR-RLK_2257

AGTGTGAAATTTTTGCAGAACAAAATATCGACTAATCATAAAATGTTGTAGTTCCTCCTTGTAACATCTATCATTGTCAAGTTCACTTTAGGATGTTGGACTTTATGTAAAAGAGGCATT

>M6-PGSC0003DMG400012929M6LRR-RLK_2318

TAACATCTATCATTGTCAAGTTCACTTTAGGATGTTGGACTTTATGTAAAAGAGGCATTCAGCTGACACCGAGAAGAAAACTATTACACCACAGGTTATTTTTGTCTTCAAATTCTTTTT

>M6-PGSC0003DMG400012929M6LRR-RLK_2379

GCTGACACCGAGAAGAAAACTATTACACCACAGGTTATTTTTGTCTTCAAATTCTTTTTTAACTAAGTGGAGAAGTCCACGACGTTAAGCTTTCTGATAGGTAAATATAAAACAAAAAAG

>M6-PGSC0003DMG400012929M6LRR-RLK_244

ATTGGAATGGAATCACATGCGATGAAAATCAGAAGGTGACTTCTATTTCTCTGTCGTCGAAGAATTTGAGTGGTTACATTCCTTCTGAAATCGGAGCACTTTCTTCTCTATCAATTCTTT

>M6-PGSC0003DMG400012929M6LRR-RLK_2440

ACTAAGTGGAGAAGTCCACGACGTTAAGCTTTCTGATAGGTAAATATAAAACAAAAAAGATAATAAGCAAGACACGAAAACGAAATTTCATCTGATTCCAAGACTTGGTAAGTCATTCCC

>M6-PGSC0003DMG400012929M6LRR-RLK_2501

AATAAGCAAGACACGAAAACGAAATTTCATCTGATTCCAAGACTTGGTAAGTCATTCCCTGTTTCCTTTAGTCTGTTTGTCTTGTTATTCAGAGGTGTCGTTTAATGTACCTATAATTTG

>M6-PGSC0003DMG400012929M6LRR-RLK_2562

TTTCCTTTAGTCTGTTTGTCTTGTTATTCAGAGGTGTCGTTTAATGTACCTATAATTTGTATTGTGGTTATAGAATCCCTGAATAGGCAGTTACTGGACTTGTGTTAGCCTTGTATTTTG

>M6-PGSC0003DMG400012929M6LRR-RLK_2623

TTGTGGTTATAGAATCCCTGAATAGGCAGTTACTGGACTTGTGTTAGCCTTGTATTTTGCTCTCTTTATTTTCAGAACCTTGCACTTCACAAAAACATTAATGTTCTTTTGAACTCCAGT

>M6-PGSC0003DMG400012929M6LRR-RLK_2684

CTCTTTATTTTCAGAACCTTGCACTTCACAAAAACATTAATGTTCTTTTGAACTCCAGTTTGTTTTTCTTAACTTCAATCTTTGTGTACAAGTCTTCTCCACATAATCCTTCCCTTTTGC

>M6-PGSC0003DMG400012929M6LRR-RLK_2745

GTTTTTCTTAACTTCAATCTTTGTGTACAAGTCTTCTCCACATAATCCTTCCCTTTTGCTCGTACTTTGGTATTGTCAATTGTGAACCAATCCTATATAAACACAAGTGGAAATGAAGAT

>M6-PGSC0003DMG400012929M6LRR-RLK_2806

GTACTTTGGTATTGTCAATTGTGAACCAATCCTATATAAACACAAGTGGAAATGAAGATTGCACCTACTTTGGGGTGGCTTCTTGAGAATGTGGTTCCTCTAGTTCATATTGGTAATTTT

>M6-PGSC0003DMG400012929M6LRR-RLK_2867

CACCTACTTTGGGGTGGCTTCTTGAGAATGTGGTTCCTCTAGTTCATATTGGTAATTTTTTGTCTCGACAAAATACCAACTGACCTATTTCTTGTTTTCCTTGACTCAGACTTCCATTGT

>M6-PGSC0003DMG400012929M6LRR-RLK_2928

GTCTCGACAAAATACCAACTGACCTATTTCTTGTTTTCCTTGACTCAGACTTCCATTGTTTCAAGCACATATTTTTGTACCAATTTCAGGATTTGAAGTGAAAAATGTTCTATTTGTTAT

>M6-PGSC0003DMG400012929M6LRR-RLK_2989

CAAGCACATATTTTTGTACCAATTTCAGGATTTGAAGTGAAAAATGTTCTATTTGTTATTCTGAAAAGCTGTCATCGATTTGGCATGAATCACTTTTTAATTTATCTTCCACAAGTCCAA

>M6-PGSC0003DMG400012929M6LRR-RLK_305

GAATTTGAGTGGTTACATTCCTTCTGAAATCGGAGCACTTTCTTCTCTATCAATTCTTTCTCTTTCCTACAACAACTTTTCAAAACCAATCCCTATTCGTCTCTTCAATGCTACTTCCCT

>M6-PGSC0003DMG400012929M6LRR-RLK_3050

TGAAAAGCTGTCATCGATTTGGCATGAATCACTTTTTAATTTATCTTCCACAAGTCCAAAAATATTTTTTGGATTTCAAAACTTGCATTTTCCGGCAAAAAATTTCTCATGGAAAAAAAC

>M6-PGSC0003DMG400012929M6LRR-RLK_3111

ATATTTTTTGGATTTCAAAACTTGCATTTTCCGGCAAAAAATTTCTCATGGAAAAAAACACATACACAACTAATTTCTCTAGAGAACTGGGTGATTTTCTTTTTATGTTGGAGCATTATC

>M6-PGSC0003DMG400012929M6LRR-RLK_3172

ATACACAACTAATTTCTCTAGAGAACTGGGTGATTTTCTTTTTATGTTGGAGCATTATCCTTGTTAATCAATCTTTTTATGTTTATTTTTTACAATTGCAGGAGGACTTGGTAATAACTT

>M6-PGSC0003DMG400012929M6LRR-RLK_3233

TGTTAATCAATCTTTTTATGTTTATTTTTTACAATTGCAGGAGGACTTGGTAATAACTTGCCACCTCTGTCATGGGCAGCTAGATTGAAGATTGCTCAAGGCACAGCTCGGGGTCTAATG

>M6-PGSC0003DMG400012929M6LRR-RLK_3294

CACCTCTGTCATGGGCAGCTAGATTGAAGATTGCTCAAGGCACAGCTCGGGGTCTAATGCACATTCATGAGTGTAATCCCAGGAAGTATGTTCATGGGAACATAAATTCAACAAAAATCC

>M6-PGSC0003DMG400012929M6LRR-RLK_3355

CATTCATGAGTGTAATCCCAGGAAGTATGTTCATGGGAACATAAATTCAACAAAAATCCTTCTTGATGATGACTTGCAAGCCTACATATCTGGTTTTGGATTGACTCGCCTTGTTTCAGG

>M6-PGSC0003DMG400012929M6LRR-RLK_3416

CTTGATGATGACTTGCAAGCCTACATATCTGGTTTTGGATTGACTCGCCTTGTTTCAGGTTGCTCCAAGCCCATAAACAGTAGTACCAAAAAGCTAAGTACAAGTCAAATCATAATAAGT

>M6-PGSC0003DMG400012929M6LRR-RLK_3477

GCTCCAAGCCCATAAACAGTAGTACCAAAAAGCTAAGTACAAGTCAAATCATAATAAGTCCTCAAAACTCAACTCCGTCTTGTACTATGTACATGGCACCTGAGGGTCGAGTGCCTGGCT

>M6-PGSC0003DMG400012929M6LRR-RLK_3538

TCAAAACTCAACTCCGTCTTGTACTATGTACATGGCACCTGAGGGTCGAGTGCCTGGCTCCAAGTTCACCCAGAAAAGCGACATCTACTCATTTGGTATGGTACTTTTGGAGATTCTGAC

>M6-PGSC0003DMG400012929M6LRR-RLK_3599

AAGTTCACCCAGAAAAGCGACATCTACTCATTTGGTATGGTACTTTTGGAGATTCTGACTGGTCAGTTGACAGATGGAGGATCGGAGGTGGAGGATGATGGGAAGGGGCTTGAGAGTTTT

>M6-PGSC0003DMG400012929M6LRR-RLK_366

CTTTCCTACAACAACTTTTCAAAACCAATCCCTATTCGTCTCTTCAATGCTACTTCCCTTCACTCCCTTGACCTCTCTAACAATGCTTTCACAGGATTCTTACCTCAACAAATCACATCC

>M6-PGSC0003DMG400012929M6LRR-RLK_3660

GTCAGTTGACAGATGGAGGATCGGAGGTGGAGGATGATGGGAAGGGGCTTGAGAGTTTTGTTAGAAAGGTTTTTAGACAAGAACGACCCTTGTCTGAGATCATAGACCCTGCTCTCTTGA

>M6-PGSC0003DMG400012929M6LRR-RLK_3721

TAGAAAGGTTTTTAGACAAGAACGACCCTTGTCTGAGATCATAGACCCTGCTCTCTTGAATGAGGTTCATGCTAAAAAGCAAGTTGTTGCAGCATTTCATATCGCTCTCAACTGCACAGA

>M6-PGSC0003DMG400012929M6LRR-RLK_3782

GAGGTTCATGCTAAAAAGCAAGTTGTTGCAGCATTTCATATCGCTCTCAACTGCACAGAACTGGATCCTGAGCTTCGGCCAAGGATGAGAACAGTTTCTGATAACCTTGATCGCATCAAA

>M6-PGSC0003DMG400012929M6LRR-RLK_3843

TGGATCCTGAGCTTCGGCCAAGGATGAGAACAGTTTCTGATAACCTTGATCGCATCAAACTGCAGTAAAAGAAACTTGATCAATGTAAAGGCTTTCAATGTTTTGCTGACCGCCTATTGT

>M6-PGSC0003DMG400012929M6LRR-RLK_3904

GCAGTAAAAGAAACTTGATCAATGTAAAGGCTTTCAATGTTTTGCTGACCGCCTATTGTAGAGCTGTTTAGTAATTGTGAATCAGAATTTTCAGTTTGTGTTGTGCCAAATCCTAGATTT

>M6-PGSC0003DMG400012929M6LRR-RLK_3965

AGCTGTTTAGTAATTGTGAATCAGAATTTTCAGTTTGTGTTGTGCCAAATCCTAGATTTGGCGTTTCTATTGTAACTCTACTAATTCAATGTTCTATTTGTTGAATCATCAATTGAAAAT

>M6-PGSC0003DMG400012929M6LRR-RLK_4026

CGTTTCTATTGTAACTCTACTAATTCAATGTTCTATTTGTTGAATCATCAATTGAAAATTGCTTGTGCTAGCGAATCTTACAAGGCTATTTGTGTTTGATGTACGGACATGATCTGAATT

>M6-PGSC0003DMG400012929M6LRR-RLK_4087

CTTGTGCTAGCGAATCTTACAAGGCTATTTGTGTTTGATGTACGGACATGATCTGAATTCCAGATAACATGCAAAGGAGCATGGTCAGACCCTGCCTATTCAAATGATGTAGTTCAATAG

>M6-PGSC0003DMG400012929M6LRR-RLK_4148

AGATAACATGCAAAGGAGCATGGTCAGACCCTGCCTATTCAAATGATGTAGTTCAATAGTTGGAAAGTTCTGTATGAATTCTTGATTGCCCATTACCCTATTCAATCTTTTGAAACTACA

>M6-PGSC0003DMG400012929M6LRR-RLK_4189

AATGATGTAGTTCAATAGTTGGAAAGTTCTGTATGAATTCTTGATTGCCCATTACCCTATTCAATCTTTTGAAACTACACTCCACCTCTATTCTCTCATTCTACCAAGTGAAGTTGCTAC

>M6-PGSC0003DMG400012929M6LRR-RLK_427

ACTCCCTTGACCTCTCTAACAATGCTTTCACAGGATTCTTACCTCAACAAATCACATCCCTTATAAACCTTAAACACCTTGATCTTTCTTCTAATTTTCTCAATGGCTCACTTCCCCAAG

>M6-PGSC0003DMG400012929M6LRR-RLK_488

TATAAACCTTAAACACCTTGATCTTTCTTCTAATTTTCTCAATGGCTCACTTCCCCAAGAGCTAACTCATCTTACACTTCTCACTGGAACTTTAAATCTTTCATACAACAGATTTACCGG

>M6-PGSC0003DMG400012929M6LRR-RLK_549

CTAACTCATCTTACACTTCTCACTGGAACTTTAAATCTTTCATACAACAGATTTACCGGCCAACTTCCGGCGAGTTATGGTAAGTTTCCTGTAACATTGAGTTTAGACCTTCGGCATAAC

>M6-PGSC0003DMG400012929M6LRR-RLK_61

TTGTTGCTGGTTAAATGATGAATGTTGTTGTAGTTGTGATCCTTAGTTTTTTCACATTTTCAATCTCTGTTATATTTTCATTGAATTCAGATGGTGTTTCTCTGTTAGCTTTGAAATCTG

>M6-PGSC0003DMG400012929M6LRR-RLK_610

AACTTCCGGCGAGTTATGGTAAGTTTCCTGTAACATTGAGTTTAGACCTTCGGCATAACAATCTGACCGGAAAGATACCTTCAATCGGGTCGTTGTTGAATCAAGGGCCTACTGCATTTT

>M6-PGSC0003DMG400012929M6LRR-RLK_671

TCTGACCGGAAAGATACCTTCAATCGGGTCGTTGTTGAATCAAGGGCCTACTGCATTTTCCGGTAACCCTTTTCTTTGTGGGTTTCCATTGGAGACTCCGTGTGCAGAACCAGAAGCTCA

>M6-PGSC0003DMG400012929M6LRR-RLK_732

GGTAACCCTTTTCTTTGTGGGTTTCCATTGGAGACTCCGTGTGCAGAACCAGAAGCTCAAAAACCACCACCGAACCCGAACCCTAATGAGGAGGAGGTGGGGTTTGTAGGAAAAGGGAAA

>M6-PGSC0003DMG400012929M6LRR-RLK_793

AACCACCACCGAACCCGAACCCTAATGAGGAGGAGGTGGGGTTTGTAGGAAAAGGGAAATCGGGAAATGGGTCAGTGGTGGTTTCTTTGATCTCCGGCGTATCGGTTGTGATTGGGGTGA

>M6-PGSC0003DMG400012929M6LRR-RLK_854

GGGAAATGGGTCAGTGGTGGTTTCTTTGATCTCCGGCGTATCGGTTGTGATTGGGGTGATGTTTGTTTCAGTGTTGGTGTTGAGGAGAAAATGGAAGTTGGATGAGGGAAAAATGGGAAA

>M6-PGSC0003DMG400012929M6LRR-RLK_915

TTTGTTTCAGTGTTGGTGTTGAGGAGAAAATGGAAGTTGGATGAGGGAAAAATGGGAAAAGAAAAGTTGGAGAAGCCAGAGGAGGGGCAAAAGGGTAAGTATGTAGTGTTAGACGAGGGG

>M6-PGSC0003DMG400012929M6LRR-RLK_976

AAAAGTTGGAGAAGCCAGAGGAGGGGCAAAAGGGTAAGTATGTAGTGTTAGACGAGGGGTTTGGGATGGAACTAGAGGATTTGTTAAGGGCTTCAGCTTATGTAGTAGGTAAGAGTAGGA

>M6-PGSC0003DMG400012955-(rev)M6LRR-RLK_0

CACCCCATTCCAACAGAGAAATACAGAGAGAAAGAGCAAAATACAAAAGAGACACCAACTCATTTGCCACTCTACCCCAAATATAAAAACCCTAATTTCCCTCTTTTTTTCTCCTTTCCC

>M6-PGSC0003DMG400012955-(rev)M6LRR-RLK_1037

TTTCCTGGAAAATTGTTGGAGGCATTTGAGAATTTGATGGTTTTGGATTTGAGGAATAATGCTATAATGGGGCAGCTTCCTTCTAACGGGTTTATGCATAATTTGAGGGTTTTAAGGCTT

>M6-PGSC0003DMG400012955-(rev)M6LRR-RLK_1098

CTATAATGGGGCAGCTTCCTTCTAACGGGTTTATGCATAATTTGAGGGTTTTAAGGCTTGGGAATAATAAGCTTTATGGATTGATTCCCGATGAGCTCTTGCAGGGAACAGGTCCATTGG

>M6-PGSC0003DMG400012955-(rev)M6LRR-RLK_1159

GAATAATAAGCTTTATGGATTGATTCCCGATGAGCTCTTGCAGGGAACAGGTCCATTGGAAGAACTGGATCTTAGTGGCAATGGATTTTCAGGTTGGGAAATGGATATCAGTTTTTTGGA

>M6-PGSC0003DMG400012955-(rev)M6LRR-RLK_122

CTCTTTCCCCACACCCCAACCCCTCTCTTACCACCCCACCTACCCTAATCTCTCTCTCTGTCTCTGAACTCTTTCTTGGATTTCCATGGTGCTCTCTCCATAGTCATCCTTAAGCTTTTC

>M6-PGSC0003DMG400012955-(rev)M6LRR-RLK_1220

GAACTGGATCTTAGTGGCAATGGATTTTCAGGTTGGGAAATGGATATCAGTTTTTTGGAATATTTACATTTCTTAGATCAGTATATATTGTTGTTGTTGAATGATCATAATCAAATACAC

>M6-PGSC0003DMG400012955-(rev)M6LRR-RLK_1281

ATTTACATTTCTTAGATCAGTATATATTGTTGTTGTTGAATGATCATAATCAAATACACCTTTTATTTGTATAGTGTTTTTGTGGATGCCTTTAACTTCAAGCCTCTTAATTTTTGGAGG

>M6-PGSC0003DMG400012955-(rev)M6LRR-RLK_1342

TTTATTTGTATAGTGTTTTTGTGGATGCCTTTAACTTCAAGCCTCTTAATTTTTGGAGGTGGACTAAAATTGGAAACTTTCTTCTGTTTTAGGCATTGAAACTTTTACAGTTATGCAAAG

>M6-PGSC0003DMG400012955-(rev)M6LRR-RLK_1586

TGTAAAATCTCGTTCTTTTTGTTTTCTTGTCAGGTTATAATTAGTTTTCTTGTGTGAGTAGTGGTTGACAATCTATATAACAGAGAATAGAAATCTTCAAGGTCTTTCAGCTTATGCCAA

>M6-PGSC0003DMG400012955-(rev)M6LRR-RLK_1647

TGGTTGACAATCTATATAACAGAGAATAGAAATCTTCAAGGTCTTTCAGCTTATGCCAAATACAGAAAAGTTTGCTTTTTTGAGTAAGTAGGTTTTTTAAAAAATGTTTTAACAAAACCA

>M6-PGSC0003DMG400012955-(rev)M6LRR-RLK_1708

ACAGAAAAGTTTGCTTTTTTGAGTAAGTAGGTTTTTTAAAAAATGTTTTAACAAAACCAGATTTTGAACAAACTTTTACCGCAAGCTTTTTTTGTTTGCCTCAACAGAACGTGAGCAGCG

>M6-PGSC0003DMG400012955-(rev)M6LRR-RLK_1769

TTTTGAACAAACTTTTACCGCAAGCTTTTTTTGTTTGCCTCAACAGAACGTGAGCAGCGTGGACTGTATAGACTAGTTTGAATTGCAAAGCATGTTCAATAGAACTTGCCCATAATAAAT

>M6-PGSC0003DMG400012955-(rev)M6LRR-RLK_183

CTCTGAACTCTTTCTTGGATTTCCATGGTGCTCTCTCCATAGTCATCCTTAAGCTTTTCTTCCTTTTACCCTTTTTCAGACTTTCCTTCTTTTTCTCAATGCCTCATTTTTAGCACTTAA

>M6-PGSC0003DMG400012955-(rev)M6LRR-RLK_1830

GACTGTATAGACTAGTTTGAATTGCAAAGCATGTTCAATAGAACTTGCCCATAATAAATAGTCAACAAAGTGTTTCCTAATTGTAGTAAATTTAATAAAAGCCCCTAATGGAATAGTTGC

>M6-PGSC0003DMG400012955-(rev)M6LRR-RLK_1891

TCAACAAAGTGTTTCCTAATTGTAGTAAATTTAATAAAAGCCCCTAATGGAATAGTTGCATGAATGATTGCATCATTAGTAGCCATAAATAATCCTTTTTGCAATGGTATAACTGTATGC

>M6-PGSC0003DMG400012955-(rev)M6LRR-RLK_1952

GAATGATTGCATCATTAGTAGCCATAAATAATCCTTTTTGCAATGGTATAACTGTATGCTCAAAGTTCATGGCAAGCTAGCTGGAGCAATATTTAGCCCAGTGTCATCAAAGGCGCTTAA

>M6-PGSC0003DMG400012955-(rev)M6LRR-RLK_2135

GTCATCAAGCTTCTAAGGCATACTTTTCCTTAAGTTTATATATTTGTAATTCATGCTTAGGACTAAAAATAAATATATTTATTTTTTCTCCCTTTTCGCCTTTTTCCATTAAAGCCCATA

>M6-PGSC0003DMG400012955-(rev)M6LRR-RLK_2257

TGTGCTTAAAGCCCAATCAAACCTTAGAGCTTTTTTACTCTTTTTGCTTTCGATAAGACTGATTTAGCCTTTATCTTGAAAAATTGAAAATGTTGAATTTTTTCTTCTGCTTCCAATAGC

>M6-PGSC0003DMG400012955-(rev)M6LRR-RLK_2318

ATTTAGCCTTTATCTTGAAAAATTGAAAATGTTGAATTTTTTCTTCTGCTTCCAATAGCCAATGTTATAGCACAATATTTGTACTTACAGTAAAATGAGTTTTATTTCATTAATGAGCTA

>M6-PGSC0003DMG400012955-(rev)M6LRR-RLK_2379

ATGTTATAGCACAATATTTGTACTTACAGTAAAATGAGTTTTATTTCATTAATGAGCTATTTGCTCTTAATATGCTCATCTTTCTATTTTGTTATCATTGCATTACACCGTCCAGGGCTT

>M6-PGSC0003DMG400012955-(rev)M6LRR-RLK_244

CCTTTTACCCTTTTTCAGACTTTCCTTCTTTTTCTCAATGCCTCATTTTTAGCACTTAACTCACATTCTCAACCCAATGACTCTCTCTCAATCTCTCTCTTTACTCCTCCTTGTACTTGT

>M6-PGSC0003DMG400012955-(rev)M6LRR-RLK_2440

TGCTCTTAATATGCTCATCTTTCTATTTTGTTATCATTGCATTACACCGTCCAGGGCTTCCCCCTTCCCTCTAGTATAGGCCGCTCATGATGACGTTGGGCCTTGGAATTGGAAAATAAT

>M6-PGSC0003DMG400012955-(rev)M6LRR-RLK_2501

CCCTTCCCTCTAGTATAGGCCGCTCATGATGACGTTGGGCCTTGGAATTGGAAAATAATTGATGACAACATTTCTTTTTTGATAAGGAATTGATGACAACATTCCTGTCTTCATTATTAT

>M6-PGSC0003DMG400012955-(rev)M6LRR-RLK_2562

ATGACAACATTTCTTTTTTGATAAGGAATTGATGACAACATTCCTGTCTTCATTATTATCAAATCAGTACTCAGATTAGAGAAAAGAATGGAGTCGGTGAGACTTGATTATTGAAGAAAA

>M6-PGSC0003DMG400012955-(rev)M6LRR-RLK_2623

AATCAGTACTCAGATTAGAGAAAAGAATGGAGTCGGTGAGACTTGATTATTGAAGAAAAGCAGAGCAAATTTACAAGGGTGATAATTTGGGTCTCTGAGCTTAAAACTCCTCGGTGACCC

>M6-PGSC0003DMG400012955-(rev)M6LRR-RLK_2684

AGAGCAAATTTACAAGGGTGATAATTTGGGTCTCTGAGCTTAAAACTCCTCGGTGACCCTCTGATAGGGAAGTTTGTTCTGTTTCATGAGCTACAGAGAGGGTTCAAAATTTCATATTTC

>M6-PGSC0003DMG400012955-(rev)M6LRR-RLK_2745

TGATAGGGAAGTTTGTTCTGTTTCATGAGCTACAGAGAGGGTTCAAAATTTCATATTTCTTTAGTTACACTTGGATATCTTTCATCATAGGACCAGAGCGTGGTTCCATTTAGAGCCTAA

>M6-PGSC0003DMG400012955-(rev)M6LRR-RLK_2806

TAGTTACACTTGGATATCTTTCATCATAGGACCAGAGCGTGGTTCCATTTAGAGCCTAAATGTGGAGATCCTCTGTTCAGAAGTTGAATGGTGCCGTTGACTGATGACTGATATTTCATT

>M6-PGSC0003DMG400012955-(rev)M6LRR-RLK_2867

GTGGAGATCCTCTGTTCAGAAGTTGAATGGTGCCGTTGACTGATGACTGATATTTCATTCTTCTCATCTTGCATTTATTCTTTGAGGAAATGACTCTCAGATCAGCTAGATAGTTTAGTA

>M6-PGSC0003DMG400012955-(rev)M6LRR-RLK_2928

TCTCATCTTGCATTTATTCTTTGAGGAAATGACTCTCAGATCAGCTAGATAGTTTAGTATACTTGTCTCACTAATATACTTTTTTGCCTTCTAATTTTTTTAGGCTATATAGTATTTCTA

>M6-PGSC0003DMG400012955-(rev)M6LRR-RLK_2989

CTTGTCTCACTAATATACTTTTTTGCCTTCTAATTTTTTTAGGCTATATAGTATTTCTACCATATTCTTTAGCTAGAAATAGTTGCACTTGGACCTGAAAGTTAAATTGGATGTTTAATA

>M6-PGSC0003DMG400012955-(rev)M6LRR-RLK_305

CACATTCTCAACCCAATGACTCTCTCTCAATCTCTCTCTTTACTCCTCCTTGTACTTGTGTTTCGGCTCAATGGGTCTTCTGCTGCAGAAGATGAAGTTCGATCACTTTTGGAGTTCAAA

>M6-PGSC0003DMG400012955-(rev)M6LRR-RLK_3050

ATATTCTTTAGCTAGAAATAGTTGCACTTGGACCTGAAAGTTAAATTGGATGTTTAATAACAATGCACATTTTTTTTTAATAATATGAGCTCTTTTTCTGTTGGACTTGTAAACTTATTC

>M6-PGSC0003DMG400012955-(rev)M6LRR-RLK_3111

AATGCACATTTTTTTTTAATAATATGAGCTCTTTTTCTGTTGGACTTGTAAACTTATTCTAAAGAGATATTTAAAGTTTAATATTTAGTTTCTACGCATGCAGGTTCAATTCCGATTGTG

>M6-PGSC0003DMG400012955-(rev)M6LRR-RLK_3172

AAGAGATATTTAAAGTTTAATATTTAGTTTCTACGCATGCAGGTTCAATTCCGATTGTGAATTCAACAAAGTTGAGAGTTCTAAATATTTCATCGAACCACCTATTAGGTTCACTGCCAT

>M6-PGSC0003DMG400012955-(rev)M6LRR-RLK_3233

TTCAACAAAGTTGAGAGTTCTAAATATTTCATCGAACCACCTATTAGGTTCACTGCCATCCTCCATAGAAAATTGTGCAGTTGTGGATTTGAGCAGAAATATGCTTGTGGATGGTATATC

>M6-PGSC0003DMG400012955-(rev)M6LRR-RLK_3294

TCCATAGAAAATTGTGCAGTTGTGGATTTGAGCAGAAATATGCTTGTGGATGGTATATCAGTTATTGAAAGCTGGGAAGTCAATTTGGAAATTATTGATTTGAGTTCAAATAGGTTAACT

>M6-PGSC0003DMG400012955-(rev)M6LRR-RLK_3355

TTATTGAAAGCTGGGAAGTCAATTTGGAAATTATTGATTTGAGTTCAAATAGGTTAACTGGAAATATTCCCAATATAACCTCTCAGTTTCAACTATTGACTTCACTTAATTTTGGAAACA

>M6-PGSC0003DMG400012955-(rev)M6LRR-RLK_3416

AAATATTCCCAATATAACCTCTCAGTTTCAACTATTGACTTCACTTAATTTTGGAAACAATTCTCTTGAGGGAACCTTACCTTCAGCACTGGGCACCTTGCCAAGACTGGTTAAACTTGA

>M6-PGSC0003DMG400012955-(rev)M6LRR-RLK_3477

TCTCTTGAGGGAACCTTACCTTCAGCACTGGGCACCTTGCCAAGACTGGTTAAACTTGATCTCAGCACCAATAAACTTGGTGGACCGATTCCATCTACATTTTTTACTTCGACGACATTG

>M6-PGSC0003DMG400012955-(rev)M6LRR-RLK_3538

TCAGCACCAATAAACTTGGTGGACCGATTCCATCTACATTTTTTACTTCGACGACATTGATGAACCTAAATATATCTGGAAATCAATTGTCAGGATCAATTCCTCTTGAAGGCTCTCATG

>M6-PGSC0003DMG400012955-(rev)M6LRR-RLK_3599

GAACCTAAATATATCTGGAAATCAATTGTCAGGATCAATTCCTCTTGAAGGCTCTCATGCTAGCGAATTAATTGTTCAATCACCATACCCTGCACTGGAGTCTCTTGATCTTTCAGAAAA

>M6-PGSC0003DMG400012955-(rev)M6LRR-RLK_366

TTCGGCTCAATGGGTCTTCTGCTGCAGAAGATGAAGTTCGATCACTTTTGGAGTTCAAAAAGGGTATTAAAAACGACCCATTAGGTAAAATCTTCAGTTCATGGAGTCAAACGGGTCTTT

>M6-PGSC0003DMG400012955-(rev)M6LRR-RLK_3660

AGCGAATTAATTGTTCAATCACCATACCCTGCACTGGAGTCTCTTGATCTTTCAGAAAACAATTTCACAGGTAATTTGTCTTCTGCCATTGGTAATCTGCGGAGGCTTCAAGTGCTAAAT

>M6-PGSC0003DMG400012955-(rev)M6LRR-RLK_3721

ATTTCACAGGTAATTTGTCTTCTGCCATTGGTAATCTGCGGAGGCTTCAAGTGCTAAATCTTGCTAAGAATCAGTTATTAGGCATGCTGCCTACTGAATTGGGTGACCTTAGAAGTTTGG

>M6-PGSC0003DMG400012955-(rev)M6LRR-RLK_3782

TGCTAAGAATCAGTTATTAGGCATGCTGCCTACTGAATTGGGTGACCTTAGAAGTTTGGAATTCCTTGATATATCTAACAACAATTTTAGTGGTATGATCCCCGAAAATCTTTCATCAAA

>M6-PGSC0003DMG400012955-(rev)M6LRR-RLK_3843

TTCCTTGATATATCTAACAACAATTTTAGTGGTATGATCCCCGAAAATCTTTCATCAAATTTGAGGGTGTTTAATGTGTCTAACAATGAATTAAGTGGTGCTATCCCTGATAACTTGAGA

>M6-PGSC0003DMG400012955-(rev)M6LRR-RLK_3904

TGAGGGTGTTTAATGTGTCTAACAATGAATTAAGTGGTGCTATCCCTGATAACTTGAGAAACTTTAATGAAAGTTCATTTCGTCCTGGAAACTCAAATTTGGCAATCCCAAGAAACTGGT

>M6-PGSC0003DMG400012955-(rev)M6LRR-RLK_3965

CTTTAATGAAAGTTCATTTCGTCCTGGAAACTCAAATTTGGCAATCCCAAGAAACTGGTTACATGATAATCATGGTGATCCTGATCAAAATAGTCAGCACCATCATAACTCGAAATCCAG

>M6-PGSC0003DMG400012955-(rev)M6LRR-RLK_4026

CATGATAATCATGGTGATCCTGATCAAAATAGTCAGCACCATCATAACTCGAAATCCAGTATCAGGGTGGCTATTATTCTTGCCTCAGTTGGAGCTGCTTTAATGATTGGTGTTGTTCTC

>M6-PGSC0003DMG400012955-(rev)M6LRR-RLK_4087

TCAGGGTGGCTATTATTCTTGCCTCAGTTGGAGCTGCTTTAATGATTGGTGTTGTTCTCCTGGCTTACCATCGACAACAGTTCCAAGATTTCCACTTACCAAGTGGATTTAACAGCCAAT

>M6-PGSC0003DMG400012955-(rev)M6LRR-RLK_4148

GGCTTACCATCGACAACAGTTCCAAGATTTCCACTTACCAAGTGGATTTAACAGCCAATCTGCTGGAAGGGATGTTAAACTCGGGAGGTTCAGCCGGCCTGGAATTTTCAAGTTCCATGG

>M6-PGSC0003DMG400012955-(rev)M6LRR-RLK_4209

GCTGGAAGGGATGTTAAACTCGGGAGGTTCAGCCGGCCTGGAATTTTCAAGTTCCATGGCTCCTCAGAACCACCACCAATTTCCTTGAGTTTCTCTAATGATCACTTGCTCACAGCGAAC

>M6-PGSC0003DMG400012955-(rev)M6LRR-RLK_427

GGGTATTAAAAACGACCCATTAGGTAAAATCTTCAGTTCATGGAGTCAAACGGGTCTTTCAAATCCATCAGCATGTCCAAAGTCGTTTCATGGTGTCGTTTGTGATGAAAACTCTGATTC

>M6-PGSC0003DMG400012955-(rev)M6LRR-RLK_4270

CCTCAGAACCACCACCAATTTCCTTGAGTTTCTCTAATGATCACTTGCTCACAGCGAACTAGAGATCTCTGTCTGGGCAAATTGAGTCTGGTACAGAAATTGATGAGCAAGTTTTTCCAG

>M6-PGSC0003DMG400012955-(rev)M6LRR-RLK_4331

GAGATCTCTGTCTGGGCAAATTGAGTCTGGTACAGAAATTGATGAGCAAGTTTTTCCAGAAGGAGTAACTGCAGTTTCAGCATCTACGCATCTTGGCACTGTGGGTAATAATCCTGCAAC

>M6-PGSC0003DMG400012955-(rev)M6LRR-RLK_4392

GGAGTAACTGCAGTTTCAGCATCTACGCATCTTGGCACTGTGGGTAATAATCCTGCAACATCTGGCCGGAGATCCTCTCCAGGCTCTCCAATAGCTTCCTCCCCTCGTTTCGTCGACACA

>M6-PGSC0003DMG400012955-(rev)M6LRR-RLK_4453

CTGGCCGGAGATCCTCTCCAGGCTCTCCAATAGCTTCCTCCCCTCGTTTCGTCGACACAGTTGAACAACCAGAGACATTGGATGTGTATTCACCGGATCGATTGGCTAGGGAATTGTTCT

>M6-PGSC0003DMG400012955-(rev)M6LRR-RLK_4514

TGAACAACCAGAGACATTGGATGTGTATTCACCGGATCGATTGGCTAGGGAATTGTTCTTCCTGGATGGTTCACTGTCATTTACTGCTGAGGAGTTATCTCGTGCCACTGCTGAAGTTCT

>M6-PGSC0003DMG400012955-(rev)M6LRR-RLK_4575

CTGGATGGTTCACTGTCATTTACTGCTGAGGAGTTATCTCGTGCCACTGCTGAAGTTCTGGGTAGAAGTAGCCATGACACATTGTATAAAGCTACTCTGAATAGTGGGCATGTTCTTACT

>M6-PGSC0003DMG400012955-(rev)M6LRR-RLK_4636

GTAGAAGTAGCCATGACACATTGTATAAAGCTACTCTGAATAGTGGGCATGTTCTTACTGTCAAGTGGTTGCGGGTTGGGTTGGTAAAAAACAAGAAAGAGTTTGCAAAGGAAGTTAAAA

>M6-PGSC0003DMG400012955-(rev)M6LRR-RLK_4697

CAAGTGGTTGCGGGTTGGGTTGGTAAAAAACAAGAAAGAGTTTGCAAAGGAAGTTAAAAAGATTGGATCTATTAGGCACCCAAATGCTGTTCCTTTACGAGCATATTACTGGGGACCTCG

>M6-PGSC0003DMG400012955-(rev)M6LRR-RLK_4758

ATTGGATCTATTAGGCACCCAAATGCTGTTCCTTTACGAGCATATTACTGGGGACCTCGGGAACAAGAAAGGCTTATTTTAGCAGACTACATACCAGGAGATAGTTTGGAGTTGCATCTC

>M6-PGSC0003DMG400012955-(rev)M6LRR-RLK_4819

AACAAGAAAGGCTTATTTTAGCAGACTACATACCAGGAGATAGTTTGGAGTTGCATCTCTATGGTAAGATTTTCTTACGAATAACTTCAACAATGATATTCCATGTCTTCCATTGTTGTT

>M6-PGSC0003DMG400012955-(rev)M6LRR-RLK_488

AATCCATCAGCATGTCCAAAGTCGTTTCATGGTGTCGTTTGTGATGAAAACTCTGATTCTGTTTTCTCCATTTCTCTTGACGGGCTTGGGCTTGTAGGTGACTTGAAGTTCTCAACTCTT

>M6-PGSC0003DMG400012955-(rev)M6LRR-RLK_4880

TGGTAAGATTTTCTTACGAATAACTTCAACAATGATATTCCATGTCTTCCATTGTTGTTTGGTTACTGTTATTAGAAAGTTCCCTTTTGCTGGTCTGGTGGGTGATTATCTTGTGTTAGG

>M6-PGSC0003DMG400012955-(rev)M6LRR-RLK_4941

GTTACTGTTATTAGAAAGTTCCCTTTTGCTGGTCTGGTGGGTGATTATCTTGTGTTAGGACCAATATTGATATCATTCGCATAAATAGTTTCTATAGCAACATAGCTTATTTCTAGTGGT

>M6-PGSC0003DMG400012955-(rev)M6LRR-RLK_5002

CAATATTGATATCATTCGCATAAATAGTTTCTATAGCAACATAGCTTATTTCTAGTGGTGGTTTTCTAATTGCTGAGATCTCTTGTGTTTTGTGGGATTGTTTGATAAAAATTATGGAGG

>M6-PGSC0003DMG400012955-(rev)M6LRR-RLK_5063

TTTTCTAATTGCTGAGATCTCTTGTGTTTTGTGGGATTGTTTGATAAAAATTATGGAGGATAAAGTTGACTTGGCCCTATTATTTATTTTTGGTTGGCCTCTCTTGCAAATGTCCCTCTT

>M6-PGSC0003DMG400012955-(rev)M6LRR-RLK_5124

AAAGTTGACTTGGCCCTATTATTTATTTTTGGTTGGCCTCTCTTGCAAATGTCCCTCTTTTACATGTCTTACAATATGCAGATAGTGATGAGATCTTATTTGGATCTGTCGTCTCTTATG

>M6-PGSC0003DMG400012955-(rev)M6LRR-RLK_5185

ACATGTCTTACAATATGCAGATAGTGATGAGATCTTATTTGGATCTGTCGTCTCTTATGAATCCAGTGGTAGAGTTGTTGATCAAGCGGAAAAGGAAAAAAATAGGACTGCAATTGTAGA

>M6-PGSC0003DMG400012955-(rev)M6LRR-RLK_5246

TCCAGTGGTAGAGTTGTTGATCAAGCGGAAAAGGAAAAAAATAGGACTGCAATTGTAGAATCTGAATGATTTTTTACAAAATTATGTTTGTTAGAATGTATGTGCAGATCTTTATGTGTC

>M6-PGSC0003DMG400012955-(rev)M6LRR-RLK_5307

CTGAATGATTTTTTACAAAATTATGTTTGTTAGAATGTATGTGCAGATCTTTATGTGTCATCTGGTAACCTTCAAAGGAGCTCCAAAAAGGTACAATAGATCTTTGGTTCACTAATTGCC

>M6-PGSC0003DMG400012955-(rev)M6LRR-RLK_5368

CTGGTAACCTTCAAAGGAGCTCCAAAAAGGTACAATAGATCTTTGGTTCACTAATTGCCGTACCCAACAGCCACACATCAATAGCCCATTAAAATTAGGGAGTAGACCTATTCTAAGATC

>M6-PGSC0003DMG400012955-(rev)M6LRR-RLK_5429

ACCCAACAGCCACACATCAATAGCCCATTAAAATTAGGGAGTAGACCTATTCTAAGATCTCGTTAACATGACCTAGAATGTAGTGATAGTTGCTCAACAATCCGCTTTAAATAGCATTTC

>M6-PGSC0003DMG400012955-(rev)M6LRR-RLK_549

TTTTCTCCATTTCTCTTGACGGGCTTGGGCTTGTAGGTGACTTGAAGTTCTCAACTCTTAGTGGGTTGAAGCAGCTCAAAATTCTTAGTCTTTCTGGTAACTCTTTTACGGGTAGGGTTG

>M6-PGSC0003DMG400012955-(rev)M6LRR-RLK_5490

GTTAACATGACCTAGAATGTAGTGATAGTTGCTCAACAATCCGCTTTAAATAGCATTTCCATTGTTTCTGAATCATTCATGATGGTAAAACTTGATAAATATGGAGAGTTGGCTTTGTCA

>M6-PGSC0003DMG400012955-(rev)M6LRR-RLK_5551

TTGTTTCTGAATCATTCATGATGGTAAAACTTGATAAATATGGAGAGTTGGCTTTGTCATAGTATCCCCTGTGCAGTGAGGACTGCTTACTTGTAGACATGTTGGGGGTTGTTGGAAACT

>M6-PGSC0003DMG400012955-(rev)M6LRR-RLK_5612

GTATCCCCTGTGCAGTGAGGACTGCTTACTTGTAGACATGTTGGGGGTTGTTGGAAACTTTTTACTTTTTGTAATTTTTTGCAGTTTTAAGTTTTTAATGAATGTTTACTTTTTACAGAG

>M6-PGSC0003DMG400012955-(rev)M6LRR-RLK_5673

TTACTTTTTGTAATTTTTTGCAGTTTTAAGTTTTTAATGAATGTTTACTTTTTACAGAGACAACCCCTCGGAGGTACTCCCCGTTATCCTTCAACCAAAGGTTAAAAGTTTCCATTGAAG

>M6-PGSC0003DMG400012955-(rev)M6LRR-RLK_5734

AACCCCTCGGAGGTACTCCCCGTTATCCTTCAACCAAAGGTTAAAAGTTTCCATTGAAGTTGCTCGGGGTTTGACATATCTGCATGAGAGAGGGCTGCCTCATGGAGACTTGAAACCTAC

>M6-PGSC0003DMG400012955-(rev)M6LRR-RLK_5795

GCTCGGGGTTTGACATATCTGCATGAGAGAGGGCTGCCTCATGGAGACTTGAAACCTACAAATATAATCTTGGTTGGTGCTGATTACAGTGTCCGCCTCACTGATTATGGTCTCCACCGC

>M6-PGSC0003DMG400012955-(rev)M6LRR-RLK_5856

ATATAATCTTGGTTGGTGCTGATTACAGTGTCCGCCTCACTGATTATGGTCTCCACCGCCTGATGACTCCAGCAGGGATTGCAGAGCAGATACTGAACCTAGGGGCTCTAGGATACCGTG

>M6-PGSC0003DMG400012955-(rev)M6LRR-RLK_5917

GATGACTCCAGCAGGGATTGCAGAGCAGATACTGAACCTAGGGGCTCTAGGATACCGTGCTCCTGAACTTGCTACTGCGACCAAACCTATCCCATCATTTAAGGCTGATGTATATGCTCT

>M6-PGSC0003DMG400012955-(rev)M6LRR-RLK_5978

CCTGAACTTGCTACTGCGACCAAACCTATCCCATCATTTAAGGCTGATGTATATGCTCTTGGTGTGATTCTAATGGAATTGCTGACAAGAAGAAGTGCTGGAGACTTAATCTCAGCACAC

>M6-PGSC0003DMG400012955-(rev)M6LRR-RLK_6039

GTGTGATTCTAATGGAATTGCTGACAAGAAGAAGTGCTGGAGACTTAATCTCAGCACACTCTGCTGCAGTTGATTTAACAGATTGGGTTAGGTTGTGTGATCAAGAAGGTAGGGGAATGG

>M6-PGSC0003DMG400012955-(rev)M6LRR-RLK_61

ATTTGCCACTCTACCCCAAATATAAAAACCCTAATTTCCCTCTTTTTTTCTCCTTTCCCAACTCTTTCCCCACACCCCAACCCCTCTCTTACCACCCCACCTACCCTAATCTCTCTCTCT

>M6-PGSC0003DMG400012955-(rev)M6LRR-RLK_610

TGGGTTGAAGCAGCTCAAAATTCTTAGTCTTTCTGGTAACTCTTTTACGGGTAGGGTTGTTCCTGCTTTGGGTTCAATGTTAACTCTTCAGCATTTGGATCTCTCTGGAAATCAGTTTTA

>M6-PGSC0003DMG400012955-(rev)M6LRR-RLK_6100

TGCTGCAGTTGATTTAACAGATTGGGTTAGGTTGTGTGATCAAGAAGGTAGGGGAATGGACTGTATCGATAGGGACATCGCAGGAGGAGAAGAACACTGCAAAGCGATGGATGATCTGCT

>M6-PGSC0003DMG400012955-(rev)M6LRR-RLK_6161

TGTATCGATAGGGACATCGCAGGAGGAGAAGAACACTGCAAAGCGATGGATGATCTGCTAGCTGTGTCGTTAAGGTGTATTCTTTCGATAAACGAAAGGCCAAACATCAGACAAGTTGTT

>M6-PGSC0003DMG400012955-(rev)M6LRR-RLK_6222

CTGTGTCGTTAAGGTGTATTCTTTCGATAAACGAAAGGCCAAACATCAGACAAGTTGTTGAAAATCTGGGTTCCATATCTGTGTGAAATTTTTGTACATGTTGTCTTGAAATTACAATTC

>M6-PGSC0003DMG400012955-(rev)M6LRR-RLK_6283

AAATCTGGGTTCCATATCTGTGTGAAATTTTTGTACATGTTGTCTTGAAATTACAATTCTTTTTTCTTTTCCTTCACATTTTTTATTATGTCCTATTGGTTGCCTTCTAATGTTGGTCAA

>M6-PGSC0003DMG400012955-(rev)M6LRR-RLK_6344

TTTTCTTTTCCTTCACATTTTTTATTATGTCCTATTGGTTGCCTTCTAATGTTGGTCAATCCTTACATATTCCCCATTTTCTTTCCCCTCCCCCCTTCAATTTTTTTGTAAAGGGATGAT

>M6-PGSC0003DMG400012955-(rev)M6LRR-RLK_6405

CTTACATATTCCCCATTTTCTTTCCCCTCCCCCCTTCAATTTTTTTGTAAAGGGATGATCAGTCCATTGATTGATTGGTTGACATTTTTCTCTAATTAGAAAAGATTGCAATTATTTATG

>M6-PGSC0003DMG400012955-(rev)M6LRR-RLK_671

CCTGCTTTGGGTTCAATGTTAACTCTTCAGCATTTGGATCTCTCTGGAAATCAGTTTTATGGAACAATTCCAGCACGAATTAATGAGCTTTGGGGTTTGAATTACCTCAATTTGTCCAAT

>M6-PGSC0003DMG400012955-(rev)M6LRR-RLK_732

GAACAATTCCAGCACGAATTAATGAGCTTTGGGGTTTGAATTACCTCAATTTGTCCAATAATAATTTTACATTTGGTTACCCAATTGGAATTAGTAATCTTCAGCAGTTGAGAGTACTGG

>M6-PGSC0003DMG400012955-(rev)M6LRR-RLK_793

TAATTTTACATTTGGTTACCCAATTGGAATTAGTAATCTTCAGCAGTTGAGAGTACTGGATTTGCATAACAATGGGATTTGGGGTGATATCGGGGAGTTGTTTTTGGAATTGAAGCGTAT

>M6-PGSC0003DMG400012955-(rev)M6LRR-RLK_854

TTGCATAACAATGGGATTTGGGGTGATATCGGGGAGTTGTTTTTGGAATTGAAGCGTATTGAGCATCTTGATTTGAGTAACAACTCGTTTTTCGGGTCTCTTCCTACGAGCCCGGAAAAT

>M6-PGSC0003DMG400012955-(rev)M6LRR-RLK_915

AGCATCTTGATTTGAGTAACAACTCGTTTTTCGGGTCTCTTCCTACGAGCCCGGAAAATGTATCACTCTCTTCGACAATTCAGGTCATGAATTTGAGTCACAACAACTTGGGTGGTGGTT

>M6-PGSC0003DMG400012955-(rev)M6LRR-RLK_976

ATCACTCTCTTCGACAATTCAGGTCATGAATTTGAGTCACAACAACTTGGGTGGTGGTTTTTTTCCTGGAAAATTGTTGGAGGCATTTGAGAATTTGATGGTTTTGGATTTGAGGAATAA

>M6-PGSC0003DMG400012992M6LRR-RLK_1037

GAAATGCAAGAAGTTGACTAATTTTCGAGTAGCCAGAAATAGAATTAGTGGTCACATACCACCAGAGATTGGAAATCTCAAAGGTCTTCAAGGACTAGATCTTTCAGCAAATCATTTGGT

>M6-PGSC0003DMG400012992M6LRR-RLK_1098

CCAGAGATTGGAAATCTCAAAGGTCTTCAAGGACTAGATCTTTCAGCAAATCATTTGGTTGGGAAGATCCCTAGGGAACTTGGAAAATTGACCTCTCTAGTTAATCTTTTTTTGCAAAAC

>M6-PGSC0003DMG400012992M6LRR-RLK_1159

GGAAGATCCCTAGGGAACTTGGAAAATTGACCTCTCTAGTTAATCTTTTTTTGCAAAACAACCAGATTTCTGGCAATATTCCTATGGAACTTGGATCAATGACAAAGTTAGAATCCCTTG

>M6-PGSC0003DMG400012992M6LRR-RLK_1220

CCAGATTTCTGGCAATATTCCTATGGAACTTGGATCAATGACAAAGTTAGAATCCCTTGATCTATCAAACAATATGTTGAATGGGTCGATCCCAACATTTATAGGAGATTACATACACTT

>M6-PGSC0003DMG400012992M6LRR-RLK_1281

CTATCAAACAATATGTTGAATGGGTCGATCCCAACATTTATAGGAGATTACATACACTTGTTTCAGTTGAACCTGAGCAACAACAAATTTGGCCAAAATATTCCAATGGAGATAGGGAAG

>M6-PGSC0003DMG400012992M6LRR-RLK_1342

TTCAGTTGAACCTGAGCAACAACAAATTTGGCCAAAATATTCCAATGGAGATAGGGAAGATAACTCAGCTTAATGTACTTGATTTGAGCAATAATCTTCTGGTTGGAGATATATCCCCTC

>M6-PGSC0003DMG400012992M6LRR-RLK_1403

AACTCAGCTTAATGTACTTGATTTGAGCAATAATCTTCTGGTTGGAGATATATCCCCTCAGTTAGCCAATTTAAAGGTCTTGGTAAACTTAAACCTTTCCCACAATGGCCTATCTGGGCG

>M6-PGSC0003DMG400012992M6LRR-RLK_1464

TTAGCCAATTTAAAGGTCTTGGTAAACTTAAACCTTTCCCACAATGGCCTATCTGGGCGCATTCCTAAAGAAGTTGAAAGTTTGACTGGTTTGCAGTATGTCGTATTGTCATACAATGAG

>M6-PGSC0003DMG400012992M6LRR-RLK_1525

TTCCTAAAGAAGTTGAAAGTTTGACTGGTTTGCAGTATGTCGTATTGTCATACAATGAGTTGGAAGGTCCAATCCCCAATAATAAAGCTTTTATGAATGCTTCATTAGAAGGTAACAAAG

>M6-PGSC0003DMG400012992M6LRR-RLK_1586

GGAAGGTCCAATCCCCAATAATAAAGCTTTTATGAATGCTTCATTAGAAGGTAACAAAGGTCTTTGCGGCAATGTAACAGGACTCCAGCCCTGTGATAGGCCATCTTCTGTGGTGAAGAA

>M6-PGSC0003DMG400012992M6LRR-RLK_1647

CTTTGCGGCAATGTAACAGGACTCCAGCCCTGTGATAGGCCATCTTCTGTGGTGAAGAAGCACTCAATGGCAAAGGGATGTAAAGTCATCCTCATCACTGTACTTTCTGTTATGGGAGCA

>M6-PGSC0003DMG400012992M6LRR-RLK_1708

ACTCAATGGCAAAGGGATGTAAAGTCATCCTCATCACTGTACTTTCTGTTATGGGAGCACTACTACTGCTCTGTGTTTTCATTAGTGTTCTTTTTATGTGTAATAAAAGAAGGAGAGTTA

>M6-PGSC0003DMG400012992M6LRR-RLK_1769

ACTACTGCTCTGTGTTTTCATTAGTGTTCTTTTTATGTGTAATAAAAGAAGGAGAGTTAGAGACGTTGAAAGAAGGGATGGTGATGGTTGGCTTTCAATATCTATGTTAGATGGAAAGGC

>M6-PGSC0003DMG400012992M6LRR-RLK_1830

GACGTTGAAAGAAGGGATGGTGATGGTTGGCTTTCAATATCTATGTTAGATGGAAAGGCATTGTACAGGGACATCTTAAATGCCACAGAAGAGTTTGATGTAAAATTTTGCATCGGGCAT

>M6-PGSC0003DMG400012992M6LRR-RLK_1891

TGTACAGGGACATCTTAAATGCCACAGAAGAGTTTGATGTAAAATTTTGCATCGGGCATGGAGGACACGGAAGCGTTTACAAGGTAAACCTTCAATCATTAGGGAATATAGCTGTGAAGA

>M6-PGSC0003DMG400012992M6LRR-RLK_1952

AGGACACGGAAGCGTTTACAAGGTAAACCTTCAATCATTAGGGAATATAGCTGTGAAGAGACTTCATTCTTCATATCAGAATACACATCCCAAAAGCTTCATGAATGAAGTAAGGGCATT

>M6-PGSC0003DMG400012992M6LRR-RLK_2257

TAGCTTACATGCACCAGGATTGTTCACCACCCATTGTTCATCGAGACATATCAAGTAGTAATGTTTTGCTTGATTCAGAGTATGAAGCTCGTGTTGCAGACTTTGGCATATCAAAGCTTC

>M6-PGSC0003DMG400012992M6LRR-RLK_2318

TGTTTTGCTTGATTCAGAGTATGAAGCTCGTGTTGCAGACTTTGGCATATCAAAGCTTCTCAAGCCAAACTCATCCAATTGCACAGCGCTTGCAGGGACATATGGTTATGTTGCACCTGG

>M6-PGSC0003DMG400012992M6LRR-RLK_2379

AAGCCAAACTCATCCAATTGCACAGCGCTTGCAGGGACATATGGTTATGTTGCACCTGGTAAGTTTAAAATATTTCATGAACTAAAGCCAAACATTCTATTGTTTATATCTCATTTACAT

>M6-PGSC0003DMG400012992M6LRR-RLK_2440

AGTTTAAAATATTTCATGAACTAAAGCCAAACATTCTATTGTTTATATCTCATTTACATGTTTATATTGTTTTGGAAATTGTATTGTTCAGAGTTTGCATATACTATGAAGGTTACTCAA

>M6-PGSC0003DMG400012992M6LRR-RLK_2501

TTATATTGTTTTGGAAATTGTATTGTTCAGAGTTTGCATATACTATGAAGGTTACTCAAATGTGTGATGTTTATAGCTTTGGAGTATTATCTTTGGAGATAATCAAAGGAAAGCATCTTG

>M6-PGSC0003DMG400012992M6LRR-RLK_2562

GTGTGATGTTTATAGCTTTGGAGTATTATCTTTGGAGATAATCAAAGGAAAGCATCTTGGGGAATACATTACCGTGCTAGCAAATTCATCTACTATAGATCATGAGCATCTTAGTGATTT

>M6-PGSC0003DMG400012992M6LRR-RLK_2623

GAATACATTACCGTGCTAGCAAATTCATCTACTATAGATCATGAGCATCTTAGTGATTTGTTGGATGAACGCCTTCCATATCCTGAAGATAGAGTAAAAGAGGTTTTTATTTTTATCATC

>M6-PGSC0003DMG400012992M6LRR-RLK_2684

TGGATGAACGCCTTCCATATCCTGAAGATAGAGTAAAAGAGGTTTTTATTTTTATCATCAAGCTAGCATGCTCTTGTTTGCTTGAAACTCCAAAATCAAGGCCAACAATGCACTTCATCT

>M6-PGSC0003DMG400012992M6LRR-RLK_305

TAGAGAGATGGGAAATTTGAAAAGCCTCACTGATTTGGATTTATCCACTAATCATTTTATTGGTCCTATTCCTAGTGAGTTGGGAAATCTGATAAATCTCACATAGATAGATTTATCTAA

>M6-PGSC0003DMG400012992M6LRR-RLK_549

AATGGTCTGTCCTTGTACGAAAATCAACTTACTGGTCCAATTCCTGCTTCCATTGGCAATTTGAGAAGCTTGCGATTTCTGTATCTCAGTGCCAACAAACTTACTGGTCCAATTCCCGCT

>M6-PGSC0003DMG400012992M6LRR-RLK_610

TGAGAAGCTTGCGATTTCTGTATCTCAGTGCCAACAAACTTACTGGTCCAATTCCCGCTTCCTTTGGCAATTTAAGAAACTTGCAATTTCTGTATCTGAGTGCCAACAAACTTTACGGTT

>M6-PGSC0003DMG400012992M6LRR-RLK_671

CTTTGGCAATTTAAGAAACTTGCAATTTCTGTATCTGAGTGCCAACAAACTTTACGGTTCTATTCCAAAAGAACTTGCATATTTGGATAACTTGGAAGAGCTGATAATAAGTGAAAATCA

>M6-PGSC0003DMG400012992M6LRR-RLK_732

ATTCCAAAAGAACTTGCATATTTGGATAACTTGGAAGAGCTGATAATAAGTGAAAATCAGTTTTCTGGTCAAATGCCTGAGCATCTTTGCCAAGGTGGGAAACTTGAGAACTTTACGGTG

>M6-PGSC0003DMG400012992M6LRR-RLK_793

TTTCTGGTCAAATGCCTGAGCATCTTTGCCAAGGTGGGAAACTTGAGAACTTTACGGTGAATAGTAATAAGTTGTCAGGGCCTATACCAAGAAGCATGTCCAAGTGCTCGAGTTTCAAAA

>M6-PGSC0003DMG400012992M6LRR-RLK_854

TAGTAATAAGTTGTCAGGGCCTATACCAAGAAGCATGTCCAAGTGCTCGAGTTTCAAAAGGGTTCGCTTTGATAACAACAGTTTTACTGGAAATTTATCTGAAGCTTTTGGTATCTATCC

>M6-PGSC0003DMG400012992M6LRR-RLK_915

GTTCGCTTTGATAACAACAGTTTTACTGGAAATTTATCTGAAGCTTTTGGTATCTATCCGGAGCTACAGTTTATTAATTTGAGCAAAAATGATTTTCATGGTGAACTCAGTAGCAACTGG

>M6-PGSC0003DMG400012992M6LRR-RLK_976

AGCTACAGTTTATTAATTTGAGCAAAAATGATTTTCATGGTGAACTCAGTAGCAACTGGGGGAAATGCAAGAAGTTGACTAATTTTCGAGTAGCCAGAAATAGAATTAGTGGTCACATAC

>M6-PGSC0003DMG400013198M6LRR-RLK_0

GATGGGATAATTATTACTGCAGCAGATTTTCAAGCACTTCAAGCTTTTAAACAAGAATTGGTTGATCCAAAAGGGTTCTTGAAAAGCTGGAATGATAGTGGATTTGGAGCTTGTTCTGGT

>M6-PGSC0003DMG400013198M6LRR-RLK_1008

CCAGCTTCTTTAGATACTTTACCAAATCTTAGTGCCTTCAATGTCTCATATAATAATCTGTCTGGTCCTGTTCCAACTCATCTCGCACGGGAATTCAACTCGAGTGCCTTTGTGGGTAAT

>M6-PGSC0003DMG400013198M6LRR-RLK_1071

GGTCCTGTTCCAACTCATCTCGCACGGGAATTCAACTCGAGTGCCTTTGTGGGTAATCTTCAGCTATGTGGTTACAGTGCATCAACCCCATGTCCAATCTCTCCAGTTAGCCCATCCCCC

>M6-PGSC0003DMG400013198M6LRR-RLK_1134

CTATGTGGTTACAGTGCATCAACCCCATGTCCAATCTCTCCAGTTAGCCCATCCCCCGAAACACCAAAAAAGCAACGTCGTAAGTTAAGTACTAAGGACATAATTCTCATAGCAGGAGGG

>M6-PGSC0003DMG400013198M6LRR-RLK_1197

CCAAAAAAGCAACGTCGTAAGTTAAGTACTAAGGACATAATTCTCATAGCAGGAGGGGCACTTCTCATAATCTTGGCTCTGCTATGTTGCATTCTCCTATGCTGCTTGATCAGGAAAAGA

>M6-PGSC0003DMG400013198M6LRR-RLK_126

CTTGGTATTAAATGTGCTCAAGGACAAGTTATTGTTATTCAGCTTCCATGGAGAGGTTTAGGTGGAAGAATTACTGAAAGAATTGGACAATTTCAGTCTCTTAGAAAACTCAGCTTACAT

>M6-PGSC0003DMG400013198M6LRR-RLK_1260

CTCATAATCTTGGCTCTGCTATGTTGCATTCTCCTATGCTGCTTGATCAGGAAAAGATCTGCAGCTGAAGCAGGGAAGGATGGTCAAGGCACGAGTAGGGTGGCTGCAGCAGCCAGAGGT

>M6-PGSC0003DMG400013198M6LRR-RLK_1323

GCTGAAGCAGGGAAGGATGGTCAAGGCACGAGTAGGGTGGCTGCAGCAGCCAGAGGTGAAAAGGGTGTTCCACCAACTGCTGGAGAAGTTGAAGCAGCAGGAGGAGGAGATACAGGAGGA

>M6-PGSC0003DMG400013198M6LRR-RLK_1386

GGTGTTCCACCAACTGCTGGAGAAGTTGAAGCAGCAGGAGGAGGAGATACAGGAGGAAAGCTTGTCCATTTTGATGGACCTATTGTGTTCACAGCAGATGATCTTCTTTGTGCAACTGCC

>M6-PGSC0003DMG400013198M6LRR-RLK_1449

GTCCATTTTGATGGACCTATTGTGTTCACAGCAGATGATCTTCTTTGTGCAACTGCCGAAATAATGGGAAAGAGCACTTATGGTACGGTGTACAAGGCCACGTTAGAGGATGGCGATCAA

>M6-PGSC0003DMG400013198M6LRR-RLK_1512

ATGGGAAAGAGCACTTATGGTACGGTGTACAAGGCCACGTTAGAGGATGGCGATCAAGTTGCTGTGAAAAGATTGAGAGAGAAGATCACAAGAGGTCAAAGGGAATTTGAGAGTGAAGTC

>M6-PGSC0003DMG400013198M6LRR-RLK_1575

GTGAAAAGATTGAGAGAGAAGATCACAAGAGGTCAAAGGGAATTTGAGAGTGAAGTCAATATTCTTGGCAAGATCAGGCATCCAAATCTTTTGGCCCTTAGAGCATATTACATGGGACCT

>M6-PGSC0003DMG400013198M6LRR-RLK_1638

CTTGGCAAGATCAGGCATCCAAATCTTTTGGCCCTTAGAGCATATTACATGGGACCTAAAGGAGAGAAACTTCTTGTTTTTGACTACATGCCTAAGGGAAGTTTGGCAACTTTTCTTCAT

>M6-PGSC0003DMG400013198M6LRR-RLK_1701

GAGAAACTTCTTGTTTTTGACTACATGCCTAAGGGAAGTTTGGCAACTTTTCTTCATGGTAAGCACAAAATCAGTTTTCCATTTCACTTTGTATTTGTGAGAACTTCAGATTATAAATCT

>M6-PGSC0003DMG400013198M6LRR-RLK_1764

CACAAAATCAGTTTTCCATTTCACTTTGTATTTGTGAGAACTTCAGATTATAAATCTTTCTGTTTTTTGCAGCTCGCAGTCCTGATACACCAATTGATTGGGCAACAAGGATGAGAATAG

>M6-PGSC0003DMG400013198M6LRR-RLK_1827

TTTTTGCAGCTCGCAGTCCTGATACACCAATTGATTGGGCAACAAGGATGAGAATAGCAAAGGGTACAGCAAGGGGATTGCTCTTCCTTCATACAAATGCCAACATCATTCATGGGAATC

>M6-PGSC0003DMG400013198M6LRR-RLK_189

GGAAGAATTACTGAAAGAATTGGACAATTTCAGTCTCTTAGAAAACTCAGCTTACATGATAATGTCATTGGTGGTTCAATTCCTTCTACTTTGGGACTCATACCTAATCTTAGAGGACTT

>M6-PGSC0003DMG400013198M6LRR-RLK_1890

GTACAGCAAGGGGATTGCTCTTCCTTCATACAAATGCCAACATCATTCATGGGAATCTTACATCGAGCAATGTTTTACTTGATGACAACACAAATGCAAAAATTGCAGATTATGGTCTTT

>M6-PGSC0003DMG400013198M6LRR-RLK_1953

CGAGCAATGTTTTACTTGATGACAACACAAATGCAAAAATTGCAGATTATGGTCTTTCGCGCCTAATGACTGCTGCTGCAAATGCAAATGTTATTGCAACTGCTGGAGCACTTGGGTATC

>M6-PGSC0003DMG400013198M6LRR-RLK_2016

TAATGACTGCTGCTGCAAATGCAAATGTTATTGCAACTGCTGGAGCACTTGGGTATCGGGCACCTGAACTTTCAAAACTCAAGAAAGCAAACACAAAGACAGATGTATACAGCCTTGGTG

>M6-PGSC0003DMG400013198M6LRR-RLK_2079

CTGAACTTTCAAAACTCAAGAAAGCAAACACAAAGACAGATGTATACAGCCTTGGTGTTATTATACTGGAACTTCTAACTGGGAAGTCACCAGGAGAAGCAATGAATGGTGTGGATTTGC

>M6-PGSC0003DMG400013198M6LRR-RLK_2142

TACTGGAACTTCTAACTGGGAAGTCACCAGGAGAAGCAATGAATGGTGTGGATTTGCCTCAATGGGTTGCATCAATTGTGAAAGAGGAGTGGACTAATGAGGTATTTGATTTAGAACTAA

>M6-PGSC0003DMG400013198M6LRR-RLK_2205

GGGTTGCATCAATTGTGAAAGAGGAGTGGACTAATGAGGTATTTGATTTAGAACTAATGAGGGATGCATCCGTTATCGGTGATGAATTATTGAATACCTTGAAATTGGCTTTGCATTGTG

>M6-PGSC0003DMG400013198M6LRR-RLK_2268

ATGCATCCGTTATCGGTGATGAATTATTGAATACCTTGAAATTGGCTTTGCATTGTGTTGATCCGTCACCATCAGCTCGACCAGAAGTTCAGCAACTTCTCCAGCAACTAGAAGAAATTA

>M6-PGSC0003DMG400013198M6LRR-RLK_2302

CTTGAAATTGGCTTTGCATTGTGTTGATCCGTCACCATCAGCTCGACCAGAAGTTCAGCAACTTCTCCAGCAACTAGAAGAAATTAGACCTGAAACAGCTACCAGTTCTGGGGACGACGG

>M6-PGSC0003DMG400013198M6LRR-RLK_252

GTCATTGGTGGTTCAATTCCTTCTACTTTGGGACTCATACCTAATCTTAGAGGACTTCAACTTTTTAACAATAGGTTATCTGGTTCTATTCCTGCTTCTCTTGGTTTATGCCCTCTTCTT

>M6-PGSC0003DMG400013198M6LRR-RLK_315

TTTAACAATAGGTTATCTGGTTCTATTCCTGCTTCTCTTGGTTTATGCCCTCTTCTTCAAACACTTGATCTCAGCAATAACTCATTCTCTGGTGCAATTCCAGACAGTCTTGTTAATTCA

>M6-PGSC0003DMG400013198M6LRR-RLK_378

CTTGATCTCAGCAATAACTCATTCTCTGGTGCAATTCCAGACAGTCTTGTTAATTCAACTAAGCTTTATAGGCTTAATCTTAGCCATAATTCTTTGTCTGGTTCAATTCCCACAAGTCTC

>M6-PGSC0003DMG400013198M6LRR-RLK_441

CTTTATAGGCTTAATCTTAGCCATAATTCTTTGTCTGGTTCAATTCCCACAAGTCTCACACAATCTCCCGCACTCATCTTTCTTGATCTCAAGTATAACAATCTTTCAGGCTCAATACCA

>M6-PGSC0003DMG400013198M6LRR-RLK_504

TCTCCCGCACTCATCTTTCTTGATCTCAAGTATAACAATCTTTCAGGCTCAATACCAGATACTTGGGATGGAAATGGAAATGGAAAAAGACTCTTCCAACTTCAGTCTCTTACACTTGAT

>M6-PGSC0003DMG400013198M6LRR-RLK_567

TGGGATGGAAATGGAAATGGAAAAAGACTCTTCCAACTTCAGTCTCTTACACTTGATCACAACTTCTTTTCTGGAAGTATTCCTGCTTCCTTAGGCAAGTTGAATGAACTTGTTGAGCTT

>M6-PGSC0003DMG400013198M6LRR-RLK_63

GATCCAAAAGGGTTCTTGAAAAGCTGGAATGATAGTGGATTTGGAGCTTGTTCTGGTGGATGGCTTGGTATTAAATGTGCTCAAGGACAAGTTATTGTTATTCAGCTTCCATGGAGAGGT

>M6-PGSC0003DMG400013198M6LRR-RLK_630

TTCTTTTCTGGAAGTATTCCTGCTTCCTTAGGCAAGTTGAATGAACTTGTTGAGCTTTCACTTAGTCATAATCAATTGACTGGAGTTATCCCAAGTCATATTGAGGGACTCTCTAGGCTT

>M6-PGSC0003DMG400013198M6LRR-RLK_693

AGTCATAATCAATTGACTGGAGTTATCCCAAGTCATATTGAGGGACTCTCTAGGCTTACAACACTTGATTTGTCTTATAATGCCATTAATGGAAGCTTGTCTGATAGCTTCTTGAATCTA

>M6-PGSC0003DMG400013198M6LRR-RLK_756

CTTGATTTGTCTTATAATGCCATTAATGGAAGCTTGTCTGATAGCTTCTTGAATCTATCCTCACTAGTGGTCTTGAACTTGGAAAGCAACCAACTTGATAACCAAATCCCAGCAGCCATA

>M6-PGSC0003DMG400013198M6LRR-RLK_819

CTAGTGGTCTTGAACTTGGAAAGCAACCAACTTGATAACCAAATCCCAGCAGCCATAATAAAATTGCAGAAACTCTCAGTTCTCAACTTGAGGAGTAACCACTTTAGTGGTGATATTCCT

>M6-PGSC0003DMG400013198M6LRR-RLK_882

TTGCAGAAACTCTCAGTTCTCAACTTGAGGAGTAACCACTTTAGTGGTGATATTCCTGTCACTATAGGAAACATCTCTACTCTTAGACAACTAGATTTAGCTCATAATAATATAAGTGGA

>M6-PGSC0003DMG400013198M6LRR-RLK_945

ATAGGAAACATCTCTACTCTTAGACAACTAGATTTAGCTCATAATAATATAAGTGGAGAAATCCCAGCTTCTTTAGATACTTTACCAAATCTTAGTGCCTTCAATGTCTCATATAATAAT

>M6-PGSC0003DMG400013383M6G-LecRK_1054

GAATTGTTCAGATAAGAATGATTGTTTAGGTTTTTTGTTTAAGTATGATGGGACTGGATATTGTGTGCTTGTTTTGGAGAAATTGTTGTATGGATATTGGTCACCAGGTACTGAGTTTGT

>M6-PGSC0003DMG400013383M6G-LecRK_1116

GTGTGCTTGTTTTGGAGAAATTGTTGTATGGATATTGGTCACCAGGTACTGAGTTTGTTATGTATTTGAGAGTTGATAGTAAAGAGAACGATATATCGAATTTCACAGGCCTGACTAGTT

>M6-PGSC0003DMG400013383M6G-LecRK_1178

TATTTGAGAGTTGATAGTAAAGAGAACGATATATCGAATTTCACAGGCCTGACTAGTTTGATGGAGACTTCATGTCCAGTGAGAATTAGCCTTCCATTTCCACCAGAAGAGTCTAAGACA

>M6-PGSC0003DMG400013383M6G-LecRK_124

ATCATTCAATTCCAAATCCCCTCCTTGGAATCCAACCCAAAACCAAATCCTACTTTCACCCAACTCCACTTTCGCTGCTGGGTTTCTTCAATCTTCCCAAAATTCCTTCAATTTCTCCAT

>M6-PGSC0003DMG400013383M6G-LecRK_1240

GGAGACTTCATGTCCAGTGAGAATTAGCCTTCCATTTCCACCAGAAGAGTCTAAGACAACCACTAGAAATATTGTGATCATTTCGATGCTTTTTGCAGCTGAGTTGATTAGTGGAGTGTT

>M6-PGSC0003DMG400013383M6G-LecRK_1302

CTAGAAATATTGTGATCATTTCGATGCTTTTTGCAGCTGAGTTGATTAGTGGAGTGTTCTTCTTTTGGGCATTTCTCAAGAAGTATATTAAGTATAGGGATATGGCTAGGACTTTTGGTC

>M6-PGSC0003DMG400013383M6G-LecRK_1364

TTTTGGGCATTTCTCAAGAAGTATATTAAGTATAGGGATATGGCTAGGACTTTTGGTCTTGAAGTCATGCCTGCGATTGGACCGAAAAGGTTTAGCTTTTCTGAGATCAAGAATGCAACT

>M6-PGSC0003DMG400013383M6G-LecRK_1426

AGTCATGCCTGCGATTGGACCGAAAAGGTTTAGCTTTTCTGAGATCAAGAATGCAACTAATGATTTCACTGATAAAATTGGGAGGGGTGGATTTGGAGATGTTTACAAAGGGAAGTTGAG

>M6-PGSC0003DMG400013383M6G-LecRK_1488

ATTTCACTGATAAAATTGGGAGGGGTGGATTTGGAGATGTTTACAAAGGGAAGTTGAGTGATGGACGTGTTGTTGCCGTGAAGTGCTTGAAGAATGTAAAAGGTGGTGATGCTGAGTTCT

>M6-PGSC0003DMG400013383M6G-LecRK_1550

GGACGTGTTGTTGCCGTGAAGTGCTTGAAGAATGTAAAAGGTGGTGATGCTGAGTTCTGGGCTGAGGTCACGATCATAGCACGTATGCACCATTTGAATTTGGTGAGGTTGTGGGGGTTT

>M6-PGSC0003DMG400013383M6G-LecRK_1612

TGAGGTCACGATCATAGCACGTATGCACCATTTGAATTTGGTGAGGTTGTGGGGGTTTTGTGCTGAGAAGGGTAGAAGGATACTTGTGTATGAGTATGTACCTAATGGTTCATTGGGTGA

>M6-PGSC0003DMG400013383M6G-LecRK_1674

CTGAGAAGGGTAGAAGGATACTTGTGTATGAGTATGTACCTAATGGTTCATTGGGTGAATTCCTCTTCCAAAAGGCACCGATCCAGTCACCAGATGGGCAGAAACCCATACTCGATTGGA

>M6-PGSC0003DMG400013383M6G-LecRK_1736

CTCTTCCAAAAGGCACCGATCCAGTCACCAGATGGGCAGAAACCCATACTCGATTGGAATATAAGGTACAGGATTGCGCTTGGTGTGGCACGAGCAATTGCCTATCTACATGAAGAGTGT

>M6-PGSC0003DMG400013383M6G-LecRK_1798

AAGGTACAGGATTGCGCTTGGTGTGGCACGAGCAATTGCCTATCTACATGAAGAGTGTTTGGAATGGGTATTACATTGTGATATCAAGCCTGAAAATATCCTCTTAGGCGACGATTTCTG

>M6-PGSC0003DMG400013383M6G-LecRK_186

ACTCCACTTTCGCTGCTGGGTTTCTTCAATCTTCCCAAAATTCCTTCAATTTCTCCATTTGGTACTACAAAATCCCAGTCAAAACCGTTGTTTGGTCAGCTAACCCAAATTCCCCACTTA

>M6-PGSC0003DMG400013383M6G-LecRK_1860

AATGGGTATTACATTGTGATATCAAGCCTGAAAATATCCTCTTAGGCGACGATTTCTGCCCAAAGGTGTCTGATTTTGGGCTAGCGAAGCTAAAGAAGAAAGAGGAGATGATGACCATGT

>M6-PGSC0003DMG400013383M6G-LecRK_1922

AAGGTGTCTGATTTTGGGCTAGCGAAGCTAAAGAAGAAAGAGGAGATGATGACCATGTCAAGATTTCGAGGCACGCCAGGATATGTAGCACCAGAATGGACCAAGGCTGATCCAATTACT

>M6-PGSC0003DMG400013383M6G-LecRK_1984

ATTTCGAGGCACGCCAGGATATGTAGCACCAGAATGGACCAAGGCTGATCCAATTACTCCAAAAGCCGATGTGTACAGCTTTGGATTGGTGCTGTTGGAGATAGTTTCAGGTACAAGGAA

>M6-PGSC0003DMG400013383M6G-LecRK_2046

AAGCCGATGTGTACAGCTTTGGATTGGTGCTGTTGGAGATAGTTTCAGGTACAAGGAACTTTGAGCATCATAACTCAAAGGTGGAAAGTGATCAATGGTTTTTCCCTGCTTGGGCATTTG

>M6-PGSC0003DMG400013383M6G-LecRK_2108

GAGCATCATAACTCAAAGGTGGAAAGTGATCAATGGTTTTTCCCTGCTTGGGCATTTGATAAGGTGTTTAAGGATATGAATGTTGACGATATTTTGGACCCTCAAATCAAACAATCGTAT

>M6-PGSC0003DMG400013383M6G-LecRK_2170

GGTGTTTAAGGATATGAATGTTGACGATATTTTGGACCCTCAAATCAAACAATCGTATGATTCAAGGGCACACTTTGACTTAGTGAACAGGATGGTGAAGACAGCAATGTGGTGCATCCA

>M6-PGSC0003DMG400013383M6G-LecRK_2232

CAAGGGCACACTTTGACTTAGTGAACAGGATGGTGAAGACAGCAATGTGGTGCATCCAAGATCAGCCAGATGCACGACCATCAATGGGGAAAGTGGCTAAGATGTTGGAAGGGACGGTGG

>M6-PGSC0003DMG400013383M6G-LecRK_2294

CAGCCAGATGCACGACCATCAATGGGGAAAGTGGCTAAGATGTTGGAAGGGACGGTGGAGATCATCGAGCCAAAGAAGCCTACCATTTTCTTCTTATCAGAGGAATAATATAAGGCTCAA

>M6-PGSC0003DMG400013383M6G-LecRK_2356

CATCGAGCCAAAGAAGCCTACCATTTTCTTCTTATCAGAGGAATAATATAAGGCTCAATCCTCTTCATAATTTCCTCAAAATGCTGTCTCAACAAGTTGCAGTAAGCATGTATTCCATGT

>M6-PGSC0003DMG400013383M6G-LecRK_2418

CTTCATAATTTCCTCAAAATGCTGTCTCAACAAGTTGCAGTAAGCATGTATTCCATGTTGTTTGAATTTGGATCATGTTGTGTTTTGTCTGTTTTTTTTTTCCAGTTTTGTAATCTCTTG

>M6-PGSC0003DMG400013383M6G-LecRK_2477

GTTTGAATTTGGATCATGTTGTGTTTTGTCTGTTTTTTTTTTCCAGTTTTGTAATCTCTTGTGTTGTAGTTGTAAAGTACATTTTGATCAATGAAGAAGCAAATGGTTGTTACATAGAAG

>M6-PGSC0003DMG400013383M6G-LecRK_248

TACTACAAAATCCCAGTCAAAACCGTTGTTTGGTCAGCTAACCCAAATTCCCCACTTAATTCCTCAGCTACCCTTTTCATTTCTTCCTCCGGTGAACTCAAGCTAACCCCATCTACATCT

>M6-PGSC0003DMG400013383M6G-LecRK_310

CTCAGCTACCCTTTTCATTTCTTCCTCCGGTGAACTCAAGCTAACCCCATCTACATCTTCCTCAGCTCCAAATTTATGGCCTTCAATTATTCGAAACACCTCCTCTGTTCTTTTCCTACA

>M6-PGSC0003DMG400013383M6G-LecRK_372

CAGCTCCAAATTTATGGCCTTCAATTATTCGAAACACCTCCTCTGTTCTTTTCCTACAAGAAGATGGAAGCTTGGTTTATGCAAACTGGAACAGTTTCTTGAACCCAACTGATACTTTTC

>M6-PGSC0003DMG400013383M6G-LecRK_434

GATGGAAGCTTGGTTTATGCAAACTGGAACAGTTTCTTGAACCCAACTGATACTTTTCTTCCGACCCAGAACATCACCGGAACAAATTTAACTTCCGGTAATGGAAAATTCCAGTTTGAT

>M6-PGSC0003DMG400013383M6G-LecRK_496

GACCCAGAACATCACCGGAACAAATTTAACTTCCGGTAATGGAAAATTCCAGTTTGATGGATCGAATTTGAATACCCTTTTCTTCGATGAAAATGATAGTTATTTCACTTTTCCCCAAAA

>M6-PGSC0003DMG400013383M6G-LecRK_558

CGAATTTGAATACCCTTTTCTTCGATGAAAATGATAGTTATTTCACTTTTCCCCAAAATGCCCTTCAGAGGTTGGAGGAAACAGGGGAAGTTACTCAAGTTAATGGAAGATTTTTTTCGT

>M6-PGSC0003DMG400013383M6G-LecRK_620

CTTCAGAGGTTGGAGGAAACAGGGGAAGTTACTCAAGTTAATGGAAGATTTTTTTCGTCGGATTTCGGGGAGAAAGGGAAACTGAGAAGGATGAAATTGGATGAAGATGGGAATATGAGA

>M6-PGSC0003DMG400013383M6G-LecRK_682

TTTCGGGGAGAAAGGGAAACTGAGAAGGATGAAATTGGATGAAGATGGGAATATGAGAATCTATAGTTTTGATCTTAGTTCAAGTTTAGCTAAAAATTGGACAATTGTTTGGGAAGCAGT

>M6-PGSC0003DMG400013383M6G-LecRK_744

ATAGTTTTGATCTTAGTTCAAGTTTAGCTAAAAATTGGACAATTGTTTGGGAAGCAGTTAATCAATTATGTACAATACATGGTACTTGTGGTACAAACTCTATATGCTTGTATGATACTT

>M6-PGSC0003DMG400013383M6G-LecRK_806

CAATTATGTACAATACATGGTACTTGTGGTACAAACTCTATATGCTTGTATGATACTTCTACAACTCAAACTTCTTGTGTTTGTCCACCTGGTTTTAGGAAAGACGCTAGTAAATCTTGT

>M6-PGSC0003DMG400013383M6G-LecRK_868

AACTCAAACTTCTTGTGTTTGTCCACCTGGTTTTAGGAAAGACGCTAGTAAATCTTGTGTGAGGAAAATTCCATTGATGACGAAGGATTCGAAGGCTAGTAAGTATTTACCTCTTGATTT

>M6-PGSC0003DMG400013383M6G-LecRK_930

GGAAAATTCCATTGATGACGAAGGATTCGAAGGCTAGTAAGTATTTACCTCTTGATTTTGTTAGTTTTACTGGTGTGGGTAACCAAACAGACCTTAAAGCTTTGAGCTTTTCGAGTTGTG

>M6-PGSC0003DMG400013383M6G-LecRK_992

AGTTTTACTGGTGTGGGTAACCAAACAGACCTTAAAGCTTTGAGCTTTTCGAGTTGTGAAAAGAATTGTTCAGATAAGAATGATTGTTTAGGTTTTTTGTTTAAGTATGATGGGACTGGA

>M6-PGSC0003DMG400013468-(rev)M6RLKantifungi(GNK)_1037

ACACTGGGTATGCTTTTGTTGTTCTTCTCTGCTGATCCCAAAGATATTGTAGGGAACCTTCCATATTGCCTCATGGCGTGGAATACAAGTTGCTGTAAAAAAGTTTGGGGAGGATGTGAT

>M6-PGSC0003DMG400013468-(rev)M6RLKantifungi(GNK)_1098

CATATTGCCTCATGGCGTGGAATACAAGTTGCTGTAAAAAAGTTTGGGGAGGATGTGATTGCTGATGAGGATAAAGTGTGAGTAACTTAAAAGTAATAGTGCTTCTATTTTTCTACTTAC

>M6-PGSC0003DMG400013468-(rev)M6RLKantifungi(GNK)_1159

CTGATGAGGATAAAGTGTGAGTAACTTAAAAGTAATAGTGCTTCTATTTTTCTACTTACAAGGGGAAAACCCGTAGTTGCTACTTCACATTGTTTAGCTAGCATCTCTCATTTATTTTTT

>M6-PGSC0003DMG400013468-(rev)M6RLKantifungi(GNK)_122

TGTACTGGCTTTTCCTTTTTTCCTAGTCTTATCTATTTGATTTACTGTTGTCATATTTCTATGTCTGTTATCTTGGAAGATTCTAGTGACAAAGGGTTTACCTAGTGTAGCATGATGATC

>M6-PGSC0003DMG400013468-(rev)M6RLKantifungi(GNK)_1220

GGGGAAAACCCGTAGTTGCTACTTCACATTGTTTAGCTAGCATCTCTCATTTATTTTTTGTGGATATTTACATTCCCTAACAGTATAGCTTCTGCAGGAGCGCATTTAGAGATGAGCTTG

>M6-PGSC0003DMG400013468-(rev)M6RLKantifungi(GNK)_1281

GGATATTTACATTCCCTAACAGTATAGCTTCTGCAGGAGCGCATTTAGAGATGAGCTTGCGTTGCTCCAGAAGATGAGACATCCTAATGTGGTGCAATTTCTTGGTGCTGTAACACAAAG

>M6-PGSC0003DMG400013468-(rev)M6RLKantifungi(GNK)_1342

TTGCTCCAGAAGATGAGACATCCTAATGTGGTGCAATTTCTTGGTGCTGTAACACAAAGTAGTCCTATGATGATAGTGACGGAGTACTTACCAAAGGTCTTACTTATTAACTCTTGACAG

>M6-PGSC0003DMG400013468-(rev)M6RLKantifungi(GNK)_1403

GTCCTATGATGATAGTGACGGAGTACTTACCAAAGGTCTTACTTATTAACTCTTGACAGTTTGCAGAATTATATTTGACAACATTTTTATACACTGTTTTCTCCCTTATAGTTCTATCCA

>M6-PGSC0003DMG400013468-(rev)M6RLKantifungi(GNK)_1464

TGCAGAATTATATTTGACAACATTTTTATACACTGTTTTCTCCCTTATAGTTCTATCCATTTTTTCAGTTTACATTCTCTTTTCTTCTTTTTGACCATATGTTTGAAACTTTTCCAACAA

>M6-PGSC0003DMG400013468-(rev)M6RLKantifungi(GNK)_1525

TTTTCAGTTTACATTCTCTTTTCTTCTTTTTGACCATATGTTTGAAACTTTTCCAACAAACCTATCACTTTGAAATGACTTTTCCATGCCGTATGTTTCCAGGGTGATCTCCGTGAATAT

>M6-PGSC0003DMG400013468-(rev)M6RLKantifungi(GNK)_1586

CTATCACTTTGAAATGACTTTTCCATGCCGTATGTTTCCAGGGTGATCTCCGTGAATATTTGAATAGAAAAGGAGCACTTAAACCAACTAAAGCTCTCAGATTTGCAATGGACATCGCGA

>M6-PGSC0003DMG400013468-(rev)M6RLKantifungi(GNK)_1647

GAATAGAAAAGGAGCACTTAAACCAACTAAAGCTCTCAGATTTGCAATGGACATCGCGAGGTTCACATCCACTCTGTGCCACTGACCAATATCATTAACTCTTTATAAATTGCAAGATTA

>M6-PGSC0003DMG400013468-(rev)M6RLKantifungi(GNK)_1708

TTCACATCCACTCTGTGCCACTGACCAATATCATTAACTCTTTATAAATTGCAAGATTAATGGTTTAGATTTTTCTTAAAAAAAACAATTTTTTCATTATGCATTACTTATAAAACTTTG

>M6-PGSC0003DMG400013468-(rev)M6RLKantifungi(GNK)_1769

GGTTTAGATTTTTCTTAAAAAAAACAATTTTTTCATTATGCATTACTTATAAAACTTTGAGTTAATGTTCAAAAATACACTTAAAAGTATTCGTTTTTTTGAGTTTACCACCTGAACTAT

>M6-PGSC0003DMG400013468-(rev)M6RLKantifungi(GNK)_183

TGTCTGTTATCTTGGAAGATTCTAGTGACAAAGGGTTTACCTAGTGTAGCATGATGATCACCTTAGAGTTCAATATTGAATTAGTAAGATTGTACTATTGGAATGGAAGCAGCAGTTGCC

>M6-PGSC0003DMG400013468-(rev)M6RLKantifungi(GNK)_1830

TTAATGTTCAAAAATACACTTAAAAGTATTCGTTTTTTTGAGTTTACCACCTGAACTATGCGAGTTTCCTACCTAAACTAGCACCAACTATTTTTCAAAATTCACCTCCTCGATCTGTTG

>M6-PGSC0003DMG400013468-(rev)M6RLKantifungi(GNK)_1891

GAGTTTCCTACCTAAACTAGCACCAACTATTTTTCAAAATTCACCTCCTCGATCTGTTGTTCCCCTTTTCCAACCTGAAATATAACCATCGTTCAGGTAAGAAAAGAGAAGAATTGATGT

>M6-PGSC0003DMG400013468-(rev)M6RLKantifungi(GNK)_1952

CCCCTTTTCCAACCTGAAATATAACCATCGTTCAGGTAAGAAAAGAGAAGAATTGATGTTGAATTGTGTTTTGATGAGTAGTTCTGGTAGAAAACTCGTACACTTGATAGCTCCGGTATG

>M6-PGSC0003DMG400013468-(rev)M6RLKantifungi(GNK)_2074

ACTCGAAAAACTGGAATACTTTAGGTGTGTTTTTGACCGTTCACTCTAAAGCTTTTTTTGGATGTAAAATATCAATGTCTTTTAGTCCTTTGATTGGAGCATTACTGCTGAGTATTAAAA

>M6-PGSC0003DMG400013468-(rev)M6RLKantifungi(GNK)_2135

ATGTAAAATATCAATGTCTTTTAGTCCTTTGATTGGAGCATTACTGCTGAGTATTAAAATTCTCATATATAATATTTGGTTTTAGGGGAATGAATTACCTGCATGAACATAAAGAAGCAA

>M6-PGSC0003DMG400013468-(rev)M6RLKantifungi(GNK)_2196

CTCATATATAATATTTGGTTTTAGGGGAATGAATTACCTGCATGAACATAAAGAAGCAATTATTCATCGTGATCTTGAGCCTTCGTAAGTTTCACTTTCGTCAGGTTACCTGAATTACAT

>M6-PGSC0003DMG400013468-(rev)M6RLKantifungi(GNK)_2257

ATTCATCGTGATCTTGAGCCTTCGTAAGTTTCACTTTCGTCAGGTTACCTGAATTACATTTTGTCTATACTAACGCCTAGCCAAGTTTTCGAGTAGCATCAATGCTTTCATATTGTGATT

>M6-PGSC0003DMG400013468-(rev)M6RLKantifungi(GNK)_2318

TGTCTATACTAACGCCTAGCCAAGTTTTCGAGTAGCATCAATGCTTTCATATTGTGATTTGAGACTCTTCATGTAATAATAGTTCACTTTTTTTTGGAGTAAGATAACAGAAATATTTTG

>M6-PGSC0003DMG400013468-(rev)M6RLKantifungi(GNK)_2379

AGACTCTTCATGTAATAATAGTTCACTTTTTTTTGGAGTAAGATAACAGAAATATTTTGCTGGATGATACTGGACATCTGAAAGTTGCTGACTTTGGAGTAAGCAAGCTACTGAAAGTTA

>M6-PGSC0003DMG400013468-(rev)M6RLKantifungi(GNK)_244

CTTAGAGTTCAATATTGAATTAGTAAGATTGTACTATTGGAATGGAAGCAGCAGTTGCCTGAGGCAAGTATGAGTAGATAAGGGATGTTCAATGGAGGAAAACTGAGGCTAGCCTCATCT

>M6-PGSC0003DMG400013468-(rev)M6RLKantifungi(GNK)_2440

GGATGATACTGGACATCTGAAAGTTGCTGACTTTGGAGTAAGCAAGCTACTGAAAGTTACAAAAAGGGTCAAAGAAGATAAGCCTTTGACATACGAAGACAGTTATCGTGAGTACTTCTT

>M6-PGSC0003DMG400013468-(rev)M6RLKantifungi(GNK)_2501

AAAAGGGTCAAAGAAGATAAGCCTTTGACATACGAAGACAGTTATCGTGAGTACTTCTTCTTTTAGATATTCTAGTCCTCTATTCCTTCTCCATTGGAAAAGATATGGTAGCTTGTTACT

>M6-PGSC0003DMG400013468-(rev)M6RLKantifungi(GNK)_2562

TTTAGATATTCTAGTCCTCTATTCCTTCTCCATTGGAAAAGATATGGTAGCTTGTTACTGTGGAAAATAATATATGCGTGAGTGTTTTCAAGTTTACTTGATAGCTCCAAAAAAAGCTTT

>M6-PGSC0003DMG400013468-(rev)M6RLKantifungi(GNK)_2623

GGAAAATAATATATGCGTGAGTGTTTTCAAGTTTACTTGATAGCTCCAAAAAAAGCTTTCTTTATTTGAGATGTTGTGATTTGATCCTGTTTAATGATATTTGAGAACCGATATTTGGAT

>M6-PGSC0003DMG400013468-(rev)M6RLKantifungi(GNK)_2684

TTATTTGAGATGTTGTGATTTGATCCTGTTTAATGATATTTGAGAACCGATATTTGGATTTCTTCTAATATTTTCATCATCTTTTTTTCTAAAAAAAAACTTTTCAATTCTTGTGCAACA

>M6-PGSC0003DMG400013468-(rev)M6RLKantifungi(GNK)_2745

CTTCTAATATTTTCATCATCTTTTTTTCTAAAAAAAAACTTTTCAATTCTTGTGCAACAAATAGGCCGATATGTAGCTCCAGAGGTTTTCAGGAATGAGGAATATGATACCAAAGTTGAT

>M6-PGSC0003DMG400013468-(rev)M6RLKantifungi(GNK)_2806

TAGGCCGATATGTAGCTCCAGAGGTTTTCAGGAATGAGGAATATGATACCAAAGTTGATGTCTTTTCATTTGCCTTGATTTTACAAGAGGTAAAATTTTGTCTTATCAGATTCTTGTTTA

>M6-PGSC0003DMG400013468-(rev)M6RLKantifungi(GNK)_2867

CTTTTCATTTGCCTTGATTTTACAAGAGGTAAAATTTTGTCTTATCAGATTCTTGTTTAAACTTTGAGGAAGAACAGTTTGTTTGCACTCTTGTGGACCATATCTTTATTTTATGTTTAA

>M6-PGSC0003DMG400013468-(rev)M6RLKantifungi(GNK)_2928

CTTTGAGGAAGAACAGTTTGTTTGCACTCTTGTGGACCATATCTTTATTTTATGTTTAAGAGTGTGTGAATGCAAGGAGTATTATGTTCCGTGTTCTTAAAAGTGAAAGCGCAAAACTGG

>M6-PGSC0003DMG400013468-(rev)M6RLKantifungi(GNK)_2989

GTGTGTGAATGCAAGGAGTATTATGTTCCGTGTTCTTAAAAGTGAAAGCGCAAAACTGGAAGTGAACACATACTTTTTTAAAATGAAATCGAAATGAGTGCTCAATCTGTAGAAAATCAG

>M6-PGSC0003DMG400013468-(rev)M6RLKantifungi(GNK)_305

AGGCAAGTATGAGTAGATAAGGGATGTTCAATGGAGGAAAACTGAGGCTAGCCTCATCTGATCACATCTTAGACTGATCTGTGGCATATTGTTCAGCTGCATAGGATTGCTTATCTTCTT

>M6-PGSC0003DMG400013468-(rev)M6RLKantifungi(GNK)_3050

GTGAACACATACTTTTTTAAAATGAAATCGAAATGAGTGCTCAATCTGTAGAAAATCAGAGATCGTGTATAACATATATGTATTTAATACTAATGGCATCAAATTACAAATAAATTACTA

>M6-PGSC0003DMG400013468-(rev)M6RLKantifungi(GNK)_3111

ATCGTGTATAACATATATGTATTTAATACTAATGGCATCAAATTACAAATAAATTACTAACTTAAGAATTCAATATATAATGATACCATATTATTCTTGCAATTCTGCTTCAGAAACATT

>M6-PGSC0003DMG400013468-(rev)M6RLKantifungi(GNK)_3355

AGATCAAATTTAACATGATATAAATATGTTTCAAAGGTCACAGAAATATACAGTGATTATCGTTTACTTGAATATTGATCTGATCTATTTATGTTGTATATCTGTGTAAGTTTGAAAGTA

>M6-PGSC0003DMG400013468-(rev)M6RLKantifungi(GNK)_3416

GTTTACTTGAATATTGATCTGATCTATTTATGTTGTATATCTGTGTAAGTTTGAAAGTAACTCCGGTTTGATGAAATTTAGTGAGTCAAACATGTGTGTCTTTTGCAGTTTCTCTTTGCA

>M6-PGSC0003DMG400013468-(rev)M6RLKantifungi(GNK)_3477

TCCGGTTTGATGAAATTTAGTGAGTCAAACATGTGTGTCTTTTGCAGTTTCTCTTTGCAGAAACAGGTGATATATTTATCTTTGTGTTGTGCTGTATTTTAGATGATTGAAGGCTATCCC

>M6-PGSC0003DMG400013468-(rev)M6RLKantifungi(GNK)_3538

AACAGGTGATATATTTATCTTTGTGTTGTGCTGTATTTTAGATGATTGAAGGCTATCCCCCTTTCCATGCGAAGGAAGAAAACGATGTACCTAAATTGTATGCTGCTAAAGAGCGCCCTC

>M6-PGSC0003DMG400013468-(rev)M6RLKantifungi(GNK)_3599

TTTCCATGCGAAGGAAGAAAACGATGTACCTAAATTGTATGCTGCTAAAGAGCGCCCTCCTTTTAAAGCCCCGGCAAAGTTTTATGCCCATGGACTGAAAGAGTAAGTTCTATATTTTAC

>M6-PGSC0003DMG400013468-(rev)M6RLKantifungi(GNK)_366

TCACATCTTAGACTGATCTGTGGCATATTGTTCAGCTGCATAGGATTGCTTATCTTCTTCTTTTGGAACCTTGGCTAACTATTTGTTGATTAGTGTGTACAATCTGATGCTTACAAGAAT

>M6-PGSC0003DMG400013468-(rev)M6RLKantifungi(GNK)_3660

TTTAAAGCCCCGGCAAAGTTTTATGCCCATGGACTGAAAGAGTAAGTTCTATATTTTACATCAGTTGCTACTATAAGTCACGATATTTTATTTTAGTATAATAAACCATGCTATTCTAGG

>M6-PGSC0003DMG400013468-(rev)M6RLKantifungi(GNK)_3721

CAGTTGCTACTATAAGTCACGATATTTTATTTTAGTATAATAAACCATGCTATTCTAGGGTAAATATCGATTGTTATAATCTGAGAATATGAAAACTCATGTCAATTAGGTGTGAGAGGA

>M6-PGSC0003DMG400013468-(rev)M6RLKantifungi(GNK)_3904

TTCTAAATAGTTACCTGGTACTTGCAACTGTGGCATAGTAAACGTACTTAAAAAGAACACAAAAAAACTAATAAGAACTGAACAGACCTACAATTATAAAATCAAACTATTCAGCATTTG

>M6-PGSC0003DMG400013468-(rev)M6RLKantifungi(GNK)_3965

AAAAAACTAATAAGAACTGAACAGACCTACAATTATAAAATCAAACTATTCAGCATTTGCGATCCTTTCCTTCCGTATAGGTGGCAGCTTCTTTCCACCTCTCTGTTGCACATAAAACAC

>M6-PGSC0003DMG400013468-(rev)M6RLKantifungi(GNK)_4026

ATCCTTTCCTTCCGTATAGGTGGCAGCTTCTTTCCACCTCTCTGTTGCACATAAAACACCTACACATCTACGTATCATGATCCCCCTTTTCTGTAGGTTGTTTTGTGTAAGGAAAGCATC

>M6-PGSC0003DMG400013468-(rev)M6RLKantifungi(GNK)_4087

ACACATCTACGTATCATGATCCCCCTTTTCTGTAGGTTGTTTTGTGTAAGGAAAGCATCCCAGGCGACATGCTATACGAAAAAAGAGGGCTTTAGACTTACCAAAGCTGTTTCTATATCC

>M6-PGSC0003DMG400013468-(rev)M6RLKantifungi(GNK)_4148

AGGCGACATGCTATACGAAAAAAGAGGGCTTTAGACTTACCAAAGCTGTTTCTATATCCCATTGGTAAAGCTGGTGGAGGGAGAAGCCCATATACGGGTTAAGCCGAGCCCAATAGCTTT

>M6-PGSC0003DMG400013468-(rev)M6RLKantifungi(GNK)_4209

TTGGTAAAGCTGGTGGAGGGAGAAGCCCATATACGGGTTAAGCCGAGCCCAATAGCTTTTGCTCAAACAATGTATCTTTGTTAAGAGATTCATTAAATATGTATACATATGAAATTTAGA

>M6-PGSC0003DMG400013468-(rev)M6RLKantifungi(GNK)_427

TTTGGAACCTTGGCTAACTATTTGTTGATTAGTGTGTACAATCTGATGCTTACAAGAATTGTGAGATGACTCATCTAATTGTTTACTGAAGTGGCCTAGTATATCTTTCTTCTCCAATAT

>M6-PGSC0003DMG400013468-(rev)M6RLKantifungi(GNK)_4270

CTCAAACAATGTATCTTTGTTAAGAGATTCATTAAATATGTATACATATGAAATTTAGAACCTAATGCATGAAGTCGTCGTTCTAAAATTTACAATCCATAAAGTTGAAATCCTGGCTCT

>M6-PGSC0003DMG400013468-(rev)M6RLKantifungi(GNK)_4331

CTAATGCATGAAGTCGTCGTTCTAAAATTTACAATCCATAAAGTTGAAATCCTGGCTCTACCTTAGTTGGCTTTAATCTTGATTCTACTTATACATAGATTAGTAGGGTTATAACTGATC

>M6-PGSC0003DMG400013468-(rev)M6RLKantifungi(GNK)_4392

CTTAGTTGGCTTTAATCTTGATTCTACTTATACATAGATTAGTAGGGTTATAACTGATCAACTGGTGTCTTTACGTAGATGGTAGCATTTAAACAAACTGGGATCTTGGAAAATGTCCTT

>M6-PGSC0003DMG400013468-(rev)M6RLKantifungi(GNK)_4453

CTGGTGTCTTTACGTAGATGGTAGCATTTAAACAAACTGGGATCTTGGAAAATGTCCTTATTTTCGTTTTATTATGTATAAATTTTACCATTTCATGATCATCAAGTTGTTGTTGTCTCC

>M6-PGSC0003DMG400013468-(rev)M6RLKantifungi(GNK)_4819

AATAGTAATATTAGGGTTGCTTTTGGTGTTCAGATTGATCGAAGAGTGTTGGAATGACAAGCCAACCGAGCGGCCAAGTTTCAAGCAAATTATCCCAAGGCTTGAGTTCATCTATAAAAA

>M6-PGSC0003DMG400013468-(rev)M6RLKantifungi(GNK)_4880

CCAACCGAGCGGCCAAGTTTCAAGCAAATTATCCCAAGGCTTGAGTTCATCTATAAAAAGTCTGACCACAACAAGCGCTGGAAGGTAAAATATCCATTAGTGGTTCTTGCACTTCAAACT

>M6-PGSC0003DMG400013468-(rev)M6RLKantifungi(GNK)_4941

CTGACCACAACAAGCGCTGGAAGGTAAAATATCCATTAGTGGTTCTTGCACTTCAAACTATCGTATTGTTTTGATGTAGTGTCTGTTCCTTTTTTAGTATGTCTTGTTATGCTTTCTTTA

>M6-PGSC0003DMG400013468-(rev)M6RLKantifungi(GNK)_5246

GTATTGGTGTTTTTGATCTTTTCACATTTTGGTCATCAATGAATCAAACTATTTGCCATTTGTTTCATCGAATAGAGACATATTAGCTCTAAATCACTCGAATTTCATGCTTTTGTGAGA

>M6-PGSC0003DMG400013468-(rev)M6RLKantifungi(GNK)_5307

GTTTCATCGAATAGAGACATATTAGCTCTAAATCACTCGAATTTCATGCTTTTGTGAGATATTGACTTCTAGTGAAAAAACAATATCCTGGCTCTGATTTGATAAGTAGCTCATCTGAGA

>M6-PGSC0003DMG400013468-(rev)M6RLKantifungi(GNK)_5368

TTGACTTCTAGTGAAAAAACAATATCCTGGCTCTGATTTGATAAGTAGCTCATCTGAGATACTATTAGTGGACGAGTTAGTTATAATTAGCTCAGATGTAATTTCTTTGTTCAGTGTTGG

>M6-PGSC0003DMG400013468-(rev)M6RLKantifungi(GNK)_5429

CTATTAGTGGACGAGTTAGTTATAATTAGCTCAGATGTAATTTCTTTGTTCAGTGTTGGTTACTTGACTCAATAATGTCCCTTCAGTCCCAAACAAGTTGGGGTCGACTATATGAATCTG

>M6-PGSC0003DMG400013468-(rev)M6RLKantifungi(GNK)_5490

ACTTGACTCAATAATGTCCCTTCAGTCCCAAACAAGTTGGGGTCGACTATATGAATCTGAGCTAATCACCACGTTCCCATTTAATTTCATCTCAGTCCAACATTATTACACAAAACCTTA

>M6-PGSC0003DMG400013468-(rev)M6RLKantifungi(GNK)_5551

CTAATCACCACGTTCCCATTTAATTTCATCTCAGTCCAACATTATTACACAAAACCTTAATCTCGCCACGATCACTGCAGTTTAGCATAACATATTATTCAGTAGATACCTAAGTGCCTC

>M6-PGSC0003DMG400013468-(rev)M6RLKantifungi(GNK)_5612

CTCGCCACGATCACTGCAGTTTAGCATAACATATTATTCAGTAGATACCTAAGTGCCTCTTTTTCACCATGAATACTTTGCTACATGCAGGTAATTAGACCATTGAACTGCTTTCAGCAT

>M6-PGSC0003DMG400013468-(rev)M6RLKantifungi(GNK)_5673

TTTCACCATGAATACTTTGCTACATGCAGGTAATTAGACCATTGAACTGCTTTCAGCATTTAGCGGCAATGTGGAAGAAAGATGGTTCAGGTTTAAGTAGTCGGAATCGTTCATCTCGGT

>M6-PGSC0003DMG400013468-(rev)M6RLKantifungi(GNK)_5734

AGCGGCAATGTGGAAGAAAGATGGTTCAGGTTTAAGTAGTCGGAATCGTTCATCTCGGTCTATGCGCCACATCTAAGTAAGTTTCTGGATAAAGTTTTTCTACTTCTCCACATTGTCTGC

>M6-PGSC0003DMG400013468-(rev)M6RLKantifungi(GNK)_5793

CTATGCGCCACATCTAAGTAAGTTTCTGGATAAAGTTTTTCTACTTCTCCACATTGTCTGCAAAAATATTCAGTTGCTCTTAGTAGTATATAGGCTGTTGGATTATTTTCTAGTTTCAAA

>M6-PGSC0003DMG400013468-(rev)M6RLKantifungi(GNK)_61

ATCCATTTTTCATTCTGCCATCATTGTTAGTTATCATTCTCCAACCTGAGACTCTGTAGTCTGTACTGGCTTTTCCTTTTTTCCTAGTCTTATCTATTTGATTTACTGTTGTCATATTTC

>M6-PGSC0003DMG400013489-(rev)M6LRR-RLK_0

AAAGAAAGAAACTTGGCATTTGGGTGTTTTGAGGTTTTTCCAGTTCTTGAAAATGCTAGTTGTGTAACGACTGAAACTTTGTCATTTGGGTGTTTGAGGTTTTTTCAGTTTCTTGAAAGA

>M6-PGSC0003DMG400013489-(rev)M6LRR-RLK_1037

CTACGATTTCCTCTCGTTATATTGAATCTTTGATGCAATGACAAACTGAAAAGTGAAAACAGTGGAAGTAATCTAATAGAGTGACTGTTTTTCCATATAATAATGTGCAGTTGTAGCTTT

>M6-PGSC0003DMG400013489-(rev)M6LRR-RLK_1098

GTGGAAGTAATCTAATAGAGTGACTGTTTTTCCATATAATAATGTGCAGTTGTAGCTTTGACGGAAATTAAGAAGGCCTTACATGACCCCTATAATGTTCTGGACAATTGGGATGTGAAT

>M6-PGSC0003DMG400013489-(rev)M6LRR-RLK_1159

CGGAAATTAAGAAGGCCTTACATGACCCCTATAATGTTCTGGACAATTGGGATGTGAATTCTGTAGACCCCTGCAGTTGGAGAATGGTCACTTGCTCCTTGGATGGCTATGTGTCTGCTC

>M6-PGSC0003DMG400013489-(rev)M6LRR-RLK_122

TAGTTGTATAAAGACTGGAACTTTATCATTTGGGTGTTTGAGCTTTTTCCAGTTCTTGAAAAGCTAGTTGTGTAAAGACTGAAACTTTATCATCTGAGTGTTTGAGGTTTTCAATCTTGA

>M6-PGSC0003DMG400013489-(rev)M6LRR-RLK_1220

TGTAGACCCCTGCAGTTGGAGAATGGTCACTTGCTCCTTGGATGGCTATGTGTCTGCTCTGTAAGTTGTTTGGATGAAAGAGTAATTTAGTAGAAACTTAGTTGGCTCATGAAAAGGCCT

>M6-PGSC0003DMG400013489-(rev)M6LRR-RLK_1281

TAAGTTGTTTGGATGAAAGAGTAATTTAGTAGAAACTTAGTTGGCTCATGAAAAGGCCTGATTTAATCCAGTAAATTTATCGCTGAATATTTTCTCTTGCATCAGAGCACTACCTAGTCA

>M6-PGSC0003DMG400013489-(rev)M6LRR-RLK_1342

TTTAATCCAGTAAATTTATCGCTGAATATTTTCTCTTGCATCAGAGCACTACCTAGTCAAAGTCTAGCTGGCACCTTATCGCCTGGAATTGGCAATCTTACTAACTTGCAATATGTGTAA

>M6-PGSC0003DMG400013489-(rev)M6LRR-RLK_1403

GTCTAGCTGGCACCTTATCGCCTGGAATTGGCAATCTTACTAACTTGCAATATGTGTAAGTTCTTGACTATATACATCTCTTAGTTGCTGCTACTTGGTTCCGTAATACCTAAGGTCACT

>M6-PGSC0003DMG400013489-(rev)M6LRR-RLK_1464

TCTTGACTATATACATCTCTTAGTTGCTGCTACTTGGTTCCGTAATACCTAAGGTCACTTCTTATATGATTTTCTTAGTGTTATTAAGCCATTTTCTCGCAAACTGTTCTTAACTTGTTC

>M6-PGSC0003DMG400013489-(rev)M6LRR-RLK_1525

TTATATGATTTTCTTAGTGTTATTAAGCCATTTTCTCGCAAACTGTTCTTAACTTGTTCATGTAGATTGTTGCAAAACAATGCCATTTCTGGTCATATTCCTGATGTGATTGGGAAGTTA

>M6-PGSC0003DMG400013489-(rev)M6LRR-RLK_1586

GTAGATTGTTGCAAAACAATGCCATTTCTGGTCATATTCCTGATGTGATTGGGAAGTTACAAAAGCTTCAGACACTTGATCTTTCCAACAATAAATTTGAAGGCGAAATACCCACTTCCT

>M6-PGSC0003DMG400013489-(rev)M6LRR-RLK_1647

AAAGCTTCAGACACTTGATCTTTCCAACAATAAATTTGAAGGCGAAATACCCACTTCCTTCGAAGACCTGAAGAATTTGAATTATTTGTGAGTTCATGAATTTTCACTTATTATTATTCA

>M6-PGSC0003DMG400013489-(rev)M6LRR-RLK_1708

GAAGACCTGAAGAATTTGAATTATTTGTGAGTTCATGAATTTTCACTTATTATTATTCACTTTCAAGACTCCAACCATATATTTCACTCTGTCAAAGATTGAATACATCATGCTACCTAT

>M6-PGSC0003DMG400013489-(rev)M6LRR-RLK_1769

TTCAAGACTCCAACCATATATTTCACTCTGTCAAAGATTGAATACATCATGCTACCTATTGACTCAAATCTTCTGTTTCGTCCTCATCCTTGTGATTGACTTAGCTGATCATTGCTTGCA

>M6-PGSC0003DMG400013489-(rev)M6LRR-RLK_183

AGCTAGTTGTGTAAAGACTGAAACTTTATCATCTGAGTGTTTGAGGTTTTCAATCTTGAAATGAGCTAGTTGTGTAAAGACTGAAACTTTGTCATTTGGGTGTTTGAGATTTTCCAATTC

>M6-PGSC0003DMG400013489-(rev)M6LRR-RLK_1830

ACTCAAATCTTCTGTTTCGTCCTCATCCTTGTGATTGACTTAGCTGATCATTGCTTGCAGGGATGTCTCTTATAATAATCTAAGTGGTACCCTTCCTAAAATATCAGCGCGAGCTTTCAA

>M6-PGSC0003DMG400013489-(rev)M6LRR-RLK_1891

GATGTCTCTTATAATAATCTAAGTGGTACCCTTCCTAAAATATCAGCGCGAGCTTTCAAGTAAGTCGGCCTTTTCTATTGCCGTAATATACTCAGTTATTTCATTTTCAAGTAGTTATCT

>M6-PGSC0003DMG400013489-(rev)M6LRR-RLK_1952

AAGTCGGCCTTTTCTATTGCCGTAATATACTCAGTTATTTCATTTTCAAGTAGTTATCTTAATATTATTGAAGTTAAACTATTGCTGTACGGCTGTACATGTAAATCTCGATGCTTAGTT

>M6-PGSC0003DMG400013489-(rev)M6LRR-RLK_2013

ATATTATTGAAGTTAAACTATTGCTGTACGGCTGTACATGTAAATCTCGATGCTTAGTTGATAAAAATCCATTATTAAAAATACATTGACAAATGCTTTTGGGTAAACTGGTACTTGAAA

>M6-PGSC0003DMG400013489-(rev)M6LRR-RLK_2074

TAAAAATCCATTATTAAAAATACATTGACAAATGCTTTTGGGTAAACTGGTACTTGAAAAGTATTTCATTAGTATTCCACACTATAGTGAGATTGACTTCCTCTTCATGGCTTTCAGAGT

>M6-PGSC0003DMG400013489-(rev)M6LRR-RLK_2135

TATTTCATTAGTATTCCACACTATAGTGAGATTGACTTCCTCTTCATGGCTTTCAGAGTGCTCGGGAATCCTTTAATTTGTGGTCAAGGTTCTGAGAATAATTGCTCAGCAGTTTATCCA

>M6-PGSC0003DMG400013489-(rev)M6LRR-RLK_2196

TCGGGAATCCTTTAATTTGTGGTCAAGGTTCTGAGAATAATTGCTCAGCAGTTTATCCAGAACCACTGTCCTTCCCACCTGATAGTCTGGCAGGTACAGATAGCATTTTCATGCTCAAGT

>M6-PGSC0003DMG400013489-(rev)M6LRR-RLK_2257

ACCACTGTCCTTCCCACCTGATAGTCTGGCAGGTACAGATAGCATTTTCATGCTCAAGTCCCTTCCCTTGCACCTGAAAATGGAAAACTGTTAAAAACATATTATACATACTGAGGAAGT

>M6-PGSC0003DMG400013489-(rev)M6LRR-RLK_2318

CTTCCCTTGCACCTGAAAATGGAAAACTGTTAAAAACATATTATACATACTGAGGAAGTTTTGCTATTTGTGTTATTGAAAATGTGTTGCATCTATTATTTTTGATATCACAGATCAATC

>M6-PGSC0003DMG400013489-(rev)M6LRR-RLK_2379

TGCTATTTGTGTTATTGAAAATGTGTTGCATCTATTATTTTTGATATCACAGATCAATCTGGTGCTGCACGCAAGAGCCGTCGTGTGGTTGTTGCGTTTGTTGCAAGTTTTGGTGCTGCT

>M6-PGSC0003DMG400013489-(rev)M6LRR-RLK_244

TGAGCTAGTTGTGTAAAGACTGAAACTTTGTCATTTGGGTGTTTGAGATTTTCCAATTCTTGAAAGCTAGTTGTGTAAAGATTGAAACTTTGTCATTTGGGTGTTTGAGGTTTGCAATCT

>M6-PGSC0003DMG400013489-(rev)M6LRR-RLK_2440

GTGCTGCACGCAAGAGCCGTCGTGTGGTTGTTGCGTTTGTTGCAAGTTTTGGTGCTGCTTTCTTGGTCATAATAGCCCTCGCTTTGCTTCTTTGGTGGCGCTACAGAAATAATCAGCAGA

>M6-PGSC0003DMG400013489-(rev)M6LRR-RLK_2501

CTTGGTCATAATAGCCCTCGCTTTGCTTCTTTGGTGGCGCTACAGAAATAATCAGCAGATCTTCTTTGATGTCAATGGTTAGTTGATTTACTTTCCCCCCCTCTTCTCTATGATAATTTG

>M6-PGSC0003DMG400013489-(rev)M6LRR-RLK_2562

TTCTTTGATGTCAATGGTTAGTTGATTTACTTTCCCCCCCTCTTCTCTATGATAATTTGAGCTTTTATTAGTTCTGCATTTCTGATTCTTTTCCTGCCACGGATATATAAGTTGAGCCAA

>M6-PGSC0003DMG400013489-(rev)M6LRR-RLK_2623

CTTTTATTAGTTCTGCATTTCTGATTCTTTTCCTGCCACGGATATATAAGTTGAGCCAAGTGAGCTTTTGCATTATTGTCTGAGTGATTGGTGTTAGGTATCATGTGAATATTTTTTAAG

>M6-PGSC0003DMG400013489-(rev)M6LRR-RLK_2684

GAGCTTTTGCATTATTGTCTGAGTGATTGGTGTTAGGTATCATGTGAATATTTTTTAAGCTAGGATGTTACAGAGTCACCTGCTTGTGGTTTGAATCTTTATTTTAATCTCTAAAAAGAT

>M6-PGSC0003DMG400013489-(rev)M6LRR-RLK_2745

AGGATGTTACAGAGTCACCTGCTTGTGGTTTGAATCTTTATTTTAATCTCTAAAAAGATTTTTGTGATTGTTGATTTTAAATTCATTTGTTTAGTTTTAGAGAGGACATGAGTGGTCAAG

>M6-PGSC0003DMG400013489-(rev)M6LRR-RLK_2806

TTGTGATTGTTGATTTTAAATTCATTTGTTTAGTTTTAGAGAGGACATGAGTGGTCAAGGAATAGGAATTGGGGATAAATTCTGTGTTTACCATATTCAATAAACCTGTTGAATATGAAG

>M6-PGSC0003DMG400013489-(rev)M6LRR-RLK_2867

ATAGGAATTGGGGATAAATTCTGTGTTTACCATATTCAATAAACCTGTTGAATATGAAGAACTTTCGAACCCCGTCAGTTAGAAAATTATGCCCTTTGGCATACTCAGTAGTTCTGGTCA

>M6-PGSC0003DMG400013489-(rev)M6LRR-RLK_2928

CTTTCGAACCCCGTCAGTTAGAAAATTATGCCCTTTGGCATACTCAGTAGTTCTGGTCAATTTTCCTAGTAAGGACTCAGAAAAGCTGTTTGAATTTTATTGTTATCGGGACAAGTCCTG

>M6-PGSC0003DMG400013489-(rev)M6LRR-RLK_2989

TTTCCTAGTAAGGACTCAGAAAAGCTGTTTGAATTTTATTGTTATCGGGACAAGTCCTGATCCACTACAGGCTACCTTAAACATTTAGTTCCAAACATTGACAGAATAATTTGTTAATCA

>M6-PGSC0003DMG400013489-(rev)M6LRR-RLK_305

GAAAGCTAGTTGTGTAAAGATTGAAACTTTGTCATTTGGGTGTTTGAGGTTTGCAATCTTGAAATGAACTAGTTGTGTAAATACAGAGACTTTGTCATTTGGGTGTTTGAGCTTTTTCCA

>M6-PGSC0003DMG400013489-(rev)M6LRR-RLK_3050

CCACTACAGGCTACCTTAAACATTTAGTTCCAAACATTGACAGAATAATTTGTTAATCACATATCTGAATAATTACACTTGCATTTTCTTGTGCAAGTTGAAGTAAATATTTCTGTAAAA

>M6-PGSC0003DMG400013489-(rev)M6LRR-RLK_3111

TATCTGAATAATTACACTTGCATTTTCTTGTGCAAGTTGAAGTAAATATTTCTGTAAAAAGGAAGATGGTGTTATGTTCACTCTCTCGGGCCACCATGCCTGGTGAATTGCCTTGCCACA

>M6-PGSC0003DMG400013489-(rev)M6LRR-RLK_3172

GAAGATGGTGTTATGTTCACTCTCTCGGGCCACCATGCCTGGTGAATTGCCTTGCCACATTCATTGCGTTGAAAGTTACTCTTGATAATCTTGTACTTTGGTTGTCTGATTCATATATGT

>M6-PGSC0003DMG400013489-(rev)M6LRR-RLK_3233

CATTGCGTTGAAAGTTACTCTTGATAATCTTGTACTTTGGTTGTCTGATTCATATATGTGGATCAGTTTTGGGACGTTTTTGCCAACCTCTATTATTTTCTCCCTGTTTCACTTTTCAAA

>M6-PGSC0003DMG400013489-(rev)M6LRR-RLK_3294

ATCAGTTTTGGGACGTTTTTGCCAACCTCTATTATTTTCTCCCTGTTTCACTTTTCAAAAGATTTTCTCCATCTGTCTCTAACTATTCCTCAAAGTTTTATCTGAGTAACGACTTCTCAA

>M6-PGSC0003DMG400013489-(rev)M6LRR-RLK_366

AAATGAACTAGTTGTGTAAATACAGAGACTTTGTCATTTGGGTGTTTGAGCTTTTTCCAGTTTCTTGAAAGAGCTAGTTATGAAAACACTGAAACTTTATCATCTGTGTGTTTGAGATCT

>M6-PGSC0003DMG400013489-(rev)M6LRR-RLK_3721

CCCCTATATGCCAAATGGAAGTGTAGCATCACGATTAAAAGGTTAGCTCTTGCCTCCATGTGTTAATACATAATACTCACTATTGATTTCAGATAATGAGTCCAACATTTCCATTTATAA

>M6-PGSC0003DMG400013489-(rev)M6LRR-RLK_3782

GTTAATACATAATACTCACTATTGATTTCAGATAATGAGTCCAACATTTCCATTTATAAAAGTTAAATATTCTCACATGGTGTTTGTTTGTTACTACTCCCTTCATTCATTTTGTTTGAC

>M6-PGSC0003DMG400013489-(rev)M6LRR-RLK_3843

GTTAAATATTCTCACATGGTGTTTGTTTGTTACTACTCCCTTCATTCATTTTGTTTGACAGATAATTGATAGGATACAGAGTTTAAAATTTTAATTTCTTTCATTTATTACTTTGTAGTA

>M6-PGSC0003DMG400013489-(rev)M6LRR-RLK_3904

ATAATTGATAGGATACAGAGTTTAAAATTTTAATTTCTTTCATTTATTACTTTGTAGTAGAATAAATTTATTAAATAGTGGTTATTATACTTGTTAAAGAGAACATCATATAAATTTTAG

>M6-PGSC0003DMG400013489-(rev)M6LRR-RLK_3965

ATAAATTTATTAAATAGTGGTTATTATACTTGTTAAAGAGAACATCATATAAATTTTAGAATGTGGATAGTATGAATTTGAGTTCTTAAAGCTGTAAAAGGTGTACGGACAAATATAATA

>M6-PGSC0003DMG400013489-(rev)M6LRR-RLK_4026

TGTGGATAGTATGAATTTGAGTTCTTAAAGCTGTAAAAGGTGTACGGACAAATATAATACTAGAATAGTGCCAAATTTTGGACCATGTCTAATTTAAAAGAAGTCATGTTAGTGAACAGT

>M6-PGSC0003DMG400013489-(rev)M6LRR-RLK_4087

AGAATAGTGCCAAATTTTGGACCATGTCTAATTTAAAAGAAGTCATGTTAGTGAACAGTCCCTACTTCCCAATTTATTGTCATTAAACATTAGTGAGTGCTCTGAAGACCAGCAACCGTT

>M6-PGSC0003DMG400013489-(rev)M6LRR-RLK_4148

CTACTTCCCAATTTATTGTCATTAAACATTAGTGAGTGCTCTGAAGACCAGCAACCGTTGTCATTGGTGAATTGAGGCCATAAGAGTCTGGTTAAATACAAGCTTAGATGTTATGACCCA

>M6-PGSC0003DMG400013489-(rev)M6LRR-RLK_4209

CATTGGTGAATTGAGGCCATAAGAGTCTGGTTAAATACAAGCTTAGATGTTATGACCCACCCTTACGTCAGTCCATCCTTGAGCCCGCGGTGGGCTGCCCACCTTCACATACTCTGTACT

>M6-PGSC0003DMG400013489-(rev)M6LRR-RLK_427

TTCTTGAAAGAGCTAGTTATGAAAACACTGAAACTTTATCATCTGTGTGTTTGAGATCTTCAATCTTGAAATGAGCAAGTTTATCAGGTGTTTGTGGAGGTGTTGAAAATGGATAAAAAG

>M6-PGSC0003DMG400013489-(rev)M6LRR-RLK_4270

CTTACGTCAGTCCATCCTTGAGCCCGCGGTGGGCTGCCCACCTTCACATACTCTGTACTGCTGTAGTTATATGATAAGACATTATCAGTTTGCTGCTGGCTGAAACCATTGTTGTGTGTT

>M6-PGSC0003DMG400013489-(rev)M6LRR-RLK_4331

TGTAGTTATATGATAAGACATTATCAGTTTGCTGCTGGCTGAAACCATTGTTGTGTGTTTCTGCATGTTCGAAAGCCAAGTTCATTTTACTTTTCTGGAACTTTATCAGTTACTTTTTAA

>M6-PGSC0003DMG400013489-(rev)M6LRR-RLK_4392

TGCATGTTCGAAAGCCAAGTTCATTTTACTTTTCTGGAACTTTATCAGTTACTTTTTAATAGTTTCCAAGTAACTGAATTTTGTAACAGTTGAAGCTGATACATATATTGGCTTGATTGA

>M6-PGSC0003DMG400013489-(rev)M6LRR-RLK_4453

GTTTCCAAGTAACTGAATTTTGTAACAGTTGAAGCTGATACATATATTGGCTTGATTGAAAATGCATTGTGGTTTCTAAAATTTGCAGATAGCATAAATGGTAAACCAGTTTTAGACTGG

>M6-PGSC0003DMG400013489-(rev)M6LRR-RLK_4514

ATGCATTGTGGTTTCTAAAATTTGCAGATAGCATAAATGGTAAACCAGTTTTAGACTGGTCAAGGCGGAAAAAGATAGCACAAGGTACAGCAAGAGGGCTAGTTTACTTGCATGAGCAGT

>M6-PGSC0003DMG400013489-(rev)M6LRR-RLK_4575

AAGGCGGAAAAAGATAGCACAAGGTACAGCAAGAGGGCTAGTTTACTTGCATGAGCAGTGTGATCCCAAAATTATCCATCGTGATGTCAAAGCCGCAAACATTCTGTTGGATGAAGAATT

>M6-PGSC0003DMG400013489-(rev)M6LRR-RLK_4636

GATCCCAAAATTATCCATCGTGATGTCAAAGCCGCAAACATTCTGTTGGATGAAGAATTTGAGGCAGTAGTTGGTGATTTTGGGTTGGCAAAGCTCTTGGATCACCACGATTCCCATGTG

>M6-PGSC0003DMG400013489-(rev)M6LRR-RLK_4697

AGGCAGTAGTTGGTGATTTTGGGTTGGCAAAGCTCTTGGATCACCACGATTCCCATGTGACAACTGCTGTTAGGGGCACCATTGGCCACATTGCTCCTGAATATTTGTCTACTGGCCAAT

>M6-PGSC0003DMG400013489-(rev)M6LRR-RLK_4758

AACTGCTGTTAGGGGCACCATTGGCCACATTGCTCCTGAATATTTGTCTACTGGCCAATCATCTGAAAAAACTGACGTTTTTGGGTTTGGAATCTTGCTACTTGAGCTGATTACTGGCCA

>M6-PGSC0003DMG400013489-(rev)M6LRR-RLK_4819

TCTGAAAAAACTGACGTTTTTGGGTTTGGAATCTTGCTACTTGAGCTGATTACTGGCCAGAAAGCTATGGACTTTGGTCGAGGGGCCAACCAGAACGGTGTTGTTATGCTTGATTGGGTG

>M6-PGSC0003DMG400013489-(rev)M6LRR-RLK_488

AATCTTGAAATGAGCAAGTTTATCAGGTGTTTGTGGAGGTGTTGAAAATGGATAAAAAGGGCATTTTTTTTAATGATAAATTATGGGGTTTTGTTAGAAAAATTACTTTGTTGAGTAGGT

>M6-PGSC0003DMG400013489-(rev)M6LRR-RLK_4880

AAGCTATGGACTTTGGTCGAGGGGCCAACCAGAACGGTGTTGTTATGCTTGATTGGGTGCGTCTCTCCTCTTTCTCTTTCTCTGGATGTTGTACCAAGTACTTTTTGAAATATGTACTGT

>M6-PGSC0003DMG400013489-(rev)M6LRR-RLK_4941

TCTCTCCTCTTTCTCTTTCTCTGGATGTTGTACCAAGTACTTTTTGAAATATGTACTGTGAGTTGTGAGAGGGAATAAGCTATACAGAGGCGGGTTATGCGTCTCTCCTCTCTCTCTTTC

>M6-PGSC0003DMG400013489-(rev)M6LRR-RLK_5002

GTTGTGAGAGGGAATAAGCTATACAGAGGCGGGTTATGCGTCTCTCCTCTCTCTCTTTCTCATAGCTGTTCTCTTTTCAGTTATTAGGAGTTGTAGGTTCAAACCTCAAATATAGCTAGC

>M6-PGSC0003DMG400013489-(rev)M6LRR-RLK_5063

ATAGCTGTTCTCTTTTCAGTTATTAGGAGTTGTAGGTTCAAACCTCAAATATAGCTAGCTAGTGGTGTGTACATGTAGTTTACAAGGCTCTTTTCCATTAGCATACTTCAACGAACTCTC

>M6-PGSC0003DMG400013489-(rev)M6LRR-RLK_5124

GTGGTGTGTACATGTAGTTTACAAGGCTCTTTTCCATTAGCATACTTCAACGAACTCTCTCGTTTAATTACTATTAATGATATGCAGGTGAGAAAGCTGCATCAAGAGGAGAAACTGAAC

>M6-PGSC0003DMG400013489-(rev)M6LRR-RLK_5185

GTTTAATTACTATTAATGATATGCAGGTGAGAAAGCTGCATCAAGAGGAGAAACTGAACCTTATGGTAGACAACAATTTGAAAAATAACTTTGACCAGATTGAGCTAGAAGAGATGGTTA

>M6-PGSC0003DMG400013489-(rev)M6LRR-RLK_5246

TATGGTAGACAACAATTTGAAAAATAACTTTGACCAGATTGAGCTAGAAGAGATGGTTAAAGTTGCCCTTCTATGCACCCATTTCAACCCTTCTCATCGCCCAAAAATGTCCGAAGCATT

>M6-PGSC0003DMG400013489-(rev)M6LRR-RLK_5307

GTTGCCCTTCTATGCACCCATTTCAACCCTTCTCATCGCCCAAAAATGTCCGAAGCATTAAGGATGTTAGAAGGCGATGGATTGGCTGAAAAATGGGAGGCATCACAAAAGCTCGAGACT

>M6-PGSC0003DMG400013489-(rev)M6LRR-RLK_5368

GGATGTTAGAAGGCGATGGATTGGCTGAAAAATGGGAGGCATCACAAAAGCTCGAGACTCCAAGAAACCGAGCTTTTGAACATACACCACAAAGATATTCCAACTACGTCGAAGACTCTT

>M6-PGSC0003DMG400013489-(rev)M6LRR-RLK_5429

AAGAAACCGAGCTTTTGAACATACACCACAAAGATATTCCAACTACGTCGAAGACTCTTCACTAGTTGAACCAATGGAACTTTCTGGACCTAGATGACATCCATTTTTCGTCCTCGATTC

>M6-PGSC0003DMG400013489-(rev)M6LRR-RLK_549

CATTTTTTTTAATGATAAATTATGGGGTTTTGTTAGAAAAATTACTTTGTTGAGTAGGTGAGATTGTAGAAATCTGTACTGGAAAGAATGTTGTATCTGTTTTGGAATGTGGGGCTGCTA

>M6-PGSC0003DMG400013489-(rev)M6LRR-RLK_5490

CTAGTTGAACCAATGGAACTTTCTGGACCTAGATGACATCCATTTTTCGTCCTCGATTCTTATCATCCTTGAGTTGCTAAAGTTGAAGTGAGAAGTATTTCATTGCTGTTGAATGTATGT

>M6-PGSC0003DMG400013489-(rev)M6LRR-RLK_5551

ATCATCCTTGAGTTGCTAAAGTTGAAGTGAGAAGTATTTCATTGCTGTTGAATGTATGTATTCATTTTGTAAAAGAAAAGGGAATAAGTCATGTGTAAGCTGAGTTCATATATGATATAT

>M6-PGSC0003DMG400013489-(rev)M6LRR-RLK_5612

TCATTTTGTAAAAGAAAAGGGAATAAGTCATGTGTAAGCTGAGTTCATATATGATATATTCCCTTGTTCTTTCAATTTTTGAAATGATATGCTATGTAATTATGATTTCATTTGTTCTTT

>M6-PGSC0003DMG400013489-(rev)M6LRR-RLK_5673

CCTTGTTCTTTCAATTTTTGAAATGATATGCTATGTAATTATGATTTCATTTGTTCTTTTTATAAGAAAGATTTTGAGATTTATGAGTAATAAAACGCCATAGGCTTGCCATTCTCATTA

>M6-PGSC0003DMG400013489-(rev)M6LRR-RLK_61

GTGTAACGACTGAAACTTTGTCATTTGGGTGTTTGAGGTTTTTTCAGTTTCTTGAAAGAGCTAGTTGTATAAAGACTGGAACTTTATCATTTGGGTGTTTGAGCTTTTTCCAGTTCTTGA

>M6-PGSC0003DMG400013489-(rev)M6LRR-RLK_610

GATTGTAGAAATCTGTACTGGAAAGAATGTTGTATCTGTTTTGGAATGTGGGGCTGCTACTGTTTTGTACATTGATGGAGAGTTCTTCTGCTACACTTTCTCCTGCTGGTGTAAACTATG

>M6-PGSC0003DMG400013489-(rev)M6LRR-RLK_671

GTTTTGTACATTGATGGAGAGTTCTTCTGCTACACTTTCTCCTGCTGGTGTAAACTATGAAGGTTAGCTTCTTTCTTGAAATTTCTCACTTGTACTGAAAATTTCCTTTTATCAAATTAA

>M6-PGSC0003DMG400013489-(rev)M6LRR-RLK_732

GGTTAGCTTCTTTCTTGAAATTTCTCACTTGTACTGAAAATTTCCTTTTATCAAATTAAGCTAAGAGTAGAAGAACTTGTTAATTCCTCTGGATCAGAGTTAGAGCCACATGGGTGCAAG

>M6-PGSC0003DMG400013489-(rev)M6LRR-RLK_976

TGAACCCCTTAGGCCTTTGCAATGTATATGAGTATGTGACTATGTGTTCTATCTTTTTTCTCTACGATTTCCTCTCGTTATATTGAATCTTTGATGCAATGACAAACTGAAAAGTGAAAA

>M6-PGSC0003DMG400013500M6LRR-RLK_0

ATGGTTGGTTCTTTACCTCTTGAAATTGGAAATTTAAAGGCTGCGACACTGATAGATCTGTCAATGAATCAATTCTCAAATGGAATTCCTATAGAAATTGGAGGATTGCAGAATCTGGTG

>M6-PGSC0003DMG400013500M6LRR-RLK_128

TTTGAGGCACAACAAGTTACAAGGATCTATACCTGACTCATTGAGCAACATGGTAGGTTTGGAATTCCTAGACCTTTCTCATAACAATATATCGGGAACCATTCCTATGTCTTTGGAGAA

>M6-PGSC0003DMG400013500M6LRR-RLK_256

ACATGAAGTATTTCAATGTTTCTGTCAACATGTTGCATGGTGAAATTCCCTCGGGGGGTCCTTTCAAGAACCTCTCGAGTCAGTTTTTCATCAACAATGAAGCATTGTGTGGTTCTTCAA

>M6-PGSC0003DMG400013500M6LRR-RLK_320

CAAGAACCTCTCGAGTCAGTTTTTCATCAACAATGAAGCATTGTGTGGTTCTTCAAGGTTTAGTGTCCCGCCATGCCCCACATCATCAAAGCATAGATCAAATAGGAAAAAAGTGCTTGT

>M6-PGSC0003DMG400013500M6LRR-RLK_384

GTCCCGCCATGCCCCACATCATCAAAGCATAGATCAAATAGGAAAAAAGTGCTTGTTCTATTTCTTATGCTGGGAGTTGCACTTGTATTTGTTCCTATTACCCTTTTGTTTGTATGGATA

>M6-PGSC0003DMG400013500M6LRR-RLK_448

TTATGCTGGGAGTTGCACTTGTATTTGTTCCTATTACCCTTTTGTTTGTATGGATAAAGCATAGAAGAGATAAGAAATACCCTCAACAAGATGATTCATTGTCTATTTCAACGAGAGGAA

>M6-PGSC0003DMG400013500M6LRR-RLK_512

AAGAGATAAGAAATACCCTCAACAAGATGATTCATTGTCTATTTCAACGAGAGGAAGAATTTCGTACTATGAATTGCTCCAAGCAACTGATTCTCTCAGCGAGAGTAATCTGATTGGTTC

>M6-PGSC0003DMG400013500M6LRR-RLK_576

TACTATGAATTGCTCCAAGCAACTGATTCTCTCAGCGAGAGTAATCTGATTGGTTCCGGAAGTTTTGGCTCTGTCTACAAAGGCATTCTCAGAAGTGGGACTGCTATTGCAGTTAAAGTG

>M6-PGSC0003DMG400013500M6LRR-RLK_64

TGAATCAATTCTCAAATGGAATTCCTATAGAAATTGGAGGATTGCAGAATCTGGTGCACCTTTCTTTGAGGCACAACAAGTTACAAGGATCTATACCTGACTCATTGAGCAACATGGTAG

>M6-PGSC0003DMG400013500M6LRR-RLK_640

TTGGCTCTGTCTACAAAGGCATTCTCAGAAGTGGGACTGCTATTGCAGTTAAAGTGTTTAATCTGCAACTGGATATGGCATTCAAGAGTTTCGATACAGAATGTGATGTTTTGCGCAACC

>M6-PGSC0003DMG400013500M6LRR-RLK_704

GCAACTGGATATGGCATTCAAGAGTTTCGATACAGAATGTGATGTTTTGCGCAACCTTCGCCATAGGAATCTCATAAAAGTCATCACTAGTTGTTCCAACATTGATTTTAAGGCTTTAGT

>M6-PGSC0003DMG400013500M6LRR-RLK_768

AGGAATCTCATAAAAGTCATCACTAGTTGTTCCAACATTGATTTTAAGGCTTTAGTGCTCGAGTATATGCCTAATGGGAGTCTTGAGAAGTATTTGTATTCTCACAACTACTTCTTGGAC

>M6-PGSC0003DMG400013500M6LRR-RLK_828

GAGTATATGCCTAATGGGAGTCTTGAGAAGTATTTGTATTCTCACAACTACTTCTTGGACATCAAGCAAAGACTAAGCATAATAATAGATGTGGCATGTGCATTGGAATATCTCCACCAT

>M6-PGSC0003DMG400013524M6RLKantifungi(GNK)_0

TTTTTTTTTTTCTTTCTTCTTTGTCACTGTCCTCTGTTCTTTATCAGAGTTCCTAAAGCATCTGCAATATAATTTTCTACAGCCATCTTGCTCAACAAATCGATTTTTCGCGATCGAAAG

>M6-PGSC0003DMG400013524M6RLKantifungi(GNK)_1037

TTTAACAAGGATATATGAGCTATATATATTACAAAGTCAAAGAGTATATTATGCCCTTTTTTCCTAATATTTTTTACATTTTCAGTGTGTACAAATCAATCTTAGTCTAATATAGGATGC

>M6-PGSC0003DMG400013524M6RLKantifungi(GNK)_1098

TCCTAATATTTTTTACATTTTCAGTGTGTACAAATCAATCTTAGTCTAATATAGGATGCTGCAGTAGTTTCAACATTCAACCATGAATATTTCTGGTTCCCTTTTTTGACTTTGTTGACA

>M6-PGSC0003DMG400013524M6RLKantifungi(GNK)_1159

CAGTAGTTTCAACATTCAACCATGAATATTTCTGGTTCCCTTTTTTGACTTTGTTGACAACATTTTGGTTTATGAAAATGAAGCTGCCTCATCATTTCTTCATTTTTTCCATATTATACA

>M6-PGSC0003DMG400013524M6RLKantifungi(GNK)_122

AAGATTTTTGAGTTTTAAGTGTCTTTTGCTTTCACTTTTTTCTTTTATGTGTTTTTCCAAAGAAACTGCTCAAATTGAGAATTCTCTGTATTCAGCTCTGAGGCTTGTGTAATTTTTGAT

>M6-PGSC0003DMG400013524M6RLKantifungi(GNK)_1220

ATTTTGGTTTATGAAAATGAAGCTGCCTCATCATTTCTTCATTTTTTCCATATTATACAATAGTTCAATAATGCTTTTAAATGCAGCGATTTGATGTTCTTATATTTGATTGATTTCACT

>M6-PGSC0003DMG400013524M6RLKantifungi(GNK)_1281

AGTTCAATAATGCTTTTAAATGCAGCGATTTGATGTTCTTATATTTGATTGATTTCACTTATTAGATCTTGCAGTCGCGTAATGGAGAAAGCAGTGACATTAGTTGCATTTAGGCTATTT

>M6-PGSC0003DMG400013524M6RLKantifungi(GNK)_1342

TTAGATCTTGCAGTCGCGTAATGGAGAAAGCAGTGACATTAGTTGCATTTAGGCTATTTGTTGTTCTGTCGATTTTACTTCTACCAAATGCATCAGAGGCAGAACCAAGATCTCAGACGG

>M6-PGSC0003DMG400013524M6RLKantifungi(GNK)_1403

TGTTCTGTCGATTTTACTTCTACCAAATGCATCAGAGGCAGAACCAAGATCTCAGACGGTCCAGATCATATGTAGTACGCAATATGAGCATAATACAACAGCTTTTGTCCCGAATTTTGT

>M6-PGSC0003DMG400013524M6RLKantifungi(GNK)_1464

CAGATCATATGTAGTACGCAATATGAGCATAATACAACAGCTTTTGTCCCGAATTTTGTTGCCACAATGGAAACAATAAGTGATCAAATGAGAACTCAAGGCTATGGAATTGCACTCACC

>M6-PGSC0003DMG400013524M6RLKantifungi(GNK)_1525

CCACAATGGAAACAATAAGTGATCAAATGAGAACTCAAGGCTATGGAATTGCACTCACCGGGAGTGGACCGGATGCTAACTATGGACTAGCACAGTGTTATGGGGATCTTTCTTTACTTG

>M6-PGSC0003DMG400013524M6RLKantifungi(GNK)_1586

GAGTGGACCGGATGCTAACTATGGACTAGCACAGTGTTATGGGGATCTTTCTTTACTTGACTGTGTCCTATGCTATGCTGAAGCTCGAACTGTCCTTCCTCAATGTTTTCCTTTCAATGG

>M6-PGSC0003DMG400013524M6RLKantifungi(GNK)_1647

TGTGTCCTATGCTATGCTGAAGCTCGAACTGTCCTTCCTCAATGTTTTCCTTTCAATGGAGGTCGAATTTATCTAGATGGATGCTTTATGCGAGCTGAGAATTACACGTTTGATCACCAG

>M6-PGSC0003DMG400013524M6RLKantifungi(GNK)_1708

GTCGAATTTATCTAGATGGATGCTTTATGCGAGCTGAGAATTACACGTTTGATCACCAGTATTTAGGACCTGAGGACAGACATGTTTGTGGAAATAGAACCAGAAAGGATTCTTTGTTCC

>M6-PGSC0003DMG400013524M6RLKantifungi(GNK)_1769

TTTAGGACCTGAGGACAGACATGTTTGTGGAAATAGAACCAGAAAGGATTCTTTGTTCCAACAAACGGCTAGGCGGGCTGTTCAGCAGGCGGTTACTAATGCACCAAACAATAATGGCTA

>M6-PGSC0003DMG400013524M6RLKantifungi(GNK)_183

GAAACTGCTCAAATTGAGAATTCTCTGTATTCAGCTCTGAGGCTTGTGTAATTTTTGATGCAATTACTATATCTTAATTTCTTCTAATATGATTAAGATATATGAAAGATGAGTTTTTGA

>M6-PGSC0003DMG400013524M6RLKantifungi(GNK)_1830

CAAACGGCTAGGCGGGCTGTTCAGCAGGCGGTTACTAATGCACCAAACAATAATGGCTATGCCCGTGCTGAATTGGCAGTTCCTGGGGCAACGAACGAGACAGCTTATGTCCTAGCTGAT

>M6-PGSC0003DMG400013524M6RLKantifungi(GNK)_1891

CCCGTGCTGAATTGGCAGTTCCTGGGGCAACGAACGAGACAGCTTATGTCCTAGCTGATTGCTGGAAAACACTGAGTGCTAACTCGTGCCGAGCTTGTTTGGAAAATGCATCTGCCTCCA

>M6-PGSC0003DMG400013524M6RLKantifungi(GNK)_1952

CTGGAAAACACTGAGTGCTAACTCGTGCCGAGCTTGTTTGGAAAATGCATCTGCCTCCATGATGGGGTGCTTACCATGGTCAGAAGGACGAGCGTTGTACACTGGATGCTTCCTGAGGTA

>M6-PGSC0003DMG400013524M6RLKantifungi(GNK)_2013

ATGGGGTGCTTACCATGGTCAGAAGGACGAGCGTTGTACACTGGATGCTTCCTGAGGTACTCGGATGTTAATTTTCTTAATGCCATTGCTAGCAGTGAAGACTCATCGCCAAATGGTAAT

>M6-PGSC0003DMG400013524M6RLKantifungi(GNK)_2074

CGGATGTTAATTTTCTTAATGCCATTGCTAGCAGTGAAGACTCATCGCCAAATGGTAATCTTTACTTTGTCTCCTTCCTTTTTCCTTTAAGTCAATCACCGAAACTGTTATCTAGACCAT

>M6-PGSC0003DMG400013524M6RLKantifungi(GNK)_2135

TTACTTTGTCTCCTTCCTTTTTCCTTTAAGTCAATCACCGAAACTGTTATCTAGACCATGTAGCTGATATATTGACGTATTTTCAGTGGTAGTGATTGTAATTGCTGTTGTTAGTGCTGT

>M6-PGSC0003DMG400013524M6RLKantifungi(GNK)_2196

AGCTGATATATTGACGTATTTTCAGTGGTAGTGATTGTAATTGCTGTTGTTAGTGCTGTGGTCGTCTTCGTACTGGGAGCTATTATCGCCTTTTATATGTGGAAGAACAAACAAATACAG

>M6-PGSC0003DMG400013524M6RLKantifungi(GNK)_2257

TCGTCTTCGTACTGGGAGCTATTATCGCCTTTTATATGTGGAAGAACAAACAAATACAGAAGAAGAGAAAAGGTAAAACATTTTGGTGATATTTCAGCTAAGGCTTTAGCTTTGAATGTC

>M6-PGSC0003DMG400013524M6RLKantifungi(GNK)_2318

GAAGAGAAAAGGTAAAACATTTTGGTGATATTTCAGCTAAGGCTTTAGCTTTGAATGTCTCCTAAAGCAACTAAGGAGTTCGAAATTTTCAGGTTCCAATGATGTGGAGAAGTTAGTGAA

>M6-PGSC0003DMG400013524M6RLKantifungi(GNK)_2379

CTAAAGCAACTAAGGAGTTCGAAATTTTCAGGTTCCAATGATGTGGAGAAGTTAGTGAAGACCCTTAACGACAGCAGCCTGAACTTCAAGTATTCAACTCTTGAGAAAGCCACCGCATCT

>M6-PGSC0003DMG400013524M6RLKantifungi(GNK)_244

AATTACTATATCTTAATTTCTTCTAATATGATTAAGATATATGAAAGATGAGTTTTTGAAGATGGGGAGTAAAGTTCTGAAAATTGGTCAATTGGGGGTATTTTAGTCTTTTTAATTATT

>M6-PGSC0003DMG400013524M6RLKantifungi(GNK)_2440

CCCTTAACGACAGCAGCCTGAACTTCAAGTATTCAACTCTTGAGAAAGCCACCGCATCTTTTGATGAGGCGAACAAGCTCGGGCAAGGTGGATTTGGCACAGTTTATAAGGTAGTGAGCT

>M6-PGSC0003DMG400013524M6RLKantifungi(GNK)_2562

AAATGCTTAGCTCGTTAATATTATTCACATAATCACATCAGAGTAAATCTATAATATGCTTATTATATTGTAGGGAGTTCTACAAGACGGGAGAGAGATAGCTGTCAAAAGGCTGTTCTT

>M6-PGSC0003DMG400013524M6RLKantifungi(GNK)_2623

ATTATATTGTAGGGAGTTCTACAAGACGGGAGAGAGATAGCTGTCAAAAGGCTGTTCTTCAACAACAAACACAGAGCTGCAGATTTTTACAATGAAGTCAATATTATTAGTAGTGTTGAG

>M6-PGSC0003DMG400013524M6RLKantifungi(GNK)_2684

ACAACAAACACAGAGCTGCAGATTTTTACAATGAAGTCAATATTATTAGTAGTGTTGAGCACAAAAATCTAGTAAGGTTATTAGGATGCAGCTGTTCTGGACCTGAAAGCCTTCTTGTAT

>M6-PGSC0003DMG400013524M6RLKantifungi(GNK)_2745

CAAAAATCTAGTAAGGTTATTAGGATGCAGCTGTTCTGGACCTGAAAGCCTTCTTGTATATGAGTTCCTCCCGAACCAGAGTCTTGATCGGTTTATCTTTGGTATGTAGAAGACTTTGTG

>M6-PGSC0003DMG400013524M6RLKantifungi(GNK)_2806

GAGTTCCTCCCGAACCAGAGTCTTGATCGGTTTATCTTTGGTATGTAGAAGACTTTGTGACTTTGACACACTCGATTCTTACTGGCATGAAATTAATAGAATTCACCCTGTATGTTGAAA

>M6-PGSC0003DMG400013524M6RLKantifungi(GNK)_2867

TTTGACACACTCGATTCTTACTGGCATGAAATTAATAGAATTCACCCTGTATGTTGAAATTGGCAGACGCTACCAAAGGCAAAACATTGAATTGGGAGAAAAGATTTGACATCATTATAG

>M6-PGSC0003DMG400013524M6RLKantifungi(GNK)_2928

GGCAGACGCTACCAAAGGCAAAACATTGAATTGGGAGAAAAGATTTGACATCATTATAGGGACTGCAGAAGGCTTGGTGTACCTCCACGAAAACAATAGAACGAGGATTATTCATAGAGA

>M6-PGSC0003DMG400013524M6RLKantifungi(GNK)_2989

ACTGCAGAAGGCTTGGTGTACCTCCACGAAAACAATAGAACGAGGATTATTCATAGAGATATAAAAGCCAGCAATATCCTGTTGGATTCGAGGCTCAGAGCAAAAATTGCTGATTTTGGG

>M6-PGSC0003DMG400013524M6RLKantifungi(GNK)_3050

TAAAAGCCAGCAATATCCTGTTGGATTCGAGGCTCAGAGCAAAAATTGCTGATTTTGGGTTGGCCAGGTCTTTCCAAGAAGACAAGAGTCACATAAGCACCGCTATAGCTGGGACTTTGT

>M6-PGSC0003DMG400013524M6RLKantifungi(GNK)_3111

GGCCAGGTCTTTCCAAGAAGACAAGAGTCACATAAGCACCGCTATAGCTGGGACTTTGTTAGTGATTCCTTACTTTTACCATATAAATCATAGCTCAAAACTTATGTTCGCGAAACATTA

>M6-PGSC0003DMG400013524M6RLKantifungi(GNK)_3172

GTGATTCCTTACTTTTACCATATAAATCATAGCTCAAAACTTATGTTCGCGAAACATTAACAGCCAGCAGTGTAATTAGTTATCTCATATGGTTAATTTTCTGTTATTTCCCTTGCAGAG

>M6-PGSC0003DMG400013524M6RLKantifungi(GNK)_3233

AGCCAGCAGTGTAATTAGTTATCTCATATGGTTAATTTTCTGTTATTTCCCTTGCAGAGGTTATATGGCACCAGAATACTTAGCTCATGGTCAATTAACTGAAAAGGCAGATGTTTACAG

>M6-PGSC0003DMG400013524M6RLKantifungi(GNK)_3294

TATATGGCACCAGAATACTTAGCTCATGGTCAATTAACTGAAAAGGCAGATGTTTACAGCTTTGGTGTTCTTTTATTGGAGATTGTTACCGGAAGGCAGAATAACAGAAGCAAAAACACT

>M6-PGSC0003DMG400013524M6RLKantifungi(GNK)_3355

TTGGTGTTCTTTTATTGGAGATTGTTACCGGAAGGCAGAATAACAGAAGCAAAAACACTGAATACTCTGACAGTTTAGTTTCCATTGTAAGTTGTGAACCAATTCTTCTAGTTACTTGAA

>M6-PGSC0003DMG400013524M6RLKantifungi(GNK)_3416

ATACTCTGACAGTTTAGTTTCCATTGTAAGTTGTGAACCAATTCTTCTAGTTACTTGAATCTATACAATTGCAGACAAGAATGTTTTTGCTCATTGATTTAACTATTGCCGTGGATGAAA

>M6-PGSC0003DMG400013524M6RLKantifungi(GNK)_3477

TATACAATTGCAGACAAGAATGTTTTTGCTCATTGATTTAACTATTGCCGTGGATGAAACAGACGTGGCAGCATTTTCAGCATGCGAGAGTAGAGGAACTGTTTGATCCAAATCTAATGC

>M6-PGSC0003DMG400013524M6RLKantifungi(GNK)_3538

GACGTGGCAGCATTTTCAGCATGCGAGAGTAGAGGAACTGTTTGATCCAAATCTAATGCTGCACAACTATCACACAAACAATGTAAAAAAGGAGGTTTTAAGAGTGGTACATGTAGGACT

>M6-PGSC0003DMG400013524M6RLKantifungi(GNK)_3599

CACAACTATCACACAAACAATGTAAAAAAGGAGGTTTTAAGAGTGGTACATGTAGGACTTTTGTGCACACAAGAGATTCCGGGATTAAGACCGTCTATGTCTAAGGCATTGCAGATGCTA

>M6-PGSC0003DMG400013524M6RLKantifungi(GNK)_366

TGTTATTATATTATCTAGCTTGTGCCAAGCTGTGATTGAGGTGGGGTCTTACTTTTTACATACTCCATGAAAATCAATGTTAGTTGGTTTTTTTGGTATAATCATCAATGCTGACCTCTC

>M6-PGSC0003DMG400013524M6RLKantifungi(GNK)_3660

TGTGCACACAAGAGATTCCGGGATTAAGACCGTCTATGTCTAAGGCATTGCAGATGCTAGTAAAGAAAGAAGAAGAGTTACCTGCACCAACTAATCCTCCCTTTGTAGATGAGAAAACTA

>M6-PGSC0003DMG400013524M6RLKantifungi(GNK)_3721

AAAGAAAGAAGAAGAGTTACCTGCACCAACTAATCCTCCCTTTGTAGATGAGAAAACTATGGAGCTTCATGACCCTTGGGATACCCCATCTCATCCCCATGGAGATGATTCAGCTTCAAT

>M6-PGSC0003DMG400013524M6RLKantifungi(GNK)_3782

GAGCTTCATGACCCTTGGGATACCCCATCTCATCCCCATGGAGATGATTCAGCTTCAATTGCAAACATTTCTCACAGCTCATTCTACCCCAGATGATGTTTCAGTTTCCTCAAACTAAGA

>M6-PGSC0003DMG400013524M6RLKantifungi(GNK)_3843

CAAACATTTCTCACAGCTCATTCTACCCCAGATGATGTTTCAGTTTCCTCAAACTAAGACATGATATAAATCTGCTAGAGAATGGATCTCGTGAATCTCAAGTATTATCTTGCCTCATGT

>M6-PGSC0003DMG400013524M6RLKantifungi(GNK)_3904

TGATATAAATCTGCTAGAGAATGGATCTCGTGAATCTCAAGTATTATCTTGCCTCATGTTCCTTTTTGTCATACTCTAGGAGCAGATGTTTGTTAGCAACAGGTTGCTTTGTCATTGTAA

>M6-PGSC0003DMG400013524M6RLKantifungi(GNK)_3965

CTTTTTGTCATACTCTAGGAGCAGATGTTTGTTAGCAACAGGTTGCTTTGTCATTGTAATGGTGATGTGCTTGCACAAATCTTGCGGGAAGTTTATCATTGTAGGTAGATACACATTATA

>M6-PGSC0003DMG400013524M6RLKantifungi(GNK)_4026

GTGATGTGCTTGCACAAATCTTGCGGGAAGTTTATCATTGTAGGTAGATACACATTATATCTTGTCATTTGTAATGGTGAAGTAAGAGATTGTTGGACATACTATCGGTAAGATTGTAGT

>M6-PGSC0003DMG400013524M6RLKantifungi(GNK)_4087

TTGTCATTTGTAATGGTGAAGTAAGAGATTGTTGGACATACTATCGGTAAGATTGTAGTACAAGATTAAATATAGTTTATCTTGATTCCAAGAATGGTTTGATTCCAACAGTGTAAAAAG

>M6-PGSC0003DMG400013524M6RLKantifungi(GNK)_4142

TAGTACAAGATTAAATATAGTTTATCTTGATTCCAAGAATGGTTTGATTCCAACAGTGTAAAAAGATCGCTTGGTCATTGATGTTTTGTATTTCTGTTATACAAATTGGTTACTGTAAAC

>M6-PGSC0003DMG400013524M6RLKantifungi(GNK)_427

ACTCCATGAAAATCAATGTTAGTTGGTTTTTTTGGTATAATCATCAATGCTGACCTCTCTTTTGTTTCTAGATGTATCACAATTCTCAACTTTCTGTCCTATTTTATACTACTATATACT

>M6-PGSC0003DMG400013524M6RLKantifungi(GNK)_488

TTGTTTCTAGATGTATCACAATTCTCAACTTTCTGTCCTATTTTATACTACTATATACTTTTCAAATTGATGAGTTTAGCTTTTATATATTGAAAGTTAGATTATTAAAATAAGTGGTAT

>M6-PGSC0003DMG400013524M6RLKantifungi(GNK)_549

TCAAATTGATGAGTTTAGCTTTTATATATTGAAAGTTAGATTATTAAAATAAGTGGTATTTAGTAACATGGAAAAAAAATCATTTTTTGGAAAGGGGGGAATTATATTGTTCAACATTCT

>M6-PGSC0003DMG400013524M6RLKantifungi(GNK)_61

CTGCAATATAATTTTCTACAGCCATCTTGCTCAACAAATCGATTTTTCGCGATCGAAAGGTAAGATTTTTGAGTTTTAAGTGTCTTTTGCTTTCACTTTTTTCTTTTATGTGTTTTTCCA

>M6-PGSC0003DMG400013524M6RLKantifungi(GNK)_610

AGTAACATGGAAAAAAAATCATTTTTTGGAAAGGGGGGAATTATATTGTTCAACATTCTGATTTAGAGTTGATATATTTTTTGTTAAAAAAATGTCATATATATACTCTTGTCGTCTAAC

>M6-PGSC0003DMG400013524M6RLKantifungi(GNK)_671

TTTAGAGTTGATATATTTTTTGTTAAAAAAATGTCATATATATACTCTTGTCGTCTAACAAATAGAGCATTGTTTACCCTTGTTAACAAATTTATATTTACTGAATTAAAAAAAAAAAGT

>M6-PGSC0003DMG400013524M6RLKantifungi(GNK)_732

ATAGAGCATTGTTTACCCTTGTTAACAAATTTATATTTACTGAATTAAAAAAAAAAAGTAATCCTAAAATTGGTCTTTTAATTTCTAAAATGTTATGCGACTTTAAAAAATTACCTCACT

>M6-PGSC0003DMG400013524M6RLKantifungi(GNK)_793

TCCTAAAATTGGTCTTTTAATTTCTAAAATGTTATGCGACTTTAAAAAATTACCTCACTCATTTTTTTACCTCTCCAATCCAACTAAAAAAATGAGTGAAATTATTTTTTAAGGCTCGTG

>M6-PGSC0003DMG400013524M6RLKantifungi(GNK)_854

TTTTTTTACCTCTCCAATCCAACTAAAAAAATGAGTGAAATTATTTTTTAAGGCTCGTGATATCTTTTTAAATTAAAAAACCAATTTTAGATTATTTTTTTAATTAAATAAATATGACTT

>M6-PGSC0003DMG400013524M6RLKantifungi(GNK)_915

ATCTTTTTAAATTAAAAAACCAATTTTAGATTATTTTTTTAATTAAATAAATATGACTTTGTTAATGAAAAGTATAAATATGCTATATTTATTAGATGATGAAGATATATATATGTCACT

>M6-PGSC0003DMG400013524M6RLKantifungi(GNK)_976

TTAATGAAAAGTATAAATATGCTATATTTATTAGATGATGAAGATATATATATGTCACTTTTTTAACAAGGATATATGAGCTATATATATTACAAAGTCAAAGAGTATATTATGCCCTTT

>M6-PGSC0003DMG400013525M6LRR-RLK_0

GTAAACTTCCATCCATCAGCAAGTTGTAGCTCATAATAAACCTCTAAATTATGTCATAAATTTCAATTCTCTTAGCTTAATTCCTTCAAATTCTTCATTAATGGGGTTTTCATCGAAATG

>M6-PGSC0003DMG400013525M6LRR-RLK_1116

GTCATATATCTTTACGGATGTCATGTTTAGTTGTGTAATCATCTTATTTAACATATGCTTGCGATGTTTCTTGGTTTCATGAGTTATTTCAATCTTATCCAATATTCAATGATTTTAATG

>M6-PGSC0003DMG400013525M6LRR-RLK_1178

GATGTTTCTTGGTTTCATGAGTTATTTCAATCTTATCCAATATTCAATGATTTTAATGTTTAATTTTGTTTTTGTATGATTTGCAATGATACTTGCTGCAAGCCTAACACTATTGGATGT

>M6-PGSC0003DMG400013525M6LRR-RLK_124

TTTGTATTTACGTTATTTTCATTCATAAATCTTTCAATTTCTGATCCAAGAGCAACAGAAGCAGCTCTAATTTGTACAAACAGAACAGCTGCACAATCCAATCGACAAGTCTATGTAGCA

>M6-PGSC0003DMG400013525M6LRR-RLK_1240

ATTTTGTTTTTGTATGATTTGCAATGATACTTGCTGCAAGCCTAACACTATTGGATGTAGAGCAAGAAAAAGTTCAACTTGTTCTGCCCAATTATTTTTTTTACTAACAAAACAGATAAC

>M6-PGSC0003DMG400013525M6LRR-RLK_1302

CAAGAAAAAGTTCAACTTGTTCTGCCCAATTATTTTTTTTACTAACAAAACAGATAACTACTTTGTCATTTGGAATGATTTGAAAAACGGCGGTTGGAAATCGAGGTGATTTCGACATTT

>M6-PGSC0003DMG400013525M6LRR-RLK_1364

TTGTCATTTGGAATGATTTGAAAAACGGCGGTTGGAAATCGAGGTGATTTCGACATTTCTTGAAAGTTGCTCTTGTTATCTGTTTAAAATGTGGTAGTTTGTATTTCTTGGATTTTAGTT

>M6-PGSC0003DMG400013525M6LRR-RLK_1426

AAAGTTGCTCTTGTTATCTGTTTAAAATGTGGTAGTTTGTATTTCTTGGATTTTAGTTCCATTCCAGGATTTAATAACTATCTAGACATTACTTGTGATGATGTTTCTTTTTGTCATCTG

>M6-PGSC0003DMG400013525M6LRR-RLK_1488

TCCAGGATTTAATAACTATCTAGACATTACTTGTGATGATGTTTCTTTTTGTCATCTGATAAGTTGAAATGGTTGACTTATATATATACATGAAATGAATGAAGAGTGAATCAGCATTGA

>M6-PGSC0003DMG400013525M6LRR-RLK_1550

GTTGAAATGGTTGACTTATATATATACATGAAATGAATGAAGAGTGAATCAGCATTGAGGCAAATGCAGACAATGAATCTTTTAGAAGCTGTGACAAAATAAACTGTTTATACAAAACTA

>M6-PGSC0003DMG400013525M6LRR-RLK_1612

AATGCAGACAATGAATCTTTTAGAAGCTGTGACAAAATAAACTGTTTATACAAAACTATGTCATTCATGTTGTCGTATTTTGATTAATTATCAGGAGGATCGAATCGTTTGGCTGTTATT

>M6-PGSC0003DMG400013525M6LRR-RLK_1674

ATTCATGTTGTCGTATTTTGATTAATTATCAGGAGGATCGAATCGTTTGGCTGTTATTCTTGCAACAACTCTGAGTATTTCTGCTTTCTTGCTATTTGTATCGGCTGTTTCATTCTTTGT

>M6-PGSC0003DMG400013525M6LRR-RLK_1736

CAACAACTCTGAGTATTTCTGCTTTCTTGCTATTTGTATCGGCTGTTTCATTCTTTGTGCGGAGGAAGGTTCTGAAACAAAAGAGAGGTACTAATTTTTAAGTCAAATCATTTTCATTCT

>M6-PGSC0003DMG400013525M6LRR-RLK_1798

AGGAAGGTTCTGAAACAAAAGAGAGGTACTAATTTTTAAGTCAAATCATTTTCATTCTTTCCGAAAAAAGAAATCTCATTTAAGTCTGATTGATCAAATGGTAATTTGACTTGAATTTGC

>M6-PGSC0003DMG400013525M6LRR-RLK_186

AGCTCTAATTTGTACAAACAGAACAGCTGCACAATCCAATCGACAAGTCTATGTAGCAACTTTTCTAGAAGCCATGGATTCCATTACGCCATTAGTGGGGCGAGACAAATTTGGAGCAGT

>M6-PGSC0003DMG400013525M6LRR-RLK_1860

GAAAAAAGAAATCTCATTTAAGTCTGATTGATCAAATGGTAATTTGACTTGAATTTGCTTCGATGATGTAGAGAAGAAGCAGCTAGGAGCTCTCATAAGAACGGTTAATAAATCAAAATT

>M6-PGSC0003DMG400013525M6LRR-RLK_1922

ATGATGTAGAGAAGAAGCAGCTAGGAGCTCTCATAAGAACGGTTAATAAATCAAAATTGAATATTTCATATGAAACTCTTGAGAAGGCGGCAAATTACTTTAACAACTCGAATAAATTGG

>M6-PGSC0003DMG400013525M6LRR-RLK_1984

ATTTCATATGAAACTCTTGAGAAGGCGGCAAATTACTTTAACAACTCGAATAAATTGGGACAAGGAGGTTCTGGTTCTGTTTACAAGGTAATTCTCTTTGAACATTATATTCATTTTTCT

>M6-PGSC0003DMG400013525M6LRR-RLK_2046

AGGAGGTTCTGGTTCTGTTTACAAGGTAATTCTCTTTGAACATTATATTCATTTTTCTTATACTAATGAGATTTGCCTTACGATAGTCGTGTTTACACATGAAATAGGGGATCTTGCCAG

>M6-PGSC0003DMG400013525M6LRR-RLK_2108

CTAATGAGATTTGCCTTACGATAGTCGTGTTTACACATGAAATAGGGGATCTTGCCAGACGGGCAGGTTGTTGCTATAAAGAGGCTCTTCTTCAATACGAGGCAGTGGGTTGATCATTTC

>M6-PGSC0003DMG400013525M6LRR-RLK_2170

GCAGGTTGTTGCTATAAAGAGGCTCTTCTTCAATACGAGGCAGTGGGTTGATCATTTCTTTAATGAGGTCAATTTGATTAGCGGCATCCATCACAAAAACTTGGTAAAGCTATTGGGTTG

>M6-PGSC0003DMG400013525M6LRR-RLK_2232

ATGAGGTCAATTTGATTAGCGGCATCCATCACAAAAACTTGGTAAAGCTATTGGGTTGCAGCATTACAGGGCCGGAAAGTCTTCTAGTGTATGAGTACGTGCCCAATCATAGTCTTGCGG

>M6-PGSC0003DMG400013525M6LRR-RLK_2294

ATTACAGGGCCGGAAAGTCTTCTAGTGTATGAGTACGTGCCCAATCATAGTCTTGCGGATTACGTCTTTGGTAAGTATCCAATCCACCTTAATTTATGAAAATTTTTAAATTGGTGTAGT

>M6-PGSC0003DMG400013525M6LRR-RLK_2356

CGTCTTTGGTAAGTATCCAATCCACCTTAATTTATGAAAATTTTTAAATTGGTGTAGTCTTATCTCGTTATGTAAATGCCTTTGTTTTGCACTGAATCAGACACAAAGAACCTTCAACCA

>M6-PGSC0003DMG400013525M6LRR-RLK_2418

TCTCGTTATGTAAATGCCTTTGTTTTGCACTGAATCAGACACAAAGAACCTTCAACCACTTACATGGAGTCAAAGATACAAAATTGTATTGGGTACTGCTGAGGGATTGGCCTATCTTCA

>M6-PGSC0003DMG400013525M6LRR-RLK_248

TTCTAGAAGCCATGGATTCCATTACGCCATTAGTGGGGCGAGACAAATTTGGAGCAGTAATTAATGGAACAGGGAATAATACAGTTTATGCGTTTGGTGAATGTATGAAGGACTTATCAC

>M6-PGSC0003DMG400013525M6LRR-RLK_2480

CATGGAGTCAAAGATACAAAATTGTATTGGGTACTGCTGAGGGATTGGCCTATCTTCATGAAGAATCGAAATTAAGGATCATCCACCGAGATATAAAGTTGAGCAATGTTCTTCTAGATG

>M6-PGSC0003DMG400013525M6LRR-RLK_2542

GAATCGAAATTAAGGATCATCCACCGAGATATAAAGTTGAGCAATGTTCTTCTAGATGAAGATTTCACGGCAAAAATAGCTGATTTTGGTCTAGCCAGGTTATTTCCTGAAGACAAGAGC

>M6-PGSC0003DMG400013525M6LRR-RLK_2604

TTTCACGGCAAAAATAGCTGATTTTGGTCTAGCCAGGTTATTTCCTGAAGACAAGAGCCACATTAGCACAGCTATTGCTGGCACTCTGTAAGTTCAATAACTTTTTCCGGTTACTTGTTA

>M6-PGSC0003DMG400013525M6LRR-RLK_2666

TTAGCACAGCTATTGCTGGCACTCTGTAAGTTCAATAACTTTTTCCGGTTACTTGTTACTTTTTCTTTAATCAATACACTAAGAAAGTGCTGAATTAATAAGTTTTGGCATTGCAGCGGT

>M6-PGSC0003DMG400013525M6LRR-RLK_2728

TTCTTTAATCAATACACTAAGAAAGTGCTGAATTAATAAGTTTTGGCATTGCAGCGGTTACATGGCTCCAGAGTACGTTGTACGTGGCATACTGACCGAAAAAGCTGATGTCTATAGTTT

>M6-PGSC0003DMG400013525M6LRR-RLK_2790

TGGCTCCAGAGTACGTTGTACGTGGCATACTGACCGAAAAAGCTGATGTCTATAGTTTTGGAGTTCTTGTTGTCGAAGTCGTGTGTGGAAAGAAGAACAATTATGTTTTCAAGAACACTA

>M6-PGSC0003DMG400013525M6LRR-RLK_2852

GTTCTTGTTGTCGAAGTCGTGTGTGGAAAGAAGAACAATTATGTTTTCAAGAACACTAATTCTCTTCTGCAGCAGGTACTTTCTTGTGATATTTTTTCTTTCCTTAACGGACATTAATAG

>M6-PGSC0003DMG400013525M6LRR-RLK_2914

TCTTCTGCAGCAGGTACTTTCTTGTGATATTTTTTCTTTCCTTAACGGACATTAATAGTATTTACTCAACTCATGTCATTGTGGTTTCTAGTTGTGGAACTTGTATGGGATGGGGAAATC

>M6-PGSC0003DMG400013525M6LRR-RLK_2976

TACTCAACTCATGTCATTGTGGTTTCTAGTTGTGGAACTTGTATGGGATGGGGAAATCTTATGAAGCAGTTGACCCTTTATTAGAAGGTAACTTCAATAAAGAGGAGGCATCTAAGCTGC

>M6-PGSC0003DMG400013525M6LRR-RLK_3038

GAAGCAGTTGACCCTTTATTAGAAGGTAACTTCAATAAAGAGGAGGCATCTAAGCTGCTGCAAGTAGGTCTTCTTTGTGTACAAGCATCAGCAGAACTACGACCCTCCATGTCAAGTGTC

>M6-PGSC0003DMG400013525M6LRR-RLK_310

AATGGAACAGGGAATAATACAGTTTATGCGTTTGGTGAATGTATGAAGGACTTATCACAAACAGATTGTAATCTCTGTTTTGCTCAGTGCAAAACTCAAATCCTACGATGCTTACCTTTT

>M6-PGSC0003DMG400013525M6LRR-RLK_3100

AGTAGGTCTTCTTTGTGTACAAGCATCAGCAGAACTACGACCCTCCATGTCAAGTGTCGTGAAAATGCTCACCGAGAACCATGAAATTCCTCTGCCAACACAGCCACCATTCCTCAATTC

>M6-PGSC0003DMG400013525M6LRR-RLK_3162

AAATGCTCACCGAGAACCATGAAATTCCTCTGCCAACACAGCCACCATTCCTCAATTCTGGTAGTTCAGAATTTAGTCCATTTAATCTGCATGGCAAAAGATTTTTCGGTCAACCAAGCT

>M6-PGSC0003DMG400013525M6LRR-RLK_3224

AGTTCAGAATTTAGTCCATTTAATCTGCATGGCAAAAGATTTTTCGGTCAACCAAGCTCCTCTACACAATCTTCTGGGAATAAATTGACGGAGAGTTGGATTGAGCCTAGATAAACTAAG

>M6-PGSC0003DMG400013525M6LRR-RLK_3286

TACACAATCTTCTGGGAATAAATTGACGGAGAGTTGGATTGAGCCTAGATAAACTAAGTTGATCATCTGGAATTGGTATTTGATTTTGTTCTCAGCAAAGGCCAACTTCTTGAAGGTGAA

>M6-PGSC0003DMG400013525M6LRR-RLK_3348

TCATCTGGAATTGGTATTTGATTTTGTTCTCAGCAAAGGCCAACTTCTTGAAGGTGAATTGTTCAGCCAACTCAAGTTTTTTCTGTGATTAACTAGTACTCACTACTCACTCTCCATTTT

>M6-PGSC0003DMG400013525M6LRR-RLK_3410

TCAGCCAACTCAAGTTTTTTCTGTGATTAACTAGTACTCACTACTCACTCTCCATTTTTTTTTACTGATATTTGTTTAAAGTCTTATTTAGTCAACTTCCTTTTGTGAATACCCCAGTGA

>M6-PGSC0003DMG400013525M6LRR-RLK_3472

TACTGATATTTGTTTAAAGTCTTATTTAGTCAACTTCCTTTTGTGAATACCCCAGTGAGCTATTTCATTTCTAATGTAAAAAAATTGTATAGTATCTGCAGCAGTTGCCATTAACTATAT

>M6-PGSC0003DMG400013525M6LRR-RLK_3625

AATTTTCATCAAGGATTATGCTTCACAATTGTCATCATCTTGTCTTAGAAGTGGCTATTCAGAATCCCCTGCCTAAAGTAACAGGCATTGGTGAAACTACACATTACAATTACATCTACA

>M6-PGSC0003DMG400013525M6LRR-RLK_372

AGATTGTAATCTCTGTTTTGCTCAGTGCAAAACTCAAATCCTACGATGCTTACCTTTTCAAAGATTAGTTACTGGTGGGAGATTATATTATGATGGGTGTTTCCTAAGGTACGATTATTA

>M6-PGSC0003DMG400013525M6LRR-RLK_434

GATTAGTTACTGGTGGGAGATTATATTATGATGGGTGTTTCCTAAGGTACGATTATTATAGATTTTTTAATGAAAGTTTGAGTTCCGTTGATAGGACGATCTGTGGGAACACGAGTTTTT

>M6-PGSC0003DMG400013525M6LRR-RLK_496

TTTTTTAATGAAAGTTTGAGTTCCGTTGATAGGACGATCTGTGGGAACACGAGTTTTTCTGGGAATCAGAGTTTGTTTAGTGAAAATGTTGGGAAATTGGCGAGGGATTTGGAGTTTGGG

>M6-PGSC0003DMG400013525M6LRR-RLK_558

GAATCAGAGTTTGTTTAGTGAAAATGTTGGGAAATTGGCGAGGGATTTGGAGTTTGGGGGTTTGAGGAATGATGGGTTTTTGACTGCAGTTGTGAGTACTGGGAATTTGAGTGTTTATGG

>M6-PGSC0003DMG400013525M6LRR-RLK_62

TCAATTCTCTTAGCTTAATTCCTTCAAATTCTTCATTAATGGGGTTTTCATCGAAATGGGTTTTTGTATTTACGTTATTTTCATTCATAAATCTTTCAATTTCTGATCCAAGAGCAACAG

>M6-PGSC0003DMG400013525M6LRR-RLK_620

TGAGGAATGATGGGTTTTTGACTGCAGTTGTGAGTACTGGGAATTTGAGTGTTTATGGGTTAGCTCAATGTTGGGAATTTGTGAATGGAAGTGCTTGTCAAAGGTGTCTTTCTGATGCAG

>M6-PGSC0003DMG400013525M6LRR-RLK_682

GCTCAATGTTGGGAATTTGTGAATGGAAGTGCTTGTCAAAGGTGTCTTTCTGATGCAGTTTTGAAGATTAGTTCTTGTTCGCCTAAGGAAGAAGGGAGGGTGTTGAATACTGGTTGTTAT

>M6-PGSC0003DMG400013525M6LRR-RLK_744

GAAGATTAGTTCTTGTTCGCCTAAGGAAGAAGGGAGGGTGTTGAATACTGGTTGTTATGTTAGGTATTCTACTCAGAGGTTCTTTAATAATTCTGCAAGTGATACTCCTCCTGCTGGAAA

>M6-PGSC0003DMG400013525M6LRR-RLK_806

GGTATTCTACTCAGAGGTTCTTTAATAATTCTGCAAGTGATACTCCTCCTGCTGGAAATGGAGGTATGTTCTTTCTTTCTTGAATGTATGAAATGAATAAATTGTTTCGTTTAGTTGTGC

>M6-PGSC0003DMG400013525M6LRR-RLK_868

GGTATGTTCTTTCTTTCTTGAATGTATGAAATGAATAAATTGTTTCGTTTAGTTGTGCGATCATTTCGTCTAAAAGTGTAAGCTATTAGAGATAGTACTTTTTGCTTGACATGTCATATC

>M6-PGSC0003DMG400013525M6LRR-RLK_930

ATTTCGTCTAAAAGTGTAAGCTATTAGAGATAGTACTTTTTGCTTGACATGTCATATCTTTACATGGTGTCATTGGGTCTGACTGTGAAGGGGTGTTAGAATGGTTAAAGTTTCACATTG

>M6-PGSC0003DMG400013644M6LRR-RLP_0

TTTCCACATAACCGAACAAGACACATTTTCTGTCTTAGATAACTCATCCTTTCTCTTCCCTTCTTCTTCTTCTCTCTGTTCTCCACATTTCACGTACACTTTCAGAAACATTGTTGATTT

>M6-PGSC0003DMG400013644M6LRR-RLP_1054

CTATTATAACTCAAAAGAGCCTATTAAAGCGTCTGAACTGTATAGTAGAGATGACAGTTACTTGAATAAGGCAAAAGCCACGTTTCTTGTGGATGGAATTGGTGTTGGGCTTCGATCTGG

>M6-PGSC0003DMG400013644M6LRR-RLP_1116

TGAATAAGGCAAAAGCCACGTTTCTTGTGGATGGAATTGGTGTTGGGCTTCGATCTGGGGATGCTTTATCAGGGCCTAAGATTGTTGATTCCACCTTAAAGCCGGCGTCAATAAGTGGTA

>M6-PGSC0003DMG400013644M6LRR-RLP_1178

GCTTTATCAGGGCCTAAGATTGTTGATTCCACCTTAAAGCCGGCGTCAATAAGTGGTATGTTACATCTATTTGCTGCACCGTATTTCTAGTGAATTTGATTATATAATTTGTTGTATTGT

>M6-PGSC0003DMG400013644M6LRR-RLP_124

TGGACAGAAAAGCCACTAAATAGTGTGAGTCAATTTCTTTCCTTAAATAATTGTTTGGAGTTTTGTTTTAATGGGAAGTTTTTTTTTTTCATTTTTATGAGTGTTTTTTCCGTGTTTTTG

>M6-PGSC0003DMG400013644M6LRR-RLP_1240

ACATCTATTTGCTGCACCGTATTTCTAGTGAATTTGATTATATAATTTGTTGTATTGTTGTGTTGACTCAAAGTTCGTATTTTTTTGATCAAGAAGACACTGGTTTCTTTGTTTGTAGGT

>M6-PGSC0003DMG400013644M6LRR-RLP_1302

TTGACTCAAAGTTCGTATTTTTTTGATCAAGAAGACACTGGTTTCTTTGTTTGTAGGTCAAGATGGTGAGAAATTAAGTTTGATTAAGCTTGCTAGTATGATTGAAGTGTCATCCAAAAA

>M6-PGSC0003DMG400013644M6LRR-RLP_1364

ATGGTGAGAAATTAAGTTTGATTAAGCTTGCTAGTATGATTGAAGTGTCATCCAAAAAAGGAAGTCGAGAACTTATCCTTCGGAATAAGTTGTCTGACCAAGTAGAATGGCTTCCAGATT

>M6-PGSC0003DMG400013644M6LRR-RLP_1426

AGTCGAGAACTTATCCTTCGGAATAAGTTGTCTGACCAAGTAGAATGGCTTCCAGATTCTATAGGGAAGCTATCAAGTTTGATCACTTTGGATCTGTCTGAGAATCGAATTACGGTCCTA

>M6-PGSC0003DMG400013644M6LRR-RLP_1488

AGGGAAGCTATCAAGTTTGATCACTTTGGATCTGTCTGAGAATCGAATTACGGTCCTACCCACCACCATAGGGGGGCTTTTATCATTGCAAAAGCTAGATTTGCACTCAAACAAAATTGT

>M6-PGSC0003DMG400013644M6LRR-RLP_1550

CCACCATAGGGGGGCTTTTATCATTGCAAAAGCTAGATTTGCACTCAAACAAAATTGTTGAACTGCCTGATTGTATTGGAGATCTTCTTAACTTGGTCTATCTTGATCTCAGTGGAAATA

>M6-PGSC0003DMG400013644M6LRR-RLP_1612

CTGCCTGATTGTATTGGAGATCTTCTTAACTTGGTCTATCTTGATCTCAGTGGAAATAACTTGAAGTTGTTGCCCGCGAGTTTTGCTAGGTTAGCCCACCTTCAAGAACTTGATTTGAGC

>M6-PGSC0003DMG400013644M6LRR-RLP_1674

GAAGTTGTTGCCCGCGAGTTTTGCTAGGTTAGCCCACCTTCAAGAACTTGATTTGAGCTCTAACATGCTATCAGTACTCCCTGAGACAATTGGCTCCTTAGTAAGTCTTAAGAAATTGAT

>M6-PGSC0003DMG400013644M6LRR-RLP_1736

ACATGCTATCAGTACTCCCTGAGACAATTGGCTCCTTAGTAAGTCTTAAGAAATTGATTGTGGAAACTAATGACCTTGAGGAACTTCCTCACACTATTGGCCAATGTACTTCGCTCAAAG

>M6-PGSC0003DMG400013644M6LRR-RLP_1798

GAAACTAATGACCTTGAGGAACTTCCTCACACTATTGGCCAATGTACTTCGCTCAAAGAGCTTCGTGTGGACTATAATCATCTTAAAGCTCTTCCTGAAGCAGTGGGACGAATTGAGTCT

>M6-PGSC0003DMG400013644M6LRR-RLP_186

TTGTTTTAATGGGAAGTTTTTTTTTTTCATTTTTATGAGTGTTTTTTCCGTGTTTTTGAAAACAGTTGTTTTCCATATAGGAAAATACGGCGAAAAATAAGCTTTACTTTGCATGCCTTC

>M6-PGSC0003DMG400013644M6LRR-RLP_1860

TCGTGTGGACTATAATCATCTTAAAGCTCTTCCTGAAGCAGTGGGACGAATTGAGTCTTTGGAGATTCTGACTGCGCGGTACAATAACATTAGGCTGTTGCCTACAACTATGTCATCCCT

>M6-PGSC0003DMG400013644M6LRR-RLP_1922

AGATTCTGACTGCGCGGTACAATAACATTAGGCTGTTGCCTACAACTATGTCATCCCTTACAAGTTTGAAAGAGCTGAATGTTAGTTTCAATGAAATCGAATCAGTGCCTGAGAGCTTGT

>M6-PGSC0003DMG400013644M6LRR-RLP_1984

AGTTTGAAAGAGCTGAATGTTAGTTTCAATGAAATCGAATCAGTGCCTGAGAGCTTGTGTTTTGCTACTTCACTGGTAAAGCTGAACATAAGCAACAATTTTGCAGACCTGAGATCACTA

>M6-PGSC0003DMG400013644M6LRR-RLP_2046

TGCTACTTCACTGGTAAAGCTGAACATAAGCAACAATTTTGCAGACCTGAGATCACTACCAAGGTCCATTGGTAACCTTGAGTTGCTTGAAGAATTGGACATGAGTAACAACCAGATACG

>M6-PGSC0003DMG400013644M6LRR-RLP_2108

GGTCCATTGGTAACCTTGAGTTGCTTGAAGAATTGGACATGAGTAACAACCAGATACGCGTTCTTCCAGATTCGTTTAGGATGCTCACAAATTTGCGGGTCCTTAAAACCGATGGAAATC

>M6-PGSC0003DMG400013644M6LRR-RLP_2170

CTTCCAGATTCGTTTAGGATGCTCACAAATTTGCGGGTCCTTAAAACCGATGGAAATCCTTTGGAAGTGCCACCAAGAAGCATCGTTGAGAAGGGAGCACAGGTTTAATTTTGTTTCGTT

>M6-PGSC0003DMG400013644M6LRR-RLP_2232

GGAAGTGCCACCAAGAAGCATCGTTGAGAAGGGAGCACAGGTTTAATTTTGTTTCGTTTAACTATTATGCTTTCTGAGTTAGTGTAGTATTTAATTATCTTAGCTCTTCTCTTGGATATA

>M6-PGSC0003DMG400013644M6LRR-RLP_2294

TATTATGCTTTCTGAGTTAGTGTAGTATTTAATTATCTTAGCTCTTCTCTTGGATATATCAATCACTATTGTTTTATTTGCTCGGATCCTCAGGCTGTTGTACAATATATGTCTGATCTG

>M6-PGSC0003DMG400013644M6LRR-RLP_2356

TCACTATTGTTTTATTTGCTCGGATCCTCAGGCTGTTGTACAATATATGTCTGATCTGGTTGCCAATAGAGAGGTAAAGGCGCAACCCTTAAAGAAGAAGAAGAAATCATGGACTCATAT

>M6-PGSC0003DMG400013644M6LRR-RLP_2418

CCAATAGAGAGGTAAAGGCGCAACCCTTAAAGAAGAAGAAGAAATCATGGACTCATATATGCTGCTTTTCAAGTTCCAACAAACGTCAGCGCAATGGCAGTATGGACTATGTTAATGCCT

>M6-PGSC0003DMG400013644M6LRR-RLP_248

CAGTTGTTTTCCATATAGGAAAATACGGCGAAAAATAAGCTTTACTTTGCATGCCTTCATGTGGGTAAAGGTTTGAATTTGCTGTCATCGTTGATTGGCGCTTTCTAGAAACTTGTAGAA

>M6-PGSC0003DMG400013644M6LRR-RLP_2480

TGCTTTTCAAGTTCCAACAAACGTCAGCGCAATGGCAGTATGGACTATGTTAATGCCTGATGTGTTCCACACAAATTTGGTATGATTATATATTTCTCCTTCTTACCAAGTTAGTCATTA

>M6-PGSC0003DMG400013644M6LRR-RLP_2542

TGTTCCACACAAATTTGGTATGATTATATATTTCTCCTTCTTACCAAGTTAGTCATTATGGTACTGAACTTTCCATATTTCGTAAATTTCGTGTTCAGTCAAATTAGTCTAGTCTATGAG

>M6-PGSC0003DMG400013644M6LRR-RLP_2604

ACTGAACTTTCCATATTTCGTAAATTTCGTGTTCAGTCAAATTAGTCTAGTCTATGAGAGGTAGTTTTAAATATATATATTGTTAATAAATGGGAATGGTGCATTATACATGGTGCTGAT

>M6-PGSC0003DMG400013644M6LRR-RLP_2666

AGTTTTAAATATATATATTGTTAATAAATGGGAATGGTGCATTATACATGGTGCTGATTAGTGTTAATTAGAATAATGAAAATGAAGGGTTGTAAGGGTAACGGCTCACATGATGTCTGT

>M6-PGSC0003DMG400013644M6LRR-RLP_2728

GTTAATTAGAATAATGAAAATGAAGGGTTGTAAGGGTAACGGCTCACATGATGTCTGTGTTATCCGTTGTAGGGAAATGAGCTGTGATTTGGCTGGACGAAAGCTGCAAAATTTCAGTTG

>M6-PGSC0003DMG400013644M6LRR-RLP_2790

TCCGTTGTAGGGAAATGAGCTGTGATTTGGCTGGACGAAAGCTGCAAAATTTCAGTTGAATTCGAACATTTTTGCCAGCGAGATTCAATCAACTTTCTTCTTGTTCCAGATAAGCTCTAA

>M6-PGSC0003DMG400013644M6LRR-RLP_2852

CGAACATTTTTGCCAGCGAGATTCAATCAACTTTCTTCTTGTTCCAGATAAGCTCTAATCCACTGCCACGTGGTTTTTTATTCTCATTTCATGTAATAATTCTTTTAATGCTCTTTCCCT

>M6-PGSC0003DMG400013644M6LRR-RLP_2914

CTGCCACGTGGTTTTTTATTCTCATTTCATGTAATAATTCTTTTAATGCTCTTTCCCTGTATTTTCCTCGATTATGATGTAAAATATCAAATGAGATGTAAAATCTTATCTTATTTTTTG

>M6-PGSC0003DMG400013644M6LRR-RLP_2947

ATAATTCTTTTAATGCTCTTTCCCTGTATTTTCCTCGATTATGATGTAAAATATCAAATGAGATGTAAAATCTTATCTTATTTTTTGACTTTCAATATATTCAATGAATTCCAGAATTTC

>M6-PGSC0003DMG400013644M6LRR-RLP_310

GGGTAAAGGTTTGAATTTGCTGTCATCGTTGATTGGCGCTTTCTAGAAACTTGTAGAAACAAAAAACTTAGCTGCGGTTTTCATTTTTCTTGATCTGCTGATTGGTTTAGATATAGCGGA

>M6-PGSC0003DMG400013644M6LRR-RLP_372

AAAACTTAGCTGCGGTTTTCATTTTTCTTGATCTGCTGATTGGTTTAGATATAGCGGAATTCGTCTTTGTTTTTTGCTTTGAATAATACATACACAACGAGGAAAAGGGTAGTCTGAGTC

>M6-PGSC0003DMG400013644M6LRR-RLP_434

GTCTTTGTTTTTTGCTTTGAATAATACATACACAACGAGGAAAAGGGTAGTCTGAGTCTAATTTTTGTGGCCGGAGTTGTATTTATGTTGAATTAAATTTATCTGGGTGTTTTTTGTTTT

>M6-PGSC0003DMG400013644M6LRR-RLP_496

TTTTGTGGCCGGAGTTGTATTTATGTTGAATTAAATTTATCTGGGTGTTTTTTGTTTTCGAGAAATTTTCAGCGGATTATTGAAAATGGGGTCGTCAGGAATGTCGATTGATGAGATTGT

>M6-PGSC0003DMG400013644M6LRR-RLP_558

AAATTTTCAGCGGATTATTGAAAATGGGGTCGTCAGGAATGTCGATTGATGAGATTGTGCAAGAAATTATGAGAATTCATGGATCTCTACCTACAAGGCCATCGATTGATGAGGTTGAAG

>M6-PGSC0003DMG400013644M6LRR-RLP_62

CTTCTTCTTCTCTCTGTTCTCCACATTTCACGTACACTTTCAGAAACATTGTTGATTTACTTTGGACAGAAAAGCCACTAAATAGTGTGAGTCAATTTCTTTCCTTAAATAATTGTTTGG

>M6-PGSC0003DMG400013644M6LRR-RLP_620

GAAATTATGAGAATTCATGGATCTCTACCTACAAGGCCATCGATTGATGAGGTTGAAGCAGCAAGAGGTTTAATTGGAAATCTGGAGAAAGAAGATCAGTTGAAAATAGAAGCTATCATG

>M6-PGSC0003DMG400013644M6LRR-RLP_682

AAGAGGTTTAATTGGAAATCTGGAGAAAGAAGATCAGTTGAAAATAGAAGCTATCATGAGGCAAAATAAAAGGAAAGGTGTACCAGAAGAGCTGTTCAAAATTTTGCAAGAGATGCAGAG

>M6-PGSC0003DMG400013644M6LRR-RLP_744

AAAATAAAAGGAAAGGTGTACCAGAAGAGCTGTTCAAAATTTTGCAAGAGATGCAGAGGAACGCTGTTCTTTTTCAGAGTAAAGAACAAAAGAGGGAGGCGTTGAAACTTTTAGATCTCG

>M6-PGSC0003DMG400013644M6LRR-RLP_806

GCTGTTCTTTTTCAGAGTAAAGAACAAAAGAGGGAGGCGTTGAAACTTTTAGATCTCGACAATATTCACAGTGTGTTCGACGATTTGATTCAGAGAGCTACGAAATGTTTGCCTTCTAAA

>M6-PGSC0003DMG400013644M6LRR-RLP_868

TATTCACAGTGTGTTCGACGATTTGATTCAGAGAGCTACGAAATGTTTGCCTTCTAAATCTCACGATAATGACCCGTCTTCTTCTTCAATTGAACCGAATGATTTATCTTTGGCAAACCA

>M6-PGSC0003DMG400013644M6LRR-RLP_930

ACGATAATGACCCGTCTTCTTCTTCAATTGAACCGAATGATTTATCTTTGGCAAACCAAGTTTCCTTCTCCGGCGGCGGCATTAATTCTCCGACAACTACAGCTGCGGCTGCTTCTTCGT

>M6-PGSC0003DMG400013644M6LRR-RLP_992

TCCTTCTCCGGCGGCGGCATTAATTCTCCGACAACTACAGCTGCGGCTGCTTCTTCGTCGAGCTATTATAACTCAAAAGAGCCTATTAAAGCGTCTGAACTGTATAGTAGAGATGACAGT

>M6-PGSC0003DMG400013682M6LRR-RLK_0

CTAAGCTTCTCTTTCCCTCACCTCCCTCCCATCTCTCTCCTCTTTGTGCTTCCCCATGCATGCCCACACAATCCCATTTCCCTAGAGATGTTGAAATTCTTTTTGCATGCCCTCTAAATC

>M6-PGSC0003DMG400013682M6LRR-RLK_1054

TGATCAAGTTCATATCATCAGTAGATCATCAATCCATTTCCCAATCAATAGGAGACTCTTTAACACTTCTTTCACACGCATCTCCCAAAACAGAGATGAACCTGACTCTGACACCATATA

>M6-PGSC0003DMG400013682M6LRR-RLK_1116

ACACTTCTTTCACACGCATCTCCCAAAACAGAGATGAACCTGACTCTGACACCATATAAAGATATGACGTTTGCGTCTAACTAAATCTCAAAGTTAACTCACGAGGAAAAAATTGTCAAA

>M6-PGSC0003DMG400013682M6LRR-RLK_1178

TATGACGTTTGCGTCTAACTAAATCTCAAAGTTAACTCACGAGGAAAAAATTGTCAAAGTCCCAATCAGCACATTCCCCATATGGCGCTTTAACCCACATTCTAACAATAGTTCTTCGTC

>M6-PGSC0003DMG400013682M6LRR-RLK_124

TATTTTCTCATTTATAATTGCCCTTTGTAAAGGGCGCGCACCCACAAGTGTGCGCACCCCTACAACTAGTCTATGCAATGTCGGTGGCTTATCGTTATAGCAACCACAATCGCCAGTATC

>M6-PGSC0003DMG400013682M6LRR-RLK_1240

CAATCAGCACATTCCCCATATGGCGCTTTAACCCACATTCTAACAATAGTTCTTCGTCATATTGACAATAGATCAAGAATTGTAGTTCATAACAATTTTTATATAGTTTTTAAATATCTA

>M6-PGSC0003DMG400013682M6LRR-RLK_1302

TGACAATAGATCAAGAATTGTAGTTCATAACAATTTTTATATAGTTTTTAAATATCTAAATTTTTTAATTTTAGAATATTGAGTTAATCTAAACATGCTCTCATTTTTGTTCATTTGGTA

>M6-PGSC0003DMG400013682M6LRR-RLK_1364

TTTTAATTTTAGAATATTGAGTTAATCTAAACATGCTCTCATTTTTGTTCATTTGGTAAAATAATACAGGTAATGATCTTTGTGATGGTCCATTATCGAAATGCACAAGTGAACCAAAAG

>M6-PGSC0003DMG400013682M6LRR-RLK_1426

AATACAGGTAATGATCTTTGTGATGGTCCATTATCGAAATGCACAAGTGAACCAAAAGTAGCTCTCTGGACAATTATATTGGTGGTTATCGCAGTTGCAGCAGCAGTAGCAGCAATTGTT

>M6-PGSC0003DMG400013682M6LRR-RLK_1488

TCTCTGGACAATTATATTGGTGGTTATCGCAGTTGCAGCAGCAGTAGCAGCAATTGTTGTTGTTATCATCATTCTTCGTCGGGGCAAGCAGACACCCGAAACTGAAACAAGACCAACACC

>M6-PGSC0003DMG400013682M6LRR-RLK_1550

TTATCATCATTCTTCGTCGGGGCAAGCAGACACCCGAAACTGAAACAAGACCAACACCTACACCCTCTGGAGCTGCAGGGGGCGCTACTAACCAAACAGGAGCGCCCTCAGCAGCAGATC

>M6-PGSC0003DMG400013682M6LRR-RLK_1612

CCCTCTGGAGCTGCAGGGGGCGCTACTAACCAAACAGGAGCGCCCTCAGCAGCAGATCTAAACAAAATGGAACAGGGCAGCAATCAAGCAATGGCTGCTCCTGCTCCTGACCAGTCTCCA

>M6-PGSC0003DMG400013682M6LRR-RLK_1674

CAAAATGGAACAGGGCAGCAATCAAGCAATGGCTGCTCCTGCTCCTGACCAGTCTCCAGAGGGGACCGGTGGACTAAATAATAATAAGAGGCCTGAAGTACAAGCAATACAAAAACTTTT

>M6-PGSC0003DMG400013682M6LRR-RLK_1736

GGACCGGTGGACTAAATAATAATAAGAGGCCTGAAGTACAAGCAATACAAAAACTTTTATTCTTAAAAGATGATATTGAAAAGTTTGATTTGCCAGATCTACTAAAGGCTTCAGCTGAAA

>M6-PGSC0003DMG400013682M6LRR-RLK_1798

TTAAAAGATGATATTGAAAAGTTTGATTTGCCAGATCTACTAAAGGCTTCAGCTGAAATATTGGGTAGCGGTGTATTTGGTTCGACTTATAAAGCTGCACTTAGCATGGGACGTGTTATG

>M6-PGSC0003DMG400013682M6LRR-RLK_186

CAACTAGTCTATGCAATGTCGGTGGCTTATCGTTATAGCAACCACAATCGCCAGTATCGTTATCACCACCACCATCTCTTAATCCTTGTCGTGTTATTATTACTAGTCGTTGTCCCGATC

>M6-PGSC0003DMG400013682M6LRR-RLK_1860

GGGTAGCGGTGTATTTGGTTCGACTTATAAAGCTGCACTTAGCATGGGACGTGTTATGGTTGTCAAAAGATTTAGGCAAATGAATAATGTAGGCAAGGAAGATTTTCATGAGCATATGAG

>M6-PGSC0003DMG400013682M6LRR-RLK_1922

TCAAAAGATTTAGGCAAATGAATAATGTAGGCAAGGAAGATTTTCATGAGCATATGAGAAGGCTAGGTAGGTTGAGTCATAAGAACTTACTTCCTGTTGTAGCTTTTTACTATAGGAAAG

>M6-PGSC0003DMG400013682M6LRR-RLK_1984

CTAGGTAGGTTGAGTCATAAGAACTTACTTCCTGTTGTAGCTTTTTACTATAGGAAAGAGGAAAAGCTTTTGGTCTCAGAGTATGTCAACAATGTCAGCCTTGCAGTTCATCTTCATGGT

>M6-PGSC0003DMG400013682M6LRR-RLK_2046

AAAGCTTTTGGTCTCAGAGTATGTCAACAATGTCAGCCTTGCAGTTCATCTTCATGGTACGTATTTCATACTTTTAAGTTTCCGCGCTAAAATTAGATCATAAAATCATGATTTGCGCTA

>M6-PGSC0003DMG400013682M6LRR-RLK_2108

ATTTCATACTTTTAAGTTTCCGCGCTAAAATTAGATCATAAAATCATGATTTGCGCTACCAGTTTATTATATTAACTCAATCTATTTTAACCTTTGTTTTATTCAATTCATTTCAAAACA

>M6-PGSC0003DMG400013682M6LRR-RLK_2170

TTTATTATATTAACTCAATCTATTTTAACCTTTGTTTTATTCAATTCATTTCAAAACAGAGTTGAGATATGCAACTCAAATCTTTTGAAGAGGGCTCATTAATTTGTTGTTCTCTCTACC

>M6-PGSC0003DMG400013682M6LRR-RLK_2232

TGAGATATGCAACTCAAATCTTTTGAAGAGGGCTCATTAATTTGTTGTTCTCTCTACCTACTCTACTAGCTAGAGTGATGATCTGACGTGTGATCATCTCATCTAAGATATTAAGTTTTT

>M6-PGSC0003DMG400013682M6LRR-RLK_2294

CTACTAGCTAGAGTGATGATCTGACGTGTGATCATCTCATCTAAGATATTAAGTTTTTACTATGATGAACTCTGTGATCATCTCATTATACATATTGTTTATTTTTTTCAGGTAATAAAT

>M6-PGSC0003DMG400013682M6LRR-RLK_2356

TGATGAACTCTGTGATCATCTCATTATACATATTGTTTATTTTTTTCAGGTAATAAATCGCGGGGGCAACCAAGTCTTGATTGGCCAACTCGATTAAAAATCGTAAAGGGAGTAGCCAAA

>M6-PGSC0003DMG400013682M6LRR-RLK_2418

GGGGCAACCAAGTCTTGATTGGCCAACTCGATTAAAAATCGTAAAGGGAGTAGCCAAAGGACTATTGTACCTCTACAACGAGCTACCAAGCTTGACAGCACCCCACGGTCACCTCAAGTC

>M6-PGSC0003DMG400013682M6LRR-RLK_248

TCACCACCACCATCTCTTAATCCTTGTCGTGTTATTATTACTAGTCGTTGTCCCGATCAAATCGGATAGTTCTGAGGCAGAAATCCTCTTTCGATTTAGCAAGTCCTTGCAGAAAAATGG

>M6-PGSC0003DMG400013682M6LRR-RLK_2480

TATTGTACCTCTACAACGAGCTACCAAGCTTGACAGCACCCCACGGTCACCTCAAGTCCTCGAACGTGCTTCTGAATGAATCCTACGAGCCCCTCCTCACGGACTATGCATTACTACCCG

>M6-PGSC0003DMG400013682M6LRR-RLK_2542

AACGTGCTTCTGAATGAATCCTACGAGCCCCTCCTCACGGACTATGCATTACTACCCGTAGTAAACCTAGAACACGCTCAAGAACACATGATTGCTTACAAATCTCCTGAATTCAAACAT

>M6-PGSC0003DMG400013682M6LRR-RLK_2604

AAACCTAGAACACGCTCAAGAACACATGATTGCTTACAAATCTCCTGAATTCAAACATAGTGGTCGTATCACACGTAAAAACGATGTGTGGACGCTTGGAATATTAATCCTCGAGATCAT

>M6-PGSC0003DMG400013682M6LRR-RLK_2666

GTCGTATCACACGTAAAAACGATGTGTGGACGCTTGGAATATTAATCCTCGAGATCATAACGGGTAAATTCCCGTCTAATTTCTTGCAACAAGGCAAGGGGAGTGACACGGATTTGGCTA

>M6-PGSC0003DMG400013682M6LRR-RLK_2728

GGTAAATTCCCGTCTAATTTCTTGCAACAAGGCAAGGGGAGTGACACGGATTTGGCTACGTGGGTAAGGTCCGTGGTAAATGAGGACATGACGGAGGTAGATGTATTCGAGAAGGAAATG

>M6-PGSC0003DMG400013682M6LRR-RLK_2790

GGTAAGGTCCGTGGTAAATGAGGACATGACGGAGGTAGATGTATTCGAGAAGGAAATGAGAGGTACGACGAATTCAGAGGGGGAAATGATGAAGTTATTGAAGATAGCTTTGGGTTGTTG

>M6-PGSC0003DMG400013682M6LRR-RLK_2852

GTACGACGAATTCAGAGGGGGAAATGATGAAGTTATTGAAGATAGCTTTGGGTTGTTGTGATCTTGATATGGAAAAGAGGTTTGATATGAAGGAAGCAATGGAAAGGATTGAGGAAGTAA

>M6-PGSC0003DMG400013682M6LRR-RLK_2914

CTTGATATGGAAAAGAGGTTTGATATGAAGGAAGCAATGGAAAGGATTGAGGAAGTAAAAGAGAGAGATGGGGATGATGATTTTTACTCTACAAGAGGATTATCTGATGATTATACTCAA

>M6-PGSC0003DMG400013682M6LRR-RLK_2976

GAGAGATGGGGATGATGATTTTTACTCTACAAGAGGATTATCTGATGATTATACTCAAGTTTCTATGAACATTTGAAGAGTGCTGGTTATACATACTCATAGTTCGTGTGTAACACTTTT

>M6-PGSC0003DMG400013682M6LRR-RLK_3038

CTATGAACATTTGAAGAGTGCTGGTTATACATACTCATAGTTCGTGTGTAACACTTTTTTATTTGTTCAAAAATCATCATGGTAAATATGGAATGGGACTTTTATTTTGTTTGATGCATG

>M6-PGSC0003DMG400013682M6LRR-RLK_310

CGGATAGTTCTGAGGCAGAAATCCTCTTTCGATTTAGCAAGTCCTTGCAGAAAAATGGTGCAACCGCTAATTGGAATACGAAAGTATTGCCTTGCGATAAAAATACAGATAAACCTAATT

>M6-PGSC0003DMG400013682M6LRR-RLK_3100

TTGTTCAAAAATCATCATGGTAAATATGGAATGGGACTTTTATTTTGTTTGATGCATGGTTGGGGGTCGTATATATATTGGTCAATGAAACCACAAAATAAATAAATATGAAAACAAAAA

>M6-PGSC0003DMG400013682M6LRR-RLK_3224

AGAGATAGAAAAAATGAAGTTAAGTGTTTTTTTGTTGTGATTCAATTGAAGCAAGGGAACATGTAAATTGTCCTTGATGAATCTAGTTTAGCTAGTTGCCTTCACTTTTTTTCTTTTCTT

>M6-PGSC0003DMG400013682M6LRR-RLK_3286

GTAAATTGTCCTTGATGAATCTAGTTTAGCTAGTTGCCTTCACTTTTTTTCTTTTCTTTTCTTTTTGAAACTTTATCTAGTTTATTATATTTGGGGCTTGTAAAAATTAAGTAAATCCAA

>M6-PGSC0003DMG400013682M6LRR-RLK_3325

TCACTTTTTTTCTTTTCTTTTCTTTTTGAAACTTTATCTAGTTTATTATATTTGGGGCTTGTAAAAATTAAGTAAATCCAAAAGCATGCAAGAACAAAGAATTATAAATCTTTCAATTGC

>M6-PGSC0003DMG400013682M6LRR-RLK_372

ACCGCTAATTGGAATACGAAAGTATTGCCTTGCGATAAAAATACAGATAAACCTAATTGGGATAATGTTATTTGCGAAAACGGATTCGTTTTCGGGTTGCAATTGGAGAGTAAAGGCTTA

>M6-PGSC0003DMG400013682M6LRR-RLK_434

TAATGTTATTTGCGAAAACGGATTCGTTTTCGGGTTGCAATTGGAGAGTAAAGGCTTAAGTGGGGCAATAGATGTGGATGCTCTTAAAGACTTGCCTAATTTCAGGACCATAAGTGTTAT

>M6-PGSC0003DMG400013682M6LRR-RLK_496

GGGCAATAGATGTGGATGCTCTTAAAGACTTGCCTAATTTCAGGACCATAAGTGTTATGAACAATAATTTTGAAGGACCTTTACCGAATTTAAGTAAACTTGCGGGGTTAAAAACGGCTT

>M6-PGSC0003DMG400013682M6LRR-RLK_558

AATAATTTTGAAGGACCTTTACCGAATTTAAGTAAACTTGCGGGGTTAAAAACGGCTTATTTCACAAATAACAAATTTTCTGGCCAAATTGACAATACTCTTTTTGAAGGGATGCATTGG

>M6-PGSC0003DMG400013682M6LRR-RLK_62

CCCACACAATCCCATTTCCCTAGAGATGTTGAAATTCTTTTTGCATGCCCTCTAAATCATAGTATTTTCTCATTTATAATTGCCCTTTGTAAAGGGCGCGCACCCACAAGTGTGCGCACC

>M6-PGSC0003DMG400013682M6LRR-RLK_620

CACAAATAACAAATTTTCTGGCCAAATTGACAATACTCTTTTTGAAGGGATGCATTGGTTAAAAAAACTTCATCTTGCTAATAATCAATTAAGTGGTAAAATCCCATCTGTTTTGGGTCA

>M6-PGSC0003DMG400013682M6LRR-RLK_682

AAAAACTTCATCTTGCTAATAATCAATTAAGTGGTAAAATCCCATCTGTTTTGGGTCAATTGCCTAAGCTTACAGAATTGAGACTTGAAAACAACAAATTTGAAGGCCAAATACCTGATT

>M6-PGSC0003DMG400013682M6LRR-RLK_744

CCTAAGCTTACAGAATTGAGACTTGAAAACAACAAATTTGAAGGCCAAATACCTGATTTTGTTCAAGAGAGACTTATGGATATGAATTTTGCTAACAACTCTCTTGAGGGTCCAATTCCT

>M6-PGSC0003DMG400013682M6LRR-RLK_806

TCAAGAGAGACTTATGGATATGAATTTTGCTAACAACTCTCTTGAGGGTCCAATTCCTCATGGCCTTACTAGTCTCAAACCTTCAGCATTTGAAGGTAACTTTTTTTTCCATGTGTAGGA

>M6-PGSC0003DMG400013682M6LRR-RLK_868

GCCTTACTAGTCTCAAACCTTCAGCATTTGAAGGTAACTTTTTTTTCCATGTGTAGGATAAATTCCATTTTCTGAAATTATTGTAACACTTATCTAGCTACGACGTACACAAATGTCTAC

>M6-PGSC0003DMG400013682M6LRR-RLK_930

TTCCATTTTCTGAAATTATTGTAACACTTATCTAGCTACGACGTACACAAATGTCTACATTTTAAAATGTAGTCATAAAGATATGTCATGTATGCCTAACTCCACCTTAGAAGCTAGCAA

>M6-PGSC0003DMG400013682M6LRR-RLK_992

TAAAATGTAGTCATAAAGATATGTCATGTATGCCTAACTCCACCTTAGAAGCTAGCAAGGATTGATCAAGTTCATATCATCAGTAGATCATCAATCCATTTCCCAATCAATAGGAGACTC

>M6-PGSC0003DMG400013766M6LRR-RLP_0

TACAATTAGCAATTTGCCTTGTAATGTACTGGATGCTGTTCTTGGATGCTTGCCTTGGAAAGATGCAGTGAGAACCAGTATCTTGTCAAAAGACTGGAGGTACAAATGGGTAACACGTCA

>M6-PGSC0003DMG400013766M6LRR-RLP_1054

CCTCAAATATTCTTTAGTACCAAGTGTGGGTGGGGGATATTTATGAACCTGTTGTACAGTTGCTACGAGCCCAACCATCCTCAAGCGGTGTTGTGAAGCTTCAAAGAGTTCAAGTGAACA

>M6-PGSC0003DMG400013766M6LRR-RLP_1178

TACTGGTCTCGAGATGGAAATGGAGATTATGAAGTTTATATTGGCATCTGCACCTATACTTGAGGAGATCTTCATTTGAAATTGTATGCGTTTCCTTTTTCGTTCGTGTAAACAAATGAT

>M6-PGSC0003DMG400013766M6LRR-RLP_1238

TGAGGAGATCTTCATTTGAAATTGTATGCGTTTCCTTTTTCGTTCGTGTAAACAAATGATCGATGAGATGAAAGAATTCCGCAGAGCATCAACCAATGTTGAATTCACATTTGAAGAAAT

>M6-PGSC0003DMG400013766M6LRR-RLP_124

CTTGATTTTAATGATGAGTTTTTCAAGTCTTTCAAACAAGATGAAGAAGCCAAGAGAATCATTTACCAAGTCCTATTAGTCCATTAAGGACCAATACTGAAGTTTAGACTCTGTAGGATT

>M6-PGSC0003DMG400013766M6LRR-RLP_186

TTACCAAGTCCTATTAGTCCATTAAGGACCAATACTGAAGTTTAGACTCTGTAGGATTACGAGTTGTCCAGGAATTCACCCTTCATGTAAGCTTAGGAAACAAATATCATTCACCTCATC

>M6-PGSC0003DMG400013766M6LRR-RLP_248

GTTGTCCAGGAATTCACCCTTCATGTAAGCTTAGGAAACAAATATCATTCACCTCATCATCTTTTTACATTCCAGCAACTAAGGTTTTTGGAACTTCAAGATTGCTTGTTCCACCCTCCA

>M6-PGSC0003DMG400013766M6LRR-RLP_372

GTTTCAAAGGGTTTGAGAAGCTTATTAATCTCGATCTTGTACGTGTCACTTTTGATCCATCAATATTTACTTATCTCATCTCTAAATGCTCGTTGCTTGAACGGTTGAGGTTGCGTTGCT

>M6-PGSC0003DMG400013766M6LRR-RLP_434

ATATTTACTTATCTCATCTCTAAATGCTCGTTGCTTGAACGGTTGAGGTTGCGTTGCTGCACAAACTTTGACATCCTTGAAATTGATGCCGCTAATCTCAAATTCTATGAGTTTATTGGA

>M6-PGSC0003DMG400013766M6LRR-RLP_496

AAACTTTGACATCCTTGAAATTGATGCCGCTAATCTCAAATTCTATGAGTTTATTGGAAAAACAAAATCCATTTCCTTCAAGAACGCTCCTATGCTTGAAAAAGTTACTGTGGCTTTCCT

>M6-PGSC0003DMG400013766M6LRR-RLP_558

CAAAATCCATTTCCTTCAAGAACGCTCCTATGCTTGAAAAAGTTACTGTGGCTTTCCTTGGACGACAACTTTTGACTGATACATCTCTTGTTTGCTCTAATTTCCCAAAGTTCTTCCATT

>M6-PGSC0003DMG400013766M6LRR-RLP_620

CGACAACTTTTGACTGATACATCTCTTGTTTGCTCTAATTTCCCAAAGTTCTTCCATTACATGCCTTCACTGCTGGATATGGATATATGCGGTACAATATTAGAGGTAAGGTACATTTGT

>M6-PGSC0003DMG400013766M6LRR-RLP_682

GCCTTCACTGCTGGATATGGATATATGCGGTACAATATTAGAGGTAAGGTACATTTGTGATATTACTATCATAACTAAATGATGAATCTGATTGGTGTCTTAAAGTAATTTTTGACTTTC

>M6-PGSC0003DMG400013766M6LRR-RLP_744

TTACTATCATAACTAAATGATGAATCTGATTGGTGTCTTAAAGTAATTTTTGACTTTCTTGTAGTACTTGATCTAGGGAGGTTTACCAGAGAGTCCCCCAACTGCTCTGAACAATATCAA

>M6-PGSC0003DMG400013766M6LRR-RLP_868

CTCACTATTTCGTCCGTGTCTTTAAGAAACGCCGAGGTGGTTTCAAGTGCAGTTTACTTGATTACAAGCTGTCCTAAATTACAAGATTTTACACTTGAATTTGTAAGCACTCTCTTTGTC

>M6-PGSC0003DMG400013766M6LRR-RLP_930

TACAAGCTGTCCTAAATTACAAGATTTTACACTTGAATTTGTAAGCACTCTCTTTGTCTTTTTTTTTTCATTAAGGTGATTGAATATGGTTTCTTATTCATCTGAACTCGCCTAACTAAC

>M6-PGSC0003DMG400013766M6LRR-RLP_992

TTTTTTCATTAAGGTGATTGAATATGGTTTCTTATTCATCTGAACTCGCCTAACTAACCGTTCCTCAAATATTCTTTAGTACCAAGTGTGGGTGGGGGATATTTATGAACCTGTTGTACA

>M6-PGSC0003DMG400013882M6G-LecRK_0

TTCATTTTGGTCCAGAGTTTAAAGTTTGAGGAACCTAAACAGAGAAGATTACCTTGTTAGAAAGCATCAGCTATGTGTCAGAAGAAAGAGTTGTTGATAGTAATTGTTTCATTTCTTGTC

>M6-PGSC0003DMG400013882M6G-LecRK_1037

ACAATGGTTCTTGAGAGATGGCACTCATGGATGTTTGAGGAAGAAAAATGAAGAGGTGTGCAACAGTGGTGAGGGATTCGTCACATTAAGTCATGTAAAGGTCCCCGACACAGGTGCAGC

>M6-PGSC0003DMG400013882M6G-LecRK_1098

AACAGTGGTGAGGGATTCGTCACATTAAGTCATGTAAAGGTCCCCGACACAGGTGCAGCTCGGATGAACAAGAGCATGGGATTGAAGGAATGTGAAGAGTTGTGTCTGAAGAACTGCTCC

>M6-PGSC0003DMG400013882M6G-LecRK_1159

GGATGAACAAGAGCATGGGATTGAAGGAATGTGAAGAGTTGTGTCTGAAGAACTGCTCCTGCACAGCCTATGCAAGTGCTAATATTAGTGCAGGAGGAAGTGGATGCATCACTTGGTATG

>M6-PGSC0003DMG400013882M6G-LecRK_122

ATTTTGTAGTTCCATAGATACCATTTCCTTTAACCAATCTCTCAAAGATGGTGATTTGTTGATCTCTTCAGGGAAATCTTTTGCTTTGGGTTTCTTTGGGAATTCCCCTGGTAAACGTTA

>M6-PGSC0003DMG400013882M6G-LecRK_1220

CACAGCCTATGCAAGTGCTAATATTAGTGCAGGAGGAAGTGGATGCATCACTTGGTATGGTGAGTTGACAGACATAAAGCAATTTATAGATGGGGCCCAAGATTTATATATCAGAGTCTC

>M6-PGSC0003DMG400013882M6G-LecRK_1281

GAGTTGACAGACATAAAGCAATTTATAGATGGGGCCCAAGATTTATATATCAGAGTCTCTGCATCTGATTTAGGTAAGGAGTTAACTTAATATATTATGTGAGGATTTAGGTAATTCAAT

>M6-PGSC0003DMG400013882M6G-LecRK_1342

CATCTGATTTAGGTAAGGAGTTAACTTAATATATTATGTGAGGATTTAGGTAATTCAATATTATGTGGAGAAGAATGTGCAAGTTGATCATTTTTGGTTAGTAGTAGTACACAGTATCTT

>M6-PGSC0003DMG400013882M6G-LecRK_1403

TATGTGGAGAAGAATGTGCAAGTTGATCATTTTTGGTTAGTAGTAGTACACAGTATCTTACCCTGGTGTTTACTGGAAGTTCTCATTTTTTGTTTGTTCAGCGCAATTCTCGAAGAATTC

>M6-PGSC0003DMG400013882M6G-LecRK_1464

CCTGGTGTTTACTGGAAGTTCTCATTTTTTGTTTGTTCAGCGCAATTCTCGAAGAATTCAAGAGGCCATAATAGGAAAAGAATGATAGGGATTCTAGTAGGGTCTGCTGCAGCAATAATT

>M6-PGSC0003DMG400013882M6G-LecRK_1525

GAGGCCATAATAGGAAAAGAATGATAGGGATTCTAGTAGGGTCTGCTGCAGCAATAATTCTTTCACTCTCTTTGGCATGTTGTTTGGTGATCAATATAAGGAGAAATGGTAAGGATATTA

>M6-PGSC0003DMG400013882M6G-LecRK_1586

TTCACTCTCTTTGGCATGTTGTTTGGTGATCAATATAAGGAGAAATGGTAAGGATATTATTCTACATCATACTTTAGTCGAATATTTTCAGTATCTGTACAGCAGCTACAATGGTGAAGC

>M6-PGSC0003DMG400013882M6G-LecRK_1647

CTACATCATACTTTAGTCGAATATTTTCAGTATCTGTACAGCAGCTACAATGGTGAAGCAACTATAGCAACCTAGACCTTCTACTACCTTTCTATATCGTGCAGCAAAAACAATAACATT

>M6-PGSC0003DMG400013882M6G-LecRK_1708

CTATAGCAACCTAGACCTTCTACTACCTTTCTATATCGTGCAGCAAAAACAATAACATTACTGCAAATTTCATTTGATCCTAATCGGTGTTTGCTGATGTCACAGATAATGAGAGAAGTG

>M6-PGSC0003DMG400013882M6G-LecRK_1769

TGCAAATTTCATTTGATCCTAATCGGTGTTTGCTGATGTCACAGATAATGAGAGAAGTGAAAGTATAGCATCCTATGACGGCATGGATGAAAGTAAACATACAGAAATCTCTATCTTTGA

>M6-PGSC0003DMG400013882M6G-LecRK_183

ATCTCTTCAGGGAAATCTTTTGCTTTGGGTTTCTTTGGGAATTCCCCTGGTAAACGTTACGTCGGAATTTGGTACAACAATGTTCCTGAACTAACTGTTGTTTGGGTTGCCAACAGAGAC

>M6-PGSC0003DMG400013882M6G-LecRK_1830

AGTATAGCATCCTATGACGGCATGGATGAAAGTAAACATACAGAAATCTCTATCTTTGATTTAACCACAATAACTAATGCCACTGATAAGTTCTCTGATGCTAACAAACTTGGTGAAGGA

>M6-PGSC0003DMG400013882M6G-LecRK_1891

TAACCACAATAACTAATGCCACTGATAAGTTCTCTGATGCTAACAAACTTGGTGAAGGAGGATTTGGTAGCGTATACAAGGTAACAACTCCCAAACATGATTGAATTCCTAACAATCAAC

>M6-PGSC0003DMG400013882M6G-LecRK_1952

ATTTGGTAGCGTATACAAGGTAACAACTCCCAAACATGATTGAATTCCTAACAATCAACATTATCTTTATAGGAGGCTACAAATAGATTTAATAATACTTGTTAAACCAAGATTTTCTCC

>M6-PGSC0003DMG400013882M6G-LecRK_2013

TATCTTTATAGGAGGCTACAAATAGATTTAATAATACTTGTTAAACCAAGATTTTCTCCCTAAAAATAAGAAGAAGAAGCTATGTTTACCAAATAATAATCTGGAATCAATATCAGGGTC

>M6-PGSC0003DMG400013882M6G-LecRK_2074

AAAAATAAGAAGAAGAAGCTATGTTTACCAAATAATAATCTGGAATCAATATCAGGGTCACCTGACAGATGGACAAGTAATAGCTGTCAAAAGACTTTCAGTAACTTCAGGGCAAGGGAC

>M6-PGSC0003DMG400013882M6G-LecRK_2135

CTGACAGATGGACAAGTAATAGCTGTCAAAAGACTTTCAGTAACTTCAGGGCAAGGGACCGAGGAGTTCAAGAATGAAGTCACACTAATTGCAAGACTTCAGCACAGAAATTTGGTGAGG

>M6-PGSC0003DMG400013882M6G-LecRK_2196

AGGAGTTCAAGAATGAAGTCACACTAATTGCAAGACTTCAGCACAGAAATTTGGTGAGGCTTTTAGGATGTTGCGTTCAACGAGGAGAGAAGATGTTGGTTTATGAATATATGCCAAACA

>M6-PGSC0003DMG400013882M6G-LecRK_2257

TTTAGGATGTTGCGTTCAACGAGGAGAGAAGATGTTGGTTTATGAATATATGCCAAACAAATCCTTGGACAGTTTTATTTTTGGTAAGGAACATCTCCAGAACCGGAATTTCTTCCATTG

>M6-PGSC0003DMG400013882M6G-LecRK_2318

TCCTTGGACAGTTTTATTTTTGGTAAGGAACATCTCCAGAACCGGAATTTCTTCCATTGACCTCCAGTCTCTACATGCTTTTACATGTTTACTGATGATTAATGATTACTGCTTTAACGT

>M6-PGSC0003DMG400013882M6G-LecRK_2379

CTCCAGTCTCTACATGCTTTTACATGTTTACTGATGATTAATGATTACTGCTTTAACGTGTGTGTGTGTCCCTAAAAGATTTCTTATGCACAAATTTTTAAACTGTGATCCAGATAAAAC

>M6-PGSC0003DMG400013882M6G-LecRK_244

TCGGAATTTGGTACAACAATGTTCCTGAACTAACTGTTGTTTGGGTTGCCAACAGAGACAATCCTGTTAATGGCACATCTGGGATTCTCACCATCGATTCAACTGGAAATCTTGTTGTAC

>M6-PGSC0003DMG400013882M6G-LecRK_2440

GTGTGTGTCCCTAAAAGATTTCTTATGCACAAATTTTTAAACTGTGATCCAGATAAAACAAAAGGGTCTTTGTTGGATTGGGGAAAACGGTTTGAAATCATACATGGGATTGCACGAGGA

>M6-PGSC0003DMG400013882M6G-LecRK_2501

AAGGGTCTTTGTTGGATTGGGGAAAACGGTTTGAAATCATACATGGGATTGCACGAGGAATGTTATACCTTCACCAAGACTCTAGACTAAGAATTATTCATAGGGATCTAAAAGCAAGCA

>M6-PGSC0003DMG400013882M6G-LecRK_2562

GTTATACCTTCACCAAGACTCTAGACTAAGAATTATTCATAGGGATCTAAAAGCAAGCAATGTTTTACTCGATGCCTCCATGCAACCAAAAATATCAGACTTTGGAATGGCCAGAATATT

>M6-PGSC0003DMG400013882M6G-LecRK_2623

GTTTTACTCGATGCCTCCATGCAACCAAAAATATCAGACTTTGGAATGGCCAGAATATTTGGAGGTGACCAAATGGAAGCAAACACAAATCGAGTTGTTGGAACATAGTAAGTATGTCAA

>M6-PGSC0003DMG400013882M6G-LecRK_2684

GAGGTGACCAAATGGAAGCAAACACAAATCGAGTTGTTGGAACATAGTAAGTATGTCAAGTTGTGTATAAACATAAATGAGTTATGTTAGATATGTGCATGTGATCTTTAAAATGATAAA

>M6-PGSC0003DMG400013882M6G-LecRK_2745

TGTGTATAAACATAAATGAGTTATGTTAGATATGTGCATGTGATCTTTAAAATGATAAATCTTGACATTGCTTATCTATTGCAGTGGTTATATGTCACCTGAATATGCAATGGTAGGGCA

>M6-PGSC0003DMG400013882M6G-LecRK_2806

TTGACATTGCTTATCTATTGCAGTGGTTATATGTCACCTGAATATGCAATGGTAGGGCACTTTTCAACAAAGTCTGATGTTTTCAGTTTCGGTGTTTTGTGTTTGGAAATTATCACTGGC

>M6-PGSC0003DMG400013882M6G-LecRK_2867

TTTCAACAAAGTCTGATGTTTTCAGTTTCGGTGTTTTGTGTTTGGAAATTATCACTGGCAGAAAGAACAACAGTCATAAGGATCAGGAGCAATCTCGACATTTAGTTGGATATGTAAGTA

>M6-PGSC0003DMG400013882M6G-LecRK_2928

AAAGAACAACAGTCATAAGGATCAGGAGCAATCTCGACATTTAGTTGGATATGTAAGTACAAACATATATATGCTGTTAAAATCGATAAATTGCATTTATTATCTTACAGTTGCCTTAGA

>M6-PGSC0003DMG400013882M6G-LecRK_2989

AACATATATATGCTGTTAAAATCGATAAATTGCATTTATTATCTTACAGTTGCCTTAGACATAAGCTGTGTCATGGATGGTAGTACTGCAAATAATAATTTGGAAACAATTCCCTTTCAG

>M6-PGSC0003DMG400013882M6G-LecRK_305

TCCTGTTAATGGCACATCTGGGATTCTCACCATCGATTCAACTGGAAATCTTGTTGTACATCTTGATGCGGAAACAAAAACTACAGCCTGGAGAACAAATGTTTCTTCAGCTACAAAACG

>M6-PGSC0003DMG400013882M6G-LecRK_3050

TAAGCTGTGTCATGGATGGTAGTACTGCAAATAATAATTTGGAAACAATTCCCTTTCAGGTTGTTAAAAGTGCTGAATAGATCGAGATTTTGTATTTCAGAATTTCAATTTTTGGTGTTA

>M6-PGSC0003DMG400013882M6G-LecRK_3111

TGTTAAAAGTGCTGAATAGATCGAGATTTTGTATTTCAGAATTTCAATTTTTGGTGTTATAGTCGCTTCTCTTATAGAGAAGAAATTGTCTGAAATTTTGATTGTGATGCATTGTGGTGT

>M6-PGSC0003DMG400013882M6G-LecRK_3172

GTCGCTTCTCTTATAGAGAAGAAATTGTCTGAAATTTTGATTGTGATGCATTGTGGTGTTTCTATTTCAGGTTTGGGATCTTTGGAAGGATGAAAAGGCATTAGATGTTGTTGATCCATT

>M6-PGSC0003DMG400013882M6G-LecRK_3233

CTATTTCAGGTTTGGGATCTTTGGAAGGATGAAAAGGCATTAGATGTTGTTGATCCATTGTTAGGTGGCTCATATGAAGCTTGTGAAGTTTTGAGATGCATCCACATTGGCCTTTTGTGT

>M6-PGSC0003DMG400013882M6G-LecRK_3294

TAGGTGGCTCATATGAAGCTTGTGAAGTTTTGAGATGCATCCACATTGGCCTTTTGTGTGTACAACCGTTTACAGATGACAGACCGACTATGTCAGAATTGGTCTTCATGCTGTGCAATG

>M6-PGSC0003DMG400013882M6G-LecRK_3355

ACAACCGTTTACAGATGACAGACCGACTATGTCAGAATTGGTCTTCATGCTGTGCAATGAAACAAATCTTCCTTTCCCTAAACAACCTAGCCTTATATTCAGATCAGAGAATCATAGTTC

>M6-PGSC0003DMG400013882M6G-LecRK_3416

ACAAATCTTCCTTTCCCTAAACAACCTAGCCTTATATTCAGATCAGAGAATCATAGTTCAGTTAAACACTCTTCATCAGCAAGTATAGCAACATCTTCTGTCAACGATATGTCTATTTCA

>M6-PGSC0003DMG400013882M6G-LecRK_3477

TTAAACACTCTTCATCAGCAAGTATAGCAACATCTTCTGTCAACGATATGTCTATTTCAACAGTTCATGGTCGCTAGAGCTTATGCTAACTATTCAAGAACTATTTTGAAAGAAATACAG

>M6-PGSC0003DMG400013882M6G-LecRK_3538

AGTTCATGGTCGCTAGAGCTTATGCTAACTATTCAAGAACTATTTTGAAAGAAATACAGAATTCCTTTTCTGTACTTTGCTTTCATATTATTAGAGAGATTTATTGGTGAGTTATACACA

>M6-PGSC0003DMG400013882M6G-LecRK_3599

TTCCTTTTCTGTACTTTGCTTTCATATTATTAGAGAGATTTATTGGTGAGTTATACACAACATTTAACATGCTAGTCCATTCTTGATTGAAACTAGTGGGGTTGGGCTCTAATAGCACTG

>M6-PGSC0003DMG400013882M6G-LecRK_366

CTTGATGCGGAAACAAAAACTACAGCCTGGAGAACAAATGTTTCTTCAGCTACAAAACGAGCAAACTCGTATACAGCTAAGCTGGAGGATTCAGGGAATTTTATATTGTTTCAAGACTCC

>M6-PGSC0003DMG400013882M6G-LecRK_3660

ATTTAACATGCTAGTCCATTCTTGATTGAAACTAGTGGGGTTGGGCTCTAATAGCACTGGAACCAAAACTGACTTCTGATTCCCCATGGCCCATGTTACAGACTTAAAGGTAGGTGGGTG

>M6-PGSC0003DMG400013882M6G-LecRK_3721

ACCAAAACTGACTTCTGATTCCCCATGGCCCATGTTACAGACTTAAAGGTAGGTGGGTGGCAAAGGATTGGTAGGAAAGAAGCATAAACACAGAGGGTTTCTATTTCATCTGTGAATTAA

>M6-PGSC0003DMG400013882M6G-LecRK_3782

AAAGGATTGGTAGGAAAGAAGCATAAACACAGAGGGTTTCTATTTCATCTGTGAATTAAATGGTGCTCTTTGTCAGCTGTATTTGATAATTGCCACACTTCATGTTTTCTTTAACAGTCT

>M6-PGSC0003DMG400013882M6G-LecRK_3843

GGTGCTCTTTGTCAGCTGTATTTGATAATTGCCACACTTCATGTTTTCTTTAACAGTCTTCTCTGTGTTCATCAATGCTCTAAAAGCAACAAAGGCTAATCCTGCTAGCCACACACAACT

>M6-PGSC0003DMG400013882M6G-LecRK_3904

TCTGTGTTCATCAATGCTCTAAAAGCAACAAAGGCTAATCCTGCTAGCCACACACAACTTTTCAGTGTTTTTATTGATTTGCAGGATAGTAATTTGGTTGTGGAGTTTTGTCAATTAATG

>M6-PGSC0003DMG400013882M6G-LecRK_3965

TCAGTGTTTTTATTGATTTGCAGGATAGTAATTTGGTTGTGGAGTTTTGTCAATTAATGAGATATTTAACTATTATTAGTTGGTTATGTTTTTCCTCAAAAATTATTGATAAAAAAATAC

>M6-PGSC0003DMG400013882M6G-LecRK_4026

ATATTTAACTATTATTAGTTGGTTATGTTTTTCCTCAAAAATTATTGATAAAAAAATACAAACATGTAAGTTATCTTATGAAAGATTTTAACTTTTGTTTCATAATTAGATCATGAGATA

>M6-PGSC0003DMG400013882M6G-LecRK_4087

ACATGTAAGTTATCTTATGAAAGATTTTAACTTTTGTTTCATAATTAGATCATGAGATAGACATGTATCAAATACAACAATAATATTATGATCATGAGTAAGAGCACTGTTGTCTCAATC

>M6-PGSC0003DMG400013882M6G-LecRK_4148

CATGTATCAAATACAACAATAATATTATGATCATGAGTAAGAGCACTGTTGTCTCAATCTTTCTGTGATGAGTATCTGTAAAAGTAATATCAACTAGTCTATTGAAAATCTTTTAGCATT

>M6-PGSC0003DMG400013882M6G-LecRK_4209

TCTGTGATGAGTATCTGTAAAAGTAATATCAACTAGTCTATTGAAAATCTTTTAGCATTATTTGCATTGAAACATTGAAGACTATATCATACTTTGGGAAGTAATGAGCACTAGATACAT

>M6-PGSC0003DMG400013882M6G-LecRK_427

CAAACTCGTATACAGCTAAGCTGGAGGATTCAGGGAATTTTATATTGTTTCAAGACTCCAAAATGGATGTTACTGAGTGGCAGAGCTTTGATTATCCTACTAATACTCTGCTTCCTTCAA

>M6-PGSC0003DMG400013882M6G-LecRK_4270

TTGCATTGAAACATTGAAGACTATATCATACTTTGGGAAGTAATGAGCACTAGATACATTTGATAAACAGATAACAGGAATACATATACACATCAGACAACTAACGAACATAAGGTGCTG

>M6-PGSC0003DMG400013882M6G-LecRK_4331

GATAAACAGATAACAGGAATACATATACACATCAGACAACTAACGAACATAAGGTGCTGTAATTGTCACCTCGTTGATCGAAGATGATGCTGTTTCGGTAGTAATCAAGTCTGAGTTAGG

>M6-PGSC0003DMG400013882M6G-LecRK_4392

ATTGTCACCTCGTTGATCGAAGATGATGCTGTTTCGGTAGTAATCAAGTCTGAGTTAGGGGTAATGAAGGGCTGCTTCGGGAAGGGAACTGTTGCTTCAATACTTAGCATGGACAGAATT

>M6-PGSC0003DMG400013882M6G-LecRK_4453

TAATGAAGGGCTGCTTCGGGAAGGGAACTGTTGCTTCAATACTTAGCATGGACAGAATTGTTGACATGGTTGGTCTGTCTGCTGCGTTGAGTTGCAAACACAATAGTCCTACTAGAATGC

>M6-PGSC0003DMG400013882M6G-LecRK_4514

TGACATGGTTGGTCTGTCTGCTGCGTTGAGTTGCAAACACAATAGTCCTACTAGAATGCAGCGTAATGCTTCCTCACTGGGACAAGAATCAACTATTGATGGATCTACCAGCTCAAAGGC

>M6-PGSC0003DMG400013882M6G-LecRK_4557

AGTCCTACTAGAATGCAGCGTAATGCTTCCTCACTGGGACAAGAATCAACTATTGATGGATCTACCAGCTCAAAGGCCTTACCATCATTCCACAACTCCCAGGCCTGAAGCAGCAGTTCA

>M6-PGSC0003DMG400013882M6G-LecRK_488

AATGGATGTTACTGAGTGGCAGAGCTTTGATTATCCTACTAATACTCTGCTTCCTTCAATGAAATATGGAATAGACAAGAGGACGGGTTTGAACCGGTTTCTAACATCCTGGAAATCGTT

>M6-PGSC0003DMG400013882M6G-LecRK_549

AAATATGGAATAGACAAGAGGACGGGTTTGAACCGGTTTCTAACATCCTGGAAATCGTTGAACGATCCAGGCACTGGAGAGTATCGTTACACAATGGAGCTCAATGGGACACCTCAAGTT

>M6-PGSC0003DMG400013882M6G-LecRK_61

AAGCATCAGCTATGTGTCAGAAGAAAGAGTTGTTGATAGTAATTGTTTCATTTCTTGTCCCATTTTGTAGTTCCATAGATACCATTTCCTTTAACCAATCTCTCAAAGATGGTGATTTGT

>M6-PGSC0003DMG400013882M6G-LecRK_610

ACGATCCAGGCACTGGAGAGTATCGTTACACAATGGAGCTCAATGGGACACCTCAAGTTTTCTTGTACAAGAACTCTAGCAGGATTTGGCGGACAGGATCCTGGACGGGTCATGGTTGGA

>M6-PGSC0003DMG400013882M6G-LecRK_671

CTTGTACAAGAACTCTAGCAGGATTTGGCGGACAGGATCCTGGACGGGTCATGGTTGGAGTGGTGTACCAGAAATGAGCCCACGATTCATCTTCAGCCTTAGTTATGTGGACAATGATAC

>M6-PGSC0003DMG400013882M6G-LecRK_732

GGTGTACCAGAAATGAGCCCACGATTCATCTTCAGCCTTAGTTATGTGGACAATGATACCGAGGTTTCCTTGACATATGGCATACATGACAGTTCAATAATCTCAAGAATGGTCCTGAAT

>M6-PGSC0003DMG400013882M6G-LecRK_793

AGGTTTCCTTGACATATGGCATACATGACAGTTCAATAATCTCAAGAATGGTCCTGAATGAATCAGGGATCGTGAACCGGTTAACTTGGCAAGAGAGTGAGCAAAAGTGGGTGCAATTTT

>M6-PGSC0003DMG400013882M6G-LecRK_854

ATCAGGGATCGTGAACCGGTTAACTTGGCAAGAGAGTGAGCAAAAGTGGGTGCAATTTTGGTCTGCCCCTAAAGACCCTTGTGACAATTATGAACATTGTGGAGCATTTAGCAATTGCAA

>M6-PGSC0003DMG400013882M6G-LecRK_915

TCTGCCCCTAAAGACCCTTGTGACAATTATGAACATTGTGGAGCATTTAGCAATTGCAATCTGTTCAATTTGGCTGAATTCGAATGCAGTTGTCTTCCAGGGTACGAGCCAAAGCTAAGC

>M6-PGSC0003DMG400013882M6G-LecRK_976

TGTTCAATTTGGCTGAATTCGAATGCAGTTGTCTTCCAGGGTACGAGCCAAAGCTAAGCCGACAATGGTTCTTGAGAGATGGCACTCATGGATGTTTGAGGAAGAAAAATGAAGAGGTGT

>M6-PGSC0003DMG400013884M6RLKWAX-EGF_0

CAAAATTTGTGATAGGCCTTCCATTTGTAATTGTATTTCTGGTGATCAAATTCAAAAGGAGGCATTTGTCGATGTATGATAAGATTGAAGGCTTTTTACAAACACAAAATAATTTCATGC

>M6-PGSC0003DMG400013884M6RLKWAX-EGF_1008

AGCCTCTTCAATCTCAACCGATTGTTCATCAGATCATGGGAAGTTCTATGACATTTTCGTCTGATTCAATGGCTTTGTTAGAAAATTCTGCTGATAATCCCGTTGAACTAGACATTTGTT

>M6-PGSC0003DMG400013884M6RLKWAX-EGF_1071

ATTCAATGGCTTTGTTAGAAAATTCTGCTGATAATCCCGTTGAACTAGACATTTGTTCTGATTGACTATAGTTGATGTAATGTTTATGTGAAACAAATATAGCCCACCCAACAATGTTTG

>M6-PGSC0003DMG400013884M6RLKWAX-EGF_1134

GACTATAGTTGATGTAATGTTTATGTGAAACAAATATAGCCCACCCAACAATGTTTGGGACTAAGGGTGTTGTTGCTATTGTTGTACTAGTTTAAAAACACACCTAGAAGCTAGCTGACC

>M6-PGSC0003DMG400013884M6RLKWAX-EGF_1193

ACTAAGGGTGTTGTTGCTATTGTTGTACTAGTTTAAAAACACACCTAGAAGCTAGCTGACCAAGTGTTCGAATACAATAGCCAAAACGCTCGTGATTACTTAATTTGCCCGTAAAATTTC

>M6-PGSC0003DMG400013884M6RLKWAX-EGF_126

GATATAATTACTCCCACATAAAGAGAATGACAAGAGGATTCAAAGAGAAATTAGGTGAGGGAGGTTATGGCAGTGTATACAAAGGAAAGCTTCAAAGTGGAAGGGATGTAGCAGTGAAGA

>M6-PGSC0003DMG400013884M6RLKWAX-EGF_378

ACTTCATGCCCAACGGATCACTCGACAAGTATATCAGCATCAGTCAAGAAGGAAGTCGTCTGTTAAGTTGGCAGAGGAAGTATGACATTATTCTTGGAGTGGCTCAAGGAATTGGGTATT

>M6-PGSC0003DMG400013884M6RLKWAX-EGF_567

ATTTCATTCCAAAGATTTCTGACTTTGGGCTTGCGAAATTATATCCAACAGATAATAGCATAGTGACTCTCACAGCTGCTCGAGGAACAATTGGTTATGTAGCTCCAGAGTTGATCAGCA

>M6-PGSC0003DMG400013884M6RLKWAX-EGF_63

ATTTGTCGATGTATGATAAGATTGAAGGCTTTTTACAAACACAAAATAATTTCATGCCAATCAGATATAATTACTCCCACATAAAGAGAATGACAAGAGGATTCAAAGAGAAATTAGGTG

>M6-PGSC0003DMG400013884M6RLKWAX-EGF_756

ACTTGAAAAGACATGAAGTTGCAAATGAAGAGAATTCCAGCCAATATTTTCCTTATTATATTTACGATAAGTTCAACAAGGGGAAGGAGATTGTGGTGGATGAAGAAGCAAATGATGATG

>M6-PGSC0003DMG400013884M6RLKWAX-EGF_819

ACGATAAGTTCAACAAGGGGAAGGAGATTGTGGTGGATGAAGAAGCAAATGATGATGAAAAGAAGATGGTTAGAAAGCTGGCTTTAGTTGCGTTATGGTGCATACAAACAAATCCGATAC

>M6-PGSC0003DMG400013884M6RLKWAX-EGF_882

AGATGGTTAGAAAGCTGGCTTTAGTTGCGTTATGGTGCATACAAACAAATCCGATACAACGCCCTTCGATGAGTAGAGTATTAGAAATGCTTGAAGGTGAAGTTGAAGTGCTAGAAGTAC

>M6-PGSC0003DMG400013885M6G-LecRK_0

GGGATGTGTACCAAAATACCATCTTCCTTCCTTCCCTCTTCCTATCAAACAAGAGAAGAAGTACACCAACAAATGTCTCAAACACCAACTCTGCAGCAAAACAAAATACTCTTTTCTTAA

>M6-PGSC0003DMG400013885M6G-LecRK_1037

GGGGTATCTGGAATTTGTGACACAAATATTTCACCAATCTGCAAATGTATGGTGGGATTCAAGCCTAAAAATCAAGTGGCCTGGGAATTGCGAGATGGGTCTGATGGTTGTGTTCGATTT

>M6-PGSC0003DMG400013885M6G-LecRK_1098

AGCCTAAAAATCAAGTGGCCTGGGAATTGCGAGATGGGTCTGATGGTTGTGTTCGATTTCATAAATTGGATTGTAAAACTGATAAATTTAACGTATTGAGGAACATGAAGTTGCCAGATA

>M6-PGSC0003DMG400013885M6G-LecRK_1159

TAAATTGGATTGTAAAACTGATAAATTTAACGTATTGAGGAACATGAAGTTGCCAGATACGACGAATTCGTTTGTGGATACAAATATGAATTTGGATGAATGTGAAGCTATGTGTATGAA

>M6-PGSC0003DMG400013885M6G-LecRK_122

TCCACAATGAGAAACACCAAAGATTACTACTTTCACCAATTCTTGTTCTTGATTTTGATTTCTCAATTTCTTCATCCAATTCTTGCTATACCCACTGATACAATCACCCCAACACAGCCT

>M6-PGSC0003DMG400013885M6G-LecRK_1220

ACGAATTCGTTTGTGGATACAAATATGAATTTGGATGAATGTGAAGCTATGTGTATGAAGAATTGTTCTTGCACGGCCTATACAAATTCGAATATTAGCGGAAGTGGGTCGGGTTGTGTG

>M6-PGSC0003DMG400013885M6G-LecRK_1281

ATTGTTCTTGCACGGCCTATACAAATTCGAATATTAGCGGAAGTGGGTCGGGTTGTGTGATTTGGAGCACCGAACTTATTGACATGCGGCAGTATACGGCGGCGGAAGGTGGCCAAGTTC

>M6-PGSC0003DMG400013885M6G-LecRK_1342

TTGGAGCACCGAACTTATTGACATGCGGCAGTATACGGCGGCGGAAGGTGGCCAAGTTCTCTACGTTAGGGTGGCTTCTTCTGACGCAGGTATCATCCTATTTCTCTACTAGCTTCGCTT

>M6-PGSC0003DMG400013885M6G-LecRK_1403

TACGTTAGGGTGGCTTCTTCTGACGCAGGTATCATCCTATTTCTCTACTAGCTTCGCTTGCCATTTCTTTTTTCGCTTAGTACAATGATAACGATTAATAATCGATAAAAAATAGTTGTA

>M6-PGSC0003DMG400013885M6G-LecRK_1464

CATTTCTTTTTTCGCTTAGTACAATGATAACGATTAATAATCGATAAAAAATAGTTGTATTAAAAAGAATTTGTACAATTAATTTTAAGTAATTTGGGGTTGAAAATTGATGGTTAGATA

>M6-PGSC0003DMG400013885M6G-LecRK_1525

AAAAAGAATTTGTACAATTAATTTTAAGTAATTTGGGGTTGAAAATTGATGGTTAGATAATCCACTTCTCTGTCAAGATCAAATTTTGGGTCACCGGGATGATTTGAGTCGACTAGGTCT

>M6-PGSC0003DMG400013885M6G-LecRK_1586

CCACTTCTCTGTCAAGATCAAATTTTGGGTCACCGGGATGATTTGAGTCGACTAGGTCTGCACGAGTGGAGTTAGTTGGAATTTTTTATTTACTTGTTAGGTTAGACGATATTTCAACAC

>M6-PGSC0003DMG400013885M6G-LecRK_1647

ACGAGTGGAGTTAGTTGGAATTTTTTATTTACTTGTTAGGTTAGACGATATTTCAACACAGAGAAGTAGAGTAAAATAGAAAGGGTGATGAAGTTTTTGGAAAATATTTTTGTGGAAGAT

>M6-PGSC0003DMG400013885M6G-LecRK_1708

AGAAGTAGAGTAAAATAGAAAGGGTGATGAAGTTTTTGGAAAATATTTTTGTGGAAGATTGGAGAGTATTTCCATGATTGAAAATATGCAATTTAATATTCTCGTTTAAGTTTATGTGTC

>M6-PGSC0003DMG400013885M6G-LecRK_183

CTCAATTTCTTCATCCAATTCTTGCTATACCCACTGATACAATCACCCCAACACAGCCTTTAACCAAAGATCAAACCTTAGTATCTTCCGATCAACACTTTGAATTAGGCTTTTTCTCTC

>M6-PGSC0003DMG400013885M6G-LecRK_2013

AAATTGAGACGAAGAAATAGTAATAATATTTTGCATATTTTTGTCGGAAGTTGATTTATCTCATTCATTTGTAAACTTGCTATTCTCTGCATTATTCCACTATATACATACTGTTCTGTT

>M6-PGSC0003DMG400013885M6G-LecRK_2074

CATTCATTTGTAAACTTGCTATTCTCTGCATTATTCCACTATATACATACTGTTCTGTTTGAATCTATTTTCATAATTTCAAAGTCGAGTTCAAGACGAAAAATCTATTTGTCCAAAAAT

>M6-PGSC0003DMG400013885M6G-LecRK_2135

AATCTATTTTCATAATTTCAAAGTCGAGTTCAAGACGAAAAATCTATTTGTCCAAAAATTGTTAGTACCAGTTTTGCATGTCATCCAATCCAAATGGACCAAATATTCCTTGAATAATAT

>M6-PGSC0003DMG400013885M6G-LecRK_2196

TTAGTACCAGTTTTGCATGTCATCCAATCCAAATGGACCAAATATTCCTTGAATAATATTTTAAATGTATTGTTTTGTCATTTTAATAAGAAGCATATTACACATTATCATAATTTTAAT

>M6-PGSC0003DMG400013885M6G-LecRK_2257

TAAATGTATTGTTTTGTCATTTTAATAAGAAGCATATTACACATTATCATAATTTTAATTTACTCTTTTCCTATTGAATTCCAAAGTTGGGAAGCATGCCGTTAGATTTGCCTCTAATAG

>M6-PGSC0003DMG400013885M6G-LecRK_2318

ACTCTTTTCCTATTGAATTCCAAAGTTGGGAAGCATGCCGTTAGATTTGCCTCTAATAGATGAAATTACAAAAATATATTTATTTTTTTGAATTTTTCTTTTCTATTTTTTTTTTTTTGG

>M6-PGSC0003DMG400013885M6G-LecRK_2379

GAAATTACAAAAATATATTTATTTTTTTGAATTTTTCTTTTCTATTTTTTTTTTTTTGGGTATTGCTATTTTTTTTAAACCACCCCCCCCCCCCCCCTCCACAAACGTCATGCAGAAATT

>M6-PGSC0003DMG400013885M6G-LecRK_244

AACCAAAGATCAAACCTTAGTATCTTCCGATCAACACTTTGAATTAGGCTTTTTCTCTCCCGGCGGTGCAAATTCAGACAAATGGTATATCGGTATTTGGTACAAAGAGATTCAAGACAG

>M6-PGSC0003DMG400013885M6G-LecRK_2440

ATTGCTATTTTTTTTAAACCACCCCCCCCCCCCCCCTCCACAAACGTCATGCAGAAATTAAATTAGAGATTTTATTTTGAAAATAAATACTCTACCCCCTTTGAACAACACGTATAGTTG

>M6-PGSC0003DMG400013885M6G-LecRK_2501

ATTAGAGATTTTATTTTGAAAATAAATACTCTACCCCCTTTGAACAACACGTATAGTTGTCTTTTAATATCCTTGGTCCCCTTTTTAAATTCCCTTTCACGTTACCTACTTTCGATCTAA

>M6-PGSC0003DMG400013885M6G-LecRK_2562

TTTTAATATCCTTGGTCCCCTTTTTAAATTCCCTTTCACGTTACCTACTTTCGATCTAATATTTTACATAATCAACTAGATTTCAAAATAAAAATAAAATTCATAATTGCTAGTTGAAAA

>M6-PGSC0003DMG400013885M6G-LecRK_2623

TTTTACATAATCAACTAGATTTCAAAATAAAAATAAAATTCATAATTGCTAGTTGAAAAAACAAGAAGTTAGTATAATAATTATTGAGTGCATTTAATCCTGAAAAGGACTTGTAAACAA

>M6-PGSC0003DMG400013885M6G-LecRK_2684

CAAGAAGTTAGTATAATAATTATTGAGTGCATTTAATCCTGAAAAGGACTTGTAAACAACCTACTTTTCAAAGTTATTCTGCTAATGAGTTTGCCAATATATTCTTTGTGGTATGCACGT

>M6-PGSC0003DMG400013885M6G-LecRK_2745

TACTTTTCAAAGTTATTCTGCTAATGAGTTTGCCAATATATTCTTTGTGGTATGCACGTTTTCAATATAATTTTTCTTTTTCTATGCTTTGTGTATTTTACTTTGAAAAATCGAAAAGTA

>M6-PGSC0003DMG400013885M6G-LecRK_2806

TCAATATAATTTTTCTTTTTCTATGCTTTGTGTATTTTACTTTGAAAAATCGAAAAGTATAAAAAACCTAACCATATCTACCAATCCCCCATCAACAATTTGCCTTCAATAGGGACTTAA

>M6-PGSC0003DMG400013885M6G-LecRK_2867

AAAAACCTAACCATATCTACCAATCCCCCATCAACAATTTGCCTTCAATAGGGACTTAATTTTATACAATTTGCTTTCAATAGGGACTTAATTTTATTTATTTACTATATTTGGACTTTC

>M6-PGSC0003DMG400013885M6G-LecRK_2928

TTATACAATTTGCTTTCAATAGGGACTTAATTTTATTTATTTACTATATTTGGACTTTCTTTTTCTCATTTGATTTGATAGGGATGATTAGTATTTGGAAAATATGTTACACGGCCAAGT

>M6-PGSC0003DMG400013885M6G-LecRK_2989

TTTCTCATTTGATTTGATAGGGATGATTAGTATTTGGAAAATATGTTACACGGCCAAGTATCTCAAGGCTCTAAAAGTAATCAGCCACCTAATTCCACCTTCTTTCCAATCACCTGTTCA

>M6-PGSC0003DMG400013885M6G-LecRK_305

GGCGGTGCAAATTCAGACAAATGGTATATCGGTATTTGGTACAAAGAGATTCAAGACAGGACTATTGTCTGGGTTGCAAACAGAGCTAAACCCCTTTCAGCTTCATCAACATCAGTCCTA

>M6-PGSC0003DMG400013885M6G-LecRK_3050

CTCAAGGCTCTAAAAGTAATCAGCCACCTAATTCCACCTTCTTTCCAATCACCTGTTCATAAATTTGAACAAACATTATCGATTTATTGTCACAATTCAGCTCGTAATAATAATAAATAA

>M6-PGSC0003DMG400013885M6G-LecRK_3111

AATTTGAACAAACATTATCGATTTATTGTCACAATTCAGCTCGTAATAATAATAAATAATCGAAATGTGCATTTGCTTTTTCTTTTGTACTTTCCTCTTTAAATCTGCATGCTTAAGAGG

>M6-PGSC0003DMG400013885M6G-LecRK_3172

GAAATGTGCATTTGCTTTTTCTTTTGTACTTTCCTCTTTAAATCTGCATGCTTAAGAGGAAACTTTTAATTATGGTGGGAAATAAAATTTCCAAATAAAGCTTATTTAGCTCTAAATTAT

>M6-PGSC0003DMG400013885M6G-LecRK_3233

ACTTTTAATTATGGTGGGAAATAAAATTTCCAAATAAAGCTTATTTAGCTCTAAATTATTTTCCATTAATTGGTCCCACAATAAATGGGTCCCACAATAAATGGGTCCCACAAAGAGATA

>M6-PGSC0003DMG400013885M6G-LecRK_3294

TCCATTAATTGGTCCCACAATAAATGGGTCCCACAATAAATGGGTCCCACAAAGAGATACTCCTTTGCATTTCAAAACGGTGACGACGCAATTTCCTGGTCGGTGTAATGGGCCCTTACA

>M6-PGSC0003DMG400013885M6G-LecRK_3355

CCTTTGCATTTCAAAACGGTGACGACGCAATTTCCTGGTCGGTGTAATGGGCCCTTACATGGCCGTCGATCTTGACTTTTATTGGGCCATCCCGGCCCAGTTGTTTTCCCTTTTTTCTCC

>M6-PGSC0003DMG400013885M6G-LecRK_3416

GCCGTCGATCTTGACTTTTATTGGGCCATCCCGGCCCAGTTGTTTTCCCTTTTTTCTCCTTTTTAGAGAAAAAAAGCATATGTTTCCTAATGTAATACAAGAAAATGATTGGACACCAAT

>M6-PGSC0003DMG400013885M6G-LecRK_3477

TTTAGAGAAAAAAAGCATATGTTTCCTAATGTAATACAAGAAAATGATTGGACACCAATTCCTTTGACACGTTTGGGAAATGGTCATATACAGGTTTGAAAAGTAAGGTTTGGTGGGGTA

>M6-PGSC0003DMG400013885M6G-LecRK_3538

CTTTGACACGTTTGGGAAATGGTCATATACAGGTTTGAAAAGTAAGGTTTGGTGGGGTAAATTAATTCAAATATTACAAAGGGATTAAGAAGTATTTATTAAGCTAATTATTTTTTTTTA

>M6-PGSC0003DMG400013885M6G-LecRK_3599

TTAATTCAAATATTACAAAGGGATTAAGAAGTATTTATTAAGCTAATTATTTTTTTTTACCAAGAATATGTCATGGATGATATTAATAGTTTTTTTTGGACACAACTTTGTGAAGTAGGT

>M6-PGSC0003DMG400013885M6G-LecRK_366

CTATTGTCTGGGTTGCAAACAGAGCTAAACCCCTTTCAGCTTCATCAACATCAGTCCTAAAAATCGCAGAAATTGGTACCCTTCTTCTCGTCGATGGCCAAACTGGAAATTCCGTCTGGT

>M6-PGSC0003DMG400013885M6G-LecRK_3660

AAGAATATGTCATGGATGATATTAATAGTTTTTTTTGGACACAACTTTGTGAAGTAGGTGAATTAGATCATGTTTAGTGACACCATTGGACTGTAGTCTTTTCAATTCAAACTTTGCTAT

>M6-PGSC0003DMG400013885M6G-LecRK_3721

ATTAGATCATGTTTAGTGACACCATTGGACTGTAGTCTTTTCAATTCAAACTTTGCTATTGTCACCGAAAATGATCTTTAACAACACTATAGGTTTTCATACATAAAGAGACATTCAAAT

>M6-PGSC0003DMG400013885M6G-LecRK_3782

TCACCGAAAATGATCTTTAACAACACTATAGGTTTTCATACATAAAGAGACATTCAAATTTGGCCTCAGCTTGCAAGTAAACACCTTAACTTTAGTATACATACCTAGACACCTCAAATT

>M6-PGSC0003DMG400013885M6G-LecRK_4087

CAAGTTTGAGTGTTTCTTTATGTATTATGCCAGTAAAAAATGATGAACTTCTTCTTGCAGAACTAAACTCTTGAATAAAATTAAGCATGATTAATTAACTCGAAAAACTTTTAATTTCAG

>M6-PGSC0003DMG400013885M6G-LecRK_4148

ACTAAACTCTTGAATAAAATTAAGCATGATTAATTAACTCGAAAAACTTTTAATTTCAGTGCAAATTGGAGGTGTGGGATCTGGAAATAGTTCTGGCAAGACAAAAATAGTTGCAATAGC

>M6-PGSC0003DMG400013885M6G-LecRK_4209

CAAATTGGAGGTGTGGGATCTGGAAATAGTTCTGGCAAGACAAAAATAGTTGCAATAGCTGCTGGGGTTACAGTTGGTATAGCTCTTGTGCTATTTGGACTAACCCTTTGCTTCTTATCA

>M6-PGSC0003DMG400013885M6G-LecRK_427

AATCGCAGAAATTGGTACCCTTCTTCTCGTCGATGGCCAAACTGGAAATTCCGTCTGGTCTTCCGAGCAAACTCCGGCGACCAACGTAGTTGCCCAACTCCTTGACTCCGGGAACTTTGT

>M6-PGSC0003DMG400013885M6G-LecRK_4270

CTGGGGTTACAGTTGGTATAGCTCTTGTGCTATTTGGACTAACCCTTTGCTTCTTATCAAAGAGAAGAAAACATCAAGGTTCAATCCGAACTAAATCCGTAAACAGAGGTATTATATGAA

>M6-PGSC0003DMG400013885M6G-LecRK_4331

GAGAAGAAAACATCAAGGTTCAATCCGAACTAAATCCGTAAACAGAGGTATTATATGAATCTATCATGATAAATAAGATTGTCTAATATAAAATGCTTCATTTTCTAATGAATGTTTCTT

>M6-PGSC0003DMG400013885M6G-LecRK_4392

TATCATGATAAATAAGATTGTCTAATATAAAATGCTTCATTTTCTAATGAATGTTTCTTATGATCCTAAACCAGGCACTAGTGAACGAAGTCAAGAACTTCTGATGAATGCAACTATTAT

>M6-PGSC0003DMG400013885M6G-LecRK_4453

GATCCTAAACCAGGCACTAGTGAACGAAGTCAAGAACTTCTGATGAATGCAACTATTATTCCAAGTAAAAGAGAATTTTCTGGTGAAACTTCCACGGAAGAGTTTGAATTGCCATTGTTT

>M6-PGSC0003DMG400013885M6G-LecRK_4514

CAAGTAAAAGAGAATTTTCTGGTGAAACTTCCACGGAAGAGTTTGAATTGCCATTGTTTGATTACAGTACCCTAGCTACGGCAACAGAAAATTTTTCTGATGCGACTAAGTTGGGCCAAG

>M6-PGSC0003DMG400013885M6G-LecRK_4575

TTACAGTACCCTAGCTACGGCAACAGAAAATTTTTCTGATGCGACTAAGTTGGGCCAAGGTGGATTTGGTTGTGTTTACAAGGTAAAATCTCAGTTATATGTAAACCATAGTGGTATAAT

>M6-PGSC0003DMG400013885M6G-LecRK_4636

GGATTTGGTTGTGTTTACAAGGTAAAATCTCAGTTATATGTAAACCATAGTGGTATAATAAAGAAGAAAAAATGATCGACAAACTCAATAATATAATCTTTTGATCAGGCAATGCTGGTT

>M6-PGSC0003DMG400013885M6G-LecRK_4697

AGAAGAAAAAATGATCGACAAACTCAATAATATAATCTTTTGATCAGGCAATGCTGGTTGGTCAAGAAGTAGCAGTCAAAAGGCTCTCGAAGAATTCCGGACAAGGAGTAGAGGAATTTA

>M6-PGSC0003DMG400013885M6G-LecRK_4758

TCAAGAAGTAGCAGTCAAAAGGCTCTCGAAGAATTCCGGACAAGGAGTAGAGGAATTTAAAAATGAGCTAAGATTAATTGCCAGGCTTCAACACAGAAATCTAGTCCGGCTTCTTGGCTG

>M6-PGSC0003DMG400013885M6G-LecRK_4819

AATGAGCTAAGATTAATTGCCAGGCTTCAACACAGAAATCTAGTCCGGCTTCTTGGCTGTTGTGTTGATATGGAAGAGAAGATGCTGATCTATGAATACCTGGAGAATAAAAGTTTAGAT

>M6-PGSC0003DMG400013885M6G-LecRK_488

TCCGAGCAAACTCCGGCGACCAACGTAGTTGCCCAACTCCTTGACTCCGGGAACTTTGTCATTCGACCGGAAAACGATGACAGAGAACAGAGCTACCTGTGGCAGAGCTTTGATTATCCA

>M6-PGSC0003DMG400013885M6G-LecRK_4880

GTGTTGATATGGAAGAGAAGATGCTGATCTATGAATACCTGGAGAATAAAAGTTTAGATTCAATTTTGTTCAGTAAGTCTCTTTCGTTGTATCGCCAAGAGAGTAGTGTTTCACATCATT

>M6-PGSC0003DMG400013885M6G-LecRK_4941

AATTTTGTTCAGTAAGTCTCTTTCGTTGTATCGCCAAGAGAGTAGTGTTTCACATCATTTACTTACAGGCTTATATAATTTGTGACTCACATTTGAATATGTTCCTCAGATAAACAGAAT

>M6-PGSC0003DMG400013885M6G-LecRK_5002

CTTACAGGCTTATATAATTTGTGACTCACATTTGAATATGTTCCTCAGATAAACAGAATAGCTCGCTGCTCAACTGGCAAAAGCGGTTCAATATTATTTGTGGTATTGCTCGGGGGCTTC

>M6-PGSC0003DMG400013885M6G-LecRK_5063

CTCGCTGCTCAACTGGCAAAAGCGGTTCAATATTATTTGTGGTATTGCTCGGGGGCTTCTATACCTTCACCAAGATTCAAGATTTAGGATTATCCACAGAGACCTTAAAGCAAGCAATAT

>M6-PGSC0003DMG400013885M6G-LecRK_5185

TGCTTGATAAGGATTTGACCCCCAAAATATCAGATTTTGGCATGGCAAGAATATTTGGCGGAGATGAGACTGAAGGAAATACAAAAAGAGTAGTTGGAACCTAGTAAGCACTCAACATAA

>M6-PGSC0003DMG400013885M6G-LecRK_5246

AGATGAGACTGAAGGAAATACAAAAAGAGTAGTTGGAACCTAGTAAGCACTCAACATAATTTCTTCACATAATTCATAGAACCTTTAAAGCTGTTACTAAATCTGTTTTTTTGTTTTGAT

>M6-PGSC0003DMG400013885M6G-LecRK_5307

TCTTCACATAATTCATAGAACCTTTAAAGCTGTTACTAAATCTGTTTTTTTGTTTTGATGTAGTGGTTACATGTCTCCGGAATATGCAATGGATGGCCTCTTCTCTGTTAAGTCCGATGT

>M6-PGSC0003DMG400013885M6G-LecRK_5368

AGTGGTTACATGTCTCCGGAATATGCAATGGATGGCCTCTTCTCTGTTAAGTCCGATGTTTTCAGCTTTGGGGTTTTGGTGTTAGAAATAGTAACCGGGAAGAAGAACAGGGGATTCTAT

>M6-PGSC0003DMG400013885M6G-LecRK_5429

TCAGCTTTGGGGTTTTGGTGTTAGAAATAGTAACCGGGAAGAAGAACAGGGGATTCTATTATCAAAATAACCAACTCAACCTTCTTGGACATGTAAGCTTCATTATTGGTAATACATTCA

>M6-PGSC0003DMG400013885M6G-LecRK_549

TTCGACCGGAAAACGATGACAGAGAACAGAGCTACCTGTGGCAGAGCTTTGATTATCCAACAAACACTTTATTACCTGGGATGAAACTTGGGTGGGATTCAAAATCCGGGATGAACAGAA

>M6-PGSC0003DMG400013885M6G-LecRK_5490

TCAAAATAACCAACTCAACCTTCTTGGACATGTAAGCTTCATTATTGGTAATACATTCATCATTTGATTCATTTCTGATAAACAAAGTAACCGATTATGCAGGCGTGGAGACTATGGAAA

>M6-PGSC0003DMG400013885M6G-LecRK_5551

ATTTGATTCATTTCTGATAAACAAAGTAACCGATTATGCAGGCGTGGAGACTATGGAAAGAAGGGAGCGGCTCAGAGCTACTAGATCCATCTTTTGGAGAATCGTTTTCTCCGTCTGAAG

>M6-PGSC0003DMG400013885M6G-LecRK_5612

AGGGAGCGGCTCAGAGCTACTAGATCCATCTTTTGGAGAATCGTTTTCTCCGTCTGAAGTAATGAGATGTATACAAGTTGGACTGTTGTGTGTCCAGGAACAAGCAGAAGACAGACCAAA

>M6-PGSC0003DMG400013885M6G-LecRK_5673

ATGAGATGTATACAAGTTGGACTGTTGTGTGTCCAGGAACAAGCAGAAGACAGACCAAACATGGCCACTGTGGTTTTGATGCTAGGTAGCGAAAGCGCGTCACTACCTCAGCCTAAAAAC

>M6-PGSC0003DMG400013885M6G-LecRK_5734

TGGCCACTGTGGTTTTGATGCTAGGTAGCGAAAGCGCGTCACTACCTCAGCCTAAAAACCCAGGATTTTGCCTTGGAAGAAGACCCGTTGATTCTGATTCATATTCGACTAATTATGAAG

>M6-PGSC0003DMG400013885M6G-LecRK_5795

AGGATTTTGCCTTGGAAGAAGACCCGTTGATTCTGATTCATATTCGACTAATTATGAAGAAACTTGCACTGTCAATCAAGTTACAGTTACCATGATAGATCCCCGGTAGTTGATGGTATA

>M6-PGSC0003DMG400013885M6G-LecRK_5856

ACTTGCACTGTCAATCAAGTTACAGTTACCATGATAGATCCCCGGTAGTTGATGGTATATATCGCCCTCAAGAAGTGTATATTACTACAAGAGGTTCACAATTTTCTCTCGTATAATTTC

>M6-PGSC0003DMG400013885M6G-LecRK_5917

TCGCCCTCAAGAAGTGTATATTACTACAAGAGGTTCACAATTTTCTCTCGTATAATTTCTCAGTTTTTGCTTTTTGGGTAAGTTTTGTGTAAGATAAAGAGTATAATTAGGTCTTTTTTT

>M6-PGSC0003DMG400013885M6G-LecRK_5978

AGTTTTTGCTTTTTGGGTAAGTTTTGTGTAAGATAAAGAGTATAATTAGGTCTTTTTTTCTCATTACATACAATTGTGAGTTCAAAAATCTGTCTTTTTTCATCATCAATGTAATAGTCC

>M6-PGSC0003DMG400013885M6G-LecRK_6025

AGGTCTTTTTTTCTCATTACATACAATTGTGAGTTCAAAAATCTGTCTTTTTTCATCATCAATGTAATAGTCCAATCATTCCTTTTCCATATCCTTGAAATCCATTGTAAATCACTGTGC

>M6-PGSC0003DMG400013885M6G-LecRK_61

TACACCAACAAATGTCTCAAACACCAACTCTGCAGCAAAACAAAATACTCTTTTCTTAAATTCCACAATGAGAAACACCAAAGATTACTACTTTCACCAATTCTTGTTCTTGATTTTGAT

>M6-PGSC0003DMG400013885M6G-LecRK_610

AAACACTTTATTACCTGGGATGAAACTTGGGTGGGATTCAAAATCCGGGATGAACAGAAACATAACATCATGGAAATCAGCAATCGATCCAGCTCCCGGAGATTACACCTTCAAGATCAA

>M6-PGSC0003DMG400013885M6G-LecRK_671

ATAACATCATGGAAATCAGCAATCGATCCAGCTCCCGGAGATTACACCTTCAAGATCAACACCAGTGGGTTCCCGGAAGTTTACTTAACAAATAAACAACAAACTATCTACAGGAGCGGC

>M6-PGSC0003DMG400013885M6G-LecRK_732

CCAGTGGGTTCCCGGAAGTTTACTTAACAAATAAACAACAAACTATCTACAGGAGCGGCGCTTGGAATGGAATTAGATTCAGTGGGGTACCGGAAATGAAGGCAAGTGAGATTATCTCAT

>M6-PGSC0003DMG400013885M6G-LecRK_793

TTGGAATGGAATTAGATTCAGTGGGGTACCGGAAATGAAGGCAAGTGAGATTATCTCATTCGAATTTCAGTTCAAATCAGATGAAATTACTTACATGTTCAAATTACACAACAAAACTTT

>M6-PGSC0003DMG400013885M6G-LecRK_854

GAATTTCAGTTCAAATCAGATGAAATTACTTACATGTTCAAATTACACAACAAAACTTTATACTCAAGATTGTCTGTGAATCATACTGGTTATCTTGAAAGATTCACTTGGATCCCAACA

>M6-PGSC0003DMG400013885M6G-LecRK_915

ACTCAAGATTGTCTGTGAATCATACTGGTTATCTTGAAAGATTCACTTGGATCCCAACAAGCAACATGTGGAATCGATTTTGGTATGCTCCAAAAGATCAATGTGACGGATACACAGAGT

>M6-PGSC0003DMG400013885M6G-LecRK_976

CAACATGTGGAATCGATTTTGGTATGCTCCAAAAGATCAATGTGACGGATACACAGAGTGTGGGGTATCTGGAATTTGTGACACAAATATTTCACCAATCTGCAAATGTATGGTGGGATT

>M6-PGSC0003DMG400013886M6G-LecRK_0

GCCTCATCAGAACCTTCTTCTTTCCTTTTGTTTTCTATACCCTTTACCATTCTCCTTTCTAAATTCCCAATTTTCTCCATTCTTGATTTGTGGCATCATCATTTCCAAATTCTTGAATGG

>M6-PGSC0003DMG400013886M6G-LecRK_1037

TGTGATAAAGATGTATGCCAATGTTTAAAAGGTTTTGTACCACGTGTTCAGGAGCAATGGCATGCTGGGAACCGAACCGGAGGATGTGTTAGGAAGACAGAATTGGAATGCAGGAGAAAT

>M6-PGSC0003DMG400013886M6G-LecRK_1098

ATGCTGGGAACCGAACCGGAGGATGTGTTAGGAAGACAGAATTGGAATGCAGGAGAAATAGCAGTGTTTCAAGGAATGATAGTTCAAAAGATGATGGATTCTCGACTATTCGGAGAGTTA

>M6-PGSC0003DMG400013886M6G-LecRK_1159

CAGTGTTTCAAGGAATGATAGTTCAAAAGATGATGGATTCTCGACTATTCGGAGAGTTAAATTACCTGACCATGCTGATGTTTCAGAAATAACCACAGACGAGTGCAAAATCAGGTGCCT

>M6-PGSC0003DMG400013886M6G-LecRK_1220

TTACCTGACCATGCTGATGTTTCAGAAATAACCACAGACGAGTGCAAAATCAGGTGCCTGAATGATTGTTCCTGCAATGCTTATGCTTATGTCAGAGGAATCAATTGTATGATATGGCGT

>M6-PGSC0003DMG400013886M6G-LecRK_1281

ATGATTGTTCCTGCAATGCTTATGCTTATGTCAGAGGAATCAATTGTATGATATGGCGTAATGATTTAGTTGATATAGAGCATTTTCAGGAAGGTGGGAACACTCTTTATGTTCGTCTTC

>M6-PGSC0003DMG400013886M6G-LecRK_1342

TGATTTAGTTGATATAGAGCATTTTCAGGAAGGTGGGAACACTCTTTATGTTCGTCTTCATCCTTCTGATATCGGTAAGCACAATACTTTCTGCTGAATTCATAAATCTCTTACCAGTTT

>M6-PGSC0003DMG400013886M6G-LecRK_1403

CCTTCTGATATCGGTAAGCACAATACTTTCTGCTGAATTCATAAATCTCTTACCAGTTTTCTTCTCATTCGTGCAATTATACAAATCTAAAACCAAAGTAGTTTGTGCTTTAGTACAAAG

>M6-PGSC0003DMG400013886M6G-LecRK_1464

TTCTCATTCGTGCAATTATACAAATCTAAAACCAAAGTAGTTTGTGCTTTAGTACAAAGATCAATTTCTTGTTTGTCTTTACTCTTTATTTGTAGCAAGCTCAAACAATAAGCAATATAT

>M6-PGSC0003DMG400013886M6G-LecRK_1525

CAATTTCTTGTTTGTCTTTACTCTTTATTTGTAGCAAGCTCAAACAATAAGCAATATATCTTTTTCTTCCTTGTGTTTGAAGAATGATTAGCCTTTCTAATTTCTTGATTTCTCCTTTGA

>M6-PGSC0003DMG400013886M6G-LecRK_1586

TTTTCTTCCTTGTGTTTGAAGAATGATTAGCCTTTCTAATTTCTTGATTTCTCCTTTGAATATCTTTTGAAACAGGTCAAAAAAAGAAGACGATCATAATTGTTGTAATATCAATTCTAG

>M6-PGSC0003DMG400013886M6G-LecRK_1647

ATCTTTTGAAACAGGTCAAAAAAAGAAGACGATCATAATTGTTGTAATATCAATTCTAGCAGCTCTGGCTTTGGTTGTTATGGTGGCCATTTGGCTAGTATGCAAGTACAGGGCCAGAAA

>M6-PGSC0003DMG400013886M6G-LecRK_1708

GCTCTGGCTTTGGTTGTTATGGTGGCCATTTGGCTAGTATGCAAGTACAGGGCCAGAAAACGAGGTAAATTTTGATGCAGTAACATTTCTGACTTGTTAGTTGCATTTATTATATGTTCC

>M6-PGSC0003DMG400013886M6G-LecRK_1769

GAGGTAAATTTTGATGCAGTAACATTTCTGACTTGTTAGTTGCATTTATTATATGTTCCATATTGTACTTATACCTTTGACTTGCAATTTTCAGAATCCAAGAGGACCAGTGAAATTCCC

>M6-PGSC0003DMG400013886M6G-LecRK_183

CATTCTGTTTAGGTGGTGACAGAGTTACACTAGGAGAGATTATTAAAGATGGTGATAACATCACTTCTAAAGGAGGGGATTTTGTGTTGGGGTTCTTTAGTCCTGCAAATACTAGTAAAA

>M6-PGSC0003DMG400013886M6G-LecRK_1830

ATTGTACTTATACCTTTGACTTGCAATTTTCAGAATCCAAGAGGACCAGTGAAATTCCCAAAAACCATTTAGTTAGGAGTGGAGAGTTCTCCACAGAATATTCTGGACCAGGCGACATCA

>M6-PGSC0003DMG400013886M6G-LecRK_1891

AAACCATTTAGTTAGGAGTGGAGAGTTCTCCACAGAATATTCTGGACCAGGCGACATCAGTGCTGAAGGGCATCAAGGAAATGGTTCAGAATTAGCATTTTTCAGCTTCAGCATGGTGGC

>M6-PGSC0003DMG400013886M6G-LecRK_1952

GCTGAAGGGCATCAAGGAAATGGTTCAGAATTAGCATTTTTCAGCTTCAGCATGGTGGCCACAGCTACTGATGACTTCTCTCTTGCAAACAAGCTTGGGCAAGGGGGATTTGGCCCCGTT

>M6-PGSC0003DMG400013886M6G-LecRK_2013

CAGCTACTGATGACTTCTCTCTTGCAAACAAGCTTGGGCAAGGGGGATTTGGCCCCGTTTATAAGGTAAATTTTCAATCTTGATAGATTATTCTGTTCCTTTAGGAAGTATATGGCCATG

>M6-PGSC0003DMG400013886M6G-LecRK_2074

TAAGGTAAATTTTCAATCTTGATAGATTATTCTGTTCCTTTAGGAAGTATATGGCCATGTTTTGGACTAAAGTTGATGTTTTTCAAATCTTTAGGGAAAGCTACCTTGTGGTCAAGAAGT

>M6-PGSC0003DMG400013886M6G-LecRK_2135

TTGGACTAAAGTTGATGTTTTTCAAATCTTTAGGGAAAGCTACCTTGTGGTCAAGAAGTTGCTGTAAAAAGGCTTTCACAAAAGTCTGGACAAGGCGATGAGGAGTTCAAGAATGAGATC

>M6-PGSC0003DMG400013886M6G-LecRK_2196

CTGTAAAAAGGCTTTCACAAAAGTCTGGACAAGGCGATGAGGAGTTCAAGAATGAGATCACTCTAATAGCAAAATTGCAACACAGAAATCTTGTTAGACTTTTGGGTTGCTGTGTTGAAG

>M6-PGSC0003DMG400013886M6G-LecRK_2257

TCTAATAGCAAAATTGCAACACAGAAATCTTGTTAGACTTTTGGGTTGCTGTGTTGAAGGAGAAGAAAAAATGCTGATTTACGAGTACATGCCCAACAAAAGCTTAGATACATTTCTATT

>M6-PGSC0003DMG400013886M6G-LecRK_2318

GAAGAAAAAATGCTGATTTACGAGTACATGCCCAACAAAAGCTTAGATACATTTCTATTTGGTAAGTTCAGGAAATTCTATCAAAATAATTTCTGCTGCTCCCACAGATGTACTATAGCA

>M6-PGSC0003DMG400013886M6G-LecRK_2379

GTAAGTTCAGGAAATTCTATCAAAATAATTTCTGCTGCTCCCACAGATGTACTATAGCAACACATCTCATTGAGATAGACACTAACAACTGTATTGGCTGTTAAACTTTGACAGATACTG

>M6-PGSC0003DMG400013886M6G-LecRK_244

CACTTCTAAAGGAGGGGATTTTGTGTTGGGGTTCTTTAGTCCTGCAAATACTAGTAAAAGGTATCTTGGTATATGGTATGTTGACGTACCAGTAAAGACATATATTTGGGTTGCCAACAG

>M6-PGSC0003DMG400013886M6G-LecRK_2440

ACATCTCATTGAGATAGACACTAACAACTGTATTGGCTGTTAAACTTTGACAGATACTGCTAGGAAATCCCAGTTAGACTGGAGAAAACGCTTCAACATTATCGAAGGAATTGCCCGAGG

>M6-PGSC0003DMG400013886M6G-LecRK_2501

AGGAAATCCCAGTTAGACTGGAGAAAACGCTTCAACATTATCGAAGGAATTGCCCGAGGATTACTGTATCTCCATAGAGATTCAAGGCTTAGAATAATCCATAGGGATCTAAAGGCTAGT

>M6-PGSC0003DMG400013886M6G-LecRK_2562

TACTGTATCTCCATAGAGATTCAAGGCTTAGAATAATCCATAGGGATCTAAAGGCTAGTAACATTCTGTTGGATGAAGAGATGACACCAAAAATTTCAGACTTTGGCATGGCCAGAATAT

>M6-PGSC0003DMG400013886M6G-LecRK_2623

CATTCTGTTGGATGAAGAGATGACACCAAAAATTTCAGACTTTGGCATGGCCAGAATATTTGGAGGAAACCAAAATGAAGCAAATACAAATAGGGTCGTAGGAACATAGTAAGTATCATT

>M6-PGSC0003DMG400013886M6G-LecRK_2684

GGAGGAAACCAAAATGAAGCAAATACAAATAGGGTCGTAGGAACATAGTAAGTATCATTTGATTTTAAATCTTTTAAGAGACTAATTTCAAAATTTCTTGTTCCTGGAGCTAAATCCATT

>M6-PGSC0003DMG400013886M6G-LecRK_2745

ATTTTAAATCTTTTAAGAGACTAATTTCAAAATTTCTTGTTCCTGGAGCTAAATCCATTTCATTTTGTACCTTGTTGCTAAACAACAGTGGATATATGGCTCCTGAGTATGCTATGGAGG

>M6-PGSC0003DMG400013886M6G-LecRK_2806

ATTTTGTACCTTGTTGCTAAACAACAGTGGATATATGGCTCCTGAGTATGCTATGGAGGGCCTGTTCTCTGGCAAGTCTGATGTCTACAGCTTCGGCATAATATTGCTTGAAATTATATG

>M6-PGSC0003DMG400013886M6G-LecRK_2867

CTGTTCTCTGGCAAGTCTGATGTCTACAGCTTCGGCATAATATTGCTTGAAATTATATGTGGACGTAGAAATACAAGCTTTCGGACAGATGAGCATTCAGGCATCATTGGATATGTAAGT

>M6-PGSC0003DMG400013886M6G-LecRK_2928

GACGTAGAAATACAAGCTTTCGGACAGATGAGCATTCAGGCATCATTGGATATGTAAGTCTGACTAATTTTCTCCAACTATTTCAATCTTTTTAGTCCAGAACAAGCTGTCTGAATAATG

>M6-PGSC0003DMG400013886M6G-LecRK_2989

GACTAATTTTCTCCAACTATTTCAATCTTTTTAGTCCAGAACAAGCTGTCTGAATAATGGGGCTAATTTCTTGTGTGATGAGCAATGACGGAGTGATCATCTTTCTCATATTTTTCTGGT

>M6-PGSC0003DMG400013886M6G-LecRK_305

TATCTTGGTATATGGTATGTTGACGTACCAGTAAAGACATATATTTGGGTTGCCAACAGGAACAATCCTGTTCATGATAAAAATGGTACCTTTTCAATTGATAAAAATGGGAATTTGGTG

>M6-PGSC0003DMG400013886M6G-LecRK_3050

GCTAATTTCTTGTGTGATGAGCAATGACGGAGTGATCATCTTTCTCATATTTTTCTGGTGGAAAATATTTCAGGCATGGGAGAAGTGGGATGAAGGTAGACCAATGGACCTAGTGGATCG

>M6-PGSC0003DMG400013886M6G-LecRK_3111

AAAATATTTCAGGCATGGGAGAAGTGGGATGAAGGTAGACCAATGGACCTAGTGGATCGTTCGATTTGGGATGGTTGTCAACACAATGAAGCATTGAGATGTATACACTTGGCATTGCTT

>M6-PGSC0003DMG400013886M6G-LecRK_3172

CGATTTGGGATGGTTGTCAACACAATGAAGCATTGAGATGTATACACTTGGCATTGCTTTGTGTTCAAGATTTGGCCGCTCACAGACCAAACATGTCTTCTGTGGTTTTGATGCTGGAAA

>M6-PGSC0003DMG400013886M6G-LecRK_3233

TGTTCAAGATTTGGCCGCTCACAGACCAAACATGTCTTCTGTGGTTTTGATGCTGGAAACTGACAATGTAAGGTTGCCCTTGCCTAGGCAACCTACCTATACTTCAATGAGAAGATCTGT

>M6-PGSC0003DMG400013886M6G-LecRK_3294

GACAATGTAAGGTTGCCCTTGCCTAGGCAACCTACCTATACTTCAATGAGAAGATCTGTGGATGAAGATATATGGCATGGGAATCAGGATTTAACATCTTCCAACAATGTCACAGTTAGT

>M6-PGSC0003DMG400013886M6G-LecRK_3355

ATGAAGATATATGGCATGGGAATCAGGATTTAACATCTTCCAACAATGTCACAGTTAGTGTACTAATAGGTAGATGATTGGTTAATGCTTGTTCCTTCTTACATACCTGCTTCATTTGAA

>M6-PGSC0003DMG400013886M6G-LecRK_3416

ACTAATAGGTAGATGATTGGTTAATGCTTGTTCCTTCTTACATACCTGCTTCATTTGAAGAGAATGGGATCAAGTAGTGTAACTATGTAAAGTTATACTTGGAACTATGTTCATAATTTT

>M6-PGSC0003DMG400013886M6G-LecRK_3477

GAATGGGATCAAGTAGTGTAACTATGTAAAGTTATACTTGGAACTATGTTCATAATTTTTTGTTAGCTAGGTAAACTTTAGCACATCATGTGTATTGTAATTAGAAGGTGGTTCAGCTCT

>M6-PGSC0003DMG400013886M6G-LecRK_3538

GTTAGCTAGGTAAACTTTAGCACATCATGTGTATTGTAATTAGAAGGTGGTTCAGCTCTGTAAGCTGAAAGAGAGATATTTCACAACCTTGTTCAATCAGTATGCTTGAAGTTTTTTTCT

>M6-PGSC0003DMG400013886M6G-LecRK_3599

AAGCTGAAAGAGAGATATTTCACAACCTTGTTCAATCAGTATGCTTGAAGTTTTTTTCTTTACTGTTTGTAGATGGAGATGCTCCACCAATCGAGCAAACCCCACTTGTCATATCTAGTT

>M6-PGSC0003DMG400013886M6G-LecRK_366

ACAATCCTGTTCATGATAAAAATGGTACCTTTTCAATTGATAAAAATGGGAATTTGGTGGTTAAAGATGGACATGGGGATTTACTATGGTCGTCTAATGTTTCTGTTAATACCACAAATT

>M6-PGSC0003DMG400013886M6G-LecRK_427

TAAAGATGGACATGGGGATTTACTATGGTCGTCTAATGTTTCTGTTAATACCACAAATTCTACGGCTTGTCTGAGAGACGAGGGAAATCTTGTCATCCTTAATAATGACAGGAATGCTAC

>M6-PGSC0003DMG400013886M6G-LecRK_488

ACGGCTTGTCTGAGAGACGAGGGAAATCTTGTCATCCTTAATAATGACAGGAATGCTACGAGGTTAAATTCTGAATTGTGGAAGAGCTTCTCGGATCCTACAGATACATTTTTGCCTGGA

>M6-PGSC0003DMG400013886M6G-LecRK_549

GGTTAAATTCTGAATTGTGGAAGAGCTTCTCGGATCCTACAGATACATTTTTGCCTGGAATGGAAGTTCTCATAGAAAGACGAGGCCAAGAGCAAAAAGTTTTCCGTTCTTGGACAAATG

>M6-PGSC0003DMG400013886M6G-LecRK_610

GGAAGTTCTCATAGAAAGACGAGGCCAAGAGCAAAAAGTTTTCCGTTCTTGGACAAATGAGAGTGACCCTTCACCTGGGAGGTACTCAATGGGAGTTGATCCTCGCGGAACACCTCAGAT

>M6-PGSC0003DMG400013886M6G-LecRK_671

AGTGACCCTTCACCTGGGAGGTACTCAATGGGAGTTGATCCTCGCGGAACACCTCAGATTGTTATTTGGGATGGACCGAATAGACGATGGAGGAGTGGACATTTTGATGGAGCGGAATTC

>M6-PGSC0003DMG400013886M6G-LecRK_732

TTATTTGGGATGGACCGAATAGACGATGGAGGAGTGGACATTTTGATGGAGCGGAATTCATTGGTGTTCCAGATGTGATTAGAACTACTTTTTTCTCTGGCTTCAGAATTCAAAACCAAG

>M6-PGSC0003DMG400013886M6G-LecRK_793

TGGTGTTCCAGATGTGATTAGAACTACTTTTTTCTCTGGCTTCAGAATTCAAAACCAAGGAGACGATAAATTGTTACTCACTTACAGTGCCTCAAATACATCTTCTTTTGTGAGGTTTCA

>M6-PGSC0003DMG400013886M6G-LecRK_854

GACGATAAATTGTTACTCACTTACAGTGCCTCAAATACATCTTCTTTTGTGAGGTTTCAGCTTACTGTGACTGGAAATGAGCTGCAACAGAGGTGGAACGAAGACCAGGGTGAATGGAAC

>M6-PGSC0003DMG400013886M6G-LecRK_915

TTACTGTGACTGGAAATGAGCTGCAACAGAGGTGGAACGAAGACCAGGGTGAATGGAACACGTTGCAATCTAGGCCTGTAGGTGGTTGTGACTTGTATAACTTCTGTGGGAATTTTGCAG

>M6-PGSC0003DMG400013886M6G-LecRK_976

GTTGCAATCTAGGCCTGTAGGTGGTTGTGACTTGTATAACTTCTGTGGGAATTTTGCAGAATGTGATAAAGATGTATGCCAATGTTTAAAAGGTTTTGTACCACGTGTTCAGGAGCAATG

>M6-PGSC0003DMG400013887M6G-LecRK_1054

TCCTTTACCTTCACAGAGACTCAAGACTATGAATTATTCATAGAGATCTGAAGGCAAGCAATATCCTGTTGGATGAATACCTAAACCCAAAAATTTCAAATTTTGGCATGGCGAGGATTT

>M6-PGSC0003DMG400013887M6G-LecRK_1116

ATCCTGTTGGATGAATACCTAAACCCAAAAATTTCAAATTTTGGCATGGCGAGGATTTTTGGAGCAATCAAGATCAGGCCAACACTATAAGAGTTGTTGGCACATAGTATGTTTGCATTG

>M6-PGSC0003DMG400013887M6G-LecRK_1178

AGCAATCAAGATCAGGCCAACACTATAAGAGTTGTTGGCACATAGTATGTTTGCATTGCTTCCTTCATTTTTTCATGGATGAACTGCCATCATTAGTCATTGATTTTCTGTGTAATATGT

>M6-PGSC0003DMG400013887M6G-LecRK_124

ATGCTCACGAAGAACTGTGTATGGCTCATACATGTTTGCAGCTGCAAATGACAAGAAAGACATTCCTGTAGCAGCTATTGCAACCACAGTCTTAATAGGCTCAATAACAGTTATCTTATG

>M6-PGSC0003DMG400013887M6G-LecRK_1240

CTTCATTTTTTCATGGATGAACTGCCATCATTAGTCATTGATTTTCTGTGTAATATGTTATGCAGTGGTTACATGGCCCCTGAATATTCAATGTACGGAAGATTCTCAGAAAAATCAGAT

>M6-PGSC0003DMG400013887M6G-LecRK_1302

CAGTGGTTACATGGCCCCTGAATATTCAATGTACGGAAGATTCTCAGAAAAATCAGATGTCTACAGCTTCAGTGTGTTGGTATTGGAAATTGTCAGTGGAAGGAAGAACTCTAGTTTTTA

>M6-PGSC0003DMG400013887M6G-LecRK_1364

ACAGCTTCAGTGTGTTGGTATTGGAAATTGTCAGTGGAAGGAAGAACTCTAGTTTTTATGATGATGAACTGACTCTACTTGCATATGTAAGTTCATTCTTGCAACTATATATGAAGCTTT

>M6-PGSC0003DMG400013887M6G-LecRK_1426

GATGAACTGACTCTACTTGCATATGTAAGTTCATTCTTGCAACTATATATGAAGCTTTAATGTTCTGCTAGCTCATCAAACCAAAAAAATAGAGGAGAAAAATTATGGAACTTCTGAAAA

>M6-PGSC0003DMG400013887M6G-LecRK_1488

TTCTGCTAGCTCATCAAACCAAAAAAATAGAGGAGAAAAATTATGGAACTTCTGAAAATAACATTTTTGCTGTTTGACAATGTCATTGTTCATTAGGCATGGAAGTTGTGGAATGAAAAC

>M6-PGSC0003DMG400013887M6G-LecRK_1550

ATTTTTGCTGTTTGACAATGTCATTGTTCATTAGGCATGGAAGTTGTGGAATGAAAACAATATCATAAAATTGATAGACCCCAAAATATTTGATTCATGCTTTGAAAAACAGATGGTGAG

>M6-PGSC0003DMG400013887M6G-LecRK_1612

TCATAAAATTGATAGACCCCAAAATATTTGATTCATGCTTTGAAAAACAGATGGTGAGATGTGTACATATTGGATTATTATGTGTTCAAGAATATGCAGAAGATAGGCCAAATGTCACCA

>M6-PGSC0003DMG400013887M6G-LecRK_1674

GTACATATTGGATTATTATGTGTTCAAGAATATGCAGAAGATAGGCCAAATGTCACCACAGTTCTGTTAATGCTCACTAGTGATATTGCTGAACTACCTACTCCTAAACAACCTGCATTT

>M6-PGSC0003DMG400013887M6G-LecRK_1736

TCTGTTAATGCTCACTAGTGATATTGCTGAACTACCTACTCCTAAACAACCTGCATTTACCGGAGGACATGCTTCACCACAGCAAGTATCTTCTAAAAGCCAAGGGTCCGTGAATGCTGA

>M6-PGSC0003DMG400013887M6G-LecRK_186

TTCCTGTAGCAGCTATTGCAACCACAGTCTTAATAGGCTCAATAACAGTTATCTTATGTGGATATCTTTTCTGGAAATTGTTGGCTAAGCACAGAGGTAATTAATCAAATAAGATTTTTG

>M6-PGSC0003DMG400013887M6G-LecRK_1860

ATAACTGTATTGGAACCACGATAGGATGCAGAGTTTTCAACAGGCAGGCAAAGATCAGTACTTTCCCTAAAATGGTAACAGTAGCATAGTATTTCTTTCTTTTGAATGTGCATAGGGTAA

>M6-PGSC0003DMG400013887M6G-LecRK_1922

TTCCCTAAAATGGTAACAGTAGCATAGTATTTCTTTCTTTTGAATGTGCATAGGGTAAAACACGCTCCTGCAATAACTCGCAAAACACACAAAAGAGGTAAAAGTGCACTAGCTATGTGA

>M6-PGSC0003DMG400013887M6G-LecRK_1984

CGCTCCTGCAATAACTCGCAAAACACACAAAAGAGGTAAAAGTGCACTAGCTATGTGATGAATTAATTACAGAGTGAGCTTGGAGAACATGAAACAGTTGATCAATTTTTAACTTTGCAT

>M6-PGSC0003DMG400013887M6G-LecRK_2046

TTAATTACAGAGTGAGCTTGGAGAACATGAAACAGTTGATCAATTTTTAACTTTGCATTATGAAAATCCTTAGAAAGAAAGTAGACAAAAGGACAAAAAGAACCGCAAATCCCCAATTAA

>M6-PGSC0003DMG400013887M6G-LecRK_2108

AAAATCCTTAGAAAGAAAGTAGACAAAAGGACAAAAAGAACCGCAAATCCCCAATTAACAAGTTCAAATAATGGCAATCAAAGACTTGAAAAATCATACCTGCTGTACTGATACACGAAA

>M6-PGSC0003DMG400013887M6G-LecRK_2170

TTCAAATAATGGCAATCAAAGACTTGAAAAATCATACCTGCTGTACTGATACACGAAACGGCGTAAGCGAAAAATAGCCACTCGTTTTACCACTGTTTTTTTTCTTAGAAAATTACAGAG

>M6-PGSC0003DMG400013887M6G-LecRK_2232

GTAAGCGAAAAATAGCCACTCGTTTTACCACTGTTTTTTTTCTTAGAAAATTACAGAGAAGTGAATTGGTGAATTAATCAATGTTGAGGAAAAGTTAACTCTGCATTATTTTAGGTCATT

>M6-PGSC0003DMG400013887M6G-LecRK_2294

GAATTGGTGAATTAATCAATGTTGAGGAAAAGTTAACTCTGCATTATTTTAGGTCATTCACATTTCACACAAGTTATTTGGGTGTTGGTTAAAGACTATAATTGTTTCATTTTTAAAATT

>M6-PGSC0003DMG400013887M6G-LecRK_248

TATCTTTTCTGGAAATTGTTGGCTAAGCACAGAGGTAATTAATCAAATAAGATTTTTGAGAAGACAAACTAGAACTTCTCTGTCTGCAAATGGTTTGATTTCTGAATGTCCACTCTAAAC

>M6-PGSC0003DMG400013887M6G-LecRK_310

GACAAACTAGAACTTCTCTGTCTGCAAATGGTTTGATTTCTGAATGTCCACTCTAAACCAGAAAGGAAGAGGAAAAATGAAGCACTCTTAAGAGAAGCATCTCCAAAATTTTACCAAGAA

>M6-PGSC0003DMG400013887M6G-LecRK_372

AAGGAAGAGGAAAAATGAAGCACTCTTAAGAGAAGCATCTCCAAAATTTTACCAAGAAGGCATGAGTAAAGATGACATCAATCAAGTCAAAATTGAAGACATCACCTTGTACATATTTGA

>M6-PGSC0003DMG400013887M6G-LecRK_434

TGAGTAAAGATGACATCAATCAAGTCAAAATTGAAGACATCACCTTGTACATATTTGATATGTTAGCGAATGCCACTGACAGTTTTCACCCAGCTAGCAAGCTGGGGCAAGGAGGCTTTG

>M6-PGSC0003DMG400013887M6G-LecRK_558

AGTCTACAAAGTAATGTTTCTTGATGCATCTCTGCAGTGTTCTGACAAATGAACACAATATAAAATAAATTAGATCAATTATCTGCAGGGACAATTGCCATATGGACAAGAAATTGTAGT

>M6-PGSC0003DMG400013887M6G-LecRK_62

AGTTACCAGTTTCAAAAAAAATAAAAATCAATTAGTGCTTTCACTTCCATGGAAAGAAAGATATGCTCACGAAGAACTGTGTATGGCTCATACATGTTTGCAGCTGCAAATGACAAGAAA

>M6-PGSC0003DMG400013887M6G-LecRK_620

AAATAAATTAGATCAATTATCTGCAGGGACAATTGCCATATGGACAAGAAATTGTAGTAAAAAGGCTTTCACAGTCTTCCGGTCAGGGGCTACAGGAGTTTATGAATGAGGTCGTTGTGA

>M6-PGSC0003DMG400013887M6G-LecRK_744

TAGACTACAACATCGTAATCTTGTTAGACTCCTTGGCTGTTGCACAGAGAGAGGGGAAAAGATGCTGGTTTATGATTTCATGCCAAATAGAAGCTTAGATGCATATGTTTTTGGTCAGTG

>M6-PGSC0003DMG400013887M6G-LecRK_806

TGCTGGTTTATGATTTCATGCCAAATAGAAGCTTAGATGCATATGTTTTTGGTCAGTGCAGAATTTTCTTTCATTGGTTTTGCCTGTCCTTCAACTTAAGTACTTCCAGAATCAAAAGAA

>M6-PGSC0003DMG400013887M6G-LecRK_868

ATTTTCTTTCATTGGTTTTGCCTGTCCTTCAACTTAAGTACTTCCAGAATCAAAAGAAAGCTGTATTTCTTTAAGGAAACAAAATCCATCATACATTTCTCAGTGATACCTTTTCAGGTT

>M6-PGSC0003DMG400013887M6G-LecRK_930

GTATTTCTTTAAGGAAACAAAATCCATCATACATTTCTCAGTGATACCTTTTCAGGTTCACACCAGGAAAAGGTCCTTGATTGGGGTAAACGAGCCATCATTATTGAAGGAACTGGTCGA

>M6-PGSC0003DMG400013898M6LRR-RLP_0

TATATATATATATGAGAACCCTTTTAATGTAAAAGCTCAAAAAAGTTGTATTATTGTTCTTGAAACCAAGTACTCTACCTATTAGACATGACAACTTCTTACAGTACCTTCCTTTACATT

>M6-PGSC0003DMG400013898M6LRR-RLP_1054

TCGCTTGAATTCAACCATTCCTTCTGAGCTAGGCCTTTGTACTAACCTCATTCTCTTGGCTCTATCAGGGAATGCCCTGCAAGGACCATTGCCTATATCCTTGTCTACAGTGACCAAGTT

>M6-PGSC0003DMG400013898M6LRR-RLP_1116

TATCAGGGAATGCCCTGCAAGGACCATTGCCTATATCCTTGTCTACAGTGACCAAGTTATCTGACTTGGGCTTGTCTGATAATTTTCTTTCTGGTGAGGTTCAATCATATCTCATTACCA

>M6-PGSC0003DMG400013898M6LRR-RLP_1178

GACTTGGGCTTGTCTGATAATTTTCTTTCTGGTGAGGTTCAATCATATCTCATTACCAATTGGACTGAGCTGACATCTTTGCAGCTTCAAAACAATTCATTTTCAGGGAAAATTCCACCT

>M6-PGSC0003DMG400013898M6LRR-RLP_124

TTGTGCTTCTTCTTTTTTCACTTCCACTCACAATCACATCATCAGCAACAACTGAAGCTGAAGCTCTTGTCAAATGGAAGATGAACTTACCTTCTACTTCTTTCTTGGATTCATGGTCCA

>M6-PGSC0003DMG400013898M6LRR-RLP_1240

GACTGAGCTGACATCTTTGCAGCTTCAAAACAATTCATTTTCAGGGAAAATTCCACCTGAAATCAGCCAGTTGAAAAACCTTGTATATCTTTACCTCTATAGTAACAATTTCACTGGTTC

>M6-PGSC0003DMG400013898M6LRR-RLP_1302

TCAGCCAGTTGAAAAACCTTGTATATCTTTACCTCTATAGTAACAATTTCACTGGTTCCATTCCCTATCAGATTGGTAAGTTGCAAAACTTGCTGGATCTAGATGTCTCTGAGAACCAGC

>M6-PGSC0003DMG400013898M6LRR-RLP_1364

CCCTATCAGATTGGTAAGTTGCAAAACTTGCTGGATCTAGATGTCTCTGAGAACCAGCTTTCAGGCACAATACCTCCAACCATTGGAAACCTGACCAATCTAAGGACGTTGCAGCTTTTT

>M6-PGSC0003DMG400013898M6LRR-RLP_1426

AGGCACAATACCTCCAACCATTGGAAACCTGACCAATCTAAGGACGTTGCAGCTTTTTCGGAACAACATCAGTGGAACCATTCCTTCTGAGATTGGCAAATTGACATCTCTTCAAACCCT

>M6-PGSC0003DMG400013898M6LRR-RLP_1488

ACAACATCAGTGGAACCATTCCTTCTGAGATTGGCAAATTGACATCTCTTCAAACCCTTGACCTCAACACTAACCGGTTTAGTGGTGAGCTGCCCGACAGCATTTCTAACCTCGGCAGTC

>M6-PGSC0003DMG400013898M6LRR-RLP_1550

CTCAACACTAACCGGTTTAGTGGTGAGCTGCCCGACAGCATTTCTAACCTCGGCAGTCTGAAGCTTCTTGAGTTCCTTTCTCTTAGTAACAACCAGTTTTCAGGTGTGATCCCTTTGTCT

>M6-PGSC0003DMG400013898M6LRR-RLP_1612

GCTTCTTGAGTTCCTTTCTCTTAGTAACAACCAGTTTTCAGGTGTGATCCCTTTGTCTATTTGCAATTCGACATCACTACAAACTCTAGTTTTCACAAGAAACAATTTCAGGGGAGAGAT

>M6-PGSC0003DMG400013898M6LRR-RLP_1674

GCAATTCGACATCACTACAAACTCTAGTTTTCACAAGAAACAATTTCAGGGGAGAGATTCCACAATGTTTGGGTAATATCAGTGGCCTCTCCATTTTGGATATGCGCTACAACAATCTTT

>M6-PGSC0003DMG400013898M6LRR-RLP_1736

CAATGTTTGGGTAATATCAGTGGCCTCTCCATTTTGGATATGCGCTACAACAATCTTTCTGGGATTCTTCCAACAAGTTTTGATATTGGATGTTCACTTAGAACCTTCAACTTTCATGGA

>M6-PGSC0003DMG400013898M6LRR-RLP_1798

GATTCTTCCAACAAGTTTTGATATTGGATGTTCACTTAGAACCTTCAACTTTCATGGAAATGAACTACAAGGGAAAATCCCTCGATCTTTAGCCAATTGTAAAGAGTTGAAAGTTCTTGA

>M6-PGSC0003DMG400013898M6LRR-RLP_186

GCTCTTGTCAAATGGAAGATGAACTTACCTTCTACTTCTTTCTTGGATTCATGGTCCATTTTGAATCTCGGAAACTTGTGCAATTGGACATCTATTGTCTGCAATGCTGGTGGAACAATT

>M6-PGSC0003DMG400013898M6LRR-RLP_2108

CTATTTCAACATTTGAAAGCCATGAAGACAACTAATCAAACAAGGAAGGCACCAATTGAAGGAAGCAACTATTACCAAGACTCGGTAGTTGTTGTAATGAAGGGACACATCTTTGAAGTT

>M6-PGSC0003DMG400013898M6LRR-RLP_2170

AAGCAACTATTACCAAGACTCGGTAGTTGTTGTAATGAAGGGACACATCTTTGAAGTTGTGAGAATCTTGTATTTATACACGAGTATGGATCTTTCAAGTAACAAATTTGAAGGACTTAT

>M6-PGSC0003DMG400013898M6LRR-RLP_248

GAATCTCGGAAACTTGTGCAATTGGACATCTATTGTCTGCAATGCTGGTGGAACAATTTCCGAAATCAATCTTTCTGATGCAGGCCTCTCAGGCACTCTTGATCAGCTTGACTTCACTTC

>M6-PGSC0003DMG400013898M6LRR-RLP_2542

TTCATATGAAGGTAATGATAGATTACATGGATTTCCAGTTTCACAAGGTTGTGGTAATGATCAGGTATCAGAGACAAATTATACCATATCTGCACTAGACGAACAAGAAAGCAATTCGGA

>M6-PGSC0003DMG400013898M6LRR-RLP_2728

CATAATATATTTCATGATCTCAACTTGAACTCCAATATGGCTAGCAAGAATCATGGAAGAGTTGGAACACAATTATTATGCAAAGTAGAAAGAAGCAATGAGGTAAAAGGCATTACAGAA

>M6-PGSC0003DMG400013898M6LRR-RLP_2790

TGGAACACAATTATTATGCAAAGTAGAAAGAAGCAATGAGGTAAAAGGCATTACAGAAGAAGAAAATAATCGCTTCTAGACAAGTTACCAACTGAGTATTCAAGTTAAGGTCTGCTGATT

>M6-PGSC0003DMG400013898M6LRR-RLP_2852

AAAATAATCGCTTCTAGACAAGTTACCAACTGAGTATTCAAGTTAAGGTCTGCTGATTTTTAGTATATAGTAATATATATTTATAGTTTCGTTACAACAACAGCTAATCGATACTTTATA

>M6-PGSC0003DMG400013898M6LRR-RLP_310

AAATCAATCTTTCTGATGCAGGCCTCTCAGGCACTCTTGATCAGCTTGACTTCACTTCATTTCCGAGTCTCACAAGTTATTCTCTCAATGGCAACAACTTCAGTGGATCGATACCATCAA

>M6-PGSC0003DMG400013898M6LRR-RLP_372

CCGAGTCTCACAAGTTATTCTCTCAATGGCAACAACTTCAGTGGATCGATACCATCAAATATTGGTAATGCTTCTATGCTCACTTTCTTGGACCTGAGCAACAATATCTTGGAGGGTGTC

>M6-PGSC0003DMG400013898M6LRR-RLP_434

TGGTAATGCTTCTATGCTCACTTTCTTGGACCTGAGCAACAATATCTTGGAGGGTGTCATACCAGAAGAAATTGGGAAGTTAACACAACTTGAATATCTCAGTTTCTATAACAACAATAT

>M6-PGSC0003DMG400013898M6LRR-RLP_496

CAGAAGAAATTGGGAAGTTAACACAACTTGAATATCTCAGTTTCTATAACAACAATATCATTGGTGTTATTCCCTACCAAATAAGTAATCTTCAGAAGGCATGGCACTTGGATCTTGGAT

>M6-PGSC0003DMG400013898M6LRR-RLP_558

GGTGTTATTCCCTACCAAATAAGTAATCTTCAGAAGGCATGGCACTTGGATCTTGGATCAAATTTTTAGAAACTCCTGATTGGTCCAAATTGAGGAATATGCCTATGTTGACATATTTGA

>M6-PGSC0003DMG400013898M6LRR-RLP_62

AAACCAAGTACTCTACCTATTAGACATGACAACTTCTTACAGTACCTTCCTTTACATTCTACTTGTGCTTCTTCTTTTTTCACTTCCACTCACAATCACATCATCAGCAACAACTGAAGC

>M6-PGSC0003DMG400013898M6LRR-RLP_620

TTTTTAGAAACTCCTGATTGGTCCAAATTGAGGAATATGCCTATGTTGACATATTTGAGCTTTGGTTACAATGAACTAACATTAGAATTCCCTGAATTTGTACTCTGTTGCCATAATCTG

>M6-PGSC0003DMG400013898M6LRR-RLP_682

TGGTTACAATGAACTAACATTAGAATTCCCTGAATTTGTACTCTGTTGCCATAATCTGACTTACCTCGATTTGTCGATAAACCATTTGAATGGTTCAATCCCAGAAACACTATTCACCAA

>M6-PGSC0003DMG400013898M6LRR-RLP_744

ACCTCGATTTGTCGATAAACCATTTGAATGGTTCAATCCCAGAAACACTATTCACCAACTTGGATAAGCTTGAACACCTCAATCTTTCATCTAATTCATTTCAAGGTTCATTGTCACCAA

>M6-PGSC0003DMG400013898M6LRR-RLP_806

GATAAGCTTGAACACCTCAATCTTTCATCTAATTCATTTCAAGGTTCATTGTCACCAAATTTTACCAAGTTGACCAAGTTAAAAGAACTTCAGCTTGGTTCCAACATGTTTTCGGGCTTA

>M6-PGSC0003DMG400013898M6LRR-RLP_868

TACCAAGTTGACCAAGTTAAAAGAACTTCAGCTTGGTTCCAACATGTTTTCGGGCTTAATTCCTGATGAAATTTCTCTGATCAGCAGTCTTGAAGTAGTTGTACTTTTCAATAATTCACT

>M6-PGSC0003DMG400013898M6LRR-RLP_930

CTGATGAAATTTCTCTGATCAGCAGTCTTGAAGTAGTTGTACTTTTCAATAATTCACTTCAAGGAAAGATTCCATCTTCTATAGGTAGACTTAGAAATCTTCAACAACTTGATCTTCGAA

>M6-PGSC0003DMG400013898M6LRR-RLP_992

GGAAAGATTCCATCTTCTATAGGTAGACTTAGAAATCTTCAACAACTTGATCTTCGAAAGAATCGCTTGAATTCAACCATTCCTTCTGAGCTAGGCCTTTGTACTAACCTCATTCTCTTG

>M6-PGSC0003DMG400013928M6OtherRLKTM_1054

TTGTTAACGGGTCAGGTCCTTCGCCACCGTCAGTATCCCCACAGCCAACGCCAATGCCATCTCAGTGGAATACGAGAAATGTGGCATTTCTAGTGGTAGGTTGTGTTGGGTCCTTAATGA

>M6-PGSC0003DMG400013928M6OtherRLKTM_1116

CAGTGGAATACGAGAAATGTGGCATTTCTAGTGGTAGGTTGTGTTGGGTCCTTAATGATGTTGAGTGTTCTTGTTATAATGTTTCTCAAGCATTGCAAGATTAGAGGATGCAGAGTACAC

>M6-PGSC0003DMG400013928M6OtherRLKTM_1178

GAGTGTTCTTGTTATAATGTTTCTCAAGCATTGCAAGATTAGAGGATGCAGAGTACACGACTCTGGCCGCCTTGATGAGGTCGGGACACCTCCACAGCAAGGCAGCCAGGCATCTCAAGT

>M6-PGSC0003DMG400013928M6OtherRLKTM_124

CTTCTAATCAAACATTAGTTTGTGCATTGATTTCCTCTTCCTTATATCCCCAAGAATCATCTCTCAATTGTACTAGTTTCCCTCAGGGTATTCAAATCCCTTTAAACCCTTCTGTTTCCT

>M6-PGSC0003DMG400013928M6OtherRLKTM_1240

CTGGCCGCCTTGATGAGGTCGGGACACCTCCACAGCAAGGCAGCCAGGCATCTCAAGTTCAAGATCAATTAGGTCCTCAGCCATCAATCTTGGAAAAAAGACCAAGTAAATTCCTTAGCA

>M6-PGSC0003DMG400013928M6OtherRLKTM_1302

GATCAATTAGGTCCTCAGCCATCAATCTTGGAAAAAAGACCAAGTAAATTCCTTAGCATGGGAAATGGGGGTCACTTAGAAGAATTTTCCTTGCAAGTGTTACTTCTAGTGACGAATAAC

>M6-PGSC0003DMG400013928M6OtherRLKTM_1364

AAATGGGGGTCACTTAGAAGAATTTTCCTTGCAAGTGTTACTTCTAGTGACGAATAACTTCTCCGAAGAGCACAAAATCGGGACAGGAAGTTTTGGATCTGTTTATCATGCTACATTAGA

>M6-PGSC0003DMG400013928M6OtherRLKTM_1426

CCGAAGAGCACAAAATCGGGACAGGAAGTTTTGGATCTGTTTATCATGCTACATTAGATGACGGTCGCGAAGTAGCCATAAAAAGGGCAGAGGCTTCAGCCTCATCTTCCTATGCTGGTG

>M6-PGSC0003DMG400013928M6OtherRLKTM_1488

GGTCGCGAAGTAGCCATAAAAAGGGCAGAGGCTTCAGCCTCATCTTCCTATGCTGGTGGTTCAAAGTATAGACAAGAGGACAAAGACAGTGCATTCCTCAATGAGCTAGACTTCTTGTCA

>M6-PGSC0003DMG400013928M6OtherRLKTM_1550

AAAGTATAGACAAGAGGACAAAGACAGTGCATTCCTCAATGAGCTAGACTTCTTGTCACGACTCAATCACAAGAACCTCGTTACGCTATTTGGCTATTGTGAAGATAACAACGAACGAGT

>M6-PGSC0003DMG400013928M6OtherRLKTM_1612

TCAATCACAAGAACCTCGTTACGCTATTTGGCTATTGTGAAGATAACAACGAACGAGTCTTGGTTTTCGAGTACATGAACAATGGCACCCTCCATGACCATCTCCACAAGCTTGAGACCT

>M6-PGSC0003DMG400013928M6OtherRLKTM_1674

GTTTTCGAGTACATGAACAATGGCACCCTCCATGACCATCTCCACAAGCTTGAGACCTCATCACTAATATCATGGACTGCTAGGATCAAGGTAGCATTGGACGCAGCCCGTGGAATCGAG

>M6-PGSC0003DMG400013928M6OtherRLKTM_1736

ACTAATATCATGGACTGCTAGGATCAAGGTAGCATTGGACGCAGCCCGTGGAATCGAGTACCTCCACGAATACGCAGTCCCAGCTGTCATCCACCGAGACATCAAGTCATCCAACATATT

>M6-PGSC0003DMG400013928M6OtherRLKTM_1798

TCCACGAATACGCAGTCCCAGCTGTCATCCACCGAGACATCAAGTCATCCAACATATTGCTCGACACCAATGGGAACGCCAAGGTGTCTGATTTTGGACTGTCCCTAATGGGACCACAAG

>M6-PGSC0003DMG400013928M6OtherRLKTM_186

CTCAATTGTACTAGTTTCCCTCAGGGTATTCAAATCCCTTTAAACCCTTCTGTTTCCTTTACTGGAATTGTGGGTGGAAATGGTTTCCTTTGTGGGTTAACCTCTTCTTCTAATTCAATC

>M6-PGSC0003DMG400013928M6OtherRLKTM_1860

GACACCAATGGGAACGCCAAGGTGTCTGATTTTGGACTGTCCCTAATGGGACCACAAGAGGACGAATCACACCTTTCTTTGCGTGCAGCGGGGACGGTGGGTTACATGGACCCTGAGTAC

>M6-PGSC0003DMG400013928M6OtherRLKTM_1922

CGAATCACACCTTTCTTTGCGTGCAGCGGGGACGGTGGGTTACATGGACCCTGAGTACTACAGATTGCAACAACTGACAACAAAAAGTGATGTGTATAGTTTTGGAGTAATGATGCTAGA

>M6-PGSC0003DMG400013928M6OtherRLKTM_1984

GATTGCAACAACTGACAACAAAAAGTGATGTGTATAGTTTTGGAGTAATGATGCTAGAGTTGTTGTCAGGTTACAAGGCGATTCACAAGAATGAGAATGGTGTACCAAGGAATGTGGTTG

>M6-PGSC0003DMG400013928M6OtherRLKTM_2046

TTGTCAGGTTACAAGGCGATTCACAAGAATGAGAATGGTGTACCAAGGAATGTGGTTGATTTTGTCGTGCCATACATAGTACAAGATGAGATTCATAGAGTGTTGGATGGTAGAGTACCA

>M6-PGSC0003DMG400013928M6OtherRLKTM_2108

TGTCGTGCCATACATAGTACAAGATGAGATTCATAGAGTGTTGGATGGTAGAGTACCACCACCAACACCATTTGAGATAGAGGCAGTGTCATATGTAGGTTATCTTGCAGCTGATTGTAC

>M6-PGSC0003DMG400013928M6OtherRLKTM_2170

CAACACCATTTGAGATAGAGGCAGTGTCATATGTAGGTTATCTTGCAGCTGATTGTACTACATTAGAAGGTAGAGATCGACCAACAATGACTGAAATTGTAAATAACCTTGAAAGAGCCT

>M6-PGSC0003DMG400013928M6OtherRLKTM_2232

TTAGAAGGTAGAGATCGACCAACAATGACTGAAATTGTAAATAACCTTGAAAGAGCCTTGCAAGCATGTTTGACATCTCCAAATTTCTCTCGATCCAACACCGATTCATCCACATAATCA

>M6-PGSC0003DMG400013928M6OtherRLKTM_2294

AGCATGTTTGACATCTCCAAATTTCTCTCGATCCAACACCGATTCATCCACATAATCATATCATTACTTGGCTTTGGATTGTGTATATTGTAATTACAACATTTTTCTTATTTGTATTTA

>M6-PGSC0003DMG400013928M6OtherRLKTM_2333

CGATTCATCCACATAATCATATCATTACTTGGCTTTGGATTGTGTATATTGTAATTACAACATTTTTCTTATTTGTATTTAATCATTTTCTCAAACTTTACACACCATTTCTTTCCTAAA

>M6-PGSC0003DMG400013928M6OtherRLKTM_248

TGGAATTGTGGGTGGAAATGGTTTCCTTTGTGGGTTAACCTCTTCTTCTAATTCAATCATGGTGTGTTGGAGGTTTTCAGATAATGGGACTGATTTGAATTATAAAAGGGTTTATATTGG

>M6-PGSC0003DMG400013928M6OtherRLKTM_310

TGTGTTGGAGGTTTTCAGATAATGGGACTGATTTGAATTATAAAAGGGTTTATATTGGCCCATTGCTTACAAATCTTGATTCTGGAAATTCCCATGTTTGTGGAATTGTTAATGGAACCA

>M6-PGSC0003DMG400013928M6OtherRLKTM_372

TTGCTTACAAATCTTGATTCTGGAAATTCCCATGTTTGTGGAATTGTTAATGGAACCAATAGGCTTCAGTGTTGGCAGTGGCATGAATTCAGTTCATCAAATAACAGTTTGATAACTTCA

>M6-PGSC0003DMG400013928M6OtherRLKTM_434

GCTTCAGTGTTGGCAGTGGCATGAATTCAGTTCATCAAATAACAGTTTGATAACTTCAAATCTTGCTGTTGGAGAAGATTTTGTTTGTGGGTTGTTGCCATCGGGTCAAATTCAATGTCT

>M6-PGSC0003DMG400013928M6OtherRLKTM_496

TTGCTGTTGGAGAAGATTTTGTTTGTGGGTTGTTGCCATCGGGTCAAATTCAATGTCTAGGAAGCTTTAGAAATGCTATTGATGTTGTTCCTTCAGGGAATTACAGTGAAATTGCAGCTG

>M6-PGSC0003DMG400013928M6OtherRLKTM_558

AGCTTTAGAAATGCTATTGATGTTGTTCCTTCAGGGAATTACAGTGAAATTGCAGCTGGTTCTCAATACGCTTGTGCCATCTCCATGAATGGTAGTTTGGCTTGTTGGGGAAATATGGTA

>M6-PGSC0003DMG400013928M6OtherRLKTM_62

TTTCATTTTTTCCCTTTCTTTCCAACCCATTTCTTCACTTTCCACTATTGCCATTTCCAAAACTTCTAATCAAACATTAGTTTGTGCATTGATTTCCTCTTCCTTATATCCCCAAGAATC

>M6-PGSC0003DMG400013928M6OtherRLKTM_620

TCAATACGCTTGTGCCATCTCCATGAATGGTAGTTTGGCTTGTTGGGGAAATATGGTAGTAGAAAAGCCTGTTGACCAATTCAAGTCTCTTGCTTTAGGTGATACTAGGGGTTGTGGTTT

>M6-PGSC0003DMG400013928M6OtherRLKTM_682

AAAAGCCTGTTGACCAATTCAAGTCTCTTGCTTTAGGTGATACTAGGGGTTGTGGTTTGAGGGTTAATGGGAAAGTGGTTTGCTGGGGTGAAAATGGTTTTAGTCTCCCTTCAACTTTGA

>M6-PGSC0003DMG400013928M6OtherRLKTM_744

GTTAATGGGAAAGTGGTTTGCTGGGGTGAAAATGGTTTTAGTCTCCCTTCAACTTTGAGTGATATATCTTTTGAAACATTAGAAGCAAAACGAGATATTTTTTGTGGCATTGCCACCTCA

>M6-PGSC0003DMG400013928M6OtherRLKTM_806

TATATCTTTTGAAACATTAGAAGCAAAACGAGATATTTTTTGTGGCATTGCCACCTCAAATTATTCATTGTTCTGTTGGGGAAATGAGATTTTCAATTCAAATCCAGTAGTCTTTGAAGA

>M6-PGSC0003DMG400013928M6OtherRLKTM_868

ATTCATTGTTCTGTTGGGGAAATGAGATTTTCAATTCAAATCCAGTAGTCTTTGAAGAAGTTGTTCCAGGACCATGTACCATGTCATGTCCTTGTGCACCTTTAGATAACTATGCTAGGT

>M6-PGSC0003DMG400013953-(rev)M6LRR-RLK_1071

CATAACAAATGCTTCCAAGCTTCAGACACTGGCACTAAATGATAACTCTCTCAAAGGAACTATTCCTAACAATTTGGGGAATCTTCGTGAGCTGCAAATACTGCTCCTACACACTAATCA

>M6-PGSC0003DMG400013953-(rev)M6LRR-RLK_1134

TCCTAACAATTTGGGGAATCTTCGTGAGCTGCAAATACTGCTCCTACACACTAATCAACTTACCAATGAACCAGGAAAGCATGCGTTGAAATTCTTCAATTCTTTGGCGAACTGTAGGAT

>M6-PGSC0003DMG400013953-(rev)M6LRR-RLK_1197

CAATGAACCAGGAAAGCATGCGTTGAAATTCTTCAATTCTTTGGCGAACTGTAGGATGTTGAGATATCTAGAAGTGGGTAACAATCCGTTGAGTGGAGTTCTGCCAAATTCTATTGGGAA

>M6-PGSC0003DMG400013953-(rev)M6LRR-RLK_1260

ATATCTAGAAGTGGGTAACAATCCGTTGAGTGGAGTTCTGCCAAATTCTATTGGGAATCTCTCATCTACTATTGAAAAAAATTATTTAGCAGATGCACACATCAATGGCCTCATCCCCAC

>M6-PGSC0003DMG400013953-(rev)M6LRR-RLK_1386

AGGCAACATGACCGGTCTTACATCCTTAAGCCTTGGCGGAAACAACTTGACAGGGAATATTCCTTCTGAGATTGGTAAGCTTAAACAACTCCAAGGACTCGATCTATCTAACAATAAATT

>M6-PGSC0003DMG400013953-(rev)M6LRR-RLK_1449

TTCTGAGATTGGTAAGCTTAAACAACTCCAAGGACTCGATCTATCTAACAATAAATTGCAAGGACATATTCCAGAAGTGGTATGCAATTTGTCTGATTTGGTTCAATTATATCTGTATGG

>M6-PGSC0003DMG400013953-(rev)M6LRR-RLK_1512

ACATATTCCAGAAGTGGTATGCAATTTGTCTGATTTGGTTCAATTATATCTGTATGGTAATGAGCTCTCTGGATTAATTCCAGAATGCTTAGGAAATCTTACCATGCTTCAACAACTTTA

>M6-PGSC0003DMG400013953-(rev)M6LRR-RLK_1575

GCTCTCTGGATTAATTCCAGAATGCTTAGGAAATCTTACCATGCTTCAACAACTTTATTTGGCTTCTAATAAATTTTCATCAAAATTTCCATTGACCCTTTGGAAAATGAGTGGTCTTCT

>M6-PGSC0003DMG400013953-(rev)M6LRR-RLK_1638

TTCTAATAAATTTTCATCAAAATTTCCATTGACCCTTTGGAAAATGAGTGGTCTTCTCTATCTAAACGTGTCACAGAATTCAATAGAGGGAGAACTTCCATCAGATATTGGAGGACTGAA

>M6-PGSC0003DMG400013953-(rev)M6LRR-RLK_1701

AAACGTGTCACAGAATTCAATAGAGGGAGAACTTCCATCAGATATTGGAGGACTGAAAGCCATTGTAGAACTATATCTTTCTGGTAACCACTTTTCAGGCATGATATCAACCAGATTCGG

>M6-PGSC0003DMG400013953-(rev)M6LRR-RLK_1764

TGTAGAACTATATCTTTCTGGTAACCACTTTTCAGGCATGATATCAACCAGATTCGGGGAACTCCAAAATCTGCAGTATCTCGACCTATCGAAAAATTCATTTTTTGGCCGAATTCCATT

>M6-PGSC0003DMG400013953-(rev)M6LRR-RLK_1827

CCAAAATCTGCAGTATCTCGACCTATCGAAAAATTCATTTTTTGGCCGAATTCCATTATCCTTTGCCAACTTGATAAACTTGGAATTCTTGAATTTGTCTTTAAATGCATTGTCAGGTAC

>M6-PGSC0003DMG400013953-(rev)M6LRR-RLK_1890

TGCCAACTTGATAAACTTGGAATTCTTGAATTTGTCTTTAAATGCATTGTCAGGTACTATTCCTAAGTCTTTGGAAAAACTCTCATACCTTAAAAGCATTAATGTTTCATTTAATGATTT

>M6-PGSC0003DMG400013953-(rev)M6LRR-RLK_315

TAGCCTCACGGGTATAATCCCTCCTTATGTTGGAAATGCCACAAAATTGATGAACATCAATTTGTCTGGGAATAGAATCAATGATAACATTCCAATGGAGATCAGTAATCTTAGCCAACT

>M6-PGSC0003DMG400013953-(rev)M6LRR-RLK_378

GTCTGGGAATAGAATCAATGATAACATTCCAATGGAGATCAGTAATCTTAGCCAACTTGCATTTTTGTCCTTGTTCAATAATCAATTAACAGGTTCCATTCCTCCAACACTGTTTAATAT

>M6-PGSC0003DMG400013953-(rev)M6LRR-RLK_441

TTTGTCCTTGTTCAATAATCAATTAACAGGTTCCATTCCTCCAACACTGTTTAATATCTCGTCACTACTTGGCATATATCTGTCATTCAATGGCCTTTCTGGTCCTCTCTTGCTAGATGA

>M6-PGSC0003DMG400013953-(rev)M6LRR-RLK_504

ACTACTTGGCATATATCTGTCATTCAATGGCCTTTCTGGTCCTCTCTTGCTAGATGAAGGGAATATTGTGTCAAATCTGAAGCATTTAAGTATGTCTAACAACCAAATTTCTGGTTGCAT

>M6-PGSC0003DMG400013953-(rev)M6LRR-RLK_567

TATTGTGTCAAATCTGAAGCATTTAAGTATGTCTAACAACCAAATTTCTGGTTGCATTCCTTCCAACATATGCCAACTCACAGAGCTCAAACTTTTGTCCATATCTTTCAACAATATAAC

>M6-PGSC0003DMG400013953-(rev)M6LRR-RLK_630

CAACATATGCCAACTCACAGAGCTCAAACTTTTGTCCATATCTTTCAACAATATAACTGGAGACATACCTAGAAATATTGGTTGTTTATCCAAACTCGAAAGGTTTTATATTGGTGAAAA

>M6-PGSC0003DMG400013953-(rev)M6LRR-RLK_693

CATACCTAGAAATATTGGTTGTTTATCCAAACTCGAAAGGTTTTATATTGGTGAAAATCCAATAGAAGGGACTATTCCCACTTCATTGGGAAATATTTCTACTCTGCAATATCTTGATTG

>M6-PGSC0003DMG400013953-(rev)M6LRR-RLK_756

AGAAGGGACTATTCCCACTTCATTGGGAAATATTTCTACTCTGCAATATCTTGATTGTGTAGACAATTGGATGAAGGGGCAAATTCCTCAAGAATTAGGGAAGCTATCAAATTTGAGGGA

>M6-PGSC0003DMG400013953-(rev)M6LRR-RLK_819

CAATTGGATGAAGGGGCAAATTCCTCAAGAATTAGGGAAGCTATCAAATTTGAGGGAATTAAGCTTTGCCAATAATTCTAATCTTATTGGTCAAATTCCAGAGGCTATTTTCAACATATC

>M6-PGSC0003DMG400013953-(rev)M6LRR-RLK_882

CTTTGCCAATAATTCTAATCTTATTGGTCAAATTCCAGAGGCTATTTTCAACATATCTTCTTTGGAAGTGATCAATTTTAGTTTGAATAACCTCTCGGGGAGAATTCCAACCACTACAGG

>M6-PGSC0003DMG400013955-(rev)M6LRR-RLP_124

TGTCACGTTCGATTTTCTTCTAAAGAAAAGATTAAAATTTCTTCCATTTTGTGCTAATTAGTGAATGTGTGTTGTGTAACACAGACTTATAAGGTTGACTCCTATTGACTTGCTCTTAAC

>M6-PGSC0003DMG400013955-(rev)M6LRR-RLP_186

GAATGTGTGTTGTGTAACACAGACTTATAAGGTTGACTCCTATTGACTTGCTCTTAACAATATAATGAAAATTTAGCAACAATTATTCTTTCTATTATGACTTTTATTTCGTTTTGTGAA

>M6-PGSC0003DMG400013955-(rev)M6LRR-RLP_248

TAATGAAAATTTAGCAACAATTATTCTTTCTATTATGACTTTTATTTCGTTTTGTGAAACACAACCCAAATAATGTTTTTTTTCTTTTTGCTACTTATTTTCCTCCTTTTAATCTGTTTT

>M6-PGSC0003DMG400013955-(rev)M6LRR-RLP_2728

TCTATGTGGAGTGCACATATTAGAGATTCCTGCTTGTGCTATCACTAATCCTGGAAAACAATCAAAGCTTAAGGAGGTTATGCTAAAAATTGTTACTCCAGTGGTTATTGCATCCTTTAT

>M6-PGSC0003DMG400013955-(rev)M6LRR-RLP_2852

TTCTTGTTCGTTTCAATTTGGATAATGAAACGACAGAAGAAAGGGAAGTCCAAAGATGTGGAAAAGATACCGGAGATGGGGACTCATCAATTAGTTTCTTATCACGAGATTCAACGAGCA

>M6-PGSC0003DMG400013955-(rev)M6LRR-RLP_310

AACCCAAATAATGTTTTTTTTCTTTTTGCTACTTATTTTCCTCCTTTTAATCTGTTTTCACATTTTATTTGTTTTAGTGCTATATATATAGTGGAGTAACTTTGGTCACTGGAAATACAA

>M6-PGSC0003DMG400013955-(rev)M6LRR-RLP_3116

TGTGAAGTGATGAGAAATGTTAGACACAGAAATCTTGTTACAGTGATTACTACATGTTCTAGTGACTATATAAGAGCCTTTGTTCTGCGATATATGCCCAATGGGAGTCTTGAGAATTGG

>M6-PGSC0003DMG400013955-(rev)M6LRR-RLP_372

TTTTATTTGTTTTAGTGCTATATATATAGTGGAGTAACTTTGGTCACTGGAAATACAACACACAAGTTAATATGGAGAAGCACATCTTCTTATTGATTCTTCTCTTTCTAATTCAATTTT

>M6-PGSC0003DMG400013955-(rev)M6LRR-RLP_434

CAAGTTAATATGGAGAAGCACATCTTCTTATTGATTCTTCTCTTTCTAATTCAATTTTCTATATCACTTGCTTCCTCTAATGAGACTGACCAACAAGCTCTGTTAGCTTTCCAAAATCTT

>M6-PGSC0003DMG400013955-(rev)M6LRR-RLP_496

ATCACTTGCTTCCTCTAATGAGACTGACCAACAAGCTCTGTTAGCTTTCCAAAATCTTATTACAAGTCCTAATCATTTTTTGGCCAATAATTGGACCAAAAATACTTCCTTTTGCACTTG

>M6-PGSC0003DMG400013955-(rev)M6LRR-RLP_62

ATTTTATTTTTATTTAAAAAAAAAAGTGAAACCACACGTGACAACGTGTGTAGTATAGTGCGTGTCACGTTCGATTTTCTTCTAAAGAAAAGATTAAAATTTCTTCCATTTTGTGCTAAT

>M6-PGSC0003DMG400013955-(rev)M6LRR-RLP_868

TCAAGTAATTTCAATGGCTTTCAATGAACTCGGTGGTGAAATGTGGAGAGGCCCATGGTATGTACCAGAACTCAGAGTCTTAAATCTCGGGAACAATAGCCTCACGGGTATAATCCCTCC

>M6-PGSC0003DMG400013955-(rev)M6LRR-RLP_930

TACCAGAACTCAGAGTCTTAAATCTCGGGAACAATAGCCTCACGGGTATAATCCCTCCTTATGTTGGAAATGCCACAAAATTGATGAACATCAATTTGTCTGGGAATAGAATCAATGATA

>M6-PGSC0003DMG400013956-(rev)M6LRR-RLK_256

CCATGGCTTATACAGAGACATTGGGCACTCTTGGATATATTGCACCAGGTAAAAGAAAATCTACTCTCATTGATTTTCTCTTATCATAATTAAGCCTCTCCAAGTTGTAGAAGTAAAAAG

>M6-PGSC0003DMG400013956-(rev)M6LRR-RLK_320

TCTCATTGATTTTCTCTTATCATAATTAAGCCTCTCCAAGTTGTAGAAGTAAAAAGAGAATTATTGTTGTATTTCAATTGAATTAACTTTTTTTTAAATCCTTTTTTAAGAATATGGCTC

>M6-PGSC0003DMG400013989-(rev)M6OtherRLKTM_0

TTGTTGATTCAAATGGAGAAAAAAGGAAATGTACTGATGCAAAAATATGATTTGAGGAGATTATTAGGTCAAGGCAACTTTGCTAAGGTTTATTACGGAAGGAATCTTGAAACGGGACAG

>M6-PGSC0003DMG400013989-(rev)M6OtherRLKTM_1040

GCAATTCATTTCCGCGAAGCCTGCCTCTTCTATCATATCCAAACTTGAGGAAGTTGGCAGGAATCTAAAGCTGGAAGTAATGAAGAAAGAAGCTGGATCTATGAGATTAGAGGGATCGGG

>M6-PGSC0003DMG400013989-(rev)M6OtherRLKTM_1105

TAAAGCTGGAAGTAATGAAGAAAGAAGCTGGATCTATGAGATTAGAGGGATCGGGTGAAGGTAGATACGGGACTTTGTCCATCGATACTGAAATATCTGAAATTACTCCGTCCTTCCACT

>M6-PGSC0003DMG400013989-(rev)M6OtherRLKTM_1170

TACGGGACTTTGTCCATCGATACTGAAATATCTGAAATTACTCCGTCCTTCCACTTGGTTGAGCTGAAAAAGTCATATGGTGATAAGGTAGAGTACCAAAAGATGCTGAAACAAGTTATA

>M6-PGSC0003DMG400013989-(rev)M6OtherRLKTM_1217

CTTCCACTTGGTTGAGCTGAAAAAGTCATATGGTGATAAGGTAGAGTACCAAAAGATGCTGAAACAAGTTATAAGACCTGCTCTGGAGGAAATTGTTTGGGCTTGGCAAGGCGAGCAGCC

>M6-PGSC0003DMG400013989-(rev)M6OtherRLKTM_130

TTAAGGTAATTGACAAAGAGAAGGTTCTTAAGGCCGGACTGATTGAACAAACGAAGAGAGAGATATCTGTTATGGCACTAGTTGAACATCCAAATGTTCTACAGCTATACGAGGTCATGG

>M6-PGSC0003DMG400013989-(rev)M6OtherRLKTM_195

TCTGTTATGGCACTAGTTGAACATCCAAATGTTCTACAGCTATACGAGGTCATGGCAACTAAATCTAAGATTTATTTAGTCATTGAACATGCCAAAGGCGGTGAGCTTTTCAAAAAGTTG

>M6-PGSC0003DMG400013989-(rev)M6OtherRLKTM_260

TAAGATTTATTTAGTCATTGAACATGCCAAAGGCGGTGAGCTTTTCAAAAAGTTGACAAAGGGGAGGCTCAAGGAAAAATTAGCTAGGAAGTACTTTCAGCAATTGATTAGCGCTGTTGA

>M6-PGSC0003DMG400013989-(rev)M6OtherRLKTM_325

GGCTCAAGGAAAAATTAGCTAGGAAGTACTTTCAGCAATTGATTAGCGCTGTTGAATGTTGCCACAGTCGAGATGTTTATCACCGTGATCTGAAACCAGAAAATGTGCTTCTAGATGAGA

>M6-PGSC0003DMG400013989-(rev)M6OtherRLKTM_390

AGTCGAGATGTTTATCACCGTGATCTGAAACCAGAAAATGTGCTTCTAGATGAGAATGGAAATCTTAAGGTATCAGACTTTGGATTAAGCGCCCTGGCTGAATCTAAGAGGCAGGATGGC

>M6-PGSC0003DMG400013989-(rev)M6OtherRLKTM_455

TAAGGTATCAGACTTTGGATTAAGCGCCCTGGCTGAATCTAAGAGGCAGGATGGCTTACTCCACACAACATGTGGAACCCCAGCTTACGTTGCACCTGAGGTGATCAGTCGAAAAGGATA

>M6-PGSC0003DMG400013989-(rev)M6OtherRLKTM_520

CAACATGTGGAACCCCAGCTTACGTTGCACCTGAGGTGATCAGTCGAAAAGGATATGACGGTGCCAAAGCTGATATCTGGTCTTGTGGGGTGATCTTATTTGTTTTGCTAGCCGGTTATC

>M6-PGSC0003DMG400013989-(rev)M6OtherRLKTM_585

AAAGCTGATATCTGGTCTTGTGGGGTGATCTTATTTGTTTTGCTAGCCGGTTATCTTCCATTCCAAGACTCAAATCTTATGGAGATTTATAGGAAGATAAAGCAAGCTGAGTTCAAATGT

>M6-PGSC0003DMG400013989-(rev)M6OtherRLKTM_65

AGGTCAAGGCAACTTTGCTAAGGTTTATTACGGAAGGAATCTTGAAACGGGACAGAGTGTAGCCATTAAGGTAATTGACAAAGAGAAGGTTCTTAAGGCCGGACTGATTGAACAAACGAA

>M6-PGSC0003DMG400013989-(rev)M6OtherRLKTM_650

AGACTCAAATCTTATGGAGATTTATAGGAAGATAAAGCAAGCTGAGTTCAAATGTCCAAATTGGTTCCCACCAGAAGTGCGGAAATTACTTTCTAAAATCCTTGATCCAAACCCTCGTAC

>M6-PGSC0003DMG400013989-(rev)M6OtherRLKTM_715

TCCCACCAGAAGTGCGGAAATTACTTTCTAAAATCCTTGATCCAAACCCTCGTACAAGGATTTCCATAGCAAAGATAAAGGAAAGCTCATGGTTCAAGAAAGGTTTGGAGTCTAGAAACG

>M6-PGSC0003DMG400013989-(rev)M6OtherRLKTM_780

ATAGCAAAGATAAAGGAAAGCTCATGGTTCAAGAAAGGTTTGGAGTCTAGAAACGCGGTAACCAAAGTAGAAGAAAAGGAAAAGTTTAGCCTAGACGCCAACGCTACTATTCAAGATTGT

>M6-PGSC0003DMG400013989-(rev)M6OtherRLKTM_845

AGTAGAAGAAAAGGAAAAGTTTAGCCTAGACGCCAACGCTACTATTCAAGATTGTACACATAACAGTATCTCTCCCTCTGAGTCAAAGCTAGAGTTGCTAAAACCGTCAAATCTAAATGC

>M6-PGSC0003DMG400013989-(rev)M6OtherRLKTM_910

GTATCTCTCCCTCTGAGTCAAAGCTAGAGTTGCTAAAACCGTCAAATCTAAATGCATTTGATATAATCTCTCTTTCAAGTGGATTTGACTTGTCCGGTTTATTCATAACAAAGGATCAGA

>M6-PGSC0003DMG400013989-(rev)M6OtherRLKTM_975

ATCTCTCTTTCAAGTGGATTTGACTTGTCCGGTTTATTCATAACAAAGGATCAGAAGGAAGACTTGCAATTCATTTCCGCGAAGCCTGCCTCTTCTATCATATCCAAACTTGAGGAAGTT

>M6-PGSC0003DMG400014060-(rev)M6RLKWAX-EGF_0

TAGAAATTTAACTCATCGTCGGACAACCTGGTTCATGGTGGCGACAGGTAACTCATCTTCTTTTTTCTTCTCTGTAACTGGCACTGCAGTTTCGTAGATACCATTGCTTCCCTTTTGCTC

>M6-PGSC0003DMG400014060-(rev)M6RLKWAX-EGF_1071

TGATGTTGGAGTTGATCGCACAAGTGAAATCTACTTTCCACATTGGCTCTATCAGCGAATTGAACTAAGTGAAGAGCTTCAACTCCCTGGAATAATGAATGAAGAGGAGAATGAATGTGC

>M6-PGSC0003DMG400014060-(rev)M6RLKWAX-EGF_1134

ACTAAGTGAAGAGCTTCAACTCCCTGGAATAATGAATGAAGAGGAGAATGAATGTGCAAGAAAGATGGTTATAGCAAGTTTATGGTGCATACAGACTGATCCCTCAAATCGACCATCGAT

>M6-PGSC0003DMG400014060-(rev)M6RLKWAX-EGF_1258

AAGGTTGTGGAAATGTTAGAAGGGAAACTAGACTCTTTACAAATGCCTCCCAAGCCTTACCTATATTCTCCCTCAAGAACTGAGGTAGATTCATCAGTACTAGAGCTGGCCTAGTCTTAA

>M6-PGSC0003DMG400014060-(rev)M6RLKWAX-EGF_126

CTTGTATTTCTTGTCTCTGTACGTATCTTCTATATGAATTGTCCAAAACACAGACTGTTCCCCAAATAGTTCATAGAACTTCCAGACTGAATTTAATTTATATTCTCTGATGGTGCATGA

>M6-PGSC0003DMG400014060-(rev)M6RLKWAX-EGF_189

AAATAGTTCATAGAACTTCCAGACTGAATTTAATTTATATTCTCTGATGGTGCATGACTTCATTTTGCTTTCCACAGGTGCAGTTTTGTTTACAGTTGGACTGTTGGTTTTACTCTTCTG

>M6-PGSC0003DMG400014060-(rev)M6RLKWAX-EGF_315

AGAGAAGATTTTGTGGTACAAATGCCTCAGGTTTTGGGAATCTAATGCTGAGGATCACCAAAAAGTTGAAGAATTCTTAAAGAACTATGGATCATATGCTCTAAACAGATACAATTATAC

>M6-PGSC0003DMG400014060-(rev)M6RLKWAX-EGF_378

AGTTGAAGAATTCTTAAAGAACTATGGATCATATGCTCTAAACAGATACAATTATACAGACATCAAAAAAATCACCAGTCACTTCAGAAGCAAACTAGGCCAAGGAGGATTTGGTAATGT

>M6-PGSC0003DMG400014060-(rev)M6RLKWAX-EGF_504

AGGAAGTTTGCGTAACGGGAGTCAAGTGGCAGTGAAGATCCTGAATAAACTAAAAGGCAGTGGTGAAGATTTCATTAACGAGGTGGCAAGTATTAGCAGGACATCGCATGTTAACATCGT

>M6-PGSC0003DMG400014060-(rev)M6RLKWAX-EGF_567

TGAAGATTTCATTAACGAGGTGGCAAGTATTAGCAGGACATCGCATGTTAACATCGTGAGCCTTGTGGGATTTTGTTTTGAGGGTCACAAAAGAGCTCTCATATATGAGTTCATGCCAAA

>M6-PGSC0003DMG400014060-(rev)M6RLKWAX-EGF_63

TTTCTTCTCTGTAACTGGCACTGCAGTTTCGTAGATACCATTGCTTCCCTTTTGCTCTTTCGACTTGTATTTCTTGTCTCTGTACGTATCTTCTATATGAATTGTCCAAAACACAGACTG

>M6-PGSC0003DMG400014060-(rev)M6RLKWAX-EGF_630

TGTGGGATTTTGTTTTGAGGGTCACAAAAGAGCTCTCATATATGAGTTCATGCCAAATGGATCTCTTGAAAAGTTTATCTATGAGGAAAGATCCAACAGTGTTCGCCAATTGGGTTGGCC

>M6-PGSC0003DMG400014060-(rev)M6RLKWAX-EGF_693

TCTTGAAAAGTTTATCTATGAGGAAAGATCCAACAGTGTTCGCCAATTGGGTTGGCCAATACTGTACAAAATCACACTAGGCATTGCTCGAGGATTGGAGTACTTGCACCGTGGTTGTAA

>M6-PGSC0003DMG400014060-(rev)M6RLKWAX-EGF_882

CTCTGACTTTGGTCTAGCCAAACTGTGCATGAAAAAGGAGAGCATTGTGTCAATGCTAGGTGCACGAGGAACAATCGGTTATATTGCTCCAGAAATTGTTTGCAGAAACTTGGGAGGTAT

>M6-PGSC0003DMG400014064M6RLKWAX-EGF_0

AATGGTTGTGTTTCTGATATCAAGAACGTGTCACTCGATCATAGTGCTTTTAAGTTTGTCAAGGAATCAAGAATTTACTTTTTGTCAAAGTGTAATGGATCGATATCTGAGGATCTTTTG

>M6-PGSC0003DMG400014064M6RLKWAX-EGF_1054

AATTGGAACCATAAGTTATCTTTTCCCACCTCATGTTCTATTTTGAGTAGACAACAATATTACTTTGTACATTCAAATGCACAAATAACAGTATGCGGATTGATAGTTATTTTCAAAGTT

>M6-PGSC0003DMG400014064M6RLKWAX-EGF_1116

CTTTGTACATTCAAATGCACAAATAACAGTATGCGGATTGATAGTTATTTTCAAAGTTATAAGAACTACTCGAGTTGTGATGCCTTTAGCTTTTAGTGTCATAAACCAAGAATTGATCGA

>M6-PGSC0003DMG400014064M6RLKWAX-EGF_1178

GAACTACTCGAGTTGTGATGCCTTTAGCTTTTAGTGTCATAAACCAAGAATTGATCGAAGCCATAGTATTCAAGCTAGTGATCTTCCTGCAAATTCATCGTTCACCTTCAGAACCAAATT

>M6-PGSC0003DMG400014064M6RLKWAX-EGF_1240

ATAGTATTCAAGCTAGTGATCTTCCTGCAAATTCATCGTTCACCTTCAGAACCAAATTCTACTGATTTGTTTGATCTATTGACTCCTAGTTTCACATTAGATTGGGAATTGTCGTTAAAT

>M6-PGSC0003DMG400014064M6RLKWAX-EGF_1302

TGATTTGTTTGATCTATTGACTCCTAGTTTCACATTAGATTGGGAATTGTCGTTAAATTATTCCAGCTGTCTTCATAGAGAAGGGCGATACCAAAGTGACATCGAAAACAACTTCTTTTA

>M6-PGSC0003DMG400014064M6RLKWAX-EGF_1364

CCAGCTGTCTTCATAGAGAAGGGCGATACCAAAGTGACATCGAAAACAACTTCTTTTACTCCAGAGGTATTGGATATAAGAAGCTTGTCTGTTGCTTGTGTATGTCTGGTTTCATTTTTG

>M6-PGSC0003DMG400014064M6RLKWAX-EGF_1426

AGAGGTATTGGATATAAGAAGCTTGTCTGTTGCTTGTGTATGTCTGGTTTCATTTTTGGAAAGGATTTAGATGTGTGTTGGTTGGTTCATTCCAAGATGTTTAATATGTCAACGTGAACG

>M6-PGSC0003DMG400014064M6RLKWAX-EGF_1488

GGATTTAGATGTGTGTTGGTTGGTTCATTCCAAGATGTTTAATATGTCAACGTGAACGGAAGTGAATTGCTTTCACTCTTATTGTGACTAATTAATCTTTCATTTCTGTGTATTTGCAGA

>M6-PGSC0003DMG400014064M6RLKWAX-EGF_1550

TGAATTGCTTTCACTCTTATTGTGACTAATTAATCTTTCATTTCTGTGTATTTGCAGAAGCAAAAAGAAATTTGGGACTGATACTAGGCACAGGTACATGTCTGTTATCTGTAATTAGAC

>M6-PGSC0003DMG400014064M6RLKWAX-EGF_1612

AAAAGAAATTTGGGACTGATACTAGGCACAGGTACATGTCTGTTATCTGTAATTAGACTTGCACAAAATCTCCTATTGCCTTGCAAATGTTGAAATTGTTGTGTTGCCTCAAAACCAGGA

>M6-PGSC0003DMG400014064M6RLKWAX-EGF_1674

ACAAAATCTCCTATTGCCTTGCAAATGTTGAAATTGTTGTGTTGCCTCAAAACCAGGATCTTTCTGGAGCATGACTTTCCTTTTTTCCCCCACATTTGCAGTTTTCTCTTGCATCGGAGT

>M6-PGSC0003DMG400014064M6RLKWAX-EGF_1798

ATTTTGCTCTTCTGCTTGAGAAAAAAGATTTTCTGGCACAAAAATATCAGGTTTTGGGAATCTATAGCTGAGGATCACCGAAATATTGAAGCATTTCTGAAGAACTTTGGGTCATATGCT

>M6-PGSC0003DMG400014064M6RLKWAX-EGF_186

GAGGATGAAAGCTTTGAGTCTGCATTACAAATGTGTAAAAATCATGTAATGGTACCAGTAGAAATGCTTGGAAATGAAGGGAGTAATCGAGTTATTGACTATCAAGTGCTACTGAGAAGG

>M6-PGSC0003DMG400014064M6RLKWAX-EGF_1860

TATAGCTGAGGATCACCGAAATATTGAAGCATTTCTGAAGAACTTTGGGTCATATGCTCCAAAAAGATACAGCTATACAGACATCAAGCGAATCACCAATCACTTTAAAAACAAACTAGG

>M6-PGSC0003DMG400014064M6RLKWAX-EGF_1922

AAAGATACAGCTATACAGACATCAAGCGAATCACCAATCACTTTAAAAACAAACTAGGTCAAGGAGGATTTGGTAATGTATACAGAGGAAGTTTGTGTAACGGGAGTCAAGTGGCAGTAA

>M6-PGSC0003DMG400014064M6RLKWAX-EGF_2046

CCTGAATGAACTAAAAGGCAGTGGTGAGATTTCATTAACGAAGTGGCAAGTATTAGCAGAACATCACATGTTAACATCGTGAGCCTTGTGGGATTTTGTTTTGAGGGTCACAAAAGAGCT

>M6-PGSC0003DMG400014064M6RLKWAX-EGF_2108

ATCACATGTTAACATCGTGAGCCTTGTGGGATTTTGTTTTGAGGGTCACAAAAGAGCTCTTTTATATGAGTTCATGCCAAATGGATCTCTTGAAAAGTTTATTTATGAGGAAATATCTGA

>M6-PGSC0003DMG400014064M6RLKWAX-EGF_2170

TATATGAGTTCATGCCAAATGGATCTCTTGAAAAGTTTATTTATGAGGAAATATCTGACGGAGTTCGCCAATTAGGCTGGCAAATATTGTATGAAATTGCATTAGGCATTGCTCGAGGAT

>M6-PGSC0003DMG400014064M6RLKWAX-EGF_248

AATGCTTGGAAATGAAGGGAGTAATCGAGTTATTGACTATCAAGTGCTACTGAGAAGGGGGTTTAGATTGAGCTGGATATCAAGTAACTGCAGTGACTGTGCAGAAAGTGGAGGACGTTG

>M6-PGSC0003DMG400014064M6RLKWAX-EGF_2666

AATGAATGAAGAGGAGAACGAATGTGCAAGAAAGATGGTGATAGCAAGTTTGTGGTGCATACAGACTGATCCTGCAAATCAACCATCAATGAGCAAGGTTGTGGAAATGTTAGAAGGGAA

>M6-PGSC0003DMG400014064M6RLKWAX-EGF_2976

TTTTTTCGATAGACTCTCTATGCCATGGAACAAAAGCGATTAATTCGTCCATCATTCCCAAGAATGTCAACAATTTTGTTACAATTCTTTCTGATTTCTTCTTCATCTCTTAGTTGAGTG

>M6-PGSC0003DMG400014064M6RLKWAX-EGF_3038

AATGTCAACAATTTTGTTACAATTCTTTCTGATTTCTTCTTCATCTCTTAGTTGAGTGTTTTATTCAGCATTGACATCAGAGCTTACTATATTGGTTCTGTCAAATGAAAAGTGTTCTCA

>M6-PGSC0003DMG400014064M6RLKWAX-EGF_310

TTAGATTGAGCTGGATATCAAGTAACTGCAGTGACTGTGCAGAAAGTGGAGGACGTTGCGGATTTGATGTTAACAATTATCAATTCAAATGTTTCTGCACTGACAGGCCTCATGCTTCGA

>M6-PGSC0003DMG400014064M6RLKWAX-EGF_372

TTTGATGTTAACAATTATCAATTCAAATGTTTCTGCACTGACAGGCCTCATGCTTCGAGTTGCCCACCTACCAGTAAGGCCCTATCTAGCAGTTAATTTTCCAGTATAATTATGCGGAAC

>M6-PGSC0003DMG400014064M6RLKWAX-EGF_434

CCCACCTACCAGTAAGGCCCTATCTAGCAGTTAATTTTCCAGTATAATTATGCGGAACCTCATCTTATGATTAGGCACCGTGCTTTAGGTTCTTTTCTTGTGATTATCAAGCGCACAAAA

>M6-PGSC0003DMG400014064M6RLKWAX-EGF_496

TCTTATGATTAGGCACCGTGCTTTAGGTTCTTTTCTTGTGATTATCAAGCGCACAAAAAAGTATCGTTTGATGATCTAGGCATCAAGATGCAGCTGCATTAAGCATTTGGATTATGTGTT

>M6-PGSC0003DMG400014064M6RLKWAX-EGF_558

ATCGTTTGATGATCTAGGCATCAAGATGCAGCTGCATTAAGCATTTGGATTATGTGTTAACCACAAAATTTTGTGCAGCCAACTTTTAAAAAAAGAAGGTAAATATAAAAATTGGTGGTA

>M6-PGSC0003DMG400014064M6RLKWAX-EGF_62

GGAATCAAGAATTTACTTTTTGTCAAAGTGTAATGGATCGATATCTGAGGATCTTTTGAAGCACAGGATTGGTTTTGGTTGTGGTGGAGAAAATGGGAATGATTGGGGTCTTGCAATGTT

>M6-PGSC0003DMG400014064M6RLKWAX-EGF_620

ACAAAATTTTGTGCAGCCAACTTTTAAAAAAAGAAGGTAAATATAAAAATTGGTGGTAGGGGAAGGGGAAATGCGGGAGGGGATTACAAGATGGGGGATCGAACACCAACAAGGTGAAAC

>M6-PGSC0003DMG400014064M6RLKWAX-EGF_744

GGTAGCCAACCAATTGAGGTACGAAGATTTCCCTGTGCAGCCAAGTTAATATTCTCGTAGCAGCTAATTACTTCTCTGATGAATGGTGTCGTGTTCTGCTTAATTACCATGCATTTCAGG

>M6-PGSC0003DMG400014064M6RLKWAX-EGF_806

GCTAATTACTTCTCTGATGAATGGTGTCGTGTTCTGCTTAATTACCATGCATTTCAGGGTTTGACCTCTTGTCCTGTTAACCCTGTGGTTATGCCAGCAGTATTTTCTGTTTGTTATTTG

>M6-PGSC0003DMG400014064M6RLKWAX-EGF_868

GACCTCTTGTCCTGTTAACCCTGTGGTTATGCCAGCAGTATTTTCTGTTTGTTATTTGTTGAAGTTTAGAGTTAGACCAATGACTGCAGCCTCCTTTTCTGCTATGTTCTCTATGTTTCA

>M6-PGSC0003DMG400014064M6RLKWAX-EGF_930

AGTTTAGAGTTAGACCAATGACTGCAGCCTCCTTTTCTGCTATGTTCTCTATGTTTCATCGGCATATCTTCAACTTAGTTCAATTAGAGAGTAAATCTTGTCCAAAAGAGTTTCGTTGTG

>M6-PGSC0003DMG400014064M6RLKWAX-EGF_992

CATATCTTCAACTTAGTTCAATTAGAGAGTAAATCTTGTCCAAAAGAGTTTCGTTGTGGAAGAATTGGAACCATAAGTTATCTTTTCCCACCTCATGTTCTATTTTGAGTAGACAACAAT

>M6-PGSC0003DMG400014281-(rev)M6LRR-RLK_0

AAGCCAAAGGAGCATCTGAGTATCAACATTGCAACCTTTGAGCAGTCTTTGTTACGGATTAACCCTACAGCTATTCTCTCAGCCACTGAGAACTTCAGCAAGAGTTACATCATTGGTGAT

>M6-PGSC0003DMG400014281-(rev)M6LRR-RLK_126

TTTGGTACTGTATACAAAGCCAAACTTCCTGAGGGCCAGAACATTGCTGTCAAGAGGCTGAACGGAGGCCACATGCACGGTGATCGCGAGTTCTTTGCTGAAATGGAAACAATTGGTAAA

>M6-PGSC0003DMG400014281-(rev)M6LRR-RLK_189

GGAGGCCACATGCACGGTGATCGCGAGTTCTTTGCTGAAATGGAAACAATTGGTAAAGTAAAACATGAAAATCTGGTGCCATTGCTTGGTTATTGTGTCTTTGCAGATGAGAGGTTCTTG

>M6-PGSC0003DMG400014281-(rev)M6LRR-RLK_252

CATGAAAATCTGGTGCCATTGCTTGGTTATTGTGTCTTTGCAGATGAGAGGTTCTTGATATATGAATACATGGAGAATGGAAGCCTTGATTTCTGGTTGAGAAACCAAGCAGATGCAGTA

>M6-PGSC0003DMG400014281-(rev)M6LRR-RLK_315

GAATACATGGAGAATGGAAGCCTTGATTTCTGGTTGAGAAACCAAGCAGATGCAGTAGAAGCACTTGACTGGCCAACTCGTTTCAAGATTTGTCTAGGGTCAGCTGTCGGACTTTCCTTT

>M6-PGSC0003DMG400014281-(rev)M6LRR-RLK_378

CTTGACTGGCCAACTCGTTTCAAGATTTGTCTAGGGTCAGCTGTCGGACTTTCCTTTCTACACCATGGTTTTGTTCCACACATCATCCACAGAGACATCAAGTCAAGCAACATACTCCTA

>M6-PGSC0003DMG400014281-(rev)M6LRR-RLK_441

CATGGTTTTGTTCCACACATCATCCACAGAGACATCAAGTCAAGCAACATACTCCTAGACAAGAACTTTGAACCACGAGTCTCTGACTTTGGCCTGGCTCGAATAATTAGTGCTTGTGAG

>M6-PGSC0003DMG400014281-(rev)M6LRR-RLK_504

AACTTTGAACCACGAGTCTCTGACTTTGGCCTGGCTCGAATAATTAGTGCTTGTGAGAGCCATGTTAGCACAATCCTTGCTGGTACATTCGGGTACATACCTCCCGAGTATGGACAAACC

>M6-PGSC0003DMG400014281-(rev)M6LRR-RLK_567

GTTAGCACAATCCTTGCTGGTACATTCGGGTACATACCTCCCGAGTATGGACAAACCATGACGGCCACAACAAAGGGTGACATCTACAGCTTTGGAGTGGTGATGTTGGAACTGGTAACA

>M6-PGSC0003DMG400014281-(rev)M6LRR-RLK_623

CATGACGGCCACAACAAAGGGTGACATCTACAGCTTTGGAGTGGTGATGTTGGAACTGGTAACAGGAAGGGCACCAACAGGACAGGCTGATGTTGAAGGAGGCAATCTTGTTGGCTGGGT

>M6-PGSC0003DMG400014281-(rev)M6LRR-RLK_63

CCTACAGCTATTCTCTCAGCCACTGAGAACTTCAGCAAGAGTTACATCATTGGTGATGGTGGTTTTGGTACTGTATACAAAGCCAAACTTCCTGAGGGCCAGAACATTGCTGTCAAGAGG

>M6-PGSC0003DMG400014463-(rev)M6LRR-RLP_0

GCTAAAGATGGGGTAATGCAATTGTGCTCACTCTCGTCTTTTTCTGTCTTGAAGTTTCTTCATCCATATTACCTCATCACTCTCACAAACATCAAATTCAAACCATTTCCCTTCTAAAAT

>M6-PGSC0003DMG400014463-(rev)M6LRR-RLP_1037

CTATCACTAATTTATTTAGAATTGAGTCATAACCACTTTTCTGGTCAGCTTGAGGATTTCAAGTCAAATTCACTATTCTATATTGGTTTAAGCAATAATCAGCTGGAAGGCCATCTTCCA

>M6-PGSC0003DMG400014463-(rev)M6LRR-RLP_1098

AGTCAAATTCACTATTCTATATTGGTTTAAGCAATAATCAGCTGGAAGGCCATCTTCCAAAGTCAATTCAAAACCTTGTGAACCTGACAACTCTTGATCTTTCATCGAATAATTTTAGTG

>M6-PGSC0003DMG400014463-(rev)M6LRR-RLP_1159

GTCAATTCAAAACCTTGTGAACCTGACAACTCTTGATCTTTCATCGAATAATTTTAGTGGCAATGTGGATGTCAGCCTCTTTTCAAACCTCAAACAACTTTCGTATCTTATTCTTTCATC

>M6-PGSC0003DMG400014463-(rev)M6LRR-RLP_122

AAGAAGTCCTTTACTATAGATCCCTCAGCCTCTTATTTTTGTCAAGACTTCGGTAAGAAGTTTTATCCAAAAACGAGTTCATGGAACATGAGCAGAGATTGCTGCTCATGGGATGGAGTT

>M6-PGSC0003DMG400014463-(rev)M6LRR-RLP_1220

AATGTGGATGTCAGCCTCTTTTCAAACCTCAAACAACTTTCGTATCTTATTCTTTCATCGAATAATTTTAGTGGCAATGTGGATTTCAGCCTCTTTTCAAACCTCAAACAGCTTTCATAT

>M6-PGSC0003DMG400014463-(rev)M6LRR-RLP_1281

ATAATTTTAGTGGCAATGTGGATTTCAGCCTCTTTTCAAACCTCAAACAGCTTTCATATCTTATTCTTTCTTCCAATAATTTTATTGGAAATGTAGATGTCAGCCTTTTTTCAGACCTCA

>M6-PGSC0003DMG400014463-(rev)M6LRR-RLP_1342

TATTCTTTCTTCCAATAATTTTATTGGAAATGTAGATGTCAGCCTTTTTTCAGACCTCAAACAGCTTTCATATCTTAGTCTTTCACAAAATCATTCCTCATTGACCAATGAAAACAAAGT

>M6-PGSC0003DMG400014463-(rev)M6LRR-RLP_1403

CAGCTTTCATATCTTAGTCTTTCACAAAATCATTCCTCATTGACCAATGAAAACAAAGTTAAGGTTACTTTGCCAGGATCTCTTTATACGTTACAATTGGCTGCATGTGAAGTGAAAGAA

>M6-PGSC0003DMG400014463-(rev)M6LRR-RLP_1464

AGGTTACTTTGCCAGGATCTCTTTATACGTTACAATTGGCTGCATGTGAAGTGAAAGAATTGGAGTTTCTAAAATCCACAAATCATCTTGTTTTCTTGGATCTGTCAAATAATAAGCTTC

>M6-PGSC0003DMG400014463-(rev)M6LRR-RLP_1525

GGAGTTTCTAAAATCCACAAATCATCTTGTTTTCTTGGATCTGTCAAATAATAAGCTTCAAGGAAGAATCCCAGATTGGGCATGGTCTAAATGGATGTTTTCATTGCAATATCTAGATCT

>M6-PGSC0003DMG400014463-(rev)M6LRR-RLP_1586

GGAAGAATCCCAGATTGGGCATGGTCTAAATGGATGTTTTCATTGCAATATCTAGATCTATCCTACAACATGTTGACAAGTGTTGATTCAATTCCTCTTCAATCACTAGATACTATTGAT

>M6-PGSC0003DMG400014463-(rev)M6LRR-RLP_1647

CCTACAACATGTTGACAAGTGTTGATTCAATTCCTCTTCAATCACTAGATACTATTGATTTACGGTCCAATTTGCTTCAAGGTTCACTACCTGTTCCACCAAATTCCACAATATTCTTCT

>M6-PGSC0003DMG400014463-(rev)M6LRR-RLP_1708

ACGGTCCAATTTGCTTCAAGGTTCACTACCTGTTCCACCAAATTCCACAATATTCTTCTTAATATCCCAAAATAATCTCACTGGAGAAATTCCTGCATCTATTTGCAATTTGATGTCACT

>M6-PGSC0003DMG400014463-(rev)M6LRR-RLP_1769

ATATCCCAAAATAATCTCACTGGAGAAATTCCTGCATCTATTTGCAATTTGATGTCACTCCAAATTTTGGATTTGGCGAGAAACAATTGAGAAATTCCACAATGTTTGGGAAATATGACT

>M6-PGSC0003DMG400014463-(rev)M6LRR-RLP_1830

AAATTTTGGATTTGGCGAGAAACAATTGAGAAATTCCACAATGTTTGGGAAATATGACTCCCCTCCGGGTTTTGGATATGCGCCACAACAATCTTTCTGGGAATATTCCACCAACTTTCA

>M6-PGSC0003DMG400014463-(rev)M6LRR-RLP_1891

CCTCCGGGTTTTGGATATGCGCCACAACAATCTTTCTGGGAATATTCCACCAACTTTCAGTAATGAAAGTATATTGAGCAGCCTCAACTTGCATGGCAATAATCTAGAGGGGAAAATTCC

>M6-PGSC0003DMG400014463-(rev)M6LRR-RLP_2074

CACATTCCCTGTGTGGTTGGGAGCTCTTCCAGATCTGCTCATTTTAAGCTTGAGATCAAACAAATTGCATGGACCAATCAGAACATCAAGTTGGAGTGAGAACATGTTTCCTAAGCTTCG

>M6-PGSC0003DMG400014463-(rev)M6LRR-RLP_2196

TCATAGATCTTTCCTACAATGCCTTCTCGGGAAACTTACCTTCGAGTTTGTTTCAACATCTTAAAACCATGAGGACAATTGATCCATCAAGGGAAGCAGCAAGACATGGTGGAAATACAT

>M6-PGSC0003DMG400014463-(rev)M6LRR-RLP_2257

TAAAACCATGAGGACAATTGATCCATCAAGGGAAGCAGCAAGACATGGTGGAAATACATATTACCAAGATTCTATTATAGTTGCAATCAAGGGATCTGAGCGTGAAATTGTCAGAATCTT

>M6-PGSC0003DMG400014463-(rev)M6LRR-RLP_2318

TACCAAGATTCTATTATAGTTGCAATCAAGGGATCTGAGCGTGAAATTGTCAGAATCTTGTATTTGTACACCGTTATCGATCTTTCAAGTAATAAATTTAGAGGGCAAATTCCAAGTATT

>M6-PGSC0003DMG400014463-(rev)M6LRR-RLP_2379

ATTTGTACACCGTTATCGATCTTTCAAGTAATAAATTTAGAGGGCAAATTCCAAGTATTATGGGGCATCTCAATGCAGTTCACATATTGAATTTATCTCATAATGCATTGCAAGGTGATA

>M6-PGSC0003DMG400014463-(rev)M6LRR-RLP_2440

GGGGCATCTCAATGCAGTTCACATATTGAATTTATCTCATAATGCATTGCAAGGTGATATACCGACATTATTCGGAGATTTATCTTCACTTGAATCATTGGACTTGTCAGGAAACAAACT

>M6-PGSC0003DMG400014463-(rev)M6LRR-RLP_2684

TATGAAGGCAATGATGGATTACGTGGATTTCCTATTTCAAAAGGCTGTGGCGATACTCATCGTCTCTAAAGAATCCACGCCCTTTTTTTAGCGTTTAGGGCCGGGCGTCATGAAACAAGA

>M6-PGSC0003DMG400014463-(rev)M6LRR-RLP_2745

GTCTCTAAAGAATCCACGCCCTTTTTTTAGCGTTTAGGGCCGGGCGTCATGAAACAAGAATTCAGAATTTTCTAAGTGTTTCGTGTGTATCAATTCATGAATTTTTGGAAATATGACTGA

>M6-PGSC0003DMG400014463-(rev)M6LRR-RLP_2799

CAAGAATTCAGAATTTTCTAAGTGTTTCGTGTGTATCAATTCATGAATTTTTGGAAATATGACTGATCGAATATTTTTGCCCAAATATTTATTGAAATCATCTGTAATGACCTAGTGAAC

>M6-PGSC0003DMG400014463-(rev)M6LRR-RLP_305

ACCATTGATTCCAATAGCAGCCTATTCCAACTCTCTCATCTCCAAAGGCTTGATCTTTCTTTAAATAACTTCTCCAATTCTCACATCTCCCCTGAATTTGGCAGGTTTTCGAGCTTGACA

>M6-PGSC0003DMG400014463-(rev)M6LRR-RLP_366

TAAATAACTTCTCCAATTCTCACATCTCCCCTGAATTTGGCAGGTTTTCGAGCTTGACACATCTTGATCTTTCAGGCTCCTATTTTTCAGGTCAAATTCCTTTTGAAATCTCTTATCTTT

>M6-PGSC0003DMG400014463-(rev)M6LRR-RLP_427

TCTTGATCTTTCAGGCTCCTATTTTTCAGGTCAAATTCCTTTTGAAATCTCTTATCTTTCCAAGTTACACTCTTTTCGTCTCTCCACAAATATTGGGTTTGTTGGCCTCCCCTCATATTT

>M6-PGSC0003DMG400014463-(rev)M6LRR-RLP_488

AAGTTACACTCTTTTCGTCTCTCCACAAATATTGGGTTTGTTGGCCTCCCCTCATATTTTGGGCCCATACAAAGACTCGCGGCCCATGATTTTAAATTGCTCCTTCAGAATTTGACCCAA

>M6-PGSC0003DMG400014463-(rev)M6LRR-RLP_61

ATCCATATTACCTCATCACTCTCACAAACATCAAATTCAAACCATTTCCCTTCTAAAATTCAAGAAGTCCTTTACTATAGATCCCTCAGCCTCTTATTTTTGTCAAGACTTCGGTAAGAA

>M6-PGSC0003DMG400014463-(rev)M6LRR-RLP_610

AAGAGAACTTGATTTTAGTTTCATAAACATCTCTTCAACCATTCCTCCGAATTTTTCTTCTCATTTCACAACTCTGAGTCTGGGATTAACAGGATTGTATGGGATAATACCTGAGCGTAT

>M6-PGSC0003DMG400014463-(rev)M6LRR-RLP_671

CATTTCACAACTCTGAGTCTGGGATTAACAGGATTGTATGGGATAATACCTGAGCGTATTTTTCACCTTCCCAACTTGAAAAAACTTGACTTATCCAGCAGTTATGAGCTCAGTGGTTAT

>M6-PGSC0003DMG400014463-(rev)M6LRR-RLP_732

TTCACCTTCCCAACTTGAAAAAACTTGACTTATCCAGCAGTTATGAGCTCAGTGGTTATTTTCCAAAGACCAAATGGAACAACAACACATCTCTTAGGAAATTAGAACTCAGTGGCGTGA

>M6-PGSC0003DMG400014463-(rev)M6LRR-RLP_793

TCCAAAGACCAAATGGAACAACAACACATCTCTTAGGAAATTAGAACTCAGTGGCGTGAATTTTTCTGATAATATACCTGAGTCTCTTGGCTATCATCTAACTTCGCTGAATTATTTTTC

>M6-PGSC0003DMG400014463-(rev)M6LRR-RLP_854

TTTTCTGATAATATACCTGAGTCTCTTGGCTATCATCTAACTTCGCTGAATTATTTTTCTCTTTCGCATTGCAAGCTTTGGGGGCCTATTCCTAAATCTCTTTCCAATCTCACTCACATA

>M6-PGSC0003DMG400014463-(rev)M6LRR-RLP_915

TTTCGCATTGCAAGCTTTGGGGGCCTATTCCTAAATCTCTTTCCAATCTCACTCACATAGAGTATTTGTACCTTGATGGTAACTTGTTGAATGGAACAATACCATTAGGGATGTTCTCCC

>M6-PGSC0003DMG400014463-(rev)M6LRR-RLP_976

GTATTTGTACCTTGATGGTAACTTGTTGAATGGAACAATACCATTAGGGATGTTCTCCCTTCTATCACTAATTTATTTAGAATTGAGTCATAACCACTTTTCTGGTCAGCTTGAGGATTT

>M6-PGSC0003DMG400014576-(rev)M6LRR-RLK_1037

GCAAAAGAAAAAAATTCACACTTTTTTGGCAAGAGATGCCCTACTGTCCCATTGTGCTCTACAGCTTTTTTGCTTATCTTTTTGACCTCACTTAGTATTCATATCTTGCAACTGTTTTGG

>M6-PGSC0003DMG400014576-(rev)M6LRR-RLK_1098

CAGCTTTTTTGCTTATCTTTTTGACCTCACTTAGTATTCATATCTTGCAACTGTTTTGGTAGGTGGATAGTGAAATATAACCTCTTGTTTTTTTTCTGGTTCAAATGGTAAAGAATCAAC

>M6-PGSC0003DMG400014576-(rev)M6LRR-RLK_1159

GGTGGATAGTGAAATATAACCTCTTGTTTTTTTTCTGGTTCAAATGGTAAAGAATCAACTTCTTGTTGAATTGCACCTGTCATTCATGTAACAGGAGACAGCTGCTTCATATGTTAAATT

>M6-PGSC0003DMG400014576-(rev)M6LRR-RLK_122

TACTTTGCTAAAGATTGAGTAGTTGTTGGTGAGTTTGGTGTATGATTTCATCACTTGTTTTGGCTTATTCATGGTAAATAGTATTCAACCTTTTGTTCTTTAAGAAATTCATAGTACTAA

>M6-PGSC0003DMG400014576-(rev)M6LRR-RLK_1220

CTTGTTGAATTGCACCTGTCATTCATGTAACAGGAGACAGCTGCTTCATATGTTAAATTTTCTCATTTTAGAATTATGAACAATTATGTGTTACTATTCCTACCATCAAAGTCTGCTATG

>M6-PGSC0003DMG400014576-(rev)M6LRR-RLK_1281

CTCATTTTAGAATTATGAACAATTATGTGTTACTATTCCTACCATCAAAGTCTGCTATGTCAACTTCAAACTCCTTGTCCTCTCTGCCCAGGTTTTTGTTCTTTTGTCTAGTTAAAGTTG

>M6-PGSC0003DMG400014576-(rev)M6LRR-RLK_1586

GTGTAAGTGTTCCATTTTATAGTAGTTTGAGTAAATGTTTTGATGGACATGAATCTTCTTGCTACAGGTAACAGAGGGCCTGGAAGAACTCCTTTAGACTGGACAACAAGGTTAAAGATT

>M6-PGSC0003DMG400014576-(rev)M6LRR-RLK_1647

CTACAGGTAACAGAGGGCCTGGAAGAACTCCTTTAGACTGGACAACAAGGTTAAAGATTGCAGCCGGAGCAGCTCGAGGGCTAGCCTTCATCCACAACTCATGCAAGTCCTTGAAACTCA

>M6-PGSC0003DMG400014576-(rev)M6LRR-RLK_1708

AGCCGGAGCAGCTCGAGGGCTAGCCTTCATCCACAACTCATGCAAGTCCTTGAAACTCACTCATGGCAACATCAAATCAACTAACATCCTCATTGACAAGAGCGGCAATGCACGTATCTC

>M6-PGSC0003DMG400014576-(rev)M6LRR-RLK_1769

CATGGCAACATCAAATCAACTAACATCCTCATTGACAAGAGCGGCAATGCACGTATCTCCGACTTCGGTCTAGCCATCTTCGCCACACCATCTTCCGTCCCAAAAACCAACGGCTACCGA

>M6-PGSC0003DMG400014576-(rev)M6LRR-RLK_183

GGCTTATTCATGGTAAATAGTATTCAACCTTTTGTTCTTTAAGAAATTCATAGTACTAAAATATTTGTGAATTACTTGTTTCTTAAAGAAAAATGTGATCTTTAGCTTTTTGTTGTAGAA

>M6-PGSC0003DMG400014576-(rev)M6LRR-RLK_1830

ACTTCGGTCTAGCCATCTTCGCCACACCATCTTCCGTCCCAAAAACCAACGGCTACCGAGCCCCTGAAGTCGCATTGGATGGTCGTAAAATAACTCAAAAGTCCGACATCTACTCATTCG

>M6-PGSC0003DMG400014576-(rev)M6LRR-RLK_1891

CCCTGAAGTCGCATTGGATGGTCGTAAAATAACTCAAAAGTCCGACATCTACTCATTCGGGGTCCTATTACTCGAGCTGCTAACCGGTAAATGCCCCTCAGTCGTGGACAACGGTGGTGG

>M6-PGSC0003DMG400014576-(rev)M6LRR-RLK_1952

GTCCTATTACTCGAGCTGCTAACCGGTAAATGCCCCTCAGTCGTGGACAACGGTGGTGGCCTCACAACCGGCTACGGAGGGGTTGTGGACTTACCGAGATGGGTGCAATCCGTGGTGAGG

>M6-PGSC0003DMG400014576-(rev)M6LRR-RLK_2013

TCACAACCGGCTACGGAGGGGTTGTGGACTTACCGAGATGGGTGCAATCCGTGGTGAGGGAAGAATGGACCGCGGAAGTGTTCGATTTGGAGTTGATGAGGTATAAGGACATTGAGGAGG

>M6-PGSC0003DMG400014576-(rev)M6LRR-RLK_2135

ATGGTGGGGCTATTGCAGATAGCTATGGCATGTACTTCAACATCACCGGATCAAAGGCCTAAAATAAATTATGTTGTGAAAATGATTGAAGAGTTGAGAGGTGTTGAAGTTTCACCTTCT

>M6-PGSC0003DMG400014576-(rev)M6LRR-RLK_2318

GCAGCTAGTCAGTAGTGAGAGTTTTTTTTTTTCTACTAGTTTGTTATTCTGAGTTATGTAAATTAACCTCTTAGTTTTAATCCTTTTATCTATAAGATATAATTTAGGAGAAGTTTTATA

>M6-PGSC0003DMG400014576-(rev)M6LRR-RLK_244

TATTTGTGAATTACTTGTTTCTTAAAGAAAAATGTGATCTTTAGCTTTTTGTTGTAGAATAGATTATTGATTGACCAATAAGGATTATAGTGGAGTTGTAAGTACTTCGTCATCATTAAT

>M6-PGSC0003DMG400014576-(rev)M6LRR-RLK_305

GATTATTGATTGACCAATAAGGATTATAGTGGAGTTGTAAGTACTTCGTCATCATTAATCTGAGGTTTCAAGTTCGAGCCTTGAGTATGGACTTGCCCTTGTTAGGTAGCGGTTTGTCCT

>M6-PGSC0003DMG400014576-(rev)M6LRR-RLK_366

GAGGTTTCAAGTTCGAGCCTTGAGTATGGACTTGCCCTTGTTAGGTAGCGGTTTGTCCTACGTGGGACTTCCTAGCGCGAACCCGTATCCGGCTCCAATGCGGTTGCCGTACACCGGATG

>M6-PGSC0003DMG400014576-(rev)M6LRR-RLK_427

GTGGGACTTCCTAGCGCGAACCCGTATCCGGCTCCAATGCGGTTGCCGTACACCGGATGGGAAAAAAGGAGAAGATTATTGATTGATTTGGTGGTAAAGTTTGGCAAGTAGTATAGCATA

>M6-PGSC0003DMG400014576-(rev)M6LRR-RLK_488

AAAAAAGGAGAAGATTATTGATTGATTTGGTGGTAAAGTTTGGCAAGTAGTATAGCATAATCACATGGGGTTTGAGTCTGGCTTTTTGTCTTTAATTGCTTTTGGTCTGCTACTGTTAAC

>M6-PGSC0003DMG400014576-(rev)M6LRR-RLK_549

CACATGGGGTTTGAGTCTGGCTTTTTGTCTTTAATTGCTTTTGGTCTGCTACTGTTAACCAGGACTAGCCTTCTGATGCAATAGGAATAAAGATGGGATGCCTCAATACTAGACAGTATT

>M6-PGSC0003DMG400014576-(rev)M6LRR-RLK_61

AGGACTGGTCTCTAATGCATATAAGAATAGAAACCCAAATTTGGATAGTTTGAAAGCATATTACTTTGCTAAAGATTGAGTAGTTGTTGGTGAGTTTGGTGTATGATTTCATCACTTGTT

>M6-PGSC0003DMG400014576-(rev)M6LRR-RLK_610

GGACTAGCCTTCTGATGCAATAGGAATAAAGATGGGATGCCTCAATACTAGACAGTATTTGCTTTAGTTAATGTTTCTATTTACATAATTGAAACTAGCCATGTGGGAATATTGAAAATC

>M6-PGSC0003DMG400014576-(rev)M6LRR-RLK_671

CTTTAGTTAATGTTTCTATTTACATAATTGAAACTAGCCATGTGGGAATATTGAAAATCTATGTTGCTCGGACTCTTTAAAAATATCGAGGGTGCGGCAACATGAGTAAGTCTTACATAG

>M6-PGSC0003DMG400014576-(rev)M6LRR-RLK_732

TGTTGCTCGGACTCTTTAAAAATATCGAGGGTGCGGCAACATGAGTAAGTCTTACATAGTCTTTTCAAGCTAAAACTATAGAATAGTATGTTACTTTATTAACAATCTTTGTCACATAAG

>M6-PGSC0003DMG400014576-(rev)M6LRR-RLK_793

TTTTCAAGCTAAAACTATAGAATAGTATGTTACTTTATTAACAATCTTTGTCACATAAGCTATTCTTGATTTTCAAATGTAGGGAACTTAAATTATTTAACAATCTTTATCATCTTATTC

>M6-PGSC0003DMG400014576-(rev)M6LRR-RLK_854

ATTCTTGATTTTCAAATGTAGGGAACTTAAATTATTTAACAATCTTTATCATCTTATTCTTGAAAAAGTTTTTAAAGGAATCTAATTTACTCATGATGGTCCTAAAGTGACAGAAGATTA

>M6-PGSC0003DMG400014576-(rev)M6LRR-RLK_915

GAAAAAGTTTTTAAAGGAATCTAATTTACTCATGATGGTCCTAAAGTGACAGAAGATTAGAGCAAAGGACTTGTTGTACCAGATGAATAGAATTATGTGTTCCCTCTGATTTTTGGGCCA

>M6-PGSC0003DMG400014576-(rev)M6LRR-RLK_976

GCAAAGGACTTGTTGTACCAGATGAATAGAATTATGTGTTCCCTCTGATTTTTGGGCCAAAGCAAAAGAAAAAAATTCACACTTTTTTGGCAAGAGATGCCCTACTGTCCCATTGTGCTC

>M6-PGSC0003DMG400014577-(rev)M6RLKWAX-EGF_0

ATGGATTTAGTTTCTTGGTTTTTCGTCTTCATTTTGTTTCATCAGAGGTGTTTTGATTCTGTTCAAGGACAGAACACACCTTATTCAAACTGTCCTCAAGAATTCCAATGTGGAAGTTCA

>M6-PGSC0003DMG400014577-(rev)M6RLKWAX-EGF_1008

GGATCATCGAAATATTGAAGCGTTTCTGAAGAACTACGGGCCATATGCTCCAATGAGATATAGTTATTCAGACATCAAAAGGATGACTAGTAGCTTCAAAAACAAACTAGGCCAAGGAGG

>M6-PGSC0003DMG400014577-(rev)M6RLKWAX-EGF_1071

TTATTCAGACATCAAAAGGATGACTAGTAGCTTCAAAAACAAACTAGGCCAAGGAGGATATGGTTATGTATACAAAGGAAGTTTACATAATGGAAATCCTGTTGCTGTGAAGGTTTTGAA

>M6-PGSC0003DMG400014577-(rev)M6RLKWAX-EGF_1134

TTATGTATACAAAGGAAGTTTACATAATGGAAATCCTGTTGCTGTGAAGGTTTTGAATGGACTAAAAGGCAGTGGTGAAGAATTCATTAATGAGGTATCGAGTATTAGCAGGACATCACA

>M6-PGSC0003DMG400014577-(rev)M6RLKWAX-EGF_1197

AAAAGGCAGTGGTGAAGAATTCATTAATGAGGTATCGAGTATTAGCAGGACATCACATGTTAACATCGTGAGCCTCGTGGGATTCTGTTTTGAGGGTCGAAAAAGAGCTCTAGTTTACGA

>M6-PGSC0003DMG400014577-(rev)M6RLKWAX-EGF_126

CTGAGTTTTCCTTTCTATAAACCCGATCAACCTGATTGCGGATTATGTAAAGTTGATTGTGATACTGCTCCACATCCGATGATCGAATTGGAAGGAGTAAAGTATCAAGTTCTTGAAAAA

>M6-PGSC0003DMG400014577-(rev)M6RLKWAX-EGF_1260

CATCGTGAGCCTCGTGGGATTCTGTTTTGAGGGTCGAAAAAGAGCTCTAGTTTACGAATTCATGCCAAATGGATCACTTGAAAAGTTTATCTATGAAGAAAGATCAGACACAGACAGAGT

>M6-PGSC0003DMG400014577-(rev)M6RLKWAX-EGF_1323

GCCAAATGGATCACTTGAAAAGTTTATCTATGAAGAAAGATCAGACACAGACAGAGTTCGTCAACTAGATTGGCAAGTACTGTATAAAATTGCATTAGGCATTGCTCGTGGATTGGAGTA

>M6-PGSC0003DMG400014577-(rev)M6RLKWAX-EGF_1386

ACTAGATTGGCAAGTACTGTATAAAATTGCATTAGGCATTGCTCGTGGATTGGAGTATTTGCATCGTGGTTGTAACACTAGAATTCTACATTTTGATATCAAGCCTCATAACATACTACT

>M6-PGSC0003DMG400014577-(rev)M6RLKWAX-EGF_1512

AGATTTCTGTCCTAAGATATCTGACTTTGGACTTGCTAAACTCTGCGTAAAAAAGGAAAGTATAGTGTCAATGTTAGGTGCACGAGGGACTATCGGTTACATTGCTCCTGAAATTGTTTG

>M6-PGSC0003DMG400014577-(rev)M6RLKWAX-EGF_1575

AGTGTCAATGTTAGGTGCACGAGGGACTATCGGTTACATTGCTCCTGAAATTGTTTGTAGAAACTTGGGAGGTGTCTCACACAAGTCTGATGTTTATAGCTATGGAATGATGGTGTTAGA

>M6-PGSC0003DMG400014577-(rev)M6RLKWAX-EGF_1638

CTTGGGAGGTGTCTCACACAAGTCTGATGTTTATAGCTATGGAATGATGGTGTTAGAGATGGTTGGAGGGAGGAAAAATGTTGTTGATCGAACGAGTGAAGTCTACTTTCCAGATTGGCT

>M6-PGSC0003DMG400014577-(rev)M6RLKWAX-EGF_1701

TGGAGGGAGGAAAAATGTTGTTGATCGAACGAGTGAAGTCTACTTTCCAGATTGGCTTTATCAACGAATTGAACTAGACGAAGAACTTCAATTAATTGGGATAATGAATGAAGAGGAGAA

>M6-PGSC0003DMG400014577-(rev)M6RLKWAX-EGF_1764

ACGAATTGAACTAGACGAAGAACTTCAATTAATTGGGATAATGAATGAAGAGGAGAAAGAATGTGCGCGAAAAATGGTGATAGTGAGTCTATGGTGCATCCAAACTGGTCCTTCAAATAG

>M6-PGSC0003DMG400014577-(rev)M6RLKWAX-EGF_1827

TGCGCGAAAAATGGTGATAGTGAGTCTATGGTGCATCCAAACTGGTCCTTCAAATAGACCATCTATGACTAAAGTTGTCGAAATGCTAGAAGGGAACTTAAACTCTTTGCAAATTCCTCC

>M6-PGSC0003DMG400014577-(rev)M6RLKWAX-EGF_189

ACTGCTCCACATCCGATGATCGAATTGGAAGGAGTAAAGTATCAAGTTCTTGAAAAACAGAATGTTTTTGTCAAAATTAAGGATCCTGTTCTTGAAAATCTCTTAAAGAAAAACAGTTGT

>M6-PGSC0003DMG400014577-(rev)M6RLKWAX-EGF_1890

TATGACTAAAGTTGTCGAAATGCTAGAAGGGAACTTAAACTCTTTGCAAATTCCTCCAACACCTTACCTATATTCTTCCTCAAGGTCAGAAGTAAAATTGATCTCATCAGTAGAAGAATC

>M6-PGSC0003DMG400014577-(rev)M6RLKWAX-EGF_1927

AACTCTTTGCAAATTCCTCCAACACCTTACCTATATTCTTCCTCAAGGTCAGAAGTAAAATTGATCTCATCAGTAGAAGAATCGAGTTCCCGATCAACTCTCTCTACCTCCACAAGGTAG

>M6-PGSC0003DMG400014577-(rev)M6RLKWAX-EGF_252

GTTTTTGTCAAAATTAAGGATCCTGTTCTTGAAAATCTCTTAAAGAAAAACAGTTGTCGATCTTTCGAAAGAAATCTATCTCTCCCAATATCTCCTATAATCACCTATAGTGTCAACCCT

>M6-PGSC0003DMG400014577-(rev)M6RLKWAX-EGF_315

TTCGAAAGAAATCTATCTCTCCCAATATCTCCTATAATCACCTATAGTGTCAACCCTTTCCTCAACTTGTTTAAATGCAATACTAGTAATGATATCTCGTACGAGGGCTTTCAAAGTTAC

>M6-PGSC0003DMG400014577-(rev)M6RLKWAX-EGF_378

AACTTGTTTAAATGCAATACTAGTAATGATATCTCGTACGAGGGCTTTCAAAGTTACAATGGTTGCGAAAGTTTCACTTTATACTATAATCATTCGTCCGTTTCTTCTATTGGATCTAAT

>M6-PGSC0003DMG400014577-(rev)M6RLKWAX-EGF_441

TGCGAAAGTTTCACTTTATACTATAATCATTCGTCCGTTTCTTCTATTGGATCTAATCTTCTTAGTAAATGTTCAATTATTCAATTGCCTATGGTGTCAGGTTATGAATCAAAATCGAAC

>M6-PGSC0003DMG400014577-(rev)M6RLKWAX-EGF_504

AGTAAATGTTCAATTATTCAATTGCCTATGGTGTCAGGTTATGAATCAAAATCGAACAATAGTAACCTGTTCGATATGTTGGCTGCTGATGTTTCTCTAGGATGGACAGTATCTCATACA

>M6-PGSC0003DMG400014577-(rev)M6RLKWAX-EGF_567

AACCTGTTCGATATGTTGGCTGCTGATGTTTCTCTAGGATGGACAGTATCTCATACATGTAATCAGTGTTATTCGAGTGGAGGACGATGTCAAGTTGATAGTAACAACAAATTTCATTGT

>M6-PGSC0003DMG400014577-(rev)M6RLKWAX-EGF_63

CAAGGACAGAACACACCTTATTCAAACTGTCCTCAAGAATTCCAATGTGGAAGTTCAGAAATTCTGAGTTTTCCTTTCTATAAACCCGATCAACCTGATTGCGGATTATGTAAAGTTGAT

>M6-PGSC0003DMG400014577-(rev)M6RLKWAX-EGF_630

CAGTGTTATTCGAGTGGAGGACGATGTCAAGTTGATAGTAACAACAAATTTCATTGTTCTGTTAGTGATCCAGGCAAAGGTATTCAAGAAGTAAATTATGCTTGATTCCTTCATCTTAGT

>M6-PGSC0003DMG400014577-(rev)M6RLKWAX-EGF_693

AGTGATCCAGGCAAAGGTATTCAAGAAGTAAATTATGCTTGATTCCTTCATCTTAGTACTCTTTATTTATTCGTTTTTCGTATTGCAGGAAAGCATCTTCGTAGAACATGGATCGTTGTT

>M6-PGSC0003DMG400014577-(rev)M6RLKWAX-EGF_756

TATTTATTCGTTTTTCGTATTGCAGGAAAGCATCTTCGTAGAACATGGATCGTTGTTTTAGCCACAGGTAACTCTTTTGTAGAAACAGTACTAGTTGCATTGTTCCGAATAGAGACTCTT

>M6-PGSC0003DMG400014577-(rev)M6RLKWAX-EGF_819

ACAGGTAACTCTTTTGTAGAAACAGTACTAGTTGCATTGTTCCGAATAGAGACTCTTCACGGTGTATGACTTCATTTTCCTCCGTATAGTTGCAGTTTTCGGTTGCATTGGAATCTTGAT

>M6-PGSC0003DMG400014577-(rev)M6RLKWAX-EGF_882

GTATGACTTCATTTTCCTCCGTATAGTTGCAGTTTTCGGTTGCATTGGAATCTTGATTTTGCTGTTCTGCTTCAGAAAAAAGATTTTCTGGCATAAGTACCTCAGGTTTTGGGAATCAGA

>M6-PGSC0003DMG400014577-(rev)M6RLKWAX-EGF_945

GTTCTGCTTCAGAAAAAAGATTTTCTGGCATAAGTACCTCAGGTTTTGGGAATCAGAAGCTGAGGATCATCGAAATATTGAAGCGTTTCTGAAGAACTACGGGCCATATGCTCCAATGAG

>M6-PGSC0003DMG400014717-(rev)M6LRR-RLP_0

CGTACAAGGATATTGAGTAGACAGTTGATGTATCAGACTCGATACTTCAAAAGAATGGAGGAGAAGACGATCTGCAGCTGCAAGTGTGCATGAAGACAATATATGAAATAACTGAGCAAA

>M6-PGSC0003DMG400014717-(rev)M6LRR-RLP_1054

AAGAGTTTAGAAAAAAATGGGAGTAGATGTGTAACACATGGATAAAGACAATAAAAAGACAATAATATTTGAGTGACTTTTCTCTTACAAAACAAAGAAAAGTGACTTTCTGAGATAATT

>M6-PGSC0003DMG400014717-(rev)M6LRR-RLP_1116

TAATATTTGAGTGACTTTTCTCTTACAAAACAAAGAAAAGTGACTTTCTGAGATAATTATCTATAGTTCAATGACCACATAAGAAATCAACTTTTGGCATCTACTACATATGAATTAGAG

>M6-PGSC0003DMG400014717-(rev)M6LRR-RLP_1178

ATAGTTCAATGACCACATAAGAAATCAACTTTTGGCATCTACTACATATGAATTAGAGATAATGTAGTTTAGAAATTTATTAATCTAAGACTGAAAATTGTTGAAACAAATATGTCAATG

>M6-PGSC0003DMG400014717-(rev)M6LRR-RLP_124

CTTTGAAACAAACAGTTCACAATTCATATATCAGAAAAATAATTTCCCAGCACGTCTAAAACAAGCAAAAATGGAAAAGAAAAACAAAACAGTCCCAAGGATAAATCTATATTGTCACAA

>M6-PGSC0003DMG400014717-(rev)M6LRR-RLP_1240

TGTAGTTTAGAAATTTATTAATCTAAGACTGAAAATTGTTGAAACAAATATGTCAATGTGTGATAACAAAACTATTTTTGCCACTTCGGTTAAGTCCTTGTTAGAATTAGTGTTGGTCTT

>M6-PGSC0003DMG400014717-(rev)M6LRR-RLP_1302

ATAACAAAACTATTTTTGCCACTTCGGTTAAGTCCTTGTTAGAATTAGTGTTGGTCTTTAATTATAATTTACGTAAAATATAAAATCCCATGAACCCGAAATAATAAACTCAAAATATGA

>M6-PGSC0003DMG400014717-(rev)M6LRR-RLP_1364

TATAATTTACGTAAAATATAAAATCCCATGAACCCGAAATAATAAACTCAAAATATGATGATGGTTTCCAGTAAAATATTCTCTTCACTTCAGTTTTTCACTCTTTTATATCTCCTTACA

>M6-PGSC0003DMG400014717-(rev)M6LRR-RLP_1426

GGTTTCCAGTAAAATATTCTCTTCACTTCAGTTTTTCACTCTTTTATATCTCCTTACAGTTACTTTTGCTTCCAATGAGGAAGCAACTGCCCTCTTGAAATGGAAAGCAACTTTCAAGAA

>M6-PGSC0003DMG400014717-(rev)M6LRR-RLP_1550

AATAATTCCTTATTGGCTTCATGGACGCCAAGTTCTAATGCATGCAGGGACTGGTATGGAGTTAAATGCTTTAATGGTAGGGTAAACACGTTGAATATTACAAGTACTAGTGTCAGTGCT

>M6-PGSC0003DMG400014717-(rev)M6LRR-RLP_1674

TCTATGCTTTTCCATTTTCATCTCTCCCTTTCTTGAGAATCTTAATCTTAGCATGAACAATTTCTCTGGCACCATTCCACCTGAGATTAGTAATCTCACTAATTTTGTCTATCTTGACTT

>M6-PGSC0003DMG400014717-(rev)M6LRR-RLP_1736

TCTCTGGCACCATTCCACCTGAGATTAGTAATCTCACTAATTTTGTCTATCTTGACTTGAACTTCAATCAGATTTCAGGAACAATCCCACCACAAATCGGATCACTAGCCAAGCGTCAGA

>M6-PGSC0003DMG400014717-(rev)M6LRR-RLP_186

AAGCAAAAATGGAAAAGAAAAACAAAACAGTCCCAAGGATAAATCTATATTGTCACAACTTAAAAAGATTTTAGCTCCTGTCTCACCACACCAGCCCTCTGCAATGTCCACAACACACGA

>M6-PGSC0003DMG400014717-(rev)M6LRR-RLP_248

AAAAGATTTTAGCTCCTGTCTCACCACACCAGCCCTCTGCAATGTCCACAACACACGAATCAGATCGATGCATAACCATCACAACGGTAAAAAGATTATGCAATCACACCTTGATCTTTG

>M6-PGSC0003DMG400014717-(rev)M6LRR-RLP_2480

TGGGGAATTTAAGAAATGTGCAAAGAATGTTTCTTGATGAAAACAATCTCACTGAAGAAATTCCTTTATCTCTGTGCAATTTAAAATCACTTGAAATTTTGTATTTGCGGAGAAACAACT

>M6-PGSC0003DMG400014717-(rev)M6LRR-RLP_2542

CCTTTATCTCTGTGCAATTTAAAATCACTTGAAATTTTGTATTTGCGGAGAAACAACTTGAAGAGAAAAGTTCCGCAATGTTTGGATAATATCAGTGGCCTCCAGGTTTTGACGATGTCA

>M6-PGSC0003DMG400014717-(rev)M6LRR-RLP_2604

GAGAAAAGTTCCGCAATGTTTGGATAATATCAGTGGCCTCCAGGTTTTGACGATGTCACGTAATAATCTTAGTGGAGAGATCCCTTCATCCATTTCCAATTTAAACTCACTACAAATTCT

>M6-PGSC0003DMG400014717-(rev)M6LRR-RLP_2666

ATAATCTTAGTGGAGAGATCCCTTCATCCATTTCCAATTTAAACTCACTACAAATTCTAGATTTGGGAAGAAATAGTCTGGAGGGAGCAATTCCAGAATGCTTTGGCAATATTAATACCC

>M6-PGSC0003DMG400014717-(rev)M6LRR-RLP_2728

TTGGGAAGAAATAGTCTGGAGGGAGCAATTCCAGAATGCTTTGGCAATATTAATACCCTTCAGGCTTTTGATATGCAGAACAACAAACTTTCTGGGACTCGTCCAACAATTTTAGCATTG

>M6-PGSC0003DMG400014717-(rev)M6LRR-RLP_2790

GGCTTTTGATATGCAGAACAACAAACTTTCTGGGACTCGTCCAACAATTTTAGCATTGGAAGTTCACTTAGAAGTTTCAATTTGCATGGCAATGAACTAGAGGGGGGAAATCCCTCGATC

>M6-PGSC0003DMG400014717-(rev)M6LRR-RLP_2976

CATGTGGTTGGGAACTTTGCTAGAGCTGAGAGTTTTAAGATTGACATCGAATAAATTGTATGGACCCATAAGATCTTCAGGGGCTGAAATCATGTTTCCTGATCTTCGAACCATAGATCT

>M6-PGSC0003DMG400014717-(rev)M6LRR-RLP_3038

GACCCATAAGATCTTCAGGGGCTGAAATCATGTTTCCTGATCTTCGAACCATAGATCTCTCTAACAATACCTTCTCGAAAGACTTACCAACGAGTTTGTTTCAACATTTGAAAGGCATGA

>M6-PGSC0003DMG400014717-(rev)M6LRR-RLP_310

GATCGATGCATAACCATCACAACGGTAAAAAGATTATGCAATCACACCTTGATCTTTGCCAACATGAGCTTAAACCTGTCAAAATTGTTGAGTGAGGCCCTCCTCTTCTGCACGATCAAC

>M6-PGSC0003DMG400014717-(rev)M6LRR-RLP_3162

AATTGATCAAACAATGATGGTACCAAGTTATGAAGGAGATGGAGATTACAAAGACTCGATAGTTGTTGTATCAAAGGGATTGAAGCTTGAAGTTGTGAGAATTTTGTCTTTGTACACAGT

>M6-PGSC0003DMG400014717-(rev)M6LRR-RLP_3224

TTGTTGTATCAAAGGGATTGAAGCTTGAAGTTGTGAGAATTTTGTCTTTGTACACAGTTATCGACCTTTCAAACAACAAATTTGAAGGACATATTCCTAGTGTTCTAGGAGATCTCATTG

>M6-PGSC0003DMG400014717-(rev)M6LRR-RLP_3596

TGGATTCCCCATTTCAAAAGGTTGTGGCAATGATCCTGTGTCAGAGACAAATTATATAGTATCTGCGCTAGCTGATCAAGAAAGCAATTCTAAATTTCTCAATGATTTTTGGAAAGCTGC

>M6-PGSC0003DMG400014717-(rev)M6LRR-RLP_372

CATGAGCTTAAACCTGTCAAAATTGTTGAGTGAGGCCCTCCTCTTCTGCACGATCAACTTCCTTTCTTAGGAACTGGACTCCCACTTGGTCTTCACATCTGCGAGGACATAAAGCAAATG

>M6-PGSC0003DMG400014717-(rev)M6LRR-RLP_434

TTTCTTAGGAACTGGACTCCCACTTGGTCTTCACATCTGCGAGGACATAAAGCAAATGAGAGTTAACCAAGTTCTTATGTAACCCTGCCGTTAGCCAAATGACTCAAATTTACCACTTCT

>M6-PGSC0003DMG400014717-(rev)M6LRR-RLP_496

TTAACCAAGTTCTTATGTAACCCTGCCGTTAGCCAAATGACTCAAATTTACCACTTCTGGCTGATGAAAACCGTTTTTCCTAATCAAAAATTCAAATTTGAATACTGCAATTCACCCAAC

>M6-PGSC0003DMG400014717-(rev)M6LRR-RLP_558

GATGAAAACCGTTTTTCCTAATCAAAAATTCAAATTTGAATACTGCAATTCACCCAACGTACACGTATCTTGATCTCTAGCTCGGCCACTGTTTCCAGTGATATGTATTTTCTGAGGGGA

>M6-PGSC0003DMG400014717-(rev)M6LRR-RLP_62

GAAGACGATCTGCAGCTGCAAGTGTGCATGAAGACAATATATGAAATAACTGAGCAAACATTCTTTGAAACAAACAGTTCACAATTCATATATCAGAAAAATAATTTCCCAGCACGTCTA

>M6-PGSC0003DMG400014717-(rev)M6LRR-RLP_620

ACGTATCTTGATCTCTAGCTCGGCCACTGTTTCCAGTGATATGTATTTTCTGAGGGGACTAATTAACGCATAGTCTAGTCACTTTAGTATTTAGTATTGTAACTCTAGTGTTACAAAGTG

>M6-PGSC0003DMG400014717-(rev)M6LRR-RLP_682

TTAACGCATAGTCTAGTCACTTTAGTATTTAGTATTGTAACTCTAGTGTTACAAAGTGTAAGTTTAGTAGTGACGATTCATGAGATTAACCTCGAAATTTTGTGATGACTGAAAATCAGA

>M6-PGSC0003DMG400014717-(rev)M6LRR-RLP_744

TTTAGTAGTGACGATTCATGAGATTAACCTCGAAATTTTGTGATGACTGAAAATCAGAGAAAGTAACACAAGATGCTGTTCTATATTATGAAAAAGATTAGAAAGATGAAACTCTACTTC

>M6-PGSC0003DMG400014717-(rev)M6LRR-RLP_806

GTAACACAAGATGCTGTTCTATATTATGAAAAAGATTAGAAAGATGAAACTCTACTTCTTTTTAAGCTTTGTATTCTAATTGATGGAATTTCTTTTCCGTATGATAGTTTTGTTTTACTT

>M6-PGSC0003DMG400014717-(rev)M6LRR-RLP_868

TAAGCTTTGTATTCTAATTGATGGAATTTCTTTTCCGTATGATAGTTTTGTTTTACTTGGAAAAAGTGGGAGTAGATGTGTAACACATTGGATAAAGACAATAAAAGGACAAAAGCACAA

>M6-PGSC0003DMG400014717-(rev)M6LRR-RLP_930

AAAGTGGGAGTAGATGTGTAACACATTGGATAAAGACAATAAAAGGACAAAAGCACAACAACCCAAATTTGACTAAAGAATGACAACATTTTCTCTTATAAAATTGAACAGACAAAGTGT

>M6-PGSC0003DMG400014717-(rev)M6LRR-RLP_992

CCAAATTTGACTAAAGAATGACAACATTTTCTCTTATAAAATTGAACAGACAAAGTGTGAATAAGAGTTTAGAAAAAAATGGGAGTAGATGTGTAACACATGGATAAAGACAATAAAAAG

>M6-PGSC0003DMG400014718-(rev)M6LRR-RLP_1152

GTGTCTGCGCTAGAAGATCAAGAAAGAAATTCTGAATTTTTCAATGATTTTTGGAAAGCAGCTCTGATGGGCTATGGCAGTCGACTATGTATTGGCTTATCCATAATATATTTCTTGATC

>M6-PGSC0003DMG400014718-(rev)M6LRR-RLP_128

AAGTGGCCTCCAGGTTCTAGTGATGTCACATAATGATCTCAGTGGAGAGATGCCTCCATCTATTTCCAATTTAACGTCACTACAAATTCTAGATTTGGGCAGAAATAATCTAGAGGGAGC

>M6-PGSC0003DMG400014718-(rev)M6LRR-RLP_192

TCCAATTTAACGTCACTACAAATTCTAGATTTGGGCAGAAATAATCTAGAGGGAGCAATTCCACAATGTTTTGGCAATAATAGTAGCCTCCAGGCATTTGATATGCAGAATAACAAACTT

>M6-PGSC0003DMG400014718-(rev)M6LRR-RLP_256

AATGTTTTGGCAATAATAGTAGCCTCCAGGCATTTGATATGCAGAATAACAAACTTTCTGGGACTCTTCCAACAAATTTTAGCATTGGAAGTTCACTTATAAGTCTCAACTTGCATGACA

>M6-PGSC0003DMG400014718-(rev)M6LRR-RLP_512

GACATCGAATAAACTGCATGGACCTATAAGATTTTCAGTGGCTGAAATCATGTTTCCTGATCTTCGAATCATAGATCTCTCTAGCAATGCCTTTTCGAAAGACTTACCAAAGAGTCTGTT

>M6-PGSC0003DMG400014718-(rev)M6LRR-RLP_576

CGAATCATAGATCTCTCTAGCAATGCCTTTTCGAAAGACTTACCAAAGAGTCTGTTTCAACAATTGAAAGGCATGAGGACTGTTGATCAAACAATGAAGACAACAAGTTATCTAGGAGGT

>M6-PGSC0003DMG400014718-(rev)M6LRR-RLP_64

TTGAAATGTTTTATATGCCGAGAAACAACTTGAAGGGAAAAGTTCCGCAATGTTTGGGTAATATAAGTGGCCTCCAGGTTCTAGTGATGTCACATAATGATCTCAGTGGAGAGATGCCTC

>M6-PGSC0003DMG400014718-(rev)M6LRR-RLP_640

TGAAAGGCATGAGGACTGTTGATCAAACAATGAAGACAACAAGTTATCTAGGAGGTAGATATAACTACCAAGACTTGGTAGTTGTTGTAACGAAGGGATTGAAGCTTGAAGTTGTGAAAA

>M6-PGSC0003DMG400014718-(rev)M6LRR-RLP_704

CTACCAAGACTTGGTAGTTGTTGTAACGAAGGGATTGAAGCTTGAAGTTGTGAAAATTTTGTCTTTGTACACAGTTATCGATCTTTCAAACAACAAATTTGAAGGACATATACCTAGTGT

>M6-PGSC0003DMG400014718-(rev)M6LRR-RLP_832

ATCTTATTGCGATTCGGGTACTTAATTTATCTCATAATGGGTTGCAAGGCTATATATCACCATCACTTGGAAGTTTATCTGTACTGGAATCATTAGACCTTTCATTTAACCGCCTTTCGG

>M6-PGSC0003DMG400014719-(rev)M6LRR-RLP_252

TTGATATGCAGAATAACAAACTTTCTGGGACTCTTCCAACAAATTTTAGCATTGGAAGTTCACTTATAAGTCTCAACTTGCATGACAATGAACTAGAGGGGGAAATCCCTCGGTCTTTGG

>M6-PGSC0003DMG400014719-(rev)M6LRR-RLP_441

TGGGAACTTTACCAGAGCTGAGAGTTCTAAGATTGACATCGAATAAACTGCATGGACCTATAAGATTTTCAGTGGCTGAAATCATGTTTCCTGATCTTCGAATCATAGATCTCTCTAGCA

>M6-PGSC0003DMG400014719-(rev)M6LRR-RLP_756

ACAACAAATTTGAAGGACATATACCTAGTGTTCTAGGAGATCTTATTGCGATTCGGGTACTTAATTTATCTCATAATGGGTTGCAAGGCTATATATCACCATCACTTGGAAGTTTATCTG

>M6-PGSC0003DMG400014736-(rev)M6LRR-RLP_120

GATCTTTCAGAAAATCAATTTTATGGTCAAATCCCACAAAGCTTTTCAAGTTTGTTCACCTTGAGCTACTTGAATCTATCTGATAACAATTTGTCAGGTATGATACCATTGAGCACTCAA

>M6-PGSC0003DMG400014738-(rev)M6LRR-RLP_458

GGATTGGTTTTACATTTCAATGGCTATAGGATTTGCACTAAGCTTTTGGGGTGTTTGTGGTTCATTGCTTTTCAAGAGATCATGGAGACATGCTTATTTTTGTTTTCTAGACCGTAGCTG

>M6-PGSC0003DMG400014739-(rev)M6LRR-RLP_0

AGTCCAAAATATTACACACCATGAAAGTGTTAGCAATAACACTATGGGTTCTCTTGTTTTTTATTTTGAAAAATACCAAGTTTGTCATTTGTGTAGGGATTTGTAGAGAAAATGAGCAAC

>M6-PGSC0003DMG400014739-(rev)M6LRR-RLP_1054

TACTTCTTTGATGGAATTGAGACTAGATGGTAATAAACTTAATGGAACTCTTCCGAAAAGTCTTGGACAACTTTCAATGCTAGAATACTTTTTCATTTCTAACAATAAGTTGGAAGGTGT

>M6-PGSC0003DMG400014739-(rev)M6LRR-RLP_1116

TTGGACAACTTTCAATGCTAGAATACTTTTTCATTTCTAACAATAAGTTGGAAGGTGTTGTAACAGAGAGTCATTTCTCTAAATTAACGCAATTACAACTCTTCTCTGCATCTAGAAATA

>M6-PGSC0003DMG400014739-(rev)M6LRR-RLP_1178

ACAGAGAGTCATTTCTCTAAATTAACGCAATTACAACTCTTCTCTGCATCTAGAAATAATCTGACTTTAAAAGTGAGTAGGAATTGGATTCCACCTTTTCAAGTCACAAACATTGCGATA

>M6-PGSC0003DMG400014739-(rev)M6LRR-RLP_1240

GACTTTAAAAGTGAGTAGGAATTGGATTCCACCTTTTCAAGTCACAAACATTGCGATAGGCGGGTGGAATATAGGCCCTTTGTTTCCAATGTGGCTCCGAACTCAGGAGATGATAATGAA

>M6-PGSC0003DMG400014739-(rev)M6LRR-RLP_1364

GACATATCAGATTGTGGAATACAAGGTGAGGTTCCAACTTGGTTTTGGAACTTATCTTCTCAAATTGTTTTTCTCAATCTTTCTCACAATCAATTTATTGGTGAGGTTCCAATCATCTCA

>M6-PGSC0003DMG400014739-(rev)M6LRR-RLP_1426

AATTGTTTTTCTCAATCTTTCTCACAATCAATTTATTGGTGAGGTTCCAATCATCTCAACAGCTTCGTGGTTAGTTGGATTGGGAGGTCCTTGGTTAATGTATTTAGGTTCCAACAATTT

>M6-PGSC0003DMG400014739-(rev)M6LRR-RLP_1488

CTTCGTGGTTAGTTGGATTGGGAGGTCCTTGGTTAATGTATTTAGGTTCCAACAATTTTAGTGGATCACTACCACGTATTTCAACGAGTATAACTGAGTTAGGTCTCTCGAACAATTTTT

>M6-PGSC0003DMG400014739-(rev)M6LRR-RLP_1550

GGATCACTACCACGTATTTCAACGAGTATAACTGAGTTAGGTCTCTCGAACAATTTTTTTTCAAAAGGTTTATCCAACTTCTTGTGTGAAGCAAAGAATGTATCTTATAATTTGGAGATC

>M6-PGSC0003DMG400014739-(rev)M6LRR-RLP_1612

AAAAGGTTTATCCAACTTCTTGTGTGAAGCAAAGAATGTATCTTATAATTTGGAGATCTTAAATCTCGGGGGAAATGATTTATCAGAAGAAATTCCTGATTGTTGGATGAATTGGCCTGA

>M6-PGSC0003DMG400014739-(rev)M6LRR-RLP_1674

ATCTCGGGGGAAATGATTTATCAGAAGAAATTCCTGATTGTTGGATGAATTGGCCTGAGTTGGAAGTTTTAATCTTGAGGGACAATAATTTGATTGGAAGCTTACCAAGATCTATGGAGG

>M6-PGSC0003DMG400014739-(rev)M6LRR-RLP_1736

GAAGTTTTAATCTTGAGGGACAATAATTTGATTGGAAGCTTACCAAGATCTATGGAGGTTTTGAGTAATTTGCTATCATTAGACTTACGAAGAAATAGACTTAACGGTCCGTTTCCTTCA

>M6-PGSC0003DMG400014739-(rev)M6LRR-RLP_1798

GAGTAATTTGCTATCATTAGACTTACGAAGAAATAGACTTAACGGTCCGTTTCCTTCATCCTTGAAAAATTGTACGAAACTGCATAAAATAGATTTAGCCGAGAATGAGTTTGCTGGGGG

>M6-PGSC0003DMG400014739-(rev)M6LRR-RLP_186

GAAAAGATTGTTGTGAATGGGATGGAGTTGTGTGCAACAATCTAACTCGTCGTGTGATTGAACTACGCATAAGTTATCTAAGGATCGATAACCTAGAGTGGTTGTCAAGTCTTTTGAGTC

>M6-PGSC0003DMG400014739-(rev)M6LRR-RLP_1860

TGAAAAATTGTACGAAACTGCATAAAATAGATTTAGCCGAGAATGAGTTTGCTGGGGGACTACCATCTTAGTTGGGAATGAGGTTTTCAACCTTGATAGTATTTATCCTTCGGTCCAATA

>M6-PGSC0003DMG400014739-(rev)M6LRR-RLP_1922

CCATCTTAGTTGGGAATGAGGTTTTCAACCTTGATAGTATTTATCCTTCGGTCCAATAAATTCAATGGTGAATTGCCTCAAGAACTTTGTCAACTCAAAGATCTTCAGATCTTAGACCTT

>M6-PGSC0003DMG400014739-(rev)M6LRR-RLP_1984

CAATGGTGAATTGCCTCAAGAACTTTGTCAACTCAAAGATCTTCAGATCTTAGACCTTGCAAACAATACATTTGTTGGAATCATACCAAGGTGTATTGGTAATTTCAGTGCAATGGTCAA

>M6-PGSC0003DMG400014739-(rev)M6LRR-RLP_2046

ACAATACATTTGTTGGAATCATACCAAGGTGTATTGGTAATTTCAGTGCAATGGTCAAAGGAAAAAAGGAAGTGGGAGATGATGATGAGTTAAATTATTCCTACTTTTTGGGAGTTTTGA

>M6-PGSC0003DMG400014739-(rev)M6LRR-RLP_248

CTACGCATAAGTTATCTAAGGATCGATAACCTAGAGTGGTTGTCAAGTCTTTTGAGTCTTGAGAACCTGGAAATGTATCATGTGGATCTCCACAAAGCAACTAACTGGCTACAGGTTATT

>M6-PGSC0003DMG400014739-(rev)M6LRR-RLP_2728

TTATTTTTGTTTTCTAGACCGTAGCTGGGAAATGTTGCTTGCAAAGTTATCAGTATGTTGAGAGATATCATTAAGGATATTGCTATCATCAAAAGTTAGTTTGTGTGCTTACATTTCTCA

>M6-PGSC0003DMG400014739-(rev)M6LRR-RLP_2790

AGATATCATTAAGGATATTGCTATCATCAAAAGTTAGTTTGTGTGCTTACATTTCTCATGTTAGCAAACATGTTGTTTCATTTTGTATTCAAATGAATGTTTCTGAAGATGTTCAAACAT

>M6-PGSC0003DMG400014739-(rev)M6LRR-RLP_310

GAACCTGGAAATGTATCATGTGGATCTCCACAAAGCAACTAACTGGCTACAGGTTATTAACATGCTTCCTTCTTTGGTTGATCTAACGTTATCCAATTGTAGTCTTCATAATATAACAAA

>M6-PGSC0003DMG400014739-(rev)M6LRR-RLP_372

TGCTTCCTTCTTTGGTTGATCTAACGTTATCCAATTGTAGTCTTCATAATATAACAAATCTACTTCATCACAATCTTTCTTCACTTGAAACCCTCGATCTTTCTAGTACTAAGTTTAATT

>M6-PGSC0003DMG400014739-(rev)M6LRR-RLP_434

CTTCATCACAATCTTTCTTCACTTGAAACCCTCGATCTTTCTAGTACTAAGTTTAATTCACCTATTCCCAATTGGGTTTTCAACCTTTCCAATCTTGTTTCTCTTAATTTGAGTGATTGT

>M6-PGSC0003DMG400014739-(rev)M6LRR-RLP_496

TATTCCCAATTGGGTTTTCAACCTTTCCAATCTTGTTTCTCTTAATTTGAGTGATTGTAATTTTACTGGCCCATTTCCCGACTTTCCTGTTAACTTGACTTCTCTCACTACCTTCAAGGC

>M6-PGSC0003DMG400014739-(rev)M6LRR-RLP_558

TTACTGGCCCATTTCCCGACTTTCCTGTTAACTTGACTTCTCTCACTACCTTCAAGGCTTTTGAAAACTCTTTCAATTGTCTTTTACCCGAATGGTTGTTTGATTTAAGTAATCTAGAAC

>M6-PGSC0003DMG400014739-(rev)M6LRR-RLP_62

ATTTTGAAAAATACCAAGTTTGTCATTTGTGTAGGGATTTGTAGAGAAAATGAGCAACGAGCATTGGAGAGCTTGAAGAAAGAAGTAGATGATCCCTTGGATATTCCCTCTTCTTGGGTT

>M6-PGSC0003DMG400014739-(rev)M6LRR-RLP_620

GAAAACTCTTTCAATTGTCTTTTACCCGAATGGTTGTTTGATTTAAGTAATCTAGAACATCTTAACCTCCGTTCTAGTGGTATTAAAGGTGCAATACCAAATAAGAGTGGAAACACAACA

>M6-PGSC0003DMG400014739-(rev)M6LRR-RLP_682

TAACCTCCGTTCTAGTGGTATTAAAGGTGCAATACCAAATAAGAGTGGAAACACAACAACACTTAAATATCTCGATCTTTTTGATAATAACCTTAACTCCACCATACCAAATTGGTTATA

>M6-PGSC0003DMG400014739-(rev)M6LRR-RLP_744

TTAAATATCTCGATCTTTTTGATAATAACCTTAACTCCACCATACCAAATTGGTTATATCAATGCAAAGATCTAGAATCACTTTATCTTACTGAAAACCGTCTTGAGGGAACAGTTTCAA

>M6-PGSC0003DMG400014739-(rev)M6LRR-RLP_806

TGCAAAGATCTAGAATCACTTTATCTTACTGAAAACCGTCTTGAGGGAACAGTTTCAAGTCTGATTTCAAATTTGAGCTCAATAACTAATATTGACCTTTCTGGTAATATGCTTACTGGA

>M6-PGSC0003DMG400014739-(rev)M6LRR-RLP_868

GATTTCAAATTTGAGCTCAATAACTAATATTGACCTTTCTGGTAATATGCTTACTGGAAAGTTACCAAATGTAATTGGAATTTTAGGGAAATTAGAATTTCTTTCTCTTTCAAAAAATCA

>M6-PGSC0003DMG400014739-(rev)M6LRR-RLP_930

TACCAAATGTAATTGGAATTTTAGGGAAATTAGAATTTCTTTCTCTTTCAAAAAATCAATTTGAGGGACAAGTATCTGAATTATTCAAAGGTAGGAGTAATTTCTTTTCAAGGGGATTGG

>M6-PGSC0003DMG400014739-(rev)M6LRR-RLP_992

GAGGGACAAGTATCTGAATTATTCAAAGGTAGGAGTAATTTCTTTTCAAGGGGATTGGGAAATACTTCTTTGATGGAATTGAGACTAGATGGTAATAAACTTAATGGAACTCTTCCGAAA

>M6-PGSC0003DMG400014740-(rev)M6LRR-RLP_0

AGGCCCTTTGTTTCCAATGTGGCTCCGAACTCAGGAGATGATAATGAATGTTGACATATCAGATTGTGGAATACAAGGTGAGGTTCCAACTTGGTTTTGGAACTTATCTTCTCAAATTGT

>M6-PGSC0003DMG400014829M6LRR-RLP_0

CAAATATCGAAATTTTCCTAAACTACTTTGCACGAAAAAACAAAGAGGAGGAGGAAGAAATAGTATTCGTAGATTACATCTGAGCTTGACAACAAATTATGCAAGATTTTGAGTGAAACA

>M6-PGSC0003DMG400014829M6LRR-RLP_1054

CAATGTTGTCAAATGCTCGAAGAGTTGATTCTTCGTGATCATAGGATGGAGGGTGGTTGGTTGTCTGCTCTTTCCTATTGTGAAAATTTGAACACCTTGAGGTTTATGTCTTGTAAGAGT

>M6-PGSC0003DMG400014829M6LRR-RLP_1116

GTCTGCTCTTTCCTATTGTGAAAATTTGAACACCTTGAGGTTTATGTCTTGTAAGAGTATTGATCATTGTGGATGGTTTGATGAAGATGTAGGATCTTGCCCAACACTTGAAAGATTACA

>M6-PGSC0003DMG400014829M6LRR-RLP_1178

ATCATTGTGGATGGTTTGATGAAGATGTAGGATCTTGCCCAACACTTGAAAGATTACATTTCGAGAAATGTCAATTAAGAAATAAGGAGAGTTTGAGAACGTTGTTTCTACTATGTCAAG

>M6-PGSC0003DMG400014829M6LRR-RLP_124

GACCTTTCATGGTGTTTCTTGTTGGCTGCTAAGAAAACAATGGCGTTTTCACCTGAAAAATCAAATTTCAGGTCACCATTGAAAGGGCCACCATCTCCTAGCTGGAATGATTTGTTGCTC

>M6-PGSC0003DMG400014829M6LRR-RLP_1240

GAGAAATGTCAATTAAGAAATAAGGAGAGTTTGAGAACGTTGTTTCTACTATGTCAAGAAGTTAGAGAGGTTATTTTCCACAACTGTTGGGGACTGGATAATGAAATGTTCAGTCTATCC

>M6-PGSC0003DMG400014829M6LRR-RLP_1302

TAGAGAGGTTATTTTCCACAACTGTTGGGGACTGGATAATGAAATGTTCAGTCTATCCACTGGTCTAAGGTATTGGTTATATTATTCTGTTCTCGTTTACTTTGCTTGCTGTTGCTGGAA

>M6-PGSC0003DMG400014829M6LRR-RLP_1364

GTCTAAGGTATTGGTTATATTATTCTGTTCTCGTTTACTTTGCTTGCTGTTGCTGGAAAATTTCGTGCTGGATTAGCATTTTGTTATGCATTTGACATACATGAACTTATTGAGTTTAGG

>M6-PGSC0003DMG400014829M6LRR-RLP_1426

TCGTGCTGGATTAGCATTTTGTTATGCATTTGACATACATGAACTTATTGAGTTTAGGTTTTATGACCTTGAAATGTAGTTCCTTATTTTTGTTAAAAATGACTTTTTAGTGTTGCTGTA

>M6-PGSC0003DMG400014829M6LRR-RLP_1488

ATGACCTTGAAATGTAGTTCCTTATTTTTGTTAAAAATGACTTTTTAGTGTTGCTGTAACCCCCATCCCCCACCCCTCGGGTATAATAGACGACCAAATGAAAGGATAGTTTGCTTGTAA

>M6-PGSC0003DMG400014829M6LRR-RLP_1550

CCATCCCCCACCCCTCGGGTATAATAGACGACCAAATGAAAGGATAGTTTGCTTGTAATAGTGCATTTTATTTTGGTCATAGTATACCCATTGATCTTGTTTTCTTGGTAAAAGGATAAG

>M6-PGSC0003DMG400014829M6LRR-RLP_1612

GCATTTTATTTTGGTCATAGTATACCCATTGATCTTGTTTTCTTGGTAAAAGGATAAGTGTGATAAATTTCCTTAGCTATTTGTGTTGGGGATAGATTGTTTTTTTAAGGCCAGTTTGTA

>M6-PGSC0003DMG400014829M6LRR-RLP_1674

ATAAATTTCCTTAGCTATTTGTGTTGGGGATAGATTGTTTTTTTAAGGCCAGTTTGTAGATGCATTTGTAATCATGCACCGTCTTAGTGTTTCAGTTATTAATGTTAAAATAGCCTTACC

>M6-PGSC0003DMG400014829M6LRR-RLP_1736

CATTTGTAATCATGCACCGTCTTAGTGTTTCAGTTATTAATGTTAAAATAGCCTTACCATTTCTCGAAGAGTATACACTTCCATCTTGCATTAGAGAAAACACATAATGTTGATGGCATG

>M6-PGSC0003DMG400014829M6LRR-RLP_1798

CTCGAAGAGTATACACTTCCATCTTGCATTAGAGAAAACACATAATGTTGATGGCATGACAATTATGTTTCAATTCATTATTGATGCTTGATTTATCAATATCATGTGTGCTATCTTGGA

>M6-PGSC0003DMG400014829M6LRR-RLP_186

AAATTTCAGGTCACCATTGAAAGGGCCACCATCTCCTAGCTGGAATGATTTGTTGCTCAAGAACAAGAAAGCACTCAATCATGTCCTTTTCACTATGCATCTTCAATCCAGGTCCAATAA

>M6-PGSC0003DMG400014829M6LRR-RLP_1860

TTATGTTTCAATTCATTATTGATGCTTGATTTATCAATATCATGTGTGCTATCTTGGATTATAAAGAGTCCAGTTTTGAGTTTTATACATTGAACTTGCTTCAGTTCTGACTGGCAATTT

>M6-PGSC0003DMG400014829M6LRR-RLP_1922

AAAGAGTCCAGTTTTGAGTTTTATACATTGAACTTGCTTCAGTTCTGACTGGCAATTTGGTATCATATTTTGTTTTGCATTCACCTGGAGCAACAGTAAAGTTGTTTCCGTGTGACCTAT

>M6-PGSC0003DMG400014829M6LRR-RLP_2046

CACCAATTTGAGCCGTGGAAGTAGCCACTAATGCTTGCATTAGGAGGGTATCCCCTTGCGTGAACAATGGATAATTTATGCACAAGGCTGCCCTTAAAGTTCACTTGAGTAAAGGGCCTG

>M6-PGSC0003DMG400014829M6LRR-RLP_2108

AACAATGGATAATTTATGCACAAGGCTGCCCTTAAAGTTCACTTGAGTAAAGGGCCTGTAATTAACTAAATATAACATTATGTTAGGAATTTGGGTTTCACAGCTTCATTTTGGTGTTAT

>M6-PGSC0003DMG400014829M6LRR-RLP_2170

TAACTAAATATAACATTATGTTAGGAATTTGGGTTTCACAGCTTCATTTTGGTGTTATTTTCAACAGGAGAGTGAAGTCTCTTTGCCTAGAAAGCTGTTCACTGCTCACAACTGAAGGCC

>M6-PGSC0003DMG400014829M6LRR-RLP_2232

AACAGGAGAGTGAAGTCTCTTTGCCTAGAAAGCTGTTCACTGCTCACAACTGAAGGCCTCGAATCTGCACTCCTTTCATGGAAGGAGATCCAGAGCCTCAAGGTGATATCATGTGGCAAT

>M6-PGSC0003DMG400014829M6LRR-RLP_2294

ATCTGCACTCCTTTCATGGAAGGAGATCCAGAGCCTCAAGGTGATATCATGTGGCAATATAAAGGATAGTGAAATCAGTCCAGCACTTTCTACCTTGTTCTCCACGCTAAAAGATTTTCA

>M6-PGSC0003DMG400014829M6LRR-RLP_2356

AGGATAGTGAAATCAGTCCAGCACTTTCTACCTTGTTCTCCACGCTAAAAGATTTTCAGTGGAGACCAGACACAAAATCTCTTCTTTCAGCTGGTCTTGTGGGAACTGGCATGCGAAAAA

>M6-PGSC0003DMG400014829M6LRR-RLP_2418

AGACCAGACACAAAATCTCTTCTTTCAGCTGGTCTTGTGGGAACTGGCATGCGAAAAAGAGGCAGTAAATTATTCAAGAAGACGTGCGACTGGAAGTCACTGCCTGGTGCATAGACTGGC

>M6-PGSC0003DMG400014829M6LRR-RLP_248

ACAAGAAAGCACTCAATCATGTCCTTTTCACTATGCATCTTCAATCCAGGTCCAATAAGCAAACCCCATCTTCATCCCCTTCTCCACAACTTCGCAAATCCCTAGATTTCACTCTTCCCT

>M6-PGSC0003DMG400014829M6LRR-RLP_2480

CAGTAAATTATTCAAGAAGACGTGCGACTGGAAGTCACTGCCTGGTGCATAGACTGGCTTCCTCATCATGCATTCAGACAATAAATTATTCAAGAAGACATGCGACTGGAAGTCACTGCC

>M6-PGSC0003DMG400014829M6LRR-RLP_2542

TCATCATGCATTCAGACAATAAATTATTCAAGAAGACATGCGACTGGAAGTCACTGCCTGGTGTATAGACTGGCTTTCTCATCATGCATTCAGACAACTCTACTAGATTTTACCTCCTGA

>M6-PGSC0003DMG400014829M6LRR-RLP_2604

GTATAGACTGGCTTTCTCATCATGCATTCAGACAACTCTACTAGATTTTACCTCCTGATTTATGCATATGAGAACTCTAACACATGAAAGAGGTCAGTGCGACACTTTGCTCCAATACTG

>M6-PGSC0003DMG400014829M6LRR-RLP_2666

TGCATATGAGAACTCTAACACATGAAAGAGGTCAGTGCGACACTTTGCTCCAATACTGTCAGCAATGAGGCCTTGATGTCATTATCCTGTATTTGTAGTCGCTTAAGTATGTAACGTAGG

>M6-PGSC0003DMG400014829M6LRR-RLP_2728

CAATGAGGCCTTGATGTCATTATCCTGTATTTGTAGTCGCTTAAGTATGTAACGTAGGAACTATAGGATCGAAGATCGGCACTTTCTCAGGGTAGAGATGGAGAAAGAGGCCTAAGTTAA

>M6-PGSC0003DMG400014829M6LRR-RLP_2790

ATAGGATCGAAGATCGGCACTTTCTCAGGGTAGAGATGGAGAAAGAGGCCTAAGTTAATGTATAGAGCTCGATGCTGTAACCTCATGAGCATTCAGCTAAAGCCATCTCGCTGTATCTTT

>M6-PGSC0003DMG400014829M6LRR-RLP_2852

TAGAGCTCGATGCTGTAACCTCATGAGCATTCAGCTAAAGCCATCTCGCTGTATCTTTGTCAAGAAGCTTAATGGTATTTCTGTATTTGGAACCCCAAAAACTCTTAAAAGGGGTTTTCC

>M6-PGSC0003DMG400014829M6LRR-RLP_2914

AGAAGCTTAATGGTATTTCTGTATTTGGAACCCCAAAAACTCTTAAAAGGGGTTTTCCTATCTACGATGTTATAAAGAATTTTGGCACCTTTTATGCCATACTACTGCCTTCCACAATTG

>M6-PGSC0003DMG400014829M6LRR-RLP_2976

TACGATGTTATAAAGAATTTTGGCACCTTTTATGCCATACTACTGCCTTCCACAATTGTGTGAAAGCATATTTGATATGAACTATACCAGCAAGCATGTCATTTGTGGAGGAGACGAGAA

>M6-PGSC0003DMG400014829M6LRR-RLP_3011

CATACTACTGCCTTCCACAATTGTGTGAAAGCATATTTGATATGAACTATACCAGCAAGCATGTCATTTGTGGAGGAGACGAGAACCAGAATAATTTTCTACATCAATGGCATCTATGTC

>M6-PGSC0003DMG400014829M6LRR-RLP_310

ACCCCATCTTCATCCCCTTCTCCACAACTTCGCAAATCCCTAGATTTCACTCTTCCCTCACTTGTATCCGATCCAATTTCTCTTCTTTCTGATCAAATACTTCTTTCCATTCTCTCTAAA

>M6-PGSC0003DMG400014829M6LRR-RLP_372

TGTATCCGATCCAATTTCTCTTCTTTCTGATCAAATACTTCTTTCCATTCTCTCTAAACTCCCTGATTCACAAAGAAATTGTAACTCTCTTGTATCTAAACGTTGGCTCAATCTCCAAGG

>M6-PGSC0003DMG400014829M6LRR-RLP_434

CTGATTCACAAAGAAATTGTAACTCTCTTGTATCTAAACGTTGGCTCAATCTCCAAGGTCGTCTGGTACGGTCGATTAAGCTTCTTGATTGGGATTTCCTTGTTTCGGGTCGGTTGTTTG

>M6-PGSC0003DMG400014829M6LRR-RLP_496

CTGGTACGGTCGATTAAGCTTCTTGATTGGGATTTCCTTGTTTCGGGTCGGTTGTTTGTTCGGTTCCCTAATCTTATCCATGTTGATTTGGTTAATGGGTCTCTCATTTCACCGGGGAAT

>M6-PGSC0003DMG400014829M6LRR-RLP_558

GTTCCCTAATCTTATCCATGTTGATTTGGTTAATGGGTCTCTCATTTCACCGGGGAATGCAGGTGTTTTTTGCTCTAACCGGTTGCTTTCTTCTCACGTGGATTGTAGCGCTGATGCGAA

>M6-PGSC0003DMG400014829M6LRR-RLP_62

GTATTCGTAGATTACATCTGAGCTTGACAACAAATTATGCAAGATTTTGAGTGAAACAACCAGACCTTTCATGGTGTTTCTTGTTGGCTGCTAAGAAAACAATGGCGTTTTCACCTGAAA

>M6-PGSC0003DMG400014829M6LRR-RLP_620

GTGTTTTTTGCTCTAACCGGTTGCTTTCTTCTCACGTGGATTGTAGCGCTGATGCGAAAGATTGGTTTCTTAAGGAAAGATTTGTGTTGCCTTCTGATGAGATTGATAAGGGGTTGAGAA

>M6-PGSC0003DMG400014829M6LRR-RLP_682

TGGTTTCTTAAGGAAAGATTTGTGTTGCCTTCTGATGAGATTGATAAGGGGTTGAGAATACTAGCAAGTGGGTGTCCTAATTTGCGTAGATTGTTGGTGGTAAATGCTAGTGAATTGGGA

>M6-PGSC0003DMG400014829M6LRR-RLP_744

AGCAAGTGGGTGTCCTAATTTGCGTAGATTGTTGGTGGTAAATGCTAGTGAATTGGGATTGTTGGGGGTAGCTGAGGAATGTCCAACTTTGCAAGTGTTGGAATTGCACAGATGTAATGA

>M6-PGSC0003DMG400014829M6LRR-RLP_806

TGGGGGTAGCTGAGGAATGTCCAACTTTGCAAGTGTTGGAATTGCACAGATGTAATGATCAGGTTCTTCGTGGTATAGCGGCCTGTCAGAACTTGCAGATATTGAGATTGATTGGAAATG

>M6-PGSC0003DMG400014829M6LRR-RLP_868

GTTCTTCGTGGTATAGCGGCCTGTCAGAACTTGCAGATATTGAGATTGATTGGAAATGTTGATGGGTTTTATAAATCTTTGGTTACTGATATTGGATTGACTATTTTAGCTCAGGGGTGT

>M6-PGSC0003DMG400014829M6LRR-RLP_930

TGGGTTTTATAAATCTTTGGTTACTGATATTGGATTGACTATTTTAGCTCAGGGGTGTAAGAGGTTGGTGAAGCTGGAGCTGAGTGGTTGTGAAGGGAGCTATGAAGGGATTAAGGCAAT

>M6-PGSC0003DMG400014829M6LRR-RLP_992

GGTTGGTGAAGCTGGAGCTGAGTGGTTGTGAAGGGAGCTATGAAGGGATTAAGGCAATAGCACAATGTTGTCAAATGCTCGAAGAGTTGATTCTTCGTGATCATAGGATGGAGGGTGGTT

>M6-PGSC0003DMG400014875-(rev)M6RLKmalectin_1054

TGATAGATTTGAACAATTTGCAGTTCCTTTTTATGTTGATTTTGTGGTGGATTCAGATGGTTCTGGTTTTATGAATATCTCAGTTGGTCCTCGGAATGATTCAAAGTCTATAAATGCTTT

>M6-PGSC0003DMG400014875-(rev)M6RLKmalectin_1116

CTGGTTTTATGAATATCTCAGTTGGTCCTCGGAATGATTCAAAGTCTATAAATGCTTTTTTGAATGGGGTGGAAATAATGGAGTTGATTAATGATCGGGTTGAGTGTAGGGGAGAAAAGA

>M6-PGSC0003DMG400014875-(rev)M6RLKmalectin_124

CACCTTCCCTGAAAAACACTTCATCAATTGTGGTTCTGATTCTCCCATCGTACTCAACCGCGGCGGCGGCAGCACCACTTTTGCCGGCGACTTGAAATTCGACCCCAACCACAGTGATTC

>M6-PGSC0003DMG400014875-(rev)M6RLKmalectin_1302

TGGTGATCTTGTGTTGTTTGAAACCAAGAAAAGCACAAACGGTTGAGAACTCGAGTTGGTGGCCAAGGATGAGTGTTAATGCTGGAAGTACAGAGGATAGGACTCCTGTAAGAACTCCAA

>M6-PGSC0003DMG400014875-(rev)M6RLKmalectin_1364

CCAAGGATGAGTGTTAATGCTGGAAGTACAGAGGATAGGACTCCTGTAAGAACTCCAATTGGTAGCACTACTCCTGATATGAACCTTGGGTTAAAGATATCTTTCGCAGAGATTTTGCAT

>M6-PGSC0003DMG400014875-(rev)M6RLKmalectin_1426

TAGCACTACTCCTGATATGAACCTTGGGTTAAAGATATCTTTCGCAGAGATTTTGCATGCTACTAATAAGTTTGATCCCAAATTTATGATTGGTGAGGGTGGTTTTGGGAAAGTTTACAA

>M6-PGSC0003DMG400014875-(rev)M6RLKmalectin_1488

CTAATAAGTTTGATCCCAAATTTATGATTGGTGAGGGTGGTTTTGGGAAAGTTTACAAAGGAACTTTACACAGTGGTGTTAAAGTGGCTGTGAAAAGAAGCGAAGCTGGACATGGGCAGG

>M6-PGSC0003DMG400014875-(rev)M6RLKmalectin_1550

ACTTTACACAGTGGTGTTAAAGTGGCTGTGAAAAGAAGCGAAGCTGGACATGGGCAGGGTCTTATGGAATTTCAAACTCAGATAATGCTCTTGTCCAAAATCCGCCATCAGCATCTTGTT

>M6-PGSC0003DMG400014875-(rev)M6RLKmalectin_1612

TATGGAATTTCAAACTCAGATAATGCTCTTGTCCAAAATCCGCCATCAGCATCTTGTTTCATTGATTGGTTATTGTGATGAACGAGATGAGATGATACTTGTTTATGAATTCATGGCGAA

>M6-PGSC0003DMG400014875-(rev)M6RLKmalectin_1674

TGATTGGTTATTGTGATGAACGAGATGAGATGATACTTGTTTATGAATTCATGGCGAAGGGAACTTTGAGAGAGCATTTGTACAGTTCGAATGAAGATCTTGGTAAATCATCTTCACGGT

>M6-PGSC0003DMG400014875-(rev)M6RLKmalectin_1736

ACTTTGAGAGAGCATTTGTACAGTTCGAATGAAGATCTTGGTAAATCATCTTCACGGTCAAAATTATCGTGGGATCAAAGGCTTCAAATTTGTATAGGTGCAGCTAATGGTCTACAGTAC

>M6-PGSC0003DMG400014875-(rev)M6RLKmalectin_1798

ATTATCGTGGGATCAAAGGCTTCAAATTTGTATAGGTGCAGCTAATGGTCTACAGTACCTTCATGCCGGTTTACCTGAGCCAACTATTCACAGGGATATTAAGTCAACAAACATACTTCT

>M6-PGSC0003DMG400014875-(rev)M6RLKmalectin_186

GCGGCGGCAGCACCACTTTTGCCGGCGACTTGAAATTCGACCCCAACCACAGTGATTCCGTTAAAGACCACACCACTACTACAGGTTTAGCTGGAATCTATCAAACCGCCAGAAAGTTCA

>M6-PGSC0003DMG400014875-(rev)M6RLKmalectin_1860

ATGCCGGTTTACCTGAGCCAACTATTCACAGGGATATTAAGTCAACAAACATACTTCTTGATGAAGATTATGTTGCTAAAGTTGCTGATTTCGGTCTCTCCAAATCAGGTCAACCTGAAG

>M6-PGSC0003DMG400014875-(rev)M6RLKmalectin_1922

GAAGATTATGTTGCTAAAGTTGCTGATTTCGGTCTCTCCAAATCAGGTCAACCTGAAGAAACACACATTGTTACTGATGTTAAAGGTAGCTTCGGTTATCTTGATCCCGAGTACCTGAAA

>M6-PGSC0003DMG400014875-(rev)M6RLKmalectin_1984

ACACATTGTTACTGATGTTAAAGGTAGCTTCGGTTATCTTGATCCCGAGTACCTGAAATCAATGCAACTAACTCAAAAGTCTGATGTCTATTCTTTTGGAGTTGTACTTTTTGAAGTACT

>M6-PGSC0003DMG400014875-(rev)M6RLKmalectin_2046

TGCAACTAACTCAAAAGTCTGATGTCTATTCTTTTGGAGTTGTACTTTTTGAAGTACTTTGTGCTAGACCAGCTGTTGATAATATGCTTCCAAGAAATCAAATGAACTTAGCTGAATGGG

>M6-PGSC0003DMG400014875-(rev)M6RLKmalectin_2108

GCTAGACCAGCTGTTGATAATATGCTTCCAAGAAATCAAATGAACTTAGCTGAATGGGGACTTTCTTGGCTAAAAGAAAACCAGATCGAAAAGATAATTGATCCATTTCTTGTAGGTAAA

>M6-PGSC0003DMG400014875-(rev)M6RLKmalectin_2170

TTCTTGGCTAAAAGAAAACCAGATCGAAAAGATAATTGATCCATTTCTTGTAGGTAAAATTAACCCAAACTCTCTGAGGAAATTCGGAGAAATAGCAGAGAAGTGTCTGCAAGAGAATGG

>M6-PGSC0003DMG400014875-(rev)M6RLKmalectin_2232

ACCCAAACTCTCTGAGGAAATTCGGAGAAATAGCAGAGAAGTGTCTGCAAGAGAATGGCACTGATAGGCCTAAGATGATGGATGTGTTGTGGGACTTGGATTACGCTCGACAGCTTCAAC

>M6-PGSC0003DMG400014875-(rev)M6RLKmalectin_2294

GATAGGCCTAAGATGATGGATGTGTTGTGGGACTTGGATTACGCTCGACAGCTTCAACACCCTACAATGCCCCAACAATCTCATGAGGATACCAACTCCGATGTTTCGTGGCAAGTGGCA

>M6-PGSC0003DMG400014875-(rev)M6RLKmalectin_2356

TACAATGCCCCAACAATCTCATGAGGATACCAACTCCGATGTTTCGTGGCAAGTGGCATTGCCTGGTATCAATCGCTTACCTTCTATAAATGTAAGCACAAGCTTTGTCAGTGTCACTGA

>M6-PGSC0003DMG400014875-(rev)M6RLKmalectin_2418

CTGGTATCAATCGCTTACCTTCTATAAATGTAAGCACAAGCTTTGTCAGTGTCACTGATAGCGAGGCTTTTTCAGAATTGAGGATCGATGAAACCAGATGATTTGTGGCCTCGTTGTTGT

>M6-PGSC0003DMG400014875-(rev)M6RLKmalectin_248

AAAGACCACACCACTACTACAGGTTTAGCTGGAATCTATCAAACCGCCAGAAAGTTCACTAAATCTGGGTCATACAATCTTCGAATTGATGAAAATGGGGTTTACATGGTACGGCTTCAT

>M6-PGSC0003DMG400014875-(rev)M6RLKmalectin_2480

GAGGCTTTTTCAGAATTGAGGATCGATGAAACCAGATGATTTGTGGCCTCGTTGTTGTTAGCATTCATCTGATGATATGTAATTTCATCTTATTCTACTTCATTCTGTCTAATGTAATTG

>M6-PGSC0003DMG400014875-(rev)M6RLKmalectin_2542

ATTCATCTGATGATATGTAATTTCATCTTATTCTACTTCATTCTGTCTAATGTAATTGTGTATCAAAACAGTGTGTAATAGTCAGTGTTCAGTTTTAATATAGGCAAATGCCTCTTGACA

>M6-PGSC0003DMG400014875-(rev)M6RLKmalectin_2602

TATCAAAACAGTGTGTAATAGTCAGTGTTCAGTTTTAATATAGGCAAATGCCTCTTGACAATGCATCTGTATTGATAGTATATAATATTTTGATAATAAAGTCTCGTGTCTATTCTCCCA

>M6-PGSC0003DMG400014875-(rev)M6RLKmalectin_310

ATCTGGGTCATACAATCTTCGAATTGATGAAAATGGGGTTTACATGGTACGGCTTCATTTCTTTCCTTTCGTTTCTCTATTCGATGCGAAATTCAGTGTTTCCGCTTCTGGGTTTTTGCT

>M6-PGSC0003DMG400014875-(rev)M6RLKmalectin_372

TTCCTTTCGTTTCTCTATTCGATGCGAAATTCAGTGTTTCCGCTTCTGGGTTTTTGCTATTGTCCAATTTTAGCATACCTAGAAACTTAATAACTTCCCCTGTTATCAAAGAGTTTCTGG

>M6-PGSC0003DMG400014875-(rev)M6RLKmalectin_434

TCCAATTTTAGCATACCTAGAAACTTAATAACTTCCCCTGTTATCAAAGAGTTTCTGGTTCCTGTACTAAAAGGTCGTAATTTGGAAATCATTTTCATACCTGATTCTTCTTTTGCATTT

>M6-PGSC0003DMG400014875-(rev)M6RLKmalectin_496

TGTACTAAAAGGTCGTAATTTGGAAATCATTTTCATACCTGATTCTTCTTTTGCATTTGTTAATGCAATAGAAGCATTTGTTACTCCTCAAAGTTTCATTCCTAACTCTAATGACACAAC

>M6-PGSC0003DMG400014875-(rev)M6RLKmalectin_558

ATGCAATAGAAGCATTTGTTACTCCTCAAAGTTTCATTCCTAACTCTAATGACACAACCCATGTCACTAGCAGCAAAATGGATTTATCCTCGAGTGTTTTGAATGTTATTCATAGGATTA

>M6-PGSC0003DMG400014875-(rev)M6RLKmalectin_62

CTCTCATTTCTCACTTTTTCTCTTCTCTTCCTCCCATTTTCCATTTCGTCATCACCTAATTACACCTTCCCTGAAAAACACTTCATCAATTGTGGTTCTGATTCTCCCATCGTACTCAAC

>M6-PGSC0003DMG400014875-(rev)M6RLKmalectin_620

GTCACTAGCAGCAAAATGGATTTATCCTCGAGTGTTTTGAATGTTATTCATAGGATTAATGTTGGGGGTTCTAATATTACCGACGATATGGATACCATGAGGAGAAAATGGGTAGCTGAT

>M6-PGSC0003DMG400014875-(rev)M6RLKmalectin_682

TGGGGGTTCTAATATTACCGACGATATGGATACCATGAGGAGAAAATGGGTAGCTGATGATGATTACTTACTTATCAAAAGATTAGCTATGAACCTTCCAATGTATAGTAAAAGGCCTAA

>M6-PGSC0003DMG400014875-(rev)M6RLKmalectin_744

ATTACTTACTTATCAAAAGATTAGCTATGAACCTTCCAATGTATAGTAAAAGGCCTAACTATGGTCCAGTAGAAGGGGCTACTCAGTATGATGCTCCTGATTTGGTTTATAAAACTGCCA

>M6-PGSC0003DMG400014875-(rev)M6RLKmalectin_806

GGTCCAGTAGAAGGGGCTACTCAGTATGATGCTCCTGATTTGGTTTATAAAACTGCCAAAGAAATGAATAGTAGTCATGTTGAGACTCAAAATGACAACTTCTTTAACATAACTTGGCGT

>M6-PGSC0003DMG400014875-(rev)M6RLKmalectin_868

AATGAATAGTAGTCATGTTGAGACTCAAAATGACAACTTCTTTAACATAACTTGGCGTTTTGAGGTGAAAAGGAATGCTACATTTTTTGTACGTCTACACTTTTGTGACATTGTTAGTGA

>M6-PGSC0003DMG400014875-(rev)M6RLKmalectin_930

AGGTGAAAAGGAATGCTACATTTTTTGTACGTCTACACTTTTGTGACATTGTTAGTGATCAGCGGAATGGTACAGTTTTCAATGCCTATATATATGGTATGTTTGGTAAGCCAATTAGTC

>M6-PGSC0003DMG400014875-(rev)M6RLKmalectin_992

CGGAATGGTACAGTTTTCAATGCCTATATATATGGTATGTTTGGTAAGCCAATTAGTCCATATGATAGATTTGAACAATTTGCAGTTCCTTTTTATGTTGATTTTGTGGTGGATTCAGAT

>M6-PGSC0003DMG400014917M6LRR-RLK_0

GTTCATTACTACTATTTTTATTTGTATCTCAATTTTGTCAAAAATAAAAAAATCCCCAACAAATGATGTGGAGGATATTCGTTTTTCTTGTTCTTTTACTCAATTCTCATACCGGATTTT

>M6-PGSC0003DMG400014917M6LRR-RLK_1054

ACTAGTGATTATTTACTTGTATTGGAAAAAGAAAGACTCTGGTGGTTGCAGCTGTACTGGAAAAGGGAAATTTGGTGGAAATGAGAAGAGAATGCTTTGTAATTTCCCATGTATTAATGG

>M6-PGSC0003DMG400014917M6LRR-RLK_1116

AAGGGAAATTTGGTGGAAATGAGAAGAGAATGCTTTGTAATTTCCCATGTATTAATGGATTTCCGAATAATGATTCGGAAGTTGAGTCTGAGAAGGGAGGTGGCGGTGGGGCTAGTGGTG

>M6-PGSC0003DMG400014917M6LRR-RLK_124

AACTTCCGACGGACTTTCACTTCTATCTCTAAAATCCGCCATGGATGATGGCGGTGGTGGTACAGTTTTCTCCGATTGGAATGAAAACGATGATACTCCTTGTACTTGGACTGGCATTTC

>M6-PGSC0003DMG400014917M6LRR-RLK_1240

TGTTAGTGGTGGGGAGGGTGATCTTGTGGCTATTGATAAAGGTTTTAGCTTTGAGCTTGATGAGCTGTTGAGGGCTTCTGCTTATGTGTTGGGAAAGAGTGGGTTGGGGATAGTGTACAA

>M6-PGSC0003DMG400014917M6LRR-RLK_1302

AGCTGTTGAGGGCTTCTGCTTATGTGTTGGGAAAGAGTGGGTTGGGGATAGTGTACAAGGTTGTGCTTGGGAACGGTATCCCAGTGGCAGTGAGGAGGTTAGGGGAAGGTGGGGAACAGA

>M6-PGSC0003DMG400014917M6LRR-RLK_1364

GTGCTTGGGAACGGTATCCCAGTGGCAGTGAGGAGGTTAGGGGAAGGTGGGGAACAGAGATATAAGGAGTTTGTTGCTGAGATTCAGGCAATCGGAAGGGTTAAGCATCCTAATGTTGTG

>M6-PGSC0003DMG400014917M6LRR-RLK_1426

TAAGGAGTTTGTTGCTGAGATTCAGGCAATCGGAAGGGTTAAGCATCCTAATGTTGTGAAGTTAAGAGCTTATTATTGGGCTCCTGATGAGAAACTTCTCATTAGTGATTTCATTTCCAA

>M6-PGSC0003DMG400014917M6LRR-RLK_1488

TAAGAGCTTATTATTGGGCTCCTGATGAGAAACTTCTCATTAGTGATTTCATTTCCAACGGCAACTTGGCTTCTGCTCTTCACGGTGAGCTTTTCATTATAGTAAAACATCTTCAGTTTT

>M6-PGSC0003DMG400014917M6LRR-RLK_1550

AACTTGGCTTCTGCTCTTCACGGTGAGCTTTTCATTATAGTAAAACATCTTCAGTTTTGTTTGTCTAGTAGATATAACCTGTTGAAAGAATCATTAACACATTCATATTTTAGATATCGG

>M6-PGSC0003DMG400014917M6LRR-RLK_1612

GTCTAGTAGATATAACCTGTTGAAAGAATCATTAACACATTCATATTTTAGATATCGGGGTATACTATATTCTTATCATTTAGTAGCATGATACACTGATGGATGTTTCGTACTTTCCAA

>M6-PGSC0003DMG400014917M6LRR-RLK_1674

TACTATATTCTTATCATTTAGTAGCATGATACACTGATGGATGTTTCGTACTTTCCAAGATCGATATGTGTAAAGTGTAATTCTTTTGCTGGATTAGTATTTTACTTATTGATCACTCTT

>M6-PGSC0003DMG400014917M6LRR-RLK_1736

GATATGTGTAAAGTGTAATTCTTTTGCTGGATTAGTATTTTACTTATTGATCACTCTTTTGAGGAGTGGAAATACAGTACAAGACTTTTGGTATTAGGTTAGTTAGGAGTTTGGGATGTT

>M6-PGSC0003DMG400014917M6LRR-RLK_1798

GGAGTGGAAATACAGTACAAGACTTTTGGTATTAGGTTAGTTAGGAGTTTGGGATGTTCTAGTATTTATTCACTAGTATAACTGTTTAATATATTCCCGATGACAAAATAATGACTTTTA

>M6-PGSC0003DMG400014917M6LRR-RLK_186

CAGTTTTCTCCGATTGGAATGAAAACGATGATACTCCTTGTACTTGGACTGGCATTTCATGTGCTAACATTTCTGGTTCTTCAGAGCCACGTGTCGTTGGGATTACTCTTTCCGGTAAGA

>M6-PGSC0003DMG400014917M6LRR-RLK_1860

TATTTATTCACTAGTATAACTGTTTAATATATTCCCGATGACAAAATAATGACTTTTAGCTGTTGTGTGTTCCTTGATGATTTTGAATAGCAATTATTTATATGCCATCTAGTGATAATT

>M6-PGSC0003DMG400014917M6LRR-RLK_1922

TTGTGTGTTCCTTGATGATTTTGAATAGCAATTATTTATATGCCATCTAGTGATAATTTTATATTGAATGTGTCCCATGTGCTTGCAATCCTTGAGTGAAGGATGAGTTTTGTTTGCGAC

>M6-PGSC0003DMG400014917M6LRR-RLK_1984

ATTGAATGTGTCCCATGTGCTTGCAATCCTTGAGTGAAGGATGAGTTTTGTTTGCGACTTGATAATTTGCGTAGGATTTTTGTTTAATTGTATCAGCTGATAGGATAGTCTTTATTATGT

>M6-PGSC0003DMG400014917M6LRR-RLK_2046

TAATTTGCGTAGGATTTTTGTTTAATTGTATCAGCTGATAGGATAGTCTTTATTATGTATCATAGATGAAAAGTTATCTGCATAAAGGGTAAACCAACAACCTAACATTTTCCATCCATT

>M6-PGSC0003DMG400014917M6LRR-RLK_2108

TAGATGAAAAGTTATCTGCATAAAGGGTAAACCAACAACCTAACATTTTCCATCCATTTCAATGGATGTTAAAGCACATAGTAGTATATTAACTTATAATGATCCAGCTTAAGTCAATTT

>M6-PGSC0003DMG400014917M6LRR-RLK_2170

TGGATGTTAAAGCACATAGTAGTATATTAACTTATAATGATCCAGCTTAAGTCAATTTGCAGACTTTAACGATTTACTGCCGATGCAATAATCATATGTTGACATCATATCGAACCTGGT

>M6-PGSC0003DMG400014917M6LRR-RLK_2232

ACTTTAACGATTTACTGCCGATGCAATAATCATATGTTGACATCATATCGAACCTGGTAGAAAGGGCATTTGGGCATGTTTTGAAGAATTTAGTCTCTCTTTTTTCATGTGACATATGGT

>M6-PGSC0003DMG400014917M6LRR-RLK_2294

AGGGCATTTGGGCATGTTTTGAAGAATTTAGTCTCTCTTTTTTCATGTGACATATGGTTGACTGCCCCGATAAACCTTGCGGAAAATGATTTATCATCAATTAGAAGCAAACCAATAAGT

>M6-PGSC0003DMG400014917M6LRR-RLK_2356

TGCCCCGATAAACCTTGCGGAAAATGATTTATCATCAATTAGAAGCAAACCAATAAGTACAAGGGGCCAAGAACTGAATTATGTCATGTACTAGGTCAATTCCTTAAGTTGTGGAGTTAC

>M6-PGSC0003DMG400014917M6LRR-RLK_2418

GGGGCCAAGAACTGAATTATGTCATGTACTAGGTCAATTCCTTAAGTTGTGGAGTTACGTATTCTGTCTCTCAAAAGAGTCTAGATATTACATTGTTTTATTATGTCTTTTAGAGATGTA

>M6-PGSC0003DMG400014917M6LRR-RLK_248

GCTAACATTTCTGGTTCTTCAGAGCCACGTGTCGTTGGGATTACTCTTTCCGGTAAGAATCTCCGGGGATACTTACCGTCTGAGCTGGGAACGTTACTCTATCTCCGGCGACTGAATCTT

>M6-PGSC0003DMG400014917M6LRR-RLK_2480

TCTGTCTCTCAAAAGAGTCTAGATATTACATTGTTTTATTATGTCTTTTAGAGATGTATTACTGATTTGCATATGCTAATATTCAAGAATTTTCGAATCGCTCCGCCTGGTGCAGTTTGA

>M6-PGSC0003DMG400014917M6LRR-RLK_2542

TGATTTGCATATGCTAATATTCAAGAATTTTCGAATCGCTCCGCCTGGTGCAGTTTGACTTTACAGATGTTTACTTTTTCGAAGCAAACTGGAATCTGCAGTATCTGGTCTGGTAACTAC

>M6-PGSC0003DMG400014917M6LRR-RLK_2604

ACAGATGTTTACTTTTTCGAAGCAAACTGGAATCTGCAGTATCTGGTCTGGTAACTACCATCTAGCATGAGCGGGACTTATACTCCCTTGAGATGGATTCTAGACTTCGAATTTTTAAAA

>M6-PGSC0003DMG400014917M6LRR-RLK_2666

TAGCATGAGCGGGACTTATACTCCCTTGAGATGGATTCTAGACTTCGAATTTTTAAAACTTAGTTTTCTGCAATTGAATTTTGCACTGTTGCTGTTGGTATAGCTGCTTGAATGTTACAT

>M6-PGSC0003DMG400014917M6LRR-RLK_2728

GTTTTCTGCAATTGAATTTTGCACTGTTGCTGTTGGTATAGCTGCTTGAATGTTACATTTTTAAAAAGAATTAGATTTTTTCCCCTGATACAAATCGTTGTTTACATTTATGAATATCTA

>M6-PGSC0003DMG400014917M6LRR-RLK_2790

AAAAAGAATTAGATTTTTTCCCCTGATACAAATCGTTGTTTACATTTATGAATATCTACTTCACTGCCTCATATACATGTCACTATGAATCTATAACACATTCGTTACCCAAGAGTAACT

>M6-PGSC0003DMG400014917M6LRR-RLK_2852

ACTGCCTCATATACATGTCACTATGAATCTATAACACATTCGTTACCCAAGAGTAACTGCTTGCTTTCTGAGACAAATATATTTGATGACCTTTGGTTCTATGGTACCTGATGTTCCCAG

>M6-PGSC0003DMG400014917M6LRR-RLK_2914

GCTTTCTGAGACAAATATATTTGATGACCTTTGGTTCTATGGTACCTGATGTTCCCAGTTTCTTAATTTACCTGTTTTTAAAATTATGTAGCTGGAATTTGACAGTAGAGCAACTTACCT

>M6-PGSC0003DMG400014917M6LRR-RLK_2976

TTAATTTACCTGTTTTTAAAATTATGTAGCTGGAATTTGACAGTAGAGCAACTTACCTTTTGCATTCTGAAGAAGTTGAGGAGTGTTTTCTTTCAGCATCTCTTTTTCCAAATTGTTTTG

>M6-PGSC0003DMG400014917M6LRR-RLK_3038

CATTCTGAAGAAGTTGAGGAGTGTTTTCTTTCAGCATCTCTTTTTCCAAATTGTTTTGCCATGCTAGATCTGTATGTATTGACGTAGTGACAGTAGTTTCATGCTCTAGCTGATTGAGAT

>M6-PGSC0003DMG400014917M6LRR-RLK_310

CCGGGGATACTTACCGTCTGAGCTGGGAACGTTACTCTATCTCCGGCGACTGAATCTTCATGGAAATAATATTTACGGTTCTATCCCTGACCCTTTATTCAATGCGACGTCGCTTCATAG

>M6-PGSC0003DMG400014917M6LRR-RLK_3100

GCTAGATCTGTATGTATTGACGTAGTGACAGTAGTTTCATGCTCTAGCTGATTGAGATTAGCCAAATTAGTGTGAACACCGATTTGCCTATTGAGCTGAAAAGTTATTAACTTGTATCTT

>M6-PGSC0003DMG400014917M6LRR-RLK_3162

CAAATTAGTGTGAACACCGATTTGCCTATTGAGCTGAAAAGTTATTAACTTGTATCTTGTTCGTCATACTGTGGTAAATGTTGTGATGACAATCTCGTTTGCCTTTATACAGGGAGAAAT

>M6-PGSC0003DMG400014917M6LRR-RLK_3224

GTCATACTGTGGTAAATGTTGTGATGACAATCTCGTTTGCCTTTATACAGGGAGAAATGGGCAACCATCACCTAGTTTAACATGGTCAACGAGACTGAAAATTGCTAAGGGAACAGCGCG

>M6-PGSC0003DMG400014917M6LRR-RLK_3286

AACCATCACCTAGTTTAACATGGTCAACGAGACTGAAAATTGCTAAGGGAACAGCGCGAGGTTTAGCCTATCTCCATGAGTGTAGCCCAAGGAAATTTGTCCATGGAGACGTCAAACCTT

>M6-PGSC0003DMG400014917M6LRR-RLK_3348

TTAGCCTATCTCCATGAGTGTAGCCCAAGGAAATTTGTCCATGGAGACGTCAAACCTTCCAATATTCTCCTTGATACTGAACTGCAGCCATATATATCTGATTTTGGCCTCAATCGATTA

>M6-PGSC0003DMG400014917M6LRR-RLK_3410

TATTCTCCTTGATACTGAACTGCAGCCATATATATCTGATTTTGGCCTCAATCGATTAATCAATATCACGGGAAACAACCCCTCCTCTTCTGGTGGATTTATGGGTGGAGCTCTTCCTTA

>M6-PGSC0003DMG400014917M6LRR-RLK_3472

ATATCACGGGAAACAACCCCTCCTCTTCTGGTGGATTTATGGGTGGAGCTCTTCCTTACTTGAAGCCTGCTCAACCAGAACGTCCCAATAACTATCGAGCACCAGAAGCTCGAATCACTG

>M6-PGSC0003DMG400014917M6LRR-RLK_3534

AAGCCTGCTCAACCAGAACGTCCCAATAACTATCGAGCACCAGAAGCTCGAATCACTGGTAACAGACCCACCCAAAAATGGGATGTGTACTCATTCGGGGTGGTCTTGCTTGAACTGCTG

>M6-PGSC0003DMG400014917M6LRR-RLK_3596

CAGACCCACCCAAAAATGGGATGTGTACTCATTCGGGGTGGTCTTGCTTGAACTGCTGACTGGAAAGTCTCCTGACCTCTCAGTTCCTACAACATCCACTTCAACGGAGGTTCCAGATCT

>M6-PGSC0003DMG400014917M6LRR-RLK_3658

GAAAGTCTCCTGACCTCTCAGTTCCTACAACATCCACTTCAACGGAGGTTCCAGATCTTGTAAGATGGGTAAGAAAGGGATTCGAGGAGCAAAATCCACTGTCGGATATGGTAGAACCAA

>M6-PGSC0003DMG400014917M6LRR-RLK_372

GAAATAATATTTACGGTTCTATCCCTGACCCTTTATTCAATGCGACGTCGCTTCATAGTATTTATTTGTATGATAATAATATCTCTGGTATACTTCCTCCTTCTGTTTGTAATCTTCCTC

>M6-PGSC0003DMG400014917M6LRR-RLK_3720

AGATGGGTAAGAAAGGGATTCGAGGAGCAAAATCCACTGTCGGATATGGTAGAACCAATGTTGCTTCAAGAAGTCCATGCTAAGAAGGAAGTACTAGCTGTATTTCATGTAGCTCTTGCA

>M6-PGSC0003DMG400014917M6LRR-RLK_3782

GCTTCAAGAAGTCCATGCTAAGAAGGAAGTACTAGCTGTATTTCATGTAGCTCTTGCATGCACCGAGGCAGATCCAGATATTCGACCAAGGATGAAAACTATCTCAGAAAACATTGAGAA

>M6-PGSC0003DMG400014917M6LRR-RLK_3844

CCGAGGCAGATCCAGATATTCGACCAAGGATGAAAACTATCTCAGAAAACATTGAGAAAGTTGGAGCTTAAACTTTTCGGTACACTCATGTCAAGGAGTTCCTGAGTTTCATGGACTTCT

>M6-PGSC0003DMG400014917M6LRR-RLK_3906

GGAGCTTAAACTTTTCGGTACACTCATGTCAAGGAGTTCCTGAGTTTCATGGACTTCTTTAATCCTTTTTTTTGCAGTAATTGCTCCTATTATTTTTTTTCTTTTAGAGGTTAATCAAAT

>M6-PGSC0003DMG400014917M6LRR-RLK_3968

TCCTTTTTTTTGCAGTAATTGCTCCTATTATTTTTTTTCTTTTAGAGGTTAATCAAATCTTTACTTAACTACAGTACAAAATACTAGTGTGCTGCAACAAAATTCCTGTCATTTTTGAGA

>M6-PGSC0003DMG400014917M6LRR-RLK_4030

ACTTAACTACAGTACAAAATACTAGTGTGCTGCAACAAAATTCCTGTCATTTTTGAGAAGTTAATTAGTTGCTTATCTCTTTAATCTATTTCTGTAATTCTGATGTTAAAACAACTTTTC

>M6-PGSC0003DMG400014917M6LRR-RLK_434

TATTTGTATGATAATAATATCTCTGGTATACTTCCTCCTTCTGTTTGTAATCTTCCTCGTTTGCAGAATTTGGATATTTCTGATAATTCACTTTCTGGAACTTTCTCTAAGGATCTACGT

>M6-PGSC0003DMG400014917M6LRR-RLK_496

GCAGAATTTGGATATTTCTGATAATTCACTTTCTGGAACTTTCTCTAAGGATCTACGTAATTGTAGGCAGTTACAGAGGCTTATTCTAGCTCGGAATAAATTCTCCGGTGAGATTCCGGT

>M6-PGSC0003DMG400014917M6LRR-RLK_558

GTAGGCAGTTACAGAGGCTTATTCTAGCTCGGAATAAATTCTCCGGTGAGATTCCGGTGGGGGTGTTCCCGGAGTTGGCGAATTTGGAACAACTTGATCTTTCGTCGAATTTGTTCAATG

>M6-PGSC0003DMG400014917M6LRR-RLK_62

ATGATGTGGAGGATATTCGTTTTTCTTGTTCTTTTACTCAATTCTCATACCGGATTTTGTCTAACTTCCGACGGACTTTCACTTCTATCTCTAAAATCCGCCATGGATGATGGCGGTGGT

>M6-PGSC0003DMG400014917M6LRR-RLK_620

GTGTTCCCGGAGTTGGCGAATTTGGAACAACTTGATCTTTCGTCGAATTTGTTCAATGGTTCGATTCCTGAAGATATTGGTGAATTGAAGTCACTATCTGGAACTTTGAATCTTTCTTTT

>M6-PGSC0003DMG400014917M6LRR-RLK_682

GATTCCTGAAGATATTGGTGAATTGAAGTCACTATCTGGAACTTTGAATCTTTCTTTTAATCATTTTAGTGGAAGAATTCCCAAATCTGTAGGTGATTTACCTTTAACTGTTAGCTTTGA

>M6-PGSC0003DMG400014917M6LRR-RLK_744

ATTTTAGTGGAAGAATTCCCAAATCTGTAGGTGATTTACCTTTAACTGTTAGCTTTGATCTAAGGAATAATAATTTATCTGGTGAAATTCCTCAGACTGGTTCTTTTGCTAATCAAGGTC

>M6-PGSC0003DMG400014917M6LRR-RLK_806

AGGAATAATAATTTATCTGGTGAAATTCCTCAGACTGGTTCTTTTGCTAATCAAGGTCCGACGGCTTTTTTGAATAATCCCTTGTTATGTGGATTTCCTTTACAAAAAAACTGTAAAAAT

>M6-PGSC0003DMG400014917M6LRR-RLK_868

GGCTTTTTTGAATAATCCCTTGTTATGTGGATTTCCTTTACAAAAAAACTGTAAAAATAGTTCAAACAATTCAACACAAGTTCAGGGTTCGTCTGATAACGAGGGAACGAATTCGCGCAA

>M6-PGSC0003DMG400014917M6LRR-RLK_930

CAAACAATTCAACACAAGTTCAGGGTTCGTCTGATAACGAGGGAACGAATTCGCGCAAGGGGTTAAAACCTGGATTCATTATATTGATATGTTTGGCTGATGCATTTGGAGTGGCATTTA

>M6-PGSC0003DMG400014917M6LRR-RLK_992

TTAAAACCTGGATTCATTATATTGATATGTTTGGCTGATGCATTTGGAGTGGCATTTATAGGACTAGTGATTATTTACTTGTATTGGAAAAAGAAAGACTCTGGTGGTTGCAGCTGTACT

>M6-PGSC0003DMG400015081M6LRR-RLK_0

CACAACAAGTTGCTAGGATCTATTCCTGACTCAGTGAGCAACATGGTAGGTTTGGAATTCCTAGACATTTCTCATAATAATATATCAGGAACCATTCCTATGTCTATGGAGAAACTTCAA

>M6-PGSC0003DMG400015081M6LRR-RLK_1024

TTTTTATTTCCAATCAATAAAGTATATTGCATCTTTAAATCAATTGTTTGACTGTAGAGTATGGACTGGATGGACTGGTGTCAATAAAATGTGATGTCTATAGTTACGGGATTATGTTGC

>M6-PGSC0003DMG400015081M6LRR-RLK_1280

ACAGAAGGAGTTAGATATCGTGGCATCAATCATGAAAGTAGCATTAGATTGTTGTGCTGAATCTCCGACAAGAAGGACAAACATGAAAGATGTTGTAGGGATGCTACAGAAGATCAAGAT

>M6-PGSC0003DMG400015081M6LRR-RLK_1390

AGATCAAGATTCAACTTCTTGCATGCTGAGCATGTCCTCTGTATCTCTTGTTCTGAATCGATCACTTTTTTTTGTTTTGTTTTAGCACTTGATTTGTGCATGATTCTGTTTTCAAAAATG

>M6-PGSC0003DMG400015081M6LRR-RLK_192
