## Supplementary material for "RLP/K enrichment sequencing; a novel method to identify receptor-like protein (*RLP*) and receptor-like kinase (*RLK*) genes": Note S5

Supplementary Note S5. Java script for calling the informative SNPs

import java.util.List;

import java.io.*;

import java.util.ArrayList;

import com.beust.jcommander.JCommander;

import com.beust.jcommander.Parameter;

public class FilterSNPs

{

static BufferedWriter writerGFF = null;

public static void main(String[] args) throws Exception {

// set up JCommander

JCommanderParams params = new JCommanderParams();

JCommander jc = new JCommander(params);

try {

jc.parse(args);

} catch (Exception e) {

jc.usage();

System.exit(1);

}

jc.setProgramName("FilterSNPs");

// help?

if (params.help) {

jc.usage();

System.exit(1);

}

// parse command line arguments

File vcfFile = params.getVCFfile();

Integer minDepth = params.getDepth();

ArrayList<String> alleles = params.getAlleleRatios();

Integer leeway = params.getLeeway();

String outputFile = params.getOutput();

ArrayList<String> resistant = new ArrayList<String>();

ArrayList<String> susceptible = new ArrayList<String>();

// check command line args

for (String al : alleles) {

String[] al_array = al.split("-");

susceptible.add(al_array[0]);

resistant.add(al_array[1]);

}

//I/O

writerGFF = new BufferedWriter(new FileWriter(outputFile));

Float lowerCutoffRes = null;

Float upperCutoffRes = null;

Float lowerCutoffSus = null;

Float upperCutoffSus = null;

// assign expected SNP ratios

for (String al : alleles) {

String[] al_array = al.split("-");

Float s = Float.parseFloat(al_array[0]);

Float r = Float.parseFloat(al_array[1]);

upperCutoffRes = r + leeway;

if ( upperCutoffRes > 100 ) {

upperCutoffRes = (float) 100;

}

lowerCutoffRes = r - leeway;

if ( lowerCutoffRes < 0 ) {

lowerCutoffRes = (float) 0;

}

upperCutoffSus = s + leeway;

if ( upperCutoffSus > 100 ) {

upperCutoffSus = (float) 100;

}

lowerCutoffSus = s - leeway;

if ( lowerCutoffSus < 0 ) {

lowerCutoffSus = (float) 0;

}

//do the filtering

System.out.println("Filtering SNPs");

filterSNPs(vcfFile, lowerCutoffRes, upperCutoffRes, upperCutoffSus, lowerCutoffSus, minDepth);

}

//clean up

writerGFF.close();

System.out.println("Pipeline complete!");

}

//------------------------------------------------------------------

//	Remove SNPs outside the expected allele ratios

private static void filterSNPs(File infile, float lowerCutoffRes, float upperCutoffRes, float upperCutoffSus, float lowerCutoffSus, int minDepth) throws IOException

{

BufferedReader reader = new BufferedReader(new FileReader(infile));

//parse the file

String line = null;

//ignore header

reader.readLine();

int lineCount = 0;

while((line = reader.readLine()) != null)

{

//ignore header lines

if(line.startsWith("#"))

continue;

lineCount++;

//	System.out.println("\nprocessing line " + lineCount);

String [] tokens = line.split("\t");

//sample input

//	#CHROM	POS	ID	REF	ALT	QUAL	FILTER	INFO	FORMAT	Sample1	Sample2

//	Contig3	133	.	T	A	.	PASS	ADP=59;WT=1;HET=1;HOM=0;NC=0	GT:GQ:SDP:DP:RD:AD:FREQ:PVAL:RBQ:ABQ:RDF:RDR:ADF:ADR	0/0:49:64:64:55:9:14.06%:1.4447E-3:37:37:47:8:9:0	0/1:35:55:55:44:11:20%:2.8065E-4:36:37:39:5:11:0

//get the variant allele frequency for both samples

//Sample1 is BS, Sample 2 is BR

//sample info is incolumns 9 and 10 (0-based)

String sample1Info = tokens[9];

String sample2Info = tokens[10];

//	System.out.println("sample1Info = " + sample1Info);

//within each of these we want the FREQ tag which is the 7th element of the string separated by ":" characters

String sample1FREQStr;

String sample2FREQStr;

try

{

sample1FREQStr = sample1Info.split(":")[6];

sample2FREQStr = sample2Info.split(":")[6];

}

catch (ArrayIndexOutOfBoundsException e)

{

continue;

}

//need to strip off the % sign at the end

sample1FREQStr = sample1FREQStr.substring(0, sample1FREQStr.indexOf("%"));

sample2FREQStr = sample2FREQStr.substring(0, sample2FREQStr.indexOf("%"));

//	System.out.println("sample1FREQStr = " + sample1FREQStr);

//now parse these

float sample1FREQ = Float.parseFloat(sample1FREQStr);

float sample2FREQ = Float.parseFloat(sample2FREQStr);

//System.out.println("sample1FREQ = " + sample1FREQ + "\tsample2FREQ = " + sample2FREQ);

//also extract the read depth for each sample

int readDepthSample1 = Integer.parseInt(sample1Info.split(":")[2]);

int readDepthSample2 = Integer.parseInt(sample2Info.split(":")[2]);

//now test our conditions

boolean depthPass = minDepth <= readDepthSample1 && minDepth <= readDepthSample2;

boolean BRPass = false;

boolean BSPass = false;

BRPass = sample2FREQ >= lowerCutoffRes && sample2FREQ <= upperCutoffRes;

BSPass = sample1FREQ >= lowerCutoffSus && sample1FREQ <= upperCutoffSus;

//if (BSPass) {

//	System.out.println("sample1FREQ = " + sample1FREQ + " passed: " + BSPass + "; sample2FREQ = " + sample2FREQ + " passed: " + BRPass);

//}

if(BSPass && BRPass && depthPass)

{

//System.out.println("SNP passes");

writerGFF.write(line + "\n");

}

}

reader.close();

}

//------------------------------------------------------------------

// Command line parameters

static class JCommanderParams {

// <vcfFile> <minDepth> <resistantAltRatio? lower:higher> <inBetween? true|false> <susceptibleAltRatio? lower:higher> <inBetween? true|false>

@Parameter

private List<String> parameters = new ArrayList<>();

@Parameter(names = { "-v", "--vcf" }, description = "VCF file to filter", required=true)

private File vcfFile;

@Parameter(names = "--debug", description = "Debug mode", hidden = true)

private boolean debug = false;

@Parameter(names = { "-d", "--depth" }, description = "Read depth" )

private Integer depth = 50;

@Parameter(names = {"-a", "--alleleRatios" }, description = "Allele ratios, given in form: susceptible-resistant. e.g. For homozygous, opposite alleles, use \"0-100 100-0\"", variableArity = true)

private List<String> allele = new ArrayList<String>();

@Parameter(names = {"-l", "--leeway"}, description = "Leeway in allele ratios")

private Integer leeway = 10;

@Parameter(names = {"-h", "--help"}, description = "Help", help = true)

private boolean help;

@Parameter(names = {"-o", "--output"}, description = "Output file")

private String output = "filteredSNPs.vcf";

//------------------------------------------------------------------

// return statements

public File getVCFfile() {

return vcfFile;

}

public Integer getDepth() {

return depth;

}

public boolean getDebug() {

return debug;

}

public ArrayList<String> getAlleleRatios() {

return (ArrayList<String>) allele;

}

public Integer getLeeway() {

return leeway;

}

public boolean help() {

return help;

}

public String getOutput() {

return output;

}

}

}
